# The Arabidopsis RNA-binding protein SGS3 is recruited to the chromatin remodeler CHR11 to promote siRNA production from protein-coding genes

**DOI:** 10.1101/2024.10.03.612444

**Authors:** Taline Elmayan, Thomas Blein, Emilie Elvira-Matelot, Ivan Le Masson, Aurélie Christ, Nathalie Bouteiller, Martin D. Crespi, Hervé Vaucheret

## Abstract

In plants, aberrant RNAs produced by endogenous genes or transgenes are normally degraded by the nuclear and cytosolic RNA quality control (RQC) pathways. Under certain biotic or abiotic stresses, RQC is impaired, and aberrant RNAs are converted into siRNAs that initiate post-transcriptional gene silencing (PTGS) in the cytosol. How aberrant RNAs are selected and brought to the cytoplasm is not known. Here we show that the RNA-binding protein SUPPRESSOR OF GENE SILENCING (SGS)3 shuttles between the cytosol and the nucleus where it associates with the ISWI-like CHROMATIN REMODELER (CHR)11. CHR11 binds to transgenes prone to trigger PTGS. Knocking down *CHR11* and its paralog *CHR17* strongly reduces transgene PTGS, suggesting that SGS3 recruitment by CHR11/17 facilitates PTGS initiation. CHR11 is also enriched at endogenous protein-coding genes (PCGs) producing nat-siRNAs and va-siRNAs under biotic or abiotic stresses, and this production is reduced in *chr11 chr17* double mutants at genome-wide level. Moreover, impairing CHR11 and CHR17 rescues the lethal phenotype caused by the massive production of siRNAs from PCGs in RQC-deficient mutants. We propose that SGS3 recruitment by CHR11/17 allows exporting RNAs to the cytosol to initiate the production of siRNAs.

**One Sentence Summary:** The plant PTGS component SGS3 is recruited by the chromatin remodeler CHR11 to select nuclear RNAs for export to the cytosol and siRNA production.

## INTRODUCTION

In plants, post-transcriptional gene silencing (PTGS) is a siRNA-based defense mechanism that eliminates exogenous RNAs in the cytosol. When PTGS targets viruses, dsRNA intermediates of viral replication or viral ssRNAs converted to dsRNA by cellular RNA-dependent RNA polymerases (RDRs) are processed by DICER-LIKE2/4 (DCL2/4) into 21-22-nt siRNAs that are loaded onto ARGONAUTE1/2 (AGO1/2) ^1^. AGO/siRNA complexes cleave viral ssRNA, thus reducing the virus title. When DCL2-dependent 22-nt siRNAs are loaded onto AGO1, AGO1 is supposed to interact with the RNA-binding protein SGS3 ^2^, which protects AGO1-associated RNA from degradation ^3^. SGS3 is present in cytosolic siRNA-bodies where RDR6 and AGO1 are also present ^4–7^, suggesting that AGO1-SGS3-protected viral RNAs are transformed into dsRNA by RDR6, thus creating an amplification loop that increases the production of siRNAs and further contributes to decreasing virus title. In the case of sense transgenes, PTGS also takes place at some loci, but the way it is initiated is not fully understood. Although sense transgenes should not produce dsRNA, it has been proposed that transgene arrangements at particular loci allow the production of dsRNA or the production of aberrant RNAs that can be transformed into dsRNA by RDR6, resulting in a form of PTGS referred to as S-PTGS. Supporting this hypothesis, an uncapped antisense RNA was detected in the well-characterized *Arabidopsis* line L1 ^8^. This line carries a *p35S:GUS-tRbcS* transgene producing a *GUS* mRNA that accumulates at high level in *ago1*, *dcl2dcl4*, *rdr6* or *sgs3* mutant backgrounds, but is degraded by PTGS in wild-type (WT) plants. An uncapped antisense RNA, which 5’ end maps in the *tRbcS*, accumulates at higher level in *rdr6* compared with WT, indicating that it is a PTGS target and not a PTGS product. It accumulates at much higher level when the nuclear and cytosolic exoribonucleases XRN3 and XRN4 are both impaired, indicating that this uncapped RNA likely is exported from the nucleus to the cytosol, and is targeted by both nuclear and cytosolic RNA Quality Control (RQC) pathway. Still, major questions remain unanswered regarding the S-PTGS initiation phase, including the way transgene aberrant RNAs are selectively exported to the cytosol. Here, we show that the ISWI-like CHROMATIN REMODELER (CHR)11 and its paralog *CHR17* contribute to siRNA production by interacting with the RNA-binding protein SGS3, which shuttles between cytosol and nucleus. We propose a model where CHR11 and SGS3 are major nuclear determinants of transgene S-PTGS. Moreover, genome-wide approaches suggest that this model generally applies to the production of siRNAs from endogenous protein-coding genes.

## RESULTS

### SGS3 interacts with CHR11, an ISWI-like chromatin remodeling protein

A yeast-two-hybrid screen was performed using an Arabidopsis cDNA library and SGS3 as a bait. Among the candidates identified in this screen, one corresponded to *CHR11* (At3g06400), which encodes one of the 42 Arabidopsis chromatin remodeling proteins ^9^. Deletion analyses revealed that, at least in yeast, the coiled coil domain of SGS3 interacts with the HAND-SANT-SLIDE domains located at the C-terminus of CHR11 (Fig. 1A and Supplementary Fig. 1).

**Fig. 1.**
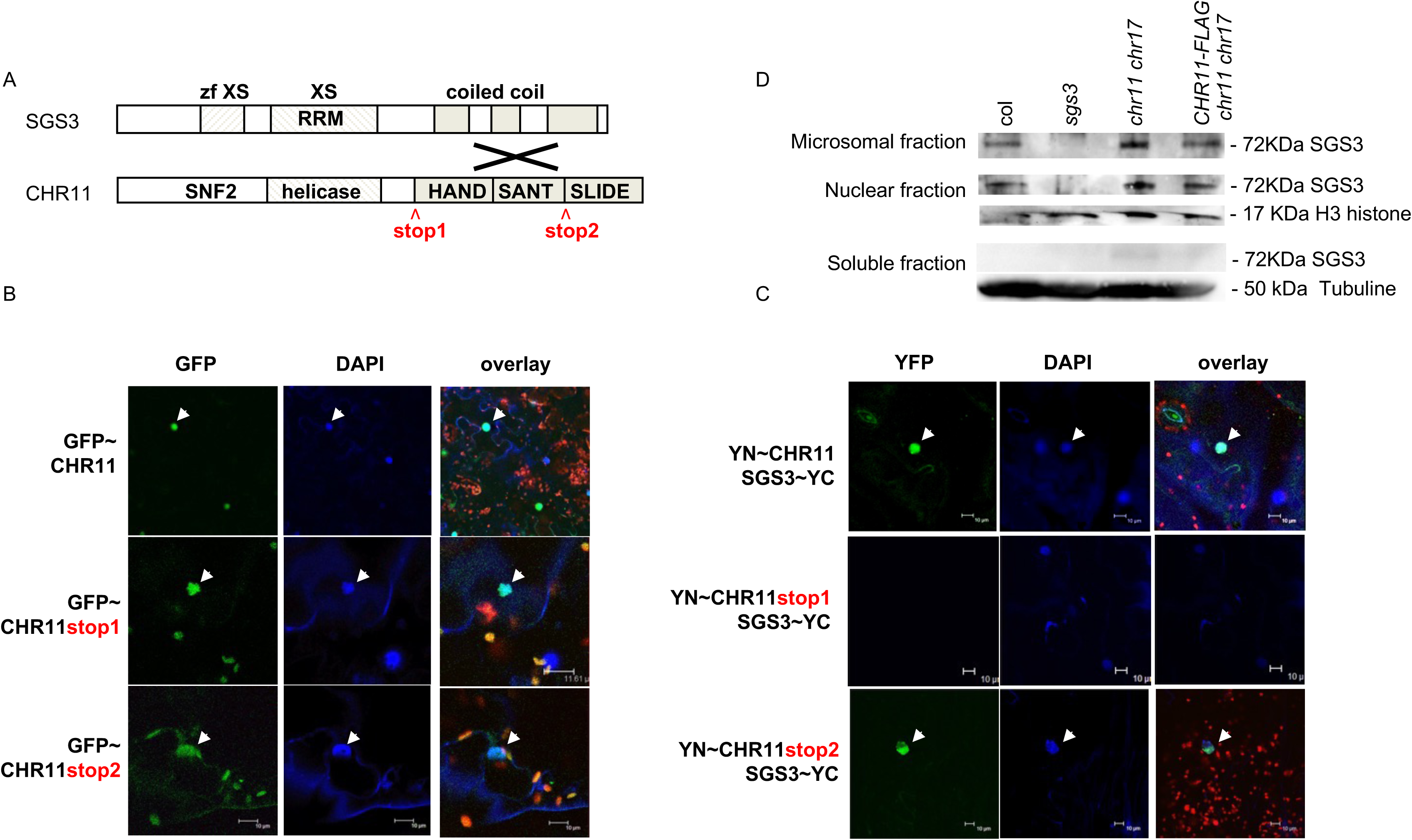
SGS3 interacts with CHR11 in the nucleus. A. SGS3 and CHR11 structures. CHR11 and SGS3 domains interacting in yeast two-hybrid are shown in grey. Sequences of the CHR11 clones retrieved in the in yeast two-hybrid screen are shown in Supplementary Fig. 1. B. Localization of CHR11 full-length and truncated forms (stop1 and stop2) in *N. benthamiana*. Channels are indicated above each column. The overlay is shown for DAPI/YFP/chlorophyll fluorescences. C. BiFC Interaction between SGS3 and CHR11 (full-length and truncated forms) in nucleus of *N. benthamiana* (pointed by arrows). Subcellular localization of reconstructed YFP was determined in the leaf epidermis for SGS3 protein in fusion with the C-terminal part of YFP (YC) and CHR11 proteins in fusion with the N-terminal part of YFP (YN). Channels are indicated above each column. The overlay is shown for DAPI/YFP/chlorophyll fluorescences. D. SGS3 is associated with fractions enriched in microsomal and nuclear compartments in wild-type Arabidopsis, in the *chr11 chr17* double mutant, and in the *chr11 chr17* double mutant complemented with the *p35S:FLAG-CHR11*construct. The *sgs3-1* null allele is used a negative control.

Confirmation of this interaction in planta was obtained by BiFC experiments performed by agro-infiltration in *N. benthamiana*. Indeed, co-infiltration of *YFP^Nter^-CHR11* and *SGS3-YFP^Cter^* generated a nuclear YFP signal similar to the GFP signal obtained when infiltrating *GFP-CHR11* (Fig. 1B and 1C). Deletion of the HAND-SANT-SLIDE domains did not abolish the nuclear localization of CHR11 (*GFP-CHR11stop1*), but abolished the interaction with SGS3 (*YFP^Nter^-CHR11stop1* + *SGS3-YFP^Cter^*), confirming the yeast-two hybrid results. Deletion of only the SLIDE domain did not abolish the nuclear localization of CHR11 (*GFP-CHR11stop2*), nor the interaction with SGS3 (*YFP^Nter^-CHR11stop2 + SGS3-YFP^Cter^*), indicating that SGS3 interacts with CHR11 through its HAND-SANT domains (Fig. 1B and C).

### SGS3 shuttles between the nucleus and the cytosol

SGS3 was reported as localizing mostly in cytosolic foci that also contain AGO1, AGO7 and RDR6, which are referred to as siRNA bodies ^4–7^. Nevertheless, one report also described SGS3 in the nucleoplasm after immuno-localization on isolated nuclei ^10^. Moreover, two recent reports revealed that truncated forms of the SGS3 protein lacking its prion-like domain, which likely makes it non-functional, localize in the nucleus ^11,12^; suggesting that SGS3 could have some function in the nucleus. To get further information on the localization of the native SGS3 protein, western blots were performed on fractionated cells (Fig. 1D). In addition to being found in the microsomal fraction ^6^, SGS3 was associated to the fractions enriched in nuclear compartments (Fig. 1D), suggesting a dual localization of SGS3 in the nucleus and cytosol of Arabidopsis cells.

To further examine the partition of SGS3 between the nucleus and the cytosol, a tagged SGS3 protein was expressed using the *SGS3* promoter to stay as close as possible to physiological levels, and introduced into *sgs3* mutants carrying the PTGS transgene locus *L1*. *L1/sgs3/pSGS3:SGS3-Venus* transformants showed complementation of *L1* PTGS but gave no detectable signal in confocal or spinning disk analysis, likely due to the very low level of expression of the *SGS3* promoter. Therefore, *SGS3-GFP* was subsequently expressed under the *RDR6* promoter, which is expressed at higher level than *SGS3* according to the ePlant software tool (http://bar.utoronto.ca/efp/cgi-bin/efpWeb.cgi) ^13^, thus avoiding the use of strong promoters such as UBQ10 or 35S. Introduction of *pRDR6:SGS3-GFP* in *L1/sgs3* plants restored *L1* PTGS as efficiently as the *pSGS3:SGS3-Venus* construct, and GFP could be detected in confocal analysis of *L1/sgs3/pRDR6:SGS3-GFP* plants. However, the GFP signal was only cytosolic, similar to that observed with *p35S:SGS3-GFP* construct (Fig. 2A), suggesting that the amount of SGS3 in the nucleus is very low, likely because it is actively exported to the cytosol.

**Fig. 2.**
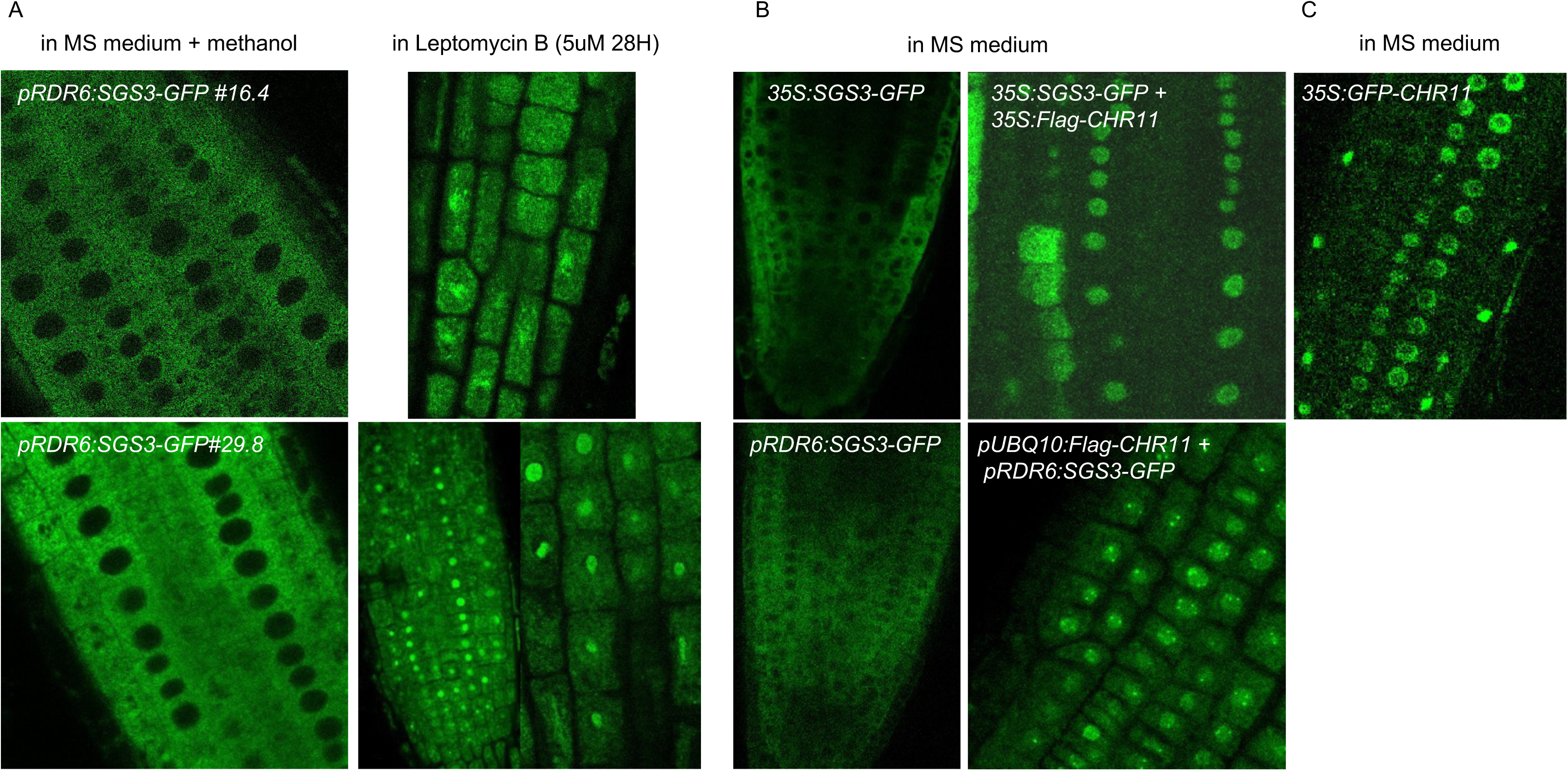
SGS3 Is a Nucleo-Cytosolic Shuttling Protein. A. SGS3-GFP relocalizes to the nucleus after inhibition of nuclear export by leptomycin B. Two independent *sgs3/pRDR6:SGS3-GFP* complemented lines were analyzed. B. *CHR11* over-expression increases retention of SGS3 in the nucleus in different combination of lines expressing *SGS3-GFP* under the *35S* or *RDR6* promoters (both complementing the *sgs3-1* mutant) and *FLAG-CHR11* under the *35S* or *UBQ10* promoters (both complementing a *chr11 chr17* double mutant). C. Nuclear localization of a control line expressing *GFP-CHR11* under the *35S* promoter.

To test this hypothesis, *L1/sgs3/pRDR6:SGS3-GFP* plants were treated with Leptomycin B, an inhibitor of nucleo-cytosolic export. After treating plants with 5 uM Leptomycin B during 28 hours, the GFP signal was mostly seen in the nucleus, but not in the same foci as H2B-RFP (Fig. 2A and Supplementary Fig. 2A), indicating that SGS3 shuttles between the nuclear and cytosolic compartments. LMB-treated *pUBQ10:GFP-ATG8A* transgenic plants ^14^ used as a control showed unaltered cytosolic GFP signal in cytosol and autophagic bodies (Supplementary Fig. 2B). Moreover, *p35S:RDR6-GFP,* a cytosolic partner of SGS3 in siRNA bodies, never show nuclear GFP signal after the same LMB treatment (Supplementary Fig. 2B).

Because SGS3 interacts with CHR11 in the nucleus, we asked whether increasing CHR11 level in the nucleus could increase the retention of SGS3 in the nucleus. To test this, *L1/sgs3/p35S:SGS3-GFP* plants were crossed with *p35S:FLAG-CHR11* plants. Whereas the GFP signal in *L1/sgs3/p35S:SGS3-GFP* plants was only cytosolic, a GFP signal was detected in the nucleus of *L1/sgs3/p35S:SGS3-GFP* x *p35S:FLAG-CHR11* plants, although in a stochastic and non-homogeneous manner (Fig. 2B and Supplementary Fig. 3). This result was confirmed by transforming *L1/sgs3/pRDR6:SGS3-GFP* plants with a *pUBQ10:FLAG-CHR11* construct. Whereas the GFP signal was only cytosolic in *L1/sgs3/pRDR6:SGS3-GFP* plants, a nuclear SGS3-GFP signal was detected in some *L1/sgs3/pRDR6:SGS3-GFP/p35S:FLAG-CHR11* transgenic plants (Fig. 2B and Supplementary Fig. 3), indicating that increasing CHR11 level increases the fraction of nuclear SGS3. Nevertheless, SGS3 import in the nucleus appears independent of CHR11 and CHR17. Indeed, western blot analysis showed that SGS3 is detected in the nucleus of *chr11 chr17* plants (Fig. 1D). Contrasting SGS3, CHR11 is never detected in the cytosol, indicating that the CHR11-SGS3 complex is purely nuclear (Fig2C), and dissociates before SGS3 re-export to the cytosol.

### The two ISWI-like chromatin remodeling factors CHR11 and CHR17 act redundantly in transgene S-PTGS

*CHROMATIN REMODELING PROTEIN 11* (*CHR11)* is a chromatin remodeling factor that belongs to the ISWI sub-family, which also comprises *CHR17*. CHR11 and CHR17 proteins share 98.6 % identity and ensure redundant functions. *chr11* or *chr17* single mutants do not exhibit any obvious developmental defects, while the *chr11 chr17* double mutant is dwarf and sterile ^15^ (Supplementary Fig. 4). Previous analyses revealed that CHR11 binds to the body of 17393 genes in Arabidopsis, and that 3222 genes are up-regulated and 4188 down-regulated in the *chr11 chr17* double mutant, which likely explains its drastic phenotype ^16,17^. It was also reported that the plant-specific ISWI complex containing CHR11 and CHR17 functions to partially derepress genes and transposable elements subjected to transcriptional gene silencing (TGS) through their function in regulating nucleosome distribution ^18^. However, no report to date has established a link between the ISWI complex and PTGS.

To determine if CHR11 and CHR17 could play a role in transgene S-PTGS by attracting SGS3 to transgene RNAs, we examined the effect of impairing *CHR11* and *CHR17* on *L1* PTGS. However, the only *chr11* and *chr17* mutants available belong to the SALK and GABI collections, which interferes with the expression of the *p35S:GUS* transgene carried by the *L1* line ^19,20^, making the use of these mutants impossible. Therefore, we transformed *L1* with a *pUBQ10:CHR11* construct and screened for transformants exhibiting a *chr11 chr17* double mutant phenotype due to co-suppression of the *CHR11* and *CHR17* endogenous copies by the *pUBQ10:CHR11* transgene. Among 50 primary transformants, 15 showed a phenotype resembling that of the *chr11 chr17* double mutant, among which two were fertile (Supplementary Fig. 5). Analysis of the progeny of these two transformants showed that *CHR11* and *CHR17* mRNA levels were reduced (Supplementary Fig. 6), and that *L1* S-PTGS was delayed compared to *L1* controls, suggesting a role for *CHR11/CHR17* in transgene S-PTGS (Fig. 3A and 3B).

**Fig. 3.**
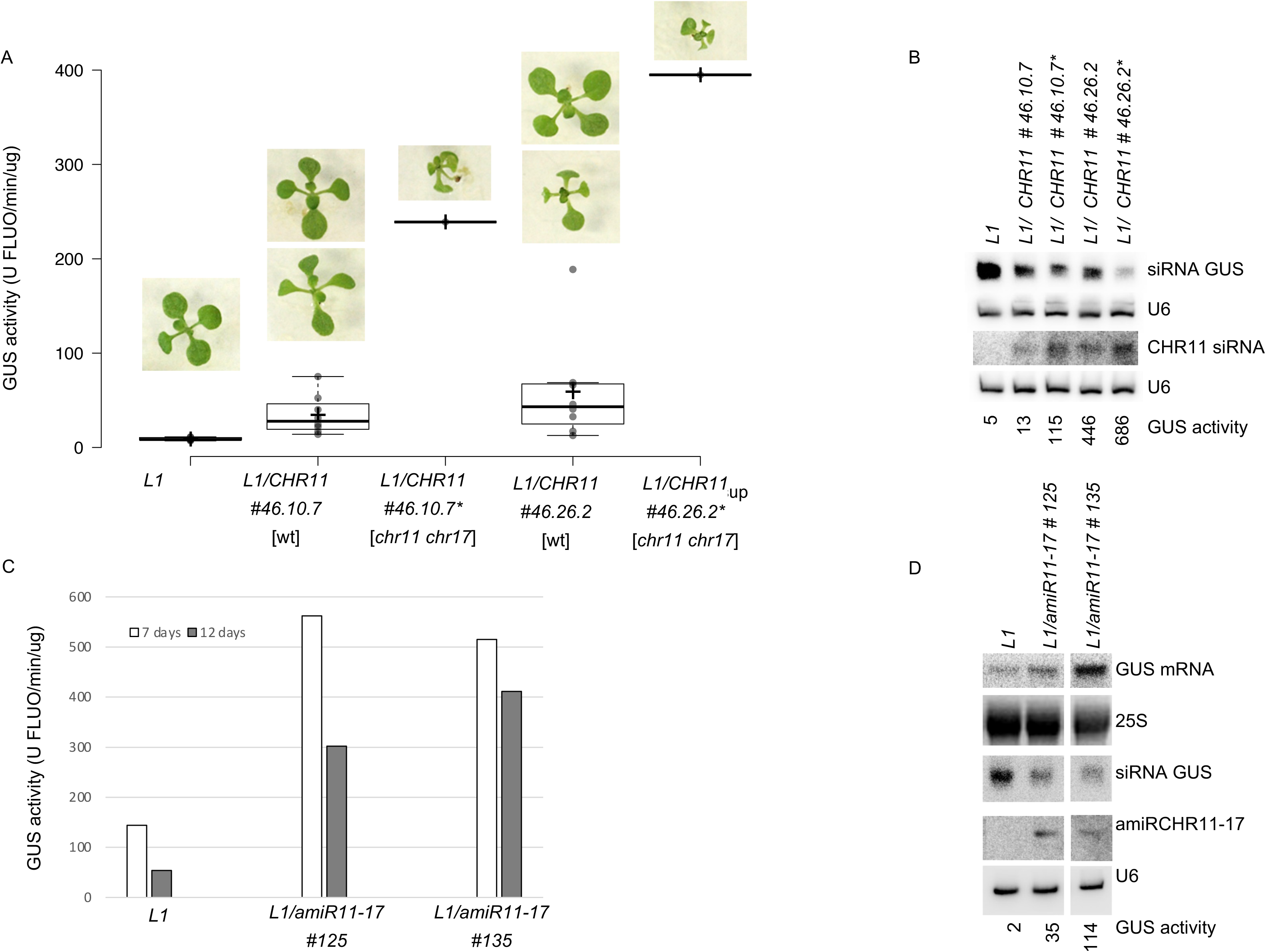
*L1* S-PTGS is delayed when partially impairing *CHR11* and *CHR17*. A. GUS activity at 12 days in the progeny of line *L1/ pUBQ10:CHR11* #46. Plants exhibiting a wild-type phenotype [wt] (#46.10.7 and #46.26.2) were harvested individually (means are indicated by a cross on box plot). Plants exhibiting a *chr11 chr17* phenotype indicative of *CHR11/CHR17* cosuppression (#46.10.7* and #46.26.2*) were pooled to form a unique sample. B. *GUS* and *CHR11* siRNA accumulation at 12 days in the progeny of line *L1/ pUBQ10:CHR11* line #46. Plants were harvested in bulk. GUS activity is given below for each sample used for RNA extraction. C. Kinetics of GUS activity in the progeny of lines *L1/ pUBQ10:amirCHR11-17* #125 and #135, which exhibit strong *CHR11/CHR17* cosuppression. At 7 and 12 days plants were harvested in bulk (results at 17, 24 and 40 days are shown in Supplementary Fig. 8). D. *GUS* mRNA, *GUS* siRNA and *amiRCHR11-17* accumulation at 18 days in the progeny of lines *L1/ pUBQ10:amirCHR11-17* lines #125 and #135 harvested in bulk. GUS activity is given below for each sample used for RNA extraction.

To test further this hypothesis, *L1* was transformed with a *pUBQ10:amiRCHR11-17* construct, which produces an artificial miRNA targeting both *CHR11* and *CHR17* mRNAs. Among 140 primary transformants, 30 showed mild to severe phenotypes resembling those of the *chr11 chr17* double mutant. The two transformants exhibiting the most severe phenotypes showed impaired *L1* S-PTGS, but these transformants were sterile (Supplementary Fig. 7). Analysis of the progeny of two fertile transformants exhibiting less severe phenotypes showed that *CHR11* and *CHR17* mRNA levels were reduced (Supplementary Fig. 6B), and that *L1 S-*PTGS was delayed compared to *L1* controls (Fig. 3C, 3D and Supplementary Fig. 8), demonstrating a role of CHR11/17 in PTGS.

A simple explanation for S-PTGS impairment in *chr11 chr17* double mutant could be that transgene transcription is reduced. To test this hypothesis, line *6b4* was transformed with the *pUBQ10:amiRCHR11-17* construct. Line *6b4* carries the same *p35S:GUS* transgene as line L1; however, unlike line *L1*, which triggers PTGS spontaneously, line *6b4* does not trigger S-PTGS spontaneously. Rather, it stably expresses the *p35S:GUS* transgene, i.e. GUS activity levels are similar in *6b4* and *6b4 rdr6* or *6b4 sgs3* plants. This is likely because the *6b4* locus produces lower amounts of aberrant RNAs than the *L1* locus. Nevertheless, the *6b4* locus triggers S-PTGS when RQC is impaired ^8,21–27^, indicating that *6b4*-derived aberrant RNAs can trigger S-PTGS when they are not degraded by the RQC machinery. Transformation of line *6b4* with the *pUBQ10:amiRCHR11-17* generated transformants that showed GUS activity levels similar to *6b4* controls (Supplementary Fig. 9), even when *CHR11* and *CHR17* mRNA levels were reduced (Supplementary Fig. 6C), indicating that impairment of *CHR11/17* does not modify the level of transcription of the *p35S:GUS* transgene. Together, these results suggest that the reason why CHR11/17 impairment affects S-PTGS is not due to changes in the level of transcription of transgene mRNA. Nevertheless, it remains possible that transgene transcription is qualitatively affected, modifying the ratio between functional mRNAs and aberrant RNAs.

### CHR11 physically interacts with the *p35S:GUS* transgene

The way CHR11/17 promotes *L1* S-PTGS could be by interacting with the *p35S:GUS* transgene locus to facilitate the addressing of *GUS* RNAs to SGS3 through the CHR11/SGS3 interaction. To test this hypothesis, *L1/pUBQ10:GFP-CHR11* and *L1/pUBQ10:Flag-CHR11* lines that complement the developmental defects of a *chr11 chr17* double mutant were selected (Supplementary Fig. 10). ChIP-qPCR experiments performed on these plants revealed an enrichment of CHR11 on the *p35S:GUS* transgene at the *L1* locus (Fig. 4).

**Fig. 4:**
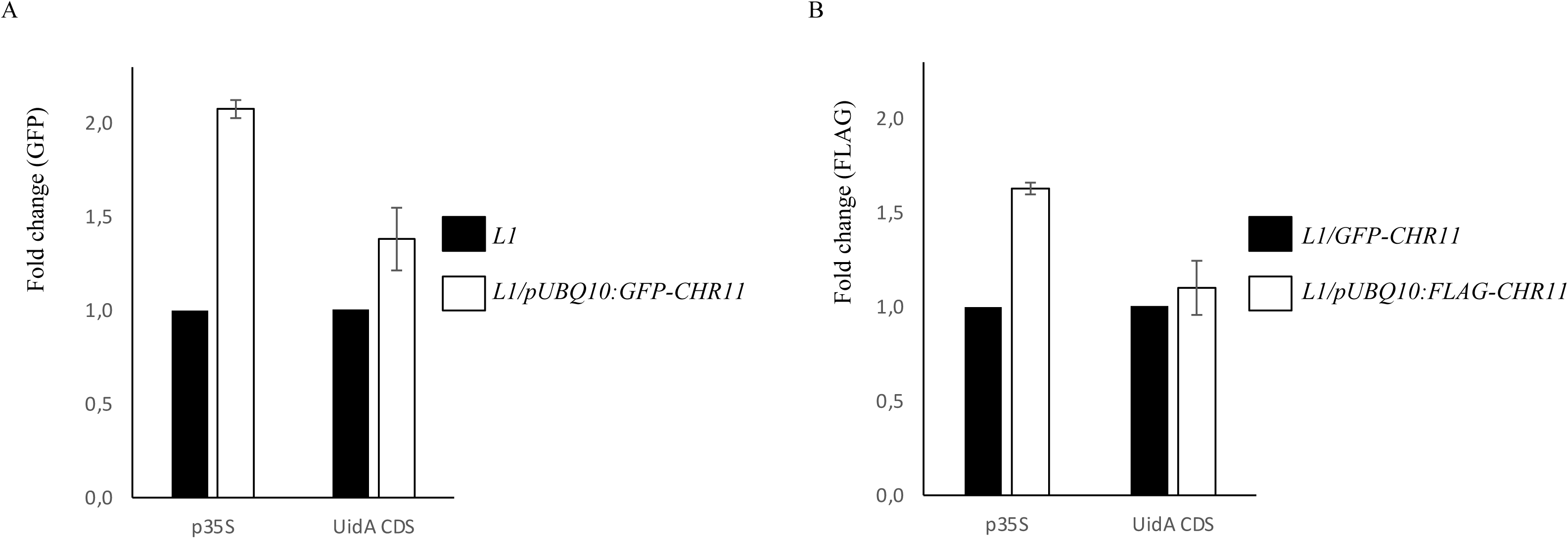
CHR11 interacts with *p35S:GUS* transgene at the *L1* locus. ChIP-qPCR analyses were performed on 15-day-old seedlings of the indicated genotypes. Graphical representation shows the fold change as the mean of two (in A) or three (in B) biological repeats. Error bars represent the standard deviation. A. ChIP was performed on the *L1/pUBQ10:GFP-CHR11* line using GFP antibodies for IP. The *L1* line was used as a control, followed by normalization to *GAPDH*. B. ChIP was performed on the *L1/pUBQ10:FLAG-CHR11* using Flag antibodies for IP. Here, the *L1/pUBQ10:GFP-CHR11,* which expressed *CHR11* at a similar level than in *L1/pUBQ10:FLAG-CHR11*, was used as a control, followed by normalization to *GAPDH*.

Because the *6b4* locus was used as a control to rule out the hypothesis that S-PTGS impairment in *chr11 chr17* double mutant is due to reduced transgene transcription (Supplementary Fig. 9), ChIP-qPCR experiments performed on *6b4/pUBQ10:GFP-CHR11* plants revealed that CHR11 also interacts with the *p35S:GUS* transgene at the *6b4* locus (Supplementary Fig. 11). In addition to validating the *6b4* locus as a control, this result supports the hypothesis that the *6b4* locus triggers S-PTGS in an RQC-deficient background through CHR11/SGS3 selection/recruitment, similar to the *L1* locus. Altogether, these results suggest that CHR11/17 generally promotes transgene S-PTGS by binding to transgene loci, thus allowing the association of transgene RNAs with SGS3 in the nucleus and their export to the cytosol to initiate S-PTGS.

### CHR11 promotes the production of siRNAs from endogenous protein-coding genes

The results presented above suggest a model in which SGS3 is recruited by CHR11 and presumably its paralog CHR17 to transgene DNA, allowing certain nascent transgene RNAs to bind SGS3, thus limiting their degradation by RQC. Moreover, SGS3 shuttling between nucleus and cytosol likely promotes the export of SGS3-bound transgene RNAs to the cytosol where they can be fully converted to dsRNA by RDR6 in siRNA bodies.

The next critical question was to assess how general is this mechanism for siRNA production from endogenous genes. To address this question, whole-genome data of CHR11 binding in wildtype plants ^17^ were mined for various categories of genes producing different types of siRNAs. In addition, small RNA sequencing was performed to determine the accumulation of endogenous siRNAs in wildtype and *chr11 chr17* plants. At first, *TAS* genes, which produce SGS3-dependent ta-siRNAs, were analyzed for CHR11 binding. Whereas 46% of the whole set of Arabidopsis genes bind CHR11 ^17^, none of the seven *TAS* genes showed binding (Fig. 6A and Supplementary Table 1). Moreover, the accumulation of ta-siRNAs was not significantly affected in *chr11 chr17* (Fig. 5) as compared to wt. This result is coherent with the fact that *TAS* RNAs become substrates for SGS3 and RDR6 only after they are cleaved by miRNA-guided AGOs in the cytosol. Therefore, the binding of CHR11 to *TAS* genes and the association of SGS3 to *TAS* RNAs in the nucleus do not appear necessary to promote the export of *TAS* RNAs.

**Fig. 5:**
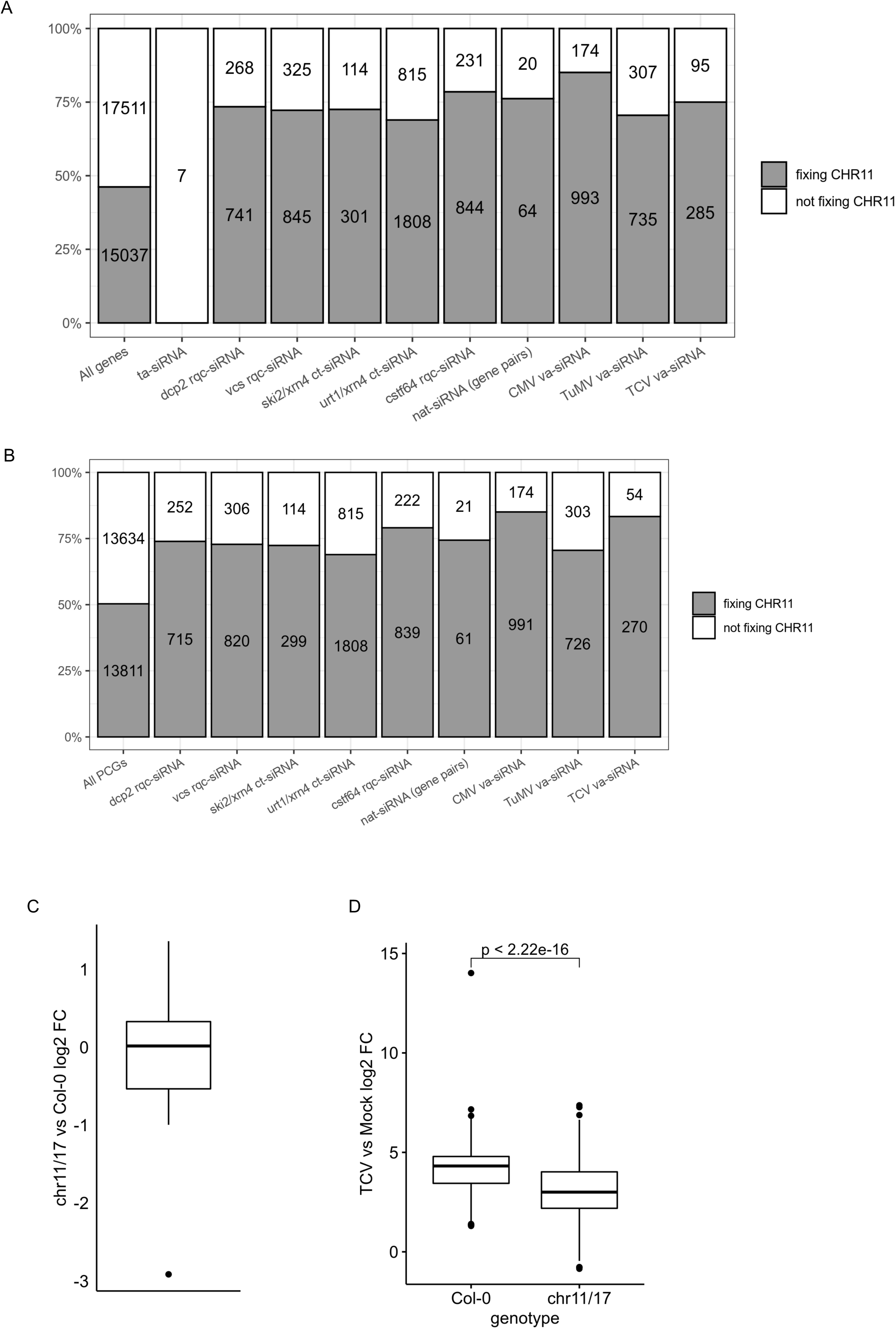
CHR11 is enriched at endogenous PCGs producing siRNAs and promotes siRNA production. A.Number and proportion of whole Arabidopsis genes (whole-genome or sub-categories producing siRNAs) fixing CHR11 or not according to 22. ta-siRNA: trans-acting siRNAs, va-siRNA: virus-activated siRNAs, nat-siRNA: natural antisense transcripts siRNAs, rqc-siRNA: endogenous siRNAs produced when RQC is impaired, ct-siRNA : endogenous siRNAs produced by coding transcripts; B. Number and proportion of Arabidopsis PCGs (whole-genome or sub-categories producing siRNAs) fixing CHR11 or not according to 22. C. Variation of 21/22nt ta-siRNA accumulation between *chr11 chr17* and Col-0. Values are log2 fold changes according to DESeq2 differential analysis. D. Accumulation of va-siRNAs upon TCV infection of Col-0 and *chr11 chr17*. Values are log2 fold changes according to DESeq2 differential analysis compared to Mock treatment. The number shown above indicates the p-values of the difference of log2 fold change determined by a Wilcoxon test.

**Fig. 6:**
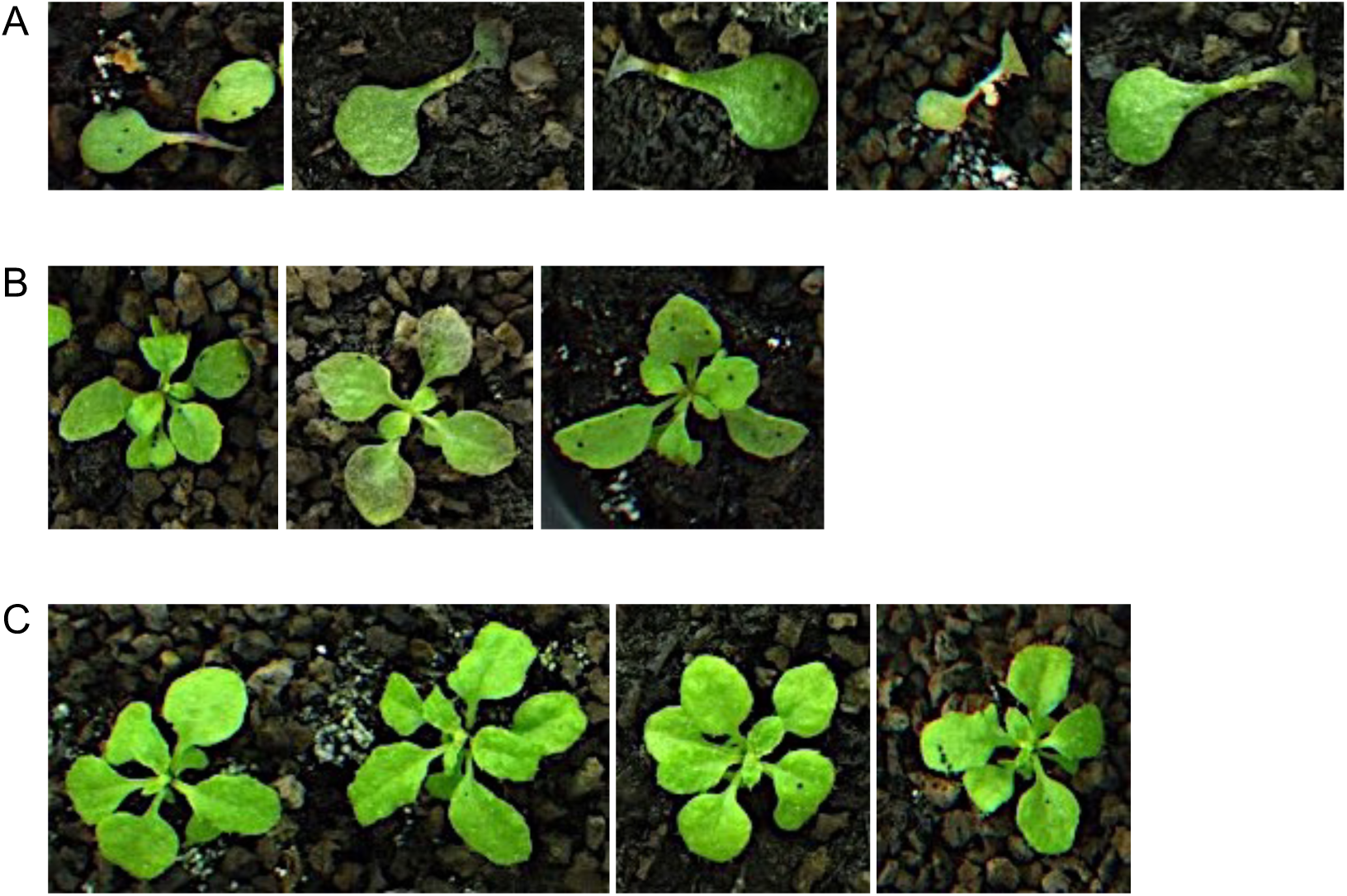
Simultaneous impairment of *CHR11 and CHR17* rescue the lethality of *ski3 xrn4 double* mutants. Pictures of representative plants of the indicated genotypes grown for 27 days in short days. All plants were grown at the same time. All pictures at the same scale. A. *ski3 xrn4 chr17* B. *ski3 xrn4 chr11 chr17* C. *chr11 chr17*

In a second time, protein-coding genes (PCGs) that produce siRNAs without the requirement of AGO/miRNA-guided RNA cleavage were examined because these PCGs could require CHR11 binding and SGS3 association in the nucleus to allow RNA export to the cytosol. Generally, PCGs do not produce siRNAs, except when RQC is impaired ^23,25,26,28–30^. Compared to the 50% of Arabidopsis PCGs that bind CHR11 ^17^, a significant enrichment was found for PCGs producing siRNAs in *dcp2* (74%), *vcs* (73%), *ski2 xrn4* (72%), *urt1 xrn4* (69%) and *cstf64* (79%) (Fig. 5 and Supplementary Table 1). In addition to RQC deficiency, specific conditions, including biotic and abiotic stresses, allow PCGs to produce siRNAs. Remarkably, a significant enrichment (74%) was also observed for PCG/NAT pairs (i.e. PCGs that overlap other protein-coding or non-protein coding genes), which, under certain biotic and abiotic stresses, produce endogenous siRNAs referred to as nat-siRNAs ^31^. Lastly, an enrichment in CHR11 binding was observed for PCGs producing endogenous siRNAs referred to as va-siRNAs, which are produced after infection with viruses such as CMV (85%), or TuMV (71%) (Fig. 5 and Supplementary Table 1). Together, these results support the relevance of CHR11 in the production of siRNAs from PCGs.

To further test whether CHR11 could promote siRNA production from PCGs, the effect of CHR11/17 impairment on endogenous siRNAs synthesis was investigated. It is likely that *chr11 chr17* mutants would not tolerate all biotic or abiotic stresses required to induce the production of nat-siRNAs. In contrast, it appeared possible to infect *chr11 chr17* double mutants with viruses. Infection with CMV was not efficient enough in our hands, but infection with TCV was very successful on the *chr11 chr17* double mutant. Small RNAseq analysis in TCV-infected wildtype plants revealed that TCV infection results in the production of va-siRNAs from PCGs that also produce va-siRNAs in CMV-and TuMV-infected plants (Supplementary Fig. 12). Moreover, an enrichment of CHR11 binding was also found for PCGs producing va-siRNAs under TCV infection (83%) (Fig. 5 and Supplementary Table 1). Comparing TCV-infected wildtype plants and TCV-infected *chr11 chr17* mutants revealed that the induction of va-siRNAs was significantly reduced in *chr11 chr17* plants (Fig. 5), further supporting the link between CHR11/17 and the production of endogenous vasiRNAs. To determine if this reduced induction of va-siRNAs was not simply due to reduced transcription of these PCGs in *chr11 chr17* double mutants, mRNAseq data of wildtype plants and *chr11 chr17* double mutants ^17^ were compared to our data. No relation or bias could be found between the variation of va-siRNAs accumulation between *chr11 chr17* and Col plants with the differential accumulation of their mRNA transcripts between the two genotypes for the full set of genes (Supplementary Fig. 13A) or for the TCV va-siRNA-producing genes (Supplementary Fig. 13B). These results rule out the possibility that the reduced induction of va-siRNAs in *chr11 chr17* double mutants is due to reduced transcription of these PCGs, similar to what was observed for transgenes (Supplementary Fig. 9).

To further examine the role of CHR11/17 in the production of siRNAs from PCGs, we investigated whether impairing CHR11 and CHR17 activity could rescue the growth of RQC-deficient mutants. Indeed, RQC-deficient mutants such as the *vcs* and *dcp2* single mutants ^25^ or the *ski2 xrn4* double mutant ^30^ die at an early stage of development, likely because of the massive production of siRNAs from PCGs. This lethal phenotype can be rescued by impairing *RDR6* or *SGS3*, which prevents the production of most of these siRNAs ^25,30^. The *chr11* and *chr17* mutations could not be combined with *dcp2* or *vcs* because *DCP2* and *CHR17* are genetically linked and so are *VCS* and *CHR11*. Therefore, *chr11* and *chr17* were combined with *ski3* and *xrn4*. The *ski3 xrn4* double mutant die at the two-cotyledons stage, whereas the *ski3 xrn4 rdr6* triple mutant is viable and indistinguishable from the *rdr6* single mutant (supplemental Fig. 14). The *chr17 ski3 xrn4* triple mutant did not grow better than *ski3 xrn4,* but the *chr11 chr17 ski3 xrn4* quadruple mutant grew as well as *chr11 chr17* (Fig. 6), indicating that impairing CHR11 and CHR17 rescues the lethal phenotype caused by the massive production of siRNAs from PCGs, thus confirming that CHR11 and CHR17 participate to the production of rogue siRNAs from endogenous gene when RQC is deficient.

## DISCUSSION

Although sense transgene PTGS (S-PTGS) was first described 30 years ago ^32,33^, the way it is initiated is still not fully understood. It is assumed that transgene aberrant RNAs are transformed into dsRNA by the RNA-dependent RNA polymerase RDR6, but how these aberrant RNAs are selected and brought to RDR6 and how they escape degradation by RNA quality control (RQC) pathways was not resolved. Here we propose a model in which the binding of CHR11 (and presumably CHR17) to transgene DNA allows the recruitment of SGS3 to nascent transgene RNAs, protecting them from a full degradation by RQC and allowing their export to the cytosol to initiate S-PTGS (Fig. 7). *CHR11* and *CHR17* domains are shared by the ISWI and SWR1 complexes involved in nucleosome sliding and H2AZ deposition at a large number of genomic loci ^17,18,34^. CHR11 and CHR17 interact with actors of the ISWI and SWR1 complexes by their SLIDE domain, which is dispensable for interaction with SGS3. Therefore, by interacting with CHR11 HAND and/or SANT domains, SGS3 appears, to form a complex that has not been described before. Of note, Luo et al ^17^ did not recovered SGS3 by IP-MS using a *pCHR11:CHR11-FLAG* transgenic line. This could be explained by the fact that the interaction between SGS3 and CHR11 was only detected by Bi-FC when a TAG epitope was fused to the N terminal part of CHR11 (this work).

**Fig. 7:**
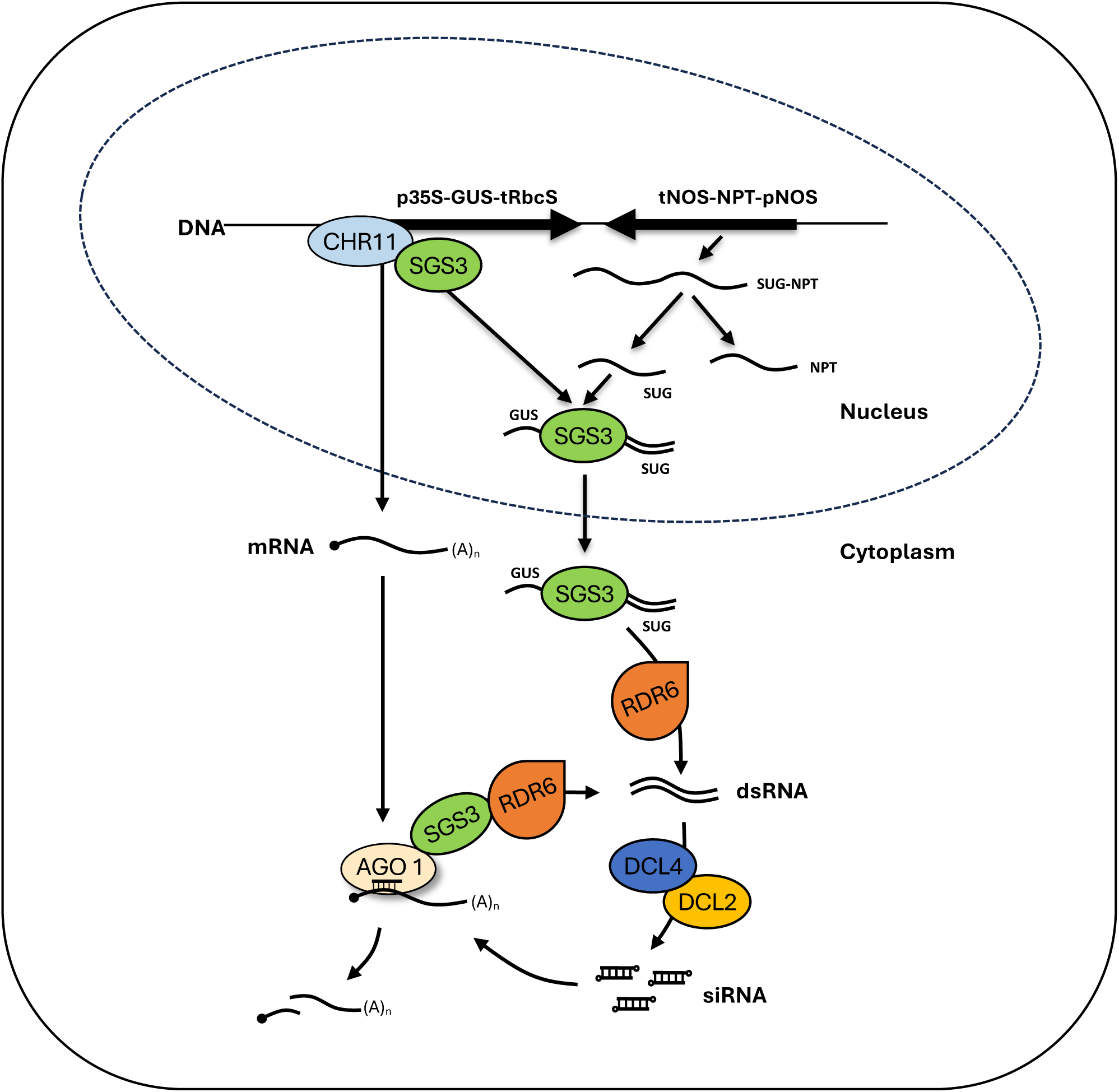
Tentative model of siRNA production via the action of nuclear CHR11 and SGS3. In the nucleus, SGS3 is recruited at CHR11-binding loci. When the genomic arrangement allows the production of sense and antisense RNAs forming dsRNA with a 5’ overhang, SGS3 binds to the dsRNA structure, which probably releases the CHR11-SGS3 interaction, allowing its export to the cytosol. In the cytosol, RNAs bound to SGS3 are converted to dsRNA by RDR6, followed by processing by DCL4 and DCL2 into 21-nt siRNAs that can execute PTGS and 22-nt siRNAs that allow PTGS amplification. The model described for the p35S-GUS transgene locus likely applies for endogenous PCG/NAT pairs.

SGS3 recruitment by CHR11/17 is reminiscent of the recruitment of the SGS3-like protein IDN2 by SWI3B, a sub-unit of the SWI/SNF complex that promotes RNA directed DNA methylation (RdDM) and transcriptional gene silencing (TGS) mediated by 24-nt siRNAs derived from transposons and repeats transcribed by the plant specific polymerases Pol IV and Pol V ^35^. Similar to SGS3, which interacts with CHR11 through its coiled-coil domain, IDN2 interacts with SWI3B through a similar domain. Whether IDN2 protects nascent Pol V-dependent RNAs against degradation by nuclear RQC is not known, but it is a reasonable hypothesis explaining the role of IDN2 in RdDM and TGS. If SGS3 plays a similar role in PTGS, it should protect aberrant RNAs in both nuclear and cytosolic compartments, and presumably travel from nucleus to cytosol with its RNA partners. SGS3 was reproducibly found in the cytosol ^5–7^ but was described only once in the nucleoplasm of isolated nuclei ^10^. However, truncated forms of the SGS3 protein lacking its prion-like domain, which likely makes it non-functional, were found in the nucleus ^11,12^; suggesting that SGS3 could have some function in the nucleus. Here we show that in fact the native SGS3 shuttles between cytosol and nucleus. When nucleo-cytosolic export is not inhibited, SGS3 is hardly detectable in the nucleus, except when CHR11 level is increased in the nucleus, suggesting that SGS3 is actively exported from the nucleus to the cytosol. This efficient export, together with the RNA binding and protecting effect of SGS3 against degradation by RQC, certainly contributes to addressing transgene RNAs to the cytosol to initiate PTGS.

Whole-genome analysis revealed that CHR11 bind to 50% of the protein-coding genes (PCGs) of the Arabidopsis genome, suggesting that half of the endogenous PCGs could have the potential to produce siRNAs through the SGS3 pathway. So far, only 5000 endogenous PCGs were shown to produce siRNAs, either when RQC is impaired by mutations ^25,29,30,36^, or when biotic or abiotic stress somehow challenge the functioning of RQC ^37,38^. However, not all cell types could be analyzed because of the dramatic developmental defects of RQC-deficient mutants and infected plants, so the number of endogenous PCGs producing siRNAs is certainly under-estimated. Nevertheless, given that the vast majority (70-85%) of the known siRNA-producing PCGs binds CHR11 (Fig. 5 and Supplementary Table 1), it is likely that CHR11 favor the attraction of SGS3 to the RNAs transcribed by these PCGs. Although the genetic requirement for the production of these PCG-derived siRNAs has not always been determined, most of them were shown to depend on RDR1 or RDR6, which generally act in concert with SGS3 to produce 21- and 22-nt siRNAs ^39^.

Given that CHR11 binds to the *p35S:GUS* transgene (Fig. 4), we propose a revised model in which S-PTGS is initiated in the nucleus owing to the interaction between CHR11 and SGS3 (Fig. 7). Consistent with the strong affinity of SGS3 to dsRNA exhibiting a 5’ overhang ^40^, we propose that SGS3 binds to dsRNA formed by the annealing of the full-length *GUS* mRNA to the uncapped antisense RNA that is complementary to the 3’ end of the *GUS* mRNA. We previously suggested that this uncapped RNA, referred to as *SUG*, results from the processing of a read-through transcript that originates in the adjacent and convergent *pNOS-NPT-tNOS* transgene because the *NOS* terminator does not efficiently terminate transcription ^8^. Following cleavage at the polyadenylation site of the *NPT* transcript, the downstream part of the read-through transcript, i.e. the uncapped *SUG* RNA, could anneal with the *GUS* mRNA. SGS3 could then bind to the *GUS/SUG* duplex in the nuclei because it has a 5’ overhang.

To explain how SGS3 and CHR11 separates to allow exporting the SGS3-dsRNA complex to the cytosol, we propose a model similar to that described for the shuttling of AGO1 ^41^. In the AGO1 model, the native AGO1 protein produced in the cytosol has an exposed NLS and a buried NES, allowing its import in the nucleus. When it binds miRNAs in the nucleus, AGO1 changes conformation, which exposes its NES, allowing the AGO1/miRNA complexes to be exported to the cytosol. We propose, that when it binds to dsRNA, SGS3 changes conformation, which disrupts the CHR11-SGS3 association. The binding of SGS3 to dsRNA and/or the dissociation of SGS3 from CHR11 may also exposes its NES, allowing the SGS3-dsRNA complex to be exported to the cytosol. Following its export to the cytosol, the SGS3-bound *GUS*/*SUG* duplex could be processed into 21- and 22-nt siRNAs by DCL4 and DCL2 to initiate PTGS, which is subsequently amplified by RDR6. *GUS* and/or *SUG* RNAs could also be separately transformed into dsRNA by RDR6 prior to DCL2/DCL4-mediated processing into 21- and 22-nt siRNAs.

This model could also explain the formation of nat-siRNAs that originate from overlapping transcription units producing natural antisense pairs of RNA (*NAT* transcripts). Indeed, the first nat-siRNAs described was 24-nt long, which is the signature of DCL3 processing in the nucleus, and derived from the overlapping region of an mRNA and its NAT, while 21-nt siRNAs, which are the signature of DCL4 processing in the cytosol, were produced from the rest of the transcripts^31^. This suggests that following cleavage by nuclear DCL3, the two annealed transcripts are protected from degradation by SGS3 and exported to the cytosol where each are transformed into dsRNA by RDR6 to produce siRNAs. Therefore, the high affinity of *NAT* pairs for CHR11/17 together with the high probability that annealed *NAT* transcripts are cleaved by a nuclear DCL likely explain how CHR11-mediated SGS3 recruitment to *NAT* transcripts may lead to producing nat-siRNAs, similar to what happens at transgene loci producing unintended antisense RNAs. Given the large overlap between the set of PCGs producing nat-siRNAs and the sets of PCGs producing siRNAs when RQC is impaired or during virus infection (Supplementary Fig. 15), nat-siRNAs, rqc-siRNAs and va-siRNAs could represent a unique class of siRNAs whose common determinant is their arrangement as PCG/NAT pairs. The complete list of *bona fide* PCG/NAT pairs is not known because the NAT partner could produce a non-conventional RNA that cannot be easily identified. Indeed, it could produce a non-coding RNA that cannot be predicted based on the genomic sequence and/or that is expressed only under particular conditions. Alternatively, it could produce uncapped or non-polyadenylated RNAs that cannot be cloned in regular RNAseq experiments. Although it is not possible to prove that all rqc-siRNAs and va-siRNAs originate from *bona fide* PCG/NAT pairs, the fact that impairing CHR11 and CHR17 rescues the lethal phenotype caused by the massive production of siRNAs from PCGs in the RQC-deficient *ski3 xrn4* double mutant (Fig. 6) strongly suggests that our model for transgene S-PTGS applies to a majority of PCG-derived siRNAs.

## METHODS

### Plant material, growth conditions, transformation, and virus inoculation

*p35S:GUS* lines *6b4* and *L1* and mutants *sgs3-1, ski3-3, xrn4-5, chr11-1 and -2, chr17-1* and -*2* have been described previously ^5,16,27,42–46^, *chr17-3* (SALK 629656) and *chr17-4* (SALK 585156) derive from the SALK T-DNA collection. The three double mutants *chr11-1 chr17-1, chr11-2 chr17-1* and *chr11-2 chr17-4* were generated and used indifferently for analyses since they share the same developmental defects. *Arabidopsis* seeds were sown *in vitro* on a nutritive medium (1.3% S-medium Duchefa, 1% Phytoblend agar) and vernalized at 4°C for at least 2 days before being transferred to soil in culture chambers. Plants were grown at 23°C, 70% humidity, 120 μE.m^−2^ lighting and 16 h light/8 h dark (long-days) or 8 h light/16 h dark (short-days) photoperiod. To generate transgene/mutant combination lines, *6b4* and *L1* were transformed by agrobacterium or crossed to the corresponding mutants/transgenic lines, and F2 progenies were genotyped to identified plants homozygous for both the transgene and the mutation(s). Transformations were performed as described previously ^5^. For virus infection, 20-day-old plants grown under short-day conditions were infected with TCV by mechanical inoculation ^47^. Plants were grown for 3 weeks, after which the total aerial parts of 4 to 12 plants were harvested.

### Cloning and constructs

*pSGS3:SGS3-GFP* was first obtained using a recombineering-based gene tagging system as described in ^13^. To ensure that all cis-regulatory sequences are included, we have used a bacterial homologous recombination system to insert Venus fluorescent gene into SGS3 genomic sequence harbored by transformation-competent bacterial artificial chromosomes (TACs) before transfer to Agrobacterium and plant transformation. cDNAs or coding sequence plus introns from ATG to stop of *SGS3* and *CHR11* were cloned in the GATEWAY™ compatible vector pDONR207 (Invitrogen) using the following primers : attB2SGS3f/ attB2SGS3R and attB1CHR11Fbis/ attB1CHR11Rbis (Supplementary Table 3). *SGS3* was transferred to the binary vectors pB7FWG2 to make the *p35S:SGS3-GFP* and in pRDR6-pGWB4 to make the *pRDR6:SGS3-GFP*. *pRDR6-pGWB4* was obtained by cloning *RDR6* promoter in pGWB4 ^48^ digested by *Hind*III after amplification by the primers pairs proSGS2F-hIII / proSGS2R-HIII.

*CHR11* was transferred to the binary vectors pB7WGF2 to make the *p35S:GFP-CHR11*, to pGWB12 to make *p35S:Flag-CHR11*, pUB-Dest ^49^ to make *pUBQ10:CHR11*, and pUBC-GFP-DEST to make *pUBQ10-GFP-CHR11*. *pUBQ10:Flag-CHR11* was obtained by cloning in pDONR207 *CHR11* cDNA amplified by attB1FlagCHR11F and attB2CHR11Rbis primers prior recombination in the compatible destination vector pUB-DEST. QuikChange™ (Stratagene, Arcueil, France) PCR-based mutagenesis was used to introduce premature stop1 and stop2 in *CHR11* cloned in pDONR207 using CHR11stop1Fbis /CHR11stop1Rbis and CHR11stop2F/CHR11stop2R pairs of primers to obtained *p35S:GFP-CHR11stop1* and *stop2*. We used the *MIR319a* precursor to engineer artificial micro RNA (amiRNA) for targeting both *CHR11* and *17* as described before ^50^. We replace the original *miR319a* with the artificial sequence *amiRCHR11-17*, 5’-TATCATGAAACGGTCGCACTA-3’, using WMD3-Web microRNA designer (http://wmd3.weigelworld.org/) to choose the best sequence for targeting both *CHR11* and *CHR17*. *AmiRCHR11-17* was synthesized and cloned in pUC57 by GenScript Corporation (New Jersey 08854-3900). Primers amiRf and amiRr were used to amplified *amiRCHR11-17* in its precursor backbone for cloning in the pDONR207 vector prior recombination in the compatible destination vector pUB-Dest to obtained *pUBQ10:amirCHR11-17*.

CDS of *RDR6* without its stop codon was cloned in *Sal*I and *Not*I of pENTR1A vector after amplification by the primers pairs RDR6f/RDR6r (Supplementary Table 3) prior recombination in the compatible destination vector pH7FWG2 to obtained *p35S:RDR6-GFP*.

The genomic sequence of *H2B* (At3g45980) under the control of 1kb of it’s own promoter have been cloned in the Gateway pGWB553 vector ^48^ with mRFP Cterminus tag, using At3g45980-F1 and At3g45980-R1 primers (Supplementary Table 3). The genomic sequence of *NUP54* (At1g24310) under the control of 1kb of it’s own promoter have been cloned in the Gateway pGWB543 vector ^48^, with CFP Cterminus tag, using At1g24310-F1 and At1g24310-R1 primers (Supplementary Table 3).

### GUS activity and Molecular analyses

GUS protein was extracted and GUS activity was quantified as described before 21 by monitoring the quantity of 4-methylumbelliferone products generated from the substrate 4- methylumbelliferyl-b-D-glucuronide (Duchefa) on a fluorometer (Thermo Scientific fluoroskan ascent).

Molecular analyses (DNA sequencing, RNA gel blot analysis) were performed as described previously ^21^. All RNA gel blot analyses were performed using 10μg of total RNA. *UidA, 25S DNA probes* and *U6* oligo probes have been described before ^21^. Oligo probe *amiRCHR11* was 32P-end-labelled to detect *amiRCHR11-17* and *CHR11* PCR fragment amplified by primers attB1CHR11Fbis/ attB2CHR11Rbis was used as probe to detect *CHR11* siRNA accumulated in co-suppressed *L1/pUBQ10:CHR11* lines (Supplementary Table 3).

For the reverse transcription, RNA used for northern blot analysis were treated with DNaseI (Invitrogen) and 1 μg of DNA-free RNA was reverse transcribed with an oligo d(T)^18^NN using the RevertAid H Minus Reverse Transcriptase (ThermoFisher, http://www.thermofisher.com/). Specific amplification of CHR11 and CHR17 were performed by using the CHR11f1 and r1, CHR17f and r primers (Supplementary Table 3), and qPCR results were normalized to GAPDH as described before ^51^.

### Yeast two-hybrid and BiFC cloning assays

*SGS3* cDNA and the C-terminal three coiled-coil domains of *SGS3* (corresponding to the amino acids 463–626) were cloned into pLex10 vector to produce fusion proteins with the *LexA* DNA-binding domain as described before ^5^. *SGS3* cDNA was used as bait in yeast two-hybrid screen of the 3-days-old etiolated *Arabidopsis* cDNA library CD4-22 ^52^. The yeast two-hybrid screen and interaction confirmation was performed as described before ^53^. Six clones corresponding to the C terminal part of *CHR11* were retrieved in this screen and were confirmed to interact with both *SGS3* and its C-terminal three coiled-coil domains in yeast two hybrid assay.

*SGS3* and *CHR11* cDNAs without their stop codon were transferred into the BiFC GATEWAY™-modified vector prior to being recombined with the N- and C-terminal parts of YFP (YN and YC) to produce p*35S:YN-CHR11*, p*35S:CHR11-YN*, p*35S:YC-CHR11*, p*35S:CHR11-YC,* p*35S:YN-SGS3*, p*35S:SGS3-YN,* p*35S:YC-SGS3* and p*35S:SGS3-YC* destination vectors as described before ^5^. The eight possible combinations were tested in BiFC, but only the combinations *YN-CHR11 / SGS3-YC* and *YN-CHR11 / YC-SGS3* gave an YFP signal. Results are shown for for the *YN-CHR11 / SGS3-YC* combination, which gave the best signal. *Nicotiana benthamiana* plants grown in the glasshouse under 13 h light, 25°C day temperature and 17 °C night temperature were used for all the agro-infiltration experiments.

The leaves were infiltrated with an overnight culture of *Agrobacterium tumefaciens* strain C58C1 that was re-suspended to an absorbance at 600 nm of 0.1 with the infiltration medium (10mM MES, pH5.6, 10mM MgCl2, 200 μM acetosyringone). For co-infiltration, equal volumes of both *Agrobacterium* cultures were mixed before infiltration. Observations were performed 72h after infiltration. Confocal microscopy was performed on an inverted TCS- SP2-AOBS spectral confocal laser-scanning microscope (Leica Microsystèmes SAS, Rueil-Malmaison, France) as previously described ^5^. Samples were excited with a 514 nm argon laser (50%) with an emission band of 520–550 nm for YFP detection and 640–700 nm for chlorophyll autofluorescence.

### Imaging

For confocal imaging, *Arabidopsis* roots were directly imaged on a Leica TCS-SP5 (Leica Microsystems) equipped with photomultiplier tube and hybrid detectors with an HCX PL APO CS 20.0 × 0.70 IMM objective. GFP was imaged with 488-nm excitation using an Argon laser.

### Leptomycin B treatment

For the nuclear export inhibition experiment, five days-old seedlings were transferred to 100ul liquid MS-medium with Leptomycin B (Sigma LMB, at 5uM final concentration corresponding to methanol at 4.7% final) and incubated 28 hours before imaging. Controls were incubated 28H in 100ul liquid MS-medium with methanol added at 4.7% final.

### Subcellular fractionations and western blot

Two grams of rosette leaves of 31 days-old plants were ground in 16 ml buffer supplemented with 1.14M sucrose (10 mM Tris–HCl pH 7.5, 5 mM MgCl^2^, 4 mM spermidine, 1 mM spermine, 1mM DTT and protease inhibitor cocktail sigma p9599). The sample was filtered through Miracloth and centrifuged at 1000 *g* for 10 min at 4°C. Supernatants recovered after the first centrifugation was supplemented with 0.15% Triton X100 and was centrifuged at 100 000g 45 min at 4°C to obtain the microsomal fractions (pellet). The first pellet was washed twice with 10 ml buffer (10 mM Tris–HCl pH 7.5, 5 mM MgCl^2^, 4 mM spermidine, 1 mM spermine, 1mM DTT and protease inhibitor cocktail sigma p9599) supplemented with 0.15% Triton X100. The last pellet corresponding to the nuclear fraction was resuspend in lysis buffer (10 mM Tris–HCl pH 7, 2% SDS, 10% glycerol and protease inhibitor cocktail sigma p9599) and sonicate on bioruptor. Protein concentration was quantified using a detergent compatible BCA kit (Bio-Rad) and 50 μg of protein were loaded on 8% SDS PAGE. Proteins were electroblotted onto nitrocellulose membranes (Amersham Hybond ECL). The membrane was blocked in 5% non-fat dry milk in 1× TBSTT (0.25% Tween-20, 0.1% Triton X100, NaCl 150 mM) for 1 h at room temperature, rinsed for 5 min in 1× TBST, and incubated with primary antibody in 5% non-fat dry milk and 1× TBST for 1 h at room temperature. The membrane was then rinsed in 1× TBST for 20 min before incubation with HRP-coupled secondary IgGs. Antigens were detected using chemiluminescence for HRP immunoblot (Amersham ECL Plus). The antibodies used were anti-SGS3 (ref B152 from Santacruz, 1/200e dilution) anti-H3 (ab1791 from Abcam, 1/3000e dilution).

### ChIP

ChIP was performed on chromatin from 2 g crosslinked in vitro plantlets 15 days after germination mainly as previously described ^54^. After 2 × 5 cycles of sonication (30 s “ON”, 30 s “OFF”, High intensity) with a Bioruptor UCD200 (Diagenode), the chromatin solution was diluted 10-fold to a final volume of 3 mL with ChIP dilution buffer before IP. To stabilize protein complexes, double crosslink was also tested using first disuccinimidyl glutarate 2 mM for 45 min at RT then formaldehyde 1% final concentration for 7 minutes essentially as described in ^55^, followed by sonication using Covaris S220 ultrasonicator 12 min with 5% duty cycle, 105W peak power and 200 cycles per burst.

For ChIP on *L1/pUBQ10:GFP-CHR11*, 30 μL of GFP-trap-M beads (gfm-20/500ul chromotech) was washed twice and resuspended in 60 μL of ChIP dilution buffer.

For ChIP on *L1*and *6b4/pUBQ10:Flag-CHR11*, 60uL of magnetic beads G was washed twice and resuspended in 60 μL of ChIP dilution buffer. 5 μL of anti-FLAG M2 clone SIGMA(F3165) were added to the beads G and incubated for at least 3 h at 4°C with gentle rotation on a wheel. After three washes the beads plus antibodies were resuspended in 100 μL of ChIP dilution buffer.

1 mL of the chromatin solution was added to the antibodies plus beads (GFP-trap-M or G beads plus anti-Flag) and was incubated overnight at 4°C with gentle rotation for GFP or Flag capture. The washing of beads was performed as described ^56^. After the last TE wash, the reverse crosslinking (5 h at 65°C) and elution were performed using IPure kit (Diagenode). The final elution was performed in 60 μL, and the chromatin was stored at −20°C until analysis.

Using Biorad-CFX-Maestro Software, the ChIP was analyzed by qPCR on 2 μL of the chromatin. Each primer pair are listed in Supplementary Table 3. The mean of the three qPCRs results (with SD < 0.4 cycle threshold) was used for each point. Glyceraldehyde-3-phosphate dehydrogenase (GAPDH)) was used as an internal reference ^57^. Results are represented as fold change: normalized expression (delta delta Cq) given by the ratio of Relative Quantity of the sample (2^(Cq control-Cq sample)^ for each identical primers with 100% of efficiency) divided by the Relative Quantity of internal reference as described before ^58^. Here control is L1 for ChIP on *L1/pUBQ10:GFP-CHR11* and *L1/ pUBQ10:GFP-CHR11* for ChIP on *L1/pUBQ10:Flag-CHR11*, sample is our target and internal reference is GAPDH. Two to four biological replicates were analyzed each time. Results show the mean and SD of the independent biological replicates.

### Small RNAseq library construction, sequencing and analysis

Small RNA (sRNA) libraries were constructed from 1µg of total RNA treated with DNaseI using the NEBNext® Multiplex Small RNA Library Prep Set for Illumina® kit (New England Biolabs) according to manufacturer’s instructions. Libraries were sequenced on a NextSeq 500 Sequencing System (Illumina) using 75-nt single-end reads. The sequencing adapters were removed using cutadapt (v2.10) and sequence matching Arabidopsis ribosomal or transfer RNA were discarded using bowtie (v1.3.1). The reads of length 21nt, 22nt and 24nt were then mapped on TAIR10 genome with the help of ShortStack (v3.8.5) without mismatch (--mismatches 0), keeping all primary multi-mapping (--bowtie_m all) and correcting for multi-mapped reads according to the uniquely mapped reads (--mmap u). For each gene annotation in Araport11 (coding genes, non-coding RNA and miRNA precursor), the accumulation of 21/22nt was counted with ShortStack. The number of sequence corresponding to each mature miRNA of Arabidopsis in miRBase (v22) was used to measure miRNA abundance. 21/22nt sRNA accumulation in each library was normalized using the median of ratio of the corresponding miRNA counts inside DESeq2 (v1.34.0). Differential 21/22nt sRNA accumulation and log2 fold changes between each genotype/condition couple were computed using DESeq2. FDR correction of the p-value was used.

## Acknowledgments

We thank Hervé Ferry and Philippe Maréchal for taking care of the plants. We thank François Roudier for discussion and advices on ChIP protocol adapted for chromatin remodeler proteins and on RIP protocol.

## Author contributions

T.E and H.V. designed the experiments. T.E., T.B, E.E.M., I.L.M., A.C. and N.B. performed the experiments. T.E, T.B., M.D.C. and H.V. analyzed the data and wrote the manuscript.

## Data Availability statement

Data supporting the findings of this work are available within the paper and its Supplementary Information files.

A reporting summary for this Article is available as a Supplementary Information file.

The datasets and plant materials generated and analyzed during the current study are available from the corresponding author upon request.

Small RNAseq data are accessible through NCBI’s Gene Expression Omnibus (GSE278529). All relevant data are available from the authors

## Supplementary Materials

Figures S1-S15

Tables S1-S2

