## supplemental-Fig-and-tables for "The Arabidopsis RNA-binding protein SGS3 is recruited to the chromatin remodeler CHR11 to promote siRNA production from protein-coding genes"

**This PDF file includes:**

Figs. S1 to S15

Tables S1 to S2

**Fig. S1. Sequence of *CHR11* retrieved in the yeast two-hybrid screen of 3-days-old etiolated *Arabidopsis* cDNA library CD4-22 [1] using SGS3 as bait.**

Interaction was observed in L40 yeast, between the C terminal part of *CHR11* fused to *GAL4* activation domain and *SGS3* or its coiled-coil C terminal domain fused to *LexA* binding domain in plex.

A. *SGS3* sequence used as bait in fusion with *LexA* binding domain. The C terminal part which contains the three coiled-coil domains of *SGS3* is in bold.

B. *CHR11* sequence. Sequence of *CHR11* C terminal part retrieved 6 times in the screen is shown in bold. E664 and E892 mutated respectively in stop codon 1 and 2 (corresponding to the two truncated forms used in BiFC) are underlined.

**A.**

MSSRAGPMSKEKNVQGGYRPEVEQLVQGLAGTRLASSQDDGGEWEVISKKNKNKPGNTSGKTWVSQNSNPPRAWGGQQQGRGSNVSGRGNNVSGRGNGNG
RGIQANISGRGRALSRKYDNNFVAPPPVSRPPLEGGWNWQARGGSAQHTAVQEFPDVEDDVDNASEEENDSDALDDSDDDLASDDYDSDVSQKSHGSRKQ
NKWFKKFFGSLDSLSIEQINEPQRQWHCPACQNGPGAIDWYNLHPLLAHARTKGARRVKLHRELAEVLEKDLQMRGASVIPCGEIYGQWKGLGEDEKDYE
IVWPPMVIIMNTRLDKDDNDKWLGMGNQELLEYFDKYEALRARHSYGPQGHRGMSVLMFESSATGYLEAERLHRELAEMGLDRIAWGQKRSMFSGGVRQL
YGFLATKQDLDIFNQHSQGKTRLKFELKSYQEMVVKELRQISEDNQQLNYFKNKLSKQNKHA**KVLEESLEIMSEKLRRTAEDNRIVRQRTKMQHEQNREE**
**MDAHDRFFMDSIKQIHERRDAKEENFEMLQQQERAKVVGQQQQNINPSSNDDCRKRAEEVSSFIEFQEKEMEEFVEEREMLIKDQEKKMEDMKKRHHEEI**
**FDLEKEFDEALEQLMYKHGLHNEDD***

**B.**

MARNSNSDEAFSSEEEEERVKDNEEEDEEELEAVARSSGSDDDEVAAADESPVSDGEAAPVEDDYEDEEDEEKAEISKREKARLKEMQKLKKQKIQEMLE
SQNASIDADMNNKGKGRLKYLLQQTELFAHFAKSDGSSSQKKAKGRGRHASKITEEEEDEEYLKEEEDGLTGSGNTRLLTQPSCIQGKMRDYQLAGLNWL
IRLYENGINGILADEMGLGKTLQTISLLAYLHEYRGINGPHMVVAPKSTLGNWMNEIRRFCPVLRAVKFLGNPEERRHIREDLLVAGKFDICVTSFEMAI
KEKTALRRFSWRYIIIDEAHRIKNENSLLSKTMRLFSTNYRLLITGTPLQNNLHELWALLNFLLPEIFSSAETFDEWFQISGENDQQEVVQQLHKVLRPF
LLRRLKSDVEKGLPPKKETILKVGMSQMQKQYYKALLQKDLEAVNAGGERKRLLNIAMQLRKCCNHPYLFQGAEPGPPYTTGDHLITNAGKMVLLDKLLP
KLKERDSRVLIFSQMTRLLDILEDYLMYRGYLYCRIDGNTGGDERDASIEAYNKPGSEKFVFLLSTRAGGLGINLATADVVILYDSDWNPQVDLQAQDRA
HRIGQKKEVQVFRFCTESAIEEKVIERAYKKLALDALVIQQGRLAEQKTVNKDELLQMVRYGAEM**VFSSKDSTITDEDIDRIIAKGEEATAELDAKMKKF**
**TEDAIQFKMDDSADFYDFDDDNKDENKLDFKKIVSDNWNDPPKRERKRNYSESEYFKQTLRQGAPAKPKEPRIPRMPQLHDFQFFNIQRLTELYEKEVRY**
**LMQTHQKNQLKDTIDVEEPEGGDPLTTEEVEEKEGLLEEGFSTWSRRDFNTFLRACEKYGRNDIKSIASEMEGKTEEEVERYAKVFKERYKELNDYDRII**
**KNIERGEARISRKDEIMKAIGKKLDRYRNPWLELKIQYGQNKGKLYNEECDRFMICMIHKLGYGNWDELKAAFRTSSVFRFDWFVKSRTSQELARRCDTL**
**IRLIEKENQEFDERERQARKEKKLAKSATPSKRPLGRQASESPSSTKKRKHLSMR***

**Fig. S2. Predominant cytosolic localization of SGS3-GFP in normal condition.**

A. Nuclear localization of SGS3 fused to GFP after inhibition of nuclear export by leptomycin B in crossed plants which expressed H2B-RFP and NUP-CFP.

B. 5uM LMB treatment did not alter the expected localization of GFP-ATG8A in autophagic bodies and cytosol, nor the cytosolic localization of RDR6-GFP.

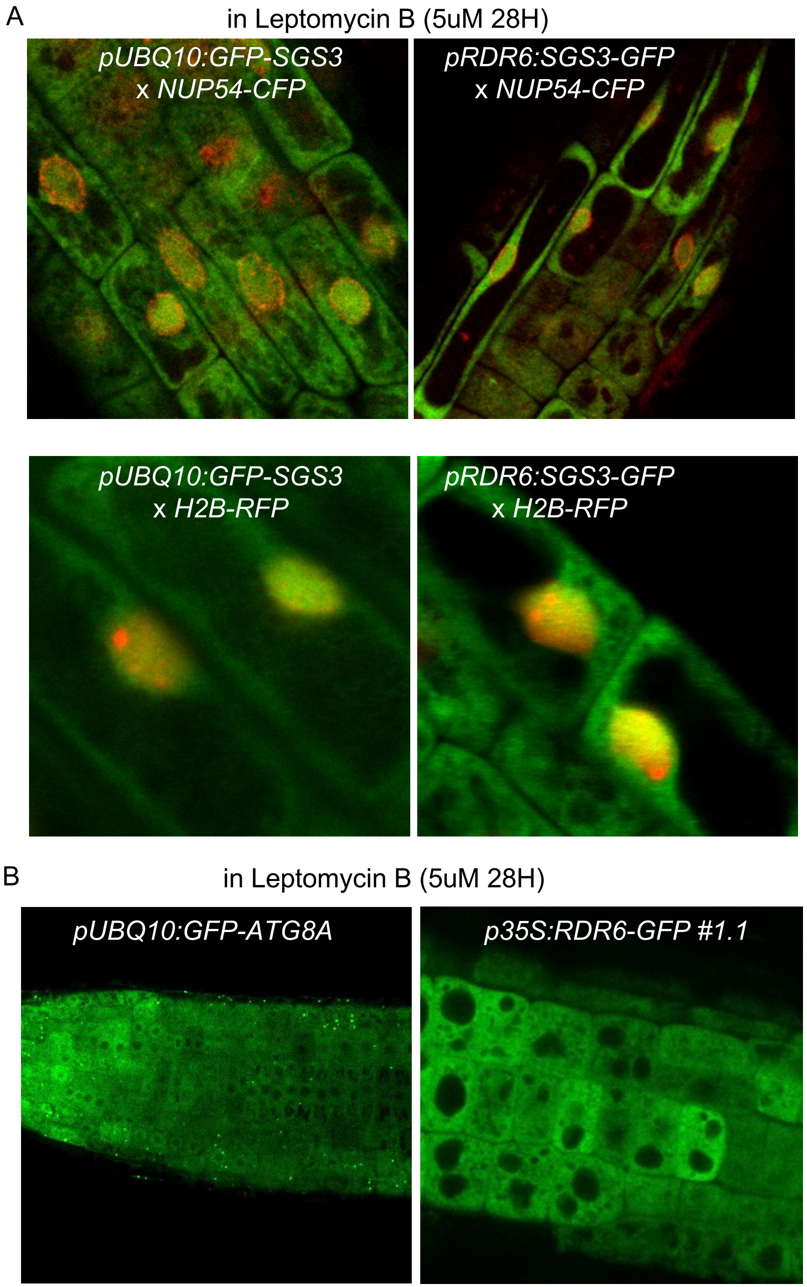
A
in Leptomycin B (5uM 28H)
pUBQ10:GFP-SGS3
x NUP54-CFP
pRDR6:SGS3-GFP
x NUP54-CFP
pUBQ10:GFP-SGS3
x H2B-RFP
pRDR6:SGS3-GFP
x H2B-RFP
B
in Leptomycin B (5uM 28H)
pUBQ10:GFP-ATG8A
p35S:RDR6-GFP #1.1

**Fig. S3.** *CHR11* over-expression increases retention of SGS3 in the nucleus in a stochastic manner.

A. Pictures of different *35S:SGS3-GFP / 35S:Flag-CHR11* plants obtained by crossing
B. Pictures of different *pRDR6:SGS3-GFP / pUBQ10:Flag-CHR11* plants obtained by crossing
C. Pictures of different *pRDR6:SGS3-GFP / pUBQ10:Flag-CHR11* plants obtained by transforming *pRDR6:SGS3-GFP* with *pUBQ10:Flag-CHR11*

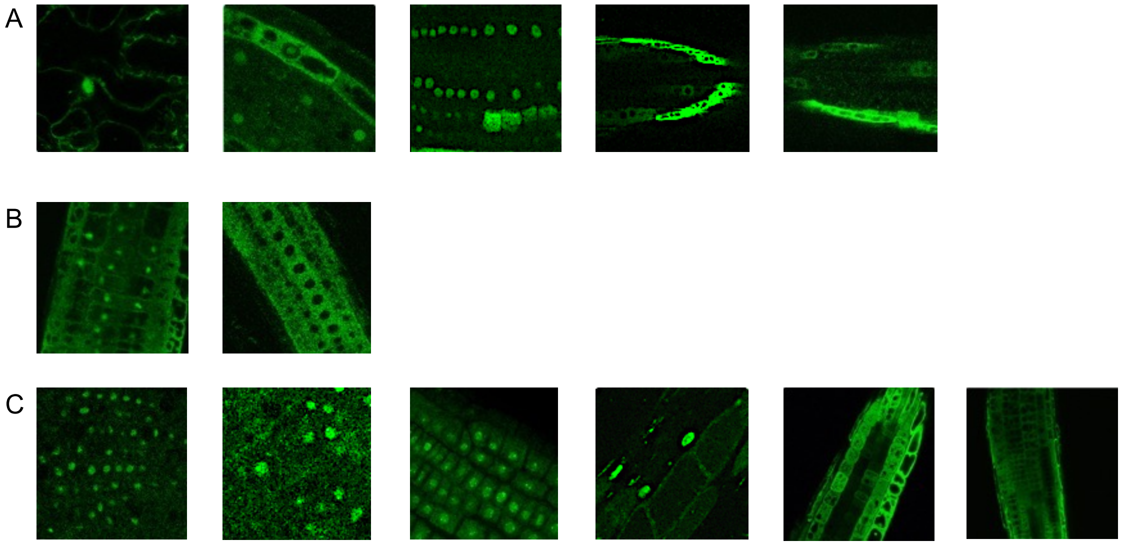

**Fig. S4. Developmental defects of the *chr11chr17* double mutant**

The *chr11chr17* double mutant is sterile and show several developmental defects: (early and terminal flowering, almost no petals, sepal fusion, very small stamen lacking pollen, defect in floral meristem maintenance, short roots, branched trichomes).

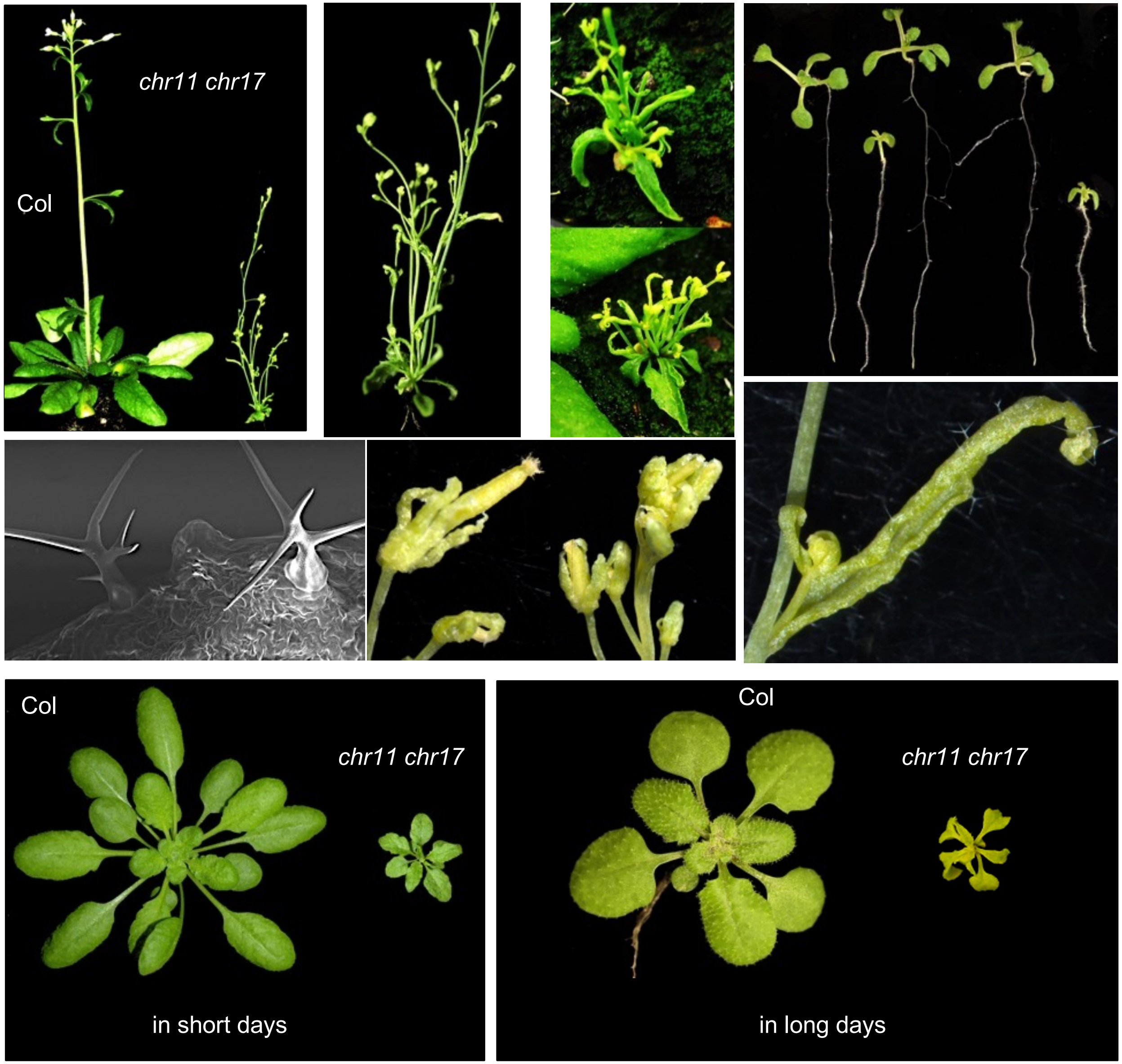

**Fig. S5. *L1* PTGS is delayed in *L1/pUBQ10:CHR11* transgenic lines exhibiting *CHR11/CHR17* cosuppression.**

A. Severity of the phenotypes of 50 *L1* primary transformants carrying the *pUBQ10:CHR11* construct.

B. Photographs of two representative fertile transgenic lines exhibiting a severe *CHR11/CHR17* silencing phenotype.

A

Phenotypes of 50 L1 primary transformants with *pUBQ10::CHR11*

| | n T1 | lines |
|---|---|---|
| letal | 1 | |
| mild phenotype | 12 | #46, #38 |
| Very mild phenotype | 2 | |
| wt phenotype | 35 | |

B

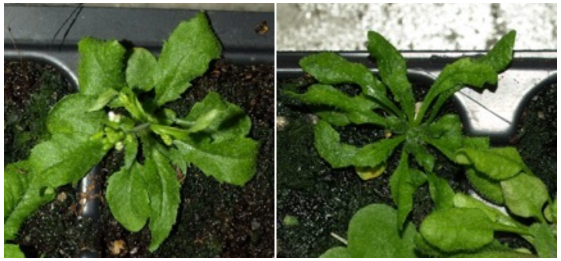

**Fig. S6. Analysis of CHR11 and CHR17 expression by RT-qPCR.**

*A. In L1, L1/ CHR11 # 46.26.2* and L1/ CHR11 # 46.10.7** co-suppressed plants analyzed in Fig3B.

*B. in L1 and L1 amiR11-17 #125* silenced plants analyzed in Fig3D,

*C. in 6b4 and 6b4 amiR11-17 #6.15** silenced plants analyzed in Fig. S7F.

Results are expressed as a fold change compared to *L1* or 6b4 ( = 1) and normalized to *GAPDH.*

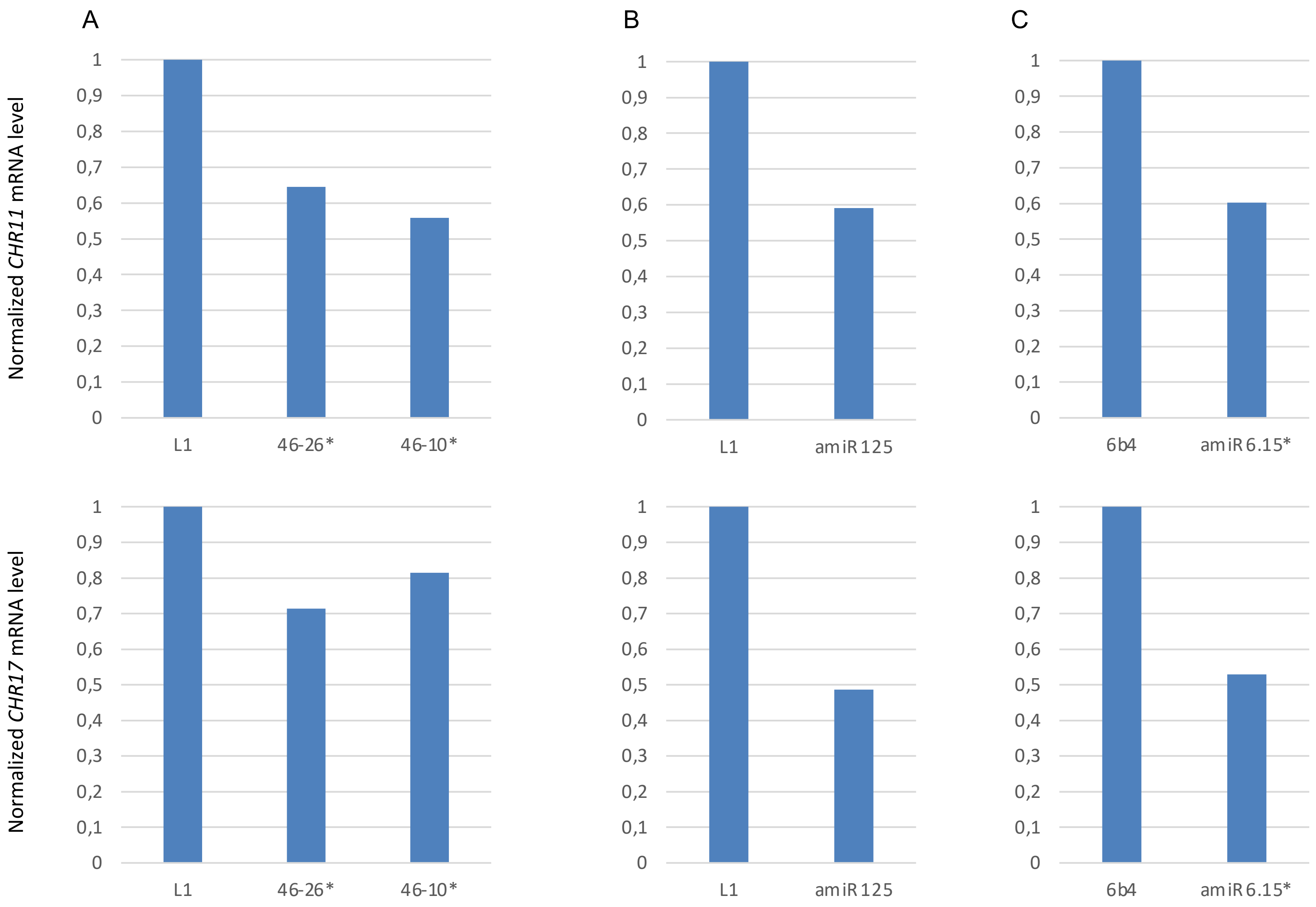

**Fig. S7. : *L1* S-PTGS is delayed in transgenic lines expressing an artificial miRNA targeting *CHR11* and *CHR17*.**

A. Severity of the phenotypes of 140 L1 primary transformants.

B. Photographs of three independent transformants exhibiting severe phenotypes.

C. GUS activity in 30 primary transformants showing a mild to severe phenotype at 46 days.

A Phenotypes of *L1* primary transformants carrying the *pUBQ10:amiRCHR11-17* construct

| | number | lines |
|---|---|---|
| letal | 11 | |
| sterile | 2 | #109, #140 |
| mild phenotype | 5 | #12, #82, #125, #135, #133 |
| very mild phenotype | 11 | #67, #76, #101, #104, #106, #111, #115, #126, #132, #136, #137, |
| wild-type phenotype | 110 | |

B

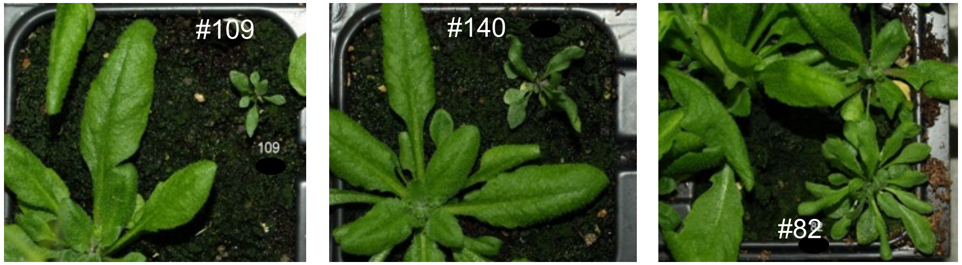


C

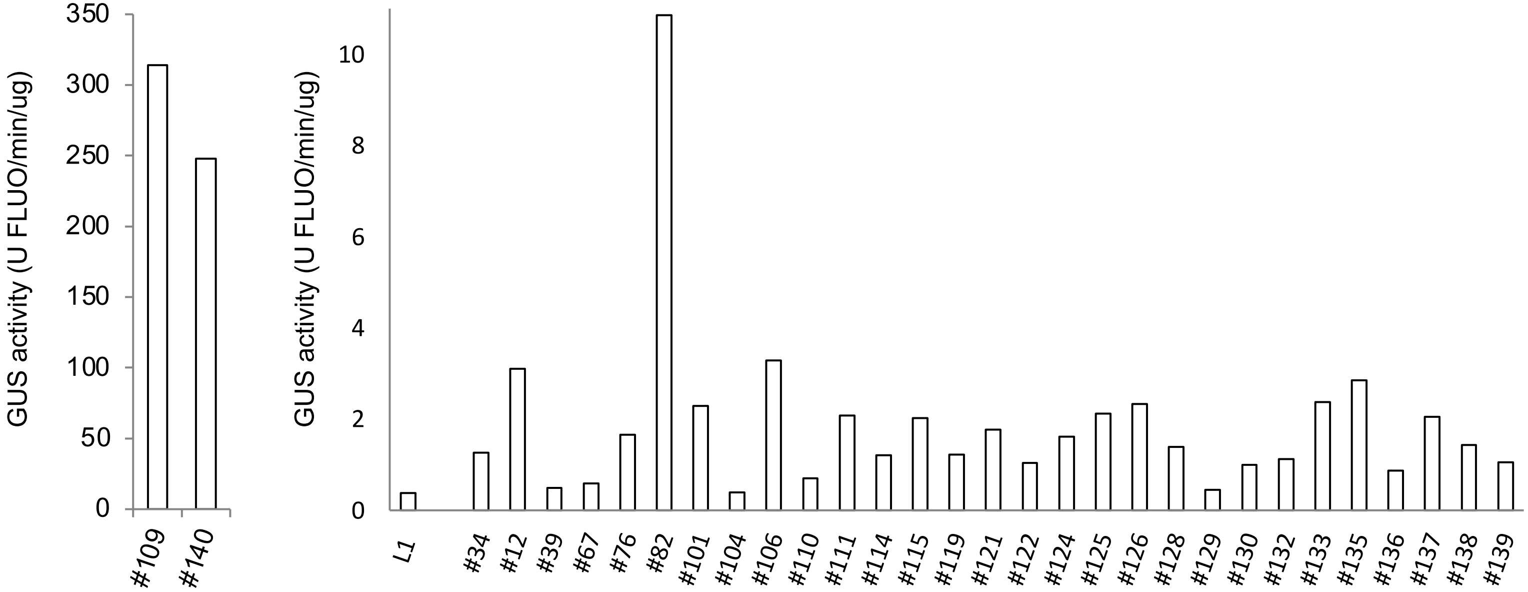

**Fig. S8. : *L1* S-PTGS is delayed in transgenic lines expressing an artificial miRNA targeting *CHR11* and *CHR17*.**

Kinetics of GUS activity at 17, 24 and 40 days in the progeny of *L1/ pUBQ10:amirCHR11-17* lines #125 and #135, which exhibit strong *CHR11/CHR17* silencing. Plants were harvested individually (means are indicated by a cross on box plot).

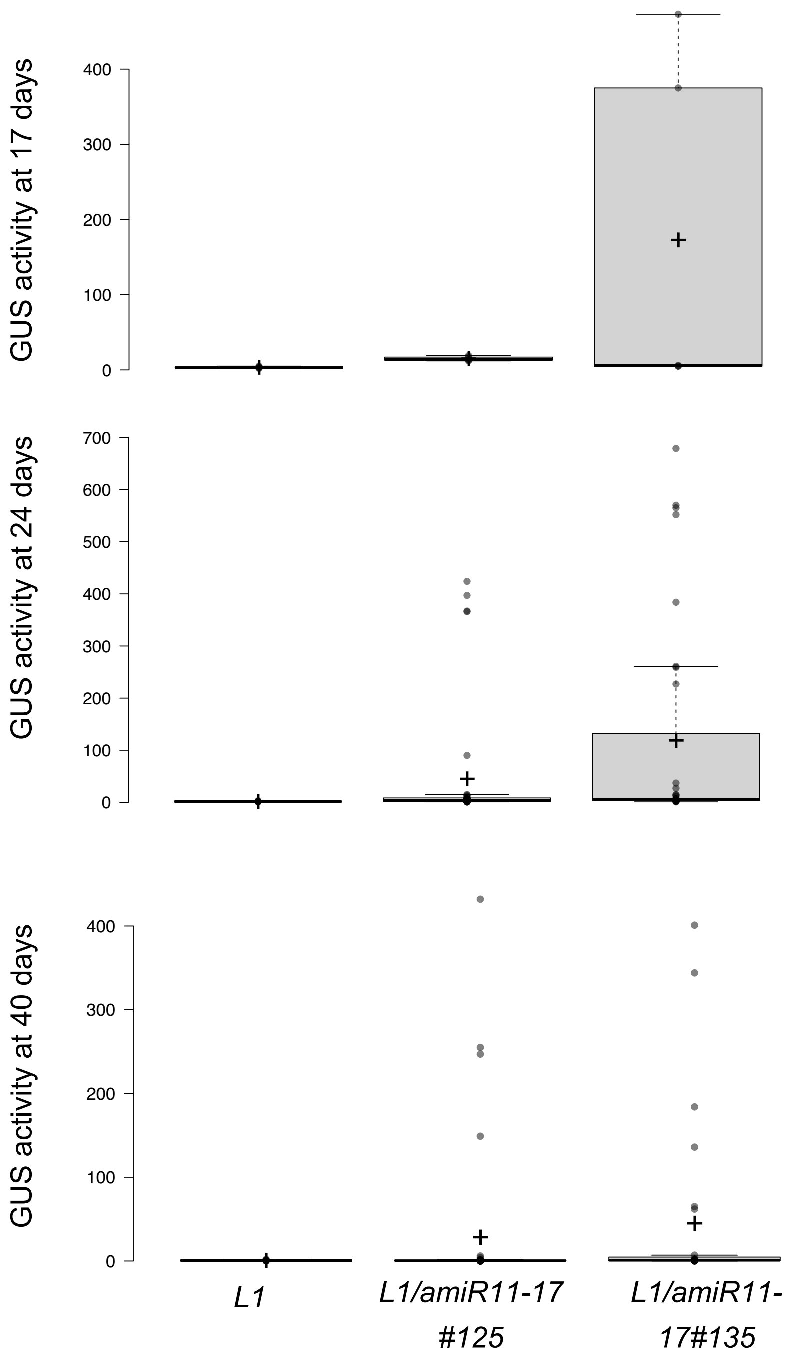

**Fig.S9. GUS activity is not affected in the *p35S:GUS* line *6b4/pUBQ10:amiRCHR11-17* transformants*.***

A. Severity of the phenotypes of 320 *6b4* primary transformants.

B. Photographs of four independent transformants exhibiting severe phenotypes.

C. Means of GUS activity in the 12 transformants exhibiting the most severe phenotype at 58 days in short days (including lines #3, #14, #56 and #59 shown in B).

D. Means of GUS activity in the 22 transformants exhibiting the most severe phenotype at 41 days in long days.

E. GUS activity at 19 days in progeny plants of *6b4/pUBQ10:amiRCHR11-17* line #6. Plants exhibiting a wild-type phenotype [wt] (#6.15), a *CHR11/CHR17* silencing phenotype [*chr11 chr17*] (#6.15*) were harvested individually (two to three plants for each genotype). Error bars correspond to standard deviation.

F. Accumulation of *amiRCHR11-17* at 19 days in plants analyzed in E.

A

Phenotypes of 320 6b4 primary transformants with *pUBQ10:amiRCHR11-17*

| | n T1 with mild or severe phenotype | % of T1 with mild or severe phenotype |
|---|---|---|
| 227 transfer in long days | 22 | 10% |
| 93 transfer in short days | 12 | 13% |

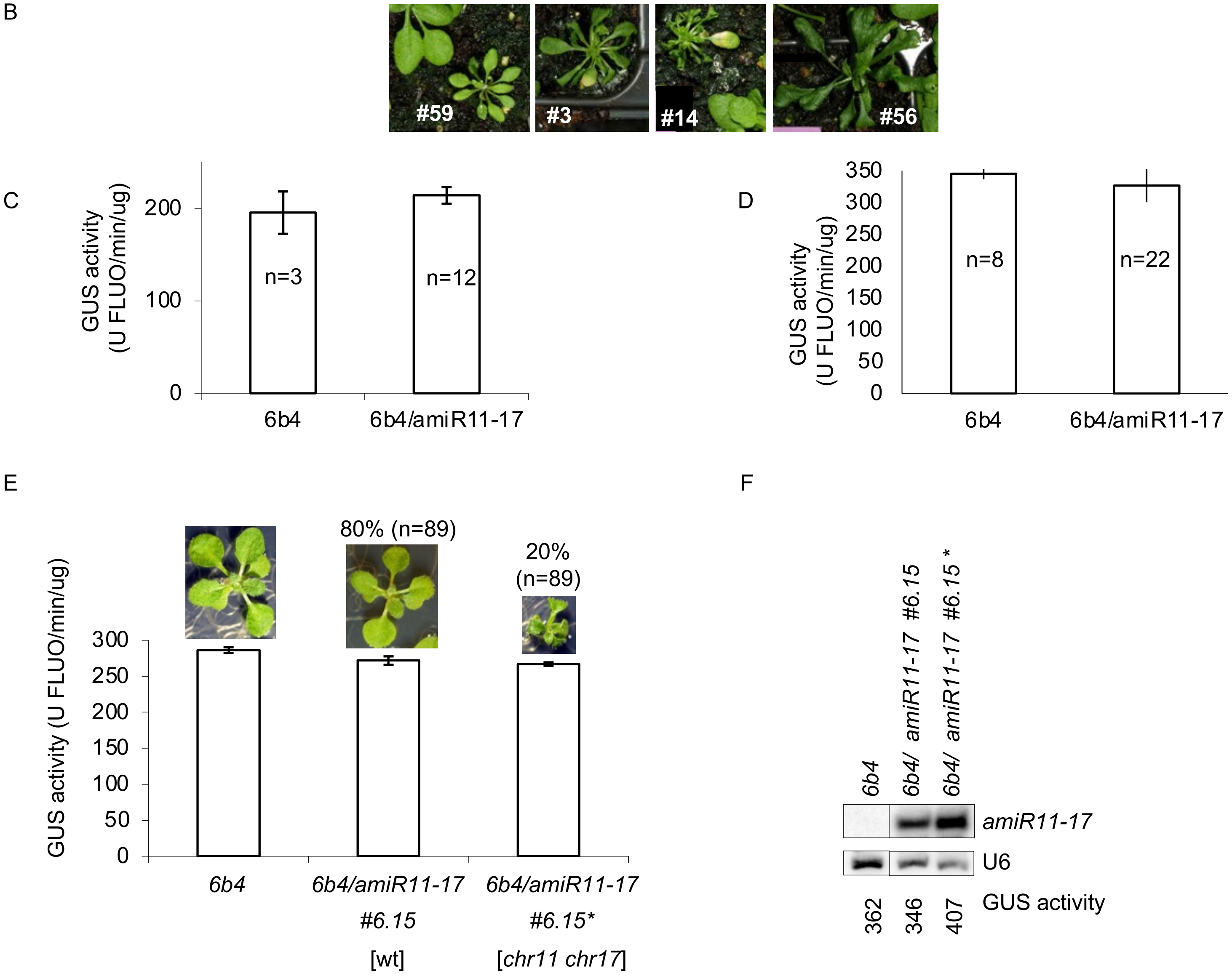

**Fig. S10. Complementation of the developmental defects of *chr11 chr17* by the *pUBQ10:GFP-CHR11* and *pUBQ10:FLAG-CHR11* constructs.**

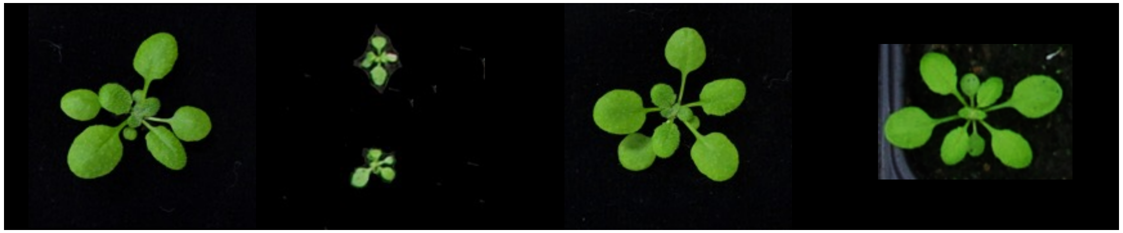

Col | *chr11 chr17* | *pUBQ10:FLAG-CHR11 chr11 chr17* | *pUBQ10:GFP-CHR11 chr11 chr17*

**Fig. S11. CHR11 interacts with *p35S:GUS* transgene at the *6b4* locus.**

ChIP-qPCR analyses were performed on 15-day-old seedlings of the indicated genotypes. ChIP was performed on the *6b4/pUBQ10:GFP-CHR11* line using GFP antibodies for IP. The *6b4* line was used as a control, followed by normalization to *GAPDH*. Graphical representation shows the fold change as the mean of four biological repeats. Error bars represent the standard deviation.

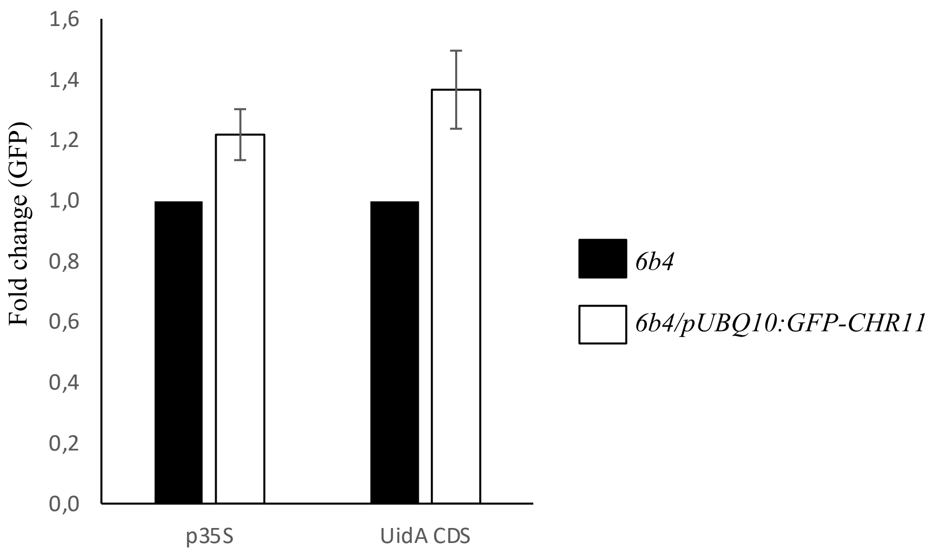

**Fig. S12. Overlap between endogenous PCGs producing va-siRNAs after, CMV, TuMV or TCV infection.**

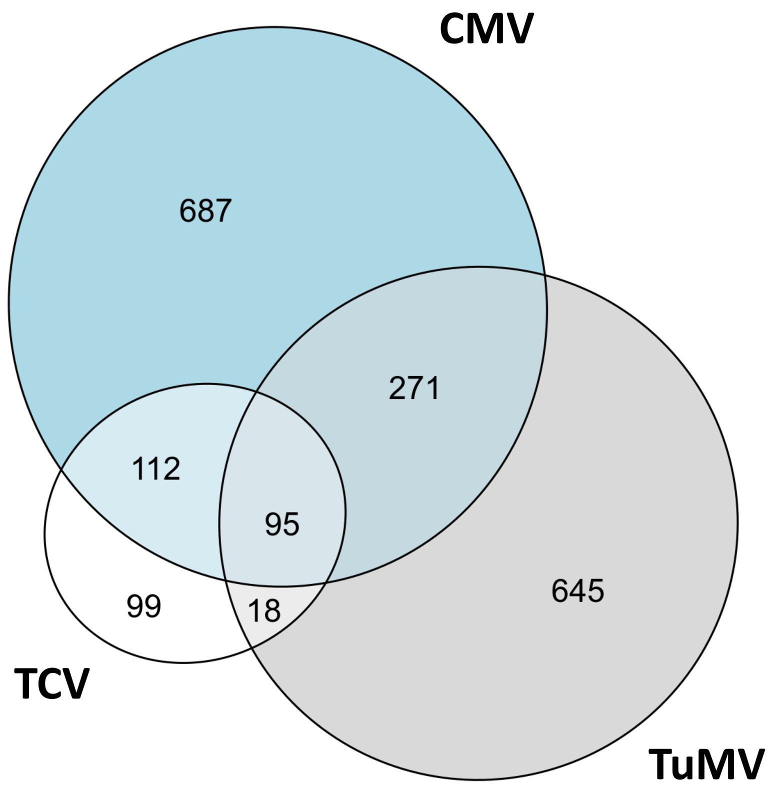

**Fig. S13. Reduced induction of va-siRNAs in *chr11 chr17* is not due to reduced transcription of the corresponding PCGs.**

Relation of the va-siRNAs induction upon TCV application in Col-0 and *chr11 chr17* double mutant according to the differential accumulation of their corresponding mRNA: either up regulated, unregulated or down regulated for all Arabidopsis genes (A) or TCV induced va-siRNA genes (B) Values are log2 fold changes according to DESeq2 differential analysis compared to Mock treatment for Col-0 (x axis) or *chr11 chr17* double mutant (y axis). Light grey points correspond to all genes while colors indicate the differentially upregulated genes (red, left), unregulated genes (dark grey center) and downregulated genes (blue, right).

A

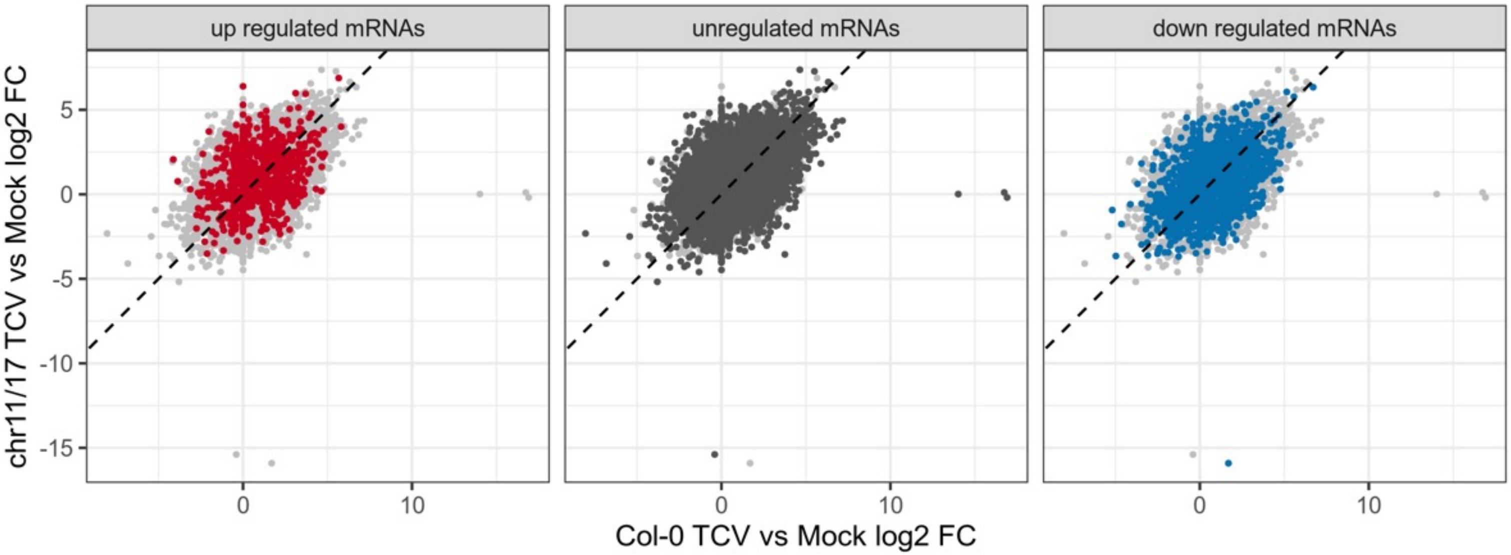


B

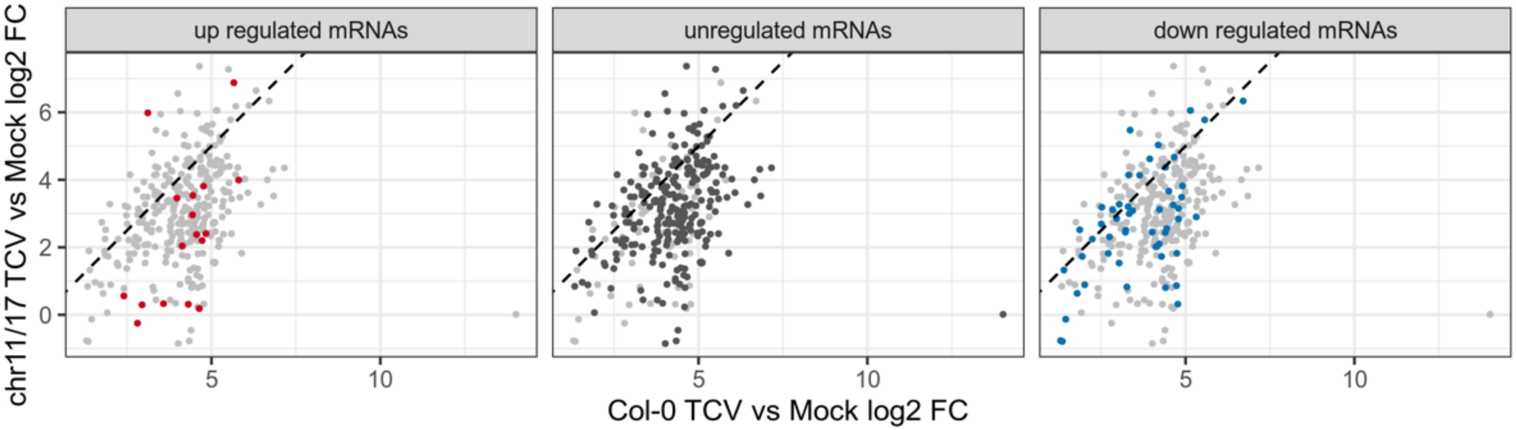

**Fig. S14. RDR6 impairment rescues the lethality of *ski3 xrn4 double* mutants.**

Pictures of the progeny of a *rdr6/rdr6 xrn4/xrn4 ski3/SKI3* plant

*A.* *rdr6/rdr6 xrn4/xrn4 ski3/SKI3* and *rdr6/rdr6 xrn4/xrn4 SKI3/SKI3* plants

*B.* *rdr6/rdr6 xrn4/xrn4 ski3/ski3* plants

A B

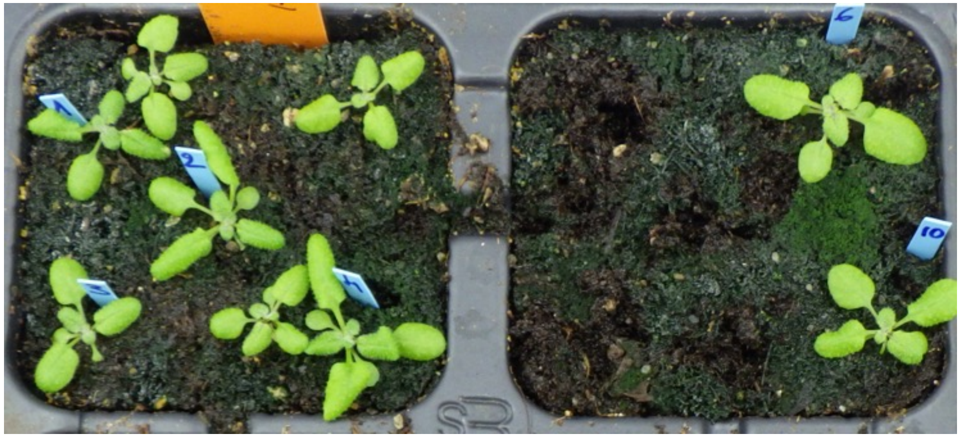

**Fig. S15.: Overlap between known PCG/NAT pairs producing nat-siRNAs and PCGs producing siRNAs when RQC is impaired by mutation or biotic stress.**

The combination matrix identifies the intersections, while the bars up to it encode the size of each intersection. Among the 84 PCG/NAT pairs , 48 generated siRNA in at least one of the other situations.

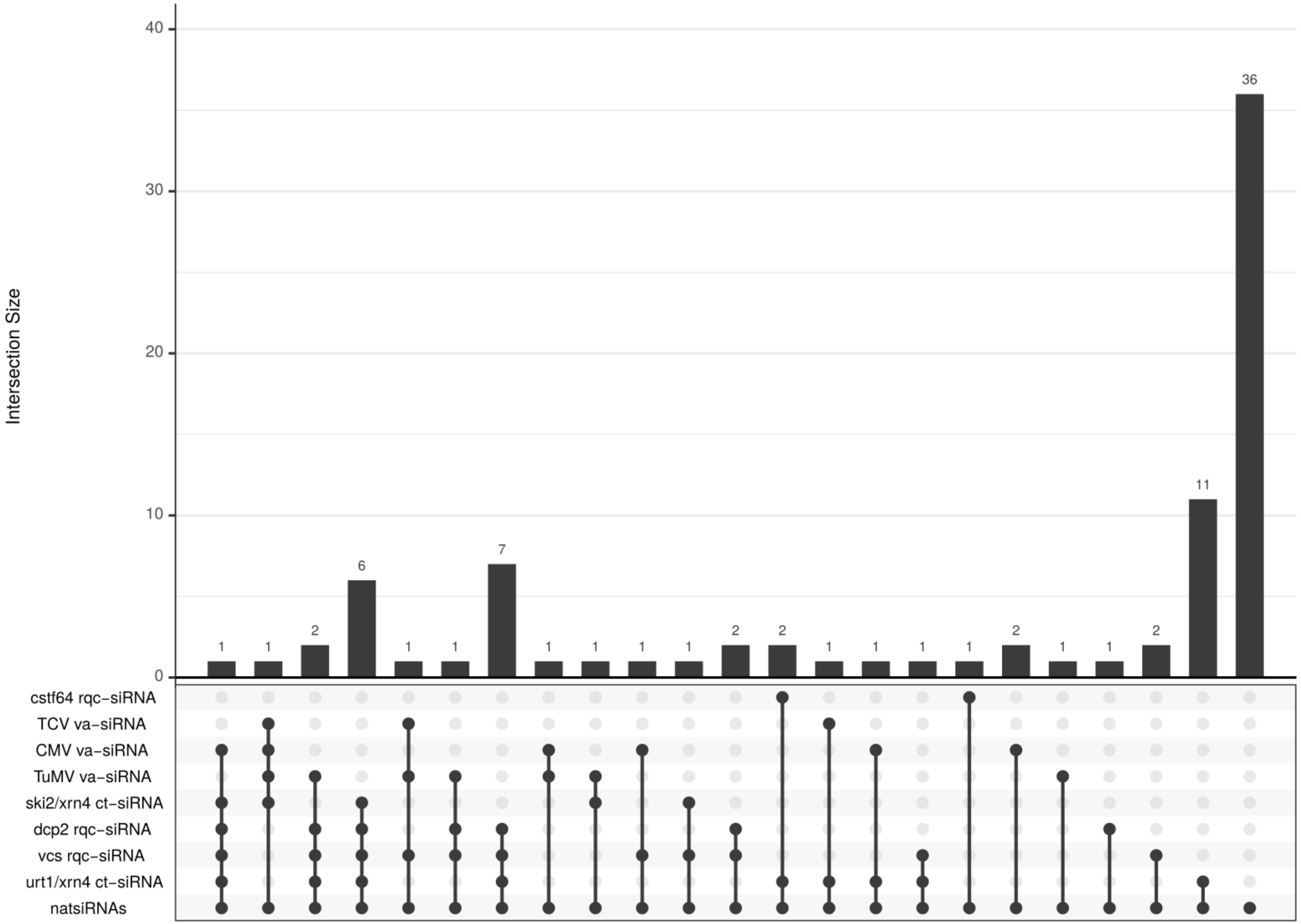

**Supplementary Table 1: CHR11 binding in siRNA-producing endogenous genes**

| | number of genes | number of genes considered | genes fixing CHR11 | genes not fixing CHR11 | % bound to CHR11 | median log2 FC in chr11/17 | median log2 FC in Col-0 | enrichment p-value |
|---|---|---|---|---|---|---|---|---|
| **A. All genes** | | | | | | | | |
| All genes | 32548 | 32548 | 15037 | 17511 | 46% | 0,88 | 0,43 | 0 |
| ta-siRNA | 7 | 7 | 0 | 7 | 0% | -1,43 | -1,54 | 9,87E-01 |
| dcp2 rqc-siRNA | 1351 | 1009 | 741 | 268 | 73% | 0,56 | 1,01 | 9,86E-72 |
| vcs rqc-siRNA | 1250 | 1170 | 845 | 325 | 72% | 0,73 | 1,12 | 7,10E-76 |
| ski2/xrn4 ct-siRNA | 441 | 415 | 301 | 114 | 73% | 0,04 | 0,70 | 2,03E-28 |
| urt1/xrn4 ct-siRNA | 2659 | 2623 | 1808 | 815 | 69% | 0,74 | 0,88 | 5,54E-133 |
| cstf64 rqc-siRNA | 1111 | 1075 | 844 | 231 | 79% | 1,32 | 1,25 | 1,95E-108 |
| nat-siRNA (gene pairs) | 84 | 84 | 64 | 20 | 76% | - | - | 5,10E-09 |
| nat-siRNA (individual genes) | 168 | 161 | 120 | 41 | 75% | 0,17 | 0,29 | 6,25E-14 |
| CMV va-siRNA | 1172 | 1167 | 993 | 174 | 85% | 1,62 | 1,88 | 3,85E-175 |
| TuMV va-siRNA | 1068 | 1042 | 735 | 307 | 71% | 1,31 | 1,45 | 3,84E-59 |
| TCV va-siRNA | 380 | 380 | 285 | 95 | 75% | 2,88 | 4,31 | 3,89E-31 |
| **B. PCGs** | | | | | | | | |
| All PCGs | 27445 | 27445 | 13811 | 13634 | 50% | 1,04 | 0,50 | 0 |
| dcp2 rqc-siRNA | 967 | 967 | 715 | 252 | 74% | 0,62 | 1,05 | 4,0E-53 |
| vcs rqc-siRNA | 1126 | 1126 | 820 | 306 | 73% | 0,77 | 1,17 | 4,1E-56 |
| ski2/xrn4 ct-siRNA | 413 | 413 | 299 | 114 | 72% | 0,04 | 0,69 | 1,1E-20 |
| urt1/xrn4 ct-siRNA | 2623 | 2623 | 1808 | 815 | 69% | 0,74 | 0,88 | 6,8E-92 |
| cstf64 rqc-siRNA | 1061 | 1061 | 839 | 222 | 79% | 1,33 | 1,26 | 4,4E-87 |
| nat-siRNA (gene pairs) | 82 | 82 | 61 | 21 | 74% | - | - | 2,4E-06 |
| nat-siRNA (individual genes) | 110 | 110 | 80 | 30 | 73% | 0,21 | 0,35 | 4,9E-07 |
| CMV va-siRNA | 1165 | 1165 | 991 | 174 | 85% | 1,63 | 1,88 | 2,0E-143 |
| TuMV va-siRNA | 1029 | 1029 | 726 | 303 | 71% | 1,33 | 1,50 | 1,4E-41 |
| TCV va-siRNA | 324 | 324 | 270 | 54 | 83% | 3,00 | 4,32 | 1,9E-36 |

**Supplementary Table 2: List of Primers used in this work.**

| name | purpose | sequence |
|---|---|---|
| SGS3Bam | SGS3 cDNA - amplification for cloning in plex10 | AT GGATCC AGT TCT AGG GCT GGt CCA AT |
| SGS3ccBam | SGS3 coiled-coil domain -amplification for cloning in plex10 | AC GGATCC AAG GTG CTT GAG GAA TCT |
| SGS3Sal | SGS3 cDNA - amplification for cloning in plex10 | GG GTCGAC TCA ATC ATC TTC ATT GTG AAG GCC |
| attB1SGS3f | SGS3 cDNA - amplification for cloning in pDONR207 | GGGG ACA AGT TTG TAC AAA AAA GCA GGC TTA ATG AGT TCT AGG GCT GGT C |
| attB2SGS3r | SGS3 cDNA - amplification for cloning in pDONR207 | GGG GAC CAC TTT GTA CAA GAA AGC TGG GTA ATC ATC TTC ATT GTG AAG GC |
| attB1SGS3KpnIf | SGS3 CDS plus introns plus KpnI- amplification for cloning in pDONR207 | GGGG ACA AGT TTG TAC AAA AAA GCA GGC TTA GGT ACC ATG AGT TCT AGG GCT GGT C |
| attB2SGS3SpeIR | SGS3 CDS plus introns plus SpeI- amplification for cloning in pDONR207 | GGGG AC CAC TTT GTA CAA GAA AGC TGG GTA ACT AGT ATC ATC TTC ATT GTG AAG GC |
| RDR6f | RDR6 cDNA - amplification for cloning in SalI / NotI in pENTR1A | GGCGTCGACATGGGGTCAGAGGGAAATATG |
| RDR6r | RDR6 cDNA - amplification for cloning in SalI / NotI in pENTR1A | GGCGCGGCCGCAAGGATCCGAGACGCTGAGCAAGAAAC |
| attB1CHR11Fbis | CHR11 cDNA - amplification for cloning in pDONR207 | GGGG ACA AGT TTG TAC AAA AAA GCA GGC TCC ATG GCG AGA AAT TCG AA |
| attB2CHR11Rbis | CHR11 cDNA - amplification for cloning in pDONR207 | GGGGACCACTTTGTACAAGAAAGCTGGGT TCA TCT CAT CGA CAG GTG CTT |
| attB1FlagCHR11F | Flag plus CHR11 cDNA- amplification for cloning in pDONR207 | GGGGACAAGTTTGTACAAAAAAGCAGGCTCC ATG GAC TAC AAG GAT GAC GAT GAC AAG GGC GCG AGA AAT TCG AAT TCC GAT GAG GC |
| attB2CHR11Rbis | Flag plus CHR11 cDNA- amplification for cloning in pDONR207 | GGGGACCACTTTGTACAAGAAAGCTGGGT TCA TCT CAT CGA CAG GTG CTT |
| CHR11stop1Fbis | CHR11 mutagenesis | GGTAAGATATGGTGCTTAGATGGTGTTCAGTTC |
| CHR11stop1Rbis | CHR11 mutagenesis | GAACTGAACACCATCTAAGCACCATATCTTACC |
| CHR11stop2F | CHR11 mutagenesis | AAA GAG CGG TAC AAG TAG CTG AAC GAC TAT G |
| CHR11stop2R | CHR11 mutagenesis | CAT AGT CGT TCA GCT ACT TGT ACC GCT CTT T |
| proSGS2F-hIII | pRDR6 amplification for cloning in pGWB4 HindIII | AAGCTTGCAAACCTGTCATATCTGTGG |
| proSGS2R-HIII | pRDR6 amplification for cloning in pGWB4 HindIII | AAGCTTTTCTCCTGAAAAAGAAACATAC |
| amiRf | amiR-CHR11-17 - amplification for cloning in pDONOR207 | GGGG ACA AGT TTG TAC AAA AAA GCA GGC T CTGCAGCCCcaaacacacgc |
| amiRr | amiR-CHR11-17 - amplification for cloning in pDONOR207 | GGGG AC CAC TTT GTA CAA GAA AGC TGG GT ATCCCCCCATGGCGATGCCTT |
| amiRCHR11 | amiRCHR11 oligoprobe for northern | TAGTGCGACCGTTTCATGATA |
| 35SLT60rev | p35S (-240bp to -50bp from +1 transcription site) -qPCR amplification for ChIP | AAGGATAGTGGGATTGTGCG |
| 35S1LT58fwd | p35S (-240bp to -50bp from +1 transcription site) -qPCR amplification for ChIP | CTACAAATGCCATCATTGCG |
| GUS1LT58fwd | UidA (230bp downstram ATG= 5' GUS) - qPCR amplification for ChIP and RIP | TCCTGTAGAAACCCCAACCC |
| GUS1LT58rev | UidA (230bp downstram ATG= 5' GUS) - qPCR amplification for ChIP and RIP | TGGCCTGCCCAACCTTT |
| GUS-central-fw | UidA (central GUS) - qPCR amplification for RIP | ctgctgtcggctttaacctc |
| GUS-central-rev | UidA (central GUS) - qPCR amplification for RIP | tgagcgtcgcagaacattac |
| At1g13440-F5 | GAPDH (180bp amplified from 950bp downstream ATG) -qPCR amplification | GGTACGACAACGAATGGGGT |
| At1g13440-R5 | GAPDH (180bp amplified from 950bp downstream ATG) -qPCR amplification | TGACTGCGCATGGAATCAGT |
| CHR11F1 | CHR11 RT-qPCR amplification | ATGTGTTTACGGATCTGTCG |
| CHR11R1 | CHR11 RT-qPCR amplification | CTAAGTTGCCATCCAATGGC |
| CHR17F | CHR17 RT-qPCR amplification | CGGGAAAGCGTAGGAAATAAG |
| CHR17R | CHR17 RT-qPCR amplification | ACGCACTACTAGGAATGTGG |
| At3g45980-F1 | H2B gene - amplification for cloning | GGGGACAAGTTTGTACAAAAAAGCAGGCTTCCAATTCGCATTTTTTCAAACTA |
| At3g45980-R1 | H2B gene - amplification for cloning | GGGGACCACTTTGTACAAGAAAGCTGGGTCAGAGCTTGTGAATTTGGTAAC |
| At1g24310-F1 | NUP54 gene - amplification for cloning | GGGGACAAGTTTGTACAAAAAAGCAGGCTGGTAATGACATAATAACTCTAAAAG |
| At1g24310-R1 | NUP54 gene - amplification for cloning | GGGGACCACTTTGTACAAGAAAGCTGGGTCAGAGCTTGTGAATTTGGTAAC |
