## Supplementary material for "The Arabidopsis RNA-binding protein SGS3 is recruited to the chromatin remodeler CHR11 to promote siRNA production from protein-coding genes": dataset1

**Dataset 1: Differential analysis of 21/22nt accumulation on Araport1 genes**

Dataset 1.1 Differential analysis of 21/22nt accumulation on Araport11 genes in uninfecte
Dataset 1.2 Differential analysis of 21/22nt accumulation on Araport11 genes in wild type
Dataset 1.3 Differential analysis of 21/22nt accumulation on Araport11 genes in *chr11 ch*
Dataset 1.4 Differential analysis of 21/22nt accumulation on Araport11 genes in *chr11 ch*
Dataset 1.5 Differential analysis of 21/22nt accumulation on Araport11 genes in *chr11 ch*
Dataset 1.6 Differential analysis of 21/22nt accumulation on Araport11 genes in *chr11 ch*

ed *chr11 chr17* relative to wild type.
infected by TCV relative to uninfected wild type.
*r17* infected by TCV relative to uninfected *chr11 chr17*.
*r17* infected by TCV relative to infected wild type.
*r17* infected by TCV relative to uninfected wild type.
*r17* uninfected by TCV relative to infected wild type.

**Dataset 1.1 Differential analysis of 21/22nt accumulation on Araport11 genes in u**

| ID | baseMean | log2FoldChange | lfcSE | stat | pvalue | padj |
|---|---|---|---|---|---|---|
| AT5G23065 | 1776,68 | -3,59 | 0,26 | -13,63 | 2,72E-42 | 5,62E-39 |
| AT5G14545 | 7868,55 | 15,69 | 1,20 | 13,13 | 2,25E-39 | 2,32E-36 |
| AT5G14550 | 7872,48 | 14,73 | 1,20 | 12,31 | 7,68E-35 | 5,29E-32 |
| AT3G44570 | 48,95 | -5,36 | 0,51 | -10,45 | 1,50E-25 | 7,75E-23 |
| AT3G44580 | 31,33 | -4,05 | 0,42 | -9,65 | 5,15E-22 | 2,13E-19 |
| AT4G23713 | 128,07 | 3,45 | 0,36 | 9,62 | 6,36E-22 | 2,19E-19 |
| AT4G23387 | 136,04 | 4,66 | 0,49 | 9,51 | 1,95E-21 | 5,77E-19 |
| AT4G03039 | 93,29 | 8,78 | 1,18 | 7,44 | 1,01E-13 | 2,17E-11 |
| AT4G03038 | 93,33 | 8,78 | 1,18 | 7,44 | 1,02E-13 | 2,17E-11 |
| AT4G04365 | 93,18 | 8,77 | 1,18 | 7,43 | 1,05E-13 | 2,17E-11 |
| AT3G51375 | 135,65 | -2,58 | 0,40 | -6,38 | 1,73E-10 | 3,24E-08 |
| AT2G01020 | 7030,11 | -2,23 | 0,35 | -6,32 | 2,70E-10 | 4,64E-08 |
| AT4G03455 | 16,01 | -8,21 | 1,31 | -6,25 | 3,99E-10 | 5,90E-08 |
| AT4G12770 | 19,67 | 4,69 | 0,75 | 6,26 | 3,85E-10 | 5,90E-08 |
| AT4G04565 | 64,98 | 2,38 | 0,38 | 6,24 | 4,32E-10 | 5,95E-08 |
| AT2G16030 | 25,88 | 3,94 | 0,63 | 6,21 | 5,21E-10 | 6,33E-08 |
| AT5G04275 | 20,42 | 7,69 | 1,24 | 6,22 | 4,93E-10 | 6,33E-08 |
| AT1G66725 | 1003,08 | 1,88 | 0,31 | 6,14 | 8,22E-10 | 9,44E-08 |
| AT4G21595 | 28,61 | 7,60 | 1,24 | 6,13 | 8,82E-10 | 9,59E-08 |
| AT5G43603 | 44,77 | 7,48 | 1,24 | 6,06 | 1,39E-09 | 1,44E-07 |
| AT5G14565 | 6609,25 | -1,56 | 0,26 | -5,91 | 3,44E-09 | 3,38E-07 |
| AT5G38540 | 8,36 | -5,02 | 0,87 | -5,79 | 6,92E-09 | 6,50E-07 |
| AT3G14210 | 12,80 | -3,52 | 0,61 | -5,75 | 8,94E-09 | 8,03E-07 |
| AT2G16586 | 20,98 | 3,75 | 0,65 | 5,74 | 9,47E-09 | 8,15E-07 |
| AT2G06555 | 20,57 | 3,51 | 0,61 | 5,71 | 1,14E-08 | 9,45E-07 |
| AT1G53683 | 15,44 | -7,30 | 1,30 | -5,61 | 1,99E-08 | 1,58E-06 |
| AT3G55734 | 132,05 | 1,71 | 0,31 | 5,55 | 2,82E-08 | 2,16E-06 |
| AT2G40805 | 280,87 | 2,22 | 0,40 | 5,53 | 3,23E-08 | 2,38E-06 |
| AT3G59884 | 100,21 | 2,38 | 0,43 | 5,49 | 4,05E-08 | 2,89E-06 |
| AT5G66045 | 69,41 | 6,92 | 1,27 | 5,45 | 5,11E-08 | 3,52E-06 |
| AT5G08717 | 14,65 | 7,02 | 1,33 | 5,26 | 1,44E-07 | 9,01E-06 |
| AT5G53740 | 20,74 | -2,55 | 0,48 | -5,27 | 1,40E-07 | 9,01E-06 |
| AT4G03445 | 45,78 | 1,62 | 0,31 | 5,27 | 1,36E-07 | 9,01E-06 |
| AT4G08691 | 15,19 | 6,58 | 1,26 | 5,23 | 1,66E-07 | 1,01E-05 |
| AT5G46845 | 66,07 | -1,42 | 0,27 | -5,19 | 2,11E-07 | 1,25E-05 |
| AT5G27606 | 14,40 | 5,01 | 0,98 | 5,11 | 3,14E-07 | 1,72E-05 |
| AT2G35940 | 16,85 | 3,52 | 0,69 | 5,11 | 3,17E-07 | 1,72E-05 |
| AT4G08755 | 25,91 | 6,34 | 1,24 | 5,12 | 3,13E-07 | 1,72E-05 |
| AT4G06075 | 405,29 | 1,63 | 0,32 | 5,06 | 4,29E-07 | 2,27E-05 |
| AT2G41310 | 487,34 | -1,38 | 0,27 | -5,05 | 4,50E-07 | 2,33E-05 |
| AT2G09250 | 487,12 | -1,38 | 0,27 | -5,04 | 4,62E-07 | 2,33E-05 |
| AT5G63715 | 27,38 | -2,08 | 0,41 | -5,01 | 5,54E-07 | 2,73E-05 |
| AT1G07051 | 29,68 | -2,72 | 0,54 | -5,00 | 5,71E-07 | 2,75E-05 |
| AT2G35945 | 15,59 | 3,41 | 0,70 | 4,91 | 9,18E-07 | 4,31E-05 |
| AT5G54075 | 15,55 | 5,26 | 1,09 | 4,83 | 1,36E-06 | 6,26E-05 |
| AT4G09135 | 26,82 | 6,04 | 1,25 | 4,82 | 1,47E-06 | 6,45E-05 |
| AT5G54070 | 15,72 | 5,27 | 1,09 | 4,82 | 1,44E-06 | 6,45E-05 |
| AT4G03580 | 13,67 | 4,16 | 0,87 | 4,77 | 1,82E-06 | 7,82E-05 |
| AT1G01183 | 317,62 | -1,66 | 0,35 | -4,76 | 1,96E-06 | 8,26E-05 |
| AT3G01845 | 219,09 | 0,92 | 0,20 | 4,65 | 3,30E-06 | 1,36E-04 |
| AT2G34430 | 28,33 | -1,94 | 0,42 | -4,64 | 3,56E-06 | 1,44E-04 |
| AT5G04455 | 10,87 | 5,92 | 1,28 | 4,61 | 4,11E-06 | 1,60E-04 |
| AT5G04445 | 10,87 | 5,92 | 1,28 | 4,61 | 4,11E-06 | 1,60E-04 |

| | | | | | | |
|---|---|---|---|---|---|---|
| AT5G52310 | 5,67 | -4,52 | 1,00 | -4,51 | 6,56E-06 | 2,51E-04 |
| AT4G00150 | 10,48 | 3,71 | 0,83 | 4,50 | 6,79E-06 | 2,55E-04 |
| AT3G08370 | 6,58 | 6,08 | 1,39 | 4,37 | 1,26E-05 | 4,64E-04 |
| AT3G43833 | 7,44 | 6,10 | 1,41 | 4,34 | 1,41E-05 | 5,12E-04 |
| AT1G51150 | 4,36 | -6,22 | 1,44 | -4,33 | 1,50E-05 | 5,36E-04 |
| AT4G07875 | 11,03 | 5,65 | 1,31 | 4,30 | 1,72E-05 | 6,04E-04 |
| AT5G43740 | 103,64 | -1,79 | 0,42 | -4,28 | 1,90E-05 | 6,56E-04 |
| AT4G00885 | 370,66 | -1,21 | 0,28 | -4,26 | 2,05E-05 | 6,96E-04 |
| AT2G16018 | 9,68 | 3,92 | 0,92 | 4,24 | 2,27E-05 | 7,56E-04 |
| AT4G23375 | 8,41 | 4,84 | 1,16 | 4,18 | 2,92E-05 | 9,57E-04 |
| AT5G10946 | 91,61 | -1,02 | 0,25 | -4,14 | 3,52E-05 | 1,14E-03 |
| AT5G10945 | 91,45 | -1,02 | 0,25 | -4,10 | 4,12E-05 | 1,31E-03 |
| AT3G02020 | 19,73 | 2,10 | 0,51 | 4,09 | 4,29E-05 | 1,34E-03 |
| AT5G41663 | 49,62 | 1,84 | 0,45 | 4,08 | 4,56E-05 | 1,41E-03 |
| AT2G05070 | 19,17 | -2,00 | 0,49 | -4,04 | 5,28E-05 | 1,61E-03 |
| AT4G08165 | 5,10 | 6,15 | 1,53 | 4,03 | 5,53E-05 | 1,66E-03 |
| AT2G39675 | 2246,76 | -1,36 | 0,34 | -4,02 | 5,73E-05 | 1,69E-03 |
| AT2G46685 | 3232,16 | -1,04 | 0,26 | -3,95 | 7,89E-05 | 2,30E-03 |
| AT4G08175 | 7,31 | -3,33 | 0,86 | -3,89 | 1,01E-04 | 2,90E-03 |
| AT4G06130 | 392,23 | -1,58 | 0,41 | -3,87 | 1,11E-04 | 3,13E-03 |
| AT1G80180 | 13,61 | 2,38 | 0,62 | 3,86 | 1,15E-04 | 3,22E-03 |
| AT1G12294 | 124,95 | -1,05 | 0,27 | -3,85 | 1,18E-04 | 3,25E-03 |
| AT1G48267 | 15177,13 | -1,28 | 0,34 | -3,82 | 1,35E-04 | 3,68E-03 |
| AT4G13575 | 426,34 | -1,51 | 0,40 | -3,81 | 1,40E-04 | 3,76E-03 |
| AT2G06420 | 4,36 | 5,55 | 1,46 | 3,79 | 1,48E-04 | 3,93E-03 |
| AT5G41905 | 1173,00 | -0,95 | 0,25 | -3,75 | 1,75E-04 | 4,58E-03 |
| AT2G45160 | 10,70 | -2,03 | 0,54 | -3,74 | 1,83E-04 | 4,73E-03 |
| AT2G01021 | 3194,91 | -1,00 | 0,27 | -3,74 | 1,87E-04 | 4,77E-03 |
| AT5G04853 | 3,91 | 5,29 | 1,42 | 3,71 | 2,04E-04 | 5,15E-03 |
| AT5G26650 | 7,57 | 4,56 | 1,23 | 3,70 | 2,12E-04 | 5,28E-03 |
| AT1G11700 | 23,46 | 1,61 | 0,44 | 3,68 | 2,34E-04 | 5,76E-03 |
| AT4G27600 | 12,18 | 4,02 | 1,10 | 3,64 | 2,68E-04 | 6,51E-03 |
| AT4G12780 | 15,31 | 2,06 | 0,57 | 3,64 | 2,76E-04 | 6,63E-03 |
| AT5G41765 | 5,08 | 4,98 | 1,37 | 3,63 | 2,81E-04 | 6,68E-03 |
| AT3G63375 | 13,10 | -1,86 | 0,52 | -3,57 | 3,57E-04 | 8,33E-03 |
| AT5G53902 | 6,85 | 4,94 | 1,38 | 3,57 | 3,62E-04 | 8,33E-03 |
| AT5G53905 | 6,85 | 4,94 | 1,38 | 3,57 | 3,62E-04 | 8,33E-03 |
| AT5G07570 | 4,01 | 5,49 | 1,56 | 3,52 | 4,31E-04 | 9,69E-03 |
| AT2G10606 | 1219,01 | -0,98 | 0,28 | -3,52 | 4,27E-04 | 9,69E-03 |
| AT5G06640 | 3,20 | 5,39 | 1,53 | 3,51 | 4,40E-04 | 9,77E-03 |
| AT1G06247 | 2,53 | -5,38 | 1,55 | -3,47 | 5,23E-04 | 1,15E-02 |
| AT5G21100 | 6,61 | 3,21 | 0,94 | 3,41 | 6,53E-04 | 1,40E-02 |
| AT1G29910 | 27,17 | -1,66 | 0,49 | -3,41 | 6,47E-04 | 1,40E-02 |
| AT2G24980 | 3,78 | 5,14 | 1,52 | 3,39 | 7,03E-04 | 1,50E-02 |
| AT3G01835 | 72,10 | 0,96 | 0,28 | 3,38 | 7,23E-04 | 1,53E-02 |
| AT2G04605 | 3,46 | 5,15 | 1,53 | 3,37 | 7,49E-04 | 1,56E-02 |
| AT3G61198 | 7,65 | 3,45 | 1,03 | 3,35 | 7,99E-04 | 1,65E-02 |
| AT1G07977 | 4,37 | -3,48 | 1,05 | -3,31 | 9,47E-04 | 1,92E-02 |
| AT1G13360 | 23,38 | 1,50 | 0,45 | 3,31 | 9,48E-04 | 1,92E-02 |
| AT3G19380 | 2,79 | 4,95 | 1,50 | 3,30 | 9,77E-04 | 1,96E-02 |
| AT3G48720 | 18,61 | 1,68 | 0,51 | 3,28 | 1,05E-03 | 2,07E-02 |
| AT3G10815 | 5,65 | 4,52 | 1,38 | 3,28 | 1,05E-03 | 2,07E-02 |
| AT4G11373 | 5,92 | 3,03 | 0,93 | 3,27 | 1,08E-03 | 2,10E-02 |
| AT5G03305 | 5,94 | 3,11 | 0,95 | 3,26 | 1,13E-03 | 2,16E-02 |
| AT3G01313 | 12,85 | 2,05 | 0,63 | 3,26 | 1,12E-03 | 2,16E-02 |
| AT1G78476 | 11,35 | 1,87 | 0,57 | 3,25 | 1,17E-03 | 2,22E-02 |

| | | | | | | |
|---|---|---|---|---|---|---|
| AT1G78478 | 10,59 | 1,90 | 0,59 | 3,23 | 1,24E-03 | 2,34E-02 |
| AT1G50440 | 6,47 | 3,17 | 0,98 | 3,22 | 1,28E-03 | 2,37E-02 |
| AT5G06320 | 5,00 | 4,86 | 1,51 | 3,22 | 1,29E-03 | 2,37E-02 |
| AT1G09817 | 10,25 | 1,95 | 0,61 | 3,20 | 1,38E-03 | 2,53E-02 |
| AT5G09505 | 92,76 | -0,93 | 0,29 | -3,19 | 1,40E-03 | 2,54E-02 |
| AT5G41750 | 8,05 | 3,51 | 1,11 | 3,18 | 1,48E-03 | 2,66E-02 |
| AT2G16020 | 3,14 | 4,83 | 1,52 | 3,17 | 1,52E-03 | 2,70E-02 |
| AT1G06273 | 2,17 | -5,59 | 1,77 | -3,15 | 1,64E-03 | 2,88E-02 |
| AT1G04907 | 10,89 | 4,26 | 1,35 | 3,15 | 1,65E-03 | 2,88E-02 |
| AT1G69797 | 12,98 | 1,91 | 0,61 | 3,14 | 1,69E-03 | 2,94E-02 |
| AT4G24415 | 10,68 | -1,66 | 0,53 | -3,13 | 1,72E-03 | 2,96E-02 |
| AT1G62930 | 53,37 | -1,08 | 0,35 | -3,11 | 1,88E-03 | 3,21E-02 |
| AT3G06345 | 8,24 | 2,38 | 0,77 | 3,10 | 1,92E-03 | 3,25E-02 |
| AT3G47350 | 23,89 | 1,86 | 0,60 | 3,10 | 1,96E-03 | 3,27E-02 |
| AT1G69792 | 2,23 | -5,61 | 1,81 | -3,10 | 1,95E-03 | 3,27E-02 |
| AT4G08335 | 4,06 | 4,48 | 1,45 | 3,08 | 2,07E-03 | 3,42E-02 |
| AT5G56070 | 3,26 | 5,03 | 1,64 | 3,06 | 2,18E-03 | 3,58E-02 |
| AT5G03552 | 58,24 | -0,92 | 0,30 | -3,05 | 2,29E-03 | 3,72E-02 |
| AT1G07633 | 6,19 | 3,06 | 1,00 | 3,04 | 2,33E-03 | 3,74E-02 |
| AT2G42540 | 2,58 | -4,90 | 1,61 | -3,05 | 2,32E-03 | 3,74E-02 |
| AT5G03553 | 58,75 | -0,92 | 0,30 | -3,04 | 2,40E-03 | 3,81E-02 |
| AT4G04570 | 4,35 | -3,38 | 1,11 | -3,03 | 2,45E-03 | 3,87E-02 |
| AT1G61520 | 19,44 | -1,55 | 0,51 | -3,02 | 2,51E-03 | 3,93E-02 |
| AT1G47497 | 2,25 | -5,45 | 1,81 | -3,01 | 2,58E-03 | 4,01E-02 |
| AT4G13495 | 869,17 | 1,00 | 0,33 | 3,01 | 2,61E-03 | 4,03E-02 |
| AT3G12120 | 5,50 | -3,12 | 1,04 | -3,00 | 2,69E-03 | 4,11E-02 |
| AT3G59940 | 11,46 | 1,71 | 0,57 | 2,98 | 2,85E-03 | 4,30E-02 |
| AT5G45307 | 438,83 | -0,94 | 0,31 | -2,98 | 2,85E-03 | 4,30E-02 |
| AT1G12290 | 131,54 | -0,79 | 0,26 | -2,98 | 2,88E-03 | 4,31E-02 |
| AT1G09070 | 2,87 | 4,59 | 1,55 | 2,96 | 3,10E-03 | 4,61E-02 |
| AT3G12630 | 8,51 | 2,46 | 0,84 | 2,94 | 3,26E-03 | 4,78E-02 |
| AT2G21330 | 7,94 | -2,10 | 0,71 | -2,94 | 3,24E-03 | 4,78E-02 |
| AT3G09275 | 11,41 | 1,70 | 0,58 | 2,94 | 3,30E-03 | 4,80E-02 |
| AT3G54590 | 2,31 | 4,94 | 1,68 | 2,93 | 3,35E-03 | 4,81E-02 |
| AT3G55512 | 2,56 | 5,23 | 1,78 | 2,93 | 3,35E-03 | 4,81E-02 |
| AT5G66570 | 20,06 | -1,70 | 0,58 | -2,92 | 3,48E-03 | 4,97E-02 |
| AT1G05780 | 4,77 | 3,27 | 1,12 | 2,91 | 3,56E-03 | 5,05E-02 |
| AT3G45780 | 10,09 | -1,67 | 0,57 | -2,91 | 3,65E-03 | 5,14E-02 |
| AT1G09310 | 8,77 | -1,83 | 0,63 | -2,90 | 3,79E-03 | 5,27E-02 |
| AT5G41740 | 6,05 | 4,22 | 1,46 | 2,89 | 3,80E-03 | 5,27E-02 |
| AT1G48410 | 43,79 | 1,14 | 0,40 | 2,89 | 3,83E-03 | 5,27E-02 |
| AT5G21940 | 4,72 | 3,03 | 1,05 | 2,88 | 3,96E-03 | 5,42E-02 |
| AT1G63070 | 3,60 | -3,15 | 1,09 | -2,88 | 3,98E-03 | 5,42E-02 |
| AT5G08925 | 2,86 | 5,07 | 1,76 | 2,88 | 4,01E-03 | 5,42E-02 |
| AT2G08665 | 53,97 | -1,15 | 0,40 | -2,86 | 4,24E-03 | 5,59E-02 |
| AT3G61470 | 27,14 | -1,28 | 0,45 | -2,86 | 4,23E-03 | 5,59E-02 |
| AT2G34420 | 53,97 | -1,15 | 0,40 | -2,86 | 4,24E-03 | 5,59E-02 |
| AT4G22410 | 2,93 | 4,22 | 1,47 | 2,86 | 4,24E-03 | 5,59E-02 |
| AT2G38325 | 156,77 | -0,67 | 0,23 | -2,85 | 4,37E-03 | 5,72E-02 |
| AT1G66783 | 14,56 | 2,08 | 0,73 | 2,84 | 4,45E-03 | 5,79E-02 |
| AT4G09735 | 2,23 | 4,87 | 1,71 | 2,84 | 4,48E-03 | 5,79E-02 |
| AT4G00140 | 2,38 | 4,66 | 1,64 | 2,84 | 4,54E-03 | 5,83E-02 |
| AT2G02840 | 4,55 | 4,27 | 1,51 | 2,83 | 4,69E-03 | 5,98E-02 |
| AT1G29930 | 124,68 | -1,09 | 0,39 | -2,80 | 5,15E-03 | 6,53E-02 |
| AT5G05505 | 4,00 | -2,84 | 1,02 | -2,79 | 5,24E-03 | 6,61E-02 |
| AT2G03445 | 309,93 | 0,73 | 0,26 | 2,79 | 5,28E-03 | 6,62E-02 |

| | | | | | | |
|---|---|---|---|---|---|---|
| AT1G10522 | 24,13 | 1,27 | 0,46 | 2,78 | 5,37E-03 | 6,69E-02 |
| AT5G10380 | 7,30 | 3,80 | 1,37 | 2,78 | 5,50E-03 | 6,80E-02 |
| AT4G08620 | 2,91 | 4,55 | 1,64 | 2,77 | 5,56E-03 | 6,84E-02 |
| AT3G10985 | 6,51 | 2,42 | 0,87 | 2,77 | 5,68E-03 | 6,94E-02 |
| AT1G63880 | 4,32 | 3,38 | 1,23 | 2,75 | 5,89E-03 | 7,16E-02 |
| AT5G27607 | 2,38 | 5,04 | 1,84 | 2,74 | 6,06E-03 | 7,32E-02 |
| AT1G04897 | 2,68 | 4,26 | 1,55 | 2,74 | 6,09E-03 | 7,32E-02 |
| AT1G54410 | 19,25 | -1,64 | 0,60 | -2,73 | 6,33E-03 | 7,56E-02 |
| AT3G43990 | 14,30 | -1,39 | 0,51 | -2,72 | 6,55E-03 | 7,79E-02 |
| AT4G20170 | 3,89 | 3,40 | 1,25 | 2,71 | 6,78E-03 | 8,01E-02 |
| AT1G20510 | 2,74 | 4,42 | 1,64 | 2,70 | 6,94E-03 | 8,15E-02 |
| AT1G62910 | 98,21 | 0,88 | 0,33 | 2,69 | 7,25E-03 | 8,46E-02 |
| AT1G31173 | 488,28 | 0,66 | 0,25 | 2,68 | 7,32E-03 | 8,50E-02 |
| AT1G29265 | 54,27 | 1,67 | 0,62 | 2,68 | 7,42E-03 | 8,57E-02 |
| AT5G25610 | 6,58 | -1,84 | 0,69 | -2,67 | 7,69E-03 | 8,83E-02 |
| AT4G08630 | 1,97 | 4,54 | 1,71 | 2,66 | 7,78E-03 | 8,88E-02 |
| AT3G04250 | 2,26 | 4,77 | 1,80 | 2,65 | 7,95E-03 | 9,03E-02 |
| AT2G40000 | 5,19 | 2,58 | 0,97 | 2,65 | 8,03E-03 | 9,07E-02 |
| AT1G18740 | 2,60 | 3,98 | 1,50 | 2,64 | 8,18E-03 | 9,09E-02 |
| AT3G25010 | 4,35 | 3,58 | 1,35 | 2,64 | 8,18E-03 | 9,09E-02 |
| AT5G24660 | 2,17 | 4,46 | 1,69 | 2,64 | 8,18E-03 | 9,09E-02 |
| AT4G31500 | 6,34 | 2,79 | 1,06 | 2,64 | 8,27E-03 | 9,14E-02 |
| AT5G02325 | 15,37 | 1,74 | 0,67 | 2,62 | 8,80E-03 | 9,63E-02 |
| AT4G26850 | 7,34 | 2,61 | 1,00 | 2,62 | 8,79E-03 | 9,63E-02 |
| AT2G15970 | 3,79 | -2,68 | 1,02 | -2,61 | 8,93E-03 | 9,71E-02 |
| AT4G29780 | 2,35 | 4,69 | 1,79 | 2,61 | 9,00E-03 | 9,74E-02 |
| AT3G13080 | 13,73 | 3,47 | 1,33 | 2,60 | 9,19E-03 | 9,90E-02 |
| AT5G60408 | 515,37 | 0,75 | 0,29 | 2,59 | 9,59E-03 | 1,03E-01 |
| AT2G16040 | 2,68 | 4,01 | 1,55 | 2,59 | 9,71E-03 | 1,03E-01 |
| AT1G35215 | 4,81 | 2,41 | 0,93 | 2,59 | 9,73E-03 | 1,03E-01 |
| AT5G26920 | 3,70 | 3,60 | 1,40 | 2,57 | 1,03E-02 | 1,09E-01 |
| AT1G64340 | 2,24 | 4,92 | 1,92 | 2,56 | 1,05E-02 | 1,10E-01 |
| AT5G08712 | 123,44 | 0,87 | 0,34 | 2,55 | 1,08E-02 | 1,13E-01 |
| AT3G23690 | 28,16 | -1,04 | 0,41 | -2,55 | 1,09E-02 | 1,13E-01 |
| AT3G57480 | 3,07 | -3,12 | 1,23 | -2,54 | 1,12E-02 | 1,16E-01 |
| AT5G54770 | 28,55 | -1,15 | 0,45 | -2,53 | 1,14E-02 | 1,17E-01 |
| AT1G08833 | 15,74 | -1,21 | 0,48 | -2,52 | 1,17E-02 | 1,18E-01 |
| AT4G17788 | 20,18 | -1,20 | 0,48 | -2,52 | 1,17E-02 | 1,18E-01 |
| AT4G32410 | 11,90 | -1,96 | 0,77 | -2,53 | 1,15E-02 | 1,18E-01 |
| AT4G32940 | 4,72 | 2,98 | 1,18 | 2,52 | 1,17E-02 | 1,18E-01 |
| AT1G22530 | 2,83 | -4,12 | 1,64 | -2,52 | 1,18E-02 | 1,19E-01 |
| AT4G21366 | 2,02 | 4,01 | 1,60 | 2,51 | 1,21E-02 | 1,19E-01 |
| AT5G15022 | 21,95 | 1,29 | 0,51 | 2,51 | 1,20E-02 | 1,19E-01 |
| AT5G07875 | 4,76 | 3,06 | 1,22 | 2,51 | 1,21E-02 | 1,19E-01 |
| AT1G67740 | 3,89 | -2,07 | 0,82 | -2,51 | 1,20E-02 | 1,19E-01 |
| AT5G05170 | 36,40 | -1,73 | 0,69 | -2,50 | 1,24E-02 | 1,21E-01 |
| AT4G04825 | 145,71 | -1,40 | 0,56 | -2,50 | 1,25E-02 | 1,22E-01 |
| AT4G30960 | 5,13 | 3,34 | 1,34 | 2,50 | 1,26E-02 | 1,22E-01 |
| AT1G23720 | 2,26 | 4,86 | 1,95 | 2,49 | 1,29E-02 | 1,25E-01 |
| AT2G01010 | 3,38 | 2,73 | 1,10 | 2,47 | 1,33E-02 | 1,28E-01 |
| AT2G28056 | 34,24 | -0,90 | 0,37 | -2,47 | 1,37E-02 | 1,31E-01 |
| AT2G22500 | 4,35 | 3,12 | 1,27 | 2,46 | 1,38E-02 | 1,31E-01 |
| AT5G38420 | 5,05 | -2,24 | 0,91 | -2,46 | 1,39E-02 | 1,32E-01 |
| AT1G07920 | 7,76 | -1,87 | 0,76 | -2,45 | 1,43E-02 | 1,35E-01 |
| AT1G20340 | 11,85 | -1,28 | 0,53 | -2,43 | 1,50E-02 | 1,41E-01 |
| AT2G18320 | 12,44 | -1,14 | 0,47 | -2,43 | 1,50E-02 | 1,41E-01 |

| | | | | | | |
|---|---|---|---|---|---|---|
| AT2G16500 | 6,48 | 3,25 | 1,34 | 2,42 | 1,53E-02 | 1,43E-01 |
| AT1G07135 | 2,46 | 3,74 | 1,55 | 2,42 | 1,57E-02 | 1,45E-01 |
| AT1G08623 | 2,21 | -3,22 | 1,33 | -2,41 | 1,58E-02 | 1,45E-01 |
| AT5G41490 | 2,24 | 3,85 | 1,60 | 2,41 | 1,58E-02 | 1,45E-01 |
| AT1G66795 | 14,00 | 1,63 | 0,68 | 2,41 | 1,60E-02 | 1,46E-01 |
| AT4G02320 | 3,77 | -2,95 | 1,22 | -2,41 | 1,61E-02 | 1,46E-01 |
| AT1G60072 | 5,42 | 3,29 | 1,37 | 2,40 | 1,64E-02 | 1,48E-01 |
| AT3G55850 | 2,66 | 3,95 | 1,65 | 2,40 | 1,64E-02 | 1,48E-01 |
| AT1G26973 | 11,41 | 1,88 | 0,79 | 2,39 | 1,69E-02 | 1,51E-01 |
| AT3G60630 | 5,65 | 2,29 | 0,96 | 2,39 | 1,68E-02 | 1,51E-01 |
| AT1G31280 | 162,06 | -1,65 | 0,69 | -2,39 | 1,71E-02 | 1,52E-01 |
| AT4G05180 | 7,70 | -1,74 | 0,73 | -2,37 | 1,76E-02 | 1,57E-01 |
| AT5G01747 | 60,67 | -0,62 | 0,26 | -2,37 | 1,78E-02 | 1,57E-01 |
| AT5G02565 | 2,69 | -2,40 | 1,03 | -2,34 | 1,92E-02 | 1,69E-01 |
| AT1G26975 | 8,83 | -1,72 | 0,73 | -2,34 | 1,95E-02 | 1,70E-01 |
| AT2G07981 | 2,13 | 4,48 | 1,92 | 2,34 | 1,95E-02 | 1,70E-01 |
| AT3G55980 | 3,63 | 2,66 | 1,14 | 2,33 | 1,96E-02 | 1,70E-01 |
| AT4G30160 | 4,90 | 3,32 | 1,43 | 2,32 | 2,05E-02 | 1,77E-01 |
| AT4G23915 | 12,58 | 1,41 | 0,61 | 2,31 | 2,06E-02 | 1,77E-01 |
| AT1G28480 | 1,91 | -3,84 | 1,66 | -2,31 | 2,06E-02 | 1,77E-01 |
| AT3G03175 | 7,33 | 1,85 | 0,81 | 2,29 | 2,23E-02 | 1,90E-01 |
| AT1G04393 | 5,00 | 2,23 | 0,97 | 2,29 | 2,23E-02 | 1,90E-01 |
| AT4G05360 | 2,15 | 3,57 | 1,56 | 2,28 | 2,24E-02 | 1,90E-01 |
| AT4G10340 | 23,35 | -1,15 | 0,50 | -2,28 | 2,28E-02 | 1,93E-01 |
| AT1G10760 | 16,93 | -1,42 | 0,63 | -2,26 | 2,35E-02 | 1,97E-01 |
| AT2G33770 | 4,11 | 3,14 | 1,39 | 2,26 | 2,36E-02 | 1,97E-01 |
| AT4G38760 | 3,63 | 2,74 | 1,21 | 2,26 | 2,37E-02 | 1,97E-01 |
| AT4G11570 | 2,16 | -3,01 | 1,34 | -2,25 | 2,43E-02 | 2,02E-01 |
| AT4G27500 | 2,93 | 3,46 | 1,54 | 2,25 | 2,45E-02 | 2,02E-01 |
| AT4G07855 | 2,93 | 3,46 | 1,54 | 2,25 | 2,45E-02 | 2,02E-01 |
| AT5G15970 | 2,85 | -2,91 | 1,29 | -2,25 | 2,46E-02 | 2,02E-01 |
| AT1G05570 | 10,12 | 1,56 | 0,70 | 2,25 | 2,47E-02 | 2,02E-01 |
| AT4G16890 | 15,81 | 2,06 | 0,92 | 2,24 | 2,51E-02 | 2,05E-01 |
| AT4G04835 | 19,24 | -1,17 | 0,52 | -2,23 | 2,55E-02 | 2,07E-01 |
| AT3G07725 | 10,80 | 1,24 | 0,56 | 2,23 | 2,58E-02 | 2,08E-01 |
| AT3G14385 | 15,96 | 1,19 | 0,53 | 2,23 | 2,57E-02 | 2,08E-01 |
| AT2G21660 | 24,65 | -1,03 | 0,46 | -2,23 | 2,59E-02 | 2,08E-01 |
| AT2G14560 | 27,90 | 1,14 | 0,51 | 2,21 | 2,69E-02 | 2,10E-01 |
| AT4G30972 | 11,83 | 1,22 | 0,55 | 2,22 | 2,67E-02 | 2,10E-01 |
| AT5G65750 | 4,51 | -3,61 | 1,63 | -2,22 | 2,68E-02 | 2,10E-01 |
| AT2G07671 | 2,57 | -2,46 | 1,11 | -2,22 | 2,66E-02 | 2,10E-01 |
| AT3G48870 | 10,82 | -3,46 | 1,56 | -2,22 | 2,65E-02 | 2,10E-01 |
| AT4G30750 | 3,95 | 3,26 | 1,47 | 2,21 | 2,68E-02 | 2,10E-01 |
| AT4G30975 | 11,87 | 1,22 | 0,55 | 2,21 | 2,73E-02 | 2,13E-01 |
| AT3G20362 | 7,16 | -1,69 | 0,77 | -2,20 | 2,79E-02 | 2,17E-01 |
| AT3G23685 | 5,28 | 3,09 | 1,41 | 2,19 | 2,82E-02 | 2,18E-01 |
| AT1G43320 | 2,75 | 3,91 | 1,79 | 2,19 | 2,88E-02 | 2,22E-01 |
| AT1G47660 | 2,01 | 3,58 | 1,64 | 2,19 | 2,88E-02 | 2,22E-01 |
| AT2G29920 | 2,43 | 4,12 | 1,89 | 2,18 | 2,91E-02 | 2,23E-01 |
| AT2G34100 | 7,49 | 1,60 | 0,74 | 2,17 | 2,97E-02 | 2,27E-01 |
| AT3G61897 | 2,76 | 5,59 | 2,57 | 2,17 | 3,00E-02 | 2,28E-01 |
| AT1G21680 | 4,80 | -2,00 | 0,92 | -2,16 | 3,05E-02 | 2,30E-01 |
| AT1G71880 | 1,96 | 3,82 | 1,77 | 2,16 | 3,05E-02 | 2,30E-01 |
| AT5G09585 | 3,20 | 2,85 | 1,32 | 2,16 | 3,05E-02 | 2,30E-01 |
| AT5G09495 | 4,86 | -1,75 | 0,81 | -2,16 | 3,08E-02 | 2,30E-01 |
| AT1G11870 | 3,10 | 2,73 | 1,26 | 2,16 | 3,10E-02 | 2,31E-01 |

| | | | | | | |
|---|---|---|---|---|---|---|
| AT2G39885 | 9,28 | 2,61 | 1,21 | 2,16 | 3,11E-02 | 2,31E-01 |
| AT3G28550 | 2,18 | 3,91 | 1,83 | 2,14 | 3,25E-02 | 2,41E-01 |
| AT3G25795 | 22,21 | 2,91 | 1,37 | 2,13 | 3,29E-02 | 2,43E-01 |
| AT5G65005 | 2,56 | -2,17 | 1,02 | -2,13 | 3,34E-02 | 2,45E-01 |
| AT1G04820 | 2,61 | -2,55 | 1,20 | -2,12 | 3,37E-02 | 2,46E-01 |
| AT1G77815 | 2,65 | 3,40 | 1,60 | 2,12 | 3,36E-02 | 2,46E-01 |
| AT5G42050 | 3,62 | 2,55 | 1,20 | 2,12 | 3,38E-02 | 2,46E-01 |
| AT3G50930 | 3,98 | 2,66 | 1,26 | 2,12 | 3,41E-02 | 2,47E-01 |
| AT3G07515 | 2,88 | 3,33 | 1,58 | 2,11 | 3,46E-02 | 2,50E-01 |
| AT5G64260 | 2,41 | 3,22 | 1,53 | 2,11 | 3,51E-02 | 2,53E-01 |
| AT5G49360 | 3,80 | 3,01 | 1,43 | 2,10 | 3,54E-02 | 2,54E-01 |
| AT2G02835 | 3,94 | 2,38 | 1,13 | 2,10 | 3,55E-02 | 2,54E-01 |
| AT3G49120 | 5,10 | 2,53 | 1,21 | 2,09 | 3,62E-02 | 2,57E-01 |
| AT5G02700 | 1,99 | 3,40 | 1,62 | 2,09 | 3,62E-02 | 2,57E-01 |
| AT1G07637 | 2,62 | 2,93 | 1,40 | 2,09 | 3,68E-02 | 2,60E-01 |
| AT1G56070 | 55,72 | -0,80 | 0,38 | -2,09 | 3,68E-02 | 2,60E-01 |
| AT3G06435 | 378,12 | 0,52 | 0,25 | 2,08 | 3,74E-02 | 2,63E-01 |
| AT3G29796 | 2,77 | 2,98 | 1,43 | 2,08 | 3,78E-02 | 2,65E-01 |
| AT3G09105 | 30,37 | 1,04 | 0,50 | 2,06 | 3,91E-02 | 2,73E-01 |
| AT3G06436 | 20,96 | 0,90 | 0,44 | 2,06 | 3,93E-02 | 2,74E-01 |
| AT1G72600 | 18,04 | -0,92 | 0,45 | -2,06 | 3,95E-02 | 2,74E-01 |
| AT5G35735 | 2,19 | 3,05 | 1,49 | 2,05 | 4,02E-02 | 2,78E-01 |
| AT2G05315 | 4,98 | 2,04 | 1,00 | 2,05 | 4,07E-02 | 2,80E-01 |
| AT1G15010 | 2,66 | 2,78 | 1,36 | 2,04 | 4,13E-02 | 2,84E-01 |
| AT1G06680 | 10,44 | -1,23 | 0,61 | -2,03 | 4,19E-02 | 2,84E-01 |
| AT4G07070 | 70,24 | -0,93 | 0,46 | -2,04 | 4,17E-02 | 2,84E-01 |
| AT5G38190 | 2,86 | 2,39 | 1,18 | 2,03 | 4,21E-02 | 2,84E-01 |
| AT4G38970 | 12,35 | -1,03 | 0,51 | -2,03 | 4,19E-02 | 2,84E-01 |
| AT5G65860 | 5,04 | -1,64 | 0,81 | -2,04 | 4,16E-02 | 2,84E-01 |
| AT1G72610 | 17,51 | -0,92 | 0,45 | -2,03 | 4,27E-02 | 2,87E-01 |
| AT2G14080 | 4,62 | 3,04 | 1,51 | 2,02 | 4,34E-02 | 2,92E-01 |
| AT1G44100 | 2,02 | 3,17 | 1,58 | 2,01 | 4,39E-02 | 2,94E-01 |
| AT3G22960 | 3,17 | -2,63 | 1,31 | -2,01 | 4,47E-02 | 2,96E-01 |
| AT3G23125 | 239,13 | 0,89 | 0,44 | 2,01 | 4,44E-02 | 2,96E-01 |
| AT4G04020 | 1,87 | -3,33 | 1,66 | -2,01 | 4,49E-02 | 2,96E-01 |
| AT2G23348 | 3,88 | 2,28 | 1,14 | 2,01 | 4,49E-02 | 2,96E-01 |
| AT5G02490 | 4,18 | 2,94 | 1,46 | 2,01 | 4,48E-02 | 2,96E-01 |
| AT4G18880 | 2,25 | 3,69 | 1,84 | 2,00 | 4,55E-02 | 2,98E-01 |
| AT4G31877 | 9,33 | -1,17 | 0,59 | -2,00 | 4,55E-02 | 2,98E-01 |
| AT2G04885 | 2,17 | 3,05 | 1,53 | 2,00 | 4,59E-02 | 3,00E-01 |
| AT5G27950 | 2,87 | -2,21 | 1,11 | -1,99 | 4,63E-02 | 3,01E-01 |
| AT5G13650 | 5,78 | -1,67 | 0,84 | -1,99 | 4,69E-02 | 3,04E-01 |
| AT2G39730 | 47,53 | -0,82 | 0,41 | -1,98 | 4,72E-02 | 3,05E-01 |
| AT3G58196 | 21,50 | 0,95 | 0,48 | 1,98 | 4,77E-02 | 3,07E-01 |
| AT2G29925 | 2,27 | 3,92 | 1,98 | 1,98 | 4,79E-02 | 3,07E-01 |
| AT1G74470 | 19,25 | -0,98 | 0,50 | -1,97 | 4,92E-02 | 3,11E-01 |
| AT2G08986 | 10,75 | 1,20 | 0,61 | 1,97 | 4,93E-02 | 3,11E-01 |
| AT2G18960 | 62,63 | -1,37 | 0,70 | -1,97 | 4,91E-02 | 3,11E-01 |
| AT1G05423 | 13,10 | -0,98 | 0,50 | -1,97 | 4,94E-02 | 3,11E-01 |
| AT4G33010 | 11,10 | -1,46 | 0,74 | -1,97 | 4,87E-02 | 3,11E-01 |
| AT5G24690 | 2,42 | -2,86 | 1,45 | -1,97 | 4,88E-02 | 3,11E-01 |
| AT1G20450 | 3,27 | -1,96 | 1,00 | -1,96 | 4,98E-02 | 3,13E-01 |
| AT4G33050 | 2,86 | 2,84 | 1,45 | 1,95 | 5,09E-02 | 3,19E-01 |
| AT3G44703 | 4,72 | -1,65 | 0,85 | -1,95 | 5,16E-02 | 3,20E-01 |
| AT1G07493 | 1,97 | -2,82 | 1,45 | -1,95 | 5,16E-02 | 3,20E-01 |
| AT2G06520 | 10,63 | -1,43 | 0,73 | -1,95 | 5,16E-02 | 3,20E-01 |

| | | | | | | |
|---|---|---|---|---|---|---|
| AT5G09000 | 2,90 | -2,03 | 1,04 | -1,95 | 5,17E-02 | 3,20E-01 |
| AT2G44490 | 17,93 | -0,94 | 0,49 | -1,93 | 5,32E-02 | 3,22E-01 |
| AT3G07790 | 2,60 | 2,97 | 1,54 | 1,93 | 5,32E-02 | 3,22E-01 |
| AT3G26740 | 15,30 | -0,96 | 0,50 | -1,94 | 5,24E-02 | 3,22E-01 |
| AT3G28180 | 8,56 | -1,89 | 0,98 | -1,93 | 5,32E-02 | 3,22E-01 |
| AT3G62290 | 5,55 | 2,71 | 1,40 | 1,94 | 5,30E-02 | 3,22E-01 |
| AT4G08593 | 1,89 | 3,49 | 1,80 | 1,93 | 5,31E-02 | 3,22E-01 |
| AT1G79750 | 4,46 | -2,78 | 1,43 | -1,94 | 5,23E-02 | 3,22E-01 |
| AT2G23347 | 3,51 | 2,20 | 1,14 | 1,94 | 5,27E-02 | 3,22E-01 |
| AT1G21202 | 4,89 | 2,22 | 1,15 | 1,93 | 5,37E-02 | 3,22E-01 |
| AT3G43160 | 4,62 | 1,72 | 0,89 | 1,93 | 5,39E-02 | 3,22E-01 |
| AT1G05637 | 4,89 | 2,22 | 1,15 | 1,93 | 5,37E-02 | 3,22E-01 |
| AT5G28823 | 1,86 | 3,66 | 1,90 | 1,93 | 5,38E-02 | 3,22E-01 |
| AT1G37130 | 39,12 | 1,08 | 0,56 | 1,93 | 5,40E-02 | 3,22E-01 |
| AT4G30210 | 1,86 | 3,60 | 1,87 | 1,92 | 5,48E-02 | 3,25E-01 |
| AT3G09162 | 2,15 | -2,80 | 1,46 | -1,91 | 5,56E-02 | 3,29E-01 |
| AT1G50200 | 4,92 | -2,75 | 1,44 | -1,91 | 5,67E-02 | 3,35E-01 |
| AT3G06835 | 2,07 | -2,83 | 1,48 | -1,90 | 5,69E-02 | 3,35E-01 |
| AT1G78020 | 1,89 | 2,86 | 1,51 | 1,90 | 5,76E-02 | 3,35E-01 |
| AT2G18570 | 4,57 | 2,08 | 1,10 | 1,90 | 5,71E-02 | 3,35E-01 |
| AT2G24103 | 3,17 | 2,83 | 1,49 | 1,90 | 5,75E-02 | 3,35E-01 |
| AT2G10940 | 5,77 | -1,76 | 0,93 | -1,90 | 5,73E-02 | 3,35E-01 |
| AT1G21200 | 5,07 | 1,94 | 1,02 | 1,89 | 5,87E-02 | 3,37E-01 |
| AT1G28070 | 4,21 | -4,16 | 2,20 | -1,89 | 5,84E-02 | 3,37E-01 |
| AT4G14160 | 2,71 | 3,21 | 1,70 | 1,89 | 5,88E-02 | 3,37E-01 |
| AT1G08380 | 7,50 | -1,35 | 0,71 | -1,89 | 5,84E-02 | 3,37E-01 |
| AT3G23680 | 9,12 | 2,68 | 1,42 | 1,89 | 5,88E-02 | 3,37E-01 |
| AT3G54660 | 2,19 | -3,22 | 1,70 | -1,89 | 5,86E-02 | 3,37E-01 |
| AT1G63980 | 3,67 | 2,87 | 1,52 | 1,88 | 5,95E-02 | 3,39E-01 |
| AT1G78570 | 2,32 | 3,04 | 1,61 | 1,88 | 5,96E-02 | 3,39E-01 |
| AT3G01290 | 2,12 | 2,89 | 1,54 | 1,88 | 5,98E-02 | 3,40E-01 |
| AT1G15690 | 11,49 | -1,19 | 0,64 | -1,87 | 6,13E-02 | 3,42E-01 |
| AT1G35710 | 7,53 | 2,75 | 1,47 | 1,88 | 6,05E-02 | 3,42E-01 |
| AT4G26730 | 2,38 | 3,23 | 1,73 | 1,87 | 6,11E-02 | 3,42E-01 |
| AT5G61440 | 4,44 | -1,42 | 0,76 | -1,88 | 6,07E-02 | 3,42E-01 |
| AT2G14610 | 3,13 | 2,39 | 1,27 | 1,87 | 6,09E-02 | 3,42E-01 |
| AT3G11010 | 3,23 | 3,08 | 1,65 | 1,87 | 6,12E-02 | 3,42E-01 |
| AT4G21640 | 1,99 | -2,37 | 1,26 | -1,87 | 6,08E-02 | 3,42E-01 |
| AT3G25020 | 3,41 | 1,87 | 1,00 | 1,87 | 6,15E-02 | 3,42E-01 |
| AT1G20620 | 38,72 | -0,79 | 0,42 | -1,87 | 6,22E-02 | 3,44E-01 |
| AT4G04205 | 2,87 | 2,03 | 1,09 | 1,87 | 6,21E-02 | 3,44E-01 |
| AT1G76490 | 3,44 | -1,96 | 1,05 | -1,86 | 6,26E-02 | 3,44E-01 |
| AT3G55800 | 2,45 | -2,08 | 1,12 | -1,86 | 6,24E-02 | 3,44E-01 |
| AT3G48057 | 47,97 | 1,04 | 0,56 | 1,86 | 6,31E-02 | 3,46E-01 |
| AT2G26530 | 1,88 | 3,22 | 1,73 | 1,86 | 6,33E-02 | 3,46E-01 |
| AT3G48200 | 2,11 | -3,02 | 1,63 | -1,85 | 6,37E-02 | 3,48E-01 |
| AT1G56510 | 2,98 | 2,92 | 1,58 | 1,85 | 6,49E-02 | 3,51E-01 |
| AT2G48060 | 4,01 | 2,77 | 1,50 | 1,85 | 6,46E-02 | 3,51E-01 |
| AT3G53460 | 6,35 | -1,55 | 0,84 | -1,84 | 6,50E-02 | 3,51E-01 |
| AT4G01800 | 8,35 | -2,47 | 1,34 | -1,85 | 6,49E-02 | 3,51E-01 |
| AT4G38770 | 7,08 | -1,07 | 0,58 | -1,84 | 6,57E-02 | 3,54E-01 |
| AT3G57710 | 4,92 | 1,58 | 0,86 | 1,84 | 6,62E-02 | 3,56E-01 |
| AT2G20410 | 6,12 | -1,37 | 0,75 | -1,83 | 6,68E-02 | 3,57E-01 |
| AT3G41761 | 2,51 | -2,63 | 1,44 | -1,83 | 6,70E-02 | 3,57E-01 |
| AT5G54390 | 4,62 | 3,01 | 1,64 | 1,83 | 6,72E-02 | 3,57E-01 |
| AT3G47470 | 25,61 | -0,82 | 0,45 | -1,83 | 6,72E-02 | 3,57E-01 |

| | | | | | | |
|---|---|---|---|---|---|---|
| AT1G50920 | 2,95 | -1,91 | 1,04 | -1,83 | 6,77E-02 | 3,59E-01 |
| AT4G00975 | 11,27 | -1,00 | 0,55 | -1,82 | 6,82E-02 | 3,60E-01 |
| AT3G15353 | 5,91 | 1,77 | 0,97 | 1,82 | 6,81E-02 | 3,60E-01 |
| AT3G18830 | 2,46 | 2,85 | 1,56 | 1,82 | 6,86E-02 | 3,61E-01 |
| AT1G55670 | 5,12 | -1,26 | 0,69 | -1,82 | 6,92E-02 | 3,63E-01 |
| AT3G28925 | 2,17 | 2,64 | 1,45 | 1,82 | 6,94E-02 | 3,63E-01 |
| AT2G04435 | 3,56 | 2,20 | 1,22 | 1,81 | 7,08E-02 | 3,65E-01 |
| AT2G39175 | 43,76 | 0,66 | 0,37 | 1,81 | 7,08E-02 | 3,65E-01 |
| AT3G29320 | 2,29 | -2,04 | 1,13 | -1,81 | 7,02E-02 | 3,65E-01 |
| AT3G25510 | 1,96 | -3,22 | 1,78 | -1,81 | 7,05E-02 | 3,65E-01 |
| AT5G01820 | 9,75 | 2,28 | 1,26 | 1,81 | 7,05E-02 | 3,65E-01 |
| AT5G17165 | 3,90 | 1,82 | 1,01 | 1,81 | 7,09E-02 | 3,65E-01 |
| AT1G32630 | 4,04 | 1,69 | 0,94 | 1,80 | 7,23E-02 | 3,69E-01 |
| AT5G28560 | 3,51 | 1,89 | 1,05 | 1,80 | 7,20E-02 | 3,69E-01 |
| AT1G32640 | 4,04 | 1,69 | 0,94 | 1,80 | 7,23E-02 | 3,69E-01 |
| AT1G52110 | 2,50 | -2,09 | 1,16 | -1,80 | 7,21E-02 | 3,69E-01 |
| AT1G29900 | 11,05 | 1,99 | 1,11 | 1,79 | 7,29E-02 | 3,69E-01 |
| AT2G04455 | 3,39 | 2,24 | 1,25 | 1,80 | 7,26E-02 | 3,69E-01 |
| AT3G54580 | 1,94 | 3,07 | 1,71 | 1,79 | 7,28E-02 | 3,69E-01 |
| AT3G47347 | 15,63 | 0,82 | 0,46 | 1,79 | 7,40E-02 | 3,73E-01 |
| AT2G45540 | 12,39 | 2,10 | 1,18 | 1,79 | 7,39E-02 | 3,73E-01 |
| AT4G17090 | 4,63 | -1,60 | 0,90 | -1,79 | 7,41E-02 | 3,73E-01 |
| AT1G21250 | 6,46 | 1,56 | 0,87 | 1,78 | 7,45E-02 | 3,74E-01 |
| AT1G13930 | 7,00 | -1,22 | 0,69 | -1,78 | 7,54E-02 | 3,76E-01 |
| AT1G07793 | 2,22 | -2,01 | 1,14 | -1,77 | 7,64E-02 | 3,76E-01 |
| AT3G59780 | 2,03 | -2,48 | 1,40 | -1,77 | 7,64E-02 | 3,76E-01 |
| AT5G01530 | 20,05 | -1,02 | 0,57 | -1,77 | 7,61E-02 | 3,76E-01 |
| AT5G20230 | 2,06 | 3,04 | 1,71 | 1,78 | 7,56E-02 | 3,76E-01 |
| AT2G15420 | 2,17 | 2,61 | 1,47 | 1,78 | 7,58E-02 | 3,76E-01 |
| AT4G05050 | 3,49 | 1,98 | 1,12 | 1,77 | 7,64E-02 | 3,76E-01 |
| AT4G35750 | 1,89 | 2,85 | 1,61 | 1,77 | 7,60E-02 | 3,76E-01 |
| AT5G67300 | 3,74 | 2,25 | 1,27 | 1,76 | 7,82E-02 | 3,83E-01 |
| AT2G39800 | 6,12 | -1,92 | 1,09 | -1,76 | 7,81E-02 | 3,83E-01 |
| AT1G47657 | 2,52 | 2,50 | 1,42 | 1,76 | 7,84E-02 | 3,83E-01 |
| AT3G56940 | 6,51 | -1,36 | 0,77 | -1,76 | 7,90E-02 | 3,85E-01 |
| AT1G53687 | 5,07 | -1,45 | 0,82 | -1,75 | 7,94E-02 | 3,86E-01 |
| AT2G26355 | 2,39 | 2,78 | 1,58 | 1,75 | 7,95E-02 | 3,86E-01 |
| AT1G15820 | 12,09 | -1,09 | 0,62 | -1,75 | 7,99E-02 | 3,87E-01 |
| AT1G77760 | 2,44 | 2,88 | 1,65 | 1,75 | 8,02E-02 | 3,87E-01 |
| AT1G74100 | 5,98 | 1,68 | 0,96 | 1,75 | 8,08E-02 | 3,88E-01 |
| AT3G44260 | 2,79 | 2,11 | 1,21 | 1,75 | 8,06E-02 | 3,88E-01 |
| AT1G74453 | 1,99 | 2,49 | 1,43 | 1,74 | 8,15E-02 | 3,91E-01 |
| AT4G30440 | 3,67 | 1,98 | 1,14 | 1,74 | 8,25E-02 | 3,95E-01 |
| AT5G39570 | 4,04 | -1,52 | 0,88 | -1,73 | 8,37E-02 | 3,99E-01 |
| AT1G47720 | 6,21 | 2,49 | 1,44 | 1,73 | 8,35E-02 | 3,99E-01 |
| AT3G50590 | 4,47 | -2,32 | 1,34 | -1,73 | 8,44E-02 | 4,00E-01 |
| AT4G12800 | 10,28 | -1,17 | 0,68 | -1,73 | 8,43E-02 | 4,00E-01 |
| AT1G19090 | 4,93 | 1,50 | 0,87 | 1,72 | 8,50E-02 | 4,02E-01 |
| AT5G01920 | 2,56 | -3,09 | 1,80 | -1,72 | 8,54E-02 | 4,02E-01 |
| AT5G54270 | 15,64 | -0,78 | 0,46 | -1,72 | 8,55E-02 | 4,02E-01 |
| AT5G39770 | 12,51 | 0,92 | 0,54 | 1,72 | 8,53E-02 | 4,02E-01 |
| AT2G16600 | 11,35 | 1,16 | 0,68 | 1,72 | 8,60E-02 | 4,03E-01 |
| AT2G21100 | 3,61 | -1,54 | 0,90 | -1,71 | 8,64E-02 | 4,04E-01 |
| AT4G31570 | 11,16 | 2,48 | 1,45 | 1,71 | 8,74E-02 | 4,04E-01 |
| AT5G19200 | 4,59 | -1,95 | 1,14 | -1,71 | 8,70E-02 | 4,04E-01 |
| AT1G62035 | 5,54 | 1,28 | 0,75 | 1,71 | 8,74E-02 | 4,04E-01 |

| | | | | | | |
|---|---|---|---|---|---|---|
| AT3G56910 | 1,94 | -2,65 | 1,55 | -1,71 | 8,74E-02 | 4,04E-01 |
| AT4G28080 | 23,67 | -1,38 | 0,81 | -1,71 | 8,74E-02 | 4,04E-01 |
| AT5G23110 | 11,58 | 2,19 | 1,28 | 1,71 | 8,79E-02 | 4,05E-01 |
| AT3G03780 | 4,16 | -1,96 | 1,16 | -1,68 | 9,21E-02 | 4,24E-01 |
| AT5G46520 | 3,26 | 2,67 | 1,59 | 1,68 | 9,26E-02 | 4,24E-01 |
| AT5G09065 | 4,44 | 1,66 | 0,99 | 1,68 | 9,24E-02 | 4,24E-01 |
| AT3G63003 | 4,44 | -1,61 | 0,96 | -1,68 | 9,39E-02 | 4,29E-01 |
| AT4G04095 | 1,94 | -2,46 | 1,47 | -1,67 | 9,44E-02 | 4,30E-01 |
| AT2G33150 | 2,47 | -1,97 | 1,18 | -1,67 | 9,45E-02 | 4,30E-01 |
| AT4G00450 | 3,39 | 2,64 | 1,58 | 1,67 | 9,53E-02 | 4,31E-01 |
| AT1G27770 | 5,67 | 2,23 | 1,34 | 1,67 | 9,53E-02 | 4,31E-01 |
| AT4G34980 | 4,56 | 1,54 | 0,92 | 1,67 | 9,54E-02 | 4,31E-01 |
| AT5G64040 | 3,44 | -1,52 | 0,91 | -1,66 | 9,63E-02 | 4,34E-01 |
| AT4G18030 | 2,74 | -2,12 | 1,28 | -1,66 | 9,65E-02 | 4,34E-01 |
| AT1G06220 | 3,15 | -2,94 | 1,77 | -1,66 | 9,69E-02 | 4,35E-01 |
| AT4G21060 | 1,98 | -1,89 | 1,14 | -1,66 | 9,78E-02 | 4,37E-01 |
| AT3G57260 | 8,35 | 1,23 | 0,74 | 1,66 | 9,77E-02 | 4,37E-01 |
| AT4G01850 | 4,27 | -2,39 | 1,45 | -1,65 | 9,89E-02 | 4,41E-01 |
| AT5G66190 | 3,89 | -1,55 | 0,94 | -1,65 | 9,91E-02 | 4,41E-01 |
| AT4G32640 | 2,41 | -2,49 | 1,51 | -1,65 | 9,96E-02 | 4,42E-01 |
| AT3G44444 | 1,95 | 2,23 | 1,35 | 1,65 | 9,95E-02 | 4,42E-01 |
| AT1G09797 | 19,74 | 1,58 | 0,96 | 1,64 | 1,02E-01 | 4,49E-01 |
| AT1G09793 | 19,74 | 1,58 | 0,96 | 1,64 | 1,02E-01 | 4,49E-01 |
| AT3G54050 | 3,78 | -1,55 | 0,95 | -1,63 | 1,02E-01 | 4,51E-01 |
| AT3G07765 | 2,13 | -1,80 | 1,11 | -1,62 | 1,05E-01 | 4,60E-01 |
| AT3G57720 | 13,95 | 1,07 | 0,66 | 1,62 | 1,05E-01 | 4,60E-01 |
| AT4G29900 | 4,72 | 1,55 | 0,96 | 1,62 | 1,05E-01 | 4,62E-01 |
| AT5G07940 | 5,66 | 2,37 | 1,47 | 1,61 | 1,07E-01 | 4,67E-01 |
| AT4G21960 | 15,77 | -0,73 | 0,45 | -1,61 | 1,07E-01 | 4,67E-01 |
| AT5G63840 | 2,51 | 2,53 | 1,58 | 1,60 | 1,09E-01 | 4,73E-01 |
| AT4G38550 | 2,97 | 2,22 | 1,38 | 1,60 | 1,10E-01 | 4,74E-01 |
| AT4G12320 | 1,93 | -2,72 | 1,70 | -1,60 | 1,10E-01 | 4,74E-01 |
| AT4G38552 | 2,97 | 2,22 | 1,38 | 1,60 | 1,10E-01 | 4,74E-01 |
| AT5G35630 | 4,18 | -1,46 | 0,91 | -1,60 | 1,11E-01 | 4,76E-01 |
| AT3G15920 | 8,54 | 1,14 | 0,71 | 1,60 | 1,11E-01 | 4,76E-01 |
| AT1G79570 | 2,57 | 2,50 | 1,57 | 1,59 | 1,11E-01 | 4,77E-01 |
| AT3G22235 | 3,04 | -1,86 | 1,17 | -1,59 | 1,11E-01 | 4,77E-01 |
| AT4G20735 | 3,19 | 2,43 | 1,53 | 1,59 | 1,12E-01 | 4,78E-01 |
| AT4G24690 | 16,56 | 1,01 | 0,64 | 1,59 | 1,12E-01 | 4,78E-01 |
| AT4G36360 | 3,63 | -2,47 | 1,56 | -1,59 | 1,12E-01 | 4,78E-01 |
| AT1G29370 | 3,57 | -1,85 | 1,17 | -1,58 | 1,13E-01 | 4,81E-01 |
| AT1G04977 | 45,78 | -0,47 | 0,30 | -1,58 | 1,14E-01 | 4,82E-01 |
| AT5G26030 | 2,16 | 2,25 | 1,43 | 1,58 | 1,14E-01 | 4,82E-01 |
| AT3G01500 | 4,00 | -1,54 | 0,98 | -1,58 | 1,14E-01 | 4,82E-01 |
| AT4G06235 | 7,69 | -1,43 | 0,90 | -1,58 | 1,14E-01 | 4,82E-01 |
| AT5G05010 | 2,08 | -2,34 | 1,48 | -1,58 | 1,15E-01 | 4,83E-01 |
| AT3G04340 | 3,02 | -2,49 | 1,58 | -1,58 | 1,15E-01 | 4,83E-01 |
| AT3G09630 | 1,89 | -2,17 | 1,38 | -1,57 | 1,16E-01 | 4,85E-01 |
| AT3G17185 | 156,12 | -0,39 | 0,25 | -1,57 | 1,17E-01 | 4,88E-01 |
| AT4G03560 | 2,24 | -1,86 | 1,19 | -1,57 | 1,17E-01 | 4,88E-01 |
| AT5G13390 | 2,05 | -2,34 | 1,50 | -1,57 | 1,17E-01 | 4,88E-01 |
| AT4G25520 | 4,54 | -1,30 | 0,83 | -1,57 | 1,17E-01 | 4,88E-01 |
| AT1G12240 | 2,62 | -2,28 | 1,46 | -1,56 | 1,18E-01 | 4,89E-01 |
| AT3G57520 | 3,92 | -2,18 | 1,39 | -1,56 | 1,18E-01 | 4,89E-01 |
| AT4G13885 | 7,77 | -0,96 | 0,61 | -1,56 | 1,18E-01 | 4,89E-01 |
| AT2G05520 | 18,30 | -0,58 | 0,37 | -1,55 | 1,20E-01 | 4,95E-01 |

| | | | | | | |
|---|---|---|---|---|---|---|
| AT1G24460 | 4,84 | -2,10 | 1,36 | -1,55 | 1,21E-01 | 4,96E-01 |
| AT2G25250 | 2,81 | 1,91 | 1,23 | 1,55 | 1,21E-01 | 4,96E-01 |
| AT2G46765 | 2,57 | 1,76 | 1,14 | 1,55 | 1,21E-01 | 4,96E-01 |
| AT1G73687 | 3456,65 | -0,40 | 0,26 | -1,55 | 1,21E-01 | 4,96E-01 |
| AT3G23920 | 2,81 | 2,41 | 1,56 | 1,55 | 1,22E-01 | 4,98E-01 |
| AT2G03855 | 12,69 | -0,80 | 0,52 | -1,54 | 1,23E-01 | 5,00E-01 |
| AT4G02520 | 4,48 | 1,48 | 0,96 | 1,54 | 1,23E-01 | 5,01E-01 |
| AT3G58193 | 8,75 | 1,13 | 0,74 | 1,54 | 1,24E-01 | 5,02E-01 |
| AT4G23170 | 6,38 | 1,41 | 0,92 | 1,54 | 1,24E-01 | 5,02E-01 |
| AT4G11600 | 2,32 | -1,62 | 1,06 | -1,54 | 1,24E-01 | 5,02E-01 |
| AT1G80440 | 3,83 | 1,84 | 1,20 | 1,53 | 1,26E-01 | 5,06E-01 |
| AT3G01345 | 6,06 | 1,94 | 1,27 | 1,53 | 1,26E-01 | 5,06E-01 |
| AT3G20310 | 2,42 | 1,93 | 1,26 | 1,53 | 1,26E-01 | 5,06E-01 |
| AT5G04305 | 5,44 | 1,26 | 0,82 | 1,53 | 1,26E-01 | 5,06E-01 |
| AT4G39364 | 2,19 | 2,39 | 1,57 | 1,53 | 1,26E-01 | 5,06E-01 |
| AT5G46110 | 7,87 | -1,11 | 0,73 | -1,52 | 1,27E-01 | 5,08E-01 |
| AT1G18745 | 15,99 | -0,77 | 0,50 | -1,52 | 1,28E-01 | 5,08E-01 |
| AT1G79920 | 8,70 | -1,58 | 1,04 | -1,52 | 1,28E-01 | 5,08E-01 |
| AT4G15258 | 2,27 | 2,22 | 1,46 | 1,52 | 1,28E-01 | 5,08E-01 |
| AT4G16860 | 6,11 | 1,54 | 1,01 | 1,52 | 1,28E-01 | 5,08E-01 |
| AT1G69572 | 2,44 | 2,05 | 1,35 | 1,52 | 1,29E-01 | 5,11E-01 |
| AT1G69570 | 2,44 | 2,05 | 1,35 | 1,52 | 1,29E-01 | 5,11E-01 |
| AT5G36228 | 1,95 | -2,01 | 1,33 | -1,51 | 1,31E-01 | 5,15E-01 |
| AT3G60190 | 1,87 | -1,94 | 1,29 | -1,51 | 1,31E-01 | 5,15E-01 |
| AT1G21750 | 2,14 | 2,85 | 1,89 | 1,51 | 1,31E-01 | 5,17E-01 |
| AT5G20720 | 3,41 | 2,41 | 1,60 | 1,51 | 1,32E-01 | 5,18E-01 |
| AT3G54890 | 13,73 | -0,90 | 0,60 | -1,50 | 1,33E-01 | 5,19E-01 |
| AT5G44800 | 4,77 | 2,09 | 1,39 | 1,50 | 1,33E-01 | 5,19E-01 |
| AT5G12150 | 4,21 | 2,51 | 1,68 | 1,50 | 1,34E-01 | 5,22E-01 |
| AT5G42820 | 2,08 | -2,60 | 1,73 | -1,50 | 1,34E-01 | 5,22E-01 |
| AT5G07350 | 9,52 | 2,15 | 1,44 | 1,49 | 1,35E-01 | 5,25E-01 |
| AT5G54250 | 1,88 | -2,74 | 1,83 | -1,49 | 1,35E-01 | 5,25E-01 |
| AT1G22610 | 11,17 | 1,70 | 1,15 | 1,49 | 1,37E-01 | 5,29E-01 |
| AT3G57410 | 3,70 | -2,59 | 1,75 | -1,49 | 1,37E-01 | 5,29E-01 |
| AT4G14910 | 7,50 | 2,07 | 1,39 | 1,49 | 1,37E-01 | 5,29E-01 |
| AT4G16950 | 9,81 | 1,28 | 0,86 | 1,49 | 1,37E-01 | 5,29E-01 |
| AT5G46510 | 5,48 | -1,36 | 0,92 | -1,48 | 1,39E-01 | 5,32E-01 |
| AT4G16900 | 2,10 | 2,34 | 1,58 | 1,48 | 1,39E-01 | 5,32E-01 |
| AT5G20730 | 2,22 | 2,33 | 1,57 | 1,48 | 1,39E-01 | 5,32E-01 |
| AT1G02470 | 5,92 | 1,58 | 1,07 | 1,48 | 1,40E-01 | 5,34E-01 |
| AT3G08940 | 8,84 | -1,09 | 0,74 | -1,47 | 1,42E-01 | 5,39E-01 |
| AT1G65440 | 2,14 | -1,96 | 1,33 | -1,47 | 1,42E-01 | 5,39E-01 |
| AT5G42390 | 8,64 | -1,27 | 0,87 | -1,46 | 1,43E-01 | 5,42E-01 |
| AT1G03410 | 2,63 | 1,82 | 1,24 | 1,47 | 1,43E-01 | 5,42E-01 |
| AT3G23325 | 69,65 | 0,93 | 0,64 | 1,46 | 1,44E-01 | 5,43E-01 |
| AT5G09590 | 2,74 | -2,48 | 1,70 | -1,46 | 1,45E-01 | 5,43E-01 |
| AT1G36730 | 3,83 | 2,20 | 1,51 | 1,46 | 1,44E-01 | 5,43E-01 |
| AT1G65350 | 2,42 | 1,90 | 1,31 | 1,46 | 1,45E-01 | 5,43E-01 |
| AT4G13577 | 2,60 | -1,64 | 1,12 | -1,46 | 1,44E-01 | 5,43E-01 |
| AT5G14800 | 5,34 | -1,92 | 1,32 | -1,46 | 1,45E-01 | 5,43E-01 |
| AT5G05485 | 5,35 | -1,54 | 1,05 | -1,46 | 1,44E-01 | 5,43E-01 |
| AT1G76010 | 2,04 | -1,89 | 1,30 | -1,45 | 1,46E-01 | 5,47E-01 |
| AT1G67450 | 1,86 | 2,11 | 1,45 | 1,45 | 1,47E-01 | 5,47E-01 |
| AT3G09840 | 22,25 | 0,93 | 0,64 | 1,45 | 1,48E-01 | 5,47E-01 |
| AT4G31480 | 6,09 | 2,18 | 1,50 | 1,45 | 1,47E-01 | 5,47E-01 |
| AT2G36530 | 5,68 | 1,56 | 1,08 | 1,45 | 1,48E-01 | 5,47E-01 |

| | | | | | | |
|---|---|---|---|---|---|---|
| AT4G13918 | 2,19 | 2,35 | 1,62 | 1,45 | 1,48E-01 | 5,47E-01 |
| AT4G35830 | 9,01 | -1,23 | 0,85 | -1,45 | 1,48E-01 | 5,47E-01 |
| AT3G48560 | 3,89 | -1,31 | 0,90 | -1,44 | 1,49E-01 | 5,49E-01 |
| AT5G54810 | 4,59 | -1,59 | 1,10 | -1,44 | 1,49E-01 | 5,49E-01 |
| AT1G04183 | 2,59 | 1,79 | 1,24 | 1,44 | 1,49E-01 | 5,49E-01 |
| AT4G00430 | 2,16 | -1,72 | 1,19 | -1,44 | 1,49E-01 | 5,49E-01 |
| AT1G08537 | 8,63 | 0,94 | 0,65 | 1,44 | 1,50E-01 | 5,50E-01 |
| AT4G13920 | 2,03 | 2,45 | 1,71 | 1,44 | 1,51E-01 | 5,52E-01 |
| AT1G26985 | 10,26 | -1,26 | 0,89 | -1,43 | 1,54E-01 | 5,54E-01 |
| AT2G25510 | 7,52 | 1,21 | 0,85 | 1,43 | 1,53E-01 | 5,54E-01 |
| AT2G46820 | 2,21 | -1,57 | 1,10 | -1,43 | 1,54E-01 | 5,54E-01 |
| AT3G44006 | 2,21 | -1,84 | 1,29 | -1,43 | 1,52E-01 | 5,54E-01 |
| AT5G43725 | 9,10 | 0,82 | 0,57 | 1,43 | 1,53E-01 | 5,54E-01 |
| AT5G43730 | 9,10 | 0,82 | 0,57 | 1,43 | 1,53E-01 | 5,54E-01 |
| AT1G12820 | 53,20 | -0,39 | 0,27 | -1,43 | 1,52E-01 | 5,54E-01 |
| AT1G70320 | 18,40 | 1,24 | 0,87 | 1,43 | 1,53E-01 | 5,54E-01 |
| AT2G04955 | 3,52 | 1,46 | 1,02 | 1,43 | 1,53E-01 | 5,54E-01 |
| AT5G45510 | 8,42 | 1,61 | 1,13 | 1,42 | 1,54E-01 | 5,54E-01 |
| AT2G40840 | 4,12 | -1,36 | 0,96 | -1,42 | 1,55E-01 | 5,56E-01 |
| AT5G60360 | 2,97 | -1,78 | 1,25 | -1,42 | 1,55E-01 | 5,56E-01 |
| AT2G38540 | 3,59 | -1,36 | 0,96 | -1,42 | 1,56E-01 | 5,56E-01 |
| AT5G19940 | 2,37 | -1,61 | 1,13 | -1,42 | 1,56E-01 | 5,56E-01 |
| AT5G03145 | 2,30 | -1,78 | 1,25 | -1,42 | 1,56E-01 | 5,56E-01 |
| AT1G74260 | 9,63 | 1,64 | 1,16 | 1,42 | 1,57E-01 | 5,56E-01 |
| AT5G63195 | 1,93 | -2,56 | 1,80 | -1,42 | 1,56E-01 | 5,56E-01 |
| AT4G16720 | 3,07 | -2,42 | 1,71 | -1,41 | 1,57E-01 | 5,58E-01 |
| AT3G56825 | 56,81 | 0,50 | 0,35 | 1,41 | 1,58E-01 | 5,59E-01 |
| AT5G06530 | 2,30 | -2,47 | 1,75 | -1,41 | 1,58E-01 | 5,59E-01 |
| AT1G09970 | 4,22 | 2,07 | 1,47 | 1,41 | 1,59E-01 | 5,63E-01 |
| AT1G05805 | 1,94 | 3,05 | 2,17 | 1,40 | 1,61E-01 | 5,64E-01 |
| AT1G51690 | 2,38 | -2,01 | 1,44 | -1,40 | 1,62E-01 | 5,64E-01 |
| AT1G62914 | 457,48 | 0,45 | 0,33 | 1,40 | 1,62E-01 | 5,64E-01 |
| AT2G30950 | 3,17 | -1,46 | 1,05 | -1,40 | 1,62E-01 | 5,64E-01 |
| AT3G62250 | 1,91 | -2,48 | 1,77 | -1,40 | 1,61E-01 | 5,64E-01 |
| AT4G14500 | 7,04 | 1,00 | 0,72 | 1,40 | 1,62E-01 | 5,64E-01 |
| AT4G27440 | 13,15 | -0,72 | 0,51 | -1,40 | 1,62E-01 | 5,64E-01 |
| AT1G12090 | 8,60 | -0,94 | 0,67 | -1,40 | 1,61E-01 | 5,64E-01 |
| AT3G25520 | 3,31 | -1,63 | 1,16 | -1,40 | 1,62E-01 | 5,64E-01 |
| AT4G02130 | 3,77 | 1,51 | 1,08 | 1,40 | 1,63E-01 | 5,65E-01 |
| AT1G61226 | 20,72 | -0,65 | 0,47 | -1,39 | 1,64E-01 | 5,68E-01 |
| AT5G20020 | 2,46 | 2,65 | 1,91 | 1,39 | 1,65E-01 | 5,68E-01 |
| AT2G46400 | 1,95 | 2,31 | 1,66 | 1,39 | 1,64E-01 | 5,68E-01 |
| AT4G36040 | 3,49 | 1,72 | 1,24 | 1,39 | 1,65E-01 | 5,68E-01 |
| AT1G06227 | 343,33 | 0,51 | 0,37 | 1,39 | 1,66E-01 | 5,69E-01 |
| AT1G76065 | 4,52 | 1,48 | 1,07 | 1,39 | 1,65E-01 | 5,69E-01 |
| AT2G08770 | 10,72 | -0,86 | 0,62 | -1,39 | 1,66E-01 | 5,69E-01 |
| AT2G42490 | 2,30 | 2,17 | 1,57 | 1,38 | 1,67E-01 | 5,71E-01 |
| AT5G38430 | 4,72 | -1,15 | 0,83 | -1,38 | 1,67E-01 | 5,72E-01 |
| AT1G17220 | 3,72 | -1,50 | 1,09 | -1,38 | 1,69E-01 | 5,75E-01 |
| AT5G57990 | 5,76 | -1,98 | 1,43 | -1,38 | 1,69E-01 | 5,75E-01 |
| AT1G52230 | 2,95 | -1,22 | 0,89 | -1,37 | 1,70E-01 | 5,76E-01 |
| AT2G47015 | 1248,43 | 0,43 | 0,31 | 1,37 | 1,70E-01 | 5,76E-01 |
| AT2G47020 | 1248,75 | 0,43 | 0,31 | 1,37 | 1,69E-01 | 5,76E-01 |
| AT3G04400 | 2,05 | -1,66 | 1,21 | -1,37 | 1,71E-01 | 5,77E-01 |
| AT4G24570 | 3,86 | 1,23 | 0,90 | 1,37 | 1,71E-01 | 5,77E-01 |
| AT1G67090 | 31,55 | -0,61 | 0,44 | -1,37 | 1,71E-01 | 5,77E-01 |

| | | | | | | |
|---|---|---|---|---|---|---|
| AT5G26340 | 2,20 | -1,82 | 1,33 | -1,37 | 1,71E-01 | 5,77E-01 |
| AT5G49440 | 5,06 | -1,12 | 0,82 | -1,37 | 1,72E-01 | 5,77E-01 |
| AT3G52430 | 2,34 | 2,42 | 1,77 | 1,36 | 1,72E-01 | 5,78E-01 |
| AT1G63130 | 240,21 | -0,37 | 0,27 | -1,36 | 1,75E-01 | 5,85E-01 |
| AT1G08857 | 2,42 | 2,74 | 2,02 | 1,36 | 1,75E-01 | 5,85E-01 |
| AT4G20830 | 3,02 | 1,61 | 1,19 | 1,35 | 1,76E-01 | 5,87E-01 |
| AT4G13510 | 4,76 | 1,25 | 0,92 | 1,35 | 1,77E-01 | 5,89E-01 |
| AT1G66970 | 4,60 | 1,42 | 1,05 | 1,35 | 1,77E-01 | 5,89E-01 |
| AT5G49980 | 4,39 | 2,01 | 1,49 | 1,35 | 1,77E-01 | 5,89E-01 |
| AT1G06237 | 336,80 | 0,51 | 0,38 | 1,34 | 1,79E-01 | 5,92E-01 |
| AT3G54920 | 2,20 | 2,04 | 1,52 | 1,35 | 1,78E-01 | 5,92E-01 |
| AT3G24480 | 4,54 | 1,69 | 1,25 | 1,34 | 1,79E-01 | 5,92E-01 |
| AT2G37160 | 3,77 | 1,25 | 0,93 | 1,34 | 1,80E-01 | 5,93E-01 |
| AT3G28540 | 2,73 | 2,11 | 1,58 | 1,34 | 1,80E-01 | 5,93E-01 |
| AT2G32240 | 2,12 | 2,21 | 1,65 | 1,34 | 1,80E-01 | 5,93E-01 |
| AT5G57560 | 1,92 | 1,95 | 1,45 | 1,34 | 1,80E-01 | 5,93E-01 |
| AT4G28570 | 3,15 | -1,26 | 0,94 | -1,34 | 1,81E-01 | 5,93E-01 |
| AT2G20990 | 4,50 | -1,55 | 1,16 | -1,33 | 1,82E-01 | 5,95E-01 |
| AT2G08855 | 3,73 | 1,25 | 0,94 | 1,33 | 1,84E-01 | 5,95E-01 |
| AT3G23326 | 69,32 | 0,85 | 0,64 | 1,33 | 1,84E-01 | 5,95E-01 |
| AT3G61630 | 1,88 | 2,51 | 1,88 | 1,33 | 1,82E-01 | 5,95E-01 |
| AT5G11740 | 4,10 | 1,37 | 1,03 | 1,33 | 1,84E-01 | 5,95E-01 |
| AT1G18260 | 1,96 | 2,27 | 1,70 | 1,33 | 1,83E-01 | 5,95E-01 |
| AT1G23310 | 12,45 | -0,82 | 0,62 | -1,33 | 1,84E-01 | 5,95E-01 |
| AT3G04470 | 2,70 | 2,08 | 1,57 | 1,33 | 1,84E-01 | 5,95E-01 |
| AT4G23650 | 3,54 | 2,06 | 1,55 | 1,33 | 1,83E-01 | 5,95E-01 |
| AT4G34890 | 2,69 | 2,19 | 1,65 | 1,33 | 1,83E-01 | 5,95E-01 |
| AT2G21390 | 23,77 | 1,78 | 1,35 | 1,32 | 1,86E-01 | 6,01E-01 |
| AT1G73870 | 2,16 | -1,80 | 1,37 | -1,32 | 1,88E-01 | 6,07E-01 |
| AT3G20820 | 3,82 | 1,45 | 1,10 | 1,31 | 1,89E-01 | 6,07E-01 |
| AT2G18280 | 2,10 | -2,47 | 1,88 | -1,31 | 1,89E-01 | 6,08E-01 |
| AT1G19960 | 1,88 | -2,25 | 1,73 | -1,30 | 1,92E-01 | 6,08E-01 |
| AT1G21270 | 2,66 | 2,39 | 1,83 | 1,30 | 1,92E-01 | 6,08E-01 |
| AT1G32720 | 2,47 | 1,52 | 1,17 | 1,30 | 1,94E-01 | 6,08E-01 |
| AT2G40140 | 4,76 | 1,69 | 1,30 | 1,30 | 1,93E-01 | 6,08E-01 |
| AT2G47980 | 2,39 | 2,12 | 1,62 | 1,31 | 1,91E-01 | 6,08E-01 |
| AT4G21280 | 3,20 | -1,36 | 1,04 | -1,31 | 1,92E-01 | 6,08E-01 |
| AT4G34150 | 1,90 | 2,10 | 1,61 | 1,31 | 1,92E-01 | 6,08E-01 |
| AT5G04560 | 3,61 | 2,01 | 1,54 | 1,30 | 1,92E-01 | 6,08E-01 |
| AT5G63190 | 2,74 | -1,82 | 1,40 | -1,30 | 1,93E-01 | 6,08E-01 |
| AT1G08115 | 40,91 | 0,54 | 0,42 | 1,30 | 1,94E-01 | 6,08E-01 |
| AT2G47940 | 2,47 | -2,32 | 1,77 | -1,31 | 1,91E-01 | 6,08E-01 |
| AT3G55320 | 2,94 | -2,03 | 1,56 | -1,30 | 1,93E-01 | 6,08E-01 |
| AT4G10710 | 3,86 | 2,14 | 1,63 | 1,31 | 1,91E-01 | 6,08E-01 |
| AT4G13505 | 4,72 | 1,19 | 0,92 | 1,30 | 1,94E-01 | 6,08E-01 |
| AT4G06110 | 22,73 | 0,65 | 0,50 | 1,30 | 1,93E-01 | 6,08E-01 |
| AT5G09515 | 9,09 | -0,84 | 0,65 | -1,30 | 1,94E-01 | 6,08E-01 |
| AT3G08530 | 12,53 | -0,83 | 0,64 | -1,30 | 1,95E-01 | 6,09E-01 |
| AT3G50820 | 7,18 | -1,05 | 0,81 | -1,30 | 1,95E-01 | 6,09E-01 |
| AT1G05993 | 11,40 | 0,73 | 0,56 | 1,29 | 1,96E-01 | 6,11E-01 |
| AT1G75040 | 3,07 | -1,27 | 0,98 | -1,29 | 1,97E-01 | 6,13E-01 |
| AT2G07629 | 5,41 | -1,10 | 0,85 | -1,29 | 1,97E-01 | 6,13E-01 |
| AT3G05700 | 20,03 | 0,95 | 0,73 | 1,29 | 1,98E-01 | 6,15E-01 |
| AT1G55673 | 3,49 | -1,11 | 0,86 | -1,29 | 1,99E-01 | 6,15E-01 |
| AT2G26780 | 3,27 | 2,01 | 1,57 | 1,28 | 2,00E-01 | 6,16E-01 |
| AT4G00360 | 2,92 | -1,28 | 1,00 | -1,28 | 2,00E-01 | 6,16E-01 |

| | | | | | | |
|---|---|---|---|---|---|---|
| AT5G20250 | 2,34 | 2,03 | 1,58 | 1,28 | 1,99E-01 | 6,16E-01 |
| AT1G14740 | 3,27 | 1,82 | 1,42 | 1,28 | 2,00E-01 | 6,16E-01 |
| AT4G31170 | 2,81 | 2,22 | 1,74 | 1,28 | 2,00E-01 | 6,17E-01 |
| AT4G39730 | 3,06 | -1,48 | 1,16 | -1,28 | 2,01E-01 | 6,17E-01 |
| AT5G01775 | 4,59 | 1,20 | 0,94 | 1,28 | 2,02E-01 | 6,17E-01 |
| AT4G37930 | 4,20 | -1,08 | 0,85 | -1,28 | 2,01E-01 | 6,17E-01 |
| AT5G42020 | 4,47 | 1,32 | 1,03 | 1,28 | 2,02E-01 | 6,17E-01 |
| AT3G13440 | 3,42 | 1,53 | 1,20 | 1,27 | 2,03E-01 | 6,18E-01 |
| AT3G26810 | 23,04 | -0,53 | 0,42 | -1,27 | 2,03E-01 | 6,18E-01 |
| AT5G25280 | 3,55 | 1,86 | 1,46 | 1,27 | 2,03E-01 | 6,18E-01 |
| AT2G05710 | 3,35 | 1,73 | 1,36 | 1,27 | 2,04E-01 | 6,20E-01 |
| AT4G25530 | 5,35 | 1,02 | 0,81 | 1,27 | 2,04E-01 | 6,20E-01 |
| AT1G09780 | 4,32 | 1,43 | 1,13 | 1,27 | 2,05E-01 | 6,20E-01 |
| AT1G49240 | 7,10 | -1,49 | 1,17 | -1,27 | 2,05E-01 | 6,21E-01 |
| AT1G48090 | 11,18 | 0,98 | 0,78 | 1,27 | 2,06E-01 | 6,21E-01 |
| AT4G32260 | 3,86 | -1,18 | 0,93 | -1,27 | 2,05E-01 | 6,21E-01 |
| AT1G31330 | 6,03 | -0,81 | 0,64 | -1,26 | 2,07E-01 | 6,24E-01 |
| AT4G32990 | 2,22 | -1,42 | 1,13 | -1,26 | 2,08E-01 | 6,25E-01 |
| AT4G00630 | 4,40 | -1,83 | 1,45 | -1,26 | 2,08E-01 | 6,25E-01 |
| AT3G44720 | 3,49 | 1,77 | 1,41 | 1,26 | 2,08E-01 | 6,25E-01 |
| AT5G57710 | 4,28 | 1,25 | 1,00 | 1,25 | 2,10E-01 | 6,26E-01 |
| AT5G18410 | 3,15 | 1,42 | 1,13 | 1,26 | 2,09E-01 | 6,26E-01 |
| AT5G45480 | 2,31 | 1,96 | 1,56 | 1,25 | 2,09E-01 | 6,26E-01 |
| AT2G26975 | 1,93 | 1,83 | 1,47 | 1,25 | 2,12E-01 | 6,29E-01 |
| AT4G20360 | 5,64 | -0,88 | 0,71 | -1,25 | 2,12E-01 | 6,29E-01 |
| AT4G20362 | 5,64 | -0,88 | 0,71 | -1,25 | 2,12E-01 | 6,29E-01 |
| AT5G42530 | 4,10 | -0,95 | 0,76 | -1,25 | 2,12E-01 | 6,29E-01 |
| AT3G19820 | 10,30 | -2,01 | 1,62 | -1,25 | 2,12E-01 | 6,29E-01 |
| AT1G55920 | 3,12 | 1,55 | 1,25 | 1,24 | 2,13E-01 | 6,31E-01 |
| AT5G15845 | 2,91 | 1,96 | 1,57 | 1,24 | 2,14E-01 | 6,31E-01 |
| AT3G23810 | 2,25 | -1,57 | 1,27 | -1,24 | 2,14E-01 | 6,31E-01 |
| AT2G36390 | 2,07 | -1,75 | 1,41 | -1,24 | 2,14E-01 | 6,31E-01 |
| AT3G06325 | 6,42 | 0,96 | 0,77 | 1,24 | 2,14E-01 | 6,31E-01 |
| AT1G01550 | 2,54 | 1,63 | 1,32 | 1,24 | 2,15E-01 | 6,31E-01 |
| AT5G04720 | 2,75 | 2,30 | 1,86 | 1,24 | 2,16E-01 | 6,33E-01 |
| AT1G07940 | 3,71 | -1,19 | 0,96 | -1,24 | 2,16E-01 | 6,33E-01 |
| AT4G28750 | 4,56 | -1,26 | 1,02 | -1,24 | 2,16E-01 | 6,33E-01 |
| AT3G29130 | 2,05 | 2,00 | 1,62 | 1,23 | 2,17E-01 | 6,34E-01 |
| AT4G19110 | 3,30 | 2,11 | 1,71 | 1,23 | 2,17E-01 | 6,35E-01 |
| AT3G52240 | 2,27 | -2,24 | 1,82 | -1,23 | 2,18E-01 | 6,37E-01 |
| AT3G04165 | 2,58 | 1,51 | 1,23 | 1,23 | 2,20E-01 | 6,38E-01 |
| AT5G17020 | 2,03 | -1,81 | 1,47 | -1,23 | 2,19E-01 | 6,38E-01 |
| AT4G37460 | 2,98 | 1,97 | 1,61 | 1,23 | 2,19E-01 | 6,38E-01 |
| AT1G04410 | 3,62 | -1,22 | 1,00 | -1,22 | 2,21E-01 | 6,42E-01 |
| AT1G52400 | 1,99 | -1,81 | 1,49 | -1,22 | 2,23E-01 | 6,42E-01 |
| AT3G05590 | 1,96 | -1,54 | 1,27 | -1,22 | 2,23E-01 | 6,42E-01 |
| AT5G01785 | 2,53 | 1,87 | 1,53 | 1,22 | 2,22E-01 | 6,42E-01 |
| AT5G13000 | 3,30 | 1,72 | 1,41 | 1,22 | 2,23E-01 | 6,42E-01 |
| AT5G07325 | 8,05 | 0,88 | 0,72 | 1,22 | 2,23E-01 | 6,42E-01 |
| AT1G55591 | 5,61 | -2,20 | 1,81 | -1,22 | 2,24E-01 | 6,43E-01 |
| AT5G61010 | 3,05 | 1,88 | 1,55 | 1,22 | 2,24E-01 | 6,43E-01 |
| AT4G20835 | 2,66 | 1,48 | 1,22 | 1,22 | 2,24E-01 | 6,43E-01 |
| AT1G30410 | 4,36 | -2,20 | 1,81 | -1,21 | 2,25E-01 | 6,44E-01 |
| AT1G65450 | 2,07 | -1,59 | 1,32 | -1,21 | 2,26E-01 | 6,44E-01 |
| AT1G80070 | 7,13 | 1,47 | 1,21 | 1,21 | 2,26E-01 | 6,44E-01 |
| AT3G55410 | 3,63 | 1,81 | 1,50 | 1,21 | 2,26E-01 | 6,44E-01 |

| | | | | | | |
|---|---|---|---|---|---|---|
| AT1G03060 | 7,63 | -1,18 | 0,98 | -1,21 | 2,26E-01 | 6,44E-01 |
| AT4G32250 | 2,07 | 2,06 | 1,70 | 1,21 | 2,26E-01 | 6,44E-01 |
| AT4G13880 | 2,00 | 2,03 | 1,68 | 1,21 | 2,27E-01 | 6,44E-01 |
| AT2G33730 | 2,24 | -1,71 | 1,41 | -1,21 | 2,27E-01 | 6,44E-01 |
| AT4G38200 | 2,17 | -2,16 | 1,79 | -1,20 | 2,28E-01 | 6,45E-01 |
| AT2G05100 | 10,97 | -0,61 | 0,50 | -1,21 | 2,28E-01 | 6,45E-01 |
| AT5G15850 | 2,49 | 1,94 | 1,61 | 1,20 | 2,28E-01 | 6,45E-01 |
| AT4G02970 | 72,42 | 0,39 | 0,32 | 1,20 | 2,29E-01 | 6,45E-01 |
| AT4G39850 | 2,19 | 2,07 | 1,72 | 1,20 | 2,29E-01 | 6,45E-01 |
| AT2G07718 | 3,36 | 1,30 | 1,09 | 1,20 | 2,32E-01 | 6,50E-01 |
| AT4G17410 | 2,55 | 2,14 | 1,79 | 1,20 | 2,32E-01 | 6,50E-01 |
| AT5G14060 | 2,66 | 1,48 | 1,24 | 1,20 | 2,32E-01 | 6,50E-01 |
| AT1G20440 | 3,88 | -1,00 | 0,84 | -1,20 | 2,32E-01 | 6,50E-01 |
| AT3G06510 | 3,56 | 1,68 | 1,41 | 1,19 | 2,34E-01 | 6,53E-01 |
| AT1G55540 | 2,72 | -2,03 | 1,71 | -1,19 | 2,34E-01 | 6,53E-01 |
| AT4G38775 | 2,66 | -1,14 | 0,96 | -1,19 | 2,35E-01 | 6,56E-01 |
| AT2G41840 | 3,13 | -1,60 | 1,35 | -1,19 | 2,35E-01 | 6,56E-01 |
| AT2G39770 | 2,97 | 1,62 | 1,37 | 1,18 | 2,36E-01 | 6,58E-01 |
| AT1G61140 | 2,12 | 1,77 | 1,50 | 1,18 | 2,37E-01 | 6,59E-01 |
| AT1G33120 | 1,97 | -1,61 | 1,37 | -1,18 | 2,40E-01 | 6,64E-01 |
| AT3G61430 | 1,94 | -2,17 | 1,84 | -1,18 | 2,39E-01 | 6,64E-01 |
| AT1G33950 | 1,88 | 1,81 | 1,54 | 1,17 | 2,40E-01 | 6,64E-01 |
| AT2G30110 | 3,88 | 1,44 | 1,22 | 1,17 | 2,40E-01 | 6,64E-01 |
| AT4G36080 | 9,35 | 1,53 | 1,30 | 1,17 | 2,41E-01 | 6,64E-01 |
| AT1G07930 | 3,92 | 1,27 | 1,08 | 1,17 | 2,42E-01 | 6,65E-01 |
| AT1G17290 | 2,38 | 1,68 | 1,43 | 1,17 | 2,42E-01 | 6,65E-01 |
| AT2G47410 | 3,41 | -2,02 | 1,73 | -1,17 | 2,43E-01 | 6,65E-01 |
| AT4G28755 | 2,23 | -1,42 | 1,22 | -1,17 | 2,43E-01 | 6,65E-01 |
| AT5G03280 | 3,28 | 1,81 | 1,55 | 1,17 | 2,43E-01 | 6,65E-01 |
| AT4G24190 | 5,76 | 0,94 | 0,80 | 1,17 | 2,42E-01 | 6,65E-01 |
| AT4G27090 | 2,08 | -1,62 | 1,39 | -1,17 | 2,43E-01 | 6,65E-01 |
| AT5G47430 | 3,26 | 1,26 | 1,08 | 1,17 | 2,43E-01 | 6,65E-01 |
| AT5G47560 | 3,13 | 1,28 | 1,10 | 1,16 | 2,44E-01 | 6,66E-01 |
| AT4G35000 | 3,13 | -1,43 | 1,23 | -1,16 | 2,46E-01 | 6,69E-01 |
| AT1G19373 | 3,55 | 1,20 | 1,04 | 1,15 | 2,48E-01 | 6,72E-01 |
| AT1G30380 | 5,72 | -1,14 | 0,99 | -1,16 | 2,48E-01 | 6,72E-01 |
| AT3G56705 | 15,48 | 0,62 | 0,54 | 1,15 | 2,49E-01 | 6,72E-01 |
| AT1G76062 | 4,01 | 1,33 | 1,15 | 1,16 | 2,48E-01 | 6,72E-01 |
| AT2G37220 | 3,40 | -1,13 | 0,98 | -1,15 | 2,48E-01 | 6,72E-01 |
| AT3G19030 | 2,04 | 1,77 | 1,53 | 1,15 | 2,49E-01 | 6,72E-01 |
| AT5G10840 | 2,05 | -1,57 | 1,36 | -1,15 | 2,49E-01 | 6,72E-01 |
| AT1G11860 | 2,12 | -1,33 | 1,18 | -1,13 | 2,59E-01 | 6,76E-01 |
| AT1G76630 | 2,14 | -2,25 | 1,98 | -1,14 | 2,56E-01 | 6,76E-01 |
| AT1G78900 | 8,95 | -0,80 | 0,71 | -1,13 | 2,58E-01 | 6,76E-01 |
| AT2G41100 | 6,69 | 0,86 | 0,76 | 1,14 | 2,56E-01 | 6,76E-01 |
| AT2G45960 | 8,71 | 0,90 | 0,79 | 1,14 | 2,56E-01 | 6,76E-01 |
| AT2G47400 | 11,67 | 0,68 | 0,60 | 1,13 | 2,58E-01 | 6,76E-01 |
| AT3G14650 | 2,33 | 1,77 | 1,57 | 1,13 | 2,59E-01 | 6,76E-01 |
| AT3G27865 | 3,38 | 1,15 | 1,02 | 1,13 | 2,58E-01 | 6,76E-01 |
| AT3G52800 | 1,93 | 1,75 | 1,54 | 1,13 | 2,58E-01 | 6,76E-01 |
| AT4G06225 | 6,57 | -1,06 | 0,93 | -1,13 | 2,59E-01 | 6,76E-01 |
| AT4G21150 | 2,92 | 1,82 | 1,60 | 1,14 | 2,55E-01 | 6,76E-01 |
| AT4G30100 | 3,56 | 1,67 | 1,48 | 1,13 | 2,59E-01 | 6,76E-01 |
| AT5G09410 | 3,14 | -2,14 | 1,87 | -1,15 | 2,52E-01 | 6,76E-01 |
| AT1G15130 | 4,06 | -1,70 | 1,50 | -1,13 | 2,56E-01 | 6,76E-01 |
| AT1G27430 | 3,11 | 1,84 | 1,63 | 1,13 | 2,59E-01 | 6,76E-01 |

| | | | | | | |
|---|---|---|---|---|---|---|
| AT1G07593 | 3,78 | -1,68 | 1,48 | -1,13 | 2,57E-01 | 6,76E-01 |
| AT1G76550 | 2,61 | 1,85 | 1,63 | 1,14 | 2,55E-01 | 6,76E-01 |
| AT2G01570 | 3,41 | 1,49 | 1,30 | 1,15 | 2,52E-01 | 6,76E-01 |
| AT2G02741 | 19,79 | 0,49 | 0,43 | 1,13 | 2,58E-01 | 6,76E-01 |
| AT2G35387 | 52,44 | -0,39 | 0,34 | -1,14 | 2,56E-01 | 6,76E-01 |
| AT2G44120 | 4,00 | -1,98 | 1,73 | -1,15 | 2,51E-01 | 6,76E-01 |
| AT3G06055 | 2,49 | 1,38 | 1,21 | 1,14 | 2,53E-01 | 6,76E-01 |
| AT4G13430 | 3,55 | 1,25 | 1,09 | 1,15 | 2,51E-01 | 6,76E-01 |
| AT4G24715 | 2,06 | -1,30 | 1,13 | -1,14 | 2,53E-01 | 6,76E-01 |
| AT5G03240 | 5,81 | -0,96 | 0,85 | -1,13 | 2,59E-01 | 6,76E-01 |
| AT5G24890 | 2,39 | 1,44 | 1,27 | 1,14 | 2,55E-01 | 6,76E-01 |
| AT5G51430 | 4,74 | -2,15 | 1,89 | -1,14 | 2,55E-01 | 6,76E-01 |
| AT4G13940 | 35,08 | -0,50 | 0,44 | -1,13 | 2,60E-01 | 6,76E-01 |
| AT1G05273 | 8,42 | 0,72 | 0,64 | 1,12 | 2,63E-01 | 6,77E-01 |
| AT2G38440 | 3,79 | 1,62 | 1,45 | 1,12 | 2,63E-01 | 6,77E-01 |
| AT3G08947 | 5,24 | 1,75 | 1,57 | 1,12 | 2,63E-01 | 6,77E-01 |
| AT4G13494 | 321,70 | 0,54 | 0,48 | 1,12 | 2,64E-01 | 6,77E-01 |
| AT4G40100 | 3,82 | -0,99 | 0,89 | -1,12 | 2,64E-01 | 6,77E-01 |
| AT5G04810 | 3,05 | -1,68 | 1,50 | -1,12 | 2,61E-01 | 6,77E-01 |
| AT1G16820 | 8,42 | 0,72 | 0,64 | 1,12 | 2,63E-01 | 6,77E-01 |
| AT3G53260 | 2,56 | 1,82 | 1,62 | 1,12 | 2,61E-01 | 6,77E-01 |
| AT4G29200 | 3,91 | 0,95 | 0,84 | 1,12 | 2,61E-01 | 6,77E-01 |
| AT5G20280 | 6,18 | -1,14 | 1,02 | -1,12 | 2,62E-01 | 6,77E-01 |
| AT5G61020 | 2,23 | -2,05 | 1,83 | -1,12 | 2,62E-01 | 6,77E-01 |
| AT4G39080 | 4,27 | -1,89 | 1,69 | -1,12 | 2,64E-01 | 6,78E-01 |
| AT2G26211 | 125,25 | 0,55 | 0,49 | 1,11 | 2,65E-01 | 6,79E-01 |
| AT5G18370 | 3,23 | -2,07 | 1,86 | -1,11 | 2,65E-01 | 6,79E-01 |
| AT5G58410 | 3,13 | -1,73 | 1,56 | -1,11 | 2,67E-01 | 6,82E-01 |
| AT4G13564 | 27,21 | 0,43 | 0,38 | 1,11 | 2,67E-01 | 6,82E-01 |
| AT2G32730 | 3,18 | 1,81 | 1,64 | 1,11 | 2,68E-01 | 6,84E-01 |
| AT5G57870 | 8,84 | 1,31 | 1,18 | 1,11 | 2,69E-01 | 6,84E-01 |
| AT1G64430 | 2,85 | -2,11 | 1,91 | -1,10 | 2,69E-01 | 6,85E-01 |
| AT4G36630 | 4,11 | -2,18 | 1,98 | -1,10 | 2,70E-01 | 6,85E-01 |
| AT1G12775 | 21,87 | 0,49 | 0,45 | 1,10 | 2,70E-01 | 6,86E-01 |
| AT1G62180 | 2,66 | 1,49 | 1,36 | 1,10 | 2,72E-01 | 6,88E-01 |
| AT3G56150 | 2,36 | 1,81 | 1,65 | 1,10 | 2,72E-01 | 6,88E-01 |
| AT1G01040 | 4,73 | 0,98 | 0,89 | 1,10 | 2,73E-01 | 6,88E-01 |
| AT3G44860 | 2,37 | 1,55 | 1,41 | 1,10 | 2,73E-01 | 6,88E-01 |
| AT2G47275 | 830,57 | -0,34 | 0,31 | -1,10 | 2,73E-01 | 6,89E-01 |
| AT1G59870 | 72,89 | 0,40 | 0,37 | 1,09 | 2,75E-01 | 6,90E-01 |
| AT2G20570 | 7,71 | 0,91 | 0,84 | 1,09 | 2,75E-01 | 6,90E-01 |
| AT2G38800 | 2,71 | 1,80 | 1,65 | 1,09 | 2,76E-01 | 6,90E-01 |
| AT2G41640 | 6,36 | 0,70 | 0,64 | 1,09 | 2,75E-01 | 6,90E-01 |
| AT3G60860 | 2,94 | 1,83 | 1,68 | 1,09 | 2,76E-01 | 6,90E-01 |
| AT4G06240 | 6,18 | -1,05 | 0,96 | -1,09 | 2,76E-01 | 6,90E-01 |
| AT5G59732 | 3,71 | 1,24 | 1,14 | 1,09 | 2,77E-01 | 6,90E-01 |
| AT3G56400 | 2,09 | 1,61 | 1,48 | 1,09 | 2,76E-01 | 6,90E-01 |
| AT5G59730 | 3,71 | 1,24 | 1,14 | 1,09 | 2,77E-01 | 6,90E-01 |
| AT3G22970 | 2,77 | 1,20 | 1,10 | 1,09 | 2,78E-01 | 6,92E-01 |
| AT4G39160 | 1,99 | 1,66 | 1,54 | 1,08 | 2,81E-01 | 6,96E-01 |
| AT5G07725 | 8,72 | 0,81 | 0,75 | 1,08 | 2,81E-01 | 6,96E-01 |
| AT1G26880 | 2,18 | -1,47 | 1,36 | -1,08 | 2,80E-01 | 6,96E-01 |
| AT3G52500 | 2,36 | 1,17 | 1,09 | 1,08 | 2,80E-01 | 6,96E-01 |
| AT1G05960 | 4,07 | -1,93 | 1,80 | -1,07 | 2,83E-01 | 6,99E-01 |
| AT1G08053 | 2,32 | -1,22 | 1,14 | -1,07 | 2,83E-01 | 6,99E-01 |
| AT1G70290 | 2,53 | 1,50 | 1,40 | 1,07 | 2,84E-01 | 6,99E-01 |

| | | | | | | |
|---|---|---|---|---|---|---|
| AT2G02750 | 20,13 | 0,46 | 0,43 | 1,08 | 2,82E-01 | 6,99E-01 |
| AT3G14310 | 2,07 | -1,66 | 1,55 | -1,07 | 2,84E-01 | 6,99E-01 |
| AT3G05985 | 2,35 | 1,68 | 1,56 | 1,07 | 2,84E-01 | 6,99E-01 |
| AT4G01010 | 2,44 | -1,25 | 1,17 | -1,07 | 2,84E-01 | 6,99E-01 |
| AT4G20850 | 8,05 | 1,15 | 1,07 | 1,07 | 2,85E-01 | 7,00E-01 |
| AT1G63740 | 1,93 | 1,39 | 1,31 | 1,07 | 2,86E-01 | 7,02E-01 |
| AT4G25880 | 1,93 | -1,97 | 1,85 | -1,07 | 2,86E-01 | 7,02E-01 |
| AT4G17330 | 4,91 | 1,67 | 1,57 | 1,06 | 2,89E-01 | 7,06E-01 |
| AT4G31180 | 4,29 | -1,15 | 1,08 | -1,06 | 2,89E-01 | 7,06E-01 |
| AT2G31890 | 2,01 | 1,89 | 1,78 | 1,06 | 2,89E-01 | 7,06E-01 |
| AT2G01008 | 28,92 | -0,39 | 0,37 | -1,05 | 2,93E-01 | 7,09E-01 |
| AT2G41430 | 6,85 | 0,82 | 0,78 | 1,06 | 2,91E-01 | 7,09E-01 |
| AT3G07870 | 5,57 | 1,59 | 1,52 | 1,05 | 2,93E-01 | 7,09E-01 |
| AT4G13570 | 36,83 | 0,39 | 0,37 | 1,05 | 2,93E-01 | 7,09E-01 |
| AT5G04415 | 4,71 | 0,99 | 0,94 | 1,05 | 2,93E-01 | 7,09E-01 |
| AT1G53310 | 3,45 | 1,37 | 1,29 | 1,06 | 2,91E-01 | 7,09E-01 |
| AT1G63770 | 7,56 | -0,91 | 0,87 | -1,05 | 2,93E-01 | 7,09E-01 |
| AT3G20690 | 3,62 | -1,01 | 0,96 | -1,05 | 2,93E-01 | 7,09E-01 |
| AT3G52930 | 3,81 | -1,01 | 0,96 | -1,05 | 2,92E-01 | 7,09E-01 |
| AT5G12860 | 2,32 | -1,91 | 1,82 | -1,05 | 2,94E-01 | 7,09E-01 |
| AT4G06320 | 7,79 | -0,64 | 0,61 | -1,05 | 2,94E-01 | 7,09E-01 |
| AT4G04340 | 2,36 | -1,71 | 1,64 | -1,05 | 2,96E-01 | 7,11E-01 |
| AT1G32270 | 1,96 | -1,48 | 1,42 | -1,04 | 2,96E-01 | 7,12E-01 |
| AT4G17340 | 2,05 | 1,77 | 1,70 | 1,04 | 2,96E-01 | 7,12E-01 |
| AT5G39500 | 6,69 | 1,58 | 1,52 | 1,04 | 2,98E-01 | 7,14E-01 |
| AT4G14504 | 4,83 | 0,83 | 0,80 | 1,04 | 3,00E-01 | 7,16E-01 |
| AT4G26255 | 3,93 | 1,33 | 1,28 | 1,04 | 3,00E-01 | 7,16E-01 |
| AT4G02770 | 8,37 | -0,63 | 0,61 | -1,04 | 3,00E-01 | 7,16E-01 |
| AT5G40395 | 2,01 | 1,31 | 1,26 | 1,04 | 2,99E-01 | 7,16E-01 |
| AT1G35501 | 18,58 | -0,51 | 0,49 | -1,03 | 3,02E-01 | 7,17E-01 |
| AT3G07855 | 4,66 | -0,77 | 0,75 | -1,03 | 3,03E-01 | 7,17E-01 |
| AT2G25660 | 3,83 | 1,66 | 1,61 | 1,03 | 3,03E-01 | 7,17E-01 |
| AT3G13724 | 5,13 | -0,81 | 0,78 | -1,03 | 3,01E-01 | 7,17E-01 |
| AT4G08850 | 2,82 | 1,57 | 1,53 | 1,03 | 3,03E-01 | 7,17E-01 |
| AT4G39260 | 6,03 | -0,70 | 0,68 | -1,03 | 3,02E-01 | 7,17E-01 |
| AT5G46270 | 2,21 | 1,78 | 1,73 | 1,03 | 3,01E-01 | 7,17E-01 |
| AT4G19120 | 2,16 | -1,17 | 1,13 | -1,03 | 3,03E-01 | 7,17E-01 |
| AT1G06243 | 5,81 | 0,69 | 0,67 | 1,03 | 3,04E-01 | 7,18E-01 |
| AT3G47910 | 2,73 | 1,57 | 1,53 | 1,03 | 3,05E-01 | 7,19E-01 |
| AT5G14270 | 2,74 | 1,57 | 1,53 | 1,02 | 3,05E-01 | 7,19E-01 |
| AT5G46430 | 2,53 | 1,34 | 1,31 | 1,02 | 3,06E-01 | 7,19E-01 |
| AT1G09937 | 6,18 | 0,78 | 0,77 | 1,02 | 3,06E-01 | 7,19E-01 |
| AT1G11910 | 5,12 | -0,98 | 0,96 | -1,02 | 3,07E-01 | 7,21E-01 |
| AT1G40104 | 7,42 | 0,81 | 0,79 | 1,02 | 3,09E-01 | 7,24E-01 |
| AT1G12620 | 2,40 | 1,24 | 1,22 | 1,02 | 3,09E-01 | 7,24E-01 |
| AT2G16485 | 2,25 | -1,54 | 1,52 | -1,02 | 3,09E-01 | 7,24E-01 |
| AT1G64583 | 1,87 | -1,20 | 1,19 | -1,01 | 3,10E-01 | 7,26E-01 |
| AT1G63005 | 3,14 | -1,15 | 1,14 | -1,01 | 3,12E-01 | 7,28E-01 |
| AT4G24413 | 2,10 | -1,10 | 1,09 | -1,01 | 3,12E-01 | 7,28E-01 |
| AT1G60950 | 5,66 | -0,81 | 0,81 | -1,01 | 3,14E-01 | 7,31E-01 |
| AT4G07975 | 9,02 | -0,94 | 0,94 | -1,01 | 3,14E-01 | 7,31E-01 |
| AT2G41410 | 4,84 | 1,13 | 1,12 | 1,01 | 3,14E-01 | 7,31E-01 |
| AT4G02760 | 2,62 | 1,49 | 1,48 | 1,00 | 3,15E-01 | 7,32E-01 |
| AT4G15560 | 2,23 | -1,23 | 1,23 | -1,00 | 3,16E-01 | 7,33E-01 |
| AT1G79040 | 10,12 | -0,74 | 0,74 | -1,00 | 3,17E-01 | 7,34E-01 |
| AT4G10800 | 6,52 | -0,69 | 0,69 | -1,00 | 3,17E-01 | 7,34E-01 |

| | | | | | | |
|---|---|---|---|---|---|---|
| AT4G00700 | 2,06 | 1,81 | 1,81 | 1,00 | 3,18E-01 | 7,34E-01 |
| AT1G05913 | 9,06 | 0,75 | 0,75 | 1,00 | 3,19E-01 | 7,35E-01 |
| AT1G31163 | 6,82 | -0,61 | 0,62 | -1,00 | 3,19E-01 | 7,35E-01 |
| AT2G37190 | 11,90 | 1,39 | 1,39 | 1,00 | 3,19E-01 | 7,35E-01 |
| AT5G46210 | 3,09 | -1,44 | 1,45 | -1,00 | 3,19E-01 | 7,35E-01 |
| AT1G62590 | 20,75 | 0,41 | 0,42 | 0,99 | 3,21E-01 | 7,37E-01 |
| AT1G69830 | 1,91 | -1,19 | 1,20 | -0,99 | 3,21E-01 | 7,37E-01 |
| AT5G26360 | 2,70 | 1,51 | 1,52 | 0,99 | 3,21E-01 | 7,37E-01 |
| AT3G07100 | 6,58 | 0,99 | 1,00 | 0,99 | 3,22E-01 | 7,38E-01 |
| AT5G42950 | 18,25 | 1,25 | 1,26 | 0,99 | 3,23E-01 | 7,40E-01 |
| AT3G01715 | 19,56 | 0,74 | 0,75 | 0,99 | 3,24E-01 | 7,40E-01 |
| AT1G61740 | 2,13 | 1,33 | 1,35 | 0,99 | 3,24E-01 | 7,40E-01 |
| AT4G18670 | 7,64 | -0,61 | 0,62 | -0,99 | 3,25E-01 | 7,41E-01 |
| AT1G21580 | 3,07 | 1,49 | 1,51 | 0,98 | 3,26E-01 | 7,42E-01 |
| AT1G49700 | 4,39 | 1,23 | 1,25 | 0,98 | 3,26E-01 | 7,42E-01 |
| AT1G32230 | 5,15 | 1,24 | 1,27 | 0,98 | 3,27E-01 | 7,43E-01 |
| AT2G36910 | 8,87 | 1,21 | 1,23 | 0,98 | 3,27E-01 | 7,44E-01 |
| AT1G58230 | 3,69 | 1,50 | 1,54 | 0,98 | 3,29E-01 | 7,44E-01 |
| AT3G46780 | 9,83 | -0,83 | 0,85 | -0,98 | 3,28E-01 | 7,44E-01 |
| AT1G61100 | 1,97 | 1,76 | 1,80 | 0,98 | 3,28E-01 | 7,44E-01 |
| AT1G61275 | 11,06 | 0,55 | 0,56 | 0,97 | 3,30E-01 | 7,45E-01 |
| AT1G32750 | 1,94 | -1,60 | 1,64 | -0,97 | 3,30E-01 | 7,45E-01 |
| AT5G22500 | 2,19 | -1,17 | 1,21 | -0,97 | 3,30E-01 | 7,45E-01 |
| AT2G04030 | 4,28 | -1,01 | 1,04 | -0,97 | 3,32E-01 | 7,45E-01 |
| AT3G02045 | 5,72 | -0,74 | 0,76 | -0,97 | 3,32E-01 | 7,45E-01 |
| AT3G14270 | 2,35 | 1,39 | 1,44 | 0,97 | 3,32E-01 | 7,45E-01 |
| AT3G44630 | 4,39 | 1,15 | 1,18 | 0,97 | 3,31E-01 | 7,45E-01 |
| AT3G05655 | 1,87 | 1,49 | 1,54 | 0,97 | 3,32E-01 | 7,45E-01 |
| AT1G70940 | 2,12 | 1,60 | 1,66 | 0,97 | 3,34E-01 | 7,48E-01 |
| AT4G11900 | 2,31 | 1,12 | 1,16 | 0,97 | 3,34E-01 | 7,48E-01 |
| AT2G31820 | 4,59 | 0,75 | 0,78 | 0,96 | 3,35E-01 | 7,50E-01 |
| AT3G07860 | 5,76 | -0,74 | 0,77 | -0,96 | 3,36E-01 | 7,50E-01 |
| AT1G06040 | 2,85 | -1,17 | 1,22 | -0,96 | 3,36E-01 | 7,50E-01 |
| AT2G20190 | 8,93 | -1,21 | 1,25 | -0,96 | 3,36E-01 | 7,50E-01 |
| AT1G49760 | 5,88 | -1,29 | 1,34 | -0,96 | 3,37E-01 | 7,50E-01 |
| AT3G24542 | 2,63 | -1,05 | 1,10 | -0,95 | 3,40E-01 | 7,52E-01 |
| AT4G05320 | 17,67 | -0,49 | 0,52 | -0,96 | 3,39E-01 | 7,52E-01 |
| AT4G16190 | 11,22 | -0,59 | 0,61 | -0,95 | 3,40E-01 | 7,52E-01 |
| AT1G22620 | 12,70 | 0,89 | 0,93 | 0,95 | 3,40E-01 | 7,52E-01 |
| AT2G25800 | 4,68 | 1,41 | 1,47 | 0,96 | 3,39E-01 | 7,52E-01 |
| AT3G23780 | 4,59 | 1,34 | 1,40 | 0,96 | 3,39E-01 | 7,52E-01 |
| AT4G16340 | 3,35 | -1,52 | 1,59 | -0,96 | 3,39E-01 | 7,52E-01 |
| AT1G80000 | 1,90 | -2,09 | 2,20 | -0,95 | 3,41E-01 | 7,53E-01 |
| AT1G12900 | 4,70 | -0,73 | 0,76 | -0,95 | 3,41E-01 | 7,53E-01 |
| AT5G14570 | 2,87 | 1,20 | 1,26 | 0,95 | 3,42E-01 | 7,54E-01 |
| AT5G11560 | 3,42 | -1,08 | 1,14 | -0,95 | 3,43E-01 | 7,54E-01 |
| AT4G06810 | 54,17 | 0,70 | 0,73 | 0,95 | 3,44E-01 | 7,55E-01 |
| AT4G06805 | 54,05 | 0,70 | 0,74 | 0,95 | 3,44E-01 | 7,55E-01 |
| AT5G40910 | 2,64 | 1,27 | 1,34 | 0,94 | 3,45E-01 | 7,56E-01 |
| AT3G08580 | 9,77 | -0,72 | 0,77 | -0,94 | 3,45E-01 | 7,56E-01 |
| AT3G24612 | 3,02 | 1,06 | 1,12 | 0,94 | 3,45E-01 | 7,56E-01 |
| AT5G13890 | 14,03 | 0,72 | 0,76 | 0,94 | 3,45E-01 | 7,56E-01 |
| AT3G03990 | 2,37 | 1,41 | 1,50 | 0,94 | 3,47E-01 | 7,57E-01 |
| AT5G40340 | 2,24 | -1,03 | 1,09 | -0,94 | 3,47E-01 | 7,57E-01 |
| AT5G14740 | 2,79 | -1,34 | 1,42 | -0,94 | 3,47E-01 | 7,57E-01 |
| AT2G01910 | 2,66 | 1,36 | 1,45 | 0,93 | 3,50E-01 | 7,59E-01 |

| | | | | | | |
|---|---|---|---|---|---|---|
| AT2G39010 | 4,44 | -0,99 | 1,06 | -0,94 | 3,48E-01 | 7,59E-01 |
| AT3G49230 | 1,94 | 1,35 | 1,45 | 0,93 | 3,51E-01 | 7,59E-01 |
| AT1G13440 | 5,54 | -0,69 | 0,74 | -0,93 | 3,50E-01 | 7,59E-01 |
| AT1G56340 | 4,49 | 0,90 | 0,97 | 0,93 | 3,52E-01 | 7,59E-01 |
| AT2G08390 | 5,34 | -0,78 | 0,84 | -0,93 | 3,50E-01 | 7,59E-01 |
| AT2G47390 | 2,13 | 1,72 | 1,84 | 0,94 | 3,49E-01 | 7,59E-01 |
| AT3G12780 | 5,73 | -0,67 | 0,72 | -0,93 | 3,50E-01 | 7,59E-01 |
| AT5G54940 | 2,86 | 0,98 | 1,06 | 0,93 | 3,51E-01 | 7,59E-01 |
| AT5G58430 | 1,95 | 1,40 | 1,50 | 0,93 | 3,51E-01 | 7,59E-01 |
| AT2G25430 | 4,01 | -1,44 | 1,56 | -0,93 | 3,54E-01 | 7,59E-01 |
| AT3G41768 | 91,21 | -0,26 | 0,28 | -0,93 | 3,54E-01 | 7,59E-01 |
| AT5G02310 | 4,95 | 0,92 | 1,00 | 0,93 | 3,54E-01 | 7,59E-01 |
| AT5G36180 | 3,47 | 0,88 | 0,95 | 0,93 | 3,52E-01 | 7,59E-01 |
| AT2G17200 | 1,89 | 1,55 | 1,67 | 0,93 | 3,53E-01 | 7,59E-01 |
| AT3G03425 | 4,95 | 0,70 | 0,76 | 0,93 | 3,54E-01 | 7,59E-01 |
| AT5G46260 | 1,90 | 1,59 | 1,71 | 0,93 | 3,54E-01 | 7,59E-01 |
| AT5G50920 | 18,45 | -0,48 | 0,52 | -0,92 | 3,55E-01 | 7,60E-01 |
| AT2G41740 | 6,26 | -1,15 | 1,24 | -0,92 | 3,57E-01 | 7,61E-01 |
| AT3G09390 | 6,61 | 0,79 | 0,86 | 0,92 | 3,56E-01 | 7,61E-01 |
| AT3G58640 | 2,56 | -1,73 | 1,87 | -0,92 | 3,57E-01 | 7,61E-01 |
| AT1G14670 | 2,24 | 1,36 | 1,48 | 0,92 | 3,57E-01 | 7,61E-01 |
| AT2G35750 | 26,17 | -0,43 | 0,47 | -0,92 | 3,58E-01 | 7,61E-01 |
| AT4G17140 | 11,35 | -0,86 | 0,94 | -0,92 | 3,58E-01 | 7,61E-01 |
| AT3G48860 | 1,96 | -2,26 | 2,47 | -0,92 | 3,59E-01 | 7,63E-01 |
| AT1G44575 | 5,52 | -0,63 | 0,69 | -0,91 | 3,62E-01 | 7,66E-01 |
| AT5G46420 | 2,51 | 1,35 | 1,48 | 0,91 | 3,62E-01 | 7,66E-01 |
| AT1G62390 | 2,10 | 1,87 | 2,05 | 0,91 | 3,62E-01 | 7,66E-01 |
| AT1G78830 | 2,12 | -1,18 | 1,29 | -0,91 | 3,62E-01 | 7,66E-01 |
| AT1G18075 | 4243,99 | -0,16 | 0,18 | -0,90 | 3,67E-01 | 7,68E-01 |
| AT1G63330 | 7,00 | -0,58 | 0,64 | -0,90 | 3,66E-01 | 7,68E-01 |
| AT3G11830 | 1,90 | 1,53 | 1,69 | 0,91 | 3,64E-01 | 7,68E-01 |
| AT3G16850 | 3,96 | -1,16 | 1,28 | -0,90 | 3,67E-01 | 7,68E-01 |
| AT4G31490 | 3,09 | -1,24 | 1,38 | -0,90 | 3,68E-01 | 7,68E-01 |
| AT5G59505 | 3,52 | 1,15 | 1,28 | 0,90 | 3,67E-01 | 7,68E-01 |
| AT5G62690 | 4,26 | -0,81 | 0,89 | -0,90 | 3,66E-01 | 7,68E-01 |
| AT1G04527 | 11,72 | -0,47 | 0,52 | -0,91 | 3,65E-01 | 7,68E-01 |
| AT1G26850 | 2,19 | -1,30 | 1,44 | -0,90 | 3,68E-01 | 7,68E-01 |
| AT2G27310 | 2,22 | 1,20 | 1,33 | 0,91 | 3,65E-01 | 7,68E-01 |
| AT2G29450 | 2,19 | 1,43 | 1,59 | 0,90 | 3,66E-01 | 7,68E-01 |
| AT3G04240 | 3,51 | -1,21 | 1,34 | -0,90 | 3,68E-01 | 7,68E-01 |
| AT5G11700 | 1,99 | 1,50 | 1,66 | 0,90 | 3,66E-01 | 7,68E-01 |
| AT4G09650 | 3,00 | -1,15 | 1,27 | -0,90 | 3,69E-01 | 7,69E-01 |
| AT4G21710 | 2,69 | 1,58 | 1,76 | 0,90 | 3,69E-01 | 7,69E-01 |
| AT5G46450 | 2,49 | 1,51 | 1,68 | 0,90 | 3,70E-01 | 7,70E-01 |
| AT1G51805 | 2,63 | -1,03 | 1,14 | -0,90 | 3,70E-01 | 7,70E-01 |
| AT5G27660 | 6,15 | 0,75 | 0,84 | 0,90 | 3,71E-01 | 7,70E-01 |
| AT4G04970 | 2,63 | 1,81 | 2,04 | 0,89 | 3,74E-01 | 7,71E-01 |
| AT4G26090 | 1,90 | 1,43 | 1,60 | 0,89 | 3,74E-01 | 7,71E-01 |
| AT4G29090 | 2,85 | 1,04 | 1,17 | 0,89 | 3,74E-01 | 7,71E-01 |
| AT4G31160 | 2,16 | 1,71 | 1,92 | 0,89 | 3,73E-01 | 7,71E-01 |
| AT5G51174 | 65,86 | 0,36 | 0,40 | 0,89 | 3,73E-01 | 7,71E-01 |
| AT1G05917 | 8,49 | 0,65 | 0,73 | 0,89 | 3,72E-01 | 7,71E-01 |
| AT2G16676 | 2,63 | -0,97 | 1,09 | -0,89 | 3,74E-01 | 7,71E-01 |
| AT5G53370 | 2,35 | 1,44 | 1,61 | 0,89 | 3,72E-01 | 7,71E-01 |
| AT3G27690 | 1,98 | -1,13 | 1,27 | -0,89 | 3,75E-01 | 7,71E-01 |
| AT2G20722 | 3,24 | 0,95 | 1,07 | 0,89 | 3,76E-01 | 7,72E-01 |

| | | | | | | |
|---|---|---|---|---|---|---|
| AT4G30190 | 7,61 | -0,92 | 1,03 | -0,89 | 3,76E-01 | 7,72E-01 |
| AT3G24929 | 2,38 | 1,09 | 1,23 | 0,88 | 3,77E-01 | 7,73E-01 |
| AT5G04930 | 5,77 | 1,18 | 1,33 | 0,88 | 3,77E-01 | 7,73E-01 |
| AT5G60640 | 2,58 | -1,03 | 1,17 | -0,88 | 3,79E-01 | 7,76E-01 |
| AT1G48920 | 4,08 | -0,81 | 0,93 | -0,88 | 3,81E-01 | 7,78E-01 |
| AT1G52780 | 3,16 | 1,24 | 1,42 | 0,87 | 3,82E-01 | 7,78E-01 |
| AT2G35260 | 3,89 | -0,79 | 0,91 | -0,88 | 3,81E-01 | 7,78E-01 |
| AT2G43110 | 4,82 | 0,84 | 0,95 | 0,88 | 3,81E-01 | 7,78E-01 |
| AT5G02575 | 6,74 | 0,78 | 0,89 | 0,87 | 3,82E-01 | 7,78E-01 |
| AT1G69550 | 1,87 | 1,58 | 1,81 | 0,87 | 3,82E-01 | 7,78E-01 |
| AT3G11940 | 3,79 | -0,91 | 1,04 | -0,87 | 3,82E-01 | 7,78E-01 |
| AT1G27595 | 2,01 | 1,55 | 1,77 | 0,87 | 3,83E-01 | 7,78E-01 |
| AT5G63800 | 1,87 | -1,01 | 1,16 | -0,87 | 3,83E-01 | 7,78E-01 |
| AT2G46255 | 72,33 | -0,34 | 0,39 | -0,87 | 3,85E-01 | 7,79E-01 |
| AT1G31540 | 3,00 | 1,30 | 1,50 | 0,87 | 3,85E-01 | 7,79E-01 |
| AT3G16640 | 7,07 | -0,77 | 0,88 | -0,87 | 3,84E-01 | 7,79E-01 |
| AT1G16880 | 2,91 | -1,19 | 1,37 | -0,86 | 3,87E-01 | 7,81E-01 |
| AT2G36350 | 2,13 | 1,62 | 1,87 | 0,87 | 3,87E-01 | 7,81E-01 |
| AT5G05660 | 1,92 | -2,19 | 2,53 | -0,87 | 3,87E-01 | 7,81E-01 |
| AT5G67030 | 1,94 | 1,40 | 1,61 | 0,87 | 3,87E-01 | 7,81E-01 |
| AT3G62420 | 1,96 | 1,70 | 1,97 | 0,86 | 3,88E-01 | 7,81E-01 |
| AT4G23140 | 2,18 | 1,32 | 1,53 | 0,86 | 3,89E-01 | 7,82E-01 |
| AT2G36880 | 2,29 | 1,24 | 1,44 | 0,86 | 3,89E-01 | 7,82E-01 |
| AT3G62700 | 3,87 | -0,85 | 0,99 | -0,86 | 3,89E-01 | 7,82E-01 |
| AT5G03940 | 2,83 | -1,03 | 1,20 | -0,86 | 3,91E-01 | 7,85E-01 |
| AT1G05817 | 1,94 | 1,12 | 1,31 | 0,85 | 3,93E-01 | 7,87E-01 |
| AT5G43760 | 1,93 | -1,20 | 1,40 | -0,85 | 3,93E-01 | 7,88E-01 |
| AT2G35050 | 3,09 | -1,17 | 1,37 | -0,85 | 3,94E-01 | 7,88E-01 |
| AT5G46760 | 2,52 | 1,25 | 1,46 | 0,85 | 3,94E-01 | 7,88E-01 |
| AT5G10450 | 3,14 | -0,93 | 1,09 | -0,85 | 3,94E-01 | 7,88E-01 |
| AT3G47342 | 15,52 | 0,45 | 0,53 | 0,85 | 3,96E-01 | 7,88E-01 |
| AT5G52070 | 6,76 | 0,88 | 1,04 | 0,85 | 3,96E-01 | 7,88E-01 |
| AT1G01960 | 2,74 | 1,28 | 1,51 | 0,85 | 3,96E-01 | 7,88E-01 |
| AT1G63360 | 4,56 | -0,67 | 0,79 | -0,85 | 3,97E-01 | 7,88E-01 |
| AT3G00980 | 2,10 | 1,47 | 1,72 | 0,85 | 3,95E-01 | 7,88E-01 |
| AT4G13810 | 2,84 | 1,21 | 1,42 | 0,85 | 3,97E-01 | 7,88E-01 |
| AT5G55565 | 4,30 | -0,77 | 0,91 | -0,85 | 3,97E-01 | 7,88E-01 |
| AT5G04140 | 18,61 | -0,57 | 0,68 | -0,84 | 3,99E-01 | 7,91E-01 |
| AT2G05455 | 4,14 | -0,71 | 0,84 | -0,84 | 3,99E-01 | 7,91E-01 |
| AT4G12290 | 3,10 | 1,14 | 1,36 | 0,84 | 4,01E-01 | 7,91E-01 |
| AT5G11670 | 3,41 | 0,88 | 1,05 | 0,84 | 4,01E-01 | 7,91E-01 |
| AT3G10745 | 164,34 | -0,28 | 0,33 | -0,84 | 4,00E-01 | 7,91E-01 |
| AT2G26080 | 10,37 | -0,72 | 0,86 | -0,84 | 4,02E-01 | 7,93E-01 |
| AT1G07890 | 4,08 | 0,79 | 0,95 | 0,84 | 4,03E-01 | 7,93E-01 |
| AT4G31210 | 1,90 | -1,54 | 1,84 | -0,84 | 4,03E-01 | 7,93E-01 |
| AT2G24270 | 3,15 | 1,06 | 1,27 | 0,84 | 4,03E-01 | 7,93E-01 |
| AT2G30520 | 8,24 | -1,01 | 1,21 | -0,83 | 4,04E-01 | 7,93E-01 |
| AT4G00730 | 3,19 | 1,40 | 1,68 | 0,83 | 4,04E-01 | 7,93E-01 |
| AT2G38230 | 6,88 | -0,52 | 0,63 | -0,83 | 4,06E-01 | 7,95E-01 |
| AT3G50950 | 4,46 | 0,83 | 1,00 | 0,83 | 4,06E-01 | 7,95E-01 |
| AT2G28550 | 2,55 | -0,84 | 1,01 | -0,83 | 4,06E-01 | 7,95E-01 |
| AT3G48050 | 3,14 | 0,98 | 1,18 | 0,83 | 4,06E-01 | 7,95E-01 |
| AT4G03260 | 2,29 | -1,14 | 1,37 | -0,83 | 4,08E-01 | 7,96E-01 |
| AT2G02950 | 6,29 | 0,57 | 0,69 | 0,83 | 4,08E-01 | 7,96E-01 |
| AT1G53640 | 2,14 | -1,15 | 1,39 | -0,83 | 4,08E-01 | 7,96E-01 |
| AT1G20015 | 7,84 | -0,59 | 0,72 | -0,82 | 4,10E-01 | 7,98E-01 |

| | | | | | | |
|---|---|---|---|---|---|---|
| AT3G13750 | 2,68 | -1,08 | 1,32 | -0,82 | 4,10E-01 | 7,98E-01 |
| AT2G42600 | 13,57 | -0,53 | 0,65 | -0,82 | 4,10E-01 | 7,98E-01 |
| AT1G06277 | 5,95 | 0,58 | 0,71 | 0,82 | 4,14E-01 | 8,00E-01 |
| AT3G14420 | 4,42 | -0,74 | 0,90 | -0,82 | 4,14E-01 | 8,00E-01 |
| AT1G79930 | 9,47 | 1,17 | 1,43 | 0,82 | 4,14E-01 | 8,00E-01 |
| AT2G07360 | 3,99 | -1,27 | 1,55 | -0,82 | 4,14E-01 | 8,00E-01 |
| AT2G22430 | 4,62 | 0,84 | 1,03 | 0,82 | 4,14E-01 | 8,00E-01 |
| AT2G25450 | 4,94 | 0,82 | 1,00 | 0,82 | 4,13E-01 | 8,00E-01 |
| AT3G46970 | 7,07 | -0,55 | 0,68 | -0,82 | 4,14E-01 | 8,00E-01 |
| AT5G13887 | 13,72 | 0,71 | 0,87 | 0,82 | 4,14E-01 | 8,00E-01 |
| AT5G09040 | 2,31 | 0,92 | 1,13 | 0,81 | 4,15E-01 | 8,01E-01 |
| AT2G46340 | 1,90 | 1,61 | 1,98 | 0,81 | 4,17E-01 | 8,03E-01 |
| AT1G74090 | 6,35 | 0,64 | 0,79 | 0,81 | 4,17E-01 | 8,03E-01 |
| AT1G72645 | 2,19 | 0,92 | 1,14 | 0,81 | 4,18E-01 | 8,03E-01 |
| AT4G22690 | 3,37 | 0,81 | 1,01 | 0,81 | 4,18E-01 | 8,03E-01 |
| AT1G18080 | 2,60 | -1,00 | 1,25 | -0,80 | 4,23E-01 | 8,05E-01 |
| AT1G34300 | 2,71 | 1,36 | 1,70 | 0,80 | 4,23E-01 | 8,05E-01 |
| AT1G35720 | 3,50 | -0,78 | 0,97 | -0,80 | 4,22E-01 | 8,05E-01 |
| AT2G25170 | 3,04 | 1,05 | 1,30 | 0,80 | 4,22E-01 | 8,05E-01 |
| AT1G13210 | 5,46 | 0,94 | 1,17 | 0,81 | 4,20E-01 | 8,05E-01 |
| AT2G27400 | 624,49 | -0,27 | 0,33 | -0,80 | 4,21E-01 | 8,05E-01 |
| AT2G32690 | 2,01 | -0,91 | 1,14 | -0,80 | 4,22E-01 | 8,05E-01 |
| AT2G33170 | 2,19 | 1,39 | 1,73 | 0,80 | 4,22E-01 | 8,05E-01 |
| AT3G59770 | 3,31 | 1,20 | 1,49 | 0,80 | 4,22E-01 | 8,05E-01 |
| AT4G04125 | 2,22 | -0,95 | 1,19 | -0,80 | 4,23E-01 | 8,05E-01 |
| AT3G16785 | 1,89 | 1,11 | 1,38 | 0,80 | 4,24E-01 | 8,06E-01 |
| AT4G13340 | 14,47 | 0,43 | 0,54 | 0,80 | 4,24E-01 | 8,06E-01 |
| AT1G08813 | 8,42 | 0,52 | 0,65 | 0,80 | 4,26E-01 | 8,07E-01 |
| AT5G08670 | 5,95 | 0,81 | 1,02 | 0,80 | 4,26E-01 | 8,07E-01 |
| AT5G13630 | 269,68 | -0,39 | 0,48 | -0,80 | 4,26E-01 | 8,07E-01 |
| AT2G41650 | 5,84 | 0,53 | 0,67 | 0,79 | 4,28E-01 | 8,07E-01 |
| AT5G56030 | 5,79 | 0,63 | 0,80 | 0,79 | 4,27E-01 | 8,07E-01 |
| AT1G35560 | 2,43 | -0,93 | 1,18 | -0,79 | 4,28E-01 | 8,07E-01 |
| AT2G36460 | 1,86 | -1,15 | 1,45 | -0,79 | 4,27E-01 | 8,07E-01 |
| AT4G03415 | 1,93 | 1,49 | 1,87 | 0,79 | 4,28E-01 | 8,07E-01 |
| AT5G66200 | 3,05 | 1,24 | 1,56 | 0,80 | 4,26E-01 | 8,07E-01 |
| AT1G69620 | 2,14 | 1,09 | 1,39 | 0,79 | 4,32E-01 | 8,11E-01 |
| AT2G22300 | 2,59 | 1,24 | 1,58 | 0,79 | 4,32E-01 | 8,11E-01 |
| AT4G32285 | 2,40 | 1,26 | 1,60 | 0,79 | 4,32E-01 | 8,11E-01 |
| AT3G13530 | 2,08 | 1,41 | 1,79 | 0,79 | 4,32E-01 | 8,11E-01 |
| AT1G55325 | 2,49 | 0,95 | 1,22 | 0,78 | 4,34E-01 | 8,13E-01 |
| AT2G26910 | 1,87 | 1,12 | 1,43 | 0,78 | 4,34E-01 | 8,13E-01 |
| AT5G40450 | 4,44 | -0,72 | 0,92 | -0,78 | 4,34E-01 | 8,13E-01 |
| AT4G18197 | 1,87 | 0,97 | 1,25 | 0,78 | 4,35E-01 | 8,13E-01 |
| AT1G05163 | 491,36 | 0,21 | 0,27 | 0,78 | 4,36E-01 | 8,13E-01 |
| AT1G15405 | 491,36 | 0,21 | 0,27 | 0,78 | 4,36E-01 | 8,13E-01 |
| AT1G49750 | 3,00 | -0,85 | 1,09 | -0,78 | 4,37E-01 | 8,15E-01 |
| AT3G54810 | 2,87 | 0,95 | 1,22 | 0,77 | 4,38E-01 | 8,16E-01 |
| AT5G54130 | 6,22 | 0,65 | 0,84 | 0,77 | 4,39E-01 | 8,16E-01 |
| AT2G43570 | 2,25 | 1,32 | 1,70 | 0,78 | 4,38E-01 | 8,16E-01 |
| AT3G51860 | 2,20 | 1,28 | 1,65 | 0,77 | 4,39E-01 | 8,16E-01 |
| AT1G20760 | 7,86 | 1,13 | 1,46 | 0,77 | 4,41E-01 | 8,17E-01 |
| AT2G20580 | 2,81 | -1,29 | 1,68 | -0,77 | 4,40E-01 | 8,17E-01 |
| AT4G19170 | 5,23 | 0,58 | 0,75 | 0,77 | 4,40E-01 | 8,17E-01 |
| AT4G36760 | 3,13 | -1,10 | 1,42 | -0,77 | 4,41E-01 | 8,17E-01 |
| AT4G03020 | 2,29 | -0,95 | 1,24 | -0,77 | 4,42E-01 | 8,17E-01 |

| | | | | | | |
|---|---|---|---|---|---|---|
| AT2G03330 | 8,86 | -0,49 | 0,63 | -0,77 | 4,42E-01 | 8,17E-01 |
| AT2G01320 | 2,02 | 1,19 | 1,56 | 0,77 | 4,43E-01 | 8,19E-01 |
| AT2G22990 | 2,03 | 0,91 | 1,20 | 0,76 | 4,48E-01 | 8,20E-01 |
| AT4G27870 | 2,09 | -1,19 | 1,57 | -0,76 | 4,48E-01 | 8,20E-01 |
| AT5G02870 | 2,41 | -0,98 | 1,30 | -0,76 | 4,49E-01 | 8,20E-01 |
| AT5G08690 | 2,64 | 1,02 | 1,34 | 0,76 | 4,47E-01 | 8,20E-01 |
| AT1G55860 | 12,82 | -0,62 | 0,82 | -0,76 | 4,48E-01 | 8,20E-01 |
| AT1G56280 | 2,59 | -0,85 | 1,12 | -0,76 | 4,48E-01 | 8,20E-01 |
| AT2G30570 | 3,97 | -0,63 | 0,83 | -0,76 | 4,48E-01 | 8,20E-01 |
| AT3G00985 | 2,03 | 1,34 | 1,76 | 0,76 | 4,47E-01 | 8,20E-01 |
| AT3G45190 | 2,88 | 1,18 | 1,55 | 0,76 | 4,46E-01 | 8,20E-01 |
| AT4G33300 | 3,93 | 0,97 | 1,28 | 0,76 | 4,48E-01 | 8,20E-01 |
| AT5G02960 | 2,13 | -1,02 | 1,33 | -0,76 | 4,45E-01 | 8,20E-01 |
| AT5G06600 | 5,36 | 1,12 | 1,47 | 0,76 | 4,47E-01 | 8,20E-01 |
| AT5G53480 | 6,73 | -0,79 | 1,04 | -0,76 | 4,50E-01 | 8,21E-01 |
| AT1G18310 | 4,64 | 0,76 | 1,01 | 0,75 | 4,52E-01 | 8,24E-01 |
| AT2G47450 | 6,02 | 0,63 | 0,84 | 0,75 | 4,53E-01 | 8,25E-01 |
| AT3G09465 | 2,48 | -0,72 | 0,95 | -0,75 | 4,53E-01 | 8,26E-01 |
| AT3G54280 | 2,37 | -1,12 | 1,50 | -0,75 | 4,55E-01 | 8,26E-01 |
| AT4G02510 | 15,59 | 0,46 | 0,62 | 0,75 | 4,54E-01 | 8,26E-01 |
| AT1G08450 | 3,29 | 1,22 | 1,63 | 0,75 | 4,54E-01 | 8,26E-01 |
| AT1G54040 | 2,21 | 0,95 | 1,27 | 0,75 | 4,55E-01 | 8,26E-01 |
| AT2G38220 | 2,26 | 0,87 | 1,17 | 0,74 | 4,57E-01 | 8,28E-01 |
| AT2G22440 | 2,40 | -0,77 | 1,04 | -0,74 | 4,59E-01 | 8,29E-01 |
| AT3G62010 | 2,16 | -0,99 | 1,33 | -0,74 | 4,59E-01 | 8,29E-01 |
| AT5G58040 | 3,29 | 1,22 | 1,64 | 0,74 | 4,59E-01 | 8,29E-01 |
| AT4G34710 | 3,32 | 1,15 | 1,55 | 0,74 | 4,58E-01 | 8,29E-01 |
| AT5G02480 | 2,67 | -1,20 | 1,63 | -0,74 | 4,61E-01 | 8,32E-01 |
| AT2G26500 | 2,52 | 1,06 | 1,44 | 0,73 | 4,63E-01 | 8,33E-01 |
| AT2G28290 | 15,57 | 0,86 | 1,18 | 0,73 | 4,64E-01 | 8,33E-01 |
| AT5G05190 | 2,00 | 1,31 | 1,79 | 0,73 | 4,64E-01 | 8,33E-01 |
| AT5G41790 | 2,54 | -0,98 | 1,33 | -0,73 | 4,63E-01 | 8,33E-01 |
| AT1G48953 | 8,24 | -0,53 | 0,73 | -0,73 | 4,64E-01 | 8,33E-01 |
| AT1G59760 | 3,05 | 1,21 | 1,65 | 0,73 | 4,63E-01 | 8,33E-01 |
| AT5G48860 | 3,76 | -0,67 | 0,92 | -0,73 | 4,64E-01 | 8,33E-01 |
| AT1G02860 | 4,18 | 1,08 | 1,49 | 0,73 | 4,68E-01 | 8,33E-01 |
| AT1G15670 | 2,03 | 0,94 | 1,32 | 0,71 | 4,76E-01 | 8,33E-01 |
| AT1G30230 | 5,95 | -0,80 | 1,12 | -0,72 | 4,72E-01 | 8,33E-01 |
| AT2G31660 | 3,59 | -1,28 | 1,78 | -0,72 | 4,73E-01 | 8,33E-01 |
| AT3G07390 | 2,06 | 0,93 | 1,31 | 0,72 | 4,74E-01 | 8,33E-01 |
| AT3G14010 | 2,74 | 1,25 | 1,74 | 0,72 | 4,73E-01 | 8,33E-01 |
| AT3G23450 | 2,19 | 0,87 | 1,22 | 0,71 | 4,76E-01 | 8,33E-01 |
| AT3G52250 | 4,06 | 1,09 | 1,51 | 0,72 | 4,70E-01 | 8,33E-01 |
| AT3G55200 | 2,62 | 1,38 | 1,93 | 0,71 | 4,75E-01 | 8,33E-01 |
| AT3G56360 | 3,83 | -0,79 | 1,11 | -0,71 | 4,75E-01 | 8,33E-01 |
| AT3G56680 | 2,66 | -1,19 | 1,67 | -0,71 | 4,76E-01 | 8,33E-01 |
| AT3G62410 | 1,88 | -0,86 | 1,21 | -0,71 | 4,77E-01 | 8,33E-01 |
| AT3G63070 | 1,86 | 1,48 | 2,05 | 0,72 | 4,69E-01 | 8,33E-01 |
| AT4G14400 | 8,78 | -0,58 | 0,80 | -0,72 | 4,73E-01 | 8,33E-01 |
| AT5G11790 | 2,58 | 0,81 | 1,11 | 0,73 | 4,67E-01 | 8,33E-01 |
| AT5G28919 | 2,64 | 0,92 | 1,27 | 0,73 | 4,68E-01 | 8,33E-01 |
| AT5G51070 | 3,71 | 0,92 | 1,29 | 0,72 | 4,72E-01 | 8,33E-01 |
| AT1G12920 | 1,87 | -1,09 | 1,51 | -0,72 | 4,72E-01 | 8,33E-01 |
| AT1G47128 | 7,64 | 0,50 | 0,69 | 0,72 | 4,73E-01 | 8,33E-01 |
| AT1G55350 | 1,89 | -1,53 | 2,15 | -0,71 | 4,76E-01 | 8,33E-01 |
| AT2G20721 | 2,89 | 0,75 | 1,04 | 0,72 | 4,71E-01 | 8,33E-01 |

| | | | | | | |
|---|---|---|---|---|---|---|
| AT2G20780 | 1,93 | 1,24 | 1,75 | 0,71 | 4,77E-01 | 8,33E-01 |
| AT2G26890 | 8,74 | -0,71 | 0,98 | -0,73 | 4,68E-01 | 8,33E-01 |
| AT3G06650 | 3,17 | -1,33 | 1,83 | -0,73 | 4,67E-01 | 8,33E-01 |
| AT3G29190 | 2,69 | -0,85 | 1,18 | -0,72 | 4,70E-01 | 8,33E-01 |
| AT3G29360 | 2,83 | 0,98 | 1,37 | 0,72 | 4,71E-01 | 8,33E-01 |
| AT4G11420 | 3,20 | 1,28 | 1,80 | 0,71 | 4,77E-01 | 8,33E-01 |
| AT4G36648 | 2,47 | 1,19 | 1,66 | 0,72 | 4,74E-01 | 8,33E-01 |
| AT5G03040 | 2,14 | -1,18 | 1,67 | -0,71 | 4,78E-01 | 8,33E-01 |
| AT5G09730 | 35,07 | -0,32 | 0,45 | -0,72 | 4,69E-01 | 8,33E-01 |
| AT5G15200 | 4,33 | -0,62 | 0,85 | -0,73 | 4,66E-01 | 8,33E-01 |
| AT5G04905 | 2,64 | 0,92 | 1,27 | 0,73 | 4,68E-01 | 8,33E-01 |
| AT5G67360 | 6,15 | 0,82 | 1,13 | 0,73 | 4,65E-01 | 8,33E-01 |
| AT1G80680 | 4,25 | 1,08 | 1,53 | 0,70 | 4,81E-01 | 8,35E-01 |
| AT2G47800 | 2,70 | -1,15 | 1,62 | -0,71 | 4,79E-01 | 8,35E-01 |
| AT5G10180 | 2,74 | 1,17 | 1,67 | 0,70 | 4,81E-01 | 8,35E-01 |
| AT5G16620 | 2,01 | -0,81 | 1,15 | -0,71 | 4,81E-01 | 8,35E-01 |
| AT1G67120 | 8,25 | -0,63 | 0,90 | -0,71 | 4,80E-01 | 8,35E-01 |
| AT3G62200 | 2,36 | 1,20 | 1,71 | 0,70 | 4,82E-01 | 8,35E-01 |
| AT5G28920 | 2,84 | 0,89 | 1,26 | 0,71 | 4,81E-01 | 8,35E-01 |
| AT5G58150 | 3,53 | 1,34 | 1,91 | 0,70 | 4,81E-01 | 8,35E-01 |
| AT5G38410 | 6,26 | -0,62 | 0,88 | -0,70 | 4,82E-01 | 8,35E-01 |
| AT3G22380 | 8,79 | 0,63 | 0,90 | 0,70 | 4,84E-01 | 8,35E-01 |
| AT4G39400 | 3,54 | 0,76 | 1,08 | 0,70 | 4,85E-01 | 8,35E-01 |
| AT1G14610 | 2,42 | 1,20 | 1,71 | 0,70 | 4,84E-01 | 8,35E-01 |
| AT3G13920 | 8,10 | -0,65 | 0,92 | -0,70 | 4,83E-01 | 8,35E-01 |
| AT4G16490 | 2,72 | 1,37 | 1,96 | 0,70 | 4,84E-01 | 8,35E-01 |
| AT5G04590 | 4,30 | 0,82 | 1,17 | 0,70 | 4,85E-01 | 8,36E-01 |
| AT1G08597 | 22,78 | 0,42 | 0,60 | 0,70 | 4,85E-01 | 8,36E-01 |
| AT1G31910 | 2,79 | -0,79 | 1,15 | -0,69 | 4,90E-01 | 8,37E-01 |
| AT1G63150 | 176,58 | -0,19 | 0,28 | -0,69 | 4,91E-01 | 8,37E-01 |
| AT1G68945 | 2,15 | 0,77 | 1,11 | 0,69 | 4,88E-01 | 8,37E-01 |
| AT3G16370 | 1,93 | -0,86 | 1,24 | -0,69 | 4,90E-01 | 8,37E-01 |
| AT3G47348 | 6,89 | 0,73 | 1,06 | 0,69 | 4,91E-01 | 8,37E-01 |
| AT1G29470 | 3,69 | 0,83 | 1,19 | 0,70 | 4,87E-01 | 8,37E-01 |
| AT2G35384 | 13,85 | 0,43 | 0,63 | 0,69 | 4,91E-01 | 8,37E-01 |
| AT3G04765 | 2,83 | -0,67 | 0,97 | -0,69 | 4,90E-01 | 8,37E-01 |
| AT3G26000 | 2,02 | 1,18 | 1,70 | 0,69 | 4,88E-01 | 8,37E-01 |
| AT3G48000 | 4,79 | -0,73 | 1,07 | -0,69 | 4,91E-01 | 8,37E-01 |
| AT4G00800 | 2,43 | 1,19 | 1,71 | 0,69 | 4,88E-01 | 8,37E-01 |
| AT5G44320 | 2,18 | 1,20 | 1,73 | 0,69 | 4,88E-01 | 8,37E-01 |
| AT1G66880 | 6,48 | 0,66 | 0,96 | 0,69 | 4,93E-01 | 8,37E-01 |
| AT2G45940 | 2,82 | -1,26 | 1,84 | -0,69 | 4,93E-01 | 8,37E-01 |
| AT1G20960 | 9,33 | -0,60 | 0,87 | -0,68 | 4,93E-01 | 8,37E-01 |
| AT1G62750 | 9,17 | -0,58 | 0,85 | -0,68 | 4,94E-01 | 8,37E-01 |
| AT2G44570 | 2,88 | -0,75 | 1,09 | -0,69 | 4,93E-01 | 8,37E-01 |
| AT5G00765 | 3,11 | -0,71 | 1,03 | -0,69 | 4,92E-01 | 8,37E-01 |
| AT1G05562 | 6,13 | -0,47 | 0,70 | -0,67 | 5,03E-01 | 8,37E-01 |
| AT1G20260 | 2,71 | -0,79 | 1,17 | -0,67 | 5,02E-01 | 8,37E-01 |
| AT1G32060 | 5,29 | -0,65 | 0,97 | -0,67 | 5,02E-01 | 8,37E-01 |
| AT1G32582 | 26,60 | 0,38 | 0,57 | 0,67 | 5,03E-01 | 8,37E-01 |
| AT1G54090 | 3,59 | 1,19 | 1,78 | 0,67 | 5,03E-01 | 8,37E-01 |
| AT2G07727 | 3,34 | -0,86 | 1,27 | -0,68 | 4,98E-01 | 8,37E-01 |
| AT2G41440 | 2,60 | -1,26 | 1,87 | -0,67 | 5,03E-01 | 8,37E-01 |
| AT4G24280 | 6,53 | -0,74 | 1,10 | -0,68 | 4,98E-01 | 8,37E-01 |
| AT5G16640 | 4,15 | 0,55 | 0,81 | 0,67 | 5,02E-01 | 8,37E-01 |
| AT5G36230 | 2,63 | 1,20 | 1,77 | 0,68 | 4,96E-01 | 8,37E-01 |

| | | | | | | |
|---|---|---|---|---|---|---|
| AT1G05560 | 6,13 | -0,47 | 0,70 | -0,67 | 5,03E-01 | 8,37E-01 |
| AT1G21590 | 2,73 | -0,91 | 1,35 | -0,67 | 5,02E-01 | 8,37E-01 |
| AT1G09027 | 2,11 | 0,76 | 1,13 | 0,67 | 5,00E-01 | 8,37E-01 |
| AT1G80250 | 2,27 | -1,30 | 1,92 | -0,68 | 4,98E-01 | 8,37E-01 |
| AT2G41720 | 1,87 | 1,18 | 1,73 | 0,68 | 4,94E-01 | 8,37E-01 |
| AT2G45930 | 2,64 | -1,26 | 1,87 | -0,68 | 4,99E-01 | 8,37E-01 |
| AT3G10650 | 3,29 | -1,25 | 1,84 | -0,68 | 4,95E-01 | 8,37E-01 |
| AT3G11410 | 4,48 | -0,74 | 1,10 | -0,68 | 4,99E-01 | 8,37E-01 |
| AT3G22790 | 2,45 | -1,26 | 1,88 | -0,67 | 5,03E-01 | 8,37E-01 |
| AT3G24180 | 1,99 | -1,01 | 1,51 | -0,67 | 5,03E-01 | 8,37E-01 |
| AT4G02380 | 7,09 | 0,55 | 0,81 | 0,68 | 4,99E-01 | 8,37E-01 |
| AT5G24900 | 2,20 | 0,85 | 1,27 | 0,67 | 5,01E-01 | 8,37E-01 |
| AT5G48380 | 2,92 | 0,83 | 1,21 | 0,68 | 4,95E-01 | 8,37E-01 |
| AT5G63020 | 8,08 | -0,43 | 0,63 | -0,68 | 4,98E-01 | 8,37E-01 |
| AT2G01270 | 3,86 | -1,22 | 1,83 | -0,67 | 5,04E-01 | 8,37E-01 |
| AT2G24790 | 3,54 | 1,10 | 1,65 | 0,67 | 5,04E-01 | 8,37E-01 |
| AT3G18290 | 9,13 | 0,83 | 1,24 | 0,67 | 5,06E-01 | 8,38E-01 |
| AT3G41762 | 25,36 | -0,28 | 0,42 | -0,67 | 5,06E-01 | 8,38E-01 |
| AT4G08125 | 3,11 | -0,58 | 0,87 | -0,67 | 5,05E-01 | 8,38E-01 |
| AT1G64140 | 5,12 | 0,87 | 1,31 | 0,66 | 5,07E-01 | 8,38E-01 |
| AT5G35210 | 2,88 | -1,12 | 1,69 | -0,66 | 5,07E-01 | 8,38E-01 |
| AT2G09610 | 9,53 | -0,39 | 0,59 | -0,66 | 5,07E-01 | 8,38E-01 |
| AT5G01915 | 2,27 | 0,88 | 1,33 | 0,66 | 5,08E-01 | 8,38E-01 |
| AT1G24706 | 2,66 | 0,98 | 1,52 | 0,65 | 5,18E-01 | 8,39E-01 |
| AT1G64500 | 2,64 | 0,73 | 1,12 | 0,65 | 5,17E-01 | 8,39E-01 |
| AT2G03890 | 2,03 | 0,86 | 1,30 | 0,66 | 5,11E-01 | 8,39E-01 |
| AT2G18790 | 2,11 | 1,07 | 1,66 | 0,64 | 5,20E-01 | 8,39E-01 |
| AT2G23350 | 29,35 | 0,77 | 1,20 | 0,65 | 5,19E-01 | 8,39E-01 |
| AT2G28000 | 11,21 | -0,54 | 0,84 | -0,64 | 5,21E-01 | 8,39E-01 |
| AT2G34208 | 2,28 | -1,26 | 1,93 | -0,65 | 5,14E-01 | 8,39E-01 |
| AT2G34410 | 2,89 | -0,87 | 1,34 | -0,65 | 5,18E-01 | 8,39E-01 |
| AT3G06355 | 6711,54 | 0,23 | 0,36 | 0,65 | 5,14E-01 | 8,39E-01 |
| AT3G53400 | 1,89 | 1,23 | 1,89 | 0,65 | 5,13E-01 | 8,39E-01 |
| AT4G34110 | 4,15 | -0,58 | 0,88 | -0,66 | 5,09E-01 | 8,39E-01 |
| AT5G10540 | 2,27 | 1,07 | 1,66 | 0,64 | 5,19E-01 | 8,39E-01 |
| AT5G44270 | 2,68 | 0,69 | 1,07 | 0,64 | 5,19E-01 | 8,39E-01 |
| AT5G48385 | 2,37 | 1,08 | 1,65 | 0,65 | 5,15E-01 | 8,39E-01 |
| AT5G49615 | 2,21 | 1,00 | 1,52 | 0,66 | 5,12E-01 | 8,39E-01 |
| AT5G64940 | 2,50 | 1,12 | 1,74 | 0,64 | 5,19E-01 | 8,39E-01 |
| AT1G04517 | 11,17 | 0,36 | 0,56 | 0,64 | 5,21E-01 | 8,39E-01 |
| AT1G12300 | 2,68 | -0,63 | 0,96 | -0,65 | 5,13E-01 | 8,39E-01 |
| AT1G13640 | 2,82 | -1,19 | 1,85 | -0,64 | 5,21E-01 | 8,39E-01 |
| AT1G29400 | 2,11 | 0,96 | 1,46 | 0,65 | 5,13E-01 | 8,39E-01 |
| AT1G55490 | 3,72 | -0,71 | 1,08 | -0,66 | 5,10E-01 | 8,39E-01 |
| AT1G58250 | 4,39 | 0,95 | 1,44 | 0,66 | 5,09E-01 | 8,39E-01 |
| AT2G05765 | 3,87 | 0,64 | 0,99 | 0,65 | 5,17E-01 | 8,39E-01 |
| AT2G23810 | 3,17 | 0,81 | 1,23 | 0,65 | 5,13E-01 | 8,39E-01 |
| AT3G01316 | 2,03 | -0,79 | 1,20 | -0,66 | 5,11E-01 | 8,39E-01 |
| AT4G13560 | 52,08 | 0,22 | 0,33 | 0,66 | 5,12E-01 | 8,39E-01 |
| AT4G35800 | 5,60 | -0,68 | 1,05 | -0,65 | 5,17E-01 | 8,39E-01 |
| AT5G13840 | 2,03 | -0,88 | 1,35 | -0,65 | 5,16E-01 | 8,39E-01 |
| AT5G26000 | 4,62 | -0,51 | 0,78 | -0,65 | 5,16E-01 | 8,39E-01 |
| AT5G28540 | 7,23 | 0,48 | 0,73 | 0,65 | 5,13E-01 | 8,39E-01 |
| AT5G66030 | 2,48 | -1,22 | 1,90 | -0,64 | 5,20E-01 | 8,39E-01 |
| AT1G30400 | 2,52 | -0,87 | 1,36 | -0,64 | 5,21E-01 | 8,39E-01 |
| AT4G26110 | 2,20 | 1,06 | 1,65 | 0,64 | 5,23E-01 | 8,39E-01 |

| | | | | | | |
|---|---|---|---|---|---|---|
| AT1G30330 | 1,89 | 0,86 | 1,34 | 0,64 | 5,22E-01 | 8,39E-01 |
| AT3G24982 | 2,19 | 0,80 | 1,25 | 0,64 | 5,22E-01 | 8,39E-01 |
| AT1G58602 | 11,97 | 0,37 | 0,59 | 0,64 | 5,25E-01 | 8,40E-01 |
| AT1G79520 | 27,20 | -0,32 | 0,50 | -0,63 | 5,26E-01 | 8,40E-01 |
| AT2G36850 | 3,05 | 1,06 | 1,67 | 0,63 | 5,25E-01 | 8,40E-01 |
| AT2G41560 | 2,99 | 0,78 | 1,23 | 0,64 | 5,24E-01 | 8,40E-01 |
| AT3G08170 | 3,83 | 0,68 | 1,07 | 0,64 | 5,25E-01 | 8,40E-01 |
| AT4G29410 | 17,21 | 0,33 | 0,52 | 0,64 | 5,25E-01 | 8,40E-01 |
| AT1G76030 | 2,84 | -0,76 | 1,21 | -0,63 | 5,27E-01 | 8,40E-01 |
| AT5G14430 | 2,60 | 1,13 | 1,79 | 0,63 | 5,28E-01 | 8,40E-01 |
| AT5G07745 | 20,93 | -0,41 | 0,66 | -0,63 | 5,29E-01 | 8,40E-01 |
| AT2G29670 | 3,81 | -0,66 | 1,05 | -0,63 | 5,28E-01 | 8,40E-01 |
| AT2G08725 | 34,05 | 0,37 | 0,58 | 0,63 | 5,26E-01 | 8,40E-01 |
| AT3G01770 | 1,96 | 1,10 | 1,74 | 0,63 | 5,29E-01 | 8,40E-01 |
| AT3G26450 | 2,27 | -0,75 | 1,20 | -0,63 | 5,28E-01 | 8,40E-01 |
| AT4G39090 | 7,03 | -0,48 | 0,76 | -0,63 | 5,28E-01 | 8,40E-01 |
| AT5G14520 | 3,02 | -1,21 | 1,92 | -0,63 | 5,29E-01 | 8,40E-01 |
| AT1G60140 | 2,14 | -0,96 | 1,53 | -0,63 | 5,30E-01 | 8,41E-01 |
| AT4G04640 | 5,86 | -0,47 | 0,74 | -0,63 | 5,30E-01 | 8,41E-01 |
| AT3G04455 | 2,46 | -0,82 | 1,31 | -0,63 | 5,31E-01 | 8,42E-01 |
| AT1G20020 | 2,67 | -0,62 | 0,99 | -0,63 | 5,32E-01 | 8,42E-01 |
| AT2G09700 | 2,16 | 0,93 | 1,49 | 0,62 | 5,33E-01 | 8,42E-01 |
| AT5G03350 | 2,45 | -0,68 | 1,09 | -0,62 | 5,33E-01 | 8,42E-01 |
| AT4G03280 | 3,09 | 0,72 | 1,16 | 0,62 | 5,35E-01 | 8,43E-01 |
| AT4G02660 | 2,17 | 1,13 | 1,82 | 0,62 | 5,34E-01 | 8,43E-01 |
| AT5G01350 | 8,44 | -0,40 | 0,64 | -0,62 | 5,34E-01 | 8,43E-01 |
| AT3G08943 | 11,39 | -0,88 | 1,41 | -0,62 | 5,35E-01 | 8,43E-01 |
| AT5G08530 | 3,76 | -0,62 | 0,99 | -0,62 | 5,36E-01 | 8,43E-01 |
| AT1G12013 | 2,08 | -0,74 | 1,19 | -0,62 | 5,37E-01 | 8,44E-01 |
| AT5G45095 | 2,04 | 0,84 | 1,37 | 0,62 | 5,38E-01 | 8,45E-01 |
| AT5G47490 | 2,21 | 1,00 | 1,62 | 0,62 | 5,38E-01 | 8,45E-01 |
| AT1G46696 | 1,91 | 0,90 | 1,46 | 0,61 | 5,39E-01 | 8,45E-01 |
| AT5G39050 | 4,52 | -0,60 | 0,98 | -0,61 | 5,39E-01 | 8,45E-01 |
| AT2G33340 | 1,98 | 1,11 | 1,82 | 0,61 | 5,40E-01 | 8,46E-01 |
| AT2G07698 | 9,74 | -0,36 | 0,59 | -0,61 | 5,41E-01 | 8,47E-01 |
| AT3G22440 | 1,87 | 0,83 | 1,36 | 0,61 | 5,41E-01 | 8,47E-01 |
| AT5G64220 | 2,50 | -1,19 | 1,94 | -0,61 | 5,42E-01 | 8,47E-01 |
| AT5G04540 | 1,96 | -1,09 | 1,78 | -0,61 | 5,42E-01 | 8,47E-01 |
| AT1G31730 | 1,97 | -1,27 | 2,10 | -0,61 | 5,44E-01 | 8,47E-01 |
| AT4G10120 | 1,86 | 1,03 | 1,69 | 0,61 | 5,44E-01 | 8,47E-01 |
| AT5G53460 | 17,48 | 0,37 | 0,62 | 0,61 | 5,44E-01 | 8,47E-01 |
| AT5G60600 | 2,43 | -0,62 | 1,02 | -0,61 | 5,43E-01 | 8,47E-01 |
| AT3G28900 | 3,87 | 0,60 | 0,98 | 0,61 | 5,44E-01 | 8,47E-01 |
| AT5G19330 | 2,99 | -0,94 | 1,56 | -0,61 | 5,44E-01 | 8,47E-01 |
| AT5G49740 | 2,42 | 0,87 | 1,44 | 0,60 | 5,45E-01 | 8,48E-01 |
| AT1G09770 | 3,11 | 0,92 | 1,53 | 0,60 | 5,49E-01 | 8,48E-01 |
| AT3G41979 | 6596,11 | 0,21 | 0,36 | 0,60 | 5,48E-01 | 8,48E-01 |
| AT4G39420 | 9,97 | -0,61 | 1,01 | -0,60 | 5,47E-01 | 8,48E-01 |
| AT1G09210 | 5,83 | -0,48 | 0,80 | -0,60 | 5,48E-01 | 8,48E-01 |
| AT2G29980 | 3,37 | -0,84 | 1,41 | -0,60 | 5,49E-01 | 8,48E-01 |
| AT2G36470 | 2,06 | 0,99 | 1,64 | 0,60 | 5,49E-01 | 8,48E-01 |
| AT3G51550 | 4,70 | 0,55 | 0,92 | 0,60 | 5,49E-01 | 8,48E-01 |
| AT5G37600 | 1,93 | 1,02 | 1,69 | 0,60 | 5,46E-01 | 8,48E-01 |
| AT5G57350 | 4,58 | 0,99 | 1,66 | 0,60 | 5,50E-01 | 8,49E-01 |
| AT2G35110 | 1,99 | -0,91 | 1,52 | -0,60 | 5,51E-01 | 8,50E-01 |
| AT2G44560 | 4,56 | -0,42 | 0,71 | -0,60 | 5,51E-01 | 8,50E-01 |

| | | | | | | |
|---|---|---|---|---|---|---|
| AT2G45180 | 2,18 | -0,71 | 1,20 | -0,59 | 5,53E-01 | 8,52E-01 |
| AT3G02090 | 3,68 | 0,77 | 1,30 | 0,59 | 5,54E-01 | 8,52E-01 |
| AT3G51050 | 3,03 | -1,13 | 1,90 | -0,59 | 5,54E-01 | 8,52E-01 |
| AT5G50290 | 244,15 | 0,21 | 0,36 | 0,59 | 5,55E-01 | 8,53E-01 |
| AT3G50370 | 9,94 | 0,62 | 1,06 | 0,59 | 5,55E-01 | 8,53E-01 |
| AT2G07806 | 2,71 | -0,73 | 1,24 | -0,59 | 5,58E-01 | 8,53E-01 |
| AT5G58465 | 4,24 | -0,50 | 0,84 | -0,59 | 5,57E-01 | 8,53E-01 |
| AT1G01320 | 16,50 | 0,44 | 0,75 | 0,59 | 5,57E-01 | 8,53E-01 |
| AT2G43410 | 2,84 | 0,85 | 1,44 | 0,59 | 5,56E-01 | 8,53E-01 |
| AT3G46740 | 3,41 | -0,72 | 1,23 | -0,59 | 5,57E-01 | 8,53E-01 |
| AT4G38220 | 1,96 | -1,13 | 1,94 | -0,58 | 5,59E-01 | 8,54E-01 |
| AT5G64420 | 4,10 | 0,89 | 1,53 | 0,58 | 5,60E-01 | 8,54E-01 |
| AT5G09670 | 2,53 | -0,77 | 1,32 | -0,58 | 5,59E-01 | 8,54E-01 |
| AT1G70770 | 12,70 | 0,70 | 1,20 | 0,58 | 5,61E-01 | 8,54E-01 |
| AT3G18217 | 147,37 | -0,14 | 0,24 | -0,58 | 5,60E-01 | 8,54E-01 |
| AT2G09555 | 2,08 | 0,69 | 1,19 | 0,58 | 5,61E-01 | 8,55E-01 |
| AT1G03130 | 1,99 | -0,64 | 1,11 | -0,58 | 5,63E-01 | 8,55E-01 |
| AT2G25490 | 3,68 | 0,60 | 1,04 | 0,58 | 5,63E-01 | 8,55E-01 |
| AT2G31141 | 64,55 | 0,17 | 0,30 | 0,58 | 5,63E-01 | 8,55E-01 |
| AT5G15400 | 2,82 | 0,90 | 1,55 | 0,58 | 5,63E-01 | 8,55E-01 |
| AT1G50480 | 9,27 | 0,63 | 1,09 | 0,58 | 5,64E-01 | 8,56E-01 |
| AT1G29920 | 50,33 | -0,25 | 0,43 | -0,58 | 5,65E-01 | 8,57E-01 |
| AT2G21920 | 4,05 | -0,46 | 0,80 | -0,57 | 5,65E-01 | 8,57E-01 |
| AT5G38850 | 479,48 | -0,12 | 0,20 | -0,57 | 5,66E-01 | 8,57E-01 |
| AT2G44410 | 2,51 | 1,16 | 2,03 | 0,57 | 5,69E-01 | 8,59E-01 |
| AT4G28760 | 2,13 | 1,02 | 1,78 | 0,57 | 5,68E-01 | 8,59E-01 |
| AT3G48115 | 1,91 | 0,82 | 1,45 | 0,57 | 5,69E-01 | 8,59E-01 |
| AT4G03295 | 3,55 | -0,49 | 0,86 | -0,57 | 5,69E-01 | 8,59E-01 |
| AT4G37910 | 5,28 | 0,71 | 1,24 | 0,57 | 5,68E-01 | 8,59E-01 |
| AT3G23000 | 7,09 | 0,73 | 1,28 | 0,57 | 5,71E-01 | 8,60E-01 |
| AT3G13330 | 3,62 | 0,77 | 1,35 | 0,57 | 5,71E-01 | 8,60E-01 |
| AT1G25440 | 6,25 | 0,64 | 1,13 | 0,57 | 5,71E-01 | 8,60E-01 |
| AT4G00570 | 3,02 | 0,99 | 1,75 | 0,56 | 5,72E-01 | 8,61E-01 |
| AT5G20290 | 5,09 | -0,52 | 0,92 | -0,56 | 5,73E-01 | 8,61E-01 |
| AT2G13560 | 2,19 | -0,86 | 1,54 | -0,56 | 5,74E-01 | 8,61E-01 |
| AT4G08920 | 2,64 | 0,95 | 1,69 | 0,56 | 5,74E-01 | 8,61E-01 |
| AT4G18700 | 2,69 | -0,75 | 1,33 | -0,56 | 5,74E-01 | 8,61E-01 |
| AT1G20860 | 3,03 | 0,60 | 1,07 | 0,56 | 5,75E-01 | 8,62E-01 |
| AT5G67420 | 2,71 | 0,65 | 1,16 | 0,56 | 5,75E-01 | 8,62E-01 |
| AT1G55450 | 2,26 | 0,78 | 1,39 | 0,56 | 5,76E-01 | 8,62E-01 |
| AT4G19395 | 418,73 | -0,12 | 0,21 | -0,56 | 5,77E-01 | 8,63E-01 |
| AT3G26440 | 2,47 | -0,67 | 1,21 | -0,56 | 5,78E-01 | 8,63E-01 |
| AT1G64385 | 1,99 | 1,05 | 1,90 | 0,56 | 5,79E-01 | 8,64E-01 |
| AT3G55610 | 4,81 | -0,74 | 1,34 | -0,56 | 5,79E-01 | 8,64E-01 |
| AT2G31880 | 6,93 | 0,58 | 1,05 | 0,55 | 5,79E-01 | 8,64E-01 |
| AT1G63750 | 2,64 | -0,73 | 1,33 | -0,55 | 5,84E-01 | 8,64E-01 |
| AT4G16660 | 2,21 | 0,84 | 1,53 | 0,55 | 5,83E-01 | 8,64E-01 |
| AT5G27720 | 3,01 | 0,59 | 1,06 | 0,55 | 5,80E-01 | 8,64E-01 |
| AT5G63145 | 2,83 | -0,59 | 1,06 | -0,55 | 5,80E-01 | 8,64E-01 |
| AT1G05850 | 2,93 | -0,70 | 1,28 | -0,55 | 5,84E-01 | 8,64E-01 |
| AT2G37710 | 4,76 | -0,47 | 0,85 | -0,55 | 5,81E-01 | 8,64E-01 |
| AT2G38170 | 3,76 | -0,59 | 1,08 | -0,55 | 5,83E-01 | 8,64E-01 |
| AT3G57730 | 5,17 | 0,45 | 0,82 | 0,55 | 5,82E-01 | 8,64E-01 |
| AT5G08180 | 2,30 | 0,58 | 1,06 | 0,55 | 5,81E-01 | 8,64E-01 |
| AT5G17760 | 1,99 | 1,12 | 2,03 | 0,55 | 5,82E-01 | 8,64E-01 |
| AT5G42080 | 4,02 | 0,68 | 1,24 | 0,55 | 5,82E-01 | 8,64E-01 |

| | | | | | | |
|---|---|---|---|---|---|---|
| AT3G14067 | 4,84 | -0,50 | 0,91 | -0,55 | 5,84E-01 | 8,64E-01 |
| AT5G58950 | 2,42 | -0,68 | 1,24 | -0,55 | 5,85E-01 | 8,64E-01 |
| AT1G18160 | 2,26 | 0,93 | 1,74 | 0,54 | 5,92E-01 | 8,67E-01 |
| AT1G72930 | 3,01 | 0,77 | 1,42 | 0,54 | 5,87E-01 | 8,67E-01 |
| AT2G30860 | 6,26 | 0,40 | 0,75 | 0,54 | 5,92E-01 | 8,67E-01 |
| AT2G47070 | 5,92 | 0,61 | 1,13 | 0,54 | 5,90E-01 | 8,67E-01 |
| AT3G25690 | 2,72 | -0,63 | 1,18 | -0,54 | 5,91E-01 | 8,67E-01 |
| AT3G07735 | 4,39 | -0,43 | 0,79 | -0,54 | 5,91E-01 | 8,67E-01 |
| AT5G13100 | 3,43 | 0,89 | 1,67 | 0,54 | 5,92E-01 | 8,67E-01 |
| AT5G60410 | 2,90 | -0,76 | 1,41 | -0,54 | 5,91E-01 | 8,67E-01 |
| AT3G63460 | 4,10 | 0,71 | 1,32 | 0,54 | 5,91E-01 | 8,67E-01 |
| AT4G04465 | 2,27 | 0,65 | 1,19 | 0,54 | 5,87E-01 | 8,67E-01 |
| AT4G40040 | 3,01 | 0,63 | 1,17 | 0,54 | 5,91E-01 | 8,67E-01 |
| AT5G08185 | 1387,13 | 0,17 | 0,32 | 0,54 | 5,89E-01 | 8,67E-01 |
| AT5G15020 | 2,14 | -0,91 | 1,69 | -0,54 | 5,92E-01 | 8,67E-01 |
| AT5G49720 | 3,89 | -0,50 | 0,92 | -0,54 | 5,89E-01 | 8,67E-01 |
| AT1G33415 | 2,72 | 0,52 | 0,98 | 0,53 | 5,99E-01 | 8,67E-01 |
| AT3G06720 | 1,89 | -1,11 | 2,11 | -0,53 | 5,98E-01 | 8,67E-01 |
| AT3G11540 | 2,22 | 1,02 | 1,94 | 0,53 | 5,98E-01 | 8,67E-01 |
| AT3G15730 | 15,00 | -0,31 | 0,58 | -0,53 | 5,97E-01 | 8,67E-01 |
| AT3G47620 | 3,09 | 0,61 | 1,14 | 0,53 | 5,94E-01 | 8,67E-01 |
| AT5G37510 | 3,25 | -0,61 | 1,14 | -0,53 | 5,96E-01 | 8,67E-01 |
| AT5G57660 | 4,42 | -0,43 | 0,82 | -0,53 | 5,96E-01 | 8,67E-01 |
| AT1G04733 | 20,09 | 0,25 | 0,47 | 0,53 | 5,98E-01 | 8,67E-01 |
| AT1G22060 | 2,54 | -0,69 | 1,31 | -0,52 | 6,00E-01 | 8,67E-01 |
| AT3G02170 | 3,18 | -0,90 | 1,71 | -0,53 | 5,96E-01 | 8,67E-01 |
| AT3G04210 | 3,60 | 0,55 | 1,05 | 0,53 | 5,98E-01 | 8,67E-01 |
| AT3G13870 | 4,03 | -0,96 | 1,80 | -0,53 | 5,95E-01 | 8,67E-01 |
| AT3G56070 | 2,75 | -0,53 | 1,01 | -0,53 | 5,97E-01 | 8,67E-01 |
| AT4G06705 | 4,08 | 0,47 | 0,89 | 0,53 | 5,93E-01 | 8,67E-01 |
| AT5G04550 | 2,33 | 0,84 | 1,61 | 0,53 | 5,99E-01 | 8,67E-01 |
| AT5G19820 | 4,02 | 0,88 | 1,67 | 0,53 | 5,99E-01 | 8,67E-01 |
| AT4G05150 | 3,81 | 0,62 | 1,17 | 0,52 | 6,00E-01 | 8,68E-01 |
| AT4G39363 | 6,83 | 0,38 | 0,73 | 0,52 | 6,01E-01 | 8,68E-01 |
| AT1G58290 | 5,16 | -0,54 | 1,03 | -0,52 | 6,02E-01 | 8,69E-01 |
| AT1G07590 | 3,38 | 0,59 | 1,14 | 0,52 | 6,02E-01 | 8,69E-01 |
| AT2G25900 | 2,31 | -0,58 | 1,11 | -0,52 | 6,05E-01 | 8,69E-01 |
| AT3G03870 | 3,66 | 0,77 | 1,49 | 0,52 | 6,05E-01 | 8,69E-01 |
| AT3G03105 | 2,11 | 0,64 | 1,24 | 0,52 | 6,03E-01 | 8,69E-01 |
| AT2G18480 | 2,00 | -0,74 | 1,44 | -0,52 | 6,04E-01 | 8,69E-01 |
| AT3G51770 | 2,18 | -0,88 | 1,69 | -0,52 | 6,05E-01 | 8,69E-01 |
| AT5G42650 | 3,24 | -0,56 | 1,08 | -0,52 | 6,04E-01 | 8,69E-01 |
| AT3G45140 | 15,78 | -0,26 | 0,51 | -0,51 | 6,07E-01 | 8,69E-01 |
| AT4G06695 | 4,20 | 0,47 | 0,91 | 0,51 | 6,07E-01 | 8,69E-01 |
| AT5G04425 | 2,72 | 0,56 | 1,08 | 0,51 | 6,07E-01 | 8,69E-01 |
| AT5G06175 | 3,94 | 0,43 | 0,83 | 0,51 | 6,07E-01 | 8,69E-01 |
| AT3G11130 | 25,04 | 0,38 | 0,74 | 0,52 | 6,06E-01 | 8,69E-01 |
| AT2G22125 | 19,76 | -0,44 | 0,86 | -0,51 | 6,08E-01 | 8,70E-01 |
| AT5G61420 | 2,01 | -0,84 | 1,64 | -0,51 | 6,09E-01 | 8,71E-01 |
| AT1G04430 | 7,26 | -0,53 | 1,05 | -0,51 | 6,13E-01 | 8,71E-01 |
| AT1G63630 | 3,43 | 0,45 | 0,88 | 0,51 | 6,12E-01 | 8,71E-01 |
| AT1G66730 | 1,91 | -0,58 | 1,15 | -0,51 | 6,13E-01 | 8,71E-01 |
| AT2G04405 | 3,40 | -0,51 | 1,01 | -0,51 | 6,10E-01 | 8,71E-01 |
| AT2G07725 | 2,14 | -0,59 | 1,15 | -0,51 | 6,11E-01 | 8,71E-01 |
| AT3G44310 | 10,52 | -0,33 | 0,66 | -0,51 | 6,13E-01 | 8,71E-01 |
| AT3G56408 | 2,59 | 0,69 | 1,38 | 0,50 | 6,14E-01 | 8,71E-01 |

| | | | | | | |
|---|---|---|---|---|---|---|
| AT5G13850 | 2,26 | -0,66 | 1,30 | -0,51 | 6,13E-01 | 8,71E-01 |
| AT5G19400 | 2,38 | 1,00 | 1,99 | 0,50 | 6,14E-01 | 8,71E-01 |
| AT1G63615 | 3,43 | 0,45 | 0,88 | 0,51 | 6,12E-01 | 8,71E-01 |
| AT1G79550 | 2,41 | 0,65 | 1,27 | 0,51 | 6,11E-01 | 8,71E-01 |
| AT1G06267 | 15,22 | 0,26 | 0,53 | 0,50 | 6,20E-01 | 8,76E-01 |
| AT2G19850 | 5,66 | 0,46 | 0,93 | 0,49 | 6,23E-01 | 8,76E-01 |
| AT5G11040 | 2,15 | 0,86 | 1,75 | 0,49 | 6,22E-01 | 8,76E-01 |
| AT5G11490 | 1,97 | -0,86 | 1,73 | -0,50 | 6,21E-01 | 8,76E-01 |
| AT5G20010 | 1,98 | -0,78 | 1,57 | -0,50 | 6,20E-01 | 8,76E-01 |
| AT5G49760 | 3,34 | -0,52 | 1,05 | -0,49 | 6,22E-01 | 8,76E-01 |
| AT1G79090 | 2,69 | 0,90 | 1,82 | 0,49 | 6,22E-01 | 8,76E-01 |
| AT3G59020 | 5,79 | 0,65 | 1,32 | 0,49 | 6,22E-01 | 8,76E-01 |
| AT3G60320 | 3,74 | 0,48 | 0,96 | 0,50 | 6,18E-01 | 8,76E-01 |
| AT3G63380 | 3,14 | 0,90 | 1,81 | 0,50 | 6,19E-01 | 8,76E-01 |
| AT5G49730 | 3,55 | -0,57 | 1,16 | -0,49 | 6,22E-01 | 8,76E-01 |
| AT5G60900 | 5,69 | 0,77 | 1,54 | 0,50 | 6,20E-01 | 8,76E-01 |
| AT2G17840 | 2,90 | -0,57 | 1,16 | -0,49 | 6,23E-01 | 8,76E-01 |
| AT3G47500 | 2,57 | -0,52 | 1,07 | -0,49 | 6,27E-01 | 8,80E-01 |
| AT3G63500 | 4,53 | 0,71 | 1,48 | 0,48 | 6,29E-01 | 8,82E-01 |
| AT5G04105 | 18,20 | 0,26 | 0,54 | 0,48 | 6,28E-01 | 8,82E-01 |
| AT3G03475 | 2,58 | -0,54 | 1,11 | -0,48 | 6,30E-01 | 8,83E-01 |
| AT2G22660 | 2,02 | -0,75 | 1,57 | -0,48 | 6,32E-01 | 8,84E-01 |
| AT1G33420 | 2,44 | 0,47 | 1,00 | 0,48 | 6,34E-01 | 8,84E-01 |
| AT1G72410 | 1,92 | 0,88 | 1,85 | 0,48 | 6,33E-01 | 8,84E-01 |
| AT3G59710 | 6,69 | 0,36 | 0,76 | 0,48 | 6,31E-01 | 8,84E-01 |
| AT3G09225 | 6,69 | 0,36 | 0,76 | 0,48 | 6,31E-01 | 8,84E-01 |
| AT4G03380 | 7,25 | 0,32 | 0,67 | 0,48 | 6,32E-01 | 8,84E-01 |
| AT5G02645 | 2,33 | 0,57 | 1,19 | 0,48 | 6,33E-01 | 8,84E-01 |
| AT5G25060 | 1,92 | 0,71 | 1,50 | 0,48 | 6,33E-01 | 8,84E-01 |
| AT5G49910 | 2,46 | -0,65 | 1,38 | -0,47 | 6,35E-01 | 8,85E-01 |
| AT3G23120 | 2,24 | 0,78 | 1,64 | 0,47 | 6,35E-01 | 8,85E-01 |
| AT2G45380 | 11,33 | -0,24 | 0,51 | -0,47 | 6,37E-01 | 8,86E-01 |
| AT1G42970 | 13,13 | -0,28 | 0,60 | -0,47 | 6,37E-01 | 8,86E-01 |
| AT5G28740 | 1,92 | -0,82 | 1,74 | -0,47 | 6,39E-01 | 8,88E-01 |
| AT2G25730 | 2,88 | 0,64 | 1,37 | 0,47 | 6,39E-01 | 8,88E-01 |
| AT3G08510 | 2,33 | -0,72 | 1,54 | -0,47 | 6,40E-01 | 8,88E-01 |
| AT1G75335 | 3,97 | 0,74 | 1,58 | 0,47 | 6,42E-01 | 8,90E-01 |
| AT1G72370 | 2,39 | 0,60 | 1,28 | 0,47 | 6,42E-01 | 8,90E-01 |
| AT1G27340 | 2,10 | 0,85 | 1,84 | 0,46 | 6,43E-01 | 8,90E-01 |
| AT4G11270 | 3,79 | -0,46 | 1,00 | -0,46 | 6,43E-01 | 8,90E-01 |
| AT5G15680 | 7,75 | 0,67 | 1,46 | 0,46 | 6,46E-01 | 8,93E-01 |
| AT1G09640 | 2,66 | -0,81 | 1,78 | -0,46 | 6,49E-01 | 8,95E-01 |
| AT1G29418 | 9,45 | -0,24 | 0,53 | -0,45 | 6,53E-01 | 8,95E-01 |
| AT3G13300 | 8,42 | -0,61 | 1,36 | -0,45 | 6,52E-01 | 8,95E-01 |
| AT4G39366 | 2,55 | -0,49 | 1,10 | -0,45 | 6,54E-01 | 8,95E-01 |
| AT5G15833 | 13,93 | -0,37 | 0,81 | -0,45 | 6,52E-01 | 8,95E-01 |
| AT5G17920 | 34,31 | -0,21 | 0,45 | -0,46 | 6,48E-01 | 8,95E-01 |
| AT1G15750 | 35,69 | -0,50 | 1,11 | -0,45 | 6,52E-01 | 8,95E-01 |
| AT1G69940 | 3,40 | 0,41 | 0,92 | 0,45 | 6,54E-01 | 8,95E-01 |
| AT1G74456 | 11,08 | 0,27 | 0,60 | 0,46 | 6,48E-01 | 8,95E-01 |
| AT2G03250 | 2,57 | -0,49 | 1,08 | -0,45 | 6,53E-01 | 8,95E-01 |
| AT3G26520 | 6,24 | -0,36 | 0,81 | -0,45 | 6,52E-01 | 8,95E-01 |
| AT3G56410 | 2,56 | 0,62 | 1,37 | 0,45 | 6,50E-01 | 8,95E-01 |
| AT4G00720 | 2,07 | -0,75 | 1,67 | -0,45 | 6,54E-01 | 8,95E-01 |
| AT4G02890 | 2,39 | -0,55 | 1,22 | -0,45 | 6,50E-01 | 8,95E-01 |
| AT4G35790 | 4,32 | -0,72 | 1,61 | -0,45 | 6,53E-01 | 8,95E-01 |

| | | | | | | |
|---|---|---|---|---|---|---|
| AT5G24810 | 2,74 | 0,70 | 1,55 | 0,45 | 6,50E-01 | 8,95E-01 |
| AT5G56970 | 2,67 | -0,45 | 1,00 | -0,45 | 6,51E-01 | 8,95E-01 |
| AT1G09877 | 23,86 | -0,22 | 0,49 | -0,45 | 6,56E-01 | 8,96E-01 |
| AT5G51090 | 2,18 | 0,53 | 1,18 | 0,45 | 6,56E-01 | 8,96E-01 |
| AT1G28420 | 2,78 | -0,68 | 1,54 | -0,44 | 6,57E-01 | 8,96E-01 |
| AT1G61190 | 2,38 | -0,63 | 1,43 | -0,44 | 6,60E-01 | 8,97E-01 |
| AT1G78080 | 3,84 | 0,48 | 1,09 | 0,44 | 6,58E-01 | 8,97E-01 |
| AT2G46430 | 2,08 | 0,74 | 1,68 | 0,44 | 6,58E-01 | 8,97E-01 |
| AT2G27860 | 4,64 | 0,43 | 0,98 | 0,44 | 6,59E-01 | 8,97E-01 |
| AT3G05675 | 9,08 | 0,35 | 0,81 | 0,44 | 6,60E-01 | 8,97E-01 |
| AT5G06165 | 4,41 | 0,39 | 0,88 | 0,44 | 6,60E-01 | 8,97E-01 |
| AT5G44340 | 14,19 | -0,33 | 0,76 | -0,44 | 6,59E-01 | 8,97E-01 |
| AT4G39350 | 3,27 | -0,47 | 1,07 | -0,44 | 6,61E-01 | 8,97E-01 |
| AT3G19420 | 2,24 | 0,83 | 1,90 | 0,44 | 6,62E-01 | 8,97E-01 |
| AT4G02555 | 3,06 | 0,50 | 1,14 | 0,44 | 6,61E-01 | 8,97E-01 |
| AT1G16635 | 19,67 | -0,24 | 0,54 | -0,44 | 6,62E-01 | 8,97E-01 |
| AT5G12370 | 4,35 | 0,67 | 1,53 | 0,44 | 6,63E-01 | 8,97E-01 |
| AT2G04375 | 5,42 | -0,35 | 0,81 | -0,43 | 6,65E-01 | 9,00E-01 |
| AT2G34250 | 2,94 | -0,48 | 1,10 | -0,43 | 6,65E-01 | 9,00E-01 |
| AT5G19120 | 2,51 | 0,46 | 1,09 | 0,43 | 6,69E-01 | 9,04E-01 |
| AT1G59980 | 3,55 | -0,41 | 0,97 | -0,42 | 6,72E-01 | 9,05E-01 |
| AT5G14780 | 2,20 | -0,48 | 1,13 | -0,42 | 6,71E-01 | 9,05E-01 |
| AT5G38990 | 2,97 | 0,53 | 1,25 | 0,42 | 6,72E-01 | 9,05E-01 |
| AT5G62000 | 10,38 | 0,28 | 0,65 | 0,43 | 6,70E-01 | 9,05E-01 |
| AT1G09800 | 20,58 | 0,20 | 0,47 | 0,42 | 6,73E-01 | 9,05E-01 |
| AT1G33240 | 1,91 | 0,65 | 1,53 | 0,42 | 6,71E-01 | 9,05E-01 |
| AT4G39800 | 2,68 | -0,48 | 1,13 | -0,42 | 6,73E-01 | 9,05E-01 |
| AT5G65380 | 2,69 | -0,67 | 1,56 | -0,43 | 6,70E-01 | 9,05E-01 |
| AT5G46470 | 3,68 | -0,53 | 1,27 | -0,42 | 6,74E-01 | 9,06E-01 |
| AT5G45470 | 2,49 | -0,68 | 1,61 | -0,42 | 6,74E-01 | 9,06E-01 |
| AT2G46460 | 2,59 | 0,57 | 1,35 | 0,42 | 6,75E-01 | 9,06E-01 |
| AT1G07250 | 2,07 | 0,53 | 1,28 | 0,41 | 6,79E-01 | 9,09E-01 |
| AT1G26558 | 2,72 | 0,49 | 1,19 | 0,41 | 6,79E-01 | 9,09E-01 |
| AT1G67840 | 1,96 | 0,64 | 1,55 | 0,41 | 6,79E-01 | 9,09E-01 |
| AT3G22231 | 4,08 | -0,34 | 0,82 | -0,42 | 6,77E-01 | 9,09E-01 |
| AT4G29060 | 2,00 | -0,60 | 1,45 | -0,41 | 6,82E-01 | 9,12E-01 |
| AT5G10572 | 2,06 | 0,52 | 1,27 | 0,41 | 6,82E-01 | 9,12E-01 |
| AT1G28270 | 2,06 | -0,52 | 1,27 | -0,41 | 6,84E-01 | 9,13E-01 |
| AT3G44480 | 4,70 | -0,41 | 1,02 | -0,41 | 6,83E-01 | 9,13E-01 |
| AT5G14040 | 2,00 | -0,71 | 1,74 | -0,41 | 6,84E-01 | 9,13E-01 |
| AT5G19140 | 6,43 | 0,27 | 0,67 | 0,41 | 6,85E-01 | 9,13E-01 |
| AT5G19150 | 6,35 | 0,28 | 0,68 | 0,41 | 6,85E-01 | 9,13E-01 |
| AT4G34100 | 3,45 | 0,47 | 1,16 | 0,40 | 6,86E-01 | 9,13E-01 |
| AT4G34071 | 2,35 | -0,48 | 1,20 | -0,40 | 6,88E-01 | 9,14E-01 |
| AT1G29940 | 2,05 | -0,74 | 1,83 | -0,40 | 6,87E-01 | 9,14E-01 |
| AT2G44540 | 4,48 | 0,33 | 0,82 | 0,40 | 6,88E-01 | 9,14E-01 |
| AT5G07735 | 1,90 | 0,47 | 1,19 | 0,40 | 6,89E-01 | 9,15E-01 |
| AT3G21805 | 21,83 | 0,27 | 0,68 | 0,40 | 6,90E-01 | 9,15E-01 |
| AT3G19170 | 4,99 | -0,41 | 1,02 | -0,40 | 6,91E-01 | 9,17E-01 |
| AT1G43910 | 2,35 | 0,74 | 1,88 | 0,40 | 6,92E-01 | 9,17E-01 |
| AT4G12650 | 2,23 | -0,49 | 1,23 | -0,40 | 6,92E-01 | 9,17E-01 |
| AT2G38195 | 4,06 | 0,46 | 1,16 | 0,39 | 6,93E-01 | 9,17E-01 |
| AT5G01155 | 4,02 | 0,37 | 0,93 | 0,39 | 6,93E-01 | 9,17E-01 |
| AT5G52530 | 3,03 | -0,60 | 1,51 | -0,40 | 6,93E-01 | 9,17E-01 |
| AT2G30920 | 2,18 | 0,47 | 1,21 | 0,39 | 6,94E-01 | 9,17E-01 |
| AT5G20950 | 3,45 | -0,47 | 1,18 | -0,39 | 6,94E-01 | 9,17E-01 |

| | | | | | | |
|---|---|---|---|---|---|---|
| AT1G05247 | 19,59 | -0,22 | 0,55 | -0,39 | 6,95E-01 | 9,18E-01 |
| AT1G62540 | 2,48 | 0,54 | 1,39 | 0,39 | 6,98E-01 | 9,19E-01 |
| AT2G29080 | 1,97 | -0,57 | 1,46 | -0,39 | 6,98E-01 | 9,19E-01 |
| AT5G18380 | 4,02 | -0,36 | 0,93 | -0,39 | 6,98E-01 | 9,19E-01 |
| AT5G61455 | 11,96 | -0,24 | 0,61 | -0,39 | 6,97E-01 | 9,19E-01 |
| AT5G60790 | 11,59 | 0,42 | 1,08 | 0,39 | 6,98E-01 | 9,19E-01 |
| AT1G02500 | 10,85 | -0,40 | 1,02 | -0,39 | 7,00E-01 | 9,19E-01 |
| AT3G26730 | 1,96 | 0,57 | 1,48 | 0,39 | 6,99E-01 | 9,19E-01 |
| AT5G52470 | 2,57 | 0,49 | 1,26 | 0,39 | 7,00E-01 | 9,19E-01 |
| AT3G18490 | 8,11 | 0,23 | 0,60 | 0,38 | 7,01E-01 | 9,20E-01 |
| AT5G09870 | 2,84 | -0,43 | 1,12 | -0,38 | 7,02E-01 | 9,20E-01 |
| AT3G04930 | 1,93 | 0,68 | 1,77 | 0,38 | 7,03E-01 | 9,20E-01 |
| AT5G02305 | 2,96 | 0,41 | 1,09 | 0,38 | 7,03E-01 | 9,20E-01 |
| AT4G38600 | 8,47 | 0,50 | 1,32 | 0,38 | 7,03E-01 | 9,20E-01 |
| AT1G79690 | 1,96 | -0,74 | 1,96 | -0,38 | 7,04E-01 | 9,21E-01 |
| AT2G07605 | 7,80 | -0,25 | 0,67 | -0,38 | 7,05E-01 | 9,21E-01 |
| AT3G17400 | 6,94 | -0,28 | 0,75 | -0,38 | 7,05E-01 | 9,21E-01 |
| AT2G16660 | 2,17 | 0,44 | 1,17 | 0,38 | 7,05E-01 | 9,21E-01 |
| AT5G24710 | 4,46 | 0,54 | 1,42 | 0,38 | 7,05E-01 | 9,21E-01 |
| AT1G14320 | 3,20 | 0,52 | 1,40 | 0,37 | 7,08E-01 | 9,22E-01 |
| AT1G27535 | 2,34 | 0,40 | 1,06 | 0,37 | 7,09E-01 | 9,22E-01 |
| AT1G32583 | 28,27 | 0,21 | 0,55 | 0,37 | 7,09E-01 | 9,22E-01 |
| AT1G74720 | 1,98 | 0,72 | 1,91 | 0,37 | 7,08E-01 | 9,22E-01 |
| AT3G49470 | 2,41 | -0,49 | 1,31 | -0,37 | 7,09E-01 | 9,22E-01 |
| AT5G09810 | 3,96 | -0,41 | 1,10 | -0,37 | 7,09E-01 | 9,22E-01 |
| AT5G65630 | 3,70 | 0,40 | 1,07 | 0,37 | 7,08E-01 | 9,22E-01 |
| AT2G48160 | 6,20 | -0,62 | 1,68 | -0,37 | 7,11E-01 | 9,23E-01 |
| AT1G05035 | 3,55 | 0,36 | 0,99 | 0,37 | 7,12E-01 | 9,24E-01 |
| AT1G54115 | 2,18 | 0,66 | 1,79 | 0,37 | 7,12E-01 | 9,24E-01 |
| AT3G13290 | 2,32 | -0,55 | 1,50 | -0,36 | 7,16E-01 | 9,26E-01 |
| AT1G65720 | 3,55 | 0,37 | 1,01 | 0,36 | 7,15E-01 | 9,26E-01 |
| AT1G76180 | 6,03 | 0,25 | 0,69 | 0,36 | 7,15E-01 | 9,26E-01 |
| AT4G25100 | 2,11 | 0,51 | 1,39 | 0,37 | 7,15E-01 | 9,26E-01 |
| AT1G07810 | 6,32 | 0,47 | 1,29 | 0,36 | 7,17E-01 | 9,27E-01 |
| AT5G61780 | 4,25 | 0,44 | 1,22 | 0,36 | 7,18E-01 | 9,27E-01 |
| AT2G45245 | 2,00 | 0,44 | 1,21 | 0,36 | 7,18E-01 | 9,27E-01 |
| AT5G40480 | 2,70 | 0,44 | 1,25 | 0,36 | 7,22E-01 | 9,30E-01 |
| AT5G62700 | 1,86 | 0,59 | 1,66 | 0,36 | 7,22E-01 | 9,30E-01 |
| AT1G09010 | 3,02 | 0,57 | 1,61 | 0,36 | 7,22E-01 | 9,30E-01 |
| AT2G05685 | 2,44 | 0,48 | 1,34 | 0,36 | 7,22E-01 | 9,30E-01 |
| AT3G09200 | 3,98 | -0,45 | 1,26 | -0,36 | 7,22E-01 | 9,30E-01 |
| AT3G03710 | 4,42 | -0,60 | 1,68 | -0,35 | 7,23E-01 | 9,30E-01 |
| AT3G61580 | 5,37 | 0,35 | 1,00 | 0,35 | 7,25E-01 | 9,31E-01 |
| AT1G66100 | 8,07 | 0,23 | 0,67 | 0,35 | 7,27E-01 | 9,32E-01 |
| AT5G10760 | 4,94 | 0,32 | 0,90 | 0,35 | 7,27E-01 | 9,32E-01 |
| AT5G43310 | 2,37 | -0,57 | 1,62 | -0,35 | 7,26E-01 | 9,32E-01 |
| AT1G30470 | 3,87 | 0,56 | 1,63 | 0,34 | 7,31E-01 | 9,32E-01 |
| AT3G05325 | 9,46 | 0,22 | 0,63 | 0,35 | 7,30E-01 | 9,32E-01 |
| AT4G05420 | 7,60 | -0,41 | 1,19 | -0,35 | 7,28E-01 | 9,32E-01 |
| AT4G18210 | 1,90 | -0,53 | 1,55 | -0,35 | 7,30E-01 | 9,32E-01 |
| AT1G19835 | 5,86 | 0,53 | 1,55 | 0,34 | 7,31E-01 | 9,32E-01 |
| AT1G31835 | 2,35 | -0,38 | 1,11 | -0,34 | 7,31E-01 | 9,32E-01 |
| AT1G64720 | 13,91 | -0,29 | 0,83 | -0,35 | 7,30E-01 | 9,32E-01 |
| AT2G25320 | 3,21 | 0,55 | 1,62 | 0,34 | 7,32E-01 | 9,32E-01 |
| AT2G42590 | 2,06 | -0,45 | 1,29 | -0,34 | 7,30E-01 | 9,32E-01 |
| AT5G05200 | 1,99 | -0,74 | 2,14 | -0,35 | 7,29E-01 | 9,32E-01 |

| | | | | | | |
|---|---|---|---|---|---|---|
| AT5G22770 | 2,48 | 0,47 | 1,34 | 0,35 | 7,27E-01 | 9,32E-01 |
| AT1G29465 | 25,62 | 0,15 | 0,44 | 0,34 | 7,32E-01 | 9,32E-01 |
| AT4G34950 | 4,16 | -0,30 | 0,87 | -0,34 | 7,33E-01 | 9,33E-01 |
| AT2G09670 | 5,26 | -0,49 | 1,43 | -0,34 | 7,34E-01 | 9,33E-01 |
| AT3G01695 | 8,72 | 0,28 | 0,83 | 0,34 | 7,34E-01 | 9,33E-01 |
| AT4G01050 | 4,58 | -0,47 | 1,38 | -0,34 | 7,34E-01 | 9,33E-01 |
| AT5G19390 | 2,78 | 0,66 | 1,95 | 0,34 | 7,35E-01 | 9,33E-01 |
| AT1G10290 | 9,22 | -0,46 | 1,37 | -0,34 | 7,36E-01 | 9,33E-01 |
| AT2G44060 | 2,21 | -0,39 | 1,17 | -0,34 | 7,37E-01 | 9,33E-01 |
| AT3G03050 | 2,11 | -0,68 | 2,03 | -0,34 | 7,36E-01 | 9,33E-01 |
| AT5G24300 | 2,35 | -0,50 | 1,49 | -0,34 | 7,37E-01 | 9,33E-01 |
| AT5G47910 | 5,04 | 0,38 | 1,13 | 0,34 | 7,35E-01 | 9,33E-01 |
| AT3G53420 | 2,26 | -0,43 | 1,29 | -0,34 | 7,38E-01 | 9,33E-01 |
| AT4G32020 | 12,23 | 0,24 | 0,73 | 0,33 | 7,39E-01 | 9,34E-01 |
| AT5G10960 | 3,18 | 0,47 | 1,40 | 0,33 | 7,39E-01 | 9,34E-01 |
| AT1G77560 | 2,14 | 0,52 | 1,55 | 0,33 | 7,40E-01 | 9,34E-01 |
| AT1G30360 | 3,38 | 0,32 | 0,98 | 0,33 | 7,41E-01 | 9,34E-01 |
| AT2G43710 | 3,74 | -0,45 | 1,36 | -0,33 | 7,41E-01 | 9,34E-01 |
| AT4G18600 | 3,50 | 0,60 | 1,81 | 0,33 | 7,41E-01 | 9,34E-01 |
| AT3G28899 | 2,18 | 0,41 | 1,24 | 0,33 | 7,43E-01 | 9,35E-01 |
| AT3G46530 | 2,60 | 0,40 | 1,21 | 0,33 | 7,43E-01 | 9,35E-01 |
| AT4G06800 | 2,19 | 0,35 | 1,06 | 0,33 | 7,42E-01 | 9,35E-01 |
| AT2G09685 | 3,15 | -0,30 | 0,93 | -0,33 | 7,44E-01 | 9,35E-01 |
| AT1G13320 | 2,17 | 0,53 | 1,62 | 0,33 | 7,44E-01 | 9,35E-01 |
| AT5G07970 | 3,91 | 0,58 | 1,78 | 0,33 | 7,44E-01 | 9,35E-01 |
| AT1G47265 | 3,77 | 0,29 | 0,91 | 0,32 | 7,48E-01 | 9,39E-01 |
| AT4G37260 | 2,49 | 0,41 | 1,28 | 0,32 | 7,49E-01 | 9,39E-01 |
| AT2G40030 | 5,36 | 0,54 | 1,70 | 0,32 | 7,50E-01 | 9,39E-01 |
| AT3G53430 | 2,55 | -0,34 | 1,06 | -0,32 | 7,50E-01 | 9,39E-01 |
| AT1G02080 | 3,21 | -0,48 | 1,52 | -0,32 | 7,51E-01 | 9,40E-01 |
| AT1G64710 | 2,31 | -0,55 | 1,75 | -0,32 | 7,52E-01 | 9,40E-01 |
| AT3G23100 | 2,14 | 0,51 | 1,64 | 0,31 | 7,54E-01 | 9,41E-01 |
| AT5G47260 | 4,17 | 0,25 | 0,79 | 0,31 | 7,53E-01 | 9,41E-01 |
| AT1G50030 | 2,19 | 0,50 | 1,59 | 0,31 | 7,54E-01 | 9,41E-01 |
| AT1G63080 | 109,45 | -0,09 | 0,27 | -0,31 | 7,53E-01 | 9,41E-01 |
| AT4G16990 | 5,03 | -0,28 | 0,89 | -0,31 | 7,54E-01 | 9,41E-01 |
| AT1G77300 | 4,17 | 0,53 | 1,71 | 0,31 | 7,55E-01 | 9,42E-01 |
| AT1G13110 | 2,32 | -0,36 | 1,17 | -0,31 | 7,56E-01 | 9,42E-01 |
| AT2G22330 | 4,57 | -0,52 | 1,67 | -0,31 | 7,56E-01 | 9,42E-01 |
| AT4G17390 | 6,32 | 0,25 | 0,81 | 0,31 | 7,58E-01 | 9,43E-01 |
| AT2G07645 | 4,48 | 0,24 | 0,77 | 0,31 | 7,58E-01 | 9,43E-01 |
| AT1G15940 | 2,10 | -0,54 | 1,78 | -0,30 | 7,62E-01 | 9,46E-01 |
| AT3G09445 | 4,99 | -0,25 | 0,82 | -0,30 | 7,62E-01 | 9,46E-01 |
| AT4G34830 | 2,44 | -0,49 | 1,63 | -0,30 | 7,62E-01 | 9,46E-01 |
| AT5G00760 | 4,31 | -0,25 | 0,81 | -0,30 | 7,62E-01 | 9,46E-01 |
| AT1G79990 | 2,18 | 0,57 | 1,90 | 0,30 | 7,63E-01 | 9,46E-01 |
| AT1G72390 | 2,40 | 0,54 | 1,81 | 0,30 | 7,67E-01 | 9,47E-01 |
| AT3G60750 | 11,99 | -0,17 | 0,57 | -0,30 | 7,66E-01 | 9,47E-01 |
| AT1G20010 | 2,01 | -0,38 | 1,29 | -0,30 | 7,66E-01 | 9,47E-01 |
| AT1G75380 | 2,78 | -0,31 | 1,02 | -0,30 | 7,64E-01 | 9,47E-01 |
| AT2G38280 | 1,89 | 0,55 | 1,84 | 0,30 | 7,65E-01 | 9,47E-01 |
| AT5G28840 | 2,50 | -0,33 | 1,11 | -0,30 | 7,67E-01 | 9,47E-01 |
| AT3G04155 | 3,42 | -0,28 | 0,94 | -0,30 | 7,68E-01 | 9,47E-01 |
| AT5G26830 | 2,03 | -0,52 | 1,75 | -0,30 | 7,67E-01 | 9,47E-01 |
| AT2G20950 | 3,17 | -0,30 | 1,01 | -0,29 | 7,69E-01 | 9,48E-01 |
| AT3G58050 | 1,93 | -0,53 | 1,79 | -0,29 | 7,69E-01 | 9,48E-01 |

| | | | | | | |
|---|---|---|---|---|---|---|
| AT3G04740 | 2,26 | 0,48 | 1,65 | 0,29 | 7,74E-01 | 9,53E-01 |
| AT3G44110 | 2,03 | -0,38 | 1,32 | -0,29 | 7,73E-01 | 9,53E-01 |
| AT1G10950 | 6,24 | 0,33 | 1,15 | 0,29 | 7,75E-01 | 9,53E-01 |
| AT3G24615 | 9,91 | -0,17 | 0,59 | -0,29 | 7,74E-01 | 9,53E-01 |
| AT1G77230 | 3,05 | -0,32 | 1,11 | -0,28 | 7,77E-01 | 9,53E-01 |
| AT3G52140 | 9,89 | 0,28 | 1,00 | 0,28 | 7,76E-01 | 9,53E-01 |
| AT4G24680 | 3,35 | -0,37 | 1,31 | -0,28 | 7,77E-01 | 9,53E-01 |
| AT1G23390 | 2,44 | -0,33 | 1,19 | -0,28 | 7,78E-01 | 9,54E-01 |
| AT4G03550 | 3,00 | -0,44 | 1,57 | -0,28 | 7,81E-01 | 9,56E-01 |
| AT5G61140 | 4,17 | 0,34 | 1,21 | 0,28 | 7,81E-01 | 9,56E-01 |
| AT1G61690 | 1,91 | -0,53 | 1,89 | -0,28 | 7,80E-01 | 9,56E-01 |
| AT2G22496 | 20,32 | -0,15 | 0,54 | -0,27 | 7,83E-01 | 9,57E-01 |
| AT3G19190 | 3,92 | 0,49 | 1,80 | 0,27 | 7,83E-01 | 9,57E-01 |
| AT4G12570 | 2,25 | 0,49 | 1,76 | 0,28 | 7,82E-01 | 9,57E-01 |
| AT3G17390 | 4,31 | -0,23 | 0,85 | -0,27 | 7,83E-01 | 9,57E-01 |
| AT2G34750 | 2,36 | -0,43 | 1,57 | -0,27 | 7,85E-01 | 9,58E-01 |
| AT5G54310 | 4,38 | -0,38 | 1,39 | -0,27 | 7,85E-01 | 9,58E-01 |
| AT4G04890 | 2,53 | -0,46 | 1,70 | -0,27 | 7,86E-01 | 9,59E-01 |
| AT5G24350 | 3,06 | 0,52 | 1,93 | 0,27 | 7,87E-01 | 9,59E-01 |
| AT3G46620 | 4,33 | 0,27 | 1,02 | 0,27 | 7,88E-01 | 9,60E-01 |
| AT3G20770 | 2,68 | -0,41 | 1,53 | -0,27 | 7,89E-01 | 9,61E-01 |
| AT1G33560 | 2,19 | 0,54 | 2,04 | 0,27 | 7,91E-01 | 9,61E-01 |
| AT5G52170 | 2,49 | 0,43 | 1,60 | 0,27 | 7,90E-01 | 9,61E-01 |
| AT3G17900 | 1,92 | 0,49 | 1,84 | 0,26 | 7,91E-01 | 9,62E-01 |
| AT3G18780 | 17,06 | 0,18 | 0,70 | 0,26 | 7,93E-01 | 9,62E-01 |
| AT4G24770 | 3,23 | -0,29 | 1,10 | -0,26 | 7,94E-01 | 9,63E-01 |
| AT3G13525 | 2,38 | -0,26 | 1,01 | -0,26 | 7,95E-01 | 9,64E-01 |
| AT5G01410 | 2,12 | -0,36 | 1,38 | -0,26 | 7,96E-01 | 9,64E-01 |
| AT3G13857 | 124,31 | 0,10 | 0,38 | 0,26 | 7,98E-01 | 9,65E-01 |
| AT3G13855 | 124,31 | 0,10 | 0,38 | 0,26 | 7,98E-01 | 9,65E-01 |
| AT5G27850 | 1,94 | -0,40 | 1,57 | -0,26 | 7,97E-01 | 9,65E-01 |
| AT2G41105 | 5,08 | 0,23 | 0,90 | 0,26 | 7,97E-01 | 9,65E-01 |
| AT3G22910 | 2,19 | -0,47 | 1,84 | -0,25 | 7,99E-01 | 9,65E-01 |
| AT1G04210 | 3,48 | -0,31 | 1,25 | -0,25 | 8,01E-01 | 9,66E-01 |
| AT1G17470 | 1,97 | -0,44 | 1,77 | -0,25 | 8,05E-01 | 9,66E-01 |
| AT1G20970 | 4,69 | -0,33 | 1,32 | -0,25 | 8,05E-01 | 9,66E-01 |
| AT1G72180 | 2,71 | -0,27 | 1,08 | -0,25 | 8,05E-01 | 9,66E-01 |
| AT2G43150 | 1,94 | -0,40 | 1,58 | -0,25 | 8,00E-01 | 9,66E-01 |
| AT2G47170 | 1,90 | 0,32 | 1,30 | 0,24 | 8,07E-01 | 9,66E-01 |
| AT3G14930 | 2,04 | -0,33 | 1,36 | -0,24 | 8,06E-01 | 9,66E-01 |
| AT4G30993 | 2,23 | 0,33 | 1,34 | 0,24 | 8,07E-01 | 9,66E-01 |
| AT4G35770 | 2,85 | 0,28 | 1,12 | 0,25 | 8,04E-01 | 9,66E-01 |
| AT5G15650 | 3,70 | -0,27 | 1,09 | -0,24 | 8,07E-01 | 9,66E-01 |
| AT5G64740 | 4,40 | -0,23 | 0,95 | -0,25 | 8,05E-01 | 9,66E-01 |
| AT5G65480 | 2,00 | 0,39 | 1,56 | 0,25 | 8,06E-01 | 9,66E-01 |
| AT1G80410 | 5,85 | 0,40 | 1,59 | 0,25 | 8,02E-01 | 9,66E-01 |
| AT2G13360 | 3,44 | -0,21 | 0,85 | -0,25 | 8,04E-01 | 9,66E-01 |
| AT2G14910 | 2,97 | -0,30 | 1,21 | -0,25 | 8,02E-01 | 9,66E-01 |
| AT2G41530 | 2,42 | -0,32 | 1,26 | -0,25 | 8,01E-01 | 9,66E-01 |
| AT3G17850 | 4,74 | 0,40 | 1,57 | 0,25 | 8,00E-01 | 9,66E-01 |
| AT4G27520 | 5,05 | -0,23 | 0,94 | -0,24 | 8,07E-01 | 9,66E-01 |
| AT1G70185 | 5,33 | -0,26 | 1,11 | -0,24 | 8,12E-01 | 9,66E-01 |
| AT3G49910 | 2,91 | -0,29 | 1,22 | -0,24 | 8,12E-01 | 9,66E-01 |
| AT5G13010 | 4,55 | -0,31 | 1,29 | -0,24 | 8,09E-01 | 9,66E-01 |
| AT5G47010 | 2,34 | -0,33 | 1,38 | -0,24 | 8,12E-01 | 9,66E-01 |
| AT2G18440 | 3,48 | -0,24 | 1,00 | -0,24 | 8,10E-01 | 9,66E-01 |

| | | | | | | |
|---|---|---|---|---|---|---|
| AT2G46500 | 2,79 | 0,42 | 1,76 | 0,24 | 8,11E-01 | 9,66E-01 |
| AT3G01120 | 2,55 | -0,39 | 1,62 | -0,24 | 8,11E-01 | 9,66E-01 |
| AT3G13405 | 1,91 | -0,32 | 1,31 | -0,24 | 8,10E-01 | 9,66E-01 |
| AT4G16920 | 1,96 | 0,35 | 1,48 | 0,24 | 8,12E-01 | 9,66E-01 |
| AT4G16983 | 2,85 | -0,24 | 0,98 | -0,24 | 8,10E-01 | 9,66E-01 |
| AT1G31580 | 10,89 | 0,12 | 0,55 | 0,23 | 8,19E-01 | 9,67E-01 |
| AT1G43170 | 3,66 | -0,24 | 1,06 | -0,23 | 8,17E-01 | 9,67E-01 |
| AT1G63230 | 3,31 | -0,20 | 0,85 | -0,23 | 8,18E-01 | 9,67E-01 |
| AT2G15530 | 40,01 | 0,10 | 0,42 | 0,23 | 8,17E-01 | 9,67E-01 |
| AT2G38040 | 4,80 | -0,28 | 1,20 | -0,23 | 8,18E-01 | 9,67E-01 |
| AT5G51630 | 1,97 | 0,42 | 1,79 | 0,24 | 8,13E-01 | 9,67E-01 |
| AT3G01310 | 2,04 | 0,47 | 2,03 | 0,23 | 8,18E-01 | 9,67E-01 |
| AT3G13460 | 10,31 | -0,15 | 0,66 | -0,23 | 8,18E-01 | 9,67E-01 |
| AT3G08165 | 3,95 | 0,23 | 1,02 | 0,23 | 8,19E-01 | 9,67E-01 |
| AT3G53740 | 2,01 | -0,49 | 2,13 | -0,23 | 8,17E-01 | 9,67E-01 |
| AT4G35090 | 2,83 | -0,23 | 1,00 | -0,23 | 8,19E-01 | 9,67E-01 |
| AT5G18360 | 2,96 | -0,37 | 1,56 | -0,23 | 8,15E-01 | 9,67E-01 |
| AT5G57110 | 2,43 | -0,35 | 1,48 | -0,24 | 8,14E-01 | 9,67E-01 |
| AT5G02880 | 3,00 | -0,29 | 1,26 | -0,23 | 8,20E-01 | 9,67E-01 |
| AT5G08525 | 4,34 | -0,23 | 1,04 | -0,23 | 8,21E-01 | 9,67E-01 |
| AT3G16140 | 1,91 | -0,30 | 1,32 | -0,23 | 8,20E-01 | 9,67E-01 |
| AT2G39681 | 1805,59 | 0,07 | 0,32 | 0,23 | 8,21E-01 | 9,67E-01 |
| AT2G43680 | 14,92 | -0,17 | 0,77 | -0,23 | 8,22E-01 | 9,67E-01 |
| AT1G29350 | 2,20 | 0,38 | 1,71 | 0,22 | 8,23E-01 | 9,68E-01 |
| AT1G36160 | 9,82 | 0,22 | 0,97 | 0,22 | 8,24E-01 | 9,69E-01 |
| AT5G39080 | 2,20 | 0,30 | 1,34 | 0,22 | 8,24E-01 | 9,69E-01 |
| AT5G26742 | 9,59 | -0,17 | 0,79 | -0,22 | 8,25E-01 | 9,69E-01 |
| AT2G15620 | 3,38 | -0,26 | 1,18 | -0,22 | 8,27E-01 | 9,71E-01 |
| AT1G14580 | 3,99 | 0,22 | 0,99 | 0,22 | 8,28E-01 | 9,71E-01 |
| AT2G43160 | 6,80 | 0,33 | 1,53 | 0,22 | 8,29E-01 | 9,71E-01 |
| AT2G46192 | 21,12 | -0,12 | 0,58 | -0,22 | 8,29E-01 | 9,71E-01 |
| AT3G59140 | 3,91 | 0,45 | 2,07 | 0,22 | 8,29E-01 | 9,71E-01 |
| AT5G24930 | 2,10 | 0,32 | 1,49 | 0,21 | 8,31E-01 | 9,72E-01 |
| AT3G48201 | 30,66 | 0,15 | 0,70 | 0,21 | 8,31E-01 | 9,72E-01 |
| AT3G61690 | 2,25 | 0,35 | 1,67 | 0,21 | 8,32E-01 | 9,72E-01 |
| AT3G20760 | 5,52 | 0,32 | 1,51 | 0,21 | 8,32E-01 | 9,72E-01 |
| AT3G51800 | 4,28 | -0,17 | 0,82 | -0,21 | 8,32E-01 | 9,72E-01 |
| AT1G20330 | 2,79 | -0,30 | 1,42 | -0,21 | 8,34E-01 | 9,72E-01 |
| AT2G03820 | 4,25 | 0,33 | 1,56 | 0,21 | 8,33E-01 | 9,72E-01 |
| AT4G27430 | 2,23 | -0,34 | 1,64 | -0,21 | 8,34E-01 | 9,73E-01 |
| AT1G10170 | 9,09 | 0,23 | 1,13 | 0,21 | 8,36E-01 | 9,74E-01 |
| AT3G18370 | 2,55 | 0,32 | 1,57 | 0,20 | 8,38E-01 | 9,76E-01 |
| AT3G04775 | 14,57 | 0,09 | 0,46 | 0,20 | 8,39E-01 | 9,76E-01 |
| AT1G05427 | 2,11 | -0,28 | 1,37 | -0,20 | 8,39E-01 | 9,76E-01 |
| AT5G12470 | 4,58 | 0,32 | 1,62 | 0,20 | 8,41E-01 | 9,77E-01 |
| AT3G05420 | 2,70 | 0,32 | 1,64 | 0,19 | 8,46E-01 | 9,79E-01 |
| AT3G23820 | 9,86 | 0,11 | 0,55 | 0,19 | 8,46E-01 | 9,79E-01 |
| AT5G47480 | 1,95 | 0,33 | 1,69 | 0,19 | 8,46E-01 | 9,79E-01 |
| AT5G48620 | 2,46 | -0,28 | 1,46 | -0,19 | 8,46E-01 | 9,79E-01 |
| AT5G62165 | 7,10 | -0,14 | 0,73 | -0,20 | 8,45E-01 | 9,79E-01 |
| AT5G65700 | 1,88 | -0,25 | 1,29 | -0,20 | 8,44E-01 | 9,79E-01 |
| AT2G17930 | 7,05 | 0,18 | 0,94 | 0,20 | 8,44E-01 | 9,79E-01 |
| AT4G01810 | 2,76 | 0,34 | 1,72 | 0,20 | 8,44E-01 | 9,79E-01 |
| AT5G53170 | 2,82 | 0,28 | 1,42 | 0,20 | 8,43E-01 | 9,79E-01 |
| AT1G07000 | 3,04 | -0,29 | 1,51 | -0,19 | 8,47E-01 | 9,79E-01 |
| AT4G02570 | 1,99 | -0,30 | 1,57 | -0,19 | 8,49E-01 | 9,80E-01 |

| | | | | | | |
|---|---|---|---|---|---|---|
| AT4G16980 | 2,88 | -0,18 | 0,97 | -0,19 | 8,49E-01 | 9,80E-01 |
| AT3G23110 | 1,95 | 0,33 | 1,74 | 0,19 | 8,49E-01 | 9,80E-01 |
| AT1G01510 | 2,79 | 0,25 | 1,74 | 0,14 | 8,87E-01 | 9,80E-01 |
| AT1G04425 | 1,91 | 0,20 | 1,22 | 0,16 | 8,72E-01 | 9,80E-01 |
| AT1G11650 | 3,30 | -0,15 | 1,11 | -0,13 | 8,94E-01 | 9,80E-01 |
| AT1G15500 | 7,05 | 0,24 | 1,55 | 0,15 | 8,79E-01 | 9,80E-01 |
| AT1G18010 | 5,80 | 0,12 | 0,72 | 0,17 | 8,65E-01 | 9,80E-01 |
| AT1G19680 | 2,21 | 0,36 | 2,04 | 0,18 | 8,59E-01 | 9,80E-01 |
| AT1G21630 | 4,05 | -0,31 | 1,82 | -0,17 | 8,66E-01 | 9,80E-01 |
| AT1G06667 | 5,78 | 0,11 | 0,79 | 0,14 | 8,90E-01 | 9,80E-01 |
| AT1G33680 | 2,02 | -0,30 | 2,10 | -0,14 | 8,85E-01 | 9,80E-01 |
| AT1G48300 | 8,14 | 0,26 | 1,39 | 0,19 | 8,53E-01 | 9,80E-01 |
| AT1G52360 | 2,02 | -0,22 | 1,51 | -0,14 | 8,85E-01 | 9,80E-01 |
| AT1G53900 | 2,01 | 0,21 | 1,18 | 0,18 | 8,60E-01 | 9,80E-01 |
| AT1G62020 | 12,54 | -0,13 | 0,95 | -0,14 | 8,89E-01 | 9,80E-01 |
| AT1G62630 | 2,10 | -0,19 | 1,16 | -0,17 | 8,69E-01 | 9,80E-01 |
| AT1G65930 | 2,39 | -0,24 | 1,44 | -0,17 | 8,68E-01 | 9,80E-01 |
| AT1G76135 | 2,70 | 0,21 | 1,19 | 0,18 | 8,61E-01 | 9,80E-01 |
| AT1G09687 | 2,04 | 0,16 | 1,19 | 0,13 | 8,95E-01 | 9,80E-01 |
| AT1G78206 | 14,29 | 0,09 | 0,53 | 0,16 | 8,70E-01 | 9,80E-01 |
| AT2G03140 | 2,19 | 0,26 | 1,62 | 0,16 | 8,71E-01 | 9,80E-01 |
| AT2G03150 | 2,08 | 0,26 | 1,71 | 0,15 | 8,78E-01 | 9,80E-01 |
| AT2G19360 | 2,70 | 0,15 | 1,11 | 0,14 | 8,90E-01 | 9,80E-01 |
| AT2G08370 | 4,11 | -0,12 | 0,82 | -0,15 | 8,81E-01 | 9,80E-01 |
| AT2G34655 | 4,12 | 0,15 | 0,96 | 0,16 | 8,72E-01 | 9,80E-01 |
| AT2G41090 | 3,05 | 0,19 | 1,05 | 0,18 | 8,58E-01 | 9,80E-01 |
| AT2G41900 | 16,69 | -0,20 | 1,15 | -0,17 | 8,65E-01 | 9,80E-01 |
| AT2G46830 | 2,43 | 0,27 | 1,85 | 0,15 | 8,83E-01 | 9,80E-01 |
| AT2G47440 | 2,19 | 0,28 | 1,51 | 0,19 | 8,51E-01 | 9,80E-01 |
| AT3G01370 | 2,22 | -0,17 | 1,20 | -0,14 | 8,86E-01 | 9,80E-01 |
| AT3G01460 | 5,55 | 0,22 | 1,60 | 0,14 | 8,89E-01 | 9,80E-01 |
| AT3G14470 | 2,64 | -0,26 | 1,93 | -0,14 | 8,92E-01 | 9,80E-01 |
| AT3G21760 | 4,32 | 0,11 | 0,81 | 0,14 | 8,92E-01 | 9,80E-01 |
| AT3G22890 | 4,47 | 0,16 | 0,95 | 0,17 | 8,63E-01 | 9,80E-01 |
| AT3G26650 | 12,31 | 0,11 | 0,66 | 0,17 | 8,65E-01 | 9,80E-01 |
| AT4G00231 | 1,99 | 0,21 | 1,51 | 0,14 | 8,89E-01 | 9,80E-01 |
| AT4G02350 | 1,95 | -0,27 | 1,94 | -0,14 | 8,91E-01 | 9,80E-01 |
| AT4G03420 | 2,29 | 0,24 | 1,77 | 0,14 | 8,91E-01 | 9,80E-01 |
| AT4G10210 | 2,08 | -0,18 | 1,30 | -0,14 | 8,89E-01 | 9,80E-01 |
| AT4G06375 | 3,09 | 0,20 | 1,33 | 0,15 | 8,79E-01 | 9,80E-01 |
| AT4G20260 | 2,30 | 0,22 | 1,17 | 0,19 | 8,53E-01 | 9,80E-01 |
| AT4G23420 | 23,63 | -0,07 | 0,48 | -0,14 | 8,91E-01 | 9,80E-01 |
| AT4G07120 | 22,20 | -0,07 | 0,52 | -0,14 | 8,89E-01 | 9,80E-01 |
| AT4G23630 | 1,89 | 0,19 | 1,27 | 0,15 | 8,82E-01 | 9,80E-01 |
| AT4G25080 | 4,26 | -0,22 | 1,17 | -0,18 | 8,54E-01 | 9,80E-01 |
| AT4G38440 | 2,75 | -0,21 | 1,47 | -0,14 | 8,87E-01 | 9,80E-01 |
| AT5G42090 | 4,37 | -0,13 | 1,00 | -0,13 | 8,94E-01 | 9,80E-01 |
| AT5G42270 | 4,20 | -0,17 | 0,94 | -0,18 | 8,54E-01 | 9,80E-01 |
| AT5G45430 | 3,88 | 0,16 | 1,22 | 0,13 | 8,94E-01 | 9,80E-01 |
| AT5G45472 | 4,93 | 0,17 | 1,03 | 0,17 | 8,66E-01 | 9,80E-01 |
| AT5G56010 | 6,62 | 0,11 | 0,84 | 0,13 | 8,94E-01 | 9,80E-01 |
| AT5G62350 | 2,26 | -0,14 | 1,08 | -0,13 | 8,95E-01 | 9,80E-01 |
| AT5G64270 | 15,19 | 0,20 | 1,23 | 0,16 | 8,71E-01 | 9,80E-01 |
| AT5G65610 | 2,72 | 0,17 | 1,19 | 0,14 | 8,85E-01 | 9,80E-01 |
| AT1G05010 | 2,06 | -0,25 | 1,48 | -0,17 | 8,67E-01 | 9,80E-01 |
| AT1G17210 | 2,32 | 0,32 | 1,91 | 0,17 | 8,65E-01 | 9,80E-01 |

| | | | | | | |
|---|---|---|---|---|---|---|
| AT1G19410 | 3,18 | -0,13 | 0,91 | -0,14 | 8,89E-01 | 9,80E-01 |
| AT1G49040 | 3,74 | -0,33 | 1,82 | -0,18 | 8,55E-01 | 9,80E-01 |
| AT1G50250 | 2,35 | -0,16 | 1,17 | -0,14 | 8,91E-01 | 9,80E-01 |
| AT1G08103 | 6,40 | 0,13 | 0,83 | 0,16 | 8,71E-01 | 9,80E-01 |
| AT1G62310 | 4,32 | -0,12 | 0,80 | -0,15 | 8,77E-01 | 9,80E-01 |
| AT1G67960 | 1,96 | 0,27 | 1,99 | 0,14 | 8,91E-01 | 9,80E-01 |
| AT1G71002 | 2,81 | -0,15 | 1,14 | -0,13 | 8,94E-01 | 9,80E-01 |
| AT2G04965 | 3,81 | -0,13 | 0,79 | -0,16 | 8,71E-01 | 9,80E-01 |
| AT2G07815 | 2,06 | -0,17 | 1,20 | -0,14 | 8,90E-01 | 9,80E-01 |
| AT2G20010 | 1,89 | -0,29 | 1,62 | -0,18 | 8,57E-01 | 9,80E-01 |
| AT2G26250 | 1,96 | -0,16 | 1,11 | -0,15 | 8,83E-01 | 9,80E-01 |
| AT2G31400 | 2,56 | 0,23 | 1,52 | 0,15 | 8,81E-01 | 9,80E-01 |
| AT2G35747 | 15,41 | -0,09 | 0,61 | -0,15 | 8,83E-01 | 9,80E-01 |
| AT2G09465 | 4,22 | 0,13 | 0,85 | 0,16 | 8,74E-01 | 9,80E-01 |
| AT2G43900 | 9,76 | -0,26 | 1,74 | -0,15 | 8,82E-01 | 9,80E-01 |
| AT3G02260 | 23,16 | -0,12 | 0,67 | -0,18 | 8,55E-01 | 9,80E-01 |
| AT3G08850 | 2,35 | 0,26 | 1,70 | 0,15 | 8,80E-01 | 9,80E-01 |
| AT3G10380 | 2,66 | 0,17 | 1,24 | 0,13 | 8,94E-01 | 9,80E-01 |
| AT3G15880 | 1,94 | -0,23 | 1,39 | -0,16 | 8,69E-01 | 9,80E-01 |
| AT3G17800 | 1,86 | -0,20 | 1,50 | -0,13 | 8,94E-01 | 9,80E-01 |
| AT3G21755 | 4,36 | 0,15 | 0,82 | 0,18 | 8,58E-01 | 9,80E-01 |
| AT3G28956 | 2,37 | -0,19 | 1,03 | -0,18 | 8,55E-01 | 9,80E-01 |
| AT3G52880 | 2,23 | -0,17 | 1,16 | -0,15 | 8,83E-01 | 9,80E-01 |
| AT3G55220 | 4,51 | -0,23 | 1,46 | -0,16 | 8,75E-01 | 9,80E-01 |
| AT3G60770 | 1,96 | 0,22 | 1,53 | 0,15 | 8,85E-01 | 9,80E-01 |
| AT3G62980 | 20,74 | 0,09 | 0,60 | 0,15 | 8,84E-01 | 9,80E-01 |
| AT3G63410 | 2,71 | 0,25 | 1,35 | 0,18 | 8,55E-01 | 9,80E-01 |
| AT4G02480 | 22,30 | -0,15 | 1,01 | -0,15 | 8,84E-01 | 9,80E-01 |
| AT4G14960 | 3,57 | -0,14 | 1,03 | -0,14 | 8,89E-01 | 9,80E-01 |
| AT4G16143 | 2,03 | 0,21 | 1,35 | 0,15 | 8,79E-01 | 9,80E-01 |
| AT4G23850 | 2,43 | 0,19 | 1,11 | 0,17 | 8,66E-01 | 9,80E-01 |
| AT4G26480 | 2,39 | -0,27 | 1,50 | -0,18 | 8,59E-01 | 9,80E-01 |
| AT4G26540 | 3,42 | 0,17 | 1,08 | 0,16 | 8,73E-01 | 9,80E-01 |
| AT4G31880 | 2,70 | -0,29 | 1,90 | -0,15 | 8,80E-01 | 9,80E-01 |
| AT4G33240 | 2,23 | 0,26 | 1,57 | 0,16 | 8,69E-01 | 9,80E-01 |
| AT4G36140 | 2,98 | 0,23 | 1,56 | 0,15 | 8,84E-01 | 9,80E-01 |
| AT5G05690 | 2,09 | 0,23 | 1,70 | 0,13 | 8,94E-01 | 9,80E-01 |
| AT5G16210 | 1,91 | -0,35 | 1,97 | -0,18 | 8,59E-01 | 9,80E-01 |
| AT5G17230 | 2,13 | 0,26 | 1,42 | 0,19 | 8,53E-01 | 9,80E-01 |
| AT5G45500 | 2,29 | 0,31 | 2,05 | 0,15 | 8,79E-01 | 9,80E-01 |
| AT5G51300 | 1,97 | 0,31 | 2,33 | 0,13 | 8,95E-01 | 9,80E-01 |
| AT5G53440 | 2,08 | -0,21 | 1,41 | -0,15 | 8,79E-01 | 9,80E-01 |
| AT5G61790 | 5,40 | 0,16 | 0,92 | 0,18 | 8,60E-01 | 9,80E-01 |
| AT1G04810 | 3,41 | -0,24 | 1,97 | -0,12 | 9,04E-01 | 9,80E-01 |
| AT1G50010 | 2,93 | 0,15 | 1,21 | 0,13 | 8,98E-01 | 9,80E-01 |
| AT3G19180 | 1,86 | -0,26 | 2,15 | -0,12 | 9,03E-01 | 9,80E-01 |
| AT3G20010 | 2,80 | -0,26 | 2,05 | -0,13 | 8,98E-01 | 9,80E-01 |
| AT3G46600 | 2,19 | 0,15 | 1,20 | 0,13 | 8,99E-01 | 9,80E-01 |
| AT3G54730 | 3,74 | 0,11 | 0,84 | 0,13 | 8,99E-01 | 9,80E-01 |
| AT3G57300 | 2,34 | 0,21 | 1,67 | 0,13 | 8,99E-01 | 9,80E-01 |
| AT4G01840 | 3,87 | -0,13 | 1,02 | -0,12 | 9,02E-01 | 9,80E-01 |
| AT4G13493 | 160,77 | 0,04 | 0,30 | 0,12 | 9,02E-01 | 9,80E-01 |
| AT5G46250 | 6,03 | -0,17 | 1,43 | -0,12 | 9,05E-01 | 9,80E-01 |
| AT1G08117 | 4,98 | 0,13 | 1,04 | 0,12 | 9,02E-01 | 9,80E-01 |
| AT1G69360 | 2,19 | -0,26 | 2,03 | -0,13 | 8,97E-01 | 9,80E-01 |
| AT3G08730 | 2,48 | -0,24 | 1,99 | -0,12 | 9,05E-01 | 9,80E-01 |

| | | | | | | |
|---|---|---|---|---|---|---|
| AT3G13070 | 1,98 | -0,26 | 2,10 | -0,12 | 9,01E-01 | 9,80E-01 |
| AT4G24520 | 3,90 | 0,22 | 1,87 | 0,12 | 9,04E-01 | 9,80E-01 |
| AT5G12120 | 2,73 | 0,22 | 1,73 | 0,13 | 9,00E-01 | 9,80E-01 |
| AT5G42930 | 2,68 | -0,13 | 1,05 | -0,12 | 9,04E-01 | 9,80E-01 |
| AT5G58550 | 1,86 | -0,26 | 2,17 | -0,12 | 9,04E-01 | 9,80E-01 |
| AT5G60450 | 7,78 | 0,08 | 0,61 | 0,13 | 8,98E-01 | 9,80E-01 |
| AT5G61310 | 4,76 | 0,10 | 0,79 | 0,12 | 9,04E-01 | 9,80E-01 |
| AT1G75660 | 6,24 | 0,18 | 1,56 | 0,12 | 9,07E-01 | 9,81E-01 |
| AT5G07290 | 1,99 | -0,29 | 2,43 | -0,12 | 9,06E-01 | 9,81E-01 |
| AT5G23630 | 2,35 | 0,18 | 1,56 | 0,12 | 9,07E-01 | 9,81E-01 |
| AT5G52882 | 2,07 | -0,24 | 2,05 | -0,12 | 9,06E-01 | 9,81E-01 |
| AT1G06148 | 2,19 | 0,15 | 1,27 | 0,12 | 9,08E-01 | 9,81E-01 |
| AT1G04080 | 1,89 | -0,17 | 1,58 | -0,11 | 9,15E-01 | 9,81E-01 |
| AT1G54270 | 2,59 | 0,11 | 1,01 | 0,11 | 9,14E-01 | 9,81E-01 |
| AT1G71220 | 2,96 | 0,16 | 1,46 | 0,11 | 9,13E-01 | 9,81E-01 |
| AT1G78040 | 2,48 | 0,13 | 1,23 | 0,11 | 9,14E-01 | 9,81E-01 |
| AT2G01422 | 2,93 | 0,15 | 1,37 | 0,11 | 9,13E-01 | 9,81E-01 |
| AT2G46020 | 3,28 | -0,13 | 1,27 | -0,11 | 9,15E-01 | 9,81E-01 |
| AT3G03380 | 4,30 | -0,15 | 1,40 | -0,11 | 9,14E-01 | 9,81E-01 |
| AT4G34670 | 4,28 | 0,15 | 1,36 | 0,11 | 9,13E-01 | 9,81E-01 |
| AT1G09570 | 2,05 | 0,17 | 1,49 | 0,11 | 9,09E-01 | 9,81E-01 |
| AT3G07160 | 3,87 | 0,14 | 1,28 | 0,11 | 9,15E-01 | 9,81E-01 |
| AT3G30390 | 2,05 | -0,13 | 1,20 | -0,11 | 9,11E-01 | 9,81E-01 |
| AT3G08210 | 3,75 | -0,10 | 0,86 | -0,11 | 9,11E-01 | 9,81E-01 |
| AT5G20700 | 23,90 | 0,05 | 0,46 | 0,11 | 9,10E-01 | 9,81E-01 |
| AT5G62190 | 2,05 | 0,19 | 1,73 | 0,11 | 9,12E-01 | 9,81E-01 |
| AT5G64840 | 5,36 | 0,13 | 1,19 | 0,11 | 9,12E-01 | 9,81E-01 |
| AT2G43375 | 6,65 | 0,07 | 0,66 | 0,10 | 9,18E-01 | 9,83E-01 |
| AT1G20690 | 2,85 | 0,11 | 1,03 | 0,10 | 9,19E-01 | 9,83E-01 |
| AT1G20691 | 2,85 | 0,11 | 1,03 | 0,10 | 9,19E-01 | 9,83E-01 |
| AT4G26530 | 4,43 | -0,09 | 0,88 | -0,10 | 9,20E-01 | 9,84E-01 |
| AT5G54440 | 2,06 | 0,16 | 1,56 | 0,10 | 9,21E-01 | 9,84E-01 |
| AT1G79000 | 2,98 | -0,22 | 2,20 | -0,10 | 9,20E-01 | 9,84E-01 |
| AT3G43300 | 11,08 | -0,10 | 1,00 | -0,10 | 9,21E-01 | 9,84E-01 |
| AT5G12250 | 2,28 | 0,12 | 1,21 | 0,10 | 9,21E-01 | 9,84E-01 |
| AT5G60390 | 16,82 | -0,05 | 0,53 | -0,10 | 9,22E-01 | 9,84E-01 |
| AT5G09660 | 2,38 | -0,10 | 1,02 | -0,10 | 9,23E-01 | 9,84E-01 |
| AT1G64790 | 7,36 | 0,08 | 0,88 | 0,10 | 9,24E-01 | 9,85E-01 |
| AT5G24990 | 2,50 | -0,11 | 1,11 | -0,09 | 9,25E-01 | 9,85E-01 |
| AT1G08347 | 16,30 | 0,04 | 0,44 | 0,09 | 9,26E-01 | 9,86E-01 |
| AT4G13770 | 2,50 | 0,12 | 1,35 | 0,09 | 9,27E-01 | 9,87E-01 |
| AT1G13980 | 8,14 | 0,13 | 1,55 | 0,08 | 9,32E-01 | 9,89E-01 |
| AT1G27320 | 2,96 | 0,13 | 1,66 | 0,08 | 9,37E-01 | 9,89E-01 |
| AT1G59610 | 3,98 | 0,12 | 1,39 | 0,09 | 9,30E-01 | 9,89E-01 |
| AT1G65960 | 3,84 | -0,10 | 1,18 | -0,09 | 9,32E-01 | 9,89E-01 |
| AT2G07890 | 2,64 | -0,09 | 1,07 | -0,08 | 9,36E-01 | 9,89E-01 |
| AT2G34110 | 1,88 | -0,12 | 1,31 | -0,09 | 9,30E-01 | 9,89E-01 |
| AT4G26542 | 2,99 | 0,08 | 1,08 | 0,08 | 9,40E-01 | 9,89E-01 |
| AT4G31980 | 2,53 | 0,09 | 1,24 | 0,08 | 9,40E-01 | 9,89E-01 |
| AT5G13530 | 2,40 | -0,09 | 1,26 | -0,08 | 9,40E-01 | 9,89E-01 |
| AT5G27807 | 30,37 | -0,03 | 0,40 | -0,07 | 9,41E-01 | 9,89E-01 |
| AT5G58140 | 8,36 | 0,09 | 1,08 | 0,08 | 9,34E-01 | 9,89E-01 |
| AT5G62162 | 10,16 | 0,06 | 0,82 | 0,07 | 9,42E-01 | 9,89E-01 |
| AT1G06190 | 5,18 | -0,12 | 1,40 | -0,09 | 9,32E-01 | 9,89E-01 |
| AT1G11640 | 3,15 | 0,13 | 1,70 | 0,08 | 9,40E-01 | 9,89E-01 |
| AT1G68560 | 3,34 | -0,09 | 1,09 | -0,08 | 9,33E-01 | 9,89E-01 |

| | | | | | | |
|---|---|---|---|---|---|---|
| AT1G77180 | 2,45 | 0,11 | 1,52 | 0,07 | 9,41E-01 | 9,89E-01 |
| AT2G35382 | 3,86 | 0,07 | 0,86 | 0,08 | 9,36E-01 | 9,89E-01 |
| AT2G43970 | 15,13 | -0,05 | 0,72 | -0,07 | 9,41E-01 | 9,89E-01 |
| AT3G03435 | 3,28 | -0,08 | 1,03 | -0,07 | 9,41E-01 | 9,89E-01 |
| AT4G16370 | 3,80 | 0,08 | 1,07 | 0,07 | 9,41E-01 | 9,89E-01 |
| AT4G27640 | 8,75 | 0,13 | 1,61 | 0,08 | 9,35E-01 | 9,89E-01 |
| AT5G04315 | 6,08 | 0,07 | 0,85 | 0,08 | 9,36E-01 | 9,89E-01 |
| AT5G04325 | 6,08 | 0,07 | 0,85 | 0,08 | 9,36E-01 | 9,89E-01 |
| AT5G43470 | 3,38 | 0,12 | 1,36 | 0,08 | 9,32E-01 | 9,89E-01 |
| AT5G58070 | 2,01 | -0,11 | 1,50 | -0,08 | 9,39E-01 | 9,89E-01 |
| AT5G58330 | 1,92 | -0,14 | 1,65 | -0,08 | 9,33E-01 | 9,89E-01 |
| AT5G61410 | 2,49 | 0,09 | 1,07 | 0,08 | 9,33E-01 | 9,89E-01 |
| AT1G68010 | 2,53 | -0,09 | 1,28 | -0,07 | 9,42E-01 | 9,89E-01 |
| AT2G18020 | 2,64 | -0,10 | 1,37 | -0,07 | 9,43E-01 | 9,89E-01 |
| AT3G22120 | 4,08 | -0,07 | 0,97 | -0,07 | 9,44E-01 | 9,89E-01 |
| AT5G02380 | 2,77 | -0,10 | 1,36 | -0,07 | 9,44E-01 | 9,89E-01 |
| AT1G60870 | 2,59 | 0,11 | 1,70 | 0,07 | 9,47E-01 | 9,90E-01 |
| AT3G58030 | 2,17 | 0,10 | 1,59 | 0,07 | 9,48E-01 | 9,90E-01 |
| AT1G07600 | 1,87 | 0,09 | 1,44 | 0,07 | 9,47E-01 | 9,90E-01 |
| AT1G15290 | 5,85 | -0,09 | 1,36 | -0,07 | 9,47E-01 | 9,90E-01 |
| AT3G62360 | 4,39 | 0,11 | 1,62 | 0,07 | 9,48E-01 | 9,90E-01 |
| AT4G14360 | 1,93 | 0,09 | 1,40 | 0,07 | 9,48E-01 | 9,90E-01 |
| AT2G04280 | 2,22 | 0,11 | 1,72 | 0,06 | 9,49E-01 | 9,90E-01 |
| AT4G19520 | 3,67 | -0,06 | 1,00 | -0,06 | 9,50E-01 | 9,91E-01 |
| AT1G21130 | 2,17 | 0,08 | 1,32 | 0,06 | 9,50E-01 | 9,91E-01 |
| AT1G04480 | 2,74 | 0,07 | 1,27 | 0,06 | 9,55E-01 | 9,91E-01 |
| AT1G06230 | 2,14 | 0,10 | 1,64 | 0,06 | 9,53E-01 | 9,91E-01 |
| AT2G04755 | 2,18 | 0,10 | 1,72 | 0,06 | 9,53E-01 | 9,91E-01 |
| AT3G22121 | 3,96 | -0,06 | 0,99 | -0,06 | 9,53E-01 | 9,91E-01 |
| AT3G58610 | 2,68 | 0,07 | 1,28 | 0,06 | 9,53E-01 | 9,91E-01 |
| AT4G18150 | 2,31 | 0,08 | 1,31 | 0,06 | 9,54E-01 | 9,91E-01 |
| AT1G18000 | 4,70 | -0,06 | 1,01 | -0,06 | 9,55E-01 | 9,91E-01 |
| AT1G05563 | 2,75 | -0,06 | 1,08 | -0,06 | 9,53E-01 | 9,91E-01 |
| AT1G68530 | 2,97 | 0,05 | 0,95 | 0,06 | 9,55E-01 | 9,91E-01 |
| AT2G28910 | 4,89 | -0,08 | 1,40 | -0,06 | 9,55E-01 | 9,91E-01 |
| AT5G02500 | 62,24 | -0,02 | 0,37 | -0,06 | 9,52E-01 | 9,91E-01 |
| AT1G50055 | 182,36 | -0,02 | 0,28 | -0,06 | 9,56E-01 | 9,91E-01 |
| AT2G09690 | 14,22 | 0,04 | 0,72 | 0,05 | 9,60E-01 | 9,93E-01 |
| AT3G04120 | 5,88 | -0,04 | 0,79 | -0,05 | 9,60E-01 | 9,93E-01 |
| AT2G35744 | 14,81 | -0,03 | 0,51 | -0,05 | 9,60E-01 | 9,93E-01 |
| AT2G46560 | 4,43 | 0,06 | 1,23 | 0,05 | 9,59E-01 | 9,93E-01 |
| AT1G07233 | 2,51 | 0,05 | 1,03 | 0,05 | 9,61E-01 | 9,94E-01 |
| AT3G06530 | 4,41 | 0,07 | 1,45 | 0,05 | 9,62E-01 | 9,94E-01 |
| AT5G35407 | 1076,14 | 0,02 | 0,37 | 0,05 | 9,61E-01 | 9,94E-01 |
| AT3G51950 | 4,09 | -0,04 | 0,97 | -0,05 | 9,63E-01 | 9,95E-01 |
| AT1G03743 | 2,10 | 0,06 | 1,28 | 0,04 | 9,65E-01 | 9,95E-01 |
| AT1G11785 | 2,48 | -0,04 | 0,97 | -0,04 | 9,64E-01 | 9,95E-01 |
| AT1G62670 | 9,93 | -0,02 | 0,56 | -0,04 | 9,65E-01 | 9,95E-01 |
| AT2G29630 | 5,51 | -0,03 | 0,75 | -0,04 | 9,66E-01 | 9,95E-01 |
| AT5G39740 | 3,11 | 0,05 | 1,24 | 0,04 | 9,67E-01 | 9,95E-01 |
| AT5G42092 | 3,68 | 0,05 | 1,11 | 0,04 | 9,67E-01 | 9,95E-01 |
| AT3G22886 | 587,34 | -0,01 | 0,25 | -0,04 | 9,69E-01 | 9,97E-01 |
| AT1G79280 | 4,52 | -0,05 | 1,28 | -0,04 | 9,70E-01 | 9,97E-01 |
| AT3G01540 | 20,21 | -0,04 | 0,99 | -0,04 | 9,70E-01 | 9,97E-01 |
| AT5G13770 | 2,99 | 0,06 | 1,78 | 0,04 | 9,71E-01 | 9,98E-01 |
| AT1G05240 | 2,51 | -0,04 | 1,05 | -0,03 | 9,73E-01 | 9,99E-01 |

| | | | | | | |
|---|---|---|---|---|---|---|
| AT1G06950 | 2,18 | 0,07 | 2,02 | 0,03 | 9,73E-01 | 9,99E-01 |
| AT1G36180 | 2,20 | 0,05 | 1,61 | 0,03 | 9,77E-01 | 1,00E+00 |
| AT1G42550 | 2,32 | 0,00 | 1,30 | 0,00 | 1,00E+00 | 1,00E+00 |
| AT1G63400 | 45,16 | 0,00 | 0,40 | 0,01 | 9,93E-01 | 1,00E+00 |
| AT1G72150 | 7,68 | 0,00 | 0,66 | -0,01 | 9,94E-01 | 1,00E+00 |
| AT2G01600 | 1,95 | -0,04 | 1,89 | -0,02 | 9,83E-01 | 1,00E+00 |
| AT2G42270 | 6,29 | 0,00 | 1,80 | 0,00 | 1,00E+00 | 1,00E+00 |
| AT2G43890 | 2,70 | 0,00 | 2,05 | 0,00 | 1,00E+00 | 1,00E+00 |
| AT3G02470 | 10,40 | 0,02 | 0,65 | 0,03 | 9,78E-01 | 1,00E+00 |
| AT3G04130 | 2,12 | 0,00 | 2,28 | 0,00 | 1,00E+00 | 1,00E+00 |
| AT4G01150 | 2,10 | 0,03 | 1,18 | 0,03 | 9,77E-01 | 1,00E+00 |
| AT4G28652 | 2,08 | 0,00 | 2,15 | 0,00 | 1,00E+00 | 1,00E+00 |
| AT4G34450 | 21,43 | -0,02 | 0,96 | -0,02 | 9,82E-01 | 1,00E+00 |
| AT5G23380 | 2,25 | 0,00 | 2,05 | 0,00 | 1,00E+00 | 1,00E+00 |
| AT5G44180 | 3,26 | 0,00 | 1,69 | 0,00 | 9,99E-01 | 1,00E+00 |
| AT5G46315 | 17,01 | 0,00 | 1,73 | 0,00 | 1,00E+00 | 1,00E+00 |
| AT5G06845 | 6,24 | 0,02 | 1,18 | 0,02 | 9,84E-01 | 1,00E+00 |
| AT5G47690 | 3,16 | -0,01 | 1,22 | -0,01 | 9,91E-01 | 1,00E+00 |
| AT5G65950 | 6,84 | 0,00 | 1,89 | 0,00 | 1,00E+00 | 1,00E+00 |
| AT1G04625 | 1,86 | 0,00 | 2,21 | 0,00 | 1,00E+00 | 1,00E+00 |
| AT1G09620 | 17,50 | -0,01 | 1,09 | -0,01 | 9,90E-01 | 1,00E+00 |
| AT1G11735 | 8,53 | -0,01 | 0,66 | -0,01 | 9,89E-01 | 1,00E+00 |
| AT1G13350 | 2,82 | 0,00 | 2,04 | 0,00 | 1,00E+00 | 1,00E+00 |
| AT1G27870 | 1,97 | -0,01 | 1,34 | -0,01 | 9,91E-01 | 1,00E+00 |
| AT1G48650 | 3,21 | 0,00 | 1,69 | 0,00 | 9,98E-01 | 1,00E+00 |
| AT1G08327 | 2,78 | 0,03 | 1,48 | 0,02 | 9,82E-01 | 1,00E+00 |
| AT1G68520 | 2,00 | -0,03 | 1,35 | -0,02 | 9,80E-01 | 1,00E+00 |
| AT1G77680 | 1,90 | 0,00 | 2,19 | 0,00 | 1,00E+00 | 1,00E+00 |
| AT2G33100 | 2,25 | 0,00 | 2,08 | 0,00 | 1,00E+00 | 1,00E+00 |
| AT2G36490 | 1,90 | 0,00 | 2,87 | 0,00 | 1,00E+00 | 1,00E+00 |
| AT2G44550 | 3,60 | 0,00 | 0,90 | 0,00 | 9,97E-01 | 1,00E+00 |
| AT3G09440 | 5,91 | 0,00 | 0,80 | -0,01 | 9,95E-01 | 1,00E+00 |
| AT3G13090 | 2,27 | 0,00 | 2,35 | 0,00 | 1,00E+00 | 1,00E+00 |
| AT3G13100 | 2,83 | 0,00 | 1,59 | 0,00 | 9,99E-01 | 1,00E+00 |
| AT3G17510 | 2,73 | -0,02 | 1,87 | -0,01 | 9,93E-01 | 1,00E+00 |
| AT3G49500 | 2,34 | 0,00 | 2,01 | 0,00 | 1,00E+00 | 1,00E+00 |
| AT3G52700 | 2,20 | 0,01 | 1,07 | 0,01 | 9,90E-01 | 1,00E+00 |
| AT3G55020 | 1,94 | 0,03 | 1,77 | 0,02 | 9,88E-01 | 1,00E+00 |
| AT3G57330 | 3,24 | 0,01 | 1,45 | 0,00 | 9,97E-01 | 1,00E+00 |
| AT4G01860 | 3,28 | 0,00 | 1,95 | 0,00 | 1,00E+00 | 1,00E+00 |
| AT4G16940 | 4,86 | 0,00 | 1,88 | 0,00 | 1,00E+00 | 1,00E+00 |
| AT4G27000 | 10,07 | 0,01 | 1,11 | 0,01 | 9,89E-01 | 1,00E+00 |
| AT4G28650 | 2,08 | 0,00 | 2,15 | 0,00 | 1,00E+00 | 1,00E+00 |
| AT4G37900 | 1,96 | 0,00 | 2,11 | 0,00 | 1,00E+00 | 1,00E+00 |
| AT4G38510 | 2,20 | -0,03 | 1,73 | -0,02 | 9,87E-01 | 1,00E+00 |
| AT5G05980 | 2,96 | 0,00 | 2,02 | 0,00 | 1,00E+00 | 1,00E+00 |
| AT5G18350 | 2,51 | 0,00 | 2,09 | 0,00 | 1,00E+00 | 1,00E+00 |
| AT5G24280 | 2,00 | 0,00 | 3,20 | 0,00 | 1,00E+00 | 1,00E+00 |
| AT5G25270 | 3,67 | 0,00 | 1,92 | 0,00 | 1,00E+00 | 1,00E+00 |
| AT5G28040 | 2,94 | 0,00 | 1,24 | 0,00 | 9,99E-01 | 1,00E+00 |
| AT5G40384 | 2,72 | -0,03 | 1,38 | -0,02 | 9,84E-01 | 1,00E+00 |
| AT5G44090 | 1,96 | 0,04 | 1,64 | 0,03 | 9,79E-01 | 1,00E+00 |
| AT5G48160 | 6,17 | 0,03 | 1,19 | 0,02 | 9,81E-01 | 1,00E+00 |
| AT5G60920 | 1,88 | 0,00 | 1,29 | 0,00 | 9,98E-01 | 1,00E+00 |
| AT1G01046 | 1,63 | 2,60 | 1,66 | 1,57 | 1,16E-01 | |
| AT1G01160 | 0,43 | 1,90 | 3,23 | 0,59 | 5,57E-01 | |

| | | | | | |
|---|---|---|---|---|---|
| AT1G01180 | 0,85 | 0,67 | 2,33 | 0,29 | 7,72E-01 |
| AT1G01220 | 0,11 | -0,26 | 4,41 | -0,06 | 9,53E-01 |
| AT1G01225 | 0,31 | -0,41 | 4,38 | -0,09 | 9,25E-01 |
| AT1G01230 | 0,37 | -1,31 | 4,39 | -0,30 | 7,65E-01 |
| AT1G01240 | 0,19 | 1,17 | 4,34 | 0,27 | 7,87E-01 |
| AT1G01260 | 0,53 | -1,43 | 2,71 | -0,53 | 5,98E-01 |
| AT1G01300 | 1,08 | 0,67 | 1,75 | 0,38 | 7,04E-01 |
| AT1G01350 | 0,29 | -0,30 | 4,40 | -0,07 | 9,46E-01 |
| AT1G01448 | 0,59 | 0,00 | 4,40 | 0,00 | 1,00E+00 |
| AT1G01480 | 0,12 | 0,00 | 4,41 | 0,00 | 1,00E+00 |
| AT1G01520 | 0,12 | 0,00 | 4,41 | 0,00 | 1,00E+00 |
| AT1G01540 | 0,22 | 0,80 | 4,35 | 0,18 | 8,55E-01 |
| AT1G01560 | 1,60 | 0,76 | 2,07 | 0,37 | 7,13E-01 |
| AT1G01630 | 0,58 | -1,29 | 3,84 | -0,34 | 7,37E-01 |
| AT1G01710 | 0,82 | -0,98 | 3,03 | -0,32 | 7,46E-01 |
| AT1G01720 | 1,37 | 0,95 | 1,54 | 0,62 | 5,37E-01 |
| AT1G04047 | 0,70 | 1,39 | 2,47 | 0,56 | 5,73E-01 |
| AT1G01730 | 0,40 | 0,00 | 4,41 | 0,00 | 1,00E+00 |
| AT1G01790 | 0,83 | -0,66 | 1,91 | -0,35 | 7,29E-01 |
| AT1G01810 | 0,12 | 0,00 | 4,41 | 0,00 | 1,00E+00 |
| AT1G01950 | 0,18 | 0,80 | 4,35 | 0,18 | 8,54E-01 |
| AT1G02010 | 0,10 | 0,24 | 4,38 | 0,05 | 9,57E-01 |
| AT1G02100 | 0,49 | 0,03 | 3,17 | 0,01 | 9,93E-01 |
| AT1G02120 | 0,36 | -0,39 | 3,48 | -0,11 | 9,11E-01 |
| AT1G02150 | 1,38 | 0,37 | 1,76 | 0,21 | 8,31E-01 |
| AT1G02205 | 1,06 | -0,30 | 1,78 | -0,17 | 8,65E-01 |
| AT1G04093 | 0,08 | 0,21 | 4,39 | 0,05 | 9,62E-01 |
| AT1G02300 | 0,34 | 0,38 | 4,38 | 0,09 | 9,30E-01 |
| AT1G02305 | 0,80 | -1,41 | 2,52 | -0,56 | 5,77E-01 |
| AT1G04103 | 0,08 | 0,00 | 4,41 | 0,00 | 1,00E+00 |
| AT1G02390 | 0,19 | 0,81 | 4,35 | 0,19 | 8,53E-01 |
| AT1G02420 | 0,52 | 0,00 | 3,96 | 0,00 | 1,00E+00 |
| AT1G02560 | 1,57 | -0,04 | 1,62 | -0,02 | 9,82E-01 |
| AT1G02690 | 0,11 | 0,88 | 4,35 | 0,20 | 8,39E-01 |
| AT1G02750 | 0,51 | 0,00 | 4,41 | 0,00 | 1,00E+00 |
| AT1G02816 | 0,38 | 0,78 | 3,38 | 0,23 | 8,19E-01 |
| AT1G02850 | 0,62 | -1,16 | 3,01 | -0,39 | 7,00E-01 |
| AT1G03160 | 0,95 | -0,99 | 2,11 | -0,47 | 6,40E-01 |
| AT1G03190 | 0,39 | 0,65 | 3,47 | 0,19 | 8,52E-01 |
| AT1G03220 | 1,11 | 1,21 | 2,13 | 0,57 | 5,69E-01 |
| AT1G03280 | 0,25 | -0,31 | 4,40 | -0,07 | 9,44E-01 |
| AT1G03290 | 0,75 | 1,88 | 2,42 | 0,78 | 4,38E-01 |
| AT1G03310 | 0,40 | -0,10 | 4,27 | -0,02 | 9,82E-01 |
| AT1G03360 | 0,35 | 0,94 | 3,50 | 0,27 | 7,89E-01 |
| AT1G03365 | 0,40 | -1,22 | 3,45 | -0,35 | 7,24E-01 |
| AT1G03370 | 0,51 | 1,76 | 3,95 | 0,44 | 6,57E-01 |
| AT1G03380 | 1,12 | 0,63 | 2,06 | 0,31 | 7,60E-01 |
| AT1G03430 | 0,37 | -0,59 | 4,36 | -0,13 | 8,93E-01 |
| AT1G03440 | 0,84 | -1,03 | 3,07 | -0,34 | 7,36E-01 |
| AT1G03545 | 0,70 | -0,82 | 2,29 | -0,36 | 7,20E-01 |
| AT1G03600 | 1,40 | 0,55 | 1,66 | 0,33 | 7,38E-01 |
| AT1G03610 | 0,72 | -1,07 | 3,12 | -0,34 | 7,30E-01 |
| AT1G03630 | 0,51 | 1,17 | 2,26 | 0,52 | 6,04E-01 |
| AT1G03687 | 0,11 | 0,88 | 4,35 | 0,20 | 8,39E-01 |
| AT1G03740 | 0,96 | 1,71 | 2,02 | 0,85 | 3,97E-01 |
| AT1G03810 | 0,77 | 2,33 | 2,15 | 1,09 | 2,78E-01 |

| | | | | | |
|---|---|---|---|---|---|
| AT1G03900 | 1,11 | 0,89 | 1,99 | 0,45 | 6,53E-01 |
| AT1G03905 | 0,24 | 0,00 | 4,41 | 0,00 | 1,00E+00 |
| AT1G03910 | 0,50 | 0,06 | 3,10 | 0,02 | 9,85E-01 |
| AT1G03930 | 0,53 | 0,46 | 2,76 | 0,16 | 8,69E-01 |
| AT1G04217 | 0,82 | -0,83 | 1,88 | -0,44 | 6,56E-01 |
| AT1G03935 | 0,16 | -0,56 | 4,37 | -0,13 | 8,98E-01 |
| AT1G04223 | 0,98 | 1,85 | 1,90 | 0,98 | 3,29E-01 |
| AT1G04227 | 0,32 | -1,34 | 3,63 | -0,37 | 7,12E-01 |
| AT1G03960 | 0,15 | -2,07 | 4,39 | -0,47 | 6,38E-01 |
| AT1G03970 | 0,36 | 1,96 | 4,30 | 0,46 | 6,49E-01 |
| AT1G04020 | 0,33 | -2,20 | 4,41 | -0,50 | 6,17E-01 |
| AT1G04130 | 0,50 | 2,13 | 2,60 | 0,82 | 4,11E-01 |
| AT1G04170 | 1,02 | -0,26 | 2,25 | -0,12 | 9,07E-01 |
| AT1G04247 | 0,30 | 0,79 | 4,34 | 0,18 | 8,55E-01 |
| AT1G04230 | 0,70 | -3,00 | 2,87 | -1,04 | 2,97E-01 |
| AT1G04295 | 0,43 | 1,32 | 2,76 | 0,48 | 6,32E-01 |
| AT1G04350 | 0,25 | 0,04 | 4,32 | 0,01 | 9,93E-01 |
| AT1G04420 | 1,20 | 1,21 | 1,91 | 0,63 | 5,26E-01 |
| AT1G04440 | 0,50 | 1,59 | 2,69 | 0,59 | 5,55E-01 |
| AT1G04273 | 0,37 | 0,12 | 3,43 | 0,04 | 9,72E-01 |
| AT1G04277 | 0,93 | -1,04 | 2,00 | -0,52 | 6,02E-01 |
| AT1G04510 | 0,67 | -0,27 | 3,07 | -0,09 | 9,29E-01 |
| AT1G04540 | 1,05 | 0,31 | 2,35 | 0,13 | 8,96E-01 |
| AT1G04690 | 0,77 | 0,56 | 2,39 | 0,23 | 8,15E-01 |
| AT1G04710 | 0,47 | 0,00 | 4,41 | 0,00 | 1,00E+00 |
| AT1G04760 | 0,36 | 0,53 | 4,37 | 0,12 | 9,04E-01 |
| AT1G04945 | 0,47 | -1,31 | 3,24 | -0,40 | 6,87E-01 |
| AT1G04970 | 0,34 | -3,14 | 4,36 | -0,72 | 4,71E-01 |
| AT1G05087 | 0,44 | -0,30 | 4,14 | -0,07 | 9,43E-01 |
| AT1G05090 | 0,12 | 0,00 | 4,41 | 0,00 | 1,00E+00 |
| AT1G05140 | 0,22 | 0,81 | 4,35 | 0,19 | 8,52E-01 |
| AT1G05200 | 1,12 | -2,31 | 1,86 | -1,24 | 2,15E-01 |
| AT1G05360 | 0,08 | 0,00 | 4,41 | 0,00 | 1,00E+00 |
| AT1G05380 | 1,42 | -0,24 | 2,18 | -0,11 | 9,13E-01 |
| AT1G05500 | 0,37 | 1,08 | 4,34 | 0,25 | 8,03E-01 |
| AT1G04397 | 0,08 | 0,00 | 4,41 | 0,00 | 1,00E+00 |
| AT1G05600 | 0,08 | 0,21 | 4,39 | 0,05 | 9,62E-01 |
| AT1G05630 | 1,84 | 1,22 | 1,85 | 0,66 | 5,10E-01 |
| AT1G04423 | 1,18 | 2,28 | 1,94 | 1,18 | 2,40E-01 |
| AT1G05790 | 0,76 | -0,30 | 2,61 | -0,11 | 9,10E-01 |
| AT1G05830 | 0,53 | 0,24 | 3,31 | 0,07 | 9,41E-01 |
| AT1G05890 | 0,68 | 0,00 | 4,41 | 0,00 | 1,00E+00 |
| AT1G05900 | 0,21 | 0,08 | 4,38 | 0,02 | 9,85E-01 |
| AT1G05910 | 0,59 | 0,00 | 3,86 | 0,00 | 1,00E+00 |
| AT1G05980 | 0,23 | 0,16 | 4,32 | 0,04 | 9,71E-01 |
| AT1G06002 | 0,95 | -0,50 | 1,95 | -0,25 | 7,99E-01 |
| AT1G06030 | 0,96 | 0,00 | 3,41 | 0,00 | 1,00E+00 |
| AT1G06290 | 0,60 | 0,09 | 2,77 | 0,03 | 9,73E-01 |
| AT1G06380 | 0,45 | -0,73 | 4,13 | -0,18 | 8,60E-01 |
| AT1G06390 | 0,40 | 0,43 | 3,43 | 0,13 | 9,00E-01 |
| AT1G06410 | 1,61 | 1,82 | 1,75 | 1,04 | 2,99E-01 |
| AT1G06530 | 0,96 | -1,12 | 2,24 | -0,50 | 6,17E-01 |
| AT1G06640 | 1,12 | 2,93 | 1,81 | 1,62 | 1,05E-01 |
| AT1G06645 | 0,50 | 0,27 | 4,04 | 0,07 | 9,47E-01 |
| AT1G06650 | 0,69 | 0,88 | 3,03 | 0,29 | 7,71E-01 |
| AT1G06670 | 1,38 | 0,54 | 2,02 | 0,27 | 7,91E-01 |

| | | | | | |
|---|---|---|---|---|---|
| AT1G06750 | 0,24 | 0,00 | 4,41 | 0,00 | 1,00E+00 |
| AT1G06780 | 0,12 | 0,27 | 4,38 | 0,06 | 9,50E-01 |
| AT1G06910 | 0,23 | -0,35 | 4,39 | -0,08 | 9,36E-01 |
| AT1G06930 | 0,08 | 0,00 | 4,41 | 0,00 | 1,00E+00 |
| AT1G07010 | 0,86 | 1,15 | 2,95 | 0,39 | 6,96E-01 |
| AT1G04537 | 0,24 | 0,00 | 4,41 | 0,00 | 1,00E+00 |
| AT1G07080 | 0,86 | -1,47 | 1,99 | -0,74 | 4,60E-01 |
| AT1G07130 | 0,28 | 0,60 | 4,37 | 0,14 | 8,91E-01 |
| AT1G07170 | 0,78 | -0,20 | 2,21 | -0,09 | 9,28E-01 |
| AT1G07180 | 0,37 | 0,32 | 3,47 | 0,09 | 9,26E-01 |
| AT1G07210 | 0,17 | 0,00 | 4,41 | 0,00 | 1,00E+00 |
| AT1G07240 | 0,59 | -0,12 | 3,30 | -0,04 | 9,70E-01 |
| AT1G07280 | 1,33 | -3,16 | 1,99 | -1,58 | 1,13E-01 |
| AT1G07320 | 1,43 | -1,09 | 1,61 | -0,68 | 4,97E-01 |
| AT1G07470 | 0,61 | 0,00 | 3,82 | 0,00 | 1,00E+00 |
| AT1G07510 | 1,74 | -0,74 | 1,65 | -0,45 | 6,52E-01 |
| AT1G07630 | 0,70 | 1,45 | 2,95 | 0,49 | 6,23E-01 |
| AT1G07705 | 0,12 | -0,26 | 4,41 | -0,06 | 9,53E-01 |
| AT1G07728 | 1,03 | 0,00 | 2,68 | 0,00 | 1,00E+00 |
| AT1G07790 | 0,25 | -0,74 | 4,35 | -0,17 | 8,66E-01 |
| AT1G07830 | 0,43 | 0,49 | 4,19 | 0,12 | 9,07E-01 |
| AT1G07990 | 0,78 | 1,03 | 3,04 | 0,34 | 7,34E-01 |
| AT1G08110 | 0,41 | -0,20 | 4,36 | -0,05 | 9,64E-01 |
| AT1G08190 | 0,34 | 0,65 | 3,55 | 0,18 | 8,55E-01 |
| AT1G08260 | 0,51 | 1,23 | 4,09 | 0,30 | 7,63E-01 |
| AT1G08360 | 1,45 | 0,13 | 1,92 | 0,07 | 9,48E-01 |
| AT1G08370 | 0,71 | -0,30 | 2,62 | -0,11 | 9,09E-01 |
| AT1G08410 | 0,87 | -1,67 | 2,54 | -0,66 | 5,11E-01 |
| AT1G08420 | 1,72 | 1,07 | 1,88 | 0,57 | 5,67E-01 |
| AT1G08460 | 0,84 | -0,08 | 1,99 | -0,04 | 9,67E-01 |
| AT1G08465 | 0,35 | 0,39 | 4,38 | 0,09 | 9,30E-01 |
| AT1G08480 | 0,59 | -0,57 | 3,22 | -0,18 | 8,60E-01 |
| AT1G08520 | 1,05 | -0,65 | 1,83 | -0,35 | 7,24E-01 |
| AT1G08530 | 0,48 | -0,27 | 4,07 | -0,07 | 9,48E-01 |
| AT1G08550 | 0,85 | -1,90 | 2,28 | -0,84 | 4,04E-01 |
| AT1G08570 | 0,49 | 0,24 | 4,10 | 0,06 | 9,54E-01 |
| AT1G08590 | 0,74 | -0,26 | 3,01 | -0,09 | 9,31E-01 |
| AT1G08600 | 0,40 | -0,51 | 3,34 | -0,15 | 8,78E-01 |
| AT1G08620 | 1,06 | 0,22 | 1,99 | 0,11 | 9,13E-01 |
| AT1G08650 | 0,89 | -1,72 | 2,89 | -0,60 | 5,51E-01 |
| AT1G08680 | 0,17 | 1,58 | 4,31 | 0,37 | 7,15E-01 |
| AT1G08710 | 0,30 | 0,00 | 4,41 | 0,00 | 1,00E+00 |
| AT1G08720 | 1,46 | -1,24 | 2,27 | -0,55 | 5,84E-01 |
| AT1G08760 | 0,68 | 0,40 | 3,05 | 0,13 | 8,96E-01 |
| AT1G08770 | 0,33 | 1,58 | 3,49 | 0,45 | 6,50E-01 |
| AT1G08830 | 1,29 | -1,69 | 2,16 | -0,78 | 4,33E-01 |
| AT1G08900 | 0,34 | 0,78 | 3,49 | 0,22 | 8,24E-01 |
| AT1G08910 | 0,43 | 1,14 | 4,16 | 0,27 | 7,84E-01 |
| AT1G08920 | 0,59 | -2,88 | 3,20 | -0,90 | 3,67E-01 |
| AT1G08930 | 1,31 | -0,10 | 1,57 | -0,07 | 9,47E-01 |
| AT1G08970 | 0,82 | -0,30 | 2,65 | -0,11 | 9,10E-01 |
| AT1G08980 | 0,91 | -0,56 | 2,25 | -0,25 | 8,02E-01 |
| AT1G09040 | 0,45 | -0,23 | 3,38 | -0,07 | 9,45E-01 |
| AT1G09240 | 1,84 | 2,86 | 2,23 | 1,28 | 2,01E-01 |
| AT1G09250 | 0,32 | -0,30 | 4,40 | -0,07 | 9,45E-01 |
| AT1G09260 | 0,62 | -4,40 | 4,32 | -1,02 | 3,08E-01 |

| | | | | | |
|---|---|---|---|---|---|
| AT1G09270 | 0,12 | 0,00 | 4,41 | 0,00 | 1,00E+00 |
| AT1G09300 | 0,36 | -0,16 | 4,39 | -0,04 | 9,70E-01 |
| AT1G09340 | 1,73 | -1,03 | 1,33 | -0,77 | 4,39E-01 |
| AT1G09390 | 0,23 | 1,10 | 4,34 | 0,25 | 8,00E-01 |
| AT1G09421 | 0,17 | -0,38 | 4,39 | -0,09 | 9,31E-01 |
| AT1G09430 | 0,79 | 1,56 | 2,39 | 0,65 | 5,15E-01 |
| AT1G09450 | 0,24 | 0,59 | 4,36 | 0,14 | 8,92E-01 |
| AT1G09490 | 0,05 | 0,18 | 4,38 | 0,04 | 9,68E-01 |
| AT1G09500 | 0,08 | 0,00 | 4,41 | 0,00 | 1,00E+00 |
| AT1G09530 | 0,63 | -2,43 | 3,71 | -0,66 | 5,12E-01 |
| AT1G09560 | 0,29 | 0,00 | 4,41 | 0,00 | 1,00E+00 |
| AT1G09590 | 0,27 | 0,38 | 4,31 | 0,09 | 9,30E-01 |
| AT1G09600 | 0,94 | 0,00 | 2,96 | 0,00 | 1,00E+00 |
| AT1G09610 | 0,36 | 0,00 | 4,41 | 0,00 | 1,00E+00 |
| AT1G09645 | 0,33 | 0,00 | 4,41 | 0,00 | 1,00E+00 |
| AT1G09665 | 0,25 | 0,00 | 4,41 | 0,00 | 1,00E+00 |
| AT1G09710 | 0,91 | 0,24 | 2,94 | 0,08 | 9,36E-01 |
| AT1G09750 | 0,82 | -0,38 | 1,76 | -0,22 | 8,29E-01 |
| AT1G09795 | 0,58 | 0,42 | 3,90 | 0,11 | 9,15E-01 |
| AT1G09815 | 0,21 | -0,33 | 4,32 | -0,08 | 9,39E-01 |
| AT1G10050 | 0,78 | -2,70 | 2,26 | -1,20 | 2,32E-01 |
| AT1G10070 | 0,24 | 0,00 | 4,41 | 0,00 | 1,00E+00 |
| AT1G10130 | 1,07 | 0,95 | 1,99 | 0,48 | 6,34E-01 |
| AT1G10150 | 1,47 | 0,39 | 1,55 | 0,25 | 7,99E-01 |
| AT1G10270 | 0,58 | -1,28 | 2,66 | -0,48 | 6,29E-01 |
| AT1G10450 | 0,33 | -0,36 | 4,39 | -0,08 | 9,34E-01 |
| AT1G10490 | 0,65 | -1,27 | 2,60 | -0,49 | 6,24E-01 |
| AT1G10570 | 0,16 | -0,21 | 4,40 | -0,05 | 9,63E-01 |
| AT1G10600 | 1,27 | 0,52 | 2,13 | 0,25 | 8,05E-01 |
| AT1G10610 | 0,90 | 0,00 | 3,45 | 0,00 | 1,00E+00 |
| AT1G10660 | 0,54 | -1,37 | 4,04 | -0,34 | 7,35E-01 |
| AT1G10670 | 0,76 | -1,52 | 3,46 | -0,44 | 6,61E-01 |
| AT1G10700 | 0,45 | 0,08 | 4,38 | 0,02 | 9,86E-01 |
| AT1G10720 | 0,48 | -0,56 | 2,70 | -0,21 | 8,36E-01 |
| AT1G10745 | 1,57 | 1,05 | 1,60 | 0,66 | 5,11E-01 |
| AT1G10790 | 0,08 | 0,00 | 4,41 | 0,00 | 1,00E+00 |
| AT1G10850 | 0,85 | 0,00 | 3,60 | 0,00 | 1,00E+00 |
| AT1G10910 | 0,08 | 0,00 | 4,41 | 0,00 | 1,00E+00 |
| AT1G10960 | 0,86 | -3,74 | 2,88 | -1,30 | 1,94E-01 |
| AT1G11020 | 0,27 | 0,13 | 4,38 | 0,03 | 9,76E-01 |
| AT1G11050 | 0,22 | 1,46 | 4,32 | 0,34 | 7,36E-01 |
| AT1G11260 | 1,39 | -1,64 | 1,59 | -1,04 | 3,01E-01 |
| AT1G11300 | 0,32 | -1,18 | 4,40 | -0,27 | 7,88E-01 |
| AT1G11303 | 0,25 | 0,00 | 4,41 | 0,00 | 1,00E+00 |
| AT1G11320 | 0,08 | 0,00 | 4,41 | 0,00 | 1,00E+00 |
| AT1G11330 | 0,64 | 0,89 | 2,96 | 0,30 | 7,64E-01 |
| AT1G11400 | 0,26 | 0,83 | 4,35 | 0,19 | 8,48E-01 |
| AT1G11410 | 0,36 | -0,12 | 3,45 | -0,03 | 9,73E-01 |
| AT1G11430 | 0,35 | -0,65 | 3,54 | -0,18 | 8,55E-01 |
| AT1G11440 | 0,54 | 0,08 | 3,07 | 0,03 | 9,79E-01 |
| AT1G11580 | 0,31 | -0,38 | 4,39 | -0,09 | 9,32E-01 |
| AT1G11590 | 0,67 | 0,12 | 2,26 | 0,05 | 9,58E-01 |
| AT1G11593 | 0,40 | 2,45 | 4,14 | 0,59 | 5,53E-01 |
| AT1G11660 | 1,14 | -0,27 | 2,20 | -0,12 | 9,04E-01 |
| AT1G11680 | 0,83 | -1,67 | 2,81 | -0,59 | 5,53E-01 |
| AT1G11710 | 0,17 | 0,00 | 4,41 | 0,00 | 1,00E+00 |

| | | | | | |
|---|---|---|---|---|---|
| AT1G11750 | 0,56 | -2,14 | 3,15 | -0,68 | 4,97E-01 |
| AT1G11790 | 0,42 | -2,19 | 4,41 | -0,50 | 6,20E-01 |
| AT1G11840 | 1,70 | 1,45 | 1,53 | 0,95 | 3,42E-01 |
| AT1G11890 | 1,50 | 0,13 | 1,81 | 0,07 | 9,45E-01 |
| AT1G11905 | 0,75 | -0,53 | 2,31 | -0,23 | 8,19E-01 |
| AT1G11930 | 0,68 | -2,06 | 2,30 | -0,90 | 3,70E-01 |
| AT1G12015 | 0,46 | -1,01 | 4,32 | -0,23 | 8,15E-01 |
| AT1G12050 | 1,22 | 1,71 | 1,71 | 1,00 | 3,16E-01 |
| AT1G12060 | 0,12 | -0,36 | 4,39 | -0,08 | 9,34E-01 |
| AT1G12140 | 0,12 | 0,00 | 4,41 | 0,00 | 1,00E+00 |
| AT1G12210 | 0,35 | -0,88 | 4,36 | -0,20 | 8,40E-01 |
| AT1G12220 | 1,65 | 0,38 | 1,30 | 0,29 | 7,72E-01 |
| AT1G12230 | 0,35 | 0,89 | 3,50 | 0,26 | 7,98E-01 |
| AT1G12250 | 0,19 | -0,57 | 4,35 | -0,13 | 8,95E-01 |
| AT1G12270 | 0,62 | -1,53 | 2,63 | -0,58 | 5,60E-01 |
| AT1G12320 | 0,82 | 3,06 | 2,75 | 1,11 | 2,65E-01 |
| AT1G12360 | 0,65 | 0,34 | 3,85 | 0,09 | 9,29E-01 |
| AT1G12410 | 0,67 | -4,47 | 3,71 | -1,20 | 2,29E-01 |
| AT1G12460 | 0,83 | -0,69 | 1,89 | -0,36 | 7,16E-01 |
| AT1G12580 | 0,43 | 1,37 | 2,80 | 0,49 | 6,24E-01 |
| AT1G12760 | 0,76 | 1,51 | 2,22 | 0,68 | 4,97E-01 |
| AT1G12810 | 0,42 | -0,05 | 2,35 | -0,02 | 9,84E-01 |
| AT1G12840 | 1,05 | -1,38 | 1,88 | -0,73 | 4,63E-01 |
| AT1G12860 | 0,23 | -1,81 | 4,34 | -0,42 | 6,76E-01 |
| AT1G12990 | 1,14 | 1,25 | 2,14 | 0,58 | 5,59E-01 |
| AT1G13040 | 0,24 | 0,60 | 4,36 | 0,14 | 8,91E-01 |
| AT1G13060 | 0,83 | -1,61 | 3,09 | -0,52 | 6,02E-01 |
| AT1G13090 | 1,29 | -0,06 | 1,78 | -0,03 | 9,72E-01 |
| AT1G13100 | 0,26 | -0,64 | 4,30 | -0,15 | 8,82E-01 |
| AT1G13180 | 0,14 | 0,31 | 4,37 | 0,07 | 9,44E-01 |
| AT1G13190 | 1,26 | 3,11 | 2,45 | 1,27 | 2,05E-01 |
| AT1G13220 | 0,56 | 0,45 | 3,84 | 0,12 | 9,08E-01 |
| AT1G13260 | 1,59 | -0,20 | 1,67 | -0,12 | 9,06E-01 |
| AT1G13270 | 0,60 | -4,23 | 3,01 | -1,41 | 1,59E-01 |
| AT1G13330 | 0,70 | 0,90 | 2,59 | 0,35 | 7,28E-01 |
| AT1G13400 | 0,23 | 0,24 | 4,39 | 0,06 | 9,56E-01 |
| AT1G13430 | 0,17 | 0,00 | 4,41 | 0,00 | 1,00E+00 |
| AT1G13460 | 0,37 | -2,55 | 4,36 | -0,58 | 5,59E-01 |
| AT1G13820 | 0,11 | -0,38 | 4,39 | -0,09 | 9,31E-01 |
| AT1G13850 | 0,13 | 0,00 | 4,41 | 0,00 | 1,00E+00 |
| AT1G13880 | 0,45 | 0,49 | 3,38 | 0,15 | 8,84E-01 |
| AT1G13990 | 0,47 | 0,43 | 3,34 | 0,13 | 8,99E-01 |
| AT1G14000 | 0,81 | -1,28 | 2,27 | -0,56 | 5,74E-01 |
| AT1G14020 | 0,41 | -0,20 | 4,25 | -0,05 | 9,62E-01 |
| AT1G05047 | 0,12 | 0,00 | 4,41 | 0,00 | 1,00E+00 |
| AT1G14110 | 0,52 | 2,34 | 3,80 | 0,62 | 5,38E-01 |
| AT1G14150 | 0,05 | 0,18 | 4,38 | 0,04 | 9,68E-01 |
| AT1G14185 | 0,05 | 0,18 | 4,38 | 0,04 | 9,68E-01 |
| AT1G14230 | 0,08 | 0,47 | 4,38 | 0,11 | 9,15E-01 |
| AT1G14250 | 0,32 | -1,98 | 4,32 | -0,46 | 6,46E-01 |
| AT1G14280 | 0,63 | 2,26 | 2,97 | 0,76 | 4,47E-01 |
| AT1G14310 | 0,25 | -0,35 | 4,40 | -0,08 | 9,37E-01 |
| AT1G14345 | 0,53 | 1,50 | 2,72 | 0,55 | 5,81E-01 |
| AT1G14350 | 0,37 | -1,13 | 4,35 | -0,26 | 7,94E-01 |
| AT1G14370 | 0,91 | 1,17 | 1,86 | 0,63 | 5,30E-01 |
| AT1G14470 | 0,37 | -3,32 | 4,36 | -0,76 | 4,46E-01 |

| | | | | | |
|---|---|---|---|---|---|
| AT1G14480 | 0,08 | 0,21 | 4,39 | 0,05 | 9,62E-01 |
| AT1G14560 | 0,21 | -1,29 | 4,40 | -0,29 | 7,69E-01 |
| AT1G14570 | 1,23 | 1,57 | 1,89 | 0,83 | 4,07E-01 |
| AT1G14650 | 1,32 | 0,48 | 2,36 | 0,20 | 8,40E-01 |
| AT1G14800 | 1,13 | 1,09 | 1,81 | 0,60 | 5,48E-01 |
| AT1G14860 | 0,08 | 0,21 | 4,39 | 0,05 | 9,62E-01 |
| AT1G14920 | 1,77 | -1,91 | 1,43 | -1,34 | 1,80E-01 |
| AT1G15030 | 0,23 | 0,58 | 4,37 | 0,13 | 8,94E-01 |
| AT1G15120 | 0,30 | 0,23 | 4,37 | 0,05 | 9,58E-01 |
| AT1G15125 | 1,84 | 1,30 | 1,69 | 0,77 | 4,41E-01 |
| AT1G15190 | 0,17 | 0,00 | 4,41 | 0,00 | 1,00E+00 |
| AT1G15240 | 0,58 | 0,23 | 2,65 | 0,09 | 9,32E-01 |
| AT1G15260 | 0,61 | -2,76 | 3,07 | -0,90 | 3,70E-01 |
| AT1G15390 | 0,48 | 0,00 | 4,41 | 0,00 | 1,00E+00 |
| AT1G15430 | 0,55 | -0,45 | 3,92 | -0,12 | 9,08E-01 |
| AT1G15650 | 0,36 | 0,00 | 4,41 | 0,00 | 1,00E+00 |
| AT1G05183 | 0,40 | -0,34 | 3,43 | -0,10 | 9,21E-01 |
| AT1G15720 | 0,33 | 0,28 | 4,37 | 0,06 | 9,49E-01 |
| AT1G15740 | 0,65 | 2,63 | 2,42 | 1,09 | 2,78E-01 |
| AT1G15810 | 0,77 | 0,69 | 1,99 | 0,35 | 7,29E-01 |
| AT1G05207 | 1,81 | -2,47 | 1,31 | -1,89 | 5,90E-02 |
| AT1G15890 | 1,49 | 0,15 | 1,76 | 0,08 | 9,33E-01 |
| AT1G15910 | 0,69 | -0,26 | 3,64 | -0,07 | 9,43E-01 |
| AT1G15920 | 0,80 | 0,00 | 2,88 | 0,00 | 1,00E+00 |
| AT1G15930 | 1,08 | -2,76 | 2,03 | -1,36 | 1,75E-01 |
| AT1G15950 | 0,56 | 0,41 | 2,66 | 0,15 | 8,78E-01 |
| AT1G15980 | 1,06 | -0,46 | 1,72 | -0,27 | 7,87E-01 |
| AT1G16060 | 0,12 | 0,27 | 4,38 | 0,06 | 9,50E-01 |
| AT1G16080 | 0,31 | -2,19 | 4,41 | -0,50 | 6,20E-01 |
| AT1G16090 | 0,22 | 0,20 | 4,38 | 0,05 | 9,63E-01 |
| AT1G16110 | 0,61 | 0,00 | 3,73 | 0,00 | 1,00E+00 |
| AT1G16180 | 1,23 | 0,55 | 1,94 | 0,28 | 7,78E-01 |
| AT1G16250 | 0,81 | 0,19 | 2,22 | 0,09 | 9,31E-01 |
| AT1G16270 | 1,59 | -0,82 | 1,65 | -0,50 | 6,17E-01 |
| AT1G16350 | 0,39 | -2,44 | 4,23 | -0,58 | 5,64E-01 |
| AT1G16400 | 0,30 | 0,19 | 4,33 | 0,04 | 9,65E-01 |
| AT1G16410 | 1,19 | -0,24 | 1,38 | -0,17 | 8,64E-01 |
| AT1G16470 | 0,19 | 1,00 | 4,33 | 0,23 | 8,17E-01 |
| AT1G16500 | 0,25 | 0,02 | 4,37 | 0,01 | 9,96E-01 |
| AT1G16560 | 0,44 | -2,15 | 3,36 | -0,64 | 5,22E-01 |
| AT1G16670 | 0,35 | 0,84 | 4,35 | 0,19 | 8,46E-01 |
| AT1G16680 | 0,85 | -2,11 | 2,40 | -0,88 | 3,80E-01 |
| AT1G16690 | 0,12 | -0,36 | 4,39 | -0,08 | 9,34E-01 |
| AT1G16700 | 0,62 | -0,17 | 3,11 | -0,05 | 9,56E-01 |
| AT1G16710 | 1,85 | -1,48 | 1,85 | -0,80 | 4,24E-01 |
| AT1G16720 | 1,64 | -1,93 | 1,88 | -1,03 | 3,03E-01 |
| AT1G16740 | 0,37 | -0,34 | 4,40 | -0,08 | 9,38E-01 |
| AT1G16810 | 0,12 | 0,00 | 4,41 | 0,00 | 1,00E+00 |
| AT1G16860 | 0,38 | -2,05 | 4,25 | -0,48 | 6,29E-01 |
| AT1G17050 | 0,20 | -2,07 | 4,39 | -0,47 | 6,37E-01 |
| AT1G17070 | 0,73 | -1,08 | 3,65 | -0,30 | 7,68E-01 |
| AT1G17100 | 0,86 | -0,69 | 1,78 | -0,39 | 6,97E-01 |
| AT1G17120 | 0,36 | 0,52 | 4,36 | 0,12 | 9,05E-01 |
| AT1G17150 | 0,40 | 0,53 | 3,40 | 0,16 | 8,77E-01 |
| AT1G17160 | 0,36 | -0,32 | 4,36 | -0,07 | 9,41E-01 |
| AT1G17200 | 0,19 | -0,02 | 4,35 | -0,01 | 9,96E-01 |

| | | | | | |
|---|---|---|---|---|---|
| AT1G17230 | 0,51 | 2,58 | 3,04 | 0,85 | 3,96E-01 |
| AT1G17255 | 1,02 | 0,41 | 2,76 | 0,15 | 8,81E-01 |
| AT1G17260 | 0,13 | 0,00 | 4,41 | 0,00 | 1,00E+00 |
| AT1G17300 | 0,61 | 0,00 | 3,73 | 0,00 | 1,00E+00 |
| AT1G17340 | 0,57 | 0,98 | 2,23 | 0,44 | 6,61E-01 |
| AT1G17360 | 1,17 | -2,19 | 2,22 | -0,99 | 3,23E-01 |
| AT1G17420 | 1,07 | 1,16 | 2,06 | 0,56 | 5,73E-01 |
| AT1G17460 | 0,31 | 0,07 | 4,38 | 0,02 | 9,87E-01 |
| AT1G17490 | 0,39 | 0,69 | 3,42 | 0,20 | 8,41E-01 |
| AT1G17500 | 0,69 | -0,43 | 2,32 | -0,19 | 8,53E-01 |
| AT1G17530 | 0,57 | 1,06 | 3,12 | 0,34 | 7,34E-01 |
| AT1G17550 | 1,05 | 0,25 | 1,92 | 0,13 | 8,97E-01 |
| AT1G17570 | 0,08 | 0,21 | 4,39 | 0,05 | 9,62E-01 |
| AT1G17580 | 1,15 | 0,81 | 2,28 | 0,36 | 7,23E-01 |
| AT1G17610 | 0,33 | 1,50 | 4,33 | 0,35 | 7,30E-01 |
| AT1G17620 | 1,43 | 1,89 | 1,72 | 1,09 | 2,74E-01 |
| AT1G17680 | 0,28 | -2,10 | 4,40 | -0,48 | 6,33E-01 |
| AT1G17745 | 0,96 | 2,28 | 1,97 | 1,15 | 2,48E-01 |
| AT1G17750 | 0,21 | -0,29 | 4,40 | -0,06 | 9,48E-01 |
| AT1G17840 | 1,51 | -0,07 | 1,61 | -0,04 | 9,67E-01 |
| AT1G17860 | 0,25 | -2,14 | 4,40 | -0,49 | 6,28E-01 |
| AT1G17970 | 0,47 | -0,18 | 4,06 | -0,04 | 9,64E-01 |
| AT1G18020 | 0,78 | 2,47 | 2,93 | 0,84 | 4,00E-01 |
| AT1G18090 | 0,88 | 0,00 | 3,64 | 0,00 | 1,00E+00 |
| AT1G18130 | 0,21 | -0,93 | 4,35 | -0,21 | 8,31E-01 |
| AT1G18170 | 0,31 | -2,19 | 4,41 | -0,50 | 6,20E-01 |
| AT1G18300 | 0,38 | 1,36 | 4,29 | 0,32 | 7,50E-01 |
| AT1G18390 | 0,88 | 1,98 | 2,07 | 0,96 | 3,39E-01 |
| AT1G18450 | 0,27 | -0,35 | 4,35 | -0,08 | 9,35E-01 |
| AT1G18460 | 0,79 | -1,89 | 1,99 | -0,95 | 3,43E-01 |
| AT1G18480 | 0,17 | 0,07 | 4,38 | 0,02 | 9,88E-01 |
| AT1G18485 | 1,73 | -2,16 | 2,28 | -0,95 | 3,43E-01 |
| AT1G18490 | 0,64 | 0,00 | 4,41 | 0,00 | 1,00E+00 |
| AT1G18500 | 0,63 | -0,74 | 3,02 | -0,24 | 8,07E-01 |
| AT1G18570 | 0,88 | 1,88 | 2,07 | 0,91 | 3,64E-01 |
| AT1G18580 | 0,40 | 0,00 | 4,41 | 0,00 | 1,00E+00 |
| AT1G18600 | 0,74 | -0,81 | 2,50 | -0,32 | 7,46E-01 |
| AT1G18620 | 0,88 | 0,70 | 2,17 | 0,32 | 7,46E-01 |
| AT1G18700 | 1,03 | -0,32 | 2,44 | -0,13 | 8,94E-01 |
| AT1G18730 | 0,47 | -2,32 | 4,11 | -0,56 | 5,72E-01 |
| AT1G18820 | 0,52 | -1,14 | 4,30 | -0,27 | 7,91E-01 |
| AT1G18870 | 0,46 | -2,40 | 4,37 | -0,55 | 5,83E-01 |
| AT1G18880 | 1,72 | 2,32 | 1,60 | 1,45 | 1,47E-01 |
| AT1G18879 | 1,60 | 2,27 | 1,68 | 1,35 | 1,78E-01 |
| AT1G18900 | 0,70 | -0,29 | 3,65 | -0,08 | 9,36E-01 |
| AT1G18950 | 0,83 | 2,29 | 2,12 | 1,08 | 2,80E-01 |
| AT1G19020 | 0,56 | 1,26 | 2,33 | 0,54 | 5,88E-01 |
| AT1G19025 | 0,65 | -0,29 | 3,13 | -0,09 | 9,27E-01 |
| AT1G19086 | 0,19 | 1,80 | 4,31 | 0,42 | 6,76E-01 |
| AT1G19110 | 0,80 | -1,30 | 2,34 | -0,56 | 5,77E-01 |
| AT1G19150 | 0,10 | 0,86 | 4,35 | 0,20 | 8,43E-01 |
| AT1G19180 | 1,37 | 3,59 | 1,89 | 1,90 | 5,70E-02 |
| AT1G19190 | 0,39 | 0,00 | 4,40 | 0,00 | 1,00E+00 |
| AT1G19290 | 0,57 | 0,00 | 3,83 | 0,00 | 1,00E+00 |
| AT1G19350 | 0,52 | 1,04 | 3,91 | 0,27 | 7,90E-01 |
| AT1G19376 | 0,56 | 2,59 | 2,99 | 0,87 | 3,87E-01 |

| | | | | | |
|---|---|---|---|---|---|
| AT1G19464 | 0,18 | 1,59 | 4,31 | 0,37 | 7,12E-01 |
| AT1G19580 | 0,28 | -2,11 | 4,40 | -0,48 | 6,32E-01 |
| AT1G05507 | 0,08 | 0,21 | 4,39 | 0,05 | 9,62E-01 |
| AT1G19710 | 0,37 | 0,00 | 4,35 | 0,00 | 1,00E+00 |
| AT1G19770 | 0,78 | 2,98 | 2,70 | 1,10 | 2,70E-01 |
| AT1G19800 | 0,29 | -1,98 | 4,38 | -0,45 | 6,51E-01 |
| AT1G19830 | 0,33 | 1,89 | 4,31 | 0,44 | 6,62E-01 |
| AT1G19850 | 0,68 | 1,34 | 2,92 | 0,46 | 6,45E-01 |
| AT1G19910 | 0,96 | 1,11 | 2,37 | 0,47 | 6,39E-01 |
| AT1G19990 | 0,28 | 1,76 | 4,30 | 0,41 | 6,82E-01 |
| AT1G20050 | 0,40 | -0,82 | 3,49 | -0,23 | 8,14E-01 |
| AT1G05523 | 0,59 | -0,09 | 2,49 | -0,04 | 9,71E-01 |
| AT1G20090 | 0,64 | -0,89 | 2,54 | -0,35 | 7,28E-01 |
| AT1G20110 | 0,46 | -1,66 | 2,70 | -0,61 | 5,39E-01 |
| AT1G20280 | 0,77 | 0,37 | 1,86 | 0,20 | 8,42E-01 |
| AT1G20300 | 0,15 | -0,57 | 4,37 | -0,13 | 8,96E-01 |
| AT1G20310 | 0,17 | 0,00 | 4,41 | 0,00 | 1,00E+00 |
| AT1G20375 | 1,50 | 2,35 | 1,77 | 1,32 | 1,85E-01 |
| AT1G20630 | 1,62 | 0,60 | 1,89 | 0,32 | 7,48E-01 |
| AT1G20640 | 1,05 | 2,02 | 2,06 | 0,98 | 3,26E-01 |
| AT1G20693 | 1,05 | 0,02 | 1,99 | 0,01 | 9,94E-01 |
| AT1G20696 | 0,69 | -0,28 | 2,22 | -0,13 | 8,99E-01 |
| AT1G20780 | 1,16 | -3,20 | 1,88 | -1,71 | 8,82E-02 |
| AT1G20823 | 0,81 | 2,44 | 2,17 | 1,12 | 2,62E-01 |
| AT1G20920 | 0,49 | -2,11 | 3,25 | -0,65 | 5,16E-01 |
| AT1G20980 | 1,44 | -1,18 | 1,83 | -0,64 | 5,21E-01 |
| AT1G21000 | 0,77 | 2,22 | 2,14 | 1,04 | 2,99E-01 |
| AT1G21050 | 0,58 | 2,04 | 2,91 | 0,70 | 4,85E-01 |
| AT1G21065 | 0,84 | 1,10 | 2,07 | 0,53 | 5,96E-01 |
| AT1G21140 | 0,08 | 0,00 | 4,41 | 0,00 | 1,00E+00 |
| AT1G21170 | 0,55 | 0,40 | 3,75 | 0,11 | 9,14E-01 |
| AT1G21210 | 0,63 | -0,37 | 3,02 | -0,12 | 9,02E-01 |
| AT1G21230 | 0,43 | 2,48 | 3,19 | 0,78 | 4,37E-01 |
| AT1G21240 | 0,38 | -0,27 | 4,31 | -0,06 | 9,49E-01 |
| AT1G21370 | 0,21 | 0,39 | 4,36 | 0,09 | 9,29E-01 |
| AT1G21430 | 0,53 | 0,00 | 4,40 | 0,00 | 1,00E+00 |
| AT1G21560 | 0,41 | 0,00 | 4,19 | 0,00 | 1,00E+00 |
| AT1G21640 | 0,89 | 0,62 | 1,96 | 0,32 | 7,53E-01 |
| AT1G21660 | 0,28 | -0,36 | 4,39 | -0,08 | 9,35E-01 |
| AT1G21670 | 0,86 | -2,07 | 3,37 | -0,62 | 5,38E-01 |
| AT1G21690 | 0,29 | 0,09 | 4,38 | 0,02 | 9,84E-01 |
| AT1G21720 | 0,34 | 0,38 | 4,38 | 0,09 | 9,30E-01 |
| AT1G21722 | 0,08 | 0,00 | 4,41 | 0,00 | 1,00E+00 |
| AT1G21780 | 0,75 | 1,05 | 2,47 | 0,42 | 6,72E-01 |
| AT1G21835 | 0,94 | 0,62 | 1,95 | 0,32 | 7,49E-01 |
| AT1G21840 | 0,12 | 0,00 | 4,41 | 0,00 | 1,00E+00 |
| AT1G21880 | 0,12 | -0,26 | 4,41 | -0,06 | 9,53E-01 |
| AT1G21910 | 0,33 | -3,33 | 4,36 | -0,76 | 4,45E-01 |
| AT1G21980 | 0,19 | 0,84 | 4,35 | 0,19 | 8,47E-01 |
| AT1G22040 | 0,05 | 0,18 | 4,38 | 0,04 | 9,68E-01 |
| AT1G22070 | 0,41 | 0,82 | 4,32 | 0,19 | 8,49E-01 |
| AT1G22190 | 1,08 | -1,59 | 1,70 | -0,93 | 3,50E-01 |
| AT1G22280 | 0,53 | 1,41 | 3,31 | 0,43 | 6,70E-01 |
| AT1G22330 | 0,11 | 0,88 | 4,35 | 0,20 | 8,39E-01 |
| AT1G22410 | 1,37 | -0,22 | 1,37 | -0,16 | 8,71E-01 |
| AT1G22500 | 0,40 | 0,00 | 4,41 | 0,00 | 1,00E+00 |

| | | | | | |
|---|---|---|---|---|---|
| AT1G22590 | 0,24 | 1,34 | 4,33 | 0,31 | 7,57E-01 |
| AT1G22640 | 0,56 | 2,16 | 3,68 | 0,59 | 5,57E-01 |
| AT1G22730 | 0,55 | 0,00 | 3,92 | 0,00 | 1,00E+00 |
| AT1G22750 | 0,35 | 1,36 | 3,46 | 0,39 | 6,94E-01 |
| AT1G22770 | 1,61 | 0,85 | 1,38 | 0,62 | 5,38E-01 |
| AT1G22780 | 0,54 | 1,04 | 3,24 | 0,32 | 7,48E-01 |
| AT1G22790 | 0,64 | -0,26 | 3,93 | -0,07 | 9,47E-01 |
| AT1G05767 | 0,23 | -1,26 | 4,37 | -0,29 | 7,72E-01 |
| AT1G22840 | 0,62 | 0,62 | 3,21 | 0,19 | 8,47E-01 |
| AT1G22860 | 0,86 | 0,44 | 2,18 | 0,20 | 8,41E-01 |
| AT1G22870 | 0,66 | -0,26 | 3,03 | -0,09 | 9,31E-01 |
| AT1G22880 | 1,11 | -1,23 | 2,99 | -0,41 | 6,79E-01 |
| AT1G22882 | 0,84 | 0,00 | 3,44 | 0,00 | 1,00E+00 |
| AT1G22930 | 1,70 | 1,81 | 1,80 | 1,00 | 3,15E-01 |
| AT1G22940 | 0,55 | -1,23 | 2,81 | -0,44 | 6,62E-01 |
| AT1G23030 | 0,27 | -2,13 | 4,39 | -0,49 | 6,27E-01 |
| AT1G23040 | 0,34 | 0,95 | 4,35 | 0,22 | 8,27E-01 |
| AT1G23050 | 1,19 | 0,54 | 2,23 | 0,24 | 8,08E-01 |
| AT1G23100 | 0,08 | 0,00 | 4,41 | 0,00 | 1,00E+00 |
| AT1G23130 | 0,08 | 0,00 | 4,41 | 0,00 | 1,00E+00 |
| AT1G23190 | 1,73 | -0,30 | 1,82 | -0,17 | 8,67E-01 |
| AT1G23230 | 1,20 | 1,19 | 1,73 | 0,69 | 4,91E-01 |
| AT1G23290 | 1,36 | -0,61 | 1,61 | -0,38 | 7,07E-01 |
| AT1G23440 | 0,65 | -2,21 | 3,09 | -0,72 | 4,74E-01 |
| AT1G23450 | 0,21 | 0,00 | 4,41 | 0,00 | 1,00E+00 |
| AT1G23490 | 1,31 | 1,21 | 1,61 | 0,75 | 4,52E-01 |
| AT1G23710 | 1,29 | 2,62 | 1,78 | 1,47 | 1,41E-01 |
| AT1G23820 | 1,10 | 0,52 | 3,02 | 0,17 | 8,64E-01 |
| AT1G05837 | 0,53 | -0,28 | 3,23 | -0,09 | 9,31E-01 |
| AT1G23870 | 0,51 | 0,82 | 3,85 | 0,21 | 8,31E-01 |
| AT1G23880 | 0,08 | 0,00 | 4,41 | 0,00 | 1,00E+00 |
| AT1G23900 | 1,85 | 0,27 | 1,82 | 0,15 | 8,82E-01 |
| AT1G23915 | 1,01 | 0,87 | 2,58 | 0,34 | 7,36E-01 |
| AT1G24145 | 0,48 | 0,77 | 3,15 | 0,25 | 8,06E-01 |
| AT1G24147 | 0,16 | -0,21 | 4,40 | -0,05 | 9,63E-01 |
| AT1G24150 | 0,48 | -0,41 | 3,99 | -0,10 | 9,18E-01 |
| AT1G24280 | 0,44 | -2,44 | 4,38 | -0,56 | 5,79E-01 |
| AT1G24340 | 0,28 | 0,28 | 3,73 | 0,07 | 9,40E-01 |
| AT1G24360 | 0,34 | 1,25 | 4,33 | 0,29 | 7,72E-01 |
| AT1G24380 | 0,38 | 0,79 | 2,79 | 0,28 | 7,76E-01 |
| AT1G24490 | 0,45 | 0,89 | 3,25 | 0,27 | 7,83E-01 |
| AT1G24530 | 0,17 | 0,07 | 4,38 | 0,02 | 9,87E-01 |
| AT1G24560 | 0,68 | 0,93 | 2,15 | 0,44 | 6,63E-01 |
| AT1G24570 | 0,08 | 0,00 | 4,41 | 0,00 | 1,00E+00 |
| AT1G25275 | 1,34 | -0,63 | 1,54 | -0,41 | 6,81E-01 |
| AT1G25280 | 0,98 | 0,55 | 2,53 | 0,22 | 8,27E-01 |
| AT1G25480 | 0,19 | 1,60 | 4,31 | 0,37 | 7,11E-01 |
| AT1G25490 | 1,64 | -1,40 | 1,30 | -1,08 | 2,79E-01 |
| AT1G25520 | 0,81 | -0,40 | 3,60 | -0,11 | 9,12E-01 |
| AT1G25550 | 1,33 | 1,90 | 1,88 | 1,01 | 3,14E-01 |
| AT1G26130 | 0,54 | -0,27 | 3,94 | -0,07 | 9,45E-01 |
| AT1G26220 | 0,37 | 1,88 | 4,31 | 0,44 | 6,63E-01 |
| AT1G26250 | 0,44 | 2,66 | 4,03 | 0,66 | 5,09E-01 |
| AT1G26460 | 0,79 | -0,29 | 2,95 | -0,10 | 9,20E-01 |
| AT1G26470 | 0,34 | 0,00 | 4,41 | 0,00 | 1,00E+00 |
| AT1G26550 | 0,37 | 1,95 | 4,30 | 0,45 | 6,49E-01 |

| | | | | | |
|---|---|---|---|---|---|
| AT1G26560 | 0,24 | -0,47 | 4,36 | -0,11 | 9,15E-01 |
| AT1G26580 | 0,53 | 1,13 | 2,70 | 0,42 | 6,77E-01 |
| AT1G26630 | 1,29 | 0,17 | 1,56 | 0,11 | 9,16E-01 |
| AT1G26650 | 1,02 | -2,19 | 2,25 | -0,97 | 3,31E-01 |
| AT1G26665 | 0,17 | 0,00 | 4,41 | 0,00 | 1,00E+00 |
| AT1G26670 | 1,04 | 0,79 | 2,37 | 0,33 | 7,38E-01 |
| AT1G26770 | 0,19 | 1,60 | 4,31 | 0,37 | 7,11E-01 |
| AT1G26830 | 0,72 | 1,98 | 2,41 | 0,82 | 4,12E-01 |
| AT1G26920 | 0,73 | 0,52 | 2,13 | 0,25 | 8,06E-01 |
| AT1G26940 | 0,05 | 0,18 | 4,38 | 0,04 | 9,68E-01 |
| AT1G27000 | 0,38 | -1,76 | 4,36 | -0,40 | 6,86E-01 |
| AT1G27060 | 0,73 | 0,00 | 3,01 | 0,00 | 1,00E+00 |
| AT1G27130 | 0,51 | -0,19 | 3,11 | -0,06 | 9,50E-01 |
| AT1G27150 | 0,43 | -0,31 | 4,40 | -0,07 | 9,45E-01 |
| AT1G27170 | 0,35 | 0,44 | 3,47 | 0,13 | 8,98E-01 |
| AT1G27180 | 0,99 | 0,00 | 2,75 | 0,00 | 1,00E+00 |
| AT1G27290 | 1,54 | 1,72 | 1,48 | 1,17 | 2,43E-01 |
| AT1G27300 | 0,40 | -0,40 | 4,22 | -0,09 | 9,25E-01 |
| AT1G27330 | 0,50 | -2,04 | 4,38 | -0,47 | 6,41E-01 |
| AT1G27360 | 0,08 | 0,00 | 4,41 | 0,00 | 1,00E+00 |
| AT1G27400 | 0,67 | 0,22 | 2,16 | 0,10 | 9,19E-01 |
| AT1G06087 | 0,12 | -0,36 | 4,39 | -0,08 | 9,34E-01 |
| AT1G27450 | 0,37 | -1,33 | 4,35 | -0,31 | 7,60E-01 |
| AT1G27460 | 0,11 | -0,26 | 4,41 | -0,06 | 9,53E-01 |
| AT1G27480 | 0,41 | -2,05 | 4,23 | -0,48 | 6,28E-01 |
| AT1G27520 | 0,14 | 0,04 | 4,38 | 0,01 | 9,93E-01 |
| AT1G06097 | 0,36 | -1,27 | 4,34 | -0,29 | 7,71E-01 |
| AT1G27570 | 1,68 | 4,14 | 1,88 | 2,20 | 2,78E-02 |
| AT1G27630 | 0,58 | 1,59 | 3,95 | 0,40 | 6,87E-01 |
| AT1G27650 | 0,97 | -0,75 | 2,16 | -0,35 | 7,28E-01 |
| AT1G27750 | 0,63 | 1,90 | 2,62 | 0,73 | 4,68E-01 |
| AT1G27752 | 1,43 | 1,71 | 2,21 | 0,77 | 4,40E-01 |
| AT1G27760 | 0,65 | 0,54 | 2,60 | 0,21 | 8,35E-01 |
| AT1G27850 | 0,84 | 0,07 | 2,31 | 0,03 | 9,78E-01 |
| AT1G27950 | 0,55 | 0,39 | 3,73 | 0,11 | 9,16E-01 |
| AT1G28005 | 0,32 | 2,33 | 4,29 | 0,54 | 5,87E-01 |
| AT1G28060 | 0,32 | -1,23 | 4,35 | -0,28 | 7,77E-01 |
| AT1G28135 | 0,08 | 0,21 | 4,39 | 0,05 | 9,62E-01 |
| AT1G28340 | 0,87 | -1,41 | 2,81 | -0,50 | 6,15E-01 |
| AT1G28350 | 1,15 | 0,66 | 2,17 | 0,30 | 7,62E-01 |
| AT1G28380 | 1,21 | 0,81 | 2,20 | 0,37 | 7,13E-01 |
| AT1G28440 | 1,72 | -0,13 | 1,33 | -0,10 | 9,20E-01 |
| AT1G06163 | 0,36 | 0,62 | 4,31 | 0,14 | 8,86E-01 |
| AT1G28490 | 0,17 | -0,29 | 4,40 | -0,07 | 9,47E-01 |
| AT1G28520 | 0,40 | 0,50 | 3,39 | 0,15 | 8,83E-01 |
| AT1G28680 | 0,55 | -0,10 | 3,23 | -0,03 | 9,75E-01 |
| AT1G28780 | 0,49 | 1,68 | 4,05 | 0,41 | 6,78E-01 |
| AT1G28760 | 0,10 | 0,24 | 4,38 | 0,05 | 9,57E-01 |
| AT1G29070 | 1,74 | -0,90 | 1,44 | -0,63 | 5,32E-01 |
| AT1G29120 | 0,19 | -0,34 | 4,40 | -0,08 | 9,38E-01 |
| AT1G29150 | 0,99 | 1,12 | 2,44 | 0,46 | 6,46E-01 |
| AT1G29200 | 0,08 | 0,00 | 4,41 | 0,00 | 1,00E+00 |
| AT1G29260 | 0,92 | 0,70 | 1,85 | 0,38 | 7,05E-01 |
| AT1G29310 | 0,68 | -2,11 | 3,06 | -0,69 | 4,91E-01 |
| AT1G29340 | 0,94 | 1,54 | 2,11 | 0,73 | 4,66E-01 |
| AT1G29380 | 0,65 | 0,00 | 4,41 | 0,00 | 1,00E+00 |

| | | | | | |
|---|---|---|---|---|---|
| AT1G06233 | 0,68 | -1,22 | 2,39 | -0,51 | 6,10E-01 |
| AT1G06253 | 1,09 | -0,95 | 1,54 | -0,62 | 5,36E-01 |
| AT1G06263 | 0,52 | -3,29 | 4,01 | -0,82 | 4,11E-01 |
| AT1G06283 | 1,64 | 1,10 | 1,30 | 0,84 | 4,01E-01 |
| AT1G29620 | 0,47 | -0,45 | 3,17 | -0,14 | 8,87E-01 |
| AT1G29630 | 0,24 | -0,26 | 4,41 | -0,06 | 9,53E-01 |
| AT1G29660 | 1,40 | -0,76 | 1,30 | -0,58 | 5,59E-01 |
| AT1G29670 | 1,73 | -1,31 | 1,39 | -0,94 | 3,45E-01 |
| AT1G29760 | 0,40 | -1,18 | 4,28 | -0,28 | 7,83E-01 |
| AT1G29770 | 0,71 | 0,00 | 4,41 | 0,00 | 1,00E+00 |
| AT1G29800 | 0,74 | -0,29 | 3,80 | -0,08 | 9,38E-01 |
| AT1G29830 | 1,13 | 0,11 | 2,00 | 0,05 | 9,57E-01 |
| AT1G29950 | 0,49 | 0,00 | 4,41 | 0,00 | 1,00E+00 |
| AT1G30010 | 0,08 | 0,21 | 4,39 | 0,05 | 9,62E-01 |
| AT1G30200 | 0,70 | -0,50 | 2,32 | -0,21 | 8,31E-01 |
| AT1G30260 | 1,16 | -0,14 | 2,21 | -0,06 | 9,49E-01 |
| AT1G30420 | 0,38 | 0,00 | 4,41 | 0,00 | 1,00E+00 |
| AT1G30430 | 0,16 | -0,56 | 4,37 | -0,13 | 8,98E-01 |
| AT1G30440 | 1,10 | -0,91 | 1,69 | -0,54 | 5,90E-01 |
| AT1G30450 | 0,86 | -1,55 | 2,14 | -0,72 | 4,69E-01 |
| AT1G30520 | 0,41 | 0,00 | 4,41 | 0,00 | 1,00E+00 |
| AT1G30530 | 0,46 | -0,48 | 3,26 | -0,15 | 8,84E-01 |
| AT1G30610 | 0,23 | 0,56 | 4,37 | 0,13 | 8,99E-01 |
| AT1G30620 | 0,56 | 1,18 | 3,27 | 0,36 | 7,18E-01 |
| AT1G30700 | 0,12 | 0,00 | 4,41 | 0,00 | 1,00E+00 |
| AT1G30720 | 0,53 | 0,00 | 4,40 | 0,00 | 1,00E+00 |
| AT1G30757 | 0,16 | 1,26 | 4,33 | 0,29 | 7,71E-01 |
| AT1G30800 | 0,20 | 0,00 | 4,41 | 0,00 | 1,00E+00 |
| AT1G30910 | 0,25 | 0,84 | 4,36 | 0,19 | 8,48E-01 |
| AT1G06393 | 0,32 | -1,96 | 4,30 | -0,46 | 6,49E-01 |
| AT1G31095 | 0,66 | 1,60 | 2,82 | 0,57 | 5,72E-01 |
| AT1G31170 | 0,22 | -0,83 | 4,36 | -0,19 | 8,49E-01 |
| AT1G31175 | 0,74 | -1,07 | 2,53 | -0,42 | 6,71E-01 |
| AT1G31190 | 0,86 | 2,62 | 2,39 | 1,10 | 2,73E-01 |
| AT1G31220 | 0,22 | 0,81 | 4,35 | 0,19 | 8,52E-01 |
| AT1G31290 | 0,15 | 0,47 | 4,37 | 0,11 | 9,15E-01 |
| AT1G31420 | 0,89 | -2,93 | 2,17 | -1,35 | 1,77E-01 |
| AT1G31480 | 0,40 | 0,00 | 4,41 | 0,00 | 1,00E+00 |
| AT1G06447 | 0,24 | 0,14 | 4,38 | 0,03 | 9,74E-01 |
| AT1G31710 | 0,82 | 3,46 | 3,45 | 1,00 | 3,15E-01 |
| AT1G31780 | 0,28 | -0,67 | 4,38 | -0,15 | 8,78E-01 |
| AT1G31800 | 0,33 | -1,58 | 3,48 | -0,45 | 6,49E-01 |
| AT1G31850 | 1,13 | -0,62 | 2,27 | -0,27 | 7,84E-01 |
| AT1G31930 | 0,12 | 0,00 | 4,41 | 0,00 | 1,00E+00 |
| AT1G31970 | 0,74 | 0,86 | 3,18 | 0,27 | 7,88E-01 |
| AT1G32050 | 0,65 | -1,11 | 2,28 | -0,49 | 6,25E-01 |
| AT1G32160 | 0,70 | -0,82 | 2,94 | -0,28 | 7,80E-01 |
| AT1G32220 | 0,21 | 1,42 | 4,32 | 0,33 | 7,42E-01 |
| AT1G32225 | 0,70 | 2,14 | 2,31 | 0,93 | 3,54E-01 |
| AT1G32320 | 0,23 | 0,23 | 4,35 | 0,05 | 9,58E-01 |
| AT1G32337 | 0,30 | -0,64 | 4,32 | -0,15 | 8,83E-01 |
| AT1G32360 | 0,41 | -2,15 | 3,40 | -0,63 | 5,26E-01 |
| AT1G06547 | 0,19 | 1,47 | 4,32 | 0,34 | 7,35E-01 |
| AT1G32385 | 0,41 | -0,71 | 2,68 | -0,26 | 7,91E-01 |
| AT1G32415 | 0,72 | 0,00 | 2,97 | 0,00 | 1,00E+00 |
| AT1G32460 | 0,41 | 0,61 | 4,32 | 0,14 | 8,88E-01 |

| | | | | | |
|---|---|---|---|---|---|
| AT1G32580 | 0,32 | 0,00 | 4,41 | 0,00 | 1,00E+00 |
| AT1G06597 | 1,64 | -2,16 | 1,35 | -1,61 | 1,08E-01 |
| AT1G32700 | 0,46 | -0,98 | 3,32 | -0,29 | 7,69E-01 |
| AT1G32810 | 0,36 | -1,25 | 3,51 | -0,36 | 7,21E-01 |
| AT1G06623 | 0,26 | 1,96 | 4,30 | 0,46 | 6,48E-01 |
| AT1G32870 | 0,59 | -0,44 | 3,88 | -0,11 | 9,10E-01 |
| AT1G32920 | 1,13 | 2,17 | 1,81 | 1,20 | 2,31E-01 |
| AT1G32928 | 0,50 | 0,60 | 4,08 | 0,15 | 8,84E-01 |
| AT1G32940 | 0,41 | 0,00 | 4,41 | 0,00 | 1,00E+00 |
| AT1G32950 | 0,62 | 0,00 | 3,75 | 0,00 | 1,00E+00 |
| AT1G32960 | 0,35 | 0,56 | 4,37 | 0,13 | 8,99E-01 |
| AT1G32990 | 0,48 | -2,59 | 4,33 | -0,60 | 5,50E-01 |
| AT1G33010 | 0,19 | 1,60 | 4,31 | 0,37 | 7,11E-01 |
| AT1G33040 | 0,24 | -0,53 | 4,37 | -0,12 | 9,03E-01 |
| AT1G33070 | 1,82 | 0,04 | 1,57 | 0,02 | 9,82E-01 |
| AT1G33110 | 0,55 | 0,00 | 3,90 | 0,00 | 1,00E+00 |
| AT1G33140 | 1,84 | -0,81 | 1,68 | -0,48 | 6,29E-01 |
| AT1G33170 | 0,44 | -2,15 | 3,36 | -0,64 | 5,22E-01 |
| AT1G33260 | 0,52 | 0,10 | 4,05 | 0,02 | 9,81E-01 |
| AT1G33360 | 0,47 | 1,26 | 4,05 | 0,31 | 7,56E-01 |
| AT1G33470 | 0,71 | 0,00 | 3,62 | 0,00 | 1,00E+00 |
| AT1G33590 | 1,70 | 0,13 | 1,24 | 0,10 | 9,17E-01 |
| AT1G33600 | 1,39 | 0,52 | 1,48 | 0,35 | 7,27E-01 |
| AT1G33610 | 0,15 | -0,35 | 4,39 | -0,08 | 9,36E-01 |
| AT1G33612 | 0,17 | -1,50 | 4,37 | -0,34 | 7,31E-01 |
| AT1G33770 | 0,49 | -1,08 | 4,02 | -0,27 | 7,87E-01 |
| AT1G33811 | 0,18 | -0,84 | 4,35 | -0,19 | 8,47E-01 |
| AT1G33910 | 0,08 | 0,21 | 4,39 | 0,05 | 9,62E-01 |
| AT1G33930 | 0,36 | -0,70 | 4,26 | -0,17 | 8,69E-01 |
| AT1G33960 | 1,02 | -0,35 | 3,23 | -0,11 | 9,14E-01 |
| AT1G33980 | 0,92 | 0,00 | 4,26 | 0,00 | 1,00E+00 |
| AT1G33990 | 0,95 | -1,57 | 1,95 | -0,80 | 4,21E-01 |
| AT1G06723 | 0,59 | -2,32 | 2,60 | -0,89 | 3,74E-01 |
| AT1G34020 | 0,78 | -1,91 | 2,49 | -0,77 | 4,44E-01 |
| AT1G34070 | 0,46 | 2,74 | 4,00 | 0,69 | 4,92E-01 |
| AT1G34130 | 1,45 | -0,85 | 1,58 | -0,53 | 5,93E-01 |
| AT1G34150 | 1,09 | 1,23 | 2,47 | 0,50 | 6,19E-01 |
| AT1G34190 | 1,14 | 1,22 | 2,01 | 0,61 | 5,44E-01 |
| AT1G34200 | 0,21 | -0,32 | 4,40 | -0,07 | 9,43E-01 |
| AT1G34210 | 0,51 | -0,26 | 4,41 | -0,06 | 9,53E-01 |
| AT1G34260 | 1,42 | 1,35 | 2,09 | 0,65 | 5,18E-01 |
| AT1G34290 | 0,25 | 1,91 | 4,30 | 0,45 | 6,56E-01 |
| AT1G06747 | 0,08 | 0,21 | 4,39 | 0,05 | 9,62E-01 |
| AT1G34355 | 0,86 | -0,35 | 2,94 | -0,12 | 9,06E-01 |
| AT1G34370 | 1,49 | -0,35 | 2,19 | -0,16 | 8,74E-01 |
| AT1G34390 | 0,68 | 1,95 | 3,63 | 0,54 | 5,92E-01 |
| AT1G34410 | 0,91 | 3,84 | 3,20 | 1,20 | 2,30E-01 |
| AT1G34420 | 0,36 | 1,22 | 4,29 | 0,28 | 7,76E-01 |
| AT1G34460 | 0,31 | 2,15 | 4,30 | 0,50 | 6,16E-01 |
| AT1G34580 | 0,53 | -3,87 | 4,32 | -0,90 | 3,70E-01 |
| AT1G34770 | 0,12 | 0,00 | 4,41 | 0,00 | 1,00E+00 |
| AT1G34844 | 0,08 | 0,00 | 4,41 | 0,00 | 1,00E+00 |
| AT1G35030 | 0,42 | -0,18 | 3,35 | -0,05 | 9,57E-01 |
| AT1G35040 | 0,23 | 1,50 | 4,31 | 0,35 | 7,29E-01 |
| AT1G35140 | 0,39 | 1,98 | 4,18 | 0,47 | 6,36E-01 |
| AT1G35160 | 0,59 | -2,80 | 2,54 | -1,10 | 2,70E-01 |

| | | | | | |
|---|---|---|---|---|---|
| AT1G06827 | 0,56 | -0,57 | 2,64 | -0,22 | 8,27E-01 |
| AT1G35220 | 0,89 | -0,26 | 2,53 | -0,10 | 9,17E-01 |
| AT1G35230 | 1,07 | 0,66 | 2,24 | 0,29 | 7,69E-01 |
| AT1G35242 | 0,19 | 0,47 | 4,36 | 0,11 | 9,14E-01 |
| AT1G35420 | 0,41 | -1,17 | 3,33 | -0,35 | 7,26E-01 |
| AT1G35460 | 0,12 | 0,00 | 4,41 | 0,00 | 1,00E+00 |
| AT1G35500 | 0,14 | 1,29 | 4,33 | 0,30 | 7,65E-01 |
| AT1G35510 | 0,12 | 0,11 | 4,38 | 0,03 | 9,80E-01 |
| AT1G35515 | 0,52 | 0,34 | 2,36 | 0,14 | 8,86E-01 |
| AT1G35516 | 0,52 | 0,34 | 2,36 | 0,14 | 8,86E-01 |
| AT1G35530 | 0,08 | 0,00 | 4,41 | 0,00 | 1,00E+00 |
| AT1G35540 | 0,95 | 1,12 | 1,82 | 0,62 | 5,36E-01 |
| AT1G35614 | 1,12 | -0,45 | 1,93 | -0,23 | 8,17E-01 |
| AT1G35620 | 0,51 | 0,74 | 3,24 | 0,23 | 8,18E-01 |
| AT1G35660 | 0,28 | -0,26 | 4,41 | -0,06 | 9,53E-01 |
| AT1G35670 | 0,21 | -0,32 | 4,40 | -0,07 | 9,43E-01 |
| AT1G35680 | 0,81 | -0,97 | 2,22 | -0,44 | 6,63E-01 |
| AT1G06927 | 0,83 | 2,90 | 2,71 | 1,07 | 2,84E-01 |
| AT1G35850 | 0,23 | 1,85 | 4,30 | 0,43 | 6,66E-01 |
| AT1G36050 | 0,26 | 0,05 | 4,38 | 0,01 | 9,91E-01 |
| AT1G06963 | 0,05 | 0,18 | 4,38 | 0,04 | 9,68E-01 |
| AT1G36320 | 0,22 | 0,15 | 4,33 | 0,03 | 9,73E-01 |
| AT1G36510 | 0,10 | 0,86 | 4,35 | 0,20 | 8,43E-01 |
| AT1G06987 | 0,58 | -0,74 | 4,35 | -0,17 | 8,64E-01 |
| AT1G36675 | 0,81 | 0,06 | 2,46 | 0,03 | 9,80E-01 |
| AT1G36745 | 0,98 | 2,76 | 2,22 | 1,25 | 2,13E-01 |
| AT1G36756 | 0,94 | 3,59 | 2,41 | 1,49 | 1,37E-01 |
| AT1G36940 | 0,22 | 1,63 | 4,31 | 0,38 | 7,05E-01 |
| AT1G36980 | 0,52 | -1,28 | 3,24 | -0,40 | 6,92E-01 |
| AT1G36992 | 0,32 | -0,30 | 4,40 | -0,07 | 9,46E-01 |
| AT1G37113 | 0,08 | 0,00 | 4,41 | 0,00 | 1,00E+00 |
| AT1G42470 | 1,71 | -1,61 | 1,97 | -0,82 | 4,14E-01 |
| AT1G42525 | 0,30 | 1,59 | 4,32 | 0,37 | 7,14E-01 |
| AT1G42540 | 1,07 | -0,31 | 2,41 | -0,13 | 8,98E-01 |
| AT1G07137 | 0,05 | 0,18 | 4,38 | 0,04 | 9,68E-01 |
| AT1G42960 | 0,66 | -3,75 | 4,29 | -0,87 | 3,82E-01 |
| AT1G42980 | 1,30 | -0,55 | 1,99 | -0,28 | 7,82E-01 |
| AT1G43040 | 0,16 | 1,26 | 4,33 | 0,29 | 7,71E-01 |
| AT1G43145 | 1,44 | 1,34 | 1,65 | 0,81 | 4,18E-01 |
| AT1G43171 | 0,72 | 0,00 | 4,41 | 0,00 | 1,00E+00 |
| AT1G43330 | 0,87 | 3,72 | 2,51 | 1,48 | 1,39E-01 |
| AT1G43670 | 0,63 | -1,28 | 2,55 | -0,50 | 6,17E-01 |
| AT1G43680 | 0,25 | 0,00 | 4,41 | 0,00 | 1,00E+00 |
| AT1G43700 | 0,72 | -0,18 | 1,84 | -0,10 | 9,20E-01 |
| AT1G43722 | 0,18 | 1,59 | 4,31 | 0,37 | 7,12E-01 |
| AT1G43770 | 1,22 | -2,29 | 1,59 | -1,44 | 1,51E-01 |
| AT1G43780 | 0,25 | -3,44 | 4,34 | -0,79 | 4,28E-01 |
| AT1G43815 | 1,28 | 3,23 | 2,01 | 1,61 | 1,08E-01 |
| AT1G43850 | 1,82 | 2,65 | 1,74 | 1,52 | 1,28E-01 |
| AT1G43890 | 0,40 | -0,29 | 4,35 | -0,07 | 9,47E-01 |
| AT1G43970 | 0,43 | 3,00 | 4,27 | 0,70 | 4,82E-01 |
| AT1G44120 | 0,95 | 1,24 | 2,73 | 0,45 | 6,49E-01 |
| AT1G44750 | 0,33 | 0,00 | 4,41 | 0,00 | 1,00E+00 |
| AT1G44780 | 0,29 | -1,98 | 4,38 | -0,45 | 6,51E-01 |
| AT1G44810 | 0,26 | -0,45 | 3,77 | -0,12 | 9,05E-01 |
| AT1G44890 | 0,08 | 0,21 | 4,39 | 0,05 | 9,62E-01 |

| | | | | | |
|---|---|---|---|---|---|
| AT1G44900 | 0,12 | 0,00 | 4,41 | 0,00 | 1,00E+00 |
| AT1G44910 | 0,50 | 0,78 | 2,29 | 0,34 | 7,32E-01 |
| AT1G44960 | 0,10 | 0,24 | 4,38 | 0,05 | 9,57E-01 |
| AT1G45000 | 0,47 | -0,49 | 2,80 | -0,17 | 8,61E-01 |
| AT1G45165 | 0,70 | 0,98 | 2,53 | 0,39 | 7,00E-01 |
| AT1G45180 | 0,30 | 0,00 | 4,41 | 0,00 | 1,00E+00 |
| AT1G45474 | 0,41 | -0,60 | 3,21 | -0,19 | 8,51E-01 |
| AT1G45688 | 0,89 | 2,62 | 2,33 | 1,13 | 2,59E-01 |
| AT1G45976 | 0,41 | 0,03 | 4,24 | 0,01 | 9,94E-01 |
| AT1G46264 | 0,10 | 0,24 | 4,38 | 0,05 | 9,57E-01 |
| AT1G46554 | 0,28 | -0,04 | 4,39 | -0,01 | 9,93E-01 |
| AT1G47210 | 0,93 | -0,40 | 2,68 | -0,15 | 8,81E-01 |
| AT1G07327 | 1,09 | 1,55 | 2,11 | 0,74 | 4,61E-01 |
| AT1G47250 | 0,35 | -1,35 | 4,39 | -0,31 | 7,58E-01 |
| AT1G47270 | 0,13 | 0,00 | 4,41 | 0,00 | 1,00E+00 |
| AT1G07357 | 0,11 | 0,88 | 4,35 | 0,20 | 8,39E-01 |
| AT1G47480 | 1,07 | -0,16 | 1,88 | -0,08 | 9,33E-01 |
| AT1G47490 | 0,30 | 0,00 | 4,41 | 0,00 | 1,00E+00 |
| AT1G47500 | 0,37 | 0,00 | 4,36 | 0,00 | 1,00E+00 |
| AT1G47530 | 1,40 | -1,38 | 1,68 | -0,82 | 4,13E-01 |
| AT1G47550 | 0,50 | 1,30 | 4,15 | 0,31 | 7,54E-01 |
| AT1G47680 | 0,12 | 0,49 | 4,38 | 0,11 | 9,11E-01 |
| AT1G47690 | 0,26 | 0,25 | 4,37 | 0,06 | 9,55E-01 |
| AT1G47770 | 0,12 | 0,49 | 4,38 | 0,11 | 9,11E-01 |
| AT1G47890 | 1,24 | 0,34 | 2,15 | 0,16 | 8,74E-01 |
| AT1G47940 | 0,36 | 2,61 | 4,28 | 0,61 | 5,42E-01 |
| AT1G47970 | 1,63 | 0,02 | 1,59 | 0,01 | 9,89E-01 |
| AT1G48050 | 0,54 | -3,61 | 4,35 | -0,83 | 4,06E-01 |
| AT1G48100 | 0,12 | -0,26 | 4,41 | -0,06 | 9,53E-01 |
| AT1G48170 | 0,32 | 0,79 | 3,59 | 0,22 | 8,27E-01 |
| AT1G48210 | 0,24 | -0,26 | 4,41 | -0,06 | 9,53E-01 |
| AT1G48230 | 0,28 | -0,26 | 4,41 | -0,06 | 9,53E-01 |
| AT1G48315 | 0,17 | 0,00 | 4,41 | 0,00 | 1,00E+00 |
| AT1G48360 | 1,34 | -0,38 | 2,30 | -0,17 | 8,69E-01 |
| AT1G48460 | 0,51 | 1,02 | 3,08 | 0,33 | 7,41E-01 |
| AT1G07487 | 0,32 | 0,26 | 3,59 | 0,07 | 9,43E-01 |
| AT1G48480 | 0,38 | -0,28 | 4,27 | -0,07 | 9,48E-01 |
| AT1G48540 | 1,33 | 1,75 | 2,00 | 0,87 | 3,82E-01 |
| AT1G48560 | 0,14 | 1,29 | 4,33 | 0,30 | 7,65E-01 |
| AT1G48600 | 0,62 | -0,90 | 2,18 | -0,41 | 6,81E-01 |
| AT1G48640 | 0,26 | 0,00 | 4,41 | 0,00 | 1,00E+00 |
| AT1G48725 | 1,45 | 2,61 | 1,63 | 1,60 | 1,09E-01 |
| AT1G48820 | 0,24 | -1,76 | 4,37 | -0,40 | 6,87E-01 |
| AT1G49010 | 0,27 | 1,33 | 4,33 | 0,31 | 7,58E-01 |
| AT1G49032 | 0,71 | 1,66 | 2,21 | 0,75 | 4,53E-01 |
| AT1G49050 | 1,80 | 1,21 | 2,05 | 0,59 | 5,54E-01 |
| AT1G49190 | 1,12 | 1,68 | 1,97 | 0,85 | 3,94E-01 |
| AT1G49200 | 0,08 | 0,21 | 4,39 | 0,05 | 9,62E-01 |
| AT1G49210 | 0,15 | 0,02 | 4,36 | 0,00 | 9,96E-01 |
| AT1G49230 | 0,38 | 0,85 | 3,43 | 0,25 | 8,05E-01 |
| AT1G49245 | 0,90 | 1,34 | 2,17 | 0,62 | 5,38E-01 |
| AT1G49300 | 0,19 | 0,91 | 4,34 | 0,21 | 8,34E-01 |
| AT1G49340 | 1,22 | 0,52 | 2,05 | 0,25 | 8,00E-01 |
| AT1G49390 | 0,08 | 0,00 | 4,41 | 0,00 | 1,00E+00 |
| AT1G49470 | 0,15 | 0,90 | 4,35 | 0,21 | 8,36E-01 |
| AT1G49510 | 0,25 | -3,44 | 4,34 | -0,79 | 4,28E-01 |

| | | | | | |
|---|---|---|---|---|---|
| AT1G49680 | 0,43 | 1,11 | 3,27 | 0,34 | 7,35E-01 |
| AT1G49720 | 0,23 | -2,48 | 4,38 | -0,57 | 5,71E-01 |
| AT1G49820 | 0,28 | -2,10 | 4,40 | -0,48 | 6,33E-01 |
| AT1G49890 | 0,23 | -0,41 | 4,38 | -0,09 | 9,26E-01 |
| AT1G49930 | 0,39 | 0,00 | 4,40 | 0,00 | 1,00E+00 |
| AT1G49975 | 1,21 | 3,32 | 2,41 | 1,37 | 1,69E-01 |
| AT1G49980 | 0,53 | -0,38 | 3,22 | -0,12 | 9,07E-01 |
| AT1G49990 | 0,80 | 0,00 | 4,37 | 0,00 | 1,00E+00 |
| AT1G50050 | 0,43 | 1,29 | 3,18 | 0,41 | 6,84E-01 |
| AT1G07613 | 1,47 | 0,04 | 1,37 | 0,03 | 9,75E-01 |
| AT1G50120 | 0,25 | -0,40 | 4,38 | -0,09 | 9,27E-01 |
| AT1G50170 | 0,08 | 0,00 | 4,41 | 0,00 | 1,00E+00 |
| AT1G50240 | 1,24 | -1,05 | 2,02 | -0,52 | 6,02E-01 |
| AT1G50270 | 0,45 | -0,72 | 3,34 | -0,22 | 8,29E-01 |
| AT1G50350 | 0,30 | 0,84 | 4,29 | 0,20 | 8,44E-01 |
| AT1G50360 | 0,23 | 0,09 | 4,34 | 0,02 | 9,84E-01 |
| AT1G50370 | 0,39 | 0,47 | 3,46 | 0,14 | 8,92E-01 |
| AT1G50380 | 0,57 | 1,23 | 3,18 | 0,39 | 6,99E-01 |
| AT1G50410 | 1,51 | 0,38 | 2,55 | 0,15 | 8,81E-01 |
| AT1G50560 | 0,41 | -0,20 | 4,34 | -0,05 | 9,63E-01 |
| AT1G50660 | 1,10 | 2,88 | 1,88 | 1,54 | 1,25E-01 |
| AT1G50730 | 0,12 | 0,00 | 4,41 | 0,00 | 1,00E+00 |
| AT1G50740 | 0,72 | -0,37 | 2,97 | -0,12 | 9,01E-01 |
| AT1G50900 | 1,85 | -1,72 | 1,38 | -1,24 | 2,14E-01 |
| AT1G51060 | 0,42 | -1,29 | 4,19 | -0,31 | 7,58E-01 |
| AT1G51100 | 0,17 | 1,58 | 4,31 | 0,37 | 7,15E-01 |
| AT1G51110 | 0,52 | -1,50 | 3,12 | -0,48 | 6,31E-01 |
| AT1G51160 | 0,67 | -2,28 | 3,05 | -0,75 | 4,55E-01 |
| AT1G51172 | 1,67 | 3,27 | 1,68 | 1,94 | 5,18E-02 |
| AT1G51200 | 1,27 | 1,01 | 2,27 | 0,45 | 6,55E-01 |
| AT1G51350 | 0,44 | 0,52 | 3,30 | 0,16 | 8,74E-01 |
| AT1G51390 | 0,24 | -0,67 | 4,38 | -0,15 | 8,78E-01 |
| AT1G07737 | 0,83 | -0,13 | 2,10 | -0,06 | 9,52E-01 |
| AT1G51402 | 1,01 | -0,16 | 1,99 | -0,08 | 9,35E-01 |
| AT1G51440 | 0,14 | 0,89 | 4,35 | 0,21 | 8,37E-01 |
| AT1G51540 | 0,08 | 0,21 | 4,39 | 0,05 | 9,62E-01 |
| AT1G51580 | 0,35 | -0,97 | 4,31 | -0,23 | 8,21E-01 |
| AT1G51600 | 0,08 | 0,00 | 4,41 | 0,00 | 1,00E+00 |
| AT1G51645 | 0,08 | 0,00 | 4,41 | 0,00 | 1,00E+00 |
| AT1G51660 | 0,75 | 0,75 | 2,69 | 0,28 | 7,80E-01 |
| AT1G51670 | 0,25 | 0,00 | 4,41 | 0,00 | 1,00E+00 |
| AT1G51720 | 0,29 | -0,39 | 4,39 | -0,09 | 9,30E-01 |
| AT1G51745 | 0,21 | 0,00 | 4,41 | 0,00 | 1,00E+00 |
| AT1G51760 | 0,56 | 0,66 | 2,60 | 0,25 | 8,01E-01 |
| AT1G51950 | 0,22 | 1,16 | 4,34 | 0,27 | 7,89E-01 |
| AT1G52080 | 0,08 | 0,00 | 4,41 | 0,00 | 1,00E+00 |
| AT1G52190 | 0,40 | -0,37 | 3,42 | -0,11 | 9,14E-01 |
| AT1G52310 | 0,73 | 0,47 | 2,56 | 0,18 | 8,54E-01 |
| AT1G52320 | 0,99 | 0,72 | 1,76 | 0,41 | 6,82E-01 |
| AT1G52380 | 0,74 | -1,58 | 2,99 | -0,53 | 5,96E-01 |
| AT1G52410 | 0,68 | -2,77 | 2,54 | -1,09 | 2,76E-01 |
| AT1G52420 | 0,99 | 1,14 | 2,86 | 0,40 | 6,90E-01 |
| AT1G52500 | 0,39 | -2,23 | 4,40 | -0,51 | 6,12E-01 |
| AT1G52600 | 0,08 | 0,00 | 4,41 | 0,00 | 1,00E+00 |
| AT1G52618 | 0,68 | 3,28 | 2,80 | 1,17 | 2,41E-01 |
| AT1G52620 | 0,78 | -0,51 | 4,32 | -0,12 | 9,06E-01 |

| | | | | | |
|---|---|---|---|---|---|
| AT1G52720 | 0,19 | 1,60 | 4,31 | 0,37 | 7,11E-01 |
| AT1G52740 | 1,01 | 1,29 | 2,02 | 0,64 | 5,23E-01 |
| AT1G52760 | 0,33 | -0,17 | 3,56 | -0,05 | 9,63E-01 |
| AT1G52870 | 0,92 | 3,35 | 2,50 | 1,34 | 1,80E-01 |
| AT1G52905 | 0,68 | 2,78 | 2,78 | 1,00 | 3,17E-01 |
| AT1G52930 | 0,12 | 0,00 | 4,41 | 0,00 | 1,00E+00 |
| AT1G52940 | 0,76 | 1,97 | 2,42 | 0,81 | 4,17E-01 |
| AT1G52950 | 0,39 | 1,03 | 3,44 | 0,30 | 7,64E-01 |
| AT1G52980 | 1,44 | -0,48 | 1,98 | -0,25 | 8,06E-01 |
| AT1G53050 | 0,64 | 0,00 | 4,41 | 0,00 | 1,00E+00 |
| AT1G53090 | 0,57 | -2,60 | 4,36 | -0,60 | 5,50E-01 |
| AT1G53165 | 0,43 | -0,18 | 2,81 | -0,06 | 9,50E-01 |
| AT1G53210 | 1,83 | 0,86 | 1,47 | 0,59 | 5,57E-01 |
| AT1G53233 | 0,99 | 1,72 | 2,25 | 0,77 | 4,44E-01 |
| AT1G53300 | 0,67 | 2,65 | 2,54 | 1,04 | 2,98E-01 |
| AT1G53330 | 0,35 | 0,00 | 4,39 | 0,00 | 1,00E+00 |
| AT1G53350 | 0,25 | 0,29 | 4,37 | 0,07 | 9,47E-01 |
| AT1G53390 | 0,38 | 1,11 | 4,34 | 0,25 | 7,99E-01 |
| AT1G53400 | 0,40 | 0,00 | 4,41 | 0,00 | 1,00E+00 |
| AT1G53430 | 1,51 | -1,79 | 1,56 | -1,15 | 2,50E-01 |
| AT1G53440 | 1,17 | -0,35 | 1,46 | -0,24 | 8,12E-01 |
| AT1G07887 | 0,84 | 0,38 | 2,91 | 0,13 | 8,96E-01 |
| AT1G53490 | 0,92 | 0,50 | 2,38 | 0,21 | 8,35E-01 |
| AT1G07893 | 0,39 | -2,16 | 4,30 | -0,50 | 6,16E-01 |
| AT1G07897 | 0,95 | 0,23 | 1,78 | 0,13 | 8,97E-01 |
| AT1G53570 | 0,96 | -1,32 | 2,43 | -0,54 | 5,88E-01 |
| AT1G53590 | 0,26 | -0,02 | 4,35 | 0,00 | 9,97E-01 |
| AT1G53655 | 1,78 | -0,63 | 1,50 | -0,42 | 6,76E-01 |
| AT1G53670 | 0,12 | 0,00 | 4,41 | 0,00 | 1,00E+00 |
| AT1G53720 | 0,26 | -2,72 | 4,37 | -0,62 | 5,33E-01 |
| AT1G53730 | 1,50 | 0,41 | 2,21 | 0,19 | 8,52E-01 |
| AT1G53840 | 1,21 | 1,48 | 1,86 | 0,80 | 4,26E-01 |
| AT1G53870 | 0,49 | 2,22 | 3,90 | 0,57 | 5,70E-01 |
| AT1G53875 | 1,66 | -0,47 | 1,22 | -0,38 | 7,01E-01 |
| AT1G53885 | 0,24 | -0,53 | 4,37 | -0,12 | 9,03E-01 |
| AT1G53890 | 1,37 | 1,44 | 1,49 | 0,97 | 3,35E-01 |
| AT1G53910 | 0,62 | 1,57 | 2,43 | 0,65 | 5,17E-01 |
| AT1G54030 | 0,19 | 1,60 | 4,31 | 0,37 | 7,11E-01 |
| AT1G54130 | 1,27 | -2,48 | 1,98 | -1,26 | 2,09E-01 |
| AT1G54150 | 0,44 | 1,07 | 3,44 | 0,31 | 7,55E-01 |
| AT1G54200 | 0,80 | 2,61 | 2,45 | 1,07 | 2,85E-01 |
| AT1G54215 | 0,08 | 0,00 | 4,41 | 0,00 | 1,00E+00 |
| AT1G07957 | 0,34 | 1,89 | 3,42 | 0,55 | 5,82E-01 |
| AT1G54310 | 0,08 | 0,00 | 4,41 | 0,00 | 1,00E+00 |
| AT1G54340 | 0,28 | 0,38 | 4,36 | 0,09 | 9,31E-01 |
| AT1G54350 | 0,37 | -0,30 | 4,34 | -0,07 | 9,45E-01 |
| AT1G54360 | 0,57 | 0,00 | 4,41 | 0,00 | 1,00E+00 |
| AT1G54420 | 1,07 | 2,18 | 2,47 | 0,88 | 3,78E-01 |
| AT1G54440 | 0,25 | -0,31 | 4,40 | -0,07 | 9,44E-01 |
| AT1G54490 | 1,36 | 1,64 | 1,87 | 0,87 | 3,82E-01 |
| AT1G54550 | 0,56 | 2,65 | 2,88 | 0,92 | 3,59E-01 |
| AT1G54680 | 0,26 | 0,00 | 4,41 | 0,00 | 1,00E+00 |
| AT1G54720 | 0,66 | 2,18 | 3,00 | 0,73 | 4,67E-01 |
| AT1G54780 | 1,22 | -0,41 | 1,50 | -0,27 | 7,84E-01 |
| AT1G54820 | 0,40 | -0,81 | 4,32 | -0,19 | 8,51E-01 |
| AT1G54920 | 1,30 | -1,14 | 2,01 | -0,56 | 5,73E-01 |

| | | | | | |
|---|---|---|---|---|---|
| AT1G55000 | 0,70 | -0,61 | 2,63 | -0,23 | 8,16E-01 |
| AT1G55020 | 0,29 | 0,35 | 4,37 | 0,08 | 9,36E-01 |
| AT1G55040 | 0,41 | -0,39 | 4,24 | -0,09 | 9,27E-01 |
| AT1G55090 | 0,81 | 0,96 | 2,95 | 0,32 | 7,46E-01 |
| AT1G55120 | 1,23 | -1,48 | 1,55 | -0,96 | 3,39E-01 |
| AT1G55150 | 0,30 | 0,00 | 4,41 | 0,00 | 1,00E+00 |
| AT1G55160 | 0,99 | 2,19 | 2,38 | 0,92 | 3,57E-01 |
| AT1G55175 | 0,08 | 0,00 | 4,41 | 0,00 | 1,00E+00 |
| AT1G55320 | 0,25 | 0,00 | 4,41 | 0,00 | 1,00E+00 |
| AT1G55335 | 0,51 | -2,40 | 3,17 | -0,76 | 4,48E-01 |
| AT1G55370 | 0,24 | 0,00 | 4,41 | 0,00 | 1,00E+00 |
| AT1G55460 | 0,32 | -0,64 | 4,37 | -0,15 | 8,83E-01 |
| AT1G55480 | 0,48 | 0,45 | 4,12 | 0,11 | 9,13E-01 |
| AT1G55500 | 0,61 | 0,46 | 2,36 | 0,20 | 8,45E-01 |
| AT1G55510 | 0,20 | -2,08 | 4,39 | -0,47 | 6,35E-01 |
| AT1G55840 | 0,33 | -0,40 | 4,38 | -0,09 | 9,27E-01 |
| AT1G55850 | 0,50 | -2,60 | 4,38 | -0,59 | 5,52E-01 |
| AT1G08073 | 0,33 | -2,09 | 4,39 | -0,48 | 6,35E-01 |
| AT1G55890 | 0,12 | -0,36 | 4,39 | -0,08 | 9,34E-01 |
| AT1G55910 | 0,98 | 0,63 | 1,97 | 0,32 | 7,49E-01 |
| AT1G55915 | 0,81 | 1,67 | 2,99 | 0,56 | 5,75E-01 |
| AT1G08077 | 0,62 | 2,11 | 2,91 | 0,73 | 4,67E-01 |
| AT1G56045 | 0,75 | -2,35 | 2,58 | -0,91 | 3,62E-01 |
| AT1G56050 | 1,07 | 0,00 | 2,82 | 0,00 | 1,00E+00 |
| AT1G56190 | 1,11 | 0,46 | 2,14 | 0,22 | 8,29E-01 |
| AT1G56200 | 1,42 | -0,95 | 1,72 | -0,55 | 5,80E-01 |
| AT1G56210 | 0,05 | 0,18 | 4,38 | 0,04 | 9,68E-01 |
| AT1G56220 | 1,48 | 0,01 | 1,49 | 0,00 | 9,97E-01 |
| AT1G56450 | 0,83 | 0,11 | 2,19 | 0,05 | 9,58E-01 |
| AT1G56460 | 0,70 | 1,02 | 2,94 | 0,35 | 7,28E-01 |
| AT1G56500 | 0,95 | -1,05 | 1,89 | -0,56 | 5,77E-01 |
| AT1G56540 | 0,56 | 0,00 | 4,41 | 0,00 | 1,00E+00 |
| AT1G56560 | 0,52 | 0,00 | 3,96 | 0,00 | 1,00E+00 |
| AT1G56570 | 0,28 | 0,38 | 4,36 | 0,09 | 9,31E-01 |
| AT1G56600 | 0,19 | -0,23 | 4,35 | -0,05 | 9,58E-01 |
| AT1G56610 | 0,17 | -3,06 | 4,35 | -0,70 | 4,82E-01 |
| AT1G56660 | 1,02 | 1,87 | 1,76 | 1,07 | 2,86E-01 |
| AT1G56700 | 0,33 | -0,84 | 3,55 | -0,24 | 8,14E-01 |
| AT1G57620 | 0,88 | -1,82 | 3,09 | -0,59 | 5,57E-01 |
| AT1G57630 | 0,41 | 0,42 | 4,36 | 0,10 | 9,24E-01 |
| AT1G57650 | 0,46 | -0,67 | 2,64 | -0,26 | 7,99E-01 |
| AT1G57720 | 1,72 | 0,41 | 1,65 | 0,25 | 8,02E-01 |
| AT1G57760 | 0,12 | 0,35 | 4,37 | 0,08 | 9,35E-01 |
| AT1G57765 | 0,15 | 0,47 | 4,37 | 0,11 | 9,14E-01 |
| AT1G58190 | 0,62 | 1,07 | 2,59 | 0,41 | 6,79E-01 |
| AT1G58215 | 0,57 | 0,00 | 4,40 | 0,00 | 1,00E+00 |
| AT1G58223 | 0,54 | -0,26 | 3,92 | -0,07 | 9,47E-01 |
| AT1G58260 | 0,12 | 0,27 | 4,38 | 0,06 | 9,50E-01 |
| AT1G58360 | 0,57 | -1,38 | 3,89 | -0,35 | 7,23E-01 |
| AT1G58525 | 0,11 | -0,26 | 4,41 | -0,06 | 9,53E-01 |
| AT1G58520 | 0,55 | -0,13 | 3,29 | -0,04 | 9,69E-01 |
| AT1G58643 | 0,42 | -0,77 | 4,36 | -0,18 | 8,60E-01 |
| AT1G58807 | 1,55 | 0,82 | 1,79 | 0,46 | 6,46E-01 |
| AT1G58808 | 1,22 | -1,90 | 1,94 | -0,98 | 3,27E-01 |
| AT1G58848 | 1,34 | 1,16 | 2,05 | 0,57 | 5,70E-01 |
| AT1G59124 | 1,27 | -0,73 | 1,68 | -0,44 | 6,62E-01 |

| | | | | | |
|---|---|---|---|---|---|
| AT1G59218 | 1,28 | 1,66 | 1,91 | 0,87 | 3,85E-01 |
| AT1G59590 | 0,59 | 1,05 | 3,21 | 0,33 | 7,44E-01 |
| AT1G59620 | 0,81 | 0,00 | 4,39 | 0,00 | 1,00E+00 |
| AT1G59650 | 0,54 | 0,23 | 3,90 | 0,06 | 9,54E-01 |
| AT1G59710 | 0,56 | -2,42 | 3,26 | -0,74 | 4,57E-01 |
| AT1G59750 | 1,67 | -2,09 | 2,60 | -0,80 | 4,21E-01 |
| AT1G59820 | 1,45 | 2,06 | 2,11 | 0,98 | 3,29E-01 |
| AT1G59890 | 0,93 | 1,57 | 2,48 | 0,63 | 5,27E-01 |
| AT1G59900 | 1,24 | 0,45 | 1,74 | 0,26 | 7,93E-01 |
| AT1G60025 | 0,05 | 0,18 | 4,38 | 0,04 | 9,68E-01 |
| AT1G60073 | 1,58 | 3,72 | 1,87 | 1,99 | 4,68E-02 |
| AT1G60170 | 1,29 | 1,89 | 2,39 | 0,79 | 4,30E-01 |
| AT1G60190 | 0,89 | 0,52 | 2,29 | 0,23 | 8,21E-01 |
| AT1G60220 | 0,08 | 0,00 | 4,41 | 0,00 | 1,00E+00 |
| AT1G60420 | 0,70 | -0,80 | 2,91 | -0,27 | 7,85E-01 |
| AT1G60545 | 1,07 | 0,00 | 3,13 | 0,00 | 1,00E+00 |
| AT1G60650 | 0,21 | 0,00 | 4,41 | 0,00 | 1,00E+00 |
| AT1G60720 | 1,62 | 4,42 | 1,90 | 2,32 | 2,01E-02 |
| AT1G60900 | 0,65 | -0,63 | 3,20 | -0,20 | 8,43E-01 |
| AT1G08353 | 0,98 | -1,55 | 1,98 | -0,78 | 4,34E-01 |
| AT1G61030 | 0,52 | -2,26 | 3,98 | -0,57 | 5,71E-01 |
| AT1G61040 | 0,54 | 0,23 | 3,90 | 0,06 | 9,54E-01 |
| AT1G61150 | 0,81 | -1,68 | 3,64 | -0,46 | 6,45E-01 |
| AT1G61170 | 0,33 | 0,43 | 4,36 | 0,10 | 9,21E-01 |
| AT1G61180 | 0,58 | -0,36 | 2,77 | -0,13 | 8,97E-01 |
| AT1G61210 | 1,07 | -1,82 | 1,81 | -1,00 | 3,16E-01 |
| AT1G61224 | 1,62 | 0,46 | 1,64 | 0,28 | 7,78E-01 |
| AT1G61250 | 0,18 | 1,20 | 4,34 | 0,28 | 7,82E-01 |
| AT1G61280 | 0,12 | 0,00 | 4,41 | 0,00 | 1,00E+00 |
| AT1G61350 | 0,76 | 3,14 | 2,28 | 1,38 | 1,69E-01 |
| AT1G61450 | 0,12 | 0,00 | 4,41 | 0,00 | 1,00E+00 |
| AT1G61470 | 0,18 | 0,93 | 4,34 | 0,21 | 8,30E-01 |
| AT1G61610 | 0,12 | 0,35 | 4,37 | 0,08 | 9,35E-01 |
| AT1G61620 | 0,37 | -2,19 | 4,41 | -0,50 | 6,20E-01 |
| AT1G61790 | 1,42 | -2,13 | 1,95 | -1,09 | 2,76E-01 |
| AT1G61800 | 0,56 | 0,00 | 4,41 | 0,00 | 1,00E+00 |
| AT1G61820 | 0,08 | 0,00 | 4,41 | 0,00 | 1,00E+00 |
| AT1G61850 | 0,80 | -1,63 | 2,35 | -0,69 | 4,89E-01 |
| AT1G61990 | 0,32 | -0,32 | 4,40 | -0,07 | 9,42E-01 |
| AT1G62170 | 0,36 | 0,53 | 4,37 | 0,12 | 9,04E-01 |
| AT1G08477 | 1,64 | 0,98 | 1,54 | 0,64 | 5,24E-01 |
| AT1G62270 | 0,29 | 2,25 | 4,29 | 0,52 | 6,00E-01 |
| AT1G08517 | 0,77 | -0,70 | 2,31 | -0,30 | 7,63E-01 |
| AT1G62380 | 1,16 | -0,66 | 1,57 | -0,42 | 6,72E-01 |
| AT1G62410 | 0,32 | -0,98 | 4,28 | -0,23 | 8,19E-01 |
| AT1G62480 | 0,12 | 0,35 | 4,37 | 0,08 | 9,35E-01 |
| AT1G62560 | 1,49 | 0,76 | 2,95 | 0,26 | 7,96E-01 |
| AT1G62570 | 0,39 | -0,18 | 4,33 | -0,04 | 9,67E-01 |
| AT1G62620 | 0,57 | -0,39 | 3,11 | -0,12 | 9,01E-01 |
| AT1G62660 | 1,32 | -1,43 | 1,93 | -0,74 | 4,58E-01 |
| AT1G62720 | 0,24 | -0,07 | 4,30 | -0,02 | 9,87E-01 |
| AT1G62790 | 0,51 | 0,00 | 4,41 | 0,00 | 1,00E+00 |
| AT1G62895 | 1,43 | -2,25 | 1,73 | -1,30 | 1,93E-01 |
| AT1G62920 | 0,97 | 0,50 | 2,82 | 0,18 | 8,60E-01 |
| AT1G62970 | 0,33 | 0,12 | 3,52 | 0,03 | 9,72E-01 |
| AT1G62975 | 0,08 | 0,00 | 4,41 | 0,00 | 1,00E+00 |

| | | | | | |
|---|---|---|---|---|---|
| AT1G63000 | 0,67 | 2,08 | 3,78 | 0,55 | 5,82E-01 |
| AT1G63110 | 0,32 | -0,29 | 4,40 | -0,07 | 9,47E-01 |
| AT1G63170 | 0,08 | 0,00 | 4,41 | 0,00 | 1,00E+00 |
| AT1G63430 | 0,74 | 0,66 | 3,93 | 0,17 | 8,66E-01 |
| AT1G63500 | 0,59 | 0,65 | 2,67 | 0,24 | 8,07E-01 |
| AT1G63540 | 0,37 | 2,60 | 4,28 | 0,61 | 5,44E-01 |
| AT1G63680 | 0,43 | 0,87 | 3,26 | 0,27 | 7,89E-01 |
| AT1G63690 | 1,03 | 0,58 | 1,87 | 0,31 | 7,55E-01 |
| AT1G63730 | 0,24 | 2,12 | 4,29 | 0,49 | 6,21E-01 |
| AT1G63830 | 0,49 | 1,68 | 4,05 | 0,41 | 6,78E-01 |
| AT1G63850 | 0,53 | -1,24 | 3,97 | -0,31 | 7,56E-01 |
| AT1G08643 | 0,79 | 2,48 | 2,93 | 0,85 | 3,97E-01 |
| AT1G63900 | 0,95 | 1,59 | 2,18 | 0,73 | 4,66E-01 |
| AT1G63940 | 0,81 | 0,38 | 1,99 | 0,19 | 8,48E-01 |
| AT1G63950 | 1,81 | 0,53 | 1,62 | 0,33 | 7,43E-01 |
| AT1G64060 | 0,86 | 1,37 | 2,41 | 0,57 | 5,69E-01 |
| AT1G64065 | 0,29 | 0,84 | 4,36 | 0,19 | 8,47E-01 |
| AT1G64090 | 0,96 | -2,90 | 2,20 | -1,32 | 1,87E-01 |
| AT1G64100 | 0,61 | -1,94 | 3,77 | -0,52 | 6,06E-01 |
| AT1G64120 | 0,13 | 0,94 | 4,35 | 0,22 | 8,29E-01 |
| AT1G64130 | 0,12 | 0,00 | 4,41 | 0,00 | 1,00E+00 |
| AT1G08657 | 0,60 | -0,85 | 2,64 | -0,32 | 7,47E-01 |
| AT1G64230 | 1,39 | -3,11 | 1,82 | -1,71 | 8,80E-02 |
| AT1G64370 | 0,17 | 0,22 | 4,32 | 0,05 | 9,59E-01 |
| AT1G64390 | 1,48 | -2,04 | 2,07 | -0,98 | 3,25E-01 |
| AT1G64520 | 0,27 | 1,31 | 4,33 | 0,30 | 7,62E-01 |
| AT1G64530 | 0,54 | -2,17 | 3,18 | -0,68 | 4,95E-01 |
| AT1G64550 | 0,64 | -0,14 | 2,68 | -0,05 | 9,59E-01 |
| AT1G64570 | 0,71 | -1,08 | 2,56 | -0,42 | 6,74E-01 |
| AT1G64572 | 0,51 | 0,00 | 4,41 | 0,00 | 1,00E+00 |
| AT1G64620 | 0,45 | 2,06 | 3,20 | 0,64 | 5,20E-01 |
| AT1G64680 | 1,48 | -2,13 | 1,60 | -1,33 | 1,85E-01 |
| AT1G64740 | 0,25 | 1,91 | 4,30 | 0,45 | 6,56E-01 |
| AT1G64770 | 1,29 | -0,68 | 1,58 | -0,43 | 6,65E-01 |
| AT1G64795 | 0,74 | 0,58 | 1,83 | 0,32 | 7,50E-01 |
| AT1G64810 | 0,44 | -0,27 | 4,36 | -0,06 | 9,50E-01 |
| AT1G64820 | 0,08 | 0,00 | 4,41 | 0,00 | 1,00E+00 |
| AT1G64860 | 0,65 | -2,76 | 3,12 | -0,88 | 3,76E-01 |
| AT1G64900 | 1,03 | -1,09 | 2,02 | -0,54 | 5,89E-01 |
| AT1G64950 | 0,34 | -0,09 | 3,46 | -0,03 | 9,79E-01 |
| AT1G64960 | 0,60 | -1,32 | 3,99 | -0,33 | 7,40E-01 |
| AT1G65010 | 1,62 | -1,92 | 1,61 | -1,20 | 2,32E-01 |
| AT1G65220 | 0,59 | 0,02 | 2,72 | 0,01 | 9,95E-01 |
| AT1G65230 | 1,04 | 0,37 | 2,17 | 0,17 | 8,64E-01 |
| AT1G65260 | 0,21 | 1,90 | 4,30 | 0,44 | 6,58E-01 |
| AT1G65270 | 0,21 | 1,18 | 4,34 | 0,27 | 7,86E-01 |
| AT1G65280 | 0,39 | 0,62 | 4,29 | 0,14 | 8,86E-01 |
| AT1G65295 | 0,05 | 0,18 | 4,38 | 0,04 | 9,68E-01 |
| AT1G65360 | 0,90 | -4,33 | 2,30 | -1,88 | 5,95E-02 |
| AT1G08753 | 0,15 | -2,07 | 4,39 | -0,47 | 6,38E-01 |
| AT1G65490 | 0,49 | 0,65 | 4,35 | 0,15 | 8,81E-01 |
| AT1G65590 | 0,23 | -0,41 | 4,38 | -0,09 | 9,26E-01 |
| AT1G65681 | 0,20 | 1,82 | 4,30 | 0,42 | 6,72E-01 |
| AT1G65730 | 0,08 | 0,00 | 4,41 | 0,00 | 1,00E+00 |
| AT1G65790 | 0,93 | 0,28 | 1,97 | 0,14 | 8,87E-01 |
| AT1G65800 | 1,17 | 0,33 | 1,59 | 0,21 | 8,33E-01 |

| | | | | | |
|---|---|---|---|---|---|
| AT1G65820 | 0,38 | 1,58 | 2,84 | 0,56 | 5,78E-01 |
| AT1G65840 | 0,61 | 0,62 | 3,84 | 0,16 | 8,72E-01 |
| AT1G08793 | 0,15 | 0,90 | 4,35 | 0,21 | 8,36E-01 |
| AT1G66090 | 0,59 | -0,40 | 3,25 | -0,12 | 9,02E-01 |
| AT1G66150 | 1,55 | -1,12 | 1,26 | -0,89 | 3,73E-01 |
| AT1G66160 | 0,48 | -1,36 | 4,10 | -0,33 | 7,40E-01 |
| AT1G66180 | 0,53 | 2,13 | 3,12 | 0,68 | 4,96E-01 |
| AT1G66230 | 0,24 | 0,00 | 4,41 | 0,00 | 1,00E+00 |
| AT1G66340 | 0,60 | -1,34 | 3,04 | -0,44 | 6,61E-01 |
| AT1G66345 | 0,47 | -0,28 | 4,08 | -0,07 | 9,46E-01 |
| AT1G66430 | 0,50 | -3,31 | 4,31 | -0,77 | 4,43E-01 |
| AT1G66465 | 0,33 | -2,14 | 4,40 | -0,49 | 6,27E-01 |
| AT1G66580 | 1,55 | -0,22 | 1,64 | -0,13 | 8,94E-01 |
| AT1G66640 | 0,21 | 0,05 | 4,34 | 0,01 | 9,92E-01 |
| AT1G66750 | 0,31 | -2,19 | 4,41 | -0,50 | 6,20E-01 |
| AT1G66820 | 0,53 | -2,87 | 4,38 | -0,66 | 5,12E-01 |
| AT1G66900 | 0,20 | -0,15 | 4,39 | -0,03 | 9,72E-01 |
| AT1G67105 | 0,86 | -0,80 | 1,70 | -0,47 | 6,40E-01 |
| AT1G67190 | 0,34 | 0,79 | 4,34 | 0,18 | 8,55E-01 |
| AT1G08923 | 0,11 | 0,88 | 4,35 | 0,20 | 8,39E-01 |
| AT1G67420 | 0,13 | 0,00 | 4,41 | 0,00 | 1,00E+00 |
| AT1G67430 | 1,82 | 0,68 | 1,37 | 0,50 | 6,20E-01 |
| AT1G08937 | 0,46 | 2,12 | 3,21 | 0,66 | 5,08E-01 |
| AT1G67470 | 0,43 | 0,11 | 3,35 | 0,03 | 9,74E-01 |
| AT1G67480 | 0,49 | 0,22 | 4,37 | 0,05 | 9,60E-01 |
| AT1G67530 | 0,54 | -0,31 | 3,96 | -0,08 | 9,38E-01 |
| AT1G67560 | 0,64 | 0,00 | 4,40 | 0,00 | 1,00E+00 |
| AT1G67620 | 0,15 | 0,82 | 4,35 | 0,19 | 8,50E-01 |
| AT1G67635 | 1,69 | 4,16 | 1,80 | 2,31 | 2,08E-02 |
| AT1G67700 | 0,46 | 2,17 | 3,90 | 0,56 | 5,79E-01 |
| AT1G67720 | 0,53 | 0,00 | 3,87 | 0,00 | 1,00E+00 |
| AT1G67792 | 0,38 | 0,00 | 4,41 | 0,00 | 1,00E+00 |
| AT1G67900 | 1,04 | -0,66 | 2,34 | -0,28 | 7,77E-01 |
| AT1G67940 | 0,66 | 0,00 | 3,04 | 0,00 | 1,00E+00 |
| AT1G08963 | 0,39 | 0,00 | 4,40 | 0,00 | 1,00E+00 |
| AT1G67980 | 0,18 | -2,46 | 4,37 | -0,56 | 5,73E-01 |
| AT1G67990 | 0,55 | -2,70 | 3,83 | -0,70 | 4,81E-01 |
| AT1G68050 | 0,24 | -1,16 | 4,40 | -0,26 | 7,92E-01 |
| AT1G68100 | 0,43 | -2,19 | 4,19 | -0,52 | 6,02E-01 |
| AT1G68110 | 0,27 | -2,13 | 4,39 | -0,49 | 6,27E-01 |
| AT1G68310 | 0,12 | -0,26 | 4,41 | -0,06 | 9,53E-01 |
| AT1G68440 | 1,32 | 2,18 | 1,73 | 1,26 | 2,08E-01 |
| AT1G68490 | 0,97 | 1,05 | 2,53 | 0,42 | 6,78E-01 |
| AT1G68570 | 0,58 | 0,62 | 2,55 | 0,24 | 8,07E-01 |
| AT1G68580 | 1,44 | 0,79 | 1,74 | 0,46 | 6,48E-01 |
| AT1G68585 | 0,27 | 0,13 | 4,38 | 0,03 | 9,76E-01 |
| AT1G68620 | 0,33 | 0,00 | 4,41 | 0,00 | 1,00E+00 |
| AT1G68650 | 0,32 | 0,00 | 4,41 | 0,00 | 1,00E+00 |
| AT1G68670 | 0,63 | 1,15 | 2,97 | 0,39 | 6,99E-01 |
| AT1G68720 | 1,50 | -1,82 | 2,10 | -0,87 | 3,84E-01 |
| AT1G68840 | 1,23 | -2,28 | 1,71 | -1,33 | 1,82E-01 |
| AT1G68890 | 1,09 | -1,11 | 2,15 | -0,51 | 6,07E-01 |
| AT1G68980 | 0,24 | 0,00 | 4,41 | 0,00 | 1,00E+00 |
| AT1G69085 | 0,08 | 0,47 | 4,38 | 0,11 | 9,15E-01 |
| AT1G69160 | 0,39 | 1,35 | 3,26 | 0,41 | 6,79E-01 |
| AT1G69170 | 0,40 | 0,00 | 4,41 | 0,00 | 1,00E+00 |

| | | | | | |
|---|---|---|---|---|---|
| AT1G69200 | 0,21 | 0,08 | 4,38 | 0,02 | 9,85E-01 |
| AT1G69252 | 0,90 | -0,07 | 2,86 | -0,02 | 9,81E-01 |
| AT1G69260 | 0,63 | 0,06 | 2,55 | 0,02 | 9,82E-01 |
| AT1G69350 | 0,78 | -0,98 | 2,51 | -0,39 | 6,95E-01 |
| AT1G69390 | 0,12 | 0,11 | 4,38 | 0,03 | 9,80E-01 |
| AT1G69410 | 0,27 | -1,97 | 4,36 | -0,45 | 6,51E-01 |
| AT1G69420 | 0,65 | -0,34 | 2,63 | -0,13 | 8,98E-01 |
| AT1G69510 | 0,10 | 0,86 | 4,35 | 0,20 | 8,43E-01 |
| AT1G69523 | 0,10 | 0,86 | 4,35 | 0,20 | 8,43E-01 |
| AT1G69526 | 0,08 | 0,00 | 4,41 | 0,00 | 1,00E+00 |
| AT1G69530 | 0,93 | 1,19 | 1,87 | 0,63 | 5,26E-01 |
| AT1G69580 | 0,26 | -0,91 | 4,38 | -0,21 | 8,36E-01 |
| AT1G69610 | 0,21 | 0,05 | 4,38 | 0,01 | 9,91E-01 |
| AT1G09087 | 0,37 | -0,78 | 3,52 | -0,22 | 8,25E-01 |
| AT1G69690 | 0,67 | 0,26 | 2,40 | 0,11 | 9,12E-01 |
| AT1G69740 | 0,99 | 2,68 | 1,89 | 1,42 | 1,56E-01 |
| AT1G69750 | 0,10 | 0,86 | 4,35 | 0,20 | 8,43E-01 |
| AT1G69870 | 0,38 | -1,44 | 4,38 | -0,33 | 7,43E-01 |
| AT1G70060 | 1,48 | 1,28 | 2,19 | 0,58 | 5,59E-01 |
| AT1G70090 | 0,99 | 1,79 | 2,08 | 0,86 | 3,90E-01 |
| AT1G70100 | 0,38 | -1,40 | 4,34 | -0,32 | 7,46E-01 |
| AT1G70160 | 1,19 | -0,77 | 1,60 | -0,48 | 6,32E-01 |
| AT1G70180 | 1,28 | -0,28 | 2,13 | -0,13 | 8,93E-01 |
| AT1G70190 | 0,40 | -0,32 | 4,40 | -0,07 | 9,42E-01 |
| AT1G70210 | 0,79 | 1,25 | 2,26 | 0,55 | 5,80E-01 |
| AT1G70230 | 0,59 | -0,13 | 2,59 | -0,05 | 9,60E-01 |
| AT1G70250 | 0,71 | -2,19 | 3,04 | -0,72 | 4,73E-01 |
| AT1G70330 | 0,33 | 0,70 | 3,53 | 0,20 | 8,43E-01 |
| AT1G70480 | 1,12 | 0,98 | 1,95 | 0,50 | 6,17E-01 |
| AT1G70510 | 0,12 | 0,00 | 4,41 | 0,00 | 1,00E+00 |
| AT1G70550 | 0,14 | 0,81 | 4,35 | 0,19 | 8,52E-01 |
| AT1G70570 | 0,36 | -2,40 | 4,37 | -0,55 | 5,83E-01 |
| AT1G70580 | 0,81 | -3,16 | 2,48 | -1,28 | 2,02E-01 |
| AT1G70620 | 1,32 | 0,60 | 2,29 | 0,26 | 7,92E-01 |
| AT1G70630 | 0,36 | -0,11 | 4,39 | -0,02 | 9,80E-01 |
| AT1G70645 | 0,05 | 0,18 | 4,38 | 0,04 | 9,68E-01 |
| AT1G70650 | 0,26 | -0,03 | 4,37 | -0,01 | 9,95E-01 |
| AT1G70660 | 0,20 | 0,00 | 4,41 | 0,00 | 1,00E+00 |
| AT1G70700 | 0,60 | 0,20 | 2,47 | 0,08 | 9,35E-01 |
| AT1G70760 | 0,35 | -0,24 | 3,49 | -0,07 | 9,45E-01 |
| AT1G70810 | 0,62 | -2,91 | 2,66 | -1,09 | 2,74E-01 |
| AT1G70820 | 0,43 | -1,00 | 2,75 | -0,36 | 7,15E-01 |
| AT1G70895 | 0,08 | 0,21 | 4,39 | 0,05 | 9,62E-01 |
| AT1G70900 | 1,03 | -0,48 | 2,08 | -0,23 | 8,18E-01 |
| AT1G70960 | 0,12 | 0,35 | 4,37 | 0,08 | 9,35E-01 |
| AT1G70980 | 0,40 | -0,38 | 4,34 | -0,09 | 9,30E-01 |
| AT1G71010 | 1,50 | 2,61 | 1,94 | 1,35 | 1,78E-01 |
| AT1G71090 | 0,35 | 0,59 | 3,56 | 0,17 | 8,68E-01 |
| AT1G71100 | 1,11 | -0,94 | 1,66 | -0,57 | 5,71E-01 |
| AT1G71110 | 0,15 | 0,47 | 4,37 | 0,11 | 9,15E-01 |
| AT1G71130 | 0,37 | 0,14 | 4,38 | 0,03 | 9,74E-01 |
| AT1G71180 | 0,42 | 0,29 | 4,30 | 0,07 | 9,46E-01 |
| AT1G71270 | 0,45 | 1,02 | 4,16 | 0,24 | 8,07E-01 |
| AT1G71300 | 0,38 | 0,00 | 4,41 | 0,00 | 1,00E+00 |
| AT1G71320 | 0,39 | 2,63 | 4,13 | 0,64 | 5,24E-01 |
| AT1G71480 | 0,80 | -0,08 | 2,96 | -0,03 | 9,79E-01 |

| | | | | | |
|---|---|---|---|---|---|
| AT1G09257 | 0,17 | 0,00 | 4,41 | 0,00 | 1,00E+00 |
| AT1G71500 | 0,95 | -0,15 | 1,92 | -0,08 | 9,40E-01 |
| AT1G71695 | 0,82 | -0,24 | 2,52 | -0,10 | 9,24E-01 |
| AT1G71720 | 0,53 | -4,40 | 3,78 | -1,17 | 2,44E-01 |
| AT1G71800 | 0,67 | 0,00 | 3,72 | 0,00 | 1,00E+00 |
| AT1G71820 | 1,53 | 0,02 | 1,42 | 0,02 | 9,86E-01 |
| AT1G71840 | 1,05 | 3,87 | 2,10 | 1,85 | 6,48E-02 |
| AT1G71900 | 0,51 | 0,00 | 4,41 | 0,00 | 1,00E+00 |
| AT1G71910 | 0,44 | 1,52 | 3,16 | 0,48 | 6,30E-01 |
| AT1G71920 | 0,73 | 1,50 | 2,31 | 0,65 | 5,18E-01 |
| AT1G71930 | 1,13 | 2,07 | 2,05 | 1,01 | 3,12E-01 |
| AT1G71970 | 0,14 | 0,31 | 4,37 | 0,07 | 9,44E-01 |
| AT1G71980 | 0,46 | 2,11 | 3,12 | 0,68 | 4,98E-01 |
| AT1G72010 | 0,42 | -0,19 | 3,30 | -0,06 | 9,55E-01 |
| AT1G72030 | 0,75 | 0,25 | 3,68 | 0,07 | 9,45E-01 |
| AT1G72050 | 0,75 | 0,00 | 3,72 | 0,00 | 1,00E+00 |
| AT1G72090 | 0,34 | -0,26 | 4,41 | -0,06 | 9,53E-01 |
| AT1G72240 | 0,60 | -0,11 | 3,91 | -0,03 | 9,77E-01 |
| AT1G72310 | 0,25 | 0,00 | 4,41 | 0,00 | 1,00E+00 |
| AT1G72330 | 0,67 | 0,06 | 2,93 | 0,02 | 9,83E-01 |
| AT1G72470 | 0,43 | 0,00 | 4,41 | 0,00 | 1,00E+00 |
| AT1G72510 | 0,57 | -1,00 | 3,11 | -0,32 | 7,48E-01 |
| AT1G72520 | 0,91 | 1,48 | 2,00 | 0,74 | 4,59E-01 |
| AT1G72530 | 0,05 | 0,18 | 4,38 | 0,04 | 9,68E-01 |
| AT1G09333 | 1,54 | 0,28 | 1,47 | 0,19 | 8,49E-01 |
| AT1G72650 | 0,79 | 0,93 | 2,40 | 0,39 | 7,00E-01 |
| AT1G72700 | 1,31 | 2,19 | 1,88 | 1,17 | 2,43E-01 |
| AT1G72710 | 1,73 | -0,09 | 1,48 | -0,06 | 9,52E-01 |
| AT1G72750 | 0,63 | 1,90 | 2,18 | 0,87 | 3,83E-01 |
| AT1G72770 | 1,51 | 1,23 | 1,69 | 0,73 | 4,64E-01 |
| AT1G72790 | 0,71 | 0,77 | 2,98 | 0,26 | 7,97E-01 |
| AT1G72855 | 0,40 | 0,00 | 4,41 | 0,00 | 1,00E+00 |
| AT1G72900 | 0,32 | -0,47 | 3,56 | -0,13 | 8,94E-01 |
| AT1G72910 | 1,29 | 2,08 | 1,81 | 1,15 | 2,52E-01 |
| AT1G72920 | 0,44 | -0,54 | 3,41 | -0,16 | 8,73E-01 |
| AT1G72940 | 0,97 | 0,00 | 4,10 | 0,00 | 1,00E+00 |
| AT1G72970 | 0,16 | 0,15 | 4,38 | 0,04 | 9,72E-01 |
| AT1G73030 | 0,82 | 2,33 | 2,38 | 0,98 | 3,28E-01 |
| AT1G73060 | 0,24 | 0,77 | 4,35 | 0,18 | 8,60E-01 |
| AT1G73080 | 1,38 | 0,91 | 1,68 | 0,54 | 5,88E-01 |
| AT1G73100 | 0,70 | 0,01 | 2,64 | 0,00 | 9,96E-01 |
| AT1G73180 | 1,25 | -0,34 | 2,17 | -0,16 | 8,75E-01 |
| AT1G73430 | 1,05 | 0,27 | 2,28 | 0,12 | 9,04E-01 |
| AT1G73450 | 1,12 | 1,36 | 2,00 | 0,68 | 4,96E-01 |
| AT1G73460 | 0,79 | 1,03 | 2,53 | 0,41 | 6,85E-01 |
| AT1G73470 | 0,23 | -0,34 | 4,40 | -0,08 | 9,38E-01 |
| AT1G73480 | 1,37 | -2,64 | 2,11 | -1,25 | 2,10E-01 |
| AT1G73600 | 1,37 | -1,57 | 1,87 | -0,84 | 4,01E-01 |
| AT1G73630 | 0,10 | 0,08 | 4,38 | 0,02 | 9,85E-01 |
| AT1G73670 | 0,46 | -1,61 | 3,35 | -0,48 | 6,31E-01 |
| AT1G73710 | 0,22 | 0,20 | 4,38 | 0,05 | 9,63E-01 |
| AT1G73750 | 0,72 | 0,00 | 4,41 | 0,00 | 1,00E+00 |
| AT1G73770 | 0,08 | 0,00 | 4,41 | 0,00 | 1,00E+00 |
| AT1G73850 | 0,17 | 0,07 | 4,38 | 0,02 | 9,88E-01 |
| AT1G73875 | 0,39 | 0,11 | 4,25 | 0,03 | 9,79E-01 |
| AT1G73885 | 0,62 | 0,38 | 3,09 | 0,12 | 9,03E-01 |

| | | | | | |
|---|---|---|---|---|---|
| AT1G73920 | 0,62 | 1,52 | 2,56 | 0,59 | 5,53E-01 |
| AT1G73970 | 0,95 | -0,88 | 2,26 | -0,39 | 6,96E-01 |
| AT1G74160 | 1,08 | -1,54 | 1,90 | -0,81 | 4,16E-01 |
| AT1G74250 | 1,42 | 2,59 | 1,84 | 1,41 | 1,59E-01 |
| AT1G74290 | 1,78 | 1,17 | 1,60 | 0,73 | 4,64E-01 |
| AT1G74360 | 1,21 | -0,30 | 2,60 | -0,11 | 9,09E-01 |
| AT1G74380 | 1,29 | 2,62 | 1,81 | 1,45 | 1,47E-01 |
| AT1G74430 | 0,38 | -0,28 | 4,32 | -0,06 | 9,49E-01 |
| AT1G74450 | 0,99 | 3,02 | 1,94 | 1,56 | 1,20E-01 |
| AT1G74580 | 0,25 | 0,09 | 4,38 | 0,02 | 9,84E-01 |
| AT1G74680 | 0,57 | 0,64 | 3,25 | 0,20 | 8,44E-01 |
| AT1G74710 | 0,60 | -0,46 | 2,58 | -0,18 | 8,58E-01 |
| AT1G74730 | 0,44 | -1,35 | 3,26 | -0,41 | 6,80E-01 |
| AT1G74750 | 0,50 | 0,00 | 4,41 | 0,00 | 1,00E+00 |
| AT1G74780 | 0,63 | -0,36 | 3,04 | -0,12 | 9,06E-01 |
| AT1G74790 | 0,51 | 0,63 | 3,95 | 0,16 | 8,74E-01 |
| AT1G74870 | 1,10 | -0,37 | 3,36 | -0,11 | 9,13E-01 |
| AT1G74930 | 0,81 | 2,55 | 2,62 | 0,97 | 3,32E-01 |
| AT1G74940 | 0,58 | 1,60 | 3,04 | 0,53 | 5,99E-01 |
| AT1G75080 | 0,21 | -0,29 | 4,40 | -0,06 | 9,48E-01 |
| AT1G09533 | 1,00 | 0,58 | 1,84 | 0,31 | 7,53E-01 |
| AT1G75163 | 0,94 | 0,83 | 1,93 | 0,43 | 6,68E-01 |
| AT1G75166 | 1,22 | 2,37 | 1,97 | 1,20 | 2,29E-01 |
| AT1G75170 | 0,12 | -0,26 | 4,41 | -0,06 | 9,53E-01 |
| AT1G09543 | 0,38 | 0,77 | 3,34 | 0,23 | 8,17E-01 |
| AT1G75190 | 0,22 | 0,26 | 4,34 | 0,06 | 9,52E-01 |
| AT1G75210 | 0,77 | -3,81 | 2,87 | -1,33 | 1,84E-01 |
| AT1G75280 | 0,22 | 1,63 | 4,31 | 0,38 | 7,05E-01 |
| AT1G75310 | 0,47 | -2,61 | 4,36 | -0,60 | 5,50E-01 |
| AT1G75350 | 0,83 | 0,10 | 2,18 | 0,05 | 9,63E-01 |
| AT1G75400 | 0,32 | 0,87 | 4,34 | 0,20 | 8,41E-01 |
| AT1G75420 | 0,31 | 0,07 | 4,38 | 0,02 | 9,87E-01 |
| AT1G75440 | 0,28 | -0,36 | 4,39 | -0,08 | 9,35E-01 |
| AT1G75460 | 0,84 | -2,31 | 2,27 | -1,02 | 3,09E-01 |
| AT1G75580 | 0,17 | -2,21 | 4,40 | -0,50 | 6,16E-01 |
| AT1G75710 | 0,31 | 0,72 | 4,29 | 0,17 | 8,68E-01 |
| AT1G75785 | 0,12 | 0,27 | 4,38 | 0,06 | 9,50E-01 |
| AT1G75800 | 1,71 | -2,12 | 1,98 | -1,07 | 2,85E-01 |
| AT1G75810 | 0,60 | -1,15 | 2,71 | -0,42 | 6,72E-01 |
| AT1G75840 | 0,75 | -0,81 | 2,27 | -0,35 | 7,23E-01 |
| AT1G75900 | 0,32 | 1,29 | 4,33 | 0,30 | 7,66E-01 |
| AT1G75950 | 1,80 | -1,71 | 1,24 | -1,38 | 1,67E-01 |
| AT1G75960 | 0,45 | 0,58 | 4,26 | 0,14 | 8,91E-01 |
| AT1G76000 | 0,58 | 0,87 | 3,02 | 0,29 | 7,74E-01 |
| AT1G76050 | 0,66 | -0,94 | 3,05 | -0,31 | 7,57E-01 |
| AT1G76140 | 0,48 | 1,09 | 4,00 | 0,27 | 7,85E-01 |
| AT1G76185 | 0,16 | 0,31 | 4,37 | 0,07 | 9,44E-01 |
| AT1G76190 | 0,11 | 0,88 | 4,35 | 0,20 | 8,39E-01 |
| AT1G76320 | 0,32 | 0,43 | 4,36 | 0,10 | 9,22E-01 |
| AT1G76350 | 0,12 | -0,36 | 4,39 | -0,08 | 9,34E-01 |
| AT1G76390 | 0,51 | 0,67 | 3,21 | 0,21 | 8,34E-01 |
| AT1G76450 | 0,08 | 0,00 | 4,41 | 0,00 | 1,00E+00 |
| AT1G76520 | 0,73 | -1,06 | 2,20 | -0,48 | 6,28E-01 |
| AT1G76580 | 0,80 | 0,64 | 3,55 | 0,18 | 8,56E-01 |
| AT1G76590 | 0,69 | -0,76 | 2,83 | -0,27 | 7,89E-01 |
| AT1G76600 | 0,49 | 1,33 | 3,21 | 0,42 | 6,78E-01 |

| | | | | | |
|---|---|---|---|---|---|
| AT1G76620 | 0,25 | -1,50 | 4,34 | -0,34 | 7,30E-01 |
| AT1G76680 | 0,14 | -0,82 | 4,37 | -0,19 | 8,50E-01 |
| AT1G76740 | 0,12 | 0,00 | 4,41 | 0,00 | 1,00E+00 |
| AT1G76780 | 1,47 | -1,06 | 1,78 | -0,60 | 5,52E-01 |
| AT1G76820 | 0,53 | -1,09 | 3,20 | -0,34 | 7,34E-01 |
| AT1G76850 | 1,15 | -0,82 | 1,88 | -0,44 | 6,61E-01 |
| AT1G76880 | 1,06 | -0,53 | 1,64 | -0,32 | 7,47E-01 |
| AT1G76900 | 0,91 | -1,25 | 2,74 | -0,46 | 6,48E-01 |
| AT1G76920 | 0,59 | -0,29 | 3,87 | -0,07 | 9,41E-01 |
| AT1G76950 | 0,83 | -1,14 | 2,78 | -0,41 | 6,82E-01 |
| AT1G76990 | 1,49 | 0,26 | 1,37 | 0,19 | 8,51E-01 |
| AT1G77080 | 0,41 | 0,36 | 3,23 | 0,11 | 9,11E-01 |
| AT1G77140 | 0,25 | 0,28 | 4,39 | 0,06 | 9,50E-01 |
| AT1G77235 | 1,77 | -0,39 | 1,60 | -0,24 | 8,09E-01 |
| AT1G77320 | 0,26 | 0,00 | 4,41 | 0,00 | 1,00E+00 |
| AT1G77350 | 0,15 | 0,90 | 4,35 | 0,21 | 8,36E-01 |
| AT1G77420 | 0,12 | 0,00 | 4,41 | 0,00 | 1,00E+00 |
| AT1G77460 | 0,94 | 0,55 | 2,70 | 0,21 | 8,37E-01 |
| AT1G77490 | 0,64 | 0,76 | 2,96 | 0,26 | 7,97E-01 |
| AT1G77500 | 0,37 | -1,09 | 4,39 | -0,25 | 8,04E-01 |
| AT1G77510 | 1,26 | 0,05 | 1,94 | 0,03 | 9,78E-01 |
| AT1G77610 | 0,53 | -0,02 | 4,34 | 0,00 | 9,97E-01 |
| AT1G77710 | 0,32 | 1,57 | 4,32 | 0,36 | 7,17E-01 |
| AT1G77740 | 1,74 | -1,32 | 1,78 | -0,74 | 4,61E-01 |
| AT1G77800 | 1,13 | 0,50 | 2,05 | 0,24 | 8,07E-01 |
| AT1G77840 | 0,31 | 0,43 | 3,62 | 0,12 | 9,05E-01 |
| AT1G77850 | 1,01 | 1,67 | 2,15 | 0,78 | 4,38E-01 |
| AT1G77890 | 0,98 | -1,98 | 2,03 | -0,97 | 3,30E-01 |
| AT1G77930 | 1,14 | -1,10 | 2,30 | -0,48 | 6,33E-01 |
| AT1G78010 | 0,90 | 1,27 | 2,30 | 0,55 | 5,83E-01 |
| AT1G78070 | 1,66 | -1,12 | 1,40 | -0,80 | 4,21E-01 |
| AT1G78100 | 0,64 | 0,23 | 2,15 | 0,11 | 9,16E-01 |
| AT1G78150 | 0,22 | 0,00 | 4,37 | 0,00 | 1,00E+00 |
| AT1G78200 | 0,54 | 0,00 | 2,60 | 0,00 | 9,99E-01 |
| AT1G78280 | 1,59 | 1,35 | 2,27 | 0,59 | 5,52E-01 |
| AT1G78300 | 0,54 | 0,63 | 3,16 | 0,20 | 8,42E-01 |
| AT1G78420 | 0,54 | 0,59 | 2,75 | 0,21 | 8,30E-01 |
| AT1G78490 | 0,05 | 0,18 | 4,38 | 0,04 | 9,68E-01 |
| AT1G78500 | 0,17 | 0,00 | 4,41 | 0,00 | 1,00E+00 |
| AT1G78600 | 0,20 | 0,40 | 4,37 | 0,09 | 9,27E-01 |
| AT1G78630 | 1,31 | 0,21 | 1,76 | 0,12 | 9,05E-01 |
| AT1G78670 | 0,52 | -0,81 | 3,14 | -0,26 | 7,96E-01 |
| AT1G78680 | 0,90 | 1,45 | 1,93 | 0,75 | 4,52E-01 |
| AT1G78700 | 0,37 | -0,34 | 4,39 | -0,08 | 9,39E-01 |
| AT1G78710 | 0,38 | 0,00 | 4,41 | 0,00 | 1,00E+00 |
| AT1G78770 | 0,21 | -0,29 | 4,40 | -0,06 | 9,48E-01 |
| AT1G78870 | 0,28 | 0,03 | 4,38 | 0,01 | 9,95E-01 |
| AT1G78890 | 0,36 | -2,09 | 4,39 | -0,47 | 6,35E-01 |
| AT1G78910 | 0,26 | 0,00 | 4,41 | 0,00 | 1,00E+00 |
| AT1G78920 | 1,61 | -0,44 | 1,70 | -0,26 | 7,97E-01 |
| AT1G79060 | 0,14 | 1,29 | 4,33 | 0,30 | 7,65E-01 |
| AT1G79103 | 0,05 | 0,18 | 4,38 | 0,04 | 9,68E-01 |
| AT1G79110 | 1,67 | 0,58 | 1,54 | 0,38 | 7,06E-01 |
| AT1G09847 | 0,44 | -0,37 | 3,12 | -0,12 | 9,07E-01 |
| AT1G79190 | 0,88 | 0,00 | 2,85 | 0,00 | 1,00E+00 |
| AT1G79230 | 0,71 | -0,27 | 2,99 | -0,09 | 9,27E-01 |

| | | | | | |
|---|---|---|---|---|---|
| AT1G79240 | 0,20 | -0,26 | 4,41 | -0,06 | 9,53E-01 |
| AT1G79270 | 1,18 | 0,04 | 1,88 | 0,02 | 9,82E-01 |
| AT1G79340 | 0,91 | -0,28 | 1,86 | -0,15 | 8,81E-01 |
| AT1G79380 | 0,91 | 0,32 | 2,92 | 0,11 | 9,12E-01 |
| AT1G79560 | 0,28 | 0,42 | 4,36 | 0,10 | 9,24E-01 |
| AT1G79590 | 0,18 | 0,15 | 4,38 | 0,03 | 9,73E-01 |
| AT1G79660 | 0,63 | 0,00 | 3,73 | 0,00 | 1,00E+00 |
| AT1G09903 | 0,08 | 0,00 | 4,41 | 0,00 | 1,00E+00 |
| AT1G79810 | 0,41 | 1,37 | 3,28 | 0,42 | 6,75E-01 |
| AT1G79840 | 0,77 | -0,23 | 3,05 | -0,08 | 9,39E-01 |
| AT1G79870 | 0,21 | 0,51 | 4,36 | 0,12 | 9,08E-01 |
| AT1G79890 | 0,21 | 0,00 | 4,41 | 0,00 | 1,00E+00 |
| AT1G79910 | 0,41 | 0,00 | 4,22 | 0,00 | 1,00E+00 |
| AT1G79940 | 1,20 | -1,26 | 1,56 | -0,81 | 4,19E-01 |
| AT1G79950 | 0,49 | -2,09 | 3,99 | -0,52 | 6,00E-01 |
| AT1G79970 | 0,27 | 0,80 | 4,35 | 0,18 | 8,54E-01 |
| AT1G79985 | 0,75 | 1,47 | 2,46 | 0,60 | 5,50E-01 |
| AT1G80010 | 1,24 | -1,29 | 1,81 | -0,71 | 4,78E-01 |
| AT1G80160 | 0,37 | -0,26 | 4,40 | -0,06 | 9,52E-01 |
| AT1G80260 | 0,59 | -2,06 | 3,02 | -0,68 | 4,96E-01 |
| AT1G80300 | 1,04 | 0,00 | 3,37 | 0,00 | 1,00E+00 |
| AT1G80310 | 0,64 | -1,03 | 2,61 | -0,39 | 6,94E-01 |
| AT1G80380 | 1,15 | -1,84 | 1,70 | -1,08 | 2,79E-01 |
| AT1G09963 | 0,96 | -1,14 | 2,61 | -0,43 | 6,64E-01 |
| AT1G80430 | 0,19 | 0,53 | 4,37 | 0,12 | 9,03E-01 |
| AT1G80510 | 0,61 | 0,62 | 3,84 | 0,16 | 8,72E-01 |
| AT1G80620 | 0,35 | -1,61 | 4,34 | -0,37 | 7,11E-01 |
| AT1G80770 | 0,54 | 0,32 | 3,20 | 0,10 | 9,21E-01 |
| AT1G80780 | 0,60 | 1,35 | 3,11 | 0,44 | 6,63E-01 |
| AT1G80790 | 0,69 | -2,03 | 2,94 | -0,69 | 4,89E-01 |
| AT1G80810 | 0,38 | -0,27 | 4,31 | -0,06 | 9,49E-01 |
| AT1G80840 | 1,09 | 3,83 | 2,38 | 1,61 | 1,08E-01 |
| AT1G80870 | 0,27 | 0,42 | 4,36 | 0,10 | 9,23E-01 |
| AT2G01070 | 0,81 | -0,26 | 2,61 | -0,10 | 9,20E-01 |
| AT2G01080 | 0,35 | 0,89 | 3,50 | 0,26 | 7,98E-01 |
| AT2G01100 | 0,12 | 0,35 | 4,37 | 0,08 | 9,35E-01 |
| AT2G01120 | 0,92 | 0,00 | 2,93 | 0,00 | 1,00E+00 |
| AT2G01190 | 0,50 | 0,64 | 3,11 | 0,21 | 8,37E-01 |
| AT2G01400 | 0,39 | -0,33 | 4,40 | -0,08 | 9,40E-01 |
| AT2G03975 | 0,34 | -0,03 | 3,53 | -0,01 | 9,93E-01 |
| AT2G01480 | 0,34 | 0,04 | 4,38 | 0,01 | 9,92E-01 |
| AT2G01620 | 0,60 | 1,22 | 2,56 | 0,47 | 6,35E-01 |
| AT2G01680 | 0,39 | 0,87 | 4,34 | 0,20 | 8,41E-01 |
| AT2G01750 | 0,24 | 0,52 | 4,31 | 0,12 | 9,04E-01 |
| AT2G01850 | 1,74 | -1,12 | 1,31 | -0,86 | 3,91E-01 |
| AT2G01860 | 0,98 | 1,76 | 2,47 | 0,71 | 4,75E-01 |
| AT2G01900 | 1,22 | 0,22 | 1,62 | 0,14 | 8,92E-01 |
| AT2G01980 | 0,63 | -0,86 | 3,12 | -0,28 | 7,82E-01 |
| AT2G02040 | 0,10 | 0,08 | 4,38 | 0,02 | 9,85E-01 |
| AT2G02050 | 0,58 | 1,27 | 2,48 | 0,51 | 6,07E-01 |
| AT2G02070 | 0,84 | -0,23 | 1,87 | -0,12 | 9,01E-01 |
| AT2G02090 | 1,22 | 1,37 | 2,40 | 0,57 | 5,69E-01 |
| AT2G02100 | 0,54 | -2,50 | 3,89 | -0,64 | 5,20E-01 |
| AT2G02440 | 0,68 | -1,86 | 2,97 | -0,63 | 5,30E-01 |
| AT2G02450 | 0,66 | 0,57 | 2,48 | 0,23 | 8,18E-01 |
| AT2G02470 | 0,18 | 0,31 | 4,38 | 0,07 | 9,44E-01 |

| | | | | | |
|---|---|---|---|---|---|
| AT2G02550 | 1,01 | 2,56 | 1,98 | 1,29 | 1,95E-01 |
| AT2G02560 | 1,80 | -0,27 | 1,57 | -0,17 | 8,61E-01 |
| AT2G02700 | 0,39 | 0,00 | 4,40 | 0,00 | 1,00E+00 |
| AT2G04295 | 0,12 | 0,11 | 4,38 | 0,03 | 9,80E-01 |
| AT2G02760 | 0,54 | -1,42 | 2,55 | -0,56 | 5,76E-01 |
| AT2G02790 | 0,39 | -1,75 | 4,24 | -0,41 | 6,80E-01 |
| AT2G02795 | 0,14 | 0,81 | 4,35 | 0,19 | 8,52E-01 |
| AT2G02810 | 0,62 | 0,63 | 3,10 | 0,20 | 8,40E-01 |
| AT2G04355 | 0,47 | 1,23 | 3,34 | 0,37 | 7,13E-01 |
| AT2G04365 | 0,97 | 2,24 | 2,32 | 0,96 | 3,35E-01 |
| AT2G02870 | 1,53 | 1,27 | 1,92 | 0,66 | 5,07E-01 |
| AT2G02955 | 0,36 | 0,12 | 4,38 | 0,03 | 9,79E-01 |
| AT2G03070 | 0,25 | 0,00 | 4,41 | 0,00 | 1,00E+00 |
| AT2G03120 | 0,64 | -1,58 | 2,59 | -0,61 | 5,42E-01 |
| AT2G03130 | 0,12 | 0,00 | 4,41 | 0,00 | 1,00E+00 |
| AT2G03270 | 0,39 | 1,58 | 2,81 | 0,56 | 5,74E-01 |
| AT2G03410 | 0,43 | -1,34 | 4,15 | -0,32 | 7,47E-01 |
| AT2G03450 | 0,32 | -0,18 | 4,40 | -0,04 | 9,67E-01 |
| AT2G03500 | 0,52 | 0,59 | 3,13 | 0,19 | 8,50E-01 |
| AT2G03520 | 0,08 | 0,00 | 4,41 | 0,00 | 1,00E+00 |
| AT2G03550 | 0,42 | -2,96 | 4,16 | -0,71 | 4,78E-01 |
| AT2G03567 | 0,47 | 0,15 | 3,28 | 0,05 | 9,63E-01 |
| AT2G03680 | 0,82 | -0,84 | 2,24 | -0,38 | 7,08E-01 |
| AT2G03710 | 0,17 | 0,00 | 4,41 | 0,00 | 1,00E+00 |
| AT2G03823 | 0,37 | 0,84 | 2,83 | 0,30 | 7,65E-01 |
| AT2G03870 | 0,41 | 0,00 | 4,19 | 0,00 | 1,00E+00 |
| AT2G03932 | 0,20 | 1,82 | 4,30 | 0,42 | 6,72E-01 |
| AT2G03931 | 0,05 | 0,18 | 4,38 | 0,04 | 9,68E-01 |
| AT2G03933 | 0,14 | 1,29 | 4,33 | 0,30 | 7,65E-01 |
| AT2G03936 | 0,18 | 0,90 | 4,35 | 0,21 | 8,36E-01 |
| AT2G03965 | 0,22 | 1,82 | 4,30 | 0,42 | 6,72E-01 |
| AT2G03980 | 0,82 | -1,20 | 2,52 | -0,48 | 6,33E-01 |
| AT2G04039 | 0,42 | 0,00 | 4,18 | 0,00 | 1,00E+00 |
| AT2G04735 | 0,15 | -0,28 | 4,35 | -0,06 | 9,50E-01 |
| AT2G04235 | 0,05 | 0,18 | 4,38 | 0,04 | 9,68E-01 |
| AT2G04270 | 1,17 | -0,39 | 2,22 | -0,17 | 8,61E-01 |
| AT2G04340 | 0,38 | 0,00 | 4,41 | 0,00 | 1,00E+00 |
| AT2G04350 | 1,50 | -2,32 | 1,61 | -1,44 | 1,51E-01 |
| AT2G04380 | 0,56 | 0,52 | 2,69 | 0,19 | 8,46E-01 |
| AT2G04390 | 0,85 | 0,98 | 2,20 | 0,45 | 6,55E-01 |
| AT2G04400 | 0,61 | -0,08 | 2,55 | -0,03 | 9,76E-01 |
| AT2G04450 | 0,21 | -0,29 | 4,40 | -0,06 | 9,48E-01 |
| AT2G04495 | 0,43 | 0,68 | 3,40 | 0,20 | 8,40E-01 |
| AT2G04550 | 0,15 | -2,07 | 4,39 | -0,47 | 6,38E-01 |
| AT2G04700 | 0,71 | 0,15 | 2,28 | 0,06 | 9,49E-01 |
| AT2G04740 | 0,38 | 0,00 | 4,41 | 0,00 | 1,00E+00 |
| AT2G04780 | 0,24 | 0,14 | 4,38 | 0,03 | 9,74E-01 |
| AT2G04795 | 0,15 | 0,90 | 4,35 | 0,21 | 8,36E-01 |
| AT2G04850 | 0,08 | 0,00 | 4,41 | 0,00 | 1,00E+00 |
| AT2G04865 | 0,25 | 0,00 | 4,41 | 0,00 | 1,00E+00 |
| AT2G04900 | 0,19 | 0,37 | 4,36 | 0,08 | 9,32E-01 |
| AT2G05060 | 0,21 | 0,00 | 4,41 | 0,00 | 1,00E+00 |
| AT2G05170 | 1,80 | 0,07 | 1,70 | 0,04 | 9,66E-01 |
| AT2G05180 | 0,08 | 0,00 | 4,41 | 0,00 | 1,00E+00 |
| AT2G05210 | 0,12 | 0,27 | 4,38 | 0,06 | 9,50E-01 |
| AT2G05320 | 0,92 | 0,06 | 1,98 | 0,03 | 9,75E-01 |

| | | | | | |
|---|---|---|---|---|---|
| AT2G05350 | 0,23 | 0,71 | 4,36 | 0,16 | 8,70E-01 |
| AT2G05380 | 0,99 | -3,02 | 1,98 | -1,53 | 1,27E-01 |
| AT2G05540 | 0,30 | 1,65 | 4,31 | 0,38 | 7,02E-01 |
| AT2G04915 | 0,39 | -1,34 | 3,37 | -0,40 | 6,91E-01 |
| AT2G04935 | 1,06 | 3,27 | 1,87 | 1,75 | 8,00E-02 |
| AT2G05635 | 0,15 | 0,82 | 4,35 | 0,19 | 8,50E-01 |
| AT2G04985 | 0,95 | 0,59 | 2,03 | 0,29 | 7,70E-01 |
| AT2G05790 | 0,44 | -0,42 | 3,24 | -0,13 | 8,97E-01 |
| AT2G05812 | 1,46 | 2,09 | 1,74 | 1,20 | 2,30E-01 |
| AT2G05830 | 0,15 | 0,47 | 4,37 | 0,11 | 9,15E-01 |
| AT2G05840 | 0,17 | -2,21 | 4,40 | -0,50 | 6,16E-01 |
| AT2G05990 | 0,47 | -1,23 | 2,39 | -0,51 | 6,08E-01 |
| AT2G05995 | 0,63 | -1,72 | 2,96 | -0,58 | 5,62E-01 |
| AT2G06025 | 0,59 | 0,78 | 2,61 | 0,30 | 7,65E-01 |
| AT2G06095 | 0,70 | -3,63 | 3,45 | -1,05 | 2,93E-01 |
| AT2G06105 | 0,23 | 1,67 | 4,31 | 0,39 | 6,99E-01 |
| AT2G06510 | 1,80 | 1,00 | 1,53 | 0,65 | 5,14E-01 |
| AT2G06845 | 0,18 | 0,82 | 4,35 | 0,19 | 8,51E-01 |
| AT2G06850 | 0,75 | -1,78 | 2,47 | -0,72 | 4,72E-01 |
| AT2G06904 | 0,38 | 2,36 | 4,29 | 0,55 | 5,82E-01 |
| AT2G06990 | 1,10 | 1,03 | 1,93 | 0,53 | 5,95E-01 |
| AT2G07050 | 0,64 | 2,44 | 2,93 | 0,83 | 4,05E-01 |
| AT2G07200 | 0,15 | 1,21 | 4,33 | 0,28 | 7,81E-01 |
| AT2G07565 | 0,91 | 0,45 | 2,20 | 0,20 | 8,38E-01 |
| AT2G07749 | 0,59 | 0,20 | 3,08 | 0,07 | 9,47E-01 |
| AT2G07672 | 0,25 | 1,91 | 4,30 | 0,45 | 6,56E-01 |
| AT2G07745 | 0,19 | -1,98 | 4,34 | -0,46 | 6,48E-01 |
| AT2G07674 | 0,81 | 1,48 | 3,68 | 0,40 | 6,88E-01 |
| AT2G07623 | 0,23 | 0,71 | 4,36 | 0,16 | 8,70E-01 |
| AT2G07678 | 0,32 | -2,10 | 4,40 | -0,48 | 6,33E-01 |
| AT2G07669 | 0,33 | -0,15 | 4,33 | -0,03 | 9,72E-01 |
| AT2G07681 | 0,75 | 2,58 | 2,14 | 1,21 | 2,27E-01 |
| AT2G07631 | 0,18 | -0,12 | 4,35 | -0,03 | 9,78E-01 |
| AT2G05215 | 0,68 | -0,52 | 2,43 | -0,22 | 8,30E-01 |
| AT2G07695 | 1,27 | 0,94 | 1,82 | 0,52 | 6,05E-01 |
| AT2G07798 | 1,27 | 0,13 | 1,62 | 0,08 | 9,35E-01 |
| AT2G07667 | 1,00 | 0,77 | 2,17 | 0,36 | 7,21E-01 |
| AT2G07638 | 1,46 | -3,51 | 1,67 | -2,10 | 3,53E-02 |
| AT2G07707 | 0,32 | -0,34 | 4,35 | -0,08 | 9,37E-01 |
| AT2G07708 | 0,57 | -1,17 | 4,32 | -0,27 | 7,87E-01 |
| AT2G07715 | 0,94 | 0,12 | 1,65 | 0,07 | 9,41E-01 |
| AT2G07728 | 0,11 | -0,38 | 4,39 | -0,09 | 9,31E-01 |
| AT2G07656 | 0,60 | 1,88 | 2,92 | 0,64 | 5,21E-01 |
| AT2G07827 | 0,36 | -0,53 | 3,51 | -0,15 | 8,79E-01 |
| AT2G07787 | 0,79 | 1,14 | 2,12 | 0,54 | 5,90E-01 |
| AT2G07659 | 0,28 | 0,82 | 4,30 | 0,19 | 8,49E-01 |
| AT2G05355 | 1,78 | -1,10 | 1,33 | -0,83 | 4,09E-01 |
| AT2G07765 | 0,12 | 0,00 | 4,41 | 0,00 | 1,00E+00 |
| AT2G07741 | 0,27 | 0,55 | 4,32 | 0,13 | 8,99E-01 |
| AT2G07680 | 0,80 | 2,24 | 2,35 | 0,95 | 3,40E-01 |
| AT2G09990 | 0,53 | -0,73 | 3,11 | -0,24 | 8,13E-01 |
| AT2G05485 | 0,27 | 1,44 | 4,33 | 0,33 | 7,39E-01 |
| AT2G10545 | 0,08 | 0,21 | 4,39 | 0,05 | 9,62E-01 |
| AT2G10556 | 0,88 | 0,90 | 2,73 | 0,33 | 7,42E-01 |
| AT2G10965 | 0,86 | 1,10 | 1,92 | 0,57 | 5,67E-01 |
| AT2G11010 | 0,65 | 2,15 | 2,93 | 0,73 | 4,63E-01 |

| | | | | | |
|---|---|---|---|---|---|
| AT2G11035 | 0,48 | -0,52 | 4,03 | -0,13 | 8,98E-01 |
| AT2G11045 | 0,08 | 0,21 | 4,39 | 0,05 | 9,62E-01 |
| AT2G11225 | 0,91 | 2,25 | 1,86 | 1,21 | 2,27E-01 |
| AT2G11620 | 0,14 | 1,29 | 4,33 | 0,30 | 7,65E-01 |
| AT2G11851 | 0,57 | 2,92 | 2,92 | 1,00 | 3,17E-01 |
| AT2G05825 | 0,69 | 1,07 | 2,22 | 0,48 | 6,32E-01 |
| AT2G13100 | 0,26 | -0,85 | 4,35 | -0,19 | 8,45E-01 |
| AT2G13290 | 0,12 | 0,11 | 4,38 | 0,03 | 9,80E-01 |
| AT2G05885 | 0,14 | 1,29 | 4,33 | 0,30 | 7,65E-01 |
| AT2G13430 | 0,44 | 0,16 | 3,23 | 0,05 | 9,61E-01 |
| AT2G13500 | 0,11 | 0,88 | 4,35 | 0,20 | 8,39E-01 |
| AT2G13540 | 0,64 | -1,34 | 2,28 | -0,59 | 5,55E-01 |
| AT2G05925 | 0,12 | 0,35 | 4,37 | 0,08 | 9,35E-01 |
| AT2G13790 | 0,49 | 1,50 | 3,29 | 0,46 | 6,48E-01 |
| AT2G13845 | 0,67 | 1,23 | 2,28 | 0,54 | 5,90E-01 |
| AT2G13960 | 0,27 | 1,23 | 4,34 | 0,28 | 7,76E-01 |
| AT2G14070 | 0,19 | -0,02 | 4,35 | -0,01 | 9,96E-01 |
| AT2G14115 | 0,49 | 0,87 | 4,35 | 0,20 | 8,42E-01 |
| AT2G14395 | 0,63 | -2,36 | 4,32 | -0,55 | 5,85E-01 |
| AT2G14440 | 0,43 | -1,06 | 4,06 | -0,26 | 7,94E-01 |
| AT2G14530 | 0,21 | 1,42 | 4,32 | 0,33 | 7,42E-01 |
| AT2G14700 | 0,64 | 3,01 | 2,88 | 1,05 | 2,96E-01 |
| AT2G14740 | 1,25 | 0,12 | 2,58 | 0,05 | 9,64E-01 |
| AT2G14750 | 1,64 | -0,31 | 1,47 | -0,21 | 8,33E-01 |
| AT2G14765 | 0,40 | 2,24 | 4,30 | 0,52 | 6,02E-01 |
| AT2G14825 | 0,08 | 0,00 | 4,41 | 0,00 | 1,00E+00 |
| AT2G14835 | 0,23 | -0,33 | 4,40 | -0,08 | 9,40E-01 |
| AT2G14850 | 0,21 | -0,18 | 4,40 | -0,04 | 9,66E-01 |
| AT2G14870 | 0,17 | -3,06 | 4,35 | -0,70 | 4,82E-01 |
| AT2G14878 | 1,23 | 0,68 | 1,50 | 0,45 | 6,53E-01 |
| AT2G14890 | 0,77 | 0,98 | 2,16 | 0,45 | 6,50E-01 |
| AT2G06305 | 0,18 | 0,93 | 4,34 | 0,21 | 8,30E-01 |
| AT2G15042 | 0,11 | -0,38 | 4,39 | -0,09 | 9,31E-01 |
| AT2G15050 | 0,64 | 0,34 | 2,76 | 0,12 | 9,01E-01 |
| AT2G15080 | 0,92 | 1,18 | 2,12 | 0,56 | 5,77E-01 |
| AT2G15090 | 1,27 | 0,38 | 2,88 | 0,13 | 8,94E-01 |
| AT2G15300 | 0,31 | -2,50 | 4,38 | -0,57 | 5,68E-01 |
| AT2G15430 | 0,16 | 0,63 | 4,36 | 0,14 | 8,85E-01 |
| AT2G15450 | 0,29 | 0,50 | 4,37 | 0,12 | 9,08E-01 |
| AT2G15470 | 0,34 | 1,97 | 4,30 | 0,46 | 6,46E-01 |
| AT2G15580 | 0,29 | 0,40 | 3,66 | 0,11 | 9,13E-01 |
| AT2G15710 | 0,40 | -0,12 | 4,24 | -0,03 | 9,77E-01 |
| AT2G15730 | 0,08 | 0,21 | 4,39 | 0,05 | 9,62E-01 |
| AT2G15900 | 0,47 | 0,11 | 2,72 | 0,04 | 9,68E-01 |
| AT2G15960 | 1,09 | 0,62 | 2,05 | 0,30 | 7,64E-01 |
| AT2G06535 | 1,38 | 3,36 | 1,95 | 1,73 | 8,42E-02 |
| AT2G16019 | 1,05 | 2,84 | 2,08 | 1,37 | 1,71E-01 |
| AT2G16280 | 0,93 | 2,02 | 2,15 | 0,94 | 3,48E-01 |
| AT2G16365 | 0,96 | -0,30 | 2,23 | -0,14 | 8,92E-01 |
| AT2G16380 | 0,44 | 0,28 | 4,21 | 0,07 | 9,46E-01 |
| AT2G16390 | 0,40 | 0,00 | 4,41 | 0,00 | 1,00E+00 |
| AT2G16630 | 0,14 | 0,31 | 4,37 | 0,07 | 9,44E-01 |
| AT2G16700 | 0,28 | -1,17 | 4,40 | -0,27 | 7,90E-01 |
| AT2G16710 | 0,72 | 0,89 | 3,73 | 0,24 | 8,11E-01 |
| AT2G16800 | 0,26 | -0,69 | 4,32 | -0,16 | 8,74E-01 |
| AT2G16950 | 1,26 | -1,45 | 2,35 | -0,62 | 5,36E-01 |

| | | | | | |
|---|---|---|---|---|---|
| AT2G16970 | 1,25 | -3,95 | 1,80 | -2,19 | 2,83E-02 |
| AT2G17033 | 0,44 | -2,15 | 4,22 | -0,51 | 6,11E-01 |
| AT2G17040 | 1,57 | 2,41 | 1,82 | 1,33 | 1,84E-01 |
| AT2G17120 | 0,23 | 1,10 | 4,34 | 0,25 | 8,00E-01 |
| AT2G17140 | 0,42 | -0,35 | 3,42 | -0,10 | 9,19E-01 |
| AT2G17170 | 0,08 | 0,00 | 4,41 | 0,00 | 1,00E+00 |
| AT2G17240 | 0,52 | -2,23 | 3,15 | -0,71 | 4,78E-01 |
| AT2G17250 | 0,25 | 0,00 | 4,41 | 0,00 | 1,00E+00 |
| AT2G17265 | 1,56 | 0,12 | 1,62 | 0,07 | 9,42E-01 |
| AT2G17270 | 0,25 | 0,02 | 4,37 | 0,01 | 9,96E-01 |
| AT2G17290 | 0,78 | -0,14 | 2,34 | -0,06 | 9,51E-01 |
| AT2G06905 | 1,12 | -2,18 | 2,07 | -1,05 | 2,94E-01 |
| AT2G17360 | 0,51 | 0,14 | 3,21 | 0,05 | 9,64E-01 |
| AT2G17390 | 0,66 | -2,12 | 3,07 | -0,69 | 4,90E-01 |
| AT2G17410 | 1,45 | 0,96 | 1,89 | 0,51 | 6,12E-01 |
| AT2G17420 | 1,05 | 2,67 | 2,05 | 1,31 | 1,92E-01 |
| AT2G17440 | 1,80 | -0,30 | 2,03 | -0,15 | 8,82E-01 |
| AT2G17520 | 1,17 | 0,25 | 1,80 | 0,14 | 8,91E-01 |
| AT2G17530 | 0,50 | 0,00 | 4,06 | 0,00 | 1,00E+00 |
| AT2G17570 | 0,42 | 0,00 | 4,22 | 0,00 | 1,00E+00 |
| AT2G17580 | 0,66 | -2,10 | 3,12 | -0,67 | 5,01E-01 |
| AT2G17705 | 0,42 | 1,97 | 3,23 | 0,61 | 5,42E-01 |
| AT2G17720 | 0,20 | 0,00 | 4,41 | 0,00 | 1,00E+00 |
| AT2G17760 | 0,65 | 0,11 | 3,01 | 0,04 | 9,70E-01 |
| AT2G17787 | 0,33 | -0,06 | 4,39 | -0,01 | 9,90E-01 |
| AT2G17790 | 0,80 | 0,41 | 2,47 | 0,17 | 8,68E-01 |
| AT2G17905 | 0,41 | 0,84 | 3,30 | 0,25 | 8,00E-01 |
| AT2G17920 | 0,15 | 0,56 | 4,37 | 0,13 | 8,99E-01 |
| AT2G17980 | 1,01 | -0,21 | 2,71 | -0,08 | 9,39E-01 |
| AT2G17990 | 0,69 | -0,30 | 2,52 | -0,12 | 9,05E-01 |
| AT2G18000 | 0,29 | 0,00 | 4,41 | 0,00 | 1,00E+00 |
| AT2G18030 | 0,68 | -0,17 | 3,07 | -0,05 | 9,57E-01 |
| AT2G18040 | 0,20 | 0,89 | 4,36 | 0,20 | 8,39E-01 |
| AT2G18090 | 1,01 | 0,63 | 1,91 | 0,33 | 7,43E-01 |
| AT2G18100 | 0,48 | 0,87 | 4,06 | 0,21 | 8,31E-01 |
| AT2G18110 | 0,62 | -1,55 | 2,98 | -0,52 | 6,02E-01 |
| AT2G18390 | 0,28 | -0,36 | 4,39 | -0,08 | 9,35E-01 |
| AT2G18420 | 0,81 | 0,00 | 3,55 | 0,00 | 1,00E+00 |
| AT2G18530 | 0,38 | -0,54 | 3,46 | -0,16 | 8,76E-01 |
| AT2G18560 | 1,54 | -0,72 | 2,34 | -0,31 | 7,59E-01 |
| AT2G18630 | 0,16 | -0,26 | 4,41 | -0,06 | 9,53E-01 |
| AT2G18670 | 1,13 | 0,48 | 1,57 | 0,30 | 7,61E-01 |
| AT2G18680 | 0,16 | -0,22 | 4,40 | -0,05 | 9,60E-01 |
| AT2G18690 | 1,29 | 3,16 | 1,89 | 1,68 | 9,38E-02 |
| AT2G18700 | 0,27 | -1,62 | 4,37 | -0,37 | 7,11E-01 |
| AT2G18730 | 0,58 | -0,66 | 2,51 | -0,26 | 7,92E-01 |
| AT2G18735 | 0,22 | -0,38 | 4,37 | -0,09 | 9,30E-01 |
| AT2G18750 | 0,67 | 0,58 | 2,07 | 0,28 | 7,80E-01 |
| AT2G18760 | 1,48 | 2,03 | 1,76 | 1,15 | 2,50E-01 |
| AT2G18850 | 0,52 | 0,00 | 4,38 | 0,00 | 1,00E+00 |
| AT2G18880 | 0,25 | 0,00 | 4,41 | 0,00 | 1,00E+00 |
| AT2G18917 | 0,08 | 0,21 | 4,39 | 0,05 | 9,62E-01 |
| AT2G07095 | 0,55 | 0,47 | 3,10 | 0,15 | 8,79E-01 |
| AT2G18938 | 0,72 | 0,14 | 2,57 | 0,06 | 9,55E-01 |
| AT2G18950 | 0,45 | 1,02 | 4,16 | 0,24 | 8,07E-01 |
| AT2G19110 | 0,15 | -0,39 | 4,39 | -0,09 | 9,29E-01 |

| | | | | | |
|---|---|---|---|---|---|
| AT2G19130 | 0,89 | -0,15 | 2,09 | -0,07 | 9,43E-01 |
| AT2G19160 | 0,39 | -0,26 | 4,41 | -0,06 | 9,53E-01 |
| AT2G19260 | 0,46 | 0,81 | 4,06 | 0,20 | 8,42E-01 |
| AT2G19280 | 0,36 | 0,00 | 4,41 | 0,00 | 1,00E+00 |
| AT2G19420 | 0,28 | 1,41 | 4,32 | 0,33 | 7,44E-01 |
| AT2G19430 | 0,24 | -1,36 | 4,35 | -0,31 | 7,54E-01 |
| AT2G19450 | 0,42 | -2,13 | 4,27 | -0,50 | 6,18E-01 |
| AT2G07205 | 0,75 | 2,53 | 3,58 | 0,71 | 4,81E-01 |
| AT2G19480 | 1,78 | -0,66 | 1,54 | -0,43 | 6,69E-01 |
| AT2G19490 | 0,36 | 0,00 | 4,41 | 0,00 | 1,00E+00 |
| AT2G19520 | 0,55 | 1,81 | 4,05 | 0,45 | 6,55E-01 |
| AT2G19540 | 0,46 | 0,22 | 2,73 | 0,08 | 9,35E-01 |
| AT2G19572 | 0,52 | 2,71 | 2,99 | 0,91 | 3,65E-01 |
| AT2G19582 | 0,25 | 1,90 | 4,30 | 0,44 | 6,59E-01 |
| AT2G19600 | 0,64 | 0,45 | 3,88 | 0,12 | 9,07E-01 |
| AT2G19640 | 0,53 | 0,00 | 3,93 | 0,00 | 1,00E+00 |
| AT2G19710 | 1,44 | 2,41 | 2,11 | 1,14 | 2,54E-01 |
| AT2G19730 | 0,57 | 0,54 | 4,04 | 0,13 | 8,94E-01 |
| AT2G19740 | 0,67 | 0,43 | 2,18 | 0,20 | 8,44E-01 |
| AT2G19810 | 0,34 | 1,74 | 3,49 | 0,50 | 6,18E-01 |
| AT2G19880 | 0,75 | -1,27 | 2,56 | -0,50 | 6,20E-01 |
| AT2G20020 | 1,07 | -0,09 | 2,17 | -0,04 | 9,68E-01 |
| AT2G20100 | 0,33 | -0,26 | 4,41 | -0,06 | 9,53E-01 |
| AT2G20110 | 0,83 | 0,00 | 4,39 | 0,00 | 1,00E+00 |
| AT2G20142 | 0,28 | -0,22 | 4,40 | -0,05 | 9,60E-01 |
| AT2G20230 | 0,26 | -0,80 | 4,37 | -0,18 | 8,55E-01 |
| AT2G20260 | 1,52 | -1,69 | 1,57 | -1,08 | 2,79E-01 |
| AT2G20340 | 0,57 | -1,85 | 3,97 | -0,47 | 6,42E-01 |
| AT2G20360 | 0,22 | -0,83 | 4,36 | -0,19 | 8,49E-01 |
| AT2G20362 | 0,08 | 0,00 | 4,41 | 0,00 | 1,00E+00 |
| AT2G20370 | 1,01 | 1,61 | 2,54 | 0,64 | 5,25E-01 |
| AT2G20420 | 0,62 | -0,39 | 2,21 | -0,18 | 8,61E-01 |
| AT2G20450 | 0,40 | -2,14 | 4,21 | -0,51 | 6,12E-01 |
| AT2G20530 | 0,21 | -1,67 | 4,36 | -0,38 | 7,01E-01 |
| AT2G20690 | 1,05 | -0,74 | 2,42 | -0,31 | 7,59E-01 |
| AT2G20710 | 0,08 | 0,00 | 4,41 | 0,00 | 1,00E+00 |
| AT2G07335 | 0,46 | -2,29 | 4,21 | -0,54 | 5,87E-01 |
| AT2G20750 | 0,59 | -2,19 | 3,86 | -0,57 | 5,71E-01 |
| AT2G20770 | 0,17 | 0,00 | 4,41 | 0,00 | 1,00E+00 |
| AT2G20820 | 0,25 | -0,35 | 4,40 | -0,08 | 9,37E-01 |
| AT2G20860 | 1,36 | -0,71 | 1,75 | -0,41 | 6,85E-01 |
| AT2G20890 | 1,04 | -1,98 | 1,81 | -1,10 | 2,73E-01 |
| AT2G20920 | 0,65 | -0,31 | 2,62 | -0,12 | 9,05E-01 |
| AT2G20960 | 0,82 | -0,40 | 2,11 | -0,19 | 8,48E-01 |
| AT2G21160 | 0,91 | 0,04 | 3,08 | 0,01 | 9,90E-01 |
| AT2G21187 | 0,89 | 1,87 | 1,88 | 0,99 | 3,20E-01 |
| AT2G21190 | 0,23 | -1,08 | 4,33 | -0,25 | 8,03E-01 |
| AT2G21270 | 0,21 | 0,00 | 4,41 | 0,00 | 1,00E+00 |
| AT2G21380 | 1,04 | -0,65 | 1,80 | -0,36 | 7,19E-01 |
| AT2G21410 | 1,46 | -0,83 | 1,68 | -0,49 | 6,24E-01 |
| AT2G21520 | 0,84 | 0,19 | 2,15 | 0,09 | 9,28E-01 |
| AT2G21530 | 0,61 | -0,92 | 2,91 | -0,32 | 7,52E-01 |
| AT2G21550 | 0,95 | -2,49 | 2,45 | -1,01 | 3,10E-01 |
| AT2G21580 | 1,04 | 0,00 | 2,91 | 0,00 | 1,00E+00 |
| AT2G21620 | 1,16 | -2,43 | 1,94 | -1,25 | 2,10E-01 |
| AT2G21630 | 0,32 | 0,00 | 4,41 | 0,00 | 1,00E+00 |

| | | | | | |
|---|---|---|---|---|---|
| AT2G21770 | 0,18 | 0,31 | 4,38 | 0,07 | 9,44E-01 |
| AT2G21790 | 0,74 | 0,44 | 2,57 | 0,17 | 8,65E-01 |
| AT2G21850 | 1,02 | -0,40 | 2,70 | -0,15 | 8,83E-01 |
| AT2G07475 | 0,24 | -0,26 | 4,41 | -0,06 | 9,53E-01 |
| AT2G21960 | 0,80 | -0,86 | 2,67 | -0,32 | 7,48E-01 |
| AT2G07495 | 0,12 | -0,26 | 4,41 | -0,06 | 9,53E-01 |
| AT2G22080 | 0,65 | -1,25 | 2,69 | -0,47 | 6,41E-01 |
| AT2G22120 | 0,32 | 0,00 | 4,41 | 0,00 | 1,00E+00 |
| AT2G22425 | 0,10 | 0,86 | 4,35 | 0,20 | 8,43E-01 |
| AT2G22450 | 0,24 | -0,02 | 4,38 | 0,00 | 9,97E-01 |
| AT2G22480 | 0,23 | -0,41 | 4,38 | -0,09 | 9,26E-01 |
| AT2G22540 | 1,53 | 1,92 | 1,68 | 1,14 | 2,55E-01 |
| AT2G22670 | 0,80 | 3,41 | 2,23 | 1,53 | 1,27E-01 |
| AT2G22680 | 0,25 | 0,28 | 4,39 | 0,06 | 9,50E-01 |
| AT2G22720 | 0,99 | -0,33 | 2,85 | -0,12 | 9,07E-01 |
| AT2G07845 | 0,43 | -0,31 | 4,18 | -0,07 | 9,40E-01 |
| AT2G22790 | 0,08 | 0,00 | 4,41 | 0,00 | 1,00E+00 |
| AT2G22840 | 1,11 | -1,07 | 2,29 | -0,47 | 6,39E-01 |
| AT2G07850 | 0,72 | -1,17 | 1,83 | -0,64 | 5,24E-01 |
| AT2G22910 | 0,32 | 0,52 | 4,38 | 0,12 | 9,06E-01 |
| AT2G23080 | 0,38 | 0,71 | 4,36 | 0,16 | 8,71E-01 |
| AT2G23120 | 0,92 | 0,04 | 2,26 | 0,02 | 9,84E-01 |
| AT2G23130 | 0,41 | -0,12 | 2,72 | -0,05 | 9,64E-01 |
| AT2G23142 | 1,46 | -5,61 | 2,36 | -2,37 | 1,76E-02 |
| AT2G23200 | 1,64 | -0,14 | 1,73 | -0,08 | 9,35E-01 |
| AT2G23310 | 0,84 | 0,22 | 3,01 | 0,07 | 9,43E-01 |
| AT2G07895 | 0,15 | -0,31 | 4,40 | -0,07 | 9,43E-01 |
| AT2G23320 | 1,39 | 1,34 | 1,82 | 0,74 | 4,59E-01 |
| AT2G23420 | 0,47 | 0,00 | 4,41 | 0,00 | 1,00E+00 |
| AT2G23460 | 1,50 | -1,19 | 1,72 | -0,69 | 4,90E-01 |
| AT2G07915 | 1,63 | 0,51 | 2,07 | 0,25 | 8,06E-01 |
| AT2G23672 | 1,50 | -1,51 | 1,56 | -0,97 | 3,33E-01 |
| AT2G23680 | 0,31 | -2,21 | 4,40 | -0,50 | 6,15E-01 |
| AT2G23740 | 0,12 | 0,27 | 4,38 | 0,06 | 9,50E-01 |
| AT2G23890 | 0,37 | -1,28 | 3,54 | -0,36 | 7,19E-01 |
| AT2G23930 | 1,70 | -1,83 | 1,61 | -1,14 | 2,56E-01 |
| AT2G23985 | 0,17 | -3,06 | 4,35 | -0,70 | 4,82E-01 |
| AT2G24090 | 0,67 | -0,89 | 2,57 | -0,34 | 7,30E-01 |
| AT2G24180 | 1,36 | -2,53 | 2,55 | -0,99 | 3,21E-01 |
| AT2G24240 | 0,31 | 0,96 | 4,34 | 0,22 | 8,24E-01 |
| AT2G24280 | 1,45 | -1,52 | 1,46 | -1,04 | 2,99E-01 |
| AT2G24300 | 0,15 | 0,47 | 4,37 | 0,11 | 9,15E-01 |
| AT2G08015 | 0,35 | 0,21 | 4,37 | 0,05 | 9,62E-01 |
| AT2G24350 | 0,50 | 0,00 | 4,05 | 0,00 | 1,00E+00 |
| AT2G24390 | 0,92 | -3,70 | 2,89 | -1,28 | 2,01E-01 |
| AT2G24380 | 0,65 | -4,01 | 3,76 | -1,07 | 2,86E-01 |
| AT2G24420 | 1,14 | -2,42 | 1,99 | -1,21 | 2,26E-01 |
| AT2G08045 | 0,23 | 0,85 | 4,34 | 0,20 | 8,45E-01 |
| AT2G08050 | 0,32 | 1,54 | 4,32 | 0,36 | 7,21E-01 |
| AT2G24540 | 0,88 | 0,18 | 1,70 | 0,10 | 9,17E-01 |
| AT2G24590 | 0,49 | 0,00 | 4,02 | 0,00 | 1,00E+00 |
| AT2G24592 | 0,83 | 0,95 | 2,21 | 0,43 | 6,66E-01 |
| AT2G24765 | 0,69 | 0,00 | 3,00 | 0,00 | 1,00E+00 |
| AT2G25000 | 0,52 | -0,28 | 3,92 | -0,07 | 9,43E-01 |
| AT2G25080 | 0,61 | -0,27 | 2,56 | -0,10 | 9,17E-01 |
| AT2G25110 | 0,19 | 0,43 | 4,36 | 0,10 | 9,21E-01 |

| | | | | | |
|---|---|---|---|---|---|
| AT2G25171 | 1,84 | 1,79 | 1,63 | 1,09 | 2,74E-01 |
| AT2G25190 | 0,14 | 0,89 | 4,35 | 0,21 | 8,37E-01 |
| AT2G25370 | 1,47 | 0,38 | 1,77 | 0,22 | 8,29E-01 |
| AT2G25410 | 1,30 | 3,74 | 2,19 | 1,71 | 8,80E-02 |
| AT2G25480 | 0,05 | 0,18 | 4,38 | 0,04 | 9,68E-01 |
| AT2G25520 | 1,73 | 2,69 | 2,02 | 1,33 | 1,82E-01 |
| AT2G25560 | 0,73 | -0,26 | 2,94 | -0,09 | 9,29E-01 |
| AT2G25580 | 0,08 | 0,00 | 4,41 | 0,00 | 1,00E+00 |
| AT2G25590 | 0,21 | -0,18 | 4,40 | -0,04 | 9,66E-01 |
| AT2G25625 | 0,15 | -0,35 | 4,39 | -0,08 | 9,36E-01 |
| AT2G25630 | 0,50 | 2,52 | 3,05 | 0,83 | 4,09E-01 |
| AT2G25650 | 0,70 | 0,00 | 3,02 | 0,00 | 1,00E+00 |
| AT2G25790 | 0,18 | 1,20 | 4,34 | 0,28 | 7,82E-01 |
| AT2G25840 | 0,59 | 1,54 | 3,90 | 0,40 | 6,93E-01 |
| AT2G25870 | 1,69 | 2,07 | 1,82 | 1,14 | 2,55E-01 |
| AT2G25930 | 1,26 | 0,78 | 1,78 | 0,44 | 6,60E-01 |
| AT2G26060 | 1,00 | 0,00 | 4,37 | 0,00 | 1,00E+00 |
| AT2G26110 | 0,61 | 2,65 | 2,86 | 0,93 | 3,53E-01 |
| AT2G26170 | 0,47 | 0,46 | 3,26 | 0,14 | 8,88E-01 |
| AT2G26270 | 0,21 | 0,00 | 4,41 | 0,00 | 1,00E+00 |
| AT2G26300 | 1,09 | -2,13 | 2,63 | -0,81 | 4,19E-01 |
| AT2G26340 | 0,57 | -0,24 | 3,17 | -0,08 | 9,39E-01 |
| AT2G26440 | 1,40 | -2,17 | 1,89 | -1,15 | 2,52E-01 |
| AT2G26510 | 0,24 | -0,02 | 4,38 | 0,00 | 9,97E-01 |
| AT2G26570 | 0,43 | -1,51 | 3,33 | -0,45 | 6,50E-01 |
| AT2G26600 | 0,45 | 0,47 | 3,26 | 0,14 | 8,86E-01 |
| AT2G26610 | 0,30 | 0,04 | 4,38 | 0,01 | 9,93E-01 |
| AT2G26640 | 0,71 | -3,59 | 3,62 | -0,99 | 3,21E-01 |
| AT2G26660 | 0,22 | 0,08 | 4,38 | 0,02 | 9,86E-01 |
| AT2G26692 | 0,08 | 0,00 | 4,41 | 0,00 | 1,00E+00 |
| AT2G26730 | 0,85 | -0,82 | 1,95 | -0,42 | 6,72E-01 |
| AT2G26920 | 0,60 | -1,28 | 3,17 | -0,41 | 6,85E-01 |
| AT2G27030 | 0,72 | 2,42 | 2,54 | 0,95 | 3,40E-01 |
| AT2G27050 | 0,99 | 1,92 | 1,89 | 1,02 | 3,10E-01 |
| AT2G27060 | 0,17 | 0,00 | 4,41 | 0,00 | 1,00E+00 |
| AT2G27080 | 0,39 | 2,15 | 4,15 | 0,52 | 6,05E-01 |
| AT2G27100 | 0,72 | 0,00 | 3,66 | 0,00 | 1,00E+00 |
| AT2G27130 | 0,30 | 1,65 | 4,31 | 0,38 | 7,02E-01 |
| AT2G27150 | 1,72 | 0,06 | 2,23 | 0,03 | 9,80E-01 |
| AT2G27200 | 0,21 | 0,39 | 4,36 | 0,09 | 9,29E-01 |
| AT2G27210 | 0,89 | 0,07 | 2,31 | 0,03 | 9,76E-01 |
| AT2G27230 | 1,31 | 0,88 | 1,73 | 0,51 | 6,13E-01 |
| AT2G27260 | 0,40 | 0,45 | 3,36 | 0,13 | 8,93E-01 |
| AT2G27320 | 0,85 | 0,53 | 2,35 | 0,23 | 8,21E-01 |
| AT2G27380 | 0,13 | 0,94 | 4,35 | 0,22 | 8,29E-01 |
| AT2G27460 | 0,28 | -2,88 | 4,36 | -0,66 | 5,09E-01 |
| AT2G27540 | 0,95 | 3,85 | 3,04 | 1,27 | 2,04E-01 |
| AT2G27570 | 0,28 | 0,00 | 4,41 | 0,00 | 1,00E+00 |
| AT2G27600 | 1,39 | 1,13 | 1,90 | 0,60 | 5,51E-01 |
| AT2G27690 | 0,16 | 0,15 | 4,38 | 0,04 | 9,72E-01 |
| AT2G27710 | 0,57 | -0,41 | 3,23 | -0,13 | 9,00E-01 |
| AT2G27720 | 1,70 | -2,94 | 1,94 | -1,51 | 1,30E-01 |
| AT2G27760 | 0,08 | 0,00 | 4,41 | 0,00 | 1,00E+00 |
| AT2G27810 | 0,54 | 0,10 | 2,71 | 0,04 | 9,71E-01 |
| AT2G27820 | 0,23 | -0,54 | 4,36 | -0,12 | 9,02E-01 |
| AT2G27900 | 1,32 | 0,90 | 1,72 | 0,52 | 6,00E-01 |

| | | | | | |
|---|---|---|---|---|---|
| AT2G28070 | 0,87 | -0,26 | 2,97 | -0,09 | 9,30E-01 |
| AT2G28120 | 0,29 | 0,00 | 4,41 | 0,00 | 1,00E+00 |
| AT2G28130 | 0,59 | 0,00 | 4,40 | 0,00 | 1,00E+00 |
| AT2G28190 | 0,42 | 1,33 | 4,29 | 0,31 | 7,57E-01 |
| AT2G28250 | 0,08 | 0,00 | 4,41 | 0,00 | 1,00E+00 |
| AT2G28310 | 0,30 | 0,14 | 4,38 | 0,03 | 9,74E-01 |
| AT2G28320 | 0,43 | 0,94 | 4,17 | 0,22 | 8,22E-01 |
| AT2G28350 | 0,96 | 0,44 | 2,15 | 0,21 | 8,37E-01 |
| AT2G28370 | 0,25 | -0,35 | 4,40 | -0,08 | 9,37E-01 |
| AT2G28460 | 0,18 | 0,25 | 4,32 | 0,06 | 9,54E-01 |
| AT2G28520 | 0,65 | 0,44 | 2,94 | 0,15 | 8,82E-01 |
| AT2G28540 | 0,87 | -2,10 | 2,47 | -0,85 | 3,94E-01 |
| AT2G28605 | 0,37 | -0,03 | 4,31 | -0,01 | 9,95E-01 |
| AT2G28660 | 0,10 | 0,86 | 4,35 | 0,20 | 8,43E-01 |
| AT2G28671 | 0,51 | 0,76 | 3,03 | 0,25 | 8,02E-01 |
| AT2G28710 | 0,24 | 0,79 | 4,36 | 0,18 | 8,56E-01 |
| AT2G28720 | 0,49 | 0,52 | 4,10 | 0,13 | 8,98E-01 |
| AT2G28730 | 1,02 | 1,39 | 1,74 | 0,80 | 4,25E-01 |
| AT2G28810 | 0,71 | 1,39 | 2,60 | 0,54 | 5,92E-01 |
| AT2G28840 | 1,76 | -0,61 | 1,61 | -0,38 | 7,06E-01 |
| AT2G28930 | 0,12 | -0,36 | 4,39 | -0,08 | 9,34E-01 |
| AT2G29020 | 0,11 | 0,88 | 4,35 | 0,20 | 8,39E-01 |
| AT2G29065 | 0,30 | 0,00 | 4,41 | 0,00 | 1,00E+00 |
| AT2G29070 | 0,15 | -0,31 | 4,40 | -0,07 | 9,43E-01 |
| AT2G29140 | 0,94 | 0,14 | 2,53 | 0,05 | 9,56E-01 |
| AT2G29180 | 0,47 | -1,28 | 4,12 | -0,31 | 7,56E-01 |
| AT2G29210 | 1,16 | 0,44 | 2,23 | 0,20 | 8,42E-01 |
| AT2G29310 | 0,46 | -0,85 | 3,30 | -0,26 | 7,96E-01 |
| AT2G29350 | 0,65 | 0,61 | 3,89 | 0,16 | 8,75E-01 |
| AT2G29360 | 0,51 | 0,00 | 4,41 | 0,00 | 1,00E+00 |
| AT2G29510 | 0,82 | 0,00 | 3,44 | 0,00 | 1,00E+00 |
| AT2G29600 | 0,47 | 2,37 | 3,13 | 0,76 | 4,50E-01 |
| AT2G29640 | 0,37 | 0,00 | 4,35 | 0,00 | 1,00E+00 |
| AT2G29660 | 0,20 | 0,00 | 4,41 | 0,00 | 1,00E+00 |
| AT2G29690 | 0,14 | -0,82 | 4,37 | -0,19 | 8,50E-01 |
| AT2G29700 | 0,24 | -0,67 | 4,38 | -0,15 | 8,78E-01 |
| AT2G29970 | 1,50 | -0,59 | 1,29 | -0,46 | 6,46E-01 |
| AT2G30020 | 0,83 | 3,37 | 3,44 | 0,98 | 3,28E-01 |
| AT2G30040 | 0,38 | 0,77 | 3,36 | 0,23 | 8,19E-01 |
| AT2G30050 | 0,66 | -0,71 | 2,51 | -0,28 | 7,79E-01 |
| AT2G30140 | 0,54 | -0,27 | 3,94 | -0,07 | 9,45E-01 |
| AT2G30150 | 0,10 | 0,24 | 4,38 | 0,05 | 9,57E-01 |
| AT2G30180 | 1,06 | -0,23 | 1,78 | -0,13 | 8,99E-01 |
| AT2G08375 | 1,24 | 1,75 | 1,79 | 0,98 | 3,27E-01 |
| AT2G30230 | 0,50 | -0,08 | 3,24 | -0,03 | 9,80E-01 |
| AT2G30362 | 0,41 | 2,28 | 4,13 | 0,55 | 5,81E-01 |
| AT2G30410 | 0,72 | -1,18 | 2,37 | -0,50 | 6,20E-01 |
| AT2G30440 | 1,74 | 2,27 | 1,53 | 1,49 | 1,37E-01 |
| AT2G30530 | 0,20 | 0,00 | 4,41 | 0,00 | 1,00E+00 |
| AT2G30550 | 1,44 | -1,35 | 1,96 | -0,69 | 4,89E-01 |
| AT2G30600 | 0,51 | -1,90 | 4,30 | -0,44 | 6,59E-01 |
| AT2G30620 | 0,85 | -0,75 | 2,22 | -0,34 | 7,37E-01 |
| AT2G30730 | 0,89 | 0,59 | 2,45 | 0,24 | 8,10E-01 |
| AT2G30780 | 0,62 | -1,23 | 3,72 | -0,33 | 7,41E-01 |
| AT2G30870 | 1,65 | 0,59 | 1,35 | 0,44 | 6,59E-01 |
| AT2G30930 | 0,49 | 0,14 | 2,66 | 0,05 | 9,58E-01 |

| | | | | | |
|---|---|---|---|---|---|
| AT2G30970 | 0,46 | -0,26 | 4,41 | -0,06 | 9,53E-01 |
| AT2G31010 | 0,95 | 0,00 | 2,48 | 0,00 | 1,00E+00 |
| AT2G31020 | 0,26 | 0,00 | 4,41 | 0,00 | 1,00E+00 |
| AT2G31200 | 0,16 | 0,74 | 4,36 | 0,17 | 8,66E-01 |
| AT2G31215 | 0,52 | 0,61 | 2,51 | 0,24 | 8,09E-01 |
| AT2G31280 | 1,47 | 0,23 | 2,27 | 0,10 | 9,19E-01 |
| AT2G31290 | 0,71 | -1,11 | 3,18 | -0,35 | 7,27E-01 |
| AT2G31350 | 0,38 | -0,26 | 4,36 | -0,06 | 9,52E-01 |
| AT2G31370 | 0,19 | -1,29 | 4,39 | -0,29 | 7,69E-01 |
| AT2G31380 | 1,06 | -1,27 | 2,10 | -0,61 | 5,43E-01 |
| AT2G08435 | 0,79 | 0,00 | 3,59 | 0,00 | 1,00E+00 |
| AT2G31560 | 0,67 | 1,40 | 2,41 | 0,58 | 5,62E-01 |
| AT2G31585 | 0,47 | -2,19 | 4,04 | -0,54 | 5,88E-01 |
| AT2G31610 | 1,80 | 1,61 | 1,71 | 0,94 | 3,46E-01 |
| AT2G31750 | 0,75 | -0,61 | 2,99 | -0,20 | 8,38E-01 |
| AT2G31790 | 0,30 | 1,61 | 3,60 | 0,45 | 6,54E-01 |
| AT2G31810 | 1,10 | 0,27 | 2,13 | 0,13 | 9,00E-01 |
| AT2G31960 | 1,68 | 0,43 | 1,63 | 0,26 | 7,93E-01 |
| AT2G31970 | 0,18 | 0,15 | 4,38 | 0,03 | 9,73E-01 |
| AT2G32010 | 0,37 | 0,37 | 4,33 | 0,09 | 9,32E-01 |
| AT2G32040 | 0,08 | 0,00 | 4,41 | 0,00 | 1,00E+00 |
| AT2G32100 | 1,15 | 2,48 | 1,97 | 1,26 | 2,08E-01 |
| AT2G32150 | 1,00 | 3,10 | 2,38 | 1,31 | 1,92E-01 |
| AT2G32250 | 0,51 | -1,19 | 3,25 | -0,37 | 7,13E-01 |
| AT2G32260 | 0,44 | 0,19 | 4,14 | 0,05 | 9,63E-01 |
| AT2G32295 | 0,53 | -0,62 | 2,68 | -0,23 | 8,16E-01 |
| AT2G32320 | 0,15 | 0,47 | 4,37 | 0,11 | 9,15E-01 |
| AT2G32440 | 0,16 | -0,21 | 4,40 | -0,05 | 9,63E-01 |
| AT2G32450 | 0,56 | 1,34 | 2,60 | 0,52 | 6,06E-01 |
| AT2G32698 | 0,12 | 0,00 | 4,41 | 0,00 | 1,00E+00 |
| AT2G32700 | 1,61 | -0,91 | 1,46 | -0,63 | 5,32E-01 |
| AT2G32800 | 1,51 | -1,05 | 1,53 | -0,69 | 4,90E-01 |
| AT2G32900 | 0,29 | -0,37 | 4,39 | -0,08 | 9,34E-01 |
| AT2G32910 | 0,68 | 0,30 | 2,99 | 0,10 | 9,19E-01 |
| AT2G33080 | 0,74 | 0,00 | 3,58 | 0,00 | 1,00E+00 |
| AT2G33160 | 0,25 | 0,00 | 4,41 | 0,00 | 1,00E+00 |
| AT2G33220 | 0,54 | -0,17 | 3,25 | -0,05 | 9,57E-01 |
| AT2G33250 | 0,76 | -1,63 | 4,37 | -0,37 | 7,10E-01 |
| AT2G33290 | 0,29 | 0,40 | 3,66 | 0,11 | 9,13E-01 |
| AT2G33390 | 0,25 | 0,28 | 4,39 | 0,06 | 9,50E-01 |
| AT2G33420 | 0,37 | 0,00 | 4,35 | 0,00 | 1,00E+00 |
| AT2G08585 | 0,32 | 0,42 | 3,64 | 0,12 | 9,08E-01 |
| AT2G33435 | 1,31 | -1,42 | 1,67 | -0,85 | 3,97E-01 |
| AT2G33450 | 0,32 | -0,72 | 4,33 | -0,17 | 8,68E-01 |
| AT2G33490 | 0,35 | -0,40 | 4,35 | -0,09 | 9,27E-01 |
| AT2G33510 | 0,26 | 0,08 | 4,38 | 0,02 | 9,86E-01 |
| AT2G33570 | 0,17 | 1,19 | 4,33 | 0,28 | 7,83E-01 |
| AT2G33610 | 0,37 | -0,32 | 4,34 | -0,07 | 9,42E-01 |
| AT2G33650 | 0,29 | -2,86 | 4,36 | -0,65 | 5,13E-01 |
| AT2G33700 | 0,56 | 1,98 | 3,95 | 0,50 | 6,15E-01 |
| AT2G33760 | 0,64 | 0,00 | 4,41 | 0,00 | 1,00E+00 |
| AT2G33810 | 0,37 | -0,75 | 4,32 | -0,17 | 8,61E-01 |
| AT2G33840 | 0,52 | -0,92 | 3,22 | -0,29 | 7,74E-01 |
| AT2G33845 | 0,47 | 0,49 | 4,10 | 0,12 | 9,06E-01 |
| AT2G33950 | 0,39 | 0,16 | 4,32 | 0,04 | 9,71E-01 |
| AT2G34120 | 0,27 | 1,93 | 4,30 | 0,45 | 6,53E-01 |

| | | | | | |
|---|---|---|---|---|---|
| AT2G34240 | 0,12 | 0,00 | 4,41 | 0,00 | 1,00E+00 |
| AT2G34357 | 1,72 | -1,03 | 1,76 | -0,59 | 5,58E-01 |
| AT2G34460 | 0,99 | 1,58 | 1,84 | 0,86 | 3,91E-01 |
| AT2G34660 | 0,69 | -0,48 | 2,60 | -0,19 | 8,53E-01 |
| AT2G34730 | 0,50 | -1,03 | 2,69 | -0,38 | 7,01E-01 |
| AT2G34770 | 0,22 | 0,08 | 4,38 | 0,02 | 9,86E-01 |
| AT2G34780 | 0,99 | 1,50 | 2,17 | 0,69 | 4,90E-01 |
| AT2G34810 | 0,16 | 0,15 | 4,38 | 0,04 | 9,72E-01 |
| AT2G34860 | 1,10 | 0,23 | 2,36 | 0,10 | 9,22E-01 |
| AT2G34970 | 0,22 | 0,00 | 4,37 | 0,00 | 1,00E+00 |
| AT2G35020 | 0,32 | 0,52 | 4,38 | 0,12 | 9,06E-01 |
| AT2G35170 | 0,10 | 0,24 | 4,38 | 0,05 | 9,57E-01 |
| AT2G35230 | 0,50 | 0,70 | 4,36 | 0,16 | 8,73E-01 |
| AT2G35330 | 0,35 | 0,98 | 4,34 | 0,23 | 8,22E-01 |
| AT2G35340 | 0,80 | 0,00 | 2,91 | 0,00 | 1,00E+00 |
| AT2G35370 | 0,43 | -0,80 | 4,31 | -0,18 | 8,53E-01 |
| AT2G35630 | 1,16 | -1,25 | 1,80 | -0,70 | 4,87E-01 |
| AT2G35635 | 0,64 | -3,36 | 4,35 | -0,77 | 4,39E-01 |
| AT2G35650 | 0,68 | -1,24 | 2,36 | -0,53 | 5,98E-01 |
| AT2G08750 | 0,08 | 0,00 | 4,41 | 0,00 | 1,00E+00 |
| AT2G35720 | 0,19 | 0,67 | 4,36 | 0,15 | 8,78E-01 |
| AT2G35760 | 0,17 | 0,07 | 4,38 | 0,02 | 9,88E-01 |
| AT2G35800 | 1,02 | -0,77 | 1,83 | -0,42 | 6,74E-01 |
| AT2G35820 | 0,71 | 0,00 | 4,41 | 0,00 | 1,00E+00 |
| AT2G35830 | 1,03 | 1,49 | 2,04 | 0,73 | 4,64E-01 |
| AT2G35840 | 0,63 | 1,14 | 2,52 | 0,45 | 6,52E-01 |
| AT2G35860 | 0,74 | 0,15 | 2,23 | 0,07 | 9,47E-01 |
| AT2G35920 | 0,77 | -1,35 | 2,51 | -0,54 | 5,90E-01 |
| AT2G35935 | 0,49 | 2,99 | 3,83 | 0,78 | 4,35E-01 |
| AT2G35960 | 0,20 | 1,44 | 4,33 | 0,33 | 7,40E-01 |
| AT2G36000 | 0,80 | 0,32 | 2,57 | 0,12 | 9,01E-01 |
| AT2G36010 | 0,59 | -1,82 | 3,09 | -0,59 | 5,56E-01 |
| AT2G36026 | 0,70 | -1,29 | 3,73 | -0,35 | 7,29E-01 |
| AT2G36090 | 0,69 | 3,50 | 3,49 | 1,00 | 3,15E-01 |
| AT2G36130 | 0,56 | 0,00 | 4,41 | 0,00 | 1,00E+00 |
| AT2G36160 | 1,81 | -0,50 | 1,89 | -0,27 | 7,90E-01 |
| AT2G36320 | 1,39 | -0,78 | 1,58 | -0,49 | 6,21E-01 |
| AT2G36330 | 0,55 | 0,00 | 4,39 | 0,00 | 1,00E+00 |
| AT2G36430 | 1,44 | 1,92 | 1,86 | 1,03 | 3,04E-01 |
| AT2G36500 | 0,54 | 0,00 | 3,19 | 0,00 | 1,00E+00 |
| AT2G36510 | 0,37 | 0,00 | 4,36 | 0,00 | 1,00E+00 |
| AT2G36570 | 0,52 | -0,29 | 4,40 | -0,06 | 9,48E-01 |
| AT2G36580 | 1,14 | 0,24 | 1,66 | 0,14 | 8,87E-01 |
| AT2G36600 | 0,66 | -3,13 | 2,21 | -1,42 | 1,57E-01 |
| AT2G36720 | 1,25 | -0,32 | 2,28 | -0,14 | 8,88E-01 |
| AT2G36792 | 0,34 | 1,94 | 4,30 | 0,45 | 6,52E-01 |
| AT2G36830 | 0,86 | -0,15 | 2,29 | -0,07 | 9,47E-01 |
| AT2G36920 | 0,12 | 0,00 | 4,41 | 0,00 | 1,00E+00 |
| AT2G36950 | 0,34 | 0,00 | 4,41 | 0,00 | 1,00E+00 |
| AT2G36960 | 0,18 | 0,82 | 4,35 | 0,19 | 8,51E-01 |
| AT2G36970 | 0,19 | 0,43 | 4,36 | 0,10 | 9,21E-01 |
| AT2G36980 | 0,05 | 0,18 | 4,38 | 0,04 | 9,68E-01 |
| AT2G37025 | 0,19 | -1,98 | 4,34 | -0,46 | 6,48E-01 |
| AT2G37050 | 0,93 | -1,22 | 2,45 | -0,50 | 6,19E-01 |
| AT2G37110 | 1,16 | 2,77 | 2,27 | 1,22 | 2,22E-01 |
| AT2G37180 | 0,08 | 0,00 | 4,41 | 0,00 | 1,00E+00 |

| | | | | | |
|---|---|---|---|---|---|
| AT2G37250 | 0,74 | -0,03 | 2,25 | -0,02 | 9,88E-01 |
| AT2G08860 | 0,31 | -0,41 | 3,62 | -0,11 | 9,10E-01 |
| AT2G37320 | 0,08 | 0,00 | 4,41 | 0,00 | 1,00E+00 |
| AT2G37330 | 0,66 | -0,26 | 3,17 | -0,08 | 9,34E-01 |
| AT2G37362 | 0,21 | 0,00 | 4,41 | 0,00 | 1,00E+00 |
| AT2G37370 | 0,40 | 2,43 | 4,29 | 0,57 | 5,72E-01 |
| AT2G37470 | 0,26 | 0,09 | 4,38 | 0,02 | 9,84E-01 |
| AT2G37480 | 1,35 | 0,65 | 2,22 | 0,30 | 7,68E-01 |
| AT2G37555 | 0,61 | -0,26 | 3,09 | -0,08 | 9,32E-01 |
| AT2G37580 | 0,12 | 0,27 | 4,38 | 0,06 | 9,50E-01 |
| AT2G37650 | 1,11 | 1,56 | 2,12 | 0,74 | 4,61E-01 |
| AT2G37690 | 0,31 | -0,31 | 4,36 | -0,07 | 9,44E-01 |
| AT2G37770 | 0,28 | -0,35 | 4,40 | -0,08 | 9,37E-01 |
| AT2G37860 | 0,83 | 1,93 | 2,21 | 0,87 | 3,83E-01 |
| AT2G37940 | 0,73 | 0,68 | 2,94 | 0,23 | 8,18E-01 |
| AT2G37970 | 0,77 | 0,81 | 2,70 | 0,30 | 7,65E-01 |
| AT2G37990 | 0,11 | -0,38 | 4,39 | -0,09 | 9,31E-01 |
| AT2G38010 | 0,53 | 0,88 | 3,09 | 0,29 | 7,75E-01 |
| AT2G38090 | 0,20 | 0,00 | 4,41 | 0,00 | 1,00E+00 |
| AT2G38120 | 0,38 | 0,35 | 2,79 | 0,13 | 8,99E-01 |
| AT2G38140 | 0,26 | -0,12 | 3,76 | -0,03 | 9,75E-01 |
| AT2G38210 | 1,49 | -0,90 | 1,75 | -0,51 | 6,08E-01 |
| AT2G38310 | 0,71 | 0,36 | 2,97 | 0,12 | 9,03E-01 |
| AT2G38365 | 0,20 | 0,00 | 4,41 | 0,00 | 1,00E+00 |
| AT2G38420 | 0,26 | 0,00 | 4,41 | 0,00 | 1,00E+00 |
| AT2G38470 | 1,30 | 1,62 | 1,82 | 0,89 | 3,72E-01 |
| AT2G38560 | 1,56 | 0,96 | 2,07 | 0,46 | 6,44E-01 |
| AT2G38590 | 0,78 | 3,37 | 2,65 | 1,27 | 2,03E-01 |
| AT2G38670 | 0,86 | -2,61 | 2,21 | -1,18 | 2,37E-01 |
| AT2G38760 | 0,08 | 0,00 | 4,41 | 0,00 | 1,00E+00 |
| AT2G38780 | 0,82 | 0,25 | 1,76 | 0,14 | 8,87E-01 |
| AT2G38820 | 0,12 | -0,26 | 4,41 | -0,06 | 9,53E-01 |
| AT2G38840 | 0,33 | -0,37 | 4,39 | -0,08 | 9,33E-01 |
| AT2G38940 | 0,25 | 0,00 | 4,41 | 0,00 | 1,00E+00 |
| AT2G38950 | 0,65 | -1,17 | 2,72 | -0,43 | 6,67E-01 |
| AT2G39190 | 0,15 | 0,47 | 4,37 | 0,11 | 9,14E-01 |
| AT2G39230 | 0,19 | 1,00 | 4,33 | 0,23 | 8,17E-01 |
| AT2G39270 | 0,08 | 0,00 | 4,41 | 0,00 | 1,00E+00 |
| AT2G39340 | 0,19 | 1,00 | 4,33 | 0,23 | 8,17E-01 |
| AT2G39400 | 0,15 | -0,39 | 4,39 | -0,09 | 9,29E-01 |
| AT2G39420 | 0,56 | -0,51 | 3,14 | -0,16 | 8,70E-01 |
| AT2G39430 | 0,52 | 0,00 | 3,96 | 0,00 | 1,00E+00 |
| AT2G39460 | 0,89 | 1,40 | 1,89 | 0,74 | 4,58E-01 |
| AT2G39470 | 0,81 | -1,23 | 2,32 | -0,53 | 5,95E-01 |
| AT2G39570 | 1,60 | 0,88 | 2,88 | 0,31 | 7,59E-01 |
| AT2G39580 | 1,16 | -2,13 | 2,30 | -0,93 | 3,53E-01 |
| AT2G39670 | 1,03 | -0,47 | 2,34 | -0,20 | 8,40E-01 |
| AT2G39725 | 0,50 | 0,00 | 4,00 | 0,00 | 1,00E+00 |
| AT2G39740 | 0,63 | 0,92 | 3,06 | 0,30 | 7,64E-01 |
| AT2G39780 | 1,17 | -0,84 | 2,27 | -0,37 | 7,11E-01 |
| AT2G39810 | 1,48 | 0,97 | 2,51 | 0,39 | 6,98E-01 |
| AT2G39840 | 0,84 | 1,64 | 2,51 | 0,65 | 5,13E-01 |
| AT2G39850 | 0,54 | -0,22 | 4,31 | -0,05 | 9,59E-01 |
| AT2G39870 | 0,43 | -0,24 | 4,27 | -0,06 | 9,54E-01 |
| AT2G39930 | 1,27 | -0,71 | 1,80 | -0,39 | 6,93E-01 |
| AT2G40060 | 0,99 | 1,66 | 2,40 | 0,69 | 4,90E-01 |

| | | | | | |
|---|---|---|---|---|---|
| AT2G40095 | 0,64 | -0,27 | 3,19 | -0,09 | 9,32E-01 |
| AT2G40130 | 0,14 | 0,13 | 4,38 | 0,03 | 9,77E-01 |
| AT2G40150 | 0,08 | 0,00 | 4,41 | 0,00 | 1,00E+00 |
| AT2G40160 | 0,42 | 0,00 | 4,19 | 0,00 | 1,00E+00 |
| AT2G40200 | 0,38 | 1,40 | 4,32 | 0,32 | 7,46E-01 |
| AT2G09120 | 0,31 | -3,66 | 4,33 | -0,84 | 3,98E-01 |
| AT2G40310 | 0,83 | 2,07 | 2,18 | 0,95 | 3,41E-01 |
| AT2G40400 | 0,82 | -0,60 | 2,85 | -0,21 | 8,34E-01 |
| AT2G40410 | 0,54 | 0,57 | 4,09 | 0,14 | 8,89E-01 |
| AT2G40435 | 0,46 | -2,04 | 4,37 | -0,47 | 6,41E-01 |
| AT2G40460 | 0,63 | -0,29 | 2,40 | -0,12 | 9,05E-01 |
| AT2G40480 | 0,31 | 0,34 | 4,38 | 0,08 | 9,38E-01 |
| AT2G40490 | 1,23 | -0,76 | 1,91 | -0,40 | 6,92E-01 |
| AT2G40540 | 0,78 | 0,06 | 2,45 | 0,02 | 9,82E-01 |
| AT2G40550 | 0,20 | 0,00 | 4,41 | 0,00 | 1,00E+00 |
| AT2G40660 | 0,08 | 0,21 | 4,39 | 0,05 | 9,62E-01 |
| AT2G40670 | 0,10 | 0,24 | 4,38 | 0,05 | 9,57E-01 |
| AT2G40760 | 0,35 | -2,19 | 4,41 | -0,50 | 6,20E-01 |
| AT2G40765 | 0,08 | 0,00 | 4,41 | 0,00 | 1,00E+00 |
| AT2G40810 | 0,08 | 0,00 | 4,41 | 0,00 | 1,00E+00 |
| AT2G40830 | 0,33 | 0,00 | 4,41 | 0,00 | 1,00E+00 |
| AT2G40880 | 1,07 | -1,54 | 2,17 | -0,71 | 4,78E-01 |
| AT2G40955 | 1,38 | 3,87 | 1,82 | 2,12 | 3,40E-02 |
| AT2G40980 | 0,41 | -2,05 | 4,23 | -0,48 | 6,28E-01 |
| AT2G41080 | 0,59 | 1,57 | 2,62 | 0,60 | 5,49E-01 |
| AT2G41110 | 0,26 | -0,03 | 3,79 | -0,01 | 9,94E-01 |
| AT2G09230 | 0,12 | 0,00 | 4,41 | 0,00 | 1,00E+00 |
| AT2G41140 | 0,37 | -0,30 | 4,34 | -0,07 | 9,45E-01 |
| AT2G41200 | 0,84 | 0,44 | 2,12 | 0,21 | 8,36E-01 |
| AT2G41210 | 0,42 | -0,34 | 4,20 | -0,08 | 9,36E-01 |
| AT2G41220 | 0,64 | 1,77 | 2,95 | 0,60 | 5,48E-01 |
| AT2G41460 | 0,13 | 0,00 | 4,41 | 0,00 | 1,00E+00 |
| AT2G41510 | 0,08 | 0,00 | 4,41 | 0,00 | 1,00E+00 |
| AT2G41520 | 0,59 | -0,37 | 3,07 | -0,12 | 9,04E-01 |
| AT2G41590 | 1,66 | -0,08 | 1,51 | -0,05 | 9,59E-01 |
| AT2G41670 | 0,12 | 0,00 | 4,41 | 0,00 | 1,00E+00 |
| AT2G41730 | 0,99 | 0,47 | 2,18 | 0,22 | 8,29E-01 |
| AT2G41820 | 0,34 | 0,00 | 4,41 | 0,00 | 1,00E+00 |
| AT2G41880 | 0,26 | -0,03 | 3,79 | -0,01 | 9,94E-01 |
| AT2G41920 | 1,42 | 4,36 | 1,90 | 2,30 | 2,16E-02 |
| AT2G41940 | 1,76 | 3,17 | 1,54 | 2,06 | 3,93E-02 |
| AT2G42130 | 0,48 | -0,49 | 3,24 | -0,15 | 8,81E-01 |
| AT2G42220 | 0,98 | -2,49 | 1,95 | -1,27 | 2,03E-01 |
| AT2G42310 | 0,61 | -0,21 | 2,56 | -0,08 | 9,34E-01 |
| AT2G42320 | 0,63 | -1,86 | 2,65 | -0,70 | 4,83E-01 |
| AT2G42430 | 0,21 | 1,90 | 4,30 | 0,44 | 6,58E-01 |
| AT2G42580 | 1,32 | -1,07 | 1,97 | -0,55 | 5,85E-01 |
| AT2G42610 | 0,29 | -1,33 | 4,39 | -0,30 | 7,62E-01 |
| AT2G42620 | 1,45 | 0,62 | 2,47 | 0,25 | 8,00E-01 |
| AT2G42700 | 0,72 | -0,28 | 3,13 | -0,09 | 9,28E-01 |
| AT2G42710 | 0,12 | -0,36 | 4,39 | -0,08 | 9,34E-01 |
| AT2G42750 | 0,57 | 0,00 | 3,85 | 0,00 | 1,00E+00 |
| AT2G42780 | 0,28 | -0,06 | 4,35 | -0,01 | 9,89E-01 |
| AT2G42890 | 0,32 | -1,26 | 4,37 | -0,29 | 7,72E-01 |
| AT2G42900 | 0,43 | 0,34 | 4,38 | 0,08 | 9,38E-01 |
| AT2G43010 | 1,28 | -1,56 | 1,81 | -0,86 | 3,87E-01 |

| | | | | | |
|---|---|---|---|---|---|
| AT2G43020 | 1,32 | 3,67 | 2,00 | 1,83 | 6,66E-02 |
| AT2G43030 | 0,58 | 0,30 | 3,11 | 0,10 | 9,24E-01 |
| AT2G43060 | 1,34 | -0,80 | 1,38 | -0,58 | 5,65E-01 |
| AT2G43080 | 0,63 | 1,12 | 3,08 | 0,36 | 7,17E-01 |
| AT2G43090 | 0,61 | -0,09 | 2,60 | -0,03 | 9,74E-01 |
| AT2G43100 | 0,56 | 0,51 | 3,29 | 0,16 | 8,76E-01 |
| AT2G43137 | 1,43 | -0,53 | 1,73 | -0,30 | 7,61E-01 |
| AT2G43138 | 1,21 | 3,44 | 1,97 | 1,74 | 8,15E-02 |
| AT2G43141 | 1,64 | 0,05 | 1,76 | 0,03 | 9,76E-01 |
| AT2G43210 | 0,63 | -2,08 | 3,05 | -0,68 | 4,96E-01 |
| AT2G43230 | 0,55 | 0,00 | 3,90 | 0,00 | 1,00E+00 |
| AT2G43250 | 0,31 | -0,40 | 4,38 | -0,09 | 9,27E-01 |
| AT2G43280 | 0,56 | 0,00 | 4,39 | 0,00 | 1,00E+00 |
| AT2G09440 | 0,08 | 0,00 | 4,41 | 0,00 | 1,00E+00 |
| AT2G43340 | 0,44 | 1,06 | 3,37 | 0,31 | 7,53E-01 |
| AT2G43400 | 0,37 | -0,13 | 4,36 | -0,03 | 9,76E-01 |
| AT2G43440 | 0,11 | 0,68 | 4,37 | 0,16 | 8,76E-01 |
| AT2G43540 | 1,53 | 2,85 | 1,83 | 1,56 | 1,19E-01 |
| AT2G43630 | 0,44 | -0,30 | 4,14 | -0,07 | 9,43E-01 |
| AT2G43650 | 0,83 | -0,49 | 2,50 | -0,20 | 8,45E-01 |
| AT2G43690 | 0,68 | 0,00 | 4,41 | 0,00 | 1,00E+00 |
| AT2G43790 | 0,39 | -1,26 | 2,82 | -0,45 | 6,55E-01 |
| AT2G43800 | 0,08 | 0,21 | 4,39 | 0,05 | 9,62E-01 |
| AT2G43820 | 0,45 | -0,36 | 3,32 | -0,11 | 9,13E-01 |
| AT2G44080 | 0,60 | 0,21 | 3,80 | 0,06 | 9,56E-01 |
| AT2G44100 | 0,65 | -0,93 | 2,57 | -0,36 | 7,17E-01 |
| AT2G44130 | 0,72 | -0,33 | 2,39 | -0,14 | 8,91E-01 |
| AT2G44160 | 0,80 | 1,90 | 2,41 | 0,79 | 4,31E-01 |
| AT2G44175 | 0,43 | 2,78 | 4,06 | 0,68 | 4,94E-01 |
| AT2G44200 | 0,72 | 1,11 | 2,57 | 0,43 | 6,68E-01 |
| AT2G44210 | 1,13 | -0,09 | 1,61 | -0,05 | 9,58E-01 |
| AT2G44230 | 0,76 | -1,25 | 2,62 | -0,48 | 6,31E-01 |
| AT2G44240 | 0,17 | 0,84 | 4,36 | 0,19 | 8,47E-01 |
| AT2G44260 | 0,20 | -0,26 | 4,41 | -0,06 | 9,53E-01 |
| AT2G44310 | 1,85 | 2,00 | 1,36 | 1,47 | 1,42E-01 |
| AT2G44350 | 0,40 | 0,75 | 3,35 | 0,22 | 8,23E-01 |
| AT2G44420 | 0,20 | 0,00 | 4,41 | 0,00 | 1,00E+00 |
| AT2G44430 | 0,96 | 1,66 | 1,84 | 0,90 | 3,67E-01 |
| AT2G44500 | 0,91 | 2,20 | 2,84 | 0,77 | 4,40E-01 |
| AT2G44520 | 0,23 | -0,35 | 4,39 | -0,08 | 9,36E-01 |
| AT2G44530 | 0,99 | 0,67 | 2,37 | 0,28 | 7,76E-01 |
| AT2G44600 | 0,10 | 0,24 | 4,38 | 0,05 | 9,57E-01 |
| AT2G44640 | 1,50 | 0,47 | 1,93 | 0,25 | 8,06E-01 |
| AT2G44670 | 1,15 | 1,13 | 1,97 | 0,58 | 5,64E-01 |
| AT2G44710 | 1,40 | 1,32 | 1,84 | 0,72 | 4,74E-01 |
| AT2G44735 | 0,42 | -0,30 | 4,17 | -0,07 | 9,43E-01 |
| AT2G44770 | 0,15 | -0,35 | 4,39 | -0,08 | 9,36E-01 |
| AT2G44800 | 0,19 | 0,27 | 4,37 | 0,06 | 9,50E-01 |
| AT2G44870 | 0,51 | -0,99 | 2,59 | -0,38 | 7,04E-01 |
| AT2G44880 | 0,91 | 0,00 | 2,87 | 0,00 | 1,00E+00 |
| AT2G44925 | 0,92 | 0,61 | 1,72 | 0,35 | 7,23E-01 |
| AT2G44940 | 0,52 | -0,37 | 2,80 | -0,13 | 8,95E-01 |
| AT2G45000 | 0,38 | 0,85 | 3,36 | 0,25 | 8,01E-01 |
| AT2G45020 | 0,36 | -2,08 | 4,32 | -0,48 | 6,31E-01 |
| AT2G45030 | 0,91 | 0,64 | 2,84 | 0,22 | 8,23E-01 |
| AT2G45140 | 0,33 | -2,04 | 4,38 | -0,47 | 6,42E-01 |

| | | | | | |
|---|---|---|---|---|---|
| AT2G45240 | 0,80 | -0,94 | 3,11 | -0,30 | 7,62E-01 |
| AT2G45250 | 0,26 | 0,00 | 4,41 | 0,00 | 1,00E+00 |
| AT2G45300 | 0,10 | 0,08 | 4,38 | 0,02 | 9,85E-01 |
| AT2G45310 | 0,67 | 0,75 | 2,50 | 0,30 | 7,63E-01 |
| AT2G45390 | 0,17 | -3,06 | 4,35 | -0,70 | 4,82E-01 |
| AT2G45440 | 0,53 | -0,12 | 3,91 | -0,03 | 9,76E-01 |
| AT2G45510 | 0,84 | 0,56 | 2,16 | 0,26 | 7,96E-01 |
| AT2G45530 | 0,98 | -0,81 | 1,95 | -0,42 | 6,77E-01 |
| AT2G09650 | 0,36 | 0,19 | 4,33 | 0,04 | 9,66E-01 |
| AT2G45590 | 0,37 | 0,44 | 4,32 | 0,10 | 9,19E-01 |
| AT2G45620 | 1,35 | -2,40 | 1,98 | -1,22 | 2,24E-01 |
| AT2G45630 | 0,80 | 0,97 | 2,59 | 0,37 | 7,09E-01 |
| AT2G45685 | 1,79 | 0,28 | 2,01 | 0,14 | 8,88E-01 |
| AT2G45710 | 0,45 | 0,21 | 4,34 | 0,05 | 9,62E-01 |
| AT2G45720 | 0,29 | -0,51 | 4,37 | -0,12 | 9,08E-01 |
| AT2G45740 | 0,35 | -1,08 | 3,54 | -0,31 | 7,60E-01 |
| AT2G45790 | 0,17 | -1,50 | 4,37 | -0,34 | 7,31E-01 |
| AT2G45810 | 0,90 | -0,24 | 2,35 | -0,10 | 9,18E-01 |
| AT2G45910 | 0,97 | 2,07 | 2,03 | 1,02 | 3,09E-01 |
| AT2G46100 | 0,37 | -0,08 | 2,75 | -0,03 | 9,77E-01 |
| AT2G46130 | 1,45 | -1,46 | 1,48 | -0,98 | 3,25E-01 |
| AT2G46170 | 0,69 | 0,05 | 3,76 | 0,01 | 9,90E-01 |
| AT2G46190 | 0,70 | 0,73 | 2,07 | 0,35 | 7,24E-01 |
| AT2G09695 | 1,56 | -1,31 | 1,43 | -0,92 | 3,59E-01 |
| AT2G09705 | 0,89 | -0,26 | 2,37 | -0,11 | 9,12E-01 |
| AT2G46220 | 1,83 | 0,34 | 1,68 | 0,20 | 8,40E-01 |
| AT2G46225 | 0,27 | -0,06 | 4,34 | -0,01 | 9,89E-01 |
| AT2G46240 | 0,70 | -2,04 | 2,60 | -0,79 | 4,32E-01 |
| AT2G46260 | 0,08 | 0,00 | 4,41 | 0,00 | 1,00E+00 |
| AT2G46270 | 0,25 | 0,00 | 4,41 | 0,00 | 1,00E+00 |
| AT2G46370 | 0,44 | 1,34 | 3,40 | 0,39 | 6,94E-01 |
| AT2G46380 | 1,50 | 0,00 | 2,39 | 0,00 | 1,00E+00 |
| AT2G46420 | 0,84 | 0,46 | 2,84 | 0,16 | 8,72E-01 |
| AT2G46440 | 1,43 | 1,69 | 1,61 | 1,05 | 2,92E-01 |
| AT2G46450 | 1,09 | 0,67 | 2,27 | 0,30 | 7,67E-01 |
| AT2G46490 | 0,75 | 1,22 | 2,89 | 0,42 | 6,74E-01 |
| AT2G46520 | 1,16 | 2,00 | 1,85 | 1,08 | 2,78E-01 |
| AT2G46570 | 0,51 | -0,35 | 4,07 | -0,09 | 9,32E-01 |
| AT2G46590 | 0,52 | 2,42 | 2,98 | 0,81 | 4,18E-01 |
| AT2G46600 | 1,11 | -0,01 | 1,97 | -0,01 | 9,95E-01 |
| AT2G46630 | 0,28 | 0,47 | 4,32 | 0,11 | 9,14E-01 |
| AT2G46650 | 0,42 | 1,34 | 3,32 | 0,40 | 6,86E-01 |
| AT2G46690 | 0,43 | 0,94 | 4,17 | 0,22 | 8,22E-01 |
| AT2G46710 | 0,30 | 0,58 | 3,64 | 0,16 | 8,74E-01 |
| AT2G46790 | 0,54 | -0,52 | 2,65 | -0,20 | 8,45E-01 |
| AT2G46800 | 1,02 | 0,83 | 1,96 | 0,42 | 6,71E-01 |
| AT2G46810 | 0,59 | -0,43 | 3,06 | -0,14 | 8,89E-01 |
| AT2G46900 | 1,50 | -4,18 | 2,09 | -2,00 | 4,54E-02 |
| AT2G46910 | 0,44 | 0,72 | 4,20 | 0,17 | 8,63E-01 |
| AT2G46990 | 0,08 | 0,00 | 4,41 | 0,00 | 1,00E+00 |
| AT2G47110 | 0,66 | 1,53 | 2,15 | 0,71 | 4,77E-01 |
| AT2G09815 | 0,23 | 1,67 | 4,31 | 0,39 | 6,99E-01 |
| AT2G47190 | 0,40 | 0,00 | 4,41 | 0,00 | 1,00E+00 |
| AT2G47220 | 0,25 | -2,14 | 4,40 | -0,49 | 6,28E-01 |
| AT2G47240 | 1,70 | -0,04 | 1,38 | -0,03 | 9,78E-01 |
| AT2G47320 | 0,28 | -1,15 | 4,35 | -0,26 | 7,92E-01 |

| | | | | | |
|---|---|---|---|---|---|
| AT2G47420 | 0,42 | -3,00 | 4,17 | -0,72 | 4,71E-01 |
| AT2G47470 | 1,77 | -0,04 | 1,68 | -0,02 | 9,81E-01 |
| AT2G47510 | 1,59 | -1,55 | 1,86 | -0,83 | 4,04E-01 |
| AT2G47610 | 1,29 | -0,26 | 1,93 | -0,13 | 8,95E-01 |
| AT2G47620 | 0,26 | -0,03 | 3,79 | -0,01 | 9,94E-01 |
| AT2G47630 | 0,42 | 0,25 | 2,79 | 0,09 | 9,29E-01 |
| AT2G47640 | 0,28 | 0,87 | 4,36 | 0,20 | 8,43E-01 |
| AT2G47690 | 0,55 | -1,37 | 4,02 | -0,34 | 7,32E-01 |
| AT2G47730 | 0,80 | 1,77 | 2,25 | 0,79 | 4,30E-01 |
| AT2G47820 | 1,37 | 0,78 | 2,59 | 0,30 | 7,62E-01 |
| AT2G47850 | 0,53 | 1,06 | 3,16 | 0,34 | 7,37E-01 |
| AT2G47910 | 0,92 | 1,70 | 2,02 | 0,84 | 3,99E-01 |
| AT2G47935 | 0,72 | -1,60 | 2,80 | -0,57 | 5,68E-01 |
| AT2G47960 | 1,39 | 1,79 | 1,93 | 0,93 | 3,52E-01 |
| AT2G47970 | 0,70 | 0,00 | 3,58 | 0,00 | 1,00E+00 |
| AT2G48010 | 0,33 | 0,56 | 4,37 | 0,13 | 8,98E-01 |
| AT2G48020 | 0,56 | 1,01 | 3,25 | 0,31 | 7,56E-01 |
| AT2G48110 | 1,17 | -0,29 | 2,23 | -0,13 | 8,96E-01 |
| AT2G48120 | 0,32 | -2,11 | 4,40 | -0,48 | 6,32E-01 |
| AT3G01040 | 0,51 | -1,51 | 3,18 | -0,47 | 6,35E-01 |
| AT3G01050 | 0,18 | -0,81 | 4,37 | -0,19 | 8,53E-01 |
| AT3G01080 | 0,17 | -0,38 | 4,39 | -0,09 | 9,31E-01 |
| AT3G01085 | 1,09 | 0,51 | 1,75 | 0,29 | 7,69E-01 |
| AT3G01150 | 0,45 | 0,86 | 3,29 | 0,26 | 7,93E-01 |
| AT3G01170 | 0,42 | -0,34 | 4,20 | -0,08 | 9,36E-01 |
| AT3G01175 | 0,08 | 0,00 | 4,41 | 0,00 | 1,00E+00 |
| AT3G01210 | 0,10 | 0,08 | 4,38 | 0,02 | 9,85E-01 |
| AT3G01280 | 1,02 | -0,20 | 1,80 | -0,11 | 9,12E-01 |
| AT3G01320 | 1,14 | 1,56 | 1,90 | 0,82 | 4,13E-01 |
| AT3G01440 | 0,31 | 0,01 | 4,31 | 0,00 | 9,99E-01 |
| AT3G01475 | 1,05 | 0,50 | 2,07 | 0,24 | 8,08E-01 |
| AT3G01480 | 1,15 | -1,20 | 1,84 | -0,65 | 5,14E-01 |
| AT3G01560 | 0,68 | -0,49 | 2,59 | -0,19 | 8,49E-01 |
| AT3G01590 | 0,40 | -0,32 | 4,40 | -0,07 | 9,42E-01 |
| AT3G01610 | 0,25 | 0,03 | 4,38 | 0,01 | 9,95E-01 |
| AT3G01660 | 0,12 | -0,26 | 4,41 | -0,06 | 9,53E-01 |
| AT3G01690 | 1,08 | 0,66 | 1,79 | 0,37 | 7,13E-01 |
| AT3G01740 | 0,28 | -1,28 | 4,38 | -0,29 | 7,70E-01 |
| AT3G01750 | 0,72 | -2,28 | 3,10 | -0,74 | 4,61E-01 |
| AT3G01780 | 1,57 | 1,54 | 1,77 | 0,87 | 3,84E-01 |
| AT3G01810 | 0,37 | -3,52 | 4,35 | -0,81 | 4,18E-01 |
| AT3G01830 | 0,94 | 1,56 | 2,25 | 0,69 | 4,89E-01 |
| AT3G01920 | 0,37 | -0,12 | 4,36 | -0,03 | 9,78E-01 |
| AT3G02010 | 0,37 | 0,00 | 4,41 | 0,00 | 1,00E+00 |
| AT3G02030 | 0,12 | 0,27 | 4,38 | 0,06 | 9,50E-01 |
| AT3G02050 | 0,37 | 0,00 | 4,36 | 0,00 | 1,00E+00 |
| AT3G02060 | 1,23 | 0,11 | 2,05 | 0,05 | 9,57E-01 |
| AT3G02065 | 0,54 | 0,00 | 3,87 | 0,00 | 1,00E+00 |
| AT3G02070 | 0,35 | 1,30 | 3,53 | 0,37 | 7,13E-01 |
| AT3G02130 | 1,18 | 0,47 | 2,30 | 0,21 | 8,37E-01 |
| AT3G02200 | 1,75 | 0,25 | 1,93 | 0,13 | 8,97E-01 |
| AT3G02280 | 0,25 | 0,28 | 4,39 | 0,06 | 9,50E-01 |
| AT3G02290 | 0,16 | -0,26 | 4,41 | -0,06 | 9,53E-01 |
| AT3G02300 | 0,16 | 1,02 | 4,34 | 0,24 | 8,13E-01 |
| AT3G02340 | 0,50 | -1,71 | 4,27 | -0,40 | 6,89E-01 |
| AT3G02350 | 0,84 | 0,18 | 2,04 | 0,09 | 9,29E-01 |

| | | | | | |
|---|---|---|---|---|---|
| AT3G02360 | 1,53 | -0,16 | 1,98 | -0,08 | 9,37E-01 |
| AT3G02370 | 0,47 | -1,38 | 3,33 | -0,41 | 6,78E-01 |
| AT3G02400 | 0,66 | 0,00 | 4,41 | 0,00 | 1,00E+00 |
| AT3G02490 | 0,33 | -0,26 | 4,41 | -0,06 | 9,53E-01 |
| AT3G02560 | 0,49 | -0,34 | 3,98 | -0,09 | 9,31E-01 |
| AT3G02610 | 0,81 | -1,73 | 2,75 | -0,63 | 5,30E-01 |
| AT3G02630 | 0,37 | -0,38 | 4,34 | -0,09 | 9,30E-01 |
| AT3G02690 | 0,66 | 0,51 | 2,64 | 0,19 | 8,47E-01 |
| AT3G02720 | 0,46 | -0,01 | 4,37 | 0,00 | 9,98E-01 |
| AT3G02830 | 1,22 | 1,78 | 2,22 | 0,80 | 4,23E-01 |
| AT3G02832 | 1,51 | 1,15 | 1,50 | 0,77 | 4,44E-01 |
| AT3G01145 | 0,57 | 1,74 | 2,47 | 0,70 | 4,82E-01 |
| AT3G02880 | 1,19 | 1,22 | 1,89 | 0,64 | 5,20E-01 |
| AT3G02900 | 0,28 | 2,16 | 4,29 | 0,50 | 6,15E-01 |
| AT3G02910 | 0,77 | 0,69 | 2,40 | 0,29 | 7,73E-01 |
| AT3G03020 | 0,43 | 0,49 | 4,19 | 0,12 | 9,07E-01 |
| AT3G03030 | 0,40 | 0,00 | 4,41 | 0,00 | 1,00E+00 |
| AT3G03040 | 0,24 | 0,59 | 4,36 | 0,14 | 8,92E-01 |
| AT3G03110 | 0,55 | 0,00 | 4,41 | 0,00 | 1,00E+00 |
| AT3G03160 | 0,47 | 0,35 | 4,03 | 0,09 | 9,30E-01 |
| AT3G03210 | 0,10 | 0,86 | 4,35 | 0,20 | 8,43E-01 |
| AT3G03320 | 0,15 | 0,02 | 4,36 | 0,00 | 9,96E-01 |
| AT3G03340 | 0,40 | -0,26 | 4,41 | -0,06 | 9,53E-01 |
| AT3G03350 | 0,24 | 0,00 | 4,41 | 0,00 | 1,00E+00 |
| AT3G03360 | 0,08 | 0,00 | 4,41 | 0,00 | 1,00E+00 |
| AT3G03610 | 0,27 | -3,27 | 4,36 | -0,75 | 4,53E-01 |
| AT3G03640 | 0,25 | -0,30 | 4,33 | -0,07 | 9,45E-01 |
| AT3G03680 | 1,60 | -0,44 | 1,58 | -0,28 | 7,81E-01 |
| AT3G03790 | 1,73 | 1,11 | 1,83 | 0,61 | 5,42E-01 |
| AT3G03860 | 0,25 | 1,90 | 4,30 | 0,44 | 6,59E-01 |
| AT3G03880 | 0,28 | 1,29 | 4,32 | 0,30 | 7,65E-01 |
| AT3G03950 | 0,32 | -2,00 | 3,60 | -0,55 | 5,79E-01 |
| AT3G03960 | 1,45 | -0,40 | 1,65 | -0,24 | 8,07E-01 |
| AT3G04140 | 0,99 | -3,22 | 2,38 | -1,36 | 1,75E-01 |
| AT3G04490 | 0,26 | 0,60 | 4,37 | 0,14 | 8,90E-01 |
| AT3G04530 | 0,08 | 0,00 | 4,41 | 0,00 | 1,00E+00 |
| AT3G04550 | 0,64 | -1,96 | 2,08 | -0,94 | 3,47E-01 |
| AT3G04790 | 1,26 | -2,76 | 2,02 | -1,36 | 1,72E-01 |
| AT3G04810 | 0,59 | 0,00 | 4,40 | 0,00 | 1,00E+00 |
| AT3G04840 | 0,80 | 0,09 | 2,40 | 0,04 | 9,70E-01 |
| AT3G04870 | 0,49 | -0,66 | 3,06 | -0,21 | 8,30E-01 |
| AT3G04910 | 0,86 | 0,98 | 1,76 | 0,55 | 5,79E-01 |
| AT3G04920 | 0,51 | -1,57 | 4,06 | -0,39 | 7,00E-01 |
| AT3G04940 | 0,44 | -2,15 | 4,22 | -0,51 | 6,11E-01 |
| AT3G01555 | 0,12 | -0,36 | 4,39 | -0,08 | 9,34E-01 |
| AT3G01575 | 0,10 | 0,86 | 4,35 | 0,20 | 8,43E-01 |
| AT3G05070 | 0,17 | -0,78 | 4,36 | -0,18 | 8,59E-01 |
| AT3G05120 | 0,72 | 0,54 | 2,21 | 0,24 | 8,08E-01 |
| AT3G05200 | 0,41 | 0,85 | 4,23 | 0,20 | 8,41E-01 |
| AT3G05220 | 1,84 | 1,79 | 1,63 | 1,10 | 2,70E-01 |
| AT3G05230 | 0,28 | -0,14 | 4,39 | -0,03 | 9,75E-01 |
| AT3G05380 | 0,92 | -1,29 | 2,41 | -0,54 | 5,92E-01 |
| AT3G05410 | 1,68 | 1,48 | 1,52 | 0,97 | 3,30E-01 |
| AT3G05430 | 0,21 | 0,00 | 4,41 | 0,00 | 1,00E+00 |
| AT3G05490 | 0,28 | -0,06 | 4,35 | -0,01 | 9,89E-01 |
| AT3G05500 | 0,55 | 0,97 | 2,69 | 0,36 | 7,19E-01 |

| | | | | | |
|---|---|---|---|---|---|
| AT3G05520 | 0,43 | 0,38 | 2,81 | 0,14 | 8,92E-01 |
| AT3G05530 | 1,78 | -0,96 | 1,62 | -0,59 | 5,54E-01 |
| AT3G05545 | 0,82 | 1,12 | 2,27 | 0,49 | 6,21E-01 |
| AT3G05560 | 0,45 | -1,21 | 3,23 | -0,37 | 7,08E-01 |
| AT3G05630 | 0,21 | 0,00 | 4,41 | 0,00 | 1,00E+00 |
| AT3G05670 | 0,41 | 0,33 | 4,24 | 0,08 | 9,39E-01 |
| AT3G05710 | 0,95 | 0,05 | 2,21 | 0,02 | 9,81E-01 |
| AT3G05725 | 0,64 | 0,67 | 2,27 | 0,30 | 7,68E-01 |
| AT3G05750 | 0,61 | -0,33 | 3,75 | -0,09 | 9,29E-01 |
| AT3G05760 | 0,68 | -0,26 | 3,76 | -0,07 | 9,44E-01 |
| AT3G05830 | 0,23 | 0,23 | 4,35 | 0,05 | 9,58E-01 |
| AT3G05840 | 0,50 | 0,00 | 4,04 | 0,00 | 1,00E+00 |
| AT3G05905 | 0,51 | -1,16 | 3,18 | -0,37 | 7,15E-01 |
| AT3G06125 | 0,60 | 1,25 | 2,29 | 0,54 | 5,86E-01 |
| AT3G06170 | 0,51 | 0,44 | 3,06 | 0,14 | 8,86E-01 |
| AT3G06180 | 0,25 | 0,00 | 4,41 | 0,00 | 1,00E+00 |
| AT3G06200 | 0,41 | 1,31 | 4,32 | 0,30 | 7,63E-01 |
| AT3G06210 | 0,20 | 0,00 | 4,41 | 0,00 | 1,00E+00 |
| AT3G06240 | 0,61 | -1,86 | 2,34 | -0,80 | 4,25E-01 |
| AT3G06380 | 0,55 | -2,05 | 3,91 | -0,52 | 6,01E-01 |
| AT3G06400 | 0,95 | 2,28 | 2,57 | 0,88 | 3,76E-01 |
| AT3G06455 | 0,58 | -0,20 | 4,35 | -0,05 | 9,64E-01 |
| AT3G06470 | 0,36 | -1,23 | 4,35 | -0,28 | 7,77E-01 |
| AT3G06500 | 1,24 | 1,78 | 2,10 | 0,85 | 3,96E-01 |
| AT3G06520 | 0,50 | 0,00 | 3,98 | 0,00 | 1,00E+00 |
| AT3G06610 | 0,15 | 0,90 | 4,35 | 0,21 | 8,36E-01 |
| AT3G06660 | 0,08 | 0,21 | 4,39 | 0,05 | 9,62E-01 |
| AT3G06730 | 0,97 | 0,40 | 3,15 | 0,13 | 8,99E-01 |
| AT3G06740 | 0,50 | 1,25 | 4,07 | 0,31 | 7,58E-01 |
| AT3G06810 | 1,67 | -1,11 | 1,93 | -0,57 | 5,67E-01 |
| AT3G06830 | 0,08 | 0,00 | 4,41 | 0,00 | 1,00E+00 |
| AT3G06860 | 1,03 | -0,24 | 1,97 | -0,12 | 9,04E-01 |
| AT3G06868 | 0,29 | -0,49 | 4,36 | -0,11 | 9,11E-01 |
| AT3G06895 | 1,19 | 2,11 | 1,89 | 1,12 | 2,63E-01 |
| AT3G06920 | 0,42 | -1,07 | 4,31 | -0,25 | 8,04E-01 |
| AT3G06980 | 0,81 | -3,52 | 2,99 | -1,18 | 2,39E-01 |
| AT3G07110 | 1,39 | -0,84 | 1,85 | -0,45 | 6,51E-01 |
| AT3G07200 | 0,08 | 0,00 | 4,41 | 0,00 | 1,00E+00 |
| AT3G07230 | 0,29 | -2,29 | 4,35 | -0,53 | 5,98E-01 |
| AT3G07270 | 0,85 | -1,20 | 2,26 | -0,53 | 5,95E-01 |
| AT3G07350 | 0,42 | -0,64 | 3,33 | -0,19 | 8,48E-01 |
| AT3G07360 | 1,00 | 2,31 | 2,39 | 0,97 | 3,33E-01 |
| AT3G07400 | 0,87 | 0,49 | 2,27 | 0,21 | 8,31E-01 |
| AT3G07430 | 0,32 | -0,62 | 4,32 | -0,14 | 8,85E-01 |
| AT3G07440 | 0,08 | 0,00 | 4,41 | 0,00 | 1,00E+00 |
| AT3G07480 | 0,45 | 0,67 | 4,28 | 0,16 | 8,76E-01 |
| AT3G07565 | 0,19 | 0,81 | 4,35 | 0,19 | 8,53E-01 |
| AT3G07610 | 0,83 | 0,84 | 2,59 | 0,33 | 7,44E-01 |
| AT3G07640 | 0,17 | 0,00 | 4,41 | 0,00 | 1,00E+00 |
| AT3G07650 | 0,97 | -0,51 | 3,46 | -0,15 | 8,82E-01 |
| AT3G07680 | 0,59 | 0,83 | 3,25 | 0,26 | 7,98E-01 |
| AT3G07690 | 0,32 | 0,00 | 4,41 | 0,00 | 1,00E+00 |
| AT3G07770 | 1,44 | 0,63 | 1,78 | 0,35 | 7,23E-01 |
| AT3G07780 | 1,15 | -0,29 | 2,12 | -0,14 | 8,91E-01 |
| AT3G07810 | 0,90 | 0,69 | 2,00 | 0,35 | 7,29E-01 |
| AT3G07880 | 0,28 | 0,43 | 4,36 | 0,10 | 9,22E-01 |

| | | | | | |
|---|---|---|---|---|---|
| AT3G07950 | 0,43 | -2,19 | 4,41 | -0,50 | 6,20E-01 |
| AT3G08010 | 0,17 | 0,22 | 4,32 | 0,05 | 9,59E-01 |
| AT3G08030 | 1,37 | -0,40 | 1,50 | -0,27 | 7,90E-01 |
| AT3G08550 | 0,31 | 1,21 | 4,34 | 0,28 | 7,80E-01 |
| AT3G08560 | 1,47 | -3,56 | 1,95 | -1,83 | 6,79E-02 |
| AT3G08600 | 0,52 | 0,84 | 3,87 | 0,22 | 8,29E-01 |
| AT3G08640 | 0,20 | -2,07 | 4,39 | -0,47 | 6,37E-01 |
| AT3G08690 | 0,22 | 0,53 | 4,36 | 0,12 | 9,03E-01 |
| AT3G08920 | 0,32 | 1,01 | 4,35 | 0,23 | 8,17E-01 |
| AT3G08930 | 0,51 | -0,13 | 3,24 | -0,04 | 9,68E-01 |
| AT3G08950 | 0,44 | 1,38 | 4,18 | 0,33 | 7,41E-01 |
| AT3G08980 | 0,33 | -0,40 | 4,38 | -0,09 | 9,27E-01 |
| AT3G09020 | 0,57 | -0,51 | 4,03 | -0,13 | 8,99E-01 |
| AT3G09030 | 0,51 | 1,15 | 2,64 | 0,43 | 6,64E-01 |
| AT3G09035 | 0,19 | 0,44 | 4,36 | 0,10 | 9,21E-01 |
| AT3G09085 | 0,40 | 0,41 | 3,43 | 0,12 | 9,05E-01 |
| AT3G02105 | 0,40 | 0,00 | 4,41 | 0,00 | 1,00E+00 |
| AT3G09180 | 0,23 | -0,41 | 4,38 | -0,09 | 9,26E-01 |
| AT3G09300 | 1,07 | -2,28 | 2,54 | -0,90 | 3,69E-01 |
| AT3G09350 | 0,51 | -3,35 | 4,35 | -0,77 | 4,41E-01 |
| AT3G09370 | 0,29 | 0,00 | 4,41 | 0,00 | 1,00E+00 |
| AT3G09430 | 1,27 | 0,00 | 2,30 | 0,00 | 1,00E+00 |
| AT3G09500 | 0,81 | -1,93 | 2,25 | -0,86 | 3,92E-01 |
| AT3G09510 | 0,52 | 1,43 | 3,06 | 0,47 | 6,40E-01 |
| AT3G09570 | 0,70 | -1,28 | 2,61 | -0,49 | 6,24E-01 |
| AT3G09600 | 0,41 | -0,35 | 3,40 | -0,10 | 9,18E-01 |
| AT3G09650 | 0,58 | 0,00 | 4,41 | 0,00 | 1,00E+00 |
| AT3G09670 | 0,48 | 0,00 | 4,41 | 0,00 | 1,00E+00 |
| AT3G09690 | 0,55 | 0,00 | 4,41 | 0,00 | 1,00E+00 |
| AT3G09810 | 0,32 | 0,80 | 4,35 | 0,18 | 8,55E-01 |
| AT3G09820 | 0,77 | -2,33 | 2,31 | -1,01 | 3,14E-01 |
| AT3G09830 | 0,46 | 0,86 | 3,23 | 0,27 | 7,90E-01 |
| AT3G09880 | 0,75 | 0,65 | 2,60 | 0,25 | 8,03E-01 |
| AT3G09890 | 0,43 | -2,19 | 4,41 | -0,50 | 6,20E-01 |
| AT3G09900 | 0,19 | -0,30 | 4,40 | -0,07 | 9,45E-01 |
| AT3G10060 | 0,36 | 1,01 | 4,35 | 0,23 | 8,16E-01 |
| AT3G10120 | 0,12 | 0,00 | 4,41 | 0,00 | 1,00E+00 |
| AT3G10190 | 0,32 | -0,37 | 4,39 | -0,09 | 9,32E-01 |
| AT3G10200 | 0,08 | 0,00 | 4,41 | 0,00 | 1,00E+00 |
| AT3G10300 | 0,68 | 1,02 | 2,78 | 0,37 | 7,14E-01 |
| AT3G10310 | 0,16 | -0,21 | 4,40 | -0,05 | 9,63E-01 |
| AT3G10340 | 0,80 | 1,06 | 2,89 | 0,37 | 7,13E-01 |
| AT3G10370 | 0,28 | 0,80 | 4,35 | 0,18 | 8,54E-01 |
| AT3G10390 | 0,49 | 0,55 | 3,26 | 0,17 | 8,66E-01 |
| AT3G10420 | 1,49 | -0,36 | 1,71 | -0,21 | 8,32E-01 |
| AT3G10480 | 0,72 | 0,00 | 3,59 | 0,00 | 1,00E+00 |
| AT3G10490 | 0,18 | -0,81 | 4,37 | -0,19 | 8,53E-01 |
| AT3G10527 | 0,27 | -1,17 | 4,40 | -0,27 | 7,91E-01 |
| AT3G10540 | 0,56 | 0,13 | 2,62 | 0,05 | 9,60E-01 |
| AT3G10570 | 0,12 | -0,36 | 4,39 | -0,08 | 9,34E-01 |
| AT3G10760 | 0,48 | 1,06 | 3,33 | 0,32 | 7,50E-01 |
| AT3G10800 | 1,54 | 1,32 | 2,05 | 0,64 | 5,21E-01 |
| AT3G10820 | 0,24 | 0,00 | 4,41 | 0,00 | 1,00E+00 |
| AT3G10860 | 0,23 | 0,85 | 4,34 | 0,20 | 8,45E-01 |
| AT3G10920 | 1,12 | -2,40 | 1,85 | -1,30 | 1,95E-01 |
| AT3G10950 | 0,19 | 1,06 | 4,34 | 0,24 | 8,07E-01 |

| | | | | | |
|---|---|---|---|---|---|
| AT3G10980 | 0,79 | 0,95 | 3,16 | 0,30 | 7,63E-01 |
| AT3G11020 | 0,05 | 0,18 | 4,38 | 0,04 | 9,68E-01 |
| AT3G11080 | 0,18 | 1,20 | 4,34 | 0,28 | 7,82E-01 |
| AT3G11090 | 0,60 | 0,89 | 3,20 | 0,28 | 7,81E-01 |
| AT3G11120 | 0,39 | 0,93 | 3,30 | 0,28 | 7,78E-01 |
| AT3G11170 | 1,02 | -0,52 | 1,56 | -0,33 | 7,40E-01 |
| AT3G11230 | 0,53 | -2,80 | 4,37 | -0,64 | 5,22E-01 |
| AT3G11250 | 0,11 | 0,88 | 4,35 | 0,20 | 8,39E-01 |
| AT3G11270 | 0,10 | 0,08 | 4,38 | 0,02 | 9,85E-01 |
| AT3G11397 | 0,44 | -2,67 | 4,12 | -0,65 | 5,18E-01 |
| AT3G11400 | 0,87 | 0,47 | 2,42 | 0,19 | 8,47E-01 |
| AT3G11420 | 0,26 | 0,38 | 4,32 | 0,09 | 9,31E-01 |
| AT3G11440 | 0,65 | 0,00 | 3,09 | 0,00 | 1,00E+00 |
| AT3G11560 | 1,70 | 1,25 | 1,93 | 0,65 | 5,17E-01 |
| AT3G02465 | 0,30 | 0,81 | 4,34 | 0,19 | 8,53E-01 |
| AT3G11630 | 0,96 | 0,18 | 1,67 | 0,11 | 9,14E-01 |
| AT3G11700 | 0,50 | 0,38 | 3,20 | 0,12 | 9,05E-01 |
| AT3G11760 | 0,23 | -0,35 | 4,39 | -0,08 | 9,36E-01 |
| AT3G11810 | 0,52 | 1,00 | 3,81 | 0,26 | 7,94E-01 |
| AT3G11900 | 0,13 | 0,00 | 4,41 | 0,00 | 1,00E+00 |
| AT3G11930 | 1,35 | -3,37 | 3,37 | -1,00 | 3,17E-01 |
| AT3G11960 | 1,04 | -1,19 | 2,18 | -0,55 | 5,83E-01 |
| AT3G12010 | 0,40 | 1,19 | 4,33 | 0,27 | 7,83E-01 |
| AT3G12020 | 0,79 | -0,25 | 2,54 | -0,10 | 9,23E-01 |
| AT3G12050 | 0,32 | 1,79 | 3,49 | 0,51 | 6,08E-01 |
| AT3G12080 | 1,08 | -0,42 | 2,62 | -0,16 | 8,74E-01 |
| AT3G12270 | 0,77 | -1,25 | 2,62 | -0,48 | 6,32E-01 |
| AT3G12290 | 0,81 | 2,31 | 2,48 | 0,93 | 3,51E-01 |
| AT3G12320 | 0,90 | -1,28 | 2,27 | -0,56 | 5,75E-01 |
| AT3G12360 | 0,71 | 0,75 | 2,50 | 0,30 | 7,63E-01 |
| AT3G12390 | 0,69 | -1,26 | 2,49 | -0,50 | 6,14E-01 |
| AT3G12480 | 0,26 | -1,24 | 4,39 | -0,28 | 7,78E-01 |
| AT3G12502 | 0,08 | 0,00 | 4,41 | 0,00 | 1,00E+00 |
| AT3G12570 | 1,24 | -2,54 | 2,77 | -0,92 | 3,60E-01 |
| AT3G12600 | 0,05 | 0,18 | 4,38 | 0,04 | 9,68E-01 |
| AT3G12640 | 0,60 | -0,55 | 2,34 | -0,23 | 8,16E-01 |
| AT3G12700 | 0,20 | 0,40 | 4,37 | 0,09 | 9,27E-01 |
| AT3G12740 | 0,52 | 2,03 | 3,26 | 0,62 | 5,33E-01 |
| AT3G12760 | 0,80 | -1,34 | 2,45 | -0,55 | 5,83E-01 |
| AT3G12810 | 1,56 | 1,22 | 1,89 | 0,64 | 5,20E-01 |
| AT3G12915 | 0,53 | 0,00 | 4,41 | 0,00 | 1,00E+00 |
| AT3G12980 | 1,82 | -1,38 | 1,93 | -0,72 | 4,73E-01 |
| AT3G13060 | 0,87 | 2,16 | 2,25 | 0,96 | 3,36E-01 |
| AT3G13062 | 1,32 | 0,46 | 1,57 | 0,29 | 7,69E-01 |
| AT3G13065 | 0,88 | 0,00 | 3,51 | 0,00 | 1,00E+00 |
| AT3G13200 | 1,18 | 0,87 | 1,73 | 0,51 | 6,12E-01 |
| AT3G13210 | 0,13 | 0,00 | 4,41 | 0,00 | 1,00E+00 |
| AT3G13340 | 1,00 | 0,01 | 1,89 | 0,00 | 9,96E-01 |
| AT3G13370 | 0,63 | 3,41 | 4,27 | 0,80 | 4,24E-01 |
| AT3G13380 | 0,67 | 1,64 | 2,46 | 0,67 | 5,05E-01 |
| AT3G13437 | 0,08 | 0,00 | 4,41 | 0,00 | 1,00E+00 |
| AT3G13445 | 0,19 | -1,25 | 4,36 | -0,29 | 7,74E-01 |
| AT3G13470 | 0,61 | 0,95 | 3,91 | 0,24 | 8,08E-01 |
| AT3G02725 | 1,24 | -0,35 | 1,80 | -0,19 | 8,48E-01 |
| AT3G13662 | 0,34 | 1,52 | 4,33 | 0,35 | 7,26E-01 |
| AT3G13690 | 0,67 | -0,32 | 3,86 | -0,08 | 9,35E-01 |

| | | | | | |
|---|---|---|---|---|---|
| AT3G13760 | 0,08 | 0,00 | 4,41 | 0,00 | 1,00E+00 |
| AT3G13770 | 0,43 | -0,40 | 4,38 | -0,09 | 9,27E-01 |
| AT3G13810 | 0,42 | 1,73 | 4,17 | 0,42 | 6,77E-01 |
| AT3G13930 | 0,81 | 0,41 | 2,11 | 0,20 | 8,44E-01 |
| AT3G13950 | 0,55 | 1,41 | 3,93 | 0,36 | 7,21E-01 |
| AT3G14020 | 0,35 | 0,00 | 4,39 | 0,00 | 1,00E+00 |
| AT3G14100 | 0,69 | 1,17 | 2,52 | 0,46 | 6,43E-01 |
| AT3G14120 | 1,41 | -0,28 | 2,25 | -0,13 | 8,99E-01 |
| AT3G14160 | 0,19 | 0,43 | 4,36 | 0,10 | 9,21E-01 |
| AT3G14220 | 0,64 | -1,31 | 3,84 | -0,34 | 7,33E-01 |
| AT3G14290 | 0,42 | -2,34 | 4,18 | -0,56 | 5,76E-01 |
| AT3G14415 | 1,72 | 0,48 | 1,43 | 0,33 | 7,38E-01 |
| AT3G14560 | 0,20 | 0,00 | 4,41 | 0,00 | 1,00E+00 |
| AT3G14595 | 1,01 | -0,79 | 2,08 | -0,38 | 7,03E-01 |
| AT3G14600 | 1,18 | 0,17 | 1,82 | 0,10 | 9,24E-01 |
| AT3G14610 | 0,28 | 0,00 | 4,41 | 0,00 | 1,00E+00 |
| AT3G14620 | 0,63 | 1,07 | 2,50 | 0,43 | 6,70E-01 |
| AT3G14640 | 0,33 | 0,00 | 4,41 | 0,00 | 1,00E+00 |
| AT3G14660 | 1,69 | 0,40 | 1,30 | 0,30 | 7,61E-01 |
| AT3G14670 | 1,07 | 1,77 | 1,88 | 0,94 | 3,48E-01 |
| AT3G14690 | 0,63 | 0,00 | 2,17 | 0,00 | 9,98E-01 |
| AT3G14720 | 0,35 | 1,29 | 4,33 | 0,30 | 7,67E-01 |
| AT3G14790 | 0,27 | -0,39 | 4,39 | -0,09 | 9,29E-01 |
| AT3G14830 | 0,30 | 0,04 | 4,38 | 0,01 | 9,93E-01 |
| AT3G14840 | 1,81 | 0,45 | 1,62 | 0,28 | 7,81E-01 |
| AT3G14870 | 0,25 | 0,17 | 4,38 | 0,04 | 9,70E-01 |
| AT3G14920 | 0,30 | 0,00 | 4,41 | 0,00 | 1,00E+00 |
| AT3G14940 | 0,68 | 0,14 | 3,61 | 0,04 | 9,70E-01 |
| AT3G14990 | 0,39 | 0,36 | 2,85 | 0,13 | 9,00E-01 |
| AT3G15000 | 0,95 | 0,43 | 2,29 | 0,19 | 8,51E-01 |
| AT3G15010 | 0,55 | -0,28 | 3,16 | -0,09 | 9,31E-01 |
| AT3G15020 | 1,02 | -0,26 | 2,59 | -0,10 | 9,19E-01 |
| AT3G15030 | 0,69 | -1,04 | 3,08 | -0,34 | 7,35E-01 |
| AT3G15090 | 0,12 | 0,00 | 4,41 | 0,00 | 1,00E+00 |
| AT3G15095 | 1,44 | -0,51 | 1,44 | -0,35 | 7,23E-01 |
| AT3G15160 | 1,08 | 0,63 | 2,32 | 0,27 | 7,87E-01 |
| AT3G15180 | 0,21 | 0,08 | 4,38 | 0,02 | 9,85E-01 |
| AT3G15190 | 1,26 | -0,73 | 1,78 | -0,41 | 6,79E-01 |
| AT3G15210 | 1,70 | 0,60 | 1,25 | 0,48 | 6,29E-01 |
| AT3G15260 | 0,39 | -1,33 | 4,34 | -0,31 | 7,59E-01 |
| AT3G15290 | 0,26 | 0,00 | 4,41 | 0,00 | 1,00E+00 |
| AT3G15350 | 1,19 | -1,29 | 2,47 | -0,52 | 6,01E-01 |
| AT3G15355 | 0,91 | 0,55 | 3,81 | 0,15 | 8,84E-01 |
| AT3G15360 | 1,20 | -0,90 | 1,59 | -0,57 | 5,71E-01 |
| AT3G15380 | 0,12 | 0,00 | 4,41 | 0,00 | 1,00E+00 |
| AT3G15450 | 0,77 | -1,32 | 2,33 | -0,57 | 5,69E-01 |
| AT3G15518 | 0,17 | 0,00 | 4,41 | 0,00 | 1,00E+00 |
| AT3G15550 | 0,17 | -2,13 | 4,40 | -0,48 | 6,28E-01 |
| AT3G03145 | 0,66 | 1,08 | 2,90 | 0,37 | 7,10E-01 |
| AT3G15635 | 0,25 | 0,75 | 4,37 | 0,17 | 8,64E-01 |
| AT3G15750 | 0,08 | 0,00 | 4,41 | 0,00 | 1,00E+00 |
| AT3G15760 | 0,84 | 0,16 | 2,59 | 0,06 | 9,51E-01 |
| AT3G15790 | 0,67 | -0,37 | 2,97 | -0,12 | 9,01E-01 |
| AT3G15820 | 0,32 | 0,43 | 4,36 | 0,10 | 9,22E-01 |
| AT3G15850 | 0,57 | -0,18 | 2,54 | -0,07 | 9,43E-01 |
| AT3G15930 | 0,12 | 0,00 | 4,41 | 0,00 | 1,00E+00 |

| | | | | | |
|---|---|---|---|---|---|
| AT3G16010 | 0,15 | 0,82 | 4,35 | 0,19 | 8,50E-01 |
| AT3G16060 | 0,47 | -0,96 | 4,10 | -0,23 | 8,15E-01 |
| AT3G16080 | 0,49 | 0,55 | 3,30 | 0,17 | 8,67E-01 |
| AT3G16130 | 0,87 | 0,05 | 1,99 | 0,03 | 9,80E-01 |
| AT3G16175 | 0,08 | 0,00 | 4,41 | 0,00 | 1,00E+00 |
| AT3G16240 | 1,39 | 0,00 | 1,44 | 0,00 | 9,98E-01 |
| AT3G16260 | 0,72 | 0,00 | 3,61 | 0,00 | 1,00E+00 |
| AT3G16270 | 0,71 | -1,61 | 2,39 | -0,67 | 5,01E-01 |
| AT3G16340 | 0,41 | 0,00 | 4,41 | 0,00 | 1,00E+00 |
| AT3G16420 | 0,28 | -0,16 | 4,40 | -0,04 | 9,70E-01 |
| AT3G16460 | 0,31 | 1,31 | 4,33 | 0,30 | 7,62E-01 |
| AT3G16525 | 0,84 | 0,21 | 1,97 | 0,11 | 9,15E-01 |
| AT3G16570 | 0,68 | 0,22 | 2,49 | 0,09 | 9,31E-01 |
| AT3G16710 | 0,43 | -1,28 | 4,26 | -0,30 | 7,64E-01 |
| AT3G16720 | 0,43 | 0,79 | 4,12 | 0,19 | 8,47E-01 |
| AT3G16740 | 0,38 | -0,54 | 3,46 | -0,16 | 8,76E-01 |
| AT3G16760 | 0,57 | 0,82 | 3,81 | 0,22 | 8,29E-01 |
| AT3G16770 | 1,33 | -1,61 | 2,30 | -0,70 | 4,82E-01 |
| AT3G16800 | 0,16 | -0,56 | 4,37 | -0,13 | 8,98E-01 |
| AT3G16830 | 1,75 | -0,51 | 1,49 | -0,34 | 7,34E-01 |
| AT3G16857 | 0,72 | -0,26 | 3,88 | -0,07 | 9,46E-01 |
| AT3G16940 | 0,78 | -0,98 | 2,97 | -0,33 | 7,42E-01 |
| AT3G17100 | 1,17 | -1,12 | 1,81 | -0,62 | 5,37E-01 |
| AT3G17160 | 0,19 | -0,37 | 4,39 | -0,08 | 9,33E-01 |
| AT3G17190 | 1,85 | 2,10 | 1,55 | 1,35 | 1,77E-01 |
| AT3G17205 | 0,35 | 1,46 | 4,32 | 0,34 | 7,35E-01 |
| AT3G17210 | 0,32 | -0,30 | 4,40 | -0,07 | 9,46E-01 |
| AT3G03465 | 0,80 | 1,17 | 2,21 | 0,53 | 5,96E-01 |
| AT3G17360 | 0,65 | -0,29 | 3,13 | -0,09 | 9,27E-01 |
| AT3G17380 | 0,28 | 0,00 | 4,41 | 0,00 | 1,00E+00 |
| AT3G17410 | 0,79 | 0,61 | 1,96 | 0,31 | 7,55E-01 |
| AT3G17470 | 0,76 | -0,39 | 3,82 | -0,10 | 9,18E-01 |
| AT3G17490 | 0,56 | 2,98 | 3,81 | 0,78 | 4,33E-01 |
| AT3G17626 | 0,20 | 0,00 | 4,41 | 0,00 | 1,00E+00 |
| AT3G17640 | 0,16 | -0,56 | 4,37 | -0,13 | 8,98E-01 |
| AT3G17650 | 0,88 | 0,26 | 2,12 | 0,12 | 9,01E-01 |
| AT3G17690 | 0,12 | 0,00 | 4,41 | 0,00 | 1,00E+00 |
| AT3G17700 | 0,35 | 1,13 | 4,33 | 0,26 | 7,94E-01 |
| AT3G17712 | 0,66 | -0,30 | 3,78 | -0,08 | 9,36E-01 |
| AT3G17750 | 1,26 | -0,51 | 1,75 | -0,29 | 7,70E-01 |
| AT3G17765 | 0,21 | 0,00 | 4,41 | 0,00 | 1,00E+00 |
| AT3G17790 | 0,83 | -1,73 | 2,57 | -0,67 | 5,02E-01 |
| AT3G03535 | 0,92 | -0,40 | 2,27 | -0,18 | 8,60E-01 |
| AT3G17810 | 0,53 | 0,42 | 2,79 | 0,15 | 8,80E-01 |
| AT3G17820 | 0,62 | -0,56 | 3,25 | -0,17 | 8,64E-01 |
| AT3G17860 | 0,36 | 1,59 | 4,32 | 0,37 | 7,13E-01 |
| AT3G17880 | 0,42 | 1,63 | 3,25 | 0,50 | 6,15E-01 |
| AT3G17890 | 0,10 | 0,08 | 4,38 | 0,02 | 9,85E-01 |
| AT3G17970 | 0,77 | -1,43 | 2,97 | -0,48 | 6,30E-01 |
| AT3G17980 | 0,35 | 0,00 | 4,39 | 0,00 | 1,00E+00 |
| AT3G18000 | 0,28 | -0,22 | 4,40 | -0,05 | 9,60E-01 |
| AT3G18020 | 0,41 | -1,33 | 4,24 | -0,31 | 7,54E-01 |
| AT3G18040 | 0,26 | -0,11 | 4,34 | -0,03 | 9,79E-01 |
| AT3G18050 | 1,52 | -2,43 | 1,53 | -1,59 | 1,12E-01 |
| AT3G18060 | 1,35 | 0,07 | 1,65 | 0,04 | 9,65E-01 |
| AT3G18070 | 0,11 | -0,26 | 4,41 | -0,06 | 9,53E-01 |

| | | | | | |
|---|---|---|---|---|---|
| AT3G18080 | 1,62 | -0,44 | 1,44 | -0,31 | 7,60E-01 |
| AT3G18090 | 1,26 | 0,00 | 3,11 | 0,00 | 1,00E+00 |
| AT3G18100 | 0,29 | 0,00 | 4,41 | 0,00 | 1,00E+00 |
| AT3G03625 | 0,40 | 0,75 | 3,35 | 0,22 | 8,23E-01 |
| AT3G18190 | 1,46 | -1,33 | 1,63 | -0,81 | 4,15E-01 |
| AT3G18240 | 0,43 | -0,92 | 4,23 | -0,22 | 8,27E-01 |
| AT3G18270 | 0,50 | -0,78 | 4,02 | -0,19 | 8,47E-01 |
| AT3G18340 | 0,19 | 1,60 | 4,31 | 0,37 | 7,11E-01 |
| AT3G18390 | 1,15 | -1,01 | 1,83 | -0,55 | 5,82E-01 |
| AT3G18440 | 0,08 | 0,00 | 4,41 | 0,00 | 1,00E+00 |
| AT3G18480 | 0,44 | 0,46 | 3,28 | 0,14 | 8,89E-01 |
| AT3G18740 | 0,73 | -1,49 | 2,25 | -0,66 | 5,07E-01 |
| AT3G18790 | 0,65 | -1,77 | 2,52 | -0,70 | 4,83E-01 |
| AT3G18800 | 0,33 | 0,00 | 4,41 | 0,00 | 1,00E+00 |
| AT3G18820 | 0,39 | -0,01 | 4,14 | 0,00 | 9,98E-01 |
| AT3G18827 | 0,57 | -1,40 | 3,09 | -0,45 | 6,52E-01 |
| AT3G18860 | 0,59 | 1,93 | 2,51 | 0,77 | 4,43E-01 |
| AT3G18890 | 0,86 | -0,21 | 2,26 | -0,09 | 9,25E-01 |
| AT3G18915 | 0,30 | 1,72 | 4,32 | 0,40 | 6,90E-01 |
| AT3G18960 | 0,08 | 0,00 | 4,41 | 0,00 | 1,00E+00 |
| AT3G18980 | 1,15 | -0,40 | 2,09 | -0,19 | 8,49E-01 |
| AT3G19050 | 0,62 | 0,00 | 3,09 | 0,00 | 1,00E+00 |
| AT3G19100 | 1,08 | 0,21 | 2,13 | 0,10 | 9,21E-01 |
| AT3G19240 | 1,33 | 1,65 | 1,67 | 0,99 | 3,23E-01 |
| AT3G19290 | 0,28 | -0,24 | 4,40 | -0,05 | 9,57E-01 |
| AT3G19390 | 0,35 | 0,00 | 4,39 | 0,00 | 1,00E+00 |
| AT3G19470 | 0,30 | 0,46 | 4,36 | 0,10 | 9,17E-01 |
| AT3G19480 | 0,52 | 1,00 | 3,23 | 0,31 | 7,57E-01 |
| AT3G19520 | 0,05 | 0,18 | 4,38 | 0,04 | 9,68E-01 |
| AT3G19540 | 1,20 | 1,92 | 2,09 | 0,92 | 3,58E-01 |
| AT3G19560 | 1,58 | 0,95 | 1,93 | 0,49 | 6,22E-01 |
| AT3G19640 | 0,86 | -0,26 | 2,56 | -0,10 | 9,18E-01 |
| AT3G19680 | 0,92 | -1,95 | 2,02 | -0,97 | 3,33E-01 |
| AT3G19760 | 1,20 | -0,53 | 1,74 | -0,30 | 7,62E-01 |
| AT3G19770 | 0,75 | -1,81 | 3,05 | -0,59 | 5,53E-01 |
| AT3G19780 | 1,34 | 0,10 | 2,30 | 0,04 | 9,65E-01 |
| AT3G19800 | 0,78 | -2,33 | 2,55 | -0,91 | 3,62E-01 |
| AT3G19860 | 0,48 | -2,14 | 4,07 | -0,52 | 6,00E-01 |
| AT3G19870 | 0,37 | 0,69 | 4,32 | 0,16 | 8,73E-01 |
| AT3G19910 | 1,61 | 1,39 | 1,63 | 0,86 | 3,92E-01 |
| AT3G19930 | 1,47 | 1,54 | 1,60 | 0,96 | 3,35E-01 |
| AT3G19960 | 1,65 | 1,60 | 1,74 | 0,92 | 3,59E-01 |
| AT3G19980 | 0,93 | 0,49 | 2,43 | 0,20 | 8,40E-01 |
| AT3G19990 | 0,32 | 0,00 | 4,41 | 0,00 | 1,00E+00 |
| AT3G20000 | 0,74 | -2,40 | 3,67 | -0,66 | 5,12E-01 |
| AT3G20030 | 0,79 | 2,08 | 2,36 | 0,88 | 3,78E-01 |
| AT3G20140 | 0,20 | 0,00 | 4,41 | 0,00 | 1,00E+00 |
| AT3G20150 | 0,54 | 0,00 | 4,41 | 0,00 | 1,00E+00 |
| AT3G20200 | 0,21 | 0,00 | 4,41 | 0,00 | 1,00E+00 |
| AT3G20410 | 0,37 | -0,03 | 4,31 | -0,01 | 9,95E-01 |
| AT3G20500 | 0,33 | 0,00 | 4,41 | 0,00 | 1,00E+00 |
| AT3G20520 | 0,48 | 0,00 | 4,41 | 0,00 | 1,00E+00 |
| AT3G20540 | 0,90 | 0,46 | 2,49 | 0,18 | 8,53E-01 |
| AT3G20600 | 0,25 | 0,28 | 4,39 | 0,06 | 9,50E-01 |
| AT3G03995 | 0,62 | 3,12 | 4,27 | 0,73 | 4,65E-01 |
| AT3G20720 | 0,42 | 1,63 | 3,39 | 0,48 | 6,31E-01 |

| | | | | | |
|---|---|---|---|---|---|
| AT3G20730 | 0,40 | 0,00 | 4,41 | 0,00 | 1,00E+00 |
| AT3G20790 | 0,23 | -2,50 | 4,38 | -0,57 | 5,68E-01 |
| AT3G20890 | 0,34 | 0,09 | 4,38 | 0,02 | 9,84E-01 |
| AT3G20950 | 1,75 | 2,66 | 1,66 | 1,60 | 1,10E-01 |
| AT3G20970 | 0,63 | -1,07 | 3,05 | -0,35 | 7,25E-01 |
| AT3G21130 | 0,35 | 1,68 | 4,32 | 0,39 | 6,97E-01 |
| AT3G21170 | 1,15 | 4,02 | 2,30 | 1,74 | 8,10E-02 |
| AT3G21175 | 0,18 | 1,20 | 4,34 | 0,28 | 7,82E-01 |
| AT3G21220 | 0,51 | 0,80 | 4,01 | 0,20 | 8,41E-01 |
| AT3G21270 | 0,29 | -1,33 | 4,39 | -0,30 | 7,62E-01 |
| AT3G21295 | 0,41 | -0,54 | 3,42 | -0,16 | 8,76E-01 |
| AT3G21310 | 0,36 | 0,00 | 4,41 | 0,00 | 1,00E+00 |
| AT3G21465 | 0,14 | -0,82 | 4,37 | -0,19 | 8,50E-01 |
| AT3G21473 | 0,48 | -0,29 | 3,92 | -0,07 | 9,42E-01 |
| AT3G21560 | 0,31 | 0,07 | 4,38 | 0,02 | 9,87E-01 |
| AT3G21690 | 0,96 | 1,69 | 1,98 | 0,85 | 3,94E-01 |
| AT3G21750 | 1,77 | 0,47 | 1,37 | 0,34 | 7,35E-01 |
| AT3G04175 | 0,99 | 1,86 | 1,98 | 0,94 | 3,47E-01 |
| AT3G21990 | 0,46 | 2,73 | 4,28 | 0,64 | 5,24E-01 |
| AT3G22072 | 0,17 | -3,06 | 4,35 | -0,70 | 4,82E-01 |
| AT3G22104 | 0,62 | 0,32 | 2,98 | 0,11 | 9,14E-01 |
| AT3G22125 | 1,40 | 0,71 | 1,58 | 0,45 | 6,52E-01 |
| AT3G22150 | 0,49 | 0,00 | 4,03 | 0,00 | 1,00E+00 |
| AT3G22200 | 0,76 | -0,08 | 2,26 | -0,04 | 9,71E-01 |
| AT3G04255 | 0,24 | -0,67 | 4,38 | -0,15 | 8,78E-01 |
| AT3G22270 | 0,79 | 0,04 | 2,55 | 0,02 | 9,86E-01 |
| AT3G22290 | 0,33 | 0,00 | 4,41 | 0,00 | 1,00E+00 |
| AT3G22330 | 0,85 | -1,17 | 2,10 | -0,56 | 5,77E-01 |
| AT3G22350 | 0,91 | 3,50 | 2,52 | 1,39 | 1,64E-01 |
| AT3G22370 | 0,72 | 1,67 | 2,57 | 0,65 | 5,15E-01 |
| AT3G22425 | 0,12 | 0,11 | 4,38 | 0,03 | 9,80E-01 |
| AT3G22430 | 0,19 | 1,22 | 4,34 | 0,28 | 7,79E-01 |
| AT3G22520 | 0,75 | -0,21 | 2,64 | -0,08 | 9,37E-01 |
| AT3G22760 | 0,51 | -0,38 | 3,20 | -0,12 | 9,06E-01 |
| AT3G22810 | 0,55 | 0,00 | 4,39 | 0,00 | 1,00E+00 |
| AT3G22845 | 0,51 | 0,00 | 4,41 | 0,00 | 1,00E+00 |
| AT3G22870 | 0,92 | 3,43 | 2,12 | 1,62 | 1,05E-01 |
| AT3G22900 | 0,24 | 0,00 | 4,41 | 0,00 | 1,00E+00 |
| AT3G22935 | 0,26 | 2,23 | 4,29 | 0,52 | 6,03E-01 |
| AT3G22940 | 1,72 | 3,44 | 1,79 | 1,92 | 5,50E-02 |
| AT3G23010 | 0,77 | -0,19 | 3,62 | -0,05 | 9,58E-01 |
| AT3G23050 | 0,64 | 0,71 | 2,59 | 0,27 | 7,84E-01 |
| AT3G04395 | 0,16 | 0,15 | 4,38 | 0,04 | 9,72E-01 |
| AT3G23250 | 0,25 | 1,09 | 4,34 | 0,25 | 8,02E-01 |
| AT3G23280 | 0,45 | 0,13 | 4,05 | 0,03 | 9,74E-01 |
| AT3G23300 | 0,75 | 1,15 | 2,20 | 0,52 | 6,01E-01 |
| AT3G23310 | 0,63 | -0,28 | 2,22 | -0,13 | 8,99E-01 |
| AT3G23380 | 0,21 | 0,59 | 4,37 | 0,14 | 8,92E-01 |
| AT3G23390 | 0,15 | 0,82 | 4,35 | 0,19 | 8,50E-01 |
| AT3G23410 | 0,36 | -1,72 | 4,32 | -0,40 | 6,92E-01 |
| AT3G23530 | 1,09 | -1,16 | 2,03 | -0,57 | 5,68E-01 |
| AT3G23580 | 0,57 | -0,91 | 2,28 | -0,40 | 6,90E-01 |
| AT3G23590 | 1,64 | -0,05 | 1,93 | -0,02 | 9,81E-01 |
| AT3G23600 | 1,16 | 2,09 | 1,97 | 1,06 | 2,88E-01 |
| AT3G23640 | 0,83 | -0,31 | 2,55 | -0,12 | 9,04E-01 |
| AT3G23750 | 0,94 | 0,14 | 1,69 | 0,09 | 9,32E-01 |

| | | | | | |
|---|---|---|---|---|---|
| AT3G23790 | 0,38 | 0,61 | 4,37 | 0,14 | 8,89E-01 |
| AT3G23800 | 0,20 | 0,00 | 4,41 | 0,00 | 1,00E+00 |
| AT3G23890 | 0,56 | -0,05 | 4,02 | -0,01 | 9,90E-01 |
| AT3G23990 | 0,73 | -2,14 | 3,75 | -0,57 | 5,69E-01 |
| AT3G24000 | 0,34 | -0,30 | 4,40 | -0,07 | 9,46E-01 |
| AT3G24050 | 0,88 | -0,84 | 1,97 | -0,43 | 6,68E-01 |
| AT3G24070 | 0,38 | 0,00 | 4,41 | 0,00 | 1,00E+00 |
| AT3G24160 | 0,45 | 0,21 | 3,41 | 0,06 | 9,50E-01 |
| AT3G24190 | 0,59 | -1,11 | 3,14 | -0,35 | 7,23E-01 |
| AT3G24255 | 0,43 | -0,49 | 2,81 | -0,17 | 8,63E-01 |
| AT3G24490 | 0,64 | 0,00 | 3,67 | 0,00 | 1,00E+00 |
| AT3G24500 | 0,22 | 1,63 | 4,31 | 0,38 | 7,05E-01 |
| AT3G24514 | 0,29 | 2,23 | 4,29 | 0,52 | 6,04E-01 |
| AT3G24520 | 0,92 | -0,10 | 2,37 | -0,04 | 9,65E-01 |
| AT3G24550 | 0,60 | 1,63 | 2,94 | 0,56 | 5,78E-01 |
| AT3G24580 | 0,47 | 0,88 | 4,05 | 0,22 | 8,28E-01 |
| AT3G24590 | 0,21 | 0,00 | 4,41 | 0,00 | 1,00E+00 |
| AT3G24830 | 0,92 | -1,69 | 1,95 | -0,87 | 3,85E-01 |
| AT3G25012 | 0,95 | 0,61 | 1,94 | 0,31 | 7,54E-01 |
| AT3G25013 | 0,18 | 1,20 | 4,34 | 0,28 | 7,82E-01 |
| AT3G25030 | 0,21 | -0,71 | 4,38 | -0,16 | 8,71E-01 |
| AT3G25040 | 0,17 | -0,78 | 4,36 | -0,18 | 8,59E-01 |
| AT3G25070 | 0,31 | 1,29 | 3,58 | 0,36 | 7,18E-01 |
| AT3G25130 | 0,57 | -3,91 | 4,32 | -0,91 | 3,65E-01 |
| AT3G25140 | 1,51 | -2,45 | 2,14 | -1,15 | 2,52E-01 |
| AT3G25250 | 0,60 | -2,02 | 2,66 | -0,76 | 4,49E-01 |
| AT3G25290 | 0,29 | -0,36 | 4,39 | -0,08 | 9,35E-01 |
| AT3G25440 | 0,23 | 0,24 | 4,39 | 0,06 | 9,56E-01 |
| AT3G25470 | 0,67 | -2,92 | 3,76 | -0,78 | 4,37E-01 |
| AT3G25490 | 0,05 | 0,18 | 4,38 | 0,04 | 9,68E-01 |
| AT3G25585 | 0,71 | -2,18 | 2,67 | -0,82 | 4,13E-01 |
| AT3G25590 | 0,75 | 2,47 | 2,39 | 1,04 | 3,00E-01 |
| AT3G25700 | 0,66 | -0,80 | 2,20 | -0,37 | 7,15E-01 |
| AT3G04955 | 0,15 | -0,39 | 4,39 | -0,09 | 9,29E-01 |
| AT3G25720 | 0,26 | 1,66 | 4,32 | 0,39 | 7,00E-01 |
| AT3G25760 | 0,49 | -2,04 | 4,38 | -0,47 | 6,42E-01 |
| AT3G25770 | 1,09 | -1,52 | 1,70 | -0,89 | 3,71E-01 |
| AT3G25780 | 0,49 | 2,29 | 2,60 | 0,88 | 3,79E-01 |
| AT3G25860 | 0,63 | -1,97 | 3,11 | -0,63 | 5,27E-01 |
| AT3G25890 | 0,89 | 0,54 | 2,20 | 0,25 | 8,04E-01 |
| AT3G25910 | 0,36 | -0,35 | 4,39 | -0,08 | 9,36E-01 |
| AT3G26020 | 0,28 | -0,42 | 4,36 | -0,10 | 9,23E-01 |
| AT3G26030 | 1,16 | -0,66 | 2,03 | -0,32 | 7,45E-01 |
| AT3G26070 | 0,46 | -0,44 | 4,20 | -0,10 | 9,17E-01 |
| AT3G26080 | 0,26 | 0,00 | 4,41 | 0,00 | 1,00E+00 |
| AT3G26085 | 0,30 | -0,35 | 4,40 | -0,08 | 9,37E-01 |
| AT3G26090 | 0,21 | 0,05 | 4,38 | 0,01 | 9,91E-01 |
| AT3G04975 | 1,19 | 0,55 | 1,80 | 0,30 | 7,61E-01 |
| AT3G04985 | 0,58 | 0,72 | 2,52 | 0,29 | 7,74E-01 |
| AT3G26370 | 0,48 | -1,27 | 2,80 | -0,45 | 6,50E-01 |
| AT3G26395 | 0,16 | -0,22 | 4,40 | -0,05 | 9,60E-01 |
| AT3G26400 | 0,95 | -0,15 | 2,36 | -0,06 | 9,49E-01 |
| AT3G26430 | 1,60 | -0,36 | 1,60 | -0,23 | 8,22E-01 |
| AT3G26420 | 1,60 | -0,36 | 1,60 | -0,23 | 8,22E-01 |
| AT3G26512 | 0,38 | 2,09 | 3,30 | 0,63 | 5,27E-01 |
| AT3G26570 | 1,54 | -0,63 | 1,81 | -0,35 | 7,26E-01 |

| | | | | | |
|---|---|---|---|---|---|
| AT3G26580 | 0,73 | -0,83 | 2,14 | -0,39 | 6,97E-01 |
| AT3G26600 | 1,07 | 1,10 | 2,05 | 0,54 | 5,92E-01 |
| AT3G26640 | 0,37 | 1,89 | 3,35 | 0,56 | 5,73E-01 |
| AT3G26710 | 1,14 | -2,04 | 2,16 | -0,94 | 3,46E-01 |
| AT3G26720 | 0,82 | -0,45 | 2,59 | -0,17 | 8,61E-01 |
| AT3G26742 | 0,18 | 0,82 | 4,35 | 0,19 | 8,51E-01 |
| AT3G26780 | 1,24 | 1,36 | 2,09 | 0,65 | 5,16E-01 |
| AT3G26782 | 0,41 | -0,32 | 4,20 | -0,08 | 9,39E-01 |
| AT3G26800 | 0,34 | 0,00 | 3,49 | 0,00 | 1,00E+00 |
| AT3G26840 | 1,06 | -0,26 | 2,45 | -0,11 | 9,15E-01 |
| AT3G27090 | 1,24 | -0,27 | 2,02 | -0,13 | 8,94E-01 |
| AT3G27160 | 0,41 | -0,85 | 3,31 | -0,26 | 7,97E-01 |
| AT3G27170 | 0,68 | -2,08 | 2,96 | -0,71 | 4,81E-01 |
| AT3G27230 | 0,45 | 0,27 | 3,38 | 0,08 | 9,37E-01 |
| AT3G27250 | 0,85 | -0,60 | 2,46 | -0,24 | 8,09E-01 |
| AT3G27260 | 1,09 | 1,28 | 2,03 | 0,63 | 5,28E-01 |
| AT3G27270 | 0,62 | -2,18 | 3,84 | -0,57 | 5,71E-01 |
| AT3G27280 | 0,60 | -1,33 | 3,09 | -0,43 | 6,67E-01 |
| AT3G27325 | 0,56 | 1,04 | 2,66 | 0,39 | 6,97E-01 |
| AT3G27329 | 0,81 | -1,28 | 1,95 | -0,66 | 5,12E-01 |
| AT3G27331 | 0,96 | 3,79 | 3,12 | 1,22 | 2,24E-01 |
| AT3G27350 | 0,51 | 0,85 | 3,96 | 0,22 | 8,29E-01 |
| AT3G27400 | 1,80 | -0,71 | 1,17 | -0,61 | 5,45E-01 |
| AT3G27430 | 0,35 | -0,40 | 3,48 | -0,11 | 9,09E-01 |
| AT3G27470 | 1,23 | 2,52 | 1,87 | 1,35 | 1,77E-01 |
| AT3G27540 | 0,29 | 0,13 | 3,65 | 0,03 | 9,73E-01 |
| AT3G27555 | 0,30 | 1,44 | 4,32 | 0,33 | 7,38E-01 |
| AT3G27670 | 1,19 | 1,31 | 2,31 | 0,57 | 5,72E-01 |
| AT3G05295 | 0,48 | 0,78 | 2,83 | 0,28 | 7,83E-01 |
| AT3G27820 | 1,03 | 1,15 | 1,95 | 0,59 | 5,55E-01 |
| AT3G27830 | 0,47 | 2,18 | 3,34 | 0,65 | 5,14E-01 |
| AT3G27850 | 1,55 | -2,23 | 1,58 | -1,41 | 1,58E-01 |
| AT3G27860 | 0,12 | 0,00 | 4,41 | 0,00 | 1,00E+00 |
| AT3G05335 | 1,73 | 0,06 | 1,26 | 0,04 | 9,64E-01 |
| AT3G05355 | 1,75 | -0,27 | 1,30 | -0,21 | 8,34E-01 |
| AT3G27870 | 0,75 | 0,46 | 2,62 | 0,18 | 8,61E-01 |
| AT3G27940 | 0,56 | 2,82 | 2,88 | 0,98 | 3,28E-01 |
| AT3G27980 | 0,34 | 0,85 | 4,27 | 0,20 | 8,42E-01 |
| AT3G27999 | 0,43 | 2,65 | 4,28 | 0,62 | 5,36E-01 |
| AT3G28040 | 1,16 | -0,62 | 2,22 | -0,28 | 7,80E-01 |
| AT3G28050 | 0,37 | -0,33 | 4,40 | -0,07 | 9,41E-01 |
| AT3G28080 | 0,70 | 1,31 | 2,20 | 0,59 | 5,53E-01 |
| AT3G28100 | 1,10 | 2,04 | 1,79 | 1,14 | 2,55E-01 |
| AT3G28170 | 0,75 | 0,00 | 3,62 | 0,00 | 1,00E+00 |
| AT3G28200 | 0,56 | -4,19 | 4,30 | -0,97 | 3,31E-01 |
| AT3G28220 | 0,69 | -3,40 | 2,88 | -1,18 | 2,39E-01 |
| AT3G28270 | 0,14 | 0,89 | 4,35 | 0,21 | 8,37E-01 |
| AT3G28290 | 1,10 | -4,54 | 2,41 | -1,89 | 5,93E-02 |
| AT3G28300 | 1,56 | -5,50 | 2,67 | -2,06 | 3,90E-02 |
| AT3G28450 | 0,78 | 0,44 | 2,44 | 0,18 | 8,57E-01 |
| AT3G28500 | 0,30 | 1,37 | 4,33 | 0,32 | 7,51E-01 |
| AT3G28510 | 0,99 | -0,31 | 2,62 | -0,12 | 9,06E-01 |
| AT3G05515 | 0,60 | 1,41 | 2,82 | 0,50 | 6,17E-01 |
| AT3G28580 | 0,77 | 1,13 | 3,45 | 0,33 | 7,43E-01 |
| AT3G28590 | 0,05 | 0,18 | 4,38 | 0,04 | 9,68E-01 |
| AT3G28650 | 0,20 | 0,40 | 4,37 | 0,09 | 9,27E-01 |

| | | | | | |
|---|---|---|---|---|---|
| AT3G28670 | 0,33 | 0,22 | 4,39 | 0,05 | 9,61E-01 |
| AT3G28690 | 0,39 | -0,92 | 4,24 | -0,22 | 8,27E-01 |
| AT3G28715 | 0,62 | -0,79 | 2,35 | -0,34 | 7,37E-01 |
| AT3G28720 | 0,91 | -0,14 | 3,26 | -0,04 | 9,67E-01 |
| AT3G28730 | 0,64 | -0,35 | 3,02 | -0,12 | 9,08E-01 |
| AT3G28870 | 0,74 | -0,61 | 3,72 | -0,16 | 8,69E-01 |
| AT3G28910 | 0,61 | 1,34 | 2,66 | 0,50 | 6,14E-01 |
| AT3G05595 | 0,35 | -0,93 | 4,38 | -0,21 | 8,32E-01 |
| AT3G28922 | 0,12 | 0,27 | 4,38 | 0,06 | 9,50E-01 |
| AT3G28958 | 0,58 | 2,01 | 3,01 | 0,67 | 5,05E-01 |
| AT3G29000 | 0,25 | 0,34 | 4,37 | 0,08 | 9,39E-01 |
| AT3G29010 | 0,38 | 0,00 | 4,41 | 0,00 | 1,00E+00 |
| AT3G29033 | 0,57 | 2,28 | 2,89 | 0,79 | 4,31E-01 |
| AT3G29080 | 1,28 | 1,51 | 1,57 | 0,96 | 3,35E-01 |
| AT3G29180 | 1,27 | 0,37 | 2,91 | 0,13 | 9,00E-01 |
| AT3G05715 | 0,40 | 0,01 | 4,27 | 0,00 | 9,98E-01 |
| AT3G29240 | 1,56 | 2,74 | 1,85 | 1,48 | 1,38E-01 |
| AT3G29270 | 0,39 | 0,00 | 4,40 | 0,00 | 1,00E+00 |
| AT3G29290 | 0,20 | -0,26 | 4,41 | -0,06 | 9,53E-01 |
| AT3G29370 | 0,05 | 0,18 | 4,38 | 0,04 | 9,68E-01 |
| AT3G29390 | 0,18 | -0,81 | 4,37 | -0,19 | 8,53E-01 |
| AT3G29450 | 0,14 | 1,29 | 4,33 | 0,30 | 7,65E-01 |
| AT3G29455 | 0,27 | 2,26 | 4,29 | 0,53 | 5,98E-01 |
| AT3G05805 | 0,31 | 1,40 | 4,33 | 0,32 | 7,46E-01 |
| AT3G29638 | 0,16 | 0,35 | 4,37 | 0,08 | 9,37E-01 |
| AT3G29670 | 0,94 | 0,19 | 1,99 | 0,10 | 9,23E-01 |
| AT3G30210 | 0,11 | -0,38 | 4,39 | -0,09 | 9,31E-01 |
| AT3G30320 | 0,40 | 0,98 | 4,31 | 0,23 | 8,20E-01 |
| AT3G30380 | 0,49 | -1,24 | 3,33 | -0,37 | 7,10E-01 |
| AT3G30383 | 0,12 | 0,00 | 4,41 | 0,00 | 1,00E+00 |
| AT3G30520 | 1,35 | 4,41 | 2,22 | 1,99 | 4,70E-02 |
| AT3G30525 | 1,14 | 3,71 | 2,63 | 1,41 | 1,59E-01 |
| AT3G30720 | 0,61 | -2,26 | 3,04 | -0,74 | 4,57E-01 |
| AT3G30770 | 0,13 | 0,94 | 4,35 | 0,22 | 8,29E-01 |
| AT3G30820 | 0,18 | 1,20 | 4,34 | 0,28 | 7,82E-01 |
| AT3G31950 | 0,59 | -0,58 | 2,16 | -0,27 | 7,87E-01 |
| AT3G32150 | 0,20 | 1,86 | 4,30 | 0,43 | 6,66E-01 |
| AT3G32280 | 0,39 | -0,37 | 4,21 | -0,09 | 9,30E-01 |
| AT3G32896 | 0,28 | 1,45 | 4,33 | 0,33 | 7,38E-01 |
| AT3G33530 | 1,04 | 1,12 | 1,94 | 0,58 | 5,65E-01 |
| AT3G42075 | 0,24 | -0,02 | 4,38 | 0,00 | 9,97E-01 |
| AT3G42170 | 0,48 | -0,35 | 3,24 | -0,11 | 9,13E-01 |
| AT3G42310 | 0,36 | 1,13 | 4,29 | 0,26 | 7,93E-01 |
| AT3G42560 | 0,10 | 0,08 | 4,38 | 0,02 | 9,85E-01 |
| AT3G42640 | 0,44 | 1,76 | 2,69 | 0,66 | 5,12E-01 |
| AT3G42870 | 1,35 | -0,10 | 1,69 | -0,06 | 9,51E-01 |
| AT3G42950 | 0,12 | 0,00 | 4,41 | 0,00 | 1,00E+00 |
| AT3G06495 | 0,51 | 1,08 | 3,12 | 0,35 | 7,29E-01 |
| AT3G43148 | 0,82 | 2,10 | 2,22 | 0,95 | 3,45E-01 |
| AT3G43150 | 0,30 | 1,89 | 3,57 | 0,53 | 5,98E-01 |
| AT3G43210 | 0,25 | 0,00 | 4,41 | 0,00 | 1,00E+00 |
| AT3G43230 | 1,49 | 0,53 | 1,93 | 0,27 | 7,85E-01 |
| AT3G43260 | 1,20 | 2,77 | 2,01 | 1,38 | 1,68E-01 |
| AT3G06565 | 0,05 | 0,18 | 4,38 | 0,04 | 9,68E-01 |
| AT3G43290 | 0,21 | 1,07 | 4,34 | 0,25 | 8,05E-01 |
| AT3G06625 | 0,13 | -0,61 | 4,37 | -0,14 | 8,89E-01 |

| | | | | | |
|---|---|---|---|---|---|
| AT3G43490 | 1,48 | 2,15 | 1,95 | 1,10 | 2,70E-01 |
| AT3G43503 | 0,30 | -0,27 | 3,66 | -0,07 | 9,41E-01 |
| AT3G06635 | 0,40 | 2,24 | 4,30 | 0,52 | 6,03E-01 |
| AT3G43670 | 0,53 | 1,46 | 2,73 | 0,53 | 5,93E-01 |
| AT3G43682 | 0,33 | 1,13 | 4,34 | 0,26 | 7,95E-01 |
| AT3G43740 | 0,28 | 0,04 | 4,32 | 0,01 | 9,92E-01 |
| AT3G43800 | 0,45 | -1,47 | 3,31 | -0,44 | 6,57E-01 |
| AT3G43837 | 0,47 | 1,85 | 4,01 | 0,46 | 6,44E-01 |
| AT3G43920 | 0,47 | 1,44 | 3,24 | 0,45 | 6,56E-01 |
| AT3G43950 | 0,37 | 1,08 | 4,34 | 0,25 | 8,03E-01 |
| AT3G44130 | 0,22 | 1,00 | 4,35 | 0,23 | 8,18E-01 |
| AT3G44150 | 0,18 | -2,46 | 4,37 | -0,56 | 5,73E-01 |
| AT3G44210 | 1,26 | 2,46 | 2,35 | 1,05 | 2,95E-01 |
| AT3G44230 | 0,25 | -0,73 | 4,35 | -0,17 | 8,67E-01 |
| AT3G44235 | 0,05 | 0,18 | 4,38 | 0,04 | 9,68E-01 |
| AT3G44300 | 1,82 | 1,86 | 1,68 | 1,10 | 2,70E-01 |
| AT3G44320 | 0,76 | -2,03 | 2,92 | -0,69 | 4,87E-01 |
| AT3G44330 | 1,72 | -0,26 | 2,28 | -0,11 | 9,09E-01 |
| AT3G44405 | 0,08 | 0,00 | 4,41 | 0,00 | 1,00E+00 |
| AT3G44430 | 1,70 | 1,62 | 1,66 | 0,98 | 3,29E-01 |
| AT3G44550 | 0,99 | 1,15 | 2,41 | 0,48 | 6,34E-01 |
| AT3G44560 | 0,14 | 0,89 | 4,35 | 0,21 | 8,37E-01 |
| AT3G44620 | 0,30 | 0,73 | 3,56 | 0,20 | 8,38E-01 |
| AT3G44670 | 1,62 | 0,54 | 1,77 | 0,31 | 7,58E-01 |
| AT3G44730 | 0,52 | -0,20 | 2,28 | -0,09 | 9,28E-01 |
| AT3G44798 | 0,43 | 0,49 | 4,19 | 0,12 | 9,07E-01 |
| AT3G44880 | 0,54 | -0,01 | 3,96 | 0,00 | 9,98E-01 |
| AT3G44900 | 0,86 | 0,00 | 2,53 | 0,00 | 1,00E+00 |
| AT3G44935 | 1,28 | 3,62 | 1,99 | 1,82 | 6,93E-02 |
| AT3G44980 | 0,39 | -0,12 | 3,45 | -0,03 | 9,73E-01 |
| AT3G45010 | 0,31 | 1,10 | 4,34 | 0,25 | 8,01E-01 |
| AT3G45400 | 0,33 | 0,49 | 4,36 | 0,11 | 9,10E-01 |
| AT3G45620 | 1,31 | -0,01 | 2,49 | 0,00 | 9,98E-01 |
| AT3G45640 | 1,79 | 2,14 | 1,61 | 1,33 | 1,82E-01 |
| AT3G45650 | 1,57 | 0,00 | 2,22 | 0,00 | 1,00E+00 |
| AT3G45660 | 1,01 | 0,00 | 2,70 | 0,00 | 1,00E+00 |
| AT3G45670 | 0,15 | 0,90 | 4,35 | 0,21 | 8,36E-01 |
| AT3G45730 | 0,15 | -0,31 | 4,40 | -0,07 | 9,43E-01 |
| AT3G45740 | 0,36 | 0,00 | 4,41 | 0,00 | 1,00E+00 |
| AT3G45830 | 1,79 | 1,03 | 1,75 | 0,59 | 5,55E-01 |
| AT3G45870 | 0,08 | 0,00 | 4,41 | 0,00 | 1,00E+00 |
| AT3G45930 | 0,32 | -2,19 | 4,38 | -0,50 | 6,17E-01 |
| AT3G45965 | 0,21 | -3,00 | 4,36 | -0,69 | 4,91E-01 |
| AT3G45970 | 0,98 | -0,41 | 2,77 | -0,15 | 8,82E-01 |
| AT3G46040 | 0,85 | -0,52 | 2,31 | -0,23 | 8,22E-01 |
| AT3G46060 | 1,03 | 0,97 | 2,09 | 0,46 | 6,42E-01 |
| AT3G46130 | 0,12 | 0,00 | 4,41 | 0,00 | 1,00E+00 |
| AT3G46220 | 0,60 | -1,24 | 2,67 | -0,46 | 6,42E-01 |
| AT3G46290 | 1,13 | -0,63 | 1,99 | -0,32 | 7,51E-01 |
| AT3G46370 | 0,25 | 0,44 | 4,30 | 0,10 | 9,19E-01 |
| AT3G46490 | 0,32 | 0,31 | 4,28 | 0,07 | 9,42E-01 |
| AT3G46520 | 0,37 | 0,00 | 4,41 | 0,00 | 1,00E+00 |
| AT3G46560 | 0,33 | 2,25 | 4,29 | 0,53 | 5,99E-01 |
| AT3G46590 | 1,40 | -1,89 | 1,76 | -1,07 | 2,83E-01 |
| AT3G07505 | 1,14 | 0,58 | 1,75 | 0,33 | 7,41E-01 |
| AT3G46640 | 1,12 | -1,93 | 2,27 | -0,85 | 3,96E-01 |

| | | | | | |
|---|---|---|---|---|---|
| AT3G46658 | 0,59 | 0,97 | 2,66 | 0,36 | 7,15E-01 |
| AT3G46668 | 0,57 | -2,67 | 3,20 | -0,83 | 4,04E-01 |
| AT3G07555 | 1,02 | 1,48 | 2,11 | 0,70 | 4,84E-01 |
| AT3G07585 | 1,61 | 0,71 | 1,49 | 0,48 | 6,34E-01 |
| AT3G46930 | 0,21 | 0,00 | 4,41 | 0,00 | 1,00E+00 |
| AT3G47060 | 0,35 | -2,19 | 4,41 | -0,50 | 6,20E-01 |
| AT3G47140 | 0,11 | 0,88 | 4,35 | 0,20 | 8,39E-01 |
| AT3G07695 | 0,33 | 0,69 | 4,35 | 0,16 | 8,73E-01 |
| AT3G47290 | 0,08 | 0,00 | 4,41 | 0,00 | 1,00E+00 |
| AT3G47295 | 0,08 | 0,21 | 4,39 | 0,05 | 9,62E-01 |
| AT3G07715 | 0,48 | -0,31 | 2,70 | -0,11 | 9,09E-01 |
| AT3G47490 | 0,22 | 1,00 | 4,35 | 0,23 | 8,18E-01 |
| AT3G47520 | 0,29 | -1,27 | 4,40 | -0,29 | 7,72E-01 |
| AT3G47550 | 0,50 | 0,07 | 3,06 | 0,02 | 9,83E-01 |
| AT3G47570 | 0,50 | -0,31 | 3,32 | -0,09 | 9,26E-01 |
| AT3G47610 | 0,61 | 1,76 | 2,52 | 0,70 | 4,85E-01 |
| AT3G47650 | 0,65 | 0,00 | 3,09 | 0,00 | 1,00E+00 |
| AT3G47660 | 0,72 | 0,00 | 2,97 | 0,00 | 1,00E+00 |
| AT3G07795 | 0,21 | -0,29 | 4,40 | -0,06 | 9,48E-01 |
| AT3G07805 | 0,89 | -3,71 | 2,73 | -1,36 | 1,74E-01 |
| AT3G47770 | 0,17 | 0,00 | 4,41 | 0,00 | 1,00E+00 |
| AT3G47800 | 1,07 | -0,57 | 1,85 | -0,31 | 7,58E-01 |
| AT3G47820 | 1,04 | 1,17 | 1,79 | 0,66 | 5,12E-01 |
| AT3G47833 | 0,22 | -0,68 | 4,34 | -0,16 | 8,76E-01 |
| AT3G47920 | 1,60 | -2,42 | 1,36 | -1,78 | 7,53E-02 |
| AT3G47930 | 0,52 | 0,00 | 3,96 | 0,00 | 1,00E+00 |
| AT3G47950 | 0,34 | 0,22 | 4,37 | 0,05 | 9,59E-01 |
| AT3G48040 | 0,17 | 0,07 | 4,38 | 0,02 | 9,88E-01 |
| AT3G48070 | 0,40 | -1,10 | 4,35 | -0,25 | 8,00E-01 |
| AT3G48110 | 0,52 | -0,37 | 3,21 | -0,11 | 9,09E-01 |
| AT3G48140 | 0,97 | 0,81 | 2,15 | 0,38 | 7,06E-01 |
| AT3G48190 | 1,19 | 0,83 | 2,55 | 0,33 | 7,45E-01 |
| AT3G48310 | 0,40 | -1,21 | 3,43 | -0,35 | 7,24E-01 |
| AT3G48350 | 1,84 | 0,39 | 2,15 | 0,18 | 8,55E-01 |
| AT3G48380 | 0,47 | 0,99 | 3,22 | 0,31 | 7,59E-01 |
| AT3G48420 | 0,42 | -1,18 | 4,29 | -0,27 | 7,84E-01 |
| AT3G48430 | 1,70 | -1,18 | 2,08 | -0,57 | 5,70E-01 |
| AT3G48460 | 0,51 | -0,49 | 4,37 | -0,11 | 9,11E-01 |
| AT3G48500 | 0,57 | -2,08 | 3,93 | -0,53 | 5,96E-01 |
| AT3G48530 | 0,79 | 1,05 | 2,16 | 0,49 | 6,27E-01 |
| AT3G48620 | 0,08 | 0,00 | 4,41 | 0,00 | 1,00E+00 |
| AT3G48680 | 0,15 | -0,35 | 4,39 | -0,08 | 9,36E-01 |
| AT3G48710 | 0,27 | 0,66 | 4,37 | 0,15 | 8,80E-01 |
| AT3G48730 | 0,57 | 0,04 | 2,76 | 0,01 | 9,89E-01 |
| AT3G48750 | 0,54 | -1,05 | 2,77 | -0,38 | 7,05E-01 |
| AT3G48760 | 0,76 | -1,25 | 2,62 | -0,48 | 6,32E-01 |
| AT3G07995 | 0,83 | 3,21 | 2,24 | 1,43 | 1,51E-01 |
| AT3G48780 | 0,82 | 0,18 | 2,68 | 0,07 | 9,47E-01 |
| AT3G48810 | 0,58 | -2,46 | 3,73 | -0,66 | 5,09E-01 |
| AT3G48890 | 0,66 | -2,21 | 2,76 | -0,80 | 4,25E-01 |
| AT3G49000 | 0,27 | 0,78 | 4,36 | 0,18 | 8,58E-01 |
| AT3G49110 | 0,08 | 0,00 | 4,41 | 0,00 | 1,00E+00 |
| AT3G49130 | 1,37 | 0,00 | 2,56 | 0,00 | 1,00E+00 |
| AT3G49220 | 0,92 | -0,52 | 2,07 | -0,25 | 8,00E-01 |
| AT3G49240 | 0,54 | -2,61 | 4,06 | -0,64 | 5,20E-01 |
| AT3G49260 | 0,64 | 0,00 | 4,41 | 0,00 | 1,00E+00 |

| | | | | | |
|---|---|---|---|---|---|
| AT3G49430 | 0,93 | -1,58 | 2,14 | -0,74 | 4,59E-01 |
| AT3G49520 | 1,37 | 3,35 | 1,82 | 1,84 | 6,63E-02 |
| AT3G49590 | 1,44 | 1,59 | 1,86 | 0,86 | 3,92E-01 |
| AT3G49670 | 1,02 | -0,79 | 2,52 | -0,31 | 7,54E-01 |
| AT3G49680 | 0,45 | -0,59 | 3,22 | -0,18 | 8,55E-01 |
| AT3G49710 | 0,32 | 0,00 | 4,41 | 0,00 | 1,00E+00 |
| AT3G49780 | 0,25 | 0,55 | 4,31 | 0,13 | 8,98E-01 |
| AT3G50000 | 0,43 | 0,48 | 4,23 | 0,11 | 9,09E-01 |
| AT3G50080 | 0,21 | -0,39 | 4,39 | -0,09 | 9,30E-01 |
| AT3G50100 | 0,05 | 0,18 | 4,38 | 0,04 | 9,68E-01 |
| AT3G50270 | 0,60 | -2,23 | 3,15 | -0,71 | 4,80E-01 |
| AT3G50350 | 0,42 | 0,43 | 3,35 | 0,13 | 8,98E-01 |
| AT3G50360 | 0,31 | -0,93 | 4,37 | -0,21 | 8,32E-01 |
| AT3G50430 | 0,16 | 1,58 | 4,31 | 0,37 | 7,14E-01 |
| AT3G50480 | 1,71 | 2,86 | 1,63 | 1,75 | 8,03E-02 |
| AT3G50530 | 0,25 | -1,08 | 4,39 | -0,25 | 8,06E-01 |
| AT3G50540 | 1,39 | -1,79 | 1,59 | -1,12 | 2,61E-01 |
| AT3G50550 | 0,51 | 0,55 | 3,24 | 0,17 | 8,66E-01 |
| AT3G50630 | 0,26 | -1,21 | 4,39 | -0,27 | 7,84E-01 |
| AT3G50700 | 0,53 | 0,59 | 2,64 | 0,22 | 8,24E-01 |
| AT3G50770 | 1,04 | 0,53 | 2,47 | 0,21 | 8,30E-01 |
| AT3G50890 | 0,25 | 1,75 | 4,31 | 0,41 | 6,84E-01 |
| AT3G50910 | 0,94 | -0,25 | 2,50 | -0,10 | 9,21E-01 |
| AT3G50970 | 0,39 | -0,33 | 4,40 | -0,08 | 9,40E-01 |
| AT3G51110 | 0,46 | 0,04 | 4,04 | 0,01 | 9,93E-01 |
| AT3G51140 | 0,97 | 0,63 | 2,21 | 0,29 | 7,74E-01 |
| AT3G51180 | 0,25 | 0,28 | 4,39 | 0,06 | 9,50E-01 |
| AT3G51260 | 0,46 | -0,26 | 3,38 | -0,08 | 9,39E-01 |
| AT3G51270 | 0,33 | 0,03 | 4,38 | 0,01 | 9,95E-01 |
| AT3G51325 | 0,36 | 0,76 | 4,36 | 0,17 | 8,62E-01 |
| AT3G51370 | 1,46 | -0,88 | 1,90 | -0,46 | 6,46E-01 |
| AT3G51420 | 0,66 | 0,25 | 2,19 | 0,12 | 9,08E-01 |
| AT3G51430 | 0,32 | 0,56 | 4,37 | 0,13 | 8,97E-01 |
| AT3G51440 | 0,80 | 0,59 | 2,30 | 0,26 | 7,97E-01 |
| AT3G51450 | 0,43 | -1,00 | 2,75 | -0,36 | 7,15E-01 |
| AT3G51460 | 0,27 | 0,30 | 4,39 | 0,07 | 9,46E-01 |
| AT3G51480 | 0,45 | 0,00 | 4,41 | 0,00 | 1,00E+00 |
| AT3G51500 | 0,44 | 0,00 | 4,41 | 0,00 | 1,00E+00 |
| AT3G51510 | 0,44 | 1,07 | 3,44 | 0,31 | 7,55E-01 |
| AT3G51540 | 0,25 | 0,00 | 4,41 | 0,00 | 1,00E+00 |
| AT3G51620 | 1,34 | -1,41 | 1,98 | -0,71 | 4,76E-01 |
| AT3G51630 | 0,87 | 0,00 | 3,34 | 0,00 | 1,00E+00 |
| AT3G51640 | 0,80 | -0,26 | 2,86 | -0,09 | 9,27E-01 |
| AT3G51650 | 1,02 | -0,38 | 2,34 | -0,16 | 8,72E-01 |
| AT3G51730 | 1,38 | -1,74 | 1,51 | -1,15 | 2,49E-01 |
| AT3G51830 | 0,25 | -0,36 | 4,39 | -0,08 | 9,35E-01 |
| AT3G51850 | 1,69 | 0,89 | 1,96 | 0,45 | 6,49E-01 |
| AT3G51870 | 1,46 | -0,26 | 2,26 | -0,11 | 9,10E-01 |
| AT3G51970 | 0,40 | 0,49 | 3,41 | 0,14 | 8,87E-01 |
| AT3G52080 | 0,33 | -0,20 | 4,39 | -0,05 | 9,64E-01 |
| AT3G52100 | 0,15 | 0,47 | 4,37 | 0,11 | 9,15E-01 |
| AT3G52105 | 0,19 | 1,17 | 4,34 | 0,27 | 7,87E-01 |
| AT3G52120 | 1,03 | 0,87 | 2,26 | 0,38 | 7,02E-01 |
| AT3G52150 | 0,83 | -0,03 | 2,63 | -0,01 | 9,91E-01 |
| AT3G52155 | 0,45 | 0,00 | 4,41 | 0,00 | 1,00E+00 |
| AT3G52200 | 0,34 | 1,39 | 4,33 | 0,32 | 7,48E-01 |

| | | | | | |
|---|---|---|---|---|---|
| AT3G52210 | 0,11 | -0,26 | 4,41 | -0,06 | 9,53E-01 |
| AT3G52230 | 0,83 | 0,78 | 3,08 | 0,25 | 7,99E-01 |
| AT3G52300 | 0,40 | -0,43 | 4,37 | -0,10 | 9,22E-01 |
| AT3G52340 | 0,38 | -0,31 | 4,35 | -0,07 | 9,43E-01 |
| AT3G52370 | 0,08 | 0,00 | 4,41 | 0,00 | 1,00E+00 |
| AT3G52380 | 1,54 | -1,84 | 2,00 | -0,92 | 3,58E-01 |
| AT3G52470 | 0,30 | 1,93 | 4,30 | 0,45 | 6,54E-01 |
| AT3G52570 | 0,29 | 1,10 | 4,34 | 0,25 | 7,99E-01 |
| AT3G52590 | 1,74 | 3,14 | 1,70 | 1,85 | 6,45E-02 |
| AT3G08275 | 0,60 | 1,78 | 3,02 | 0,59 | 5,56E-01 |
| AT3G52610 | 0,30 | 0,00 | 4,41 | 0,00 | 1,00E+00 |
| AT3G52680 | 0,59 | 2,13 | 2,49 | 0,86 | 3,91E-01 |
| AT3G52710 | 0,88 | 0,67 | 2,00 | 0,33 | 7,38E-01 |
| AT3G08285 | 1,59 | 2,52 | 1,65 | 1,53 | 1,27E-01 |
| AT3G08295 | 0,73 | 3,41 | 3,34 | 1,02 | 3,08E-01 |
| AT3G52830 | 1,16 | 0,82 | 1,67 | 0,49 | 6,25E-01 |
| AT3G52840 | 0,82 | -2,75 | 2,53 | -1,09 | 2,76E-01 |
| AT3G52850 | 1,76 | -1,56 | 1,38 | -1,13 | 2,58E-01 |
| AT3G52950 | 0,45 | -0,26 | 3,35 | -0,08 | 9,38E-01 |
| AT3G52960 | 0,80 | -0,07 | 1,84 | -0,04 | 9,72E-01 |
| AT3G52990 | 1,11 | -0,75 | 1,51 | -0,49 | 6,22E-01 |
| AT3G53000 | 0,31 | 0,19 | 4,38 | 0,04 | 9,65E-01 |
| AT3G53030 | 0,57 | 0,73 | 4,00 | 0,18 | 8,56E-01 |
| AT3G53050 | 0,05 | 0,18 | 4,38 | 0,04 | 9,68E-01 |
| AT3G53065 | 0,19 | 1,00 | 4,33 | 0,23 | 8,17E-01 |
| AT3G53090 | 0,36 | -0,95 | 4,37 | -0,22 | 8,28E-01 |
| AT3G53110 | 0,40 | 0,80 | 3,38 | 0,24 | 8,14E-01 |
| AT3G53180 | 0,20 | 1,10 | 4,35 | 0,25 | 8,00E-01 |
| AT3G53190 | 0,21 | 0,08 | 4,38 | 0,02 | 9,85E-01 |
| AT3G53230 | 1,14 | -0,23 | 2,88 | -0,08 | 9,37E-01 |
| AT3G53240 | 0,55 | -0,37 | 3,95 | -0,09 | 9,26E-01 |
| AT3G53305 | 0,72 | 1,56 | 2,12 | 0,74 | 4,60E-01 |
| AT3G53360 | 0,08 | 0,00 | 4,41 | 0,00 | 1,00E+00 |
| AT3G53390 | 0,27 | 1,01 | 4,34 | 0,23 | 8,17E-01 |
| AT3G53470 | 0,77 | -2,06 | 2,88 | -0,72 | 4,75E-01 |
| AT3G53520 | 0,83 | -1,12 | 2,59 | -0,43 | 6,66E-01 |
| AT3G53670 | 0,90 | 0,47 | 2,14 | 0,22 | 8,26E-01 |
| AT3G53700 | 0,94 | -0,85 | 1,86 | -0,46 | 6,47E-01 |
| AT3G53870 | 0,80 | -1,05 | 2,51 | -0,42 | 6,76E-01 |
| AT3G54220 | 0,50 | -1,15 | 3,27 | -0,35 | 7,25E-01 |
| AT3G54230 | 1,41 | 1,32 | 2,15 | 0,61 | 5,40E-01 |
| AT3G54366 | 1,11 | 2,38 | 2,08 | 1,15 | 2,52E-01 |
| AT3G54420 | 0,17 | -0,29 | 4,40 | -0,07 | 9,47E-01 |
| AT3G54540 | 1,42 | 0,58 | 2,06 | 0,28 | 7,78E-01 |
| AT3G54600 | 0,51 | 0,35 | 4,00 | 0,09 | 9,31E-01 |
| AT3G54610 | 0,47 | -2,05 | 4,03 | -0,51 | 6,12E-01 |
| AT3G54625 | 0,68 | 0,00 | 4,40 | 0,00 | 1,00E+00 |
| AT3G54680 | 0,75 | 1,18 | 2,92 | 0,40 | 6,86E-01 |
| AT3G54800 | 0,59 | 1,81 | 2,57 | 0,71 | 4,80E-01 |
| AT3G54820 | 0,15 | 0,47 | 4,37 | 0,11 | 9,14E-01 |
| AT3G54850 | 0,81 | -0,92 | 2,56 | -0,36 | 7,20E-01 |
| AT3G54980 | 0,41 | -0,20 | 4,25 | -0,05 | 9,62E-01 |
| AT3G55030 | 0,39 | -1,65 | 4,23 | -0,39 | 6,96E-01 |
| AT3G55250 | 0,63 | -1,05 | 2,27 | -0,46 | 6,44E-01 |
| AT3G55260 | 0,43 | 2,50 | 4,00 | 0,62 | 5,33E-01 |
| AT3G55270 | 1,69 | -0,26 | 2,08 | -0,13 | 9,00E-01 |

| | | | | | |
|---|---|---|---|---|---|
| AT3G55280 | 0,61 | 0,19 | 3,68 | 0,05 | 9,59E-01 |
| AT3G55370 | 0,12 | 0,27 | 4,38 | 0,06 | 9,50E-01 |
| AT3G55420 | 0,14 | 0,81 | 4,35 | 0,19 | 8,52E-01 |
| AT3G55440 | 1,26 | 1,13 | 1,75 | 0,65 | 5,18E-01 |
| AT3G55450 | 0,15 | -0,35 | 4,39 | -0,08 | 9,36E-01 |
| AT3G55460 | 1,02 | -0,84 | 1,91 | -0,44 | 6,59E-01 |
| AT3G55485 | 0,83 | 0,53 | 2,21 | 0,24 | 8,12E-01 |
| AT3G08485 | 0,83 | 0,53 | 2,21 | 0,24 | 8,12E-01 |
| AT3G55520 | 0,40 | 0,00 | 4,41 | 0,00 | 1,00E+00 |
| AT3G55620 | 0,54 | 1,04 | 2,66 | 0,39 | 6,95E-01 |
| AT3G55630 | 0,42 | 0,00 | 4,41 | 0,00 | 1,00E+00 |
| AT3G55677 | 0,24 | 0,77 | 4,35 | 0,18 | 8,60E-01 |
| AT3G55760 | 0,99 | -3,67 | 2,08 | -1,77 | 7,69E-02 |
| AT3G55770 | 0,98 | -0,37 | 1,97 | -0,19 | 8,51E-01 |
| AT3G55780 | 0,67 | 0,00 | 3,70 | 0,00 | 1,00E+00 |
| AT3G55830 | 1,72 | -0,51 | 1,77 | -0,29 | 7,73E-01 |
| AT3G55880 | 0,30 | 1,61 | 3,60 | 0,45 | 6,54E-01 |
| AT3G55940 | 0,10 | 0,24 | 4,38 | 0,05 | 9,57E-01 |
| AT3G56130 | 0,33 | -0,92 | 4,33 | -0,21 | 8,31E-01 |
| AT3G56140 | 0,74 | -0,65 | 2,47 | -0,26 | 7,93E-01 |
| AT3G56180 | 0,21 | 0,08 | 4,38 | 0,02 | 9,85E-01 |
| AT3G56310 | 0,76 | 0,85 | 2,44 | 0,35 | 7,28E-01 |
| AT3G56330 | 0,52 | 0,00 | 3,94 | 0,00 | 1,00E+00 |
| AT3G56390 | 0,30 | 0,00 | 4,41 | 0,00 | 1,00E+00 |
| AT3G56450 | 0,39 | -2,19 | 4,41 | -0,50 | 6,20E-01 |
| AT3G08665 | 0,76 | -0,12 | 2,97 | -0,04 | 9,68E-01 |
| AT3G56470 | 0,12 | 0,00 | 4,41 | 0,00 | 1,00E+00 |
| AT3G56490 | 0,11 | -0,26 | 4,41 | -0,06 | 9,53E-01 |
| AT3G56590 | 0,57 | 0,66 | 3,01 | 0,22 | 8,27E-01 |
| AT3G56640 | 0,59 | 0,28 | 3,16 | 0,09 | 9,30E-01 |
| AT3G56650 | 0,53 | -2,33 | 3,97 | -0,59 | 5,58E-01 |
| AT3G56720 | 0,17 | -0,58 | 4,38 | -0,13 | 8,94E-01 |
| AT3G56740 | 0,49 | 0,63 | 4,04 | 0,16 | 8,76E-01 |
| AT3G56780 | 0,32 | 0,04 | 4,38 | 0,01 | 9,92E-01 |
| AT3G56840 | 0,64 | 0,66 | 2,56 | 0,26 | 7,96E-01 |
| AT3G56880 | 1,69 | -0,44 | 1,32 | -0,33 | 7,40E-01 |
| AT3G08785 | 1,51 | -2,86 | 1,83 | -1,56 | 1,18E-01 |
| AT3G56960 | 0,60 | 0,00 | 4,40 | 0,00 | 1,00E+00 |
| AT3G57040 | 1,37 | -0,71 | 1,49 | -0,48 | 6,31E-01 |
| AT3G57062 | 1,47 | 1,20 | 1,94 | 0,62 | 5,37E-01 |
| AT3G57070 | 0,24 | 0,98 | 4,28 | 0,23 | 8,19E-01 |
| AT3G57080 | 0,21 | 0,00 | 4,41 | 0,00 | 1,00E+00 |
| AT3G57090 | 0,59 | 0,85 | 2,24 | 0,38 | 7,05E-01 |
| AT3G08845 | 0,12 | -0,36 | 4,39 | -0,08 | 9,34E-01 |
| AT3G57230 | 0,38 | 0,01 | 2,88 | 0,00 | 9,98E-01 |
| AT3G57340 | 0,42 | -0,12 | 3,43 | -0,03 | 9,73E-01 |
| AT3G57390 | 0,11 | -0,26 | 4,41 | -0,06 | 9,53E-01 |
| AT3G57400 | 1,77 | 1,96 | 1,58 | 1,24 | 2,13E-01 |
| AT3G57450 | 1,58 | 1,80 | 1,65 | 1,09 | 2,77E-01 |
| AT3G57550 | 0,50 | 1,05 | 3,21 | 0,33 | 7,44E-01 |
| AT3G57570 | 0,40 | -1,47 | 4,22 | -0,35 | 7,28E-01 |
| AT3G57587 | 0,73 | -0,73 | 2,02 | -0,36 | 7,17E-01 |
| AT3G57645 | 1,71 | 3,09 | 1,88 | 1,64 | 1,01E-01 |
| AT3G57650 | 0,53 | 1,58 | 2,63 | 0,60 | 5,48E-01 |
| AT3G57660 | 1,78 | 0,48 | 1,97 | 0,24 | 8,09E-01 |
| AT3G57700 | 1,50 | 0,73 | 1,51 | 0,48 | 6,31E-01 |

| | | | | | |
|---|---|---|---|---|---|
| AT3G08975 | 0,42 | 0,00 | 4,18 | 0,00 | 1,00E+00 |
| AT3G57760 | 0,08 | 0,00 | 4,41 | 0,00 | 1,00E+00 |
| AT3G57770 | 1,17 | 0,80 | 1,57 | 0,51 | 6,07E-01 |
| AT3G57810 | 0,21 | -0,71 | 4,38 | -0,16 | 8,71E-01 |
| AT3G57830 | 0,31 | -0,68 | 4,35 | -0,16 | 8,76E-01 |
| AT3G57890 | 1,04 | -2,13 | 1,84 | -1,16 | 2,47E-01 |
| AT3G57940 | 0,20 | 1,10 | 4,35 | 0,25 | 8,00E-01 |
| AT3G57970 | 0,17 | 0,00 | 4,41 | 0,00 | 1,00E+00 |
| AT3G58020 | 0,08 | 0,00 | 4,41 | 0,00 | 1,00E+00 |
| AT3G58110 | 1,07 | 0,00 | 3,34 | 0,00 | 1,00E+00 |
| AT3G58160 | 0,08 | 0,00 | 4,41 | 0,00 | 1,00E+00 |
| AT3G58490 | 0,30 | 1,75 | 4,31 | 0,41 | 6,85E-01 |
| AT3G58510 | 0,99 | -0,20 | 1,93 | -0,10 | 9,19E-01 |
| AT3G58530 | 0,46 | 0,01 | 4,02 | 0,00 | 9,97E-01 |
| AT3G58570 | 0,71 | 0,27 | 2,56 | 0,10 | 9,17E-01 |
| AT3G58680 | 0,61 | -0,30 | 2,58 | -0,12 | 9,08E-01 |
| AT3G58690 | 0,58 | -0,26 | 3,16 | -0,08 | 9,34E-01 |
| AT3G58700 | 0,20 | 0,00 | 4,41 | 0,00 | 1,00E+00 |
| AT3G58795 | 0,23 | -1,08 | 4,33 | -0,25 | 8,03E-01 |
| AT3G59030 | 0,64 | 0,00 | 3,75 | 0,00 | 1,00E+00 |
| AT3G59068 | 1,78 | 1,97 | 1,62 | 1,21 | 2,24E-01 |
| AT3G59080 | 0,19 | 0,27 | 4,37 | 0,06 | 9,50E-01 |
| AT3G59090 | 0,80 | -0,29 | 2,56 | -0,11 | 9,10E-01 |
| AT3G59100 | 0,79 | -0,26 | 2,92 | -0,09 | 9,28E-01 |
| AT3G59110 | 0,41 | 0,29 | 4,22 | 0,07 | 9,46E-01 |
| AT3G59210 | 0,24 | 0,79 | 4,36 | 0,18 | 8,56E-01 |
| AT3G59290 | 1,04 | 0,55 | 1,87 | 0,29 | 7,68E-01 |
| AT3G59300 | 0,19 | 0,91 | 4,34 | 0,21 | 8,34E-01 |
| AT3G59350 | 0,86 | -1,80 | 2,51 | -0,72 | 4,74E-01 |
| AT3G59380 | 0,24 | 0,79 | 4,36 | 0,18 | 8,56E-01 |
| AT3G59410 | 1,72 | 0,47 | 1,52 | 0,31 | 7,59E-01 |
| AT3G59500 | 0,47 | -0,87 | 3,36 | -0,26 | 7,95E-01 |
| AT3G59520 | 0,08 | 0,00 | 4,41 | 0,00 | 1,00E+00 |
| AT3G59530 | 0,53 | 0,00 | 4,41 | 0,00 | 1,00E+00 |
| AT3G59670 | 0,30 | -0,15 | 4,35 | -0,03 | 9,72E-01 |
| AT3G59700 | 0,10 | 0,86 | 4,35 | 0,20 | 8,43E-01 |
| AT3G59810 | 0,46 | -2,04 | 4,37 | -0,47 | 6,41E-01 |
| AT3G59920 | 0,37 | 1,58 | 4,32 | 0,37 | 7,15E-01 |
| AT3G59970 | 1,71 | 1,00 | 1,41 | 0,71 | 4,78E-01 |
| AT3G60080 | 1,12 | 2,81 | 1,95 | 1,44 | 1,50E-01 |
| AT3G60110 | 0,48 | 0,31 | 4,39 | 0,07 | 9,44E-01 |
| AT3G60130 | 0,63 | -2,04 | 3,24 | -0,63 | 5,29E-01 |
| AT3G60140 | 0,58 | -2,16 | 3,86 | -0,56 | 5,77E-01 |
| AT3G60240 | 1,46 | 0,04 | 1,45 | 0,03 | 9,76E-01 |
| AT3G60245 | 0,58 | 0,34 | 2,38 | 0,14 | 8,85E-01 |
| AT3G60290 | 0,27 | -0,28 | 4,36 | -0,06 | 9,50E-01 |
| AT3G60300 | 0,58 | -0,43 | 2,67 | -0,16 | 8,72E-01 |
| AT3G60310 | 0,49 | -0,84 | 3,98 | -0,21 | 8,33E-01 |
| AT3G60410 | 0,24 | 0,00 | 4,41 | 0,00 | 1,00E+00 |
| AT3G60415 | 1,73 | -0,26 | 2,08 | -0,13 | 9,00E-01 |
| AT3G60420 | 0,55 | 0,00 | 3,91 | 0,00 | 1,00E+00 |
| AT3G60440 | 0,83 | -2,19 | 3,53 | -0,62 | 5,36E-01 |
| AT3G60450 | 1,00 | 0,00 | 2,79 | 0,00 | 1,00E+00 |
| AT3G60500 | 0,28 | -0,22 | 4,40 | -0,05 | 9,60E-01 |
| AT3G60530 | 0,83 | -0,36 | 1,84 | -0,20 | 8,43E-01 |
| AT3G60600 | 1,58 | -0,74 | 1,96 | -0,38 | 7,05E-01 |

| | | | | | |
|---|---|---|---|---|---|
| AT3G60690 | 1,35 | 2,32 | 1,88 | 1,23 | 2,19E-01 |
| AT3G60910 | 0,08 | 0,47 | 4,38 | 0,11 | 9,15E-01 |
| AT3G60950 | 1,11 | 0,49 | 1,83 | 0,27 | 7,90E-01 |
| AT3G60960 | 1,54 | -0,14 | 1,54 | -0,09 | 9,26E-01 |
| AT3G09515 | 1,27 | 0,51 | 1,87 | 0,27 | 7,86E-01 |
| AT3G60961 | 1,33 | -1,34 | 2,27 | -0,59 | 5,56E-01 |
| AT3G09525 | 0,12 | 0,00 | 4,41 | 0,00 | 1,00E+00 |
| AT3G60970 | 0,12 | 0,00 | 4,41 | 0,00 | 1,00E+00 |
| AT3G61030 | 0,61 | -0,88 | 1,94 | -0,45 | 6,51E-01 |
| AT3G61050 | 1,51 | -1,29 | 1,51 | -0,85 | 3,93E-01 |
| AT3G61080 | 0,12 | 0,00 | 4,41 | 0,00 | 1,00E+00 |
| AT3G61110 | 1,40 | -1,13 | 1,76 | -0,64 | 5,21E-01 |
| AT3G61130 | 1,07 | 0,01 | 2,42 | 0,01 | 9,95E-01 |
| AT3G61140 | 0,73 | 0,02 | 2,10 | 0,01 | 9,93E-01 |
| AT3G61150 | 0,68 | -0,40 | 2,92 | -0,14 | 8,91E-01 |
| AT3G61180 | 0,94 | -0,69 | 2,45 | -0,28 | 7,77E-01 |
| AT3G61240 | 1,00 | 1,04 | 1,96 | 0,53 | 5,96E-01 |
| AT3G61280 | 0,41 | 0,63 | 3,40 | 0,19 | 8,53E-01 |
| AT3G61320 | 0,08 | 0,21 | 4,39 | 0,05 | 9,62E-01 |
| AT3G61440 | 1,04 | 0,28 | 2,44 | 0,12 | 9,08E-01 |
| AT3G61480 | 0,44 | -2,15 | 3,36 | -0,64 | 5,22E-01 |
| AT3G61490 | 0,55 | 0,00 | 4,41 | 0,00 | 1,00E+00 |
| AT3G61540 | 0,29 | -0,46 | 4,35 | -0,10 | 9,17E-01 |
| AT3G61550 | 0,37 | -0,01 | 3,50 | 0,00 | 9,99E-01 |
| AT3G61600 | 1,22 | -1,09 | 2,03 | -0,54 | 5,91E-01 |
| AT3G61670 | 0,29 | -0,12 | 4,39 | -0,03 | 9,79E-01 |
| AT3G61700 | 1,46 | 0,88 | 1,51 | 0,58 | 5,62E-01 |
| AT3G61755 | 0,16 | 0,63 | 4,36 | 0,14 | 8,85E-01 |
| AT3G61810 | 0,05 | 0,18 | 4,38 | 0,04 | 9,68E-01 |
| AT3G61850 | 0,15 | 0,47 | 4,37 | 0,11 | 9,14E-01 |
| AT3G61870 | 1,16 | -0,91 | 1,48 | -0,62 | 5,38E-01 |
| AT3G61950 | 0,11 | -0,26 | 4,41 | -0,06 | 9,53E-01 |
| AT3G62030 | 0,98 | -1,08 | 2,03 | -0,53 | 5,94E-01 |
| AT3G62130 | 0,39 | 1,08 | 4,33 | 0,25 | 8,02E-01 |
| AT3G62190 | 0,24 | -0,68 | 4,38 | -0,16 | 8,76E-01 |
| AT3G62245 | 0,42 | 2,26 | 4,04 | 0,56 | 5,76E-01 |
| AT3G62300 | 0,93 | -2,89 | 3,42 | -0,85 | 3,98E-01 |
| AT3G62370 | 0,90 | -0,96 | 2,19 | -0,44 | 6,60E-01 |
| AT3G62530 | 0,72 | 0,37 | 2,44 | 0,15 | 8,78E-01 |
| AT3G62550 | 0,70 | -3,02 | 3,69 | -0,82 | 4,13E-01 |
| AT3G62560 | 0,61 | 0,94 | 3,14 | 0,30 | 7,66E-01 |
| AT3G62580 | 0,64 | 1,61 | 2,22 | 0,72 | 4,70E-01 |
| AT3G09815 | 0,08 | 0,00 | 4,41 | 0,00 | 1,00E+00 |
| AT3G09835 | 1,22 | -0,60 | 1,54 | -0,39 | 6,94E-01 |
| AT3G62620 | 0,29 | -0,15 | 4,35 | -0,03 | 9,72E-01 |
| AT3G62650 | 1,34 | 0,48 | 1,86 | 0,26 | 7,95E-01 |
| AT3G62660 | 1,25 | 0,33 | 1,62 | 0,20 | 8,40E-01 |
| AT3G62720 | 0,40 | 1,52 | 4,11 | 0,37 | 7,12E-01 |
| AT3G62750 | 0,68 | 0,76 | 2,41 | 0,32 | 7,53E-01 |
| AT3G62820 | 0,26 | 0,97 | 4,34 | 0,22 | 8,24E-01 |
| AT3G62830 | 0,86 | 0,19 | 1,90 | 0,10 | 9,20E-01 |
| AT3G62890 | 0,20 | 0,87 | 4,35 | 0,20 | 8,41E-01 |
| AT3G63000 | 0,33 | -0,38 | 4,39 | -0,09 | 9,32E-01 |
| AT3G63010 | 0,38 | 1,39 | 3,47 | 0,40 | 6,88E-01 |
| AT3G63080 | 0,53 | 0,97 | 2,73 | 0,35 | 7,24E-01 |
| AT3G63130 | 0,43 | -0,92 | 4,12 | -0,22 | 8,24E-01 |

| | | | | | |
|---|---|---|---|---|---|
| AT3G63170 | 0,32 | -0,18 | 4,40 | -0,04 | 9,67E-01 |
| AT3G63215 | 0,08 | 0,21 | 4,39 | 0,05 | 9,62E-01 |
| AT3G63300 | 0,72 | 0,00 | 2,97 | 0,00 | 1,00E+00 |
| AT3G63370 | 0,57 | -0,36 | 3,21 | -0,11 | 9,11E-01 |
| AT3G63400 | 0,95 | -0,74 | 2,21 | -0,34 | 7,37E-01 |
| AT3G63420 | 0,46 | 0,00 | 4,41 | 0,00 | 1,00E+00 |
| AT3G63470 | 0,42 | 0,00 | 4,24 | 0,00 | 1,00E+00 |
| AT3G63490 | 1,82 | 0,20 | 1,95 | 0,10 | 9,17E-01 |
| AT3G09995 | 0,45 | -2,05 | 4,10 | -0,50 | 6,17E-01 |
| AT3G63520 | 0,87 | -0,59 | 2,32 | -0,26 | 7,98E-01 |
| AT4G00030 | 0,38 | -0,92 | 4,33 | -0,21 | 8,32E-01 |
| AT4G00060 | 1,77 | 0,37 | 1,72 | 0,22 | 8,27E-01 |
| AT4G00100 | 1,39 | -2,09 | 1,90 | -1,10 | 2,73E-01 |
| AT4G00163 | 0,65 | -0,66 | 2,53 | -0,26 | 7,95E-01 |
| AT4G00230 | 0,69 | 0,00 | 4,40 | 0,00 | 1,00E+00 |
| AT4G00300 | 1,08 | 0,67 | 1,66 | 0,40 | 6,87E-01 |
| AT4G00340 | 0,31 | 1,06 | 4,34 | 0,24 | 8,08E-01 |
| AT4G00440 | 0,33 | -0,27 | 4,41 | -0,06 | 9,52E-01 |
| AT4G00490 | 0,44 | -0,42 | 4,39 | -0,10 | 9,24E-01 |
| AT4G00580 | 0,48 | 1,61 | 3,95 | 0,41 | 6,83E-01 |
| AT4G00660 | 1,18 | -0,37 | 2,16 | -0,17 | 8,63E-01 |
| AT4G00710 | 0,48 | -2,37 | 4,39 | -0,54 | 5,89E-01 |
| AT4G00750 | 0,96 | 0,17 | 1,76 | 0,09 | 9,25E-01 |
| AT4G00820 | 0,05 | 0,18 | 4,38 | 0,04 | 9,68E-01 |
| AT4G00830 | 0,92 | 0,51 | 2,07 | 0,25 | 8,06E-01 |
| AT4G00889 | 0,23 | 2,01 | 4,30 | 0,47 | 6,40E-01 |
| AT4G00930 | 0,17 | 0,00 | 4,41 | 0,00 | 1,00E+00 |
| AT4G00953 | 0,67 | -0,50 | 2,50 | -0,20 | 8,42E-01 |
| AT4G00955 | 0,42 | 0,17 | 2,75 | 0,06 | 9,51E-01 |
| AT4G00960 | 1,30 | 2,13 | 2,00 | 1,07 | 2,86E-01 |
| AT4G00970 | 0,33 | 2,10 | 4,30 | 0,49 | 6,26E-01 |
| AT4G00990 | 0,61 | -0,32 | 4,01 | -0,08 | 9,37E-01 |
| AT4G01020 | 0,37 | -0,55 | 4,36 | -0,13 | 9,00E-01 |
| AT4G01026 | 0,56 | 0,86 | 2,68 | 0,32 | 7,48E-01 |
| AT4G01040 | 0,12 | -0,26 | 4,41 | -0,06 | 9,53E-01 |
| AT4G01100 | 0,48 | 0,57 | 3,17 | 0,18 | 8,58E-01 |
| AT4G01130 | 0,11 | 0,88 | 4,35 | 0,20 | 8,39E-01 |
| AT4G01280 | 0,59 | -0,20 | 3,16 | -0,06 | 9,51E-01 |
| AT4G01320 | 0,90 | -0,26 | 2,33 | -0,11 | 9,10E-01 |
| AT4G01330 | 0,80 | 1,61 | 2,47 | 0,65 | 5,14E-01 |
| AT4G01340 | 1,48 | -2,91 | 1,52 | -1,92 | 5,47E-02 |
| AT4G01360 | 0,89 | 2,00 | 2,88 | 0,69 | 4,88E-01 |
| AT4G01370 | 0,27 | 1,20 | 4,33 | 0,28 | 7,82E-01 |
| AT4G01460 | 0,10 | 0,08 | 4,38 | 0,02 | 9,85E-01 |
| AT4G01570 | 0,24 | -1,29 | 4,39 | -0,29 | 7,69E-01 |
| AT4G03905 | 0,55 | 1,81 | 3,89 | 0,46 | 6,42E-01 |
| AT4G03925 | 0,76 | -0,01 | 2,16 | -0,01 | 9,96E-01 |
| AT4G01610 | 0,62 | 2,19 | 2,46 | 0,89 | 3,74E-01 |
| AT4G01690 | 1,14 | -0,42 | 1,68 | -0,25 | 8,04E-01 |
| AT4G01720 | 0,44 | -0,24 | 4,17 | -0,06 | 9,54E-01 |
| AT4G01995 | 0,08 | 0,00 | 4,41 | 0,00 | 1,00E+00 |
| AT4G02000 | 0,69 | -0,35 | 2,00 | -0,18 | 8,59E-01 |
| AT4G02010 | 0,14 | 0,04 | 4,38 | 0,01 | 9,93E-01 |
| AT4G02020 | 0,55 | -0,10 | 3,23 | -0,03 | 9,75E-01 |
| AT4G02030 | 0,75 | 0,00 | 2,92 | 0,00 | 1,00E+00 |
| AT4G02070 | 0,12 | 0,00 | 4,41 | 0,00 | 1,00E+00 |

| | | | | | |
|---|---|---|---|---|---|
| AT4G02080 | 0,57 | -2,01 | 2,67 | -0,75 | 4,51E-01 |
| AT4G02100 | 0,08 | 0,21 | 4,39 | 0,05 | 9,62E-01 |
| AT4G02110 | 0,21 | 0,00 | 4,41 | 0,00 | 1,00E+00 |
| AT4G02200 | 0,55 | 1,16 | 3,85 | 0,30 | 7,64E-01 |
| AT4G02220 | 0,29 | 0,29 | 4,37 | 0,07 | 9,47E-01 |
| AT4G02260 | 0,56 | 0,34 | 2,98 | 0,11 | 9,09E-01 |
| AT4G02330 | 0,22 | -0,64 | 4,32 | -0,15 | 8,82E-01 |
| AT4G02340 | 1,02 | 0,67 | 2,24 | 0,30 | 7,64E-01 |
| AT4G02370 | 0,72 | -0,99 | 2,98 | -0,33 | 7,39E-01 |
| AT4G02390 | 0,10 | 0,08 | 4,38 | 0,02 | 9,85E-01 |
| AT4G02400 | 0,19 | 0,43 | 4,36 | 0,10 | 9,21E-01 |
| AT4G02440 | 0,90 | 1,59 | 1,92 | 0,83 | 4,09E-01 |
| AT4G04135 | 0,05 | 0,18 | 4,38 | 0,04 | 9,68E-01 |
| AT4G02500 | 1,37 | -0,88 | 2,87 | -0,31 | 7,59E-01 |
| AT4G04195 | 0,38 | 1,91 | 3,35 | 0,57 | 5,69E-01 |
| AT4G02540 | 0,29 | -0,82 | 3,65 | -0,23 | 8,22E-01 |
| AT4G02580 | 1,19 | -0,78 | 2,18 | -0,36 | 7,22E-01 |
| AT4G02600 | 0,66 | -0,92 | 2,26 | -0,41 | 6,85E-01 |
| AT4G02610 | 0,31 | -0,40 | 4,37 | -0,09 | 9,27E-01 |
| AT4G02650 | 0,24 | 0,00 | 4,41 | 0,00 | 1,00E+00 |
| AT4G02700 | 1,40 | 0,59 | 1,68 | 0,35 | 7,27E-01 |
| AT4G02725 | 0,69 | -2,71 | 2,98 | -0,91 | 3,63E-01 |
| AT4G02790 | 0,08 | 0,00 | 4,41 | 0,00 | 1,00E+00 |
| AT4G02880 | 0,95 | 1,12 | 2,22 | 0,50 | 6,14E-01 |
| AT4G02920 | 1,02 | -0,81 | 2,36 | -0,34 | 7,31E-01 |
| AT4G02940 | 0,12 | 0,49 | 4,38 | 0,11 | 9,11E-01 |
| AT4G02990 | 0,69 | 1,75 | 2,52 | 0,70 | 4,87E-01 |
| AT4G03000 | 0,64 | -1,20 | 2,99 | -0,40 | 6,87E-01 |
| AT4G04425 | 0,08 | 0,00 | 4,41 | 0,00 | 1,00E+00 |
| AT4G03100 | 0,38 | 0,15 | 4,38 | 0,03 | 9,73E-01 |
| AT4G03120 | 0,23 | -0,34 | 4,40 | -0,08 | 9,38E-01 |
| AT4G03180 | 0,27 | -1,59 | 4,38 | -0,36 | 7,17E-01 |
| AT4G03200 | 0,82 | -3,03 | 2,47 | -1,23 | 2,19E-01 |
| AT4G03205 | 0,48 | -0,38 | 3,27 | -0,12 | 9,08E-01 |
| AT4G03210 | 0,53 | 0,00 | 4,40 | 0,00 | 1,00E+00 |
| AT4G03285 | 0,16 | 0,15 | 4,38 | 0,04 | 9,72E-01 |
| AT4G03410 | 0,58 | 1,02 | 3,76 | 0,27 | 7,87E-01 |
| AT4G03500 | 0,47 | 15,90 | 4,36 | 3,65 | 2,65E-04 |
| AT4G03830 | 0,61 | 0,85 | 2,53 | 0,34 | 7,37E-01 |
| AT4G03930 | 0,65 | -0,58 | 3,12 | -0,19 | 8,53E-01 |
| AT4G03940 | 0,17 | -0,29 | 4,40 | -0,07 | 9,47E-01 |
| AT4G03945 | 0,92 | 1,81 | 2,30 | 0,79 | 4,29E-01 |
| AT4G04110 | 0,65 | -0,42 | 4,36 | -0,10 | 9,23E-01 |
| AT4G04180 | 0,08 | 0,21 | 4,39 | 0,05 | 9,62E-01 |
| AT4G04210 | 0,39 | 1,31 | 2,86 | 0,46 | 6,47E-01 |
| AT4G04220 | 0,89 | 0,87 | 2,21 | 0,39 | 6,95E-01 |
| AT4G04795 | 0,87 | 3,38 | 2,47 | 1,37 | 1,71E-01 |
| AT4G04315 | 0,10 | 0,08 | 4,38 | 0,02 | 9,85E-01 |
| AT4G04350 | 0,42 | 0,32 | 3,32 | 0,10 | 9,24E-01 |
| AT4G04450 | 0,17 | -1,50 | 4,37 | -0,34 | 7,31E-01 |
| AT4G04510 | 0,17 | 0,07 | 4,38 | 0,02 | 9,88E-01 |
| AT4G04540 | 0,26 | -0,11 | 4,34 | -0,03 | 9,79E-01 |
| AT4G04610 | 0,73 | -1,00 | 2,07 | -0,48 | 6,30E-01 |
| AT4G04670 | 0,48 | 0,00 | 4,41 | 0,00 | 1,00E+00 |
| AT4G04750 | 0,90 | 0,35 | 1,74 | 0,20 | 8,42E-01 |
| AT4G04760 | 1,18 | 0,96 | 1,93 | 0,50 | 6,20E-01 |

| | | | | | |
|---|---|---|---|---|---|
| AT4G04800 | 0,52 | -0,38 | 3,91 | -0,10 | 9,23E-01 |
| AT4G04840 | 1,00 | 0,40 | 1,89 | 0,21 | 8,32E-01 |
| AT4G04850 | 0,79 | -1,26 | 2,01 | -0,63 | 5,30E-01 |
| AT4G04860 | 0,28 | -0,44 | 3,73 | -0,12 | 9,07E-01 |
| AT4G04920 | 1,70 | 1,03 | 1,92 | 0,54 | 5,93E-01 |
| AT4G04925 | 1,79 | 2,36 | 1,60 | 1,48 | 1,39E-01 |
| AT4G04960 | 0,39 | 1,31 | 2,85 | 0,46 | 6,44E-01 |
| AT4G05020 | 0,40 | 0,95 | 4,28 | 0,22 | 8,25E-01 |
| AT4G05080 | 0,23 | 1,85 | 4,30 | 0,43 | 6,66E-01 |
| AT4G05105 | 0,17 | 0,00 | 4,41 | 0,00 | 1,00E+00 |
| AT4G05160 | 0,46 | 0,19 | 4,12 | 0,05 | 9,64E-01 |
| AT4G05612 | 0,12 | 0,00 | 4,41 | 0,00 | 1,00E+00 |
| AT4G05632 | 0,08 | 0,21 | 4,39 | 0,05 | 9,62E-01 |
| AT4G06479 | 0,10 | 0,86 | 4,35 | 0,20 | 8,43E-01 |
| AT4G06598 | 0,74 | -0,28 | 3,57 | -0,08 | 9,38E-01 |
| AT4G06688 | 0,68 | 1,47 | 2,45 | 0,60 | 5,49E-01 |
| AT4G07390 | 0,27 | 1,28 | 3,72 | 0,34 | 7,31E-01 |
| AT4G07455 | 0,24 | 0,00 | 4,41 | 0,00 | 1,00E+00 |
| AT4G07825 | 1,05 | -0,41 | 2,70 | -0,15 | 8,80E-01 |
| AT4G07932 | 0,51 | -0,10 | 3,96 | -0,02 | 9,80E-01 |
| AT4G07965 | 0,15 | -0,35 | 4,39 | -0,08 | 9,36E-01 |
| AT4G08025 | 0,16 | 0,41 | 4,37 | 0,09 | 9,25E-01 |
| AT4G05305 | 0,12 | 1,17 | 4,34 | 0,27 | 7,88E-01 |
| AT4G08097 | 0,66 | 0,48 | 2,42 | 0,20 | 8,43E-01 |
| AT4G08180 | 1,01 | -2,97 | 2,14 | -1,39 | 1,65E-01 |
| AT4G08267 | 0,63 | -0,91 | 4,35 | -0,21 | 8,35E-01 |
| AT4G08270 | 0,17 | -0,38 | 4,39 | -0,09 | 9,31E-01 |
| AT4G08320 | 0,31 | 1,78 | 4,30 | 0,41 | 6,79E-01 |
| AT4G08350 | 0,65 | 0,94 | 2,54 | 0,37 | 7,11E-01 |
| AT4G08395 | 0,29 | 1,95 | 4,30 | 0,45 | 6,50E-01 |
| AT4G05455 | 0,68 | -1,03 | 2,36 | -0,44 | 6,63E-01 |
| AT4G08450 | 0,11 | 0,68 | 4,37 | 0,16 | 8,76E-01 |
| AT4G08455 | 0,63 | -0,27 | 4,41 | -0,06 | 9,50E-01 |
| AT4G08510 | 0,33 | -1,29 | 4,40 | -0,29 | 7,70E-01 |
| AT4G08545 | 1,36 | 3,62 | 1,76 | 2,05 | 4,02E-02 |
| AT4G08560 | 0,64 | -2,46 | 4,37 | -0,56 | 5,73E-01 |
| AT4G08780 | 0,43 | -0,35 | 4,20 | -0,08 | 9,34E-01 |
| AT4G05605 | 0,39 | 1,71 | 4,31 | 0,40 | 6,91E-01 |
| AT4G08930 | 0,71 | -1,27 | 2,53 | -0,50 | 6,16E-01 |
| AT4G08950 | 0,37 | 1,41 | 4,31 | 0,33 | 7,43E-01 |
| AT4G08980 | 0,65 | 0,38 | 2,93 | 0,13 | 8,96E-01 |
| AT4G08990 | 0,69 | 3,08 | 3,64 | 0,85 | 3,97E-01 |
| AT4G09000 | 1,74 | 1,28 | 1,37 | 0,94 | 3,49E-01 |
| AT4G09020 | 1,04 | -0,46 | 1,84 | -0,25 | 8,01E-01 |
| AT4G09030 | 0,16 | 0,41 | 4,37 | 0,09 | 9,25E-01 |
| AT4G09150 | 0,62 | 0,20 | 3,05 | 0,07 | 9,48E-01 |
| AT4G09160 | 0,36 | 0,00 | 4,41 | 0,00 | 1,00E+00 |
| AT4G09250 | 0,12 | -0,26 | 4,41 | -0,06 | 9,53E-01 |
| AT4G09310 | 0,21 | -2,19 | 4,41 | -0,50 | 6,20E-01 |
| AT4G05705 | 0,10 | 0,86 | 4,35 | 0,20 | 8,43E-01 |
| AT4G09320 | 1,06 | 0,10 | 1,53 | 0,07 | 9,48E-01 |
| AT4G09360 | 0,71 | 1,85 | 2,36 | 0,78 | 4,33E-01 |
| AT4G09450 | 1,44 | -4,21 | 1,81 | -2,32 | 2,02E-02 |
| AT4G09460 | 0,37 | -0,31 | 4,35 | -0,07 | 9,43E-01 |
| AT4G09545 | 0,31 | 1,07 | 4,28 | 0,25 | 8,02E-01 |
| AT4G09570 | 0,54 | 0,66 | 2,30 | 0,29 | 7,73E-01 |

| | | | | | |
|---|---|---|---|---|---|
| AT4G09630 | 0,87 | 1,80 | 2,08 | 0,87 | 3,86E-01 |
| AT4G09660 | 0,30 | 2,09 | 4,29 | 0,49 | 6,26E-01 |
| AT4G09680 | 0,45 | 0,00 | 4,07 | 0,00 | 1,00E+00 |
| AT4G09690 | 1,50 | -1,23 | 1,46 | -0,84 | 3,99E-01 |
| AT4G09730 | 0,51 | -1,97 | 3,13 | -0,63 | 5,30E-01 |
| AT4G09800 | 0,41 | -0,53 | 4,26 | -0,12 | 9,01E-01 |
| AT4G09820 | 0,31 | -1,68 | 4,34 | -0,39 | 6,98E-01 |
| AT4G09830 | 0,41 | -1,10 | 3,42 | -0,32 | 7,46E-01 |
| AT4G09850 | 0,21 | 0,34 | 4,37 | 0,08 | 9,37E-01 |
| AT4G05745 | 0,45 | 1,36 | 3,16 | 0,43 | 6,68E-01 |
| AT4G09890 | 0,18 | 0,09 | 4,38 | 0,02 | 9,84E-01 |
| AT4G10060 | 0,79 | 1,26 | 2,22 | 0,57 | 5,70E-01 |
| AT4G10170 | 0,08 | 0,00 | 4,41 | 0,00 | 1,00E+00 |
| AT4G10180 | 0,21 | -0,29 | 4,40 | -0,06 | 9,48E-01 |
| AT4G10465 | 0,60 | 1,58 | 2,54 | 0,62 | 5,36E-01 |
| AT4G10610 | 1,30 | 0,06 | 2,22 | 0,03 | 9,78E-01 |
| AT4G10613 | 0,77 | 1,05 | 1,99 | 0,53 | 5,96E-01 |
| AT4G10620 | 0,24 | 0,00 | 4,41 | 0,00 | 1,00E+00 |
| AT4G10660 | 0,11 | 0,88 | 4,35 | 0,20 | 8,39E-01 |
| AT4G10840 | 1,51 | 0,86 | 2,14 | 0,40 | 6,88E-01 |
| AT4G11000 | 0,26 | 0,03 | 4,38 | 0,01 | 9,95E-01 |
| AT4G11070 | 0,37 | -1,01 | 4,34 | -0,23 | 8,16E-01 |
| AT4G11100 | 0,32 | 0,80 | 4,34 | 0,18 | 8,55E-01 |
| AT4G11110 | 0,51 | -1,84 | 2,60 | -0,71 | 4,78E-01 |
| AT4G11130 | 0,54 | 0,00 | 3,97 | 0,00 | 1,00E+00 |
| AT4G11150 | 1,03 | 0,06 | 1,55 | 0,04 | 9,72E-01 |
| AT4G11160 | 1,61 | -0,52 | 1,77 | -0,30 | 7,67E-01 |
| AT4G11170 | 0,50 | -0,94 | 3,03 | -0,31 | 7,58E-01 |
| AT4G11240 | 1,11 | -0,26 | 2,20 | -0,12 | 9,05E-01 |
| AT4G11280 | 0,67 | 0,37 | 3,05 | 0,12 | 9,03E-01 |
| AT4G11330 | 0,53 | -1,08 | 3,18 | -0,34 | 7,34E-01 |
| AT4G11350 | 0,19 | 0,47 | 4,36 | 0,11 | 9,14E-01 |
| AT4G11360 | 0,31 | -3,20 | 4,34 | -0,74 | 4,60E-01 |
| AT4G11380 | 1,75 | -2,96 | 1,57 | -1,88 | 5,97E-02 |
| AT4G05885 | 0,80 | -4,01 | 2,64 | -1,52 | 1,28E-01 |
| AT4G05900 | 1,70 | 1,29 | 1,53 | 0,84 | 3,99E-01 |
| AT4G11660 | 0,57 | 1,64 | 3,23 | 0,51 | 6,11E-01 |
| AT4G11720 | 0,60 | -1,93 | 2,97 | -0,65 | 5,16E-01 |
| AT4G11740 | 0,35 | 1,34 | 4,33 | 0,31 | 7,58E-01 |
| AT4G11880 | 0,28 | -0,06 | 4,35 | -0,01 | 9,89E-01 |
| AT4G11890 | 0,62 | -1,34 | 3,13 | -0,43 | 6,68E-01 |
| AT4G11910 | 0,12 | 0,00 | 4,41 | 0,00 | 1,00E+00 |
| AT4G12020 | 1,23 | 0,39 | 2,67 | 0,15 | 8,84E-01 |
| AT4G12030 | 0,68 | -0,33 | 3,59 | -0,09 | 9,28E-01 |
| AT4G12040 | 1,45 | -0,35 | 1,84 | -0,19 | 8,48E-01 |
| AT4G12070 | 0,19 | -0,02 | 4,35 | -0,01 | 9,96E-01 |
| AT4G12240 | 0,26 | 0,13 | 4,38 | 0,03 | 9,77E-01 |
| AT4G05985 | 0,10 | 0,08 | 4,38 | 0,02 | 9,85E-01 |
| AT4G12390 | 0,50 | 0,33 | 4,34 | 0,08 | 9,40E-01 |
| AT4G12600 | 0,92 | -1,38 | 3,06 | -0,45 | 6,53E-01 |
| AT4G12640 | 0,26 | -0,45 | 3,79 | -0,12 | 9,06E-01 |
| AT4G12680 | 0,21 | 0,00 | 4,41 | 0,00 | 1,00E+00 |
| AT4G12690 | 0,12 | -0,36 | 4,39 | -0,08 | 9,34E-01 |
| AT4G12720 | 1,54 | 2,99 | 1,75 | 1,71 | 8,82E-02 |
| AT4G12830 | 0,34 | -0,40 | 4,33 | -0,09 | 9,27E-01 |
| AT4G12900 | 0,05 | 0,18 | 4,38 | 0,04 | 9,68E-01 |

| | | | | | |
|---|---|---|---|---|---|
| AT4G05995 | 0,98 | 3,59 | 2,42 | 1,48 | 1,38E-01 |
| AT4G13010 | 0,63 | -1,50 | 3,56 | -0,42 | 6,74E-01 |
| AT4G13020 | 0,54 | -0,24 | 3,19 | -0,08 | 9,39E-01 |
| AT4G06010 | 0,28 | 0,60 | 4,37 | 0,14 | 8,91E-01 |
| AT4G06015 | 0,10 | 0,24 | 4,38 | 0,05 | 9,57E-01 |
| AT4G13245 | 0,48 | 0,59 | 3,18 | 0,18 | 8,54E-01 |
| AT4G13270 | 0,50 | -1,10 | 3,24 | -0,34 | 7,33E-01 |
| AT4G13390 | 1,53 | 4,29 | 1,87 | 2,29 | 2,18E-02 |
| AT4G13440 | 0,58 | -0,15 | 4,04 | -0,04 | 9,70E-01 |
| AT4G13530 | 0,95 | -1,28 | 2,02 | -0,64 | 5,25E-01 |
| AT4G13572 | 0,31 | -2,06 | 4,39 | -0,47 | 6,39E-01 |
| AT4G13610 | 0,37 | 0,72 | 2,81 | 0,26 | 7,98E-01 |
| AT4G13615 | 0,92 | -0,19 | 2,01 | -0,09 | 9,25E-01 |
| AT4G13670 | 0,92 | 0,00 | 3,31 | 0,00 | 1,00E+00 |
| AT4G13730 | 0,24 | 0,48 | 4,37 | 0,11 | 9,12E-01 |
| AT4G13750 | 0,39 | -0,20 | 3,33 | -0,06 | 9,52E-01 |
| AT4G13760 | 0,54 | 1,63 | 3,14 | 0,52 | 6,05E-01 |
| AT4G13830 | 0,70 | 1,47 | 2,88 | 0,51 | 6,11E-01 |
| AT4G13850 | 0,81 | -1,15 | 2,02 | -0,57 | 5,69E-01 |
| AT4G13860 | 0,43 | -0,28 | 4,19 | -0,07 | 9,46E-01 |
| AT4G13992 | 1,30 | 0,34 | 1,80 | 0,19 | 8,50E-01 |
| AT4G14000 | 0,12 | 1,17 | 4,34 | 0,27 | 7,88E-01 |
| AT4G14140 | 0,81 | 0,62 | 1,91 | 0,33 | 7,45E-01 |
| AT4G14220 | 0,46 | -0,34 | 4,40 | -0,08 | 9,38E-01 |
| AT4G14270 | 0,92 | 1,71 | 1,83 | 0,93 | 3,51E-01 |
| AT4G14300 | 0,77 | -2,56 | 3,54 | -0,72 | 4,70E-01 |
| AT4G14330 | 0,14 | 0,04 | 4,38 | 0,01 | 9,93E-01 |
| AT4G14540 | 0,71 | -0,08 | 2,34 | -0,03 | 9,72E-01 |
| AT4G14548 | 1,46 | 3,32 | 1,83 | 1,81 | 7,02E-02 |
| AT4G14560 | 0,12 | 0,59 | 4,36 | 0,14 | 8,92E-01 |
| AT4G14570 | 0,70 | -0,37 | 2,91 | -0,13 | 8,99E-01 |
| AT4G14630 | 0,43 | 1,83 | 3,95 | 0,46 | 6,44E-01 |
| AT4G14700 | 0,12 | 0,00 | 4,41 | 0,00 | 1,00E+00 |
| AT4G06245 | 1,52 | -5,43 | 2,08 | -2,62 | 8,87E-03 |
| AT4G06255 | 0,39 | 0,87 | 4,34 | 0,20 | 8,41E-01 |
| AT4G06260 | 0,42 | 1,16 | 3,29 | 0,35 | 7,24E-01 |
| AT4G14760 | 0,91 | -1,43 | 2,49 | -0,57 | 5,66E-01 |
| AT4G14800 | 0,34 | -0,26 | 4,41 | -0,06 | 9,53E-01 |
| AT4G14810 | 0,45 | -0,12 | 3,34 | -0,03 | 9,72E-01 |
| AT4G14850 | 0,15 | 0,82 | 4,35 | 0,19 | 8,50E-01 |
| AT4G14870 | 0,48 | 1,08 | 2,60 | 0,41 | 6,79E-01 |
| AT4G14890 | 0,73 | 0,77 | 2,17 | 0,36 | 7,22E-01 |
| AT4G14950 | 0,54 | 0,18 | 2,62 | 0,07 | 9,46E-01 |
| AT4G15000 | 1,11 | -2,33 | 1,71 | -1,36 | 1,73E-01 |
| AT4G15020 | 0,86 | -1,17 | 2,49 | -0,47 | 6,39E-01 |
| AT4G15030 | 0,61 | -0,57 | 2,30 | -0,25 | 8,06E-01 |
| AT4G15090 | 0,08 | 0,00 | 4,41 | 0,00 | 1,00E+00 |
| AT4G15096 | 1,15 | 1,93 | 1,98 | 0,97 | 3,30E-01 |
| AT4G15165 | 0,40 | -0,12 | 4,39 | -0,03 | 9,78E-01 |
| AT4G15180 | 1,84 | 1,11 | 1,67 | 0,66 | 5,07E-01 |
| AT4G15233 | 0,90 | -0,26 | 2,53 | -0,10 | 9,18E-01 |
| AT4G15245 | 0,49 | 1,37 | 2,57 | 0,53 | 5,94E-01 |
| AT4G15260 | 0,57 | 1,37 | 3,02 | 0,45 | 6,50E-01 |
| AT4G15280 | 1,02 | 0,88 | 2,18 | 0,40 | 6,87E-01 |
| AT4G15415 | 0,24 | 0,34 | 4,37 | 0,08 | 9,37E-01 |
| AT4G15420 | 1,24 | -0,32 | 2,33 | -0,14 | 8,90E-01 |

| | | | | | |
|---|---|---|---|---|---|
| AT4G15440 | 0,99 | 0,84 | 2,24 | 0,37 | 7,10E-01 |
| AT4G15450 | 0,24 | 0,77 | 4,35 | 0,18 | 8,60E-01 |
| AT4G15470 | 1,75 | 2,11 | 1,60 | 1,32 | 1,88E-01 |
| AT4G15475 | 0,96 | 0,00 | 4,14 | 0,00 | 1,00E+00 |
| AT4G15540 | 0,39 | 1,48 | 3,45 | 0,43 | 6,68E-01 |
| AT4G15545 | 1,17 | -0,35 | 1,61 | -0,21 | 8,30E-01 |
| AT4G15660 | 0,10 | 0,08 | 4,38 | 0,02 | 9,85E-01 |
| AT4G15700 | 0,08 | 0,21 | 4,39 | 0,05 | 9,62E-01 |
| AT4G15720 | 0,08 | 0,00 | 4,41 | 0,00 | 1,00E+00 |
| AT4G15770 | 0,18 | 0,31 | 4,38 | 0,07 | 9,44E-01 |
| AT4G15775 | 0,24 | -2,39 | 4,34 | -0,55 | 5,82E-01 |
| AT4G15800 | 0,76 | 1,24 | 2,18 | 0,57 | 5,68E-01 |
| AT4G15820 | 0,10 | 0,24 | 4,38 | 0,05 | 9,57E-01 |
| AT4G15840 | 0,73 | -0,45 | 2,13 | -0,21 | 8,32E-01 |
| AT4G15850 | 0,38 | 0,00 | 4,41 | 0,00 | 1,00E+00 |
| AT4G15880 | 0,12 | 0,27 | 4,38 | 0,06 | 9,50E-01 |
| AT4G15940 | 0,10 | 0,86 | 4,35 | 0,20 | 8,43E-01 |
| AT4G16008 | 0,08 | 0,00 | 4,41 | 0,00 | 1,00E+00 |
| AT4G16015 | 0,19 | 0,43 | 4,36 | 0,10 | 9,21E-01 |
| AT4G16130 | 1,05 | -0,55 | 2,25 | -0,24 | 8,09E-01 |
| AT4G16140 | 0,34 | 0,71 | 4,31 | 0,16 | 8,69E-01 |
| AT4G16150 | 0,50 | 0,69 | 4,13 | 0,17 | 8,67E-01 |
| AT4G16310 | 1,01 | 2,00 | 2,04 | 0,98 | 3,29E-01 |
| AT4G16360 | 0,61 | -2,49 | 2,61 | -0,95 | 3,42E-01 |
| AT4G16380 | 0,81 | 2,16 | 2,49 | 0,87 | 3,86E-01 |
| AT4G16390 | 0,67 | -0,02 | 2,34 | -0,01 | 9,94E-01 |
| AT4G16400 | 1,31 | -0,02 | 2,12 | -0,01 | 9,92E-01 |
| AT4G16430 | 0,28 | -0,24 | 4,40 | -0,05 | 9,57E-01 |
| AT4G16480 | 0,65 | -0,30 | 3,65 | -0,08 | 9,35E-01 |
| AT4G16510 | 0,53 | -3,42 | 4,01 | -0,85 | 3,93E-01 |
| AT4G16590 | 0,08 | 0,00 | 4,41 | 0,00 | 1,00E+00 |
| AT4G16650 | 0,41 | -3,04 | 4,22 | -0,72 | 4,72E-01 |
| AT4G16670 | 0,65 | -1,28 | 2,38 | -0,54 | 5,92E-01 |
| AT4G16710 | 0,63 | 0,49 | 2,60 | 0,19 | 8,51E-01 |
| AT4G16780 | 0,60 | -0,55 | 3,93 | -0,14 | 8,90E-01 |
| AT4G16830 | 0,78 | -1,45 | 3,02 | -0,48 | 6,31E-01 |
| AT4G16845 | 0,11 | 0,88 | 4,35 | 0,20 | 8,39E-01 |
| AT4G16850 | 0,29 | 0,00 | 4,41 | 0,00 | 1,00E+00 |
| AT4G16855 | 0,17 | 0,00 | 4,41 | 0,00 | 1,00E+00 |
| AT4G16955 | 0,78 | 0,25 | 3,11 | 0,08 | 9,37E-01 |
| AT4G17010 | 0,55 | -3,81 | 4,33 | -0,88 | 3,79E-01 |
| AT4G17080 | 0,57 | 0,00 | 4,40 | 0,00 | 1,00E+00 |
| AT4G17085 | 0,42 | 0,00 | 4,41 | 0,00 | 1,00E+00 |
| AT4G06665 | 1,22 | -1,50 | 1,75 | -0,86 | 3,90E-01 |
| AT4G17098 | 0,61 | -1,73 | 2,43 | -0,71 | 4,78E-01 |
| AT4G17245 | 0,66 | 0,86 | 2,60 | 0,33 | 7,40E-01 |
| AT4G17250 | 0,41 | -0,38 | 4,39 | -0,09 | 9,31E-01 |
| AT4G17270 | 0,64 | 1,84 | 2,98 | 0,62 | 5,37E-01 |
| AT4G17300 | 0,71 | 1,19 | 2,22 | 0,54 | 5,91E-01 |
| AT4G17380 | 0,44 | 0,42 | 2,80 | 0,15 | 8,82E-01 |
| AT4G17500 | 0,96 | 2,97 | 2,18 | 1,36 | 1,74E-01 |
| AT4G17520 | 1,02 | -1,21 | 2,56 | -0,47 | 6,38E-01 |
| AT4G17600 | 1,35 | -1,41 | 1,50 | -0,94 | 3,49E-01 |
| AT4G17616 | 0,78 | 0,00 | 3,52 | 0,00 | 1,00E+00 |
| AT4G17620 | 1,00 | 0,23 | 2,53 | 0,09 | 9,26E-01 |
| AT4G17650 | 0,32 | -2,12 | 4,40 | -0,48 | 6,30E-01 |

| | | | | | |
|---|---|---|---|---|---|
| AT4G17770 | 1,15 | -0,51 | 1,85 | -0,27 | 7,85E-01 |
| AT4G17810 | 0,14 | -2,43 | 4,37 | -0,55 | 5,79E-01 |
| AT4G17830 | 0,83 | 0,07 | 2,55 | 0,03 | 9,77E-01 |
| AT4G17870 | 1,03 | 2,09 | 2,26 | 0,93 | 3,55E-01 |
| AT4G17900 | 0,47 | -0,55 | 2,80 | -0,20 | 8,45E-01 |
| AT4G18070 | 1,02 | -0,71 | 2,15 | -0,33 | 7,41E-01 |
| AT4G18160 | 0,40 | 0,15 | 4,37 | 0,04 | 9,72E-01 |
| AT4G18195 | 0,22 | 0,90 | 4,35 | 0,21 | 8,36E-01 |
| AT4G18205 | 1,32 | 0,22 | 1,41 | 0,15 | 8,78E-01 |
| AT4G18220 | 1,16 | 0,11 | 1,72 | 0,06 | 9,50E-01 |
| AT4G18240 | 1,25 | -0,29 | 1,88 | -0,15 | 8,78E-01 |
| AT4G18370 | 0,87 | -0,99 | 2,41 | -0,41 | 6,81E-01 |
| AT4G18380 | 0,33 | -2,04 | 4,38 | -0,47 | 6,42E-01 |
| AT4G06795 | 0,56 | -0,40 | 3,11 | -0,13 | 8,98E-01 |
| AT4G18530 | 0,35 | -0,93 | 4,38 | -0,21 | 8,32E-01 |
| AT4G18570 | 0,37 | -0,16 | 2,90 | -0,06 | 9,56E-01 |
| AT4G18596 | 1,78 | -0,27 | 2,06 | -0,13 | 8,96E-01 |
| AT4G18710 | 1,19 | -0,18 | 2,00 | -0,09 | 9,28E-01 |
| AT4G18725 | 0,19 | 1,00 | 4,33 | 0,23 | 8,17E-01 |
| AT4G18730 | 1,33 | -0,51 | 1,96 | -0,26 | 7,96E-01 |
| AT4G18750 | 0,50 | 0,43 | 4,10 | 0,10 | 9,17E-01 |
| AT4G18820 | 1,31 | -1,53 | 2,10 | -0,73 | 4,65E-01 |
| AT4G18830 | 0,19 | 0,84 | 4,35 | 0,19 | 8,47E-01 |
| AT4G18890 | 0,44 | 2,41 | 4,00 | 0,60 | 5,47E-01 |
| AT4G18905 | 0,35 | 0,39 | 4,38 | 0,09 | 9,30E-01 |
| AT4G18930 | 0,31 | -0,57 | 4,33 | -0,13 | 8,96E-01 |
| AT4G18950 | 1,00 | 0,43 | 1,98 | 0,22 | 8,29E-01 |
| AT4G19020 | 0,17 | -0,29 | 4,40 | -0,07 | 9,47E-01 |
| AT4G06865 | 0,10 | 0,08 | 4,38 | 0,02 | 9,85E-01 |
| AT4G19090 | 1,32 | -0,08 | 1,78 | -0,05 | 9,63E-01 |
| AT4G19100 | 1,38 | 2,10 | 2,02 | 1,04 | 2,98E-01 |
| AT4G19190 | 0,39 | -1,20 | 4,26 | -0,28 | 7,78E-01 |
| AT4G19200 | 0,86 | -0,10 | 2,14 | -0,04 | 9,64E-01 |
| AT4G19210 | 1,66 | -1,13 | 1,72 | -0,66 | 5,08E-01 |
| AT4G19450 | 0,65 | 0,23 | 4,39 | 0,05 | 9,59E-01 |
| AT4G19490 | 0,82 | 0,26 | 2,82 | 0,09 | 9,28E-01 |
| AT4G19500 | 0,89 | 0,39 | 2,36 | 0,17 | 8,68E-01 |
| AT4G19510 | 1,30 | 0,48 | 2,11 | 0,23 | 8,21E-01 |
| AT4G19530 | 1,16 | 0,95 | 1,84 | 0,52 | 6,05E-01 |
| AT4G19540 | 0,68 | -0,17 | 3,04 | -0,05 | 9,57E-01 |
| AT4G19580 | 0,44 | -0,30 | 4,14 | -0,07 | 9,43E-01 |
| AT4G19600 | 0,21 | -2,83 | 4,36 | -0,65 | 5,16E-01 |
| AT4G19610 | 0,37 | -0,74 | 3,49 | -0,21 | 8,33E-01 |
| AT4G19650 | 0,19 | -0,37 | 4,39 | -0,08 | 9,33E-01 |
| AT4G19710 | 1,47 | -1,24 | 1,80 | -0,69 | 4,92E-01 |
| AT4G19960 | 0,50 | 1,10 | 3,22 | 0,34 | 7,33E-01 |
| AT4G20110 | 0,83 | 1,21 | 2,28 | 0,53 | 5,96E-01 |
| AT4G20140 | 0,84 | 0,00 | 4,30 | 0,00 | 1,00E+00 |
| AT4G06950 | 0,12 | 0,11 | 4,38 | 0,03 | 9,80E-01 |
| AT4G20270 | 0,28 | 0,43 | 4,36 | 0,10 | 9,22E-01 |
| AT4G20300 | 0,68 | -0,34 | 2,42 | -0,14 | 8,87E-01 |
| AT4G20310 | 0,08 | 0,00 | 4,41 | 0,00 | 1,00E+00 |
| AT4G20400 | 1,23 | -2,32 | 2,26 | -1,03 | 3,04E-01 |
| AT4G20720 | 0,34 | -0,30 | 4,40 | -0,07 | 9,46E-01 |
| AT4G20840 | 1,22 | 0,00 | 2,63 | 0,00 | 1,00E+00 |
| AT4G20860 | 0,52 | 1,03 | 3,13 | 0,33 | 7,41E-01 |

| | | | | | |
|---|---|---|---|---|---|
| AT4G20890 | 0,68 | -0,43 | 2,58 | -0,16 | 8,69E-01 |
| AT4G20940 | 0,56 | 0,00 | 4,41 | 0,00 | 1,00E+00 |
| AT4G20960 | 0,48 | 1,63 | 2,69 | 0,61 | 5,44E-01 |
| AT4G20980 | 1,36 | -0,84 | 1,91 | -0,44 | 6,60E-01 |
| AT4G21105 | 0,54 | -0,25 | 4,35 | -0,06 | 9,55E-01 |
| AT4G21160 | 0,15 | -0,57 | 4,37 | -0,13 | 8,96E-01 |
| AT4G21180 | 0,08 | 0,00 | 4,41 | 0,00 | 1,00E+00 |
| AT4G21210 | 0,33 | -0,54 | 4,36 | -0,12 | 9,01E-01 |
| AT4G21300 | 0,05 | 0,18 | 4,38 | 0,04 | 9,68E-01 |
| AT4G21323 | 0,38 | 0,00 | 4,41 | 0,00 | 1,00E+00 |
| AT4G21450 | 0,15 | -0,26 | 4,41 | -0,06 | 9,53E-01 |
| AT4G21470 | 0,32 | -1,23 | 4,39 | -0,28 | 7,79E-01 |
| AT4G21515 | 0,05 | 0,18 | 4,38 | 0,04 | 9,68E-01 |
| AT4G21540 | 0,68 | 0,60 | 2,95 | 0,20 | 8,38E-01 |
| AT4G21550 | 0,37 | -0,34 | 4,40 | -0,08 | 9,38E-01 |
| AT4G21620 | 0,72 | -0,12 | 2,58 | -0,05 | 9,64E-01 |
| AT4G21670 | 1,62 | 0,19 | 1,33 | 0,14 | 8,89E-01 |
| AT4G21740 | 0,47 | -0,68 | 2,37 | -0,29 | 7,75E-01 |
| AT4G21750 | 0,81 | -2,19 | 2,90 | -0,75 | 4,51E-01 |
| AT4G21790 | 0,64 | 0,51 | 2,59 | 0,20 | 8,45E-01 |
| AT4G21800 | 0,53 | 0,87 | 3,87 | 0,23 | 8,21E-01 |
| AT4G07030 | 0,38 | 0,00 | 4,41 | 0,00 | 1,00E+00 |
| AT4G21940 | 0,15 | -0,31 | 4,40 | -0,07 | 9,43E-01 |
| AT4G22010 | 0,22 | -1,15 | 4,40 | -0,26 | 7,94E-01 |
| AT4G22050 | 0,20 | 1,44 | 4,33 | 0,33 | 7,40E-01 |
| AT4G07040 | 0,12 | 0,49 | 4,38 | 0,11 | 9,11E-01 |
| AT4G22130 | 0,44 | 1,60 | 2,73 | 0,59 | 5,57E-01 |
| AT4G22270 | 0,20 | 0,00 | 4,41 | 0,00 | 1,00E+00 |
| AT4G07075 | 1,23 | 1,69 | 1,75 | 0,97 | 3,33E-01 |
| AT4G07080 | 1,23 | 1,69 | 1,75 | 0,97 | 3,33E-01 |
| AT4G22330 | 0,17 | 0,00 | 4,41 | 0,00 | 1,00E+00 |
| AT4G22467 | 0,12 | 0,11 | 4,38 | 0,03 | 9,80E-01 |
| AT4G22475 | 0,49 | 1,74 | 3,82 | 0,46 | 6,48E-01 |
| AT4G22670 | 0,32 | 0,43 | 4,32 | 0,10 | 9,20E-01 |
| AT4G22710 | 1,76 | 1,54 | 1,70 | 0,90 | 3,66E-01 |
| AT4G22720 | 0,28 | -0,22 | 4,40 | -0,05 | 9,60E-01 |
| AT4G22860 | 0,08 | 0,00 | 4,41 | 0,00 | 1,00E+00 |
| AT4G22890 | 1,50 | -1,29 | 1,56 | -0,83 | 4,07E-01 |
| AT4G22910 | 0,08 | 0,00 | 4,41 | 0,00 | 1,00E+00 |
| AT4G22960 | 0,12 | 0,11 | 4,38 | 0,03 | 9,80E-01 |
| AT4G23030 | 0,17 | 0,07 | 4,38 | 0,02 | 9,88E-01 |
| AT4G23050 | 0,25 | 0,00 | 4,41 | 0,00 | 1,00E+00 |
| AT4G23060 | 0,21 | 0,00 | 4,41 | 0,00 | 1,00E+00 |
| AT4G23150 | 0,15 | -0,28 | 4,35 | -0,06 | 9,50E-01 |
| AT4G23160 | 0,75 | 1,58 | 4,32 | 0,37 | 7,15E-01 |
| AT4G23180 | 1,57 | 1,70 | 1,62 | 1,05 | 2,94E-01 |
| AT4G23270 | 0,94 | -0,52 | 2,17 | -0,24 | 8,11E-01 |
| AT4G23400 | 0,56 | -0,53 | 2,16 | -0,24 | 8,07E-01 |
| AT4G23440 | 0,12 | 0,00 | 4,41 | 0,00 | 1,00E+00 |
| AT4G23470 | 0,84 | 2,09 | 2,05 | 1,02 | 3,09E-01 |
| AT4G23530 | 1,11 | 0,88 | 2,17 | 0,41 | 6,85E-01 |
| AT4G23570 | 0,44 | -0,72 | 3,32 | -0,22 | 8,28E-01 |
| AT4G23600 | 1,02 | -3,17 | 2,60 | -1,22 | 2,23E-01 |
| AT4G23740 | 0,40 | 0,22 | 4,39 | 0,05 | 9,59E-01 |
| AT4G23860 | 0,25 | 0,28 | 4,39 | 0,06 | 9,50E-01 |
| AT4G07190 | 0,45 | -1,02 | 3,37 | -0,30 | 7,63E-01 |

| | | | | | |
|---|---|---|---|---|---|
| AT4G23895 | 0,24 | -0,53 | 4,37 | -0,12 | 9,03E-01 |
| AT4G23940 | 1,06 | -0,29 | 2,41 | -0,12 | 9,04E-01 |
| AT4G23950 | 0,16 | -0,26 | 4,41 | -0,06 | 9,53E-01 |
| AT4G23980 | 1,47 | -0,21 | 1,64 | -0,13 | 8,98E-01 |
| AT4G24000 | 0,08 | 0,00 | 4,41 | 0,00 | 1,00E+00 |
| AT4G24010 | 0,33 | -2,04 | 4,38 | -0,47 | 6,42E-01 |
| AT4G24020 | 1,49 | 1,41 | 1,72 | 0,82 | 4,12E-01 |
| AT4G24060 | 0,12 | 0,00 | 4,41 | 0,00 | 1,00E+00 |
| AT4G24090 | 0,53 | -4,57 | 4,30 | -1,06 | 2,87E-01 |
| AT4G24120 | 1,68 | -0,26 | 2,37 | -0,11 | 9,12E-01 |
| AT4G24130 | 0,24 | 0,00 | 4,41 | 0,00 | 1,00E+00 |
| AT4G24160 | 0,15 | 0,47 | 4,37 | 0,11 | 9,15E-01 |
| AT4G24175 | 0,21 | 0,00 | 4,41 | 0,00 | 1,00E+00 |
| AT4G24220 | 1,52 | -1,12 | 1,46 | -0,77 | 4,43E-01 |
| AT4G24290 | 0,72 | -0,52 | 2,51 | -0,21 | 8,37E-01 |
| AT4G24350 | 0,25 | 0,55 | 4,31 | 0,13 | 8,98E-01 |
| AT4G24440 | 0,43 | -1,24 | 4,26 | -0,29 | 7,71E-01 |
| AT4G24460 | 0,18 | -2,46 | 4,37 | -0,56 | 5,73E-01 |
| AT4G24470 | 0,57 | -0,26 | 3,94 | -0,07 | 9,47E-01 |
| AT4G24550 | 0,38 | -1,47 | 3,44 | -0,43 | 6,69E-01 |
| AT4G24790 | 0,10 | 0,24 | 4,38 | 0,05 | 9,57E-01 |
| AT4G24800 | 0,98 | -0,16 | 2,73 | -0,06 | 9,54E-01 |
| AT4G24805 | 0,88 | 0,18 | 2,36 | 0,07 | 9,40E-01 |
| AT4G24880 | 0,08 | 0,21 | 4,39 | 0,05 | 9,62E-01 |
| AT4G07295 | 0,21 | -0,52 | 4,36 | -0,12 | 9,05E-01 |
| AT4G25030 | 0,40 | -0,18 | 2,73 | -0,06 | 9,49E-01 |
| AT4G25050 | 1,37 | -0,98 | 1,62 | -0,61 | 5,45E-01 |
| AT4G25070 | 0,53 | 0,00 | 4,39 | 0,00 | 1,00E+00 |
| AT4G25120 | 0,50 | -1,99 | 3,27 | -0,61 | 5,44E-01 |
| AT4G25170 | 1,05 | -0,55 | 1,94 | -0,28 | 7,76E-01 |
| AT4G25210 | 0,75 | 2,45 | 2,79 | 0,88 | 3,80E-01 |
| AT4G25230 | 0,77 | 0,53 | 2,20 | 0,24 | 8,10E-01 |
| AT4G25360 | 0,25 | -0,33 | 4,40 | -0,08 | 9,40E-01 |
| AT4G25370 | 0,59 | -2,42 | 3,89 | -0,62 | 5,34E-01 |
| AT4G25390 | 0,17 | 1,19 | 4,33 | 0,28 | 7,83E-01 |
| AT4G25400 | 0,31 | 2,22 | 4,29 | 0,52 | 6,05E-01 |
| AT4G25500 | 1,12 | 0,89 | 2,05 | 0,43 | 6,64E-01 |
| AT4G25515 | 0,32 | -0,38 | 4,39 | -0,09 | 9,32E-01 |
| AT4G25550 | 0,38 | 0,00 | 4,41 | 0,00 | 1,00E+00 |
| AT4G25610 | 0,54 | -0,26 | 3,25 | -0,08 | 9,36E-01 |
| AT4G25630 | 1,57 | 2,37 | 1,91 | 1,24 | 2,15E-01 |
| AT4G25631 | 1,33 | 3,01 | 2,11 | 1,42 | 1,55E-01 |
| AT4G25660 | 0,81 | -1,42 | 2,34 | -0,61 | 5,44E-01 |
| AT4G25810 | 0,12 | 0,00 | 4,41 | 0,00 | 1,00E+00 |
| AT4G25835 | 0,21 | -1,29 | 4,40 | -0,29 | 7,69E-01 |
| AT4G25900 | 0,36 | -0,26 | 4,36 | -0,06 | 9,52E-01 |
| AT4G25910 | 0,69 | 0,00 | 3,61 | 0,00 | 1,00E+00 |
| AT4G25960 | 1,53 | -0,87 | 1,39 | -0,63 | 5,31E-01 |
| AT4G25970 | 1,00 | 1,07 | 1,92 | 0,56 | 5,77E-01 |
| AT4G26000 | 0,96 | 0,54 | 2,52 | 0,21 | 8,31E-01 |
| AT4G26120 | 0,24 | 0,00 | 4,41 | 0,00 | 1,00E+00 |
| AT4G26170 | 0,08 | 0,00 | 4,41 | 0,00 | 1,00E+00 |
| AT4G26210 | 0,36 | 0,00 | 4,41 | 0,00 | 1,00E+00 |
| AT4G26230 | 0,82 | 0,32 | 2,65 | 0,12 | 9,05E-01 |
| AT4G26450 | 0,45 | -0,10 | 3,30 | -0,03 | 9,77E-01 |
| AT4G26455 | 0,54 | 0,00 | 3,87 | 0,00 | 1,00E+00 |

| | | | | | |
|---|---|---|---|---|---|
| AT4G26510 | 0,72 | -0,58 | 2,53 | -0,23 | 8,19E-01 |
| AT4G26520 | 1,37 | -0,21 | 2,25 | -0,09 | 9,26E-01 |
| AT4G26650 | 0,56 | -0,37 | 2,37 | -0,16 | 8,76E-01 |
| AT4G26670 | 0,33 | -1,63 | 4,32 | -0,38 | 7,05E-01 |
| AT4G26680 | 0,40 | 0,42 | 4,18 | 0,10 | 9,19E-01 |
| AT4G26700 | 0,28 | -0,40 | 4,37 | -0,09 | 9,28E-01 |
| AT4G26790 | 0,42 | 2,35 | 4,03 | 0,58 | 5,59E-01 |
| AT4G26800 | 0,12 | 0,35 | 4,37 | 0,08 | 9,35E-01 |
| AT4G26840 | 0,82 | 0,23 | 2,22 | 0,10 | 9,19E-01 |
| AT4G26870 | 0,92 | 0,74 | 2,29 | 0,32 | 7,48E-01 |
| AT4G26970 | 1,79 | 0,02 | 1,78 | 0,01 | 9,89E-01 |
| AT4G27010 | 0,79 | 1,51 | 2,12 | 0,71 | 4,77E-01 |
| AT4G27030 | 0,63 | -1,99 | 2,60 | -0,76 | 4,45E-01 |
| AT4G07775 | 0,58 | 2,40 | 2,90 | 0,83 | 4,08E-01 |
| AT4G27080 | 1,09 | -0,10 | 2,22 | -0,04 | 9,65E-01 |
| AT4G27260 | 0,26 | 0,00 | 4,41 | 0,00 | 1,00E+00 |
| AT4G27290 | 0,45 | 0,00 | 4,41 | 0,00 | 1,00E+00 |
| AT4G27310 | 0,93 | 0,49 | 1,92 | 0,25 | 8,00E-01 |
| AT4G27380 | 0,21 | -3,00 | 4,36 | -0,69 | 4,91E-01 |
| AT4G27395 | 0,73 | -4,51 | 2,78 | -1,62 | 1,05E-01 |
| AT4G27652 | 0,57 | 2,83 | 3,64 | 0,78 | 4,38E-01 |
| AT4G27654 | 0,56 | 1,83 | 2,59 | 0,71 | 4,81E-01 |
| AT4G27657 | 0,12 | 0,11 | 4,38 | 0,03 | 9,80E-01 |
| AT4G27680 | 0,93 | 0,99 | 2,19 | 0,45 | 6,50E-01 |
| AT4G27710 | 0,52 | -1,45 | 3,24 | -0,45 | 6,55E-01 |
| AT4G27720 | 1,38 | 1,98 | 1,79 | 1,11 | 2,67E-01 |
| AT4G27740 | 0,52 | 2,97 | 3,74 | 0,79 | 4,28E-01 |
| AT4G27750 | 0,69 | -2,12 | 4,36 | -0,49 | 6,28E-01 |
| AT4G27760 | 0,28 | 0,00 | 4,41 | 0,00 | 1,00E+00 |
| AT4G27765 | 0,37 | 0,00 | 4,35 | 0,00 | 1,00E+00 |
| AT4G27780 | 0,50 | 0,00 | 4,01 | 0,00 | 1,00E+00 |
| AT4G27860 | 0,21 | 0,70 | 4,35 | 0,16 | 8,72E-01 |
| AT4G27880 | 0,08 | 0,00 | 4,41 | 0,00 | 1,00E+00 |
| AT4G27910 | 0,46 | 0,00 | 4,14 | 0,00 | 1,00E+00 |
| AT4G27940 | 1,14 | -0,34 | 2,37 | -0,15 | 8,84E-01 |
| AT4G28010 | 0,17 | -0,38 | 4,39 | -0,09 | 9,31E-01 |
| AT4G28210 | 0,08 | 0,21 | 4,39 | 0,05 | 9,62E-01 |
| AT4G28220 | 0,27 | 1,33 | 4,33 | 0,31 | 7,58E-01 |
| AT4G28260 | 0,80 | -0,26 | 3,03 | -0,09 | 9,31E-01 |
| AT4G28300 | 0,21 | -0,29 | 4,40 | -0,06 | 9,48E-01 |
| AT4G28362 | 0,21 | 0,00 | 4,41 | 0,00 | 1,00E+00 |
| AT4G28400 | 0,26 | 0,81 | 4,36 | 0,19 | 8,53E-01 |
| AT4G28440 | 0,16 | 0,39 | 4,37 | 0,09 | 9,29E-01 |
| AT4G28490 | 0,74 | 1,38 | 3,63 | 0,38 | 7,05E-01 |
| AT4G28600 | 1,03 | -0,70 | 2,24 | -0,31 | 7,56E-01 |
| AT4G28660 | 0,48 | -1,51 | 4,02 | -0,37 | 7,08E-01 |
| AT4G28706 | 0,39 | -0,71 | 4,25 | -0,17 | 8,67E-01 |
| AT4G28710 | 0,41 | 0,47 | 4,23 | 0,11 | 9,11E-01 |
| AT4G28740 | 0,90 | 0,22 | 2,46 | 0,09 | 9,28E-01 |
| AT4G28780 | 0,15 | 0,90 | 4,35 | 0,21 | 8,36E-01 |
| AT4G28860 | 0,76 | -1,75 | 3,15 | -0,56 | 5,78E-01 |
| AT4G28870 | 0,12 | 1,17 | 4,34 | 0,27 | 7,88E-01 |
| AT4G28880 | 1,20 | 0,69 | 1,75 | 0,39 | 6,95E-01 |
| AT4G28980 | 0,73 | -1,32 | 2,59 | -0,51 | 6,09E-01 |
| AT4G29020 | 0,17 | 0,07 | 4,38 | 0,02 | 9,88E-01 |
| AT4G29035 | 0,22 | 0,90 | 4,35 | 0,21 | 8,36E-01 |

| | | | | | |
|---|---|---|---|---|---|
| AT4G29040 | 1,10 | 2,44 | 2,03 | 1,20 | 2,30E-01 |
| AT4G29070 | 0,33 | 0,85 | 4,36 | 0,20 | 8,45E-01 |
| AT4G29110 | 0,63 | 2,56 | 2,84 | 0,90 | 3,67E-01 |
| AT4G29120 | 0,68 | 0,88 | 2,89 | 0,30 | 7,61E-01 |
| AT4G29160 | 0,87 | 0,28 | 2,03 | 0,14 | 8,90E-01 |
| AT4G29170 | 0,92 | 0,00 | 4,41 | 0,00 | 1,00E+00 |
| AT4G29180 | 0,96 | 0,00 | 2,77 | 0,00 | 1,00E+00 |
| AT4G29210 | 0,68 | -1,40 | 2,64 | -0,53 | 5,97E-01 |
| AT4G29350 | 0,47 | 0,60 | 4,06 | 0,15 | 8,82E-01 |
| AT4G29380 | 1,74 | -1,29 | 1,93 | -0,67 | 5,06E-01 |
| AT4G29400 | 0,21 | 0,08 | 4,38 | 0,02 | 9,85E-01 |
| AT4G29510 | 0,71 | -0,26 | 2,36 | -0,11 | 9,14E-01 |
| AT4G29740 | 1,04 | -0,94 | 1,91 | -0,49 | 6,22E-01 |
| AT4G29750 | 0,66 | 0,17 | 2,59 | 0,06 | 9,49E-01 |
| AT4G29790 | 1,30 | -2,11 | 2,14 | -0,99 | 3,24E-01 |
| AT4G08245 | 0,54 | 2,16 | 3,10 | 0,70 | 4,86E-01 |
| AT4G08255 | 0,51 | 2,30 | 4,30 | 0,53 | 5,93E-01 |
| AT4G29950 | 0,57 | 1,45 | 2,63 | 0,55 | 5,83E-01 |
| AT4G30010 | 0,55 | 0,56 | 3,24 | 0,17 | 8,63E-01 |
| AT4G30020 | 1,12 | -3,06 | 1,99 | -1,54 | 1,25E-01 |
| AT4G30130 | 0,24 | 0,45 | 4,38 | 0,10 | 9,18E-01 |
| AT4G30150 | 1,08 | -0,52 | 2,23 | -0,23 | 8,16E-01 |
| AT4G30310 | 0,60 | -2,09 | 3,75 | -0,56 | 5,76E-01 |
| AT4G30350 | 0,36 | 1,64 | 4,31 | 0,38 | 7,04E-01 |
| AT4G30400 | 0,15 | 0,82 | 4,35 | 0,19 | 8,50E-01 |
| AT4G08285 | 0,08 | 0,00 | 4,41 | 0,00 | 1,00E+00 |
| AT4G30480 | 0,31 | -2,06 | 4,39 | -0,47 | 6,39E-01 |
| AT4G30530 | 1,06 | -0,30 | 1,57 | -0,19 | 8,51E-01 |
| AT4G30650 | 0,58 | -0,97 | 2,60 | -0,37 | 7,09E-01 |
| AT4G30660 | 0,47 | -1,45 | 2,67 | -0,54 | 5,87E-01 |
| AT4G30760 | 1,76 | 3,43 | 1,67 | 2,05 | 4,05E-02 |
| AT4G30780 | 0,23 | 0,24 | 4,39 | 0,06 | 9,56E-01 |
| AT4G30800 | 0,17 | -3,06 | 4,35 | -0,70 | 4,82E-01 |
| AT4G30940 | 0,31 | 1,06 | 4,34 | 0,24 | 8,08E-01 |
| AT4G30990 | 1,34 | -1,14 | 2,07 | -0,55 | 5,83E-01 |
| AT4G08425 | 1,45 | 0,16 | 1,40 | 0,11 | 9,11E-01 |
| AT4G31080 | 0,70 | 0,51 | 2,95 | 0,17 | 8,62E-01 |
| AT4G31115 | 0,40 | -2,11 | 4,27 | -0,49 | 6,22E-01 |
| AT4G31300 | 0,37 | 0,69 | 4,29 | 0,16 | 8,72E-01 |
| AT4G31340 | 0,63 | -3,05 | 3,80 | -0,80 | 4,23E-01 |
| AT4G31390 | 0,47 | -0,49 | 2,80 | -0,17 | 8,61E-01 |
| AT4G31430 | 0,50 | -0,26 | 4,04 | -0,06 | 9,48E-01 |
| AT4G31530 | 0,77 | -0,16 | 2,47 | -0,06 | 9,49E-01 |
| AT4G31580 | 0,97 | 1,87 | 2,09 | 0,89 | 3,72E-01 |
| AT4G31710 | 0,50 | 0,00 | 4,01 | 0,00 | 1,00E+00 |
| AT4G31715 | 0,08 | 0,00 | 4,41 | 0,00 | 1,00E+00 |
| AT4G31780 | 1,36 | 1,41 | 2,87 | 0,49 | 6,23E-01 |
| AT4G31790 | 0,26 | 0,07 | 4,34 | 0,02 | 9,87E-01 |
| AT4G31800 | 1,69 | 3,61 | 1,80 | 2,00 | 4,50E-02 |
| AT4G31805 | 0,98 | 0,00 | 3,43 | 0,00 | 1,00E+00 |
| AT4G31850 | 0,54 | 1,16 | 3,29 | 0,35 | 7,24E-01 |
| AT4G31910 | 0,49 | 0,65 | 2,60 | 0,25 | 8,03E-01 |
| AT4G08615 | 0,91 | -0,30 | 2,09 | -0,15 | 8,84E-01 |
| AT4G08625 | 1,54 | 0,27 | 1,38 | 0,20 | 8,43E-01 |
| AT4G31985 | 0,37 | 0,00 | 4,41 | 0,00 | 1,00E+00 |
| AT4G32010 | 1,47 | -0,31 | 2,21 | -0,14 | 8,89E-01 |

| | | | | | |
|---|---|---|---|---|---|
| AT4G32030 | 0,47 | -0,48 | 4,24 | -0,11 | 9,09E-01 |
| AT4G32040 | 0,84 | -0,34 | 3,58 | -0,09 | 9,25E-01 |
| AT4G32060 | 1,61 | 0,10 | 1,32 | 0,08 | 9,39E-01 |
| AT4G32160 | 0,93 | -0,36 | 2,00 | -0,18 | 8,57E-01 |
| AT4G32190 | 0,86 | -0,35 | 2,84 | -0,12 | 9,02E-01 |
| AT4G32200 | 0,65 | -2,24 | 3,69 | -0,61 | 5,44E-01 |
| AT4G32265 | 1,11 | -2,75 | 1,65 | -1,67 | 9,59E-02 |
| AT4G32300 | 0,70 | 1,89 | 2,18 | 0,87 | 3,86E-01 |
| AT4G32330 | 0,80 | 2,47 | 2,13 | 1,16 | 2,48E-01 |
| AT4G32360 | 0,05 | 0,18 | 4,38 | 0,04 | 9,68E-01 |
| AT4G32390 | 1,34 | 0,50 | 1,88 | 0,27 | 7,88E-01 |
| AT4G32430 | 0,12 | 0,11 | 4,38 | 0,03 | 9,80E-01 |
| AT4G32551 | 0,49 | 0,04 | 2,74 | 0,01 | 9,89E-01 |
| AT4G32600 | 0,53 | 1,64 | 3,30 | 0,50 | 6,20E-01 |
| AT4G32620 | 1,61 | 0,43 | 2,27 | 0,19 | 8,51E-01 |
| AT4G32630 | 0,68 | 0,00 | 4,41 | 0,00 | 1,00E+00 |
| AT4G32700 | 0,67 | 0,00 | 3,73 | 0,00 | 1,00E+00 |
| AT4G32720 | 0,62 | -1,09 | 3,09 | -0,35 | 7,25E-01 |
| AT4G32730 | 0,45 | 0,00 | 4,07 | 0,00 | 1,00E+00 |
| AT4G32760 | 1,73 | 0,22 | 1,86 | 0,12 | 9,04E-01 |
| AT4G32770 | 0,32 | 0,74 | 4,36 | 0,17 | 8,65E-01 |
| AT4G32800 | 0,31 | 0,59 | 4,37 | 0,13 | 8,93E-01 |
| AT4G32850 | 1,20 | -1,78 | 1,86 | -0,96 | 3,38E-01 |
| AT4G32910 | 0,98 | -3,70 | 2,11 | -1,75 | 7,99E-02 |
| AT4G32930 | 0,91 | 0,24 | 2,53 | 0,10 | 9,24E-01 |
| AT4G33000 | 0,14 | 1,29 | 4,33 | 0,30 | 7,65E-01 |
| AT4G33030 | 0,99 | -2,00 | 2,67 | -0,75 | 4,54E-01 |
| AT4G33060 | 0,15 | -0,35 | 4,39 | -0,08 | 9,36E-01 |
| AT4G33180 | 0,05 | 0,18 | 4,38 | 0,04 | 9,68E-01 |
| AT4G33220 | 0,80 | -0,04 | 3,13 | -0,01 | 9,89E-01 |
| AT4G33290 | 1,50 | 3,65 | 1,70 | 2,14 | 3,21E-02 |
| AT4G08785 | 0,69 | 0,72 | 3,79 | 0,19 | 8,50E-01 |
| AT4G33310 | 0,49 | 0,65 | 4,35 | 0,15 | 8,81E-01 |
| AT4G33380 | 0,54 | -0,58 | 3,34 | -0,17 | 8,63E-01 |
| AT4G33410 | 0,32 | -0,25 | 4,41 | -0,06 | 9,55E-01 |
| AT4G33510 | 1,23 | -0,19 | 1,48 | -0,13 | 8,97E-01 |
| AT4G33520 | 1,37 | 0,41 | 2,37 | 0,17 | 8,62E-01 |
| AT4G33540 | 0,41 | 0,96 | 4,26 | 0,23 | 8,21E-01 |
| AT4G33565 | 0,80 | 2,68 | 2,68 | 1,00 | 3,16E-01 |
| AT4G33580 | 1,02 | 0,36 | 3,11 | 0,12 | 9,08E-01 |
| AT4G33620 | 0,33 | -0,38 | 4,39 | -0,09 | 9,32E-01 |
| AT4G33630 | 1,15 | -0,19 | 2,13 | -0,09 | 9,29E-01 |
| AT4G33650 | 1,75 | -0,34 | 1,75 | -0,20 | 8,45E-01 |
| AT4G33666 | 0,24 | 0,59 | 4,36 | 0,14 | 8,92E-01 |
| AT4G33690 | 0,48 | 1,91 | 3,98 | 0,48 | 6,32E-01 |
| AT4G33890 | 1,49 | -0,33 | 1,89 | -0,17 | 8,63E-01 |
| AT4G33920 | 0,65 | 2,92 | 3,54 | 0,82 | 4,09E-01 |
| AT4G33960 | 0,45 | -4,17 | 4,31 | -0,97 | 3,33E-01 |
| AT4G34030 | 0,40 | -1,00 | 3,35 | -0,30 | 7,64E-01 |
| AT4G34035 | 0,32 | 0,00 | 4,41 | 0,00 | 1,00E+00 |
| AT4G34050 | 1,48 | 2,51 | 1,63 | 1,54 | 1,24E-01 |
| AT4G34060 | 0,12 | 0,00 | 4,41 | 0,00 | 1,00E+00 |
| AT4G34120 | 1,07 | -0,80 | 1,83 | -0,44 | 6,62E-01 |
| AT4G34160 | 0,73 | 1,20 | 2,44 | 0,49 | 6,23E-01 |
| AT4G34240 | 0,30 | 0,19 | 4,33 | 0,04 | 9,65E-01 |
| AT4G34260 | 1,45 | -0,37 | 2,23 | -0,17 | 8,68E-01 |

| | | | | | |
|---|---|---|---|---|---|
| AT4G34280 | 0,96 | 0,00 | 2,89 | 0,00 | 1,00E+00 |
| AT4G34290 | 0,10 | 0,08 | 4,38 | 0,02 | 9,85E-01 |
| AT4G34310 | 0,63 | -0,35 | 4,39 | -0,08 | 9,36E-01 |
| AT4G34330 | 0,08 | 0,00 | 4,41 | 0,00 | 1,00E+00 |
| AT4G34370 | 0,50 | -2,04 | 3,95 | -0,52 | 6,06E-01 |
| AT4G34390 | 1,80 | 2,81 | 1,75 | 1,61 | 1,08E-01 |
| AT4G34400 | 0,59 | 0,00 | 3,82 | 0,00 | 1,00E+00 |
| AT4G34410 | 0,77 | 3,34 | 2,74 | 1,22 | 2,22E-01 |
| AT4G34430 | 1,22 | -2,33 | 2,33 | -1,00 | 3,17E-01 |
| AT4G34440 | 0,20 | 0,00 | 4,41 | 0,00 | 1,00E+00 |
| AT4G34640 | 0,93 | 0,00 | 2,92 | 0,00 | 1,00E+00 |
| AT4G34680 | 0,14 | 0,89 | 4,35 | 0,21 | 8,37E-01 |
| AT4G34700 | 0,83 | 0,72 | 2,50 | 0,29 | 7,73E-01 |
| AT4G34870 | 1,35 | 1,71 | 1,74 | 0,99 | 3,25E-01 |
| AT4G34881 | 0,53 | -0,42 | 2,63 | -0,16 | 8,73E-01 |
| AT4G34910 | 0,61 | -0,34 | 3,14 | -0,11 | 9,14E-01 |
| AT4G34920 | 0,49 | 0,98 | 3,23 | 0,30 | 7,61E-01 |
| AT4G35040 | 0,31 | -0,38 | 4,39 | -0,09 | 9,32E-01 |
| AT4G35100 | 1,16 | -2,06 | 2,46 | -0,83 | 4,04E-01 |
| AT4G35310 | 1,84 | -0,63 | 1,39 | -0,46 | 6,49E-01 |
| AT4G35370 | 0,13 | 0,00 | 4,41 | 0,00 | 1,00E+00 |
| AT4G35450 | 0,43 | -0,71 | 3,33 | -0,21 | 8,30E-01 |
| AT4G35460 | 0,66 | 0,00 | 3,04 | 0,00 | 1,00E+00 |
| AT4G35470 | 0,67 | 0,55 | 2,59 | 0,21 | 8,33E-01 |
| AT4G35500 | 0,79 | 0,00 | 2,89 | 0,00 | 1,00E+00 |
| AT4G35560 | 0,72 | -1,96 | 2,98 | -0,66 | 5,10E-01 |
| AT4G35600 | 1,15 | 0,42 | 2,05 | 0,20 | 8,39E-01 |
| AT4G35630 | 0,33 | -0,63 | 3,61 | -0,17 | 8,62E-01 |
| AT4G35850 | 0,77 | -2,25 | 2,26 | -1,00 | 3,19E-01 |
| AT4G35870 | 0,76 | 0,67 | 2,56 | 0,26 | 7,94E-01 |
| AT4G35880 | 0,17 | 0,00 | 4,41 | 0,00 | 1,00E+00 |
| AT4G35890 | 1,08 | 1,34 | 2,14 | 0,63 | 5,30E-01 |
| AT4G35940 | 1,33 | 0,42 | 1,87 | 0,23 | 8,21E-01 |
| AT4G36032 | 1,21 | 2,53 | 2,15 | 1,18 | 2,40E-01 |
| AT4G36052 | 0,12 | 0,00 | 4,41 | 0,00 | 1,00E+00 |
| AT4G36130 | 1,27 | -2,20 | 2,17 | -1,01 | 3,11E-01 |
| AT4G36150 | 1,48 | -2,83 | 2,64 | -1,07 | 2,84E-01 |
| AT4G36190 | 0,22 | 0,60 | 4,30 | 0,14 | 8,89E-01 |
| AT4G36195 | 0,86 | 0,33 | 2,12 | 0,15 | 8,77E-01 |
| AT4G36210 | 1,25 | 2,29 | 1,82 | 1,25 | 2,09E-01 |
| AT4G36250 | 0,69 | 0,12 | 2,43 | 0,05 | 9,60E-01 |
| AT4G36540 | 0,24 | 0,00 | 4,41 | 0,00 | 1,00E+00 |
| AT4G36710 | 1,27 | -1,81 | 2,48 | -0,73 | 4,65E-01 |
| AT4G36720 | 0,82 | 0,00 | 2,91 | 0,00 | 1,00E+00 |
| AT4G09375 | 0,14 | -0,82 | 4,37 | -0,19 | 8,50E-01 |
| AT4G36810 | 0,84 | -0,44 | 2,45 | -0,18 | 8,59E-01 |
| AT4G36870 | 0,25 | 0,44 | 4,30 | 0,10 | 9,19E-01 |
| AT4G36920 | 0,14 | -0,82 | 4,37 | -0,19 | 8,50E-01 |
| AT4G36925 | 1,06 | -0,41 | 1,75 | -0,24 | 8,13E-01 |
| AT4G36990 | 0,59 | -0,37 | 3,09 | -0,12 | 9,06E-01 |
| AT4G37000 | 0,23 | 0,24 | 4,39 | 0,06 | 9,56E-01 |
| AT4G37020 | 0,15 | -0,26 | 4,41 | -0,06 | 9,53E-01 |
| AT4G37080 | 0,19 | -1,14 | 4,40 | -0,26 | 7,95E-01 |
| AT4G37130 | 0,62 | 0,00 | 3,73 | 0,00 | 1,00E+00 |
| AT4G37180 | 0,47 | 0,36 | 4,08 | 0,09 | 9,29E-01 |
| AT4G37190 | 0,32 | -2,12 | 4,40 | -0,48 | 6,29E-01 |

| | | | | | |
|---|---|---|---|---|---|
| AT4G37210 | 0,15 | -2,07 | 4,39 | -0,47 | 6,38E-01 |
| AT4G37300 | 1,24 | 0,82 | 2,17 | 0,38 | 7,04E-01 |
| AT4G37430 | 0,86 | -1,12 | 3,44 | -0,33 | 7,44E-01 |
| AT4G37440 | 0,34 | 0,53 | 4,37 | 0,12 | 9,03E-01 |
| AT4G37550 | 0,88 | 0,81 | 2,16 | 0,38 | 7,06E-01 |
| AT4G37560 | 0,42 | 0,00 | 4,41 | 0,00 | 1,00E+00 |
| AT4G37650 | 1,47 | 2,01 | 1,81 | 1,11 | 2,68E-01 |
| AT4G37680 | 0,49 | 1,39 | 3,18 | 0,44 | 6,62E-01 |
| AT4G37730 | 0,75 | 1,23 | 2,06 | 0,60 | 5,51E-01 |
| AT4G37760 | 0,63 | -0,42 | 2,20 | -0,19 | 8,48E-01 |
| AT4G37790 | 0,15 | 0,02 | 4,36 | 0,00 | 9,96E-01 |
| AT4G37980 | 0,67 | 0,17 | 2,02 | 0,08 | 9,34E-01 |
| AT4G38040 | 0,20 | 0,00 | 4,41 | 0,00 | 1,00E+00 |
| AT4G38170 | 0,97 | -0,10 | 2,18 | -0,04 | 9,65E-01 |
| AT4G09685 | 0,46 | 0,00 | 4,06 | 0,00 | 1,00E+00 |
| AT4G38215 | 0,75 | 0,00 | 4,39 | 0,00 | 1,00E+00 |
| AT4G38225 | 0,78 | 1,99 | 2,49 | 0,80 | 4,24E-01 |
| AT4G38250 | 1,25 | -0,71 | 1,72 | -0,41 | 6,81E-01 |
| AT4G38270 | 0,51 | -2,02 | 3,15 | -0,64 | 5,21E-01 |
| AT4G38290 | 0,11 | -0,26 | 4,41 | -0,06 | 9,53E-01 |
| AT4G38320 | 0,19 | -0,40 | 4,39 | -0,09 | 9,28E-01 |
| AT4G38330 | 0,12 | 0,00 | 4,41 | 0,00 | 1,00E+00 |
| AT4G38360 | 0,15 | -0,35 | 4,39 | -0,08 | 9,36E-01 |
| AT4G38430 | 0,81 | -0,39 | 2,79 | -0,14 | 8,89E-01 |
| AT4G38460 | 1,41 | 2,29 | 1,80 | 1,27 | 2,03E-01 |
| AT4G38500 | 0,68 | -0,26 | 3,61 | -0,07 | 9,42E-01 |
| AT4G38540 | 0,10 | 0,24 | 4,38 | 0,05 | 9,57E-01 |
| AT4G38560 | 1,22 | -0,29 | 2,59 | -0,11 | 9,09E-01 |
| AT4G38580 | 0,50 | -2,49 | 4,37 | -0,57 | 5,68E-01 |
| AT4G38620 | 0,24 | -1,51 | 4,38 | -0,34 | 7,31E-01 |
| AT4G38790 | 0,69 | -3,05 | 3,02 | -1,01 | 3,12E-01 |
| AT4G38860 | 0,38 | 1,72 | 4,32 | 0,40 | 6,91E-01 |
| AT4G38880 | 0,39 | 0,00 | 4,40 | 0,00 | 1,00E+00 |
| AT4G38920 | 0,99 | -0,51 | 2,17 | -0,24 | 8,13E-01 |
| AT4G38930 | 0,41 | -0,31 | 4,40 | -0,07 | 9,43E-01 |
| AT4G39020 | 0,55 | 1,52 | 4,33 | 0,35 | 7,26E-01 |
| AT4G39030 | 0,29 | 0,00 | 4,41 | 0,00 | 1,00E+00 |
| AT4G39050 | 1,56 | 0,06 | 1,55 | 0,04 | 9,69E-01 |
| AT4G39200 | 1,73 | -1,99 | 2,04 | -0,98 | 3,29E-01 |
| AT4G39210 | 1,07 | -3,49 | 2,27 | -1,53 | 1,25E-01 |
| AT4G39270 | 1,16 | 0,24 | 1,61 | 0,15 | 8,81E-01 |
| AT4G39330 | 1,70 | 0,52 | 1,39 | 0,38 | 7,06E-01 |
| AT4G39361 | 1,80 | 1,82 | 1,71 | 1,06 | 2,88E-01 |
| AT4G39620 | 0,18 | 0,15 | 4,38 | 0,03 | 9,73E-01 |
| AT4G39640 | 0,76 | 0,20 | 2,55 | 0,08 | 9,38E-01 |
| AT4G39670 | 0,55 | 1,47 | 3,17 | 0,46 | 6,42E-01 |
| AT4G39840 | 1,07 | 2,64 | 1,95 | 1,36 | 1,75E-01 |
| AT4G39860 | 0,51 | 0,80 | 2,29 | 0,35 | 7,25E-01 |
| AT4G39870 | 0,35 | -1,23 | 4,39 | -0,28 | 7,79E-01 |
| AT4G39940 | 0,36 | -0,16 | 4,39 | -0,04 | 9,70E-01 |
| AT4G39950 | 1,14 | -1,32 | 1,77 | -0,75 | 4,55E-01 |
| AT4G39952 | 0,28 | 0,00 | 4,41 | 0,00 | 1,00E+00 |
| AT4G39980 | 0,50 | 1,21 | 3,17 | 0,38 | 7,02E-01 |
| AT4G40063 | 0,73 | -2,19 | 3,07 | -0,71 | 4,76E-01 |
| AT5G00730 | 1,64 | -1,17 | 1,51 | -0,78 | 4,36E-01 |
| AT5G01170 | 0,10 | 0,08 | 4,38 | 0,02 | 9,85E-01 |

| | | | | | |
|---|---|---|---|---|---|
| AT5G01210 | 1,09 | 1,13 | 1,72 | 0,66 | 5,11E-01 |
| AT5G01240 | 0,22 | -0,15 | 4,35 | -0,03 | 9,72E-01 |
| AT5G01260 | 0,54 | -0,34 | 2,42 | -0,14 | 8,88E-01 |
| AT5G01290 | 0,28 | -0,38 | 4,39 | -0,09 | 9,30E-01 |
| AT5G01460 | 0,28 | -0,16 | 4,40 | -0,04 | 9,70E-01 |
| AT5G01542 | 0,46 | 0,40 | 4,10 | 0,10 | 9,21E-01 |
| AT5G01590 | 0,94 | -2,13 | 2,19 | -0,97 | 3,33E-01 |
| AT5G01595 | 0,81 | 1,30 | 2,49 | 0,52 | 6,01E-01 |
| AT5G01690 | 0,08 | 0,00 | 4,41 | 0,00 | 1,00E+00 |
| AT5G01732 | 0,29 | 0,00 | 4,41 | 0,00 | 1,00E+00 |
| AT5G01740 | 0,15 | 1,21 | 4,33 | 0,28 | 7,81E-01 |
| AT5G01750 | 1,69 | 1,12 | 1,75 | 0,64 | 5,21E-01 |
| AT5G01760 | 0,79 | 0,00 | 2,88 | 0,00 | 1,00E+00 |
| AT5G01800 | 0,74 | -0,30 | 2,58 | -0,12 | 9,08E-01 |
| AT5G01810 | 0,55 | 0,00 | 3,91 | 0,00 | 1,00E+00 |
| AT5G01850 | 0,24 | -0,76 | 4,37 | -0,17 | 8,62E-01 |
| AT5G01960 | 1,82 | -0,51 | 2,20 | -0,23 | 8,18E-01 |
| AT5G01980 | 1,71 | 0,79 | 1,96 | 0,40 | 6,89E-01 |
| AT5G02035 | 0,47 | -2,04 | 4,02 | -0,51 | 6,11E-01 |
| AT5G02040 | 0,25 | 0,00 | 4,41 | 0,00 | 1,00E+00 |
| AT5G02100 | 0,70 | -1,57 | 3,79 | -0,41 | 6,78E-01 |
| AT5G02120 | 0,39 | -1,55 | 4,32 | -0,36 | 7,19E-01 |
| AT5G00820 | 1,31 | -5,19 | 2,53 | -2,05 | 4,05E-02 |
| AT5G02160 | 1,30 | -2,45 | 1,49 | -1,64 | 1,00E-01 |
| AT5G02240 | 0,32 | -1,26 | 4,40 | -0,29 | 7,75E-01 |
| AT5G02250 | 1,33 | -1,37 | 2,37 | -0,58 | 5,64E-01 |
| AT5G02280 | 0,11 | -0,26 | 4,41 | -0,06 | 9,53E-01 |
| AT5G02420 | 0,12 | -0,26 | 4,41 | -0,06 | 9,53E-01 |
| AT5G02450 | 0,59 | -1,75 | 3,16 | -0,55 | 5,80E-01 |
| AT5G02620 | 0,32 | 0,48 | 4,37 | 0,11 | 9,12E-01 |
| AT5G02630 | 0,30 | 0,00 | 4,41 | 0,00 | 1,00E+00 |
| AT5G02650 | 0,40 | 0,00 | 4,41 | 0,00 | 1,00E+00 |
| AT5G02760 | 0,40 | -1,99 | 3,46 | -0,57 | 5,66E-01 |
| AT5G02790 | 0,39 | 2,10 | 3,41 | 0,61 | 5,39E-01 |
| AT5G02840 | 0,40 | -0,20 | 4,22 | -0,05 | 9,63E-01 |
| AT5G02850 | 0,54 | -0,26 | 3,17 | -0,08 | 9,34E-01 |
| AT5G02860 | 1,11 | -0,05 | 2,09 | -0,03 | 9,79E-01 |
| AT5G02950 | 0,61 | -3,02 | 3,84 | -0,78 | 4,33E-01 |
| AT5G02995 | 0,13 | -0,61 | 4,37 | -0,14 | 8,89E-01 |
| AT5G03020 | 0,28 | 0,00 | 4,41 | 0,00 | 1,00E+00 |
| AT5G03160 | 0,68 | 1,53 | 2,45 | 0,63 | 5,32E-01 |
| AT5G03190 | 0,92 | 1,23 | 2,06 | 0,60 | 5,51E-01 |
| AT5G03290 | 1,24 | 1,29 | 1,74 | 0,74 | 4,58E-01 |
| AT5G03300 | 0,81 | -0,13 | 2,38 | -0,06 | 9,56E-01 |
| AT5G03330 | 0,61 | -0,39 | 3,15 | -0,12 | 9,01E-01 |
| AT5G03355 | 0,46 | 0,45 | 3,30 | 0,14 | 8,93E-01 |
| AT5G03415 | 0,36 | 0,41 | 4,36 | 0,09 | 9,24E-01 |
| AT5G03420 | 0,63 | 0,86 | 2,61 | 0,33 | 7,43E-01 |
| AT5G03455 | 0,25 | 0,00 | 4,41 | 0,00 | 1,00E+00 |
| AT5G03460 | 0,54 | -0,36 | 3,18 | -0,11 | 9,10E-01 |
| AT5G03470 | 0,40 | -0,26 | 4,41 | -0,06 | 9,53E-01 |
| AT5G03490 | 0,11 | -0,38 | 4,39 | -0,09 | 9,31E-01 |
| AT5G03540 | 1,05 | -0,08 | 2,88 | -0,03 | 9,77E-01 |
| AT5G03650 | 0,91 | -0,23 | 1,72 | -0,14 | 8,92E-01 |
| AT5G03905 | 0,30 | 0,31 | 4,39 | 0,07 | 9,44E-01 |
| AT5G04040 | 1,27 | -1,39 | 2,07 | -0,67 | 5,03E-01 |

| | | | | | |
|---|---|---|---|---|---|
| AT5G04050 | 0,10 | 0,08 | 4,38 | 0,02 | 9,85E-01 |
| AT5G04060 | 0,37 | 0,14 | 4,38 | 0,03 | 9,74E-01 |
| AT5G04170 | 0,76 | -0,14 | 2,56 | -0,05 | 9,57E-01 |
| AT5G04190 | 0,13 | 0,94 | 4,35 | 0,22 | 8,29E-01 |
| AT5G04240 | 0,88 | -2,36 | 3,39 | -0,70 | 4,86E-01 |
| AT5G04280 | 1,25 | 0,65 | 1,96 | 0,33 | 7,40E-01 |
| AT5G04290 | 0,66 | 0,46 | 3,23 | 0,14 | 8,87E-01 |
| AT5G04360 | 0,45 | -2,06 | 4,17 | -0,49 | 6,22E-01 |
| AT5G00960 | 0,96 | 2,62 | 2,16 | 1,21 | 2,26E-01 |
| AT5G04410 | 1,77 | -0,72 | 2,15 | -0,34 | 7,36E-01 |
| AT5G04460 | 0,56 | -0,26 | 3,20 | -0,08 | 9,36E-01 |
| AT5G04490 | 0,46 | 1,68 | 3,31 | 0,51 | 6,11E-01 |
| AT5G04510 | 0,58 | -1,36 | 2,73 | -0,50 | 6,18E-01 |
| AT5G04600 | 0,71 | -0,76 | 2,20 | -0,35 | 7,28E-01 |
| AT5G04620 | 0,43 | 0,00 | 4,41 | 0,00 | 1,00E+00 |
| AT5G04630 | 0,57 | 0,00 | 3,84 | 0,00 | 1,00E+00 |
| AT5G04660 | 0,24 | -0,68 | 4,38 | -0,16 | 8,76E-01 |
| AT5G00985 | 0,28 | 0,00 | 4,41 | 0,00 | 1,00E+00 |
| AT5G04800 | 0,67 | 0,21 | 2,21 | 0,10 | 9,23E-01 |
| AT5G04885 | 0,27 | 1,28 | 3,72 | 0,34 | 7,31E-01 |
| AT5G04910 | 0,42 | 0,83 | 4,13 | 0,20 | 8,41E-01 |
| AT5G05070 | 0,40 | -0,05 | 4,32 | -0,01 | 9,90E-01 |
| AT5G05090 | 0,35 | 1,41 | 3,53 | 0,40 | 6,89E-01 |
| AT5G05160 | 0,72 | -0,39 | 2,92 | -0,13 | 8,93E-01 |
| AT5G05210 | 0,28 | 1,05 | 4,34 | 0,24 | 8,09E-01 |
| AT5G05220 | 0,08 | 0,00 | 4,41 | 0,00 | 1,00E+00 |
| AT5G05230 | 0,08 | 0,00 | 4,41 | 0,00 | 1,00E+00 |
| AT5G05310 | 0,18 | 0,93 | 4,34 | 0,21 | 8,30E-01 |
| AT5G05440 | 0,16 | -0,26 | 4,41 | -0,06 | 9,53E-01 |
| AT5G05490 | 0,55 | 0,00 | 4,41 | 0,00 | 1,00E+00 |
| AT5G05560 | 1,14 | 1,06 | 1,92 | 0,55 | 5,80E-01 |
| AT5G05570 | 0,62 | -0,28 | 4,40 | -0,06 | 9,49E-01 |
| AT5G05600 | 0,31 | -1,73 | 4,33 | -0,40 | 6,89E-01 |
| AT5G05630 | 0,08 | 0,21 | 4,39 | 0,05 | 9,62E-01 |
| AT5G05680 | 1,56 | -0,31 | 2,09 | -0,15 | 8,80E-01 |
| AT5G05710 | 0,20 | 0,00 | 4,41 | 0,00 | 1,00E+00 |
| AT5G05750 | 1,85 | -0,32 | 2,34 | -0,14 | 8,90E-01 |
| AT5G05780 | 0,96 | -2,64 | 2,54 | -1,04 | 2,98E-01 |
| AT5G05950 | 0,83 | 1,17 | 2,48 | 0,47 | 6,37E-01 |
| AT5G05970 | 0,30 | -0,06 | 4,37 | -0,01 | 9,88E-01 |
| AT5G05985 | 0,55 | 0,00 | 3,90 | 0,00 | 1,00E+00 |
| AT5G06000 | 0,23 | 0,42 | 4,31 | 0,10 | 9,23E-01 |
| AT5G06050 | 0,39 | -2,12 | 4,31 | -0,49 | 6,22E-01 |
| AT5G06100 | 0,21 | -0,29 | 4,40 | -0,06 | 9,48E-01 |
| AT5G06120 | 1,17 | -2,21 | 2,59 | -0,85 | 3,94E-01 |
| AT5G06170 | 0,31 | 0,90 | 4,28 | 0,21 | 8,34E-01 |
| AT5G06220 | 0,84 | -1,21 | 2,62 | -0,46 | 6,44E-01 |
| AT5G06265 | 0,26 | 0,51 | 4,31 | 0,12 | 9,07E-01 |
| AT5G06290 | 1,37 | -3,02 | 1,79 | -1,69 | 9,16E-02 |
| AT5G06310 | 0,33 | 1,39 | 4,33 | 0,32 | 7,49E-01 |
| AT5G01255 | 0,70 | 1,82 | 2,81 | 0,65 | 5,17E-01 |
| AT5G06350 | 0,36 | -0,09 | 3,46 | -0,02 | 9,80E-01 |
| AT5G06360 | 0,40 | -0,05 | 4,32 | -0,01 | 9,90E-01 |
| AT5G06380 | 0,08 | 0,00 | 4,41 | 0,00 | 1,00E+00 |
| AT5G06390 | 0,14 | 0,81 | 4,35 | 0,19 | 8,52E-01 |
| AT5G06460 | 1,60 | 0,49 | 2,08 | 0,24 | 8,13E-01 |

| | | | | | |
|---|---|---|---|---|---|
| AT5G06645 | 0,58 | 3,33 | 4,27 | 0,78 | 4,35E-01 |
| AT5G06660 | 0,96 | 1,08 | 3,59 | 0,30 | 7,63E-01 |
| AT5G06680 | 0,39 | 0,50 | 4,33 | 0,12 | 9,07E-01 |
| AT5G06700 | 0,91 | -1,16 | 1,68 | -0,69 | 4,89E-01 |
| AT5G06811 | 0,11 | -0,26 | 4,41 | -0,06 | 9,53E-01 |
| AT5G06820 | 0,12 | 0,00 | 4,41 | 0,00 | 1,00E+00 |
| AT5G06860 | 0,22 | 0,15 | 4,33 | 0,03 | 9,73E-01 |
| AT5G06870 | 0,67 | 2,34 | 2,78 | 0,84 | 4,00E-01 |
| AT5G06950 | 0,61 | -0,26 | 3,18 | -0,08 | 9,34E-01 |
| AT5G06980 | 0,16 | -0,22 | 4,40 | -0,05 | 9,60E-01 |
| AT5G07090 | 1,54 | -1,53 | 1,30 | -1,18 | 2,40E-01 |
| AT5G07110 | 0,85 | 0,00 | 3,46 | 0,00 | 1,00E+00 |
| AT5G01315 | 0,64 | -0,33 | 3,05 | -0,11 | 9,14E-01 |
| AT5G07130 | 0,33 | -2,09 | 4,39 | -0,48 | 6,35E-01 |
| AT5G07140 | 1,24 | 0,00 | 2,66 | 0,00 | 1,00E+00 |
| AT5G07210 | 1,09 | -0,84 | 1,79 | -0,47 | 6,40E-01 |
| AT5G07240 | 1,19 | -1,35 | 2,26 | -0,60 | 5,50E-01 |
| AT5G07270 | 0,15 | 0,82 | 4,35 | 0,19 | 8,50E-01 |
| AT5G07340 | 0,98 | 0,24 | 2,34 | 0,10 | 9,18E-01 |
| AT5G07410 | 1,62 | 3,38 | 1,83 | 1,85 | 6,40E-02 |
| AT5G07430 | 0,50 | 0,81 | 3,90 | 0,21 | 8,36E-01 |
| AT5G07440 | 0,36 | -2,89 | 4,37 | -0,66 | 5,08E-01 |
| AT5G07580 | 0,75 | 1,10 | 2,23 | 0,49 | 6,21E-01 |
| AT5G07610 | 0,48 | 2,08 | 4,02 | 0,52 | 6,05E-01 |
| AT5G07760 | 0,08 | 0,00 | 4,41 | 0,00 | 1,00E+00 |
| AT5G07770 | 0,16 | -0,21 | 4,40 | -0,05 | 9,63E-01 |
| AT5G07790 | 0,58 | -0,28 | 4,41 | -0,06 | 9,50E-01 |
| AT5G07830 | 0,86 | 0,16 | 2,53 | 0,06 | 9,49E-01 |
| AT5G07860 | 0,58 | -0,26 | 3,96 | -0,07 | 9,47E-01 |
| AT5G07870 | 0,49 | 0,00 | 4,41 | 0,00 | 1,00E+00 |
| AT5G07960 | 0,12 | -0,26 | 4,41 | -0,06 | 9,53E-01 |
| AT5G08050 | 0,48 | -0,56 | 4,20 | -0,13 | 8,94E-01 |
| AT5G08060 | 0,10 | 0,08 | 4,38 | 0,02 | 9,85E-01 |
| AT5G08070 | 0,35 | 0,00 | 4,39 | 0,00 | 1,00E+00 |
| AT5G08139 | 0,37 | 2,13 | 4,28 | 0,50 | 6,19E-01 |
| AT5G08190 | 0,29 | 0,00 | 4,41 | 0,00 | 1,00E+00 |
| AT5G08260 | 0,90 | -1,86 | 2,44 | -0,76 | 4,45E-01 |
| AT5G01545 | 0,37 | -3,32 | 4,36 | -0,76 | 4,46E-01 |
| AT5G08290 | 0,74 | 0,27 | 2,12 | 0,13 | 8,99E-01 |
| AT5G08300 | 0,94 | 2,18 | 2,37 | 0,92 | 3,56E-01 |
| AT5G08330 | 1,56 | 1,54 | 1,80 | 0,85 | 3,93E-01 |
| AT5G08335 | 0,18 | 0,82 | 4,35 | 0,19 | 8,51E-01 |
| AT5G08340 | 0,58 | -0,26 | 4,05 | -0,06 | 9,48E-01 |
| AT5G08390 | 0,26 | 0,25 | 4,37 | 0,06 | 9,55E-01 |
| AT5G08430 | 0,57 | 0,00 | 4,41 | 0,00 | 1,00E+00 |
| AT5G08440 | 0,59 | -0,26 | 3,96 | -0,07 | 9,47E-01 |
| AT5G08470 | 0,81 | -0,14 | 2,86 | -0,05 | 9,62E-01 |
| AT5G08510 | 0,51 | -1,22 | 3,32 | -0,37 | 7,13E-01 |
| AT5G08540 | 0,41 | -0,24 | 3,39 | -0,07 | 9,44E-01 |
| AT5G01555 | 0,36 | 0,70 | 4,36 | 0,16 | 8,72E-01 |
| AT5G08570 | 1,20 | -1,92 | 1,55 | -1,24 | 2,13E-01 |
| AT5G08590 | 0,63 | -1,10 | 2,64 | -0,41 | 6,79E-01 |
| AT5G08600 | 0,28 | 0,00 | 4,41 | 0,00 | 1,00E+00 |
| AT5G08610 | 1,71 | -1,43 | 1,73 | -0,82 | 4,09E-01 |
| AT5G08620 | 1,12 | -1,85 | 1,80 | -1,03 | 3,05E-01 |
| AT5G08660 | 0,33 | 0,15 | 4,38 | 0,04 | 9,72E-01 |

| | | | | | |
|---|---|---|---|---|---|
| AT5G08680 | 1,77 | -0,78 | 1,64 | -0,47 | 6,36E-01 |
| AT5G08695 | 0,20 | 0,00 | 4,41 | 0,00 | 1,00E+00 |
| AT5G08750 | 0,11 | -0,38 | 4,39 | -0,09 | 9,31E-01 |
| AT5G08760 | 0,55 | -0,80 | 3,86 | -0,21 | 8,35E-01 |
| AT5G08780 | 0,49 | -0,26 | 3,26 | -0,08 | 9,36E-01 |
| AT5G09220 | 0,82 | -0,90 | 2,89 | -0,31 | 7,56E-01 |
| AT5G09400 | 1,22 | 0,05 | 2,14 | 0,02 | 9,80E-01 |
| AT5G09440 | 0,77 | 1,76 | 2,35 | 0,75 | 4,55E-01 |
| AT5G09445 | 0,26 | 0,00 | 4,41 | 0,00 | 1,00E+00 |
| AT5G09450 | 1,25 | -0,70 | 2,20 | -0,32 | 7,50E-01 |
| AT5G09600 | 0,62 | -1,12 | 3,12 | -0,36 | 7,20E-01 |
| AT5G09760 | 0,14 | 0,81 | 4,35 | 0,19 | 8,52E-01 |
| AT5G10010 | 0,47 | 0,18 | 3,37 | 0,05 | 9,58E-01 |
| AT5G10020 | 0,41 | 0,00 | 4,41 | 0,00 | 1,00E+00 |
| AT5G10100 | 0,16 | 0,31 | 4,37 | 0,07 | 9,44E-01 |
| AT5G10190 | 0,41 | -0,35 | 3,40 | -0,10 | 9,18E-01 |
| AT5G10200 | 0,81 | 0,00 | 3,51 | 0,00 | 1,00E+00 |
| AT5G10340 | 1,40 | 0,36 | 2,01 | 0,18 | 8,57E-01 |
| AT5G10370 | 1,36 | -1,17 | 1,86 | -0,63 | 5,30E-01 |
| AT5G10410 | 0,05 | 0,18 | 4,38 | 0,04 | 9,68E-01 |
| AT5G10460 | 0,08 | 0,00 | 4,41 | 0,00 | 1,00E+00 |
| AT5G01765 | 1,15 | 0,26 | 1,99 | 0,13 | 8,97E-01 |
| AT5G10600 | 0,40 | 1,03 | 4,28 | 0,24 | 8,09E-01 |
| AT5G10630 | 0,28 | 0,87 | 4,35 | 0,20 | 8,42E-01 |
| AT5G10660 | 0,46 | 0,17 | 2,75 | 0,06 | 9,50E-01 |
| AT5G10695 | 0,85 | 1,17 | 1,99 | 0,59 | 5,57E-01 |
| AT5G10745 | 0,35 | 1,67 | 3,48 | 0,48 | 6,30E-01 |
| AT5G10750 | 1,12 | -0,48 | 2,21 | -0,22 | 8,27E-01 |
| AT5G10780 | 0,76 | -0,56 | 2,56 | -0,22 | 8,27E-01 |
| AT5G10790 | 0,93 | 0,88 | 2,15 | 0,41 | 6,84E-01 |
| AT5G10920 | 0,91 | -0,05 | 2,27 | -0,02 | 9,82E-01 |
| AT5G11060 | 1,02 | 0,98 | 1,65 | 0,59 | 5,53E-01 |
| AT5G11090 | 0,49 | -0,01 | 2,73 | 0,00 | 9,96E-01 |
| AT5G11110 | 0,50 | 1,04 | 4,00 | 0,26 | 7,96E-01 |
| AT5G11170 | 1,42 | -1,24 | 2,09 | -0,59 | 5,52E-01 |
| AT5G11200 | 0,33 | -0,36 | 4,39 | -0,08 | 9,34E-01 |
| AT5G11280 | 0,20 | 0,53 | 4,37 | 0,12 | 9,04E-01 |
| AT5G11340 | 0,28 | -0,32 | 4,40 | -0,07 | 9,43E-01 |
| AT5G11350 | 0,82 | -2,25 | 3,02 | -0,75 | 4,55E-01 |
| AT5G11390 | 0,30 | 0,00 | 4,41 | 0,00 | 1,00E+00 |
| AT5G11420 | 1,15 | 1,00 | 1,92 | 0,52 | 6,05E-01 |
| AT5G11430 | 0,70 | -0,76 | 2,27 | -0,34 | 7,36E-01 |
| AT5G11500 | 0,94 | -1,62 | 1,96 | -0,82 | 4,10E-01 |
| AT5G11510 | 0,81 | -0,26 | 2,54 | -0,10 | 9,18E-01 |
| AT5G11530 | 0,81 | -0,90 | 2,43 | -0,37 | 7,13E-01 |
| AT5G11580 | 0,12 | -0,36 | 4,39 | -0,08 | 9,34E-01 |
| AT5G11650 | 0,38 | 1,25 | 3,44 | 0,36 | 7,17E-01 |
| AT5G11680 | 0,19 | 0,43 | 4,36 | 0,10 | 9,21E-01 |
| AT5G11710 | 1,24 | -0,06 | 1,66 | -0,04 | 9,71E-01 |
| AT5G11720 | 1,68 | -0,22 | 2,28 | -0,10 | 9,24E-01 |
| AT5G11730 | 0,89 | 0,90 | 1,97 | 0,46 | 6,46E-01 |
| AT5G11760 | 0,22 | 0,14 | 4,38 | 0,03 | 9,75E-01 |
| AT5G11890 | 0,82 | 1,01 | 2,13 | 0,47 | 6,35E-01 |
| AT5G11920 | 0,12 | -0,36 | 4,39 | -0,08 | 9,34E-01 |
| AT5G11977 | 0,12 | 0,00 | 4,41 | 0,00 | 1,00E+00 |
| AT5G12090 | 0,79 | 1,19 | 2,18 | 0,55 | 5,86E-01 |

| | | | | | |
|---|---|---|---|---|---|
| AT5G12100 | 0,44 | 0,00 | 4,41 | 0,00 | 1,00E+00 |
| AT5G12110 | 1,52 | -0,82 | 1,71 | -0,48 | 6,31E-01 |
| AT5G12210 | 0,65 | 0,14 | 3,76 | 0,04 | 9,71E-01 |
| AT5G12260 | 0,32 | -1,26 | 4,40 | -0,29 | 7,75E-01 |
| AT5G12350 | 0,49 | -0,28 | 4,21 | -0,07 | 9,47E-01 |
| AT5G12400 | 0,93 | 0,62 | 2,91 | 0,21 | 8,32E-01 |
| AT5G12850 | 0,97 | 1,36 | 1,95 | 0,70 | 4,84E-01 |
| AT5G12950 | 0,53 | 1,36 | 2,50 | 0,54 | 5,87E-01 |
| AT5G12960 | 0,27 | -2,13 | 4,39 | -0,49 | 6,27E-01 |
| AT5G12970 | 0,24 | 0,00 | 4,41 | 0,00 | 1,00E+00 |
| AT5G13030 | 0,96 | -1,23 | 1,78 | -0,69 | 4,91E-01 |
| AT5G13180 | 1,85 | 1,19 | 1,54 | 0,77 | 4,41E-01 |
| AT5G13200 | 1,26 | 2,07 | 1,75 | 1,18 | 2,36E-01 |
| AT5G13210 | 0,35 | 0,44 | 3,47 | 0,13 | 8,98E-01 |
| AT5G13260 | 0,92 | -2,77 | 2,17 | -1,28 | 2,01E-01 |
| AT5G13280 | 0,45 | -0,17 | 4,19 | -0,04 | 9,68E-01 |
| AT5G13310 | 0,31 | -0,39 | 4,39 | -0,09 | 9,29E-01 |
| AT5G13320 | 1,27 | 2,06 | 1,98 | 1,04 | 2,98E-01 |
| AT5G13330 | 0,37 | 0,00 | 4,41 | 0,00 | 1,00E+00 |
| AT5G13370 | 0,68 | -1,71 | 3,15 | -0,54 | 5,87E-01 |
| AT5G13380 | 0,54 | -2,39 | 4,39 | -0,54 | 5,86E-01 |
| AT5G13500 | 0,32 | 1,08 | 4,34 | 0,25 | 8,03E-01 |
| AT5G13510 | 0,48 | -1,97 | 3,33 | -0,59 | 5,53E-01 |
| AT5G13580 | 0,29 | 0,00 | 4,41 | 0,00 | 1,00E+00 |
| AT5G13640 | 0,84 | -1,07 | 3,09 | -0,35 | 7,28E-01 |
| AT5G13690 | 0,61 | -0,35 | 3,19 | -0,11 | 9,13E-01 |
| AT5G13720 | 0,21 | -1,55 | 4,38 | -0,35 | 7,24E-01 |
| AT5G13740 | 0,19 | 0,43 | 4,36 | 0,10 | 9,21E-01 |
| AT5G13750 | 0,26 | 0,00 | 4,41 | 0,00 | 1,00E+00 |
| AT5G13760 | 0,26 | 0,79 | 4,35 | 0,18 | 8,55E-01 |
| AT5G13810 | 0,93 | 0,88 | 1,92 | 0,46 | 6,46E-01 |
| AT5G13820 | 0,28 | -0,32 | 4,40 | -0,07 | 9,41E-01 |
| AT5G13930 | 1,07 | -2,32 | 2,37 | -0,98 | 3,27E-01 |
| AT5G13980 | 0,97 | -1,43 | 2,29 | -0,62 | 5,32E-01 |
| AT5G14030 | 0,98 | 0,46 | 2,56 | 0,18 | 8,56E-01 |
| AT5G14050 | 0,50 | 0,87 | 3,20 | 0,27 | 7,85E-01 |
| AT5G14120 | 1,26 | 0,91 | 1,66 | 0,55 | 5,85E-01 |
| AT5G14200 | 1,42 | 0,94 | 1,58 | 0,60 | 5,51E-01 |
| AT5G14230 | 0,39 | 1,92 | 3,39 | 0,57 | 5,70E-01 |
| AT5G14260 | 0,68 | -0,26 | 2,52 | -0,10 | 9,19E-01 |
| AT5G14280 | 0,43 | 0,00 | 4,40 | 0,00 | 1,00E+00 |
| AT5G14285 | 0,21 | 0,00 | 4,41 | 0,00 | 1,00E+00 |
| AT5G14480 | 0,25 | -1,08 | 4,39 | -0,25 | 8,06E-01 |
| AT5G14610 | 0,64 | 1,56 | 3,22 | 0,48 | 6,29E-01 |
| AT5G14670 | 0,08 | 0,00 | 4,41 | 0,00 | 1,00E+00 |
| AT5G14880 | 0,37 | 0,48 | 3,48 | 0,14 | 8,90E-01 |
| AT5G14890 | 0,99 | 0,00 | 4,15 | 0,00 | 1,00E+00 |
| AT5G14910 | 0,77 | -0,45 | 2,61 | -0,17 | 8,64E-01 |
| AT5G14930 | 1,80 | 0,99 | 2,31 | 0,43 | 6,68E-01 |
| AT5G14970 | 0,18 | 0,09 | 4,38 | 0,02 | 9,84E-01 |
| AT5G14990 | 0,42 | 0,00 | 4,24 | 0,00 | 1,00E+00 |
| AT5G15010 | 0,38 | -3,01 | 4,32 | -0,70 | 4,85E-01 |
| AT5G02295 | 1,40 | 1,55 | 1,50 | 1,04 | 3,00E-01 |
| AT5G15080 | 0,92 | -2,04 | 2,96 | -0,69 | 4,91E-01 |
| AT5G15230 | 0,74 | -2,47 | 2,92 | -0,85 | 3,98E-01 |
| AT5G15240 | 0,05 | 0,18 | 4,38 | 0,04 | 9,68E-01 |

| | | | | | |
|---|---|---|---|---|---|
| AT5G15270 | 0,61 | -0,12 | 3,17 | -0,04 | 9,71E-01 |
| AT5G15280 | 0,54 | 0,00 | 3,92 | 0,00 | 1,00E+00 |
| AT5G15320 | 0,21 | -0,95 | 4,38 | -0,22 | 8,29E-01 |
| AT5G15330 | 0,11 | -0,26 | 4,41 | -0,06 | 9,53E-01 |
| AT5G15530 | 0,12 | 0,00 | 4,41 | 0,00 | 1,00E+00 |
| AT5G15570 | 0,44 | 0,00 | 4,41 | 0,00 | 1,00E+00 |
| AT5G15610 | 0,89 | -0,35 | 1,88 | -0,19 | 8,51E-01 |
| AT5G15685 | 0,72 | 0,34 | 1,94 | 0,18 | 8,60E-01 |
| AT5G15710 | 0,74 | 0,44 | 3,67 | 0,12 | 9,05E-01 |
| AT5G15730 | 0,59 | 0,47 | 2,67 | 0,18 | 8,60E-01 |
| AT5G15750 | 0,54 | 1,24 | 3,87 | 0,32 | 7,48E-01 |
| AT5G15820 | 0,22 | 0,08 | 4,38 | 0,02 | 9,86E-01 |
| AT5G15930 | 0,32 | -0,32 | 4,40 | -0,07 | 9,42E-01 |
| AT5G15950 | 1,23 | 0,00 | 2,32 | 0,00 | 1,00E+00 |
| AT5G15960 | 0,51 | -3,96 | 4,32 | -0,92 | 3,60E-01 |
| AT5G16010 | 0,36 | 0,52 | 4,36 | 0,12 | 9,05E-01 |
| AT5G16030 | 0,58 | -1,81 | 3,19 | -0,57 | 5,71E-01 |
| AT5G16120 | 0,30 | -0,26 | 4,41 | -0,06 | 9,53E-01 |
| AT5G16130 | 1,11 | 0,68 | 2,27 | 0,30 | 7,64E-01 |
| AT5G16150 | 1,33 | -0,97 | 1,48 | -0,66 | 5,11E-01 |
| AT5G16260 | 0,11 | 0,88 | 4,35 | 0,20 | 8,39E-01 |
| AT5G16270 | 1,71 | 0,68 | 1,72 | 0,40 | 6,93E-01 |
| AT5G16290 | 0,43 | -1,49 | 4,30 | -0,35 | 7,29E-01 |
| AT5G16300 | 1,23 | 0,63 | 1,85 | 0,34 | 7,32E-01 |
| AT5G16440 | 0,67 | 0,56 | 2,64 | 0,21 | 8,32E-01 |
| AT5G16470 | 0,69 | -0,12 | 2,56 | -0,05 | 9,62E-01 |
| AT5G16540 | 0,32 | 0,00 | 4,41 | 0,00 | 1,00E+00 |
| AT5G16550 | 0,57 | -0,65 | 2,73 | -0,24 | 8,13E-01 |
| AT5G02555 | 0,05 | 0,18 | 4,38 | 0,04 | 9,68E-01 |
| AT5G16590 | 0,87 | -0,04 | 2,28 | -0,02 | 9,85E-01 |
| AT5G16630 | 0,92 | -0,27 | 2,80 | -0,10 | 9,24E-01 |
| AT5G16715 | 0,74 | -2,55 | 2,44 | -1,04 | 2,96E-01 |
| AT5G16730 | 1,05 | -0,67 | 1,84 | -0,37 | 7,14E-01 |
| AT5G16760 | 1,05 | 1,55 | 2,22 | 0,70 | 4,86E-01 |
| AT5G16820 | 0,94 | -0,26 | 2,47 | -0,11 | 9,15E-01 |
| AT5G16880 | 1,27 | -1,24 | 1,67 | -0,74 | 4,57E-01 |
| AT5G16900 | 0,33 | 0,00 | 4,41 | 0,00 | 1,00E+00 |
| AT5G16910 | 0,98 | 0,00 | 2,73 | 0,00 | 1,00E+00 |
| AT5G17167 | 0,20 | 1,10 | 4,35 | 0,25 | 8,00E-01 |
| AT5G17170 | 0,33 | -0,56 | 4,35 | -0,13 | 8,98E-01 |
| AT5G17190 | 0,51 | 1,95 | 3,10 | 0,63 | 5,28E-01 |
| AT5G17270 | 0,61 | 0,00 | 4,41 | 0,00 | 1,00E+00 |
| AT5G17280 | 0,29 | -0,70 | 4,38 | -0,16 | 8,73E-01 |
| AT5G17305 | 0,08 | 0,00 | 4,41 | 0,00 | 1,00E+00 |
| AT5G17440 | 0,24 | -0,34 | 4,40 | -0,08 | 9,38E-01 |
| AT5G17530 | 0,59 | -1,57 | 3,12 | -0,50 | 6,14E-01 |
| AT5G17580 | 0,17 | 0,00 | 4,41 | 0,00 | 1,00E+00 |
| AT5G17630 | 0,33 | -0,39 | 4,39 | -0,09 | 9,30E-01 |
| AT5G17670 | 0,39 | 0,00 | 4,40 | 0,00 | 1,00E+00 |
| AT5G17680 | 0,30 | 0,00 | 4,41 | 0,00 | 1,00E+00 |
| AT5G17850 | 0,18 | 1,62 | 4,31 | 0,38 | 7,08E-01 |
| AT5G17870 | 0,86 | -1,41 | 2,15 | -0,65 | 5,14E-01 |
| AT5G17890 | 1,77 | 1,03 | 1,98 | 0,52 | 6,02E-01 |
| AT5G17900 | 0,30 | 0,81 | 4,35 | 0,19 | 8,53E-01 |
| AT5G17910 | 1,47 | -1,42 | 1,82 | -0,78 | 4,36E-01 |
| AT5G17980 | 0,38 | 0,03 | 4,28 | 0,01 | 9,95E-01 |

| | | | | | |
|---|---|---|---|---|---|
| AT5G17990 | 0,55 | -1,16 | 4,05 | -0,29 | 7,75E-01 |
| AT5G18040 | 0,71 | 0,74 | 2,53 | 0,29 | 7,71E-01 |
| AT5G18065 | 0,30 | -0,27 | 3,66 | -0,07 | 9,41E-01 |
| AT5G18070 | 0,62 | 0,53 | 2,64 | 0,20 | 8,40E-01 |
| AT5G18085 | 0,24 | 0,00 | 4,41 | 0,00 | 1,00E+00 |
| AT5G18100 | 0,33 | -0,54 | 4,36 | -0,12 | 9,01E-01 |
| AT5G18130 | 1,11 | 0,79 | 2,37 | 0,33 | 7,39E-01 |
| AT5G18140 | 1,06 | 0,40 | 2,68 | 0,15 | 8,82E-01 |
| AT5G18170 | 0,16 | 0,35 | 4,37 | 0,08 | 9,37E-01 |
| AT5G18200 | 0,08 | 0,00 | 4,41 | 0,00 | 1,00E+00 |
| AT5G18245 | 1,46 | 4,64 | 2,69 | 1,72 | 8,53E-02 |
| AT5G18255 | 1,25 | -2,75 | 1,57 | -1,75 | 7,94E-02 |
| AT5G18310 | 0,76 | 0,76 | 2,27 | 0,33 | 7,38E-01 |
| AT5G18390 | 0,33 | -0,38 | 4,39 | -0,09 | 9,31E-01 |
| AT5G18470 | 1,32 | 1,45 | 1,94 | 0,75 | 4,54E-01 |
| AT5G18500 | 0,37 | -0,55 | 4,36 | -0,13 | 9,00E-01 |
| AT5G18520 | 1,46 | 0,56 | 1,58 | 0,36 | 7,22E-01 |
| AT5G18580 | 0,90 | -0,61 | 2,10 | -0,29 | 7,71E-01 |
| AT5G18610 | 1,43 | 0,25 | 2,16 | 0,12 | 9,08E-01 |
| AT5G18630 | 0,28 | 1,31 | 4,32 | 0,30 | 7,62E-01 |
| AT5G18636 | 0,58 | 1,00 | 2,31 | 0,43 | 6,65E-01 |
| AT5G18650 | 0,68 | -0,02 | 3,02 | -0,01 | 9,94E-01 |
| AT5G18670 | 0,21 | 0,00 | 4,41 | 0,00 | 1,00E+00 |
| AT5G18750 | 0,77 | 0,42 | 3,45 | 0,12 | 9,03E-01 |
| AT5G18830 | 0,51 | -0,42 | 3,27 | -0,13 | 8,99E-01 |
| AT5G18890 | 0,98 | -3,13 | 4,28 | -0,73 | 4,64E-01 |
| AT5G18937 | 0,54 | 3,31 | 4,27 | 0,78 | 4,37E-01 |
| AT5G18940 | 1,00 | -0,36 | 2,85 | -0,13 | 8,99E-01 |
| AT5G18960 | 0,35 | 0,48 | 3,50 | 0,14 | 8,92E-01 |
| AT5G19000 | 0,44 | 0,46 | 3,41 | 0,14 | 8,92E-01 |
| AT5G19090 | 0,78 | 2,87 | 2,42 | 1,19 | 2,35E-01 |
| AT5G19180 | 0,77 | 0,78 | 2,20 | 0,35 | 7,23E-01 |
| AT5G19190 | 0,37 | 0,60 | 4,36 | 0,14 | 8,90E-01 |
| AT5G19210 | 0,49 | 1,74 | 3,29 | 0,53 | 5,97E-01 |
| AT5G19221 | 0,47 | 0,15 | 4,38 | 0,03 | 9,74E-01 |
| AT5G19240 | 0,65 | 0,03 | 2,02 | 0,01 | 9,90E-01 |
| AT5G19250 | 0,98 | -1,11 | 2,59 | -0,43 | 6,70E-01 |
| AT5G19290 | 0,41 | 0,99 | 3,36 | 0,29 | 7,68E-01 |
| AT5G19440 | 0,36 | -0,26 | 4,41 | -0,06 | 9,53E-01 |
| AT5G19500 | 1,08 | 0,49 | 2,46 | 0,20 | 8,43E-01 |
| AT5G19540 | 0,77 | -1,55 | 1,97 | -0,79 | 4,30E-01 |
| AT5G19550 | 0,90 | 0,60 | 2,03 | 0,29 | 7,69E-01 |
| AT5G19570 | 0,52 | 0,23 | 3,99 | 0,06 | 9,54E-01 |
| AT5G19620 | 0,50 | 0,35 | 3,99 | 0,09 | 9,29E-01 |
| AT5G19680 | 0,19 | -1,25 | 4,36 | -0,29 | 7,74E-01 |
| AT5G19690 | 1,74 | 1,85 | 1,51 | 1,22 | 2,21E-01 |
| AT5G19750 | 0,55 | -2,05 | 4,39 | -0,47 | 6,40E-01 |
| AT5G19780 | 1,35 | -2,25 | 1,71 | -1,31 | 1,89E-01 |
| AT5G19990 | 0,77 | -0,47 | 2,29 | -0,21 | 8,36E-01 |
| AT5G20080 | 0,48 | -0,03 | 3,31 | -0,01 | 9,93E-01 |
| AT5G20120 | 0,55 | 0,00 | 4,39 | 0,00 | 1,00E+00 |
| AT5G20130 | 0,68 | -2,29 | 3,78 | -0,61 | 5,45E-01 |
| AT5G20150 | 0,56 | 0,41 | 2,67 | 0,15 | 8,78E-01 |
| AT5G20190 | 1,21 | 0,95 | 2,00 | 0,47 | 6,35E-01 |
| AT5G20200 | 1,58 | -0,27 | 2,09 | -0,13 | 8,96E-01 |
| AT5G20220 | 0,21 | -0,29 | 4,40 | -0,06 | 9,48E-01 |

| | | | | | |
|---|---|---|---|---|---|
| AT5G20350 | 1,36 | -0,50 | 1,78 | -0,28 | 7,79E-01 |
| AT5G20410 | 0,63 | 0,00 | 3,73 | 0,00 | 1,00E+00 |
| AT5G20480 | 0,15 | 0,82 | 4,35 | 0,19 | 8,50E-01 |
| AT5G20500 | 0,33 | 1,03 | 4,34 | 0,24 | 8,12E-01 |
| AT5G20590 | 0,85 | -0,97 | 2,20 | -0,44 | 6,61E-01 |
| AT5G20610 | 1,39 | 0,52 | 1,98 | 0,26 | 7,92E-01 |
| AT5G20660 | 1,27 | 0,89 | 2,02 | 0,44 | 6,58E-01 |
| AT5G20680 | 0,31 | -0,68 | 4,35 | -0,16 | 8,76E-01 |
| AT5G20710 | 0,12 | 0,00 | 4,41 | 0,00 | 1,00E+00 |
| AT5G20900 | 0,45 | 0,55 | 3,28 | 0,17 | 8,66E-01 |
| AT5G03255 | 0,31 | -0,40 | 4,38 | -0,09 | 9,27E-01 |
| AT5G20960 | 1,02 | 1,57 | 2,62 | 0,60 | 5,49E-01 |
| AT5G21020 | 1,50 | 0,29 | 1,39 | 0,21 | 8,36E-01 |
| AT5G21090 | 1,39 | 2,97 | 1,93 | 1,54 | 1,23E-01 |
| AT5G21105 | 1,68 | 1,82 | 1,70 | 1,07 | 2,85E-01 |
| AT5G21150 | 0,12 | 0,00 | 4,41 | 0,00 | 1,00E+00 |
| AT5G21222 | 1,01 | 0,14 | 2,81 | 0,05 | 9,60E-01 |
| AT5G21326 | 0,90 | 0,27 | 2,86 | 0,10 | 9,24E-01 |
| AT5G21482 | 0,24 | 1,34 | 4,33 | 0,31 | 7,57E-01 |
| AT5G21930 | 1,60 | -0,49 | 1,56 | -0,32 | 7,52E-01 |
| AT5G22000 | 1,27 | 0,02 | 1,96 | 0,01 | 9,91E-01 |
| AT5G22050 | 0,36 | 0,06 | 4,34 | 0,01 | 9,89E-01 |
| AT5G22070 | 0,57 | 1,12 | 2,97 | 0,38 | 7,06E-01 |
| AT5G22300 | 0,30 | 0,53 | 3,65 | 0,14 | 8,86E-01 |
| AT5G22315 | 0,20 | 0,40 | 4,37 | 0,09 | 9,27E-01 |
| AT5G22340 | 0,95 | -1,46 | 2,27 | -0,64 | 5,20E-01 |
| AT5G22420 | 0,42 | 0,00 | 4,22 | 0,00 | 1,00E+00 |
| AT5G22580 | 0,46 | -1,37 | 4,36 | -0,31 | 7,53E-01 |
| AT5G22590 | 1,03 | 3,95 | 2,52 | 1,57 | 1,17E-01 |
| AT5G22630 | 1,52 | -0,42 | 1,72 | -0,24 | 8,08E-01 |
| AT5G22640 | 1,19 | -1,21 | 1,87 | -0,65 | 5,17E-01 |
| AT5G22650 | 0,66 | 0,00 | 2,32 | 0,00 | 9,99E-01 |
| AT5G22690 | 0,12 | 0,00 | 4,41 | 0,00 | 1,00E+00 |
| AT5G22760 | 1,25 | -0,71 | 2,06 | -0,34 | 7,30E-01 |
| AT5G22794 | 0,08 | 0,00 | 4,41 | 0,00 | 1,00E+00 |
| AT5G22830 | 0,08 | 0,21 | 4,39 | 0,05 | 9,62E-01 |
| AT5G22950 | 0,12 | 0,11 | 4,38 | 0,03 | 9,80E-01 |
| AT5G23010 | 0,84 | 1,13 | 2,04 | 0,55 | 5,80E-01 |
| AT5G23070 | 0,19 | 0,50 | 4,37 | 0,12 | 9,08E-01 |
| AT5G03575 | 0,26 | -2,72 | 4,37 | -0,62 | 5,33E-01 |
| AT5G23120 | 0,65 | -1,00 | 2,17 | -0,46 | 6,46E-01 |
| AT5G23140 | 0,95 | -0,35 | 2,42 | -0,15 | 8,84E-01 |
| AT5G23150 | 1,41 | 1,12 | 2,24 | 0,50 | 6,16E-01 |
| AT5G23210 | 0,16 | 0,15 | 4,38 | 0,04 | 9,72E-01 |
| AT5G23280 | 1,01 | 1,45 | 2,53 | 0,57 | 5,67E-01 |
| AT5G23320 | 0,08 | 0,00 | 4,41 | 0,00 | 1,00E+00 |
| AT5G23330 | 0,43 | 0,00 | 4,41 | 0,00 | 1,00E+00 |
| AT5G23340 | 1,40 | -0,26 | 2,12 | -0,12 | 9,02E-01 |
| AT5G23490 | 0,49 | 1,08 | 2,69 | 0,40 | 6,88E-01 |
| AT5G23540 | 0,96 | 0,78 | 1,83 | 0,42 | 6,73E-01 |
| AT5G23570 | 0,58 | -1,38 | 2,25 | -0,61 | 5,40E-01 |
| AT5G23575 | 1,00 | 1,87 | 1,88 | 0,99 | 3,22E-01 |
| AT5G23600 | 0,08 | 0,00 | 4,41 | 0,00 | 1,00E+00 |
| AT5G23670 | 0,72 | -0,21 | 3,11 | -0,07 | 9,46E-01 |
| AT5G23720 | 0,51 | -1,22 | 3,32 | -0,37 | 7,13E-01 |
| AT5G23730 | 0,26 | 0,03 | 4,38 | 0,01 | 9,95E-01 |

| | | | | | |
|---|---|---|---|---|---|
| AT5G23890 | 0,99 | -0,21 | 2,85 | -0,07 | 9,41E-01 |
| AT5G23950 | 0,35 | 0,84 | 4,35 | 0,19 | 8,46E-01 |
| AT5G24020 | 0,16 | 0,15 | 4,38 | 0,04 | 9,72E-01 |
| AT5G24200 | 1,34 | 2,61 | 1,68 | 1,55 | 1,20E-01 |
| AT5G24210 | 0,96 | -0,72 | 2,15 | -0,33 | 7,39E-01 |
| AT5G24360 | 0,40 | -1,29 | 3,41 | -0,38 | 7,06E-01 |
| AT5G24380 | 0,41 | -1,10 | 3,42 | -0,32 | 7,46E-01 |
| AT5G24450 | 0,69 | 1,84 | 3,67 | 0,50 | 6,17E-01 |
| AT5G24460 | 0,88 | -1,19 | 2,01 | -0,59 | 5,55E-01 |
| AT5G24490 | 1,61 | -1,33 | 1,89 | -0,70 | 4,81E-01 |
| AT5G24530 | 1,77 | 0,54 | 1,37 | 0,39 | 6,94E-01 |
| AT5G24610 | 0,55 | -2,13 | 2,64 | -0,81 | 4,18E-01 |
| AT5G24830 | 0,08 | 0,00 | 4,41 | 0,00 | 1,00E+00 |
| AT5G24980 | 0,43 | -1,68 | 3,36 | -0,50 | 6,18E-01 |
| AT5G25040 | 0,67 | 3,04 | 2,35 | 1,29 | 1,96E-01 |
| AT5G25120 | 0,51 | 1,65 | 3,14 | 0,53 | 5,99E-01 |
| AT5G03925 | 0,17 | 1,58 | 4,31 | 0,37 | 7,15E-01 |
| AT5G25130 | 1,42 | 1,70 | 1,59 | 1,07 | 2,85E-01 |
| AT5G25140 | 0,28 | -0,35 | 4,40 | -0,08 | 9,37E-01 |
| AT5G25150 | 0,27 | 0,84 | 4,35 | 0,19 | 8,47E-01 |
| AT5G25170 | 0,33 | 0,18 | 4,37 | 0,04 | 9,67E-01 |
| AT5G25210 | 0,42 | -0,18 | 3,35 | -0,05 | 9,57E-01 |
| AT5G25220 | 1,05 | 2,61 | 2,07 | 1,26 | 2,07E-01 |
| AT5G25230 | 1,66 | 0,59 | 1,95 | 0,30 | 7,65E-01 |
| AT5G25250 | 1,43 | 3,80 | 1,91 | 1,99 | 4,68E-02 |
| AT5G25260 | 1,02 | -2,30 | 2,83 | -0,81 | 4,17E-01 |
| AT5G25340 | 0,46 | 0,00 | 4,14 | 0,00 | 1,00E+00 |
| AT5G25460 | 1,53 | -1,63 | 1,79 | -0,91 | 3,62E-01 |
| AT5G25520 | 1,48 | 0,00 | 2,61 | 0,00 | 1,00E+00 |
| AT5G25560 | 0,53 | -0,26 | 3,26 | -0,08 | 9,36E-01 |
| AT5G25580 | 0,12 | 0,00 | 4,41 | 0,00 | 1,00E+00 |
| AT5G25630 | 0,47 | -0,85 | 3,22 | -0,26 | 7,92E-01 |
| AT5G25754 | 1,48 | 0,38 | 1,45 | 0,26 | 7,92E-01 |
| AT5G04145 | 0,89 | -1,72 | 2,27 | -0,76 | 4,48E-01 |
| AT5G25900 | 0,33 | -2,19 | 4,41 | -0,50 | 6,20E-01 |
| AT5G25910 | 0,60 | 0,00 | 3,79 | 0,00 | 1,00E+00 |
| AT5G25930 | 0,70 | 2,19 | 2,45 | 0,90 | 3,71E-01 |
| AT5G25980 | 1,32 | 0,92 | 1,60 | 0,58 | 5,63E-01 |
| AT5G26038 | 1,13 | 2,09 | 2,10 | 1,00 | 3,20E-01 |
| AT5G26146 | 0,89 | -0,73 | 2,16 | -0,34 | 7,37E-01 |
| AT5G26147 | 0,89 | -0,73 | 2,16 | -0,34 | 7,37E-01 |
| AT5G26200 | 0,12 | 0,00 | 4,41 | 0,00 | 1,00E+00 |
| AT5G26240 | 0,84 | -2,10 | 2,13 | -0,99 | 3,24E-01 |
| AT5G26270 | 0,26 | -0,06 | 4,33 | -0,01 | 9,89E-01 |
| AT5G26290 | 0,79 | -0,74 | 2,06 | -0,36 | 7,19E-01 |
| AT5G26570 | 1,51 | -1,88 | 1,85 | -1,02 | 3,09E-01 |
| AT5G26740 | 0,58 | -1,20 | 2,67 | -0,45 | 6,53E-01 |
| AT5G26731 | 0,38 | 1,07 | 4,34 | 0,25 | 8,04E-01 |
| AT5G26710 | 0,73 | -0,26 | 2,64 | -0,10 | 9,21E-01 |
| AT5G26670 | 0,26 | -2,46 | 4,37 | -0,56 | 5,74E-01 |
| AT5G26610 | 0,16 | 1,58 | 4,31 | 0,37 | 7,14E-01 |
| AT5G26600 | 0,82 | -0,81 | 2,57 | -0,32 | 7,53E-01 |
| AT5G26780 | 0,39 | -0,14 | 2,86 | -0,05 | 9,61E-01 |
| AT5G26820 | 0,42 | 0,00 | 4,41 | 0,00 | 1,00E+00 |
| AT5G26850 | 1,21 | -3,14 | 2,33 | -1,34 | 1,79E-01 |
| AT5G26860 | 1,16 | 0,27 | 1,96 | 0,14 | 8,89E-01 |

| | | | | | |
|---|---|---|---|---|---|
| AT5G26930 | 0,33 | 0,00 | 4,41 | 0,00 | 1,00E+00 |
| AT5G26940 | 0,66 | 0,00 | 3,80 | 0,00 | 1,00E+00 |
| AT5G26990 | 0,23 | 0,24 | 4,39 | 0,06 | 9,56E-01 |
| AT5G27120 | 0,91 | -2,46 | 3,53 | -0,70 | 4,86E-01 |
| AT5G27150 | 0,57 | 0,90 | 3,04 | 0,30 | 7,68E-01 |
| AT5G27170 | 0,25 | 1,56 | 4,32 | 0,36 | 7,19E-01 |
| AT5G27230 | 0,08 | 0,21 | 4,39 | 0,05 | 9,62E-01 |
| AT5G27240 | 0,05 | 0,18 | 4,38 | 0,04 | 9,68E-01 |
| AT5G27270 | 0,17 | -0,78 | 4,36 | -0,18 | 8,59E-01 |
| AT5G27280 | 0,34 | -0,48 | 3,57 | -0,14 | 8,92E-01 |
| AT5G27300 | 0,08 | 0,21 | 4,39 | 0,05 | 9,62E-01 |
| AT5G27320 | 0,08 | 0,21 | 4,39 | 0,05 | 9,62E-01 |
| AT5G27340 | 0,54 | 2,76 | 3,69 | 0,75 | 4,54E-01 |
| AT5G27350 | 0,64 | 0,17 | 2,43 | 0,07 | 9,43E-01 |
| AT5G27410 | 0,42 | 0,83 | 4,11 | 0,20 | 8,40E-01 |
| AT5G27420 | 1,44 | 0,50 | 1,84 | 0,27 | 7,87E-01 |
| AT5G27470 | 0,51 | 0,56 | 3,16 | 0,18 | 8,60E-01 |
| AT5G27530 | 0,32 | -0,26 | 4,30 | -0,06 | 9,52E-01 |
| AT5G27540 | 0,68 | -0,41 | 3,58 | -0,12 | 9,08E-01 |
| AT5G27600 | 0,18 | 0,80 | 4,35 | 0,18 | 8,54E-01 |
| AT5G27610 | 0,08 | 0,21 | 4,39 | 0,05 | 9,62E-01 |
| AT5G27630 | 0,17 | 1,19 | 4,33 | 0,28 | 7,83E-01 |
| AT5G27650 | 0,94 | -0,04 | 2,82 | -0,02 | 9,88E-01 |
| AT5G27710 | 0,67 | 1,55 | 2,47 | 0,63 | 5,30E-01 |
| AT5G27730 | 0,69 | 0,78 | 2,04 | 0,38 | 7,02E-01 |
| AT5G27760 | 0,27 | 0,30 | 4,39 | 0,07 | 9,46E-01 |
| AT5G27770 | 0,86 | -0,34 | 2,47 | -0,14 | 8,89E-01 |
| AT5G27800 | 0,16 | 1,16 | 4,34 | 0,27 | 7,89E-01 |
| AT5G27830 | 0,23 | -0,34 | 4,40 | -0,08 | 9,38E-01 |
| AT5G27890 | 0,40 | 2,06 | 4,30 | 0,48 | 6,33E-01 |
| AT5G27893 | 1,07 | -1,29 | 1,62 | -0,80 | 4,23E-01 |
| AT5G27970 | 1,51 | 0,49 | 2,26 | 0,22 | 8,28E-01 |
| AT5G28010 | 1,26 | -0,44 | 1,47 | -0,30 | 7,66E-01 |
| AT5G04625 | 0,10 | 0,86 | 4,35 | 0,20 | 8,43E-01 |
| AT5G04665 | 0,11 | 0,88 | 4,35 | 0,20 | 8,39E-01 |
| AT5G28090 | 0,29 | 1,38 | 4,33 | 0,32 | 7,50E-01 |
| AT5G04675 | 0,19 | 1,60 | 4,31 | 0,37 | 7,11E-01 |
| AT5G28320 | 0,89 | 0,00 | 2,81 | 0,00 | 1,00E+00 |
| AT5G28350 | 0,24 | -0,09 | 4,39 | -0,02 | 9,84E-01 |
| AT5G28410 | 0,36 | -1,62 | 4,34 | -0,37 | 7,08E-01 |
| AT5G28463 | 0,37 | 0,95 | 4,26 | 0,22 | 8,23E-01 |
| AT5G28500 | 0,70 | -0,16 | 2,50 | -0,07 | 9,47E-01 |
| AT5G04815 | 0,05 | 0,18 | 4,38 | 0,04 | 9,68E-01 |
| AT5G28810 | 0,66 | 2,24 | 2,77 | 0,81 | 4,19E-01 |
| AT5G28830 | 0,05 | 0,18 | 4,38 | 0,04 | 9,68E-01 |
| AT5G28900 | 0,40 | -1,27 | 4,32 | -0,29 | 7,69E-01 |
| AT5G29624 | 0,12 | 0,00 | 4,41 | 0,00 | 1,00E+00 |
| AT5G30341 | 0,12 | 0,49 | 4,38 | 0,11 | 9,11E-01 |
| AT5G04955 | 1,06 | 4,29 | 3,80 | 1,13 | 2,58E-01 |
| AT5G32440 | 1,33 | 2,41 | 1,89 | 1,28 | 2,02E-01 |
| AT5G32613 | 0,26 | 1,70 | 4,31 | 0,39 | 6,94E-01 |
| AT5G33290 | 0,62 | 0,07 | 3,96 | 0,02 | 9,86E-01 |
| AT5G33320 | 0,61 | 0,03 | 2,32 | 0,01 | 9,91E-01 |
| AT5G33390 | 0,38 | -3,19 | 4,36 | -0,73 | 4,64E-01 |
| AT5G34869 | 0,19 | 0,44 | 4,36 | 0,10 | 9,21E-01 |
| AT5G34871 | 0,12 | -0,26 | 4,41 | -0,06 | 9,53E-01 |

| | | | | | |
|---|---|---|---|---|---|
| AT5G34870 | 0,38 | -0,54 | 3,46 | -0,16 | 8,76E-01 |
| AT5G35160 | 1,68 | 0,34 | 1,55 | 0,22 | 8,27E-01 |
| AT5G35170 | 0,47 | -0,68 | 2,37 | -0,29 | 7,75E-01 |
| AT5G35180 | 0,52 | -0,28 | 2,66 | -0,11 | 9,15E-01 |
| AT5G35190 | 0,84 | 2,62 | 2,73 | 0,96 | 3,36E-01 |
| AT5G35300 | 0,23 | 1,56 | 4,31 | 0,36 | 7,18E-01 |
| AT5G35320 | 0,21 | 0,08 | 4,38 | 0,02 | 9,85E-01 |
| AT5G35360 | 1,57 | 0,90 | 1,84 | 0,49 | 6,25E-01 |
| AT5G35430 | 1,24 | -1,27 | 2,43 | -0,52 | 6,00E-01 |
| AT5G35450 | 0,40 | 0,00 | 4,41 | 0,00 | 1,00E+00 |
| AT5G05195 | 0,23 | 1,19 | 4,34 | 0,27 | 7,84E-01 |
| AT5G35490 | 1,40 | 3,89 | 2,25 | 1,72 | 8,48E-02 |
| AT5G35730 | 0,82 | -2,07 | 2,83 | -0,73 | 4,64E-01 |
| AT5G35840 | 1,35 | 0,10 | 2,28 | 0,04 | 9,66E-01 |
| AT5G35917 | 0,29 | 1,96 | 4,30 | 0,45 | 6,49E-01 |
| AT5G35920 | 1,48 | 2,80 | 1,74 | 1,61 | 1,07E-01 |
| AT5G35926 | 1,15 | 0,64 | 1,79 | 0,36 | 7,22E-01 |
| AT5G35980 | 1,15 | 0,38 | 1,78 | 0,22 | 8,29E-01 |
| AT5G36001 | 0,12 | 0,00 | 4,41 | 0,00 | 1,00E+00 |
| AT5G36080 | 1,49 | 3,61 | 1,84 | 1,97 | 4,92E-02 |
| AT5G36290 | 0,47 | 0,81 | 2,63 | 0,31 | 7,59E-01 |
| AT5G36300 | 0,87 | 1,71 | 2,32 | 0,73 | 4,63E-01 |
| AT5G36740 | 0,08 | 0,00 | 4,41 | 0,00 | 1,00E+00 |
| AT5G36870 | 1,07 | -1,30 | 1,77 | -0,74 | 4,62E-01 |
| AT5G36900 | 0,54 | 1,30 | 3,11 | 0,42 | 6,75E-01 |
| AT5G36940 | 0,51 | -0,34 | 3,96 | -0,09 | 9,32E-01 |
| AT5G37020 | 0,62 | -0,25 | 2,54 | -0,10 | 9,23E-01 |
| AT5G37070 | 0,43 | -2,80 | 4,37 | -0,64 | 5,22E-01 |
| AT5G37130 | 0,66 | 1,45 | 2,68 | 0,54 | 5,88E-01 |
| AT5G37150 | 0,98 | 2,72 | 2,18 | 1,25 | 2,13E-01 |
| AT5G37540 | 0,17 | 0,00 | 4,41 | 0,00 | 1,00E+00 |
| AT5G37630 | 0,94 | -0,28 | 2,44 | -0,11 | 9,09E-01 |
| AT5G37760 | 0,15 | -0,31 | 4,40 | -0,07 | 9,43E-01 |
| AT5G37793 | 0,51 | -1,33 | 2,53 | -0,52 | 6,01E-01 |
| AT5G37850 | 0,30 | 0,00 | 4,41 | 0,00 | 1,00E+00 |
| AT5G37950 | 0,11 | 0,88 | 4,35 | 0,20 | 8,39E-01 |
| AT5G38155 | 0,12 | 0,11 | 4,38 | 0,03 | 9,80E-01 |
| AT5G38210 | 1,00 | 3,17 | 2,13 | 1,49 | 1,37E-01 |
| AT5G38380 | 0,15 | -0,57 | 4,37 | -0,13 | 8,96E-01 |
| AT5G38470 | 0,73 | 0,18 | 3,72 | 0,05 | 9,61E-01 |
| AT5G38480 | 1,20 | 0,18 | 1,86 | 0,10 | 9,23E-01 |
| AT5G38560 | 0,34 | 1,18 | 3,49 | 0,34 | 7,35E-01 |
| AT5G38910 | 0,16 | 1,02 | 4,34 | 0,24 | 8,13E-01 |
| AT5G38980 | 0,57 | 1,18 | 2,66 | 0,44 | 6,58E-01 |
| AT5G39000 | 0,70 | 0,29 | 4,39 | 0,07 | 9,48E-01 |
| AT5G39030 | 0,81 | -0,16 | 2,93 | -0,05 | 9,57E-01 |
| AT5G39040 | 0,29 | -2,19 | 4,41 | -0,50 | 6,20E-01 |
| AT5G39250 | 0,34 | 0,41 | 4,37 | 0,09 | 9,25E-01 |
| AT5G39320 | 0,82 | 0,00 | 4,40 | 0,00 | 1,00E+00 |
| AT5G39350 | 0,21 | -0,37 | 4,36 | -0,09 | 9,32E-01 |
| AT5G39510 | 0,77 | 2,42 | 2,32 | 1,05 | 2,95E-01 |
| AT5G39590 | 0,82 | 1,82 | 2,37 | 0,77 | 4,42E-01 |
| AT5G39660 | 0,40 | -0,26 | 4,41 | -0,06 | 9,53E-01 |
| AT5G39670 | 0,40 | 1,28 | 4,17 | 0,31 | 7,59E-01 |
| AT5G39690 | 1,03 | 0,64 | 2,08 | 0,31 | 7,60E-01 |
| AT5G39710 | 0,25 | 0,00 | 4,41 | 0,00 | 1,00E+00 |

| | | | | | |
|---|---|---|---|---|---|
| AT5G39730 | 0,19 | 1,06 | 4,34 | 0,24 | 8,07E-01 |
| AT5G39775 | 0,53 | 1,03 | 4,11 | 0,25 | 8,02E-01 |
| AT5G39850 | 0,08 | 0,00 | 4,41 | 0,00 | 1,00E+00 |
| AT5G39960 | 0,21 | 0,00 | 4,41 | 0,00 | 1,00E+00 |
| AT5G39990 | 0,08 | 0,00 | 4,41 | 0,00 | 1,00E+00 |
| AT5G40060 | 1,33 | -0,92 | 2,58 | -0,36 | 7,22E-01 |
| AT5G40140 | 0,31 | 0,91 | 3,62 | 0,25 | 8,02E-01 |
| AT5G40200 | 0,62 | -0,41 | 2,71 | -0,15 | 8,80E-01 |
| AT5G40380 | 0,46 | 1,29 | 3,91 | 0,33 | 7,41E-01 |
| AT5G40400 | 1,55 | 0,91 | 1,34 | 0,68 | 4,96E-01 |
| AT5G40650 | 0,74 | -0,33 | 2,92 | -0,11 | 9,09E-01 |
| AT5G40670 | 1,01 | -1,26 | 2,21 | -0,57 | 5,69E-01 |
| AT5G40680 | 0,17 | 0,00 | 4,41 | 0,00 | 1,00E+00 |
| AT5G40690 | 0,62 | -0,09 | 3,74 | -0,02 | 9,80E-01 |
| AT5G40720 | 0,32 | 1,97 | 4,30 | 0,46 | 6,47E-01 |
| AT5G40760 | 1,16 | 0,57 | 2,15 | 0,27 | 7,91E-01 |
| AT5G40780 | 1,36 | 0,38 | 1,78 | 0,21 | 8,33E-01 |
| AT5G40810 | 1,32 | -2,04 | 1,70 | -1,20 | 2,30E-01 |
| AT5G40850 | 0,08 | 0,47 | 4,38 | 0,11 | 9,15E-01 |
| AT5G40870 | 0,14 | 0,81 | 4,35 | 0,19 | 8,52E-01 |
| AT5G40880 | 0,51 | -0,26 | 4,03 | -0,06 | 9,49E-01 |
| AT5G40900 | 0,15 | 0,02 | 4,36 | 0,00 | 9,96E-01 |
| AT5G41010 | 0,28 | 1,12 | 4,34 | 0,26 | 7,97E-01 |
| AT5G06115 | 0,20 | 0,89 | 4,36 | 0,20 | 8,39E-01 |
| AT5G41070 | 0,20 | 0,40 | 4,37 | 0,09 | 9,27E-01 |
| AT5G41100 | 0,28 | 1,13 | 4,33 | 0,26 | 7,94E-01 |
| AT5G41110 | 0,22 | 0,53 | 4,36 | 0,12 | 9,03E-01 |
| AT5G41140 | 0,59 | -1,33 | 3,84 | -0,35 | 7,30E-01 |
| AT5G41260 | 0,61 | 0,90 | 2,34 | 0,39 | 6,99E-01 |
| AT5G41360 | 0,66 | -1,05 | 3,72 | -0,28 | 7,77E-01 |
| AT5G41370 | 0,15 | 0,90 | 4,35 | 0,21 | 8,36E-01 |
| AT5G41410 | 1,13 | 1,49 | 1,93 | 0,78 | 4,38E-01 |
| AT5G41471 | 1,79 | -0,74 | 1,63 | -0,46 | 6,49E-01 |
| AT5G41500 | 1,05 | 2,72 | 2,04 | 1,33 | 1,83E-01 |
| AT5G41510 | 1,76 | 3,62 | 1,77 | 2,05 | 4,07E-02 |
| AT5G41660 | 0,71 | 2,34 | 2,90 | 0,81 | 4,19E-01 |
| AT5G41700 | 1,24 | -2,47 | 1,85 | -1,33 | 1,83E-01 |
| AT5G41761 | 0,83 | 3,18 | 2,28 | 1,39 | 1,63E-01 |
| AT5G41770 | 0,61 | -1,10 | 2,58 | -0,42 | 6,71E-01 |
| AT5G41860 | 0,77 | 0,23 | 4,39 | 0,05 | 9,59E-01 |
| AT5G41920 | 0,61 | 0,98 | 4,33 | 0,23 | 8,20E-01 |
| AT5G41940 | 0,98 | -2,01 | 2,08 | -0,97 | 3,33E-01 |
| AT5G41950 | 1,46 | -0,30 | 2,22 | -0,13 | 8,94E-01 |
| AT5G41960 | 0,15 | -2,07 | 4,39 | -0,47 | 6,38E-01 |
| AT5G41970 | 0,85 | 0,00 | 3,48 | 0,00 | 1,00E+00 |
| AT5G42010 | 0,60 | 0,44 | 3,19 | 0,14 | 8,90E-01 |
| AT5G42220 | 0,70 | 1,57 | 2,37 | 0,66 | 5,07E-01 |
| AT5G42250 | 0,33 | 0,00 | 4,41 | 0,00 | 1,00E+00 |
| AT5G42260 | 0,28 | 2,34 | 4,29 | 0,55 | 5,85E-01 |
| AT5G42290 | 0,08 | 0,00 | 4,41 | 0,00 | 1,00E+00 |
| AT5G42310 | 0,50 | 0,85 | 4,09 | 0,21 | 8,35E-01 |
| AT5G42350 | 0,37 | -0,24 | 4,39 | -0,05 | 9,57E-01 |
| AT5G42420 | 0,74 | 0,00 | 3,71 | 0,00 | 1,00E+00 |
| AT5G42460 | 0,33 | 0,00 | 4,41 | 0,00 | 1,00E+00 |
| AT5G42470 | 0,22 | -0,68 | 4,34 | -0,16 | 8,76E-01 |
| AT5G42480 | 0,45 | 0,00 | 4,41 | 0,00 | 1,00E+00 |

| | | | | | |
|---|---|---|---|---|---|
| AT5G42504 | 0,17 | -1,50 | 4,37 | -0,34 | 7,31E-01 |
| AT5G42520 | 0,71 | -1,47 | 2,34 | -0,63 | 5,31E-01 |
| AT5G42540 | 1,02 | -0,26 | 2,53 | -0,10 | 9,17E-01 |
| AT5G42620 | 0,80 | -0,66 | 2,37 | -0,28 | 7,81E-01 |
| AT5G42640 | 0,08 | 0,21 | 4,39 | 0,05 | 9,62E-01 |
| AT5G42670 | 0,16 | 1,26 | 4,33 | 0,29 | 7,71E-01 |
| AT5G42680 | 0,80 | 1,30 | 3,17 | 0,41 | 6,81E-01 |
| AT5G42720 | 0,19 | 0,27 | 4,37 | 0,06 | 9,50E-01 |
| AT5G42740 | 1,16 | -0,92 | 2,86 | -0,32 | 7,47E-01 |
| AT5G42760 | 0,12 | 0,27 | 4,38 | 0,06 | 9,50E-01 |
| AT5G06395 | 1,55 | -2,13 | 2,09 | -1,02 | 3,08E-01 |
| AT5G42825 | 0,29 | -2,17 | 4,41 | -0,49 | 6,23E-01 |
| AT5G42895 | 1,80 | 1,54 | 1,58 | 0,97 | 3,31E-01 |
| AT5G42965 | 0,19 | -1,14 | 4,40 | -0,26 | 7,95E-01 |
| AT5G42980 | 0,59 | -0,69 | 2,62 | -0,26 | 7,91E-01 |
| AT5G43100 | 0,78 | -0,16 | 2,25 | -0,07 | 9,45E-01 |
| AT5G43150 | 0,61 | -0,13 | 3,13 | -0,04 | 9,66E-01 |
| AT5G06505 | 0,36 | 0,05 | 4,35 | 0,01 | 9,90E-01 |
| AT5G43460 | 1,01 | -1,24 | 2,21 | -0,56 | 5,74E-01 |
| AT5G43500 | 1,47 | -2,54 | 1,58 | -1,61 | 1,07E-01 |
| AT5G43525 | 0,69 | 3,09 | 2,74 | 1,13 | 2,60E-01 |
| AT5G43560 | 0,55 | 0,82 | 3,17 | 0,26 | 7,96E-01 |
| AT5G43610 | 1,12 | -0,28 | 2,12 | -0,13 | 8,93E-01 |
| AT5G43620 | 0,36 | -1,28 | 4,34 | -0,29 | 7,69E-01 |
| AT5G43700 | 0,52 | -2,87 | 3,14 | -0,91 | 3,60E-01 |
| AT5G43810 | 1,04 | 0,55 | 2,77 | 0,20 | 8,44E-01 |
| AT5G43930 | 0,73 | 0,50 | 2,59 | 0,19 | 8,48E-01 |
| AT5G43940 | 1,17 | 1,84 | 2,02 | 0,91 | 3,63E-01 |
| AT5G44020 | 1,03 | -0,18 | 1,98 | -0,09 | 9,26E-01 |
| AT5G44050 | 1,57 | -3,77 | 1,98 | -1,91 | 5,65E-02 |
| AT5G44060 | 0,47 | 0,75 | 2,74 | 0,27 | 7,84E-01 |
| AT5G44070 | 1,32 | 0,62 | 1,58 | 0,39 | 6,94E-01 |
| AT5G44130 | 0,80 | -3,89 | 2,79 | -1,39 | 1,63E-01 |
| AT5G44190 | 1,58 | -0,90 | 1,58 | -0,57 | 5,68E-01 |
| AT5G44230 | 0,12 | 0,00 | 4,41 | 0,00 | 1,00E+00 |
| AT5G44240 | 1,02 | -1,56 | 1,99 | -0,78 | 4,33E-01 |
| AT5G06565 | 0,63 | -0,39 | 2,55 | -0,15 | 8,79E-01 |
| AT5G44286 | 0,85 | 0,80 | 2,16 | 0,37 | 7,12E-01 |
| AT5G44530 | 1,17 | -2,55 | 2,73 | -0,93 | 3,50E-01 |
| AT5G44574 | 0,91 | 0,21 | 3,02 | 0,07 | 9,45E-01 |
| AT5G44580 | 0,52 | -0,55 | 3,97 | -0,14 | 8,90E-01 |
| AT5G44585 | 1,38 | -0,29 | 1,66 | -0,18 | 8,60E-01 |
| AT5G06605 | 1,31 | -0,81 | 1,72 | -0,47 | 6,36E-01 |
| AT5G44640 | 1,59 | 2,09 | 1,66 | 1,26 | 2,08E-01 |
| AT5G44650 | 0,46 | -2,91 | 4,06 | -0,72 | 4,74E-01 |
| AT5G44720 | 0,16 | 0,41 | 4,37 | 0,09 | 9,25E-01 |
| AT5G44750 | 0,74 | -1,19 | 2,35 | -0,50 | 6,14E-01 |
| AT5G44760 | 0,36 | 0,00 | 4,41 | 0,00 | 1,00E+00 |
| AT5G44770 | 0,22 | 1,19 | 4,34 | 0,28 | 7,83E-01 |
| AT5G44870 | 1,81 | 0,61 | 1,34 | 0,46 | 6,48E-01 |
| AT5G45000 | 0,38 | -1,52 | 4,32 | -0,35 | 7,24E-01 |
| AT5G06695 | 0,59 | -2,19 | 3,16 | -0,69 | 4,89E-01 |
| AT5G45030 | 0,59 | -2,19 | 3,16 | -0,69 | 4,89E-01 |
| AT5G45060 | 1,56 | -1,07 | 1,73 | -0,62 | 5,36E-01 |
| AT5G45100 | 1,72 | 1,45 | 2,01 | 0,72 | 4,70E-01 |
| AT5G45110 | 1,72 | 1,99 | 1,63 | 1,22 | 2,22E-01 |

| | | | | | |
|---|---|---|---|---|---|
| AT5G45130 | 0,78 | -1,18 | 3,09 | -0,38 | 7,02E-01 |
| AT5G45150 | 0,08 | 0,00 | 4,41 | 0,00 | 1,00E+00 |
| AT5G45240 | 0,12 | 0,00 | 4,41 | 0,00 | 1,00E+00 |
| AT5G45260 | 0,75 | -0,28 | 3,00 | -0,09 | 9,26E-01 |
| AT5G45390 | 0,62 | -0,40 | 2,21 | -0,18 | 8,55E-01 |
| AT5G45400 | 0,76 | -1,19 | 2,64 | -0,45 | 6,53E-01 |
| AT5G45450 | 0,08 | 0,00 | 4,41 | 0,00 | 1,00E+00 |
| AT5G45475 | 0,74 | -0,38 | 2,14 | -0,18 | 8,60E-01 |
| AT5G45490 | 1,25 | 0,65 | 1,68 | 0,38 | 7,01E-01 |
| AT5G45570 | 0,68 | 2,09 | 2,36 | 0,89 | 3,76E-01 |
| AT5G45620 | 0,59 | 1,24 | 3,10 | 0,40 | 6,88E-01 |
| AT5G45670 | 0,10 | 0,86 | 4,35 | 0,20 | 8,43E-01 |
| AT5G45680 | 0,26 | 0,60 | 4,37 | 0,14 | 8,90E-01 |
| AT5G45710 | 0,55 | -0,35 | 3,87 | -0,09 | 9,28E-01 |
| AT5G45750 | 0,19 | 1,60 | 4,31 | 0,37 | 7,11E-01 |
| AT5G45770 | 0,12 | 0,00 | 4,41 | 0,00 | 1,00E+00 |
| AT5G45920 | 1,21 | -3,47 | 1,82 | -1,90 | 5,71E-02 |
| AT5G46070 | 1,78 | 0,58 | 2,03 | 0,29 | 7,75E-01 |
| AT5G46330 | 0,47 | 0,35 | 4,03 | 0,09 | 9,30E-01 |
| AT5G46410 | 0,44 | 15,41 | 4,38 | 3,52 | 4,34E-04 |
| AT5G06825 | 0,67 | 3,34 | 2,97 | 1,13 | 2,61E-01 |
| AT5G46460 | 0,90 | 0,00 | 3,38 | 0,00 | 1,00E+00 |
| AT5G46490 | 1,46 | 2,20 | 1,76 | 1,25 | 2,10E-01 |
| AT5G46500 | 1,11 | -0,36 | 2,60 | -0,14 | 8,91E-01 |
| AT5G06835 | 1,13 | -5,40 | 2,42 | -2,23 | 2,55E-02 |
| AT5G46630 | 1,04 | -2,10 | 2,07 | -1,01 | 3,11E-01 |
| AT5G46640 | 1,47 | -2,19 | 2,24 | -0,98 | 3,28E-01 |
| AT5G46660 | 1,02 | 4,08 | 3,97 | 1,03 | 3,05E-01 |
| AT5G46670 | 0,74 | 2,69 | 2,64 | 1,02 | 3,09E-01 |
| AT5G46730 | 0,08 | 0,21 | 4,39 | 0,05 | 9,62E-01 |
| AT5G46780 | 0,35 | -0,28 | 4,35 | -0,07 | 9,48E-01 |
| AT5G46825 | 1,18 | -0,06 | 1,38 | -0,05 | 9,63E-01 |
| AT5G46840 | 0,62 | -1,46 | 3,13 | -0,47 | 6,41E-01 |
| AT5G46850 | 0,29 | -0,39 | 4,39 | -0,09 | 9,30E-01 |
| AT5G47020 | 1,57 | 1,11 | 1,86 | 0,60 | 5,50E-01 |
| AT5G47030 | 0,63 | -1,22 | 3,82 | -0,32 | 7,49E-01 |
| AT5G47120 | 1,06 | 3,11 | 2,04 | 1,53 | 1,26E-01 |
| AT5G47130 | 0,25 | -0,51 | 4,36 | -0,12 | 9,07E-01 |
| AT5G47140 | 0,26 | -0,02 | 4,35 | 0,00 | 9,97E-01 |
| AT5G47200 | 0,34 | -1,17 | 4,40 | -0,27 | 7,90E-01 |
| AT5G47230 | 0,27 | 1,92 | 4,31 | 0,45 | 6,56E-01 |
| AT5G47240 | 0,14 | 1,29 | 4,33 | 0,30 | 7,65E-01 |
| AT5G47280 | 1,15 | 1,17 | 1,73 | 0,68 | 4,98E-01 |
| AT5G47300 | 1,47 | 4,04 | 1,78 | 2,27 | 2,32E-02 |
| AT5G47320 | 0,19 | 1,17 | 4,34 | 0,27 | 7,87E-01 |
| AT5G47390 | 0,64 | 0,32 | 2,09 | 0,15 | 8,78E-01 |
| AT5G47420 | 0,50 | 0,00 | 4,05 | 0,00 | 1,00E+00 |
| AT5G47435 | 1,48 | -0,34 | 2,20 | -0,15 | 8,79E-01 |
| AT5G47640 | 0,70 | -1,85 | 2,19 | -0,85 | 3,97E-01 |
| AT5G47660 | 0,08 | 0,21 | 4,39 | 0,05 | 9,62E-01 |
| AT5G47720 | 0,16 | -0,22 | 4,40 | -0,05 | 9,60E-01 |
| AT5G47770 | 0,39 | 0,42 | 4,25 | 0,10 | 9,20E-01 |
| AT5G47780 | 0,52 | 0,22 | 2,69 | 0,08 | 9,34E-01 |
| AT5G47790 | 0,28 | -2,10 | 4,39 | -0,48 | 6,33E-01 |
| AT5G47800 | 0,24 | 0,00 | 4,41 | 0,00 | 1,00E+00 |
| AT5G47820 | 0,70 | -0,37 | 3,74 | -0,10 | 9,22E-01 |

| | | | | | |
|---|---|---|---|---|---|
| AT5G47840 | 0,45 | 1,07 | 2,65 | 0,40 | 6,86E-01 |
| AT5G47860 | 0,53 | -0,78 | 3,92 | -0,20 | 8,43E-01 |
| AT5G47890 | 1,07 | -2,91 | 2,14 | -1,36 | 1,74E-01 |
| AT5G48190 | 1,46 | -1,72 | 1,58 | -1,09 | 2,77E-01 |
| AT5G48220 | 0,16 | -0,26 | 4,41 | -0,06 | 9,53E-01 |
| AT5G48290 | 0,36 | 0,00 | 4,41 | 0,00 | 1,00E+00 |
| AT5G48300 | 1,23 | -0,94 | 1,85 | -0,51 | 6,13E-01 |
| AT5G48340 | 0,30 | 0,00 | 4,41 | 0,00 | 1,00E+00 |
| AT5G48360 | 0,08 | 0,21 | 4,39 | 0,05 | 9,62E-01 |
| AT5G48390 | 0,69 | 0,00 | 3,61 | 0,00 | 1,00E+00 |
| AT5G48400 | 0,56 | 0,00 | 3,88 | 0,00 | 1,00E+00 |
| AT5G48410 | 0,12 | 0,00 | 4,41 | 0,00 | 1,00E+00 |
| AT5G07095 | 1,59 | -0,55 | 1,38 | -0,40 | 6,92E-01 |
| AT5G07105 | 0,86 | 1,45 | 2,04 | 0,71 | 4,77E-01 |
| AT5G48545 | 0,30 | 0,58 | 3,64 | 0,16 | 8,74E-01 |
| AT5G48720 | 0,16 | -0,21 | 4,40 | -0,05 | 9,63E-01 |
| AT5G48730 | 0,12 | 0,00 | 4,41 | 0,00 | 1,00E+00 |
| AT5G48775 | 0,08 | 0,21 | 4,39 | 0,05 | 9,62E-01 |
| AT5G48800 | 0,90 | -0,38 | 3,00 | -0,13 | 8,99E-01 |
| AT5G48900 | 0,13 | -0,61 | 4,37 | -0,14 | 8,89E-01 |
| AT5G49010 | 1,17 | -0,84 | 1,78 | -0,47 | 6,37E-01 |
| AT5G49020 | 0,22 | -0,15 | 4,35 | -0,03 | 9,72E-01 |
| AT5G49060 | 0,17 | -0,29 | 4,40 | -0,07 | 9,47E-01 |
| AT5G49160 | 0,93 | 1,15 | 2,59 | 0,44 | 6,57E-01 |
| AT5G49215 | 0,08 | 0,21 | 4,39 | 0,05 | 9,62E-01 |
| AT5G49250 | 0,66 | 3,00 | 2,73 | 1,10 | 2,72E-01 |
| AT5G49260 | 0,08 | 0,21 | 4,39 | 0,05 | 9,62E-01 |
| AT5G49450 | 0,19 | -0,34 | 4,40 | -0,08 | 9,38E-01 |
| AT5G49460 | 0,62 | -0,35 | 3,08 | -0,11 | 9,10E-01 |
| AT5G49470 | 0,29 | -0,39 | 4,39 | -0,09 | 9,30E-01 |
| AT5G49480 | 1,15 | 0,46 | 2,40 | 0,19 | 8,49E-01 |
| AT5G49510 | 0,19 | 0,47 | 4,37 | 0,11 | 9,15E-01 |
| AT5G49520 | 0,12 | 0,27 | 4,38 | 0,06 | 9,50E-01 |
| AT5G49530 | 0,46 | 0,19 | 4,12 | 0,05 | 9,64E-01 |
| AT5G49590 | 0,86 | 0,24 | 2,66 | 0,09 | 9,28E-01 |
| AT5G49640 | 0,08 | 0,00 | 4,41 | 0,00 | 1,00E+00 |
| AT5G49650 | 0,61 | -0,95 | 2,30 | -0,41 | 6,78E-01 |
| AT5G49810 | 1,47 | -3,27 | 2,02 | -1,62 | 1,05E-01 |
| AT5G49840 | 0,10 | 0,24 | 4,38 | 0,05 | 9,57E-01 |
| AT5G49945 | 0,51 | -2,05 | 3,90 | -0,53 | 5,99E-01 |
| AT5G49970 | 0,29 | 0,49 | 4,36 | 0,11 | 9,11E-01 |
| AT5G50020 | 0,47 | 0,00 | 4,40 | 0,00 | 1,00E+00 |
| AT5G50160 | 1,06 | 1,17 | 2,29 | 0,51 | 6,09E-01 |
| AT5G50170 | 0,08 | 0,00 | 4,41 | 0,00 | 1,00E+00 |
| AT5G50200 | 0,08 | 0,21 | 4,39 | 0,05 | 9,62E-01 |
| AT5G07605 | 0,25 | 1,35 | 4,33 | 0,31 | 7,56E-01 |
| AT5G50310 | 0,21 | -0,29 | 4,40 | -0,06 | 9,48E-01 |
| AT5G50580 | 0,65 | 0,00 | 3,72 | 0,00 | 1,00E+00 |
| AT5G50680 | 1,09 | 2,06 | 2,36 | 0,87 | 3,83E-01 |
| AT5G50717 | 0,11 | 0,88 | 4,35 | 0,20 | 8,39E-01 |
| AT5G50780 | 0,59 | -2,43 | 4,38 | -0,55 | 5,80E-01 |
| AT5G50850 | 0,82 | -0,50 | 2,42 | -0,21 | 8,37E-01 |
| AT5G50950 | 1,52 | -0,27 | 1,24 | -0,22 | 8,27E-01 |
| AT5G50960 | 0,50 | -2,04 | 3,95 | -0,52 | 6,06E-01 |
| AT5G51010 | 0,08 | 0,00 | 4,41 | 0,00 | 1,00E+00 |
| AT5G51050 | 0,20 | -0,26 | 4,41 | -0,06 | 9,53E-01 |

| | | | | | |
|---|---|---|---|---|---|
| AT5G51055 | 0,34 | -0,04 | 3,43 | -0,01 | 9,91E-01 |
| AT5G51120 | 0,08 | 0,00 | 4,41 | 0,00 | 1,00E+00 |
| AT5G51150 | 0,28 | -0,36 | 4,39 | -0,08 | 9,35E-01 |
| AT5G51180 | 0,38 | 0,01 | 2,88 | 0,00 | 9,98E-01 |
| AT5G51200 | 1,04 | -0,78 | 2,22 | -0,35 | 7,24E-01 |
| AT5G51230 | 0,21 | -0,52 | 4,36 | -0,12 | 9,05E-01 |
| AT5G51280 | 0,48 | 1,58 | 2,64 | 0,60 | 5,51E-01 |
| AT5G51340 | 0,39 | -0,82 | 3,43 | -0,24 | 8,11E-01 |
| AT5G51450 | 0,53 | -2,86 | 3,94 | -0,73 | 4,68E-01 |
| AT5G51560 | 0,28 | -0,16 | 4,40 | -0,04 | 9,70E-01 |
| AT5G51570 | 0,40 | 0,62 | 3,36 | 0,18 | 8,55E-01 |
| AT5G51660 | 1,44 | 0,21 | 1,89 | 0,11 | 9,13E-01 |
| AT5G51670 | 0,12 | -0,26 | 4,41 | -0,06 | 9,53E-01 |
| AT5G51680 | 0,64 | 0,54 | 2,67 | 0,20 | 8,39E-01 |
| AT5G51750 | 0,75 | 0,17 | 3,59 | 0,05 | 9,63E-01 |
| AT5G51760 | 1,35 | -0,58 | 1,66 | -0,35 | 7,27E-01 |
| AT5G51830 | 0,27 | 1,20 | 4,33 | 0,28 | 7,82E-01 |
| AT5G51960 | 0,11 | 0,88 | 4,35 | 0,20 | 8,39E-01 |
| AT5G51970 | 1,04 | 2,56 | 1,82 | 1,40 | 1,61E-01 |
| AT5G52050 | 0,61 | 2,52 | 2,49 | 1,01 | 3,11E-01 |
| AT5G52115 | 1,12 | 0,75 | 2,39 | 0,31 | 7,54E-01 |
| AT5G52240 | 1,16 | -0,42 | 2,08 | -0,20 | 8,39E-01 |
| AT5G52280 | 0,34 | -0,03 | 4,37 | -0,01 | 9,94E-01 |
| AT5G52471 | 1,40 | 0,22 | 1,68 | 0,13 | 8,96E-01 |
| AT5G52510 | 0,69 | -2,08 | 3,06 | -0,68 | 4,95E-01 |
| AT5G52520 | 0,23 | 0,81 | 4,35 | 0,19 | 8,52E-01 |
| AT5G52580 | 0,50 | 1,63 | 3,22 | 0,51 | 6,12E-01 |
| AT5G52630 | 0,08 | 0,00 | 4,41 | 0,00 | 1,00E+00 |
| AT5G52700 | 0,14 | 1,29 | 4,33 | 0,30 | 7,65E-01 |
| AT5G52750 | 0,33 | 1,36 | 4,33 | 0,31 | 7,53E-01 |
| AT5G52760 | 0,52 | 2,26 | 2,56 | 0,88 | 3,77E-01 |
| AT5G52920 | 1,32 | -0,71 | 2,14 | -0,33 | 7,39E-01 |
| AT5G07945 | 0,52 | 2,11 | 3,92 | 0,54 | 5,91E-01 |
| AT5G52940 | 0,11 | -0,38 | 4,39 | -0,09 | 9,31E-01 |
| AT5G53020 | 0,27 | -0,06 | 4,34 | -0,01 | 9,89E-01 |
| AT5G53030 | 0,24 | -0,67 | 4,38 | -0,15 | 8,78E-01 |
| AT5G53048 | 0,25 | -0,53 | 4,37 | -0,12 | 9,03E-01 |
| AT5G53060 | 0,48 | 0,43 | 4,01 | 0,11 | 9,14E-01 |
| AT5G53080 | 0,12 | 0,00 | 4,41 | 0,00 | 1,00E+00 |
| AT5G53110 | 0,31 | -3,20 | 4,34 | -0,74 | 4,60E-01 |
| AT5G53120 | 0,82 | 1,47 | 2,57 | 0,57 | 5,68E-01 |
| AT5G53140 | 1,45 | 0,63 | 1,91 | 0,33 | 7,42E-01 |
| AT5G53150 | 0,95 | -0,38 | 2,28 | -0,17 | 8,69E-01 |
| AT5G53160 | 1,14 | 2,90 | 1,86 | 1,56 | 1,18E-01 |
| AT5G53220 | 0,48 | 0,77 | 3,14 | 0,24 | 8,07E-01 |
| AT5G53280 | 0,44 | -0,36 | 4,18 | -0,09 | 9,31E-01 |
| AT5G53350 | 0,54 | -2,06 | 3,15 | -0,66 | 5,12E-01 |
| AT5G53360 | 0,08 | 0,00 | 4,41 | 0,00 | 1,00E+00 |
| AT5G53410 | 1,13 | -1,29 | 1,64 | -0,79 | 4,30E-01 |
| AT5G53420 | 0,48 | 0,52 | 3,32 | 0,16 | 8,75E-01 |
| AT5G53430 | 1,00 | -0,69 | 1,98 | -0,35 | 7,28E-01 |
| AT5G53450 | 0,71 | -0,35 | 2,63 | -0,13 | 8,93E-01 |
| AT5G53550 | 0,99 | 0,80 | 2,38 | 0,34 | 7,35E-01 |
| AT5G53560 | 1,33 | 1,19 | 1,70 | 0,70 | 4,83E-01 |
| AT5G53590 | 0,48 | -2,36 | 4,06 | -0,58 | 5,62E-01 |
| AT5G53620 | 1,09 | -0,45 | 1,95 | -0,23 | 8,17E-01 |

| | | | | | |
|---|---|---|---|---|---|
| AT5G53860 | 0,25 | 1,59 | 3,76 | 0,42 | 6,73E-01 |
| AT5G53890 | 0,71 | 0,11 | 2,49 | 0,04 | 9,66E-01 |
| AT5G54035 | 0,15 | 1,42 | 4,32 | 0,33 | 7,43E-01 |
| AT5G54043 | 0,22 | 1,46 | 4,32 | 0,34 | 7,36E-01 |
| AT5G54080 | 0,99 | 0,50 | 2,30 | 0,22 | 8,27E-01 |
| AT5G54110 | 0,33 | 0,00 | 4,41 | 0,00 | 1,00E+00 |
| AT5G54140 | 0,24 | 0,00 | 4,41 | 0,00 | 1,00E+00 |
| AT5G54145 | 0,12 | 1,17 | 4,34 | 0,27 | 7,88E-01 |
| AT5G54160 | 0,46 | -1,65 | 4,09 | -0,40 | 6,86E-01 |
| AT5G54180 | 0,12 | 0,59 | 4,36 | 0,14 | 8,92E-01 |
| AT5G54290 | 0,66 | 0,17 | 2,64 | 0,06 | 9,49E-01 |
| AT5G54500 | 0,31 | -1,02 | 4,32 | -0,24 | 8,14E-01 |
| AT5G54540 | 0,58 | -0,13 | 2,69 | -0,05 | 9,60E-01 |
| AT5G54600 | 0,87 | -1,08 | 2,06 | -0,53 | 6,00E-01 |
| AT5G08225 | 0,36 | 0,22 | 4,39 | 0,05 | 9,60E-01 |
| AT5G54640 | 0,05 | 0,18 | 4,38 | 0,04 | 9,68E-01 |
| AT5G54680 | 0,72 | 0,31 | 2,50 | 0,12 | 9,02E-01 |
| AT5G54750 | 0,55 | -0,83 | 3,92 | -0,21 | 8,33E-01 |
| AT5G54760 | 0,49 | -0,50 | 2,63 | -0,19 | 8,49E-01 |
| AT5G54800 | 0,08 | 0,47 | 4,38 | 0,11 | 9,15E-01 |
| AT5G54830 | 0,76 | -0,36 | 2,88 | -0,12 | 9,01E-01 |
| AT5G54855 | 0,18 | -2,46 | 4,37 | -0,56 | 5,73E-01 |
| AT5G54860 | 0,26 | 0,00 | 4,41 | 0,00 | 1,00E+00 |
| AT5G54900 | 0,82 | -1,06 | 2,51 | -0,42 | 6,71E-01 |
| AT5G54920 | 0,83 | 0,00 | 4,39 | 0,00 | 1,00E+00 |
| AT5G55060 | 0,14 | 0,81 | 4,35 | 0,19 | 8,52E-01 |
| AT5G55070 | 1,01 | 0,06 | 2,02 | 0,03 | 9,75E-01 |
| AT5G55190 | 0,80 | -0,31 | 2,40 | -0,13 | 8,98E-01 |
| AT5G55220 | 0,88 | -1,68 | 2,45 | -0,68 | 4,94E-01 |
| AT5G55230 | 0,95 | 0,92 | 2,04 | 0,45 | 6,51E-01 |
| AT5G55450 | 0,62 | -1,45 | 3,09 | -0,47 | 6,39E-01 |
| AT5G55460 | 0,37 | 0,18 | 3,40 | 0,05 | 9,57E-01 |
| AT5G55500 | 0,57 | -0,34 | 3,19 | -0,11 | 9,15E-01 |
| AT5G55530 | 0,30 | 0,08 | 4,38 | 0,02 | 9,86E-01 |
| AT5G55580 | 0,70 | -1,57 | 3,90 | -0,40 | 6,88E-01 |
| AT5G55620 | 0,43 | 0,96 | 4,21 | 0,23 | 8,19E-01 |
| AT5G55630 | 0,12 | -0,26 | 4,41 | -0,06 | 9,53E-01 |
| AT5G55660 | 0,90 | -1,49 | 2,06 | -0,73 | 4,68E-01 |
| AT5G55840 | 0,22 | 0,83 | 4,35 | 0,19 | 8,49E-01 |
| AT5G55850 | 0,24 | 1,18 | 4,34 | 0,27 | 7,85E-01 |
| AT5G55900 | 0,10 | 0,86 | 4,35 | 0,20 | 8,43E-01 |
| AT5G55930 | 0,70 | 0,04 | 3,00 | 0,01 | 9,88E-01 |
| AT5G55960 | 0,59 | -1,04 | 2,98 | -0,35 | 7,27E-01 |
| AT5G55990 | 0,33 | 0,06 | 4,38 | 0,01 | 9,89E-01 |
| AT5G56020 | 0,32 | 0,00 | 4,41 | 0,00 | 1,00E+00 |
| AT5G56040 | 0,73 | -0,11 | 2,45 | -0,05 | 9,63E-01 |
| AT5G56060 | 0,67 | 0,18 | 4,34 | 0,04 | 9,67E-01 |
| AT5G56140 | 0,98 | 0,71 | 1,99 | 0,36 | 7,22E-01 |
| AT5G56160 | 0,49 | -3,54 | 4,34 | -0,82 | 4,15E-01 |
| AT5G56170 | 0,53 | 0,71 | 4,33 | 0,16 | 8,70E-01 |
| AT5G56270 | 0,76 | 0,00 | 2,94 | 0,00 | 1,00E+00 |
| AT5G56290 | 0,52 | 0,19 | 2,36 | 0,08 | 9,34E-01 |
| AT5G56340 | 0,44 | -2,15 | 3,36 | -0,64 | 5,22E-01 |
| AT5G08495 | 0,43 | 0,26 | 4,13 | 0,06 | 9,49E-01 |
| AT5G56460 | 0,74 | -2,44 | 3,87 | -0,63 | 5,29E-01 |
| AT5G56500 | 0,32 | -0,18 | 4,40 | -0,04 | 9,67E-01 |

| | | | | | |
|---|---|---|---|---|---|
| AT5G56590 | 0,24 | 0,14 | 4,38 | 0,03 | 9,74E-01 |
| AT5G56630 | 0,71 | -0,59 | 2,59 | -0,23 | 8,20E-01 |
| AT5G56650 | 0,64 | 1,26 | 2,32 | 0,54 | 5,87E-01 |
| AT5G56680 | 1,22 | -1,67 | 1,92 | -0,87 | 3,82E-01 |
| AT5G56720 | 0,12 | -0,36 | 4,39 | -0,08 | 9,34E-01 |
| AT5G56870 | 0,19 | 0,84 | 4,35 | 0,19 | 8,47E-01 |
| AT5G56910 | 0,15 | -0,35 | 4,39 | -0,08 | 9,36E-01 |
| AT5G56920 | 0,65 | 2,72 | 2,97 | 0,92 | 3,59E-01 |
| AT5G56940 | 0,08 | 0,47 | 4,38 | 0,11 | 9,15E-01 |
| AT5G56950 | 0,44 | -2,86 | 3,28 | -0,87 | 3,82E-01 |
| AT5G57015 | 0,32 | 1,20 | 4,33 | 0,28 | 7,82E-01 |
| AT5G57020 | 1,25 | -0,23 | 1,74 | -0,13 | 8,96E-01 |
| AT5G57030 | 0,86 | -2,95 | 2,43 | -1,21 | 2,25E-01 |
| AT5G57035 | 1,04 | -1,55 | 2,65 | -0,58 | 5,59E-01 |
| AT5G57070 | 0,42 | -0,15 | 4,27 | -0,04 | 9,71E-01 |
| AT5G57150 | 0,25 | 0,00 | 4,41 | 0,00 | 1,00E+00 |
| AT5G57180 | 1,05 | -1,44 | 1,76 | -0,82 | 4,12E-01 |
| AT5G57320 | 0,08 | 0,00 | 4,41 | 0,00 | 1,00E+00 |
| AT5G57330 | 0,49 | -0,27 | 2,81 | -0,10 | 9,22E-01 |
| AT5G57340 | 0,20 | 1,44 | 4,33 | 0,33 | 7,40E-01 |
| AT5G57360 | 0,96 | 0,60 | 1,93 | 0,31 | 7,55E-01 |
| AT5G57370 | 0,31 | 0,19 | 4,38 | 0,04 | 9,65E-01 |
| AT5G57460 | 1,07 | 0,20 | 1,90 | 0,11 | 9,14E-01 |
| AT5G57520 | 0,31 | 2,33 | 4,29 | 0,54 | 5,87E-01 |
| AT5G57610 | 0,89 | -0,22 | 1,91 | -0,12 | 9,07E-01 |
| AT5G57655 | 1,67 | 0,39 | 1,57 | 0,24 | 8,07E-01 |
| AT5G57735 | 0,19 | 1,30 | 4,33 | 0,30 | 7,64E-01 |
| AT5G57800 | 0,61 | 0,17 | 2,47 | 0,07 | 9,45E-01 |
| AT5G08805 | 1,03 | 2,11 | 2,37 | 0,89 | 3,73E-01 |
| AT5G57880 | 0,48 | -2,09 | 4,12 | -0,51 | 6,11E-01 |
| AT5G57930 | 1,66 | -2,21 | 2,13 | -1,04 | 2,99E-01 |
| AT5G57940 | 0,35 | -0,35 | 3,57 | -0,10 | 9,23E-01 |
| AT5G58000 | 0,08 | 0,00 | 4,41 | 0,00 | 1,00E+00 |
| AT5G58020 | 0,39 | -3,94 | 4,32 | -0,91 | 3,62E-01 |
| AT5G58060 | 0,59 | 0,78 | 2,59 | 0,30 | 7,63E-01 |
| AT5G58100 | 0,97 | 0,38 | 1,77 | 0,21 | 8,31E-01 |
| AT5G58110 | 0,27 | 0,14 | 4,33 | 0,03 | 9,75E-01 |
| AT5G58120 | 1,53 | -0,18 | 1,38 | -0,13 | 8,93E-01 |
| AT5G08820 | 0,19 | -0,40 | 4,39 | -0,09 | 9,28E-01 |
| AT5G58160 | 1,77 | 0,00 | 2,67 | 0,00 | 1,00E+00 |
| AT5G58200 | 0,40 | -2,42 | 4,36 | -0,55 | 5,79E-01 |
| AT5G58230 | 0,77 | 0,24 | 2,55 | 0,09 | 9,26E-01 |
| AT5G58250 | 0,32 | 0,06 | 4,33 | 0,01 | 9,88E-01 |
| AT5G58270 | 0,43 | -1,00 | 4,24 | -0,24 | 8,13E-01 |
| AT5G58290 | 1,43 | -0,43 | 1,57 | -0,28 | 7,82E-01 |
| AT5G58300 | 0,99 | -0,36 | 2,41 | -0,15 | 8,81E-01 |
| AT5G58320 | 0,24 | 0,00 | 4,41 | 0,00 | 1,00E+00 |
| AT5G58350 | 1,12 | 0,96 | 2,07 | 0,46 | 6,42E-01 |
| AT5G58370 | 0,29 | -0,38 | 4,39 | -0,09 | 9,30E-01 |
| AT5G58375 | 0,48 | 0,08 | 4,38 | 0,02 | 9,86E-01 |
| AT5G58420 | 1,59 | -0,66 | 1,81 | -0,36 | 7,16E-01 |
| AT5G58450 | 0,68 | -0,36 | 3,00 | -0,12 | 9,05E-01 |
| AT5G58495 | 0,12 | 0,00 | 4,41 | 0,00 | 1,00E+00 |
| AT5G58520 | 0,39 | 0,88 | 4,35 | 0,20 | 8,40E-01 |
| AT5G58540 | 1,83 | -0,35 | 2,02 | -0,17 | 8,62E-01 |
| AT5G58560 | 0,47 | 0,94 | 3,24 | 0,29 | 7,73E-01 |

| | | | | | |
|---|---|---|---|---|---|
| AT5G08855 | 0,43 | 1,05 | 4,15 | 0,25 | 8,01E-01 |
| AT5G58620 | 0,34 | 0,00 | 4,41 | 0,00 | 1,00E+00 |
| AT5G58640 | 0,71 | -1,07 | 3,02 | -0,36 | 7,22E-01 |
| AT5G58710 | 0,66 | -1,79 | 3,03 | -0,59 | 5,54E-01 |
| AT5G58990 | 0,29 | -0,15 | 4,35 | -0,03 | 9,72E-01 |
| AT5G59160 | 0,32 | -1,28 | 4,38 | -0,29 | 7,70E-01 |
| AT5G59170 | 0,26 | 0,00 | 4,41 | 0,00 | 1,00E+00 |
| AT5G59250 | 0,50 | -2,07 | 3,22 | -0,64 | 5,20E-01 |
| AT5G59350 | 0,35 | -0,40 | 4,38 | -0,09 | 9,27E-01 |
| AT5G59420 | 1,43 | -0,18 | 1,87 | -0,09 | 9,24E-01 |
| AT5G59430 | 0,64 | -0,61 | 2,52 | -0,24 | 8,09E-01 |
| AT5G59450 | 0,61 | 0,63 | 3,87 | 0,16 | 8,70E-01 |
| AT5G59610 | 0,51 | -0,28 | 4,05 | -0,07 | 9,45E-01 |
| AT5G59660 | 0,05 | 0,18 | 4,38 | 0,04 | 9,68E-01 |
| AT5G59670 | 0,28 | 0,72 | 3,61 | 0,20 | 8,42E-01 |
| AT5G59680 | 0,64 | 0,00 | 4,41 | 0,00 | 1,00E+00 |
| AT5G59690 | 0,64 | 1,11 | 2,95 | 0,37 | 7,08E-01 |
| AT5G59820 | 0,42 | -0,19 | 3,30 | -0,06 | 9,55E-01 |
| AT5G59830 | 0,85 | -2,36 | 2,98 | -0,79 | 4,29E-01 |
| AT5G59860 | 0,43 | 0,00 | 4,41 | 0,00 | 1,00E+00 |
| AT5G59880 | 0,87 | -1,37 | 2,56 | -0,53 | 5,93E-01 |
| AT5G59910 | 0,91 | 3,21 | 2,15 | 1,50 | 1,35E-01 |
| AT5G59920 | 0,58 | 0,00 | 3,80 | 0,00 | 1,00E+00 |
| AT5G59950 | 1,16 | -0,83 | 1,93 | -0,43 | 6,68E-01 |
| AT5G59980 | 0,39 | 0,00 | 4,40 | 0,00 | 1,00E+00 |
| AT5G60040 | 0,79 | -0,25 | 2,54 | -0,10 | 9,23E-01 |
| AT5G60110 | 0,25 | 0,00 | 4,41 | 0,00 | 1,00E+00 |
| AT5G60120 | 0,67 | -0,87 | 2,85 | -0,30 | 7,61E-01 |
| AT5G60170 | 1,68 | -2,22 | 1,94 | -1,15 | 2,52E-01 |
| AT5G60270 | 0,48 | 0,00 | 4,41 | 0,00 | 1,00E+00 |
| AT5G60280 | 0,76 | 0,35 | 3,56 | 0,10 | 9,21E-01 |
| AT5G60300 | 0,52 | -0,27 | 3,96 | -0,07 | 9,46E-01 |
| AT5G60320 | 0,25 | 0,00 | 4,41 | 0,00 | 1,00E+00 |
| AT5G60550 | 0,49 | -0,94 | 3,24 | -0,29 | 7,73E-01 |
| AT5G60560 | 0,60 | 2,38 | 3,01 | 0,79 | 4,29E-01 |
| AT5G60570 | 0,88 | 2,19 | 2,11 | 1,04 | 2,99E-01 |
| AT5G60610 | 0,72 | 0,00 | 3,66 | 0,00 | 1,00E+00 |
| AT5G60620 | 0,40 | -0,45 | 3,51 | -0,13 | 8,98E-01 |
| AT5G60680 | 1,59 | 2,18 | 1,88 | 1,16 | 2,44E-01 |
| AT5G60780 | 1,02 | -0,31 | 2,35 | -0,13 | 8,95E-01 |
| AT5G60820 | 0,45 | -0,18 | 4,17 | -0,04 | 9,65E-01 |
| AT5G60850 | 1,63 | -0,97 | 1,93 | -0,50 | 6,16E-01 |
| AT5G60890 | 0,24 | 0,00 | 4,41 | 0,00 | 1,00E+00 |
| AT5G60945 | 1,24 | 0,17 | 1,49 | 0,11 | 9,10E-01 |
| AT5G60963 | 0,50 | -0,35 | 2,70 | -0,13 | 8,96E-01 |
| AT5G60964 | 0,93 | 1,30 | 3,31 | 0,39 | 6,94E-01 |
| AT5G60980 | 1,13 | 0,51 | 2,24 | 0,23 | 8,19E-01 |
| AT5G61190 | 0,65 | -2,92 | 2,55 | -1,14 | 2,53E-01 |
| AT5G61200 | 0,25 | 0,00 | 4,41 | 0,00 | 1,00E+00 |
| AT5G61210 | 1,18 | 0,85 | 1,87 | 0,46 | 6,49E-01 |
| AT5G61230 | 1,00 | -2,02 | 2,06 | -0,98 | 3,26E-01 |
| AT5G61240 | 0,29 | -1,28 | 4,35 | -0,30 | 7,68E-01 |
| AT5G61380 | 0,78 | -1,32 | 2,52 | -0,53 | 5,99E-01 |
| AT5G61400 | 0,78 | 0,00 | 3,52 | 0,00 | 1,00E+00 |
| AT5G61470 | 0,39 | 2,44 | 3,28 | 0,74 | 4,57E-01 |
| AT5G61490 | 0,17 | 0,00 | 4,41 | 0,00 | 1,00E+00 |

| | | | | | |
|---|---|---|---|---|---|
| AT5G61550 | 0,85 | 0,00 | 3,43 | 0,00 | 1,00E+00 |
| AT5G61570 | 0,19 | -0,02 | 4,35 | -0,01 | 9,96E-01 |
| AT5G61580 | 0,26 | 0,03 | 4,38 | 0,01 | 9,95E-01 |
| AT5G61950 | 0,32 | 0,00 | 4,41 | 0,00 | 1,00E+00 |
| AT5G61970 | 0,49 | 0,49 | 3,34 | 0,15 | 8,83E-01 |
| AT5G61980 | 0,55 | 0,00 | 4,41 | 0,00 | 1,00E+00 |
| AT5G62030 | 0,41 | 0,86 | 4,16 | 0,21 | 8,36E-01 |
| AT5G62270 | 0,20 | 0,87 | 4,35 | 0,20 | 8,41E-01 |
| AT5G62290 | 0,13 | 0,94 | 4,35 | 0,22 | 8,29E-01 |
| AT5G62360 | 0,27 | 0,84 | 4,35 | 0,19 | 8,47E-01 |
| AT5G62460 | 0,44 | -0,30 | 4,22 | -0,07 | 9,43E-01 |
| AT5G62500 | 0,05 | 0,18 | 4,38 | 0,04 | 9,68E-01 |
| AT5G62520 | 0,28 | -2,10 | 4,40 | -0,48 | 6,33E-01 |
| AT5G62560 | 0,37 | 0,32 | 3,47 | 0,09 | 9,26E-01 |
| AT5G62570 | 0,48 | 1,58 | 2,65 | 0,60 | 5,51E-01 |
| AT5G62575 | 0,21 | 0,00 | 4,41 | 0,00 | 1,00E+00 |
| AT5G62620 | 0,35 | 1,06 | 4,34 | 0,24 | 8,08E-01 |
| AT5G62670 | 1,85 | -1,33 | 1,91 | -0,70 | 4,87E-01 |
| AT5G62710 | 0,23 | -0,41 | 4,38 | -0,09 | 9,26E-01 |
| AT5G62720 | 0,31 | 0,55 | 4,38 | 0,13 | 8,99E-01 |
| AT5G62740 | 0,43 | -1,66 | 4,13 | -0,40 | 6,87E-01 |
| AT5G62770 | 0,31 | -0,41 | 4,38 | -0,09 | 9,25E-01 |
| AT5G62890 | 1,11 | 0,58 | 2,11 | 0,28 | 7,83E-01 |
| AT5G62910 | 0,36 | -1,25 | 4,40 | -0,28 | 7,77E-01 |
| AT5G62970 | 0,35 | 0,52 | 3,50 | 0,15 | 8,83E-01 |
| AT5G63010 | 0,41 | 0,47 | 3,35 | 0,14 | 8,88E-01 |
| AT5G09120 | 1,28 | -0,02 | 1,63 | -0,01 | 9,89E-01 |
| AT5G63050 | 0,55 | -1,36 | 4,07 | -0,33 | 7,39E-01 |
| AT5G63130 | 0,64 | -1,83 | 3,06 | -0,60 | 5,49E-01 |
| AT5G63200 | 0,75 | 0,36 | 3,56 | 0,10 | 9,20E-01 |
| AT5G63260 | 0,08 | 0,00 | 4,41 | 0,00 | 1,00E+00 |
| AT5G63280 | 0,12 | 0,00 | 4,41 | 0,00 | 1,00E+00 |
| AT5G09150 | 0,24 | 0,00 | 4,41 | 0,00 | 1,00E+00 |
| AT5G63300 | 0,47 | 0,00 | 4,41 | 0,00 | 1,00E+00 |
| AT5G63340 | 0,26 | 1,70 | 4,31 | 0,39 | 6,94E-01 |
| AT5G63420 | 0,69 | -2,49 | 3,11 | -0,80 | 4,23E-01 |
| AT5G63480 | 0,12 | 1,17 | 4,34 | 0,27 | 7,88E-01 |
| AT5G63510 | 0,37 | -0,24 | 4,35 | -0,06 | 9,55E-01 |
| AT5G63570 | 1,81 | -2,71 | 1,69 | -1,60 | 1,10E-01 |
| AT5G63640 | 0,36 | -2,10 | 4,40 | -0,48 | 6,33E-01 |
| AT5G63680 | 0,63 | 0,50 | 3,19 | 0,16 | 8,76E-01 |
| AT5G63770 | 0,64 | -0,50 | 3,02 | -0,17 | 8,69E-01 |
| AT5G63780 | 0,69 | 0,51 | 2,20 | 0,23 | 8,17E-01 |
| AT5G63810 | 0,68 | -2,05 | 3,07 | -0,67 | 5,04E-01 |
| AT5G63940 | 0,43 | 1,97 | 3,25 | 0,61 | 5,45E-01 |
| AT5G63960 | 0,45 | -0,26 | 4,41 | -0,06 | 9,53E-01 |
| AT5G63980 | 0,08 | 0,00 | 4,41 | 0,00 | 1,00E+00 |
| AT5G64030 | 1,58 | -2,56 | 1,52 | -1,69 | 9,09E-02 |
| AT5G64090 | 0,72 | -0,33 | 2,57 | -0,13 | 8,98E-01 |
| AT5G64160 | 0,20 | -1,20 | 4,38 | -0,27 | 7,85E-01 |
| AT5G64190 | 0,08 | 0,00 | 4,41 | 0,00 | 1,00E+00 |
| AT5G64240 | 0,28 | 0,60 | 4,37 | 0,14 | 8,91E-01 |
| AT5G64300 | 1,43 | 0,03 | 1,57 | 0,02 | 9,85E-01 |
| AT5G64310 | 0,80 | 2,35 | 2,72 | 0,86 | 3,87E-01 |
| AT5G64330 | 1,11 | -2,01 | 1,88 | -1,07 | 2,85E-01 |
| AT5G64370 | 0,51 | -0,39 | 3,98 | -0,10 | 9,22E-01 |

| | | | | | |
|---|---|---|---|---|---|
| AT5G64380 | 0,10 | 0,08 | 4,38 | 0,02 | 9,85E-01 |
| AT5G64400 | 0,52 | -1,58 | 4,14 | -0,38 | 7,03E-01 |
| AT5G64440 | 0,42 | 1,07 | 3,29 | 0,32 | 7,45E-01 |
| AT5G64500 | 0,82 | -1,81 | 2,23 | -0,81 | 4,16E-01 |
| AT5G64505 | 0,26 | 0,00 | 4,41 | 0,00 | 1,00E+00 |
| AT5G64572 | 1,22 | 1,89 | 1,91 | 0,99 | 3,23E-01 |
| AT5G64690 | 0,11 | 0,88 | 4,35 | 0,20 | 8,39E-01 |
| AT5G64750 | 0,12 | 0,00 | 4,41 | 0,00 | 1,00E+00 |
| AT5G64813 | 0,60 | 0,40 | 2,63 | 0,15 | 8,79E-01 |
| AT5G09285 | 0,45 | 0,00 | 4,07 | 0,00 | 1,00E+00 |
| AT5G64950 | 0,48 | 0,00 | 4,41 | 0,00 | 1,00E+00 |
| AT5G64960 | 0,59 | -1,16 | 3,22 | -0,36 | 7,19E-01 |
| AT5G65010 | 1,00 | -1,85 | 1,76 | -1,05 | 2,94E-01 |
| AT5G65015 | 0,39 | -0,33 | 4,40 | -0,08 | 9,40E-01 |
| AT5G65210 | 0,26 | 0,00 | 4,41 | 0,00 | 1,00E+00 |
| AT5G65220 | 1,17 | -1,95 | 1,96 | -0,99 | 3,21E-01 |
| AT5G65230 | 0,08 | 0,00 | 4,41 | 0,00 | 1,00E+00 |
| AT5G65270 | 0,10 | 0,08 | 4,38 | 0,02 | 9,85E-01 |
| AT5G65290 | 0,47 | 0,00 | 4,40 | 0,00 | 1,00E+00 |
| AT5G65400 | 0,17 | -2,21 | 4,40 | -0,50 | 6,16E-01 |
| AT5G65440 | 0,58 | 0,49 | 3,08 | 0,16 | 8,73E-01 |
| AT5G65470 | 0,30 | 1,16 | 4,33 | 0,27 | 7,90E-01 |
| AT5G65510 | 0,12 | 0,11 | 4,38 | 0,03 | 9,80E-01 |
| AT5G65570 | 0,25 | 0,00 | 4,41 | 0,00 | 1,00E+00 |
| AT5G65620 | 1,40 | 1,53 | 1,81 | 0,85 | 3,98E-01 |
| AT5G65640 | 0,08 | 0,00 | 4,41 | 0,00 | 1,00E+00 |
| AT5G65670 | 0,91 | -0,65 | 2,01 | -0,32 | 7,47E-01 |
| AT5G65710 | 0,77 | 1,93 | 2,12 | 0,91 | 3,63E-01 |
| AT5G65720 | 1,65 | -2,45 | 1,54 | -1,59 | 1,11E-01 |
| AT5G65760 | 0,55 | 1,87 | 2,60 | 0,72 | 4,71E-01 |
| AT5G65770 | 1,30 | -0,48 | 1,87 | -0,26 | 7,98E-01 |
| AT5G65780 | 1,41 | -0,86 | 1,79 | -0,48 | 6,29E-01 |
| AT5G65890 | 0,05 | 0,18 | 4,38 | 0,04 | 9,68E-01 |
| AT5G65960 | 1,37 | -1,29 | 2,17 | -0,59 | 5,53E-01 |
| AT5G65990 | 0,70 | -0,43 | 1,99 | -0,22 | 8,27E-01 |
| AT5G66052 | 0,53 | -0,34 | 4,08 | -0,08 | 9,34E-01 |
| AT5G66070 | 0,10 | 0,08 | 4,38 | 0,02 | 9,85E-01 |
| AT5G66100 | 0,24 | 0,79 | 4,36 | 0,18 | 8,56E-01 |
| AT5G66110 | 0,30 | 0,00 | 4,41 | 0,00 | 1,00E+00 |
| AT5G66160 | 0,31 | 0,51 | 4,38 | 0,12 | 9,07E-01 |
| AT5G66170 | 0,33 | 0,00 | 4,41 | 0,00 | 1,00E+00 |
| AT5G66240 | 1,02 | 0,90 | 2,36 | 0,38 | 7,03E-01 |
| AT5G66360 | 0,60 | 0,00 | 3,87 | 0,00 | 1,00E+00 |
| AT5G66450 | 0,53 | 1,36 | 3,16 | 0,43 | 6,67E-01 |
| AT5G66560 | 1,38 | 0,20 | 1,86 | 0,11 | 9,14E-01 |
| AT5G66562 | 1,20 | 0,75 | 1,63 | 0,46 | 6,44E-01 |
| AT5G66564 | 1,37 | 1,16 | 1,44 | 0,80 | 4,23E-01 |
| AT5G66568 | 0,20 | 1,10 | 4,35 | 0,25 | 8,00E-01 |
| AT5G66760 | 1,08 | -2,07 | 2,04 | -1,01 | 3,10E-01 |
| AT5G66770 | 0,43 | -0,86 | 4,37 | -0,20 | 8,43E-01 |
| AT5G66790 | 0,74 | -2,54 | 2,19 | -1,16 | 2,45E-01 |
| AT5G66810 | 0,67 | -0,29 | 2,64 | -0,11 | 9,13E-01 |
| AT5G66930 | 0,12 | -0,36 | 4,39 | -0,08 | 9,34E-01 |
| AT5G66950 | 1,10 | -0,84 | 1,52 | -0,55 | 5,83E-01 |
| AT5G67070 | 0,51 | 0,63 | 3,89 | 0,16 | 8,71E-01 |
| AT5G67080 | 0,30 | 0,31 | 4,39 | 0,07 | 9,44E-01 |

| | | | | | |
|---|---|---|---|---|---|
| AT5G67170 | 0,08 | 0,00 | 4,41 | 0,00 | 1,00E+00 |
| AT5G67260 | 0,65 | 1,88 | 3,90 | 0,48 | 6,29E-01 |
| AT5G67320 | 0,96 | -1,12 | 2,72 | -0,41 | 6,81E-01 |
| AT5G67330 | 1,26 | -0,18 | 1,92 | -0,10 | 9,24E-01 |
| AT5G67340 | 0,24 | -0,26 | 4,41 | -0,06 | 9,53E-01 |
| AT5G67385 | 0,80 | 1,34 | 2,18 | 0,62 | 5,38E-01 |
| AT5G67455 | 0,10 | 0,08 | 4,38 | 0,02 | 9,85E-01 |
| AT5G67470 | 1,09 | 1,76 | 2,09 | 0,84 | 4,01E-01 |
| AT5G67488 | 1,20 | 0,62 | 1,85 | 0,33 | 7,38E-01 |
| AT5G67500 | 0,21 | -0,35 | 4,40 | -0,08 | 9,37E-01 |
| AT5G67530 | 0,08 | 0,00 | 4,41 | 0,00 | 1,00E+00 |
| AT5G67560 | 0,70 | 0,30 | 3,70 | 0,08 | 9,36E-01 |
| AT5G67570 | 0,48 | 0,70 | 4,16 | 0,17 | 8,67E-01 |
| AT5G67590 | 0,32 | -0,13 | 4,35 | -0,03 | 9,77E-01 |
| AT1G01020 | 0,15 | 0,82 | 4,35 | 0,19 | 8,50E-01 |
| AT1G03993 | 0,11 | -0,26 | 4,41 | -0,06 | 9,53E-01 |
| AT1G01050 | 0,66 | 0,00 | 3,04 | 0,00 | 1,00E+00 |
| AT1G01060 | 0,69 | -0,35 | 2,57 | -0,14 | 8,91E-01 |
| AT1G01090 | 1,61 | -0,71 | 1,56 | -0,46 | 6,48E-01 |
| AT1G01100 | 1,47 | -1,32 | 1,55 | -0,85 | 3,93E-01 |
| AT1G01120 | 1,43 | 0,81 | 1,44 | 0,57 | 5,72E-01 |
| AT1G01130 | 0,29 | 0,54 | 4,37 | 0,12 | 9,01E-01 |
| AT1G01140 | 0,54 | 2,33 | 2,51 | 0,93 | 3,54E-01 |
| AT1G01170 | 0,58 | 2,01 | 3,01 | 0,67 | 5,05E-01 |
| AT1G01440 | 0,35 | 0,64 | 4,37 | 0,15 | 8,83E-01 |
| AT1G01450 | 0,59 | 0,00 | 4,40 | 0,00 | 1,00E+00 |
| AT1G01470 | 1,29 | -0,01 | 1,41 | -0,01 | 9,93E-01 |
| AT1G01490 | 0,15 | -0,57 | 4,37 | -0,13 | 8,96E-01 |
| AT1G01530 | 0,08 | 0,47 | 4,38 | 0,11 | 9,15E-01 |
| AT1G01610 | 0,73 | -1,76 | 2,28 | -0,77 | 4,39E-01 |
| AT1G01620 | 1,23 | -0,50 | 1,86 | -0,27 | 7,87E-01 |
| AT1G01650 | 0,59 | 1,90 | 2,89 | 0,66 | 5,12E-01 |
| AT1G04027 | 0,46 | 1,89 | 3,10 | 0,61 | 5,42E-01 |
| AT1G01680 | 0,72 | 1,07 | 2,54 | 0,42 | 6,74E-01 |
| AT1G01725 | 1,09 | 0,15 | 1,84 | 0,08 | 9,33E-01 |
| AT1G04063 | 0,34 | -0,03 | 4,31 | -0,01 | 9,95E-01 |
| AT1G01820 | 0,86 | -1,13 | 2,22 | -0,51 | 6,12E-01 |
| AT1G01830 | 1,22 | -0,03 | 2,00 | -0,01 | 9,89E-01 |
| AT1G01880 | 1,08 | -4,13 | 2,24 | -1,84 | 6,52E-02 |
| AT1G01890 | 1,04 | -4,05 | 2,59 | -1,57 | 1,17E-01 |
| AT1G01930 | 0,16 | -0,26 | 4,41 | -0,06 | 9,53E-01 |
| AT1G01990 | 0,21 | -0,32 | 4,40 | -0,07 | 9,43E-01 |
| AT1G02020 | 0,45 | 0,00 | 4,14 | 0,00 | 1,00E+00 |
| AT1G02090 | 0,84 | -0,11 | 2,89 | -0,04 | 9,70E-01 |
| AT1G02140 | 0,21 | -0,95 | 4,38 | -0,22 | 8,29E-01 |
| AT1G02260 | 0,54 | -0,26 | 3,30 | -0,08 | 9,38E-01 |
| AT1G02280 | 0,61 | -1,28 | 3,13 | -0,41 | 6,83E-01 |
| AT1G02310 | 0,21 | 0,00 | 4,41 | 0,00 | 1,00E+00 |
| AT1G04117 | 0,17 | -0,29 | 4,40 | -0,07 | 9,47E-01 |
| AT1G02430 | 0,17 | -0,33 | 4,40 | -0,08 | 9,40E-01 |
| AT1G02450 | 0,20 | -2,08 | 4,39 | -0,47 | 6,35E-01 |
| AT1G02610 | 0,33 | -0,31 | 4,35 | -0,07 | 9,44E-01 |
| AT1G02660 | 0,36 | 0,23 | 4,37 | 0,05 | 9,57E-01 |
| AT1G02770 | 0,41 | 0,00 | 4,41 | 0,00 | 1,00E+00 |
| AT1G02780 | 1,83 | -0,13 | 1,20 | -0,11 | 9,16E-01 |
| AT1G02870 | 0,35 | -2,14 | 4,39 | -0,49 | 6,26E-01 |

| | | | | | |
|---|---|---|---|---|---|
| AT1G02890 | 1,79 | 0,58 | 1,74 | 0,33 | 7,40E-01 |
| AT1G02910 | 0,70 | -2,16 | 2,61 | -0,83 | 4,07E-01 |
| AT1G02920 | 0,41 | 1,70 | 3,26 | 0,52 | 6,02E-01 |
| AT1G02930 | 0,49 | 0,84 | 3,30 | 0,25 | 7,99E-01 |
| AT1G02960 | 0,08 | 0,00 | 4,41 | 0,00 | 1,00E+00 |
| AT1G03000 | 0,70 | 0,03 | 2,69 | 0,01 | 9,91E-01 |
| AT1G03040 | 0,62 | -0,58 | 2,69 | -0,22 | 8,30E-01 |
| AT1G03080 | 1,73 | 0,49 | 1,57 | 0,31 | 7,53E-01 |
| AT1G03140 | 0,90 | 1,35 | 1,84 | 0,73 | 4,63E-01 |
| AT1G03240 | 1,51 | 0,19 | 1,93 | 0,10 | 9,20E-01 |
| AT1G03300 | 0,12 | 0,00 | 4,41 | 0,00 | 1,00E+00 |
| AT1G03400 | 0,49 | 0,46 | 2,80 | 0,17 | 8,68E-01 |
| AT1G03445 | 0,28 | -0,26 | 4,41 | -0,06 | 9,53E-01 |
| AT1G03475 | 0,69 | -0,80 | 2,14 | -0,38 | 7,07E-01 |
| AT1G03520 | 0,48 | -0,03 | 4,06 | -0,01 | 9,94E-01 |
| AT1G03530 | 0,41 | 0,00 | 4,41 | 0,00 | 1,00E+00 |
| AT1G03550 | 0,70 | -0,82 | 2,29 | -0,36 | 7,20E-01 |
| AT1G03590 | 0,41 | -0,20 | 4,25 | -0,05 | 9,62E-01 |
| AT1G03660 | 0,52 | 0,54 | 2,75 | 0,20 | 8,44E-01 |
| AT1G03680 | 1,73 | -0,94 | 1,27 | -0,74 | 4,60E-01 |
| AT1G03730 | 1,35 | -1,97 | 2,33 | -0,85 | 3,97E-01 |
| AT1G03746 | 0,46 | -0,33 | 4,02 | -0,08 | 9,34E-01 |
| AT1G03760 | 0,08 | 0,21 | 4,39 | 0,05 | 9,62E-01 |
| AT1G03830 | 0,65 | -1,80 | 2,55 | -0,71 | 4,80E-01 |
| AT1G03870 | 0,58 | -2,22 | 3,18 | -0,70 | 4,84E-01 |
| AT1G03940 | 1,20 | -1,91 | 1,70 | -1,12 | 2,62E-01 |
| AT1G04233 | 0,08 | 0,00 | 4,41 | 0,00 | 1,00E+00 |
| AT1G03980 | 0,11 | 0,88 | 4,35 | 0,20 | 8,39E-01 |
| AT1G04000 | 0,55 | 1,17 | 3,02 | 0,39 | 6,98E-01 |
| AT1G04010 | 0,05 | 0,18 | 4,38 | 0,04 | 9,68E-01 |
| AT1G04050 | 0,08 | 0,21 | 4,39 | 0,05 | 9,62E-01 |
| AT1G04070 | 0,16 | -0,21 | 4,40 | -0,05 | 9,63E-01 |
| AT1G04110 | 0,35 | 0,00 | 4,39 | 0,00 | 1,00E+00 |
| AT1G04120 | 1,63 | 1,77 | 1,77 | 1,00 | 3,16E-01 |
| AT1G04140 | 1,54 | 0,55 | 1,93 | 0,28 | 7,77E-01 |
| AT1G04190 | 0,13 | -0,61 | 4,37 | -0,14 | 8,89E-01 |
| AT1G04220 | 0,35 | 0,81 | 4,35 | 0,19 | 8,53E-01 |
| AT1G04240 | 0,91 | -2,54 | 2,12 | -1,20 | 2,31E-01 |
| AT1G04260 | 0,47 | -0,52 | 2,75 | -0,19 | 8,51E-01 |
| AT1G04270 | 1,04 | -0,87 | 2,04 | -0,43 | 6,70E-01 |
| AT1G04280 | 0,49 | -0,93 | 3,24 | -0,29 | 7,74E-01 |
| AT1G04290 | 0,43 | 1,32 | 2,76 | 0,48 | 6,32E-01 |
| AT1G04300 | 0,83 | 0,44 | 3,08 | 0,14 | 8,87E-01 |
| AT1G04380 | 0,20 | 0,00 | 4,41 | 0,00 | 1,00E+00 |
| AT1G04390 | 0,19 | 0,43 | 4,36 | 0,10 | 9,21E-01 |
| AT1G04400 | 0,89 | -1,14 | 2,22 | -0,51 | 6,09E-01 |
| AT1G04263 | 1,35 | -4,53 | 2,37 | -1,91 | 5,64E-02 |
| AT1G04530 | 0,42 | 0,89 | 3,32 | 0,27 | 7,90E-01 |
| AT1G04570 | 0,11 | -0,38 | 4,39 | -0,09 | 9,31E-01 |
| AT1G04630 | 0,66 | 0,00 | 3,81 | 0,00 | 1,00E+00 |
| AT1G04680 | 0,74 | -1,05 | 2,97 | -0,35 | 7,25E-01 |
| AT1G04730 | 0,74 | -1,13 | 2,58 | -0,44 | 6,61E-01 |
| AT1G04750 | 0,67 | -1,15 | 2,18 | -0,53 | 5,98E-01 |
| AT1G04770 | 0,89 | 0,98 | 2,19 | 0,45 | 6,55E-01 |
| AT1G04780 | 0,86 | -0,04 | 2,03 | -0,02 | 9,84E-01 |
| AT1G04840 | 0,63 | 0,00 | 3,80 | 0,00 | 1,00E+00 |

| | | | | | |
|---|---|---|---|---|---|
| AT1G04850 | 0,86 | 0,86 | 2,80 | 0,31 | 7,60E-01 |
| AT1G04860 | 0,57 | -0,20 | 4,01 | -0,05 | 9,60E-01 |
| AT1G04870 | 0,84 | -2,69 | 3,05 | -0,88 | 3,79E-01 |
| AT1G04880 | 0,21 | -0,33 | 4,32 | -0,08 | 9,39E-01 |
| AT1G04910 | 0,78 | 0,20 | 2,90 | 0,07 | 9,46E-01 |
| AT1G04950 | 0,50 | -0,31 | 3,32 | -0,09 | 9,26E-01 |
| AT1G04960 | 0,36 | -1,29 | 4,40 | -0,29 | 7,70E-01 |
| AT1G04980 | 0,36 | 0,00 | 4,41 | 0,00 | 1,00E+00 |
| AT1G04985 | 0,26 | 0,83 | 4,35 | 0,19 | 8,48E-01 |
| AT1G04990 | 0,84 | -1,96 | 2,52 | -0,78 | 4,36E-01 |
| AT1G05030 | 1,78 | 0,78 | 1,50 | 0,52 | 6,01E-01 |
| AT1G05120 | 0,53 | 0,00 | 4,40 | 0,00 | 1,00E+00 |
| AT1G05150 | 0,85 | -0,36 | 2,20 | -0,16 | 8,71E-01 |
| AT1G05160 | 0,59 | 0,00 | 4,40 | 0,00 | 1,00E+00 |
| AT1G05170 | 0,47 | -1,71 | 4,37 | -0,39 | 6,95E-01 |
| AT1G05190 | 0,78 | 0,79 | 2,40 | 0,33 | 7,41E-01 |
| AT1G05205 | 0,08 | 0,00 | 4,41 | 0,00 | 1,00E+00 |
| AT1G05230 | 0,45 | 0,00 | 4,14 | 0,00 | 1,00E+00 |
| AT1G05300 | 0,18 | 0,90 | 4,35 | 0,21 | 8,36E-01 |
| AT1G05350 | 0,28 | -0,22 | 4,40 | -0,05 | 9,60E-01 |
| AT1G05520 | 0,61 | 0,32 | 3,10 | 0,10 | 9,17E-01 |
| AT1G05575 | 0,16 | 1,16 | 4,34 | 0,27 | 7,89E-01 |
| AT1G05640 | 0,20 | 0,40 | 4,37 | 0,09 | 9,27E-01 |
| AT1G05670 | 0,15 | -0,26 | 4,41 | -0,06 | 9,53E-01 |
| AT1G05720 | 0,13 | -0,61 | 4,37 | -0,14 | 8,89E-01 |
| AT1G05783 | 0,96 | -0,04 | 1,84 | -0,02 | 9,81E-01 |
| AT1G05840 | 0,75 | 1,74 | 2,19 | 0,79 | 4,27E-01 |
| AT1G05870 | 0,42 | 1,29 | 4,17 | 0,31 | 7,57E-01 |
| AT1G05950 | 0,12 | 0,00 | 4,41 | 0,00 | 1,00E+00 |
| AT1G05970 | 0,08 | 0,00 | 4,41 | 0,00 | 1,00E+00 |
| AT1G06000 | 0,95 | -0,50 | 1,95 | -0,25 | 7,99E-01 |
| AT1G06130 | 0,62 | -2,11 | 3,12 | -0,67 | 5,00E-01 |
| AT1G06150 | 1,38 | 0,80 | 1,99 | 0,40 | 6,89E-01 |
| AT1G06250 | 0,72 | 0,00 | 4,41 | 0,00 | 1,00E+00 |
| AT1G06265 | 0,08 | 0,00 | 4,41 | 0,00 | 1,00E+00 |
| AT1G06360 | 0,33 | -1,45 | 4,30 | -0,34 | 7,35E-01 |
| AT1G06400 | 0,21 | 1,07 | 4,34 | 0,25 | 8,05E-01 |
| AT1G06430 | 1,82 | -2,69 | 1,40 | -1,91 | 5,56E-02 |
| AT1G06440 | 0,58 | 0,88 | 2,71 | 0,32 | 7,46E-01 |
| AT1G06460 | 1,79 | -0,86 | 1,47 | -0,58 | 5,59E-01 |
| AT1G06540 | 1,04 | -1,14 | 2,24 | -0,51 | 6,12E-01 |
| AT1G06570 | 0,97 | -3,06 | 2,56 | -1,19 | 2,32E-01 |
| AT1G06580 | 1,59 | -1,63 | 1,28 | -1,27 | 2,04E-01 |
| AT1G04487 | 0,39 | 0,17 | 4,24 | 0,04 | 9,68E-01 |
| AT1G06660 | 0,50 | 0,00 | 4,05 | 0,00 | 1,00E+00 |
| AT1G06690 | 0,36 | -2,10 | 4,40 | -0,48 | 6,33E-01 |
| AT1G06700 | 1,69 | -1,16 | 1,73 | -0,67 | 5,02E-01 |
| AT1G06710 | 0,26 | 0,00 | 4,41 | 0,00 | 1,00E+00 |
| AT1G06720 | 0,55 | -0,97 | 4,00 | -0,24 | 8,09E-01 |
| AT1G06760 | 1,45 | 0,12 | 1,45 | 0,08 | 9,33E-01 |
| AT1G06810 | 0,08 | 0,00 | 4,41 | 0,00 | 1,00E+00 |
| AT1G06840 | 0,17 | -0,58 | 4,38 | -0,13 | 8,94E-01 |
| AT1G06890 | 0,40 | -0,46 | 2,84 | -0,16 | 8,70E-01 |
| AT1G06900 | 1,35 | -1,30 | 1,70 | -0,77 | 4,44E-01 |
| AT1G07020 | 0,55 | -2,19 | 4,41 | -0,50 | 6,20E-01 |
| AT1G07050 | 0,42 | -3,49 | 4,11 | -0,85 | 3,96E-01 |

| | | | | | |
|---|---|---|---|---|---|
| AT1G07090 | 0,43 | 0,60 | 4,32 | 0,14 | 8,89E-01 |
| AT1G07110 | 1,24 | 0,00 | 2,52 | 0,00 | 1,00E+00 |
| AT1G07128 | 0,28 | 0,60 | 4,37 | 0,14 | 8,91E-01 |
| AT1G07140 | 1,81 | -0,33 | 1,68 | -0,20 | 8,43E-01 |
| AT1G07200 | 1,82 | 1,48 | 1,82 | 0,81 | 4,18E-01 |
| AT1G07220 | 0,25 | -3,44 | 4,34 | -0,79 | 4,28E-01 |
| AT1G07260 | 0,48 | 0,24 | 4,37 | 0,06 | 9,56E-01 |
| AT1G04573 | 0,66 | 0,00 | 4,41 | 0,00 | 1,00E+00 |
| AT1G07310 | 0,69 | -2,15 | 2,62 | -0,82 | 4,12E-01 |
| AT1G07350 | 0,45 | -0,26 | 3,36 | -0,08 | 9,38E-01 |
| AT1G07360 | 0,57 | -1,09 | 3,06 | -0,36 | 7,22E-01 |
| AT1G07430 | 0,32 | -2,12 | 4,40 | -0,48 | 6,30E-01 |
| AT1G07440 | 0,75 | 0,97 | 2,58 | 0,37 | 7,08E-01 |
| AT1G07480 | 0,80 | 0,05 | 2,28 | 0,02 | 9,82E-01 |
| AT1G07500 | 0,08 | 0,21 | 4,39 | 0,05 | 9,62E-01 |
| AT1G07520 | 1,20 | -2,45 | 2,23 | -1,10 | 2,72E-01 |
| AT1G07530 | 1,00 | -0,32 | 1,95 | -0,17 | 8,69E-01 |
| AT1G07640 | 0,24 | -0,42 | 4,36 | -0,10 | 9,23E-01 |
| AT1G07670 | 1,74 | -1,24 | 1,43 | -0,87 | 3,85E-01 |
| AT1G07720 | 1,26 | 0,10 | 1,48 | 0,07 | 9,45E-01 |
| AT1G07725 | 1,03 | 0,00 | 2,68 | 0,00 | 1,00E+00 |
| AT1G07740 | 0,29 | 0,00 | 4,41 | 0,00 | 1,00E+00 |
| AT1G07730 | 0,29 | 0,00 | 4,41 | 0,00 | 1,00E+00 |
| AT1G07750 | 0,25 | -3,44 | 4,34 | -0,79 | 4,28E-01 |
| AT1G07770 | 0,87 | 0,43 | 1,70 | 0,25 | 8,02E-01 |
| AT1G07870 | 0,12 | -0,26 | 4,41 | -0,06 | 9,53E-01 |
| AT1G07970 | 0,62 | 0,00 | 3,73 | 0,00 | 1,00E+00 |
| AT1G08040 | 0,40 | -1,19 | 4,36 | -0,27 | 7,84E-01 |
| AT1G08050 | 0,33 | -0,36 | 4,39 | -0,08 | 9,34E-01 |
| AT1G08060 | 1,18 | 0,37 | 2,24 | 0,17 | 8,69E-01 |
| AT1G08125 | 0,32 | 0,04 | 4,38 | 0,01 | 9,92E-01 |
| AT1G08200 | 1,82 | 0,43 | 1,56 | 0,28 | 7,82E-01 |
| AT1G08510 | 1,46 | -0,97 | 1,41 | -0,69 | 4,92E-01 |
| AT1G08580 | 0,24 | 0,00 | 4,41 | 0,00 | 1,00E+00 |
| AT1G08592 | 0,74 | -0,26 | 3,01 | -0,09 | 9,31E-01 |
| AT1G08610 | 0,28 | 0,02 | 4,38 | 0,01 | 9,96E-01 |
| AT1G08640 | 0,47 | -0,01 | 4,00 | 0,00 | 9,98E-01 |
| AT1G08645 | 0,64 | -1,61 | 3,71 | -0,43 | 6,65E-01 |
| AT1G08660 | 0,10 | 0,86 | 4,35 | 0,20 | 8,43E-01 |
| AT1G08700 | 1,22 | -1,12 | 2,31 | -0,49 | 6,27E-01 |
| AT1G08800 | 0,74 | -1,09 | 3,00 | -0,36 | 7,17E-01 |
| AT1G08820 | 0,89 | 2,24 | 2,62 | 0,85 | 3,93E-01 |
| AT1G08880 | 0,12 | 0,00 | 4,41 | 0,00 | 1,00E+00 |
| AT1G09020 | 0,78 | -0,20 | 2,29 | -0,09 | 9,29E-01 |
| AT1G09060 | 1,04 | -0,62 | 2,40 | -0,26 | 7,96E-01 |
| AT1G09080 | 0,40 | 0,77 | 4,31 | 0,18 | 8,59E-01 |
| AT1G09130 | 0,46 | -0,81 | 3,20 | -0,25 | 8,00E-01 |
| AT1G09160 | 0,40 | -0,96 | 3,41 | -0,28 | 7,79E-01 |
| AT1G09190 | 0,17 | 0,00 | 4,41 | 0,00 | 1,00E+00 |
| AT1G09195 | 0,70 | -0,07 | 2,99 | -0,02 | 9,81E-01 |
| AT1G09200 | 0,34 | 2,35 | 4,29 | 0,55 | 5,84E-01 |
| AT1G09230 | 0,23 | 0,24 | 4,39 | 0,06 | 9,56E-01 |
| AT1G09280 | 1,11 | -1,61 | 1,70 | -0,95 | 3,43E-01 |
| AT1G09420 | 0,88 | 0,42 | 3,41 | 0,12 | 9,02E-01 |
| AT1G09440 | 0,21 | -2,83 | 4,36 | -0,65 | 5,16E-01 |
| AT1G04707 | 0,36 | 0,00 | 4,41 | 0,00 | 1,00E+00 |

| | | | | | |
|---|---|---|---|---|---|
| AT1G09630 | 0,08 | 0,00 | 4,41 | 0,00 | 1,00E+00 |
| AT1G04717 | 0,54 | 0,00 | 3,89 | 0,00 | 1,00E+00 |
| AT1G09680 | 1,19 | -0,26 | 2,70 | -0,10 | 9,23E-01 |
| AT1G09690 | 1,78 | -0,22 | 1,44 | -0,15 | 8,78E-01 |
| AT1G09730 | 0,83 | -1,32 | 2,82 | -0,47 | 6,40E-01 |
| AT1G09760 | 0,17 | -0,33 | 4,40 | -0,08 | 9,40E-01 |
| AT1G09810 | 0,42 | -0,07 | 2,81 | -0,02 | 9,80E-01 |
| AT1G09880 | 0,33 | 0,00 | 4,41 | 0,00 | 1,00E+00 |
| AT1G09910 | 0,78 | -0,30 | 2,51 | -0,12 | 9,05E-01 |
| AT1G10020 | 0,10 | 0,08 | 4,38 | 0,02 | 9,85E-01 |
| AT1G10090 | 0,57 | -1,70 | 2,65 | -0,64 | 5,21E-01 |
| AT1G10155 | 1,47 | 0,39 | 1,55 | 0,25 | 7,99E-01 |
| AT1G10180 | 0,61 | -0,66 | 2,64 | -0,25 | 8,02E-01 |
| AT1G10190 | 0,94 | 1,01 | 2,55 | 0,40 | 6,92E-01 |
| AT1G10200 | 0,57 | 0,15 | 3,89 | 0,04 | 9,70E-01 |
| AT1G10310 | 0,42 | 0,00 | 4,41 | 0,00 | 1,00E+00 |
| AT1G10320 | 0,47 | -0,30 | 4,04 | -0,08 | 9,40E-01 |
| AT1G10340 | 1,00 | 0,87 | 2,29 | 0,38 | 7,05E-01 |
| AT1G10350 | 0,52 | 0,00 | 3,96 | 0,00 | 1,00E+00 |
| AT1G10390 | 1,30 | 1,53 | 1,99 | 0,77 | 4,41E-01 |
| AT1G10400 | 0,08 | 0,00 | 4,41 | 0,00 | 1,00E+00 |
| AT1G10410 | 0,96 | 1,41 | 1,96 | 0,72 | 4,71E-01 |
| AT1G10430 | 0,87 | 0,67 | 2,08 | 0,32 | 7,46E-01 |
| AT1G10470 | 1,21 | 0,21 | 1,74 | 0,12 | 9,03E-01 |
| AT1G10500 | 1,39 | -2,26 | 2,50 | -0,90 | 3,67E-01 |
| AT1G10580 | 0,57 | -2,99 | 3,88 | -0,77 | 4,41E-01 |
| AT1G10586 | 1,03 | 1,61 | 1,96 | 0,82 | 4,11E-01 |
| AT1G10590 | 0,69 | 0,80 | 3,05 | 0,26 | 7,93E-01 |
| AT1G10630 | 0,65 | -1,17 | 2,74 | -0,43 | 6,71E-01 |
| AT1G10640 | 0,05 | 0,18 | 4,38 | 0,04 | 9,68E-01 |
| AT1G10680 | 0,08 | 0,00 | 4,41 | 0,00 | 1,00E+00 |
| AT1G10840 | 0,88 | 0,42 | 1,98 | 0,21 | 8,31E-01 |
| AT1G10870 | 1,11 | -0,47 | 2,08 | -0,23 | 8,20E-01 |
| AT1G10900 | 1,03 | -1,37 | 2,93 | -0,47 | 6,41E-01 |
| AT1G10920 | 0,82 | 0,00 | 4,41 | 0,00 | 1,00E+00 |
| AT1G10930 | 0,74 | -0,38 | 3,02 | -0,12 | 9,01E-01 |
| AT1G10940 | 0,48 | 0,84 | 3,33 | 0,25 | 8,01E-01 |
| AT1G04833 | 0,22 | 1,46 | 4,32 | 0,34 | 7,36E-01 |
| AT1G11100 | 0,45 | 0,76 | 3,40 | 0,22 | 8,23E-01 |
| AT1G11210 | 0,56 | 1,61 | 2,49 | 0,65 | 5,17E-01 |
| AT1G11240 | 0,05 | 0,18 | 4,38 | 0,04 | 9,68E-01 |
| AT1G11270 | 0,83 | 3,44 | 2,22 | 1,55 | 1,21E-01 |
| AT1G11280 | 0,45 | -0,04 | 2,77 | -0,01 | 9,89E-01 |
| AT1G11290 | 0,63 | 0,00 | 3,73 | 0,00 | 1,00E+00 |
| AT1G11310 | 1,46 | 0,82 | 1,72 | 0,48 | 6,35E-01 |
| AT1G11340 | 0,08 | 0,00 | 4,41 | 0,00 | 1,00E+00 |
| AT1G11350 | 0,52 | 0,41 | 3,27 | 0,13 | 9,00E-01 |
| AT1G11360 | 0,33 | 0,00 | 4,33 | 0,00 | 1,00E+00 |
| AT1G11480 | 0,70 | 0,39 | 2,25 | 0,17 | 8,63E-01 |
| AT1G11630 | 0,35 | 0,00 | 4,39 | 0,00 | 1,00E+00 |
| AT1G11820 | 0,25 | -0,80 | 4,37 | -0,18 | 8,54E-01 |
| AT1G12000 | 0,52 | 1,65 | 3,06 | 0,54 | 5,91E-01 |
| AT1G12020 | 0,63 | 1,51 | 2,12 | 0,71 | 4,78E-01 |
| AT1G12120 | 0,70 | 0,04 | 2,61 | 0,02 | 9,88E-01 |
| AT1G12211 | 0,63 | -0,98 | 3,17 | -0,31 | 7,57E-01 |
| AT1G12280 | 1,78 | -0,53 | 1,37 | -0,39 | 6,99E-01 |

| | | | | | |
|---|---|---|---|---|---|
| AT1G12310 | 0,63 | 2,12 | 3,12 | 0,68 | 4,97E-01 |
| AT1G12380 | 0,49 | 1,68 | 4,05 | 0,41 | 6,78E-01 |
| AT1G12440 | 1,21 | 1,11 | 1,78 | 0,63 | 5,31E-01 |
| AT1G12500 | 0,25 | 0,29 | 4,37 | 0,07 | 9,47E-01 |
| AT1G12520 | 0,51 | -2,29 | 4,37 | -0,52 | 6,01E-01 |
| AT1G12590 | 0,78 | -0,39 | 1,68 | -0,23 | 8,17E-01 |
| AT1G12610 | 0,19 | 0,84 | 4,35 | 0,19 | 8,47E-01 |
| AT1G12615 | 1,59 | 4,55 | 2,02 | 2,26 | 2,40E-02 |
| AT1G12640 | 0,58 | 0,00 | 4,41 | 0,00 | 1,00E+00 |
| AT1G12680 | 1,43 | 0,82 | 1,61 | 0,51 | 6,12E-01 |
| AT1G12700 | 0,22 | 0,80 | 4,35 | 0,18 | 8,55E-01 |
| AT1G12780 | 0,71 | 0,84 | 2,27 | 0,37 | 7,12E-01 |
| AT1G12800 | 0,97 | -2,00 | 2,53 | -0,79 | 4,31E-01 |
| AT1G12910 | 1,07 | -1,26 | 2,23 | -0,56 | 5,72E-01 |
| AT1G13020 | 0,59 | -1,24 | 2,71 | -0,46 | 6,48E-01 |
| AT1G13120 | 0,23 | -0,38 | 4,39 | -0,09 | 9,32E-01 |
| AT1G13170 | 0,46 | -0,35 | 3,29 | -0,11 | 9,16E-01 |
| AT1G13200 | 0,66 | 0,00 | 3,71 | 0,00 | 1,00E+00 |
| AT1G13245 | 0,44 | -2,34 | 4,09 | -0,57 | 5,68E-01 |
| AT1G13310 | 0,35 | 0,00 | 4,39 | 0,00 | 1,00E+00 |
| AT1G13340 | 0,95 | 1,68 | 2,22 | 0,76 | 4,50E-01 |
| AT1G13390 | 0,17 | 0,00 | 4,41 | 0,00 | 1,00E+00 |
| AT1G13450 | 0,76 | 0,60 | 2,33 | 0,26 | 7,96E-01 |
| AT1G13470 | 1,76 | 1,64 | 1,65 | 1,00 | 3,19E-01 |
| AT1G13520 | 0,54 | -0,39 | 4,39 | -0,09 | 9,30E-01 |
| AT1G13560 | 0,43 | -0,40 | 4,38 | -0,09 | 9,27E-01 |
| AT1G13825 | 0,76 | 0,48 | 3,63 | 0,13 | 8,96E-01 |
| AT1G13860 | 0,10 | 0,24 | 4,38 | 0,05 | 9,57E-01 |
| AT1G13870 | 0,12 | 0,49 | 4,38 | 0,11 | 9,11E-01 |
| AT1G13900 | 0,21 | -0,35 | 4,40 | -0,08 | 9,37E-01 |
| AT1G13940 | 0,86 | 1,22 | 2,47 | 0,49 | 6,21E-01 |
| AT1G14010 | 0,36 | 1,01 | 4,35 | 0,23 | 8,16E-01 |
| AT1G14030 | 0,62 | -1,81 | 3,16 | -0,57 | 5,67E-01 |
| AT1G14290 | 0,26 | 0,00 | 4,41 | 0,00 | 1,00E+00 |
| AT1G14300 | 0,22 | -1,49 | 4,36 | -0,34 | 7,33E-01 |
| AT1G14330 | 0,59 | -0,58 | 2,68 | -0,22 | 8,28E-01 |
| AT1G14360 | 0,18 | 0,31 | 4,38 | 0,07 | 9,44E-01 |
| AT1G14400 | 0,66 | -1,01 | 3,14 | -0,32 | 7,48E-01 |
| AT1G14450 | 0,11 | 0,68 | 4,37 | 0,16 | 8,76E-01 |
| AT1G14455 | 0,14 | -2,43 | 4,37 | -0,55 | 5,79E-01 |
| AT1G14518 | 0,84 | 3,14 | 2,29 | 1,37 | 1,70E-01 |
| AT1G14590 | 0,42 | 0,00 | 4,24 | 0,00 | 1,00E+00 |
| AT1G14690 | 0,78 | -2,06 | 2,35 | -0,87 | 3,82E-01 |
| AT1G14710 | 0,51 | -2,05 | 3,92 | -0,52 | 6,02E-01 |
| AT1G14720 | 0,26 | -2,46 | 4,37 | -0,56 | 5,74E-01 |
| AT1G14790 | 1,42 | 1,34 | 2,27 | 0,59 | 5,54E-01 |
| AT1G14810 | 1,02 | 0,43 | 2,28 | 0,19 | 8,52E-01 |
| AT1G14830 | 0,90 | 1,31 | 2,82 | 0,46 | 6,42E-01 |
| AT1G14840 | 0,51 | -0,82 | 2,25 | -0,36 | 7,17E-01 |
| AT1G14850 | 1,26 | 1,06 | 1,95 | 0,54 | 5,87E-01 |
| AT1G14880 | 1,19 | 2,42 | 1,89 | 1,28 | 2,00E-01 |
| AT1G14900 | 0,38 | -3,11 | 4,34 | -0,72 | 4,74E-01 |
| AT1G14910 | 0,80 | -0,80 | 2,55 | -0,32 | 7,52E-01 |
| AT1G14980 | 0,57 | -2,08 | 3,13 | -0,66 | 5,07E-01 |
| AT1G15002 | 1,82 | 0,10 | 1,29 | 0,08 | 9,36E-01 |
| AT1G05137 | 0,28 | -0,67 | 4,38 | -0,15 | 8,78E-01 |

| | | | | | |
|---|---|---|---|---|---|
| AT1G15020 | 0,98 | 0,00 | 2,74 | 0,00 | 1,00E+00 |
| AT1G15040 | 0,21 | 0,39 | 4,36 | 0,09 | 9,29E-01 |
| AT1G15100 | 1,26 | 1,17 | 2,51 | 0,47 | 6,40E-01 |
| AT1G15140 | 0,76 | -0,28 | 2,19 | -0,13 | 8,99E-01 |
| AT1G15175 | 0,27 | -0,41 | 4,38 | -0,09 | 9,25E-01 |
| AT1G15210 | 0,17 | 0,00 | 4,41 | 0,00 | 1,00E+00 |
| AT1G15220 | 0,11 | -0,38 | 4,39 | -0,09 | 9,31E-01 |
| AT1G15340 | 0,21 | 0,15 | 4,33 | 0,03 | 9,72E-01 |
| AT1G15350 | 1,25 | 0,26 | 2,22 | 0,12 | 9,07E-01 |
| AT1G15400 | 0,43 | 1,61 | 2,77 | 0,58 | 5,62E-01 |
| AT1G15440 | 0,87 | -1,73 | 2,27 | -0,76 | 4,47E-01 |
| AT1G15520 | 0,46 | 1,19 | 3,24 | 0,37 | 7,13E-01 |
| AT1G15530 | 1,02 | -2,03 | 2,41 | -0,84 | 4,00E-01 |
| AT1G15550 | 0,24 | 0,00 | 4,41 | 0,00 | 1,00E+00 |
| AT1G15700 | 0,40 | -0,34 | 3,43 | -0,10 | 9,21E-01 |
| AT1G05187 | 0,24 | 0,00 | 4,41 | 0,00 | 1,00E+00 |
| AT1G15730 | 0,44 | 2,14 | 4,06 | 0,53 | 5,98E-01 |
| AT1G15780 | 1,01 | -0,89 | 1,89 | -0,47 | 6,36E-01 |
| AT1G15790 | 1,05 | 1,18 | 2,87 | 0,41 | 6,82E-01 |
| AT1G05203 | 0,26 | 0,00 | 4,41 | 0,00 | 1,00E+00 |
| AT1G15880 | 0,20 | 0,40 | 4,37 | 0,09 | 9,27E-01 |
| AT1G15970 | 0,11 | 0,88 | 4,35 | 0,20 | 8,39E-01 |
| AT1G16010 | 0,99 | -1,28 | 2,43 | -0,52 | 6,00E-01 |
| AT1G16070 | 0,40 | 0,22 | 4,39 | 0,05 | 9,59E-01 |
| AT1G16210 | 0,60 | -1,03 | 3,10 | -0,33 | 7,40E-01 |
| AT1G16240 | 0,80 | -1,79 | 2,42 | -0,74 | 4,61E-01 |
| AT1G16260 | 1,63 | 0,43 | 1,87 | 0,23 | 8,20E-01 |
| AT1G16460 | 0,57 | 0,34 | 2,35 | 0,15 | 8,84E-01 |
| AT1G16570 | 0,28 | 0,60 | 4,37 | 0,14 | 8,91E-01 |
| AT1G16610 | 1,02 | 1,23 | 3,13 | 0,39 | 6,94E-01 |
| AT1G16750 | 0,49 | -0,35 | 3,22 | -0,11 | 9,13E-01 |
| AT1G16780 | 0,12 | 0,00 | 4,41 | 0,00 | 1,00E+00 |
| AT1G16800 | 1,44 | 0,57 | 2,43 | 0,23 | 8,16E-01 |
| AT1G16870 | 0,95 | 0,00 | 3,45 | 0,00 | 1,00E+00 |
| AT1G16890 | 1,30 | -0,07 | 1,57 | -0,04 | 9,66E-01 |
| AT1G16900 | 0,65 | 0,00 | 3,78 | 0,00 | 1,00E+00 |
| AT1G16920 | 0,22 | 0,79 | 4,35 | 0,18 | 8,56E-01 |
| AT1G16970 | 0,37 | -0,81 | 3,51 | -0,23 | 8,18E-01 |
| AT1G17080 | 0,60 | 0,24 | 3,24 | 0,08 | 9,40E-01 |
| AT1G17110 | 0,57 | -2,04 | 2,66 | -0,76 | 4,45E-01 |
| AT1G17140 | 0,95 | -2,39 | 3,49 | -0,69 | 4,93E-01 |
| AT1G17147 | 0,12 | 1,17 | 4,34 | 0,27 | 7,88E-01 |
| AT1G17232 | 0,51 | 2,58 | 3,04 | 0,85 | 3,96E-01 |
| AT1G17240 | 0,05 | 0,18 | 4,38 | 0,04 | 9,68E-01 |
| AT1G17250 | 0,99 | -0,26 | 2,74 | -0,10 | 9,24E-01 |
| AT1G17370 | 0,81 | -1,05 | 2,59 | -0,40 | 6,86E-01 |
| AT1G17380 | 0,50 | 2,22 | 2,60 | 0,86 | 3,92E-01 |
| AT1G05313 | 0,32 | 1,75 | 4,31 | 0,41 | 6,84E-01 |
| AT1G17440 | 0,46 | 1,90 | 3,28 | 0,58 | 5,62E-01 |
| AT1G17450 | 1,22 | -0,21 | 1,98 | -0,10 | 9,17E-01 |
| AT1G17590 | 0,46 | -0,98 | 3,32 | -0,29 | 7,69E-01 |
| AT1G17600 | 1,00 | 1,70 | 2,10 | 0,81 | 4,19E-01 |
| AT1G17665 | 0,20 | -2,08 | 4,39 | -0,47 | 6,36E-01 |
| AT1G17670 | 0,37 | -1,37 | 4,35 | -0,32 | 7,52E-01 |
| AT1G17690 | 0,25 | 0,00 | 4,41 | 0,00 | 1,00E+00 |
| AT1G17720 | 0,66 | 0,82 | 2,24 | 0,37 | 7,13E-01 |

| | | | | | |
|---|---|---|---|---|---|
| AT1G17744 | 0,16 | -0,26 | 4,41 | -0,06 | 9,53E-01 |
| AT1G17790 | 0,63 | -1,35 | 3,18 | -0,43 | 6,71E-01 |
| AT1G05333 | 0,71 | 0,00 | 3,62 | 0,00 | 1,00E+00 |
| AT1G17820 | 0,71 | 0,00 | 3,62 | 0,00 | 1,00E+00 |
| AT1G17850 | 0,36 | -0,33 | 4,40 | -0,08 | 9,39E-01 |
| AT1G05347 | 0,21 | -2,16 | 4,40 | -0,49 | 6,25E-01 |
| AT1G17870 | 0,39 | -0,40 | 4,31 | -0,09 | 9,26E-01 |
| AT1G17880 | 0,59 | -2,56 | 3,92 | -0,65 | 5,13E-01 |
| AT1G17890 | 0,42 | -0,26 | 4,41 | -0,06 | 9,53E-01 |
| AT1G17930 | 0,20 | 0,00 | 4,41 | 0,00 | 1,00E+00 |
| AT1G17980 | 1,06 | -0,95 | 2,05 | -0,46 | 6,45E-01 |
| AT1G17990 | 0,53 | 1,06 | 3,16 | 0,34 | 7,37E-01 |
| AT1G18050 | 0,60 | -0,44 | 2,47 | -0,18 | 8,59E-01 |
| AT1G18060 | 0,60 | -0,10 | 2,35 | -0,04 | 9,67E-01 |
| AT1G18070 | 0,60 | -2,14 | 2,60 | -0,82 | 4,10E-01 |
| AT1G18100 | 0,36 | 0,00 | 4,41 | 0,00 | 1,00E+00 |
| AT1G18150 | 1,19 | -3,76 | 2,22 | -1,70 | 8,99E-02 |
| AT1G18210 | 0,61 | 0,60 | 2,17 | 0,28 | 7,82E-01 |
| AT1G18330 | 0,22 | 0,08 | 4,38 | 0,02 | 9,86E-01 |
| AT1G18360 | 1,25 | -1,01 | 1,67 | -0,60 | 5,46E-01 |
| AT1G18382 | 0,44 | 1,79 | 2,72 | 0,66 | 5,11E-01 |
| AT1G18470 | 0,48 | 0,10 | 2,60 | 0,04 | 9,70E-01 |
| AT1G18540 | 1,15 | 0,02 | 2,09 | 0,01 | 9,92E-01 |
| AT1G18720 | 0,23 | 0,85 | 4,34 | 0,20 | 8,45E-01 |
| AT1G05433 | 0,53 | 2,93 | 3,00 | 0,98 | 3,29E-01 |
| AT1G18840 | 0,43 | 0,44 | 3,39 | 0,13 | 8,96E-01 |
| AT1G18890 | 0,28 | -0,36 | 4,39 | -0,08 | 9,35E-01 |
| AT1G18990 | 0,45 | -0,26 | 4,24 | -0,06 | 9,52E-01 |
| AT1G19000 | 0,86 | -0,17 | 1,67 | -0,10 | 9,18E-01 |
| AT1G19130 | 0,49 | -1,35 | 2,74 | -0,49 | 6,23E-01 |
| AT1G19140 | 0,44 | -0,26 | 4,40 | -0,06 | 9,54E-01 |
| AT1G19210 | 0,10 | 0,86 | 4,35 | 0,20 | 8,43E-01 |
| AT1G19260 | 0,97 | 3,36 | 2,51 | 1,34 | 1,80E-01 |
| AT1G19300 | 0,88 | 0,78 | 2,47 | 0,32 | 7,51E-01 |
| AT1G19360 | 0,50 | -0,26 | 4,04 | -0,06 | 9,48E-01 |
| AT1G19370 | 0,41 | -1,34 | 4,32 | -0,31 | 7,57E-01 |
| AT1G19400 | 1,69 | 1,38 | 1,42 | 0,97 | 3,32E-01 |
| AT1G19430 | 0,39 | -0,90 | 4,25 | -0,21 | 8,32E-01 |
| AT1G19450 | 0,33 | -0,33 | 4,34 | -0,08 | 9,40E-01 |
| AT1G19570 | 0,89 | -0,49 | 1,88 | -0,26 | 7,94E-01 |
| AT1G19640 | 0,14 | -2,43 | 4,37 | -0,55 | 5,79E-01 |
| AT1G19660 | 0,55 | 1,01 | 2,58 | 0,39 | 6,95E-01 |
| AT1G19670 | 1,59 | 0,56 | 1,75 | 0,32 | 7,50E-01 |
| AT1G19700 | 0,25 | -1,50 | 4,33 | -0,34 | 7,30E-01 |
| AT1G19715 | 1,55 | -0,16 | 2,30 | -0,07 | 9,45E-01 |
| AT1G19720 | 1,66 | 0,55 | 1,76 | 0,31 | 7,57E-01 |
| AT1G19780 | 0,50 | 1,58 | 4,03 | 0,39 | 6,95E-01 |
| AT1G19860 | 0,15 | 0,47 | 4,37 | 0,11 | 9,15E-01 |
| AT1G19870 | 0,80 | -0,70 | 2,55 | -0,27 | 7,85E-01 |
| AT1G19880 | 0,41 | 2,09 | 3,36 | 0,62 | 5,34E-01 |
| AT1G19920 | 1,00 | -0,28 | 2,45 | -0,11 | 9,09E-01 |
| AT1G20040 | 0,34 | -1,35 | 4,36 | -0,31 | 7,57E-01 |
| AT1G20070 | 0,59 | -0,09 | 2,49 | -0,04 | 9,71E-01 |
| AT1G20100 | 1,77 | 0,33 | 1,88 | 0,18 | 8,61E-01 |
| AT1G20160 | 0,33 | 0,00 | 4,41 | 0,00 | 1,00E+00 |
| AT1G20190 | 0,64 | -2,60 | 3,16 | -0,82 | 4,11E-01 |

| | | | | | |
|---|---|---|---|---|---|
| AT1G20200 | 0,33 | -0,05 | 3,54 | -0,01 | 9,88E-01 |
| AT1G20320 | 0,05 | 0,18 | 4,38 | 0,04 | 9,68E-01 |
| AT1G20400 | 1,62 | 2,97 | 1,67 | 1,78 | 7,45E-02 |
| AT1G20430 | 0,89 | -2,66 | 2,91 | -0,91 | 3,62E-01 |
| AT1G20490 | 0,48 | -2,37 | 4,39 | -0,54 | 5,89E-01 |
| AT1G20560 | 0,25 | 0,10 | 4,38 | 0,02 | 9,82E-01 |
| AT1G20570 | 0,08 | 0,00 | 4,41 | 0,00 | 1,00E+00 |
| AT1G20650 | 0,41 | 0,69 | 4,35 | 0,16 | 8,73E-01 |
| AT1G20670 | 0,31 | -0,39 | 4,39 | -0,09 | 9,29E-01 |
| AT1G20770 | 0,16 | -0,26 | 4,41 | -0,06 | 9,53E-01 |
| AT1G20840 | 1,09 | -0,27 | 1,77 | -0,15 | 8,80E-01 |
| AT1G20880 | 0,25 | -2,73 | 4,36 | -0,63 | 5,32E-01 |
| AT1G20890 | 0,48 | -0,30 | 4,06 | -0,07 | 9,41E-01 |
| AT1G20950 | 0,69 | -0,97 | 2,58 | -0,37 | 7,08E-01 |
| AT1G21080 | 0,61 | 2,15 | 2,49 | 0,86 | 3,88E-01 |
| AT1G05627 | 0,05 | 0,18 | 4,38 | 0,04 | 9,68E-01 |
| AT1G21160 | 0,26 | 1,00 | 4,35 | 0,23 | 8,19E-01 |
| AT1G21190 | 0,45 | -0,15 | 4,11 | -0,04 | 9,70E-01 |
| AT1G21310 | 0,44 | -0,30 | 2,79 | -0,11 | 9,13E-01 |
| AT1G21380 | 1,01 | 1,70 | 2,27 | 0,75 | 4,55E-01 |
| AT1G21390 | 0,08 | 0,00 | 4,41 | 0,00 | 1,00E+00 |
| AT1G21440 | 1,10 | -0,01 | 2,64 | 0,00 | 9,97E-01 |
| AT1G21500 | 0,51 | 0,84 | 3,02 | 0,28 | 7,80E-01 |
| AT1G21520 | 0,17 | -0,33 | 4,40 | -0,08 | 9,40E-01 |
| AT1G21550 | 0,17 | -0,29 | 4,40 | -0,07 | 9,47E-01 |
| AT1G21600 | 0,77 | 2,33 | 2,12 | 1,10 | 2,72E-01 |
| AT1G21650 | 0,46 | -1,11 | 3,32 | -0,33 | 7,38E-01 |
| AT1G21770 | 0,66 | 1,15 | 2,55 | 0,45 | 6,52E-01 |
| AT1G21890 | 0,36 | 2,30 | 4,29 | 0,54 | 5,92E-01 |
| AT1G21900 | 0,26 | 0,07 | 4,34 | 0,02 | 9,87E-01 |
| AT1G21920 | 0,41 | -1,04 | 3,34 | -0,31 | 7,55E-01 |
| AT1G21928 | 0,29 | 0,69 | 4,35 | 0,16 | 8,74E-01 |
| AT1G22180 | 0,30 | 1,89 | 3,57 | 0,53 | 5,98E-01 |
| AT1G22200 | 0,54 | -2,04 | 3,89 | -0,52 | 6,01E-01 |
| AT1G22300 | 0,58 | 0,14 | 2,53 | 0,05 | 9,56E-01 |
| AT1G22360 | 0,63 | -1,21 | 2,69 | -0,45 | 6,53E-01 |
| AT1G22400 | 0,23 | 0,47 | 4,36 | 0,11 | 9,14E-01 |
| AT1G22550 | 0,19 | 1,17 | 4,34 | 0,27 | 7,87E-01 |
| AT1G22570 | 0,46 | -1,30 | 4,08 | -0,32 | 7,50E-01 |
| AT1G22710 | 0,50 | 1,29 | 2,61 | 0,50 | 6,20E-01 |
| AT1G22850 | 1,38 | -4,18 | 1,88 | -2,22 | 2,64E-02 |
| AT1G23020 | 0,20 | 0,00 | 4,41 | 0,00 | 1,00E+00 |
| AT1G23052 | 1,19 | 0,54 | 2,23 | 0,24 | 8,08E-01 |
| AT1G23080 | 0,61 | 1,89 | 3,27 | 0,58 | 5,64E-01 |
| AT1G05793 | 0,08 | 0,21 | 4,39 | 0,05 | 9,62E-01 |
| AT1G23205 | 0,35 | 0,57 | 4,33 | 0,13 | 8,96E-01 |
| AT1G23260 | 0,22 | 0,11 | 4,38 | 0,02 | 9,80E-01 |
| AT1G23300 | 1,57 | -0,34 | 1,51 | -0,22 | 8,23E-01 |
| AT1G23330 | 0,10 | 0,86 | 4,35 | 0,20 | 8,43E-01 |
| AT1G23650 | 0,08 | 0,00 | 4,41 | 0,00 | 1,00E+00 |
| AT1G23670 | 0,22 | 1,98 | 4,30 | 0,46 | 6,45E-01 |
| AT1G23740 | 1,33 | -2,04 | 1,78 | -1,15 | 2,51E-01 |
| AT1G23780 | 0,31 | -0,38 | 4,39 | -0,09 | 9,32E-01 |
| AT1G23830 | 0,53 | -0,28 | 3,23 | -0,09 | 9,31E-01 |
| AT1G23840 | 0,48 | -0,37 | 3,23 | -0,12 | 9,08E-01 |
| AT1G23850 | 0,35 | 0,21 | 4,39 | 0,05 | 9,61E-01 |

| | | | | | |
|---|---|---|---|---|---|
| AT1G23960 | 0,23 | 1,03 | 4,35 | 0,24 | 8,12E-01 |
| AT1G24040 | 0,10 | 0,24 | 4,38 | 0,05 | 9,57E-01 |
| AT1G24100 | 1,00 | 1,70 | 1,97 | 0,86 | 3,88E-01 |
| AT1G24120 | 0,29 | 0,28 | 4,31 | 0,06 | 9,48E-01 |
| AT1G24140 | 0,74 | -0,26 | 3,07 | -0,09 | 9,32E-01 |
| AT1G24160 | 0,84 | 0,00 | 3,44 | 0,00 | 1,00E+00 |
| AT1G24190 | 0,84 | -2,65 | 2,58 | -1,03 | 3,04E-01 |
| AT1G24267 | 0,48 | -1,23 | 3,27 | -0,38 | 7,07E-01 |
| AT1G24290 | 0,17 | -0,33 | 4,40 | -0,08 | 9,40E-01 |
| AT1G24300 | 1,68 | -0,05 | 1,76 | -0,03 | 9,77E-01 |
| AT1G24350 | 0,14 | -0,82 | 4,37 | -0,19 | 8,50E-01 |
| AT1G24510 | 1,77 | 1,07 | 1,46 | 0,74 | 4,62E-01 |
| AT1G05907 | 0,77 | 1,88 | 2,38 | 0,79 | 4,31E-01 |
| AT1G24764 | 0,53 | -0,35 | 3,25 | -0,11 | 9,14E-01 |
| AT1G24822 | 0,94 | 0,00 | 4,40 | 0,00 | 1,00E+00 |
| AT1G25220 | 0,36 | 0,00 | 4,41 | 0,00 | 1,00E+00 |
| AT1G25350 | 1,21 | -0,60 | 1,87 | -0,32 | 7,48E-01 |
| AT1G25370 | 0,12 | 0,00 | 4,41 | 0,00 | 1,00E+00 |
| AT1G25390 | 0,51 | 0,76 | 3,10 | 0,25 | 8,05E-01 |
| AT1G25400 | 0,63 | 1,14 | 2,90 | 0,39 | 6,94E-01 |
| AT1G25500 | 0,25 | 0,59 | 4,36 | 0,14 | 8,92E-01 |
| AT1G25540 | 0,94 | 1,54 | 2,34 | 0,66 | 5,11E-01 |
| AT1G25560 | 0,90 | 1,50 | 1,95 | 0,77 | 4,41E-01 |
| AT1G25570 | 0,49 | 0,00 | 4,41 | 0,00 | 1,00E+00 |
| AT1G26110 | 1,79 | -0,71 | 1,53 | -0,46 | 6,45E-01 |
| AT1G26150 | 0,88 | 0,66 | 2,28 | 0,29 | 7,72E-01 |
| AT1G26160 | 0,24 | 0,59 | 4,36 | 0,14 | 8,92E-01 |
| AT1G26170 | 0,54 | -0,06 | 3,16 | -0,02 | 9,86E-01 |
| AT1G26208 | 0,43 | 1,59 | 4,33 | 0,37 | 7,13E-01 |
| AT1G26218 | 0,37 | 1,88 | 4,31 | 0,44 | 6,63E-01 |
| AT1G26233 | 0,08 | 0,21 | 4,39 | 0,05 | 9,62E-01 |
| AT1G26240 | 0,88 | 3,61 | 2,55 | 1,42 | 1,56E-01 |
| AT1G26255 | 0,44 | 2,66 | 4,03 | 0,66 | 5,09E-01 |
| AT1G26270 | 1,28 | 0,48 | 2,23 | 0,21 | 8,30E-01 |
| AT1G26380 | 0,46 | 0,23 | 4,35 | 0,05 | 9,58E-01 |
| AT1G26390 | 0,25 | 0,02 | 4,37 | 0,01 | 9,96E-01 |
| AT1G26420 | 0,40 | 0,01 | 4,27 | 0,00 | 9,98E-01 |
| AT1G26690 | 0,33 | -0,36 | 4,39 | -0,08 | 9,34E-01 |
| AT1G26761 | 0,85 | -0,29 | 4,40 | -0,07 | 9,48E-01 |
| AT1G26800 | 0,42 | 0,70 | 2,78 | 0,25 | 8,01E-01 |
| AT1G06033 | 1,64 | -2,04 | 1,46 | -1,39 | 1,63E-01 |
| AT1G26930 | 0,36 | 0,85 | 4,33 | 0,20 | 8,45E-01 |
| AT1G27008 | 0,08 | 0,00 | 4,41 | 0,00 | 1,00E+00 |
| AT1G27090 | 1,85 | -0,53 | 1,23 | -0,43 | 6,69E-01 |
| AT1G27100 | 1,46 | 3,00 | 1,73 | 1,74 | 8,21E-02 |
| AT1G27190 | 0,77 | 1,17 | 2,23 | 0,53 | 5,99E-01 |
| AT1G27200 | 0,57 | 0,67 | 2,68 | 0,25 | 8,04E-01 |
| AT1G27210 | 0,26 | 1,41 | 4,33 | 0,33 | 7,44E-01 |
| AT1G27350 | 0,20 | 0,34 | 4,37 | 0,08 | 9,39E-01 |
| AT1G27370 | 0,51 | 0,43 | 4,12 | 0,10 | 9,18E-01 |
| AT1G27385 | 0,43 | -2,20 | 4,41 | -0,50 | 6,18E-01 |
| AT1G27440 | 0,24 | -0,34 | 4,40 | -0,08 | 9,38E-01 |
| AT1G27500 | 0,29 | 0,00 | 4,41 | 0,00 | 1,00E+00 |
| AT1G27510 | 0,44 | 0,78 | 3,26 | 0,24 | 8,11E-01 |
| AT1G06093 | 0,29 | -0,71 | 4,38 | -0,16 | 8,70E-01 |
| AT1G27600 | 1,12 | -3,71 | 2,13 | -1,74 | 8,21E-02 |

| | | | | | |
|---|---|---|---|---|---|
| AT1G27700 | 0,50 | 0,00 | 4,41 | 0,00 | 1,00E+00 |
| AT1G27720 | 0,50 | 0,00 | 4,00 | 0,00 | 1,00E+00 |
| AT1G27730 | 1,60 | 0,21 | 1,26 | 0,17 | 8,67E-01 |
| AT1G27900 | 0,40 | 2,20 | 3,26 | 0,67 | 5,00E-01 |
| AT1G27930 | 0,53 | -1,27 | 3,14 | -0,40 | 6,86E-01 |
| AT1G28000 | 0,35 | 2,46 | 4,28 | 0,57 | 5,65E-01 |
| AT1G28120 | 0,73 | 0,00 | 2,99 | 0,00 | 1,00E+00 |
| AT1G28140 | 1,15 | -2,00 | 2,04 | -0,98 | 3,25E-01 |
| AT1G28150 | 0,60 | -0,27 | 3,11 | -0,09 | 9,30E-01 |
| AT1G28190 | 0,15 | -0,39 | 4,39 | -0,09 | 9,29E-01 |
| AT1G28200 | 0,24 | -1,87 | 4,35 | -0,43 | 6,68E-01 |
| AT1G28230 | 0,78 | 0,80 | 2,15 | 0,37 | 7,09E-01 |
| AT1G28240 | 0,33 | -0,16 | 4,35 | -0,04 | 9,70E-01 |
| AT1G28260 | 1,27 | -0,49 | 2,09 | -0,23 | 8,15E-01 |
| AT1G28280 | 1,03 | 2,64 | 1,99 | 1,33 | 1,84E-01 |
| AT1G28290 | 0,27 | 1,23 | 4,34 | 0,28 | 7,76E-01 |
| AT1G28310 | 0,66 | 0,97 | 2,03 | 0,48 | 6,31E-01 |
| AT1G28330 | 0,60 | -1,55 | 3,10 | -0,50 | 6,18E-01 |
| AT1G28370 | 0,92 | 2,10 | 2,22 | 0,95 | 3,44E-01 |
| AT1G28390 | 0,25 | 0,00 | 4,41 | 0,00 | 1,00E+00 |
| AT1G28400 | 0,35 | -0,59 | 3,38 | -0,18 | 8,61E-01 |
| AT1G28570 | 0,12 | -0,26 | 4,41 | -0,06 | 9,53E-01 |
| AT1G28580 | 0,48 | -0,29 | 3,97 | -0,07 | 9,42E-01 |
| AT1G28600 | 1,10 | -1,01 | 1,83 | -0,55 | 5,82E-01 |
| AT1G28630 | 0,79 | -0,91 | 2,60 | -0,35 | 7,27E-01 |
| AT1G28660 | 0,08 | 0,00 | 4,41 | 0,00 | 1,00E+00 |
| AT1G28685 | 0,55 | -0,10 | 3,23 | -0,03 | 9,75E-01 |
| AT1G28765 | 0,17 | 0,00 | 4,41 | 0,00 | 1,00E+00 |
| AT1G28960 | 0,56 | -0,18 | 3,99 | -0,05 | 9,64E-01 |
| AT1G29050 | 0,14 | 1,29 | 4,33 | 0,30 | 7,65E-01 |
| AT1G29071 | 1,34 | 1,71 | 1,59 | 1,08 | 2,82E-01 |
| AT1G29170 | 0,29 | 0,85 | 4,35 | 0,20 | 8,44E-01 |
| AT1G29240 | 0,32 | 0,00 | 4,41 | 0,00 | 1,00E+00 |
| AT1G29300 | 0,08 | 0,47 | 4,38 | 0,11 | 9,15E-01 |
| AT1G29390 | 0,82 | -0,28 | 2,29 | -0,12 | 9,03E-01 |
| AT1G29395 | 0,39 | -2,43 | 3,37 | -0,72 | 4,72E-01 |
| AT1G29530 | 0,36 | -0,81 | 3,52 | -0,23 | 8,17E-01 |
| AT1G29690 | 0,27 | 0,15 | 4,34 | 0,04 | 9,72E-01 |
| AT1G29700 | 0,34 | -3,34 | 4,34 | -0,77 | 4,42E-01 |
| AT1G29720 | 0,71 | -1,96 | 2,60 | -0,75 | 4,51E-01 |
| AT1G29750 | 0,26 | 0,32 | 4,37 | 0,07 | 9,42E-01 |
| AT1G29880 | 1,19 | 2,39 | 1,99 | 1,20 | 2,31E-01 |
| AT1G29970 | 0,83 | 0,44 | 2,56 | 0,17 | 8,65E-01 |
| AT1G30000 | 0,84 | 0,92 | 2,69 | 0,34 | 7,33E-01 |
| AT1G30060 | 0,17 | 1,58 | 4,31 | 0,37 | 7,15E-01 |
| AT1G30070 | 0,15 | 0,02 | 4,36 | 0,00 | 9,96E-01 |
| AT1G30090 | 0,16 | 0,15 | 4,38 | 0,04 | 9,72E-01 |
| AT1G30120 | 1,11 | 1,67 | 1,97 | 0,85 | 3,97E-01 |
| AT1G30210 | 0,29 | 0,09 | 4,38 | 0,02 | 9,84E-01 |
| AT1G30250 | 0,60 | 1,55 | 2,49 | 0,62 | 5,34E-01 |
| AT1G30370 | 0,55 | -0,26 | 3,20 | -0,08 | 9,34E-01 |
| AT1G30490 | 1,77 | -2,19 | 2,15 | -1,02 | 3,09E-01 |
| AT1G30500 | 0,64 | 0,18 | 3,89 | 0,05 | 9,62E-01 |
| AT1G06347 | 0,94 | 0,13 | 2,04 | 0,06 | 9,51E-01 |
| AT1G30540 | 0,42 | -0,63 | 4,17 | -0,15 | 8,79E-01 |
| AT1G30580 | 1,07 | -1,55 | 2,04 | -0,76 | 4,47E-01 |

| | | | | | |
|---|---|---|---|---|---|
| AT1G30590 | 0,66 | 3,15 | 2,79 | 1,13 | 2,59E-01 |
| AT1G30755 | 0,62 | 1,48 | 2,63 | 0,56 | 5,73E-01 |
| AT1G30780 | 0,83 | -0,16 | 2,06 | -0,08 | 9,37E-01 |
| AT1G30810 | 0,83 | -1,92 | 1,94 | -0,99 | 3,21E-01 |
| AT1G30880 | 0,08 | 0,00 | 4,41 | 0,00 | 1,00E+00 |
| AT1G30960 | 0,37 | 0,00 | 4,41 | 0,00 | 1,00E+00 |
| AT1G06407 | 0,34 | 1,30 | 3,57 | 0,37 | 7,15E-01 |
| AT1G31130 | 1,47 | 1,87 | 1,73 | 1,09 | 2,78E-01 |
| AT1G31160 | 0,42 | 0,00 | 4,19 | 0,00 | 1,00E+00 |
| AT1G31180 | 0,17 | -0,33 | 4,40 | -0,08 | 9,40E-01 |
| AT1G31230 | 1,31 | 0,55 | 1,64 | 0,34 | 7,36E-01 |
| AT1G31240 | 0,22 | 0,60 | 4,30 | 0,14 | 8,89E-01 |
| AT1G31350 | 0,45 | -0,36 | 3,32 | -0,11 | 9,13E-01 |
| AT1G31410 | 0,08 | 0,00 | 4,41 | 0,00 | 1,00E+00 |
| AT1G31470 | 0,15 | -0,31 | 4,40 | -0,07 | 9,43E-01 |
| AT1G31550 | 0,81 | -2,65 | 2,59 | -1,02 | 3,05E-01 |
| AT1G31600 | 0,81 | -1,89 | 2,84 | -0,66 | 5,07E-01 |
| AT1G31640 | 0,23 | 1,10 | 4,34 | 0,25 | 8,00E-01 |
| AT1G31650 | 0,25 | 0,35 | 4,37 | 0,08 | 9,37E-01 |
| AT1G31660 | 0,51 | -2,19 | 4,41 | -0,50 | 6,20E-01 |
| AT1G31740 | 0,22 | 0,80 | 4,35 | 0,18 | 8,55E-01 |
| AT1G31810 | 0,57 | -0,39 | 2,37 | -0,17 | 8,69E-01 |
| AT1G31812 | 0,29 | -0,15 | 4,35 | -0,03 | 9,72E-01 |
| AT1G31814 | 0,37 | 0,00 | 4,41 | 0,00 | 1,00E+00 |
| AT1G31820 | 0,94 | 0,01 | 2,84 | 0,00 | 9,96E-01 |
| AT1G31830 | 0,66 | -0,39 | 2,69 | -0,15 | 8,84E-01 |
| AT1G06453 | 1,48 | -0,55 | 1,66 | -0,33 | 7,42E-01 |
| AT1G31860 | 1,08 | 1,80 | 1,82 | 0,99 | 3,24E-01 |
| AT1G06463 | 0,27 | 1,06 | 4,34 | 0,25 | 8,06E-01 |
| AT1G31920 | 0,52 | 0,52 | 2,70 | 0,19 | 8,48E-01 |
| AT1G31940 | 0,35 | 0,28 | 4,39 | 0,06 | 9,49E-01 |
| AT1G06473 | 0,38 | -1,57 | 4,35 | -0,36 | 7,19E-01 |
| AT1G31960 | 1,81 | 0,03 | 1,28 | 0,03 | 9,78E-01 |
| AT1G32000 | 0,35 | 1,46 | 4,32 | 0,34 | 7,35E-01 |
| AT1G32010 | 1,81 | 2,19 | 1,56 | 1,40 | 1,62E-01 |
| AT1G32070 | 0,34 | 0,00 | 4,41 | 0,00 | 1,00E+00 |
| AT1G32080 | 0,85 | 0,16 | 1,90 | 0,08 | 9,35E-01 |
| AT1G32130 | 1,27 | -1,90 | 1,99 | -0,95 | 3,40E-01 |
| AT1G32200 | 0,75 | -1,18 | 1,99 | -0,59 | 5,53E-01 |
| AT1G32340 | 0,12 | -0,36 | 4,39 | -0,08 | 9,34E-01 |
| AT1G06537 | 1,54 | 0,63 | 2,02 | 0,31 | 7,56E-01 |
| AT1G32400 | 0,47 | 1,91 | 2,68 | 0,71 | 4,77E-01 |
| AT1G32450 | 0,27 | 0,54 | 4,36 | 0,12 | 9,02E-01 |
| AT1G06573 | 0,41 | 0,61 | 4,32 | 0,14 | 8,88E-01 |
| AT1G32470 | 0,50 | 0,38 | 2,60 | 0,15 | 8,83E-01 |
| AT1G32490 | 0,70 | -0,10 | 2,27 | -0,04 | 9,66E-01 |
| AT1G32500 | 0,52 | -0,08 | 2,74 | -0,03 | 9,76E-01 |
| AT1G32530 | 0,53 | 0,46 | 2,76 | 0,16 | 8,69E-01 |
| AT1G06577 | 0,16 | -0,56 | 4,37 | -0,13 | 8,98E-01 |
| AT1G32713 | 1,03 | 2,20 | 2,24 | 0,98 | 3,25E-01 |
| AT1G32790 | 1,57 | 0,59 | 2,15 | 0,28 | 7,82E-01 |
| AT1G32860 | 0,05 | 0,18 | 4,38 | 0,04 | 9,68E-01 |
| AT1G32880 | 0,26 | 0,00 | 4,41 | 0,00 | 1,00E+00 |
| AT1G32900 | 0,61 | 2,23 | 3,07 | 0,73 | 4,66E-01 |
| AT1G32970 | 0,17 | 0,00 | 4,41 | 0,00 | 1,00E+00 |
| AT1G33050 | 0,95 | -0,41 | 2,10 | -0,19 | 8,47E-01 |

| | | | | | |
|---|---|---|---|---|---|
| AT1G06643 | 0,36 | -2,89 | 4,37 | -0,66 | 5,08E-01 |
| AT1G33250 | 0,90 | 1,19 | 2,31 | 0,52 | 6,06E-01 |
| AT1G33270 | 0,24 | 0,00 | 4,41 | 0,00 | 1,00E+00 |
| AT1G06673 | 0,47 | 1,17 | 3,21 | 0,36 | 7,15E-01 |
| AT1G33390 | 1,05 | -1,22 | 2,50 | -0,49 | 6,27E-01 |
| AT1G33410 | 1,37 | 1,59 | 1,76 | 0,90 | 3,68E-01 |
| AT1G33430 | 0,08 | 0,00 | 4,41 | 0,00 | 1,00E+00 |
| AT1G33520 | 0,47 | -1,36 | 3,26 | -0,42 | 6,77E-01 |
| AT1G33615 | 0,17 | -1,50 | 4,37 | -0,34 | 7,31E-01 |
| AT1G33700 | 0,88 | 0,70 | 2,24 | 0,31 | 7,54E-01 |
| AT1G33720 | 1,30 | 0,32 | 1,89 | 0,17 | 8,65E-01 |
| AT1G33780 | 0,08 | 0,47 | 4,38 | 0,11 | 9,15E-01 |
| AT1G33870 | 0,60 | -1,17 | 2,53 | -0,46 | 6,44E-01 |
| AT1G33970 | 0,67 | 0,22 | 2,64 | 0,08 | 9,34E-01 |
| AT1G34030 | 0,44 | -1,70 | 2,83 | -0,60 | 5,47E-01 |
| AT1G34042 | 0,29 | 1,66 | 4,32 | 0,39 | 7,00E-01 |
| AT1G34110 | 0,91 | 1,17 | 1,86 | 0,63 | 5,30E-01 |
| AT1G34170 | 0,20 | -0,22 | 4,40 | -0,05 | 9,61E-01 |
| AT1G34220 | 1,01 | 0,42 | 2,99 | 0,14 | 8,89E-01 |
| AT1G34270 | 0,12 | 0,00 | 4,41 | 0,00 | 1,00E+00 |
| AT1G34310 | 0,97 | 2,60 | 2,57 | 1,01 | 3,11E-01 |
| AT1G34315 | 0,23 | -0,36 | 4,39 | -0,08 | 9,36E-01 |
| AT1G34380 | 0,08 | 0,00 | 4,41 | 0,00 | 1,00E+00 |
| AT1G34440 | 1,54 | 1,37 | 1,53 | 0,90 | 3,70E-01 |
| AT1G34550 | 0,63 | 0,71 | 2,46 | 0,29 | 7,72E-01 |
| AT1G34750 | 0,23 | 1,31 | 4,33 | 0,30 | 7,62E-01 |
| AT1G34780 | 0,68 | -0,26 | 2,99 | -0,09 | 9,30E-01 |
| AT1G35150 | 0,16 | 1,05 | 4,35 | 0,24 | 8,10E-01 |
| AT1G35240 | 1,26 | 1,90 | 1,85 | 1,03 | 3,04E-01 |
| AT1G35250 | 0,19 | 1,47 | 4,32 | 0,34 | 7,35E-01 |
| AT1G35320 | 0,59 | 2,83 | 2,88 | 0,98 | 3,25E-01 |
| AT1G35350 | 0,96 | 2,11 | 1,92 | 1,10 | 2,71E-01 |
| AT1G06883 | 0,19 | 1,80 | 4,31 | 0,42 | 6,76E-01 |
| AT1G06897 | 0,91 | -0,89 | 3,27 | -0,27 | 7,85E-01 |
| AT1G06903 | 0,10 | 0,86 | 4,35 | 0,20 | 8,43E-01 |
| AT1G35520 | 1,01 | 3,66 | 2,43 | 1,51 | 1,32E-01 |
| AT1G35545 | 0,38 | 1,14 | 3,32 | 0,34 | 7,31E-01 |
| AT1G35580 | 1,07 | 1,21 | 1,77 | 0,68 | 4,93E-01 |
| AT1G35610 | 0,22 | -0,38 | 4,37 | -0,09 | 9,30E-01 |
| AT1G35625 | 0,08 | 0,00 | 4,41 | 0,00 | 1,00E+00 |
| AT1G35730 | 0,17 | 0,00 | 4,41 | 0,00 | 1,00E+00 |
| AT1G35820 | 1,40 | 4,58 | 2,78 | 1,65 | 9,94E-02 |
| AT1G35895 | 0,55 | -0,63 | 3,21 | -0,20 | 8,44E-01 |
| AT1G36070 | 0,25 | 0,00 | 4,41 | 0,00 | 1,00E+00 |
| AT1G36085 | 0,68 | 3,64 | 4,26 | 0,85 | 3,93E-01 |
| AT1G36230 | 0,12 | 1,17 | 4,34 | 0,27 | 7,88E-01 |
| AT1G36325 | 1,41 | -0,02 | 1,50 | -0,01 | 9,91E-01 |
| AT1G36370 | 0,71 | 0,06 | 2,42 | 0,02 | 9,82E-01 |
| AT1G36390 | 0,17 | -2,21 | 4,40 | -0,50 | 6,16E-01 |
| AT1G36622 | 0,40 | -0,19 | 3,43 | -0,06 | 9,56E-01 |
| AT1G36990 | 1,02 | 0,31 | 2,00 | 0,15 | 8,78E-01 |
| AT1G07013 | 0,15 | 0,02 | 4,36 | 0,00 | 9,96E-01 |
| AT1G40087 | 0,19 | 1,60 | 4,31 | 0,37 | 7,11E-01 |
| AT1G41830 | 0,28 | 1,66 | 4,31 | 0,39 | 6,99E-01 |
| AT1G41920 | 0,51 | -0,05 | 3,24 | -0,02 | 9,88E-01 |
| AT1G42440 | 0,46 | 1,44 | 3,36 | 0,43 | 6,68E-01 |

| | | | | | |
|---|---|---|---|---|---|
| AT1G42560 | 0,05 | 0,18 | 4,38 | 0,04 | 9,68E-01 |
| AT1G42990 | 1,44 | 3,13 | 1,72 | 1,82 | 6,85E-02 |
| AT1G43190 | 0,38 | 0,48 | 4,38 | 0,11 | 9,13E-01 |
| AT1G43560 | 0,22 | -0,15 | 4,35 | -0,03 | 9,72E-01 |
| AT1G43580 | 0,21 | -0,95 | 4,38 | -0,22 | 8,29E-01 |
| AT1G43620 | 0,44 | 0,77 | 4,23 | 0,18 | 8,56E-01 |
| AT1G43665 | 0,88 | 2,48 | 3,18 | 0,78 | 4,37E-01 |
| AT1G43710 | 0,50 | 0,66 | 3,26 | 0,20 | 8,40E-01 |
| AT1G43730 | 0,40 | 1,02 | 4,22 | 0,24 | 8,10E-01 |
| AT1G43760 | 0,66 | -0,88 | 2,42 | -0,37 | 7,15E-01 |
| AT1G43825 | 0,81 | 2,64 | 2,69 | 0,98 | 3,26E-01 |
| AT1G43860 | 1,69 | -0,19 | 1,79 | -0,10 | 9,17E-01 |
| AT1G43950 | 0,46 | 1,95 | 4,02 | 0,48 | 6,28E-01 |
| AT1G44000 | 0,08 | 0,00 | 4,41 | 0,00 | 1,00E+00 |
| AT1G44030 | 0,25 | 0,55 | 4,31 | 0,13 | 8,99E-01 |
| AT1G44080 | 1,35 | -2,56 | 1,80 | -1,42 | 1,56E-01 |
| AT1G44085 | 0,65 | 3,14 | 2,81 | 1,12 | 2,64E-01 |
| AT1G44090 | 0,05 | 0,18 | 4,38 | 0,04 | 9,68E-01 |
| AT1G07243 | 1,82 | 2,84 | 1,60 | 1,77 | 7,62E-02 |
| AT1G44125 | 0,91 | 0,92 | 2,75 | 0,34 | 7,36E-01 |
| AT1G44170 | 0,30 | 2,09 | 4,29 | 0,49 | 6,26E-01 |
| AT1G44350 | 0,41 | 0,03 | 4,24 | 0,01 | 9,94E-01 |
| AT1G44446 | 1,72 | -0,11 | 1,57 | -0,07 | 9,46E-01 |
| AT1G44542 | 0,40 | 1,47 | 4,33 | 0,34 | 7,33E-01 |
| AT1G44770 | 0,41 | 1,21 | 3,34 | 0,36 | 7,17E-01 |
| AT1G44790 | 0,21 | 1,51 | 4,32 | 0,35 | 7,27E-01 |
| AT1G44920 | 0,44 | 0,54 | 3,33 | 0,16 | 8,72E-01 |
| AT1G44980 | 0,59 | -0,15 | 2,62 | -0,06 | 9,54E-01 |
| AT1G45063 | 0,43 | -3,83 | 4,33 | -0,89 | 3,75E-01 |
| AT1G45145 | 0,72 | 1,28 | 3,14 | 0,41 | 6,85E-01 |
| AT1G45150 | 0,25 | 0,35 | 4,37 | 0,08 | 9,37E-01 |
| AT1G45160 | 1,73 | -1,60 | 1,86 | -0,86 | 3,89E-01 |
| AT1G45163 | 0,70 | 0,98 | 2,53 | 0,39 | 7,00E-01 |
| AT1G45207 | 0,37 | 0,00 | 4,41 | 0,00 | 1,00E+00 |
| AT1G07323 | 0,55 | 1,73 | 2,70 | 0,64 | 5,22E-01 |
| AT1G47200 | 0,16 | 1,02 | 4,34 | 0,24 | 8,13E-01 |
| AT1G47240 | 0,43 | -0,31 | 4,18 | -0,07 | 9,40E-01 |
| AT1G07333 | 0,81 | 2,45 | 3,22 | 0,76 | 4,47E-01 |
| AT1G47260 | 1,01 | 0,26 | 1,57 | 0,17 | 8,68E-01 |
| AT1G47280 | 1,31 | 3,16 | 1,99 | 1,59 | 1,13E-01 |
| AT1G07347 | 0,18 | 1,59 | 4,31 | 0,37 | 7,12E-01 |
| AT1G47380 | 0,18 | -0,81 | 4,37 | -0,19 | 8,53E-01 |
| AT1G47420 | 0,71 | 1,25 | 2,24 | 0,56 | 5,77E-01 |
| AT1G07383 | 0,55 | 0,16 | 2,59 | 0,06 | 9,51E-01 |
| AT1G07393 | 1,67 | -1,51 | 1,48 | -1,02 | 3,05E-01 |
| AT1G47640 | 0,35 | 0,89 | 3,50 | 0,26 | 7,98E-01 |
| AT1G47670 | 0,05 | 0,18 | 4,38 | 0,04 | 9,68E-01 |
| AT1G47765 | 0,55 | 0,83 | 2,33 | 0,36 | 7,23E-01 |
| AT1G47900 | 0,45 | -2,74 | 4,16 | -0,66 | 5,11E-01 |
| AT1G47960 | 1,43 | -0,92 | 1,37 | -0,67 | 5,00E-01 |
| AT1G48000 | 0,50 | -0,26 | 4,33 | -0,06 | 9,53E-01 |
| AT1G48030 | 1,26 | 1,52 | 2,22 | 0,69 | 4,93E-01 |
| AT1G48040 | 0,63 | 0,00 | 4,41 | 0,00 | 1,00E+00 |
| AT1G48110 | 0,42 | -1,47 | 4,38 | -0,34 | 7,37E-01 |
| AT1G48120 | 0,82 | 0,18 | 2,46 | 0,07 | 9,43E-01 |
| AT1G48160 | 0,15 | 0,47 | 4,37 | 0,11 | 9,15E-01 |

| | | | | | |
|---|---|---|---|---|---|
| AT1G48285 | 0,80 | 0,92 | 2,28 | 0,40 | 6,87E-01 |
| AT1G48320 | 0,17 | 0,00 | 4,41 | 0,00 | 1,00E+00 |
| AT1G48380 | 0,10 | 0,08 | 4,38 | 0,02 | 9,85E-01 |
| AT1G48405 | 0,82 | 0,00 | 3,43 | 0,00 | 1,00E+00 |
| AT1G48430 | 0,15 | -0,35 | 4,39 | -0,08 | 9,36E-01 |
| AT1G48450 | 1,45 | -0,53 | 1,94 | -0,27 | 7,86E-01 |
| AT1G48550 | 1,08 | -0,98 | 2,71 | -0,36 | 7,17E-01 |
| AT1G48570 | 0,26 | -0,82 | 4,36 | -0,19 | 8,50E-01 |
| AT1G48580 | 0,08 | 0,00 | 4,41 | 0,00 | 1,00E+00 |
| AT1G48620 | 0,46 | 1,07 | 3,14 | 0,34 | 7,33E-01 |
| AT1G48630 | 1,25 | -1,71 | 2,01 | -0,85 | 3,94E-01 |
| AT1G48730 | 0,75 | 1,66 | 2,15 | 0,77 | 4,40E-01 |
| AT1G48742 | 0,46 | -2,74 | 3,14 | -0,87 | 3,82E-01 |
| AT1G48760 | 1,82 | -2,40 | 2,08 | -1,15 | 2,50E-01 |
| AT1G48790 | 0,44 | 0,44 | 3,26 | 0,13 | 8,94E-01 |
| AT1G48850 | 0,39 | 0,17 | 4,37 | 0,04 | 9,70E-01 |
| AT1G48900 | 0,23 | -1,56 | 4,37 | -0,36 | 7,21E-01 |
| AT1G48930 | 0,71 | 0,00 | 2,96 | 0,00 | 1,00E+00 |
| AT1G48970 | 1,37 | -0,83 | 1,88 | -0,44 | 6,58E-01 |
| AT1G49005 | 0,38 | 0,13 | 4,29 | 0,03 | 9,75E-01 |
| AT1G49140 | 0,12 | 0,11 | 4,38 | 0,03 | 9,80E-01 |
| AT1G49180 | 0,08 | 0,00 | 4,41 | 0,00 | 1,00E+00 |
| AT1G49205 | 0,08 | 0,21 | 4,39 | 0,05 | 9,62E-01 |
| AT1G49225 | 0,48 | 1,59 | 3,19 | 0,50 | 6,20E-01 |
| AT1G49270 | 0,10 | 0,86 | 4,35 | 0,20 | 8,43E-01 |
| AT1G49490 | 0,32 | -0,30 | 4,40 | -0,07 | 9,45E-01 |
| AT1G49500 | 1,83 | 0,20 | 1,22 | 0,16 | 8,72E-01 |
| AT1G49540 | 0,83 | -1,69 | 2,50 | -0,68 | 4,98E-01 |
| AT1G49600 | 0,82 | 0,44 | 2,21 | 0,20 | 8,43E-01 |
| AT1G49670 | 0,26 | 1,93 | 4,30 | 0,45 | 6,54E-01 |
| AT1G49710 | 0,34 | 1,58 | 3,47 | 0,46 | 6,49E-01 |
| AT1G49780 | 1,22 | 0,13 | 1,60 | 0,08 | 9,35E-01 |
| AT1G49850 | 0,39 | -0,33 | 4,40 | -0,08 | 9,40E-01 |
| AT1G49970 | 0,66 | 0,06 | 2,14 | 0,03 | 9,78E-01 |
| AT1G07603 | 0,61 | 0,44 | 3,26 | 0,14 | 8,92E-01 |
| AT1G50020 | 0,58 | -1,55 | 4,00 | -0,39 | 6,99E-01 |
| AT1G50140 | 0,20 | -2,08 | 4,39 | -0,47 | 6,36E-01 |
| AT1G50190 | 0,32 | -0,32 | 4,40 | -0,07 | 9,42E-01 |
| AT1G50260 | 0,32 | -1,26 | 4,37 | -0,29 | 7,73E-01 |
| AT1G50280 | 0,61 | -0,33 | 2,23 | -0,15 | 8,82E-01 |
| AT1G50320 | 0,25 | -0,80 | 4,37 | -0,18 | 8,54E-01 |
| AT1G07627 | 0,18 | 0,93 | 4,34 | 0,21 | 8,30E-01 |
| AT1G50420 | 0,49 | 0,00 | 4,41 | 0,00 | 1,00E+00 |
| AT1G50430 | 1,02 | 1,55 | 1,95 | 0,80 | 4,25E-01 |
| AT1G50450 | 0,63 | -1,35 | 3,02 | -0,45 | 6,54E-01 |
| AT1G50500 | 0,57 | -2,27 | 3,13 | -0,73 | 4,68E-01 |
| AT1G07647 | 0,08 | 0,00 | 4,41 | 0,00 | 1,00E+00 |
| AT1G50570 | 0,65 | 0,75 | 2,63 | 0,29 | 7,74E-01 |
| AT1G50600 | 0,88 | 0,77 | 2,10 | 0,37 | 7,12E-01 |
| AT1G50640 | 1,08 | 0,14 | 2,02 | 0,07 | 9,43E-01 |
| AT1G50659 | 0,43 | 2,28 | 3,20 | 0,71 | 4,76E-01 |
| AT1G50750 | 0,08 | 0,21 | 4,39 | 0,05 | 9,62E-01 |
| AT1G50790 | 0,12 | 0,00 | 4,41 | 0,00 | 1,00E+00 |
| AT1G50820 | 0,50 | 0,00 | 4,04 | 0,00 | 1,00E+00 |
| AT1G50910 | 1,39 | 0,51 | 1,66 | 0,31 | 7,57E-01 |
| AT1G50980 | 0,39 | -0,33 | 4,40 | -0,08 | 9,40E-01 |

| | | | | | |
|---|---|---|---|---|---|
| AT1G51080 | 0,73 | 1,49 | 2,59 | 0,58 | 5,65E-01 |
| AT1G07733 | 0,24 | -0,67 | 4,38 | -0,15 | 8,78E-01 |
| AT1G51400 | 0,93 | -0,18 | 2,00 | -0,09 | 9,27E-01 |
| AT1G51480 | 0,33 | 0,00 | 4,41 | 0,00 | 1,00E+00 |
| AT1G51500 | 1,21 | -0,94 | 1,83 | -0,51 | 6,09E-01 |
| AT1G51510 | 0,91 | 1,23 | 2,13 | 0,58 | 5,63E-01 |
| AT1G51560 | 0,45 | -0,30 | 3,38 | -0,09 | 9,29E-01 |
| AT1G51570 | 1,72 | -0,11 | 1,82 | -0,06 | 9,50E-01 |
| AT1G51590 | 0,26 | 1,66 | 4,32 | 0,39 | 7,00E-01 |
| AT1G51630 | 0,52 | 0,89 | 3,05 | 0,29 | 7,71E-01 |
| AT1G51680 | 0,80 | 2,19 | 2,14 | 1,02 | 3,06E-01 |
| AT1G07763 | 0,56 | -3,98 | 4,32 | -0,92 | 3,57E-01 |
| AT1G51710 | 0,45 | -2,09 | 4,17 | -0,50 | 6,17E-01 |
| AT1G51730 | 0,47 | -1,38 | 3,23 | -0,43 | 6,70E-01 |
| AT1G51790 | 0,25 | 0,28 | 4,39 | 0,06 | 9,50E-01 |
| AT1G51890 | 0,23 | -0,33 | 4,40 | -0,08 | 9,40E-01 |
| AT1G51940 | 1,08 | -0,84 | 2,54 | -0,33 | 7,41E-01 |
| AT1G51980 | 1,85 | 0,00 | 1,42 | 0,00 | 9,99E-01 |
| AT1G52000 | 0,46 | -2,70 | 4,04 | -0,67 | 5,03E-01 |
| AT1G52040 | 0,54 | -3,58 | 3,91 | -0,92 | 3,59E-01 |
| AT1G52150 | 1,30 | 0,14 | 2,51 | 0,06 | 9,55E-01 |
| AT1G52160 | 0,61 | 1,75 | 3,09 | 0,57 | 5,70E-01 |
| AT1G52180 | 1,00 | -1,49 | 1,82 | -0,82 | 4,13E-01 |
| AT1G52185 | 0,32 | -2,21 | 4,33 | -0,51 | 6,10E-01 |
| AT1G52200 | 0,25 | 0,00 | 4,41 | 0,00 | 1,00E+00 |
| AT1G52220 | 0,76 | -1,25 | 2,13 | -0,59 | 5,58E-01 |
| AT1G52290 | 0,41 | 0,01 | 4,28 | 0,00 | 9,98E-01 |
| AT1G52300 | 0,35 | -0,16 | 4,34 | -0,04 | 9,71E-01 |
| AT1G52340 | 0,08 | 0,21 | 4,39 | 0,05 | 9,62E-01 |
| AT1G52342 | 0,45 | 0,00 | 4,41 | 0,00 | 1,00E+00 |
| AT1G52510 | 0,47 | -0,72 | 4,24 | -0,17 | 8,65E-01 |
| AT1G52570 | 1,59 | 0,00 | 2,45 | 0,00 | 1,00E+00 |
| AT1G52750 | 0,31 | -0,39 | 4,39 | -0,09 | 9,29E-01 |
| AT1G52880 | 0,40 | -1,27 | 3,48 | -0,37 | 7,14E-01 |
| AT1G52890 | 0,42 | -0,29 | 4,21 | -0,07 | 9,45E-01 |
| AT1G52970 | 1,13 | -2,64 | 1,84 | -1,43 | 1,52E-01 |
| AT1G52990 | 0,41 | 2,84 | 4,28 | 0,66 | 5,07E-01 |
| AT1G53000 | 0,49 | -0,37 | 4,07 | -0,09 | 9,28E-01 |
| AT1G53025 | 0,43 | 0,87 | 3,37 | 0,26 | 7,96E-01 |
| AT1G53035 | 0,12 | 0,00 | 4,41 | 0,00 | 1,00E+00 |
| AT1G53170 | 0,65 | -0,24 | 2,44 | -0,10 | 9,21E-01 |
| AT1G53200 | 0,24 | 0,00 | 4,41 | 0,00 | 1,00E+00 |
| AT1G53230 | 1,22 | 1,89 | 2,18 | 0,86 | 3,87E-01 |
| AT1G53240 | 1,16 | -0,43 | 1,69 | -0,25 | 8,01E-01 |
| AT1G53265 | 0,53 | -2,88 | 3,99 | -0,72 | 4,71E-01 |
| AT1G53320 | 0,35 | 0,39 | 4,38 | 0,09 | 9,30E-01 |
| AT1G53450 | 0,86 | -0,47 | 1,77 | -0,26 | 7,92E-01 |
| AT1G53480 | 0,84 | 0,38 | 2,91 | 0,13 | 8,96E-01 |
| AT1G53500 | 0,50 | -0,96 | 3,27 | -0,29 | 7,69E-01 |
| AT1G53510 | 0,76 | -0,74 | 2,45 | -0,30 | 7,62E-01 |
| AT1G53541 | 0,64 | 1,20 | 2,48 | 0,49 | 6,27E-01 |
| AT1G53580 | 0,12 | 0,27 | 4,38 | 0,06 | 9,50E-01 |
| AT1G53710 | 0,67 | -2,19 | 3,05 | -0,72 | 4,73E-01 |
| AT1G53750 | 1,03 | 0,17 | 1,99 | 0,08 | 9,33E-01 |
| AT1G53760 | 0,08 | 0,00 | 4,41 | 0,00 | 1,00E+00 |
| AT1G53770 | 0,33 | 0,00 | 4,41 | 0,00 | 1,00E+00 |

| | | | | | |
|---|---|---|---|---|---|
| AT1G53780 | 1,40 | 0,59 | 2,14 | 0,27 | 7,84E-01 |
| AT1G54050 | 0,12 | 0,27 | 4,38 | 0,06 | 9,50E-01 |
| AT1G54080 | 1,16 | -1,76 | 2,15 | -0,82 | 4,13E-01 |
| AT1G54100 | 1,08 | 0,00 | 1,95 | 0,00 | 9,98E-01 |
| AT1G54140 | 0,15 | 0,47 | 4,37 | 0,11 | 9,15E-01 |
| AT1G54170 | 0,82 | -3,17 | 2,14 | -1,48 | 1,38E-01 |
| AT1G54320 | 0,67 | -1,39 | 2,38 | -0,59 | 5,58E-01 |
| AT1G54370 | 0,38 | -0,26 | 4,36 | -0,06 | 9,52E-01 |
| AT1G54500 | 0,85 | 1,79 | 2,05 | 0,87 | 3,84E-01 |
| AT1G54590 | 0,10 | 0,08 | 4,38 | 0,02 | 9,85E-01 |
| AT1G54630 | 0,35 | -0,88 | 4,36 | -0,20 | 8,40E-01 |
| AT1G54690 | 0,48 | 0,92 | 2,77 | 0,33 | 7,38E-01 |
| AT1G54775 | 0,57 | -0,66 | 3,74 | -0,18 | 8,59E-01 |
| AT1G54840 | 0,49 | -0,38 | 4,39 | -0,09 | 9,31E-01 |
| AT1G54950 | 0,44 | 0,61 | 3,96 | 0,15 | 8,78E-01 |
| AT1G55152 | 0,78 | 2,10 | 2,45 | 0,86 | 3,92E-01 |
| AT1G55180 | 0,78 | -0,32 | 2,91 | -0,11 | 9,13E-01 |
| AT1G55200 | 0,08 | 0,00 | 4,41 | 0,00 | 1,00E+00 |
| AT1G08043 | 0,30 | 0,00 | 4,41 | 0,00 | 1,00E+00 |
| AT1G55270 | 1,00 | 0,00 | 3,28 | 0,00 | 1,00E+00 |
| AT1G55300 | 0,10 | 0,24 | 4,38 | 0,05 | 9,57E-01 |
| AT1G55330 | 0,51 | -2,40 | 3,17 | -0,76 | 4,48E-01 |
| AT1G55360 | 0,31 | 0,81 | 4,35 | 0,19 | 8,52E-01 |
| AT1G55530 | 0,83 | -1,01 | 2,23 | -0,45 | 6,50E-01 |
| AT1G55535 | 0,40 | -2,12 | 4,28 | -0,50 | 6,19E-01 |
| AT1G55590 | 0,61 | -0,80 | 2,68 | -0,30 | 7,65E-01 |
| AT1G55610 | 0,12 | 0,27 | 4,38 | 0,06 | 9,50E-01 |
| AT1G55620 | 0,57 | 0,77 | 3,10 | 0,25 | 8,05E-01 |
| AT1G55675 | 0,68 | 1,01 | 3,74 | 0,27 | 7,87E-01 |
| AT1G55680 | 1,44 | -1,35 | 2,23 | -0,60 | 5,47E-01 |
| AT1G55690 | 0,12 | 0,00 | 4,41 | 0,00 | 1,00E+00 |
| AT1G55830 | 0,23 | 0,81 | 4,35 | 0,19 | 8,52E-01 |
| AT1G55880 | 0,36 | 0,00 | 4,41 | 0,00 | 1,00E+00 |
| AT1G55950 | 0,93 | 2,79 | 2,13 | 1,31 | 1,91E-01 |
| AT1G55960 | 0,25 | -0,81 | 4,36 | -0,19 | 8,52E-01 |
| AT1G55970 | 0,17 | 0,00 | 4,41 | 0,00 | 1,00E+00 |
| AT1G56010 | 0,42 | -0,26 | 4,18 | -0,06 | 9,50E-01 |
| AT1G56100 | 0,86 | -2,19 | 2,77 | -0,79 | 4,29E-01 |
| AT1G56110 | 1,35 | -0,28 | 1,82 | -0,16 | 8,77E-01 |
| AT1G56120 | 0,47 | -1,14 | 4,04 | -0,28 | 7,78E-01 |
| AT1G56140 | 0,44 | 1,18 | 3,29 | 0,36 | 7,20E-01 |
| AT1G56145 | 0,80 | 0,00 | 2,87 | 0,00 | 1,00E+00 |
| AT1G56165 | 0,20 | 0,00 | 4,41 | 0,00 | 1,00E+00 |
| AT1G56230 | 0,19 | -0,58 | 4,38 | -0,13 | 8,95E-01 |
| AT1G08107 | 0,31 | 1,09 | 3,54 | 0,31 | 7,59E-01 |
| AT1G08113 | 0,55 | -1,59 | 3,12 | -0,51 | 6,11E-01 |
| AT1G08123 | 0,56 | 1,38 | 2,47 | 0,56 | 5,75E-01 |
| AT1G56300 | 0,85 | 0,20 | 2,81 | 0,07 | 9,44E-01 |
| AT1G56330 | 0,78 | 0,81 | 2,39 | 0,34 | 7,35E-01 |
| AT1G56345 | 0,19 | 1,00 | 4,33 | 0,23 | 8,17E-01 |
| AT1G56440 | 0,27 | 0,54 | 4,36 | 0,12 | 9,02E-01 |
| AT1G56520 | 1,21 | 1,98 | 2,30 | 0,86 | 3,89E-01 |
| AT1G56550 | 0,17 | -2,21 | 4,40 | -0,50 | 6,16E-01 |
| AT1G56580 | 0,24 | 0,59 | 4,36 | 0,14 | 8,92E-01 |
| AT1G56630 | 1,84 | -6,07 | 2,10 | -2,89 | 3,85E-03 |
| AT1G56650 | 1,01 | 0,00 | 4,04 | 0,00 | 1,00E+00 |

| | | | | | |
|---|---|---|---|---|---|
| AT1G57680 | 1,45 | -0,31 | 1,98 | -0,16 | 8,75E-01 |
| AT1G57775 | 0,38 | 1,14 | 3,32 | 0,34 | 7,31E-01 |
| AT1G57777 | 0,31 | 1,31 | 3,61 | 0,36 | 7,16E-01 |
| AT1G57860 | 0,30 | -2,51 | 4,38 | -0,57 | 5,67E-01 |
| AT1G57870 | 0,40 | 0,17 | 4,26 | 0,04 | 9,68E-01 |
| AT1G57943 | 0,08 | 0,00 | 4,41 | 0,00 | 1,00E+00 |
| AT1G57990 | 0,77 | 1,50 | 2,55 | 0,59 | 5,55E-01 |
| AT1G58030 | 0,73 | -0,32 | 2,57 | -0,12 | 9,02E-01 |
| AT1G58060 | 0,60 | 0,90 | 3,13 | 0,29 | 7,74E-01 |
| AT1G58080 | 0,29 | -0,35 | 4,40 | -0,08 | 9,37E-01 |
| AT1G58100 | 1,20 | -0,40 | 2,05 | -0,20 | 8,43E-01 |
| AT1G58110 | 0,53 | -1,96 | 3,96 | -0,50 | 6,20E-01 |
| AT1G58200 | 0,16 | 1,02 | 4,34 | 0,24 | 8,13E-01 |
| AT1G58220 | 0,54 | -0,26 | 3,92 | -0,07 | 9,47E-01 |
| AT1G58300 | 1,33 | -1,71 | 1,53 | -1,12 | 2,64E-01 |
| AT1G58350 | 0,80 | 1,66 | 2,57 | 0,65 | 5,19E-01 |
| AT1G58390 | 0,70 | 1,20 | 2,87 | 0,42 | 6,77E-01 |
| AT1G58440 | 0,58 | -0,69 | 2,90 | -0,24 | 8,11E-01 |
| AT1G58684 | 0,49 | -0,72 | 4,29 | -0,17 | 8,67E-01 |
| AT1G58725 | 0,33 | 0,63 | 4,32 | 0,15 | 8,84E-01 |
| AT1G58766 | 0,16 | 0,15 | 4,38 | 0,04 | 9,72E-01 |
| AT1G59030 | 0,44 | -0,12 | 4,21 | -0,03 | 9,77E-01 |
| AT1G59077 | 0,16 | -0,56 | 4,37 | -0,13 | 8,98E-01 |
| AT1G59406 | 0,14 | 0,89 | 4,35 | 0,21 | 8,37E-01 |
| AT1G59453 | 0,25 | 1,56 | 4,32 | 0,36 | 7,17E-01 |
| AT1G59720 | 0,61 | 0,33 | 2,74 | 0,12 | 9,05E-01 |
| AT1G59780 | 0,52 | 0,48 | 3,12 | 0,15 | 8,77E-01 |
| AT1G59835 | 1,47 | -3,84 | 1,74 | -2,21 | 2,71E-02 |
| AT1G59910 | 1,37 | 2,71 | 2,05 | 1,32 | 1,86E-01 |
| AT1G59920 | 0,42 | 1,31 | 3,22 | 0,41 | 6,84E-01 |
| AT1G59930 | 1,11 | 3,24 | 2,34 | 1,39 | 1,65E-01 |
| AT1G59990 | 1,13 | -0,30 | 2,24 | -0,13 | 8,94E-01 |
| AT1G60000 | 0,38 | 0,07 | 3,51 | 0,02 | 9,84E-01 |
| AT1G60070 | 1,35 | -0,29 | 2,26 | -0,13 | 8,98E-01 |
| AT1G60160 | 0,43 | 1,15 | 3,22 | 0,36 | 7,22E-01 |
| AT1G60200 | 0,68 | 1,94 | 3,16 | 0,61 | 5,39E-01 |
| AT1G60230 | 0,48 | 0,51 | 4,05 | 0,13 | 9,00E-01 |
| AT1G60260 | 0,77 | -0,16 | 2,47 | -0,06 | 9,48E-01 |
| AT1G08317 | 0,08 | 0,00 | 4,41 | 0,00 | 1,00E+00 |
| AT1G60430 | 0,45 | 0,00 | 4,07 | 0,00 | 1,00E+00 |
| AT1G60490 | 0,25 | -0,80 | 4,37 | -0,18 | 8,54E-01 |
| AT1G60550 | 0,38 | 0,00 | 4,41 | 0,00 | 1,00E+00 |
| AT1G60560 | 0,17 | 0,00 | 4,41 | 0,00 | 1,00E+00 |
| AT1G60590 | 0,17 | -3,06 | 4,35 | -0,70 | 4,82E-01 |
| AT1G60630 | 0,08 | 0,00 | 4,41 | 0,00 | 1,00E+00 |
| AT1G60640 | 0,12 | -0,26 | 4,41 | -0,06 | 9,53E-01 |
| AT1G60700 | 0,48 | 0,00 | 4,41 | 0,00 | 1,00E+00 |
| AT1G60710 | 0,41 | 0,00 | 4,41 | 0,00 | 1,00E+00 |
| AT1G60770 | 1,20 | 0,00 | 2,33 | 0,00 | 1,00E+00 |
| AT1G60780 | 0,33 | 1,59 | 4,32 | 0,37 | 7,13E-01 |
| AT1G60800 | 0,56 | -3,23 | 3,08 | -1,05 | 2,94E-01 |
| AT1G60810 | 0,12 | 1,17 | 4,34 | 0,27 | 7,88E-01 |
| AT1G60850 | 0,11 | -0,26 | 4,41 | -0,06 | 9,53E-01 |
| AT1G60890 | 0,24 | -2,09 | 4,39 | -0,48 | 6,34E-01 |
| AT1G60930 | 0,24 | 0,00 | 4,41 | 0,00 | 1,00E+00 |
| AT1G60940 | 0,93 | -0,56 | 2,21 | -0,25 | 7,99E-01 |

| | | | | | |
|---|---|---|---|---|---|
| AT1G60960 | 1,57 | 0,17 | 1,80 | 0,10 | 9,23E-01 |
| AT1G60990 | 0,24 | 0,00 | 4,41 | 0,00 | 1,00E+00 |
| AT1G60995 | 0,35 | 1,46 | 4,32 | 0,34 | 7,35E-01 |
| AT1G61010 | 0,89 | 0,47 | 2,41 | 0,19 | 8,46E-01 |
| AT1G61200 | 1,13 | -0,42 | 1,47 | -0,28 | 7,76E-01 |
| AT1G61260 | 0,47 | 2,02 | 3,21 | 0,63 | 5,28E-01 |
| AT1G61300 | 1,06 | -1,24 | 2,23 | -0,56 | 5,76E-01 |
| AT1G61310 | 0,33 | 0,79 | 4,30 | 0,18 | 8,55E-01 |
| AT1G61360 | 0,15 | 0,47 | 4,37 | 0,11 | 9,15E-01 |
| AT1G61370 | 0,17 | 0,07 | 4,38 | 0,02 | 9,87E-01 |
| AT1G61380 | 0,34 | 1,15 | 4,33 | 0,26 | 7,91E-01 |
| AT1G61420 | 0,12 | 0,00 | 4,41 | 0,00 | 1,00E+00 |
| AT1G61560 | 0,37 | 0,22 | 4,39 | 0,05 | 9,59E-01 |
| AT1G08417 | 0,75 | 2,22 | 2,77 | 0,80 | 4,23E-01 |
| AT1G61660 | 0,65 | 0,84 | 2,93 | 0,29 | 7,73E-01 |
| AT1G61730 | 1,18 | -3,24 | 2,27 | -1,43 | 1,53E-01 |
| AT1G61890 | 1,56 | 3,30 | 1,80 | 1,84 | 6,62E-02 |
| AT1G61900 | 0,26 | 0,08 | 4,32 | 0,02 | 9,85E-01 |
| AT1G62030 | 1,53 | 2,42 | 1,70 | 1,42 | 1,55E-01 |
| AT1G62085 | 0,60 | 0,89 | 3,20 | 0,28 | 7,81E-01 |
| AT1G62300 | 0,27 | 1,11 | 4,34 | 0,25 | 7,99E-01 |
| AT1G62422 | 0,51 | -0,11 | 2,68 | -0,04 | 9,66E-01 |
| AT1G62610 | 0,76 | 0,09 | 2,47 | 0,04 | 9,71E-01 |
| AT1G62680 | 1,24 | -0,65 | 1,89 | -0,34 | 7,33E-01 |
| AT1G62760 | 0,29 | 1,20 | 4,33 | 0,28 | 7,83E-01 |
| AT1G62780 | 0,49 | 0,06 | 3,17 | 0,02 | 9,85E-01 |
| AT1G62810 | 0,97 | -0,32 | 2,18 | -0,15 | 8,82E-01 |
| AT1G62915 | 0,64 | -0,47 | 2,84 | -0,16 | 8,69E-01 |
| AT1G62960 | 0,36 | -0,95 | 4,37 | -0,22 | 8,28E-01 |
| AT1G63010 | 0,16 | -0,56 | 4,37 | -0,13 | 8,98E-01 |
| AT1G63020 | 0,44 | 0,00 | 4,41 | 0,00 | 1,00E+00 |
| AT1G63050 | 0,34 | 0,09 | 4,38 | 0,02 | 9,84E-01 |
| AT1G63090 | 1,34 | 0,10 | 1,57 | 0,07 | 9,48E-01 |
| AT1G63250 | 0,13 | 0,00 | 4,41 | 0,00 | 1,00E+00 |
| AT1G63290 | 0,12 | -0,26 | 4,41 | -0,06 | 9,53E-01 |
| AT1G63320 | 0,20 | 1,86 | 4,30 | 0,43 | 6,66E-01 |
| AT1G63410 | 0,62 | -0,01 | 2,59 | 0,00 | 9,98E-01 |
| AT1G63490 | 0,53 | 0,85 | 3,23 | 0,26 | 7,93E-01 |
| AT1G63660 | 0,92 | 0,10 | 2,24 | 0,04 | 9,65E-01 |
| AT1G63670 | 0,18 | 0,31 | 4,38 | 0,07 | 9,44E-01 |
| AT1G63700 | 1,36 | 0,72 | 1,68 | 0,43 | 6,67E-01 |
| AT1G63720 | 0,19 | 0,50 | 4,37 | 0,12 | 9,08E-01 |
| AT1G08627 | 1,18 | -3,83 | 1,96 | -1,95 | 5,10E-02 |
| AT1G63800 | 0,47 | -0,25 | 4,33 | -0,06 | 9,53E-01 |
| AT1G63810 | 0,31 | -2,52 | 4,38 | -0,58 | 5,65E-01 |
| AT1G63840 | 0,40 | -0,32 | 4,40 | -0,07 | 9,42E-01 |
| AT1G63860 | 1,09 | 0,36 | 2,01 | 0,18 | 8,58E-01 |
| AT1G63870 | 0,58 | -0,33 | 3,11 | -0,11 | 9,16E-01 |
| AT1G63970 | 0,51 | -0,51 | 3,28 | -0,16 | 8,76E-01 |
| AT1G64080 | 0,15 | 0,82 | 4,35 | 0,19 | 8,50E-01 |
| AT1G64190 | 1,11 | -0,26 | 1,82 | -0,14 | 8,87E-01 |
| AT1G64200 | 0,78 | -2,67 | 4,37 | -0,61 | 5,42E-01 |
| AT1G64280 | 0,86 | 0,53 | 2,62 | 0,20 | 8,40E-01 |
| AT1G64380 | 0,37 | 0,69 | 4,29 | 0,16 | 8,72E-01 |
| AT1G64460 | 0,72 | -0,33 | 2,57 | -0,13 | 8,98E-01 |
| AT1G64480 | 0,18 | 0,93 | 4,34 | 0,21 | 8,30E-01 |

| | | | | | |
|---|---|---|---|---|---|
| AT1G64510 | 0,32 | 0,02 | 4,34 | 0,00 | 9,97E-01 |
| AT1G64580 | 0,24 | -1,51 | 4,38 | -0,34 | 7,31E-01 |
| AT1G64610 | 0,62 | 0,07 | 2,65 | 0,03 | 9,80E-01 |
| AT1G64640 | 0,11 | -0,38 | 4,39 | -0,09 | 9,31E-01 |
| AT1G64650 | 0,59 | 2,42 | 2,43 | 1,00 | 3,20E-01 |
| AT1G64670 | 0,10 | 0,24 | 4,38 | 0,05 | 9,57E-01 |
| AT1G64750 | 0,36 | -2,40 | 4,37 | -0,55 | 5,83E-01 |
| AT1G64970 | 0,63 | -0,33 | 2,62 | -0,13 | 8,99E-01 |
| AT1G64980 | 1,26 | -0,64 | 1,54 | -0,41 | 6,79E-01 |
| AT1G65040 | 0,28 | 0,81 | 3,62 | 0,22 | 8,23E-01 |
| AT1G65150 | 0,20 | -2,08 | 4,39 | -0,47 | 6,35E-01 |
| AT1G65190 | 0,43 | -1,94 | 4,10 | -0,47 | 6,36E-01 |
| AT1G08747 | 0,81 | -1,11 | 2,94 | -0,38 | 7,05E-01 |
| AT1G65290 | 0,77 | -0,07 | 2,27 | -0,03 | 9,74E-01 |
| AT1G65320 | 0,39 | -1,31 | 4,35 | -0,30 | 7,63E-01 |
| AT1G65346 | 0,34 | 1,77 | 4,31 | 0,41 | 6,82E-01 |
| AT1G65420 | 0,15 | -2,07 | 4,39 | -0,47 | 6,38E-01 |
| AT1G65430 | 0,52 | -0,80 | 3,17 | -0,25 | 8,02E-01 |
| AT1G65520 | 0,29 | -0,69 | 4,33 | -0,16 | 8,73E-01 |
| AT1G65540 | 1,25 | -2,01 | 2,12 | -0,95 | 3,43E-01 |
| AT1G65550 | 0,21 | -2,19 | 4,41 | -0,50 | 6,20E-01 |
| AT1G65580 | 0,46 | -0,59 | 2,68 | -0,22 | 8,24E-01 |
| AT1G65690 | 0,12 | 0,00 | 4,41 | 0,00 | 1,00E+00 |
| AT1G65950 | 0,28 | 0,00 | 4,41 | 0,00 | 1,00E+00 |
| AT1G08797 | 0,63 | -0,26 | 4,41 | -0,06 | 9,53E-01 |
| AT1G65980 | 1,27 | -0,31 | 2,34 | -0,13 | 8,95E-01 |
| AT1G65985 | 0,33 | -0,59 | 4,34 | -0,14 | 8,92E-01 |
| AT1G65990 | 1,24 | 3,75 | 1,99 | 1,89 | 5,92E-02 |
| AT1G66200 | 1,38 | 0,50 | 1,43 | 0,35 | 7,27E-01 |
| AT1G66235 | 0,12 | 0,11 | 4,38 | 0,03 | 9,80E-01 |
| AT1G66240 | 0,12 | -0,26 | 4,41 | -0,06 | 9,53E-01 |
| AT1G66260 | 1,28 | 1,05 | 1,73 | 0,61 | 5,43E-01 |
| AT1G66290 | 0,49 | -3,17 | 3,17 | -1,00 | 3,17E-01 |
| AT1G66330 | 0,52 | -0,69 | 2,65 | -0,26 | 7,96E-01 |
| AT1G66410 | 1,68 | 0,36 | 1,54 | 0,23 | 8,15E-01 |
| AT1G66460 | 0,25 | -2,17 | 4,41 | -0,49 | 6,22E-01 |
| AT1G66490 | 0,90 | -2,60 | 2,43 | -1,07 | 2,85E-01 |
| AT1G66500 | 0,25 | 0,00 | 4,41 | 0,00 | 1,00E+00 |
| AT1G66570 | 1,67 | 0,59 | 1,43 | 0,42 | 6,76E-01 |
| AT1G66660 | 0,44 | 0,48 | 4,16 | 0,12 | 9,08E-01 |
| AT1G66670 | 1,37 | 1,92 | 1,80 | 1,06 | 2,87E-01 |
| AT1G66680 | 0,59 | 0,00 | 4,41 | 0,00 | 1,00E+00 |
| AT1G66700 | 0,46 | 0,07 | 4,38 | 0,02 | 9,87E-01 |
| AT1G66830 | 0,68 | -1,35 | 3,81 | -0,35 | 7,23E-01 |
| AT1G66910 | 0,65 | 0,45 | 2,33 | 0,19 | 8,45E-01 |
| AT1G66940 | 0,34 | 1,59 | 4,30 | 0,37 | 7,12E-01 |
| AT1G66980 | 1,51 | -0,55 | 1,52 | -0,36 | 7,17E-01 |
| AT1G67000 | 0,91 | 0,61 | 1,82 | 0,33 | 7,39E-01 |
| AT1G67050 | 0,28 | 0,72 | 3,61 | 0,20 | 8,42E-01 |
| AT1G67080 | 0,36 | -0,33 | 4,40 | -0,08 | 9,40E-01 |
| AT1G67140 | 0,95 | 0,48 | 2,61 | 0,18 | 8,54E-01 |
| AT1G67230 | 0,62 | -0,25 | 2,54 | -0,10 | 9,23E-01 |
| AT1G67235 | 0,29 | 2,32 | 4,29 | 0,54 | 5,89E-01 |
| AT1G67250 | 0,34 | -0,81 | 4,36 | -0,19 | 8,53E-01 |
| AT1G67280 | 1,72 | -1,94 | 1,60 | -1,22 | 2,24E-01 |
| AT1G67300 | 0,37 | -0,34 | 4,36 | -0,08 | 9,38E-01 |

| | | | | | |
|---|---|---|---|---|---|
| AT1G67310 | 0,33 | -0,36 | 4,39 | -0,08 | 9,34E-01 |
| AT1G67325 | 0,61 | -0,35 | 3,08 | -0,11 | 9,10E-01 |
| AT1G67360 | 0,73 | -1,04 | 2,76 | -0,38 | 7,07E-01 |
| AT1G67481 | 1,30 | 2,12 | 1,95 | 1,09 | 2,76E-01 |
| AT1G67490 | 0,57 | 0,88 | 3,15 | 0,28 | 7,79E-01 |
| AT1G67500 | 0,52 | -0,35 | 4,07 | -0,08 | 9,32E-01 |
| AT1G67510 | 0,38 | -0,16 | 4,26 | -0,04 | 9,71E-01 |
| AT1G67580 | 1,21 | 0,71 | 2,02 | 0,35 | 7,27E-01 |
| AT1G67590 | 0,17 | 0,07 | 4,38 | 0,02 | 9,88E-01 |
| AT1G67680 | 0,43 | -0,38 | 4,39 | -0,09 | 9,30E-01 |
| AT1G67785 | 0,15 | -0,57 | 4,37 | -0,13 | 8,96E-01 |
| AT1G67790 | 0,26 | 0,00 | 4,41 | 0,00 | 1,00E+00 |
| AT1G67860 | 0,58 | 0,49 | 3,10 | 0,16 | 8,75E-01 |
| AT1G67865 | 0,24 | -1,16 | 4,40 | -0,26 | 7,92E-01 |
| AT1G67870 | 0,89 | 0,59 | 1,75 | 0,34 | 7,35E-01 |
| AT1G67930 | 1,11 | 0,58 | 1,81 | 0,32 | 7,49E-01 |
| AT1G67950 | 0,39 | 0,00 | 4,40 | 0,00 | 1,00E+00 |
| AT1G68060 | 0,56 | -1,24 | 3,22 | -0,39 | 7,00E-01 |
| AT1G68140 | 0,87 | 1,80 | 2,14 | 0,84 | 3,99E-01 |
| AT1G68325 | 0,05 | 0,18 | 4,38 | 0,04 | 9,68E-01 |
| AT1G68370 | 0,32 | -2,12 | 4,40 | -0,48 | 6,29E-01 |
| AT1G68550 | 0,87 | 1,57 | 1,98 | 0,79 | 4,29E-01 |
| AT1G08997 | 0,58 | -0,92 | 3,76 | -0,24 | 8,07E-01 |
| AT1G68590 | 1,41 | 1,94 | 1,85 | 1,05 | 2,95E-01 |
| AT1G68660 | 0,71 | -1,80 | 2,22 | -0,81 | 4,19E-01 |
| AT1G68710 | 0,46 | 0,00 | 4,41 | 0,00 | 1,00E+00 |
| AT1G68830 | 0,57 | -1,17 | 2,66 | -0,44 | 6,59E-01 |
| AT1G68910 | 0,08 | 0,00 | 4,41 | 0,00 | 1,00E+00 |
| AT1G69020 | 0,30 | 0,00 | 4,41 | 0,00 | 1,00E+00 |
| AT1G69070 | 0,46 | -0,29 | 3,39 | -0,08 | 9,33E-01 |
| AT1G69220 | 1,28 | -1,12 | 2,19 | -0,51 | 6,08E-01 |
| AT1G69270 | 0,78 | -1,29 | 3,15 | -0,41 | 6,81E-01 |
| AT1G69280 | 0,21 | -2,19 | 4,41 | -0,50 | 6,20E-01 |
| AT1G69290 | 0,15 | -0,31 | 4,40 | -0,07 | 9,43E-01 |
| AT1G69310 | 0,25 | -0,31 | 4,40 | -0,07 | 9,44E-01 |
| AT1G69450 | 0,38 | 1,34 | 4,30 | 0,31 | 7,55E-01 |
| AT1G69480 | 0,55 | 0,00 | 4,39 | 0,00 | 1,00E+00 |
| AT1G69630 | 0,59 | 0,00 | 4,41 | 0,00 | 1,00E+00 |
| AT1G69640 | 0,90 | -0,44 | 2,38 | -0,19 | 8,53E-01 |
| AT1G69730 | 0,73 | 2,71 | 2,30 | 1,18 | 2,40E-01 |
| AT1G69760 | 0,47 | 1,36 | 3,10 | 0,44 | 6,61E-01 |
| AT1G69770 | 1,05 | -0,33 | 2,45 | -0,13 | 8,94E-01 |
| AT1G69795 | 1,39 | -4,59 | 2,37 | -1,94 | 5,27E-02 |
| AT1G69840 | 0,65 | 1,91 | 3,00 | 0,64 | 5,24E-01 |
| AT1G69850 | 0,22 | 1,16 | 4,34 | 0,27 | 7,89E-01 |
| AT1G69890 | 1,49 | 2,14 | 1,84 | 1,16 | 2,45E-01 |
| AT1G69960 | 0,51 | -1,32 | 3,24 | -0,41 | 6,83E-01 |
| AT1G70000 | 0,98 | -1,25 | 2,23 | -0,56 | 5,74E-01 |
| AT1G70070 | 1,07 | -2,41 | 2,05 | -1,18 | 2,40E-01 |
| AT1G70200 | 0,37 | 0,06 | 4,32 | 0,01 | 9,88E-01 |
| AT1G70280 | 0,17 | 0,00 | 4,41 | 0,00 | 1,00E+00 |
| AT1G70300 | 0,38 | -0,32 | 4,33 | -0,07 | 9,41E-01 |
| AT1G70310 | 0,89 | -3,34 | 3,04 | -1,10 | 2,72E-01 |
| AT1G70340 | 0,34 | -0,48 | 3,58 | -0,13 | 8,94E-01 |
| AT1G70370 | 0,57 | -0,02 | 2,55 | -0,01 | 9,94E-01 |
| AT1G70410 | 0,70 | -0,42 | 2,56 | -0,17 | 8,69E-01 |

| | | | | | |
|---|---|---|---|---|---|
| AT1G70420 | 0,44 | -1,62 | 4,03 | -0,40 | 6,88E-01 |
| AT1G70490 | 1,85 | 0,17 | 1,36 | 0,12 | 9,02E-01 |
| AT1G70520 | 0,53 | -3,12 | 4,36 | -0,71 | 4,75E-01 |
| AT1G70530 | 1,72 | 0,31 | 2,02 | 0,16 | 8,77E-01 |
| AT1G70600 | 0,97 | -1,47 | 2,98 | -0,49 | 6,23E-01 |
| AT1G70730 | 0,91 | -2,21 | 2,47 | -0,89 | 3,72E-01 |
| AT1G70740 | 0,10 | 0,86 | 4,35 | 0,20 | 8,43E-01 |
| AT1G70750 | 0,44 | 0,00 | 4,41 | 0,00 | 1,00E+00 |
| AT1G70780 | 0,73 | 0,68 | 2,79 | 0,24 | 8,07E-01 |
| AT1G70830 | 0,27 | 0,78 | 4,36 | 0,18 | 8,58E-01 |
| AT1G70890 | 1,35 | -1,95 | 1,50 | -1,30 | 1,93E-01 |
| AT1G71020 | 0,94 | 0,00 | 2,49 | 0,00 | 1,00E+00 |
| AT1G71030 | 0,60 | -0,94 | 2,41 | -0,39 | 6,98E-01 |
| AT1G71040 | 0,83 | 1,70 | 2,10 | 0,81 | 4,18E-01 |
| AT1G71060 | 0,73 | 0,00 | 3,54 | 0,00 | 1,00E+00 |
| AT1G71070 | 0,49 | -2,07 | 3,22 | -0,64 | 5,20E-01 |
| AT1G71080 | 0,90 | -0,17 | 3,00 | -0,06 | 9,54E-01 |
| AT1G71160 | 0,69 | 0,00 | 3,65 | 0,00 | 1,00E+00 |
| AT1G71190 | 0,19 | 0,47 | 4,36 | 0,11 | 9,14E-01 |
| AT1G71210 | 0,73 | 0,28 | 2,97 | 0,09 | 9,25E-01 |
| AT1G71310 | 0,27 | -2,52 | 4,38 | -0,58 | 5,65E-01 |
| AT1G71330 | 0,30 | 0,00 | 4,41 | 0,00 | 1,00E+00 |
| AT1G71340 | 0,92 | 2,23 | 2,30 | 0,97 | 3,33E-01 |
| AT1G71350 | 0,21 | 0,00 | 4,41 | 0,00 | 1,00E+00 |
| AT1G71410 | 1,45 | 1,25 | 2,06 | 0,61 | 5,44E-01 |
| AT1G09253 | 0,32 | -0,10 | 4,39 | -0,02 | 9,81E-01 |
| AT1G71490 | 0,42 | -0,26 | 4,25 | -0,06 | 9,51E-01 |
| AT1G71710 | 0,44 | -1,09 | 2,83 | -0,39 | 6,99E-01 |
| AT1G71770 | 0,40 | 1,05 | 4,14 | 0,25 | 8,00E-01 |
| AT1G71870 | 0,88 | 2,17 | 2,76 | 0,79 | 4,31E-01 |
| AT1G71960 | 0,26 | -0,45 | 3,79 | -0,12 | 9,06E-01 |
| AT1G72020 | 0,56 | 0,41 | 2,62 | 0,16 | 8,76E-01 |
| AT1G72040 | 0,83 | 0,12 | 2,18 | 0,05 | 9,57E-01 |
| AT1G72160 | 0,64 | 0,03 | 2,18 | 0,01 | 9,88E-01 |
| AT1G72190 | 0,52 | -2,32 | 3,94 | -0,59 | 5,55E-01 |
| AT1G72270 | 0,12 | -0,36 | 4,39 | -0,08 | 9,34E-01 |
| AT1G72300 | 0,54 | -0,26 | 3,17 | -0,08 | 9,34E-01 |
| AT1G72320 | 0,39 | 0,47 | 3,46 | 0,14 | 8,92E-01 |
| AT1G72440 | 1,79 | -0,45 | 1,91 | -0,24 | 8,12E-01 |
| AT1G72450 | 1,12 | 0,09 | 1,59 | 0,05 | 9,57E-01 |
| AT1G72480 | 0,35 | -2,19 | 4,41 | -0,50 | 6,20E-01 |
| AT1G72500 | 0,46 | -0,34 | 4,40 | -0,08 | 9,38E-01 |
| AT1G72550 | 0,79 | -2,04 | 2,54 | -0,80 | 4,22E-01 |
| AT1G72560 | 1,09 | -1,24 | 2,25 | -0,55 | 5,83E-01 |
| AT1G09327 | 0,20 | -2,07 | 4,39 | -0,47 | 6,37E-01 |
| AT1G72670 | 0,20 | 0,00 | 4,41 | 0,00 | 1,00E+00 |
| AT1G72720 | 0,35 | 0,00 | 4,39 | 0,00 | 1,00E+00 |
| AT1G72730 | 0,32 | 0,00 | 4,41 | 0,00 | 1,00E+00 |
| AT1G72760 | 0,65 | 0,00 | 4,41 | 0,00 | 1,00E+00 |
| AT1G72840 | 0,41 | -0,26 | 4,41 | -0,06 | 9,53E-01 |
| AT1G72860 | 0,44 | 0,00 | 4,41 | 0,00 | 1,00E+00 |
| AT1G73010 | 0,40 | 0,00 | 4,41 | 0,00 | 1,00E+00 |
| AT1G09363 | 0,41 | 2,02 | 3,27 | 0,62 | 5,37E-01 |
| AT1G73110 | 0,63 | -3,58 | 3,82 | -0,94 | 3,48E-01 |
| AT1G73170 | 1,76 | 2,13 | 1,96 | 1,09 | 2,76E-01 |
| AT1G73200 | 0,73 | -0,33 | 2,32 | -0,14 | 8,88E-01 |

| | | | | | |
|---|---|---|---|---|---|
| AT1G73230 | 0,17 | 0,00 | 4,41 | 0,00 | 1,00E+00 |
| AT1G73260 | 0,63 | 0,00 | 3,09 | 0,00 | 1,00E+00 |
| AT1G73370 | 0,52 | 0,00 | 4,41 | 0,00 | 1,00E+00 |
| AT1G73390 | 0,25 | 0,27 | 4,37 | 0,06 | 9,52E-01 |
| AT1G73490 | 0,37 | -0,38 | 4,39 | -0,09 | 9,31E-01 |
| AT1G73500 | 0,92 | -0,58 | 1,93 | -0,30 | 7,64E-01 |
| AT1G73510 | 0,45 | 0,00 | 4,14 | 0,00 | 1,00E+00 |
| AT1G73530 | 0,46 | 0,89 | 4,17 | 0,21 | 8,32E-01 |
| AT1G73540 | 0,81 | 1,74 | 2,15 | 0,81 | 4,20E-01 |
| AT1G73650 | 0,96 | -2,38 | 2,48 | -0,96 | 3,36E-01 |
| AT1G73655 | 0,74 | 0,53 | 3,80 | 0,14 | 8,88E-01 |
| AT1G73660 | 1,58 | 0,76 | 1,99 | 0,38 | 7,04E-01 |
| AT1G73680 | 0,85 | 0,12 | 2,06 | 0,06 | 9,54E-01 |
| AT1G73730 | 0,17 | 0,00 | 4,41 | 0,00 | 1,00E+00 |
| AT1G73760 | 0,39 | 1,24 | 4,26 | 0,29 | 7,71E-01 |
| AT1G73805 | 1,81 | 2,97 | 1,77 | 1,68 | 9,38E-02 |
| AT1G73930 | 0,25 | 0,00 | 4,41 | 0,00 | 1,00E+00 |
| AT1G73950 | 0,45 | -2,11 | 4,40 | -0,48 | 6,32E-01 |
| AT1G73960 | 0,29 | -0,37 | 4,39 | -0,08 | 9,34E-01 |
| AT1G73980 | 0,67 | 1,59 | 2,56 | 0,62 | 5,35E-01 |
| AT1G74050 | 0,43 | 0,55 | 3,31 | 0,17 | 8,68E-01 |
| AT1G74060 | 0,72 | -0,35 | 4,30 | -0,08 | 9,35E-01 |
| AT1G74070 | 0,42 | -1,00 | 4,15 | -0,24 | 8,09E-01 |
| AT1G74120 | 0,82 | 0,70 | 2,85 | 0,24 | 8,07E-01 |
| AT1G74170 | 0,18 | 0,09 | 4,38 | 0,02 | 9,84E-01 |
| AT1G74340 | 0,31 | -2,19 | 4,41 | -0,50 | 6,20E-01 |
| AT1G74370 | 0,53 | 0,81 | 3,92 | 0,21 | 8,36E-01 |
| AT1G74390 | 0,95 | -2,09 | 2,81 | -0,74 | 4,57E-01 |
| AT1G74420 | 0,45 | 0,00 | 4,41 | 0,00 | 1,00E+00 |
| AT1G74440 | 0,80 | 0,92 | 2,96 | 0,31 | 7,57E-01 |
| AT1G09467 | 0,50 | -2,21 | 4,13 | -0,54 | 5,92E-01 |
| AT1G74560 | 0,37 | 0,97 | 3,37 | 0,29 | 7,73E-01 |
| AT1G74690 | 0,95 | -1,73 | 2,24 | -0,77 | 4,39E-01 |
| AT1G74770 | 0,42 | 0,00 | 4,24 | 0,00 | 1,00E+00 |
| AT1G74800 | 0,56 | 1,34 | 3,15 | 0,42 | 6,71E-01 |
| AT1G74830 | 0,12 | 0,00 | 4,41 | 0,00 | 1,00E+00 |
| AT1G74850 | 0,94 | -0,11 | 2,18 | -0,05 | 9,61E-01 |
| AT1G74880 | 0,11 | -0,26 | 4,41 | -0,06 | 9,53E-01 |
| AT1G74920 | 0,86 | -0,41 | 2,57 | -0,16 | 8,74E-01 |
| AT1G74950 | 0,28 | 1,17 | 4,34 | 0,27 | 7,88E-01 |
| AT1G74960 | 1,51 | -0,31 | 2,83 | -0,11 | 9,13E-01 |
| AT1G74970 | 0,46 | -1,27 | 4,33 | -0,29 | 7,70E-01 |
| AT1G75010 | 0,20 | 1,86 | 4,30 | 0,43 | 6,66E-01 |
| AT1G75100 | 0,60 | 0,84 | 2,24 | 0,38 | 7,07E-01 |
| AT1G75130 | 0,12 | -0,26 | 4,41 | -0,06 | 9,53E-01 |
| AT1G75180 | 0,38 | 0,77 | 3,34 | 0,23 | 8,17E-01 |
| AT1G75200 | 0,15 | -0,26 | 4,41 | -0,06 | 9,53E-01 |
| AT1G75220 | 0,55 | 1,21 | 3,17 | 0,38 | 7,03E-01 |
| AT1G75370 | 1,12 | -0,93 | 1,59 | -0,59 | 5,57E-01 |
| AT1G75410 | 0,90 | -1,44 | 2,51 | -0,57 | 5,68E-01 |
| AT1G75450 | 0,11 | -0,38 | 4,39 | -0,09 | 9,31E-01 |
| AT1G75500 | 0,97 | -0,20 | 1,82 | -0,11 | 9,12E-01 |
| AT1G75680 | 0,99 | -0,60 | 1,52 | -0,39 | 6,94E-01 |
| AT1G75690 | 0,84 | -0,72 | 1,87 | -0,38 | 7,02E-01 |
| AT1G75750 | 1,13 | -3,52 | 2,32 | -1,52 | 1,28E-01 |
| AT1G75780 | 0,25 | -0,55 | 4,36 | -0,13 | 8,99E-01 |

| | | | | | |
|---|---|---|---|---|---|
| AT1G75820 | 1,17 | 0,29 | 2,40 | 0,12 | 9,03E-01 |
| AT1G75860 | 0,67 | -0,51 | 2,37 | -0,22 | 8,29E-01 |
| AT1G75945 | 0,28 | -0,26 | 4,41 | -0,06 | 9,53E-01 |
| AT1G76060 | 0,20 | 0,40 | 4,37 | 0,09 | 9,27E-01 |
| AT1G76080 | 1,03 | -0,25 | 1,56 | -0,16 | 8,74E-01 |
| AT1G76090 | 1,72 | 0,73 | 1,52 | 0,48 | 6,32E-01 |
| AT1G76100 | 1,22 | 0,65 | 1,96 | 0,33 | 7,40E-01 |
| AT1G76120 | 0,15 | -0,39 | 4,39 | -0,09 | 9,29E-01 |
| AT1G76150 | 0,72 | 1,13 | 2,92 | 0,39 | 7,00E-01 |
| AT1G76160 | 0,88 | -0,25 | 2,05 | -0,12 | 9,01E-01 |
| AT1G76270 | 0,08 | 0,00 | 4,41 | 0,00 | 1,00E+00 |
| AT1G76280 | 0,12 | -0,26 | 4,41 | -0,06 | 9,53E-01 |
| AT1G09627 | 0,45 | -0,22 | 4,10 | -0,05 | 9,57E-01 |
| AT1G76400 | 0,57 | 0,93 | 3,08 | 0,30 | 7,63E-01 |
| AT1G76460 | 0,53 | -1,56 | 3,23 | -0,48 | 6,29E-01 |
| AT1G76640 | 0,08 | 0,00 | 4,41 | 0,00 | 1,00E+00 |
| AT1G76650 | 0,42 | 1,53 | 3,28 | 0,47 | 6,41E-01 |
| AT1G76660 | 0,13 | -0,61 | 4,37 | -0,14 | 8,89E-01 |
| AT1G76670 | 1,15 | -0,39 | 2,22 | -0,18 | 8,60E-01 |
| AT1G76700 | 0,39 | -2,19 | 4,41 | -0,50 | 6,20E-01 |
| AT1G76810 | 1,64 | -0,05 | 1,26 | -0,04 | 9,67E-01 |
| AT1G76878 | 1,06 | -0,53 | 1,64 | -0,32 | 7,47E-01 |
| AT1G76890 | 0,74 | 1,39 | 1,95 | 0,71 | 4,75E-01 |
| AT1G76955 | 0,30 | 0,00 | 4,41 | 0,00 | 1,00E+00 |
| AT1G76960 | 0,40 | 0,30 | 3,41 | 0,09 | 9,29E-01 |
| AT1G76965 | 0,25 | 0,00 | 4,41 | 0,00 | 1,00E+00 |
| AT1G76970 | 0,43 | -0,23 | 3,28 | -0,07 | 9,45E-01 |
| AT1G76980 | 0,73 | 0,88 | 3,09 | 0,29 | 7,75E-01 |
| AT1G77030 | 0,32 | 1,57 | 4,32 | 0,36 | 7,17E-01 |
| AT1G77060 | 0,12 | 0,27 | 4,38 | 0,06 | 9,50E-01 |
| AT1G77100 | 0,82 | 2,27 | 3,63 | 0,63 | 5,32E-01 |
| AT1G77145 | 0,08 | 0,00 | 4,41 | 0,00 | 1,00E+00 |
| AT1G77170 | 0,63 | -0,33 | 3,07 | -0,11 | 9,15E-01 |
| AT1G77240 | 1,26 | -0,30 | 2,17 | -0,14 | 8,92E-01 |
| AT1G77250 | 0,55 | 0,00 | 3,85 | 0,00 | 1,00E+00 |
| AT1G77260 | 0,57 | 0,25 | 3,89 | 0,07 | 9,48E-01 |
| AT1G77480 | 0,33 | -3,33 | 4,36 | -0,76 | 4,45E-01 |
| AT1G77580 | 0,18 | 0,80 | 4,35 | 0,18 | 8,54E-01 |
| AT1G77600 | 0,35 | 0,32 | 4,36 | 0,07 | 9,42E-01 |
| AT1G77660 | 0,33 | -0,38 | 4,39 | -0,09 | 9,32E-01 |
| AT1G77810 | 0,58 | 0,67 | 3,77 | 0,18 | 8,59E-01 |
| AT1G77932 | 0,47 | 0,00 | 4,41 | 0,00 | 1,00E+00 |
| AT1G77940 | 0,55 | -0,02 | 2,61 | -0,01 | 9,94E-01 |
| AT1G77960 | 1,37 | -0,06 | 1,57 | -0,04 | 9,69E-01 |
| AT1G77980 | 0,77 | 0,99 | 2,00 | 0,50 | 6,19E-01 |
| AT1G77990 | 0,37 | 1,46 | 4,32 | 0,34 | 7,36E-01 |
| AT1G78060 | 0,43 | -1,31 | 4,33 | -0,30 | 7,63E-01 |
| AT1G78210 | 0,32 | 1,06 | 4,34 | 0,24 | 8,07E-01 |
| AT1G78240 | 0,51 | -2,02 | 2,76 | -0,73 | 4,64E-01 |
| AT1G78370 | 1,22 | 2,00 | 2,18 | 0,92 | 3,59E-01 |
| AT1G78380 | 1,48 | -0,86 | 1,49 | -0,58 | 5,61E-01 |
| AT1G78460 | 0,60 | -2,19 | 2,57 | -0,86 | 3,92E-01 |
| AT1G78480 | 0,26 | 0,08 | 4,38 | 0,02 | 9,86E-01 |
| AT1G78510 | 0,69 | -1,86 | 3,75 | -0,50 | 6,19E-01 |
| AT1G78580 | 0,70 | -1,79 | 2,28 | -0,78 | 4,32E-01 |
| AT1G78800 | 0,21 | -0,32 | 4,40 | -0,07 | 9,43E-01 |

| | | | | | |
|---|---|---|---|---|---|
| AT1G78820 | 0,54 | -1,33 | 2,25 | -0,59 | 5,55E-01 |
| AT1G78850 | 0,94 | -2,46 | 1,89 | -1,31 | 1,92E-01 |
| AT1G78865 | 0,13 | 0,94 | 4,35 | 0,22 | 8,29E-01 |
| AT1G78880 | 0,88 | -1,47 | 2,83 | -0,52 | 6,04E-01 |
| AT1G78915 | 0,18 | 0,25 | 4,32 | 0,06 | 9,54E-01 |
| AT1G78990 | 0,38 | 0,00 | 4,41 | 0,00 | 1,00E+00 |
| AT1G78995 | 0,24 | -0,68 | 4,38 | -0,16 | 8,76E-01 |
| AT1G79020 | 0,25 | 0,00 | 4,41 | 0,00 | 1,00E+00 |
| AT1G79030 | 0,53 | -0,13 | 3,96 | -0,03 | 9,73E-01 |
| AT1G79050 | 0,58 | -0,87 | 3,16 | -0,27 | 7,84E-01 |
| AT1G79080 | 1,23 | -2,21 | 2,39 | -0,92 | 3,56E-01 |
| AT1G79120 | 0,10 | 0,86 | 4,35 | 0,20 | 8,43E-01 |
| AT1G79150 | 0,35 | -1,64 | 4,38 | -0,37 | 7,09E-01 |
| AT1G79350 | 0,49 | 0,49 | 2,78 | 0,18 | 8,60E-01 |
| AT1G79430 | 0,17 | 0,07 | 4,38 | 0,02 | 9,88E-01 |
| AT1G79440 | 0,39 | -0,51 | 3,54 | -0,14 | 8,86E-01 |
| AT1G09867 | 1,21 | 0,76 | 2,11 | 0,36 | 7,19E-01 |
| AT1G09883 | 0,79 | 1,61 | 2,84 | 0,57 | 5,71E-01 |
| AT1G79540 | 0,53 | -0,30 | 3,22 | -0,09 | 9,25E-01 |
| AT1G79600 | 1,43 | 0,33 | 1,97 | 0,17 | 8,68E-01 |
| AT1G79640 | 0,40 | -0,32 | 4,40 | -0,07 | 9,42E-01 |
| AT1G79670 | 0,90 | -0,62 | 2,66 | -0,23 | 8,16E-01 |
| AT1G79700 | 0,08 | 0,00 | 4,41 | 0,00 | 1,00E+00 |
| AT1G79720 | 0,47 | -2,11 | 3,33 | -0,64 | 5,25E-01 |
| AT1G79730 | 0,63 | 1,04 | 3,85 | 0,27 | 7,87E-01 |
| AT1G79740 | 0,25 | 0,00 | 4,41 | 0,00 | 1,00E+00 |
| AT1G79790 | 0,05 | 0,18 | 4,38 | 0,04 | 9,68E-01 |
| AT1G79830 | 1,06 | 0,48 | 1,77 | 0,27 | 7,85E-01 |
| AT1G79880 | 0,11 | -0,26 | 4,41 | -0,06 | 9,53E-01 |
| AT1G79900 | 0,42 | 0,00 | 4,24 | 0,00 | 1,00E+00 |
| AT1G80030 | 1,07 | -2,78 | 2,22 | -1,25 | 2,09E-01 |
| AT1G80040 | 0,61 | -1,26 | 2,62 | -0,48 | 6,31E-01 |
| AT1G80080 | 0,50 | 0,36 | 3,18 | 0,11 | 9,10E-01 |
| AT1G80325 | 1,08 | -1,36 | 2,05 | -0,66 | 5,06E-01 |
| AT1G80350 | 0,38 | 0,00 | 4,41 | 0,00 | 1,00E+00 |
| AT1G80420 | 0,34 | -0,04 | 4,33 | -0,01 | 9,92E-01 |
| AT1G80460 | 0,24 | -0,02 | 4,38 | 0,00 | 9,97E-01 |
| AT1G80480 | 0,56 | 1,12 | 3,15 | 0,36 | 7,22E-01 |
| AT1G80490 | 1,23 | -0,19 | 1,89 | -0,10 | 9,20E-01 |
| AT1G80530 | 0,68 | -2,30 | 2,98 | -0,77 | 4,39E-01 |
| AT1G80600 | 1,40 | -1,94 | 1,92 | -1,01 | 3,12E-01 |
| AT1G80610 | 0,35 | 1,82 | 3,41 | 0,53 | 5,94E-01 |
| AT1G80630 | 0,28 | -0,36 | 4,39 | -0,08 | 9,35E-01 |
| AT1G80660 | 1,14 | -0,26 | 2,24 | -0,12 | 9,07E-01 |
| AT1G80670 | 0,39 | 0,00 | 4,40 | 0,00 | 1,00E+00 |
| AT1G80760 | 0,62 | -0,20 | 3,78 | -0,05 | 9,59E-01 |
| AT1G80830 | 0,27 | 1,01 | 4,34 | 0,23 | 8,17E-01 |
| AT1G80850 | 0,20 | 1,82 | 4,30 | 0,42 | 6,72E-01 |
| AT1G80920 | 0,74 | 2,14 | 2,15 | 1,00 | 3,19E-01 |
| AT1G80930 | 0,69 | 1,33 | 2,43 | 0,55 | 5,84E-01 |
| AT1G80950 | 0,35 | 1,03 | 4,34 | 0,24 | 8,12E-01 |
| AT2G01023 | 0,35 | -0,75 | 2,88 | -0,26 | 7,94E-01 |
| AT2G01050 | 0,45 | 1,84 | 4,30 | 0,43 | 6,69E-01 |
| AT2G01060 | 0,57 | 1,91 | 2,55 | 0,75 | 4,53E-01 |
| AT2G01110 | 0,90 | 0,16 | 2,54 | 0,06 | 9,51E-01 |
| AT2G01130 | 1,10 | -1,81 | 2,19 | -0,83 | 4,08E-01 |

| | | | | | |
|---|---|---|---|---|---|
| AT2G01140 | 1,55 | 2,51 | 1,90 | 1,32 | 1,87E-01 |
| AT2G01180 | 0,86 | 0,74 | 1,78 | 0,42 | 6,77E-01 |
| AT2G01250 | 0,52 | -2,32 | 3,97 | -0,58 | 5,59E-01 |
| AT2G01275 | 0,19 | 1,06 | 4,34 | 0,24 | 8,07E-01 |
| AT2G01290 | 0,18 | 0,15 | 4,38 | 0,03 | 9,73E-01 |
| AT2G01300 | 0,08 | 0,00 | 4,41 | 0,00 | 1,00E+00 |
| AT2G01310 | 0,05 | 0,18 | 4,38 | 0,04 | 9,68E-01 |
| AT2G01350 | 0,12 | 1,17 | 4,34 | 0,27 | 7,88E-01 |
| AT2G01410 | 0,34 | -0,03 | 3,53 | -0,01 | 9,93E-01 |
| AT2G01420 | 1,24 | -0,19 | 1,48 | -0,13 | 9,00E-01 |
| AT2G01440 | 1,25 | 0,09 | 2,55 | 0,03 | 9,73E-01 |
| AT2G01450 | 1,15 | 2,74 | 1,77 | 1,55 | 1,22E-01 |
| AT2G01505 | 0,30 | 1,23 | 4,33 | 0,28 | 7,76E-01 |
| AT2G01630 | 0,87 | -0,31 | 2,92 | -0,11 | 9,16E-01 |
| AT2G01650 | 1,03 | 1,39 | 2,19 | 0,63 | 5,26E-01 |
| AT2G01670 | 0,39 | 0,26 | 3,31 | 0,08 | 9,38E-01 |
| AT2G01690 | 0,55 | 1,27 | 3,80 | 0,33 | 7,39E-01 |
| AT2G01720 | 1,57 | 0,26 | 2,40 | 0,11 | 9,13E-01 |
| AT2G01755 | 0,14 | 0,89 | 4,35 | 0,21 | 8,37E-01 |
| AT2G01820 | 1,26 | -2,17 | 2,53 | -0,86 | 3,91E-01 |
| AT2G01830 | 0,91 | -0,59 | 1,91 | -0,31 | 7,56E-01 |
| AT2G04125 | 0,12 | 0,11 | 4,38 | 0,03 | 9,80E-01 |
| AT2G01870 | 0,33 | -0,39 | 4,39 | -0,09 | 9,30E-01 |
| AT2G01918 | 1,01 | 0,06 | 1,91 | 0,03 | 9,77E-01 |
| AT2G01930 | 0,16 | -0,26 | 4,41 | -0,06 | 9,53E-01 |
| AT2G01970 | 1,49 | -0,31 | 1,63 | -0,19 | 8,48E-01 |
| AT2G01990 | 0,25 | 1,45 | 4,33 | 0,34 | 7,37E-01 |
| AT2G02080 | 1,03 | -0,72 | 2,49 | -0,29 | 7,73E-01 |
| AT2G02103 | 0,70 | 0,60 | 2,40 | 0,25 | 8,01E-01 |
| AT2G02148 | 0,42 | 0,00 | 4,18 | 0,00 | 1,00E+00 |
| AT2G02150 | 0,39 | -0,40 | 4,31 | -0,09 | 9,26E-01 |
| AT2G02160 | 0,60 | 0,05 | 2,53 | 0,02 | 9,85E-01 |
| AT2G02220 | 0,80 | -0,53 | 2,47 | -0,21 | 8,31E-01 |
| AT2G02230 | 0,17 | -0,33 | 4,40 | -0,08 | 9,40E-01 |
| AT2G02340 | 1,23 | -1,17 | 1,96 | -0,60 | 5,51E-01 |
| AT2G02360 | 0,16 | 0,15 | 4,38 | 0,04 | 9,72E-01 |
| AT2G02400 | 0,84 | 2,95 | 2,22 | 1,33 | 1,84E-01 |
| AT2G02520 | 0,08 | 0,47 | 4,38 | 0,11 | 9,15E-01 |
| AT2G02570 | 0,25 | 1,56 | 4,32 | 0,36 | 7,17E-01 |
| AT2G02635 | 0,17 | -0,52 | 4,36 | -0,12 | 9,04E-01 |
| AT2G02695 | 0,58 | 2,82 | 2,93 | 0,96 | 3,37E-01 |
| AT2G02710 | 0,66 | 1,08 | 2,51 | 0,43 | 6,67E-01 |
| AT2G02780 | 0,28 | -0,12 | 4,29 | -0,03 | 9,78E-01 |
| AT2G02800 | 0,78 | -0,44 | 2,41 | -0,18 | 8,56E-01 |
| AT2G04345 | 0,62 | 2,38 | 2,58 | 0,92 | 3,55E-01 |
| AT2G02930 | 0,49 | 1,33 | 3,21 | 0,42 | 6,78E-01 |
| AT2G03050 | 0,25 | -0,27 | 4,40 | -0,06 | 9,51E-01 |
| AT2G03090 | 0,08 | 0,00 | 4,41 | 0,00 | 1,00E+00 |
| AT2G03240 | 0,24 | 0,00 | 4,41 | 0,00 | 1,00E+00 |
| AT2G03340 | 0,45 | 0,55 | 3,32 | 0,17 | 8,69E-01 |
| AT2G03390 | 0,61 | -1,63 | 2,32 | -0,70 | 4,84E-01 |
| AT2G03420 | 0,43 | -1,34 | 4,15 | -0,32 | 7,47E-01 |
| AT2G03430 | 0,52 | 0,00 | 4,41 | 0,00 | 1,00E+00 |
| AT2G03440 | 0,35 | 1,52 | 4,32 | 0,35 | 7,24E-01 |
| AT2G03460 | 0,08 | 0,00 | 4,41 | 0,00 | 1,00E+00 |
| AT2G03470 | 1,22 | 1,19 | 2,42 | 0,49 | 6,22E-01 |

| | | | | | |
|---|---|---|---|---|---|
| AT2G03490 | 0,08 | 0,00 | 4,41 | 0,00 | 1,00E+00 |
| AT2G03620 | 0,33 | 0,00 | 4,41 | 0,00 | 1,00E+00 |
| AT2G03640 | 0,13 | 0,00 | 4,41 | 0,00 | 1,00E+00 |
| AT2G03690 | 0,23 | 0,24 | 4,39 | 0,06 | 9,56E-01 |
| AT2G03730 | 0,29 | 0,07 | 4,34 | 0,02 | 9,88E-01 |
| AT2G03750 | 1,29 | 0,17 | 1,42 | 0,12 | 9,07E-01 |
| AT2G03760 | 0,93 | 1,95 | 3,46 | 0,56 | 5,73E-01 |
| AT2G03770 | 0,15 | -0,35 | 4,39 | -0,08 | 9,36E-01 |
| AT2G03821 | 1,77 | 0,90 | 1,26 | 0,72 | 4,73E-01 |
| AT2G03930 | 1,60 | 0,76 | 1,33 | 0,57 | 5,67E-01 |
| AT2G04655 | 0,14 | 1,29 | 4,33 | 0,30 | 7,65E-01 |
| AT2G04031 | 0,15 | 1,21 | 4,33 | 0,28 | 7,81E-01 |
| AT2G04037 | 0,16 | 0,74 | 4,36 | 0,17 | 8,66E-01 |
| AT2G04050 | 0,12 | 0,00 | 4,41 | 0,00 | 1,00E+00 |
| AT2G04060 | 1,72 | 4,46 | 2,07 | 2,15 | 3,14E-02 |
| AT2G04062 | 0,70 | 3,16 | 2,84 | 1,11 | 2,67E-01 |
| AT2G04063 | 0,69 | 3,08 | 2,69 | 1,15 | 2,51E-01 |
| AT2G04066 | 0,42 | 0,96 | 4,35 | 0,22 | 8,25E-01 |
| AT2G04070 | 0,33 | 0,00 | 4,41 | 0,00 | 1,00E+00 |
| AT2G04080 | 0,61 | 1,12 | 2,57 | 0,44 | 6,63E-01 |
| AT2G04160 | 0,43 | 2,15 | 4,02 | 0,54 | 5,92E-01 |
| AT2G04230 | 1,09 | -1,36 | 2,25 | -0,60 | 5,46E-01 |
| AT2G04360 | 0,70 | 0,51 | 2,56 | 0,20 | 8,43E-01 |
| AT2G04378 | 0,56 | 0,52 | 2,69 | 0,19 | 8,46E-01 |
| AT2G04520 | 0,26 | -0,64 | 4,30 | -0,15 | 8,82E-01 |
| AT2G04620 | 0,17 | 0,00 | 4,41 | 0,00 | 1,00E+00 |
| AT2G04630 | 0,19 | 0,67 | 4,36 | 0,15 | 8,78E-01 |
| AT2G04842 | 0,88 | -0,15 | 3,08 | -0,05 | 9,62E-01 |
| AT2G04870 | 0,55 | 0,69 | 2,57 | 0,27 | 7,88E-01 |
| AT2G04890 | 0,11 | -0,38 | 4,39 | -0,09 | 9,31E-01 |
| AT2G05120 | 0,91 | -1,33 | 2,26 | -0,59 | 5,56E-01 |
| AT2G05220 | 0,38 | 0,23 | 4,39 | 0,05 | 9,58E-01 |
| AT2G05250 | 0,41 | 0,00 | 4,41 | 0,00 | 1,00E+00 |
| AT2G04895 | 0,17 | 1,19 | 4,33 | 0,28 | 7,83E-01 |
| AT2G05752 | 0,90 | 2,79 | 2,27 | 1,23 | 2,19E-01 |
| AT2G04995 | 1,03 | 0,84 | 2,01 | 0,42 | 6,75E-01 |
| AT2G05810 | 1,10 | 1,75 | 1,93 | 0,91 | 3,65E-01 |
| AT2G05914 | 0,51 | 1,12 | 4,31 | 0,26 | 7,95E-01 |
| AT2G05920 | 1,42 | 0,25 | 1,55 | 0,16 | 8,73E-01 |
| AT2G05940 | 0,97 | 2,10 | 1,92 | 1,09 | 2,75E-01 |
| AT2G06000 | 0,52 | 0,41 | 2,66 | 0,15 | 8,78E-01 |
| AT2G06040 | 0,24 | -1,18 | 4,39 | -0,27 | 7,87E-01 |
| AT2G06050 | 0,61 | 2,20 | 2,99 | 0,74 | 4,61E-01 |
| AT2G06530 | 0,61 | -0,34 | 3,25 | -0,11 | 9,16E-01 |
| AT2G06541 | 0,49 | 1,75 | 4,08 | 0,43 | 6,68E-01 |
| AT2G06645 | 0,15 | -0,28 | 4,35 | -0,06 | 9,50E-01 |
| AT2G06675 | 0,19 | 1,60 | 4,31 | 0,37 | 7,11E-01 |
| AT2G06906 | 0,12 | -0,26 | 4,41 | -0,06 | 9,53E-01 |
| AT2G06925 | 0,35 | -0,72 | 3,46 | -0,21 | 8,36E-01 |
| AT2G06960 | 0,68 | 2,49 | 2,72 | 0,91 | 3,60E-01 |
| AT2G07020 | 0,35 | 2,39 | 4,28 | 0,56 | 5,77E-01 |
| AT2G07120 | 0,12 | 0,35 | 4,37 | 0,08 | 9,35E-01 |
| AT2G07180 | 1,49 | -0,13 | 1,91 | -0,07 | 9,45E-01 |
| AT2G07240 | 1,04 | 0,44 | 1,76 | 0,25 | 8,01E-01 |
| AT2G07560 | 0,12 | -0,36 | 4,39 | -0,08 | 9,34E-01 |
| AT2G07771 | 0,58 | 0,70 | 2,62 | 0,27 | 7,89E-01 |

| | | | | | |
|---|---|---|---|---|---|
| AT2G07777 | 0,89 | -1,43 | 2,00 | -0,72 | 4,74E-01 |
| AT2G07779 | 0,28 | 0,47 | 4,37 | 0,11 | 9,14E-01 |
| AT2G07621 | 0,12 | 0,11 | 4,38 | 0,03 | 9,80E-01 |
| AT2G07774 | 0,25 | -0,05 | 4,30 | -0,01 | 9,92E-01 |
| AT2G07687 | 1,11 | -1,09 | 1,94 | -0,56 | 5,75E-01 |
| AT2G05205 | 0,68 | -0,52 | 2,43 | -0,22 | 8,30E-01 |
| AT2G07599 | 0,90 | -0,20 | 2,05 | -0,10 | 9,22E-01 |
| AT2G07706 | 1,10 | -0,90 | 2,12 | -0,42 | 6,72E-01 |
| AT2G07641 | 0,45 | 0,01 | 2,79 | 0,00 | 9,98E-01 |
| AT2G07719 | 0,62 | -1,22 | 2,63 | -0,46 | 6,43E-01 |
| AT2G07756 | 1,02 | 0,00 | 1,74 | 0,00 | 9,99E-01 |
| AT2G07734 | 0,71 | -0,21 | 2,82 | -0,07 | 9,41E-01 |
| AT2G07658 | 0,65 | 0,25 | 2,51 | 0,10 | 9,19E-01 |
| AT2G07662 | 1,06 | -0,52 | 2,08 | -0,25 | 8,05E-01 |
| AT2G07835 | 0,78 | -0,28 | 2,94 | -0,09 | 9,25E-01 |
| AT2G07690 | 0,71 | 0,51 | 2,57 | 0,20 | 8,43E-01 |
| AT2G07760 | 0,23 | -1,08 | 4,38 | -0,25 | 8,06E-01 |
| AT2G09795 | 0,34 | -0,30 | 4,40 | -0,07 | 9,46E-01 |
| AT2G05465 | 0,27 | 1,44 | 4,33 | 0,33 | 7,39E-01 |
| AT2G10557 | 1,21 | 0,00 | 3,79 | 0,00 | 1,00E+00 |
| AT2G10608 | 0,36 | -0,82 | 4,37 | -0,19 | 8,51E-01 |
| AT2G05565 | 0,43 | 2,50 | 4,00 | 0,62 | 5,33E-01 |
| AT2G10625 | 0,25 | 1,94 | 4,30 | 0,45 | 6,52E-01 |
| AT2G10950 | 0,47 | 2,73 | 3,90 | 0,70 | 4,84E-01 |
| AT2G10955 | 0,26 | 1,22 | 4,31 | 0,28 | 7,77E-01 |
| AT2G11000 | 0,08 | 0,47 | 4,38 | 0,11 | 9,15E-01 |
| AT2G11852 | 0,26 | 1,39 | 4,33 | 0,32 | 7,48E-01 |
| AT2G11910 | 1,43 | -0,98 | 1,66 | -0,59 | 5,57E-01 |
| AT2G12400 | 0,28 | -0,42 | 4,36 | -0,10 | 9,23E-01 |
| AT2G12405 | 0,18 | 0,25 | 4,32 | 0,06 | 9,54E-01 |
| AT2G05705 | 0,21 | 0,69 | 4,36 | 0,16 | 8,75E-01 |
| AT2G12461 | 0,12 | 0,00 | 4,41 | 0,00 | 1,00E+00 |
| AT2G05835 | 0,29 | 0,07 | 4,32 | 0,02 | 9,86E-01 |
| AT2G12945 | 0,05 | 0,18 | 4,38 | 0,04 | 9,68E-01 |
| AT2G13370 | 1,20 | 0,01 | 2,40 | 0,01 | 9,95E-01 |
| AT2G13450 | 1,51 | -1,74 | 1,50 | -1,16 | 2,46E-01 |
| AT2G13550 | 0,22 | 0,91 | 4,34 | 0,21 | 8,34E-01 |
| AT2G13640 | 0,11 | -0,26 | 4,41 | -0,06 | 9,53E-01 |
| AT2G13650 | 0,22 | 0,81 | 4,35 | 0,19 | 8,52E-01 |
| AT2G13820 | 1,49 | 4,41 | 2,72 | 1,62 | 1,04E-01 |
| AT2G14100 | 0,15 | 0,90 | 4,35 | 0,21 | 8,36E-01 |
| AT2G14120 | 0,98 | 1,90 | 2,08 | 0,91 | 3,62E-01 |
| AT2G14170 | 1,22 | -0,71 | 1,96 | -0,36 | 7,17E-01 |
| AT2G14260 | 0,26 | 0,09 | 4,38 | 0,02 | 9,84E-01 |
| AT2G14265 | 1,52 | 0,49 | 1,54 | 0,32 | 7,51E-01 |
| AT2G14288 | 0,20 | 1,36 | 4,32 | 0,31 | 7,54E-01 |
| AT2G14510 | 0,15 | 0,90 | 4,35 | 0,21 | 8,36E-01 |
| AT2G14540 | 0,12 | -0,26 | 4,41 | -0,06 | 9,53E-01 |
| AT2G14670 | 1,21 | 1,28 | 2,28 | 0,56 | 5,73E-01 |
| AT2G14720 | 1,05 | 1,78 | 2,36 | 0,76 | 4,49E-01 |
| AT2G14755 | 0,32 | 2,42 | 4,28 | 0,56 | 5,72E-01 |
| AT2G15000 | 0,23 | 0,24 | 4,39 | 0,06 | 9,56E-01 |
| AT2G15020 | 0,51 | -2,18 | 3,26 | -0,67 | 5,03E-01 |
| AT2G06355 | 0,23 | 0,09 | 4,34 | 0,02 | 9,84E-01 |
| AT2G15110 | 0,63 | 1,23 | 2,94 | 0,42 | 6,76E-01 |
| AT2G15290 | 0,58 | -2,04 | 3,81 | -0,53 | 5,93E-01 |

| | | | | | |
|---|---|---|---|---|---|
| AT2G15292 | 1,13 | -0,14 | 1,70 | -0,08 | 9,34E-01 |
| AT2G15320 | 1,25 | 1,55 | 1,54 | 1,00 | 3,16E-01 |
| AT2G15390 | 0,37 | 0,52 | 4,36 | 0,12 | 9,05E-01 |
| AT2G15555 | 1,03 | -0,78 | 1,67 | -0,47 | 6,40E-01 |
| AT2G15560 | 0,05 | 0,18 | 4,38 | 0,04 | 9,68E-01 |
| AT2G15790 | 0,41 | 0,59 | 4,19 | 0,14 | 8,89E-01 |
| AT2G15860 | 0,36 | 0,47 | 4,32 | 0,11 | 9,14E-01 |
| AT2G15880 | 1,49 | -1,13 | 1,88 | -0,60 | 5,47E-01 |
| AT2G15890 | 1,14 | 1,05 | 1,69 | 0,62 | 5,33E-01 |
| AT2G06515 | 0,69 | -3,29 | 3,64 | -0,90 | 3,67E-01 |
| AT2G15980 | 0,29 | -1,55 | 3,64 | -0,43 | 6,71E-01 |
| AT2G16016 | 1,06 | 1,22 | 2,06 | 0,59 | 5,53E-01 |
| AT2G06575 | 0,75 | 2,17 | 2,03 | 1,07 | 2,85E-01 |
| AT2G16070 | 0,64 | -0,90 | 3,83 | -0,23 | 8,15E-01 |
| AT2G16090 | 0,55 | 1,14 | 2,55 | 0,45 | 6,56E-01 |
| AT2G16370 | 0,11 | -0,26 | 4,41 | -0,06 | 9,53E-01 |
| AT2G16400 | 1,00 | -0,03 | 1,88 | -0,01 | 9,89E-01 |
| AT2G16405 | 0,20 | -2,08 | 4,39 | -0,47 | 6,35E-01 |
| AT2G16440 | 1,04 | 0,39 | 3,61 | 0,11 | 9,15E-01 |
| AT2G16450 | 0,33 | 0,00 | 4,41 | 0,00 | 1,00E+00 |
| AT2G16490 | 1,37 | -1,17 | 2,12 | -0,55 | 5,82E-01 |
| AT2G16505 | 0,85 | 0,85 | 2,11 | 0,40 | 6,86E-01 |
| AT2G16510 | 1,63 | -1,27 | 1,68 | -0,76 | 4,48E-01 |
| AT2G16620 | 0,18 | 1,20 | 4,34 | 0,28 | 7,82E-01 |
| AT2G16640 | 1,05 | 1,28 | 2,21 | 0,58 | 5,63E-01 |
| AT2G16650 | 0,37 | 0,00 | 4,41 | 0,00 | 1,00E+00 |
| AT2G16870 | 1,12 | 0,44 | 2,14 | 0,21 | 8,36E-01 |
| AT2G16880 | 0,51 | 0,00 | 4,41 | 0,00 | 1,00E+00 |
| AT2G16920 | 0,25 | -2,19 | 4,41 | -0,50 | 6,20E-01 |
| AT2G16940 | 0,88 | 1,02 | 2,18 | 0,47 | 6,41E-01 |
| AT2G17020 | 0,78 | -0,41 | 2,14 | -0,19 | 8,50E-01 |
| AT2G17110 | 0,49 | 0,24 | 2,69 | 0,09 | 9,29E-01 |
| AT2G17130 | 0,45 | -0,36 | 4,24 | -0,09 | 9,32E-01 |
| AT2G17190 | 0,94 | 0,26 | 2,35 | 0,11 | 9,12E-01 |
| AT2G17210 | 0,23 | -0,35 | 4,39 | -0,08 | 9,36E-01 |
| AT2G17220 | 0,39 | -0,36 | 4,39 | -0,08 | 9,35E-01 |
| AT2G17230 | 0,92 | 1,08 | 2,43 | 0,44 | 6,57E-01 |
| AT2G17300 | 0,17 | -0,33 | 4,40 | -0,08 | 9,40E-01 |
| AT2G06935 | 0,05 | 0,18 | 4,38 | 0,04 | 9,68E-01 |
| AT2G17340 | 0,86 | -0,67 | 2,34 | -0,29 | 7,74E-01 |
| AT2G17350 | 0,34 | -1,35 | 4,34 | -0,31 | 7,56E-01 |
| AT2G17450 | 0,63 | 1,30 | 2,32 | 0,56 | 5,76E-01 |
| AT2G17470 | 0,10 | 0,24 | 4,38 | 0,05 | 9,57E-01 |
| AT2G17510 | 1,71 | 0,05 | 1,44 | 0,04 | 9,70E-01 |
| AT2G17550 | 0,71 | 0,46 | 2,51 | 0,19 | 8,53E-01 |
| AT2G17560 | 0,46 | -0,36 | 4,39 | -0,08 | 9,36E-01 |
| AT2G17700 | 0,43 | -1,29 | 4,11 | -0,31 | 7,53E-01 |
| AT2G17820 | 0,96 | -1,26 | 2,06 | -0,61 | 5,41E-01 |
| AT2G17870 | 0,45 | 0,57 | 4,14 | 0,14 | 8,90E-01 |
| AT2G18130 | 0,22 | 1,46 | 4,32 | 0,34 | 7,36E-01 |
| AT2G18160 | 1,66 | 3,19 | 1,77 | 1,81 | 7,09E-02 |
| AT2G18170 | 0,50 | -0,31 | 3,32 | -0,09 | 9,26E-01 |
| AT2G18193 | 1,29 | -0,29 | 2,33 | -0,12 | 9,02E-01 |
| AT2G18290 | 0,19 | 0,44 | 4,36 | 0,10 | 9,21E-01 |
| AT2G18300 | 1,33 | -1,53 | 1,99 | -0,77 | 4,42E-01 |
| AT2G18310 | 0,43 | -0,40 | 4,34 | -0,09 | 9,26E-01 |

| | | | | | |
|---|---|---|---|---|---|
| AT2G18520 | 0,25 | 0,00 | 4,41 | 0,00 | 1,00E+00 |
| AT2G18600 | 0,24 | 0,00 | 4,41 | 0,00 | 1,00E+00 |
| AT2G18660 | 0,89 | 1,18 | 2,10 | 0,56 | 5,74E-01 |
| AT2G18685 | 0,08 | 0,00 | 4,41 | 0,00 | 1,00E+00 |
| AT2G18900 | 1,01 | 0,86 | 2,06 | 0,42 | 6,78E-01 |
| AT2G18915 | 0,29 | 0,05 | 4,38 | 0,01 | 9,91E-01 |
| AT2G18940 | 0,31 | -0,32 | 3,59 | -0,09 | 9,28E-01 |
| AT2G19120 | 0,71 | -2,30 | 3,60 | -0,64 | 5,22E-01 |
| AT2G07135 | 0,16 | 1,58 | 4,31 | 0,37 | 7,14E-01 |
| AT2G07165 | 0,21 | 0,28 | 4,37 | 0,06 | 9,49E-01 |
| AT2G19290 | 0,05 | 0,18 | 4,38 | 0,04 | 9,68E-01 |
| AT2G19310 | 0,12 | 0,35 | 4,37 | 0,08 | 9,35E-01 |
| AT2G19340 | 0,43 | -0,37 | 4,40 | -0,08 | 9,33E-01 |
| AT2G19365 | 0,81 | -0,47 | 1,97 | -0,24 | 8,12E-01 |
| AT2G19390 | 0,76 | 2,13 | 2,34 | 0,91 | 3,61E-01 |
| AT2G07195 | 0,51 | 1,91 | 2,59 | 0,74 | 4,61E-01 |
| AT2G19400 | 0,15 | 0,02 | 4,36 | 0,00 | 9,96E-01 |
| AT2G19470 | 0,28 | 0,66 | 4,35 | 0,15 | 8,79E-01 |
| AT2G19560 | 0,20 | -0,26 | 4,41 | -0,06 | 9,53E-01 |
| AT2G19570 | 0,52 | 2,71 | 2,99 | 0,91 | 3,65E-01 |
| AT2G19580 | 0,25 | 1,90 | 4,30 | 0,44 | 6,59E-01 |
| AT2G19760 | 0,25 | 1,73 | 4,31 | 0,40 | 6,88E-01 |
| AT2G19780 | 0,19 | -0,02 | 4,35 | -0,01 | 9,96E-01 |
| AT2G19910 | 0,11 | -0,26 | 4,41 | -0,06 | 9,53E-01 |
| AT2G20000 | 0,44 | -1,26 | 3,36 | -0,37 | 7,08E-01 |
| AT2G20050 | 0,77 | 1,08 | 2,41 | 0,45 | 6,53E-01 |
| AT2G20120 | 0,38 | 0,15 | 4,38 | 0,03 | 9,73E-01 |
| AT2G20180 | 0,53 | -0,64 | 2,57 | -0,25 | 8,02E-01 |
| AT2G20240 | 1,31 | -0,49 | 2,66 | -0,19 | 8,53E-01 |
| AT2G20270 | 0,73 | -0,67 | 3,07 | -0,22 | 8,26E-01 |
| AT2G20280 | 0,90 | -0,37 | 2,82 | -0,13 | 8,96E-01 |
| AT2G20300 | 0,57 | 0,43 | 2,62 | 0,16 | 8,70E-01 |
| AT2G20310 | 0,15 | -0,31 | 4,40 | -0,07 | 9,43E-01 |
| AT2G20380 | 0,08 | 0,00 | 4,41 | 0,00 | 1,00E+00 |
| AT2G20562 | 0,11 | 0,88 | 4,35 | 0,20 | 8,39E-01 |
| AT2G20610 | 0,62 | 0,90 | 2,48 | 0,36 | 7,17E-01 |
| AT2G20630 | 0,57 | -0,79 | 2,52 | -0,31 | 7,53E-01 |
| AT2G20670 | 0,99 | -2,91 | 2,17 | -1,34 | 1,79E-01 |
| AT2G07325 | 0,08 | 0,00 | 4,41 | 0,00 | 1,00E+00 |
| AT2G20720 | 0,50 | -2,22 | 3,30 | -0,67 | 5,01E-01 |
| AT2G20723 | 1,21 | 0,60 | 1,55 | 0,39 | 6,98E-01 |
| AT2G20725 | 0,17 | 0,00 | 4,41 | 0,00 | 1,00E+00 |
| AT2G20760 | 1,00 | -0,54 | 2,18 | -0,25 | 8,06E-01 |
| AT2G20790 | 0,65 | -0,47 | 3,78 | -0,12 | 9,02E-01 |
| AT2G20825 | 0,24 | 0,00 | 4,41 | 0,00 | 1,00E+00 |
| AT2G20830 | 0,42 | 2,10 | 3,22 | 0,65 | 5,14E-01 |
| AT2G20850 | 0,80 | -1,56 | 3,45 | -0,45 | 6,51E-01 |
| AT2G20900 | 0,70 | -0,15 | 3,10 | -0,05 | 9,63E-01 |
| AT2G21030 | 1,78 | -0,09 | 1,42 | -0,07 | 9,47E-01 |
| AT2G21060 | 0,37 | 0,61 | 4,29 | 0,14 | 8,87E-01 |
| AT2G21090 | 0,12 | 0,00 | 4,41 | 0,00 | 1,00E+00 |
| AT2G21120 | 1,14 | 2,19 | 1,86 | 1,18 | 2,38E-01 |
| AT2G21170 | 1,20 | -1,43 | 1,61 | -0,89 | 3,76E-01 |
| AT2G21185 | 0,86 | 1,76 | 1,88 | 0,93 | 3,52E-01 |
| AT2G21188 | 0,47 | -1,27 | 4,28 | -0,30 | 7,66E-01 |
| AT2G21210 | 1,12 | 1,51 | 2,15 | 0,70 | 4,83E-01 |

| | | | | | |
|---|---|---|---|---|---|
| AT2G21240 | 0,62 | -0,58 | 2,27 | -0,26 | 7,97E-01 |
| AT2G21280 | 0,38 | 0,00 | 4,41 | 0,00 | 1,00E+00 |
| AT2G21300 | 0,99 | -0,49 | 2,36 | -0,21 | 8,36E-01 |
| AT2G21350 | 0,05 | 0,18 | 4,38 | 0,04 | 9,68E-01 |
| AT2G21370 | 0,82 | 0,59 | 2,02 | 0,29 | 7,72E-01 |
| AT2G21440 | 0,17 | 0,00 | 4,41 | 0,00 | 1,00E+00 |
| AT2G21480 | 0,25 | 0,00 | 4,41 | 0,00 | 1,00E+00 |
| AT2G21500 | 0,51 | 1,52 | 3,07 | 0,50 | 6,20E-01 |
| AT2G21560 | 0,27 | 2,30 | 4,29 | 0,54 | 5,92E-01 |
| AT2G21870 | 1,17 | -1,76 | 2,14 | -0,82 | 4,11E-01 |
| AT2G21950 | 0,24 | -0,26 | 4,41 | -0,06 | 9,53E-01 |
| AT2G21970 | 0,61 | -3,98 | 3,85 | -1,03 | 3,02E-01 |
| AT2G22010 | 0,65 | 0,84 | 2,49 | 0,34 | 7,35E-01 |
| AT2G22090 | 0,34 | -0,66 | 4,34 | -0,15 | 8,79E-01 |
| AT2G22140 | 0,05 | 0,18 | 4,38 | 0,04 | 9,68E-01 |
| AT2G22160 | 0,35 | 2,17 | 4,29 | 0,51 | 6,13E-01 |
| AT2G22240 | 0,21 | -0,37 | 4,36 | -0,09 | 9,32E-01 |
| AT2G22250 | 0,26 | 0,00 | 4,41 | 0,00 | 1,00E+00 |
| AT2G07555 | 0,37 | 1,58 | 4,21 | 0,38 | 7,07E-01 |
| AT2G22400 | 0,55 | 0,52 | 3,83 | 0,14 | 8,91E-01 |
| AT2G22470 | 0,19 | -0,40 | 4,39 | -0,09 | 9,28E-01 |
| AT2G22490 | 1,17 | 0,00 | 2,55 | 0,00 | 1,00E+00 |
| AT2G22740 | 0,43 | -0,31 | 4,18 | -0,07 | 9,40E-01 |
| AT2G22850 | 0,76 | -0,91 | 1,81 | -0,50 | 6,16E-01 |
| AT2G22890 | 0,53 | 2,29 | 3,01 | 0,76 | 4,47E-01 |
| AT2G22940 | 0,87 | 3,38 | 2,58 | 1,31 | 1,90E-01 |
| AT2G23070 | 0,53 | 0,20 | 2,15 | 0,09 | 9,26E-01 |
| AT2G23090 | 0,40 | -0,29 | 4,35 | -0,07 | 9,47E-01 |
| AT2G23100 | 0,16 | -0,26 | 4,41 | -0,06 | 9,53E-01 |
| AT2G23140 | 0,38 | 1,16 | 4,24 | 0,27 | 7,85E-01 |
| AT2G23210 | 0,71 | -0,39 | 2,95 | -0,13 | 8,95E-01 |
| AT2G23290 | 0,64 | 0,51 | 2,59 | 0,20 | 8,44E-01 |
| AT2G23390 | 0,32 | -1,23 | 4,35 | -0,28 | 7,77E-01 |
| AT2G23430 | 0,47 | 1,91 | 3,34 | 0,57 | 5,67E-01 |
| AT2G23450 | 0,62 | -0,39 | 2,22 | -0,17 | 8,62E-01 |
| AT2G23520 | 0,43 | 0,88 | 3,37 | 0,26 | 7,94E-01 |
| AT2G07910 | 1,63 | 0,51 | 2,07 | 0,25 | 8,06E-01 |
| AT2G23550 | 1,59 | 0,45 | 2,08 | 0,21 | 8,30E-01 |
| AT2G23600 | 1,45 | -0,51 | 1,83 | -0,28 | 7,80E-01 |
| AT2G23610 | 0,63 | 1,54 | 2,95 | 0,52 | 6,02E-01 |
| AT2G23670 | 1,50 | -1,51 | 1,56 | -0,97 | 3,33E-01 |
| AT2G23760 | 0,34 | 0,19 | 4,34 | 0,04 | 9,64E-01 |
| AT2G23790 | 0,79 | -0,46 | 3,01 | -0,15 | 8,78E-01 |
| AT2G23840 | 0,70 | -1,71 | 2,99 | -0,57 | 5,68E-01 |
| AT2G23987 | 0,17 | -3,06 | 4,35 | -0,70 | 4,82E-01 |
| AT2G24120 | 0,81 | 0,33 | 2,07 | 0,16 | 8,74E-01 |
| AT2G24150 | 0,79 | -0,54 | 2,26 | -0,24 | 8,10E-01 |
| AT2G24170 | 0,17 | -0,29 | 4,40 | -0,07 | 9,47E-01 |
| AT2G24200 | 1,48 | 1,25 | 1,68 | 0,74 | 4,58E-01 |
| AT2G24230 | 0,59 | 1,04 | 2,47 | 0,42 | 6,75E-01 |
| AT2G24330 | 0,33 | -0,33 | 4,34 | -0,08 | 9,40E-01 |
| AT2G24340 | 1,46 | 2,99 | 1,74 | 1,72 | 8,54E-02 |
| AT2G24360 | 1,19 | 0,69 | 1,90 | 0,37 | 7,15E-01 |
| AT2G24500 | 1,13 | -1,09 | 2,14 | -0,51 | 6,09E-01 |
| AT2G08055 | 0,33 | 0,70 | 4,35 | 0,16 | 8,72E-01 |
| AT2G24545 | 1,16 | 0,24 | 1,61 | 0,15 | 8,81E-01 |

| | | | | | |
|---|---|---|---|---|---|
| AT2G24550 | 1,50 | 1,75 | 1,69 | 1,04 | 3,00E-01 |
| AT2G24600 | 1,31 | 1,62 | 2,31 | 0,70 | 4,84E-01 |
| AT2G24640 | 0,24 | 0,00 | 4,41 | 0,00 | 1,00E+00 |
| AT2G24710 | 0,19 | -0,34 | 4,40 | -0,08 | 9,38E-01 |
| AT2G24755 | 0,71 | 1,00 | 2,62 | 0,38 | 7,01E-01 |
| AT2G24780 | 0,92 | -1,74 | 2,23 | -0,78 | 4,36E-01 |
| AT2G24850 | 1,45 | 1,19 | 1,82 | 0,65 | 5,13E-01 |
| AT2G24870 | 1,09 | 0,06 | 2,22 | 0,03 | 9,77E-01 |
| AT2G24880 | 1,30 | -1,71 | 1,47 | -1,16 | 2,45E-01 |
| AT2G25011 | 1,24 | 0,42 | 1,74 | 0,24 | 8,09E-01 |
| AT2G25050 | 0,47 | -2,61 | 3,17 | -0,82 | 4,10E-01 |
| AT2G25070 | 0,45 | 0,67 | 2,73 | 0,24 | 8,07E-01 |
| AT2G08105 | 0,74 | -3,31 | 2,37 | -1,40 | 1,62E-01 |
| AT2G25180 | 0,76 | -2,37 | 3,02 | -0,78 | 4,33E-01 |
| AT2G25200 | 0,69 | -1,21 | 3,77 | -0,32 | 7,49E-01 |
| AT2G25300 | 0,15 | -2,07 | 4,39 | -0,47 | 6,38E-01 |
| AT2G25310 | 0,20 | 0,40 | 4,37 | 0,09 | 9,27E-01 |
| AT2G25570 | 0,45 | 0,31 | 4,39 | 0,07 | 9,44E-01 |
| AT2G25610 | 0,53 | 0,54 | 3,16 | 0,17 | 8,65E-01 |
| AT2G25670 | 0,93 | -0,36 | 2,28 | -0,16 | 8,75E-01 |
| AT2G25760 | 0,93 | -0,38 | 2,92 | -0,13 | 8,96E-01 |
| AT2G25850 | 0,16 | 0,15 | 4,38 | 0,04 | 9,72E-01 |
| AT2G25910 | 0,91 | -0,30 | 2,26 | -0,13 | 8,95E-01 |
| AT2G25970 | 1,54 | 0,95 | 1,94 | 0,49 | 6,26E-01 |
| AT2G26030 | 0,31 | 0,42 | 3,59 | 0,12 | 9,06E-01 |
| AT2G26070 | 0,39 | 0,00 | 4,40 | 0,00 | 1,00E+00 |
| AT2G26140 | 1,61 | -1,33 | 1,65 | -0,80 | 4,22E-01 |
| AT2G26190 | 1,38 | 1,65 | 1,84 | 0,89 | 3,72E-01 |
| AT2G26280 | 1,11 | 1,67 | 2,42 | 0,69 | 4,90E-01 |
| AT2G26430 | 0,38 | -0,32 | 4,33 | -0,07 | 9,41E-01 |
| AT2G26540 | 0,42 | 0,48 | 4,35 | 0,11 | 9,12E-01 |
| AT2G26560 | 1,67 | 2,25 | 1,63 | 1,38 | 1,67E-01 |
| AT2G26580 | 0,26 | -0,91 | 4,38 | -0,21 | 8,36E-01 |
| AT2G26590 | 0,25 | -0,05 | 4,35 | -0,01 | 9,90E-01 |
| AT2G26620 | 0,18 | 1,20 | 4,34 | 0,28 | 7,82E-01 |
| AT2G26680 | 0,33 | 0,00 | 4,41 | 0,00 | 1,00E+00 |
| AT2G26690 | 0,68 | -0,52 | 2,62 | -0,20 | 8,42E-01 |
| AT2G26740 | 0,33 | 0,49 | 4,36 | 0,11 | 9,10E-01 |
| AT2G08175 | 1,14 | 0,67 | 1,73 | 0,38 | 7,00E-01 |
| AT2G26790 | 0,21 | 0,00 | 4,41 | 0,00 | 1,00E+00 |
| AT2G26800 | 0,41 | -1,25 | 3,45 | -0,36 | 7,18E-01 |
| AT2G26840 | 0,08 | 0,00 | 4,41 | 0,00 | 1,00E+00 |
| AT2G26900 | 1,34 | -1,18 | 1,63 | -0,73 | 4,67E-01 |
| AT2G26930 | 0,69 | -2,71 | 2,98 | -0,91 | 3,63E-01 |
| AT2G26980 | 0,61 | -0,74 | 2,28 | -0,32 | 7,47E-01 |
| AT2G26990 | 0,56 | -1,29 | 3,20 | -0,40 | 6,87E-01 |
| AT2G27020 | 0,74 | -1,36 | 2,53 | -0,54 | 5,92E-01 |
| AT2G27040 | 0,44 | -0,10 | 3,33 | -0,03 | 9,75E-01 |
| AT2G27070 | 0,56 | -3,40 | 3,01 | -1,13 | 2,59E-01 |
| AT2G27090 | 0,77 | 0,41 | 3,12 | 0,13 | 8,96E-01 |
| AT2G27170 | 0,61 | -1,77 | 2,47 | -0,72 | 4,74E-01 |
| AT2G27190 | 0,39 | -0,71 | 4,25 | -0,17 | 8,67E-01 |
| AT2G27290 | 0,57 | -0,66 | 2,65 | -0,25 | 8,05E-01 |
| AT2G27330 | 0,10 | 0,86 | 4,35 | 0,20 | 8,43E-01 |
| AT2G27350 | 0,23 | 0,81 | 4,35 | 0,19 | 8,52E-01 |
| AT2G27385 | 1,03 | -1,03 | 1,87 | -0,55 | 5,82E-01 |

| | | | | | |
|---|---|---|---|---|---|
| AT2G27500 | 0,69 | 2,71 | 2,46 | 1,10 | 2,69E-01 |
| AT2G27510 | 0,44 | -0,57 | 4,21 | -0,13 | 8,93E-01 |
| AT2G27530 | 0,92 | -0,15 | 1,88 | -0,08 | 9,35E-01 |
| AT2G27580 | 0,43 | 0,48 | 4,23 | 0,11 | 9,09E-01 |
| AT2G27660 | 0,41 | -0,34 | 4,40 | -0,08 | 9,39E-01 |
| AT2G27680 | 0,72 | -0,66 | 2,02 | -0,33 | 7,42E-01 |
| AT2G27770 | 0,38 | 1,84 | 4,30 | 0,43 | 6,68E-01 |
| AT2G27950 | 1,06 | 0,90 | 2,10 | 0,43 | 6,70E-01 |
| AT2G28080 | 0,87 | 3,23 | 2,69 | 1,20 | 2,31E-01 |
| AT2G28170 | 0,05 | 0,18 | 4,38 | 0,04 | 9,68E-01 |
| AT2G28360 | 1,42 | 0,28 | 1,53 | 0,19 | 8,53E-01 |
| AT2G28380 | 0,74 | 0,43 | 2,97 | 0,14 | 8,86E-01 |
| AT2G28390 | 0,41 | 0,00 | 4,41 | 0,00 | 1,00E+00 |
| AT2G28400 | 0,81 | -2,56 | 2,49 | -1,03 | 3,04E-01 |
| AT2G28450 | 1,48 | 0,41 | 2,08 | 0,20 | 8,42E-01 |
| AT2G28470 | 0,87 | 2,82 | 2,67 | 1,06 | 2,90E-01 |
| AT2G28480 | 0,12 | -0,36 | 4,39 | -0,08 | 9,34E-01 |
| AT2G28620 | 1,12 | 2,83 | 2,12 | 1,33 | 1,82E-01 |
| AT2G08275 | 1,12 | 2,83 | 2,12 | 1,33 | 1,82E-01 |
| AT2G28630 | 0,37 | 0,04 | 4,28 | 0,01 | 9,92E-01 |
| AT2G28640 | 0,62 | -0,02 | 2,89 | -0,01 | 9,95E-01 |
| AT2G28670 | 0,51 | 0,76 | 3,03 | 0,25 | 8,02E-01 |
| AT2G28740 | 0,44 | 1,23 | 4,15 | 0,29 | 7,68E-01 |
| AT2G28800 | 0,62 | 1,55 | 2,89 | 0,53 | 5,93E-01 |
| AT2G08300 | 0,20 | 0,40 | 4,37 | 0,09 | 9,27E-01 |
| AT2G28880 | 0,33 | 0,00 | 4,41 | 0,00 | 1,00E+00 |
| AT2G28890 | 0,77 | 1,51 | 2,20 | 0,68 | 4,94E-01 |
| AT2G28900 | 0,81 | -0,49 | 2,37 | -0,21 | 8,35E-01 |
| AT2G28950 | 0,45 | -0,13 | 4,12 | -0,03 | 9,75E-01 |
| AT2G29030 | 0,32 | 1,48 | 4,32 | 0,34 | 7,31E-01 |
| AT2G29040 | 0,08 | 0,21 | 4,39 | 0,05 | 9,62E-01 |
| AT2G29110 | 0,41 | -0,40 | 4,17 | -0,10 | 9,23E-01 |
| AT2G29120 | 1,16 | -1,27 | 2,34 | -0,54 | 5,86E-01 |
| AT2G29170 | 0,18 | 0,90 | 4,35 | 0,21 | 8,36E-01 |
| AT2G08330 | 0,26 | 0,00 | 4,41 | 0,00 | 1,00E+00 |
| AT2G29190 | 0,61 | 0,58 | 2,71 | 0,21 | 8,32E-01 |
| AT2G29220 | 0,39 | 0,00 | 4,40 | 0,00 | 1,00E+00 |
| AT2G29390 | 0,71 | -0,88 | 2,54 | -0,35 | 7,27E-01 |
| AT2G29400 | 0,98 | 1,23 | 1,98 | 0,62 | 5,34E-01 |
| AT2G29550 | 0,75 | 1,55 | 2,30 | 0,67 | 5,02E-01 |
| AT2G29570 | 0,14 | 0,81 | 4,35 | 0,19 | 8,52E-01 |
| AT2G29605 | 0,65 | 2,23 | 2,80 | 0,79 | 4,27E-01 |
| AT2G29650 | 0,77 | -2,68 | 2,52 | -1,06 | 2,89E-01 |
| AT2G30250 | 0,71 | -0,28 | 2,38 | -0,12 | 9,06E-01 |
| AT2G30260 | 0,44 | -1,15 | 3,32 | -0,35 | 7,29E-01 |
| AT2G30270 | 0,18 | 0,35 | 4,37 | 0,08 | 9,36E-01 |
| AT2G30360 | 0,37 | 1,70 | 4,31 | 0,39 | 6,94E-01 |
| AT2G30390 | 0,49 | -1,26 | 3,33 | -0,38 | 7,04E-01 |
| AT2G30470 | 0,73 | -1,39 | 2,27 | -0,61 | 5,40E-01 |
| AT2G30490 | 0,91 | -0,23 | 3,08 | -0,07 | 9,40E-01 |
| AT2G30500 | 1,56 | -1,64 | 2,01 | -0,81 | 4,16E-01 |
| AT2G30505 | 0,35 | -1,45 | 4,35 | -0,33 | 7,38E-01 |
| AT2G30700 | 0,36 | -1,21 | 4,35 | -0,28 | 7,82E-01 |
| AT2G30710 | 0,35 | -0,40 | 4,38 | -0,09 | 9,27E-01 |
| AT2G30720 | 0,49 | -0,69 | 3,28 | -0,21 | 8,33E-01 |
| AT2G30790 | 0,21 | 0,08 | 4,38 | 0,02 | 9,85E-01 |

| | | | | | |
|---|---|---|---|---|---|
| AT2G30800 | 1,27 | -0,14 | 2,04 | -0,07 | 9,45E-01 |
| AT2G30850 | 0,41 | -2,62 | 4,36 | -0,60 | 5,48E-01 |
| AT2G30880 | 0,34 | 0,05 | 4,38 | 0,01 | 9,91E-01 |
| AT2G08420 | 0,54 | -3,59 | 3,99 | -0,90 | 3,69E-01 |
| AT2G30980 | 0,41 | -0,35 | 4,39 | -0,08 | 9,36E-01 |
| AT2G31040 | 0,19 | -0,58 | 4,38 | -0,13 | 8,95E-01 |
| AT2G31060 | 0,49 | -0,26 | 4,41 | -0,06 | 9,53E-01 |
| AT2G31070 | 0,79 | 2,53 | 2,26 | 1,12 | 2,62E-01 |
| AT2G31130 | 0,17 | 0,84 | 4,36 | 0,19 | 8,47E-01 |
| AT2G31170 | 0,45 | -2,04 | 4,36 | -0,47 | 6,40E-01 |
| AT2G31240 | 0,25 | 0,00 | 4,41 | 0,00 | 1,00E+00 |
| AT2G31250 | 0,39 | 0,00 | 4,40 | 0,00 | 1,00E+00 |
| AT2G31260 | 0,53 | 1,56 | 2,55 | 0,61 | 5,40E-01 |
| AT2G08430 | 0,08 | 0,00 | 4,41 | 0,00 | 1,00E+00 |
| AT2G31360 | 1,30 | -0,86 | 1,93 | -0,45 | 6,55E-01 |
| AT2G31390 | 0,86 | -0,26 | 2,99 | -0,09 | 9,30E-01 |
| AT2G31410 | 0,37 | 0,00 | 4,41 | 0,00 | 1,00E+00 |
| AT2G31440 | 0,44 | 0,00 | 4,41 | 0,00 | 1,00E+00 |
| AT2G31510 | 0,63 | -2,18 | 3,02 | -0,72 | 4,71E-01 |
| AT2G31570 | 0,14 | 0,04 | 4,38 | 0,01 | 9,93E-01 |
| AT2G31670 | 0,73 | -2,34 | 2,53 | -0,92 | 3,56E-01 |
| AT2G31730 | 0,53 | -2,06 | 4,36 | -0,47 | 6,36E-01 |
| AT2G31740 | 0,52 | -2,14 | 3,27 | -0,65 | 5,14E-01 |
| AT2G31751 | 0,97 | -0,78 | 2,43 | -0,32 | 7,49E-01 |
| AT2G31800 | 0,85 | 1,13 | 2,11 | 0,54 | 5,92E-01 |
| AT2G31830 | 0,34 | -0,26 | 4,41 | -0,06 | 9,53E-01 |
| AT2G31865 | 0,37 | 0,00 | 4,41 | 0,00 | 1,00E+00 |
| AT2G31870 | 0,27 | 1,29 | 3,74 | 0,35 | 7,30E-01 |
| AT2G08475 | 0,81 | 0,44 | 2,43 | 0,18 | 8,55E-01 |
| AT2G31990 | 0,31 | 0,55 | 4,36 | 0,13 | 9,00E-01 |
| AT2G32000 | 0,55 | 0,86 | 2,65 | 0,32 | 7,47E-01 |
| AT2G32030 | 0,22 | 0,80 | 4,35 | 0,18 | 8,55E-01 |
| AT2G32050 | 0,08 | 0,00 | 4,41 | 0,00 | 1,00E+00 |
| AT2G32060 | 0,34 | 1,59 | 4,32 | 0,37 | 7,14E-01 |
| AT2G32070 | 0,64 | -0,50 | 2,62 | -0,19 | 8,49E-01 |
| AT2G32080 | 0,57 | -0,56 | 2,61 | -0,22 | 8,29E-01 |
| AT2G32130 | 0,28 | -0,35 | 4,40 | -0,08 | 9,37E-01 |
| AT2G32230 | 0,70 | 0,00 | 2,98 | 0,00 | 1,00E+00 |
| AT2G32235 | 0,12 | 0,00 | 4,41 | 0,00 | 1,00E+00 |
| AT2G32390 | 0,36 | 0,00 | 4,41 | 0,00 | 1,00E+00 |
| AT2G32400 | 0,29 | 0,00 | 4,41 | 0,00 | 1,00E+00 |
| AT2G32415 | 0,42 | 0,00 | 4,41 | 0,00 | 1,00E+00 |
| AT2G32480 | 0,87 | -1,66 | 2,24 | -0,74 | 4,59E-01 |
| AT2G32520 | 0,35 | -0,83 | 4,36 | -0,19 | 8,49E-01 |
| AT2G32600 | 0,08 | 0,00 | 4,41 | 0,00 | 1,00E+00 |
| AT2G32680 | 1,55 | 1,94 | 1,76 | 1,10 | 2,71E-01 |
| AT2G32810 | 0,70 | -0,78 | 2,34 | -0,33 | 7,40E-01 |
| AT2G32920 | 1,12 | 0,03 | 1,81 | 0,02 | 9,87E-01 |
| AT2G32940 | 0,12 | 0,00 | 4,41 | 0,00 | 1,00E+00 |
| AT2G33040 | 0,74 | 0,98 | 2,29 | 0,43 | 6,69E-01 |
| AT2G33051 | 0,64 | -2,05 | 3,90 | -0,53 | 6,00E-01 |
| AT2G33070 | 0,08 | 0,00 | 4,41 | 0,00 | 1,00E+00 |
| AT2G33110 | 0,40 | 0,00 | 4,41 | 0,00 | 1,00E+00 |
| AT2G33120 | 0,77 | 2,17 | 2,10 | 1,03 | 3,02E-01 |
| AT2G33175 | 0,67 | 1,77 | 2,34 | 0,76 | 4,49E-01 |
| AT2G33210 | 0,70 | 1,30 | 2,49 | 0,52 | 6,02E-01 |

| | | | | | |
|---|---|---|---|---|---|
| AT2G33260 | 0,24 | 0,00 | 4,41 | 0,00 | 1,00E+00 |
| AT2G33320 | 0,54 | 1,60 | 2,57 | 0,62 | 5,35E-01 |
| AT2G33370 | 0,46 | 0,70 | 2,71 | 0,26 | 7,96E-01 |
| AT2G33385 | 0,08 | 0,21 | 4,39 | 0,05 | 9,62E-01 |
| AT2G33460 | 0,28 | -0,32 | 4,40 | -0,07 | 9,43E-01 |
| AT2G33470 | 0,39 | 0,85 | 3,35 | 0,25 | 7,99E-01 |
| AT2G33500 | 0,51 | -2,43 | 4,38 | -0,55 | 5,80E-01 |
| AT2G33530 | 0,99 | 0,02 | 2,03 | 0,01 | 9,91E-01 |
| AT2G33540 | 0,96 | 1,55 | 1,84 | 0,84 | 3,99E-01 |
| AT2G33550 | 0,30 | 1,44 | 4,32 | 0,33 | 7,38E-01 |
| AT2G33580 | 0,75 | 1,47 | 2,18 | 0,67 | 5,00E-01 |
| AT2G33630 | 0,50 | -0,53 | 3,26 | -0,16 | 8,71E-01 |
| AT2G33655 | 0,29 | -2,86 | 4,36 | -0,65 | 5,13E-01 |
| AT2G33800 | 0,66 | -1,82 | 2,54 | -0,72 | 4,75E-01 |
| AT2G33815 | 0,37 | -0,75 | 4,32 | -0,17 | 8,61E-01 |
| AT2G33830 | 1,76 | -1,95 | 1,33 | -1,46 | 1,44E-01 |
| AT2G33860 | 0,43 | 0,94 | 4,17 | 0,22 | 8,22E-01 |
| AT2G33980 | 0,47 | 0,13 | 3,31 | 0,04 | 9,68E-01 |
| AT2G34040 | 1,26 | -1,23 | 1,61 | -0,76 | 4,46E-01 |
| AT2G34070 | 0,35 | 0,30 | 3,54 | 0,08 | 9,33E-01 |
| AT2G34090 | 0,11 | 0,88 | 4,35 | 0,20 | 8,39E-01 |
| AT2G34290 | 0,45 | 2,39 | 3,97 | 0,60 | 5,47E-01 |
| AT2G34300 | 0,72 | -0,34 | 4,40 | -0,08 | 9,37E-01 |
| AT2G34450 | 0,60 | 0,00 | 3,87 | 0,00 | 1,00E+00 |
| AT2G34470 | 0,95 | 0,20 | 2,22 | 0,09 | 9,28E-01 |
| AT2G34480 | 1,66 | 1,26 | 1,52 | 0,83 | 4,08E-01 |
| AT2G34490 | 0,49 | 1,83 | 4,06 | 0,45 | 6,52E-01 |
| AT2G34510 | 0,33 | -2,80 | 4,37 | -0,64 | 5,21E-01 |
| AT2G34520 | 0,36 | 0,00 | 4,41 | 0,00 | 1,00E+00 |
| AT2G34585 | 1,42 | 2,77 | 1,72 | 1,61 | 1,08E-01 |
| AT2G34680 | 1,27 | 1,25 | 2,16 | 0,58 | 5,65E-01 |
| AT2G34710 | 1,10 | -0,78 | 2,58 | -0,30 | 7,64E-01 |
| AT2G34790 | 0,43 | 1,44 | 4,09 | 0,35 | 7,25E-01 |
| AT2G34920 | 0,05 | 0,18 | 4,38 | 0,04 | 9,68E-01 |
| AT2G34930 | 0,44 | 1,25 | 3,40 | 0,37 | 7,14E-01 |
| AT2G35030 | 0,48 | -2,18 | 4,05 | -0,54 | 5,90E-01 |
| AT2G35040 | 0,75 | -2,35 | 2,98 | -0,79 | 4,31E-01 |
| AT2G35060 | 1,08 | -0,32 | 1,63 | -0,19 | 8,46E-01 |
| AT2G35120 | 0,10 | 0,86 | 4,35 | 0,20 | 8,43E-01 |
| AT2G35250 | 1,05 | -1,30 | 1,77 | -0,73 | 4,65E-01 |
| AT2G35360 | 0,45 | -0,17 | 4,19 | -0,04 | 9,68E-01 |
| AT2G35390 | 0,25 | 0,00 | 4,41 | 0,00 | 1,00E+00 |
| AT2G35420 | 0,17 | -0,33 | 4,40 | -0,08 | 9,40E-01 |
| AT2G35490 | 0,67 | 1,11 | 2,12 | 0,53 | 6,00E-01 |
| AT2G35510 | 0,57 | 0,45 | 3,27 | 0,14 | 8,91E-01 |
| AT2G35610 | 0,17 | -0,29 | 4,40 | -0,07 | 9,47E-01 |
| AT2G35680 | 0,73 | 0,29 | 2,42 | 0,12 | 9,04E-01 |
| AT2G35730 | 0,35 | 0,00 | 4,39 | 0,00 | 1,00E+00 |
| AT2G35742 | 0,34 | 1,11 | 3,43 | 0,32 | 7,46E-01 |
| AT2G08755 | 0,29 | 0,53 | 4,36 | 0,12 | 9,04E-01 |
| AT2G08765 | 0,51 | -0,70 | 3,11 | -0,23 | 8,22E-01 |
| AT2G35859 | 0,71 | -0,37 | 2,67 | -0,14 | 8,89E-01 |
| AT2G35880 | 0,24 | 0,00 | 4,41 | 0,00 | 1,00E+00 |
| AT2G35930 | 0,76 | 2,94 | 2,75 | 1,07 | 2,84E-01 |
| AT2G36030 | 0,62 | -1,78 | 4,30 | -0,41 | 6,80E-01 |
| AT2G36050 | 0,69 | 0,87 | 2,87 | 0,30 | 7,63E-01 |

| | | | | | |
|---|---|---|---|---|---|
| AT2G36060 | 0,48 | -0,38 | 2,77 | -0,14 | 8,90E-01 |
| AT2G36070 | 0,34 | 0,09 | 4,38 | 0,02 | 9,84E-01 |
| AT2G36080 | 0,43 | 0,11 | 3,30 | 0,03 | 9,73E-01 |
| AT2G36180 | 0,76 | 3,06 | 3,29 | 0,93 | 3,53E-01 |
| AT2G36200 | 0,33 | 0,57 | 4,37 | 0,13 | 8,97E-01 |
| AT2G36230 | 0,20 | 1,25 | 4,33 | 0,29 | 7,72E-01 |
| AT2G36250 | 0,55 | -0,14 | 3,11 | -0,05 | 9,63E-01 |
| AT2G36300 | 0,44 | 1,37 | 3,29 | 0,42 | 6,77E-01 |
| AT2G36360 | 0,43 | -0,12 | 3,23 | -0,04 | 9,69E-01 |
| AT2G36400 | 0,83 | 0,68 | 3,05 | 0,22 | 8,23E-01 |
| AT2G36480 | 1,71 | 0,63 | 1,98 | 0,32 | 7,51E-01 |
| AT2G36620 | 0,25 | 0,09 | 4,38 | 0,02 | 9,84E-01 |
| AT2G36680 | 0,35 | -1,24 | 4,36 | -0,28 | 7,76E-01 |
| AT2G36710 | 0,27 | 0,55 | 4,32 | 0,13 | 8,99E-01 |
| AT2G36750 | 0,21 | 0,00 | 4,41 | 0,00 | 1,00E+00 |
| AT2G36790 | 0,34 | 1,94 | 4,30 | 0,45 | 6,52E-01 |
| AT2G36800 | 0,85 | -0,26 | 2,48 | -0,11 | 9,16E-01 |
| AT2G36810 | 1,10 | -0,59 | 2,17 | -0,27 | 7,87E-01 |
| AT2G36835 | 0,30 | 0,41 | 3,59 | 0,11 | 9,10E-01 |
| AT2G36895 | 0,35 | 0,00 | 4,39 | 0,00 | 1,00E+00 |
| AT2G36900 | 0,55 | -0,42 | 3,83 | -0,11 | 9,13E-01 |
| AT2G36930 | 0,54 | 0,23 | 3,90 | 0,06 | 9,54E-01 |
| AT2G36990 | 1,19 | -0,37 | 2,68 | -0,14 | 8,92E-01 |
| AT2G37035 | 0,78 | -0,31 | 2,43 | -0,13 | 8,97E-01 |
| AT2G37040 | 1,58 | 1,28 | 1,84 | 0,70 | 4,84E-01 |
| AT2G37080 | 0,12 | 0,00 | 4,41 | 0,00 | 1,00E+00 |
| AT2G37130 | 0,68 | 1,65 | 3,02 | 0,55 | 5,84E-01 |
| AT2G37150 | 0,75 | 0,00 | 3,52 | 0,00 | 1,00E+00 |
| AT2G37170 | 0,90 | -0,27 | 2,49 | -0,11 | 9,12E-01 |
| AT2G37200 | 0,31 | 0,43 | 3,62 | 0,12 | 9,05E-01 |
| AT2G37230 | 0,56 | -0,16 | 3,18 | -0,05 | 9,60E-01 |
| AT2G37270 | 1,75 | -1,60 | 1,61 | -0,99 | 3,21E-01 |
| AT2G37310 | 0,77 | 0,00 | 3,61 | 0,00 | 1,00E+00 |
| AT2G37340 | 1,12 | -2,12 | 2,39 | -0,89 | 3,75E-01 |
| AT2G37360 | 0,21 | 0,00 | 4,41 | 0,00 | 1,00E+00 |
| AT2G37400 | 0,25 | 0,00 | 4,41 | 0,00 | 1,00E+00 |
| AT2G37410 | 0,19 | -0,34 | 4,40 | -0,08 | 9,38E-01 |
| AT2G37520 | 1,03 | -0,17 | 1,84 | -0,09 | 9,28E-01 |
| AT2G37540 | 0,69 | 0,00 | 3,01 | 0,00 | 1,00E+00 |
| AT2G37550 | 1,38 | 0,61 | 2,06 | 0,30 | 7,68E-01 |
| AT2G37570 | 0,98 | -0,61 | 1,90 | -0,32 | 7,49E-01 |
| AT2G37630 | 0,98 | 0,43 | 2,10 | 0,20 | 8,39E-01 |
| AT2G37660 | 1,35 | -0,46 | 1,50 | -0,31 | 7,59E-01 |
| AT2G37678 | 0,57 | -1,74 | 4,33 | -0,40 | 6,88E-01 |
| AT2G08895 | 1,00 | 2,38 | 2,22 | 1,07 | 2,84E-01 |
| AT2G08900 | 0,86 | -0,25 | 1,87 | -0,13 | 8,93E-01 |
| AT2G08910 | 1,52 | 1,31 | 1,43 | 0,92 | 3,60E-01 |
| AT2G08915 | 0,73 | 0,80 | 2,75 | 0,29 | 7,70E-01 |
| AT2G38000 | 0,32 | 1,03 | 4,35 | 0,24 | 8,12E-01 |
| AT2G38020 | 1,29 | -1,57 | 2,30 | -0,68 | 4,95E-01 |
| AT2G38160 | 0,87 | 0,00 | 3,37 | 0,00 | 1,00E+00 |
| AT2G38270 | 0,22 | 0,60 | 4,30 | 0,14 | 8,89E-01 |
| AT2G38290 | 0,66 | -0,38 | 2,99 | -0,13 | 9,00E-01 |
| AT2G38340 | 0,17 | 0,07 | 4,38 | 0,02 | 9,87E-01 |
| AT2G38360 | 0,39 | 1,58 | 3,31 | 0,48 | 6,33E-01 |
| AT2G38410 | 0,35 | -1,24 | 3,47 | -0,36 | 7,20E-01 |

| | | | | | |
|---|---|---|---|---|---|
| AT2G38480 | 0,17 | -0,29 | 4,40 | -0,07 | 9,47E-01 |
| AT2G38610 | 0,83 | -1,34 | 2,58 | -0,52 | 6,02E-01 |
| AT2G38650 | 1,16 | -0,91 | 2,03 | -0,45 | 6,54E-01 |
| AT2G38680 | 0,08 | 0,21 | 4,39 | 0,05 | 9,62E-01 |
| AT2G38750 | 0,05 | 0,18 | 4,38 | 0,04 | 9,68E-01 |
| AT2G38770 | 1,53 | -0,24 | 1,77 | -0,13 | 8,93E-01 |
| AT2G38790 | 0,55 | 0,89 | 2,34 | 0,38 | 7,02E-01 |
| AT2G38860 | 0,41 | 0,64 | 3,42 | 0,19 | 8,51E-01 |
| AT2G39000 | 1,06 | -2,32 | 3,00 | -0,77 | 4,41E-01 |
| AT2G39080 | 0,20 | 0,00 | 4,41 | 0,00 | 1,00E+00 |
| AT2G39130 | 0,08 | 0,21 | 4,39 | 0,05 | 9,62E-01 |
| AT2G39160 | 0,24 | -1,28 | 4,38 | -0,29 | 7,70E-01 |
| AT2G39170 | 0,19 | 0,37 | 4,36 | 0,08 | 9,32E-01 |
| AT2G09010 | 0,08 | 0,21 | 4,39 | 0,05 | 9,62E-01 |
| AT2G39210 | 0,91 | 0,00 | 2,88 | 0,00 | 1,00E+00 |
| AT2G39260 | 1,36 | -0,61 | 1,56 | -0,39 | 6,99E-01 |
| AT2G39350 | 0,49 | 0,22 | 3,25 | 0,07 | 9,47E-01 |
| AT2G39360 | 0,80 | 0,00 | 3,58 | 0,00 | 1,00E+00 |
| AT2G09045 | 0,52 | 0,00 | 3,96 | 0,00 | 1,00E+00 |
| AT2G39450 | 0,29 | 0,00 | 4,41 | 0,00 | 1,00E+00 |
| AT2G39480 | 1,05 | 2,23 | 2,10 | 1,06 | 2,88E-01 |
| AT2G39650 | 0,49 | 1,84 | 4,00 | 0,46 | 6,46E-01 |
| AT2G39710 | 0,60 | 0,24 | 3,23 | 0,07 | 9,41E-01 |
| AT2G39720 | 1,30 | 0,04 | 2,51 | 0,01 | 9,88E-01 |
| AT2G39750 | 1,24 | -0,90 | 1,97 | -0,45 | 6,50E-01 |
| AT2G09080 | 0,50 | -0,31 | 3,32 | -0,09 | 9,26E-01 |
| AT2G39805 | 0,30 | 0,00 | 4,41 | 0,00 | 1,00E+00 |
| AT2G39920 | 0,18 | -1,25 | 4,37 | -0,29 | 7,74E-01 |
| AT2G09090 | 0,82 | -2,47 | 2,91 | -0,85 | 3,96E-01 |
| AT2G39940 | 1,84 | 0,38 | 2,11 | 0,18 | 8,58E-01 |
| AT2G39950 | 0,75 | -1,94 | 2,61 | -0,74 | 4,58E-01 |
| AT2G39960 | 0,25 | -2,95 | 4,37 | -0,68 | 5,00E-01 |
| AT2G39990 | 1,46 | -0,64 | 2,62 | -0,24 | 8,07E-01 |
| AT2G40070 | 0,52 | 0,00 | 3,96 | 0,00 | 1,00E+00 |
| AT2G40080 | 0,23 | 0,81 | 4,35 | 0,19 | 8,52E-01 |
| AT2G40165 | 0,38 | 0,00 | 4,41 | 0,00 | 1,00E+00 |
| AT2G40205 | 0,27 | 1,11 | 4,34 | 0,25 | 7,99E-01 |
| AT2G40270 | 1,34 | 1,52 | 1,71 | 0,89 | 3,74E-01 |
| AT2G40280 | 0,63 | -3,05 | 3,04 | -1,01 | 3,15E-01 |
| AT2G40350 | 0,12 | 0,00 | 4,41 | 0,00 | 1,00E+00 |
| AT2G40360 | 0,28 | 1,80 | 4,30 | 0,42 | 6,75E-01 |
| AT2G40430 | 0,45 | -0,16 | 4,17 | -0,04 | 9,69E-01 |
| AT2G40475 | 0,89 | 3,14 | 2,21 | 1,42 | 1,55E-01 |
| AT2G40590 | 0,50 | -1,60 | 2,66 | -0,60 | 5,47E-01 |
| AT2G40600 | 0,11 | -0,38 | 4,39 | -0,09 | 9,31E-01 |
| AT2G40610 | 0,37 | -2,34 | 4,35 | -0,54 | 5,91E-01 |
| AT2G40650 | 0,77 | 2,08 | 2,24 | 0,93 | 3,53E-01 |
| AT2G40730 | 0,34 | 0,70 | 3,42 | 0,21 | 8,37E-01 |
| AT2G40750 | 0,46 | -3,40 | 4,14 | -0,82 | 4,11E-01 |
| AT2G40770 | 0,51 | -1,55 | 3,98 | -0,39 | 6,97E-01 |
| AT2G40815 | 0,24 | 0,00 | 4,41 | 0,00 | 1,00E+00 |
| AT2G40860 | 0,30 | 0,00 | 4,41 | 0,00 | 1,00E+00 |
| AT2G40890 | 0,53 | -0,57 | 3,92 | -0,15 | 8,84E-01 |
| AT2G40920 | 0,35 | -1,33 | 4,39 | -0,30 | 7,61E-01 |
| AT2G40925 | 0,65 | 0,00 | 3,74 | 0,00 | 1,00E+00 |
| AT2G40930 | 1,74 | -2,16 | 2,11 | -1,02 | 3,06E-01 |

| | | | | | |
|---|---|---|---|---|---|
| AT2G40950 | 0,74 | -0,06 | 3,60 | -0,02 | 9,86E-01 |
| AT2G40970 | 0,42 | 0,00 | 4,41 | 0,00 | 1,00E+00 |
| AT2G09225 | 0,24 | -0,68 | 4,38 | -0,16 | 8,76E-01 |
| AT2G41120 | 0,21 | 0,00 | 4,41 | 0,00 | 1,00E+00 |
| AT2G41160 | 0,39 | -0,33 | 4,40 | -0,08 | 9,40E-01 |
| AT2G41250 | 0,87 | 2,41 | 2,37 | 1,02 | 3,10E-01 |
| AT2G41420 | 0,11 | 0,88 | 4,35 | 0,20 | 8,39E-01 |
| AT2G41475 | 0,71 | 0,00 | 2,57 | 0,00 | 9,99E-01 |
| AT2G41500 | 0,96 | -1,30 | 2,29 | -0,57 | 5,70E-01 |
| AT2G41600 | 0,58 | -0,46 | 2,65 | -0,17 | 8,62E-01 |
| AT2G41620 | 1,68 | -1,24 | 2,08 | -0,59 | 5,52E-01 |
| AT2G41616 | 0,33 | 0,00 | 4,41 | 0,00 | 1,00E+00 |
| AT2G41630 | 0,92 | 0,33 | 2,83 | 0,12 | 9,08E-01 |
| AT2G41680 | 0,32 | 0,84 | 4,34 | 0,19 | 8,47E-01 |
| AT2G41700 | 1,62 | 0,89 | 2,06 | 0,43 | 6,66E-01 |
| AT2G41770 | 0,40 | 0,62 | 4,30 | 0,14 | 8,86E-01 |
| AT2G41790 | 0,58 | 0,18 | 2,60 | 0,07 | 9,44E-01 |
| AT2G41830 | 0,40 | -2,14 | 4,23 | -0,51 | 6,13E-01 |
| AT2G41835 | 0,70 | 0,51 | 3,76 | 0,14 | 8,92E-01 |
| AT2G41870 | 0,29 | -0,30 | 4,40 | -0,07 | 9,46E-01 |
| AT2G41890 | 0,43 | 1,38 | 4,16 | 0,33 | 7,39E-01 |
| AT2G41945 | 0,60 | 0,36 | 3,98 | 0,09 | 9,28E-01 |
| AT2G41960 | 0,92 | -1,22 | 2,04 | -0,60 | 5,49E-01 |
| AT2G42010 | 0,72 | -0,09 | 1,96 | -0,05 | 9,63E-01 |
| AT2G42020 | 0,48 | -1,61 | 4,31 | -0,37 | 7,08E-01 |
| AT2G42070 | 0,23 | 1,31 | 4,33 | 0,30 | 7,62E-01 |
| AT2G42230 | 0,42 | -0,26 | 4,18 | -0,06 | 9,50E-01 |
| AT2G42280 | 0,33 | 0,00 | 4,41 | 0,00 | 1,00E+00 |
| AT2G42365 | 0,35 | 0,00 | 4,39 | 0,00 | 1,00E+00 |
| AT2G42400 | 1,01 | -1,08 | 2,23 | -0,49 | 6,27E-01 |
| AT2G42500 | 0,82 | 0,74 | 2,30 | 0,32 | 7,48E-01 |
| AT2G42530 | 0,98 | -2,72 | 1,92 | -1,42 | 1,56E-01 |
| AT2G42690 | 1,34 | -0,26 | 1,42 | -0,18 | 8,53E-01 |
| AT2G42760 | 0,47 | 1,49 | 2,60 | 0,57 | 5,67E-01 |
| AT2G42790 | 0,53 | -1,14 | 2,75 | -0,42 | 6,78E-01 |
| AT2G42800 | 0,18 | 1,59 | 4,31 | 0,37 | 7,12E-01 |
| AT2G42810 | 0,84 | -1,00 | 2,06 | -0,48 | 6,29E-01 |
| AT2G42840 | 0,22 | 1,63 | 4,31 | 0,38 | 7,05E-01 |
| AT2G42865 | 0,08 | 0,00 | 4,41 | 0,00 | 1,00E+00 |
| AT2G42870 | 0,10 | 0,24 | 4,38 | 0,05 | 9,57E-01 |
| AT2G42880 | 0,41 | 1,13 | 3,46 | 0,33 | 7,44E-01 |
| AT2G43180 | 0,70 | 0,00 | 3,01 | 0,00 | 1,00E+00 |
| AT2G43260 | 0,20 | 0,00 | 4,41 | 0,00 | 1,00E+00 |
| AT2G43290 | 0,50 | 0,89 | 3,21 | 0,28 | 7,81E-01 |
| AT2G43320 | 0,22 | 0,11 | 4,38 | 0,02 | 9,80E-01 |
| AT2G43360 | 0,33 | -2,12 | 4,40 | -0,48 | 6,30E-01 |
| AT2G43420 | 0,11 | -0,38 | 4,39 | -0,09 | 9,31E-01 |
| AT2G43430 | 0,59 | -1,00 | 2,56 | -0,39 | 6,97E-01 |
| AT2G43460 | 0,41 | -0,10 | 3,37 | -0,03 | 9,76E-01 |
| AT2G43490 | 0,44 | -0,30 | 4,14 | -0,07 | 9,43E-01 |
| AT2G43560 | 0,25 | -0,40 | 4,38 | -0,09 | 9,27E-01 |
| AT2G43750 | 1,46 | 1,00 | 1,73 | 0,58 | 5,64E-01 |
| AT2G43865 | 0,15 | 0,90 | 4,35 | 0,21 | 8,36E-01 |
| AT2G43910 | 0,26 | -0,45 | 3,77 | -0,12 | 9,05E-01 |
| AT2G43940 | 0,08 | 0,00 | 4,41 | 0,00 | 1,00E+00 |
| AT2G43945 | 0,47 | -2,68 | 4,37 | -0,61 | 5,40E-01 |

| | | | | | |
|---|---|---|---|---|---|
| AT2G43950 | 0,24 | 0,00 | 4,41 | 0,00 | 1,00E+00 |
| AT2G44090 | 0,69 | -1,18 | 2,72 | -0,43 | 6,64E-01 |
| AT2G44140 | 0,91 | -0,08 | 2,15 | -0,04 | 9,72E-01 |
| AT2G44180 | 0,46 | 1,64 | 3,26 | 0,50 | 6,15E-01 |
| AT2G44255 | 0,20 | -0,26 | 4,41 | -0,06 | 9,53E-01 |
| AT2G44360 | 0,63 | 15,84 | 4,34 | 3,65 | 2,63E-04 |
| AT2G44610 | 0,48 | -0,56 | 4,20 | -0,13 | 8,94E-01 |
| AT2G44660 | 0,35 | 0,42 | 3,52 | 0,12 | 9,05E-01 |
| AT2G44760 | 0,32 | 0,42 | 3,61 | 0,12 | 9,07E-01 |
| AT2G44798 | 0,36 | 0,77 | 4,36 | 0,18 | 8,60E-01 |
| AT2G44950 | 0,77 | -1,72 | 2,33 | -0,74 | 4,61E-01 |
| AT2G45070 | 0,80 | 1,29 | 2,05 | 0,63 | 5,31E-01 |
| AT2G45220 | 0,54 | 0,00 | 4,41 | 0,00 | 1,00E+00 |
| AT2G45315 | 0,67 | 0,75 | 2,50 | 0,30 | 7,63E-01 |
| AT2G45460 | 0,20 | 1,52 | 4,31 | 0,35 | 7,25E-01 |
| AT2G45470 | 1,65 | 1,03 | 1,90 | 0,54 | 5,86E-01 |
| AT2G45520 | 0,22 | 0,15 | 4,33 | 0,03 | 9,73E-01 |
| AT2G09640 | 1,68 | 0,74 | 2,13 | 0,34 | 7,30E-01 |
| AT2G45560 | 0,47 | 0,00 | 4,40 | 0,00 | 1,00E+00 |
| AT2G45660 | 0,55 | 0,86 | 2,29 | 0,38 | 7,05E-01 |
| AT2G45670 | 0,95 | -2,28 | 2,45 | -0,93 | 3,54E-01 |
| AT2G45680 | 1,79 | 0,28 | 2,01 | 0,14 | 8,88E-01 |
| AT2G45700 | 0,38 | -1,40 | 4,34 | -0,32 | 7,46E-01 |
| AT2G45820 | 0,98 | -1,14 | 1,63 | -0,70 | 4,84E-01 |
| AT2G45970 | 0,56 | 0,86 | 2,68 | 0,32 | 7,48E-01 |
| AT2G45980 | 0,45 | 0,40 | 4,13 | 0,10 | 9,23E-01 |
| AT2G46030 | 0,57 | 0,00 | 4,41 | 0,00 | 1,00E+00 |
| AT2G46040 | 0,20 | 0,00 | 4,41 | 0,00 | 1,00E+00 |
| AT2G46060 | 0,49 | -3,54 | 4,34 | -0,82 | 4,15E-01 |
| AT2G46080 | 0,51 | 0,88 | 3,24 | 0,27 | 7,86E-01 |
| AT2G46180 | 0,35 | 0,88 | 4,36 | 0,20 | 8,41E-01 |
| AT2G46280 | 1,16 | -3,12 | 1,92 | -1,62 | 1,05E-01 |
| AT2G46330 | 0,93 | -1,06 | 1,91 | -0,56 | 5,79E-01 |
| AT2G46390 | 0,46 | 0,00 | 4,15 | 0,00 | 1,00E+00 |
| AT2G46505 | 0,89 | 0,47 | 2,41 | 0,19 | 8,46E-01 |
| AT2G46510 | 1,07 | 2,06 | 2,04 | 1,01 | 3,13E-01 |
| AT2G46540 | 0,12 | 0,49 | 4,38 | 0,11 | 9,11E-01 |
| AT2G46550 | 0,57 | -0,25 | 2,75 | -0,09 | 9,27E-01 |
| AT2G46572 | 0,51 | -0,35 | 4,07 | -0,09 | 9,32E-01 |
| AT2G46610 | 0,85 | 0,65 | 2,01 | 0,32 | 7,45E-01 |
| AT2G46620 | 1,45 | 1,41 | 1,87 | 0,75 | 4,53E-01 |
| AT2G46680 | 0,94 | 0,14 | 2,33 | 0,06 | 9,52E-01 |
| AT2G46840 | 0,08 | 0,00 | 4,41 | 0,00 | 1,00E+00 |
| AT2G46920 | 0,44 | -0,39 | 4,09 | -0,10 | 9,24E-01 |
| AT2G46980 | 0,24 | 0,00 | 4,41 | 0,00 | 1,00E+00 |
| AT2G47000 | 1,78 | -0,14 | 2,30 | -0,06 | 9,51E-01 |
| AT2G47090 | 0,25 | 0,00 | 4,41 | 0,00 | 1,00E+00 |
| AT2G47180 | 1,41 | 0,10 | 1,75 | 0,06 | 9,56E-01 |
| AT2G47250 | 1,15 | -2,24 | 2,47 | -0,91 | 3,64E-01 |
| AT2G47330 | 0,60 | -0,05 | 3,14 | -0,01 | 9,88E-01 |
| AT2G47600 | 0,61 | 0,45 | 2,60 | 0,17 | 8,61E-01 |
| AT2G47650 | 1,46 | -1,12 | 1,87 | -0,60 | 5,48E-01 |
| AT2G47680 | 0,89 | 0,89 | 2,16 | 0,41 | 6,81E-01 |
| AT2G47700 | 0,93 | 1,61 | 2,45 | 0,66 | 5,10E-01 |
| AT2G47840 | 0,52 | -1,30 | 3,25 | -0,40 | 6,88E-01 |
| AT2G47990 | 0,88 | 0,00 | 2,81 | 0,00 | 1,00E+00 |

| | | | | | |
|---|---|---|---|---|---|
| AT2G48000 | 1,00 | 0,00 | 3,48 | 0,00 | 1,00E+00 |
| AT2G48030 | 0,73 | -0,50 | 2,50 | -0,20 | 8,42E-01 |
| AT2G48130 | 0,91 | -0,08 | 1,84 | -0,04 | 9,65E-01 |
| AT3G01090 | 0,63 | 1,12 | 2,18 | 0,52 | 6,06E-01 |
| AT3G01100 | 0,90 | 0,83 | 2,27 | 0,37 | 7,15E-01 |
| AT3G01130 | 0,21 | 0,08 | 4,38 | 0,02 | 9,85E-01 |
| AT3G01160 | 0,24 | -1,28 | 4,38 | -0,29 | 7,70E-01 |
| AT3G01180 | 0,84 | -0,43 | 1,88 | -0,23 | 8,18E-01 |
| AT3G01300 | 0,08 | 0,00 | 4,41 | 0,00 | 1,00E+00 |
| AT3G01340 | 0,50 | -0,82 | 3,24 | -0,25 | 8,00E-01 |
| AT3G01380 | 1,09 | -1,13 | 2,13 | -0,53 | 5,94E-01 |
| AT3G01390 | 0,38 | -1,57 | 4,35 | -0,36 | 7,19E-01 |
| AT3G01415 | 0,40 | 0,65 | 4,32 | 0,15 | 8,80E-01 |
| AT3G01420 | 0,31 | -0,40 | 4,38 | -0,09 | 9,27E-01 |
| AT3G01430 | 0,79 | 1,00 | 3,01 | 0,33 | 7,39E-01 |
| AT3G01470 | 1,49 | 0,33 | 1,81 | 0,18 | 8,56E-01 |
| AT3G01550 | 0,10 | 0,08 | 4,38 | 0,02 | 9,85E-01 |
| AT3G01720 | 0,80 | 0,65 | 2,94 | 0,22 | 8,24E-01 |
| AT3G01820 | 0,81 | -3,04 | 3,65 | -0,83 | 4,05E-01 |
| AT3G01840 | 0,88 | 0,67 | 2,29 | 0,29 | 7,71E-01 |
| AT3G01910 | 0,46 | -1,39 | 4,13 | -0,34 | 7,37E-01 |
| AT3G01930 | 0,62 | -2,04 | 3,04 | -0,67 | 5,01E-01 |
| AT3G01980 | 0,16 | -0,22 | 4,40 | -0,05 | 9,60E-01 |
| AT3G02040 | 0,68 | 0,82 | 2,50 | 0,33 | 7,44E-01 |
| AT3G02080 | 1,20 | -1,06 | 2,16 | -0,49 | 6,23E-01 |
| AT3G02190 | 0,44 | -0,21 | 3,34 | -0,06 | 9,50E-01 |
| AT3G02210 | 0,52 | 0,36 | 4,05 | 0,09 | 9,30E-01 |
| AT3G02250 | 0,34 | -1,56 | 4,34 | -0,36 | 7,19E-01 |
| AT3G02380 | 0,91 | 0,43 | 2,40 | 0,18 | 8,58E-01 |
| AT3G02410 | 0,66 | 0,00 | 4,41 | 0,00 | 1,00E+00 |
| AT3G02450 | 0,68 | 0,15 | 2,32 | 0,06 | 9,49E-01 |
| AT3G02520 | 1,11 | -1,27 | 1,62 | -0,78 | 4,34E-01 |
| AT3G02530 | 0,67 | -0,04 | 2,65 | -0,02 | 9,87E-01 |
| AT3G02540 | 1,81 | 0,47 | 1,76 | 0,27 | 7,91E-01 |
| AT3G02700 | 0,39 | -1,22 | 4,23 | -0,29 | 7,73E-01 |
| AT3G02730 | 0,92 | -1,97 | 2,83 | -0,70 | 4,86E-01 |
| AT3G02750 | 1,16 | -0,89 | 1,97 | -0,45 | 6,50E-01 |
| AT3G02760 | 0,94 | -0,21 | 2,85 | -0,07 | 9,42E-01 |
| AT3G01205 | 0,80 | 0,89 | 2,35 | 0,38 | 7,04E-01 |
| AT3G03090 | 0,29 | -3,05 | 4,36 | -0,70 | 4,85E-01 |
| AT3G03150 | 0,22 | 0,26 | 4,34 | 0,06 | 9,52E-01 |
| AT3G03220 | 0,16 | 1,58 | 4,31 | 0,37 | 7,14E-01 |
| AT3G03250 | 1,00 | 1,59 | 1,84 | 0,87 | 3,86E-01 |
| AT3G03300 | 0,33 | 0,00 | 4,41 | 0,00 | 1,00E+00 |
| AT3G03305 | 0,32 | -0,32 | 4,40 | -0,07 | 9,42E-01 |
| AT3G03310 | 0,29 | 0,00 | 4,41 | 0,00 | 1,00E+00 |
| AT3G03330 | 0,30 | 0,01 | 4,37 | 0,00 | 9,97E-01 |
| AT3G03405 | 0,16 | 1,58 | 4,31 | 0,37 | 7,14E-01 |
| AT3G03460 | 0,17 | 0,00 | 4,41 | 0,00 | 1,00E+00 |
| AT3G03470 | 1,68 | 2,50 | 1,86 | 1,34 | 1,79E-01 |
| AT3G03480 | 0,08 | 0,00 | 4,41 | 0,00 | 1,00E+00 |
| AT3G03550 | 0,10 | 0,08 | 4,38 | 0,02 | 9,85E-01 |
| AT3G03560 | 0,61 | -0,36 | 3,06 | -0,12 | 9,06E-01 |
| AT3G03580 | 0,26 | 0,00 | 4,41 | 0,00 | 1,00E+00 |
| AT3G03590 | 0,08 | 0,00 | 4,41 | 0,00 | 1,00E+00 |
| AT3G03600 | 0,30 | -0,23 | 4,31 | -0,05 | 9,58E-01 |

| | | | | | |
|---|---|---|---|---|---|
| AT3G03720 | 0,26 | 0,84 | 4,34 | 0,19 | 8,47E-01 |
| AT3G03770 | 0,99 | -0,31 | 3,20 | -0,10 | 9,22E-01 |
| AT3G03890 | 0,79 | 0,89 | 2,49 | 0,36 | 7,21E-01 |
| AT3G03920 | 0,42 | -1,58 | 3,28 | -0,48 | 6,30E-01 |
| AT3G03940 | 0,41 | 1,01 | 2,75 | 0,37 | 7,12E-01 |
| AT3G04030 | 0,25 | 2,14 | 4,29 | 0,50 | 6,19E-01 |
| AT3G04080 | 1,01 | -1,09 | 2,17 | -0,50 | 6,16E-01 |
| AT3G04150 | 0,26 | 0,00 | 4,41 | 0,00 | 1,00E+00 |
| AT3G04200 | 0,62 | 0,00 | 4,40 | 0,00 | 1,00E+00 |
| AT3G04220 | 0,33 | 0,10 | 4,38 | 0,02 | 9,81E-01 |
| AT3G04260 | 0,61 | -2,17 | 4,34 | -0,50 | 6,18E-01 |
| AT3G04290 | 0,84 | -1,86 | 3,10 | -0,60 | 5,48E-01 |
| AT3G04350 | 0,35 | 2,17 | 4,29 | 0,51 | 6,13E-01 |
| AT3G04480 | 0,84 | 1,08 | 2,38 | 0,45 | 6,50E-01 |
| AT3G04500 | 0,20 | 0,40 | 4,37 | 0,09 | 9,27E-01 |
| AT3G04580 | 1,03 | -1,26 | 2,22 | -0,57 | 5,72E-01 |
| AT3G04640 | 0,85 | 2,21 | 2,08 | 1,07 | 2,86E-01 |
| AT3G04690 | 0,22 | -0,90 | 4,35 | -0,21 | 8,36E-01 |
| AT3G04720 | 0,50 | 2,09 | 3,05 | 0,69 | 4,93E-01 |
| AT3G04730 | 1,02 | -0,29 | 1,64 | -0,18 | 8,61E-01 |
| AT3G04750 | 0,57 | 0,48 | 2,69 | 0,18 | 8,60E-01 |
| AT3G04760 | 0,89 | 1,85 | 2,05 | 0,90 | 3,67E-01 |
| AT3G04860 | 0,12 | 0,00 | 4,41 | 0,00 | 1,00E+00 |
| AT3G04980 | 0,46 | 0,00 | 4,41 | 0,00 | 1,00E+00 |
| AT3G05030 | 0,28 | 0,80 | 4,35 | 0,18 | 8,54E-01 |
| AT3G01585 | 0,10 | 0,86 | 4,35 | 0,20 | 8,43E-01 |
| AT3G05040 | 1,58 | -0,91 | 1,72 | -0,53 | 5,95E-01 |
| AT3G05050 | 0,38 | -0,26 | 4,41 | -0,06 | 9,53E-01 |
| AT3G05060 | 0,60 | -0,41 | 3,76 | -0,11 | 9,13E-01 |
| AT3G05090 | 0,52 | 1,77 | 2,58 | 0,68 | 4,94E-01 |
| AT3G05165 | 0,83 | 0,66 | 2,05 | 0,32 | 7,47E-01 |
| AT3G05240 | 0,13 | 0,00 | 4,41 | 0,00 | 1,00E+00 |
| AT3G05270 | 0,58 | -0,32 | 2,70 | -0,12 | 9,07E-01 |
| AT3G05280 | 0,59 | -2,19 | 3,86 | -0,57 | 5,71E-01 |
| AT3G05540 | 1,04 | -1,30 | 1,65 | -0,79 | 4,29E-01 |
| AT3G05650 | 0,98 | 1,40 | 2,32 | 0,60 | 5,46E-01 |
| AT3G05660 | 0,79 | 0,27 | 2,83 | 0,09 | 9,25E-01 |
| AT3G05680 | 1,68 | -0,48 | 1,70 | -0,28 | 7,78E-01 |
| AT3G05690 | 0,11 | 0,88 | 4,35 | 0,20 | 8,39E-01 |
| AT3G05770 | 0,46 | 1,90 | 3,28 | 0,58 | 5,62E-01 |
| AT3G05810 | 0,21 | -0,95 | 4,38 | -0,22 | 8,29E-01 |
| AT3G05850 | 0,90 | 0,90 | 2,55 | 0,35 | 7,24E-01 |
| AT3G05880 | 0,55 | -2,65 | 3,12 | -0,85 | 3,95E-01 |
| AT3G05900 | 0,51 | -1,16 | 3,18 | -0,37 | 7,15E-01 |
| AT3G05910 | 0,68 | -2,43 | 2,53 | -0,96 | 3,37E-01 |
| AT3G05932 | 0,27 | -0,38 | 4,39 | -0,09 | 9,31E-01 |
| AT3G05970 | 0,71 | -0,59 | 2,14 | -0,27 | 7,84E-01 |
| AT3G06030 | 0,29 | 0,00 | 4,41 | 0,00 | 1,00E+00 |
| AT3G06050 | 0,18 | 0,25 | 4,32 | 0,06 | 9,54E-01 |
| AT3G06060 | 0,17 | 0,07 | 4,38 | 0,02 | 9,88E-01 |
| AT3G06080 | 1,38 | 0,49 | 1,49 | 0,33 | 7,44E-01 |
| AT3G06150 | 0,98 | -1,40 | 1,58 | -0,89 | 3,76E-01 |
| AT3G06190 | 0,71 | 1,49 | 3,10 | 0,48 | 6,31E-01 |
| AT3G06250 | 0,74 | -0,69 | 2,92 | -0,23 | 8,14E-01 |
| AT3G06290 | 0,72 | 1,11 | 2,49 | 0,44 | 6,57E-01 |
| AT3G06310 | 0,08 | 0,00 | 4,41 | 0,00 | 1,00E+00 |

| | | | | | |
|---|---|---|---|---|---|
| AT3G06350 | 0,64 | 0,00 | 3,10 | 0,00 | 1,00E+00 |
| AT3G06430 | 0,12 | 0,00 | 4,41 | 0,00 | 1,00E+00 |
| AT3G06440 | 0,12 | 0,00 | 4,41 | 0,00 | 1,00E+00 |
| AT3G06450 | 0,61 | 0,88 | 2,69 | 0,33 | 7,44E-01 |
| AT3G06480 | 1,05 | -0,02 | 2,06 | -0,01 | 9,94E-01 |
| AT3G06483 | 0,45 | 0,87 | 2,69 | 0,32 | 7,47E-01 |
| AT3G06540 | 0,08 | 0,00 | 4,41 | 0,00 | 1,00E+00 |
| AT3G06550 | 0,32 | -0,13 | 4,35 | -0,03 | 9,76E-01 |
| AT3G06580 | 0,15 | -0,31 | 4,40 | -0,07 | 9,43E-01 |
| AT3G06590 | 0,40 | 0,32 | 4,37 | 0,07 | 9,41E-01 |
| AT3G06620 | 0,12 | 0,00 | 4,41 | 0,00 | 1,00E+00 |
| AT3G06665 | 0,25 | 2,14 | 4,29 | 0,50 | 6,19E-01 |
| AT3G06670 | 1,42 | -0,34 | 1,89 | -0,18 | 8,56E-01 |
| AT3G06680 | 0,28 | 1,08 | 4,35 | 0,25 | 8,03E-01 |
| AT3G06690 | 0,20 | 0,40 | 4,37 | 0,09 | 9,27E-01 |
| AT3G06700 | 0,57 | 0,32 | 2,71 | 0,12 | 9,06E-01 |
| AT3G06790 | 0,16 | -0,26 | 4,41 | -0,06 | 9,53E-01 |
| AT3G06820 | 0,08 | 0,00 | 4,41 | 0,00 | 1,00E+00 |
| AT3G06870 | 0,21 | 0,69 | 4,36 | 0,16 | 8,75E-01 |
| AT3G06900 | 1,19 | 2,11 | 1,89 | 1,12 | 2,63E-01 |
| AT3G06930 | 0,40 | -0,40 | 4,22 | -0,09 | 9,25E-01 |
| AT3G06960 | 0,89 | -0,74 | 2,34 | -0,31 | 7,53E-01 |
| AT3G07010 | 0,41 | -0,59 | 4,25 | -0,14 | 8,89E-01 |
| AT3G07020 | 0,54 | 0,00 | 3,87 | 0,00 | 1,00E+00 |
| AT3G07040 | 1,04 | -0,35 | 2,36 | -0,15 | 8,82E-01 |
| AT3G07050 | 0,53 | 0,60 | 3,17 | 0,19 | 8,50E-01 |
| AT3G07115 | 0,08 | 0,21 | 4,39 | 0,05 | 9,62E-01 |
| AT3G07180 | 0,12 | 0,00 | 4,41 | 0,00 | 1,00E+00 |
| AT3G07215 | 0,55 | 1,69 | 2,51 | 0,67 | 5,01E-01 |
| AT3G07273 | 0,08 | 0,00 | 4,41 | 0,00 | 1,00E+00 |
| AT3G07290 | 0,69 | 0,00 | 3,59 | 0,00 | 1,00E+00 |
| AT3G07300 | 0,29 | 0,00 | 4,41 | 0,00 | 1,00E+00 |
| AT3G07330 | 0,98 | 1,97 | 2,50 | 0,79 | 4,30E-01 |
| AT3G07380 | 1,22 | 0,71 | 1,47 | 0,48 | 6,28E-01 |
| AT3G07460 | 0,63 | -0,50 | 2,46 | -0,20 | 8,40E-01 |
| AT3G07470 | 0,58 | -0,61 | 3,23 | -0,19 | 8,49E-01 |
| AT3G07520 | 0,51 | 1,43 | 3,21 | 0,45 | 6,56E-01 |
| AT3G07540 | 0,08 | 0,00 | 4,41 | 0,00 | 1,00E+00 |
| AT3G07570 | 1,83 | -0,54 | 1,42 | -0,38 | 7,01E-01 |
| AT3G07660 | 1,40 | 1,23 | 2,02 | 0,61 | 5,42E-01 |
| AT3G07700 | 0,94 | -0,49 | 1,76 | -0,28 | 7,82E-01 |
| AT3G07800 | 0,53 | 0,46 | 2,76 | 0,17 | 8,69E-01 |
| AT3G07890 | 0,34 | 0,79 | 4,34 | 0,18 | 8,55E-01 |
| AT3G08570 | 1,10 | 0,00 | 2,36 | 0,00 | 1,00E+00 |
| AT3G08590 | 1,52 | -1,72 | 1,58 | -1,09 | 2,76E-01 |
| AT3G08610 | 0,59 | -1,17 | 3,25 | -0,36 | 7,18E-01 |
| AT3G08650 | 0,24 | 0,45 | 4,38 | 0,10 | 9,18E-01 |
| AT3G08720 | 1,37 | 1,54 | 2,05 | 0,75 | 4,51E-01 |
| AT3G08740 | 1,74 | 0,77 | 1,63 | 0,47 | 6,35E-01 |
| AT3G08760 | 0,70 | 0,30 | 2,29 | 0,13 | 8,94E-01 |
| AT3G08840 | 0,67 | -0,30 | 3,12 | -0,10 | 9,23E-01 |
| AT3G08870 | 0,75 | -2,04 | 3,52 | -0,58 | 5,63E-01 |
| AT3G08960 | 0,38 | 1,53 | 4,31 | 0,35 | 7,23E-01 |
| AT3G09040 | 0,17 | 0,00 | 4,41 | 0,00 | 1,00E+00 |
| AT3G09070 | 1,31 | 0,18 | 1,93 | 0,09 | 9,28E-01 |
| AT3G09090 | 1,29 | -1,37 | 2,07 | -0,66 | 5,09E-01 |

| | | | | | |
|---|---|---|---|---|---|
| AT3G09190 | 0,55 | -0,01 | 3,79 | 0,00 | 9,97E-01 |
| AT3G09210 | 0,27 | -2,46 | 4,37 | -0,56 | 5,74E-01 |
| AT3G09270 | 0,53 | 0,08 | 2,71 | 0,03 | 9,78E-01 |
| AT3G09320 | 0,85 | 0,79 | 2,45 | 0,32 | 7,46E-01 |
| AT3G09410 | 0,82 | 0,00 | 2,55 | 0,00 | 1,00E+00 |
| AT3G09470 | 0,82 | -0,38 | 3,55 | -0,11 | 9,15E-01 |
| AT3G09560 | 0,97 | -1,52 | 2,27 | -0,67 | 5,02E-01 |
| AT3G09580 | 0,48 | 0,80 | 3,41 | 0,23 | 8,15E-01 |
| AT3G09770 | 0,38 | 0,23 | 4,39 | 0,05 | 9,58E-01 |
| AT3G09780 | 0,23 | 0,24 | 4,39 | 0,06 | 9,56E-01 |
| AT3G09850 | 1,65 | -1,56 | 1,83 | -0,86 | 3,92E-01 |
| AT3G09870 | 0,08 | 0,00 | 4,41 | 0,00 | 1,00E+00 |
| AT3G09920 | 0,16 | -0,21 | 4,40 | -0,05 | 9,63E-01 |
| AT3G10010 | 0,35 | 0,28 | 4,39 | 0,06 | 9,49E-01 |
| AT3G10020 | 0,27 | -1,59 | 4,38 | -0,36 | 7,16E-01 |
| AT3G10030 | 0,88 | -3,72 | 2,85 | -1,31 | 1,92E-01 |
| AT3G10050 | 0,25 | -1,25 | 4,40 | -0,28 | 7,77E-01 |
| AT3G10070 | 0,46 | 0,56 | 4,28 | 0,13 | 8,97E-01 |
| AT3G10130 | 0,21 | -0,29 | 4,40 | -0,06 | 9,48E-01 |
| AT3G10160 | 0,36 | 0,78 | 3,44 | 0,23 | 8,21E-01 |
| AT3G10230 | 1,13 | 1,13 | 2,30 | 0,49 | 6,22E-01 |
| AT3G10250 | 0,08 | 0,00 | 4,41 | 0,00 | 1,00E+00 |
| AT3G10270 | 0,56 | -1,74 | 3,10 | -0,56 | 5,74E-01 |
| AT3G10320 | 0,12 | 0,00 | 4,41 | 0,00 | 1,00E+00 |
| AT3G10360 | 0,63 | -0,26 | 3,83 | -0,07 | 9,45E-01 |
| AT3G10405 | 0,68 | -0,26 | 4,41 | -0,06 | 9,53E-01 |
| AT3G10410 | 1,05 | -1,34 | 1,82 | -0,74 | 4,61E-01 |
| AT3G10520 | 0,15 | 0,02 | 4,36 | 0,00 | 9,96E-01 |
| AT3G10525 | 0,39 | -1,30 | 3,48 | -0,37 | 7,10E-01 |
| AT3G10630 | 0,99 | 0,86 | 2,08 | 0,41 | 6,80E-01 |
| AT3G10640 | 0,33 | -0,16 | 4,35 | -0,04 | 9,70E-01 |
| AT3G10660 | 0,27 | -0,41 | 4,38 | -0,09 | 9,25E-01 |
| AT3G10690 | 0,77 | 0,00 | 3,54 | 0,00 | 1,00E+00 |
| AT3G10720 | 0,36 | -0,27 | 4,29 | -0,06 | 9,50E-01 |
| AT3G10730 | 0,58 | -2,08 | 3,04 | -0,68 | 4,94E-01 |
| AT3G10770 | 0,74 | -1,38 | 2,57 | -0,54 | 5,91E-01 |
| AT3G10810 | 0,46 | -0,29 | 4,26 | -0,07 | 9,46E-01 |
| AT3G10840 | 0,41 | 0,00 | 4,41 | 0,00 | 1,00E+00 |
| AT3G10850 | 0,36 | 1,39 | 4,33 | 0,32 | 7,49E-01 |
| AT3G10900 | 0,31 | 2,13 | 4,30 | 0,50 | 6,20E-01 |
| AT3G10915 | 0,24 | 0,60 | 4,36 | 0,14 | 8,91E-01 |
| AT3G10930 | 0,61 | 2,43 | 2,46 | 0,99 | 3,24E-01 |
| AT3G10960 | 1,23 | 2,97 | 1,74 | 1,70 | 8,86E-02 |
| AT3G11030 | 0,56 | 0,07 | 3,83 | 0,02 | 9,85E-01 |
| AT3G11100 | 0,47 | -1,34 | 4,06 | -0,33 | 7,42E-01 |
| AT3G11280 | 0,31 | -0,69 | 4,38 | -0,16 | 8,74E-01 |
| AT3G11290 | 0,30 | 0,00 | 4,37 | 0,00 | 9,99E-01 |
| AT3G11310 | 0,31 | 2,15 | 4,29 | 0,50 | 6,17E-01 |
| AT3G11320 | 0,17 | -0,33 | 4,40 | -0,08 | 9,40E-01 |
| AT3G11330 | 0,97 | 0,00 | 2,75 | 0,00 | 1,00E+00 |
| AT3G11435 | 0,23 | -3,23 | 4,35 | -0,74 | 4,58E-01 |
| AT3G11450 | 1,01 | 1,77 | 2,00 | 0,88 | 3,77E-01 |
| AT3G11510 | 1,34 | -0,81 | 1,80 | -0,45 | 6,53E-01 |
| AT3G02435 | 1,34 | 2,18 | 1,83 | 1,19 | 2,33E-01 |
| AT3G02445 | 0,88 | 1,33 | 3,33 | 0,40 | 6,89E-01 |
| AT3G02455 | 0,99 | 0,61 | 2,96 | 0,21 | 8,37E-01 |

| | | | | | |
|---|---|---|---|---|---|
| AT3G11570 | 0,77 | 0,84 | 3,10 | 0,27 | 7,87E-01 |
| AT3G11580 | 0,26 | 1,36 | 4,32 | 0,31 | 7,54E-01 |
| AT3G11650 | 0,48 | 2,25 | 3,16 | 0,71 | 4,76E-01 |
| AT3G11660 | 1,33 | 3,16 | 3,34 | 0,95 | 3,45E-01 |
| AT3G11670 | 1,77 | -1,32 | 1,75 | -0,75 | 4,51E-01 |
| AT3G11680 | 0,22 | 1,16 | 4,34 | 0,27 | 7,89E-01 |
| AT3G11710 | 1,08 | 0,26 | 2,03 | 0,13 | 8,98E-01 |
| AT3G11730 | 0,25 | 0,00 | 4,41 | 0,00 | 1,00E+00 |
| AT3G11770 | 0,28 | 0,02 | 4,38 | 0,01 | 9,96E-01 |
| AT3G11780 | 0,21 | 0,12 | 4,38 | 0,03 | 9,78E-01 |
| AT3G11820 | 1,51 | 0,94 | 1,90 | 0,49 | 6,21E-01 |
| AT3G11840 | 0,47 | 0,48 | 4,14 | 0,12 | 9,07E-01 |
| AT3G11910 | 1,81 | -3,36 | 1,66 | -2,03 | 4,25E-02 |
| AT3G11964 | 0,92 | -0,08 | 1,89 | -0,04 | 9,66E-01 |
| AT3G12030 | 0,12 | 0,00 | 4,41 | 0,00 | 1,00E+00 |
| AT3G12130 | 0,80 | -1,04 | 2,63 | -0,40 | 6,92E-01 |
| AT3G12140 | 0,25 | 0,00 | 4,34 | 0,00 | 1,00E+00 |
| AT3G12145 | 0,15 | 0,90 | 4,35 | 0,21 | 8,36E-01 |
| AT3G12200 | 0,53 | 1,16 | 3,90 | 0,30 | 7,65E-01 |
| AT3G12260 | 0,32 | 0,24 | 4,34 | 0,05 | 9,56E-01 |
| AT3G12280 | 1,55 | -0,12 | 1,69 | -0,07 | 9,42E-01 |
| AT3G12400 | 0,71 | 0,73 | 1,97 | 0,37 | 7,12E-01 |
| AT3G12490 | 0,47 | 0,51 | 2,39 | 0,21 | 8,31E-01 |
| AT3G12500 | 0,08 | 0,00 | 4,41 | 0,00 | 1,00E+00 |
| AT3G12580 | 0,43 | 0,00 | 4,41 | 0,00 | 1,00E+00 |
| AT3G12587 | 0,28 | -0,16 | 4,40 | -0,04 | 9,70E-01 |
| AT3G12590 | 0,82 | -0,26 | 2,58 | -0,10 | 9,19E-01 |
| AT3G12610 | 0,59 | 0,38 | 2,30 | 0,16 | 8,70E-01 |
| AT3G12670 | 1,09 | -1,25 | 2,55 | -0,49 | 6,25E-01 |
| AT3G12680 | 0,32 | 0,56 | 4,37 | 0,13 | 8,97E-01 |
| AT3G12690 | 0,57 | 0,00 | 4,41 | 0,00 | 1,00E+00 |
| AT3G12750 | 0,45 | 0,00 | 4,07 | 0,00 | 1,00E+00 |
| AT3G12770 | 0,67 | -0,37 | 2,97 | -0,12 | 9,01E-01 |
| AT3G12800 | 0,96 | 0,84 | 1,79 | 0,47 | 6,39E-01 |
| AT3G12820 | 0,21 | 0,00 | 4,41 | 0,00 | 1,00E+00 |
| AT3G12920 | 1,14 | 3,01 | 2,71 | 1,11 | 2,67E-01 |
| AT3G12940 | 0,28 | 0,99 | 4,35 | 0,23 | 8,20E-01 |
| AT3G12950 | 0,48 | 0,00 | 4,41 | 0,00 | 1,00E+00 |
| AT3G12965 | 0,28 | 1,08 | 4,35 | 0,25 | 8,03E-01 |
| AT3G13000 | 0,24 | 0,00 | 4,41 | 0,00 | 1,00E+00 |
| AT3G13010 | 0,29 | 0,00 | 4,41 | 0,00 | 1,00E+00 |
| AT3G13020 | 0,33 | 0,00 | 4,41 | 0,00 | 1,00E+00 |
| AT3G13030 | 0,97 | -2,19 | 2,48 | -0,88 | 3,78E-01 |
| AT3G13040 | 0,58 | 0,19 | 2,62 | 0,07 | 9,43E-01 |
| AT3G13061 | 0,23 | 1,25 | 4,34 | 0,29 | 7,73E-01 |
| AT3G13110 | 1,06 | -1,78 | 2,12 | -0,84 | 4,00E-01 |
| AT3G13120 | 0,60 | -2,30 | 3,04 | -0,75 | 4,51E-01 |
| AT3G13150 | 0,37 | 0,00 | 4,41 | 0,00 | 1,00E+00 |
| AT3G13160 | 0,08 | 0,00 | 4,41 | 0,00 | 1,00E+00 |
| AT3G13220 | 0,37 | 0,00 | 4,41 | 0,00 | 1,00E+00 |
| AT3G13222 | 0,48 | 0,00 | 4,41 | 0,00 | 1,00E+00 |
| AT3G13225 | 0,32 | -0,36 | 4,39 | -0,08 | 9,35E-01 |
| AT3G13226 | 0,08 | 0,00 | 4,41 | 0,00 | 1,00E+00 |
| AT3G13235 | 0,44 | 0,43 | 2,80 | 0,15 | 8,79E-01 |
| AT3G13310 | 0,25 | -0,55 | 4,35 | -0,13 | 8,99E-01 |
| AT3G13390 | 0,05 | 0,18 | 4,38 | 0,04 | 9,68E-01 |

| | | | | | |
|---|---|---|---|---|---|
| AT3G13404 | 0,46 | 0,96 | 4,02 | 0,24 | 8,11E-01 |
| AT3G13410 | 1,64 | 0,23 | 2,04 | 0,11 | 9,09E-01 |
| AT3G13450 | 0,75 | 0,00 | 2,92 | 0,00 | 1,00E+00 |
| AT3G13480 | 0,20 | 0,53 | 4,37 | 0,12 | 9,04E-01 |
| AT3G13490 | 0,57 | -1,26 | 3,16 | -0,40 | 6,90E-01 |
| AT3G13570 | 0,44 | -2,93 | 4,20 | -0,70 | 4,85E-01 |
| AT3G13720 | 0,96 | 1,59 | 1,69 | 0,94 | 3,48E-01 |
| AT3G13772 | 0,72 | -0,27 | 2,93 | -0,09 | 9,27E-01 |
| AT3G13860 | 0,89 | -0,52 | 2,13 | -0,24 | 8,07E-01 |
| AT3G13990 | 1,05 | -2,69 | 2,34 | -1,15 | 2,50E-01 |
| AT3G14050 | 1,74 | 1,38 | 1,50 | 0,92 | 3,57E-01 |
| AT3G14070 | 0,23 | -0,40 | 4,38 | -0,09 | 9,27E-01 |
| AT3G14075 | 0,26 | -1,24 | 4,39 | -0,28 | 7,78E-01 |
| AT3G14090 | 0,68 | -0,12 | 3,14 | -0,04 | 9,69E-01 |
| AT3G14110 | 0,61 | -1,26 | 2,63 | -0,48 | 6,31E-01 |
| AT3G14172 | 1,38 | -1,30 | 2,07 | -0,63 | 5,29E-01 |
| AT3G14180 | 0,40 | 0,00 | 4,41 | 0,00 | 1,00E+00 |
| AT3G14205 | 0,51 | -0,01 | 2,79 | 0,00 | 9,97E-01 |
| AT3G14230 | 0,40 | -0,40 | 4,22 | -0,09 | 9,25E-01 |
| AT3G14240 | 1,34 | -1,02 | 1,77 | -0,58 | 5,63E-01 |
| AT3G14330 | 0,56 | -2,19 | 3,12 | -0,70 | 4,83E-01 |
| AT3G14350 | 1,44 | 0,60 | 1,56 | 0,38 | 7,01E-01 |
| AT3G14400 | 1,24 | 0,00 | 3,33 | 0,00 | 1,00E+00 |
| AT3G14430 | 0,25 | -2,19 | 4,41 | -0,50 | 6,20E-01 |
| AT3G14440 | 0,33 | 1,03 | 4,34 | 0,24 | 8,12E-01 |
| AT3G14460 | 0,50 | 0,38 | 3,99 | 0,10 | 9,24E-01 |
| AT3G14590 | 0,14 | 0,81 | 4,35 | 0,19 | 8,52E-01 |
| AT3G14810 | 0,79 | -1,65 | 2,46 | -0,67 | 5,03E-01 |
| AT3G14900 | 0,17 | 0,00 | 4,41 | 0,00 | 1,00E+00 |
| AT3G15110 | 0,15 | 0,02 | 4,36 | 0,00 | 9,96E-01 |
| AT3G15120 | 0,82 | 0,94 | 2,51 | 0,37 | 7,10E-01 |
| AT3G15150 | 0,08 | 0,21 | 4,39 | 0,05 | 9,62E-01 |
| AT3G15220 | 0,77 | -1,34 | 2,54 | -0,53 | 5,98E-01 |
| AT3G15354 | 0,91 | -0,50 | 2,26 | -0,22 | 8,23E-01 |
| AT3G15356 | 0,36 | -0,26 | 4,41 | -0,06 | 9,53E-01 |
| AT3G15470 | 0,41 | 0,32 | 2,80 | 0,11 | 9,09E-01 |
| AT3G03125 | 0,17 | 0,00 | 4,41 | 0,00 | 1,00E+00 |
| AT3G15530 | 1,09 | 2,67 | 2,41 | 1,11 | 2,68E-01 |
| AT3G15570 | 0,83 | 1,75 | 2,65 | 0,66 | 5,09E-01 |
| AT3G15590 | 0,41 | -0,01 | 4,30 | 0,00 | 9,97E-01 |
| AT3G15630 | 0,46 | 2,53 | 3,16 | 0,80 | 4,24E-01 |
| AT3G15720 | 0,20 | 1,25 | 4,33 | 0,29 | 7,72E-01 |
| AT3G15810 | 0,58 | 1,68 | 2,42 | 0,69 | 4,88E-01 |
| AT3G15840 | 1,45 | -0,31 | 1,30 | -0,23 | 8,15E-01 |
| AT3G15900 | 0,29 | -0,15 | 4,35 | -0,03 | 9,72E-01 |
| AT3G15940 | 1,43 | -2,31 | 1,99 | -1,17 | 2,44E-01 |
| AT3G15970 | 0,63 | 0,73 | 2,64 | 0,28 | 7,82E-01 |
| AT3G15980 | 1,59 | 1,23 | 1,56 | 0,79 | 4,32E-01 |
| AT3G16000 | 0,51 | -0,27 | 3,12 | -0,09 | 9,32E-01 |
| AT3G16110 | 0,21 | 0,00 | 4,41 | 0,00 | 1,00E+00 |
| AT3G16190 | 0,21 | 0,00 | 4,41 | 0,00 | 1,00E+00 |
| AT3G03235 | 0,34 | 0,00 | 4,41 | 0,00 | 1,00E+00 |
| AT3G16290 | 0,84 | -0,15 | 1,90 | -0,08 | 9,39E-01 |
| AT3G16310 | 0,55 | -0,31 | 3,87 | -0,08 | 9,36E-01 |
| AT3G16500 | 0,17 | 0,07 | 4,38 | 0,02 | 9,87E-01 |
| AT3G16510 | 0,65 | 1,52 | 3,01 | 0,50 | 6,15E-01 |

| | | | | | |
|---|---|---|---|---|---|
| AT3G16520 | 0,76 | 0,93 | 2,06 | 0,45 | 6,51E-01 |
| AT3G16530 | 0,27 | 1,01 | 4,34 | 0,23 | 8,17E-01 |
| AT3G16560 | 0,66 | 0,45 | 2,58 | 0,17 | 8,62E-01 |
| AT3G16630 | 1,01 | 1,10 | 2,01 | 0,55 | 5,84E-01 |
| AT3G16820 | 1,66 | 2,29 | 1,45 | 1,58 | 1,15E-01 |
| AT3G16840 | 0,19 | 0,84 | 4,35 | 0,19 | 8,47E-01 |
| AT3G16860 | 0,47 | 1,58 | 3,97 | 0,40 | 6,90E-01 |
| AT3G16870 | 0,39 | 0,00 | 4,40 | 0,00 | 1,00E+00 |
| AT3G16890 | 0,19 | -0,70 | 4,37 | -0,16 | 8,72E-01 |
| AT3G17020 | 1,21 | -0,23 | 2,13 | -0,11 | 9,13E-01 |
| AT3G03415 | 1,34 | 0,47 | 1,75 | 0,27 | 7,86E-01 |
| AT3G03445 | 0,77 | 0,50 | 2,65 | 0,19 | 8,50E-01 |
| AT3G17170 | 0,31 | -2,40 | 4,36 | -0,55 | 5,82E-01 |
| AT3G17240 | 1,76 | -0,26 | 2,07 | -0,13 | 8,98E-01 |
| AT3G17320 | 0,12 | 0,59 | 4,36 | 0,14 | 8,92E-01 |
| AT3G17330 | 0,20 | -0,26 | 4,41 | -0,06 | 9,53E-01 |
| AT3G17420 | 0,17 | -2,13 | 4,40 | -0,48 | 6,28E-01 |
| AT3G17440 | 0,08 | 0,00 | 4,41 | 0,00 | 1,00E+00 |
| AT3G17450 | 0,60 | -0,36 | 3,05 | -0,12 | 9,07E-01 |
| AT3G17655 | 0,19 | -0,30 | 4,40 | -0,07 | 9,45E-01 |
| AT3G17740 | 0,37 | -0,34 | 4,36 | -0,08 | 9,38E-01 |
| AT3G17760 | 0,21 | 0,00 | 4,41 | 0,00 | 1,00E+00 |
| AT3G17770 | 1,51 | 1,03 | 1,63 | 0,63 | 5,28E-01 |
| AT3G03585 | 0,05 | 0,18 | 4,38 | 0,04 | 9,68E-01 |
| AT3G17940 | 0,81 | -0,80 | 2,58 | -0,31 | 7,55E-01 |
| AT3G18030 | 0,52 | 0,36 | 4,05 | 0,09 | 9,30E-01 |
| AT3G18035 | 1,23 | -0,25 | 1,53 | -0,16 | 8,71E-01 |
| AT3G18110 | 1,16 | 0,00 | 2,66 | 0,00 | 1,00E+00 |
| AT3G18130 | 0,88 | 0,53 | 2,48 | 0,21 | 8,32E-01 |
| AT3G18140 | 0,30 | 1,44 | 4,33 | 0,33 | 7,40E-01 |
| AT3G18250 | 0,55 | 0,29 | 4,39 | 0,07 | 9,47E-01 |
| AT3G18380 | 0,25 | 0,00 | 4,41 | 0,00 | 1,00E+00 |
| AT3G18420 | 0,20 | 0,87 | 4,35 | 0,20 | 8,41E-01 |
| AT3G18535 | 0,67 | 2,47 | 2,39 | 1,03 | 3,01E-01 |
| AT3G18600 | 0,25 | 0,00 | 4,41 | 0,00 | 1,00E+00 |
| AT3G18610 | 0,55 | 0,00 | 4,39 | 0,00 | 1,00E+00 |
| AT3G18630 | 0,20 | 0,00 | 4,41 | 0,00 | 1,00E+00 |
| AT3G18700 | 1,19 | 2,51 | 2,08 | 1,21 | 2,27E-01 |
| AT3G18750 | 0,17 | 0,00 | 4,41 | 0,00 | 1,00E+00 |
| AT3G18930 | 0,90 | 2,58 | 2,17 | 1,19 | 2,34E-01 |
| AT3G18970 | 0,08 | 0,00 | 4,41 | 0,00 | 1,00E+00 |
| AT3G19000 | 0,24 | 0,65 | 4,37 | 0,15 | 8,83E-01 |
| AT3G19010 | 1,81 | -0,23 | 1,53 | -0,15 | 8,81E-01 |
| AT3G19020 | 0,54 | 0,81 | 3,18 | 0,25 | 7,99E-01 |
| AT3G19150 | 0,16 | 0,39 | 4,37 | 0,09 | 9,29E-01 |
| AT3G03815 | 0,25 | 0,00 | 4,41 | 0,00 | 1,00E+00 |
| AT3G19300 | 0,12 | -0,26 | 4,41 | -0,06 | 9,53E-01 |
| AT3G19340 | 0,12 | 0,11 | 4,38 | 0,03 | 9,80E-01 |
| AT3G19430 | 0,25 | 0,00 | 4,41 | 0,00 | 1,00E+00 |
| AT3G19490 | 0,72 | 0,54 | 2,58 | 0,21 | 8,34E-01 |
| AT3G19510 | 0,12 | 0,27 | 4,38 | 0,06 | 9,50E-01 |
| AT3G19515 | 0,59 | -1,12 | 3,19 | -0,35 | 7,26E-01 |
| AT3G19553 | 0,42 | 1,40 | 4,10 | 0,34 | 7,33E-01 |
| AT3G19580 | 0,72 | 2,03 | 2,46 | 0,83 | 4,09E-01 |
| AT3G19620 | 0,32 | 0,00 | 4,41 | 0,00 | 1,00E+00 |
| AT3G19630 | 0,39 | 0,00 | 4,40 | 0,00 | 1,00E+00 |

| | | | | | |
|---|---|---|---|---|---|
| AT3G19710 | 0,69 | 1,58 | 2,49 | 0,64 | 5,25E-01 |
| AT3G19720 | 1,17 | 1,16 | 2,51 | 0,46 | 6,43E-01 |
| AT3G19740 | 0,59 | -0,09 | 3,18 | -0,03 | 9,76E-01 |
| AT3G19790 | 0,71 | 1,55 | 2,03 | 0,76 | 4,45E-01 |
| AT3G19810 | 0,70 | -3,06 | 4,36 | -0,70 | 4,83E-01 |
| AT3G19830 | 0,67 | 2,52 | 2,93 | 0,86 | 3,90E-01 |
| AT3G19850 | 0,50 | 0,78 | 3,15 | 0,25 | 8,05E-01 |
| AT3G19880 | 0,35 | 1,30 | 3,44 | 0,38 | 7,05E-01 |
| AT3G19895 | 0,23 | 1,67 | 4,31 | 0,39 | 6,99E-01 |
| AT3G19900 | 0,17 | 0,00 | 4,41 | 0,00 | 1,00E+00 |
| AT3G20015 | 0,62 | 1,38 | 3,06 | 0,45 | 6,53E-01 |
| AT3G20050 | 0,74 | -1,34 | 2,51 | -0,53 | 5,95E-01 |
| AT3G20060 | 0,33 | -0,17 | 4,35 | -0,04 | 9,69E-01 |
| AT3G20230 | 0,35 | 0,78 | 4,36 | 0,18 | 8,59E-01 |
| AT3G20250 | 1,61 | -0,21 | 1,97 | -0,11 | 9,13E-01 |
| AT3G20290 | 0,63 | 1,57 | 3,04 | 0,52 | 6,05E-01 |
| AT3G20320 | 0,51 | 0,37 | 2,72 | 0,14 | 8,91E-01 |
| AT3G20390 | 0,62 | -0,31 | 2,10 | -0,15 | 8,82E-01 |
| AT3G20470 | 0,38 | -0,43 | 4,36 | -0,10 | 9,22E-01 |
| AT3G20620 | 0,68 | 0,00 | 3,67 | 0,00 | 1,00E+00 |
| AT3G20630 | 1,20 | 0,32 | 1,86 | 0,17 | 8,62E-01 |
| AT3G20655 | 0,30 | 0,00 | 4,41 | 0,00 | 1,00E+00 |
| AT3G20700 | 1,05 | 3,96 | 2,38 | 1,66 | 9,65E-02 |
| AT3G03985 | 0,54 | 3,20 | 4,27 | 0,75 | 4,54E-01 |
| AT3G20830 | 0,60 | -1,07 | 3,22 | -0,33 | 7,39E-01 |
| AT3G21055 | 1,34 | -1,60 | 1,61 | -0,99 | 3,22E-01 |
| AT3G21070 | 0,23 | 0,81 | 4,35 | 0,19 | 8,52E-01 |
| AT3G21140 | 0,25 | -0,39 | 4,39 | -0,09 | 9,29E-01 |
| AT3G21160 | 0,11 | -0,26 | 4,41 | -0,06 | 9,53E-01 |
| AT3G21215 | 0,11 | -0,38 | 4,39 | -0,09 | 9,31E-01 |
| AT3G21300 | 0,80 | 0,60 | 2,33 | 0,26 | 7,96E-01 |
| AT3G21430 | 0,61 | 1,61 | 3,13 | 0,51 | 6,07E-01 |
| AT3G21540 | 0,20 | -2,07 | 4,39 | -0,47 | 6,37E-01 |
| AT3G21570 | 0,87 | -0,13 | 2,73 | -0,05 | 9,61E-01 |
| AT3G21630 | 0,12 | 0,11 | 4,38 | 0,03 | 9,80E-01 |
| AT3G21650 | 0,33 | 0,00 | 4,41 | 0,00 | 1,00E+00 |
| AT3G21670 | 1,42 | 0,06 | 2,52 | 0,02 | 9,81E-01 |
| AT3G21740 | 0,24 | -0,34 | 4,40 | -0,08 | 9,38E-01 |
| AT3G21790 | 0,96 | 0,92 | 2,13 | 0,43 | 6,64E-01 |
| AT3G21865 | 0,48 | 1,35 | 2,71 | 0,50 | 6,20E-01 |
| AT3G21870 | 1,78 | -1,56 | 1,50 | -1,04 | 2,97E-01 |
| AT3G22070 | 0,17 | -3,06 | 4,35 | -0,70 | 4,82E-01 |
| AT3G22110 | 0,73 | 0,72 | 2,51 | 0,29 | 7,74E-01 |
| AT3G22142 | 1,40 | 0,71 | 1,58 | 0,45 | 6,52E-01 |
| AT3G22170 | 1,10 | 1,25 | 2,13 | 0,59 | 5,58E-01 |
| AT3G22190 | 0,78 | 0,27 | 3,01 | 0,09 | 9,28E-01 |
| AT3G22210 | 0,48 | -0,98 | 3,32 | -0,30 | 7,67E-01 |
| AT3G22220 | 0,36 | -1,20 | 3,54 | -0,34 | 7,36E-01 |
| AT3G22230 | 0,95 | 1,40 | 1,99 | 0,70 | 4,82E-01 |
| AT3G22240 | 1,25 | 2,06 | 1,84 | 1,12 | 2,63E-01 |
| AT3G22320 | 0,41 | -0,01 | 4,30 | 0,00 | 9,97E-01 |
| AT3G22421 | 0,96 | 2,44 | 2,66 | 0,92 | 3,58E-01 |
| AT3G22470 | 0,59 | -0,50 | 2,66 | -0,19 | 8,50E-01 |
| AT3G22530 | 0,38 | -2,31 | 4,37 | -0,53 | 5,97E-01 |
| AT3G22550 | 0,60 | 0,31 | 3,13 | 0,10 | 9,21E-01 |
| AT3G22580 | 0,15 | 0,02 | 4,36 | 0,00 | 9,96E-01 |

| | | | | | |
|---|---|---|---|---|---|
| AT3G22600 | 0,17 | 0,00 | 4,41 | 0,00 | 1,00E+00 |
| AT3G22630 | 0,68 | 1,29 | 2,25 | 0,57 | 5,66E-01 |
| AT3G22660 | 0,44 | -0,13 | 4,39 | -0,03 | 9,77E-01 |
| AT3G22690 | 0,29 | -0,76 | 4,38 | -0,17 | 8,63E-01 |
| AT3G22710 | 1,14 | 1,31 | 1,97 | 0,66 | 5,06E-01 |
| AT3G22720 | 0,08 | 0,21 | 4,39 | 0,05 | 9,62E-01 |
| AT3G22730 | 1,61 | 2,18 | 1,54 | 1,42 | 1,56E-01 |
| AT3G22770 | 0,68 | -0,72 | 3,20 | -0,23 | 8,21E-01 |
| AT3G22950 | 0,15 | -0,57 | 4,37 | -0,13 | 8,96E-01 |
| AT3G22980 | 0,74 | -0,45 | 2,54 | -0,18 | 8,58E-01 |
| AT3G23020 | 0,53 | 0,00 | 3,90 | 0,00 | 1,00E+00 |
| AT3G23030 | 0,69 | -1,00 | 2,05 | -0,49 | 6,24E-01 |
| AT3G23080 | 0,08 | 0,00 | 4,41 | 0,00 | 1,00E+00 |
| AT3G23170 | 0,50 | 1,60 | 2,65 | 0,60 | 5,46E-01 |
| AT3G23172 | 0,24 | 0,94 | 4,35 | 0,22 | 8,29E-01 |
| AT3G23400 | 0,58 | 0,35 | 3,24 | 0,11 | 9,14E-01 |
| AT3G23420 | 0,32 | 1,84 | 4,30 | 0,43 | 6,69E-01 |
| AT3G23490 | 0,26 | 1,68 | 4,31 | 0,39 | 6,96E-01 |
| AT3G23570 | 0,32 | -0,29 | 4,40 | -0,07 | 9,47E-01 |
| AT3G23633 | 0,12 | 1,17 | 4,34 | 0,27 | 7,88E-01 |
| AT3G23650 | 0,55 | 2,76 | 2,89 | 0,96 | 3,39E-01 |
| AT3G04585 | 0,72 | 3,09 | 2,77 | 1,11 | 2,65E-01 |
| AT3G23660 | 0,75 | -0,37 | 2,99 | -0,12 | 9,01E-01 |
| AT3G23700 | 0,24 | 0,00 | 4,33 | 0,00 | 9,99E-01 |
| AT3G23740 | 0,08 | 0,00 | 4,41 | 0,00 | 1,00E+00 |
| AT3G23830 | 0,27 | -2,24 | 4,40 | -0,51 | 6,10E-01 |
| AT3G23900 | 0,77 | 0,41 | 2,83 | 0,14 | 8,86E-01 |
| AT3G23970 | 0,08 | 0,00 | 4,41 | 0,00 | 1,00E+00 |
| AT3G23980 | 0,76 | 0,83 | 2,50 | 0,33 | 7,41E-01 |
| AT3G24080 | 0,22 | 0,20 | 4,38 | 0,05 | 9,63E-01 |
| AT3G24120 | 0,83 | -0,37 | 3,56 | -0,10 | 9,18E-01 |
| AT3G24140 | 0,26 | 0,57 | 4,31 | 0,13 | 8,94E-01 |
| AT3G24170 | 1,33 | 1,03 | 1,98 | 0,52 | 6,02E-01 |
| AT3G24200 | 0,11 | -0,26 | 4,41 | -0,06 | 9,53E-01 |
| AT3G24260 | 0,15 | 1,21 | 4,33 | 0,28 | 7,81E-01 |
| AT3G24270 | 0,91 | 2,44 | 2,31 | 1,06 | 2,91E-01 |
| AT3G24320 | 0,63 | -0,23 | 3,14 | -0,07 | 9,42E-01 |
| AT3G24430 | 0,54 | -0,94 | 2,65 | -0,35 | 7,23E-01 |
| AT3G24440 | 0,50 | -0,26 | 3,29 | -0,08 | 9,36E-01 |
| AT3G24465 | 0,32 | -0,39 | 3,59 | -0,11 | 9,13E-01 |
| AT3G24503 | 1,13 | 0,73 | 1,82 | 0,40 | 6,89E-01 |
| AT3G24518 | 0,96 | 0,21 | 2,37 | 0,09 | 9,28E-01 |
| AT3G24614 | 0,96 | 1,70 | 1,91 | 0,89 | 3,75E-01 |
| AT3G24670 | 0,20 | -0,26 | 4,41 | -0,06 | 9,53E-01 |
| AT3G24870 | 1,16 | -1,28 | 1,94 | -0,66 | 5,09E-01 |
| AT3G24880 | 0,67 | 1,70 | 3,05 | 0,56 | 5,78E-01 |
| AT3G24900 | 0,51 | 0,95 | 3,14 | 0,30 | 7,63E-01 |
| AT3G25090 | 0,87 | 2,46 | 1,97 | 1,25 | 2,12E-01 |
| AT3G25150 | 1,03 | -0,39 | 2,65 | -0,15 | 8,83E-01 |
| AT3G25180 | 0,15 | 0,02 | 4,36 | 0,00 | 9,96E-01 |
| AT3G25460 | 1,48 | -0,30 | 1,46 | -0,21 | 8,36E-01 |
| AT3G25500 | 1,85 | 1,85 | 1,66 | 1,11 | 2,65E-01 |
| AT3G25530 | 0,19 | 1,22 | 4,34 | 0,28 | 7,79E-01 |
| AT3G25560 | 0,41 | -0,31 | 4,40 | -0,07 | 9,43E-01 |
| AT3G25610 | 1,05 | -2,00 | 2,40 | -0,84 | 4,03E-01 |
| AT3G25660 | 0,47 | -2,04 | 4,02 | -0,51 | 6,11E-01 |

| | | | | | |
|---|---|---|---|---|---|
| AT3G04935 | 0,15 | -0,39 | 4,39 | -0,09 | 9,29E-01 |
| AT3G25800 | 1,14 | -3,18 | 2,01 | -1,58 | 1,14E-01 |
| AT3G25840 | 1,32 | -0,27 | 1,77 | -0,15 | 8,80E-01 |
| AT3G25855 | 1,70 | -5,02 | 1,93 | -2,60 | 9,26E-03 |
| AT3G25870 | 0,65 | -0,38 | 2,39 | -0,16 | 8,75E-01 |
| AT3G25880 | 1,38 | 0,00 | 2,71 | 0,00 | 1,00E+00 |
| AT3G25882 | 0,50 | 0,28 | 4,07 | 0,07 | 9,45E-01 |
| AT3G25920 | 0,58 | 0,06 | 2,31 | 0,03 | 9,78E-01 |
| AT3G25990 | 0,75 | 0,00 | 3,54 | 0,00 | 1,00E+00 |
| AT3G26170 | 1,29 | -0,22 | 1,61 | -0,14 | 8,90E-01 |
| AT3G26180 | 0,36 | 1,27 | 4,33 | 0,29 | 7,70E-01 |
| AT3G26210 | 1,30 | 2,75 | 2,06 | 1,33 | 1,82E-01 |
| AT3G26220 | 0,49 | 1,89 | 4,02 | 0,47 | 6,39E-01 |
| AT3G26230 | 0,79 | 0,92 | 2,44 | 0,38 | 7,06E-01 |
| AT3G26280 | 1,16 | -0,33 | 1,65 | -0,20 | 8,43E-01 |
| AT3G26290 | 0,78 | -0,44 | 1,81 | -0,24 | 8,07E-01 |
| AT3G26300 | 0,36 | -2,09 | 4,39 | -0,47 | 6,35E-01 |
| AT3G26340 | 0,62 | -1,50 | 3,82 | -0,39 | 6,95E-01 |
| AT3G26410 | 0,31 | -2,06 | 4,39 | -0,47 | 6,39E-01 |
| AT3G26500 | 1,32 | 3,57 | 1,92 | 1,86 | 6,31E-02 |
| AT3G26510 | 0,66 | 1,62 | 2,12 | 0,77 | 4,44E-01 |
| AT3G26550 | 0,20 | 0,00 | 4,41 | 0,00 | 1,00E+00 |
| AT3G26560 | 0,40 | 0,90 | 3,32 | 0,27 | 7,86E-01 |
| AT3G26590 | 0,41 | 0,75 | 4,35 | 0,17 | 8,63E-01 |
| AT3G26616 | 0,23 | 1,31 | 4,33 | 0,30 | 7,62E-01 |
| AT3G26630 | 0,69 | -0,36 | 2,57 | -0,14 | 8,89E-01 |
| AT3G26670 | 0,54 | 0,75 | 3,34 | 0,22 | 8,22E-01 |
| AT3G26744 | 0,39 | -2,12 | 4,31 | -0,49 | 6,22E-01 |
| AT3G26812 | 0,17 | 1,19 | 4,33 | 0,28 | 7,83E-01 |
| AT3G26819 | 0,05 | 0,18 | 4,38 | 0,04 | 9,68E-01 |
| AT3G26860 | 0,30 | -3,42 | 4,34 | -0,79 | 4,31E-01 |
| AT3G26870 | 1,55 | -2,64 | 1,83 | -1,44 | 1,49E-01 |
| AT3G26890 | 0,29 | 1,10 | 4,34 | 0,25 | 8,01E-01 |
| AT3G26932 | 0,90 | 1,96 | 2,29 | 0,86 | 3,91E-01 |
| AT3G26935 | 0,27 | 0,54 | 4,36 | 0,12 | 9,02E-01 |
| AT3G26980 | 0,14 | 0,13 | 4,38 | 0,03 | 9,77E-01 |
| AT3G27010 | 0,67 | -2,37 | 3,68 | -0,64 | 5,21E-01 |
| AT3G27020 | 0,51 | 1,30 | 2,39 | 0,55 | 5,85E-01 |
| AT3G27040 | 1,43 | 1,19 | 1,65 | 0,72 | 4,69E-01 |
| AT3G27060 | 0,90 | 0,59 | 2,32 | 0,25 | 7,99E-01 |
| AT3G27100 | 0,74 | 0,00 | 4,38 | 0,00 | 1,00E+00 |
| AT3G27240 | 0,37 | 1,89 | 3,35 | 0,56 | 5,73E-01 |
| AT3G27300 | 0,49 | -2,07 | 3,28 | -0,63 | 5,27E-01 |
| AT3G27380 | 0,45 | -2,04 | 4,36 | -0,47 | 6,40E-01 |
| AT3G05235 | 0,99 | 0,94 | 2,29 | 0,41 | 6,80E-01 |
| AT3G05245 | 0,33 | 0,47 | 4,33 | 0,11 | 9,13E-01 |
| AT3G27530 | 0,76 | 0,77 | 2,71 | 0,28 | 7,76E-01 |
| AT3G27600 | 1,14 | 3,49 | 2,25 | 1,55 | 1,20E-01 |
| AT3G27610 | 0,33 | 0,22 | 4,39 | 0,05 | 9,61E-01 |
| AT3G27680 | 0,86 | 1,13 | 2,17 | 0,52 | 6,02E-01 |
| AT3G27700 | 1,39 | 1,02 | 2,00 | 0,51 | 6,09E-01 |
| AT3G27740 | 0,52 | -2,19 | 4,08 | -0,54 | 5,91E-01 |
| AT3G27770 | 0,43 | -0,53 | 2,85 | -0,18 | 8,54E-01 |
| AT3G27906 | 1,35 | 1,53 | 1,67 | 0,92 | 3,57E-01 |
| AT3G27925 | 0,65 | -1,87 | 3,00 | -0,62 | 5,32E-01 |
| AT3G27930 | 0,20 | 1,25 | 4,33 | 0,29 | 7,72E-01 |

| | | | | | |
|---|---|---|---|---|---|
| AT3G27990 | 0,64 | 0,71 | 2,35 | 0,30 | 7,62E-01 |
| AT3G28340 | 0,22 | 1,60 | 4,32 | 0,37 | 7,11E-01 |
| AT3G28380 | 0,08 | 0,00 | 4,41 | 0,00 | 1,00E+00 |
| AT3G28410 | 0,14 | 0,89 | 4,35 | 0,21 | 8,37E-01 |
| AT3G28674 | 0,28 | 1,12 | 4,34 | 0,26 | 7,97E-01 |
| AT3G28695 | 0,43 | 1,93 | 4,13 | 0,47 | 6,41E-01 |
| AT3G28710 | 0,99 | 0,36 | 1,72 | 0,21 | 8,33E-01 |
| AT3G28780 | 0,61 | 0,61 | 3,05 | 0,20 | 8,41E-01 |
| AT3G28860 | 1,33 | -1,45 | 2,41 | -0,60 | 5,46E-01 |
| AT3G28890 | 0,43 | -2,27 | 4,40 | -0,52 | 6,05E-01 |
| AT3G28920 | 1,35 | 0,42 | 1,79 | 0,23 | 8,16E-01 |
| AT3G28930 | 0,77 | 1,61 | 2,90 | 0,56 | 5,78E-01 |
| AT3G28940 | 1,30 | 1,21 | 1,68 | 0,72 | 4,72E-01 |
| AT3G28950 | 0,97 | -0,52 | 1,83 | -0,29 | 7,75E-01 |
| AT3G28960 | 0,30 | 1,63 | 4,31 | 0,38 | 7,05E-01 |
| AT3G28970 | 0,52 | -0,34 | 4,06 | -0,08 | 9,33E-01 |
| AT3G29030 | 0,33 | 0,71 | 4,27 | 0,17 | 8,68E-01 |
| AT3G05665 | 0,80 | 3,55 | 2,60 | 1,37 | 1,72E-01 |
| AT3G29110 | 0,53 | -0,49 | 2,49 | -0,20 | 8,44E-01 |
| AT3G29160 | 0,59 | -1,62 | 4,31 | -0,38 | 7,07E-01 |
| AT3G29200 | 0,59 | -0,32 | 3,16 | -0,10 | 9,20E-01 |
| AT3G29350 | 0,17 | 0,07 | 4,38 | 0,02 | 9,88E-01 |
| AT3G29400 | 0,67 | 1,89 | 2,52 | 0,75 | 4,53E-01 |
| AT3G29570 | 0,38 | 1,07 | 4,34 | 0,25 | 8,04E-01 |
| AT3G05745 | 0,14 | 0,31 | 4,37 | 0,07 | 9,44E-01 |
| AT3G29633 | 0,14 | -2,43 | 4,37 | -0,55 | 5,79E-01 |
| AT3G29639 | 0,16 | 1,16 | 4,34 | 0,27 | 7,89E-01 |
| AT3G05895 | 0,08 | 0,21 | 4,39 | 0,05 | 9,62E-01 |
| AT3G29680 | 0,25 | 1,19 | 4,34 | 0,27 | 7,85E-01 |
| AT3G29720 | 0,40 | -0,38 | 4,34 | -0,09 | 9,30E-01 |
| AT3G29760 | 0,67 | -0,30 | 3,12 | -0,10 | 9,23E-01 |
| AT3G29785 | 0,22 | 1,48 | 4,32 | 0,34 | 7,32E-01 |
| AT3G30160 | 0,58 | 2,03 | 2,90 | 0,70 | 4,84E-01 |
| AT3G30200 | 0,12 | 0,27 | 4,38 | 0,06 | 9,50E-01 |
| AT3G30220 | 0,08 | 0,21 | 4,39 | 0,05 | 9,62E-01 |
| AT3G30230 | 0,54 | 1,63 | 3,88 | 0,42 | 6,74E-01 |
| AT3G30300 | 0,53 | 0,45 | 4,38 | 0,10 | 9,18E-01 |
| AT3G30840 | 0,21 | -0,32 | 4,40 | -0,07 | 9,43E-01 |
| AT3G32190 | 0,34 | 0,78 | 3,49 | 0,22 | 8,24E-01 |
| AT3G32420 | 0,10 | 0,08 | 4,38 | 0,02 | 9,85E-01 |
| AT3G32904 | 0,65 | 1,22 | 3,10 | 0,39 | 6,93E-01 |
| AT3G33528 | 0,61 | -0,94 | 2,48 | -0,38 | 7,06E-01 |
| AT3G42050 | 1,44 | 0,14 | 2,09 | 0,07 | 9,47E-01 |
| AT3G42060 | 0,32 | 0,83 | 4,30 | 0,19 | 8,48E-01 |
| AT3G42140 | 0,14 | 0,89 | 4,35 | 0,21 | 8,37E-01 |
| AT3G42155 | 0,34 | 1,63 | 4,32 | 0,38 | 7,05E-01 |
| AT3G42550 | 0,12 | 0,49 | 4,38 | 0,11 | 9,11E-01 |
| AT3G42723 | 0,12 | 0,11 | 4,38 | 0,03 | 9,80E-01 |
| AT3G42790 | 0,59 | 1,70 | 3,15 | 0,54 | 5,91E-01 |
| AT3G43153 | 0,99 | 1,67 | 2,55 | 0,66 | 5,12E-01 |
| AT3G43190 | 0,26 | 0,60 | 4,37 | 0,14 | 8,90E-01 |
| AT3G43220 | 0,79 | -2,04 | 2,94 | -0,70 | 4,87E-01 |
| AT3G43240 | 0,86 | -0,25 | 2,84 | -0,09 | 9,30E-01 |
| AT3G43270 | 0,68 | -1,55 | 2,25 | -0,69 | 4,92E-01 |
| AT3G43280 | 0,11 | 0,88 | 4,35 | 0,20 | 8,39E-01 |
| AT3G43291 | 0,65 | 1,24 | 2,37 | 0,52 | 6,02E-01 |

| | | | | | |
|---|---|---|---|---|---|
| AT3G06595 | 0,10 | 0,08 | 4,38 | 0,02 | 9,85E-01 |
| AT3G43432 | 0,16 | 1,05 | 4,35 | 0,24 | 8,10E-01 |
| AT3G43470 | 0,45 | 2,56 | 3,90 | 0,66 | 5,11E-01 |
| AT3G43500 | 0,45 | 2,86 | 3,91 | 0,73 | 4,65E-01 |
| AT3G43600 | 0,98 | -4,27 | 2,15 | -1,99 | 4,67E-02 |
| AT3G43610 | 0,74 | 0,81 | 2,61 | 0,31 | 7,56E-01 |
| AT3G06645 | 0,23 | 0,23 | 4,35 | 0,05 | 9,58E-01 |
| AT3G06655 | 1,06 | 2,69 | 2,04 | 1,31 | 1,89E-01 |
| AT3G43720 | 0,87 | -4,78 | 2,67 | -1,79 | 7,39E-02 |
| AT3G43810 | 0,51 | -0,40 | 3,06 | -0,13 | 8,96E-01 |
| AT3G43880 | 0,10 | 0,24 | 4,38 | 0,05 | 9,57E-01 |
| AT3G43930 | 0,08 | 0,00 | 4,41 | 0,00 | 1,00E+00 |
| AT3G43940 | 0,53 | 2,22 | 3,83 | 0,58 | 5,62E-01 |
| AT3G44010 | 1,00 | 0,27 | 1,87 | 0,15 | 8,83E-01 |
| AT3G44070 | 0,37 | -1,17 | 3,34 | -0,35 | 7,27E-01 |
| AT3G44160 | 0,43 | -0,92 | 4,12 | -0,22 | 8,24E-01 |
| AT3G44265 | 0,40 | 1,99 | 4,26 | 0,47 | 6,40E-01 |
| AT3G44340 | 1,56 | 1,53 | 1,74 | 0,88 | 3,80E-01 |
| AT3G44400 | 0,82 | 1,57 | 2,74 | 0,57 | 5,66E-01 |
| AT3G44435 | 0,70 | 1,68 | 3,77 | 0,45 | 6,56E-01 |
| AT3G44450 | 0,56 | 1,01 | 3,01 | 0,34 | 7,36E-01 |
| AT3G44460 | 0,11 | 0,88 | 4,35 | 0,20 | 8,39E-01 |
| AT3G44635 | 1,31 | 0,00 | 3,05 | 0,00 | 1,00E+00 |
| AT3G44680 | 0,12 | -0,26 | 4,41 | -0,06 | 9,53E-01 |
| AT3G44753 | 0,14 | 0,81 | 4,35 | 0,19 | 8,52E-01 |
| AT3G44780 | 0,10 | 0,86 | 4,35 | 0,20 | 8,43E-01 |
| AT3G44820 | 0,38 | -0,38 | 4,39 | -0,09 | 9,31E-01 |
| AT3G44990 | 0,60 | -3,71 | 3,61 | -1,03 | 3,04E-01 |
| AT3G45030 | 0,91 | -1,76 | 2,42 | -0,73 | 4,67E-01 |
| AT3G45040 | 0,29 | 1,95 | 4,30 | 0,45 | 6,50E-01 |
| AT3G45060 | 0,08 | 0,00 | 4,41 | 0,00 | 1,00E+00 |
| AT3G45090 | 1,49 | -2,02 | 2,10 | -0,96 | 3,36E-01 |
| AT3G06955 | 0,21 | 1,90 | 4,30 | 0,44 | 6,58E-01 |
| AT3G45130 | 0,82 | 1,25 | 2,20 | 0,57 | 5,71E-01 |
| AT3G45180 | 1,00 | -0,93 | 1,78 | -0,52 | 6,02E-01 |
| AT3G07065 | 0,05 | 0,18 | 4,38 | 0,04 | 9,68E-01 |
| AT3G45240 | 0,25 | 0,00 | 4,41 | 0,00 | 1,00E+00 |
| AT3G07095 | 0,20 | -2,08 | 4,39 | -0,47 | 6,35E-01 |
| AT3G45260 | 1,01 | -0,38 | 1,91 | -0,20 | 8,41E-01 |
| AT3G45300 | 0,17 | -1,50 | 4,37 | -0,34 | 7,31E-01 |
| AT3G45430 | 0,20 | 0,00 | 4,41 | 0,00 | 1,00E+00 |
| AT3G45443 | 0,08 | 0,21 | 4,39 | 0,05 | 9,62E-01 |
| AT3G45600 | 0,92 | -1,89 | 1,94 | -0,98 | 3,29E-01 |
| AT3G45630 | 0,80 | 1,25 | 2,30 | 0,54 | 5,87E-01 |
| AT3G45770 | 0,14 | 0,89 | 4,35 | 0,21 | 8,37E-01 |
| AT3G45820 | 0,59 | -0,05 | 2,88 | -0,02 | 9,86E-01 |
| AT3G45850 | 1,07 | -0,03 | 2,52 | -0,01 | 9,91E-01 |
| AT3G45860 | 0,95 | 1,22 | 2,13 | 0,57 | 5,66E-01 |
| AT3G45910 | 1,00 | 1,38 | 2,43 | 0,57 | 5,71E-01 |
| AT3G45990 | 0,46 | -1,34 | 3,33 | -0,40 | 6,88E-01 |
| AT3G46000 | 0,30 | -0,27 | 3,66 | -0,07 | 9,41E-01 |
| AT3G46030 | 0,35 | 1,55 | 4,33 | 0,36 | 7,20E-01 |
| AT3G46090 | 0,72 | 2,80 | 2,31 | 1,21 | 2,27E-01 |
| AT3G46100 | 0,51 | -1,99 | 3,91 | -0,51 | 6,12E-01 |
| AT3G46210 | 0,56 | 2,18 | 3,84 | 0,57 | 5,71E-01 |
| AT3G07375 | 0,21 | -2,19 | 4,41 | -0,50 | 6,20E-01 |

| | | | | | |
|---|---|---|---|---|---|
| AT3G46320 | 0,61 | 15,98 | 4,34 | 3,68 | 2,35E-04 |
| AT3G46360 | 0,82 | -2,80 | 2,69 | -1,04 | 2,98E-01 |
| AT3G46390 | 0,25 | 0,57 | 4,36 | 0,13 | 8,97E-01 |
| AT3G46440 | 0,45 | 0,00 | 4,41 | 0,00 | 1,00E+00 |
| AT3G46450 | 0,53 | 0,21 | 3,92 | 0,05 | 9,57E-01 |
| AT3G46470 | 0,38 | 1,59 | 4,25 | 0,37 | 7,09E-01 |
| AT3G46510 | 1,83 | 0,47 | 1,38 | 0,34 | 7,33E-01 |
| AT3G46540 | 0,31 | -0,93 | 4,37 | -0,21 | 8,32E-01 |
| AT3G46550 | 1,58 | 2,10 | 1,47 | 1,43 | 1,53E-01 |
| AT3G46610 | 1,41 | 0,74 | 1,66 | 0,45 | 6,53E-01 |
| AT3G46630 | 0,81 | -2,10 | 2,14 | -0,98 | 3,26E-01 |
| AT3G46650 | 0,58 | -2,09 | 3,20 | -0,65 | 5,14E-01 |
| AT3G46660 | 0,59 | 0,97 | 2,66 | 0,36 | 7,15E-01 |
| AT3G46670 | 0,57 | -2,67 | 3,20 | -0,83 | 4,04E-01 |
| AT3G46690 | 0,12 | 0,00 | 4,41 | 0,00 | 1,00E+00 |
| AT3G46730 | 0,12 | 0,00 | 4,41 | 0,00 | 1,00E+00 |
| AT3G46790 | 0,53 | 0,00 | 4,40 | 0,00 | 1,00E+00 |
| AT3G46845 | 1,29 | 1,88 | 1,85 | 1,01 | 3,11E-01 |
| AT3G46920 | 0,08 | 0,00 | 4,41 | 0,00 | 1,00E+00 |
| AT3G46960 | 0,92 | 1,20 | 2,35 | 0,51 | 6,11E-01 |
| AT3G47000 | 0,33 | 0,15 | 4,38 | 0,04 | 9,72E-01 |
| AT3G47070 | 0,67 | -0,66 | 2,54 | -0,26 | 7,94E-01 |
| AT3G47090 | 0,54 | -0,40 | 3,83 | -0,11 | 9,16E-01 |
| AT3G47160 | 0,74 | -0,05 | 2,37 | -0,02 | 9,83E-01 |
| AT3G47250 | 1,84 | 1,77 | 1,61 | 1,10 | 2,72E-01 |
| AT3G47370 | 0,35 | 0,66 | 4,35 | 0,15 | 8,79E-01 |
| AT3G47420 | 0,51 | 1,15 | 2,64 | 0,43 | 6,64E-01 |
| AT3G47480 | 0,70 | 2,53 | 2,90 | 0,87 | 3,83E-01 |
| AT3G47670 | 0,97 | 0,26 | 1,99 | 0,13 | 8,97E-01 |
| AT3G47675 | 0,21 | 0,00 | 4,41 | 0,00 | 1,00E+00 |
| AT3G47680 | 0,08 | 0,00 | 4,41 | 0,00 | 1,00E+00 |
| AT3G47700 | 0,43 | 1,32 | 2,76 | 0,48 | 6,32E-01 |
| AT3G47730 | 0,93 | -3,71 | 2,71 | -1,37 | 1,71E-01 |
| AT3G47810 | 0,27 | -0,41 | 4,38 | -0,09 | 9,25E-01 |
| AT3G47860 | 0,23 | 0,94 | 4,35 | 0,22 | 8,29E-01 |
| AT3G47890 | 1,54 | -2,47 | 1,62 | -1,53 | 1,27E-01 |
| AT3G47940 | 0,35 | 0,04 | 4,38 | 0,01 | 9,93E-01 |
| AT3G47960 | 1,01 | 1,55 | 1,85 | 0,84 | 4,02E-01 |
| AT3G47990 | 0,32 | -0,38 | 4,39 | -0,09 | 9,32E-01 |
| AT3G48060 | 1,66 | 1,54 | 2,02 | 0,76 | 4,46E-01 |
| AT3G48080 | 1,04 | 0,51 | 2,21 | 0,23 | 8,17E-01 |
| AT3G48090 | 1,08 | 0,59 | 2,82 | 0,21 | 8,36E-01 |
| AT3G48100 | 0,85 | -0,36 | 4,36 | -0,08 | 9,34E-01 |
| AT3G48131 | 0,31 | 1,76 | 4,31 | 0,41 | 6,83E-01 |
| AT3G48150 | 0,55 | 0,00 | 3,90 | 0,00 | 1,00E+00 |
| AT3G48170 | 0,90 | -1,51 | 2,59 | -0,58 | 5,60E-01 |
| AT3G48195 | 0,48 | 0,77 | 3,15 | 0,25 | 8,06E-01 |
| AT3G48360 | 1,03 | -2,19 | 2,97 | -0,74 | 4,62E-01 |
| AT3G48520 | 0,38 | 0,00 | 4,41 | 0,00 | 1,00E+00 |
| AT3G48570 | 0,28 | 1,13 | 4,33 | 0,26 | 7,94E-01 |
| AT3G48640 | 0,25 | 0,09 | 4,38 | 0,02 | 9,84E-01 |
| AT3G48690 | 0,79 | -1,28 | 2,61 | -0,49 | 6,23E-01 |
| AT3G48740 | 0,68 | 0,85 | 2,17 | 0,39 | 6,95E-01 |
| AT3G48770 | 1,49 | 2,67 | 1,67 | 1,60 | 1,10E-01 |
| AT3G08005 | 0,05 | 0,18 | 4,38 | 0,04 | 9,68E-01 |
| AT3G48840 | 0,12 | -0,36 | 4,39 | -0,08 | 9,34E-01 |

| | | | | | |
|---|---|---|---|---|---|
| AT3G48930 | 0,53 | 0,50 | 2,73 | 0,18 | 8,55E-01 |
| AT3G48990 | 0,58 | -2,16 | 3,11 | -0,69 | 4,87E-01 |
| AT3G49010 | 1,04 | -0,58 | 1,72 | -0,34 | 7,35E-01 |
| AT3G49055 | 0,42 | 0,00 | 4,41 | 0,00 | 1,00E+00 |
| AT3G49060 | 0,44 | 1,16 | 3,18 | 0,37 | 7,15E-01 |
| AT3G49070 | 1,57 | -5,68 | 2,95 | -1,93 | 5,41E-02 |
| AT3G49140 | 0,93 | -1,60 | 2,08 | -0,77 | 4,42E-01 |
| AT3G49160 | 0,11 | -0,26 | 4,41 | -0,06 | 9,53E-01 |
| AT3G49250 | 0,54 | -2,61 | 4,06 | -0,64 | 5,20E-01 |
| AT3G49350 | 0,36 | -1,26 | 4,37 | -0,29 | 7,73E-01 |
| AT3G49490 | 1,14 | 0,49 | 2,28 | 0,22 | 8,28E-01 |
| AT3G49530 | 1,18 | -0,26 | 2,06 | -0,13 | 9,00E-01 |
| AT3G49600 | 1,57 | -0,34 | 2,14 | -0,16 | 8,72E-01 |
| AT3G49601 | 0,27 | 1,71 | 4,31 | 0,40 | 6,92E-01 |
| AT3G49650 | 0,08 | 0,00 | 4,41 | 0,00 | 1,00E+00 |
| AT3G49720 | 1,09 | 2,98 | 2,26 | 1,32 | 1,87E-01 |
| AT3G49790 | 0,38 | 2,09 | 3,30 | 0,63 | 5,27E-01 |
| AT3G49800 | 1,37 | 1,00 | 1,65 | 0,60 | 5,47E-01 |
| AT3G49870 | 0,08 | 0,00 | 4,41 | 0,00 | 1,00E+00 |
| AT3G49980 | 0,21 | -0,37 | 4,36 | -0,09 | 9,32E-01 |
| AT3G49990 | 0,57 | 0,14 | 3,80 | 0,04 | 9,70E-01 |
| AT3G50060 | 0,43 | 0,62 | 2,74 | 0,23 | 8,20E-01 |
| AT3G50110 | 0,18 | 0,82 | 4,35 | 0,19 | 8,51E-01 |
| AT3G50120 | 0,78 | -2,62 | 2,10 | -1,25 | 2,12E-01 |
| AT3G50240 | 0,71 | 0,00 | 4,41 | 0,00 | 1,00E+00 |
| AT3G50320 | 1,17 | 3,10 | 2,08 | 1,49 | 1,35E-01 |
| AT3G50380 | 1,07 | 0,16 | 2,54 | 0,06 | 9,50E-01 |
| AT3G50500 | 1,35 | -0,90 | 1,88 | -0,48 | 6,32E-01 |
| AT3G50670 | 0,33 | 0,36 | 4,37 | 0,08 | 9,35E-01 |
| AT3G50690 | 0,85 | -0,99 | 1,72 | -0,58 | 5,63E-01 |
| AT3G50740 | 0,53 | -0,54 | 3,13 | -0,17 | 8,62E-01 |
| AT3G50750 | 1,00 | 2,08 | 2,08 | 1,00 | 3,18E-01 |
| AT3G50780 | 0,08 | 0,00 | 4,41 | 0,00 | 1,00E+00 |
| AT3G50790 | 0,29 | 0,28 | 4,37 | 0,06 | 9,48E-01 |
| AT3G50808 | 0,10 | 0,86 | 4,35 | 0,20 | 8,43E-01 |
| AT3G50825 | 0,41 | 2,73 | 4,28 | 0,64 | 5,23E-01 |
| AT3G50920 | 0,48 | -2,09 | 4,09 | -0,51 | 6,08E-01 |
| AT3G51000 | 0,52 | 1,52 | 3,20 | 0,48 | 6,34E-01 |
| AT3G51120 | 1,56 | -0,26 | 2,09 | -0,13 | 9,00E-01 |
| AT3G51135 | 0,51 | 1,27 | 2,71 | 0,47 | 6,39E-01 |
| AT3G51160 | 0,58 | -0,87 | 3,12 | -0,28 | 7,80E-01 |
| AT3G51265 | 0,13 | 0,94 | 4,35 | 0,22 | 8,29E-01 |
| AT3G51310 | 0,51 | -1,29 | 3,25 | -0,40 | 6,92E-01 |
| AT3G51580 | 0,25 | -0,31 | 4,40 | -0,07 | 9,44E-01 |
| AT3G51600 | 1,48 | -1,71 | 1,40 | -1,22 | 2,23E-01 |
| AT3G51660 | 0,43 | -0,06 | 4,13 | -0,01 | 9,89E-01 |
| AT3G51790 | 0,60 | 0,00 | 3,76 | 0,00 | 1,00E+00 |
| AT3G51820 | 0,59 | -0,30 | 3,92 | -0,08 | 9,38E-01 |
| AT3G51840 | 0,33 | 0,32 | 4,37 | 0,07 | 9,42E-01 |
| AT3G51880 | 0,48 | -0,38 | 3,27 | -0,12 | 9,08E-01 |
| AT3G51890 | 1,41 | 0,57 | 2,82 | 0,20 | 8,39E-01 |
| AT3G51895 | 0,82 | -2,40 | 2,85 | -0,84 | 4,00E-01 |
| AT3G51920 | 0,46 | 0,29 | 2,71 | 0,11 | 9,14E-01 |
| AT3G08215 | 0,08 | 0,00 | 4,41 | 0,00 | 1,00E+00 |
| AT3G51980 | 0,42 | -1,58 | 3,28 | -0,48 | 6,30E-01 |
| AT3G52060 | 0,46 | -1,10 | 2,78 | -0,40 | 6,91E-01 |

| | | | | | |
|---|---|---|---|---|---|
| AT3G52072 | 0,22 | 1,16 | 4,34 | 0,27 | 7,89E-01 |
| AT3G08230 | 0,12 | 0,27 | 4,38 | 0,06 | 9,50E-01 |
| AT3G52110 | 0,12 | -0,26 | 4,41 | -0,06 | 9,53E-01 |
| AT3G52180 | 0,66 | 0,87 | 2,65 | 0,33 | 7,43E-01 |
| AT3G52190 | 0,13 | 0,00 | 4,41 | 0,00 | 1,00E+00 |
| AT3G52220 | 0,11 | -0,26 | 4,41 | -0,06 | 9,53E-01 |
| AT3G52260 | 0,64 | 0,00 | 3,67 | 0,00 | 1,00E+00 |
| AT3G52400 | 1,57 | 1,72 | 1,52 | 1,13 | 2,58E-01 |
| AT3G52480 | 0,53 | 2,56 | 2,97 | 0,86 | 3,89E-01 |
| AT3G52560 | 0,47 | -0,97 | 3,36 | -0,29 | 7,73E-01 |
| AT3G52640 | 0,21 | 0,00 | 4,41 | 0,00 | 1,00E+00 |
| AT3G52730 | 0,69 | -1,86 | 3,75 | -0,50 | 6,19E-01 |
| AT3G52740 | 0,17 | 0,07 | 4,38 | 0,02 | 9,88E-01 |
| AT3G52870 | 0,19 | 1,60 | 4,31 | 0,37 | 7,11E-01 |
| AT3G53015 | 1,03 | -1,30 | 2,34 | -0,56 | 5,78E-01 |
| AT3G53016 | 0,52 | -0,16 | 4,00 | -0,04 | 9,68E-01 |
| AT3G53100 | 0,69 | 0,36 | 3,07 | 0,12 | 9,05E-01 |
| AT3G53380 | 0,40 | 1,92 | 4,16 | 0,46 | 6,45E-01 |
| AT3G53410 | 0,47 | 0,00 | 4,41 | 0,00 | 1,00E+00 |
| AT3G53500 | 0,57 | -0,83 | 2,72 | -0,31 | 7,59E-01 |
| AT3G53510 | 0,25 | 0,00 | 4,41 | 0,00 | 1,00E+00 |
| AT3G53540 | 1,08 | -2,19 | 2,41 | -0,91 | 3,65E-01 |
| AT3G53560 | 0,23 | 0,81 | 4,35 | 0,19 | 8,52E-01 |
| AT3G53570 | 0,21 | 0,00 | 4,41 | 0,00 | 1,00E+00 |
| AT3G53610 | 0,53 | 1,94 | 3,12 | 0,62 | 5,35E-01 |
| AT3G53710 | 0,37 | -0,15 | 3,45 | -0,04 | 9,66E-01 |
| AT3G53730 | 0,67 | 2,19 | 2,43 | 0,90 | 3,68E-01 |
| AT3G53810 | 0,44 | 1,37 | 4,27 | 0,32 | 7,49E-01 |
| AT3G53890 | 0,52 | -2,14 | 3,27 | -0,65 | 5,14E-01 |
| AT3G53910 | 1,01 | 3,23 | 2,12 | 1,52 | 1,28E-01 |
| AT3G53920 | 0,24 | 0,00 | 4,41 | 0,00 | 1,00E+00 |
| AT3G53960 | 0,98 | -1,10 | 2,60 | -0,42 | 6,72E-01 |
| AT3G53990 | 1,83 | -2,59 | 1,39 | -1,86 | 6,30E-02 |
| AT3G54100 | 0,47 | 0,48 | 3,28 | 0,15 | 8,84E-01 |
| AT3G54120 | 0,20 | 0,00 | 4,41 | 0,00 | 1,00E+00 |
| AT3G54140 | 0,70 | 1,64 | 2,64 | 0,62 | 5,33E-01 |
| AT3G54210 | 1,01 | -0,52 | 1,51 | -0,34 | 7,31E-01 |
| AT3G54290 | 0,42 | 0,15 | 4,28 | 0,03 | 9,72E-01 |
| AT3G54350 | 1,02 | 1,23 | 2,09 | 0,59 | 5,54E-01 |
| AT3G54360 | 1,14 | -1,96 | 2,00 | -0,98 | 3,25E-01 |
| AT3G54400 | 0,32 | -2,12 | 4,40 | -0,48 | 6,30E-01 |
| AT3G54440 | 0,61 | 0,21 | 2,13 | 0,10 | 9,20E-01 |
| AT3G54460 | 0,97 | -0,32 | 2,86 | -0,11 | 9,12E-01 |
| AT3G54470 | 0,44 | -2,15 | 3,36 | -0,64 | 5,22E-01 |
| AT3G54500 | 1,37 | -0,22 | 2,28 | -0,10 | 9,22E-01 |
| AT3G54620 | 0,24 | 0,00 | 4,41 | 0,00 | 1,00E+00 |
| AT3G54630 | 0,68 | 0,00 | 4,40 | 0,00 | 1,00E+00 |
| AT3G54640 | 0,22 | 0,81 | 4,35 | 0,19 | 8,52E-01 |
| AT3G54650 | 0,62 | 1,11 | 3,20 | 0,35 | 7,29E-01 |
| AT3G54750 | 0,48 | 0,00 | 4,41 | 0,00 | 1,00E+00 |
| AT3G54760 | 1,04 | 0,81 | 2,05 | 0,40 | 6,92E-01 |
| AT3G54790 | 1,31 | -1,19 | 1,86 | -0,64 | 5,21E-01 |
| AT3G54860 | 0,55 | 1,19 | 3,03 | 0,39 | 6,95E-01 |
| AT3G54900 | 0,95 | -1,67 | 1,95 | -0,86 | 3,92E-01 |
| AT3G54925 | 1,76 | -0,26 | 1,56 | -0,17 | 8,68E-01 |
| AT3G54960 | 0,90 | 0,49 | 1,93 | 0,25 | 8,01E-01 |

| | | | | | |
|---|---|---|---|---|---|
| AT3G55040 | 0,21 | -2,67 | 4,37 | -0,61 | 5,40E-01 |
| AT3G55050 | 0,45 | 0,89 | 3,31 | 0,27 | 7,88E-01 |
| AT3G55060 | 0,14 | 0,89 | 4,35 | 0,21 | 8,37E-01 |
| AT3G55130 | 0,51 | -0,88 | 3,01 | -0,29 | 7,69E-01 |
| AT3G55160 | 0,95 | -1,91 | 1,88 | -1,02 | 3,09E-01 |
| AT3G55252 | 0,66 | 0,15 | 2,18 | 0,07 | 9,45E-01 |
| AT3G55254 | 0,80 | -0,30 | 2,16 | -0,14 | 8,88E-01 |
| AT3G55330 | 0,44 | -0,30 | 2,79 | -0,11 | 9,13E-01 |
| AT3G55360 | 0,60 | -0,35 | 2,46 | -0,14 | 8,86E-01 |
| AT3G55400 | 0,18 | -2,46 | 4,37 | -0,56 | 5,74E-01 |
| AT3G55430 | 0,77 | -1,09 | 1,93 | -0,57 | 5,72E-01 |
| AT3G55480 | 1,64 | -0,53 | 1,95 | -0,27 | 7,84E-01 |
| AT3G55840 | 1,41 | -0,85 | 1,88 | -0,45 | 6,51E-01 |
| AT3G55860 | 0,83 | 0,02 | 1,94 | 0,01 | 9,91E-01 |
| AT3G55935 | 0,46 | 1,15 | 2,70 | 0,43 | 6,70E-01 |
| AT3G55950 | 0,63 | 2,60 | 2,42 | 1,07 | 2,83E-01 |
| AT3G55960 | 0,64 | 0,18 | 2,63 | 0,07 | 9,46E-01 |
| AT3G56040 | 0,77 | -0,72 | 2,20 | -0,33 | 7,45E-01 |
| AT3G56050 | 0,97 | -0,11 | 2,02 | -0,05 | 9,56E-01 |
| AT3G56085 | 0,12 | 0,11 | 4,38 | 0,03 | 9,80E-01 |
| AT3G56090 | 0,34 | -1,48 | 3,48 | -0,43 | 6,70E-01 |
| AT3G56110 | 0,41 | 0,50 | 3,33 | 0,15 | 8,80E-01 |
| AT3G56190 | 1,16 | 0,12 | 1,71 | 0,07 | 9,43E-01 |
| AT3G56210 | 0,45 | 0,35 | 4,37 | 0,08 | 9,36E-01 |
| AT3G56240 | 0,35 | 0,95 | 2,83 | 0,34 | 7,37E-01 |
| AT3G56290 | 1,19 | -0,16 | 2,36 | -0,07 | 9,47E-01 |
| AT3G56320 | 0,58 | -0,36 | 3,92 | -0,09 | 9,27E-01 |
| AT3G56340 | 0,44 | -0,10 | 3,33 | -0,03 | 9,75E-01 |
| AT3G56370 | 0,84 | 0,14 | 2,43 | 0,06 | 9,55E-01 |
| AT3G56460 | 1,07 | 1,31 | 2,10 | 0,62 | 5,33E-01 |
| AT3G56480 | 0,39 | 0,36 | 2,85 | 0,13 | 9,00E-01 |
| AT3G56670 | 0,38 | 1,35 | 4,32 | 0,31 | 7,54E-01 |
| AT3G56690 | 0,43 | -0,53 | 4,15 | -0,13 | 8,99E-01 |
| AT3G56790 | 0,33 | 0,22 | 4,39 | 0,05 | 9,61E-01 |
| AT3G56800 | 1,39 | 2,03 | 1,59 | 1,28 | 2,02E-01 |
| AT3G56820 | 0,25 | 0,00 | 4,41 | 0,00 | 1,00E+00 |
| AT3G56830 | 0,17 | 0,00 | 4,41 | 0,00 | 1,00E+00 |
| AT3G56850 | 0,48 | 0,84 | 3,33 | 0,25 | 8,01E-01 |
| AT3G56860 | 0,49 | 0,82 | 2,67 | 0,31 | 7,59E-01 |
| AT3G56950 | 0,79 | -1,70 | 2,16 | -0,79 | 4,31E-01 |
| AT3G56970 | 0,12 | 0,00 | 4,41 | 0,00 | 1,00E+00 |
| AT3G56990 | 0,46 | -2,87 | 3,19 | -0,90 | 3,69E-01 |
| AT3G57020 | 0,42 | -0,77 | 4,36 | -0,18 | 8,60E-01 |
| AT3G57030 | 0,05 | 0,18 | 4,38 | 0,04 | 9,68E-01 |
| AT3G08825 | 0,72 | -0,85 | 2,91 | -0,29 | 7,70E-01 |
| AT3G57050 | 0,52 | -0,09 | 2,64 | -0,04 | 9,71E-01 |
| AT3G08855 | 0,12 | -0,36 | 4,39 | -0,08 | 9,34E-01 |
| AT3G57140 | 0,48 | 0,41 | 3,18 | 0,13 | 8,97E-01 |
| AT3G57240 | 0,24 | 0,00 | 4,41 | 0,00 | 1,00E+00 |
| AT3G57290 | 1,20 | 1,19 | 2,23 | 0,53 | 5,93E-01 |
| AT3G57350 | 0,44 | -0,34 | 3,37 | -0,10 | 9,21E-01 |
| AT3G57420 | 0,74 | 0,60 | 2,26 | 0,27 | 7,89E-01 |
| AT3G57460 | 0,24 | 1,90 | 4,30 | 0,44 | 6,58E-01 |
| AT3G08935 | 1,61 | -3,29 | 1,76 | -1,87 | 6,12E-02 |
| AT3G57470 | 0,21 | -1,55 | 4,38 | -0,35 | 7,24E-01 |
| AT3G57530 | 1,08 | 1,84 | 1,97 | 0,93 | 3,51E-01 |

| | | | | | |
|---|---|---|---|---|---|
| AT3G57540 | 0,08 | 0,21 | 4,39 | 0,05 | 9,62E-01 |
| AT3G57580 | 0,45 | 1,18 | 3,24 | 0,36 | 7,16E-01 |
| AT3G57610 | 1,05 | -0,36 | 4,39 | -0,08 | 9,35E-01 |
| AT3G57640 | 0,11 | 0,88 | 4,35 | 0,20 | 8,39E-01 |
| AT3G57765 | 1,26 | 0,75 | 1,52 | 0,50 | 6,20E-01 |
| AT3G57780 | 0,43 | -2,20 | 4,41 | -0,50 | 6,18E-01 |
| AT3G57790 | 0,44 | 0,22 | 4,39 | 0,05 | 9,59E-01 |
| AT3G57800 | 0,37 | 2,28 | 4,29 | 0,53 | 5,96E-01 |
| AT3G57870 | 0,53 | 0,00 | 4,41 | 0,00 | 1,00E+00 |
| AT3G57880 | 0,23 | 0,85 | 4,34 | 0,20 | 8,45E-01 |
| AT3G57930 | 0,30 | 0,41 | 4,37 | 0,09 | 9,26E-01 |
| AT3G57965 | 0,28 | -2,10 | 4,40 | -0,48 | 6,33E-01 |
| AT3G57980 | 0,74 | 2,16 | 2,79 | 0,77 | 4,40E-01 |
| AT3G57990 | 0,40 | 1,61 | 3,33 | 0,48 | 6,29E-01 |
| AT3G58100 | 1,03 | 0,00 | 3,44 | 0,00 | 1,00E+00 |
| AT3G58120 | 0,15 | -2,07 | 4,39 | -0,47 | 6,38E-01 |
| AT3G58140 | 0,42 | 0,85 | 3,30 | 0,26 | 7,96E-01 |
| AT3G58170 | 0,42 | -1,96 | 4,25 | -0,46 | 6,45E-01 |
| AT3G58210 | 0,64 | 0,00 | 4,41 | 0,00 | 1,00E+00 |
| AT3G58270 | 0,99 | 0,23 | 2,64 | 0,09 | 9,30E-01 |
| AT3G58500 | 1,34 | -0,12 | 1,59 | -0,08 | 9,39E-01 |
| AT3G58560 | 0,23 | 1,19 | 4,34 | 0,27 | 7,84E-01 |
| AT3G58580 | 0,38 | -1,30 | 4,36 | -0,30 | 7,66E-01 |
| AT3G58670 | 0,12 | -0,36 | 4,39 | -0,08 | 9,34E-01 |
| AT3G58730 | 1,29 | -1,28 | 1,53 | -0,83 | 4,04E-01 |
| AT3G58750 | 0,61 | -2,53 | 3,86 | -0,66 | 5,12E-01 |
| AT3G58790 | 0,34 | -0,45 | 4,32 | -0,10 | 9,17E-01 |
| AT3G58970 | 1,53 | 0,00 | 2,57 | 0,00 | 1,00E+00 |
| AT3G58990 | 0,12 | 0,27 | 4,38 | 0,06 | 9,50E-01 |
| AT3G59040 | 1,65 | -0,26 | 2,09 | -0,13 | 9,00E-01 |
| AT3G59050 | 0,52 | 1,12 | 3,13 | 0,36 | 7,21E-01 |
| AT3G59060 | 1,47 | 1,92 | 1,66 | 1,16 | 2,48E-01 |
| AT3G59070 | 1,78 | 1,97 | 1,62 | 1,21 | 2,24E-01 |
| AT3G59120 | 0,10 | 0,08 | 4,38 | 0,02 | 9,85E-01 |
| AT3G59280 | 0,32 | 0,52 | 4,38 | 0,12 | 9,06E-01 |
| AT3G59360 | 0,61 | 0,96 | 2,98 | 0,32 | 7,48E-01 |
| AT3G59400 | 1,33 | 0,52 | 1,53 | 0,34 | 7,36E-01 |
| AT3G59540 | 1,10 | -2,16 | 1,91 | -1,13 | 2,59E-01 |
| AT3G59570 | 0,57 | -0,22 | 3,00 | -0,07 | 9,41E-01 |
| AT3G59660 | 1,15 | 1,62 | 1,99 | 0,81 | 4,16E-01 |
| AT3G59760 | 0,20 | -0,96 | 4,35 | -0,22 | 8,26E-01 |
| AT3G59765 | 1,81 | -3,28 | 1,54 | -2,14 | 3,27E-02 |
| AT3G59840 | 0,15 | 0,82 | 4,35 | 0,19 | 8,50E-01 |
| AT3G59910 | 0,48 | -0,34 | 4,11 | -0,08 | 9,35E-01 |
| AT3G59950 | 1,22 | 1,09 | 2,37 | 0,46 | 6,44E-01 |
| AT3G59980 | 0,35 | 0,48 | 3,50 | 0,14 | 8,92E-01 |
| AT3G60030 | 1,14 | -1,84 | 2,22 | -0,83 | 4,09E-01 |
| AT3G09335 | 0,23 | -0,54 | 4,36 | -0,12 | 9,02E-01 |
| AT3G60050 | 0,12 | 0,00 | 4,41 | 0,00 | 1,00E+00 |
| AT3G60070 | 0,35 | -2,14 | 4,39 | -0,49 | 6,26E-01 |
| AT3G60160 | 1,27 | 1,55 | 2,04 | 0,76 | 4,46E-01 |
| AT3G60176 | 0,85 | 1,83 | 2,68 | 0,68 | 4,94E-01 |
| AT3G60220 | 0,15 | 1,21 | 4,33 | 0,28 | 7,81E-01 |
| AT3G60250 | 1,02 | -1,24 | 2,21 | -0,56 | 5,73E-01 |
| AT3G60350 | 1,30 | -2,10 | 2,15 | -0,97 | 3,30E-01 |
| AT3G60380 | 0,49 | 0,04 | 3,96 | 0,01 | 9,91E-01 |

| | | | | | |
|---|---|---|---|---|---|
| AT3G60540 | 0,08 | 0,21 | 4,39 | 0,05 | 9,62E-01 |
| AT3G60670 | 0,12 | 0,27 | 4,38 | 0,06 | 9,50E-01 |
| AT3G60740 | 0,66 | 2,37 | 2,40 | 0,99 | 3,24E-01 |
| AT3G60760 | 0,12 | 0,00 | 4,41 | 0,00 | 1,00E+00 |
| AT3G60820 | 0,15 | 0,02 | 4,36 | 0,00 | 9,96E-01 |
| AT3G60990 | 0,73 | -1,23 | 2,60 | -0,47 | 6,37E-01 |
| AT3G61070 | 0,50 | 0,43 | 4,10 | 0,10 | 9,17E-01 |
| AT3G09565 | 0,50 | -0,39 | 3,91 | -0,10 | 9,20E-01 |
| AT3G61190 | 1,32 | 1,32 | 1,85 | 0,71 | 4,75E-01 |
| AT3G61200 | 0,69 | -0,74 | 2,51 | -0,30 | 7,67E-01 |
| AT3G61260 | 0,83 | -1,61 | 2,39 | -0,68 | 4,99E-01 |
| AT3G61310 | 0,38 | 0,00 | 4,41 | 0,00 | 1,00E+00 |
| AT3G61380 | 0,18 | 0,02 | 4,37 | 0,00 | 9,96E-01 |
| AT3G61420 | 0,37 | -0,34 | 4,40 | -0,08 | 9,38E-01 |
| AT3G61460 | 0,83 | -0,49 | 1,72 | -0,28 | 7,76E-01 |
| AT3G61530 | 0,25 | -2,14 | 4,40 | -0,49 | 6,28E-01 |
| AT3G61640 | 0,38 | 1,89 | 3,32 | 0,57 | 5,68E-01 |
| AT3G61740 | 0,78 | -0,26 | 2,57 | -0,10 | 9,19E-01 |
| AT3G61750 | 0,24 | 0,00 | 4,41 | 0,00 | 1,00E+00 |
| AT3G61780 | 1,45 | 1,12 | 1,89 | 0,59 | 5,55E-01 |
| AT3G61800 | 0,23 | 1,75 | 4,31 | 0,41 | 6,84E-01 |
| AT3G61820 | 1,36 | -2,02 | 1,49 | -1,36 | 1,75E-01 |
| AT3G61860 | 0,39 | 0,42 | 4,36 | 0,10 | 9,23E-01 |
| AT3G61890 | 0,44 | -0,21 | 3,45 | -0,06 | 9,52E-01 |
| AT3G61960 | 1,02 | -0,85 | 1,89 | -0,45 | 6,53E-01 |
| AT3G61990 | 0,15 | 0,47 | 4,37 | 0,11 | 9,14E-01 |
| AT3G62070 | 0,12 | 1,17 | 4,34 | 0,27 | 7,88E-01 |
| AT3G62120 | 1,00 | -1,12 | 2,15 | -0,52 | 6,01E-01 |
| AT3G62240 | 0,93 | 1,12 | 2,92 | 0,38 | 7,01E-01 |
| AT3G62260 | 0,41 | 0,28 | 4,16 | 0,07 | 9,47E-01 |
| AT3G62310 | 0,33 | -3,19 | 4,35 | -0,73 | 4,64E-01 |
| AT3G62330 | 0,73 | -0,21 | 3,03 | -0,07 | 9,45E-01 |
| AT3G62470 | 0,24 | 0,00 | 4,41 | 0,00 | 1,00E+00 |
| AT3G62540 | 0,28 | 0,00 | 4,41 | 0,00 | 1,00E+00 |
| AT3G09795 | 0,21 | 1,55 | 4,32 | 0,36 | 7,19E-01 |
| AT3G62590 | 0,64 | 1,03 | 2,99 | 0,34 | 7,31E-01 |
| AT3G62600 | 0,95 | 1,22 | 2,01 | 0,61 | 5,43E-01 |
| AT3G09825 | 1,22 | -0,60 | 1,54 | -0,39 | 6,94E-01 |
| AT3G09845 | 0,08 | 0,21 | 4,39 | 0,05 | 9,62E-01 |
| AT3G62630 | 0,91 | 1,92 | 2,32 | 0,83 | 4,09E-01 |
| AT3G62800 | 0,37 | 0,00 | 4,41 | 0,00 | 1,00E+00 |
| AT3G62840 | 0,66 | -0,14 | 3,16 | -0,04 | 9,65E-01 |
| AT3G62860 | 0,81 | 0,00 | 3,59 | 0,00 | 1,00E+00 |
| AT3G62870 | 1,22 | -2,08 | 2,05 | -1,01 | 3,10E-01 |
| AT3G62900 | 0,94 | 0,27 | 2,34 | 0,11 | 9,09E-01 |
| AT3G62910 | 0,49 | 0,59 | 4,06 | 0,14 | 8,85E-01 |
| AT3G62940 | 0,31 | 0,14 | 4,33 | 0,03 | 9,75E-01 |
| AT3G63052 | 0,42 | 0,85 | 3,30 | 0,26 | 7,96E-01 |
| AT3G63110 | 0,56 | 0,14 | 2,54 | 0,06 | 9,56E-01 |
| AT3G63140 | 0,93 | -0,33 | 1,84 | -0,18 | 8,58E-01 |
| AT3G63150 | 0,90 | 1,33 | 2,49 | 0,54 | 5,93E-01 |
| AT3G63160 | 1,15 | -0,62 | 1,73 | -0,36 | 7,20E-01 |
| AT3G63180 | 1,24 | -2,12 | 1,95 | -1,09 | 2,77E-01 |
| AT3G63190 | 0,25 | 0,55 | 4,31 | 0,13 | 8,99E-01 |
| AT3G63210 | 0,47 | 1,52 | 4,24 | 0,36 | 7,20E-01 |
| AT3G63310 | 1,42 | -0,45 | 2,14 | -0,21 | 8,32E-01 |

| | | | | | |
|---|---|---|---|---|---|
| AT3G63340 | 1,13 | -0,38 | 2,64 | -0,14 | 8,86E-01 |
| AT3G63390 | 0,24 | 1,13 | 4,34 | 0,26 | 7,95E-01 |
| AT3G63540 | 1,40 | -0,23 | 1,54 | -0,15 | 8,79E-01 |
| AT4G00005 | 0,19 | -0,58 | 4,38 | -0,13 | 8,95E-01 |
| AT4G00020 | 0,79 | -0,68 | 2,11 | -0,32 | 7,46E-01 |
| AT4G00070 | 0,43 | -0,92 | 4,23 | -0,22 | 8,27E-01 |
| AT4G00110 | 1,14 | -0,22 | 2,48 | -0,09 | 9,29E-01 |
| AT4G00120 | 1,32 | 1,39 | 1,59 | 0,88 | 3,81E-01 |
| AT4G00165 | 0,65 | -0,66 | 2,53 | -0,26 | 7,95E-01 |
| AT4G00180 | 0,45 | -2,04 | 4,36 | -0,47 | 6,40E-01 |
| AT4G00280 | 0,25 | 0,00 | 4,41 | 0,00 | 1,00E+00 |
| AT4G00290 | 0,50 | -0,78 | 2,66 | -0,29 | 7,71E-01 |
| AT4G00330 | 0,80 | 0,92 | 2,15 | 0,43 | 6,70E-01 |
| AT4G03585 | 1,51 | -0,70 | 1,44 | -0,49 | 6,27E-01 |
| AT4G00370 | 0,63 | -3,80 | 2,57 | -1,48 | 1,39E-01 |
| AT4G00400 | 0,42 | -0,19 | 2,70 | -0,07 | 9,44E-01 |
| AT4G00500 | 0,24 | -1,87 | 4,35 | -0,43 | 6,68E-01 |
| AT4G00540 | 0,54 | 1,08 | 2,45 | 0,44 | 6,60E-01 |
| AT4G00550 | 0,15 | -0,35 | 4,39 | -0,08 | 9,36E-01 |
| AT4G00560 | 0,40 | -0,36 | 4,39 | -0,08 | 9,35E-01 |
| AT4G00620 | 0,40 | 0,04 | 4,21 | 0,01 | 9,91E-01 |
| AT4G00740 | 0,31 | 0,95 | 4,35 | 0,22 | 8,27E-01 |
| AT4G00810 | 0,25 | 0,00 | 4,41 | 0,00 | 1,00E+00 |
| AT4G00890 | 0,23 | 2,01 | 4,30 | 0,47 | 6,40E-01 |
| AT4G00900 | 1,09 | -2,46 | 2,11 | -1,17 | 2,43E-01 |
| AT4G00980 | 0,33 | -0,38 | 4,39 | -0,09 | 9,31E-01 |
| AT4G00985 | 0,26 | -2,46 | 4,37 | -0,56 | 5,74E-01 |
| AT4G01000 | 0,50 | 0,16 | 3,96 | 0,04 | 9,67E-01 |
| AT4G01030 | 0,51 | 1,18 | 3,08 | 0,38 | 7,02E-01 |
| AT4G01037 | 0,21 | -1,51 | 4,33 | -0,35 | 7,28E-01 |
| AT4G01070 | 1,43 | -0,36 | 1,58 | -0,23 | 8,21E-01 |
| AT4G01090 | 0,44 | 0,42 | 3,29 | 0,13 | 8,98E-01 |
| AT4G01110 | 0,08 | 0,00 | 4,41 | 0,00 | 1,00E+00 |
| AT4G01120 | 0,18 | -2,46 | 4,37 | -0,56 | 5,73E-01 |
| AT4G01180 | 0,08 | 0,00 | 4,41 | 0,00 | 1,00E+00 |
| AT4G01190 | 0,20 | 0,00 | 4,41 | 0,00 | 1,00E+00 |
| AT4G01200 | 0,46 | 0,00 | 4,41 | 0,00 | 1,00E+00 |
| AT4G01210 | 1,50 | 1,63 | 1,96 | 0,83 | 4,05E-01 |
| AT4G01240 | 0,94 | 2,74 | 2,04 | 1,34 | 1,81E-01 |
| AT4G01290 | 1,74 | -0,16 | 1,55 | -0,10 | 9,18E-01 |
| AT4G01310 | 0,66 | 1,06 | 3,02 | 0,35 | 7,24E-01 |
| AT4G01395 | 0,71 | 0,80 | 2,21 | 0,36 | 7,17E-01 |
| AT4G01533 | 1,62 | 3,02 | 1,92 | 1,57 | 1,16E-01 |
| AT4G01540 | 0,20 | 0,53 | 4,37 | 0,12 | 9,04E-01 |
| AT4G01550 | 0,48 | -3,11 | 4,34 | -0,72 | 4,74E-01 |
| AT4G01700 | 0,60 | -1,49 | 3,19 | -0,47 | 6,40E-01 |
| AT4G01870 | 0,45 | -2,06 | 4,17 | -0,49 | 6,22E-01 |
| AT4G01950 | 0,49 | 1,38 | 2,63 | 0,53 | 5,99E-01 |
| AT4G01985 | 0,41 | -1,85 | 4,33 | -0,43 | 6,68E-01 |
| AT4G02195 | 0,71 | -0,50 | 2,62 | -0,19 | 8,50E-01 |
| AT4G02210 | 0,65 | 1,95 | 2,88 | 0,68 | 5,00E-01 |
| AT4G02230 | 0,28 | -0,36 | 4,39 | -0,08 | 9,35E-01 |
| AT4G02405 | 0,05 | 0,18 | 4,38 | 0,04 | 9,68E-01 |
| AT4G02410 | 0,84 | 0,52 | 2,95 | 0,18 | 8,59E-01 |
| AT4G02420 | 1,44 | 1,89 | 1,71 | 1,10 | 2,71E-01 |
| AT4G02450 | 0,79 | 1,48 | 2,38 | 0,62 | 5,34E-01 |

| | | | | | |
|---|---|---|---|---|---|
| AT4G02460 | 0,19 | -0,40 | 4,39 | -0,09 | 9,28E-01 |
| AT4G02530 | 0,43 | -0,39 | 3,23 | -0,12 | 9,04E-01 |
| AT4G04185 | 0,38 | 1,91 | 3,35 | 0,57 | 5,69E-01 |
| AT4G02550 | 0,12 | 0,27 | 4,38 | 0,06 | 9,50E-01 |
| AT4G02560 | 0,57 | -1,16 | 2,75 | -0,42 | 6,74E-01 |
| AT4G02590 | 0,27 | -2,13 | 4,39 | -0,49 | 6,27E-01 |
| AT4G02620 | 0,23 | -2,48 | 4,38 | -0,57 | 5,71E-01 |
| AT4G02680 | 0,66 | -0,36 | 3,88 | -0,09 | 9,27E-01 |
| AT4G02710 | 0,71 | -0,25 | 3,02 | -0,08 | 9,35E-01 |
| AT4G02715 | 1,63 | -1,32 | 2,08 | -0,64 | 5,24E-01 |
| AT4G02735 | 0,41 | 0,00 | 4,41 | 0,00 | 1,00E+00 |
| AT4G02740 | 0,60 | 1,03 | 3,07 | 0,34 | 7,37E-01 |
| AT4G02750 | 0,98 | 0,00 | 2,41 | 0,00 | 1,00E+00 |
| AT4G02910 | 0,98 | 0,08 | 2,32 | 0,03 | 9,72E-01 |
| AT4G02930 | 0,46 | 0,00 | 4,41 | 0,00 | 1,00E+00 |
| AT4G03030 | 1,74 | 0,40 | 1,94 | 0,21 | 8,37E-01 |
| AT4G03080 | 1,69 | -0,21 | 1,57 | -0,14 | 8,91E-01 |
| AT4G03090 | 0,91 | -0,98 | 1,68 | -0,58 | 5,59E-01 |
| AT4G03110 | 0,81 | -0,28 | 2,98 | -0,09 | 9,25E-01 |
| AT4G03130 | 0,08 | 0,00 | 4,41 | 0,00 | 1,00E+00 |
| AT4G03150 | 0,26 | 0,00 | 4,41 | 0,00 | 1,00E+00 |
| AT4G03190 | 0,84 | -2,02 | 2,32 | -0,87 | 3,84E-01 |
| AT4G03230 | 0,47 | 0,18 | 4,03 | 0,05 | 9,64E-01 |
| AT4G03270 | 0,81 | 0,00 | 2,91 | 0,00 | 1,00E+00 |
| AT4G04475 | 0,42 | 0,78 | 3,30 | 0,24 | 8,13E-01 |
| AT4G04485 | 0,66 | -0,09 | 2,48 | -0,03 | 9,72E-01 |
| AT4G03340 | 0,12 | 0,00 | 4,41 | 0,00 | 1,00E+00 |
| AT4G03390 | 0,56 | 0,59 | 2,75 | 0,21 | 8,30E-01 |
| AT4G03400 | 0,95 | 2,50 | 2,16 | 1,16 | 2,46E-01 |
| AT4G03430 | 1,44 | 2,37 | 1,68 | 1,41 | 1,58E-01 |
| AT4G03435 | 0,08 | 0,21 | 4,39 | 0,05 | 9,62E-01 |
| AT4G03438 | 0,26 | 1,17 | 4,34 | 0,27 | 7,87E-01 |
| AT4G03450 | 0,60 | 1,24 | 2,66 | 0,47 | 6,42E-01 |
| AT4G03490 | 0,42 | 0,87 | 4,30 | 0,20 | 8,39E-01 |
| AT4G03505 | 0,25 | 0,00 | 4,41 | 0,00 | 1,00E+00 |
| AT4G03520 | 1,13 | -1,89 | 1,85 | -1,03 | 3,05E-01 |
| AT4G03570 | 0,98 | -0,09 | 1,52 | -0,06 | 9,53E-01 |
| AT4G03590 | 1,16 | 2,77 | 2,20 | 1,26 | 2,08E-01 |
| AT4G03728 | 0,69 | 1,17 | 3,62 | 0,32 | 7,46E-01 |
| AT4G03740 | 1,37 | 3,09 | 1,93 | 1,61 | 1,08E-01 |
| AT4G04675 | 1,07 | -1,18 | 1,93 | -0,61 | 5,40E-01 |
| AT4G04030 | 0,82 | -0,97 | 2,25 | -0,43 | 6,66E-01 |
| AT4G04221 | 1,00 | 1,82 | 2,15 | 0,85 | 3,97E-01 |
| AT4G04805 | 1,38 | 4,21 | 2,19 | 1,92 | 5,47E-02 |
| AT4G04370 | 1,03 | -0,26 | 2,27 | -0,12 | 9,08E-01 |
| AT4G04470 | 0,08 | 0,21 | 4,39 | 0,05 | 9,62E-01 |
| AT4G04580 | 0,11 | -0,38 | 4,39 | -0,09 | 9,31E-01 |
| AT4G04614 | 0,51 | 0,00 | 4,41 | 0,00 | 1,00E+00 |
| AT4G04632 | 0,44 | 1,47 | 4,09 | 0,36 | 7,19E-01 |
| AT4G04650 | 0,82 | 2,54 | 2,23 | 1,14 | 2,56E-01 |
| AT4G04710 | 0,47 | 0,48 | 3,28 | 0,15 | 8,84E-01 |
| AT4G04720 | 1,28 | -3,72 | 2,05 | -1,82 | 6,94E-02 |
| AT4G04740 | 0,87 | 0,35 | 2,00 | 0,17 | 8,62E-01 |
| AT4G04745 | 1,61 | 0,02 | 1,36 | 0,01 | 9,89E-01 |
| AT4G04770 | 1,41 | 0,21 | 1,90 | 0,11 | 9,11E-01 |
| AT4G04910 | 0,75 | -0,57 | 2,13 | -0,27 | 7,88E-01 |

| | | | | | |
|---|---|---|---|---|---|
| AT4G04950 | 0,83 | -3,90 | 3,62 | -1,08 | 2,81E-01 |
| AT4G04957 | 0,91 | 3,57 | 2,55 | 1,40 | 1,62E-01 |
| AT4G04980 | 0,11 | 0,88 | 4,35 | 0,20 | 8,39E-01 |
| AT4G05070 | 1,30 | -1,38 | 1,56 | -0,88 | 3,78E-01 |
| AT4G05110 | 0,17 | 0,00 | 4,41 | 0,00 | 1,00E+00 |
| AT4G05085 | 0,05 | 0,18 | 4,38 | 0,04 | 9,68E-01 |
| AT4G05330 | 0,44 | -0,05 | 4,14 | -0,01 | 9,90E-01 |
| AT4G05410 | 0,51 | -1,21 | 4,12 | -0,29 | 7,69E-01 |
| AT4G05470 | 0,12 | -0,26 | 4,41 | -0,06 | 9,53E-01 |
| AT4G05631 | 0,41 | -0,36 | 3,43 | -0,11 | 9,15E-01 |
| AT4G06526 | 0,56 | -0,29 | 3,86 | -0,08 | 9,40E-01 |
| AT4G06639 | 0,21 | 0,59 | 4,37 | 0,14 | 8,92E-01 |
| AT4G06599 | 0,16 | 0,74 | 4,36 | 0,17 | 8,66E-01 |
| AT4G07350 | 0,08 | 0,21 | 4,39 | 0,05 | 9,62E-01 |
| AT4G07400 | 0,28 | -0,54 | 4,36 | -0,12 | 9,01E-01 |
| AT4G07410 | 1,28 | 1,31 | 2,24 | 0,59 | 5,58E-01 |
| AT4G07524 | 0,19 | 0,44 | 4,36 | 0,10 | 9,21E-01 |
| AT4G07526 | 0,83 | -3,18 | 2,36 | -1,35 | 1,78E-01 |
| AT4G05225 | 0,74 | -0,92 | 2,86 | -0,32 | 7,47E-01 |
| AT4G07990 | 0,20 | 0,40 | 4,37 | 0,09 | 9,27E-01 |
| AT4G08230 | 0,30 | 2,06 | 4,30 | 0,48 | 6,32E-01 |
| AT4G08310 | 0,85 | 0,92 | 1,94 | 0,48 | 6,35E-01 |
| AT4G05415 | 0,95 | 0,00 | 2,79 | 0,00 | 1,00E+00 |
| AT4G08370 | 0,46 | 1,59 | 2,68 | 0,59 | 5,52E-01 |
| AT4G08400 | 0,54 | 2,97 | 2,93 | 1,02 | 3,09E-01 |
| AT4G08410 | 1,29 | 4,01 | 2,22 | 1,81 | 7,09E-02 |
| AT4G05535 | 0,95 | -1,97 | 2,42 | -0,81 | 4,16E-01 |
| AT4G08460 | 0,62 | -0,90 | 2,65 | -0,34 | 7,35E-01 |
| AT4G08470 | 1,82 | -0,32 | 2,26 | -0,14 | 8,88E-01 |
| AT4G08480 | 0,67 | 1,49 | 3,08 | 0,48 | 6,28E-01 |
| AT4G08485 | 0,63 | -3,79 | 3,67 | -1,03 | 3,01E-01 |
| AT4G08590 | 0,56 | 0,00 | 4,41 | 0,00 | 1,00E+00 |
| AT4G08760 | 1,19 | 3,28 | 2,16 | 1,52 | 1,28E-01 |
| AT4G08810 | 0,77 | -1,24 | 2,70 | -0,46 | 6,46E-01 |
| AT4G08870 | 1,44 | -4,18 | 1,70 | -2,46 | 1,39E-02 |
| AT4G08874 | 0,12 | 0,11 | 4,38 | 0,03 | 9,80E-01 |
| AT4G08940 | 0,46 | 0,71 | 3,36 | 0,21 | 8,33E-01 |
| AT4G09010 | 1,11 | 0,14 | 1,85 | 0,08 | 9,39E-01 |
| AT4G09012 | 0,55 | -0,21 | 3,24 | -0,07 | 9,48E-01 |
| AT4G09035 | 0,16 | 0,41 | 4,37 | 0,09 | 9,25E-01 |
| AT4G09040 | 0,47 | 0,43 | 4,37 | 0,10 | 9,21E-01 |
| AT4G09340 | 0,44 | -0,37 | 3,32 | -0,11 | 9,12E-01 |
| AT4G09350 | 0,26 | -0,91 | 4,38 | -0,21 | 8,36E-01 |
| AT4G09490 | 0,39 | -3,94 | 4,33 | -0,91 | 3,62E-01 |
| AT4G09510 | 0,38 | -2,06 | 3,36 | -0,61 | 5,40E-01 |
| AT4G09620 | 0,40 | 0,00 | 4,41 | 0,00 | 1,00E+00 |
| AT4G09640 | 0,98 | -0,95 | 1,69 | -0,56 | 5,74E-01 |
| AT4G05740 | 1,41 | -0,51 | 1,38 | -0,37 | 7,10E-01 |
| AT4G09900 | 0,13 | -0,61 | 4,37 | -0,14 | 8,89E-01 |
| AT4G09960 | 0,46 | -1,37 | 4,36 | -0,31 | 7,53E-01 |
| AT4G09980 | 0,57 | 0,50 | 2,68 | 0,19 | 8,51E-01 |
| AT4G10070 | 1,18 | -0,35 | 2,36 | -0,15 | 8,81E-01 |
| AT4G10080 | 1,04 | -0,92 | 2,02 | -0,46 | 6,48E-01 |
| AT4G10130 | 0,55 | -0,40 | 3,86 | -0,10 | 9,17E-01 |
| AT4G10320 | 1,43 | 1,15 | 1,51 | 0,76 | 4,46E-01 |
| AT4G10400 | 0,39 | 0,00 | 4,40 | 0,00 | 1,00E+00 |

| | | | | | |
|---|---|---|---|---|---|
| AT4G10420 | 1,07 | 0,00 | 2,79 | 0,00 | 1,00E+00 |
| AT4G10440 | 0,08 | 0,00 | 4,41 | 0,00 | 1,00E+00 |
| AT4G10480 | 0,41 | -0,98 | 4,35 | -0,23 | 8,21E-01 |
| AT4G10540 | 0,12 | 0,00 | 4,41 | 0,00 | 1,00E+00 |
| AT4G10550 | 0,52 | 0,00 | 3,96 | 0,00 | 1,00E+00 |
| AT4G10570 | 1,14 | 1,16 | 1,88 | 0,62 | 5,35E-01 |
| AT4G10590 | 1,40 | -0,35 | 2,19 | -0,16 | 8,74E-01 |
| AT4G05800 | 1,37 | 3,32 | 1,91 | 1,74 | 8,22E-02 |
| AT4G10700 | 0,54 | 3,12 | 3,69 | 0,85 | 3,97E-01 |
| AT4G10730 | 0,28 | 0,00 | 4,41 | 0,00 | 1,00E+00 |
| AT4G10740 | 1,27 | 4,19 | 2,21 | 1,90 | 5,78E-02 |
| AT4G10760 | 0,19 | 1,60 | 4,31 | 0,37 | 7,11E-01 |
| AT4G10770 | 0,10 | 0,08 | 4,38 | 0,02 | 9,85E-01 |
| AT4G10790 | 1,30 | -1,92 | 1,53 | -1,25 | 2,10E-01 |
| AT4G10920 | 0,27 | 1,29 | 3,74 | 0,35 | 7,30E-01 |
| AT4G10960 | 0,05 | 0,18 | 4,38 | 0,04 | 9,68E-01 |
| AT4G10970 | 0,10 | 0,08 | 4,38 | 0,02 | 9,85E-01 |
| AT4G11010 | 0,17 | -0,33 | 4,40 | -0,08 | 9,40E-01 |
| AT4G11050 | 0,37 | 0,18 | 3,40 | 0,05 | 9,57E-01 |
| AT4G11080 | 0,05 | 0,18 | 4,38 | 0,04 | 9,68E-01 |
| AT4G05855 | 1,27 | -2,29 | 2,20 | -1,04 | 2,98E-01 |
| AT4G11216 | 0,26 | 0,00 | 4,41 | 0,00 | 1,00E+00 |
| AT4G11220 | 0,29 | 0,14 | 4,33 | 0,03 | 9,74E-01 |
| AT4G11260 | 0,66 | 0,39 | 4,35 | 0,09 | 9,28E-01 |
| AT4G11300 | 0,45 | 0,05 | 4,17 | 0,01 | 9,91E-01 |
| AT4G11340 | 1,18 | -0,95 | 1,49 | -0,63 | 5,26E-01 |
| AT4G11410 | 0,24 | 0,00 | 4,41 | 0,00 | 1,00E+00 |
| AT4G05880 | 0,80 | -4,01 | 2,64 | -1,52 | 1,28E-01 |
| AT4G11650 | 0,43 | 0,00 | 4,41 | 0,00 | 1,00E+00 |
| AT4G11670 | 0,44 | 0,42 | 2,80 | 0,15 | 8,82E-01 |
| AT4G11800 | 1,18 | 1,07 | 1,70 | 0,63 | 5,30E-01 |
| AT4G05925 | 1,28 | 1,00 | 1,85 | 0,54 | 5,89E-01 |
| AT4G11820 | 0,25 | -3,44 | 4,34 | -0,79 | 4,28E-01 |
| AT4G11830 | 1,70 | -2,93 | 2,09 | -1,40 | 1,62E-01 |
| AT4G11840 | 1,34 | -0,80 | 1,61 | -0,50 | 6,20E-01 |
| AT4G11850 | 1,56 | -0,13 | 1,61 | -0,08 | 9,33E-01 |
| AT4G11860 | 0,25 | 0,00 | 4,41 | 0,00 | 1,00E+00 |
| AT4G11920 | 0,40 | 1,74 | 4,13 | 0,42 | 6,74E-01 |
| AT4G05930 | 0,37 | 1,58 | 4,22 | 0,37 | 7,08E-01 |
| AT4G12010 | 0,59 | -0,29 | 3,76 | -0,08 | 9,39E-01 |
| AT4G05980 | 0,40 | 0,22 | 4,39 | 0,05 | 9,59E-01 |
| AT4G12300 | 1,00 | 1,08 | 2,19 | 0,49 | 6,23E-01 |
| AT4G12310 | 0,40 | 0,11 | 2,85 | 0,04 | 9,69E-01 |
| AT4G12330 | 0,81 | -3,17 | 2,57 | -1,23 | 2,18E-01 |
| AT4G12420 | 1,53 | 1,73 | 2,03 | 0,86 | 3,92E-01 |
| AT4G12590 | 0,64 | 0,92 | 2,45 | 0,37 | 7,08E-01 |
| AT4G12710 | 0,37 | 0,00 | 4,41 | 0,00 | 1,00E+00 |
| AT4G12730 | 0,13 | -0,61 | 4,37 | -0,14 | 8,89E-01 |
| AT4G12750 | 1,05 | -1,22 | 2,23 | -0,55 | 5,85E-01 |
| AT4G12790 | 0,53 | 1,76 | 3,28 | 0,54 | 5,92E-01 |
| AT4G12820 | 0,35 | -4,04 | 4,32 | -0,93 | 3,50E-01 |
| AT4G12880 | 0,35 | -4,04 | 4,32 | -0,93 | 3,50E-01 |
| AT4G12980 | 0,57 | 0,88 | 2,60 | 0,34 | 7,36E-01 |
| AT4G13150 | 0,55 | 0,00 | 4,41 | 0,00 | 1,00E+00 |
| AT4G13250 | 1,54 | 0,62 | 2,06 | 0,30 | 7,65E-01 |
| AT4G13350 | 0,30 | 1,39 | 4,33 | 0,32 | 7,49E-01 |

| | | | | | |
|---|---|---|---|---|---|
| AT4G13395 | 0,15 | 1,21 | 4,33 | 0,28 | 7,81E-01 |
| AT4G13420 | 0,12 | 0,00 | 4,41 | 0,00 | 1,00E+00 |
| AT4G13550 | 0,15 | -0,31 | 4,40 | -0,07 | 9,43E-01 |
| AT4G13590 | 0,56 | 0,84 | 2,61 | 0,32 | 7,47E-01 |
| AT4G13630 | 0,80 | 0,00 | 3,54 | 0,00 | 1,00E+00 |
| AT4G13640 | 0,21 | 0,00 | 4,41 | 0,00 | 1,00E+00 |
| AT4G13780 | 1,18 | -1,04 | 1,99 | -0,52 | 6,02E-01 |
| AT4G13800 | 0,45 | -0,34 | 3,31 | -0,10 | 9,18E-01 |
| AT4G13820 | 0,22 | 0,83 | 4,35 | 0,19 | 8,49E-01 |
| AT4G13840 | 1,76 | -1,29 | 1,47 | -0,88 | 3,79E-01 |
| AT4G13870 | 0,08 | 0,00 | 4,41 | 0,00 | 1,00E+00 |
| AT4G13930 | 1,56 | 1,06 | 1,76 | 0,60 | 5,47E-01 |
| AT4G13950 | 0,08 | 0,00 | 4,41 | 0,00 | 1,00E+00 |
| AT4G13970 | 0,50 | -0,41 | 3,93 | -0,10 | 9,17E-01 |
| AT4G14010 | 0,10 | 0,86 | 4,35 | 0,20 | 8,43E-01 |
| AT4G14030 | 1,74 | -1,20 | 1,57 | -0,76 | 4,46E-01 |
| AT4G14040 | 0,70 | 0,69 | 2,62 | 0,26 | 7,93E-01 |
| AT4G14050 | 0,14 | 0,13 | 4,38 | 0,03 | 9,77E-01 |
| AT4G14070 | 0,95 | -0,39 | 2,72 | -0,14 | 8,87E-01 |
| AT4G14090 | 0,47 | 1,37 | 2,78 | 0,49 | 6,24E-01 |
| AT4G14110 | 0,26 | 0,00 | 4,41 | 0,00 | 1,00E+00 |
| AT4G14200 | 0,21 | 0,36 | 4,37 | 0,08 | 9,34E-01 |
| AT4G14210 | 0,70 | 0,65 | 2,06 | 0,32 | 7,53E-01 |
| AT4G14230 | 0,73 | -0,39 | 3,06 | -0,13 | 8,99E-01 |
| AT4G14240 | 0,36 | -1,73 | 4,35 | -0,40 | 6,91E-01 |
| AT4G14340 | 0,26 | 0,00 | 4,41 | 0,00 | 1,00E+00 |
| AT4G14342 | 0,72 | 0,00 | 4,41 | 0,00 | 1,00E+00 |
| AT4G14350 | 0,16 | 0,39 | 4,37 | 0,09 | 9,29E-01 |
| AT4G14365 | 1,73 | 1,90 | 1,79 | 1,06 | 2,90E-01 |
| AT4G14370 | 0,47 | 0,60 | 2,80 | 0,22 | 8,29E-01 |
| AT4G14385 | 0,08 | 0,00 | 4,41 | 0,00 | 1,00E+00 |
| AT4G14420 | 0,43 | 0,00 | 4,41 | 0,00 | 1,00E+00 |
| AT4G14430 | 1,12 | -1,89 | 1,71 | -1,11 | 2,68E-01 |
| AT4G14440 | 0,19 | 1,47 | 4,32 | 0,34 | 7,35E-01 |
| AT4G14510 | 1,36 | 0,08 | 1,99 | 0,04 | 9,69E-01 |
| AT4G14680 | 0,20 | 0,00 | 4,41 | 0,00 | 1,00E+00 |
| AT4G14710 | 0,27 | 0,24 | 4,32 | 0,06 | 9,55E-01 |
| AT4G14740 | 0,35 | -1,61 | 4,34 | -0,37 | 7,11E-01 |
| AT4G14746 | 0,16 | 0,63 | 4,36 | 0,14 | 8,85E-01 |
| AT4G14790 | 0,23 | 0,24 | 4,39 | 0,06 | 9,56E-01 |
| AT4G14811 | 0,45 | -0,12 | 3,34 | -0,03 | 9,72E-01 |
| AT4G14880 | 1,32 | 1,21 | 1,60 | 0,75 | 4,51E-01 |
| AT4G14900 | 1,30 | 1,11 | 1,79 | 0,62 | 5,34E-01 |
| AT4G14905 | 0,68 | -0,30 | 2,50 | -0,12 | 9,05E-01 |
| AT4G14920 | 0,44 | -0,30 | 4,22 | -0,07 | 9,43E-01 |
| AT4G14990 | 0,87 | -0,08 | 2,03 | -0,04 | 9,68E-01 |
| AT4G15050 | 0,52 | 2,12 | 3,79 | 0,56 | 5,75E-01 |
| AT4G15053 | 0,24 | 1,60 | 4,31 | 0,37 | 7,10E-01 |
| AT4G15080 | 0,43 | 0,60 | 4,16 | 0,14 | 8,85E-01 |
| AT4G15200 | 0,36 | 0,00 | 4,41 | 0,00 | 1,00E+00 |
| AT4G15210 | 0,12 | 0,00 | 4,41 | 0,00 | 1,00E+00 |
| AT4G15242 | 0,72 | 1,97 | 2,05 | 0,96 | 3,36E-01 |
| AT4G06305 | 0,44 | -1,62 | 4,03 | -0,40 | 6,88E-01 |
| AT4G06310 | 0,35 | 0,13 | 4,38 | 0,03 | 9,76E-01 |
| AT4G06315 | 1,00 | 4,06 | 2,99 | 1,36 | 1,75E-01 |
| AT4G15270 | 0,91 | -0,23 | 2,29 | -0,10 | 9,21E-01 |

| | | | | | |
|---|---|---|---|---|---|
| AT4G15490 | 0,92 | 1,57 | 2,09 | 0,75 | 4,53E-01 |
| AT4G15530 | 1,47 | 1,01 | 1,93 | 0,53 | 5,99E-01 |
| AT4G15550 | 0,31 | -0,40 | 4,37 | -0,09 | 9,27E-01 |
| AT4G15810 | 0,14 | 0,89 | 4,35 | 0,21 | 8,37E-01 |
| AT4G06340 | 0,10 | 0,24 | 4,38 | 0,05 | 9,57E-01 |
| AT4G15920 | 0,36 | 0,26 | 4,30 | 0,06 | 9,52E-01 |
| AT4G15955 | 0,12 | 0,00 | 4,41 | 0,00 | 1,00E+00 |
| AT4G16070 | 0,52 | -0,05 | 2,72 | -0,02 | 9,85E-01 |
| AT4G16141 | 0,51 | -1,08 | 4,04 | -0,27 | 7,89E-01 |
| AT4G16144 | 1,36 | -0,72 | 2,11 | -0,34 | 7,35E-01 |
| AT4G16155 | 0,30 | 0,14 | 4,38 | 0,03 | 9,74E-01 |
| AT4G16180 | 1,78 | 0,05 | 2,39 | 0,02 | 9,83E-01 |
| AT4G16215 | 0,40 | 2,44 | 3,25 | 0,75 | 4,52E-01 |
| AT4G16250 | 0,73 | -3,86 | 2,94 | -1,31 | 1,89E-01 |
| AT4G16280 | 0,45 | -0,36 | 3,32 | -0,11 | 9,13E-01 |
| AT4G16330 | 0,88 | 1,29 | 2,89 | 0,45 | 6,55E-01 |
| AT4G16410 | 1,15 | -0,10 | 2,16 | -0,04 | 9,64E-01 |
| AT4G16470 | 0,11 | -0,26 | 4,41 | -0,06 | 9,53E-01 |
| AT4G16500 | 1,23 | 0,72 | 1,64 | 0,44 | 6,58E-01 |
| AT4G16515 | 0,18 | -2,46 | 4,37 | -0,56 | 5,73E-01 |
| AT4G16520 | 0,32 | 1,15 | 4,34 | 0,27 | 7,90E-01 |
| AT4G16540 | 0,20 | 0,40 | 4,37 | 0,09 | 9,27E-01 |
| AT4G16580 | 0,33 | -0,17 | 3,56 | -0,05 | 9,63E-01 |
| AT4G16630 | 0,66 | -1,04 | 3,78 | -0,27 | 7,84E-01 |
| AT4G16680 | 0,40 | 0,00 | 4,41 | 0,00 | 1,00E+00 |
| AT4G16690 | 0,88 | 2,38 | 2,10 | 1,13 | 2,57E-01 |
| AT4G16750 | 0,15 | -0,57 | 4,37 | -0,13 | 8,96E-01 |
| AT4G16760 | 1,69 | -0,80 | 1,62 | -0,49 | 6,23E-01 |
| AT4G16790 | 0,35 | 0,55 | 3,52 | 0,16 | 8,75E-01 |
| AT4G16880 | 1,01 | 1,38 | 2,12 | 0,65 | 5,16E-01 |
| AT4G16892 | 1,06 | 1,33 | 2,47 | 0,54 | 5,89E-01 |
| AT4G16957 | 0,81 | 0,00 | 3,51 | 0,00 | 1,00E+00 |
| AT4G16960 | 0,92 | 0,00 | 4,16 | 0,00 | 1,00E+00 |
| AT4G17040 | 0,44 | -2,93 | 4,20 | -0,70 | 4,85E-01 |
| AT4G17060 | 0,64 | 0,00 | 3,73 | 0,00 | 1,00E+00 |
| AT4G17070 | 0,40 | 0,45 | 3,36 | 0,13 | 8,93E-01 |
| AT4G17100 | 0,77 | -2,24 | 2,22 | -1,01 | 3,13E-01 |
| AT4G17190 | 0,27 | 1,33 | 4,33 | 0,31 | 7,58E-01 |
| AT4G17230 | 1,13 | 2,50 | 1,84 | 1,36 | 1,75E-01 |
| AT4G17243 | 0,66 | 0,86 | 2,60 | 0,33 | 7,40E-01 |
| AT4G17280 | 0,46 | -0,31 | 3,35 | -0,09 | 9,25E-01 |
| AT4G17430 | 0,16 | 1,26 | 4,33 | 0,29 | 7,71E-01 |
| AT4G17486 | 0,51 | 0,00 | 4,41 | 0,00 | 1,00E+00 |
| AT4G17490 | 0,05 | 0,18 | 4,38 | 0,04 | 9,68E-01 |
| AT4G17510 | 0,15 | 0,47 | 4,37 | 0,11 | 9,14E-01 |
| AT4G17550 | 0,46 | -1,28 | 4,16 | -0,31 | 7,59E-01 |
| AT4G17570 | 0,79 | 2,08 | 2,43 | 0,86 | 3,90E-01 |
| AT4G17590 | 0,76 | 1,98 | 2,76 | 0,72 | 4,73E-01 |
| AT4G17610 | 0,22 | 0,80 | 4,35 | 0,18 | 8,55E-01 |
| AT4G17670 | 0,33 | 0,00 | 4,41 | 0,00 | 1,00E+00 |
| AT4G17720 | 0,25 | -0,81 | 4,36 | -0,19 | 8,52E-01 |
| AT4G17740 | 0,24 | -0,53 | 4,37 | -0,12 | 9,03E-01 |
| AT4G17840 | 0,82 | 1,25 | 2,34 | 0,54 | 5,92E-01 |
| AT4G17880 | 0,51 | -1,11 | 3,23 | -0,34 | 7,31E-01 |
| AT4G17950 | 0,50 | -3,00 | 4,08 | -0,74 | 4,62E-01 |
| AT4G18010 | 1,46 | 2,58 | 1,94 | 1,33 | 1,85E-01 |

| | | | | | |
|---|---|---|---|---|---|
| AT4G18020 | 0,33 | -0,27 | 4,41 | -0,06 | 9,52E-01 |
| AT4G18040 | 0,22 | -0,15 | 4,35 | -0,03 | 9,72E-01 |
| AT4G18100 | 0,24 | 0,52 | 4,31 | 0,12 | 9,04E-01 |
| AT4G18110 | 0,16 | -0,26 | 4,41 | -0,06 | 9,53E-01 |
| AT4G18130 | 0,68 | -0,38 | 2,93 | -0,13 | 8,97E-01 |
| AT4G18140 | 0,37 | -0,13 | 4,36 | -0,03 | 9,77E-01 |
| AT4G18215 | 1,61 | 0,19 | 1,58 | 0,12 | 9,03E-01 |
| AT4G18230 | 0,15 | -0,35 | 4,39 | -0,08 | 9,36E-01 |
| AT4G18250 | 0,30 | 0,00 | 4,41 | 0,00 | 1,00E+00 |
| AT4G18253 | 0,68 | 0,15 | 2,56 | 0,06 | 9,54E-01 |
| AT4G18255 | 0,35 | 0,00 | 4,39 | 0,00 | 1,00E+00 |
| AT4G18390 | 1,03 | -2,51 | 2,06 | -1,22 | 2,22E-01 |
| AT4G18440 | 0,69 | 1,61 | 3,54 | 0,46 | 6,48E-01 |
| AT4G18480 | 1,72 | -1,13 | 1,46 | -0,77 | 4,39E-01 |
| AT4G18520 | 0,18 | 0,15 | 4,38 | 0,03 | 9,73E-01 |
| AT4G18580 | 0,08 | 0,00 | 4,41 | 0,00 | 1,00E+00 |
| AT4G18790 | 0,08 | 0,00 | 4,41 | 0,00 | 1,00E+00 |
| AT4G18810 | 0,53 | -0,80 | 2,71 | -0,30 | 7,67E-01 |
| AT4G18840 | 0,38 | -0,18 | 3,44 | -0,05 | 9,59E-01 |
| AT4G19006 | 0,25 | -0,40 | 4,38 | -0,09 | 9,27E-01 |
| AT4G19040 | 0,87 | 0,06 | 3,35 | 0,02 | 9,86E-01 |
| AT4G19045 | 0,12 | 0,00 | 4,41 | 0,00 | 1,00E+00 |
| AT4G19050 | 0,78 | -2,15 | 2,85 | -0,75 | 4,51E-01 |
| AT4G19060 | 0,46 | 0,43 | 4,05 | 0,10 | 9,16E-01 |
| AT4G19080 | 1,21 | -1,26 | 1,91 | -0,66 | 5,11E-01 |
| AT4G19095 | 0,58 | 0,84 | 2,58 | 0,33 | 7,44E-01 |
| AT4G06870 | 0,16 | 1,16 | 4,34 | 0,27 | 7,89E-01 |
| AT4G19180 | 0,15 | -0,31 | 4,40 | -0,07 | 9,43E-01 |
| AT4G19220 | 0,37 | 0,00 | 4,36 | 0,00 | 1,00E+00 |
| AT4G19240 | 0,33 | 0,03 | 4,38 | 0,01 | 9,95E-01 |
| AT4G19410 | 1,78 | -0,09 | 1,59 | -0,06 | 9,55E-01 |
| AT4G19420 | 0,68 | 1,29 | 2,43 | 0,53 | 5,95E-01 |
| AT4G19440 | 0,20 | -0,26 | 4,41 | -0,06 | 9,53E-01 |
| AT4G19640 | 0,35 | 1,57 | 3,43 | 0,46 | 6,48E-01 |
| AT4G19660 | 1,11 | 0,88 | 1,98 | 0,44 | 6,57E-01 |
| AT4G19670 | 0,77 | 0,00 | 3,62 | 0,00 | 1,00E+00 |
| AT4G06915 | 0,34 | 0,78 | 4,34 | 0,18 | 8,58E-01 |
| AT4G19700 | 1,20 | 1,74 | 2,04 | 0,85 | 3,94E-01 |
| AT4G19720 | 0,28 | -0,77 | 4,32 | -0,18 | 8,58E-01 |
| AT4G19860 | 1,15 | -0,25 | 1,87 | -0,13 | 8,95E-01 |
| AT4G19870 | 0,10 | 0,24 | 4,38 | 0,05 | 9,57E-01 |
| AT4G19880 | 0,52 | -0,17 | 2,62 | -0,06 | 9,49E-01 |
| AT4G19890 | 0,56 | -0,37 | 3,16 | -0,12 | 9,07E-01 |
| AT4G19900 | 0,17 | 0,00 | 4,41 | 0,00 | 1,00E+00 |
| AT4G19985 | 0,13 | 0,00 | 4,41 | 0,00 | 1,00E+00 |
| AT4G20095 | 0,45 | -4,17 | 4,31 | -0,97 | 3,33E-01 |
| AT4G20150 | 0,58 | -1,86 | 2,60 | -0,72 | 4,73E-01 |
| AT4G06945 | 1,00 | 0,81 | 2,87 | 0,28 | 7,79E-01 |
| AT4G20210 | 0,49 | 0,73 | 2,56 | 0,29 | 7,74E-01 |
| AT4G20330 | 0,48 | -1,27 | 2,77 | -0,46 | 6,47E-01 |
| AT4G20410 | 1,25 | -0,94 | 2,00 | -0,47 | 6,38E-01 |
| AT4G20440 | 0,64 | 0,73 | 2,19 | 0,33 | 7,40E-01 |
| AT4G20690 | 0,21 | 0,00 | 4,41 | 0,00 | 1,00E+00 |
| AT4G20880 | 0,74 | 1,64 | 1,91 | 0,86 | 3,91E-01 |
| AT4G20910 | 0,27 | -0,39 | 4,39 | -0,09 | 9,29E-01 |
| AT4G20953 | 0,51 | 2,30 | 3,21 | 0,72 | 4,72E-01 |

| | | | | | |
|---|---|---|---|---|---|
| AT4G21100 | 1,08 | -1,36 | 1,81 | -0,75 | 4,53E-01 |
| AT4G21140 | 0,20 | 0,00 | 4,41 | 0,00 | 1,00E+00 |
| AT4G21170 | 0,28 | 0,69 | 4,29 | 0,16 | 8,72E-01 |
| AT4G06985 | 0,08 | 0,00 | 4,41 | 0,00 | 1,00E+00 |
| AT4G21190 | 0,08 | 0,00 | 4,41 | 0,00 | 1,00E+00 |
| AT4G21350 | 0,08 | 0,00 | 4,41 | 0,00 | 1,00E+00 |
| AT4G21380 | 1,64 | 1,92 | 1,78 | 1,08 | 2,82E-01 |
| AT4G21400 | 0,16 | -0,21 | 4,40 | -0,05 | 9,63E-01 |
| AT4G21430 | 0,22 | 0,26 | 4,34 | 0,06 | 9,52E-01 |
| AT4G21510 | 0,05 | 0,18 | 4,38 | 0,04 | 9,68E-01 |
| AT4G21520 | 0,22 | 0,08 | 4,38 | 0,02 | 9,86E-01 |
| AT4G21560 | 0,38 | 0,41 | 4,37 | 0,09 | 9,25E-01 |
| AT4G21570 | 1,42 | 0,44 | 1,95 | 0,23 | 8,22E-01 |
| AT4G21660 | 1,13 | 1,45 | 2,25 | 0,64 | 5,20E-01 |
| AT4G21810 | 1,02 | 0,42 | 2,14 | 0,19 | 8,46E-01 |
| AT4G21860 | 0,50 | -2,21 | 3,35 | -0,66 | 5,10E-01 |
| AT4G21890 | 0,12 | 0,00 | 4,41 | 0,00 | 1,00E+00 |
| AT4G21910 | 0,54 | -0,36 | 4,33 | -0,08 | 9,33E-01 |
| AT4G21990 | 0,81 | 1,00 | 2,58 | 0,39 | 6,97E-01 |
| AT4G22120 | 0,12 | 0,00 | 4,41 | 0,00 | 1,00E+00 |
| AT4G22150 | 0,22 | -1,49 | 4,36 | -0,34 | 7,33E-01 |
| AT4G22190 | 0,98 | -1,26 | 1,91 | -0,66 | 5,08E-01 |
| AT4G22200 | 0,21 | 0,28 | 4,37 | 0,06 | 9,49E-01 |
| AT4G22240 | 1,83 | -0,69 | 1,29 | -0,53 | 5,94E-01 |
| AT4G22250 | 0,16 | 0,15 | 4,38 | 0,04 | 9,72E-01 |
| AT4G22260 | 0,18 | -0,12 | 4,35 | -0,03 | 9,78E-01 |
| AT4G22265 | 0,20 | 0,00 | 4,41 | 0,00 | 1,00E+00 |
| AT4G22285 | 0,20 | -1,20 | 4,38 | -0,27 | 7,85E-01 |
| AT4G22290 | 0,23 | 1,67 | 4,31 | 0,39 | 6,99E-01 |
| AT4G22350 | 0,69 | 0,73 | 3,65 | 0,20 | 8,42E-01 |
| AT4G22380 | 0,51 | 2,13 | 2,60 | 0,82 | 4,15E-01 |
| AT4G22470 | 0,12 | 0,11 | 4,38 | 0,03 | 9,80E-01 |
| AT4G22485 | 0,49 | 1,74 | 3,82 | 0,46 | 6,48E-01 |
| AT4G22550 | 0,26 | -0,45 | 3,79 | -0,12 | 9,06E-01 |
| AT4G22570 | 0,24 | -0,34 | 4,40 | -0,08 | 9,38E-01 |
| AT4G22745 | 0,42 | 0,00 | 4,19 | 0,00 | 1,00E+00 |
| AT4G22750 | 1,03 | -1,57 | 2,31 | -0,68 | 4,98E-01 |
| AT4G22755 | 0,47 | 0,00 | 4,41 | 0,00 | 1,00E+00 |
| AT4G22770 | 1,29 | -0,45 | 1,50 | -0,30 | 7,64E-01 |
| AT4G22780 | 0,24 | 0,25 | 4,37 | 0,06 | 9,53E-01 |
| AT4G22970 | 0,52 | 0,16 | 3,90 | 0,04 | 9,68E-01 |
| AT4G22980 | 0,47 | 1,39 | 3,95 | 0,35 | 7,25E-01 |
| AT4G23010 | 0,83 | -0,32 | 2,48 | -0,13 | 8,98E-01 |
| AT4G23040 | 0,28 | -2,11 | 4,37 | -0,48 | 6,30E-01 |
| AT4G23100 | 1,83 | -0,16 | 1,36 | -0,12 | 9,06E-01 |
| AT4G23130 | 0,55 | -0,35 | 3,99 | -0,09 | 9,31E-01 |
| AT4G23190 | 1,18 | 2,27 | 1,96 | 1,16 | 2,46E-01 |
| AT4G23205 | 0,08 | 0,00 | 4,41 | 0,00 | 1,00E+00 |
| AT4G23200 | 0,43 | 0,00 | 4,40 | 0,00 | 1,00E+00 |
| AT4G23210 | 0,21 | -0,32 | 4,40 | -0,07 | 9,43E-01 |
| AT4G23220 | 0,76 | 1,10 | 2,43 | 0,45 | 6,51E-01 |
| AT4G23240 | 0,37 | 0,00 | 4,41 | 0,00 | 1,00E+00 |
| AT4G23250 | 0,45 | 1,36 | 3,94 | 0,34 | 7,30E-01 |
| AT4G23260 | 0,19 | -0,40 | 4,39 | -0,09 | 9,28E-01 |
| AT4G23290 | 0,10 | 0,86 | 4,35 | 0,20 | 8,43E-01 |
| AT4G23320 | 0,08 | 0,00 | 4,41 | 0,00 | 1,00E+00 |

| | | | | | |
|---|---|---|---|---|---|
| AT4G23460 | 1,64 | 0,19 | 1,75 | 0,11 | 9,15E-01 |
| AT4G23490 | 0,15 | -0,57 | 4,37 | -0,13 | 8,96E-01 |
| AT4G07130 | 0,46 | -0,10 | 4,19 | -0,03 | 9,80E-01 |
| AT4G23540 | 0,67 | -1,47 | 3,14 | -0,47 | 6,40E-01 |
| AT4G23560 | 0,10 | 0,86 | 4,35 | 0,20 | 8,43E-01 |
| AT4G07135 | 1,25 | 0,95 | 2,12 | 0,45 | 6,53E-01 |
| AT4G07145 | 0,96 | 0,00 | 2,79 | 0,00 | 1,00E+00 |
| AT4G23620 | 0,31 | 0,48 | 4,37 | 0,11 | 9,12E-01 |
| AT4G23640 | 0,77 | 0,37 | 2,40 | 0,15 | 8,78E-01 |
| AT4G23670 | 1,50 | -1,31 | 1,44 | -0,91 | 3,61E-01 |
| AT4G23700 | 0,29 | 0,00 | 4,41 | 0,00 | 1,00E+00 |
| AT4G23770 | 0,22 | 0,07 | 4,34 | 0,02 | 9,87E-01 |
| AT4G23810 | 0,35 | 0,39 | 4,38 | 0,09 | 9,30E-01 |
| AT4G23820 | 0,63 | 1,06 | 2,19 | 0,49 | 6,27E-01 |
| AT4G23880 | 0,47 | 1,54 | 3,10 | 0,50 | 6,18E-01 |
| AT4G23890 | 0,66 | -1,79 | 2,65 | -0,67 | 5,01E-01 |
| AT4G24180 | 0,37 | 0,00 | 4,41 | 0,00 | 1,00E+00 |
| AT4G24200 | 0,70 | 0,63 | 3,01 | 0,21 | 8,34E-01 |
| AT4G07250 | 0,17 | -3,06 | 4,35 | -0,70 | 4,82E-01 |
| AT4G24300 | 0,45 | 2,70 | 4,28 | 0,63 | 5,29E-01 |
| AT4G24320 | 0,33 | -2,16 | 4,40 | -0,49 | 6,23E-01 |
| AT4G24330 | 0,84 | 0,73 | 1,84 | 0,40 | 6,90E-01 |
| AT4G07255 | 0,10 | 0,86 | 4,35 | 0,20 | 8,43E-01 |
| AT4G24390 | 0,29 | 0,35 | 4,37 | 0,08 | 9,35E-01 |
| AT4G24490 | 0,72 | -0,36 | 2,54 | -0,14 | 8,87E-01 |
| AT4G24530 | 0,08 | 0,00 | 4,41 | 0,00 | 1,00E+00 |
| AT4G24560 | 1,21 | -1,39 | 2,13 | -0,65 | 5,15E-01 |
| AT4G24580 | 0,46 | 0,35 | 4,14 | 0,09 | 9,32E-01 |
| AT4G24610 | 0,67 | 0,00 | 3,05 | 0,00 | 1,00E+00 |
| AT4G24620 | 0,42 | -0,16 | 4,19 | -0,04 | 9,70E-01 |
| AT4G24660 | 1,04 | -0,03 | 2,03 | -0,01 | 9,90E-01 |
| AT4G24670 | 0,08 | 0,21 | 4,39 | 0,05 | 9,62E-01 |
| AT4G24730 | 0,45 | 0,00 | 4,14 | 0,00 | 1,00E+00 |
| AT4G24740 | 0,63 | 0,49 | 3,14 | 0,16 | 8,76E-01 |
| AT4G24750 | 0,28 | 0,00 | 4,41 | 0,00 | 1,00E+00 |
| AT4G24780 | 1,27 | 2,80 | 2,15 | 1,30 | 1,92E-01 |
| AT4G07285 | 0,27 | -1,39 | 4,30 | -0,32 | 7,47E-01 |
| AT4G24820 | 1,43 | 1,05 | 1,82 | 0,58 | 5,65E-01 |
| AT4G24830 | 0,25 | 0,55 | 4,31 | 0,13 | 8,99E-01 |
| AT4G24840 | 0,53 | 0,31 | 3,07 | 0,10 | 9,18E-01 |
| AT4G24920 | 0,86 | 0,00 | 2,88 | 0,00 | 1,00E+00 |
| AT4G24990 | 0,31 | 1,06 | 4,34 | 0,24 | 8,08E-01 |
| AT4G25090 | 0,29 | 0,00 | 4,41 | 0,00 | 1,00E+00 |
| AT4G25270 | 0,45 | 0,00 | 4,14 | 0,00 | 1,00E+00 |
| AT4G25280 | 0,71 | 0,00 | 4,41 | 0,00 | 1,00E+00 |
| AT4G25290 | 0,52 | -0,38 | 3,91 | -0,10 | 9,23E-01 |
| AT4G25340 | 0,19 | -1,29 | 4,39 | -0,29 | 7,69E-01 |
| AT4G25435 | 0,88 | -4,09 | 2,69 | -1,52 | 1,29E-01 |
| AT4G25540 | 0,13 | 0,00 | 4,41 | 0,00 | 1,00E+00 |
| AT4G25570 | 0,43 | -1,12 | 4,15 | -0,27 | 7,88E-01 |
| AT4G25640 | 1,48 | 0,32 | 1,79 | 0,18 | 8,58E-01 |
| AT4G25650 | 1,11 | -1,87 | 1,64 | -1,15 | 2,52E-01 |
| AT4G25670 | 0,79 | -0,26 | 2,96 | -0,09 | 9,29E-01 |
| AT4G25690 | 0,56 | -0,48 | 3,30 | -0,15 | 8,84E-01 |
| AT4G25730 | 1,71 | -0,29 | 1,93 | -0,15 | 8,80E-01 |
| AT4G25740 | 0,38 | 0,48 | 4,38 | 0,11 | 9,13E-01 |

| | | | | | |
|---|---|---|---|---|---|
| AT4G25870 | 0,26 | 0,00 | 4,41 | 0,00 | 1,00E+00 |
| AT4G25940 | 0,39 | -2,19 | 4,41 | -0,50 | 6,20E-01 |
| AT4G25990 | 0,26 | 1,22 | 4,34 | 0,28 | 7,79E-01 |
| AT4G26080 | 0,87 | -2,46 | 2,46 | -1,00 | 3,17E-01 |
| AT4G26095 | 1,81 | 1,42 | 1,62 | 0,88 | 3,79E-01 |
| AT4G26100 | 0,81 | 0,37 | 2,13 | 0,17 | 8,61E-01 |
| AT4G26130 | 1,45 | -2,40 | 1,64 | -1,46 | 1,44E-01 |
| AT4G26140 | 0,21 | 0,08 | 4,38 | 0,02 | 9,85E-01 |
| AT4G26190 | 0,25 | 1,09 | 4,34 | 0,25 | 8,02E-01 |
| AT4G26300 | 0,69 | -1,65 | 2,66 | -0,62 | 5,36E-01 |
| AT4G07565 | 0,79 | 0,88 | 1,90 | 0,46 | 6,44E-01 |
| AT4G07625 | 0,79 | -1,19 | 2,57 | -0,46 | 6,44E-01 |
| AT4G07635 | 0,12 | 0,11 | 4,38 | 0,03 | 9,80E-01 |
| AT4G26370 | 0,48 | -1,50 | 4,02 | -0,37 | 7,10E-01 |
| AT4G26375 | 0,40 | 0,00 | 4,41 | 0,00 | 1,00E+00 |
| AT4G26400 | 0,60 | -0,26 | 4,00 | -0,07 | 9,48E-01 |
| AT4G26620 | 0,28 | -1,26 | 4,37 | -0,29 | 7,73E-01 |
| AT4G26630 | 0,72 | 1,77 | 2,46 | 0,72 | 4,70E-01 |
| AT4G26640 | 1,00 | -1,27 | 2,39 | -0,53 | 5,95E-01 |
| AT4G26690 | 1,51 | 1,30 | 1,68 | 0,77 | 4,40E-01 |
| AT4G26750 | 0,43 | -1,36 | 4,22 | -0,32 | 7,47E-01 |
| AT4G26760 | 0,64 | -1,16 | 3,17 | -0,37 | 7,14E-01 |
| AT4G26795 | 0,42 | 2,35 | 4,03 | 0,58 | 5,59E-01 |
| AT4G26860 | 0,37 | -0,17 | 4,34 | -0,04 | 9,68E-01 |
| AT4G26910 | 1,33 | -1,92 | 1,90 | -1,01 | 3,11E-01 |
| AT4G26940 | 0,43 | -1,29 | 4,11 | -0,31 | 7,53E-01 |
| AT4G26980 | 0,78 | -0,26 | 2,57 | -0,10 | 9,19E-01 |
| AT4G27020 | 0,48 | 0,00 | 3,39 | 0,00 | 9,99E-01 |
| AT4G07765 | 0,66 | 2,76 | 2,77 | 1,00 | 3,18E-01 |
| AT4G27060 | 1,71 | 1,38 | 1,83 | 0,76 | 4,50E-01 |
| AT4G27130 | 0,28 | 1,81 | 3,65 | 0,50 | 6,19E-01 |
| AT4G27270 | 0,66 | 0,09 | 3,91 | 0,02 | 9,81E-01 |
| AT4G27280 | 0,84 | 3,22 | 2,74 | 1,17 | 2,41E-01 |
| AT4G27300 | 0,83 | -0,32 | 2,37 | -0,14 | 8,91E-01 |
| AT4G27320 | 1,19 | -0,99 | 2,26 | -0,44 | 6,62E-01 |
| AT4G27340 | 0,21 | 0,12 | 4,38 | 0,03 | 9,78E-01 |
| AT4G27410 | 0,98 | 0,00 | 2,50 | 0,00 | 1,00E+00 |
| AT4G27450 | 0,72 | 0,05 | 2,35 | 0,02 | 9,82E-01 |
| AT4G27480 | 0,65 | 0,00 | 3,67 | 0,00 | 1,00E+00 |
| AT4G27650 | 0,39 | -1,66 | 3,45 | -0,48 | 6,30E-01 |
| AT4G27700 | 0,47 | 0,95 | 3,29 | 0,29 | 7,73E-01 |
| AT4G27730 | 0,37 | 0,00 | 4,41 | 0,00 | 1,00E+00 |
| AT4G27800 | 1,09 | -1,94 | 2,00 | -0,97 | 3,31E-01 |
| AT4G27830 | 0,84 | 1,12 | 1,95 | 0,57 | 5,67E-01 |
| AT4G27840 | 0,43 | 0,51 | 4,36 | 0,12 | 9,07E-01 |
| AT4G27885 | 0,16 | 1,26 | 4,33 | 0,29 | 7,71E-01 |
| AT4G27960 | 0,38 | 2,35 | 4,22 | 0,56 | 5,78E-01 |
| AT4G28030 | 0,45 | -0,44 | 2,80 | -0,16 | 8,74E-01 |
| AT4G28050 | 0,43 | -2,04 | 4,38 | -0,47 | 6,42E-01 |
| AT4G28060 | 0,25 | 0,00 | 4,41 | 0,00 | 1,00E+00 |
| AT4G28180 | 0,69 | 2,11 | 2,51 | 0,84 | 4,00E-01 |
| AT4G28200 | 0,59 | -2,30 | 2,70 | -0,85 | 3,94E-01 |
| AT4G28230 | 0,33 | 0,00 | 4,41 | 0,00 | 1,00E+00 |
| AT4G28240 | 0,66 | 0,31 | 2,13 | 0,15 | 8,85E-01 |
| AT4G28320 | 0,20 | 0,00 | 4,41 | 0,00 | 1,00E+00 |
| AT4G28390 | 0,33 | 0,00 | 4,41 | 0,00 | 1,00E+00 |

| | | | | | |
|---|---|---|---|---|---|
| AT4G28450 | 0,61 | 0,00 | 4,41 | 0,00 | 1,00E+00 |
| AT4G28470 | 0,62 | 0,42 | 3,18 | 0,13 | 8,95E-01 |
| AT4G28610 | 0,93 | 1,17 | 2,65 | 0,44 | 6,59E-01 |
| AT4G28630 | 0,12 | 0,00 | 4,41 | 0,00 | 1,00E+00 |
| AT4G28770 | 0,20 | 0,89 | 4,36 | 0,20 | 8,39E-01 |
| AT4G28910 | 1,06 | -2,97 | 2,48 | -1,20 | 2,30E-01 |
| AT4G29010 | 1,63 | 0,02 | 1,57 | 0,01 | 9,90E-01 |
| AT4G29050 | 0,13 | -0,61 | 4,37 | -0,14 | 8,89E-01 |
| AT4G29080 | 0,28 | 0,66 | 4,35 | 0,15 | 8,79E-01 |
| AT4G29130 | 1,04 | -3,10 | 1,88 | -1,65 | 9,88E-02 |
| AT4G29190 | 1,79 | -1,68 | 1,48 | -1,14 | 2,55E-01 |
| AT4G29220 | 0,63 | -1,04 | 3,07 | -0,34 | 7,35E-01 |
| AT4G29260 | 0,48 | 2,83 | 3,94 | 0,72 | 4,73E-01 |
| AT4G29300 | 1,28 | -0,59 | 1,46 | -0,41 | 6,84E-01 |
| AT4G29305 | 0,76 | 2,34 | 2,30 | 1,02 | 3,09E-01 |
| AT4G08155 | 1,40 | 2,55 | 1,67 | 1,53 | 1,27E-01 |
| AT4G29440 | 0,66 | 1,24 | 2,98 | 0,42 | 6,78E-01 |
| AT4G29520 | 0,77 | 0,27 | 2,63 | 0,10 | 9,19E-01 |
| AT4G29654 | 1,07 | 3,34 | 2,57 | 1,30 | 1,94E-01 |
| AT4G29670 | 0,65 | 0,45 | 2,31 | 0,20 | 8,45E-01 |
| AT4G29680 | 0,85 | -0,60 | 3,07 | -0,20 | 8,45E-01 |
| AT4G29720 | 0,17 | -2,13 | 4,40 | -0,48 | 6,28E-01 |
| AT4G29735 | 0,17 | -2,21 | 4,40 | -0,50 | 6,16E-01 |
| AT4G29810 | 0,27 | 0,57 | 4,37 | 0,13 | 8,97E-01 |
| AT4G29820 | 0,24 | 0,59 | 4,36 | 0,14 | 8,92E-01 |
| AT4G29840 | 0,58 | -0,88 | 2,31 | -0,38 | 7,04E-01 |
| AT4G29880 | 0,21 | 0,50 | 4,37 | 0,11 | 9,09E-01 |
| AT4G29905 | 1,17 | 1,61 | 1,76 | 0,92 | 3,60E-01 |
| AT4G29910 | 0,08 | 0,21 | 4,39 | 0,05 | 9,62E-01 |
| AT4G29920 | 0,05 | 0,18 | 4,38 | 0,04 | 9,68E-01 |
| AT4G29940 | 0,28 | 0,00 | 4,41 | 0,00 | 1,00E+00 |
| AT4G29960 | 0,50 | 0,00 | 4,00 | 0,00 | 1,00E+00 |
| AT4G29990 | 0,08 | 0,00 | 4,41 | 0,00 | 1,00E+00 |
| AT4G30000 | 0,38 | 0,00 | 4,41 | 0,00 | 1,00E+00 |
| AT4G30030 | 0,45 | 0,72 | 4,35 | 0,17 | 8,68E-01 |
| AT4G30060 | 0,08 | 0,00 | 4,41 | 0,00 | 1,00E+00 |
| AT4G30080 | 0,31 | 0,81 | 4,35 | 0,19 | 8,52E-01 |
| AT4G30110 | 1,05 | 0,00 | 3,27 | 0,00 | 1,00E+00 |
| AT4G30200 | 1,47 | -0,22 | 1,74 | -0,13 | 8,97E-01 |
| AT4G30240 | 0,30 | 0,00 | 4,37 | 0,00 | 9,99E-01 |
| AT4G30260 | 0,47 | -1,42 | 3,24 | -0,44 | 6,62E-01 |
| AT4G30270 | 1,68 | 0,83 | 1,51 | 0,55 | 5,83E-01 |
| AT4G30280 | 0,12 | 0,49 | 4,38 | 0,11 | 9,11E-01 |
| AT4G30360 | 0,21 | 1,90 | 4,30 | 0,44 | 6,58E-01 |
| AT4G30410 | 0,12 | 0,27 | 4,38 | 0,06 | 9,50E-01 |
| AT4G30430 | 0,08 | 0,00 | 4,41 | 0,00 | 1,00E+00 |
| AT4G30560 | 0,74 | 0,00 | 3,59 | 0,00 | 1,00E+00 |
| AT4G30580 | 0,92 | 0,00 | 3,50 | 0,00 | 1,00E+00 |
| AT4G30600 | 1,60 | 1,45 | 1,66 | 0,87 | 3,82E-01 |
| AT4G30620 | 0,23 | 0,24 | 4,39 | 0,06 | 9,56E-01 |
| AT4G30700 | 0,08 | 0,00 | 4,41 | 0,00 | 1,00E+00 |
| AT4G30710 | 0,12 | 0,00 | 4,41 | 0,00 | 1,00E+00 |
| AT4G30720 | 1,29 | 0,94 | 1,90 | 0,49 | 6,23E-01 |
| AT4G30740 | 1,03 | 3,84 | 2,34 | 1,64 | 1,01E-01 |
| AT4G30790 | 0,99 | 1,97 | 2,24 | 0,88 | 3,79E-01 |
| AT4G30890 | 0,60 | 0,52 | 3,04 | 0,17 | 8,65E-01 |

| | | | | | |
|---|---|---|---|---|---|
| AT4G30920 | 1,19 | -1,81 | 1,83 | -0,99 | 3,24E-01 |
| AT4G30950 | 1,64 | -1,84 | 1,56 | -1,18 | 2,38E-01 |
| AT4G31020 | 0,29 | 0,00 | 4,41 | 0,00 | 1,00E+00 |
| AT4G31120 | 0,87 | -1,02 | 2,58 | -0,40 | 6,92E-01 |
| AT4G31130 | 0,59 | -0,90 | 3,15 | -0,29 | 7,74E-01 |
| AT4G31420 | 0,28 | 0,02 | 4,38 | 0,01 | 9,96E-01 |
| AT4G31450 | 0,46 | -3,40 | 4,14 | -0,82 | 4,11E-01 |
| AT4G31540 | 0,43 | 0,13 | 4,13 | 0,03 | 9,74E-01 |
| AT4G31550 | 1,02 | 0,07 | 1,87 | 0,04 | 9,70E-01 |
| AT4G31590 | 0,63 | 1,14 | 2,62 | 0,43 | 6,65E-01 |
| AT4G31700 | 1,83 | 0,52 | 1,42 | 0,37 | 7,11E-01 |
| AT4G31810 | 0,08 | 0,00 | 4,41 | 0,00 | 1,00E+00 |
| AT4G31820 | 0,43 | -0,51 | 2,85 | -0,18 | 8,59E-01 |
| AT4G31990 | 1,81 | -0,77 | 1,79 | -0,43 | 6,69E-01 |
| AT4G32070 | 0,58 | -0,26 | 3,96 | -0,07 | 9,47E-01 |
| AT4G32120 | 0,15 | 0,47 | 4,37 | 0,11 | 9,15E-01 |
| AT4G32130 | 0,18 | 1,20 | 4,34 | 0,28 | 7,82E-01 |
| AT4G32150 | 1,17 | 1,26 | 1,78 | 0,70 | 4,81E-01 |
| AT4G32180 | 1,00 | -3,43 | 2,58 | -1,33 | 1,84E-01 |
| AT4G32208 | 0,37 | 0,00 | 4,36 | 0,00 | 1,00E+00 |
| AT4G32340 | 1,02 | 1,74 | 1,85 | 0,94 | 3,47E-01 |
| AT4G32420 | 0,78 | -1,41 | 2,58 | -0,55 | 5,84E-01 |
| AT4G32460 | 0,31 | -0,89 | 4,38 | -0,20 | 8,39E-01 |
| AT4G32470 | 0,29 | 1,27 | 4,33 | 0,29 | 7,69E-01 |
| AT4G32510 | 0,56 | 0,00 | 3,84 | 0,00 | 1,00E+00 |
| AT4G32520 | 1,00 | -0,97 | 1,96 | -0,49 | 6,22E-01 |
| AT4G32530 | 0,89 | -1,02 | 2,56 | -0,40 | 6,90E-01 |
| AT4G32670 | 0,15 | -0,39 | 4,39 | -0,09 | 9,29E-01 |
| AT4G32750 | 0,18 | -1,13 | 4,39 | -0,26 | 7,96E-01 |
| AT4G32820 | 0,83 | -0,40 | 2,79 | -0,14 | 8,85E-01 |
| AT4G32880 | 1,09 | 0,95 | 2,12 | 0,45 | 6,55E-01 |
| AT4G32900 | 0,12 | -0,26 | 4,41 | -0,06 | 9,53E-01 |
| AT4G32920 | 0,70 | 0,88 | 2,87 | 0,31 | 7,60E-01 |
| AT4G33040 | 0,37 | -1,17 | 4,39 | -0,27 | 7,90E-01 |
| AT4G33090 | 0,64 | 0,14 | 2,98 | 0,05 | 9,63E-01 |
| AT4G33110 | 0,17 | 0,66 | 4,36 | 0,15 | 8,79E-01 |
| AT4G33150 | 0,65 | 0,00 | 3,73 | 0,00 | 1,00E+00 |
| AT4G33200 | 0,29 | 0,00 | 4,41 | 0,00 | 1,00E+00 |
| AT4G33210 | 1,11 | -0,26 | 2,20 | -0,12 | 9,05E-01 |
| AT4G33250 | 0,89 | 0,07 | 2,08 | 0,03 | 9,74E-01 |
| AT4G33260 | 0,12 | 0,00 | 4,41 | 0,00 | 1,00E+00 |
| AT4G33270 | 0,12 | 0,49 | 4,38 | 0,11 | 9,11E-01 |
| AT4G33400 | 0,33 | -2,09 | 4,39 | -0,48 | 6,35E-01 |
| AT4G33430 | 1,19 | 1,19 | 1,98 | 0,60 | 5,47E-01 |
| AT4G33470 | 0,67 | -0,40 | 3,00 | -0,13 | 8,95E-01 |
| AT4G33495 | 0,29 | 0,00 | 4,41 | 0,00 | 1,00E+00 |
| AT4G33500 | 0,96 | -2,01 | 2,53 | -0,79 | 4,27E-01 |
| AT4G33530 | 1,44 | -1,31 | 1,71 | -0,77 | 4,43E-01 |
| AT4G33585 | 1,20 | 0,12 | 3,02 | 0,04 | 9,69E-01 |
| AT4G33625 | 0,16 | 0,35 | 4,37 | 0,08 | 9,37E-01 |
| AT4G33670 | 0,76 | 0,83 | 2,20 | 0,38 | 7,05E-01 |
| AT4G33680 | 0,20 | -0,15 | 4,39 | -0,03 | 9,72E-01 |
| AT4G33700 | 0,28 | -1,06 | 4,33 | -0,25 | 8,06E-01 |
| AT4G33790 | 0,16 | -0,21 | 4,40 | -0,05 | 9,63E-01 |
| AT4G33860 | 0,68 | 0,00 | 3,71 | 0,00 | 1,00E+00 |
| AT4G33865 | 0,81 | -0,67 | 2,13 | -0,31 | 7,55E-01 |

| | | | | | |
|---|---|---|---|---|---|
| AT4G33910 | 0,26 | 0,00 | 4,41 | 0,00 | 1,00E+00 |
| AT4G34020 | 0,41 | -3,29 | 4,36 | -0,75 | 4,50E-01 |
| AT4G34040 | 0,82 | 0,83 | 2,37 | 0,35 | 7,27E-01 |
| AT4G34070 | 0,72 | -2,50 | 2,66 | -0,94 | 3,49E-01 |
| AT4G34090 | 0,71 | -2,04 | 4,38 | -0,47 | 6,42E-01 |
| AT4G34131 | 0,52 | 0,00 | 3,96 | 0,00 | 1,00E+00 |
| AT4G34180 | 0,68 | -1,16 | 2,93 | -0,40 | 6,91E-01 |
| AT4G34190 | 1,52 | 1,80 | 1,77 | 1,02 | 3,07E-01 |
| AT4G34200 | 1,37 | 0,19 | 1,88 | 0,10 | 9,20E-01 |
| AT4G34220 | 1,59 | 0,90 | 1,54 | 0,58 | 5,59E-01 |
| AT4G34332 | 0,08 | 0,00 | 4,41 | 0,00 | 1,00E+00 |
| AT4G34340 | 0,12 | 0,00 | 4,41 | 0,00 | 1,00E+00 |
| AT4G34350 | 1,55 | -0,01 | 1,65 | 0,00 | 9,97E-01 |
| AT4G08985 | 0,45 | 2,58 | 3,16 | 0,81 | 4,15E-01 |
| AT4G34460 | 0,42 | 1,30 | 3,27 | 0,40 | 6,90E-01 |
| AT4G34480 | 1,19 | 0,64 | 1,91 | 0,33 | 7,40E-01 |
| AT4G34490 | 1,39 | -0,51 | 1,72 | -0,29 | 7,69E-01 |
| AT4G34610 | 0,37 | 0,50 | 4,36 | 0,11 | 9,09E-01 |
| AT4G34620 | 1,51 | -0,67 | 1,93 | -0,35 | 7,29E-01 |
| AT4G34630 | 0,16 | 0,74 | 4,36 | 0,17 | 8,66E-01 |
| AT4G34690 | 1,55 | 3,08 | 1,88 | 1,64 | 1,02E-01 |
| AT4G34720 | 1,83 | -1,28 | 1,25 | -1,02 | 3,08E-01 |
| AT4G34730 | 0,55 | 0,72 | 2,60 | 0,28 | 7,82E-01 |
| AT4G34740 | 0,69 | -1,61 | 2,89 | -0,56 | 5,77E-01 |
| AT4G34860 | 0,08 | 0,00 | 4,41 | 0,00 | 1,00E+00 |
| AT4G34900 | 0,65 | 0,00 | 3,69 | 0,00 | 1,00E+00 |
| AT4G35060 | 0,12 | 0,00 | 4,41 | 0,00 | 1,00E+00 |
| AT4G35140 | 0,36 | -1,27 | 4,34 | -0,29 | 7,71E-01 |
| AT4G35180 | 0,66 | 0,12 | 3,01 | 0,04 | 9,69E-01 |
| AT4G35230 | 1,11 | 2,63 | 2,00 | 1,32 | 1,88E-01 |
| AT4G35240 | 1,02 | -0,38 | 2,21 | -0,17 | 8,64E-01 |
| AT4G35260 | 0,53 | -0,26 | 3,05 | -0,09 | 9,32E-01 |
| AT4G35300 | 1,43 | -0,67 | 2,02 | -0,33 | 7,39E-01 |
| AT4G35320 | 0,51 | 2,57 | 3,09 | 0,83 | 4,04E-01 |
| AT4G35360 | 0,54 | -2,11 | 4,03 | -0,52 | 6,01E-01 |
| AT4G35440 | 0,24 | 1,13 | 4,34 | 0,26 | 7,95E-01 |
| AT4G35570 | 0,29 | -0,12 | 4,39 | -0,03 | 9,79E-01 |
| AT4G35580 | 0,56 | -0,33 | 3,09 | -0,11 | 9,14E-01 |
| AT4G35780 | 0,15 | -0,31 | 4,40 | -0,07 | 9,43E-01 |
| AT4G35785 | 0,84 | -0,26 | 2,94 | -0,09 | 9,29E-01 |
| AT4G35920 | 0,70 | 0,00 | 4,41 | 0,00 | 1,00E+00 |
| AT4G36010 | 0,74 | -0,77 | 2,64 | -0,29 | 7,71E-01 |
| AT4G36030 | 1,21 | 2,53 | 2,15 | 1,18 | 2,40E-01 |
| AT4G36050 | 0,52 | 0,00 | 4,41 | 0,00 | 1,00E+00 |
| AT4G36070 | 0,08 | 0,00 | 4,41 | 0,00 | 1,00E+00 |
| AT4G36220 | 1,38 | 0,50 | 1,54 | 0,32 | 7,47E-01 |
| AT4G36270 | 0,14 | 0,31 | 4,37 | 0,07 | 9,44E-01 |
| AT4G36280 | 0,08 | 0,00 | 4,41 | 0,00 | 1,00E+00 |
| AT4G36380 | 0,24 | -2,39 | 4,34 | -0,55 | 5,82E-01 |
| AT4G36390 | 0,60 | 1,24 | 2,57 | 0,48 | 6,30E-01 |
| AT4G36460 | 0,50 | 1,36 | 3,07 | 0,44 | 6,57E-01 |
| AT4G36500 | 0,97 | -0,04 | 2,16 | -0,02 | 9,85E-01 |
| AT4G36520 | 1,04 | 1,34 | 1,76 | 0,76 | 4,47E-01 |
| AT4G36550 | 0,53 | 1,00 | 3,18 | 0,31 | 7,54E-01 |
| AT4G36640 | 1,54 | 0,00 | 2,28 | 0,00 | 1,00E+00 |
| AT4G36670 | 0,44 | 0,65 | 3,31 | 0,20 | 8,45E-01 |

| | | | | | |
|---|---|---|---|---|---|
| AT4G36690 | 0,56 | 0,50 | 3,27 | 0,15 | 8,77E-01 |
| AT4G36730 | 0,51 | -0,84 | 3,10 | -0,27 | 7,87E-01 |
| AT4G36770 | 0,41 | 0,01 | 4,19 | 0,00 | 9,98E-01 |
| AT4G36800 | 0,60 | -1,20 | 3,07 | -0,39 | 6,96E-01 |
| AT4G36808 | 0,88 | -0,16 | 2,43 | -0,07 | 9,47E-01 |
| AT4G36860 | 0,41 | -2,03 | 4,26 | -0,48 | 6,33E-01 |
| AT4G36890 | 0,60 | 0,07 | 3,96 | 0,02 | 9,85E-01 |
| AT4G36970 | 0,18 | 0,31 | 4,38 | 0,07 | 9,44E-01 |
| AT4G36980 | 0,30 | 0,00 | 4,41 | 0,00 | 1,00E+00 |
| AT4G37100 | 0,62 | 0,54 | 3,10 | 0,18 | 8,61E-01 |
| AT4G37270 | 0,72 | -0,53 | 2,56 | -0,21 | 8,37E-01 |
| AT4G37280 | 0,63 | 0,00 | 3,73 | 0,00 | 1,00E+00 |
| AT4G37370 | 0,86 | 0,34 | 2,67 | 0,13 | 8,99E-01 |
| AT4G37432 | 0,86 | -1,12 | 3,44 | -0,33 | 7,44E-01 |
| AT4G37470 | 1,39 | -0,09 | 1,55 | -0,06 | 9,55E-01 |
| AT4G37510 | 0,72 | -0,14 | 2,47 | -0,06 | 9,55E-01 |
| AT4G37540 | 0,41 | 0,63 | 4,32 | 0,15 | 8,84E-01 |
| AT4G37553 | 0,42 | 0,00 | 4,41 | 0,00 | 1,00E+00 |
| AT4G37590 | 0,76 | 0,68 | 2,49 | 0,27 | 7,84E-01 |
| AT4G37610 | 0,33 | -0,63 | 3,61 | -0,17 | 8,62E-01 |
| AT4G37640 | 1,71 | 1,10 | 1,71 | 0,64 | 5,20E-01 |
| AT4G37740 | 0,08 | 0,21 | 4,39 | 0,05 | 9,62E-01 |
| AT4G37800 | 0,23 | 1,67 | 4,31 | 0,39 | 6,99E-01 |
| AT4G37830 | 0,60 | 0,13 | 2,61 | 0,05 | 9,60E-01 |
| AT4G37850 | 0,24 | 0,00 | 4,41 | 0,00 | 1,00E+00 |
| AT4G37870 | 0,70 | -0,61 | 2,11 | -0,29 | 7,71E-01 |
| AT4G37920 | 0,32 | -0,34 | 4,40 | -0,08 | 9,39E-01 |
| AT4G38010 | 0,08 | 0,00 | 4,41 | 0,00 | 1,00E+00 |
| AT4G38050 | 0,35 | -2,19 | 4,41 | -0,50 | 6,20E-01 |
| AT4G38120 | 0,89 | 0,40 | 2,34 | 0,17 | 8,64E-01 |
| AT4G38130 | 0,71 | -0,11 | 1,90 | -0,06 | 9,53E-01 |
| AT4G38180 | 0,60 | -0,38 | 2,35 | -0,16 | 8,72E-01 |
| AT4G38210 | 0,75 | 0,00 | 4,39 | 0,00 | 1,00E+00 |
| AT4G38230 | 0,40 | 0,43 | 3,43 | 0,13 | 9,00E-01 |
| AT4G38240 | 0,62 | 0,00 | 3,75 | 0,00 | 1,00E+00 |
| AT4G09695 | 1,25 | -0,71 | 1,72 | -0,41 | 6,81E-01 |
| AT4G38300 | 0,68 | 0,00 | 3,67 | 0,00 | 1,00E+00 |
| AT4G38350 | 1,14 | -1,45 | 1,95 | -0,74 | 4,56E-01 |
| AT4G38380 | 0,43 | -2,19 | 4,19 | -0,52 | 6,02E-01 |
| AT4G38520 | 0,55 | 1,55 | 2,30 | 0,67 | 5,01E-01 |
| AT4G38545 | 0,17 | 0,66 | 4,36 | 0,15 | 8,79E-01 |
| AT4G38570 | 0,81 | -0,16 | 2,93 | -0,05 | 9,57E-01 |
| AT4G38630 | 0,74 | -1,22 | 2,07 | -0,59 | 5,55E-01 |
| AT4G38640 | 0,43 | 0,56 | 3,38 | 0,17 | 8,69E-01 |
| AT4G38660 | 0,37 | 0,83 | 4,36 | 0,19 | 8,48E-01 |
| AT4G38670 | 0,49 | -2,07 | 3,23 | -0,64 | 5,22E-01 |
| AT4G38680 | 1,18 | 0,90 | 1,56 | 0,58 | 5,63E-01 |
| AT4G38700 | 0,34 | 1,59 | 4,30 | 0,37 | 7,12E-01 |
| AT4G38710 | 1,00 | 0,22 | 1,54 | 0,15 | 8,84E-01 |
| AT4G38730 | 0,38 | 0,00 | 4,41 | 0,00 | 1,00E+00 |
| AT4G38740 | 1,15 | 2,17 | 1,92 | 1,13 | 2,58E-01 |
| AT4G38780 | 1,10 | 0,66 | 2,63 | 0,25 | 8,03E-01 |
| AT4G38800 | 0,65 | 0,08 | 2,49 | 0,03 | 9,75E-01 |
| AT4G38810 | 0,68 | -1,41 | 2,53 | -0,56 | 5,76E-01 |
| AT4G38870 | 0,21 | -0,32 | 4,40 | -0,07 | 9,43E-01 |
| AT4G38890 | 0,48 | 0,77 | 3,25 | 0,24 | 8,13E-01 |

| | | | | | |
|---|---|---|---|---|---|
| AT4G38900 | 0,33 | 0,03 | 4,38 | 0,01 | 9,95E-01 |
| AT4G38932 | 0,58 | -0,32 | 2,74 | -0,12 | 9,06E-01 |
| AT4G38950 | 1,10 | -0,26 | 2,24 | -0,12 | 9,07E-01 |
| AT4G39100 | 0,33 | 0,00 | 4,41 | 0,00 | 1,00E+00 |
| AT4G39120 | 0,19 | -0,23 | 4,40 | -0,05 | 9,58E-01 |
| AT4G39140 | 0,65 | 0,93 | 3,86 | 0,24 | 8,09E-01 |
| AT4G39180 | 0,48 | 0,12 | 4,38 | 0,03 | 9,79E-01 |
| AT4G39190 | 0,69 | 3,16 | 2,37 | 1,33 | 1,82E-01 |
| AT4G39195 | 0,79 | -0,26 | 3,03 | -0,09 | 9,31E-01 |
| AT4G39240 | 1,33 | -0,26 | 2,41 | -0,11 | 9,14E-01 |
| AT4G39280 | 1,06 | 1,13 | 1,63 | 0,69 | 4,89E-01 |
| AT4G39460 | 0,85 | -2,04 | 2,51 | -0,81 | 4,18E-01 |
| AT4G39500 | 0,20 | 0,00 | 4,41 | 0,00 | 1,00E+00 |
| AT4G39510 | 0,43 | -2,64 | 4,37 | -0,60 | 5,45E-01 |
| AT4G39520 | 1,02 | -0,59 | 1,58 | -0,37 | 7,09E-01 |
| AT4G39550 | 0,17 | 0,00 | 4,41 | 0,00 | 1,00E+00 |
| AT4G39560 | 0,59 | 0,00 | 3,86 | 0,00 | 1,00E+00 |
| AT4G39570 | 0,55 | -2,06 | 3,81 | -0,54 | 5,90E-01 |
| AT4G39580 | 0,15 | -0,39 | 4,39 | -0,09 | 9,29E-01 |
| AT4G39590 | 0,08 | 0,00 | 4,41 | 0,00 | 1,00E+00 |
| AT4G39630 | 0,05 | 0,18 | 4,38 | 0,04 | 9,68E-01 |
| AT4G39680 | 0,30 | 1,93 | 4,30 | 0,45 | 6,54E-01 |
| AT4G39710 | 0,56 | 1,26 | 3,88 | 0,33 | 7,45E-01 |
| AT4G39780 | 0,21 | -0,95 | 4,38 | -0,22 | 8,29E-01 |
| AT4G39820 | 0,46 | -0,29 | 4,12 | -0,07 | 9,44E-01 |
| AT4G39838 | 1,07 | 2,64 | 1,95 | 1,36 | 1,75E-01 |
| AT4G39865 | 0,25 | 0,00 | 4,41 | 0,00 | 1,00E+00 |
| AT4G39910 | 0,28 | -0,90 | 4,37 | -0,21 | 8,37E-01 |
| AT4G40000 | 0,43 | -2,19 | 4,19 | -0,52 | 6,02E-01 |
| AT4G40020 | 0,75 | 0,00 | 2,91 | 0,00 | 1,00E+00 |
| AT4G40030 | 1,32 | 0,67 | 1,84 | 0,36 | 7,16E-01 |
| AT4G40045 | 0,12 | 0,00 | 4,41 | 0,00 | 1,00E+00 |
| AT4G40050 | 1,65 | 1,51 | 1,82 | 0,83 | 4,04E-01 |
| AT4G40060 | 1,22 | 2,52 | 1,80 | 1,40 | 1,61E-01 |
| AT4G40065 | 1,04 | -0,52 | 2,01 | -0,26 | 7,97E-01 |
| AT5G01010 | 1,03 | -1,28 | 2,49 | -0,51 | 6,09E-01 |
| AT5G01020 | 1,61 | 1,70 | 1,80 | 0,95 | 3,44E-01 |
| AT5G01100 | 0,27 | 1,01 | 4,34 | 0,23 | 8,17E-01 |
| AT5G01215 | 1,09 | 1,13 | 1,72 | 0,66 | 5,11E-01 |
| AT5G01220 | 0,12 | 0,00 | 4,41 | 0,00 | 1,00E+00 |
| AT5G01270 | 0,34 | -0,26 | 4,41 | -0,06 | 9,53E-01 |
| AT5G01365 | 0,66 | -2,04 | 4,37 | -0,47 | 6,41E-01 |
| AT5G01400 | 1,70 | -0,24 | 2,17 | -0,11 | 9,12E-01 |
| AT5G01540 | 0,46 | 0,40 | 4,10 | 0,10 | 9,21E-01 |
| AT5G01560 | 1,57 | 1,03 | 1,93 | 0,54 | 5,91E-01 |
| AT5G01600 | 1,52 | 2,83 | 1,74 | 1,62 | 1,04E-01 |
| AT5G01630 | 1,03 | 0,25 | 2,74 | 0,09 | 9,26E-01 |
| AT5G01700 | 0,62 | 0,00 | 4,41 | 0,00 | 1,00E+00 |
| AT5G01710 | 1,65 | 1,29 | 1,70 | 0,75 | 4,51E-01 |
| AT5G01720 | 0,49 | 0,83 | 3,96 | 0,21 | 8,35E-01 |
| AT5G01730 | 0,29 | 0,00 | 4,41 | 0,00 | 1,00E+00 |
| AT5G01770 | 0,54 | 0,00 | 3,87 | 0,00 | 1,00E+00 |
| AT5G01780 | 0,19 | 0,91 | 4,34 | 0,21 | 8,34E-01 |
| AT5G01890 | 0,64 | 1,15 | 2,62 | 0,44 | 6,61E-01 |
| AT5G01930 | 1,04 | 0,00 | 3,22 | 0,00 | 1,00E+00 |
| AT5G01950 | 0,59 | -0,12 | 2,65 | -0,05 | 9,64E-01 |

| | | | | | |
|---|---|---|---|---|---|
| AT5G01970 | 1,20 | -0,31 | 2,29 | -0,13 | 8,93E-01 |
| AT5G01990 | 1,12 | -0,27 | 2,35 | -0,11 | 9,09E-01 |
| AT5G02130 | 0,08 | 0,21 | 4,39 | 0,05 | 9,62E-01 |
| AT5G02150 | 1,74 | -4,19 | 1,79 | -2,34 | 1,93E-02 |
| AT5G02290 | 0,48 | 0,92 | 2,77 | 0,33 | 7,38E-01 |
| AT5G02502 | 0,33 | -0,17 | 4,35 | -0,04 | 9,69E-01 |
| AT5G02530 | 0,37 | -1,38 | 4,39 | -0,32 | 7,53E-01 |
| AT5G02670 | 0,44 | 0,52 | 3,30 | 0,16 | 8,74E-01 |
| AT5G02770 | 0,75 | 1,91 | 2,36 | 0,81 | 4,17E-01 |
| AT5G02800 | 0,65 | -0,23 | 3,07 | -0,08 | 9,40E-01 |
| AT5G02810 | 1,73 | -0,51 | 1,82 | -0,28 | 7,78E-01 |
| AT5G02830 | 0,85 | 1,17 | 1,91 | 0,61 | 5,40E-01 |
| AT5G02890 | 0,18 | -0,12 | 4,35 | -0,03 | 9,78E-01 |
| AT5G02940 | 0,88 | 2,21 | 2,42 | 0,91 | 3,61E-01 |
| AT5G03140 | 0,48 | 0,09 | 2,68 | 0,03 | 9,72E-01 |
| AT5G03170 | 0,32 | -2,10 | 4,40 | -0,48 | 6,33E-01 |
| AT5G03204 | 0,29 | -0,15 | 4,35 | -0,03 | 9,72E-01 |
| AT5G03210 | 0,71 | 1,28 | 2,36 | 0,54 | 5,87E-01 |
| AT5G03340 | 1,55 | 0,29 | 1,80 | 0,16 | 8,71E-01 |
| AT5G03380 | 0,84 | 0,85 | 2,28 | 0,37 | 7,08E-01 |
| AT5G03400 | 1,62 | 2,61 | 1,66 | 1,58 | 1,15E-01 |
| AT5G03430 | 0,65 | 1,43 | 3,08 | 0,46 | 6,44E-01 |
| AT5G03450 | 0,39 | -1,18 | 4,40 | -0,27 | 7,88E-01 |
| AT5G03555 | 1,08 | -0,34 | 2,17 | -0,16 | 8,75E-01 |
| AT5G03560 | 0,43 | 0,79 | 4,12 | 0,19 | 8,47E-01 |
| AT5G03630 | 0,55 | -1,81 | 3,04 | -0,59 | 5,52E-01 |
| AT5G03700 | 0,11 | -0,26 | 4,41 | -0,06 | 9,53E-01 |
| AT5G03730 | 0,62 | -2,49 | 2,57 | -0,97 | 3,33E-01 |
| AT5G03760 | 0,44 | -1,62 | 4,06 | -0,40 | 6,90E-01 |
| AT5G03800 | 0,30 | 0,00 | 4,41 | 0,00 | 1,00E+00 |
| AT5G03850 | 0,83 | 0,09 | 2,98 | 0,03 | 9,76E-01 |
| AT5G03880 | 0,57 | -0,42 | 3,10 | -0,13 | 8,93E-01 |
| AT5G03910 | 0,83 | -1,70 | 2,42 | -0,70 | 4,82E-01 |
| AT5G03970 | 0,19 | -0,37 | 4,39 | -0,08 | 9,33E-01 |
| AT5G04020 | 0,78 | -0,53 | 1,92 | -0,27 | 7,83E-01 |
| AT5G04130 | 0,05 | 0,18 | 4,38 | 0,04 | 9,68E-01 |
| AT5G04267 | 0,16 | 1,10 | 4,35 | 0,25 | 8,00E-01 |
| AT5G04340 | 1,00 | -1,72 | 1,86 | -0,93 | 3,53E-01 |
| AT5G00955 | 1,17 | 2,82 | 1,88 | 1,50 | 1,34E-01 |
| AT5G04420 | 0,49 | 0,00 | 4,03 | 0,00 | 1,00E+00 |
| AT5G04430 | 1,54 | -0,63 | 1,84 | -0,34 | 7,30E-01 |
| AT5G04470 | 0,16 | 0,15 | 4,38 | 0,04 | 9,72E-01 |
| AT5G04480 | 0,41 | 1,39 | 3,29 | 0,42 | 6,73E-01 |
| AT5G04670 | 1,32 | -1,10 | 2,01 | -0,55 | 5,84E-01 |
| AT5G04700 | 0,30 | -0,26 | 4,41 | -0,06 | 9,53E-01 |
| AT5G04710 | 0,12 | 0,00 | 4,41 | 0,00 | 1,00E+00 |
| AT5G04730 | 1,06 | -0,37 | 2,32 | -0,16 | 8,75E-01 |
| AT5G04760 | 0,48 | 0,00 | 4,41 | 0,00 | 1,00E+00 |
| AT5G04830 | 0,60 | 0,64 | 3,17 | 0,20 | 8,40E-01 |
| AT5G04850 | 0,17 | -0,33 | 4,40 | -0,08 | 9,40E-01 |
| AT5G04860 | 0,46 | 0,42 | 4,21 | 0,10 | 9,20E-01 |
| AT5G04870 | 0,78 | 0,13 | 2,21 | 0,06 | 9,54E-01 |
| AT5G04940 | 0,64 | 0,42 | 3,01 | 0,14 | 8,90E-01 |
| AT5G04950 | 0,11 | 0,88 | 4,35 | 0,20 | 8,39E-01 |
| AT5G04990 | 0,74 | -0,31 | 3,53 | -0,09 | 9,31E-01 |
| AT5G05060 | 0,37 | -3,32 | 4,36 | -0,76 | 4,46E-01 |

| | | | | | |
|---|---|---|---|---|---|
| AT5G05080 | 0,77 | -0,78 | 4,32 | -0,18 | 8,57E-01 |
| AT5G05085 | 0,35 | 1,41 | 3,53 | 0,40 | 6,89E-01 |
| AT5G05100 | 0,72 | -2,28 | 3,03 | -0,75 | 4,52E-01 |
| AT5G01005 | 0,75 | -0,40 | 3,02 | -0,13 | 8,94E-01 |
| AT5G05260 | 0,23 | 0,92 | 4,34 | 0,21 | 8,33E-01 |
| AT5G01055 | 0,10 | 0,86 | 4,35 | 0,20 | 8,43E-01 |
| AT5G05480 | 0,31 | 0,43 | 3,62 | 0,12 | 9,05E-01 |
| AT5G05670 | 0,17 | -0,38 | 4,39 | -0,09 | 9,31E-01 |
| AT5G05700 | 0,69 | -0,29 | 3,05 | -0,09 | 9,25E-01 |
| AT5G05730 | 0,58 | -1,19 | 3,28 | -0,36 | 7,17E-01 |
| AT5G01095 | 0,95 | 0,00 | 4,41 | 0,00 | 1,00E+00 |
| AT5G05800 | 0,36 | 0,00 | 4,41 | 0,00 | 1,00E+00 |
| AT5G06043 | 0,95 | -2,36 | 2,47 | -0,96 | 3,39E-01 |
| AT5G06060 | 0,53 | -0,34 | 3,16 | -0,11 | 9,13E-01 |
| AT5G06110 | 0,69 | -0,21 | 3,01 | -0,07 | 9,46E-01 |
| AT5G06140 | 0,36 | 0,00 | 4,40 | 0,00 | 1,00E+00 |
| AT5G06160 | 0,43 | -0,12 | 2,77 | -0,04 | 9,67E-01 |
| AT5G01165 | 0,96 | 0,91 | 1,86 | 0,49 | 6,25E-01 |
| AT5G01195 | 1,58 | 0,43 | 1,32 | 0,32 | 7,47E-01 |
| AT5G06230 | 0,49 | 0,00 | 3,96 | 0,00 | 1,00E+00 |
| AT5G06260 | 0,46 | -0,75 | 3,31 | -0,23 | 8,22E-01 |
| AT5G06330 | 0,08 | 0,21 | 4,39 | 0,05 | 9,62E-01 |
| AT5G06410 | 0,25 | 0,00 | 4,41 | 0,00 | 1,00E+00 |
| AT5G06430 | 0,51 | 0,00 | 4,41 | 0,00 | 1,00E+00 |
| AT5G06440 | 0,83 | -2,03 | 3,60 | -0,57 | 5,72E-01 |
| AT5G06520 | 0,44 | 0,00 | 4,41 | 0,00 | 1,00E+00 |
| AT5G06540 | 0,48 | 0,00 | 4,41 | 0,00 | 1,00E+00 |
| AT5G06560 | 0,78 | 0,08 | 2,29 | 0,04 | 9,72E-01 |
| AT5G06630 | 1,85 | 4,41 | 1,73 | 2,55 | 1,08E-02 |
| AT5G06670 | 0,24 | 0,00 | 4,33 | 0,00 | 9,99E-01 |
| AT5G06690 | 0,35 | 0,52 | 3,50 | 0,15 | 8,83E-01 |
| AT5G06750 | 0,11 | -0,38 | 4,39 | -0,09 | 9,31E-01 |
| AT5G06770 | 0,36 | 0,06 | 4,34 | 0,01 | 9,89E-01 |
| AT5G06865 | 0,22 | 0,15 | 4,33 | 0,03 | 9,73E-01 |
| AT5G06970 | 0,87 | 0,02 | 2,05 | 0,01 | 9,92E-01 |
| AT5G01295 | 0,28 | 0,00 | 4,41 | 0,00 | 1,00E+00 |
| AT5G07120 | 0,92 | 1,31 | 2,44 | 0,54 | 5,91E-01 |
| AT5G07220 | 0,71 | 0,95 | 2,20 | 0,43 | 6,65E-01 |
| AT5G07250 | 0,62 | -3,36 | 2,98 | -1,13 | 2,59E-01 |
| AT5G07360 | 0,50 | 0,36 | 3,20 | 0,11 | 9,11E-01 |
| AT5G07590 | 0,15 | -0,35 | 4,39 | -0,08 | 9,36E-01 |
| AT5G07620 | 0,13 | -0,61 | 4,37 | -0,14 | 8,89E-01 |
| AT5G07675 | 0,20 | 1,36 | 4,32 | 0,31 | 7,54E-01 |
| AT5G07740 | 1,77 | 0,38 | 1,53 | 0,25 | 8,02E-01 |
| AT5G07810 | 0,26 | 0,00 | 4,41 | 0,00 | 1,00E+00 |
| AT5G07950 | 0,39 | 0,00 | 4,40 | 0,00 | 1,00E+00 |
| AT5G07980 | 1,38 | -0,30 | 2,09 | -0,14 | 8,87E-01 |
| AT5G08040 | 0,14 | -2,43 | 4,37 | -0,55 | 5,79E-01 |
| AT5G08075 | 0,48 | 1,11 | 4,05 | 0,28 | 7,83E-01 |
| AT5G08110 | 0,94 | 0,00 | 2,50 | 0,00 | 1,00E+00 |
| AT5G08130 | 1,09 | 1,80 | 1,97 | 0,91 | 3,61E-01 |
| AT5G08160 | 1,05 | -0,02 | 2,14 | -0,01 | 9,91E-01 |
| AT5G08170 | 0,71 | -0,15 | 3,68 | -0,04 | 9,68E-01 |
| AT5G01495 | 0,69 | 1,14 | 1,97 | 0,58 | 5,61E-01 |
| AT5G01505 | 1,02 | 0,81 | 1,55 | 0,52 | 6,03E-01 |
| AT5G01515 | 0,11 | -0,26 | 4,41 | -0,06 | 9,53E-01 |

| | | | | | |
|---|---|---|---|---|---|
| AT5G08230 | 0,62 | 0,93 | 3,14 | 0,30 | 7,66E-01 |
| AT5G08280 | 1,65 | -2,98 | 1,54 | -1,94 | 5,24E-02 |
| AT5G08305 | 0,42 | -0,29 | 4,21 | -0,07 | 9,45E-01 |
| AT5G08315 | 0,44 | -0,26 | 4,41 | -0,06 | 9,53E-01 |
| AT5G08350 | 0,26 | 0,00 | 4,41 | 0,00 | 1,00E+00 |
| AT5G08450 | 0,66 | -0,02 | 2,76 | -0,01 | 9,95E-01 |
| AT5G08520 | 1,35 | -2,02 | 1,44 | -1,41 | 1,60E-01 |
| AT5G08550 | 1,83 | 0,20 | 1,73 | 0,12 | 9,07E-01 |
| AT5G08560 | 0,55 | 0,94 | 2,67 | 0,35 | 7,25E-01 |
| AT5G08650 | 1,18 | -1,34 | 1,87 | -0,72 | 4,72E-01 |
| AT5G08770 | 1,04 | -0,39 | 1,90 | -0,20 | 8,38E-01 |
| AT5G08790 | 1,51 | 1,91 | 1,71 | 1,12 | 2,65E-01 |
| AT5G09330 | 1,10 | 1,39 | 1,74 | 0,80 | 4,23E-01 |
| AT5G09443 | 0,26 | 0,00 | 4,41 | 0,00 | 1,00E+00 |
| AT5G09460 | 0,93 | -0,48 | 2,07 | -0,23 | 8,18E-01 |
| AT5G09510 | 0,57 | 0,31 | 4,02 | 0,08 | 9,39E-01 |
| AT5G09620 | 0,88 | 2,51 | 2,34 | 1,08 | 2,82E-01 |
| AT5G09650 | 0,62 | -0,01 | 2,59 | 0,00 | 9,98E-01 |
| AT5G09820 | 0,63 | 1,19 | 3,12 | 0,38 | 7,02E-01 |
| AT5G09850 | 0,67 | 1,61 | 3,05 | 0,53 | 5,97E-01 |
| AT5G09880 | 0,62 | 1,25 | 2,96 | 0,42 | 6,72E-01 |
| AT5G09890 | 0,52 | 0,76 | 3,93 | 0,19 | 8,46E-01 |
| AT5G09900 | 1,74 | 0,68 | 1,75 | 0,39 | 7,00E-01 |
| AT5G09930 | 0,08 | 0,00 | 4,41 | 0,00 | 1,00E+00 |
| AT5G10030 | 0,52 | -0,27 | 4,01 | -0,07 | 9,47E-01 |
| AT5G10235 | 0,28 | -2,11 | 4,40 | -0,48 | 6,32E-01 |
| AT5G10290 | 0,76 | -0,02 | 2,44 | -0,01 | 9,94E-01 |
| AT5G10330 | 0,57 | 0,11 | 2,69 | 0,04 | 9,68E-01 |
| AT5G10350 | 0,66 | 0,00 | 4,38 | 0,00 | 1,00E+00 |
| AT5G10360 | 1,44 | -0,11 | 1,50 | -0,07 | 9,43E-01 |
| AT5G10390 | 0,27 | 1,05 | 4,35 | 0,24 | 8,10E-01 |
| AT5G10400 | 0,23 | 2,01 | 4,30 | 0,47 | 6,40E-01 |
| AT5G10470 | 0,57 | 0,54 | 3,27 | 0,17 | 8,68E-01 |
| AT5G10490 | 0,10 | 0,08 | 4,38 | 0,02 | 9,85E-01 |
| AT5G10550 | 0,86 | 0,13 | 2,36 | 0,05 | 9,57E-01 |
| AT5G10605 | 0,65 | 1,40 | 3,11 | 0,45 | 6,53E-01 |
| AT5G10650 | 0,25 | 0,00 | 4,41 | 0,00 | 1,00E+00 |
| AT5G10690 | 0,38 | 1,09 | 4,34 | 0,25 | 8,01E-01 |
| AT5G01815 | 0,80 | 0,98 | 2,00 | 0,49 | 6,24E-01 |
| AT5G10700 | 0,56 | 1,80 | 3,15 | 0,57 | 5,68E-01 |
| AT5G10730 | 0,23 | 0,23 | 4,35 | 0,05 | 9,58E-01 |
| AT5G10770 | 0,94 | 2,28 | 2,25 | 1,01 | 3,13E-01 |
| AT5G10800 | 0,30 | 0,00 | 4,41 | 0,00 | 1,00E+00 |
| AT5G10820 | 0,63 | 0,00 | 4,41 | 0,00 | 1,00E+00 |
| AT5G10860 | 1,76 | 0,13 | 1,59 | 0,08 | 9,36E-01 |
| AT5G10930 | 1,14 | 0,30 | 1,93 | 0,15 | 8,78E-01 |
| AT5G10940 | 0,77 | -0,26 | 3,68 | -0,07 | 9,43E-01 |
| AT5G10965 | 0,75 | -0,32 | 2,53 | -0,13 | 9,00E-01 |
| AT5G10980 | 0,55 | -0,44 | 3,83 | -0,12 | 9,08E-01 |
| AT5G11000 | 0,45 | 0,99 | 4,20 | 0,24 | 8,14E-01 |
| AT5G11070 | 1,00 | 2,71 | 2,05 | 1,32 | 1,85E-01 |
| AT5G11180 | 0,81 | -1,28 | 2,20 | -0,58 | 5,61E-01 |
| AT5G11210 | 1,06 | 0,00 | 3,28 | 0,00 | 1,00E+00 |
| AT5G11250 | 1,58 | 0,05 | 1,58 | 0,03 | 9,74E-01 |
| AT5G11270 | 0,22 | 0,14 | 4,38 | 0,03 | 9,75E-01 |
| AT5G11330 | 1,01 | 0,86 | 1,89 | 0,46 | 6,48E-01 |

| | | | | | |
|---|---|---|---|---|---|
| AT5G11480 | 0,46 | -1,42 | 3,15 | -0,45 | 6,52E-01 |
| AT5G11520 | 1,24 | -0,79 | 1,88 | -0,42 | 6,73E-01 |
| AT5G11550 | 0,28 | 0,03 | 4,38 | 0,01 | 9,95E-01 |
| AT5G11770 | 0,54 | -0,51 | 2,64 | -0,19 | 8,46E-01 |
| AT5G11850 | 0,25 | 0,29 | 4,37 | 0,07 | 9,47E-01 |
| AT5G11860 | 0,12 | 0,00 | 4,41 | 0,00 | 1,00E+00 |
| AT5G11910 | 0,27 | -0,38 | 4,39 | -0,09 | 9,31E-01 |
| AT5G11960 | 0,36 | 0,00 | 4,41 | 0,00 | 1,00E+00 |
| AT5G12010 | 0,59 | -0,75 | 3,14 | -0,24 | 8,12E-01 |
| AT5G12080 | 0,25 | 0,00 | 4,41 | 0,00 | 1,00E+00 |
| AT5G12140 | 0,22 | 0,79 | 4,35 | 0,18 | 8,56E-01 |
| AT5G12200 | 0,44 | -1,90 | 3,29 | -0,58 | 5,63E-01 |
| AT5G12310 | 0,42 | 1,63 | 3,32 | 0,49 | 6,24E-01 |
| AT5G12390 | 0,11 | -0,26 | 4,41 | -0,06 | 9,53E-01 |
| AT5G12430 | 1,28 | 0,00 | 2,52 | 0,00 | 1,00E+00 |
| AT5G12440 | 0,88 | 0,00 | 2,83 | 0,00 | 1,00E+00 |
| AT5G12480 | 0,34 | 0,22 | 4,34 | 0,05 | 9,59E-01 |
| AT5G12890 | 0,77 | 0,00 | 3,73 | 0,00 | 1,00E+00 |
| AT5G12900 | 0,23 | 0,71 | 4,36 | 0,16 | 8,70E-01 |
| AT5G13020 | 0,81 | -1,92 | 2,30 | -0,83 | 4,05E-01 |
| AT5G13050 | 0,42 | 0,47 | 4,19 | 0,11 | 9,11E-01 |
| AT5G13090 | 0,10 | 0,24 | 4,38 | 0,05 | 9,57E-01 |
| AT5G13120 | 0,17 | -1,50 | 4,37 | -0,34 | 7,31E-01 |
| AT5G13140 | 0,23 | -1,08 | 4,33 | -0,25 | 8,03E-01 |
| AT5G13170 | 0,48 | 0,00 | 4,41 | 0,00 | 1,00E+00 |
| AT5G13190 | 0,99 | 1,47 | 1,89 | 0,77 | 4,39E-01 |
| AT5G02055 | 0,48 | -0,48 | 2,65 | -0,18 | 8,56E-01 |
| AT5G13225 | 0,48 | -0,48 | 2,65 | -0,18 | 8,56E-01 |
| AT5G13270 | 0,40 | -0,40 | 4,22 | -0,09 | 9,25E-01 |
| AT5G13300 | 0,77 | 0,18 | 2,53 | 0,07 | 9,44E-01 |
| AT5G13400 | 0,43 | -0,34 | 4,40 | -0,08 | 9,38E-01 |
| AT5G13430 | 0,54 | 0,00 | 3,87 | 0,00 | 1,00E+00 |
| AT5G13440 | 0,29 | 1,81 | 4,30 | 0,42 | 6,73E-01 |
| AT5G13450 | 0,96 | 0,67 | 2,00 | 0,33 | 7,40E-01 |
| AT5G13550 | 0,46 | -0,31 | 4,06 | -0,08 | 9,38E-01 |
| AT5G13560 | 1,05 | -1,94 | 2,20 | -0,88 | 3,78E-01 |
| AT5G13590 | 1,32 | -0,40 | 2,15 | -0,19 | 8,51E-01 |
| AT5G13680 | 1,03 | 0,64 | 1,92 | 0,33 | 7,40E-01 |
| AT5G13710 | 0,59 | 0,91 | 3,09 | 0,29 | 7,69E-01 |
| AT5G13780 | 0,62 | 0,33 | 3,13 | 0,11 | 9,15E-01 |
| AT5G13800 | 0,74 | 0,34 | 2,35 | 0,15 | 8,83E-01 |
| AT5G13825 | 0,39 | -2,48 | 4,20 | -0,59 | 5,54E-01 |
| AT5G13950 | 1,63 | -0,48 | 1,64 | -0,29 | 7,72E-01 |
| AT5G13990 | 0,12 | 0,11 | 4,38 | 0,03 | 9,80E-01 |
| AT5G14100 | 0,11 | -0,26 | 4,41 | -0,06 | 9,53E-01 |
| AT5G14170 | 1,07 | -0,31 | 2,37 | -0,13 | 8,95E-01 |
| AT5G14210 | 0,50 | -0,60 | 3,31 | -0,18 | 8,57E-01 |
| AT5G14240 | 0,57 | 0,28 | 2,71 | 0,10 | 9,19E-01 |
| AT5G14320 | 0,97 | -0,41 | 1,69 | -0,24 | 8,10E-01 |
| AT5G14390 | 0,52 | 0,79 | 4,13 | 0,19 | 8,48E-01 |
| AT5G14420 | 0,18 | 0,15 | 4,38 | 0,03 | 9,73E-01 |
| AT5G14530 | 0,86 | 0,40 | 2,91 | 0,14 | 8,90E-01 |
| AT5G14540 | 0,54 | -0,07 | 3,98 | -0,02 | 9,87E-01 |
| AT5G02245 | 0,10 | 0,86 | 4,35 | 0,20 | 8,43E-01 |
| AT5G14580 | 1,26 | 0,19 | 2,79 | 0,07 | 9,45E-01 |
| AT5G14590 | 0,98 | -1,41 | 1,97 | -0,71 | 4,75E-01 |

| | | | | | |
|---|---|---|---|---|---|
| AT5G14700 | 0,24 | 1,90 | 4,30 | 0,44 | 6,58E-01 |
| AT5G14720 | 0,96 | 1,23 | 1,97 | 0,62 | 5,33E-01 |
| AT5G14770 | 0,20 | -2,08 | 4,39 | -0,47 | 6,35E-01 |
| AT5G14820 | 0,29 | -0,12 | 4,39 | -0,03 | 9,79E-01 |
| AT5G14940 | 1,18 | 0,61 | 2,40 | 0,25 | 7,99E-01 |
| AT5G14950 | 1,52 | 0,14 | 1,59 | 0,09 | 9,29E-01 |
| AT5G15050 | 0,10 | 0,08 | 4,38 | 0,02 | 9,85E-01 |
| AT5G15210 | 0,28 | -0,26 | 4,41 | -0,06 | 9,53E-01 |
| AT5G15250 | 0,05 | 0,18 | 4,38 | 0,04 | 9,68E-01 |
| AT5G15360 | 0,52 | 1,56 | 2,52 | 0,62 | 5,37E-01 |
| AT5G15410 | 1,41 | -2,12 | 1,90 | -1,12 | 2,63E-01 |
| AT5G15420 | 1,66 | -1,24 | 1,39 | -0,89 | 3,73E-01 |
| AT5G15450 | 1,23 | -0,18 | 1,86 | -0,10 | 9,21E-01 |
| AT5G15460 | 0,08 | 0,00 | 4,41 | 0,00 | 1,00E+00 |
| AT5G15470 | 0,21 | -1,29 | 4,40 | -0,29 | 7,69E-01 |
| AT5G15490 | 0,40 | 0,45 | 3,41 | 0,13 | 8,95E-01 |
| AT5G15537 | 0,21 | 0,00 | 4,41 | 0,00 | 1,00E+00 |
| AT5G15540 | 1,01 | -0,33 | 2,10 | -0,16 | 8,73E-01 |
| AT5G15580 | 0,70 | -2,17 | 2,51 | -0,87 | 3,86E-01 |
| AT5G15690 | 0,53 | 0,43 | 2,66 | 0,16 | 8,71E-01 |
| AT5G15700 | 0,61 | 0,69 | 2,30 | 0,30 | 7,66E-01 |
| AT5G15740 | 0,28 | 1,45 | 4,33 | 0,33 | 7,38E-01 |
| AT5G15780 | 0,22 | 0,15 | 4,33 | 0,03 | 9,73E-01 |
| AT5G15800 | 0,08 | 0,47 | 4,38 | 0,11 | 9,15E-01 |
| AT5G15802 | 0,08 | 0,21 | 4,39 | 0,05 | 9,62E-01 |
| AT5G15860 | 0,19 | -0,37 | 4,39 | -0,08 | 9,33E-01 |
| AT5G16050 | 0,47 | -0,32 | 2,70 | -0,12 | 9,05E-01 |
| AT5G16070 | 0,93 | -1,37 | 2,27 | -0,60 | 5,46E-01 |
| AT5G16110 | 1,09 | -0,12 | 1,94 | -0,06 | 9,51E-01 |
| AT5G16200 | 0,36 | 0,00 | 4,41 | 0,00 | 1,00E+00 |
| AT5G16220 | 0,18 | 1,59 | 4,31 | 0,37 | 7,12E-01 |
| AT5G16280 | 0,67 | 0,00 | 3,00 | 0,00 | 1,00E+00 |
| AT5G16360 | 0,08 | 0,00 | 4,41 | 0,00 | 1,00E+00 |
| AT5G16370 | 1,57 | -0,22 | 1,81 | -0,12 | 9,04E-01 |
| AT5G16380 | 0,25 | -1,25 | 4,40 | -0,28 | 7,77E-01 |
| AT5G16390 | 0,61 | -2,14 | 3,75 | -0,57 | 5,68E-01 |
| AT5G16480 | 0,56 | -0,39 | 4,39 | -0,09 | 9,30E-01 |
| AT5G16490 | 0,08 | 0,21 | 4,39 | 0,05 | 9,62E-01 |
| AT5G16520 | 0,47 | -0,52 | 4,20 | -0,12 | 9,01E-01 |
| AT5G16560 | 0,42 | -0,90 | 4,31 | -0,21 | 8,34E-01 |
| AT5G16610 | 0,08 | 0,00 | 4,41 | 0,00 | 1,00E+00 |
| AT5G16660 | 0,23 | 0,43 | 4,36 | 0,10 | 9,22E-01 |
| AT5G16680 | 0,78 | 1,59 | 2,21 | 0,72 | 4,71E-01 |
| AT5G16740 | 0,42 | -0,26 | 4,41 | -0,06 | 9,53E-01 |
| AT5G16750 | 0,89 | 0,24 | 2,14 | 0,11 | 9,12E-01 |
| AT5G16780 | 0,54 | 2,12 | 3,89 | 0,55 | 5,85E-01 |
| AT5G16800 | 0,21 | 0,08 | 4,38 | 0,02 | 9,85E-01 |
| AT5G16830 | 0,10 | 0,08 | 4,38 | 0,02 | 9,85E-01 |
| AT5G16980 | 0,08 | 0,00 | 4,41 | 0,00 | 1,00E+00 |
| AT5G16990 | 0,20 | -0,15 | 4,39 | -0,03 | 9,72E-01 |
| AT5G17000 | 0,20 | 0,40 | 4,37 | 0,09 | 9,27E-01 |
| AT5G17010 | 0,24 | 0,54 | 4,37 | 0,12 | 9,01E-01 |
| AT5G02655 | 0,55 | -0,63 | 2,42 | -0,26 | 7,94E-01 |
| AT5G02665 | 1,25 | 0,68 | 2,19 | 0,31 | 7,58E-01 |
| AT5G02675 | 0,37 | 0,29 | 4,32 | 0,07 | 9,46E-01 |
| AT5G17290 | 0,65 | -0,04 | 3,10 | -0,01 | 9,89E-01 |

| | | | | | |
|---|---|---|---|---|---|
| AT5G17300 | 0,69 | -0,39 | 3,05 | -0,13 | 8,99E-01 |
| AT5G17320 | 0,38 | -1,20 | 4,38 | -0,27 | 7,84E-01 |
| AT5G17380 | 0,80 | -0,37 | 2,54 | -0,15 | 8,84E-01 |
| AT5G17420 | 0,40 | -0,26 | 4,41 | -0,06 | 9,53E-01 |
| AT5G17460 | 0,52 | -4,59 | 4,30 | -1,07 | 2,85E-01 |
| AT5G17520 | 0,17 | -3,06 | 4,35 | -0,70 | 4,82E-01 |
| AT5G17550 | 0,08 | 0,00 | 4,41 | 0,00 | 1,00E+00 |
| AT5G17600 | 0,90 | 2,22 | 2,15 | 1,04 | 3,00E-01 |
| AT5G17620 | 0,42 | -0,26 | 4,41 | -0,06 | 9,53E-01 |
| AT5G17640 | 0,35 | 0,86 | 4,35 | 0,20 | 8,43E-01 |
| AT5G17690 | 0,20 | -0,26 | 4,41 | -0,06 | 9,53E-01 |
| AT5G17710 | 1,85 | 0,69 | 1,62 | 0,43 | 6,67E-01 |
| AT5G17770 | 0,49 | 1,65 | 3,18 | 0,52 | 6,05E-01 |
| AT5G17860 | 0,63 | 2,08 | 2,82 | 0,74 | 4,60E-01 |
| AT5G17880 | 1,61 | 0,36 | 2,14 | 0,17 | 8,66E-01 |
| AT5G17930 | 0,48 | 0,00 | 4,41 | 0,00 | 1,00E+00 |
| AT5G18050 | 0,24 | 0,00 | 4,41 | 0,00 | 1,00E+00 |
| AT5G18110 | 0,39 | 0,07 | 4,34 | 0,02 | 9,87E-01 |
| AT5G18190 | 0,63 | -1,21 | 2,69 | -0,45 | 6,53E-01 |
| AT5G18230 | 0,99 | -0,19 | 2,89 | -0,07 | 9,47E-01 |
| AT5G18240 | 1,40 | 4,57 | 2,74 | 1,67 | 9,52E-02 |
| AT5G18260 | 0,49 | 0,00 | 4,41 | 0,00 | 1,00E+00 |
| AT5G18280 | 0,49 | 0,46 | 3,18 | 0,15 | 8,84E-01 |
| AT5G18290 | 0,36 | 0,00 | 4,41 | 0,00 | 1,00E+00 |
| AT5G18340 | 0,58 | -0,36 | 4,02 | -0,09 | 9,29E-01 |
| AT5G18400 | 0,32 | 1,29 | 4,33 | 0,30 | 7,65E-01 |
| AT5G02895 | 0,88 | 1,11 | 2,75 | 0,41 | 6,85E-01 |
| AT5G18475 | 0,21 | 0,00 | 4,41 | 0,00 | 1,00E+00 |
| AT5G18480 | 0,62 | -1,86 | 3,02 | -0,62 | 5,37E-01 |
| AT5G18490 | 1,80 | 0,60 | 1,93 | 0,31 | 7,55E-01 |
| AT5G18525 | 1,70 | -0,26 | 2,09 | -0,13 | 9,00E-01 |
| AT5G18570 | 0,43 | 1,13 | 4,02 | 0,28 | 7,78E-01 |
| AT5G18590 | 0,48 | 1,08 | 3,26 | 0,33 | 7,39E-01 |
| AT5G18620 | 0,19 | 0,91 | 4,34 | 0,21 | 8,34E-01 |
| AT5G18700 | 0,89 | -0,19 | 2,86 | -0,07 | 9,48E-01 |
| AT5G02935 | 0,05 | 0,18 | 4,38 | 0,04 | 9,68E-01 |
| AT5G19010 | 0,59 | 0,84 | 2,52 | 0,33 | 7,40E-01 |
| AT5G19130 | 0,62 | -0,23 | 2,68 | -0,09 | 9,31E-01 |
| AT5G19220 | 0,51 | 0,09 | 4,38 | 0,02 | 9,83E-01 |
| AT5G19360 | 0,72 | 0,00 | 4,41 | 0,00 | 1,00E+00 |
| AT5G19410 | 0,62 | 0,00 | 3,05 | 0,00 | 1,00E+00 |
| AT5G19420 | 1,09 | 1,04 | 2,00 | 0,52 | 6,04E-01 |
| AT5G19450 | 0,87 | 0,26 | 1,94 | 0,13 | 8,93E-01 |
| AT5G19485 | 0,20 | 0,00 | 4,41 | 0,00 | 1,00E+00 |
| AT5G19510 | 1,83 | -0,12 | 1,61 | -0,07 | 9,40E-01 |
| AT5G19590 | 0,42 | 2,08 | 3,20 | 0,65 | 5,16E-01 |
| AT5G19660 | 0,36 | 0,80 | 4,34 | 0,18 | 8,54E-01 |
| AT5G19760 | 0,90 | -0,06 | 2,57 | -0,02 | 9,80E-01 |
| AT5G19770 | 1,75 | 0,97 | 1,91 | 0,51 | 6,10E-01 |
| AT5G19850 | 0,75 | 0,51 | 3,70 | 0,14 | 8,90E-01 |
| AT5G19860 | 0,51 | -0,28 | 4,05 | -0,07 | 9,45E-01 |
| AT5G19900 | 0,73 | 1,60 | 3,04 | 0,53 | 5,99E-01 |
| AT5G19920 | 0,37 | 0,00 | 4,41 | 0,00 | 1,00E+00 |
| AT5G19930 | 0,12 | 0,27 | 4,38 | 0,06 | 9,50E-01 |
| AT5G19950 | 0,27 | 0,80 | 4,35 | 0,18 | 8,54E-01 |
| AT5G19980 | 0,33 | 0,12 | 4,36 | 0,03 | 9,78E-01 |

| | | | | | |
|---|---|---|---|---|---|
| AT5G20040 | 0,12 | 0,00 | 4,41 | 0,00 | 1,00E+00 |
| AT5G20160 | 0,08 | 0,21 | 4,39 | 0,05 | 9,62E-01 |
| AT5G20170 | 0,19 | 0,81 | 4,35 | 0,19 | 8,53E-01 |
| AT5G20225 | 0,17 | 0,00 | 4,41 | 0,00 | 1,00E+00 |
| AT5G20260 | 0,52 | 0,00 | 4,41 | 0,00 | 1,00E+00 |
| AT5G20270 | 0,14 | 0,81 | 4,35 | 0,19 | 8,52E-01 |
| AT5G20300 | 1,26 | 0,42 | 2,56 | 0,16 | 8,69E-01 |
| AT5G20320 | 1,21 | 0,00 | 2,64 | 0,00 | 1,00E+00 |
| AT5G20360 | 0,79 | -0,81 | 2,13 | -0,38 | 7,03E-01 |
| AT5G20380 | 0,45 | -3,29 | 4,35 | -0,76 | 4,49E-01 |
| AT5G20490 | 1,59 | -0,87 | 1,73 | -0,50 | 6,16E-01 |
| AT5G20520 | 1,55 | -0,91 | 1,62 | -0,56 | 5,73E-01 |
| AT5G20570 | 0,58 | -0,32 | 3,17 | -0,10 | 9,19E-01 |
| AT5G20600 | 1,15 | 0,00 | 2,34 | 0,00 | 1,00E+00 |
| AT5G20620 | 0,96 | 0,54 | 2,52 | 0,21 | 8,31E-01 |
| AT5G20630 | 1,03 | -0,28 | 1,85 | -0,15 | 8,78E-01 |
| AT5G20650 | 0,87 | 0,87 | 3,07 | 0,28 | 7,78E-01 |
| AT5G20820 | 0,08 | 0,21 | 4,39 | 0,05 | 9,62E-01 |
| AT5G20830 | 0,97 | -0,62 | 2,51 | -0,25 | 8,03E-01 |
| AT5G20840 | 1,18 | 1,89 | 1,71 | 1,10 | 2,70E-01 |
| AT5G20890 | 1,15 | 1,06 | 2,27 | 0,47 | 6,41E-01 |
| AT5G20920 | 0,64 | -0,49 | 3,22 | -0,15 | 8,78E-01 |
| AT5G20940 | 0,27 | -0,40 | 4,38 | -0,09 | 9,28E-01 |
| AT5G03265 | 0,23 | 0,56 | 4,38 | 0,13 | 8,98E-01 |
| AT5G20980 | 0,81 | 0,45 | 2,93 | 0,16 | 8,77E-01 |
| AT5G20990 | 1,65 | 1,72 | 1,94 | 0,89 | 3,75E-01 |
| AT5G21040 | 0,32 | -0,25 | 4,41 | -0,06 | 9,55E-01 |
| AT5G21160 | 0,41 | 1,54 | 4,31 | 0,36 | 7,20E-01 |
| AT5G21274 | 0,23 | 0,43 | 4,36 | 0,10 | 9,22E-01 |
| AT5G21920 | 0,36 | -0,78 | 4,36 | -0,18 | 8,58E-01 |
| AT5G21950 | 0,08 | 0,00 | 4,41 | 0,00 | 1,00E+00 |
| AT5G22010 | 0,59 | -0,37 | 3,09 | -0,12 | 9,06E-01 |
| AT5G22020 | 0,55 | -1,39 | 3,90 | -0,36 | 7,21E-01 |
| AT5G22030 | 0,76 | 0,04 | 3,45 | 0,01 | 9,91E-01 |
| AT5G22060 | 1,34 | 1,69 | 2,31 | 0,73 | 4,64E-01 |
| AT5G22080 | 0,35 | 1,89 | 3,47 | 0,55 | 5,86E-01 |
| AT5G22090 | 0,53 | -0,02 | 3,31 | -0,01 | 9,95E-01 |
| AT5G22250 | 1,15 | 3,35 | 2,04 | 1,64 | 1,00E-01 |
| AT5G22350 | 0,24 | 1,35 | 4,32 | 0,31 | 7,54E-01 |
| AT5G22360 | 0,12 | 0,00 | 4,41 | 0,00 | 1,00E+00 |
| AT5G22390 | 0,49 | -2,89 | 3,19 | -0,91 | 3,65E-01 |
| AT5G22440 | 0,73 | 0,51 | 3,03 | 0,17 | 8,66E-01 |
| AT5G22450 | 0,87 | 1,32 | 1,91 | 0,69 | 4,90E-01 |
| AT5G22505 | 1,58 | -0,60 | 1,52 | -0,39 | 6,96E-01 |
| AT5G22510 | 0,44 | 1,38 | 4,18 | 0,33 | 7,41E-01 |
| AT5G03425 | 0,62 | -3,97 | 3,59 | -1,10 | 2,70E-01 |
| AT5G22540 | 0,22 | 0,53 | 4,36 | 0,12 | 9,03E-01 |
| AT5G22570 | 0,26 | 0,03 | 4,38 | 0,01 | 9,95E-01 |
| AT5G22700 | 0,44 | -0,27 | 4,17 | -0,06 | 9,48E-01 |
| AT5G22740 | 0,58 | -0,02 | 3,92 | 0,00 | 9,96E-01 |
| AT5G22750 | 0,55 | 0,00 | 4,39 | 0,00 | 1,00E+00 |
| AT5G03475 | 1,21 | 0,22 | 1,73 | 0,13 | 8,97E-01 |
| AT5G22780 | 1,47 | 1,80 | 1,70 | 1,06 | 2,91E-01 |
| AT5G22800 | 1,71 | -1,97 | 1,35 | -1,46 | 1,45E-01 |
| AT5G22850 | 0,23 | -0,36 | 4,39 | -0,08 | 9,36E-01 |
| AT5G22860 | 0,35 | -0,07 | 3,48 | -0,02 | 9,84E-01 |

| | | | | | |
|---|---|---|---|---|---|
| AT5G23040 | 0,63 | 0,43 | 2,40 | 0,18 | 8,58E-01 |
| AT5G23050 | 0,45 | 0,00 | 4,14 | 0,00 | 1,00E+00 |
| AT5G23060 | 1,11 | -0,83 | 1,67 | -0,50 | 6,17E-01 |
| AT5G23080 | 1,07 | -0,13 | 2,71 | -0,05 | 9,63E-01 |
| AT5G23240 | 0,89 | 0,14 | 2,13 | 0,06 | 9,48E-01 |
| AT5G23300 | 0,32 | -0,10 | 4,39 | -0,02 | 9,81E-01 |
| AT5G23390 | 0,47 | -1,35 | 4,01 | -0,34 | 7,37E-01 |
| AT5G23405 | 0,14 | 0,13 | 4,38 | 0,03 | 9,77E-01 |
| AT5G23430 | 0,90 | 1,90 | 2,16 | 0,88 | 3,78E-01 |
| AT5G23440 | 0,16 | -0,56 | 4,37 | -0,13 | 8,98E-01 |
| AT5G23450 | 0,83 | 0,50 | 2,95 | 0,17 | 8,66E-01 |
| AT5G23580 | 0,83 | -0,48 | 2,32 | -0,21 | 8,35E-01 |
| AT5G23590 | 0,61 | 0,00 | 3,75 | 0,00 | 1,00E+00 |
| AT5G23610 | 1,31 | -1,43 | 2,13 | -0,67 | 5,03E-01 |
| AT5G23660 | 0,79 | 2,68 | 3,60 | 0,74 | 4,57E-01 |
| AT5G23740 | 0,24 | 1,90 | 4,30 | 0,44 | 6,58E-01 |
| AT5G23900 | 0,57 | -1,70 | 2,72 | -0,62 | 5,32E-01 |
| AT5G23940 | 0,08 | 0,21 | 4,39 | 0,05 | 9,62E-01 |
| AT5G23990 | 0,29 | 0,54 | 4,32 | 0,12 | 9,01E-01 |
| AT5G24120 | 0,77 | 0,24 | 2,55 | 0,09 | 9,26E-01 |
| AT5G24140 | 0,08 | 0,21 | 4,39 | 0,05 | 9,62E-01 |
| AT5G24150 | 0,21 | 0,00 | 4,41 | 0,00 | 1,00E+00 |
| AT5G24160 | 0,65 | 0,99 | 2,93 | 0,34 | 7,36E-01 |
| AT5G24240 | 1,01 | -0,64 | 1,65 | -0,39 | 6,97E-01 |
| AT5G24260 | 1,06 | 1,26 | 2,20 | 0,57 | 5,68E-01 |
| AT5G24270 | 1,64 | 0,09 | 1,34 | 0,07 | 9,47E-01 |
| AT5G24320 | 0,20 | -1,20 | 4,38 | -0,27 | 7,85E-01 |
| AT5G24355 | 0,08 | 0,00 | 4,41 | 0,00 | 1,00E+00 |
| AT5G24430 | 0,56 | -0,55 | 3,92 | -0,14 | 8,88E-01 |
| AT5G24470 | 1,29 | -0,47 | 2,59 | -0,18 | 8,57E-01 |
| AT5G24480 | 0,23 | 0,43 | 4,36 | 0,10 | 9,22E-01 |
| AT5G24500 | 0,24 | -0,02 | 4,38 | 0,00 | 9,97E-01 |
| AT5G24520 | 0,95 | 1,26 | 2,09 | 0,60 | 5,47E-01 |
| AT5G24590 | 1,51 | 3,79 | 1,78 | 2,12 | 3,38E-02 |
| AT5G24655 | 0,37 | 1,81 | 4,32 | 0,42 | 6,75E-01 |
| AT5G24735 | 0,57 | 0,92 | 3,25 | 0,28 | 7,78E-01 |
| AT5G24740 | 1,40 | -0,69 | 1,73 | -0,40 | 6,89E-01 |
| AT5G24770 | 0,62 | 0,16 | 2,21 | 0,07 | 9,43E-01 |
| AT5G24780 | 0,08 | 0,21 | 4,39 | 0,05 | 9,62E-01 |
| AT5G24940 | 0,08 | 0,00 | 4,41 | 0,00 | 1,00E+00 |
| AT5G24960 | 0,36 | 2,33 | 4,29 | 0,54 | 5,87E-01 |
| AT5G25000 | 0,16 | 0,41 | 4,37 | 0,09 | 9,25E-01 |
| AT5G25070 | 1,26 | 0,53 | 1,99 | 0,27 | 7,90E-01 |
| AT5G25100 | 1,32 | -1,93 | 1,59 | -1,21 | 2,25E-01 |
| AT5G25200 | 0,60 | 1,70 | 2,52 | 0,67 | 5,00E-01 |
| AT5G25265 | 0,67 | -1,17 | 3,01 | -0,39 | 6,98E-01 |
| AT5G25350 | 1,15 | -1,23 | 2,21 | -0,56 | 5,77E-01 |
| AT5G25370 | 0,71 | -1,81 | 2,71 | -0,67 | 5,03E-01 |
| AT5G03985 | 0,37 | -3,32 | 4,36 | -0,76 | 4,46E-01 |
| AT5G25480 | 0,27 | -2,46 | 4,37 | -0,56 | 5,74E-01 |
| AT5G25510 | 0,49 | -1,66 | 3,22 | -0,51 | 6,07E-01 |
| AT5G25530 | 1,06 | 0,00 | 3,35 | 0,00 | 1,00E+00 |
| AT5G25756 | 0,35 | 2,67 | 4,28 | 0,62 | 5,32E-01 |
| AT5G25757 | 0,87 | -2,37 | 1,95 | -1,21 | 2,25E-01 |
| AT5G25770 | 0,28 | 0,00 | 4,41 | 0,00 | 1,00E+00 |
| AT5G04085 | 0,59 | 3,20 | 4,27 | 0,75 | 4,54E-01 |

| | | | | | |
|---|---|---|---|---|---|
| AT5G25780 | 1,13 | -2,71 | 1,89 | -1,43 | 1,52E-01 |
| AT5G25820 | 0,43 | 0,34 | 4,38 | 0,08 | 9,38E-01 |
| AT5G25830 | 0,05 | 0,18 | 4,38 | 0,04 | 9,68E-01 |
| AT5G25920 | 0,08 | 0,00 | 4,41 | 0,00 | 1,00E+00 |
| AT5G26170 | 0,17 | -0,29 | 4,40 | -0,07 | 9,47E-01 |
| AT5G26210 | 0,43 | -0,40 | 4,22 | -0,09 | 9,25E-01 |
| AT5G26230 | 0,08 | 0,47 | 4,38 | 0,11 | 9,15E-01 |
| AT5G26680 | 0,15 | 0,82 | 4,35 | 0,19 | 8,50E-01 |
| AT5G04245 | 1,57 | 4,27 | 2,16 | 1,97 | 4,84E-02 |
| AT5G26640 | 0,17 | 0,00 | 4,41 | 0,00 | 1,00E+00 |
| AT5G26622 | 0,46 | 1,25 | 4,32 | 0,29 | 7,73E-01 |
| AT5G04255 | 0,21 | 0,08 | 4,38 | 0,02 | 9,85E-01 |
| AT5G26760 | 0,42 | 0,00 | 4,19 | 0,00 | 1,00E+00 |
| AT5G26770 | 1,06 | -0,46 | 1,75 | -0,26 | 7,91E-01 |
| AT5G26880 | 0,63 | -1,21 | 3,13 | -0,39 | 6,99E-01 |
| AT5G26910 | 0,66 | -1,70 | 2,61 | -0,65 | 5,15E-01 |
| AT5G26960 | 0,27 | 0,66 | 4,29 | 0,15 | 8,77E-01 |
| AT5G26970 | 0,84 | -0,80 | 1,72 | -0,47 | 6,41E-01 |
| AT5G27030 | 1,03 | 1,78 | 2,02 | 0,88 | 3,79E-01 |
| AT5G27100 | 0,25 | 0,00 | 4,41 | 0,00 | 1,00E+00 |
| AT5G27238 | 0,05 | 0,18 | 4,38 | 0,04 | 9,68E-01 |
| AT5G04375 | 0,24 | -0,02 | 4,38 | 0,00 | 9,97E-01 |
| AT5G27330 | 0,05 | 0,18 | 4,38 | 0,04 | 9,68E-01 |
| AT5G27380 | 0,63 | -2,47 | 3,65 | -0,68 | 4,99E-01 |
| AT5G27550 | 0,34 | -0,30 | 4,40 | -0,07 | 9,46E-01 |
| AT5G27640 | 1,34 | -0,07 | 1,95 | -0,03 | 9,73E-01 |
| AT5G27700 | 0,49 | -0,23 | 3,32 | -0,07 | 9,44E-01 |
| AT5G27715 | 0,29 | 1,56 | 4,32 | 0,36 | 7,19E-01 |
| AT5G04435 | 0,39 | 0,32 | 4,17 | 0,08 | 9,38E-01 |
| AT5G27790 | 0,75 | 3,49 | 3,45 | 1,01 | 3,12E-01 |
| AT5G27840 | 0,72 | -0,29 | 2,93 | -0,10 | 9,21E-01 |
| AT5G27870 | 1,24 | 0,22 | 2,15 | 0,10 | 9,19E-01 |
| AT5G27920 | 0,08 | 0,00 | 4,41 | 0,00 | 1,00E+00 |
| AT5G27960 | 0,74 | 3,22 | 2,62 | 1,23 | 2,19E-01 |
| AT5G28020 | 0,34 | -0,91 | 3,56 | -0,26 | 7,98E-01 |
| AT5G28030 | 0,66 | -0,26 | 3,03 | -0,09 | 9,31E-01 |
| AT5G28050 | 0,28 | 0,42 | 4,36 | 0,10 | 9,24E-01 |
| AT5G04635 | 0,50 | -0,19 | 3,15 | -0,06 | 9,53E-01 |
| AT5G28060 | 0,65 | -0,42 | 2,24 | -0,19 | 8,51E-01 |
| AT5G04685 | 0,19 | 1,60 | 4,31 | 0,37 | 7,11E-01 |
| AT5G28190 | 0,36 | 1,60 | 4,32 | 0,37 | 7,11E-01 |
| AT5G28220 | 0,30 | 0,84 | 4,29 | 0,20 | 8,44E-01 |
| AT5G28235 | 0,34 | 2,58 | 4,28 | 0,60 | 5,47E-01 |
| AT5G28300 | 1,51 | -0,72 | 1,37 | -0,53 | 5,98E-01 |
| AT5G28310 | 0,08 | 0,00 | 4,41 | 0,00 | 1,00E+00 |
| AT5G28400 | 1,06 | -0,45 | 1,96 | -0,23 | 8,20E-01 |
| AT5G28442 | 0,36 | 2,10 | 3,34 | 0,63 | 5,31E-01 |
| AT5G28450 | 0,85 | 0,78 | 2,17 | 0,36 | 7,18E-01 |
| AT5G28530 | 0,22 | 1,19 | 4,34 | 0,28 | 7,83E-01 |
| AT5G28550 | 0,21 | 1,18 | 4,34 | 0,27 | 7,86E-01 |
| AT5G28640 | 0,08 | 0,21 | 4,39 | 0,05 | 9,62E-01 |
| AT5G28750 | 0,65 | 0,33 | 2,15 | 0,15 | 8,78E-01 |
| AT5G28770 | 0,50 | 0,16 | 3,96 | 0,04 | 9,67E-01 |
| AT5G28824 | 1,47 | 1,35 | 1,84 | 0,73 | 4,63E-01 |
| AT5G28850 | 0,48 | 0,94 | 3,24 | 0,29 | 7,72E-01 |
| AT5G28910 | 0,82 | 0,30 | 2,43 | 0,12 | 9,02E-01 |

| | | | | | |
|---|---|---|---|---|---|
| AT5G04925 | 0,18 | -0,84 | 4,35 | -0,19 | 8,47E-01 |
| AT5G30510 | 1,83 | -0,76 | 1,46 | -0,52 | 6,02E-01 |
| AT5G32470 | 0,85 | 0,50 | 2,14 | 0,23 | 8,15E-01 |
| AT5G32590 | 0,68 | 1,13 | 2,15 | 0,52 | 6,00E-01 |
| AT5G33210 | 0,08 | 0,21 | 4,39 | 0,05 | 9,62E-01 |
| AT5G33439 | 0,11 | 0,88 | 4,35 | 0,20 | 8,39E-01 |
| AT5G34850 | 1,64 | -0,62 | 1,47 | -0,43 | 6,70E-01 |
| AT5G35100 | 0,30 | -0,03 | 3,65 | -0,01 | 9,93E-01 |
| AT5G35200 | 0,52 | 0,42 | 3,13 | 0,13 | 8,94E-01 |
| AT5G35220 | 0,25 | 0,00 | 4,41 | 0,00 | 1,00E+00 |
| AT5G35230 | 0,96 | 3,52 | 2,57 | 1,37 | 1,71E-01 |
| AT5G35370 | 0,70 | -0,61 | 2,02 | -0,30 | 7,63E-01 |
| AT5G35475 | 0,36 | 0,79 | 4,32 | 0,18 | 8,55E-01 |
| AT5G35480 | 0,65 | 2,58 | 3,01 | 0,86 | 3,91E-01 |
| AT5G35530 | 0,88 | -0,37 | 2,48 | -0,15 | 8,82E-01 |
| AT5G35560 | 0,64 | -1,19 | 2,33 | -0,51 | 6,11E-01 |
| AT5G35570 | 0,29 | 0,00 | 4,41 | 0,00 | 1,00E+00 |
| AT5G35590 | 0,12 | 0,49 | 4,38 | 0,11 | 9,11E-01 |
| AT5G35603 | 0,59 | 1,51 | 4,06 | 0,37 | 7,09E-01 |
| AT5G35604 | 0,45 | 1,57 | 3,93 | 0,40 | 6,90E-01 |
| AT5G35620 | 0,26 | -0,39 | 4,37 | -0,09 | 9,29E-01 |
| AT5G35690 | 0,37 | -0,26 | 4,38 | -0,06 | 9,52E-01 |
| AT5G35700 | 0,37 | -0,37 | 4,33 | -0,09 | 9,31E-01 |
| AT5G35750 | 1,20 | -1,36 | 2,10 | -0,65 | 5,17E-01 |
| AT5G35753 | 0,77 | -1,22 | 2,34 | -0,52 | 6,02E-01 |
| AT5G35790 | 0,34 | -1,38 | 4,36 | -0,32 | 7,51E-01 |
| AT5G35810 | 0,15 | 0,47 | 4,37 | 0,11 | 9,14E-01 |
| AT5G35890 | 0,19 | 1,17 | 4,34 | 0,27 | 7,87E-01 |
| AT5G35910 | 0,15 | -0,26 | 4,41 | -0,06 | 9,53E-01 |
| AT5G35913 | 0,08 | 0,00 | 4,41 | 0,00 | 1,00E+00 |
| AT5G35970 | 1,60 | -0,75 | 1,26 | -0,60 | 5,50E-01 |
| AT5G35995 | 0,29 | -0,34 | 4,40 | -0,08 | 9,38E-01 |
| AT5G36002 | 0,12 | 0,00 | 4,41 | 0,00 | 1,00E+00 |
| AT5G36120 | 0,05 | 0,18 | 4,38 | 0,04 | 9,68E-01 |
| AT5G36150 | 0,29 | -1,45 | 4,32 | -0,34 | 7,37E-01 |
| AT5G36160 | 0,29 | 0,09 | 4,38 | 0,02 | 9,84E-01 |
| AT5G36200 | 1,52 | 2,70 | 1,87 | 1,44 | 1,49E-01 |
| AT5G36210 | 1,11 | -3,17 | 1,95 | -1,62 | 1,04E-01 |
| AT5G36220 | 0,80 | 0,32 | 2,21 | 0,15 | 8,85E-01 |
| AT5G36700 | 0,58 | -1,24 | 2,71 | -0,46 | 6,48E-01 |
| AT5G36790 | 0,17 | -0,58 | 4,38 | -0,13 | 8,94E-01 |
| AT5G36880 | 0,93 | -0,38 | 2,38 | -0,16 | 8,73E-01 |
| AT5G36930 | 1,17 | 0,26 | 2,09 | 0,12 | 9,01E-01 |
| AT5G37260 | 0,78 | 0,00 | 3,44 | 0,00 | 1,00E+00 |
| AT5G37310 | 0,77 | -0,29 | 3,48 | -0,08 | 9,32E-01 |
| AT5G37370 | 0,59 | -2,19 | 3,78 | -0,58 | 5,63E-01 |
| AT5G37480 | 0,13 | 0,94 | 4,35 | 0,22 | 8,29E-01 |
| AT5G37590 | 0,66 | 0,00 | 3,69 | 0,00 | 1,00E+00 |
| AT5G37720 | 0,46 | -0,26 | 3,38 | -0,08 | 9,39E-01 |
| AT5G37770 | 1,11 | 2,71 | 2,66 | 1,02 | 3,07E-01 |
| AT5G37780 | 0,80 | 0,68 | 2,09 | 0,32 | 7,47E-01 |
| AT5G05545 | 0,31 | 2,22 | 4,29 | 0,52 | 6,05E-01 |
| AT5G37830 | 1,35 | 0,11 | 2,13 | 0,05 | 9,57E-01 |
| AT5G37890 | 0,36 | 0,00 | 4,41 | 0,00 | 1,00E+00 |
| AT5G05565 | 0,45 | 1,40 | 4,10 | 0,34 | 7,32E-01 |
| AT5G38020 | 0,42 | -3,73 | 4,33 | -0,86 | 3,89E-01 |

| | | | | | |
|---|---|---|---|---|---|
| AT5G38080 | 0,08 | 0,00 | 4,41 | 0,00 | 1,00E+00 |
| AT5G38140 | 0,10 | 0,86 | 4,35 | 0,20 | 8,43E-01 |
| AT5G38212 | 1,00 | 3,17 | 2,13 | 1,49 | 1,37E-01 |
| AT5G38240 | 0,41 | 0,00 | 4,41 | 0,00 | 1,00E+00 |
| AT5G38280 | 0,74 | -1,77 | 3,56 | -0,50 | 6,20E-01 |
| AT5G38450 | 0,46 | 0,53 | 4,09 | 0,13 | 8,97E-01 |
| AT5G05755 | 0,05 | 0,18 | 4,38 | 0,04 | 9,68E-01 |
| AT5G38510 | 0,76 | -1,85 | 2,49 | -0,74 | 4,57E-01 |
| AT5G38640 | 0,50 | -0,48 | 2,72 | -0,18 | 8,58E-01 |
| AT5G38660 | 1,59 | -0,19 | 1,76 | -0,11 | 9,14E-01 |
| AT5G38650 | 1,00 | -0,08 | 2,34 | -0,03 | 9,73E-01 |
| AT5G38830 | 0,49 | -2,98 | 4,37 | -0,68 | 4,96E-01 |
| AT5G39130 | 0,12 | 0,27 | 4,38 | 0,06 | 9,50E-01 |
| AT5G39160 | 0,34 | 2,37 | 4,28 | 0,55 | 5,80E-01 |
| AT5G05895 | 0,62 | 1,64 | 2,90 | 0,57 | 5,72E-01 |
| AT5G39480 | 0,08 | 0,21 | 4,39 | 0,05 | 9,62E-01 |
| AT5G39520 | 0,33 | -0,26 | 4,41 | -0,06 | 9,53E-01 |
| AT5G39530 | 0,77 | -3,78 | 2,90 | -1,31 | 1,92E-01 |
| AT5G39600 | 0,66 | -2,72 | 2,26 | -1,20 | 2,28E-01 |
| AT5G39610 | 0,52 | -0,41 | 3,94 | -0,10 | 9,18E-01 |
| AT5G05995 | 0,40 | 1,28 | 4,17 | 0,31 | 7,59E-01 |
| AT5G39680 | 0,42 | 0,00 | 4,41 | 0,00 | 1,00E+00 |
| AT5G39693 | 0,43 | -0,32 | 3,20 | -0,10 | 9,20E-01 |
| AT5G39920 | 0,25 | -2,07 | 4,37 | -0,47 | 6,35E-01 |
| AT5G39950 | 0,95 | 0,07 | 2,45 | 0,03 | 9,78E-01 |
| AT5G40170 | 0,68 | 0,81 | 2,21 | 0,37 | 7,14E-01 |
| AT5G40240 | 0,14 | 0,13 | 4,38 | 0,03 | 9,77E-01 |
| AT5G40370 | 0,37 | 0,00 | 3,40 | 0,00 | 9,99E-01 |
| AT5G40470 | 0,27 | -0,39 | 4,39 | -0,09 | 9,29E-01 |
| AT5G40580 | 0,60 | 0,00 | 4,40 | 0,00 | 1,00E+00 |
| AT5G40660 | 0,29 | 0,00 | 4,41 | 0,00 | 1,00E+00 |
| AT5G40740 | 0,85 | 0,79 | 2,45 | 0,32 | 7,46E-01 |
| AT5G40770 | 1,47 | -1,92 | 1,89 | -1,02 | 3,10E-01 |
| AT5G40820 | 0,95 | 0,54 | 2,41 | 0,22 | 8,23E-01 |
| AT5G40830 | 0,63 | 0,61 | 2,66 | 0,23 | 8,19E-01 |
| AT5G40890 | 1,74 | -0,61 | 1,53 | -0,40 | 6,88E-01 |
| AT5G40940 | 0,56 | 0,00 | 3,88 | 0,00 | 1,00E+00 |
| AT5G41020 | 0,73 | 1,36 | 2,59 | 0,52 | 6,00E-01 |
| AT5G41130 | 0,21 | 0,00 | 4,41 | 0,00 | 1,00E+00 |
| AT5G41150 | 0,33 | 0,00 | 4,41 | 0,00 | 1,00E+00 |
| AT5G41190 | 0,83 | 0,69 | 2,96 | 0,23 | 8,14E-01 |
| AT5G41330 | 0,40 | -0,37 | 3,42 | -0,11 | 9,14E-01 |
| AT5G41400 | 0,65 | 2,34 | 2,60 | 0,90 | 3,68E-01 |
| AT5G41460 | 0,17 | -3,06 | 4,35 | -0,70 | 4,82E-01 |
| AT5G41520 | 0,58 | 1,33 | 2,72 | 0,49 | 6,24E-01 |
| AT5G41670 | 0,43 | 1,74 | 2,77 | 0,63 | 5,29E-01 |
| AT5G06235 | 0,77 | 0,23 | 4,39 | 0,05 | 9,59E-01 |
| AT5G41870 | 0,38 | -0,60 | 4,29 | -0,14 | 8,89E-01 |
| AT5G41990 | 1,56 | -1,16 | 1,99 | -0,58 | 5,60E-01 |
| AT5G42100 | 1,07 | -0,29 | 1,55 | -0,19 | 8,51E-01 |
| AT5G42110 | 0,47 | 1,97 | 3,21 | 0,61 | 5,40E-01 |
| AT5G42130 | 0,21 | -0,32 | 4,40 | -0,07 | 9,43E-01 |
| AT5G42140 | 0,12 | 0,11 | 4,38 | 0,03 | 9,80E-01 |
| AT5G42190 | 0,65 | 0,09 | 2,51 | 0,04 | 9,71E-01 |
| AT5G42280 | 0,14 | 0,13 | 4,38 | 0,03 | 9,77E-01 |
| AT5G42300 | 0,48 | 1,91 | 3,89 | 0,49 | 6,23E-01 |

| | | | | | |
|---|---|---|---|---|---|
| AT5G42360 | 1,47 | 0,51 | 1,94 | 0,26 | 7,91E-01 |
| AT5G42380 | 1,28 | 2,35 | 2,09 | 1,12 | 2,63E-01 |
| AT5G42400 | 0,21 | 0,23 | 4,37 | 0,05 | 9,58E-01 |
| AT5G06305 | 1,02 | -0,23 | 1,57 | -0,15 | 8,82E-01 |
| AT5G42490 | 0,56 | -0,26 | 3,93 | -0,07 | 9,47E-01 |
| AT5G42567 | 0,58 | 3,40 | 4,27 | 0,80 | 4,25E-01 |
| AT5G42570 | 0,15 | -2,07 | 4,39 | -0,47 | 6,38E-01 |
| AT5G42660 | 0,12 | 0,11 | 4,38 | 0,03 | 9,80E-01 |
| AT5G42750 | 1,69 | 2,25 | 1,50 | 1,50 | 1,34E-01 |
| AT5G42790 | 0,93 | 0,05 | 1,92 | 0,03 | 9,80E-01 |
| AT5G42800 | 0,38 | 0,00 | 4,41 | 0,00 | 1,00E+00 |
| AT5G42850 | 1,15 | -1,34 | 2,23 | -0,60 | 5,46E-01 |
| AT5G42870 | 0,81 | 1,51 | 2,34 | 0,64 | 5,20E-01 |
| AT5G42890 | 0,08 | 0,21 | 4,39 | 0,05 | 9,62E-01 |
| AT5G42900 | 0,25 | 0,00 | 4,41 | 0,00 | 1,00E+00 |
| AT5G42905 | 0,52 | 1,90 | 3,79 | 0,50 | 6,17E-01 |
| AT5G42940 | 0,81 | -1,14 | 2,64 | -0,43 | 6,67E-01 |
| AT5G42955 | 0,65 | -4,01 | 3,78 | -1,06 | 2,89E-01 |
| AT5G42957 | 0,83 | 0,76 | 2,21 | 0,34 | 7,32E-01 |
| AT5G42990 | 0,17 | 0,00 | 4,41 | 0,00 | 1,00E+00 |
| AT5G43010 | 0,13 | -0,61 | 4,37 | -0,14 | 8,89E-01 |
| AT5G43020 | 0,41 | 0,86 | 4,16 | 0,21 | 8,36E-01 |
| AT5G43130 | 1,30 | -0,22 | 2,36 | -0,09 | 9,25E-01 |
| AT5G43190 | 1,41 | 0,33 | 1,71 | 0,19 | 8,48E-01 |
| AT5G43260 | 0,52 | 0,74 | 2,68 | 0,27 | 7,84E-01 |
| AT5G43270 | 0,76 | 0,73 | 3,06 | 0,24 | 8,13E-01 |
| AT5G43285 | 0,35 | -2,75 | 4,31 | -0,64 | 5,24E-01 |
| AT5G43320 | 0,66 | 0,45 | 2,67 | 0,17 | 8,66E-01 |
| AT5G43440 | 0,40 | 0,11 | 4,34 | 0,03 | 9,79E-01 |
| AT5G43670 | 0,40 | 0,17 | 4,17 | 0,04 | 9,68E-01 |
| AT5G43710 | 0,21 | 0,00 | 4,41 | 0,00 | 1,00E+00 |
| AT5G43745 | 0,36 | -2,09 | 4,39 | -0,47 | 6,35E-01 |
| AT5G43750 | 0,47 | 0,99 | 4,34 | 0,23 | 8,19E-01 |
| AT5G43780 | 0,63 | 2,24 | 2,64 | 0,85 | 3,96E-01 |
| AT5G43790 | 0,45 | -0,97 | 4,28 | -0,23 | 8,22E-01 |
| AT5G43830 | 0,91 | -1,24 | 1,92 | -0,65 | 5,16E-01 |
| AT5G43850 | 0,29 | -1,55 | 3,64 | -0,43 | 6,71E-01 |
| AT5G43880 | 0,19 | 0,56 | 4,30 | 0,13 | 8,97E-01 |
| AT5G43900 | 1,55 | -0,25 | 1,87 | -0,13 | 8,94E-01 |
| AT5G43950 | 0,65 | -0,47 | 2,69 | -0,18 | 8,61E-01 |
| AT5G43960 | 0,54 | -1,11 | 3,17 | -0,35 | 7,26E-01 |
| AT5G44000 | 0,95 | 1,22 | 2,29 | 0,53 | 5,94E-01 |
| AT5G44005 | 0,16 | -0,21 | 4,40 | -0,05 | 9,63E-01 |
| AT5G44100 | 0,66 | -0,24 | 2,61 | -0,09 | 9,26E-01 |
| AT5G44250 | 0,12 | 0,00 | 4,41 | 0,00 | 1,00E+00 |
| AT5G44260 | 0,22 | -0,90 | 4,35 | -0,21 | 8,36E-01 |
| AT5G44283 | 0,91 | -0,53 | 2,44 | -0,22 | 8,29E-01 |
| AT5G44290 | 1,25 | -0,02 | 1,91 | -0,01 | 9,93E-01 |
| AT5G44306 | 0,93 | 2,12 | 2,86 | 0,74 | 4,59E-01 |
| AT5G44495 | 1,23 | 0,01 | 1,35 | 0,01 | 9,92E-01 |
| AT5G44510 | 0,61 | 0,00 | 4,41 | 0,00 | 1,00E+00 |
| AT5G44568 | 0,48 | -0,52 | 2,77 | -0,19 | 8,51E-01 |
| AT5G06595 | 1,38 | -0,29 | 1,66 | -0,18 | 8,60E-01 |
| AT5G44670 | 0,13 | 0,00 | 4,41 | 0,00 | 1,00E+00 |
| AT5G44680 | 0,48 | -3,22 | 4,31 | -0,75 | 4,54E-01 |
| AT5G44785 | 0,10 | 0,08 | 4,38 | 0,02 | 9,85E-01 |

| | | | | | |
|---|---|---|---|---|---|
| AT5G44790 | 1,35 | 0,28 | 1,85 | 0,15 | 8,78E-01 |
| AT5G44820 | 0,08 | 0,00 | 4,41 | 0,00 | 1,00E+00 |
| AT5G44840 | 0,14 | -2,43 | 4,37 | -0,55 | 5,79E-01 |
| AT5G45010 | 0,58 | -0,46 | 3,17 | -0,15 | 8,84E-01 |
| AT5G45040 | 0,08 | 0,00 | 4,41 | 0,00 | 1,00E+00 |
| AT5G45050 | 0,86 | 0,01 | 2,18 | 0,00 | 9,98E-01 |
| AT5G45170 | 0,11 | 0,88 | 4,35 | 0,20 | 8,39E-01 |
| AT5G45180 | 0,66 | -3,58 | 3,89 | -0,92 | 3,58E-01 |
| AT5G45230 | 0,97 | 0,90 | 1,80 | 0,50 | 6,18E-01 |
| AT5G45250 | 0,93 | 0,92 | 2,17 | 0,42 | 6,73E-01 |
| AT5G45275 | 0,48 | -0,37 | 4,40 | -0,08 | 9,34E-01 |
| AT5G45290 | 1,09 | -0,30 | 2,25 | -0,13 | 8,95E-01 |
| AT5G45340 | 0,20 | 1,44 | 4,33 | 0,33 | 7,40E-01 |
| AT5G45360 | 0,08 | 0,00 | 4,41 | 0,00 | 1,00E+00 |
| AT5G45410 | 0,46 | -2,19 | 4,09 | -0,53 | 5,93E-01 |
| AT5G45420 | 0,27 | -0,06 | 4,34 | -0,01 | 9,89E-01 |
| AT5G45440 | 0,74 | 0,00 | 3,53 | 0,00 | 1,00E+00 |
| AT5G45460 | 0,18 | 0,31 | 4,38 | 0,07 | 9,44E-01 |
| AT5G45573 | 0,61 | 2,76 | 2,84 | 0,97 | 3,30E-01 |
| AT5G45630 | 0,05 | 0,18 | 4,38 | 0,04 | 9,68E-01 |
| AT5G45650 | 0,61 | 1,73 | 3,20 | 0,54 | 5,89E-01 |
| AT5G45775 | 0,86 | -0,65 | 2,03 | -0,32 | 7,47E-01 |
| AT5G45800 | 0,52 | 0,34 | 2,36 | 0,14 | 8,86E-01 |
| AT5G45820 | 1,25 | 0,39 | 1,52 | 0,26 | 7,96E-01 |
| AT5G46020 | 1,24 | -1,66 | 2,36 | -0,70 | 4,82E-01 |
| AT5G46040 | 0,11 | -0,38 | 4,39 | -0,09 | 9,31E-01 |
| AT5G46050 | 0,37 | -0,34 | 4,40 | -0,08 | 9,38E-01 |
| AT5G46280 | 0,57 | 0,74 | 2,97 | 0,25 | 8,03E-01 |
| AT5G46290 | 0,63 | -2,13 | 3,13 | -0,68 | 4,96E-01 |
| AT5G46340 | 0,61 | -3,05 | 3,93 | -0,77 | 4,38E-01 |
| AT5G46395 | 0,11 | 0,88 | 4,35 | 0,20 | 8,39E-01 |
| AT5G46580 | 0,75 | -1,57 | 2,28 | -0,69 | 4,91E-01 |
| AT5G46620 | 0,64 | -2,50 | 2,98 | -0,84 | 4,03E-01 |
| AT5G46750 | 0,66 | 0,43 | 3,80 | 0,11 | 9,09E-01 |
| AT5G46790 | 0,53 | 0,19 | 3,04 | 0,06 | 9,49E-01 |
| AT5G46800 | 0,83 | -1,37 | 2,17 | -0,63 | 5,28E-01 |
| AT5G46860 | 0,46 | -0,68 | 3,99 | -0,17 | 8,65E-01 |
| AT5G46970 | 0,65 | 0,16 | 2,47 | 0,07 | 9,47E-01 |
| AT5G47040 | 1,34 | 1,88 | 1,78 | 1,05 | 2,93E-01 |
| AT5G47070 | 0,97 | 2,65 | 2,31 | 1,15 | 2,50E-01 |
| AT5G47080 | 0,13 | 0,00 | 4,41 | 0,00 | 1,00E+00 |
| AT5G47110 | 1,25 | 1,52 | 1,72 | 0,89 | 3,76E-01 |
| AT5G47190 | 0,93 | -1,56 | 2,06 | -0,76 | 4,49E-01 |
| AT5G47210 | 1,48 | 1,18 | 1,92 | 0,62 | 5,37E-01 |
| AT5G47220 | 0,78 | 1,24 | 2,58 | 0,48 | 6,30E-01 |
| AT5G47250 | 0,28 | -0,75 | 4,32 | -0,17 | 8,63E-01 |
| AT5G47370 | 0,28 | 0,00 | 4,41 | 0,00 | 1,00E+00 |
| AT5G47460 | 0,15 | -0,35 | 4,39 | -0,08 | 9,36E-01 |
| AT5G47570 | 0,30 | 0,03 | 4,35 | 0,01 | 9,94E-01 |
| AT5G47580 | 0,64 | 0,00 | 3,71 | 0,00 | 1,00E+00 |
| AT5G47650 | 0,20 | 0,00 | 4,41 | 0,00 | 1,00E+00 |
| AT5G47700 | 0,72 | -1,32 | 2,16 | -0,61 | 5,41E-01 |
| AT5G47730 | 0,16 | 0,15 | 4,38 | 0,04 | 9,72E-01 |
| AT5G47750 | 1,50 | 0,32 | 2,42 | 0,13 | 8,96E-01 |
| AT5G47760 | 0,12 | 0,11 | 4,38 | 0,03 | 9,80E-01 |
| AT5G47810 | 0,55 | -1,95 | 2,71 | -0,72 | 4,73E-01 |

| | | | | | |
|---|---|---|---|---|---|
| AT5G47830 | 0,54 | -2,49 | 4,38 | -0,57 | 5,69E-01 |
| AT5G47850 | 0,49 | 0,88 | 4,02 | 0,22 | 8,26E-01 |
| AT5G47880 | 0,55 | 0,47 | 3,21 | 0,15 | 8,83E-01 |
| AT5G47930 | 0,53 | 1,39 | 2,57 | 0,54 | 5,88E-01 |
| AT5G48000 | 0,12 | 0,49 | 4,38 | 0,11 | 9,11E-01 |
| AT5G07055 | 0,58 | -0,53 | 3,00 | -0,18 | 8,60E-01 |
| AT5G48120 | 1,18 | -0,04 | 1,96 | -0,02 | 9,83E-01 |
| AT5G48150 | 0,85 | -2,41 | 3,59 | -0,67 | 5,02E-01 |
| AT5G48180 | 0,98 | 1,01 | 2,04 | 0,49 | 6,22E-01 |
| AT5G48375 | 0,26 | 0,00 | 4,41 | 0,00 | 1,00E+00 |
| AT5G48412 | 0,64 | 0,36 | 2,53 | 0,14 | 8,88E-01 |
| AT5G48440 | 0,43 | 0,48 | 4,23 | 0,11 | 9,09E-01 |
| AT5G48540 | 0,08 | 0,00 | 4,41 | 0,00 | 1,00E+00 |
| AT5G48580 | 0,41 | -0,32 | 4,20 | -0,08 | 9,39E-01 |
| AT5G48640 | 0,19 | -0,02 | 4,35 | -0,01 | 9,96E-01 |
| AT5G48657 | 0,40 | 1,76 | 3,29 | 0,53 | 5,93E-01 |
| AT5G07265 | 0,12 | 0,00 | 4,41 | 0,00 | 1,00E+00 |
| AT5G48770 | 0,08 | 0,21 | 4,39 | 0,05 | 9,62E-01 |
| AT5G07275 | 0,71 | 0,00 | 4,41 | 0,00 | 1,00E+00 |
| AT5G48810 | 0,80 | 0,31 | 2,38 | 0,13 | 8,95E-01 |
| AT5G48830 | 0,28 | -0,36 | 4,39 | -0,08 | 9,35E-01 |
| AT5G48880 | 0,58 | -0,17 | 4,36 | -0,04 | 9,69E-01 |
| AT5G48910 | 0,76 | 0,86 | 2,18 | 0,39 | 6,93E-01 |
| AT5G48930 | 1,24 | -0,56 | 1,78 | -0,32 | 7,52E-01 |
| AT5G49030 | 1,79 | -2,22 | 1,58 | -1,41 | 1,60E-01 |
| AT5G49100 | 1,11 | 1,53 | 1,85 | 0,83 | 4,09E-01 |
| AT5G49300 | 0,36 | 0,00 | 4,41 | 0,00 | 1,00E+00 |
| AT5G49350 | 0,08 | 0,00 | 4,41 | 0,00 | 1,00E+00 |
| AT5G49420 | 0,35 | 0,32 | 4,33 | 0,07 | 9,40E-01 |
| AT5G49430 | 1,30 | 1,04 | 1,76 | 0,59 | 5,55E-01 |
| AT5G49540 | 0,90 | -2,23 | 2,57 | -0,87 | 3,87E-01 |
| AT5G49550 | 0,27 | 0,13 | 4,38 | 0,03 | 9,76E-01 |
| AT5G49555 | 0,40 | -0,60 | 4,33 | -0,14 | 8,90E-01 |
| AT5G49570 | 0,11 | -0,26 | 4,41 | -0,06 | 9,53E-01 |
| AT5G49580 | 0,46 | -1,63 | 3,98 | -0,41 | 6,82E-01 |
| AT5G49630 | 0,19 | 1,60 | 4,31 | 0,37 | 7,11E-01 |
| AT5G49645 | 0,08 | 0,00 | 4,41 | 0,00 | 1,00E+00 |
| AT5G49660 | 0,65 | 0,06 | 2,54 | 0,03 | 9,80E-01 |
| AT5G49665 | 0,50 | -0,33 | 4,05 | -0,08 | 9,34E-01 |
| AT5G49830 | 1,27 | -0,05 | 1,92 | -0,03 | 9,80E-01 |
| AT5G49890 | 0,22 | 0,06 | 4,34 | 0,01 | 9,89E-01 |
| AT5G49900 | 0,41 | -0,30 | 4,22 | -0,07 | 9,43E-01 |
| AT5G49930 | 0,74 | -1,31 | 2,21 | -0,59 | 5,55E-01 |
| AT5G49950 | 0,25 | -0,57 | 4,38 | -0,13 | 8,97E-01 |
| AT5G50000 | 0,84 | -0,38 | 1,90 | -0,20 | 8,43E-01 |
| AT5G50150 | 0,10 | 0,24 | 4,38 | 0,05 | 9,57E-01 |
| AT5G50220 | 0,28 | -0,16 | 4,40 | -0,04 | 9,70E-01 |
| AT5G50320 | 0,77 | 0,52 | 3,03 | 0,17 | 8,63E-01 |
| AT5G50340 | 0,25 | 0,00 | 4,41 | 0,00 | 1,00E+00 |
| AT5G50350 | 0,29 | -0,15 | 4,35 | -0,03 | 9,72E-01 |
| AT5G50380 | 0,97 | 1,21 | 2,09 | 0,58 | 5,62E-01 |
| AT5G50390 | 0,17 | 0,00 | 4,41 | 0,00 | 1,00E+00 |
| AT5G50400 | 0,54 | 0,49 | 3,97 | 0,12 | 9,02E-01 |
| AT5G50460 | 0,36 | -1,29 | 4,40 | -0,29 | 7,70E-01 |
| AT5G50480 | 0,53 | -3,46 | 3,78 | -0,92 | 3,60E-01 |
| AT5G50570 | 0,31 | 0,14 | 4,33 | 0,03 | 9,75E-01 |

| | | | | | |
|---|---|---|---|---|---|
| AT5G50630 | 0,05 | 0,18 | 4,38 | 0,04 | 9,68E-01 |
| AT5G50640 | 0,24 | 0,00 | 4,41 | 0,00 | 1,00E+00 |
| AT5G50670 | 0,93 | 0,62 | 2,38 | 0,26 | 7,95E-01 |
| AT5G50860 | 0,49 | 0,00 | 3,97 | 0,00 | 1,00E+00 |
| AT5G50870 | 0,08 | 0,00 | 4,41 | 0,00 | 1,00E+00 |
| AT5G50900 | 0,83 | -0,18 | 2,47 | -0,07 | 9,42E-01 |
| AT5G51000 | 0,61 | 1,68 | 2,94 | 0,57 | 5,68E-01 |
| AT5G51020 | 0,37 | -0,37 | 4,33 | -0,09 | 9,31E-01 |
| AT5G51110 | 0,78 | -1,47 | 2,24 | -0,65 | 5,13E-01 |
| AT5G51170 | 0,12 | 0,00 | 4,41 | 0,00 | 1,00E+00 |
| AT5G51195 | 0,57 | -0,37 | 2,48 | -0,15 | 8,81E-01 |
| AT5G51220 | 0,23 | 1,25 | 4,34 | 0,29 | 7,73E-01 |
| AT5G51290 | 0,61 | -0,34 | 3,11 | -0,11 | 9,14E-01 |
| AT5G51390 | 0,29 | 0,00 | 4,41 | 0,00 | 1,00E+00 |
| AT5G51400 | 1,21 | -1,08 | 1,60 | -0,67 | 5,00E-01 |
| AT5G51550 | 1,02 | 1,36 | 1,82 | 0,75 | 4,55E-01 |
| AT5G51640 | 0,39 | 0,48 | 3,42 | 0,14 | 8,89E-01 |
| AT5G51650 | 0,32 | 1,79 | 3,49 | 0,51 | 6,08E-01 |
| AT5G51690 | 0,72 | 0,54 | 2,58 | 0,21 | 8,35E-01 |
| AT5G51710 | 0,10 | 0,86 | 4,35 | 0,20 | 8,43E-01 |
| AT5G51820 | 0,69 | -2,42 | 2,55 | -0,95 | 3,42E-01 |
| AT5G52060 | 0,54 | 0,26 | 2,64 | 0,10 | 9,23E-01 |
| AT5G52090 | 0,73 | 1,80 | 2,12 | 0,85 | 3,96E-01 |
| AT5G52120 | 0,47 | -1,71 | 4,37 | -0,39 | 6,95E-01 |
| AT5G52180 | 0,61 | -0,39 | 3,15 | -0,12 | 9,01E-01 |
| AT5G52190 | 1,80 | 0,05 | 1,92 | 0,02 | 9,81E-01 |
| AT5G52320 | 1,77 | -0,50 | 1,34 | -0,37 | 7,11E-01 |
| AT5G52330 | 0,15 | 0,02 | 4,36 | 0,00 | 9,96E-01 |
| AT5G52370 | 0,37 | -0,31 | 4,35 | -0,07 | 9,43E-01 |
| AT5G52450 | 0,44 | -0,36 | 3,31 | -0,11 | 9,13E-01 |
| AT5G52540 | 1,60 | 0,77 | 1,70 | 0,46 | 6,49E-01 |
| AT5G52550 | 0,25 | -0,39 | 4,39 | -0,09 | 9,29E-01 |
| AT5G52650 | 0,64 | -1,31 | 3,69 | -0,36 | 7,22E-01 |
| AT5G52780 | 0,46 | -0,26 | 3,20 | -0,08 | 9,35E-01 |
| AT5G52797 | 0,48 | 1,97 | 3,19 | 0,62 | 5,37E-01 |
| AT5G53000 | 0,29 | -0,12 | 4,39 | -0,03 | 9,79E-01 |
| AT5G53010 | 0,50 | -1,25 | 3,38 | -0,37 | 7,12E-01 |
| AT5G53050 | 0,49 | -0,84 | 3,11 | -0,27 | 7,88E-01 |
| AT5G53130 | 0,72 | 0,91 | 2,92 | 0,31 | 7,56E-01 |
| AT5G07955 | 0,08 | 0,00 | 4,41 | 0,00 | 1,00E+00 |
| AT5G53300 | 0,89 | -0,41 | 2,76 | -0,15 | 8,82E-01 |
| AT5G53320 | 0,58 | 0,00 | 3,11 | 0,00 | 1,00E+00 |
| AT5G07975 | 0,08 | 0,00 | 4,41 | 0,00 | 1,00E+00 |
| AT5G53490 | 0,65 | 1,33 | 2,15 | 0,62 | 5,38E-01 |
| AT5G53500 | 0,52 | 0,63 | 3,09 | 0,20 | 8,38E-01 |
| AT5G53530 | 0,37 | -1,37 | 4,35 | -0,32 | 7,52E-01 |
| AT5G53570 | 0,72 | -0,42 | 2,51 | -0,17 | 8,67E-01 |
| AT5G53580 | 0,36 | -0,48 | 3,57 | -0,13 | 8,94E-01 |
| AT5G53730 | 0,14 | 0,81 | 4,35 | 0,19 | 8,52E-01 |
| AT5G53850 | 0,47 | -2,61 | 4,36 | -0,60 | 5,50E-01 |
| AT5G53870 | 0,19 | 1,00 | 4,33 | 0,23 | 8,17E-01 |
| AT5G53900 | 0,79 | 0,57 | 2,25 | 0,25 | 8,01E-01 |
| AT5G54030 | 0,27 | 2,26 | 4,29 | 0,53 | 5,98E-01 |
| AT5G54040 | 0,27 | 1,98 | 4,30 | 0,46 | 6,46E-01 |
| AT5G54050 | 0,38 | 1,64 | 4,33 | 0,38 | 7,05E-01 |
| AT5G54060 | 0,12 | 0,00 | 4,41 | 0,00 | 1,00E+00 |

| | | | | | |
|---|---|---|---|---|---|
| AT5G54090 | 1,00 | -2,13 | 2,44 | -0,87 | 3,83E-01 |
| AT5G54100 | 0,65 | -0,34 | 3,03 | -0,11 | 9,09E-01 |
| AT5G54170 | 0,64 | 0,40 | 2,27 | 0,18 | 8,61E-01 |
| AT5G54280 | 1,29 | 1,51 | 1,72 | 0,88 | 3,80E-01 |
| AT5G54380 | 1,11 | 1,05 | 1,97 | 0,53 | 5,95E-01 |
| AT5G54410 | 0,82 | 0,96 | 2,00 | 0,48 | 6,32E-01 |
| AT5G54430 | 1,12 | 0,48 | 2,55 | 0,19 | 8,52E-01 |
| AT5G54460 | 1,09 | 3,36 | 3,22 | 1,04 | 2,97E-01 |
| AT5G54470 | 0,35 | 1,57 | 3,43 | 0,46 | 6,48E-01 |
| AT5G54510 | 0,08 | 0,00 | 4,41 | 0,00 | 1,00E+00 |
| AT5G54610 | 0,49 | 0,24 | 3,91 | 0,06 | 9,50E-01 |
| AT5G54620 | 0,12 | 0,00 | 4,41 | 0,00 | 1,00E+00 |
| AT5G54630 | 0,29 | -1,63 | 4,31 | -0,38 | 7,05E-01 |
| AT5G54650 | 0,47 | 0,00 | 4,41 | 0,00 | 1,00E+00 |
| AT5G54690 | 0,39 | 0,00 | 4,40 | 0,00 | 1,00E+00 |
| AT5G54700 | 0,89 | 1,91 | 2,18 | 0,88 | 3,80E-01 |
| AT5G54710 | 0,79 | 2,23 | 2,39 | 0,93 | 3,50E-01 |
| AT5G54720 | 0,73 | 2,18 | 2,25 | 0,97 | 3,33E-01 |
| AT5G54780 | 1,16 | -0,48 | 2,00 | -0,24 | 8,10E-01 |
| AT5G54930 | 0,40 | -2,14 | 4,23 | -0,51 | 6,13E-01 |
| AT5G54960 | 0,98 | -1,14 | 2,08 | -0,55 | 5,83E-01 |
| AT5G55040 | 0,74 | 0,60 | 2,15 | 0,28 | 7,81E-01 |
| AT5G55055 | 0,05 | 0,18 | 4,38 | 0,04 | 9,68E-01 |
| AT5G55100 | 1,12 | 0,00 | 2,62 | 0,00 | 1,00E+00 |
| AT5G55130 | 0,31 | 1,29 | 3,58 | 0,36 | 7,18E-01 |
| AT5G55150 | 1,21 | 0,00 | 2,64 | 0,00 | 1,00E+00 |
| AT5G55160 | 0,37 | -0,26 | 4,41 | -0,06 | 9,53E-01 |
| AT5G55210 | 0,08 | 0,21 | 4,39 | 0,05 | 9,62E-01 |
| AT5G55250 | 0,40 | -0,04 | 4,28 | -0,01 | 9,92E-01 |
| AT5G55280 | 1,13 | -1,23 | 2,17 | -0,57 | 5,70E-01 |
| AT5G55290 | 0,30 | 1,44 | 4,32 | 0,33 | 7,38E-01 |
| AT5G55300 | 0,70 | -2,51 | 2,31 | -1,09 | 2,77E-01 |
| AT5G55340 | 0,19 | 0,37 | 4,36 | 0,08 | 9,32E-01 |
| AT5G55390 | 0,36 | 2,10 | 4,30 | 0,49 | 6,25E-01 |
| AT5G08275 | 0,59 | -2,19 | 3,86 | -0,57 | 5,72E-01 |
| AT5G08285 | 0,30 | -0,87 | 4,34 | -0,20 | 8,41E-01 |
| AT5G55510 | 0,05 | 0,18 | 4,38 | 0,04 | 9,68E-01 |
| AT5G55540 | 1,63 | -1,39 | 1,78 | -0,78 | 4,36E-01 |
| AT5G55550 | 0,34 | 0,00 | 4,41 | 0,00 | 1,00E+00 |
| AT5G55570 | 1,30 | -5,37 | 2,96 | -1,81 | 7,01E-02 |
| AT5G55600 | 0,31 | -0,46 | 3,66 | -0,13 | 9,00E-01 |
| AT5G55670 | 0,30 | 1,59 | 4,32 | 0,37 | 7,14E-01 |
| AT5G55710 | 0,92 | -1,18 | 2,57 | -0,46 | 6,46E-01 |
| AT5G55740 | 0,27 | 0,78 | 4,36 | 0,18 | 8,58E-01 |
| AT5G55790 | 0,33 | 0,63 | 4,36 | 0,14 | 8,85E-01 |
| AT5G55835 | 0,61 | 0,30 | 2,31 | 0,13 | 8,98E-01 |
| AT5G55910 | 1,51 | -1,95 | 1,55 | -1,26 | 2,07E-01 |
| AT5G55920 | 0,81 | 0,68 | 2,52 | 0,27 | 7,89E-01 |
| AT5G55940 | 0,50 | 0,00 | 3,97 | 0,00 | 1,00E+00 |
| AT5G56000 | 1,26 | -0,23 | 1,72 | -0,13 | 8,94E-01 |
| AT5G56180 | 0,23 | 0,24 | 4,39 | 0,06 | 9,56E-01 |
| AT5G56240 | 1,23 | 0,89 | 2,36 | 0,38 | 7,06E-01 |
| AT5G56250 | 0,87 | 0,93 | 2,85 | 0,33 | 7,45E-01 |
| AT5G56260 | 0,25 | -0,57 | 4,38 | -0,13 | 8,97E-01 |
| AT5G56280 | 0,08 | 0,21 | 4,39 | 0,05 | 9,62E-01 |
| AT5G56350 | 0,73 | -2,77 | 2,04 | -1,36 | 1,75E-01 |

| | | | | | |
|---|---|---|---|---|---|
| AT5G56360 | 0,87 | -3,02 | 2,83 | -1,07 | 2,85E-01 |
| AT5G56410 | 0,30 | 0,14 | 4,38 | 0,03 | 9,74E-01 |
| AT5G56450 | 0,20 | -0,15 | 4,39 | -0,03 | 9,72E-01 |
| AT5G56710 | 0,79 | 0,00 | 3,49 | 0,00 | 1,00E+00 |
| AT5G56730 | 0,65 | -1,08 | 2,63 | -0,41 | 6,82E-01 |
| AT5G56760 | 0,72 | -1,33 | 2,80 | -0,48 | 6,34E-01 |
| AT5G56860 | 0,23 | 1,67 | 4,31 | 0,39 | 6,99E-01 |
| AT5G56890 | 0,16 | -0,22 | 4,40 | -0,05 | 9,60E-01 |
| AT5G56900 | 1,69 | 1,54 | 1,83 | 0,84 | 4,01E-01 |
| AT5G56980 | 0,28 | -0,26 | 4,41 | -0,06 | 9,53E-01 |
| AT5G57040 | 0,80 | 0,84 | 2,17 | 0,39 | 6,99E-01 |
| AT5G57160 | 0,81 | -1,24 | 2,58 | -0,48 | 6,30E-01 |
| AT5G57170 | 0,08 | 0,21 | 4,39 | 0,05 | 9,62E-01 |
| AT5G57250 | 0,54 | -0,27 | 3,97 | -0,07 | 9,45E-01 |
| AT5G57270 | 0,45 | -1,45 | 4,38 | -0,33 | 7,41E-01 |
| AT5G57290 | 0,69 | 0,09 | 2,47 | 0,03 | 9,72E-01 |
| AT5G57345 | 0,70 | 0,46 | 2,25 | 0,21 | 8,37E-01 |
| AT5G57580 | 0,90 | -1,50 | 2,52 | -0,60 | 5,51E-01 |
| AT5G57630 | 0,49 | 0,12 | 3,35 | 0,03 | 9,73E-01 |
| AT5G57670 | 0,16 | 1,16 | 4,34 | 0,27 | 7,89E-01 |
| AT5G57780 | 0,36 | 1,22 | 4,33 | 0,28 | 7,78E-01 |
| AT5G57785 | 0,56 | 1,83 | 2,59 | 0,71 | 4,81E-01 |
| AT5G57790 | 0,05 | 0,18 | 4,38 | 0,04 | 9,68E-01 |
| AT5G57850 | 0,51 | -2,17 | 4,34 | -0,50 | 6,18E-01 |
| AT5G57900 | 0,27 | -1,08 | 4,38 | -0,25 | 8,05E-01 |
| AT5G57960 | 0,12 | 0,59 | 4,36 | 0,14 | 8,92E-01 |
| AT5G58030 | 0,11 | -0,26 | 4,41 | -0,06 | 9,53E-01 |
| AT5G58050 | 0,17 | 0,00 | 4,41 | 0,00 | 1,00E+00 |
| AT5G58090 | 0,76 | 0,56 | 2,46 | 0,23 | 8,19E-01 |
| AT5G58130 | 0,84 | -0,29 | 2,81 | -0,10 | 9,17E-01 |
| AT5G58170 | 0,90 | 0,00 | 3,33 | 0,00 | 1,00E+00 |
| AT5G58220 | 0,12 | 0,35 | 4,37 | 0,08 | 9,35E-01 |
| AT5G58260 | 0,45 | -1,32 | 3,36 | -0,39 | 6,94E-01 |
| AT5G58380 | 0,93 | 0,39 | 2,23 | 0,17 | 8,61E-01 |
| AT5G58440 | 0,43 | -1,22 | 2,83 | -0,43 | 6,65E-01 |
| AT5G58570 | 0,61 | 1,21 | 3,90 | 0,31 | 7,57E-01 |
| AT5G08860 | 0,08 | 0,21 | 4,39 | 0,05 | 9,62E-01 |
| AT5G58600 | 0,29 | 0,00 | 4,41 | 0,00 | 1,00E+00 |
| AT5G58650 | 0,16 | 0,39 | 4,37 | 0,09 | 9,29E-01 |
| AT5G58670 | 0,69 | -2,14 | 2,29 | -0,93 | 3,50E-01 |
| AT5G58720 | 0,64 | 0,17 | 2,61 | 0,07 | 9,48E-01 |
| AT5G58730 | 0,50 | -0,26 | 4,04 | -0,06 | 9,48E-01 |
| AT5G58870 | 0,47 | 0,40 | 3,35 | 0,12 | 9,04E-01 |
| AT5G58900 | 0,28 | 0,00 | 4,41 | 0,00 | 1,00E+00 |
| AT5G58930 | 1,32 | -1,17 | 2,13 | -0,55 | 5,83E-01 |
| AT5G59030 | 0,49 | 0,00 | 3,97 | 0,00 | 1,00E+00 |
| AT5G59130 | 0,21 | 0,00 | 4,41 | 0,00 | 1,00E+00 |
| AT5G59220 | 0,62 | 0,00 | 3,83 | 0,00 | 1,00E+00 |
| AT5G59290 | 0,24 | -0,02 | 4,38 | 0,00 | 9,97E-01 |
| AT5G08920 | 0,49 | -0,05 | 4,06 | -0,01 | 9,91E-01 |
| AT5G59480 | 0,17 | 0,00 | 4,41 | 0,00 | 1,00E+00 |
| AT5G59550 | 1,55 | 2,28 | 1,81 | 1,26 | 2,08E-01 |
| AT5G59662 | 0,05 | 0,18 | 4,38 | 0,04 | 9,68E-01 |
| AT5G59700 | 0,33 | -0,26 | 4,41 | -0,06 | 9,53E-01 |
| AT5G59770 | 0,46 | -0,84 | 3,26 | -0,26 | 7,97E-01 |
| AT5G59780 | 1,19 | 0,30 | 2,14 | 0,14 | 8,88E-01 |

| | | | | | |
|---|---|---|---|---|---|
| AT5G59840 | 0,17 | -0,78 | 4,36 | -0,18 | 8,59E-01 |
| AT5G59850 | 0,69 | -1,11 | 2,55 | -0,44 | 6,63E-01 |
| AT5G59870 | 0,19 | 0,81 | 4,36 | 0,19 | 8,52E-01 |
| AT5G59900 | 0,16 | -0,56 | 4,37 | -0,13 | 8,98E-01 |
| AT5G59960 | 0,11 | -0,26 | 4,41 | -0,06 | 9,53E-01 |
| AT5G60050 | 0,15 | -2,06 | 4,39 | -0,47 | 6,38E-01 |
| AT5G60100 | 0,30 | -0,75 | 4,33 | -0,17 | 8,63E-01 |
| AT5G60160 | 0,49 | 0,49 | 4,05 | 0,12 | 9,05E-01 |
| AT5G60180 | 0,21 | 0,00 | 4,41 | 0,00 | 1,00E+00 |
| AT5G60210 | 0,29 | -1,33 | 4,39 | -0,30 | 7,62E-01 |
| AT5G60340 | 0,36 | 0,53 | 4,37 | 0,12 | 9,04E-01 |
| AT5G60400 | 0,12 | 0,00 | 4,41 | 0,00 | 1,00E+00 |
| AT5G60460 | 0,27 | 0,38 | 4,31 | 0,09 | 9,30E-01 |
| AT5G60670 | 0,64 | -0,77 | 2,34 | -0,33 | 7,40E-01 |
| AT5G60700 | 1,32 | 0,00 | 2,48 | 0,00 | 1,00E+00 |
| AT5G60710 | 1,48 | -0,72 | 1,69 | -0,43 | 6,70E-01 |
| AT5G60800 | 0,66 | -1,32 | 3,12 | -0,42 | 6,73E-01 |
| AT5G60840 | 0,12 | 1,17 | 4,34 | 0,27 | 7,88E-01 |
| AT5G60930 | 0,24 | -0,34 | 4,40 | -0,08 | 9,38E-01 |
| AT5G60960 | 0,29 | -0,30 | 4,40 | -0,07 | 9,46E-01 |
| AT5G60990 | 0,16 | 0,15 | 4,38 | 0,04 | 9,72E-01 |
| AT5G61040 | 0,88 | 0,37 | 4,32 | 0,09 | 9,32E-01 |
| AT5G61060 | 0,37 | 0,55 | 3,47 | 0,16 | 8,75E-01 |
| AT5G61130 | 0,17 | 0,00 | 4,41 | 0,00 | 1,00E+00 |
| AT5G61150 | 0,86 | 0,73 | 2,60 | 0,28 | 7,79E-01 |
| AT5G61250 | 0,23 | -0,34 | 4,40 | -0,08 | 9,38E-01 |
| AT5G61390 | 0,43 | 0,00 | 4,41 | 0,00 | 1,00E+00 |
| AT5G09005 | 1,51 | -0,63 | 1,28 | -0,49 | 6,23E-01 |
| AT5G61460 | 0,21 | 0,00 | 4,41 | 0,00 | 1,00E+00 |
| AT5G61480 | 0,83 | 0,28 | 2,20 | 0,13 | 8,97E-01 |
| AT5G61500 | 0,76 | 0,49 | 2,93 | 0,17 | 8,66E-01 |
| AT5G61510 | 0,56 | -1,11 | 3,34 | -0,33 | 7,39E-01 |
| AT5G61520 | 0,33 | 0,26 | 4,37 | 0,06 | 9,52E-01 |
| AT5G61590 | 1,28 | -0,61 | 1,63 | -0,37 | 7,09E-01 |
| AT5G61600 | 0,57 | 1,66 | 2,51 | 0,66 | 5,08E-01 |
| AT5G61800 | 0,12 | 0,00 | 4,41 | 0,00 | 1,00E+00 |
| AT5G61810 | 0,48 | 0,84 | 3,33 | 0,25 | 8,01E-01 |
| AT5G61820 | 1,07 | 0,80 | 2,43 | 0,33 | 7,42E-01 |
| AT5G61840 | 0,89 | 1,51 | 2,24 | 0,68 | 4,98E-01 |
| AT5G61910 | 1,60 | 1,44 | 1,67 | 0,86 | 3,90E-01 |
| AT5G61900 | 1,09 | 1,12 | 1,85 | 0,60 | 5,47E-01 |
| AT5G61960 | 0,94 | -0,53 | 2,59 | -0,20 | 8,39E-01 |
| AT5G61990 | 0,30 | 0,00 | 4,41 | 0,00 | 1,00E+00 |
| AT5G62090 | 0,32 | -1,73 | 4,34 | -0,40 | 6,90E-01 |
| AT5G62110 | 0,34 | 1,37 | 4,32 | 0,32 | 7,50E-01 |
| AT5G62130 | 0,45 | 1,56 | 3,13 | 0,50 | 6,20E-01 |
| AT5G62140 | 1,15 | 0,74 | 1,60 | 0,46 | 6,45E-01 |
| AT5G62180 | 0,08 | 0,00 | 4,41 | 0,00 | 1,00E+00 |
| AT5G62220 | 1,34 | 1,74 | 1,62 | 1,07 | 2,83E-01 |
| AT5G62300 | 1,07 | -2,72 | 1,95 | -1,39 | 1,65E-01 |
| AT5G62430 | 0,29 | 0,28 | 4,37 | 0,06 | 9,48E-01 |
| AT5G62470 | 0,30 | -0,20 | 4,27 | -0,05 | 9,63E-01 |
| AT5G62530 | 0,56 | -2,44 | 3,07 | -0,79 | 4,27E-01 |
| AT5G62600 | 0,50 | -0,26 | 4,04 | -0,06 | 9,48E-01 |
| AT5G62610 | 0,31 | 1,03 | 4,34 | 0,24 | 8,13E-01 |
| AT5G62630 | 0,37 | -2,68 | 4,38 | -0,61 | 5,41E-01 |

| | | | | | |
|---|---|---|---|---|---|
| AT5G62650 | 0,39 | -2,86 | 4,35 | -0,66 | 5,12E-01 |
| AT5G62680 | 0,33 | 0,06 | 4,38 | 0,01 | 9,88E-01 |
| AT5G62790 | 0,60 | 0,25 | 2,43 | 0,10 | 9,17E-01 |
| AT5G63080 | 0,23 | -1,08 | 4,38 | -0,25 | 8,06E-01 |
| AT5G63100 | 0,29 | -0,35 | 4,40 | -0,08 | 9,37E-01 |
| AT5G63110 | 0,58 | -1,43 | 3,10 | -0,46 | 6,44E-01 |
| AT5G63120 | 0,65 | 0,47 | 3,13 | 0,15 | 8,79E-01 |
| AT5G63160 | 0,51 | 2,68 | 3,05 | 0,88 | 3,79E-01 |
| AT5G63180 | 0,25 | -1,50 | 4,33 | -0,34 | 7,30E-01 |
| AT5G63290 | 0,56 | 0,00 | 4,41 | 0,00 | 1,00E+00 |
| AT5G63310 | 0,47 | -2,02 | 4,39 | -0,46 | 6,45E-01 |
| AT5G63320 | 0,52 | 2,90 | 3,82 | 0,76 | 4,48E-01 |
| AT5G63370 | 1,26 | -0,50 | 1,93 | -0,26 | 7,96E-01 |
| AT5G63400 | 0,30 | -2,59 | 4,37 | -0,59 | 5,53E-01 |
| AT5G63410 | 0,46 | 2,15 | 2,66 | 0,81 | 4,18E-01 |
| AT5G63470 | 0,44 | 2,46 | 3,98 | 0,62 | 5,36E-01 |
| AT5G63610 | 0,51 | -0,26 | 4,02 | -0,07 | 9,48E-01 |
| AT5G63625 | 1,28 | -0,51 | 1,67 | -0,30 | 7,60E-01 |
| AT5G63760 | 0,90 | 1,49 | 2,40 | 0,62 | 5,35E-01 |
| AT5G63790 | 0,54 | 0,00 | 4,41 | 0,00 | 1,00E+00 |
| AT5G63860 | 0,75 | -0,52 | 2,28 | -0,23 | 8,19E-01 |
| AT5G63870 | 0,08 | 0,00 | 4,41 | 0,00 | 1,00E+00 |
| AT5G63890 | 0,39 | -2,64 | 4,27 | -0,62 | 5,37E-01 |
| AT5G63910 | 0,66 | -0,17 | 2,99 | -0,06 | 9,55E-01 |
| AT5G64020 | 0,23 | -0,34 | 4,40 | -0,08 | 9,38E-01 |
| AT5G64050 | 1,12 | -1,17 | 1,72 | -0,68 | 4,95E-01 |
| AT5G64070 | 1,53 | 1,30 | 1,67 | 0,78 | 4,37E-01 |
| AT5G64130 | 0,35 | -0,65 | 3,54 | -0,18 | 8,55E-01 |
| AT5G64140 | 1,06 | 1,91 | 1,74 | 1,09 | 2,74E-01 |
| AT5G64170 | 0,64 | 1,60 | 3,14 | 0,51 | 6,10E-01 |
| AT5G64200 | 0,28 | 0,03 | 4,38 | 0,01 | 9,95E-01 |
| AT5G64280 | 1,39 | 1,21 | 1,85 | 0,65 | 5,13E-01 |
| AT5G64290 | 0,42 | -1,19 | 4,18 | -0,29 | 7,75E-01 |
| AT5G64340 | 1,19 | 0,56 | 1,80 | 0,31 | 7,54E-01 |
| AT5G64350 | 0,41 | -1,84 | 4,24 | -0,43 | 6,64E-01 |
| AT5G64360 | 0,62 | -0,31 | 3,76 | -0,08 | 9,34E-01 |
| AT5G64410 | 0,79 | 0,15 | 2,16 | 0,07 | 9,44E-01 |
| AT5G64430 | 0,96 | -0,59 | 2,25 | -0,26 | 7,94E-01 |
| AT5G64460 | 0,08 | 0,21 | 4,39 | 0,05 | 9,62E-01 |
| AT5G64550 | 0,25 | 0,00 | 4,41 | 0,00 | 1,00E+00 |
| AT5G64570 | 1,49 | 0,51 | 1,65 | 0,31 | 7,59E-01 |
| AT5G64580 | 0,77 | -0,30 | 2,70 | -0,11 | 9,11E-01 |
| AT5G64610 | 0,28 | 1,08 | 4,34 | 0,25 | 8,04E-01 |
| AT5G64660 | 0,37 | 0,63 | 3,39 | 0,19 | 8,53E-01 |
| AT5G64670 | 0,17 | -2,13 | 4,40 | -0,48 | 6,28E-01 |
| AT5G64760 | 0,83 | 0,68 | 3,05 | 0,22 | 8,23E-01 |
| AT5G64830 | 0,77 | -1,19 | 2,60 | -0,46 | 6,46E-01 |
| AT5G64860 | 0,28 | 0,43 | 4,36 | 0,10 | 9,21E-01 |
| AT5G64870 | 0,67 | 2,80 | 2,77 | 1,01 | 3,12E-01 |
| AT5G64920 | 1,21 | 0,84 | 1,99 | 0,42 | 6,74E-01 |
| AT5G64930 | 0,39 | 0,00 | 4,40 | 0,00 | 1,00E+00 |
| AT5G64980 | 0,12 | 0,00 | 4,41 | 0,00 | 1,00E+00 |
| AT5G65110 | 0,46 | -0,98 | 3,21 | -0,30 | 7,61E-01 |
| AT5G65200 | 0,16 | 0,15 | 4,38 | 0,04 | 9,72E-01 |
| AT5G65240 | 0,27 | 1,28 | 3,72 | 0,34 | 7,31E-01 |
| AT5G65310 | 1,18 | -0,27 | 2,04 | -0,13 | 8,93E-01 |

| | | | | | |
|---|---|---|---|---|---|
| AT5G65360 | 0,52 | 1,39 | 3,13 | 0,44 | 6,58E-01 |
| AT5G65430 | 0,62 | 1,48 | 2,63 | 0,56 | 5,73E-01 |
| AT5G65560 | 0,37 | 0,51 | 4,37 | 0,12 | 9,07E-01 |
| AT5G65685 | 0,97 | -0,28 | 2,04 | -0,14 | 8,89E-01 |
| AT5G65840 | 0,17 | 0,07 | 4,38 | 0,02 | 9,87E-01 |
| AT5G65910 | 1,14 | -2,04 | 3,33 | -0,61 | 5,40E-01 |
| AT5G65930 | 0,99 | 0,21 | 2,22 | 0,10 | 9,23E-01 |
| AT5G09535 | 0,24 | 0,00 | 4,41 | 0,00 | 1,00E+00 |
| AT5G65970 | 0,45 | 0,00 | 4,41 | 0,00 | 1,00E+00 |
| AT5G66055 | 0,63 | -2,10 | 2,60 | -0,81 | 4,19E-01 |
| AT5G66120 | 0,85 | 0,58 | 2,15 | 0,27 | 7,87E-01 |
| AT5G66210 | 1,05 | 1,89 | 2,17 | 0,87 | 3,84E-01 |
| AT5G66300 | 0,13 | 0,94 | 4,35 | 0,22 | 8,29E-01 |
| AT5G66310 | 0,12 | 0,00 | 4,41 | 0,00 | 1,00E+00 |
| AT5G66320 | 0,48 | 0,89 | 3,35 | 0,26 | 7,91E-01 |
| AT5G66330 | 0,11 | 0,68 | 4,37 | 0,16 | 8,76E-01 |
| AT5G66400 | 0,42 | -3,85 | 4,19 | -0,92 | 3,57E-01 |
| AT5G66440 | 0,33 | 0,22 | 4,39 | 0,05 | 9,61E-01 |
| AT5G66460 | 0,57 | -0,38 | 4,39 | -0,09 | 9,31E-01 |
| AT5G66500 | 0,15 | -0,31 | 4,40 | -0,07 | 9,43E-01 |
| AT5G66530 | 0,97 | -2,82 | 2,52 | -1,12 | 2,64E-01 |
| AT5G66540 | 0,21 | 0,39 | 4,36 | 0,09 | 9,29E-01 |
| AT5G66558 | 1,38 | 0,20 | 1,86 | 0,11 | 9,14E-01 |
| AT5G66595 | 0,74 | 1,80 | 2,32 | 0,78 | 4,37E-01 |
| AT5G66607 | 0,08 | 0,21 | 4,39 | 0,05 | 9,62E-01 |
| AT5G66650 | 0,26 | 0,00 | 4,41 | 0,00 | 1,00E+00 |
| AT5G66675 | 0,51 | 2,61 | 3,03 | 0,86 | 3,90E-01 |
| AT5G66680 | 0,76 | 1,68 | 3,69 | 0,45 | 6,50E-01 |
| AT5G66720 | 0,41 | -0,27 | 4,41 | -0,06 | 9,51E-01 |
| AT5G66730 | 0,31 | -0,35 | 4,35 | -0,08 | 9,35E-01 |
| AT5G66850 | 0,24 | -2,09 | 4,39 | -0,48 | 6,34E-01 |
| AT5G66890 | 0,75 | 0,00 | 3,56 | 0,00 | 1,00E+00 |
| AT5G66900 | 1,52 | 0,72 | 1,88 | 0,38 | 7,03E-01 |
| AT5G66910 | 0,38 | 1,44 | 3,39 | 0,43 | 6,71E-01 |
| AT5G67120 | 0,79 | 2,82 | 2,68 | 1,05 | 2,93E-01 |
| AT5G67150 | 0,35 | 0,81 | 4,35 | 0,19 | 8,53E-01 |
| AT5G67190 | 0,91 | 0,63 | 2,47 | 0,25 | 7,99E-01 |
| AT5G67220 | 0,60 | -0,03 | 2,54 | -0,01 | 9,91E-01 |
| AT5G67240 | 1,06 | -1,25 | 1,94 | -0,64 | 5,20E-01 |
| AT5G67245 | 0,45 | 0,31 | 4,39 | 0,07 | 9,44E-01 |
| AT5G67250 | 1,03 | 2,88 | 2,00 | 1,44 | 1,49E-01 |
| AT5G67280 | 0,58 | 0,40 | 2,58 | 0,16 | 8,76E-01 |
| AT5G67290 | 0,82 | 1,57 | 2,45 | 0,64 | 5,21E-01 |
| AT5G09855 | 0,69 | 1,64 | 2,44 | 0,67 | 5,02E-01 |
| AT5G67310 | 0,78 | -2,09 | 2,93 | -0,72 | 4,74E-01 |
| AT5G67350 | 0,75 | 1,19 | 2,49 | 0,48 | 6,32E-01 |
| AT5G67380 | 0,30 | 1,22 | 4,34 | 0,28 | 7,79E-01 |
| AT5G67460 | 0,10 | 0,08 | 4,38 | 0,02 | 9,85E-01 |
| AT5G67480 | 1,04 | 0,35 | 2,61 | 0,13 | 8,95E-01 |
| AT5G67490 | 1,16 | 0,65 | 1,85 | 0,35 | 7,26E-01 |
| AT5G67580 | 0,39 | -0,72 | 2,81 | -0,26 | 7,98E-01 |
| AT5G67610 | 0,37 | 0,00 | 4,41 | 0,00 | 1,00E+00 |

ıninfected *chr11 chr17* relative to wild type.

| type |
|---|
| miRNA_precursor |
| miRNA_precursor |
| protein_coding |
| protein_coding |
| protein_coding |
| miRNA_precursor |
| miRNA_precursor |
| miRNA_precursor |
| ncRNA |
| miRNA_precursor |
| miRNA_precursor |
| ribosomal_rna |
| miRNA_precursor |
| protein_coding |
| ncRNA |
| protein_coding |
| miRNA_precursor |
| miRNA_precursor |
| miRNA_precursor |
| miRNA_precursor |
| miRNA_precursor |
| protein_coding |
| protein_coding |
| protein_coding |
| protein_coding |
| miRNA_precursor |
| miRNA_precursor |
| miRNA_precursor |
| miRNA_precursor |
| miRNA_precursor |
| miRNA_precursor |
| protein_coding |
| miRNA_precursor |
| protein_coding |
| miRNA_precursor |
| protein_coding |
| protein_coding |
| small_nucle(ol)ar_rna |
| miRNA_precursor |
| protein_coding |
| ncRNA |
| miRNA_precursor |
| miRNA_precursor |
| ncRNA |
| small_nucle(ol)ar_rna |
| small_nucle(ol)ar_rna |
| protein_coding |
| protein_coding |
| miRNA_precursor |
| ncRNA |
| protein_coding |
| small_nucle(ol)ar_rna |
| ncRNA |

protein_coding
protein_coding
ncRNA
protein_coding
protein_coding
small_nucle(ol)ar_rna
protein_coding
miRNA_precursor
protein_coding
miRNA_precursor
protein_coding
miRNA_precursor
protein_coding
miRNA_precursor
protein_coding
small_nucle(ol)ar_rna
ncRNA
miRNA_precursor
small_nucle(ol)ar_rna
miRNA_precursor
protein_coding
miRNA_precursor
miRNA_precursor
protein_coding
protein_coding
miRNA_precursor
protein_coding
protein_coding
protein_coding
protein_coding
protein_coding
protein_coding
protein_coding
protein_coding
miRNA_precursor
small_nucle(ol)ar_rna
protein_coding
protein_coding
miRNA_precursor
protein_coding
small_nucle(ol)ar_rna
protein_coding
protein_coding
protein_coding
ncRNA
ncRNA
ncRNA
ncRNA
protein_coding
protein_coding
protein_coding
protein_coding
protein_coding
miRNA_precursor
small_nucle(ol)ar_rna
protein_coding

miRNA_precursor
protein_coding
protein_coding
miRNA_precursor
ncRNA
protein_coding
protein_coding
small_nucle(ol)ar_rna
small_nucle(ol)ar_rna
miRNA_precursor
miRNA_precursor
protein_coding
miRNA_precursor
protein_coding
miRNA_precursor
miRNA_precursor
protein_coding
miRNA_precursor
miRNA_precursor
protein_coding
protein_coding
protein_coding
protein_coding
protein_coding
ncRNA
protein_coding
protein_coding
miRNA_precursor
protein_coding
protein_coding
protein_coding
protein_coding
ncRNA
protein_coding
miRNA_precursor
protein_coding
protein_coding
protein_coding
protein_coding
protein_coding
protein_coding
protein_coding
protein_coding
miRNA_precursor
ncRNA
protein_coding
protein_coding
protein_coding
miRNA_precursor
miRNA_precursor
miRNA_precursor
protein_coding
protein_coding
protein_coding
miRNA_precursor
miRNA_precursor

protein_coding
protein_coding
protein_coding
protein_coding
protein_coding
protein_coding
small_nucle(ol)ar_rna
protein_coding
protein_coding
protein_coding
protein_coding
protein_coding
miRNA_precursor
miRNA_precursor
protein_coding
protein_coding
protein_coding
protein_coding
protein_coding
protein_coding
protein_coding
protein_coding
small_nucle(ol)ar_rna
protein_coding
protein_coding
protein_coding
protein_coding
miRNA_precursor
protein_coding
protein_coding
protein_coding
protein_coding
miRNA_precursor
protein_coding
protein_coding
protein_coding
ncRNA
miRNA_precursor
protein_coding
protein_coding
protein_coding
protein_coding
ncRNA
miRNA_precursor
protein_coding
protein_coding
miRNA_precursor
protein_coding
protein_coding
ribosomal_rna
miRNA_precursor
protein_coding
protein_coding
protein_coding
protein_coding
protein_coding

protein_coding
protein_coding
ncRNA
protein_coding
miRNA_precursor
protein_coding
miRNA_precursor
protein_coding
miRNA_precursor
protein_coding
protein_coding
protein_coding
miRNA_precursor
small_nucle(ol)ar_rna
miRNA_precursor
protein_coding
protein_coding
protein_coding
pre_trna
protein_coding
small_nucle(ol)ar_rna
miRNA_precursor
protein_coding
protein_coding
protein_coding
protein_coding
protein_coding
protein_coding
protein_coding
ncRNA
protein_coding
protein_coding
protein_coding
miRNA_precursor
small_nucle(ol)ar_rna
miRNA_precursor
protein_coding
protein_coding
miRNA_precursor
protein_coding
protein_coding
protein_coding
protein_coding
ncRNA
protein_coding
protein_coding
protein_coding
protein_coding
protein_coding
protein_coding
miRNA_precursor
protein_coding
protein_coding
small_nucle(ol)ar_rna
small_nucle(ol)ar_rna
protein_coding

miRNA_precursor
protein_coding
ncRNA
protein_coding
protein_coding
protein_coding
protein_coding
protein_coding
miRNA_precursor
protein_coding
protein_coding
protein_coding
protein_coding
protein_coding
ncRNA
protein_coding
protein_coding
protein_coding
ncRNA
protein_coding
protein_coding
protein_coding
pre_trna
protein_coding
protein_coding
ncRNA
protein_coding
protein_coding
protein_coding
protein_coding
protein_coding
protein_coding
protein_coding
miRNA_precursor
protein_coding
protein_coding
protein_coding
protein_coding
miRNA_precursor
ncRNA
protein_coding
protein_coding
protein_coding
small_nucle(ol)ar_rna
protein_coding
protein_coding
protein_coding
protein_coding
small_nucle(ol)ar_rna
protein_coding
protein_coding
protein_coding
protein_coding
protein_coding
ncRNA
protein_coding

small_nucle(ol)ar_rna
protein_coding
protein_coding
protein_coding
protein_coding
protein_coding
protein_coding
protein_coding
miRNA_precursor
miRNA_precursor
protein_coding
miRNA_precursor
protein_coding
protein_coding
protein_coding
protein_coding
protein_coding
ncRNA
protein_coding
protein_coding
miRNA_precursor
protein_coding
protein_coding
protein_coding
protein_coding
protein_coding
protein_coding
protein_coding
protein_coding
protein_coding
protein_coding
protein_coding
protein_coding
protein_coding
protein_coding
protein_coding
protein_coding
protein_coding
protein_coding
protein_coding
small_nucle(ol)ar_rna
protein_coding
protein_coding
miRNA_precursor
protein_coding
protein_coding
protein_coding
protein_coding
protein_coding
protein_coding
protein_coding
protein_coding
protein_coding
ncRNA
protein_coding
protein_coding

protein_coding
ncRNA
protein_coding
protein_coding
protein_coding
protein_coding
ncRNA
miRNA_precursor
protein_coding
protein_coding
protein_coding
protein_coding
protein_coding
protein_coding
protein_coding
protein_coding
protein_coding
small_nucle(ol)ar_rna
protein_coding
small_nucle(ol)ar_rna
protein_coding
protein_coding
protein_coding
protein_coding
ncRNA
protein_coding
protein_coding
protein_coding
protein_coding
protein_coding
protein_coding
protein_coding
protein_coding
protein_coding
protein_coding
miRNA_precursor
ncRNA
protein_coding
protein_coding
protein_coding
protein_coding
small_nucle(ol)ar_rna
protein_coding
protein_coding
protein_coding
protein_coding
protein_coding
protein_coding
protein_coding
protein_coding
protein_coding
protein_coding
protein_coding
protein_coding
protein_coding
miRNA_precursor

protein_coding
protein_coding
protein_coding
protein_coding
protein_coding
ncRNA
pre_trna
miRNA_precursor
protein_coding
protein_coding
protein_coding
protein_coding
protein_coding
protein_coding
protein_coding
protein_coding
protein_coding
protein_coding
protein_coding
protein_coding
miRNA_precursor
ncRNA
ncRNA
protein_coding
ncRNA
protein_coding
protein_coding
protein_coding
protein_coding
protein_coding
protein_coding
protein_coding
ncRNA
protein_coding
protein_coding
protein_coding
protein_coding
protein_coding
protein_coding
protein_coding
protein_coding
ncRNA
protein_coding
protein_coding
ncRNA
protein_coding
protein_coding
protein_coding
ncRNA
protein_coding
protein_coding
protein_coding
protein_coding
protein_coding
protein_coding
protein_coding

protein_coding
protein_coding
protein_coding
miRNA_precursor
protein_coding
miRNA_precursor
protein_coding
small_nucle(ol)ar_rna
protein_coding
protein_coding
protein_coding
protein_coding
protein_coding
small_nucle(ol)ar_rna
small_nucle(ol)ar_rna
protein_coding
ncRNA
protein_coding
small_nucle(ol)ar_rna
protein_coding
ncRNA
protein_coding
protein_coding
protein_coding
protein_coding
protein_coding
protein_coding
protein_coding
protein_coding
protein_coding
protein_coding
protein_coding
protein_coding
protein_coding
protein_coding
protein_coding
protein_coding
protein_coding
protein_coding
protein_coding
protein_coding
protein_coding
protein_coding
protein_coding
protein_coding
protein_coding
protein_coding
protein_coding
protein_coding
protein_coding
ncRNA
protein_coding
protein_coding
protein_coding
protein_coding
protein_coding

ncRNA
protein_coding
protein_coding
protein_coding
ncRNA
protein_coding
ncRNA
protein_coding
miRNA_precursor
protein_coding
protein_coding
protein_coding
ncRNA
protein_coding
protein_coding
protein_coding
small_nucle(ol)ar_rna
protein_coding
protein_coding
protein_coding
protein_coding
protein_coding
miRNA_precursor
protein_coding
ncRNA
protein_coding
small_nucle(ol)ar_rna
protein_coding
protein_coding
protein_coding
protein_coding
protein_coding
protein_coding
protein_coding
protein_coding
protein_coding
protein_coding
protein_coding
protein_coding
miRNA_precursor
protein_coding
protein_coding
protein_coding
ncRNA
protein_coding
small_nucle(ol)ar_rna
protein_coding
protein_coding
protein_coding
protein_coding
protein_coding
miRNA_precursor
protein_coding
protein_coding
protein_coding
protein_coding

protein_coding
protein_coding
protein_coding
protein_coding
ncRNA
protein_coding
protein_coding
protein_coding
protein_coding
small_nucle(ol)ar_rna
protein_coding
protein_coding
protein_coding
protein_coding
protein_coding
protein_coding
protein_coding
protein_coding
miRNA_precursor
miRNA_precursor
protein_coding
protein_coding
protein_coding
protein_coding
protein_coding
protein_coding
protein_coding
protein_coding
protein_coding
protein_coding
protein_coding
protein_coding
protein_coding
protein_coding
protein_coding
protein_coding
protein_coding
protein_coding
protein_coding
protein_coding
small_nucle(ol)ar_rna
protein_coding
protein_coding
protein_coding
ncRNA
miRNA_precursor
small_nucle(ol)ar_rna
protein_coding
protein_coding
small_nucle(ol)ar_rna
protein_coding
protein_coding
protein_coding
protein_coding
protein_coding
protein_coding

protein_coding
protein_coding
protein_coding
protein_coding
ncRNA
protein_coding
protein_coding
protein_coding
protein_coding
protein_coding
protein_coding
protein_coding
protein_coding
protein_coding
protein_coding
protein_coding
protein_coding
protein_coding
protein_coding
protein_coding
protein_coding
protein_coding
protein_coding
protein_coding
protein_coding
ncRNA
protein_coding
protein_coding
protein_coding
ncRNA
protein_coding
protein_coding
ncRNA
protein_coding
protein_coding
protein_coding
protein_coding
protein_coding
protein_coding
protein_coding
small_nucle(ol)ar_rna
protein_coding
protein_coding
protein_coding
protein_coding
protein_coding
small_nucle(ol)ar_rna
protein_coding
ncRNA
miRNA_precursor
protein_coding
protein_coding
protein_coding
protein_coding
protein_coding
protein_coding

protein_coding
protein_coding
protein_coding
protein_coding
protein_coding
protein_coding
protein_coding
protein_coding
protein_coding
protein_coding
protein_coding
protein_coding
protein_coding
protein_coding
protein_coding
protein_coding
protein_coding
protein_coding
protein_coding
protein_coding
protein_coding
protein_coding
protein_coding
protein_coding
protein_coding
protein_coding
protein_coding
protein_coding
protein_coding
protein_coding
protein_coding
protein_coding
protein_coding
protein_coding
small_nucle(ol)ar_rna
protein_coding
small_nucle(ol)ar_rna
miRNA_precursor
protein_coding
protein_coding
protein_coding
protein_coding
protein_coding
protein_coding
protein_coding
protein_coding
protein_coding
protein_coding
small_nucle(ol)ar_rna
protein_coding
ncRNA
protein_coding
protein_coding
protein_coding
protein_coding
protein_coding

small_nucle(ol)ar_rna
protein_coding
protein_coding
miRNA_precursor
small_nucle(ol)ar_rna
protein_coding
ncRNA
protein_coding
protein_coding
protein_coding
protein_coding
protein_coding
protein_coding
small_nucle(ol)ar_rna
protein_coding
protein_coding
miRNA_precursor
protein_coding
protein_coding
protein_coding
protein_coding
protein_coding
protein_coding
protein_coding
protein_coding
miRNA_precursor
protein_coding
protein_coding
miRNA_precursor
protein_coding
protein_coding
protein_coding
protein_coding
protein_coding
protein_coding
protein_coding
protein_coding
protein_coding
miRNA_precursor
protein_coding
protein_coding
protein_coding
protein_coding
protein_coding
small_nucle(ol)ar_rna
ncRNA
protein_coding
protein_coding
protein_coding
protein_coding
small_nucle(ol)ar_rna
protein_coding
protein_coding
protein_coding
miRNA_precursor
protein_coding

protein_coding
protein_coding
miRNA_precursor
protein_coding
protein_coding
protein_coding
protein_coding
protein_coding
protein_coding
protein_coding
protein_coding
protein_coding
protein_coding
protein_coding
miRNA_precursor
protein_coding
protein_coding
protein_coding
protein_coding
protein_coding
small_nucle(ol)ar_rna
protein_coding
protein_coding
protein_coding
protein_coding
miRNA_precursor
ncRNA
protein_coding
small_nucle(ol)ar_rna
miRNA_precursor
ncRNA
protein_coding
miRNA_precursor
protein_coding
protein_coding
protein_coding
protein_coding
small_nucle(ol)ar_rna
protein_coding
protein_coding
protein_coding
ncRNA
protein_coding
protein_coding
protein_coding
protein_coding
protein_coding
miRNA_precursor
protein_coding
protein_coding
ncRNA
protein_coding
protein_coding
protein_coding
protein_coding
protein_coding

protein_coding
ncRNA
protein_coding
protein_coding
protein_coding
protein_coding
protein_coding
protein_coding
protein_coding
protein_coding
small_nucle(ol)ar_rna
protein_coding
protein_coding
protein_coding
protein_coding
protein_coding
protein_coding
protein_coding
protein_coding
protein_coding
small_nucle(ol)ar_rna
protein_coding
protein_coding
protein_coding
small_nucle(ol)ar_rna
protein_coding
protein_coding
ncRNA
protein_coding
protein_coding
protein_coding
protein_coding
protein_coding
protein_coding
protein_coding
protein_coding
protein_coding
protein_coding
protein_coding
protein_coding
protein_coding
protein_coding
protein_coding
protein_coding
protein_coding
protein_coding
ncRNA
ncRNA
protein_coding
protein_coding
small_nucle(ol)ar_rna
protein_coding
protein_coding
protein_coding
protein_coding
protein_coding

protein_coding
protein_coding
protein_coding
protein_coding
small_nucle(ol)ar_rna
protein_coding
protein_coding
protein_coding
protein_coding
protein_coding
ribosomal_rna
protein_coding
protein_coding
protein_coding
small_nucle(ol)ar_rna
protein_coding
protein_coding
protein_coding
protein_coding
protein_coding
protein_coding
protein_coding
protein_coding
protein_coding
protein_coding
protein_coding
protein_coding
protein_coding
miRNA_precursor
protein_coding
protein_coding
protein_coding
protein_coding
miRNA_precursor
protein_coding
small_nucle(ol)ar_rna
protein_coding
protein_coding
protein_coding
protein_coding
protein_coding
protein_coding
protein_coding
protein_coding
protein_coding
protein_coding
protein_coding
protein_coding
protein_coding
protein_coding
small_nucle(ol)ar_rna
small_nucle(ol)ar_rna
protein_coding
protein_coding
protein_coding
small_nucle(ol)ar_rna

protein_coding
protein_coding
protein_coding
protein_coding
protein_coding
protein_coding
protein_coding
protein_coding
small_nucle(ol)ar_rna
protein_coding
protein_coding
protein_coding
protein_coding
miRNA_precursor
protein_coding
protein_coding
protein_coding
protein_coding
protein_coding
protein_coding
protein_coding
protein_coding
protein_coding
protein_coding
protein_coding
ncRNA
protein_coding
protein_coding
protein_coding
protein_coding
small_nucle(ol)ar_rna
protein_coding
protein_coding
protein_coding
ncRNA
protein_coding
protein_coding
protein_coding
miRNA_precursor
protein_coding
protein_coding
miRNA_precursor
protein_coding
protein_coding
protein_coding
protein_coding
protein_coding
protein_coding
protein_coding
protein_coding
protein_coding
protein_coding
protein_coding
protein_coding
protein_coding
small_nucle(ol)ar_rna

protein_coding
protein_coding
small_nucle(ol)ar_rna
protein_coding
protein_coding
protein_coding
protein_coding
protein_coding
protein_coding
miRNA_precursor
ncRNA
protein_coding
protein_coding
protein_coding
protein_coding
protein_coding
protein_coding
protein_coding
protein_coding
protein_coding
ncRNA
protein_coding
protein_coding
protein_coding
miRNA_precursor
protein_coding
protein_coding
miRNA_precursor
protein_coding
protein_coding
protein_coding
protein_coding
protein_coding
protein_coding
protein_coding
protein_coding
protein_coding
protein_coding
protein_coding
protein_coding
protein_coding
protein_coding
protein_coding
protein_coding
ncRNA
ncRNA
protein_coding
protein_coding
protein_coding
protein_coding
protein_coding
protein_coding
protein_coding
protein_coding
protein_coding
protein_coding

protein_coding
protein_coding
protein_coding
protein_coding
protein_coding
protein_coding
protein_coding
protein_coding
protein_coding
small_nucle(ol)ar_rna
protein_coding
protein_coding
protein_coding
protein_coding
protein_coding
protein_coding
protein_coding
ncRNA
protein_coding
protein_coding
protein_coding
protein_coding
protein_coding
protein_coding
protein_coding
protein_coding
protein_coding
protein_coding
protein_coding
protein_coding
protein_coding
protein_coding
protein_coding
protein_coding
protein_coding
protein_coding
protein_coding
protein_coding
protein_coding
protein_coding
protein_coding
protein_coding
protein_coding
protein_coding
protein_coding
protein_coding
protein_coding
protein_coding
protein_coding
protein_coding
protein_coding
protein_coding
protein_coding
protein_coding
protein_coding
small_nucle(ol)ar_rna

protein_coding
protein_coding
protein_coding
protein_coding
protein_coding
protein_coding
ncRNA
protein_coding
protein_coding
protein_coding
ncRNA
protein_coding
protein_coding
protein_coding
protein_coding
protein_coding
protein_coding
protein_coding
protein_coding
protein_coding
protein_coding
protein_coding
protein_coding
protein_coding
protein_coding
protein_coding
protein_coding
miRNA_precursor
protein_coding
protein_coding
protein_coding
protein_coding
small_nucle(ol)ar_rna
protein_coding
small_nucle(ol)ar_rna
miRNA_precursor
protein_coding
protein_coding
protein_coding
protein_coding
protein_coding
protein_coding
protein_coding
protein_coding
protein_coding
small_nucle(ol)ar_rna
ncRNA
protein_coding
protein_coding
miRNA_precursor
protein_coding
protein_coding
protein_coding
protein_coding
protein_coding
protein_coding

protein_coding
protein_coding
small_nucle(ol)ar_rna
pre_trna
protein_coding
protein_coding
protein_coding
protein_coding
protein_coding
protein_coding
protein_coding
protein_coding
protein_coding
protein_coding
protein_coding
protein_coding
protein_coding
protein_coding
small_nucle(ol)ar_rna
protein_coding
protein_coding
ncRNA
miRNA_precursor
protein_coding
protein_coding
protein_coding
protein_coding
protein_coding
protein_coding
miRNA_precursor
protein_coding
ncRNA
protein_coding
protein_coding
protein_coding
protein_coding
protein_coding
ncRNA
protein_coding
small_nucle(ol)ar_rna
protein_coding
protein_coding
protein_coding
protein_coding
protein_coding
small_nucle(ol)ar_rna
protein_coding
small_nucle(ol)ar_rna
protein_coding
protein_coding
protein_coding
protein_coding
protein_coding
protein_coding
protein_coding
protein_coding

protein_coding
protein_coding
protein_coding
protein_coding
protein_coding
protein_coding
ncRNA
protein_coding
protein_coding
protein_coding
ncRNA
protein_coding
small_nucle(ol)ar_rna
protein_coding
protein_coding
protein_coding
protein_coding
protein_coding
protein_coding
ncRNA
protein_coding
small_nucle(ol)ar_rna
protein_coding
protein_coding
protein_coding
protein_coding
protein_coding
protein_coding
small_nucle(ol)ar_rna
protein_coding
protein_coding
protein_coding
protein_coding
protein_coding
protein_coding
protein_coding
protein_coding
protein_coding
protein_coding
protein_coding
protein_coding
protein_coding
protein_coding
protein_coding
protein_coding
protein_coding
ribosomal_rna
protein_coding
protein_coding
protein_coding
protein_coding
protein_coding
protein_coding
protein_coding
protein_coding
protein_coding

protein_coding
protein_coding
protein_coding
protein_coding
protein_coding
protein_coding
miRNA_precursor
protein_coding
protein_coding
protein_coding
protein_coding
protein_coding
protein_coding
protein_coding
miRNA_precursor
ncRNA
protein_coding
protein_coding
protein_coding
protein_coding
protein_coding
protein_coding
protein_coding
protein_coding
protein_coding
protein_coding
ncRNA
small_nucle(ol)ar_rna
protein_coding
protein_coding
protein_coding
protein_coding
protein_coding
protein_coding
protein_coding
protein_coding
protein_coding
protein_coding
protein_coding
protein_coding
miRNA_precursor
protein_coding
protein_coding
protein_coding
protein_coding
protein_coding
protein_coding
protein_coding
pre_trna
protein_coding
protein_coding
protein_coding
protein_coding
protein_coding
protein_coding
protein_coding

protein_coding
protein_coding
protein_coding
protein_coding
protein_coding
protein_coding
protein_coding
small_nucle(ol)ar_rna
protein_coding
protein_coding
protein_coding
small_nucle(ol)ar_rna
protein_coding
miRNA_precursor
protein_coding
protein_coding
ncRNA
protein_coding
protein_coding
protein_coding
protein_coding
protein_coding
protein_coding
small_nucle(ol)ar_rna
protein_coding
protein_coding
protein_coding
protein_coding
protein_coding
small_nucle(ol)ar_rna
protein_coding
protein_coding
protein_coding
small_nucle(ol)ar_rna
protein_coding
protein_coding
protein_coding
protein_coding
ncRNA
protein_coding
protein_coding
protein_coding
protein_coding
ncRNA
small_nucle(ol)ar_rna
miRNA_precursor
protein_coding
protein_coding
protein_coding
protein_coding
protein_coding
protein_coding
ncRNA
protein_coding
protein_coding
ncRNA

protein_coding
protein_coding
protein_coding
protein_coding
small_nucle(ol)ar_rna
protein_coding
protein_coding
protein_coding
protein_coding
protein_coding
protein_coding
protein_coding
protein_coding
protein_coding
protein_coding
protein_coding
protein_coding
protein_coding
protein_coding
small_nucle(ol)ar_rna
ncRNA
protein_coding
protein_coding
protein_coding
protein_coding
small_nucle(ol)ar_rna
protein_coding
ncRNA
protein_coding
protein_coding
protein_coding
protein_coding
protein_coding
protein_coding
protein_coding
protein_coding
protein_coding
protein_coding
protein_coding
protein_coding
protein_coding
protein_coding
protein_coding
protein_coding
small_nucle(ol)ar_rna
miRNA_precursor
protein_coding
protein_coding
protein_coding
small_nucle(ol)ar_rna
protein_coding
protein_coding
protein_coding
protein_coding
protein_coding
protein_coding

protein_coding
protein_coding
small_nucle(ol)ar_rna
protein_coding
protein_coding
protein_coding
protein_coding
protein_coding
protein_coding
protein_coding
ncRNA
protein_coding
protein_coding
protein_coding
small_nucle(ol)ar_rna
ncRNA
protein_coding
small_nucle(ol)ar_rna
protein_coding
protein_coding
protein_coding
protein_coding
protein_coding
protein_coding
protein_coding
protein_coding
protein_coding
protein_coding
protein_coding
protein_coding
protein_coding
protein_coding
ncRNA
protein_coding
protein_coding
protein_coding
small_nucle(ol)ar_rna
protein_coding
protein_coding
protein_coding
protein_coding
protein_coding
protein_coding
ncRNA
protein_coding
protein_coding
small_nucle(ol)ar_rna
small_nucle(ol)ar_rna
protein_coding
protein_coding
protein_coding
protein_coding
ncRNA
protein_coding
protein_coding
protein_coding

small_nucle(ol)ar_rna
protein_coding
protein_coding
protein_coding
small_nucle(ol)ar_rna
protein_coding
protein_coding
protein_coding
protein_coding
protein_coding
protein_coding
protein_coding
small_nucle(ol)ar_rna
protein_coding
protein_coding
small_nucle(ol)ar_rna
protein_coding
protein_coding
protein_coding
protein_coding
protein_coding
protein_coding
protein_coding
protein_coding
protein_coding
protein_coding
protein_coding
protein_coding
protein_coding
protein_coding
protein_coding
protein_coding
protein_coding
protein_coding
protein_coding
ncRNA
protein_coding
protein_coding
protein_coding
ncRNA
protein_coding
protein_coding
protein_coding
protein_coding
protein_coding
protein_coding
protein_coding
small_nucle(ol)ar_rna
protein_coding
protein_coding
protein_coding
protein_coding
protein_coding
protein_coding
protein_coding
protein_coding

protein_coding
protein_coding
protein_coding
miRNA_precursor
small_nucle(ol)ar_rna
protein_coding
protein_coding
protein_coding
protein_coding
protein_coding
protein_coding
protein_coding
protein_coding
protein_coding
protein_coding
pre_trna
protein_coding
protein_coding
protein_coding
protein_coding
protein_coding
ncRNA
small_nucle(ol)ar_rna
protein_coding
protein_coding
protein_coding
protein_coding
protein_coding
protein_coding
protein_coding
protein_coding
protein_coding
protein_coding
protein_coding
protein_coding
protein_coding
protein_coding
protein_coding
protein_coding
protein_coding
ncRNA
protein_coding
ncRNA
protein_coding
small_nucle(ol)ar_rna
protein_coding
protein_coding
protein_coding
protein_coding
protein_coding
protein_coding
protein_coding
small_nucle(ol)ar_rna
protein_coding
protein_coding
protein_coding

protein_coding
protein_coding
protein_coding
small_nucle(ol)ar_rna
protein_coding
protein_coding
protein_coding
protein_coding
protein_coding
protein_coding
protein_coding
miRNA_precursor
protein_coding
protein_coding
protein_coding
protein_coding
protein_coding
protein_coding
protein_coding
protein_coding
protein_coding
protein_coding
protein_coding
protein_coding
protein_coding
protein_coding
small_nucle(ol)ar_rna
protein_coding
protein_coding
small_nucle(ol)ar_rna
protein_coding
protein_coding
protein_coding
protein_coding
protein_coding
protein_coding
protein_coding
protein_coding
protein_coding
protein_coding
protein_coding
protein_coding
protein_coding
protein_coding
protein_coding
protein_coding
protein_coding
protein_coding
protein_coding
protein_coding
protein_coding
ncRNA
protein_coding
protein_coding
protein_coding
ncRNA

protein_coding
protein_coding
miRNA_precursor
protein_coding
protein_coding
protein_coding
protein_coding
protein_coding
protein_coding
protein_coding
protein_coding
protein_coding
protein_coding
small_nucle(ol)ar_rna
protein_coding
protein_coding
protein_coding
protein_coding
protein_coding
small_nucle(ol)ar_rna
protein_coding
ncRNA
protein_coding
protein_coding
protein_coding
protein_coding
protein_coding
protein_coding
protein_coding
protein_coding
ncRNA
protein_coding
protein_coding
miRNA_precursor
protein_coding
protein_coding
protein_coding
protein_coding
protein_coding
protein_coding
protein_coding
protein_coding
ncRNA
small_nucle(ol)ar_rna
protein_coding
protein_coding
protein_coding
protein_coding
protein_coding
protein_coding
protein_coding
protein_coding
protein_coding
protein_coding
protein_coding
protein_coding

protein_coding
protein_coding
protein_coding
ncRNA
protein_coding
protein_coding
protein_coding
protein_coding
protein_coding
small_nucle(ol)ar_rna
protein_coding
protein_coding
protein_coding
protein_coding
protein_coding
protein_coding
protein_coding
miRNA_precursor
miRNA_precursor
miRNA_precursor
protein_coding
protein_coding
protein_coding
small_nucle(ol)ar_rna
protein_coding
protein_coding
protein_coding
protein_coding
protein_coding
protein_coding
protein_coding
protein_coding
protein_coding
protein_coding
protein_coding
protein_coding
protein_coding
protein_coding
protein_coding
small_nucle(ol)ar_rna
protein_coding
protein_coding
small_nucle(ol)ar_rna
protein_coding
protein_coding
protein_coding
protein_coding
protein_coding
protein_coding
ncRNA
protein_coding
protein_coding
protein_coding
protein_coding
protein_coding
protein_coding

protein_coding
protein_coding
protein_coding
ncRNA
protein_coding
protein_coding
miRNA_precursor
small_nucle(ol)ar_rna
protein_coding
protein_coding
protein_coding
protein_coding
small_nucle(ol)ar_rna
small_nucle(ol)ar_rna
protein_coding
protein_coding
protein_coding
protein_coding
protein_coding
protein_coding
ncRNA
protein_coding
protein_coding
protein_coding
protein_coding
protein_coding
protein_coding
protein_coding
protein_coding
protein_coding
protein_coding
protein_coding
protein_coding
protein_coding
protein_coding
protein_coding
protein_coding
protein_coding
protein_coding
protein_coding
protein_coding
protein_coding
protein_coding
protein_coding
protein_coding
protein_coding
protein_coding
protein_coding
protein_coding
protein_coding
protein_coding
protein_coding
miRNA_precursor
protein_coding
small_nucle(ol)ar_rna
protein_coding
protein_coding

protein_coding
protein_coding
protein_coding
protein_coding
protein_coding
protein_coding
protein_coding
protein_coding
protein_coding
protein_coding
protein_coding
protein_coding
protein_coding
protein_coding
protein_coding
protein_coding
ncRNA
protein_coding
protein_coding
protein_coding
protein_coding
protein_coding
protein_coding
small_nucle(ol)ar_rna
protein_coding
protein_coding
protein_coding
ncRNA
protein_coding
ncRNA
protein_coding
protein_coding
protein_coding
protein_coding
protein_coding
protein_coding
protein_coding
protein_coding
protein_coding
small_nucle(ol)ar_rna
protein_coding
protein_coding
protein_coding
protein_coding
protein_coding
ncRNA
protein_coding
ncRNA
protein_coding
protein_coding
miRNA_precursor
protein_coding
miRNA_precursor
protein_coding
pre_trna
protein_coding

protein_coding
small_nucle(ol)ar_rna
protein_coding
ncRNA
protein_coding
protein_coding
small_nucle(ol)ar_rna
ncRNA
protein_coding
protein_coding
protein_coding
protein_coding
protein_coding
protein_coding
protein_coding
protein_coding
protein_coding
protein_coding
protein_coding
protein_coding
protein_coding
protein_coding
protein_coding
protein_coding
protein_coding
protein_coding
protein_coding
ncRNA
ncRNA
protein_coding
protein_coding
protein_coding
small_nucle(ol)ar_rna
protein_coding
protein_coding
protein_coding
ncRNA
small_nucle(ol)ar_rna
protein_coding
small_nucle(ol)ar_rna
protein_coding
ncRNA
protein_coding
miRNA_precursor
protein_coding
small_nucle(ol)ar_rna
protein_coding
protein_coding
protein_coding
protein_coding
ncRNA
miRNA_precursor
protein_coding
protein_coding
protein_coding
protein_coding

protein_coding
protein_coding
protein_coding
protein_coding
protein_coding
protein_coding
protein_coding
protein_coding
protein_coding
protein_coding
protein_coding
ncRNA
protein_coding
protein_coding
protein_coding
small_nucle(ol)ar_rna
ncRNA
protein_coding
protein_coding
protein_coding
protein_coding
miRNA_precursor
protein_coding
protein_coding
protein_coding
small_nucle(ol)ar_rna
protein_coding
protein_coding
protein_coding
protein_coding
protein_coding
protein_coding
protein_coding
protein_coding
protein_coding
protein_coding
protein_coding
protein_coding
protein_coding
protein_coding
protein_coding
protein_coding
protein_coding
protein_coding
protein_coding
protein_coding
protein_coding
protein_coding
protein_coding
protein_coding
miRNA_precursor
protein_coding
protein_coding
protein_coding
miRNA_precursor
protein_coding

protein_coding
protein_coding
protein_coding
protein_coding
protein_coding
protein_coding
protein_coding
protein_coding
ncRNA
protein_coding
protein_coding
protein_coding
protein_coding
protein_coding
protein_coding
protein_coding
ncRNA
protein_coding
protein_coding
protein_coding
protein_coding
protein_coding
protein_coding
protein_coding
protein_coding
protein_coding
ncRNA
protein_coding
protein_coding
ncRNA
protein_coding
protein_coding
protein_coding
protein_coding
protein_coding
protein_coding
protein_coding
protein_coding
protein_coding
protein_coding
protein_coding
protein_coding
protein_coding
protein_coding
protein_coding
protein_coding
protein_coding
protein_coding
protein_coding
ncRNA
protein_coding
protein_coding
protein_coding
protein_coding
protein_coding
protein_coding

protein_coding
protein_coding
protein_coding
protein_coding
small_nucle(ol)ar_rna
small_nucle(ol)ar_rna
small_nucle(ol)ar_rna
small_nucle(ol)ar_rna
protein_coding
protein_coding
protein_coding
protein_coding
protein_coding
ncRNA
protein_coding
ncRNA
protein_coding
protein_coding
protein_coding
small_nucle(ol)ar_rna
small_nucle(ol)ar_rna
protein_coding
protein_coding
protein_coding
protein_coding
protein_coding
protein_coding
protein_coding
protein_coding
protein_coding
protein_coding
protein_coding
protein_coding
protein_coding
protein_coding
ncRNA
protein_coding
protein_coding
miRNA_precursor
protein_coding
protein_coding
protein_coding
protein_coding
protein_coding
pre_trna
ncRNA
protein_coding
protein_coding
protein_coding
protein_coding
protein_coding
protein_coding
protein_coding
protein_coding
protein_coding
protein_coding

protein_coding
protein_coding
protein_coding
protein_coding
protein_coding
ncRNA
protein_coding
protein_coding
protein_coding
protein_coding
protein_coding
protein_coding
protein_coding
protein_coding
protein_coding
protein_coding
protein_coding
protein_coding
ncRNA
protein_coding
protein_coding
protein_coding
protein_coding
protein_coding
protein_coding
protein_coding
protein_coding
protein_coding
protein_coding
protein_coding
protein_coding
protein_coding
protein_coding
protein_coding
protein_coding
protein_coding
protein_coding
protein_coding
protein_coding
protein_coding
protein_coding
protein_coding
protein_coding
protein_coding
protein_coding
protein_coding
protein_coding
protein_coding
protein_coding
protein_coding
protein_coding
protein_coding
protein_coding
protein_coding
protein_coding
protein_coding

protein_coding
protein_coding
protein_coding
protein_coding
ncRNA
protein_coding
protein_coding
protein_coding
protein_coding
protein_coding
protein_coding
protein_coding
protein_coding
protein_coding
protein_coding
protein_coding
protein_coding
protein_coding
protein_coding
protein_coding
protein_coding
protein_coding
protein_coding
protein_coding
protein_coding
protein_coding
protein_coding
protein_coding
protein_coding
protein_coding
protein_coding
protein_coding
protein_coding
protein_coding
protein_coding
protein_coding
protein_coding
protein_coding
protein_coding
protein_coding
protein_coding
protein_coding
protein_coding
protein_coding
protein_coding
protein_coding
protein_coding
protein_coding
protein_coding
protein_coding
protein_coding
protein_coding
protein_coding
protein_coding
protein_coding
protein_coding

protein_coding
protein_coding
protein_coding
protein_coding
protein_coding
protein_coding
small_nucle(ol)ar_rna
protein_coding
protein_coding
protein_coding
protein_coding
protein_coding
protein_coding
protein_coding
protein_coding
protein_coding
protein_coding
protein_coding
protein_coding
protein_coding
protein_coding
protein_coding
protein_coding
protein_coding
protein_coding
protein_coding
protein_coding
protein_coding
protein_coding
protein_coding
protein_coding
protein_coding
protein_coding
protein_coding
protein_coding
protein_coding
protein_coding
protein_coding
protein_coding
pre_trna
protein_coding
protein_coding
protein_coding
protein_coding
ncRNA
protein_coding
protein_coding
protein_coding
protein_coding
protein_coding
protein_coding
protein_coding
protein_coding
protein_coding
protein_coding
protein_coding

protein_coding
protein_coding
protein_coding
protein_coding
protein_coding
protein_coding
protein_coding
protein_coding
protein_coding
protein_coding
protein_coding
protein_coding
protein_coding
protein_coding
protein_coding
pre_trna
ncRNA
protein_coding
protein_coding
protein_coding
ncRNA
protein_coding
protein_coding
protein_coding
protein_coding
protein_coding
protein_coding
protein_coding
protein_coding
protein_coding
protein_coding
protein_coding
protein_coding
protein_coding
protein_coding
protein_coding
protein_coding
protein_coding
protein_coding
protein_coding
protein_coding
protein_coding
protein_coding
protein_coding
protein_coding
protein_coding
protein_coding
protein_coding
protein_coding
protein_coding
protein_coding
protein_coding
protein_coding
protein_coding
protein_coding
protein_coding

protein_coding
ncRNA
protein_coding
protein_coding
protein_coding
protein_coding
protein_coding
protein_coding
protein_coding
protein_coding
protein_coding
protein_coding
pre_trna
protein_coding
protein_coding
protein_coding
protein_coding
protein_coding
protein_coding
protein_coding
protein_coding
protein_coding
protein_coding
protein_coding
protein_coding
protein_coding
protein_coding
protein_coding
protein_coding
protein_coding
protein_coding
protein_coding
protein_coding
protein_coding
protein_coding
protein_coding
protein_coding
protein_coding
protein_coding
protein_coding
pre_trna
protein_coding
protein_coding
miRNA_precursor
protein_coding
protein_coding
protein_coding
protein_coding
protein_coding
protein_coding
protein_coding
protein_coding
protein_coding
protein_coding
protein_coding
small_nucle(ol)ar_rna

miRNA_precursor
protein_coding
ncRNA
protein_coding
protein_coding
protein_coding
protein_coding
protein_coding
protein_coding
protein_coding
protein_coding
ncRNA
protein_coding
protein_coding
protein_coding
protein_coding
protein_coding
miRNA_precursor
protein_coding
protein_coding
protein_coding
protein_coding
protein_coding
protein_coding
protein_coding
protein_coding
protein_coding
protein_coding
protein_coding
protein_coding
protein_coding
protein_coding
protein_coding
protein_coding
protein_coding
protein_coding
protein_coding
protein_coding
protein_coding
protein_coding
protein_coding
protein_coding
protein_coding
protein_coding
protein_coding
protein_coding
protein_coding
protein_coding
protein_coding
protein_coding
protein_coding
protein_coding
protein_coding
protein_coding
protein_coding
protein_coding

protein_coding
protein_coding
protein_coding
protein_coding
protein_coding
protein_coding
protein_coding
ncRNA
protein_coding
protein_coding
protein_coding
protein_coding
protein_coding
protein_coding
protein_coding
protein_coding
protein_coding
protein_coding
protein_coding
protein_coding
protein_coding
protein_coding
protein_coding
protein_coding
protein_coding
protein_coding
protein_coding
protein_coding
ncRNA
protein_coding
protein_coding
protein_coding
protein_coding
protein_coding
protein_coding
protein_coding
protein_coding
protein_coding
protein_coding
protein_coding
protein_coding
protein_coding
protein_coding
protein_coding
protein_coding
protein_coding
protein_coding
protein_coding
protein_coding
protein_coding
protein_coding
protein_coding
protein_coding
protein_coding
protein_coding
protein_coding

protein_coding
protein_coding
protein_coding
protein_coding
protein_coding
protein_coding
protein_coding
protein_coding
protein_coding
protein_coding
protein_coding
protein_coding
protein_coding
protein_coding
protein_coding
protein_coding
protein_coding
protein_coding
protein_coding
protein_coding
protein_coding
small_nucle(ol)ar_rna
protein_coding
protein_coding
protein_coding
protein_coding
ncRNA
protein_coding
protein_coding
protein_coding
protein_coding
protein_coding
protein_coding
protein_coding
protein_coding
protein_coding
protein_coding
protein_coding
protein_coding
protein_coding
protein_coding
protein_coding
ncRNA
protein_coding
protein_coding
protein_coding
pre_trna
protein_coding
protein_coding
protein_coding
protein_coding
protein_coding
protein_coding
protein_coding
protein_coding
protein_coding

small_nucle(ol)ar_rna
small_nucle(ol)ar_rna
small_nucle(ol)ar_rna
small_nucle(ol)ar_rna
protein_coding
protein_coding
protein_coding
protein_coding
protein_coding
protein_coding
protein_coding
protein_coding
protein_coding
protein_coding
protein_coding
protein_coding
protein_coding
pre_trna
protein_coding
protein_coding
protein_coding
protein_coding
protein_coding
protein_coding
protein_coding
protein_coding
protein_coding
protein_coding
protein_coding
ncRNA
protein_coding
protein_coding
protein_coding
protein_coding
protein_coding
protein_coding
protein_coding
protein_coding
ncRNA
protein_coding
protein_coding
protein_coding
protein_coding
protein_coding
protein_coding
protein_coding
protein_coding
protein_coding
protein_coding
protein_coding
protein_coding
protein_coding
small_nucle(ol)ar_rna
small_nucle(ol)ar_rna
protein_coding
protein_coding

protein_coding
miRNA_precursor
protein_coding
protein_coding
miRNA_precursor
protein_coding
protein_coding
protein_coding
protein_coding
protein_coding
protein_coding
protein_coding
protein_coding
protein_coding
protein_coding
protein_coding
protein_coding
protein_coding
protein_coding
protein_coding
protein_coding
protein_coding
protein_coding
protein_coding
protein_coding
protein_coding
protein_coding
protein_coding
protein_coding
protein_coding
protein_coding
protein_coding
ncRNA
protein_coding
protein_coding
protein_coding
protein_coding
protein_coding
protein_coding
protein_coding
protein_coding
protein_coding
ncRNA
protein_coding
protein_coding
protein_coding
protein_coding
protein_coding
protein_coding
protein_coding
protein_coding
ncRNA
protein_coding
protein_coding
protein_coding
protein_coding

ncRNA
protein_coding
protein_coding
protein_coding
protein_coding
protein_coding
protein_coding
protein_coding
protein_coding
protein_coding
protein_coding
protein_coding
protein_coding
protein_coding
protein_coding
protein_coding
protein_coding
ncRNA
protein_coding
protein_coding
ncRNA
protein_coding
protein_coding
ncRNA
protein_coding
protein_coding
protein_coding
protein_coding
protein_coding
ncRNA
protein_coding
protein_coding
protein_coding
protein_coding
miRNA_precursor
protein_coding
protein_coding
protein_coding
protein_coding
protein_coding
protein_coding
protein_coding
protein_coding
protein_coding
protein_coding
protein_coding
protein_coding
protein_coding
protein_coding
protein_coding
protein_coding
protein_coding
protein_coding
protein_coding
protein_coding
protein_coding

protein_coding
protein_coding
protein_coding
protein_coding
protein_coding
protein_coding
protein_coding
protein_coding
protein_coding
protein_coding
ncRNA
protein_coding
ncRNA
protein_coding
protein_coding
miRNA_precursor
protein_coding
protein_coding
protein_coding
protein_coding
protein_coding
protein_coding
protein_coding
protein_coding
protein_coding
protein_coding
protein_coding
protein_coding
protein_coding
protein_coding
protein_coding
protein_coding
ncRNA
protein_coding
protein_coding
small_nucle(ol)ar_rna
protein_coding
protein_coding
protein_coding
protein_coding
protein_coding
protein_coding
protein_coding
protein_coding
protein_coding
protein_coding
protein_coding
protein_coding
protein_coding
protein_coding
protein_coding
protein_coding
protein_coding
protein_coding
protein_coding
protein_coding

protein_coding
protein_coding
protein_coding
protein_coding
protein_coding
protein_coding
protein_coding
protein_coding
protein_coding
miRNA_precursor
protein_coding
protein_coding
protein_coding
protein_coding
protein_coding
protein_coding
protein_coding
protein_coding
protein_coding
protein_coding
protein_coding
protein_coding
protein_coding
protein_coding
protein_coding
protein_coding
protein_coding
protein_coding
protein_coding
protein_coding
protein_coding
protein_coding
ncRNA
protein_coding
protein_coding
protein_coding
protein_coding
protein_coding
ncRNA
protein_coding
protein_coding
protein_coding
protein_coding
protein_coding
protein_coding
protein_coding
protein_coding
protein_coding
protein_coding
protein_coding
protein_coding
protein_coding
protein_coding
protein_coding
protein_coding
protein_coding

protein_coding
protein_coding
protein_coding
protein_coding
protein_coding
protein_coding
protein_coding
protein_coding
protein_coding
protein_coding
protein_coding
protein_coding
protein_coding
ncRNA
protein_coding
protein_coding
protein_coding
protein_coding
protein_coding
protein_coding
protein_coding
ncRNA
protein_coding
ncRNA
small_nucle(ol)ar_rna
protein_coding
protein_coding
protein_coding
protein_coding
protein_coding
protein_coding
protein_coding
protein_coding
protein_coding
protein_coding
protein_coding
protein_coding
protein_coding
protein_coding
protein_coding
protein_coding
protein_coding
small_nucle(ol)ar_rna
protein_coding
protein_coding
protein_coding
protein_coding
protein_coding
protein_coding
protein_coding
protein_coding
protein_coding
protein_coding
protein_coding
protein_coding
protein_coding

protein_coding
protein_coding
protein_coding
protein_coding
protein_coding
protein_coding
protein_coding
protein_coding
protein_coding
protein_coding
protein_coding
protein_coding
protein_coding
protein_coding
protein_coding
protein_coding
protein_coding
ncRNA
protein_coding
protein_coding
protein_coding
ncRNA
protein_coding
protein_coding
protein_coding
protein_coding
protein_coding
protein_coding
protein_coding
protein_coding
protein_coding
protein_coding
protein_coding
protein_coding
protein_coding
protein_coding
protein_coding
protein_coding
protein_coding
protein_coding
protein_coding
protein_coding
protein_coding
protein_coding
protein_coding
protein_coding
protein_coding
protein_coding
protein_coding
protein_coding
protein_coding
protein_coding
protein_coding
ncRNA
protein_coding
protein_coding

protein_coding
protein_coding
protein_coding
protein_coding
protein_coding
protein_coding
protein_coding
protein_coding
protein_coding
miRNA_precursor
miRNA_precursor
protein_coding
protein_coding
protein_coding
protein_coding
ncRNA
protein_coding
protein_coding
protein_coding
small_nucle(ol)ar_rna
protein_coding
protein_coding
protein_coding
protein_coding
protein_coding
protein_coding
miRNA_precursor
protein_coding
protein_coding
protein_coding
protein_coding
protein_coding
protein_coding
protein_coding
protein_coding
protein_coding
protein_coding
protein_coding
protein_coding
protein_coding
ncRNA
protein_coding
ncRNA
protein_coding
protein_coding
protein_coding
protein_coding
protein_coding
protein_coding
protein_coding
protein_coding
protein_coding
protein_coding
protein_coding
protein_coding
protein_coding

protein_coding
protein_coding
protein_coding
protein_coding
protein_coding
protein_coding
protein_coding
protein_coding
protein_coding
protein_coding
protein_coding
small_nucle(ol)ar_rna
protein_coding
protein_coding
protein_coding
protein_coding
protein_coding
protein_coding
protein_coding
pre_trna
protein_coding
ncRNA
protein_coding
protein_coding
protein_coding
protein_coding
protein_coding
protein_coding
protein_coding
ncRNA
protein_coding
protein_coding
protein_coding
protein_coding
protein_coding
protein_coding
protein_coding
protein_coding
protein_coding
protein_coding
protein_coding
protein_coding
protein_coding
protein_coding
protein_coding
protein_coding
protein_coding
protein_coding
protein_coding
ncRNA
protein_coding
protein_coding
protein_coding
protein_coding
protein_coding
protein_coding

protein_coding
protein_coding
miRNA_precursor
protein_coding
protein_coding
protein_coding
protein_coding
protein_coding
protein_coding
protein_coding
protein_coding
protein_coding
protein_coding
protein_coding
protein_coding
protein_coding
protein_coding
ncRNA
protein_coding
ncRNA
protein_coding
protein_coding
small_nucle(ol)ar_rna
protein_coding
protein_coding
protein_coding
protein_coding
protein_coding
protein_coding
protein_coding
protein_coding
ncRNA
protein_coding
protein_coding
ncRNA
protein_coding
protein_coding
protein_coding
protein_coding
protein_coding
protein_coding
protein_coding
protein_coding
protein_coding
protein_coding
protein_coding
protein_coding
protein_coding
protein_coding
protein_coding
protein_coding
protein_coding
protein_coding
protein_coding
protein_coding
protein_coding

protein_coding
ncRNA
protein_coding
protein_coding
protein_coding
protein_coding
protein_coding
protein_coding
protein_coding
protein_coding
protein_coding
protein_coding
protein_coding
small_nucle(ol)ar_rna
protein_coding
protein_coding
protein_coding
protein_coding
protein_coding
protein_coding
protein_coding
protein_coding
protein_coding
protein_coding
protein_coding
protein_coding
protein_coding
protein_coding
protein_coding
protein_coding
protein_coding
protein_coding
protein_coding
protein_coding
protein_coding
miRNA_precursor
protein_coding
protein_coding
protein_coding
protein_coding
protein_coding
protein_coding
protein_coding
protein_coding
protein_coding
protein_coding
protein_coding
protein_coding
protein_coding
protein_coding
protein_coding
protein_coding
protein_coding
protein_coding
protein_coding
protein_coding

ncRNA
protein_coding
protein_coding
protein_coding
protein_coding
protein_coding
protein_coding
protein_coding
protein_coding
protein_coding
protein_coding
protein_coding
protein_coding
protein_coding
protein_coding
protein_coding
protein_coding
protein_coding
protein_coding
protein_coding
protein_coding
protein_coding
protein_coding
protein_coding
small_nucle(ol)ar_rna
protein_coding
protein_coding
protein_coding
protein_coding
protein_coding
protein_coding
ncRNA
protein_coding
protein_coding
protein_coding
protein_coding
protein_coding
protein_coding
protein_coding
protein_coding
protein_coding
protein_coding
protein_coding
protein_coding
protein_coding
protein_coding
protein_coding
protein_coding
protein_coding
protein_coding
protein_coding
protein_coding
protein_coding
protein_coding
protein_coding
protein_coding

protein_coding
protein_coding
protein_coding
protein_coding
protein_coding
protein_coding
protein_coding
protein_coding
protein_coding
protein_coding
protein_coding
protein_coding
protein_coding
protein_coding
protein_coding
protein_coding
protein_coding
protein_coding
protein_coding
protein_coding
small_nucle(ol)ar_rna
small_nucle(ol)ar_rna
small_nucle(ol)ar_rna
protein_coding
ncRNA
protein_coding
protein_coding
protein_coding
protein_coding
protein_coding
protein_coding
protein_coding
protein_coding
protein_coding
protein_coding
protein_coding
protein_coding
protein_coding
protein_coding
protein_coding
protein_coding
protein_coding
protein_coding
pre_trna
protein_coding
protein_coding
protein_coding
protein_coding
protein_coding
protein_coding
protein_coding
protein_coding
protein_coding
protein_coding
protein_coding
protein_coding

protein_coding
protein_coding
protein_coding
protein_coding
protein_coding
protein_coding
protein_coding
protein_coding
protein_coding
protein_coding
protein_coding
protein_coding
protein_coding
miRNA_precursor
protein_coding
protein_coding
protein_coding
protein_coding
protein_coding
protein_coding
protein_coding
protein_coding
protein_coding
protein_coding
protein_coding
protein_coding
protein_coding
protein_coding
protein_coding
protein_coding
protein_coding
protein_coding
protein_coding
protein_coding
protein_coding
protein_coding
protein_coding
protein_coding
protein_coding
protein_coding
protein_coding
protein_coding
protein_coding
protein_coding
protein_coding
protein_coding
protein_coding
protein_coding
protein_coding
protein_coding
protein_coding
protein_coding
protein_coding
ncRNA
protein_coding
protein_coding

pre_trna
protein_coding
protein_coding
protein_coding
protein_coding
protein_coding
protein_coding
ncRNA
protein_coding
protein_coding
protein_coding
protein_coding
protein_coding
protein_coding
protein_coding
protein_coding
protein_coding
protein_coding
protein_coding
protein_coding
protein_coding
protein_coding
protein_coding
ncRNA
pre_trna
protein_coding
protein_coding
protein_coding
protein_coding
protein_coding
protein_coding
protein_coding
protein_coding
protein_coding
protein_coding
protein_coding
protein_coding
protein_coding
protein_coding
ncRNA
protein_coding
protein_coding
protein_coding
protein_coding
protein_coding
protein_coding
protein_coding
protein_coding
protein_coding
protein_coding
protein_coding
protein_coding
protein_coding
protein_coding
protein_coding
protein_coding

protein_coding
protein_coding
protein_coding
ncRNA
protein_coding
protein_coding
protein_coding
protein_coding
ncRNA
ncRNA
protein_coding
protein_coding
protein_coding
protein_coding
protein_coding
protein_coding
protein_coding
protein_coding
protein_coding
protein_coding
protein_coding
protein_coding
protein_coding
protein_coding
protein_coding
protein_coding
protein_coding
protein_coding
protein_coding
protein_coding
protein_coding
protein_coding
protein_coding
ncRNA
protein_coding
protein_coding
protein_coding
protein_coding
protein_coding
protein_coding
protein_coding
protein_coding
protein_coding
protein_coding
protein_coding
protein_coding
protein_coding
protein_coding
protein_coding
protein_coding
protein_coding
protein_coding
protein_coding
protein_coding
protein_coding
protein_coding

protein_coding
protein_coding
protein_coding
ncRNA
miRNA_precursor
protein_coding
ncRNA
protein_coding
ncRNA
protein_coding
protein_coding
protein_coding
ncRNA
protein_coding
protein_coding
protein_coding
protein_coding
protein_coding
protein_coding
protein_coding
protein_coding
protein_coding
protein_coding
protein_coding
protein_coding
protein_coding
pre_trna
protein_coding
protein_coding
protein_coding
protein_coding
protein_coding
protein_coding
ncRNA
protein_coding
protein_coding
protein_coding
protein_coding
protein_coding
protein_coding
protein_coding
protein_coding
protein_coding
protein_coding
protein_coding
protein_coding
pre_trna
pre_trna
protein_coding
protein_coding
protein_coding
ncRNA
protein_coding
protein_coding
protein_coding
protein_coding

protein_coding
protein_coding
protein_coding
protein_coding
protein_coding
ncRNA
protein_coding
protein_coding
ncRNA
protein_coding
protein_coding
protein_coding
ncRNA
protein_coding
protein_coding
protein_coding
protein_coding
ncRNA
protein_coding
protein_coding
protein_coding
protein_coding
protein_coding
protein_coding
protein_coding
protein_coding
protein_coding
protein_coding
protein_coding
ncRNA
protein_coding
ncRNA
protein_coding
protein_coding
protein_coding
protein_coding
protein_coding
protein_coding
protein_coding
protein_coding
protein_coding
protein_coding
protein_coding
protein_coding
protein_coding
ncRNA
protein_coding
protein_coding
protein_coding
protein_coding
protein_coding
protein_coding
protein_coding
protein_coding
protein_coding
protein_coding

protein_coding
protein_coding
protein_coding
protein_coding
protein_coding
protein_coding
protein_coding
protein_coding
protein_coding
protein_coding
protein_coding
ncRNA
protein_coding
protein_coding
protein_coding
protein_coding
protein_coding
protein_coding
protein_coding
protein_coding
protein_coding
protein_coding
protein_coding
protein_coding
protein_coding
protein_coding
protein_coding
protein_coding
protein_coding
protein_coding
protein_coding
protein_coding
protein_coding
protein_coding
protein_coding
protein_coding
protein_coding
protein_coding
protein_coding
protein_coding
protein_coding
protein_coding
protein_coding
protein_coding
protein_coding
protein_coding
ncRNA
protein_coding
protein_coding
protein_coding
protein_coding
ncRNA
ncRNA
protein_coding
protein_coding
protein_coding

protein_coding
protein_coding
protein_coding
protein_coding
protein_coding
protein_coding
protein_coding
miRNA_precursor
protein_coding
protein_coding
protein_coding
protein_coding
ncRNA
ncRNA
protein_coding
protein_coding
protein_coding
protein_coding
protein_coding
protein_coding
protein_coding
protein_coding
protein_coding
protein_coding
protein_coding
protein_coding
protein_coding
protein_coding
protein_coding
protein_coding
protein_coding
protein_coding
protein_coding
protein_coding
protein_coding
protein_coding
ncRNA
protein_coding
protein_coding
protein_coding
protein_coding
protein_coding
protein_coding
protein_coding
protein_coding
ncRNA
protein_coding
protein_coding
protein_coding
protein_coding
protein_coding
protein_coding
protein_coding
protein_coding
protein_coding
protein_coding

protein_coding
protein_coding
protein_coding
ncRNA
protein_coding
ncRNA
protein_coding
protein_coding
protein_coding
protein_coding
protein_coding
protein_coding
protein_coding
protein_coding
protein_coding
ncRNA
protein_coding
protein_coding
ncRNA
protein_coding
protein_coding
protein_coding
protein_coding
protein_coding
protein_coding
protein_coding
miRNA_precursor
protein_coding
protein_coding
protein_coding
miRNA_precursor
ncRNA
protein_coding
protein_coding
protein_coding
protein_coding
protein_coding
protein_coding
protein_coding
protein_coding
protein_coding
protein_coding
miRNA_precursor
protein_coding
protein_coding
pre_trna
protein_coding
ncRNA
ncRNA
protein_coding
protein_coding
ncRNA
protein_coding
protein_coding
protein_coding
protein_coding

miRNA_precursor
protein_coding
protein_coding
protein_coding
protein_coding
protein_coding
protein_coding
protein_coding
protein_coding
protein_coding
protein_coding
protein_coding
protein_coding
protein_coding
protein_coding
protein_coding
protein_coding
protein_coding
protein_coding
protein_coding
protein_coding
protein_coding
protein_coding
protein_coding
protein_coding
protein_coding
protein_coding
protein_coding
protein_coding
ncRNA
protein_coding
protein_coding
protein_coding
protein_coding
protein_coding
protein_coding
protein_coding
protein_coding
protein_coding
protein_coding
protein_coding
protein_coding
protein_coding
protein_coding
protein_coding
protein_coding
protein_coding
protein_coding
protein_coding
protein_coding
protein_coding
protein_coding
protein_coding
protein_coding
protein_coding
protein_coding

protein_coding
protein_coding
protein_coding
protein_coding
protein_coding
protein_coding
protein_coding
protein_coding
protein_coding
protein_coding
protein_coding
protein_coding
protein_coding
protein_coding
protein_coding
protein_coding
protein_coding
pre_trna
protein_coding
protein_coding
protein_coding
protein_coding
protein_coding
protein_coding
protein_coding
protein_coding
protein_coding
protein_coding
protein_coding
protein_coding
protein_coding
protein_coding
protein_coding
protein_coding
protein_coding
protein_coding
protein_coding
protein_coding
protein_coding
protein_coding
protein_coding
protein_coding
pre_trna
small_nucle(ol)ar_rna
protein_coding
ncRNA
protein_coding
protein_coding
protein_coding
protein_coding
protein_coding
protein_coding
protein_coding
protein_coding
protein_coding
protein_coding

protein_coding
protein_coding
protein_coding
protein_coding
protein_coding
protein_coding
protein_coding
protein_coding
protein_coding
protein_coding
ncRNA
protein_coding
ncRNA
protein_coding
protein_coding
protein_coding
protein_coding
protein_coding
protein_coding
protein_coding
protein_coding
protein_coding
protein_coding
protein_coding
protein_coding
protein_coding
protein_coding
protein_coding
protein_coding
miRNA_precursor
protein_coding
protein_coding
protein_coding
protein_coding
protein_coding
protein_coding
protein_coding
protein_coding
protein_coding
protein_coding
protein_coding
small_nucle(ol)ar_rna
protein_coding
protein_coding
protein_coding
protein_coding
protein_coding
protein_coding
pre_trna
protein_coding
protein_coding
protein_coding
protein_coding
protein_coding
pre_trna
protein_coding

protein_coding
protein_coding
protein_coding
protein_coding
protein_coding
protein_coding
protein_coding
protein_coding
protein_coding
protein_coding
protein_coding
protein_coding
protein_coding
protein_coding
protein_coding
protein_coding
protein_coding
protein_coding
protein_coding
ncRNA
protein_coding
protein_coding
protein_coding
protein_coding
protein_coding
protein_coding
protein_coding
protein_coding
protein_coding
protein_coding
protein_coding
protein_coding
protein_coding
protein_coding
protein_coding
protein_coding
protein_coding
protein_coding
protein_coding
protein_coding
pre_trna
protein_coding
protein_coding
pre_trna
protein_coding
ncRNA
protein_coding
protein_coding
protein_coding
protein_coding
protein_coding
protein_coding
protein_coding
protein_coding
protein_coding
protein_coding

protein_coding
small_nucle(ol)ar_rna
protein_coding
protein_coding
ncRNA
protein_coding
protein_coding
protein_coding
ncRNA
protein_coding
protein_coding
protein_coding
protein_coding
protein_coding
protein_coding
protein_coding
protein_coding
protein_coding
protein_coding
protein_coding
protein_coding
protein_coding
protein_coding
protein_coding
protein_coding
protein_coding
protein_coding
protein_coding
protein_coding
protein_coding
protein_coding
protein_coding
protein_coding
protein_coding
protein_coding
protein_coding
protein_coding
protein_coding
protein_coding
protein_coding
protein_coding
protein_coding
protein_coding
protein_coding
protein_coding
protein_coding
protein_coding
protein_coding
protein_coding
protein_coding
protein_coding
protein_coding
protein_coding
protein_coding
protein_coding
protein_coding

protein_coding
protein_coding
protein_coding
protein_coding
protein_coding
miRNA_precursor
protein_coding
protein_coding
protein_coding
protein_coding
protein_coding
protein_coding
protein_coding
protein_coding
protein_coding
protein_coding
protein_coding
protein_coding
protein_coding
protein_coding
protein_coding
protein_coding
protein_coding
protein_coding
protein_coding
protein_coding
ncRNA
protein_coding
protein_coding
protein_coding
protein_coding
protein_coding
protein_coding
protein_coding
protein_coding
protein_coding
protein_coding
protein_coding
protein_coding
protein_coding
protein_coding
protein_coding
protein_coding
protein_coding
protein_coding
protein_coding
protein_coding
protein_coding
protein_coding
protein_coding
protein_coding
protein_coding
protein_coding
protein_coding
protein_coding
protein_coding

protein_coding
protein_coding
protein_coding
protein_coding
protein_coding
protein_coding
small_nucle(ol)ar_rna
small_nucle(ol)ar_rna
small_nucle(ol)ar_rna
protein_coding
protein_coding
protein_coding
protein_coding
ncRNA
protein_coding
protein_coding
protein_coding
protein_coding
protein_coding
protein_coding
protein_coding
protein_coding
protein_coding
protein_coding
protein_coding
protein_coding
protein_coding
protein_coding
protein_coding
protein_coding
protein_coding
protein_coding
protein_coding
protein_coding
protein_coding
protein_coding
protein_coding
protein_coding
protein_coding
protein_coding
protein_coding
protein_coding
protein_coding
protein_coding
protein_coding
protein_coding
protein_coding
protein_coding
protein_coding
protein_coding
protein_coding
protein_coding
protein_coding
pre_trna
protein_coding
protein_coding

protein_coding
protein_coding
protein_coding
protein_coding
pre_trna
protein_coding
protein_coding
protein_coding
ncRNA
protein_coding
protein_coding
protein_coding
ncRNA
protein_coding
protein_coding
protein_coding
protein_coding
protein_coding
protein_coding
protein_coding
protein_coding
protein_coding
protein_coding
small_nucle(ol)ar_rna
small_nucle(ol)ar_rna
protein_coding
protein_coding
protein_coding
protein_coding
protein_coding
protein_coding
protein_coding
protein_coding
protein_coding
protein_coding
protein_coding
protein_coding
protein_coding
protein_coding
protein_coding
protein_coding
protein_coding
protein_coding
protein_coding
protein_coding
protein_coding
protein_coding
protein_coding
protein_coding
protein_coding
protein_coding
small_nucle(ol)ar_rna
protein_coding
protein_coding
protein_coding
protein_coding

protein_coding
protein_coding
protein_coding
protein_coding
protein_coding
protein_coding
protein_coding
protein_coding
protein_coding
protein_coding
protein_coding
protein_coding
protein_coding
protein_coding
protein_coding
protein_coding
protein_coding
protein_coding
protein_coding
protein_coding
protein_coding
protein_coding
protein_coding
protein_coding
protein_coding
protein_coding
protein_coding
protein_coding
protein_coding
protein_coding
protein_coding
protein_coding
protein_coding
protein_coding
protein_coding
protein_coding
protein_coding
protein_coding
protein_coding
protein_coding
protein_coding
protein_coding
protein_coding
protein_coding
protein_coding
protein_coding
protein_coding
protein_coding
protein_coding
protein_coding
protein_coding
protein_coding
protein_coding
protein_coding
protein_coding
protein_coding

protein_coding
protein_coding
protein_coding
protein_coding
protein_coding
protein_coding
protein_coding
protein_coding
protein_coding
protein_coding
ncRNA
small_nucle(ol)ar_rna
protein_coding
protein_coding
protein_coding
protein_coding
protein_coding
protein_coding
protein_coding
protein_coding
protein_coding
protein_coding
protein_coding
protein_coding
protein_coding
protein_coding
protein_coding
protein_coding
protein_coding
protein_coding
protein_coding
protein_coding
protein_coding
protein_coding
protein_coding
protein_coding
protein_coding
protein_coding
protein_coding
protein_coding
protein_coding
protein_coding
protein_coding
protein_coding
ncRNA
ncRNA
protein_coding
protein_coding
protein_coding
protein_coding
protein_coding
protein_coding
protein_coding
protein_coding
protein_coding
protein_coding

protein_coding
protein_coding
protein_coding
protein_coding
protein_coding
protein_coding
protein_coding
protein_coding
protein_coding
protein_coding
protein_coding
protein_coding
ncRNA
ncRNA
protein_coding
protein_coding
protein_coding
protein_coding
protein_coding
protein_coding
protein_coding
protein_coding
protein_coding
protein_coding
protein_coding
protein_coding
protein_coding
protein_coding
protein_coding
protein_coding
protein_coding
protein_coding
protein_coding
protein_coding
protein_coding
protein_coding
protein_coding
protein_coding
protein_coding
protein_coding
protein_coding
protein_coding
protein_coding
protein_coding
protein_coding
protein_coding
protein_coding
protein_coding
protein_coding
protein_coding
protein_coding
protein_coding
protein_coding
protein_coding
protein_coding
protein_coding

protein_coding
protein_coding
protein_coding
protein_coding
protein_coding
protein_coding
protein_coding
protein_coding
protein_coding
protein_coding
protein_coding
protein_coding
protein_coding
protein_coding
protein_coding
protein_coding
ncRNA
protein_coding
protein_coding
protein_coding
protein_coding
protein_coding
protein_coding
protein_coding
protein_coding
protein_coding
protein_coding
protein_coding
protein_coding
protein_coding
protein_coding
protein_coding
protein_coding
protein_coding
protein_coding
protein_coding
protein_coding
protein_coding
protein_coding
protein_coding
protein_coding
protein_coding
protein_coding
protein_coding
protein_coding
protein_coding
protein_coding
protein_coding
protein_coding
protein_coding
protein_coding
protein_coding
protein_coding
protein_coding
protein_coding
protein_coding

protein_coding
protein_coding
protein_coding
protein_coding
protein_coding
protein_coding
protein_coding
protein_coding
protein_coding
protein_coding
protein_coding
protein_coding
protein_coding
protein_coding
ncRNA
protein_coding
protein_coding
protein_coding
protein_coding
protein_coding
protein_coding
protein_coding
protein_coding
protein_coding
protein_coding
protein_coding
protein_coding
protein_coding
protein_coding
protein_coding
protein_coding
protein_coding
ncRNA
protein_coding
protein_coding
protein_coding
protein_coding
protein_coding
protein_coding
protein_coding
protein_coding
protein_coding
protein_coding
protein_coding
protein_coding
protein_coding
protein_coding
protein_coding
protein_coding
protein_coding
protein_coding
protein_coding
protein_coding
small_nucle(ol)ar_rna
protein_coding
protein_coding

protein_coding
protein_coding
protein_coding
protein_coding
protein_coding
protein_coding
protein_coding
protein_coding
protein_coding
protein_coding
protein_coding
protein_coding
protein_coding
protein_coding
protein_coding
protein_coding
protein_coding
protein_coding
protein_coding
protein_coding
protein_coding
protein_coding
protein_coding
protein_coding
protein_coding
protein_coding
protein_coding
protein_coding
protein_coding
protein_coding
protein_coding
protein_coding
protein_coding
protein_coding
protein_coding
protein_coding
protein_coding
protein_coding
protein_coding
protein_coding
protein_coding
protein_coding
protein_coding
protein_coding
protein_coding
protein_coding
protein_coding
protein_coding
ncRNA
protein_coding
protein_coding
protein_coding
protein_coding
protein_coding
protein_coding
protein_coding

protein_coding
protein_coding
protein_coding
protein_coding
protein_coding
protein_coding
protein_coding
protein_coding
protein_coding
protein_coding
protein_coding
protein_coding
protein_coding
protein_coding
protein_coding
protein_coding
protein_coding
protein_coding
protein_coding
protein_coding
protein_coding
protein_coding
protein_coding
protein_coding
protein_coding
protein_coding
protein_coding
ncRNA
protein_coding
protein_coding
protein_coding
protein_coding
protein_coding
protein_coding
protein_coding
protein_coding
protein_coding
protein_coding
protein_coding
protein_coding
protein_coding
protein_coding
ncRNA
protein_coding
protein_coding
protein_coding
protein_coding
protein_coding
protein_coding
protein_coding
protein_coding
protein_coding
protein_coding
protein_coding
protein_coding
protein_coding

protein_coding
protein_coding
protein_coding
miRNA_precursor
protein_coding
protein_coding
protein_coding
protein_coding
protein_coding
protein_coding
protein_coding
protein_coding
protein_coding
protein_coding
protein_coding
miRNA_precursor
protein_coding
protein_coding
protein_coding
protein_coding
protein_coding
protein_coding
protein_coding
protein_coding
protein_coding
protein_coding
protein_coding
protein_coding
protein_coding
protein_coding
protein_coding
protein_coding
protein_coding
protein_coding
protein_coding
protein_coding
protein_coding
protein_coding
protein_coding
protein_coding
protein_coding
protein_coding
protein_coding
protein_coding
protein_coding
protein_coding
protein_coding
protein_coding
protein_coding
protein_coding
protein_coding
protein_coding
protein_coding
protein_coding
ncRNA
protein_coding

protein_coding
protein_coding
protein_coding
protein_coding
protein_coding
protein_coding
protein_coding
protein_coding
protein_coding
protein_coding
protein_coding
protein_coding
protein_coding
protein_coding
protein_coding
protein_coding
protein_coding
small_nucle(ol)ar_rna
protein_coding
ncRNA
protein_coding
protein_coding
protein_coding
protein_coding
ncRNA
protein_coding
protein_coding
protein_coding
protein_coding
protein_coding
protein_coding
protein_coding
protein_coding
protein_coding
protein_coding
protein_coding
protein_coding
protein_coding
protein_coding
protein_coding
protein_coding
protein_coding
ncRNA
protein_coding
protein_coding
protein_coding
protein_coding
protein_coding
protein_coding
protein_coding
protein_coding
protein_coding
protein_coding
protein_coding
protein_coding
protein_coding

protein_coding
protein_coding
protein_coding
protein_coding
protein_coding
protein_coding
protein_coding
protein_coding
protein_coding
protein_coding
protein_coding
protein_coding
protein_coding
protein_coding
protein_coding
protein_coding
protein_coding
protein_coding
protein_coding
protein_coding
protein_coding
protein_coding
protein_coding
protein_coding
protein_coding
protein_coding
protein_coding
protein_coding
protein_coding
protein_coding
protein_coding
protein_coding
protein_coding
ncRNA
protein_coding
protein_coding
protein_coding
protein_coding
protein_coding
protein_coding
protein_coding
protein_coding
protein_coding
protein_coding
protein_coding
protein_coding
protein_coding
ncRNA
small_nucle(ol)ar_rna
protein_coding
protein_coding
protein_coding
protein_coding
protein_coding
protein_coding
protein_coding

protein_coding
protein_coding
protein_coding
protein_coding
protein_coding
protein_coding
protein_coding
protein_coding
protein_coding
protein_coding
protein_coding
protein_coding
protein_coding
protein_coding
protein_coding
protein_coding
protein_coding
protein_coding
protein_coding
protein_coding
protein_coding
protein_coding
protein_coding
protein_coding
protein_coding
protein_coding
pre_trna
protein_coding
ncRNA
protein_coding
protein_coding
protein_coding
protein_coding
small_nucle(ol)ar_rna
small_nucle(ol)ar_rna
protein_coding
protein_coding
protein_coding
protein_coding
protein_coding
protein_coding
protein_coding
protein_coding
protein_coding
protein_coding
protein_coding
protein_coding
protein_coding
protein_coding
protein_coding
protein_coding
protein_coding
miRNA_precursor
protein_coding
protein_coding
protein_coding

protein_coding
protein_coding
protein_coding
protein_coding
protein_coding
protein_coding
protein_coding
ncRNA
protein_coding
protein_coding
protein_coding
protein_coding
protein_coding
protein_coding
protein_coding
small_nucle(ol)ar_rna
protein_coding
protein_coding
protein_coding
protein_coding
protein_coding
protein_coding
protein_coding
ncRNA
protein_coding
protein_coding
protein_coding
protein_coding
protein_coding
protein_coding
protein_coding
protein_coding
protein_coding
protein_coding
protein_coding
protein_coding
protein_coding
protein_coding
protein_coding
protein_coding
protein_coding
protein_coding
protein_coding
protein_coding
protein_coding
protein_coding
protein_coding
ncRNA
protein_coding
protein_coding
protein_coding
protein_coding
protein_coding
ncRNA
protein_coding
ncRNA

protein_coding
protein_coding
ncRNA
protein_coding
protein_coding
protein_coding
protein_coding
protein_coding
protein_coding
protein_coding
protein_coding
protein_coding
protein_coding
protein_coding
protein_coding
protein_coding
protein_coding
protein_coding
protein_coding
protein_coding
protein_coding
protein_coding
protein_coding
protein_coding
protein_coding
ncRNA
protein_coding
protein_coding
protein_coding
protein_coding
protein_coding
protein_coding
protein_coding
protein_coding
protein_coding
protein_coding
protein_coding
protein_coding
protein_coding
protein_coding
protein_coding
protein_coding
pre_trna
protein_coding
protein_coding
protein_coding
protein_coding
protein_coding
protein_coding
protein_coding
protein_coding
protein_coding
protein_coding
protein_coding
ncRNA
protein_coding

ncRNA
ncRNA
ncRNA
ncRNA
protein_coding
protein_coding
protein_coding
miRNA_precursor
protein_coding
protein_coding
small_nucle(ol)ar_rna
protein_coding
protein_coding
protein_coding
protein_coding
protein_coding
protein_coding
protein_coding
ncRNA
ncRNA
protein_coding
protein_coding
protein_coding
protein_coding
protein_coding
protein_coding
protein_coding
protein_coding
protein_coding
protein_coding
protein_coding
protein_coding
protein_coding
protein_coding
protein_coding
protein_coding
protein_coding
protein_coding
protein_coding
protein_coding
protein_coding
protein_coding
protein_coding
protein_coding
protein_coding
protein_coding
ncRNA
protein_coding
protein_coding
protein_coding
protein_coding
protein_coding
protein_coding
protein_coding
protein_coding
protein_coding

protein_coding
protein_coding
protein_coding
protein_coding
protein_coding
protein_coding
protein_coding
protein_coding
protein_coding
protein_coding
protein_coding
protein_coding
protein_coding
protein_coding
protein_coding
protein_coding
protein_coding
protein_coding
protein_coding
protein_coding
protein_coding
protein_coding
protein_coding
protein_coding
protein_coding
protein_coding
protein_coding
protein_coding
protein_coding
protein_coding
protein_coding
protein_coding
protein_coding
protein_coding
protein_coding
protein_coding
protein_coding
protein_coding
protein_coding
protein_coding
protein_coding
protein_coding
protein_coding
protein_coding
protein_coding
protein_coding
protein_coding
protein_coding
protein_coding
protein_coding
protein_coding
protein_coding
protein_coding
protein_coding
protein_coding
protein_coding

protein_coding
protein_coding
protein_coding
protein_coding
protein_coding
protein_coding
protein_coding
protein_coding
protein_coding
small_nucle(ol)ar_rna
protein_coding
protein_coding
protein_coding
miRNA_precursor
ncRNA
protein_coding
protein_coding
protein_coding
protein_coding
protein_coding
protein_coding
protein_coding
protein_coding
protein_coding
protein_coding
protein_coding
protein_coding
protein_coding
protein_coding
protein_coding
protein_coding
protein_coding
protein_coding
protein_coding
protein_coding
protein_coding
protein_coding
protein_coding
protein_coding
protein_coding
protein_coding
ncRNA
protein_coding
protein_coding
protein_coding
protein_coding
ncRNA
protein_coding
protein_coding
protein_coding
protein_coding
protein_coding
protein_coding
protein_coding
protein_coding
protein_coding

protein_coding
protein_coding
protein_coding
protein_coding
protein_coding
protein_coding
ncRNA
small_nucle(ol)ar_rna
protein_coding
protein_coding
protein_coding
protein_coding
protein_coding
protein_coding
protein_coding
protein_coding
protein_coding
protein_coding
protein_coding
protein_coding
protein_coding
protein_coding
protein_coding
protein_coding
protein_coding
ncRNA
protein_coding
protein_coding
protein_coding
protein_coding
protein_coding
protein_coding
protein_coding
protein_coding
protein_coding
protein_coding
ncRNA
protein_coding
protein_coding
protein_coding
protein_coding
protein_coding
protein_coding
ncRNA
protein_coding
protein_coding
protein_coding
protein_coding
protein_coding
protein_coding
protein_coding
protein_coding
small_nucle(ol)ar_rna
protein_coding
protein_coding
protein_coding

ncRNA
protein_coding
protein_coding
protein_coding
protein_coding
protein_coding
protein_coding
protein_coding
protein_coding
protein_coding
protein_coding
protein_coding
protein_coding
protein_coding
protein_coding
protein_coding
protein_coding
protein_coding
ncRNA
protein_coding
ncRNA
protein_coding
protein_coding
protein_coding
protein_coding
protein_coding
protein_coding
protein_coding
protein_coding
protein_coding
protein_coding
protein_coding
protein_coding
protein_coding
protein_coding
protein_coding
protein_coding
protein_coding
protein_coding
protein_coding
protein_coding
protein_coding
protein_coding
protein_coding
protein_coding
protein_coding
protein_coding
protein_coding
protein_coding
protein_coding
protein_coding
protein_coding
protein_coding
protein_coding
protein_coding
protein_coding

protein_coding
protein_coding
protein_coding
protein_coding
miRNA_precursor
protein_coding
ncRNA
protein_coding
protein_coding
protein_coding
protein_coding
protein_coding
protein_coding
protein_coding
protein_coding
protein_coding
protein_coding
protein_coding
protein_coding
protein_coding
protein_coding
protein_coding
protein_coding
protein_coding
protein_coding
protein_coding
protein_coding
pre_trna
protein_coding
protein_coding
protein_coding
protein_coding
protein_coding
protein_coding
protein_coding
pre_trna
protein_coding
protein_coding
protein_coding
protein_coding
protein_coding
protein_coding
ncRNA
ncRNA
protein_coding
protein_coding
protein_coding
protein_coding
protein_coding
protein_coding
protein_coding
protein_coding
protein_coding
protein_coding
protein_coding
protein_coding

protein_coding
protein_coding
protein_coding
protein_coding
protein_coding
protein_coding
protein_coding
protein_coding
ncRNA
protein_coding
protein_coding
protein_coding
protein_coding
protein_coding
protein_coding
protein_coding
protein_coding
protein_coding
protein_coding
protein_coding
protein_coding
protein_coding
protein_coding
protein_coding
protein_coding
protein_coding
protein_coding
protein_coding
protein_coding
protein_coding
protein_coding
protein_coding
protein_coding
protein_coding
protein_coding
protein_coding
protein_coding
protein_coding
protein_coding
protein_coding
protein_coding
protein_coding
protein_coding
protein_coding
protein_coding
ncRNA
small_nucle(ol)ar_rna
protein_coding
protein_coding
protein_coding
protein_coding
protein_coding
protein_coding
protein_coding
protein_coding
protein_coding

protein_coding
protein_coding
protein_coding
protein_coding
protein_coding
protein_coding
protein_coding
protein_coding
protein_coding
protein_coding
protein_coding
protein_coding
ncRNA
protein_coding
ncRNA
protein_coding
protein_coding
protein_coding
protein_coding
protein_coding
protein_coding
protein_coding
protein_coding
protein_coding
protein_coding
protein_coding
protein_coding
protein_coding
ncRNA
protein_coding
protein_coding
protein_coding
protein_coding
protein_coding
protein_coding
pre_trna
protein_coding
protein_coding
protein_coding
protein_coding
protein_coding
protein_coding
protein_coding
protein_coding
protein_coding
protein_coding
ncRNA
ncRNA
protein_coding
protein_coding
protein_coding
protein_coding
protein_coding
protein_coding
protein_coding
protein_coding

protein_coding
protein_coding
protein_coding
protein_coding
protein_coding
protein_coding
protein_coding
protein_coding
protein_coding
miRNA_precursor
protein_coding
protein_coding
protein_coding
protein_coding
protein_coding
protein_coding
protein_coding
protein_coding
protein_coding
protein_coding
protein_coding
protein_coding
miRNA_precursor
protein_coding
protein_coding
protein_coding
protein_coding
protein_coding
protein_coding
protein_coding
miRNA_precursor
protein_coding
protein_coding
protein_coding
protein_coding
protein_coding
protein_coding
ncRNA
protein_coding
protein_coding
protein_coding
protein_coding
protein_coding
protein_coding
protein_coding
protein_coding
protein_coding
protein_coding
protein_coding
ncRNA
protein_coding
protein_coding
protein_coding
protein_coding
protein_coding
protein_coding

protein_coding
protein_coding
protein_coding
protein_coding
protein_coding
protein_coding
protein_coding
protein_coding
protein_coding
ncRNA
protein_coding
protein_coding
protein_coding
protein_coding
protein_coding
protein_coding
protein_coding
protein_coding
protein_coding
protein_coding
protein_coding
protein_coding
protein_coding
protein_coding
protein_coding
protein_coding
protein_coding
protein_coding
protein_coding
protein_coding
protein_coding
protein_coding
protein_coding
protein_coding
ncRNA
ncRNA
protein_coding
protein_coding
protein_coding
protein_coding
protein_coding
protein_coding
protein_coding
protein_coding
protein_coding
protein_coding
protein_coding
ncRNA
protein_coding
protein_coding
protein_coding
protein_coding
protein_coding
protein_coding
protein_coding
protein_coding

ncRNA
protein_coding
protein_coding
small_nucle(ol)ar_rna
small_nucle(ol)ar_rna
small_nucle(ol)ar_rna
protein_coding
protein_coding
protein_coding
protein_coding
protein_coding
protein_coding
protein_coding
protein_coding
protein_coding
protein_coding
protein_coding
protein_coding
protein_coding
protein_coding
protein_coding
protein_coding
protein_coding
protein_coding
protein_coding
protein_coding
protein_coding
protein_coding
ncRNA
protein_coding
protein_coding
protein_coding
protein_coding
small_nucle(ol)ar_rna
small_nucle(ol)ar_rna
miRNA_precursor
protein_coding
protein_coding
protein_coding
protein_coding
protein_coding
protein_coding
protein_coding
protein_coding
protein_coding
protein_coding
protein_coding
protein_coding
protein_coding
protein_coding
protein_coding
protein_coding
protein_coding
protein_coding
protein_coding
protein_coding

protein_coding
protein_coding
protein_coding
protein_coding
protein_coding
protein_coding
protein_coding
protein_coding
protein_coding
protein_coding
pre_trna
protein_coding
protein_coding
protein_coding
protein_coding
protein_coding
protein_coding
protein_coding
protein_coding
protein_coding
protein_coding
protein_coding
protein_coding
protein_coding
protein_coding
protein_coding
protein_coding
protein_coding
protein_coding
protein_coding
protein_coding
protein_coding
protein_coding
protein_coding
protein_coding
protein_coding
protein_coding
protein_coding
protein_coding
protein_coding
protein_coding
protein_coding
protein_coding
ncRNA
ncRNA
protein_coding
protein_coding
protein_coding
protein_coding
protein_coding
protein_coding
protein_coding
protein_coding
protein_coding
protein_coding
protein_coding

protein_coding
protein_coding
protein_coding
protein_coding
protein_coding
protein_coding
protein_coding
protein_coding
protein_coding
protein_coding
protein_coding
protein_coding
protein_coding
ncRNA
protein_coding
protein_coding
protein_coding
protein_coding
pre_trna
protein_coding
protein_coding
protein_coding
protein_coding
protein_coding
protein_coding
protein_coding
protein_coding
protein_coding
miRNA_precursor
protein_coding
protein_coding
protein_coding
protein_coding
protein_coding
protein_coding
protein_coding
protein_coding
protein_coding
protein_coding
protein_coding
protein_coding
protein_coding
protein_coding
protein_coding
protein_coding
protein_coding
protein_coding
protein_coding
ncRNA
protein_coding
protein_coding
protein_coding
protein_coding
protein_coding
protein_coding
protein_coding

protein_coding
protein_coding
protein_coding
protein_coding
protein_coding
protein_coding
protein_coding
protein_coding
protein_coding
protein_coding
protein_coding
protein_coding
protein_coding
protein_coding
protein_coding
protein_coding
protein_coding
protein_coding
protein_coding
protein_coding
protein_coding
ncRNA
protein_coding
protein_coding
protein_coding
ncRNA
protein_coding
protein_coding
small_nucle(ol)ar_rna
miRNA_precursor
protein_coding
protein_coding
protein_coding
protein_coding
protein_coding
protein_coding
protein_coding
protein_coding
protein_coding
protein_coding
protein_coding
protein_coding
protein_coding
protein_coding
protein_coding
protein_coding
protein_coding
protein_coding
protein_coding
protein_coding
protein_coding
protein_coding
protein_coding
protein_coding
protein_coding
ncRNA

protein_coding
protein_coding
protein_coding
protein_coding
protein_coding
protein_coding
protein_coding
protein_coding
protein_coding
protein_coding
protein_coding
protein_coding
protein_coding
protein_coding
protein_coding
protein_coding
protein_coding
protein_coding
protein_coding
protein_coding
protein_coding
protein_coding
protein_coding
protein_coding
ncRNA
protein_coding
protein_coding
protein_coding
protein_coding
protein_coding
protein_coding
protein_coding
protein_coding
protein_coding
protein_coding
protein_coding
protein_coding
protein_coding
protein_coding
protein_coding
protein_coding
small_nucle(ol)ar_rna
protein_coding
protein_coding
protein_coding
protein_coding
protein_coding
protein_coding
protein_coding
protein_coding
protein_coding
protein_coding
protein_coding
protein_coding
protein_coding
protein_coding

protein_coding
protein_coding
protein_coding
protein_coding
protein_coding
protein_coding
protein_coding
protein_coding
protein_coding
protein_coding
protein_coding
protein_coding
protein_coding
ncRNA
protein_coding
protein_coding
protein_coding
protein_coding
protein_coding
pre_trna
protein_coding
protein_coding
protein_coding
protein_coding
protein_coding
protein_coding
protein_coding
protein_coding
protein_coding
miRNA_precursor
protein_coding
protein_coding
protein_coding
protein_coding
protein_coding
protein_coding
protein_coding
protein_coding
protein_coding
protein_coding
pre_trna
protein_coding
protein_coding
protein_coding
protein_coding
protein_coding
protein_coding
protein_coding
protein_coding
protein_coding
protein_coding
protein_coding
protein_coding
protein_coding
protein_coding
protein_coding

protein_coding
protein_coding
protein_coding
protein_coding
protein_coding
protein_coding
protein_coding
protein_coding
protein_coding
protein_coding
protein_coding
protein_coding
protein_coding
protein_coding
protein_coding
ncRNA
ncRNA
protein_coding
protein_coding
protein_coding
protein_coding
protein_coding
protein_coding
protein_coding
protein_coding
ncRNA
protein_coding
protein_coding
protein_coding
protein_coding
protein_coding
protein_coding
protein_coding
protein_coding
protein_coding
small_nucle(ol)ar_rna
protein_coding
protein_coding
protein_coding
protein_coding
protein_coding
protein_coding
protein_coding
protein_coding
protein_coding
protein_coding
protein_coding
protein_coding
protein_coding
protein_coding
protein_coding
protein_coding
small_nucle(ol)ar_rna
small_nucle(ol)ar_rna
protein_coding
protein_coding

protein_coding
protein_coding
protein_coding
protein_coding
protein_coding
protein_coding
pre_trna
protein_coding
protein_coding
protein_coding
protein_coding
protein_coding
protein_coding
protein_coding
protein_coding
protein_coding
protein_coding
protein_coding
protein_coding
protein_coding
protein_coding
protein_coding
protein_coding
protein_coding
protein_coding
protein_coding
protein_coding
protein_coding
protein_coding
protein_coding
protein_coding
ncRNA
protein_coding
protein_coding
protein_coding
protein_coding
protein_coding
protein_coding
protein_coding
protein_coding
protein_coding
protein_coding
protein_coding
protein_coding
protein_coding
protein_coding
protein_coding
protein_coding
protein_coding
pre_trna
protein_coding
protein_coding
protein_coding
protein_coding
protein_coding
protein_coding

protein_coding
protein_coding
protein_coding
protein_coding
protein_coding
protein_coding
protein_coding
protein_coding
protein_coding
protein_coding
protein_coding
protein_coding
protein_coding
protein_coding
protein_coding
protein_coding
protein_coding
protein_coding
protein_coding
protein_coding
protein_coding
protein_coding
protein_coding
protein_coding
protein_coding
protein_coding
protein_coding
protein_coding
protein_coding
protein_coding
protein_coding
protein_coding
protein_coding
ncRNA
ncRNA
protein_coding
protein_coding
protein_coding
protein_coding
protein_coding
protein_coding
protein_coding
protein_coding
protein_coding
ncRNA
protein_coding
protein_coding
protein_coding
protein_coding
protein_coding
protein_coding
protein_coding
protein_coding
protein_coding
protein_coding
protein_coding

protein_coding
protein_coding
protein_coding
protein_coding
protein_coding
protein_coding
protein_coding
protein_coding
protein_coding
protein_coding
protein_coding
protein_coding
protein_coding
protein_coding
ncRNA
ncRNA
protein_coding
protein_coding
protein_coding
protein_coding
protein_coding
protein_coding
protein_coding
protein_coding
protein_coding
protein_coding
protein_coding
protein_coding
protein_coding
protein_coding
protein_coding
protein_coding
protein_coding
protein_coding
protein_coding
protein_coding
protein_coding
protein_coding
protein_coding
protein_coding
protein_coding
protein_coding
small_nucle(ol)ar_rna
protein_coding
protein_coding
protein_coding
protein_coding
protein_coding
protein_coding
protein_coding
protein_coding
protein_coding
protein_coding
protein_coding
ncRNA
protein_coding

protein_coding
protein_coding
protein_coding
protein_coding
protein_coding
ncRNA
protein_coding
ncRNA
protein_coding
ncRNA
protein_coding
protein_coding
protein_coding
protein_coding
protein_coding
protein_coding
protein_coding
protein_coding
miRNA_precursor
protein_coding
protein_coding
protein_coding
ncRNA
protein_coding
protein_coding
protein_coding
protein_coding
protein_coding
protein_coding
protein_coding
protein_coding
protein_coding
protein_coding
protein_coding
protein_coding
protein_coding
protein_coding
protein_coding
protein_coding
protein_coding
protein_coding
protein_coding
protein_coding
protein_coding
protein_coding
protein_coding
protein_coding
protein_coding
protein_coding
protein_coding
protein_coding
protein_coding
protein_coding
protein_coding
protein_coding
protein_coding

protein_coding
protein_coding
protein_coding
protein_coding
protein_coding
protein_coding
protein_coding
protein_coding
ncRNA
protein_coding
protein_coding
protein_coding
protein_coding
protein_coding
protein_coding
protein_coding
protein_coding
ncRNA
protein_coding
protein_coding
protein_coding
protein_coding
protein_coding
protein_coding
protein_coding
protein_coding
protein_coding
protein_coding
protein_coding
protein_coding
protein_coding
protein_coding
protein_coding
protein_coding
protein_coding
protein_coding
protein_coding
protein_coding
protein_coding
protein_coding
pre_trna
protein_coding
protein_coding
protein_coding
protein_coding
protein_coding
protein_coding
protein_coding
protein_coding
protein_coding
ncRNA
protein_coding
protein_coding
protein_coding
protein_coding
protein_coding

protein_coding
protein_coding
protein_coding
protein_coding
protein_coding
protein_coding
protein_coding
protein_coding
protein_coding
protein_coding
protein_coding
protein_coding
ncRNA
protein_coding
protein_coding
protein_coding
protein_coding
protein_coding
protein_coding
protein_coding
protein_coding
protein_coding
protein_coding
protein_coding
protein_coding
protein_coding
protein_coding
protein_coding
protein_coding
protein_coding
protein_coding
protein_coding
protein_coding
protein_coding
protein_coding
protein_coding
protein_coding
ncRNA
protein_coding
protein_coding
protein_coding
protein_coding
protein_coding
protein_coding
protein_coding
protein_coding
protein_coding
protein_coding
protein_coding
ncRNA
protein_coding
protein_coding
protein_coding
protein_coding
protein_coding
protein_coding

protein_coding
protein_coding
protein_coding
protein_coding
protein_coding
protein_coding
protein_coding
protein_coding
protein_coding
protein_coding
protein_coding
protein_coding
protein_coding
protein_coding
protein_coding
protein_coding
protein_coding
protein_coding
protein_coding
protein_coding
protein_coding
ncRNA
protein_coding
protein_coding
protein_coding
protein_coding
protein_coding
protein_coding
protein_coding
protein_coding
protein_coding
protein_coding
protein_coding
protein_coding
protein_coding
protein_coding
protein_coding
protein_coding
protein_coding
protein_coding
protein_coding
protein_coding
protein_coding
protein_coding
protein_coding
protein_coding
protein_coding
protein_coding
protein_coding
protein_coding
protein_coding
protein_coding
protein_coding
protein_coding
miRNA_precursor
protein_coding

protein_coding
protein_coding
protein_coding
protein_coding
protein_coding
protein_coding
protein_coding
protein_coding
protein_coding
protein_coding
protein_coding
protein_coding
protein_coding
protein_coding
protein_coding
protein_coding
protein_coding
protein_coding
protein_coding
protein_coding
protein_coding
protein_coding
protein_coding
protein_coding
protein_coding
protein_coding
protein_coding
protein_coding
protein_coding
protein_coding
protein_coding
protein_coding
protein_coding
protein_coding
protein_coding
protein_coding
protein_coding
protein_coding
protein_coding
protein_coding
protein_coding
protein_coding
protein_coding
protein_coding
protein_coding
protein_coding
protein_coding
protein_coding
protein_coding
protein_coding
protein_coding
protein_coding
small_nucle(ol)ar_rna
protein_coding
protein_coding
protein_coding

protein_coding
protein_coding
protein_coding
protein_coding
protein_coding
protein_coding
protein_coding
protein_coding
protein_coding
protein_coding
protein_coding
protein_coding
protein_coding
protein_coding
protein_coding
protein_coding
protein_coding
protein_coding
protein_coding
protein_coding
protein_coding
protein_coding
protein_coding
protein_coding
protein_coding
protein_coding
protein_coding
protein_coding
ncRNA
protein_coding
protein_coding
protein_coding
protein_coding
protein_coding
protein_coding
protein_coding
protein_coding
protein_coding
protein_coding
protein_coding
protein_coding
protein_coding
protein_coding
protein_coding
protein_coding
protein_coding
protein_coding
protein_coding
protein_coding
protein_coding
protein_coding
protein_coding
protein_coding
protein_coding
protein_coding
protein_coding

protein_coding
protein_coding
protein_coding
protein_coding
pre_trna
protein_coding
protein_coding
protein_coding
protein_coding
protein_coding
ncRNA
ncRNA
protein_coding
protein_coding
protein_coding
protein_coding
protein_coding
protein_coding
protein_coding
protein_coding
protein_coding
protein_coding
protein_coding
protein_coding
protein_coding
protein_coding
protein_coding
protein_coding
protein_coding
protein_coding
protein_coding
protein_coding
protein_coding
protein_coding
ncRNA
protein_coding
protein_coding
protein_coding
protein_coding
protein_coding
protein_coding
protein_coding
protein_coding
protein_coding
protein_coding
protein_coding
protein_coding
protein_coding
protein_coding
protein_coding
protein_coding
protein_coding
protein_coding
protein_coding
protein_coding
protein_coding

protein_coding
protein_coding
protein_coding
protein_coding
protein_coding
protein_coding
protein_coding
protein_coding
protein_coding
protein_coding
ncRNA
protein_coding
protein_coding
protein_coding
protein_coding
protein_coding
protein_coding
protein_coding
protein_coding
protein_coding
protein_coding
protein_coding
protein_coding
protein_coding
pre_trna
protein_coding
protein_coding
protein_coding
protein_coding
protein_coding
protein_coding
protein_coding
protein_coding
protein_coding
protein_coding
protein_coding
protein_coding
protein_coding
protein_coding
ncRNA
protein_coding
protein_coding
protein_coding
protein_coding
protein_coding
protein_coding
protein_coding
protein_coding
protein_coding
protein_coding
protein_coding
protein_coding
protein_coding
protein_coding
protein_coding
protein_coding

protein_coding
protein_coding
protein_coding
protein_coding
protein_coding
protein_coding
protein_coding
protein_coding
protein_coding
protein_coding
protein_coding
protein_coding
protein_coding
protein_coding
protein_coding
protein_coding
ncRNA
protein_coding
protein_coding
protein_coding
protein_coding
protein_coding
protein_coding
protein_coding
protein_coding
protein_coding
protein_coding
protein_coding
protein_coding
protein_coding
protein_coding
protein_coding
protein_coding
miRNA_precursor
protein_coding
protein_coding
protein_coding
protein_coding
miRNA_precursor
ncRNA
miRNA_precursor
protein_coding
protein_coding
protein_coding
protein_coding
protein_coding
protein_coding
protein_coding
protein_coding
protein_coding
protein_coding
protein_coding
protein_coding
protein_coding
protein_coding
protein_coding

protein_coding
protein_coding
protein_coding
protein_coding
protein_coding
protein_coding
protein_coding
protein_coding
protein_coding
protein_coding
protein_coding
protein_coding
protein_coding
protein_coding
protein_coding
protein_coding
protein_coding
protein_coding
protein_coding
protein_coding
protein_coding
protein_coding
protein_coding
protein_coding
protein_coding
protein_coding
protein_coding
protein_coding
protein_coding
protein_coding
protein_coding
protein_coding
protein_coding
ncRNA
ncRNA
protein_coding
ncRNA
protein_coding
protein_coding
protein_coding
protein_coding
protein_coding
ncRNA
protein_coding
protein_coding
protein_coding
protein_coding
protein_coding
miRNA_precursor
protein_coding
protein_coding
protein_coding
protein_coding
protein_coding
protein_coding
ncRNA

protein_coding
protein_coding
protein_coding
protein_coding
protein_coding
protein_coding
protein_coding
protein_coding
protein_coding
protein_coding
ncRNA
protein_coding
protein_coding
protein_coding
protein_coding
protein_coding
protein_coding
protein_coding
protein_coding
protein_coding
protein_coding
protein_coding
protein_coding
protein_coding
protein_coding
protein_coding
protein_coding
protein_coding
protein_coding
protein_coding
protein_coding
protein_coding
protein_coding
protein_coding
protein_coding
protein_coding
pre_trna
protein_coding
protein_coding
protein_coding
protein_coding
protein_coding
protein_coding
protein_coding
protein_coding
protein_coding
protein_coding
protein_coding
protein_coding
protein_coding
protein_coding
protein_coding
protein_coding
protein_coding
protein_coding
protein_coding

protein_coding
protein_coding
protein_coding
protein_coding
protein_coding
protein_coding
protein_coding
protein_coding
protein_coding
protein_coding
protein_coding
protein_coding
protein_coding
protein_coding
protein_coding
protein_coding
protein_coding
protein_coding
protein_coding
protein_coding
protein_coding
protein_coding
protein_coding
small_nucle(ol)ar_rna
protein_coding
protein_coding
protein_coding
protein_coding
protein_coding
protein_coding
protein_coding
protein_coding
small_nucle(ol)ar_rna
protein_coding
protein_coding
protein_coding
protein_coding
protein_coding
protein_coding
protein_coding
protein_coding
protein_coding
protein_coding
protein_coding
protein_coding
protein_coding
protein_coding
protein_coding
protein_coding
protein_coding
protein_coding
protein_coding
protein_coding
protein_coding
protein_coding
protein_coding

protein_coding
protein_coding
protein_coding
protein_coding
protein_coding
protein_coding
protein_coding
protein_coding
protein_coding
protein_coding
ncRNA
protein_coding
protein_coding
protein_coding
protein_coding
protein_coding
protein_coding
ncRNA
protein_coding
protein_coding
protein_coding
protein_coding
protein_coding
protein_coding
protein_coding
protein_coding
protein_coding
protein_coding
protein_coding
protein_coding
protein_coding
protein_coding
protein_coding
protein_coding
protein_coding
protein_coding
small_nucle(ol)ar_rna
small_nucle(ol)ar_rna
protein_coding
protein_coding
protein_coding
protein_coding
miRNA_precursor
protein_coding
protein_coding
protein_coding
protein_coding
protein_coding
protein_coding
protein_coding
protein_coding
ncRNA
protein_coding
protein_coding
protein_coding
protein_coding

protein_coding
protein_coding
protein_coding
protein_coding
protein_coding
protein_coding
protein_coding
ncRNA
protein_coding
protein_coding
protein_coding
protein_coding
protein_coding
protein_coding
protein_coding
protein_coding
protein_coding
protein_coding
protein_coding
protein_coding
small_nucle(ol)ar_rna
protein_coding
protein_coding
protein_coding
small_nucle(ol)ar_rna
protein_coding
protein_coding
protein_coding
protein_coding
protein_coding
protein_coding
protein_coding
protein_coding
protein_coding
protein_coding
protein_coding
protein_coding
protein_coding
protein_coding
protein_coding
protein_coding
protein_coding
protein_coding
protein_coding
protein_coding
protein_coding
protein_coding
protein_coding
protein_coding
protein_coding
protein_coding
protein_coding
protein_coding
protein_coding
protein_coding
protein_coding

protein_coding
protein_coding
protein_coding
protein_coding
protein_coding
protein_coding
protein_coding
protein_coding
protein_coding
protein_coding
protein_coding
protein_coding
small_nucle(ol)ar_rna
small_nucle(ol)ar_rna
protein_coding
protein_coding
protein_coding
ncRNA
protein_coding
protein_coding
protein_coding
protein_coding
protein_coding
protein_coding
protein_coding
protein_coding
protein_coding
protein_coding
protein_coding
protein_coding
protein_coding
protein_coding
protein_coding
protein_coding
protein_coding
protein_coding
protein_coding
protein_coding
protein_coding
protein_coding
protein_coding
protein_coding
protein_coding
protein_coding
protein_coding
ncRNA
protein_coding
protein_coding
protein_coding
miRNA_precursor
protein_coding
protein_coding
protein_coding
protein_coding
protein_coding
protein_coding

pre_trna
protein_coding
protein_coding
protein_coding
protein_coding
protein_coding
protein_coding
protein_coding
protein_coding
protein_coding
protein_coding
protein_coding
protein_coding
protein_coding
protein_coding
protein_coding
protein_coding
protein_coding
protein_coding
protein_coding
protein_coding
protein_coding
protein_coding
small_nucle(ol)ar_rna
protein_coding
protein_coding
protein_coding
protein_coding
protein_coding
protein_coding
protein_coding
protein_coding
ncRNA
protein_coding
protein_coding
protein_coding
ncRNA
protein_coding
protein_coding
protein_coding
protein_coding
protein_coding
protein_coding
protein_coding
protein_coding
protein_coding
protein_coding
protein_coding
protein_coding
protein_coding
protein_coding
protein_coding
protein_coding
protein_coding
protein_coding
protein_coding

protein_coding
protein_coding
protein_coding
protein_coding
protein_coding
protein_coding
protein_coding
protein_coding
protein_coding
protein_coding
protein_coding
protein_coding
protein_coding
protein_coding
ncRNA
protein_coding
protein_coding
protein_coding
protein_coding
protein_coding
protein_coding
protein_coding
protein_coding
protein_coding
protein_coding
protein_coding
protein_coding
protein_coding
protein_coding
protein_coding
protein_coding
protein_coding
protein_coding
protein_coding
protein_coding
protein_coding
protein_coding
protein_coding
protein_coding
protein_coding
protein_coding
protein_coding
protein_coding
protein_coding
protein_coding
protein_coding
protein_coding
protein_coding
protein_coding
protein_coding
protein_coding
protein_coding
protein_coding
small_nucle(ol)ar_rna
protein_coding
protein_coding

protein_coding
protein_coding
protein_coding
protein_coding
protein_coding
protein_coding
protein_coding
protein_coding
protein_coding
protein_coding
protein_coding
protein_coding
protein_coding
protein_coding
protein_coding
protein_coding
protein_coding
protein_coding
protein_coding
protein_coding
protein_coding
protein_coding
protein_coding
protein_coding
protein_coding
protein_coding
ncRNA
protein_coding
small_nucle(ol)ar_rna
protein_coding
protein_coding
protein_coding
protein_coding
protein_coding
protein_coding
protein_coding
protein_coding
protein_coding
ncRNA
protein_coding
protein_coding
protein_coding
protein_coding
protein_coding
protein_coding
protein_coding
protein_coding
protein_coding
protein_coding
protein_coding
protein_coding
protein_coding
pre_trna
protein_coding
protein_coding
protein_coding

ncRNA
protein_coding
protein_coding
protein_coding
protein_coding
protein_coding
protein_coding
protein_coding
protein_coding
protein_coding
protein_coding
protein_coding
protein_coding
protein_coding
protein_coding
protein_coding
protein_coding
protein_coding
protein_coding
protein_coding
protein_coding
protein_coding
protein_coding
protein_coding
protein_coding
protein_coding
protein_coding
protein_coding
protein_coding
protein_coding
protein_coding
protein_coding
protein_coding
protein_coding
protein_coding
protein_coding
protein_coding
protein_coding
protein_coding
protein_coding
protein_coding
protein_coding
protein_coding
protein_coding
pre_trna
protein_coding
protein_coding
protein_coding
protein_coding
protein_coding
protein_coding
protein_coding
protein_coding
protein_coding
protein_coding
protein_coding

protein_coding
protein_coding
protein_coding
protein_coding
protein_coding
protein_coding
protein_coding
protein_coding
protein_coding
protein_coding
protein_coding
protein_coding
protein_coding
protein_coding
protein_coding
protein_coding
protein_coding
protein_coding
protein_coding
protein_coding
protein_coding
protein_coding
protein_coding
protein_coding
protein_coding
protein_coding
ncRNA
protein_coding
protein_coding
protein_coding
protein_coding
protein_coding
ncRNA
protein_coding
protein_coding
protein_coding
protein_coding
protein_coding
protein_coding
protein_coding
protein_coding
protein_coding
protein_coding
protein_coding
protein_coding
protein_coding
protein_coding
protein_coding
protein_coding
protein_coding
protein_coding
protein_coding
protein_coding
protein_coding
protein_coding
protein_coding

protein_coding
protein_coding
protein_coding
protein_coding
pre_trna
ncRNA
protein_coding
protein_coding
protein_coding
ncRNA
protein_coding
protein_coding
protein_coding
pre_trna
protein_coding
protein_coding
protein_coding
protein_coding
protein_coding
protein_coding
protein_coding
protein_coding
protein_coding
protein_coding
protein_coding
protein_coding
protein_coding
protein_coding
protein_coding
protein_coding
protein_coding
protein_coding
protein_coding
protein_coding
protein_coding
protein_coding
protein_coding
protein_coding
protein_coding
protein_coding
protein_coding
protein_coding
protein_coding
protein_coding
protein_coding
protein_coding
small_nucle(ol)ar_rna
small_nucle(ol)ar_rna
pre_trna
protein_coding
protein_coding
protein_coding
protein_coding
protein_coding
protein_coding
protein_coding
protein_coding

protein_coding
protein_coding
protein_coding
protein_coding
protein_coding
protein_coding
pre_trna
protein_coding
ncRNA
protein_coding
protein_coding
protein_coding
protein_coding
protein_coding
protein_coding
ncRNA
protein_coding
protein_coding
protein_coding
protein_coding
protein_coding
protein_coding
protein_coding
protein_coding
protein_coding
protein_coding
protein_coding
protein_coding
protein_coding
protein_coding
protein_coding
protein_coding
miRNA_precursor
protein_coding
protein_coding
ncRNA
protein_coding
protein_coding
protein_coding
pre_trna
protein_coding
protein_coding
protein_coding
protein_coding
protein_coding
protein_coding
protein_coding
protein_coding
ncRNA
protein_coding
protein_coding
protein_coding
protein_coding
protein_coding
protein_coding
protein_coding

protein_coding
protein_coding
protein_coding
protein_coding
protein_coding
protein_coding
protein_coding
protein_coding
protein_coding
protein_coding
protein_coding
protein_coding
protein_coding
protein_coding
protein_coding
protein_coding
protein_coding
protein_coding
protein_coding
protein_coding
protein_coding
small_nucle(ol)ar_rna
protein_coding
protein_coding
protein_coding
protein_coding
ncRNA
protein_coding
protein_coding
protein_coding
protein_coding
protein_coding
protein_coding
protein_coding
protein_coding
protein_coding
protein_coding
protein_coding
protein_coding
protein_coding
protein_coding
protein_coding
protein_coding
protein_coding
protein_coding
protein_coding
small_nucle(ol)ar_rna
protein_coding
protein_coding
protein_coding
protein_coding
protein_coding
protein_coding
protein_coding
protein_coding
protein_coding

protein_coding
protein_coding
protein_coding
protein_coding
protein_coding
protein_coding
protein_coding
protein_coding
protein_coding
protein_coding
protein_coding
protein_coding
protein_coding
protein_coding
protein_coding
protein_coding
protein_coding
protein_coding
protein_coding
protein_coding
protein_coding
protein_coding
protein_coding
protein_coding
protein_coding
protein_coding
protein_coding
protein_coding
protein_coding
protein_coding
protein_coding
protein_coding
protein_coding
protein_coding
ncRNA
protein_coding
protein_coding
protein_coding
protein_coding
protein_coding
protein_coding
protein_coding
protein_coding
ncRNA
protein_coding
protein_coding
protein_coding
protein_coding
protein_coding
protein_coding
protein_coding
protein_coding
protein_coding
protein_coding
protein_coding
protein_coding

protein_coding
protein_coding
ncRNA
protein_coding
protein_coding
protein_coding
protein_coding
ncRNA
protein_coding
protein_coding
protein_coding
protein_coding
protein_coding
protein_coding
protein_coding
protein_coding
protein_coding
protein_coding
protein_coding
protein_coding
protein_coding
protein_coding
protein_coding
protein_coding
protein_coding
protein_coding
protein_coding
protein_coding
protein_coding
protein_coding
protein_coding
protein_coding
protein_coding
protein_coding
ncRNA
protein_coding
protein_coding
protein_coding
protein_coding
protein_coding
protein_coding
protein_coding
protein_coding
protein_coding
protein_coding
protein_coding
protein_coding
protein_coding
protein_coding
protein_coding
protein_coding
protein_coding
protein_coding
protein_coding
protein_coding
ncRNA

protein_coding
ncRNA
protein_coding
protein_coding
protein_coding
protein_coding
protein_coding
protein_coding
protein_coding
protein_coding
protein_coding
protein_coding
protein_coding
protein_coding
protein_coding
protein_coding
protein_coding
protein_coding
protein_coding
protein_coding
protein_coding
protein_coding
protein_coding
protein_coding
protein_coding
protein_coding
protein_coding
protein_coding
protein_coding
protein_coding
protein_coding
protein_coding
protein_coding
protein_coding
protein_coding
protein_coding
protein_coding
ncRNA
protein_coding
protein_coding
protein_coding
protein_coding
protein_coding
protein_coding
protein_coding
protein_coding
protein_coding
protein_coding
protein_coding
protein_coding
protein_coding
protein_coding
protein_coding
protein_coding
protein_coding
protein_coding

protein_coding
protein_coding
protein_coding
protein_coding
protein_coding
pre_trna
protein_coding
protein_coding
protein_coding
protein_coding
protein_coding
protein_coding
protein_coding
protein_coding
protein_coding
protein_coding
protein_coding
protein_coding
protein_coding
protein_coding
protein_coding
protein_coding
protein_coding
protein_coding
protein_coding
protein_coding
protein_coding
protein_coding
protein_coding
protein_coding
protein_coding
protein_coding
protein_coding
protein_coding
protein_coding
protein_coding
protein_coding
protein_coding
protein_coding
protein_coding
ncRNA
protein_coding
protein_coding
protein_coding
protein_coding
protein_coding
protein_coding
protein_coding
protein_coding
protein_coding
protein_coding
protein_coding
protein_coding
protein_coding
ncRNA
small_nucle(ol)ar_rna

protein_coding
protein_coding
protein_coding
protein_coding
ncRNA
protein_coding
protein_coding
protein_coding
protein_coding
protein_coding
protein_coding
protein_coding
protein_coding
protein_coding
protein_coding
ncRNA
protein_coding
protein_coding
protein_coding
ncRNA
protein_coding
protein_coding
protein_coding
protein_coding
protein_coding
protein_coding
protein_coding
protein_coding
protein_coding
protein_coding
protein_coding
protein_coding
protein_coding
protein_coding
protein_coding
protein_coding
protein_coding
protein_coding
protein_coding
protein_coding
protein_coding
protein_coding
ncRNA
protein_coding
protein_coding
protein_coding
protein_coding
ncRNA
protein_coding
protein_coding
protein_coding
protein_coding
protein_coding
pre_trna
protein_coding
protein_coding

protein_coding
protein_coding
ncRNA
protein_coding
protein_coding
ncRNA
protein_coding
protein_coding
protein_coding
protein_coding
protein_coding
protein_coding
protein_coding
protein_coding
protein_coding
protein_coding
protein_coding
protein_coding
protein_coding
protein_coding
ncRNA
protein_coding
protein_coding
protein_coding
small_nucle(ol)ar_rna
protein_coding
protein_coding
protein_coding
protein_coding
protein_coding
protein_coding
protein_coding
protein_coding
protein_coding
protein_coding
protein_coding
protein_coding
protein_coding
protein_coding
protein_coding
protein_coding
protein_coding
protein_coding
protein_coding
protein_coding
protein_coding
protein_coding
protein_coding
protein_coding
protein_coding
protein_coding
pre_trna
protein_coding
protein_coding
protein_coding
protein_coding

protein_coding
protein_coding
protein_coding
protein_coding
protein_coding
protein_coding
protein_coding
protein_coding
protein_coding
protein_coding
protein_coding
protein_coding
protein_coding
protein_coding
protein_coding
miRNA_precursor
protein_coding
protein_coding
protein_coding
protein_coding
protein_coding
protein_coding
protein_coding
protein_coding
protein_coding
protein_coding
protein_coding
protein_coding
protein_coding
protein_coding
protein_coding
protein_coding
protein_coding
protein_coding
protein_coding
protein_coding
protein_coding
protein_coding
protein_coding
protein_coding
protein_coding
protein_coding
ncRNA
protein_coding
ncRNA
protein_coding
protein_coding
protein_coding
protein_coding
protein_coding
protein_coding
protein_coding
protein_coding
protein_coding
protein_coding
protein_coding

protein_coding
protein_coding
protein_coding
protein_coding
protein_coding
protein_coding
protein_coding
protein_coding
protein_coding
protein_coding
protein_coding
protein_coding
small_nucle(ol)ar_rna
protein_coding
protein_coding
protein_coding
protein_coding
protein_coding
protein_coding
protein_coding
protein_coding
protein_coding
protein_coding
protein_coding
protein_coding
protein_coding
protein_coding
protein_coding
ncRNA
ncRNA
small_nucle(ol)ar_rna
protein_coding
protein_coding
protein_coding
protein_coding
protein_coding
protein_coding
protein_coding
protein_coding
protein_coding
small_nucle(ol)ar_rna
protein_coding
protein_coding
protein_coding
protein_coding
protein_coding
protein_coding
protein_coding
protein_coding
protein_coding
protein_coding
protein_coding
protein_coding
protein_coding
ncRNA
protein_coding

protein_coding
protein_coding
protein_coding
protein_coding
protein_coding
protein_coding
protein_coding
protein_coding
protein_coding
protein_coding
protein_coding
protein_coding
protein_coding
protein_coding
protein_coding
protein_coding
protein_coding
protein_coding
protein_coding
protein_coding
protein_coding
protein_coding
protein_coding
protein_coding
protein_coding
protein_coding
ncRNA
protein_coding
protein_coding
protein_coding
small_nucle(ol)ar_rna
protein_coding
protein_coding
protein_coding
protein_coding
protein_coding
protein_coding
protein_coding
protein_coding
protein_coding
protein_coding
protein_coding
protein_coding
protein_coding
protein_coding
protein_coding
protein_coding
protein_coding
protein_coding
protein_coding
protein_coding
protein_coding
protein_coding
ncRNA
protein_coding
protein_coding

protein_coding
protein_coding
protein_coding
protein_coding
protein_coding
protein_coding
ncRNA
protein_coding
protein_coding
protein_coding
protein_coding
protein_coding
protein_coding
protein_coding
protein_coding
protein_coding
protein_coding
protein_coding
protein_coding
protein_coding
protein_coding
protein_coding
protein_coding
protein_coding
protein_coding
protein_coding
small_nucle(ol)ar_rna
protein_coding
small_nucle(ol)ar_rna
protein_coding
protein_coding
ncRNA
protein_coding
protein_coding
protein_coding
protein_coding
protein_coding
protein_coding
protein_coding
protein_coding
ncRNA
protein_coding
protein_coding
ncRNA
protein_coding
protein_coding
protein_coding
protein_coding
ncRNA
miRNA_precursor
protein_coding
protein_coding
protein_coding
protein_coding
protein_coding
protein_coding

miRNA_precursor
protein_coding
protein_coding
small_nucle(ol)ar_rna
protein_coding
protein_coding
protein_coding
protein_coding
ncRNA
protein_coding
protein_coding
protein_coding
protein_coding
protein_coding
protein_coding
protein_coding
protein_coding
protein_coding
protein_coding
protein_coding
protein_coding
protein_coding
protein_coding
protein_coding
protein_coding
protein_coding
protein_coding
protein_coding
protein_coding
protein_coding
protein_coding
protein_coding
ncRNA
ncRNA
miRNA_precursor
protein_coding
ncRNA
protein_coding
protein_coding
protein_coding
protein_coding
protein_coding
protein_coding
protein_coding
protein_coding
protein_coding
protein_coding
protein_coding
protein_coding
protein_coding
protein_coding
ncRNA
protein_coding
protein_coding
protein_coding
protein_coding

protein_coding
protein_coding
protein_coding
protein_coding
protein_coding
protein_coding
protein_coding
protein_coding
protein_coding
protein_coding
protein_coding
protein_coding
protein_coding
protein_coding
protein_coding
protein_coding
protein_coding
protein_coding
miRNA_precursor
ncRNA
protein_coding
protein_coding
protein_coding
protein_coding
protein_coding
protein_coding
protein_coding
protein_coding
protein_coding
protein_coding
protein_coding
protein_coding
protein_coding
protein_coding
ncRNA
protein_coding
protein_coding
ncRNA
protein_coding
protein_coding
ncRNA
protein_coding
protein_coding
ncRNA
small_nucle(ol)ar_rna
protein_coding
protein_coding
protein_coding
protein_coding
protein_coding
protein_coding
protein_coding
protein_coding
protein_coding
protein_coding
protein_coding

protein_coding
protein_coding
protein_coding
protein_coding
protein_coding
protein_coding
protein_coding
protein_coding
protein_coding
protein_coding
protein_coding
protein_coding
miRNA_precursor
protein_coding
protein_coding
protein_coding
protein_coding
protein_coding
protein_coding
protein_coding
protein_coding
protein_coding
protein_coding
protein_coding
protein_coding
protein_coding
protein_coding
protein_coding
protein_coding
protein_coding
protein_coding
protein_coding
protein_coding
protein_coding
ncRNA
protein_coding
protein_coding
protein_coding
protein_coding
protein_coding
protein_coding
ncRNA
protein_coding
protein_coding
protein_coding
protein_coding
ncRNA
protein_coding
protein_coding
protein_coding
ncRNA
protein_coding
protein_coding
protein_coding
protein_coding
protein_coding

protein_coding
ncRNA
protein_coding
protein_coding
protein_coding
protein_coding
protein_coding
protein_coding
protein_coding
protein_coding
protein_coding
ncRNA
protein_coding
protein_coding
protein_coding
protein_coding
protein_coding
protein_coding
protein_coding
protein_coding
protein_coding
protein_coding
protein_coding
miRNA_precursor
protein_coding
protein_coding
protein_coding
protein_coding
protein_coding
protein_coding
protein_coding
protein_coding
protein_coding
protein_coding
protein_coding
protein_coding
protein_coding
protein_coding
protein_coding
protein_coding
protein_coding
protein_coding
protein_coding
protein_coding
protein_coding
protein_coding
protein_coding
protein_coding
protein_coding
protein_coding
protein_coding
protein_coding
protein_coding
protein_coding
protein_coding
protein_coding
protein_coding

protein_coding
protein_coding
protein_coding
protein_coding
protein_coding
protein_coding
protein_coding
protein_coding
protein_coding
protein_coding
protein_coding
protein_coding
protein_coding
protein_coding
protein_coding
protein_coding
protein_coding
protein_coding
ncRNA
protein_coding
protein_coding
protein_coding
protein_coding
protein_coding
protein_coding
protein_coding
protein_coding
protein_coding
protein_coding
protein_coding
protein_coding
protein_coding
protein_coding
protein_coding
protein_coding
protein_coding
protein_coding
protein_coding
protein_coding
protein_coding
protein_coding
protein_coding
ncRNA
protein_coding
small_nucle(ol)ar_rna
small_nucle(ol)ar_rna
small_nucle(ol)ar_rna
protein_coding
protein_coding
protein_coding
protein_coding
protein_coding
protein_coding
protein_coding
protein_coding
protein_coding

protein_coding
protein_coding
protein_coding
protein_coding
protein_coding
protein_coding
protein_coding
protein_coding
protein_coding
protein_coding
protein_coding
protein_coding
protein_coding
protein_coding
protein_coding
protein_coding
protein_coding
protein_coding
protein_coding
protein_coding
protein_coding
protein_coding
protein_coding
protein_coding
protein_coding
protein_coding
protein_coding
protein_coding
protein_coding
protein_coding
protein_coding
protein_coding
protein_coding
protein_coding
protein_coding
protein_coding
protein_coding
protein_coding
ncRNA
protein_coding
protein_coding
protein_coding
protein_coding
protein_coding
protein_coding
protein_coding
protein_coding
protein_coding
protein_coding
protein_coding
protein_coding
protein_coding
protein_coding
protein_coding
protein_coding
protein_coding

protein_coding
protein_coding
protein_coding
protein_coding
protein_coding
protein_coding
protein_coding
protein_coding
protein_coding
protein_coding
protein_coding
protein_coding
protein_coding
ncRNA
protein_coding
protein_coding
protein_coding
protein_coding
protein_coding
protein_coding
protein_coding
protein_coding
protein_coding
protein_coding
protein_coding
protein_coding
protein_coding
protein_coding
protein_coding
protein_coding
protein_coding
protein_coding
protein_coding
protein_coding
protein_coding
protein_coding
protein_coding
protein_coding
protein_coding
protein_coding
protein_coding
protein_coding
ncRNA
protein_coding
protein_coding
protein_coding
protein_coding
protein_coding
protein_coding
protein_coding
protein_coding
protein_coding
protein_coding
protein_coding
protein_coding
protein_coding

protein_coding
protein_coding
protein_coding
protein_coding
protein_coding
protein_coding
protein_coding
protein_coding
protein_coding
protein_coding
protein_coding
protein_coding
ncRNA
protein_coding
protein_coding
protein_coding
protein_coding
protein_coding
protein_coding
protein_coding
protein_coding
protein_coding
protein_coding
protein_coding
ncRNA
protein_coding
protein_coding
protein_coding
protein_coding
protein_coding
protein_coding
protein_coding
protein_coding
protein_coding
protein_coding
protein_coding
protein_coding
protein_coding
protein_coding
protein_coding
protein_coding
protein_coding
protein_coding
protein_coding
protein_coding
protein_coding
protein_coding
protein_coding
protein_coding
protein_coding
protein_coding
protein_coding
protein_coding
protein_coding
protein_coding
protein_coding

protein_coding
protein_coding
protein_coding
miRNA_precursor
protein_coding
protein_coding
protein_coding
protein_coding
protein_coding
protein_coding
protein_coding
protein_coding
protein_coding
protein_coding
protein_coding
protein_coding
protein_coding
protein_coding
protein_coding
protein_coding
protein_coding
protein_coding
ncRNA
protein_coding
protein_coding
protein_coding
protein_coding
protein_coding
protein_coding
protein_coding
protein_coding
protein_coding
protein_coding
protein_coding
protein_coding
protein_coding
protein_coding
protein_coding
protein_coding
protein_coding
protein_coding
protein_coding
miRNA_precursor
protein_coding
protein_coding
protein_coding
protein_coding
protein_coding
protein_coding
protein_coding
protein_coding
protein_coding
protein_coding
protein_coding
protein_coding
protein_coding

| |
|---|
| protein_coding |
| protein_coding |
| protein_coding |
| protein_coding |
| protein_coding |
| protein_coding |
| protein_coding |
| protein_coding |
| protein_coding |
| protein_coding |
| protein_coding |
| protein_coding |
| protein_coding |
| protein_coding |
| protein_coding |
| protein_coding |
| protein_coding |
| protein_coding |
| protein_coding |
| protein_coding |
| protein_coding |
| protein_coding |
| protein_coding |
| protein_coding |
| protein_coding |
| ncRNA |
| protein_coding |
| protein_coding |
| protein_coding |
| protein_coding |
| protein_coding |
| protein_coding |
| protein_coding |
| protein_coding |
| protein_coding |
| protein_coding |
| protein_coding |
| protein_coding |
| protein_coding |
| protein_coding |
| protein_coding |
| protein_coding |
| protein_coding |
| protein_coding |
| ncRNA |
| protein_coding |
| protein_coding |
| protein_coding |
| protein_coding |
| protein_coding |
| protein_coding |
| protein_coding |
| ncRNA |
| protein_coding |
| protein_coding |
| protein_coding |

protein_coding
protein_coding
protein_coding
protein_coding
protein_coding
protein_coding
protein_coding
protein_coding
protein_coding
protein_coding
protein_coding
protein_coding
protein_coding
protein_coding
protein_coding
protein_coding
protein_coding
protein_coding
protein_coding
protein_coding
protein_coding
protein_coding
protein_coding
protein_coding
protein_coding
protein_coding
protein_coding
protein_coding
protein_coding
protein_coding
small_nucle(ol)ar_rna
protein_coding
protein_coding
protein_coding
protein_coding
protein_coding
protein_coding
protein_coding
protein_coding
protein_coding
protein_coding
protein_coding
protein_coding
protein_coding
protein_coding
protein_coding
protein_coding
protein_coding
protein_coding
protein_coding
protein_coding
protein_coding
protein_coding
protein_coding
protein_coding
protein_coding

protein_coding
protein_coding
protein_coding
protein_coding
protein_coding
protein_coding
protein_coding
protein_coding
protein_coding
protein_coding
protein_coding
protein_coding
ncRNA
protein_coding
protein_coding
protein_coding
protein_coding
protein_coding
protein_coding
protein_coding
protein_coding
ncRNA
protein_coding
protein_coding
protein_coding
protein_coding
protein_coding
protein_coding
protein_coding
protein_coding
protein_coding
protein_coding
protein_coding
protein_coding
protein_coding
protein_coding
protein_coding
protein_coding
protein_coding
protein_coding
protein_coding
protein_coding
protein_coding
protein_coding
protein_coding
protein_coding
protein_coding
protein_coding
protein_coding
protein_coding
protein_coding
protein_coding
protein_coding
protein_coding
protein_coding
protein_coding

protein_coding
protein_coding
ncRNA
protein_coding
protein_coding
protein_coding
protein_coding
protein_coding
protein_coding
protein_coding
protein_coding
protein_coding
protein_coding
protein_coding
protein_coding
protein_coding
small_nucle(ol)ar_rna
small_nucle(ol)ar_rna
protein_coding
protein_coding
protein_coding
protein_coding
protein_coding
protein_coding
protein_coding
protein_coding
protein_coding
protein_coding
protein_coding
protein_coding
protein_coding
protein_coding
protein_coding
ncRNA
protein_coding
protein_coding
protein_coding
protein_coding
protein_coding
protein_coding
protein_coding
protein_coding
protein_coding
protein_coding
protein_coding
protein_coding
protein_coding
protein_coding
protein_coding
protein_coding
protein_coding
protein_coding
protein_coding
protein_coding
protein_coding
protein_coding

protein_coding
protein_coding
protein_coding
protein_coding
protein_coding
protein_coding
protein_coding
protein_coding
protein_coding
protein_coding
protein_coding
protein_coding
protein_coding
protein_coding
protein_coding
protein_coding
protein_coding
protein_coding
protein_coding
protein_coding
protein_coding
ncRNA
protein_coding
protein_coding
protein_coding
protein_coding
protein_coding
protein_coding
protein_coding
protein_coding
protein_coding
protein_coding
protein_coding
protein_coding
protein_coding
protein_coding
protein_coding
protein_coding
protein_coding
protein_coding
protein_coding
protein_coding
protein_coding
protein_coding
ncRNA
protein_coding
protein_coding
protein_coding
protein_coding
protein_coding
protein_coding
protein_coding
protein_coding
protein_coding
protein_coding
protein_coding

pre_trna
protein_coding
protein_coding
protein_coding
protein_coding
protein_coding
protein_coding
protein_coding
protein_coding
protein_coding
ncRNA
protein_coding
protein_coding
protein_coding
protein_coding
protein_coding
protein_coding
protein_coding
protein_coding
protein_coding
protein_coding
protein_coding
protein_coding
protein_coding
protein_coding
protein_coding
protein_coding
protein_coding
protein_coding
protein_coding
protein_coding
protein_coding
protein_coding
ncRNA
protein_coding
ncRNA
protein_coding
ncRNA
protein_coding
protein_coding
protein_coding
protein_coding
protein_coding
protein_coding
protein_coding
protein_coding
protein_coding
protein_coding
protein_coding
protein_coding
protein_coding
protein_coding
protein_coding
protein_coding
protein_coding
protein_coding

protein_coding
protein_coding
protein_coding
protein_coding
protein_coding
ncRNA
protein_coding
protein_coding
protein_coding
protein_coding
pre_trna
protein_coding
protein_coding
protein_coding
protein_coding
protein_coding
protein_coding
ncRNA
ncRNA
protein_coding
protein_coding
small_nucle(ol)ar_rna
protein_coding
protein_coding
protein_coding
protein_coding
ncRNA
protein_coding
protein_coding
protein_coding
ncRNA
protein_coding
ncRNA
protein_coding
protein_coding
protein_coding
protein_coding
protein_coding
protein_coding
protein_coding
protein_coding
protein_coding
protein_coding
protein_coding
protein_coding
protein_coding
protein_coding
protein_coding
protein_coding
protein_coding
protein_coding
protein_coding
protein_coding
protein_coding
ncRNA
protein_coding
protein_coding

ncRNA
protein_coding
protein_coding
ncRNA
protein_coding
protein_coding
protein_coding
protein_coding
protein_coding
ncRNA
protein_coding
protein_coding
ncRNA
protein_coding
protein_coding
protein_coding
protein_coding
protein_coding
protein_coding
protein_coding
protein_coding
protein_coding
protein_coding
protein_coding
protein_coding
protein_coding
protein_coding
protein_coding
protein_coding
protein_coding
protein_coding
protein_coding
protein_coding
protein_coding
protein_coding
protein_coding
protein_coding
protein_coding
ncRNA
protein_coding
protein_coding
protein_coding
protein_coding
protein_coding
protein_coding
protein_coding
protein_coding
protein_coding
protein_coding
protein_coding
protein_coding
protein_coding
protein_coding
protein_coding
pre_trna

protein_coding
protein_coding
protein_coding
protein_coding
protein_coding
protein_coding
protein_coding
protein_coding
ncRNA
ncRNA
protein_coding
protein_coding
protein_coding
protein_coding
protein_coding
miRNA_precursor
protein_coding
protein_coding
protein_coding
protein_coding
protein_coding
protein_coding
protein_coding
protein_coding
protein_coding
protein_coding
protein_coding
protein_coding
protein_coding
protein_coding
protein_coding
protein_coding
protein_coding
protein_coding
protein_coding
protein_coding
protein_coding
protein_coding
ncRNA
protein_coding
small_nucle(ol)ar_rna
protein_coding
protein_coding
protein_coding
protein_coding
protein_coding
protein_coding
protein_coding
protein_coding
protein_coding
protein_coding
protein_coding
protein_coding
protein_coding
ncRNA
protein_coding

protein_coding
protein_coding
protein_coding
protein_coding
protein_coding
protein_coding
protein_coding
protein_coding
protein_coding
protein_coding
protein_coding
protein_coding
protein_coding
protein_coding
protein_coding
protein_coding
protein_coding
protein_coding
ncRNA
protein_coding
protein_coding
protein_coding
protein_coding
protein_coding
protein_coding
protein_coding
protein_coding
protein_coding
protein_coding
protein_coding
protein_coding
protein_coding
protein_coding
protein_coding
protein_coding
protein_coding
miRNA_precursor
protein_coding
protein_coding
protein_coding
protein_coding
protein_coding
protein_coding
protein_coding
protein_coding
protein_coding
protein_coding
protein_coding
protein_coding
protein_coding
protein_coding
protein_coding
protein_coding
protein_coding
ncRNA
ncRNA

protein_coding
protein_coding
protein_coding
protein_coding
ncRNA
protein_coding
protein_coding
protein_coding
protein_coding
miRNA_precursor
protein_coding
protein_coding
ncRNA
protein_coding
protein_coding
protein_coding
protein_coding
protein_coding
protein_coding
protein_coding
protein_coding
protein_coding
protein_coding
protein_coding
protein_coding
protein_coding
protein_coding
protein_coding
protein_coding
protein_coding
protein_coding
protein_coding
protein_coding
protein_coding
protein_coding
protein_coding
protein_coding
protein_coding
protein_coding
ncRNA
protein_coding
protein_coding
protein_coding
protein_coding
protein_coding
protein_coding
protein_coding
protein_coding
protein_coding
protein_coding
protein_coding
protein_coding
protein_coding
protein_coding
protein_coding
protein_coding
protein_coding

protein_coding
protein_coding
protein_coding
protein_coding
protein_coding
protein_coding
protein_coding
protein_coding
protein_coding
protein_coding
protein_coding
protein_coding
protein_coding
protein_coding
protein_coding
protein_coding
protein_coding
protein_coding
miRNA_precursor
protein_coding
protein_coding
protein_coding
protein_coding
protein_coding
ncRNA
protein_coding
protein_coding
protein_coding
protein_coding
pre_trna
protein_coding
protein_coding
protein_coding
protein_coding
ncRNA
protein_coding
protein_coding
protein_coding
protein_coding
protein_coding
protein_coding
protein_coding
protein_coding
protein_coding
protein_coding
protein_coding
protein_coding
protein_coding
protein_coding
protein_coding
protein_coding
protein_coding
protein_coding
protein_coding
protein_coding
protein_coding

protein_coding
pre_trna
protein_coding
ncRNA
protein_coding
protein_coding
protein_coding
protein_coding
protein_coding
protein_coding
protein_coding
protein_coding
protein_coding
ncRNA
protein_coding
protein_coding
protein_coding
protein_coding
protein_coding
protein_coding
protein_coding
protein_coding
protein_coding
ncRNA
protein_coding
protein_coding
protein_coding
protein_coding
ncRNA
protein_coding
protein_coding
protein_coding
protein_coding
protein_coding
protein_coding
protein_coding
protein_coding
protein_coding
protein_coding
protein_coding
protein_coding
protein_coding
protein_coding
protein_coding
protein_coding
protein_coding
protein_coding
protein_coding
protein_coding
protein_coding
ncRNA
protein_coding
protein_coding
protein_coding
protein_coding
protein_coding

protein_coding
protein_coding
protein_coding
protein_coding
protein_coding
protein_coding
protein_coding
protein_coding
protein_coding
protein_coding
protein_coding
protein_coding
protein_coding
protein_coding
ncRNA
protein_coding
protein_coding
protein_coding
protein_coding
protein_coding
protein_coding
protein_coding
protein_coding
protein_coding
protein_coding
protein_coding
protein_coding
protein_coding
protein_coding
protein_coding
protein_coding
protein_coding
protein_coding
protein_coding
protein_coding
protein_coding
protein_coding
protein_coding
protein_coding
protein_coding
protein_coding
protein_coding
protein_coding
protein_coding
protein_coding
protein_coding
protein_coding
protein_coding
small_nucle(ol)ar_rna
small_nucle(ol)ar_rna
small_nucle(ol)ar_rna
ncRNA
protein_coding
protein_coding
protein_coding
protein_coding

protein_coding
protein_coding
protein_coding
protein_coding
protein_coding
protein_coding
protein_coding
protein_coding
protein_coding
protein_coding
protein_coding
protein_coding
protein_coding
protein_coding
protein_coding
protein_coding
protein_coding
protein_coding
protein_coding
protein_coding
protein_coding
protein_coding
protein_coding
protein_coding
protein_coding
protein_coding
protein_coding
protein_coding
protein_coding
protein_coding
protein_coding
protein_coding
protein_coding
protein_coding
protein_coding
protein_coding
protein_coding
protein_coding
protein_coding
protein_coding
protein_coding
protein_coding
protein_coding
protein_coding
ncRNA
small_nucle(ol)ar_rna
small_nucle(ol)ar_rna
ncRNA
protein_coding
protein_coding
protein_coding
protein_coding
protein_coding
protein_coding
protein_coding
protein_coding

protein_coding
protein_coding
protein_coding
protein_coding
protein_coding
protein_coding
protein_coding
protein_coding
protein_coding
protein_coding
protein_coding
protein_coding
protein_coding
ncRNA
protein_coding
protein_coding
protein_coding
protein_coding
ncRNA
protein_coding
protein_coding
protein_coding
protein_coding
protein_coding
protein_coding
miRNA_precursor
protein_coding
protein_coding
ncRNA
protein_coding
protein_coding
protein_coding
protein_coding
protein_coding
protein_coding
protein_coding
protein_coding
protein_coding
protein_coding
protein_coding
protein_coding
protein_coding
protein_coding
protein_coding
protein_coding
protein_coding
protein_coding
protein_coding
protein_coding
protein_coding
protein_coding
protein_coding
protein_coding
protein_coding
protein_coding
protein_coding

protein_coding
protein_coding
ncRNA
protein_coding
protein_coding
protein_coding
protein_coding
protein_coding
protein_coding
protein_coding
protein_coding
miRNA_precursor
protein_coding
protein_coding
protein_coding
protein_coding
protein_coding
protein_coding
protein_coding
protein_coding
protein_coding
protein_coding
protein_coding
protein_coding
pre_trna
protein_coding
protein_coding
protein_coding
ncRNA
protein_coding
protein_coding
protein_coding
protein_coding
protein_coding
protein_coding
protein_coding
protein_coding
protein_coding
protein_coding
protein_coding
protein_coding
protein_coding
protein_coding
protein_coding
protein_coding
protein_coding
protein_coding
protein_coding
protein_coding
protein_coding
protein_coding
protein_coding
protein_coding
protein_coding
protein_coding
protein_coding
protein_coding

protein_coding
protein_coding
protein_coding
protein_coding
ncRNA
protein_coding
protein_coding
protein_coding
protein_coding
ncRNA
protein_coding
protein_coding
protein_coding
ncRNA
protein_coding
protein_coding
protein_coding
ncRNA
protein_coding
protein_coding
protein_coding
protein_coding
protein_coding
protein_coding
protein_coding
protein_coding
protein_coding
protein_coding
protein_coding
protein_coding
protein_coding
protein_coding
protein_coding
protein_coding
protein_coding
protein_coding
protein_coding
protein_coding
ncRNA
protein_coding
protein_coding
protein_coding
protein_coding
protein_coding
protein_coding
protein_coding
protein_coding
protein_coding
protein_coding
protein_coding
protein_coding
protein_coding
protein_coding
protein_coding
protein_coding
protein_coding

protein_coding
protein_coding
protein_coding
protein_coding
protein_coding
protein_coding
protein_coding
protein_coding
protein_coding
protein_coding
protein_coding
protein_coding
protein_coding
protein_coding
protein_coding
protein_coding
protein_coding
protein_coding
protein_coding
protein_coding
protein_coding
protein_coding
protein_coding
protein_coding
protein_coding
protein_coding
protein_coding
protein_coding
protein_coding
protein_coding
protein_coding
protein_coding
protein_coding
protein_coding
protein_coding
protein_coding
protein_coding
protein_coding
ncRNA
protein_coding
protein_coding
protein_coding
protein_coding
protein_coding
protein_coding
protein_coding
protein_coding
protein_coding
protein_coding
protein_coding
protein_coding
protein_coding
protein_coding
protein_coding
protein_coding
protein_coding

protein_coding
protein_coding
protein_coding
protein_coding
protein_coding
protein_coding
protein_coding
protein_coding
protein_coding
protein_coding
protein_coding
protein_coding
protein_coding
protein_coding
protein_coding
protein_coding
protein_coding
protein_coding
protein_coding
protein_coding
protein_coding
protein_coding
protein_coding
protein_coding
protein_coding
ncRNA
protein_coding
protein_coding
protein_coding
protein_coding
protein_coding
protein_coding
protein_coding
protein_coding
protein_coding
protein_coding
protein_coding
protein_coding
protein_coding
protein_coding
protein_coding
protein_coding
protein_coding
protein_coding
protein_coding
ncRNA
protein_coding
protein_coding
protein_coding
protein_coding
protein_coding
protein_coding
protein_coding
protein_coding
protein_coding
protein_coding

protein_coding
protein_coding
protein_coding
protein_coding
protein_coding
protein_coding
protein_coding
protein_coding
protein_coding
protein_coding
protein_coding
pre_trna
protein_coding
protein_coding
protein_coding
protein_coding
protein_coding
protein_coding
protein_coding
small_nucle(ol)ar_rna
protein_coding
protein_coding
protein_coding
protein_coding
protein_coding
protein_coding
pre_trna
protein_coding
ncRNA
protein_coding
protein_coding
protein_coding
protein_coding
protein_coding
protein_coding
protein_coding
protein_coding
protein_coding
protein_coding
protein_coding
protein_coding
protein_coding
protein_coding
protein_coding
protein_coding
protein_coding
protein_coding
protein_coding
protein_coding
protein_coding
protein_coding
protein_coding
protein_coding
protein_coding
protein_coding
protein_coding

protein_coding
protein_coding
protein_coding
protein_coding
protein_coding
protein_coding
protein_coding
protein_coding
protein_coding
protein_coding
protein_coding
protein_coding
protein_coding
protein_coding
protein_coding
protein_coding
protein_coding
protein_coding
protein_coding
protein_coding
protein_coding
protein_coding
protein_coding
protein_coding
protein_coding
protein_coding
protein_coding
protein_coding
protein_coding
protein_coding
protein_coding
protein_coding
protein_coding
protein_coding
protein_coding
protein_coding
protein_coding
protein_coding
protein_coding
protein_coding
protein_coding
protein_coding
protein_coding
protein_coding
protein_coding
protein_coding
protein_coding
protein_coding
protein_coding
protein_coding
miRNA_precursor
protein_coding
protein_coding
small_nucle(ol)ar_rna
small_nucle(ol)ar_rna
ncRNA

protein_coding
protein_coding
protein_coding
protein_coding
protein_coding
protein_coding
protein_coding
protein_coding
protein_coding
protein_coding
protein_coding
protein_coding
protein_coding
protein_coding
protein_coding
protein_coding
protein_coding
protein_coding
protein_coding
protein_coding
protein_coding
protein_coding
protein_coding
protein_coding
protein_coding
protein_coding
protein_coding
protein_coding
protein_coding
protein_coding
protein_coding
protein_coding
protein_coding
protein_coding
protein_coding
protein_coding
protein_coding
protein_coding
ncRNA
protein_coding
protein_coding
protein_coding
protein_coding
protein_coding
ncRNA
protein_coding
protein_coding
protein_coding
protein_coding
protein_coding
protein_coding
protein_coding
protein_coding
protein_coding
protein_coding
protein_coding

protein_coding
protein_coding
protein_coding
protein_coding
protein_coding
protein_coding
protein_coding
protein_coding
protein_coding
protein_coding
protein_coding
protein_coding
protein_coding
protein_coding
protein_coding
protein_coding
protein_coding
protein_coding
protein_coding
protein_coding
protein_coding
protein_coding
protein_coding
protein_coding
protein_coding
protein_coding
protein_coding
protein_coding
protein_coding
protein_coding
protein_coding
protein_coding
protein_coding
protein_coding
protein_coding
protein_coding
ncRNA
protein_coding
protein_coding
protein_coding
protein_coding
protein_coding
protein_coding
protein_coding
protein_coding
protein_coding
protein_coding
protein_coding
protein_coding
protein_coding
protein_coding
miRNA_precursor
protein_coding
protein_coding
protein_coding
protein_coding

protein_coding
protein_coding
protein_coding
protein_coding
protein_coding
protein_coding
protein_coding
protein_coding
protein_coding
protein_coding
small_nucle(ol)ar_rna
small_nucle(ol)ar_rna
protein_coding
protein_coding
protein_coding
protein_coding
protein_coding
protein_coding
protein_coding
protein_coding
protein_coding
protein_coding
protein_coding
small_nucle(ol)ar_rna
protein_coding
protein_coding
protein_coding
protein_coding
protein_coding
protein_coding
protein_coding
protein_coding
protein_coding
protein_coding
protein_coding
protein_coding
protein_coding
protein_coding
protein_coding
protein_coding
protein_coding
protein_coding
protein_coding
protein_coding
protein_coding
ncRNA
protein_coding
protein_coding
protein_coding
protein_coding
protein_coding
protein_coding
protein_coding
protein_coding
protein_coding
protein_coding

protein_coding
protein_coding
protein_coding
protein_coding
protein_coding
protein_coding
protein_coding
protein_coding
protein_coding
protein_coding
protein_coding
protein_coding
protein_coding
protein_coding
protein_coding
protein_coding
protein_coding
protein_coding
protein_coding
protein_coding
protein_coding
pre_trna
protein_coding
ncRNA
protein_coding
protein_coding
protein_coding
protein_coding
protein_coding
protein_coding
protein_coding
protein_coding
protein_coding
protein_coding
protein_coding
protein_coding
protein_coding
protein_coding
protein_coding
protein_coding
protein_coding
protein_coding
protein_coding
protein_coding
protein_coding
protein_coding
protein_coding
protein_coding
protein_coding
protein_coding
protein_coding
protein_coding
protein_coding
protein_coding
protein_coding
protein_coding

protein_coding
protein_coding
protein_coding
protein_coding
protein_coding
protein_coding
protein_coding
protein_coding
protein_coding
protein_coding
protein_coding
protein_coding
protein_coding
protein_coding
protein_coding
protein_coding
protein_coding
protein_coding
protein_coding
protein_coding
protein_coding
ncRNA
protein_coding
protein_coding
protein_coding
protein_coding
protein_coding
protein_coding
protein_coding
protein_coding
protein_coding
protein_coding
protein_coding
protein_coding
protein_coding
protein_coding
protein_coding
protein_coding
protein_coding
protein_coding
protein_coding
ncRNA
small_nucle(ol)ar_rna
protein_coding
protein_coding
protein_coding
protein_coding
protein_coding
protein_coding
protein_coding
protein_coding
protein_coding
protein_coding
protein_coding
protein_coding
protein_coding

ncRNA
protein_coding
protein_coding
protein_coding
protein_coding
protein_coding
protein_coding
protein_coding
protein_coding
protein_coding
protein_coding
protein_coding
protein_coding
protein_coding
protein_coding
protein_coding
protein_coding
protein_coding
protein_coding
protein_coding
protein_coding
protein_coding
protein_coding
protein_coding
protein_coding
protein_coding
protein_coding
protein_coding
miRNA_precursor
miRNA_precursor
protein_coding
protein_coding
protein_coding
protein_coding
protein_coding
protein_coding
protein_coding
protein_coding
protein_coding
protein_coding
protein_coding
protein_coding
protein_coding
protein_coding
ncRNA
ncRNA
protein_coding
protein_coding
protein_coding
protein_coding
protein_coding
protein_coding
protein_coding
protein_coding
protein_coding
protein_coding

ncRNA
protein_coding
protein_coding
protein_coding
protein_coding
pre_trna
protein_coding
protein_coding
protein_coding
protein_coding
protein_coding
protein_coding
protein_coding
protein_coding
protein_coding
protein_coding
protein_coding
ncRNA
protein_coding
protein_coding
protein_coding
protein_coding
protein_coding
protein_coding
ncRNA
protein_coding
protein_coding
ncRNA
protein_coding
protein_coding
protein_coding
protein_coding
protein_coding
protein_coding
protein_coding
protein_coding
protein_coding
protein_coding
protein_coding
protein_coding
protein_coding
protein_coding
protein_coding
protein_coding
protein_coding
protein_coding
protein_coding
protein_coding
protein_coding
protein_coding
protein_coding
protein_coding
protein_coding
protein_coding
protein_coding
protein_coding

miRNA_precursor
protein_coding
protein_coding
protein_coding
protein_coding
protein_coding
ncRNA
ncRNA
protein_coding
protein_coding
protein_coding
protein_coding
protein_coding
protein_coding
protein_coding
protein_coding
protein_coding
protein_coding
protein_coding
protein_coding
protein_coding
protein_coding
protein_coding
protein_coding
protein_coding
protein_coding
protein_coding
protein_coding
protein_coding
protein_coding
protein_coding
protein_coding
ncRNA
protein_coding
protein_coding
miRNA_precursor
protein_coding
miRNA_precursor
protein_coding
protein_coding
protein_coding
protein_coding
protein_coding
protein_coding
protein_coding
protein_coding
protein_coding
protein_coding
protein_coding
protein_coding
protein_coding
protein_coding
protein_coding
protein_coding
protein_coding
ncRNA

protein_coding
protein_coding
protein_coding
protein_coding
protein_coding
protein_coding
protein_coding
protein_coding
protein_coding
protein_coding
protein_coding
protein_coding
protein_coding
protein_coding
protein_coding
protein_coding
protein_coding
protein_coding
protein_coding
protein_coding
protein_coding
protein_coding
protein_coding
protein_coding
protein_coding
protein_coding
protein_coding
protein_coding
protein_coding
protein_coding
protein_coding
protein_coding
protein_coding
protein_coding
protein_coding
protein_coding
protein_coding
protein_coding
protein_coding
protein_coding
protein_coding
protein_coding
protein_coding
protein_coding
protein_coding
protein_coding
protein_coding
protein_coding
protein_coding
protein_coding
protein_coding
protein_coding
protein_coding
protein_coding
ncRNA
protein_coding

protein_coding
protein_coding
protein_coding
protein_coding
protein_coding
protein_coding
protein_coding
protein_coding
protein_coding
protein_coding
protein_coding
protein_coding
protein_coding
protein_coding
protein_coding
protein_coding
protein_coding
protein_coding
protein_coding
protein_coding
protein_coding
protein_coding
protein_coding
protein_coding
protein_coding
protein_coding
protein_coding
protein_coding
protein_coding
protein_coding
protein_coding
protein_coding
protein_coding
protein_coding
protein_coding
small_nucle(ol)ar_rna
protein_coding
protein_coding
protein_coding
pre_trna
protein_coding
pre_trna
protein_coding
protein_coding
protein_coding
protein_coding
protein_coding
protein_coding
protein_coding
protein_coding
protein_coding
protein_coding
protein_coding
ncRNA
protein_coding
protein_coding

ncRNA
ncRNA
protein_coding
protein_coding
protein_coding
protein_coding
protein_coding
protein_coding
protein_coding
protein_coding
protein_coding
protein_coding
protein_coding
protein_coding
miRNA_precursor
miRNA_precursor
protein_coding
protein_coding
protein_coding
protein_coding
protein_coding
protein_coding
protein_coding
protein_coding
protein_coding
protein_coding
protein_coding
protein_coding
protein_coding
protein_coding
protein_coding
protein_coding
protein_coding
protein_coding
protein_coding
protein_coding
protein_coding
protein_coding
protein_coding
protein_coding
protein_coding
protein_coding
protein_coding
protein_coding
protein_coding
protein_coding
protein_coding
protein_coding
protein_coding
protein_coding
protein_coding
protein_coding
protein_coding
protein_coding
protein_coding
protein_coding

protein_coding
protein_coding
protein_coding
protein_coding
protein_coding
protein_coding
protein_coding
protein_coding
protein_coding
protein_coding
protein_coding
protein_coding
protein_coding
protein_coding
protein_coding
protein_coding
protein_coding
protein_coding
protein_coding
pre_trna
protein_coding
protein_coding
protein_coding
protein_coding
protein_coding
protein_coding
protein_coding
protein_coding
protein_coding
protein_coding
protein_coding
protein_coding
protein_coding
protein_coding
protein_coding
protein_coding
protein_coding
protein_coding
protein_coding
protein_coding
protein_coding
protein_coding
protein_coding
protein_coding
ncRNA
protein_coding
ncRNA
protein_coding
protein_coding
protein_coding
protein_coding
protein_coding
protein_coding
ncRNA
protein_coding
protein_coding

protein_coding
protein_coding
protein_coding
protein_coding
small_nucle(ol)ar_rna
protein_coding
protein_coding
protein_coding
protein_coding
protein_coding
protein_coding
ncRNA
protein_coding
protein_coding
protein_coding
protein_coding
protein_coding
protein_coding
protein_coding
protein_coding
protein_coding
protein_coding
protein_coding
protein_coding
protein_coding
protein_coding
protein_coding
protein_coding
protein_coding
protein_coding
protein_coding
protein_coding
protein_coding
protein_coding
protein_coding
protein_coding
protein_coding
protein_coding
protein_coding
protein_coding
protein_coding
ncRNA
protein_coding
protein_coding
protein_coding
protein_coding
protein_coding
ncRNA
protein_coding
protein_coding
protein_coding
ncRNA
protein_coding
protein_coding
protein_coding
protein_coding

protein_coding
protein_coding
protein_coding
protein_coding
protein_coding
protein_coding
protein_coding
ncRNA
protein_coding
protein_coding
protein_coding
protein_coding
protein_coding
protein_coding
protein_coding
protein_coding
protein_coding
protein_coding
protein_coding
protein_coding
protein_coding
protein_coding
protein_coding
protein_coding
protein_coding
protein_coding
protein_coding
protein_coding
protein_coding
protein_coding
protein_coding
protein_coding
protein_coding
protein_coding
ncRNA
protein_coding
protein_coding
ncRNA
ncRNA
protein_coding
protein_coding
protein_coding
protein_coding
protein_coding
protein_coding
protein_coding
protein_coding
protein_coding
protein_coding
protein_coding
protein_coding
protein_coding
protein_coding
protein_coding
protein_coding
protein_coding

protein_coding
protein_coding
protein_coding
protein_coding
protein_coding
protein_coding
protein_coding
protein_coding
protein_coding
protein_coding
protein_coding
protein_coding
protein_coding
ncRNA
protein_coding
protein_coding
protein_coding
protein_coding
protein_coding
protein_coding
protein_coding
protein_coding
protein_coding
protein_coding
protein_coding
protein_coding
pre_trna
protein_coding
protein_coding
protein_coding
protein_coding
protein_coding
protein_coding
protein_coding
protein_coding
protein_coding
protein_coding
protein_coding
protein_coding
protein_coding
protein_coding
protein_coding
ncRNA
protein_coding
protein_coding
protein_coding
protein_coding
protein_coding
protein_coding
protein_coding
protein_coding
protein_coding
protein_coding
protein_coding
protein_coding
protein_coding

protein_coding
protein_coding
small_nucle(ol)ar_rna
protein_coding
protein_coding
protein_coding
protein_coding
protein_coding
protein_coding
protein_coding
protein_coding
protein_coding
protein_coding
protein_coding
protein_coding
protein_coding
protein_coding
protein_coding
protein_coding
protein_coding
protein_coding
protein_coding
protein_coding
protein_coding
small_nucle(ol)ar_rna
small_nucle(ol)ar_rna
protein_coding
protein_coding
protein_coding
protein_coding
pre_trna
miRNA_precursor
protein_coding
protein_coding
protein_coding
protein_coding
protein_coding
protein_coding
protein_coding
protein_coding
ncRNA
protein_coding
ncRNA
ncRNA
protein_coding
protein_coding
protein_coding
protein_coding
protein_coding
protein_coding
protein_coding
protein_coding
protein_coding
protein_coding
protein_coding
protein_coding

protein_coding
protein_coding
protein_coding
protein_coding
protein_coding
ncRNA
protein_coding
protein_coding
protein_coding
protein_coding
protein_coding
protein_coding
protein_coding
protein_coding
protein_coding
protein_coding
protein_coding
protein_coding
ncRNA
protein_coding
protein_coding
protein_coding
miRNA_precursor
protein_coding
protein_coding
protein_coding
ncRNA
protein_coding
protein_coding
protein_coding
protein_coding
protein_coding
protein_coding
protein_coding
protein_coding
protein_coding
protein_coding
protein_coding
protein_coding
protein_coding
protein_coding
protein_coding
protein_coding
protein_coding
protein_coding
protein_coding
protein_coding
ncRNA
protein_coding
protein_coding
protein_coding
protein_coding
protein_coding
protein_coding
protein_coding
protein_coding

protein_coding
protein_coding
protein_coding
protein_coding
protein_coding
protein_coding
protein_coding
miRNA_precursor
protein_coding
protein_coding
protein_coding
protein_coding
protein_coding
protein_coding
protein_coding
protein_coding
protein_coding
protein_coding
protein_coding
protein_coding
miRNA_precursor
pre_trna
protein_coding
protein_coding
protein_coding
protein_coding
protein_coding
ncRNA
protein_coding
protein_coding
protein_coding
miRNA_precursor
protein_coding
protein_coding
protein_coding
protein_coding
protein_coding
protein_coding
miRNA_precursor
protein_coding
ncRNA
protein_coding
protein_coding
protein_coding
protein_coding
protein_coding
protein_coding
protein_coding
protein_coding
protein_coding
protein_coding
protein_coding
protein_coding
protein_coding
protein_coding
protein_coding

protein_coding
protein_coding
protein_coding
protein_coding
protein_coding
protein_coding
protein_coding
protein_coding
protein_coding
protein_coding
protein_coding
protein_coding
protein_coding
protein_coding
protein_coding
protein_coding
protein_coding
protein_coding
protein_coding
protein_coding
protein_coding
protein_coding
protein_coding
protein_coding
protein_coding
protein_coding
protein_coding
protein_coding
protein_coding
protein_coding
protein_coding
protein_coding
protein_coding
protein_coding
protein_coding
protein_coding
protein_coding
protein_coding
protein_coding
protein_coding
miRNA_precursor
protein_coding
protein_coding
protein_coding
protein_coding
protein_coding
protein_coding
protein_coding
protein_coding
protein_coding
protein_coding
ncRNA
small_nucle(ol)ar_rna
small_nucle(ol)ar_rna
small_nucle(ol)ar_rna
protein_coding

protein_coding
protein_coding
protein_coding
protein_coding
ncRNA
protein_coding
protein_coding
protein_coding
protein_coding
protein_coding
protein_coding
protein_coding
protein_coding
protein_coding
protein_coding
protein_coding
protein_coding
protein_coding
protein_coding
protein_coding
protein_coding
protein_coding
protein_coding
protein_coding
protein_coding
protein_coding
protein_coding
protein_coding
protein_coding
protein_coding
ncRNA
protein_coding
protein_coding
protein_coding
protein_coding
protein_coding
protein_coding
protein_coding
protein_coding
protein_coding
protein_coding
protein_coding
protein_coding
protein_coding
protein_coding
protein_coding
protein_coding
protein_coding
protein_coding
protein_coding
protein_coding
protein_coding
protein_coding
protein_coding
protein_coding
protein_coding

protein_coding
protein_coding
protein_coding
protein_coding
protein_coding
protein_coding
protein_coding
protein_coding
protein_coding
protein_coding
pre_trna
protein_coding
protein_coding
protein_coding
protein_coding
protein_coding
protein_coding
protein_coding
protein_coding
protein_coding
protein_coding
protein_coding
protein_coding
protein_coding
protein_coding
protein_coding
ncRNA
protein_coding
protein_coding
protein_coding
protein_coding
protein_coding
protein_coding
protein_coding
protein_coding
protein_coding
miRNA_precursor
protein_coding
protein_coding
protein_coding
protein_coding
protein_coding
protein_coding
protein_coding
protein_coding
protein_coding
protein_coding
ncRNA
protein_coding
protein_coding
protein_coding
protein_coding
protein_coding
protein_coding
protein_coding
protein_coding

protein_coding
protein_coding
protein_coding
ncRNA
protein_coding
protein_coding
protein_coding
protein_coding
protein_coding
protein_coding
protein_coding
protein_coding
protein_coding
protein_coding
protein_coding
protein_coding
protein_coding
protein_coding
protein_coding
protein_coding
protein_coding
protein_coding
protein_coding
protein_coding
protein_coding
protein_coding
pre_trna
protein_coding
protein_coding
protein_coding
protein_coding
protein_coding
protein_coding
protein_coding
protein_coding
protein_coding
protein_coding
protein_coding
protein_coding
protein_coding
protein_coding
protein_coding
protein_coding
protein_coding
protein_coding
protein_coding
protein_coding
ncRNA
protein_coding
protein_coding
protein_coding
protein_coding
protein_coding
protein_coding
protein_coding
protein_coding

protein_coding
protein_coding
ncRNA
protein_coding
protein_coding
ncRNA
small_nucle(ol)ar_rna
protein_coding
protein_coding
protein_coding
protein_coding
protein_coding
protein_coding
protein_coding
protein_coding
protein_coding
protein_coding
protein_coding
ncRNA
protein_coding
protein_coding
protein_coding
ncRNA
protein_coding
protein_coding
protein_coding
protein_coding
protein_coding
protein_coding
protein_coding
protein_coding
protein_coding
protein_coding
protein_coding
protein_coding
protein_coding
ncRNA
protein_coding
protein_coding
protein_coding
protein_coding
protein_coding
protein_coding
protein_coding
protein_coding
protein_coding
protein_coding
pre_trna
protein_coding
protein_coding
protein_coding
protein_coding
protein_coding
protein_coding
protein_coding
protein_coding

protein_coding
protein_coding
protein_coding
protein_coding
ncRNA
protein_coding
protein_coding
protein_coding
protein_coding
protein_coding
miRNA_precursor
small_nucle(ol)ar_rna
small_nucle(ol)ar_rna
protein_coding
pre_trna
protein_coding
protein_coding
protein_coding
protein_coding
protein_coding
protein_coding
protein_coding
ncRNA
protein_coding
protein_coding
protein_coding
protein_coding
protein_coding
ncRNA
protein_coding
protein_coding
protein_coding
protein_coding
protein_coding
protein_coding
protein_coding
protein_coding
protein_coding
protein_coding
protein_coding
protein_coding
protein_coding
protein_coding
protein_coding
protein_coding
pre_trna
protein_coding
protein_coding
protein_coding
protein_coding
protein_coding
protein_coding
protein_coding
protein_coding
protein_coding
protein_coding

protein_coding
protein_coding
protein_coding
protein_coding
protein_coding
protein_coding
protein_coding
protein_coding
protein_coding
protein_coding
protein_coding
protein_coding
protein_coding
protein_coding
protein_coding
small_nucle(ol)ar_rna
protein_coding
protein_coding
protein_coding
protein_coding
protein_coding
protein_coding
protein_coding
protein_coding
protein_coding
protein_coding
protein_coding
protein_coding
protein_coding
protein_coding
protein_coding
protein_coding
protein_coding
protein_coding
protein_coding
protein_coding
protein_coding
protein_coding
protein_coding
protein_coding
protein_coding
protein_coding
protein_coding
protein_coding
protein_coding
protein_coding
protein_coding
protein_coding
protein_coding
protein_coding
protein_coding
protein_coding
protein_coding
protein_coding
protein_coding
protein_coding

protein_coding
protein_coding
protein_coding
protein_coding
protein_coding
protein_coding
protein_coding
protein_coding
protein_coding
protein_coding
protein_coding
protein_coding
protein_coding
protein_coding
protein_coding
protein_coding
protein_coding
protein_coding
protein_coding
protein_coding
protein_coding
protein_coding
protein_coding
protein_coding
protein_coding
protein_coding
protein_coding
protein_coding
protein_coding
protein_coding
protein_coding
protein_coding
protein_coding
protein_coding
protein_coding
protein_coding
protein_coding
protein_coding
protein_coding
protein_coding
protein_coding
protein_coding
protein_coding
protein_coding
protein_coding
protein_coding
protein_coding
protein_coding
protein_coding
protein_coding
protein_coding
protein_coding
protein_coding
protein_coding
protein_coding
protein_coding

protein_coding
protein_coding
protein_coding
protein_coding
protein_coding
protein_coding
protein_coding
protein_coding
protein_coding
protein_coding
ncRNA
protein_coding
protein_coding
ncRNA
protein_coding
protein_coding
protein_coding
protein_coding
protein_coding
protein_coding
protein_coding
protein_coding
protein_coding
protein_coding
protein_coding
protein_coding
protein_coding
protein_coding
protein_coding
protein_coding
protein_coding
protein_coding
protein_coding
protein_coding
protein_coding
protein_coding
protein_coding
protein_coding
protein_coding
protein_coding
protein_coding
protein_coding
protein_coding
protein_coding
protein_coding
protein_coding
protein_coding
protein_coding
protein_coding
protein_coding
protein_coding
protein_coding
protein_coding
protein_coding
protein_coding
protein_coding

protein_coding
protein_coding
protein_coding
protein_coding
ncRNA
protein_coding
protein_coding
protein_coding
protein_coding
protein_coding
protein_coding
protein_coding
protein_coding
ncRNA
protein_coding
protein_coding
protein_coding
ncRNA
protein_coding
protein_coding
protein_coding
protein_coding
protein_coding
protein_coding
protein_coding
protein_coding
protein_coding
protein_coding
protein_coding
protein_coding
protein_coding
protein_coding
protein_coding
protein_coding
protein_coding
ncRNA
protein_coding
protein_coding
protein_coding
protein_coding
ncRNA
protein_coding
protein_coding
protein_coding
protein_coding
protein_coding
protein_coding
protein_coding
protein_coding
protein_coding
protein_coding
protein_coding
protein_coding
protein_coding
protein_coding
protein_coding

protein_coding
ncRNA
protein_coding
protein_coding
protein_coding
protein_coding
protein_coding
protein_coding
pre_trna
protein_coding
protein_coding
protein_coding
protein_coding
protein_coding
protein_coding
protein_coding
protein_coding
protein_coding
protein_coding
protein_coding
protein_coding
protein_coding
protein_coding
protein_coding
protein_coding
ncRNA
pre_trna
protein_coding
protein_coding
protein_coding
protein_coding
protein_coding
protein_coding
protein_coding
ncRNA
protein_coding
protein_coding
protein_coding
ncRNA
protein_coding
protein_coding
pre_trna
protein_coding
protein_coding
protein_coding
protein_coding
protein_coding
protein_coding
protein_coding
protein_coding
protein_coding
protein_coding
protein_coding
protein_coding
protein_coding
protein_coding

protein_coding
protein_coding
protein_coding
protein_coding
protein_coding
protein_coding
protein_coding
protein_coding
protein_coding
protein_coding
protein_coding
protein_coding
protein_coding
protein_coding
protein_coding
protein_coding
protein_coding
protein_coding
protein_coding
protein_coding
protein_coding
protein_coding
protein_coding
protein_coding
protein_coding
protein_coding
protein_coding
protein_coding
protein_coding
protein_coding
protein_coding
protein_coding
protein_coding
protein_coding
protein_coding
protein_coding
protein_coding
protein_coding
ncRNA
protein_coding
protein_coding
protein_coding
protein_coding
protein_coding
protein_coding
protein_coding
protein_coding
protein_coding
protein_coding
protein_coding
protein_coding
protein_coding
protein_coding
protein_coding
protein_coding
protein_coding

protein_coding
protein_coding
protein_coding
ncRNA
protein_coding
ncRNA
protein_coding
protein_coding
protein_coding
protein_coding
ncRNA
protein_coding
protein_coding
protein_coding
protein_coding
protein_coding
protein_coding
small_nucle(ol)ar_rna
small_nucle(ol)ar_rna
protein_coding
protein_coding
protein_coding
protein_coding
protein_coding
protein_coding
protein_coding
protein_coding
protein_coding
protein_coding
protein_coding
protein_coding
protein_coding
protein_coding
ncRNA
protein_coding
ncRNA
protein_coding
protein_coding
protein_coding
protein_coding
protein_coding
protein_coding
pre_trna
protein_coding
protein_coding
protein_coding
protein_coding
protein_coding
pre_trna
protein_coding
protein_coding
protein_coding
protein_coding
small_nucle(ol)ar_rna
small_nucle(ol)ar_rna
small_nucle(ol)ar_rna

protein_coding
protein_coding
protein_coding
protein_coding
protein_coding
protein_coding
protein_coding
protein_coding
protein_coding
protein_coding
protein_coding
protein_coding
protein_coding
ncRNA
protein_coding
protein_coding
protein_coding
protein_coding
protein_coding
protein_coding
protein_coding
protein_coding
protein_coding
protein_coding
protein_coding
pre_trna
protein_coding
protein_coding
protein_coding
protein_coding
protein_coding
protein_coding
protein_coding
protein_coding
protein_coding
protein_coding
protein_coding
protein_coding
ncRNA
protein_coding
protein_coding
protein_coding
protein_coding
protein_coding
protein_coding
protein_coding
protein_coding
ncRNA
protein_coding
protein_coding
protein_coding
protein_coding
protein_coding
protein_coding
protein_coding
protein_coding

protein_coding
protein_coding
protein_coding
protein_coding
protein_coding
protein_coding
protein_coding
protein_coding
protein_coding
protein_coding
protein_coding
protein_coding
protein_coding
protein_coding
protein_coding
protein_coding
protein_coding
protein_coding
protein_coding
protein_coding
protein_coding
protein_coding
protein_coding
protein_coding
protein_coding
protein_coding
ncRNA
small_nucle(ol)ar_rna
protein_coding
protein_coding
protein_coding
protein_coding
protein_coding
protein_coding
protein_coding
protein_coding
protein_coding
protein_coding
protein_coding
protein_coding
protein_coding
protein_coding
protein_coding
protein_coding
protein_coding
protein_coding
protein_coding
protein_coding
protein_coding
protein_coding
protein_coding
protein_coding
protein_coding
small_nucle(ol)ar_rna
protein_coding
protein_coding

protein_coding
protein_coding
protein_coding
protein_coding
protein_coding
protein_coding
protein_coding
protein_coding
protein_coding
protein_coding
protein_coding
protein_coding
protein_coding
protein_coding
protein_coding
protein_coding
protein_coding
protein_coding
protein_coding
protein_coding
protein_coding
protein_coding
protein_coding
protein_coding
protein_coding
protein_coding
protein_coding
protein_coding
protein_coding
protein_coding
protein_coding
protein_coding
protein_coding
protein_coding
protein_coding
protein_coding
protein_coding
protein_coding
protein_coding
protein_coding
protein_coding
protein_coding
protein_coding
protein_coding
protein_coding
protein_coding
protein_coding
protein_coding
protein_coding
protein_coding
protein_coding
protein_coding
small_nucle(ol)ar_rna
small_nucle(ol)ar_rna
small_nucle(ol)ar_rna
protein_coding

protein_coding
protein_coding
protein_coding
protein_coding
protein_coding
protein_coding
protein_coding
protein_coding
protein_coding
protein_coding
protein_coding
protein_coding
protein_coding
protein_coding
protein_coding
protein_coding
protein_coding
protein_coding
protein_coding
protein_coding
protein_coding
protein_coding
protein_coding
protein_coding
protein_coding
protein_coding
ncRNA
protein_coding
protein_coding
protein_coding
protein_coding
protein_coding
protein_coding
protein_coding
protein_coding
ncRNA
protein_coding
protein_coding
protein_coding
protein_coding
protein_coding
protein_coding
protein_coding
protein_coding
protein_coding
protein_coding
protein_coding
protein_coding
protein_coding
protein_coding
protein_coding
protein_coding
protein_coding
protein_coding
protein_coding
protein_coding

protein_coding
protein_coding
protein_coding
ncRNA
protein_coding
protein_coding
protein_coding
protein_coding
protein_coding
protein_coding
protein_coding
protein_coding
protein_coding
protein_coding
protein_coding
protein_coding
protein_coding
protein_coding
protein_coding
protein_coding
protein_coding
protein_coding
protein_coding
miRNA_precursor
protein_coding
protein_coding
protein_coding
protein_coding
protein_coding
protein_coding
protein_coding
protein_coding
protein_coding
protein_coding
protein_coding
protein_coding
protein_coding
protein_coding
protein_coding
protein_coding
protein_coding
protein_coding
protein_coding
protein_coding
protein_coding
miRNA_precursor
protein_coding
protein_coding
protein_coding
protein_coding
protein_coding
miRNA_precursor
protein_coding
protein_coding
protein_coding
protein_coding

protein_coding
protein_coding
protein_coding
protein_coding
protein_coding
protein_coding
protein_coding
protein_coding
protein_coding
protein_coding
protein_coding
protein_coding
protein_coding
protein_coding
protein_coding
protein_coding
protein_coding
protein_coding
protein_coding
protein_coding
protein_coding
protein_coding
protein_coding
protein_coding
protein_coding
protein_coding
protein_coding
protein_coding
protein_coding
protein_coding
protein_coding
protein_coding
protein_coding
protein_coding
protein_coding
ncRNA
protein_coding
protein_coding
protein_coding
protein_coding
protein_coding
protein_coding
protein_coding
protein_coding
protein_coding
protein_coding
protein_coding
protein_coding
ncRNA
protein_coding
protein_coding
protein_coding
protein_coding
protein_coding
protein_coding
small_nucle(ol)ar_rna

protein_coding
protein_coding
protein_coding
protein_coding
protein_coding
protein_coding
protein_coding
protein_coding
ncRNA
protein_coding
protein_coding
miRNA_precursor
protein_coding
protein_coding
protein_coding
protein_coding
protein_coding
protein_coding
protein_coding
protein_coding
protein_coding
ncRNA
protein_coding
protein_coding
protein_coding
protein_coding
protein_coding
pre_trna
ncRNA
protein_coding
protein_coding
protein_coding
protein_coding
protein_coding
protein_coding
protein_coding
protein_coding
ncRNA
protein_coding
ncRNA
protein_coding
protein_coding
protein_coding
protein_coding
protein_coding
protein_coding
protein_coding
protein_coding
protein_coding
protein_coding
protein_coding
protein_coding
protein_coding
ncRNA
protein_coding
protein_coding

ncRNA
protein_coding
protein_coding
protein_coding
protein_coding
ncRNA
protein_coding
protein_coding
protein_coding
protein_coding
protein_coding
protein_coding
protein_coding
protein_coding
protein_coding
protein_coding
protein_coding
protein_coding
protein_coding
protein_coding
protein_coding
protein_coding
protein_coding
protein_coding
protein_coding
protein_coding
protein_coding
protein_coding
protein_coding
protein_coding
protein_coding
protein_coding
ncRNA
protein_coding
protein_coding
protein_coding
protein_coding
protein_coding
protein_coding
protein_coding
protein_coding
protein_coding
protein_coding
protein_coding
protein_coding
protein_coding
protein_coding
protein_coding
protein_coding
protein_coding
protein_coding
ncRNA
protein_coding
protein_coding
ncRNA
protein_coding

protein_coding
protein_coding
ncRNA
protein_coding
protein_coding
protein_coding
ncRNA
protein_coding
protein_coding
protein_coding
protein_coding
protein_coding
protein_coding
protein_coding
ncRNA
protein_coding
protein_coding
protein_coding
protein_coding
protein_coding
ncRNA
protein_coding
miRNA_precursor
protein_coding
protein_coding
protein_coding
protein_coding
protein_coding
protein_coding
protein_coding
protein_coding
protein_coding
protein_coding
protein_coding
protein_coding
protein_coding
protein_coding
protein_coding
protein_coding
protein_coding
protein_coding
protein_coding
protein_coding
protein_coding
protein_coding
protein_coding
ncRNA
protein_coding
protein_coding
protein_coding
protein_coding
protein_coding
protein_coding
protein_coding
protein_coding
protein_coding

protein_coding
protein_coding
protein_coding
small_nucle(ol)ar_rna
protein_coding
protein_coding
protein_coding
protein_coding
protein_coding
protein_coding
protein_coding
protein_coding
protein_coding
protein_coding
protein_coding
protein_coding
protein_coding
protein_coding
protein_coding
protein_coding
protein_coding
protein_coding
protein_coding
protein_coding
protein_coding
protein_coding
protein_coding
protein_coding
protein_coding
protein_coding
protein_coding
protein_coding
protein_coding
protein_coding
protein_coding
protein_coding
protein_coding
protein_coding
protein_coding
protein_coding
protein_coding
protein_coding
protein_coding
protein_coding
protein_coding
protein_coding
pre_trna
protein_coding
protein_coding
protein_coding
protein_coding
protein_coding
ncRNA
protein_coding
protein_coding
protein_coding

protein_coding
protein_coding
protein_coding
protein_coding
protein_coding
protein_coding
protein_coding
protein_coding
protein_coding
protein_coding
protein_coding
protein_coding
protein_coding
protein_coding
protein_coding
protein_coding
protein_coding
protein_coding
protein_coding
protein_coding
protein_coding
protein_coding
protein_coding
protein_coding
protein_coding
protein_coding
protein_coding
protein_coding
protein_coding
protein_coding
protein_coding
protein_coding
protein_coding
protein_coding
protein_coding
protein_coding
protein_coding
protein_coding
protein_coding
protein_coding
protein_coding
protein_coding
protein_coding
protein_coding
protein_coding
protein_coding
protein_coding
protein_coding
protein_coding
protein_coding
protein_coding
protein_coding
protein_coding
protein_coding
protein_coding
protein_coding

protein_coding
protein_coding
protein_coding
protein_coding
protein_coding
miRNA_precursor
protein_coding
protein_coding
protein_coding
protein_coding
ncRNA
protein_coding
protein_coding
protein_coding
protein_coding
protein_coding
ncRNA
protein_coding
ncRNA
protein_coding
protein_coding
protein_coding
protein_coding
protein_coding
protein_coding
protein_coding
protein_coding
protein_coding
protein_coding
protein_coding
protein_coding
protein_coding
protein_coding
protein_coding
protein_coding
protein_coding
protein_coding
protein_coding
protein_coding
protein_coding
protein_coding
protein_coding
protein_coding
protein_coding
protein_coding
protein_coding
protein_coding
protein_coding
protein_coding
protein_coding
protein_coding
protein_coding
protein_coding
protein_coding
protein_coding
protein_coding

protein_coding
protein_coding
protein_coding
protein_coding
protein_coding
protein_coding
protein_coding
protein_coding
protein_coding
protein_coding
protein_coding
protein_coding
protein_coding
protein_coding
protein_coding
protein_coding
protein_coding
protein_coding
protein_coding
protein_coding
protein_coding
protein_coding
protein_coding
protein_coding
protein_coding
protein_coding
protein_coding
protein_coding
protein_coding
protein_coding
protein_coding
protein_coding
protein_coding
protein_coding
miRNA_precursor
protein_coding
protein_coding
protein_coding
protein_coding
ncRNA
protein_coding
protein_coding
ncRNA
protein_coding
protein_coding
protein_coding
protein_coding
protein_coding
protein_coding
protein_coding
protein_coding
protein_coding
protein_coding
protein_coding
protein_coding
protein_coding

protein_coding
protein_coding
protein_coding
protein_coding
protein_coding
protein_coding
protein_coding
protein_coding
protein_coding
protein_coding
protein_coding
protein_coding
protein_coding
protein_coding
protein_coding
protein_coding
protein_coding
protein_coding
protein_coding
protein_coding
protein_coding
protein_coding
pre_trna
protein_coding
protein_coding
protein_coding
protein_coding
protein_coding
protein_coding
protein_coding
protein_coding
protein_coding
protein_coding
protein_coding
ncRNA
ncRNA
protein_coding
protein_coding
protein_coding
protein_coding
protein_coding
protein_coding
protein_coding
protein_coding
protein_coding
miRNA_precursor
protein_coding
protein_coding
protein_coding
protein_coding
protein_coding
protein_coding
protein_coding
protein_coding
protein_coding
protein_coding

protein_coding
protein_coding
protein_coding
protein_coding
protein_coding
protein_coding
protein_coding
protein_coding
protein_coding
protein_coding
protein_coding
protein_coding
protein_coding
protein_coding
protein_coding
protein_coding
protein_coding
protein_coding
protein_coding
protein_coding
protein_coding
protein_coding
protein_coding
protein_coding
protein_coding
protein_coding
protein_coding
protein_coding
protein_coding
protein_coding
protein_coding
protein_coding
protein_coding
protein_coding
protein_coding
protein_coding
small_nucle(ol)ar_rna
protein_coding
protein_coding
protein_coding
protein_coding
protein_coding
protein_coding
protein_coding
protein_coding
protein_coding
protein_coding
protein_coding
protein_coding
ncRNA
protein_coding
protein_coding
ncRNA
protein_coding
protein_coding
protein_coding

protein_coding
protein_coding
protein_coding
protein_coding
protein_coding
protein_coding
protein_coding
protein_coding
protein_coding
protein_coding
protein_coding
protein_coding
protein_coding
protein_coding
protein_coding
protein_coding
protein_coding
protein_coding
protein_coding
protein_coding
protein_coding
protein_coding
protein_coding
protein_coding
protein_coding
protein_coding
protein_coding
small_nucle(ol)ar_rna
protein_coding
protein_coding
protein_coding
protein_coding
protein_coding
protein_coding
protein_coding
protein_coding
protein_coding
protein_coding
protein_coding
protein_coding
protein_coding
protein_coding
protein_coding
protein_coding
protein_coding
protein_coding
protein_coding
protein_coding
protein_coding
protein_coding
protein_coding
protein_coding
protein_coding
protein_coding
protein_coding
protein_coding

protein_coding
protein_coding
protein_coding
protein_coding
protein_coding
protein_coding
protein_coding
protein_coding
protein_coding
protein_coding
protein_coding
protein_coding
protein_coding
protein_coding
protein_coding
protein_coding
protein_coding
protein_coding
protein_coding
protein_coding
protein_coding
protein_coding
protein_coding
protein_coding
protein_coding
protein_coding
protein_coding
protein_coding
protein_coding
protein_coding
protein_coding
protein_coding
protein_coding
protein_coding
protein_coding
protein_coding
protein_coding
protein_coding
protein_coding
protein_coding
protein_coding
protein_coding
protein_coding
protein_coding
protein_coding
protein_coding
protein_coding
protein_coding
protein_coding
protein_coding
protein_coding
protein_coding
protein_coding
protein_coding
protein_coding
protein_coding

protein_coding
protein_coding
protein_coding
protein_coding
protein_coding
protein_coding
protein_coding
ncRNA
protein_coding
protein_coding
protein_coding
protein_coding
protein_coding
protein_coding
protein_coding
protein_coding
protein_coding
protein_coding
protein_coding
protein_coding
protein_coding
protein_coding
ncRNA
protein_coding
protein_coding
protein_coding
protein_coding
protein_coding
protein_coding
protein_coding
protein_coding
protein_coding
protein_coding
protein_coding
protein_coding
protein_coding
protein_coding
protein_coding
protein_coding
protein_coding
protein_coding
protein_coding
protein_coding
ncRNA
protein_coding
protein_coding
protein_coding
protein_coding
protein_coding
protein_coding
protein_coding
protein_coding

**Dataset 1.2 Differential analysis of 21/22nt accumulation on Araport11 genes in v**

| ID | baseMean | log2FoldChange | lfcSE | stat | pvalue | padj |
|---|---|---|---|---|---|---|
| AT5G38850 | 479,48 | -3,36 | 0,23 | -14,44 | 2,71E-47 | 6,25E-44 |
| AT5G14565 | 6609,25 | -3,83 | 0,27 | -14,37 | 8,31E-47 | 9,59E-44 |
| AT2G26211 | 125,25 | 6,10 | 0,45 | 13,65 | 2,00E-42 | 1,54E-39 |
| AT5G13630 | 269,68 | 5,57 | 0,44 | 12,61 | 1,95E-36 | 1,12E-33 |
| AT5G14545 | 7868,55 | 14,98 | 1,20 | 12,53 | 5,14E-36 | 2,37E-33 |
| AT5G08185 | 1387,13 | 3,86 | 0,31 | 12,26 | 1,51E-34 | 5,83E-32 |
| AT5G14550 | 7872,48 | 14,02 | 1,20 | 11,72 | 1,06E-31 | 3,50E-29 |
| AT1G63130 | 240,21 | -3,36 | 0,31 | -10,74 | 6,73E-27 | 1,94E-24 |
| AT1G63150 | 176,58 | -3,63 | 0,34 | -10,58 | 3,84E-26 | 9,85E-24 |
| AT3G17185 | 156,12 | -3,18 | 0,31 | -10,37 | 3,39E-25 | 7,12E-23 |
| AT3G23325 | 69,65 | 6,08 | 0,59 | 10,38 | 3,17E-25 | 7,12E-23 |
| AT3G23326 | 69,32 | 6,07 | 0,59 | 10,31 | 6,18E-25 | 1,19E-22 |
| AT4G00885 | 370,66 | -3,11 | 0,30 | -10,29 | 8,19E-25 | 1,45E-22 |
| AT1G31280 | 162,06 | 5,66 | 0,57 | 9,94 | 2,67E-23 | 4,40E-21 |
| AT3G59884 | 100,21 | 4,23 | 0,43 | 9,81 | 1,06E-22 | 1,64E-20 |
| AT1G18075 | 4243,99 | -1,67 | 0,18 | -9,44 | 3,55E-21 | 5,13E-19 |
| AT1G63080 | 109,45 | -3,19 | 0,36 | -8,94 | 3,81E-19 | 5,17E-17 |
| AT5G10946 | 91,61 | -2,74 | 0,31 | -8,87 | 7,25E-19 | 9,29E-17 |
| AT4G06810 | 54,17 | 5,90 | 0,67 | 8,85 | 8,78E-19 | 1,07E-16 |
| AT4G06805 | 54,05 | 5,89 | 0,67 | 8,83 | 1,05E-18 | 1,22E-16 |
| AT5G10945 | 91,45 | -2,73 | 0,31 | -8,81 | 1,22E-18 | 1,34E-16 |
| AT2G47275 | 830,57 | 2,73 | 0,31 | 8,79 | 1,50E-18 | 1,57E-16 |
| AT1G62914 | 457,48 | -3,02 | 0,35 | -8,75 | 2,19E-18 | 2,20E-16 |
| AT2G10606 | 1219,01 | 2,28 | 0,28 | 8,23 | 1,83E-16 | 1,76E-14 |
| AT1G29265 | 54,27 | 4,93 | 0,60 | 8,20 | 2,34E-16 | 2,16E-14 |
| AT1G56070 | 55,72 | 2,81 | 0,35 | 8,16 | 3,44E-16 | 3,06E-14 |
| AT1G12820 | 53,20 | -3,49 | 0,44 | -7,96 | 1,68E-15 | 1,44E-13 |
| AT4G06130 | 392,23 | -3,27 | 0,42 | -7,78 | 7,17E-15 | 5,91E-13 |
| AT5G66045 | 69,41 | 9,62 | 1,27 | 7,59 | 3,19E-14 | 2,54E-12 |
| AT5G03553 | 58,75 | -2,97 | 0,39 | -7,58 | 3,47E-14 | 2,66E-12 |
| AT4G13575 | 426,34 | -3,08 | 0,41 | -7,58 | 3,58E-14 | 2,66E-12 |
| AT3G01845 | 219,09 | -1,79 | 0,24 | -7,57 | 3,85E-14 | 2,78E-12 |
| AT5G03552 | 58,24 | -2,95 | 0,39 | -7,56 | 3,97E-14 | 2,78E-12 |
| AT1G04977 | 45,78 | -3,59 | 0,48 | -7,48 | 7,28E-14 | 4,94E-12 |
| AT5G43740 | 103,64 | -3,45 | 0,46 | -7,45 | 9,08E-14 | 5,99E-12 |
| AT1G62910 | 98,21 | -3,20 | 0,44 | -7,29 | 3,04E-13 | 1,95E-11 |
| AT2G46255 | 72,33 | -3,50 | 0,49 | -7,15 | 8,64E-13 | 5,37E-11 |
| AT5G02500 | 62,24 | 2,55 | 0,36 | 7,15 | 8,83E-13 | 5,37E-11 |
| AT1G62930 | 53,37 | -3,11 | 0,44 | -7,10 | 1,21E-12 | 7,00E-11 |
| AT2G18960 | 62,63 | 4,08 | 0,57 | 7,11 | 1,18E-12 | 7,00E-11 |
| AT3G48201 | 30,66 | 4,46 | 0,64 | 6,98 | 2,93E-12 | 1,65E-10 |
| AT2G15530 | 40,01 | 2,75 | 0,40 | 6,86 | 7,13E-12 | 3,92E-10 |
| AT1G37130 | 39,12 | 3,70 | 0,54 | 6,84 | 7,93E-12 | 4,26E-10 |
| AT1G15750 | 35,69 | 6,11 | 0,91 | 6,70 | 2,13E-11 | 1,12E-09 |
| AT2G01020 | 7030,11 | -2,34 | 0,35 | -6,62 | 3,53E-11 | 1,81E-09 |
| AT5G05170 | 36,40 | 3,70 | 0,56 | 6,62 | 3,63E-11 | 1,82E-09 |
| AT4G07070 | 70,24 | -3,51 | 0,54 | -6,54 | 6,34E-11 | 3,11E-09 |
| AT4G04565 | 64,98 | 2,57 | 0,39 | 6,52 | 7,05E-11 | 3,39E-09 |
| AT5G63715 | 27,38 | -8,08 | 1,24 | -6,51 | 7,53E-11 | 3,55E-09 |
| AT1G63400 | 45,16 | -8,05 | 1,24 | -6,48 | 9,33E-11 | 4,31E-09 |
| AT1G48267 | 15177,13 | -2,13 | 0,34 | -6,34 | 2,28E-10 | 1,03E-08 |
| AT3G11130 | 25,04 | 4,34 | 0,69 | 6,30 | 3,03E-10 | 1,34E-08 |
| AT2G23350 | 29,35 | 6,81 | 1,09 | 6,26 | 3,81E-10 | 1,66E-08 |

| | | | | | | |
|---|---|---|---|---|---|---|
| AT4G09135 | 26,82 | 7,72 | 1,25 | 6,16 | 7,32E-10 | 3,13E-08 |
| AT5G43603 | 44,77 | 7,60 | 1,24 | 6,12 | 9,39E-10 | 3,94E-08 |
| AT1G50055 | 182,36 | -1,87 | 0,31 | -6,06 | 1,37E-09 | 5,63E-08 |
| AT5G46315 | 17,01 | 7,58 | 1,27 | 5,96 | 2,51E-09 | 1,02E-07 |
| AT3G01540 | 20,21 | 5,11 | 0,86 | 5,95 | 2,71E-09 | 1,08E-07 |
| AT2G22125 | 19,76 | 4,44 | 0,75 | 5,91 | 3,49E-09 | 1,37E-07 |
| AT4G34450 | 21,43 | 4,96 | 0,84 | 5,89 | 3,77E-09 | 1,44E-07 |
| AT3G23690 | 28,16 | -5,43 | 0,92 | -5,89 | 3,85E-09 | 1,44E-07 |
| AT4G08755 | 25,91 | 7,33 | 1,24 | 5,89 | 3,88E-09 | 1,44E-07 |
| AT1G01320 | 16,50 | 4,01 | 0,69 | 5,83 | 5,56E-09 | 2,04E-07 |
| AT3G05700 | 20,03 | 4,07 | 0,71 | 5,75 | 9,03E-09 | 3,26E-07 |
| AT1G70320 | 18,40 | 4,65 | 0,82 | 5,66 | 1,50E-08 | 5,32E-07 |
| AT2G21390 | 23,77 | 7,16 | 1,27 | 5,66 | 1,54E-08 | 5,39E-07 |
| AT3G01715 | 19,56 | 4,05 | 0,72 | 5,60 | 2,11E-08 | 7,27E-07 |
| AT1G09797 | 19,74 | 5,05 | 0,92 | 5,49 | 3,92E-08 | 1,31E-06 |
| AT1G09793 | 19,74 | 5,05 | 0,92 | 5,49 | 3,92E-08 | 1,31E-06 |
| AT1G62590 | 20,75 | -6,82 | 1,25 | -5,47 | 4,41E-08 | 1,45E-06 |
| AT1G09620 | 17,50 | 5,16 | 0,95 | 5,44 | 5,20E-08 | 1,69E-06 |
| AT5G46845 | 66,07 | -1,67 | 0,31 | -5,40 | 6,77E-08 | 2,17E-06 |
| AT1G73687 | 3456,65 | -1,41 | 0,26 | -5,38 | 7,46E-08 | 2,36E-06 |
| AT4G04825 | 145,71 | -3,09 | 0,58 | -5,31 | 1,07E-07 | 3,29E-06 |
| AT5G07745 | 20,93 | 3,19 | 0,60 | 5,31 | 1,07E-07 | 3,29E-06 |
| AT3G09840 | 22,25 | 3,26 | 0,62 | 5,22 | 1,77E-07 | 5,38E-06 |
| AT3G51375 | 135,65 | 1,93 | 0,37 | 5,17 | 2,38E-07 | 7,08E-06 |
| AT5G07350 | 9,52 | 6,84 | 1,32 | 5,17 | 2,39E-07 | 7,08E-06 |
| AT4G02480 | 22,30 | 4,68 | 0,91 | 5,12 | 3,10E-07 | 9,06E-06 |
| AT1G06227 | 343,33 | -1,93 | 0,38 | -5,10 | 3,41E-07 | 9,83E-06 |
| AT5G17920 | 34,31 | 2,21 | 0,44 | 5,07 | 4,06E-07 | 1,14E-05 |
| AT2G43900 | 9,76 | 6,71 | 1,32 | 5,07 | 4,05E-07 | 1,14E-05 |
| AT5G64270 | 15,19 | 5,46 | 1,08 | 5,05 | 4,32E-07 | 1,20E-05 |
| AT1G06237 | 336,80 | -1,95 | 0,39 | -5,02 | 5,30E-07 | 1,46E-05 |
| AT3G02260 | 23,16 | 3,10 | 0,62 | 5,00 | 5,71E-07 | 1,55E-05 |
| AT1G59870 | 72,89 | 1,83 | 0,37 | 4,94 | 7,80E-07 | 2,09E-05 |
| AT2G28290 | 15,57 | 5,29 | 1,08 | 4,91 | 9,34E-07 | 2,48E-05 |
| AT3G18780 | 17,06 | 3,22 | 0,66 | 4,88 | 1,06E-06 | 2,77E-05 |
| AT4G31570 | 11,16 | 6,65 | 1,36 | 4,88 | 1,08E-06 | 2,79E-05 |
| AT4G07875 | 11,03 | 6,44 | 1,32 | 4,86 | 1,15E-06 | 2,94E-05 |
| AT5G42950 | 18,25 | 5,71 | 1,19 | 4,81 | 1,51E-06 | 3,84E-05 |
| AT5G53460 | 17,48 | 2,85 | 0,59 | 4,80 | 1,58E-06 | 3,93E-05 |
| AT3G23680 | 9,12 | 6,46 | 1,35 | 4,80 | 1,58E-06 | 3,93E-05 |
| AT5G04275 | 20,42 | 6,01 | 1,26 | 4,76 | 1,93E-06 | 4,75E-05 |
| AT4G28080 | 23,67 | 3,31 | 0,70 | 4,76 | 1,97E-06 | 4,79E-05 |
| AT2G01008 | 28,92 | -2,39 | 0,50 | -4,75 | 2,02E-06 | 4,81E-05 |
| AT1G29920 | 50,33 | 2,01 | 0,42 | 4,75 | 2,00E-06 | 4,81E-05 |
| AT1G70770 | 12,70 | 5,14 | 1,08 | 4,74 | 2,10E-06 | 4,95E-05 |
| AT2G41900 | 16,69 | 4,79 | 1,01 | 4,73 | 2,20E-06 | 5,14E-05 |
| AT1G05163 | 491,36 | -1,29 | 0,28 | -4,70 | 2,59E-06 | 5,91E-05 |
| AT1G15405 | 491,36 | -1,29 | 0,28 | -4,70 | 2,59E-06 | 5,91E-05 |
| AT1G26973 | 11,41 | 3,67 | 0,78 | 4,68 | 2,81E-06 | 6,36E-05 |
| AT4G36080 | 9,35 | 5,60 | 1,20 | 4,65 | 3,31E-06 | 7,42E-05 |
| AT5G53740 | 20,74 | -2,77 | 0,60 | -4,63 | 3,71E-06 | 8,22E-05 |
| AT3G19820 | 10,30 | 5,14 | 1,12 | 4,57 | 4,84E-06 | 1,06E-04 |
| AT5G15833 | 13,93 | 3,35 | 0,74 | 4,54 | 5,59E-06 | 1,22E-04 |
| AT2G39675 | 2246,76 | -1,54 | 0,34 | -4,53 | 5,89E-06 | 1,27E-04 |
| AT4G16890 | 15,81 | 4,12 | 0,91 | 4,53 | 5,96E-06 | 1,27E-04 |
| AT5G60790 | 11,59 | 4,45 | 0,99 | 4,50 | 6,93E-06 | 1,47E-04 |

| | | | | | | |
|---|---|---|---|---|---|---|
| AT4G23713 | 128,07 | -3,31 | 0,74 | -4,49 | 7,10E-06 | 1,49E-04 |
| AT3G52140 | 9,89 | 4,13 | 0,92 | 4,49 | 7,18E-06 | 1,49E-04 |
| AT3G50370 | 9,94 | 4,38 | 0,98 | 4,47 | 7,74E-06 | 1,59E-04 |
| AT3G48870 | 10,82 | 4,64 | 1,04 | 4,46 | 8,03E-06 | 1,64E-04 |
| AT5G44340 | 14,19 | 3,12 | 0,70 | 4,46 | 8,13E-06 | 1,65E-04 |
| AT3G62980 | 20,74 | 2,58 | 0,58 | 4,46 | 8,30E-06 | 1,67E-04 |
| AT1G22610 | 11,17 | 4,90 | 1,10 | 4,45 | 8,40E-06 | 1,67E-04 |
| AT1G04907 | 10,89 | 6,01 | 1,35 | 4,45 | 8,61E-06 | 1,70E-04 |
| AT3G13080 | 13,73 | 5,81 | 1,31 | 4,43 | 9,32E-06 | 1,82E-04 |
| AT1G29900 | 11,05 | 4,76 | 1,08 | 4,41 | 1,02E-05 | 1,98E-04 |
| AT5G08717 | 14,65 | 6,00 | 1,36 | 4,41 | 1,03E-05 | 1,98E-04 |
| AT5G23110 | 11,58 | 5,40 | 1,23 | 4,38 | 1,17E-05 | 2,21E-04 |
| AT5G39500 | 6,69 | 5,99 | 1,37 | 4,38 | 1,17E-05 | 2,21E-04 |
| AT1G22620 | 12,70 | 3,90 | 0,89 | 4,38 | 1,18E-05 | 2,21E-04 |
| AT2G47020 | 1248,75 | 1,37 | 0,31 | 4,38 | 1,19E-05 | 2,22E-04 |
| AT2G47015 | 1248,43 | 1,37 | 0,31 | 4,37 | 1,22E-05 | 2,22E-04 |
| AT1G08597 | 22,78 | -4,05 | 0,93 | -4,38 | 1,21E-05 | 2,22E-04 |
| AT3G10745 | 164,34 | 1,45 | 0,33 | 4,38 | 1,21E-05 | 2,22E-04 |
| AT3G43300 | 11,08 | 3,93 | 0,90 | 4,37 | 1,24E-05 | 2,24E-04 |
| AT5G10380 | 7,30 | 5,89 | 1,35 | 4,35 | 1,35E-05 | 2,40E-04 |
| AT3G08943 | 11,39 | 4,97 | 1,14 | 4,35 | 1,34E-05 | 2,40E-04 |
| AT1G64720 | 13,91 | 3,30 | 0,76 | 4,33 | 1,47E-05 | 2,60E-04 |
| AT2G28000 | 11,21 | 3,23 | 0,75 | 4,31 | 1,63E-05 | 2,83E-04 |
| AT5G60408 | 515,37 | -1,29 | 0,30 | -4,31 | 1,62E-05 | 2,83E-04 |
| AT4G13940 | 35,08 | 1,85 | 0,43 | 4,30 | 1,67E-05 | 2,88E-04 |
| AT4G27600 | 12,18 | 4,77 | 1,11 | 4,29 | 1,81E-05 | 3,10E-04 |
| AT2G37190 | 11,90 | 5,53 | 1,30 | 4,27 | 1,97E-05 | 3,34E-04 |
| AT2G41310 | 487,34 | -1,18 | 0,28 | -4,27 | 2,00E-05 | 3,36E-04 |
| AT2G09250 | 487,12 | -1,18 | 0,28 | -4,26 | 2,02E-05 | 3,37E-04 |
| AT1G12775 | 21,87 | -2,89 | 0,68 | -4,25 | 2,10E-05 | 3,49E-04 |
| AT5G57870 | 8,84 | 4,74 | 1,11 | 4,25 | 2,12E-05 | 3,50E-04 |
| AT3G55734 | 132,05 | 1,34 | 0,32 | 4,23 | 2,33E-05 | 3,78E-04 |
| AT2G43970 | 15,13 | 2,84 | 0,67 | 4,23 | 2,31E-05 | 3,78E-04 |
| AT5G65950 | 6,84 | 6,32 | 1,50 | 4,21 | 2,55E-05 | 4,12E-04 |
| AT1G79930 | 9,47 | 5,52 | 1,31 | 4,20 | 2,63E-05 | 4,22E-04 |
| AT5G50290 | 244,15 | -1,57 | 0,38 | -4,19 | 2,85E-05 | 4,53E-04 |
| AT4G27640 | 8,75 | 5,78 | 1,39 | 4,17 | 3,00E-05 | 4,74E-04 |
| AT3G18290 | 9,13 | 4,79 | 1,15 | 4,17 | 3,09E-05 | 4,85E-04 |
| AT3G13857 | 124,31 | 1,56 | 0,38 | 4,12 | 3,72E-05 | 5,72E-04 |
| AT3G13855 | 124,31 | 1,56 | 0,38 | 4,12 | 3,72E-05 | 5,72E-04 |
| AT1G74260 | 9,63 | 4,59 | 1,11 | 4,13 | 3,68E-05 | 5,72E-04 |
| AT2G43680 | 14,92 | 2,96 | 0,72 | 4,12 | 3,79E-05 | 5,79E-04 |
| AT4G27000 | 10,07 | 4,07 | 0,99 | 4,11 | 3,92E-05 | 5,96E-04 |
| AT2G20190 | 8,93 | 4,11 | 1,01 | 4,05 | 5,06E-05 | 7,63E-04 |
| AT1G10170 | 9,09 | 4,14 | 1,03 | 4,03 | 5,47E-05 | 8,20E-04 |
| AT1G02500 | 10,85 | 3,69 | 0,92 | 4,03 | 5,52E-05 | 8,21E-04 |
| AT3G47350 | 23,89 | 2,49 | 0,62 | 4,03 | 5,61E-05 | 8,31E-04 |
| AT5G04140 | 18,61 | 2,52 | 0,63 | 4,02 | 5,77E-05 | 8,48E-04 |
| AT1G15500 | 7,05 | 5,29 | 1,32 | 4,01 | 6,18E-05 | 9,03E-04 |
| AT5G15680 | 7,75 | 5,23 | 1,31 | 4,00 | 6,31E-05 | 9,16E-04 |
| AT2G43160 | 6,80 | 5,26 | 1,32 | 3,99 | 6,48E-05 | 9,34E-04 |
| AT4G17140 | 11,35 | 3,22 | 0,81 | 3,98 | 6,89E-05 | 9,88E-04 |
| AT1G10290 | 9,22 | 4,66 | 1,17 | 3,97 | 7,24E-05 | 1,03E-03 |
| AT1G48300 | 8,14 | 4,91 | 1,24 | 3,96 | 7,61E-05 | 1,08E-03 |
| AT3G41768 | 91,21 | -1,22 | 0,31 | -3,95 | 7,67E-05 | 1,08E-03 |
| AT2G28056 | 34,24 | -1,69 | 0,43 | -3,95 | 7,71E-05 | 1,08E-03 |

| | | | | | | |
|---|---|---|---|---|---|---|
| AT2G01021 | 3194,91 | -1,06 | 0,27 | -3,95 | 7,93E-05 | 1,10E-03 |
| AT1G08537 | 8,63 | -5,28 | 1,34 | -3,94 | 8,14E-05 | 1,13E-03 |
| AT2G42270 | 6,29 | 5,50 | 1,40 | 3,93 | 8,54E-05 | 1,17E-03 |
| AT2G36910 | 8,87 | 4,59 | 1,17 | 3,92 | 8,80E-05 | 1,20E-03 |
| AT5G07940 | 5,66 | 5,50 | 1,40 | 3,92 | 8,90E-05 | 1,21E-03 |
| AT2G45540 | 12,39 | 4,52 | 1,16 | 3,91 | 9,28E-05 | 1,25E-03 |
| AT2G39885 | 9,28 | 4,69 | 1,20 | 3,91 | 9,34E-05 | 1,25E-03 |
| AT5G04455 | 10,87 | 5,18 | 1,33 | 3,90 | 9,64E-05 | 1,28E-03 |
| AT5G04445 | 10,87 | 5,18 | 1,33 | 3,90 | 9,64E-05 | 1,28E-03 |
| AT4G32020 | 12,23 | 2,72 | 0,70 | 3,89 | 9,82E-05 | 1,30E-03 |
| AT4G07975 | 9,02 | 3,12 | 0,80 | 3,89 | 9,91E-05 | 1,30E-03 |
| AT1G47720 | 6,21 | 5,35 | 1,38 | 3,89 | 1,01E-04 | 1,32E-03 |
| AT1G62020 | 12,54 | 3,39 | 0,88 | 3,87 | 1,11E-04 | 1,42E-03 |
| AT3G23685 | 5,28 | 5,35 | 1,38 | 3,87 | 1,11E-04 | 1,42E-03 |
| AT3G20760 | 5,52 | 5,09 | 1,32 | 3,87 | 1,11E-04 | 1,42E-03 |
| AT1G75660 | 6,24 | 5,15 | 1,34 | 3,85 | 1,17E-04 | 1,49E-03 |
| AT1G13980 | 8,14 | 5,10 | 1,33 | 3,84 | 1,22E-04 | 1,55E-03 |
| AT3G26810 | 23,04 | -1,99 | 0,52 | -3,84 | 1,23E-04 | 1,55E-03 |
| AT1G62670 | 9,93 | -4,97 | 1,29 | -3,84 | 1,24E-04 | 1,56E-03 |
| AT4G13560 | 52,08 | 1,30 | 0,34 | 3,83 | 1,28E-04 | 1,59E-03 |
| AT3G08947 | 5,24 | 5,53 | 1,44 | 3,83 | 1,29E-04 | 1,60E-03 |
| AT4G16940 | 4,86 | 5,72 | 1,50 | 3,81 | 1,37E-04 | 1,69E-03 |
| AT4G14910 | 7,50 | 5,10 | 1,34 | 3,79 | 1,48E-04 | 1,82E-03 |
| AT2G25660 | 3,83 | 5,59 | 1,47 | 3,79 | 1,50E-04 | 1,84E-03 |
| AT1G36160 | 9,82 | 3,43 | 0,91 | 3,79 | 1,52E-04 | 1,84E-03 |
| AT5G62165 | 7,10 | -5,18 | 1,37 | -3,78 | 1,59E-04 | 1,92E-03 |
| AT4G31480 | 6,09 | 5,36 | 1,42 | 3,77 | 1,66E-04 | 2,00E-03 |
| AT1G77300 | 4,17 | 5,37 | 1,43 | 3,75 | 1,77E-04 | 2,12E-03 |
| AT3G14385 | 15,96 | 2,05 | 0,55 | 3,73 | 1,92E-04 | 2,29E-03 |
| AT2G40030 | 5,36 | 5,25 | 1,41 | 3,73 | 1,94E-04 | 2,29E-03 |
| AT5G09505 | 92,76 | -1,17 | 0,31 | -3,73 | 1,94E-04 | 2,29E-03 |
| AT4G39420 | 9,97 | 3,33 | 0,90 | 3,72 | 2,01E-04 | 2,36E-03 |
| AT3G43990 | 14,30 | -4,18 | 1,13 | -3,72 | 2,03E-04 | 2,37E-03 |
| AT1G50480 | 9,27 | 3,74 | 1,02 | 3,68 | 2,35E-04 | 2,72E-03 |
| AT4G01800 | 8,35 | 3,36 | 0,91 | 3,68 | 2,36E-04 | 2,72E-03 |
| AT1G55591 | 5,61 | 5,12 | 1,40 | 3,67 | 2,42E-04 | 2,78E-03 |
| AT3G22886 | 587,34 | -0,93 | 0,25 | -3,66 | 2,52E-04 | 2,86E-03 |
| AT2G30520 | 8,24 | 3,64 | 1,00 | 3,66 | 2,51E-04 | 2,86E-03 |
| AT4G20850 | 8,05 | 3,79 | 1,04 | 3,66 | 2,53E-04 | 2,86E-03 |
| AT1G48090 | 11,18 | 2,81 | 0,77 | 3,66 | 2,57E-04 | 2,89E-03 |
| AT1G29930 | 124,68 | 1,40 | 0,38 | 3,65 | 2,60E-04 | 2,92E-03 |
| AT4G24520 | 3,90 | 5,84 | 1,60 | 3,65 | 2,62E-04 | 2,92E-03 |
| AT4G19395 | 418,73 | 0,77 | 0,21 | 3,63 | 2,81E-04 | 3,12E-03 |
| AT1G19835 | 5,86 | 4,99 | 1,38 | 3,62 | 2,97E-04 | 3,28E-03 |
| AT1G79920 | 8,70 | 3,00 | 0,83 | 3,61 | 3,04E-04 | 3,34E-03 |
| AT4G38600 | 8,47 | 4,39 | 1,22 | 3,60 | 3,12E-04 | 3,42E-03 |
| AT1G20760 | 7,86 | 4,87 | 1,36 | 3,59 | 3,36E-04 | 3,66E-03 |
| AT4G13564 | 27,21 | 1,41 | 0,39 | 3,58 | 3,43E-04 | 3,70E-03 |
| AT4G23650 | 3,54 | 5,24 | 1,46 | 3,58 | 3,42E-04 | 3,70E-03 |
| AT2G22496 | 20,32 | 1,89 | 0,53 | 3,57 | 3,53E-04 | 3,79E-03 |
| AT4G13570 | 36,83 | 1,35 | 0,38 | 3,57 | 3,58E-04 | 3,83E-03 |
| AT5G13100 | 3,43 | 5,32 | 1,49 | 3,56 | 3,65E-04 | 3,86E-03 |
| AT2G48060 | 4,01 | 5,21 | 1,46 | 3,56 | 3,65E-04 | 3,86E-03 |
| AT5G54070 | 15,72 | 4,02 | 1,13 | 3,56 | 3,69E-04 | 3,89E-03 |
| AT1G80410 | 5,85 | 4,96 | 1,39 | 3,56 | 3,76E-04 | 3,94E-03 |
| AT3G06435 | 378,12 | -0,91 | 0,26 | -3,55 | 3,92E-04 | 4,09E-03 |

| | | | | | | |
|---|---|---|---|---|---|---|
| AT3G23000 | 7,09 | 4,23 | 1,19 | 3,54 | 3,99E-04 | 4,15E-03 |
| AT1G21630 | 4,05 | 5,25 | 1,49 | 3,54 | 4,05E-04 | 4,19E-03 |
| AT1G49760 | 5,88 | 3,76 | 1,06 | 3,53 | 4,11E-04 | 4,23E-03 |
| AT2G02741 | 19,79 | -2,12 | 0,60 | -3,53 | 4,18E-04 | 4,29E-03 |
| AT1G36730 | 3,83 | 5,05 | 1,44 | 3,52 | 4,37E-04 | 4,46E-03 |
| AT5G04560 | 3,61 | 5,11 | 1,45 | 3,51 | 4,41E-04 | 4,48E-03 |
| AT5G54075 | 15,55 | 3,95 | 1,13 | 3,51 | 4,45E-04 | 4,50E-03 |
| AT3G23780 | 4,59 | 4,59 | 1,31 | 3,50 | 4,68E-04 | 4,71E-03 |
| AT5G35407 | 1076,14 | 1,29 | 0,37 | 3,50 | 4,72E-04 | 4,73E-03 |
| AT1G80070 | 7,13 | 4,10 | 1,18 | 3,48 | 4,97E-04 | 4,97E-03 |
| AT4G24690 | 16,56 | 2,24 | 0,64 | 3,47 | 5,13E-04 | 5,10E-03 |
| AT4G05420 | 7,60 | 3,60 | 1,04 | 3,47 | 5,26E-04 | 5,21E-03 |
| AT2G26890 | 8,74 | 2,98 | 0,86 | 3,46 | 5,33E-04 | 5,25E-03 |
| AT5G41740 | 6,05 | 5,11 | 1,48 | 3,46 | 5,40E-04 | 5,31E-03 |
| AT2G02750 | 20,13 | -2,03 | 0,59 | -3,46 | 5,45E-04 | 5,33E-03 |
| AT3G07870 | 5,57 | 4,98 | 1,45 | 3,44 | 5,74E-04 | 5,59E-03 |
| AT1G07810 | 6,32 | 4,00 | 1,18 | 3,40 | 6,83E-04 | 6,60E-03 |
| AT1G63980 | 3,67 | 5,09 | 1,50 | 3,40 | 6,84E-04 | 6,60E-03 |
| AT1G23310 | 12,45 | 1,94 | 0,57 | 3,38 | 7,33E-04 | 7,05E-03 |
| AT2G35750 | 26,17 | -1,85 | 0,55 | -3,36 | 7,89E-04 | 7,56E-03 |
| AT3G01460 | 5,55 | 4,72 | 1,41 | 3,35 | 8,03E-04 | 7,58E-03 |
| AT5G43725 | 9,10 | -4,31 | 1,28 | -3,35 | 8,04E-04 | 7,58E-03 |
| AT5G43730 | 9,10 | -4,31 | 1,28 | -3,35 | 8,04E-04 | 7,58E-03 |
| AT5G57350 | 4,58 | 4,97 | 1,48 | 3,35 | 8,03E-04 | 7,58E-03 |
| AT1G15290 | 5,85 | 4,03 | 1,21 | 3,34 | 8,29E-04 | 7,74E-03 |
| AT1G59760 | 3,05 | 4,96 | 1,48 | 3,34 | 8,28E-04 | 7,74E-03 |
| AT5G04105 | 18,20 | 1,80 | 0,54 | 3,34 | 8,42E-04 | 7,84E-03 |
| AT4G01860 | 3,28 | 5,33 | 1,60 | 3,34 | 8,47E-04 | 7,85E-03 |
| AT5G51430 | 4,74 | 5,03 | 1,51 | 3,34 | 8,51E-04 | 7,86E-03 |
| AT1G64140 | 5,12 | 4,09 | 1,23 | 3,33 | 8,70E-04 | 7,97E-03 |
| AT5G45307 | 438,83 | 1,04 | 0,31 | 3,33 | 8,67E-04 | 7,97E-03 |
| AT5G06600 | 5,36 | 4,57 | 1,37 | 3,33 | 8,75E-04 | 7,98E-03 |
| AT5G26920 | 3,70 | 4,73 | 1,42 | 3,33 | 8,83E-04 | 8,03E-03 |
| AT1G35501 | 18,58 | -2,00 | 0,60 | -3,31 | 9,25E-04 | 8,28E-03 |
| AT1G54090 | 3,59 | 5,32 | 1,61 | 3,31 | 9,25E-04 | 8,28E-03 |
| AT2G16500 | 6,48 | 4,48 | 1,35 | 3,31 | 9,24E-04 | 8,28E-03 |
| AT5G25270 | 3,67 | 5,17 | 1,56 | 3,31 | 9,21E-04 | 8,28E-03 |
| AT5G61010 | 3,05 | 4,84 | 1,46 | 3,30 | 9,52E-04 | 8,48E-03 |
| AT3G62290 | 5,55 | 4,59 | 1,39 | 3,30 | 9,63E-04 | 8,52E-03 |
| AT5G06845 | 6,24 | 3,55 | 1,07 | 3,30 | 9,66E-04 | 8,52E-03 |
| AT4G37910 | 5,28 | 3,80 | 1,15 | 3,30 | 9,67E-04 | 8,52E-03 |
| AT5G12470 | 4,58 | 4,79 | 1,45 | 3,30 | 9,74E-04 | 8,54E-03 |
| AT5G12370 | 4,35 | 4,55 | 1,39 | 3,28 | 1,05E-03 | 9,16E-03 |
| AT3G03870 | 3,66 | 4,44 | 1,36 | 3,27 | 1,06E-03 | 9,26E-03 |
| AT2G45160 | 10,70 | -2,13 | 0,65 | -3,27 | 1,07E-03 | 9,26E-03 |
| AT5G20720 | 3,41 | 5,05 | 1,55 | 3,27 | 1,09E-03 | 9,39E-03 |
| AT1G10760 | 16,93 | 1,85 | 0,57 | 3,25 | 1,15E-03 | 9,93E-03 |
| AT5G02490 | 4,18 | 4,72 | 1,46 | 3,24 | 1,18E-03 | 1,01E-02 |
| AT4G10710 | 3,86 | 5,07 | 1,56 | 3,24 | 1,19E-03 | 1,01E-02 |
| AT3G46780 | 9,83 | 2,51 | 0,77 | 3,24 | 1,19E-03 | 1,02E-02 |
| AT2G25800 | 4,68 | 4,54 | 1,40 | 3,23 | 1,22E-03 | 1,03E-02 |
| AT5G52310 | 5,67 | -3,69 | 1,15 | -3,22 | 1,27E-03 | 1,08E-02 |
| AT1G48410 | 43,79 | 1,33 | 0,42 | 3,21 | 1,31E-03 | 1,11E-02 |
| AT3G24480 | 4,54 | 3,94 | 1,23 | 3,20 | 1,38E-03 | 1,16E-02 |
| AT5G12150 | 4,21 | 5,22 | 1,63 | 3,20 | 1,38E-03 | 1,16E-02 |
| AT3G03710 | 4,42 | 4,41 | 1,38 | 3,20 | 1,40E-03 | 1,16E-02 |

| | | | | | | |
|---|---|---|---|---|---|---|
| AT5G54770 | 28,55 | -1,62 | 0,51 | -3,19 | 1,41E-03 | 1,17E-02 |
| AT3G57260 | 8,35 | 2,41 | 0,75 | 3,19 | 1,43E-03 | 1,18E-02 |
| AT1G10950 | 6,24 | 3,45 | 1,08 | 3,18 | 1,46E-03 | 1,20E-02 |
| AT4G26850 | 7,34 | 3,26 | 1,03 | 3,18 | 1,46E-03 | 1,20E-02 |
| AT1G12294 | 124,95 | 0,86 | 0,27 | 3,18 | 1,49E-03 | 1,22E-02 |
| AT2G44540 | 4,48 | -4,49 | 1,42 | -3,17 | 1,51E-03 | 1,23E-02 |
| AT3G55410 | 3,63 | 4,59 | 1,45 | 3,17 | 1,55E-03 | 1,25E-02 |
| AT5G38540 | 8,36 | -2,30 | 0,73 | -3,17 | 1,55E-03 | 1,25E-02 |
| AT3G25010 | 4,35 | 4,34 | 1,37 | 3,16 | 1,58E-03 | 1,27E-02 |
| AT3G55220 | 4,51 | 3,91 | 1,24 | 3,15 | 1,64E-03 | 1,32E-02 |
| AT1G05423 | 13,10 | -1,98 | 0,63 | -3,15 | 1,66E-03 | 1,33E-02 |
| AT1G06267 | 15,22 | -2,30 | 0,73 | -3,14 | 1,67E-03 | 1,33E-02 |
| AT5G41750 | 8,05 | 3,58 | 1,14 | 3,14 | 1,68E-03 | 1,34E-02 |
| AT4G13340 | 14,47 | 1,72 | 0,55 | 3,14 | 1,71E-03 | 1,35E-02 |
| AT1G12290 | 131,54 | 0,83 | 0,26 | 3,14 | 1,71E-03 | 1,35E-02 |
| AT2G36530 | 5,68 | 3,36 | 1,07 | 3,14 | 1,71E-03 | 1,35E-02 |
| AT4G27440 | 13,15 | -2,04 | 0,65 | -3,13 | 1,75E-03 | 1,36E-02 |
| AT4G06110 | 22,73 | 1,59 | 0,51 | 3,13 | 1,75E-03 | 1,36E-02 |
| AT4G16950 | 9,81 | 2,71 | 0,87 | 3,13 | 1,74E-03 | 1,36E-02 |
| AT1G04430 | 7,26 | 2,94 | 0,94 | 3,13 | 1,77E-03 | 1,37E-02 |
| AT2G09670 | 5,26 | 3,89 | 1,24 | 3,12 | 1,79E-03 | 1,39E-02 |
| AT3G59020 | 5,79 | 3,88 | 1,24 | 3,12 | 1,80E-03 | 1,39E-02 |
| AT4G38770 | 7,08 | -2,81 | 0,90 | -3,12 | 1,81E-03 | 1,39E-02 |
| AT2G17930 | 7,05 | 2,77 | 0,89 | 3,11 | 1,86E-03 | 1,43E-02 |
| AT4G21710 | 2,69 | 5,09 | 1,64 | 3,11 | 1,87E-03 | 1,43E-02 |
| AT5G63020 | 8,08 | -3,21 | 1,03 | -3,11 | 1,87E-03 | 1,43E-02 |
| AT1G49040 | 3,74 | 4,72 | 1,52 | 3,11 | 1,89E-03 | 1,44E-02 |
| AT1G32230 | 5,15 | 3,81 | 1,23 | 3,10 | 1,94E-03 | 1,47E-02 |
| AT4G06075 | 405,29 | -1,03 | 0,33 | -3,09 | 1,98E-03 | 1,49E-02 |
| AT5G54310 | 4,38 | 3,80 | 1,23 | 3,09 | 1,98E-03 | 1,49E-02 |
| AT5G58040 | 3,29 | 4,60 | 1,49 | 3,09 | 2,00E-03 | 1,49E-02 |
| AT3G55610 | 4,81 | 3,56 | 1,15 | 3,09 | 2,00E-03 | 1,49E-02 |
| AT3G21805 | 21,83 | -2,53 | 0,82 | -3,09 | 2,02E-03 | 1,50E-02 |
| AT1G18745 | 15,99 | -1,88 | 0,61 | -3,09 | 2,01E-03 | 1,50E-02 |
| AT4G30160 | 4,90 | 4,49 | 1,46 | 3,09 | 2,03E-03 | 1,50E-02 |
| AT5G08670 | 5,95 | 3,06 | 0,99 | 3,08 | 2,05E-03 | 1,51E-02 |
| AT5G44800 | 4,77 | 4,22 | 1,37 | 3,08 | 2,09E-03 | 1,53E-02 |
| AT4G31880 | 2,70 | 4,84 | 1,57 | 3,08 | 2,10E-03 | 1,54E-02 |
| AT4G34670 | 4,28 | 3,77 | 1,23 | 3,07 | 2,16E-03 | 1,58E-02 |
| AT1G20960 | 9,33 | 2,44 | 0,80 | 3,06 | 2,18E-03 | 1,59E-02 |
| AT4G00570 | 3,02 | 4,78 | 1,56 | 3,06 | 2,20E-03 | 1,60E-02 |
| AT1G67120 | 8,25 | 2,48 | 0,81 | 3,06 | 2,22E-03 | 1,60E-02 |
| AT5G15845 | 2,91 | 4,60 | 1,50 | 3,06 | 2,22E-03 | 1,60E-02 |
| AT3G62360 | 4,39 | 4,32 | 1,41 | 3,05 | 2,26E-03 | 1,62E-02 |
| AT2G22330 | 4,57 | 4,35 | 1,43 | 3,05 | 2,30E-03 | 1,64E-02 |
| AT3G05675 | 9,08 | 2,41 | 0,79 | 3,05 | 2,29E-03 | 1,64E-02 |
| AT1G75335 | 3,97 | 4,44 | 1,46 | 3,04 | 2,34E-03 | 1,67E-02 |
| AT1G29465 | 25,62 | -1,62 | 0,53 | -3,03 | 2,41E-03 | 1,70E-02 |
| AT1G14610 | 2,42 | 4,72 | 1,55 | 3,04 | 2,40E-03 | 1,70E-02 |
| AT2G03820 | 4,25 | 4,25 | 1,40 | 3,04 | 2,40E-03 | 1,70E-02 |
| AT2G31660 | 3,59 | 4,34 | 1,43 | 3,03 | 2,47E-03 | 1,74E-02 |
| AT2G42600 | 13,57 | 1,87 | 0,62 | 3,03 | 2,48E-03 | 1,74E-02 |
| AT5G25280 | 3,55 | 4,32 | 1,43 | 3,03 | 2,49E-03 | 1,74E-02 |
| AT3G19190 | 3,92 | 4,67 | 1,55 | 3,01 | 2,58E-03 | 1,80E-02 |
| AT1G35710 | 7,53 | 4,40 | 1,46 | 3,01 | 2,65E-03 | 1,84E-02 |
| AT3G17850 | 4,74 | 4,25 | 1,41 | 3,00 | 2,68E-03 | 1,86E-02 |

| | | | | | | |
|---|---|---|---|---|---|---|
| AT5G20730 | 2,22 | 4,60 | 1,54 | 3,00 | 2,73E-03 | 1,89E-02 |
| AT3G13920 | 8,10 | 2,56 | 0,86 | 2,99 | 2,76E-03 | 1,90E-02 |
| AT1G04810 | 3,41 | 4,94 | 1,65 | 2,99 | 2,82E-03 | 1,94E-02 |
| AT3G54730 | 3,74 | -4,24 | 1,42 | -2,98 | 2,86E-03 | 1,95E-02 |
| AT5G58140 | 8,36 | 3,04 | 1,02 | 2,98 | 2,86E-03 | 1,95E-02 |
| AT2G07360 | 3,99 | 3,74 | 1,25 | 2,98 | 2,87E-03 | 1,95E-02 |
| AT4G00730 | 3,19 | 4,63 | 1,56 | 2,98 | 2,93E-03 | 1,99E-02 |
| AT3G14470 | 2,64 | 4,72 | 1,59 | 2,97 | 2,96E-03 | 2,00E-02 |
| AT2G05520 | 18,30 | -1,42 | 0,48 | -2,97 | 3,01E-03 | 2,03E-02 |
| AT2G01270 | 3,86 | 4,31 | 1,45 | 2,96 | 3,06E-03 | 2,06E-02 |
| AT2G48160 | 6,20 | 4,30 | 1,46 | 2,95 | 3,16E-03 | 2,12E-02 |
| AT4G00450 | 3,39 | 4,60 | 1,56 | 2,95 | 3,18E-03 | 2,13E-02 |
| AT3G63380 | 3,14 | 4,76 | 1,62 | 2,95 | 3,20E-03 | 2,13E-02 |
| AT5G57990 | 5,76 | 3,24 | 1,10 | 2,95 | 3,20E-03 | 2,13E-02 |
| AT1G30470 | 3,87 | 4,38 | 1,49 | 2,94 | 3,25E-03 | 2,15E-02 |
| AT5G63840 | 2,51 | 4,56 | 1,55 | 2,94 | 3,28E-03 | 2,16E-02 |
| AT1G76550 | 2,61 | 4,56 | 1,55 | 2,94 | 3,28E-03 | 2,16E-02 |
| AT3G63460 | 4,10 | 3,66 | 1,25 | 2,94 | 3,31E-03 | 2,18E-02 |
| AT2G44120 | 4,00 | 3,83 | 1,31 | 2,93 | 3,36E-03 | 2,20E-02 |
| AT4G24415 | 10,68 | -1,89 | 0,64 | -2,93 | 3,39E-03 | 2,22E-02 |
| AT2G32730 | 3,18 | 4,56 | 1,56 | 2,93 | 3,43E-03 | 2,24E-02 |
| AT5G64420 | 4,10 | 4,16 | 1,43 | 2,92 | 3,52E-03 | 2,29E-02 |
| AT1G09010 | 3,02 | 4,29 | 1,47 | 2,91 | 3,65E-03 | 2,36E-02 |
| AT3G63500 | 4,53 | 3,92 | 1,35 | 2,90 | 3,74E-03 | 2,42E-02 |
| AT1G34300 | 2,71 | 4,54 | 1,57 | 2,89 | 3,81E-03 | 2,45E-02 |
| AT2G38440 | 3,79 | 4,08 | 1,41 | 2,89 | 3,80E-03 | 2,45E-02 |
| AT5G14520 | 3,02 | 4,55 | 1,57 | 2,89 | 3,82E-03 | 2,45E-02 |
| AT2G03855 | 12,69 | -1,91 | 0,66 | -2,89 | 3,87E-03 | 2,47E-02 |
| AT3G04470 | 2,70 | 4,39 | 1,52 | 2,88 | 3,94E-03 | 2,51E-02 |
| AT1G18010 | 5,80 | -3,83 | 1,34 | -2,87 | 4,16E-03 | 2,64E-02 |
| AT3G01695 | 8,72 | 2,32 | 0,81 | 2,86 | 4,19E-03 | 2,64E-02 |
| AT5G08712 | 123,44 | 1,00 | 0,35 | 2,87 | 4,15E-03 | 2,64E-02 |
| AT2G46400 | 1,95 | 4,64 | 1,62 | 2,86 | 4,17E-03 | 2,64E-02 |
| AT3G08730 | 2,48 | 4,82 | 1,68 | 2,86 | 4,19E-03 | 2,64E-02 |
| AT1G58250 | 4,39 | 3,88 | 1,35 | 2,86 | 4,22E-03 | 2,64E-02 |
| AT1G79090 | 2,69 | 4,70 | 1,64 | 2,86 | 4,26E-03 | 2,67E-02 |
| AT2G05100 | 10,97 | -1,93 | 0,68 | -2,86 | 4,28E-03 | 2,67E-02 |
| AT5G05980 | 2,96 | 4,84 | 1,69 | 2,86 | 4,29E-03 | 2,67E-02 |
| AT2G40140 | 4,76 | 3,65 | 1,28 | 2,85 | 4,44E-03 | 2,70E-02 |
| AT2G47070 | 5,92 | 3,09 | 1,08 | 2,85 | 4,36E-03 | 2,70E-02 |
| AT4G27500 | 2,93 | 4,44 | 1,56 | 2,85 | 4,44E-03 | 2,70E-02 |
| AT1G11640 | 3,15 | 4,28 | 1,50 | 2,85 | 4,39E-03 | 2,70E-02 |
| AT1G13350 | 2,82 | 4,87 | 1,71 | 2,85 | 4,40E-03 | 2,70E-02 |
| AT4G18600 | 3,50 | 4,46 | 1,57 | 2,85 | 4,38E-03 | 2,70E-02 |
| AT4G07855 | 2,93 | 4,44 | 1,56 | 2,85 | 4,44E-03 | 2,70E-02 |
| AT4G37460 | 2,98 | 4,41 | 1,55 | 2,85 | 4,43E-03 | 2,70E-02 |
| AT5G19390 | 2,78 | 4,89 | 1,72 | 2,84 | 4,47E-03 | 2,71E-02 |
| AT4G16900 | 2,10 | 4,41 | 1,55 | 2,84 | 4,58E-03 | 2,78E-02 |
| AT5G49980 | 4,39 | 4,15 | 1,47 | 2,83 | 4,62E-03 | 2,79E-02 |
| AT3G10650 | 3,29 | 4,23 | 1,49 | 2,83 | 4,65E-03 | 2,80E-02 |
| AT4G17330 | 4,91 | 4,31 | 1,53 | 2,83 | 4,71E-03 | 2,83E-02 |
| AT4G39080 | 4,27 | 3,64 | 1,29 | 2,82 | 4,80E-03 | 2,86E-02 |
| AT5G02575 | 6,74 | 2,50 | 0,89 | 2,82 | 4,80E-03 | 2,86E-02 |
| AT1G05960 | 4,07 | 3,94 | 1,40 | 2,82 | 4,82E-03 | 2,86E-02 |
| AT4G36630 | 4,11 | 4,57 | 1,62 | 2,82 | 4,83E-03 | 2,86E-02 |
| AT5G18370 | 3,23 | 4,20 | 1,49 | 2,82 | 4,80E-03 | 2,86E-02 |

| | | | | | | |
|---|---|---|---|---|---|---|
| AT5G46450 | 2,49 | 4,44 | 1,58 | 2,82 | 4,85E-03 | 2,87E-02 |
| AT1G14740 | 3,27 | 3,93 | 1,40 | 2,81 | 4,92E-03 | 2,90E-02 |
| AT1G62750 | 9,17 | 2,24 | 0,80 | 2,81 | 4,99E-03 | 2,94E-02 |
| AT3G06650 | 3,17 | 4,28 | 1,53 | 2,80 | 5,03E-03 | 2,96E-02 |
| AT4G39850 | 2,19 | 4,66 | 1,66 | 2,80 | 5,06E-03 | 2,96E-02 |
| AT4G34710 | 3,32 | 4,13 | 1,47 | 2,80 | 5,07E-03 | 2,96E-02 |
| AT1G72410 | 1,92 | 4,74 | 1,69 | 2,80 | 5,09E-03 | 2,97E-02 |
| AT5G07970 | 3,91 | 4,29 | 1,53 | 2,80 | 5,16E-03 | 3,00E-02 |
| AT3G19420 | 2,24 | 4,77 | 1,71 | 2,80 | 5,17E-03 | 3,00E-02 |
| AT1G30410 | 4,36 | 4,01 | 1,43 | 2,79 | 5,20E-03 | 3,01E-02 |
| AT2G46830 | 2,43 | 4,50 | 1,62 | 2,78 | 5,40E-03 | 3,11E-02 |
| AT3G07790 | 2,60 | 4,31 | 1,55 | 2,78 | 5,39E-03 | 3,11E-02 |
| AT5G24350 | 3,06 | 4,70 | 1,69 | 2,78 | 5,42E-03 | 3,11E-02 |
| AT1G08380 | 7,50 | -2,50 | 0,90 | -2,77 | 5,53E-03 | 3,17E-02 |
| AT4G28760 | 2,13 | 4,48 | 1,62 | 2,77 | 5,62E-03 | 3,20E-02 |
| AT5G46520 | 3,26 | 4,37 | 1,58 | 2,77 | 5,62E-03 | 3,20E-02 |
| AT2G18320 | 12,44 | -1,56 | 0,56 | -2,76 | 5,69E-03 | 3,24E-02 |
| AT1G80680 | 4,25 | 4,02 | 1,45 | 2,76 | 5,72E-03 | 3,24E-02 |
| AT5G24710 | 4,46 | 3,70 | 1,34 | 2,76 | 5,75E-03 | 3,25E-02 |
| AT1G25440 | 6,25 | 3,03 | 1,10 | 2,75 | 5,91E-03 | 3,34E-02 |
| AT2G25430 | 4,01 | 3,41 | 1,24 | 2,75 | 6,02E-03 | 3,39E-02 |
| AT5G06320 | 5,00 | 4,31 | 1,57 | 2,74 | 6,07E-03 | 3,41E-02 |
| AT3G44580 | 31,33 | -0,98 | 0,36 | -2,74 | 6,10E-03 | 3,42E-02 |
| AT2G14560 | 27,90 | 1,45 | 0,53 | 2,73 | 6,30E-03 | 3,52E-02 |
| AT4G09650 | 3,00 | -4,64 | 1,70 | -2,73 | 6,34E-03 | 3,53E-02 |
| AT1G63005 | 3,14 | -4,35 | 1,60 | -2,73 | 6,36E-03 | 3,53E-02 |
| AT2G03445 | 309,93 | -0,74 | 0,27 | -2,73 | 6,42E-03 | 3,56E-02 |
| AT1G49700 | 4,39 | 3,33 | 1,22 | 2,73 | 6,42E-03 | 3,56E-02 |
| AT5G51174 | 65,86 | -1,17 | 0,43 | -2,72 | 6,56E-03 | 3,62E-02 |
| AT3G13300 | 8,42 | 3,34 | 1,23 | 2,72 | 6,61E-03 | 3,64E-02 |
| AT1G27430 | 3,11 | 4,27 | 1,57 | 2,71 | 6,65E-03 | 3,65E-02 |
| AT3G26650 | 12,31 | 1,79 | 0,66 | 2,71 | 6,73E-03 | 3,69E-02 |
| AT2G33340 | 1,98 | 4,60 | 1,70 | 2,71 | 6,74E-03 | 3,69E-02 |
| AT4G01050 | 4,58 | 3,33 | 1,23 | 2,70 | 6,84E-03 | 3,73E-02 |
| AT3G17900 | 1,92 | 4,41 | 1,63 | 2,70 | 6,96E-03 | 3,79E-02 |
| AT2G31880 | 6,93 | 2,76 | 1,02 | 2,69 | 7,08E-03 | 3,83E-02 |
| AT5G23380 | 2,25 | 4,67 | 1,73 | 2,69 | 7,08E-03 | 3,83E-02 |
| AT4G06235 | 7,69 | -3,05 | 1,13 | -2,69 | 7,07E-03 | 3,83E-02 |
| AT1G64430 | 2,85 | 4,19 | 1,56 | 2,69 | 7,11E-03 | 3,83E-02 |
| AT3G20010 | 2,80 | 4,67 | 1,74 | 2,69 | 7,14E-03 | 3,84E-02 |
| AT3G52250 | 4,06 | 3,87 | 1,44 | 2,69 | 7,16E-03 | 3,84E-02 |
| AT1G27340 | 2,10 | 4,49 | 1,67 | 2,69 | 7,19E-03 | 3,85E-02 |
| AT4G01840 | 3,87 | -4,11 | 1,53 | -2,68 | 7,26E-03 | 3,88E-02 |
| AT1G06190 | 5,18 | 3,37 | 1,26 | 2,68 | 7,30E-03 | 3,88E-02 |
| AT4G31500 | 6,34 | 2,95 | 1,10 | 2,68 | 7,29E-03 | 3,88E-02 |
| AT3G41762 | 25,36 | -1,31 | 0,49 | -2,68 | 7,33E-03 | 3,89E-02 |
| AT2G44570 | 2,88 | -4,19 | 1,57 | -2,67 | 7,48E-03 | 3,94E-02 |
| AT4G00800 | 2,43 | 4,22 | 1,58 | 2,67 | 7,48E-03 | 3,94E-02 |
| AT5G45500 | 2,29 | 4,92 | 1,84 | 2,67 | 7,49E-03 | 3,94E-02 |
| AT5G13000 | 3,30 | 3,69 | 1,39 | 2,66 | 7,80E-03 | 4,10E-02 |
| AT4G19110 | 3,30 | 4,44 | 1,67 | 2,66 | 7,82E-03 | 4,10E-02 |
| AT5G48160 | 6,17 | 2,96 | 1,11 | 2,66 | 7,93E-03 | 4,15E-02 |
| AT3G60860 | 2,94 | 4,28 | 1,62 | 2,65 | 8,07E-03 | 4,20E-02 |
| AT5G65750 | 4,51 | 3,11 | 1,18 | 2,65 | 8,05E-03 | 4,20E-02 |
| AT1G49240 | 7,10 | 2,59 | 0,98 | 2,65 | 8,13E-03 | 4,23E-02 |
| AT2G41650 | 5,84 | -3,55 | 1,34 | -2,65 | 8,15E-03 | 4,23E-02 |

| | | | | | | |
|---|---|---|---|---|---|---|
| AT5G41663 | 49,62 | -1,43 | 0,54 | -2,65 | 8,17E-03 | 4,23E-02 |
| AT1G70185 | 5,33 | 2,71 | 1,03 | 2,64 | 8,21E-03 | 4,24E-02 |
| AT5G44180 | 3,26 | 3,93 | 1,49 | 2,64 | 8,25E-03 | 4,25E-02 |
| AT1G79570 | 2,57 | 4,14 | 1,57 | 2,64 | 8,28E-03 | 4,25E-02 |
| AT5G46270 | 2,21 | 4,37 | 1,66 | 2,63 | 8,42E-03 | 4,32E-02 |
| AT5G60900 | 5,69 | 3,80 | 1,44 | 2,63 | 8,48E-03 | 4,34E-02 |
| AT1G05570 | 10,12 | 1,92 | 0,73 | 2,63 | 8,52E-03 | 4,35E-02 |
| AT1G53683 | 15,44 | 1,50 | 0,57 | 2,63 | 8,66E-03 | 4,41E-02 |
| AT1G60072 | 5,42 | 3,70 | 1,41 | 2,62 | 8,72E-03 | 4,43E-02 |
| AT5G19400 | 2,38 | 4,80 | 1,83 | 2,62 | 8,73E-03 | 4,43E-02 |
| AT3G06530 | 4,41 | 3,43 | 1,31 | 2,62 | 8,82E-03 | 4,45E-02 |
| AT3G28540 | 2,73 | 4,06 | 1,55 | 2,62 | 8,86E-03 | 4,45E-02 |
| AT4G30100 | 3,56 | 3,80 | 1,45 | 2,62 | 8,83E-03 | 4,45E-02 |
| AT5G14430 | 2,60 | 4,32 | 1,65 | 2,62 | 8,85E-03 | 4,45E-02 |
| AT5G19820 | 4,02 | 4,11 | 1,57 | 2,62 | 8,87E-03 | 4,45E-02 |
| AT5G13010 | 4,55 | 3,09 | 1,18 | 2,61 | 8,96E-03 | 4,48E-02 |
| AT5G15850 | 2,49 | 4,10 | 1,57 | 2,61 | 8,95E-03 | 4,48E-02 |
| AT2G35747 | 15,41 | -1,94 | 0,74 | -2,61 | 9,00E-03 | 4,48E-02 |
| AT1G02470 | 5,92 | 2,81 | 1,08 | 2,61 | 9,10E-03 | 4,52E-02 |
| AT3G57520 | 3,92 | 2,85 | 1,09 | 2,61 | 9,09E-03 | 4,52E-02 |
| AT3G55850 | 2,66 | 4,39 | 1,68 | 2,61 | 9,12E-03 | 4,52E-02 |
| AT4G32410 | 11,90 | 1,75 | 0,67 | 2,60 | 9,21E-03 | 4,55E-02 |
| AT2G45930 | 2,64 | 3,98 | 1,53 | 2,60 | 9,27E-03 | 4,57E-02 |
| AT2G41740 | 6,26 | 2,79 | 1,07 | 2,60 | 9,32E-03 | 4,58E-02 |
| AT3G26740 | 15,30 | -1,50 | 0,58 | -2,60 | 9,35E-03 | 4,59E-02 |
| AT5G14800 | 5,34 | 2,83 | 1,09 | 2,60 | 9,39E-03 | 4,60E-02 |
| AT2G33100 | 2,25 | 4,59 | 1,77 | 2,59 | 9,51E-03 | 4,65E-02 |
| AT5G49360 | 3,80 | 3,78 | 1,46 | 2,59 | 9,54E-03 | 4,66E-02 |
| AT3G57410 | 3,70 | 3,49 | 1,35 | 2,59 | 9,71E-03 | 4,73E-02 |
| AT3G05420 | 2,70 | 3,84 | 1,49 | 2,58 | 9,75E-03 | 4,74E-02 |
| AT5G61780 | 4,25 | 3,02 | 1,17 | 2,58 | 9,80E-03 | 4,75E-02 |
| AT1G53310 | 3,45 | 3,29 | 1,28 | 2,57 | 1,01E-02 | 4,88E-02 |
| AT4G16340 | 3,35 | 3,33 | 1,29 | 2,57 | 1,01E-02 | 4,88E-02 |
| AT1G01510 | 2,79 | 4,06 | 1,58 | 2,57 | 1,03E-02 | 4,96E-02 |
| AT1G79000 | 2,98 | 4,93 | 1,92 | 2,57 | 1,03E-02 | 4,96E-02 |
| AT1G55860 | 12,82 | 1,99 | 0,78 | 2,56 | 1,04E-02 | 4,97E-02 |
| AT3G57660 | 1,78 | 4,54 | 1,77 | 2,56 | 1,04E-02 | 4,98E-02 |
| AT5G24990 | 2,50 | -4,07 | 1,59 | -2,56 | 1,05E-02 | 5,01E-02 |
| AT5G64220 | 2,50 | 4,18 | 1,64 | 2,56 | 1,05E-02 | 5,01E-02 |
| AT4G30190 | 7,61 | 2,38 | 0,93 | 2,56 | 1,05E-02 | 5,01E-02 |
| AT3G28956 | 2,37 | -3,91 | 1,53 | -2,56 | 1,06E-02 | 5,02E-02 |
| AT2G05455 | 4,14 | -3,62 | 1,42 | -2,56 | 1,06E-02 | 5,03E-02 |
| AT1G48953 | 8,24 | -2,39 | 0,94 | -2,55 | 1,07E-02 | 5,05E-02 |
| AT3G07100 | 6,58 | 2,56 | 1,00 | 2,55 | 1,07E-02 | 5,05E-02 |
| AT5G18350 | 2,51 | 4,54 | 1,78 | 2,55 | 1,09E-02 | 5,11E-02 |
| AT1G13640 | 2,82 | 3,79 | 1,49 | 2,55 | 1,09E-02 | 5,13E-02 |
| AT3G52430 | 2,34 | 4,43 | 1,74 | 2,54 | 1,10E-02 | 5,17E-02 |
| AT3G61580 | 5,37 | 2,46 | 0,97 | 2,54 | 1,11E-02 | 5,19E-02 |
| AT1G33680 | 2,02 | 4,63 | 1,83 | 2,54 | 1,12E-02 | 5,20E-02 |
| AT3G48350 | 1,84 | 4,91 | 1,94 | 2,54 | 1,12E-02 | 5,20E-02 |
| AT1G79280 | 4,52 | 2,97 | 1,17 | 2,54 | 1,11E-02 | 5,20E-02 |
| AT2G30110 | 3,88 | 3,09 | 1,22 | 2,54 | 1,12E-02 | 5,20E-02 |
| AT3G45190 | 2,88 | 3,77 | 1,49 | 2,53 | 1,13E-02 | 5,23E-02 |
| AT5G01747 | 60,67 | -0,74 | 0,29 | -2,53 | 1,13E-02 | 5,24E-02 |
| AT3G47910 | 2,73 | 3,78 | 1,49 | 2,53 | 1,14E-02 | 5,25E-02 |
| AT1G08833 | 15,74 | -1,38 | 0,55 | -2,53 | 1,14E-02 | 5,27E-02 |

| | | | | | | |
|---|---|---|---|---|---|---|
| AT1G58290 | 5,16 | 2,39 | 0,94 | 2,52 | 1,16E-02 | 5,33E-02 |
| AT4G06225 | 6,57 | -3,09 | 1,22 | -2,52 | 1,17E-02 | 5,35E-02 |
| AT4G11420 | 3,20 | 4,25 | 1,69 | 2,52 | 1,17E-02 | 5,35E-02 |
| AT1G43910 | 2,35 | 4,23 | 1,68 | 2,52 | 1,18E-02 | 5,40E-02 |
| AT5G11040 | 2,15 | 4,09 | 1,63 | 2,52 | 1,19E-02 | 5,42E-02 |
| AT4G00630 | 4,40 | 2,80 | 1,12 | 2,51 | 1,19E-02 | 5,42E-02 |
| AT4G35800 | 5,60 | 2,41 | 0,96 | 2,51 | 1,19E-02 | 5,42E-02 |
| AT3G03380 | 4,30 | 3,21 | 1,28 | 2,51 | 1,20E-02 | 5,44E-02 |
| AT5G14270 | 2,74 | 3,76 | 1,50 | 2,51 | 1,21E-02 | 5,47E-02 |
| AT1G24706 | 2,66 | 3,63 | 1,45 | 2,50 | 1,23E-02 | 5,55E-02 |
| AT5G46420 | 2,51 | 3,59 | 1,44 | 2,50 | 1,24E-02 | 5,59E-02 |
| AT1G79990 | 2,18 | 4,29 | 1,72 | 2,50 | 1,24E-02 | 5,60E-02 |
| AT4G14160 | 2,71 | 4,30 | 1,72 | 2,50 | 1,25E-02 | 5,60E-02 |
| AT3G51050 | 3,03 | 4,05 | 1,63 | 2,49 | 1,27E-02 | 5,69E-02 |
| AT1G17210 | 2,32 | 4,19 | 1,68 | 2,49 | 1,27E-02 | 5,69E-02 |
| AT2G02840 | 4,55 | 3,86 | 1,55 | 2,49 | 1,29E-02 | 5,77E-02 |
| AT1G07977 | 4,37 | -3,03 | 1,22 | -2,48 | 1,30E-02 | 5,78E-02 |
| AT5G46070 | 1,78 | 4,63 | 1,86 | 2,48 | 1,30E-02 | 5,78E-02 |
| AT2G36480 | 1,71 | 4,40 | 1,78 | 2,48 | 1,33E-02 | 5,89E-02 |
| AT2G36850 | 3,05 | 3,93 | 1,59 | 2,47 | 1,36E-02 | 6,02E-02 |
| AT5G58150 | 3,53 | 4,45 | 1,80 | 2,47 | 1,36E-02 | 6,02E-02 |
| AT1G30230 | 5,95 | 2,48 | 1,00 | 2,46 | 1,37E-02 | 6,05E-02 |
| AT4G06240 | 6,18 | -3,07 | 1,25 | -2,46 | 1,38E-02 | 6,05E-02 |
| AT4G04890 | 2,53 | 3,69 | 1,50 | 2,46 | 1,37E-02 | 6,05E-02 |
| AT5G54390 | 4,62 | 4,10 | 1,67 | 2,45 | 1,41E-02 | 6,20E-02 |
| AT1G77680 | 1,90 | 4,65 | 1,90 | 2,45 | 1,42E-02 | 6,22E-02 |
| AT3G11540 | 2,22 | 4,44 | 1,81 | 2,45 | 1,43E-02 | 6,25E-02 |
| AT5G20700 | 23,90 | 1,15 | 0,47 | 2,45 | 1,44E-02 | 6,28E-02 |
| AT2G27860 | 4,64 | 2,36 | 0,97 | 2,44 | 1,46E-02 | 6,34E-02 |
| AT2G27400 | 624,49 | -0,81 | 0,33 | -2,44 | 1,46E-02 | 6,35E-02 |
| AT2G24103 | 3,17 | 3,69 | 1,51 | 2,44 | 1,47E-02 | 6,38E-02 |
| AT2G43890 | 2,70 | 4,28 | 1,76 | 2,44 | 1,48E-02 | 6,40E-02 |
| AT2G26780 | 3,27 | 3,78 | 1,55 | 2,43 | 1,50E-02 | 6,46E-02 |
| AT5G20280 | 6,18 | 2,18 | 0,89 | 2,43 | 1,50E-02 | 6,48E-02 |
| AT3G15730 | 15,00 | 1,41 | 0,58 | 2,43 | 1,50E-02 | 6,48E-02 |
| AT1G64710 | 2,31 | 3,78 | 1,56 | 2,43 | 1,51E-02 | 6,51E-02 |
| AT5G09410 | 3,14 | 3,69 | 1,52 | 2,43 | 1,52E-02 | 6,51E-02 |
| AT5G12120 | 2,73 | 3,82 | 1,57 | 2,43 | 1,52E-02 | 6,52E-02 |
| AT3G02090 | 3,68 | 3,06 | 1,26 | 2,42 | 1,53E-02 | 6,55E-02 |
| AT1G78570 | 2,32 | 4,00 | 1,65 | 2,42 | 1,54E-02 | 6,56E-02 |
| AT1G80250 | 2,27 | 3,93 | 1,62 | 2,42 | 1,54E-02 | 6,56E-02 |
| AT2G46500 | 2,79 | 3,94 | 1,63 | 2,42 | 1,55E-02 | 6,61E-02 |
| AT5G46470 | 3,68 | 2,76 | 1,14 | 2,42 | 1,57E-02 | 6,64E-02 |
| AT5G47260 | 4,17 | -3,38 | 1,40 | -2,42 | 1,57E-02 | 6,64E-02 |
| AT3G13870 | 4,03 | 3,82 | 1,58 | 2,42 | 1,57E-02 | 6,64E-02 |
| AT3G22790 | 2,45 | 3,69 | 1,53 | 2,42 | 1,57E-02 | 6,64E-02 |
| AT5G36230 | 2,63 | 3,99 | 1,66 | 2,41 | 1,60E-02 | 6,74E-02 |
| AT1G26985 | 10,26 | 1,93 | 0,80 | 2,41 | 1,60E-02 | 6,75E-02 |
| AT3G13330 | 3,62 | 3,14 | 1,30 | 2,41 | 1,61E-02 | 6,76E-02 |
| AT3G57300 | 2,34 | 3,63 | 1,51 | 2,40 | 1,62E-02 | 6,79E-02 |
| AT4G01810 | 2,76 | 3,79 | 1,58 | 2,40 | 1,62E-02 | 6,79E-02 |
| AT5G46250 | 6,03 | 3,14 | 1,31 | 2,40 | 1,63E-02 | 6,81E-02 |
| AT5G65380 | 2,69 | 3,24 | 1,35 | 2,40 | 1,63E-02 | 6,81E-02 |
| AT1G09310 | 8,77 | -1,73 | 0,72 | -2,40 | 1,66E-02 | 6,88E-02 |
| AT5G04930 | 5,77 | 3,14 | 1,31 | 2,40 | 1,66E-02 | 6,88E-02 |
| AT5G17890 | 1,77 | 4,38 | 1,83 | 2,39 | 1,68E-02 | 6,94E-02 |

| | | | | | | |
|---|---|---|---|---|---|---|
| AT1G74720 | 1,98 | 4,18 | 1,76 | 2,38 | 1,72E-02 | 7,10E-02 |
| AT5G67360 | 6,15 | 2,66 | 1,12 | 2,38 | 1,72E-02 | 7,12E-02 |
| AT5G52882 | 2,07 | 4,21 | 1,77 | 2,38 | 1,74E-02 | 7,19E-02 |
| AT2G26080 | 10,37 | 1,92 | 0,81 | 2,37 | 1,75E-02 | 7,20E-02 |
| AT3G01835 | 72,10 | -0,77 | 0,33 | -2,37 | 1,76E-02 | 7,20E-02 |
| AT4G18670 | 7,64 | -1,99 | 0,84 | -2,38 | 1,75E-02 | 7,20E-02 |
| AT1G05562 | 6,13 | -2,38 | 1,00 | -2,37 | 1,77E-02 | 7,23E-02 |
| AT1G05560 | 6,13 | -2,38 | 1,00 | -2,37 | 1,77E-02 | 7,23E-02 |
| AT5G62190 | 2,05 | 3,71 | 1,57 | 2,37 | 1,78E-02 | 7,26E-02 |
| AT5G44320 | 2,18 | 3,85 | 1,63 | 2,37 | 1,79E-02 | 7,28E-02 |
| AT1G63770 | 7,56 | 1,88 | 0,80 | 2,37 | 1,80E-02 | 7,32E-02 |
| AT1G59610 | 3,98 | 3,05 | 1,29 | 2,36 | 1,82E-02 | 7,36E-02 |
| AT2G08770 | 10,72 | -1,72 | 0,73 | -2,36 | 1,82E-02 | 7,36E-02 |
| AT4G14504 | 4,83 | -3,33 | 1,41 | -2,36 | 1,82E-02 | 7,37E-02 |
| AT3G08530 | 12,53 | 1,45 | 0,61 | 2,36 | 1,83E-02 | 7,37E-02 |
| AT1G31173 | 488,28 | 0,59 | 0,25 | 2,36 | 1,84E-02 | 7,43E-02 |
| AT1G06220 | 3,15 | 3,26 | 1,38 | 2,35 | 1,86E-02 | 7,46E-02 |
| AT3G49500 | 2,34 | 4,05 | 1,73 | 2,35 | 1,89E-02 | 7,57E-02 |
| AT5G66030 | 2,48 | 3,73 | 1,59 | 2,35 | 1,89E-02 | 7,57E-02 |
| AT1G48650 | 3,21 | 3,54 | 1,51 | 2,34 | 1,92E-02 | 7,67E-02 |
| AT1G04210 | 3,48 | 2,68 | 1,15 | 2,34 | 1,94E-02 | 7,75E-02 |
| AT1G11700 | 23,46 | -1,47 | 0,63 | -2,34 | 1,95E-02 | 7,75E-02 |
| AT3G13460 | 10,31 | 1,53 | 0,65 | 2,34 | 1,95E-02 | 7,75E-02 |
| AT3G22380 | 8,79 | 2,10 | 0,90 | 2,33 | 1,96E-02 | 7,78E-02 |
| AT4G31160 | 2,16 | 4,32 | 1,85 | 2,33 | 1,98E-02 | 7,86E-02 |
| AT1G09770 | 3,11 | 3,40 | 1,46 | 2,33 | 2,00E-02 | 7,92E-02 |
| AT3G09200 | 3,98 | 2,70 | 1,16 | 2,32 | 2,02E-02 | 7,96E-02 |
| AT3G25500 | 1,85 | 3,78 | 1,63 | 2,32 | 2,03E-02 | 7,99E-02 |
| AT1G03060 | 7,63 | 2,02 | 0,87 | 2,32 | 2,03E-02 | 8,01E-02 |
| AT5G64940 | 2,50 | 3,88 | 1,67 | 2,32 | 2,04E-02 | 8,03E-02 |
| AT1G72390 | 2,40 | 3,76 | 1,62 | 2,31 | 2,07E-02 | 8,11E-02 |
| AT4G24280 | 6,53 | 2,33 | 1,01 | 2,31 | 2,07E-02 | 8,11E-02 |
| AT1G79040 | 10,12 | -2,01 | 0,87 | -2,31 | 2,08E-02 | 8,11E-02 |
| AT3G44630 | 4,39 | 2,73 | 1,18 | 2,31 | 2,08E-02 | 8,11E-02 |
| AT3G17510 | 2,73 | 3,92 | 1,70 | 2,31 | 2,08E-02 | 8,11E-02 |
| AT4G02380 | 7,09 | 1,90 | 0,82 | 2,31 | 2,08E-02 | 8,11E-02 |
| AT3G20690 | 3,62 | -3,19 | 1,38 | -2,31 | 2,09E-02 | 8,14E-02 |
| AT1G76135 | 2,70 | -3,79 | 1,65 | -2,30 | 2,15E-02 | 8,27E-02 |
| AT2G40805 | 280,87 | -0,97 | 0,42 | -2,30 | 2,15E-02 | 8,27E-02 |
| AT3G61630 | 1,88 | 4,29 | 1,86 | 2,30 | 2,13E-02 | 8,27E-02 |
| AT4G25530 | 5,35 | -3,16 | 1,37 | -2,30 | 2,15E-02 | 8,27E-02 |
| AT1G01183 | 317,62 | -0,81 | 0,35 | -2,30 | 2,14E-02 | 8,27E-02 |
| AT1G33560 | 2,19 | 4,28 | 1,86 | 2,29 | 2,18E-02 | 8,36E-02 |
| AT2G25730 | 2,88 | 3,03 | 1,32 | 2,29 | 2,18E-02 | 8,36E-02 |
| AT1G28480 | 1,91 | -4,13 | 1,80 | -2,29 | 2,20E-02 | 8,42E-02 |
| AT5G51070 | 3,71 | 2,89 | 1,26 | 2,29 | 2,23E-02 | 8,53E-02 |
| AT1G70940 | 2,12 | 3,71 | 1,63 | 2,28 | 2,25E-02 | 8,60E-02 |
| AT2G24790 | 3,54 | 3,64 | 1,60 | 2,28 | 2,25E-02 | 8,60E-02 |
| AT3G07855 | 4,66 | -2,49 | 1,09 | -2,28 | 2,27E-02 | 8,64E-02 |
| AT3G55200 | 2,62 | 4,26 | 1,87 | 2,28 | 2,28E-02 | 8,68E-02 |
| AT5G26742 | 9,59 | 1,77 | 0,78 | 2,28 | 2,29E-02 | 8,68E-02 |
| AT1G09970 | 4,22 | 3,37 | 1,48 | 2,27 | 2,30E-02 | 8,69E-02 |
| AT2G20570 | 7,71 | 1,95 | 0,86 | 2,27 | 2,30E-02 | 8,69E-02 |
| AT4G04835 | 19,24 | 1,15 | 0,51 | 2,27 | 2,29E-02 | 8,69E-02 |
| AT1G20970 | 4,69 | 2,81 | 1,24 | 2,26 | 2,36E-02 | 8,91E-02 |
| AT5G07875 | 4,76 | 2,88 | 1,27 | 2,26 | 2,37E-02 | 8,92E-02 |

| | | | | | | |
|---|---|---|---|---|---|---|
| AT3G01316 | 2,03 | -3,68 | 1,63 | -2,26 | 2,38E-02 | 8,94E-02 |
| AT1G14320 | 3,20 | 3,01 | 1,34 | 2,25 | 2,42E-02 | 9,05E-02 |
| AT1G09780 | 4,32 | 2,59 | 1,15 | 2,25 | 2,41E-02 | 9,05E-02 |
| AT4G28652 | 2,08 | 4,22 | 1,87 | 2,25 | 2,43E-02 | 9,05E-02 |
| AT2G29980 | 3,37 | 2,82 | 1,25 | 2,25 | 2,43E-02 | 9,05E-02 |
| AT4G28650 | 2,08 | 4,22 | 1,87 | 2,25 | 2,43E-02 | 9,05E-02 |
| AT3G02045 | 5,72 | -2,53 | 1,12 | -2,25 | 2,45E-02 | 9,09E-02 |
| AT5G09515 | 9,09 | -1,76 | 0,78 | -2,25 | 2,45E-02 | 9,09E-02 |
| AT5G16210 | 1,91 | 3,91 | 1,74 | 2,25 | 2,45E-02 | 9,10E-02 |
| AT3G60415 | 1,73 | 4,05 | 1,80 | 2,25 | 2,46E-02 | 9,11E-02 |
| AT1G21250 | 6,46 | 2,05 | 0,91 | 2,25 | 2,47E-02 | 9,15E-02 |
| AT4G04020 | 1,87 | -4,16 | 1,85 | -2,24 | 2,48E-02 | 9,16E-02 |
| AT1G55540 | 2,72 | 3,10 | 1,38 | 2,24 | 2,49E-02 | 9,17E-02 |
| AT1G53687 | 5,07 | -2,28 | 1,02 | -2,24 | 2,51E-02 | 9,17E-02 |
| AT5G05190 | 2,00 | 3,82 | 1,70 | 2,24 | 2,50E-02 | 9,17E-02 |
| AT1G12300 | 2,68 | -3,32 | 1,48 | -2,24 | 2,50E-02 | 9,17E-02 |
| AT1G27770 | 5,67 | 3,05 | 1,36 | 2,24 | 2,50E-02 | 9,17E-02 |
| AT5G05505 | 4,00 | -2,73 | 1,22 | -2,24 | 2,51E-02 | 9,17E-02 |
| AT2G43710 | 3,74 | 2,79 | 1,25 | 2,24 | 2,52E-02 | 9,21E-02 |
| AT3G07860 | 5,76 | -2,52 | 1,13 | -2,24 | 2,53E-02 | 9,22E-02 |
| AT5G61140 | 4,17 | 2,59 | 1,16 | 2,24 | 2,54E-02 | 9,25E-02 |
| AT1G64790 | 7,36 | 1,94 | 0,87 | 2,23 | 2,55E-02 | 9,26E-02 |
| AT4G01850 | 4,27 | 2,44 | 1,09 | 2,23 | 2,55E-02 | 9,26E-02 |
| AT1G08420 | 1,72 | 3,91 | 1,75 | 2,23 | 2,58E-02 | 9,27E-02 |
| AT1G21270 | 2,66 | 4,07 | 1,83 | 2,23 | 2,58E-02 | 9,27E-02 |
| AT2G41640 | 6,36 | -2,65 | 1,19 | -2,23 | 2,59E-02 | 9,27E-02 |
| AT3G06355 | 6711,54 | 0,79 | 0,36 | 2,23 | 2,59E-02 | 9,27E-02 |
| AT1G69360 | 2,19 | 3,91 | 1,75 | 2,23 | 2,58E-02 | 9,27E-02 |
| AT1G77760 | 2,44 | 3,77 | 1,69 | 2,23 | 2,56E-02 | 9,27E-02 |
| AT3G63003 | 4,44 | -3,02 | 1,35 | -2,23 | 2,59E-02 | 9,27E-02 |
| AT5G42080 | 4,02 | 2,71 | 1,21 | 2,23 | 2,57E-02 | 9,27E-02 |
| AT5G55565 | 4,30 | -2,94 | 1,32 | -2,23 | 2,58E-02 | 9,27E-02 |
| AT5G03280 | 3,28 | 3,45 | 1,55 | 2,23 | 2,60E-02 | 9,29E-02 |
| AT2G35387 | 52,44 | -0,83 | 0,37 | -2,22 | 2,63E-02 | 9,37E-02 |
| AT4G02660 | 2,17 | 3,81 | 1,72 | 2,22 | 2,65E-02 | 9,41E-02 |
| AT4G36140 | 2,98 | 3,16 | 1,42 | 2,22 | 2,65E-02 | 9,41E-02 |
| AT2G41560 | 2,99 | 2,68 | 1,21 | 2,22 | 2,65E-02 | 9,42E-02 |
| AT2G01570 | 3,41 | 2,89 | 1,31 | 2,21 | 2,69E-02 | 9,55E-02 |
| AT4G25080 | 4,26 | 2,43 | 1,10 | 2,21 | 2,72E-02 | 9,62E-02 |
| AT5G42020 | 4,47 | 2,33 | 1,05 | 2,21 | 2,72E-02 | 9,62E-02 |
| AT3G41979 | 6596,11 | 0,79 | 0,36 | 2,20 | 2,77E-02 | 9,78E-02 |
| AT5G01820 | 9,75 | 2,85 | 1,30 | 2,20 | 2,78E-02 | 9,78E-02 |
| AT5G18525 | 1,70 | 4,01 | 1,82 | 2,20 | 2,78E-02 | 9,78E-02 |
| AT3G41761 | 2,51 | -3,76 | 1,71 | -2,20 | 2,79E-02 | 9,78E-02 |
| AT5G61455 | 11,96 | 1,34 | 0,61 | 2,20 | 2,79E-02 | 9,78E-02 |
| AT4G38440 | 2,75 | 2,94 | 1,34 | 2,20 | 2,80E-02 | 9,80E-02 |
| AT4G31877 | 9,33 | -1,50 | 0,68 | -2,20 | 2,80E-02 | 9,80E-02 |
| AT2G47980 | 2,39 | 3,59 | 1,63 | 2,19 | 2,82E-02 | 9,81E-02 |
| AT5G27807 | 30,37 | -1,03 | 0,47 | -2,19 | 2,84E-02 | 9,81E-02 |
| AT1G18310 | 4,64 | -3,34 | 1,52 | -2,19 | 2,83E-02 | 9,81E-02 |
| AT3G08580 | 9,77 | 1,61 | 0,73 | 2,19 | 2,82E-02 | 9,81E-02 |
| AT3G20770 | 2,68 | 3,01 | 1,37 | 2,19 | 2,84E-02 | 9,81E-02 |
| AT3G08165 | 3,95 | -3,38 | 1,54 | -2,20 | 2,81E-02 | 9,81E-02 |
| AT4G34890 | 2,69 | 3,65 | 1,66 | 2,19 | 2,82E-02 | 9,81E-02 |
| AT5G17710 | 1,85 | 3,35 | 1,53 | 2,19 | 2,83E-02 | 9,81E-02 |
| AT1G69792 | 2,23 | -3,97 | 1,81 | -2,19 | 2,84E-02 | 9,81E-02 |

| | | | | | | |
|---|---|---|---|---|---|---|
| AT2G07645 | 4,48 | -2,75 | 1,25 | -2,19 | 2,85E-02 | 9,83E-02 |
| AT1G27320 | 2,96 | 3,30 | 1,51 | 2,19 | 2,87E-02 | 9,85E-02 |
| AT2G38040 | 4,80 | 2,48 | 1,13 | 2,19 | 2,87E-02 | 9,85E-02 |
| AT3G04740 | 2,26 | 3,40 | 1,55 | 2,19 | 2,87E-02 | 9,85E-02 |
| AT1G16635 | 19,67 | -1,36 | 0,62 | -2,19 | 2,88E-02 | 9,85E-02 |
| AT4G21150 | 2,92 | 3,49 | 1,60 | 2,19 | 2,88E-02 | 9,85E-02 |
| AT4G32940 | 4,72 | 2,72 | 1,25 | 2,18 | 2,91E-02 | 9,92E-02 |
| AT3G50590 | 4,47 | 2,37 | 1,09 | 2,18 | 2,92E-02 | 9,96E-02 |
| AT5G02310 | 4,95 | 2,20 | 1,01 | 2,18 | 2,93E-02 | 9,96E-02 |
| AT3G44720 | 3,49 | 3,09 | 1,42 | 2,18 | 2,93E-02 | 9,97E-02 |
| AT3G50820 | 7,18 | -2,11 | 0,97 | -2,17 | 2,97E-02 | 1,01E-01 |
| AT1G18740 | 2,60 | 3,50 | 1,61 | 2,17 | 2,99E-02 | 1,01E-01 |
| AT2G46685 | 3232,16 | -0,57 | 0,26 | -2,17 | 2,98E-02 | 1,01E-01 |
| AT5G25610 | 6,58 | -1,74 | 0,80 | -2,17 | 2,98E-02 | 1,01E-01 |
| AT1G18260 | 1,96 | 3,71 | 1,71 | 2,17 | 3,02E-02 | 1,02E-01 |
| AT1G58230 | 3,69 | 3,30 | 1,52 | 2,17 | 3,03E-02 | 1,02E-01 |
| AT4G38775 | 2,66 | -3,17 | 1,47 | -2,17 | 3,04E-02 | 1,02E-01 |
| AT1G31730 | 1,97 | 3,93 | 1,82 | 2,16 | 3,08E-02 | 1,03E-01 |
| AT3G60630 | 5,65 | 2,18 | 1,01 | 2,16 | 3,08E-02 | 1,03E-01 |
| AT4G27060 | 1,71 | 3,78 | 1,76 | 2,15 | 3,13E-02 | 1,05E-01 |
| AT5G05690 | 2,09 | 3,42 | 1,59 | 2,15 | 3,16E-02 | 1,06E-01 |
| AT2G36350 | 2,13 | 3,91 | 1,82 | 2,15 | 3,17E-02 | 1,06E-01 |
| AT5G17760 | 1,99 | 4,11 | 1,91 | 2,15 | 3,17E-02 | 1,06E-01 |
| AT3G62200 | 2,36 | 3,57 | 1,66 | 2,15 | 3,18E-02 | 1,06E-01 |
| AT4G02350 | 1,95 | 3,84 | 1,79 | 2,14 | 3,21E-02 | 1,07E-01 |
| AT5G14930 | 1,80 | 4,66 | 2,18 | 2,14 | 3,22E-02 | 1,07E-01 |
| AT2G46460 | 2,59 | -3,80 | 1,77 | -2,14 | 3,25E-02 | 1,08E-01 |
| AT2G33770 | 4,11 | 3,11 | 1,46 | 2,14 | 3,25E-02 | 1,08E-01 |
| AT3G10815 | 5,65 | 3,28 | 1,53 | 2,14 | 3,26E-02 | 1,08E-01 |
| AT4G35750 | 1,89 | 3,56 | 1,67 | 2,14 | 3,27E-02 | 1,08E-01 |
| AT4G16490 | 2,72 | 4,00 | 1,87 | 2,13 | 3,28E-02 | 1,08E-01 |
| AT1G13320 | 2,17 | 3,24 | 1,52 | 2,13 | 3,31E-02 | 1,09E-01 |
| AT1G21580 | 3,07 | 3,21 | 1,51 | 2,13 | 3,31E-02 | 1,09E-01 |
| AT5G14740 | 2,79 | 2,58 | 1,21 | 2,13 | 3,32E-02 | 1,09E-01 |
| AT1G05247 | 19,59 | -1,34 | 0,63 | -2,13 | 3,34E-02 | 1,10E-01 |
| AT5G66200 | 3,05 | 3,26 | 1,53 | 2,13 | 3,36E-02 | 1,10E-01 |
| AT3G14010 | 2,74 | 3,61 | 1,70 | 2,12 | 3,37E-02 | 1,10E-01 |
| AT5G35210 | 2,88 | 3,01 | 1,42 | 2,12 | 3,37E-02 | 1,10E-01 |
| AT2G45940 | 2,82 | 3,28 | 1,55 | 2,12 | 3,37E-02 | 1,10E-01 |
| AT5G39740 | 3,11 | 2,51 | 1,18 | 2,12 | 3,38E-02 | 1,10E-01 |
| AT1G61520 | 19,44 | -1,16 | 0,55 | -2,12 | 3,39E-02 | 1,10E-01 |
| AT1G08450 | 3,29 | 3,38 | 1,59 | 2,12 | 3,40E-02 | 1,11E-01 |
| AT1G50200 | 4,92 | 2,27 | 1,07 | 2,12 | 3,42E-02 | 1,11E-01 |
| AT4G10800 | 6,52 | -2,06 | 0,97 | -2,11 | 3,45E-02 | 1,12E-01 |
| AT3G56680 | 2,66 | 2,97 | 1,41 | 2,11 | 3,49E-02 | 1,13E-01 |
| AT5G17165 | 3,90 | -3,24 | 1,53 | -2,11 | 3,49E-02 | 1,13E-01 |
| AT1G05630 | 1,84 | 3,72 | 1,77 | 2,11 | 3,51E-02 | 1,13E-01 |
| AT3G04130 | 2,12 | 4,26 | 2,02 | 2,11 | 3,51E-02 | 1,13E-01 |
| AT1G29470 | 3,69 | 2,49 | 1,18 | 2,11 | 3,52E-02 | 1,13E-01 |
| AT3G54920 | 2,20 | 3,23 | 1,54 | 2,10 | 3,56E-02 | 1,14E-01 |
| AT4G38510 | 2,20 | 3,37 | 1,60 | 2,10 | 3,56E-02 | 1,14E-01 |
| AT2G23348 | 3,88 | 2,49 | 1,18 | 2,10 | 3,57E-02 | 1,14E-01 |
| AT5G01980 | 1,71 | 3,80 | 1,82 | 2,09 | 3,63E-02 | 1,16E-01 |
| AT4G21640 | 1,99 | -3,32 | 1,59 | -2,09 | 3,64E-02 | 1,16E-01 |
| AT4G36040 | 3,49 | 2,64 | 1,27 | 2,09 | 3,67E-02 | 1,17E-01 |
| AT2G02950 | 6,29 | -2,07 | 0,99 | -2,09 | 3,69E-02 | 1,17E-01 |

| | | | | | | |
|---|---|---|---|---|---|---|
| AT2G21100 | 3,61 | -2,45 | 1,17 | -2,09 | 3,70E-02 | 1,18E-01 |
| AT3G52590 | 1,74 | 3,66 | 1,76 | 2,08 | 3,72E-02 | 1,18E-01 |
| AT4G08920 | 2,64 | 3,40 | 1,63 | 2,08 | 3,73E-02 | 1,18E-01 |
| AT3G26000 | 2,02 | 3,46 | 1,66 | 2,08 | 3,74E-02 | 1,18E-01 |
| AT2G15620 | 3,38 | 2,33 | 1,12 | 2,08 | 3,75E-02 | 1,18E-01 |
| AT2G39730 | 47,53 | 0,85 | 0,41 | 2,08 | 3,75E-02 | 1,18E-01 |
| AT1G66795 | 14,00 | 1,51 | 0,73 | 2,08 | 3,76E-02 | 1,19E-01 |
| AT2G46560 | 4,43 | 2,42 | 1,17 | 2,08 | 3,79E-02 | 1,19E-01 |
| AT5G15400 | 2,82 | 3,12 | 1,51 | 2,07 | 3,81E-02 | 1,20E-01 |
| AT2G22430 | 4,62 | 2,16 | 1,04 | 2,07 | 3,83E-02 | 1,20E-01 |
| AT1G69550 | 1,87 | 3,65 | 1,76 | 2,07 | 3,84E-02 | 1,20E-01 |
| AT5G62162 | 10,16 | 1,70 | 0,82 | 2,07 | 3,85E-02 | 1,20E-01 |
| AT4G05180 | 7,70 | -1,71 | 0,83 | -2,07 | 3,85E-02 | 1,20E-01 |
| AT1G67960 | 1,96 | 3,74 | 1,81 | 2,07 | 3,87E-02 | 1,21E-01 |
| AT1G56510 | 2,98 | 3,35 | 1,62 | 2,06 | 3,90E-02 | 1,21E-01 |
| AT3G03790 | 1,73 | 3,59 | 1,74 | 2,07 | 3,89E-02 | 1,21E-01 |
| AT5G02565 | 2,69 | -2,75 | 1,33 | -2,06 | 3,89E-02 | 1,21E-01 |
| AT1G61100 | 1,97 | 3,69 | 1,79 | 2,06 | 3,90E-02 | 1,21E-01 |
| AT4G35790 | 4,32 | 2,92 | 1,41 | 2,06 | 3,91E-02 | 1,21E-01 |
| AT1G04625 | 1,86 | 4,04 | 1,96 | 2,06 | 3,91E-02 | 1,21E-01 |
| AT4G31170 | 2,81 | 3,59 | 1,74 | 2,06 | 3,92E-02 | 1,21E-01 |
| AT4G02510 | 15,59 | 1,31 | 0,63 | 2,06 | 3,93E-02 | 1,22E-01 |
| AT2G14080 | 4,62 | 3,22 | 1,56 | 2,06 | 3,95E-02 | 1,22E-01 |
| AT1G13210 | 5,46 | 2,42 | 1,17 | 2,06 | 3,94E-02 | 1,22E-01 |
| AT3G61690 | 2,25 | 3,31 | 1,61 | 2,06 | 3,97E-02 | 1,22E-01 |
| AT3G58640 | 2,56 | 3,13 | 1,52 | 2,06 | 3,97E-02 | 1,22E-01 |
| AT3G13530 | 2,08 | 3,58 | 1,74 | 2,06 | 3,98E-02 | 1,22E-01 |
| AT2G46020 | 3,28 | 2,47 | 1,20 | 2,05 | 3,99E-02 | 1,22E-01 |
| AT1G51150 | 4,36 | -2,14 | 1,04 | -2,05 | 4,00E-02 | 1,22E-01 |
| AT5G15022 | 21,95 | -1,32 | 0,64 | -2,05 | 4,01E-02 | 1,23E-01 |
| AT1G63070 | 3,60 | -2,75 | 1,34 | -2,05 | 4,02E-02 | 1,23E-01 |
| AT4G37900 | 1,96 | 3,80 | 1,85 | 2,05 | 4,03E-02 | 1,23E-01 |
| AT1G27595 | 2,01 | 3,55 | 1,74 | 2,04 | 4,11E-02 | 1,25E-01 |
| AT5G10960 | 3,18 | 2,75 | 1,35 | 2,03 | 4,19E-02 | 1,27E-01 |
| AT5G47480 | 1,95 | 3,18 | 1,57 | 2,03 | 4,22E-02 | 1,28E-01 |
| AT3G17240 | 1,76 | 3,68 | 1,81 | 2,03 | 4,24E-02 | 1,29E-01 |
| AT3G16850 | 3,96 | 2,28 | 1,12 | 2,03 | 4,26E-02 | 1,29E-01 |
| AT1G01550 | 2,54 | 2,72 | 1,34 | 2,03 | 4,28E-02 | 1,30E-01 |
| AT2G44410 | 2,51 | 3,91 | 1,93 | 2,02 | 4,32E-02 | 1,30E-01 |
| AT3G19380 | 2,79 | 3,32 | 1,64 | 2,02 | 4,32E-02 | 1,30E-01 |
| AT5G19330 | 2,99 | 2,77 | 1,37 | 2,02 | 4,31E-02 | 1,30E-01 |
| AT1G79750 | 4,46 | 2,17 | 1,08 | 2,02 | 4,33E-02 | 1,30E-01 |
| AT2G03150 | 2,08 | 3,26 | 1,61 | 2,02 | 4,34E-02 | 1,30E-01 |
| AT3G18490 | 8,11 | -1,66 | 0,82 | -2,02 | 4,34E-02 | 1,30E-01 |
| AT3G23920 | 2,81 | 3,27 | 1,63 | 2,01 | 4,40E-02 | 1,32E-01 |
| AT5G04810 | 3,05 | 2,39 | 1,19 | 2,01 | 4,40E-02 | 1,32E-01 |
| AT5G62000 | 10,38 | 1,34 | 0,67 | 2,01 | 4,43E-02 | 1,33E-01 |
| AT2G42490 | 2,30 | 3,21 | 1,60 | 2,01 | 4,44E-02 | 1,33E-01 |
| AT1G61275 | 11,06 | -1,55 | 0,77 | -2,01 | 4,47E-02 | 1,33E-01 |
| AT5G58540 | 1,83 | 3,65 | 1,82 | 2,01 | 4,47E-02 | 1,33E-01 |
| AT4G13885 | 7,77 | -1,47 | 0,73 | -2,00 | 4,53E-02 | 1,35E-01 |
| AT2G47400 | 11,67 | 1,26 | 0,63 | 2,00 | 4,60E-02 | 1,37E-01 |
| AT5G09900 | 1,74 | 3,33 | 1,67 | 1,99 | 4,62E-02 | 1,37E-01 |
| AT4G10120 | 1,86 | 3,27 | 1,65 | 1,98 | 4,72E-02 | 1,39E-01 |
| AT4G32990 | 2,22 | -2,99 | 1,50 | -1,98 | 4,72E-02 | 1,39E-01 |
| AT2G47000 | 1,78 | 4,17 | 2,10 | 1,99 | 4,71E-02 | 1,39E-01 |

| | | | | | | |
|---|---|---|---|---|---|---|
| AT4G16720 | 3,07 | 2,61 | 1,32 | 1,99 | 4,71E-02 | 1,39E-01 |
| AT2G22300 | 2,59 | 3,11 | 1,57 | 1,98 | 4,76E-02 | 1,40E-01 |
| AT5G10540 | 2,27 | 3,22 | 1,63 | 1,98 | 4,77E-02 | 1,40E-01 |
| AT4G02970 | 72,42 | 0,66 | 0,33 | 1,98 | 4,80E-02 | 1,41E-01 |
| AT3G13070 | 1,98 | 3,66 | 1,85 | 1,98 | 4,80E-02 | 1,41E-01 |
| AT1G31330 | 6,03 | -1,59 | 0,81 | -1,98 | 4,82E-02 | 1,41E-01 |
| AT2G41840 | 3,13 | 2,32 | 1,18 | 1,97 | 4,84E-02 | 1,42E-01 |
| AT3G53740 | 2,01 | 3,86 | 1,95 | 1,97 | 4,84E-02 | 1,42E-01 |
| AT2G22500 | 4,35 | 2,63 | 1,33 | 1,97 | 4,85E-02 | 1,42E-01 |
| AT5G47490 | 2,21 | 3,11 | 1,58 | 1,97 | 4,87E-02 | 1,42E-01 |
| AT3G57330 | 3,24 | 2,69 | 1,36 | 1,97 | 4,87E-02 | 1,42E-01 |
| AT5G04415 | 4,71 | 1,91 | 0,97 | 1,97 | 4,89E-02 | 1,42E-01 |
| AT2G25320 | 3,21 | 3,07 | 1,56 | 1,97 | 4,89E-02 | 1,42E-01 |
| AT3G46740 | 3,41 | 2,24 | 1,14 | 1,97 | 4,88E-02 | 1,42E-01 |
| AT5G19770 | 1,75 | 3,61 | 1,83 | 1,97 | 4,92E-02 | 1,43E-01 |
| AT3G18217 | 147,37 | 0,48 | 0,25 | 1,96 | 4,97E-02 | 1,44E-01 |
| AT3G51550 | 4,70 | 1,84 | 0,94 | 1,96 | 5,01E-02 | 1,45E-01 |
| AT4G33300 | 3,93 | 2,51 | 1,28 | 1,96 | 5,05E-02 | 1,46E-01 |
| AT3G63070 | 1,86 | 3,89 | 1,99 | 1,95 | 5,06E-02 | 1,46E-01 |
| AT2G39800 | 6,12 | 1,80 | 0,92 | 1,95 | 5,12E-02 | 1,47E-01 |
| AT4G12800 | 10,28 | -1,48 | 0,76 | -1,95 | 5,15E-02 | 1,48E-01 |
| AT4G17410 | 2,55 | 3,51 | 1,80 | 1,95 | 5,15E-02 | 1,48E-01 |
| AT2G17840 | 2,90 | -3,08 | 1,58 | -1,95 | 5,16E-02 | 1,48E-01 |
| AT3G61198 | 7,65 | 2,17 | 1,11 | 1,95 | 5,17E-02 | 1,48E-01 |
| AT2G36160 | 1,81 | 3,29 | 1,70 | 1,94 | 5,20E-02 | 1,49E-01 |
| AT4G24680 | 3,35 | 2,37 | 1,22 | 1,94 | 5,21E-02 | 1,49E-01 |
| AT3G48000 | 4,79 | 1,94 | 1,00 | 1,94 | 5,22E-02 | 1,49E-01 |
| AT4G26255 | 3,93 | 2,50 | 1,29 | 1,94 | 5,26E-02 | 1,50E-01 |
| AT3G58050 | 1,93 | 3,13 | 1,62 | 1,94 | 5,29E-02 | 1,50E-01 |
| AT5G07735 | 1,90 | -3,17 | 1,64 | -1,94 | 5,28E-02 | 1,50E-01 |
| AT5G54130 | 6,22 | 1,66 | 0,86 | 1,94 | 5,29E-02 | 1,50E-01 |
| AT2G38280 | 1,89 | 3,39 | 1,75 | 1,94 | 5,29E-02 | 1,50E-01 |
| AT4G02760 | 2,62 | 2,88 | 1,49 | 1,93 | 5,31E-02 | 1,50E-01 |
| AT5G64840 | 5,36 | 2,24 | 1,16 | 1,93 | 5,40E-02 | 1,53E-01 |
| AT1G01960 | 2,74 | 2,90 | 1,51 | 1,93 | 5,40E-02 | 1,53E-01 |
| AT2G03330 | 8,86 | -1,48 | 0,77 | -1,92 | 5,46E-02 | 1,54E-01 |
| AT4G18880 | 2,25 | 3,68 | 1,91 | 1,92 | 5,48E-02 | 1,55E-01 |
| AT1G09640 | 2,66 | 3,07 | 1,60 | 1,92 | 5,53E-02 | 1,55E-01 |
| AT3G07765 | 2,13 | -2,91 | 1,52 | -1,92 | 5,54E-02 | 1,55E-01 |
| AT1G07135 | 2,46 | 3,22 | 1,68 | 1,92 | 5,52E-02 | 1,55E-01 |
| AT3G55020 | 1,94 | 3,11 | 1,62 | 1,92 | 5,53E-02 | 1,55E-01 |
| AT5G64740 | 4,40 | 1,77 | 0,92 | 1,91 | 5,57E-02 | 1,56E-01 |
| AT5G47910 | 5,04 | 2,14 | 1,12 | 1,91 | 5,58E-02 | 1,56E-01 |
| AT5G10180 | 2,74 | 3,13 | 1,64 | 1,91 | 5,63E-02 | 1,57E-01 |
| AT2G42540 | 2,58 | -3,08 | 1,62 | -1,91 | 5,64E-02 | 1,58E-01 |
| AT3G02540 | 1,81 | 3,17 | 1,66 | 1,91 | 5,64E-02 | 1,58E-01 |
| AT1G62390 | 2,10 | 3,85 | 2,02 | 1,90 | 5,73E-02 | 1,59E-01 |
| AT1G67740 | 3,89 | -1,91 | 1,00 | -1,90 | 5,72E-02 | 1,59E-01 |
| AT2G07671 | 2,57 | -2,57 | 1,35 | -1,90 | 5,72E-02 | 1,59E-01 |
| AT3G14650 | 2,33 | 3,02 | 1,59 | 1,90 | 5,75E-02 | 1,60E-01 |
| AT2G25900 | 2,31 | -3,00 | 1,58 | -1,90 | 5,79E-02 | 1,61E-01 |
| AT1G61140 | 2,12 | 2,89 | 1,53 | 1,89 | 5,84E-02 | 1,62E-01 |
| AT4G33050 | 2,86 | 2,89 | 1,53 | 1,89 | 5,87E-02 | 1,63E-01 |
| AT5G30510 | 1,83 | -3,44 | 1,82 | -1,89 | 5,90E-02 | 1,63E-01 |
| AT4G34071 | 2,35 | -3,12 | 1,65 | -1,89 | 5,92E-02 | 1,63E-01 |
| AT2G40930 | 1,74 | 3,45 | 1,83 | 1,89 | 5,92E-02 | 1,63E-01 |

| | | | | | | |
|---|---|---|---|---|---|---|
| AT1G05240 | 2,51 | -2,83 | 1,50 | -1,88 | 5,96E-02 | 1,64E-01 |
| AT4G32760 | 1,73 | 3,27 | 1,74 | 1,88 | 5,97E-02 | 1,64E-01 |
| AT5G45510 | 8,42 | 2,21 | 1,18 | 1,88 | 5,98E-02 | 1,64E-01 |
| AT3G53260 | 2,56 | 3,08 | 1,64 | 1,88 | 5,98E-02 | 1,64E-01 |
| AT5G03040 | 2,14 | 2,67 | 1,42 | 1,88 | 6,00E-02 | 1,64E-01 |
| AT1G13930 | 7,00 | -1,50 | 0,80 | -1,88 | 6,03E-02 | 1,65E-01 |
| AT2G17440 | 1,80 | 3,42 | 1,82 | 1,88 | 6,03E-02 | 1,65E-01 |
| AT5G22500 | 2,19 | -3,07 | 1,63 | -1,88 | 6,03E-02 | 1,65E-01 |
| AT1G09570 | 2,05 | 2,68 | 1,43 | 1,88 | 6,06E-02 | 1,65E-01 |
| AT1G43850 | 1,82 | 3,38 | 1,80 | 1,88 | 6,07E-02 | 1,65E-01 |
| AT5G60410 | 2,90 | 2,37 | 1,26 | 1,87 | 6,10E-02 | 1,66E-01 |
| AT3G63410 | 2,71 | 2,45 | 1,31 | 1,87 | 6,10E-02 | 1,66E-01 |
| AT5G23630 | 2,35 | 2,79 | 1,49 | 1,87 | 6,09E-02 | 1,66E-01 |
| AT3G06720 | 1,89 | 3,42 | 1,83 | 1,87 | 6,13E-02 | 1,66E-01 |
| AT4G03030 | 1,74 | 3,44 | 1,84 | 1,87 | 6,13E-02 | 1,66E-01 |
| AT2G47390 | 2,13 | 3,44 | 1,84 | 1,87 | 6,16E-02 | 1,67E-01 |
| AT5G26830 | 2,03 | 2,96 | 1,59 | 1,87 | 6,20E-02 | 1,67E-01 |
| AT1G64583 | 1,87 | -3,01 | 1,61 | -1,87 | 6,20E-02 | 1,67E-01 |
| AT3G13525 | 2,38 | -2,81 | 1,51 | -1,86 | 6,24E-02 | 1,68E-01 |
| AT3G03175 | 7,33 | 1,62 | 0,87 | 1,86 | 6,24E-02 | 1,68E-01 |
| AT5G18360 | 2,96 | 2,63 | 1,41 | 1,86 | 6,25E-02 | 1,68E-01 |
| AT5G46260 | 1,90 | 3,20 | 1,71 | 1,86 | 6,24E-02 | 1,68E-01 |
| AT2G26355 | 2,39 | 3,13 | 1,68 | 1,86 | 6,28E-02 | 1,68E-01 |
| AT5G04720 | 2,75 | 3,51 | 1,89 | 1,86 | 6,29E-02 | 1,68E-01 |
| AT1G07000 | 3,04 | 2,60 | 1,40 | 1,86 | 6,29E-02 | 1,68E-01 |
| AT3G44480 | 4,70 | 1,81 | 0,97 | 1,86 | 6,28E-02 | 1,68E-01 |
| AT5G18380 | 4,02 | 1,69 | 0,91 | 1,86 | 6,29E-02 | 1,68E-01 |
| AT2G15970 | 3,79 | -2,12 | 1,14 | -1,86 | 6,32E-02 | 1,69E-01 |
| AT4G34390 | 1,80 | 3,37 | 1,82 | 1,86 | 6,35E-02 | 1,69E-01 |
| AT2G39770 | 2,97 | 2,59 | 1,40 | 1,85 | 6,38E-02 | 1,70E-01 |
| AT1G55670 | 5,12 | -1,55 | 0,84 | -1,85 | 6,38E-02 | 1,70E-01 |
| AT3G04930 | 1,93 | 3,15 | 1,70 | 1,85 | 6,43E-02 | 1,70E-01 |
| AT4G23420 | 23,63 | 0,90 | 0,49 | 1,85 | 6,43E-02 | 1,70E-01 |
| AT1G20340 | 11,85 | -1,09 | 0,59 | -1,85 | 6,41E-02 | 1,70E-01 |
| AT1G51570 | 1,72 | 3,16 | 1,71 | 1,85 | 6,42E-02 | 1,70E-01 |
| AT2G10940 | 5,77 | -1,96 | 1,06 | -1,85 | 6,41E-02 | 1,70E-01 |
| AT5G04550 | 2,33 | 2,95 | 1,59 | 1,85 | 6,43E-02 | 1,70E-01 |
| AT5G04315 | 6,08 | 1,57 | 0,85 | 1,85 | 6,45E-02 | 1,70E-01 |
| AT5G04325 | 6,08 | 1,57 | 0,85 | 1,85 | 6,45E-02 | 1,70E-01 |
| AT5G20020 | 2,46 | 3,60 | 1,95 | 1,85 | 6,48E-02 | 1,71E-01 |
| AT2G47800 | 2,70 | 2,63 | 1,43 | 1,84 | 6,53E-02 | 1,71E-01 |
| AT1G04897 | 2,68 | 3,14 | 1,70 | 1,84 | 6,53E-02 | 1,71E-01 |
| AT5G27950 | 2,87 | -2,53 | 1,38 | -1,84 | 6,58E-02 | 1,73E-01 |
| AT5G53480 | 6,73 | 1,80 | 0,98 | 1,84 | 6,64E-02 | 1,74E-01 |
| AT1G07200 | 1,82 | 3,29 | 1,80 | 1,83 | 6,70E-02 | 1,75E-01 |
| AT3G51800 | 4,28 | -2,17 | 1,18 | -1,83 | 6,70E-02 | 1,75E-01 |
| AT5G53170 | 2,82 | 2,51 | 1,37 | 1,83 | 6,70E-02 | 1,75E-01 |
| AT1G21750 | 2,14 | 3,56 | 1,94 | 1,83 | 6,72E-02 | 1,75E-01 |
| AT4G16370 | 3,80 | 1,92 | 1,05 | 1,83 | 6,73E-02 | 1,75E-01 |
| AT2G06520 | 10,63 | -1,46 | 0,80 | -1,82 | 6,81E-02 | 1,77E-01 |
| AT3G56150 | 2,36 | 3,04 | 1,67 | 1,82 | 6,85E-02 | 1,78E-01 |
| AT2G45180 | 2,18 | -2,97 | 1,63 | -1,82 | 6,88E-02 | 1,79E-01 |
| AT1G29350 | 2,20 | 2,94 | 1,62 | 1,82 | 6,91E-02 | 1,79E-01 |
| AT4G34830 | 2,44 | 2,67 | 1,47 | 1,82 | 6,93E-02 | 1,80E-01 |
| AT2G41440 | 2,60 | 2,88 | 1,59 | 1,81 | 6,96E-02 | 1,80E-01 |
| AT3G11830 | 1,90 | 3,06 | 1,69 | 1,81 | 6,96E-02 | 1,80E-01 |

| | | | | | | |
|---|---|---|---|---|---|---|
| AT4G02130 | 3,77 | 2,04 | 1,12 | 1,81 | 6,98E-02 | 1,80E-01 |
| AT5G16640 | 4,15 | -2,32 | 1,29 | -1,80 | 7,12E-02 | 1,84E-01 |
| AT2G02835 | 3,94 | 2,17 | 1,20 | 1,80 | 7,14E-02 | 1,84E-01 |
| AT4G03039 | 93,29 | 2,44 | 1,36 | 1,80 | 7,16E-02 | 1,84E-01 |
| AT4G03038 | 93,33 | 2,44 | 1,36 | 1,80 | 7,17E-02 | 1,84E-01 |
| AT4G04365 | 93,18 | 2,44 | 1,36 | 1,80 | 7,17E-02 | 1,84E-01 |
| AT4G38220 | 1,96 | 2,96 | 1,65 | 1,80 | 7,20E-02 | 1,85E-01 |
| AT1G20510 | 2,74 | 3,19 | 1,78 | 1,80 | 7,24E-02 | 1,85E-01 |
| AT1G66783 | 14,56 | 1,42 | 0,79 | 1,80 | 7,24E-02 | 1,85E-01 |
| AT4G26090 | 1,90 | 2,89 | 1,61 | 1,80 | 7,25E-02 | 1,85E-01 |
| AT4G30960 | 5,13 | 2,62 | 1,46 | 1,79 | 7,28E-02 | 1,85E-01 |
| AT4G16860 | 6,11 | 1,91 | 1,07 | 1,79 | 7,28E-02 | 1,85E-01 |
| AT4G26095 | 1,81 | 2,91 | 1,62 | 1,79 | 7,28E-02 | 1,85E-01 |
| AT1G24460 | 4,84 | 1,97 | 1,10 | 1,79 | 7,30E-02 | 1,86E-01 |
| AT2G05710 | 3,35 | 2,51 | 1,40 | 1,79 | 7,36E-02 | 1,86E-01 |
| AT4G38200 | 2,17 | 2,60 | 1,45 | 1,79 | 7,35E-02 | 1,86E-01 |
| AT1G73170 | 1,76 | 3,53 | 1,97 | 1,79 | 7,34E-02 | 1,86E-01 |
| AT4G06320 | 7,79 | -1,34 | 0,75 | -1,79 | 7,33E-02 | 1,86E-01 |
| AT5G11700 | 1,99 | 2,99 | 1,67 | 1,79 | 7,35E-02 | 1,86E-01 |
| AT2G39940 | 1,84 | 3,58 | 2,00 | 1,79 | 7,37E-02 | 1,86E-01 |
| AT2G27720 | 1,70 | 2,88 | 1,61 | 1,79 | 7,42E-02 | 1,87E-01 |
| AT2G36490 | 1,90 | 4,74 | 2,66 | 1,78 | 7,43E-02 | 1,87E-01 |
| AT4G31180 | 4,29 | 1,76 | 0,99 | 1,78 | 7,45E-02 | 1,87E-01 |
| AT4G08125 | 3,11 | -2,33 | 1,31 | -1,78 | 7,44E-02 | 1,87E-01 |
| AT3G04455 | 2,46 | -3,05 | 1,71 | -1,78 | 7,46E-02 | 1,87E-01 |
| AT5G14040 | 2,00 | 2,76 | 1,56 | 1,78 | 7,58E-02 | 1,90E-01 |
| AT1G36180 | 2,20 | 2,73 | 1,54 | 1,77 | 7,59E-02 | 1,91E-01 |
| AT2G23347 | 3,51 | 2,13 | 1,20 | 1,77 | 7,60E-02 | 1,91E-01 |
| AT3G50930 | 3,98 | 2,37 | 1,34 | 1,77 | 7,65E-02 | 1,91E-01 |
| AT2G18790 | 2,11 | 2,92 | 1,65 | 1,77 | 7,66E-02 | 1,92E-01 |
| AT5G39570 | 4,04 | -1,89 | 1,07 | -1,77 | 7,68E-02 | 1,92E-01 |
| AT4G36760 | 3,13 | 2,21 | 1,25 | 1,77 | 7,72E-02 | 1,93E-01 |
| AT4G00700 | 2,06 | 3,22 | 1,82 | 1,76 | 7,79E-02 | 1,94E-01 |
| AT4G03415 | 1,93 | 3,28 | 1,86 | 1,76 | 7,80E-02 | 1,94E-01 |
| AT5G26360 | 2,70 | 2,73 | 1,55 | 1,76 | 7,82E-02 | 1,94E-01 |
| AT1G15130 | 4,06 | 2,10 | 1,20 | 1,76 | 7,84E-02 | 1,95E-01 |
| AT5G01920 | 2,56 | 2,54 | 1,44 | 1,76 | 7,86E-02 | 1,95E-01 |
| AT5G20230 | 2,06 | 3,19 | 1,81 | 1,76 | 7,89E-02 | 1,96E-01 |
| AT3G14210 | 12,80 | -0,98 | 0,56 | -1,76 | 7,91E-02 | 1,96E-01 |
| AT3G06345 | 8,24 | 1,48 | 0,84 | 1,75 | 7,94E-02 | 1,96E-01 |
| AT4G26110 | 2,20 | 2,87 | 1,64 | 1,75 | 7,94E-02 | 1,96E-01 |
| AT1G08327 | 2,78 | 2,52 | 1,44 | 1,75 | 7,93E-02 | 1,96E-01 |
| AT2G31890 | 2,01 | 3,13 | 1,80 | 1,74 | 8,14E-02 | 2,01E-01 |
| AT4G27430 | 2,23 | 2,64 | 1,52 | 1,74 | 8,16E-02 | 2,01E-01 |
| AT3G13090 | 2,27 | 3,70 | 2,13 | 1,74 | 8,25E-02 | 2,03E-01 |
| AT5G67030 | 1,94 | 2,81 | 1,62 | 1,73 | 8,33E-02 | 2,05E-01 |
| AT1G54040 | 2,21 | -2,95 | 1,71 | -1,73 | 8,35E-02 | 2,05E-01 |
| AT4G32285 | 2,40 | 2,78 | 1,61 | 1,73 | 8,38E-02 | 2,05E-01 |
| AT1G29940 | 2,05 | 2,98 | 1,72 | 1,73 | 8,37E-02 | 2,05E-01 |
| AT4G02770 | 8,37 | -1,26 | 0,73 | -1,73 | 8,41E-02 | 2,06E-01 |
| AT4G34980 | 4,56 | 1,69 | 0,98 | 1,73 | 8,43E-02 | 2,06E-01 |
| AT4G15470 | 1,75 | 2,85 | 1,65 | 1,73 | 8,44E-02 | 2,06E-01 |
| AT2G26500 | 2,52 | 2,50 | 1,45 | 1,72 | 8,51E-02 | 2,07E-01 |
| AT3G09465 | 2,48 | -2,32 | 1,35 | -1,72 | 8,51E-02 | 2,07E-01 |
| AT2G45685 | 1,79 | 3,21 | 1,87 | 1,72 | 8,53E-02 | 2,08E-01 |
| AT2G45680 | 1,79 | 3,21 | 1,87 | 1,72 | 8,53E-02 | 2,08E-01 |

| | | | | | | |
|---|---|---|---|---|---|---|
| AT3G03425 | 4,95 | -2,17 | 1,26 | -1,72 | 8,55E-02 | 2,08E-01 |
| AT5G05660 | 1,92 | 3,90 | 2,27 | 1,71 | 8,65E-02 | 2,10E-01 |
| AT1G06230 | 2,14 | 2,67 | 1,56 | 1,71 | 8,67E-02 | 2,10E-01 |
| AT5G49615 | 2,21 | -3,26 | 1,90 | -1,71 | 8,70E-02 | 2,11E-01 |
| AT3G01500 | 4,00 | -1,95 | 1,14 | -1,71 | 8,71E-02 | 2,11E-01 |
| AT2G45960 | 8,71 | 1,41 | 0,83 | 1,71 | 8,74E-02 | 2,11E-01 |
| AT1G73805 | 1,81 | 3,19 | 1,87 | 1,71 | 8,77E-02 | 2,12E-01 |
| AT5G28560 | 3,51 | -2,66 | 1,56 | -1,71 | 8,81E-02 | 2,12E-01 |
| AT1G54115 | 2,18 | 2,95 | 1,73 | 1,70 | 8,85E-02 | 2,13E-01 |
| AT5G07290 | 1,99 | 3,77 | 2,21 | 1,70 | 8,86E-02 | 2,13E-01 |
| AT1G53655 | 1,78 | -3,15 | 1,85 | -1,70 | 8,87E-02 | 2,13E-01 |
| AT5G61020 | 2,23 | 2,53 | 1,49 | 1,70 | 8,93E-02 | 2,14E-01 |
| AT4G30210 | 1,86 | 3,33 | 1,97 | 1,69 | 9,05E-02 | 2,17E-01 |
| AT5G42390 | 8,64 | 1,38 | 0,81 | 1,69 | 9,08E-02 | 2,18E-01 |
| AT3G29360 | 2,83 | 2,34 | 1,38 | 1,69 | 9,10E-02 | 2,18E-01 |
| AT2G46430 | 2,08 | 2,77 | 1,64 | 1,69 | 9,15E-02 | 2,19E-01 |
| AT1G48920 | 4,08 | 1,49 | 0,88 | 1,69 | 9,20E-02 | 2,20E-01 |
| AT1G60140 | 2,14 | 2,30 | 1,37 | 1,69 | 9,20E-02 | 2,20E-01 |
| AT3G47348 | 6,89 | -2,28 | 1,35 | -1,68 | 9,22E-02 | 2,20E-01 |
| AT1G54410 | 19,25 | -1,06 | 0,63 | -1,68 | 9,28E-02 | 2,21E-01 |
| AT3G24929 | 2,38 | -2,80 | 1,67 | -1,68 | 9,39E-02 | 2,23E-01 |
| AT3G47347 | 15,63 | -0,97 | 0,58 | -1,68 | 9,38E-02 | 2,23E-01 |
| AT4G24570 | 3,86 | -2,45 | 1,46 | -1,68 | 9,37E-02 | 2,23E-01 |
| AT4G39200 | 1,73 | 2,88 | 1,72 | 1,68 | 9,38E-02 | 2,23E-01 |
| AT3G18830 | 2,46 | 2,86 | 1,71 | 1,67 | 9,40E-02 | 2,23E-01 |
| AT1G17470 | 1,97 | 2,78 | 1,66 | 1,67 | 9,41E-02 | 2,23E-01 |
| AT1G15940 | 2,10 | 2,74 | 1,64 | 1,67 | 9,47E-02 | 2,24E-01 |
| AT1G31580 | 10,89 | -1,10 | 0,66 | -1,67 | 9,48E-02 | 2,24E-01 |
| AT1G78206 | 14,29 | 0,91 | 0,55 | 1,67 | 9,51E-02 | 2,24E-01 |
| AT3G17400 | 6,94 | -1,53 | 0,91 | -1,67 | 9,51E-02 | 2,24E-01 |
| AT5G28740 | 1,92 | 2,53 | 1,52 | 1,67 | 9,51E-02 | 2,24E-01 |
| AT2G34655 | 4,12 | -2,25 | 1,35 | -1,66 | 9,62E-02 | 2,26E-01 |
| AT2G38325 | 156,77 | -0,41 | 0,24 | -1,66 | 9,62E-02 | 2,26E-01 |
| AT5G64040 | 3,44 | -1,81 | 1,09 | -1,66 | 9,61E-02 | 2,26E-01 |
| AT2G28910 | 4,89 | 2,26 | 1,36 | 1,66 | 9,66E-02 | 2,27E-01 |
| AT5G05485 | 5,35 | -1,95 | 1,17 | -1,66 | 9,67E-02 | 2,27E-01 |
| AT5G53370 | 2,35 | 2,72 | 1,63 | 1,66 | 9,68E-02 | 2,27E-01 |
| AT2G16030 | 25,88 | -1,95 | 1,18 | -1,66 | 9,73E-02 | 2,27E-01 |
| AT1G78020 | 1,89 | 2,66 | 1,61 | 1,65 | 9,82E-02 | 2,29E-01 |
| AT5G23065 | 1776,68 | 0,43 | 0,26 | 1,65 | 9,82E-02 | 2,29E-01 |
| AT5G65610 | 2,72 | -2,72 | 1,65 | -1,65 | 9,86E-02 | 2,30E-01 |
| AT3G08850 | 2,35 | 2,66 | 1,61 | 1,65 | 9,86E-02 | 2,30E-01 |
| AT4G33010 | 11,10 | 1,15 | 0,70 | 1,65 | 9,90E-02 | 2,30E-01 |
| AT5G45430 | 3,88 | 1,98 | 1,20 | 1,65 | 1,00E-01 | 2,32E-01 |
| AT3G24982 | 2,19 | -2,77 | 1,68 | -1,65 | 9,99E-02 | 2,32E-01 |
| AT5G51300 | 1,97 | 3,72 | 2,26 | 1,64 | 1,00E-01 | 2,33E-01 |
| AT1G32720 | 2,47 | -2,69 | 1,64 | -1,64 | 1,01E-01 | 2,34E-01 |
| AT2G27150 | 1,72 | 3,47 | 2,11 | 1,64 | 1,01E-01 | 2,34E-01 |
| AT5G48385 | 2,37 | 2,71 | 1,65 | 1,64 | 1,01E-01 | 2,34E-01 |
| AT3G19170 | 4,99 | -2,34 | 1,43 | -1,64 | 1,01E-01 | 2,34E-01 |
| AT2G41105 | 5,08 | -2,14 | 1,31 | -1,64 | 1,01E-01 | 2,34E-01 |
| AT3G58610 | 2,68 | 2,04 | 1,25 | 1,64 | 1,02E-01 | 2,35E-01 |
| AT1G31540 | 3,00 | 2,48 | 1,52 | 1,63 | 1,03E-01 | 2,36E-01 |
| AT4G03295 | 3,55 | -1,90 | 1,16 | -1,63 | 1,03E-01 | 2,36E-01 |
| AT5G07325 | 8,05 | 1,25 | 0,77 | 1,63 | 1,03E-01 | 2,36E-01 |
| AT3G44300 | 1,82 | 2,85 | 1,75 | 1,63 | 1,04E-01 | 2,37E-01 |

| | | | | | | |
|---|---|---|---|---|---|---|
| AT3G58196 | 21,50 | -0,94 | 0,58 | -1,63 | 1,04E-01 | 2,37E-01 |
| AT3G11010 | 3,23 | 2,91 | 1,79 | 1,63 | 1,04E-01 | 2,37E-01 |
| AT3G19030 | 2,04 | 2,59 | 1,59 | 1,63 | 1,04E-01 | 2,37E-01 |
| AT3G60320 | 3,74 | 1,60 | 0,98 | 1,63 | 1,03E-01 | 2,37E-01 |
| AT3G62420 | 1,96 | 3,20 | 1,96 | 1,63 | 1,04E-01 | 2,37E-01 |
| AT4G32260 | 3,86 | -1,87 | 1,15 | -1,63 | 1,04E-01 | 2,37E-01 |
| AT5G66190 | 3,89 | -1,82 | 1,12 | -1,62 | 1,05E-01 | 2,39E-01 |
| AT1G15125 | 1,84 | 2,76 | 1,70 | 1,62 | 1,05E-01 | 2,40E-01 |
| AT2G31610 | 1,80 | 2,80 | 1,73 | 1,62 | 1,06E-01 | 2,40E-01 |
| AT3G44430 | 1,70 | 2,73 | 1,69 | 1,62 | 1,06E-01 | 2,40E-01 |
| AT3G49120 | 5,10 | 2,12 | 1,31 | 1,62 | 1,06E-01 | 2,40E-01 |
| AT5G13770 | 2,99 | 2,71 | 1,68 | 1,62 | 1,06E-01 | 2,40E-01 |
| AT5G42050 | 3,62 | 2,07 | 1,28 | 1,62 | 1,06E-01 | 2,40E-01 |
| AT5G42270 | 4,20 | -2,12 | 1,31 | -1,62 | 1,05E-01 | 2,40E-01 |
| AT3G47250 | 1,84 | 2,68 | 1,65 | 1,62 | 1,06E-01 | 2,40E-01 |
| AT5G46210 | 3,09 | 2,03 | 1,26 | 1,62 | 1,06E-01 | 2,40E-01 |
| AT3G44570 | 48,95 | -0,62 | 0,39 | -1,61 | 1,07E-01 | 2,42E-01 |
| AT1G79690 | 1,96 | 2,84 | 1,77 | 1,61 | 1,08E-01 | 2,42E-01 |
| AT5G66570 | 20,06 | -0,97 | 0,61 | -1,61 | 1,08E-01 | 2,42E-01 |
| AT2G36880 | 2,29 | 2,36 | 1,46 | 1,61 | 1,08E-01 | 2,42E-01 |
| AT3G22910 | 2,19 | 2,65 | 1,64 | 1,61 | 1,08E-01 | 2,42E-01 |
| AT3G08210 | 3,75 | -1,96 | 1,22 | -1,61 | 1,08E-01 | 2,42E-01 |
| AT5G27660 | 6,15 | 1,41 | 0,88 | 1,61 | 1,08E-01 | 2,42E-01 |
| AT1G26975 | 8,83 | 1,08 | 0,67 | 1,61 | 1,08E-01 | 2,43E-01 |
| AT3G53400 | 1,89 | 3,00 | 1,87 | 1,60 | 1,09E-01 | 2,43E-01 |
| AT5G40910 | 2,64 | 2,21 | 1,38 | 1,60 | 1,09E-01 | 2,44E-01 |
| AT2G16586 | 20,98 | 1,18 | 0,74 | 1,60 | 1,09E-01 | 2,45E-01 |
| AT1G04080 | 1,89 | 2,39 | 1,49 | 1,60 | 1,10E-01 | 2,45E-01 |
| AT4G12570 | 2,25 | 2,74 | 1,71 | 1,60 | 1,10E-01 | 2,45E-01 |
| AT5G15970 | 2,85 | -2,26 | 1,41 | -1,60 | 1,10E-01 | 2,45E-01 |
| AT5G08530 | 3,76 | -2,07 | 1,30 | -1,60 | 1,10E-01 | 2,45E-01 |
| AT5G16270 | 1,71 | 2,68 | 1,68 | 1,59 | 1,11E-01 | 2,47E-01 |
| AT5G58550 | 1,86 | 3,14 | 1,97 | 1,59 | 1,11E-01 | 2,48E-01 |
| AT2G25520 | 1,73 | 3,30 | 2,08 | 1,59 | 1,12E-01 | 2,49E-01 |
| AT1G08115 | 40,91 | 0,69 | 0,43 | 1,59 | 1,12E-01 | 2,49E-01 |
| AT5G15650 | 3,70 | 1,70 | 1,07 | 1,59 | 1,12E-01 | 2,49E-01 |
| AT5G06175 | 3,94 | -1,85 | 1,17 | -1,59 | 1,13E-01 | 2,49E-01 |
| AT2G03821 | 1,77 | -2,67 | 1,69 | -1,58 | 1,13E-01 | 2,51E-01 |
| AT4G21366 | 2,02 | 2,82 | 1,78 | 1,58 | 1,14E-01 | 2,51E-01 |
| AT1G63230 | 3,31 | -1,83 | 1,16 | -1,58 | 1,14E-01 | 2,52E-01 |
| AT1G71880 | 1,96 | 3,01 | 1,91 | 1,58 | 1,14E-01 | 2,52E-01 |
| AT2G34208 | 2,28 | 2,67 | 1,69 | 1,58 | 1,14E-01 | 2,52E-01 |
| AT4G15180 | 1,84 | 2,65 | 1,68 | 1,58 | 1,15E-01 | 2,52E-01 |
| AT5G11740 | 4,10 | 1,73 | 1,09 | 1,58 | 1,15E-01 | 2,52E-01 |
| AT1G05993 | 11,40 | -1,15 | 0,73 | -1,58 | 1,14E-01 | 2,52E-01 |
| AT2G20010 | 1,89 | 2,42 | 1,54 | 1,58 | 1,15E-01 | 2,52E-01 |
| AT5G54810 | 4,59 | 1,53 | 0,97 | 1,58 | 1,15E-01 | 2,52E-01 |
| AT3G48050 | 3,14 | 1,93 | 1,22 | 1,58 | 1,15E-01 | 2,52E-01 |
| AT1G30490 | 1,77 | 2,97 | 1,89 | 1,57 | 1,16E-01 | 2,54E-01 |
| AT5G28040 | 2,94 | 1,91 | 1,22 | 1,57 | 1,17E-01 | 2,55E-01 |
| AT2G24270 | 3,15 | 2,04 | 1,30 | 1,57 | 1,18E-01 | 2,57E-01 |
| AT5G17230 | 2,13 | 2,18 | 1,39 | 1,57 | 1,17E-01 | 2,57E-01 |
| AT1G02080 | 3,21 | 2,25 | 1,44 | 1,56 | 1,18E-01 | 2,58E-01 |
| AT2G39681 | 1805,59 | -0,51 | 0,32 | -1,56 | 1,19E-01 | 2,59E-01 |
| AT3G07160 | 3,87 | 1,97 | 1,26 | 1,56 | 1,19E-01 | 2,59E-01 |
| AT1G50010 | 2,93 | 1,88 | 1,20 | 1,56 | 1,19E-01 | 2,59E-01 |

| | | | | | | |
|---|---|---|---|---|---|---|
| AT3G29320 | 2,29 | -2,13 | 1,37 | -1,56 | 1,20E-01 | 2,60E-01 |
| AT3G48430 | 1,70 | 2,96 | 1,90 | 1,56 | 1,19E-01 | 2,60E-01 |
| AT2G20410 | 6,12 | -1,38 | 0,89 | -1,55 | 1,20E-01 | 2,60E-01 |
| AT3G13724 | 5,13 | -1,47 | 0,94 | -1,55 | 1,20E-01 | 2,60E-01 |
| AT5G57110 | 2,43 | 2,19 | 1,41 | 1,55 | 1,20E-01 | 2,60E-01 |
| AT3G08740 | 1,74 | 2,50 | 1,61 | 1,55 | 1,21E-01 | 2,61E-01 |
| AT1G52360 | 2,02 | 2,23 | 1,44 | 1,55 | 1,21E-01 | 2,62E-01 |
| AT4G29060 | 2,00 | -2,78 | 1,79 | -1,55 | 1,21E-01 | 2,62E-01 |
| AT5G60450 | 7,78 | -1,19 | 0,77 | -1,55 | 1,22E-01 | 2,62E-01 |
| AT1G08623 | 2,21 | -2,19 | 1,41 | -1,55 | 1,22E-01 | 2,62E-01 |
| AT3G57720 | 13,95 | 1,08 | 0,70 | 1,55 | 1,22E-01 | 2,62E-01 |
| AT1G31163 | 6,82 | -1,16 | 0,75 | -1,55 | 1,22E-01 | 2,62E-01 |
| AT2G03140 | 2,19 | 2,44 | 1,58 | 1,55 | 1,22E-01 | 2,63E-01 |
| AT1G60870 | 2,59 | 2,49 | 1,61 | 1,54 | 1,23E-01 | 2,63E-01 |
| AT2G41100 | 6,69 | -1,73 | 1,12 | -1,54 | 1,23E-01 | 2,63E-01 |
| AT5G63800 | 1,87 | -2,44 | 1,58 | -1,54 | 1,23E-01 | 2,63E-01 |
| AT5G48860 | 3,76 | -1,85 | 1,20 | -1,54 | 1,23E-01 | 2,64E-01 |
| AT1G13110 | 2,32 | -2,35 | 1,52 | -1,54 | 1,24E-01 | 2,64E-01 |
| AT3G27865 | 3,38 | -2,37 | 1,54 | -1,54 | 1,24E-01 | 2,65E-01 |
| AT3G04120 | 5,88 | 1,23 | 0,80 | 1,54 | 1,24E-01 | 2,66E-01 |
| AT3G51860 | 2,20 | 2,57 | 1,67 | 1,54 | 1,25E-01 | 2,66E-01 |
| AT5G04540 | 1,96 | 2,38 | 1,55 | 1,53 | 1,25E-01 | 2,67E-01 |
| AT5G09495 | 4,86 | -1,42 | 0,93 | -1,53 | 1,25E-01 | 2,67E-01 |
| AT5G65860 | 5,04 | -1,42 | 0,93 | -1,53 | 1,26E-01 | 2,68E-01 |
| AT4G36360 | 3,63 | 1,94 | 1,27 | 1,53 | 1,26E-01 | 2,68E-01 |
| AT3G50950 | 4,46 | 1,59 | 1,04 | 1,53 | 1,26E-01 | 2,68E-01 |
| AT2G44560 | 4,56 | -1,41 | 0,93 | -1,53 | 1,27E-01 | 2,69E-01 |
| AT5G04590 | 4,30 | 1,83 | 1,20 | 1,52 | 1,28E-01 | 2,70E-01 |
| AT1G07793 | 2,22 | -2,17 | 1,42 | -1,52 | 1,28E-01 | 2,71E-01 |
| AT1G60950 | 5,66 | -1,43 | 0,94 | -1,52 | 1,29E-01 | 2,73E-01 |
| AT4G18596 | 1,78 | 2,86 | 1,88 | 1,52 | 1,29E-01 | 2,73E-01 |
| AT3G14067 | 4,84 | 1,35 | 0,89 | 1,52 | 1,30E-01 | 2,73E-01 |
| AT2G44550 | 3,60 | -1,82 | 1,20 | -1,51 | 1,31E-01 | 2,75E-01 |
| AT5G42530 | 4,10 | -1,44 | 0,95 | -1,51 | 1,31E-01 | 2,76E-01 |
| AT1G30380 | 5,72 | -1,68 | 1,11 | -1,51 | 1,31E-01 | 2,76E-01 |
| AT3G03050 | 2,11 | 2,79 | 1,85 | 1,51 | 1,31E-01 | 2,76E-01 |
| AT3G09105 | 30,37 | -0,86 | 0,57 | -1,51 | 1,31E-01 | 2,76E-01 |
| AT2G34750 | 2,36 | 2,30 | 1,52 | 1,51 | 1,31E-01 | 2,76E-01 |
| AT1G76630 | 2,14 | 2,59 | 1,72 | 1,51 | 1,32E-01 | 2,76E-01 |
| AT3G03780 | 4,16 | 1,51 | 1,00 | 1,51 | 1,32E-01 | 2,76E-01 |
| AT3G05325 | 9,46 | -1,19 | 0,79 | -1,51 | 1,32E-01 | 2,76E-01 |
| AT5G39050 | 4,52 | -1,87 | 1,24 | -1,51 | 1,32E-01 | 2,76E-01 |
| AT3G56825 | 56,81 | -0,59 | 0,39 | -1,51 | 1,32E-01 | 2,76E-01 |
| AT1G23190 | 1,73 | 2,58 | 1,71 | 1,51 | 1,32E-01 | 2,76E-01 |
| AT3G03990 | 2,37 | 2,30 | 1,53 | 1,50 | 1,32E-01 | 2,76E-01 |
| AT3G59770 | 3,31 | 2,31 | 1,54 | 1,50 | 1,33E-01 | 2,77E-01 |
| AT4G13493 | 160,77 | 0,46 | 0,30 | 1,50 | 1,34E-01 | 2,78E-01 |
| AT4G20360 | 5,64 | -1,28 | 0,86 | -1,50 | 1,34E-01 | 2,78E-01 |
| AT4G20362 | 5,64 | -1,28 | 0,86 | -1,50 | 1,34E-01 | 2,78E-01 |
| AT1G78080 | 3,84 | 1,66 | 1,11 | 1,50 | 1,34E-01 | 2,78E-01 |
| AT2G01600 | 1,95 | 2,65 | 1,77 | 1,50 | 1,34E-01 | 2,79E-01 |
| AT2G01320 | 2,02 | 2,36 | 1,58 | 1,50 | 1,35E-01 | 2,79E-01 |
| AT3G43160 | 4,62 | -2,19 | 1,46 | -1,49 | 1,35E-01 | 2,80E-01 |
| AT2G35744 | 14,81 | -0,88 | 0,59 | -1,49 | 1,35E-01 | 2,80E-01 |
| AT1G14670 | 2,24 | 2,26 | 1,52 | 1,49 | 1,36E-01 | 2,80E-01 |
| AT2G46192 | 21,12 | -0,94 | 0,63 | -1,49 | 1,36E-01 | 2,81E-01 |

| | | | | | | |
|---|---|---|---|---|---|---|
| AT3G01770 | 1,96 | 2,60 | 1,75 | 1,49 | 1,37E-01 | 2,82E-01 |
| AT4G11600 | 2,32 | -2,05 | 1,38 | -1,49 | 1,37E-01 | 2,82E-01 |
| AT1G28070 | 4,21 | 2,80 | 1,88 | 1,49 | 1,37E-01 | 2,82E-01 |
| AT4G31210 | 1,90 | 2,44 | 1,64 | 1,49 | 1,37E-01 | 2,82E-01 |
| AT2G03250 | 2,57 | -2,16 | 1,45 | -1,49 | 1,37E-01 | 2,82E-01 |
| AT3G02470 | 10,40 | 1,00 | 0,68 | 1,48 | 1,38E-01 | 2,83E-01 |
| AT1G08347 | 16,30 | -0,78 | 0,53 | -1,48 | 1,38E-01 | 2,83E-01 |
| AT3G10380 | 2,66 | 1,83 | 1,24 | 1,48 | 1,38E-01 | 2,84E-01 |
| AT5G54440 | 2,06 | 2,27 | 1,53 | 1,48 | 1,39E-01 | 2,85E-01 |
| AT4G07120 | 22,20 | 0,78 | 0,53 | 1,47 | 1,40E-01 | 2,88E-01 |
| AT3G05530 | 1,78 | 2,24 | 1,52 | 1,47 | 1,41E-01 | 2,88E-01 |
| AT4G29200 | 3,91 | -1,92 | 1,31 | -1,47 | 1,42E-01 | 2,90E-01 |
| AT3G01313 | 12,85 | -1,32 | 0,90 | -1,47 | 1,42E-01 | 2,91E-01 |
| AT5G01400 | 1,70 | 3,00 | 2,05 | 1,47 | 1,43E-01 | 2,91E-01 |
| AT3G05355 | 1,75 | -2,43 | 1,66 | -1,47 | 1,43E-01 | 2,91E-01 |
| AT1G58602 | 11,97 | 0,91 | 0,62 | 1,46 | 1,43E-01 | 2,92E-01 |
| AT1G09937 | 6,18 | 1,19 | 0,81 | 1,46 | 1,44E-01 | 2,93E-01 |
| AT3G23100 | 2,14 | 2,36 | 1,62 | 1,45 | 1,46E-01 | 2,97E-01 |
| AT4G39160 | 1,99 | 2,31 | 1,59 | 1,45 | 1,46E-01 | 2,98E-01 |
| AT4G23140 | 2,18 | 2,30 | 1,59 | 1,45 | 1,47E-01 | 2,98E-01 |
| AT3G21870 | 1,78 | -2,65 | 1,83 | -1,45 | 1,47E-01 | 2,98E-01 |
| AT4G29900 | 4,72 | 1,50 | 1,04 | 1,45 | 1,47E-01 | 2,98E-01 |
| AT5G22770 | 2,48 | 1,94 | 1,34 | 1,45 | 1,47E-01 | 2,98E-01 |
| AT1G18485 | 1,73 | 2,95 | 2,03 | 1,45 | 1,48E-01 | 2,98E-01 |
| AT2G01850 | 1,74 | -2,33 | 1,61 | -1,45 | 1,48E-01 | 2,99E-01 |
| AT1G66970 | 4,60 | 1,62 | 1,12 | 1,44 | 1,49E-01 | 3,02E-01 |
| AT1G07930 | 3,92 | 1,63 | 1,13 | 1,44 | 1,50E-01 | 3,02E-01 |
| AT3G11940 | 3,79 | 1,43 | 0,99 | 1,44 | 1,50E-01 | 3,02E-01 |
| AT3G14270 | 2,35 | 2,13 | 1,48 | 1,44 | 1,50E-01 | 3,02E-01 |
| AT3G47342 | 15,52 | -0,91 | 0,64 | -1,44 | 1,50E-01 | 3,03E-01 |
| AT1G78900 | 8,95 | 1,01 | 0,70 | 1,44 | 1,51E-01 | 3,04E-01 |
| AT1G55325 | 2,49 | 1,81 | 1,26 | 1,44 | 1,51E-01 | 3,04E-01 |
| AT5G45095 | 2,04 | -2,54 | 1,77 | -1,43 | 1,52E-01 | 3,04E-01 |
| AT1G70290 | 2,53 | 2,09 | 1,46 | 1,43 | 1,52E-01 | 3,04E-01 |
| AT3G02200 | 1,75 | 2,72 | 1,89 | 1,43 | 1,52E-01 | 3,04E-01 |
| AT1G49050 | 1,80 | 2,92 | 2,04 | 1,43 | 1,52E-01 | 3,04E-01 |
| AT4G22690 | 3,37 | 1,51 | 1,05 | 1,43 | 1,52E-01 | 3,04E-01 |
| AT1G71220 | 2,96 | 2,06 | 1,44 | 1,43 | 1,54E-01 | 3,07E-01 |
| AT4G40100 | 3,82 | -1,58 | 1,11 | -1,42 | 1,54E-01 | 3,08E-01 |
| AT1G57720 | 1,72 | 2,32 | 1,63 | 1,42 | 1,55E-01 | 3,09E-01 |
| AT2G31400 | 2,56 | 2,13 | 1,50 | 1,42 | 1,55E-01 | 3,09E-01 |
| AT4G24413 | 2,10 | -2,03 | 1,43 | -1,42 | 1,55E-01 | 3,09E-01 |
| AT1G08053 | 2,32 | -2,09 | 1,47 | -1,42 | 1,56E-01 | 3,10E-01 |
| AT1G29418 | 9,45 | -0,92 | 0,65 | -1,42 | 1,57E-01 | 3,12E-01 |
| AT1G42970 | 13,13 | 0,87 | 0,62 | 1,41 | 1,58E-01 | 3,13E-01 |
| AT4G13430 | 3,55 | 1,62 | 1,15 | 1,41 | 1,58E-01 | 3,13E-01 |
| AT5G08550 | 1,83 | 2,36 | 1,67 | 1,41 | 1,58E-01 | 3,13E-01 |
| AT3G52850 | 1,76 | -2,36 | 1,67 | -1,41 | 1,59E-01 | 3,14E-01 |
| AT4G39350 | 3,27 | 1,48 | 1,05 | 1,41 | 1,59E-01 | 3,14E-01 |
| AT5G09065 | 4,44 | 1,50 | 1,07 | 1,41 | 1,59E-01 | 3,14E-01 |
| AT1G04480 | 2,74 | 1,77 | 1,26 | 1,41 | 1,60E-01 | 3,14E-01 |
| AT4G31490 | 3,09 | 1,76 | 1,25 | 1,41 | 1,59E-01 | 3,14E-01 |
| AT5G58160 | 1,77 | 3,50 | 2,49 | 1,41 | 1,59E-01 | 3,14E-01 |
| AT1G77180 | 2,45 | 2,09 | 1,48 | 1,41 | 1,60E-01 | 3,14E-01 |
| AT1G75800 | 1,71 | 2,46 | 1,75 | 1,40 | 1,61E-01 | 3,16E-01 |
| AT4G28570 | 3,15 | -1,58 | 1,13 | -1,40 | 1,60E-01 | 3,16E-01 |

| | | | | | | |
|---|---|---|---|---|---|---|
| AT4G03550 | 3,00 | 2,08 | 1,49 | 1,40 | 1,61E-01 | 3,17E-01 |
| AT3G01310 | 2,04 | 2,76 | 1,97 | 1,40 | 1,61E-01 | 3,17E-01 |
| AT1G63880 | 4,32 | 1,92 | 1,37 | 1,40 | 1,62E-01 | 3,17E-01 |
| AT1G23900 | 1,85 | 2,45 | 1,76 | 1,39 | 1,63E-01 | 3,20E-01 |
| AT5G42820 | 2,08 | 1,94 | 1,39 | 1,39 | 1,63E-01 | 3,20E-01 |
| AT3G20362 | 7,16 | -1,15 | 0,82 | -1,39 | 1,64E-01 | 3,21E-01 |
| AT3G52240 | 2,27 | 2,08 | 1,49 | 1,39 | 1,65E-01 | 3,21E-01 |
| AT3G59140 | 3,91 | 2,77 | 1,99 | 1,39 | 1,64E-01 | 3,21E-01 |
| AT4G35830 | 9,01 | 1,13 | 0,81 | 1,39 | 1,65E-01 | 3,22E-01 |
| AT2G04030 | 4,28 | 1,36 | 0,98 | 1,39 | 1,65E-01 | 3,22E-01 |
| AT2G26910 | 1,87 | 2,05 | 1,47 | 1,39 | 1,65E-01 | 3,23E-01 |
| AT5G20290 | 5,09 | 1,26 | 0,91 | 1,38 | 1,66E-01 | 3,24E-01 |
| AT4G08470 | 1,82 | 2,94 | 2,13 | 1,38 | 1,67E-01 | 3,24E-01 |
| AT4G33240 | 2,23 | 2,13 | 1,55 | 1,38 | 1,69E-01 | 3,28E-01 |
| AT3G12980 | 1,82 | 2,46 | 1,78 | 1,38 | 1,69E-01 | 3,28E-01 |
| AT5G02810 | 1,73 | 2,35 | 1,71 | 1,37 | 1,69E-01 | 3,28E-01 |
| AT3G14930 | 2,04 | 1,81 | 1,32 | 1,37 | 1,70E-01 | 3,29E-01 |
| AT4G00150 | 10,48 | 1,32 | 0,96 | 1,37 | 1,70E-01 | 3,29E-01 |
| AT5G25060 | 1,92 | 2,08 | 1,51 | 1,37 | 1,70E-01 | 3,29E-01 |
| AT2G02560 | 1,80 | 2,06 | 1,50 | 1,37 | 1,70E-01 | 3,29E-01 |
| AT5G21940 | 4,72 | 1,61 | 1,17 | 1,37 | 1,70E-01 | 3,29E-01 |
| AT3G19180 | 1,86 | 2,74 | 2,00 | 1,37 | 1,71E-01 | 3,30E-01 |
| AT5G47430 | 3,26 | 1,56 | 1,14 | 1,37 | 1,72E-01 | 3,31E-01 |
| AT4G14500 | 7,04 | -1,31 | 0,96 | -1,37 | 1,72E-01 | 3,32E-01 |
| AT2G18280 | 2,10 | 2,11 | 1,55 | 1,36 | 1,72E-01 | 3,32E-01 |
| AT3G44260 | 2,79 | -2,28 | 1,67 | -1,36 | 1,72E-01 | 3,32E-01 |
| AT1G66725 | 1003,08 | -0,42 | 0,31 | -1,36 | 1,74E-01 | 3,33E-01 |
| AT3G06510 | 3,56 | 2,02 | 1,48 | 1,36 | 1,74E-01 | 3,33E-01 |
| AT5G46760 | 2,52 | 2,06 | 1,52 | 1,36 | 1,74E-01 | 3,33E-01 |
| AT1G20450 | 3,27 | -1,55 | 1,14 | -1,36 | 1,73E-01 | 3,33E-01 |
| AT1G79520 | 27,20 | -0,72 | 0,53 | -1,36 | 1,73E-01 | 3,33E-01 |
| AT5G37600 | 1,93 | 2,33 | 1,71 | 1,36 | 1,74E-01 | 3,33E-01 |
| AT5G11490 | 1,97 | 2,09 | 1,54 | 1,36 | 1,74E-01 | 3,33E-01 |
| AT3G22231 | 4,08 | -1,46 | 1,07 | -1,36 | 1,74E-01 | 3,33E-01 |
| AT5G40384 | 2,72 | 1,85 | 1,36 | 1,36 | 1,74E-01 | 3,33E-01 |
| AT5G58465 | 4,24 | -1,40 | 1,03 | -1,36 | 1,75E-01 | 3,34E-01 |
| AT3G60770 | 1,96 | 2,04 | 1,51 | 1,36 | 1,75E-01 | 3,34E-01 |
| AT2G19850 | 5,66 | -1,51 | 1,12 | -1,35 | 1,76E-01 | 3,35E-01 |
| AT3G48057 | 47,97 | 0,78 | 0,57 | 1,35 | 1,76E-01 | 3,35E-01 |
| AT3G23450 | 2,19 | -2,27 | 1,68 | -1,35 | 1,77E-01 | 3,36E-01 |
| AT2G25510 | 7,52 | 1,21 | 0,90 | 1,35 | 1,77E-01 | 3,37E-01 |
| AT3G23110 | 1,95 | 2,31 | 1,71 | 1,35 | 1,78E-01 | 3,38E-01 |
| AT1G02890 | 1,79 | 2,32 | 1,73 | 1,35 | 1,78E-01 | 3,39E-01 |
| AT4G19410 | 1,78 | 2,08 | 1,55 | 1,34 | 1,79E-01 | 3,40E-01 |
| AT3G48860 | 1,96 | 3,01 | 2,24 | 1,34 | 1,80E-01 | 3,40E-01 |
| AT4G04570 | 4,35 | -1,46 | 1,09 | -1,34 | 1,80E-01 | 3,40E-01 |
| AT1G32060 | 5,29 | -1,51 | 1,13 | -1,34 | 1,81E-01 | 3,41E-01 |
| AT5G24930 | 2,10 | 1,99 | 1,48 | 1,34 | 1,81E-01 | 3,41E-01 |
| AT5G51630 | 1,97 | 2,34 | 1,75 | 1,34 | 1,81E-01 | 3,41E-01 |
| AT1G56630 | 1,84 | -2,70 | 2,02 | -1,34 | 1,81E-01 | 3,41E-01 |
| AT1G61690 | 1,91 | 2,38 | 1,78 | 1,34 | 1,81E-01 | 3,41E-01 |
| AT3G04340 | 3,02 | 1,73 | 1,29 | 1,34 | 1,81E-01 | 3,41E-01 |
| AT3G25510 | 1,96 | 1,94 | 1,45 | 1,34 | 1,81E-01 | 3,41E-01 |
| AT5G03145 | 2,30 | -2,01 | 1,51 | -1,34 | 1,82E-01 | 3,42E-01 |
| AT1G48760 | 1,82 | 2,51 | 1,88 | 1,33 | 1,83E-01 | 3,43E-01 |
| AT5G19510 | 1,83 | 2,07 | 1,55 | 1,33 | 1,83E-01 | 3,43E-01 |

| | | | | | | |
|---|---|---|---|---|---|---|
| AT1G06667 | 5,78 | -1,32 | 0,99 | -1,33 | 1,85E-01 | 3,46E-01 |
| AT3G49910 | 2,91 | 1,59 | 1,20 | 1,33 | 1,85E-01 | 3,46E-01 |
| AT5G67300 | 3,74 | 1,83 | 1,38 | 1,33 | 1,85E-01 | 3,46E-01 |
| AT1G52110 | 2,50 | -1,75 | 1,32 | -1,33 | 1,85E-01 | 3,46E-01 |
| AT4G25100 | 2,11 | -2,35 | 1,77 | -1,33 | 1,85E-01 | 3,46E-01 |
| AT5G58430 | 1,95 | 2,07 | 1,56 | 1,33 | 1,84E-01 | 3,46E-01 |
| AT3G01290 | 2,12 | 2,21 | 1,67 | 1,32 | 1,85E-01 | 3,46E-01 |
| AT2G33170 | 2,19 | 2,36 | 1,78 | 1,32 | 1,86E-01 | 3,47E-01 |
| AT2G07605 | 7,80 | -1,05 | 0,80 | -1,32 | 1,87E-01 | 3,48E-01 |
| AT3G25520 | 3,31 | 1,37 | 1,04 | 1,32 | 1,88E-01 | 3,50E-01 |
| AT3G18370 | 2,55 | 2,05 | 1,56 | 1,31 | 1,89E-01 | 3,52E-01 |
| AT3G14840 | 1,81 | 2,11 | 1,61 | 1,31 | 1,91E-01 | 3,55E-01 |
| AT5G49910 | 2,46 | 1,72 | 1,32 | 1,31 | 1,91E-01 | 3,55E-01 |
| AT5G52170 | 2,49 | 2,09 | 1,60 | 1,31 | 1,91E-01 | 3,55E-01 |
| AT1G68945 | 2,15 | -2,07 | 1,58 | -1,30 | 1,92E-01 | 3,56E-01 |
| AT2G05170 | 1,80 | 2,16 | 1,66 | 1,30 | 1,92E-01 | 3,57E-01 |
| AT4G27870 | 2,09 | 1,84 | 1,41 | 1,30 | 1,93E-01 | 3,58E-01 |
| AT4G03380 | 7,25 | -1,10 | 0,84 | -1,30 | 1,93E-01 | 3,58E-01 |
| AT3G16785 | 1,89 | 1,88 | 1,45 | 1,30 | 1,94E-01 | 3,58E-01 |
| AT1G27870 | 1,97 | -2,27 | 1,75 | -1,30 | 1,94E-01 | 3,58E-01 |
| AT2G08665 | 53,97 | -0,54 | 0,41 | -1,30 | 1,95E-01 | 3,59E-01 |
| AT5G08690 | 2,64 | 1,81 | 1,39 | 1,30 | 1,94E-01 | 3,59E-01 |
| AT2G34420 | 53,97 | -0,54 | 0,41 | -1,30 | 1,95E-01 | 3,59E-01 |
| AT3G26520 | 6,24 | -1,25 | 0,97 | -1,30 | 1,94E-01 | 3,59E-01 |
| AT2G20721 | 2,89 | -1,99 | 1,54 | -1,30 | 1,95E-01 | 3,59E-01 |
| AT1G40104 | 7,42 | -1,36 | 1,05 | -1,29 | 1,96E-01 | 3,60E-01 |
| AT5G62700 | 1,86 | 2,14 | 1,66 | 1,29 | 1,96E-01 | 3,60E-01 |
| AT1G06680 | 10,44 | -0,85 | 0,65 | -1,29 | 1,96E-01 | 3,60E-01 |
| AT5G47690 | 3,16 | 1,58 | 1,22 | 1,29 | 1,97E-01 | 3,61E-01 |
| AT5G02480 | 2,67 | 1,92 | 1,49 | 1,29 | 1,97E-01 | 3,62E-01 |
| AT1G09027 | 2,11 | -2,07 | 1,60 | -1,29 | 1,98E-01 | 3,62E-01 |
| AT2G43410 | 2,84 | 1,91 | 1,48 | 1,29 | 1,98E-01 | 3,62E-01 |
| AT4G29380 | 1,74 | 2,25 | 1,75 | 1,29 | 1,98E-01 | 3,62E-01 |
| AT3G45830 | 1,79 | 2,29 | 1,78 | 1,29 | 1,98E-01 | 3,62E-01 |
| AT3G56360 | 3,83 | 1,39 | 1,08 | 1,29 | 1,99E-01 | 3,63E-01 |
| AT4G29090 | 2,85 | -2,09 | 1,63 | -1,29 | 1,99E-01 | 3,63E-01 |
| AT5G62350 | 2,26 | -1,86 | 1,45 | -1,28 | 1,99E-01 | 3,63E-01 |
| AT5G09660 | 2,38 | -1,81 | 1,41 | -1,28 | 2,00E-01 | 3,64E-01 |
| AT2G47940 | 2,47 | 1,86 | 1,45 | 1,28 | 2,01E-01 | 3,65E-01 |
| AT1G08857 | 2,42 | 2,74 | 2,14 | 1,28 | 2,01E-01 | 3,65E-01 |
| AT1G06430 | 1,82 | -1,96 | 1,53 | -1,28 | 2,02E-01 | 3,67E-01 |
| AT2G46220 | 1,83 | 2,11 | 1,66 | 1,27 | 2,04E-01 | 3,69E-01 |
| AT1G74100 | 5,98 | 1,31 | 1,03 | 1,27 | 2,03E-01 | 3,69E-01 |
| AT5G03940 | 2,83 | 1,42 | 1,12 | 1,27 | 2,03E-01 | 3,69E-01 |
| AT5G41905 | 1173,00 | -0,32 | 0,25 | -1,27 | 2,03E-01 | 3,69E-01 |
| AT4G03560 | 2,24 | -1,82 | 1,43 | -1,27 | 2,04E-01 | 3,69E-01 |
| AT5G60390 | 16,82 | 0,70 | 0,55 | 1,27 | 2,04E-01 | 3,69E-01 |
| AT1G11735 | 8,53 | 0,87 | 0,68 | 1,27 | 2,04E-01 | 3,69E-01 |
| AT1G07051 | 29,68 | 0,63 | 0,50 | 1,27 | 2,05E-01 | 3,70E-01 |
| AT2G07698 | 9,74 | -0,86 | 0,68 | -1,27 | 2,05E-01 | 3,70E-01 |
| AT3G28180 | 8,56 | -1,28 | 1,01 | -1,27 | 2,05E-01 | 3,70E-01 |
| AT3G46620 | 4,33 | 1,33 | 1,05 | 1,27 | 2,05E-01 | 3,70E-01 |
| AT2G09690 | 14,22 | -1,00 | 0,79 | -1,27 | 2,06E-01 | 3,71E-01 |
| AT2G20780 | 1,93 | 2,27 | 1,79 | 1,26 | 2,07E-01 | 3,72E-01 |
| AT3G56070 | 2,75 | -1,75 | 1,39 | -1,26 | 2,07E-01 | 3,72E-01 |
| AT5G09870 | 2,84 | 1,39 | 1,10 | 1,26 | 2,07E-01 | 3,72E-01 |

| | | | | | | |
|---|---|---|---|---|---|---|
| AT1G12620 | 2,40 | -2,11 | 1,67 | -1,26 | 2,07E-01 | 3,72E-01 |
| AT5G61440 | 4,44 | -1,09 | 0,86 | -1,26 | 2,09E-01 | 3,74E-01 |
| AT1G63630 | 3,43 | -1,49 | 1,19 | -1,26 | 2,09E-01 | 3,75E-01 |
| AT1G63615 | 3,43 | -1,49 | 1,19 | -1,26 | 2,09E-01 | 3,75E-01 |
| AT1G29400 | 2,11 | 1,89 | 1,51 | 1,26 | 2,09E-01 | 3,75E-01 |
| AT1G31960 | 1,81 | -2,13 | 1,70 | -1,25 | 2,10E-01 | 3,75E-01 |
| AT2G35110 | 1,99 | 1,81 | 1,45 | 1,25 | 2,11E-01 | 3,77E-01 |
| AT2G28550 | 2,55 | -1,58 | 1,26 | -1,25 | 2,11E-01 | 3,77E-01 |
| AT1G32270 | 1,96 | -2,00 | 1,61 | -1,24 | 2,14E-01 | 3,81E-01 |
| AT2G46820 | 2,21 | -1,62 | 1,31 | -1,24 | 2,14E-01 | 3,82E-01 |
| AT4G21060 | 1,98 | -1,64 | 1,32 | -1,24 | 2,15E-01 | 3,82E-01 |
| AT5G27606 | 14,40 | -1,90 | 1,53 | -1,24 | 2,14E-01 | 3,82E-01 |
| AT3G11410 | 4,48 | 1,33 | 1,07 | 1,24 | 2,15E-01 | 3,82E-01 |
| AT4G11380 | 1,75 | -2,09 | 1,69 | -1,24 | 2,16E-01 | 3,84E-01 |
| AT2G09685 | 3,15 | -1,49 | 1,21 | -1,24 | 2,16E-01 | 3,84E-01 |
| AT1G63360 | 4,56 | -1,16 | 0,94 | -1,23 | 2,18E-01 | 3,87E-01 |
| AT3G09390 | 6,61 | 1,12 | 0,91 | 1,23 | 2,18E-01 | 3,87E-01 |
| AT1G06950 | 2,18 | 2,42 | 1,97 | 1,23 | 2,19E-01 | 3,88E-01 |
| AT4G39366 | 2,55 | -1,75 | 1,42 | -1,23 | 2,20E-01 | 3,89E-01 |
| AT3G04765 | 2,83 | -1,50 | 1,22 | -1,23 | 2,20E-01 | 3,89E-01 |
| AT4G13880 | 2,00 | 2,18 | 1,78 | 1,23 | 2,21E-01 | 3,90E-01 |
| AT1G04410 | 3,62 | -1,41 | 1,15 | -1,22 | 2,21E-01 | 3,90E-01 |
| AT1G09070 | 2,87 | 2,25 | 1,84 | 1,22 | 2,22E-01 | 3,90E-01 |
| AT3G63490 | 1,82 | 2,31 | 1,89 | 1,22 | 2,21E-01 | 3,90E-01 |
| AT3G29190 | 2,69 | -1,82 | 1,49 | -1,22 | 2,21E-01 | 3,90E-01 |
| AT4G18700 | 2,69 | 1,53 | 1,25 | 1,22 | 2,22E-01 | 3,90E-01 |
| AT5G57560 | 1,92 | 1,89 | 1,55 | 1,22 | 2,22E-01 | 3,90E-01 |
| AT1G72370 | 2,39 | 1,60 | 1,31 | 1,22 | 2,22E-01 | 3,91E-01 |
| AT1G22930 | 1,70 | 2,30 | 1,89 | 1,22 | 2,23E-01 | 3,91E-01 |
| AT2G47450 | 6,02 | 1,09 | 0,89 | 1,22 | 2,23E-01 | 3,91E-01 |
| AT1G03680 | 1,73 | -1,92 | 1,57 | -1,22 | 2,23E-01 | 3,91E-01 |
| AT5G18410 | 3,15 | 1,46 | 1,20 | 1,22 | 2,24E-01 | 3,92E-01 |
| AT5G54270 | 15,64 | -0,61 | 0,50 | -1,21 | 2,24E-01 | 3,93E-01 |
| AT2G30570 | 3,97 | -1,22 | 1,01 | -1,22 | 2,24E-01 | 3,93E-01 |
| AT2G21030 | 1,78 | -2,11 | 1,74 | -1,21 | 2,25E-01 | 3,94E-01 |
| AT2G25450 | 4,94 | 1,28 | 1,06 | 1,21 | 2,25E-01 | 3,94E-01 |
| AT3G01120 | 2,55 | 1,91 | 1,57 | 1,21 | 2,26E-01 | 3,94E-01 |
| AT4G08850 | 2,82 | 1,94 | 1,60 | 1,21 | 2,26E-01 | 3,94E-01 |
| AT2G32690 | 2,01 | -1,73 | 1,43 | -1,21 | 2,26E-01 | 3,94E-01 |
| AT3G27400 | 1,80 | -1,81 | 1,50 | -1,21 | 2,26E-01 | 3,94E-01 |
| AT3G04240 | 3,51 | 1,50 | 1,24 | 1,21 | 2,26E-01 | 3,94E-01 |
| AT4G32250 | 2,07 | 2,18 | 1,80 | 1,21 | 2,26E-01 | 3,94E-01 |
| AT5G58410 | 3,13 | 1,58 | 1,31 | 1,21 | 2,28E-01 | 3,96E-01 |
| AT3G21755 | 4,36 | -1,28 | 1,06 | -1,21 | 2,28E-01 | 3,96E-01 |
| AT3G21760 | 4,32 | -1,28 | 1,06 | -1,20 | 2,29E-01 | 3,97E-01 |
| AT3G22890 | 4,47 | 1,19 | 0,99 | 1,20 | 2,29E-01 | 3,97E-01 |
| AT1G52400 | 1,99 | -2,08 | 1,73 | -1,20 | 2,30E-01 | 3,98E-01 |
| AT5G47010 | 2,34 | 1,62 | 1,35 | 1,20 | 2,30E-01 | 3,99E-01 |
| AT5G17020 | 2,03 | 1,54 | 1,28 | 1,20 | 2,31E-01 | 4,00E-01 |
| AT1G07593 | 3,78 | 1,64 | 1,37 | 1,20 | 2,31E-01 | 4,00E-01 |
| AT3G48720 | 18,61 | -0,76 | 0,64 | -1,20 | 2,32E-01 | 4,00E-01 |
| AT5G07570 | 4,01 | 2,23 | 1,86 | 1,20 | 2,32E-01 | 4,00E-01 |
| AT5G02325 | 15,37 | -0,97 | 0,81 | -1,19 | 2,32E-01 | 4,01E-01 |
| AT5G43310 | 2,37 | 1,82 | 1,52 | 1,19 | 2,32E-01 | 4,01E-01 |
| AT1G66880 | 6,48 | 1,21 | 1,01 | 1,19 | 2,33E-01 | 4,01E-01 |
| AT2G20990 | 4,50 | 1,27 | 1,07 | 1,19 | 2,33E-01 | 4,01E-01 |

| | | | | | | |
|---|---|---|---|---|---|---|
| AT1G33140 | 1,84 | 1,87 | 1,58 | 1,19 | 2,35E-01 | 4,03E-01 |
| AT2G34430 | 28,33 | -0,50 | 0,42 | -1,19 | 2,35E-01 | 4,03E-01 |
| AT4G16660 | 2,21 | 1,87 | 1,58 | 1,19 | 2,35E-01 | 4,03E-01 |
| AT4G02320 | 3,77 | 1,14 | 0,96 | 1,19 | 2,34E-01 | 4,03E-01 |
| AT5G05200 | 1,99 | 2,38 | 2,00 | 1,19 | 2,35E-01 | 4,03E-01 |
| AT4G16920 | 1,96 | 1,77 | 1,49 | 1,19 | 2,36E-01 | 4,04E-01 |
| AT1G74470 | 19,25 | 0,59 | 0,50 | 1,18 | 2,36E-01 | 4,04E-01 |
| AT1G56340 | 4,49 | 1,23 | 1,03 | 1,18 | 2,36E-01 | 4,04E-01 |
| AT2G22440 | 2,40 | -1,52 | 1,29 | -1,18 | 2,36E-01 | 4,04E-01 |
| AT2G42590 | 2,06 | -1,99 | 1,68 | -1,18 | 2,37E-01 | 4,05E-01 |
| AT1G19960 | 1,88 | -2,21 | 1,87 | -1,18 | 2,37E-01 | 4,05E-01 |
| AT3G55830 | 1,72 | 2,00 | 1,70 | 1,18 | 2,38E-01 | 4,06E-01 |
| AT1G52230 | 2,95 | -1,22 | 1,04 | -1,18 | 2,39E-01 | 4,07E-01 |
| AT5G61790 | 5,40 | 1,11 | 0,95 | 1,18 | 2,39E-01 | 4,08E-01 |
| AT4G38550 | 2,97 | 1,75 | 1,49 | 1,17 | 2,41E-01 | 4,09E-01 |
| AT4G38552 | 2,97 | 1,75 | 1,49 | 1,17 | 2,41E-01 | 4,09E-01 |
| AT3G55800 | 2,45 | -1,44 | 1,23 | -1,17 | 2,42E-01 | 4,11E-01 |
| AT1G22060 | 2,54 | 1,50 | 1,28 | 1,17 | 2,42E-01 | 4,11E-01 |
| AT1G69797 | 12,98 | 0,80 | 0,68 | 1,17 | 2,43E-01 | 4,12E-01 |
| AT5G41765 | 5,08 | 1,87 | 1,60 | 1,17 | 2,43E-01 | 4,12E-01 |
| AT2G35050 | 3,09 | 1,49 | 1,28 | 1,17 | 2,44E-01 | 4,13E-01 |
| AT2G09610 | 9,53 | -0,77 | 0,66 | -1,16 | 2,44E-01 | 4,14E-01 |
| AT1G06243 | 5,81 | -1,06 | 0,91 | -1,16 | 2,45E-01 | 4,14E-01 |
| AT2G01422 | 2,93 | -1,92 | 1,65 | -1,16 | 2,45E-01 | 4,15E-01 |
| AT5G36228 | 1,95 | -1,74 | 1,50 | -1,16 | 2,46E-01 | 4,15E-01 |
| AT5G35735 | 2,19 | 1,94 | 1,67 | 1,16 | 2,47E-01 | 4,16E-01 |
| AT5G40890 | 1,74 | 1,69 | 1,45 | 1,16 | 2,46E-01 | 4,16E-01 |
| AT1G09877 | 23,86 | -0,61 | 0,53 | -1,16 | 2,47E-01 | 4,16E-01 |
| AT4G25880 | 1,93 | 1,82 | 1,57 | 1,16 | 2,47E-01 | 4,17E-01 |
| AT4G15258 | 2,27 | 1,81 | 1,57 | 1,16 | 2,48E-01 | 4,17E-01 |
| AT5G52320 | 1,77 | -1,91 | 1,65 | -1,16 | 2,48E-01 | 4,17E-01 |
| AT1G31835 | 2,35 | -1,67 | 1,45 | -1,15 | 2,49E-01 | 4,19E-01 |
| AT4G39730 | 3,06 | -1,52 | 1,32 | -1,15 | 2,50E-01 | 4,19E-01 |
| AT1G07890 | 4,08 | 1,16 | 1,01 | 1,15 | 2,51E-01 | 4,21E-01 |
| AT4G11373 | 5,92 | -1,71 | 1,49 | -1,15 | 2,51E-01 | 4,21E-01 |
| AT3G15920 | 8,54 | 0,90 | 0,79 | 1,15 | 2,51E-01 | 4,21E-01 |
| AT1G07600 | 1,87 | -2,04 | 1,78 | -1,15 | 2,51E-01 | 4,21E-01 |
| AT5G57710 | 4,28 | 1,25 | 1,09 | 1,15 | 2,52E-01 | 4,21E-01 |
| AT4G21960 | 15,77 | -0,58 | 0,50 | -1,14 | 2,52E-01 | 4,22E-01 |
| AT1G06277 | 5,95 | -1,03 | 0,91 | -1,14 | 2,54E-01 | 4,25E-01 |
| AT2G38195 | 4,06 | -1,64 | 1,44 | -1,14 | 2,55E-01 | 4,25E-01 |
| AT1G64385 | 1,99 | 2,18 | 1,91 | 1,14 | 2,56E-01 | 4,27E-01 |
| AT2G34100 | 7,49 | -1,11 | 0,98 | -1,14 | 2,56E-01 | 4,27E-01 |
| AT2G38800 | 2,71 | 2,01 | 1,77 | 1,13 | 2,57E-01 | 4,27E-01 |
| AT3G06325 | 6,42 | -1,13 | 1,00 | -1,13 | 2,57E-01 | 4,27E-01 |
| AT2G20722 | 3,24 | -1,66 | 1,46 | -1,13 | 2,57E-01 | 4,27E-01 |
| AT4G11830 | 1,70 | 2,08 | 1,83 | 1,13 | 2,57E-01 | 4,27E-01 |
| AT5G56030 | 5,79 | 0,98 | 0,86 | 1,13 | 2,58E-01 | 4,28E-01 |
| AT3G51950 | 4,09 | 1,13 | 1,00 | 1,13 | 2,58E-01 | 4,28E-01 |
| AT1G49750 | 3,00 | -1,47 | 1,30 | -1,13 | 2,58E-01 | 4,29E-01 |
| AT1G06148 | 2,19 | -1,89 | 1,68 | -1,13 | 2,59E-01 | 4,29E-01 |
| AT1G33420 | 2,44 | -1,52 | 1,35 | -1,13 | 2,59E-01 | 4,30E-01 |
| AT1G55490 | 3,72 | 1,19 | 1,06 | 1,13 | 2,60E-01 | 4,30E-01 |
| AT5G38410 | 6,26 | -1,10 | 0,98 | -1,13 | 2,60E-01 | 4,30E-01 |
| AT3G62250 | 1,91 | 1,64 | 1,45 | 1,13 | 2,60E-01 | 4,30E-01 |
| AT5G62670 | 1,85 | 1,99 | 1,77 | 1,13 | 2,60E-01 | 4,30E-01 |

| | | | | | | |
|---|---|---|---|---|---|---|
| AT5G08610 | 1,71 | 1,74 | 1,55 | 1,12 | 2,62E-01 | 4,33E-01 |
| AT4G18210 | 1,90 | -2,05 | 1,83 | -1,12 | 2,62E-01 | 4,33E-01 |
| AT1G07250 | 2,07 | 1,48 | 1,32 | 1,12 | 2,64E-01 | 4,35E-01 |
| AT4G14360 | 1,93 | 1,57 | 1,41 | 1,12 | 2,64E-01 | 4,35E-01 |
| AT4G37640 | 1,71 | 1,99 | 1,78 | 1,12 | 2,64E-01 | 4,35E-01 |
| AT1G74090 | 6,35 | 0,94 | 0,85 | 1,11 | 2,66E-01 | 4,37E-01 |
| AT2G37220 | 3,40 | -1,25 | 1,13 | -1,11 | 2,66E-01 | 4,38E-01 |
| AT3G08170 | 3,83 | 1,25 | 1,13 | 1,11 | 2,68E-01 | 4,40E-01 |
| AT3G05335 | 1,73 | -1,85 | 1,68 | -1,11 | 2,69E-01 | 4,40E-01 |
| AT1G07493 | 1,97 | -1,67 | 1,51 | -1,11 | 2,68E-01 | 4,40E-01 |
| AT5G58330 | 1,92 | 1,79 | 1,62 | 1,11 | 2,69E-01 | 4,40E-01 |
| AT5G08925 | 2,86 | 2,24 | 2,03 | 1,11 | 2,69E-01 | 4,40E-01 |
| AT1G62630 | 2,10 | -1,61 | 1,46 | -1,10 | 2,70E-01 | 4,41E-01 |
| AT3G05985 | 2,35 | 1,82 | 1,65 | 1,10 | 2,70E-01 | 4,41E-01 |
| AT5G02700 | 1,99 | 1,98 | 1,80 | 1,10 | 2,70E-01 | 4,41E-01 |
| AT1G19680 | 2,21 | 2,22 | 2,02 | 1,10 | 2,71E-01 | 4,43E-01 |
| AT2G47410 | 3,41 | 1,64 | 1,49 | 1,10 | 2,72E-01 | 4,44E-01 |
| AT1G43170 | 3,66 | 1,18 | 1,07 | 1,10 | 2,72E-01 | 4,44E-01 |
| AT4G34100 | 3,45 | 1,31 | 1,20 | 1,10 | 2,73E-01 | 4,44E-01 |
| AT5G65480 | 2,00 | 1,74 | 1,59 | 1,10 | 2,73E-01 | 4,44E-01 |
| AT1G12920 | 1,87 | 1,54 | 1,41 | 1,10 | 2,73E-01 | 4,44E-01 |
| AT5G20950 | 3,45 | 1,29 | 1,17 | 1,10 | 2,73E-01 | 4,44E-01 |
| AT5G49740 | 2,42 | 1,64 | 1,50 | 1,09 | 2,74E-01 | 4,44E-01 |
| AT5G56970 | 2,67 | -1,37 | 1,25 | -1,10 | 2,73E-01 | 4,44E-01 |
| AT2G07815 | 2,06 | -1,63 | 1,49 | -1,09 | 2,74E-01 | 4,44E-01 |
| AT4G00720 | 2,07 | 1,73 | 1,58 | 1,09 | 2,75E-01 | 4,45E-01 |
| AT1G20010 | 2,01 | 1,39 | 1,27 | 1,09 | 2,77E-01 | 4,48E-01 |
| AT5G36180 | 3,47 | -1,36 | 1,25 | -1,09 | 2,78E-01 | 4,49E-01 |
| AT5G07725 | 8,72 | -0,99 | 0,91 | -1,09 | 2,78E-01 | 4,49E-01 |
| AT2G26250 | 1,96 | -1,59 | 1,46 | -1,09 | 2,78E-01 | 4,49E-01 |
| AT3G16640 | 7,07 | 0,95 | 0,88 | 1,09 | 2,78E-01 | 4,49E-01 |
| AT5G61310 | 4,76 | -1,09 | 1,00 | -1,09 | 2,78E-01 | 4,49E-01 |
| AT1G33590 | 1,70 | -1,79 | 1,66 | -1,08 | 2,79E-01 | 4,49E-01 |
| AT4G13770 | 2,50 | -1,87 | 1,72 | -1,08 | 2,79E-01 | 4,49E-01 |
| AT3G07735 | 4,39 | -1,03 | 0,95 | -1,08 | 2,79E-01 | 4,50E-01 |
| AT5G19690 | 1,74 | 1,81 | 1,68 | 1,08 | 2,80E-01 | 4,52E-01 |
| AT1G63950 | 1,81 | -2,13 | 1,98 | -1,08 | 2,81E-01 | 4,52E-01 |
| AT4G20260 | 2,30 | -1,58 | 1,47 | -1,08 | 2,81E-01 | 4,52E-01 |
| AT1G50030 | 2,19 | 1,74 | 1,62 | 1,08 | 2,81E-01 | 4,52E-01 |
| AT2G41410 | 4,84 | 1,29 | 1,19 | 1,08 | 2,81E-01 | 4,52E-01 |
| AT1G11650 | 3,30 | 1,21 | 1,12 | 1,08 | 2,82E-01 | 4,53E-01 |
| AT1G47657 | 2,52 | -1,97 | 1,83 | -1,08 | 2,82E-01 | 4,53E-01 |
| AT2G05685 | 2,44 | -1,83 | 1,70 | -1,07 | 2,83E-01 | 4,54E-01 |
| AT5G09730 | 35,07 | -0,52 | 0,48 | -1,07 | 2,83E-01 | 4,54E-01 |
| AT4G29410 | 17,21 | -0,63 | 0,59 | -1,07 | 2,84E-01 | 4,55E-01 |
| AT2G35382 | 3,86 | -1,13 | 1,06 | -1,07 | 2,84E-01 | 4,55E-01 |
| AT3G14310 | 2,07 | -1,89 | 1,76 | -1,07 | 2,85E-01 | 4,55E-01 |
| AT5G20250 | 2,34 | 1,85 | 1,74 | 1,07 | 2,85E-01 | 4,55E-01 |
| AT5G45470 | 2,49 | -2,02 | 1,89 | -1,07 | 2,85E-01 | 4,55E-01 |
| AT5G45110 | 1,72 | 1,89 | 1,77 | 1,07 | 2,86E-01 | 4,56E-01 |
| AT1G09817 | 10,25 | -0,87 | 0,81 | -1,07 | 2,86E-01 | 4,56E-01 |
| AT4G13918 | 2,19 | 1,90 | 1,78 | 1,07 | 2,86E-01 | 4,56E-01 |
| AT1G11870 | 3,10 | 1,53 | 1,44 | 1,07 | 2,86E-01 | 4,56E-01 |
| AT1G19373 | 3,55 | -1,54 | 1,45 | -1,06 | 2,88E-01 | 4,58E-01 |
| AT2G20580 | 2,81 | 1,69 | 1,59 | 1,06 | 2,88E-01 | 4,58E-01 |
| AT3G15210 | 1,70 | -1,78 | 1,67 | -1,06 | 2,88E-01 | 4,58E-01 |

| | | | | | | |
|---|---|---|---|---|---|---|
| AT4G36648 | 2,47 | 1,83 | 1,73 | 1,06 | 2,90E-01 | 4,60E-01 |
| AT5G11560 | 3,42 | 1,14 | 1,08 | 1,06 | 2,90E-01 | 4,60E-01 |
| AT3G12630 | 8,51 | 1,02 | 0,97 | 1,06 | 2,91E-01 | 4,60E-01 |
| AT1G68520 | 2,00 | 1,44 | 1,36 | 1,06 | 2,91E-01 | 4,60E-01 |
| AT3G02170 | 3,18 | 1,70 | 1,61 | 1,06 | 2,91E-01 | 4,60E-01 |
| AT1G21202 | 4,89 | 1,31 | 1,25 | 1,05 | 2,93E-01 | 4,63E-01 |
| AT1G05637 | 4,89 | 1,31 | 1,25 | 1,05 | 2,93E-01 | 4,63E-01 |
| AT2G14610 | 3,13 | 1,51 | 1,44 | 1,05 | 2,93E-01 | 4,63E-01 |
| AT4G40040 | 3,01 | -1,69 | 1,61 | -1,05 | 2,93E-01 | 4,63E-01 |
| AT1G47128 | 7,64 | 0,79 | 0,76 | 1,05 | 2,94E-01 | 4,64E-01 |
| AT5G50920 | 18,45 | 0,56 | 0,53 | 1,04 | 2,97E-01 | 4,68E-01 |
| AT5G53440 | 2,08 | 1,47 | 1,41 | 1,04 | 2,97E-01 | 4,68E-01 |
| AT1G02860 | 4,18 | 1,61 | 1,55 | 1,04 | 2,97E-01 | 4,68E-01 |
| AT1G21590 | 2,73 | 1,35 | 1,30 | 1,04 | 2,98E-01 | 4,68E-01 |
| AT1G22530 | 2,83 | 1,26 | 1,21 | 1,04 | 2,97E-01 | 4,68E-01 |
| AT1G11860 | 2,12 | -1,42 | 1,37 | -1,04 | 2,98E-01 | 4,68E-01 |
| AT3G45640 | 1,79 | 1,81 | 1,74 | 1,04 | 2,98E-01 | 4,68E-01 |
| AT3G54660 | 2,19 | 1,38 | 1,33 | 1,04 | 2,98E-01 | 4,68E-01 |
| AT5G19200 | 4,59 | 1,04 | 1,00 | 1,04 | 2,99E-01 | 4,69E-01 |
| AT3G51770 | 2,18 | 1,64 | 1,58 | 1,04 | 2,99E-01 | 4,69E-01 |
| AT4G03420 | 2,29 | 1,84 | 1,77 | 1,04 | 3,00E-01 | 4,69E-01 |
| AT1G79550 | 2,41 | 1,38 | 1,33 | 1,03 | 3,01E-01 | 4,72E-01 |
| AT4G39260 | 6,03 | -0,81 | 0,79 | -1,03 | 3,02E-01 | 4,73E-01 |
| AT5G20010 | 1,98 | 1,54 | 1,50 | 1,03 | 3,04E-01 | 4,75E-01 |
| AT2G17200 | 1,89 | 1,86 | 1,81 | 1,03 | 3,04E-01 | 4,75E-01 |
| AT3G24180 | 1,99 | 1,44 | 1,40 | 1,03 | 3,04E-01 | 4,75E-01 |
| AT1G05207 | 1,81 | -1,43 | 1,40 | -1,02 | 3,05E-01 | 4,75E-01 |
| AT1G20015 | 7,84 | -0,82 | 0,80 | -1,02 | 3,06E-01 | 4,75E-01 |
| AT2G44490 | 17,93 | 0,50 | 0,49 | 1,03 | 3,05E-01 | 4,75E-01 |
| AT3G13750 | 2,68 | 1,28 | 1,25 | 1,02 | 3,06E-01 | 4,75E-01 |
| AT3G44330 | 1,72 | 2,27 | 2,21 | 1,03 | 3,05E-01 | 4,75E-01 |
| AT5G65005 | 2,56 | -1,11 | 1,09 | -1,03 | 3,05E-01 | 4,75E-01 |
| AT3G23120 | 2,24 | 1,76 | 1,72 | 1,02 | 3,06E-01 | 4,75E-01 |
| AT4G31990 | 1,81 | 1,76 | 1,72 | 1,02 | 3,07E-01 | 4,76E-01 |
| AT4G02570 | 1,99 | 1,56 | 1,54 | 1,02 | 3,09E-01 | 4,79E-01 |
| AT5G52530 | 3,03 | 1,51 | 1,48 | 1,02 | 3,09E-01 | 4,79E-01 |
| AT5G56010 | 6,62 | 0,88 | 0,87 | 1,02 | 3,10E-01 | 4,79E-01 |
| AT1G80180 | 13,61 | -0,87 | 0,85 | -1,01 | 3,11E-01 | 4,81E-01 |
| AT2G46340 | 1,90 | 2,12 | 2,09 | 1,01 | 3,11E-01 | 4,81E-01 |
| AT1G07140 | 1,81 | 1,66 | 1,64 | 1,01 | 3,11E-01 | 4,81E-01 |
| AT1G70530 | 1,72 | 2,02 | 2,00 | 1,01 | 3,13E-01 | 4,84E-01 |
| AT1G09240 | 1,84 | 2,41 | 2,39 | 1,01 | 3,14E-01 | 4,84E-01 |
| AT3G14420 | 4,42 | -1,04 | 1,04 | -1,01 | 3,14E-01 | 4,84E-01 |
| AT1G47497 | 2,25 | -1,67 | 1,66 | -1,01 | 3,15E-01 | 4,85E-01 |
| AT3G59780 | 2,03 | -1,51 | 1,50 | -1,00 | 3,16E-01 | 4,87E-01 |
| AT4G02520 | 4,48 | 1,04 | 1,04 | 1,00 | 3,17E-01 | 4,88E-01 |
| AT5G09040 | 2,31 | -1,50 | 1,50 | -1,00 | 3,17E-01 | 4,88E-01 |
| AT2G33830 | 1,76 | -1,47 | 1,47 | -1,00 | 3,18E-01 | 4,88E-01 |
| AT2G08390 | 5,34 | -0,95 | 0,95 | -1,00 | 3,18E-01 | 4,89E-01 |
| AT1G20620 | 38,72 | -0,44 | 0,44 | -1,00 | 3,19E-01 | 4,90E-01 |
| AT1G77560 | 2,14 | 1,59 | 1,60 | 1,00 | 3,20E-01 | 4,90E-01 |
| AT4G06705 | 4,08 | -1,12 | 1,13 | -1,00 | 3,19E-01 | 4,90E-01 |
| AT4G25730 | 1,71 | 1,87 | 1,88 | 1,00 | 3,19E-01 | 4,90E-01 |
| AT1G77740 | 1,74 | 1,63 | 1,64 | 0,99 | 3,21E-01 | 4,90E-01 |
| AT3G54280 | 2,37 | 1,38 | 1,40 | 0,99 | 3,21E-01 | 4,90E-01 |
| AT5G49760 | 3,34 | 1,05 | 1,06 | 0,99 | 3,21E-01 | 4,90E-01 |

| | | | | | | |
|---|---|---|---|---|---|---|
| AT3G46530 | 2,60 | 1,25 | 1,26 | 0,99 | 3,21E-01 | 4,90E-01 |
| AT2G38230 | 6,88 | -0,72 | 0,73 | -0,99 | 3,22E-01 | 4,91E-01 |
| AT5G65630 | 3,70 | 1,12 | 1,13 | 0,99 | 3,22E-01 | 4,92E-01 |
| AT5G09590 | 2,74 | 1,41 | 1,43 | 0,99 | 3,23E-01 | 4,92E-01 |
| AT1G62180 | 2,66 | 1,47 | 1,49 | 0,99 | 3,23E-01 | 4,92E-01 |
| AT3G17390 | 4,31 | -1,01 | 1,02 | -0,99 | 3,23E-01 | 4,92E-01 |
| AT3G12120 | 5,50 | 0,79 | 0,80 | 0,99 | 3,24E-01 | 4,93E-01 |
| AT4G01010 | 2,44 | -1,30 | 1,31 | -0,99 | 3,24E-01 | 4,93E-01 |
| AT1G17290 | 2,38 | 1,52 | 1,55 | 0,98 | 3,25E-01 | 4,94E-01 |
| AT1G27535 | 2,34 | -1,43 | 1,45 | -0,98 | 3,25E-01 | 4,94E-01 |
| AT1G32010 | 1,81 | -1,90 | 1,94 | -0,98 | 3,26E-01 | 4,94E-01 |
| AT1G54270 | 2,59 | -1,36 | 1,39 | -0,98 | 3,26E-01 | 4,95E-01 |
| AT2G20950 | 3,17 | -1,24 | 1,26 | -0,98 | 3,27E-01 | 4,96E-01 |
| AT2G30440 | 1,74 | 1,63 | 1,66 | 0,98 | 3,29E-01 | 4,98E-01 |
| AT3G57730 | 5,17 | 0,86 | 0,88 | 0,98 | 3,29E-01 | 4,98E-01 |
| AT3G49470 | 2,41 | -1,55 | 1,59 | -0,98 | 3,29E-01 | 4,98E-01 |
| AT5G07740 | 1,77 | 1,53 | 1,57 | 0,98 | 3,29E-01 | 4,98E-01 |
| AT5G40340 | 2,24 | -1,26 | 1,29 | -0,97 | 3,32E-01 | 5,01E-01 |
| AT1G09210 | 5,83 | 0,79 | 0,81 | 0,97 | 3,32E-01 | 5,01E-01 |
| AT5G10760 | 4,94 | 0,93 | 0,96 | 0,97 | 3,32E-01 | 5,01E-01 |
| AT1G17220 | 3,72 | 0,98 | 1,01 | 0,97 | 3,34E-01 | 5,03E-01 |
| AT1G29670 | 1,73 | -1,52 | 1,57 | -0,97 | 3,34E-01 | 5,03E-01 |
| AT1G43320 | 2,75 | 1,98 | 2,05 | 0,97 | 3,34E-01 | 5,03E-01 |
| AT3G11670 | 1,77 | 1,58 | 1,64 | 0,97 | 3,34E-01 | 5,03E-01 |
| AT5G67420 | 2,71 | 1,18 | 1,22 | 0,97 | 3,34E-01 | 5,03E-01 |
| AT3G15880 | 1,94 | 1,34 | 1,39 | 0,96 | 3,36E-01 | 5,06E-01 |
| AT2G22660 | 2,02 | 1,44 | 1,51 | 0,96 | 3,39E-01 | 5,10E-01 |
| AT5G02305 | 2,96 | -1,33 | 1,40 | -0,95 | 3,41E-01 | 5,12E-01 |
| AT1G18160 | 2,26 | 1,74 | 1,83 | 0,95 | 3,42E-01 | 5,12E-01 |
| AT4G16180 | 1,78 | 2,24 | 2,36 | 0,95 | 3,42E-01 | 5,12E-01 |
| AT5G38430 | 4,72 | 0,77 | 0,81 | 0,95 | 3,42E-01 | 5,12E-01 |
| AT5G39770 | 12,51 | -0,62 | 0,65 | -0,95 | 3,42E-01 | 5,12E-01 |
| AT3G59710 | 6,69 | -0,86 | 0,90 | -0,95 | 3,44E-01 | 5,14E-01 |
| AT3G09225 | 6,69 | -0,86 | 0,90 | -0,95 | 3,44E-01 | 5,14E-01 |
| AT4G19520 | 3,67 | 0,99 | 1,04 | 0,95 | 3,43E-01 | 5,14E-01 |
| AT3G58193 | 8,75 | -0,85 | 0,89 | -0,95 | 3,44E-01 | 5,14E-01 |
| AT3G53420 | 2,26 | -1,45 | 1,53 | -0,95 | 3,44E-01 | 5,14E-01 |
| AT1G31910 | 2,79 | 1,06 | 1,13 | 0,94 | 3,46E-01 | 5,16E-01 |
| AT3G59940 | 11,46 | -0,70 | 0,74 | -0,94 | 3,46E-01 | 5,16E-01 |
| AT5G28540 | 7,23 | 0,74 | 0,78 | 0,94 | 3,46E-01 | 5,16E-01 |
| AT5G40480 | 2,70 | 1,24 | 1,31 | 0,94 | 3,47E-01 | 5,16E-01 |
| AT1G78478 | 10,59 | -0,72 | 0,77 | -0,94 | 3,47E-01 | 5,16E-01 |
| AT3G09275 | 11,41 | -0,70 | 0,75 | -0,94 | 3,47E-01 | 5,16E-01 |
| AT4G00430 | 2,16 | -1,23 | 1,31 | -0,94 | 3,47E-01 | 5,16E-01 |
| AT4G34110 | 4,15 | 0,84 | 0,89 | 0,94 | 3,48E-01 | 5,18E-01 |
| AT1G15820 | 12,09 | -0,62 | 0,66 | -0,94 | 3,49E-01 | 5,19E-01 |
| AT2G04435 | 3,56 | 1,27 | 1,36 | 0,93 | 3,52E-01 | 5,21E-01 |
| AT1G44575 | 5,52 | -0,74 | 0,80 | -0,93 | 3,52E-01 | 5,22E-01 |
| AT5G01960 | 1,82 | 1,98 | 2,13 | 0,93 | 3,52E-01 | 5,22E-01 |
| AT4G14365 | 1,73 | 1,79 | 1,93 | 0,93 | 3,54E-01 | 5,24E-01 |
| AT4G33650 | 1,75 | 1,59 | 1,72 | 0,93 | 3,55E-01 | 5,25E-01 |
| AT1G30360 | 3,38 | 0,96 | 1,04 | 0,92 | 3,56E-01 | 5,26E-01 |
| AT5G01915 | 2,27 | -1,60 | 1,74 | -0,92 | 3,58E-01 | 5,29E-01 |
| AT1G04425 | 1,91 | -1,42 | 1,55 | -0,92 | 3,58E-01 | 5,29E-01 |
| AT1G05035 | 3,55 | -1,11 | 1,20 | -0,92 | 3,58E-01 | 5,29E-01 |
| AT3G28900 | 3,87 | 0,95 | 1,04 | 0,92 | 3,60E-01 | 5,31E-01 |

| | | | | | | |
|---|---|---|---|---|---|---|
| AT1G05917 | 8,49 | -0,78 | 0,86 | -0,91 | 3,61E-01 | 5,32E-01 |
| AT1G20100 | 1,77 | 1,74 | 1,91 | 0,91 | 3,62E-01 | 5,33E-01 |
| AT1G08103 | 6,40 | -0,88 | 0,96 | -0,91 | 3,62E-01 | 5,33E-01 |
| AT5G45100 | 1,72 | 1,91 | 2,10 | 0,91 | 3,62E-01 | 5,33E-01 |
| AT5G09670 | 2,53 | 1,17 | 1,28 | 0,91 | 3,63E-01 | 5,34E-01 |
| AT5G35630 | 4,18 | -0,90 | 0,99 | -0,91 | 3,64E-01 | 5,34E-01 |
| AT2G40840 | 4,12 | 0,83 | 0,91 | 0,91 | 3,64E-01 | 5,35E-01 |
| AT1G21200 | 5,07 | 1,02 | 1,13 | 0,90 | 3,66E-01 | 5,36E-01 |
| AT5G63190 | 2,74 | 1,16 | 1,28 | 0,90 | 3,66E-01 | 5,36E-01 |
| AT1G05817 | 1,94 | 1,27 | 1,41 | 0,90 | 3,66E-01 | 5,36E-01 |
| AT2G04955 | 3,52 | 1,02 | 1,12 | 0,90 | 3,67E-01 | 5,36E-01 |
| AT3G24615 | 9,91 | -0,61 | 0,67 | -0,90 | 3,67E-01 | 5,36E-01 |
| AT5G42895 | 1,80 | 1,50 | 1,67 | 0,90 | 3,67E-01 | 5,37E-01 |
| AT5G10450 | 3,14 | 0,96 | 1,07 | 0,90 | 3,68E-01 | 5,38E-01 |
| AT4G13505 | 4,72 | 0,91 | 1,02 | 0,90 | 3,69E-01 | 5,39E-01 |
| AT4G25520 | 4,54 | -0,81 | 0,90 | -0,90 | 3,69E-01 | 5,39E-01 |
| AT1G15670 | 2,03 | 1,26 | 1,40 | 0,90 | 3,70E-01 | 5,40E-01 |
| AT5G38420 | 5,05 | 0,72 | 0,81 | 0,89 | 3,73E-01 | 5,44E-01 |
| AT1G53210 | 1,83 | -1,63 | 1,83 | -0,89 | 3,74E-01 | 5,44E-01 |
| AT1G05030 | 1,78 | -1,66 | 1,87 | -0,89 | 3,74E-01 | 5,44E-01 |
| AT3G57710 | 4,92 | 0,86 | 0,97 | 0,89 | 3,74E-01 | 5,44E-01 |
| AT2G04755 | 2,18 | 1,54 | 1,74 | 0,89 | 3,75E-01 | 5,45E-01 |
| AT4G13510 | 4,76 | 0,91 | 1,02 | 0,89 | 3,75E-01 | 5,45E-01 |
| AT1G20690 | 2,85 | -1,13 | 1,29 | -0,88 | 3,81E-01 | 5,52E-01 |
| AT2G26975 | 1,93 | 1,40 | 1,60 | 0,88 | 3,80E-01 | 5,52E-01 |
| AT5G13180 | 1,85 | 1,44 | 1,64 | 0,88 | 3,80E-01 | 5,52E-01 |
| AT1G20691 | 2,85 | -1,13 | 1,29 | -0,88 | 3,81E-01 | 5,52E-01 |
| AT3G44110 | 2,03 | 1,16 | 1,33 | 0,88 | 3,81E-01 | 5,52E-01 |
| AT5G28840 | 2,50 | 0,99 | 1,13 | 0,88 | 3,81E-01 | 5,52E-01 |
| AT5G49730 | 3,55 | -1,20 | 1,36 | -0,88 | 3,80E-01 | 5,52E-01 |
| AT2G04280 | 2,22 | 1,52 | 1,74 | 0,87 | 3,83E-01 | 5,53E-01 |
| AT2G33730 | 2,24 | 1,09 | 1,26 | 0,87 | 3,84E-01 | 5,54E-01 |
| AT3G22120 | 4,08 | 0,87 | 1,00 | 0,87 | 3,84E-01 | 5,54E-01 |
| AT1G11785 | 2,48 | -1,07 | 1,23 | -0,87 | 3,85E-01 | 5,56E-01 |
| AT5G54250 | 1,88 | 1,36 | 1,57 | 0,87 | 3,86E-01 | 5,57E-01 |
| AT4G23170 | 6,38 | 0,86 | 0,99 | 0,86 | 3,87E-01 | 5,58E-01 |
| AT5G03350 | 2,45 | -1,13 | 1,30 | -0,87 | 3,87E-01 | 5,58E-01 |
| AT2G07718 | 3,36 | 1,02 | 1,18 | 0,86 | 3,88E-01 | 5,59E-01 |
| AT2G18480 | 2,00 | -1,38 | 1,60 | -0,86 | 3,89E-01 | 5,60E-01 |
| AT1G72645 | 2,19 | -1,28 | 1,49 | -0,85 | 3,93E-01 | 5,66E-01 |
| AT5G27850 | 1,94 | 1,33 | 1,55 | 0,85 | 3,94E-01 | 5,66E-01 |
| AT1G63750 | 2,64 | 1,11 | 1,31 | 0,85 | 3,95E-01 | 5,66E-01 |
| AT3G27690 | 1,98 | -1,22 | 1,44 | -0,85 | 3,95E-01 | 5,66E-01 |
| AT2G05070 | 19,17 | -0,42 | 0,49 | -0,85 | 3,94E-01 | 5,66E-01 |
| AT4G13840 | 1,76 | -1,39 | 1,63 | -0,85 | 3,94E-01 | 5,66E-01 |
| AT5G14780 | 2,20 | -1,24 | 1,46 | -0,85 | 3,96E-01 | 5,67E-01 |
| AT4G03260 | 2,29 | 1,10 | 1,30 | 0,85 | 3,96E-01 | 5,67E-01 |
| AT2G04965 | 3,81 | -0,81 | 0,96 | -0,85 | 3,96E-01 | 5,67E-01 |
| AT4G32640 | 2,41 | 1,09 | 1,29 | 0,85 | 3,97E-01 | 5,68E-01 |
| AT1G05563 | 2,75 | -1,12 | 1,33 | -0,85 | 3,97E-01 | 5,68E-01 |
| AT4G27090 | 2,08 | -1,30 | 1,54 | -0,85 | 3,97E-01 | 5,68E-01 |
| AT3G00985 | 2,03 | -1,77 | 2,10 | -0,84 | 3,99E-01 | 5,69E-01 |
| AT1G66730 | 1,91 | -1,24 | 1,47 | -0,84 | 3,99E-01 | 5,70E-01 |
| AT3G13440 | 3,42 | 1,11 | 1,32 | 0,84 | 4,00E-01 | 5,70E-01 |
| AT1G51805 | 2,63 | -1,12 | 1,33 | -0,84 | 4,00E-01 | 5,70E-01 |
| AT5G52470 | 2,57 | 1,12 | 1,33 | 0,84 | 4,00E-01 | 5,70E-01 |

| | | | | | | |
|---|---|---|---|---|---|---|
| AT4G04640 | 5,86 | -0,71 | 0,84 | -0,84 | 4,01E-01 | 5,70E-01 |
| AT5G13890 | 14,03 | 0,67 | 0,80 | 0,84 | 4,02E-01 | 5,72E-01 |
| AT1G72150 | 7,68 | -0,66 | 0,79 | -0,84 | 4,03E-01 | 5,73E-01 |
| AT1G14580 | 3,99 | -0,97 | 1,16 | -0,84 | 4,04E-01 | 5,73E-01 |
| AT3G00980 | 2,10 | -1,73 | 2,07 | -0,84 | 4,03E-01 | 5,73E-01 |
| AT2G33150 | 2,47 | -1,06 | 1,28 | -0,83 | 4,05E-01 | 5,74E-01 |
| AT1G80000 | 1,90 | 1,69 | 2,04 | 0,83 | 4,06E-01 | 5,75E-01 |
| AT5G38990 | 2,97 | 1,11 | 1,33 | 0,83 | 4,07E-01 | 5,76E-01 |
| AT4G35310 | 1,84 | -1,42 | 1,71 | -0,83 | 4,07E-01 | 5,76E-01 |
| AT2G43150 | 1,94 | 1,30 | 1,57 | 0,83 | 4,09E-01 | 5,78E-01 |
| AT5G10860 | 1,76 | 1,34 | 1,62 | 0,82 | 4,10E-01 | 5,80E-01 |
| AT4G37930 | 4,20 | -0,79 | 0,96 | -0,82 | 4,11E-01 | 5,81E-01 |
| AT1G29370 | 3,57 | 0,86 | 1,06 | 0,82 | 4,13E-01 | 5,83E-01 |
| AT5G02880 | 3,00 | 1,06 | 1,29 | 0,82 | 4,14E-01 | 5,84E-01 |
| AT5G09585 | 3,20 | -1,43 | 1,75 | -0,82 | 4,14E-01 | 5,84E-01 |
| AT1G12013 | 2,08 | -1,13 | 1,38 | -0,82 | 4,15E-01 | 5,84E-01 |
| AT4G39330 | 1,70 | 1,18 | 1,45 | 0,82 | 4,15E-01 | 5,84E-01 |
| AT4G39400 | 3,54 | 0,95 | 1,16 | 0,81 | 4,15E-01 | 5,85E-01 |
| AT3G61470 | 27,14 | -0,37 | 0,46 | -0,81 | 4,17E-01 | 5,87E-01 |
| AT5G59732 | 3,71 | 1,00 | 1,23 | 0,81 | 4,18E-01 | 5,87E-01 |
| AT5G59730 | 3,71 | 1,00 | 1,23 | 0,81 | 4,18E-01 | 5,87E-01 |
| AT3G47620 | 3,09 | 0,98 | 1,21 | 0,81 | 4,18E-01 | 5,87E-01 |
| AT5G05750 | 1,85 | 1,84 | 2,28 | 0,81 | 4,19E-01 | 5,88E-01 |
| AT3G01345 | 6,06 | -1,37 | 1,69 | -0,81 | 4,19E-01 | 5,88E-01 |
| AT4G19170 | 5,23 | -0,75 | 0,93 | -0,81 | 4,19E-01 | 5,88E-01 |
| AT3G23125 | 239,13 | 0,36 | 0,45 | 0,81 | 4,20E-01 | 5,88E-01 |
| AT4G16980 | 2,88 | -0,94 | 1,17 | -0,81 | 4,20E-01 | 5,88E-01 |
| AT3G04775 | 14,57 | -0,43 | 0,53 | -0,81 | 4,21E-01 | 5,89E-01 |
| AT1G33415 | 2,72 | -1,00 | 1,25 | -0,80 | 4,22E-01 | 5,90E-01 |
| AT4G16990 | 5,03 | 0,74 | 0,92 | 0,80 | 4,22E-01 | 5,90E-01 |
| AT2G41530 | 2,42 | 1,03 | 1,28 | 0,80 | 4,22E-01 | 5,90E-01 |
| AT1G05427 | 2,11 | -1,27 | 1,59 | -0,80 | 4,25E-01 | 5,92E-01 |
| AT3G25020 | 3,41 | -1,13 | 1,41 | -0,80 | 4,25E-01 | 5,92E-01 |
| AT4G16983 | 2,85 | -0,93 | 1,17 | -0,80 | 4,25E-01 | 5,92E-01 |
| AT4G00360 | 2,92 | -0,88 | 1,11 | -0,80 | 4,26E-01 | 5,94E-01 |
| AT2G37710 | 4,76 | -0,81 | 1,02 | -0,79 | 4,28E-01 | 5,96E-01 |
| AT1G07590 | 3,38 | -1,25 | 1,58 | -0,79 | 4,29E-01 | 5,98E-01 |
| AT5G22800 | 1,71 | -1,14 | 1,45 | -0,79 | 4,32E-01 | 6,01E-01 |
| AT5G19940 | 2,37 | -0,96 | 1,23 | -0,78 | 4,35E-01 | 6,04E-01 |
| AT1G09690 | 1,78 | 1,14 | 1,46 | 0,78 | 4,35E-01 | 6,04E-01 |
| AT1G19410 | 3,18 | -0,87 | 1,12 | -0,78 | 4,35E-01 | 6,04E-01 |
| AT1G46696 | 1,91 | -1,42 | 1,81 | -0,78 | 4,35E-01 | 6,04E-01 |
| AT1G03130 | 1,99 | -1,03 | 1,32 | -0,78 | 4,36E-01 | 6,05E-01 |
| AT4G00231 | 1,99 | 1,22 | 1,58 | 0,77 | 4,40E-01 | 6,08E-01 |
| AT4G06695 | 4,20 | -0,85 | 1,10 | -0,77 | 4,40E-01 | 6,08E-01 |
| AT4G31980 | 2,53 | -1,15 | 1,49 | -0,77 | 4,40E-01 | 6,08E-01 |
| AT5G11670 | 3,41 | 0,89 | 1,15 | 0,77 | 4,40E-01 | 6,08E-01 |
| AT3G54925 | 1,76 | -1,39 | 1,80 | -0,77 | 4,40E-01 | 6,08E-01 |
| AT2G41940 | 1,76 | 1,39 | 1,80 | 0,77 | 4,40E-01 | 6,08E-01 |
| AT2G06555 | 20,57 | -0,69 | 0,90 | -0,77 | 4,41E-01 | 6,09E-01 |
| AT4G00975 | 11,27 | 0,43 | 0,55 | 0,77 | 4,42E-01 | 6,09E-01 |
| AT2G16676 | 2,63 | -0,96 | 1,25 | -0,77 | 4,42E-01 | 6,09E-01 |
| AT5G15020 | 2,14 | 1,24 | 1,62 | 0,77 | 4,42E-01 | 6,09E-01 |
| AT1G19090 | 4,93 | -0,87 | 1,14 | -0,76 | 4,45E-01 | 6,13E-01 |
| AT3G52880 | 2,23 | -1,14 | 1,50 | -0,76 | 4,47E-01 | 6,15E-01 |
| AT5G46510 | 5,48 | 0,67 | 0,88 | 0,76 | 4,48E-01 | 6,16E-01 |

| | | | | | | |
|---|---|---|---|---|---|---|
| AT1G05780 | 4,77 | 0,99 | 1,30 | 0,76 | 4,49E-01 | 6,17E-01 |
| AT1G05273 | 8,42 | 0,55 | 0,72 | 0,76 | 4,49E-01 | 6,17E-01 |
| AT1G16820 | 8,42 | 0,55 | 0,72 | 0,76 | 4,49E-01 | 6,17E-01 |
| AT3G43833 | 7,44 | 1,38 | 1,83 | 0,76 | 4,50E-01 | 6,18E-01 |
| AT4G08335 | 4,06 | 1,39 | 1,84 | 0,75 | 4,51E-01 | 6,18E-01 |
| AT1G72610 | 17,51 | -0,37 | 0,49 | -0,75 | 4,51E-01 | 6,18E-01 |
| AT4G16143 | 2,03 | 1,07 | 1,42 | 0,75 | 4,51E-01 | 6,18E-01 |
| AT4G08175 | 7,31 | -0,58 | 0,77 | -0,75 | 4,51E-01 | 6,18E-01 |
| AT5G04853 | 3,91 | 1,38 | 1,84 | 0,75 | 4,53E-01 | 6,19E-01 |
| AT5G63195 | 1,93 | 1,21 | 1,63 | 0,74 | 4,56E-01 | 6,24E-01 |
| AT1G55673 | 3,49 | -0,71 | 0,95 | -0,74 | 4,57E-01 | 6,24E-01 |
| AT2G43110 | 4,82 | 0,77 | 1,03 | 0,74 | 4,57E-01 | 6,24E-01 |
| AT3G06436 | 20,96 | -0,39 | 0,53 | -0,74 | 4,57E-01 | 6,24E-01 |
| AT4G17788 | 20,18 | 0,35 | 0,48 | 0,74 | 4,58E-01 | 6,25E-01 |
| AT3G22121 | 3,96 | 0,76 | 1,03 | 0,74 | 4,59E-01 | 6,26E-01 |
| AT4G18150 | 2,31 | -1,14 | 1,54 | -0,74 | 4,59E-01 | 6,26E-01 |
| AT5G28920 | 2,84 | 1,01 | 1,36 | 0,74 | 4,59E-01 | 6,26E-01 |
| AT3G29796 | 2,77 | -1,36 | 1,84 | -0,74 | 4,60E-01 | 6,26E-01 |
| AT3G03475 | 2,58 | -0,98 | 1,33 | -0,74 | 4,61E-01 | 6,27E-01 |
| AT5G48380 | 2,92 | 0,97 | 1,32 | 0,74 | 4,62E-01 | 6,29E-01 |
| AT5G15200 | 4,33 | 0,64 | 0,88 | 0,73 | 4,63E-01 | 6,29E-01 |
| AT2G27310 | 2,22 | 1,07 | 1,46 | 0,73 | 4,63E-01 | 6,29E-01 |
| AT5G06165 | 4,41 | -0,78 | 1,06 | -0,73 | 4,63E-01 | 6,29E-01 |
| AT1G16880 | 2,91 | 0,96 | 1,32 | 0,73 | 4,64E-01 | 6,29E-01 |
| AT1G61226 | 20,72 | -0,36 | 0,50 | -0,73 | 4,65E-01 | 6,30E-01 |
| AT4G12290 | 3,10 | 1,11 | 1,52 | 0,73 | 4,65E-01 | 6,30E-01 |
| AT1G70490 | 1,85 | 1,03 | 1,41 | 0,73 | 4,67E-01 | 6,32E-01 |
| AT3G62010 | 2,16 | 0,95 | 1,30 | 0,73 | 4,68E-01 | 6,33E-01 |
| AT4G14400 | 8,78 | 0,59 | 0,81 | 0,73 | 4,68E-01 | 6,33E-01 |
| AT5G24890 | 2,39 | 1,01 | 1,39 | 0,73 | 4,68E-01 | 6,33E-01 |
| AT2G16020 | 3,14 | 1,38 | 1,91 | 0,72 | 4,70E-01 | 6,35E-01 |
| AT1G05913 | 9,06 | -0,62 | 0,86 | -0,72 | 4,70E-01 | 6,35E-01 |
| AT2G05765 | 3,87 | -0,86 | 1,18 | -0,72 | 4,70E-01 | 6,35E-01 |
| AT2G43570 | 2,25 | 1,34 | 1,86 | 0,72 | 4,71E-01 | 6,35E-01 |
| AT1G62540 | 2,48 | -1,18 | 1,64 | -0,72 | 4,72E-01 | 6,36E-01 |
| AT2G47440 | 2,19 | 1,12 | 1,56 | 0,72 | 4,72E-01 | 6,36E-01 |
| AT1G28270 | 2,06 | -1,04 | 1,45 | -0,72 | 4,74E-01 | 6,38E-01 |
| AT2G31820 | 4,59 | -0,70 | 0,98 | -0,71 | 4,75E-01 | 6,40E-01 |
| AT2G04455 | 3,39 | 1,01 | 1,42 | 0,71 | 4,76E-01 | 6,41E-01 |
| AT1G16710 | 1,85 | 1,27 | 1,78 | 0,71 | 4,77E-01 | 6,41E-01 |
| AT1G73870 | 2,16 | 0,90 | 1,27 | 0,71 | 4,79E-01 | 6,43E-01 |
| AT5G06530 | 2,30 | 1,04 | 1,48 | 0,70 | 4,82E-01 | 6,47E-01 |
| AT5G63145 | 2,83 | -0,86 | 1,22 | -0,70 | 4,83E-01 | 6,47E-01 |
| AT5G60600 | 2,43 | -0,87 | 1,24 | -0,70 | 4,84E-01 | 6,48E-01 |
| AT4G39800 | 2,68 | 0,80 | 1,15 | 0,70 | 4,85E-01 | 6,50E-01 |
| AT1G11910 | 5,12 | -0,73 | 1,05 | -0,69 | 4,88E-01 | 6,53E-01 |
| AT2G32240 | 2,12 | 1,31 | 1,89 | 0,69 | 4,87E-01 | 6,53E-01 |
| AT4G34150 | 1,90 | 1,28 | 1,86 | 0,69 | 4,90E-01 | 6,55E-01 |
| AT4G08620 | 2,91 | 1,37 | 1,99 | 0,69 | 4,92E-01 | 6,57E-01 |
| AT5G13887 | 13,72 | 0,62 | 0,90 | 0,69 | 4,92E-01 | 6,58E-01 |
| AT3G17190 | 1,85 | -1,32 | 1,92 | -0,69 | 4,93E-01 | 6,58E-01 |
| AT1G05850 | 2,93 | -0,98 | 1,43 | -0,69 | 4,93E-01 | 6,58E-01 |
| AT1G78476 | 11,35 | -0,50 | 0,73 | -0,68 | 4,94E-01 | 6,59E-01 |
| AT5G42092 | 3,68 | 0,80 | 1,17 | 0,68 | 4,94E-01 | 6,59E-01 |
| AT2G35260 | 3,89 | -0,70 | 1,03 | -0,68 | 4,96E-01 | 6,60E-01 |
| AT3G09630 | 1,89 | -0,98 | 1,44 | -0,68 | 4,95E-01 | 6,60E-01 |

| | | | | | | |
|---|---|---|---|---|---|---|
| AT4G30760 | 1,76 | 1,36 | 1,99 | 0,68 | 4,95E-01 | 6,60E-01 |
| AT3G09440 | 5,91 | 0,58 | 0,85 | 0,68 | 4,96E-01 | 6,60E-01 |
| AT2G16485 | 2,25 | 0,99 | 1,45 | 0,68 | 4,97E-01 | 6,60E-01 |
| AT4G06800 | 2,19 | -0,90 | 1,33 | -0,68 | 4,98E-01 | 6,61E-01 |
| AT1G72440 | 1,79 | 1,28 | 1,89 | 0,68 | 4,99E-01 | 6,62E-01 |
| AT2G30920 | 2,18 | -1,04 | 1,54 | -0,68 | 4,98E-01 | 6,62E-01 |
| AT3G54590 | 2,31 | 1,38 | 2,04 | 0,68 | 4,99E-01 | 6,62E-01 |
| AT5G37510 | 3,25 | 0,79 | 1,16 | 0,68 | 4,99E-01 | 6,62E-01 |
| AT4G12770 | 19,67 | 0,64 | 0,95 | 0,67 | 5,00E-01 | 6,62E-01 |
| AT3G52500 | 2,36 | -0,98 | 1,46 | -0,67 | 5,00E-01 | 6,62E-01 |
| AT1G53900 | 2,01 | -0,96 | 1,43 | -0,67 | 5,01E-01 | 6,63E-01 |
| AT1G26110 | 1,79 | 1,02 | 1,52 | 0,67 | 5,03E-01 | 6,65E-01 |
| AT3G26450 | 2,27 | -0,91 | 1,35 | -0,67 | 5,03E-01 | 6,65E-01 |
| AT2G01910 | 2,66 | -1,14 | 1,72 | -0,66 | 5,07E-01 | 6,69E-01 |
| AT4G13920 | 2,03 | 1,30 | 1,95 | 0,66 | 5,07E-01 | 6,69E-01 |
| AT4G26480 | 2,39 | 1,01 | 1,53 | 0,66 | 5,08E-01 | 6,70E-01 |
| AT3G24542 | 2,63 | -0,80 | 1,22 | -0,66 | 5,10E-01 | 6,72E-01 |
| AT3G23810 | 2,25 | -0,89 | 1,36 | -0,66 | 5,10E-01 | 6,72E-01 |
| AT5G01350 | 8,44 | 0,45 | 0,68 | 0,66 | 5,11E-01 | 6,73E-01 |
| AT2G14910 | 2,97 | 0,82 | 1,25 | 0,66 | 5,11E-01 | 6,73E-01 |
| AT3G10985 | 6,51 | 0,68 | 1,04 | 0,65 | 5,13E-01 | 6,74E-01 |
| AT3G44006 | 2,21 | -0,88 | 1,34 | -0,65 | 5,13E-01 | 6,74E-01 |
| AT2G45380 | 11,33 | -0,38 | 0,57 | -0,66 | 5,12E-01 | 6,74E-01 |
| AT3G60190 | 1,87 | -0,89 | 1,36 | -0,65 | 5,13E-01 | 6,74E-01 |
| AT1G71002 | 2,81 | 0,78 | 1,19 | 0,65 | 5,14E-01 | 6,74E-01 |
| AT4G17340 | 2,05 | 1,25 | 1,92 | 0,65 | 5,16E-01 | 6,77E-01 |
| AT4G26540 | 3,42 | 0,74 | 1,14 | 0,65 | 5,17E-01 | 6,78E-01 |
| AT4G06375 | 3,09 | 0,89 | 1,38 | 0,65 | 5,18E-01 | 6,79E-01 |
| AT4G09735 | 2,23 | 1,31 | 2,02 | 0,65 | 5,18E-01 | 6,79E-01 |
| AT1G12900 | 4,70 | -0,55 | 0,86 | -0,65 | 5,19E-01 | 6,79E-01 |
| AT4G13810 | 2,84 | -1,17 | 1,81 | -0,64 | 5,19E-01 | 6,79E-01 |
| AT4G27520 | 5,05 | -0,68 | 1,06 | -0,64 | 5,19E-01 | 6,79E-01 |
| AT1G52780 | 3,16 | 1,02 | 1,60 | 0,64 | 5,22E-01 | 6,82E-01 |
| AT1G72930 | 3,01 | -1,02 | 1,60 | -0,64 | 5,24E-01 | 6,82E-01 |
| AT2G18570 | 4,57 | -0,90 | 1,41 | -0,64 | 5,22E-01 | 6,82E-01 |
| AT5G65700 | 1,88 | 0,85 | 1,34 | 0,64 | 5,23E-01 | 6,82E-01 |
| AT1G33240 | 1,91 | 1,03 | 1,61 | 0,64 | 5,23E-01 | 6,82E-01 |
| AT2G35940 | 16,85 | 0,53 | 0,84 | 0,64 | 5,23E-01 | 6,82E-01 |
| AT2G36470 | 2,06 | -1,25 | 1,97 | -0,64 | 5,23E-01 | 6,82E-01 |
| AT3G53460 | 6,35 | 0,52 | 0,81 | 0,64 | 5,23E-01 | 6,82E-01 |
| AT1G35720 | 3,50 | 0,63 | 0,98 | 0,64 | 5,24E-01 | 6,82E-01 |
| AT4G14960 | 3,57 | 0,69 | 1,08 | 0,64 | 5,25E-01 | 6,82E-01 |
| AT5G59505 | 3,52 | -0,94 | 1,48 | -0,64 | 5,25E-01 | 6,82E-01 |
| AT3G56400 | 2,09 | 1,04 | 1,64 | 0,63 | 5,28E-01 | 6,85E-01 |
| AT2G28840 | 1,76 | 1,01 | 1,61 | 0,63 | 5,28E-01 | 6,85E-01 |
| AT1G32750 | 1,94 | 0,99 | 1,58 | 0,63 | 5,29E-01 | 6,87E-01 |
| AT2G19360 | 2,70 | -0,81 | 1,29 | -0,63 | 5,30E-01 | 6,87E-01 |
| AT2G21330 | 7,94 | 0,42 | 0,66 | 0,63 | 5,30E-01 | 6,87E-01 |
| AT5G10840 | 2,05 | 0,81 | 1,28 | 0,63 | 5,31E-01 | 6,87E-01 |
| AT3G55980 | 3,63 | 0,83 | 1,33 | 0,63 | 5,31E-01 | 6,88E-01 |
| AT4G20170 | 3,89 | 0,94 | 1,51 | 0,62 | 5,34E-01 | 6,90E-01 |
| AT1G69940 | 3,40 | -0,71 | 1,14 | -0,62 | 5,34E-01 | 6,90E-01 |
| AT3G28550 | 2,18 | 1,32 | 2,12 | 0,62 | 5,33E-01 | 6,90E-01 |
| AT5G43470 | 3,38 | 0,88 | 1,43 | 0,62 | 5,36E-01 | 6,92E-01 |
| AT5G61420 | 2,01 | 1,00 | 1,61 | 0,62 | 5,36E-01 | 6,92E-01 |
| AT5G11790 | 2,58 | -0,91 | 1,48 | -0,62 | 5,36E-01 | 6,92E-01 |

| | | | | | | |
|---|---|---|---|---|---|---|
| AT3G58030 | 2,17 | 1,02 | 1,65 | 0,62 | 5,37E-01 | 6,92E-01 |
| AT1G72600 | 18,04 | -0,29 | 0,48 | -0,62 | 5,38E-01 | 6,93E-01 |
| AT3G57400 | 1,77 | 1,08 | 1,76 | 0,62 | 5,38E-01 | 6,93E-01 |
| AT2G21660 | 24,65 | -0,30 | 0,48 | -0,62 | 5,38E-01 | 6,93E-01 |
| AT4G17090 | 4,63 | -0,58 | 0,94 | -0,61 | 5,39E-01 | 6,94E-01 |
| AT3G59410 | 1,72 | 0,98 | 1,59 | 0,61 | 5,40E-01 | 6,94E-01 |
| AT3G50480 | 1,71 | 1,14 | 1,87 | 0,61 | 5,41E-01 | 6,95E-01 |
| AT4G18197 | 1,87 | -0,97 | 1,59 | -0,61 | 5,41E-01 | 6,95E-01 |
| AT2G36390 | 2,07 | -0,91 | 1,49 | -0,61 | 5,42E-01 | 6,96E-01 |
| AT1G12280 | 1,78 | 0,85 | 1,40 | 0,61 | 5,42E-01 | 6,96E-01 |
| AT4G29190 | 1,79 | 0,84 | 1,39 | 0,61 | 5,43E-01 | 6,96E-01 |
| AT1G51690 | 2,38 | 0,78 | 1,29 | 0,61 | 5,43E-01 | 6,96E-01 |
| AT4G24190 | 5,76 | 0,55 | 0,90 | 0,61 | 5,44E-01 | 6,97E-01 |
| AT1G08200 | 1,82 | 0,99 | 1,63 | 0,60 | 5,46E-01 | 6,99E-01 |
| AT1G35560 | 2,43 | 0,71 | 1,17 | 0,60 | 5,46E-01 | 6,99E-01 |
| AT1G13470 | 1,76 | 1,08 | 1,80 | 0,60 | 5,47E-01 | 6,99E-01 |
| AT2G16600 | 11,35 | 0,45 | 0,76 | 0,60 | 5,49E-01 | 7,02E-01 |
| AT2G25170 | 3,04 | 0,84 | 1,41 | 0,60 | 5,51E-01 | 7,03E-01 |
| AT2G04885 | 2,17 | -1,14 | 1,91 | -0,60 | 5,52E-01 | 7,04E-01 |
| AT1G06460 | 1,79 | 0,87 | 1,46 | 0,60 | 5,51E-01 | 7,04E-01 |
| AT1G32582 | 26,60 | 0,35 | 0,59 | 0,59 | 5,55E-01 | 7,06E-01 |
| AT5G13530 | 2,40 | 0,78 | 1,32 | 0,59 | 5,55E-01 | 7,06E-01 |
| AT1G68530 | 2,97 | -0,67 | 1,13 | -0,59 | 5,55E-01 | 7,06E-01 |
| AT1G68560 | 3,34 | -0,77 | 1,31 | -0,59 | 5,55E-01 | 7,06E-01 |
| AT2G41720 | 1,87 | 1,09 | 1,84 | 0,59 | 5,55E-01 | 7,06E-01 |
| AT5G02645 | 2,33 | 0,76 | 1,29 | 0,59 | 5,54E-01 | 7,06E-01 |
| AT2G43375 | 6,65 | -0,45 | 0,76 | -0,59 | 5,56E-01 | 7,06E-01 |
| AT4G23100 | 1,83 | 0,84 | 1,42 | 0,59 | 5,56E-01 | 7,06E-01 |
| AT1G61190 | 2,38 | 0,85 | 1,45 | 0,59 | 5,57E-01 | 7,07E-01 |
| AT3G12780 | 5,73 | -0,48 | 0,82 | -0,59 | 5,57E-01 | 7,07E-01 |
| AT5G09000 | 2,90 | -0,62 | 1,06 | -0,59 | 5,59E-01 | 7,08E-01 |
| AT4G04970 | 2,63 | 1,28 | 2,20 | 0,58 | 5,59E-01 | 7,09E-01 |
| AT1G42550 | 2,32 | 0,79 | 1,36 | 0,58 | 5,61E-01 | 7,11E-01 |
| AT1G21680 | 4,80 | 0,49 | 0,85 | 0,58 | 5,63E-01 | 7,12E-01 |
| AT1G29070 | 1,74 | 0,82 | 1,43 | 0,57 | 5,67E-01 | 7,12E-01 |
| AT2G22990 | 2,03 | 0,76 | 1,32 | 0,58 | 5,64E-01 | 7,12E-01 |
| AT3G22960 | 3,17 | 0,64 | 1,11 | 0,57 | 5,67E-01 | 7,12E-01 |
| AT5G45472 | 4,93 | -0,68 | 1,19 | -0,57 | 5,67E-01 | 7,12E-01 |
| AT1G07670 | 1,74 | 0,80 | 1,40 | 0,57 | 5,66E-01 | 7,12E-01 |
| AT1G12090 | 8,60 | -0,41 | 0,71 | -0,58 | 5,64E-01 | 7,12E-01 |
| AT1G26850 | 2,19 | 0,80 | 1,40 | 0,57 | 5,67E-01 | 7,12E-01 |
| AT1G62035 | 5,54 | 0,49 | 0,85 | 0,57 | 5,67E-01 | 7,12E-01 |
| AT1G76090 | 1,72 | 0,93 | 1,61 | 0,58 | 5,65E-01 | 7,12E-01 |
| AT2G26530 | 1,88 | -1,19 | 2,08 | -0,57 | 5,67E-01 | 7,12E-01 |
| AT2G09465 | 4,22 | -0,57 | 1,00 | -0,57 | 5,66E-01 | 7,12E-01 |
| AT3G47500 | 2,57 | -0,71 | 1,24 | -0,57 | 5,67E-01 | 7,12E-01 |
| AT3G56910 | 1,94 | -0,89 | 1,56 | -0,57 | 5,67E-01 | 7,12E-01 |
| AT4G04465 | 2,27 | -0,88 | 1,53 | -0,57 | 5,66E-01 | 7,12E-01 |
| AT4G22240 | 1,83 | 0,75 | 1,30 | 0,57 | 5,66E-01 | 7,12E-01 |
| AT5G44090 | 1,96 | 0,97 | 1,69 | 0,57 | 5,66E-01 | 7,12E-01 |
| AT5G47560 | 3,13 | 0,70 | 1,22 | 0,58 | 5,64E-01 | 7,12E-01 |
| AT3G11560 | 1,70 | 1,20 | 2,10 | 0,57 | 5,68E-01 | 7,12E-01 |
| AT3G62410 | 1,88 | -0,79 | 1,38 | -0,57 | 5,68E-01 | 7,12E-01 |
| AT4G00060 | 1,77 | 1,03 | 1,81 | 0,57 | 5,69E-01 | 7,12E-01 |
| AT1G45160 | 1,73 | 1,01 | 1,79 | 0,57 | 5,71E-01 | 7,14E-01 |
| AT1G75950 | 1,80 | -0,75 | 1,33 | -0,57 | 5,72E-01 | 7,15E-01 |

| | | | | | | |
|---|---|---|---|---|---|---|
| AT1G35215 | 4,81 | -0,78 | 1,37 | -0,57 | 5,72E-01 | 7,15E-01 |
| AT5G57660 | 4,42 | 0,49 | 0,86 | 0,56 | 5,73E-01 | 7,16E-01 |
| AT1G65930 | 2,39 | 0,83 | 1,49 | 0,56 | 5,74E-01 | 7,17E-01 |
| AT4G35000 | 3,13 | -0,74 | 1,31 | -0,56 | 5,74E-01 | 7,17E-01 |
| AT4G05900 | 1,70 | -1,04 | 1,85 | -0,56 | 5,75E-01 | 7,17E-01 |
| AT1G26880 | 2,18 | -0,81 | 1,45 | -0,56 | 5,76E-01 | 7,18E-01 |
| AT2G29080 | 1,97 | 0,83 | 1,48 | 0,56 | 5,76E-01 | 7,18E-01 |
| AT3G52700 | 2,20 | -0,73 | 1,32 | -0,55 | 5,80E-01 | 7,22E-01 |
| AT3G07390 | 2,06 | -0,90 | 1,63 | -0,55 | 5,81E-01 | 7,23E-01 |
| AT3G24612 | 3,02 | 0,69 | 1,25 | 0,55 | 5,81E-01 | 7,23E-01 |
| AT4G03280 | 3,09 | 0,69 | 1,26 | 0,55 | 5,82E-01 | 7,24E-01 |
| AT5G19120 | 2,51 | 0,65 | 1,17 | 0,55 | 5,82E-01 | 7,24E-01 |
| AT5G02150 | 1,74 | -0,91 | 1,66 | -0,55 | 5,83E-01 | 7,24E-01 |
| AT1G77230 | 3,05 | 0,63 | 1,16 | 0,55 | 5,85E-01 | 7,25E-01 |
| AT1G80440 | 3,83 | -0,85 | 1,57 | -0,55 | 5,86E-01 | 7,25E-01 |
| AT2G08986 | 10,75 | -0,42 | 0,77 | -0,55 | 5,86E-01 | 7,25E-01 |
| AT2G38540 | 3,59 | -0,56 | 1,02 | -0,55 | 5,85E-01 | 7,25E-01 |
| AT3G46600 | 2,19 | 0,70 | 1,29 | 0,55 | 5,85E-01 | 7,25E-01 |
| AT1G18000 | 4,70 | -0,61 | 1,11 | -0,55 | 5,86E-01 | 7,25E-01 |
| AT3G07570 | 1,83 | 0,78 | 1,44 | 0,54 | 5,86E-01 | 7,26E-01 |
| AT3G04155 | 3,42 | -0,59 | 1,09 | -0,54 | 5,87E-01 | 7,26E-01 |
| AT4G30972 | 11,83 | -0,37 | 0,68 | -0,54 | 5,88E-01 | 7,26E-01 |
| AT5G39080 | 2,20 | -0,87 | 1,61 | -0,54 | 5,87E-01 | 7,26E-01 |
| AT4G30975 | 11,87 | -0,37 | 0,68 | -0,54 | 5,89E-01 | 7,28E-01 |
| AT2G38220 | 2,26 | -0,76 | 1,41 | -0,54 | 5,90E-01 | 7,29E-01 |
| AT5G08680 | 1,77 | 0,88 | 1,63 | 0,54 | 5,92E-01 | 7,30E-01 |
| AT5G51090 | 2,18 | -0,77 | 1,44 | -0,54 | 5,92E-01 | 7,30E-01 |
| AT3G54810 | 2,87 | 0,72 | 1,35 | 0,53 | 5,94E-01 | 7,31E-01 |
| AT3G59068 | 1,78 | 0,96 | 1,80 | 0,53 | 5,94E-01 | 7,31E-01 |
| AT2G45245 | 2,00 | 0,70 | 1,31 | 0,53 | 5,94E-01 | 7,31E-01 |
| AT3G59070 | 1,78 | 0,96 | 1,80 | 0,53 | 5,94E-01 | 7,31E-01 |
| AT1G42470 | 1,71 | 1,01 | 1,90 | 0,53 | 5,96E-01 | 7,31E-01 |
| AT5G28919 | 2,64 | 0,74 | 1,40 | 0,53 | 5,96E-01 | 7,31E-01 |
| AT5G46110 | 7,87 | 0,39 | 0,73 | 0,53 | 5,96E-01 | 7,31E-01 |
| AT1G13440 | 5,54 | -0,43 | 0,82 | -0,53 | 5,96E-01 | 7,31E-01 |
| AT5G04905 | 2,64 | 0,74 | 1,40 | 0,53 | 5,96E-01 | 7,31E-01 |
| AT5G45480 | 2,31 | 0,92 | 1,73 | 0,53 | 5,96E-01 | 7,31E-01 |
| AT1G13360 | 23,38 | 0,28 | 0,53 | 0,53 | 5,98E-01 | 7,33E-01 |
| AT3G05220 | 1,84 | -1,01 | 1,91 | -0,53 | 5,99E-01 | 7,34E-01 |
| AT3G13405 | 1,91 | -0,79 | 1,52 | -0,52 | 6,00E-01 | 7,35E-01 |
| AT3G17800 | 1,86 | 0,81 | 1,55 | 0,52 | 6,00E-01 | 7,35E-01 |
| AT1G55350 | 1,89 | 1,08 | 2,07 | 0,52 | 6,02E-01 | 7,37E-01 |
| AT1G11840 | 1,70 | 0,89 | 1,72 | 0,52 | 6,04E-01 | 7,38E-01 |
| AT2G07890 | 2,64 | -0,64 | 1,24 | -0,52 | 6,04E-01 | 7,38E-01 |
| AT4G13577 | 2,60 | -0,60 | 1,17 | -0,52 | 6,06E-01 | 7,40E-01 |
| AT2G18440 | 3,48 | 0,54 | 1,06 | 0,51 | 6,08E-01 | 7,42E-01 |
| AT1G53640 | 2,14 | 0,70 | 1,37 | 0,51 | 6,09E-01 | 7,43E-01 |
| AT4G38760 | 3,63 | 0,72 | 1,42 | 0,51 | 6,10E-01 | 7,44E-01 |
| AT4G13494 | 321,70 | 0,25 | 0,48 | 0,51 | 6,11E-01 | 7,44E-01 |
| AT4G16190 | 11,22 | 0,33 | 0,64 | 0,51 | 6,11E-01 | 7,44E-01 |
| AT5G42650 | 3,24 | -0,61 | 1,20 | -0,51 | 6,11E-01 | 7,44E-01 |
| AT1G67280 | 1,72 | 0,78 | 1,54 | 0,51 | 6,12E-01 | 7,44E-01 |
| AT2G40000 | 5,19 | 0,59 | 1,16 | 0,51 | 6,12E-01 | 7,44E-01 |
| AT3G25690 | 2,72 | 0,62 | 1,22 | 0,51 | 6,13E-01 | 7,45E-01 |
| AT5G41471 | 1,79 | -0,92 | 1,82 | -0,50 | 6,14E-01 | 7,46E-01 |
| AT5G46430 | 2,53 | -0,83 | 1,67 | -0,50 | 6,17E-01 | 7,49E-01 |

| | | | | | | |
|---|---|---|---|---|---|---|
| AT3G15353 | 5,91 | -0,59 | 1,18 | -0,50 | 6,19E-01 | 7,52E-01 |
| AT2G29670 | 3,81 | 0,53 | 1,08 | 0,49 | 6,21E-01 | 7,53E-01 |
| AT1G33070 | 1,82 | -0,90 | 1,83 | -0,49 | 6,23E-01 | 7,55E-01 |
| AT5G04425 | 2,72 | -0,65 | 1,33 | -0,49 | 6,24E-01 | 7,56E-01 |
| AT4G14030 | 1,74 | 0,74 | 1,52 | 0,49 | 6,25E-01 | 7,57E-01 |
| AT5G01410 | 2,12 | -0,77 | 1,59 | -0,49 | 6,26E-01 | 7,57E-01 |
| AT3G20820 | 3,82 | -0,65 | 1,35 | -0,49 | 6,27E-01 | 7,57E-01 |
| AT1G03080 | 1,73 | 0,82 | 1,69 | 0,49 | 6,27E-01 | 7,57E-01 |
| AT5G49030 | 1,79 | 0,70 | 1,44 | 0,49 | 6,26E-01 | 7,57E-01 |
| AT4G23387 | 136,04 | -0,28 | 0,57 | -0,49 | 6,27E-01 | 7,58E-01 |
| AT5G01155 | 4,02 | -0,53 | 1,10 | -0,48 | 6,30E-01 | 7,60E-01 |
| AT5G44270 | 2,68 | -0,64 | 1,32 | -0,48 | 6,29E-01 | 7,60E-01 |
| AT3G59970 | 1,71 | -0,85 | 1,78 | -0,48 | 6,31E-01 | 7,61E-01 |
| AT1G07920 | 7,76 | 0,35 | 0,73 | 0,48 | 6,32E-01 | 7,62E-01 |
| AT3G26440 | 2,47 | -0,64 | 1,33 | -0,48 | 6,32E-01 | 7,62E-01 |
| AT2G04405 | 3,40 | -0,55 | 1,15 | -0,48 | 6,34E-01 | 7,63E-01 |
| AT3G48200 | 2,11 | 0,66 | 1,39 | 0,48 | 6,35E-01 | 7,64E-01 |
| AT1G20330 | 2,79 | 0,70 | 1,47 | 0,47 | 6,35E-01 | 7,64E-01 |
| AT2G31141 | 64,55 | -0,16 | 0,33 | -0,47 | 6,36E-01 | 7,64E-01 |
| AT1G28440 | 1,72 | 0,65 | 1,39 | 0,47 | 6,39E-01 | 7,65E-01 |
| AT2G30950 | 3,17 | 0,48 | 1,02 | 0,47 | 6,38E-01 | 7,65E-01 |
| AT4G35770 | 2,85 | -0,61 | 1,29 | -0,47 | 6,37E-01 | 7,65E-01 |
| AT4G39364 | 2,19 | 0,84 | 1,79 | 0,47 | 6,39E-01 | 7,65E-01 |
| AT1G20440 | 3,88 | -0,43 | 0,92 | -0,47 | 6,39E-01 | 7,65E-01 |
| AT1G55450 | 2,26 | -0,79 | 1,68 | -0,47 | 6,37E-01 | 7,65E-01 |
| AT2G08725 | 34,05 | -0,29 | 0,61 | -0,47 | 6,37E-01 | 7,65E-01 |
| AT4G12650 | 2,23 | 0,60 | 1,28 | 0,47 | 6,38E-01 | 7,65E-01 |
| AT5G24530 | 1,77 | 0,70 | 1,49 | 0,47 | 6,42E-01 | 7,68E-01 |
| AT5G60640 | 2,58 | -0,59 | 1,27 | -0,46 | 6,42E-01 | 7,68E-01 |
| AT5G49440 | 5,06 | -0,41 | 0,88 | -0,46 | 6,44E-01 | 7,70E-01 |
| AT1G50920 | 2,95 | -0,49 | 1,07 | -0,46 | 6,46E-01 | 7,71E-01 |
| AT3G60750 | 11,99 | 0,28 | 0,61 | 0,46 | 6,46E-01 | 7,71E-01 |
| AT5G13850 | 2,26 | 0,61 | 1,34 | 0,46 | 6,46E-01 | 7,71E-01 |
| AT5G14570 | 2,87 | 0,67 | 1,45 | 0,46 | 6,46E-01 | 7,71E-01 |
| AT1G14920 | 1,77 | -0,67 | 1,47 | -0,45 | 6,50E-01 | 7,73E-01 |
| AT2G30860 | 6,26 | 0,37 | 0,82 | 0,46 | 6,49E-01 | 7,73E-01 |
| AT2G46765 | 2,57 | 0,60 | 1,32 | 0,45 | 6,50E-01 | 7,73E-01 |
| AT4G09000 | 1,74 | 0,70 | 1,53 | 0,46 | 6,48E-01 | 7,73E-01 |
| AT1G67090 | 31,55 | 0,21 | 0,46 | 0,46 | 6,49E-01 | 7,73E-01 |
| AT1G65720 | 3,55 | 0,51 | 1,14 | 0,45 | 6,51E-01 | 7,74E-01 |
| AT3G08940 | 8,84 | -0,35 | 0,78 | -0,45 | 6,51E-01 | 7,74E-01 |
| AT2G47170 | 1,90 | -0,71 | 1,59 | -0,45 | 6,54E-01 | 7,77E-01 |
| AT3G44703 | 4,72 | -0,39 | 0,87 | -0,45 | 6,55E-01 | 7,78E-01 |
| AT2G13560 | 2,19 | -0,74 | 1,65 | -0,45 | 6,56E-01 | 7,78E-01 |
| AT4G34950 | 4,16 | 0,41 | 0,92 | 0,44 | 6,59E-01 | 7,82E-01 |
| AT2G06510 | 1,80 | 0,74 | 1,68 | 0,44 | 6,60E-01 | 7,82E-01 |
| AT3G23820 | 9,86 | 0,26 | 0,60 | 0,44 | 6,60E-01 | 7,82E-01 |
| AT4G11570 | 2,16 | -0,55 | 1,25 | -0,44 | 6,62E-01 | 7,84E-01 |
| AT4G21280 | 3,20 | -0,48 | 1,09 | -0,44 | 6,62E-01 | 7,84E-01 |
| AT1G72180 | 2,71 | 0,50 | 1,15 | 0,43 | 6,64E-01 | 7,85E-01 |
| AT4G20735 | 3,19 | -0,82 | 1,91 | -0,43 | 6,66E-01 | 7,88E-01 |
| AT4G26530 | 4,43 | 0,41 | 0,96 | 0,43 | 6,68E-01 | 7,90E-01 |
| AT4G31700 | 1,83 | -0,71 | 1,66 | -0,43 | 6,68E-01 | 7,90E-01 |
| AT1G18080 | 2,60 | 0,54 | 1,25 | 0,43 | 6,69E-01 | 7,90E-01 |
| AT5G60920 | 1,88 | -0,67 | 1,57 | -0,43 | 6,70E-01 | 7,90E-01 |
| AT3G61430 | 1,94 | 0,74 | 1,73 | 0,43 | 6,70E-01 | 7,91E-01 |

| | | | | | | |
|---|---|---|---|---|---|---|
| AT3G49230 | 1,94 | -0,73 | 1,74 | -0,42 | 6,74E-01 | 7,94E-01 |
| AT4G04125 | 2,22 | -0,55 | 1,32 | -0,42 | 6,75E-01 | 7,95E-01 |
| AT1G30330 | 1,89 | -0,67 | 1,62 | -0,42 | 6,77E-01 | 7,97E-01 |
| AT2G25490 | 3,68 | -0,52 | 1,24 | -0,42 | 6,78E-01 | 7,98E-01 |
| AT1G44446 | 1,72 | 0,68 | 1,65 | 0,41 | 6,79E-01 | 7,99E-01 |
| AT5G09810 | 3,96 | 0,47 | 1,14 | 0,41 | 6,81E-01 | 8,00E-01 |
| AT1G04527 | 11,72 | -0,24 | 0,57 | -0,41 | 6,81E-01 | 8,00E-01 |
| AT5G26340 | 2,20 | 0,52 | 1,27 | 0,41 | 6,81E-01 | 8,00E-01 |
| AT1G72710 | 1,73 | 0,64 | 1,56 | 0,41 | 6,83E-01 | 8,01E-01 |
| AT1G76490 | 3,44 | 0,41 | 0,99 | 0,41 | 6,83E-01 | 8,01E-01 |
| AT2G07806 | 2,71 | 0,51 | 1,27 | 0,40 | 6,87E-01 | 8,05E-01 |
| AT1G03743 | 2,10 | -0,60 | 1,48 | -0,40 | 6,87E-01 | 8,05E-01 |
| AT2G47240 | 1,70 | -0,65 | 1,62 | -0,40 | 6,88E-01 | 8,05E-01 |
| AT3G44310 | 10,52 | -0,29 | 0,72 | -0,40 | 6,89E-01 | 8,07E-01 |
| AT5G52190 | 1,80 | 0,80 | 2,00 | 0,40 | 6,89E-01 | 8,07E-01 |
| AT3G54050 | 3,78 | 0,37 | 0,93 | 0,40 | 6,90E-01 | 8,07E-01 |
| AT4G39090 | 7,03 | -0,33 | 0,84 | -0,40 | 6,90E-01 | 8,07E-01 |
| AT1G06247 | 2,53 | -0,47 | 1,19 | -0,40 | 6,92E-01 | 8,09E-01 |
| AT1G04733 | 20,09 | -0,21 | 0,53 | -0,40 | 6,92E-01 | 8,09E-01 |
| AT5G60360 | 2,97 | -0,50 | 1,28 | -0,39 | 6,96E-01 | 8,13E-01 |
| AT2G16660 | 2,17 | 0,51 | 1,30 | 0,39 | 6,97E-01 | 8,13E-01 |
| AT5G42090 | 4,37 | 0,42 | 1,08 | 0,39 | 6,97E-01 | 8,13E-01 |
| AT5G58950 | 2,42 | -0,54 | 1,41 | -0,38 | 7,00E-01 | 8,16E-01 |
| AT1G55920 | 3,12 | 0,54 | 1,41 | 0,38 | 7,01E-01 | 8,17E-01 |
| AT4G39361 | 1,80 | -0,78 | 2,03 | -0,38 | 7,02E-01 | 8,17E-01 |
| AT1G69620 | 2,14 | 0,59 | 1,54 | 0,38 | 7,02E-01 | 8,17E-01 |
| AT2G34410 | 2,89 | 0,52 | 1,37 | 0,38 | 7,03E-01 | 8,17E-01 |
| AT3G16140 | 1,91 | 0,53 | 1,39 | 0,38 | 7,03E-01 | 8,17E-01 |
| AT1G09340 | 1,73 | 0,51 | 1,34 | 0,38 | 7,04E-01 | 8,18E-01 |
| AT5G49720 | 3,89 | -0,39 | 1,02 | -0,38 | 7,06E-01 | 8,20E-01 |
| AT1G62310 | 4,32 | -0,34 | 0,90 | -0,38 | 7,07E-01 | 8,21E-01 |
| AT4G12320 | 1,93 | 0,54 | 1,45 | 0,38 | 7,08E-01 | 8,21E-01 |
| AT5G18490 | 1,80 | 0,75 | 2,01 | 0,38 | 7,08E-01 | 8,21E-01 |
| AT3G59765 | 1,81 | -0,51 | 1,37 | -0,37 | 7,09E-01 | 8,21E-01 |
| AT5G01530 | 20,05 | -0,22 | 0,59 | -0,37 | 7,10E-01 | 8,22E-01 |
| AT1G04393 | 5,00 | 0,43 | 1,15 | 0,37 | 7,10E-01 | 8,23E-01 |
| AT3G30390 | 2,05 | 0,48 | 1,30 | 0,37 | 7,11E-01 | 8,23E-01 |
| AT3G53430 | 2,55 | 0,42 | 1,13 | 0,37 | 7,13E-01 | 8,25E-01 |
| AT3G07515 | 2,88 | 0,69 | 1,88 | 0,37 | 7,14E-01 | 8,26E-01 |
| AT4G13495 | 869,17 | -0,12 | 0,34 | -0,36 | 7,16E-01 | 8,28E-01 |
| AT2G07629 | 5,41 | 0,31 | 0,86 | 0,36 | 7,17E-01 | 8,28E-01 |
| AT3G02020 | 19,73 | -0,22 | 0,62 | -0,35 | 7,23E-01 | 8,34E-01 |
| AT3G13290 | 2,32 | 0,55 | 1,56 | 0,35 | 7,24E-01 | 8,35E-01 |
| AT4G20830 | 3,02 | 0,50 | 1,41 | 0,35 | 7,23E-01 | 8,35E-01 |
| AT5G43760 | 1,93 | -0,53 | 1,51 | -0,35 | 7,24E-01 | 8,35E-01 |
| AT3G06055 | 2,49 | -0,54 | 1,53 | -0,35 | 7,24E-01 | 8,35E-01 |
| AT1G33120 | 1,97 | -0,51 | 1,45 | -0,35 | 7,26E-01 | 8,36E-01 |
| AT3G01370 | 2,22 | -0,49 | 1,39 | -0,35 | 7,26E-01 | 8,36E-01 |
| AT3G16830 | 1,75 | 0,54 | 1,55 | 0,35 | 7,27E-01 | 8,37E-01 |
| AT1G20260 | 2,71 | -0,44 | 1,28 | -0,35 | 7,28E-01 | 8,37E-01 |
| AT1G77235 | 1,77 | -0,60 | 1,74 | -0,35 | 7,29E-01 | 8,38E-01 |
| AT5G01775 | 4,59 | -0,39 | 1,12 | -0,34 | 7,31E-01 | 8,40E-01 |
| AT1G24510 | 1,77 | 0,56 | 1,64 | 0,34 | 7,32E-01 | 8,40E-01 |
| AT2G09555 | 2,08 | -0,49 | 1,41 | -0,34 | 7,31E-01 | 8,40E-01 |
| AT3G28925 | 2,17 | 0,59 | 1,75 | 0,34 | 7,34E-01 | 8,41E-01 |
| AT1G08117 | 4,98 | -0,39 | 1,14 | -0,34 | 7,34E-01 | 8,41E-01 |

| | | | | | | |
|---|---|---|---|---|---|---|
| AT5G24690 | 2,42 | 0,41 | 1,20 | 0,34 | 7,34E-01 | 8,41E-01 |
| AT3G56940 | 6,51 | 0,26 | 0,77 | 0,34 | 7,35E-01 | 8,41E-01 |
| AT4G05320 | 17,67 | -0,19 | 0,56 | -0,34 | 7,35E-01 | 8,42E-01 |
| AT2G07727 | 3,34 | 0,42 | 1,28 | 0,33 | 7,40E-01 | 8,47E-01 |
| AT1G76062 | 4,01 | -0,45 | 1,36 | -0,33 | 7,41E-01 | 8,47E-01 |
| AT1G61740 | 2,13 | 0,50 | 1,51 | 0,33 | 7,41E-01 | 8,47E-01 |
| AT5G26000 | 4,62 | -0,29 | 0,87 | -0,33 | 7,42E-01 | 8,47E-01 |
| AT3G08510 | 2,33 | 0,52 | 1,57 | 0,33 | 7,42E-01 | 8,48E-01 |
| AT5G04305 | 5,44 | 0,32 | 0,98 | 0,33 | 7,43E-01 | 8,48E-01 |
| AT3G45780 | 10,09 | 0,18 | 0,56 | 0,32 | 7,46E-01 | 8,51E-01 |
| AT5G48620 | 2,46 | 0,50 | 1,53 | 0,32 | 7,46E-01 | 8,51E-01 |
| AT1G66100 | 8,07 | -0,24 | 0,74 | -0,32 | 7,47E-01 | 8,51E-01 |
| AT2G34250 | 2,94 | 0,38 | 1,17 | 0,32 | 7,48E-01 | 8,52E-01 |
| AT2G44310 | 1,85 | 0,54 | 1,67 | 0,32 | 7,48E-01 | 8,52E-01 |
| AT4G37260 | 2,49 | 0,45 | 1,41 | 0,32 | 7,48E-01 | 8,52E-01 |
| AT1G32583 | 28,27 | 0,18 | 0,57 | 0,32 | 7,50E-01 | 8,53E-01 |
| AT2G03890 | 2,03 | 0,47 | 1,46 | 0,32 | 7,50E-01 | 8,53E-01 |
| AT1G30400 | 2,52 | 0,44 | 1,39 | 0,32 | 7,51E-01 | 8,53E-01 |
| AT5G08180 | 2,30 | 0,38 | 1,19 | 0,32 | 7,51E-01 | 8,53E-01 |
| AT1G09800 | 20,58 | -0,17 | 0,53 | -0,32 | 7,52E-01 | 8,53E-01 |
| AT1G56280 | 2,59 | 0,36 | 1,15 | 0,31 | 7,53E-01 | 8,54E-01 |
| AT3G04165 | 2,58 | 0,44 | 1,40 | 0,31 | 7,55E-01 | 8,56E-01 |
| AT4G04925 | 1,79 | 0,58 | 1,86 | 0,31 | 7,55E-01 | 8,56E-01 |
| AT4G08691 | 15,19 | -0,53 | 1,72 | -0,31 | 7,56E-01 | 8,57E-01 |
| AT2G34357 | 1,72 | 0,54 | 1,75 | 0,31 | 7,57E-01 | 8,57E-01 |
| AT1G05805 | 1,94 | -0,76 | 2,45 | -0,31 | 7,58E-01 | 8,58E-01 |
| AT1G05010 | 2,06 | -0,50 | 1,62 | -0,31 | 7,59E-01 | 8,59E-01 |
| AT2G38170 | 3,76 | 0,35 | 1,13 | 0,31 | 7,59E-01 | 8,59E-01 |
| AT1G09687 | 2,04 | -0,42 | 1,40 | -0,30 | 7,61E-01 | 8,60E-01 |
| AT5G52070 | 6,76 | 0,33 | 1,10 | 0,30 | 7,63E-01 | 8,62E-01 |
| AT5G53902 | 6,85 | -0,54 | 1,81 | -0,30 | 7,65E-01 | 8,62E-01 |
| AT3G03435 | 3,28 | -0,36 | 1,20 | -0,30 | 7,64E-01 | 8,62E-01 |
| AT3G20310 | 2,42 | 0,45 | 1,49 | 0,30 | 7,65E-01 | 8,62E-01 |
| AT3G29130 | 2,05 | 0,56 | 1,87 | 0,30 | 7,64E-01 | 8,62E-01 |
| AT5G53905 | 6,85 | -0,54 | 1,81 | -0,30 | 7,65E-01 | 8,62E-01 |
| AT5G24900 | 2,20 | -0,45 | 1,50 | -0,30 | 7,66E-01 | 8,63E-01 |
| AT5G03240 | 5,81 | 0,26 | 0,88 | 0,30 | 7,67E-01 | 8,64E-01 |
| AT1G10522 | 24,13 | 0,15 | 0,51 | 0,30 | 7,68E-01 | 8,64E-01 |
| AT2G07725 | 2,14 | -0,38 | 1,29 | -0,29 | 7,69E-01 | 8,65E-01 |
| AT1G63740 | 1,93 | -0,48 | 1,64 | -0,29 | 7,71E-01 | 8,67E-01 |
| AT5G24300 | 2,35 | 0,46 | 1,56 | 0,29 | 7,71E-01 | 8,67E-01 |
| AT4G04095 | 1,94 | 0,38 | 1,33 | 0,29 | 7,72E-01 | 8,67E-01 |
| AT1G49500 | 1,83 | -0,41 | 1,43 | -0,29 | 7,74E-01 | 8,69E-01 |
| AT4G03445 | 45,78 | 0,10 | 0,36 | 0,29 | 7,75E-01 | 8,69E-01 |
| AT4G03020 | 2,29 | 0,36 | 1,28 | 0,28 | 7,76E-01 | 8,70E-01 |
| AT5G14060 | 2,66 | 0,40 | 1,42 | 0,28 | 7,77E-01 | 8,71E-01 |
| AT2G04375 | 5,42 | 0,24 | 0,87 | 0,28 | 7,78E-01 | 8,71E-01 |
| AT2G16040 | 2,68 | -0,54 | 1,93 | -0,28 | 7,78E-01 | 8,71E-01 |
| AT1G07633 | 6,19 | 0,35 | 1,23 | 0,28 | 7,78E-01 | 8,71E-01 |
| AT3G48115 | 1,91 | 0,48 | 1,70 | 0,28 | 7,78E-01 | 8,71E-01 |
| AT1G50440 | 6,47 | 0,34 | 1,22 | 0,28 | 7,83E-01 | 8,74E-01 |
| AT1G75040 | 3,07 | 0,27 | 0,99 | 0,28 | 7,83E-01 | 8,74E-01 |
| AT1G76030 | 2,84 | 0,34 | 1,25 | 0,28 | 7,83E-01 | 8,74E-01 |
| AT2G29630 | 5,51 | 0,23 | 0,84 | 0,28 | 7,83E-01 | 8,74E-01 |
| AT3G54890 | 13,73 | -0,17 | 0,62 | -0,28 | 7,83E-01 | 8,74E-01 |
| AT5G54940 | 2,86 | -0,36 | 1,31 | -0,27 | 7,83E-01 | 8,74E-01 |

| | | | | | | |
|---|---|---|---|---|---|---|
| AT3G22970 | 2,77 | -0,36 | 1,33 | -0,27 | 7,84E-01 | 8,74E-01 |
| AT3G45140 | 15,78 | -0,15 | 0,56 | -0,27 | 7,84E-01 | 8,74E-01 |
| AT3G28899 | 2,18 | -0,38 | 1,40 | -0,27 | 7,86E-01 | 8,75E-01 |
| AT1G06040 | 2,85 | -0,35 | 1,29 | -0,27 | 7,86E-01 | 8,75E-01 |
| AT4G17390 | 6,32 | 0,24 | 0,88 | 0,27 | 7,86E-01 | 8,75E-01 |
| AT5G26030 | 2,16 | 0,46 | 1,72 | 0,27 | 7,89E-01 | 8,77E-01 |
| AT5G41490 | 2,24 | -0,53 | 1,97 | -0,27 | 7,88E-01 | 8,77E-01 |
| AT5G02380 | 2,77 | -0,41 | 1,53 | -0,27 | 7,90E-01 | 8,78E-01 |
| AT4G26970 | 1,79 | 0,50 | 1,88 | 0,27 | 7,90E-01 | 8,78E-01 |
| AT4G02555 | 3,06 | 0,33 | 1,26 | 0,27 | 7,91E-01 | 8,78E-01 |
| AT1G77815 | 2,65 | 0,48 | 1,83 | 0,26 | 7,91E-01 | 8,78E-01 |
| AT3G54580 | 1,94 | -0,54 | 2,05 | -0,27 | 7,91E-01 | 8,78E-01 |
| AT3G19010 | 1,81 | 0,42 | 1,60 | 0,26 | 7,92E-01 | 8,78E-01 |
| AT5G24810 | 2,74 | 0,46 | 1,76 | 0,26 | 7,93E-01 | 8,79E-01 |
| AT3G07725 | 10,80 | -0,17 | 0,67 | -0,26 | 7,94E-01 | 8,79E-01 |
| AT4G08593 | 1,89 | -0,55 | 2,13 | -0,26 | 7,96E-01 | 8,81E-01 |
| AT5G41510 | 1,76 | -0,54 | 2,11 | -0,26 | 7,96E-01 | 8,81E-01 |
| AT1G74453 | 1,99 | 0,44 | 1,71 | 0,26 | 7,97E-01 | 8,81E-01 |
| AT3G22940 | 1,72 | -0,54 | 2,12 | -0,26 | 7,98E-01 | 8,82E-01 |
| AT1G76180 | 6,03 | -0,21 | 0,81 | -0,26 | 7,98E-01 | 8,82E-01 |
| AT4G03455 | 16,01 | 0,15 | 0,61 | 0,25 | 8,01E-01 | 8,84E-01 |
| AT5G40450 | 4,44 | -0,25 | 1,00 | -0,25 | 8,01E-01 | 8,84E-01 |
| AT2G37270 | 1,75 | 0,39 | 1,57 | 0,25 | 8,02E-01 | 8,84E-01 |
| AT4G35090 | 2,83 | -0,28 | 1,13 | -0,25 | 8,02E-01 | 8,85E-01 |
| AT1G01040 | 4,73 | 0,26 | 1,03 | 0,25 | 8,03E-01 | 8,85E-01 |
| AT4G23375 | 8,41 | -0,41 | 1,65 | -0,25 | 8,04E-01 | 8,85E-01 |
| AT5G42930 | 2,68 | -0,29 | 1,18 | -0,25 | 8,06E-01 | 8,87E-01 |
| AT1G29910 | 27,17 | -0,12 | 0,49 | -0,24 | 8,07E-01 | 8,88E-01 |
| AT5G64260 | 2,41 | 0,47 | 1,91 | 0,24 | 8,07E-01 | 8,88E-01 |
| AT1G28420 | 2,78 | 0,39 | 1,61 | 0,24 | 8,08E-01 | 8,88E-01 |
| AT5G08525 | 4,34 | -0,27 | 1,13 | -0,24 | 8,08E-01 | 8,88E-01 |
| AT5G02960 | 2,13 | 0,33 | 1,35 | 0,24 | 8,09E-01 | 8,89E-01 |
| AT4G23630 | 1,89 | 0,33 | 1,39 | 0,24 | 8,10E-01 | 8,89E-01 |
| AT3G46510 | 1,83 | 0,37 | 1,55 | 0,24 | 8,11E-01 | 8,90E-01 |
| AT4G04340 | 2,36 | 0,38 | 1,60 | 0,24 | 8,12E-01 | 8,90E-01 |
| AT3G56705 | 15,48 | -0,15 | 0,61 | -0,24 | 8,13E-01 | 8,91E-01 |
| AT5G01785 | 2,53 | -0,43 | 1,80 | -0,24 | 8,13E-01 | 8,91E-01 |
| AT2G13360 | 3,44 | -0,23 | 0,96 | -0,24 | 8,14E-01 | 8,91E-01 |
| AT2G39010 | 4,44 | 0,25 | 1,08 | 0,24 | 8,14E-01 | 8,91E-01 |
| AT1G63330 | 7,00 | -0,16 | 0,69 | -0,23 | 8,18E-01 | 8,93E-01 |
| AT3G26730 | 1,96 | 0,40 | 1,71 | 0,23 | 8,17E-01 | 8,93E-01 |
| AT5G27720 | 3,01 | -0,29 | 1,25 | -0,23 | 8,18E-01 | 8,93E-01 |
| AT4G34720 | 1,83 | -0,31 | 1,35 | -0,23 | 8,17E-01 | 8,93E-01 |
| AT4G30750 | 3,95 | 0,42 | 1,85 | 0,23 | 8,19E-01 | 8,95E-01 |
| AT1G15690 | 11,49 | -0,15 | 0,66 | -0,23 | 8,20E-01 | 8,95E-01 |
| AT1G67450 | 1,86 | -0,42 | 1,83 | -0,23 | 8,20E-01 | 8,95E-01 |
| AT1G23390 | 2,44 | -0,30 | 1,33 | -0,22 | 8,22E-01 | 8,97E-01 |
| AT4G01290 | 1,74 | 0,37 | 1,65 | 0,22 | 8,22E-01 | 8,97E-01 |
| AT4G15560 | 2,23 | -0,29 | 1,29 | -0,22 | 8,23E-01 | 8,97E-01 |
| AT4G10340 | 23,35 | 0,11 | 0,51 | 0,22 | 8,24E-01 | 8,97E-01 |
| AT1G06273 | 2,17 | -0,32 | 1,44 | -0,22 | 8,26E-01 | 8,98E-01 |
| AT1G78040 | 2,48 | -0,31 | 1,41 | -0,22 | 8,28E-01 | 8,98E-01 |
| AT2G01010 | 3,38 | -0,32 | 1,49 | -0,22 | 8,28E-01 | 8,98E-01 |
| AT2G39175 | 43,76 | 0,09 | 0,41 | 0,22 | 8,25E-01 | 8,98E-01 |
| AT3G04400 | 2,05 | -0,27 | 1,24 | -0,22 | 8,26E-01 | 8,98E-01 |
| AT3G03105 | 2,11 | 0,30 | 1,38 | 0,22 | 8,27E-01 | 8,98E-01 |

| | | | | | | |
|---|---|---|---|---|---|---|
| AT3G20950 | 1,75 | 0,43 | 1,99 | 0,22 | 8,29E-01 | 8,98E-01 |
| AT4G05360 | 2,15 | 0,42 | 1,94 | 0,22 | 8,29E-01 | 8,98E-01 |
| AT5G04410 | 1,77 | -0,49 | 2,24 | -0,22 | 8,26E-01 | 8,98E-01 |
| AT1G07243 | 1,82 | 0,42 | 1,95 | 0,22 | 8,28E-01 | 8,98E-01 |
| AT1G76010 | 2,04 | -0,28 | 1,30 | -0,22 | 8,29E-01 | 8,98E-01 |
| AT3G22235 | 3,04 | -0,25 | 1,17 | -0,22 | 8,28E-01 | 8,98E-01 |
| AT4G22410 | 2,93 | 0,41 | 1,87 | 0,22 | 8,29E-01 | 8,98E-01 |
| AT4G26730 | 2,38 | 0,43 | 2,03 | 0,21 | 8,31E-01 | 8,99E-01 |
| AT1G44100 | 2,02 | 0,41 | 1,94 | 0,21 | 8,31E-01 | 8,99E-01 |
| AT2G29450 | 2,19 | 0,37 | 1,77 | 0,21 | 8,33E-01 | 8,99E-01 |
| AT5G12860 | 2,32 | 0,37 | 1,74 | 0,21 | 8,32E-01 | 8,99E-01 |
| AT5G40395 | 2,01 | 0,31 | 1,49 | 0,21 | 8,33E-01 | 8,99E-01 |
| AT1G69572 | 2,44 | 0,33 | 1,58 | 0,21 | 8,34E-01 | 9,00E-01 |
| AT1G69570 | 2,44 | 0,33 | 1,58 | 0,21 | 8,34E-01 | 9,00E-01 |
| AT1G64500 | 2,64 | 0,26 | 1,25 | 0,21 | 8,36E-01 | 9,01E-01 |
| AT5G02870 | 2,41 | 0,27 | 1,32 | 0,21 | 8,36E-01 | 9,01E-01 |
| AT3G14050 | 1,74 | -0,38 | 1,86 | -0,21 | 8,36E-01 | 9,01E-01 |
| AT1G12240 | 2,62 | 0,27 | 1,34 | 0,20 | 8,39E-01 | 9,01E-01 |
| AT1G15010 | 2,66 | -0,36 | 1,78 | -0,20 | 8,39E-01 | 9,01E-01 |
| AT1G20020 | 2,67 | 0,21 | 1,05 | 0,20 | 8,39E-01 | 9,01E-01 |
| AT3G52800 | 1,93 | 0,37 | 1,81 | 0,20 | 8,39E-01 | 9,01E-01 |
| AT4G11270 | 3,79 | -0,22 | 1,10 | -0,20 | 8,38E-01 | 9,01E-01 |
| AT5G05010 | 2,08 | 0,29 | 1,39 | 0,20 | 8,38E-01 | 9,01E-01 |
| AT5G26650 | 7,57 | -0,35 | 1,70 | -0,20 | 8,38E-01 | 9,01E-01 |
| AT1G78830 | 2,12 | 0,27 | 1,30 | 0,21 | 8,37E-01 | 9,01E-01 |
| AT5G00760 | 4,31 | 0,18 | 0,89 | 0,20 | 8,41E-01 | 9,03E-01 |
| AT1G18880 | 1,72 | 0,38 | 1,94 | 0,20 | 8,43E-01 | 9,04E-01 |
| AT1G32630 | 4,04 | -0,23 | 1,17 | -0,20 | 8,44E-01 | 9,05E-01 |
| AT1G32640 | 4,04 | -0,23 | 1,17 | -0,20 | 8,44E-01 | 9,05E-01 |
| AT2G19480 | 1,78 | -0,34 | 1,72 | -0,20 | 8,45E-01 | 9,05E-01 |
| AT3G05590 | 1,96 | 0,24 | 1,25 | 0,19 | 8,46E-01 | 9,06E-01 |
| AT3G44860 | 2,37 | 0,33 | 1,70 | 0,19 | 8,47E-01 | 9,06E-01 |
| AT3G57480 | 3,07 | 0,20 | 1,02 | 0,19 | 8,47E-01 | 9,06E-01 |
| AT2G35945 | 15,59 | 0,17 | 0,88 | 0,19 | 8,49E-01 | 9,07E-01 |
| AT3G53990 | 1,83 | -0,24 | 1,30 | -0,19 | 8,52E-01 | 9,10E-01 |
| AT4G19120 | 2,16 | 0,22 | 1,15 | 0,19 | 8,52E-01 | 9,10E-01 |
| AT1G47660 | 2,01 | 0,36 | 1,96 | 0,19 | 8,53E-01 | 9,10E-01 |
| AT3G21750 | 1,77 | -0,29 | 1,60 | -0,18 | 8,54E-01 | 9,11E-01 |
| AT1G26558 | 2,72 | 0,23 | 1,31 | 0,18 | 8,58E-01 | 9,14E-01 |
| AT1G67430 | 1,82 | 0,28 | 1,54 | 0,18 | 8,58E-01 | 9,14E-01 |
| AT2G25250 | 2,81 | 0,26 | 1,47 | 0,18 | 8,58E-01 | 9,14E-01 |
| AT4G11900 | 2,31 | 0,24 | 1,35 | 0,18 | 8,59E-01 | 9,14E-01 |
| AT5G00765 | 3,11 | 0,19 | 1,09 | 0,18 | 8,59E-01 | 9,14E-01 |
| AT1G59980 | 3,55 | -0,18 | 1,05 | -0,17 | 8,62E-01 | 9,18E-01 |
| AT2G16018 | 9,68 | 0,21 | 1,24 | 0,17 | 8,63E-01 | 9,18E-01 |
| AT5G38190 | 2,86 | -0,27 | 1,54 | -0,17 | 8,63E-01 | 9,18E-01 |
| AT3G63375 | 13,10 | -0,09 | 0,51 | -0,17 | 8,66E-01 | 9,20E-01 |
| AT4G01150 | 2,10 | 0,22 | 1,31 | 0,17 | 8,66E-01 | 9,20E-01 |
| AT2G34110 | 1,88 | -0,25 | 1,48 | -0,17 | 8,68E-01 | 9,20E-01 |
| AT1G75380 | 2,78 | -0,19 | 1,14 | -0,17 | 8,68E-01 | 9,20E-01 |
| AT2G15420 | 2,17 | -0,31 | 1,86 | -0,17 | 8,68E-01 | 9,20E-01 |
| AT4G18480 | 1,72 | -0,26 | 1,56 | -0,17 | 8,67E-01 | 9,20E-01 |
| AT5G13840 | 2,03 | 0,23 | 1,41 | 0,16 | 8,69E-01 | 9,21E-01 |
| AT4G12780 | 15,31 | 0,11 | 0,69 | 0,16 | 8,70E-01 | 9,21E-01 |
| AT1G76065 | 4,52 | -0,20 | 1,25 | -0,16 | 8,72E-01 | 9,23E-01 |
| AT3G13100 | 2,83 | 0,28 | 1,76 | 0,16 | 8,73E-01 | 9,24E-01 |

| | | | | | | |
|---|---|---|---|---|---|---|
| AT2G41430 | 6,85 | -0,14 | 0,89 | -0,16 | 8,73E-01 | 9,24E-01 |
| AT4G23850 | 2,43 | 0,20 | 1,25 | 0,16 | 8,76E-01 | 9,26E-01 |
| AT5G63570 | 1,81 | 0,24 | 1,55 | 0,16 | 8,77E-01 | 9,26E-01 |
| AT1G65450 | 2,07 | 0,20 | 1,30 | 0,15 | 8,78E-01 | 9,27E-01 |
| AT4G28755 | 2,23 | -0,19 | 1,25 | -0,15 | 8,78E-01 | 9,27E-01 |
| AT1G08813 | 8,42 | 0,11 | 0,72 | 0,15 | 8,78E-01 | 9,27E-01 |
| AT1G21130 | 2,17 | -0,22 | 1,47 | -0,15 | 8,78E-01 | 9,27E-01 |
| AT2G21920 | 4,05 | -0,13 | 0,88 | -0,15 | 8,79E-01 | 9,27E-01 |
| AT1G68010 | 2,53 | -0,21 | 1,42 | -0,15 | 8,82E-01 | 9,29E-01 |
| AT2G23930 | 1,70 | 0,23 | 1,56 | 0,15 | 8,84E-01 | 9,31E-01 |
| AT3G48560 | 3,89 | -0,14 | 0,94 | -0,14 | 8,85E-01 | 9,32E-01 |
| AT3G62700 | 3,87 | 0,15 | 1,04 | 0,14 | 8,87E-01 | 9,34E-01 |
| AT4G30993 | 2,23 | -0,22 | 1,58 | -0,14 | 8,88E-01 | 9,34E-01 |
| AT5G10572 | 2,06 | -0,20 | 1,47 | -0,14 | 8,89E-01 | 9,35E-01 |
| AT2G37160 | 3,77 | -0,16 | 1,15 | -0,14 | 8,90E-01 | 9,35E-01 |
| AT2G08855 | 3,73 | -0,16 | 1,16 | -0,14 | 8,90E-01 | 9,35E-01 |
| AT3G04210 | 3,60 | 0,16 | 1,19 | 0,14 | 8,92E-01 | 9,36E-01 |
| AT3G16370 | 1,93 | 0,17 | 1,29 | 0,13 | 8,95E-01 | 9,40E-01 |
| AT4G22710 | 1,76 | 0,24 | 1,87 | 0,13 | 8,97E-01 | 9,41E-01 |
| AT3G05655 | 1,87 | 0,21 | 1,76 | 0,12 | 9,04E-01 | 9,48E-01 |
| AT1G04820 | 2,61 | 0,12 | 1,09 | 0,11 | 9,11E-01 | 9,55E-01 |
| AT4G26542 | 2,99 | 0,13 | 1,20 | 0,11 | 9,14E-01 | 9,58E-01 |
| AT3G56408 | 2,59 | -0,17 | 1,67 | -0,10 | 9,17E-01 | 9,60E-01 |
| AT4G24770 | 3,23 | -0,12 | 1,19 | -0,10 | 9,17E-01 | 9,60E-01 |
| AT2G18020 | 2,64 | 0,16 | 1,53 | 0,10 | 9,18E-01 | 9,61E-01 |
| AT1G47265 | 3,77 | -0,10 | 1,02 | -0,10 | 9,20E-01 | 9,61E-01 |
| AT4G38970 | 12,35 | -0,05 | 0,54 | -0,10 | 9,19E-01 | 9,61E-01 |
| AT4G05150 | 3,81 | 0,13 | 1,31 | 0,10 | 9,21E-01 | 9,62E-01 |
| AT5G44870 | 1,81 | 0,15 | 1,53 | 0,10 | 9,21E-01 | 9,62E-01 |
| AT2G23810 | 3,17 | -0,14 | 1,46 | -0,10 | 9,22E-01 | 9,62E-01 |
| AT3G09162 | 2,15 | 0,13 | 1,36 | 0,10 | 9,22E-01 | 9,62E-01 |
| AT1G20860 | 3,03 | -0,12 | 1,24 | -0,10 | 9,23E-01 | 9,63E-01 |
| AT5G19140 | 6,43 | 0,07 | 0,77 | 0,09 | 9,27E-01 | 9,63E-01 |
| AT5G41790 | 2,54 | -0,13 | 1,39 | -0,09 | 9,25E-01 | 9,63E-01 |
| AT1G07637 | 2,62 | -0,16 | 1,76 | -0,09 | 9,27E-01 | 9,63E-01 |
| AT1G51980 | 1,85 | -0,15 | 1,58 | -0,09 | 9,26E-01 | 9,63E-01 |
| AT3G11910 | 1,81 | 0,13 | 1,37 | 0,09 | 9,27E-01 | 9,63E-01 |
| AT3G06835 | 2,07 | 0,12 | 1,33 | 0,09 | 9,25E-01 | 9,63E-01 |
| AT3G56410 | 2,56 | -0,16 | 1,66 | -0,10 | 9,24E-01 | 9,63E-01 |
| AT4G05050 | 3,49 | 0,12 | 1,32 | 0,09 | 9,26E-01 | 9,63E-01 |
| AT5G19150 | 6,35 | 0,07 | 0,77 | 0,09 | 9,27E-01 | 9,63E-01 |
| AT2G41090 | 3,05 | -0,11 | 1,18 | -0,09 | 9,29E-01 | 9,64E-01 |
| AT1G50250 | 2,35 | 0,12 | 1,31 | 0,09 | 9,29E-01 | 9,64E-01 |
| AT1G07940 | 3,71 | -0,09 | 1,01 | -0,09 | 9,29E-01 | 9,64E-01 |
| AT5G62690 | 4,26 | -0,08 | 0,96 | -0,09 | 9,31E-01 | 9,64E-01 |
| AT4G28750 | 4,56 | -0,09 | 1,05 | -0,09 | 9,30E-01 | 9,64E-01 |
| AT3G55320 | 2,94 | 0,12 | 1,45 | 0,08 | 9,33E-01 | 9,66E-01 |
| AT2G08370 | 4,11 | -0,08 | 0,94 | -0,08 | 9,34E-01 | 9,67E-01 |
| AT4G18030 | 2,74 | -0,10 | 1,24 | -0,08 | 9,35E-01 | 9,68E-01 |
| AT5G03305 | 5,94 | -0,10 | 1,25 | -0,08 | 9,36E-01 | 9,69E-01 |
| AT5G12250 | 2,28 | 0,11 | 1,35 | 0,08 | 9,38E-01 | 9,70E-01 |
| AT2G05315 | 4,98 | 0,09 | 1,18 | 0,08 | 9,40E-01 | 9,70E-01 |
| AT3G09445 | 4,99 | -0,07 | 0,90 | -0,08 | 9,39E-01 | 9,70E-01 |
| AT4G04205 | 2,87 | 0,10 | 1,35 | 0,08 | 9,40E-01 | 9,70E-01 |
| AT4G10210 | 2,08 | -0,11 | 1,44 | -0,08 | 9,40E-01 | 9,70E-01 |
| AT4G23915 | 12,58 | -0,06 | 0,73 | -0,08 | 9,39E-01 | 9,70E-01 |

| | | | | | | |
|---|---|---|---|---|---|---|
| AT2G36460 | 1,86 | 0,11 | 1,48 | 0,07 | 9,41E-01 | 9,70E-01 |
| AT4G02890 | 2,39 | 0,10 | 1,32 | 0,07 | 9,42E-01 | 9,71E-01 |
| AT5G13390 | 2,05 | -0,11 | 1,47 | -0,07 | 9,43E-01 | 9,71E-01 |
| AT1G15002 | 1,82 | -0,10 | 1,47 | -0,07 | 9,44E-01 | 9,72E-01 |
| AT1G74456 | 11,08 | -0,05 | 0,66 | -0,07 | 9,44E-01 | 9,72E-01 |
| AT3G47470 | 25,61 | 0,03 | 0,46 | 0,07 | 9,44E-01 | 9,72E-01 |
| AT5G13650 | 5,78 | 0,06 | 0,83 | 0,07 | 9,46E-01 | 9,73E-01 |
| AT4G20835 | 2,66 | -0,10 | 1,55 | -0,07 | 9,47E-01 | 9,74E-01 |
| AT4G39363 | 6,83 | 0,05 | 0,82 | 0,06 | 9,49E-01 | 9,75E-01 |
| AT1G65960 | 3,84 | -0,08 | 1,32 | -0,06 | 9,52E-01 | 9,77E-01 |
| AT1G27090 | 1,85 | 0,08 | 1,34 | 0,06 | 9,51E-01 | 9,77E-01 |
| AT2G09700 | 2,16 | -0,10 | 1,75 | -0,06 | 9,53E-01 | 9,78E-01 |
| AT3G14415 | 1,72 | 0,09 | 1,59 | 0,06 | 9,56E-01 | 9,80E-01 |
| AT2G17510 | 1,71 | 0,09 | 1,63 | 0,05 | 9,57E-01 | 9,81E-01 |
| AT1G67840 | 1,96 | 0,09 | 1,78 | 0,05 | 9,60E-01 | 9,84E-01 |
| AT5G61410 | 2,49 | -0,06 | 1,23 | -0,05 | 9,60E-01 | 9,84E-01 |
| AT2G05355 | 1,78 | 0,07 | 1,38 | 0,05 | 9,61E-01 | 9,84E-01 |
| AT5G21100 | 6,61 | -0,06 | 1,25 | -0,05 | 9,61E-01 | 9,84E-01 |
| AT4G24715 | 2,06 | 0,06 | 1,16 | 0,05 | 9,62E-01 | 9,84E-01 |
| AT1G02780 | 1,83 | 0,06 | 1,33 | 0,05 | 9,63E-01 | 9,85E-01 |
| AT2G35384 | 13,85 | 0,03 | 0,67 | 0,05 | 9,64E-01 | 9,85E-01 |
| AT5G58070 | 2,01 | -0,07 | 1,61 | -0,04 | 9,65E-01 | 9,86E-01 |
| AT1G04183 | 2,59 | 0,06 | 1,49 | 0,04 | 9,69E-01 | 9,89E-01 |
| AT3G22440 | 1,87 | 0,06 | 1,64 | 0,04 | 9,70E-01 | 9,90E-01 |
| AT1G69830 | 1,91 | 0,05 | 1,24 | 0,04 | 9,70E-01 | 9,90E-01 |
| AT2G44060 | 2,21 | 0,04 | 1,29 | 0,03 | 9,74E-01 | 9,93E-01 |
| AT5G16620 | 2,01 | -0,04 | 1,25 | -0,03 | 9,74E-01 | 9,93E-01 |
| AT1G03410 | 2,63 | 0,05 | 1,49 | 0,03 | 9,75E-01 | 9,93E-01 |
| AT3G25855 | 1,70 | -0,05 | 1,65 | -0,03 | 9,77E-01 | 9,95E-01 |
| AT3G52930 | 3,81 | -0,03 | 1,01 | -0,03 | 9,77E-01 | 9,95E-01 |
| AT4G30440 | 3,67 | -0,04 | 1,38 | -0,03 | 9,77E-01 | 9,95E-01 |
| AT4G03580 | 13,67 | 0,03 | 1,15 | 0,03 | 9,78E-01 | 9,95E-01 |
| AT1G33950 | 1,88 | -0,05 | 1,80 | -0,03 | 9,79E-01 | 9,95E-01 |
| AT1G65440 | 2,14 | -0,03 | 1,29 | -0,03 | 9,80E-01 | 9,96E-01 |
| AT3G44444 | 1,95 | -0,04 | 1,63 | -0,02 | 9,81E-01 | 9,96E-01 |
| AT3G46970 | 7,07 | -0,02 | 0,73 | -0,02 | 9,81E-01 | 9,96E-01 |
| AT1G74290 | 1,78 | -0,04 | 1,82 | -0,02 | 9,82E-01 | 9,96E-01 |
| AT1G04517 | 11,17 | 0,01 | 0,64 | 0,02 | 9,82E-01 | 9,97E-01 |
| AT1G07510 | 1,74 | -0,02 | 1,76 | -0,01 | 9,93E-01 | 1,00E+00 |
| AT1G23720 | 2,26 | 0,00 | 2,27 | 0,00 | 1,00E+00 | 1,00E+00 |
| AT1G07233 | 2,51 | 0,01 | 1,14 | 0,01 | 9,90E-01 | 1,00E+00 |
| AT1G50900 | 1,85 | -0,02 | 1,37 | -0,02 | 9,88E-01 | 1,00E+00 |
| AT1G64340 | 2,24 | 0,00 | 2,24 | 0,00 | 1,00E+00 | 1,00E+00 |
| AT2G06420 | 4,36 | 0,00 | 1,87 | 0,00 | 1,00E+00 | 1,00E+00 |
| AT2G07981 | 2,13 | 0,00 | 2,23 | 0,00 | 1,00E+00 | 1,00E+00 |
| AT2G24980 | 3,78 | 0,00 | 1,91 | 0,00 | 1,00E+00 | 1,00E+00 |
| AT2G25171 | 1,84 | -0,01 | 1,89 | 0,00 | 9,97E-01 | 1,00E+00 |
| AT2G29920 | 2,43 | 0,00 | 2,20 | 0,00 | 1,00E+00 | 1,00E+00 |
| AT2G47470 | 1,77 | -0,02 | 1,86 | -0,01 | 9,90E-01 | 1,00E+00 |
| AT3G04250 | 2,26 | 0,00 | 2,13 | 0,00 | 1,00E+00 | 1,00E+00 |
| AT3G55512 | 2,56 | 0,00 | 2,12 | 0,00 | 1,00E+00 | 1,00E+00 |
| AT3G57645 | 1,71 | 0,00 | 2,17 | 0,00 | 1,00E+00 | 1,00E+00 |
| AT3G61897 | 2,76 | 0,00 | 2,83 | 0,00 | 1,00E+00 | 1,00E+00 |
| AT4G29780 | 2,35 | 0,00 | 2,13 | 0,00 | 1,00E+00 | 1,00E+00 |
| AT5G27607 | 2,38 | 0,00 | 2,17 | 0,00 | 1,00E+00 | 1,00E+00 |
| AT5G56070 | 3,26 | 0,00 | 2,00 | 0,00 | 1,00E+00 | 1,00E+00 |

| | | | | | | |
|---|---|---|---|---|---|---|
| AT1G65350 | 2,42 | -0,01 | 1,60 | 0,00 | 9,97E-01 | 1,00E+00 |
| AT2G04605 | 3,46 | 0,00 | 1,92 | 0,00 | 1,00E+00 | 1,00E+00 |
| AT2G04060 | 1,72 | 0,00 | 2,36 | 0,00 | 1,00E+00 | 1,00E+00 |
| AT2G29925 | 2,27 | 0,00 | 2,28 | 0,00 | 1,00E+00 | 1,00E+00 |
| AT3G25795 | 22,21 | 0,00 | 1,74 | 0,00 | 1,00E+00 | 1,00E+00 |
| AT3G08370 | 6,58 | 0,00 | 1,82 | 0,00 | 1,00E+00 | 1,00E+00 |
| AT4G00140 | 2,38 | 0,00 | 2,00 | 0,00 | 1,00E+00 | 1,00E+00 |
| AT4G08630 | 1,97 | 0,00 | 2,05 | 0,00 | 1,00E+00 | 1,00E+00 |
| AT4G21595 | 28,61 | 0,00 | 1,71 | 0,00 | 1,00E+00 | 1,00E+00 |
| AT4G08165 | 5,10 | 0,00 | 1,92 | 0,00 | 1,00E+00 | 1,00E+00 |
| AT5G06630 | 1,85 | 0,00 | 2,07 | 0,00 | 1,00E+00 | 1,00E+00 |
| AT5G06640 | 3,20 | 0,00 | 1,92 | 0,00 | 1,00E+00 | 1,00E+00 |
| AT5G24280 | 2,00 | 0,00 | 3,20 | 0,00 | 1,00E+00 | 1,00E+00 |
| AT5G24660 | 2,17 | 0,00 | 2,04 | 0,00 | 1,00E+00 | 1,00E+00 |
| AT5G28823 | 1,86 | 0,00 | 2,20 | 0,00 | 1,00E+00 | 1,00E+00 |
| AT1G01046 | 1,63 | 1,72 | 1,84 | 0,94 | 3,48E-01 | |
| AT1G01160 | 0,43 | 1,26 | 3,34 | 0,38 | 7,05E-01 | |
| AT1G01180 | 0,85 | -0,45 | 2,48 | -0,18 | 8,56E-01 | |
| AT1G01220 | 0,11 | 0,00 | 4,41 | 0,00 | 1,00E+00 | |
| AT1G01225 | 0,31 | 1,74 | 4,38 | 0,40 | 6,91E-01 | |
| AT1G01230 | 0,37 | 0,82 | 4,38 | 0,19 | 8,52E-01 | |
| AT1G01240 | 0,19 | 0,00 | 4,41 | 0,00 | 1,00E+00 | |
| AT1G01260 | 0,53 | -0,43 | 2,78 | -0,15 | 8,77E-01 | |
| AT1G01300 | 1,08 | -1,98 | 2,06 | -0,96 | 3,36E-01 | |
| AT1G01350 | 0,29 | 1,47 | 4,40 | 0,33 | 7,39E-01 | |
| AT1G01448 | 0,59 | 0,00 | 4,40 | 0,00 | 1,00E+00 | |
| AT1G01480 | 0,12 | 0,00 | 4,41 | 0,00 | 1,00E+00 | |
| AT1G01520 | 0,12 | 0,00 | 4,41 | 0,00 | 1,00E+00 | |
| AT1G01540 | 0,22 | 0,00 | 4,41 | 0,00 | 1,00E+00 | |
| AT1G01560 | 1,60 | 3,31 | 1,97 | 1,68 | 9,23E-02 | |
| AT1G01630 | 0,58 | 1,60 | 3,78 | 0,42 | 6,73E-01 | |
| AT1G01710 | 0,82 | 0,18 | 3,08 | 0,06 | 9,54E-01 | |
| AT1G01720 | 1,37 | 1,03 | 1,69 | 0,61 | 5,41E-01 | |
| AT1G04047 | 0,70 | -0,47 | 2,67 | -0,17 | 8,61E-01 | |
| AT1G01730 | 0,40 | 2,14 | 4,38 | 0,49 | 6,25E-01 | |
| AT1G01790 | 0,83 | 0,33 | 1,98 | 0,17 | 8,68E-01 | |
| AT1G01810 | 0,12 | 0,00 | 4,41 | 0,00 | 1,00E+00 | |
| AT1G01950 | 0,18 | 0,00 | 4,41 | 0,00 | 1,00E+00 | |
| AT1G02010 | 0,10 | 0,00 | 4,41 | 0,00 | 1,00E+00 | |
| AT1G02100 | 0,49 | -0,46 | 3,24 | -0,14 | 8,88E-01 | |
| AT1G02120 | 0,36 | -0,46 | 3,55 | -0,13 | 8,97E-01 | |
| AT1G02150 | 1,38 | 1,11 | 1,84 | 0,60 | 5,47E-01 | |
| AT1G02205 | 1,06 | -2,22 | 2,05 | -1,08 | 2,80E-01 | |
| AT1G04093 | 0,08 | 0,00 | 4,41 | 0,00 | 1,00E+00 | |
| AT1G02300 | 0,34 | 1,38 | 4,41 | 0,31 | 7,54E-01 | |
| AT1G02305 | 0,80 | 0,56 | 2,50 | 0,22 | 8,24E-01 | |
| AT1G04103 | 0,08 | 0,00 | 4,41 | 0,00 | 1,00E+00 | |
| AT1G02390 | 0,19 | 0,00 | 4,41 | 0,00 | 1,00E+00 | |
| AT1G02420 | 0,52 | 1,37 | 3,96 | 0,34 | 7,30E-01 | |
| AT1G02560 | 1,57 | 1,06 | 1,65 | 0,64 | 5,23E-01 | |
| AT1G02690 | 0,11 | 0,00 | 4,41 | 0,00 | 1,00E+00 | |
| AT1G02750 | 0,51 | 2,84 | 4,33 | 0,66 | 5,11E-01 | |
| AT1G02816 | 0,38 | 1,30 | 3,44 | 0,38 | 7,06E-01 | |
| AT1G02850 | 0,62 | -0,45 | 3,02 | -0,15 | 8,82E-01 | |
| AT1G03160 | 0,95 | -0,18 | 2,19 | -0,08 | 9,33E-01 | |
| AT1G03190 | 0,39 | 1,38 | 3,53 | 0,39 | 6,96E-01 | |

| | | | | | |
|---|---|---|---|---|---|
| AT1G03220 | 1,11 | 1,54 | 2,25 | 0,69 | 4,92E-01 |
| AT1G03280 | 0,25 | 0,00 | 4,41 | 0,00 | 1,00E+00 |
| AT1G03290 | 0,75 | 0,00 | 2,61 | 0,00 | 1,00E+00 |
| AT1G03310 | 0,40 | 1,79 | 4,26 | 0,42 | 6,75E-01 |
| AT1G03360 | 0,35 | 1,36 | 3,58 | 0,38 | 7,05E-01 |
| AT1G03365 | 0,40 | 0,47 | 3,45 | 0,14 | 8,92E-01 |
| AT1G03370 | 0,51 | 0,00 | 4,05 | 0,00 | 1,00E+00 |
| AT1G03380 | 1,12 | 1,31 | 2,13 | 0,61 | 5,39E-01 |
| AT1G03430 | 0,37 | 0,62 | 4,37 | 0,14 | 8,86E-01 |
| AT1G03440 | 0,84 | 0,13 | 3,10 | 0,04 | 9,66E-01 |
| AT1G03545 | 0,70 | 0,80 | 2,29 | 0,35 | 7,25E-01 |
| AT1G03600 | 1,40 | -0,91 | 1,88 | -0,48 | 6,28E-01 |
| AT1G03610 | 0,72 | -1,84 | 3,22 | -0,57 | 5,68E-01 |
| AT1G03630 | 0,51 | 0,45 | 2,47 | 0,18 | 8,55E-01 |
| AT1G03687 | 0,11 | 0,00 | 4,41 | 0,00 | 1,00E+00 |
| AT1G03740 | 0,96 | 0,30 | 2,21 | 0,13 | 8,93E-01 |
| AT1G03810 | 0,77 | 2,88 | 2,23 | 1,29 | 1,96E-01 |
| AT1G03900 | 1,11 | 0,64 | 2,16 | 0,30 | 7,66E-01 |
| AT1G03905 | 0,24 | 1,96 | 4,39 | 0,45 | 6,55E-01 |
| AT1G03910 | 0,50 | 0,85 | 3,13 | 0,27 | 7,85E-01 |
| AT1G03930 | 0,53 | 2,46 | 2,75 | 0,90 | 3,70E-01 |
| AT1G04217 | 0,82 | -0,15 | 1,99 | -0,07 | 9,40E-01 |
| AT1G03935 | 0,16 | -0,55 | 4,41 | -0,12 | 9,01E-01 |
| AT1G04223 | 0,98 | 0,78 | 2,12 | 0,37 | 7,12E-01 |
| AT1G04227 | 0,32 | -1,32 | 3,67 | -0,36 | 7,20E-01 |
| AT1G03960 | 0,15 | -0,43 | 4,39 | -0,10 | 9,23E-01 |
| AT1G03970 | 0,36 | 0,00 | 4,41 | 0,00 | 1,00E+00 |
| AT1G04020 | 0,33 | 0,94 | 4,39 | 0,21 | 8,31E-01 |
| AT1G04130 | 0,50 | 1,38 | 2,79 | 0,50 | 6,20E-01 |
| AT1G04170 | 1,02 | 3,32 | 2,05 | 1,62 | 1,05E-01 |
| AT1G04247 | 0,30 | 1,20 | 4,39 | 0,27 | 7,85E-01 |
| AT1G04230 | 0,70 | -2,89 | 2,93 | -0,99 | 3,24E-01 |
| AT1G04295 | 0,43 | 1,33 | 2,87 | 0,47 | 6,42E-01 |
| AT1G04350 | 0,25 | -0,40 | 4,37 | -0,09 | 9,27E-01 |
| AT1G04420 | 1,20 | 2,86 | 1,91 | 1,50 | 1,34E-01 |
| AT1G04440 | 0,50 | 2,15 | 2,78 | 0,77 | 4,39E-01 |
| AT1G04273 | 0,37 | 0,53 | 3,51 | 0,15 | 8,80E-01 |
| AT1G04277 | 0,93 | 0,18 | 2,03 | 0,09 | 9,31E-01 |
| AT1G04510 | 0,67 | 2,55 | 2,99 | 0,85 | 3,94E-01 |
| AT1G04540 | 1,05 | 3,25 | 2,26 | 1,44 | 1,51E-01 |
| AT1G04690 | 0,77 | 0,54 | 2,51 | 0,21 | 8,31E-01 |
| AT1G04710 | 0,47 | 1,94 | 4,39 | 0,44 | 6,59E-01 |
| AT1G04760 | 0,36 | 0,00 | 4,41 | 0,00 | 1,00E+00 |
| AT1G04945 | 0,47 | 1,16 | 3,20 | 0,36 | 7,18E-01 |
| AT1G04970 | 0,34 | -1,50 | 4,36 | -0,34 | 7,31E-01 |
| AT1G05087 | 0,44 | 0,00 | 4,14 | 0,00 | 1,00E+00 |
| AT1G05090 | 0,12 | 0,00 | 4,41 | 0,00 | 1,00E+00 |
| AT1G05140 | 0,22 | 1,37 | 4,41 | 0,31 | 7,56E-01 |
| AT1G05200 | 1,12 | -0,63 | 1,88 | -0,33 | 7,38E-01 |
| AT1G05360 | 0,08 | 0,00 | 4,41 | 0,00 | 1,00E+00 |
| AT1G05380 | 1,42 | 1,79 | 2,14 | 0,84 | 4,03E-01 |
| AT1G05500 | 0,37 | 0,00 | 4,41 | 0,00 | 1,00E+00 |
| AT1G04397 | 0,08 | 0,00 | 4,41 | 0,00 | 1,00E+00 |
| AT1G05600 | 0,08 | 0,00 | 4,41 | 0,00 | 1,00E+00 |
| AT1G04423 | 1,18 | 1,10 | 2,11 | 0,52 | 6,00E-01 |
| AT1G05790 | 0,76 | 2,17 | 2,57 | 0,85 | 3,98E-01 |

| | | | | | |
|---|---|---|---|---|---|
| AT1G05830 | 0,53 | 0,59 | 3,38 | 0,17 | 8,62E-01 |
| AT1G05890 | 0,68 | 2,99 | 4,32 | 0,69 | 4,90E-01 |
| AT1G05900 | 0,21 | 1,25 | 4,39 | 0,29 | 7,75E-01 |
| AT1G05910 | 0,59 | 2,86 | 3,77 | 0,76 | 4,49E-01 |
| AT1G05980 | 0,23 | -0,40 | 4,38 | -0,09 | 9,28E-01 |
| AT1G06002 | 0,95 | 0,71 | 1,99 | 0,35 | 7,23E-01 |
| AT1G06030 | 0,96 | 2,57 | 3,33 | 0,77 | 4,40E-01 |
| AT1G06290 | 0,60 | 1,30 | 2,82 | 0,46 | 6,43E-01 |
| AT1G06380 | 0,45 | -0,45 | 4,20 | -0,11 | 9,15E-01 |
| AT1G06390 | 0,40 | 2,14 | 3,44 | 0,62 | 5,34E-01 |
| AT1G06410 | 1,61 | 1,92 | 1,87 | 1,03 | 3,04E-01 |
| AT1G06530 | 0,96 | 2,65 | 2,02 | 1,31 | 1,89E-01 |
| AT1G06640 | 1,12 | 2,27 | 2,02 | 1,12 | 2,62E-01 |
| AT1G06645 | 0,50 | 1,48 | 4,06 | 0,36 | 7,15E-01 |
| AT1G06650 | 0,69 | 2,42 | 3,03 | 0,80 | 4,24E-01 |
| AT1G06670 | 1,38 | 2,56 | 1,99 | 1,29 | 1,98E-01 |
| AT1G06750 | 0,24 | 1,31 | 4,40 | 0,30 | 7,65E-01 |
| AT1G06780 | 0,12 | 0,00 | 4,41 | 0,00 | 1,00E+00 |
| AT1G06910 | 0,23 | 1,38 | 4,41 | 0,31 | 7,55E-01 |
| AT1G06930 | 0,08 | 0,00 | 4,41 | 0,00 | 1,00E+00 |
| AT1G07010 | 0,86 | 2,85 | 2,96 | 0,96 | 3,36E-01 |
| AT1G04537 | 0,24 | 1,96 | 4,39 | 0,45 | 6,55E-01 |
| AT1G07080 | 0,86 | -1,35 | 2,14 | -0,63 | 5,28E-01 |
| AT1G07130 | 0,28 | 0,00 | 4,41 | 0,00 | 1,00E+00 |
| AT1G07170 | 0,78 | 0,53 | 2,29 | 0,23 | 8,17E-01 |
| AT1G07180 | 0,37 | 0,42 | 3,57 | 0,12 | 9,06E-01 |
| AT1G07210 | 0,17 | 0,00 | 4,41 | 0,00 | 1,00E+00 |
| AT1G07240 | 0,59 | 1,40 | 3,32 | 0,42 | 6,73E-01 |
| AT1G07280 | 1,33 | 1,31 | 1,77 | 0,74 | 4,59E-01 |
| AT1G07320 | 1,43 | 1,26 | 1,52 | 0,83 | 4,06E-01 |
| AT1G07470 | 0,61 | 2,93 | 3,73 | 0,79 | 4,32E-01 |
| AT1G07630 | 0,70 | 2,37 | 2,98 | 0,80 | 4,26E-01 |
| AT1G07705 | 0,12 | 0,00 | 4,41 | 0,00 | 1,00E+00 |
| AT1G07728 | 1,03 | 3,80 | 2,48 | 1,53 | 1,26E-01 |
| AT1G07790 | 0,25 | -0,90 | 4,38 | -0,21 | 8,37E-01 |
| AT1G07830 | 0,43 | 0,00 | 4,23 | 0,00 | 1,00E+00 |
| AT1G07990 | 0,78 | 1,60 | 3,08 | 0,52 | 6,04E-01 |
| AT1G08110 | 0,41 | 2,09 | 4,34 | 0,48 | 6,30E-01 |
| AT1G08190 | 0,34 | -0,54 | 3,66 | -0,15 | 8,82E-01 |
| AT1G08260 | 0,51 | 0,00 | 4,17 | 0,00 | 1,00E+00 |
| AT1G08360 | 1,45 | 1,91 | 1,90 | 1,01 | 3,15E-01 |
| AT1G08370 | 0,71 | 2,16 | 2,57 | 0,84 | 4,01E-01 |
| AT1G08410 | 0,87 | 0,54 | 2,52 | 0,22 | 8,30E-01 |
| AT1G08460 | 0,84 | -0,34 | 2,20 | -0,16 | 8,76E-01 |
| AT1G08465 | 0,35 | 1,38 | 4,41 | 0,31 | 7,54E-01 |
| AT1G08480 | 0,59 | -0,46 | 3,25 | -0,14 | 8,88E-01 |
| AT1G08520 | 1,05 | 1,48 | 1,79 | 0,83 | 4,06E-01 |
| AT1G08530 | 0,48 | 0,00 | 4,07 | 0,00 | 1,00E+00 |
| AT1G08550 | 0,85 | -0,96 | 2,34 | -0,41 | 6,82E-01 |
| AT1G08570 | 0,49 | 0,95 | 4,11 | 0,23 | 8,16E-01 |
| AT1G08590 | 0,74 | 1,38 | 3,01 | 0,46 | 6,47E-01 |
| AT1G08600 | 0,40 | -0,52 | 3,41 | -0,15 | 8,80E-01 |
| AT1G08620 | 1,06 | 2,28 | 1,94 | 1,18 | 2,39E-01 |
| AT1G08650 | 0,89 | -0,63 | 2,92 | -0,22 | 8,28E-01 |
| AT1G08680 | 0,17 | 0,00 | 4,41 | 0,00 | 1,00E+00 |
| AT1G08710 | 0,30 | 1,44 | 4,40 | 0,33 | 7,43E-01 |

| | | | | | |
|---|---|---|---|---|---|
| AT1G08720 | 1,46 | 2,04 | 2,15 | 0,95 | 3,42E-01 |
| AT1G08760 | 0,68 | 1,41 | 3,08 | 0,46 | 6,47E-01 |
| AT1G08770 | 0,33 | 1,34 | 3,59 | 0,37 | 7,09E-01 |
| AT1G08830 | 1,29 | 0,14 | 2,13 | 0,07 | 9,47E-01 |
| AT1G08900 | 0,34 | 1,31 | 3,55 | 0,37 | 7,12E-01 |
| AT1G08910 | 0,43 | 2,16 | 4,21 | 0,51 | 6,08E-01 |
| AT1G08920 | 0,59 | 0,79 | 3,12 | 0,25 | 7,99E-01 |
| AT1G08930 | 1,31 | 0,58 | 1,67 | 0,35 | 7,30E-01 |
| AT1G08970 | 0,82 | 0,31 | 2,73 | 0,11 | 9,10E-01 |
| AT1G08980 | 0,91 | 2,40 | 2,10 | 1,14 | 2,53E-01 |
| AT1G09040 | 0,45 | 2,20 | 3,34 | 0,66 | 5,09E-01 |
| AT1G09250 | 0,32 | 2,37 | 4,36 | 0,54 | 5,86E-01 |
| AT1G09260 | 0,62 | -2,76 | 4,32 | -0,64 | 5,23E-01 |
| AT1G09270 | 0,12 | 0,00 | 4,41 | 0,00 | 1,00E+00 |
| AT1G09300 | 0,36 | 1,83 | 4,38 | 0,42 | 6,76E-01 |
| AT1G09390 | 0,23 | 0,00 | 4,41 | 0,00 | 1,00E+00 |
| AT1G09421 | 0,17 | 0,00 | 4,41 | 0,00 | 1,00E+00 |
| AT1G09430 | 0,79 | 1,80 | 2,49 | 0,72 | 4,70E-01 |
| AT1G09450 | 0,24 | 1,38 | 4,41 | 0,31 | 7,54E-01 |
| AT1G09490 | 0,05 | 0,00 | 4,41 | 0,00 | 1,00E+00 |
| AT1G09500 | 0,08 | 0,00 | 4,41 | 0,00 | 1,00E+00 |
| AT1G09530 | 0,63 | 0,28 | 3,68 | 0,08 | 9,39E-01 |
| AT1G09560 | 0,29 | 1,38 | 4,41 | 0,31 | 7,54E-01 |
| AT1G09590 | 0,27 | -0,42 | 4,39 | -0,10 | 9,24E-01 |
| AT1G09600 | 0,94 | 2,89 | 2,84 | 1,02 | 3,08E-01 |
| AT1G09610 | 0,36 | 1,85 | 4,39 | 0,42 | 6,73E-01 |
| AT1G09645 | 0,33 | 2,12 | 4,37 | 0,49 | 6,27E-01 |
| AT1G09665 | 0,25 | 1,84 | 4,38 | 0,42 | 6,75E-01 |
| AT1G09710 | 0,91 | 3,12 | 2,86 | 1,09 | 2,75E-01 |
| AT1G09750 | 0,82 | -1,13 | 2,03 | -0,56 | 5,78E-01 |
| AT1G09795 | 0,58 | 0,69 | 3,97 | 0,17 | 8,61E-01 |
| AT1G09815 | 0,21 | -0,74 | 4,37 | -0,17 | 8,65E-01 |
| AT1G10050 | 0,78 | -0,40 | 2,24 | -0,18 | 8,57E-01 |
| AT1G10070 | 0,24 | 1,38 | 4,41 | 0,31 | 7,54E-01 |
| AT1G10130 | 1,07 | 1,51 | 2,07 | 0,73 | 4,66E-01 |
| AT1G10150 | 1,47 | 1,05 | 1,65 | 0,64 | 5,23E-01 |
| AT1G10270 | 0,58 | -0,48 | 2,72 | -0,18 | 8,61E-01 |
| AT1G10450 | 0,33 | 1,27 | 4,39 | 0,29 | 7,72E-01 |
| AT1G10490 | 0,65 | 1,19 | 2,55 | 0,47 | 6,42E-01 |
| AT1G10570 | 0,16 | 0,00 | 4,41 | 0,00 | 1,00E+00 |
| AT1G10600 | 1,27 | 2,78 | 2,05 | 1,36 | 1,75E-01 |
| AT1G10610 | 0,90 | 3,37 | 3,32 | 1,01 | 3,11E-01 |
| AT1G10660 | 0,54 | 1,50 | 3,97 | 0,38 | 7,05E-01 |
| AT1G10670 | 0,76 | -2,40 | 3,56 | -0,68 | 4,99E-01 |
| AT1G10700 | 0,45 | 1,33 | 4,40 | 0,30 | 7,63E-01 |
| AT1G10720 | 0,48 | 0,36 | 2,77 | 0,13 | 8,98E-01 |
| AT1G10745 | 1,57 | -0,14 | 1,79 | -0,08 | 9,38E-01 |
| AT1G10790 | 0,08 | 0,00 | 4,41 | 0,00 | 1,00E+00 |
| AT1G10850 | 0,85 | 3,22 | 3,48 | 0,93 | 3,55E-01 |
| AT1G10910 | 0,08 | 0,00 | 4,41 | 0,00 | 1,00E+00 |
| AT1G10960 | 0,86 | -0,45 | 2,80 | -0,16 | 8,73E-01 |
| AT1G11020 | 0,27 | 1,40 | 4,40 | 0,32 | 7,51E-01 |
| AT1G11050 | 0,22 | 0,00 | 4,41 | 0,00 | 1,00E+00 |
| AT1G11260 | 1,39 | -1,01 | 1,72 | -0,59 | 5,57E-01 |
| AT1G11300 | 0,32 | -0,50 | 4,40 | -0,11 | 9,09E-01 |
| AT1G11303 | 0,25 | 0,00 | 4,41 | 0,00 | 1,00E+00 |

| | | | | | |
|---|---|---|---|---|---|
| AT1G11320 | 0,08 | 0,00 | 4,41 | 0,00 | 1,00E+00 |
| AT1G11330 | 0,64 | 1,76 | 3,02 | 0,58 | 5,60E-01 |
| AT1G11400 | 0,26 | 1,36 | 4,40 | 0,31 | 7,58E-01 |
| AT1G11410 | 0,36 | -0,54 | 3,53 | -0,15 | 8,78E-01 |
| AT1G11430 | 0,35 | -1,23 | 3,62 | -0,34 | 7,33E-01 |
| AT1G11440 | 0,54 | 0,42 | 3,10 | 0,14 | 8,92E-01 |
| AT1G11580 | 0,31 | 1,34 | 4,40 | 0,31 | 7,60E-01 |
| AT1G11590 | 0,67 | -2,09 | 2,49 | -0,84 | 4,01E-01 |
| AT1G11593 | 0,40 | 0,00 | 4,26 | 0,00 | 1,00E+00 |
| AT1G11660 | 1,14 | 3,59 | 1,97 | 1,82 | 6,84E-02 |
| AT1G11680 | 0,83 | 0,84 | 2,80 | 0,30 | 7,65E-01 |
| AT1G11710 | 0,17 | 0,00 | 4,41 | 0,00 | 1,00E+00 |
| AT1G11750 | 0,56 | -2,14 | 3,21 | -0,67 | 5,05E-01 |
| AT1G11790 | 0,42 | 0,79 | 4,38 | 0,18 | 8,56E-01 |
| AT1G11890 | 1,50 | 0,78 | 1,90 | 0,41 | 6,82E-01 |
| AT1G11905 | 0,75 | 1,71 | 2,29 | 0,75 | 4,55E-01 |
| AT1G11930 | 0,68 | 0,89 | 2,20 | 0,41 | 6,85E-01 |
| AT1G12015 | 0,46 | -1,32 | 4,38 | -0,30 | 7,63E-01 |
| AT1G12050 | 1,22 | 2,59 | 1,75 | 1,48 | 1,39E-01 |
| AT1G12060 | 0,12 | 0,00 | 4,41 | 0,00 | 1,00E+00 |
| AT1G12140 | 0,12 | 0,00 | 4,41 | 0,00 | 1,00E+00 |
| AT1G12210 | 0,35 | 0,35 | 4,39 | 0,08 | 9,37E-01 |
| AT1G12220 | 1,65 | -1,06 | 1,56 | -0,68 | 4,98E-01 |
| AT1G12230 | 0,35 | 1,38 | 3,57 | 0,39 | 6,99E-01 |
| AT1G12250 | 0,19 | -0,55 | 4,38 | -0,13 | 9,00E-01 |
| AT1G12270 | 0,62 | -0,62 | 2,69 | -0,23 | 8,18E-01 |
| AT1G12320 | 0,82 | 0,00 | 2,96 | 0,00 | 1,00E+00 |
| AT1G12360 | 0,65 | 2,51 | 3,81 | 0,66 | 5,10E-01 |
| AT1G12410 | 0,67 | -1,86 | 3,71 | -0,50 | 6,16E-01 |
| AT1G12460 | 0,83 | -1,19 | 2,10 | -0,57 | 5,71E-01 |
| AT1G12580 | 0,43 | 1,35 | 2,91 | 0,46 | 6,42E-01 |
| AT1G12760 | 0,76 | 2,91 | 2,24 | 1,30 | 1,94E-01 |
| AT1G12810 | 0,42 | -1,29 | 2,53 | -0,51 | 6,11E-01 |
| AT1G12840 | 1,05 | 0,32 | 1,90 | 0,17 | 8,68E-01 |
| AT1G12860 | 0,23 | -0,99 | 4,36 | -0,23 | 8,21E-01 |
| AT1G12990 | 1,14 | 2,14 | 2,24 | 0,96 | 3,39E-01 |
| AT1G13040 | 0,24 | 0,00 | 4,41 | 0,00 | 1,00E+00 |
| AT1G13060 | 0,83 | 0,13 | 3,09 | 0,04 | 9,66E-01 |
| AT1G13090 | 1,29 | 1,44 | 1,78 | 0,81 | 4,21E-01 |
| AT1G13100 | 0,26 | -1,28 | 4,37 | -0,29 | 7,69E-01 |
| AT1G13180 | 0,14 | 0,00 | 4,41 | 0,00 | 1,00E+00 |
| AT1G13190 | 1,26 | 0,00 | 2,69 | 0,00 | 1,00E+00 |
| AT1G13220 | 0,56 | 0,00 | 3,88 | 0,00 | 1,00E+00 |
| AT1G13260 | 1,59 | 0,01 | 1,80 | 0,00 | 9,97E-01 |
| AT1G13270 | 0,60 | -2,59 | 3,01 | -0,86 | 3,89E-01 |
| AT1G13330 | 0,70 | 2,14 | 2,64 | 0,81 | 4,16E-01 |
| AT1G13400 | 0,23 | 1,38 | 4,41 | 0,31 | 7,54E-01 |
| AT1G13430 | 0,17 | 0,00 | 4,41 | 0,00 | 1,00E+00 |
| AT1G13460 | 0,37 | -0,06 | 4,35 | -0,01 | 9,89E-01 |
| AT1G13820 | 0,11 | 0,00 | 4,41 | 0,00 | 1,00E+00 |
| AT1G13850 | 0,13 | 0,00 | 4,41 | 0,00 | 1,00E+00 |
| AT1G13880 | 0,45 | -0,51 | 3,48 | -0,15 | 8,84E-01 |
| AT1G13990 | 0,47 | 2,04 | 3,35 | 0,61 | 5,43E-01 |
| AT1G14000 | 0,81 | -0,44 | 2,33 | -0,19 | 8,50E-01 |
| AT1G14020 | 0,41 | 2,14 | 4,22 | 0,51 | 6,12E-01 |
| AT1G05047 | 0,12 | 0,00 | 4,41 | 0,00 | 1,00E+00 |

| | | | | | |
|---|---|---|---|---|---|
| AT1G14110 | 0,52 | 0,00 | 3,92 | 0,00 | 1,00E+00 |
| AT1G14150 | 0,05 | 0,00 | 4,41 | 0,00 | 1,00E+00 |
| AT1G14185 | 0,05 | 0,00 | 4,41 | 0,00 | 1,00E+00 |
| AT1G14230 | 0,08 | 0,00 | 4,41 | 0,00 | 1,00E+00 |
| AT1G14250 | 0,32 | -1,72 | 4,35 | -0,40 | 6,92E-01 |
| AT1G14280 | 0,63 | 2,18 | 3,09 | 0,70 | 4,81E-01 |
| AT1G14310 | 0,25 | 1,29 | 4,39 | 0,29 | 7,69E-01 |
| AT1G14345 | 0,53 | 2,13 | 2,80 | 0,76 | 4,47E-01 |
| AT1G14350 | 0,37 | 0,61 | 4,34 | 0,14 | 8,88E-01 |
| AT1G14370 | 0,91 | 0,52 | 2,06 | 0,25 | 7,99E-01 |
| AT1G14470 | 0,37 | -1,68 | 4,36 | -0,38 | 7,01E-01 |
| AT1G14480 | 0,08 | 0,00 | 4,41 | 0,00 | 1,00E+00 |
| AT1G14560 | 0,21 | -0,54 | 4,41 | -0,12 | 9,02E-01 |
| AT1G14570 | 1,23 | 3,33 | 1,88 | 1,77 | 7,64E-02 |
| AT1G14650 | 1,32 | 2,21 | 2,37 | 0,93 | 3,50E-01 |
| AT1G14800 | 1,13 | 1,38 | 1,89 | 0,73 | 4,67E-01 |
| AT1G14860 | 0,08 | 0,00 | 4,41 | 0,00 | 1,00E+00 |
| AT1G15030 | 0,23 | 0,00 | 4,41 | 0,00 | 1,00E+00 |
| AT1G15120 | 0,30 | 1,38 | 4,41 | 0,31 | 7,54E-01 |
| AT1G15190 | 0,17 | 0,00 | 4,41 | 0,00 | 1,00E+00 |
| AT1G15240 | 0,58 | -0,46 | 2,77 | -0,17 | 8,69E-01 |
| AT1G15260 | 0,61 | -1,07 | 3,08 | -0,35 | 7,29E-01 |
| AT1G15390 | 0,48 | 1,38 | 4,41 | 0,31 | 7,54E-01 |
| AT1G15430 | 0,55 | 1,61 | 3,90 | 0,41 | 6,80E-01 |
| AT1G15650 | 0,36 | 2,37 | 4,36 | 0,54 | 5,86E-01 |
| AT1G05183 | 0,40 | 2,10 | 3,40 | 0,62 | 5,37E-01 |
| AT1G15720 | 0,33 | 1,56 | 4,39 | 0,36 | 7,22E-01 |
| AT1G15740 | 0,65 | 2,13 | 2,57 | 0,83 | 4,07E-01 |
| AT1G15810 | 0,77 | 1,21 | 2,13 | 0,57 | 5,71E-01 |
| AT1G15890 | 1,49 | 1,23 | 1,79 | 0,69 | 4,91E-01 |
| AT1G15910 | 0,69 | 3,55 | 3,51 | 1,01 | 3,12E-01 |
| AT1G15920 | 0,80 | 3,31 | 2,72 | 1,22 | 2,24E-01 |
| AT1G15930 | 1,08 | -0,21 | 1,93 | -0,11 | 9,11E-01 |
| AT1G15950 | 0,56 | 2,56 | 2,64 | 0,97 | 3,32E-01 |
| AT1G15980 | 1,06 | -1,56 | 1,99 | -0,78 | 4,33E-01 |
| AT1G16060 | 0,12 | 0,00 | 4,41 | 0,00 | 1,00E+00 |
| AT1G16080 | 0,31 | -0,54 | 4,41 | -0,12 | 9,02E-01 |
| AT1G16090 | 0,22 | 0,00 | 4,41 | 0,00 | 1,00E+00 |
| AT1G16110 | 0,61 | 1,36 | 3,73 | 0,36 | 7,16E-01 |
| AT1G16180 | 1,23 | 1,36 | 2,03 | 0,67 | 5,05E-01 |
| AT1G16250 | 0,81 | 0,34 | 2,37 | 0,14 | 8,85E-01 |
| AT1G16270 | 1,59 | 1,68 | 1,54 | 1,10 | 2,73E-01 |
| AT1G16350 | 0,39 | -1,76 | 4,23 | -0,42 | 6,78E-01 |
| AT1G16400 | 0,30 | 0,49 | 4,40 | 0,11 | 9,12E-01 |
| AT1G16410 | 1,19 | -0,88 | 1,65 | -0,53 | 5,94E-01 |
| AT1G16470 | 0,19 | 0,00 | 4,40 | 0,00 | 1,00E+00 |
| AT1G16500 | 0,25 | 1,24 | 4,38 | 0,28 | 7,78E-01 |
| AT1G16560 | 0,44 | 1,30 | 3,31 | 0,39 | 6,95E-01 |
| AT1G16670 | 0,35 | 2,13 | 4,37 | 0,49 | 6,26E-01 |
| AT1G16680 | 0,85 | -0,97 | 2,45 | -0,40 | 6,92E-01 |
| AT1G16690 | 0,12 | 0,00 | 4,41 | 0,00 | 1,00E+00 |
| AT1G16700 | 0,62 | 0,87 | 3,15 | 0,28 | 7,81E-01 |
| AT1G16720 | 1,64 | 1,54 | 1,63 | 0,94 | 3,45E-01 |
| AT1G16740 | 0,37 | 1,76 | 4,38 | 0,40 | 6,87E-01 |
| AT1G16810 | 0,12 | 0,00 | 4,41 | 0,00 | 1,00E+00 |
| AT1G16860 | 0,38 | -1,25 | 4,27 | -0,29 | 7,70E-01 |

| | | | | | |
|---|---|---|---|---|---|
| AT1G17050 | 0,20 | -0,43 | 4,39 | -0,10 | 9,22E-01 |
| AT1G17070 | 0,73 | -0,40 | 3,65 | -0,11 | 9,13E-01 |
| AT1G17100 | 0,86 | -0,09 | 1,90 | -0,05 | 9,63E-01 |
| AT1G17120 | 0,36 | 1,70 | 4,38 | 0,39 | 6,97E-01 |
| AT1G17150 | 0,40 | -0,52 | 3,51 | -0,15 | 8,83E-01 |
| AT1G17160 | 0,36 | 1,37 | 4,37 | 0,31 | 7,54E-01 |
| AT1G17200 | 0,19 | -0,54 | 4,41 | -0,12 | 9,02E-01 |
| AT1G17230 | 0,51 | 0,00 | 3,21 | 0,00 | 1,00E+00 |
| AT1G17255 | 1,02 | 1,38 | 2,82 | 0,49 | 6,25E-01 |
| AT1G17260 | 0,13 | 0,00 | 4,41 | 0,00 | 1,00E+00 |
| AT1G17300 | 0,61 | 1,36 | 3,73 | 0,36 | 7,16E-01 |
| AT1G17340 | 0,57 | 1,35 | 2,37 | 0,57 | 5,69E-01 |
| AT1G17360 | 1,17 | 0,22 | 2,17 | 0,10 | 9,18E-01 |
| AT1G17420 | 1,07 | 0,70 | 2,24 | 0,31 | 7,55E-01 |
| AT1G17460 | 0,31 | 1,49 | 4,39 | 0,34 | 7,35E-01 |
| AT1G17490 | 0,39 | -0,67 | 3,55 | -0,19 | 8,49E-01 |
| AT1G17500 | 0,69 | 1,31 | 2,32 | 0,56 | 5,73E-01 |
| AT1G17530 | 0,57 | 1,38 | 3,22 | 0,43 | 6,68E-01 |
| AT1G17550 | 1,05 | 2,29 | 1,91 | 1,20 | 2,30E-01 |
| AT1G17570 | 0,08 | 0,00 | 4,41 | 0,00 | 1,00E+00 |
| AT1G17580 | 1,15 | 3,32 | 2,18 | 1,52 | 1,29E-01 |
| AT1G17610 | 0,33 | 0,00 | 4,41 | 0,00 | 1,00E+00 |
| AT1G17620 | 1,43 | 0,45 | 1,95 | 0,23 | 8,17E-01 |
| AT1G17680 | 0,28 | -0,46 | 4,40 | -0,10 | 9,17E-01 |
| AT1G17745 | 0,96 | 2,60 | 2,10 | 1,24 | 2,15E-01 |
| AT1G17750 | 0,21 | 0,00 | 4,41 | 0,00 | 1,00E+00 |
| AT1G17840 | 1,51 | -1,06 | 1,87 | -0,57 | 5,69E-01 |
| AT1G17860 | 0,25 | -0,49 | 4,40 | -0,11 | 9,11E-01 |
| AT1G17970 | 0,47 | 0,27 | 4,12 | 0,07 | 9,48E-01 |
| AT1G18020 | 0,78 | 1,99 | 3,08 | 0,65 | 5,18E-01 |
| AT1G18090 | 0,88 | 2,35 | 3,58 | 0,66 | 5,12E-01 |
| AT1G18130 | 0,21 | -0,58 | 4,38 | -0,13 | 8,95E-01 |
| AT1G18170 | 0,31 | -0,54 | 4,41 | -0,12 | 9,02E-01 |
| AT1G18300 | 0,38 | 1,53 | 4,35 | 0,35 | 7,25E-01 |
| AT1G18390 | 0,88 | 2,94 | 2,12 | 1,39 | 1,65E-01 |
| AT1G18450 | 0,27 | -0,40 | 4,38 | -0,09 | 9,27E-01 |
| AT1G18460 | 0,79 | -1,13 | 2,07 | -0,54 | 5,87E-01 |
| AT1G18480 | 0,17 | 1,32 | 4,40 | 0,30 | 7,64E-01 |
| AT1G18490 | 0,64 | 1,35 | 4,40 | 0,31 | 7,59E-01 |
| AT1G18500 | 0,63 | 0,14 | 3,06 | 0,05 | 9,63E-01 |
| AT1G18570 | 0,88 | 2,35 | 2,19 | 1,07 | 2,84E-01 |
| AT1G18580 | 0,40 | 0,00 | 4,41 | 0,00 | 1,00E+00 |
| AT1G18600 | 0,74 | 1,32 | 2,50 | 0,53 | 5,98E-01 |
| AT1G18620 | 0,88 | 0,29 | 2,29 | 0,13 | 8,98E-01 |
| AT1G18700 | 1,03 | 3,93 | 2,24 | 1,75 | 7,97E-02 |
| AT1G18730 | 0,47 | -2,54 | 4,15 | -0,61 | 5,40E-01 |
| AT1G18820 | 0,52 | -1,54 | 4,35 | -0,35 | 7,23E-01 |
| AT1G18870 | 0,46 | -0,76 | 4,37 | -0,17 | 8,62E-01 |
| AT1G18879 | 1,60 | -0,60 | 2,01 | -0,30 | 7,65E-01 |
| AT1G18900 | 0,70 | 0,00 | 3,66 | 0,00 | 1,00E+00 |
| AT1G18950 | 0,83 | 1,34 | 2,33 | 0,57 | 5,66E-01 |
| AT1G19020 | 0,56 | 2,19 | 2,42 | 0,91 | 3,64E-01 |
| AT1G19025 | 0,65 | 3,27 | 3,00 | 1,09 | 2,75E-01 |
| AT1G19086 | 0,19 | 0,00 | 4,41 | 0,00 | 1,00E+00 |
| AT1G19110 | 0,80 | 1,83 | 2,19 | 0,84 | 4,04E-01 |
| AT1G19150 | 0,10 | 0,00 | 4,41 | 0,00 | 1,00E+00 |

| | | | | | |
|---|---|---|---|---|---|
| AT1G19180 | 1,37 | 2,75 | 2,04 | 1,35 | 1,77E-01 |
| AT1G19190 | 0,39 | 1,76 | 4,38 | 0,40 | 6,87E-01 |
| AT1G19290 | 0,57 | 1,30 | 3,83 | 0,34 | 7,34E-01 |
| AT1G19350 | 0,52 | 1,75 | 3,97 | 0,44 | 6,59E-01 |
| AT1G19376 | 0,56 | 0,00 | 3,17 | 0,00 | 1,00E+00 |
| AT1G19464 | 0,18 | 0,00 | 4,41 | 0,00 | 1,00E+00 |
| AT1G19580 | 0,28 | 0,50 | 4,40 | 0,11 | 9,10E-01 |
| AT1G05507 | 0,08 | 0,00 | 4,41 | 0,00 | 1,00E+00 |
| AT1G19710 | 0,37 | 2,13 | 4,32 | 0,49 | 6,22E-01 |
| AT1G19770 | 0,78 | 0,00 | 2,90 | 0,00 | 1,00E+00 |
| AT1G19800 | 0,29 | 0,43 | 4,37 | 0,10 | 9,22E-01 |
| AT1G19830 | 0,33 | 0,00 | 4,41 | 0,00 | 1,00E+00 |
| AT1G19850 | 0,68 | 0,00 | 3,03 | 0,00 | 1,00E+00 |
| AT1G19910 | 0,96 | 1,29 | 2,48 | 0,52 | 6,01E-01 |
| AT1G19990 | 0,28 | 0,00 | 4,41 | 0,00 | 1,00E+00 |
| AT1G20050 | 0,40 | 0,32 | 3,50 | 0,09 | 9,28E-01 |
| AT1G05523 | 0,59 | 0,43 | 2,57 | 0,17 | 8,67E-01 |
| AT1G20090 | 0,64 | 0,14 | 2,61 | 0,06 | 9,56E-01 |
| AT1G20110 | 0,46 | -1,72 | 2,76 | -0,62 | 5,34E-01 |
| AT1G20280 | 0,77 | -0,77 | 2,13 | -0,36 | 7,19E-01 |
| AT1G20300 | 0,15 | -0,43 | 4,39 | -0,10 | 9,23E-01 |
| AT1G20310 | 0,17 | 0,00 | 4,41 | 0,00 | 1,00E+00 |
| AT1G20375 | 1,50 | 4,08 | 1,76 | 2,31 | 2,06E-02 |
| AT1G20630 | 1,62 | 1,92 | 1,90 | 1,01 | 3,12E-01 |
| AT1G20640 | 1,05 | 3,37 | 2,08 | 1,62 | 1,06E-01 |
| AT1G20693 | 1,05 | -1,49 | 2,26 | -0,66 | 5,10E-01 |
| AT1G20696 | 0,69 | 0,38 | 2,32 | 0,16 | 8,69E-01 |
| AT1G20780 | 1,16 | -1,55 | 1,87 | -0,83 | 4,07E-01 |
| AT1G20823 | 0,81 | -0,52 | 2,44 | -0,21 | 8,32E-01 |
| AT1G20920 | 0,49 | 1,22 | 3,19 | 0,38 | 7,02E-01 |
| AT1G20980 | 1,44 | -0,15 | 1,89 | -0,08 | 9,37E-01 |
| AT1G21000 | 0,77 | 1,37 | 2,36 | 0,58 | 5,62E-01 |
| AT1G21050 | 0,58 | -0,46 | 3,10 | -0,15 | 8,82E-01 |
| AT1G21065 | 0,84 | -0,59 | 2,29 | -0,26 | 7,98E-01 |
| AT1G21140 | 0,08 | 0,00 | 4,41 | 0,00 | 1,00E+00 |
| AT1G21170 | 0,55 | -0,42 | 3,83 | -0,11 | 9,14E-01 |
| AT1G21210 | 0,63 | 1,34 | 3,04 | 0,44 | 6,59E-01 |
| AT1G21230 | 0,43 | 0,00 | 3,36 | 0,00 | 1,00E+00 |
| AT1G21240 | 0,38 | 1,41 | 4,31 | 0,33 | 7,42E-01 |
| AT1G21370 | 0,21 | 0,00 | 4,41 | 0,00 | 1,00E+00 |
| AT1G21430 | 0,53 | 1,22 | 4,39 | 0,28 | 7,81E-01 |
| AT1G21560 | 0,41 | 2,11 | 4,16 | 0,51 | 6,12E-01 |
| AT1G21640 | 0,89 | 1,15 | 2,06 | 0,56 | 5,77E-01 |
| AT1G21660 | 0,28 | 1,39 | 4,41 | 0,32 | 7,52E-01 |
| AT1G21670 | 0,86 | 2,26 | 3,26 | 0,69 | 4,88E-01 |
| AT1G21690 | 0,29 | 1,26 | 4,39 | 0,29 | 7,74E-01 |
| AT1G21720 | 0,34 | 1,38 | 4,41 | 0,31 | 7,54E-01 |
| AT1G21722 | 0,08 | 0,00 | 4,41 | 0,00 | 1,00E+00 |
| AT1G21780 | 0,75 | 1,19 | 2,62 | 0,45 | 6,50E-01 |
| AT1G21835 | 0,94 | 0,04 | 2,14 | 0,02 | 9,87E-01 |
| AT1G21840 | 0,12 | 0,00 | 4,41 | 0,00 | 1,00E+00 |
| AT1G21880 | 0,12 | 0,00 | 4,41 | 0,00 | 1,00E+00 |
| AT1G21910 | 0,33 | -1,69 | 4,36 | -0,39 | 6,99E-01 |
| AT1G21980 | 0,19 | 0,00 | 4,41 | 0,00 | 1,00E+00 |
| AT1G22040 | 0,05 | 0,00 | 4,41 | 0,00 | 1,00E+00 |
| AT1G22070 | 0,41 | 1,38 | 4,37 | 0,32 | 7,52E-01 |

| | | | | | |
|---|---|---|---|---|---|
| AT1G22190 | 1,08 | 0,10 | 1,72 | 0,06 | 9,53E-01 |
| AT1G22280 | 0,53 | 2,37 | 3,36 | 0,71 | 4,80E-01 |
| AT1G22330 | 0,11 | 0,00 | 4,41 | 0,00 | 1,00E+00 |
| AT1G22410 | 1,37 | -0,19 | 1,53 | -0,12 | 9,02E-01 |
| AT1G22500 | 0,40 | 0,00 | 4,41 | 0,00 | 1,00E+00 |
| AT1G22590 | 0,24 | 0,00 | 4,41 | 0,00 | 1,00E+00 |
| AT1G22640 | 0,56 | 0,00 | 3,82 | 0,00 | 1,00E+00 |
| AT1G22730 | 0,55 | 2,74 | 3,84 | 0,71 | 4,75E-01 |
| AT1G22750 | 0,35 | 1,27 | 3,54 | 0,36 | 7,19E-01 |
| AT1G22770 | 1,61 | 1,17 | 1,46 | 0,80 | 4,25E-01 |
| AT1G22780 | 0,54 | 2,20 | 3,29 | 0,67 | 5,02E-01 |
| AT1G22790 | 0,64 | 2,80 | 3,84 | 0,73 | 4,66E-01 |
| AT1G05767 | 0,23 | -0,45 | 4,39 | -0,10 | 9,18E-01 |
| AT1G22840 | 0,62 | -0,46 | 3,33 | -0,14 | 8,91E-01 |
| AT1G22860 | 0,86 | 3,25 | 2,10 | 1,55 | 1,21E-01 |
| AT1G22870 | 0,66 | 1,38 | 3,03 | 0,46 | 6,49E-01 |
| AT1G22880 | 1,11 | 3,47 | 2,81 | 1,23 | 2,17E-01 |
| AT1G22882 | 0,84 | 1,38 | 3,44 | 0,40 | 6,88E-01 |
| AT1G22940 | 0,55 | 1,31 | 2,75 | 0,48 | 6,33E-01 |
| AT1G23030 | 0,27 | 0,45 | 4,38 | 0,10 | 9,19E-01 |
| AT1G23040 | 0,34 | 0,00 | 4,41 | 0,00 | 1,00E+00 |
| AT1G23050 | 1,19 | -1,07 | 2,47 | -0,43 | 6,64E-01 |
| AT1G23100 | 0,08 | 0,00 | 4,41 | 0,00 | 1,00E+00 |
| AT1G23130 | 0,08 | 0,00 | 4,41 | 0,00 | 1,00E+00 |
| AT1G23230 | 1,20 | 1,19 | 1,92 | 0,62 | 5,36E-01 |
| AT1G23290 | 1,36 | -1,32 | 1,90 | -0,70 | 4,85E-01 |
| AT1G23440 | 0,65 | 0,39 | 3,09 | 0,13 | 9,00E-01 |
| AT1G23450 | 0,21 | 1,38 | 4,41 | 0,31 | 7,54E-01 |
| AT1G23490 | 1,31 | 0,36 | 1,83 | 0,20 | 8,44E-01 |
| AT1G23710 | 1,29 | 0,39 | 2,10 | 0,19 | 8,51E-01 |
| AT1G23820 | 1,10 | 3,21 | 2,93 | 1,10 | 2,73E-01 |
| AT1G05837 | 0,53 | 0,29 | 3,29 | 0,09 | 9,29E-01 |
| AT1G23870 | 0,51 | 0,00 | 3,93 | 0,00 | 1,00E+00 |
| AT1G23880 | 0,08 | 0,00 | 4,41 | 0,00 | 1,00E+00 |
| AT1G23915 | 1,01 | 0,00 | 2,69 | 0,00 | 1,00E+00 |
| AT1G24145 | 0,48 | 1,27 | 3,22 | 0,39 | 6,93E-01 |
| AT1G24147 | 0,16 | 0,00 | 4,41 | 0,00 | 1,00E+00 |
| AT1G24150 | 0,48 | 1,75 | 3,99 | 0,44 | 6,60E-01 |
| AT1G24280 | 0,44 | -0,79 | 4,38 | -0,18 | 8,56E-01 |
| AT1G24340 | 0,28 | -0,51 | 3,81 | -0,13 | 8,93E-01 |
| AT1G24360 | 0,34 | 1,38 | 4,41 | 0,31 | 7,54E-01 |
| AT1G24380 | 0,38 | -0,42 | 2,94 | -0,14 | 8,87E-01 |
| AT1G24490 | 0,45 | 1,40 | 3,32 | 0,42 | 6,73E-01 |
| AT1G24530 | 0,17 | 0,00 | 4,41 | 0,00 | 1,00E+00 |
| AT1G24560 | 0,68 | 0,40 | 2,35 | 0,17 | 8,64E-01 |
| AT1G24570 | 0,08 | 0,00 | 4,41 | 0,00 | 1,00E+00 |
| AT1G25275 | 1,34 | -1,10 | 1,78 | -0,62 | 5,37E-01 |
| AT1G25280 | 0,98 | 3,47 | 2,39 | 1,45 | 1,47E-01 |
| AT1G25480 | 0,19 | 0,00 | 4,41 | 0,00 | 1,00E+00 |
| AT1G25490 | 1,64 | -0,76 | 1,40 | -0,54 | 5,90E-01 |
| AT1G25520 | 0,81 | 1,08 | 3,58 | 0,30 | 7,62E-01 |
| AT1G25550 | 1,33 | -0,24 | 2,18 | -0,11 | 9,13E-01 |
| AT1G26130 | 0,54 | 1,34 | 3,93 | 0,34 | 7,33E-01 |
| AT1G26220 | 0,37 | 0,00 | 4,41 | 0,00 | 1,00E+00 |
| AT1G26250 | 0,44 | 0,00 | 4,17 | 0,00 | 1,00E+00 |
| AT1G26460 | 0,79 | 1,96 | 2,89 | 0,68 | 4,97E-01 |

| | | | | | |
|---|---|---|---|---|---|
| AT1G26470 | 0,34 | 1,38 | 4,41 | 0,31 | 7,54E-01 |
| AT1G26550 | 0,37 | 1,34 | 4,40 | 0,30 | 7,61E-01 |
| AT1G26560 | 0,24 | -0,52 | 4,40 | -0,12 | 9,06E-01 |
| AT1G26580 | 0,53 | 2,18 | 2,75 | 0,79 | 4,28E-01 |
| AT1G26630 | 1,29 | -0,25 | 1,76 | -0,14 | 8,87E-01 |
| AT1G26650 | 1,02 | 0,78 | 2,15 | 0,36 | 7,17E-01 |
| AT1G26665 | 0,17 | 0,00 | 4,41 | 0,00 | 1,00E+00 |
| AT1G26670 | 1,04 | 3,17 | 2,33 | 1,36 | 1,74E-01 |
| AT1G26770 | 0,19 | 0,00 | 4,41 | 0,00 | 1,00E+00 |
| AT1G26830 | 0,72 | 1,29 | 2,56 | 0,50 | 6,16E-01 |
| AT1G26920 | 0,73 | -0,39 | 2,36 | -0,17 | 8,67E-01 |
| AT1G26940 | 0,05 | 0,00 | 4,41 | 0,00 | 1,00E+00 |
| AT1G27000 | 0,38 | -0,07 | 4,37 | -0,02 | 9,87E-01 |
| AT1G27060 | 0,73 | 2,16 | 2,96 | 0,73 | 4,66E-01 |
| AT1G27130 | 0,51 | -1,37 | 3,23 | -0,42 | 6,71E-01 |
| AT1G27150 | 0,43 | 2,15 | 4,36 | 0,49 | 6,23E-01 |
| AT1G27170 | 0,35 | 1,35 | 3,53 | 0,38 | 7,01E-01 |
| AT1G27180 | 0,99 | 2,70 | 2,63 | 1,02 | 3,06E-01 |
| AT1G27290 | 1,54 | 0,82 | 1,65 | 0,50 | 6,18E-01 |
| AT1G27300 | 0,40 | 2,13 | 4,20 | 0,51 | 6,13E-01 |
| AT1G27330 | 0,50 | 1,01 | 4,34 | 0,23 | 8,17E-01 |
| AT1G27360 | 0,08 | 0,00 | 4,41 | 0,00 | 1,00E+00 |
| AT1G27400 | 0,67 | 1,15 | 2,22 | 0,52 | 6,05E-01 |
| AT1G06087 | 0,12 | 0,00 | 4,41 | 0,00 | 1,00E+00 |
| AT1G27450 | 0,37 | -0,98 | 4,38 | -0,22 | 8,23E-01 |
| AT1G27460 | 0,11 | 0,00 | 4,41 | 0,00 | 1,00E+00 |
| AT1G27480 | 0,41 | 0,93 | 4,21 | 0,22 | 8,26E-01 |
| AT1G27520 | 0,14 | 0,00 | 4,41 | 0,00 | 1,00E+00 |
| AT1G06097 | 0,36 | -0,09 | 4,37 | -0,02 | 9,84E-01 |
| AT1G27570 | 1,68 | 0,00 | 2,20 | 0,00 | 1,00E+00 |
| AT1G27630 | 0,58 | 2,25 | 4,00 | 0,56 | 5,74E-01 |
| AT1G27650 | 0,97 | 1,38 | 2,10 | 0,65 | 5,13E-01 |
| AT1G27750 | 0,63 | 2,32 | 2,72 | 0,85 | 3,94E-01 |
| AT1G27752 | 1,43 | 1,38 | 2,40 | 0,57 | 5,66E-01 |
| AT1G27760 | 0,65 | 2,17 | 2,60 | 0,83 | 4,04E-01 |
| AT1G27850 | 0,84 | 2,19 | 2,26 | 0,97 | 3,32E-01 |
| AT1G27950 | 0,55 | -1,17 | 3,84 | -0,30 | 7,61E-01 |
| AT1G28005 | 0,32 | 0,00 | 4,41 | 0,00 | 1,00E+00 |
| AT1G28060 | 0,32 | -0,41 | 4,38 | -0,09 | 9,26E-01 |
| AT1G28135 | 0,08 | 0,00 | 4,41 | 0,00 | 1,00E+00 |
| AT1G28340 | 0,87 | -0,73 | 2,81 | -0,26 | 7,94E-01 |
| AT1G28350 | 1,15 | 2,85 | 2,13 | 1,34 | 1,81E-01 |
| AT1G28380 | 1,21 | 1,57 | 2,28 | 0,69 | 4,92E-01 |
| AT1G06163 | 0,36 | 0,40 | 4,40 | 0,09 | 9,27E-01 |
| AT1G28490 | 0,17 | 0,00 | 4,41 | 0,00 | 1,00E+00 |
| AT1G28520 | 0,40 | 1,38 | 3,43 | 0,40 | 6,88E-01 |
| AT1G28680 | 0,55 | 0,49 | 3,30 | 0,15 | 8,83E-01 |
| AT1G28780 | 0,49 | 0,00 | 4,15 | 0,00 | 1,00E+00 |
| AT1G28760 | 0,10 | 0,00 | 4,41 | 0,00 | 1,00E+00 |
| AT1G29120 | 0,19 | 0,00 | 4,41 | 0,00 | 1,00E+00 |
| AT1G29150 | 0,99 | 3,38 | 2,39 | 1,41 | 1,58E-01 |
| AT1G29200 | 0,08 | 0,00 | 4,41 | 0,00 | 1,00E+00 |
| AT1G29260 | 0,92 | 1,93 | 1,89 | 1,02 | 3,08E-01 |
| AT1G29310 | 0,68 | 0,97 | 3,02 | 0,32 | 7,49E-01 |
| AT1G29340 | 0,94 | 0,43 | 2,27 | 0,19 | 8,50E-01 |
| AT1G29380 | 0,65 | 2,32 | 4,35 | 0,53 | 5,94E-01 |

| | | | | | |
|---|---|---|---|---|---|
| AT1G06233 | 0,68 | -0,76 | 2,49 | -0,31 | 7,59E-01 |
| AT1G06253 | 1,09 | -1,65 | 1,79 | -0,92 | 3,57E-01 |
| AT1G06263 | 0,52 | -2,53 | 4,02 | -0,63 | 5,29E-01 |
| AT1G06283 | 1,64 | -1,55 | 1,73 | -0,89 | 3,71E-01 |
| AT1G29620 | 0,47 | -1,40 | 3,28 | -0,43 | 6,69E-01 |
| AT1G29630 | 0,24 | 0,00 | 4,41 | 0,00 | 1,00E+00 |
| AT1G29660 | 1,40 | -1,36 | 1,58 | -0,86 | 3,90E-01 |
| AT1G29760 | 0,40 | 0,54 | 4,29 | 0,13 | 8,99E-01 |
| AT1G29770 | 0,71 | 1,38 | 4,41 | 0,31 | 7,54E-01 |
| AT1G29800 | 0,74 | 3,88 | 3,67 | 1,06 | 2,90E-01 |
| AT1G29830 | 1,13 | 2,04 | 1,99 | 1,02 | 3,06E-01 |
| AT1G29950 | 0,49 | 2,71 | 4,34 | 0,62 | 5,32E-01 |
| AT1G30010 | 0,08 | 0,00 | 4,41 | 0,00 | 1,00E+00 |
| AT1G30200 | 0,70 | 1,46 | 2,31 | 0,63 | 5,27E-01 |
| AT1G30260 | 1,16 | -0,32 | 2,36 | -0,14 | 8,92E-01 |
| AT1G30420 | 0,38 | 2,22 | 4,37 | 0,51 | 6,12E-01 |
| AT1G30430 | 0,16 | -0,55 | 4,41 | -0,12 | 9,01E-01 |
| AT1G30440 | 1,10 | -0,38 | 1,81 | -0,21 | 8,33E-01 |
| AT1G30450 | 0,86 | -0,42 | 2,16 | -0,19 | 8,48E-01 |
| AT1G30520 | 0,41 | 2,59 | 4,35 | 0,60 | 5,52E-01 |
| AT1G30530 | 0,46 | -1,40 | 3,36 | -0,42 | 6,76E-01 |
| AT1G30610 | 0,23 | 0,00 | 4,41 | 0,00 | 1,00E+00 |
| AT1G30620 | 0,56 | 2,06 | 3,31 | 0,62 | 5,33E-01 |
| AT1G30700 | 0,12 | 0,00 | 4,41 | 0,00 | 1,00E+00 |
| AT1G30720 | 0,53 | 1,22 | 4,39 | 0,28 | 7,81E-01 |
| AT1G30757 | 0,16 | 0,00 | 4,41 | 0,00 | 1,00E+00 |
| AT1G30800 | 0,20 | 1,33 | 4,40 | 0,30 | 7,63E-01 |
| AT1G30910 | 0,25 | 0,00 | 4,41 | 0,00 | 1,00E+00 |
| AT1G06393 | 0,32 | -1,60 | 4,34 | -0,37 | 7,12E-01 |
| AT1G31095 | 0,66 | 0,96 | 2,96 | 0,33 | 7,45E-01 |
| AT1G31170 | 0,22 | -0,44 | 4,39 | -0,10 | 9,20E-01 |
| AT1G31175 | 0,74 | 1,71 | 2,43 | 0,70 | 4,81E-01 |
| AT1G31190 | 0,86 | 1,38 | 2,62 | 0,53 | 5,98E-01 |
| AT1G31220 | 0,22 | 1,37 | 4,41 | 0,31 | 7,56E-01 |
| AT1G31290 | 0,15 | 0,00 | 4,41 | 0,00 | 1,00E+00 |
| AT1G31420 | 0,89 | -0,45 | 2,12 | -0,21 | 8,31E-01 |
| AT1G31480 | 0,40 | 0,00 | 4,41 | 0,00 | 1,00E+00 |
| AT1G06447 | 0,24 | 1,61 | 4,39 | 0,37 | 7,14E-01 |
| AT1G31710 | 0,82 | 0,00 | 3,62 | 0,00 | 1,00E+00 |
| AT1G31780 | 0,28 | -0,51 | 4,40 | -0,11 | 9,09E-01 |
| AT1G31800 | 0,33 | -1,22 | 3,52 | -0,35 | 7,28E-01 |
| AT1G31850 | 1,13 | 2,68 | 2,17 | 1,24 | 2,16E-01 |
| AT1G31930 | 0,12 | 0,00 | 4,41 | 0,00 | 1,00E+00 |
| AT1G31970 | 0,74 | 2,83 | 3,18 | 0,89 | 3,72E-01 |
| AT1G32050 | 0,65 | -0,64 | 2,41 | -0,27 | 7,90E-01 |
| AT1G32160 | 0,70 | -0,87 | 3,01 | -0,29 | 7,72E-01 |
| AT1G32220 | 0,21 | 0,00 | 4,41 | 0,00 | 1,00E+00 |
| AT1G32225 | 0,70 | 0,43 | 2,51 | 0,17 | 8,63E-01 |
| AT1G32320 | 0,23 | 0,00 | 4,38 | 0,00 | 1,00E+00 |
| AT1G32337 | 0,30 | -0,99 | 4,38 | -0,23 | 8,20E-01 |
| AT1G32360 | 0,41 | -1,47 | 3,40 | -0,43 | 6,64E-01 |
| AT1G06547 | 0,19 | 0,00 | 4,41 | 0,00 | 1,00E+00 |
| AT1G32385 | 0,41 | -1,58 | 2,81 | -0,56 | 5,75E-01 |
| AT1G32415 | 0,72 | 3,29 | 2,82 | 1,17 | 2,44E-01 |
| AT1G32460 | 0,41 | -0,50 | 4,40 | -0,11 | 9,09E-01 |
| AT1G32580 | 0,32 | 1,79 | 4,38 | 0,41 | 6,83E-01 |

| | | | | | |
|---|---|---|---|---|---|
| AT1G06597 | 1,64 | -1,95 | 1,51 | -1,29 | 1,96E-01 |
| AT1G32700 | 0,46 | 0,30 | 3,33 | 0,09 | 9,28E-01 |
| AT1G32810 | 0,36 | 0,43 | 3,52 | 0,12 | 9,03E-01 |
| AT1G06623 | 0,26 | 0,00 | 4,41 | 0,00 | 1,00E+00 |
| AT1G32870 | 0,59 | 1,62 | 3,85 | 0,42 | 6,73E-01 |
| AT1G32920 | 1,13 | 0,45 | 2,04 | 0,22 | 8,25E-01 |
| AT1G32928 | 0,50 | 2,11 | 4,09 | 0,52 | 6,07E-01 |
| AT1G32940 | 0,41 | 1,82 | 4,38 | 0,42 | 6,77E-01 |
| AT1G32950 | 0,62 | 1,34 | 3,75 | 0,36 | 7,21E-01 |
| AT1G32960 | 0,35 | 0,00 | 4,41 | 0,00 | 1,00E+00 |
| AT1G32990 | 0,48 | -1,81 | 4,35 | -0,42 | 6,76E-01 |
| AT1G33010 | 0,19 | 0,00 | 4,41 | 0,00 | 1,00E+00 |
| AT1G33040 | 0,24 | -0,50 | 4,40 | -0,11 | 9,10E-01 |
| AT1G33110 | 0,55 | 2,59 | 3,83 | 0,68 | 4,98E-01 |
| AT1G33170 | 0,44 | 1,27 | 3,32 | 0,38 | 7,02E-01 |
| AT1G33260 | 0,52 | 1,26 | 4,06 | 0,31 | 7,57E-01 |
| AT1G33360 | 0,47 | 0,00 | 4,14 | 0,00 | 1,00E+00 |
| AT1G33470 | 0,71 | 3,14 | 3,51 | 0,90 | 3,71E-01 |
| AT1G33600 | 1,39 | -0,13 | 1,73 | -0,07 | 9,41E-01 |
| AT1G33610 | 0,15 | 0,00 | 4,41 | 0,00 | 1,00E+00 |
| AT1G33612 | 0,17 | -0,82 | 4,37 | -0,19 | 8,51E-01 |
| AT1G33770 | 0,49 | 0,17 | 4,04 | 0,04 | 9,67E-01 |
| AT1G33811 | 0,18 | -0,42 | 4,39 | -0,10 | 9,23E-01 |
| AT1G33910 | 0,08 | 0,00 | 4,41 | 0,00 | 1,00E+00 |
| AT1G33930 | 0,36 | -1,65 | 4,35 | -0,38 | 7,04E-01 |
| AT1G33960 | 1,02 | 3,33 | 3,13 | 1,06 | 2,88E-01 |
| AT1G33980 | 0,92 | 2,30 | 4,19 | 0,55 | 5,83E-01 |
| AT1G33990 | 0,95 | -0,07 | 1,98 | -0,03 | 9,73E-01 |
| AT1G06723 | 0,59 | 0,39 | 2,53 | 0,15 | 8,77E-01 |
| AT1G34020 | 0,78 | -0,24 | 2,48 | -0,10 | 9,23E-01 |
| AT1G34070 | 0,46 | 0,00 | 4,14 | 0,00 | 1,00E+00 |
| AT1G34130 | 1,45 | -0,18 | 1,74 | -0,11 | 9,16E-01 |
| AT1G34150 | 1,09 | 1,59 | 2,54 | 0,63 | 5,31E-01 |
| AT1G34190 | 1,14 | 2,73 | 2,03 | 1,34 | 1,79E-01 |
| AT1G34200 | 0,21 | 0,00 | 4,41 | 0,00 | 1,00E+00 |
| AT1G34210 | 0,51 | 2,04 | 4,39 | 0,47 | 6,42E-01 |
| AT1G34260 | 1,42 | 2,23 | 2,20 | 1,01 | 3,10E-01 |
| AT1G34290 | 0,25 | 0,00 | 4,41 | 0,00 | 1,00E+00 |
| AT1G06747 | 0,08 | 0,00 | 4,41 | 0,00 | 1,00E+00 |
| AT1G34355 | 0,86 | 3,47 | 2,80 | 1,24 | 2,14E-01 |
| AT1G34370 | 1,49 | 3,24 | 2,04 | 1,59 | 1,12E-01 |
| AT1G34390 | 0,68 | -0,44 | 3,77 | -0,12 | 9,08E-01 |
| AT1G34410 | 0,91 | 0,00 | 3,39 | 0,00 | 1,00E+00 |
| AT1G34420 | 0,36 | 0,00 | 4,36 | 0,00 | 1,00E+00 |
| AT1G34460 | 0,31 | 0,00 | 4,41 | 0,00 | 1,00E+00 |
| AT1G34580 | 0,53 | -2,23 | 4,32 | -0,52 | 6,06E-01 |
| AT1G34770 | 0,12 | 0,00 | 4,41 | 0,00 | 1,00E+00 |
| AT1G34844 | 0,08 | 0,00 | 4,41 | 0,00 | 1,00E+00 |
| AT1G35030 | 0,42 | -0,54 | 3,44 | -0,16 | 8,74E-01 |
| AT1G35040 | 0,23 | 0,00 | 4,41 | 0,00 | 1,00E+00 |
| AT1G35140 | 0,39 | 1,35 | 4,29 | 0,32 | 7,52E-01 |
| AT1G35160 | 0,59 | -2,04 | 2,57 | -0,80 | 4,27E-01 |
| AT1G06827 | 0,56 | -0,51 | 2,73 | -0,19 | 8,53E-01 |
| AT1G35220 | 0,89 | 2,73 | 2,40 | 1,13 | 2,56E-01 |
| AT1G35230 | 1,07 | 3,05 | 2,15 | 1,42 | 1,56E-01 |
| AT1G35242 | 0,19 | 0,00 | 4,41 | 0,00 | 1,00E+00 |

| | | | | | |
|---|---|---|---|---|---|
| AT1G35420 | 0,41 | -1,29 | 3,38 | -0,38 | 7,03E-01 |
| AT1G35460 | 0,12 | 0,00 | 4,41 | 0,00 | 1,00E+00 |
| AT1G35500 | 0,14 | 0,00 | 4,41 | 0,00 | 1,00E+00 |
| AT1G35510 | 0,12 | 0,00 | 4,41 | 0,00 | 1,00E+00 |
| AT1G35515 | 0,52 | 0,41 | 2,50 | 0,16 | 8,69E-01 |
| AT1G35516 | 0,52 | 0,41 | 2,50 | 0,16 | 8,69E-01 |
| AT1G35530 | 0,08 | 0,00 | 4,41 | 0,00 | 1,00E+00 |
| AT1G35540 | 0,95 | -0,33 | 2,09 | -0,16 | 8,74E-01 |
| AT1G35614 | 1,12 | -1,33 | 2,15 | -0,62 | 5,36E-01 |
| AT1G35620 | 0,51 | -0,42 | 3,37 | -0,12 | 9,02E-01 |
| AT1G35660 | 0,28 | 0,00 | 4,41 | 0,00 | 1,00E+00 |
| AT1G35670 | 0,21 | 0,00 | 4,41 | 0,00 | 1,00E+00 |
| AT1G35680 | 0,81 | 1,62 | 2,11 | 0,77 | 4,42E-01 |
| AT1G06927 | 0,83 | 0,00 | 2,92 | 0,00 | 1,00E+00 |
| AT1G35850 | 0,23 | 0,00 | 4,41 | 0,00 | 1,00E+00 |
| AT1G36050 | 0,26 | 0,00 | 4,41 | 0,00 | 1,00E+00 |
| AT1G06963 | 0,05 | 0,00 | 4,41 | 0,00 | 1,00E+00 |
| AT1G36320 | 0,22 | -0,48 | 4,40 | -0,11 | 9,14E-01 |
| AT1G36510 | 0,10 | 0,00 | 4,41 | 0,00 | 1,00E+00 |
| AT1G06987 | 0,58 | -0,67 | 4,39 | -0,15 | 8,78E-01 |
| AT1G36675 | 0,81 | -0,09 | 2,58 | -0,03 | 9,72E-01 |
| AT1G36745 | 0,98 | 0,00 | 2,46 | 0,00 | 1,00E+00 |
| AT1G36756 | 0,94 | 0,00 | 2,66 | 0,00 | 1,00E+00 |
| AT1G36940 | 0,22 | 0,00 | 4,41 | 0,00 | 1,00E+00 |
| AT1G36980 | 0,52 | -0,61 | 3,25 | -0,19 | 8,52E-01 |
| AT1G36992 | 0,32 | 2,00 | 4,38 | 0,46 | 6,48E-01 |
| AT1G37113 | 0,08 | 0,00 | 4,41 | 0,00 | 1,00E+00 |
| AT1G42525 | 0,30 | 0,00 | 4,41 | 0,00 | 1,00E+00 |
| AT1G42540 | 1,07 | 2,89 | 2,28 | 1,27 | 2,06E-01 |
| AT1G07137 | 0,05 | 0,00 | 4,41 | 0,00 | 1,00E+00 |
| AT1G42960 | 0,66 | -3,06 | 4,29 | -0,71 | 4,76E-01 |
| AT1G42980 | 1,30 | 1,52 | 1,92 | 0,79 | 4,29E-01 |
| AT1G43040 | 0,16 | 0,00 | 4,41 | 0,00 | 1,00E+00 |
| AT1G43145 | 1,44 | -1,41 | 1,98 | -0,71 | 4,77E-01 |
| AT1G43171 | 0,72 | 3,54 | 4,30 | 0,82 | 4,11E-01 |
| AT1G43330 | 0,87 | 0,00 | 2,75 | 0,00 | 1,00E+00 |
| AT1G43670 | 0,63 | -0,30 | 2,58 | -0,11 | 9,09E-01 |
| AT1G43680 | 0,25 | 1,84 | 4,38 | 0,42 | 6,75E-01 |
| AT1G43700 | 0,72 | 0,13 | 2,00 | 0,07 | 9,47E-01 |
| AT1G43722 | 0,18 | 0,00 | 4,41 | 0,00 | 1,00E+00 |
| AT1G43770 | 1,22 | -0,98 | 1,66 | -0,59 | 5,54E-01 |
| AT1G43780 | 0,25 | -1,80 | 4,34 | -0,41 | 6,78E-01 |
| AT1G43815 | 1,28 | 0,42 | 2,31 | 0,18 | 8,56E-01 |
| AT1G43890 | 0,40 | 2,30 | 4,32 | 0,53 | 5,93E-01 |
| AT1G43970 | 0,43 | 0,00 | 4,41 | 0,00 | 1,00E+00 |
| AT1G44120 | 0,95 | 0,00 | 2,85 | 0,00 | 1,00E+00 |
| AT1G44750 | 0,33 | 0,00 | 4,41 | 0,00 | 1,00E+00 |
| AT1G44780 | 0,29 | 0,43 | 4,37 | 0,10 | 9,22E-01 |
| AT1G44810 | 0,26 | -0,51 | 3,81 | -0,13 | 8,94E-01 |
| AT1G44890 | 0,08 | 0,00 | 4,41 | 0,00 | 1,00E+00 |
| AT1G44900 | 0,12 | 0,00 | 4,41 | 0,00 | 1,00E+00 |
| AT1G44910 | 0,50 | 0,53 | 2,46 | 0,22 | 8,29E-01 |
| AT1G44960 | 0,10 | 0,00 | 4,41 | 0,00 | 1,00E+00 |
| AT1G45000 | 0,47 | 1,23 | 2,81 | 0,44 | 6,63E-01 |
| AT1G45165 | 0,70 | 1,32 | 2,63 | 0,50 | 6,14E-01 |
| AT1G45180 | 0,30 | 1,38 | 4,41 | 0,31 | 7,54E-01 |

| | | | | | |
|---|---|---|---|---|---|
| AT1G45474 | 0,41 | -1,74 | 3,33 | -0,52 | 6,01E-01 |
| AT1G45688 | 0,89 | 1,32 | 2,53 | 0,52 | 6,01E-01 |
| AT1G45976 | 0,41 | 1,05 | 4,24 | 0,25 | 8,04E-01 |
| AT1G46264 | 0,10 | 0,00 | 4,41 | 0,00 | 1,00E+00 |
| AT1G46554 | 0,28 | 1,28 | 4,39 | 0,29 | 7,71E-01 |
| AT1G47210 | 0,93 | 3,08 | 2,59 | 1,19 | 2,36E-01 |
| AT1G07327 | 1,09 | -1,58 | 2,38 | -0,66 | 5,06E-01 |
| AT1G47250 | 0,35 | -0,54 | 4,41 | -0,12 | 9,02E-01 |
| AT1G47270 | 0,13 | 0,00 | 4,41 | 0,00 | 1,00E+00 |
| AT1G07357 | 0,11 | 0,00 | 4,41 | 0,00 | 1,00E+00 |
| AT1G47480 | 1,07 | -1,72 | 2,11 | -0,81 | 4,16E-01 |
| AT1G47490 | 0,30 | 1,38 | 4,41 | 0,31 | 7,54E-01 |
| AT1G47500 | 0,37 | 2,11 | 4,32 | 0,49 | 6,26E-01 |
| AT1G47530 | 1,40 | 1,26 | 1,56 | 0,81 | 4,20E-01 |
| AT1G47550 | 0,50 | 2,08 | 4,20 | 0,49 | 6,21E-01 |
| AT1G47680 | 0,12 | 0,00 | 4,41 | 0,00 | 1,00E+00 |
| AT1G47690 | 0,26 | 0,00 | 4,41 | 0,00 | 1,00E+00 |
| AT1G47770 | 0,12 | 0,00 | 4,41 | 0,00 | 1,00E+00 |
| AT1G47890 | 1,24 | 3,81 | 1,98 | 1,92 | 5,45E-02 |
| AT1G47940 | 0,36 | 0,00 | 4,41 | 0,00 | 1,00E+00 |
| AT1G47970 | 1,63 | 1,65 | 1,59 | 1,03 | 3,01E-01 |
| AT1G48050 | 0,54 | -1,06 | 4,34 | -0,24 | 8,07E-01 |
| AT1G48100 | 0,12 | 0,00 | 4,41 | 0,00 | 1,00E+00 |
| AT1G48170 | 0,32 | 1,27 | 3,64 | 0,35 | 7,27E-01 |
| AT1G48210 | 0,24 | 0,00 | 4,41 | 0,00 | 1,00E+00 |
| AT1G48230 | 0,28 | 1,38 | 4,41 | 0,31 | 7,54E-01 |
| AT1G48315 | 0,17 | 0,00 | 4,41 | 0,00 | 1,00E+00 |
| AT1G48360 | 1,34 | 3,93 | 2,12 | 1,86 | 6,35E-02 |
| AT1G48460 | 0,51 | -0,54 | 3,23 | -0,17 | 8,66E-01 |
| AT1G07487 | 0,32 | -0,53 | 3,68 | -0,14 | 8,85E-01 |
| AT1G48480 | 0,38 | 0,10 | 4,31 | 0,02 | 9,82E-01 |
| AT1G48540 | 1,33 | 2,86 | 2,05 | 1,39 | 1,63E-01 |
| AT1G48560 | 0,14 | 0,00 | 4,41 | 0,00 | 1,00E+00 |
| AT1G48600 | 0,62 | -0,40 | 2,27 | -0,18 | 8,61E-01 |
| AT1G48640 | 0,26 | 1,38 | 4,41 | 0,31 | 7,54E-01 |
| AT1G48725 | 1,45 | -1,32 | 1,99 | -0,66 | 5,06E-01 |
| AT1G48820 | 0,24 | -1,08 | 4,37 | -0,25 | 8,04E-01 |
| AT1G49010 | 0,27 | 0,00 | 4,41 | 0,00 | 1,00E+00 |
| AT1G49032 | 0,71 | 2,54 | 2,27 | 1,12 | 2,64E-01 |
| AT1G49190 | 1,12 | -1,65 | 2,26 | -0,73 | 4,66E-01 |
| AT1G49200 | 0,08 | 0,00 | 4,41 | 0,00 | 1,00E+00 |
| AT1G49210 | 0,15 | 0,00 | 4,39 | 0,00 | 1,00E+00 |
| AT1G49230 | 0,38 | 2,11 | 3,46 | 0,61 | 5,43E-01 |
| AT1G49245 | 0,90 | 1,59 | 2,29 | 0,69 | 4,88E-01 |
| AT1G49300 | 0,19 | 0,00 | 4,41 | 0,00 | 1,00E+00 |
| AT1G49340 | 1,22 | 2,21 | 2,04 | 1,08 | 2,80E-01 |
| AT1G49390 | 0,08 | 0,00 | 4,41 | 0,00 | 1,00E+00 |
| AT1G49470 | 0,15 | 0,00 | 4,41 | 0,00 | 1,00E+00 |
| AT1G49510 | 0,25 | -1,80 | 4,34 | -0,41 | 6,78E-01 |
| AT1G49680 | 0,43 | -0,54 | 3,41 | -0,16 | 8,74E-01 |
| AT1G49720 | 0,23 | -0,84 | 4,38 | -0,19 | 8,48E-01 |
| AT1G49820 | 0,28 | -0,46 | 4,40 | -0,10 | 9,17E-01 |
| AT1G49890 | 0,23 | 1,27 | 4,39 | 0,29 | 7,72E-01 |
| AT1G49930 | 0,39 | 1,79 | 4,37 | 0,41 | 6,81E-01 |
| AT1G49975 | 1,21 | 2,05 | 2,61 | 0,79 | 4,32E-01 |
| AT1G49980 | 0,53 | 2,92 | 3,13 | 0,93 | 3,52E-01 |

| | | | | | |
|---|---|---|---|---|---|
| AT1G49990 | 0,80 | 0,00 | 4,37 | 0,00 | 1,00E+00 |
| AT1G50050 | 0,43 | 0,52 | 3,34 | 0,16 | 8,76E-01 |
| AT1G07613 | 1,47 | -3,02 | 1,76 | -1,71 | 8,65E-02 |
| AT1G50120 | 0,25 | 1,24 | 4,38 | 0,28 | 7,77E-01 |
| AT1G50170 | 0,08 | 0,00 | 4,41 | 0,00 | 1,00E+00 |
| AT1G50240 | 1,24 | 2,14 | 1,86 | 1,15 | 2,48E-01 |
| AT1G50270 | 0,45 | 1,18 | 3,34 | 0,35 | 7,24E-01 |
| AT1G50350 | 0,30 | -0,66 | 4,39 | -0,15 | 8,80E-01 |
| AT1G50360 | 0,23 | -0,50 | 4,40 | -0,11 | 9,10E-01 |
| AT1G50370 | 0,39 | 1,90 | 3,49 | 0,55 | 5,85E-01 |
| AT1G50380 | 0,57 | 1,31 | 3,26 | 0,40 | 6,88E-01 |
| AT1G50410 | 1,51 | 3,32 | 2,40 | 1,38 | 1,67E-01 |
| AT1G50560 | 0,41 | 2,20 | 4,31 | 0,51 | 6,10E-01 |
| AT1G50660 | 1,10 | 2,22 | 2,09 | 1,06 | 2,88E-01 |
| AT1G50730 | 0,12 | 0,00 | 4,41 | 0,00 | 1,00E+00 |
| AT1G50740 | 0,72 | 2,09 | 2,93 | 0,71 | 4,76E-01 |
| AT1G51060 | 0,42 | -1,93 | 4,26 | -0,45 | 6,51E-01 |
| AT1G51100 | 0,17 | 0,00 | 4,41 | 0,00 | 1,00E+00 |
| AT1G51110 | 0,52 | -0,52 | 3,17 | -0,16 | 8,70E-01 |
| AT1G51160 | 0,67 | 1,63 | 2,95 | 0,55 | 5,81E-01 |
| AT1G51172 | 1,67 | 0,39 | 2,03 | 0,19 | 8,48E-01 |
| AT1G51200 | 1,27 | 0,92 | 2,39 | 0,38 | 7,02E-01 |
| AT1G51350 | 0,44 | 1,38 | 3,34 | 0,41 | 6,80E-01 |
| AT1G51390 | 0,24 | -0,56 | 4,41 | -0,13 | 8,99E-01 |
| AT1G07737 | 0,83 | -2,50 | 2,36 | -1,06 | 2,90E-01 |
| AT1G51402 | 1,01 | -2,34 | 2,25 | -1,04 | 2,98E-01 |
| AT1G51440 | 0,14 | 0,00 | 4,41 | 0,00 | 1,00E+00 |
| AT1G51540 | 0,08 | 0,00 | 4,41 | 0,00 | 1,00E+00 |
| AT1G51580 | 0,35 | -1,16 | 4,35 | -0,27 | 7,90E-01 |
| AT1G51600 | 0,08 | 0,00 | 4,41 | 0,00 | 1,00E+00 |
| AT1G51645 | 0,08 | 0,00 | 4,41 | 0,00 | 1,00E+00 |
| AT1G51660 | 0,75 | 2,96 | 2,67 | 1,11 | 2,68E-01 |
| AT1G51670 | 0,25 | 1,34 | 4,40 | 0,30 | 7,61E-01 |
| AT1G51720 | 0,29 | 1,25 | 4,39 | 0,29 | 7,75E-01 |
| AT1G51745 | 0,21 | 0,00 | 4,41 | 0,00 | 1,00E+00 |
| AT1G51760 | 0,56 | 0,43 | 2,75 | 0,16 | 8,76E-01 |
| AT1G51950 | 0,22 | 0,00 | 4,41 | 0,00 | 1,00E+00 |
| AT1G52080 | 0,08 | 0,00 | 4,41 | 0,00 | 1,00E+00 |
| AT1G52190 | 0,40 | 2,11 | 3,39 | 0,62 | 5,35E-01 |
| AT1G52310 | 0,73 | 2,17 | 2,56 | 0,84 | 3,98E-01 |
| AT1G52320 | 0,99 | -0,44 | 2,02 | -0,22 | 8,27E-01 |
| AT1G52380 | 0,74 | 1,08 | 2,94 | 0,37 | 7,13E-01 |
| AT1G52410 | 0,68 | -1,13 | 2,54 | -0,45 | 6,56E-01 |
| AT1G52420 | 0,99 | 2,17 | 2,90 | 0,75 | 4,54E-01 |
| AT1G52500 | 0,39 | 0,37 | 4,40 | 0,08 | 9,33E-01 |
| AT1G52600 | 0,08 | 0,00 | 4,41 | 0,00 | 1,00E+00 |
| AT1G52618 | 0,68 | 0,00 | 3,01 | 0,00 | 1,00E+00 |
| AT1G52620 | 0,78 | 0,00 | 4,33 | 0,00 | 1,00E+00 |
| AT1G52720 | 0,19 | 0,00 | 4,41 | 0,00 | 1,00E+00 |
| AT1G52740 | 1,01 | 0,29 | 2,27 | 0,13 | 8,97E-01 |
| AT1G52760 | 0,33 | 0,31 | 3,60 | 0,09 | 9,32E-01 |
| AT1G52870 | 0,92 | 0,00 | 2,73 | 0,00 | 1,00E+00 |
| AT1G52905 | 0,68 | 0,00 | 2,97 | 0,00 | 1,00E+00 |
| AT1G52930 | 0,12 | 0,00 | 4,41 | 0,00 | 1,00E+00 |
| AT1G52940 | 0,76 | 2,02 | 2,56 | 0,79 | 4,31E-01 |
| AT1G52950 | 0,39 | -0,54 | 3,57 | -0,15 | 8,79E-01 |

| | | | | | |
|---|---|---|---|---|---|
| AT1G52980 | 1,44 | 2,71 | 1,83 | 1,48 | 1,38E-01 |
| AT1G53050 | 0,64 | 3,08 | 4,32 | 0,71 | 4,76E-01 |
| AT1G53090 | 0,57 | -0,27 | 4,34 | -0,06 | 9,50E-01 |
| AT1G53165 | 0,43 | 0,51 | 2,87 | 0,18 | 8,60E-01 |
| AT1G53233 | 0,99 | 1,41 | 2,42 | 0,58 | 5,59E-01 |
| AT1G53300 | 0,67 | 2,25 | 2,69 | 0,84 | 4,03E-01 |
| AT1G53330 | 0,35 | 1,58 | 4,36 | 0,36 | 7,18E-01 |
| AT1G53350 | 0,25 | 0,00 | 4,41 | 0,00 | 1,00E+00 |
| AT1G53390 | 0,38 | 1,57 | 4,39 | 0,36 | 7,20E-01 |
| AT1G53400 | 0,40 | 0,00 | 4,41 | 0,00 | 1,00E+00 |
| AT1G53430 | 1,51 | -0,49 | 1,60 | -0,31 | 7,57E-01 |
| AT1G53440 | 1,17 | -0,57 | 1,70 | -0,33 | 7,40E-01 |
| AT1G07887 | 0,84 | -1,00 | 3,06 | -0,33 | 7,44E-01 |
| AT1G53490 | 0,92 | -0,94 | 2,58 | -0,36 | 7,15E-01 |
| AT1G07893 | 0,39 | -1,62 | 4,32 | -0,37 | 7,08E-01 |
| AT1G07897 | 0,95 | 0,34 | 1,90 | 0,18 | 8,57E-01 |
| AT1G53570 | 0,96 | 2,60 | 2,24 | 1,16 | 2,47E-01 |
| AT1G53590 | 0,26 | -0,49 | 4,40 | -0,11 | 9,12E-01 |
| AT1G53670 | 0,12 | 0,00 | 4,41 | 0,00 | 1,00E+00 |
| AT1G53720 | 0,26 | -1,08 | 4,37 | -0,25 | 8,04E-01 |
| AT1G53730 | 1,50 | 2,63 | 2,16 | 1,21 | 2,25E-01 |
| AT1G53840 | 1,21 | -0,40 | 2,15 | -0,18 | 8,54E-01 |
| AT1G53870 | 0,49 | 0,00 | 4,02 | 0,00 | 1,00E+00 |
| AT1G53875 | 1,66 | -2,36 | 1,65 | -1,44 | 1,51E-01 |
| AT1G53885 | 0,24 | -0,50 | 4,40 | -0,11 | 9,10E-01 |
| AT1G53890 | 1,37 | -1,79 | 1,86 | -0,96 | 3,37E-01 |
| AT1G53910 | 0,62 | -0,54 | 2,64 | -0,21 | 8,37E-01 |
| AT1G54030 | 0,19 | 0,00 | 4,41 | 0,00 | 1,00E+00 |
| AT1G54130 | 1,27 | 0,54 | 1,86 | 0,29 | 7,72E-01 |
| AT1G54150 | 0,44 | 1,99 | 3,49 | 0,57 | 5,70E-01 |
| AT1G54200 | 0,80 | 2,38 | 2,57 | 0,93 | 3,54E-01 |
| AT1G54215 | 0,08 | 0,00 | 4,41 | 0,00 | 1,00E+00 |
| AT1G07957 | 0,34 | 0,00 | 3,55 | 0,00 | 1,00E+00 |
| AT1G54310 | 0,08 | 0,00 | 4,41 | 0,00 | 1,00E+00 |
| AT1G54340 | 0,28 | 1,48 | 4,40 | 0,34 | 7,36E-01 |
| AT1G54350 | 0,37 | 1,38 | 4,35 | 0,32 | 7,51E-01 |
| AT1G54360 | 0,57 | 2,85 | 4,33 | 0,66 | 5,11E-01 |
| AT1G54420 | 1,07 | -1,26 | 2,72 | -0,47 | 6,42E-01 |
| AT1G54440 | 0,25 | 0,00 | 4,41 | 0,00 | 1,00E+00 |
| AT1G54490 | 1,36 | 3,72 | 1,84 | 2,02 | 4,30E-02 |
| AT1G54550 | 0,56 | 0,00 | 3,07 | 0,00 | 1,00E+00 |
| AT1G54680 | 0,26 | 1,38 | 4,41 | 0,31 | 7,54E-01 |
| AT1G54720 | 0,66 | -0,51 | 3,19 | -0,16 | 8,73E-01 |
| AT1G54780 | 1,22 | -1,95 | 1,84 | -1,06 | 2,90E-01 |
| AT1G54820 | 0,40 | 0,36 | 4,36 | 0,08 | 9,34E-01 |
| AT1G54920 | 1,30 | 2,36 | 1,79 | 1,32 | 1,88E-01 |
| AT1G55000 | 0,70 | -0,73 | 2,68 | -0,27 | 7,85E-01 |
| AT1G55020 | 0,29 | 1,24 | 4,38 | 0,28 | 7,77E-01 |
| AT1G55040 | 0,41 | 1,70 | 4,23 | 0,40 | 6,88E-01 |
| AT1G55090 | 0,81 | 2,26 | 2,99 | 0,76 | 4,50E-01 |
| AT1G55120 | 1,23 | -1,36 | 1,75 | -0,78 | 4,37E-01 |
| AT1G55150 | 0,30 | 1,38 | 4,41 | 0,31 | 7,54E-01 |
| AT1G55160 | 0,99 | 2,06 | 2,54 | 0,81 | 4,18E-01 |
| AT1G55175 | 0,08 | 0,00 | 4,41 | 0,00 | 1,00E+00 |
| AT1G55320 | 0,25 | 1,26 | 4,39 | 0,29 | 7,74E-01 |
| AT1G55335 | 0,51 | -0,43 | 3,18 | -0,13 | 8,93E-01 |

| | | | | | |
|---|---|---|---|---|---|
| AT1G55370 | 0,24 | 1,96 | 4,39 | 0,45 | 6,55E-01 |
| AT1G55460 | 0,32 | -0,54 | 4,41 | -0,12 | 9,02E-01 |
| AT1G55480 | 0,48 | 2,07 | 4,14 | 0,50 | 6,17E-01 |
| AT1G55500 | 0,61 | 2,88 | 2,30 | 1,25 | 2,10E-01 |
| AT1G55510 | 0,20 | -0,44 | 4,39 | -0,10 | 9,20E-01 |
| AT1G55840 | 0,33 | 1,66 | 4,37 | 0,38 | 7,05E-01 |
| AT1G55850 | 0,50 | -0,96 | 4,38 | -0,22 | 8,26E-01 |
| AT1G08073 | 0,33 | 0,43 | 4,38 | 0,10 | 9,21E-01 |
| AT1G55890 | 0,12 | 0,00 | 4,41 | 0,00 | 1,00E+00 |
| AT1G55910 | 0,98 | 2,32 | 1,96 | 1,18 | 2,38E-01 |
| AT1G55915 | 0,81 | 2,36 | 3,04 | 0,78 | 4,36E-01 |
| AT1G08077 | 0,62 | 0,00 | 3,08 | 0,00 | 1,00E+00 |
| AT1G56045 | 0,75 | -1,38 | 2,63 | -0,53 | 6,00E-01 |
| AT1G56050 | 1,07 | 3,23 | 2,67 | 1,21 | 2,26E-01 |
| AT1G56190 | 1,11 | 1,08 | 2,26 | 0,48 | 6,31E-01 |
| AT1G56200 | 1,42 | 0,43 | 1,75 | 0,24 | 8,08E-01 |
| AT1G56210 | 0,05 | 0,00 | 4,41 | 0,00 | 1,00E+00 |
| AT1G56220 | 1,48 | -0,18 | 1,65 | -0,11 | 9,13E-01 |
| AT1G56450 | 0,83 | 0,28 | 2,34 | 0,12 | 9,06E-01 |
| AT1G56460 | 0,70 | 0,00 | 3,03 | 0,00 | 1,00E+00 |
| AT1G56500 | 0,95 | -0,26 | 1,98 | -0,13 | 8,96E-01 |
| AT1G56540 | 0,56 | 1,98 | 4,38 | 0,45 | 6,50E-01 |
| AT1G56560 | 0,52 | 1,38 | 3,96 | 0,35 | 7,28E-01 |
| AT1G56570 | 0,28 | 1,48 | 4,40 | 0,34 | 7,36E-01 |
| AT1G56600 | 0,19 | -0,43 | 4,39 | -0,10 | 9,22E-01 |
| AT1G56610 | 0,17 | -1,42 | 4,35 | -0,33 | 7,45E-01 |
| AT1G56660 | 1,02 | 0,49 | 2,06 | 0,24 | 8,12E-01 |
| AT1G56700 | 0,33 | -1,28 | 3,62 | -0,35 | 7,23E-01 |
| AT1G57620 | 0,88 | -0,48 | 3,10 | -0,16 | 8,76E-01 |
| AT1G57630 | 0,41 | 1,23 | 4,37 | 0,28 | 7,78E-01 |
| AT1G57650 | 0,46 | -1,63 | 2,77 | -0,59 | 5,56E-01 |
| AT1G57760 | 0,12 | 0,00 | 4,41 | 0,00 | 1,00E+00 |
| AT1G57765 | 0,15 | 0,00 | 4,41 | 0,00 | 1,00E+00 |
| AT1G58190 | 0,62 | 2,92 | 2,57 | 1,14 | 2,56E-01 |
| AT1G58215 | 0,57 | 1,23 | 4,39 | 0,28 | 7,80E-01 |
| AT1G58223 | 0,54 | 2,61 | 3,85 | 0,68 | 4,98E-01 |
| AT1G58260 | 0,12 | 0,00 | 4,41 | 0,00 | 1,00E+00 |
| AT1G58360 | 0,57 | 0,02 | 3,92 | 0,01 | 9,95E-01 |
| AT1G58525 | 0,11 | 0,00 | 4,41 | 0,00 | 1,00E+00 |
| AT1G58520 | 0,55 | 0,43 | 3,37 | 0,13 | 8,97E-01 |
| AT1G58643 | 0,42 | 1,25 | 4,34 | 0,29 | 7,73E-01 |
| AT1G58807 | 1,55 | 2,88 | 1,75 | 1,65 | 9,88E-02 |
| AT1G58808 | 1,22 | 0,95 | 1,78 | 0,54 | 5,93E-01 |
| AT1G58848 | 1,34 | 3,10 | 2,02 | 1,54 | 1,25E-01 |
| AT1G59124 | 1,27 | 1,53 | 1,59 | 0,97 | 3,34E-01 |
| AT1G59218 | 1,28 | 1,91 | 2,02 | 0,94 | 3,46E-01 |
| AT1G59590 | 0,59 | 2,58 | 3,22 | 0,80 | 4,23E-01 |
| AT1G59620 | 0,81 | 2,50 | 4,36 | 0,57 | 5,66E-01 |
| AT1G59650 | 0,54 | 1,29 | 3,93 | 0,33 | 7,43E-01 |
| AT1G59710 | 0,56 | -0,78 | 3,26 | -0,24 | 8,11E-01 |
| AT1G59750 | 1,67 | 2,72 | 2,41 | 1,13 | 2,59E-01 |
| AT1G59820 | 1,45 | 3,75 | 2,10 | 1,79 | 7,40E-02 |
| AT1G59890 | 0,93 | 1,38 | 2,64 | 0,52 | 6,01E-01 |
| AT1G59900 | 1,24 | -1,64 | 2,03 | -0,81 | 4,20E-01 |
| AT1G60025 | 0,05 | 0,00 | 4,41 | 0,00 | 1,00E+00 |
| AT1G60073 | 1,58 | 0,00 | 2,18 | 0,00 | 1,00E+00 |

| | | | | | |
|---|---|---|---|---|---|
| AT1G60170 | 1,29 | 2,60 | 2,47 | 1,05 | 2,92E-01 |
| AT1G60190 | 0,89 | 2,65 | 2,23 | 1,19 | 2,35E-01 |
| AT1G60220 | 0,08 | 0,00 | 4,41 | 0,00 | 1,00E+00 |
| AT1G60420 | 0,70 | -0,46 | 2,96 | -0,16 | 8,75E-01 |
| AT1G60545 | 1,07 | 3,10 | 3,01 | 1,03 | 3,03E-01 |
| AT1G60650 | 0,21 | 1,27 | 4,39 | 0,29 | 7,72E-01 |
| AT1G60720 | 1,62 | 1,38 | 2,21 | 0,62 | 5,33E-01 |
| AT1G60900 | 0,65 | 0,12 | 3,26 | 0,04 | 9,70E-01 |
| AT1G08353 | 0,98 | 0,72 | 1,89 | 0,38 | 7,02E-01 |
| AT1G61030 | 0,52 | 0,30 | 3,97 | 0,07 | 9,41E-01 |
| AT1G61040 | 0,54 | 1,29 | 3,93 | 0,33 | 7,43E-01 |
| AT1G61150 | 0,81 | 1,09 | 3,58 | 0,30 | 7,61E-01 |
| AT1G61170 | 0,33 | 0,00 | 4,41 | 0,00 | 1,00E+00 |
| AT1G61180 | 0,58 | 1,15 | 2,78 | 0,41 | 6,81E-01 |
| AT1G61210 | 1,07 | 0,66 | 1,77 | 0,37 | 7,08E-01 |
| AT1G61224 | 1,62 | 0,91 | 1,72 | 0,53 | 5,97E-01 |
| AT1G61250 | 0,18 | 0,00 | 4,41 | 0,00 | 1,00E+00 |
| AT1G61280 | 0,12 | 0,00 | 4,41 | 0,00 | 1,00E+00 |
| AT1G61350 | 0,76 | 1,38 | 2,53 | 0,55 | 5,86E-01 |
| AT1G61450 | 0,12 | 0,00 | 4,41 | 0,00 | 1,00E+00 |
| AT1G61470 | 0,18 | 0,00 | 4,40 | 0,00 | 1,00E+00 |
| AT1G61610 | 0,12 | 0,00 | 4,41 | 0,00 | 1,00E+00 |
| AT1G61620 | 0,37 | 0,84 | 4,38 | 0,19 | 8,48E-01 |
| AT1G61790 | 1,42 | 2,21 | 1,68 | 1,31 | 1,89E-01 |
| AT1G61800 | 0,56 | 1,37 | 4,41 | 0,31 | 7,56E-01 |
| AT1G61820 | 0,08 | 0,00 | 4,41 | 0,00 | 1,00E+00 |
| AT1G61850 | 0,80 | 0,51 | 2,34 | 0,22 | 8,27E-01 |
| AT1G61990 | 0,32 | 0,00 | 4,41 | 0,00 | 1,00E+00 |
| AT1G62170 | 0,36 | 0,00 | 4,41 | 0,00 | 1,00E+00 |
| AT1G08477 | 1,64 | 0,45 | 1,79 | 0,25 | 7,99E-01 |
| AT1G62270 | 0,29 | 0,00 | 4,41 | 0,00 | 1,00E+00 |
| AT1G08517 | 0,77 | -1,11 | 2,49 | -0,45 | 6,54E-01 |
| AT1G62380 | 1,16 | -2,41 | 1,86 | -1,30 | 1,95E-01 |
| AT1G62410 | 0,32 | -1,58 | 4,35 | -0,36 | 7,15E-01 |
| AT1G62480 | 0,12 | 0,00 | 4,41 | 0,00 | 1,00E+00 |
| AT1G62560 | 1,49 | 0,09 | 3,04 | 0,03 | 9,76E-01 |
| AT1G62570 | 0,39 | -0,54 | 4,40 | -0,12 | 9,01E-01 |
| AT1G62620 | 0,57 | 2,56 | 3,07 | 0,83 | 4,04E-01 |
| AT1G62660 | 1,32 | 1,43 | 1,79 | 0,80 | 4,24E-01 |
| AT1G62720 | 0,24 | -0,85 | 4,38 | -0,19 | 8,47E-01 |
| AT1G62790 | 0,51 | 2,84 | 4,33 | 0,66 | 5,11E-01 |
| AT1G62895 | 1,43 | -0,72 | 1,73 | -0,42 | 6,77E-01 |
| AT1G62920 | 0,97 | -1,57 | 3,01 | -0,52 | 6,01E-01 |
| AT1G62970 | 0,33 | -0,47 | 3,61 | -0,13 | 8,96E-01 |
| AT1G62975 | 0,08 | 0,00 | 4,41 | 0,00 | 1,00E+00 |
| AT1G63000 | 0,67 | 1,38 | 3,91 | 0,35 | 7,24E-01 |
| AT1G63110 | 0,32 | 1,36 | 4,40 | 0,31 | 7,58E-01 |
| AT1G63170 | 0,08 | 0,00 | 4,41 | 0,00 | 1,00E+00 |
| AT1G63430 | 0,74 | 1,45 | 3,96 | 0,37 | 7,13E-01 |
| AT1G63500 | 0,59 | 0,44 | 2,80 | 0,16 | 8,75E-01 |
| AT1G63540 | 0,37 | 0,00 | 4,41 | 0,00 | 1,00E+00 |
| AT1G63680 | 0,43 | 1,38 | 3,33 | 0,41 | 6,79E-01 |
| AT1G63690 | 1,03 | 2,37 | 1,85 | 1,28 | 2,00E-01 |
| AT1G63730 | 0,24 | 0,00 | 4,41 | 0,00 | 1,00E+00 |
| AT1G63830 | 0,49 | 0,00 | 4,15 | 0,00 | 1,00E+00 |
| AT1G63850 | 0,53 | 0,92 | 3,97 | 0,23 | 8,17E-01 |

| | | | | | |
|---|---|---|---|---|---|
| AT1G08643 | 0,79 | 0,00 | 3,11 | 0,00 | 1,00E+00 |
| AT1G63900 | 0,95 | 3,17 | 2,18 | 1,45 | 1,46E-01 |
| AT1G63940 | 0,81 | 1,81 | 2,02 | 0,90 | 3,69E-01 |
| AT1G64060 | 0,86 | 1,38 | 2,56 | 0,54 | 5,90E-01 |
| AT1G64065 | 0,29 | 1,31 | 4,40 | 0,30 | 7,67E-01 |
| AT1G64090 | 0,96 | 1,11 | 2,04 | 0,54 | 5,88E-01 |
| AT1G64100 | 0,61 | 1,22 | 3,74 | 0,33 | 7,45E-01 |
| AT1G64120 | 0,13 | 0,00 | 4,41 | 0,00 | 1,00E+00 |
| AT1G64130 | 0,12 | 0,00 | 4,41 | 0,00 | 1,00E+00 |
| AT1G08657 | 0,60 | -0,34 | 2,72 | -0,13 | 8,99E-01 |
| AT1G64230 | 1,39 | -0,09 | 1,64 | -0,05 | 9,56E-01 |
| AT1G64370 | 0,17 | -0,43 | 4,39 | -0,10 | 9,21E-01 |
| AT1G64390 | 1,48 | 0,07 | 2,02 | 0,04 | 9,71E-01 |
| AT1G64520 | 0,27 | 0,00 | 4,41 | 0,00 | 1,00E+00 |
| AT1G64530 | 0,54 | 1,26 | 3,14 | 0,40 | 6,88E-01 |
| AT1G64550 | 0,64 | 0,88 | 2,74 | 0,32 | 7,49E-01 |
| AT1G64570 | 0,71 | 0,64 | 2,54 | 0,25 | 8,00E-01 |
| AT1G64572 | 0,51 | 2,52 | 4,35 | 0,58 | 5,63E-01 |
| AT1G64620 | 0,45 | 1,32 | 3,33 | 0,40 | 6,92E-01 |
| AT1G64680 | 1,48 | -1,61 | 1,75 | -0,92 | 3,57E-01 |
| AT1G64740 | 0,25 | 0,00 | 4,41 | 0,00 | 1,00E+00 |
| AT1G64770 | 1,29 | -0,06 | 1,66 | -0,04 | 9,69E-01 |
| AT1G64795 | 0,74 | -0,29 | 2,10 | -0,14 | 8,92E-01 |
| AT1G64810 | 0,44 | 0,29 | 4,39 | 0,07 | 9,46E-01 |
| AT1G64820 | 0,08 | 0,00 | 4,41 | 0,00 | 1,00E+00 |
| AT1G64860 | 0,65 | -1,04 | 3,13 | -0,33 | 7,40E-01 |
| AT1G64900 | 1,03 | 0,78 | 1,99 | 0,39 | 6,97E-01 |
| AT1G64950 | 0,34 | -0,45 | 3,53 | -0,13 | 8,99E-01 |
| AT1G64960 | 0,60 | 0,72 | 3,98 | 0,18 | 8,57E-01 |
| AT1G65010 | 1,62 | -0,45 | 1,64 | -0,28 | 7,82E-01 |
| AT1G65220 | 0,59 | 0,29 | 2,78 | 0,10 | 9,17E-01 |
| AT1G65230 | 1,04 | 2,55 | 2,12 | 1,20 | 2,28E-01 |
| AT1G65260 | 0,21 | 0,00 | 4,41 | 0,00 | 1,00E+00 |
| AT1G65270 | 0,21 | 0,00 | 4,41 | 0,00 | 1,00E+00 |
| AT1G65280 | 0,39 | 1,27 | 4,31 | 0,29 | 7,69E-01 |
| AT1G65295 | 0,05 | 0,00 | 4,41 | 0,00 | 1,00E+00 |
| AT1G65360 | 0,90 | -0,44 | 2,17 | -0,20 | 8,38E-01 |
| AT1G08753 | 0,15 | -0,43 | 4,39 | -0,10 | 9,23E-01 |
| AT1G65490 | 0,49 | 2,75 | 4,34 | 0,63 | 5,27E-01 |
| AT1G65590 | 0,23 | 1,27 | 4,39 | 0,29 | 7,72E-01 |
| AT1G65681 | 0,20 | 0,00 | 4,41 | 0,00 | 1,00E+00 |
| AT1G65730 | 0,08 | 0,00 | 4,41 | 0,00 | 1,00E+00 |
| AT1G65790 | 0,93 | 1,66 | 2,00 | 0,83 | 4,06E-01 |
| AT1G65800 | 1,17 | -0,49 | 1,85 | -0,26 | 7,91E-01 |
| AT1G65820 | 0,38 | 1,38 | 2,97 | 0,46 | 6,43E-01 |
| AT1G65840 | 0,61 | 1,46 | 3,87 | 0,38 | 7,06E-01 |
| AT1G08793 | 0,15 | 0,00 | 4,41 | 0,00 | 1,00E+00 |
| AT1G66090 | 0,59 | 0,73 | 3,30 | 0,22 | 8,24E-01 |
| AT1G66150 | 1,55 | -0,43 | 1,38 | -0,31 | 7,54E-01 |
| AT1G66160 | 0,48 | -0,68 | 4,10 | -0,17 | 8,69E-01 |
| AT1G66180 | 0,53 | 1,88 | 3,24 | 0,58 | 5,63E-01 |
| AT1G66230 | 0,24 | 1,96 | 4,39 | 0,45 | 6,55E-01 |
| AT1G66340 | 0,60 | 0,30 | 3,04 | 0,10 | 9,21E-01 |
| AT1G66345 | 0,47 | 0,00 | 4,08 | 0,00 | 1,00E+00 |
| AT1G66430 | 0,50 | -2,63 | 4,31 | -0,61 | 5,42E-01 |
| AT1G66465 | 0,33 | -0,50 | 4,40 | -0,11 | 9,10E-01 |

| | | | | | |
|---|---|---|---|---|---|
| AT1G66580 | 1,55 | 1,08 | 1,68 | 0,64 | 5,22E-01 |
| AT1G66640 | 0,21 | -0,47 | 4,40 | -0,11 | 9,15E-01 |
| AT1G66750 | 0,31 | -0,54 | 4,41 | -0,12 | 9,02E-01 |
| AT1G66820 | 0,53 | -0,40 | 4,36 | -0,09 | 9,26E-01 |
| AT1G66900 | 0,20 | 0,00 | 4,41 | 0,00 | 1,00E+00 |
| AT1G67105 | 0,86 | -2,35 | 1,96 | -1,20 | 2,30E-01 |
| AT1G67190 | 0,34 | 1,36 | 4,41 | 0,31 | 7,57E-01 |
| AT1G08923 | 0,11 | 0,00 | 4,41 | 0,00 | 1,00E+00 |
| AT1G67420 | 0,13 | 0,00 | 4,41 | 0,00 | 1,00E+00 |
| AT1G08937 | 0,46 | 0,00 | 3,36 | 0,00 | 1,00E+00 |
| AT1G67470 | 0,43 | 2,17 | 3,35 | 0,65 | 5,17E-01 |
| AT1G67480 | 0,49 | 1,75 | 4,38 | 0,40 | 6,89E-01 |
| AT1G67530 | 0,54 | 2,89 | 3,88 | 0,75 | 4,56E-01 |
| AT1G67560 | 0,64 | 2,45 | 4,34 | 0,56 | 5,73E-01 |
| AT1G67620 | 0,15 | 0,00 | 4,41 | 0,00 | 1,00E+00 |
| AT1G67635 | 1,69 | 0,00 | 2,13 | 0,00 | 1,00E+00 |
| AT1G67700 | 0,46 | 1,26 | 4,03 | 0,31 | 7,54E-01 |
| AT1G67720 | 0,53 | 2,09 | 3,82 | 0,55 | 5,84E-01 |
| AT1G67792 | 0,38 | 2,22 | 4,37 | 0,51 | 6,12E-01 |
| AT1G67900 | 1,04 | 0,14 | 2,36 | 0,06 | 9,54E-01 |
| AT1G67940 | 0,66 | 3,29 | 2,90 | 1,14 | 2,56E-01 |
| AT1G08963 | 0,39 | 1,76 | 4,38 | 0,40 | 6,87E-01 |
| AT1G67980 | 0,18 | -0,82 | 4,37 | -0,19 | 8,51E-01 |
| AT1G67990 | 0,55 | 0,71 | 3,79 | 0,19 | 8,52E-01 |
| AT1G68050 | 0,24 | -0,48 | 4,40 | -0,11 | 9,13E-01 |
| AT1G68100 | 0,43 | 0,42 | 4,19 | 0,10 | 9,21E-01 |
| AT1G68110 | 0,27 | 0,45 | 4,38 | 0,10 | 9,19E-01 |
| AT1G68310 | 0,12 | 0,00 | 4,41 | 0,00 | 1,00E+00 |
| AT1G68440 | 1,32 | 2,31 | 1,83 | 1,26 | 2,07E-01 |
| AT1G68490 | 0,97 | 2,82 | 2,52 | 1,12 | 2,64E-01 |
| AT1G68570 | 0,58 | -0,54 | 2,70 | -0,20 | 8,40E-01 |
| AT1G68580 | 1,44 | 2,24 | 1,75 | 1,28 | 2,02E-01 |
| AT1G68585 | 0,27 | 1,40 | 4,40 | 0,32 | 7,51E-01 |
| AT1G68620 | 0,33 | 1,33 | 4,40 | 0,30 | 7,63E-01 |
| AT1G68650 | 0,32 | 1,90 | 4,39 | 0,43 | 6,65E-01 |
| AT1G68670 | 0,63 | 0,00 | 3,07 | 0,00 | 1,00E+00 |
| AT1G68720 | 1,50 | -1,21 | 2,22 | -0,54 | 5,86E-01 |
| AT1G68840 | 1,23 | 0,02 | 1,62 | 0,01 | 9,92E-01 |
| AT1G68890 | 1,09 | 1,77 | 2,02 | 0,88 | 3,81E-01 |
| AT1G68980 | 0,24 | 1,38 | 4,41 | 0,31 | 7,54E-01 |
| AT1G69085 | 0,08 | 0,00 | 4,41 | 0,00 | 1,00E+00 |
| AT1G69160 | 0,39 | -0,47 | 3,41 | -0,14 | 8,89E-01 |
| AT1G69170 | 0,40 | 0,00 | 4,41 | 0,00 | 1,00E+00 |
| AT1G69200 | 0,21 | 1,25 | 4,39 | 0,29 | 7,75E-01 |
| AT1G69252 | 0,90 | 0,00 | 2,97 | 0,00 | 9,99E-01 |
| AT1G69260 | 0,63 | -0,65 | 2,74 | -0,24 | 8,12E-01 |
| AT1G69350 | 0,78 | 1,23 | 2,49 | 0,49 | 6,21E-01 |
| AT1G69390 | 0,12 | 0,00 | 4,41 | 0,00 | 1,00E+00 |
| AT1G69410 | 0,27 | -1,25 | 4,37 | -0,29 | 7,74E-01 |
| AT1G69420 | 0,65 | 3,21 | 2,49 | 1,29 | 1,98E-01 |
| AT1G69510 | 0,10 | 0,00 | 4,41 | 0,00 | 1,00E+00 |
| AT1G69523 | 0,10 | 0,00 | 4,41 | 0,00 | 1,00E+00 |
| AT1G69526 | 0,08 | 0,00 | 4,41 | 0,00 | 1,00E+00 |
| AT1G69530 | 0,93 | 1,07 | 2,01 | 0,53 | 5,94E-01 |
| AT1G69580 | 0,26 | -0,56 | 4,41 | -0,13 | 9,00E-01 |
| AT1G69610 | 0,21 | 0,00 | 4,41 | 0,00 | 1,00E+00 |

| | | | | | |
|---|---|---|---|---|---|
| AT1G09087 | 0,37 | 0,31 | 3,55 | 0,09 | 9,30E-01 |
| AT1G69690 | 0,67 | -1,45 | 2,59 | -0,56 | 5,74E-01 |
| AT1G69740 | 0,99 | 2,21 | 2,10 | 1,05 | 2,93E-01 |
| AT1G69750 | 0,10 | 0,00 | 4,41 | 0,00 | 1,00E+00 |
| AT1G69870 | 0,38 | -0,65 | 4,40 | -0,15 | 8,83E-01 |
| AT1G70060 | 1,48 | 3,30 | 2,15 | 1,54 | 1,24E-01 |
| AT1G70090 | 0,99 | 1,34 | 2,25 | 0,60 | 5,51E-01 |
| AT1G70100 | 0,38 | -0,72 | 4,34 | -0,17 | 8,67E-01 |
| AT1G70160 | 1,19 | 0,17 | 1,68 | 0,10 | 9,21E-01 |
| AT1G70180 | 1,28 | 3,54 | 1,91 | 1,85 | 6,37E-02 |
| AT1G70190 | 0,40 | 0,00 | 4,41 | 0,00 | 1,00E+00 |
| AT1G70210 | 0,79 | -1,34 | 2,50 | -0,53 | 5,93E-01 |
| AT1G70230 | 0,59 | -0,38 | 2,75 | -0,14 | 8,90E-01 |
| AT1G70250 | 0,71 | 1,90 | 2,93 | 0,65 | 5,18E-01 |
| AT1G70330 | 0,33 | -0,52 | 3,65 | -0,14 | 8,87E-01 |
| AT1G70480 | 1,12 | 3,22 | 1,89 | 1,71 | 8,79E-02 |
| AT1G70510 | 0,12 | 0,00 | 4,41 | 0,00 | 1,00E+00 |
| AT1G70550 | 0,14 | 0,00 | 4,41 | 0,00 | 1,00E+00 |
| AT1G70570 | 0,36 | 0,10 | 4,35 | 0,02 | 9,82E-01 |
| AT1G70580 | 0,81 | 0,36 | 2,40 | 0,15 | 8,80E-01 |
| AT1G70620 | 1,32 | 3,06 | 2,19 | 1,40 | 1,61E-01 |
| AT1G70630 | 0,36 | 2,09 | 4,37 | 0,48 | 6,32E-01 |
| AT1G70645 | 0,05 | 0,00 | 4,41 | 0,00 | 1,00E+00 |
| AT1G70650 | 0,26 | 1,25 | 4,39 | 0,29 | 7,75E-01 |
| AT1G70660 | 0,20 | 1,33 | 4,40 | 0,30 | 7,63E-01 |
| AT1G70700 | 0,60 | -1,67 | 2,67 | -0,62 | 5,32E-01 |
| AT1G70760 | 0,35 | -0,43 | 3,53 | -0,12 | 9,03E-01 |
| AT1G70810 | 0,62 | 0,63 | 2,58 | 0,24 | 8,07E-01 |
| AT1G70820 | 0,43 | -0,54 | 2,82 | -0,19 | 8,47E-01 |
| AT1G70895 | 0,08 | 0,00 | 4,41 | 0,00 | 1,00E+00 |
| AT1G70900 | 1,03 | 1,55 | 2,03 | 0,76 | 4,47E-01 |
| AT1G70960 | 0,12 | 0,00 | 4,41 | 0,00 | 1,00E+00 |
| AT1G70980 | 0,40 | -0,40 | 4,38 | -0,09 | 9,28E-01 |
| AT1G71010 | 1,50 | 3,23 | 2,01 | 1,61 | 1,07E-01 |
| AT1G71090 | 0,35 | 1,38 | 3,61 | 0,38 | 7,02E-01 |
| AT1G71100 | 1,11 | -1,37 | 1,88 | -0,73 | 4,67E-01 |
| AT1G71110 | 0,15 | 0,00 | 4,41 | 0,00 | 1,00E+00 |
| AT1G71130 | 0,37 | 0,00 | 4,41 | 0,00 | 1,00E+00 |
| AT1G71180 | 0,42 | 1,45 | 4,32 | 0,34 | 7,37E-01 |
| AT1G71270 | 0,45 | 0,00 | 4,23 | 0,00 | 1,00E+00 |
| AT1G71300 | 0,38 | 2,10 | 4,36 | 0,48 | 6,31E-01 |
| AT1G71320 | 0,39 | 0,00 | 4,26 | 0,00 | 1,00E+00 |
| AT1G71480 | 0,80 | 2,01 | 2,94 | 0,68 | 4,94E-01 |
| AT1G09257 | 0,17 | 0,00 | 4,41 | 0,00 | 1,00E+00 |
| AT1G71500 | 0,95 | 1,19 | 1,96 | 0,61 | 5,43E-01 |
| AT1G71695 | 0,82 | 1,54 | 2,52 | 0,61 | 5,41E-01 |
| AT1G71720 | 0,53 | -2,76 | 3,78 | -0,73 | 4,65E-01 |
| AT1G71800 | 0,67 | 2,89 | 3,62 | 0,80 | 4,25E-01 |
| AT1G71820 | 1,53 | -0,37 | 1,67 | -0,22 | 8,27E-01 |
| AT1G71840 | 1,05 | 1,38 | 2,38 | 0,58 | 5,62E-01 |
| AT1G71900 | 0,51 | 2,52 | 4,35 | 0,58 | 5,63E-01 |
| AT1G71910 | 0,44 | -0,54 | 3,32 | -0,16 | 8,70E-01 |
| AT1G71920 | 0,73 | -1,05 | 2,55 | -0,41 | 6,81E-01 |
| AT1G71930 | 1,13 | -0,48 | 2,30 | -0,21 | 8,33E-01 |
| AT1G71970 | 0,14 | 0,00 | 4,41 | 0,00 | 1,00E+00 |
| AT1G71980 | 0,46 | 0,00 | 3,27 | 0,00 | 1,00E+00 |

| | | | | | |
|---|---|---|---|---|---|
| AT1G72010 | 0,42 | -0,54 | 3,38 | -0,16 | 8,72E-01 |
| AT1G72030 | 0,75 | 1,80 | 3,67 | 0,49 | 6,23E-01 |
| AT1G72050 | 0,75 | 3,05 | 3,62 | 0,84 | 4,00E-01 |
| AT1G72090 | 0,34 | 1,38 | 4,41 | 0,31 | 7,54E-01 |
| AT1G72240 | 0,60 | 1,99 | 3,91 | 0,51 | 6,11E-01 |
| AT1G72310 | 0,25 | 1,39 | 4,41 | 0,32 | 7,52E-01 |
| AT1G72330 | 0,67 | 2,48 | 2,90 | 0,85 | 3,93E-01 |
| AT1G72470 | 0,43 | 1,90 | 4,39 | 0,43 | 6,65E-01 |
| AT1G72510 | 0,57 | -2,11 | 3,23 | -0,65 | 5,14E-01 |
| AT1G72520 | 0,91 | 0,42 | 2,25 | 0,19 | 8,53E-01 |
| AT1G72530 | 0,05 | 0,00 | 4,41 | 0,00 | 1,00E+00 |
| AT1G09333 | 1,54 | -3,08 | 1,85 | -1,66 | 9,64E-02 |
| AT1G72650 | 0,79 | 1,34 | 2,51 | 0,54 | 5,93E-01 |
| AT1G72700 | 1,31 | 1,38 | 2,14 | 0,65 | 5,18E-01 |
| AT1G72750 | 0,63 | 2,17 | 2,32 | 0,93 | 3,50E-01 |
| AT1G72770 | 1,51 | 1,87 | 1,78 | 1,05 | 2,95E-01 |
| AT1G72790 | 0,71 | -0,54 | 3,11 | -0,17 | 8,61E-01 |
| AT1G72855 | 0,40 | 0,00 | 4,41 | 0,00 | 1,00E+00 |
| AT1G72900 | 0,32 | -0,50 | 3,62 | -0,14 | 8,90E-01 |
| AT1G72910 | 1,29 | 1,03 | 2,04 | 0,50 | 6,14E-01 |
| AT1G72920 | 0,44 | 1,04 | 3,43 | 0,30 | 7,62E-01 |
| AT1G72940 | 0,97 | 0,00 | 4,10 | 0,00 | 1,00E+00 |
| AT1G72970 | 0,16 | 0,00 | 4,41 | 0,00 | 1,00E+00 |
| AT1G73030 | 0,82 | 3,12 | 2,43 | 1,29 | 1,98E-01 |
| AT1G73060 | 0,24 | 1,45 | 4,40 | 0,33 | 7,41E-01 |
| AT1G73080 | 1,38 | 2,23 | 1,71 | 1,30 | 1,93E-01 |
| AT1G73100 | 0,70 | 1,53 | 2,65 | 0,58 | 5,63E-01 |
| AT1G73180 | 1,25 | 3,67 | 1,97 | 1,87 | 6,16E-02 |
| AT1G73430 | 1,05 | 1,50 | 2,34 | 0,64 | 5,21E-01 |
| AT1G73450 | 1,12 | 2,87 | 2,00 | 1,43 | 1,53E-01 |
| AT1G73460 | 0,79 | 2,52 | 2,54 | 0,99 | 3,21E-01 |
| AT1G73470 | 0,23 | 1,38 | 4,41 | 0,31 | 7,54E-01 |
| AT1G73480 | 1,37 | 0,42 | 1,98 | 0,21 | 8,31E-01 |
| AT1G73600 | 1,37 | -1,89 | 2,07 | -0,92 | 3,60E-01 |
| AT1G73630 | 0,10 | 0,00 | 4,41 | 0,00 | 1,00E+00 |
| AT1G73670 | 0,46 | 0,01 | 3,34 | 0,00 | 9,99E-01 |
| AT1G73710 | 0,22 | 0,00 | 4,41 | 0,00 | 1,00E+00 |
| AT1G73750 | 0,72 | 0,00 | 4,41 | 0,00 | 1,00E+00 |
| AT1G73770 | 0,08 | 0,00 | 4,41 | 0,00 | 1,00E+00 |
| AT1G73850 | 0,17 | 1,32 | 4,40 | 0,30 | 7,64E-01 |
| AT1G73875 | 0,39 | 1,43 | 4,28 | 0,34 | 7,38E-01 |
| AT1G73885 | 0,62 | -0,50 | 3,19 | -0,16 | 8,76E-01 |
| AT1G73920 | 0,62 | 2,89 | 2,57 | 1,12 | 2,61E-01 |
| AT1G73970 | 0,95 | -2,63 | 2,46 | -1,07 | 2,84E-01 |
| AT1G74160 | 1,08 | 1,93 | 1,74 | 1,11 | 2,68E-01 |
| AT1G74250 | 1,42 | 2,20 | 2,05 | 1,07 | 2,83E-01 |
| AT1G74360 | 1,21 | 1,90 | 2,56 | 0,74 | 4,57E-01 |
| AT1G74380 | 1,29 | 3,39 | 1,87 | 1,82 | 6,92E-02 |
| AT1G74430 | 0,38 | 0,18 | 4,38 | 0,04 | 9,67E-01 |
| AT1G74450 | 0,99 | 1,38 | 2,23 | 0,62 | 5,36E-01 |
| AT1G74580 | 0,25 | 1,26 | 4,39 | 0,29 | 7,75E-01 |
| AT1G74680 | 0,57 | 3,04 | 3,19 | 0,95 | 3,41E-01 |
| AT1G74710 | 0,60 | -0,29 | 2,70 | -0,11 | 9,13E-01 |
| AT1G74730 | 0,44 | -0,06 | 3,28 | -0,02 | 9,86E-01 |
| AT1G74750 | 0,50 | 2,72 | 4,34 | 0,63 | 5,30E-01 |
| AT1G74780 | 0,63 | 1,36 | 3,06 | 0,44 | 6,57E-01 |

| | | | | | |
|---|---|---|---|---|---|
| AT1G74790 | 0,51 | 2,04 | 3,96 | 0,51 | 6,07E-01 |
| AT1G74870 | 1,10 | 1,35 | 3,38 | 0,40 | 6,89E-01 |
| AT1G74930 | 0,81 | -0,59 | 2,85 | -0,21 | 8,36E-01 |
| AT1G74940 | 0,58 | 0,00 | 3,18 | 0,00 | 1,00E+00 |
| AT1G75080 | 0,21 | 0,00 | 4,41 | 0,00 | 1,00E+00 |
| AT1G09533 | 1,00 | 0,37 | 2,04 | 0,18 | 8,56E-01 |
| AT1G75163 | 0,94 | -1,10 | 2,18 | -0,50 | 6,14E-01 |
| AT1G75166 | 1,22 | 1,16 | 2,21 | 0,53 | 5,98E-01 |
| AT1G75170 | 0,12 | 0,00 | 4,41 | 0,00 | 1,00E+00 |
| AT1G09543 | 0,38 | -0,46 | 3,45 | -0,13 | 8,94E-01 |
| AT1G75190 | 0,22 | -0,54 | 4,41 | -0,12 | 9,03E-01 |
| AT1G75210 | 0,77 | -2,23 | 2,85 | -0,78 | 4,33E-01 |
| AT1G75280 | 0,22 | 0,00 | 4,41 | 0,00 | 1,00E+00 |
| AT1G75310 | 0,47 | 0,49 | 4,34 | 0,11 | 9,10E-01 |
| AT1G75350 | 0,83 | 0,44 | 2,31 | 0,19 | 8,50E-01 |
| AT1G75400 | 0,32 | 0,00 | 4,41 | 0,00 | 1,00E+00 |
| AT1G75420 | 0,31 | 1,49 | 4,39 | 0,34 | 7,35E-01 |
| AT1G75440 | 0,28 | 1,39 | 4,41 | 0,32 | 7,52E-01 |
| AT1G75460 | 0,84 | 0,86 | 2,11 | 0,41 | 6,83E-01 |
| AT1G75580 | 0,17 | -0,57 | 4,40 | -0,13 | 8,98E-01 |
| AT1G75710 | 0,31 | -0,43 | 4,39 | -0,10 | 9,22E-01 |
| AT1G75785 | 0,12 | 0,00 | 4,41 | 0,00 | 1,00E+00 |
| AT1G75810 | 0,60 | 1,83 | 2,59 | 0,71 | 4,80E-01 |
| AT1G75840 | 0,75 | 1,08 | 2,25 | 0,48 | 6,31E-01 |
| AT1G75900 | 0,32 | 1,38 | 4,41 | 0,31 | 7,55E-01 |
| AT1G75960 | 0,45 | 2,07 | 4,27 | 0,48 | 6,28E-01 |
| AT1G76000 | 0,58 | 0,00 | 3,10 | 0,00 | 1,00E+00 |
| AT1G76050 | 0,66 | 1,50 | 3,02 | 0,50 | 6,19E-01 |
| AT1G76140 | 0,48 | 0,00 | 4,07 | 0,00 | 1,00E+00 |
| AT1G76185 | 0,16 | 0,00 | 4,41 | 0,00 | 1,00E+00 |
| AT1G76190 | 0,11 | 0,00 | 4,41 | 0,00 | 1,00E+00 |
| AT1G76320 | 0,32 | 0,00 | 4,41 | 0,00 | 1,00E+00 |
| AT1G76350 | 0,12 | 0,00 | 4,41 | 0,00 | 1,00E+00 |
| AT1G76390 | 0,51 | 1,38 | 3,27 | 0,42 | 6,73E-01 |
| AT1G76450 | 0,08 | 0,00 | 4,41 | 0,00 | 1,00E+00 |
| AT1G76520 | 0,73 | 1,00 | 2,15 | 0,46 | 6,42E-01 |
| AT1G76580 | 0,80 | 2,44 | 3,54 | 0,69 | 4,90E-01 |
| AT1G76590 | 0,69 | -0,74 | 2,91 | -0,25 | 8,00E-01 |
| AT1G76600 | 0,49 | 0,00 | 3,32 | 0,00 | 1,00E+00 |
| AT1G76620 | 0,25 | -1,09 | 4,37 | -0,25 | 8,02E-01 |
| AT1G76680 | 0,14 | -0,48 | 4,40 | -0,11 | 9,13E-01 |
| AT1G76740 | 0,12 | 0,00 | 4,41 | 0,00 | 1,00E+00 |
| AT1G76780 | 1,47 | -0,27 | 1,89 | -0,14 | 8,87E-01 |
| AT1G76820 | 0,53 | 0,48 | 3,19 | 0,15 | 8,81E-01 |
| AT1G76850 | 1,15 | 1,67 | 1,79 | 0,93 | 3,51E-01 |
| AT1G76880 | 1,06 | -0,84 | 1,86 | -0,45 | 6,51E-01 |
| AT1G76900 | 0,91 | 0,37 | 2,72 | 0,14 | 8,92E-01 |
| AT1G76920 | 0,59 | 2,29 | 3,82 | 0,60 | 5,49E-01 |
| AT1G76950 | 0,83 | 1,85 | 2,69 | 0,69 | 4,91E-01 |
| AT1G76990 | 1,49 | -0,48 | 1,61 | -0,30 | 7,63E-01 |
| AT1G77080 | 0,41 | -0,80 | 3,36 | -0,24 | 8,11E-01 |
| AT1G77140 | 0,25 | 1,43 | 4,40 | 0,32 | 7,46E-01 |
| AT1G77320 | 0,26 | 1,38 | 4,41 | 0,31 | 7,54E-01 |
| AT1G77350 | 0,15 | 0,00 | 4,41 | 0,00 | 1,00E+00 |
| AT1G77420 | 0,12 | 0,00 | 4,41 | 0,00 | 1,00E+00 |
| AT1G77460 | 0,94 | 3,09 | 2,66 | 1,16 | 2,47E-01 |

| | | | | | |
|---|---|---|---|---|---|
| AT1G77490 | 0,64 | 1,19 | 3,06 | 0,39 | 6,97E-01 |
| AT1G77500 | 0,37 | 0,55 | 4,39 | 0,13 | 9,00E-01 |
| AT1G77510 | 1,26 | 2,69 | 1,84 | 1,46 | 1,43E-01 |
| AT1G77610 | 0,53 | 1,71 | 4,35 | 0,39 | 6,93E-01 |
| AT1G77710 | 0,32 | 1,38 | 4,41 | 0,31 | 7,54E-01 |
| AT1G77800 | 1,13 | 3,21 | 1,93 | 1,67 | 9,54E-02 |
| AT1G77840 | 0,31 | 1,38 | 3,68 | 0,37 | 7,08E-01 |
| AT1G77850 | 1,01 | 0,32 | 2,36 | 0,14 | 8,91E-01 |
| AT1G77890 | 0,98 | 1,96 | 1,79 | 1,09 | 2,75E-01 |
| AT1G77930 | 1,14 | 0,99 | 2,24 | 0,44 | 6,58E-01 |
| AT1G78010 | 0,90 | 3,09 | 2,28 | 1,35 | 1,76E-01 |
| AT1G78070 | 1,66 | -1,30 | 1,60 | -0,82 | 4,14E-01 |
| AT1G78100 | 0,64 | -0,38 | 2,35 | -0,16 | 8,73E-01 |
| AT1G78150 | 0,22 | 1,27 | 4,39 | 0,29 | 7,72E-01 |
| AT1G78200 | 0,54 | 1,26 | 2,62 | 0,48 | 6,31E-01 |
| AT1G78280 | 1,59 | 2,22 | 2,35 | 0,95 | 3,45E-01 |
| AT1G78300 | 0,54 | -0,45 | 3,28 | -0,14 | 8,90E-01 |
| AT1G78420 | 0,54 | 2,13 | 2,75 | 0,78 | 4,38E-01 |
| AT1G78490 | 0,05 | 0,00 | 4,41 | 0,00 | 1,00E+00 |
| AT1G78500 | 0,17 | 0,00 | 4,41 | 0,00 | 1,00E+00 |
| AT1G78600 | 0,20 | 1,40 | 4,41 | 0,32 | 7,51E-01 |
| AT1G78630 | 1,31 | 1,44 | 1,78 | 0,81 | 4,19E-01 |
| AT1G78670 | 0,52 | 0,66 | 3,16 | 0,21 | 8,34E-01 |
| AT1G78680 | 0,90 | 0,39 | 2,18 | 0,18 | 8,57E-01 |
| AT1G78700 | 0,37 | 1,56 | 4,38 | 0,36 | 7,22E-01 |
| AT1G78710 | 0,38 | 2,48 | 4,35 | 0,57 | 5,69E-01 |
| AT1G78770 | 0,21 | 0,00 | 4,41 | 0,00 | 1,00E+00 |
| AT1G78870 | 0,28 | 1,34 | 4,40 | 0,30 | 7,61E-01 |
| AT1G78890 | 0,36 | 0,97 | 4,36 | 0,22 | 8,24E-01 |
| AT1G78910 | 0,26 | 1,38 | 4,41 | 0,31 | 7,54E-01 |
| AT1G78920 | 1,61 | 0,98 | 1,73 | 0,57 | 5,70E-01 |
| AT1G79060 | 0,14 | 0,00 | 4,41 | 0,00 | 1,00E+00 |
| AT1G79103 | 0,05 | 0,00 | 4,41 | 0,00 | 1,00E+00 |
| AT1G79110 | 1,67 | 0,52 | 1,69 | 0,31 | 7,57E-01 |
| AT1G09847 | 0,44 | -1,71 | 3,26 | -0,52 | 6,00E-01 |
| AT1G79190 | 0,88 | 3,69 | 2,67 | 1,38 | 1,67E-01 |
| AT1G79230 | 0,71 | 2,50 | 2,90 | 0,86 | 3,88E-01 |
| AT1G79240 | 0,20 | 0,00 | 4,41 | 0,00 | 1,00E+00 |
| AT1G79270 | 1,18 | 1,90 | 1,86 | 1,02 | 3,09E-01 |
| AT1G79340 | 0,91 | -2,33 | 2,13 | -1,09 | 2,75E-01 |
| AT1G79380 | 0,91 | 2,17 | 2,88 | 0,75 | 4,51E-01 |
| AT1G79560 | 0,28 | 1,34 | 4,40 | 0,30 | 7,61E-01 |
| AT1G79590 | 0,18 | 0,00 | 4,41 | 0,00 | 1,00E+00 |
| AT1G79660 | 0,63 | 3,27 | 3,61 | 0,91 | 3,65E-01 |
| AT1G09903 | 0,08 | 0,00 | 4,41 | 0,00 | 1,00E+00 |
| AT1G79810 | 0,41 | 1,31 | 3,37 | 0,39 | 6,97E-01 |
| AT1G79840 | 0,77 | 2,19 | 3,01 | 0,73 | 4,67E-01 |
| AT1G79870 | 0,21 | 0,00 | 4,41 | 0,00 | 1,00E+00 |
| AT1G79890 | 0,21 | 1,27 | 4,39 | 0,29 | 7,72E-01 |
| AT1G79910 | 0,41 | 2,13 | 4,19 | 0,51 | 6,10E-01 |
| AT1G79940 | 1,20 | -0,06 | 1,61 | -0,04 | 9,70E-01 |
| AT1G79950 | 0,49 | 1,62 | 3,94 | 0,41 | 6,80E-01 |
| AT1G79970 | 0,27 | 1,30 | 4,40 | 0,29 | 7,68E-01 |
| AT1G79985 | 0,75 | 1,19 | 2,60 | 0,46 | 6,47E-01 |
| AT1G80010 | 1,24 | 0,93 | 1,77 | 0,52 | 6,00E-01 |
| AT1G80160 | 0,37 | 0,00 | 4,41 | 0,00 | 1,00E+00 |

| | | | | | |
|---|---|---|---|---|---|
| AT1G80260 | 0,59 | 0,00 | 3,01 | 0,00 | 1,00E+00 |
| AT1G80300 | 1,04 | 3,40 | 3,23 | 1,05 | 2,93E-01 |
| AT1G80310 | 0,64 | -0,29 | 2,65 | -0,11 | 9,14E-01 |
| AT1G80380 | 1,15 | -0,28 | 1,71 | -0,16 | 8,72E-01 |
| AT1G09963 | 0,96 | 2,39 | 2,45 | 0,98 | 3,28E-01 |
| AT1G80430 | 0,19 | 0,00 | 4,41 | 0,00 | 1,00E+00 |
| AT1G80510 | 0,61 | 1,46 | 3,87 | 0,38 | 7,06E-01 |
| AT1G80620 | 0,35 | -1,47 | 4,36 | -0,34 | 7,36E-01 |
| AT1G80770 | 0,54 | 1,02 | 3,26 | 0,31 | 7,54E-01 |
| AT1G80780 | 0,60 | 1,27 | 3,22 | 0,39 | 6,93E-01 |
| AT1G80790 | 0,69 | 0,50 | 2,92 | 0,17 | 8,65E-01 |
| AT1G80810 | 0,38 | 1,41 | 4,31 | 0,33 | 7,42E-01 |
| AT1G80840 | 1,09 | 0,00 | 2,63 | 0,00 | 1,00E+00 |
| AT1G80870 | 0,27 | 1,42 | 4,40 | 0,32 | 7,46E-01 |
| AT2G01070 | 0,81 | 2,70 | 2,49 | 1,08 | 2,79E-01 |
| AT2G01080 | 0,35 | 1,38 | 3,57 | 0,39 | 6,99E-01 |
| AT2G01100 | 0,12 | 0,00 | 4,41 | 0,00 | 1,00E+00 |
| AT2G01120 | 0,92 | 2,34 | 2,87 | 0,82 | 4,15E-01 |
| AT2G01190 | 0,50 | -0,52 | 3,23 | -0,16 | 8,73E-01 |
| AT2G01400 | 0,39 | 1,83 | 4,37 | 0,42 | 6,75E-01 |
| AT2G03975 | 0,34 | 0,44 | 3,60 | 0,12 | 9,02E-01 |
| AT2G01480 | 0,34 | 2,16 | 4,37 | 0,49 | 6,21E-01 |
| AT2G01620 | 0,60 | 2,48 | 2,59 | 0,96 | 3,39E-01 |
| AT2G01680 | 0,39 | 1,82 | 4,38 | 0,42 | 6,78E-01 |
| AT2G01750 | 0,24 | -0,47 | 4,40 | -0,11 | 9,15E-01 |
| AT2G01860 | 0,98 | 1,38 | 2,64 | 0,52 | 6,02E-01 |
| AT2G01900 | 1,22 | -1,90 | 1,92 | -0,99 | 3,22E-01 |
| AT2G01980 | 0,63 | 0,56 | 3,15 | 0,18 | 8,58E-01 |
| AT2G02040 | 0,10 | 0,00 | 4,41 | 0,00 | 1,00E+00 |
| AT2G02050 | 0,58 | 0,30 | 2,68 | 0,11 | 9,11E-01 |
| AT2G02070 | 0,84 | 0,10 | 2,03 | 0,05 | 9,60E-01 |
| AT2G02090 | 1,22 | 3,38 | 2,37 | 1,43 | 1,54E-01 |
| AT2G02100 | 0,54 | -1,74 | 3,90 | -0,45 | 6,55E-01 |
| AT2G02440 | 0,68 | -1,11 | 3,02 | -0,37 | 7,14E-01 |
| AT2G02450 | 0,66 | -1,48 | 2,69 | -0,55 | 5,82E-01 |
| AT2G02470 | 0,18 | 1,38 | 4,41 | 0,31 | 7,54E-01 |
| AT2G02550 | 1,01 | -0,46 | 2,27 | -0,20 | 8,40E-01 |
| AT2G02700 | 0,39 | 1,76 | 4,38 | 0,40 | 6,87E-01 |
| AT2G04295 | 0,12 | 0,00 | 4,41 | 0,00 | 1,00E+00 |
| AT2G02760 | 0,54 | -2,18 | 2,67 | -0,82 | 4,13E-01 |
| AT2G02790 | 0,39 | -0,11 | 4,24 | -0,03 | 9,79E-01 |
| AT2G02795 | 0,14 | 0,00 | 4,41 | 0,00 | 1,00E+00 |
| AT2G02810 | 0,62 | 1,38 | 3,15 | 0,44 | 6,62E-01 |
| AT2G04355 | 0,47 | 2,08 | 3,38 | 0,61 | 5,39E-01 |
| AT2G04365 | 0,97 | -0,51 | 2,57 | -0,20 | 8,42E-01 |
| AT2G02870 | 1,53 | 1,21 | 2,07 | 0,58 | 5,60E-01 |
| AT2G02955 | 0,36 | 1,82 | 4,38 | 0,41 | 6,79E-01 |
| AT2G03070 | 0,25 | 1,34 | 4,40 | 0,30 | 7,61E-01 |
| AT2G03120 | 0,64 | -0,18 | 2,61 | -0,07 | 9,45E-01 |
| AT2G03130 | 0,12 | 0,00 | 4,41 | 0,00 | 1,00E+00 |
| AT2G03270 | 0,39 | 1,32 | 2,94 | 0,45 | 6,54E-01 |
| AT2G03410 | 0,43 | -0,54 | 4,17 | -0,13 | 8,96E-01 |
| AT2G03450 | 0,32 | 2,21 | 4,37 | 0,51 | 6,13E-01 |
| AT2G03500 | 0,52 | 0,24 | 3,27 | 0,07 | 9,41E-01 |
| AT2G03520 | 0,08 | 0,00 | 4,41 | 0,00 | 1,00E+00 |
| AT2G03550 | 0,42 | -2,16 | 4,18 | -0,52 | 6,06E-01 |

| | | | | | |
|---|---|---|---|---|---|
| AT2G03567 | 0,47 | -0,92 | 3,41 | -0,27 | 7,87E-01 |
| AT2G03680 | 0,82 | 0,66 | 2,26 | 0,29 | 7,69E-01 |
| AT2G03710 | 0,17 | 0,00 | 4,41 | 0,00 | 1,00E+00 |
| AT2G03823 | 0,37 | -0,51 | 2,98 | -0,17 | 8,65E-01 |
| AT2G03870 | 0,41 | 2,11 | 4,16 | 0,51 | 6,12E-01 |
| AT2G03932 | 0,20 | 0,00 | 4,41 | 0,00 | 1,00E+00 |
| AT2G03931 | 0,05 | 0,00 | 4,41 | 0,00 | 1,00E+00 |
| AT2G03933 | 0,14 | 0,00 | 4,41 | 0,00 | 1,00E+00 |
| AT2G03936 | 0,18 | 0,00 | 4,41 | 0,00 | 1,00E+00 |
| AT2G03965 | 0,22 | 0,00 | 4,41 | 0,00 | 1,00E+00 |
| AT2G03980 | 0,82 | 0,06 | 2,56 | 0,03 | 9,80E-01 |
| AT2G04039 | 0,42 | 2,13 | 4,14 | 0,51 | 6,07E-01 |
| AT2G04735 | 0,15 | -0,43 | 4,39 | -0,10 | 9,22E-01 |
| AT2G04235 | 0,05 | 0,00 | 4,41 | 0,00 | 1,00E+00 |
| AT2G04270 | 1,17 | -0,11 | 2,35 | -0,04 | 9,64E-01 |
| AT2G04340 | 0,38 | 2,11 | 4,37 | 0,48 | 6,29E-01 |
| AT2G04350 | 1,50 | -0,20 | 1,51 | -0,13 | 8,94E-01 |
| AT2G04380 | 0,56 | 1,26 | 2,77 | 0,45 | 6,50E-01 |
| AT2G04390 | 0,85 | 2,16 | 2,25 | 0,96 | 3,36E-01 |
| AT2G04400 | 0,61 | 0,31 | 2,64 | 0,12 | 9,07E-01 |
| AT2G04450 | 0,21 | 0,00 | 4,41 | 0,00 | 1,00E+00 |
| AT2G04495 | 0,43 | 1,38 | 3,46 | 0,40 | 6,90E-01 |
| AT2G04550 | 0,15 | -0,43 | 4,39 | -0,10 | 9,23E-01 |
| AT2G04700 | 0,71 | 0,43 | 2,40 | 0,18 | 8,56E-01 |
| AT2G04740 | 0,38 | 2,22 | 4,37 | 0,51 | 6,12E-01 |
| AT2G04780 | 0,24 | 1,61 | 4,39 | 0,37 | 7,14E-01 |
| AT2G04795 | 0,15 | 0,00 | 4,41 | 0,00 | 1,00E+00 |
| AT2G04850 | 0,08 | 0,00 | 4,41 | 0,00 | 1,00E+00 |
| AT2G04865 | 0,25 | 0,00 | 4,41 | 0,00 | 1,00E+00 |
| AT2G04900 | 0,19 | 0,00 | 4,40 | 0,00 | 1,00E+00 |
| AT2G05060 | 0,21 | 1,27 | 4,39 | 0,29 | 7,72E-01 |
| AT2G05180 | 0,08 | 0,00 | 4,41 | 0,00 | 1,00E+00 |
| AT2G05210 | 0,12 | 0,00 | 4,41 | 0,00 | 1,00E+00 |
| AT2G05320 | 0,92 | -1,73 | 2,22 | -0,78 | 4,35E-01 |
| AT2G05350 | 0,23 | 0,00 | 4,41 | 0,00 | 1,00E+00 |
| AT2G05380 | 0,99 | 0,01 | 1,80 | 0,01 | 9,93E-01 |
| AT2G05540 | 0,30 | 1,38 | 4,41 | 0,31 | 7,54E-01 |
| AT2G04915 | 0,39 | -1,83 | 3,45 | -0,53 | 5,95E-01 |
| AT2G04935 | 1,06 | 2,12 | 2,07 | 1,02 | 3,06E-01 |
| AT2G05635 | 0,15 | 0,00 | 4,41 | 0,00 | 1,00E+00 |
| AT2G04985 | 0,95 | -1,62 | 2,29 | -0,71 | 4,78E-01 |
| AT2G05790 | 0,44 | 0,42 | 3,27 | 0,13 | 8,97E-01 |
| AT2G05812 | 1,46 | 1,79 | 1,89 | 0,95 | 3,44E-01 |
| AT2G05830 | 0,15 | 0,00 | 4,41 | 0,00 | 1,00E+00 |
| AT2G05840 | 0,17 | -0,57 | 4,40 | -0,13 | 8,98E-01 |
| AT2G05990 | 0,47 | -0,34 | 2,47 | -0,14 | 8,91E-01 |
| AT2G05995 | 0,63 | -1,38 | 3,05 | -0,45 | 6,51E-01 |
| AT2G06025 | 0,59 | 0,86 | 2,72 | 0,32 | 7,52E-01 |
| AT2G06095 | 0,70 | -2,93 | 3,45 | -0,85 | 3,96E-01 |
| AT2G06105 | 0,23 | 0,00 | 4,41 | 0,00 | 1,00E+00 |
| AT2G06845 | 0,18 | 0,00 | 4,41 | 0,00 | 1,00E+00 |
| AT2G06850 | 0,75 | -1,28 | 2,56 | -0,50 | 6,17E-01 |
| AT2G06904 | 0,38 | 0,00 | 4,41 | 0,00 | 1,00E+00 |
| AT2G06990 | 1,10 | -0,48 | 2,18 | -0,22 | 8,26E-01 |
| AT2G07050 | 0,64 | 0,00 | 3,10 | 0,00 | 1,00E+00 |
| AT2G07200 | 0,15 | 0,00 | 4,41 | 0,00 | 1,00E+00 |

| | | | | | |
|---|---|---|---|---|---|
| AT2G07565 | 0,91 | -1,07 | 2,45 | -0,43 | 6,64E-01 |
| AT2G07749 | 0,59 | -1,60 | 3,24 | -0,49 | 6,21E-01 |
| AT2G07672 | 0,25 | 0,00 | 4,41 | 0,00 | 1,00E+00 |
| AT2G07745 | 0,19 | -1,18 | 4,36 | -0,27 | 7,86E-01 |
| AT2G07674 | 0,81 | 1,38 | 3,79 | 0,36 | 7,16E-01 |
| AT2G07623 | 0,23 | 0,00 | 4,41 | 0,00 | 1,00E+00 |
| AT2G07678 | 0,32 | 0,95 | 4,37 | 0,22 | 8,27E-01 |
| AT2G07669 | 0,33 | 0,32 | 4,37 | 0,07 | 9,42E-01 |
| AT2G07681 | 0,75 | 2,19 | 2,32 | 0,94 | 3,45E-01 |
| AT2G07631 | 0,18 | -0,51 | 4,40 | -0,12 | 9,07E-01 |
| AT2G05215 | 0,68 | -1,96 | 2,61 | -0,75 | 4,53E-01 |
| AT2G07695 | 1,27 | -1,25 | 2,10 | -0,59 | 5,53E-01 |
| AT2G07798 | 1,27 | -1,17 | 1,92 | -0,61 | 5,40E-01 |
| AT2G07667 | 1,00 | -0,73 | 2,38 | -0,31 | 7,58E-01 |
| AT2G07638 | 1,46 | -1,01 | 1,59 | -0,64 | 5,24E-01 |
| AT2G07707 | 0,32 | -0,54 | 4,41 | -0,12 | 9,02E-01 |
| AT2G07708 | 0,57 | -0,49 | 4,33 | -0,11 | 9,11E-01 |
| AT2G07715 | 0,94 | -0,07 | 1,83 | -0,04 | 9,70E-01 |
| AT2G07728 | 0,11 | 0,00 | 4,41 | 0,00 | 1,00E+00 |
| AT2G07656 | 0,60 | -0,59 | 3,11 | -0,19 | 8,50E-01 |
| AT2G07827 | 0,36 | 0,33 | 3,55 | 0,09 | 9,27E-01 |
| AT2G07787 | 0,79 | 0,41 | 2,34 | 0,18 | 8,60E-01 |
| AT2G07659 | 0,28 | -0,47 | 4,40 | -0,11 | 9,14E-01 |
| AT2G07765 | 0,12 | 0,00 | 4,41 | 0,00 | 1,00E+00 |
| AT2G07741 | 0,27 | -0,54 | 4,41 | -0,12 | 9,02E-01 |
| AT2G07680 | 0,80 | -0,51 | 2,59 | -0,20 | 8,43E-01 |
| AT2G09990 | 0,53 | -1,19 | 3,18 | -0,37 | 7,09E-01 |
| AT2G05485 | 0,27 | 0,00 | 4,41 | 0,00 | 1,00E+00 |
| AT2G10545 | 0,08 | 0,00 | 4,41 | 0,00 | 1,00E+00 |
| AT2G10556 | 0,88 | 0,00 | 2,83 | 0,00 | 1,00E+00 |
| AT2G10965 | 0,86 | -1,32 | 2,20 | -0,60 | 5,48E-01 |
| AT2G11010 | 0,65 | 1,35 | 3,09 | 0,44 | 6,61E-01 |
| AT2G11035 | 0,48 | 0,86 | 4,05 | 0,21 | 8,31E-01 |
| AT2G11045 | 0,08 | 0,00 | 4,41 | 0,00 | 1,00E+00 |
| AT2G11225 | 0,91 | 2,86 | 1,95 | 1,47 | 1,43E-01 |
| AT2G11620 | 0,14 | 0,00 | 4,41 | 0,00 | 1,00E+00 |
| AT2G11851 | 0,57 | 0,00 | 3,11 | 0,00 | 1,00E+00 |
| AT2G05825 | 0,69 | 1,40 | 2,35 | 0,59 | 5,52E-01 |
| AT2G13100 | 0,26 | 0,43 | 4,37 | 0,10 | 9,22E-01 |
| AT2G13290 | 0,12 | 0,00 | 4,41 | 0,00 | 1,00E+00 |
| AT2G05885 | 0,14 | 0,00 | 4,41 | 0,00 | 1,00E+00 |
| AT2G13430 | 0,44 | -0,54 | 3,32 | -0,16 | 8,70E-01 |
| AT2G13500 | 0,11 | 0,00 | 4,41 | 0,00 | 1,00E+00 |
| AT2G13540 | 0,64 | -0,82 | 2,38 | -0,34 | 7,32E-01 |
| AT2G05925 | 0,12 | 0,00 | 4,41 | 0,00 | 1,00E+00 |
| AT2G13790 | 0,49 | 1,43 | 3,39 | 0,42 | 6,74E-01 |
| AT2G13845 | 0,67 | 1,38 | 2,43 | 0,57 | 5,70E-01 |
| AT2G13960 | 0,27 | 1,38 | 4,41 | 0,31 | 7,54E-01 |
| AT2G14070 | 0,19 | -0,54 | 4,41 | -0,12 | 9,02E-01 |
| AT2G14115 | 0,49 | 1,80 | 4,37 | 0,41 | 6,81E-01 |
| AT2G14395 | 0,63 | -1,41 | 4,34 | -0,33 | 7,45E-01 |
| AT2G14440 | 0,43 | -1,66 | 4,13 | -0,40 | 6,88E-01 |
| AT2G14530 | 0,21 | 0,00 | 4,41 | 0,00 | 1,00E+00 |
| AT2G14700 | 0,64 | 0,00 | 3,08 | 0,00 | 1,00E+00 |
| AT2G14740 | 1,25 | 1,33 | 2,62 | 0,51 | 6,12E-01 |
| AT2G14750 | 1,64 | 0,90 | 1,51 | 0,60 | 5,52E-01 |

| | | | | | |
|---|---|---|---|---|---|
| AT2G14765 | 0,40 | 0,00 | 4,41 | 0,00 | 1,00E+00 |
| AT2G14825 | 0,08 | 0,00 | 4,41 | 0,00 | 1,00E+00 |
| AT2G14835 | 0,23 | 0,00 | 4,41 | 0,00 | 1,00E+00 |
| AT2G14850 | 0,21 | 0,00 | 4,41 | 0,00 | 1,00E+00 |
| AT2G14870 | 0,17 | -1,42 | 4,35 | -0,33 | 7,45E-01 |
| AT2G14878 | 1,23 | 0,10 | 1,74 | 0,06 | 9,54E-01 |
| AT2G14890 | 0,77 | 1,29 | 2,30 | 0,56 | 5,75E-01 |
| AT2G06305 | 0,18 | 0,00 | 4,40 | 0,00 | 1,00E+00 |
| AT2G15042 | 0,11 | 0,00 | 4,41 | 0,00 | 1,00E+00 |
| AT2G15050 | 0,64 | -1,71 | 2,96 | -0,58 | 5,62E-01 |
| AT2G15080 | 0,92 | 1,06 | 2,24 | 0,47 | 6,36E-01 |
| AT2G15090 | 1,27 | -1,94 | 3,07 | -0,63 | 5,28E-01 |
| AT2G15300 | 0,31 | 0,07 | 4,37 | 0,02 | 9,87E-01 |
| AT2G15430 | 0,16 | 0,00 | 4,41 | 0,00 | 1,00E+00 |
| AT2G15450 | 0,29 | 0,00 | 4,40 | 0,00 | 1,00E+00 |
| AT2G15470 | 0,34 | 0,00 | 4,41 | 0,00 | 1,00E+00 |
| AT2G15580 | 0,29 | -0,51 | 3,75 | -0,14 | 8,91E-01 |
| AT2G15710 | 0,40 | -1,58 | 4,35 | -0,36 | 7,17E-01 |
| AT2G15730 | 0,08 | 0,00 | 4,41 | 0,00 | 1,00E+00 |
| AT2G15900 | 0,47 | 0,18 | 2,81 | 0,07 | 9,48E-01 |
| AT2G15960 | 1,09 | 3,28 | 1,91 | 1,72 | 8,63E-02 |
| AT2G06535 | 1,38 | 0,00 | 2,24 | 0,00 | 1,00E+00 |
| AT2G16019 | 1,05 | -0,50 | 2,36 | -0,21 | 8,33E-01 |
| AT2G16280 | 0,93 | -1,01 | 2,42 | -0,42 | 6,76E-01 |
| AT2G16365 | 0,96 | -0,27 | 2,39 | -0,12 | 9,08E-01 |
| AT2G16380 | 0,44 | 0,95 | 4,26 | 0,22 | 8,24E-01 |
| AT2G16390 | 0,40 | 0,00 | 4,41 | 0,00 | 1,00E+00 |
| AT2G16630 | 0,14 | 0,00 | 4,41 | 0,00 | 1,00E+00 |
| AT2G16700 | 0,28 | -0,49 | 4,40 | -0,11 | 9,11E-01 |
| AT2G16710 | 0,72 | 0,00 | 3,80 | 0,00 | 1,00E+00 |
| AT2G16800 | 0,26 | -1,08 | 4,37 | -0,25 | 8,05E-01 |
| AT2G16950 | 1,26 | 0,39 | 2,36 | 0,17 | 8,67E-01 |
| AT2G16970 | 1,25 | -0,82 | 1,61 | -0,51 | 6,10E-01 |
| AT2G17033 | 0,44 | 1,08 | 4,21 | 0,26 | 7,98E-01 |
| AT2G17040 | 1,57 | 3,69 | 1,84 | 2,01 | 4,47E-02 |
| AT2G17120 | 0,23 | 0,00 | 4,41 | 0,00 | 1,00E+00 |
| AT2G17140 | 0,42 | -0,59 | 3,48 | -0,17 | 8,66E-01 |
| AT2G17170 | 0,08 | 0,00 | 4,41 | 0,00 | 1,00E+00 |
| AT2G17240 | 0,52 | -0,77 | 3,15 | -0,25 | 8,06E-01 |
| AT2G17250 | 0,25 | 1,38 | 4,41 | 0,31 | 7,54E-01 |
| AT2G17265 | 1,56 | 1,13 | 1,69 | 0,67 | 5,03E-01 |
| AT2G17270 | 0,25 | 1,24 | 4,38 | 0,28 | 7,78E-01 |
| AT2G17290 | 0,78 | 1,41 | 2,33 | 0,60 | 5,47E-01 |
| AT2G06905 | 1,12 | 1,92 | 1,88 | 1,02 | 3,06E-01 |
| AT2G17360 | 0,51 | -0,13 | 3,34 | -0,04 | 9,70E-01 |
| AT2G17390 | 0,66 | 1,68 | 2,97 | 0,57 | 5,71E-01 |
| AT2G17410 | 1,45 | 4,06 | 1,75 | 2,33 | 2,00E-02 |
| AT2G17420 | 1,05 | 2,66 | 2,17 | 1,22 | 2,21E-01 |
| AT2G17520 | 1,17 | -0,32 | 2,02 | -0,16 | 8,75E-01 |
| AT2G17530 | 0,50 | 2,25 | 4,02 | 0,56 | 5,75E-01 |
| AT2G17570 | 0,42 | 2,15 | 4,19 | 0,51 | 6,08E-01 |
| AT2G17580 | 0,66 | 1,79 | 3,03 | 0,59 | 5,54E-01 |
| AT2G17705 | 0,42 | 1,27 | 3,35 | 0,38 | 7,04E-01 |
| AT2G17720 | 0,20 | 1,38 | 4,41 | 0,31 | 7,54E-01 |
| AT2G17760 | 0,65 | 2,72 | 2,96 | 0,92 | 3,57E-01 |
| AT2G17787 | 0,33 | 1,38 | 4,41 | 0,31 | 7,55E-01 |

| | | | | | |
|---|---|---|---|---|---|
| AT2G17790 | 0,80 | 0,52 | 2,59 | 0,20 | 8,40E-01 |
| AT2G17905 | 0,41 | 0,40 | 3,44 | 0,12 | 9,06E-01 |
| AT2G17920 | 0,15 | 0,00 | 4,41 | 0,00 | 1,00E+00 |
| AT2G17980 | 1,01 | 1,38 | 2,72 | 0,51 | 6,12E-01 |
| AT2G17990 | 0,69 | 1,03 | 2,53 | 0,41 | 6,83E-01 |
| AT2G18000 | 0,29 | 1,35 | 4,40 | 0,31 | 7,59E-01 |
| AT2G18030 | 0,68 | 1,38 | 3,09 | 0,45 | 6,56E-01 |
| AT2G18040 | 0,20 | 0,00 | 4,41 | 0,00 | 1,00E+00 |
| AT2G18090 | 1,01 | 1,20 | 2,05 | 0,58 | 5,59E-01 |
| AT2G18100 | 0,48 | 0,00 | 4,11 | 0,00 | 1,00E+00 |
| AT2G18110 | 0,62 | -1,68 | 3,03 | -0,56 | 5,78E-01 |
| AT2G18390 | 0,28 | 1,39 | 4,41 | 0,32 | 7,52E-01 |
| AT2G18420 | 0,81 | 3,18 | 3,43 | 0,93 | 3,55E-01 |
| AT2G18530 | 0,38 | -0,54 | 3,52 | -0,15 | 8,77E-01 |
| AT2G18560 | 1,54 | -1,02 | 2,48 | -0,41 | 6,81E-01 |
| AT2G18630 | 0,16 | 0,00 | 4,41 | 0,00 | 1,00E+00 |
| AT2G18670 | 1,13 | -1,12 | 1,90 | -0,59 | 5,55E-01 |
| AT2G18680 | 0,16 | 0,00 | 4,41 | 0,00 | 1,00E+00 |
| AT2G18690 | 1,29 | 2,22 | 2,11 | 1,05 | 2,92E-01 |
| AT2G18700 | 0,27 | -0,88 | 4,38 | -0,20 | 8,41E-01 |
| AT2G18730 | 0,58 | -1,32 | 2,64 | -0,50 | 6,17E-01 |
| AT2G18735 | 0,22 | -0,48 | 4,40 | -0,11 | 9,14E-01 |
| AT2G18750 | 0,67 | 1,26 | 2,18 | 0,58 | 5,63E-01 |
| AT2G18760 | 1,48 | 3,17 | 1,81 | 1,75 | 7,95E-02 |
| AT2G18850 | 0,52 | 1,77 | 4,35 | 0,41 | 6,83E-01 |
| AT2G18880 | 0,25 | 0,00 | 4,41 | 0,00 | 1,00E+00 |
| AT2G18917 | 0,08 | 0,00 | 4,41 | 0,00 | 1,00E+00 |
| AT2G07095 | 0,55 | 0,00 | 3,16 | 0,00 | 1,00E+00 |
| AT2G18938 | 0,72 | 1,62 | 2,59 | 0,63 | 5,31E-01 |
| AT2G18950 | 0,45 | 0,00 | 4,23 | 0,00 | 1,00E+00 |
| AT2G19110 | 0,15 | 0,00 | 4,41 | 0,00 | 1,00E+00 |
| AT2G19130 | 0,89 | 0,63 | 2,17 | 0,29 | 7,73E-01 |
| AT2G19160 | 0,39 | 0,00 | 4,41 | 0,00 | 1,00E+00 |
| AT2G19260 | 0,46 | 1,56 | 4,09 | 0,38 | 7,03E-01 |
| AT2G19280 | 0,36 | 0,00 | 4,41 | 0,00 | 1,00E+00 |
| AT2G19420 | 0,28 | 0,00 | 4,41 | 0,00 | 1,00E+00 |
| AT2G19430 | 0,24 | -1,35 | 4,38 | -0,31 | 7,58E-01 |
| AT2G19450 | 0,42 | -1,49 | 4,29 | -0,35 | 7,29E-01 |
| AT2G07205 | 0,75 | 1,34 | 3,72 | 0,36 | 7,20E-01 |
| AT2G19490 | 0,36 | 0,00 | 4,41 | 0,00 | 1,00E+00 |
| AT2G19520 | 0,55 | 1,93 | 4,10 | 0,47 | 6,38E-01 |
| AT2G19540 | 0,46 | 0,51 | 2,84 | 0,18 | 8,58E-01 |
| AT2G19572 | 0,52 | 0,00 | 3,17 | 0,00 | 1,00E+00 |
| AT2G19582 | 0,25 | 0,00 | 4,41 | 0,00 | 1,00E+00 |
| AT2G19600 | 0,64 | 0,42 | 3,97 | 0,11 | 9,16E-01 |
| AT2G19640 | 0,53 | 2,51 | 3,87 | 0,65 | 5,17E-01 |
| AT2G19710 | 1,44 | 4,02 | 2,11 | 1,91 | 5,62E-02 |
| AT2G19730 | 0,57 | -1,30 | 4,16 | -0,31 | 7,55E-01 |
| AT2G19740 | 0,67 | -0,33 | 2,40 | -0,14 | 8,91E-01 |
| AT2G19810 | 0,34 | 1,31 | 3,60 | 0,36 | 7,16E-01 |
| AT2G19880 | 0,75 | 1,31 | 2,54 | 0,52 | 6,07E-01 |
| AT2G20020 | 1,07 | 1,33 | 2,19 | 0,61 | 5,43E-01 |
| AT2G20100 | 0,33 | 1,38 | 4,41 | 0,31 | 7,54E-01 |
| AT2G20110 | 0,83 | 0,00 | 4,39 | 0,00 | 1,00E+00 |
| AT2G20142 | 0,28 | 1,34 | 4,40 | 0,30 | 7,61E-01 |
| AT2G20230 | 0,26 | 0,50 | 4,40 | 0,11 | 9,10E-01 |

| | | | | | |
|---|---|---|---|---|---|
| AT2G20260 | 1,52 | 0,58 | 1,50 | 0,39 | 6,98E-01 |
| AT2G20340 | 0,57 | -1,31 | 3,99 | -0,33 | 7,42E-01 |
| AT2G20360 | 0,22 | -0,44 | 4,39 | -0,10 | 9,20E-01 |
| AT2G20362 | 0,08 | 0,00 | 4,41 | 0,00 | 1,00E+00 |
| AT2G20370 | 1,01 | 3,44 | 2,52 | 1,37 | 1,72E-01 |
| AT2G20420 | 0,62 | -0,41 | 2,33 | -0,17 | 8,61E-01 |
| AT2G20450 | 0,40 | 0,43 | 4,21 | 0,10 | 9,19E-01 |
| AT2G20530 | 0,21 | -0,94 | 4,37 | -0,21 | 8,30E-01 |
| AT2G20690 | 1,05 | -1,81 | 2,58 | -0,70 | 4,82E-01 |
| AT2G20710 | 0,08 | 0,00 | 4,41 | 0,00 | 1,00E+00 |
| AT2G07335 | 0,46 | 1,23 | 4,17 | 0,29 | 7,69E-01 |
| AT2G20750 | 0,59 | 1,86 | 3,78 | 0,49 | 6,23E-01 |
| AT2G20770 | 0,17 | 0,00 | 4,41 | 0,00 | 1,00E+00 |
| AT2G20820 | 0,25 | 1,29 | 4,39 | 0,29 | 7,69E-01 |
| AT2G20860 | 1,36 | 1,25 | 1,73 | 0,72 | 4,69E-01 |
| AT2G20890 | 1,04 | -0,91 | 1,89 | -0,48 | 6,29E-01 |
| AT2G20920 | 0,65 | 2,89 | 2,50 | 1,16 | 2,48E-01 |
| AT2G20960 | 0,82 | 1,07 | 2,13 | 0,50 | 6,15E-01 |
| AT2G21160 | 0,91 | 0,62 | 3,17 | 0,20 | 8,44E-01 |
| AT2G21187 | 0,89 | 1,19 | 2,07 | 0,57 | 5,65E-01 |
| AT2G21190 | 0,23 | -1,17 | 4,37 | -0,27 | 7,88E-01 |
| AT2G21270 | 0,21 | 0,00 | 4,41 | 0,00 | 1,00E+00 |
| AT2G21380 | 1,04 | 0,32 | 1,89 | 0,17 | 8,67E-01 |
| AT2G21410 | 1,46 | 2,33 | 1,54 | 1,52 | 1,29E-01 |
| AT2G21520 | 0,84 | 0,35 | 2,28 | 0,15 | 8,78E-01 |
| AT2G21530 | 0,61 | -1,87 | 3,04 | -0,61 | 5,39E-01 |
| AT2G21550 | 0,95 | -3,37 | 2,59 | -1,30 | 1,95E-01 |
| AT2G21580 | 1,04 | 2,88 | 2,78 | 1,03 | 3,01E-01 |
| AT2G21620 | 1,16 | -1,47 | 2,04 | -0,72 | 4,71E-01 |
| AT2G21630 | 0,32 | 1,79 | 4,38 | 0,41 | 6,83E-01 |
| AT2G21770 | 0,18 | 1,38 | 4,41 | 0,31 | 7,54E-01 |
| AT2G21790 | 0,74 | 2,77 | 2,53 | 1,09 | 2,74E-01 |
| AT2G21850 | 1,02 | 3,49 | 2,57 | 1,36 | 1,75E-01 |
| AT2G07475 | 0,24 | 0,00 | 4,41 | 0,00 | 1,00E+00 |
| AT2G21960 | 0,80 | 2,42 | 2,57 | 0,94 | 3,45E-01 |
| AT2G07495 | 0,12 | 0,00 | 4,41 | 0,00 | 1,00E+00 |
| AT2G22080 | 0,65 | 0,38 | 2,69 | 0,14 | 8,87E-01 |
| AT2G22120 | 0,32 | 1,79 | 4,38 | 0,41 | 6,83E-01 |
| AT2G22425 | 0,10 | 0,00 | 4,41 | 0,00 | 1,00E+00 |
| AT2G22450 | 0,24 | 1,36 | 4,40 | 0,31 | 7,57E-01 |
| AT2G22480 | 0,23 | 1,27 | 4,39 | 0,29 | 7,72E-01 |
| AT2G22540 | 1,53 | 2,95 | 1,71 | 1,72 | 8,48E-02 |
| AT2G22670 | 0,80 | 1,37 | 2,49 | 0,55 | 5,81E-01 |
| AT2G22680 | 0,25 | 1,43 | 4,40 | 0,32 | 7,46E-01 |
| AT2G22720 | 0,99 | 3,34 | 2,71 | 1,23 | 2,18E-01 |
| AT2G07845 | 0,43 | 0,00 | 4,18 | 0,00 | 1,00E+00 |
| AT2G22790 | 0,08 | 0,00 | 4,41 | 0,00 | 1,00E+00 |
| AT2G22840 | 1,11 | 0,52 | 2,27 | 0,23 | 8,17E-01 |
| AT2G07850 | 0,72 | -2,36 | 2,05 | -1,15 | 2,49E-01 |
| AT2G22910 | 0,32 | 1,38 | 4,41 | 0,31 | 7,54E-01 |
| AT2G23080 | 0,38 | 1,26 | 4,39 | 0,29 | 7,74E-01 |
| AT2G23120 | 0,92 | 1,06 | 2,33 | 0,45 | 6,51E-01 |
| AT2G23130 | 0,41 | -1,29 | 2,86 | -0,45 | 6,52E-01 |
| AT2G23142 | 1,46 | -3,01 | 2,36 | -1,27 | 2,03E-01 |
| AT2G23200 | 1,64 | 2,01 | 1,67 | 1,20 | 2,31E-01 |
| AT2G23310 | 0,84 | 3,49 | 2,91 | 1,20 | 2,30E-01 |

| | | | | | |
|---|---|---|---|---|---|
| AT2G07895 | 0,15 | 0,00 | 4,41 | 0,00 | 1,00E+00 |
| AT2G23320 | 1,39 | 1,89 | 1,91 | 0,99 | 3,22E-01 |
| AT2G23420 | 0,47 | 2,10 | 4,36 | 0,48 | 6,31E-01 |
| AT2G23460 | 1,50 | 1,41 | 1,67 | 0,84 | 4,00E-01 |
| AT2G07915 | 1,63 | -3,07 | 2,35 | -1,30 | 1,93E-01 |
| AT2G23672 | 1,50 | -1,19 | 1,73 | -0,69 | 4,93E-01 |
| AT2G23680 | 0,31 | -0,57 | 4,40 | -0,13 | 8,97E-01 |
| AT2G23740 | 0,12 | 0,00 | 4,41 | 0,00 | 1,00E+00 |
| AT2G23890 | 0,37 | 0,35 | 3,54 | 0,10 | 9,22E-01 |
| AT2G23985 | 0,17 | -1,42 | 4,35 | -0,33 | 7,45E-01 |
| AT2G24090 | 0,67 | -0,14 | 2,62 | -0,05 | 9,57E-01 |
| AT2G24180 | 1,36 | -1,94 | 2,58 | -0,75 | 4,53E-01 |
| AT2G24240 | 0,31 | 0,00 | 4,40 | 0,00 | 1,00E+00 |
| AT2G24280 | 1,45 | -1,01 | 1,62 | -0,62 | 5,34E-01 |
| AT2G24300 | 0,15 | 0,00 | 4,41 | 0,00 | 1,00E+00 |
| AT2G08015 | 0,35 | 0,00 | 4,41 | 0,00 | 1,00E+00 |
| AT2G24350 | 0,50 | 2,75 | 3,97 | 0,69 | 4,89E-01 |
| AT2G24390 | 0,92 | -0,36 | 2,84 | -0,13 | 8,98E-01 |
| AT2G24380 | 0,65 | -0,94 | 3,73 | -0,25 | 8,02E-01 |
| AT2G24420 | 1,14 | -1,72 | 2,10 | -0,82 | 4,12E-01 |
| AT2G08045 | 0,23 | 0,00 | 4,41 | 0,00 | 1,00E+00 |
| AT2G08050 | 0,32 | 0,00 | 4,41 | 0,00 | 1,00E+00 |
| AT2G24540 | 0,88 | 1,12 | 1,78 | 0,63 | 5,26E-01 |
| AT2G24590 | 0,49 | 2,50 | 3,96 | 0,63 | 5,27E-01 |
| AT2G24592 | 0,83 | 2,18 | 2,26 | 0,96 | 3,36E-01 |
| AT2G24765 | 0,69 | 2,15 | 2,95 | 0,73 | 4,66E-01 |
| AT2G25000 | 0,52 | 0,00 | 3,92 | 0,00 | 1,00E+00 |
| AT2G25080 | 0,61 | -0,57 | 2,71 | -0,21 | 8,33E-01 |
| AT2G25110 | 0,19 | 0,00 | 4,41 | 0,00 | 1,00E+00 |
| AT2G25190 | 0,14 | 0,00 | 4,41 | 0,00 | 1,00E+00 |
| AT2G25370 | 1,47 | -2,02 | 2,10 | -0,96 | 3,35E-01 |
| AT2G25410 | 1,30 | 0,00 | 2,46 | 0,00 | 1,00E+00 |
| AT2G25480 | 0,05 | 0,00 | 4,41 | 0,00 | 1,00E+00 |
| AT2G25560 | 0,73 | 1,38 | 2,94 | 0,47 | 6,38E-01 |
| AT2G25580 | 0,08 | 0,00 | 4,41 | 0,00 | 1,00E+00 |
| AT2G25590 | 0,21 | 0,00 | 4,41 | 0,00 | 1,00E+00 |
| AT2G25625 | 0,15 | 0,00 | 4,41 | 0,00 | 1,00E+00 |
| AT2G25630 | 0,50 | 0,00 | 3,23 | 0,00 | 1,00E+00 |
| AT2G25650 | 0,70 | 2,13 | 2,96 | 0,72 | 4,72E-01 |
| AT2G25790 | 0,18 | 0,00 | 4,41 | 0,00 | 1,00E+00 |
| AT2G25840 | 0,59 | 1,89 | 3,97 | 0,47 | 6,35E-01 |
| AT2G25870 | 1,69 | 3,72 | 1,82 | 2,04 | 4,10E-02 |
| AT2G25930 | 1,26 | 2,01 | 1,83 | 1,10 | 2,70E-01 |
| AT2G26060 | 1,00 | 0,72 | 4,34 | 0,17 | 8,68E-01 |
| AT2G26110 | 0,61 | 1,36 | 3,05 | 0,45 | 6,54E-01 |
| AT2G26170 | 0,47 | 1,91 | 3,28 | 0,58 | 5,59E-01 |
| AT2G26270 | 0,21 | 1,27 | 4,39 | 0,29 | 7,72E-01 |
| AT2G26300 | 1,09 | 0,46 | 2,63 | 0,17 | 8,62E-01 |
| AT2G26340 | 0,57 | 2,70 | 3,08 | 0,87 | 3,82E-01 |
| AT2G26440 | 1,40 | -2,02 | 1,97 | -1,03 | 3,05E-01 |
| AT2G26510 | 0,24 | 1,36 | 4,40 | 0,31 | 7,57E-01 |
| AT2G26570 | 0,43 | 0,09 | 3,32 | 0,03 | 9,79E-01 |
| AT2G26600 | 0,45 | 1,38 | 3,31 | 0,42 | 6,77E-01 |
| AT2G26610 | 0,30 | 1,30 | 4,40 | 0,30 | 7,68E-01 |
| AT2G26640 | 0,71 | -1,95 | 3,62 | -0,54 | 5,90E-01 |
| AT2G26660 | 0,22 | 1,33 | 4,40 | 0,30 | 7,63E-01 |

| | | | | | |
|---|---|---|---|---|---|
| AT2G26692 | 0,08 | 0,00 | 4,41 | 0,00 | 1,00E+00 |
| AT2G26730 | 0,85 | -2,50 | 2,18 | -1,15 | 2,51E-01 |
| AT2G26920 | 0,60 | 0,54 | 3,16 | 0,17 | 8,64E-01 |
| AT2G27030 | 0,72 | 2,35 | 2,67 | 0,88 | 3,78E-01 |
| AT2G27050 | 0,99 | 2,27 | 2,05 | 1,11 | 2,67E-01 |
| AT2G27060 | 0,17 | 0,00 | 4,41 | 0,00 | 1,00E+00 |
| AT2G27080 | 0,39 | 0,00 | 4,26 | 0,00 | 1,00E+00 |
| AT2G27100 | 0,72 | 2,87 | 3,56 | 0,81 | 4,20E-01 |
| AT2G27130 | 0,30 | 1,38 | 4,41 | 0,31 | 7,54E-01 |
| AT2G27200 | 0,21 | 0,00 | 4,41 | 0,00 | 1,00E+00 |
| AT2G27210 | 0,89 | -1,02 | 2,47 | -0,41 | 6,79E-01 |
| AT2G27230 | 1,31 | 2,90 | 1,70 | 1,71 | 8,82E-02 |
| AT2G27260 | 0,40 | 1,34 | 3,42 | 0,39 | 6,96E-01 |
| AT2G27320 | 0,85 | -0,43 | 2,50 | -0,17 | 8,65E-01 |
| AT2G27380 | 0,13 | 0,00 | 4,41 | 0,00 | 1,00E+00 |
| AT2G27460 | 0,28 | -1,24 | 4,36 | -0,28 | 7,77E-01 |
| AT2G27540 | 0,95 | 0,00 | 3,24 | 0,00 | 1,00E+00 |
| AT2G27570 | 0,28 | 1,84 | 4,39 | 0,42 | 6,74E-01 |
| AT2G27600 | 1,39 | 1,24 | 2,07 | 0,60 | 5,49E-01 |
| AT2G27690 | 0,16 | 0,00 | 4,41 | 0,00 | 1,00E+00 |
| AT2G27710 | 0,57 | 1,15 | 3,25 | 0,36 | 7,22E-01 |
| AT2G27760 | 0,08 | 0,00 | 4,41 | 0,00 | 1,00E+00 |
| AT2G27810 | 0,54 | 0,62 | 2,79 | 0,22 | 8,24E-01 |
| AT2G27820 | 0,23 | -0,54 | 4,41 | -0,12 | 9,02E-01 |
| AT2G27900 | 1,32 | 2,59 | 1,71 | 1,51 | 1,31E-01 |
| AT2G28070 | 0,87 | 2,97 | 2,85 | 1,04 | 2,97E-01 |
| AT2G28120 | 0,29 | 1,32 | 4,40 | 0,30 | 7,64E-01 |
| AT2G28130 | 0,59 | 0,00 | 4,40 | 0,00 | 1,00E+00 |
| AT2G28190 | 0,42 | 1,96 | 4,35 | 0,45 | 6,52E-01 |
| AT2G28250 | 0,08 | 0,00 | 4,41 | 0,00 | 1,00E+00 |
| AT2G28310 | 0,30 | 1,61 | 4,39 | 0,37 | 7,14E-01 |
| AT2G28320 | 0,43 | 0,00 | 4,23 | 0,00 | 1,00E+00 |
| AT2G28350 | 0,96 | 2,62 | 2,13 | 1,23 | 2,18E-01 |
| AT2G28370 | 0,25 | 1,29 | 4,39 | 0,29 | 7,69E-01 |
| AT2G28460 | 0,18 | -0,43 | 4,39 | -0,10 | 9,21E-01 |
| AT2G28520 | 0,65 | 1,73 | 2,97 | 0,58 | 5,61E-01 |
| AT2G28540 | 0,87 | 2,31 | 2,30 | 1,00 | 3,15E-01 |
| AT2G28605 | 0,37 | 0,19 | 4,38 | 0,04 | 9,66E-01 |
| AT2G28660 | 0,10 | 0,00 | 4,41 | 0,00 | 1,00E+00 |
| AT2G28671 | 0,51 | -0,46 | 3,16 | -0,15 | 8,85E-01 |
| AT2G28710 | 0,24 | 0,00 | 4,41 | 0,00 | 1,00E+00 |
| AT2G28720 | 0,49 | -0,19 | 4,21 | -0,05 | 9,64E-01 |
| AT2G28730 | 1,02 | 2,27 | 1,80 | 1,26 | 2,09E-01 |
| AT2G28810 | 0,71 | 2,13 | 2,67 | 0,80 | 4,25E-01 |
| AT2G28930 | 0,12 | 0,00 | 4,41 | 0,00 | 1,00E+00 |
| AT2G29020 | 0,11 | 0,00 | 4,41 | 0,00 | 1,00E+00 |
| AT2G29065 | 0,30 | 1,38 | 4,41 | 0,31 | 7,54E-01 |
| AT2G29070 | 0,15 | 0,00 | 4,41 | 0,00 | 1,00E+00 |
| AT2G29140 | 0,94 | 0,30 | 2,62 | 0,12 | 9,08E-01 |
| AT2G29180 | 0,47 | 0,38 | 4,12 | 0,09 | 9,27E-01 |
| AT2G29210 | 1,16 | 3,55 | 2,13 | 1,67 | 9,54E-02 |
| AT2G29310 | 0,46 | 0,98 | 3,28 | 0,30 | 7,66E-01 |
| AT2G29350 | 0,65 | 0,00 | 3,95 | 0,00 | 1,00E+00 |
| AT2G29360 | 0,51 | 2,52 | 4,35 | 0,58 | 5,63E-01 |
| AT2G29510 | 0,82 | 1,81 | 3,40 | 0,53 | 5,94E-01 |
| AT2G29600 | 0,47 | 0,00 | 3,29 | 0,00 | 1,00E+00 |

| | | | | | |
|---|---|---|---|---|---|
| AT2G29640 | 0,37 | 2,13 | 4,32 | 0,49 | 6,22E-01 |
| AT2G29660 | 0,20 | 1,38 | 4,41 | 0,31 | 7,54E-01 |
| AT2G29690 | 0,14 | -0,48 | 4,40 | -0,11 | 9,13E-01 |
| AT2G29700 | 0,24 | -0,56 | 4,41 | -0,13 | 8,99E-01 |
| AT2G29970 | 1,50 | -2,95 | 1,67 | -1,76 | 7,84E-02 |
| AT2G30020 | 0,83 | 0,00 | 3,61 | 0,00 | 1,00E+00 |
| AT2G30040 | 0,38 | 1,32 | 3,43 | 0,39 | 6,99E-01 |
| AT2G30050 | 0,66 | 0,38 | 2,54 | 0,15 | 8,80E-01 |
| AT2G30140 | 0,54 | 1,34 | 3,93 | 0,34 | 7,33E-01 |
| AT2G30150 | 0,10 | 0,00 | 4,41 | 0,00 | 1,00E+00 |
| AT2G30180 | 1,06 | 0,51 | 1,85 | 0,27 | 7,85E-01 |
| AT2G08375 | 1,24 | 0,07 | 2,02 | 0,03 | 9,72E-01 |
| AT2G30230 | 0,50 | -0,52 | 3,31 | -0,16 | 8,75E-01 |
| AT2G30362 | 0,41 | 0,00 | 4,25 | 0,00 | 1,00E+00 |
| AT2G30410 | 0,72 | -0,13 | 2,42 | -0,06 | 9,56E-01 |
| AT2G30530 | 0,20 | 1,38 | 4,41 | 0,31 | 7,54E-01 |
| AT2G30550 | 1,44 | 1,68 | 1,85 | 0,91 | 3,64E-01 |
| AT2G30600 | 0,51 | -0,06 | 4,30 | -0,01 | 9,89E-01 |
| AT2G30620 | 0,85 | -2,00 | 2,39 | -0,84 | 4,03E-01 |
| AT2G30730 | 0,89 | 0,33 | 2,60 | 0,13 | 9,00E-01 |
| AT2G30780 | 0,62 | -0,93 | 3,77 | -0,25 | 8,05E-01 |
| AT2G30870 | 1,65 | -1,32 | 1,74 | -0,76 | 4,50E-01 |
| AT2G30930 | 0,49 | 1,28 | 2,70 | 0,47 | 6,36E-01 |
| AT2G30970 | 0,46 | 2,03 | 4,38 | 0,46 | 6,42E-01 |
| AT2G31010 | 0,95 | 3,21 | 2,31 | 1,39 | 1,65E-01 |
| AT2G31020 | 0,26 | 1,38 | 4,41 | 0,31 | 7,54E-01 |
| AT2G31200 | 0,16 | 0,00 | 4,41 | 0,00 | 1,00E+00 |
| AT2G31215 | 0,52 | -0,83 | 2,69 | -0,31 | 7,58E-01 |
| AT2G31280 | 1,47 | 3,12 | 2,15 | 1,45 | 1,47E-01 |
| AT2G31290 | 0,71 | 1,81 | 3,11 | 0,58 | 5,61E-01 |
| AT2G31350 | 0,38 | 1,38 | 4,36 | 0,32 | 7,52E-01 |
| AT2G31370 | 0,19 | -0,52 | 4,40 | -0,12 | 9,06E-01 |
| AT2G31380 | 1,06 | 2,63 | 1,93 | 1,37 | 1,72E-01 |
| AT2G08435 | 0,79 | 2,38 | 3,52 | 0,67 | 5,00E-01 |
| AT2G31560 | 0,67 | 0,39 | 2,61 | 0,15 | 8,83E-01 |
| AT2G31585 | 0,47 | 0,28 | 4,01 | 0,07 | 9,44E-01 |
| AT2G31750 | 0,75 | -1,03 | 3,07 | -0,34 | 7,36E-01 |
| AT2G31790 | 0,30 | 0,00 | 3,71 | 0,00 | 1,00E+00 |
| AT2G31810 | 1,10 | 2,47 | 2,07 | 1,19 | 2,34E-01 |
| AT2G31960 | 1,68 | 1,73 | 1,66 | 1,04 | 2,97E-01 |
| AT2G31970 | 0,18 | 0,00 | 4,41 | 0,00 | 1,00E+00 |
| AT2G32010 | 0,37 | -0,46 | 4,40 | -0,10 | 9,16E-01 |
| AT2G32040 | 0,08 | 0,00 | 4,41 | 0,00 | 1,00E+00 |
| AT2G32100 | 1,15 | -1,26 | 2,27 | -0,55 | 5,80E-01 |
| AT2G32150 | 1,00 | 2,17 | 2,55 | 0,85 | 3,95E-01 |
| AT2G32250 | 0,51 | 0,96 | 3,22 | 0,30 | 7,65E-01 |
| AT2G32260 | 0,44 | 1,38 | 4,16 | 0,33 | 7,40E-01 |
| AT2G32295 | 0,53 | -0,26 | 2,78 | -0,09 | 9,26E-01 |
| AT2G32320 | 0,15 | 0,00 | 4,41 | 0,00 | 1,00E+00 |
| AT2G32440 | 0,16 | 0,00 | 4,41 | 0,00 | 1,00E+00 |
| AT2G32450 | 0,56 | 1,38 | 2,73 | 0,51 | 6,13E-01 |
| AT2G32698 | 0,12 | 0,00 | 4,41 | 0,00 | 1,00E+00 |
| AT2G32700 | 1,61 | 0,30 | 1,52 | 0,20 | 8,43E-01 |
| AT2G32800 | 1,51 | 0,83 | 1,50 | 0,56 | 5,79E-01 |
| AT2G32900 | 0,29 | 1,27 | 4,39 | 0,29 | 7,72E-01 |
| AT2G32910 | 0,68 | 1,16 | 3,04 | 0,38 | 7,02E-01 |

| | | | | | |
|---|---|---|---|---|---|
| AT2G33080 | 0,74 | 1,37 | 3,58 | 0,38 | 7,01E-01 |
| AT2G33160 | 0,25 | 1,34 | 4,40 | 0,30 | 7,61E-01 |
| AT2G33220 | 0,54 | 2,36 | 3,20 | 0,74 | 4,61E-01 |
| AT2G33250 | 0,76 | -0,95 | 4,37 | -0,22 | 8,29E-01 |
| AT2G33290 | 0,29 | -0,51 | 3,75 | -0,14 | 8,91E-01 |
| AT2G33390 | 0,25 | 1,43 | 4,40 | 0,32 | 7,46E-01 |
| AT2G33420 | 0,37 | 2,13 | 4,32 | 0,49 | 6,22E-01 |
| AT2G08585 | 0,32 | -0,55 | 3,73 | -0,15 | 8,83E-01 |
| AT2G33435 | 1,31 | 0,69 | 1,61 | 0,43 | 6,68E-01 |
| AT2G33450 | 0,32 | -0,89 | 4,38 | -0,20 | 8,39E-01 |
| AT2G33490 | 0,35 | 1,30 | 4,37 | 0,30 | 7,66E-01 |
| AT2G33510 | 0,26 | 1,33 | 4,40 | 0,30 | 7,63E-01 |
| AT2G33570 | 0,17 | 0,00 | 4,41 | 0,00 | 1,00E+00 |
| AT2G33610 | 0,37 | 1,27 | 4,33 | 0,29 | 7,68E-01 |
| AT2G33650 | 0,29 | -1,22 | 4,36 | -0,28 | 7,80E-01 |
| AT2G33700 | 0,56 | 1,98 | 4,05 | 0,49 | 6,24E-01 |
| AT2G33760 | 0,64 | 2,36 | 4,36 | 0,54 | 5,88E-01 |
| AT2G33810 | 0,37 | 0,37 | 4,34 | 0,09 | 9,32E-01 |
| AT2G33840 | 0,52 | 1,02 | 3,22 | 0,32 | 7,51E-01 |
| AT2G33845 | 0,47 | 0,00 | 4,13 | 0,00 | 1,00E+00 |
| AT2G33950 | 0,39 | 2,16 | 4,32 | 0,50 | 6,17E-01 |
| AT2G34120 | 0,27 | 0,00 | 4,41 | 0,00 | 1,00E+00 |
| AT2G34240 | 0,12 | 0,00 | 4,41 | 0,00 | 1,00E+00 |
| AT2G34460 | 0,99 | 1,28 | 2,04 | 0,63 | 5,29E-01 |
| AT2G34660 | 0,69 | -0,34 | 2,72 | -0,13 | 9,00E-01 |
| AT2G34730 | 0,50 | -0,16 | 2,75 | -0,06 | 9,53E-01 |
| AT2G34770 | 0,22 | 1,33 | 4,40 | 0,30 | 7,63E-01 |
| AT2G34780 | 0,99 | 2,87 | 2,19 | 1,31 | 1,90E-01 |
| AT2G34810 | 0,16 | 0,00 | 4,41 | 0,00 | 1,00E+00 |
| AT2G34860 | 1,10 | 3,96 | 2,19 | 1,81 | 7,09E-02 |
| AT2G34970 | 0,22 | 1,27 | 4,39 | 0,29 | 7,72E-01 |
| AT2G35020 | 0,32 | 1,45 | 4,40 | 0,33 | 7,41E-01 |
| AT2G35170 | 0,10 | 0,00 | 4,41 | 0,00 | 1,00E+00 |
| AT2G35230 | 0,50 | 1,74 | 4,37 | 0,40 | 6,90E-01 |
| AT2G35330 | 0,35 | 2,02 | 4,38 | 0,46 | 6,45E-01 |
| AT2G35340 | 0,80 | 3,33 | 2,76 | 1,21 | 2,27E-01 |
| AT2G35370 | 0,43 | 0,70 | 4,33 | 0,16 | 8,72E-01 |
| AT2G35630 | 1,16 | 0,76 | 1,77 | 0,43 | 6,65E-01 |
| AT2G35635 | 0,64 | 0,55 | 4,28 | 0,13 | 8,97E-01 |
| AT2G35650 | 0,68 | 1,95 | 2,23 | 0,88 | 3,81E-01 |
| AT2G08750 | 0,08 | 0,00 | 4,41 | 0,00 | 1,00E+00 |
| AT2G35720 | 0,19 | 0,00 | 4,41 | 0,00 | 1,00E+00 |
| AT2G35760 | 0,17 | 1,32 | 4,40 | 0,30 | 7,64E-01 |
| AT2G35800 | 1,02 | 0,98 | 1,82 | 0,54 | 5,88E-01 |
| AT2G35820 | 0,71 | 1,38 | 4,41 | 0,31 | 7,54E-01 |
| AT2G35830 | 1,03 | 2,11 | 2,13 | 0,99 | 3,21E-01 |
| AT2G35840 | 0,63 | 2,55 | 2,54 | 1,00 | 3,16E-01 |
| AT2G35860 | 0,74 | -0,37 | 2,43 | -0,15 | 8,79E-01 |
| AT2G35920 | 0,77 | 1,21 | 2,48 | 0,49 | 6,26E-01 |
| AT2G35935 | 0,49 | 0,00 | 3,98 | 0,00 | 1,00E+00 |
| AT2G35960 | 0,20 | 0,00 | 4,41 | 0,00 | 1,00E+00 |
| AT2G36000 | 0,80 | 0,45 | 2,70 | 0,17 | 8,68E-01 |
| AT2G36010 | 0,59 | 0,82 | 3,01 | 0,27 | 7,86E-01 |
| AT2G36026 | 0,70 | -1,64 | 3,80 | -0,43 | 6,66E-01 |
| AT2G36090 | 0,69 | 0,00 | 3,66 | 0,00 | 1,00E+00 |
| AT2G36130 | 0,56 | 1,98 | 4,38 | 0,45 | 6,50E-01 |

| | | | | | |
|---|---|---|---|---|---|
| AT2G36320 | 1,39 | -1,52 | 1,86 | -0,82 | 4,14E-01 |
| AT2G36330 | 0,55 | 0,00 | 4,39 | 0,00 | 1,00E+00 |
| AT2G36430 | 1,44 | 0,22 | 2,11 | 0,10 | 9,18E-01 |
| AT2G36500 | 0,54 | 2,87 | 3,08 | 0,93 | 3,53E-01 |
| AT2G36510 | 0,37 | 2,11 | 4,32 | 0,49 | 6,26E-01 |
| AT2G36570 | 0,52 | 3,01 | 4,32 | 0,70 | 4,87E-01 |
| AT2G36580 | 1,14 | 0,81 | 1,77 | 0,46 | 6,49E-01 |
| AT2G36600 | 0,66 | -1,49 | 2,21 | -0,68 | 4,99E-01 |
| AT2G36720 | 1,25 | 3,80 | 2,07 | 1,83 | 6,67E-02 |
| AT2G36792 | 0,34 | 0,00 | 4,41 | 0,00 | 1,00E+00 |
| AT2G36830 | 0,86 | 2,28 | 2,21 | 1,03 | 3,02E-01 |
| AT2G36920 | 0,12 | 0,00 | 4,41 | 0,00 | 1,00E+00 |
| AT2G36950 | 0,34 | 1,38 | 4,41 | 0,31 | 7,54E-01 |
| AT2G36960 | 0,18 | 0,00 | 4,41 | 0,00 | 1,00E+00 |
| AT2G36970 | 0,19 | 0,00 | 4,41 | 0,00 | 1,00E+00 |
| AT2G36980 | 0,05 | 0,00 | 4,41 | 0,00 | 1,00E+00 |
| AT2G37025 | 0,19 | -1,18 | 4,36 | -0,27 | 7,86E-01 |
| AT2G37050 | 0,93 | 2,21 | 2,31 | 0,96 | 3,39E-01 |
| AT2G37110 | 1,16 | 1,31 | 2,48 | 0,53 | 5,97E-01 |
| AT2G37180 | 0,08 | 0,00 | 4,41 | 0,00 | 1,00E+00 |
| AT2G37250 | 0,74 | 0,35 | 2,37 | 0,15 | 8,81E-01 |
| AT2G08860 | 0,31 | -0,55 | 3,67 | -0,15 | 8,82E-01 |
| AT2G37320 | 0,08 | 0,00 | 4,41 | 0,00 | 1,00E+00 |
| AT2G37330 | 0,66 | 2,41 | 3,10 | 0,77 | 4,38E-01 |
| AT2G37362 | 0,21 | 0,00 | 4,41 | 0,00 | 1,00E+00 |
| AT2G37370 | 0,40 | 0,00 | 4,41 | 0,00 | 1,00E+00 |
| AT2G37470 | 0,26 | 0,00 | 4,41 | 0,00 | 1,00E+00 |
| AT2G37480 | 1,35 | 1,63 | 2,27 | 0,72 | 4,72E-01 |
| AT2G37555 | 0,61 | 1,38 | 3,09 | 0,45 | 6,56E-01 |
| AT2G37580 | 0,12 | 0,00 | 4,41 | 0,00 | 1,00E+00 |
| AT2G37650 | 1,11 | 3,19 | 2,12 | 1,50 | 1,33E-01 |
| AT2G37690 | 0,31 | 0,45 | 4,40 | 0,10 | 9,18E-01 |
| AT2G37770 | 0,28 | 0,00 | 4,41 | 0,00 | 1,00E+00 |
| AT2G37860 | 0,83 | 3,04 | 2,25 | 1,35 | 1,78E-01 |
| AT2G37940 | 0,73 | 1,71 | 2,98 | 0,57 | 5,66E-01 |
| AT2G37970 | 0,77 | -0,46 | 2,83 | -0,16 | 8,71E-01 |
| AT2G37990 | 0,11 | 0,00 | 4,41 | 0,00 | 1,00E+00 |
| AT2G38010 | 0,53 | 1,41 | 3,17 | 0,45 | 6,56E-01 |
| AT2G38090 | 0,20 | 1,33 | 4,40 | 0,30 | 7,63E-01 |
| AT2G38120 | 0,38 | -0,46 | 2,91 | -0,16 | 8,74E-01 |
| AT2G38140 | 0,26 | -0,48 | 3,83 | -0,13 | 9,00E-01 |
| AT2G38210 | 1,49 | -2,19 | 2,02 | -1,09 | 2,78E-01 |
| AT2G38310 | 0,71 | 2,05 | 2,97 | 0,69 | 4,91E-01 |
| AT2G38365 | 0,20 | 1,38 | 4,41 | 0,31 | 7,54E-01 |
| AT2G38420 | 0,26 | 1,38 | 4,41 | 0,31 | 7,54E-01 |
| AT2G38470 | 1,30 | 2,20 | 1,90 | 1,16 | 2,47E-01 |
| AT2G38560 | 1,56 | 3,21 | 2,02 | 1,59 | 1,12E-01 |
| AT2G38590 | 0,78 | 0,00 | 2,87 | 0,00 | 1,00E+00 |
| AT2G38670 | 0,86 | -0,18 | 2,15 | -0,08 | 9,35E-01 |
| AT2G38760 | 0,08 | 0,00 | 4,41 | 0,00 | 1,00E+00 |
| AT2G38780 | 0,82 | 0,49 | 1,93 | 0,25 | 8,01E-01 |
| AT2G38820 | 0,12 | 0,00 | 4,41 | 0,00 | 1,00E+00 |
| AT2G38840 | 0,33 | 1,66 | 4,38 | 0,38 | 7,05E-01 |
| AT2G38940 | 0,25 | 1,30 | 4,40 | 0,30 | 7,67E-01 |
| AT2G38950 | 0,65 | 2,11 | 2,58 | 0,82 | 4,15E-01 |
| AT2G39190 | 0,15 | 0,00 | 4,41 | 0,00 | 1,00E+00 |

| | | | | | |
|---|---|---|---|---|---|
| AT2G39230 | 0,19 | 0,00 | 4,40 | 0,00 | 1,00E+00 |
| AT2G39270 | 0,08 | 0,00 | 4,41 | 0,00 | 1,00E+00 |
| AT2G39340 | 0,19 | 0,00 | 4,40 | 0,00 | 1,00E+00 |
| AT2G39400 | 0,15 | 0,00 | 4,41 | 0,00 | 1,00E+00 |
| AT2G39420 | 0,56 | 1,34 | 3,13 | 0,43 | 6,69E-01 |
| AT2G39430 | 0,52 | 1,38 | 3,96 | 0,35 | 7,27E-01 |
| AT2G39460 | 0,89 | 0,39 | 2,13 | 0,18 | 8,54E-01 |
| AT2G39470 | 0,81 | 0,46 | 2,31 | 0,20 | 8,43E-01 |
| AT2G39570 | 1,60 | 2,58 | 2,87 | 0,90 | 3,69E-01 |
| AT2G39580 | 1,16 | 2,95 | 2,06 | 1,43 | 1,51E-01 |
| AT2G39670 | 1,03 | -0,55 | 2,42 | -0,23 | 8,20E-01 |
| AT2G39725 | 0,50 | 2,34 | 3,95 | 0,59 | 5,55E-01 |
| AT2G39740 | 0,63 | 2,09 | 3,08 | 0,68 | 4,97E-01 |
| AT2G39780 | 1,17 | 1,52 | 2,19 | 0,69 | 4,89E-01 |
| AT2G39810 | 1,48 | 4,03 | 2,41 | 1,67 | 9,43E-02 |
| AT2G39840 | 0,84 | 2,15 | 2,61 | 0,82 | 4,10E-01 |
| AT2G39850 | 0,54 | -1,21 | 4,38 | -0,28 | 7,83E-01 |
| AT2G39870 | 0,43 | -0,03 | 4,34 | -0,01 | 9,94E-01 |
| AT2G39930 | 1,27 | 1,85 | 1,69 | 1,09 | 2,74E-01 |
| AT2G40060 | 0,99 | 1,37 | 2,58 | 0,53 | 5,94E-01 |
| AT2G40095 | 0,64 | 2,17 | 3,16 | 0,69 | 4,92E-01 |
| AT2G40130 | 0,14 | 0,00 | 4,41 | 0,00 | 1,00E+00 |
| AT2G40150 | 0,08 | 0,00 | 4,41 | 0,00 | 1,00E+00 |
| AT2G40160 | 0,42 | 2,17 | 4,16 | 0,52 | 6,03E-01 |
| AT2G40200 | 0,38 | 1,38 | 4,41 | 0,31 | 7,54E-01 |
| AT2G09120 | 0,31 | -2,02 | 4,33 | -0,47 | 6,41E-01 |
| AT2G40310 | 0,83 | -0,52 | 2,43 | -0,21 | 8,30E-01 |
| AT2G40400 | 0,82 | 1,46 | 2,81 | 0,52 | 6,04E-01 |
| AT2G40410 | 0,54 | 2,93 | 4,05 | 0,72 | 4,69E-01 |
| AT2G40435 | 0,46 | 0,93 | 4,35 | 0,21 | 8,30E-01 |
| AT2G40460 | 0,63 | -1,62 | 2,57 | -0,63 | 5,28E-01 |
| AT2G40480 | 0,31 | 1,38 | 4,41 | 0,31 | 7,54E-01 |
| AT2G40490 | 1,23 | -0,58 | 2,01 | -0,29 | 7,74E-01 |
| AT2G40540 | 0,78 | -0,07 | 2,55 | -0,03 | 9,80E-01 |
| AT2G40550 | 0,20 | 1,38 | 4,41 | 0,31 | 7,54E-01 |
| AT2G40660 | 0,08 | 0,00 | 4,41 | 0,00 | 1,00E+00 |
| AT2G40670 | 0,10 | 0,00 | 4,41 | 0,00 | 1,00E+00 |
| AT2G40760 | 0,35 | -0,54 | 4,41 | -0,12 | 9,02E-01 |
| AT2G40765 | 0,08 | 0,00 | 4,41 | 0,00 | 1,00E+00 |
| AT2G40810 | 0,08 | 0,00 | 4,41 | 0,00 | 1,00E+00 |
| AT2G40830 | 0,33 | 1,38 | 4,41 | 0,31 | 7,54E-01 |
| AT2G40880 | 1,07 | 0,23 | 2,17 | 0,10 | 9,17E-01 |
| AT2G40955 | 1,38 | 1,38 | 2,15 | 0,64 | 5,20E-01 |
| AT2G40980 | 0,41 | 0,93 | 4,21 | 0,22 | 8,26E-01 |
| AT2G41080 | 0,59 | 2,40 | 2,68 | 0,89 | 3,72E-01 |
| AT2G41110 | 0,26 | -0,51 | 3,85 | -0,13 | 8,95E-01 |
| AT2G09230 | 0,12 | 0,00 | 4,41 | 0,00 | 1,00E+00 |
| AT2G41140 | 0,37 | 1,38 | 4,35 | 0,32 | 7,51E-01 |
| AT2G41200 | 0,84 | 0,95 | 2,24 | 0,42 | 6,71E-01 |
| AT2G41210 | 0,42 | 1,55 | 4,19 | 0,37 | 7,11E-01 |
| AT2G41220 | 0,64 | 1,40 | 3,08 | 0,45 | 6,51E-01 |
| AT2G41460 | 0,13 | 0,00 | 4,41 | 0,00 | 1,00E+00 |
| AT2G41510 | 0,08 | 0,00 | 4,41 | 0,00 | 1,00E+00 |
| AT2G41520 | 0,59 | 2,48 | 3,02 | 0,82 | 4,12E-01 |
| AT2G41590 | 1,66 | 0,20 | 1,65 | 0,12 | 9,04E-01 |
| AT2G41670 | 0,12 | 0,00 | 4,41 | 0,00 | 1,00E+00 |

| | | | | | |
|---|---|---|---|---|---|
| AT2G41730 | 0,99 | 3,40 | 2,06 | 1,65 | 9,91E-02 |
| AT2G41820 | 0,34 | 1,38 | 4,41 | 0,31 | 7,54E-01 |
| AT2G41880 | 0,26 | -0,51 | 3,85 | -0,13 | 8,95E-01 |
| AT2G41920 | 1,42 | 1,36 | 2,20 | 0,62 | 5,37E-01 |
| AT2G42130 | 0,48 | -0,08 | 3,31 | -0,02 | 9,81E-01 |
| AT2G42220 | 0,98 | -1,45 | 2,03 | -0,71 | 4,75E-01 |
| AT2G42310 | 0,61 | 0,44 | 2,65 | 0,16 | 8,69E-01 |
| AT2G42320 | 0,63 | -0,43 | 2,69 | -0,16 | 8,74E-01 |
| AT2G42430 | 0,21 | 0,00 | 4,41 | 0,00 | 1,00E+00 |
| AT2G42580 | 1,32 | 1,06 | 1,93 | 0,55 | 5,84E-01 |
| AT2G42610 | 0,29 | -0,57 | 4,40 | -0,13 | 8,97E-01 |
| AT2G42620 | 1,45 | 0,97 | 2,56 | 0,38 | 7,04E-01 |
| AT2G42700 | 0,72 | 2,17 | 3,09 | 0,70 | 4,82E-01 |
| AT2G42710 | 0,12 | 0,00 | 4,41 | 0,00 | 1,00E+00 |
| AT2G42750 | 0,57 | 1,33 | 3,84 | 0,35 | 7,29E-01 |
| AT2G42780 | 0,28 | -0,54 | 4,41 | -0,12 | 9,02E-01 |
| AT2G42890 | 0,32 | -0,45 | 4,39 | -0,10 | 9,19E-01 |
| AT2G42900 | 0,43 | 2,05 | 4,36 | 0,47 | 6,39E-01 |
| AT2G43010 | 1,28 | 0,81 | 1,70 | 0,48 | 6,33E-01 |
| AT2G43020 | 1,32 | 1,38 | 2,29 | 0,60 | 5,48E-01 |
| AT2G43030 | 0,58 | -0,21 | 3,21 | -0,07 | 9,47E-01 |
| AT2G43060 | 1,34 | -2,09 | 1,70 | -1,23 | 2,20E-01 |
| AT2G43080 | 0,63 | 1,38 | 3,18 | 0,43 | 6,65E-01 |
| AT2G43090 | 0,61 | -0,33 | 2,76 | -0,12 | 9,05E-01 |
| AT2G43100 | 0,56 | 0,32 | 3,40 | 0,09 | 9,26E-01 |
| AT2G43137 | 1,43 | -0,72 | 1,94 | -0,37 | 7,12E-01 |
| AT2G43138 | 1,21 | 1,38 | 2,26 | 0,61 | 5,42E-01 |
| AT2G43141 | 1,64 | 0,24 | 1,86 | 0,13 | 8,99E-01 |
| AT2G43210 | 0,63 | 0,53 | 3,05 | 0,17 | 8,63E-01 |
| AT2G43230 | 0,55 | 2,59 | 3,83 | 0,68 | 4,98E-01 |
| AT2G43250 | 0,31 | 1,29 | 4,39 | 0,29 | 7,70E-01 |
| AT2G43280 | 0,56 | 2,27 | 4,37 | 0,52 | 6,03E-01 |
| AT2G09440 | 0,08 | 0,00 | 4,41 | 0,00 | 1,00E+00 |
| AT2G43340 | 0,44 | 2,29 | 3,40 | 0,67 | 5,00E-01 |
| AT2G43400 | 0,37 | 1,38 | 4,38 | 0,31 | 7,53E-01 |
| AT2G43440 | 0,11 | 0,00 | 4,41 | 0,00 | 1,00E+00 |
| AT2G43540 | 1,53 | 1,10 | 2,09 | 0,52 | 6,00E-01 |
| AT2G43630 | 0,44 | 0,00 | 4,14 | 0,00 | 1,00E+00 |
| AT2G43650 | 0,83 | -0,04 | 2,60 | -0,02 | 9,87E-01 |
| AT2G43690 | 0,68 | 1,34 | 4,40 | 0,31 | 7,60E-01 |
| AT2G43790 | 0,39 | -1,30 | 2,89 | -0,45 | 6,53E-01 |
| AT2G43800 | 0,08 | 0,00 | 4,41 | 0,00 | 1,00E+00 |
| AT2G43820 | 0,45 | 2,17 | 3,28 | 0,66 | 5,09E-01 |
| AT2G44080 | 0,60 | 3,31 | 3,71 | 0,89 | 3,73E-01 |
| AT2G44100 | 0,65 | -1,35 | 2,65 | -0,51 | 6,10E-01 |
| AT2G44130 | 0,72 | -1,73 | 2,56 | -0,68 | 4,99E-01 |
| AT2G44160 | 0,80 | 2,50 | 2,49 | 1,00 | 3,15E-01 |
| AT2G44175 | 0,43 | 0,00 | 4,20 | 0,00 | 1,00E+00 |
| AT2G44200 | 0,72 | 1,38 | 2,69 | 0,51 | 6,09E-01 |
| AT2G44210 | 1,13 | 0,34 | 1,73 | 0,20 | 8,42E-01 |
| AT2G44230 | 0,76 | -2,26 | 2,75 | -0,82 | 4,10E-01 |
| AT2G44240 | 0,17 | 0,00 | 4,41 | 0,00 | 1,00E+00 |
| AT2G44260 | 0,20 | 0,00 | 4,41 | 0,00 | 1,00E+00 |
| AT2G44350 | 0,40 | -0,58 | 3,47 | -0,17 | 8,67E-01 |
| AT2G44420 | 0,20 | 1,38 | 4,41 | 0,31 | 7,54E-01 |
| AT2G44430 | 0,96 | -0,10 | 2,08 | -0,05 | 9,62E-01 |

| | | | | | |
|---|---|---|---|---|---|
| AT2G44500 | 0,91 | 0,00 | 3,02 | 0,00 | 1,00E+00 |
| AT2G44520 | 0,23 | 1,38 | 4,41 | 0,31 | 7,55E-01 |
| AT2G44530 | 0,99 | 1,38 | 2,45 | 0,56 | 5,73E-01 |
| AT2G44600 | 0,10 | 0,00 | 4,41 | 0,00 | 1,00E+00 |
| AT2G44640 | 1,50 | 3,98 | 1,75 | 2,27 | 2,33E-02 |
| AT2G44670 | 1,15 | 1,14 | 2,09 | 0,55 | 5,83E-01 |
| AT2G44710 | 1,40 | 3,76 | 1,76 | 2,14 | 3,27E-02 |
| AT2G44735 | 0,42 | 1,46 | 4,17 | 0,35 | 7,26E-01 |
| AT2G44770 | 0,15 | 0,00 | 4,41 | 0,00 | 1,00E+00 |
| AT2G44800 | 0,19 | 0,00 | 4,40 | 0,00 | 1,00E+00 |
| AT2G44870 | 0,51 | -0,82 | 2,67 | -0,31 | 7,59E-01 |
| AT2G44880 | 0,91 | 2,60 | 2,78 | 0,94 | 3,49E-01 |
| AT2G44925 | 0,92 | 0,41 | 1,92 | 0,21 | 8,32E-01 |
| AT2G44940 | 0,52 | 1,35 | 2,79 | 0,48 | 6,29E-01 |
| AT2G45000 | 0,38 | 0,53 | 3,49 | 0,15 | 8,79E-01 |
| AT2G45020 | 0,36 | -1,64 | 4,35 | -0,38 | 7,06E-01 |
| AT2G45030 | 0,91 | 2,11 | 2,85 | 0,74 | 4,60E-01 |
| AT2G45140 | 0,33 | 0,51 | 4,37 | 0,12 | 9,07E-01 |
| AT2G45240 | 0,80 | 0,45 | 3,12 | 0,14 | 8,86E-01 |
| AT2G45250 | 0,26 | 1,38 | 4,41 | 0,31 | 7,54E-01 |
| AT2G45300 | 0,10 | 0,00 | 4,41 | 0,00 | 1,00E+00 |
| AT2G45310 | 0,67 | 0,45 | 2,65 | 0,17 | 8,67E-01 |
| AT2G45390 | 0,17 | -1,42 | 4,35 | -0,33 | 7,45E-01 |
| AT2G45440 | 0,53 | 2,49 | 3,87 | 0,64 | 5,21E-01 |
| AT2G45510 | 0,84 | 1,53 | 2,21 | 0,69 | 4,89E-01 |
| AT2G45530 | 0,98 | 0,03 | 2,05 | 0,02 | 9,87E-01 |
| AT2G09650 | 0,36 | -0,54 | 4,41 | -0,12 | 9,02E-01 |
| AT2G45590 | 0,37 | 1,37 | 4,34 | 0,32 | 7,52E-01 |
| AT2G45620 | 1,35 | 1,13 | 1,74 | 0,65 | 5,17E-01 |
| AT2G45630 | 0,80 | 2,04 | 2,65 | 0,77 | 4,42E-01 |
| AT2G45710 | 0,45 | 1,44 | 4,35 | 0,33 | 7,41E-01 |
| AT2G45720 | 0,29 | -0,54 | 4,41 | -0,12 | 9,02E-01 |
| AT2G45740 | 0,35 | -1,22 | 3,58 | -0,34 | 7,34E-01 |
| AT2G45790 | 0,17 | -0,82 | 4,37 | -0,19 | 8,51E-01 |
| AT2G45810 | 0,90 | -1,34 | 2,51 | -0,53 | 5,95E-01 |
| AT2G45910 | 0,97 | 2,69 | 2,13 | 1,27 | 2,06E-01 |
| AT2G46100 | 0,37 | -1,22 | 2,89 | -0,42 | 6,73E-01 |
| AT2G46130 | 1,45 | -1,20 | 1,62 | -0,74 | 4,62E-01 |
| AT2G46170 | 0,69 | 3,00 | 3,70 | 0,81 | 4,18E-01 |
| AT2G46190 | 0,70 | -1,42 | 2,31 | -0,62 | 5,38E-01 |
| AT2G09695 | 1,56 | -0,21 | 1,48 | -0,14 | 8,86E-01 |
| AT2G09705 | 0,89 | -2,35 | 2,59 | -0,91 | 3,63E-01 |
| AT2G46225 | 0,27 | -0,42 | 4,39 | -0,10 | 9,24E-01 |
| AT2G46240 | 0,70 | -1,33 | 2,61 | -0,51 | 6,11E-01 |
| AT2G46260 | 0,08 | 0,00 | 4,41 | 0,00 | 1,00E+00 |
| AT2G46270 | 0,25 | 1,26 | 4,39 | 0,29 | 7,74E-01 |
| AT2G46370 | 0,44 | 2,15 | 3,47 | 0,62 | 5,35E-01 |
| AT2G46380 | 1,50 | 4,20 | 2,15 | 1,95 | 5,08E-02 |
| AT2G46420 | 0,84 | 0,24 | 2,96 | 0,08 | 9,35E-01 |
| AT2G46440 | 1,43 | 2,63 | 1,66 | 1,59 | 1,13E-01 |
| AT2G46450 | 1,09 | 2,94 | 2,23 | 1,32 | 1,88E-01 |
| AT2G46490 | 0,75 | 0,00 | 3,01 | 0,00 | 1,00E+00 |
| AT2G46520 | 1,16 | 2,26 | 2,01 | 1,12 | 2,63E-01 |
| AT2G46570 | 0,51 | 1,38 | 4,08 | 0,34 | 7,36E-01 |
| AT2G46590 | 0,52 | 0,00 | 3,15 | 0,00 | 1,00E+00 |
| AT2G46600 | 1,11 | 2,73 | 1,88 | 1,45 | 1,48E-01 |

| | | | | | |
|---|---|---|---|---|---|
| AT2G46630 | 0,28 | -0,50 | 4,40 | -0,11 | 9,10E-01 |
| AT2G46650 | 0,42 | 1,30 | 3,41 | 0,38 | 7,03E-01 |
| AT2G46690 | 0,43 | 0,00 | 4,23 | 0,00 | 1,00E+00 |
| AT2G46710 | 0,30 | -0,59 | 3,75 | -0,16 | 8,75E-01 |
| AT2G46790 | 0,54 | 0,29 | 2,70 | 0,11 | 9,15E-01 |
| AT2G46800 | 1,02 | 1,64 | 2,02 | 0,81 | 4,18E-01 |
| AT2G46810 | 0,59 | -1,87 | 3,20 | -0,58 | 5,60E-01 |
| AT2G46900 | 1,50 | 0,30 | 1,87 | 0,16 | 8,71E-01 |
| AT2G46910 | 0,44 | -0,98 | 4,32 | -0,23 | 8,21E-01 |
| AT2G46990 | 0,08 | 0,00 | 4,41 | 0,00 | 1,00E+00 |
| AT2G47110 | 0,66 | 0,43 | 2,37 | 0,18 | 8,55E-01 |
| AT2G09815 | 0,23 | 0,00 | 4,41 | 0,00 | 1,00E+00 |
| AT2G47190 | 0,40 | 0,00 | 4,41 | 0,00 | 1,00E+00 |
| AT2G47220 | 0,25 | -0,49 | 4,40 | -0,11 | 9,11E-01 |
| AT2G47320 | 0,28 | -1,07 | 4,37 | -0,25 | 8,06E-01 |
| AT2G47420 | 0,42 | -1,36 | 4,17 | -0,33 | 7,43E-01 |
| AT2G47510 | 1,59 | 1,65 | 1,68 | 0,98 | 3,26E-01 |
| AT2G47610 | 1,29 | 1,33 | 1,91 | 0,70 | 4,87E-01 |
| AT2G47620 | 0,26 | -0,51 | 3,85 | -0,13 | 8,95E-01 |
| AT2G47630 | 0,42 | 0,52 | 2,90 | 0,18 | 8,57E-01 |
| AT2G47640 | 0,28 | 1,30 | 4,40 | 0,29 | 7,68E-01 |
| AT2G47690 | 0,55 | 0,90 | 3,99 | 0,23 | 8,22E-01 |
| AT2G47730 | 0,80 | 0,30 | 2,42 | 0,12 | 9,02E-01 |
| AT2G47820 | 1,37 | 3,61 | 2,49 | 1,45 | 1,47E-01 |
| AT2G47850 | 0,53 | 2,52 | 3,17 | 0,79 | 4,27E-01 |
| AT2G47910 | 0,92 | 0,42 | 2,28 | 0,18 | 8,54E-01 |
| AT2G47935 | 0,72 | 0,72 | 2,80 | 0,26 | 7,96E-01 |
| AT2G47960 | 1,39 | 3,63 | 1,91 | 1,90 | 5,70E-02 |
| AT2G47970 | 0,70 | 2,48 | 3,51 | 0,71 | 4,79E-01 |
| AT2G48010 | 0,33 | 0,00 | 4,41 | 0,00 | 1,00E+00 |
| AT2G48020 | 0,56 | 2,28 | 3,28 | 0,69 | 4,87E-01 |
| AT2G48110 | 1,17 | 2,77 | 2,09 | 1,32 | 1,86E-01 |
| AT2G48120 | 0,32 | 0,49 | 4,40 | 0,11 | 9,10E-01 |
| AT3G01040 | 0,51 | 0,19 | 3,19 | 0,06 | 9,52E-01 |
| AT3G01050 | 0,18 | -0,47 | 4,40 | -0,11 | 9,14E-01 |
| AT3G01080 | 0,17 | 0,00 | 4,41 | 0,00 | 1,00E+00 |
| AT3G01085 | 1,09 | -0,63 | 2,04 | -0,31 | 7,56E-01 |
| AT3G01150 | 0,45 | 2,16 | 3,31 | 0,65 | 5,13E-01 |
| AT3G01170 | 0,42 | 1,55 | 4,19 | 0,37 | 7,11E-01 |
| AT3G01175 | 0,08 | 0,00 | 4,41 | 0,00 | 1,00E+00 |
| AT3G01210 | 0,10 | 0,00 | 4,41 | 0,00 | 1,00E+00 |
| AT3G01280 | 1,02 | 1,65 | 1,78 | 0,93 | 3,54E-01 |
| AT3G01320 | 1,14 | 3,32 | 1,88 | 1,76 | 7,84E-02 |
| AT3G01440 | 0,31 | -0,77 | 4,39 | -0,18 | 8,60E-01 |
| AT3G01475 | 1,05 | 3,29 | 1,95 | 1,69 | 9,10E-02 |
| AT3G01480 | 1,15 | 0,30 | 1,85 | 0,16 | 8,69E-01 |
| AT3G01560 | 0,68 | 1,21 | 2,60 | 0,46 | 6,42E-01 |
| AT3G01590 | 0,40 | 0,00 | 4,41 | 0,00 | 1,00E+00 |
| AT3G01610 | 0,25 | 0,00 | 4,41 | 0,00 | 1,00E+00 |
| AT3G01660 | 0,12 | 0,00 | 4,41 | 0,00 | 1,00E+00 |
| AT3G01690 | 1,08 | 0,09 | 1,96 | 0,04 | 9,65E-01 |
| AT3G01740 | 0,28 | -0,49 | 4,40 | -0,11 | 9,12E-01 |
| AT3G01750 | 0,72 | 1,81 | 2,98 | 0,61 | 5,44E-01 |
| AT3G01780 | 1,57 | 3,83 | 1,71 | 2,24 | 2,49E-02 |
| AT3G01810 | 0,37 | -1,88 | 4,35 | -0,43 | 6,65E-01 |
| AT3G01830 | 0,94 | 2,80 | 2,29 | 1,22 | 2,22E-01 |

| | | | | | |
|---|---|---|---|---|---|
| AT3G01920 | 0,37 | 0,95 | 4,36 | 0,22 | 8,28E-01 |
| AT3G02010 | 0,37 | 2,18 | 4,37 | 0,50 | 6,18E-01 |
| AT3G02030 | 0,12 | 0,00 | 4,41 | 0,00 | 1,00E+00 |
| AT3G02050 | 0,37 | 2,11 | 4,32 | 0,49 | 6,26E-01 |
| AT3G02060 | 1,23 | 0,52 | 2,12 | 0,25 | 8,05E-01 |
| AT3G02065 | 0,54 | 2,22 | 3,83 | 0,58 | 5,62E-01 |
| AT3G02070 | 0,35 | 1,38 | 3,62 | 0,38 | 7,03E-01 |
| AT3G02130 | 1,18 | 2,73 | 2,28 | 1,20 | 2,31E-01 |
| AT3G02280 | 0,25 | 1,43 | 4,40 | 0,32 | 7,46E-01 |
| AT3G02290 | 0,16 | 0,00 | 4,41 | 0,00 | 1,00E+00 |
| AT3G02300 | 0,16 | 0,00 | 4,41 | 0,00 | 1,00E+00 |
| AT3G02340 | 0,50 | -1,93 | 4,33 | -0,45 | 6,56E-01 |
| AT3G02350 | 0,84 | 0,04 | 2,23 | 0,02 | 9,85E-01 |
| AT3G02360 | 1,53 | 3,29 | 1,84 | 1,79 | 7,34E-02 |
| AT3G02370 | 0,47 | -0,45 | 3,36 | -0,13 | 8,93E-01 |
| AT3G02400 | 0,66 | 3,63 | 4,30 | 0,85 | 3,98E-01 |
| AT3G02490 | 0,33 | 1,38 | 4,41 | 0,31 | 7,54E-01 |
| AT3G02560 | 0,49 | 1,38 | 3,99 | 0,35 | 7,30E-01 |
| AT3G02610 | 0,81 | -2,30 | 2,84 | -0,81 | 4,19E-01 |
| AT3G02630 | 0,37 | 2,21 | 4,33 | 0,51 | 6,09E-01 |
| AT3G02690 | 0,66 | 2,90 | 2,56 | 1,13 | 2,58E-01 |
| AT3G02720 | 0,46 | 2,06 | 4,37 | 0,47 | 6,38E-01 |
| AT3G02830 | 1,22 | 2,60 | 2,30 | 1,13 | 2,58E-01 |
| AT3G02832 | 1,51 | -0,11 | 1,78 | -0,06 | 9,49E-01 |
| AT3G01145 | 0,57 | -0,47 | 2,69 | -0,18 | 8,61E-01 |
| AT3G02880 | 1,19 | 3,87 | 1,79 | 2,16 | 3,11E-02 |
| AT3G02900 | 0,28 | 0,00 | 4,41 | 0,00 | 1,00E+00 |
| AT3G02910 | 0,77 | 0,65 | 2,51 | 0,26 | 7,96E-01 |
| AT3G03020 | 0,43 | 0,00 | 4,23 | 0,00 | 1,00E+00 |
| AT3G03030 | 0,40 | 0,00 | 4,41 | 0,00 | 1,00E+00 |
| AT3G03040 | 0,24 | 1,38 | 4,41 | 0,31 | 7,54E-01 |
| AT3G03110 | 0,55 | 1,97 | 4,39 | 0,45 | 6,54E-01 |
| AT3G03160 | 0,47 | 1,33 | 4,08 | 0,33 | 7,44E-01 |
| AT3G03210 | 0,10 | 0,00 | 4,41 | 0,00 | 1,00E+00 |
| AT3G03320 | 0,15 | 0,00 | 4,39 | 0,00 | 1,00E+00 |
| AT3G03340 | 0,40 | 0,00 | 4,41 | 0,00 | 1,00E+00 |
| AT3G03350 | 0,24 | 1,37 | 4,41 | 0,31 | 7,56E-01 |
| AT3G03360 | 0,08 | 0,00 | 4,41 | 0,00 | 1,00E+00 |
| AT3G03610 | 0,27 | -1,63 | 4,36 | -0,37 | 7,09E-01 |
| AT3G03640 | 0,25 | -0,64 | 4,37 | -0,15 | 8,83E-01 |
| AT3G03680 | 1,60 | -0,27 | 1,69 | -0,16 | 8,75E-01 |
| AT3G03860 | 0,25 | 0,00 | 4,41 | 0,00 | 1,00E+00 |
| AT3G03880 | 0,28 | 0,00 | 4,41 | 0,00 | 1,00E+00 |
| AT3G03950 | 0,32 | -1,32 | 3,60 | -0,37 | 7,15E-01 |
| AT3G03960 | 1,45 | 0,52 | 1,73 | 0,30 | 7,63E-01 |
| AT3G04140 | 0,99 | 0,61 | 2,26 | 0,27 | 7,86E-01 |
| AT3G04490 | 0,26 | 1,38 | 4,41 | 0,31 | 7,54E-01 |
| AT3G04530 | 0,08 | 0,00 | 4,41 | 0,00 | 1,00E+00 |
| AT3G04550 | 0,64 | -2,37 | 2,22 | -1,07 | 2,86E-01 |
| AT3G04790 | 1,26 | -0,50 | 1,97 | -0,26 | 7,99E-01 |
| AT3G04810 | 0,59 | 0,00 | 4,40 | 0,00 | 1,00E+00 |
| AT3G04840 | 0,80 | 1,91 | 2,37 | 0,81 | 4,20E-01 |
| AT3G04870 | 0,49 | -0,71 | 3,16 | -0,23 | 8,21E-01 |
| AT3G04910 | 0,86 | -0,36 | 2,06 | -0,18 | 8,61E-01 |
| AT3G04920 | 0,51 | 0,76 | 4,02 | 0,19 | 8,51E-01 |
| AT3G04940 | 0,44 | 1,08 | 4,21 | 0,26 | 7,98E-01 |

| | | | | | |
|---|---|---|---|---|---|
| AT3G01555 | 0,12 | 0,00 | 4,41 | 0,00 | 1,00E+00 |
| AT3G01575 | 0,10 | 0,00 | 4,41 | 0,00 | 1,00E+00 |
| AT3G05070 | 0,17 | -0,48 | 4,39 | -0,11 | 9,13E-01 |
| AT3G05120 | 0,72 | 0,29 | 2,37 | 0,12 | 9,01E-01 |
| AT3G05200 | 0,41 | 1,61 | 4,25 | 0,38 | 7,06E-01 |
| AT3G05230 | 0,28 | 0,00 | 4,41 | 0,00 | 1,00E+00 |
| AT3G05380 | 0,92 | 2,01 | 2,28 | 0,88 | 3,79E-01 |
| AT3G05410 | 1,68 | 0,89 | 1,68 | 0,53 | 5,95E-01 |
| AT3G05430 | 0,21 | 0,00 | 4,41 | 0,00 | 1,00E+00 |
| AT3G05490 | 0,28 | -0,54 | 4,41 | -0,12 | 9,02E-01 |
| AT3G05500 | 0,55 | 2,16 | 2,74 | 0,79 | 4,30E-01 |
| AT3G05520 | 0,43 | 0,49 | 2,93 | 0,17 | 8,67E-01 |
| AT3G05545 | 0,82 | 0,30 | 2,42 | 0,12 | 9,02E-01 |
| AT3G05560 | 0,45 | -1,67 | 3,29 | -0,51 | 6,11E-01 |
| AT3G05630 | 0,21 | 0,00 | 4,41 | 0,00 | 1,00E+00 |
| AT3G05670 | 0,41 | 1,55 | 4,27 | 0,36 | 7,16E-01 |
| AT3G05710 | 0,95 | 2,24 | 2,18 | 1,03 | 3,04E-01 |
| AT3G05725 | 0,64 | -1,64 | 2,50 | -0,65 | 5,13E-01 |
| AT3G05750 | 0,61 | 0,00 | 3,76 | 0,00 | 1,00E+00 |
| AT3G05760 | 0,68 | 0,00 | 3,76 | 0,00 | 1,00E+00 |
| AT3G05830 | 0,23 | 0,00 | 4,38 | 0,00 | 1,00E+00 |
| AT3G05840 | 0,50 | 2,87 | 3,95 | 0,73 | 4,68E-01 |
| AT3G05905 | 0,51 | -0,54 | 3,22 | -0,17 | 8,67E-01 |
| AT3G06125 | 0,60 | 2,26 | 2,37 | 0,96 | 3,39E-01 |
| AT3G06170 | 0,51 | -0,54 | 3,18 | -0,17 | 8,64E-01 |
| AT3G06180 | 0,25 | 0,00 | 4,41 | 0,00 | 1,00E+00 |
| AT3G06200 | 0,41 | 1,38 | 4,41 | 0,31 | 7,54E-01 |
| AT3G06210 | 0,20 | 1,38 | 4,41 | 0,31 | 7,54E-01 |
| AT3G06240 | 0,61 | 0,54 | 2,30 | 0,23 | 8,15E-01 |
| AT3G06380 | 0,55 | 0,60 | 3,90 | 0,15 | 8,78E-01 |
| AT3G06400 | 0,95 | 1,34 | 2,71 | 0,50 | 6,20E-01 |
| AT3G06455 | 0,58 | 1,33 | 4,34 | 0,31 | 7,60E-01 |
| AT3G06470 | 0,36 | -0,40 | 4,38 | -0,09 | 9,27E-01 |
| AT3G06500 | 1,24 | 1,51 | 2,25 | 0,67 | 5,02E-01 |
| AT3G06520 | 0,50 | 2,16 | 3,95 | 0,55 | 5,85E-01 |
| AT3G06610 | 0,15 | 0,00 | 4,41 | 0,00 | 1,00E+00 |
| AT3G06660 | 0,08 | 0,00 | 4,41 | 0,00 | 1,00E+00 |
| AT3G06730 | 0,97 | 2,57 | 3,11 | 0,83 | 4,09E-01 |
| AT3G06740 | 0,50 | 1,98 | 4,14 | 0,48 | 6,32E-01 |
| AT3G06810 | 1,67 | 3,27 | 1,63 | 2,01 | 4,42E-02 |
| AT3G06830 | 0,08 | 0,00 | 4,41 | 0,00 | 1,00E+00 |
| AT3G06860 | 1,03 | 0,96 | 2,01 | 0,48 | 6,34E-01 |
| AT3G06868 | 0,29 | 0,59 | 4,39 | 0,13 | 8,93E-01 |
| AT3G06895 | 1,19 | 1,79 | 2,03 | 0,88 | 3,78E-01 |
| AT3G06920 | 0,42 | 0,38 | 4,33 | 0,09 | 9,29E-01 |
| AT3G06980 | 0,81 | -0,47 | 2,95 | -0,16 | 8,73E-01 |
| AT3G07110 | 1,39 | 0,98 | 1,82 | 0,54 | 5,91E-01 |
| AT3G07200 | 0,08 | 0,00 | 4,41 | 0,00 | 1,00E+00 |
| AT3G07230 | 0,29 | -1,59 | 4,35 | -0,37 | 7,14E-01 |
| AT3G07270 | 0,85 | 2,37 | 2,08 | 1,14 | 2,54E-01 |
| AT3G07350 | 0,42 | 0,53 | 3,36 | 0,16 | 8,75E-01 |
| AT3G07360 | 1,00 | 1,89 | 2,56 | 0,74 | 4,59E-01 |
| AT3G07400 | 0,87 | 3,38 | 2,16 | 1,57 | 1,17E-01 |
| AT3G07430 | 0,32 | -1,10 | 4,39 | -0,25 | 8,01E-01 |
| AT3G07440 | 0,08 | 0,00 | 4,41 | 0,00 | 1,00E+00 |
| AT3G07480 | 0,45 | 1,81 | 4,30 | 0,42 | 6,74E-01 |

| | | | | | |
|---|---|---|---|---|---|
| AT3G07565 | 0,19 | 0,00 | 4,41 | 0,00 | 1,00E+00 |
| AT3G07610 | 0,83 | 2,21 | 2,60 | 0,85 | 3,96E-01 |
| AT3G07640 | 0,17 | 0,00 | 4,41 | 0,00 | 1,00E+00 |
| AT3G07650 | 0,97 | 1,88 | 3,42 | 0,55 | 5,83E-01 |
| AT3G07680 | 0,59 | 2,74 | 3,23 | 0,85 | 3,97E-01 |
| AT3G07690 | 0,32 | 2,14 | 4,37 | 0,49 | 6,25E-01 |
| AT3G07770 | 1,44 | 2,96 | 1,71 | 1,73 | 8,32E-02 |
| AT3G07780 | 1,15 | 0,51 | 2,23 | 0,23 | 8,21E-01 |
| AT3G07810 | 0,90 | 1,96 | 2,04 | 0,96 | 3,37E-01 |
| AT3G07880 | 0,28 | 0,00 | 4,41 | 0,00 | 1,00E+00 |
| AT3G07950 | 0,43 | -0,54 | 4,41 | -0,12 | 9,02E-01 |
| AT3G08010 | 0,17 | -0,43 | 4,39 | -0,10 | 9,21E-01 |
| AT3G08030 | 1,37 | 0,09 | 1,59 | 0,06 | 9,53E-01 |
| AT3G08550 | 0,31 | 1,33 | 4,40 | 0,30 | 7,62E-01 |
| AT3G08560 | 1,47 | 0,82 | 1,72 | 0,48 | 6,34E-01 |
| AT3G08600 | 0,52 | 0,00 | 3,93 | 0,00 | 1,00E+00 |
| AT3G08640 | 0,20 | -0,43 | 4,39 | -0,10 | 9,22E-01 |
| AT3G08690 | 0,22 | 1,33 | 4,40 | 0,30 | 7,62E-01 |
| AT3G08920 | 0,32 | 1,24 | 4,39 | 0,28 | 7,77E-01 |
| AT3G08930 | 0,51 | 0,38 | 3,32 | 0,11 | 9,09E-01 |
| AT3G08950 | 0,44 | 2,41 | 4,22 | 0,57 | 5,67E-01 |
| AT3G08980 | 0,33 | 1,66 | 4,37 | 0,38 | 7,05E-01 |
| AT3G09020 | 0,57 | -0,45 | 4,07 | -0,11 | 9,11E-01 |
| AT3G09030 | 0,51 | 2,08 | 2,69 | 0,77 | 4,41E-01 |
| AT3G09035 | 0,19 | 0,00 | 4,41 | 0,00 | 1,00E+00 |
| AT3G09085 | 0,40 | 1,32 | 3,49 | 0,38 | 7,05E-01 |
| AT3G02105 | 0,40 | 0,00 | 4,41 | 0,00 | 1,00E+00 |
| AT3G09180 | 0,23 | 1,27 | 4,39 | 0,29 | 7,72E-01 |
| AT3G09300 | 1,07 | 1,25 | 2,38 | 0,53 | 5,99E-01 |
| AT3G09350 | 0,51 | -0,19 | 4,33 | -0,04 | 9,66E-01 |
| AT3G09370 | 0,29 | 1,38 | 4,41 | 0,31 | 7,54E-01 |
| AT3G09430 | 1,27 | 2,86 | 2,14 | 1,34 | 1,81E-01 |
| AT3G09500 | 0,81 | 1,24 | 2,12 | 0,58 | 5,60E-01 |
| AT3G09510 | 0,52 | -0,54 | 3,23 | -0,17 | 8,66E-01 |
| AT3G09570 | 0,70 | 1,11 | 2,58 | 0,43 | 6,66E-01 |
| AT3G09600 | 0,41 | 2,16 | 3,37 | 0,64 | 5,22E-01 |
| AT3G09650 | 0,58 | 1,36 | 4,40 | 0,31 | 7,58E-01 |
| AT3G09670 | 0,48 | 1,38 | 4,41 | 0,31 | 7,54E-01 |
| AT3G09690 | 0,55 | 2,98 | 4,33 | 0,69 | 4,92E-01 |
| AT3G09810 | 0,32 | 1,24 | 4,38 | 0,28 | 7,77E-01 |
| AT3G09820 | 0,77 | -2,31 | 2,42 | -0,96 | 3,39E-01 |
| AT3G09830 | 0,46 | -0,54 | 3,37 | -0,16 | 8,72E-01 |
| AT3G09880 | 0,75 | 2,16 | 2,61 | 0,83 | 4,08E-01 |
| AT3G09890 | 0,43 | 0,32 | 4,39 | 0,07 | 9,42E-01 |
| AT3G09900 | 0,19 | 0,00 | 4,41 | 0,00 | 1,00E+00 |
| AT3G10060 | 0,36 | 1,38 | 4,41 | 0,31 | 7,54E-01 |
| AT3G10120 | 0,12 | 0,00 | 4,41 | 0,00 | 1,00E+00 |
| AT3G10190 | 0,32 | 0,00 | 4,41 | 0,00 | 1,00E+00 |
| AT3G10200 | 0,08 | 0,00 | 4,41 | 0,00 | 1,00E+00 |
| AT3G10300 | 0,68 | -1,14 | 2,97 | -0,38 | 7,01E-01 |
| AT3G10310 | 0,16 | 0,00 | 4,41 | 0,00 | 1,00E+00 |
| AT3G10340 | 0,80 | -1,38 | 3,08 | -0,45 | 6,53E-01 |
| AT3G10370 | 0,28 | 0,00 | 4,41 | 0,00 | 1,00E+00 |
| AT3G10390 | 0,49 | 2,22 | 3,25 | 0,68 | 4,95E-01 |
| AT3G10420 | 1,49 | 1,07 | 1,73 | 0,62 | 5,36E-01 |
| AT3G10480 | 0,72 | 1,89 | 3,57 | 0,53 | 5,96E-01 |

| | | | | | |
|---|---|---|---|---|---|
| AT3G10490 | 0,18 | -0,47 | 4,40 | -0,11 | 9,14E-01 |
| AT3G10527 | 0,27 | -0,49 | 4,40 | -0,11 | 9,12E-01 |
| AT3G10540 | 0,56 | 1,23 | 2,64 | 0,46 | 6,42E-01 |
| AT3G10570 | 0,12 | 0,00 | 4,41 | 0,00 | 1,00E+00 |
| AT3G10760 | 0,48 | 2,30 | 3,36 | 0,69 | 4,93E-01 |
| AT3G10800 | 1,54 | 3,22 | 2,02 | 1,59 | 1,11E-01 |
| AT3G10820 | 0,24 | 1,96 | 4,39 | 0,45 | 6,55E-01 |
| AT3G10860 | 0,23 | 0,00 | 4,41 | 0,00 | 1,00E+00 |
| AT3G10920 | 1,12 | -1,12 | 1,91 | -0,59 | 5,58E-01 |
| AT3G10950 | 0,19 | 0,00 | 4,41 | 0,00 | 1,00E+00 |
| AT3G10980 | 0,79 | 3,08 | 3,12 | 0,99 | 3,24E-01 |
| AT3G11020 | 0,05 | 0,00 | 4,41 | 0,00 | 1,00E+00 |
| AT3G11080 | 0,18 | 0,00 | 4,41 | 0,00 | 1,00E+00 |
| AT3G11090 | 0,60 | 1,38 | 3,27 | 0,42 | 6,73E-01 |
| AT3G11120 | 0,39 | 0,53 | 3,43 | 0,15 | 8,78E-01 |
| AT3G11170 | 1,02 | -0,61 | 1,78 | -0,35 | 7,30E-01 |
| AT3G11230 | 0,53 | 0,20 | 4,34 | 0,05 | 9,63E-01 |
| AT3G11250 | 0,11 | 0,00 | 4,41 | 0,00 | 1,00E+00 |
| AT3G11270 | 0,10 | 0,00 | 4,41 | 0,00 | 1,00E+00 |
| AT3G11397 | 0,44 | -0,95 | 4,11 | -0,23 | 8,17E-01 |
| AT3G11400 | 0,87 | 3,56 | 2,31 | 1,54 | 1,23E-01 |
| AT3G11420 | 0,26 | -0,47 | 4,40 | -0,11 | 9,15E-01 |
| AT3G11440 | 0,65 | 2,23 | 3,04 | 0,73 | 4,63E-01 |
| AT3G02465 | 0,30 | 1,36 | 4,41 | 0,31 | 7,57E-01 |
| AT3G11630 | 0,96 | -1,63 | 1,96 | -0,83 | 4,06E-01 |
| AT3G11700 | 0,50 | -0,53 | 3,31 | -0,16 | 8,73E-01 |
| AT3G11760 | 0,23 | 1,38 | 4,41 | 0,31 | 7,55E-01 |
| AT3G11810 | 0,52 | 1,62 | 3,84 | 0,42 | 6,74E-01 |
| AT3G11900 | 0,13 | 0,00 | 4,41 | 0,00 | 1,00E+00 |
| AT3G11930 | 1,35 | -1,20 | 3,36 | -0,36 | 7,21E-01 |
| AT3G11960 | 1,04 | 1,76 | 2,07 | 0,85 | 3,95E-01 |
| AT3G12010 | 0,40 | 1,87 | 4,38 | 0,43 | 6,69E-01 |
| AT3G12020 | 0,79 | 3,30 | 2,38 | 1,38 | 1,67E-01 |
| AT3G12050 | 0,32 | 0,00 | 3,61 | 0,00 | 1,00E+00 |
| AT3G12080 | 1,08 | 3,09 | 2,54 | 1,22 | 2,24E-01 |
| AT3G12270 | 0,77 | -0,69 | 2,69 | -0,26 | 7,98E-01 |
| AT3G12290 | 0,81 | 2,92 | 2,54 | 1,15 | 2,51E-01 |
| AT3G12320 | 0,90 | 2,58 | 2,09 | 1,23 | 2,17E-01 |
| AT3G12360 | 0,71 | 1,34 | 2,57 | 0,52 | 6,01E-01 |
| AT3G12390 | 0,69 | 1,67 | 2,46 | 0,68 | 4,95E-01 |
| AT3G12480 | 0,26 | -0,48 | 4,40 | -0,11 | 9,13E-01 |
| AT3G12502 | 0,08 | 0,00 | 4,41 | 0,00 | 1,00E+00 |
| AT3G12570 | 1,24 | -1,86 | 2,77 | -0,67 | 5,02E-01 |
| AT3G12600 | 0,05 | 0,00 | 4,41 | 0,00 | 1,00E+00 |
| AT3G12640 | 0,60 | 1,13 | 2,36 | 0,48 | 6,30E-01 |
| AT3G12700 | 0,20 | 1,40 | 4,41 | 0,32 | 7,51E-01 |
| AT3G12740 | 0,52 | 1,38 | 3,40 | 0,41 | 6,85E-01 |
| AT3G12760 | 0,80 | 1,51 | 2,39 | 0,63 | 5,29E-01 |
| AT3G12810 | 1,56 | 3,88 | 1,79 | 2,16 | 3,06E-02 |
| AT3G12915 | 0,53 | 1,85 | 4,38 | 0,42 | 6,73E-01 |
| AT3G13060 | 0,87 | 2,49 | 2,31 | 1,08 | 2,80E-01 |
| AT3G13062 | 1,32 | 1,27 | 1,66 | 0,76 | 4,45E-01 |
| AT3G13065 | 0,88 | 2,82 | 3,41 | 0,83 | 4,09E-01 |
| AT3G13200 | 1,18 | 0,11 | 1,95 | 0,06 | 9,55E-01 |
| AT3G13210 | 0,13 | 0,00 | 4,41 | 0,00 | 1,00E+00 |
| AT3G13340 | 1,00 | -0,41 | 2,09 | -0,19 | 8,47E-01 |

| | | | | | |
|---|---|---|---|---|---|
| AT3G13370 | 0,63 | 0,00 | 4,41 | 0,00 | 1,00E+00 |
| AT3G13380 | 0,67 | 1,36 | 2,61 | 0,52 | 6,01E-01 |
| AT3G13437 | 0,08 | 0,00 | 4,41 | 0,00 | 1,00E+00 |
| AT3G13445 | 0,19 | -0,42 | 4,39 | -0,10 | 9,23E-01 |
| AT3G13470 | 0,61 | 2,09 | 3,93 | 0,53 | 5,96E-01 |
| AT3G02725 | 1,24 | 0,70 | 1,85 | 0,38 | 7,04E-01 |
| AT3G13662 | 0,34 | 1,34 | 4,40 | 0,30 | 7,61E-01 |
| AT3G13690 | 0,67 | 1,20 | 3,87 | 0,31 | 7,57E-01 |
| AT3G13760 | 0,08 | 0,00 | 4,41 | 0,00 | 1,00E+00 |
| AT3G13770 | 0,43 | 2,13 | 4,38 | 0,49 | 6,26E-01 |
| AT3G13810 | 0,42 | 1,55 | 4,24 | 0,36 | 7,15E-01 |
| AT3G13930 | 0,81 | 2,38 | 2,08 | 1,14 | 2,53E-01 |
| AT3G13950 | 0,55 | 1,38 | 4,02 | 0,34 | 7,32E-01 |
| AT3G14020 | 0,35 | 1,58 | 4,36 | 0,36 | 7,18E-01 |
| AT3G14100 | 0,69 | -0,47 | 2,71 | -0,17 | 8,63E-01 |
| AT3G14120 | 1,41 | 4,17 | 2,00 | 2,08 | 3,77E-02 |
| AT3G14160 | 0,19 | 0,00 | 4,41 | 0,00 | 1,00E+00 |
| AT3G14220 | 0,64 | 2,26 | 3,74 | 0,60 | 5,46E-01 |
| AT3G14290 | 0,42 | 0,27 | 4,18 | 0,06 | 9,49E-01 |
| AT3G14560 | 0,20 | 1,38 | 4,41 | 0,31 | 7,54E-01 |
| AT3G14595 | 1,01 | 0,11 | 2,13 | 0,05 | 9,59E-01 |
| AT3G14600 | 1,18 | 1,68 | 1,83 | 0,92 | 3,59E-01 |
| AT3G14610 | 0,28 | 1,84 | 4,39 | 0,42 | 6,74E-01 |
| AT3G14620 | 0,63 | 0,59 | 2,65 | 0,22 | 8,23E-01 |
| AT3G14640 | 0,33 | 1,79 | 4,38 | 0,41 | 6,83E-01 |
| AT3G14660 | 1,69 | -0,34 | 1,58 | -0,22 | 8,29E-01 |
| AT3G14670 | 1,07 | 0,05 | 2,12 | 0,02 | 9,81E-01 |
| AT3G14690 | 0,63 | -1,46 | 2,37 | -0,61 | 5,39E-01 |
| AT3G14720 | 0,35 | 1,58 | 4,39 | 0,36 | 7,19E-01 |
| AT3G14790 | 0,27 | 1,84 | 4,38 | 0,42 | 6,75E-01 |
| AT3G14830 | 0,30 | 1,30 | 4,40 | 0,30 | 7,68E-01 |
| AT3G14870 | 0,25 | 0,00 | 4,41 | 0,00 | 1,00E+00 |
| AT3G14920 | 0,30 | 1,38 | 4,41 | 0,31 | 7,54E-01 |
| AT3G14940 | 0,68 | 1,49 | 3,64 | 0,41 | 6,83E-01 |
| AT3G14990 | 0,39 | -0,54 | 2,97 | -0,18 | 8,54E-01 |
| AT3G15000 | 0,95 | 1,10 | 2,39 | 0,46 | 6,46E-01 |
| AT3G15010 | 0,55 | -0,44 | 3,21 | -0,14 | 8,90E-01 |
| AT3G15020 | 1,02 | 3,36 | 2,41 | 1,40 | 1,62E-01 |
| AT3G15030 | 0,69 | 2,20 | 2,97 | 0,74 | 4,60E-01 |
| AT3G15090 | 0,12 | 0,00 | 4,41 | 0,00 | 1,00E+00 |
| AT3G15095 | 1,44 | 0,39 | 1,53 | 0,26 | 7,97E-01 |
| AT3G15160 | 1,08 | 2,88 | 2,24 | 1,29 | 1,99E-01 |
| AT3G15180 | 0,21 | 1,25 | 4,39 | 0,29 | 7,75E-01 |
| AT3G15190 | 1,26 | 1,55 | 1,69 | 0,92 | 3,59E-01 |
| AT3G15260 | 0,39 | -0,54 | 4,35 | -0,13 | 9,00E-01 |
| AT3G15290 | 0,26 | 1,38 | 4,41 | 0,31 | 7,54E-01 |
| AT3G15350 | 1,19 | 3,05 | 2,27 | 1,34 | 1,80E-01 |
| AT3G15355 | 0,91 | 2,25 | 3,82 | 0,59 | 5,56E-01 |
| AT3G15360 | 1,20 | -0,61 | 1,75 | -0,35 | 7,29E-01 |
| AT3G15380 | 0,12 | 0,00 | 4,41 | 0,00 | 1,00E+00 |
| AT3G15450 | 0,77 | 2,00 | 2,22 | 0,90 | 3,67E-01 |
| AT3G15518 | 0,17 | 0,00 | 4,41 | 0,00 | 1,00E+00 |
| AT3G15550 | 0,17 | -0,49 | 4,40 | -0,11 | 9,11E-01 |
| AT3G03145 | 0,66 | 0,44 | 3,02 | 0,15 | 8,83E-01 |
| AT3G15635 | 0,25 | 0,00 | 4,41 | 0,00 | 1,00E+00 |
| AT3G15750 | 0,08 | 0,00 | 4,41 | 0,00 | 1,00E+00 |

| | | | | | |
|---|---|---|---|---|---|
| AT3G15760 | 0,84 | 0,34 | 2,71 | 0,13 | 9,00E-01 |
| AT3G15790 | 0,67 | 1,35 | 2,99 | 0,45 | 6,52E-01 |
| AT3G15820 | 0,32 | 0,00 | 4,41 | 0,00 | 1,00E+00 |
| AT3G15850 | 0,57 | -1,76 | 2,72 | -0,65 | 5,19E-01 |
| AT3G15930 | 0,12 | 0,00 | 4,41 | 0,00 | 1,00E+00 |
| AT3G16010 | 0,15 | 0,00 | 4,41 | 0,00 | 1,00E+00 |
| AT3G16060 | 0,47 | 0,16 | 4,11 | 0,04 | 9,68E-01 |
| AT3G16080 | 0,49 | 2,19 | 3,31 | 0,66 | 5,08E-01 |
| AT3G16130 | 0,87 | 0,43 | 2,14 | 0,20 | 8,42E-01 |
| AT3G16175 | 0,08 | 0,00 | 4,41 | 0,00 | 1,00E+00 |
| AT3G16240 | 1,39 | -0,23 | 1,60 | -0,14 | 8,87E-01 |
| AT3G16260 | 0,72 | 2,68 | 3,53 | 0,76 | 4,48E-01 |
| AT3G16270 | 0,71 | -2,07 | 2,50 | -0,83 | 4,08E-01 |
| AT3G16340 | 0,41 | 1,89 | 4,39 | 0,43 | 6,67E-01 |
| AT3G16420 | 0,28 | 2,12 | 4,38 | 0,48 | 6,28E-01 |
| AT3G16460 | 0,31 | 1,37 | 4,41 | 0,31 | 7,57E-01 |
| AT3G16525 | 0,84 | -1,07 | 2,23 | -0,48 | 6,31E-01 |
| AT3G16570 | 0,68 | 0,43 | 2,55 | 0,17 | 8,66E-01 |
| AT3G16710 | 0,43 | 0,11 | 4,29 | 0,03 | 9,80E-01 |
| AT3G16720 | 0,43 | 1,42 | 4,17 | 0,34 | 7,33E-01 |
| AT3G16740 | 0,38 | -0,54 | 3,52 | -0,15 | 8,77E-01 |
| AT3G16760 | 0,57 | 0,00 | 3,87 | 0,00 | 1,00E+00 |
| AT3G16770 | 1,33 | -2,86 | 2,48 | -1,15 | 2,49E-01 |
| AT3G16800 | 0,16 | -0,55 | 4,41 | -0,12 | 9,01E-01 |
| AT3G16857 | 0,72 | 2,90 | 3,78 | 0,77 | 4,44E-01 |
| AT3G16940 | 0,78 | 0,77 | 2,97 | 0,26 | 7,94E-01 |
| AT3G17100 | 1,17 | 0,49 | 1,80 | 0,27 | 7,84E-01 |
| AT3G17160 | 0,19 | 0,00 | 4,41 | 0,00 | 1,00E+00 |
| AT3G17205 | 0,35 | 1,38 | 4,41 | 0,31 | 7,54E-01 |
| AT3G17210 | 0,32 | 2,00 | 4,38 | 0,46 | 6,48E-01 |
| AT3G03465 | 0,80 | -1,08 | 2,45 | -0,44 | 6,60E-01 |
| AT3G17360 | 0,65 | 3,27 | 3,00 | 1,09 | 2,75E-01 |
| AT3G17380 | 0,28 | 2,07 | 4,38 | 0,47 | 6,37E-01 |
| AT3G17410 | 0,79 | 1,20 | 2,05 | 0,59 | 5,57E-01 |
| AT3G17470 | 0,76 | 2,68 | 3,75 | 0,71 | 4,75E-01 |
| AT3G17490 | 0,56 | 0,00 | 3,96 | 0,00 | 1,00E+00 |
| AT3G17626 | 0,20 | 1,38 | 4,41 | 0,31 | 7,54E-01 |
| AT3G17640 | 0,16 | -0,55 | 4,41 | -0,12 | 9,01E-01 |
| AT3G17650 | 0,88 | -0,57 | 2,28 | -0,25 | 8,01E-01 |
| AT3G17690 | 0,12 | 0,00 | 4,41 | 0,00 | 1,00E+00 |
| AT3G17700 | 0,35 | 0,00 | 4,41 | 0,00 | 1,00E+00 |
| AT3G17712 | 0,66 | 3,37 | 3,66 | 0,92 | 3,58E-01 |
| AT3G17750 | 1,26 | 0,92 | 1,77 | 0,52 | 6,03E-01 |
| AT3G17765 | 0,21 | 1,38 | 4,41 | 0,31 | 7,54E-01 |
| AT3G17790 | 0,83 | 0,85 | 2,51 | 0,34 | 7,36E-01 |
| AT3G03535 | 0,92 | 1,16 | 2,27 | 0,51 | 6,09E-01 |
| AT3G17810 | 0,53 | 0,40 | 2,91 | 0,14 | 8,91E-01 |
| AT3G17820 | 0,62 | 2,11 | 3,19 | 0,66 | 5,09E-01 |
| AT3G17860 | 0,36 | 0,00 | 4,41 | 0,00 | 1,00E+00 |
| AT3G17880 | 0,42 | 0,00 | 3,37 | 0,00 | 1,00E+00 |
| AT3G17890 | 0,10 | 0,00 | 4,41 | 0,00 | 1,00E+00 |
| AT3G17970 | 0,77 | -0,40 | 2,99 | -0,13 | 8,93E-01 |
| AT3G17980 | 0,35 | 1,58 | 4,36 | 0,36 | 7,18E-01 |
| AT3G18000 | 0,28 | 1,34 | 4,40 | 0,30 | 7,61E-01 |
| AT3G18020 | 0,41 | 0,71 | 4,24 | 0,17 | 8,67E-01 |
| AT3G18040 | 0,26 | -0,41 | 4,39 | -0,09 | 9,25E-01 |

| | | | | | |
|---|---|---|---|---|---|
| AT3G18050 | 1,52 | -0,32 | 1,51 | -0,21 | 8,31E-01 |
| AT3G18060 | 1,35 | 1,94 | 1,62 | 1,20 | 2,32E-01 |
| AT3G18070 | 0,11 | 0,00 | 4,41 | 0,00 | 1,00E+00 |
| AT3G18080 | 1,62 | 0,40 | 1,54 | 0,26 | 7,96E-01 |
| AT3G18090 | 1,26 | 2,15 | 3,06 | 0,70 | 4,83E-01 |
| AT3G18100 | 0,29 | 0,00 | 4,41 | 0,00 | 1,00E+00 |
| AT3G03625 | 0,40 | -0,58 | 3,47 | -0,17 | 8,67E-01 |
| AT3G18190 | 1,46 | 0,90 | 1,60 | 0,56 | 5,75E-01 |
| AT3G18240 | 0,43 | 0,32 | 4,23 | 0,08 | 9,39E-01 |
| AT3G18270 | 0,50 | 0,71 | 4,02 | 0,18 | 8,59E-01 |
| AT3G18340 | 0,19 | 0,00 | 4,41 | 0,00 | 1,00E+00 |
| AT3G18390 | 1,15 | 0,04 | 1,89 | 0,02 | 9,84E-01 |
| AT3G18440 | 0,08 | 0,00 | 4,41 | 0,00 | 1,00E+00 |
| AT3G18480 | 0,44 | 1,32 | 3,33 | 0,40 | 6,91E-01 |
| AT3G18740 | 0,73 | -1,54 | 2,34 | -0,66 | 5,09E-01 |
| AT3G18790 | 0,65 | 0,73 | 2,49 | 0,30 | 7,68E-01 |
| AT3G18800 | 0,33 | 1,79 | 4,38 | 0,41 | 6,83E-01 |
| AT3G18820 | 0,39 | -0,77 | 4,21 | -0,18 | 8,54E-01 |
| AT3G18827 | 0,57 | 0,10 | 3,11 | 0,03 | 9,75E-01 |
| AT3G18860 | 0,59 | 1,38 | 2,70 | 0,51 | 6,09E-01 |
| AT3G18890 | 0,86 | 2,23 | 2,16 | 1,04 | 3,01E-01 |
| AT3G18915 | 0,30 | 0,00 | 4,41 | 0,00 | 1,00E+00 |
| AT3G18960 | 0,08 | 0,00 | 4,41 | 0,00 | 1,00E+00 |
| AT3G18980 | 1,15 | 2,68 | 1,98 | 1,35 | 1,77E-01 |
| AT3G19050 | 0,62 | 3,26 | 2,95 | 1,11 | 2,69E-01 |
| AT3G19100 | 1,08 | 3,20 | 2,01 | 1,59 | 1,11E-01 |
| AT3G19240 | 1,33 | 2,27 | 1,75 | 1,30 | 1,93E-01 |
| AT3G19290 | 0,28 | 1,91 | 4,39 | 0,44 | 6,63E-01 |
| AT3G19390 | 0,35 | 1,58 | 4,36 | 0,36 | 7,18E-01 |
| AT3G19470 | 0,30 | 1,93 | 4,39 | 0,44 | 6,61E-01 |
| AT3G19480 | 0,52 | 1,31 | 3,31 | 0,39 | 6,93E-01 |
| AT3G19520 | 0,05 | 0,00 | 4,41 | 0,00 | 1,00E+00 |
| AT3G19540 | 1,20 | 3,64 | 2,08 | 1,75 | 8,10E-02 |
| AT3G19560 | 1,58 | 0,86 | 2,03 | 0,42 | 6,73E-01 |
| AT3G19640 | 0,86 | 2,66 | 2,44 | 1,09 | 2,77E-01 |
| AT3G19680 | 0,92 | -0,21 | 2,01 | -0,11 | 9,16E-01 |
| AT3G19760 | 1,20 | 0,03 | 1,83 | 0,02 | 9,88E-01 |
| AT3G19770 | 0,75 | 0,67 | 3,02 | 0,22 | 8,25E-01 |
| AT3G19780 | 1,34 | 2,61 | 2,26 | 1,16 | 2,47E-01 |
| AT3G19800 | 0,78 | 1,02 | 2,48 | 0,41 | 6,81E-01 |
| AT3G19860 | 0,48 | 1,35 | 4,03 | 0,34 | 7,37E-01 |
| AT3G19870 | 0,37 | 0,00 | 4,37 | 0,00 | 1,00E+00 |
| AT3G19910 | 1,61 | 1,14 | 1,83 | 0,62 | 5,33E-01 |
| AT3G19930 | 1,47 | 2,68 | 1,63 | 1,64 | 1,01E-01 |
| AT3G19960 | 1,65 | 2,39 | 1,80 | 1,33 | 1,85E-01 |
| AT3G19980 | 0,93 | 2,88 | 2,36 | 1,22 | 2,23E-01 |
| AT3G19990 | 0,32 | 1,90 | 4,39 | 0,43 | 6,65E-01 |
| AT3G20000 | 0,74 | 1,05 | 3,61 | 0,29 | 7,70E-01 |
| AT3G20030 | 0,79 | -0,50 | 2,59 | -0,19 | 8,47E-01 |
| AT3G20140 | 0,20 | 1,33 | 4,40 | 0,30 | 7,63E-01 |
| AT3G20150 | 0,54 | 3,24 | 4,31 | 0,75 | 4,52E-01 |
| AT3G20200 | 0,21 | 0,00 | 4,41 | 0,00 | 1,00E+00 |
| AT3G20410 | 0,37 | 0,19 | 4,38 | 0,04 | 9,66E-01 |
| AT3G20500 | 0,33 | 2,12 | 4,37 | 0,49 | 6,27E-01 |
| AT3G20520 | 0,48 | 1,38 | 4,41 | 0,31 | 7,54E-01 |
| AT3G20540 | 0,90 | 3,46 | 2,38 | 1,45 | 1,47E-01 |

| | | | | | |
|---|---|---|---|---|---|
| AT3G20600 | 0,25 | 1,43 | 4,40 | 0,32 | 7,46E-01 |
| AT3G03995 | 0,62 | 0,00 | 4,41 | 0,00 | 1,00E+00 |
| AT3G20720 | 0,42 | 1,38 | 3,51 | 0,39 | 6,94E-01 |
| AT3G20730 | 0,40 | 0,00 | 4,41 | 0,00 | 1,00E+00 |
| AT3G20790 | 0,23 | -0,86 | 4,38 | -0,20 | 8,45E-01 |
| AT3G20890 | 0,34 | 1,69 | 4,38 | 0,39 | 6,99E-01 |
| AT3G20970 | 0,63 | 0,49 | 3,03 | 0,16 | 8,72E-01 |
| AT3G21130 | 0,35 | 0,00 | 4,41 | 0,00 | 1,00E+00 |
| AT3G21170 | 1,15 | 0,00 | 2,56 | 0,00 | 1,00E+00 |
| AT3G21175 | 0,18 | 0,00 | 4,41 | 0,00 | 1,00E+00 |
| AT3G21220 | 0,51 | 1,00 | 4,07 | 0,24 | 8,06E-01 |
| AT3G21270 | 0,29 | -0,57 | 4,40 | -0,13 | 8,97E-01 |
| AT3G21295 | 0,41 | 0,37 | 3,47 | 0,11 | 9,14E-01 |
| AT3G21310 | 0,36 | 0,00 | 4,41 | 0,00 | 1,00E+00 |
| AT3G21465 | 0,14 | -0,48 | 4,40 | -0,11 | 9,13E-01 |
| AT3G21473 | 0,48 | -1,48 | 4,01 | -0,37 | 7,12E-01 |
| AT3G21560 | 0,31 | 1,49 | 4,39 | 0,34 | 7,35E-01 |
| AT3G21690 | 0,96 | -0,53 | 2,24 | -0,24 | 8,12E-01 |
| AT3G04175 | 0,99 | 1,30 | 2,14 | 0,61 | 5,43E-01 |
| AT3G21990 | 0,46 | 0,00 | 4,41 | 0,00 | 1,00E+00 |
| AT3G22072 | 0,17 | -1,42 | 4,35 | -0,33 | 7,45E-01 |
| AT3G22104 | 0,62 | 2,51 | 2,97 | 0,84 | 3,98E-01 |
| AT3G22125 | 1,40 | -2,30 | 1,93 | -1,19 | 2,33E-01 |
| AT3G22150 | 0,49 | 2,58 | 3,96 | 0,65 | 5,15E-01 |
| AT3G22200 | 0,76 | 0,42 | 2,38 | 0,18 | 8,61E-01 |
| AT3G04255 | 0,24 | -0,56 | 4,41 | -0,13 | 8,99E-01 |
| AT3G22270 | 0,79 | 1,57 | 2,55 | 0,61 | 5,40E-01 |
| AT3G22290 | 0,33 | 1,79 | 4,38 | 0,41 | 6,83E-01 |
| AT3G22330 | 0,85 | -0,54 | 2,19 | -0,25 | 8,06E-01 |
| AT3G22350 | 0,91 | 0,00 | 2,75 | 0,00 | 1,00E+00 |
| AT3G22370 | 0,72 | 2,90 | 2,59 | 1,12 | 2,64E-01 |
| AT3G22425 | 0,12 | 0,00 | 4,41 | 0,00 | 1,00E+00 |
| AT3G22430 | 0,19 | 0,00 | 4,41 | 0,00 | 1,00E+00 |
| AT3G22520 | 0,75 | 1,31 | 2,67 | 0,49 | 6,24E-01 |
| AT3G22760 | 0,51 | 0,50 | 3,25 | 0,15 | 8,78E-01 |
| AT3G22810 | 0,55 | 0,00 | 4,39 | 0,00 | 1,00E+00 |
| AT3G22845 | 0,51 | 2,84 | 4,33 | 0,66 | 5,11E-01 |
| AT3G22870 | 0,92 | 1,35 | 2,38 | 0,57 | 5,69E-01 |
| AT3G22900 | 0,24 | 1,38 | 4,41 | 0,31 | 7,54E-01 |
| AT3G22935 | 0,26 | 0,00 | 4,41 | 0,00 | 1,00E+00 |
| AT3G23010 | 0,77 | 3,61 | 3,50 | 1,03 | 3,03E-01 |
| AT3G23050 | 0,64 | 0,51 | 2,74 | 0,18 | 8,54E-01 |
| AT3G04395 | 0,16 | 0,00 | 4,41 | 0,00 | 1,00E+00 |
| AT3G23250 | 0,25 | 0,00 | 4,41 | 0,00 | 1,00E+00 |
| AT3G23280 | 0,45 | -1,02 | 4,14 | -0,25 | 8,05E-01 |
| AT3G23300 | 0,75 | 2,90 | 2,20 | 1,32 | 1,88E-01 |
| AT3G23310 | 0,63 | -0,54 | 2,36 | -0,23 | 8,20E-01 |
| AT3G23380 | 0,21 | 0,00 | 4,41 | 0,00 | 1,00E+00 |
| AT3G23390 | 0,15 | 0,00 | 4,41 | 0,00 | 1,00E+00 |
| AT3G23410 | 0,36 | -1,57 | 4,36 | -0,36 | 7,19E-01 |
| AT3G23530 | 1,09 | 1,65 | 1,90 | 0,87 | 3,85E-01 |
| AT3G23580 | 0,57 | -0,45 | 2,39 | -0,19 | 8,52E-01 |
| AT3G23590 | 1,64 | 1,12 | 1,97 | 0,57 | 5,69E-01 |
| AT3G23600 | 1,16 | 2,74 | 2,06 | 1,34 | 1,82E-01 |
| AT3G23640 | 0,83 | 2,38 | 2,47 | 0,97 | 3,34E-01 |
| AT3G23750 | 0,94 | 1,05 | 1,77 | 0,60 | 5,51E-01 |

| | | | | | |
|---|---|---|---|---|---|
| AT3G23790 | 0,38 | 1,74 | 4,38 | 0,40 | 6,91E-01 |
| AT3G23800 | 0,20 | 1,33 | 4,40 | 0,30 | 7,63E-01 |
| AT3G23890 | 0,56 | 2,40 | 3,99 | 0,60 | 5,48E-01 |
| AT3G23990 | 0,73 | 1,89 | 3,67 | 0,52 | 6,06E-01 |
| AT3G24000 | 0,34 | 1,46 | 4,40 | 0,33 | 7,39E-01 |
| AT3G24050 | 0,88 | -0,96 | 2,15 | -0,45 | 6,56E-01 |
| AT3G24070 | 0,38 | 1,38 | 4,41 | 0,31 | 7,54E-01 |
| AT3G24160 | 0,45 | -1,21 | 3,53 | -0,34 | 7,32E-01 |
| AT3G24190 | 0,59 | 0,53 | 3,14 | 0,17 | 8,66E-01 |
| AT3G24255 | 0,43 | 0,42 | 2,87 | 0,15 | 8,85E-01 |
| AT3G24490 | 0,64 | 1,38 | 3,67 | 0,38 | 7,07E-01 |
| AT3G24500 | 0,22 | 0,00 | 4,41 | 0,00 | 1,00E+00 |
| AT3G24514 | 0,29 | 0,00 | 4,41 | 0,00 | 1,00E+00 |
| AT3G24520 | 0,92 | -0,33 | 2,49 | -0,13 | 8,96E-01 |
| AT3G24550 | 0,60 | 0,00 | 3,07 | 0,00 | 1,00E+00 |
| AT3G24580 | 0,47 | 0,00 | 4,11 | 0,00 | 1,00E+00 |
| AT3G24590 | 0,21 | 1,27 | 4,39 | 0,29 | 7,72E-01 |
| AT3G24830 | 0,92 | -2,28 | 2,09 | -1,09 | 2,75E-01 |
| AT3G25012 | 0,95 | 2,19 | 1,94 | 1,13 | 2,58E-01 |
| AT3G25013 | 0,18 | 0,00 | 4,41 | 0,00 | 1,00E+00 |
| AT3G25030 | 0,21 | -0,48 | 4,40 | -0,11 | 9,13E-01 |
| AT3G25040 | 0,17 | -0,48 | 4,39 | -0,11 | 9,13E-01 |
| AT3G25070 | 0,31 | 0,00 | 3,68 | 0,00 | 1,00E+00 |
| AT3G25130 | 0,57 | -1,30 | 4,32 | -0,30 | 7,63E-01 |
| AT3G25140 | 1,51 | 0,03 | 2,10 | 0,01 | 9,89E-01 |
| AT3G25250 | 0,60 | -1,34 | 2,66 | -0,50 | 6,16E-01 |
| AT3G25290 | 0,29 | 1,62 | 4,38 | 0,37 | 7,11E-01 |
| AT3G25440 | 0,23 | 1,38 | 4,41 | 0,31 | 7,54E-01 |
| AT3G25470 | 0,67 | 0,38 | 3,72 | 0,10 | 9,20E-01 |
| AT3G25490 | 0,05 | 0,00 | 4,41 | 0,00 | 1,00E+00 |
| AT3G25585 | 0,71 | 0,40 | 2,61 | 0,15 | 8,80E-01 |
| AT3G25590 | 0,75 | 1,30 | 2,57 | 0,51 | 6,13E-01 |
| AT3G25700 | 0,66 | -0,41 | 2,26 | -0,18 | 8,54E-01 |
| AT3G04955 | 0,15 | 0,00 | 4,41 | 0,00 | 1,00E+00 |
| AT3G25720 | 0,26 | 0,00 | 4,41 | 0,00 | 1,00E+00 |
| AT3G25760 | 0,49 | 1,75 | 4,32 | 0,41 | 6,85E-01 |
| AT3G25770 | 1,09 | -2,11 | 1,96 | -1,08 | 2,81E-01 |
| AT3G25780 | 0,49 | 1,32 | 2,77 | 0,48 | 6,33E-01 |
| AT3G25860 | 0,63 | -2,40 | 3,20 | -0,75 | 4,52E-01 |
| AT3G25890 | 0,89 | 1,01 | 2,32 | 0,44 | 6,63E-01 |
| AT3G25910 | 0,36 | 0,00 | 4,41 | 0,00 | 1,00E+00 |
| AT3G26020 | 0,28 | -0,54 | 4,41 | -0,12 | 9,02E-01 |
| AT3G26030 | 1,16 | 1,81 | 1,99 | 0,91 | 3,63E-01 |
| AT3G26070 | 0,46 | 1,33 | 4,20 | 0,32 | 7,52E-01 |
| AT3G26080 | 0,26 | 1,38 | 4,41 | 0,31 | 7,54E-01 |
| AT3G26085 | 0,30 | 1,58 | 4,39 | 0,36 | 7,19E-01 |
| AT3G26090 | 0,21 | 0,00 | 4,41 | 0,00 | 1,00E+00 |
| AT3G04975 | 1,19 | -2,35 | 2,11 | -1,12 | 2,64E-01 |
| AT3G04985 | 0,58 | -1,22 | 2,72 | -0,45 | 6,52E-01 |
| AT3G26370 | 0,48 | 1,22 | 2,77 | 0,44 | 6,58E-01 |
| AT3G26395 | 0,16 | 0,00 | 4,41 | 0,00 | 1,00E+00 |
| AT3G26400 | 0,95 | 1,93 | 2,33 | 0,83 | 4,07E-01 |
| AT3G26430 | 1,60 | 1,95 | 1,52 | 1,28 | 2,00E-01 |
| AT3G26420 | 1,60 | 1,95 | 1,52 | 1,28 | 2,00E-01 |
| AT3G26512 | 0,38 | 0,00 | 3,44 | 0,00 | 1,00E+00 |
| AT3G26570 | 1,54 | 0,30 | 1,88 | 0,16 | 8,73E-01 |

| | | | | | |
|---|---|---|---|---|---|
| AT3G26580 | 0,73 | -1,64 | 2,30 | -0,71 | 4,76E-01 |
| AT3G26600 | 1,07 | 1,35 | 2,20 | 0,61 | 5,39E-01 |
| AT3G26640 | 0,37 | 1,38 | 3,48 | 0,40 | 6,92E-01 |
| AT3G26710 | 1,14 | 2,77 | 1,93 | 1,44 | 1,51E-01 |
| AT3G26720 | 0,82 | 0,79 | 2,61 | 0,30 | 7,63E-01 |
| AT3G26742 | 0,18 | 0,00 | 4,41 | 0,00 | 1,00E+00 |
| AT3G26780 | 1,24 | 2,63 | 2,14 | 1,23 | 2,18E-01 |
| AT3G26782 | 0,41 | 1,38 | 4,21 | 0,33 | 7,43E-01 |
| AT3G26800 | 0,34 | 0,43 | 3,53 | 0,12 | 9,03E-01 |
| AT3G26840 | 1,06 | 2,78 | 2,32 | 1,20 | 2,31E-01 |
| AT3G27090 | 1,24 | 1,78 | 2,02 | 0,88 | 3,78E-01 |
| AT3G27160 | 0,41 | -1,29 | 3,37 | -0,38 | 7,03E-01 |
| AT3G27170 | 0,68 | 0,52 | 2,96 | 0,18 | 8,61E-01 |
| AT3G27230 | 0,45 | 2,12 | 3,39 | 0,63 | 5,31E-01 |
| AT3G27250 | 0,85 | 0,53 | 2,48 | 0,21 | 8,30E-01 |
| AT3G27260 | 1,09 | 1,04 | 2,15 | 0,48 | 6,28E-01 |
| AT3G27270 | 0,62 | 0,00 | 3,83 | 0,00 | 9,99E-01 |
| AT3G27280 | 0,60 | 0,65 | 3,08 | 0,21 | 8,32E-01 |
| AT3G27325 | 0,56 | 2,13 | 2,71 | 0,79 | 4,30E-01 |
| AT3G27329 | 0,81 | -2,42 | 2,15 | -1,12 | 2,61E-01 |
| AT3G27331 | 0,96 | 0,00 | 3,31 | 0,00 | 1,00E+00 |
| AT3G27350 | 0,51 | 0,00 | 4,01 | 0,00 | 1,00E+00 |
| AT3G27430 | 0,35 | -1,24 | 3,58 | -0,35 | 7,29E-01 |
| AT3G27470 | 1,23 | 1,38 | 2,14 | 0,64 | 5,19E-01 |
| AT3G27540 | 0,29 | -0,46 | 3,73 | -0,12 | 9,01E-01 |
| AT3G27555 | 0,30 | 1,38 | 4,41 | 0,31 | 7,54E-01 |
| AT3G27670 | 1,19 | 3,17 | 2,28 | 1,39 | 1,65E-01 |
| AT3G05295 | 0,48 | 0,41 | 2,97 | 0,14 | 8,90E-01 |
| AT3G27820 | 1,03 | -0,54 | 2,19 | -0,25 | 8,04E-01 |
| AT3G27830 | 0,47 | 1,33 | 3,47 | 0,38 | 7,02E-01 |
| AT3G27850 | 1,55 | -1,38 | 1,66 | -0,83 | 4,08E-01 |
| AT3G27860 | 0,12 | 0,00 | 4,41 | 0,00 | 1,00E+00 |
| AT3G27870 | 0,75 | 2,62 | 2,59 | 1,01 | 3,12E-01 |
| AT3G27940 | 0,56 | 0,00 | 3,07 | 0,00 | 1,00E+00 |
| AT3G27980 | 0,34 | -0,80 | 4,38 | -0,18 | 8,56E-01 |
| AT3G27999 | 0,43 | 0,00 | 4,41 | 0,00 | 1,00E+00 |
| AT3G28040 | 1,16 | 1,70 | 2,17 | 0,78 | 4,34E-01 |
| AT3G28050 | 0,37 | 0,00 | 4,41 | 0,00 | 1,00E+00 |
| AT3G28080 | 0,70 | 1,31 | 2,37 | 0,55 | 5,79E-01 |
| AT3G28100 | 1,10 | -0,07 | 2,03 | -0,03 | 9,73E-01 |
| AT3G28170 | 0,75 | 2,94 | 3,53 | 0,83 | 4,04E-01 |
| AT3G28200 | 0,56 | -2,55 | 4,30 | -0,59 | 5,54E-01 |
| AT3G28220 | 0,69 | -3,12 | 2,93 | -1,07 | 2,86E-01 |
| AT3G28270 | 0,14 | 0,00 | 4,41 | 0,00 | 1,00E+00 |
| AT3G28290 | 1,10 | -3,86 | 2,41 | -1,60 | 1,09E-01 |
| AT3G28300 | 1,56 | -2,46 | 2,62 | -0,94 | 3,48E-01 |
| AT3G28450 | 0,78 | -1,51 | 2,65 | -0,57 | 5,69E-01 |
| AT3G28500 | 0,30 | 0,00 | 4,41 | 0,00 | 1,00E+00 |
| AT3G28510 | 0,99 | 1,47 | 2,63 | 0,56 | 5,76E-01 |
| AT3G05515 | 0,60 | -1,26 | 3,03 | -0,42 | 6,76E-01 |
| AT3G28580 | 0,77 | 0,00 | 3,54 | 0,00 | 1,00E+00 |
| AT3G28590 | 0,05 | 0,00 | 4,41 | 0,00 | 1,00E+00 |
| AT3G28650 | 0,20 | 1,40 | 4,41 | 0,32 | 7,51E-01 |
| AT3G28670 | 0,33 | 0,00 | 4,41 | 0,00 | 1,00E+00 |
| AT3G28690 | 0,39 | -0,48 | 4,27 | -0,11 | 9,11E-01 |
| AT3G28715 | 0,62 | 0,98 | 2,33 | 0,42 | 6,74E-01 |

| | | | | | |
|---|---|---|---|---|---|
| AT3G28720 | 0,91 | 1,89 | 3,26 | 0,58 | 5,63E-01 |
| AT3G28730 | 0,64 | 2,86 | 2,94 | 0,97 | 3,30E-01 |
| AT3G28870 | 0,74 | -0,63 | 3,75 | -0,17 | 8,66E-01 |
| AT3G28910 | 0,61 | 2,18 | 2,72 | 0,80 | 4,24E-01 |
| AT3G05595 | 0,35 | -0,54 | 4,41 | -0,12 | 9,02E-01 |
| AT3G28922 | 0,12 | 0,00 | 4,41 | 0,00 | 1,00E+00 |
| AT3G28958 | 0,58 | 0,00 | 3,16 | 0,00 | 1,00E+00 |
| AT3G29000 | 0,25 | 0,00 | 4,41 | 0,00 | 1,00E+00 |
| AT3G29010 | 0,38 | 2,10 | 4,36 | 0,48 | 6,31E-01 |
| AT3G29033 | 0,57 | 0,00 | 3,06 | 0,00 | 1,00E+00 |
| AT3G29080 | 1,28 | -0,33 | 1,92 | -0,17 | 8,63E-01 |
| AT3G29180 | 1,27 | 3,71 | 2,76 | 1,34 | 1,79E-01 |
| AT3G05715 | 0,40 | -0,64 | 4,33 | -0,15 | 8,83E-01 |
| AT3G29240 | 1,56 | 3,55 | 1,89 | 1,87 | 6,12E-02 |
| AT3G29270 | 0,39 | 1,79 | 4,37 | 0,41 | 6,81E-01 |
| AT3G29290 | 0,20 | 0,00 | 4,41 | 0,00 | 1,00E+00 |
| AT3G29370 | 0,05 | 0,00 | 4,41 | 0,00 | 1,00E+00 |
| AT3G29390 | 0,18 | -0,47 | 4,40 | -0,11 | 9,14E-01 |
| AT3G29450 | 0,14 | 0,00 | 4,41 | 0,00 | 1,00E+00 |
| AT3G29455 | 0,27 | 0,00 | 4,41 | 0,00 | 1,00E+00 |
| AT3G05805 | 0,31 | 0,00 | 4,41 | 0,00 | 1,00E+00 |
| AT3G29638 | 0,16 | 0,00 | 4,41 | 0,00 | 1,00E+00 |
| AT3G29670 | 0,94 | -2,51 | 2,27 | -1,10 | 2,69E-01 |
| AT3G30210 | 0,11 | 0,00 | 4,41 | 0,00 | 1,00E+00 |
| AT3G30320 | 0,40 | -0,53 | 4,41 | -0,12 | 9,04E-01 |
| AT3G30380 | 0,49 | 1,34 | 3,28 | 0,41 | 6,84E-01 |
| AT3G30383 | 0,12 | 0,00 | 4,41 | 0,00 | 1,00E+00 |
| AT3G30520 | 1,35 | 0,00 | 2,50 | 0,00 | 1,00E+00 |
| AT3G30525 | 1,14 | 0,00 | 2,86 | 0,00 | 1,00E+00 |
| AT3G30720 | 0,61 | -1,93 | 3,09 | -0,63 | 5,31E-01 |
| AT3G30770 | 0,13 | 0,00 | 4,41 | 0,00 | 1,00E+00 |
| AT3G30820 | 0,18 | 0,00 | 4,41 | 0,00 | 1,00E+00 |
| AT3G31950 | 0,59 | -1,73 | 2,34 | -0,74 | 4,61E-01 |
| AT3G32150 | 0,20 | 0,00 | 4,41 | 0,00 | 1,00E+00 |
| AT3G32280 | 0,39 | -1,36 | 4,29 | -0,32 | 7,52E-01 |
| AT3G32896 | 0,28 | 1,37 | 4,41 | 0,31 | 7,55E-01 |
| AT3G33530 | 1,04 | 1,70 | 2,02 | 0,84 | 3,99E-01 |
| AT3G42075 | 0,24 | 1,36 | 4,40 | 0,31 | 7,57E-01 |
| AT3G42170 | 0,48 | 2,09 | 3,20 | 0,65 | 5,14E-01 |
| AT3G42310 | 0,36 | -0,49 | 4,40 | -0,11 | 9,12E-01 |
| AT3G42560 | 0,10 | 0,00 | 4,41 | 0,00 | 1,00E+00 |
| AT3G42640 | 0,44 | 1,33 | 2,83 | 0,47 | 6,37E-01 |
| AT3G42870 | 1,35 | -2,74 | 2,00 | -1,37 | 1,70E-01 |
| AT3G42950 | 0,12 | 0,00 | 4,41 | 0,00 | 1,00E+00 |
| AT3G06495 | 0,51 | -0,45 | 3,25 | -0,14 | 8,91E-01 |
| AT3G43148 | 0,82 | -0,54 | 2,47 | -0,22 | 8,26E-01 |
| AT3G43150 | 0,30 | 0,00 | 3,70 | 0,00 | 1,00E+00 |
| AT3G43210 | 0,25 | 1,34 | 4,40 | 0,30 | 7,61E-01 |
| AT3G43230 | 1,49 | 2,86 | 1,84 | 1,55 | 1,21E-01 |
| AT3G43260 | 1,20 | -0,47 | 2,30 | -0,21 | 8,37E-01 |
| AT3G06565 | 0,05 | 0,00 | 4,41 | 0,00 | 1,00E+00 |
| AT3G43290 | 0,21 | 0,00 | 4,41 | 0,00 | 1,00E+00 |
| AT3G06625 | 0,13 | -0,43 | 4,39 | -0,10 | 9,23E-01 |
| AT3G43490 | 1,48 | 0,32 | 2,24 | 0,14 | 8,88E-01 |
| AT3G43503 | 0,30 | -0,42 | 3,70 | -0,11 | 9,09E-01 |
| AT3G06635 | 0,40 | 0,00 | 4,41 | 0,00 | 1,00E+00 |

| | | | | | |
|---|---|---|---|---|---|
| AT3G43670 | 0,53 | 2,37 | 2,79 | 0,85 | 3,96E-01 |
| AT3G43682 | 0,33 | 0,00 | 4,41 | 0,00 | 1,00E+00 |
| AT3G43740 | 0,28 | -0,68 | 4,39 | -0,15 | 8,77E-01 |
| AT3G43800 | 0,45 | 0,24 | 3,33 | 0,07 | 9,43E-01 |
| AT3G43837 | 0,47 | 1,55 | 4,10 | 0,38 | 7,06E-01 |
| AT3G43920 | 0,47 | 2,19 | 3,31 | 0,66 | 5,08E-01 |
| AT3G43950 | 0,37 | 0,00 | 4,41 | 0,00 | 1,00E+00 |
| AT3G44130 | 0,22 | 0,00 | 4,41 | 0,00 | 1,00E+00 |
| AT3G44150 | 0,18 | -0,82 | 4,37 | -0,19 | 8,51E-01 |
| AT3G44210 | 1,26 | -1,32 | 2,61 | -0,51 | 6,12E-01 |
| AT3G44230 | 0,25 | -0,90 | 4,38 | -0,21 | 8,37E-01 |
| AT3G44235 | 0,05 | 0,00 | 4,41 | 0,00 | 1,00E+00 |
| AT3G44320 | 0,76 | 1,30 | 2,86 | 0,45 | 6,50E-01 |
| AT3G44405 | 0,08 | 0,00 | 4,41 | 0,00 | 1,00E+00 |
| AT3G44550 | 0,99 | -0,56 | 2,61 | -0,21 | 8,30E-01 |
| AT3G44560 | 0,14 | 0,00 | 4,41 | 0,00 | 1,00E+00 |
| AT3G44620 | 0,30 | -0,42 | 3,68 | -0,11 | 9,09E-01 |
| AT3G44670 | 1,62 | 2,22 | 1,76 | 1,27 | 2,06E-01 |
| AT3G44730 | 0,52 | -1,34 | 2,45 | -0,55 | 5,83E-01 |
| AT3G44798 | 0,43 | 0,00 | 4,23 | 0,00 | 1,00E+00 |
| AT3G44880 | 0,54 | 0,52 | 4,03 | 0,13 | 8,97E-01 |
| AT3G44900 | 0,86 | 2,86 | 2,39 | 1,20 | 2,32E-01 |
| AT3G44935 | 1,28 | 0,00 | 2,29 | 0,00 | 1,00E+00 |
| AT3G44980 | 0,39 | 0,62 | 3,51 | 0,18 | 8,59E-01 |
| AT3G45010 | 0,31 | 1,53 | 4,39 | 0,35 | 7,28E-01 |
| AT3G45400 | 0,33 | 1,43 | 4,40 | 0,32 | 7,45E-01 |
| AT3G45620 | 1,31 | 0,83 | 2,58 | 0,32 | 7,48E-01 |
| AT3G45650 | 1,57 | 3,80 | 1,97 | 1,92 | 5,43E-02 |
| AT3G45660 | 1,01 | 3,81 | 2,51 | 1,52 | 1,28E-01 |
| AT3G45670 | 0,15 | 0,00 | 4,41 | 0,00 | 1,00E+00 |
| AT3G45730 | 0,15 | 0,00 | 4,41 | 0,00 | 1,00E+00 |
| AT3G45740 | 0,36 | 1,85 | 4,39 | 0,42 | 6,73E-01 |
| AT3G45870 | 0,08 | 0,00 | 4,41 | 0,00 | 1,00E+00 |
| AT3G45930 | 0,32 | 0,36 | 4,37 | 0,08 | 9,34E-01 |
| AT3G45965 | 0,21 | -1,36 | 4,36 | -0,31 | 7,55E-01 |
| AT3G45970 | 0,98 | 2,65 | 2,67 | 0,99 | 3,21E-01 |
| AT3G46040 | 0,85 | 1,24 | 2,26 | 0,55 | 5,83E-01 |
| AT3G46060 | 1,03 | 1,75 | 2,16 | 0,81 | 4,17E-01 |
| AT3G46130 | 0,12 | 0,00 | 4,41 | 0,00 | 1,00E+00 |
| AT3G46220 | 0,60 | 1,68 | 2,60 | 0,65 | 5,18E-01 |
| AT3G46290 | 1,13 | 1,27 | 1,99 | 0,64 | 5,22E-01 |
| AT3G46370 | 0,25 | -0,70 | 4,39 | -0,16 | 8,74E-01 |
| AT3G46490 | 0,32 | -0,80 | 4,37 | -0,18 | 8,54E-01 |
| AT3G46520 | 0,37 | 1,80 | 4,38 | 0,41 | 6,81E-01 |
| AT3G46560 | 0,33 | 0,00 | 4,41 | 0,00 | 1,00E+00 |
| AT3G46590 | 1,40 | -0,43 | 1,80 | -0,24 | 8,13E-01 |
| AT3G07505 | 1,14 | -0,07 | 1,97 | -0,03 | 9,73E-01 |
| AT3G46640 | 1,12 | 1,39 | 2,10 | 0,66 | 5,07E-01 |
| AT3G46658 | 0,59 | 1,38 | 2,76 | 0,50 | 6,17E-01 |
| AT3G46668 | 0,57 | -1,10 | 3,22 | -0,34 | 7,33E-01 |
| AT3G07555 | 1,02 | 0,53 | 2,34 | 0,23 | 8,21E-01 |
| AT3G07585 | 1,61 | -0,63 | 1,75 | -0,36 | 7,19E-01 |
| AT3G46930 | 0,21 | 0,00 | 4,41 | 0,00 | 1,00E+00 |
| AT3G47060 | 0,35 | -0,54 | 4,41 | -0,12 | 9,02E-01 |
| AT3G47140 | 0,11 | 0,00 | 4,41 | 0,00 | 1,00E+00 |
| AT3G07695 | 0,33 | 0,00 | 4,41 | 0,00 | 1,00E+00 |

| | | | | | |
|---|---|---|---|---|---|
| AT3G47290 | 0,08 | 0,00 | 4,41 | 0,00 | 1,00E+00 |
| AT3G47295 | 0,08 | 0,00 | 4,41 | 0,00 | 1,00E+00 |
| AT3G07715 | 0,48 | -0,07 | 2,79 | -0,02 | 9,80E-01 |
| AT3G47490 | 0,22 | 0,00 | 4,41 | 0,00 | 1,00E+00 |
| AT3G47520 | 0,29 | -0,54 | 4,41 | -0,12 | 9,02E-01 |
| AT3G47550 | 0,50 | -0,91 | 3,17 | -0,29 | 7,76E-01 |
| AT3G47570 | 0,50 | 2,19 | 3,29 | 0,67 | 5,05E-01 |
| AT3G47610 | 0,61 | 2,58 | 2,58 | 1,00 | 3,18E-01 |
| AT3G47650 | 0,65 | 2,23 | 3,04 | 0,73 | 4,63E-01 |
| AT3G47660 | 0,72 | 3,29 | 2,82 | 1,17 | 2,44E-01 |
| AT3G07795 | 0,21 | 0,00 | 4,41 | 0,00 | 1,00E+00 |
| AT3G07805 | 0,89 | -2,06 | 2,73 | -0,76 | 4,49E-01 |
| AT3G47770 | 0,17 | 0,00 | 4,41 | 0,00 | 1,00E+00 |
| AT3G47800 | 1,07 | 0,06 | 1,94 | 0,03 | 9,74E-01 |
| AT3G47820 | 1,04 | 1,07 | 1,93 | 0,55 | 5,79E-01 |
| AT3G47833 | 0,22 | -0,42 | 4,37 | -0,10 | 9,24E-01 |
| AT3G47920 | 1,60 | -3,02 | 1,68 | -1,80 | 7,16E-02 |
| AT3G47930 | 0,52 | 1,38 | 3,96 | 0,35 | 7,27E-01 |
| AT3G47950 | 0,34 | 2,10 | 4,37 | 0,48 | 6,31E-01 |
| AT3G48040 | 0,17 | 1,32 | 4,40 | 0,30 | 7,64E-01 |
| AT3G48070 | 0,40 | 0,54 | 4,35 | 0,12 | 9,01E-01 |
| AT3G48110 | 0,52 | 2,51 | 3,15 | 0,80 | 4,26E-01 |
| AT3G48140 | 0,97 | 1,00 | 2,30 | 0,43 | 6,65E-01 |
| AT3G48190 | 1,19 | 3,73 | 2,44 | 1,53 | 1,27E-01 |
| AT3G48310 | 0,40 | 0,43 | 3,43 | 0,13 | 9,00E-01 |
| AT3G48380 | 0,47 | 1,38 | 3,31 | 0,42 | 6,77E-01 |
| AT3G48420 | 0,42 | 0,36 | 4,30 | 0,08 | 9,34E-01 |
| AT3G48460 | 0,51 | 1,69 | 4,35 | 0,39 | 6,97E-01 |
| AT3G48500 | 0,57 | -1,57 | 3,95 | -0,40 | 6,91E-01 |
| AT3G48530 | 0,79 | 1,31 | 2,26 | 0,58 | 5,64E-01 |
| AT3G48620 | 0,08 | 0,00 | 4,41 | 0,00 | 1,00E+00 |
| AT3G48680 | 0,15 | 0,00 | 4,41 | 0,00 | 1,00E+00 |
| AT3G48710 | 0,27 | 0,00 | 4,41 | 0,00 | 1,00E+00 |
| AT3G48730 | 0,57 | 1,51 | 2,79 | 0,54 | 5,89E-01 |
| AT3G48750 | 0,54 | 1,35 | 2,74 | 0,49 | 6,22E-01 |
| AT3G48760 | 0,76 | 1,10 | 2,57 | 0,43 | 6,70E-01 |
| AT3G07995 | 0,83 | 1,32 | 2,47 | 0,54 | 5,92E-01 |
| AT3G48780 | 0,82 | -1,11 | 2,82 | -0,39 | 6,94E-01 |
| AT3G48810 | 0,58 | -0,82 | 3,73 | -0,22 | 8,26E-01 |
| AT3G48890 | 0,66 | 0,24 | 2,72 | 0,09 | 9,30E-01 |
| AT3G49000 | 0,27 | 0,00 | 4,41 | 0,00 | 1,00E+00 |
| AT3G49110 | 0,08 | 0,00 | 4,41 | 0,00 | 1,00E+00 |
| AT3G49130 | 1,37 | 3,90 | 2,35 | 1,66 | 9,70E-02 |
| AT3G49220 | 0,92 | 1,88 | 2,01 | 0,93 | 3,51E-01 |
| AT3G49240 | 0,54 | -2,71 | 4,10 | -0,66 | 5,08E-01 |
| AT3G49260 | 0,64 | 3,08 | 4,32 | 0,71 | 4,76E-01 |
| AT3G49430 | 0,93 | 0,56 | 2,13 | 0,26 | 7,94E-01 |
| AT3G49520 | 1,37 | 2,61 | 1,97 | 1,33 | 1,85E-01 |
| AT3G49590 | 1,44 | 4,17 | 1,79 | 2,34 | 1,95E-02 |
| AT3G49670 | 1,02 | 3,09 | 2,37 | 1,30 | 1,92E-01 |
| AT3G49680 | 0,45 | -1,28 | 3,31 | -0,39 | 7,00E-01 |
| AT3G49710 | 0,32 | 1,90 | 4,39 | 0,43 | 6,65E-01 |
| AT3G49780 | 0,25 | -0,47 | 4,40 | -0,11 | 9,15E-01 |
| AT3G50000 | 0,43 | 0,00 | 4,27 | 0,00 | 1,00E+00 |
| AT3G50080 | 0,21 | 0,00 | 4,41 | 0,00 | 1,00E+00 |
| AT3G50100 | 0,05 | 0,00 | 4,41 | 0,00 | 1,00E+00 |

| | | | | | |
|---|---|---|---|---|---|
| AT3G50270 | 0,60 | 1,17 | 3,11 | 0,38 | 7,07E-01 |
| AT3G50350 | 0,42 | 0,54 | 3,46 | 0,16 | 8,76E-01 |
| AT3G50360 | 0,31 | -0,45 | 4,40 | -0,10 | 9,18E-01 |
| AT3G50430 | 0,16 | 0,00 | 4,41 | 0,00 | 1,00E+00 |
| AT3G50530 | 0,25 | -0,54 | 4,41 | -0,12 | 9,02E-01 |
| AT3G50540 | 1,39 | -1,05 | 1,71 | -0,61 | 5,39E-01 |
| AT3G50550 | 0,51 | 2,73 | 3,20 | 0,86 | 3,92E-01 |
| AT3G50630 | 0,26 | -0,49 | 4,40 | -0,11 | 9,12E-01 |
| AT3G50700 | 0,53 | 0,34 | 2,78 | 0,12 | 9,02E-01 |
| AT3G50770 | 1,04 | 1,99 | 2,49 | 0,80 | 4,26E-01 |
| AT3G50890 | 0,25 | 0,00 | 4,41 | 0,00 | 1,00E+00 |
| AT3G50910 | 0,94 | 2,87 | 2,36 | 1,21 | 2,25E-01 |
| AT3G50970 | 0,39 | 1,83 | 4,37 | 0,42 | 6,75E-01 |
| AT3G51110 | 0,46 | 2,17 | 4,02 | 0,54 | 5,90E-01 |
| AT3G51140 | 0,97 | 2,50 | 2,18 | 1,15 | 2,51E-01 |
| AT3G51180 | 0,25 | 1,43 | 4,40 | 0,32 | 7,46E-01 |
| AT3G51260 | 0,46 | 2,41 | 3,32 | 0,73 | 4,67E-01 |
| AT3G51270 | 0,33 | 1,24 | 4,38 | 0,28 | 7,78E-01 |
| AT3G51325 | 0,36 | 1,38 | 4,41 | 0,31 | 7,54E-01 |
| AT3G51370 | 1,46 | 1,98 | 1,82 | 1,09 | 2,77E-01 |
| AT3G51420 | 0,66 | 0,31 | 2,32 | 0,13 | 8,94E-01 |
| AT3G51430 | 0,32 | 1,40 | 4,40 | 0,32 | 7,50E-01 |
| AT3G51440 | 0,80 | 2,28 | 2,28 | 1,00 | 3,17E-01 |
| AT3G51450 | 0,43 | -0,54 | 2,82 | -0,19 | 8,47E-01 |
| AT3G51460 | 0,27 | 1,38 | 4,41 | 0,31 | 7,54E-01 |
| AT3G51480 | 0,45 | 2,54 | 4,35 | 0,58 | 5,60E-01 |
| AT3G51500 | 0,44 | 0,00 | 4,41 | 0,00 | 1,00E+00 |
| AT3G51510 | 0,44 | 1,99 | 3,49 | 0,57 | 5,70E-01 |
| AT3G51540 | 0,25 | 1,34 | 4,40 | 0,30 | 7,61E-01 |
| AT3G51620 | 1,34 | 1,23 | 1,92 | 0,64 | 5,22E-01 |
| AT3G51630 | 0,87 | 2,47 | 3,26 | 0,76 | 4,50E-01 |
| AT3G51640 | 0,80 | 1,38 | 2,86 | 0,48 | 6,30E-01 |
| AT3G51650 | 1,02 | 3,42 | 2,19 | 1,56 | 1,18E-01 |
| AT3G51730 | 1,38 | -0,99 | 1,62 | -0,61 | 5,39E-01 |
| AT3G51830 | 0,25 | 0,00 | 4,41 | 0,00 | 1,00E+00 |
| AT3G51850 | 1,69 | 4,42 | 1,80 | 2,45 | 1,43E-02 |
| AT3G51870 | 1,46 | 4,32 | 2,00 | 2,16 | 3,04E-02 |
| AT3G51970 | 0,40 | 1,31 | 3,46 | 0,38 | 7,05E-01 |
| AT3G52080 | 0,33 | 1,41 | 4,40 | 0,32 | 7,48E-01 |
| AT3G52100 | 0,15 | 0,00 | 4,41 | 0,00 | 1,00E+00 |
| AT3G52105 | 0,19 | 0,00 | 4,41 | 0,00 | 1,00E+00 |
| AT3G52120 | 1,03 | 3,76 | 2,16 | 1,74 | 8,14E-02 |
| AT3G52150 | 0,83 | -2,38 | 2,83 | -0,84 | 4,01E-01 |
| AT3G52155 | 0,45 | 2,52 | 4,35 | 0,58 | 5,63E-01 |
| AT3G52200 | 0,34 | 1,46 | 4,40 | 0,33 | 7,40E-01 |
| AT3G52210 | 0,11 | 0,00 | 4,41 | 0,00 | 1,00E+00 |
| AT3G52230 | 0,83 | 3,61 | 3,01 | 1,20 | 2,30E-01 |
| AT3G52300 | 0,40 | 1,23 | 4,38 | 0,28 | 7,80E-01 |
| AT3G52340 | 0,38 | -0,62 | 4,40 | -0,14 | 8,87E-01 |
| AT3G52370 | 0,08 | 0,00 | 4,41 | 0,00 | 1,00E+00 |
| AT3G52380 | 1,54 | 1,05 | 1,86 | 0,57 | 5,71E-01 |
| AT3G52470 | 0,30 | 0,00 | 4,41 | 0,00 | 1,00E+00 |
| AT3G52570 | 0,29 | 0,00 | 4,41 | 0,00 | 1,00E+00 |
| AT3G08275 | 0,60 | 1,38 | 3,16 | 0,44 | 6,63E-01 |
| AT3G52610 | 0,30 | 1,38 | 4,41 | 0,31 | 7,54E-01 |
| AT3G52680 | 0,59 | 2,13 | 2,61 | 0,81 | 4,16E-01 |

| | | | | | |
|---|---|---|---|---|---|
| AT3G52710 | 0,88 | 0,38 | 2,19 | 0,17 | 8,64E-01 |
| AT3G08285 | 1,59 | 1,90 | 1,77 | 1,08 | 2,82E-01 |
| AT3G08295 | 0,73 | 0,00 | 3,52 | 0,00 | 1,00E+00 |
| AT3G52830 | 1,16 | -1,96 | 1,99 | -0,98 | 3,25E-01 |
| AT3G52840 | 0,82 | -1,99 | 2,59 | -0,77 | 4,43E-01 |
| AT3G52950 | 0,45 | 2,23 | 3,30 | 0,68 | 4,99E-01 |
| AT3G52960 | 0,80 | 0,56 | 1,97 | 0,29 | 7,75E-01 |
| AT3G52990 | 1,11 | -1,35 | 1,81 | -0,75 | 4,54E-01 |
| AT3G53000 | 0,31 | 1,38 | 4,41 | 0,31 | 7,54E-01 |
| AT3G53030 | 0,57 | 1,91 | 4,03 | 0,47 | 6,35E-01 |
| AT3G53050 | 0,05 | 0,00 | 4,41 | 0,00 | 1,00E+00 |
| AT3G53065 | 0,19 | 0,00 | 4,40 | 0,00 | 1,00E+00 |
| AT3G53090 | 0,36 | 0,29 | 4,39 | 0,07 | 9,47E-01 |
| AT3G53110 | 0,40 | 2,16 | 3,41 | 0,63 | 5,27E-01 |
| AT3G53180 | 0,20 | 0,00 | 4,41 | 0,00 | 1,00E+00 |
| AT3G53190 | 0,21 | 1,25 | 4,39 | 0,29 | 7,75E-01 |
| AT3G53230 | 1,14 | 3,62 | 2,72 | 1,33 | 1,83E-01 |
| AT3G53240 | 0,55 | 2,84 | 3,89 | 0,73 | 4,65E-01 |
| AT3G53305 | 0,72 | -0,43 | 2,36 | -0,18 | 8,55E-01 |
| AT3G53360 | 0,08 | 0,00 | 4,41 | 0,00 | 1,00E+00 |
| AT3G53390 | 0,27 | 1,35 | 4,40 | 0,31 | 7,59E-01 |
| AT3G53470 | 0,77 | 0,77 | 2,79 | 0,28 | 7,83E-01 |
| AT3G53520 | 0,83 | 1,82 | 2,49 | 0,73 | 4,64E-01 |
| AT3G53670 | 0,90 | 1,59 | 2,18 | 0,73 | 4,68E-01 |
| AT3G53700 | 0,94 | -0,90 | 2,03 | -0,44 | 6,58E-01 |
| AT3G53870 | 0,80 | -0,99 | 2,65 | -0,37 | 7,09E-01 |
| AT3G54220 | 0,50 | 0,34 | 3,29 | 0,10 | 9,18E-01 |
| AT3G54230 | 1,41 | 3,11 | 2,15 | 1,45 | 1,48E-01 |
| AT3G54366 | 1,11 | -0,52 | 2,36 | -0,22 | 8,25E-01 |
| AT3G54420 | 0,17 | 0,00 | 4,41 | 0,00 | 1,00E+00 |
| AT3G54540 | 1,42 | 2,71 | 1,99 | 1,36 | 1,73E-01 |
| AT3G54600 | 0,51 | -0,69 | 4,09 | -0,17 | 8,66E-01 |
| AT3G54610 | 0,47 | -0,40 | 4,03 | -0,10 | 9,20E-01 |
| AT3G54625 | 0,68 | 1,42 | 4,39 | 0,32 | 7,46E-01 |
| AT3G54680 | 0,75 | 2,74 | 2,92 | 0,94 | 3,47E-01 |
| AT3G54800 | 0,59 | -0,54 | 2,78 | -0,20 | 8,45E-01 |
| AT3G54820 | 0,15 | 0,00 | 4,41 | 0,00 | 1,00E+00 |
| AT3G54850 | 0,81 | 2,13 | 2,48 | 0,86 | 3,90E-01 |
| AT3G54980 | 0,41 | 2,14 | 4,22 | 0,51 | 6,12E-01 |
| AT3G55030 | 0,39 | -0,97 | 4,23 | -0,23 | 8,20E-01 |
| AT3G55250 | 0,63 | -0,53 | 2,38 | -0,22 | 8,25E-01 |
| AT3G55260 | 0,43 | 1,35 | 4,13 | 0,33 | 7,43E-01 |
| AT3G55270 | 1,69 | 4,12 | 1,79 | 2,29 | 2,18E-02 |
| AT3G55280 | 0,61 | -1,07 | 3,78 | -0,28 | 7,77E-01 |
| AT3G55370 | 0,12 | 0,00 | 4,41 | 0,00 | 1,00E+00 |
| AT3G55420 | 0,14 | 0,00 | 4,41 | 0,00 | 1,00E+00 |
| AT3G55440 | 1,26 | -0,05 | 2,00 | -0,02 | 9,82E-01 |
| AT3G55450 | 0,15 | 0,00 | 4,41 | 0,00 | 1,00E+00 |
| AT3G55460 | 1,02 | -0,63 | 2,05 | -0,31 | 7,57E-01 |
| AT3G55485 | 0,83 | -0,73 | 2,44 | -0,30 | 7,64E-01 |
| AT3G08485 | 0,83 | -0,73 | 2,44 | -0,30 | 7,64E-01 |
| AT3G55520 | 0,40 | 0,00 | 4,41 | 0,00 | 1,00E+00 |
| AT3G55620 | 0,54 | 2,16 | 2,71 | 0,80 | 4,26E-01 |
| AT3G55630 | 0,42 | 1,38 | 4,41 | 0,31 | 7,54E-01 |
| AT3G55677 | 0,24 | 1,45 | 4,40 | 0,33 | 7,41E-01 |
| AT3G55760 | 0,99 | 0,53 | 1,90 | 0,28 | 7,79E-01 |

| | | | | | |
|---|---|---|---|---|---|
| AT3G55770 | 0,98 | 1,53 | 1,93 | 0,79 | 4,29E-01 |
| AT3G55780 | 0,67 | 3,27 | 3,59 | 0,91 | 3,63E-01 |
| AT3G55880 | 0,30 | 0,00 | 3,71 | 0,00 | 1,00E+00 |
| AT3G55940 | 0,10 | 0,00 | 4,41 | 0,00 | 1,00E+00 |
| AT3G56130 | 0,33 | -1,08 | 4,39 | -0,25 | 8,06E-01 |
| AT3G56140 | 0,74 | 0,34 | 2,55 | 0,13 | 8,93E-01 |
| AT3G56180 | 0,21 | 1,25 | 4,39 | 0,29 | 7,75E-01 |
| AT3G56310 | 0,76 | 1,30 | 2,52 | 0,52 | 6,05E-01 |
| AT3G56330 | 0,52 | 1,34 | 3,93 | 0,34 | 7,33E-01 |
| AT3G56390 | 0,30 | 1,38 | 4,41 | 0,31 | 7,54E-01 |
| AT3G56450 | 0,39 | 0,64 | 4,39 | 0,14 | 8,85E-01 |
| AT3G08665 | 0,76 | 1,38 | 2,99 | 0,46 | 6,45E-01 |
| AT3G56470 | 0,12 | 0,00 | 4,41 | 0,00 | 1,00E+00 |
| AT3G56490 | 0,11 | 0,00 | 4,41 | 0,00 | 1,00E+00 |
| AT3G56590 | 0,57 | 0,32 | 3,08 | 0,10 | 9,18E-01 |
| AT3G56640 | 0,59 | 1,38 | 3,19 | 0,43 | 6,66E-01 |
| AT3G56650 | 0,53 | -0,23 | 3,98 | -0,06 | 9,55E-01 |
| AT3G56720 | 0,17 | -0,46 | 4,40 | -0,10 | 9,17E-01 |
| AT3G56740 | 0,49 | 1,64 | 4,08 | 0,40 | 6,87E-01 |
| AT3G56780 | 0,32 | 1,87 | 4,39 | 0,43 | 6,70E-01 |
| AT3G56840 | 0,64 | 2,14 | 2,59 | 0,83 | 4,09E-01 |
| AT3G56880 | 1,69 | -0,12 | 1,47 | -0,08 | 9,34E-01 |
| AT3G08785 | 1,51 | -0,80 | 1,81 | -0,44 | 6,59E-01 |
| AT3G56960 | 0,60 | 2,31 | 4,34 | 0,53 | 5,94E-01 |
| AT3G57040 | 1,37 | -1,03 | 1,70 | -0,60 | 5,45E-01 |
| AT3G57062 | 1,47 | 3,51 | 1,88 | 1,87 | 6,21E-02 |
| AT3G57070 | 0,24 | -0,44 | 4,39 | -0,10 | 9,21E-01 |
| AT3G57080 | 0,21 | 1,38 | 4,41 | 0,31 | 7,54E-01 |
| AT3G57090 | 0,59 | 0,39 | 2,42 | 0,16 | 8,70E-01 |
| AT3G08845 | 0,12 | 0,00 | 4,41 | 0,00 | 1,00E+00 |
| AT3G57230 | 0,38 | 0,42 | 2,98 | 0,14 | 8,88E-01 |
| AT3G57340 | 0,42 | 0,43 | 3,50 | 0,12 | 9,01E-01 |
| AT3G57390 | 0,11 | 0,00 | 4,41 | 0,00 | 1,00E+00 |
| AT3G57450 | 1,58 | 2,68 | 1,70 | 1,58 | 1,15E-01 |
| AT3G57550 | 0,50 | 1,39 | 3,30 | 0,42 | 6,74E-01 |
| AT3G57570 | 0,40 | 0,10 | 4,20 | 0,02 | 9,81E-01 |
| AT3G57587 | 0,73 | -1,23 | 2,23 | -0,55 | 5,79E-01 |
| AT3G57650 | 0,53 | 0,00 | 2,79 | 0,00 | 1,00E+00 |
| AT3G57700 | 1,50 | 0,31 | 1,74 | 0,18 | 8,61E-01 |
| AT3G08975 | 0,42 | 2,13 | 4,14 | 0,51 | 6,07E-01 |
| AT3G57760 | 0,08 | 0,00 | 4,41 | 0,00 | 1,00E+00 |
| AT3G57770 | 1,17 | -0,01 | 1,77 | -0,01 | 9,95E-01 |
| AT3G57810 | 0,21 | -0,48 | 4,40 | -0,11 | 9,13E-01 |
| AT3G57830 | 0,31 | -0,68 | 4,39 | -0,16 | 8,77E-01 |
| AT3G57890 | 1,04 | -0,36 | 1,85 | -0,19 | 8,48E-01 |
| AT3G57940 | 0,20 | 0,00 | 4,41 | 0,00 | 1,00E+00 |
| AT3G57970 | 0,17 | 0,00 | 4,41 | 0,00 | 1,00E+00 |
| AT3G58020 | 0,08 | 0,00 | 4,41 | 0,00 | 1,00E+00 |
| AT3G58110 | 1,07 | 1,38 | 3,34 | 0,41 | 6,80E-01 |
| AT3G58160 | 0,08 | 0,00 | 4,41 | 0,00 | 1,00E+00 |
| AT3G58490 | 0,30 | 0,00 | 4,41 | 0,00 | 1,00E+00 |
| AT3G58510 | 0,99 | 0,42 | 2,05 | 0,20 | 8,38E-01 |
| AT3G58530 | 0,46 | 2,48 | 4,01 | 0,62 | 5,36E-01 |
| AT3G58570 | 0,71 | 0,53 | 2,66 | 0,20 | 8,42E-01 |
| AT3G58680 | 0,61 | 0,34 | 2,65 | 0,13 | 8,99E-01 |
| AT3G58690 | 0,58 | 2,55 | 3,07 | 0,83 | 4,07E-01 |

| | | | | | |
|---|---|---|---|---|---|
| AT3G58700 | 0,20 | 1,33 | 4,40 | 0,30 | 7,63E-01 |
| AT3G58795 | 0,23 | -1,17 | 4,37 | -0,27 | 7,88E-01 |
| AT3G59030 | 0,64 | 1,93 | 3,73 | 0,52 | 6,04E-01 |
| AT3G59080 | 0,19 | 0,00 | 4,40 | 0,00 | 1,00E+00 |
| AT3G59090 | 0,80 | 0,25 | 2,65 | 0,10 | 9,24E-01 |
| AT3G59100 | 0,79 | 3,64 | 2,74 | 1,33 | 1,85E-01 |
| AT3G59110 | 0,41 | -0,72 | 4,30 | -0,17 | 8,68E-01 |
| AT3G59210 | 0,24 | 0,00 | 4,41 | 0,00 | 1,00E+00 |
| AT3G59290 | 1,04 | 1,09 | 1,98 | 0,55 | 5,82E-01 |
| AT3G59300 | 0,19 | 0,00 | 4,41 | 0,00 | 1,00E+00 |
| AT3G59350 | 0,86 | 1,75 | 2,40 | 0,73 | 4,68E-01 |
| AT3G59380 | 0,24 | 0,00 | 4,41 | 0,00 | 1,00E+00 |
| AT3G59500 | 0,47 | -1,72 | 3,46 | -0,50 | 6,19E-01 |
| AT3G59520 | 0,08 | 0,00 | 4,41 | 0,00 | 1,00E+00 |
| AT3G59530 | 0,53 | 3,14 | 4,32 | 0,73 | 4,68E-01 |
| AT3G59670 | 0,30 | 0,42 | 4,41 | 0,09 | 9,25E-01 |
| AT3G59700 | 0,10 | 0,00 | 4,41 | 0,00 | 1,00E+00 |
| AT3G59810 | 0,46 | 0,93 | 4,35 | 0,21 | 8,31E-01 |
| AT3G59920 | 0,37 | 1,31 | 4,40 | 0,30 | 7,66E-01 |
| AT3G60080 | 1,12 | 2,78 | 2,07 | 1,34 | 1,80E-01 |
| AT3G60110 | 0,48 | 1,39 | 4,41 | 0,32 | 7,52E-01 |
| AT3G60130 | 0,63 | -0,40 | 3,24 | -0,12 | 9,02E-01 |
| AT3G60140 | 0,58 | -1,48 | 3,86 | -0,38 | 7,02E-01 |
| AT3G60240 | 1,46 | -0,34 | 1,64 | -0,21 | 8,34E-01 |
| AT3G60245 | 0,58 | 1,18 | 2,48 | 0,47 | 6,35E-01 |
| AT3G60290 | 0,27 | -0,54 | 4,41 | -0,12 | 9,02E-01 |
| AT3G60300 | 0,58 | 0,22 | 2,75 | 0,08 | 9,35E-01 |
| AT3G60310 | 0,49 | -0,63 | 4,03 | -0,16 | 8,76E-01 |
| AT3G60410 | 0,24 | 1,37 | 4,41 | 0,31 | 7,56E-01 |
| AT3G60420 | 0,55 | 2,22 | 3,88 | 0,57 | 5,67E-01 |
| AT3G60440 | 0,83 | 2,30 | 3,41 | 0,67 | 5,00E-01 |
| AT3G60450 | 1,00 | 2,28 | 2,73 | 0,83 | 4,05E-01 |
| AT3G60500 | 0,28 | 1,34 | 4,40 | 0,30 | 7,61E-01 |
| AT3G60530 | 0,83 | -2,52 | 2,12 | -1,19 | 2,33E-01 |
| AT3G60600 | 1,58 | 2,19 | 1,84 | 1,19 | 2,34E-01 |
| AT3G60690 | 1,35 | 2,09 | 2,02 | 1,03 | 3,02E-01 |
| AT3G60910 | 0,08 | 0,00 | 4,41 | 0,00 | 1,00E+00 |
| AT3G60950 | 1,11 | -2,36 | 2,13 | -1,11 | 2,66E-01 |
| AT3G60960 | 1,54 | 0,25 | 1,67 | 0,15 | 8,82E-01 |
| AT3G09515 | 1,27 | 1,12 | 1,96 | 0,57 | 5,69E-01 |
| AT3G60961 | 1,33 | 0,33 | 2,28 | 0,14 | 8,87E-01 |
| AT3G09525 | 0,12 | 0,00 | 4,41 | 0,00 | 1,00E+00 |
| AT3G60970 | 0,12 | 0,00 | 4,41 | 0,00 | 1,00E+00 |
| AT3G61030 | 0,61 | -2,05 | 2,14 | -0,96 | 3,38E-01 |
| AT3G61050 | 1,51 | -1,05 | 1,68 | -0,62 | 5,35E-01 |
| AT3G61080 | 0,12 | 0,00 | 4,41 | 0,00 | 1,00E+00 |
| AT3G61110 | 1,40 | -0,47 | 1,84 | -0,26 | 7,97E-01 |
| AT3G61130 | 1,07 | 2,75 | 2,33 | 1,18 | 2,39E-01 |
| AT3G61140 | 0,73 | 1,95 | 2,09 | 0,93 | 3,51E-01 |
| AT3G61150 | 0,68 | 2,46 | 2,87 | 0,86 | 3,90E-01 |
| AT3G61180 | 0,94 | 0,23 | 2,51 | 0,09 | 9,26E-01 |
| AT3G61240 | 1,00 | 1,91 | 2,03 | 0,94 | 3,47E-01 |
| AT3G61280 | 0,41 | 1,30 | 3,44 | 0,38 | 7,04E-01 |
| AT3G61320 | 0,08 | 0,00 | 4,41 | 0,00 | 1,00E+00 |
| AT3G61440 | 1,04 | 1,75 | 2,45 | 0,71 | 4,75E-01 |
| AT3G61480 | 0,44 | 1,27 | 3,32 | 0,38 | 7,02E-01 |

| | | | | | |
|---|---|---|---|---|---|
| AT3G61490 | 0,55 | 2,98 | 4,33 | 0,69 | 4,92E-01 |
| AT3G61540 | 0,29 | -0,54 | 4,41 | -0,12 | 9,02E-01 |
| AT3G61550 | 0,37 | -0,55 | 3,57 | -0,15 | 8,78E-01 |
| AT3G61600 | 1,22 | 2,04 | 1,85 | 1,10 | 2,70E-01 |
| AT3G61670 | 0,29 | 1,38 | 4,41 | 0,31 | 7,54E-01 |
| AT3G61700 | 1,46 | 0,04 | 1,69 | 0,02 | 9,82E-01 |
| AT3G61755 | 0,16 | 0,00 | 4,41 | 0,00 | 1,00E+00 |
| AT3G61810 | 0,05 | 0,00 | 4,41 | 0,00 | 1,00E+00 |
| AT3G61850 | 0,15 | 0,00 | 4,41 | 0,00 | 1,00E+00 |
| AT3G61870 | 1,16 | -0,72 | 1,65 | -0,43 | 6,64E-01 |
| AT3G61950 | 0,11 | 0,00 | 4,41 | 0,00 | 1,00E+00 |
| AT3G62030 | 0,98 | -0,33 | 2,14 | -0,16 | 8,76E-01 |
| AT3G62130 | 0,39 | 1,82 | 4,37 | 0,42 | 6,77E-01 |
| AT3G62190 | 0,24 | -0,48 | 4,40 | -0,11 | 9,13E-01 |
| AT3G62245 | 0,42 | 0,00 | 4,17 | 0,00 | 1,00E+00 |
| AT3G62300 | 0,93 | -0,05 | 3,38 | -0,01 | 9,88E-01 |
| AT3G62370 | 0,90 | -1,45 | 2,32 | -0,63 | 5,32E-01 |
| AT3G62530 | 0,72 | -0,45 | 2,56 | -0,18 | 8,59E-01 |
| AT3G62550 | 0,70 | -2,24 | 3,71 | -0,60 | 5,46E-01 |
| AT3G62560 | 0,61 | 2,91 | 3,11 | 0,93 | 3,50E-01 |
| AT3G62580 | 0,64 | 2,22 | 2,34 | 0,95 | 3,44E-01 |
| AT3G09815 | 0,08 | 0,00 | 4,41 | 0,00 | 1,00E+00 |
| AT3G09835 | 1,22 | -3,06 | 1,87 | -1,63 | 1,03E-01 |
| AT3G62620 | 0,29 | -0,54 | 4,41 | -0,12 | 9,02E-01 |
| AT3G62650 | 1,34 | 2,37 | 1,83 | 1,30 | 1,95E-01 |
| AT3G62660 | 1,25 | -0,99 | 1,87 | -0,53 | 5,98E-01 |
| AT3G62720 | 0,40 | 0,00 | 4,20 | 0,00 | 1,00E+00 |
| AT3G62750 | 0,68 | 1,73 | 2,46 | 0,70 | 4,82E-01 |
| AT3G62820 | 0,26 | 1,29 | 4,39 | 0,29 | 7,70E-01 |
| AT3G62830 | 0,86 | 0,33 | 2,06 | 0,16 | 8,72E-01 |
| AT3G62890 | 0,20 | 0,00 | 4,41 | 0,00 | 1,00E+00 |
| AT3G63000 | 0,33 | 1,26 | 4,39 | 0,29 | 7,74E-01 |
| AT3G63010 | 0,38 | 1,35 | 3,57 | 0,38 | 7,05E-01 |
| AT3G63080 | 0,53 | 2,41 | 2,76 | 0,87 | 3,83E-01 |
| AT3G63130 | 0,43 | 0,36 | 4,13 | 0,09 | 9,30E-01 |
| AT3G63170 | 0,32 | 2,21 | 4,37 | 0,51 | 6,13E-01 |
| AT3G63215 | 0,08 | 0,00 | 4,41 | 0,00 | 1,00E+00 |
| AT3G63300 | 0,72 | 3,29 | 2,82 | 1,17 | 2,44E-01 |
| AT3G63370 | 0,57 | 2,70 | 3,13 | 0,86 | 3,88E-01 |
| AT3G63400 | 0,95 | 0,59 | 2,25 | 0,26 | 7,93E-01 |
| AT3G63420 | 0,46 | 1,71 | 4,38 | 0,39 | 6,95E-01 |
| AT3G63470 | 0,42 | 2,32 | 4,20 | 0,55 | 5,80E-01 |
| AT3G09995 | 0,45 | 0,54 | 4,10 | 0,13 | 8,95E-01 |
| AT3G63520 | 0,87 | 1,23 | 2,30 | 0,53 | 5,93E-01 |
| AT4G00030 | 0,38 | -1,04 | 4,37 | -0,24 | 8,13E-01 |
| AT4G00100 | 1,39 | -0,17 | 1,86 | -0,09 | 9,28E-01 |
| AT4G00163 | 0,65 | -0,02 | 2,63 | -0,01 | 9,94E-01 |
| AT4G00230 | 0,69 | 2,59 | 4,33 | 0,60 | 5,50E-01 |
| AT4G00300 | 1,08 | -0,12 | 1,91 | -0,06 | 9,51E-01 |
| AT4G00340 | 0,31 | 1,38 | 4,41 | 0,31 | 7,54E-01 |
| AT4G00440 | 0,33 | 1,38 | 4,41 | 0,31 | 7,54E-01 |
| AT4G00490 | 0,44 | 0,00 | 4,39 | 0,00 | 1,00E+00 |
| AT4G00580 | 0,48 | 1,29 | 4,05 | 0,32 | 7,50E-01 |
| AT4G00660 | 1,18 | 2,04 | 2,12 | 0,96 | 3,36E-01 |
| AT4G00710 | 0,48 | 0,23 | 4,39 | 0,05 | 9,58E-01 |
| AT4G00750 | 0,96 | 0,44 | 1,92 | 0,23 | 8,19E-01 |

| | | | | | |
|---|---|---|---|---|---|
| AT4G00820 | 0,05 | 0,00 | 4,41 | 0,00 | 1,00E+00 |
| AT4G00830 | 0,92 | 1,01 | 2,15 | 0,47 | 6,40E-01 |
| AT4G00889 | 0,23 | 0,00 | 4,41 | 0,00 | 1,00E+00 |
| AT4G00930 | 0,17 | 0,00 | 4,41 | 0,00 | 1,00E+00 |
| AT4G00953 | 0,67 | -0,37 | 2,62 | -0,14 | 8,87E-01 |
| AT4G00955 | 0,42 | -1,23 | 2,91 | -0,42 | 6,72E-01 |
| AT4G00960 | 1,30 | -0,41 | 2,26 | -0,18 | 8,57E-01 |
| AT4G00970 | 0,33 | 0,00 | 4,41 | 0,00 | 1,00E+00 |
| AT4G00990 | 0,61 | 2,89 | 3,93 | 0,73 | 4,63E-01 |
| AT4G01020 | 0,37 | 0,53 | 4,39 | 0,12 | 9,03E-01 |
| AT4G01026 | 0,56 | 2,07 | 2,72 | 0,76 | 4,46E-01 |
| AT4G01040 | 0,12 | 0,00 | 4,41 | 0,00 | 1,00E+00 |
| AT4G01100 | 0,48 | -0,54 | 3,28 | -0,17 | 8,68E-01 |
| AT4G01130 | 0,11 | 0,00 | 4,41 | 0,00 | 1,00E+00 |
| AT4G01280 | 0,59 | -0,06 | 3,25 | -0,02 | 9,85E-01 |
| AT4G01320 | 0,90 | 2,87 | 2,18 | 1,32 | 1,88E-01 |
| AT4G01330 | 0,80 | 0,47 | 2,66 | 0,18 | 8,59E-01 |
| AT4G01340 | 1,48 | -2,81 | 1,73 | -1,62 | 1,05E-01 |
| AT4G01360 | 0,89 | 1,39 | 3,04 | 0,46 | 6,48E-01 |
| AT4G01370 | 0,27 | 0,00 | 4,41 | 0,00 | 1,00E+00 |
| AT4G01460 | 0,10 | 0,00 | 4,41 | 0,00 | 1,00E+00 |
| AT4G01570 | 0,24 | -0,52 | 4,40 | -0,12 | 9,06E-01 |
| AT4G03905 | 0,55 | 0,00 | 4,00 | 0,00 | 1,00E+00 |
| AT4G03925 | 0,76 | -0,32 | 2,35 | -0,14 | 8,92E-01 |
| AT4G01610 | 0,62 | 1,36 | 2,64 | 0,51 | 6,08E-01 |
| AT4G01690 | 1,14 | 1,22 | 1,67 | 0,73 | 4,64E-01 |
| AT4G01720 | 0,44 | 2,50 | 4,12 | 0,61 | 5,44E-01 |
| AT4G01995 | 0,08 | 0,00 | 4,41 | 0,00 | 1,00E+00 |
| AT4G02000 | 0,69 | -2,05 | 2,24 | -0,91 | 3,61E-01 |
| AT4G02010 | 0,14 | 0,00 | 4,41 | 0,00 | 1,00E+00 |
| AT4G02020 | 0,55 | 0,49 | 3,30 | 0,15 | 8,83E-01 |
| AT4G02030 | 0,75 | 3,22 | 2,77 | 1,16 | 2,46E-01 |
| AT4G02070 | 0,12 | 0,00 | 4,41 | 0,00 | 1,00E+00 |
| AT4G02080 | 0,57 | -0,42 | 2,65 | -0,16 | 8,74E-01 |
| AT4G02100 | 0,08 | 0,00 | 4,41 | 0,00 | 1,00E+00 |
| AT4G02110 | 0,21 | 0,00 | 4,41 | 0,00 | 1,00E+00 |
| AT4G02200 | 0,55 | 0,00 | 3,93 | 0,00 | 1,00E+00 |
| AT4G02220 | 0,29 | 1,27 | 4,39 | 0,29 | 7,72E-01 |
| AT4G02260 | 0,56 | 0,83 | 3,02 | 0,28 | 7,83E-01 |
| AT4G02330 | 0,22 | -0,89 | 4,37 | -0,20 | 8,38E-01 |
| AT4G02340 | 1,02 | 1,60 | 2,30 | 0,69 | 4,87E-01 |
| AT4G02370 | 0,72 | -1,84 | 3,10 | -0,59 | 5,52E-01 |
| AT4G02390 | 0,10 | 0,00 | 4,41 | 0,00 | 1,00E+00 |
| AT4G02400 | 0,19 | 0,00 | 4,41 | 0,00 | 1,00E+00 |
| AT4G02440 | 0,90 | 1,21 | 2,11 | 0,57 | 5,66E-01 |
| AT4G04135 | 0,05 | 0,00 | 4,41 | 0,00 | 1,00E+00 |
| AT4G02500 | 1,37 | 0,02 | 2,88 | 0,01 | 9,94E-01 |
| AT4G04195 | 0,38 | 0,00 | 3,48 | 0,00 | 1,00E+00 |
| AT4G02540 | 0,29 | -1,27 | 3,71 | -0,34 | 7,33E-01 |
| AT4G02580 | 1,19 | 1,22 | 2,14 | 0,57 | 5,70E-01 |
| AT4G02600 | 0,66 | 0,32 | 2,31 | 0,14 | 8,88E-01 |
| AT4G02610 | 0,31 | -0,50 | 4,40 | -0,11 | 9,10E-01 |
| AT4G02650 | 0,24 | 1,37 | 4,41 | 0,31 | 7,56E-01 |
| AT4G02700 | 1,40 | -0,06 | 1,88 | -0,03 | 9,73E-01 |
| AT4G02725 | 0,69 | -0,64 | 2,96 | -0,22 | 8,28E-01 |
| AT4G02790 | 0,08 | 0,00 | 4,41 | 0,00 | 1,00E+00 |

| | | | | | |
|---|---|---|---|---|---|
| AT4G02880 | 0,95 | 3,08 | 2,21 | 1,39 | 1,63E-01 |
| AT4G02920 | 1,02 | -0,14 | 2,46 | -0,06 | 9,55E-01 |
| AT4G02940 | 0,12 | 0,00 | 4,41 | 0,00 | 1,00E+00 |
| AT4G02990 | 0,69 | 2,68 | 2,58 | 1,04 | 2,98E-01 |
| AT4G03000 | 0,64 | 0,43 | 2,99 | 0,14 | 8,85E-01 |
| AT4G04425 | 0,08 | 0,00 | 4,41 | 0,00 | 1,00E+00 |
| AT4G03100 | 0,38 | 2,11 | 4,36 | 0,48 | 6,29E-01 |
| AT4G03120 | 0,23 | 1,38 | 4,41 | 0,31 | 7,54E-01 |
| AT4G03180 | 0,27 | -0,91 | 4,38 | -0,21 | 8,35E-01 |
| AT4G03200 | 0,82 | -0,43 | 2,47 | -0,17 | 8,62E-01 |
| AT4G03205 | 0,48 | 2,50 | 3,22 | 0,78 | 4,38E-01 |
| AT4G03210 | 0,53 | 1,22 | 4,39 | 0,28 | 7,81E-01 |
| AT4G03285 | 0,16 | 0,00 | 4,41 | 0,00 | 1,00E+00 |
| AT4G03410 | 0,58 | 0,51 | 3,86 | 0,13 | 8,94E-01 |
| AT4G03500 | 0,47 | 16,91 | 4,39 | 3,85 | 1,18E-04 |
| AT4G03830 | 0,61 | 0,96 | 2,62 | 0,37 | 7,15E-01 |
| AT4G03930 | 0,65 | -0,47 | 3,15 | -0,15 | 8,82E-01 |
| AT4G03940 | 0,17 | 0,00 | 4,41 | 0,00 | 1,00E+00 |
| AT4G03945 | 0,92 | -0,54 | 2,53 | -0,21 | 8,30E-01 |
| AT4G04110 | 0,65 | 1,32 | 4,34 | 0,30 | 7,61E-01 |
| AT4G04180 | 0,08 | 0,00 | 4,41 | 0,00 | 1,00E+00 |
| AT4G04210 | 0,39 | 1,36 | 2,97 | 0,46 | 6,48E-01 |
| AT4G04220 | 0,89 | 3,40 | 2,12 | 1,60 | 1,09E-01 |
| AT4G04795 | 0,87 | 0,00 | 2,70 | 0,00 | 1,00E+00 |
| AT4G04315 | 0,10 | 0,00 | 4,41 | 0,00 | 1,00E+00 |
| AT4G04350 | 0,42 | 0,49 | 3,40 | 0,15 | 8,84E-01 |
| AT4G04450 | 0,17 | -0,82 | 4,37 | -0,19 | 8,51E-01 |
| AT4G04510 | 0,17 | 1,32 | 4,40 | 0,30 | 7,64E-01 |
| AT4G04540 | 0,26 | -0,41 | 4,39 | -0,09 | 9,25E-01 |
| AT4G04610 | 0,73 | -0,76 | 2,22 | -0,35 | 7,30E-01 |
| AT4G04670 | 0,48 | 1,38 | 4,41 | 0,31 | 7,54E-01 |
| AT4G04750 | 0,90 | -1,77 | 2,03 | -0,87 | 3,85E-01 |
| AT4G04760 | 1,18 | -0,34 | 2,17 | -0,16 | 8,76E-01 |
| AT4G04800 | 0,52 | 1,26 | 3,91 | 0,32 | 7,47E-01 |
| AT4G04840 | 1,00 | -0,44 | 2,11 | -0,21 | 8,34E-01 |
| AT4G04850 | 0,79 | 0,81 | 1,99 | 0,41 | 6,82E-01 |
| AT4G04860 | 0,28 | -0,48 | 3,79 | -0,13 | 8,98E-01 |
| AT4G04920 | 1,70 | 3,55 | 1,82 | 1,95 | 5,09E-02 |
| AT4G04960 | 0,39 | 1,38 | 2,97 | 0,47 | 6,42E-01 |
| AT4G05020 | 0,40 | 1,37 | 4,35 | 0,32 | 7,52E-01 |
| AT4G05080 | 0,23 | 0,00 | 4,41 | 0,00 | 1,00E+00 |
| AT4G05105 | 0,17 | 0,00 | 4,41 | 0,00 | 1,00E+00 |
| AT4G05160 | 0,46 | 0,00 | 4,15 | 0,00 | 1,00E+00 |
| AT4G05612 | 0,12 | 0,00 | 4,41 | 0,00 | 1,00E+00 |
| AT4G05632 | 0,08 | 0,00 | 4,41 | 0,00 | 1,00E+00 |
| AT4G06479 | 0,10 | 0,00 | 4,41 | 0,00 | 1,00E+00 |
| AT4G06598 | 0,74 | 3,74 | 3,43 | 1,09 | 2,74E-01 |
| AT4G06688 | 0,68 | 2,20 | 2,52 | 0,87 | 3,84E-01 |
| AT4G07390 | 0,27 | 0,00 | 3,81 | 0,00 | 1,00E+00 |
| AT4G07455 | 0,24 | 1,96 | 4,39 | 0,45 | 6,55E-01 |
| AT4G07825 | 1,05 | 1,64 | 2,66 | 0,61 | 5,39E-01 |
| AT4G07932 | 0,51 | -1,85 | 4,08 | -0,45 | 6,51E-01 |
| AT4G07965 | 0,15 | 0,00 | 4,41 | 0,00 | 1,00E+00 |
| AT4G08025 | 0,16 | 0,00 | 4,41 | 0,00 | 1,00E+00 |
| AT4G05305 | 0,12 | 0,00 | 4,41 | 0,00 | 1,00E+00 |
| AT4G08097 | 0,66 | -1,59 | 2,63 | -0,60 | 5,45E-01 |

| | | | | | |
|---|---|---|---|---|---|
| AT4G08180 | 1,01 | -0,43 | 2,07 | -0,21 | 8,34E-01 |
| AT4G08267 | 0,63 | 0,00 | 4,35 | 0,00 | 1,00E+00 |
| AT4G08270 | 0,17 | 0,00 | 4,41 | 0,00 | 1,00E+00 |
| AT4G08320 | 0,31 | 0,00 | 4,40 | 0,00 | 1,00E+00 |
| AT4G08350 | 0,65 | -0,46 | 2,72 | -0,17 | 8,64E-01 |
| AT4G08395 | 0,29 | 0,00 | 4,41 | 0,00 | 1,00E+00 |
| AT4G05455 | 0,68 | -2,38 | 2,53 | -0,94 | 3,46E-01 |
| AT4G08450 | 0,11 | 0,00 | 4,41 | 0,00 | 1,00E+00 |
| AT4G08455 | 0,63 | 0,00 | 4,41 | 0,00 | 1,00E+00 |
| AT4G08510 | 0,33 | -0,54 | 4,41 | -0,12 | 9,02E-01 |
| AT4G08545 | 1,36 | 1,38 | 2,09 | 0,66 | 5,09E-01 |
| AT4G08560 | 0,64 | 2,18 | 4,27 | 0,51 | 6,10E-01 |
| AT4G08780 | 0,43 | 2,53 | 4,15 | 0,61 | 5,43E-01 |
| AT4G05605 | 0,39 | 1,38 | 4,41 | 0,31 | 7,54E-01 |
| AT4G08930 | 0,71 | 0,43 | 2,55 | 0,17 | 8,65E-01 |
| AT4G08950 | 0,37 | 1,29 | 4,37 | 0,29 | 7,69E-01 |
| AT4G08980 | 0,65 | -0,54 | 3,05 | -0,18 | 8,58E-01 |
| AT4G08990 | 0,69 | 0,00 | 3,80 | 0,00 | 1,00E+00 |
| AT4G09020 | 1,04 | -2,10 | 2,07 | -1,01 | 3,10E-01 |
| AT4G09030 | 0,16 | 0,00 | 4,41 | 0,00 | 1,00E+00 |
| AT4G09150 | 0,62 | 2,13 | 3,05 | 0,70 | 4,85E-01 |
| AT4G09160 | 0,36 | 2,79 | 4,33 | 0,64 | 5,19E-01 |
| AT4G09250 | 0,12 | 0,00 | 4,41 | 0,00 | 1,00E+00 |
| AT4G09310 | 0,21 | -0,54 | 4,41 | -0,12 | 9,02E-01 |
| AT4G05705 | 0,10 | 0,00 | 4,41 | 0,00 | 1,00E+00 |
| AT4G09320 | 1,06 | -1,05 | 1,83 | -0,57 | 5,68E-01 |
| AT4G09360 | 0,71 | -0,46 | 2,57 | -0,18 | 8,59E-01 |
| AT4G09450 | 1,44 | -1,70 | 1,71 | -1,00 | 3,19E-01 |
| AT4G09460 | 0,37 | 1,29 | 4,34 | 0,30 | 7,67E-01 |
| AT4G09545 | 0,31 | -0,41 | 4,38 | -0,09 | 9,26E-01 |
| AT4G09570 | 0,54 | 1,38 | 2,39 | 0,58 | 5,65E-01 |
| AT4G09630 | 0,87 | 1,16 | 2,24 | 0,52 | 6,04E-01 |
| AT4G09660 | 0,30 | 0,00 | 4,41 | 0,00 | 1,00E+00 |
| AT4G09680 | 0,45 | 2,11 | 4,03 | 0,52 | 6,01E-01 |
| AT4G09690 | 1,50 | -2,55 | 1,81 | -1,41 | 1,58E-01 |
| AT4G09730 | 0,51 | -0,48 | 3,15 | -0,15 | 8,79E-01 |
| AT4G09800 | 0,41 | 1,15 | 4,27 | 0,27 | 7,88E-01 |
| AT4G09820 | 0,31 | -1,66 | 4,36 | -0,38 | 7,03E-01 |
| AT4G09830 | 0,41 | 0,54 | 3,42 | 0,16 | 8,75E-01 |
| AT4G09850 | 0,21 | 0,00 | 4,41 | 0,00 | 1,00E+00 |
| AT4G05745 | 0,45 | 1,33 | 3,26 | 0,41 | 6,85E-01 |
| AT4G09890 | 0,18 | 0,00 | 4,41 | 0,00 | 1,00E+00 |
| AT4G10060 | 0,79 | 1,91 | 2,32 | 0,82 | 4,09E-01 |
| AT4G10170 | 0,08 | 0,00 | 4,41 | 0,00 | 1,00E+00 |
| AT4G10180 | 0,21 | 0,00 | 4,41 | 0,00 | 1,00E+00 |
| AT4G10465 | 0,60 | -0,57 | 2,75 | -0,21 | 8,35E-01 |
| AT4G10610 | 1,30 | 1,92 | 2,20 | 0,87 | 3,83E-01 |
| AT4G10613 | 0,77 | -1,31 | 2,25 | -0,58 | 5,60E-01 |
| AT4G10620 | 0,24 | 1,96 | 4,39 | 0,45 | 6,55E-01 |
| AT4G10660 | 0,11 | 0,00 | 4,41 | 0,00 | 1,00E+00 |
| AT4G10840 | 1,51 | 3,90 | 2,03 | 1,92 | 5,47E-02 |
| AT4G11000 | 0,26 | 1,29 | 4,39 | 0,29 | 7,69E-01 |
| AT4G11070 | 0,37 | 0,76 | 4,35 | 0,17 | 8,61E-01 |
| AT4G11100 | 0,32 | 1,30 | 4,40 | 0,30 | 7,68E-01 |
| AT4G11110 | 0,51 | -1,94 | 2,67 | -0,72 | 4,69E-01 |
| AT4G11130 | 0,54 | 2,88 | 3,89 | 0,74 | 4,59E-01 |

| | | | | | |
|---|---|---|---|---|---|
| AT4G11150 | 1,03 | -2,42 | 1,88 | -1,28 | 2,00E-01 |
| AT4G11160 | 1,61 | 1,36 | 1,74 | 0,78 | 4,34E-01 |
| AT4G11170 | 0,50 | -0,64 | 3,12 | -0,21 | 8,38E-01 |
| AT4G11240 | 1,11 | 3,27 | 2,00 | 1,64 | 1,02E-01 |
| AT4G11280 | 0,67 | 0,79 | 3,11 | 0,25 | 7,99E-01 |
| AT4G11330 | 0,53 | 1,07 | 3,14 | 0,34 | 7,34E-01 |
| AT4G11350 | 0,19 | 0,00 | 4,41 | 0,00 | 1,00E+00 |
| AT4G11360 | 0,31 | -1,56 | 4,34 | -0,36 | 7,19E-01 |
| AT4G05885 | 0,80 | -3,23 | 2,67 | -1,21 | 2,27E-01 |
| AT4G11660 | 0,57 | 2,00 | 3,30 | 0,61 | 5,44E-01 |
| AT4G11720 | 0,60 | -0,20 | 2,98 | -0,07 | 9,47E-01 |
| AT4G11740 | 0,35 | 1,34 | 4,40 | 0,30 | 7,62E-01 |
| AT4G11880 | 0,28 | -0,54 | 4,41 | -0,12 | 9,02E-01 |
| AT4G11890 | 0,62 | -0,58 | 3,15 | -0,18 | 8,54E-01 |
| AT4G11910 | 0,12 | 0,00 | 4,41 | 0,00 | 1,00E+00 |
| AT4G12020 | 1,23 | 2,05 | 2,68 | 0,76 | 4,44E-01 |
| AT4G12030 | 0,68 | 0,00 | 3,60 | 0,00 | 1,00E+00 |
| AT4G12040 | 1,45 | 0,01 | 1,96 | 0,01 | 9,95E-01 |
| AT4G12070 | 0,19 | -0,54 | 4,41 | -0,12 | 9,02E-01 |
| AT4G12240 | 0,26 | 0,00 | 4,41 | 0,00 | 1,00E+00 |
| AT4G05985 | 0,10 | 0,00 | 4,41 | 0,00 | 1,00E+00 |
| AT4G12390 | 0,50 | 1,10 | 4,37 | 0,25 | 8,01E-01 |
| AT4G12600 | 0,92 | 0,28 | 3,06 | 0,09 | 9,27E-01 |
| AT4G12640 | 0,26 | -0,51 | 3,83 | -0,13 | 8,94E-01 |
| AT4G12680 | 0,21 | 0,00 | 4,41 | 0,00 | 1,00E+00 |
| AT4G12690 | 0,12 | 0,00 | 4,41 | 0,00 | 1,00E+00 |
| AT4G12720 | 1,54 | 2,86 | 1,88 | 1,52 | 1,29E-01 |
| AT4G12830 | 0,34 | -0,82 | 4,38 | -0,19 | 8,51E-01 |
| AT4G12900 | 0,05 | 0,00 | 4,41 | 0,00 | 1,00E+00 |
| AT4G05995 | 0,98 | 0,00 | 2,66 | 0,00 | 1,00E+00 |
| AT4G13010 | 0,63 | -1,96 | 3,63 | -0,54 | 5,90E-01 |
| AT4G13020 | 0,54 | -0,43 | 3,24 | -0,13 | 8,94E-01 |
| AT4G06010 | 0,28 | 0,00 | 4,41 | 0,00 | 1,00E+00 |
| AT4G06015 | 0,10 | 0,00 | 4,41 | 0,00 | 1,00E+00 |
| AT4G13245 | 0,48 | 0,38 | 3,30 | 0,11 | 9,09E-01 |
| AT4G13270 | 0,50 | -0,43 | 3,24 | -0,13 | 8,96E-01 |
| AT4G13390 | 1,53 | 1,38 | 2,19 | 0,63 | 5,29E-01 |
| AT4G13440 | 0,58 | 0,73 | 4,08 | 0,18 | 8,58E-01 |
| AT4G13530 | 0,95 | -0,64 | 2,12 | -0,30 | 7,63E-01 |
| AT4G13572 | 0,31 | 1,36 | 4,35 | 0,31 | 7,55E-01 |
| AT4G13610 | 0,37 | -0,46 | 2,95 | -0,16 | 8,75E-01 |
| AT4G13615 | 0,92 | 1,37 | 2,01 | 0,68 | 4,95E-01 |
| AT4G13670 | 0,92 | 1,38 | 3,31 | 0,42 | 6,77E-01 |
| AT4G13730 | 0,24 | 0,00 | 4,40 | 0,00 | 1,00E+00 |
| AT4G13750 | 0,39 | -0,98 | 3,43 | -0,29 | 7,75E-01 |
| AT4G13760 | 0,54 | 0,00 | 3,27 | 0,00 | 1,00E+00 |
| AT4G13830 | 0,70 | 0,42 | 2,99 | 0,14 | 8,87E-01 |
| AT4G13850 | 0,81 | -1,35 | 2,20 | -0,61 | 5,40E-01 |
| AT4G13860 | 0,43 | 0,00 | 4,19 | 0,00 | 1,00E+00 |
| AT4G13992 | 1,30 | -1,96 | 2,08 | -0,94 | 3,45E-01 |
| AT4G14000 | 0,12 | 0,00 | 4,41 | 0,00 | 1,00E+00 |
| AT4G14140 | 0,81 | -0,74 | 2,19 | -0,34 | 7,34E-01 |
| AT4G14220 | 0,46 | 2,44 | 4,35 | 0,56 | 5,75E-01 |
| AT4G14270 | 0,92 | 1,67 | 1,97 | 0,85 | 3,97E-01 |
| AT4G14300 | 0,77 | 1,13 | 3,46 | 0,33 | 7,44E-01 |
| AT4G14330 | 0,14 | 0,00 | 4,41 | 0,00 | 1,00E+00 |

| | | | | | |
|---|---|---|---|---|---|
| AT4G14540 | 0,71 | -2,08 | 2,55 | -0,81 | 4,16E-01 |
| AT4G14548 | 1,46 | 0,42 | 2,16 | 0,19 | 8,46E-01 |
| AT4G14560 | 0,12 | 0,00 | 4,41 | 0,00 | 1,00E+00 |
| AT4G14570 | 0,70 | 2,45 | 2,85 | 0,86 | 3,90E-01 |
| AT4G14630 | 0,43 | -0,41 | 4,09 | -0,10 | 9,21E-01 |
| AT4G14700 | 0,12 | 0,00 | 4,41 | 0,00 | 1,00E+00 |
| AT4G06245 | 1,52 | -2,85 | 2,06 | -1,38 | 1,67E-01 |
| AT4G06255 | 0,39 | 1,82 | 4,38 | 0,42 | 6,78E-01 |
| AT4G06260 | 0,42 | 2,12 | 3,33 | 0,64 | 5,25E-01 |
| AT4G14760 | 0,91 | 2,09 | 2,36 | 0,89 | 3,76E-01 |
| AT4G14800 | 0,34 | 1,38 | 4,41 | 0,31 | 7,54E-01 |
| AT4G14810 | 0,45 | 0,50 | 3,42 | 0,15 | 8,83E-01 |
| AT4G14850 | 0,15 | 0,00 | 4,41 | 0,00 | 1,00E+00 |
| AT4G14870 | 0,48 | 0,53 | 2,77 | 0,19 | 8,49E-01 |
| AT4G14890 | 0,73 | 0,67 | 2,32 | 0,29 | 7,71E-01 |
| AT4G14950 | 0,54 | 0,62 | 2,69 | 0,23 | 8,18E-01 |
| AT4G15000 | 1,11 | -0,57 | 1,72 | -0,33 | 7,43E-01 |
| AT4G15020 | 0,86 | 0,38 | 2,50 | 0,15 | 8,80E-01 |
| AT4G15030 | 0,61 | -0,44 | 2,44 | -0,18 | 8,56E-01 |
| AT4G15090 | 0,08 | 0,00 | 4,41 | 0,00 | 1,00E+00 |
| AT4G15096 | 1,15 | -1,71 | 2,28 | -0,75 | 4,53E-01 |
| AT4G15165 | 0,40 | 2,56 | 4,35 | 0,59 | 5,56E-01 |
| AT4G15233 | 0,90 | 2,73 | 2,41 | 1,14 | 2,56E-01 |
| AT4G15245 | 0,49 | -0,42 | 2,76 | -0,15 | 8,80E-01 |
| AT4G15260 | 0,57 | 2,48 | 3,05 | 0,81 | 4,17E-01 |
| AT4G15280 | 1,02 | 1,15 | 2,31 | 0,50 | 6,19E-01 |
| AT4G15415 | 0,24 | 1,30 | 4,40 | 0,30 | 7,68E-01 |
| AT4G15420 | 1,24 | 3,67 | 2,13 | 1,72 | 8,49E-02 |
| AT4G15440 | 0,99 | 1,24 | 2,33 | 0,53 | 5,96E-01 |
| AT4G15450 | 0,24 | 1,45 | 4,40 | 0,33 | 7,41E-01 |
| AT4G15475 | 0,96 | 3,94 | 4,01 | 0,98 | 3,26E-01 |
| AT4G15540 | 0,39 | 1,38 | 3,56 | 0,39 | 6,99E-01 |
| AT4G15545 | 1,17 | -0,07 | 1,79 | -0,04 | 9,70E-01 |
| AT4G15660 | 0,10 | 0,00 | 4,41 | 0,00 | 1,00E+00 |
| AT4G15700 | 0,08 | 0,00 | 4,41 | 0,00 | 1,00E+00 |
| AT4G15720 | 0,08 | 0,00 | 4,41 | 0,00 | 1,00E+00 |
| AT4G15770 | 0,18 | 1,38 | 4,41 | 0,31 | 7,54E-01 |
| AT4G15775 | 0,24 | -1,71 | 4,34 | -0,39 | 6,93E-01 |
| AT4G15800 | 0,76 | 3,20 | 2,15 | 1,49 | 1,36E-01 |
| AT4G15820 | 0,10 | 0,00 | 4,41 | 0,00 | 1,00E+00 |
| AT4G15840 | 0,73 | 1,92 | 2,05 | 0,94 | 3,49E-01 |
| AT4G15850 | 0,38 | 2,11 | 4,37 | 0,48 | 6,29E-01 |
| AT4G15880 | 0,12 | 0,00 | 4,41 | 0,00 | 1,00E+00 |
| AT4G15940 | 0,10 | 0,00 | 4,41 | 0,00 | 1,00E+00 |
| AT4G16008 | 0,08 | 0,00 | 4,41 | 0,00 | 1,00E+00 |
| AT4G16015 | 0,19 | 0,00 | 4,41 | 0,00 | 1,00E+00 |
| AT4G16130 | 1,05 | 1,75 | 2,23 | 0,79 | 4,31E-01 |
| AT4G16140 | 0,34 | -0,52 | 4,40 | -0,12 | 9,06E-01 |
| AT4G16150 | 0,50 | 1,49 | 4,17 | 0,36 | 7,20E-01 |
| AT4G16310 | 1,01 | 1,43 | 2,24 | 0,64 | 5,24E-01 |
| AT4G16360 | 0,61 | -0,85 | 2,62 | -0,32 | 7,46E-01 |
| AT4G16380 | 0,81 | 2,72 | 2,57 | 1,06 | 2,90E-01 |
| AT4G16390 | 0,67 | 1,40 | 2,37 | 0,59 | 5,55E-01 |
| AT4G16400 | 1,31 | 2,85 | 2,00 | 1,42 | 1,55E-01 |
| AT4G16430 | 0,28 | 1,91 | 4,39 | 0,44 | 6,63E-01 |
| AT4G16480 | 0,65 | 0,00 | 3,66 | 0,00 | 1,00E+00 |

| | | | | | |
|---|---|---|---|---|---|
| AT4G16510 | 0,53 | -0,84 | 4,01 | -0,21 | 8,34E-01 |
| AT4G16590 | 0,08 | 0,00 | 4,41 | 0,00 | 1,00E+00 |
| AT4G16650 | 0,41 | -0,45 | 4,22 | -0,11 | 9,14E-01 |
| AT4G16670 | 0,65 | 1,97 | 2,27 | 0,87 | 3,86E-01 |
| AT4G16710 | 0,63 | 1,31 | 2,65 | 0,50 | 6,20E-01 |
| AT4G16780 | 0,60 | 0,42 | 3,95 | 0,11 | 9,15E-01 |
| AT4G16830 | 0,78 | -0,39 | 3,07 | -0,13 | 8,98E-01 |
| AT4G16845 | 0,11 | 0,00 | 4,41 | 0,00 | 1,00E+00 |
| AT4G16850 | 0,29 | 1,38 | 4,41 | 0,31 | 7,54E-01 |
| AT4G16855 | 0,17 | 0,00 | 4,41 | 0,00 | 1,00E+00 |
| AT4G16955 | 0,78 | 3,60 | 3,02 | 1,19 | 2,33E-01 |
| AT4G17010 | 0,55 | -2,17 | 4,33 | -0,50 | 6,17E-01 |
| AT4G17080 | 0,57 | 2,09 | 4,37 | 0,48 | 6,34E-01 |
| AT4G17085 | 0,42 | 2,02 | 4,37 | 0,46 | 6,44E-01 |
| AT4G06665 | 1,22 | -0,83 | 1,86 | -0,44 | 6,58E-01 |
| AT4G17098 | 0,61 | -2,47 | 2,56 | -0,97 | 3,34E-01 |
| AT4G17245 | 0,66 | 2,15 | 2,64 | 0,81 | 4,16E-01 |
| AT4G17250 | 0,41 | 1,72 | 4,38 | 0,39 | 6,94E-01 |
| AT4G17270 | 0,64 | 2,90 | 3,01 | 0,96 | 3,37E-01 |
| AT4G17300 | 0,71 | 0,96 | 2,35 | 0,41 | 6,82E-01 |
| AT4G17380 | 0,44 | 2,09 | 2,82 | 0,74 | 4,58E-01 |
| AT4G17500 | 0,96 | 1,38 | 2,44 | 0,57 | 5,72E-01 |
| AT4G17520 | 1,02 | -0,10 | 2,61 | -0,04 | 9,71E-01 |
| AT4G17600 | 1,35 | -1,63 | 1,76 | -0,92 | 3,56E-01 |
| AT4G17616 | 0,78 | 2,76 | 3,44 | 0,80 | 4,22E-01 |
| AT4G17620 | 1,00 | 2,20 | 2,51 | 0,88 | 3,80E-01 |
| AT4G17650 | 0,32 | 0,48 | 4,40 | 0,11 | 9,13E-01 |
| AT4G17770 | 1,15 | 0,38 | 1,94 | 0,19 | 8,46E-01 |
| AT4G17810 | 0,14 | -0,78 | 4,37 | -0,18 | 8,58E-01 |
| AT4G17830 | 0,83 | 0,35 | 2,65 | 0,13 | 8,95E-01 |
| AT4G17870 | 1,03 | -0,50 | 2,50 | -0,20 | 8,40E-01 |
| AT4G17900 | 0,47 | 0,28 | 2,85 | 0,10 | 9,23E-01 |
| AT4G18070 | 1,02 | -1,01 | 2,34 | -0,43 | 6,66E-01 |
| AT4G18160 | 0,40 | 2,57 | 4,33 | 0,59 | 5,53E-01 |
| AT4G18195 | 0,22 | 1,35 | 4,40 | 0,31 | 7,59E-01 |
| AT4G18205 | 1,32 | -1,76 | 1,79 | -0,98 | 3,25E-01 |
| AT4G18220 | 1,16 | 0,32 | 1,85 | 0,17 | 8,64E-01 |
| AT4G18240 | 1,25 | 0,28 | 2,00 | 0,14 | 8,90E-01 |
| AT4G18370 | 0,87 | 1,75 | 2,35 | 0,75 | 4,54E-01 |
| AT4G18380 | 0,33 | 0,50 | 4,37 | 0,12 | 9,08E-01 |
| AT4G06795 | 0,56 | 2,47 | 3,06 | 0,81 | 4,19E-01 |
| AT4G18530 | 0,35 | -0,54 | 4,41 | -0,12 | 9,02E-01 |
| AT4G18570 | 0,37 | 0,37 | 2,98 | 0,12 | 9,01E-01 |
| AT4G18710 | 1,19 | 1,40 | 2,01 | 0,70 | 4,86E-01 |
| AT4G18725 | 0,19 | 0,00 | 4,40 | 0,00 | 1,00E+00 |
| AT4G18730 | 1,33 | 3,16 | 1,79 | 1,77 | 7,66E-02 |
| AT4G18750 | 0,50 | 1,38 | 4,14 | 0,33 | 7,39E-01 |
| AT4G18820 | 1,31 | 1,24 | 1,97 | 0,63 | 5,31E-01 |
| AT4G18830 | 0,19 | 0,00 | 4,41 | 0,00 | 1,00E+00 |
| AT4G18890 | 0,44 | 1,37 | 4,13 | 0,33 | 7,40E-01 |
| AT4G18905 | 0,35 | 1,38 | 4,41 | 0,31 | 7,54E-01 |
| AT4G18930 | 0,31 | -0,93 | 4,38 | -0,21 | 8,31E-01 |
| AT4G18950 | 1,00 | -0,34 | 2,22 | -0,15 | 8,77E-01 |
| AT4G19020 | 0,17 | 0,00 | 4,41 | 0,00 | 1,00E+00 |
| AT4G06865 | 0,10 | 0,00 | 4,41 | 0,00 | 1,00E+00 |
| AT4G19090 | 1,32 | 1,28 | 1,81 | 0,71 | 4,79E-01 |

| | | | | | |
|---|---|---|---|---|---|
| AT4G19100 | 1,38 | -1,77 | 2,32 | -0,76 | 4,45E-01 |
| AT4G19190 | 0,39 | -0,51 | 4,26 | -0,12 | 9,04E-01 |
| AT4G19200 | 0,86 | 0,40 | 2,24 | 0,18 | 8,59E-01 |
| AT4G19210 | 1,66 | 1,18 | 1,60 | 0,74 | 4,59E-01 |
| AT4G19450 | 0,65 | 0,00 | 4,41 | 0,00 | 1,00E+00 |
| AT4G19490 | 0,82 | 2,87 | 2,78 | 1,03 | 3,02E-01 |
| AT4G19500 | 0,89 | 2,83 | 2,33 | 1,22 | 2,23E-01 |
| AT4G19510 | 1,30 | 2,85 | 2,04 | 1,39 | 1,63E-01 |
| AT4G19530 | 1,16 | 1,74 | 1,91 | 0,91 | 3,61E-01 |
| AT4G19540 | 0,68 | 1,38 | 3,06 | 0,45 | 6,52E-01 |
| AT4G19580 | 0,44 | 0,00 | 4,14 | 0,00 | 1,00E+00 |
| AT4G19600 | 0,21 | -1,19 | 4,36 | -0,27 | 7,85E-01 |
| AT4G19610 | 0,37 | 0,42 | 3,52 | 0,12 | 9,04E-01 |
| AT4G19650 | 0,19 | 0,00 | 4,41 | 0,00 | 1,00E+00 |
| AT4G19710 | 1,47 | 1,81 | 1,67 | 1,09 | 2,77E-01 |
| AT4G19960 | 0,50 | 1,36 | 3,31 | 0,41 | 6,81E-01 |
| AT4G20110 | 0,83 | 2,39 | 2,34 | 1,02 | 3,05E-01 |
| AT4G20140 | 0,84 | 0,00 | 4,30 | 0,00 | 1,00E+00 |
| AT4G06950 | 0,12 | 0,00 | 4,41 | 0,00 | 1,00E+00 |
| AT4G20270 | 0,28 | 0,00 | 4,41 | 0,00 | 1,00E+00 |
| AT4G20300 | 0,68 | 0,00 | 2,49 | 0,00 | 1,00E+00 |
| AT4G20310 | 0,08 | 0,00 | 4,41 | 0,00 | 1,00E+00 |
| AT4G20400 | 1,23 | 1,23 | 2,11 | 0,58 | 5,62E-01 |
| AT4G20720 | 0,34 | 1,46 | 4,40 | 0,33 | 7,39E-01 |
| AT4G20840 | 1,22 | 3,89 | 2,42 | 1,61 | 1,08E-01 |
| AT4G20860 | 0,52 | 1,27 | 3,21 | 0,39 | 6,93E-01 |
| AT4G20890 | 0,68 | -0,11 | 2,67 | -0,04 | 9,66E-01 |
| AT4G20940 | 0,56 | 1,98 | 4,38 | 0,45 | 6,50E-01 |
| AT4G20960 | 0,48 | 1,38 | 2,84 | 0,49 | 6,28E-01 |
| AT4G20980 | 1,36 | -0,15 | 1,99 | -0,08 | 9,40E-01 |
| AT4G21105 | 0,54 | 1,72 | 4,35 | 0,40 | 6,92E-01 |
| AT4G21160 | 0,15 | -0,43 | 4,39 | -0,10 | 9,23E-01 |
| AT4G21180 | 0,08 | 0,00 | 4,41 | 0,00 | 1,00E+00 |
| AT4G21210 | 0,33 | -0,54 | 4,41 | -0,12 | 9,02E-01 |
| AT4G21300 | 0,05 | 0,00 | 4,41 | 0,00 | 1,00E+00 |
| AT4G21323 | 0,38 | 2,11 | 4,37 | 0,48 | 6,29E-01 |
| AT4G21450 | 0,15 | 0,00 | 4,41 | 0,00 | 1,00E+00 |
| AT4G21470 | 0,32 | -0,49 | 4,40 | -0,11 | 9,12E-01 |
| AT4G21515 | 0,05 | 0,00 | 4,41 | 0,00 | 1,00E+00 |
| AT4G21540 | 0,68 | 0,00 | 3,00 | 0,00 | 1,00E+00 |
| AT4G21550 | 0,37 | 1,76 | 4,38 | 0,40 | 6,87E-01 |
| AT4G21620 | 0,72 | 0,94 | 2,62 | 0,36 | 7,19E-01 |
| AT4G21670 | 1,62 | -0,11 | 1,51 | -0,07 | 9,43E-01 |
| AT4G21740 | 0,47 | -0,34 | 2,49 | -0,14 | 8,92E-01 |
| AT4G21750 | 0,81 | 1,49 | 2,82 | 0,53 | 5,98E-01 |
| AT4G21790 | 0,64 | 0,98 | 2,68 | 0,37 | 7,15E-01 |
| AT4G21800 | 0,53 | 2,48 | 3,87 | 0,64 | 5,21E-01 |
| AT4G07030 | 0,38 | 1,38 | 4,41 | 0,31 | 7,54E-01 |
| AT4G21940 | 0,15 | 0,00 | 4,41 | 0,00 | 1,00E+00 |
| AT4G22010 | 0,22 | -0,47 | 4,40 | -0,11 | 9,15E-01 |
| AT4G22050 | 0,20 | 0,00 | 4,41 | 0,00 | 1,00E+00 |
| AT4G07040 | 0,12 | 0,00 | 4,41 | 0,00 | 1,00E+00 |
| AT4G22130 | 0,44 | 1,38 | 2,88 | 0,48 | 6,32E-01 |
| AT4G22270 | 0,20 | 1,38 | 4,41 | 0,31 | 7,54E-01 |
| AT4G07075 | 1,23 | 0,38 | 2,05 | 0,19 | 8,52E-01 |
| AT4G07080 | 1,23 | 0,38 | 2,05 | 0,19 | 8,52E-01 |

| | | | | | |
|---|---|---|---|---|---|
| AT4G22330 | 0,17 | 0,00 | 4,41 | 0,00 | 1,00E+00 |
| AT4G22467 | 0,12 | 0,00 | 4,41 | 0,00 | 1,00E+00 |
| AT4G22475 | 0,49 | -0,40 | 3,96 | -0,10 | 9,20E-01 |
| AT4G22670 | 0,32 | -0,48 | 4,40 | -0,11 | 9,13E-01 |
| AT4G22720 | 0,28 | 1,34 | 4,40 | 0,30 | 7,61E-01 |
| AT4G22860 | 0,08 | 0,00 | 4,41 | 0,00 | 1,00E+00 |
| AT4G22890 | 1,50 | -0,87 | 1,68 | -0,52 | 6,07E-01 |
| AT4G22910 | 0,08 | 0,00 | 4,41 | 0,00 | 1,00E+00 |
| AT4G22960 | 0,12 | 0,00 | 4,41 | 0,00 | 1,00E+00 |
| AT4G23030 | 0,17 | 1,32 | 4,40 | 0,30 | 7,64E-01 |
| AT4G23050 | 0,25 | 1,30 | 4,40 | 0,30 | 7,67E-01 |
| AT4G23060 | 0,21 | 1,27 | 4,39 | 0,29 | 7,72E-01 |
| AT4G23150 | 0,15 | -0,43 | 4,39 | -0,10 | 9,22E-01 |
| AT4G23160 | 0,75 | 3,22 | 4,32 | 0,75 | 4,55E-01 |
| AT4G23180 | 1,57 | 2,21 | 1,71 | 1,30 | 1,95E-01 |
| AT4G23270 | 0,94 | 0,43 | 2,26 | 0,19 | 8,50E-01 |
| AT4G23400 | 0,56 | -1,68 | 2,35 | -0,72 | 4,73E-01 |
| AT4G23440 | 0,12 | 0,00 | 4,41 | 0,00 | 1,00E+00 |
| AT4G23470 | 0,84 | 1,38 | 2,28 | 0,61 | 5,45E-01 |
| AT4G23530 | 1,11 | 3,74 | 2,06 | 1,82 | 6,90E-02 |
| AT4G23570 | 0,44 | 0,21 | 3,37 | 0,06 | 9,51E-01 |
| AT4G23600 | 1,02 | -2,31 | 2,65 | -0,87 | 3,83E-01 |
| AT4G23740 | 0,40 | 0,00 | 4,41 | 0,00 | 1,00E+00 |
| AT4G23860 | 0,25 | 1,43 | 4,40 | 0,32 | 7,46E-01 |
| AT4G07190 | 0,45 | -0,35 | 3,43 | -0,10 | 9,20E-01 |
| AT4G23895 | 0,24 | -0,50 | 4,40 | -0,11 | 9,10E-01 |
| AT4G23940 | 1,06 | 2,72 | 2,29 | 1,19 | 2,35E-01 |
| AT4G23950 | 0,16 | 0,00 | 4,41 | 0,00 | 1,00E+00 |
| AT4G23980 | 1,47 | 1,44 | 1,63 | 0,88 | 3,77E-01 |
| AT4G24000 | 0,08 | 0,00 | 4,41 | 0,00 | 1,00E+00 |
| AT4G24010 | 0,33 | 0,50 | 4,37 | 0,12 | 9,08E-01 |
| AT4G24020 | 1,49 | 2,92 | 1,72 | 1,69 | 9,08E-02 |
| AT4G24060 | 0,12 | 0,00 | 4,41 | 0,00 | 1,00E+00 |
| AT4G24090 | 0,53 | -2,93 | 4,30 | -0,68 | 4,95E-01 |
| AT4G24120 | 1,68 | 1,38 | 2,37 | 0,58 | 5,61E-01 |
| AT4G24130 | 0,24 | 1,96 | 4,39 | 0,45 | 6,55E-01 |
| AT4G24160 | 0,15 | 0,00 | 4,41 | 0,00 | 1,00E+00 |
| AT4G24175 | 0,21 | 1,38 | 4,41 | 0,31 | 7,54E-01 |
| AT4G24220 | 1,52 | -0,15 | 1,54 | -0,10 | 9,23E-01 |
| AT4G24290 | 0,72 | 0,88 | 2,54 | 0,35 | 7,29E-01 |
| AT4G24350 | 0,25 | -0,47 | 4,40 | -0,11 | 9,15E-01 |
| AT4G24440 | 0,43 | 0,89 | 4,26 | 0,21 | 8,34E-01 |
| AT4G24460 | 0,18 | -0,82 | 4,37 | -0,19 | 8,51E-01 |
| AT4G24470 | 0,57 | 1,38 | 3,94 | 0,35 | 7,27E-01 |
| AT4G24550 | 0,38 | 0,23 | 3,45 | 0,07 | 9,46E-01 |
| AT4G24790 | 0,10 | 0,00 | 4,41 | 0,00 | 1,00E+00 |
| AT4G24800 | 0,98 | 3,09 | 2,65 | 1,17 | 2,44E-01 |
| AT4G24805 | 0,88 | -0,01 | 2,48 | -0,01 | 9,96E-01 |
| AT4G24880 | 0,08 | 0,00 | 4,41 | 0,00 | 1,00E+00 |
| AT4G07295 | 0,21 | -0,45 | 4,39 | -0,10 | 9,19E-01 |
| AT4G25030 | 0,40 | -1,27 | 2,87 | -0,44 | 6,57E-01 |
| AT4G25050 | 1,37 | 0,60 | 1,62 | 0,37 | 7,10E-01 |
| AT4G25070 | 0,53 | 1,16 | 4,38 | 0,27 | 7,90E-01 |
| AT4G25120 | 0,50 | 0,68 | 3,24 | 0,21 | 8,34E-01 |
| AT4G25170 | 1,05 | 0,64 | 1,98 | 0,33 | 7,45E-01 |
| AT4G25210 | 0,75 | 2,56 | 2,89 | 0,89 | 3,75E-01 |

| | | | | | |
|---|---|---|---|---|---|
| AT4G25230 | 0,77 | -0,62 | 2,38 | -0,26 | 7,96E-01 |
| AT4G25360 | 0,25 | 0,00 | 4,41 | 0,00 | 1,00E+00 |
| AT4G25370 | 0,59 | 0,90 | 3,85 | 0,23 | 8,15E-01 |
| AT4G25390 | 0,17 | 0,00 | 4,41 | 0,00 | 1,00E+00 |
| AT4G25400 | 0,31 | 0,00 | 4,41 | 0,00 | 1,00E+00 |
| AT4G25500 | 1,12 | 1,36 | 2,16 | 0,63 | 5,30E-01 |
| AT4G25515 | 0,32 | 1,45 | 4,40 | 0,33 | 7,42E-01 |
| AT4G25550 | 0,38 | 2,11 | 4,37 | 0,48 | 6,29E-01 |
| AT4G25610 | 0,54 | 2,30 | 3,19 | 0,72 | 4,72E-01 |
| AT4G25630 | 1,57 | 1,95 | 2,04 | 0,95 | 3,40E-01 |
| AT4G25631 | 1,33 | 2,88 | 2,22 | 1,30 | 1,95E-01 |
| AT4G25660 | 0,81 | 0,73 | 2,32 | 0,31 | 7,53E-01 |
| AT4G25810 | 0,12 | 0,00 | 4,41 | 0,00 | 1,00E+00 |
| AT4G25835 | 0,21 | -0,54 | 4,41 | -0,12 | 9,02E-01 |
| AT4G25900 | 0,36 | 0,42 | 4,41 | 0,10 | 9,24E-01 |
| AT4G25910 | 0,69 | 2,80 | 3,52 | 0,79 | 4,27E-01 |
| AT4G25960 | 1,53 | -1,15 | 1,62 | -0,71 | 4,78E-01 |
| AT4G25970 | 1,00 | 1,80 | 2,01 | 0,89 | 3,71E-01 |
| AT4G26000 | 0,96 | 2,71 | 2,47 | 1,10 | 2,72E-01 |
| AT4G26120 | 0,24 | 1,96 | 4,39 | 0,45 | 6,55E-01 |
| AT4G26170 | 0,08 | 0,00 | 4,41 | 0,00 | 1,00E+00 |
| AT4G26210 | 0,36 | 0,00 | 4,41 | 0,00 | 1,00E+00 |
| AT4G26230 | 0,82 | 2,91 | 2,56 | 1,14 | 2,56E-01 |
| AT4G26450 | 0,45 | 0,47 | 3,37 | 0,14 | 8,89E-01 |
| AT4G26455 | 0,54 | 2,10 | 3,82 | 0,55 | 5,82E-01 |
| AT4G26510 | 0,72 | 0,99 | 2,51 | 0,40 | 6,92E-01 |
| AT4G26520 | 1,37 | 4,27 | 2,01 | 2,12 | 3,38E-02 |
| AT4G26650 | 0,56 | 1,25 | 2,36 | 0,53 | 5,97E-01 |
| AT4G26670 | 0,33 | -1,28 | 4,35 | -0,29 | 7,69E-01 |
| AT4G26680 | 0,40 | 0,00 | 4,23 | 0,00 | 1,00E+00 |
| AT4G26700 | 0,28 | -0,50 | 4,40 | -0,11 | 9,10E-01 |
| AT4G26790 | 0,42 | 0,00 | 4,15 | 0,00 | 1,00E+00 |
| AT4G26800 | 0,12 | 0,00 | 4,41 | 0,00 | 1,00E+00 |
| AT4G26840 | 0,82 | 1,38 | 2,26 | 0,61 | 5,41E-01 |
| AT4G26870 | 0,92 | 2,83 | 2,27 | 1,24 | 2,13E-01 |
| AT4G27010 | 0,79 | 1,26 | 2,29 | 0,55 | 5,82E-01 |
| AT4G27030 | 0,63 | -1,31 | 2,60 | -0,50 | 6,15E-01 |
| AT4G07775 | 0,58 | 0,00 | 3,08 | 0,00 | 1,00E+00 |
| AT4G27080 | 1,09 | 1,77 | 2,20 | 0,80 | 4,21E-01 |
| AT4G27260 | 0,26 | 1,38 | 4,41 | 0,31 | 7,54E-01 |
| AT4G27290 | 0,45 | 2,55 | 4,36 | 0,58 | 5,59E-01 |
| AT4G27310 | 0,93 | -0,67 | 2,17 | -0,31 | 7,57E-01 |
| AT4G27380 | 0,21 | -1,36 | 4,36 | -0,31 | 7,55E-01 |
| AT4G27395 | 0,73 | -2,87 | 2,78 | -1,03 | 3,02E-01 |
| AT4G27652 | 0,57 | 0,00 | 3,80 | 0,00 | 1,00E+00 |
| AT4G27654 | 0,56 | 0,00 | 2,77 | 0,00 | 1,00E+00 |
| AT4G27657 | 0,12 | 0,00 | 4,41 | 0,00 | 1,00E+00 |
| AT4G27680 | 0,93 | 2,39 | 2,22 | 1,08 | 2,81E-01 |
| AT4G27710 | 0,52 | -1,50 | 3,30 | -0,45 | 6,49E-01 |
| AT4G27720 | 1,38 | 1,58 | 1,96 | 0,80 | 4,22E-01 |
| AT4G27740 | 0,52 | 0,00 | 3,89 | 0,00 | 1,00E+00 |
| AT4G27750 | 0,69 | 0,72 | 4,33 | 0,17 | 8,69E-01 |
| AT4G27760 | 0,28 | 1,84 | 4,39 | 0,42 | 6,74E-01 |
| AT4G27765 | 0,37 | 2,13 | 4,32 | 0,49 | 6,22E-01 |
| AT4G27780 | 0,50 | 2,51 | 3,95 | 0,64 | 5,24E-01 |
| AT4G27860 | 0,21 | 0,00 | 4,41 | 0,00 | 1,00E+00 |

| | | | | | |
|---|---|---|---|---|---|
| AT4G27880 | 0,08 | 0,00 | 4,41 | 0,00 | 1,00E+00 |
| AT4G27910 | 0,46 | 2,20 | 4,10 | 0,54 | 5,91E-01 |
| AT4G27940 | 1,14 | 3,52 | 2,20 | 1,60 | 1,09E-01 |
| AT4G28010 | 0,17 | 0,00 | 4,41 | 0,00 | 1,00E+00 |
| AT4G28210 | 0,08 | 0,00 | 4,41 | 0,00 | 1,00E+00 |
| AT4G28220 | 0,27 | 0,00 | 4,41 | 0,00 | 1,00E+00 |
| AT4G28260 | 0,80 | 1,38 | 3,03 | 0,46 | 6,49E-01 |
| AT4G28300 | 0,21 | 0,00 | 4,41 | 0,00 | 1,00E+00 |
| AT4G28362 | 0,21 | 1,27 | 4,39 | 0,29 | 7,72E-01 |
| AT4G28400 | 0,26 | 1,30 | 4,40 | 0,30 | 7,67E-01 |
| AT4G28440 | 0,16 | 0,00 | 4,41 | 0,00 | 1,00E+00 |
| AT4G28490 | 0,74 | 0,00 | 3,72 | 0,00 | 1,00E+00 |
| AT4G28600 | 1,03 | 0,97 | 2,26 | 0,43 | 6,67E-01 |
| AT4G28660 | 0,48 | -0,58 | 4,05 | -0,14 | 8,87E-01 |
| AT4G28706 | 0,39 | -1,43 | 4,33 | -0,33 | 7,40E-01 |
| AT4G28710 | 0,41 | 1,50 | 4,26 | 0,35 | 7,24E-01 |
| AT4G28740 | 0,90 | -0,14 | 2,63 | -0,05 | 9,57E-01 |
| AT4G28780 | 0,15 | 0,00 | 4,41 | 0,00 | 1,00E+00 |
| AT4G28860 | 0,76 | 1,56 | 3,03 | 0,52 | 6,06E-01 |
| AT4G28870 | 0,12 | 0,00 | 4,41 | 0,00 | 1,00E+00 |
| AT4G28880 | 1,20 | 2,95 | 1,69 | 1,75 | 8,04E-02 |
| AT4G28980 | 0,73 | 0,39 | 2,61 | 0,15 | 8,81E-01 |
| AT4G29020 | 0,17 | 1,32 | 4,40 | 0,30 | 7,64E-01 |
| AT4G29035 | 0,22 | 1,35 | 4,40 | 0,31 | 7,59E-01 |
| AT4G29040 | 1,10 | 1,38 | 2,29 | 0,60 | 5,46E-01 |
| AT4G29070 | 0,33 | 1,28 | 4,39 | 0,29 | 7,71E-01 |
| AT4G29110 | 0,63 | 1,44 | 3,01 | 0,48 | 6,34E-01 |
| AT4G29120 | 0,68 | 0,00 | 2,98 | 0,00 | 1,00E+00 |
| AT4G29160 | 0,87 | 1,78 | 2,04 | 0,87 | 3,82E-01 |
| AT4G29170 | 0,92 | 3,05 | 4,32 | 0,71 | 4,80E-01 |
| AT4G29180 | 0,96 | 3,62 | 2,59 | 1,40 | 1,62E-01 |
| AT4G29210 | 0,68 | 1,12 | 2,58 | 0,43 | 6,65E-01 |
| AT4G29350 | 0,47 | -0,54 | 4,15 | -0,13 | 8,96E-01 |
| AT4G29400 | 0,21 | 1,25 | 4,39 | 0,29 | 7,75E-01 |
| AT4G29510 | 0,71 | 1,18 | 2,40 | 0,49 | 6,22E-01 |
| AT4G29740 | 1,04 | -0,35 | 2,01 | -0,17 | 8,62E-01 |
| AT4G29750 | 0,66 | 0,91 | 2,66 | 0,34 | 7,32E-01 |
| AT4G29790 | 1,30 | 1,97 | 1,97 | 1,00 | 3,17E-01 |
| AT4G08245 | 0,54 | 1,38 | 3,25 | 0,42 | 6,72E-01 |
| AT4G08255 | 0,51 | 0,00 | 4,41 | 0,00 | 1,00E+00 |
| AT4G29950 | 0,57 | 2,13 | 2,71 | 0,79 | 4,31E-01 |
| AT4G30010 | 0,55 | -0,54 | 3,36 | -0,16 | 8,71E-01 |
| AT4G30020 | 1,12 | 0,06 | 1,82 | 0,04 | 9,72E-01 |
| AT4G30130 | 0,24 | 0,00 | 4,40 | 0,00 | 1,00E+00 |
| AT4G30150 | 1,08 | 1,85 | 2,16 | 0,85 | 3,93E-01 |
| AT4G30310 | 0,60 | 0,72 | 3,71 | 0,19 | 8,47E-01 |
| AT4G30350 | 0,36 | 0,00 | 4,41 | 0,00 | 1,00E+00 |
| AT4G30400 | 0,15 | 0,00 | 4,41 | 0,00 | 1,00E+00 |
| AT4G08285 | 0,08 | 0,00 | 4,41 | 0,00 | 1,00E+00 |
| AT4G30480 | 0,31 | 1,36 | 4,35 | 0,31 | 7,55E-01 |
| AT4G30530 | 1,06 | -2,74 | 1,90 | -1,44 | 1,49E-01 |
| AT4G30650 | 0,58 | -2,09 | 2,75 | -0,76 | 4,47E-01 |
| AT4G30660 | 0,47 | -0,72 | 2,72 | -0,26 | 7,92E-01 |
| AT4G30780 | 0,23 | 1,38 | 4,41 | 0,31 | 7,54E-01 |
| AT4G30800 | 0,17 | -1,42 | 4,35 | -0,33 | 7,45E-01 |
| AT4G30940 | 0,31 | 1,38 | 4,41 | 0,31 | 7,54E-01 |

| | | | | | |
|---|---|---|---|---|---|
| AT4G30990 | 1,34 | 2,62 | 1,85 | 1,41 | 1,57E-01 |
| AT4G08425 | 1,45 | -1,04 | 1,75 | -0,59 | 5,54E-01 |
| AT4G31080 | 0,70 | 2,07 | 2,96 | 0,70 | 4,84E-01 |
| AT4G31115 | 0,40 | 1,45 | 4,23 | 0,34 | 7,31E-01 |
| AT4G31300 | 0,37 | -0,68 | 4,39 | -0,16 | 8,77E-01 |
| AT4G31340 | 0,63 | 0,08 | 3,77 | 0,02 | 9,84E-01 |
| AT4G31390 | 0,47 | 1,23 | 2,81 | 0,44 | 6,63E-01 |
| AT4G31430 | 0,50 | 2,01 | 4,02 | 0,50 | 6,17E-01 |
| AT4G31530 | 0,77 | 0,47 | 2,54 | 0,18 | 8,54E-01 |
| AT4G31580 | 0,97 | 2,27 | 2,23 | 1,02 | 3,08E-01 |
| AT4G31710 | 0,50 | 2,51 | 3,95 | 0,64 | 5,24E-01 |
| AT4G31715 | 0,08 | 0,00 | 4,41 | 0,00 | 1,00E+00 |
| AT4G31780 | 1,36 | 4,14 | 2,82 | 1,47 | 1,41E-01 |
| AT4G31790 | 0,26 | -0,51 | 4,40 | -0,12 | 9,08E-01 |
| AT4G31800 | 1,69 | 3,22 | 1,91 | 1,69 | 9,19E-02 |
| AT4G31805 | 0,98 | 3,14 | 3,31 | 0,95 | 3,44E-01 |
| AT4G31850 | 0,54 | 2,49 | 3,32 | 0,75 | 4,53E-01 |
| AT4G31910 | 0,49 | -1,23 | 2,79 | -0,44 | 6,58E-01 |
| AT4G08615 | 0,91 | -1,35 | 2,33 | -0,58 | 5,63E-01 |
| AT4G08625 | 1,54 | -0,67 | 1,69 | -0,39 | 6,93E-01 |
| AT4G31985 | 0,37 | 1,97 | 4,38 | 0,45 | 6,53E-01 |
| AT4G32010 | 1,47 | 4,23 | 1,98 | 2,14 | 3,25E-02 |
| AT4G32030 | 0,47 | 0,78 | 4,26 | 0,18 | 8,55E-01 |
| AT4G32040 | 0,84 | 1,25 | 3,57 | 0,35 | 7,27E-01 |
| AT4G32060 | 1,61 | -1,78 | 1,69 | -1,05 | 2,92E-01 |
| AT4G32160 | 0,93 | -1,43 | 2,18 | -0,65 | 5,13E-01 |
| AT4G32190 | 0,86 | 1,37 | 2,86 | 0,48 | 6,32E-01 |
| AT4G32200 | 0,65 | 0,36 | 3,69 | 0,10 | 9,22E-01 |
| AT4G32265 | 1,11 | -2,20 | 1,79 | -1,22 | 2,21E-01 |
| AT4G32300 | 0,70 | 2,24 | 2,32 | 0,97 | 3,34E-01 |
| AT4G32330 | 0,80 | 2,35 | 2,29 | 1,03 | 3,04E-01 |
| AT4G32360 | 0,05 | 0,00 | 4,41 | 0,00 | 1,00E+00 |
| AT4G32390 | 1,34 | 0,52 | 2,00 | 0,26 | 7,97E-01 |
| AT4G32430 | 0,12 | 0,00 | 4,41 | 0,00 | 1,00E+00 |
| AT4G32551 | 0,49 | 0,42 | 2,84 | 0,15 | 8,83E-01 |
| AT4G32600 | 0,53 | 1,35 | 3,41 | 0,40 | 6,92E-01 |
| AT4G32620 | 1,61 | 3,19 | 2,16 | 1,48 | 1,40E-01 |
| AT4G32630 | 0,68 | 2,84 | 4,33 | 0,66 | 5,12E-01 |
| AT4G32700 | 0,67 | 2,42 | 3,67 | 0,66 | 5,09E-01 |
| AT4G32720 | 0,62 | 1,18 | 3,04 | 0,39 | 6,99E-01 |
| AT4G32730 | 0,45 | 2,10 | 4,03 | 0,52 | 6,03E-01 |
| AT4G32770 | 0,32 | 1,36 | 4,40 | 0,31 | 7,58E-01 |
| AT4G32800 | 0,31 | 1,30 | 4,40 | 0,30 | 7,68E-01 |
| AT4G32850 | 1,20 | 0,41 | 1,84 | 0,22 | 8,23E-01 |
| AT4G32910 | 0,98 | -1,10 | 2,11 | -0,52 | 6,03E-01 |
| AT4G32930 | 0,91 | 3,40 | 2,43 | 1,40 | 1,61E-01 |
| AT4G33000 | 0,14 | 0,00 | 4,41 | 0,00 | 1,00E+00 |
| AT4G33030 | 0,99 | 1,60 | 2,58 | 0,62 | 5,35E-01 |
| AT4G33060 | 0,15 | 0,00 | 4,41 | 0,00 | 1,00E+00 |
| AT4G33180 | 0,05 | 0,00 | 4,41 | 0,00 | 1,00E+00 |
| AT4G33220 | 0,80 | -1,87 | 3,28 | -0,57 | 5,68E-01 |
| AT4G33290 | 1,50 | 2,86 | 1,86 | 1,53 | 1,25E-01 |
| AT4G08785 | 0,69 | 2,84 | 3,78 | 0,75 | 4,53E-01 |
| AT4G33310 | 0,49 | 2,75 | 4,34 | 0,63 | 5,27E-01 |
| AT4G33380 | 0,54 | 0,34 | 3,39 | 0,10 | 9,21E-01 |
| AT4G33410 | 0,32 | 1,38 | 4,41 | 0,31 | 7,54E-01 |

| | | | | | |
|---|---|---|---|---|---|
| AT4G33510 | 1,23 | -0,13 | 1,68 | -0,08 | 9,37E-01 |
| AT4G33520 | 1,37 | 2,72 | 2,33 | 1,17 | 2,42E-01 |
| AT4G33540 | 0,41 | 0,00 | 4,32 | 0,00 | 1,00E+00 |
| AT4G33565 | 0,80 | 1,40 | 2,88 | 0,49 | 6,26E-01 |
| AT4G33580 | 1,02 | 1,22 | 3,15 | 0,39 | 6,99E-01 |
| AT4G33620 | 0,33 | 1,26 | 4,39 | 0,29 | 7,74E-01 |
| AT4G33630 | 1,15 | 2,65 | 2,03 | 1,30 | 1,92E-01 |
| AT4G33666 | 0,24 | 1,38 | 4,41 | 0,31 | 7,54E-01 |
| AT4G33690 | 0,48 | 2,18 | 4,06 | 0,54 | 5,91E-01 |
| AT4G33890 | 1,49 | 2,43 | 1,79 | 1,36 | 1,74E-01 |
| AT4G33920 | 0,65 | 0,00 | 3,70 | 0,00 | 1,00E+00 |
| AT4G33960 | 0,45 | -2,53 | 4,31 | -0,59 | 5,57E-01 |
| AT4G34030 | 0,40 | -1,45 | 3,44 | -0,42 | 6,73E-01 |
| AT4G34035 | 0,32 | 1,90 | 4,39 | 0,43 | 6,65E-01 |
| AT4G34050 | 1,48 | -1,27 | 1,99 | -0,64 | 5,24E-01 |
| AT4G34060 | 0,12 | 0,00 | 4,41 | 0,00 | 1,00E+00 |
| AT4G34120 | 1,07 | 0,81 | 1,83 | 0,45 | 6,56E-01 |
| AT4G34160 | 0,73 | 1,37 | 2,57 | 0,53 | 5,93E-01 |
| AT4G34240 | 0,30 | 0,49 | 4,40 | 0,11 | 9,12E-01 |
| AT4G34260 | 1,45 | 4,23 | 2,02 | 2,10 | 3,60E-02 |
| AT4G34280 | 0,96 | 3,29 | 2,74 | 1,20 | 2,30E-01 |
| AT4G34290 | 0,10 | 0,00 | 4,41 | 0,00 | 1,00E+00 |
| AT4G34310 | 0,63 | 2,50 | 4,35 | 0,57 | 5,65E-01 |
| AT4G34330 | 0,08 | 0,00 | 4,41 | 0,00 | 1,00E+00 |
| AT4G34370 | 0,50 | 0,99 | 3,92 | 0,25 | 8,00E-01 |
| AT4G34400 | 0,59 | 2,72 | 3,74 | 0,73 | 4,67E-01 |
| AT4G34410 | 0,77 | 0,00 | 2,95 | 0,00 | 1,00E+00 |
| AT4G34430 | 1,22 | 1,52 | 2,20 | 0,69 | 4,91E-01 |
| AT4G34440 | 0,20 | 1,38 | 4,41 | 0,31 | 7,54E-01 |
| AT4G34640 | 0,93 | 2,77 | 2,81 | 0,98 | 3,26E-01 |
| AT4G34680 | 0,14 | 0,00 | 4,41 | 0,00 | 1,00E+00 |
| AT4G34700 | 0,83 | 1,13 | 2,59 | 0,44 | 6,62E-01 |
| AT4G34870 | 1,35 | 1,96 | 1,85 | 1,06 | 2,89E-01 |
| AT4G34881 | 0,53 | -0,34 | 2,75 | -0,12 | 9,03E-01 |
| AT4G34910 | 0,61 | 1,30 | 3,14 | 0,41 | 6,79E-01 |
| AT4G34920 | 0,49 | 1,83 | 3,28 | 0,56 | 5,76E-01 |
| AT4G35040 | 0,31 | 1,34 | 4,40 | 0,31 | 7,60E-01 |
| AT4G35100 | 1,16 | -0,08 | 2,46 | -0,03 | 9,74E-01 |
| AT4G35370 | 0,13 | 0,00 | 4,41 | 0,00 | 1,00E+00 |
| AT4G35450 | 0,43 | -0,15 | 3,41 | -0,04 | 9,65E-01 |
| AT4G35460 | 0,66 | 3,29 | 2,90 | 1,14 | 2,56E-01 |
| AT4G35470 | 0,67 | 0,50 | 2,74 | 0,18 | 8,55E-01 |
| AT4G35500 | 0,79 | 3,30 | 2,74 | 1,20 | 2,29E-01 |
| AT4G35560 | 0,72 | 0,50 | 2,95 | 0,17 | 8,67E-01 |
| AT4G35600 | 1,15 | 3,71 | 1,93 | 1,92 | 5,44E-02 |
| AT4G35630 | 0,33 | 0,44 | 3,64 | 0,12 | 9,05E-01 |
| AT4G35850 | 0,77 | 0,24 | 2,22 | 0,11 | 9,12E-01 |
| AT4G35870 | 0,76 | 2,06 | 2,56 | 0,80 | 4,21E-01 |
| AT4G35880 | 0,17 | 0,00 | 4,41 | 0,00 | 1,00E+00 |
| AT4G35890 | 1,08 | 3,03 | 2,14 | 1,42 | 1,56E-01 |
| AT4G35940 | 1,33 | 3,10 | 1,77 | 1,75 | 8,05E-02 |
| AT4G36032 | 1,21 | 1,91 | 2,28 | 0,84 | 4,02E-01 |
| AT4G36052 | 0,12 | 0,00 | 4,41 | 0,00 | 1,00E+00 |
| AT4G36130 | 1,27 | 2,82 | 1,92 | 1,47 | 1,41E-01 |
| AT4G36150 | 1,48 | 1,17 | 2,52 | 0,47 | 6,41E-01 |
| AT4G36190 | 0,22 | -0,43 | 4,39 | -0,10 | 9,21E-01 |

| | | | | | |
|---|---|---|---|---|---|
| AT4G36195 | 0,86 | -0,68 | 2,33 | -0,29 | 7,70E-01 |
| AT4G36210 | 1,25 | 3,51 | 1,85 | 1,90 | 5,72E-02 |
| AT4G36250 | 0,69 | -0,36 | 2,57 | -0,14 | 8,89E-01 |
| AT4G36540 | 0,24 | 1,38 | 4,41 | 0,31 | 7,54E-01 |
| AT4G36710 | 1,27 | -1,09 | 2,55 | -0,43 | 6,69E-01 |
| AT4G36720 | 0,82 | 2,55 | 2,82 | 0,90 | 3,67E-01 |
| AT4G09375 | 0,14 | -0,48 | 4,40 | -0,11 | 9,13E-01 |
| AT4G36810 | 0,84 | -0,11 | 2,55 | -0,04 | 9,67E-01 |
| AT4G36870 | 0,25 | -0,70 | 4,39 | -0,16 | 8,74E-01 |
| AT4G36920 | 0,14 | -0,48 | 4,40 | -0,11 | 9,13E-01 |
| AT4G36925 | 1,06 | 0,26 | 1,85 | 0,14 | 8,87E-01 |
| AT4G36990 | 0,59 | 1,35 | 3,11 | 0,44 | 6,64E-01 |
| AT4G37000 | 0,23 | 1,38 | 4,41 | 0,31 | 7,54E-01 |
| AT4G37020 | 0,15 | 0,00 | 4,41 | 0,00 | 1,00E+00 |
| AT4G37080 | 0,19 | -0,47 | 4,40 | -0,11 | 9,16E-01 |
| AT4G37130 | 0,62 | 2,27 | 3,68 | 0,61 | 5,39E-01 |
| AT4G37180 | 0,47 | -0,57 | 4,17 | -0,14 | 8,92E-01 |
| AT4G37190 | 0,32 | 0,48 | 4,40 | 0,11 | 9,13E-01 |
| AT4G37210 | 0,15 | -0,43 | 4,39 | -0,10 | 9,23E-01 |
| AT4G37300 | 1,24 | 3,08 | 2,12 | 1,46 | 1,45E-01 |
| AT4G37430 | 0,86 | -0,40 | 3,45 | -0,11 | 9,09E-01 |
| AT4G37440 | 0,34 | 1,99 | 4,38 | 0,45 | 6,50E-01 |
| AT4G37550 | 0,88 | 1,54 | 2,22 | 0,70 | 4,86E-01 |
| AT4G37560 | 0,42 | 1,73 | 4,38 | 0,40 | 6,93E-01 |
| AT4G37650 | 1,47 | 0,68 | 2,02 | 0,34 | 7,36E-01 |
| AT4G37680 | 0,49 | 1,38 | 3,29 | 0,42 | 6,75E-01 |
| AT4G37730 | 0,75 | 0,35 | 2,30 | 0,15 | 8,78E-01 |
| AT4G37760 | 0,63 | -0,77 | 2,39 | -0,32 | 7,48E-01 |
| AT4G37790 | 0,15 | 0,00 | 4,39 | 0,00 | 1,00E+00 |
| AT4G37980 | 0,67 | -1,74 | 2,26 | -0,77 | 4,42E-01 |
| AT4G38040 | 0,20 | 1,38 | 4,41 | 0,31 | 7,54E-01 |
| AT4G38170 | 0,97 | 1,46 | 2,19 | 0,67 | 5,04E-01 |
| AT4G09685 | 0,46 | 2,13 | 4,02 | 0,53 | 5,96E-01 |
| AT4G38215 | 0,75 | 1,62 | 4,36 | 0,37 | 7,10E-01 |
| AT4G38225 | 0,78 | 2,41 | 2,60 | 0,93 | 3,53E-01 |
| AT4G38250 | 1,25 | 1,08 | 1,72 | 0,63 | 5,30E-01 |
| AT4G38270 | 0,51 | -1,27 | 3,16 | -0,40 | 6,89E-01 |
| AT4G38290 | 0,11 | 0,00 | 4,41 | 0,00 | 1,00E+00 |
| AT4G38320 | 0,19 | 0,00 | 4,41 | 0,00 | 1,00E+00 |
| AT4G38330 | 0,12 | 0,00 | 4,41 | 0,00 | 1,00E+00 |
| AT4G38360 | 0,15 | 0,00 | 4,41 | 0,00 | 1,00E+00 |
| AT4G38430 | 0,81 | 2,47 | 2,73 | 0,91 | 3,65E-01 |
| AT4G38460 | 1,41 | 2,37 | 1,89 | 1,26 | 2,09E-01 |
| AT4G38500 | 0,68 | 0,00 | 3,61 | 0,00 | 1,00E+00 |
| AT4G38540 | 0,10 | 0,00 | 4,41 | 0,00 | 1,00E+00 |
| AT4G38560 | 1,22 | 3,85 | 2,39 | 1,61 | 1,07E-01 |
| AT4G38580 | 0,50 | 1,44 | 4,31 | 0,33 | 7,38E-01 |
| AT4G38620 | 0,24 | -0,83 | 4,38 | -0,19 | 8,50E-01 |
| AT4G38790 | 0,69 | -0,53 | 3,00 | -0,18 | 8,60E-01 |
| AT4G38860 | 0,38 | 0,00 | 4,41 | 0,00 | 1,00E+00 |
| AT4G38880 | 0,39 | 1,79 | 4,37 | 0,41 | 6,81E-01 |
| AT4G38920 | 0,99 | 0,45 | 2,23 | 0,20 | 8,39E-01 |
| AT4G38930 | 0,41 | 2,00 | 4,38 | 0,46 | 6,48E-01 |
| AT4G39020 | 0,55 | 0,00 | 4,41 | 0,00 | 1,00E+00 |
| AT4G39030 | 0,29 | 1,74 | 4,38 | 0,40 | 6,90E-01 |
| AT4G39050 | 1,56 | -0,27 | 1,77 | -0,15 | 8,78E-01 |

| | | | | | |
|---|---|---|---|---|---|
| AT4G39210 | 1,07 | 0,18 | 2,14 | 0,08 | 9,33E-01 |
| AT4G39270 | 1,16 | 1,15 | 1,68 | 0,69 | 4,93E-01 |
| AT4G39620 | 0,18 | 0,00 | 4,41 | 0,00 | 1,00E+00 |
| AT4G39640 | 0,76 | 0,56 | 2,62 | 0,21 | 8,31E-01 |
| AT4G39670 | 0,55 | 0,00 | 3,29 | 0,00 | 1,00E+00 |
| AT4G39840 | 1,07 | 2,80 | 2,06 | 1,36 | 1,74E-01 |
| AT4G39860 | 0,51 | 0,53 | 2,46 | 0,22 | 8,29E-01 |
| AT4G39870 | 0,35 | -0,50 | 4,40 | -0,11 | 9,09E-01 |
| AT4G39940 | 0,36 | 1,83 | 4,38 | 0,42 | 6,76E-01 |
| AT4G39950 | 1,14 | -0,33 | 1,85 | -0,18 | 8,59E-01 |
| AT4G39952 | 0,28 | 1,84 | 4,39 | 0,42 | 6,74E-01 |
| AT4G39980 | 0,50 | 1,86 | 3,23 | 0,58 | 5,64E-01 |
| AT4G40063 | 0,73 | 0,99 | 3,04 | 0,33 | 7,45E-01 |
| AT5G00730 | 1,64 | -2,76 | 1,85 | -1,50 | 1,35E-01 |
| AT5G01170 | 0,10 | 0,00 | 4,41 | 0,00 | 1,00E+00 |
| AT5G01210 | 1,09 | -0,38 | 2,01 | -0,19 | 8,52E-01 |
| AT5G01240 | 0,22 | -0,53 | 4,41 | -0,12 | 9,05E-01 |
| AT5G01260 | 0,54 | 1,41 | 2,40 | 0,59 | 5,56E-01 |
| AT5G01290 | 0,28 | 0,00 | 4,41 | 0,00 | 1,00E+00 |
| AT5G01460 | 0,28 | 2,12 | 4,38 | 0,48 | 6,28E-01 |
| AT5G01542 | 0,46 | 1,70 | 4,10 | 0,41 | 6,80E-01 |
| AT5G01590 | 0,94 | -0,55 | 2,20 | -0,25 | 8,04E-01 |
| AT5G01595 | 0,81 | 1,32 | 2,61 | 0,51 | 6,12E-01 |
| AT5G01690 | 0,08 | 0,00 | 4,41 | 0,00 | 1,00E+00 |
| AT5G01732 | 0,29 | 1,32 | 4,40 | 0,30 | 7,64E-01 |
| AT5G01740 | 0,15 | 0,00 | 4,41 | 0,00 | 1,00E+00 |
| AT5G01750 | 1,69 | -2,01 | 2,07 | -0,97 | 3,32E-01 |
| AT5G01760 | 0,79 | 3,42 | 2,72 | 1,26 | 2,09E-01 |
| AT5G01800 | 0,74 | 2,89 | 2,45 | 1,18 | 2,38E-01 |
| AT5G01810 | 0,55 | 2,22 | 3,88 | 0,57 | 5,67E-01 |
| AT5G01850 | 0,24 | -0,46 | 4,40 | -0,10 | 9,17E-01 |
| AT5G02035 | 0,47 | -0,40 | 4,02 | -0,10 | 9,20E-01 |
| AT5G02040 | 0,25 | 1,30 | 4,40 | 0,30 | 7,67E-01 |
| AT5G02100 | 0,70 | -0,89 | 3,79 | -0,24 | 8,14E-01 |
| AT5G02120 | 0,39 | -1,63 | 4,36 | -0,37 | 7,09E-01 |
| AT5G00820 | 1,31 | -1,88 | 2,48 | -0,76 | 4,49E-01 |
| AT5G02160 | 1,30 | -2,57 | 1,73 | -1,49 | 1,37E-01 |
| AT5G02240 | 0,32 | 0,39 | 4,40 | 0,09 | 9,29E-01 |
| AT5G02250 | 1,33 | 1,78 | 2,26 | 0,79 | 4,31E-01 |
| AT5G02280 | 0,11 | 0,00 | 4,41 | 0,00 | 1,00E+00 |
| AT5G02420 | 0,12 | 0,00 | 4,41 | 0,00 | 1,00E+00 |
| AT5G02450 | 0,59 | 0,26 | 3,17 | 0,08 | 9,35E-01 |
| AT5G02620 | 0,32 | 1,78 | 4,38 | 0,41 | 6,84E-01 |
| AT5G02630 | 0,30 | 1,38 | 4,41 | 0,31 | 7,54E-01 |
| AT5G02650 | 0,40 | 0,00 | 4,41 | 0,00 | 1,00E+00 |
| AT5G02760 | 0,40 | -0,34 | 3,46 | -0,10 | 9,21E-01 |
| AT5G02790 | 0,39 | 1,38 | 3,55 | 0,39 | 6,98E-01 |
| AT5G02840 | 0,40 | 1,38 | 4,23 | 0,33 | 7,44E-01 |
| AT5G02850 | 0,54 | 2,12 | 3,12 | 0,68 | 4,96E-01 |
| AT5G02860 | 1,11 | 1,21 | 2,14 | 0,56 | 5,72E-01 |
| AT5G02950 | 0,61 | -1,37 | 3,84 | -0,36 | 7,21E-01 |
| AT5G02995 | 0,13 | -0,43 | 4,39 | -0,10 | 9,23E-01 |
| AT5G03020 | 0,28 | 1,84 | 4,39 | 0,42 | 6,74E-01 |
| AT5G03160 | 0,68 | 2,50 | 2,48 | 1,01 | 3,14E-01 |
| AT5G03190 | 0,92 | 2,87 | 2,06 | 1,39 | 1,64E-01 |
| AT5G03290 | 1,24 | 2,39 | 1,78 | 1,34 | 1,80E-01 |

| | | | | | |
|---|---|---|---|---|---|
| AT5G03300 | 0,81 | -1,24 | 2,55 | -0,49 | 6,27E-01 |
| AT5G03330 | 0,61 | 2,95 | 3,07 | 0,96 | 3,37E-01 |
| AT5G03355 | 0,46 | 1,34 | 3,35 | 0,40 | 6,89E-01 |
| AT5G03415 | 0,36 | 1,50 | 4,39 | 0,34 | 7,32E-01 |
| AT5G03420 | 0,63 | 1,31 | 2,69 | 0,49 | 6,25E-01 |
| AT5G03455 | 0,25 | 1,39 | 4,41 | 0,32 | 7,52E-01 |
| AT5G03460 | 0,54 | 3,01 | 3,09 | 0,97 | 3,31E-01 |
| AT5G03470 | 0,40 | 2,32 | 4,36 | 0,53 | 5,95E-01 |
| AT5G03490 | 0,11 | 0,00 | 4,41 | 0,00 | 1,00E+00 |
| AT5G03540 | 1,05 | 2,82 | 2,80 | 1,01 | 3,14E-01 |
| AT5G03650 | 0,91 | 0,17 | 1,88 | 0,09 | 9,28E-01 |
| AT5G03905 | 0,30 | 1,39 | 4,41 | 0,32 | 7,52E-01 |
| AT5G04040 | 1,27 | -0,64 | 2,13 | -0,30 | 7,65E-01 |
| AT5G04050 | 0,10 | 0,00 | 4,41 | 0,00 | 1,00E+00 |
| AT5G04060 | 0,37 | 0,00 | 4,41 | 0,00 | 1,00E+00 |
| AT5G04170 | 0,76 | 1,59 | 2,57 | 0,62 | 5,37E-01 |
| AT5G04190 | 0,13 | 0,00 | 4,41 | 0,00 | 1,00E+00 |
| AT5G04240 | 0,88 | -0,72 | 3,39 | -0,21 | 8,33E-01 |
| AT5G04280 | 1,25 | 0,53 | 2,15 | 0,25 | 8,05E-01 |
| AT5G04290 | 0,66 | 1,37 | 3,28 | 0,42 | 6,77E-01 |
| AT5G04360 | 0,45 | 1,33 | 4,15 | 0,32 | 7,48E-01 |
| AT5G00960 | 0,96 | 0,00 | 2,41 | 0,00 | 1,00E+00 |
| AT5G04460 | 0,56 | 1,38 | 3,20 | 0,43 | 6,67E-01 |
| AT5G04490 | 0,46 | 1,38 | 3,44 | 0,40 | 6,88E-01 |
| AT5G04510 | 0,58 | 1,01 | 2,69 | 0,38 | 7,08E-01 |
| AT5G04600 | 0,71 | 0,86 | 2,19 | 0,39 | 6,94E-01 |
| AT5G04620 | 0,43 | 2,00 | 4,37 | 0,46 | 6,47E-01 |
| AT5G04630 | 0,57 | 1,31 | 3,84 | 0,34 | 7,33E-01 |
| AT5G04660 | 0,24 | -0,48 | 4,40 | -0,11 | 9,13E-01 |
| AT5G00985 | 0,28 | 2,07 | 4,38 | 0,47 | 6,37E-01 |
| AT5G04800 | 0,67 | 1,58 | 2,24 | 0,71 | 4,81E-01 |
| AT5G04885 | 0,27 | 0,00 | 3,81 | 0,00 | 1,00E+00 |
| AT5G04910 | 0,42 | 1,72 | 4,15 | 0,41 | 6,79E-01 |
| AT5G05070 | 0,40 | 1,94 | 4,32 | 0,45 | 6,53E-01 |
| AT5G05090 | 0,35 | 1,32 | 3,61 | 0,37 | 7,14E-01 |
| AT5G05160 | 0,72 | 2,87 | 2,85 | 1,01 | 3,13E-01 |
| AT5G05210 | 0,28 | 0,00 | 4,41 | 0,00 | 1,00E+00 |
| AT5G05220 | 0,08 | 0,00 | 4,41 | 0,00 | 1,00E+00 |
| AT5G05230 | 0,08 | 0,00 | 4,41 | 0,00 | 1,00E+00 |
| AT5G05310 | 0,18 | 0,00 | 4,40 | 0,00 | 1,00E+00 |
| AT5G05440 | 0,16 | 0,00 | 4,41 | 0,00 | 1,00E+00 |
| AT5G05490 | 0,55 | 2,54 | 4,34 | 0,58 | 5,59E-01 |
| AT5G05560 | 1,14 | 1,90 | 2,00 | 0,95 | 3,43E-01 |
| AT5G05570 | 0,62 | 0,00 | 4,41 | 0,00 | 1,00E+00 |
| AT5G05600 | 0,31 | -1,59 | 4,35 | -0,37 | 7,15E-01 |
| AT5G05630 | 0,08 | 0,00 | 4,41 | 0,00 | 1,00E+00 |
| AT5G05680 | 1,56 | 4,10 | 1,84 | 2,22 | 2,62E-02 |
| AT5G05710 | 0,20 | 1,33 | 4,40 | 0,30 | 7,63E-01 |
| AT5G05780 | 0,96 | -1,89 | 2,56 | -0,74 | 4,60E-01 |
| AT5G05950 | 0,83 | 2,90 | 2,47 | 1,17 | 2,40E-01 |
| AT5G05970 | 0,30 | 1,22 | 4,38 | 0,28 | 7,80E-01 |
| AT5G05985 | 0,55 | 2,59 | 3,83 | 0,68 | 4,98E-01 |
| AT5G06000 | 0,23 | -0,40 | 4,38 | -0,09 | 9,27E-01 |
| AT5G06050 | 0,39 | 1,08 | 4,30 | 0,25 | 8,02E-01 |
| AT5G06100 | 0,21 | 0,00 | 4,41 | 0,00 | 1,00E+00 |
| AT5G06120 | 1,17 | 2,15 | 2,44 | 0,88 | 3,78E-01 |

| | | | | | |
|---|---|---|---|---|---|
| AT5G06170 | 0,31 | -0,66 | 4,39 | -0,15 | 8,81E-01 |
| AT5G06220 | 0,84 | 1,86 | 2,52 | 0,74 | 4,61E-01 |
| AT5G06265 | 0,26 | -0,48 | 4,40 | -0,11 | 9,13E-01 |
| AT5G06290 | 1,37 | -0,34 | 1,67 | -0,20 | 8,39E-01 |
| AT5G06310 | 0,33 | 0,00 | 4,41 | 0,00 | 1,00E+00 |
| AT5G01255 | 0,70 | 1,34 | 2,93 | 0,46 | 6,47E-01 |
| AT5G06350 | 0,36 | -0,54 | 3,55 | -0,15 | 8,78E-01 |
| AT5G06360 | 0,40 | 1,94 | 4,32 | 0,45 | 6,53E-01 |
| AT5G06380 | 0,08 | 0,00 | 4,41 | 0,00 | 1,00E+00 |
| AT5G06390 | 0,14 | 0,00 | 4,41 | 0,00 | 1,00E+00 |
| AT5G06460 | 1,60 | 3,94 | 1,91 | 2,06 | 3,91E-02 |
| AT5G06645 | 0,58 | 0,00 | 4,41 | 0,00 | 1,00E+00 |
| AT5G06660 | 0,96 | 0,00 | 3,67 | 0,00 | 1,00E+00 |
| AT5G06680 | 0,39 | 2,18 | 4,34 | 0,50 | 6,14E-01 |
| AT5G06700 | 0,91 | -0,45 | 1,81 | -0,25 | 8,03E-01 |
| AT5G06811 | 0,11 | 0,00 | 4,41 | 0,00 | 1,00E+00 |
| AT5G06820 | 0,12 | 0,00 | 4,41 | 0,00 | 1,00E+00 |
| AT5G06860 | 0,22 | -0,48 | 4,40 | -0,11 | 9,14E-01 |
| AT5G06870 | 0,67 | 1,68 | 2,90 | 0,58 | 5,62E-01 |
| AT5G06950 | 0,61 | 2,82 | 3,07 | 0,92 | 3,58E-01 |
| AT5G06980 | 0,16 | 0,00 | 4,41 | 0,00 | 1,00E+00 |
| AT5G07090 | 1,54 | 0,12 | 1,30 | 0,09 | 9,25E-01 |
| AT5G07110 | 0,85 | 2,52 | 3,38 | 0,74 | 4,57E-01 |
| AT5G01315 | 0,64 | 2,67 | 2,98 | 0,90 | 3,70E-01 |
| AT5G07130 | 0,33 | 0,43 | 4,38 | 0,10 | 9,21E-01 |
| AT5G07140 | 1,24 | 3,46 | 2,48 | 1,39 | 1,64E-01 |
| AT5G07210 | 1,09 | -1,92 | 2,02 | -0,95 | 3,40E-01 |
| AT5G07240 | 1,19 | 1,28 | 2,16 | 0,59 | 5,55E-01 |
| AT5G07270 | 0,15 | 0,00 | 4,41 | 0,00 | 1,00E+00 |
| AT5G07340 | 0,98 | -0,44 | 2,47 | -0,18 | 8,58E-01 |
| AT5G07410 | 1,62 | -0,53 | 2,15 | -0,25 | 8,05E-01 |
| AT5G07430 | 0,50 | 0,30 | 3,98 | 0,08 | 9,40E-01 |
| AT5G07440 | 0,36 | -0,29 | 4,37 | -0,07 | 9,47E-01 |
| AT5G07580 | 0,75 | 1,36 | 2,36 | 0,57 | 5,65E-01 |
| AT5G07610 | 0,48 | 0,00 | 4,14 | 0,00 | 1,00E+00 |
| AT5G07760 | 0,08 | 0,00 | 4,41 | 0,00 | 1,00E+00 |
| AT5G07770 | 0,16 | 0,00 | 4,41 | 0,00 | 1,00E+00 |
| AT5G07790 | 0,58 | 0,00 | 4,41 | 0,00 | 1,00E+00 |
| AT5G07830 | 0,86 | 0,58 | 2,64 | 0,22 | 8,27E-01 |
| AT5G07860 | 0,58 | 2,87 | 3,87 | 0,74 | 4,58E-01 |
| AT5G07870 | 0,49 | 2,54 | 4,35 | 0,58 | 5,60E-01 |
| AT5G07960 | 0,12 | 0,00 | 4,41 | 0,00 | 1,00E+00 |
| AT5G08050 | 0,48 | -0,59 | 4,24 | -0,14 | 8,89E-01 |
| AT5G08060 | 0,10 | 0,00 | 4,41 | 0,00 | 1,00E+00 |
| AT5G08070 | 0,35 | 1,58 | 4,36 | 0,36 | 7,18E-01 |
| AT5G08139 | 0,37 | 0,00 | 4,39 | 0,00 | 1,00E+00 |
| AT5G08190 | 0,29 | 1,35 | 4,40 | 0,31 | 7,59E-01 |
| AT5G08260 | 0,90 | 2,05 | 2,29 | 0,90 | 3,69E-01 |
| AT5G01545 | 0,37 | -1,68 | 4,36 | -0,38 | 7,01E-01 |
| AT5G08290 | 0,74 | 0,04 | 2,30 | 0,02 | 9,86E-01 |
| AT5G08300 | 0,94 | 2,64 | 2,46 | 1,07 | 2,85E-01 |
| AT5G08330 | 1,56 | 3,19 | 1,79 | 1,78 | 7,50E-02 |
| AT5G08335 | 0,18 | 0,00 | 4,41 | 0,00 | 1,00E+00 |
| AT5G08340 | 0,58 | 2,27 | 4,01 | 0,57 | 5,70E-01 |
| AT5G08390 | 0,26 | 0,00 | 4,41 | 0,00 | 1,00E+00 |
| AT5G08430 | 0,57 | 2,20 | 4,37 | 0,50 | 6,14E-01 |

| | | | | | |
|---|---|---|---|---|---|
| AT5G08440 | 0,59 | 2,73 | 3,88 | 0,70 | 4,81E-01 |
| AT5G08470 | 0,81 | 2,61 | 2,79 | 0,93 | 3,51E-01 |
| AT5G08510 | 0,51 | 1,84 | 3,23 | 0,57 | 5,69E-01 |
| AT5G08540 | 0,41 | -0,59 | 3,46 | -0,17 | 8,64E-01 |
| AT5G01555 | 0,36 | 1,36 | 4,41 | 0,31 | 7,57E-01 |
| AT5G08570 | 1,20 | -0,61 | 1,59 | -0,38 | 7,03E-01 |
| AT5G08590 | 0,63 | -0,53 | 2,72 | -0,19 | 8,46E-01 |
| AT5G08600 | 0,28 | 2,07 | 4,38 | 0,47 | 6,37E-01 |
| AT5G08620 | 1,12 | 0,42 | 1,77 | 0,24 | 8,13E-01 |
| AT5G08660 | 0,33 | 2,20 | 4,37 | 0,50 | 6,16E-01 |
| AT5G08695 | 0,20 | 1,33 | 4,40 | 0,30 | 7,63E-01 |
| AT5G08750 | 0,11 | 0,00 | 4,41 | 0,00 | 1,00E+00 |
| AT5G08760 | 0,55 | -0,38 | 3,91 | -0,10 | 9,23E-01 |
| AT5G08780 | 0,49 | 2,18 | 3,22 | 0,68 | 4,98E-01 |
| AT5G09220 | 0,82 | -2,16 | 3,03 | -0,71 | 4,77E-01 |
| AT5G09400 | 1,22 | 1,97 | 2,11 | 0,93 | 3,50E-01 |
| AT5G09440 | 0,77 | 1,70 | 2,45 | 0,69 | 4,88E-01 |
| AT5G09445 | 0,26 | 1,38 | 4,41 | 0,31 | 7,54E-01 |
| AT5G09450 | 1,25 | 2,44 | 2,03 | 1,20 | 2,29E-01 |
| AT5G09600 | 0,62 | 0,52 | 3,12 | 0,17 | 8,67E-01 |
| AT5G09760 | 0,14 | 0,00 | 4,41 | 0,00 | 1,00E+00 |
| AT5G10010 | 0,47 | 1,20 | 3,40 | 0,35 | 7,24E-01 |
| AT5G10020 | 0,41 | 2,03 | 4,38 | 0,46 | 6,43E-01 |
| AT5G10100 | 0,16 | 0,00 | 4,41 | 0,00 | 1,00E+00 |
| AT5G10190 | 0,41 | 2,16 | 3,37 | 0,64 | 5,22E-01 |
| AT5G10200 | 0,81 | 2,96 | 3,41 | 0,87 | 3,86E-01 |
| AT5G10340 | 1,40 | 0,55 | 2,11 | 0,26 | 7,94E-01 |
| AT5G10370 | 1,36 | -0,35 | 1,94 | -0,18 | 8,56E-01 |
| AT5G10410 | 0,05 | 0,00 | 4,41 | 0,00 | 1,00E+00 |
| AT5G10460 | 0,08 | 0,00 | 4,41 | 0,00 | 1,00E+00 |
| AT5G01765 | 1,15 | 1,54 | 2,04 | 0,75 | 4,51E-01 |
| AT5G10600 | 0,40 | 1,64 | 4,33 | 0,38 | 7,04E-01 |
| AT5G10630 | 0,28 | 0,00 | 4,41 | 0,00 | 1,00E+00 |
| AT5G10660 | 0,46 | 0,50 | 2,84 | 0,18 | 8,60E-01 |
| AT5G10695 | 0,85 | -1,42 | 2,26 | -0,63 | 5,32E-01 |
| AT5G10745 | 0,35 | 0,00 | 3,60 | 0,00 | 1,00E+00 |
| AT5G10750 | 1,12 | 1,00 | 2,23 | 0,45 | 6,53E-01 |
| AT5G10780 | 0,76 | 0,78 | 2,58 | 0,30 | 7,63E-01 |
| AT5G10790 | 0,93 | 3,40 | 2,07 | 1,65 | 9,98E-02 |
| AT5G10920 | 0,91 | 1,69 | 2,25 | 0,75 | 4,52E-01 |
| AT5G11060 | 1,02 | 0,40 | 1,87 | 0,21 | 8,32E-01 |
| AT5G11090 | 0,49 | 0,30 | 2,81 | 0,11 | 9,14E-01 |
| AT5G11110 | 0,50 | 1,38 | 4,07 | 0,34 | 7,35E-01 |
| AT5G11170 | 1,42 | 1,76 | 2,00 | 0,88 | 3,80E-01 |
| AT5G11200 | 0,33 | 1,27 | 4,39 | 0,29 | 7,72E-01 |
| AT5G11280 | 0,20 | 0,00 | 4,41 | 0,00 | 1,00E+00 |
| AT5G11340 | 0,28 | 1,38 | 4,41 | 0,31 | 7,54E-01 |
| AT5G11350 | 0,82 | 2,10 | 2,88 | 0,73 | 4,67E-01 |
| AT5G11390 | 0,30 | 1,38 | 4,41 | 0,31 | 7,54E-01 |
| AT5G11420 | 1,15 | 0,82 | 2,08 | 0,39 | 6,93E-01 |
| AT5G11430 | 0,70 | 0,66 | 2,31 | 0,28 | 7,76E-01 |
| AT5G11500 | 0,94 | -0,76 | 2,04 | -0,37 | 7,10E-01 |
| AT5G11510 | 0,81 | 2,25 | 2,48 | 0,91 | 3,64E-01 |
| AT5G11530 | 0,81 | 1,28 | 2,42 | 0,53 | 5,98E-01 |
| AT5G11580 | 0,12 | 0,00 | 4,41 | 0,00 | 1,00E+00 |
| AT5G11650 | 0,38 | 1,38 | 3,53 | 0,39 | 6,96E-01 |

| | | | | | |
|---|---|---|---|---|---|
| AT5G11680 | 0,19 | 0,00 | 4,41 | 0,00 | 1,00E+00 |
| AT5G11710 | 1,24 | 1,05 | 1,72 | 0,61 | 5,41E-01 |
| AT5G11720 | 1,68 | 4,39 | 2,04 | 2,15 | 3,13E-02 |
| AT5G11730 | 0,89 | 1,87 | 2,03 | 0,92 | 3,57E-01 |
| AT5G11760 | 0,22 | 1,39 | 4,41 | 0,32 | 7,52E-01 |
| AT5G11890 | 0,82 | 1,66 | 2,23 | 0,74 | 4,56E-01 |
| AT5G11920 | 0,12 | 0,00 | 4,41 | 0,00 | 1,00E+00 |
| AT5G11977 | 0,12 | 0,00 | 4,41 | 0,00 | 1,00E+00 |
| AT5G12090 | 0,79 | -1,66 | 2,44 | -0,68 | 4,96E-01 |
| AT5G12100 | 0,44 | 0,00 | 4,41 | 0,00 | 1,00E+00 |
| AT5G12110 | 1,52 | 1,57 | 1,65 | 0,95 | 3,41E-01 |
| AT5G12210 | 0,65 | 2,32 | 3,74 | 0,62 | 5,36E-01 |
| AT5G12260 | 0,32 | 0,39 | 4,40 | 0,09 | 9,29E-01 |
| AT5G12350 | 0,49 | 2,41 | 4,16 | 0,58 | 5,63E-01 |
| AT5G12400 | 0,93 | 2,07 | 2,94 | 0,70 | 4,81E-01 |
| AT5G12850 | 0,97 | 1,93 | 2,03 | 0,95 | 3,43E-01 |
| AT5G12950 | 0,53 | -0,52 | 2,69 | -0,19 | 8,47E-01 |
| AT5G12960 | 0,27 | 0,45 | 4,38 | 0,10 | 9,19E-01 |
| AT5G12970 | 0,24 | 1,96 | 4,39 | 0,45 | 6,55E-01 |
| AT5G13030 | 0,96 | -2,76 | 2,02 | -1,37 | 1,71E-01 |
| AT5G13200 | 1,26 | 1,15 | 1,96 | 0,59 | 5,57E-01 |
| AT5G13210 | 0,35 | 1,35 | 3,53 | 0,38 | 7,01E-01 |
| AT5G13260 | 0,92 | 0,09 | 2,07 | 0,05 | 9,63E-01 |
| AT5G13280 | 0,45 | 2,56 | 4,15 | 0,62 | 5,36E-01 |
| AT5G13310 | 0,31 | 1,31 | 4,40 | 0,30 | 7,65E-01 |
| AT5G13320 | 1,27 | 2,24 | 2,13 | 1,05 | 2,93E-01 |
| AT5G13330 | 0,37 | 1,38 | 4,41 | 0,31 | 7,54E-01 |
| AT5G13370 | 0,68 | -0,89 | 3,19 | -0,28 | 7,81E-01 |
| AT5G13380 | 0,54 | -0,75 | 4,39 | -0,17 | 8,65E-01 |
| AT5G13500 | 0,32 | 1,31 | 4,40 | 0,30 | 7,65E-01 |
| AT5G13510 | 0,48 | -0,33 | 3,33 | -0,10 | 9,21E-01 |
| AT5G13580 | 0,29 | 0,00 | 4,41 | 0,00 | 1,00E+00 |
| AT5G13640 | 0,84 | 2,30 | 2,98 | 0,77 | 4,41E-01 |
| AT5G13690 | 0,61 | 2,76 | 3,11 | 0,89 | 3,74E-01 |
| AT5G13720 | 0,21 | -0,87 | 4,38 | -0,20 | 8,43E-01 |
| AT5G13740 | 0,19 | 0,00 | 4,41 | 0,00 | 1,00E+00 |
| AT5G13750 | 0,26 | 1,38 | 4,41 | 0,31 | 7,54E-01 |
| AT5G13760 | 0,26 | 1,38 | 4,41 | 0,31 | 7,54E-01 |
| AT5G13810 | 0,93 | 1,55 | 1,99 | 0,78 | 4,38E-01 |
| AT5G13820 | 0,28 | 0,00 | 4,41 | 0,00 | 1,00E+00 |
| AT5G13930 | 1,07 | 1,51 | 2,25 | 0,67 | 5,03E-01 |
| AT5G13980 | 0,97 | 0,96 | 2,25 | 0,43 | 6,70E-01 |
| AT5G14030 | 0,98 | 2,58 | 2,53 | 1,02 | 3,09E-01 |
| AT5G14050 | 0,50 | 0,00 | 3,27 | 0,00 | 1,00E+00 |
| AT5G14120 | 1,26 | 1,85 | 1,70 | 1,09 | 2,77E-01 |
| AT5G14200 | 1,42 | 0,39 | 1,79 | 0,22 | 8,25E-01 |
| AT5G14230 | 0,39 | 0,00 | 3,52 | 0,00 | 1,00E+00 |
| AT5G14260 | 0,68 | 1,22 | 2,54 | 0,48 | 6,32E-01 |
| AT5G14280 | 0,43 | 1,87 | 4,38 | 0,43 | 6,70E-01 |
| AT5G14285 | 0,21 | 0,00 | 4,41 | 0,00 | 1,00E+00 |
| AT5G14480 | 0,25 | -0,54 | 4,41 | -0,12 | 9,02E-01 |
| AT5G14610 | 0,64 | 2,31 | 3,29 | 0,70 | 4,83E-01 |
| AT5G14670 | 0,08 | 0,00 | 4,41 | 0,00 | 1,00E+00 |
| AT5G14880 | 0,37 | 1,38 | 3,53 | 0,39 | 6,96E-01 |
| AT5G14890 | 0,99 | 0,00 | 4,15 | 0,00 | 1,00E+00 |
| AT5G14910 | 0,77 | 0,26 | 2,68 | 0,10 | 9,24E-01 |

| | | | | | |
|---|---|---|---|---|---|
| AT5G14970 | 0,18 | 0,00 | 4,41 | 0,00 | 1,00E+00 |
| AT5G14990 | 0,42 | 2,32 | 4,20 | 0,55 | 5,80E-01 |
| AT5G15010 | 0,38 | -2,27 | 4,33 | -0,52 | 6,01E-01 |
| AT5G02295 | 1,40 | -0,98 | 1,85 | -0,53 | 5,99E-01 |
| AT5G15080 | 0,92 | 2,47 | 2,81 | 0,88 | 3,79E-01 |
| AT5G15230 | 0,74 | 1,28 | 2,84 | 0,45 | 6,53E-01 |
| AT5G15240 | 0,05 | 0,00 | 4,41 | 0,00 | 1,00E+00 |
| AT5G15270 | 0,61 | -0,54 | 3,24 | -0,17 | 8,67E-01 |
| AT5G15280 | 0,54 | 2,67 | 3,84 | 0,70 | 4,87E-01 |
| AT5G15320 | 0,21 | -0,50 | 4,40 | -0,11 | 9,09E-01 |
| AT5G15330 | 0,11 | 0,00 | 4,41 | 0,00 | 1,00E+00 |
| AT5G15530 | 0,12 | 0,00 | 4,41 | 0,00 | 1,00E+00 |
| AT5G15570 | 0,44 | 0,00 | 4,41 | 0,00 | 1,00E+00 |
| AT5G15610 | 0,89 | 1,13 | 1,89 | 0,60 | 5,50E-01 |
| AT5G15685 | 0,72 | -0,31 | 2,17 | -0,14 | 8,88E-01 |
| AT5G15710 | 0,74 | 0,00 | 3,70 | 0,00 | 1,00E+00 |
| AT5G15730 | 0,59 | 1,35 | 2,72 | 0,49 | 6,21E-01 |
| AT5G15750 | 0,54 | 1,13 | 3,95 | 0,29 | 7,75E-01 |
| AT5G15820 | 0,22 | 1,33 | 4,40 | 0,30 | 7,63E-01 |
| AT5G15930 | 0,32 | 1,38 | 4,41 | 0,31 | 7,54E-01 |
| AT5G15950 | 1,23 | 3,23 | 2,13 | 1,52 | 1,29E-01 |
| AT5G15960 | 0,51 | -2,32 | 4,32 | -0,54 | 5,92E-01 |
| AT5G16010 | 0,36 | 1,70 | 4,38 | 0,39 | 6,97E-01 |
| AT5G16030 | 0,58 | -2,58 | 3,29 | -0,78 | 4,33E-01 |
| AT5G16120 | 0,30 | 1,38 | 4,41 | 0,31 | 7,54E-01 |
| AT5G16130 | 1,11 | 3,18 | 2,19 | 1,46 | 1,45E-01 |
| AT5G16150 | 1,33 | -0,88 | 1,66 | -0,53 | 5,95E-01 |
| AT5G16260 | 0,11 | 0,00 | 4,41 | 0,00 | 1,00E+00 |
| AT5G16290 | 0,43 | -1,39 | 4,35 | -0,32 | 7,50E-01 |
| AT5G16300 | 1,23 | 1,20 | 1,97 | 0,61 | 5,44E-01 |
| AT5G16440 | 0,67 | 3,00 | 2,57 | 1,17 | 2,43E-01 |
| AT5G16470 | 0,69 | 1,50 | 2,58 | 0,58 | 5,60E-01 |
| AT5G16540 | 0,32 | 2,14 | 4,37 | 0,49 | 6,25E-01 |
| AT5G16550 | 0,57 | 1,23 | 2,71 | 0,46 | 6,49E-01 |
| AT5G02555 | 0,05 | 0,00 | 4,41 | 0,00 | 1,00E+00 |
| AT5G16590 | 0,87 | 0,96 | 2,34 | 0,41 | 6,82E-01 |
| AT5G16630 | 0,92 | 1,35 | 2,79 | 0,48 | 6,28E-01 |
| AT5G16715 | 0,74 | 0,49 | 2,39 | 0,20 | 8,39E-01 |
| AT5G16730 | 1,05 | -1,08 | 2,06 | -0,52 | 6,02E-01 |
| AT5G16760 | 1,05 | 2,10 | 2,33 | 0,90 | 3,66E-01 |
| AT5G16820 | 0,94 | 2,65 | 2,35 | 1,13 | 2,60E-01 |
| AT5G16880 | 1,27 | 0,50 | 1,68 | 0,29 | 7,68E-01 |
| AT5G16900 | 0,33 | 1,37 | 4,41 | 0,31 | 7,56E-01 |
| AT5G16910 | 0,98 | 2,64 | 2,62 | 1,01 | 3,15E-01 |
| AT5G17167 | 0,20 | 0,00 | 4,41 | 0,00 | 1,00E+00 |
| AT5G17170 | 0,33 | 0,82 | 4,36 | 0,19 | 8,51E-01 |
| AT5G17190 | 0,51 | 2,19 | 3,21 | 0,68 | 4,95E-01 |
| AT5G17270 | 0,61 | 3,06 | 4,32 | 0,71 | 4,79E-01 |
| AT5G17280 | 0,29 | -0,51 | 4,40 | -0,12 | 9,08E-01 |
| AT5G17305 | 0,08 | 0,00 | 4,41 | 0,00 | 1,00E+00 |
| AT5G17440 | 0,24 | 1,57 | 4,39 | 0,36 | 7,20E-01 |
| AT5G17530 | 0,59 | 0,97 | 3,10 | 0,31 | 7,54E-01 |
| AT5G17580 | 0,17 | 0,00 | 4,41 | 0,00 | 1,00E+00 |
| AT5G17630 | 0,33 | 1,18 | 4,38 | 0,27 | 7,89E-01 |
| AT5G17670 | 0,39 | 1,76 | 4,38 | 0,40 | 6,87E-01 |
| AT5G17680 | 0,30 | 1,38 | 4,41 | 0,31 | 7,54E-01 |

| | | | | | |
|---|---|---|---|---|---|
| AT5G17850 | 0,18 | 0,00 | 4,41 | 0,00 | 1,00E+00 |
| AT5G17870 | 0,86 | -1,34 | 2,27 | -0,59 | 5,56E-01 |
| AT5G17900 | 0,30 | 1,28 | 4,39 | 0,29 | 7,70E-01 |
| AT5G17910 | 1,47 | 0,41 | 1,82 | 0,22 | 8,23E-01 |
| AT5G17980 | 0,38 | 1,29 | 4,30 | 0,30 | 7,64E-01 |
| AT5G17990 | 0,55 | 1,17 | 4,03 | 0,29 | 7,71E-01 |
| AT5G18040 | 0,71 | 1,38 | 2,61 | 0,53 | 5,97E-01 |
| AT5G18065 | 0,30 | -0,42 | 3,70 | -0,11 | 9,09E-01 |
| AT5G18070 | 0,62 | 2,50 | 2,59 | 0,97 | 3,35E-01 |
| AT5G18085 | 0,24 | 1,96 | 4,39 | 0,45 | 6,55E-01 |
| AT5G18100 | 0,33 | -0,54 | 4,41 | -0,12 | 9,02E-01 |
| AT5G18130 | 1,11 | 2,34 | 2,39 | 0,98 | 3,27E-01 |
| AT5G18140 | 1,06 | 3,09 | 2,65 | 1,17 | 2,43E-01 |
| AT5G18170 | 0,16 | 0,00 | 4,41 | 0,00 | 1,00E+00 |
| AT5G18200 | 0,08 | 0,00 | 4,41 | 0,00 | 1,00E+00 |
| AT5G18245 | 1,46 | 0,00 | 2,93 | 0,00 | 1,00E+00 |
| AT5G18255 | 1,25 | -0,63 | 1,54 | -0,41 | 6,82E-01 |
| AT5G18310 | 0,76 | -1,61 | 2,50 | -0,64 | 5,20E-01 |
| AT5G18390 | 0,33 | 1,72 | 4,38 | 0,39 | 6,94E-01 |
| AT5G18470 | 1,32 | 2,61 | 1,99 | 1,32 | 1,88E-01 |
| AT5G18500 | 0,37 | 0,53 | 4,39 | 0,12 | 9,03E-01 |
| AT5G18520 | 1,46 | 0,57 | 1,78 | 0,32 | 7,49E-01 |
| AT5G18580 | 0,90 | 0,26 | 2,20 | 0,12 | 9,07E-01 |
| AT5G18610 | 1,43 | 4,41 | 1,97 | 2,24 | 2,49E-02 |
| AT5G18630 | 0,28 | 1,38 | 4,41 | 0,31 | 7,54E-01 |
| AT5G18636 | 0,58 | 1,31 | 2,44 | 0,54 | 5,91E-01 |
| AT5G18650 | 0,68 | 1,30 | 3,05 | 0,43 | 6,70E-01 |
| AT5G18670 | 0,21 | 0,00 | 4,41 | 0,00 | 1,00E+00 |
| AT5G18750 | 0,77 | 0,00 | 3,50 | 0,00 | 1,00E+00 |
| AT5G18830 | 0,51 | -0,53 | 3,32 | -0,16 | 8,74E-01 |
| AT5G18890 | 0,98 | -0,01 | 4,24 | 0,00 | 9,99E-01 |
| AT5G18937 | 0,54 | 0,00 | 4,41 | 0,00 | 1,00E+00 |
| AT5G18940 | 1,00 | 3,41 | 2,72 | 1,25 | 2,10E-01 |
| AT5G18960 | 0,35 | 1,32 | 3,55 | 0,37 | 7,10E-01 |
| AT5G19000 | 0,44 | 2,53 | 3,39 | 0,75 | 4,56E-01 |
| AT5G19090 | 0,78 | 2,43 | 2,56 | 0,95 | 3,44E-01 |
| AT5G19180 | 0,77 | 0,53 | 2,38 | 0,22 | 8,24E-01 |
| AT5G19190 | 0,37 | 1,81 | 4,37 | 0,41 | 6,80E-01 |
| AT5G19210 | 0,49 | 2,19 | 3,38 | 0,65 | 5,18E-01 |
| AT5G19221 | 0,47 | 2,28 | 4,35 | 0,52 | 6,01E-01 |
| AT5G19240 | 0,65 | -1,72 | 2,25 | -0,76 | 4,46E-01 |
| AT5G19250 | 0,98 | -1,40 | 2,72 | -0,51 | 6,07E-01 |
| AT5G19290 | 0,41 | 1,27 | 3,41 | 0,37 | 7,10E-01 |
| AT5G19440 | 0,36 | 1,38 | 4,41 | 0,31 | 7,54E-01 |
| AT5G19500 | 1,08 | 3,44 | 2,34 | 1,47 | 1,42E-01 |
| AT5G19540 | 0,77 | -0,36 | 2,03 | -0,18 | 8,60E-01 |
| AT5G19550 | 0,90 | 0,05 | 2,23 | 0,02 | 9,81E-01 |
| AT5G19570 | 0,52 | 1,49 | 4,00 | 0,37 | 7,11E-01 |
| AT5G19620 | 0,50 | -0,69 | 4,09 | -0,17 | 8,65E-01 |
| AT5G19680 | 0,19 | -0,42 | 4,39 | -0,10 | 9,23E-01 |
| AT5G19750 | 0,55 | -0,41 | 4,39 | -0,09 | 9,26E-01 |
| AT5G19780 | 1,35 | -0,80 | 1,74 | -0,46 | 6,48E-01 |
| AT5G19990 | 0,77 | 2,40 | 2,16 | 1,11 | 2,66E-01 |
| AT5G20080 | 0,48 | 0,92 | 3,35 | 0,27 | 7,84E-01 |
| AT5G20120 | 0,55 | 0,00 | 4,39 | 0,00 | 1,00E+00 |
| AT5G20130 | 0,68 | 1,47 | 3,72 | 0,40 | 6,92E-01 |

| | | | | | |
|---|---|---|---|---|---|
| AT5G20150 | 0,56 | 2,11 | 2,68 | 0,79 | 4,32E-01 |
| AT5G20190 | 1,21 | 1,08 | 2,11 | 0,51 | 6,08E-01 |
| AT5G20200 | 1,58 | 3,27 | 1,88 | 1,74 | 8,19E-02 |
| AT5G20220 | 0,21 | 0,00 | 4,41 | 0,00 | 1,00E+00 |
| AT5G20350 | 1,36 | 0,59 | 1,87 | 0,31 | 7,54E-01 |
| AT5G20410 | 0,63 | 3,23 | 3,61 | 0,90 | 3,71E-01 |
| AT5G20480 | 0,15 | 0,00 | 4,41 | 0,00 | 1,00E+00 |
| AT5G20500 | 0,33 | 1,36 | 4,40 | 0,31 | 7,58E-01 |
| AT5G20590 | 0,85 | 0,79 | 2,19 | 0,36 | 7,17E-01 |
| AT5G20610 | 1,39 | 3,55 | 1,83 | 1,94 | 5,18E-02 |
| AT5G20660 | 1,27 | 3,68 | 1,92 | 1,92 | 5,53E-02 |
| AT5G20680 | 0,31 | -0,68 | 4,39 | -0,16 | 8,77E-01 |
| AT5G20710 | 0,12 | 0,00 | 4,41 | 0,00 | 1,00E+00 |
| AT5G20900 | 0,45 | 0,54 | 3,39 | 0,16 | 8,74E-01 |
| AT5G03255 | 0,31 | 1,29 | 4,39 | 0,29 | 7,70E-01 |
| AT5G20960 | 1,02 | 0,00 | 2,77 | 0,00 | 1,00E+00 |
| AT5G21020 | 1,50 | -1,31 | 1,76 | -0,74 | 4,56E-01 |
| AT5G21090 | 1,39 | 2,87 | 2,03 | 1,41 | 1,58E-01 |
| AT5G21105 | 1,68 | 0,97 | 1,96 | 0,49 | 6,21E-01 |
| AT5G21150 | 0,12 | 0,00 | 4,41 | 0,00 | 1,00E+00 |
| AT5G21222 | 1,01 | 3,32 | 2,71 | 1,23 | 2,20E-01 |
| AT5G21326 | 0,90 | 1,38 | 2,89 | 0,48 | 6,33E-01 |
| AT5G21482 | 0,24 | 0,00 | 4,41 | 0,00 | 1,00E+00 |
| AT5G21930 | 1,60 | 0,27 | 1,64 | 0,17 | 8,67E-01 |
| AT5G22000 | 1,27 | 2,85 | 1,84 | 1,55 | 1,21E-01 |
| AT5G22050 | 0,36 | -0,54 | 4,41 | -0,12 | 9,02E-01 |
| AT5G22070 | 0,57 | -0,63 | 3,13 | -0,20 | 8,41E-01 |
| AT5G22300 | 0,30 | -0,51 | 3,75 | -0,14 | 8,91E-01 |
| AT5G22315 | 0,20 | 1,40 | 4,41 | 0,32 | 7,51E-01 |
| AT5G22340 | 0,95 | 1,04 | 2,18 | 0,48 | 6,32E-01 |
| AT5G22420 | 0,42 | 2,15 | 4,19 | 0,51 | 6,08E-01 |
| AT5G22580 | 0,46 | -0,76 | 4,37 | -0,17 | 8,63E-01 |
| AT5G22590 | 1,03 | 1,38 | 2,76 | 0,50 | 6,17E-01 |
| AT5G22630 | 1,52 | 2,16 | 1,62 | 1,33 | 1,84E-01 |
| AT5G22640 | 1,19 | 0,42 | 1,87 | 0,23 | 8,21E-01 |
| AT5G22650 | 0,66 | -1,81 | 2,54 | -0,71 | 4,76E-01 |
| AT5G22690 | 0,12 | 0,00 | 4,41 | 0,00 | 1,00E+00 |
| AT5G22760 | 1,25 | 2,55 | 1,91 | 1,33 | 1,83E-01 |
| AT5G22794 | 0,08 | 0,00 | 4,41 | 0,00 | 1,00E+00 |
| AT5G22830 | 0,08 | 0,00 | 4,41 | 0,00 | 1,00E+00 |
| AT5G22950 | 0,12 | 0,00 | 4,41 | 0,00 | 1,00E+00 |
| AT5G23010 | 0,84 | 0,42 | 2,25 | 0,19 | 8,52E-01 |
| AT5G23070 | 0,19 | 0,00 | 4,41 | 0,00 | 1,00E+00 |
| AT5G03575 | 0,26 | -1,08 | 4,37 | -0,25 | 8,04E-01 |
| AT5G23120 | 0,65 | -0,43 | 2,30 | -0,19 | 8,52E-01 |
| AT5G23140 | 0,95 | 3,62 | 2,26 | 1,60 | 1,09E-01 |
| AT5G23150 | 1,41 | 3,31 | 2,20 | 1,50 | 1,33E-01 |
| AT5G23210 | 0,16 | 0,00 | 4,41 | 0,00 | 1,00E+00 |
| AT5G23280 | 1,01 | 3,71 | 2,49 | 1,49 | 1,36E-01 |
| AT5G23320 | 0,08 | 0,00 | 4,41 | 0,00 | 1,00E+00 |
| AT5G23330 | 0,43 | 2,00 | 4,37 | 0,46 | 6,47E-01 |
| AT5G23340 | 1,40 | 3,87 | 1,86 | 2,08 | 3,74E-02 |
| AT5G23490 | 0,49 | 2,08 | 2,74 | 0,76 | 4,48E-01 |
| AT5G23540 | 0,96 | 1,04 | 1,96 | 0,53 | 5,96E-01 |
| AT5G23570 | 0,58 | -2,07 | 2,38 | -0,87 | 3,83E-01 |
| AT5G23575 | 1,00 | 3,54 | 1,88 | 1,89 | 5,94E-02 |

| | | | | | |
|---|---|---|---|---|---|
| AT5G23600 | 0,08 | 0,00 | 4,41 | 0,00 | 1,00E+00 |
| AT5G23670 | 0,72 | 2,20 | 3,07 | 0,72 | 4,73E-01 |
| AT5G23720 | 0,51 | 1,84 | 3,23 | 0,57 | 5,69E-01 |
| AT5G23730 | 0,26 | 1,29 | 4,39 | 0,29 | 7,69E-01 |
| AT5G23890 | 0,99 | 3,05 | 2,77 | 1,10 | 2,70E-01 |
| AT5G23950 | 0,35 | 2,13 | 4,37 | 0,49 | 6,26E-01 |
| AT5G24020 | 0,16 | 0,00 | 4,41 | 0,00 | 1,00E+00 |
| AT5G24200 | 1,34 | 3,28 | 1,75 | 1,87 | 6,14E-02 |
| AT5G24210 | 0,96 | 1,35 | 2,13 | 0,64 | 5,25E-01 |
| AT5G24360 | 0,40 | 0,35 | 3,41 | 0,10 | 9,19E-01 |
| AT5G24380 | 0,41 | 0,54 | 3,42 | 0,16 | 8,75E-01 |
| AT5G24450 | 0,69 | 0,00 | 3,79 | 0,00 | 1,00E+00 |
| AT5G24460 | 0,88 | 0,40 | 1,98 | 0,20 | 8,41E-01 |
| AT5G24490 | 1,61 | -1,80 | 2,12 | -0,85 | 3,95E-01 |
| AT5G24610 | 0,55 | -0,91 | 2,67 | -0,34 | 7,35E-01 |
| AT5G24830 | 0,08 | 0,00 | 4,41 | 0,00 | 1,00E+00 |
| AT5G24980 | 0,43 | -1,43 | 3,41 | -0,42 | 6,75E-01 |
| AT5G25040 | 0,67 | 1,38 | 2,59 | 0,53 | 5,94E-01 |
| AT5G25120 | 0,51 | -0,53 | 3,31 | -0,16 | 8,74E-01 |
| AT5G03925 | 0,17 | 0,00 | 4,41 | 0,00 | 1,00E+00 |
| AT5G25130 | 1,42 | 0,82 | 1,79 | 0,46 | 6,46E-01 |
| AT5G25140 | 0,28 | 0,00 | 4,41 | 0,00 | 1,00E+00 |
| AT5G25150 | 0,27 | 1,46 | 4,40 | 0,33 | 7,40E-01 |
| AT5G25170 | 0,33 | 0,00 | 4,41 | 0,00 | 1,00E+00 |
| AT5G25210 | 0,42 | -0,54 | 3,44 | -0,16 | 8,74E-01 |
| AT5G25220 | 1,05 | 0,00 | 2,33 | 0,00 | 1,00E+00 |
| AT5G25230 | 1,66 | 3,65 | 1,78 | 2,06 | 3,97E-02 |
| AT5G25250 | 1,43 | 1,38 | 2,22 | 0,62 | 5,34E-01 |
| AT5G25260 | 1,02 | 0,31 | 2,83 | 0,11 | 9,14E-01 |
| AT5G25340 | 0,46 | 2,20 | 4,10 | 0,54 | 5,91E-01 |
| AT5G25460 | 1,53 | -2,34 | 2,01 | -1,16 | 2,44E-01 |
| AT5G25520 | 1,48 | 3,23 | 2,45 | 1,32 | 1,87E-01 |
| AT5G25560 | 0,53 | 2,76 | 3,16 | 0,87 | 3,83E-01 |
| AT5G25580 | 0,12 | 0,00 | 4,41 | 0,00 | 1,00E+00 |
| AT5G25630 | 0,47 | 0,43 | 3,24 | 0,13 | 8,95E-01 |
| AT5G25754 | 1,48 | 0,29 | 1,62 | 0,18 | 8,60E-01 |
| AT5G04145 | 0,89 | 1,15 | 2,20 | 0,53 | 6,00E-01 |
| AT5G25900 | 0,33 | 0,42 | 4,41 | 0,09 | 9,25E-01 |
| AT5G25910 | 0,60 | 1,38 | 3,79 | 0,36 | 7,16E-01 |
| AT5G25930 | 0,70 | 1,38 | 2,65 | 0,52 | 6,03E-01 |
| AT5G25980 | 1,32 | -2,03 | 1,94 | -1,04 | 2,97E-01 |
| AT5G26038 | 1,13 | 0,38 | 2,30 | 0,17 | 8,67E-01 |
| AT5G26146 | 0,89 | -1,85 | 2,35 | -0,79 | 4,31E-01 |
| AT5G26147 | 0,89 | -1,85 | 2,35 | -0,79 | 4,31E-01 |
| AT5G26200 | 0,12 | 0,00 | 4,41 | 0,00 | 1,00E+00 |
| AT5G26240 | 0,84 | 0,82 | 2,06 | 0,40 | 6,91E-01 |
| AT5G26270 | 0,26 | -0,65 | 4,39 | -0,15 | 8,83E-01 |
| AT5G26290 | 0,79 | -0,14 | 2,16 | -0,06 | 9,50E-01 |
| AT5G26570 | 1,51 | 0,81 | 1,69 | 0,48 | 6,31E-01 |
| AT5G26740 | 0,58 | -1,31 | 2,72 | -0,48 | 6,31E-01 |
| AT5G26731 | 0,38 | 0,00 | 4,41 | 0,00 | 1,00E+00 |
| AT5G26710 | 0,73 | 2,39 | 2,56 | 0,93 | 3,51E-01 |
| AT5G26670 | 0,26 | 0,23 | 4,36 | 0,05 | 9,58E-01 |
| AT5G26610 | 0,16 | 0,00 | 4,41 | 0,00 | 1,00E+00 |
| AT5G26600 | 0,82 | 0,24 | 2,62 | 0,09 | 9,28E-01 |
| AT5G26780 | 0,39 | 0,42 | 2,96 | 0,14 | 8,87E-01 |

| | | | | | |
|---|---|---|---|---|---|
| AT5G26820 | 0,42 | 2,12 | 4,37 | 0,48 | 6,28E-01 |
| AT5G26850 | 1,21 | 1,62 | 2,12 | 0,76 | 4,45E-01 |
| AT5G26860 | 1,16 | 1,20 | 2,04 | 0,59 | 5,56E-01 |
| AT5G26930 | 0,33 | 2,19 | 4,38 | 0,50 | 6,17E-01 |
| AT5G26940 | 0,66 | 3,20 | 3,69 | 0,87 | 3,87E-01 |
| AT5G26990 | 0,23 | 1,38 | 4,41 | 0,31 | 7,54E-01 |
| AT5G27120 | 0,91 | 1,37 | 3,46 | 0,40 | 6,92E-01 |
| AT5G27150 | 0,57 | 1,40 | 3,12 | 0,45 | 6,55E-01 |
| AT5G27170 | 0,25 | 0,00 | 4,41 | 0,00 | 1,00E+00 |
| AT5G27230 | 0,08 | 0,00 | 4,41 | 0,00 | 1,00E+00 |
| AT5G27240 | 0,05 | 0,00 | 4,41 | 0,00 | 1,00E+00 |
| AT5G27270 | 0,17 | -0,48 | 4,39 | -0,11 | 9,13E-01 |
| AT5G27280 | 0,34 | 0,29 | 3,61 | 0,08 | 9,37E-01 |
| AT5G27300 | 0,08 | 0,00 | 4,41 | 0,00 | 1,00E+00 |
| AT5G27320 | 0,08 | 0,00 | 4,41 | 0,00 | 1,00E+00 |
| AT5G27340 | 0,54 | 0,00 | 3,84 | 0,00 | 1,00E+00 |
| AT5G27350 | 0,64 | -1,74 | 2,64 | -0,66 | 5,09E-01 |
| AT5G27410 | 0,42 | 2,09 | 4,13 | 0,51 | 6,12E-01 |
| AT5G27420 | 1,44 | 1,78 | 1,88 | 0,95 | 3,44E-01 |
| AT5G27470 | 0,51 | 1,32 | 3,20 | 0,41 | 6,80E-01 |
| AT5G27530 | 0,32 | -1,15 | 4,38 | -0,26 | 7,93E-01 |
| AT5G27540 | 0,68 | 2,68 | 3,54 | 0,76 | 4,50E-01 |
| AT5G27600 | 0,18 | 0,00 | 4,41 | 0,00 | 1,00E+00 |
| AT5G27610 | 0,08 | 0,00 | 4,41 | 0,00 | 1,00E+00 |
| AT5G27630 | 0,17 | 0,00 | 4,41 | 0,00 | 1,00E+00 |
| AT5G27650 | 0,94 | 2,89 | 2,74 | 1,05 | 2,92E-01 |
| AT5G27710 | 0,67 | 1,31 | 2,61 | 0,50 | 6,16E-01 |
| AT5G27730 | 0,69 | -1,31 | 2,29 | -0,57 | 5,69E-01 |
| AT5G27760 | 0,27 | 1,38 | 4,41 | 0,31 | 7,54E-01 |
| AT5G27770 | 0,86 | 2,80 | 2,38 | 1,18 | 2,40E-01 |
| AT5G27800 | 0,16 | 0,00 | 4,41 | 0,00 | 1,00E+00 |
| AT5G27830 | 0,23 | 1,38 | 4,41 | 0,31 | 7,54E-01 |
| AT5G27890 | 0,40 | 0,00 | 4,41 | 0,00 | 1,00E+00 |
| AT5G27893 | 1,07 | -0,95 | 1,80 | -0,53 | 5,98E-01 |
| AT5G27970 | 1,51 | 0,88 | 2,38 | 0,37 | 7,12E-01 |
| AT5G28010 | 1,26 | -0,80 | 1,67 | -0,48 | 6,34E-01 |
| AT5G04625 | 0,10 | 0,00 | 4,41 | 0,00 | 1,00E+00 |
| AT5G04665 | 0,11 | 0,00 | 4,41 | 0,00 | 1,00E+00 |
| AT5G28090 | 0,29 | 0,00 | 4,41 | 0,00 | 1,00E+00 |
| AT5G04675 | 0,19 | 0,00 | 4,41 | 0,00 | 1,00E+00 |
| AT5G28320 | 0,89 | 2,15 | 2,75 | 0,78 | 4,35E-01 |
| AT5G28350 | 0,24 | 1,29 | 4,40 | 0,29 | 7,69E-01 |
| AT5G28410 | 0,36 | 0,02 | 4,35 | 0,00 | 9,97E-01 |
| AT5G28463 | 0,37 | -0,49 | 4,36 | -0,11 | 9,10E-01 |
| AT5G28500 | 0,70 | -0,61 | 2,67 | -0,23 | 8,20E-01 |
| AT5G04815 | 0,05 | 0,00 | 4,41 | 0,00 | 1,00E+00 |
| AT5G28810 | 0,66 | -0,54 | 2,98 | -0,18 | 8,55E-01 |
| AT5G28830 | 0,05 | 0,00 | 4,41 | 0,00 | 1,00E+00 |
| AT5G28900 | 0,40 | 0,64 | 4,32 | 0,15 | 8,82E-01 |
| AT5G29624 | 0,12 | 0,00 | 4,41 | 0,00 | 1,00E+00 |
| AT5G30341 | 0,12 | 0,00 | 4,41 | 0,00 | 1,00E+00 |
| AT5G04955 | 1,06 | 0,00 | 3,96 | 0,00 | 1,00E+00 |
| AT5G32440 | 1,33 | 3,22 | 1,95 | 1,65 | 9,94E-02 |
| AT5G32613 | 0,26 | 0,00 | 4,41 | 0,00 | 1,00E+00 |
| AT5G33290 | 0,62 | 3,51 | 3,88 | 0,90 | 3,66E-01 |
| AT5G33320 | 0,61 | 1,31 | 2,36 | 0,56 | 5,77E-01 |

| | | | | | |
|---|---|---|---|---|---|
| AT5G33390 | 0,38 | -1,55 | 4,36 | -0,36 | 7,22E-01 |
| AT5G34869 | 0,19 | 0,00 | 4,41 | 0,00 | 1,00E+00 |
| AT5G34871 | 0,12 | 0,00 | 4,41 | 0,00 | 1,00E+00 |
| AT5G34870 | 0,38 | -0,54 | 3,52 | -0,15 | 8,77E-01 |
| AT5G35160 | 1,68 | 2,15 | 1,53 | 1,40 | 1,62E-01 |
| AT5G35170 | 0,47 | -0,34 | 2,49 | -0,14 | 8,92E-01 |
| AT5G35180 | 0,52 | 0,29 | 2,76 | 0,10 | 9,17E-01 |
| AT5G35190 | 0,84 | 0,00 | 2,92 | 0,00 | 1,00E+00 |
| AT5G35300 | 0,23 | 0,00 | 4,41 | 0,00 | 1,00E+00 |
| AT5G35320 | 0,21 | 1,25 | 4,39 | 0,29 | 7,75E-01 |
| AT5G35360 | 1,57 | 3,76 | 1,73 | 2,16 | 3,04E-02 |
| AT5G35430 | 1,24 | 2,99 | 2,25 | 1,33 | 1,84E-01 |
| AT5G35450 | 0,40 | 0,00 | 4,41 | 0,00 | 1,00E+00 |
| AT5G05195 | 0,23 | 0,00 | 4,41 | 0,00 | 1,00E+00 |
| AT5G35490 | 1,40 | 0,00 | 2,51 | 0,00 | 1,00E+00 |
| AT5G35730 | 0,82 | 2,15 | 2,70 | 0,80 | 4,24E-01 |
| AT5G35840 | 1,35 | 0,33 | 2,41 | 0,14 | 8,90E-01 |
| AT5G35917 | 0,29 | 0,00 | 4,41 | 0,00 | 1,00E+00 |
| AT5G35920 | 1,48 | -0,54 | 2,07 | -0,26 | 7,93E-01 |
| AT5G35926 | 1,15 | 0,28 | 1,99 | 0,14 | 8,88E-01 |
| AT5G35980 | 1,15 | 0,15 | 1,95 | 0,08 | 9,39E-01 |
| AT5G36001 | 0,12 | 0,00 | 4,41 | 0,00 | 1,00E+00 |
| AT5G36080 | 1,49 | 0,00 | 2,15 | 0,00 | 1,00E+00 |
| AT5G36290 | 0,47 | 0,51 | 2,77 | 0,19 | 8,53E-01 |
| AT5G36300 | 0,87 | -0,50 | 2,55 | -0,19 | 8,46E-01 |
| AT5G36740 | 0,08 | 0,00 | 4,41 | 0,00 | 1,00E+00 |
| AT5G36870 | 1,07 | -2,17 | 2,03 | -1,07 | 2,86E-01 |
| AT5G36900 | 0,54 | 0,00 | 3,22 | 0,00 | 1,00E+00 |
| AT5G36940 | 0,51 | 0,00 | 3,97 | 0,00 | 1,00E+00 |
| AT5G37020 | 0,62 | -1,31 | 2,70 | -0,49 | 6,26E-01 |
| AT5G37070 | 0,43 | 0,20 | 4,34 | 0,05 | 9,63E-01 |
| AT5G37130 | 0,66 | 2,27 | 2,76 | 0,82 | 4,11E-01 |
| AT5G37150 | 0,98 | 0,42 | 2,45 | 0,17 | 8,65E-01 |
| AT5G37540 | 0,17 | 0,00 | 4,41 | 0,00 | 1,00E+00 |
| AT5G37630 | 0,94 | 2,09 | 2,35 | 0,89 | 3,74E-01 |
| AT5G37760 | 0,15 | 0,00 | 4,41 | 0,00 | 1,00E+00 |
| AT5G37793 | 0,51 | -2,02 | 2,64 | -0,76 | 4,45E-01 |
| AT5G37850 | 0,30 | 1,38 | 4,41 | 0,31 | 7,54E-01 |
| AT5G37950 | 0,11 | 0,00 | 4,41 | 0,00 | 1,00E+00 |
| AT5G38155 | 0,12 | 0,00 | 4,41 | 0,00 | 1,00E+00 |
| AT5G38210 | 1,00 | 0,00 | 2,40 | 0,00 | 1,00E+00 |
| AT5G38380 | 0,15 | -0,43 | 4,39 | -0,10 | 9,23E-01 |
| AT5G38470 | 0,73 | 2,77 | 3,67 | 0,75 | 4,51E-01 |
| AT5G38480 | 1,20 | 2,35 | 1,80 | 1,31 | 1,90E-01 |
| AT5G38560 | 0,34 | 1,36 | 3,58 | 0,38 | 7,05E-01 |
| AT5G38910 | 0,16 | 0,00 | 4,41 | 0,00 | 1,00E+00 |
| AT5G38980 | 0,57 | 0,44 | 2,84 | 0,15 | 8,78E-01 |
| AT5G39000 | 0,70 | 3,15 | 4,31 | 0,73 | 4,65E-01 |
| AT5G39030 | 0,81 | 2,12 | 2,90 | 0,73 | 4,66E-01 |
| AT5G39040 | 0,29 | -0,54 | 4,41 | -0,12 | 9,02E-01 |
| AT5G39250 | 0,34 | 1,68 | 4,38 | 0,38 | 7,01E-01 |
| AT5G39320 | 0,82 | 2,67 | 4,33 | 0,62 | 5,37E-01 |
| AT5G39350 | 0,21 | -0,54 | 4,41 | -0,12 | 9,02E-01 |
| AT5G39510 | 0,77 | 1,34 | 2,52 | 0,53 | 5,95E-01 |
| AT5G39590 | 0,82 | 1,27 | 2,54 | 0,50 | 6,18E-01 |
| AT5G39660 | 0,40 | 0,00 | 4,41 | 0,00 | 1,00E+00 |

| | | | | | |
|---|---|---|---|---|---|
| AT5G39670 | 0,40 | 0,00 | 4,25 | 0,00 | 1,00E+00 |
| AT5G39690 | 1,03 | 0,19 | 2,22 | 0,08 | 9,32E-01 |
| AT5G39710 | 0,25 | 1,34 | 4,40 | 0,30 | 7,61E-01 |
| AT5G39730 | 0,19 | 0,00 | 4,41 | 0,00 | 1,00E+00 |
| AT5G39775 | 0,53 | 0,55 | 4,22 | 0,13 | 8,97E-01 |
| AT5G39850 | 0,08 | 0,00 | 4,41 | 0,00 | 1,00E+00 |
| AT5G39960 | 0,21 | 1,38 | 4,41 | 0,31 | 7,54E-01 |
| AT5G39990 | 0,08 | 0,00 | 4,41 | 0,00 | 1,00E+00 |
| AT5G40060 | 1,33 | 0,89 | 2,58 | 0,34 | 7,30E-01 |
| AT5G40140 | 0,31 | 1,38 | 3,70 | 0,37 | 7,09E-01 |
| AT5G40200 | 0,62 | -0,36 | 2,84 | -0,13 | 8,99E-01 |
| AT5G40380 | 0,46 | -0,61 | 4,05 | -0,15 | 8,80E-01 |
| AT5G40400 | 1,55 | 0,24 | 1,55 | 0,16 | 8,75E-01 |
| AT5G40650 | 0,74 | 3,66 | 2,77 | 1,32 | 1,87E-01 |
| AT5G40670 | 1,01 | 0,68 | 2,18 | 0,31 | 7,53E-01 |
| AT5G40680 | 0,17 | 0,00 | 4,41 | 0,00 | 1,00E+00 |
| AT5G40690 | 0,62 | 1,36 | 3,77 | 0,36 | 7,18E-01 |
| AT5G40720 | 0,32 | 0,00 | 4,41 | 0,00 | 1,00E+00 |
| AT5G40760 | 1,16 | 3,31 | 2,04 | 1,62 | 1,05E-01 |
| AT5G40780 | 1,36 | 2,61 | 1,71 | 1,52 | 1,27E-01 |
| AT5G40810 | 1,32 | -0,65 | 1,76 | -0,37 | 7,13E-01 |
| AT5G40850 | 0,08 | 0,00 | 4,41 | 0,00 | 1,00E+00 |
| AT5G40870 | 0,14 | 0,00 | 4,41 | 0,00 | 1,00E+00 |
| AT5G40880 | 0,51 | 1,37 | 4,03 | 0,34 | 7,34E-01 |
| AT5G40900 | 0,15 | 0,00 | 4,39 | 0,00 | 1,00E+00 |
| AT5G41010 | 0,28 | 0,00 | 4,41 | 0,00 | 1,00E+00 |
| AT5G06115 | 0,20 | 0,00 | 4,41 | 0,00 | 1,00E+00 |
| AT5G41070 | 0,20 | 1,40 | 4,41 | 0,32 | 7,51E-01 |
| AT5G41100 | 0,28 | 1,38 | 4,41 | 0,31 | 7,54E-01 |
| AT5G41110 | 0,22 | 1,33 | 4,40 | 0,30 | 7,62E-01 |
| AT5G41140 | 0,59 | 1,88 | 3,78 | 0,50 | 6,19E-01 |
| AT5G41260 | 0,61 | 2,90 | 2,31 | 1,26 | 2,09E-01 |
| AT5G41360 | 0,66 | 1,36 | 3,67 | 0,37 | 7,10E-01 |
| AT5G41370 | 0,15 | 0,00 | 4,41 | 0,00 | 1,00E+00 |
| AT5G41410 | 1,13 | 1,87 | 2,03 | 0,92 | 3,58E-01 |
| AT5G41500 | 1,05 | -0,54 | 2,32 | -0,23 | 8,16E-01 |
| AT5G41660 | 0,71 | -0,51 | 3,10 | -0,16 | 8,69E-01 |
| AT5G41700 | 1,24 | -0,50 | 1,81 | -0,28 | 7,82E-01 |
| AT5G41761 | 0,83 | 1,29 | 2,49 | 0,52 | 6,05E-01 |
| AT5G41770 | 0,61 | -1,41 | 2,65 | -0,53 | 5,95E-01 |
| AT5G41860 | 0,77 | 0,00 | 4,41 | 0,00 | 1,00E+00 |
| AT5G41920 | 0,61 | 1,25 | 4,38 | 0,29 | 7,75E-01 |
| AT5G41940 | 0,98 | 0,49 | 2,00 | 0,24 | 8,07E-01 |
| AT5G41950 | 1,46 | 2,72 | 2,11 | 1,29 | 1,97E-01 |
| AT5G41960 | 0,15 | -0,43 | 4,39 | -0,10 | 9,23E-01 |
| AT5G41970 | 0,85 | 3,53 | 3,34 | 1,06 | 2,90E-01 |
| AT5G42010 | 0,60 | 1,39 | 3,24 | 0,43 | 6,68E-01 |
| AT5G42220 | 0,70 | 2,06 | 2,44 | 0,84 | 3,99E-01 |
| AT5G42250 | 0,33 | 1,37 | 4,41 | 0,31 | 7,56E-01 |
| AT5G42260 | 0,28 | 0,00 | 4,41 | 0,00 | 1,00E+00 |
| AT5G42290 | 0,08 | 0,00 | 4,41 | 0,00 | 1,00E+00 |
| AT5G42310 | 0,50 | 1,23 | 4,17 | 0,30 | 7,68E-01 |
| AT5G42350 | 0,37 | 2,10 | 4,37 | 0,48 | 6,30E-01 |
| AT5G42420 | 0,74 | 2,55 | 3,64 | 0,70 | 4,83E-01 |
| AT5G42460 | 0,33 | 1,85 | 4,39 | 0,42 | 6,73E-01 |
| AT5G42470 | 0,22 | -0,42 | 4,37 | -0,10 | 9,24E-01 |

| | | | | | |
|---|---|---|---|---|---|
| AT5G42480 | 0,45 | 1,85 | 4,39 | 0,42 | 6,73E-01 |
| AT5G42504 | 0,17 | -0,82 | 4,37 | -0,19 | 8,51E-01 |
| AT5G42520 | 0,71 | 0,42 | 2,34 | 0,18 | 8,57E-01 |
| AT5G42540 | 1,02 | 2,77 | 2,40 | 1,15 | 2,48E-01 |
| AT5G42620 | 0,80 | 0,76 | 2,39 | 0,32 | 7,50E-01 |
| AT5G42640 | 0,08 | 0,00 | 4,41 | 0,00 | 1,00E+00 |
| AT5G42670 | 0,16 | 0,00 | 4,41 | 0,00 | 1,00E+00 |
| AT5G42680 | 0,80 | 0,41 | 3,31 | 0,12 | 9,01E-01 |
| AT5G42720 | 0,19 | 0,00 | 4,40 | 0,00 | 1,00E+00 |
| AT5G42740 | 1,16 | -1,03 | 2,91 | -0,35 | 7,24E-01 |
| AT5G42760 | 0,12 | 0,00 | 4,41 | 0,00 | 1,00E+00 |
| AT5G06395 | 1,55 | 3,21 | 1,81 | 1,77 | 7,66E-02 |
| AT5G42825 | 0,29 | 0,44 | 4,41 | 0,10 | 9,21E-01 |
| AT5G42965 | 0,19 | -0,47 | 4,40 | -0,11 | 9,16E-01 |
| AT5G42980 | 0,59 | -0,61 | 2,75 | -0,22 | 8,25E-01 |
| AT5G43100 | 0,78 | 1,17 | 2,30 | 0,51 | 6,09E-01 |
| AT5G43150 | 0,61 | 2,52 | 3,08 | 0,82 | 4,13E-01 |
| AT5G06505 | 0,36 | 0,43 | 4,41 | 0,10 | 9,22E-01 |
| AT5G43460 | 1,01 | 2,59 | 1,99 | 1,30 | 1,94E-01 |
| AT5G43500 | 1,47 | -2,80 | 1,80 | -1,55 | 1,21E-01 |
| AT5G43525 | 0,69 | 0,00 | 2,95 | 0,00 | 1,00E+00 |
| AT5G43560 | 0,55 | 0,97 | 3,28 | 0,30 | 7,66E-01 |
| AT5G43610 | 1,12 | -0,31 | 2,23 | -0,14 | 8,90E-01 |
| AT5G43620 | 0,36 | -0,47 | 4,36 | -0,11 | 9,13E-01 |
| AT5G43700 | 0,52 | 0,18 | 3,10 | 0,06 | 9,52E-01 |
| AT5G43810 | 1,04 | 2,27 | 2,79 | 0,82 | 4,15E-01 |
| AT5G43930 | 0,73 | 1,81 | 2,62 | 0,69 | 4,89E-01 |
| AT5G43940 | 1,17 | 1,35 | 2,17 | 0,62 | 5,35E-01 |
| AT5G44020 | 1,03 | -0,59 | 2,12 | -0,28 | 7,82E-01 |
| AT5G44050 | 1,57 | -4,28 | 2,10 | -2,04 | 4,15E-02 |
| AT5G44060 | 0,47 | 0,42 | 2,89 | 0,14 | 8,85E-01 |
| AT5G44070 | 1,32 | 0,10 | 1,81 | 0,05 | 9,57E-01 |
| AT5G44130 | 0,80 | -2,25 | 2,79 | -0,81 | 4,20E-01 |
| AT5G44190 | 1,58 | -2,47 | 1,91 | -1,29 | 1,97E-01 |
| AT5G44230 | 0,12 | 0,00 | 4,41 | 0,00 | 1,00E+00 |
| AT5G44240 | 1,02 | 1,42 | 1,82 | 0,78 | 4,36E-01 |
| AT5G06565 | 0,63 | -1,85 | 2,73 | -0,68 | 4,98E-01 |
| AT5G44286 | 0,85 | 0,25 | 2,31 | 0,11 | 9,15E-01 |
| AT5G44530 | 1,17 | 0,98 | 2,66 | 0,37 | 7,12E-01 |
| AT5G44574 | 0,91 | -0,30 | 3,15 | -0,09 | 9,25E-01 |
| AT5G44580 | 0,52 | 1,73 | 3,93 | 0,44 | 6,60E-01 |
| AT5G44585 | 1,38 | 1,77 | 1,63 | 1,09 | 2,76E-01 |
| AT5G06605 | 1,31 | 1,80 | 1,62 | 1,11 | 2,66E-01 |
| AT5G44640 | 1,59 | -1,67 | 2,01 | -0,83 | 4,06E-01 |
| AT5G44650 | 0,46 | -0,31 | 4,06 | -0,08 | 9,40E-01 |
| AT5G44720 | 0,16 | 0,00 | 4,41 | 0,00 | 1,00E+00 |
| AT5G44750 | 0,74 | 1,41 | 2,29 | 0,62 | 5,36E-01 |
| AT5G44760 | 0,36 | 0,00 | 4,41 | 0,00 | 1,00E+00 |
| AT5G44770 | 0,22 | 0,00 | 4,41 | 0,00 | 1,00E+00 |
| AT5G45000 | 0,38 | 0,65 | 4,31 | 0,15 | 8,81E-01 |
| AT5G06695 | 0,59 | 1,42 | 3,09 | 0,46 | 6,47E-01 |
| AT5G45030 | 0,59 | 1,42 | 3,09 | 0,46 | 6,47E-01 |
| AT5G45060 | 1,56 | 0,60 | 1,74 | 0,35 | 7,28E-01 |
| AT5G45130 | 0,78 | 0,39 | 3,09 | 0,12 | 9,01E-01 |
| AT5G45150 | 0,08 | 0,00 | 4,41 | 0,00 | 1,00E+00 |
| AT5G45240 | 0,12 | 0,00 | 4,41 | 0,00 | 1,00E+00 |

| | | | | | |
|---|---|---|---|---|---|
| AT5G45260 | 0,75 | 2,95 | 2,88 | 1,02 | 3,06E-01 |
| AT5G45390 | 0,62 | -0,68 | 2,35 | -0,29 | 7,71E-01 |
| AT5G45400 | 0,76 | 2,01 | 2,51 | 0,80 | 4,22E-01 |
| AT5G45450 | 0,08 | 0,00 | 4,41 | 0,00 | 1,00E+00 |
| AT5G45475 | 0,74 | -0,80 | 2,31 | -0,35 | 7,28E-01 |
| AT5G45490 | 1,25 | -0,18 | 1,90 | -0,09 | 9,26E-01 |
| AT5G45570 | 0,68 | -0,46 | 2,59 | -0,18 | 8,58E-01 |
| AT5G45620 | 0,59 | 0,00 | 3,21 | 0,00 | 1,00E+00 |
| AT5G45670 | 0,10 | 0,00 | 4,41 | 0,00 | 1,00E+00 |
| AT5G45680 | 0,26 | 1,38 | 4,41 | 0,31 | 7,54E-01 |
| AT5G45710 | 0,55 | 2,59 | 3,81 | 0,68 | 4,97E-01 |
| AT5G45750 | 0,19 | 0,00 | 4,41 | 0,00 | 1,00E+00 |
| AT5G45770 | 0,12 | 0,00 | 4,41 | 0,00 | 1,00E+00 |
| AT5G45920 | 1,21 | -3,68 | 1,92 | -1,92 | 5,45E-02 |
| AT5G46330 | 0,47 | 1,33 | 4,08 | 0,33 | 7,44E-01 |
| AT5G46410 | 0,44 | 16,74 | 4,39 | 3,81 | 1,38E-04 |
| AT5G06825 | 0,67 | 1,30 | 3,15 | 0,41 | 6,78E-01 |
| AT5G46460 | 0,90 | 1,83 | 3,35 | 0,55 | 5,85E-01 |
| AT5G46490 | 1,46 | 3,41 | 1,79 | 1,91 | 5,64E-02 |
| AT5G46500 | 1,11 | 1,36 | 2,62 | 0,52 | 6,04E-01 |
| AT5G06835 | 1,13 | -2,79 | 2,42 | -1,16 | 2,48E-01 |
| AT5G46630 | 1,04 | 0,60 | 2,00 | 0,30 | 7,66E-01 |
| AT5G46640 | 1,47 | 3,47 | 1,96 | 1,77 | 7,67E-02 |
| AT5G46660 | 1,02 | 0,00 | 4,13 | 0,00 | 1,00E+00 |
| AT5G46670 | 0,74 | -0,46 | 2,87 | -0,16 | 8,73E-01 |
| AT5G46730 | 0,08 | 0,00 | 4,41 | 0,00 | 1,00E+00 |
| AT5G46780 | 0,35 | -0,55 | 4,41 | -0,13 | 9,00E-01 |
| AT5G46825 | 1,18 | -1,68 | 1,76 | -0,96 | 3,39E-01 |
| AT5G46840 | 0,62 | 0,48 | 3,12 | 0,15 | 8,77E-01 |
| AT5G46850 | 0,29 | 1,25 | 4,39 | 0,29 | 7,75E-01 |
| AT5G47020 | 1,57 | 2,66 | 1,87 | 1,42 | 1,55E-01 |
| AT5G47030 | 0,63 | 1,38 | 3,81 | 0,36 | 7,17E-01 |
| AT5G47120 | 1,06 | 0,00 | 2,31 | 0,00 | 1,00E+00 |
| AT5G47130 | 0,25 | -0,45 | 4,39 | -0,10 | 9,18E-01 |
| AT5G47140 | 0,26 | -0,49 | 4,40 | -0,11 | 9,12E-01 |
| AT5G47200 | 0,34 | -0,54 | 4,41 | -0,12 | 9,02E-01 |
| AT5G47230 | 0,27 | 0,00 | 4,41 | 0,00 | 1,00E+00 |
| AT5G47240 | 0,14 | 0,00 | 4,41 | 0,00 | 1,00E+00 |
| AT5G47280 | 1,15 | -1,88 | 2,05 | -0,91 | 3,61E-01 |
| AT5G47300 | 1,47 | 1,38 | 2,11 | 0,65 | 5,13E-01 |
| AT5G47320 | 0,19 | 0,00 | 4,41 | 0,00 | 1,00E+00 |
| AT5G47390 | 0,64 | 1,19 | 2,17 | 0,55 | 5,83E-01 |
| AT5G47420 | 0,50 | 2,75 | 3,97 | 0,69 | 4,89E-01 |
| AT5G47435 | 1,48 | 4,28 | 1,96 | 2,18 | 2,91E-02 |
| AT5G47640 | 0,70 | -0,16 | 2,20 | -0,07 | 9,42E-01 |
| AT5G47660 | 0,08 | 0,00 | 4,41 | 0,00 | 1,00E+00 |
| AT5G47720 | 0,16 | 0,00 | 4,41 | 0,00 | 1,00E+00 |
| AT5G47770 | 0,39 | -0,54 | 4,33 | -0,13 | 9,00E-01 |
| AT5G47780 | 0,52 | 0,43 | 2,78 | 0,15 | 8,77E-01 |
| AT5G47790 | 0,28 | -0,46 | 4,39 | -0,10 | 9,18E-01 |
| AT5G47800 | 0,24 | 1,37 | 4,41 | 0,31 | 7,56E-01 |
| AT5G47820 | 0,70 | 2,92 | 3,68 | 0,80 | 4,26E-01 |
| AT5G47840 | 0,45 | 0,45 | 2,81 | 0,16 | 8,74E-01 |
| AT5G47860 | 0,53 | -0,54 | 3,95 | -0,14 | 8,90E-01 |
| AT5G47890 | 1,07 | 1,19 | 1,97 | 0,60 | 5,48E-01 |
| AT5G48190 | 1,46 | -2,57 | 1,87 | -1,37 | 1,71E-01 |

| | | | | | |
|---|---|---|---|---|---|
| AT5G48220 | 0,16 | 0,00 | 4,41 | 0,00 | 1,00E+00 |
| AT5G48290 | 0,36 | 2,37 | 4,36 | 0,54 | 5,86E-01 |
| AT5G48300 | 1,23 | -1,36 | 2,00 | -0,68 | 4,95E-01 |
| AT5G48340 | 0,30 | 1,38 | 4,41 | 0,31 | 7,54E-01 |
| AT5G48360 | 0,08 | 0,00 | 4,41 | 0,00 | 1,00E+00 |
| AT5G48390 | 0,69 | 2,80 | 3,52 | 0,79 | 4,27E-01 |
| AT5G48400 | 0,56 | 1,38 | 3,88 | 0,36 | 7,22E-01 |
| AT5G48410 | 0,12 | 0,00 | 4,41 | 0,00 | 1,00E+00 |
| AT5G07095 | 1,59 | 0,45 | 1,44 | 0,31 | 7,55E-01 |
| AT5G07105 | 0,86 | 1,64 | 2,16 | 0,76 | 4,48E-01 |
| AT5G48545 | 0,30 | -0,59 | 3,75 | -0,16 | 8,75E-01 |
| AT5G48720 | 0,16 | 0,00 | 4,41 | 0,00 | 1,00E+00 |
| AT5G48730 | 0,12 | 0,00 | 4,41 | 0,00 | 1,00E+00 |
| AT5G48775 | 0,08 | 0,00 | 4,41 | 0,00 | 1,00E+00 |
| AT5G48800 | 0,90 | 1,88 | 2,99 | 0,63 | 5,29E-01 |
| AT5G48900 | 0,13 | -0,43 | 4,39 | -0,10 | 9,23E-01 |
| AT5G49010 | 1,17 | -0,34 | 1,88 | -0,18 | 8,55E-01 |
| AT5G49020 | 0,22 | -0,53 | 4,41 | -0,12 | 9,05E-01 |
| AT5G49060 | 0,17 | 0,00 | 4,41 | 0,00 | 1,00E+00 |
| AT5G49160 | 0,93 | 2,86 | 2,58 | 1,11 | 2,67E-01 |
| AT5G49215 | 0,08 | 0,00 | 4,41 | 0,00 | 1,00E+00 |
| AT5G49250 | 0,66 | 0,00 | 2,93 | 0,00 | 1,00E+00 |
| AT5G49260 | 0,08 | 0,00 | 4,41 | 0,00 | 1,00E+00 |
| AT5G49450 | 0,19 | 0,00 | 4,41 | 0,00 | 1,00E+00 |
| AT5G49460 | 0,62 | 2,71 | 3,00 | 0,90 | 3,66E-01 |
| AT5G49470 | 0,29 | 1,25 | 4,39 | 0,29 | 7,75E-01 |
| AT5G49480 | 1,15 | 3,97 | 2,28 | 1,74 | 8,19E-02 |
| AT5G49510 | 0,19 | 0,00 | 4,41 | 0,00 | 1,00E+00 |
| AT5G49520 | 0,12 | 0,00 | 4,41 | 0,00 | 1,00E+00 |
| AT5G49530 | 0,46 | 0,00 | 4,15 | 0,00 | 1,00E+00 |
| AT5G49590 | 0,86 | -0,81 | 2,82 | -0,29 | 7,73E-01 |
| AT5G49640 | 0,08 | 0,00 | 4,41 | 0,00 | 1,00E+00 |
| AT5G49650 | 0,61 | -0,50 | 2,38 | -0,21 | 8,35E-01 |
| AT5G49810 | 1,47 | -0,16 | 1,88 | -0,09 | 9,30E-01 |
| AT5G49840 | 0,10 | 0,00 | 4,41 | 0,00 | 1,00E+00 |
| AT5G49945 | 0,51 | -0,41 | 3,90 | -0,10 | 9,16E-01 |
| AT5G49970 | 0,29 | 0,00 | 4,41 | 0,00 | 1,00E+00 |
| AT5G50020 | 0,47 | 2,55 | 4,35 | 0,59 | 5,57E-01 |
| AT5G50160 | 1,06 | 0,43 | 2,50 | 0,17 | 8,64E-01 |
| AT5G50170 | 0,08 | 0,00 | 4,41 | 0,00 | 1,00E+00 |
| AT5G50200 | 0,08 | 0,00 | 4,41 | 0,00 | 1,00E+00 |
| AT5G07605 | 0,25 | 0,00 | 4,41 | 0,00 | 1,00E+00 |
| AT5G50310 | 0,21 | 0,00 | 4,41 | 0,00 | 1,00E+00 |
| AT5G50580 | 0,65 | 1,81 | 3,69 | 0,49 | 6,23E-01 |
| AT5G50680 | 1,09 | 1,32 | 2,54 | 0,52 | 6,02E-01 |
| AT5G50717 | 0,11 | 0,00 | 4,41 | 0,00 | 1,00E+00 |
| AT5G50780 | 0,59 | 0,59 | 4,36 | 0,13 | 8,93E-01 |
| AT5G50850 | 0,82 | -0,54 | 2,51 | -0,22 | 8,28E-01 |
| AT5G50950 | 1,52 | -1,07 | 1,54 | -0,70 | 4,86E-01 |
| AT5G50960 | 0,50 | 0,99 | 3,92 | 0,25 | 8,00E-01 |
| AT5G51010 | 0,08 | 0,00 | 4,41 | 0,00 | 1,00E+00 |
| AT5G51050 | 0,20 | 0,00 | 4,41 | 0,00 | 1,00E+00 |
| AT5G51055 | 0,34 | -1,31 | 3,56 | -0,37 | 7,13E-01 |
| AT5G51120 | 0,08 | 0,00 | 4,41 | 0,00 | 1,00E+00 |
| AT5G51150 | 0,28 | 1,39 | 4,41 | 0,32 | 7,52E-01 |
| AT5G51180 | 0,38 | 0,42 | 2,98 | 0,14 | 8,88E-01 |

| | | | | | |
|---|---|---|---|---|---|
| AT5G51200 | 1,04 | -0,33 | 2,29 | -0,15 | 8,85E-01 |
| AT5G51230 | 0,21 | -0,45 | 4,39 | -0,10 | 9,19E-01 |
| AT5G51280 | 0,48 | 1,38 | 2,80 | 0,49 | 6,22E-01 |
| AT5G51340 | 0,39 | 0,57 | 3,46 | 0,17 | 8,68E-01 |
| AT5G51450 | 0,53 | -0,20 | 3,93 | -0,05 | 9,59E-01 |
| AT5G51560 | 0,28 | 2,12 | 4,38 | 0,48 | 6,28E-01 |
| AT5G51570 | 0,40 | -0,57 | 3,48 | -0,16 | 8,69E-01 |
| AT5G51660 | 1,44 | 2,86 | 1,84 | 1,56 | 1,20E-01 |
| AT5G51670 | 0,12 | 0,00 | 4,41 | 0,00 | 1,00E+00 |
| AT5G51680 | 0,64 | 2,68 | 2,62 | 1,03 | 3,05E-01 |
| AT5G51750 | 0,75 | -0,39 | 3,66 | -0,11 | 9,16E-01 |
| AT5G51760 | 1,35 | 0,44 | 1,73 | 0,25 | 7,99E-01 |
| AT5G51830 | 0,27 | 0,00 | 4,41 | 0,00 | 1,00E+00 |
| AT5G51960 | 0,11 | 0,00 | 4,41 | 0,00 | 1,00E+00 |
| AT5G51970 | 1,04 | 2,17 | 2,02 | 1,07 | 2,83E-01 |
| AT5G52050 | 0,61 | 1,35 | 2,68 | 0,50 | 6,15E-01 |
| AT5G52115 | 1,12 | -2,40 | 2,64 | -0,91 | 3,62E-01 |
| AT5G52240 | 1,16 | 0,60 | 2,15 | 0,28 | 7,79E-01 |
| AT5G52280 | 0,34 | 1,70 | 4,38 | 0,39 | 6,98E-01 |
| AT5G52471 | 1,40 | -0,60 | 1,90 | -0,32 | 7,51E-01 |
| AT5G52510 | 0,69 | -1,41 | 3,06 | -0,46 | 6,46E-01 |
| AT5G52520 | 0,23 | 1,29 | 4,40 | 0,29 | 7,69E-01 |
| AT5G52580 | 0,50 | 0,00 | 3,35 | 0,00 | 1,00E+00 |
| AT5G52630 | 0,08 | 0,00 | 4,41 | 0,00 | 1,00E+00 |
| AT5G52700 | 0,14 | 0,00 | 4,41 | 0,00 | 1,00E+00 |
| AT5G52750 | 0,33 | 0,00 | 4,41 | 0,00 | 1,00E+00 |
| AT5G52760 | 0,52 | 1,36 | 2,75 | 0,50 | 6,20E-01 |
| AT5G52920 | 1,32 | 0,48 | 2,18 | 0,22 | 8,24E-01 |
| AT5G07945 | 0,52 | 0,00 | 4,04 | 0,00 | 1,00E+00 |
| AT5G52940 | 0,11 | 0,00 | 4,41 | 0,00 | 1,00E+00 |
| AT5G53020 | 0,27 | -0,42 | 4,39 | -0,10 | 9,24E-01 |
| AT5G53030 | 0,24 | -0,56 | 4,41 | -0,13 | 8,99E-01 |
| AT5G53048 | 0,25 | -0,50 | 4,40 | -0,11 | 9,10E-01 |
| AT5G53060 | 0,48 | 0,00 | 4,05 | 0,00 | 1,00E+00 |
| AT5G53080 | 0,12 | 0,00 | 4,41 | 0,00 | 1,00E+00 |
| AT5G53110 | 0,31 | -1,56 | 4,34 | -0,36 | 7,19E-01 |
| AT5G53120 | 0,82 | 2,29 | 2,65 | 0,86 | 3,88E-01 |
| AT5G53140 | 1,45 | 2,59 | 1,88 | 1,38 | 1,69E-01 |
| AT5G53150 | 0,95 | 2,39 | 2,15 | 1,11 | 2,65E-01 |
| AT5G53160 | 1,14 | 1,38 | 2,15 | 0,64 | 5,21E-01 |
| AT5G53220 | 0,48 | 1,26 | 3,20 | 0,39 | 6,94E-01 |
| AT5G53280 | 0,44 | 2,60 | 4,14 | 0,63 | 5,30E-01 |
| AT5G53350 | 0,54 | -0,78 | 3,18 | -0,25 | 8,06E-01 |
| AT5G53360 | 0,08 | 0,00 | 4,41 | 0,00 | 1,00E+00 |
| AT5G53410 | 1,13 | -2,54 | 1,88 | -1,35 | 1,77E-01 |
| AT5G53420 | 0,48 | 2,37 | 3,31 | 0,72 | 4,74E-01 |
| AT5G53430 | 1,00 | 0,98 | 2,00 | 0,49 | 6,24E-01 |
| AT5G53450 | 0,71 | 2,02 | 2,57 | 0,79 | 4,32E-01 |
| AT5G53550 | 0,99 | 1,96 | 2,43 | 0,80 | 4,21E-01 |
| AT5G53560 | 1,33 | 0,43 | 1,92 | 0,22 | 8,25E-01 |
| AT5G53590 | 0,48 | 0,24 | 4,06 | 0,06 | 9,52E-01 |
| AT5G53620 | 1,09 | 2,26 | 1,83 | 1,24 | 2,16E-01 |
| AT5G53860 | 0,25 | 0,00 | 3,87 | 0,00 | 1,00E+00 |
| AT5G53890 | 0,71 | -0,48 | 2,60 | -0,18 | 8,55E-01 |
| AT5G54035 | 0,15 | 0,00 | 4,41 | 0,00 | 1,00E+00 |
| AT5G54043 | 0,22 | 0,00 | 4,41 | 0,00 | 1,00E+00 |

| | | | | | |
|---|---|---|---|---|---|
| AT5G54080 | 0,99 | 2,21 | 2,31 | 0,96 | 3,38E-01 |
| AT5G54110 | 0,33 | 1,38 | 4,41 | 0,31 | 7,54E-01 |
| AT5G54140 | 0,24 | 1,38 | 4,41 | 0,31 | 7,54E-01 |
| AT5G54145 | 0,12 | 0,00 | 4,41 | 0,00 | 1,00E+00 |
| AT5G54160 | 0,46 | -1,62 | 4,13 | -0,39 | 6,95E-01 |
| AT5G54180 | 0,12 | 0,00 | 4,41 | 0,00 | 1,00E+00 |
| AT5G54290 | 0,66 | 1,20 | 2,68 | 0,45 | 6,55E-01 |
| AT5G54500 | 0,31 | -1,22 | 4,36 | -0,28 | 7,79E-01 |
| AT5G54540 | 0,58 | 0,41 | 2,79 | 0,15 | 8,83E-01 |
| AT5G54600 | 0,87 | 0,58 | 2,05 | 0,28 | 7,76E-01 |
| AT5G08225 | 0,36 | 0,00 | 4,41 | 0,00 | 1,00E+00 |
| AT5G54640 | 0,05 | 0,00 | 4,41 | 0,00 | 1,00E+00 |
| AT5G54680 | 0,72 | 1,16 | 2,55 | 0,45 | 6,50E-01 |
| AT5G54750 | 0,55 | -0,65 | 3,95 | -0,16 | 8,70E-01 |
| AT5G54760 | 0,49 | -1,66 | 2,78 | -0,60 | 5,51E-01 |
| AT5G54800 | 0,08 | 0,00 | 4,41 | 0,00 | 1,00E+00 |
| AT5G54830 | 0,76 | 1,36 | 2,90 | 0,47 | 6,40E-01 |
| AT5G54855 | 0,18 | -0,82 | 4,37 | -0,19 | 8,51E-01 |
| AT5G54860 | 0,26 | 1,38 | 4,41 | 0,31 | 7,54E-01 |
| AT5G54900 | 0,82 | 1,73 | 2,41 | 0,72 | 4,73E-01 |
| AT5G54920 | 0,83 | 0,00 | 4,39 | 0,00 | 1,00E+00 |
| AT5G55060 | 0,14 | 0,00 | 4,41 | 0,00 | 1,00E+00 |
| AT5G55070 | 1,01 | 1,37 | 2,07 | 0,66 | 5,10E-01 |
| AT5G55190 | 0,80 | 0,94 | 2,43 | 0,39 | 6,98E-01 |
| AT5G55220 | 0,88 | -0,81 | 2,54 | -0,32 | 7,50E-01 |
| AT5G55230 | 0,95 | 1,99 | 2,08 | 0,96 | 3,39E-01 |
| AT5G55450 | 0,62 | 1,43 | 3,03 | 0,47 | 6,38E-01 |
| AT5G55460 | 0,37 | 0,43 | 3,44 | 0,12 | 9,01E-01 |
| AT5G55500 | 0,57 | 1,29 | 3,19 | 0,41 | 6,85E-01 |
| AT5G55530 | 0,30 | 1,33 | 4,40 | 0,30 | 7,63E-01 |
| AT5G55580 | 0,70 | 0,20 | 3,90 | 0,05 | 9,58E-01 |
| AT5G55620 | 0,43 | -0,54 | 4,31 | -0,13 | 9,00E-01 |
| AT5G55630 | 0,12 | 0,00 | 4,41 | 0,00 | 1,00E+00 |
| AT5G55660 | 0,90 | 0,18 | 2,06 | 0,09 | 9,30E-01 |
| AT5G55840 | 0,22 | 0,00 | 4,41 | 0,00 | 1,00E+00 |
| AT5G55850 | 0,24 | 0,00 | 4,41 | 0,00 | 1,00E+00 |
| AT5G55900 | 0,10 | 0,00 | 4,41 | 0,00 | 1,00E+00 |
| AT5G55930 | 0,70 | 1,81 | 3,01 | 0,60 | 5,48E-01 |
| AT5G55960 | 0,59 | -1,90 | 3,09 | -0,62 | 5,38E-01 |
| AT5G55990 | 0,33 | 1,25 | 4,39 | 0,28 | 7,76E-01 |
| AT5G56020 | 0,32 | 2,14 | 4,37 | 0,49 | 6,25E-01 |
| AT5G56040 | 0,73 | 0,43 | 2,57 | 0,17 | 8,65E-01 |
| AT5G56060 | 0,67 | 0,00 | 4,38 | 0,00 | 1,00E+00 |
| AT5G56140 | 0,98 | 1,90 | 2,03 | 0,94 | 3,49E-01 |
| AT5G56160 | 0,49 | -0,93 | 4,34 | -0,22 | 8,30E-01 |
| AT5G56170 | 0,53 | 1,12 | 4,38 | 0,26 | 7,98E-01 |
| AT5G56270 | 0,76 | 3,32 | 2,79 | 1,19 | 2,35E-01 |
| AT5G56290 | 0,52 | 0,31 | 2,50 | 0,12 | 9,02E-01 |
| AT5G56340 | 0,44 | 1,27 | 3,32 | 0,38 | 7,02E-01 |
| AT5G08495 | 0,43 | 0,18 | 4,22 | 0,04 | 9,65E-01 |
| AT5G56460 | 0,74 | -2,77 | 3,93 | -0,70 | 4,81E-01 |
| AT5G56500 | 0,32 | 2,21 | 4,37 | 0,51 | 6,13E-01 |
| AT5G56590 | 0,24 | 1,61 | 4,39 | 0,37 | 7,14E-01 |
| AT5G56630 | 0,71 | 1,73 | 2,52 | 0,69 | 4,91E-01 |
| AT5G56650 | 0,64 | 2,76 | 2,33 | 1,18 | 2,38E-01 |
| AT5G56680 | 1,22 | 1,04 | 1,85 | 0,56 | 5,73E-01 |

| | | | | | |
|---|---|---|---|---|---|
| AT5G56720 | 0,12 | 0,00 | 4,41 | 0,00 | 1,00E+00 |
| AT5G56870 | 0,19 | 0,00 | 4,41 | 0,00 | 1,00E+00 |
| AT5G56910 | 0,15 | 0,00 | 4,41 | 0,00 | 1,00E+00 |
| AT5G56920 | 0,65 | 1,38 | 3,16 | 0,44 | 6,62E-01 |
| AT5G56940 | 0,08 | 0,00 | 4,41 | 0,00 | 1,00E+00 |
| AT5G56950 | 0,44 | -0,32 | 3,26 | -0,10 | 9,22E-01 |
| AT5G57015 | 0,32 | 1,44 | 4,40 | 0,33 | 7,44E-01 |
| AT5G57020 | 1,25 | 0,45 | 1,88 | 0,24 | 8,09E-01 |
| AT5G57030 | 0,86 | -0,41 | 2,41 | -0,17 | 8,66E-01 |
| AT5G57035 | 1,04 | 1,84 | 2,55 | 0,72 | 4,71E-01 |
| AT5G57070 | 0,42 | 2,24 | 4,25 | 0,53 | 5,98E-01 |
| AT5G57150 | 0,25 | 1,38 | 4,41 | 0,31 | 7,54E-01 |
| AT5G57180 | 1,05 | -0,70 | 1,84 | -0,38 | 7,02E-01 |
| AT5G57320 | 0,08 | 0,00 | 4,41 | 0,00 | 1,00E+00 |
| AT5G57330 | 0,49 | -0,15 | 2,90 | -0,05 | 9,59E-01 |
| AT5G57340 | 0,20 | 0,00 | 4,41 | 0,00 | 1,00E+00 |
| AT5G57360 | 0,96 | 1,30 | 2,04 | 0,64 | 5,25E-01 |
| AT5G57370 | 0,31 | 1,38 | 4,41 | 0,31 | 7,54E-01 |
| AT5G57460 | 1,07 | 1,95 | 1,88 | 1,04 | 2,99E-01 |
| AT5G57520 | 0,31 | 0,00 | 4,41 | 0,00 | 1,00E+00 |
| AT5G57610 | 0,89 | 0,64 | 1,99 | 0,32 | 7,47E-01 |
| AT5G57655 | 1,67 | -0,20 | 1,81 | -0,11 | 9,12E-01 |
| AT5G57735 | 0,19 | 0,00 | 4,41 | 0,00 | 1,00E+00 |
| AT5G57800 | 0,61 | 0,50 | 2,59 | 0,19 | 8,47E-01 |
| AT5G08805 | 1,03 | 1,31 | 2,55 | 0,51 | 6,08E-01 |
| AT5G57880 | 0,48 | 2,16 | 4,02 | 0,54 | 5,92E-01 |
| AT5G57930 | 1,66 | 2,71 | 1,88 | 1,45 | 1,48E-01 |
| AT5G57940 | 0,35 | -0,57 | 3,62 | -0,16 | 8,74E-01 |
| AT5G58000 | 0,08 | 0,00 | 4,41 | 0,00 | 1,00E+00 |
| AT5G58020 | 0,39 | -2,30 | 4,32 | -0,53 | 5,95E-01 |
| AT5G58060 | 0,59 | 2,52 | 2,60 | 0,97 | 3,32E-01 |
| AT5G58100 | 0,97 | 1,11 | 1,86 | 0,59 | 5,53E-01 |
| AT5G58110 | 0,27 | -0,47 | 4,40 | -0,11 | 9,15E-01 |
| AT5G58120 | 1,53 | -0,22 | 1,61 | -0,13 | 8,93E-01 |
| AT5G08820 | 0,19 | 0,00 | 4,41 | 0,00 | 1,00E+00 |
| AT5G58200 | 0,40 | -1,71 | 4,36 | -0,39 | 6,95E-01 |
| AT5G58230 | 0,77 | 2,17 | 2,55 | 0,85 | 3,94E-01 |
| AT5G58250 | 0,32 | -0,61 | 4,39 | -0,14 | 8,90E-01 |
| AT5G58270 | 0,43 | 0,35 | 4,25 | 0,08 | 9,34E-01 |
| AT5G58290 | 1,43 | -1,71 | 1,89 | -0,91 | 3,65E-01 |
| AT5G58300 | 0,99 | 3,63 | 2,25 | 1,61 | 1,07E-01 |
| AT5G58320 | 0,24 | 1,31 | 4,40 | 0,30 | 7,65E-01 |
| AT5G58350 | 1,12 | 2,28 | 2,12 | 1,08 | 2,81E-01 |
| AT5G58370 | 0,29 | 1,71 | 4,38 | 0,39 | 6,95E-01 |
| AT5G58375 | 0,48 | 2,22 | 4,36 | 0,51 | 6,10E-01 |
| AT5G58420 | 1,59 | 0,72 | 1,82 | 0,40 | 6,91E-01 |
| AT5G58450 | 0,68 | 2,09 | 2,96 | 0,70 | 4,81E-01 |
| AT5G58495 | 0,12 | 0,00 | 4,41 | 0,00 | 1,00E+00 |
| AT5G58520 | 0,39 | 1,95 | 4,39 | 0,44 | 6,57E-01 |
| AT5G58560 | 0,47 | 1,27 | 3,31 | 0,38 | 7,02E-01 |
| AT5G08855 | 0,43 | 0,53 | 4,27 | 0,12 | 9,01E-01 |
| AT5G58620 | 0,34 | 1,49 | 4,39 | 0,34 | 7,34E-01 |
| AT5G58640 | 0,71 | 1,51 | 2,94 | 0,51 | 6,08E-01 |
| AT5G58710 | 0,66 | -2,07 | 3,12 | -0,66 | 5,07E-01 |
| AT5G58990 | 0,29 | -0,54 | 4,41 | -0,12 | 9,02E-01 |
| AT5G59160 | 0,32 | -0,49 | 4,40 | -0,11 | 9,11E-01 |

| | | | | | |
|---|---|---|---|---|---|
| AT5G59170 | 0,26 | 1,38 | 4,41 | 0,31 | 7,54E-01 |
| AT5G59250 | 0,50 | -0,07 | 3,20 | -0,02 | 9,84E-01 |
| AT5G59350 | 0,35 | 1,78 | 4,38 | 0,41 | 6,84E-01 |
| AT5G59420 | 1,43 | 0,33 | 2,01 | 0,16 | 8,69E-01 |
| AT5G59430 | 0,64 | -1,32 | 2,66 | -0,50 | 6,20E-01 |
| AT5G59450 | 0,61 | 2,00 | 3,90 | 0,51 | 6,08E-01 |
| AT5G59610 | 0,51 | 1,38 | 4,05 | 0,34 | 7,33E-01 |
| AT5G59660 | 0,05 | 0,00 | 4,41 | 0,00 | 1,00E+00 |
| AT5G59670 | 0,28 | -0,42 | 3,72 | -0,11 | 9,09E-01 |
| AT5G59680 | 0,64 | 2,97 | 4,33 | 0,69 | 4,93E-01 |
| AT5G59690 | 0,64 | 1,53 | 3,04 | 0,50 | 6,15E-01 |
| AT5G59820 | 0,42 | -0,54 | 3,38 | -0,16 | 8,72E-01 |
| AT5G59830 | 0,85 | 2,02 | 2,84 | 0,71 | 4,78E-01 |
| AT5G59860 | 0,43 | 2,00 | 4,37 | 0,46 | 6,47E-01 |
| AT5G59880 | 0,87 | -0,33 | 2,59 | -0,13 | 9,00E-01 |
| AT5G59910 | 0,91 | 1,81 | 2,35 | 0,77 | 4,39E-01 |
| AT5G59920 | 0,58 | 2,28 | 3,76 | 0,61 | 5,44E-01 |
| AT5G59950 | 1,16 | 1,42 | 1,91 | 0,74 | 4,58E-01 |
| AT5G59980 | 0,39 | 1,76 | 4,38 | 0,40 | 6,87E-01 |
| AT5G60040 | 0,79 | 3,30 | 2,38 | 1,38 | 1,67E-01 |
| AT5G60110 | 0,25 | 1,34 | 4,40 | 0,30 | 7,61E-01 |
| AT5G60120 | 0,67 | 0,86 | 2,86 | 0,30 | 7,65E-01 |
| AT5G60170 | 1,68 | 0,98 | 1,81 | 0,54 | 5,86E-01 |
| AT5G60270 | 0,48 | 0,00 | 4,41 | 0,00 | 1,00E+00 |
| AT5G60280 | 0,76 | 0,00 | 3,59 | 0,00 | 1,00E+00 |
| AT5G60300 | 0,52 | 0,00 | 3,96 | 0,00 | 1,00E+00 |
| AT5G60320 | 0,25 | 1,38 | 4,41 | 0,31 | 7,54E-01 |
| AT5G60550 | 0,49 | 0,44 | 3,27 | 0,13 | 8,94E-01 |
| AT5G60560 | 0,60 | 0,00 | 3,19 | 0,00 | 1,00E+00 |
| AT5G60570 | 0,88 | 3,39 | 2,14 | 1,58 | 1,13E-01 |
| AT5G60610 | 0,72 | 3,09 | 3,55 | 0,87 | 3,84E-01 |
| AT5G60620 | 0,40 | 0,42 | 3,56 | 0,12 | 9,07E-01 |
| AT5G60680 | 1,59 | 2,96 | 1,95 | 1,52 | 1,29E-01 |
| AT5G60780 | 1,02 | 3,74 | 2,15 | 1,74 | 8,27E-02 |
| AT5G60820 | 0,45 | 1,87 | 4,16 | 0,45 | 6,53E-01 |
| AT5G60850 | 1,63 | 2,43 | 1,75 | 1,39 | 1,64E-01 |
| AT5G60890 | 0,24 | 1,96 | 4,39 | 0,45 | 6,55E-01 |
| AT5G60945 | 1,24 | 0,03 | 1,66 | 0,02 | 9,86E-01 |
| AT5G60963 | 0,50 | 0,49 | 2,75 | 0,18 | 8,59E-01 |
| AT5G60964 | 0,93 | -0,25 | 3,43 | -0,07 | 9,42E-01 |
| AT5G60980 | 1,13 | 2,69 | 2,19 | 1,23 | 2,19E-01 |
| AT5G61190 | 0,65 | 0,45 | 2,49 | 0,18 | 8,56E-01 |
| AT5G61200 | 0,25 | 1,84 | 4,38 | 0,42 | 6,75E-01 |
| AT5G61210 | 1,18 | 1,27 | 2,02 | 0,63 | 5,30E-01 |
| AT5G61230 | 1,00 | -2,07 | 2,16 | -0,96 | 3,38E-01 |
| AT5G61240 | 0,29 | -0,97 | 4,38 | -0,22 | 8,25E-01 |
| AT5G61380 | 0,78 | 0,99 | 2,50 | 0,40 | 6,92E-01 |
| AT5G61400 | 0,78 | 2,32 | 3,45 | 0,67 | 5,01E-01 |
| AT5G61470 | 0,39 | 0,00 | 3,43 | 0,00 | 1,00E+00 |
| AT5G61490 | 0,17 | 0,00 | 4,41 | 0,00 | 1,00E+00 |
| AT5G61550 | 0,85 | 1,93 | 3,41 | 0,57 | 5,71E-01 |
| AT5G61570 | 0,19 | -0,54 | 4,41 | -0,12 | 9,02E-01 |
| AT5G61580 | 0,26 | 1,29 | 4,39 | 0,29 | 7,69E-01 |
| AT5G61950 | 0,32 | 1,79 | 4,38 | 0,41 | 6,83E-01 |
| AT5G61970 | 0,49 | 0,52 | 3,44 | 0,15 | 8,79E-01 |
| AT5G61980 | 0,55 | 2,54 | 4,34 | 0,58 | 5,59E-01 |

| | | | | | |
|---|---|---|---|---|---|
| AT5G62030 | 0,41 | 1,24 | 4,18 | 0,30 | 7,67E-01 |
| AT5G62270 | 0,20 | 0,00 | 4,41 | 0,00 | 1,00E+00 |
| AT5G62290 | 0,13 | 0,00 | 4,41 | 0,00 | 1,00E+00 |
| AT5G62360 | 0,27 | 1,46 | 4,40 | 0,33 | 7,40E-01 |
| AT5G62460 | 0,44 | 1,97 | 4,20 | 0,47 | 6,39E-01 |
| AT5G62500 | 0,05 | 0,00 | 4,41 | 0,00 | 1,00E+00 |
| AT5G62520 | 0,28 | -0,46 | 4,40 | -0,10 | 9,17E-01 |
| AT5G62560 | 0,37 | 0,42 | 3,57 | 0,12 | 9,06E-01 |
| AT5G62570 | 0,48 | 1,28 | 2,77 | 0,46 | 6,43E-01 |
| AT5G62575 | 0,21 | 0,00 | 4,41 | 0,00 | 1,00E+00 |
| AT5G62620 | 0,35 | 1,60 | 4,39 | 0,36 | 7,16E-01 |
| AT5G62710 | 0,23 | 1,27 | 4,39 | 0,29 | 7,72E-01 |
| AT5G62720 | 0,31 | 0,00 | 4,41 | 0,00 | 1,00E+00 |
| AT5G62740 | 0,43 | -0,98 | 4,13 | -0,24 | 8,12E-01 |
| AT5G62770 | 0,31 | 1,25 | 4,39 | 0,29 | 7,75E-01 |
| AT5G62890 | 1,11 | 0,54 | 2,24 | 0,24 | 8,08E-01 |
| AT5G62910 | 0,36 | 0,39 | 4,40 | 0,09 | 9,30E-01 |
| AT5G62970 | 0,35 | -0,44 | 3,61 | -0,12 | 9,03E-01 |
| AT5G63010 | 0,41 | 1,38 | 3,40 | 0,41 | 6,85E-01 |
| AT5G09120 | 1,28 | -2,92 | 1,96 | -1,48 | 1,38E-01 |
| AT5G63050 | 0,55 | 1,78 | 4,03 | 0,44 | 6,58E-01 |
| AT5G63130 | 0,64 | -0,23 | 3,05 | -0,07 | 9,41E-01 |
| AT5G63200 | 0,75 | 2,62 | 3,51 | 0,75 | 4,55E-01 |
| AT5G63260 | 0,08 | 0,00 | 4,41 | 0,00 | 1,00E+00 |
| AT5G63280 | 0,12 | 0,00 | 4,41 | 0,00 | 1,00E+00 |
| AT5G09150 | 0,24 | 1,96 | 4,39 | 0,45 | 6,55E-01 |
| AT5G63300 | 0,47 | 1,99 | 4,38 | 0,45 | 6,50E-01 |
| AT5G63340 | 0,26 | 0,00 | 4,41 | 0,00 | 1,00E+00 |
| AT5G63420 | 0,69 | 0,70 | 3,03 | 0,23 | 8,17E-01 |
| AT5G63480 | 0,12 | 0,00 | 4,41 | 0,00 | 1,00E+00 |
| AT5G63510 | 0,37 | 1,36 | 4,35 | 0,31 | 7,54E-01 |
| AT5G63640 | 0,36 | 0,40 | 4,38 | 0,09 | 9,28E-01 |
| AT5G63680 | 0,63 | 1,99 | 3,22 | 0,62 | 5,35E-01 |
| AT5G63770 | 0,64 | 0,03 | 3,05 | 0,01 | 9,91E-01 |
| AT5G63780 | 0,69 | 1,17 | 2,31 | 0,51 | 6,13E-01 |
| AT5G63810 | 0,68 | 0,24 | 3,01 | 0,08 | 9,37E-01 |
| AT5G63940 | 0,43 | 1,38 | 3,39 | 0,41 | 6,84E-01 |
| AT5G63960 | 0,45 | 2,98 | 4,32 | 0,69 | 4,90E-01 |
| AT5G63980 | 0,08 | 0,00 | 4,41 | 0,00 | 1,00E+00 |
| AT5G64030 | 1,58 | 0,17 | 1,35 | 0,12 | 9,01E-01 |
| AT5G64090 | 0,72 | 2,13 | 2,53 | 0,84 | 4,00E-01 |
| AT5G64160 | 0,20 | -0,44 | 4,39 | -0,10 | 9,19E-01 |
| AT5G64190 | 0,08 | 0,00 | 4,41 | 0,00 | 1,00E+00 |
| AT5G64240 | 0,28 | 0,00 | 4,41 | 0,00 | 1,00E+00 |
| AT5G64300 | 1,43 | -0,66 | 1,82 | -0,36 | 7,18E-01 |
| AT5G64310 | 0,80 | 1,27 | 2,86 | 0,45 | 6,56E-01 |
| AT5G64330 | 1,11 | 0,07 | 1,84 | 0,04 | 9,72E-01 |
| AT5G64370 | 0,51 | 1,32 | 3,99 | 0,33 | 7,41E-01 |
| AT5G64380 | 0,10 | 0,00 | 4,41 | 0,00 | 1,00E+00 |
| AT5G64400 | 0,52 | -1,69 | 4,18 | -0,40 | 6,86E-01 |
| AT5G64440 | 0,42 | -0,55 | 3,43 | -0,16 | 8,73E-01 |
| AT5G64500 | 0,82 | 1,54 | 2,08 | 0,74 | 4,59E-01 |
| AT5G64505 | 0,26 | 1,38 | 4,41 | 0,31 | 7,54E-01 |
| AT5G64572 | 1,22 | 3,66 | 1,90 | 1,93 | 5,41E-02 |
| AT5G64690 | 0,11 | 0,00 | 4,41 | 0,00 | 1,00E+00 |
| AT5G64750 | 0,12 | 0,00 | 4,41 | 0,00 | 1,00E+00 |

| | | | | | |
|---|---|---|---|---|---|
| AT5G64813 | 0,60 | 2,50 | 2,62 | 0,95 | 3,41E-01 |
| AT5G09285 | 0,45 | 2,11 | 4,03 | 0,52 | 6,01E-01 |
| AT5G64950 | 0,48 | 0,00 | 4,41 | 0,00 | 1,00E+00 |
| AT5G64960 | 0,59 | 0,48 | 3,22 | 0,15 | 8,83E-01 |
| AT5G65010 | 1,00 | 0,41 | 1,66 | 0,25 | 8,03E-01 |
| AT5G65015 | 0,39 | 0,00 | 4,41 | 0,00 | 1,00E+00 |
| AT5G65210 | 0,26 | 1,38 | 4,41 | 0,31 | 7,54E-01 |
| AT5G65220 | 1,17 | -2,26 | 2,15 | -1,05 | 2,92E-01 |
| AT5G65230 | 0,08 | 0,00 | 4,41 | 0,00 | 1,00E+00 |
| AT5G65270 | 0,10 | 0,00 | 4,41 | 0,00 | 1,00E+00 |
| AT5G65290 | 0,47 | 2,43 | 4,35 | 0,56 | 5,76E-01 |
| AT5G65400 | 0,17 | -0,57 | 4,40 | -0,13 | 8,98E-01 |
| AT5G65440 | 0,58 | 2,43 | 3,05 | 0,80 | 4,24E-01 |
| AT5G65470 | 0,30 | 0,00 | 4,41 | 0,00 | 1,00E+00 |
| AT5G65510 | 0,12 | 0,00 | 4,41 | 0,00 | 1,00E+00 |
| AT5G65570 | 0,25 | 1,30 | 4,40 | 0,30 | 7,67E-01 |
| AT5G65620 | 1,40 | 2,99 | 1,81 | 1,65 | 9,93E-02 |
| AT5G65640 | 0,08 | 0,00 | 4,41 | 0,00 | 1,00E+00 |
| AT5G65670 | 0,91 | -0,53 | 2,13 | -0,25 | 8,03E-01 |
| AT5G65710 | 0,77 | 1,38 | 2,34 | 0,59 | 5,55E-01 |
| AT5G65720 | 1,65 | -0,75 | 1,56 | -0,48 | 6,31E-01 |
| AT5G65760 | 0,55 | 1,37 | 2,77 | 0,49 | 6,21E-01 |
| AT5G65770 | 1,30 | 0,70 | 1,90 | 0,37 | 7,12E-01 |
| AT5G65780 | 1,41 | 0,41 | 1,82 | 0,23 | 8,21E-01 |
| AT5G65890 | 0,05 | 0,00 | 4,41 | 0,00 | 1,00E+00 |
| AT5G65960 | 1,37 | 1,15 | 2,10 | 0,55 | 5,84E-01 |
| AT5G65990 | 0,70 | -1,25 | 2,15 | -0,58 | 5,61E-01 |
| AT5G66052 | 0,53 | 2,99 | 4,00 | 0,75 | 4,54E-01 |
| AT5G66070 | 0,10 | 0,00 | 4,41 | 0,00 | 1,00E+00 |
| AT5G66100 | 0,24 | 0,00 | 4,41 | 0,00 | 1,00E+00 |
| AT5G66110 | 0,30 | 1,38 | 4,41 | 0,31 | 7,54E-01 |
| AT5G66160 | 0,31 | 1,42 | 4,40 | 0,32 | 7,47E-01 |
| AT5G66170 | 0,33 | 1,85 | 4,39 | 0,42 | 6,73E-01 |
| AT5G66240 | 1,02 | 2,96 | 2,35 | 1,26 | 2,08E-01 |
| AT5G66360 | 0,60 | 1,97 | 3,85 | 0,51 | 6,09E-01 |
| AT5G66450 | 0,53 | 2,17 | 3,21 | 0,68 | 4,99E-01 |
| AT5G66560 | 1,38 | 2,81 | 1,79 | 1,57 | 1,17E-01 |
| AT5G66562 | 1,20 | 0,48 | 1,80 | 0,26 | 7,92E-01 |
| AT5G66564 | 1,37 | -0,98 | 1,82 | -0,54 | 5,92E-01 |
| AT5G66568 | 0,20 | 0,00 | 4,41 | 0,00 | 1,00E+00 |
| AT5G66760 | 1,08 | 1,50 | 1,81 | 0,83 | 4,08E-01 |
| AT5G66770 | 0,43 | -0,54 | 4,41 | -0,12 | 9,02E-01 |
| AT5G66790 | 0,74 | 0,46 | 2,11 | 0,22 | 8,28E-01 |
| AT5G66810 | 0,67 | 2,91 | 2,51 | 1,16 | 2,47E-01 |
| AT5G66930 | 0,12 | 0,00 | 4,41 | 0,00 | 1,00E+00 |
| AT5G66950 | 1,10 | -1,57 | 1,82 | -0,86 | 3,88E-01 |
| AT5G67070 | 0,51 | 0,51 | 3,97 | 0,13 | 8,99E-01 |
| AT5G67080 | 0,30 | 1,39 | 4,41 | 0,32 | 7,52E-01 |
| AT5G67170 | 0,08 | 0,00 | 4,41 | 0,00 | 1,00E+00 |
| AT5G67260 | 0,65 | 2,25 | 3,98 | 0,56 | 5,72E-01 |
| AT5G67320 | 0,96 | -0,42 | 2,73 | -0,15 | 8,77E-01 |
| AT5G67330 | 1,26 | 2,10 | 1,90 | 1,11 | 2,67E-01 |
| AT5G67340 | 0,24 | 0,00 | 4,41 | 0,00 | 1,00E+00 |
| AT5G67385 | 0,80 | 0,30 | 2,41 | 0,13 | 9,00E-01 |
| AT5G67455 | 0,10 | 0,00 | 4,41 | 0,00 | 1,00E+00 |
| AT5G67470 | 1,09 | 2,89 | 2,13 | 1,35 | 1,76E-01 |

| | | | | | |
|---|---|---|---|---|---|
| AT5G67488 | 1,20 | 0,34 | 2,01 | 0,17 | 8,68E-01 |
| AT5G67500 | 0,21 | 0,00 | 4,41 | 0,00 | 1,00E+00 |
| AT5G67530 | 0,08 | 0,00 | 4,41 | 0,00 | 1,00E+00 |
| AT5G67560 | 0,70 | 0,00 | 3,75 | 0,00 | 1,00E+00 |
| AT5G67570 | 0,48 | 2,49 | 4,15 | 0,60 | 5,48E-01 |
| AT5G67590 | 0,32 | 0,37 | 4,40 | 0,08 | 9,33E-01 |
| AT1G01020 | 0,15 | 0,00 | 4,41 | 0,00 | 1,00E+00 |
| AT1G03993 | 0,11 | 0,00 | 4,41 | 0,00 | 1,00E+00 |
| AT1G01050 | 0,66 | 3,29 | 2,90 | 1,14 | 2,56E-01 |
| AT1G01060 | 0,69 | 2,90 | 2,46 | 1,18 | 2,39E-01 |
| AT1G01090 | 1,61 | 1,58 | 1,49 | 1,06 | 2,87E-01 |
| AT1G01100 | 1,47 | 1,13 | 1,48 | 0,77 | 4,42E-01 |
| AT1G01120 | 1,43 | -0,94 | 1,76 | -0,53 | 5,95E-01 |
| AT1G01130 | 0,29 | 1,23 | 4,38 | 0,28 | 7,78E-01 |
| AT1G01140 | 0,54 | 1,32 | 2,70 | 0,49 | 6,23E-01 |
| AT1G01170 | 0,58 | 0,00 | 3,16 | 0,00 | 1,00E+00 |
| AT1G01440 | 0,35 | 1,38 | 4,41 | 0,31 | 7,54E-01 |
| AT1G01450 | 0,59 | 0,00 | 4,40 | 0,00 | 1,00E+00 |
| AT1G01470 | 1,29 | -0,55 | 1,67 | -0,33 | 7,42E-01 |
| AT1G01490 | 0,15 | -0,43 | 4,39 | -0,10 | 9,23E-01 |
| AT1G01530 | 0,08 | 0,00 | 4,41 | 0,00 | 1,00E+00 |
| AT1G01610 | 0,73 | -2,78 | 2,44 | -1,14 | 2,55E-01 |
| AT1G01620 | 1,23 | -0,98 | 2,05 | -0,48 | 6,31E-01 |
| AT1G01650 | 0,59 | 0,00 | 3,04 | 0,00 | 1,00E+00 |
| AT1G04027 | 0,46 | 0,00 | 3,25 | 0,00 | 1,00E+00 |
| AT1G01680 | 0,72 | 2,10 | 2,58 | 0,81 | 4,16E-01 |
| AT1G01725 | 1,09 | -1,74 | 2,10 | -0,83 | 4,08E-01 |
| AT1G04063 | 0,34 | -0,91 | 4,38 | -0,21 | 8,35E-01 |
| AT1G01820 | 0,86 | 1,13 | 2,14 | 0,53 | 5,96E-01 |
| AT1G01830 | 1,22 | -0,08 | 2,18 | -0,04 | 9,70E-01 |
| AT1G01880 | 1,08 | -3,41 | 2,25 | -1,52 | 1,29E-01 |
| AT1G01890 | 1,04 | -3,41 | 2,60 | -1,31 | 1,90E-01 |
| AT1G01930 | 0,16 | 0,00 | 4,41 | 0,00 | 1,00E+00 |
| AT1G01990 | 0,21 | 0,00 | 4,41 | 0,00 | 1,00E+00 |
| AT1G02020 | 0,45 | 2,57 | 4,08 | 0,63 | 5,29E-01 |
| AT1G02090 | 0,84 | 2,03 | 2,87 | 0,71 | 4,80E-01 |
| AT1G02140 | 0,21 | -0,50 | 4,40 | -0,11 | 9,09E-01 |
| AT1G02260 | 0,54 | 2,88 | 3,19 | 0,90 | 3,66E-01 |
| AT1G02280 | 0,61 | -0,60 | 3,13 | -0,19 | 8,49E-01 |
| AT1G02310 | 0,21 | 1,38 | 4,41 | 0,31 | 7,54E-01 |
| AT1G04117 | 0,17 | 0,00 | 4,41 | 0,00 | 1,00E+00 |
| AT1G02430 | 0,17 | 0,00 | 4,41 | 0,00 | 1,00E+00 |
| AT1G02450 | 0,20 | -0,44 | 4,39 | -0,10 | 9,20E-01 |
| AT1G02610 | 0,33 | 0,82 | 4,35 | 0,19 | 8,51E-01 |
| AT1G02660 | 0,36 | 2,48 | 4,36 | 0,57 | 5,70E-01 |
| AT1G02770 | 0,41 | 2,70 | 4,34 | 0,62 | 5,33E-01 |
| AT1G02870 | 0,35 | 1,03 | 4,36 | 0,24 | 8,13E-01 |
| AT1G02910 | 0,70 | 2,27 | 2,44 | 0,93 | 3,53E-01 |
| AT1G02920 | 0,41 | -0,45 | 3,43 | -0,13 | 8,96E-01 |
| AT1G02930 | 0,49 | 2,64 | 3,30 | 0,80 | 4,23E-01 |
| AT1G02960 | 0,08 | 0,00 | 4,41 | 0,00 | 1,00E+00 |
| AT1G03000 | 0,70 | 0,45 | 2,78 | 0,16 | 8,72E-01 |
| AT1G03040 | 0,62 | 0,57 | 2,73 | 0,21 | 8,34E-01 |
| AT1G03140 | 0,90 | 1,35 | 2,02 | 0,67 | 5,05E-01 |
| AT1G03240 | 1,51 | -1,39 | 2,16 | -0,64 | 5,20E-01 |
| AT1G03300 | 0,12 | 0,00 | 4,41 | 0,00 | 1,00E+00 |

| | | | | | |
|---|---|---|---|---|---|
| AT1G03400 | 0,49 | 2,41 | 2,79 | 0,87 | 3,86E-01 |
| AT1G03445 | 0,28 | 0,00 | 4,41 | 0,00 | 1,00E+00 |
| AT1G03475 | 0,69 | -0,72 | 2,27 | -0,32 | 7,51E-01 |
| AT1G03520 | 0,48 | 1,92 | 4,06 | 0,47 | 6,36E-01 |
| AT1G03530 | 0,41 | 2,20 | 4,37 | 0,50 | 6,16E-01 |
| AT1G03550 | 0,70 | 0,80 | 2,29 | 0,35 | 7,25E-01 |
| AT1G03590 | 0,41 | 2,14 | 4,22 | 0,51 | 6,12E-01 |
| AT1G03660 | 0,52 | 2,17 | 2,75 | 0,79 | 4,31E-01 |
| AT1G03730 | 1,35 | -3,11 | 2,50 | -1,24 | 2,13E-01 |
| AT1G03746 | 0,46 | -0,23 | 4,08 | -0,06 | 9,56E-01 |
| AT1G03760 | 0,08 | 0,00 | 4,41 | 0,00 | 1,00E+00 |
| AT1G03830 | 0,65 | 0,78 | 2,53 | 0,31 | 7,59E-01 |
| AT1G03870 | 0,58 | 0,02 | 3,17 | 0,01 | 9,95E-01 |
| AT1G03940 | 1,20 | -1,58 | 1,87 | -0,85 | 3,96E-01 |
| AT1G04233 | 0,08 | 0,00 | 4,41 | 0,00 | 1,00E+00 |
| AT1G03980 | 0,11 | 0,00 | 4,41 | 0,00 | 1,00E+00 |
| AT1G04000 | 0,55 | -0,39 | 3,17 | -0,12 | 9,02E-01 |
| AT1G04010 | 0,05 | 0,00 | 4,41 | 0,00 | 1,00E+00 |
| AT1G04050 | 0,08 | 0,00 | 4,41 | 0,00 | 1,00E+00 |
| AT1G04070 | 0,16 | 0,00 | 4,41 | 0,00 | 1,00E+00 |
| AT1G04110 | 0,35 | 1,58 | 4,36 | 0,36 | 7,18E-01 |
| AT1G04120 | 1,63 | 2,85 | 1,83 | 1,56 | 1,19E-01 |
| AT1G04140 | 1,54 | 3,82 | 1,75 | 2,19 | 2,86E-02 |
| AT1G04190 | 0,13 | -0,43 | 4,39 | -0,10 | 9,23E-01 |
| AT1G04220 | 0,35 | 1,32 | 4,40 | 0,30 | 7,63E-01 |
| AT1G04240 | 0,91 | -2,08 | 2,25 | -0,93 | 3,54E-01 |
| AT1G04260 | 0,47 | 0,38 | 2,82 | 0,13 | 8,94E-01 |
| AT1G04270 | 1,04 | -1,12 | 2,22 | -0,50 | 6,15E-01 |
| AT1G04280 | 0,49 | -1,77 | 3,33 | -0,53 | 5,96E-01 |
| AT1G04290 | 0,43 | 1,33 | 2,87 | 0,47 | 6,42E-01 |
| AT1G04300 | 0,83 | 3,43 | 3,01 | 1,14 | 2,54E-01 |
| AT1G04380 | 0,20 | 1,33 | 4,40 | 0,30 | 7,63E-01 |
| AT1G04390 | 0,19 | 0,00 | 4,41 | 0,00 | 1,00E+00 |
| AT1G04400 | 0,89 | 0,89 | 2,20 | 0,41 | 6,84E-01 |
| AT1G04263 | 1,35 | -1,93 | 2,37 | -0,81 | 4,17E-01 |
| AT1G04530 | 0,42 | 1,30 | 3,36 | 0,39 | 6,98E-01 |
| AT1G04570 | 0,11 | 0,00 | 4,41 | 0,00 | 1,00E+00 |
| AT1G04630 | 0,66 | 2,13 | 3,77 | 0,57 | 5,72E-01 |
| AT1G04680 | 0,74 | -2,37 | 3,10 | -0,76 | 4,45E-01 |
| AT1G04730 | 0,74 | 1,98 | 2,45 | 0,81 | 4,18E-01 |
| AT1G04750 | 0,67 | -0,22 | 2,25 | -0,10 | 9,23E-01 |
| AT1G04770 | 0,89 | 1,00 | 2,33 | 0,43 | 6,69E-01 |
| AT1G04780 | 0,86 | 0,06 | 2,15 | 0,03 | 9,79E-01 |
| AT1G04840 | 0,63 | 2,29 | 3,75 | 0,61 | 5,42E-01 |
| AT1G04850 | 0,86 | 2,12 | 2,82 | 0,75 | 4,53E-01 |
| AT1G04860 | 0,57 | 2,63 | 3,97 | 0,66 | 5,07E-01 |
| AT1G04870 | 0,84 | 0,07 | 3,00 | 0,02 | 9,83E-01 |
| AT1G04880 | 0,21 | -0,74 | 4,37 | -0,17 | 8,65E-01 |
| AT1G04910 | 0,78 | 1,53 | 2,91 | 0,53 | 5,98E-01 |
| AT1G04950 | 0,50 | 2,60 | 3,26 | 0,80 | 4,25E-01 |
| AT1G04960 | 0,36 | 1,14 | 4,37 | 0,26 | 7,94E-01 |
| AT1G04980 | 0,36 | 2,37 | 4,36 | 0,54 | 5,86E-01 |
| AT1G04985 | 0,26 | 1,36 | 4,40 | 0,31 | 7,58E-01 |
| AT1G04990 | 0,84 | 0,61 | 2,45 | 0,25 | 8,04E-01 |
| AT1G05120 | 0,53 | 2,11 | 4,38 | 0,48 | 6,29E-01 |
| AT1G05150 | 0,85 | 2,06 | 2,14 | 0,96 | 3,36E-01 |

| | | | | | |
|---|---|---|---|---|---|
| AT1G05160 | 0,59 | 0,00 | 4,40 | 0,00 | 1,00E+00 |
| AT1G05170 | 0,47 | 0,40 | 4,35 | 0,09 | 9,26E-01 |
| AT1G05190 | 0,78 | 1,24 | 2,48 | 0,50 | 6,18E-01 |
| AT1G05205 | 0,08 | 0,00 | 4,41 | 0,00 | 1,00E+00 |
| AT1G05230 | 0,45 | 2,57 | 4,08 | 0,63 | 5,29E-01 |
| AT1G05300 | 0,18 | 0,00 | 4,41 | 0,00 | 1,00E+00 |
| AT1G05350 | 0,28 | 1,34 | 4,40 | 0,30 | 7,61E-01 |
| AT1G05520 | 0,61 | 2,62 | 3,08 | 0,85 | 3,94E-01 |
| AT1G05575 | 0,16 | 0,00 | 4,41 | 0,00 | 1,00E+00 |
| AT1G05640 | 0,20 | 1,40 | 4,41 | 0,32 | 7,51E-01 |
| AT1G05670 | 0,15 | 0,00 | 4,41 | 0,00 | 1,00E+00 |
| AT1G05720 | 0,13 | -0,43 | 4,39 | -0,10 | 9,23E-01 |
| AT1G05783 | 0,96 | -0,26 | 2,06 | -0,12 | 9,01E-01 |
| AT1G05840 | 0,75 | 2,90 | 2,22 | 1,31 | 1,92E-01 |
| AT1G05870 | 0,42 | 1,78 | 4,21 | 0,42 | 6,72E-01 |
| AT1G05950 | 0,12 | 0,00 | 4,41 | 0,00 | 1,00E+00 |
| AT1G05970 | 0,08 | 0,00 | 4,41 | 0,00 | 1,00E+00 |
| AT1G06000 | 0,95 | 0,71 | 1,99 | 0,35 | 7,23E-01 |
| AT1G06130 | 0,62 | 2,06 | 3,01 | 0,68 | 4,94E-01 |
| AT1G06150 | 1,38 | 3,48 | 1,93 | 1,81 | 7,10E-02 |
| AT1G06250 | 0,72 | 3,54 | 4,30 | 0,82 | 4,11E-01 |
| AT1G06265 | 0,08 | 0,00 | 4,41 | 0,00 | 1,00E+00 |
| AT1G06360 | 0,33 | -1,69 | 4,35 | -0,39 | 6,98E-01 |
| AT1G06400 | 0,21 | 0,00 | 4,41 | 0,00 | 1,00E+00 |
| AT1G06440 | 0,58 | 2,20 | 2,75 | 0,80 | 4,23E-01 |
| AT1G06540 | 1,04 | 2,65 | 2,02 | 1,31 | 1,89E-01 |
| AT1G06570 | 0,97 | 0,34 | 2,50 | 0,14 | 8,93E-01 |
| AT1G06580 | 1,59 | -2,62 | 1,65 | -1,59 | 1,11E-01 |
| AT1G04487 | 0,39 | 1,38 | 4,26 | 0,32 | 7,46E-01 |
| AT1G06660 | 0,50 | 2,75 | 3,97 | 0,69 | 4,89E-01 |
| AT1G06690 | 0,36 | 0,40 | 4,38 | 0,09 | 9,28E-01 |
| AT1G06700 | 1,69 | 1,48 | 1,65 | 0,90 | 3,70E-01 |
| AT1G06710 | 0,26 | 1,38 | 4,41 | 0,31 | 7,54E-01 |
| AT1G06720 | 0,55 | 1,00 | 3,97 | 0,25 | 8,01E-01 |
| AT1G06760 | 1,45 | 1,10 | 1,49 | 0,74 | 4,60E-01 |
| AT1G06810 | 0,08 | 0,00 | 4,41 | 0,00 | 1,00E+00 |
| AT1G06840 | 0,17 | -0,46 | 4,40 | -0,10 | 9,17E-01 |
| AT1G06890 | 0,40 | -1,31 | 2,96 | -0,44 | 6,58E-01 |
| AT1G06900 | 1,35 | 0,40 | 1,70 | 0,24 | 8,14E-01 |
| AT1G07020 | 0,55 | 1,82 | 4,34 | 0,42 | 6,74E-01 |
| AT1G07050 | 0,42 | -1,84 | 4,11 | -0,45 | 6,54E-01 |
| AT1G07090 | 0,43 | 1,94 | 4,35 | 0,45 | 6,55E-01 |
| AT1G07110 | 1,24 | 3,88 | 2,31 | 1,68 | 9,29E-02 |
| AT1G07128 | 0,28 | 0,00 | 4,41 | 0,00 | 1,00E+00 |
| AT1G07220 | 0,25 | -1,80 | 4,34 | -0,41 | 6,78E-01 |
| AT1G07260 | 0,48 | 2,01 | 4,37 | 0,46 | 6,45E-01 |
| AT1G04573 | 0,66 | 3,35 | 4,31 | 0,78 | 4,37E-01 |
| AT1G07310 | 0,69 | -0,14 | 2,61 | -0,05 | 9,57E-01 |
| AT1G07350 | 0,45 | 2,23 | 3,31 | 0,67 | 5,01E-01 |
| AT1G07360 | 0,57 | -1,99 | 3,18 | -0,63 | 5,32E-01 |
| AT1G07430 | 0,32 | 0,48 | 4,40 | 0,11 | 9,13E-01 |
| AT1G07440 | 0,75 | 2,11 | 2,63 | 0,80 | 4,23E-01 |
| AT1G07480 | 0,80 | 0,99 | 2,34 | 0,42 | 6,71E-01 |
| AT1G07500 | 0,08 | 0,00 | 4,41 | 0,00 | 1,00E+00 |
| AT1G07520 | 1,20 | 0,89 | 2,13 | 0,42 | 6,76E-01 |
| AT1G07530 | 1,00 | 1,97 | 1,84 | 1,07 | 2,86E-01 |

| | | | | | |
|---|---|---|---|---|---|
| AT1G07640 | 0,24 | -0,46 | 4,40 | -0,10 | 9,17E-01 |
| AT1G07720 | 1,26 | -2,46 | 1,83 | -1,35 | 1,78E-01 |
| AT1G07725 | 1,03 | 3,80 | 2,48 | 1,53 | 1,26E-01 |
| AT1G07740 | 0,29 | 1,38 | 4,41 | 0,31 | 7,54E-01 |
| AT1G07730 | 0,29 | 1,38 | 4,41 | 0,31 | 7,54E-01 |
| AT1G07750 | 0,25 | -1,80 | 4,34 | -0,41 | 6,78E-01 |
| AT1G07770 | 0,87 | 0,53 | 1,88 | 0,28 | 7,77E-01 |
| AT1G07870 | 0,12 | 0,00 | 4,41 | 0,00 | 1,00E+00 |
| AT1G07970 | 0,62 | 2,49 | 3,66 | 0,68 | 4,96E-01 |
| AT1G08040 | 0,40 | 0,36 | 4,37 | 0,08 | 9,35E-01 |
| AT1G08050 | 0,33 | 1,27 | 4,39 | 0,29 | 7,72E-01 |
| AT1G08060 | 1,18 | 2,82 | 2,21 | 1,28 | 2,01E-01 |
| AT1G08125 | 0,32 | 1,87 | 4,39 | 0,43 | 6,70E-01 |
| AT1G08510 | 1,46 | -1,86 | 1,75 | -1,06 | 2,87E-01 |
| AT1G08580 | 0,24 | 1,96 | 4,39 | 0,45 | 6,55E-01 |
| AT1G08592 | 0,74 | 1,38 | 3,01 | 0,46 | 6,47E-01 |
| AT1G08610 | 0,28 | 1,34 | 4,40 | 0,30 | 7,61E-01 |
| AT1G08640 | 0,47 | 2,16 | 4,00 | 0,54 | 5,89E-01 |
| AT1G08645 | 0,64 | 0,22 | 3,69 | 0,06 | 9,53E-01 |
| AT1G08660 | 0,10 | 0,00 | 4,41 | 0,00 | 1,00E+00 |
| AT1G08700 | 1,22 | 0,52 | 2,31 | 0,22 | 8,23E-01 |
| AT1G08800 | 0,74 | 1,12 | 2,97 | 0,38 | 7,06E-01 |
| AT1G08820 | 0,89 | 2,76 | 2,69 | 1,02 | 3,06E-01 |
| AT1G08880 | 0,12 | 0,00 | 4,41 | 0,00 | 1,00E+00 |
| AT1G09020 | 0,78 | 1,08 | 2,33 | 0,46 | 6,45E-01 |
| AT1G09060 | 1,04 | 1,37 | 2,39 | 0,57 | 5,66E-01 |
| AT1G09080 | 0,40 | 1,38 | 4,36 | 0,32 | 7,51E-01 |
| AT1G09130 | 0,46 | -1,44 | 3,29 | -0,44 | 6,63E-01 |
| AT1G09160 | 0,40 | -1,40 | 3,48 | -0,40 | 6,87E-01 |
| AT1G09190 | 0,17 | 0,00 | 4,41 | 0,00 | 1,00E+00 |
| AT1G09195 | 0,70 | 2,54 | 2,95 | 0,86 | 3,89E-01 |
| AT1G09200 | 0,34 | 0,00 | 4,41 | 0,00 | 1,00E+00 |
| AT1G09230 | 0,23 | 1,38 | 4,41 | 0,31 | 7,54E-01 |
| AT1G09280 | 1,11 | -0,48 | 1,78 | -0,27 | 7,86E-01 |
| AT1G09420 | 0,88 | 0,00 | 3,46 | 0,00 | 1,00E+00 |
| AT1G09440 | 0,21 | -1,19 | 4,36 | -0,27 | 7,85E-01 |
| AT1G04707 | 0,36 | 1,85 | 4,39 | 0,42 | 6,73E-01 |
| AT1G09630 | 0,08 | 0,00 | 4,41 | 0,00 | 1,00E+00 |
| AT1G04717 | 0,54 | 2,30 | 3,85 | 0,60 | 5,49E-01 |
| AT1G09680 | 1,19 | 2,05 | 2,67 | 0,77 | 4,42E-01 |
| AT1G09730 | 0,83 | -0,54 | 2,84 | -0,19 | 8,48E-01 |
| AT1G09760 | 0,17 | 0,00 | 4,41 | 0,00 | 1,00E+00 |
| AT1G09810 | 0,42 | 0,42 | 2,91 | 0,14 | 8,86E-01 |
| AT1G09880 | 0,33 | 2,19 | 4,38 | 0,50 | 6,17E-01 |
| AT1G09910 | 0,78 | 3,23 | 2,36 | 1,37 | 1,70E-01 |
| AT1G10020 | 0,10 | 0,00 | 4,41 | 0,00 | 1,00E+00 |
| AT1G10090 | 0,57 | -0,47 | 2,69 | -0,17 | 8,63E-01 |
| AT1G10155 | 1,47 | 1,05 | 1,65 | 0,64 | 5,23E-01 |
| AT1G10180 | 0,61 | 2,10 | 2,56 | 0,82 | 4,12E-01 |
| AT1G10190 | 0,94 | -1,90 | 2,78 | -0,68 | 4,94E-01 |
| AT1G10200 | 0,57 | 1,80 | 3,90 | 0,46 | 6,45E-01 |
| AT1G10310 | 0,42 | 2,12 | 4,37 | 0,48 | 6,28E-01 |
| AT1G10320 | 0,47 | 0,00 | 4,05 | 0,00 | 1,00E+00 |
| AT1G10340 | 1,00 | 3,56 | 2,20 | 1,62 | 1,06E-01 |
| AT1G10350 | 0,52 | 1,38 | 3,96 | 0,35 | 7,27E-01 |
| AT1G10390 | 1,30 | 2,05 | 2,13 | 0,96 | 3,37E-01 |

| | | | | | |
|---|---|---|---|---|---|
| AT1G10400 | 0,08 | 0,00 | 4,41 | 0,00 | 1,00E+00 |
| AT1G10410 | 0,96 | 1,14 | 2,15 | 0,53 | 5,95E-01 |
| AT1G10430 | 0,87 | -0,07 | 2,30 | -0,03 | 9,76E-01 |
| AT1G10470 | 1,21 | 0,63 | 1,83 | 0,34 | 7,32E-01 |
| AT1G10500 | 1,39 | 0,35 | 2,42 | 0,14 | 8,87E-01 |
| AT1G10580 | 0,57 | -1,35 | 3,88 | -0,35 | 7,28E-01 |
| AT1G10586 | 1,03 | -1,27 | 2,25 | -0,56 | 5,73E-01 |
| AT1G10590 | 0,69 | 2,72 | 3,05 | 0,89 | 3,71E-01 |
| AT1G10630 | 0,65 | 1,95 | 2,61 | 0,75 | 4,55E-01 |
| AT1G10640 | 0,05 | 0,00 | 4,41 | 0,00 | 1,00E+00 |
| AT1G10680 | 0,08 | 0,00 | 4,41 | 0,00 | 1,00E+00 |
| AT1G10840 | 0,88 | 1,94 | 2,00 | 0,97 | 3,32E-01 |
| AT1G10870 | 1,11 | 2,38 | 1,95 | 1,22 | 2,23E-01 |
| AT1G10900 | 1,03 | 2,57 | 2,76 | 0,93 | 3,52E-01 |
| AT1G10920 | 0,82 | 2,31 | 4,36 | 0,53 | 5,96E-01 |
| AT1G10930 | 0,74 | 2,74 | 2,95 | 0,93 | 3,54E-01 |
| AT1G10940 | 0,48 | 2,43 | 3,34 | 0,73 | 4,68E-01 |
| AT1G04833 | 0,22 | 0,00 | 4,41 | 0,00 | 1,00E+00 |
| AT1G11100 | 0,45 | 1,34 | 3,46 | 0,39 | 6,98E-01 |
| AT1G11210 | 0,56 | -0,54 | 2,69 | -0,20 | 8,41E-01 |
| AT1G11240 | 0,05 | 0,00 | 4,41 | 0,00 | 1,00E+00 |
| AT1G11270 | 0,83 | 1,36 | 2,47 | 0,55 | 5,81E-01 |
| AT1G11280 | 0,45 | 1,20 | 2,82 | 0,43 | 6,70E-01 |
| AT1G11290 | 0,63 | 2,18 | 3,70 | 0,59 | 5,55E-01 |
| AT1G11310 | 1,46 | 2,23 | 1,75 | 1,28 | 2,01E-01 |
| AT1G11340 | 0,08 | 0,00 | 4,41 | 0,00 | 1,00E+00 |
| AT1G11350 | 0,52 | 1,81 | 3,31 | 0,55 | 5,85E-01 |
| AT1G11360 | 0,33 | 0,44 | 4,38 | 0,10 | 9,21E-01 |
| AT1G11480 | 0,70 | 1,55 | 2,29 | 0,68 | 4,97E-01 |
| AT1G11630 | 0,35 | 1,58 | 4,36 | 0,36 | 7,18E-01 |
| AT1G11820 | 0,25 | -0,52 | 4,40 | -0,12 | 9,06E-01 |
| AT1G12000 | 0,52 | 0,00 | 3,19 | 0,00 | 1,00E+00 |
| AT1G12020 | 0,63 | 0,53 | 2,35 | 0,22 | 8,22E-01 |
| AT1G12120 | 0,70 | 1,61 | 2,62 | 0,61 | 5,39E-01 |
| AT1G12211 | 0,63 | 0,19 | 3,20 | 0,06 | 9,52E-01 |
| AT1G12310 | 0,63 | 0,00 | 3,28 | 0,00 | 1,00E+00 |
| AT1G12380 | 0,49 | 0,00 | 4,15 | 0,00 | 1,00E+00 |
| AT1G12440 | 1,21 | 2,42 | 1,80 | 1,35 | 1,78E-01 |
| AT1G12500 | 0,25 | 0,00 | 4,41 | 0,00 | 1,00E+00 |
| AT1G12520 | 0,51 | 1,09 | 4,33 | 0,25 | 8,01E-01 |
| AT1G12590 | 0,78 | -1,07 | 1,94 | -0,55 | 5,82E-01 |
| AT1G12610 | 0,19 | 0,00 | 4,41 | 0,00 | 1,00E+00 |
| AT1G12615 | 1,59 | 0,00 | 2,32 | 0,00 | 1,00E+00 |
| AT1G12640 | 0,58 | 1,74 | 4,38 | 0,40 | 6,92E-01 |
| AT1G12680 | 1,43 | -1,44 | 1,96 | -0,73 | 4,63E-01 |
| AT1G12700 | 0,22 | 0,00 | 4,41 | 0,00 | 1,00E+00 |
| AT1G12780 | 0,71 | 0,48 | 2,46 | 0,20 | 8,45E-01 |
| AT1G12800 | 0,97 | 1,56 | 2,39 | 0,65 | 5,15E-01 |
| AT1G12910 | 1,07 | 1,22 | 2,16 | 0,56 | 5,73E-01 |
| AT1G13020 | 0,59 | 1,68 | 2,61 | 0,64 | 5,21E-01 |
| AT1G13120 | 0,23 | 0,00 | 4,41 | 0,00 | 1,00E+00 |
| AT1G13170 | 0,46 | 2,16 | 3,26 | 0,66 | 5,08E-01 |
| AT1G13200 | 0,66 | 3,20 | 3,60 | 0,89 | 3,75E-01 |
| AT1G13245 | 0,44 | -0,76 | 4,08 | -0,19 | 8,52E-01 |
| AT1G13310 | 0,35 | 1,58 | 4,36 | 0,36 | 7,18E-01 |
| AT1G13340 | 0,95 | 2,36 | 2,33 | 1,02 | 3,10E-01 |

| | | | | | |
|---|---|---|---|---|---|
| AT1G13390 | 0,17 | 0,00 | 4,41 | 0,00 | 1,00E+00 |
| AT1G13450 | 0,76 | 2,31 | 2,31 | 1,00 | 3,18E-01 |
| AT1G13520 | 0,54 | 1,87 | 4,38 | 0,43 | 6,69E-01 |
| AT1G13560 | 0,43 | 2,13 | 4,38 | 0,49 | 6,26E-01 |
| AT1G13825 | 0,76 | 0,00 | 3,67 | 0,00 | 1,00E+00 |
| AT1G13860 | 0,10 | 0,00 | 4,41 | 0,00 | 1,00E+00 |
| AT1G13870 | 0,12 | 0,00 | 4,41 | 0,00 | 1,00E+00 |
| AT1G13900 | 0,21 | 0,00 | 4,41 | 0,00 | 1,00E+00 |
| AT1G13940 | 0,86 | 2,06 | 2,57 | 0,80 | 4,22E-01 |
| AT1G14010 | 0,36 | 1,38 | 4,41 | 0,31 | 7,54E-01 |
| AT1G14030 | 0,62 | -0,08 | 3,15 | -0,03 | 9,79E-01 |
| AT1G14290 | 0,26 | 1,38 | 4,41 | 0,31 | 7,54E-01 |
| AT1G14300 | 0,22 | -0,74 | 4,37 | -0,17 | 8,66E-01 |
| AT1G14330 | 0,59 | 0,19 | 2,73 | 0,07 | 9,44E-01 |
| AT1G14360 | 0,18 | 1,38 | 4,41 | 0,31 | 7,54E-01 |
| AT1G14400 | 0,66 | -0,12 | 3,17 | -0,04 | 9,70E-01 |
| AT1G14450 | 0,11 | 0,00 | 4,41 | 0,00 | 1,00E+00 |
| AT1G14455 | 0,14 | -0,78 | 4,37 | -0,18 | 8,58E-01 |
| AT1G14518 | 0,84 | 1,45 | 2,51 | 0,58 | 5,64E-01 |
| AT1G14590 | 0,42 | 2,32 | 4,20 | 0,55 | 5,80E-01 |
| AT1G14690 | 0,78 | 1,13 | 2,19 | 0,52 | 6,05E-01 |
| AT1G14710 | 0,51 | -0,41 | 3,92 | -0,10 | 9,17E-01 |
| AT1G14720 | 0,26 | 0,23 | 4,36 | 0,05 | 9,58E-01 |
| AT1G14790 | 1,42 | 1,89 | 2,37 | 0,80 | 4,24E-01 |
| AT1G14810 | 1,02 | 3,34 | 2,16 | 1,55 | 1,21E-01 |
| AT1G14830 | 0,90 | 1,82 | 2,89 | 0,63 | 5,29E-01 |
| AT1G14840 | 0,51 | -1,70 | 2,40 | -0,71 | 4,80E-01 |
| AT1G14850 | 1,26 | 3,64 | 1,85 | 1,97 | 4,91E-02 |
| AT1G14880 | 1,19 | 0,41 | 2,18 | 0,19 | 8,51E-01 |
| AT1G14900 | 0,38 | -0,51 | 4,34 | -0,12 | 9,07E-01 |
| AT1G14910 | 0,80 | -0,26 | 2,63 | -0,10 | 9,22E-01 |
| AT1G14980 | 0,57 | 1,27 | 3,08 | 0,41 | 6,80E-01 |
| AT1G05137 | 0,28 | -0,51 | 4,40 | -0,11 | 9,09E-01 |
| AT1G15020 | 0,98 | 2,56 | 2,65 | 0,96 | 3,35E-01 |
| AT1G15040 | 0,21 | 0,00 | 4,41 | 0,00 | 1,00E+00 |
| AT1G15100 | 1,26 | 1,98 | 2,56 | 0,77 | 4,40E-01 |
| AT1G15140 | 0,76 | 0,50 | 2,23 | 0,23 | 8,22E-01 |
| AT1G15175 | 0,27 | 1,26 | 4,39 | 0,29 | 7,74E-01 |
| AT1G15210 | 0,17 | 0,00 | 4,41 | 0,00 | 1,00E+00 |
| AT1G15220 | 0,11 | 0,00 | 4,41 | 0,00 | 1,00E+00 |
| AT1G15340 | 0,21 | -0,47 | 4,40 | -0,11 | 9,15E-01 |
| AT1G15350 | 1,25 | 2,89 | 2,16 | 1,34 | 1,80E-01 |
| AT1G15400 | 0,43 | 1,33 | 2,90 | 0,46 | 6,46E-01 |
| AT1G15440 | 0,87 | 1,67 | 2,13 | 0,78 | 4,33E-01 |
| AT1G15520 | 0,46 | 1,38 | 3,34 | 0,41 | 6,80E-01 |
| AT1G15530 | 1,02 | 2,25 | 2,25 | 1,00 | 3,16E-01 |
| AT1G15550 | 0,24 | 1,96 | 4,39 | 0,45 | 6,55E-01 |
| AT1G15700 | 0,40 | 2,10 | 3,40 | 0,62 | 5,37E-01 |
| AT1G05187 | 0,24 | 1,38 | 4,41 | 0,31 | 7,54E-01 |
| AT1G15730 | 0,44 | 1,36 | 4,18 | 0,33 | 7,44E-01 |
| AT1G15780 | 1,01 | -1,19 | 2,09 | -0,57 | 5,71E-01 |
| AT1G15790 | 1,05 | 3,37 | 2,83 | 1,19 | 2,35E-01 |
| AT1G05203 | 0,26 | 1,38 | 4,41 | 0,31 | 7,54E-01 |
| AT1G15880 | 0,20 | 1,40 | 4,41 | 0,32 | 7,51E-01 |
| AT1G15970 | 0,11 | 0,00 | 4,41 | 0,00 | 1,00E+00 |
| AT1G16010 | 0,99 | 0,43 | 2,46 | 0,17 | 8,62E-01 |

| | | | | | |
|---|---|---|---|---|---|
| AT1G16070 | 0,40 | 0,00 | 4,41 | 0,00 | 1,00E+00 |
| AT1G16210 | 0,60 | 1,82 | 3,02 | 0,60 | 5,46E-01 |
| AT1G16240 | 0,80 | -1,51 | 2,49 | -0,61 | 5,44E-01 |
| AT1G16260 | 1,63 | 3,16 | 1,78 | 1,77 | 7,60E-02 |
| AT1G16460 | 0,57 | 0,42 | 2,49 | 0,17 | 8,67E-01 |
| AT1G16570 | 0,28 | 0,00 | 4,41 | 0,00 | 1,00E+00 |
| AT1G16610 | 1,02 | 3,03 | 3,11 | 0,97 | 3,30E-01 |
| AT1G16750 | 0,49 | 2,13 | 3,18 | 0,67 | 5,02E-01 |
| AT1G16780 | 0,12 | 0,00 | 4,41 | 0,00 | 1,00E+00 |
| AT1G16800 | 1,44 | 2,78 | 2,42 | 1,15 | 2,49E-01 |
| AT1G16870 | 0,95 | 3,80 | 3,30 | 1,15 | 2,49E-01 |
| AT1G16890 | 1,30 | 0,33 | 1,69 | 0,20 | 8,44E-01 |
| AT1G16900 | 0,65 | 1,84 | 3,76 | 0,49 | 6,23E-01 |
| AT1G16920 | 0,22 | 0,00 | 4,41 | 0,00 | 1,00E+00 |
| AT1G16970 | 0,37 | 0,44 | 3,54 | 0,12 | 9,02E-01 |
| AT1G17080 | 0,60 | 0,89 | 3,28 | 0,27 | 7,87E-01 |
| AT1G17110 | 0,57 | -0,52 | 2,67 | -0,19 | 8,46E-01 |
| AT1G17140 | 0,95 | 1,96 | 3,37 | 0,58 | 5,61E-01 |
| AT1G17147 | 0,12 | 0,00 | 4,41 | 0,00 | 1,00E+00 |
| AT1G17232 | 0,51 | 0,00 | 3,21 | 0,00 | 1,00E+00 |
| AT1G17240 | 0,05 | 0,00 | 4,41 | 0,00 | 1,00E+00 |
| AT1G17250 | 0,99 | 1,38 | 2,74 | 0,50 | 6,14E-01 |
| AT1G17370 | 0,81 | -0,92 | 2,71 | -0,34 | 7,33E-01 |
| AT1G17380 | 0,50 | 1,30 | 2,76 | 0,47 | 6,37E-01 |
| AT1G05313 | 0,32 | 0,00 | 4,41 | 0,00 | 1,00E+00 |
| AT1G17440 | 0,46 | 1,38 | 3,42 | 0,40 | 6,87E-01 |
| AT1G17450 | 1,22 | 1,17 | 2,02 | 0,58 | 5,63E-01 |
| AT1G17590 | 0,46 | 0,30 | 3,33 | 0,09 | 9,28E-01 |
| AT1G17600 | 1,00 | 1,38 | 2,22 | 0,62 | 5,34E-01 |
| AT1G17665 | 0,20 | -0,44 | 4,39 | -0,10 | 9,21E-01 |
| AT1G17670 | 0,37 | 0,59 | 4,36 | 0,14 | 8,92E-01 |
| AT1G17690 | 0,25 | 1,30 | 4,40 | 0,30 | 7,67E-01 |
| AT1G17720 | 0,66 | 0,49 | 2,43 | 0,20 | 8,41E-01 |
| AT1G17744 | 0,16 | 0,00 | 4,41 | 0,00 | 1,00E+00 |
| AT1G17790 | 0,63 | 1,88 | 3,06 | 0,61 | 5,39E-01 |
| AT1G05333 | 0,71 | 3,14 | 3,51 | 0,90 | 3,71E-01 |
| AT1G17820 | 0,71 | 3,14 | 3,51 | 0,90 | 3,71E-01 |
| AT1G17850 | 0,36 | 2,44 | 4,36 | 0,56 | 5,76E-01 |
| AT1G05347 | 0,21 | -0,51 | 4,40 | -0,12 | 9,07E-01 |
| AT1G17870 | 0,39 | 1,80 | 4,31 | 0,42 | 6,76E-01 |
| AT1G17880 | 0,59 | 1,83 | 3,81 | 0,48 | 6,31E-01 |
| AT1G17890 | 0,42 | 1,92 | 4,39 | 0,44 | 6,62E-01 |
| AT1G17930 | 0,20 | 1,38 | 4,41 | 0,31 | 7,54E-01 |
| AT1G17980 | 1,06 | -1,71 | 2,29 | -0,75 | 4,56E-01 |
| AT1G17990 | 0,53 | 2,52 | 3,17 | 0,79 | 4,27E-01 |
| AT1G18050 | 0,60 | -1,29 | 2,61 | -0,50 | 6,19E-01 |
| AT1G18060 | 0,60 | 1,39 | 2,38 | 0,58 | 5,60E-01 |
| AT1G18070 | 0,60 | -0,50 | 2,59 | -0,19 | 8,47E-01 |
| AT1G18100 | 0,36 | 0,00 | 4,41 | 0,00 | 1,00E+00 |
| AT1G18150 | 1,19 | 0,35 | 2,05 | 0,17 | 8,66E-01 |
| AT1G18210 | 0,61 | -0,35 | 2,40 | -0,14 | 8,85E-01 |
| AT1G18330 | 0,22 | 1,33 | 4,40 | 0,30 | 7,63E-01 |
| AT1G18360 | 1,25 | 0,66 | 1,67 | 0,40 | 6,92E-01 |
| AT1G18382 | 0,44 | 1,38 | 2,88 | 0,48 | 6,32E-01 |
| AT1G18470 | 0,48 | -1,28 | 2,76 | -0,47 | 6,41E-01 |
| AT1G18540 | 1,15 | 0,53 | 2,18 | 0,24 | 8,09E-01 |

| | | | | | |
|---|---|---|---|---|---|
| AT1G18720 | 0,23 | 0,00 | 4,41 | 0,00 | 1,00E+00 |
| AT1G05433 | 0,53 | 0,00 | 3,19 | 0,00 | 1,00E+00 |
| AT1G18840 | 0,43 | 1,98 | 3,41 | 0,58 | 5,60E-01 |
| AT1G18890 | 0,28 | 1,39 | 4,41 | 0,32 | 7,52E-01 |
| AT1G18990 | 0,45 | 2,26 | 4,21 | 0,54 | 5,90E-01 |
| AT1G19000 | 0,86 | 0,13 | 1,85 | 0,07 | 9,43E-01 |
| AT1G19130 | 0,49 | -1,67 | 2,82 | -0,59 | 5,54E-01 |
| AT1G19140 | 0,44 | 2,35 | 4,36 | 0,54 | 5,90E-01 |
| AT1G19210 | 0,10 | 0,00 | 4,41 | 0,00 | 1,00E+00 |
| AT1G19260 | 0,97 | 0,00 | 2,74 | 0,00 | 1,00E+00 |
| AT1G19300 | 0,88 | -0,54 | 2,63 | -0,21 | 8,36E-01 |
| AT1G19360 | 0,50 | 2,01 | 4,02 | 0,50 | 6,17E-01 |
| AT1G19370 | 0,41 | 0,35 | 4,33 | 0,08 | 9,36E-01 |
| AT1G19400 | 1,69 | 0,55 | 1,70 | 0,33 | 7,44E-01 |
| AT1G19430 | 0,39 | 0,77 | 4,26 | 0,18 | 8,56E-01 |
| AT1G19450 | 0,33 | 0,50 | 4,37 | 0,11 | 9,10E-01 |
| AT1G19570 | 0,89 | -0,10 | 2,03 | -0,05 | 9,61E-01 |
| AT1G19640 | 0,14 | -0,78 | 4,37 | -0,18 | 8,58E-01 |
| AT1G19660 | 0,55 | 0,39 | 2,75 | 0,14 | 8,88E-01 |
| AT1G19670 | 1,59 | -0,82 | 2,01 | -0,41 | 6,81E-01 |
| AT1G19700 | 0,25 | -1,23 | 4,36 | -0,28 | 7,79E-01 |
| AT1G19715 | 1,55 | 3,41 | 2,14 | 1,59 | 1,11E-01 |
| AT1G19720 | 1,66 | 2,73 | 1,74 | 1,57 | 1,17E-01 |
| AT1G19780 | 0,50 | 2,20 | 4,07 | 0,54 | 5,89E-01 |
| AT1G19860 | 0,15 | 0,00 | 4,41 | 0,00 | 1,00E+00 |
| AT1G19870 | 0,80 | 0,17 | 2,60 | 0,07 | 9,48E-01 |
| AT1G19880 | 0,41 | 1,29 | 3,48 | 0,37 | 7,12E-01 |
| AT1G19920 | 1,00 | 3,66 | 2,25 | 1,63 | 1,04E-01 |
| AT1G20040 | 0,34 | -0,84 | 4,38 | -0,19 | 8,49E-01 |
| AT1G20070 | 0,59 | 0,43 | 2,57 | 0,17 | 8,67E-01 |
| AT1G20160 | 0,33 | 1,37 | 4,41 | 0,31 | 7,56E-01 |
| AT1G20190 | 0,64 | -2,75 | 3,21 | -0,86 | 3,91E-01 |
| AT1G20200 | 0,33 | -0,54 | 3,61 | -0,15 | 8,82E-01 |
| AT1G20320 | 0,05 | 0,00 | 4,41 | 0,00 | 1,00E+00 |
| AT1G20400 | 1,62 | 1,11 | 1,87 | 0,59 | 5,52E-01 |
| AT1G20430 | 0,89 | -0,13 | 2,89 | -0,05 | 9,63E-01 |
| AT1G20490 | 0,48 | 0,23 | 4,39 | 0,05 | 9,58E-01 |
| AT1G20560 | 0,25 | 0,00 | 4,41 | 0,00 | 1,00E+00 |
| AT1G20570 | 0,08 | 0,00 | 4,41 | 0,00 | 1,00E+00 |
| AT1G20650 | 0,41 | 1,82 | 4,38 | 0,42 | 6,77E-01 |
| AT1G20670 | 0,31 | 1,85 | 4,39 | 0,42 | 6,74E-01 |
| AT1G20770 | 0,16 | 0,00 | 4,41 | 0,00 | 1,00E+00 |
| AT1G20840 | 1,09 | -0,38 | 1,98 | -0,19 | 8,50E-01 |
| AT1G20880 | 0,25 | -1,09 | 4,36 | -0,25 | 8,03E-01 |
| AT1G20890 | 0,48 | 0,00 | 4,06 | 0,00 | 1,00E+00 |
| AT1G20950 | 0,69 | 1,24 | 2,56 | 0,49 | 6,28E-01 |
| AT1G21080 | 0,61 | 2,25 | 2,63 | 0,86 | 3,92E-01 |
| AT1G05627 | 0,05 | 0,00 | 4,41 | 0,00 | 1,00E+00 |
| AT1G21160 | 0,26 | 0,00 | 4,41 | 0,00 | 1,00E+00 |
| AT1G21190 | 0,45 | -0,68 | 4,17 | -0,16 | 8,71E-01 |
| AT1G21310 | 0,44 | 0,52 | 2,84 | 0,18 | 8,56E-01 |
| AT1G21380 | 1,01 | 2,91 | 2,31 | 1,26 | 2,07E-01 |
| AT1G21390 | 0,08 | 0,00 | 4,41 | 0,00 | 1,00E+00 |
| AT1G21440 | 1,10 | 1,30 | 2,66 | 0,49 | 6,24E-01 |
| AT1G21500 | 0,51 | 0,10 | 3,15 | 0,03 | 9,76E-01 |
| AT1G21520 | 0,17 | 0,00 | 4,41 | 0,00 | 1,00E+00 |

| | | | | | |
|---|---|---|---|---|---|
| AT1G21550 | 0,17 | 0,00 | 4,41 | 0,00 | 1,00E+00 |
| AT1G21600 | 0,77 | 2,15 | 2,28 | 0,94 | 3,45E-01 |
| AT1G21650 | 0,46 | 0,33 | 3,33 | 0,10 | 9,20E-01 |
| AT1G21770 | 0,66 | 1,13 | 2,70 | 0,42 | 6,76E-01 |
| AT1G21890 | 0,36 | 0,00 | 4,41 | 0,00 | 1,00E+00 |
| AT1G21900 | 0,26 | -0,51 | 4,40 | -0,12 | 9,08E-01 |
| AT1G21920 | 0,41 | 0,20 | 3,37 | 0,06 | 9,53E-01 |
| AT1G21928 | 0,29 | 1,38 | 4,41 | 0,31 | 7,54E-01 |
| AT1G22180 | 0,30 | 0,00 | 3,70 | 0,00 | 1,00E+00 |
| AT1G22200 | 0,54 | 1,01 | 3,86 | 0,26 | 7,93E-01 |
| AT1G22300 | 0,58 | -1,41 | 2,70 | -0,52 | 6,03E-01 |
| AT1G22360 | 0,63 | -0,42 | 2,74 | -0,15 | 8,79E-01 |
| AT1G22400 | 0,23 | 0,00 | 4,41 | 0,00 | 1,00E+00 |
| AT1G22550 | 0,19 | 0,00 | 4,41 | 0,00 | 1,00E+00 |
| AT1G22570 | 0,46 | -0,55 | 4,09 | -0,13 | 8,94E-01 |
| AT1G22710 | 0,50 | -0,50 | 2,79 | -0,18 | 8,58E-01 |
| AT1G22850 | 1,38 | -0,78 | 1,80 | -0,44 | 6,63E-01 |
| AT1G23020 | 0,20 | 1,33 | 4,40 | 0,30 | 7,63E-01 |
| AT1G23052 | 1,19 | -1,07 | 2,47 | -0,43 | 6,64E-01 |
| AT1G23080 | 0,61 | 2,45 | 3,34 | 0,73 | 4,63E-01 |
| AT1G05793 | 0,08 | 0,00 | 4,41 | 0,00 | 1,00E+00 |
| AT1G23205 | 0,35 | 0,99 | 4,36 | 0,23 | 8,21E-01 |
| AT1G23260 | 0,22 | 0,00 | 4,41 | 0,00 | 1,00E+00 |
| AT1G23300 | 1,57 | -0,54 | 1,74 | -0,31 | 7,55E-01 |
| AT1G23330 | 0,10 | 0,00 | 4,41 | 0,00 | 1,00E+00 |
| AT1G23650 | 0,08 | 0,00 | 4,41 | 0,00 | 1,00E+00 |
| AT1G23670 | 0,22 | 0,00 | 4,41 | 0,00 | 1,00E+00 |
| AT1G23740 | 1,33 | -0,89 | 1,85 | -0,48 | 6,31E-01 |
| AT1G23780 | 0,31 | 1,34 | 4,40 | 0,31 | 7,60E-01 |
| AT1G23830 | 0,53 | 0,29 | 3,29 | 0,09 | 9,29E-01 |
| AT1G23840 | 0,48 | 2,09 | 3,19 | 0,66 | 5,12E-01 |
| AT1G23850 | 0,35 | 1,38 | 4,41 | 0,31 | 7,54E-01 |
| AT1G23960 | 0,23 | 0,00 | 4,41 | 0,00 | 1,00E+00 |
| AT1G24040 | 0,10 | 0,00 | 4,41 | 0,00 | 1,00E+00 |
| AT1G24100 | 1,00 | -0,53 | 2,23 | -0,24 | 8,13E-01 |
| AT1G24120 | 0,29 | -0,40 | 4,38 | -0,09 | 9,28E-01 |
| AT1G24140 | 0,74 | 3,48 | 2,92 | 1,19 | 2,33E-01 |
| AT1G24160 | 0,84 | 1,38 | 3,44 | 0,40 | 6,88E-01 |
| AT1G24190 | 0,84 | -1,76 | 2,61 | -0,67 | 5,01E-01 |
| AT1G24267 | 0,48 | 0,44 | 3,27 | 0,14 | 8,93E-01 |
| AT1G24290 | 0,17 | 0,00 | 4,41 | 0,00 | 1,00E+00 |
| AT1G24300 | 1,68 | 1,49 | 1,76 | 0,85 | 3,97E-01 |
| AT1G24350 | 0,14 | -0,48 | 4,40 | -0,11 | 9,13E-01 |
| AT1G05907 | 0,77 | 0,54 | 2,61 | 0,21 | 8,37E-01 |
| AT1G24764 | 0,53 | -0,79 | 3,32 | -0,24 | 8,11E-01 |
| AT1G24822 | 0,94 | 3,26 | 4,30 | 0,76 | 4,49E-01 |
| AT1G25220 | 0,36 | 0,00 | 4,41 | 0,00 | 1,00E+00 |
| AT1G25350 | 1,21 | 1,57 | 1,80 | 0,87 | 3,85E-01 |
| AT1G25370 | 0,12 | 0,00 | 4,41 | 0,00 | 1,00E+00 |
| AT1G25390 | 0,51 | 1,29 | 3,17 | 0,41 | 6,83E-01 |
| AT1G25400 | 0,63 | 1,36 | 3,00 | 0,46 | 6,49E-01 |
| AT1G25500 | 0,25 | 0,00 | 4,41 | 0,00 | 1,00E+00 |
| AT1G25540 | 0,94 | 1,38 | 2,51 | 0,55 | 5,83E-01 |
| AT1G25560 | 0,90 | -0,03 | 2,23 | -0,01 | 9,90E-01 |
| AT1G25570 | 0,49 | 3,21 | 4,31 | 0,74 | 4,57E-01 |
| AT1G26150 | 0,88 | 1,28 | 2,38 | 0,54 | 5,91E-01 |

| | | | | | |
|---|---|---|---|---|---|
| AT1G26160 | 0,24 | 1,38 | 4,41 | 0,31 | 7,54E-01 |
| AT1G26170 | 0,54 | -0,48 | 3,23 | -0,15 | 8,81E-01 |
| AT1G26208 | 0,43 | 0,00 | 4,41 | 0,00 | 1,00E+00 |
| AT1G26218 | 0,37 | 0,00 | 4,41 | 0,00 | 1,00E+00 |
| AT1G26233 | 0,08 | 0,00 | 4,41 | 0,00 | 1,00E+00 |
| AT1G26240 | 0,88 | 0,00 | 2,78 | 0,00 | 1,00E+00 |
| AT1G26255 | 0,44 | 0,00 | 4,17 | 0,00 | 1,00E+00 |
| AT1G26270 | 1,28 | 4,20 | 2,07 | 2,03 | 4,23E-02 |
| AT1G26380 | 0,46 | 1,44 | 4,35 | 0,33 | 7,40E-01 |
| AT1G26390 | 0,25 | 1,24 | 4,38 | 0,28 | 7,78E-01 |
| AT1G26420 | 0,40 | -0,64 | 4,33 | -0,15 | 8,83E-01 |
| AT1G26690 | 0,33 | 1,27 | 4,39 | 0,29 | 7,72E-01 |
| AT1G26761 | 0,85 | 0,00 | 4,40 | 0,00 | 1,00E+00 |
| AT1G26800 | 0,42 | 0,50 | 2,92 | 0,17 | 8,66E-01 |
| AT1G06033 | 1,64 | -1,80 | 1,65 | -1,09 | 2,75E-01 |
| AT1G26930 | 0,36 | 0,00 | 4,40 | 0,00 | 1,00E+00 |
| AT1G27008 | 0,08 | 0,00 | 4,41 | 0,00 | 1,00E+00 |
| AT1G27100 | 1,46 | -0,56 | 2,06 | -0,27 | 7,87E-01 |
| AT1G27190 | 0,77 | 2,14 | 2,30 | 0,93 | 3,52E-01 |
| AT1G27200 | 0,57 | 0,54 | 2,82 | 0,19 | 8,49E-01 |
| AT1G27210 | 0,26 | 0,00 | 4,41 | 0,00 | 1,00E+00 |
| AT1G27350 | 0,20 | 1,30 | 4,40 | 0,29 | 7,68E-01 |
| AT1G27370 | 0,51 | 2,48 | 4,11 | 0,60 | 5,46E-01 |
| AT1G27385 | 0,43 | 0,73 | 4,38 | 0,17 | 8,68E-01 |
| AT1G27440 | 0,24 | 1,57 | 4,39 | 0,36 | 7,20E-01 |
| AT1G27500 | 0,29 | 1,35 | 4,40 | 0,31 | 7,59E-01 |
| AT1G27510 | 0,44 | 1,28 | 3,32 | 0,38 | 7,01E-01 |
| AT1G06093 | 0,29 | 0,48 | 4,40 | 0,11 | 9,14E-01 |
| AT1G27600 | 1,12 | 1,07 | 1,90 | 0,56 | 5,73E-01 |
| AT1G27700 | 0,50 | 2,72 | 4,34 | 0,63 | 5,30E-01 |
| AT1G27720 | 0,50 | 2,34 | 3,95 | 0,59 | 5,55E-01 |
| AT1G27730 | 1,60 | -1,84 | 1,63 | -1,13 | 2,60E-01 |
| AT1G27900 | 0,40 | 0,00 | 3,41 | 0,00 | 1,00E+00 |
| AT1G27930 | 0,53 | 0,60 | 3,15 | 0,19 | 8,50E-01 |
| AT1G28000 | 0,35 | 0,00 | 4,41 | 0,00 | 1,00E+00 |
| AT1G28120 | 0,73 | 2,20 | 2,95 | 0,75 | 4,56E-01 |
| AT1G28140 | 1,15 | 0,64 | 1,93 | 0,33 | 7,42E-01 |
| AT1G28150 | 0,60 | 1,38 | 3,11 | 0,44 | 6,58E-01 |
| AT1G28190 | 0,15 | 0,00 | 4,41 | 0,00 | 1,00E+00 |
| AT1G28200 | 0,24 | -1,07 | 4,37 | -0,24 | 8,07E-01 |
| AT1G28230 | 0,78 | -0,75 | 2,36 | -0,32 | 7,52E-01 |
| AT1G28240 | 0,33 | -0,54 | 4,41 | -0,12 | 9,02E-01 |
| AT1G28260 | 1,27 | 0,33 | 2,15 | 0,15 | 8,78E-01 |
| AT1G28280 | 1,03 | 1,95 | 2,19 | 0,89 | 3,73E-01 |
| AT1G28290 | 0,27 | 1,38 | 4,41 | 0,31 | 7,54E-01 |
| AT1G28310 | 0,66 | -1,24 | 2,28 | -0,54 | 5,86E-01 |
| AT1G28330 | 0,60 | -0,81 | 3,16 | -0,26 | 7,97E-01 |
| AT1G28370 | 0,92 | -0,44 | 2,47 | -0,18 | 8,60E-01 |
| AT1G28390 | 0,25 | 1,84 | 4,38 | 0,42 | 6,75E-01 |
| AT1G28400 | 0,35 | -1,21 | 3,46 | -0,35 | 7,28E-01 |
| AT1G28570 | 0,12 | 0,00 | 4,41 | 0,00 | 1,00E+00 |
| AT1G28580 | 0,48 | 1,34 | 3,98 | 0,34 | 7,36E-01 |
| AT1G28600 | 1,10 | 1,36 | 1,79 | 0,76 | 4,50E-01 |
| AT1G28630 | 0,79 | -2,70 | 2,78 | -0,97 | 3,31E-01 |
| AT1G28660 | 0,08 | 0,00 | 4,41 | 0,00 | 1,00E+00 |
| AT1G28685 | 0,55 | 0,49 | 3,30 | 0,15 | 8,83E-01 |

| | | | | | |
|---|---|---|---|---|---|
| AT1G28765 | 0,17 | 0,00 | 4,41 | 0,00 | 1,00E+00 |
| AT1G28960 | 0,56 | 1,38 | 4,00 | 0,34 | 7,30E-01 |
| AT1G29050 | 0,14 | 0,00 | 4,41 | 0,00 | 1,00E+00 |
| AT1G29071 | 1,34 | -0,44 | 1,90 | -0,23 | 8,18E-01 |
| AT1G29170 | 0,29 | 0,00 | 4,41 | 0,00 | 1,00E+00 |
| AT1G29240 | 0,32 | 1,90 | 4,39 | 0,43 | 6,65E-01 |
| AT1G29300 | 0,08 | 0,00 | 4,41 | 0,00 | 1,00E+00 |
| AT1G29390 | 0,82 | 1,28 | 2,28 | 0,56 | 5,74E-01 |
| AT1G29395 | 0,39 | -1,64 | 3,40 | -0,48 | 6,29E-01 |
| AT1G29530 | 0,36 | 0,33 | 3,53 | 0,09 | 9,25E-01 |
| AT1G29690 | 0,27 | -0,54 | 4,41 | -0,12 | 9,02E-01 |
| AT1G29700 | 0,34 | -0,78 | 4,34 | -0,18 | 8,56E-01 |
| AT1G29720 | 0,71 | 0,79 | 2,53 | 0,31 | 7,56E-01 |
| AT1G29750 | 0,26 | 1,38 | 4,41 | 0,31 | 7,54E-01 |
| AT1G29880 | 1,19 | 2,32 | 2,16 | 1,07 | 2,82E-01 |
| AT1G29970 | 0,83 | 2,62 | 2,52 | 1,04 | 2,98E-01 |
| AT1G30000 | 0,84 | 2,66 | 2,67 | 1,00 | 3,19E-01 |
| AT1G30060 | 0,17 | 0,00 | 4,41 | 0,00 | 1,00E+00 |
| AT1G30070 | 0,15 | 0,00 | 4,39 | 0,00 | 1,00E+00 |
| AT1G30090 | 0,16 | 0,00 | 4,41 | 0,00 | 1,00E+00 |
| AT1G30120 | 1,11 | 2,18 | 2,12 | 1,03 | 3,02E-01 |
| AT1G30210 | 0,29 | 1,69 | 4,38 | 0,39 | 6,99E-01 |
| AT1G30250 | 0,60 | 0,45 | 2,70 | 0,17 | 8,68E-01 |
| AT1G30370 | 0,55 | 1,38 | 3,20 | 0,43 | 6,67E-01 |
| AT1G30500 | 0,64 | 1,42 | 3,89 | 0,37 | 7,15E-01 |
| AT1G06347 | 0,94 | 0,16 | 2,17 | 0,07 | 9,43E-01 |
| AT1G30540 | 0,42 | 1,26 | 4,15 | 0,30 | 7,61E-01 |
| AT1G30580 | 1,07 | 0,10 | 2,06 | 0,05 | 9,62E-01 |
| AT1G30590 | 0,66 | 1,33 | 2,98 | 0,45 | 6,56E-01 |
| AT1G30755 | 0,62 | 2,86 | 2,65 | 1,08 | 2,81E-01 |
| AT1G30780 | 0,83 | -1,05 | 2,29 | -0,46 | 6,46E-01 |
| AT1G30810 | 0,83 | 0,86 | 1,85 | 0,46 | 6,43E-01 |
| AT1G30880 | 0,08 | 0,00 | 4,41 | 0,00 | 1,00E+00 |
| AT1G30960 | 0,37 | 2,12 | 4,37 | 0,48 | 6,28E-01 |
| AT1G06407 | 0,34 | 1,38 | 3,66 | 0,38 | 7,06E-01 |
| AT1G31130 | 1,47 | 0,50 | 2,03 | 0,25 | 8,05E-01 |
| AT1G31160 | 0,42 | 2,17 | 4,16 | 0,52 | 6,03E-01 |
| AT1G31180 | 0,17 | 0,00 | 4,41 | 0,00 | 1,00E+00 |
| AT1G31230 | 1,31 | -0,11 | 1,85 | -0,06 | 9,52E-01 |
| AT1G31240 | 0,22 | -0,43 | 4,39 | -0,10 | 9,21E-01 |
| AT1G31350 | 0,45 | 2,17 | 3,28 | 0,66 | 5,09E-01 |
| AT1G31410 | 0,08 | 0,00 | 4,41 | 0,00 | 1,00E+00 |
| AT1G31470 | 0,15 | 0,00 | 4,41 | 0,00 | 1,00E+00 |
| AT1G31550 | 0,81 | -1,01 | 2,58 | -0,39 | 6,95E-01 |
| AT1G31600 | 0,81 | -0,73 | 2,86 | -0,26 | 7,98E-01 |
| AT1G31640 | 0,23 | 0,00 | 4,41 | 0,00 | 1,00E+00 |
| AT1G31650 | 0,25 | 1,24 | 4,38 | 0,28 | 7,77E-01 |
| AT1G31660 | 0,51 | 1,50 | 4,36 | 0,34 | 7,31E-01 |
| AT1G31740 | 0,22 | 0,00 | 4,41 | 0,00 | 1,00E+00 |
| AT1G31810 | 0,57 | 1,31 | 2,38 | 0,55 | 5,82E-01 |
| AT1G31812 | 0,29 | -0,54 | 4,41 | -0,12 | 9,02E-01 |
| AT1G31814 | 0,37 | 1,29 | 4,39 | 0,29 | 7,70E-01 |
| AT1G31820 | 0,94 | 2,11 | 2,84 | 0,74 | 4,58E-01 |
| AT1G31830 | 0,66 | 0,48 | 2,74 | 0,17 | 8,62E-01 |
| AT1G06453 | 1,48 | -3,05 | 1,97 | -1,55 | 1,22E-01 |
| AT1G31860 | 1,08 | 1,40 | 2,01 | 0,70 | 4,87E-01 |

| | | | | | |
|---|---|---|---|---|---|
| AT1G06463 | 0,27 | 0,00 | 4,41 | 0,00 | 1,00E+00 |
| AT1G31920 | 0,52 | 2,13 | 2,70 | 0,79 | 4,31E-01 |
| AT1G31940 | 0,35 | 2,30 | 4,37 | 0,53 | 5,98E-01 |
| AT1G06473 | 0,38 | -1,29 | 4,37 | -0,29 | 7,68E-01 |
| AT1G32000 | 0,35 | 1,38 | 4,41 | 0,31 | 7,54E-01 |
| AT1G32070 | 0,34 | 1,38 | 4,41 | 0,31 | 7,54E-01 |
| AT1G32080 | 0,85 | 0,51 | 2,03 | 0,25 | 8,03E-01 |
| AT1G32130 | 1,27 | 0,86 | 1,91 | 0,45 | 6,52E-01 |
| AT1G32200 | 0,75 | -1,46 | 2,18 | -0,67 | 5,04E-01 |
| AT1G32340 | 0,12 | 0,00 | 4,41 | 0,00 | 1,00E+00 |
| AT1G06537 | 1,54 | 3,07 | 1,91 | 1,60 | 1,09E-01 |
| AT1G32400 | 0,47 | 1,38 | 2,85 | 0,49 | 6,27E-01 |
| AT1G32450 | 0,27 | 1,31 | 4,40 | 0,30 | 7,66E-01 |
| AT1G06573 | 0,41 | -0,50 | 4,40 | -0,11 | 9,09E-01 |
| AT1G32470 | 0,50 | -1,13 | 2,77 | -0,41 | 6,83E-01 |
| AT1G32490 | 0,70 | 0,43 | 2,39 | 0,18 | 8,58E-01 |
| AT1G32500 | 0,52 | 1,60 | 2,74 | 0,58 | 5,60E-01 |
| AT1G32530 | 0,53 | 2,46 | 2,75 | 0,90 | 3,70E-01 |
| AT1G06577 | 0,16 | -0,55 | 4,41 | -0,12 | 9,01E-01 |
| AT1G32713 | 1,03 | -0,49 | 2,48 | -0,20 | 8,43E-01 |
| AT1G32790 | 1,57 | 1,38 | 2,22 | 0,62 | 5,34E-01 |
| AT1G32860 | 0,05 | 0,00 | 4,41 | 0,00 | 1,00E+00 |
| AT1G32880 | 0,26 | 1,38 | 4,41 | 0,31 | 7,54E-01 |
| AT1G32900 | 0,61 | 1,69 | 3,17 | 0,53 | 5,94E-01 |
| AT1G32970 | 0,17 | 0,00 | 4,41 | 0,00 | 1,00E+00 |
| AT1G33050 | 0,95 | 1,90 | 2,03 | 0,93 | 3,51E-01 |
| AT1G06643 | 0,36 | -0,29 | 4,37 | -0,07 | 9,47E-01 |
| AT1G33250 | 0,90 | 1,36 | 2,45 | 0,56 | 5,77E-01 |
| AT1G33270 | 0,24 | 1,37 | 4,41 | 0,31 | 7,56E-01 |
| AT1G06673 | 0,47 | -0,43 | 3,36 | -0,13 | 8,98E-01 |
| AT1G33390 | 1,05 | 1,77 | 2,41 | 0,73 | 4,64E-01 |
| AT1G33410 | 1,37 | 3,58 | 1,73 | 2,07 | 3,80E-02 |
| AT1G33430 | 0,08 | 0,00 | 4,41 | 0,00 | 1,00E+00 |
| AT1G33520 | 0,47 | 0,36 | 3,28 | 0,11 | 9,12E-01 |
| AT1G33615 | 0,17 | -0,82 | 4,37 | -0,19 | 8,51E-01 |
| AT1G33700 | 0,88 | 2,14 | 2,25 | 0,95 | 3,43E-01 |
| AT1G33720 | 1,30 | 0,88 | 1,97 | 0,44 | 6,56E-01 |
| AT1G33780 | 0,08 | 0,00 | 4,41 | 0,00 | 1,00E+00 |
| AT1G33870 | 0,60 | -1,60 | 2,64 | -0,60 | 5,45E-01 |
| AT1G33970 | 0,67 | 1,56 | 2,67 | 0,58 | 5,59E-01 |
| AT1G34030 | 0,44 | -0,59 | 2,85 | -0,21 | 8,35E-01 |
| AT1G34042 | 0,29 | 0,00 | 4,41 | 0,00 | 1,00E+00 |
| AT1G34110 | 0,91 | 0,52 | 2,06 | 0,25 | 7,99E-01 |
| AT1G34170 | 0,20 | 0,00 | 4,41 | 0,00 | 1,00E+00 |
| AT1G34220 | 1,01 | 1,25 | 3,01 | 0,42 | 6,78E-01 |
| AT1G34270 | 0,12 | 0,00 | 4,41 | 0,00 | 1,00E+00 |
| AT1G34310 | 0,97 | -0,54 | 2,80 | -0,19 | 8,46E-01 |
| AT1G34315 | 0,23 | 0,00 | 4,41 | 0,00 | 1,00E+00 |
| AT1G34380 | 0,08 | 0,00 | 4,41 | 0,00 | 1,00E+00 |
| AT1G34440 | 1,54 | -2,14 | 1,90 | -1,13 | 2,60E-01 |
| AT1G34550 | 0,63 | 0,30 | 2,59 | 0,12 | 9,07E-01 |
| AT1G34750 | 0,23 | 0,00 | 4,41 | 0,00 | 1,00E+00 |
| AT1G34780 | 0,68 | 1,38 | 2,99 | 0,46 | 6,44E-01 |
| AT1G35150 | 0,16 | 0,00 | 4,41 | 0,00 | 1,00E+00 |
| AT1G35240 | 1,26 | -0,60 | 2,14 | -0,28 | 7,78E-01 |
| AT1G35250 | 0,19 | 0,00 | 4,41 | 0,00 | 1,00E+00 |

| | | | | | |
|---|---|---|---|---|---|
| AT1G35320 | 0,59 | 0,00 | 3,07 | 0,00 | 1,00E+00 |
| AT1G35350 | 0,96 | 0,44 | 2,21 | 0,20 | 8,43E-01 |
| AT1G06883 | 0,19 | 0,00 | 4,41 | 0,00 | 1,00E+00 |
| AT1G06897 | 0,91 | -2,62 | 3,41 | -0,77 | 4,42E-01 |
| AT1G06903 | 0,10 | 0,00 | 4,41 | 0,00 | 1,00E+00 |
| AT1G35520 | 1,01 | 0,00 | 2,67 | 0,00 | 1,00E+00 |
| AT1G35545 | 0,38 | -0,54 | 3,46 | -0,16 | 8,75E-01 |
| AT1G35580 | 1,07 | 1,24 | 1,95 | 0,64 | 5,24E-01 |
| AT1G35610 | 0,22 | -0,48 | 4,40 | -0,11 | 9,14E-01 |
| AT1G35625 | 0,08 | 0,00 | 4,41 | 0,00 | 1,00E+00 |
| AT1G35730 | 0,17 | 0,00 | 4,41 | 0,00 | 1,00E+00 |
| AT1G35820 | 1,40 | 0,00 | 3,01 | 0,00 | 1,00E+00 |
| AT1G35895 | 0,55 | -0,89 | 3,27 | -0,27 | 7,86E-01 |
| AT1G36070 | 0,25 | 1,34 | 4,40 | 0,30 | 7,61E-01 |
| AT1G36085 | 0,68 | 0,00 | 4,41 | 0,00 | 1,00E+00 |
| AT1G36230 | 0,12 | 0,00 | 4,41 | 0,00 | 1,00E+00 |
| AT1G36325 | 1,41 | 0,60 | 1,58 | 0,38 | 7,05E-01 |
| AT1G36370 | 0,71 | 0,31 | 2,49 | 0,12 | 9,01E-01 |
| AT1G36390 | 0,17 | -0,57 | 4,40 | -0,13 | 8,98E-01 |
| AT1G36622 | 0,40 | 0,54 | 3,48 | 0,15 | 8,77E-01 |
| AT1G36990 | 1,02 | 2,02 | 1,99 | 1,01 | 3,11E-01 |
| AT1G07013 | 0,15 | 0,00 | 4,39 | 0,00 | 1,00E+00 |
| AT1G40087 | 0,19 | 0,00 | 4,41 | 0,00 | 1,00E+00 |
| AT1G41830 | 0,28 | 0,00 | 4,41 | 0,00 | 1,00E+00 |
| AT1G41920 | 0,51 | -0,56 | 3,32 | -0,17 | 8,65E-01 |
| AT1G42440 | 0,46 | 1,88 | 3,43 | 0,55 | 5,85E-01 |
| AT1G42560 | 0,05 | 0,00 | 4,41 | 0,00 | 1,00E+00 |
| AT1G42990 | 1,44 | 2,76 | 1,87 | 1,47 | 1,41E-01 |
| AT1G43190 | 0,38 | 1,60 | 4,39 | 0,36 | 7,16E-01 |
| AT1G43560 | 0,22 | -0,53 | 4,41 | -0,12 | 9,05E-01 |
| AT1G43580 | 0,21 | -0,50 | 4,40 | -0,11 | 9,09E-01 |
| AT1G43620 | 0,44 | 1,73 | 4,27 | 0,41 | 6,85E-01 |
| AT1G43665 | 0,88 | 0,00 | 3,33 | 0,00 | 1,00E+00 |
| AT1G43710 | 0,50 | 1,38 | 3,31 | 0,42 | 6,77E-01 |
| AT1G43730 | 0,40 | -0,51 | 4,33 | -0,12 | 9,06E-01 |
| AT1G43760 | 0,66 | -2,01 | 2,57 | -0,78 | 4,34E-01 |
| AT1G43825 | 0,81 | 2,68 | 2,78 | 0,96 | 3,36E-01 |
| AT1G43860 | 1,69 | 1,32 | 1,80 | 0,74 | 4,62E-01 |
| AT1G43950 | 0,46 | 1,28 | 4,10 | 0,31 | 7,55E-01 |
| AT1G44000 | 0,08 | 0,00 | 4,41 | 0,00 | 1,00E+00 |
| AT1G44030 | 0,25 | -0,47 | 4,40 | -0,11 | 9,14E-01 |
| AT1G44080 | 1,35 | -1,69 | 1,90 | -0,89 | 3,75E-01 |
| AT1G44085 | 0,65 | 0,00 | 3,01 | 0,00 | 1,00E+00 |
| AT1G44090 | 0,05 | 0,00 | 4,41 | 0,00 | 1,00E+00 |
| AT1G44125 | 0,91 | 0,00 | 2,84 | 0,00 | 1,00E+00 |
| AT1G44170 | 0,30 | 0,00 | 4,41 | 0,00 | 1,00E+00 |
| AT1G44350 | 0,41 | 1,05 | 4,24 | 0,25 | 8,04E-01 |
| AT1G44542 | 0,40 | 1,85 | 4,39 | 0,42 | 6,74E-01 |
| AT1G44770 | 0,41 | 1,29 | 3,42 | 0,38 | 7,06E-01 |
| AT1G44790 | 0,21 | 0,00 | 4,41 | 0,00 | 1,00E+00 |
| AT1G44920 | 0,44 | 0,42 | 3,44 | 0,12 | 9,03E-01 |
| AT1G44980 | 0,59 | -1,68 | 2,79 | -0,60 | 5,48E-01 |
| AT1G45063 | 0,43 | -2,19 | 4,33 | -0,51 | 6,12E-01 |
| AT1G45145 | 0,72 | 1,38 | 3,25 | 0,42 | 6,71E-01 |
| AT1G45150 | 0,25 | 1,24 | 4,38 | 0,28 | 7,77E-01 |
| AT1G45163 | 0,70 | 1,32 | 2,63 | 0,50 | 6,14E-01 |

| | | | | | |
|---|---|---|---|---|---|
| AT1G45207 | 0,37 | 2,72 | 4,34 | 0,63 | 5,31E-01 |
| AT1G07323 | 0,55 | 2,08 | 2,79 | 0,75 | 4,56E-01 |
| AT1G47200 | 0,16 | 0,00 | 4,41 | 0,00 | 1,00E+00 |
| AT1G47240 | 0,43 | 0,00 | 4,18 | 0,00 | 1,00E+00 |
| AT1G07333 | 0,81 | -0,64 | 3,40 | -0,19 | 8,50E-01 |
| AT1G47260 | 1,01 | 0,09 | 1,80 | 0,05 | 9,61E-01 |
| AT1G47280 | 1,31 | 3,75 | 2,04 | 1,85 | 6,50E-02 |
| AT1G07347 | 0,18 | 0,00 | 4,41 | 0,00 | 1,00E+00 |
| AT1G47380 | 0,18 | -0,47 | 4,40 | -0,11 | 9,14E-01 |
| AT1G47420 | 0,71 | 1,35 | 2,37 | 0,57 | 5,70E-01 |
| AT1G07383 | 0,55 | -0,37 | 2,75 | -0,13 | 8,94E-01 |
| AT1G07393 | 1,67 | -1,47 | 1,65 | -0,89 | 3,73E-01 |
| AT1G47640 | 0,35 | 1,38 | 3,57 | 0,39 | 6,99E-01 |
| AT1G47670 | 0,05 | 0,00 | 4,41 | 0,00 | 1,00E+00 |
| AT1G47765 | 0,55 | 1,22 | 2,44 | 0,50 | 6,18E-01 |
| AT1G47900 | 0,45 | 0,59 | 4,12 | 0,14 | 8,86E-01 |
| AT1G47960 | 1,43 | -1,09 | 1,57 | -0,69 | 4,88E-01 |
| AT1G48000 | 0,50 | 0,00 | 4,35 | 0,00 | 1,00E+00 |
| AT1G48030 | 1,26 | 1,37 | 2,40 | 0,57 | 5,66E-01 |
| AT1G48040 | 0,63 | 0,00 | 4,41 | 0,00 | 1,00E+00 |
| AT1G48110 | 0,42 | -0,67 | 4,39 | -0,15 | 8,78E-01 |
| AT1G48120 | 0,82 | 2,14 | 2,45 | 0,88 | 3,81E-01 |
| AT1G48160 | 0,15 | 0,00 | 4,41 | 0,00 | 1,00E+00 |
| AT1G48285 | 0,80 | 1,47 | 2,39 | 0,61 | 5,39E-01 |
| AT1G48320 | 0,17 | 0,00 | 4,41 | 0,00 | 1,00E+00 |
| AT1G48380 | 0,10 | 0,00 | 4,41 | 0,00 | 1,00E+00 |
| AT1G48405 | 0,82 | 1,38 | 3,43 | 0,40 | 6,88E-01 |
| AT1G48430 | 0,15 | 0,00 | 4,41 | 0,00 | 1,00E+00 |
| AT1G48450 | 1,45 | 2,22 | 1,87 | 1,19 | 2,35E-01 |
| AT1G48550 | 1,08 | 0,84 | 2,70 | 0,31 | 7,55E-01 |
| AT1G48570 | 0,26 | -0,45 | 4,39 | -0,10 | 9,19E-01 |
| AT1G48580 | 0,08 | 0,00 | 4,41 | 0,00 | 1,00E+00 |
| AT1G48620 | 0,46 | 0,58 | 3,28 | 0,18 | 8,60E-01 |
| AT1G48630 | 1,25 | 1,41 | 1,83 | 0,77 | 4,40E-01 |
| AT1G48730 | 0,75 | -1,23 | 2,42 | -0,51 | 6,11E-01 |
| AT1G48742 | 0,46 | -1,95 | 3,17 | -0,61 | 5,39E-01 |
| AT1G48790 | 0,44 | 1,35 | 3,33 | 0,40 | 6,86E-01 |
| AT1G48850 | 0,39 | 2,17 | 4,36 | 0,50 | 6,18E-01 |
| AT1G48900 | 0,23 | -0,86 | 4,38 | -0,20 | 8,44E-01 |
| AT1G48930 | 0,71 | 2,90 | 2,84 | 1,02 | 3,07E-01 |
| AT1G48970 | 1,37 | -0,88 | 2,04 | -0,43 | 6,66E-01 |
| AT1G49005 | 0,38 | -0,97 | 4,38 | -0,22 | 8,25E-01 |
| AT1G49140 | 0,12 | 0,00 | 4,41 | 0,00 | 1,00E+00 |
| AT1G49180 | 0,08 | 0,00 | 4,41 | 0,00 | 1,00E+00 |
| AT1G49205 | 0,08 | 0,00 | 4,41 | 0,00 | 1,00E+00 |
| AT1G49225 | 0,48 | 2,09 | 3,25 | 0,64 | 5,21E-01 |
| AT1G49270 | 0,10 | 0,00 | 4,41 | 0,00 | 1,00E+00 |
| AT1G49490 | 0,32 | 2,37 | 4,36 | 0,54 | 5,86E-01 |
| AT1G49540 | 0,83 | -1,52 | 2,56 | -0,59 | 5,52E-01 |
| AT1G49600 | 0,82 | 2,90 | 2,16 | 1,34 | 1,79E-01 |
| AT1G49670 | 0,26 | 0,00 | 4,41 | 0,00 | 1,00E+00 |
| AT1G49710 | 0,34 | 0,00 | 3,58 | 0,00 | 1,00E+00 |
| AT1G49780 | 1,22 | -0,84 | 1,90 | -0,44 | 6,60E-01 |
| AT1G49850 | 0,39 | 0,00 | 4,41 | 0,00 | 1,00E+00 |
| AT1G49970 | 0,66 | -0,39 | 2,32 | -0,17 | 8,68E-01 |
| AT1G07603 | 0,61 | 2,84 | 3,21 | 0,88 | 3,78E-01 |

| | | | | | |
|---|---|---|---|---|---|
| AT1G50020 | 0,58 | 0,96 | 3,96 | 0,24 | 8,08E-01 |
| AT1G50140 | 0,20 | -0,44 | 4,39 | -0,10 | 9,21E-01 |
| AT1G50190 | 0,32 | 1,38 | 4,41 | 0,31 | 7,54E-01 |
| AT1G50260 | 0,32 | 0,70 | 4,37 | 0,16 | 8,73E-01 |
| AT1G50280 | 0,61 | -0,53 | 2,40 | -0,22 | 8,24E-01 |
| AT1G50320 | 0,25 | -0,52 | 4,40 | -0,12 | 9,06E-01 |
| AT1G07627 | 0,18 | 0,00 | 4,40 | 0,00 | 1,00E+00 |
| AT1G50420 | 0,49 | 2,71 | 4,34 | 0,62 | 5,32E-01 |
| AT1G50430 | 1,02 | 2,74 | 1,99 | 1,38 | 1,69E-01 |
| AT1G50450 | 0,63 | 1,50 | 2,97 | 0,51 | 6,13E-01 |
| AT1G50500 | 0,57 | -0,31 | 3,14 | -0,10 | 9,22E-01 |
| AT1G07647 | 0,08 | 0,00 | 4,41 | 0,00 | 1,00E+00 |
| AT1G50570 | 0,65 | 1,19 | 2,73 | 0,44 | 6,63E-01 |
| AT1G50600 | 0,88 | 0,41 | 2,29 | 0,18 | 8,57E-01 |
| AT1G50640 | 1,08 | 2,22 | 2,00 | 1,11 | 2,67E-01 |
| AT1G50659 | 0,43 | 0,00 | 3,35 | 0,00 | 1,00E+00 |
| AT1G50750 | 0,08 | 0,00 | 4,41 | 0,00 | 1,00E+00 |
| AT1G50790 | 0,12 | 0,00 | 4,41 | 0,00 | 1,00E+00 |
| AT1G50820 | 0,50 | 2,87 | 3,95 | 0,73 | 4,68E-01 |
| AT1G50910 | 1,39 | 1,74 | 1,68 | 1,03 | 3,01E-01 |
| AT1G50980 | 0,39 | 1,83 | 4,37 | 0,42 | 6,75E-01 |
| AT1G51080 | 0,73 | 2,71 | 2,63 | 1,03 | 3,03E-01 |
| AT1G07733 | 0,24 | -0,56 | 4,41 | -0,13 | 8,99E-01 |
| AT1G51400 | 0,93 | -2,68 | 2,27 | -1,18 | 2,37E-01 |
| AT1G51480 | 0,33 | 2,19 | 4,38 | 0,50 | 6,17E-01 |
| AT1G51500 | 1,21 | -1,92 | 2,07 | -0,93 | 3,54E-01 |
| AT1G51510 | 0,91 | 1,19 | 2,28 | 0,52 | 6,02E-01 |
| AT1G51560 | 0,45 | 2,36 | 3,33 | 0,71 | 4,79E-01 |
| AT1G51590 | 0,26 | 0,00 | 4,41 | 0,00 | 1,00E+00 |
| AT1G51630 | 0,52 | -0,39 | 3,19 | -0,12 | 9,03E-01 |
| AT1G51680 | 0,80 | 2,12 | 2,27 | 0,93 | 3,51E-01 |
| AT1G07763 | 0,56 | -1,38 | 4,32 | -0,32 | 7,50E-01 |
| AT1G51710 | 0,45 | 1,29 | 4,13 | 0,31 | 7,55E-01 |
| AT1G51730 | 0,47 | -0,23 | 3,24 | -0,07 | 9,43E-01 |
| AT1G51790 | 0,25 | 1,43 | 4,40 | 0,32 | 7,46E-01 |
| AT1G51890 | 0,23 | 0,00 | 4,41 | 0,00 | 1,00E+00 |
| AT1G51940 | 1,08 | -0,76 | 2,68 | -0,28 | 7,77E-01 |
| AT1G52000 | 0,46 | -1,07 | 4,04 | -0,26 | 7,91E-01 |
| AT1G52040 | 0,54 | -0,16 | 3,87 | -0,04 | 9,66E-01 |
| AT1G52150 | 1,30 | 2,94 | 2,42 | 1,21 | 2,25E-01 |
| AT1G52160 | 0,61 | 1,96 | 3,20 | 0,61 | 5,40E-01 |
| AT1G52180 | 1,00 | -2,02 | 1,97 | -1,02 | 3,06E-01 |
| AT1G52185 | 0,32 | -1,69 | 4,34 | -0,39 | 6,98E-01 |
| AT1G52200 | 0,25 | 0,00 | 4,41 | 0,00 | 1,00E+00 |
| AT1G52220 | 0,76 | -0,91 | 2,26 | -0,41 | 6,85E-01 |
| AT1G52290 | 0,41 | 1,34 | 4,31 | 0,31 | 7,55E-01 |
| AT1G52300 | 0,35 | -0,53 | 4,41 | -0,12 | 9,04E-01 |
| AT1G52340 | 0,08 | 0,00 | 4,41 | 0,00 | 1,00E+00 |
| AT1G52342 | 0,45 | 1,37 | 4,41 | 0,31 | 7,55E-01 |
| AT1G52510 | 0,47 | 0,67 | 4,26 | 0,16 | 8,75E-01 |
| AT1G52570 | 1,59 | 4,47 | 2,20 | 2,03 | 4,23E-02 |
| AT1G52750 | 0,31 | 1,31 | 4,40 | 0,30 | 7,65E-01 |
| AT1G52880 | 0,40 | 0,36 | 3,48 | 0,10 | 9,17E-01 |
| AT1G52890 | 0,42 | 1,32 | 4,21 | 0,31 | 7,55E-01 |
| AT1G52970 | 1,13 | -1,51 | 1,88 | -0,81 | 4,21E-01 |
| AT1G52990 | 0,41 | 0,00 | 4,41 | 0,00 | 1,00E+00 |

| | | | | | |
|---|---|---|---|---|---|
| AT1G53000 | 0,49 | 2,83 | 4,02 | 0,70 | 4,82E-01 |
| AT1G53025 | 0,43 | 2,34 | 3,38 | 0,69 | 4,89E-01 |
| AT1G53035 | 0,12 | 0,00 | 4,41 | 0,00 | 1,00E+00 |
| AT1G53170 | 0,65 | -1,88 | 2,63 | -0,72 | 4,74E-01 |
| AT1G53200 | 0,24 | 1,37 | 4,41 | 0,31 | 7,56E-01 |
| AT1G53230 | 1,22 | 2,05 | 2,28 | 0,90 | 3,68E-01 |
| AT1G53240 | 1,16 | -2,41 | 1,98 | -1,22 | 2,23E-01 |
| AT1G53265 | 0,53 | -2,13 | 4,00 | -0,53 | 5,95E-01 |
| AT1G53320 | 0,35 | 1,38 | 4,41 | 0,31 | 7,54E-01 |
| AT1G53450 | 0,86 | -0,28 | 1,95 | -0,14 | 8,87E-01 |
| AT1G53480 | 0,84 | -1,00 | 3,06 | -0,33 | 7,44E-01 |
| AT1G53500 | 0,50 | -1,70 | 3,35 | -0,51 | 6,13E-01 |
| AT1G53510 | 0,76 | 0,68 | 2,48 | 0,27 | 7,84E-01 |
| AT1G53541 | 0,64 | 2,07 | 2,53 | 0,82 | 4,13E-01 |
| AT1G53580 | 0,12 | 0,00 | 4,41 | 0,00 | 1,00E+00 |
| AT1G53710 | 0,67 | 1,63 | 2,95 | 0,55 | 5,81E-01 |
| AT1G53750 | 1,03 | 1,88 | 1,98 | 0,95 | 3,43E-01 |
| AT1G53760 | 0,08 | 0,00 | 4,41 | 0,00 | 1,00E+00 |
| AT1G53770 | 0,33 | 1,79 | 4,38 | 0,41 | 6,83E-01 |
| AT1G53780 | 1,40 | 2,64 | 2,08 | 1,27 | 2,04E-01 |
| AT1G54050 | 0,12 | 0,00 | 4,41 | 0,00 | 1,00E+00 |
| AT1G54080 | 1,16 | 0,87 | 2,08 | 0,42 | 6,74E-01 |
| AT1G54100 | 1,08 | 0,05 | 2,06 | 0,03 | 9,79E-01 |
| AT1G54140 | 0,15 | 0,00 | 4,41 | 0,00 | 1,00E+00 |
| AT1G54170 | 0,82 | -0,99 | 2,11 | -0,47 | 6,37E-01 |
| AT1G54320 | 0,67 | -1,79 | 2,51 | -0,71 | 4,76E-01 |
| AT1G54370 | 0,38 | 1,38 | 4,36 | 0,32 | 7,52E-01 |
| AT1G54500 | 0,85 | 0,99 | 2,26 | 0,44 | 6,61E-01 |
| AT1G54590 | 0,10 | 0,00 | 4,41 | 0,00 | 1,00E+00 |
| AT1G54630 | 0,35 | 0,35 | 4,39 | 0,08 | 9,37E-01 |
| AT1G54690 | 0,48 | 2,28 | 2,80 | 0,81 | 4,15E-01 |
| AT1G54775 | 0,57 | -1,59 | 3,83 | -0,42 | 6,78E-01 |
| AT1G54840 | 0,49 | 2,61 | 4,33 | 0,60 | 5,47E-01 |
| AT1G54950 | 0,44 | -0,75 | 4,08 | -0,18 | 8,54E-01 |
| AT1G55152 | 0,78 | 1,38 | 2,65 | 0,52 | 6,02E-01 |
| AT1G55180 | 0,78 | 2,45 | 2,85 | 0,86 | 3,91E-01 |
| AT1G55200 | 0,08 | 0,00 | 4,41 | 0,00 | 1,00E+00 |
| AT1G08043 | 0,30 | 1,38 | 4,41 | 0,31 | 7,54E-01 |
| AT1G55270 | 1,00 | 3,33 | 3,15 | 1,06 | 2,91E-01 |
| AT1G55300 | 0,10 | 0,00 | 4,41 | 0,00 | 1,00E+00 |
| AT1G55330 | 0,51 | -0,43 | 3,18 | -0,13 | 8,93E-01 |
| AT1G55360 | 0,31 | 1,53 | 4,39 | 0,35 | 7,27E-01 |
| AT1G55530 | 0,83 | 0,71 | 2,24 | 0,32 | 7,51E-01 |
| AT1G55535 | 0,40 | 0,95 | 4,25 | 0,22 | 8,23E-01 |
| AT1G55590 | 0,61 | 1,29 | 2,66 | 0,49 | 6,26E-01 |
| AT1G55610 | 0,12 | 0,00 | 4,41 | 0,00 | 1,00E+00 |
| AT1G55620 | 0,57 | -0,43 | 3,24 | -0,13 | 8,95E-01 |
| AT1G55675 | 0,68 | 3,35 | 3,69 | 0,91 | 3,64E-01 |
| AT1G55680 | 1,44 | 0,77 | 2,21 | 0,35 | 7,26E-01 |
| AT1G55690 | 0,12 | 0,00 | 4,41 | 0,00 | 1,00E+00 |
| AT1G55830 | 0,23 | 0,00 | 4,41 | 0,00 | 1,00E+00 |
| AT1G55880 | 0,36 | 0,00 | 4,41 | 0,00 | 1,00E+00 |
| AT1G55950 | 0,93 | 0,00 | 2,39 | 0,00 | 1,00E+00 |
| AT1G55960 | 0,25 | 0,43 | 4,38 | 0,10 | 9,22E-01 |
| AT1G55970 | 0,17 | 0,00 | 4,41 | 0,00 | 1,00E+00 |
| AT1G56010 | 0,42 | 1,39 | 4,18 | 0,33 | 7,40E-01 |

| | | | | | |
|---|---|---|---|---|---|
| AT1G56100 | 0,86 | -1,55 | 2,84 | -0,55 | 5,85E-01 |
| AT1G56110 | 1,35 | 1,13 | 1,82 | 0,62 | 5,35E-01 |
| AT1G56120 | 0,47 | -0,12 | 4,06 | -0,03 | 9,76E-01 |
| AT1G56140 | 0,44 | 1,45 | 3,38 | 0,43 | 6,68E-01 |
| AT1G56145 | 0,80 | 3,32 | 2,72 | 1,22 | 2,22E-01 |
| AT1G56165 | 0,20 | 1,33 | 4,40 | 0,30 | 7,63E-01 |
| AT1G56230 | 0,19 | -0,47 | 4,40 | -0,11 | 9,15E-01 |
| AT1G08107 | 0,31 | -0,43 | 3,67 | -0,12 | 9,06E-01 |
| AT1G08113 | 0,55 | -2,15 | 3,21 | -0,67 | 5,03E-01 |
| AT1G08123 | 0,56 | -0,52 | 2,67 | -0,19 | 8,47E-01 |
| AT1G56300 | 0,85 | 0,85 | 2,85 | 0,30 | 7,65E-01 |
| AT1G56330 | 0,78 | 1,54 | 2,46 | 0,63 | 5,30E-01 |
| AT1G56345 | 0,19 | 0,00 | 4,40 | 0,00 | 1,00E+00 |
| AT1G56440 | 0,27 | 1,31 | 4,40 | 0,30 | 7,66E-01 |
| AT1G56520 | 1,21 | 3,15 | 2,32 | 1,36 | 1,74E-01 |
| AT1G56550 | 0,17 | -0,57 | 4,40 | -0,13 | 8,98E-01 |
| AT1G56580 | 0,24 | 1,38 | 4,41 | 0,31 | 7,54E-01 |
| AT1G56650 | 1,01 | 0,00 | 4,04 | 0,00 | 1,00E+00 |
| AT1G57680 | 1,45 | -0,35 | 2,12 | -0,17 | 8,68E-01 |
| AT1G57775 | 0,38 | -0,54 | 3,46 | -0,16 | 8,75E-01 |
| AT1G57777 | 0,31 | 0,00 | 3,70 | 0,00 | 1,00E+00 |
| AT1G57860 | 0,30 | 0,10 | 4,38 | 0,02 | 9,83E-01 |
| AT1G57870 | 0,40 | -0,54 | 4,33 | -0,13 | 9,00E-01 |
| AT1G57943 | 0,08 | 0,00 | 4,41 | 0,00 | 1,00E+00 |
| AT1G57990 | 0,77 | 2,47 | 2,59 | 0,95 | 3,42E-01 |
| AT1G58030 | 0,73 | 1,06 | 2,60 | 0,41 | 6,84E-01 |
| AT1G58060 | 0,60 | 1,90 | 3,19 | 0,60 | 5,50E-01 |
| AT1G58080 | 0,29 | 1,29 | 4,39 | 0,29 | 7,69E-01 |
| AT1G58100 | 1,20 | 2,85 | 1,89 | 1,51 | 1,31E-01 |
| AT1G58110 | 0,53 | 1,22 | 3,93 | 0,31 | 7,55E-01 |
| AT1G58200 | 0,16 | 0,00 | 4,41 | 0,00 | 1,00E+00 |
| AT1G58220 | 0,54 | 2,61 | 3,85 | 0,68 | 4,98E-01 |
| AT1G58300 | 1,33 | -1,42 | 1,72 | -0,82 | 4,10E-01 |
| AT1G58350 | 0,80 | 3,00 | 2,59 | 1,16 | 2,48E-01 |
| AT1G58390 | 0,70 | -0,88 | 3,06 | -0,29 | 7,74E-01 |
| AT1G58440 | 0,58 | -1,87 | 3,04 | -0,62 | 5,38E-01 |
| AT1G58684 | 0,49 | -1,56 | 4,36 | -0,36 | 7,21E-01 |
| AT1G58725 | 0,33 | -0,54 | 4,41 | -0,12 | 9,02E-01 |
| AT1G58766 | 0,16 | 0,00 | 4,41 | 0,00 | 1,00E+00 |
| AT1G59030 | 0,44 | -1,16 | 4,30 | -0,27 | 7,87E-01 |
| AT1G59077 | 0,16 | -0,55 | 4,41 | -0,12 | 9,01E-01 |
| AT1G59406 | 0,14 | 0,00 | 4,41 | 0,00 | 1,00E+00 |
| AT1G59453 | 0,25 | 0,00 | 4,41 | 0,00 | 1,00E+00 |
| AT1G59720 | 0,61 | 1,45 | 2,77 | 0,52 | 6,02E-01 |
| AT1G59780 | 0,52 | 0,70 | 3,20 | 0,22 | 8,28E-01 |
| AT1G59835 | 1,47 | -1,24 | 1,62 | -0,76 | 4,45E-01 |
| AT1G59910 | 1,37 | 2,65 | 2,19 | 1,21 | 2,25E-01 |
| AT1G59920 | 0,42 | -0,54 | 3,38 | -0,16 | 8,72E-01 |
| AT1G59930 | 1,11 | 0,00 | 2,57 | 0,00 | 1,00E+00 |
| AT1G59990 | 1,13 | 1,71 | 2,19 | 0,78 | 4,34E-01 |
| AT1G60000 | 0,38 | 0,32 | 3,58 | 0,09 | 9,29E-01 |
| AT1G60070 | 1,35 | 2,97 | 2,13 | 1,40 | 1,62E-01 |
| AT1G60160 | 0,43 | 1,28 | 3,31 | 0,39 | 6,98E-01 |
| AT1G60200 | 0,68 | 2,86 | 3,20 | 0,89 | 3,72E-01 |
| AT1G60230 | 0,48 | 2,77 | 4,01 | 0,69 | 4,89E-01 |
| AT1G60260 | 0,77 | 0,46 | 2,53 | 0,18 | 8,55E-01 |

| | | | | | |
|---|---|---|---|---|---|
| AT1G08317 | 0,08 | 0,00 | 4,41 | 0,00 | 1,00E+00 |
| AT1G60430 | 0,45 | 2,11 | 4,03 | 0,52 | 6,01E-01 |
| AT1G60490 | 0,25 | -0,52 | 4,40 | -0,12 | 9,06E-01 |
| AT1G60550 | 0,38 | 2,11 | 4,37 | 0,48 | 6,29E-01 |
| AT1G60560 | 0,17 | 0,00 | 4,41 | 0,00 | 1,00E+00 |
| AT1G60590 | 0,17 | -1,42 | 4,35 | -0,33 | 7,45E-01 |
| AT1G60630 | 0,08 | 0,00 | 4,41 | 0,00 | 1,00E+00 |
| AT1G60640 | 0,12 | 0,00 | 4,41 | 0,00 | 1,00E+00 |
| AT1G60700 | 0,48 | 1,38 | 4,41 | 0,31 | 7,54E-01 |
| AT1G60710 | 0,41 | 2,78 | 4,34 | 0,64 | 5,22E-01 |
| AT1G60770 | 1,20 | 3,56 | 2,11 | 1,68 | 9,21E-02 |
| AT1G60780 | 0,33 | 0,00 | 4,41 | 0,00 | 1,00E+00 |
| AT1G60800 | 0,56 | -0,19 | 3,04 | -0,06 | 9,51E-01 |
| AT1G60810 | 0,12 | 0,00 | 4,41 | 0,00 | 1,00E+00 |
| AT1G60850 | 0,11 | 0,00 | 4,41 | 0,00 | 1,00E+00 |
| AT1G60890 | 0,24 | -0,45 | 4,39 | -0,10 | 9,18E-01 |
| AT1G60930 | 0,24 | 1,31 | 4,40 | 0,30 | 7,65E-01 |
| AT1G60940 | 0,93 | 1,14 | 2,22 | 0,51 | 6,07E-01 |
| AT1G60960 | 1,57 | 2,73 | 1,70 | 1,60 | 1,09E-01 |
| AT1G60990 | 0,24 | 1,37 | 4,41 | 0,31 | 7,56E-01 |
| AT1G60995 | 0,35 | 1,38 | 4,41 | 0,31 | 7,54E-01 |
| AT1G61010 | 0,89 | 3,75 | 2,28 | 1,64 | 1,01E-01 |
| AT1G61200 | 1,13 | -1,73 | 1,81 | -0,96 | 3,39E-01 |
| AT1G61260 | 0,47 | 1,38 | 3,35 | 0,41 | 6,81E-01 |
| AT1G61300 | 1,06 | -0,08 | 2,28 | -0,03 | 9,72E-01 |
| AT1G61310 | 0,33 | -0,51 | 4,40 | -0,12 | 9,08E-01 |
| AT1G61360 | 0,15 | 0,00 | 4,41 | 0,00 | 1,00E+00 |
| AT1G61370 | 0,17 | 0,00 | 4,41 | 0,00 | 1,00E+00 |
| AT1G61380 | 0,34 | 0,00 | 4,41 | 0,00 | 1,00E+00 |
| AT1G61420 | 0,12 | 0,00 | 4,41 | 0,00 | 1,00E+00 |
| AT1G61560 | 0,37 | 0,00 | 4,41 | 0,00 | 1,00E+00 |
| AT1G08417 | 0,75 | 2,98 | 2,82 | 1,06 | 2,91E-01 |
| AT1G61660 | 0,65 | -0,60 | 3,07 | -0,20 | 8,44E-01 |
| AT1G61730 | 1,18 | 0,88 | 2,11 | 0,42 | 6,77E-01 |
| AT1G61890 | 1,56 | 1,84 | 2,02 | 0,91 | 3,62E-01 |
| AT1G61900 | 0,26 | -0,40 | 4,37 | -0,09 | 9,27E-01 |
| AT1G62030 | 1,53 | -1,42 | 2,05 | -0,69 | 4,89E-01 |
| AT1G62085 | 0,60 | 1,32 | 3,26 | 0,40 | 6,87E-01 |
| AT1G62300 | 0,27 | 1,38 | 4,41 | 0,31 | 7,54E-01 |
| AT1G62422 | 0,51 | 0,36 | 2,76 | 0,13 | 8,97E-01 |
| AT1G62610 | 0,76 | 3,19 | 2,37 | 1,35 | 1,78E-01 |
| AT1G62680 | 1,24 | -0,90 | 2,01 | -0,44 | 6,56E-01 |
| AT1G62760 | 0,29 | 0,00 | 4,41 | 0,00 | 1,00E+00 |
| AT1G62780 | 0,49 | -0,15 | 3,27 | -0,04 | 9,65E-01 |
| AT1G62810 | 0,97 | 1,77 | 2,13 | 0,83 | 4,06E-01 |
| AT1G62915 | 0,64 | -1,81 | 2,98 | -0,61 | 5,42E-01 |
| AT1G62960 | 0,36 | 0,29 | 4,39 | 0,07 | 9,47E-01 |
| AT1G63010 | 0,16 | -0,55 | 4,41 | -0,12 | 9,01E-01 |
| AT1G63020 | 0,44 | 0,00 | 4,41 | 0,00 | 1,00E+00 |
| AT1G63050 | 0,34 | 1,69 | 4,38 | 0,39 | 6,99E-01 |
| AT1G63090 | 1,34 | -0,25 | 1,81 | -0,14 | 8,89E-01 |
| AT1G63250 | 0,13 | 0,00 | 4,41 | 0,00 | 1,00E+00 |
| AT1G63290 | 0,12 | 0,00 | 4,41 | 0,00 | 1,00E+00 |
| AT1G63320 | 0,20 | 0,00 | 4,41 | 0,00 | 1,00E+00 |
| AT1G63410 | 0,62 | 0,43 | 2,65 | 0,16 | 8,70E-01 |
| AT1G63490 | 0,53 | 1,38 | 3,30 | 0,42 | 6,76E-01 |

| | | | | | |
|---|---|---|---|---|---|
| AT1G63660 | 0,92 | 1,96 | 2,22 | 0,88 | 3,77E-01 |
| AT1G63670 | 0,18 | 1,38 | 4,41 | 0,31 | 7,54E-01 |
| AT1G63700 | 1,36 | 1,95 | 1,73 | 1,13 | 2,59E-01 |
| AT1G63720 | 0,19 | 0,00 | 4,41 | 0,00 | 1,00E+00 |
| AT1G08627 | 1,18 | -2,08 | 1,98 | -1,05 | 2,94E-01 |
| AT1G63800 | 0,47 | 0,93 | 4,35 | 0,21 | 8,31E-01 |
| AT1G63810 | 0,31 | 0,09 | 4,38 | 0,02 | 9,84E-01 |
| AT1G63840 | 0,40 | 0,00 | 4,41 | 0,00 | 1,00E+00 |
| AT1G63860 | 1,09 | 2,56 | 1,95 | 1,31 | 1,89E-01 |
| AT1G63870 | 0,58 | 0,49 | 3,16 | 0,15 | 8,77E-01 |
| AT1G63970 | 0,51 | 0,40 | 3,33 | 0,12 | 9,03E-01 |
| AT1G64080 | 0,15 | 0,00 | 4,41 | 0,00 | 1,00E+00 |
| AT1G64190 | 1,11 | 1,06 | 1,85 | 0,57 | 5,65E-01 |
| AT1G64200 | 0,78 | -1,03 | 4,37 | -0,23 | 8,14E-01 |
| AT1G64280 | 0,86 | 2,17 | 2,63 | 0,83 | 4,09E-01 |
| AT1G64380 | 0,37 | -0,68 | 4,39 | -0,16 | 8,77E-01 |
| AT1G64460 | 0,72 | 2,13 | 2,53 | 0,84 | 4,00E-01 |
| AT1G64480 | 0,18 | 0,00 | 4,40 | 0,00 | 1,00E+00 |
| AT1G64510 | 0,32 | -0,54 | 4,41 | -0,12 | 9,02E-01 |
| AT1G64580 | 0,24 | -0,83 | 4,38 | -0,19 | 8,50E-01 |
| AT1G64610 | 0,62 | 1,91 | 2,62 | 0,73 | 4,67E-01 |
| AT1G64640 | 0,11 | 0,00 | 4,41 | 0,00 | 1,00E+00 |
| AT1G64650 | 0,59 | 1,36 | 2,63 | 0,52 | 6,04E-01 |
| AT1G64670 | 0,10 | 0,00 | 4,41 | 0,00 | 1,00E+00 |
| AT1G64750 | 0,36 | 0,10 | 4,35 | 0,02 | 9,82E-01 |
| AT1G64970 | 0,63 | 0,43 | 2,67 | 0,16 | 8,71E-01 |
| AT1G64980 | 1,26 | -0,04 | 1,65 | -0,03 | 9,80E-01 |
| AT1G65040 | 0,28 | -0,42 | 3,74 | -0,11 | 9,09E-01 |
| AT1G65150 | 0,20 | -0,44 | 4,39 | -0,10 | 9,20E-01 |
| AT1G65190 | 0,43 | -1,57 | 4,13 | -0,38 | 7,04E-01 |
| AT1G08747 | 0,81 | 1,66 | 2,88 | 0,58 | 5,65E-01 |
| AT1G65290 | 0,77 | 0,47 | 2,37 | 0,20 | 8,41E-01 |
| AT1G65320 | 0,39 | -0,92 | 4,38 | -0,21 | 8,34E-01 |
| AT1G65346 | 0,34 | 0,00 | 4,41 | 0,00 | 1,00E+00 |
| AT1G65420 | 0,15 | -0,43 | 4,39 | -0,10 | 9,23E-01 |
| AT1G65430 | 0,52 | 0,58 | 3,19 | 0,18 | 8,55E-01 |
| AT1G65520 | 0,29 | -1,03 | 4,38 | -0,23 | 8,14E-01 |
| AT1G65540 | 1,25 | -0,39 | 2,10 | -0,18 | 8,54E-01 |
| AT1G65550 | 0,21 | -0,54 | 4,41 | -0,12 | 9,02E-01 |
| AT1G65580 | 0,46 | -0,26 | 2,78 | -0,09 | 9,25E-01 |
| AT1G65690 | 0,12 | 0,00 | 4,41 | 0,00 | 1,00E+00 |
| AT1G65950 | 0,28 | 1,84 | 4,39 | 0,42 | 6,74E-01 |
| AT1G08797 | 0,63 | 0,00 | 4,41 | 0,00 | 1,00E+00 |
| AT1G65980 | 1,27 | 3,92 | 2,11 | 1,86 | 6,32E-02 |
| AT1G65985 | 0,33 | 0,43 | 4,36 | 0,10 | 9,22E-01 |
| AT1G65990 | 1,24 | 0,00 | 2,28 | 0,00 | 1,00E+00 |
| AT1G66200 | 1,38 | 0,95 | 1,52 | 0,62 | 5,34E-01 |
| AT1G66235 | 0,12 | 0,00 | 4,41 | 0,00 | 1,00E+00 |
| AT1G66240 | 0,12 | 0,00 | 4,41 | 0,00 | 1,00E+00 |
| AT1G66260 | 1,28 | 0,55 | 1,97 | 0,28 | 7,81E-01 |
| AT1G66290 | 0,49 | -2,47 | 3,18 | -0,78 | 4,38E-01 |
| AT1G66330 | 0,52 | 0,09 | 2,68 | 0,03 | 9,74E-01 |
| AT1G66410 | 1,68 | 0,72 | 1,63 | 0,44 | 6,60E-01 |
| AT1G66460 | 0,25 | -0,53 | 4,41 | -0,12 | 9,04E-01 |
| AT1G66490 | 0,90 | 0,72 | 2,31 | 0,31 | 7,54E-01 |
| AT1G66500 | 0,25 | 1,84 | 4,38 | 0,42 | 6,75E-01 |

| | | | | | |
|---|---|---|---|---|---|
| AT1G66570 | 1,67 | 0,67 | 1,54 | 0,44 | 6,63E-01 |
| AT1G66660 | 0,44 | 1,38 | 4,20 | 0,33 | 7,42E-01 |
| AT1G66670 | 1,37 | 0,32 | 2,09 | 0,15 | 8,78E-01 |
| AT1G66680 | 0,59 | 2,81 | 4,33 | 0,65 | 5,16E-01 |
| AT1G66700 | 0,46 | 0,00 | 4,41 | 0,00 | 1,00E+00 |
| AT1G66830 | 0,68 | 0,16 | 3,81 | 0,04 | 9,67E-01 |
| AT1G66910 | 0,65 | 2,90 | 2,27 | 1,28 | 2,01E-01 |
| AT1G66940 | 0,34 | 1,31 | 4,40 | 0,30 | 7,66E-01 |
| AT1G66980 | 1,51 | 0,64 | 1,56 | 0,41 | 6,82E-01 |
| AT1G67000 | 0,91 | -2,06 | 2,12 | -0,97 | 3,31E-01 |
| AT1G67050 | 0,28 | -0,42 | 3,72 | -0,11 | 9,09E-01 |
| AT1G67080 | 0,36 | 2,06 | 4,37 | 0,47 | 6,38E-01 |
| AT1G67140 | 0,95 | 3,12 | 2,53 | 1,23 | 2,18E-01 |
| AT1G67230 | 0,62 | -1,31 | 2,70 | -0,49 | 6,26E-01 |
| AT1G67235 | 0,29 | 0,00 | 4,41 | 0,00 | 1,00E+00 |
| AT1G67250 | 0,34 | 0,16 | 4,38 | 0,04 | 9,71E-01 |
| AT1G67300 | 0,37 | 1,38 | 4,37 | 0,32 | 7,52E-01 |
| AT1G67310 | 0,33 | 1,27 | 4,39 | 0,29 | 7,72E-01 |
| AT1G67325 | 0,61 | 1,29 | 3,08 | 0,42 | 6,76E-01 |
| AT1G67360 | 0,73 | -1,21 | 2,88 | -0,42 | 6,74E-01 |
| AT1G67481 | 1,30 | 3,40 | 1,97 | 1,72 | 8,51E-02 |
| AT1G67490 | 0,57 | 2,51 | 3,15 | 0,80 | 4,25E-01 |
| AT1G67500 | 0,52 | 2,53 | 4,02 | 0,63 | 5,29E-01 |
| AT1G67510 | 0,38 | -0,53 | 4,31 | -0,12 | 9,03E-01 |
| AT1G67580 | 1,21 | 2,27 | 2,03 | 1,12 | 2,64E-01 |
| AT1G67590 | 0,17 | 1,32 | 4,40 | 0,30 | 7,64E-01 |
| AT1G67680 | 0,43 | 1,91 | 4,38 | 0,44 | 6,63E-01 |
| AT1G67785 | 0,15 | -0,43 | 4,39 | -0,10 | 9,23E-01 |
| AT1G67790 | 0,26 | 1,38 | 4,41 | 0,31 | 7,54E-01 |
| AT1G67860 | 0,58 | 1,35 | 3,16 | 0,43 | 6,68E-01 |
| AT1G67865 | 0,24 | -0,48 | 4,40 | -0,11 | 9,13E-01 |
| AT1G67870 | 0,89 | -2,02 | 2,06 | -0,98 | 3,28E-01 |
| AT1G67930 | 1,11 | 1,94 | 1,84 | 1,05 | 2,92E-01 |
| AT1G67950 | 0,39 | 1,76 | 4,38 | 0,40 | 6,87E-01 |
| AT1G68060 | 0,56 | -1,26 | 3,29 | -0,38 | 7,03E-01 |
| AT1G68140 | 0,87 | 1,34 | 2,32 | 0,58 | 5,63E-01 |
| AT1G68325 | 0,05 | 0,00 | 4,41 | 0,00 | 1,00E+00 |
| AT1G68370 | 0,32 | 0,48 | 4,40 | 0,11 | 9,13E-01 |
| AT1G68550 | 0,87 | -0,44 | 2,23 | -0,19 | 8,46E-01 |
| AT1G08997 | 0,58 | -0,81 | 3,80 | -0,21 | 8,32E-01 |
| AT1G68590 | 1,41 | 3,68 | 1,84 | 2,00 | 4,59E-02 |
| AT1G68660 | 0,71 | -1,01 | 2,27 | -0,44 | 6,57E-01 |
| AT1G68710 | 0,46 | 1,75 | 4,38 | 0,40 | 6,89E-01 |
| AT1G68830 | 0,57 | -0,26 | 2,73 | -0,09 | 9,25E-01 |
| AT1G68910 | 0,08 | 0,00 | 4,41 | 0,00 | 1,00E+00 |
| AT1G69020 | 0,30 | 1,38 | 4,41 | 0,31 | 7,54E-01 |
| AT1G69070 | 0,46 | 2,55 | 3,32 | 0,77 | 4,43E-01 |
| AT1G69220 | 1,28 | 2,88 | 1,95 | 1,48 | 1,40E-01 |
| AT1G69270 | 0,78 | 1,66 | 3,08 | 0,54 | 5,90E-01 |
| AT1G69280 | 0,21 | -0,54 | 4,41 | -0,12 | 9,02E-01 |
| AT1G69290 | 0,15 | 0,00 | 4,41 | 0,00 | 1,00E+00 |
| AT1G69310 | 0,25 | 0,00 | 4,41 | 0,00 | 1,00E+00 |
| AT1G69450 | 0,38 | 1,95 | 4,36 | 0,45 | 6,54E-01 |
| AT1G69480 | 0,55 | 0,00 | 4,39 | 0,00 | 1,00E+00 |
| AT1G69630 | 0,59 | 3,10 | 4,32 | 0,72 | 4,73E-01 |
| AT1G69640 | 0,90 | -1,31 | 2,51 | -0,52 | 6,02E-01 |

| | | | | | |
|---|---|---|---|---|---|
| AT1G69730 | 0,73 | 1,36 | 2,52 | 0,54 | 5,91E-01 |
| AT1G69760 | 0,47 | 0,44 | 3,24 | 0,13 | 8,93E-01 |
| AT1G69770 | 1,05 | 3,62 | 2,27 | 1,59 | 1,11E-01 |
| AT1G69795 | 1,39 | -2,95 | 2,37 | -1,24 | 2,13E-01 |
| AT1G69840 | 0,65 | 1,31 | 3,13 | 0,42 | 6,76E-01 |
| AT1G69850 | 0,22 | 0,00 | 4,41 | 0,00 | 1,00E+00 |
| AT1G69890 | 1,49 | -0,54 | 2,14 | -0,25 | 7,99E-01 |
| AT1G69960 | 0,51 | 0,64 | 3,23 | 0,20 | 8,43E-01 |
| AT1G70000 | 0,98 | 1,18 | 2,16 | 0,55 | 5,86E-01 |
| AT1G70070 | 1,07 | -1,58 | 2,14 | -0,74 | 4,60E-01 |
| AT1G70200 | 0,37 | 0,50 | 4,37 | 0,11 | 9,09E-01 |
| AT1G70280 | 0,17 | 0,00 | 4,41 | 0,00 | 1,00E+00 |
| AT1G70300 | 0,38 | 1,52 | 4,32 | 0,35 | 7,26E-01 |
| AT1G70310 | 0,89 | -0,75 | 3,02 | -0,25 | 8,03E-01 |
| AT1G70340 | 0,34 | 0,44 | 3,62 | 0,12 | 9,04E-01 |
| AT1G70370 | 0,57 | -1,71 | 2,74 | -0,62 | 5,33E-01 |
| AT1G70410 | 0,70 | 0,74 | 2,60 | 0,28 | 7,76E-01 |
| AT1G70420 | 0,44 | -1,66 | 4,07 | -0,41 | 6,83E-01 |
| AT1G70520 | 0,53 | -1,48 | 4,37 | -0,34 | 7,35E-01 |
| AT1G70600 | 0,97 | 1,31 | 2,93 | 0,45 | 6,55E-01 |
| AT1G70730 | 0,91 | 0,16 | 2,39 | 0,07 | 9,48E-01 |
| AT1G70740 | 0,10 | 0,00 | 4,41 | 0,00 | 1,00E+00 |
| AT1G70750 | 0,44 | 0,00 | 4,41 | 0,00 | 1,00E+00 |
| AT1G70780 | 0,73 | -0,54 | 2,93 | -0,19 | 8,52E-01 |
| AT1G70830 | 0,27 | 0,00 | 4,41 | 0,00 | 1,00E+00 |
| AT1G70890 | 1,35 | -2,28 | 1,72 | -1,32 | 1,85E-01 |
| AT1G71020 | 0,94 | 3,27 | 2,31 | 1,42 | 1,56E-01 |
| AT1G71030 | 0,60 | -0,69 | 2,48 | -0,28 | 7,81E-01 |
| AT1G71040 | 0,83 | 2,91 | 2,14 | 1,36 | 1,74E-01 |
| AT1G71060 | 0,73 | 1,38 | 3,54 | 0,39 | 6,97E-01 |
| AT1G71070 | 0,49 | 1,26 | 3,16 | 0,40 | 6,90E-01 |
| AT1G71080 | 0,90 | 2,81 | 2,91 | 0,97 | 3,33E-01 |
| AT1G71160 | 0,69 | 1,83 | 3,62 | 0,50 | 6,14E-01 |
| AT1G71190 | 0,19 | 0,00 | 4,41 | 0,00 | 1,00E+00 |
| AT1G71210 | 0,73 | 2,43 | 2,95 | 0,82 | 4,11E-01 |
| AT1G71310 | 0,27 | -0,88 | 4,38 | -0,20 | 8,41E-01 |
| AT1G71330 | 0,30 | 1,38 | 4,41 | 0,31 | 7,54E-01 |
| AT1G71340 | 0,92 | 1,71 | 2,46 | 0,69 | 4,88E-01 |
| AT1G71350 | 0,21 | 0,00 | 4,41 | 0,00 | 1,00E+00 |
| AT1G71410 | 1,45 | 4,35 | 1,95 | 2,23 | 2,59E-02 |
| AT1G09253 | 0,32 | 1,38 | 4,41 | 0,31 | 7,54E-01 |
| AT1G71490 | 0,42 | 1,38 | 4,25 | 0,32 | 7,46E-01 |
| AT1G71710 | 0,44 | -0,33 | 2,91 | -0,11 | 9,11E-01 |
| AT1G71770 | 0,40 | 0,00 | 4,22 | 0,00 | 1,00E+00 |
| AT1G71870 | 0,88 | 0,00 | 2,95 | 0,00 | 1,00E+00 |
| AT1G71960 | 0,26 | -0,51 | 3,83 | -0,13 | 8,94E-01 |
| AT1G72020 | 0,56 | 0,83 | 2,71 | 0,31 | 7,60E-01 |
| AT1G72040 | 0,83 | 0,36 | 2,32 | 0,15 | 8,77E-01 |
| AT1G72160 | 0,64 | -0,29 | 2,34 | -0,12 | 9,01E-01 |
| AT1G72190 | 0,52 | -0,68 | 3,94 | -0,17 | 8,62E-01 |
| AT1G72270 | 0,12 | 0,00 | 4,41 | 0,00 | 1,00E+00 |
| AT1G72300 | 0,54 | 2,12 | 3,12 | 0,68 | 4,96E-01 |
| AT1G72320 | 0,39 | 1,90 | 3,49 | 0,55 | 5,85E-01 |
| AT1G72450 | 1,12 | -0,15 | 1,81 | -0,09 | 9,32E-01 |
| AT1G72480 | 0,35 | -0,54 | 4,41 | -0,12 | 9,02E-01 |
| AT1G72500 | 0,46 | 2,44 | 4,35 | 0,56 | 5,75E-01 |

| | | | | | |
|---|---|---|---|---|---|
| AT1G72550 | 0,79 | 1,01 | 2,41 | 0,42 | 6,75E-01 |
| AT1G72560 | 1,09 | -0,52 | 2,29 | -0,23 | 8,22E-01 |
| AT1G09327 | 0,20 | -0,43 | 4,39 | -0,10 | 9,22E-01 |
| AT1G72670 | 0,20 | 1,33 | 4,40 | 0,30 | 7,63E-01 |
| AT1G72720 | 0,35 | 1,58 | 4,36 | 0,36 | 7,18E-01 |
| AT1G72730 | 0,32 | 1,90 | 4,39 | 0,43 | 6,65E-01 |
| AT1G72760 | 0,65 | 0,00 | 4,41 | 0,00 | 1,00E+00 |
| AT1G72840 | 0,41 | 0,00 | 4,41 | 0,00 | 1,00E+00 |
| AT1G72860 | 0,44 | 0,00 | 4,41 | 0,00 | 1,00E+00 |
| AT1G73010 | 0,40 | 0,00 | 4,41 | 0,00 | 1,00E+00 |
| AT1G09363 | 0,41 | 1,35 | 3,40 | 0,40 | 6,91E-01 |
| AT1G73110 | 0,63 | -1,94 | 3,82 | -0,51 | 6,11E-01 |
| AT1G73200 | 0,73 | 1,95 | 2,24 | 0,87 | 3,82E-01 |
| AT1G73230 | 0,17 | 0,00 | 4,41 | 0,00 | 1,00E+00 |
| AT1G73260 | 0,63 | 2,91 | 2,98 | 0,98 | 3,28E-01 |
| AT1G73370 | 0,52 | 0,00 | 4,41 | 0,00 | 1,00E+00 |
| AT1G73390 | 0,25 | 0,00 | 4,41 | 0,00 | 1,00E+00 |
| AT1G73490 | 0,37 | 1,72 | 4,38 | 0,39 | 6,94E-01 |
| AT1G73500 | 0,92 | 0,66 | 1,97 | 0,34 | 7,37E-01 |
| AT1G73510 | 0,45 | 2,57 | 4,08 | 0,63 | 5,29E-01 |
| AT1G73530 | 0,46 | -0,93 | 4,30 | -0,22 | 8,28E-01 |
| AT1G73540 | 0,81 | 0,40 | 2,39 | 0,17 | 8,68E-01 |
| AT1G73650 | 0,96 | -0,70 | 2,50 | -0,28 | 7,78E-01 |
| AT1G73655 | 0,74 | 1,38 | 3,84 | 0,36 | 7,19E-01 |
| AT1G73660 | 1,58 | 2,63 | 1,98 | 1,33 | 1,84E-01 |
| AT1G73680 | 0,85 | -1,21 | 2,26 | -0,54 | 5,92E-01 |
| AT1G73730 | 0,17 | 0,00 | 4,41 | 0,00 | 1,00E+00 |
| AT1G73760 | 0,39 | 1,65 | 4,31 | 0,38 | 7,02E-01 |
| AT1G73930 | 0,25 | 1,39 | 4,41 | 0,32 | 7,52E-01 |
| AT1G73950 | 0,45 | 0,37 | 4,38 | 0,09 | 9,32E-01 |
| AT1G73960 | 0,29 | 1,27 | 4,39 | 0,29 | 7,72E-01 |
| AT1G73980 | 0,67 | 1,38 | 2,72 | 0,51 | 6,12E-01 |
| AT1G74050 | 0,43 | 1,34 | 3,35 | 0,40 | 6,88E-01 |
| AT1G74060 | 0,72 | 0,84 | 4,32 | 0,19 | 8,46E-01 |
| AT1G74070 | 0,42 | -1,91 | 4,23 | -0,45 | 6,52E-01 |
| AT1G74120 | 0,82 | 1,52 | 2,89 | 0,53 | 5,98E-01 |
| AT1G74170 | 0,18 | 0,00 | 4,41 | 0,00 | 1,00E+00 |
| AT1G74340 | 0,31 | -0,54 | 4,41 | -0,12 | 9,02E-01 |
| AT1G74370 | 0,53 | 1,42 | 3,98 | 0,36 | 7,22E-01 |
| AT1G74390 | 0,95 | 2,26 | 2,66 | 0,85 | 3,95E-01 |
| AT1G74420 | 0,45 | 2,55 | 4,36 | 0,58 | 5,59E-01 |
| AT1G74440 | 0,80 | 0,32 | 3,09 | 0,10 | 9,17E-01 |
| AT1G09467 | 0,50 | -1,53 | 4,13 | -0,37 | 7,11E-01 |
| AT1G74560 | 0,37 | -0,61 | 3,51 | -0,17 | 8,62E-01 |
| AT1G74690 | 0,95 | 0,84 | 2,15 | 0,39 | 6,97E-01 |
| AT1G74770 | 0,42 | 2,32 | 4,20 | 0,55 | 5,80E-01 |
| AT1G74800 | 0,56 | 0,00 | 3,26 | 0,00 | 1,00E+00 |
| AT1G74830 | 0,12 | 0,00 | 4,41 | 0,00 | 1,00E+00 |
| AT1G74850 | 0,94 | 2,48 | 2,13 | 1,16 | 2,44E-01 |
| AT1G74880 | 0,11 | 0,00 | 4,41 | 0,00 | 1,00E+00 |
| AT1G74920 | 0,86 | 0,33 | 2,64 | 0,13 | 9,00E-01 |
| AT1G74950 | 0,28 | 1,34 | 4,40 | 0,30 | 7,61E-01 |
| AT1G74960 | 1,51 | 3,88 | 2,66 | 1,46 | 1,44E-01 |
| AT1G74970 | 0,46 | 0,44 | 4,34 | 0,10 | 9,20E-01 |
| AT1G75010 | 0,20 | 0,00 | 4,41 | 0,00 | 1,00E+00 |
| AT1G75100 | 0,60 | 0,34 | 2,40 | 0,14 | 8,87E-01 |

| | | | | | |
|---|---|---|---|---|---|
| AT1G75130 | 0,12 | 0,00 | 4,41 | 0,00 | 1,00E+00 |
| AT1G75180 | 0,38 | -0,46 | 3,45 | -0,13 | 8,94E-01 |
| AT1G75200 | 0,15 | 0,00 | 4,41 | 0,00 | 1,00E+00 |
| AT1G75220 | 0,55 | 0,00 | 3,27 | 0,00 | 1,00E+00 |
| AT1G75370 | 1,12 | 0,62 | 1,59 | 0,39 | 6,98E-01 |
| AT1G75410 | 0,90 | -0,36 | 2,56 | -0,14 | 8,87E-01 |
| AT1G75450 | 0,11 | 0,00 | 4,41 | 0,00 | 1,00E+00 |
| AT1G75500 | 0,97 | 0,05 | 1,93 | 0,02 | 9,81E-01 |
| AT1G75680 | 0,99 | -1,63 | 1,84 | -0,88 | 3,77E-01 |
| AT1G75690 | 0,84 | 0,51 | 1,90 | 0,27 | 7,89E-01 |
| AT1G75750 | 1,13 | 0,83 | 2,13 | 0,39 | 6,98E-01 |
| AT1G75780 | 0,25 | -0,54 | 4,41 | -0,12 | 9,02E-01 |
| AT1G75820 | 1,17 | 2,95 | 2,31 | 1,28 | 2,01E-01 |
| AT1G75860 | 0,67 | 1,57 | 2,35 | 0,67 | 5,03E-01 |
| AT1G75945 | 0,28 | 1,38 | 4,41 | 0,31 | 7,54E-01 |
| AT1G76060 | 0,20 | 1,40 | 4,41 | 0,32 | 7,51E-01 |
| AT1G76080 | 1,03 | -2,39 | 1,88 | -1,28 | 2,02E-01 |
| AT1G76100 | 1,22 | 1,18 | 2,03 | 0,58 | 5,61E-01 |
| AT1G76120 | 0,15 | 0,00 | 4,41 | 0,00 | 1,00E+00 |
| AT1G76150 | 0,72 | 0,00 | 3,03 | 0,00 | 1,00E+00 |
| AT1G76160 | 0,88 | 1,30 | 2,04 | 0,63 | 5,26E-01 |
| AT1G76270 | 0,08 | 0,00 | 4,41 | 0,00 | 1,00E+00 |
| AT1G76280 | 0,12 | 0,00 | 4,41 | 0,00 | 1,00E+00 |
| AT1G09627 | 0,45 | 1,38 | 4,10 | 0,34 | 7,37E-01 |
| AT1G76400 | 0,57 | 1,38 | 3,17 | 0,44 | 6,63E-01 |
| AT1G76460 | 0,53 | 0,66 | 3,22 | 0,21 | 8,37E-01 |
| AT1G76640 | 0,08 | 0,00 | 4,41 | 0,00 | 1,00E+00 |
| AT1G76650 | 0,42 | -0,51 | 3,44 | -0,15 | 8,82E-01 |
| AT1G76660 | 0,13 | -0,43 | 4,39 | -0,10 | 9,23E-01 |
| AT1G76670 | 1,15 | 2,49 | 2,09 | 1,19 | 2,34E-01 |
| AT1G76700 | 0,39 | 0,64 | 4,39 | 0,14 | 8,85E-01 |
| AT1G76810 | 1,64 | -2,08 | 1,62 | -1,28 | 1,99E-01 |
| AT1G76878 | 1,06 | -0,84 | 1,86 | -0,45 | 6,51E-01 |
| AT1G76890 | 0,74 | 1,32 | 2,12 | 0,62 | 5,35E-01 |
| AT1G76955 | 0,30 | 1,38 | 4,41 | 0,31 | 7,54E-01 |
| AT1G76960 | 0,40 | 0,43 | 3,48 | 0,12 | 9,01E-01 |
| AT1G76965 | 0,25 | 1,26 | 4,39 | 0,29 | 7,74E-01 |
| AT1G76970 | 0,43 | -0,97 | 3,37 | -0,29 | 7,73E-01 |
| AT1G76980 | 0,73 | 1,23 | 3,11 | 0,39 | 6,93E-01 |
| AT1G77030 | 0,32 | 1,38 | 4,41 | 0,31 | 7,54E-01 |
| AT1G77060 | 0,12 | 0,00 | 4,41 | 0,00 | 1,00E+00 |
| AT1G77100 | 0,82 | 0,00 | 3,78 | 0,00 | 1,00E+00 |
| AT1G77145 | 0,08 | 0,00 | 4,41 | 0,00 | 1,00E+00 |
| AT1G77170 | 0,63 | 2,19 | 3,04 | 0,72 | 4,72E-01 |
| AT1G77240 | 1,26 | 3,24 | 1,98 | 1,64 | 1,01E-01 |
| AT1G77250 | 0,55 | 2,16 | 3,82 | 0,57 | 5,72E-01 |
| AT1G77260 | 0,57 | 1,72 | 3,90 | 0,44 | 6,60E-01 |
| AT1G77480 | 0,33 | -1,69 | 4,36 | -0,39 | 6,99E-01 |
| AT1G77580 | 0,18 | 0,00 | 4,41 | 0,00 | 1,00E+00 |
| AT1G77600 | 0,35 | 1,36 | 4,40 | 0,31 | 7,58E-01 |
| AT1G77660 | 0,33 | 1,26 | 4,39 | 0,29 | 7,74E-01 |
| AT1G77810 | 0,58 | 0,91 | 3,82 | 0,24 | 8,11E-01 |
| AT1G77932 | 0,47 | 1,94 | 4,39 | 0,44 | 6,59E-01 |
| AT1G77940 | 0,55 | 0,42 | 2,72 | 0,15 | 8,78E-01 |
| AT1G77960 | 1,37 | 0,50 | 1,68 | 0,30 | 7,65E-01 |
| AT1G77980 | 0,77 | -0,44 | 2,22 | -0,20 | 8,43E-01 |

| | | | | | |
|---|---|---|---|---|---|
| AT1G77990 | 0,37 | 0,00 | 4,41 | 0,00 | 1,00E+00 |
| AT1G78060 | 0,43 | -1,39 | 4,37 | -0,32 | 7,50E-01 |
| AT1G78210 | 0,32 | 1,38 | 4,41 | 0,31 | 7,54E-01 |
| AT1G78240 | 0,51 | 0,44 | 2,72 | 0,16 | 8,73E-01 |
| AT1G78370 | 1,22 | 2,82 | 2,25 | 1,26 | 2,09E-01 |
| AT1G78380 | 1,48 | -0,99 | 1,71 | -0,58 | 5,64E-01 |
| AT1G78460 | 0,60 | 0,14 | 2,52 | 0,05 | 9,57E-01 |
| AT1G78480 | 0,26 | 1,33 | 4,40 | 0,30 | 7,63E-01 |
| AT1G78510 | 0,69 | 0,31 | 3,70 | 0,08 | 9,33E-01 |
| AT1G78580 | 0,70 | 0,53 | 2,26 | 0,23 | 8,16E-01 |
| AT1G78800 | 0,21 | 0,00 | 4,41 | 0,00 | 1,00E+00 |
| AT1G78820 | 0,54 | -0,79 | 2,38 | -0,33 | 7,40E-01 |
| AT1G78850 | 0,94 | -2,05 | 1,97 | -1,04 | 2,96E-01 |
| AT1G78865 | 0,13 | 0,00 | 4,41 | 0,00 | 1,00E+00 |
| AT1G78880 | 0,88 | 0,01 | 2,86 | 0,00 | 9,98E-01 |
| AT1G78915 | 0,18 | -0,43 | 4,39 | -0,10 | 9,21E-01 |
| AT1G78990 | 0,38 | 2,48 | 4,35 | 0,57 | 5,69E-01 |
| AT1G78995 | 0,24 | -0,48 | 4,40 | -0,11 | 9,13E-01 |
| AT1G79020 | 0,25 | 1,84 | 4,38 | 0,42 | 6,75E-01 |
| AT1G79030 | 0,53 | 1,32 | 3,98 | 0,33 | 7,41E-01 |
| AT1G79050 | 0,58 | 1,70 | 3,12 | 0,54 | 5,86E-01 |
| AT1G79080 | 1,23 | 1,22 | 2,33 | 0,52 | 6,00E-01 |
| AT1G79120 | 0,10 | 0,00 | 4,41 | 0,00 | 1,00E+00 |
| AT1G79150 | 0,35 | -0,96 | 4,38 | -0,22 | 8,27E-01 |
| AT1G79350 | 0,49 | 2,31 | 2,78 | 0,83 | 4,06E-01 |
| AT1G79430 | 0,17 | 1,32 | 4,40 | 0,30 | 7,64E-01 |
| AT1G79440 | 0,39 | 0,42 | 3,59 | 0,12 | 9,07E-01 |
| AT1G09867 | 1,21 | -2,31 | 2,38 | -0,97 | 3,33E-01 |
| AT1G09883 | 0,79 | 1,27 | 2,96 | 0,43 | 6,69E-01 |
| AT1G79540 | 0,53 | 2,37 | 3,17 | 0,75 | 4,54E-01 |
| AT1G79600 | 1,43 | 2,64 | 1,90 | 1,39 | 1,65E-01 |
| AT1G79640 | 0,40 | 0,00 | 4,41 | 0,00 | 1,00E+00 |
| AT1G79670 | 0,90 | 0,36 | 2,72 | 0,13 | 8,96E-01 |
| AT1G79700 | 0,08 | 0,00 | 4,41 | 0,00 | 1,00E+00 |
| AT1G79720 | 0,47 | -1,40 | 3,34 | -0,42 | 6,74E-01 |
| AT1G79730 | 0,63 | 2,02 | 3,90 | 0,52 | 6,04E-01 |
| AT1G79740 | 0,25 | 1,84 | 4,38 | 0,42 | 6,75E-01 |
| AT1G79790 | 0,05 | 0,00 | 4,41 | 0,00 | 1,00E+00 |
| AT1G79830 | 1,06 | 0,12 | 1,93 | 0,06 | 9,51E-01 |
| AT1G79880 | 0,11 | 0,00 | 4,41 | 0,00 | 1,00E+00 |
| AT1G79900 | 0,42 | 2,32 | 4,20 | 0,55 | 5,80E-01 |
| AT1G80030 | 1,07 | 0,07 | 2,11 | 0,03 | 9,74E-01 |
| AT1G80040 | 0,61 | 2,12 | 2,54 | 0,83 | 4,04E-01 |
| AT1G80080 | 0,50 | 2,19 | 3,19 | 0,69 | 4,93E-01 |
| AT1G80325 | 1,08 | 0,34 | 2,07 | 0,16 | 8,71E-01 |
| AT1G80350 | 0,38 | 2,48 | 4,35 | 0,57 | 5,69E-01 |
| AT1G80420 | 0,34 | -0,48 | 4,38 | -0,11 | 9,12E-01 |
| AT1G80460 | 0,24 | 1,36 | 4,40 | 0,31 | 7,57E-01 |
| AT1G80480 | 0,56 | 1,36 | 3,25 | 0,42 | 6,75E-01 |
| AT1G80490 | 1,23 | 2,49 | 1,82 | 1,36 | 1,73E-01 |
| AT1G80530 | 0,68 | -1,49 | 3,01 | -0,49 | 6,22E-01 |
| AT1G80600 | 1,40 | 0,30 | 1,84 | 0,16 | 8,71E-01 |
| AT1G80610 | 0,35 | 0,00 | 3,54 | 0,00 | 1,00E+00 |
| AT1G80630 | 0,28 | 1,39 | 4,41 | 0,32 | 7,52E-01 |
| AT1G80660 | 1,14 | 3,62 | 2,01 | 1,80 | 7,22E-02 |
| AT1G80670 | 0,39 | 1,76 | 4,38 | 0,40 | 6,87E-01 |

| | | | | | |
|---|---|---|---|---|---|
| AT1G80760 | 0,62 | 2,63 | 3,72 | 0,71 | 4,80E-01 |
| AT1G80830 | 0,27 | 1,35 | 4,40 | 0,31 | 7,59E-01 |
| AT1G80850 | 0,20 | 0,00 | 4,41 | 0,00 | 1,00E+00 |
| AT1G80920 | 0,74 | 2,16 | 2,30 | 0,94 | 3,47E-01 |
| AT1G80930 | 0,69 | 0,41 | 2,58 | 0,16 | 8,73E-01 |
| AT1G80950 | 0,35 | 1,38 | 4,41 | 0,31 | 7,53E-01 |
| AT2G01023 | 0,35 | -1,29 | 2,98 | -0,43 | 6,65E-01 |
| AT2G01050 | 0,45 | 0,00 | 4,41 | 0,00 | 1,00E+00 |
| AT2G01060 | 0,57 | 0,00 | 2,73 | 0,00 | 1,00E+00 |
| AT2G01110 | 0,90 | 2,55 | 2,50 | 1,02 | 3,07E-01 |
| AT2G01130 | 1,10 | 1,66 | 1,99 | 0,83 | 4,04E-01 |
| AT2G01140 | 1,55 | 2,40 | 2,06 | 1,16 | 2,44E-01 |
| AT2G01180 | 0,86 | -0,50 | 2,07 | -0,24 | 8,10E-01 |
| AT2G01250 | 0,52 | 0,29 | 3,97 | 0,07 | 9,43E-01 |
| AT2G01275 | 0,19 | 0,00 | 4,41 | 0,00 | 1,00E+00 |
| AT2G01290 | 0,18 | 0,00 | 4,41 | 0,00 | 1,00E+00 |
| AT2G01300 | 0,08 | 0,00 | 4,41 | 0,00 | 1,00E+00 |
| AT2G01310 | 0,05 | 0,00 | 4,41 | 0,00 | 1,00E+00 |
| AT2G01350 | 0,12 | 0,00 | 4,41 | 0,00 | 1,00E+00 |
| AT2G01410 | 0,34 | 0,44 | 3,60 | 0,12 | 9,02E-01 |
| AT2G01420 | 1,24 | -1,75 | 1,83 | -0,96 | 3,40E-01 |
| AT2G01440 | 1,25 | 2,71 | 2,50 | 1,08 | 2,79E-01 |
| AT2G01450 | 1,15 | 2,87 | 1,89 | 1,52 | 1,29E-01 |
| AT2G01505 | 0,30 | 0,00 | 4,41 | 0,00 | 1,00E+00 |
| AT2G01630 | 0,87 | 3,40 | 2,77 | 1,23 | 2,20E-01 |
| AT2G01650 | 1,03 | 2,85 | 2,21 | 1,29 | 1,96E-01 |
| AT2G01670 | 0,39 | -0,49 | 3,41 | -0,14 | 8,86E-01 |
| AT2G01690 | 0,55 | 1,97 | 3,84 | 0,51 | 6,08E-01 |
| AT2G01720 | 1,57 | 2,29 | 2,38 | 0,96 | 3,36E-01 |
| AT2G01755 | 0,14 | 0,00 | 4,41 | 0,00 | 1,00E+00 |
| AT2G01820 | 1,26 | 1,71 | 2,41 | 0,71 | 4,78E-01 |
| AT2G01830 | 0,91 | -0,77 | 2,10 | -0,37 | 7,13E-01 |
| AT2G04125 | 0,12 | 0,00 | 4,41 | 0,00 | 1,00E+00 |
| AT2G01870 | 0,33 | 1,18 | 4,38 | 0,27 | 7,89E-01 |
| AT2G01918 | 1,01 | -1,28 | 2,18 | -0,59 | 5,57E-01 |
| AT2G01930 | 0,16 | 0,00 | 4,41 | 0,00 | 1,00E+00 |
| AT2G01970 | 1,49 | 0,50 | 1,71 | 0,29 | 7,71E-01 |
| AT2G01990 | 0,25 | 0,00 | 4,41 | 0,00 | 1,00E+00 |
| AT2G02080 | 1,03 | -0,99 | 2,61 | -0,38 | 7,05E-01 |
| AT2G02103 | 0,70 | -1,48 | 2,61 | -0,57 | 5,71E-01 |
| AT2G02148 | 0,42 | 2,13 | 4,14 | 0,51 | 6,07E-01 |
| AT2G02150 | 0,39 | 1,80 | 4,31 | 0,42 | 6,76E-01 |
| AT2G02160 | 0,60 | 0,43 | 2,64 | 0,16 | 8,70E-01 |
| AT2G02220 | 0,80 | -0,19 | 2,57 | -0,07 | 9,42E-01 |
| AT2G02230 | 0,17 | 0,00 | 4,41 | 0,00 | 1,00E+00 |
| AT2G02340 | 1,23 | -0,26 | 2,02 | -0,13 | 8,97E-01 |
| AT2G02360 | 0,16 | 0,00 | 4,41 | 0,00 | 1,00E+00 |
| AT2G02400 | 0,84 | 1,34 | 2,45 | 0,55 | 5,85E-01 |
| AT2G02520 | 0,08 | 0,00 | 4,41 | 0,00 | 1,00E+00 |
| AT2G02570 | 0,25 | 0,00 | 4,41 | 0,00 | 1,00E+00 |
| AT2G02635 | 0,17 | -0,44 | 4,39 | -0,10 | 9,19E-01 |
| AT2G02695 | 0,58 | 0,00 | 3,12 | 0,00 | 1,00E+00 |
| AT2G02710 | 0,66 | 2,17 | 2,56 | 0,85 | 3,96E-01 |
| AT2G02780 | 0,28 | -0,95 | 4,37 | -0,22 | 8,27E-01 |
| AT2G02800 | 0,78 | -0,99 | 2,52 | -0,39 | 6,94E-01 |
| AT2G04345 | 0,62 | 2,03 | 2,71 | 0,75 | 4,55E-01 |

| | | | | | |
|---|---|---|---|---|---|
| AT2G02930 | 0,49 | 0,00 | 3,32 | 0,00 | 1,00E+00 |
| AT2G03050 | 0,25 | 0,00 | 4,41 | 0,00 | 1,00E+00 |
| AT2G03090 | 0,08 | 0,00 | 4,41 | 0,00 | 1,00E+00 |
| AT2G03240 | 0,24 | 1,38 | 4,41 | 0,31 | 7,54E-01 |
| AT2G03340 | 0,45 | 2,24 | 3,31 | 0,68 | 4,98E-01 |
| AT2G03390 | 0,61 | -0,70 | 2,41 | -0,29 | 7,70E-01 |
| AT2G03420 | 0,43 | -0,54 | 4,17 | -0,13 | 8,96E-01 |
| AT2G03430 | 0,52 | 1,96 | 4,37 | 0,45 | 6,53E-01 |
| AT2G03440 | 0,35 | 1,34 | 4,40 | 0,30 | 7,62E-01 |
| AT2G03460 | 0,08 | 0,00 | 4,41 | 0,00 | 1,00E+00 |
| AT2G03470 | 1,22 | 3,01 | 2,41 | 1,25 | 2,11E-01 |
| AT2G03490 | 0,08 | 0,00 | 4,41 | 0,00 | 1,00E+00 |
| AT2G03620 | 0,33 | 1,85 | 4,39 | 0,42 | 6,73E-01 |
| AT2G03640 | 0,13 | 0,00 | 4,41 | 0,00 | 1,00E+00 |
| AT2G03690 | 0,23 | 1,38 | 4,41 | 0,31 | 7,54E-01 |
| AT2G03730 | 0,29 | -0,51 | 4,40 | -0,12 | 9,08E-01 |
| AT2G03750 | 1,29 | -1,60 | 1,79 | -0,89 | 3,73E-01 |
| AT2G03760 | 0,93 | 2,20 | 3,50 | 0,63 | 5,29E-01 |
| AT2G03770 | 0,15 | 0,00 | 4,41 | 0,00 | 1,00E+00 |
| AT2G03930 | 1,60 | -2,58 | 1,74 | -1,48 | 1,38E-01 |
| AT2G04655 | 0,14 | 0,00 | 4,41 | 0,00 | 1,00E+00 |
| AT2G04031 | 0,15 | 0,00 | 4,41 | 0,00 | 1,00E+00 |
| AT2G04037 | 0,16 | 0,00 | 4,41 | 0,00 | 1,00E+00 |
| AT2G04050 | 0,12 | 0,00 | 4,41 | 0,00 | 1,00E+00 |
| AT2G04062 | 0,70 | 0,00 | 3,04 | 0,00 | 1,00E+00 |
| AT2G04063 | 0,69 | 0,00 | 2,90 | 0,00 | 1,00E+00 |
| AT2G04066 | 0,42 | 1,50 | 4,39 | 0,34 | 7,32E-01 |
| AT2G04070 | 0,33 | 2,19 | 4,38 | 0,50 | 6,17E-01 |
| AT2G04080 | 0,61 | 2,18 | 2,63 | 0,83 | 4,07E-01 |
| AT2G04160 | 0,43 | 0,00 | 4,14 | 0,00 | 1,00E+00 |
| AT2G04230 | 1,09 | 1,06 | 2,19 | 0,48 | 6,28E-01 |
| AT2G04360 | 0,70 | 3,33 | 2,45 | 1,36 | 1,74E-01 |
| AT2G04378 | 0,56 | 1,26 | 2,77 | 0,45 | 6,50E-01 |
| AT2G04520 | 0,26 | -1,28 | 4,37 | -0,29 | 7,69E-01 |
| AT2G04620 | 0,17 | 0,00 | 4,41 | 0,00 | 1,00E+00 |
| AT2G04630 | 0,19 | 0,00 | 4,41 | 0,00 | 1,00E+00 |
| AT2G04842 | 0,88 | -0,63 | 3,16 | -0,20 | 8,41E-01 |
| AT2G04870 | 0,55 | 0,14 | 2,74 | 0,05 | 9,58E-01 |
| AT2G04890 | 0,11 | 0,00 | 4,41 | 0,00 | 1,00E+00 |
| AT2G05120 | 0,91 | 1,18 | 2,22 | 0,53 | 5,94E-01 |
| AT2G05220 | 0,38 | 1,48 | 4,40 | 0,34 | 7,36E-01 |
| AT2G05250 | 0,41 | 1,89 | 4,39 | 0,43 | 6,67E-01 |
| AT2G04895 | 0,17 | 0,00 | 4,41 | 0,00 | 1,00E+00 |
| AT2G05752 | 0,90 | 1,38 | 2,51 | 0,55 | 5,83E-01 |
| AT2G04995 | 1,03 | -1,60 | 2,27 | -0,70 | 4,83E-01 |
| AT2G05810 | 1,10 | 1,66 | 2,06 | 0,80 | 4,21E-01 |
| AT2G05914 | 0,51 | 0,00 | 4,39 | 0,00 | 1,00E+00 |
| AT2G05920 | 1,42 | -1,84 | 1,87 | -0,98 | 3,26E-01 |
| AT2G05940 | 0,97 | 0,46 | 2,21 | 0,21 | 8,35E-01 |
| AT2G06000 | 0,52 | 2,08 | 2,68 | 0,78 | 4,37E-01 |
| AT2G06040 | 0,24 | -0,47 | 4,40 | -0,11 | 9,14E-01 |
| AT2G06050 | 0,61 | -0,49 | 3,18 | -0,15 | 8,77E-01 |
| AT2G06530 | 0,61 | -1,41 | 3,36 | -0,42 | 6,73E-01 |
| AT2G06541 | 0,49 | 0,00 | 4,19 | 0,00 | 1,00E+00 |
| AT2G06645 | 0,15 | -0,43 | 4,39 | -0,10 | 9,22E-01 |
| AT2G06675 | 0,19 | 0,00 | 4,41 | 0,00 | 1,00E+00 |

| | | | | | |
|---|---|---|---|---|---|
| AT2G06906 | 0,12 | 0,00 | 4,41 | 0,00 | 1,00E+00 |
| AT2G06925 | 0,35 | -1,36 | 3,55 | -0,38 | 7,01E-01 |
| AT2G06960 | 0,68 | -0,44 | 2,94 | -0,15 | 8,82E-01 |
| AT2G07020 | 0,35 | 0,00 | 4,41 | 0,00 | 1,00E+00 |
| AT2G07120 | 0,12 | 0,00 | 4,41 | 0,00 | 1,00E+00 |
| AT2G07180 | 1,49 | 1,59 | 1,90 | 0,84 | 4,03E-01 |
| AT2G07240 | 1,04 | -2,17 | 2,06 | -1,05 | 2,93E-01 |
| AT2G07560 | 0,12 | 0,00 | 4,41 | 0,00 | 1,00E+00 |
| AT2G07771 | 0,58 | 1,26 | 2,71 | 0,46 | 6,43E-01 |
| AT2G07777 | 0,89 | -0,70 | 2,10 | -0,33 | 7,39E-01 |
| AT2G07779 | 0,28 | 1,67 | 4,38 | 0,38 | 7,03E-01 |
| AT2G07621 | 0,12 | 0,00 | 4,41 | 0,00 | 1,00E+00 |
| AT2G07774 | 0,25 | -0,85 | 4,38 | -0,19 | 8,47E-01 |
| AT2G07687 | 1,11 | -0,75 | 2,05 | -0,37 | 7,15E-01 |
| AT2G05205 | 0,68 | -1,96 | 2,61 | -0,75 | 4,53E-01 |
| AT2G07599 | 0,90 | -0,99 | 2,28 | -0,43 | 6,64E-01 |
| AT2G07706 | 1,10 | -3,08 | 2,37 | -1,30 | 1,93E-01 |
| AT2G07641 | 0,45 | 0,42 | 2,88 | 0,14 | 8,85E-01 |
| AT2G07719 | 0,62 | -0,35 | 2,69 | -0,13 | 8,97E-01 |
| AT2G07756 | 1,02 | -1,08 | 1,99 | -0,54 | 5,89E-01 |
| AT2G07734 | 0,71 | 0,61 | 2,85 | 0,21 | 8,30E-01 |
| AT2G07658 | 0,65 | 0,44 | 2,63 | 0,17 | 8,68E-01 |
| AT2G07662 | 1,06 | -2,19 | 2,30 | -0,95 | 3,41E-01 |
| AT2G07835 | 0,78 | -0,39 | 2,99 | -0,13 | 8,97E-01 |
| AT2G07690 | 0,71 | 1,93 | 2,60 | 0,74 | 4,58E-01 |
| AT2G07760 | 0,23 | -0,40 | 4,38 | -0,09 | 9,28E-01 |
| AT2G09795 | 0,34 | 1,46 | 4,40 | 0,33 | 7,39E-01 |
| AT2G05465 | 0,27 | 0,00 | 4,41 | 0,00 | 1,00E+00 |
| AT2G10557 | 1,21 | 0,00 | 3,79 | 0,00 | 1,00E+00 |
| AT2G10608 | 0,36 | 0,97 | 4,37 | 0,22 | 8,25E-01 |
| AT2G05565 | 0,43 | 1,35 | 4,13 | 0,33 | 7,43E-01 |
| AT2G10625 | 0,25 | 0,00 | 4,41 | 0,00 | 1,00E+00 |
| AT2G10950 | 0,47 | 0,00 | 4,04 | 0,00 | 1,00E+00 |
| AT2G10955 | 0,26 | 0,00 | 4,39 | 0,00 | 1,00E+00 |
| AT2G11000 | 0,08 | 0,00 | 4,41 | 0,00 | 1,00E+00 |
| AT2G11852 | 0,26 | 0,00 | 4,41 | 0,00 | 1,00E+00 |
| AT2G11910 | 1,43 | -0,28 | 1,81 | -0,15 | 8,77E-01 |
| AT2G12400 | 0,28 | -0,54 | 4,41 | -0,12 | 9,02E-01 |
| AT2G12405 | 0,18 | -0,43 | 4,39 | -0,10 | 9,21E-01 |
| AT2G05705 | 0,21 | 1,38 | 4,41 | 0,31 | 7,54E-01 |
| AT2G12461 | 0,12 | 0,00 | 4,41 | 0,00 | 1,00E+00 |
| AT2G05835 | 0,29 | -0,40 | 4,38 | -0,09 | 9,28E-01 |
| AT2G12945 | 0,05 | 0,00 | 4,41 | 0,00 | 1,00E+00 |
| AT2G13370 | 1,20 | 1,74 | 2,42 | 0,72 | 4,71E-01 |
| AT2G13450 | 1,51 | -1,10 | 1,62 | -0,68 | 4,95E-01 |
| AT2G13550 | 0,22 | 0,00 | 4,41 | 0,00 | 1,00E+00 |
| AT2G13640 | 0,11 | 0,00 | 4,41 | 0,00 | 1,00E+00 |
| AT2G13650 | 0,22 | 1,37 | 4,41 | 0,31 | 7,56E-01 |
| AT2G13820 | 1,49 | 0,00 | 2,95 | 0,00 | 1,00E+00 |
| AT2G14100 | 0,15 | 0,00 | 4,41 | 0,00 | 1,00E+00 |
| AT2G14120 | 0,98 | 2,10 | 2,23 | 0,94 | 3,47E-01 |
| AT2G14170 | 1,22 | -2,13 | 2,18 | -0,98 | 3,28E-01 |
| AT2G14260 | 0,26 | 0,00 | 4,41 | 0,00 | 1,00E+00 |
| AT2G14265 | 1,52 | -1,71 | 1,90 | -0,90 | 3,70E-01 |
| AT2G14288 | 0,20 | 0,00 | 4,41 | 0,00 | 1,00E+00 |
| AT2G14510 | 0,15 | 0,00 | 4,41 | 0,00 | 1,00E+00 |

| | | | | | |
|---|---|---|---|---|---|
| AT2G14540 | 0,12 | 0,00 | 4,41 | 0,00 | 1,00E+00 |
| AT2G14670 | 1,21 | -2,19 | 2,54 | -0,86 | 3,89E-01 |
| AT2G14720 | 1,05 | 2,85 | 2,39 | 1,19 | 2,33E-01 |
| AT2G14755 | 0,32 | 0,00 | 4,41 | 0,00 | 1,00E+00 |
| AT2G15000 | 0,23 | 1,38 | 4,41 | 0,31 | 7,54E-01 |
| AT2G15020 | 0,51 | -0,21 | 3,24 | -0,07 | 9,48E-01 |
| AT2G06355 | 0,23 | -0,50 | 4,40 | -0,11 | 9,10E-01 |
| AT2G15110 | 0,63 | -0,54 | 3,09 | -0,18 | 8,60E-01 |
| AT2G15290 | 0,58 | 1,02 | 3,78 | 0,27 | 7,87E-01 |
| AT2G15292 | 1,13 | 0,38 | 1,86 | 0,20 | 8,39E-01 |
| AT2G15320 | 1,25 | -0,26 | 1,89 | -0,14 | 8,91E-01 |
| AT2G15390 | 0,37 | 1,47 | 4,40 | 0,33 | 7,39E-01 |
| AT2G15555 | 1,03 | -0,19 | 1,83 | -0,11 | 9,15E-01 |
| AT2G15560 | 0,05 | 0,00 | 4,41 | 0,00 | 1,00E+00 |
| AT2G15790 | 0,41 | 1,24 | 4,21 | 0,29 | 7,68E-01 |
| AT2G15860 | 0,36 | -0,47 | 4,40 | -0,11 | 9,16E-01 |
| AT2G15880 | 1,49 | 2,09 | 1,71 | 1,22 | 2,21E-01 |
| AT2G15890 | 1,14 | 1,33 | 1,84 | 0,72 | 4,71E-01 |
| AT2G06515 | 0,69 | -1,54 | 3,66 | -0,42 | 6,73E-01 |
| AT2G15980 | 0,29 | -1,22 | 3,68 | -0,33 | 7,40E-01 |
| AT2G16016 | 1,06 | 0,47 | 2,22 | 0,21 | 8,31E-01 |
| AT2G06575 | 0,75 | 0,45 | 2,30 | 0,19 | 8,46E-01 |
| AT2G16070 | 0,64 | 0,97 | 3,84 | 0,25 | 8,00E-01 |
| AT2G16090 | 0,55 | 2,08 | 2,61 | 0,80 | 4,26E-01 |
| AT2G16370 | 0,11 | 0,00 | 4,41 | 0,00 | 1,00E+00 |
| AT2G16400 | 1,00 | -0,39 | 2,10 | -0,19 | 8,53E-01 |
| AT2G16405 | 0,20 | -0,44 | 4,39 | -0,10 | 9,20E-01 |
| AT2G16440 | 1,04 | 3,31 | 3,52 | 0,94 | 3,47E-01 |
| AT2G16450 | 0,33 | 1,85 | 4,39 | 0,42 | 6,73E-01 |
| AT2G16490 | 1,37 | -1,61 | 2,30 | -0,70 | 4,84E-01 |
| AT2G16505 | 0,85 | -1,25 | 2,36 | -0,53 | 5,95E-01 |
| AT2G16510 | 1,63 | -1,07 | 1,87 | -0,57 | 5,66E-01 |
| AT2G16620 | 0,18 | 0,00 | 4,41 | 0,00 | 1,00E+00 |
| AT2G16640 | 1,05 | 2,17 | 2,30 | 0,94 | 3,45E-01 |
| AT2G16650 | 0,37 | 0,00 | 4,41 | 0,00 | 1,00E+00 |
| AT2G16870 | 1,12 | 3,29 | 2,04 | 1,61 | 1,07E-01 |
| AT2G16880 | 0,51 | 1,98 | 4,39 | 0,45 | 6,52E-01 |
| AT2G16920 | 0,25 | -0,54 | 4,41 | -0,12 | 9,02E-01 |
| AT2G16940 | 0,88 | 2,13 | 2,25 | 0,95 | 3,44E-01 |
| AT2G17020 | 0,78 | 1,32 | 2,14 | 0,62 | 5,37E-01 |
| AT2G17110 | 0,49 | 0,15 | 2,78 | 0,05 | 9,57E-01 |
| AT2G17130 | 0,45 | 2,52 | 4,20 | 0,60 | 5,48E-01 |
| AT2G17190 | 0,94 | -0,48 | 2,55 | -0,19 | 8,50E-01 |
| AT2G17210 | 0,23 | 1,38 | 4,41 | 0,31 | 7,55E-01 |
| AT2G17220 | 0,39 | 2,60 | 4,35 | 0,60 | 5,49E-01 |
| AT2G17230 | 0,92 | -0,43 | 2,62 | -0,16 | 8,70E-01 |
| AT2G17300 | 0,17 | 0,00 | 4,41 | 0,00 | 1,00E+00 |
| AT2G06935 | 0,05 | 0,00 | 4,41 | 0,00 | 1,00E+00 |
| AT2G17340 | 0,86 | 1,14 | 2,32 | 0,49 | 6,22E-01 |
| AT2G17350 | 0,34 | -1,19 | 4,36 | -0,27 | 7,85E-01 |
| AT2G17450 | 0,63 | 2,66 | 2,34 | 1,14 | 2,55E-01 |
| AT2G17470 | 0,10 | 0,00 | 4,41 | 0,00 | 1,00E+00 |
| AT2G17550 | 0,71 | 1,38 | 2,59 | 0,53 | 5,94E-01 |
| AT2G17560 | 0,46 | 2,11 | 4,37 | 0,48 | 6,29E-01 |
| AT2G17700 | 0,43 | 0,35 | 4,11 | 0,09 | 9,32E-01 |
| AT2G17820 | 0,96 | -0,27 | 2,16 | -0,13 | 8,99E-01 |

| | | | | | |
|---|---|---|---|---|---|
| AT2G17870 | 0,45 | 2,24 | 4,15 | 0,54 | 5,88E-01 |
| AT2G18130 | 0,22 | 0,00 | 4,41 | 0,00 | 1,00E+00 |
| AT2G18160 | 1,66 | 3,62 | 1,83 | 1,98 | 4,81E-02 |
| AT2G18170 | 0,50 | 2,60 | 3,26 | 0,80 | 4,25E-01 |
| AT2G18193 | 1,29 | 3,96 | 2,10 | 1,88 | 5,96E-02 |
| AT2G18290 | 0,19 | 0,00 | 4,41 | 0,00 | 1,00E+00 |
| AT2G18300 | 1,33 | -0,04 | 2,00 | -0,02 | 9,85E-01 |
| AT2G18310 | 0,43 | 0,00 | 4,36 | 0,00 | 1,00E+00 |
| AT2G18520 | 0,25 | 1,84 | 4,38 | 0,42 | 6,75E-01 |
| AT2G18600 | 0,24 | 1,96 | 4,39 | 0,45 | 6,55E-01 |
| AT2G18660 | 0,89 | 0,37 | 2,33 | 0,16 | 8,75E-01 |
| AT2G18685 | 0,08 | 0,00 | 4,41 | 0,00 | 1,00E+00 |
| AT2G18900 | 1,01 | 3,51 | 1,99 | 1,76 | 7,82E-02 |
| AT2G18915 | 0,29 | 1,24 | 4,39 | 0,28 | 7,77E-01 |
| AT2G18940 | 0,31 | -0,42 | 3,62 | -0,12 | 9,08E-01 |
| AT2G19120 | 0,71 | 1,77 | 3,51 | 0,51 | 6,13E-01 |
| AT2G07135 | 0,16 | 0,00 | 4,41 | 0,00 | 1,00E+00 |
| AT2G07165 | 0,21 | 0,00 | 4,41 | 0,00 | 1,00E+00 |
| AT2G19290 | 0,05 | 0,00 | 4,41 | 0,00 | 1,00E+00 |
| AT2G19310 | 0,12 | 0,00 | 4,41 | 0,00 | 1,00E+00 |
| AT2G19340 | 0,43 | 0,00 | 4,40 | 0,00 | 1,00E+00 |
| AT2G19365 | 0,81 | -1,23 | 2,18 | -0,57 | 5,72E-01 |
| AT2G19390 | 0,76 | 1,38 | 2,54 | 0,54 | 5,87E-01 |
| AT2G07195 | 0,51 | 1,38 | 2,77 | 0,50 | 6,19E-01 |
| AT2G19400 | 0,15 | 0,00 | 4,39 | 0,00 | 1,00E+00 |
| AT2G19470 | 0,28 | 1,37 | 4,41 | 0,31 | 7,56E-01 |
| AT2G19560 | 0,20 | 0,00 | 4,41 | 0,00 | 1,00E+00 |
| AT2G19570 | 0,52 | 0,00 | 3,17 | 0,00 | 1,00E+00 |
| AT2G19580 | 0,25 | 0,00 | 4,41 | 0,00 | 1,00E+00 |
| AT2G19760 | 0,25 | 0,00 | 4,41 | 0,00 | 1,00E+00 |
| AT2G19780 | 0,19 | -0,54 | 4,41 | -0,12 | 9,02E-01 |
| AT2G19910 | 0,11 | 0,00 | 4,41 | 0,00 | 1,00E+00 |
| AT2G20000 | 0,44 | 0,43 | 3,38 | 0,13 | 8,98E-01 |
| AT2G20050 | 0,77 | 1,34 | 2,52 | 0,53 | 5,94E-01 |
| AT2G20120 | 0,38 | 2,11 | 4,36 | 0,48 | 6,29E-01 |
| AT2G20180 | 0,53 | -1,32 | 2,69 | -0,49 | 6,24E-01 |
| AT2G20240 | 1,31 | 1,00 | 2,69 | 0,37 | 7,12E-01 |
| AT2G20270 | 0,73 | 0,10 | 3,12 | 0,03 | 9,76E-01 |
| AT2G20280 | 0,90 | 2,68 | 2,73 | 0,98 | 3,25E-01 |
| AT2G20300 | 0,57 | -0,40 | 2,75 | -0,15 | 8,85E-01 |
| AT2G20310 | 0,15 | 0,00 | 4,41 | 0,00 | 1,00E+00 |
| AT2G20380 | 0,08 | 0,00 | 4,41 | 0,00 | 1,00E+00 |
| AT2G20562 | 0,11 | 0,00 | 4,41 | 0,00 | 1,00E+00 |
| AT2G20610 | 0,62 | 0,88 | 2,56 | 0,35 | 7,30E-01 |
| AT2G20630 | 0,57 | -0,60 | 2,62 | -0,23 | 8,19E-01 |
| AT2G20670 | 0,99 | 1,93 | 1,93 | 1,00 | 3,17E-01 |
| AT2G07325 | 0,08 | 0,00 | 4,41 | 0,00 | 1,00E+00 |
| AT2G20720 | 0,50 | 1,23 | 3,25 | 0,38 | 7,05E-01 |
| AT2G20723 | 1,21 | -1,12 | 1,90 | -0,59 | 5,55E-01 |
| AT2G20725 | 0,17 | 0,00 | 4,41 | 0,00 | 1,00E+00 |
| AT2G20760 | 1,00 | -0,06 | 2,26 | -0,03 | 9,80E-01 |
| AT2G20790 | 0,65 | 1,40 | 3,78 | 0,37 | 7,12E-01 |
| AT2G20825 | 0,24 | 1,31 | 4,40 | 0,30 | 7,65E-01 |
| AT2G20830 | 0,42 | 1,28 | 3,35 | 0,38 | 7,02E-01 |
| AT2G20850 | 0,80 | 0,65 | 3,45 | 0,19 | 8,50E-01 |
| AT2G20900 | 0,70 | 3,22 | 2,98 | 1,08 | 2,81E-01 |

| | | | | | |
|---|---|---|---|---|---|
| AT2G21060 | 0,37 | -0,40 | 4,38 | -0,09 | 9,28E-01 |
| AT2G21090 | 0,12 | 0,00 | 4,41 | 0,00 | 1,00E+00 |
| AT2G21120 | 1,14 | 3,52 | 1,88 | 1,87 | 6,09E-02 |
| AT2G21170 | 1,20 | 0,12 | 1,63 | 0,08 | 9,39E-01 |
| AT2G21185 | 0,86 | 1,20 | 2,08 | 0,58 | 5,65E-01 |
| AT2G21188 | 0,47 | 1,19 | 4,25 | 0,28 | 7,80E-01 |
| AT2G21210 | 1,12 | 3,16 | 2,15 | 1,47 | 1,42E-01 |
| AT2G21240 | 0,62 | -1,49 | 2,42 | -0,61 | 5,39E-01 |
| AT2G21280 | 0,38 | 2,48 | 4,35 | 0,57 | 5,69E-01 |
| AT2G21300 | 0,99 | 1,21 | 2,37 | 0,51 | 6,10E-01 |
| AT2G21350 | 0,05 | 0,00 | 4,41 | 0,00 | 1,00E+00 |
| AT2G21370 | 0,82 | 1,56 | 2,09 | 0,75 | 4,55E-01 |
| AT2G21440 | 0,17 | 0,00 | 4,41 | 0,00 | 1,00E+00 |
| AT2G21480 | 0,25 | 1,34 | 4,40 | 0,30 | 7,61E-01 |
| AT2G21500 | 0,51 | 1,26 | 3,16 | 0,40 | 6,90E-01 |
| AT2G21560 | 0,27 | 0,00 | 4,41 | 0,00 | 1,00E+00 |
| AT2G21870 | 1,17 | 0,99 | 2,03 | 0,49 | 6,25E-01 |
| AT2G21950 | 0,24 | 0,00 | 4,41 | 0,00 | 1,00E+00 |
| AT2G21970 | 0,61 | -1,42 | 3,85 | -0,37 | 7,12E-01 |
| AT2G22010 | 0,65 | 0,55 | 2,62 | 0,21 | 8,33E-01 |
| AT2G22090 | 0,34 | 0,38 | 4,36 | 0,09 | 9,31E-01 |
| AT2G22140 | 0,05 | 0,00 | 4,41 | 0,00 | 1,00E+00 |
| AT2G22160 | 0,35 | 0,00 | 4,41 | 0,00 | 1,00E+00 |
| AT2G22240 | 0,21 | -0,54 | 4,41 | -0,12 | 9,02E-01 |
| AT2G22250 | 0,26 | 1,38 | 4,41 | 0,31 | 7,54E-01 |
| AT2G07555 | 0,37 | -0,41 | 4,34 | -0,10 | 9,24E-01 |
| AT2G22400 | 0,55 | 2,15 | 3,83 | 0,56 | 5,74E-01 |
| AT2G22470 | 0,19 | 0,00 | 4,41 | 0,00 | 1,00E+00 |
| AT2G22490 | 1,17 | 4,06 | 2,33 | 1,74 | 8,13E-02 |
| AT2G22740 | 0,43 | 0,00 | 4,18 | 0,00 | 1,00E+00 |
| AT2G22850 | 0,76 | -2,37 | 2,05 | -1,16 | 2,48E-01 |
| AT2G22890 | 0,53 | 2,15 | 3,13 | 0,69 | 4,92E-01 |
| AT2G22940 | 0,87 | 0,00 | 2,80 | 0,00 | 1,00E+00 |
| AT2G23070 | 0,53 | -1,28 | 2,36 | -0,54 | 5,87E-01 |
| AT2G23090 | 0,40 | 2,30 | 4,32 | 0,53 | 5,93E-01 |
| AT2G23100 | 0,16 | 0,00 | 4,41 | 0,00 | 1,00E+00 |
| AT2G23140 | 0,38 | 0,00 | 4,32 | 0,00 | 1,00E+00 |
| AT2G23210 | 0,71 | 1,85 | 2,94 | 0,63 | 5,29E-01 |
| AT2G23290 | 0,64 | 0,21 | 2,72 | 0,08 | 9,38E-01 |
| AT2G23390 | 0,32 | -0,41 | 4,38 | -0,09 | 9,26E-01 |
| AT2G23430 | 0,47 | 1,31 | 3,45 | 0,38 | 7,05E-01 |
| AT2G23450 | 0,62 | 0,73 | 2,27 | 0,32 | 7,49E-01 |
| AT2G23520 | 0,43 | 2,37 | 3,39 | 0,70 | 4,83E-01 |
| AT2G07910 | 1,63 | -3,07 | 2,35 | -1,30 | 1,93E-01 |
| AT2G23550 | 1,59 | -3,06 | 2,37 | -1,29 | 1,96E-01 |
| AT2G23600 | 1,45 | 1,34 | 1,80 | 0,74 | 4,59E-01 |
| AT2G23610 | 0,63 | 1,07 | 3,07 | 0,35 | 7,28E-01 |
| AT2G23670 | 1,50 | -1,19 | 1,73 | -0,69 | 4,93E-01 |
| AT2G23760 | 0,34 | 0,39 | 4,40 | 0,09 | 9,29E-01 |
| AT2G23790 | 0,79 | -0,41 | 3,05 | -0,13 | 8,93E-01 |
| AT2G23840 | 0,70 | -0,02 | 2,99 | -0,01 | 9,94E-01 |
| AT2G23987 | 0,17 | -1,42 | 4,35 | -0,33 | 7,45E-01 |
| AT2G24120 | 0,81 | -0,76 | 2,27 | -0,33 | 7,38E-01 |
| AT2G24150 | 0,79 | 2,00 | 2,19 | 0,91 | 3,62E-01 |
| AT2G24170 | 0,17 | 0,00 | 4,41 | 0,00 | 1,00E+00 |
| AT2G24200 | 1,48 | -0,24 | 1,99 | -0,12 | 9,03E-01 |

| | | | | | |
|---|---|---|---|---|---|
| AT2G24230 | 0,59 | 0,19 | 2,67 | 0,07 | 9,42E-01 |
| AT2G24330 | 0,33 | 0,50 | 4,37 | 0,11 | 9,10E-01 |
| AT2G24340 | 1,46 | 0,43 | 2,06 | 0,21 | 8,34E-01 |
| AT2G24360 | 1,19 | 1,48 | 1,98 | 0,75 | 4,53E-01 |
| AT2G24500 | 1,13 | 0,11 | 2,18 | 0,05 | 9,58E-01 |
| AT2G08055 | 0,33 | 1,35 | 4,40 | 0,31 | 7,59E-01 |
| AT2G24545 | 1,16 | 1,15 | 1,68 | 0,69 | 4,93E-01 |
| AT2G24550 | 1,50 | 0,34 | 1,99 | 0,17 | 8,63E-01 |
| AT2G24600 | 1,31 | 1,38 | 2,48 | 0,56 | 5,79E-01 |
| AT2G24640 | 0,24 | 1,37 | 4,41 | 0,31 | 7,56E-01 |
| AT2G24710 | 0,19 | 0,00 | 4,41 | 0,00 | 1,00E+00 |
| AT2G24755 | 0,71 | 2,06 | 2,68 | 0,77 | 4,41E-01 |
| AT2G24780 | 0,92 | -0,87 | 2,28 | -0,38 | 7,01E-01 |
| AT2G24850 | 1,45 | 2,40 | 1,85 | 1,29 | 1,96E-01 |
| AT2G24870 | 1,09 | -0,89 | 2,43 | -0,36 | 7,16E-01 |
| AT2G24880 | 1,30 | -1,26 | 1,64 | -0,77 | 4,44E-01 |
| AT2G25011 | 1,24 | -1,03 | 2,00 | -0,51 | 6,07E-01 |
| AT2G25050 | 0,47 | -2,25 | 3,21 | -0,70 | 4,84E-01 |
| AT2G25070 | 0,45 | 0,48 | 2,88 | 0,17 | 8,68E-01 |
| AT2G08105 | 0,74 | -1,55 | 2,39 | -0,65 | 5,15E-01 |
| AT2G25180 | 0,76 | -2,14 | 3,06 | -0,70 | 4,85E-01 |
| AT2G25200 | 0,69 | -1,92 | 3,85 | -0,50 | 6,18E-01 |
| AT2G25300 | 0,15 | -0,43 | 4,39 | -0,10 | 9,23E-01 |
| AT2G25310 | 0,20 | 1,40 | 4,41 | 0,32 | 7,51E-01 |
| AT2G25570 | 0,45 | 1,38 | 4,41 | 0,31 | 7,54E-01 |
| AT2G25610 | 0,53 | 2,49 | 3,13 | 0,80 | 4,26E-01 |
| AT2G25670 | 0,93 | -0,46 | 2,44 | -0,19 | 8,51E-01 |
| AT2G25760 | 0,93 | 1,97 | 2,85 | 0,69 | 4,89E-01 |
| AT2G25850 | 0,16 | 0,00 | 4,41 | 0,00 | 1,00E+00 |
| AT2G25910 | 0,91 | -1,39 | 2,49 | -0,56 | 5,77E-01 |
| AT2G25970 | 1,54 | 2,87 | 1,92 | 1,50 | 1,34E-01 |
| AT2G26030 | 0,31 | -0,44 | 3,69 | -0,12 | 9,06E-01 |
| AT2G26070 | 0,39 | 1,79 | 4,37 | 0,41 | 6,81E-01 |
| AT2G26140 | 1,61 | 0,99 | 1,58 | 0,63 | 5,32E-01 |
| AT2G26190 | 1,38 | 2,21 | 1,92 | 1,15 | 2,50E-01 |
| AT2G26280 | 1,11 | 3,55 | 2,40 | 1,48 | 1,39E-01 |
| AT2G26430 | 0,38 | 1,52 | 4,32 | 0,35 | 7,26E-01 |
| AT2G26540 | 0,42 | 2,45 | 4,33 | 0,56 | 5,73E-01 |
| AT2G26560 | 1,67 | 1,18 | 1,88 | 0,63 | 5,31E-01 |
| AT2G26580 | 0,26 | -0,56 | 4,41 | -0,13 | 9,00E-01 |
| AT2G26590 | 0,25 | -0,50 | 4,40 | -0,11 | 9,10E-01 |
| AT2G26620 | 0,18 | 0,00 | 4,41 | 0,00 | 1,00E+00 |
| AT2G26680 | 0,33 | 1,85 | 4,39 | 0,42 | 6,73E-01 |
| AT2G26690 | 0,68 | 0,41 | 2,69 | 0,15 | 8,79E-01 |
| AT2G26740 | 0,33 | 1,43 | 4,40 | 0,32 | 7,45E-01 |
| AT2G08175 | 1,14 | -0,02 | 1,95 | -0,01 | 9,90E-01 |
| AT2G26790 | 0,21 | 0,00 | 4,41 | 0,00 | 1,00E+00 |
| AT2G26800 | 0,41 | 0,36 | 3,46 | 0,10 | 9,17E-01 |
| AT2G26840 | 0,08 | 0,00 | 4,41 | 0,00 | 1,00E+00 |
| AT2G26900 | 1,34 | -1,13 | 1,77 | -0,64 | 5,22E-01 |
| AT2G26930 | 0,69 | -0,64 | 2,96 | -0,22 | 8,28E-01 |
| AT2G26980 | 0,61 | -0,42 | 2,41 | -0,17 | 8,63E-01 |
| AT2G26990 | 0,56 | 0,56 | 3,19 | 0,17 | 8,62E-01 |
| AT2G27020 | 0,74 | -1,57 | 2,60 | -0,60 | 5,47E-01 |
| AT2G27040 | 0,44 | 0,54 | 3,39 | 0,16 | 8,73E-01 |
| AT2G27070 | 0,56 | -2,71 | 3,01 | -0,90 | 3,69E-01 |

| | | | | | |
|---|---|---|---|---|---|
| AT2G27090 | 0,77 | 1,25 | 3,14 | 0,40 | 6,90E-01 |
| AT2G27170 | 0,61 | -2,13 | 2,59 | -0,82 | 4,11E-01 |
| AT2G27190 | 0,39 | -1,43 | 4,33 | -0,33 | 7,40E-01 |
| AT2G27290 | 0,57 | -1,27 | 2,76 | -0,46 | 6,45E-01 |
| AT2G27330 | 0,10 | 0,00 | 4,41 | 0,00 | 1,00E+00 |
| AT2G27350 | 0,23 | 0,00 | 4,41 | 0,00 | 1,00E+00 |
| AT2G27385 | 1,03 | -1,71 | 2,09 | -0,82 | 4,15E-01 |
| AT2G27500 | 0,69 | 2,12 | 2,60 | 0,81 | 4,15E-01 |
| AT2G27510 | 0,44 | 0,36 | 4,25 | 0,08 | 9,32E-01 |
| AT2G27530 | 0,92 | 0,88 | 1,93 | 0,45 | 6,50E-01 |
| AT2G27580 | 0,43 | 0,00 | 4,27 | 0,00 | 1,00E+00 |
| AT2G27660 | 0,41 | 2,57 | 4,35 | 0,59 | 5,55E-01 |
| AT2G27680 | 0,72 | 0,59 | 2,07 | 0,29 | 7,74E-01 |
| AT2G27770 | 0,38 | 0,00 | 4,41 | 0,00 | 1,00E+00 |
| AT2G27950 | 1,06 | 3,42 | 2,02 | 1,69 | 9,09E-02 |
| AT2G28080 | 0,87 | 1,38 | 2,91 | 0,47 | 6,36E-01 |
| AT2G28170 | 0,05 | 0,00 | 4,41 | 0,00 | 1,00E+00 |
| AT2G28360 | 1,42 | 2,38 | 1,48 | 1,60 | 1,09E-01 |
| AT2G28380 | 0,74 | 0,00 | 3,02 | 0,00 | 1,00E+00 |
| AT2G28390 | 0,41 | 2,60 | 4,35 | 0,60 | 5,50E-01 |
| AT2G28400 | 0,81 | -0,24 | 2,47 | -0,10 | 9,24E-01 |
| AT2G28450 | 1,48 | 4,15 | 1,89 | 2,20 | 2,76E-02 |
| AT2G28470 | 0,87 | -0,49 | 2,89 | -0,17 | 8,64E-01 |
| AT2G28480 | 0,12 | 0,00 | 4,41 | 0,00 | 1,00E+00 |
| AT2G28620 | 1,12 | 0,00 | 2,38 | 0,00 | 1,00E+00 |
| AT2G08275 | 1,12 | 0,00 | 2,38 | 0,00 | 1,00E+00 |
| AT2G28630 | 0,37 | -0,54 | 4,34 | -0,13 | 9,00E-01 |
| AT2G28640 | 0,62 | -0,44 | 3,01 | -0,15 | 8,84E-01 |
| AT2G28670 | 0,51 | -0,46 | 3,16 | -0,15 | 8,85E-01 |
| AT2G28740 | 0,44 | 0,00 | 4,23 | 0,00 | 1,00E+00 |
| AT2G28800 | 0,62 | -0,91 | 3,09 | -0,30 | 7,67E-01 |
| AT2G08300 | 0,20 | 1,40 | 4,41 | 0,32 | 7,51E-01 |
| AT2G28880 | 0,33 | 1,33 | 4,40 | 0,30 | 7,62E-01 |
| AT2G28890 | 0,77 | 2,14 | 2,30 | 0,93 | 3,52E-01 |
| AT2G28900 | 0,81 | -1,42 | 2,59 | -0,55 | 5,82E-01 |
| AT2G28950 | 0,45 | 1,04 | 4,13 | 0,25 | 8,01E-01 |
| AT2G29030 | 0,32 | 1,38 | 4,41 | 0,31 | 7,54E-01 |
| AT2G29040 | 0,08 | 0,00 | 4,41 | 0,00 | 1,00E+00 |
| AT2G29110 | 0,41 | 1,67 | 4,16 | 0,40 | 6,88E-01 |
| AT2G29120 | 1,16 | 0,37 | 2,34 | 0,16 | 8,74E-01 |
| AT2G29170 | 0,18 | 0,00 | 4,41 | 0,00 | 1,00E+00 |
| AT2G08330 | 0,26 | 1,38 | 4,41 | 0,31 | 7,54E-01 |
| AT2G29190 | 0,61 | 2,33 | 2,69 | 0,87 | 3,86E-01 |
| AT2G29220 | 0,39 | 1,79 | 4,37 | 0,41 | 6,81E-01 |
| AT2G29390 | 0,71 | 0,39 | 2,58 | 0,15 | 8,80E-01 |
| AT2G29400 | 0,98 | 2,87 | 1,98 | 1,45 | 1,48E-01 |
| AT2G29550 | 0,75 | -0,54 | 2,53 | -0,22 | 8,29E-01 |
| AT2G29570 | 0,14 | 0,00 | 4,41 | 0,00 | 1,00E+00 |
| AT2G29605 | 0,65 | -0,54 | 3,01 | -0,18 | 8,56E-01 |
| AT2G29650 | 0,77 | -0,10 | 2,50 | -0,04 | 9,68E-01 |
| AT2G30250 | 0,71 | 1,91 | 2,31 | 0,83 | 4,08E-01 |
| AT2G30260 | 0,44 | 0,53 | 3,33 | 0,16 | 8,72E-01 |
| AT2G30270 | 0,18 | 0,00 | 4,41 | 0,00 | 1,00E+00 |
| AT2G30360 | 0,37 | 0,00 | 4,41 | 0,00 | 1,00E+00 |
| AT2G30390 | 0,49 | 1,81 | 3,28 | 0,55 | 5,82E-01 |
| AT2G30470 | 0,73 | -0,30 | 2,32 | -0,13 | 8,97E-01 |

| | | | | | |
|---|---|---|---|---|---|
| AT2G30490 | 0,91 | 0,54 | 3,12 | 0,17 | 8,63E-01 |
| AT2G30500 | 1,56 | 0,81 | 1,93 | 0,42 | 6,74E-01 |
| AT2G30505 | 0,35 | -0,64 | 4,38 | -0,15 | 8,84E-01 |
| AT2G30700 | 0,36 | -0,51 | 4,36 | -0,12 | 9,07E-01 |
| AT2G30710 | 0,35 | 1,78 | 4,38 | 0,41 | 6,84E-01 |
| AT2G30720 | 0,49 | 0,93 | 3,29 | 0,28 | 7,78E-01 |
| AT2G30790 | 0,21 | 1,25 | 4,39 | 0,29 | 7,75E-01 |
| AT2G30800 | 1,27 | 3,04 | 1,90 | 1,60 | 1,10E-01 |
| AT2G30850 | 0,41 | 0,34 | 4,34 | 0,08 | 9,37E-01 |
| AT2G30880 | 0,34 | 1,53 | 4,39 | 0,35 | 7,27E-01 |
| AT2G08420 | 0,54 | -0,99 | 3,99 | -0,25 | 8,05E-01 |
| AT2G30980 | 0,41 | 2,47 | 4,36 | 0,57 | 5,70E-01 |
| AT2G31040 | 0,19 | -0,47 | 4,40 | -0,11 | 9,15E-01 |
| AT2G31060 | 0,49 | 0,00 | 4,41 | 0,00 | 1,00E+00 |
| AT2G31070 | 0,79 | 1,34 | 2,47 | 0,54 | 5,86E-01 |
| AT2G31130 | 0,17 | 0,00 | 4,41 | 0,00 | 1,00E+00 |
| AT2G31170 | 0,45 | -1,23 | 4,38 | -0,28 | 7,79E-01 |
| AT2G31240 | 0,25 | 1,34 | 4,40 | 0,30 | 7,61E-01 |
| AT2G31250 | 0,39 | 1,76 | 4,38 | 0,40 | 6,87E-01 |
| AT2G31260 | 0,53 | -0,61 | 2,76 | -0,22 | 8,24E-01 |
| AT2G08430 | 0,08 | 0,00 | 4,41 | 0,00 | 1,00E+00 |
| AT2G31360 | 1,30 | -0,02 | 1,99 | -0,01 | 9,93E-01 |
| AT2G31390 | 0,86 | 2,77 | 2,88 | 0,96 | 3,36E-01 |
| AT2G31410 | 0,37 | 2,72 | 4,34 | 0,63 | 5,31E-01 |
| AT2G31440 | 0,44 | 0,00 | 4,41 | 0,00 | 1,00E+00 |
| AT2G31510 | 0,63 | -1,25 | 3,08 | -0,41 | 6,84E-01 |
| AT2G31570 | 0,14 | 0,00 | 4,41 | 0,00 | 1,00E+00 |
| AT2G31670 | 0,73 | -1,59 | 2,56 | -0,62 | 5,34E-01 |
| AT2G31730 | 0,53 | 0,04 | 4,35 | 0,01 | 9,93E-01 |
| AT2G31740 | 0,52 | 1,90 | 3,17 | 0,60 | 5,49E-01 |
| AT2G31751 | 0,97 | -0,22 | 2,48 | -0,09 | 9,31E-01 |
| AT2G31800 | 0,85 | 1,35 | 2,25 | 0,60 | 5,47E-01 |
| AT2G31830 | 0,34 | 1,38 | 4,41 | 0,31 | 7,54E-01 |
| AT2G31865 | 0,37 | 1,31 | 4,40 | 0,30 | 7,65E-01 |
| AT2G31870 | 0,27 | 0,00 | 3,83 | 0,00 | 1,00E+00 |
| AT2G08475 | 0,81 | 1,05 | 2,50 | 0,42 | 6,73E-01 |
| AT2G31990 | 0,31 | 1,29 | 4,40 | 0,29 | 7,69E-01 |
| AT2G32000 | 0,55 | 2,89 | 2,61 | 1,10 | 2,69E-01 |
| AT2G32030 | 0,22 | 0,00 | 4,41 | 0,00 | 1,00E+00 |
| AT2G32050 | 0,08 | 0,00 | 4,41 | 0,00 | 1,00E+00 |
| AT2G32060 | 0,34 | 0,00 | 4,41 | 0,00 | 1,00E+00 |
| AT2G32070 | 0,64 | -0,54 | 2,69 | -0,20 | 8,40E-01 |
| AT2G32080 | 0,57 | 0,34 | 2,68 | 0,13 | 9,00E-01 |
| AT2G32130 | 0,28 | 0,00 | 4,41 | 0,00 | 1,00E+00 |
| AT2G32230 | 0,70 | 3,22 | 2,83 | 1,14 | 2,56E-01 |
| AT2G32235 | 0,12 | 0,00 | 4,41 | 0,00 | 1,00E+00 |
| AT2G32390 | 0,36 | 0,00 | 4,41 | 0,00 | 1,00E+00 |
| AT2G32400 | 0,29 | 0,00 | 4,41 | 0,00 | 1,00E+00 |
| AT2G32415 | 0,42 | 2,54 | 4,35 | 0,58 | 5,60E-01 |
| AT2G32480 | 0,87 | 0,30 | 2,22 | 0,13 | 8,93E-01 |
| AT2G32520 | 0,35 | -0,44 | 4,39 | -0,10 | 9,20E-01 |
| AT2G32600 | 0,08 | 0,00 | 4,41 | 0,00 | 1,00E+00 |
| AT2G32680 | 1,55 | 1,08 | 2,00 | 0,54 | 5,89E-01 |
| AT2G32810 | 0,70 | -2,10 | 2,51 | -0,84 | 4,02E-01 |
| AT2G32920 | 1,12 | 1,15 | 1,86 | 0,62 | 5,35E-01 |
| AT2G32940 | 0,12 | 0,00 | 4,41 | 0,00 | 1,00E+00 |

| | | | | | |
|---|---|---|---|---|---|
| AT2G33040 | 0,74 | -1,02 | 2,50 | -0,41 | 6,84E-01 |
| AT2G33051 | 0,64 | 1,30 | 3,86 | 0,34 | 7,36E-01 |
| AT2G33070 | 0,08 | 0,00 | 4,41 | 0,00 | 1,00E+00 |
| AT2G33110 | 0,40 | 0,00 | 4,41 | 0,00 | 1,00E+00 |
| AT2G33120 | 0,77 | 2,16 | 2,26 | 0,95 | 3,40E-01 |
| AT2G33175 | 0,67 | 0,53 | 2,55 | 0,21 | 8,36E-01 |
| AT2G33210 | 0,70 | 2,17 | 2,55 | 0,85 | 3,96E-01 |
| AT2G33260 | 0,24 | 1,31 | 4,40 | 0,30 | 7,65E-01 |
| AT2G33320 | 0,54 | 1,34 | 2,70 | 0,50 | 6,20E-01 |
| AT2G33370 | 0,46 | 0,52 | 2,86 | 0,18 | 8,56E-01 |
| AT2G33385 | 0,08 | 0,00 | 4,41 | 0,00 | 1,00E+00 |
| AT2G33460 | 0,28 | 1,38 | 4,41 | 0,31 | 7,54E-01 |
| AT2G33470 | 0,39 | 1,31 | 3,43 | 0,38 | 7,03E-01 |
| AT2G33500 | 0,51 | 1,49 | 4,32 | 0,34 | 7,31E-01 |
| AT2G33530 | 0,99 | 2,47 | 1,96 | 1,26 | 2,07E-01 |
| AT2G33540 | 0,96 | 1,35 | 2,03 | 0,67 | 5,04E-01 |
| AT2G33550 | 0,30 | 1,38 | 4,41 | 0,31 | 7,54E-01 |
| AT2G33580 | 0,75 | 1,01 | 2,34 | 0,43 | 6,68E-01 |
| AT2G33630 | 0,50 | 1,90 | 3,23 | 0,59 | 5,56E-01 |
| AT2G33655 | 0,29 | -1,22 | 4,36 | -0,28 | 7,80E-01 |
| AT2G33800 | 0,66 | -0,56 | 2,56 | -0,22 | 8,27E-01 |
| AT2G33815 | 0,37 | 0,37 | 4,34 | 0,09 | 9,32E-01 |
| AT2G33860 | 0,43 | 0,00 | 4,23 | 0,00 | 1,00E+00 |
| AT2G33980 | 0,47 | 2,21 | 3,31 | 0,67 | 5,04E-01 |
| AT2G34040 | 1,26 | -1,06 | 1,81 | -0,58 | 5,59E-01 |
| AT2G34070 | 0,35 | -0,45 | 3,63 | -0,12 | 9,02E-01 |
| AT2G34090 | 0,11 | 0,00 | 4,41 | 0,00 | 1,00E+00 |
| AT2G34290 | 0,45 | 0,00 | 4,10 | 0,00 | 1,00E+00 |
| AT2G34300 | 0,72 | 1,29 | 4,39 | 0,29 | 7,69E-01 |
| AT2G34450 | 0,60 | 1,97 | 3,85 | 0,51 | 6,09E-01 |
| AT2G34470 | 0,95 | 2,88 | 2,13 | 1,35 | 1,76E-01 |
| AT2G34480 | 1,66 | 1,26 | 1,65 | 0,76 | 4,46E-01 |
| AT2G34490 | 0,49 | 0,00 | 4,17 | 0,00 | 1,00E+00 |
| AT2G34510 | 0,33 | -0,19 | 4,37 | -0,04 | 9,66E-01 |
| AT2G34520 | 0,36 | 2,79 | 4,33 | 0,64 | 5,19E-01 |
| AT2G34585 | 1,42 | 2,21 | 1,95 | 1,13 | 2,58E-01 |
| AT2G34680 | 1,27 | 1,34 | 2,31 | 0,58 | 5,62E-01 |
| AT2G34710 | 1,10 | 2,48 | 2,47 | 1,00 | 3,15E-01 |
| AT2G34790 | 0,43 | 2,14 | 4,16 | 0,51 | 6,07E-01 |
| AT2G34920 | 0,05 | 0,00 | 4,41 | 0,00 | 1,00E+00 |
| AT2G34930 | 0,44 | 0,44 | 3,52 | 0,12 | 9,02E-01 |
| AT2G35030 | 0,48 | 0,99 | 4,03 | 0,25 | 8,06E-01 |
| AT2G35040 | 0,75 | -0,87 | 3,01 | -0,29 | 7,72E-01 |
| AT2G35060 | 1,08 | -1,39 | 1,87 | -0,74 | 4,57E-01 |
| AT2G35120 | 0,10 | 0,00 | 4,41 | 0,00 | 1,00E+00 |
| AT2G35250 | 1,05 | -2,42 | 1,99 | -1,22 | 2,23E-01 |
| AT2G35360 | 0,45 | 2,56 | 4,15 | 0,62 | 5,36E-01 |
| AT2G35390 | 0,25 | 1,26 | 4,39 | 0,29 | 7,74E-01 |
| AT2G35420 | 0,17 | 0,00 | 4,41 | 0,00 | 1,00E+00 |
| AT2G35490 | 0,67 | 0,53 | 2,33 | 0,23 | 8,20E-01 |
| AT2G35510 | 0,57 | 3,05 | 3,21 | 0,95 | 3,43E-01 |
| AT2G35610 | 0,17 | 0,00 | 4,41 | 0,00 | 1,00E+00 |
| AT2G35680 | 0,73 | 0,30 | 2,53 | 0,12 | 9,06E-01 |
| AT2G35730 | 0,35 | 1,58 | 4,36 | 0,36 | 7,18E-01 |
| AT2G35742 | 0,34 | -0,43 | 3,57 | -0,12 | 9,03E-01 |
| AT2G08755 | 0,29 | 0,00 | 4,41 | 0,00 | 1,00E+00 |

| | | | | | |
|---|---|---|---|---|---|
| AT2G08765 | 0,51 | -0,58 | 3,22 | -0,18 | 8,56E-01 |
| AT2G35859 | 0,71 | -0,38 | 2,80 | -0,14 | 8,92E-01 |
| AT2G35880 | 0,24 | 1,38 | 4,41 | 0,31 | 7,54E-01 |
| AT2G35930 | 0,76 | 0,00 | 2,95 | 0,00 | 1,00E+00 |
| AT2G36030 | 0,62 | -1,87 | 4,34 | -0,43 | 6,66E-01 |
| AT2G36050 | 0,69 | 0,19 | 2,99 | 0,06 | 9,50E-01 |
| AT2G36060 | 0,48 | 1,25 | 2,79 | 0,45 | 6,55E-01 |
| AT2G36070 | 0,34 | 1,69 | 4,38 | 0,39 | 6,99E-01 |
| AT2G36080 | 0,43 | 0,58 | 3,37 | 0,17 | 8,63E-01 |
| AT2G36180 | 0,76 | 0,00 | 3,47 | 0,00 | 1,00E+00 |
| AT2G36200 | 0,33 | 1,26 | 4,39 | 0,29 | 7,75E-01 |
| AT2G36230 | 0,20 | 0,00 | 4,41 | 0,00 | 1,00E+00 |
| AT2G36250 | 0,55 | -0,41 | 3,24 | -0,13 | 8,99E-01 |
| AT2G36300 | 0,44 | 1,76 | 3,36 | 0,52 | 6,01E-01 |
| AT2G36360 | 0,43 | -1,36 | 3,36 | -0,40 | 6,86E-01 |
| AT2G36400 | 0,83 | 1,41 | 3,10 | 0,45 | 6,49E-01 |
| AT2G36620 | 0,25 | 1,26 | 4,39 | 0,29 | 7,75E-01 |
| AT2G36680 | 0,35 | 0,89 | 4,36 | 0,20 | 8,38E-01 |
| AT2G36710 | 0,27 | -0,54 | 4,41 | -0,12 | 9,02E-01 |
| AT2G36750 | 0,21 | 0,00 | 4,41 | 0,00 | 1,00E+00 |
| AT2G36790 | 0,34 | 0,00 | 4,41 | 0,00 | 1,00E+00 |
| AT2G36800 | 0,85 | 2,18 | 2,43 | 0,90 | 3,70E-01 |
| AT2G36810 | 1,10 | 1,31 | 2,14 | 0,61 | 5,41E-01 |
| AT2G36835 | 0,30 | -0,43 | 3,68 | -0,12 | 9,08E-01 |
| AT2G36895 | 0,35 | 1,58 | 4,36 | 0,36 | 7,18E-01 |
| AT2G36900 | 0,55 | 2,01 | 3,82 | 0,53 | 5,98E-01 |
| AT2G36930 | 0,54 | 1,29 | 3,93 | 0,33 | 7,43E-01 |
| AT2G36990 | 1,19 | 4,36 | 2,50 | 1,74 | 8,13E-02 |
| AT2G37035 | 0,78 | -0,02 | 2,55 | -0,01 | 9,93E-01 |
| AT2G37040 | 1,58 | 3,22 | 1,80 | 1,79 | 7,42E-02 |
| AT2G37080 | 0,12 | 0,00 | 4,41 | 0,00 | 1,00E+00 |
| AT2G37130 | 0,68 | 1,41 | 3,15 | 0,45 | 6,54E-01 |
| AT2G37150 | 0,75 | 2,56 | 3,46 | 0,74 | 4,58E-01 |
| AT2G37170 | 0,90 | 2,14 | 2,42 | 0,88 | 3,77E-01 |
| AT2G37200 | 0,31 | 1,38 | 3,68 | 0,37 | 7,08E-01 |
| AT2G37230 | 0,56 | 0,42 | 3,26 | 0,13 | 8,98E-01 |
| AT2G37310 | 0,77 | 1,97 | 3,59 | 0,55 | 5,84E-01 |
| AT2G37340 | 1,12 | 2,91 | 2,17 | 1,34 | 1,81E-01 |
| AT2G37360 | 0,21 | 0,00 | 4,41 | 0,00 | 1,00E+00 |
| AT2G37400 | 0,25 | 1,30 | 4,40 | 0,30 | 7,67E-01 |
| AT2G37410 | 0,19 | 0,00 | 4,41 | 0,00 | 1,00E+00 |
| AT2G37520 | 1,03 | -0,43 | 2,00 | -0,21 | 8,31E-01 |
| AT2G37540 | 0,69 | 2,19 | 2,96 | 0,74 | 4,59E-01 |
| AT2G37550 | 1,38 | 2,67 | 2,00 | 1,34 | 1,81E-01 |
| AT2G37570 | 0,98 | 0,50 | 1,97 | 0,25 | 8,00E-01 |
| AT2G37630 | 0,98 | 3,51 | 1,99 | 1,76 | 7,82E-02 |
| AT2G37660 | 1,35 | -1,54 | 1,80 | -0,85 | 3,94E-01 |
| AT2G37678 | 0,57 | -0,72 | 4,34 | -0,17 | 8,69E-01 |
| AT2G08895 | 1,00 | 0,00 | 2,45 | 0,00 | 1,00E+00 |
| AT2G08900 | 0,86 | -2,03 | 2,11 | -0,96 | 3,37E-01 |
| AT2G08910 | 1,52 | 0,37 | 1,63 | 0,23 | 8,19E-01 |
| AT2G08915 | 0,73 | -0,48 | 2,88 | -0,17 | 8,67E-01 |
| AT2G38000 | 0,32 | 1,29 | 4,39 | 0,29 | 7,69E-01 |
| AT2G38020 | 1,29 | 2,12 | 2,10 | 1,01 | 3,13E-01 |
| AT2G38160 | 0,87 | 3,77 | 3,22 | 1,17 | 2,41E-01 |
| AT2G38270 | 0,22 | -0,43 | 4,39 | -0,10 | 9,21E-01 |

| | | | | | |
|---|---|---|---|---|---|
| AT2G38290 | 0,66 | 2,16 | 2,95 | 0,73 | 4,65E-01 |
| AT2G38340 | 0,17 | 0,00 | 4,41 | 0,00 | 1,00E+00 |
| AT2G38360 | 0,39 | 1,26 | 3,40 | 0,37 | 7,11E-01 |
| AT2G38410 | 0,35 | 0,43 | 3,48 | 0,12 | 9,02E-01 |
| AT2G38480 | 0,17 | 0,00 | 4,41 | 0,00 | 1,00E+00 |
| AT2G38610 | 0,83 | 0,29 | 2,58 | 0,11 | 9,10E-01 |
| AT2G38650 | 1,16 | -1,93 | 2,21 | -0,87 | 3,84E-01 |
| AT2G38680 | 0,08 | 0,00 | 4,41 | 0,00 | 1,00E+00 |
| AT2G38750 | 0,05 | 0,00 | 4,41 | 0,00 | 1,00E+00 |
| AT2G38770 | 1,53 | 1,95 | 1,71 | 1,14 | 2,54E-01 |
| AT2G38790 | 0,55 | 0,41 | 2,52 | 0,16 | 8,72E-01 |
| AT2G38860 | 0,41 | 1,38 | 3,47 | 0,40 | 6,91E-01 |
| AT2G39000 | 1,06 | 1,17 | 2,95 | 0,40 | 6,91E-01 |
| AT2G39080 | 0,20 | 1,33 | 4,40 | 0,30 | 7,63E-01 |
| AT2G39130 | 0,08 | 0,00 | 4,41 | 0,00 | 1,00E+00 |
| AT2G39160 | 0,24 | -0,48 | 4,40 | -0,11 | 9,13E-01 |
| AT2G39170 | 0,19 | 0,00 | 4,40 | 0,00 | 1,00E+00 |
| AT2G09010 | 0,08 | 0,00 | 4,41 | 0,00 | 1,00E+00 |
| AT2G39210 | 0,91 | 2,74 | 2,77 | 0,99 | 3,23E-01 |
| AT2G39260 | 1,36 | -0,26 | 1,69 | -0,15 | 8,77E-01 |
| AT2G39350 | 0,49 | 2,14 | 3,25 | 0,66 | 5,11E-01 |
| AT2G39360 | 0,80 | 3,40 | 3,45 | 0,98 | 3,25E-01 |
| AT2G09045 | 0,52 | 1,38 | 3,96 | 0,35 | 7,27E-01 |
| AT2G39450 | 0,29 | 1,29 | 4,39 | 0,29 | 7,69E-01 |
| AT2G39480 | 1,05 | 2,89 | 2,18 | 1,33 | 1,84E-01 |
| AT2G39650 | 0,49 | 0,00 | 4,11 | 0,00 | 1,00E+00 |
| AT2G39710 | 0,60 | 0,48 | 3,33 | 0,15 | 8,85E-01 |
| AT2G39720 | 1,30 | 3,55 | 2,40 | 1,48 | 1,38E-01 |
| AT2G39750 | 1,24 | 1,60 | 1,87 | 0,86 | 3,91E-01 |
| AT2G09080 | 0,50 | 2,19 | 3,29 | 0,67 | 5,05E-01 |
| AT2G39805 | 0,30 | 1,38 | 4,41 | 0,31 | 7,54E-01 |
| AT2G39920 | 0,18 | -0,47 | 4,39 | -0,11 | 9,14E-01 |
| AT2G09090 | 0,82 | 1,32 | 2,82 | 0,47 | 6,38E-01 |
| AT2G39950 | 0,75 | 0,39 | 2,57 | 0,15 | 8,80E-01 |
| AT2G39960 | 0,25 | -1,31 | 4,37 | -0,30 | 7,65E-01 |
| AT2G39990 | 1,46 | 0,60 | 2,65 | 0,23 | 8,22E-01 |
| AT2G40070 | 0,52 | 1,37 | 3,96 | 0,34 | 7,30E-01 |
| AT2G40080 | 0,23 | 0,00 | 4,41 | 0,00 | 1,00E+00 |
| AT2G40165 | 0,38 | 2,11 | 4,37 | 0,48 | 6,29E-01 |
| AT2G40205 | 0,27 | 1,38 | 4,41 | 0,31 | 7,54E-01 |
| AT2G40270 | 1,34 | 1,37 | 1,91 | 0,72 | 4,73E-01 |
| AT2G40280 | 0,63 | 0,03 | 2,99 | 0,01 | 9,91E-01 |
| AT2G40350 | 0,12 | 0,00 | 4,41 | 0,00 | 1,00E+00 |
| AT2G40360 | 0,28 | 0,00 | 4,41 | 0,00 | 1,00E+00 |
| AT2G40430 | 0,45 | 1,14 | 4,20 | 0,27 | 7,85E-01 |
| AT2G40475 | 0,89 | 0,00 | 2,47 | 0,00 | 1,00E+00 |
| AT2G40590 | 0,50 | 0,53 | 2,66 | 0,20 | 8,44E-01 |
| AT2G40600 | 0,11 | 0,00 | 4,41 | 0,00 | 1,00E+00 |
| AT2G40610 | 0,37 | -1,63 | 4,36 | -0,37 | 7,09E-01 |
| AT2G40650 | 0,77 | -0,54 | 2,48 | -0,22 | 8,26E-01 |
| AT2G40730 | 0,34 | -0,43 | 3,54 | -0,12 | 9,02E-01 |
| AT2G40750 | 0,46 | -0,80 | 4,14 | -0,19 | 8,46E-01 |
| AT2G40770 | 0,51 | -0,26 | 3,99 | -0,06 | 9,49E-01 |
| AT2G40815 | 0,24 | 1,38 | 4,41 | 0,31 | 7,54E-01 |
| AT2G40860 | 0,30 | 1,38 | 4,41 | 0,31 | 7,54E-01 |
| AT2G40890 | 0,53 | -0,63 | 3,96 | -0,16 | 8,74E-01 |

| | | | | | |
|---|---|---|---|---|---|
| AT2G40920 | 0,35 | -0,54 | 4,41 | -0,12 | 9,02E-01 |
| AT2G40925 | 0,65 | 2,11 | 3,69 | 0,57 | 5,68E-01 |
| AT2G40950 | 0,74 | 1,37 | 3,62 | 0,38 | 7,05E-01 |
| AT2G40970 | 0,42 | 2,51 | 4,35 | 0,58 | 5,63E-01 |
| AT2G09225 | 0,24 | -0,48 | 4,40 | -0,11 | 9,13E-01 |
| AT2G41120 | 0,21 | 0,00 | 4,41 | 0,00 | 1,00E+00 |
| AT2G41160 | 0,39 | 0,00 | 4,41 | 0,00 | 1,00E+00 |
| AT2G41250 | 0,87 | 1,38 | 2,59 | 0,53 | 5,94E-01 |
| AT2G41420 | 0,11 | 0,00 | 4,41 | 0,00 | 1,00E+00 |
| AT2G41475 | 0,71 | 1,18 | 2,62 | 0,45 | 6,52E-01 |
| AT2G41500 | 0,96 | 2,27 | 2,13 | 1,06 | 2,88E-01 |
| AT2G41600 | 0,58 | 0,74 | 2,68 | 0,28 | 7,81E-01 |
| AT2G41620 | 1,68 | 3,06 | 1,83 | 1,67 | 9,43E-02 |
| AT2G41616 | 0,33 | 1,37 | 4,41 | 0,31 | 7,56E-01 |
| AT2G41630 | 0,92 | 3,22 | 2,72 | 1,18 | 2,37E-01 |
| AT2G41680 | 0,32 | 0,00 | 4,41 | 0,00 | 1,00E+00 |
| AT2G41700 | 1,62 | -1,17 | 2,30 | -0,51 | 6,09E-01 |
| AT2G41770 | 0,40 | 1,54 | 4,32 | 0,36 | 7,22E-01 |
| AT2G41790 | 0,58 | 0,41 | 2,71 | 0,15 | 8,80E-01 |
| AT2G41830 | 0,40 | 0,46 | 4,23 | 0,11 | 9,13E-01 |
| AT2G41835 | 0,70 | 0,00 | 3,80 | 0,00 | 1,00E+00 |
| AT2G41870 | 0,29 | 1,47 | 4,40 | 0,33 | 7,39E-01 |
| AT2G41890 | 0,43 | 1,52 | 4,23 | 0,36 | 7,19E-01 |
| AT2G41945 | 0,60 | 1,31 | 4,00 | 0,33 | 7,44E-01 |
| AT2G41960 | 0,92 | -0,34 | 2,10 | -0,16 | 8,73E-01 |
| AT2G42010 | 0,72 | -0,45 | 2,17 | -0,21 | 8,35E-01 |
| AT2G42020 | 0,48 | -1,69 | 4,35 | -0,39 | 6,98E-01 |
| AT2G42070 | 0,23 | 0,00 | 4,41 | 0,00 | 1,00E+00 |
| AT2G42230 | 0,42 | 1,39 | 4,18 | 0,33 | 7,40E-01 |
| AT2G42280 | 0,33 | 1,79 | 4,38 | 0,41 | 6,83E-01 |
| AT2G42365 | 0,35 | 1,58 | 4,36 | 0,36 | 7,18E-01 |
| AT2G42400 | 1,01 | 1,80 | 2,10 | 0,86 | 3,92E-01 |
| AT2G42500 | 0,82 | 1,34 | 2,41 | 0,55 | 5,80E-01 |
| AT2G42530 | 0,98 | -2,31 | 2,05 | -1,12 | 2,61E-01 |
| AT2G42690 | 1,34 | -1,34 | 1,76 | -0,76 | 4,46E-01 |
| AT2G42760 | 0,47 | -0,52 | 2,79 | -0,19 | 8,53E-01 |
| AT2G42790 | 0,53 | 1,39 | 2,69 | 0,52 | 6,04E-01 |
| AT2G42800 | 0,18 | 0,00 | 4,41 | 0,00 | 1,00E+00 |
| AT2G42810 | 0,84 | 1,34 | 1,97 | 0,68 | 4,95E-01 |
| AT2G42840 | 0,22 | 0,00 | 4,41 | 0,00 | 1,00E+00 |
| AT2G42865 | 0,08 | 0,00 | 4,41 | 0,00 | 1,00E+00 |
| AT2G42870 | 0,10 | 0,00 | 4,41 | 0,00 | 1,00E+00 |
| AT2G42880 | 0,41 | 1,30 | 3,52 | 0,37 | 7,12E-01 |
| AT2G43180 | 0,70 | 3,32 | 2,87 | 1,16 | 2,47E-01 |
| AT2G43260 | 0,20 | 1,38 | 4,41 | 0,31 | 7,54E-01 |
| AT2G43290 | 0,50 | 0,00 | 3,29 | 0,00 | 1,00E+00 |
| AT2G43320 | 0,22 | 0,00 | 4,41 | 0,00 | 1,00E+00 |
| AT2G43360 | 0,33 | 0,44 | 4,39 | 0,10 | 9,21E-01 |
| AT2G43420 | 0,11 | 0,00 | 4,41 | 0,00 | 1,00E+00 |
| AT2G43430 | 0,59 | 0,24 | 2,61 | 0,09 | 9,27E-01 |
| AT2G43460 | 0,41 | 0,35 | 3,44 | 0,10 | 9,18E-01 |
| AT2G43490 | 0,44 | 0,00 | 4,14 | 0,00 | 1,00E+00 |
| AT2G43560 | 0,25 | 1,24 | 4,38 | 0,28 | 7,77E-01 |
| AT2G43750 | 1,46 | 2,70 | 1,72 | 1,57 | 1,16E-01 |
| AT2G43865 | 0,15 | 0,00 | 4,41 | 0,00 | 1,00E+00 |
| AT2G43910 | 0,26 | -0,51 | 3,81 | -0,13 | 8,94E-01 |

| | | | | | |
|---|---|---|---|---|---|
| AT2G43940 | 0,08 | 0,00 | 4,41 | 0,00 | 1,00E+00 |
| AT2G43945 | 0,47 | 0,40 | 4,35 | 0,09 | 9,27E-01 |
| AT2G43950 | 0,24 | 1,37 | 4,41 | 0,31 | 7,56E-01 |
| AT2G44090 | 0,69 | 2,16 | 2,58 | 0,84 | 4,02E-01 |
| AT2G44140 | 0,91 | -1,97 | 2,36 | -0,84 | 4,03E-01 |
| AT2G44180 | 0,46 | 1,28 | 3,35 | 0,38 | 7,02E-01 |
| AT2G44255 | 0,20 | 0,00 | 4,41 | 0,00 | 1,00E+00 |
| AT2G44360 | 0,63 | 0,00 | 4,41 | 0,00 | 1,00E+00 |
| AT2G44610 | 0,48 | -0,59 | 4,24 | -0,14 | 8,89E-01 |
| AT2G44660 | 0,35 | 1,40 | 3,58 | 0,39 | 6,96E-01 |
| AT2G44760 | 0,32 | 1,31 | 3,66 | 0,36 | 7,20E-01 |
| AT2G44798 | 0,36 | 1,29 | 4,40 | 0,29 | 7,68E-01 |
| AT2G44950 | 0,77 | 1,41 | 2,17 | 0,65 | 5,16E-01 |
| AT2G45070 | 0,80 | -0,89 | 2,31 | -0,39 | 6,98E-01 |
| AT2G45220 | 0,54 | 1,28 | 4,39 | 0,29 | 7,71E-01 |
| AT2G45315 | 0,67 | 0,45 | 2,65 | 0,17 | 8,67E-01 |
| AT2G45460 | 0,20 | 0,00 | 4,41 | 0,00 | 1,00E+00 |
| AT2G45470 | 1,65 | 1,33 | 1,96 | 0,68 | 4,97E-01 |
| AT2G45520 | 0,22 | -0,48 | 4,40 | -0,11 | 9,14E-01 |
| AT2G09640 | 1,68 | 1,29 | 2,23 | 0,58 | 5,63E-01 |
| AT2G45560 | 0,47 | 2,55 | 4,35 | 0,59 | 5,57E-01 |
| AT2G45660 | 0,55 | 0,43 | 2,45 | 0,18 | 8,59E-01 |
| AT2G45670 | 0,95 | 1,20 | 2,38 | 0,51 | 6,14E-01 |
| AT2G45700 | 0,38 | -0,72 | 4,34 | -0,17 | 8,67E-01 |
| AT2G45820 | 0,98 | -1,64 | 1,90 | -0,86 | 3,88E-01 |
| AT2G45970 | 0,56 | 2,07 | 2,72 | 0,76 | 4,46E-01 |
| AT2G45980 | 0,45 | 1,52 | 4,17 | 0,36 | 7,16E-01 |
| AT2G46030 | 0,57 | 1,90 | 4,38 | 0,43 | 6,65E-01 |
| AT2G46040 | 0,20 | 1,33 | 4,40 | 0,30 | 7,63E-01 |
| AT2G46060 | 0,49 | -0,93 | 4,34 | -0,22 | 8,30E-01 |
| AT2G46080 | 0,51 | 2,46 | 3,24 | 0,76 | 4,47E-01 |
| AT2G46180 | 0,35 | 1,38 | 4,41 | 0,31 | 7,54E-01 |
| AT2G46280 | 1,16 | 0,01 | 1,74 | 0,01 | 9,96E-01 |
| AT2G46330 | 0,93 | -2,44 | 2,13 | -1,15 | 2,51E-01 |
| AT2G46390 | 0,46 | 2,39 | 4,10 | 0,58 | 5,59E-01 |
| AT2G46505 | 0,89 | 3,75 | 2,28 | 1,64 | 1,01E-01 |
| AT2G46510 | 1,07 | 2,05 | 2,14 | 0,96 | 3,39E-01 |
| AT2G46540 | 0,12 | 0,00 | 4,41 | 0,00 | 1,00E+00 |
| AT2G46550 | 0,57 | 1,31 | 2,75 | 0,48 | 6,34E-01 |
| AT2G46572 | 0,51 | 1,38 | 4,08 | 0,34 | 7,36E-01 |
| AT2G46610 | 0,85 | 2,31 | 2,01 | 1,15 | 2,50E-01 |
| AT2G46620 | 1,45 | 3,87 | 1,80 | 2,15 | 3,14E-02 |
| AT2G46680 | 0,94 | 0,31 | 2,38 | 0,13 | 8,97E-01 |
| AT2G46840 | 0,08 | 0,00 | 4,41 | 0,00 | 1,00E+00 |
| AT2G46920 | 0,44 | 1,32 | 4,10 | 0,32 | 7,48E-01 |
| AT2G46980 | 0,24 | 1,96 | 4,39 | 0,45 | 6,55E-01 |
| AT2G47090 | 0,25 | 0,00 | 4,41 | 0,00 | 1,00E+00 |
| AT2G47180 | 1,41 | 1,94 | 1,73 | 1,12 | 2,61E-01 |
| AT2G47250 | 1,15 | 1,22 | 2,40 | 0,51 | 6,10E-01 |
| AT2G47330 | 0,60 | -0,55 | 3,22 | -0,17 | 8,65E-01 |
| AT2G47600 | 0,61 | 2,25 | 2,61 | 0,86 | 3,90E-01 |
| AT2G47650 | 1,46 | 1,80 | 1,70 | 1,06 | 2,89E-01 |
| AT2G47680 | 0,89 | 3,39 | 2,08 | 1,63 | 1,03E-01 |
| AT2G47700 | 0,93 | 3,11 | 2,45 | 1,27 | 2,04E-01 |
| AT2G47840 | 0,52 | -0,84 | 3,33 | -0,25 | 8,00E-01 |
| AT2G47990 | 0,88 | 3,59 | 2,63 | 1,36 | 1,73E-01 |

| | | | | | |
|---|---|---|---|---|---|
| AT2G48000 | 1,00 | 2,02 | 3,46 | 0,59 | 5,58E-01 |
| AT2G48030 | 0,73 | -0,54 | 2,58 | -0,21 | 8,33E-01 |
| AT2G48130 | 0,91 | -1,06 | 2,08 | -0,51 | 6,09E-01 |
| AT3G01090 | 0,63 | 0,38 | 2,37 | 0,16 | 8,72E-01 |
| AT3G01100 | 0,90 | 3,55 | 2,16 | 1,64 | 1,00E-01 |
| AT3G01130 | 0,21 | 1,25 | 4,39 | 0,29 | 7,75E-01 |
| AT3G01160 | 0,24 | -0,48 | 4,40 | -0,11 | 9,13E-01 |
| AT3G01180 | 0,84 | 0,67 | 1,95 | 0,34 | 7,31E-01 |
| AT3G01300 | 0,08 | 0,00 | 4,41 | 0,00 | 1,00E+00 |
| AT3G01340 | 0,50 | -0,49 | 3,29 | -0,15 | 8,82E-01 |
| AT3G01380 | 1,09 | 2,11 | 1,98 | 1,07 | 2,85E-01 |
| AT3G01390 | 0,38 | -1,29 | 4,37 | -0,29 | 7,68E-01 |
| AT3G01415 | 0,40 | -0,54 | 4,41 | -0,12 | 9,02E-01 |
| AT3G01420 | 0,31 | 1,29 | 4,39 | 0,29 | 7,70E-01 |
| AT3G01430 | 0,79 | 2,85 | 2,99 | 0,95 | 3,40E-01 |
| AT3G01470 | 1,49 | 2,55 | 1,75 | 1,45 | 1,47E-01 |
| AT3G01550 | 0,10 | 0,00 | 4,41 | 0,00 | 1,00E+00 |
| AT3G01720 | 0,80 | 1,39 | 3,00 | 0,46 | 6,42E-01 |
| AT3G01820 | 0,81 | 0,51 | 3,58 | 0,14 | 8,86E-01 |
| AT3G01840 | 0,88 | 2,69 | 2,24 | 1,20 | 2,31E-01 |
| AT3G01910 | 0,46 | -0,61 | 4,14 | -0,15 | 8,83E-01 |
| AT3G01930 | 0,62 | 1,70 | 2,95 | 0,57 | 5,66E-01 |
| AT3G01980 | 0,16 | 0,00 | 4,41 | 0,00 | 1,00E+00 |
| AT3G02040 | 0,68 | 2,87 | 2,49 | 1,15 | 2,49E-01 |
| AT3G02080 | 1,20 | 1,52 | 2,12 | 0,72 | 4,71E-01 |
| AT3G02190 | 0,44 | 0,57 | 3,38 | 0,17 | 8,66E-01 |
| AT3G02210 | 0,52 | 1,30 | 4,08 | 0,32 | 7,49E-01 |
| AT3G02250 | 0,34 | -1,28 | 4,37 | -0,29 | 7,70E-01 |
| AT3G02380 | 0,91 | 3,41 | 2,29 | 1,48 | 1,38E-01 |
| AT3G02410 | 0,66 | 3,63 | 4,30 | 0,85 | 3,98E-01 |
| AT3G02450 | 0,68 | 1,82 | 2,32 | 0,79 | 4,32E-01 |
| AT3G02520 | 1,11 | -0,63 | 1,74 | -0,36 | 7,18E-01 |
| AT3G02530 | 0,67 | -0,10 | 2,78 | -0,04 | 9,72E-01 |
| AT3G02700 | 0,39 | 0,84 | 4,23 | 0,20 | 8,42E-01 |
| AT3G02730 | 0,92 | 1,80 | 2,67 | 0,67 | 5,00E-01 |
| AT3G02750 | 1,16 | 1,66 | 1,87 | 0,89 | 3,74E-01 |
| AT3G02760 | 0,94 | 2,93 | 2,74 | 1,07 | 2,85E-01 |
| AT3G01205 | 0,80 | 0,60 | 2,48 | 0,24 | 8,07E-01 |
| AT3G03090 | 0,29 | -0,44 | 4,36 | -0,10 | 9,19E-01 |
| AT3G03150 | 0,22 | -0,54 | 4,41 | -0,12 | 9,03E-01 |
| AT3G03220 | 0,16 | 0,00 | 4,41 | 0,00 | 1,00E+00 |
| AT3G03250 | 1,00 | 0,46 | 2,11 | 0,22 | 8,29E-01 |
| AT3G03300 | 0,33 | 0,00 | 4,41 | 0,00 | 1,00E+00 |
| AT3G03305 | 0,32 | 1,38 | 4,41 | 0,31 | 7,54E-01 |
| AT3G03310 | 0,29 | 1,35 | 4,40 | 0,31 | 7,59E-01 |
| AT3G03330 | 0,30 | 1,77 | 4,38 | 0,40 | 6,86E-01 |
| AT3G03405 | 0,16 | 0,00 | 4,41 | 0,00 | 1,00E+00 |
| AT3G03460 | 0,17 | 0,00 | 4,41 | 0,00 | 1,00E+00 |
| AT3G03470 | 1,68 | 2,20 | 2,06 | 1,07 | 2,86E-01 |
| AT3G03480 | 0,08 | 0,00 | 4,41 | 0,00 | 1,00E+00 |
| AT3G03550 | 0,10 | 0,00 | 4,41 | 0,00 | 1,00E+00 |
| AT3G03560 | 0,61 | 1,27 | 3,06 | 0,42 | 6,78E-01 |
| AT3G03580 | 0,26 | 1,38 | 4,41 | 0,31 | 7,54E-01 |
| AT3G03590 | 0,08 | 0,00 | 4,41 | 0,00 | 1,00E+00 |
| AT3G03600 | 0,30 | -0,78 | 4,37 | -0,18 | 8,59E-01 |
| AT3G03720 | 0,26 | 0,00 | 4,41 | 0,00 | 1,00E+00 |

| | | | | | |
|---|---|---|---|---|---|
| AT3G03770 | 0,99 | 0,00 | 3,21 | 0,00 | 1,00E+00 |
| AT3G03890 | 0,79 | 2,89 | 2,46 | 1,18 | 2,40E-01 |
| AT3G03920 | 0,42 | -0,70 | 3,32 | -0,21 | 8,33E-01 |
| AT3G03940 | 0,41 | -0,60 | 2,92 | -0,20 | 8,38E-01 |
| AT3G04030 | 0,25 | 0,00 | 4,41 | 0,00 | 1,00E+00 |
| AT3G04080 | 1,01 | 1,76 | 2,05 | 0,86 | 3,91E-01 |
| AT3G04150 | 0,26 | 1,38 | 4,41 | 0,31 | 7,54E-01 |
| AT3G04200 | 0,62 | 1,27 | 4,39 | 0,29 | 7,72E-01 |
| AT3G04220 | 0,33 | 0,00 | 4,41 | 0,00 | 1,00E+00 |
| AT3G04260 | 0,61 | -0,14 | 4,33 | -0,03 | 9,73E-01 |
| AT3G04290 | 0,84 | -0,52 | 3,11 | -0,17 | 8,67E-01 |
| AT3G04350 | 0,35 | 0,00 | 4,41 | 0,00 | 1,00E+00 |
| AT3G04480 | 0,84 | 1,26 | 2,45 | 0,51 | 6,08E-01 |
| AT3G04500 | 0,20 | 1,40 | 4,41 | 0,32 | 7,51E-01 |
| AT3G04580 | 1,03 | 1,22 | 2,15 | 0,57 | 5,71E-01 |
| AT3G04640 | 0,85 | 0,49 | 2,35 | 0,21 | 8,36E-01 |
| AT3G04690 | 0,22 | -0,84 | 4,38 | -0,19 | 8,47E-01 |
| AT3G04720 | 0,50 | 0,00 | 3,20 | 0,00 | 1,00E+00 |
| AT3G04730 | 1,02 | 0,45 | 1,74 | 0,26 | 7,97E-01 |
| AT3G04750 | 0,57 | 2,63 | 2,66 | 0,99 | 3,22E-01 |
| AT3G04760 | 0,89 | 1,38 | 2,27 | 0,61 | 5,43E-01 |
| AT3G04860 | 0,12 | 0,00 | 4,41 | 0,00 | 1,00E+00 |
| AT3G04980 | 0,46 | 1,42 | 4,40 | 0,32 | 7,47E-01 |
| AT3G05030 | 0,28 | 0,00 | 4,41 | 0,00 | 1,00E+00 |
| AT3G01585 | 0,10 | 0,00 | 4,41 | 0,00 | 1,00E+00 |
| AT3G05040 | 1,58 | 2,28 | 1,60 | 1,42 | 1,55E-01 |
| AT3G05050 | 0,38 | 1,38 | 4,41 | 0,31 | 7,54E-01 |
| AT3G05060 | 0,60 | 2,04 | 3,76 | 0,54 | 5,87E-01 |
| AT3G05090 | 0,52 | 1,27 | 2,71 | 0,47 | 6,39E-01 |
| AT3G05165 | 0,83 | 1,40 | 2,16 | 0,65 | 5,18E-01 |
| AT3G05240 | 0,13 | 0,00 | 4,41 | 0,00 | 1,00E+00 |
| AT3G05270 | 0,58 | 2,93 | 2,58 | 1,13 | 2,57E-01 |
| AT3G05280 | 0,59 | 1,86 | 3,78 | 0,49 | 6,23E-01 |
| AT3G05540 | 1,04 | -1,83 | 1,89 | -0,97 | 3,31E-01 |
| AT3G05650 | 0,98 | 2,11 | 2,39 | 0,88 | 3,77E-01 |
| AT3G05660 | 0,79 | 1,38 | 2,89 | 0,48 | 6,33E-01 |
| AT3G05680 | 1,68 | 0,82 | 1,74 | 0,47 | 6,39E-01 |
| AT3G05690 | 0,11 | 0,00 | 4,41 | 0,00 | 1,00E+00 |
| AT3G05770 | 0,46 | 1,38 | 3,42 | 0,40 | 6,87E-01 |
| AT3G05810 | 0,21 | -0,50 | 4,40 | -0,11 | 9,09E-01 |
| AT3G05850 | 0,90 | 3,57 | 2,46 | 1,45 | 1,47E-01 |
| AT3G05880 | 0,55 | -1,05 | 3,12 | -0,34 | 7,37E-01 |
| AT3G05900 | 0,51 | -0,54 | 3,22 | -0,17 | 8,67E-01 |
| AT3G05910 | 0,68 | -2,67 | 2,59 | -1,03 | 3,04E-01 |
| AT3G05932 | 0,27 | 1,34 | 4,40 | 0,30 | 7,61E-01 |
| AT3G05970 | 0,71 | -1,01 | 2,32 | -0,43 | 6,65E-01 |
| AT3G06030 | 0,29 | 0,00 | 4,41 | 0,00 | 1,00E+00 |
| AT3G06050 | 0,18 | -0,43 | 4,39 | -0,10 | 9,21E-01 |
| AT3G06060 | 0,17 | 1,32 | 4,40 | 0,30 | 7,64E-01 |
| AT3G06080 | 1,38 | 0,31 | 1,65 | 0,19 | 8,52E-01 |
| AT3G06150 | 0,98 | -0,78 | 1,70 | -0,46 | 6,49E-01 |
| AT3G06190 | 0,71 | 2,70 | 3,13 | 0,86 | 3,87E-01 |
| AT3G06250 | 0,74 | 0,93 | 2,93 | 0,32 | 7,51E-01 |
| AT3G06290 | 0,72 | 1,29 | 2,58 | 0,50 | 6,18E-01 |
| AT3G06310 | 0,08 | 0,00 | 4,41 | 0,00 | 1,00E+00 |
| AT3G06350 | 0,64 | 2,21 | 3,05 | 0,73 | 4,68E-01 |

| | | | | | |
|---|---|---|---|---|---|
| AT3G06430 | 0,12 | 0,00 | 4,41 | 0,00 | 1,00E+00 |
| AT3G06440 | 0,12 | 0,00 | 4,41 | 0,00 | 1,00E+00 |
| AT3G06450 | 0,61 | 2,29 | 2,72 | 0,84 | 4,01E-01 |
| AT3G06480 | 1,05 | 2,51 | 1,98 | 1,27 | 2,04E-01 |
| AT3G06483 | 0,45 | 0,54 | 2,85 | 0,19 | 8,51E-01 |
| AT3G06540 | 0,08 | 0,00 | 4,41 | 0,00 | 1,00E+00 |
| AT3G06550 | 0,32 | -0,54 | 4,41 | -0,12 | 9,02E-01 |
| AT3G06580 | 0,15 | 0,00 | 4,41 | 0,00 | 1,00E+00 |
| AT3G06590 | 0,40 | 1,82 | 4,38 | 0,42 | 6,78E-01 |
| AT3G06620 | 0,12 | 0,00 | 4,41 | 0,00 | 1,00E+00 |
| AT3G06665 | 0,25 | 0,00 | 4,41 | 0,00 | 1,00E+00 |
| AT3G06670 | 1,42 | 2,68 | 1,70 | 1,57 | 1,15E-01 |
| AT3G06680 | 0,28 | 1,35 | 4,40 | 0,31 | 7,59E-01 |
| AT3G06690 | 0,20 | 1,40 | 4,41 | 0,32 | 7,51E-01 |
| AT3G06700 | 0,57 | 1,31 | 2,76 | 0,47 | 6,35E-01 |
| AT3G06790 | 0,16 | 0,00 | 4,41 | 0,00 | 1,00E+00 |
| AT3G06820 | 0,08 | 0,00 | 4,41 | 0,00 | 1,00E+00 |
| AT3G06870 | 0,21 | 1,38 | 4,41 | 0,31 | 7,54E-01 |
| AT3G06900 | 1,19 | 1,79 | 2,03 | 0,88 | 3,78E-01 |
| AT3G06930 | 0,40 | 2,13 | 4,20 | 0,51 | 6,13E-01 |
| AT3G06960 | 0,89 | -2,44 | 2,55 | -0,96 | 3,39E-01 |
| AT3G07010 | 0,41 | 0,90 | 4,26 | 0,21 | 8,32E-01 |
| AT3G07020 | 0,54 | 2,10 | 3,82 | 0,55 | 5,82E-01 |
| AT3G07040 | 1,04 | 2,57 | 2,28 | 1,13 | 2,59E-01 |
| AT3G07050 | 0,53 | 0,62 | 3,28 | 0,19 | 8,49E-01 |
| AT3G07115 | 0,08 | 0,00 | 4,41 | 0,00 | 1,00E+00 |
| AT3G07180 | 0,12 | 0,00 | 4,41 | 0,00 | 1,00E+00 |
| AT3G07215 | 0,55 | 0,43 | 2,71 | 0,16 | 8,74E-01 |
| AT3G07273 | 0,08 | 0,00 | 4,41 | 0,00 | 1,00E+00 |
| AT3G07290 | 0,69 | 1,32 | 3,58 | 0,37 | 7,13E-01 |
| AT3G07300 | 0,29 | 1,85 | 4,39 | 0,42 | 6,73E-01 |
| AT3G07330 | 0,98 | 1,38 | 2,68 | 0,51 | 6,07E-01 |
| AT3G07380 | 1,22 | -0,45 | 1,73 | -0,26 | 7,95E-01 |
| AT3G07460 | 0,63 | -0,90 | 2,64 | -0,34 | 7,33E-01 |
| AT3G07470 | 0,58 | 1,09 | 3,22 | 0,34 | 7,35E-01 |
| AT3G07520 | 0,51 | 1,38 | 3,33 | 0,41 | 6,79E-01 |
| AT3G07540 | 0,08 | 0,00 | 4,41 | 0,00 | 1,00E+00 |
| AT3G07660 | 1,40 | 3,77 | 1,95 | 1,93 | 5,34E-02 |
| AT3G07700 | 0,94 | 0,59 | 1,85 | 0,32 | 7,51E-01 |
| AT3G07800 | 0,53 | 2,32 | 2,77 | 0,84 | 4,03E-01 |
| AT3G07890 | 0,34 | 1,36 | 4,41 | 0,31 | 7,57E-01 |
| AT3G08570 | 1,10 | 3,24 | 2,18 | 1,49 | 1,37E-01 |
| AT3G08590 | 1,52 | 0,28 | 1,57 | 0,18 | 8,56E-01 |
| AT3G08610 | 0,59 | 1,59 | 3,18 | 0,50 | 6,17E-01 |
| AT3G08650 | 0,24 | 0,00 | 4,40 | 0,00 | 1,00E+00 |
| AT3G08720 | 1,37 | 2,22 | 2,17 | 1,02 | 3,06E-01 |
| AT3G08760 | 0,70 | 0,34 | 2,43 | 0,14 | 8,89E-01 |
| AT3G08840 | 0,67 | 3,14 | 3,00 | 1,04 | 2,96E-01 |
| AT3G08870 | 0,75 | -0,39 | 3,52 | -0,11 | 9,11E-01 |
| AT3G08960 | 0,38 | 1,31 | 4,38 | 0,30 | 7,65E-01 |
| AT3G09040 | 0,17 | 0,00 | 4,41 | 0,00 | 1,00E+00 |
| AT3G09070 | 1,31 | -1,99 | 2,20 | -0,91 | 3,65E-01 |
| AT3G09090 | 1,29 | 0,93 | 2,04 | 0,46 | 6,49E-01 |
| AT3G09190 | 0,55 | 2,45 | 3,77 | 0,65 | 5,15E-01 |
| AT3G09210 | 0,27 | 0,23 | 4,36 | 0,05 | 9,58E-01 |
| AT3G09270 | 0,53 | 0,44 | 2,81 | 0,16 | 8,77E-01 |

| | | | | | |
|---|---|---|---|---|---|
| AT3G09320 | 0,85 | 1,38 | 2,54 | 0,54 | 5,87E-01 |
| AT3G09410 | 0,82 | 2,87 | 2,41 | 1,19 | 2,34E-01 |
| AT3G09470 | 0,82 | 1,34 | 3,56 | 0,37 | 7,08E-01 |
| AT3G09560 | 0,97 | 0,27 | 2,28 | 0,12 | 9,05E-01 |
| AT3G09580 | 0,48 | 2,39 | 3,41 | 0,70 | 4,84E-01 |
| AT3G09770 | 0,38 | 1,48 | 4,40 | 0,34 | 7,36E-01 |
| AT3G09780 | 0,23 | 1,38 | 4,41 | 0,31 | 7,54E-01 |
| AT3G09850 | 1,65 | 1,94 | 1,66 | 1,16 | 2,44E-01 |
| AT3G09870 | 0,08 | 0,00 | 4,41 | 0,00 | 1,00E+00 |
| AT3G09920 | 0,16 | 0,00 | 4,41 | 0,00 | 1,00E+00 |
| AT3G10010 | 0,35 | 2,30 | 4,37 | 0,53 | 5,98E-01 |
| AT3G10020 | 0,27 | -0,91 | 4,38 | -0,21 | 8,35E-01 |
| AT3G10030 | 0,88 | -1,11 | 2,85 | -0,39 | 6,98E-01 |
| AT3G10050 | 0,25 | -0,54 | 4,41 | -0,12 | 9,02E-01 |
| AT3G10070 | 0,46 | 2,43 | 4,28 | 0,57 | 5,70E-01 |
| AT3G10130 | 0,21 | 0,00 | 4,41 | 0,00 | 1,00E+00 |
| AT3G10160 | 0,36 | 1,26 | 3,49 | 0,36 | 7,18E-01 |
| AT3G10230 | 1,13 | 1,36 | 2,42 | 0,56 | 5,73E-01 |
| AT3G10250 | 0,08 | 0,00 | 4,41 | 0,00 | 1,00E+00 |
| AT3G10270 | 0,56 | -0,68 | 3,14 | -0,22 | 8,28E-01 |
| AT3G10320 | 0,12 | 0,00 | 4,41 | 0,00 | 1,00E+00 |
| AT3G10360 | 0,63 | 2,44 | 3,77 | 0,65 | 5,19E-01 |
| AT3G10405 | 0,68 | 3,18 | 4,32 | 0,74 | 4,61E-01 |
| AT3G10410 | 1,05 | -0,28 | 1,89 | -0,15 | 8,81E-01 |
| AT3G10520 | 0,15 | 0,00 | 4,39 | 0,00 | 1,00E+00 |
| AT3G10525 | 0,39 | 0,42 | 3,50 | 0,12 | 9,04E-01 |
| AT3G10630 | 0,99 | 2,12 | 2,13 | 1,00 | 3,19E-01 |
| AT3G10640 | 0,33 | -0,54 | 4,41 | -0,12 | 9,02E-01 |
| AT3G10660 | 0,27 | 1,26 | 4,39 | 0,29 | 7,74E-01 |
| AT3G10690 | 0,77 | 2,55 | 3,46 | 0,74 | 4,62E-01 |
| AT3G10720 | 0,36 | -1,35 | 4,38 | -0,31 | 7,57E-01 |
| AT3G10730 | 0,58 | 0,34 | 3,03 | 0,11 | 9,10E-01 |
| AT3G10770 | 0,74 | 0,26 | 2,57 | 0,10 | 9,21E-01 |
| AT3G10810 | 0,46 | 2,18 | 4,24 | 0,52 | 6,06E-01 |
| AT3G10840 | 0,41 | 2,70 | 4,34 | 0,62 | 5,33E-01 |
| AT3G10850 | 0,36 | 1,38 | 4,41 | 0,31 | 7,54E-01 |
| AT3G10900 | 0,31 | 0,00 | 4,41 | 0,00 | 1,00E+00 |
| AT3G10915 | 0,24 | 0,00 | 4,41 | 0,00 | 1,00E+00 |
| AT3G10930 | 0,61 | 1,41 | 2,66 | 0,53 | 5,96E-01 |
| AT3G10960 | 1,23 | 2,23 | 1,97 | 1,13 | 2,58E-01 |
| AT3G11030 | 0,56 | 2,60 | 3,79 | 0,69 | 4,93E-01 |
| AT3G11100 | 0,47 | -0,54 | 4,08 | -0,13 | 8,94E-01 |
| AT3G11280 | 0,31 | 0,60 | 4,39 | 0,14 | 8,92E-01 |
| AT3G11290 | 0,30 | 1,27 | 4,39 | 0,29 | 7,72E-01 |
| AT3G11310 | 0,31 | 0,00 | 4,41 | 0,00 | 1,00E+00 |
| AT3G11320 | 0,17 | 0,00 | 4,41 | 0,00 | 1,00E+00 |
| AT3G11330 | 0,97 | 2,63 | 2,65 | 0,99 | 3,21E-01 |
| AT3G11435 | 0,23 | -1,59 | 4,35 | -0,36 | 7,15E-01 |
| AT3G11450 | 1,01 | 3,34 | 2,01 | 1,67 | 9,56E-02 |
| AT3G11510 | 1,34 | 0,17 | 1,86 | 0,09 | 9,28E-01 |
| AT3G02435 | 1,34 | -0,33 | 2,15 | -0,15 | 8,79E-01 |
| AT3G02445 | 0,88 | -1,58 | 3,51 | -0,45 | 6,53E-01 |
| AT3G02455 | 0,99 | -0,55 | 3,09 | -0,18 | 8,58E-01 |
| AT3G11570 | 0,77 | 3,17 | 3,04 | 1,04 | 2,98E-01 |
| AT3G11580 | 0,26 | 0,00 | 4,41 | 0,00 | 1,00E+00 |
| AT3G11650 | 0,48 | 1,38 | 3,32 | 0,42 | 6,77E-01 |

| | | | | | |
|---|---|---|---|---|---|
| AT3G11660 | 1,33 | 0,00 | 3,52 | 0,00 | 1,00E+00 |
| AT3G11680 | 0,22 | 0,00 | 4,41 | 0,00 | 1,00E+00 |
| AT3G11710 | 1,08 | 0,74 | 2,13 | 0,35 | 7,29E-01 |
| AT3G11730 | 0,25 | 1,34 | 4,40 | 0,30 | 7,61E-01 |
| AT3G11770 | 0,28 | 1,34 | 4,40 | 0,30 | 7,61E-01 |
| AT3G11780 | 0,21 | 0,00 | 4,41 | 0,00 | 1,00E+00 |
| AT3G11820 | 1,51 | 2,12 | 1,93 | 1,09 | 2,74E-01 |
| AT3G11840 | 0,47 | 0,00 | 4,17 | 0,00 | 1,00E+00 |
| AT3G11964 | 0,92 | 1,24 | 1,92 | 0,65 | 5,17E-01 |
| AT3G12030 | 0,12 | 0,00 | 4,41 | 0,00 | 1,00E+00 |
| AT3G12130 | 0,80 | 2,03 | 2,54 | 0,80 | 4,25E-01 |
| AT3G12140 | 0,25 | -0,54 | 4,41 | -0,12 | 9,02E-01 |
| AT3G12145 | 0,15 | 0,00 | 4,41 | 0,00 | 1,00E+00 |
| AT3G12200 | 0,53 | 1,39 | 3,98 | 0,35 | 7,27E-01 |
| AT3G12260 | 0,32 | -0,54 | 4,41 | -0,12 | 9,02E-01 |
| AT3G12280 | 1,55 | 1,25 | 1,72 | 0,72 | 4,68E-01 |
| AT3G12400 | 0,71 | 1,30 | 2,11 | 0,62 | 5,38E-01 |
| AT3G12490 | 0,47 | 0,43 | 2,51 | 0,17 | 8,63E-01 |
| AT3G12500 | 0,08 | 0,00 | 4,41 | 0,00 | 1,00E+00 |
| AT3G12580 | 0,43 | 1,90 | 4,39 | 0,43 | 6,65E-01 |
| AT3G12587 | 0,28 | 2,12 | 4,38 | 0,48 | 6,28E-01 |
| AT3G12590 | 0,82 | 3,36 | 2,40 | 1,40 | 1,62E-01 |
| AT3G12610 | 0,59 | 1,48 | 2,36 | 0,62 | 5,32E-01 |
| AT3G12670 | 1,09 | -0,40 | 2,57 | -0,16 | 8,76E-01 |
| AT3G12680 | 0,32 | 1,40 | 4,40 | 0,32 | 7,50E-01 |
| AT3G12690 | 0,57 | 0,00 | 4,41 | 0,00 | 1,00E+00 |
| AT3G12750 | 0,45 | 2,10 | 4,03 | 0,52 | 6,03E-01 |
| AT3G12770 | 0,67 | 3,23 | 2,87 | 1,12 | 2,61E-01 |
| AT3G12800 | 0,96 | -0,51 | 2,08 | -0,25 | 8,06E-01 |
| AT3G12820 | 0,21 | 1,38 | 4,41 | 0,31 | 7,54E-01 |
| AT3G12920 | 1,14 | 2,11 | 2,86 | 0,74 | 4,60E-01 |
| AT3G12940 | 0,28 | 0,00 | 4,41 | 0,00 | 1,00E+00 |
| AT3G12950 | 0,48 | 1,38 | 4,41 | 0,31 | 7,54E-01 |
| AT3G12965 | 0,28 | 1,35 | 4,40 | 0,31 | 7,59E-01 |
| AT3G13000 | 0,24 | 1,96 | 4,39 | 0,45 | 6,55E-01 |
| AT3G13010 | 0,29 | 1,32 | 4,40 | 0,30 | 7,64E-01 |
| AT3G13020 | 0,33 | 1,71 | 4,38 | 0,39 | 6,97E-01 |
| AT3G13030 | 0,97 | 2,60 | 2,28 | 1,14 | 2,55E-01 |
| AT3G13040 | 0,58 | -0,40 | 2,78 | -0,14 | 8,86E-01 |
| AT3G13061 | 0,23 | 0,00 | 4,41 | 0,00 | 1,00E+00 |
| AT3G13110 | 1,06 | 0,98 | 2,03 | 0,48 | 6,30E-01 |
| AT3G13120 | 0,60 | -0,23 | 3,03 | -0,08 | 9,39E-01 |
| AT3G13150 | 0,37 | 1,75 | 4,38 | 0,40 | 6,89E-01 |
| AT3G13160 | 0,08 | 0,00 | 4,41 | 0,00 | 1,00E+00 |
| AT3G13220 | 0,37 | 1,35 | 4,40 | 0,31 | 7,59E-01 |
| AT3G13222 | 0,48 | 0,00 | 4,41 | 0,00 | 1,00E+00 |
| AT3G13225 | 0,32 | 0,00 | 4,41 | 0,00 | 1,00E+00 |
| AT3G13226 | 0,08 | 0,00 | 4,41 | 0,00 | 1,00E+00 |
| AT3G13235 | 0,44 | 0,44 | 2,92 | 0,15 | 8,81E-01 |
| AT3G13310 | 0,25 | -0,43 | 4,39 | -0,10 | 9,23E-01 |
| AT3G13390 | 0,05 | 0,00 | 4,41 | 0,00 | 1,00E+00 |
| AT3G13404 | 0,46 | 0,47 | 4,11 | 0,11 | 9,09E-01 |
| AT3G13410 | 1,64 | -0,02 | 2,16 | -0,01 | 9,91E-01 |
| AT3G13450 | 0,75 | 3,22 | 2,77 | 1,16 | 2,46E-01 |
| AT3G13480 | 0,20 | 0,00 | 4,41 | 0,00 | 1,00E+00 |
| AT3G13490 | 0,57 | 1,09 | 3,13 | 0,35 | 7,28E-01 |

| | | | | | |
|---|---|---|---|---|---|
| AT3G13570 | 0,44 | 0,32 | 4,18 | 0,08 | 9,39E-01 |
| AT3G13720 | 0,96 | -0,34 | 2,02 | -0,17 | 8,65E-01 |
| AT3G13772 | 0,72 | 1,38 | 2,94 | 0,47 | 6,38E-01 |
| AT3G13860 | 0,89 | 1,59 | 2,11 | 0,75 | 4,52E-01 |
| AT3G13990 | 1,05 | 2,14 | 2,12 | 1,01 | 3,13E-01 |
| AT3G14070 | 0,23 | 0,00 | 4,41 | 0,00 | 1,00E+00 |
| AT3G14075 | 0,26 | -0,48 | 4,40 | -0,11 | 9,13E-01 |
| AT3G14090 | 0,68 | 1,17 | 3,16 | 0,37 | 7,11E-01 |
| AT3G14110 | 0,61 | 1,67 | 2,59 | 0,64 | 5,19E-01 |
| AT3G14172 | 1,38 | 2,76 | 1,83 | 1,50 | 1,33E-01 |
| AT3G14180 | 0,40 | 0,00 | 4,41 | 0,00 | 1,00E+00 |
| AT3G14205 | 0,51 | 1,17 | 2,83 | 0,41 | 6,80E-01 |
| AT3G14230 | 0,40 | 2,13 | 4,20 | 0,51 | 6,13E-01 |
| AT3G14240 | 1,34 | 1,65 | 1,65 | 0,99 | 3,20E-01 |
| AT3G14330 | 0,56 | 1,19 | 3,07 | 0,39 | 6,97E-01 |
| AT3G14350 | 1,44 | 1,91 | 1,58 | 1,21 | 2,28E-01 |
| AT3G14400 | 1,24 | 2,77 | 3,23 | 0,86 | 3,91E-01 |
| AT3G14430 | 0,25 | -0,54 | 4,41 | -0,12 | 9,02E-01 |
| AT3G14440 | 0,33 | 0,00 | 4,41 | 0,00 | 1,00E+00 |
| AT3G14460 | 0,50 | 1,73 | 3,99 | 0,43 | 6,66E-01 |
| AT3G14590 | 0,14 | 0,00 | 4,41 | 0,00 | 1,00E+00 |
| AT3G14810 | 0,79 | -0,01 | 2,46 | 0,00 | 9,97E-01 |
| AT3G14900 | 0,17 | 0,00 | 4,41 | 0,00 | 1,00E+00 |
| AT3G15110 | 0,15 | 0,00 | 4,39 | 0,00 | 1,00E+00 |
| AT3G15120 | 0,82 | 3,12 | 2,46 | 1,27 | 2,06E-01 |
| AT3G15150 | 0,08 | 0,00 | 4,41 | 0,00 | 1,00E+00 |
| AT3G15220 | 0,77 | 2,16 | 2,46 | 0,88 | 3,79E-01 |
| AT3G15354 | 0,91 | 0,36 | 2,32 | 0,15 | 8,78E-01 |
| AT3G15356 | 0,36 | 2,30 | 4,37 | 0,53 | 5,99E-01 |
| AT3G15470 | 0,41 | 0,42 | 2,92 | 0,14 | 8,86E-01 |
| AT3G03125 | 0,17 | 0,00 | 4,41 | 0,00 | 1,00E+00 |
| AT3G15530 | 1,09 | 1,37 | 2,63 | 0,52 | 6,03E-01 |
| AT3G15570 | 0,83 | 0,42 | 2,81 | 0,15 | 8,81E-01 |
| AT3G15590 | 0,41 | 1,98 | 4,29 | 0,46 | 6,44E-01 |
| AT3G15630 | 0,46 | 0,00 | 3,33 | 0,00 | 1,00E+00 |
| AT3G15720 | 0,20 | 0,00 | 4,41 | 0,00 | 1,00E+00 |
| AT3G15810 | 0,58 | 0,49 | 2,64 | 0,19 | 8,52E-01 |
| AT3G15840 | 1,45 | 0,35 | 1,40 | 0,25 | 8,00E-01 |
| AT3G15900 | 0,29 | -0,54 | 4,41 | -0,12 | 9,02E-01 |
| AT3G15940 | 1,43 | 0,22 | 1,92 | 0,12 | 9,07E-01 |
| AT3G15970 | 0,63 | 1,38 | 2,71 | 0,51 | 6,11E-01 |
| AT3G15980 | 1,59 | 1,56 | 1,65 | 0,95 | 3,43E-01 |
| AT3G16000 | 0,51 | -1,74 | 3,26 | -0,53 | 5,94E-01 |
| AT3G16110 | 0,21 | 0,00 | 4,41 | 0,00 | 1,00E+00 |
| AT3G16190 | 0,21 | 1,27 | 4,39 | 0,29 | 7,72E-01 |
| AT3G03235 | 0,34 | 1,38 | 4,41 | 0,31 | 7,54E-01 |
| AT3G16290 | 0,84 | -1,62 | 2,13 | -0,76 | 4,47E-01 |
| AT3G16310 | 0,55 | 1,49 | 3,86 | 0,39 | 7,00E-01 |
| AT3G16500 | 0,17 | 0,00 | 4,41 | 0,00 | 1,00E+00 |
| AT3G16510 | 0,65 | 1,39 | 3,14 | 0,44 | 6,59E-01 |
| AT3G16520 | 0,76 | 0,05 | 2,28 | 0,02 | 9,81E-01 |
| AT3G16530 | 0,27 | 1,35 | 4,40 | 0,31 | 7,59E-01 |
| AT3G16560 | 0,66 | 2,28 | 2,58 | 0,88 | 3,78E-01 |
| AT3G16630 | 1,01 | 1,03 | 2,20 | 0,47 | 6,38E-01 |
| AT3G16820 | 1,66 | 0,90 | 1,66 | 0,54 | 5,87E-01 |
| AT3G16840 | 0,19 | 0,00 | 4,41 | 0,00 | 1,00E+00 |

| | | | | | |
|---|---|---|---|---|---|
| AT3G16860 | 0,47 | 1,07 | 4,06 | 0,26 | 7,92E-01 |
| AT3G16870 | 0,39 | 1,76 | 4,38 | 0,40 | 6,87E-01 |
| AT3G16890 | 0,19 | -0,45 | 4,39 | -0,10 | 9,19E-01 |
| AT3G17020 | 1,21 | 0,94 | 2,16 | 0,43 | 6,65E-01 |
| AT3G03415 | 1,34 | 0,58 | 1,86 | 0,31 | 7,55E-01 |
| AT3G03445 | 0,77 | 2,10 | 2,63 | 0,80 | 4,26E-01 |
| AT3G17170 | 0,31 | -1,72 | 4,36 | -0,40 | 6,93E-01 |
| AT3G17320 | 0,12 | 0,00 | 4,41 | 0,00 | 1,00E+00 |
| AT3G17330 | 0,20 | 0,00 | 4,41 | 0,00 | 1,00E+00 |
| AT3G17420 | 0,17 | -0,49 | 4,40 | -0,11 | 9,11E-01 |
| AT3G17440 | 0,08 | 0,00 | 4,41 | 0,00 | 1,00E+00 |
| AT3G17450 | 0,60 | 2,53 | 2,98 | 0,85 | 3,96E-01 |
| AT3G17655 | 0,19 | 0,00 | 4,41 | 0,00 | 1,00E+00 |
| AT3G17740 | 0,37 | 1,38 | 4,37 | 0,32 | 7,52E-01 |
| AT3G17760 | 0,21 | 1,38 | 4,41 | 0,31 | 7,54E-01 |
| AT3G17770 | 1,51 | 2,19 | 1,67 | 1,31 | 1,90E-01 |
| AT3G03585 | 0,05 | 0,00 | 4,41 | 0,00 | 1,00E+00 |
| AT3G17940 | 0,81 | 1,48 | 2,51 | 0,59 | 5,56E-01 |
| AT3G18030 | 0,52 | 1,30 | 4,08 | 0,32 | 7,49E-01 |
| AT3G18035 | 1,23 | -0,63 | 1,72 | -0,37 | 7,13E-01 |
| AT3G18110 | 1,16 | 2,64 | 2,55 | 1,04 | 3,00E-01 |
| AT3G18130 | 0,88 | 2,58 | 2,44 | 1,06 | 2,91E-01 |
| AT3G18140 | 0,30 | 1,29 | 4,40 | 0,29 | 7,68E-01 |
| AT3G18250 | 0,55 | 3,04 | 4,32 | 0,70 | 4,82E-01 |
| AT3G18380 | 0,25 | 1,34 | 4,40 | 0,30 | 7,61E-01 |
| AT3G18420 | 0,20 | 0,00 | 4,41 | 0,00 | 1,00E+00 |
| AT3G18535 | 0,67 | 1,38 | 2,61 | 0,53 | 5,97E-01 |
| AT3G18600 | 0,25 | 1,84 | 4,38 | 0,42 | 6,75E-01 |
| AT3G18610 | 0,55 | 0,00 | 4,39 | 0,00 | 1,00E+00 |
| AT3G18630 | 0,20 | 1,33 | 4,40 | 0,30 | 7,63E-01 |
| AT3G18700 | 1,19 | 0,00 | 2,33 | 0,00 | 1,00E+00 |
| AT3G18750 | 0,17 | 0,00 | 4,41 | 0,00 | 1,00E+00 |
| AT3G18930 | 0,90 | 0,39 | 2,43 | 0,16 | 8,71E-01 |
| AT3G18970 | 0,08 | 0,00 | 4,41 | 0,00 | 1,00E+00 |
| AT3G19000 | 0,24 | 0,00 | 4,41 | 0,00 | 1,00E+00 |
| AT3G19020 | 0,54 | 2,56 | 3,18 | 0,80 | 4,21E-01 |
| AT3G19150 | 0,16 | 0,00 | 4,41 | 0,00 | 1,00E+00 |
| AT3G03815 | 0,25 | 1,84 | 4,38 | 0,42 | 6,75E-01 |
| AT3G19300 | 0,12 | 0,00 | 4,41 | 0,00 | 1,00E+00 |
| AT3G19340 | 0,12 | 0,00 | 4,41 | 0,00 | 1,00E+00 |
| AT3G19430 | 0,25 | 1,84 | 4,38 | 0,42 | 6,75E-01 |
| AT3G19490 | 0,72 | 1,38 | 2,63 | 0,53 | 5,99E-01 |
| AT3G19510 | 0,12 | 0,00 | 4,41 | 0,00 | 1,00E+00 |
| AT3G19515 | 0,59 | 0,52 | 3,19 | 0,16 | 8,69E-01 |
| AT3G19553 | 0,42 | 2,11 | 4,14 | 0,51 | 6,10E-01 |
| AT3G19580 | 0,72 | 1,36 | 2,63 | 0,52 | 6,05E-01 |
| AT3G19620 | 0,32 | 2,14 | 4,37 | 0,49 | 6,25E-01 |
| AT3G19630 | 0,39 | 1,76 | 4,38 | 0,40 | 6,87E-01 |
| AT3G19710 | 0,69 | 1,33 | 2,63 | 0,51 | 6,13E-01 |
| AT3G19720 | 1,17 | 0,00 | 2,64 | 0,00 | 1,00E+00 |
| AT3G19740 | 0,59 | 0,46 | 3,25 | 0,14 | 8,87E-01 |
| AT3G19790 | 0,71 | 2,86 | 2,06 | 1,39 | 1,65E-01 |
| AT3G19810 | 0,70 | -0,18 | 4,32 | -0,04 | 9,66E-01 |
| AT3G19830 | 0,67 | 0,00 | 3,11 | 0,00 | 1,00E+00 |
| AT3G19850 | 0,50 | 2,48 | 3,16 | 0,79 | 4,32E-01 |
| AT3G19880 | 0,35 | 0,00 | 3,54 | 0,00 | 1,00E+00 |

| | | | | | |
|---|---|---|---|---|---|
| AT3G19895 | 0,23 | 0,00 | 4,41 | 0,00 | 1,00E+00 |
| AT3G19900 | 0,17 | 0,00 | 4,41 | 0,00 | 1,00E+00 |
| AT3G20015 | 0,62 | 2,94 | 3,07 | 0,96 | 3,39E-01 |
| AT3G20050 | 0,74 | 0,19 | 2,54 | 0,07 | 9,42E-01 |
| AT3G20060 | 0,33 | -0,54 | 4,41 | -0,12 | 9,02E-01 |
| AT3G20230 | 0,35 | 0,00 | 4,41 | 0,00 | 1,00E+00 |
| AT3G20250 | 1,61 | 2,94 | 1,89 | 1,55 | 1,20E-01 |
| AT3G20290 | 0,63 | 0,00 | 3,17 | 0,00 | 1,00E+00 |
| AT3G20320 | 0,51 | 0,38 | 2,82 | 0,13 | 8,93E-01 |
| AT3G20390 | 0,62 | -1,80 | 2,32 | -0,78 | 4,37E-01 |
| AT3G20470 | 0,38 | 0,33 | 4,40 | 0,07 | 9,41E-01 |
| AT3G20620 | 0,68 | 1,91 | 3,64 | 0,52 | 6,00E-01 |
| AT3G20630 | 1,20 | 2,55 | 1,81 | 1,41 | 1,58E-01 |
| AT3G20655 | 0,30 | 1,38 | 4,41 | 0,31 | 7,54E-01 |
| AT3G20700 | 1,05 | 0,00 | 2,63 | 0,00 | 1,00E+00 |
| AT3G03985 | 0,54 | 0,00 | 4,41 | 0,00 | 1,00E+00 |
| AT3G20830 | 0,60 | -1,57 | 3,30 | -0,48 | 6,34E-01 |
| AT3G21055 | 1,34 | -0,31 | 1,65 | -0,19 | 8,49E-01 |
| AT3G21070 | 0,23 | 0,00 | 4,41 | 0,00 | 1,00E+00 |
| AT3G21140 | 0,25 | 0,00 | 4,41 | 0,00 | 1,00E+00 |
| AT3G21160 | 0,11 | 0,00 | 4,41 | 0,00 | 1,00E+00 |
| AT3G21215 | 0,11 | 0,00 | 4,41 | 0,00 | 1,00E+00 |
| AT3G21300 | 0,80 | 1,87 | 2,36 | 0,79 | 4,27E-01 |
| AT3G21430 | 0,61 | 1,36 | 3,26 | 0,42 | 6,76E-01 |
| AT3G21540 | 0,20 | -0,43 | 4,39 | -0,10 | 9,22E-01 |
| AT3G21570 | 0,87 | -1,31 | 2,92 | -0,45 | 6,54E-01 |
| AT3G21630 | 0,12 | 0,00 | 4,41 | 0,00 | 1,00E+00 |
| AT3G21650 | 0,33 | 1,37 | 4,41 | 0,31 | 7,56E-01 |
| AT3G21670 | 1,42 | 2,68 | 2,47 | 1,08 | 2,78E-01 |
| AT3G21740 | 0,24 | 1,57 | 4,39 | 0,36 | 7,20E-01 |
| AT3G21790 | 0,96 | 0,37 | 2,34 | 0,16 | 8,76E-01 |
| AT3G21865 | 0,48 | 1,29 | 2,81 | 0,46 | 6,46E-01 |
| AT3G22070 | 0,17 | -1,42 | 4,35 | -0,33 | 7,45E-01 |
| AT3G22110 | 0,73 | 1,59 | 2,56 | 0,62 | 5,35E-01 |
| AT3G22142 | 1,40 | -2,30 | 1,93 | -1,19 | 2,33E-01 |
| AT3G22170 | 1,10 | 2,32 | 2,20 | 1,05 | 2,93E-01 |
| AT3G22190 | 0,78 | 2,97 | 2,92 | 1,02 | 3,09E-01 |
| AT3G22210 | 0,48 | -0,44 | 3,37 | -0,13 | 8,96E-01 |
| AT3G22220 | 0,36 | 0,45 | 3,54 | 0,13 | 9,00E-01 |
| AT3G22230 | 0,95 | 2,86 | 2,01 | 1,42 | 1,55E-01 |
| AT3G22240 | 1,25 | 0,41 | 2,13 | 0,19 | 8,49E-01 |
| AT3G22320 | 0,41 | 1,98 | 4,29 | 0,46 | 6,44E-01 |
| AT3G22421 | 0,96 | 0,00 | 2,85 | 0,00 | 1,00E+00 |
| AT3G22470 | 0,59 | 0,43 | 2,73 | 0,16 | 8,74E-01 |
| AT3G22530 | 0,38 | 0,34 | 4,37 | 0,08 | 9,37E-01 |
| AT3G22550 | 0,60 | 1,45 | 3,16 | 0,46 | 6,45E-01 |
| AT3G22580 | 0,15 | 0,00 | 4,39 | 0,00 | 1,00E+00 |
| AT3G22600 | 0,17 | 0,00 | 4,41 | 0,00 | 1,00E+00 |
| AT3G22630 | 0,68 | 2,54 | 2,30 | 1,11 | 2,68E-01 |
| AT3G22660 | 0,44 | 2,58 | 4,35 | 0,59 | 5,52E-01 |
| AT3G22690 | 0,29 | -0,51 | 4,40 | -0,12 | 9,08E-01 |
| AT3G22710 | 1,14 | 2,57 | 1,99 | 1,29 | 1,97E-01 |
| AT3G22720 | 0,08 | 0,00 | 4,41 | 0,00 | 1,00E+00 |
| AT3G22730 | 1,61 | -1,81 | 1,91 | -0,94 | 3,45E-01 |
| AT3G22770 | 0,68 | -1,69 | 3,31 | -0,51 | 6,09E-01 |
| AT3G22950 | 0,15 | -0,43 | 4,39 | -0,10 | 9,23E-01 |

| | | | | | |
|---|---|---|---|---|---|
| AT3G22980 | 0,74 | 0,77 | 2,56 | 0,30 | 7,64E-01 |
| AT3G23020 | 0,53 | 2,51 | 3,84 | 0,65 | 5,14E-01 |
| AT3G23030 | 0,69 | 0,29 | 2,10 | 0,14 | 8,90E-01 |
| AT3G23080 | 0,08 | 0,00 | 4,41 | 0,00 | 1,00E+00 |
| AT3G23170 | 0,50 | 1,32 | 2,77 | 0,48 | 6,34E-01 |
| AT3G23172 | 0,24 | 0,00 | 4,41 | 0,00 | 1,00E+00 |
| AT3G23400 | 0,58 | 0,61 | 3,32 | 0,18 | 8,54E-01 |
| AT3G23420 | 0,32 | 0,00 | 4,41 | 0,00 | 1,00E+00 |
| AT3G23490 | 0,26 | 0,00 | 4,41 | 0,00 | 1,00E+00 |
| AT3G23570 | 0,32 | 1,36 | 4,40 | 0,31 | 7,58E-01 |
| AT3G23633 | 0,12 | 0,00 | 4,41 | 0,00 | 1,00E+00 |
| AT3G23650 | 0,55 | 0,00 | 3,08 | 0,00 | 1,00E+00 |
| AT3G04585 | 0,72 | 0,00 | 2,97 | 0,00 | 1,00E+00 |
| AT3G23660 | 0,75 | 2,65 | 2,93 | 0,91 | 3,65E-01 |
| AT3G23700 | 0,24 | -0,40 | 4,38 | -0,09 | 9,26E-01 |
| AT3G23740 | 0,08 | 0,00 | 4,41 | 0,00 | 1,00E+00 |
| AT3G23830 | 0,27 | -0,60 | 4,40 | -0,14 | 8,91E-01 |
| AT3G23900 | 0,77 | 1,40 | 2,90 | 0,48 | 6,29E-01 |
| AT3G23970 | 0,08 | 0,00 | 4,41 | 0,00 | 1,00E+00 |
| AT3G23980 | 0,76 | 1,77 | 2,56 | 0,69 | 4,90E-01 |
| AT3G24080 | 0,22 | 0,00 | 4,41 | 0,00 | 1,00E+00 |
| AT3G24120 | 0,83 | 1,21 | 3,55 | 0,34 | 7,32E-01 |
| AT3G24140 | 0,26 | -0,47 | 4,40 | -0,11 | 9,15E-01 |
| AT3G24170 | 1,33 | 2,87 | 1,95 | 1,47 | 1,42E-01 |
| AT3G24200 | 0,11 | 0,00 | 4,41 | 0,00 | 1,00E+00 |
| AT3G24260 | 0,15 | 0,00 | 4,41 | 0,00 | 1,00E+00 |
| AT3G24270 | 0,91 | 0,00 | 2,54 | 0,00 | 1,00E+00 |
| AT3G24320 | 0,63 | 1,93 | 3,12 | 0,62 | 5,36E-01 |
| AT3G24430 | 0,54 | -0,25 | 2,73 | -0,09 | 9,28E-01 |
| AT3G24440 | 0,50 | 2,49 | 3,22 | 0,77 | 4,39E-01 |
| AT3G24465 | 0,32 | -0,97 | 3,67 | -0,27 | 7,91E-01 |
| AT3G24503 | 1,13 | 0,75 | 1,93 | 0,39 | 6,96E-01 |
| AT3G24518 | 0,96 | -0,14 | 2,49 | -0,05 | 9,56E-01 |
| AT3G24614 | 0,96 | 1,79 | 2,03 | 0,88 | 3,80E-01 |
| AT3G24670 | 0,20 | 0,00 | 4,41 | 0,00 | 1,00E+00 |
| AT3G24870 | 1,16 | -1,56 | 2,12 | -0,74 | 4,61E-01 |
| AT3G24880 | 0,67 | 2,97 | 3,07 | 0,97 | 3,33E-01 |
| AT3G24900 | 0,51 | 1,38 | 3,22 | 0,43 | 6,69E-01 |
| AT3G25090 | 0,87 | 0,43 | 2,26 | 0,19 | 8,51E-01 |
| AT3G25150 | 1,03 | 2,86 | 2,57 | 1,12 | 2,65E-01 |
| AT3G25180 | 0,15 | 0,00 | 4,39 | 0,00 | 1,00E+00 |
| AT3G25460 | 1,48 | -3,12 | 1,83 | -1,71 | 8,72E-02 |
| AT3G25530 | 0,19 | 0,00 | 4,41 | 0,00 | 1,00E+00 |
| AT3G25560 | 0,41 | 2,00 | 4,38 | 0,46 | 6,48E-01 |
| AT3G25610 | 1,05 | 1,36 | 2,23 | 0,61 | 5,41E-01 |
| AT3G25660 | 0,47 | -0,40 | 4,02 | -0,10 | 9,20E-01 |
| AT3G04935 | 0,15 | 0,00 | 4,41 | 0,00 | 1,00E+00 |
| AT3G25800 | 1,14 | -0,74 | 1,94 | -0,38 | 7,02E-01 |
| AT3G25840 | 1,32 | 1,78 | 1,73 | 1,03 | 3,04E-01 |
| AT3G25870 | 0,65 | -1,32 | 2,55 | -0,52 | 6,06E-01 |
| AT3G25880 | 1,38 | 4,13 | 2,50 | 1,66 | 9,78E-02 |
| AT3G25882 | 0,50 | 1,63 | 4,07 | 0,40 | 6,89E-01 |
| AT3G25920 | 0,58 | 1,28 | 2,36 | 0,54 | 5,87E-01 |
| AT3G25990 | 0,75 | 3,69 | 3,40 | 1,08 | 2,78E-01 |
| AT3G26170 | 1,29 | -1,25 | 1,92 | -0,65 | 5,15E-01 |
| AT3G26180 | 0,36 | 1,29 | 4,39 | 0,29 | 7,69E-01 |

| | | | | | |
|---|---|---|---|---|---|
| AT3G26210 | 1,30 | 2,16 | 2,25 | 0,96 | 3,36E-01 |
| AT3G26220 | 0,49 | 1,29 | 4,10 | 0,32 | 7,52E-01 |
| AT3G26230 | 0,79 | 1,43 | 2,51 | 0,57 | 5,67E-01 |
| AT3G26280 | 1,16 | 0,63 | 1,71 | 0,37 | 7,11E-01 |
| AT3G26290 | 0,78 | -0,95 | 2,05 | -0,46 | 6,44E-01 |
| AT3G26300 | 0,36 | 0,97 | 4,36 | 0,22 | 8,24E-01 |
| AT3G26340 | 0,62 | 1,89 | 3,71 | 0,51 | 6,11E-01 |
| AT3G26410 | 0,31 | 1,36 | 4,35 | 0,31 | 7,55E-01 |
| AT3G26500 | 1,32 | 0,00 | 2,22 | 0,00 | 1,00E+00 |
| AT3G26510 | 0,66 | 0,43 | 2,33 | 0,19 | 8,52E-01 |
| AT3G26550 | 0,20 | 1,38 | 4,41 | 0,31 | 7,54E-01 |
| AT3G26560 | 0,40 | -0,54 | 3,45 | -0,16 | 8,76E-01 |
| AT3G26590 | 0,41 | 2,21 | 4,37 | 0,51 | 6,13E-01 |
| AT3G26616 | 0,23 | 0,00 | 4,41 | 0,00 | 1,00E+00 |
| AT3G26630 | 0,69 | 3,20 | 2,44 | 1,31 | 1,89E-01 |
| AT3G26670 | 0,54 | 2,61 | 3,33 | 0,78 | 4,33E-01 |
| AT3G26744 | 0,39 | 1,08 | 4,30 | 0,25 | 8,02E-01 |
| AT3G26812 | 0,17 | 0,00 | 4,41 | 0,00 | 1,00E+00 |
| AT3G26819 | 0,05 | 0,00 | 4,41 | 0,00 | 1,00E+00 |
| AT3G26860 | 0,30 | -1,78 | 4,34 | -0,41 | 6,82E-01 |
| AT3G26870 | 1,55 | -3,79 | 2,04 | -1,85 | 6,36E-02 |
| AT3G26890 | 0,29 | 1,37 | 4,41 | 0,31 | 7,55E-01 |
| AT3G26932 | 0,90 | -0,43 | 2,52 | -0,17 | 8,65E-01 |
| AT3G26935 | 0,27 | 1,31 | 4,40 | 0,30 | 7,66E-01 |
| AT3G26980 | 0,14 | 0,00 | 4,41 | 0,00 | 1,00E+00 |
| AT3G27010 | 0,67 | 0,11 | 3,66 | 0,03 | 9,75E-01 |
| AT3G27020 | 0,51 | 2,15 | 2,46 | 0,87 | 3,83E-01 |
| AT3G27040 | 1,43 | -2,41 | 2,00 | -1,21 | 2,27E-01 |
| AT3G27060 | 0,90 | -1,40 | 2,53 | -0,55 | 5,82E-01 |
| AT3G27100 | 0,74 | 2,22 | 4,33 | 0,51 | 6,09E-01 |
| AT3G27240 | 0,37 | 1,38 | 3,48 | 0,40 | 6,92E-01 |
| AT3G27300 | 0,49 | 0,17 | 3,28 | 0,05 | 9,58E-01 |
| AT3G27380 | 0,45 | -1,23 | 4,38 | -0,28 | 7,79E-01 |
| AT3G05235 | 0,99 | -1,07 | 2,53 | -0,42 | 6,72E-01 |
| AT3G05245 | 0,33 | -0,54 | 4,41 | -0,12 | 9,02E-01 |
| AT3G27530 | 0,76 | 2,29 | 2,72 | 0,84 | 4,00E-01 |
| AT3G27600 | 1,14 | 2,52 | 2,40 | 1,05 | 2,92E-01 |
| AT3G27610 | 0,33 | 0,00 | 4,41 | 0,00 | 1,00E+00 |
| AT3G27680 | 0,86 | -0,48 | 2,41 | -0,20 | 8,41E-01 |
| AT3G27700 | 1,39 | 0,50 | 2,23 | 0,23 | 8,22E-01 |
| AT3G27740 | 0,52 | 1,21 | 4,04 | 0,30 | 7,65E-01 |
| AT3G27770 | 0,43 | 0,32 | 2,91 | 0,11 | 9,12E-01 |
| AT3G27906 | 1,35 | 0,92 | 1,85 | 0,50 | 6,19E-01 |
| AT3G27925 | 0,65 | -1,87 | 3,07 | -0,61 | 5,42E-01 |
| AT3G27930 | 0,20 | 0,00 | 4,41 | 0,00 | 1,00E+00 |
| AT3G27990 | 0,64 | -0,37 | 2,57 | -0,14 | 8,87E-01 |
| AT3G28340 | 0,22 | 0,00 | 4,41 | 0,00 | 1,00E+00 |
| AT3G28380 | 0,08 | 0,00 | 4,41 | 0,00 | 1,00E+00 |
| AT3G28410 | 0,14 | 0,00 | 4,41 | 0,00 | 1,00E+00 |
| AT3G28674 | 0,28 | 0,00 | 4,41 | 0,00 | 1,00E+00 |
| AT3G28695 | 0,43 | 1,45 | 4,24 | 0,34 | 7,32E-01 |
| AT3G28710 | 0,99 | 0,29 | 1,89 | 0,15 | 8,78E-01 |
| AT3G28780 | 0,61 | -1,22 | 3,21 | -0,38 | 7,03E-01 |
| AT3G28860 | 1,33 | 1,05 | 2,31 | 0,45 | 6,50E-01 |
| AT3G28890 | 0,43 | 0,31 | 4,39 | 0,07 | 9,44E-01 |
| AT3G28920 | 1,35 | 0,26 | 1,96 | 0,13 | 8,93E-01 |

| | | | | | |
|---|---|---|---|---|---|
| AT3G28930 | 0,77 | 0,00 | 3,05 | 0,00 | 1,00E+00 |
| AT3G28940 | 1,30 | 0,47 | 1,92 | 0,24 | 8,08E-01 |
| AT3G28950 | 0,97 | -0,16 | 1,98 | -0,08 | 9,34E-01 |
| AT3G28960 | 0,30 | 0,00 | 4,41 | 0,00 | 1,00E+00 |
| AT3G28970 | 0,52 | 1,29 | 4,06 | 0,32 | 7,50E-01 |
| AT3G29030 | 0,33 | -0,80 | 4,38 | -0,18 | 8,54E-01 |
| AT3G05665 | 0,80 | 1,32 | 2,81 | 0,47 | 6,39E-01 |
| AT3G29110 | 0,53 | -0,71 | 2,63 | -0,27 | 7,86E-01 |
| AT3G29160 | 0,59 | -0,40 | 4,34 | -0,09 | 9,26E-01 |
| AT3G29200 | 0,59 | 2,19 | 3,13 | 0,70 | 4,85E-01 |
| AT3G29350 | 0,17 | 1,32 | 4,40 | 0,30 | 7,64E-01 |
| AT3G29400 | 0,67 | 2,05 | 2,67 | 0,77 | 4,42E-01 |
| AT3G29570 | 0,38 | 0,00 | 4,41 | 0,00 | 1,00E+00 |
| AT3G05745 | 0,14 | 0,00 | 4,41 | 0,00 | 1,00E+00 |
| AT3G29633 | 0,14 | -0,78 | 4,37 | -0,18 | 8,58E-01 |
| AT3G29639 | 0,16 | 0,00 | 4,41 | 0,00 | 1,00E+00 |
| AT3G05895 | 0,08 | 0,00 | 4,41 | 0,00 | 1,00E+00 |
| AT3G29680 | 0,25 | 1,36 | 4,41 | 0,31 | 7,57E-01 |
| AT3G29720 | 0,40 | -0,40 | 4,38 | -0,09 | 9,28E-01 |
| AT3G29760 | 0,67 | 3,14 | 3,00 | 1,04 | 2,96E-01 |
| AT3G29785 | 0,22 | 0,00 | 4,41 | 0,00 | 1,00E+00 |
| AT3G30160 | 0,58 | -0,54 | 3,10 | -0,18 | 8,60E-01 |
| AT3G30200 | 0,12 | 0,00 | 4,41 | 0,00 | 1,00E+00 |
| AT3G30220 | 0,08 | 0,00 | 4,41 | 0,00 | 1,00E+00 |
| AT3G30230 | 0,54 | 0,00 | 3,99 | 0,00 | 1,00E+00 |
| AT3G30300 | 0,53 | 2,15 | 4,38 | 0,49 | 6,24E-01 |
| AT3G30840 | 0,21 | 0,00 | 4,41 | 0,00 | 1,00E+00 |
| AT3G32190 | 0,34 | 1,31 | 3,55 | 0,37 | 7,12E-01 |
| AT3G32420 | 0,10 | 0,00 | 4,41 | 0,00 | 1,00E+00 |
| AT3G32904 | 0,65 | 0,73 | 3,22 | 0,23 | 8,21E-01 |
| AT3G33528 | 0,61 | -1,35 | 2,59 | -0,52 | 6,03E-01 |
| AT3G42050 | 1,44 | 3,23 | 1,97 | 1,64 | 1,02E-01 |
| AT3G42060 | 0,32 | -0,49 | 4,40 | -0,11 | 9,11E-01 |
| AT3G42140 | 0,14 | 0,00 | 4,41 | 0,00 | 1,00E+00 |
| AT3G42155 | 0,34 | 1,28 | 4,39 | 0,29 | 7,70E-01 |
| AT3G42550 | 0,12 | 0,00 | 4,41 | 0,00 | 1,00E+00 |
| AT3G42723 | 0,12 | 0,00 | 4,41 | 0,00 | 1,00E+00 |
| AT3G42790 | 0,59 | 1,87 | 3,26 | 0,58 | 5,65E-01 |
| AT3G43153 | 0,99 | 0,00 | 2,70 | 0,00 | 1,00E+00 |
| AT3G43190 | 0,26 | 1,38 | 4,41 | 0,31 | 7,54E-01 |
| AT3G43220 | 0,79 | 0,06 | 2,92 | 0,02 | 9,84E-01 |
| AT3G43240 | 0,86 | 2,93 | 2,76 | 1,06 | 2,89E-01 |
| AT3G43270 | 0,68 | -2,75 | 2,43 | -1,13 | 2,58E-01 |
| AT3G43280 | 0,11 | 0,00 | 4,41 | 0,00 | 1,00E+00 |
| AT3G43291 | 0,65 | 0,08 | 2,57 | 0,03 | 9,76E-01 |
| AT3G06595 | 0,10 | 0,00 | 4,41 | 0,00 | 1,00E+00 |
| AT3G43432 | 0,16 | 0,00 | 4,41 | 0,00 | 1,00E+00 |
| AT3G43470 | 0,45 | 0,00 | 4,03 | 0,00 | 1,00E+00 |
| AT3G43500 | 0,45 | 0,00 | 4,06 | 0,00 | 1,00E+00 |
| AT3G43600 | 0,98 | -0,19 | 1,98 | -0,09 | 9,25E-01 |
| AT3G43610 | 0,74 | 2,92 | 2,56 | 1,14 | 2,55E-01 |
| AT3G06645 | 0,23 | 0,00 | 4,38 | 0,00 | 1,00E+00 |
| AT3G06655 | 1,06 | -0,50 | 2,33 | -0,22 | 8,29E-01 |
| AT3G43720 | 0,87 | -1,40 | 2,62 | -0,54 | 5,92E-01 |
| AT3G43810 | 0,51 | -0,71 | 3,20 | -0,22 | 8,25E-01 |
| AT3G43880 | 0,10 | 0,00 | 4,41 | 0,00 | 1,00E+00 |

| | | | | | |
|---|---|---|---|---|---|
| AT3G43930 | 0,08 | 0,00 | 4,41 | 0,00 | 1,00E+00 |
| AT3G43940 | 0,53 | 0,00 | 3,96 | 0,00 | 1,00E+00 |
| AT3G44010 | 1,00 | 0,26 | 2,04 | 0,13 | 8,98E-01 |
| AT3G44070 | 0,37 | -1,61 | 3,40 | -0,47 | 6,37E-01 |
| AT3G44160 | 0,43 | 0,36 | 4,13 | 0,09 | 9,30E-01 |
| AT3G44265 | 0,40 | 0,00 | 4,37 | 0,00 | 1,00E+00 |
| AT3G44340 | 1,56 | 1,82 | 1,87 | 0,98 | 3,28E-01 |
| AT3G44400 | 0,82 | 1,98 | 2,80 | 0,71 | 4,80E-01 |
| AT3G44435 | 0,70 | 1,80 | 3,83 | 0,47 | 6,39E-01 |
| AT3G44450 | 0,56 | 0,38 | 3,11 | 0,12 | 9,02E-01 |
| AT3G44460 | 0,11 | 0,00 | 4,41 | 0,00 | 1,00E+00 |
| AT3G44635 | 1,31 | 3,23 | 2,92 | 1,11 | 2,69E-01 |
| AT3G44680 | 0,12 | 0,00 | 4,41 | 0,00 | 1,00E+00 |
| AT3G44753 | 0,14 | 0,00 | 4,41 | 0,00 | 1,00E+00 |
| AT3G44780 | 0,10 | 0,00 | 4,41 | 0,00 | 1,00E+00 |
| AT3G44820 | 0,38 | 1,69 | 4,38 | 0,39 | 7,00E-01 |
| AT3G44990 | 0,60 | -2,97 | 3,62 | -0,82 | 4,12E-01 |
| AT3G45030 | 0,91 | -0,61 | 2,47 | -0,25 | 8,05E-01 |
| AT3G45040 | 0,29 | 0,00 | 4,41 | 0,00 | 1,00E+00 |
| AT3G45060 | 0,08 | 0,00 | 4,41 | 0,00 | 1,00E+00 |
| AT3G45090 | 1,49 | 2,14 | 1,84 | 1,16 | 2,45E-01 |
| AT3G06955 | 0,21 | 0,00 | 4,41 | 0,00 | 1,00E+00 |
| AT3G45130 | 0,82 | -1,70 | 2,47 | -0,69 | 4,91E-01 |
| AT3G45180 | 1,00 | -0,31 | 1,88 | -0,17 | 8,67E-01 |
| AT3G07065 | 0,05 | 0,00 | 4,41 | 0,00 | 1,00E+00 |
| AT3G45240 | 0,25 | 1,30 | 4,40 | 0,30 | 7,67E-01 |
| AT3G07095 | 0,20 | -0,44 | 4,39 | -0,10 | 9,20E-01 |
| AT3G45260 | 1,01 | 1,60 | 1,86 | 0,86 | 3,92E-01 |
| AT3G45300 | 0,17 | -0,82 | 4,37 | -0,19 | 8,51E-01 |
| AT3G45430 | 0,20 | 1,38 | 4,41 | 0,31 | 7,54E-01 |
| AT3G45443 | 0,08 | 0,00 | 4,41 | 0,00 | 1,00E+00 |
| AT3G45600 | 0,92 | -0,26 | 1,96 | -0,13 | 8,94E-01 |
| AT3G45630 | 0,80 | 2,87 | 2,29 | 1,25 | 2,11E-01 |
| AT3G45770 | 0,14 | 0,00 | 4,41 | 0,00 | 1,00E+00 |
| AT3G45820 | 0,59 | -1,64 | 3,05 | -0,54 | 5,90E-01 |
| AT3G45850 | 1,07 | 2,42 | 2,49 | 0,97 | 3,30E-01 |
| AT3G45860 | 0,95 | 3,51 | 2,07 | 1,69 | 9,07E-02 |
| AT3G45910 | 1,00 | -0,68 | 2,65 | -0,26 | 7,97E-01 |
| AT3G45990 | 0,46 | -1,64 | 3,40 | -0,48 | 6,30E-01 |
| AT3G46000 | 0,30 | -0,42 | 3,70 | -0,11 | 9,09E-01 |
| AT3G46030 | 0,35 | 1,29 | 4,40 | 0,29 | 7,69E-01 |
| AT3G46090 | 0,72 | 1,38 | 2,55 | 0,54 | 5,89E-01 |
| AT3G46100 | 0,51 | -2,40 | 3,97 | -0,60 | 5,46E-01 |
| AT3G46210 | 0,56 | 0,00 | 3,97 | 0,00 | 1,00E+00 |
| AT3G07375 | 0,21 | -0,54 | 4,41 | -0,12 | 9,02E-01 |
| AT3G46320 | 0,61 | 0,00 | 4,41 | 0,00 | 1,00E+00 |
| AT3G46360 | 0,82 | -3,38 | 2,79 | -1,21 | 2,27E-01 |
| AT3G46390 | 0,25 | 0,00 | 4,41 | 0,00 | 1,00E+00 |
| AT3G46440 | 0,45 | 2,54 | 4,35 | 0,58 | 5,60E-01 |
| AT3G46450 | 0,53 | 1,64 | 3,94 | 0,42 | 6,76E-01 |
| AT3G46470 | 0,38 | 0,00 | 4,34 | 0,00 | 1,00E+00 |
| AT3G46540 | 0,31 | -0,45 | 4,40 | -0,10 | 9,18E-01 |
| AT3G46550 | 1,58 | 0,75 | 1,74 | 0,43 | 6,67E-01 |
| AT3G46610 | 1,41 | 1,07 | 1,78 | 0,60 | 5,47E-01 |
| AT3G46630 | 0,81 | -2,18 | 2,22 | -0,98 | 3,26E-01 |
| AT3G46650 | 0,58 | 1,31 | 3,16 | 0,42 | 6,77E-01 |

| | | | | | |
|---|---|---|---|---|---|
| AT3G46660 | 0,59 | 1,38 | 2,76 | 0,50 | 6,17E-01 |
| AT3G46670 | 0,57 | -1,10 | 3,22 | -0,34 | 7,33E-01 |
| AT3G46690 | 0,12 | 0,00 | 4,41 | 0,00 | 1,00E+00 |
| AT3G46730 | 0,12 | 0,00 | 4,41 | 0,00 | 1,00E+00 |
| AT3G46790 | 0,53 | 1,22 | 4,39 | 0,28 | 7,81E-01 |
| AT3G46845 | 1,29 | 1,43 | 2,05 | 0,70 | 4,87E-01 |
| AT3G46920 | 0,08 | 0,00 | 4,41 | 0,00 | 1,00E+00 |
| AT3G46960 | 0,92 | 2,93 | 2,36 | 1,24 | 2,15E-01 |
| AT3G47000 | 0,33 | 2,20 | 4,37 | 0,50 | 6,16E-01 |
| AT3G47070 | 0,67 | -0,48 | 2,63 | -0,18 | 8,57E-01 |
| AT3G47090 | 0,54 | 1,66 | 3,82 | 0,44 | 6,63E-01 |
| AT3G47160 | 0,74 | -2,11 | 2,58 | -0,82 | 4,14E-01 |
| AT3G47370 | 0,35 | 1,78 | 4,38 | 0,41 | 6,84E-01 |
| AT3G47420 | 0,51 | 2,08 | 2,69 | 0,77 | 4,41E-01 |
| AT3G47480 | 0,70 | 1,54 | 3,05 | 0,51 | 6,13E-01 |
| AT3G47670 | 0,97 | 0,40 | 2,15 | 0,19 | 8,51E-01 |
| AT3G47675 | 0,21 | 1,27 | 4,39 | 0,29 | 7,72E-01 |
| AT3G47680 | 0,08 | 0,00 | 4,41 | 0,00 | 1,00E+00 |
| AT3G47700 | 0,43 | 1,33 | 2,87 | 0,47 | 6,42E-01 |
| AT3G47730 | 0,93 | -2,07 | 2,71 | -0,76 | 4,45E-01 |
| AT3G47810 | 0,27 | 1,26 | 4,39 | 0,29 | 7,74E-01 |
| AT3G47860 | 0,23 | 1,38 | 4,41 | 0,31 | 7,54E-01 |
| AT3G47890 | 1,54 | 0,32 | 1,52 | 0,21 | 8,32E-01 |
| AT3G47940 | 0,35 | 1,55 | 4,39 | 0,35 | 7,24E-01 |
| AT3G47960 | 1,01 | 1,21 | 2,03 | 0,60 | 5,50E-01 |
| AT3G47990 | 0,32 | 1,45 | 4,40 | 0,33 | 7,42E-01 |
| AT3G48060 | 1,66 | 3,32 | 2,00 | 1,66 | 9,77E-02 |
| AT3G48080 | 1,04 | 3,02 | 2,11 | 1,43 | 1,54E-01 |
| AT3G48090 | 1,08 | 2,31 | 2,80 | 0,83 | 4,08E-01 |
| AT3G48100 | 0,85 | 3,47 | 4,27 | 0,81 | 4,16E-01 |
| AT3G48131 | 0,31 | 0,00 | 4,41 | 0,00 | 1,00E+00 |
| AT3G48150 | 0,55 | 2,59 | 3,83 | 0,68 | 4,98E-01 |
| AT3G48170 | 0,90 | 0,58 | 2,56 | 0,23 | 8,21E-01 |
| AT3G48195 | 0,48 | 1,27 | 3,22 | 0,39 | 6,93E-01 |
| AT3G48360 | 1,03 | 2,40 | 2,82 | 0,85 | 3,96E-01 |
| AT3G48520 | 0,38 | 1,38 | 4,41 | 0,31 | 7,54E-01 |
| AT3G48570 | 0,28 | 1,38 | 4,41 | 0,31 | 7,54E-01 |
| AT3G48640 | 0,25 | 1,26 | 4,39 | 0,29 | 7,75E-01 |
| AT3G48690 | 0,79 | 2,10 | 2,48 | 0,85 | 3,95E-01 |
| AT3G48740 | 0,68 | 0,42 | 2,37 | 0,18 | 8,60E-01 |
| AT3G48770 | 1,49 | 1,77 | 1,83 | 0,97 | 3,34E-01 |
| AT3G08005 | 0,05 | 0,00 | 4,41 | 0,00 | 1,00E+00 |
| AT3G48840 | 0,12 | 0,00 | 4,41 | 0,00 | 1,00E+00 |
| AT3G48930 | 0,53 | 2,28 | 2,73 | 0,84 | 4,03E-01 |
| AT3G48990 | 0,58 | 1,15 | 3,04 | 0,38 | 7,05E-01 |
| AT3G49010 | 1,04 | -0,07 | 1,88 | -0,04 | 9,69E-01 |
| AT3G49055 | 0,42 | 2,11 | 4,36 | 0,48 | 6,29E-01 |
| AT3G49060 | 0,44 | 0,48 | 3,31 | 0,15 | 8,84E-01 |
| AT3G49070 | 1,57 | -1,68 | 2,84 | -0,59 | 5,53E-01 |
| AT3G49140 | 0,93 | 0,54 | 2,03 | 0,26 | 7,91E-01 |
| AT3G49160 | 0,11 | 0,00 | 4,41 | 0,00 | 1,00E+00 |
| AT3G49250 | 0,54 | -2,71 | 4,10 | -0,66 | 5,08E-01 |
| AT3G49350 | 0,36 | 0,43 | 4,38 | 0,10 | 9,21E-01 |
| AT3G49490 | 1,14 | 3,29 | 2,17 | 1,51 | 1,30E-01 |
| AT3G49530 | 1,18 | 1,17 | 2,08 | 0,56 | 5,74E-01 |
| AT3G49600 | 1,57 | 4,08 | 1,92 | 2,13 | 3,35E-02 |

| | | | | | |
|---|---|---|---|---|---|
| AT3G49601 | 0,27 | 0,00 | 4,41 | 0,00 | 1,00E+00 |
| AT3G49650 | 0,08 | 0,00 | 4,41 | 0,00 | 1,00E+00 |
| AT3G49720 | 1,09 | 0,29 | 2,48 | 0,12 | 9,08E-01 |
| AT3G49790 | 0,38 | 0,00 | 3,44 | 0,00 | 1,00E+00 |
| AT3G49800 | 1,37 | 1,85 | 1,74 | 1,07 | 2,86E-01 |
| AT3G49870 | 0,08 | 0,00 | 4,41 | 0,00 | 1,00E+00 |
| AT3G49980 | 0,21 | -0,54 | 4,41 | -0,12 | 9,02E-01 |
| AT3G49990 | 0,57 | 0,00 | 3,83 | 0,00 | 1,00E+00 |
| AT3G50060 | 0,43 | 0,36 | 2,86 | 0,13 | 9,00E-01 |
| AT3G50110 | 0,18 | 0,00 | 4,41 | 0,00 | 1,00E+00 |
| AT3G50120 | 0,78 | -2,87 | 2,18 | -1,31 | 1,89E-01 |
| AT3G50240 | 0,71 | 1,38 | 4,41 | 0,31 | 7,54E-01 |
| AT3G50320 | 1,17 | -0,52 | 2,36 | -0,22 | 8,25E-01 |
| AT3G50380 | 1,07 | 2,45 | 2,49 | 0,98 | 3,25E-01 |
| AT3G50500 | 1,35 | -1,74 | 2,12 | -0,82 | 4,11E-01 |
| AT3G50670 | 0,33 | 0,00 | 4,41 | 0,00 | 1,00E+00 |
| AT3G50690 | 0,85 | -1,50 | 1,97 | -0,76 | 4,45E-01 |
| AT3G50740 | 0,53 | -1,28 | 3,23 | -0,40 | 6,92E-01 |
| AT3G50750 | 1,00 | 1,58 | 2,24 | 0,70 | 4,81E-01 |
| AT3G50780 | 0,08 | 0,00 | 4,41 | 0,00 | 1,00E+00 |
| AT3G50790 | 0,29 | 1,23 | 4,38 | 0,28 | 7,79E-01 |
| AT3G50808 | 0,10 | 0,00 | 4,41 | 0,00 | 1,00E+00 |
| AT3G50825 | 0,41 | 0,00 | 4,41 | 0,00 | 1,00E+00 |
| AT3G50920 | 0,48 | 1,98 | 4,00 | 0,49 | 6,21E-01 |
| AT3G51000 | 0,52 | 1,38 | 3,32 | 0,42 | 6,78E-01 |
| AT3G51120 | 1,56 | 3,25 | 1,88 | 1,73 | 8,38E-02 |
| AT3G51135 | 0,51 | 2,39 | 2,75 | 0,87 | 3,86E-01 |
| AT3G51160 | 0,58 | 1,46 | 3,05 | 0,48 | 6,32E-01 |
| AT3G51265 | 0,13 | 0,00 | 4,41 | 0,00 | 1,00E+00 |
| AT3G51310 | 0,51 | 0,36 | 3,25 | 0,11 | 9,13E-01 |
| AT3G51580 | 0,25 | 0,00 | 4,41 | 0,00 | 1,00E+00 |
| AT3G51600 | 1,48 | -1,15 | 1,53 | -0,75 | 4,54E-01 |
| AT3G51660 | 0,43 | 0,28 | 4,20 | 0,07 | 9,46E-01 |
| AT3G51790 | 0,60 | 1,38 | 3,76 | 0,37 | 7,13E-01 |
| AT3G51820 | 0,59 | 1,38 | 3,92 | 0,35 | 7,25E-01 |
| AT3G51840 | 0,33 | 1,24 | 4,38 | 0,28 | 7,78E-01 |
| AT3G51880 | 0,48 | 2,50 | 3,22 | 0,78 | 4,38E-01 |
| AT3G51890 | 1,41 | 3,86 | 2,69 | 1,43 | 1,51E-01 |
| AT3G51895 | 0,82 | 0,20 | 2,85 | 0,07 | 9,43E-01 |
| AT3G51920 | 0,46 | 0,35 | 2,80 | 0,13 | 9,00E-01 |
| AT3G08215 | 0,08 | 0,00 | 4,41 | 0,00 | 1,00E+00 |
| AT3G51980 | 0,42 | -0,70 | 3,32 | -0,21 | 8,33E-01 |
| AT3G52060 | 0,46 | -0,25 | 2,83 | -0,09 | 9,29E-01 |
| AT3G52072 | 0,22 | 0,00 | 4,41 | 0,00 | 1,00E+00 |
| AT3G08230 | 0,12 | 0,00 | 4,41 | 0,00 | 1,00E+00 |
| AT3G52110 | 0,12 | 0,00 | 4,41 | 0,00 | 1,00E+00 |
| AT3G52180 | 0,66 | 0,49 | 2,82 | 0,17 | 8,63E-01 |
| AT3G52190 | 0,13 | 0,00 | 4,41 | 0,00 | 1,00E+00 |
| AT3G52220 | 0,11 | 0,00 | 4,41 | 0,00 | 1,00E+00 |
| AT3G52260 | 0,64 | 1,38 | 3,67 | 0,38 | 7,07E-01 |
| AT3G52400 | 1,57 | 1,33 | 1,69 | 0,79 | 4,30E-01 |
| AT3G52480 | 0,53 | 0,00 | 3,15 | 0,00 | 1,00E+00 |
| AT3G52560 | 0,47 | -0,12 | 3,41 | -0,04 | 9,71E-01 |
| AT3G52640 | 0,21 | 0,00 | 4,41 | 0,00 | 1,00E+00 |
| AT3G52730 | 0,69 | 0,31 | 3,70 | 0,08 | 9,33E-01 |
| AT3G52740 | 0,17 | 1,32 | 4,40 | 0,30 | 7,64E-01 |

| | | | | | |
|---|---|---|---|---|---|
| AT3G52870 | 0,19 | 0,00 | 4,41 | 0,00 | 1,00E+00 |
| AT3G53015 | 1,03 | 0,44 | 2,35 | 0,19 | 8,50E-01 |
| AT3G53016 | 0,52 | 0,33 | 4,04 | 0,08 | 9,36E-01 |
| AT3G53100 | 0,69 | 1,67 | 3,09 | 0,54 | 5,89E-01 |
| AT3G53380 | 0,40 | 1,35 | 4,27 | 0,32 | 7,52E-01 |
| AT3G53410 | 0,47 | 1,99 | 4,38 | 0,45 | 6,50E-01 |
| AT3G53500 | 0,57 | 0,42 | 2,76 | 0,15 | 8,78E-01 |
| AT3G53510 | 0,25 | 0,00 | 4,41 | 0,00 | 1,00E+00 |
| AT3G53540 | 1,08 | 1,59 | 2,32 | 0,69 | 4,91E-01 |
| AT3G53560 | 0,23 | 1,29 | 4,40 | 0,29 | 7,69E-01 |
| AT3G53570 | 0,21 | 1,27 | 4,39 | 0,29 | 7,72E-01 |
| AT3G53610 | 0,53 | 0,00 | 3,27 | 0,00 | 1,00E+00 |
| AT3G53710 | 0,37 | -0,57 | 3,51 | -0,16 | 8,72E-01 |
| AT3G53730 | 0,67 | 1,38 | 2,63 | 0,52 | 6,00E-01 |
| AT3G53810 | 0,44 | 1,31 | 4,34 | 0,30 | 7,63E-01 |
| AT3G53890 | 0,52 | 1,90 | 3,17 | 0,60 | 5,49E-01 |
| AT3G53910 | 1,01 | 0,00 | 2,39 | 0,00 | 1,00E+00 |
| AT3G53920 | 0,24 | 1,31 | 4,40 | 0,30 | 7,65E-01 |
| AT3G53960 | 0,98 | 1,60 | 2,51 | 0,64 | 5,24E-01 |
| AT3G54100 | 0,47 | 0,00 | 3,33 | 0,00 | 1,00E+00 |
| AT3G54120 | 0,20 | 1,33 | 4,40 | 0,30 | 7,63E-01 |
| AT3G54140 | 0,70 | 2,09 | 2,72 | 0,77 | 4,41E-01 |
| AT3G54210 | 1,01 | -2,70 | 1,84 | -1,47 | 1,42E-01 |
| AT3G54290 | 0,42 | 2,75 | 4,24 | 0,65 | 5,16E-01 |
| AT3G54350 | 1,02 | 1,38 | 2,25 | 0,61 | 5,39E-01 |
| AT3G54360 | 1,14 | 0,16 | 2,00 | 0,08 | 9,36E-01 |
| AT3G54400 | 0,32 | 0,49 | 4,40 | 0,11 | 9,12E-01 |
| AT3G54440 | 0,61 | 0,54 | 2,27 | 0,24 | 8,12E-01 |
| AT3G54460 | 0,97 | 2,83 | 2,76 | 1,03 | 3,05E-01 |
| AT3G54470 | 0,44 | 1,27 | 3,32 | 0,38 | 7,02E-01 |
| AT3G54500 | 1,37 | 3,84 | 2,06 | 1,86 | 6,28E-02 |
| AT3G54620 | 0,24 | 1,38 | 4,41 | 0,31 | 7,54E-01 |
| AT3G54630 | 0,68 | 1,42 | 4,39 | 0,32 | 7,46E-01 |
| AT3G54640 | 0,22 | 1,37 | 4,41 | 0,31 | 7,56E-01 |
| AT3G54650 | 0,62 | 2,13 | 3,24 | 0,66 | 5,10E-01 |
| AT3G54750 | 0,48 | 1,32 | 4,40 | 0,30 | 7,65E-01 |
| AT3G54760 | 1,04 | 1,68 | 2,13 | 0,79 | 4,29E-01 |
| AT3G54790 | 1,31 | 1,70 | 1,69 | 1,00 | 3,16E-01 |
| AT3G54860 | 0,55 | 1,31 | 3,12 | 0,42 | 6,75E-01 |
| AT3G54900 | 0,95 | -0,78 | 2,05 | -0,38 | 7,02E-01 |
| AT3G54960 | 0,90 | -0,37 | 2,17 | -0,17 | 8,66E-01 |
| AT3G55040 | 0,21 | -1,03 | 4,37 | -0,24 | 8,13E-01 |
| AT3G55050 | 0,45 | 0,00 | 3,38 | 0,00 | 1,00E+00 |
| AT3G55060 | 0,14 | 0,00 | 4,41 | 0,00 | 1,00E+00 |
| AT3G55130 | 0,51 | -2,07 | 3,14 | -0,66 | 5,10E-01 |
| AT3G55160 | 0,95 | 0,26 | 1,85 | 0,14 | 8,87E-01 |
| AT3G55252 | 0,66 | 0,35 | 2,33 | 0,15 | 8,80E-01 |
| AT3G55254 | 0,80 | -1,45 | 2,33 | -0,62 | 5,33E-01 |
| AT3G55330 | 0,44 | 0,52 | 2,84 | 0,18 | 8,56E-01 |
| AT3G55360 | 0,60 | -1,32 | 2,62 | -0,50 | 6,14E-01 |
| AT3G55400 | 0,18 | -0,82 | 4,37 | -0,19 | 8,52E-01 |
| AT3G55430 | 0,77 | -0,31 | 2,03 | -0,15 | 8,80E-01 |
| AT3G55480 | 1,64 | 2,82 | 1,74 | 1,62 | 1,06E-01 |
| AT3G55840 | 1,41 | 1,48 | 1,82 | 0,81 | 4,18E-01 |
| AT3G55860 | 0,83 | -2,05 | 2,19 | -0,93 | 3,51E-01 |
| AT3G55935 | 0,46 | 2,09 | 2,75 | 0,76 | 4,48E-01 |

| | | | | | |
|---|---|---|---|---|---|
| AT3G55950 | 0,63 | 1,38 | 2,64 | 0,52 | 6,01E-01 |
| AT3G55960 | 0,64 | 0,43 | 2,72 | 0,16 | 8,73E-01 |
| AT3G56040 | 0,77 | 0,53 | 2,25 | 0,23 | 8,15E-01 |
| AT3G56050 | 0,97 | 0,26 | 2,13 | 0,12 | 9,02E-01 |
| AT3G56085 | 0,12 | 0,00 | 4,41 | 0,00 | 1,00E+00 |
| AT3G56090 | 0,34 | -1,61 | 3,52 | -0,46 | 6,47E-01 |
| AT3G56110 | 0,41 | 0,43 | 3,42 | 0,13 | 8,99E-01 |
| AT3G56190 | 1,16 | -0,96 | 1,98 | -0,48 | 6,29E-01 |
| AT3G56210 | 0,45 | 2,49 | 4,36 | 0,57 | 5,68E-01 |
| AT3G56240 | 0,35 | -0,46 | 2,99 | -0,15 | 8,78E-01 |
| AT3G56290 | 1,19 | 3,97 | 2,17 | 1,83 | 6,66E-02 |
| AT3G56320 | 0,58 | 2,71 | 3,87 | 0,70 | 4,85E-01 |
| AT3G56340 | 0,44 | 0,54 | 3,39 | 0,16 | 8,73E-01 |
| AT3G56370 | 0,84 | 3,20 | 2,32 | 1,38 | 1,67E-01 |
| AT3G56460 | 1,07 | 1,94 | 2,20 | 0,88 | 3,77E-01 |
| AT3G56480 | 0,39 | -0,54 | 2,97 | -0,18 | 8,54E-01 |
| AT3G56670 | 0,38 | 1,29 | 4,38 | 0,30 | 7,68E-01 |
| AT3G56690 | 0,43 | -0,69 | 4,19 | -0,16 | 8,69E-01 |
| AT3G56790 | 0,33 | 0,00 | 4,41 | 0,00 | 1,00E+00 |
| AT3G56800 | 1,39 | 0,67 | 1,82 | 0,37 | 7,13E-01 |
| AT3G56820 | 0,25 | 1,84 | 4,38 | 0,42 | 6,75E-01 |
| AT3G56830 | 0,17 | 0,00 | 4,41 | 0,00 | 1,00E+00 |
| AT3G56850 | 0,48 | 2,43 | 3,34 | 0,73 | 4,68E-01 |
| AT3G56860 | 0,49 | -0,54 | 2,83 | -0,19 | 8,47E-01 |
| AT3G56950 | 0,79 | -1,09 | 2,27 | -0,48 | 6,31E-01 |
| AT3G56970 | 0,12 | 0,00 | 4,41 | 0,00 | 1,00E+00 |
| AT3G56990 | 0,46 | -2,10 | 3,21 | -0,65 | 5,14E-01 |
| AT3G57020 | 0,42 | 1,25 | 4,34 | 0,29 | 7,73E-01 |
| AT3G57030 | 0,05 | 0,00 | 4,41 | 0,00 | 1,00E+00 |
| AT3G08825 | 0,72 | -1,86 | 3,04 | -0,61 | 5,41E-01 |
| AT3G57050 | 0,52 | 0,47 | 2,73 | 0,17 | 8,64E-01 |
| AT3G08855 | 0,12 | 0,00 | 4,41 | 0,00 | 1,00E+00 |
| AT3G57140 | 0,48 | 1,30 | 3,23 | 0,40 | 6,88E-01 |
| AT3G57240 | 0,24 | 1,96 | 4,39 | 0,45 | 6,55E-01 |
| AT3G57290 | 1,20 | 0,36 | 2,42 | 0,15 | 8,82E-01 |
| AT3G57350 | 0,44 | 2,09 | 3,33 | 0,63 | 5,30E-01 |
| AT3G57420 | 0,74 | 1,24 | 2,38 | 0,52 | 6,02E-01 |
| AT3G57460 | 0,24 | 0,00 | 4,41 | 0,00 | 1,00E+00 |
| AT3G08935 | 1,61 | -1,00 | 1,67 | -0,60 | 5,48E-01 |
| AT3G57470 | 0,21 | -0,87 | 4,38 | -0,20 | 8,43E-01 |
| AT3G57530 | 1,08 | 1,21 | 2,17 | 0,56 | 5,77E-01 |
| AT3G57540 | 0,08 | 0,00 | 4,41 | 0,00 | 1,00E+00 |
| AT3G57580 | 0,45 | 0,54 | 3,38 | 0,16 | 8,73E-01 |
| AT3G57610 | 1,05 | 4,02 | 4,29 | 0,94 | 3,49E-01 |
| AT3G57640 | 0,11 | 0,00 | 4,41 | 0,00 | 1,00E+00 |
| AT3G57765 | 1,26 | -0,04 | 1,73 | -0,02 | 9,81E-01 |
| AT3G57780 | 0,43 | 0,73 | 4,38 | 0,17 | 8,68E-01 |
| AT3G57790 | 0,44 | 0,00 | 4,41 | 0,00 | 1,00E+00 |
| AT3G57800 | 0,37 | 0,00 | 4,41 | 0,00 | 1,00E+00 |
| AT3G57870 | 0,53 | 3,14 | 4,32 | 0,73 | 4,68E-01 |
| AT3G57880 | 0,23 | 0,00 | 4,41 | 0,00 | 1,00E+00 |
| AT3G57930 | 0,30 | 1,47 | 4,40 | 0,33 | 7,39E-01 |
| AT3G57965 | 0,28 | -0,46 | 4,40 | -0,10 | 9,17E-01 |
| AT3G57980 | 0,74 | 0,00 | 2,96 | 0,00 | 1,00E+00 |
| AT3G57990 | 0,40 | 1,31 | 3,43 | 0,38 | 7,02E-01 |
| AT3G58100 | 1,03 | 1,38 | 3,44 | 0,40 | 6,88E-01 |

| | | | | | |
|---|---|---|---|---|---|
| AT3G58120 | 0,15 | -0,43 | 4,39 | -0,10 | 9,23E-01 |
| AT3G58140 | 0,42 | 0,54 | 3,42 | 0,16 | 8,75E-01 |
| AT3G58170 | 0,42 | -0,44 | 4,27 | -0,10 | 9,17E-01 |
| AT3G58210 | 0,64 | 3,08 | 4,32 | 0,71 | 4,76E-01 |
| AT3G58270 | 0,99 | 0,38 | 2,74 | 0,14 | 8,90E-01 |
| AT3G58500 | 1,34 | -0,70 | 1,87 | -0,37 | 7,09E-01 |
| AT3G58560 | 0,23 | 0,00 | 4,41 | 0,00 | 1,00E+00 |
| AT3G58580 | 0,38 | -0,54 | 4,37 | -0,12 | 9,01E-01 |
| AT3G58670 | 0,12 | 0,00 | 4,41 | 0,00 | 1,00E+00 |
| AT3G58730 | 1,29 | -1,36 | 1,78 | -0,76 | 4,45E-01 |
| AT3G58750 | 0,61 | 0,74 | 3,82 | 0,19 | 8,46E-01 |
| AT3G58790 | 0,34 | -0,86 | 4,38 | -0,20 | 8,44E-01 |
| AT3G58970 | 1,53 | 3,53 | 2,38 | 1,48 | 1,38E-01 |
| AT3G58990 | 0,12 | 0,00 | 4,41 | 0,00 | 1,00E+00 |
| AT3G59040 | 1,65 | 4,13 | 1,80 | 2,29 | 2,21E-02 |
| AT3G59050 | 0,52 | 1,53 | 3,22 | 0,47 | 6,35E-01 |
| AT3G59060 | 1,47 | 2,67 | 1,71 | 1,56 | 1,19E-01 |
| AT3G59120 | 0,10 | 0,00 | 4,41 | 0,00 | 1,00E+00 |
| AT3G59280 | 0,32 | 1,38 | 4,41 | 0,31 | 7,54E-01 |
| AT3G59360 | 0,61 | 0,00 | 3,07 | 0,00 | 1,00E+00 |
| AT3G59400 | 1,33 | 0,89 | 1,64 | 0,55 | 5,85E-01 |
| AT3G59540 | 1,10 | 0,62 | 1,81 | 0,34 | 7,32E-01 |
| AT3G59570 | 0,57 | -1,13 | 3,12 | -0,36 | 7,18E-01 |
| AT3G59660 | 1,15 | 2,28 | 2,12 | 1,08 | 2,81E-01 |
| AT3G59760 | 0,20 | -0,84 | 4,38 | -0,19 | 8,47E-01 |
| AT3G59840 | 0,15 | 0,00 | 4,41 | 0,00 | 1,00E+00 |
| AT3G59910 | 0,48 | 2,71 | 4,05 | 0,67 | 5,04E-01 |
| AT3G59950 | 1,22 | 0,72 | 2,49 | 0,29 | 7,72E-01 |
| AT3G59980 | 0,35 | 1,32 | 3,55 | 0,37 | 7,10E-01 |
| AT3G60030 | 1,14 | 1,80 | 2,04 | 0,88 | 3,76E-01 |
| AT3G09335 | 0,23 | -0,54 | 4,41 | -0,12 | 9,02E-01 |
| AT3G60050 | 0,12 | 0,00 | 4,41 | 0,00 | 1,00E+00 |
| AT3G60070 | 0,35 | 1,03 | 4,36 | 0,24 | 8,13E-01 |
| AT3G60160 | 1,27 | 2,25 | 2,16 | 1,04 | 2,97E-01 |
| AT3G60176 | 0,85 | -0,57 | 2,88 | -0,20 | 8,42E-01 |
| AT3G60220 | 0,15 | 0,00 | 4,41 | 0,00 | 1,00E+00 |
| AT3G60250 | 1,02 | 2,85 | 1,98 | 1,44 | 1,51E-01 |
| AT3G60350 | 1,30 | 3,03 | 1,89 | 1,60 | 1,10E-01 |
| AT3G60380 | 0,49 | 2,52 | 3,95 | 0,64 | 5,23E-01 |
| AT3G60540 | 0,08 | 0,00 | 4,41 | 0,00 | 1,00E+00 |
| AT3G60670 | 0,12 | 0,00 | 4,41 | 0,00 | 1,00E+00 |
| AT3G60740 | 0,66 | 1,83 | 2,56 | 0,71 | 4,75E-01 |
| AT3G60760 | 0,12 | 0,00 | 4,41 | 0,00 | 1,00E+00 |
| AT3G60820 | 0,15 | 0,00 | 4,39 | 0,00 | 1,00E+00 |
| AT3G60990 | 0,73 | -1,01 | 2,72 | -0,37 | 7,10E-01 |
| AT3G61070 | 0,50 | 1,38 | 4,14 | 0,33 | 7,39E-01 |
| AT3G09565 | 0,50 | 0,44 | 3,93 | 0,11 | 9,11E-01 |
| AT3G61190 | 1,32 | 1,66 | 1,97 | 0,84 | 4,01E-01 |
| AT3G61200 | 0,69 | -0,94 | 2,57 | -0,36 | 7,15E-01 |
| AT3G61260 | 0,83 | -1,97 | 2,52 | -0,78 | 4,35E-01 |
| AT3G61310 | 0,38 | 2,11 | 4,37 | 0,48 | 6,29E-01 |
| AT3G61380 | 0,18 | 0,00 | 4,41 | 0,00 | 1,00E+00 |
| AT3G61420 | 0,37 | 1,76 | 4,38 | 0,40 | 6,87E-01 |
| AT3G61460 | 0,83 | -2,38 | 2,00 | -1,19 | 2,33E-01 |
| AT3G61530 | 0,25 | -0,49 | 4,40 | -0,11 | 9,11E-01 |
| AT3G61640 | 0,38 | 0,00 | 3,45 | 0,00 | 1,00E+00 |

| | | | | | |
|---|---|---|---|---|---|
| AT3G61740 | 0,78 | 2,34 | 2,49 | 0,94 | 3,46E-01 |
| AT3G61750 | 0,24 | 1,96 | 4,39 | 0,45 | 6,55E-01 |
| AT3G61780 | 1,45 | 3,27 | 1,83 | 1,78 | 7,44E-02 |
| AT3G61800 | 0,23 | 0,00 | 4,41 | 0,00 | 1,00E+00 |
| AT3G61820 | 1,36 | -1,56 | 1,62 | -0,96 | 3,35E-01 |
| AT3G61860 | 0,39 | 2,11 | 4,36 | 0,48 | 6,28E-01 |
| AT3G61890 | 0,44 | -1,38 | 3,56 | -0,39 | 6,98E-01 |
| AT3G61960 | 1,02 | 0,78 | 1,88 | 0,41 | 6,79E-01 |
| AT3G61990 | 0,15 | 0,00 | 4,41 | 0,00 | 1,00E+00 |
| AT3G62070 | 0,12 | 0,00 | 4,41 | 0,00 | 1,00E+00 |
| AT3G62120 | 1,00 | -0,37 | 2,18 | -0,17 | 8,64E-01 |
| AT3G62240 | 0,93 | 2,02 | 3,00 | 0,67 | 5,01E-01 |
| AT3G62260 | 0,41 | 1,23 | 4,17 | 0,29 | 7,68E-01 |
| AT3G62310 | 0,33 | -1,55 | 4,35 | -0,36 | 7,22E-01 |
| AT3G62330 | 0,73 | 2,00 | 2,98 | 0,67 | 5,02E-01 |
| AT3G62470 | 0,24 | 1,37 | 4,41 | 0,31 | 7,56E-01 |
| AT3G62540 | 0,28 | 1,84 | 4,39 | 0,42 | 6,74E-01 |
| AT3G09795 | 0,21 | 0,00 | 4,41 | 0,00 | 1,00E+00 |
| AT3G62590 | 0,64 | 0,00 | 3,09 | 0,00 | 1,00E+00 |
| AT3G62600 | 0,95 | 1,84 | 2,09 | 0,88 | 3,79E-01 |
| AT3G09825 | 1,22 | -3,06 | 1,87 | -1,63 | 1,03E-01 |
| AT3G09845 | 0,08 | 0,00 | 4,41 | 0,00 | 1,00E+00 |
| AT3G62630 | 0,91 | 3,20 | 2,34 | 1,37 | 1,71E-01 |
| AT3G62800 | 0,37 | 1,38 | 4,41 | 0,31 | 7,54E-01 |
| AT3G62840 | 0,66 | 2,90 | 3,07 | 0,94 | 3,46E-01 |
| AT3G62860 | 0,81 | 3,13 | 3,48 | 0,90 | 3,69E-01 |
| AT3G62870 | 1,22 | 1,54 | 1,83 | 0,84 | 4,02E-01 |
| AT3G62900 | 0,94 | -0,44 | 2,48 | -0,18 | 8,58E-01 |
| AT3G62910 | 0,49 | 2,85 | 4,02 | 0,71 | 4,78E-01 |
| AT3G62940 | 0,31 | 0,44 | 4,39 | 0,10 | 9,20E-01 |
| AT3G63052 | 0,42 | 0,54 | 3,42 | 0,16 | 8,75E-01 |
| AT3G63110 | 0,56 | -1,56 | 2,73 | -0,57 | 5,66E-01 |
| AT3G63140 | 0,93 | -1,33 | 2,10 | -0,63 | 5,26E-01 |
| AT3G63150 | 0,90 | 2,56 | 2,52 | 1,02 | 3,09E-01 |
| AT3G63160 | 1,15 | -3,10 | 2,04 | -1,52 | 1,28E-01 |
| AT3G63180 | 1,24 | 1,88 | 1,75 | 1,07 | 2,83E-01 |
| AT3G63190 | 0,25 | -0,47 | 4,40 | -0,11 | 9,14E-01 |
| AT3G63210 | 0,47 | -0,85 | 4,38 | -0,19 | 8,46E-01 |
| AT3G63310 | 1,42 | 2,51 | 1,98 | 1,27 | 2,04E-01 |
| AT3G63340 | 1,13 | 2,82 | 2,57 | 1,10 | 2,72E-01 |
| AT3G63390 | 0,24 | 0,00 | 4,41 | 0,00 | 1,00E+00 |
| AT3G63540 | 1,40 | -0,31 | 1,73 | -0,18 | 8,57E-01 |
| AT4G00005 | 0,19 | -0,47 | 4,40 | -0,11 | 9,15E-01 |
| AT4G00020 | 0,79 | -1,06 | 2,31 | -0,46 | 6,46E-01 |
| AT4G00070 | 0,43 | 0,32 | 4,23 | 0,08 | 9,39E-01 |
| AT4G00110 | 1,14 | 2,90 | 2,36 | 1,23 | 2,20E-01 |
| AT4G00120 | 1,32 | -1,16 | 1,95 | -0,59 | 5,53E-01 |
| AT4G00165 | 0,65 | -0,02 | 2,63 | -0,01 | 9,94E-01 |
| AT4G00180 | 0,45 | -1,23 | 4,38 | -0,28 | 7,79E-01 |
| AT4G00280 | 0,25 | 1,84 | 4,38 | 0,42 | 6,75E-01 |
| AT4G00290 | 0,50 | -0,38 | 2,75 | -0,14 | 8,91E-01 |
| AT4G00330 | 0,80 | 0,43 | 2,35 | 0,18 | 8,53E-01 |
| AT4G03585 | 1,51 | -0,80 | 1,61 | -0,49 | 6,21E-01 |
| AT4G00370 | 0,63 | -1,19 | 2,57 | -0,47 | 6,42E-01 |
| AT4G00400 | 0,42 | -1,35 | 2,85 | -0,47 | 6,36E-01 |
| AT4G00500 | 0,24 | -1,07 | 4,37 | -0,24 | 8,07E-01 |

| | | | | | |
|---|---|---|---|---|---|
| AT4G00540 | 0,54 | -1,24 | 2,67 | -0,46 | 6,42E-01 |
| AT4G00550 | 0,15 | 0,00 | 4,41 | 0,00 | 1,00E+00 |
| AT4G00560 | 0,40 | 0,00 | 4,41 | 0,00 | 1,00E+00 |
| AT4G00620 | 0,40 | 0,30 | 4,26 | 0,07 | 9,43E-01 |
| AT4G00740 | 0,31 | 1,38 | 4,41 | 0,31 | 7,54E-01 |
| AT4G00810 | 0,25 | 1,30 | 4,40 | 0,30 | 7,67E-01 |
| AT4G00890 | 0,23 | 0,00 | 4,41 | 0,00 | 1,00E+00 |
| AT4G00900 | 1,09 | -0,82 | 2,11 | -0,39 | 6,97E-01 |
| AT4G00980 | 0,33 | 1,72 | 4,38 | 0,39 | 6,94E-01 |
| AT4G00985 | 0,26 | 0,23 | 4,36 | 0,05 | 9,58E-01 |
| AT4G01000 | 0,50 | 0,00 | 3,99 | 0,00 | 1,00E+00 |
| AT4G01030 | 0,51 | 1,32 | 3,18 | 0,41 | 6,79E-01 |
| AT4G01037 | 0,21 | -1,22 | 4,35 | -0,28 | 7,80E-01 |
| AT4G01070 | 1,43 | -0,73 | 1,82 | -0,40 | 6,88E-01 |
| AT4G01090 | 0,44 | 1,39 | 3,36 | 0,41 | 6,80E-01 |
| AT4G01110 | 0,08 | 0,00 | 4,41 | 0,00 | 1,00E+00 |
| AT4G01120 | 0,18 | -0,82 | 4,37 | -0,19 | 8,51E-01 |
| AT4G01180 | 0,08 | 0,00 | 4,41 | 0,00 | 1,00E+00 |
| AT4G01190 | 0,20 | 1,33 | 4,40 | 0,30 | 7,63E-01 |
| AT4G01200 | 0,46 | 2,63 | 4,34 | 0,61 | 5,44E-01 |
| AT4G01210 | 1,50 | 3,53 | 1,93 | 1,83 | 6,71E-02 |
| AT4G01240 | 0,94 | 2,89 | 2,15 | 1,34 | 1,79E-01 |
| AT4G01310 | 0,66 | 1,96 | 3,06 | 0,64 | 5,22E-01 |
| AT4G01395 | 0,71 | 1,97 | 2,25 | 0,88 | 3,81E-01 |
| AT4G01533 | 1,62 | 1,65 | 2,11 | 0,78 | 4,33E-01 |
| AT4G01540 | 0,20 | 0,00 | 4,41 | 0,00 | 1,00E+00 |
| AT4G01550 | 0,48 | -1,47 | 4,34 | -0,34 | 7,35E-01 |
| AT4G01700 | 0,60 | -0,32 | 3,20 | -0,10 | 9,21E-01 |
| AT4G01870 | 0,45 | 1,33 | 4,15 | 0,32 | 7,48E-01 |
| AT4G01950 | 0,49 | 0,48 | 2,82 | 0,17 | 8,65E-01 |
| AT4G01985 | 0,41 | -1,60 | 4,35 | -0,37 | 7,13E-01 |
| AT4G02195 | 0,71 | 1,89 | 2,56 | 0,74 | 4,60E-01 |
| AT4G02210 | 0,65 | 1,25 | 2,99 | 0,42 | 6,76E-01 |
| AT4G02230 | 0,28 | 1,39 | 4,41 | 0,32 | 7,52E-01 |
| AT4G02405 | 0,05 | 0,00 | 4,41 | 0,00 | 1,00E+00 |
| AT4G02410 | 0,84 | 3,18 | 2,87 | 1,11 | 2,69E-01 |
| AT4G02420 | 1,44 | -0,33 | 2,04 | -0,16 | 8,73E-01 |
| AT4G02450 | 0,79 | -0,67 | 2,60 | -0,26 | 7,98E-01 |
| AT4G02460 | 0,19 | 0,00 | 4,41 | 0,00 | 1,00E+00 |
| AT4G02530 | 0,43 | -1,66 | 3,36 | -0,49 | 6,22E-01 |
| AT4G04185 | 0,38 | 0,00 | 3,48 | 0,00 | 1,00E+00 |
| AT4G02550 | 0,12 | 0,00 | 4,41 | 0,00 | 1,00E+00 |
| AT4G02560 | 0,57 | 1,58 | 2,67 | 0,59 | 5,54E-01 |
| AT4G02590 | 0,27 | 0,45 | 4,38 | 0,10 | 9,19E-01 |
| AT4G02620 | 0,23 | -0,84 | 4,38 | -0,19 | 8,48E-01 |
| AT4G02680 | 0,66 | 2,69 | 3,82 | 0,70 | 4,81E-01 |
| AT4G02710 | 0,71 | 2,50 | 2,94 | 0,85 | 3,96E-01 |
| AT4G02715 | 1,63 | -0,32 | 2,13 | -0,15 | 8,82E-01 |
| AT4G02735 | 0,41 | 2,70 | 4,34 | 0,62 | 5,33E-01 |
| AT4G02740 | 0,60 | 1,81 | 3,12 | 0,58 | 5,61E-01 |
| AT4G02750 | 0,98 | 2,86 | 2,26 | 1,27 | 2,06E-01 |
| AT4G02910 | 0,98 | -0,25 | 2,45 | -0,10 | 9,20E-01 |
| AT4G02930 | 0,46 | 2,12 | 4,37 | 0,49 | 6,27E-01 |
| AT4G03080 | 1,69 | 0,97 | 1,62 | 0,60 | 5,49E-01 |
| AT4G03090 | 0,91 | -2,61 | 1,95 | -1,34 | 1,80E-01 |
| AT4G03110 | 0,81 | 2,53 | 2,90 | 0,87 | 3,83E-01 |

| | | | | | |
|---|---|---|---|---|---|
| AT4G03130 | 0,08 | 0,00 | 4,41 | 0,00 | 1,00E+00 |
| AT4G03150 | 0,26 | 1,38 | 4,41 | 0,31 | 7,54E-01 |
| AT4G03190 | 0,84 | 1,24 | 2,16 | 0,57 | 5,67E-01 |
| AT4G03230 | 0,47 | 2,88 | 3,97 | 0,72 | 4,68E-01 |
| AT4G03270 | 0,81 | 2,65 | 2,81 | 0,94 | 3,47E-01 |
| AT4G04475 | 0,42 | 0,43 | 3,39 | 0,13 | 8,99E-01 |
| AT4G04485 | 0,66 | 0,39 | 2,59 | 0,15 | 8,80E-01 |
| AT4G03340 | 0,12 | 0,00 | 4,41 | 0,00 | 1,00E+00 |
| AT4G03390 | 0,56 | 2,22 | 2,74 | 0,81 | 4,19E-01 |
| AT4G03400 | 0,95 | 1,35 | 2,37 | 0,57 | 5,68E-01 |
| AT4G03430 | 1,44 | 3,20 | 1,74 | 1,84 | 6,59E-02 |
| AT4G03435 | 0,08 | 0,00 | 4,41 | 0,00 | 1,00E+00 |
| AT4G03438 | 0,26 | 1,38 | 4,41 | 0,31 | 7,54E-01 |
| AT4G03450 | 0,60 | 2,91 | 2,66 | 1,10 | 2,73E-01 |
| AT4G03490 | 0,42 | 2,25 | 4,32 | 0,52 | 6,02E-01 |
| AT4G03505 | 0,25 | 1,84 | 4,38 | 0,42 | 6,75E-01 |
| AT4G03520 | 1,13 | -1,40 | 1,97 | -0,71 | 4,79E-01 |
| AT4G03570 | 0,98 | -1,44 | 1,85 | -0,78 | 4,34E-01 |
| AT4G03590 | 1,16 | -0,55 | 2,47 | -0,22 | 8,24E-01 |
| AT4G03728 | 0,69 | -0,66 | 3,76 | -0,18 | 8,61E-01 |
| AT4G03740 | 1,37 | -0,54 | 2,23 | -0,24 | 8,08E-01 |
| AT4G04675 | 1,07 | -0,54 | 2,03 | -0,26 | 7,92E-01 |
| AT4G04030 | 0,82 | -2,84 | 2,48 | -1,15 | 2,52E-01 |
| AT4G04221 | 1,00 | 3,44 | 2,14 | 1,61 | 1,07E-01 |
| AT4G04805 | 1,38 | 0,00 | 2,47 | 0,00 | 1,00E+00 |
| AT4G04370 | 1,03 | 3,25 | 2,07 | 1,57 | 1,17E-01 |
| AT4G04470 | 0,08 | 0,00 | 4,41 | 0,00 | 1,00E+00 |
| AT4G04580 | 0,11 | 0,00 | 4,41 | 0,00 | 1,00E+00 |
| AT4G04614 | 0,51 | 2,51 | 4,35 | 0,58 | 5,63E-01 |
| AT4G04632 | 0,44 | 0,00 | 4,18 | 0,00 | 1,00E+00 |
| AT4G04650 | 0,82 | 0,48 | 2,49 | 0,19 | 8,46E-01 |
| AT4G04710 | 0,47 | 0,00 | 3,33 | 0,00 | 1,00E+00 |
| AT4G04720 | 1,28 | -1,17 | 2,03 | -0,58 | 5,64E-01 |
| AT4G04740 | 0,87 | 0,42 | 2,15 | 0,20 | 8,44E-01 |
| AT4G04745 | 1,61 | -1,22 | 1,65 | -0,74 | 4,61E-01 |
| AT4G04770 | 1,41 | 0,24 | 2,01 | 0,12 | 9,05E-01 |
| AT4G04910 | 0,75 | 0,33 | 2,20 | 0,15 | 8,82E-01 |
| AT4G04950 | 0,83 | -0,51 | 3,58 | -0,14 | 8,87E-01 |
| AT4G04957 | 0,91 | 0,00 | 2,78 | 0,00 | 1,00E+00 |
| AT4G04980 | 0,11 | 0,00 | 4,41 | 0,00 | 1,00E+00 |
| AT4G05070 | 1,30 | -0,29 | 1,62 | -0,18 | 8,59E-01 |
| AT4G05110 | 0,17 | 0,00 | 4,41 | 0,00 | 1,00E+00 |
| AT4G05085 | 0,05 | 0,00 | 4,41 | 0,00 | 1,00E+00 |
| AT4G05330 | 0,44 | 1,76 | 4,13 | 0,43 | 6,70E-01 |
| AT4G05410 | 0,51 | 1,29 | 4,09 | 0,32 | 7,52E-01 |
| AT4G05470 | 0,12 | 0,00 | 4,41 | 0,00 | 1,00E+00 |
| AT4G05631 | 0,41 | 0,54 | 3,46 | 0,15 | 8,77E-01 |
| AT4G06526 | 0,56 | -0,40 | 3,96 | -0,10 | 9,19E-01 |
| AT4G06639 | 0,21 | 0,00 | 4,41 | 0,00 | 1,00E+00 |
| AT4G06599 | 0,16 | 0,00 | 4,41 | 0,00 | 1,00E+00 |
| AT4G07350 | 0,08 | 0,00 | 4,41 | 0,00 | 1,00E+00 |
| AT4G07400 | 0,28 | -0,54 | 4,41 | -0,12 | 9,02E-01 |
| AT4G07410 | 1,28 | 3,82 | 2,17 | 1,77 | 7,75E-02 |
| AT4G07524 | 0,19 | 0,00 | 4,41 | 0,00 | 1,00E+00 |
| AT4G07526 | 0,83 | -0,37 | 2,28 | -0,16 | 8,70E-01 |
| AT4G05225 | 0,74 | -2,61 | 3,03 | -0,86 | 3,89E-01 |

| | | | | | |
|---|---|---|---|---|---|
| AT4G07990 | 0,20 | 1,40 | 4,41 | 0,32 | 7,51E-01 |
| AT4G08230 | 0,30 | 0,00 | 4,41 | 0,00 | 1,00E+00 |
| AT4G08310 | 0,85 | 2,43 | 1,95 | 1,24 | 2,14E-01 |
| AT4G05415 | 0,95 | 2,71 | 2,68 | 1,01 | 3,12E-01 |
| AT4G08370 | 0,46 | 1,37 | 2,82 | 0,49 | 6,27E-01 |
| AT4G08400 | 0,54 | 0,00 | 3,12 | 0,00 | 1,00E+00 |
| AT4G08410 | 1,29 | 0,00 | 2,49 | 0,00 | 1,00E+00 |
| AT4G05535 | 0,95 | 0,22 | 2,40 | 0,09 | 9,26E-01 |
| AT4G08460 | 0,62 | 1,51 | 2,61 | 0,58 | 5,64E-01 |
| AT4G08480 | 0,67 | 2,40 | 3,13 | 0,77 | 4,43E-01 |
| AT4G08485 | 0,63 | -3,11 | 3,67 | -0,85 | 3,96E-01 |
| AT4G08590 | 0,56 | 2,06 | 4,38 | 0,47 | 6,38E-01 |
| AT4G08760 | 1,19 | -0,44 | 2,44 | -0,18 | 8,58E-01 |
| AT4G08810 | 0,77 | 2,01 | 2,57 | 0,78 | 4,33E-01 |
| AT4G08870 | 1,44 | -1,12 | 1,56 | -0,71 | 4,75E-01 |
| AT4G08874 | 0,12 | 0,00 | 4,41 | 0,00 | 1,00E+00 |
| AT4G08940 | 0,46 | 1,38 | 3,42 | 0,40 | 6,87E-01 |
| AT4G09010 | 1,11 | 1,13 | 1,91 | 0,59 | 5,55E-01 |
| AT4G09012 | 0,55 | -0,44 | 3,29 | -0,13 | 8,95E-01 |
| AT4G09035 | 0,16 | 0,00 | 4,41 | 0,00 | 1,00E+00 |
| AT4G09040 | 0,47 | 2,19 | 4,36 | 0,50 | 6,16E-01 |
| AT4G09340 | 0,44 | 2,17 | 3,30 | 0,66 | 5,11E-01 |
| AT4G09350 | 0,26 | -0,56 | 4,41 | -0,13 | 9,00E-01 |
| AT4G09490 | 0,39 | -2,30 | 4,33 | -0,53 | 5,95E-01 |
| AT4G09510 | 0,38 | -0,39 | 3,37 | -0,12 | 9,08E-01 |
| AT4G09620 | 0,40 | 0,00 | 4,41 | 0,00 | 1,00E+00 |
| AT4G09640 | 0,98 | -0,88 | 1,84 | -0,48 | 6,33E-01 |
| AT4G05740 | 1,41 | -1,04 | 1,63 | -0,64 | 5,25E-01 |
| AT4G09900 | 0,13 | -0,43 | 4,39 | -0,10 | 9,23E-01 |
| AT4G09960 | 0,46 | -0,76 | 4,37 | -0,17 | 8,63E-01 |
| AT4G09980 | 0,57 | 2,26 | 2,69 | 0,84 | 4,01E-01 |
| AT4G10070 | 1,18 | 3,53 | 2,19 | 1,61 | 1,08E-01 |
| AT4G10080 | 1,04 | 0,30 | 2,05 | 0,15 | 8,84E-01 |
| AT4G10130 | 0,55 | 2,78 | 3,82 | 0,73 | 4,66E-01 |
| AT4G10320 | 1,43 | 0,47 | 1,76 | 0,27 | 7,89E-01 |
| AT4G10400 | 0,39 | 1,76 | 4,38 | 0,40 | 6,87E-01 |
| AT4G10420 | 1,07 | 3,67 | 2,61 | 1,41 | 1,59E-01 |
| AT4G10440 | 0,08 | 0,00 | 4,41 | 0,00 | 1,00E+00 |
| AT4G10480 | 0,41 | -0,66 | 4,37 | -0,15 | 8,80E-01 |
| AT4G10540 | 0,12 | 0,00 | 4,41 | 0,00 | 1,00E+00 |
| AT4G10550 | 0,52 | 1,38 | 3,96 | 0,35 | 7,28E-01 |
| AT4G10570 | 1,14 | 1,53 | 1,96 | 0,78 | 4,36E-01 |
| AT4G10590 | 1,40 | 1,74 | 2,14 | 0,81 | 4,15E-01 |
| AT4G05800 | 1,37 | 1,83 | 2,12 | 0,86 | 3,89E-01 |
| AT4G10700 | 0,54 | 0,00 | 3,85 | 0,00 | 1,00E+00 |
| AT4G10730 | 0,28 | 1,84 | 4,39 | 0,42 | 6,74E-01 |
| AT4G10740 | 1,27 | 0,00 | 2,48 | 0,00 | 1,00E+00 |
| AT4G10760 | 0,19 | 0,00 | 4,41 | 0,00 | 1,00E+00 |
| AT4G10770 | 0,10 | 0,00 | 4,41 | 0,00 | 1,00E+00 |
| AT4G10790 | 1,30 | -2,72 | 1,74 | -1,56 | 1,18E-01 |
| AT4G10920 | 0,27 | 0,00 | 3,83 | 0,00 | 1,00E+00 |
| AT4G10960 | 0,05 | 0,00 | 4,41 | 0,00 | 1,00E+00 |
| AT4G10970 | 0,10 | 0,00 | 4,41 | 0,00 | 1,00E+00 |
| AT4G11010 | 0,17 | 0,00 | 4,41 | 0,00 | 1,00E+00 |
| AT4G11050 | 0,37 | 0,43 | 3,44 | 0,12 | 9,01E-01 |
| AT4G11080 | 0,05 | 0,00 | 4,41 | 0,00 | 1,00E+00 |

| | | | | | |
|---|---|---|---|---|---|
| AT4G05855 | 1,27 | 1,35 | 2,02 | 0,67 | 5,04E-01 |
| AT4G11216 | 0,26 | 1,38 | 4,41 | 0,31 | 7,54E-01 |
| AT4G11220 | 0,29 | 0,49 | 4,40 | 0,11 | 9,12E-01 |
| AT4G11260 | 0,66 | 2,00 | 4,35 | 0,46 | 6,45E-01 |
| AT4G11300 | 0,45 | 1,91 | 4,17 | 0,46 | 6,47E-01 |
| AT4G11340 | 1,18 | -1,60 | 1,76 | -0,91 | 3,64E-01 |
| AT4G11410 | 0,24 | 1,37 | 4,41 | 0,31 | 7,56E-01 |
| AT4G05880 | 0,80 | -3,23 | 2,67 | -1,21 | 2,27E-01 |
| AT4G11650 | 0,43 | 2,00 | 4,37 | 0,46 | 6,47E-01 |
| AT4G11670 | 0,44 | 2,09 | 2,82 | 0,74 | 4,58E-01 |
| AT4G11800 | 1,18 | -0,20 | 2,00 | -0,10 | 9,21E-01 |
| AT4G05925 | 1,28 | 0,28 | 2,02 | 0,14 | 8,92E-01 |
| AT4G11820 | 0,25 | -1,80 | 4,34 | -0,41 | 6,78E-01 |
| AT4G11840 | 1,34 | -0,27 | 1,76 | -0,15 | 8,77E-01 |
| AT4G11850 | 1,56 | 0,20 | 1,73 | 0,12 | 9,08E-01 |
| AT4G11860 | 0,25 | 1,38 | 4,41 | 0,31 | 7,54E-01 |
| AT4G11920 | 0,40 | 0,00 | 4,22 | 0,00 | 1,00E+00 |
| AT4G05930 | 0,37 | 0,00 | 4,30 | 0,00 | 1,00E+00 |
| AT4G12010 | 0,59 | 0,00 | 3,77 | 0,00 | 1,00E+00 |
| AT4G05980 | 0,40 | 0,00 | 4,41 | 0,00 | 1,00E+00 |
| AT4G12300 | 1,00 | 0,52 | 2,40 | 0,21 | 8,30E-01 |
| AT4G12310 | 0,40 | 0,43 | 2,94 | 0,15 | 8,83E-01 |
| AT4G12330 | 0,81 | -3,25 | 2,63 | -1,24 | 2,17E-01 |
| AT4G12420 | 1,53 | 4,39 | 1,96 | 2,24 | 2,54E-02 |
| AT4G12590 | 0,64 | -0,28 | 2,67 | -0,11 | 9,16E-01 |
| AT4G12710 | 0,37 | 1,97 | 4,38 | 0,45 | 6,53E-01 |
| AT4G12730 | 0,13 | -0,43 | 4,39 | -0,10 | 9,23E-01 |
| AT4G12750 | 1,05 | 2,29 | 2,04 | 1,13 | 2,60E-01 |
| AT4G12790 | 0,53 | 1,27 | 3,38 | 0,37 | 7,08E-01 |
| AT4G12820 | 0,35 | -2,40 | 4,32 | -0,55 | 5,79E-01 |
| AT4G12880 | 0,35 | -2,40 | 4,32 | -0,55 | 5,79E-01 |
| AT4G12980 | 0,57 | -0,40 | 2,77 | -0,14 | 8,86E-01 |
| AT4G13150 | 0,55 | 2,54 | 4,35 | 0,58 | 5,60E-01 |
| AT4G13250 | 1,54 | 2,48 | 2,04 | 1,22 | 2,24E-01 |
| AT4G13350 | 0,30 | 0,00 | 4,41 | 0,00 | 1,00E+00 |
| AT4G13395 | 0,15 | 0,00 | 4,41 | 0,00 | 1,00E+00 |
| AT4G13420 | 0,12 | 0,00 | 4,41 | 0,00 | 1,00E+00 |
| AT4G13550 | 0,15 | 0,00 | 4,41 | 0,00 | 1,00E+00 |
| AT4G13590 | 0,56 | 1,18 | 2,72 | 0,43 | 6,65E-01 |
| AT4G13630 | 0,80 | 1,36 | 3,54 | 0,38 | 7,01E-01 |
| AT4G13640 | 0,21 | 0,00 | 4,41 | 0,00 | 1,00E+00 |
| AT4G13780 | 1,18 | 1,62 | 1,89 | 0,85 | 3,93E-01 |
| AT4G13800 | 0,45 | 2,13 | 3,28 | 0,65 | 5,16E-01 |
| AT4G13820 | 0,22 | 0,00 | 4,41 | 0,00 | 1,00E+00 |
| AT4G13870 | 0,08 | 0,00 | 4,41 | 0,00 | 1,00E+00 |
| AT4G13930 | 1,56 | 2,68 | 1,76 | 1,53 | 1,26E-01 |
| AT4G13950 | 0,08 | 0,00 | 4,41 | 0,00 | 1,00E+00 |
| AT4G13970 | 0,50 | 1,64 | 3,92 | 0,42 | 6,75E-01 |
| AT4G14010 | 0,10 | 0,00 | 4,41 | 0,00 | 1,00E+00 |
| AT4G14040 | 0,70 | 3,24 | 2,53 | 1,28 | 2,00E-01 |
| AT4G14050 | 0,14 | 0,00 | 4,41 | 0,00 | 1,00E+00 |
| AT4G14070 | 0,95 | 2,41 | 2,65 | 0,91 | 3,65E-01 |
| AT4G14090 | 0,47 | 1,94 | 2,87 | 0,68 | 4,99E-01 |
| AT4G14110 | 0,26 | 1,38 | 4,41 | 0,31 | 7,54E-01 |
| AT4G14200 | 0,21 | 0,00 | 4,41 | 0,00 | 1,00E+00 |
| AT4G14210 | 0,70 | 1,81 | 2,11 | 0,86 | 3,92E-01 |

| | | | | | |
|---|---|---|---|---|---|
| AT4G14230 | 0,73 | 2,05 | 3,01 | 0,68 | 4,96E-01 |
| AT4G14240 | 0,36 | 0,28 | 4,36 | 0,06 | 9,50E-01 |
| AT4G14340 | 0,26 | 1,38 | 4,41 | 0,31 | 7,54E-01 |
| AT4G14342 | 0,72 | 0,00 | 4,41 | 0,00 | 1,00E+00 |
| AT4G14350 | 0,16 | 0,00 | 4,41 | 0,00 | 1,00E+00 |
| AT4G14370 | 0,47 | 0,42 | 2,93 | 0,14 | 8,85E-01 |
| AT4G14385 | 0,08 | 0,00 | 4,41 | 0,00 | 1,00E+00 |
| AT4G14420 | 0,43 | 2,00 | 4,37 | 0,46 | 6,47E-01 |
| AT4G14430 | 1,12 | -2,38 | 1,87 | -1,27 | 2,04E-01 |
| AT4G14440 | 0,19 | 0,00 | 4,41 | 0,00 | 1,00E+00 |
| AT4G14510 | 1,36 | 0,31 | 2,11 | 0,15 | 8,85E-01 |
| AT4G14680 | 0,20 | 1,33 | 4,40 | 0,30 | 7,63E-01 |
| AT4G14710 | 0,27 | -0,41 | 4,39 | -0,09 | 9,26E-01 |
| AT4G14740 | 0,35 | -1,47 | 4,36 | -0,34 | 7,36E-01 |
| AT4G14746 | 0,16 | 0,00 | 4,41 | 0,00 | 1,00E+00 |
| AT4G14790 | 0,23 | 1,38 | 4,41 | 0,31 | 7,54E-01 |
| AT4G14811 | 0,45 | 0,50 | 3,42 | 0,15 | 8,83E-01 |
| AT4G14880 | 1,32 | 1,17 | 1,73 | 0,68 | 4,99E-01 |
| AT4G14900 | 1,30 | 2,53 | 1,81 | 1,40 | 1,61E-01 |
| AT4G14905 | 0,68 | 1,75 | 2,47 | 0,71 | 4,80E-01 |
| AT4G14920 | 0,44 | 1,97 | 4,20 | 0,47 | 6,39E-01 |
| AT4G14990 | 0,87 | 1,89 | 1,99 | 0,95 | 3,42E-01 |
| AT4G15050 | 0,52 | 0,00 | 3,90 | 0,00 | 1,00E+00 |
| AT4G15053 | 0,24 | 0,00 | 4,41 | 0,00 | 1,00E+00 |
| AT4G15080 | 0,43 | 1,73 | 4,18 | 0,41 | 6,78E-01 |
| AT4G15200 | 0,36 | 0,00 | 4,41 | 0,00 | 1,00E+00 |
| AT4G15210 | 0,12 | 0,00 | 4,41 | 0,00 | 1,00E+00 |
| AT4G15242 | 0,72 | 0,49 | 2,31 | 0,21 | 8,34E-01 |
| AT4G06305 | 0,44 | -1,66 | 4,07 | -0,41 | 6,83E-01 |
| AT4G06310 | 0,35 | 2,11 | 4,37 | 0,48 | 6,29E-01 |
| AT4G06315 | 1,00 | 1,38 | 3,20 | 0,43 | 6,66E-01 |
| AT4G15270 | 0,91 | 2,19 | 2,20 | 0,99 | 3,20E-01 |
| AT4G15490 | 0,92 | 1,36 | 2,27 | 0,60 | 5,50E-01 |
| AT4G15530 | 1,47 | 2,09 | 1,97 | 1,06 | 2,90E-01 |
| AT4G15550 | 0,31 | -0,50 | 4,40 | -0,11 | 9,10E-01 |
| AT4G15810 | 0,14 | 0,00 | 4,41 | 0,00 | 1,00E+00 |
| AT4G06340 | 0,10 | 0,00 | 4,41 | 0,00 | 1,00E+00 |
| AT4G15920 | 0,36 | -0,54 | 4,37 | -0,12 | 9,01E-01 |
| AT4G15955 | 0,12 | 0,00 | 4,41 | 0,00 | 1,00E+00 |
| AT4G16070 | 0,52 | 0,48 | 2,80 | 0,17 | 8,64E-01 |
| AT4G16141 | 0,51 | 2,51 | 3,94 | 0,64 | 5,24E-01 |
| AT4G16144 | 1,36 | -2,27 | 2,33 | -0,97 | 3,31E-01 |
| AT4G16155 | 0,30 | 1,61 | 4,39 | 0,37 | 7,14E-01 |
| AT4G16215 | 0,40 | 0,00 | 3,41 | 0,00 | 1,00E+00 |
| AT4G16250 | 0,73 | -0,49 | 2,89 | -0,17 | 8,65E-01 |
| AT4G16280 | 0,45 | 2,17 | 3,28 | 0,66 | 5,09E-01 |
| AT4G16330 | 0,88 | 0,00 | 3,01 | 0,00 | 1,00E+00 |
| AT4G16410 | 1,15 | 2,57 | 2,05 | 1,25 | 2,11E-01 |
| AT4G16470 | 0,11 | 0,00 | 4,41 | 0,00 | 1,00E+00 |
| AT4G16500 | 1,23 | 1,22 | 1,73 | 0,71 | 4,78E-01 |
| AT4G16515 | 0,18 | -0,82 | 4,37 | -0,19 | 8,51E-01 |
| AT4G16520 | 0,32 | 1,35 | 4,40 | 0,31 | 7,59E-01 |
| AT4G16540 | 0,20 | 1,40 | 4,41 | 0,32 | 7,51E-01 |
| AT4G16580 | 0,33 | 0,31 | 3,60 | 0,09 | 9,32E-01 |
| AT4G16630 | 0,66 | 2,09 | 3,70 | 0,56 | 5,73E-01 |
| AT4G16680 | 0,40 | 0,00 | 4,41 | 0,00 | 1,00E+00 |

| | | | | | |
|---|---|---|---|---|---|
| AT4G16690 | 0,88 | 1,39 | 2,33 | 0,60 | 5,50E-01 |
| AT4G16750 | 0,15 | -0,43 | 4,39 | -0,10 | 9,23E-01 |
| AT4G16760 | 1,69 | 1,38 | 1,56 | 0,89 | 3,75E-01 |
| AT4G16790 | 0,35 | 1,34 | 3,56 | 0,38 | 7,06E-01 |
| AT4G16880 | 1,01 | 1,18 | 2,26 | 0,52 | 6,01E-01 |
| AT4G16892 | 1,06 | 1,87 | 2,56 | 0,73 | 4,64E-01 |
| AT4G16957 | 0,81 | 1,95 | 3,49 | 0,56 | 5,77E-01 |
| AT4G16960 | 0,92 | 0,00 | 4,16 | 0,00 | 1,00E+00 |
| AT4G17040 | 0,44 | 0,32 | 4,18 | 0,08 | 9,39E-01 |
| AT4G17060 | 0,64 | 2,69 | 3,65 | 0,74 | 4,62E-01 |
| AT4G17070 | 0,40 | 1,34 | 3,42 | 0,39 | 6,96E-01 |
| AT4G17100 | 0,77 | -2,93 | 2,35 | -1,25 | 2,12E-01 |
| AT4G17190 | 0,27 | 0,00 | 4,41 | 0,00 | 1,00E+00 |
| AT4G17230 | 1,13 | 0,42 | 2,15 | 0,19 | 8,46E-01 |
| AT4G17243 | 0,66 | 2,15 | 2,64 | 0,81 | 4,16E-01 |
| AT4G17280 | 0,46 | 2,17 | 3,32 | 0,65 | 5,13E-01 |
| AT4G17430 | 0,16 | 0,00 | 4,41 | 0,00 | 1,00E+00 |
| AT4G17486 | 0,51 | 2,84 | 4,33 | 0,66 | 5,11E-01 |
| AT4G17490 | 0,05 | 0,00 | 4,41 | 0,00 | 1,00E+00 |
| AT4G17510 | 0,15 | 0,00 | 4,41 | 0,00 | 1,00E+00 |
| AT4G17550 | 0,46 | 0,22 | 4,17 | 0,05 | 9,58E-01 |
| AT4G17570 | 0,79 | 2,87 | 2,49 | 1,15 | 2,48E-01 |
| AT4G17590 | 0,76 | 0,00 | 2,92 | 0,00 | 1,00E+00 |
| AT4G17610 | 0,22 | 0,00 | 4,41 | 0,00 | 1,00E+00 |
| AT4G17670 | 0,33 | 1,33 | 4,40 | 0,30 | 7,62E-01 |
| AT4G17720 | 0,25 | 0,43 | 4,38 | 0,10 | 9,22E-01 |
| AT4G17740 | 0,24 | -0,50 | 4,40 | -0,11 | 9,10E-01 |
| AT4G17840 | 0,82 | 1,37 | 2,48 | 0,55 | 5,80E-01 |
| AT4G17880 | 0,51 | -0,43 | 3,23 | -0,13 | 8,94E-01 |
| AT4G17950 | 0,50 | 0,37 | 4,04 | 0,09 | 9,27E-01 |
| AT4G18010 | 1,46 | 0,00 | 2,21 | 0,00 | 1,00E+00 |
| AT4G18020 | 0,33 | 1,38 | 4,41 | 0,31 | 7,54E-01 |
| AT4G18040 | 0,22 | -0,53 | 4,41 | -0,12 | 9,05E-01 |
| AT4G18100 | 0,24 | -0,47 | 4,40 | -0,11 | 9,15E-01 |
| AT4G18110 | 0,16 | 0,00 | 4,41 | 0,00 | 1,00E+00 |
| AT4G18130 | 0,68 | 2,49 | 2,87 | 0,87 | 3,87E-01 |
| AT4G18140 | 0,37 | 0,94 | 4,36 | 0,22 | 8,29E-01 |
| AT4G18215 | 1,61 | -0,14 | 1,73 | -0,08 | 9,36E-01 |
| AT4G18230 | 0,15 | 0,00 | 4,41 | 0,00 | 1,00E+00 |
| AT4G18250 | 0,30 | 1,38 | 4,41 | 0,31 | 7,54E-01 |
| AT4G18253 | 0,68 | 2,91 | 2,48 | 1,18 | 2,39E-01 |
| AT4G18255 | 0,35 | 1,58 | 4,36 | 0,36 | 7,18E-01 |
| AT4G18390 | 1,03 | -2,01 | 2,17 | -0,93 | 3,54E-01 |
| AT4G18440 | 0,69 | -0,52 | 3,67 | -0,14 | 8,87E-01 |
| AT4G18520 | 0,18 | 0,00 | 4,41 | 0,00 | 1,00E+00 |
| AT4G18580 | 0,08 | 0,00 | 4,41 | 0,00 | 1,00E+00 |
| AT4G18790 | 0,08 | 0,00 | 4,41 | 0,00 | 1,00E+00 |
| AT4G18810 | 0,53 | -0,66 | 2,84 | -0,23 | 8,17E-01 |
| AT4G18840 | 0,38 | -0,54 | 3,51 | -0,15 | 8,77E-01 |
| AT4G19006 | 0,25 | 1,24 | 4,38 | 0,28 | 7,77E-01 |
| AT4G19040 | 0,87 | 0,00 | 3,38 | 0,00 | 1,00E+00 |
| AT4G19045 | 0,12 | 0,00 | 4,41 | 0,00 | 1,00E+00 |
| AT4G19050 | 0,78 | 1,59 | 2,75 | 0,58 | 5,64E-01 |
| AT4G19060 | 0,46 | 1,45 | 4,08 | 0,35 | 7,23E-01 |
| AT4G19080 | 1,21 | 0,35 | 1,94 | 0,18 | 8,55E-01 |
| AT4G19095 | 0,58 | 1,13 | 2,69 | 0,42 | 6,76E-01 |

| | | | | | |
|---|---|---|---|---|---|
| AT4G06870 | 0,16 | 0,00 | 4,41 | 0,00 | 1,00E+00 |
| AT4G19180 | 0,15 | 0,00 | 4,41 | 0,00 | 1,00E+00 |
| AT4G19220 | 0,37 | 2,11 | 4,32 | 0,49 | 6,26E-01 |
| AT4G19240 | 0,33 | 1,24 | 4,38 | 0,28 | 7,78E-01 |
| AT4G19420 | 0,68 | 0,47 | 2,63 | 0,18 | 8,59E-01 |
| AT4G19440 | 0,20 | 0,00 | 4,41 | 0,00 | 1,00E+00 |
| AT4G19640 | 0,35 | 1,28 | 3,53 | 0,36 | 7,16E-01 |
| AT4G19660 | 1,11 | 2,33 | 2,00 | 1,17 | 2,42E-01 |
| AT4G19670 | 0,77 | 2,73 | 3,53 | 0,77 | 4,40E-01 |
| AT4G06915 | 0,34 | 0,00 | 4,41 | 0,00 | 1,00E+00 |
| AT4G19700 | 1,20 | 0,30 | 2,21 | 0,13 | 8,93E-01 |
| AT4G19720 | 0,28 | -1,10 | 4,37 | -0,25 | 8,01E-01 |
| AT4G19860 | 1,15 | 1,33 | 1,89 | 0,70 | 4,81E-01 |
| AT4G19870 | 0,10 | 0,00 | 4,41 | 0,00 | 1,00E+00 |
| AT4G19880 | 0,52 | 0,33 | 2,71 | 0,12 | 9,03E-01 |
| AT4G19890 | 0,56 | 2,56 | 3,10 | 0,83 | 4,08E-01 |
| AT4G19900 | 0,17 | 0,00 | 4,41 | 0,00 | 1,00E+00 |
| AT4G19985 | 0,13 | 0,00 | 4,41 | 0,00 | 1,00E+00 |
| AT4G20095 | 0,45 | -2,53 | 4,31 | -0,59 | 5,57E-01 |
| AT4G20150 | 0,58 | -0,94 | 2,68 | -0,35 | 7,27E-01 |
| AT4G06945 | 1,00 | 2,59 | 2,88 | 0,90 | 3,67E-01 |
| AT4G20210 | 0,49 | -1,27 | 2,76 | -0,46 | 6,44E-01 |
| AT4G20330 | 0,48 | 1,21 | 2,74 | 0,44 | 6,58E-01 |
| AT4G20410 | 1,25 | 1,86 | 1,89 | 0,99 | 3,24E-01 |
| AT4G20440 | 0,64 | 0,53 | 2,37 | 0,22 | 8,23E-01 |
| AT4G20690 | 0,21 | 0,00 | 4,41 | 0,00 | 1,00E+00 |
| AT4G20880 | 0,74 | 1,31 | 2,11 | 0,62 | 5,34E-01 |
| AT4G20910 | 0,27 | 1,84 | 4,38 | 0,42 | 6,75E-01 |
| AT4G20953 | 0,51 | 0,00 | 3,36 | 0,00 | 1,00E+00 |
| AT4G21100 | 1,08 | 0,93 | 1,78 | 0,52 | 6,01E-01 |
| AT4G21140 | 0,20 | 1,33 | 4,40 | 0,30 | 7,63E-01 |
| AT4G21170 | 0,28 | -0,65 | 4,39 | -0,15 | 8,83E-01 |
| AT4G06985 | 0,08 | 0,00 | 4,41 | 0,00 | 1,00E+00 |
| AT4G21190 | 0,08 | 0,00 | 4,41 | 0,00 | 1,00E+00 |
| AT4G21350 | 0,08 | 0,00 | 4,41 | 0,00 | 1,00E+00 |
| AT4G21380 | 1,64 | 2,18 | 1,95 | 1,12 | 2,65E-01 |
| AT4G21400 | 0,16 | 0,00 | 4,41 | 0,00 | 1,00E+00 |
| AT4G21430 | 0,22 | -0,54 | 4,41 | -0,12 | 9,03E-01 |
| AT4G21510 | 0,05 | 0,00 | 4,41 | 0,00 | 1,00E+00 |
| AT4G21520 | 0,22 | 1,33 | 4,40 | 0,30 | 7,63E-01 |
| AT4G21560 | 0,38 | 2,11 | 4,37 | 0,48 | 6,29E-01 |
| AT4G21570 | 1,42 | -0,56 | 2,18 | -0,26 | 7,96E-01 |
| AT4G21660 | 1,13 | 3,17 | 2,24 | 1,42 | 1,56E-01 |
| AT4G21810 | 1,02 | 1,93 | 2,15 | 0,90 | 3,70E-01 |
| AT4G21860 | 0,50 | -0,63 | 3,36 | -0,19 | 8,52E-01 |
| AT4G21890 | 0,12 | 0,00 | 4,41 | 0,00 | 1,00E+00 |
| AT4G21910 | 0,54 | -0,80 | 4,38 | -0,18 | 8,56E-01 |
| AT4G21990 | 0,81 | 3,23 | 2,54 | 1,27 | 2,03E-01 |
| AT4G22120 | 0,12 | 0,00 | 4,41 | 0,00 | 1,00E+00 |
| AT4G22150 | 0,22 | -0,74 | 4,37 | -0,17 | 8,66E-01 |
| AT4G22190 | 0,98 | 1,04 | 1,81 | 0,58 | 5,64E-01 |
| AT4G22200 | 0,21 | 0,00 | 4,41 | 0,00 | 1,00E+00 |
| AT4G22250 | 0,16 | 0,00 | 4,41 | 0,00 | 1,00E+00 |
| AT4G22260 | 0,18 | -0,51 | 4,40 | -0,12 | 9,07E-01 |
| AT4G22265 | 0,20 | 1,38 | 4,41 | 0,31 | 7,54E-01 |
| AT4G22285 | 0,20 | -0,44 | 4,39 | -0,10 | 9,19E-01 |

| | | | | | |
|---|---|---|---|---|---|
| AT4G22290 | 0,23 | 0,00 | 4,41 | 0,00 | 1,00E+00 |
| AT4G22350 | 0,69 | 1,40 | 3,68 | 0,38 | 7,04E-01 |
| AT4G22380 | 0,51 | 1,38 | 2,79 | 0,49 | 6,21E-01 |
| AT4G22470 | 0,12 | 0,00 | 4,41 | 0,00 | 1,00E+00 |
| AT4G22485 | 0,49 | -0,40 | 3,96 | -0,10 | 9,20E-01 |
| AT4G22550 | 0,26 | -0,51 | 3,83 | -0,13 | 8,94E-01 |
| AT4G22570 | 0,24 | 1,57 | 4,39 | 0,36 | 7,20E-01 |
| AT4G22745 | 0,42 | 2,17 | 4,16 | 0,52 | 6,03E-01 |
| AT4G22750 | 1,03 | 0,28 | 2,32 | 0,12 | 9,05E-01 |
| AT4G22755 | 0,47 | 1,99 | 4,38 | 0,45 | 6,50E-01 |
| AT4G22770 | 1,29 | -1,02 | 1,78 | -0,57 | 5,66E-01 |
| AT4G22780 | 0,24 | 1,26 | 4,39 | 0,29 | 7,75E-01 |
| AT4G22970 | 0,52 | 0,00 | 3,92 | 0,00 | 1,00E+00 |
| AT4G22980 | 0,47 | 0,00 | 4,04 | 0,00 | 1,00E+00 |
| AT4G23010 | 0,83 | 3,58 | 2,31 | 1,55 | 1,20E-01 |
| AT4G23040 | 0,28 | -1,39 | 4,37 | -0,32 | 7,51E-01 |
| AT4G23130 | 0,55 | 1,38 | 4,01 | 0,34 | 7,31E-01 |
| AT4G23190 | 1,18 | 2,87 | 2,05 | 1,40 | 1,61E-01 |
| AT4G23205 | 0,08 | 0,00 | 4,41 | 0,00 | 1,00E+00 |
| AT4G23200 | 0,43 | 1,87 | 4,38 | 0,43 | 6,70E-01 |
| AT4G23210 | 0,21 | 0,00 | 4,41 | 0,00 | 1,00E+00 |
| AT4G23220 | 0,76 | 1,28 | 2,52 | 0,51 | 6,12E-01 |
| AT4G23240 | 0,37 | 1,38 | 4,41 | 0,31 | 7,54E-01 |
| AT4G23250 | 0,45 | 0,00 | 4,02 | 0,00 | 1,00E+00 |
| AT4G23260 | 0,19 | 0,00 | 4,41 | 0,00 | 1,00E+00 |
| AT4G23290 | 0,10 | 0,00 | 4,41 | 0,00 | 1,00E+00 |
| AT4G23320 | 0,08 | 0,00 | 4,41 | 0,00 | 1,00E+00 |
| AT4G23460 | 1,64 | 3,13 | 1,63 | 1,92 | 5,44E-02 |
| AT4G23490 | 0,15 | -0,43 | 4,39 | -0,10 | 9,23E-01 |
| AT4G07130 | 0,46 | 2,24 | 4,17 | 0,54 | 5,91E-01 |
| AT4G23540 | 0,67 | 1,83 | 3,03 | 0,60 | 5,46E-01 |
| AT4G23560 | 0,10 | 0,00 | 4,41 | 0,00 | 1,00E+00 |
| AT4G07135 | 1,25 | 3,79 | 2,01 | 1,88 | 6,02E-02 |
| AT4G07145 | 0,96 | 3,34 | 2,63 | 1,27 | 2,03E-01 |
| AT4G23620 | 0,31 | 1,26 | 4,39 | 0,29 | 7,74E-01 |
| AT4G23640 | 0,77 | 0,63 | 2,50 | 0,25 | 8,00E-01 |
| AT4G23670 | 1,50 | -2,67 | 1,77 | -1,51 | 1,31E-01 |
| AT4G23700 | 0,29 | 1,85 | 4,39 | 0,42 | 6,73E-01 |
| AT4G23770 | 0,22 | -0,48 | 4,40 | -0,11 | 9,13E-01 |
| AT4G23810 | 0,35 | 1,38 | 4,41 | 0,31 | 7,54E-01 |
| AT4G23820 | 0,63 | 1,20 | 2,33 | 0,52 | 6,05E-01 |
| AT4G23880 | 0,47 | 0,51 | 3,27 | 0,16 | 8,75E-01 |
| AT4G23890 | 0,66 | -0,07 | 2,66 | -0,03 | 9,80E-01 |
| AT4G24180 | 0,37 | 1,75 | 4,38 | 0,40 | 6,89E-01 |
| AT4G24200 | 0,70 | 2,66 | 3,01 | 0,88 | 3,76E-01 |
| AT4G07250 | 0,17 | -1,42 | 4,35 | -0,33 | 7,45E-01 |
| AT4G24300 | 0,45 | 0,00 | 4,41 | 0,00 | 1,00E+00 |
| AT4G24320 | 0,33 | -0,52 | 4,40 | -0,12 | 9,06E-01 |
| AT4G24330 | 0,84 | -0,34 | 2,11 | -0,16 | 8,72E-01 |
| AT4G07255 | 0,10 | 0,00 | 4,41 | 0,00 | 1,00E+00 |
| AT4G24390 | 0,29 | 1,30 | 4,40 | 0,30 | 7,67E-01 |
| AT4G24490 | 0,72 | 1,34 | 2,54 | 0,53 | 5,97E-01 |
| AT4G24530 | 0,08 | 0,00 | 4,41 | 0,00 | 1,00E+00 |
| AT4G24560 | 1,21 | 0,45 | 2,08 | 0,22 | 8,27E-01 |
| AT4G24580 | 0,46 | 2,68 | 4,12 | 0,65 | 5,15E-01 |
| AT4G24610 | 0,67 | 2,92 | 2,93 | 0,99 | 3,20E-01 |

| | | | | | |
|---|---|---|---|---|---|
| AT4G24620 | 0,42 | 0,28 | 4,22 | 0,07 | 9,47E-01 |
| AT4G24660 | 1,04 | -1,44 | 2,27 | -0,64 | 5,25E-01 |
| AT4G24670 | 0,08 | 0,00 | 4,41 | 0,00 | 1,00E+00 |
| AT4G24730 | 0,45 | 2,57 | 4,08 | 0,63 | 5,29E-01 |
| AT4G24740 | 0,63 | 2,55 | 3,12 | 0,82 | 4,14E-01 |
| AT4G24750 | 0,28 | 1,84 | 4,39 | 0,42 | 6,74E-01 |
| AT4G24780 | 1,27 | 0,60 | 2,37 | 0,25 | 8,00E-01 |
| AT4G07285 | 0,27 | -1,46 | 4,34 | -0,34 | 7,36E-01 |
| AT4G24820 | 1,43 | 1,93 | 1,86 | 1,04 | 3,00E-01 |
| AT4G24830 | 0,25 | -0,47 | 4,40 | -0,11 | 9,14E-01 |
| AT4G24840 | 0,53 | -0,06 | 3,17 | -0,02 | 9,85E-01 |
| AT4G24920 | 0,86 | 2,72 | 2,77 | 0,98 | 3,26E-01 |
| AT4G24990 | 0,31 | 1,38 | 4,41 | 0,31 | 7,54E-01 |
| AT4G25090 | 0,29 | 0,00 | 4,41 | 0,00 | 1,00E+00 |
| AT4G25270 | 0,45 | 2,57 | 4,08 | 0,63 | 5,29E-01 |
| AT4G25280 | 0,71 | 1,38 | 4,41 | 0,31 | 7,54E-01 |
| AT4G25290 | 0,52 | 1,26 | 3,91 | 0,32 | 7,47E-01 |
| AT4G25340 | 0,19 | -0,52 | 4,40 | -0,12 | 9,06E-01 |
| AT4G25435 | 0,88 | -0,36 | 2,59 | -0,14 | 8,89E-01 |
| AT4G25540 | 0,13 | 0,00 | 4,41 | 0,00 | 1,00E+00 |
| AT4G25570 | 0,43 | -0,23 | 4,17 | -0,06 | 9,56E-01 |
| AT4G25640 | 1,48 | 2,56 | 1,72 | 1,49 | 1,37E-01 |
| AT4G25650 | 1,11 | -0,64 | 1,71 | -0,38 | 7,07E-01 |
| AT4G25670 | 0,79 | 2,89 | 2,84 | 1,02 | 3,09E-01 |
| AT4G25690 | 0,56 | 1,78 | 3,26 | 0,55 | 5,84E-01 |
| AT4G25740 | 0,38 | 1,60 | 4,39 | 0,36 | 7,16E-01 |
| AT4G25870 | 0,26 | 1,38 | 4,41 | 0,31 | 7,54E-01 |
| AT4G25940 | 0,39 | 0,32 | 4,39 | 0,07 | 9,42E-01 |
| AT4G25990 | 0,26 | 1,38 | 4,41 | 0,31 | 7,54E-01 |
| AT4G26080 | 0,87 | -1,41 | 2,52 | -0,56 | 5,76E-01 |
| AT4G26100 | 0,81 | 1,10 | 2,20 | 0,50 | 6,18E-01 |
| AT4G26130 | 1,45 | -3,25 | 1,85 | -1,76 | 7,86E-02 |
| AT4G26140 | 0,21 | 1,25 | 4,39 | 0,29 | 7,75E-01 |
| AT4G26190 | 0,25 | 0,00 | 4,41 | 0,00 | 1,00E+00 |
| AT4G26300 | 0,69 | 0,75 | 2,60 | 0,29 | 7,72E-01 |
| AT4G07565 | 0,79 | -1,76 | 2,19 | -0,81 | 4,20E-01 |
| AT4G07625 | 0,79 | -0,67 | 2,66 | -0,25 | 8,01E-01 |
| AT4G07635 | 0,12 | 0,00 | 4,41 | 0,00 | 1,00E+00 |
| AT4G26370 | 0,48 | -0,78 | 4,05 | -0,19 | 8,48E-01 |
| AT4G26375 | 0,40 | 0,00 | 4,41 | 0,00 | 1,00E+00 |
| AT4G26400 | 0,60 | 2,34 | 3,95 | 0,59 | 5,53E-01 |
| AT4G26620 | 0,28 | -0,44 | 4,39 | -0,10 | 9,21E-01 |
| AT4G26630 | 0,72 | 1,35 | 2,61 | 0,52 | 6,06E-01 |
| AT4G26640 | 1,00 | -1,32 | 2,47 | -0,53 | 5,93E-01 |
| AT4G26690 | 1,51 | 0,39 | 1,92 | 0,20 | 8,40E-01 |
| AT4G26750 | 0,43 | -0,68 | 4,22 | -0,16 | 8,72E-01 |
| AT4G26760 | 0,64 | 0,42 | 3,18 | 0,13 | 8,95E-01 |
| AT4G26795 | 0,42 | 0,00 | 4,15 | 0,00 | 1,00E+00 |
| AT4G26860 | 0,37 | -0,54 | 4,41 | -0,12 | 9,02E-01 |
| AT4G26910 | 1,33 | 2,00 | 1,62 | 1,24 | 2,17E-01 |
| AT4G26940 | 0,43 | 0,35 | 4,11 | 0,09 | 9,32E-01 |
| AT4G26980 | 0,78 | 2,34 | 2,49 | 0,94 | 3,46E-01 |
| AT4G27020 | 0,48 | 0,42 | 3,48 | 0,12 | 9,04E-01 |
| AT4G07765 | 0,66 | 0,00 | 2,96 | 0,00 | 1,00E+00 |
| AT4G27130 | 0,28 | 0,00 | 3,77 | 0,00 | 1,00E+00 |
| AT4G27270 | 0,66 | 1,33 | 3,94 | 0,34 | 7,35E-01 |

| | | | | | |
|---|---|---|---|---|---|
| AT4G27280 | 0,84 | 1,48 | 2,93 | 0,51 | 6,12E-01 |
| AT4G27300 | 0,83 | -0,06 | 2,44 | -0,02 | 9,80E-01 |
| AT4G27320 | 1,19 | 1,56 | 2,20 | 0,71 | 4,78E-01 |
| AT4G27340 | 0,21 | 0,00 | 4,41 | 0,00 | 1,00E+00 |
| AT4G27410 | 0,98 | 3,32 | 2,31 | 1,44 | 1,51E-01 |
| AT4G27450 | 0,72 | 1,50 | 2,38 | 0,63 | 5,27E-01 |
| AT4G27480 | 0,65 | 1,34 | 3,66 | 0,37 | 7,14E-01 |
| AT4G27650 | 0,39 | -0,02 | 3,45 | -0,01 | 9,96E-01 |
| AT4G27700 | 0,47 | 2,36 | 3,31 | 0,71 | 4,75E-01 |
| AT4G27730 | 0,37 | 1,80 | 4,38 | 0,41 | 6,81E-01 |
| AT4G27800 | 1,09 | 0,29 | 1,90 | 0,15 | 8,78E-01 |
| AT4G27830 | 0,84 | 1,36 | 2,10 | 0,65 | 5,15E-01 |
| AT4G27840 | 0,43 | 2,29 | 4,37 | 0,52 | 6,01E-01 |
| AT4G27885 | 0,16 | 0,00 | 4,41 | 0,00 | 1,00E+00 |
| AT4G27960 | 0,38 | 0,00 | 4,35 | 0,00 | 1,00E+00 |
| AT4G28030 | 0,45 | 0,43 | 2,87 | 0,15 | 8,79E-01 |
| AT4G28050 | 0,43 | -0,40 | 4,38 | -0,09 | 9,28E-01 |
| AT4G28060 | 0,25 | 1,84 | 4,38 | 0,42 | 6,75E-01 |
| AT4G28180 | 0,69 | 1,37 | 2,70 | 0,51 | 6,12E-01 |
| AT4G28200 | 0,59 | -0,69 | 2,70 | -0,25 | 7,99E-01 |
| AT4G28230 | 0,33 | 1,38 | 4,41 | 0,31 | 7,54E-01 |
| AT4G28240 | 0,66 | -0,39 | 2,32 | -0,17 | 8,66E-01 |
| AT4G28320 | 0,20 | 1,33 | 4,40 | 0,30 | 7,63E-01 |
| AT4G28390 | 0,33 | 1,37 | 4,41 | 0,31 | 7,56E-01 |
| AT4G28450 | 0,61 | 3,53 | 4,30 | 0,82 | 4,11E-01 |
| AT4G28470 | 0,62 | 2,55 | 3,16 | 0,81 | 4,19E-01 |
| AT4G28610 | 0,93 | 2,70 | 2,66 | 1,02 | 3,10E-01 |
| AT4G28630 | 0,12 | 0,00 | 4,41 | 0,00 | 1,00E+00 |
| AT4G28770 | 0,20 | 0,00 | 4,41 | 0,00 | 1,00E+00 |
| AT4G28910 | 1,06 | 1,26 | 2,32 | 0,54 | 5,87E-01 |
| AT4G29010 | 1,63 | -1,46 | 1,84 | -0,79 | 4,27E-01 |
| AT4G29050 | 0,13 | -0,43 | 4,39 | -0,10 | 9,23E-01 |
| AT4G29080 | 0,28 | 1,37 | 4,41 | 0,31 | 7,56E-01 |
| AT4G29130 | 1,04 | -0,34 | 1,77 | -0,19 | 8,48E-01 |
| AT4G29220 | 0,63 | 0,21 | 3,10 | 0,07 | 9,46E-01 |
| AT4G29260 | 0,48 | 0,00 | 4,08 | 0,00 | 1,00E+00 |
| AT4G29300 | 1,28 | -1,85 | 1,74 | -1,06 | 2,88E-01 |
| AT4G29305 | 0,76 | 0,40 | 2,53 | 0,16 | 8,74E-01 |
| AT4G08155 | 1,40 | 1,87 | 1,83 | 1,02 | 3,06E-01 |
| AT4G29440 | 0,66 | 2,19 | 3,01 | 0,72 | 4,68E-01 |
| AT4G29520 | 0,77 | 1,73 | 2,64 | 0,66 | 5,12E-01 |
| AT4G29654 | 1,07 | 0,00 | 2,80 | 0,00 | 1,00E+00 |
| AT4G29670 | 0,65 | 1,25 | 2,38 | 0,53 | 5,99E-01 |
| AT4G29680 | 0,85 | 1,68 | 3,01 | 0,56 | 5,77E-01 |
| AT4G29720 | 0,17 | -0,49 | 4,40 | -0,11 | 9,11E-01 |
| AT4G29735 | 0,17 | -0,57 | 4,40 | -0,13 | 8,98E-01 |
| AT4G29810 | 0,27 | 0,00 | 4,41 | 0,00 | 1,00E+00 |
| AT4G29820 | 0,24 | 1,38 | 4,41 | 0,31 | 7,54E-01 |
| AT4G29840 | 0,58 | 0,25 | 2,38 | 0,11 | 9,16E-01 |
| AT4G29880 | 0,21 | 0,00 | 4,41 | 0,00 | 1,00E+00 |
| AT4G29905 | 1,17 | -0,43 | 2,01 | -0,22 | 8,29E-01 |
| AT4G29910 | 0,08 | 0,00 | 4,41 | 0,00 | 1,00E+00 |
| AT4G29920 | 0,05 | 0,00 | 4,41 | 0,00 | 1,00E+00 |
| AT4G29940 | 0,28 | 1,84 | 4,39 | 0,42 | 6,74E-01 |
| AT4G29960 | 0,50 | 2,34 | 3,95 | 0,59 | 5,55E-01 |
| AT4G29990 | 0,08 | 0,00 | 4,41 | 0,00 | 1,00E+00 |

| | | | | | |
|---|---|---|---|---|---|
| AT4G30000 | 0,38 | 2,48 | 4,35 | 0,57 | 5,69E-01 |
| AT4G30030 | 0,45 | 1,24 | 4,38 | 0,28 | 7,77E-01 |
| AT4G30060 | 0,08 | 0,00 | 4,41 | 0,00 | 1,00E+00 |
| AT4G30080 | 0,31 | 1,53 | 4,39 | 0,35 | 7,27E-01 |
| AT4G30110 | 1,05 | 2,14 | 3,23 | 0,66 | 5,08E-01 |
| AT4G30200 | 1,47 | 1,00 | 1,79 | 0,56 | 5,76E-01 |
| AT4G30240 | 0,30 | 1,27 | 4,39 | 0,29 | 7,72E-01 |
| AT4G30260 | 0,47 | 0,30 | 3,26 | 0,09 | 9,28E-01 |
| AT4G30270 | 1,68 | -0,32 | 1,80 | -0,18 | 8,59E-01 |
| AT4G30280 | 0,12 | 0,00 | 4,41 | 0,00 | 1,00E+00 |
| AT4G30360 | 0,21 | 0,00 | 4,41 | 0,00 | 1,00E+00 |
| AT4G30410 | 0,12 | 0,00 | 4,41 | 0,00 | 1,00E+00 |
| AT4G30430 | 0,08 | 0,00 | 4,41 | 0,00 | 1,00E+00 |
| AT4G30560 | 0,74 | 3,05 | 3,49 | 0,88 | 3,81E-01 |
| AT4G30580 | 0,92 | 2,03 | 3,47 | 0,58 | 5,59E-01 |
| AT4G30600 | 1,60 | 1,66 | 1,75 | 0,95 | 3,41E-01 |
| AT4G30620 | 0,23 | 1,38 | 4,41 | 0,31 | 7,54E-01 |
| AT4G30700 | 0,08 | 0,00 | 4,41 | 0,00 | 1,00E+00 |
| AT4G30710 | 0,12 | 0,00 | 4,41 | 0,00 | 1,00E+00 |
| AT4G30720 | 1,29 | 3,23 | 1,84 | 1,76 | 7,90E-02 |
| AT4G30740 | 1,03 | 0,00 | 2,60 | 0,00 | 1,00E+00 |
| AT4G30790 | 0,99 | 1,38 | 2,44 | 0,56 | 5,72E-01 |
| AT4G30890 | 0,60 | 2,47 | 3,02 | 0,82 | 4,14E-01 |
| AT4G30920 | 1,19 | -0,53 | 1,87 | -0,28 | 7,78E-01 |
| AT4G30950 | 1,64 | -0,17 | 1,58 | -0,11 | 9,14E-01 |
| AT4G31020 | 0,29 | 1,74 | 4,38 | 0,40 | 6,90E-01 |
| AT4G31120 | 0,87 | 2,26 | 2,46 | 0,92 | 3,59E-01 |
| AT4G31130 | 0,59 | -1,36 | 3,25 | -0,42 | 6,76E-01 |
| AT4G31420 | 0,28 | 1,34 | 4,40 | 0,30 | 7,61E-01 |
| AT4G31450 | 0,46 | -0,80 | 4,14 | -0,19 | 8,46E-01 |
| AT4G31540 | 0,43 | -1,08 | 4,22 | -0,26 | 7,99E-01 |
| AT4G31550 | 1,02 | 1,63 | 1,87 | 0,87 | 3,85E-01 |
| AT4G31590 | 0,63 | 1,38 | 2,74 | 0,50 | 6,15E-01 |
| AT4G31810 | 0,08 | 0,00 | 4,41 | 0,00 | 1,00E+00 |
| AT4G31820 | 0,43 | 0,41 | 2,91 | 0,14 | 8,89E-01 |
| AT4G32070 | 0,58 | 2,87 | 3,87 | 0,74 | 4,58E-01 |
| AT4G32120 | 0,15 | 0,00 | 4,41 | 0,00 | 1,00E+00 |
| AT4G32130 | 0,18 | 0,00 | 4,41 | 0,00 | 1,00E+00 |
| AT4G32150 | 1,17 | 2,26 | 1,82 | 1,24 | 2,14E-01 |
| AT4G32180 | 1,00 | -0,01 | 2,53 | 0,00 | 9,98E-01 |
| AT4G32208 | 0,37 | 2,11 | 4,32 | 0,49 | 6,26E-01 |
| AT4G32340 | 1,02 | 1,40 | 2,02 | 0,70 | 4,86E-01 |
| AT4G32420 | 0,78 | 0,23 | 2,58 | 0,09 | 9,29E-01 |
| AT4G32460 | 0,31 | -0,54 | 4,41 | -0,12 | 9,02E-01 |
| AT4G32470 | 0,29 | 1,29 | 4,40 | 0,29 | 7,68E-01 |
| AT4G32510 | 0,56 | 1,38 | 3,84 | 0,36 | 7,20E-01 |
| AT4G32520 | 1,00 | 0,22 | 2,02 | 0,11 | 9,14E-01 |
| AT4G32530 | 0,89 | 1,50 | 2,47 | 0,61 | 5,44E-01 |
| AT4G32670 | 0,15 | 0,00 | 4,41 | 0,00 | 1,00E+00 |
| AT4G32750 | 0,18 | -0,45 | 4,39 | -0,10 | 9,18E-01 |
| AT4G32820 | 0,83 | 1,82 | 2,78 | 0,65 | 5,14E-01 |
| AT4G32880 | 1,09 | 3,41 | 2,03 | 1,68 | 9,28E-02 |
| AT4G32900 | 0,12 | 0,00 | 4,41 | 0,00 | 1,00E+00 |
| AT4G32920 | 0,70 | 1,43 | 2,95 | 0,48 | 6,28E-01 |
| AT4G33040 | 0,37 | 0,49 | 4,40 | 0,11 | 9,11E-01 |
| AT4G33090 | 0,64 | -1,40 | 3,14 | -0,45 | 6,56E-01 |

| | | | | | |
|---|---|---|---|---|---|
| AT4G33110 | 0,17 | 0,00 | 4,41 | 0,00 | 1,00E+00 |
| AT4G33150 | 0,65 | 1,38 | 3,73 | 0,37 | 7,12E-01 |
| AT4G33200 | 0,29 | 1,35 | 4,40 | 0,31 | 7,59E-01 |
| AT4G33210 | 1,11 | 3,27 | 2,00 | 1,64 | 1,02E-01 |
| AT4G33250 | 0,89 | 1,83 | 2,06 | 0,89 | 3,75E-01 |
| AT4G33260 | 0,12 | 0,00 | 4,41 | 0,00 | 1,00E+00 |
| AT4G33270 | 0,12 | 0,00 | 4,41 | 0,00 | 1,00E+00 |
| AT4G33400 | 0,33 | 0,43 | 4,38 | 0,10 | 9,21E-01 |
| AT4G33430 | 1,19 | 1,19 | 2,13 | 0,56 | 5,78E-01 |
| AT4G33470 | 0,67 | 1,84 | 2,99 | 0,61 | 5,40E-01 |
| AT4G33495 | 0,29 | 1,35 | 4,40 | 0,31 | 7,59E-01 |
| AT4G33500 | 0,96 | -2,07 | 2,60 | -0,80 | 4,25E-01 |
| AT4G33530 | 1,44 | -0,22 | 1,80 | -0,12 | 9,05E-01 |
| AT4G33585 | 1,20 | 4,19 | 2,87 | 1,46 | 1,44E-01 |
| AT4G33625 | 0,16 | 0,00 | 4,41 | 0,00 | 1,00E+00 |
| AT4G33670 | 0,76 | 3,21 | 2,15 | 1,50 | 1,34E-01 |
| AT4G33680 | 0,20 | 0,00 | 4,41 | 0,00 | 1,00E+00 |
| AT4G33700 | 0,28 | -1,10 | 4,38 | -0,25 | 8,02E-01 |
| AT4G33790 | 0,16 | 0,00 | 4,41 | 0,00 | 1,00E+00 |
| AT4G33860 | 0,68 | 2,76 | 3,62 | 0,76 | 4,46E-01 |
| AT4G33865 | 0,81 | -0,33 | 2,26 | -0,14 | 8,85E-01 |
| AT4G33910 | 0,26 | 1,38 | 4,41 | 0,31 | 7,54E-01 |
| AT4G34020 | 0,41 | -1,65 | 4,36 | -0,38 | 7,06E-01 |
| AT4G34040 | 0,82 | 1,36 | 2,48 | 0,55 | 5,84E-01 |
| AT4G34070 | 0,72 | -0,09 | 2,62 | -0,03 | 9,72E-01 |
| AT4G34090 | 0,71 | -0,40 | 4,38 | -0,09 | 9,28E-01 |
| AT4G34131 | 0,52 | 1,38 | 3,96 | 0,35 | 7,28E-01 |
| AT4G34180 | 0,68 | -0,99 | 2,96 | -0,33 | 7,39E-01 |
| AT4G34190 | 1,52 | 2,76 | 1,84 | 1,50 | 1,34E-01 |
| AT4G34200 | 1,37 | 1,79 | 1,89 | 0,95 | 3,42E-01 |
| AT4G34220 | 1,59 | 0,33 | 1,73 | 0,19 | 8,51E-01 |
| AT4G34332 | 0,08 | 0,00 | 4,41 | 0,00 | 1,00E+00 |
| AT4G34340 | 0,12 | 0,00 | 4,41 | 0,00 | 1,00E+00 |
| AT4G34350 | 1,55 | 0,48 | 1,83 | 0,26 | 7,93E-01 |
| AT4G08985 | 0,45 | 0,00 | 3,33 | 0,00 | 1,00E+00 |
| AT4G34460 | 0,42 | 1,28 | 3,34 | 0,38 | 7,03E-01 |
| AT4G34480 | 1,19 | 1,82 | 1,96 | 0,93 | 3,53E-01 |
| AT4G34490 | 1,39 | 0,97 | 1,74 | 0,55 | 5,79E-01 |
| AT4G34610 | 0,37 | 2,20 | 4,37 | 0,50 | 6,14E-01 |
| AT4G34620 | 1,51 | 0,19 | 1,97 | 0,09 | 9,25E-01 |
| AT4G34630 | 0,16 | 0,00 | 4,41 | 0,00 | 1,00E+00 |
| AT4G34690 | 1,55 | -0,54 | 2,19 | -0,25 | 8,04E-01 |
| AT4G34730 | 0,55 | 0,91 | 2,67 | 0,34 | 7,34E-01 |
| AT4G34740 | 0,69 | 0,79 | 2,85 | 0,28 | 7,81E-01 |
| AT4G34860 | 0,08 | 0,00 | 4,41 | 0,00 | 1,00E+00 |
| AT4G34900 | 0,65 | 2,89 | 3,60 | 0,80 | 4,23E-01 |
| AT4G35060 | 0,12 | 0,00 | 4,41 | 0,00 | 1,00E+00 |
| AT4G35140 | 0,36 | -0,09 | 4,37 | -0,02 | 9,84E-01 |
| AT4G35180 | 0,66 | 2,08 | 2,99 | 0,70 | 4,86E-01 |
| AT4G35230 | 1,11 | 3,00 | 2,08 | 1,44 | 1,50E-01 |
| AT4G35240 | 1,02 | 2,60 | 2,14 | 1,22 | 2,24E-01 |
| AT4G35260 | 0,53 | 0,21 | 3,13 | 0,07 | 9,47E-01 |
| AT4G35300 | 1,43 | -0,12 | 2,09 | -0,06 | 9,54E-01 |
| AT4G35320 | 0,51 | 1,35 | 3,25 | 0,42 | 6,77E-01 |
| AT4G35360 | 0,54 | 0,68 | 3,97 | 0,17 | 8,63E-01 |
| AT4G35440 | 0,24 | 0,00 | 4,41 | 0,00 | 1,00E+00 |

| | | | | | |
|---|---|---|---|---|---|
| AT4G35570 | 0,29 | 1,38 | 4,41 | 0,31 | 7,54E-01 |
| AT4G35580 | 0,56 | 0,90 | 3,09 | 0,29 | 7,71E-01 |
| AT4G35780 | 0,15 | 0,00 | 4,41 | 0,00 | 1,00E+00 |
| AT4G35785 | 0,84 | 1,38 | 2,94 | 0,47 | 6,39E-01 |
| AT4G35920 | 0,70 | 3,60 | 4,30 | 0,84 | 4,02E-01 |
| AT4G36010 | 0,74 | -0,91 | 2,78 | -0,33 | 7,44E-01 |
| AT4G36030 | 1,21 | 1,91 | 2,28 | 0,84 | 4,02E-01 |
| AT4G36050 | 0,52 | 0,00 | 4,41 | 0,00 | 1,00E+00 |
| AT4G36070 | 0,08 | 0,00 | 4,41 | 0,00 | 1,00E+00 |
| AT4G36220 | 1,38 | 0,70 | 1,67 | 0,42 | 6,74E-01 |
| AT4G36270 | 0,14 | 0,00 | 4,41 | 0,00 | 1,00E+00 |
| AT4G36280 | 0,08 | 0,00 | 4,41 | 0,00 | 1,00E+00 |
| AT4G36380 | 0,24 | -1,71 | 4,34 | -0,39 | 6,93E-01 |
| AT4G36390 | 0,60 | 2,40 | 2,61 | 0,92 | 3,58E-01 |
| AT4G36460 | 0,50 | -0,45 | 3,23 | -0,14 | 8,90E-01 |
| AT4G36500 | 0,97 | 0,63 | 2,25 | 0,28 | 7,78E-01 |
| AT4G36520 | 1,04 | 0,53 | 2,01 | 0,26 | 7,91E-01 |
| AT4G36550 | 0,53 | 1,38 | 3,27 | 0,42 | 6,73E-01 |
| AT4G36640 | 1,54 | 3,84 | 2,04 | 1,88 | 5,99E-02 |
| AT4G36670 | 0,44 | -0,54 | 3,43 | -0,16 | 8,74E-01 |
| AT4G36690 | 0,56 | 0,53 | 3,36 | 0,16 | 8,75E-01 |
| AT4G36730 | 0,51 | -0,53 | 3,18 | -0,17 | 8,67E-01 |
| AT4G36770 | 0,41 | 1,65 | 4,19 | 0,39 | 6,95E-01 |
| AT4G36800 | 0,60 | -0,36 | 3,09 | -0,12 | 9,06E-01 |
| AT4G36808 | 0,88 | -0,15 | 2,55 | -0,06 | 9,52E-01 |
| AT4G36860 | 0,41 | -0,10 | 4,27 | -0,02 | 9,82E-01 |
| AT4G36890 | 0,60 | 3,18 | 3,89 | 0,82 | 4,14E-01 |
| AT4G36970 | 0,18 | 1,38 | 4,41 | 0,31 | 7,54E-01 |
| AT4G36980 | 0,30 | 1,38 | 4,41 | 0,31 | 7,54E-01 |
| AT4G37100 | 0,62 | 2,76 | 3,04 | 0,91 | 3,65E-01 |
| AT4G37270 | 0,72 | 0,34 | 2,60 | 0,13 | 8,97E-01 |
| AT4G37280 | 0,63 | 3,27 | 3,61 | 0,91 | 3,65E-01 |
| AT4G37370 | 0,86 | 2,90 | 2,58 | 1,12 | 2,62E-01 |
| AT4G37432 | 0,86 | -0,40 | 3,45 | -0,11 | 9,09E-01 |
| AT4G37470 | 1,39 | -1,33 | 1,79 | -0,74 | 4,57E-01 |
| AT4G37510 | 0,72 | 0,92 | 2,53 | 0,37 | 7,15E-01 |
| AT4G37540 | 0,41 | 0,44 | 4,41 | 0,10 | 9,21E-01 |
| AT4G37553 | 0,42 | 1,73 | 4,38 | 0,40 | 6,93E-01 |
| AT4G37590 | 0,76 | -0,54 | 2,65 | -0,21 | 8,37E-01 |
| AT4G37610 | 0,33 | 0,44 | 3,64 | 0,12 | 9,05E-01 |
| AT4G37740 | 0,08 | 0,00 | 4,41 | 0,00 | 1,00E+00 |
| AT4G37800 | 0,23 | 0,00 | 4,41 | 0,00 | 1,00E+00 |
| AT4G37830 | 0,60 | 1,60 | 2,63 | 0,61 | 5,44E-01 |
| AT4G37850 | 0,24 | 1,96 | 4,39 | 0,45 | 6,55E-01 |
| AT4G37870 | 0,70 | -0,66 | 2,28 | -0,29 | 7,72E-01 |
| AT4G37920 | 0,32 | 1,38 | 4,41 | 0,31 | 7,54E-01 |
| AT4G38010 | 0,08 | 0,00 | 4,41 | 0,00 | 1,00E+00 |
| AT4G38050 | 0,35 | -0,54 | 4,41 | -0,12 | 9,02E-01 |
| AT4G38120 | 0,89 | 2,70 | 2,32 | 1,16 | 2,45E-01 |
| AT4G38130 | 0,71 | -0,28 | 2,10 | -0,13 | 8,94E-01 |
| AT4G38180 | 0,60 | 1,24 | 2,34 | 0,53 | 5,96E-01 |
| AT4G38210 | 0,75 | 1,62 | 4,36 | 0,37 | 7,10E-01 |
| AT4G38230 | 0,40 | 2,14 | 3,44 | 0,62 | 5,34E-01 |
| AT4G38240 | 0,62 | 1,34 | 3,75 | 0,36 | 7,21E-01 |
| AT4G09695 | 1,25 | 1,08 | 1,72 | 0,63 | 5,30E-01 |
| AT4G38300 | 0,68 | 1,91 | 3,64 | 0,52 | 6,00E-01 |

| | | | | | |
|---|---|---|---|---|---|
| AT4G38350 | 1,14 | 0,05 | 1,97 | 0,02 | 9,80E-01 |
| AT4G38380 | 0,43 | 0,42 | 4,19 | 0,10 | 9,21E-01 |
| AT4G38520 | 0,55 | 2,20 | 2,41 | 0,91 | 3,61E-01 |
| AT4G38545 | 0,17 | 0,00 | 4,41 | 0,00 | 1,00E+00 |
| AT4G38570 | 0,81 | 2,12 | 2,90 | 0,73 | 4,66E-01 |
| AT4G38630 | 0,74 | 0,41 | 2,06 | 0,20 | 8,42E-01 |
| AT4G38640 | 0,43 | 1,87 | 3,40 | 0,55 | 5,81E-01 |
| AT4G38660 | 0,37 | 1,88 | 4,38 | 0,43 | 6,67E-01 |
| AT4G38670 | 0,49 | 1,30 | 3,18 | 0,41 | 6,83E-01 |
| AT4G38680 | 1,18 | -0,68 | 1,90 | -0,36 | 7,21E-01 |
| AT4G38700 | 0,34 | 1,31 | 4,40 | 0,30 | 7,66E-01 |
| AT4G38710 | 1,00 | -1,12 | 1,83 | -0,62 | 5,38E-01 |
| AT4G38730 | 0,38 | 2,11 | 4,37 | 0,48 | 6,29E-01 |
| AT4G38740 | 1,15 | 1,49 | 2,07 | 0,72 | 4,70E-01 |
| AT4G38780 | 1,10 | 0,00 | 2,70 | 0,00 | 1,00E+00 |
| AT4G38800 | 0,65 | -0,26 | 2,64 | -0,10 | 9,21E-01 |
| AT4G38810 | 0,68 | -0,77 | 2,59 | -0,30 | 7,66E-01 |
| AT4G38870 | 0,21 | 0,00 | 4,41 | 0,00 | 1,00E+00 |
| AT4G38890 | 0,48 | 0,51 | 3,36 | 0,15 | 8,80E-01 |
| AT4G38900 | 0,33 | 1,66 | 4,38 | 0,38 | 7,04E-01 |
| AT4G38932 | 0,58 | 1,51 | 2,72 | 0,56 | 5,78E-01 |
| AT4G38950 | 1,10 | 3,59 | 2,02 | 1,78 | 7,55E-02 |
| AT4G39100 | 0,33 | 1,30 | 4,40 | 0,30 | 7,67E-01 |
| AT4G39120 | 0,19 | 0,00 | 4,41 | 0,00 | 1,00E+00 |
| AT4G39140 | 0,65 | 2,35 | 3,87 | 0,61 | 5,43E-01 |
| AT4G39180 | 0,48 | 0,00 | 4,41 | 0,00 | 1,00E+00 |
| AT4G39190 | 0,69 | 1,38 | 2,61 | 0,53 | 5,98E-01 |
| AT4G39195 | 0,79 | 1,38 | 3,03 | 0,45 | 6,49E-01 |
| AT4G39240 | 1,33 | 2,11 | 2,33 | 0,90 | 3,66E-01 |
| AT4G39280 | 1,06 | -0,75 | 1,97 | -0,38 | 7,04E-01 |
| AT4G39460 | 0,85 | 1,79 | 2,39 | 0,75 | 4,56E-01 |
| AT4G39500 | 0,20 | 1,33 | 4,40 | 0,30 | 7,63E-01 |
| AT4G39510 | 0,43 | -0,15 | 4,35 | -0,04 | 9,72E-01 |
| AT4G39520 | 1,02 | 0,73 | 1,62 | 0,45 | 6,52E-01 |
| AT4G39550 | 0,17 | 0,00 | 4,41 | 0,00 | 1,00E+00 |
| AT4G39560 | 0,59 | 2,86 | 3,77 | 0,76 | 4,49E-01 |
| AT4G39570 | 0,55 | -0,41 | 3,81 | -0,11 | 9,13E-01 |
| AT4G39580 | 0,15 | 0,00 | 4,41 | 0,00 | 1,00E+00 |
| AT4G39590 | 0,08 | 0,00 | 4,41 | 0,00 | 1,00E+00 |
| AT4G39630 | 0,05 | 0,00 | 4,41 | 0,00 | 1,00E+00 |
| AT4G39680 | 0,30 | 0,00 | 4,41 | 0,00 | 1,00E+00 |
| AT4G39710 | 0,56 | 1,15 | 3,96 | 0,29 | 7,72E-01 |
| AT4G39780 | 0,21 | -0,50 | 4,40 | -0,11 | 9,09E-01 |
| AT4G39820 | 0,46 | 1,32 | 4,12 | 0,32 | 7,49E-01 |
| AT4G39838 | 1,07 | 2,80 | 2,06 | 1,36 | 1,74E-01 |
| AT4G39865 | 0,25 | 1,84 | 4,38 | 0,42 | 6,75E-01 |
| AT4G39910 | 0,28 | -0,61 | 4,40 | -0,14 | 8,89E-01 |
| AT4G40000 | 0,43 | 0,51 | 4,18 | 0,12 | 9,02E-01 |
| AT4G40020 | 0,75 | 3,24 | 2,76 | 1,17 | 2,40E-01 |
| AT4G40030 | 1,32 | 1,32 | 1,92 | 0,69 | 4,91E-01 |
| AT4G40045 | 0,12 | 0,00 | 4,41 | 0,00 | 1,00E+00 |
| AT4G40050 | 1,65 | 3,55 | 1,78 | 2,00 | 4,56E-02 |
| AT4G40060 | 1,22 | 0,38 | 2,12 | 0,18 | 8,57E-01 |
| AT4G40065 | 1,04 | 1,67 | 1,98 | 0,84 | 3,99E-01 |
| AT5G01010 | 1,03 | 1,39 | 2,42 | 0,57 | 5,67E-01 |
| AT5G01020 | 1,61 | 3,52 | 1,77 | 1,98 | 4,72E-02 |

| | | | | | |
|---|---|---|---|---|---|
| AT5G01100 | 0,27 | 1,35 | 4,40 | 0,31 | 7,59E-01 |
| AT5G01215 | 1,09 | -0,38 | 2,01 | -0,19 | 8,52E-01 |
| AT5G01220 | 0,12 | 0,00 | 4,41 | 0,00 | 1,00E+00 |
| AT5G01270 | 0,34 | 1,38 | 4,41 | 0,31 | 7,54E-01 |
| AT5G01365 | 0,66 | -0,40 | 4,37 | -0,09 | 9,27E-01 |
| AT5G01540 | 0,46 | 1,70 | 4,10 | 0,41 | 6,80E-01 |
| AT5G01560 | 1,57 | 3,57 | 1,83 | 1,95 | 5,08E-02 |
| AT5G01600 | 1,52 | 3,29 | 1,82 | 1,81 | 7,04E-02 |
| AT5G01630 | 1,03 | 2,42 | 2,72 | 0,89 | 3,74E-01 |
| AT5G01700 | 0,62 | 2,49 | 4,36 | 0,57 | 5,68E-01 |
| AT5G01710 | 1,65 | 2,80 | 1,70 | 1,65 | 9,99E-02 |
| AT5G01720 | 0,49 | 0,00 | 4,02 | 0,00 | 1,00E+00 |
| AT5G01730 | 0,29 | 1,32 | 4,40 | 0,30 | 7,64E-01 |
| AT5G01770 | 0,54 | 2,49 | 3,81 | 0,65 | 5,14E-01 |
| AT5G01780 | 0,19 | 0,00 | 4,41 | 0,00 | 1,00E+00 |
| AT5G01890 | 0,64 | 1,38 | 2,74 | 0,50 | 6,15E-01 |
| AT5G01930 | 1,04 | 4,06 | 3,04 | 1,33 | 1,83E-01 |
| AT5G01950 | 0,59 | 1,60 | 2,64 | 0,61 | 5,44E-01 |
| AT5G01970 | 1,20 | 2,16 | 2,24 | 0,96 | 3,35E-01 |
| AT5G01990 | 1,12 | 3,84 | 2,13 | 1,80 | 7,11E-02 |
| AT5G02130 | 0,08 | 0,00 | 4,41 | 0,00 | 1,00E+00 |
| AT5G02290 | 0,48 | 2,28 | 2,80 | 0,81 | 4,15E-01 |
| AT5G02502 | 0,33 | -0,54 | 4,41 | -0,12 | 9,02E-01 |
| AT5G02530 | 0,37 | 0,30 | 4,39 | 0,07 | 9,46E-01 |
| AT5G02670 | 0,44 | 1,38 | 3,34 | 0,41 | 6,80E-01 |
| AT5G02770 | 0,75 | 2,45 | 2,41 | 1,02 | 3,10E-01 |
| AT5G02800 | 0,65 | 3,20 | 2,94 | 1,09 | 2,77E-01 |
| AT5G02830 | 0,85 | 0,54 | 2,15 | 0,25 | 8,02E-01 |
| AT5G02890 | 0,18 | -0,51 | 4,40 | -0,12 | 9,07E-01 |
| AT5G02940 | 0,88 | 1,38 | 2,62 | 0,53 | 5,99E-01 |
| AT5G03140 | 0,48 | 0,43 | 2,78 | 0,16 | 8,76E-01 |
| AT5G03170 | 0,32 | 0,95 | 4,37 | 0,22 | 8,27E-01 |
| AT5G03204 | 0,29 | -0,54 | 4,41 | -0,12 | 9,02E-01 |
| AT5G03210 | 0,71 | -0,39 | 2,56 | -0,15 | 8,78E-01 |
| AT5G03340 | 1,55 | 0,39 | 2,00 | 0,19 | 8,47E-01 |
| AT5G03380 | 0,84 | -0,39 | 2,46 | -0,16 | 8,75E-01 |
| AT5G03400 | 1,62 | -0,45 | 2,00 | -0,22 | 8,22E-01 |
| AT5G03430 | 0,65 | 2,33 | 3,15 | 0,74 | 4,60E-01 |
| AT5G03450 | 0,39 | 1,13 | 4,38 | 0,26 | 7,96E-01 |
| AT5G03555 | 1,08 | 1,87 | 2,08 | 0,90 | 3,69E-01 |
| AT5G03560 | 0,43 | 1,42 | 4,17 | 0,34 | 7,33E-01 |
| AT5G03630 | 0,55 | -1,93 | 3,08 | -0,63 | 5,31E-01 |
| AT5G03700 | 0,11 | 0,00 | 4,41 | 0,00 | 1,00E+00 |
| AT5G03730 | 0,62 | -0,86 | 2,57 | -0,33 | 7,40E-01 |
| AT5G03760 | 0,44 | -0,57 | 4,09 | -0,14 | 8,89E-01 |
| AT5G03800 | 0,30 | 1,38 | 4,41 | 0,31 | 7,54E-01 |
| AT5G03850 | 0,83 | 1,98 | 2,95 | 0,67 | 5,03E-01 |
| AT5G03880 | 0,57 | -1,53 | 3,23 | -0,48 | 6,35E-01 |
| AT5G03910 | 0,83 | -0,20 | 2,40 | -0,08 | 9,32E-01 |
| AT5G03970 | 0,19 | 0,00 | 4,41 | 0,00 | 1,00E+00 |
| AT5G04020 | 0,78 | 0,51 | 1,99 | 0,26 | 7,97E-01 |
| AT5G04130 | 0,05 | 0,00 | 4,41 | 0,00 | 1,00E+00 |
| AT5G04267 | 0,16 | 0,00 | 4,41 | 0,00 | 1,00E+00 |
| AT5G04340 | 1,00 | -0,79 | 1,94 | -0,41 | 6,85E-01 |
| AT5G00955 | 1,17 | 1,36 | 2,15 | 0,63 | 5,26E-01 |
| AT5G04420 | 0,49 | 2,58 | 3,96 | 0,65 | 5,15E-01 |

| | | | | | |
|---|---|---|---|---|---|
| AT5G04430 | 1,54 | 0,54 | 1,91 | 0,29 | 7,75E-01 |
| AT5G04470 | 0,16 | 0,00 | 4,41 | 0,00 | 1,00E+00 |
| AT5G04480 | 0,41 | 1,31 | 3,38 | 0,39 | 6,98E-01 |
| AT5G04670 | 1,32 | -0,20 | 2,05 | -0,10 | 9,23E-01 |
| AT5G04700 | 0,30 | 1,38 | 4,41 | 0,31 | 7,54E-01 |
| AT5G04710 | 0,12 | 0,00 | 4,41 | 0,00 | 1,00E+00 |
| AT5G04730 | 1,06 | 3,21 | 2,18 | 1,47 | 1,42E-01 |
| AT5G04760 | 0,48 | 1,38 | 4,41 | 0,31 | 7,54E-01 |
| AT5G04830 | 0,60 | 1,38 | 3,22 | 0,43 | 6,68E-01 |
| AT5G04850 | 0,17 | 0,00 | 4,41 | 0,00 | 1,00E+00 |
| AT5G04860 | 0,46 | 2,84 | 4,18 | 0,68 | 4,97E-01 |
| AT5G04870 | 0,78 | 1,02 | 2,31 | 0,44 | 6,58E-01 |
| AT5G04940 | 0,64 | 2,90 | 2,99 | 0,97 | 3,31E-01 |
| AT5G04950 | 0,11 | 0,00 | 4,41 | 0,00 | 1,00E+00 |
| AT5G04990 | 0,74 | 3,55 | 3,40 | 1,04 | 2,97E-01 |
| AT5G05060 | 0,37 | -1,68 | 4,36 | -0,38 | 7,01E-01 |
| AT5G05080 | 0,77 | 0,82 | 4,32 | 0,19 | 8,50E-01 |
| AT5G05085 | 0,35 | 1,32 | 3,61 | 0,37 | 7,14E-01 |
| AT5G05100 | 0,72 | 1,83 | 2,92 | 0,63 | 5,31E-01 |
| AT5G01005 | 0,75 | 2,16 | 3,01 | 0,72 | 4,72E-01 |
| AT5G05260 | 0,23 | 0,00 | 4,41 | 0,00 | 1,00E+00 |
| AT5G01055 | 0,10 | 0,00 | 4,41 | 0,00 | 1,00E+00 |
| AT5G05480 | 0,31 | 1,38 | 3,68 | 0,37 | 7,08E-01 |
| AT5G05670 | 0,17 | 0,00 | 4,41 | 0,00 | 1,00E+00 |
| AT5G05700 | 0,69 | 1,43 | 3,04 | 0,47 | 6,38E-01 |
| AT5G05730 | 0,58 | 1,99 | 3,17 | 0,63 | 5,30E-01 |
| AT5G01095 | 0,95 | 0,00 | 4,41 | 0,00 | 1,00E+00 |
| AT5G05800 | 0,36 | 0,00 | 4,41 | 0,00 | 1,00E+00 |
| AT5G06043 | 0,95 | -2,31 | 2,58 | -0,89 | 3,71E-01 |
| AT5G06060 | 0,53 | 0,08 | 3,24 | 0,03 | 9,79E-01 |
| AT5G06110 | 0,69 | 2,90 | 2,94 | 0,99 | 3,23E-01 |
| AT5G06140 | 0,36 | 1,76 | 4,38 | 0,40 | 6,88E-01 |
| AT5G06160 | 0,43 | 0,33 | 2,84 | 0,12 | 9,06E-01 |
| AT5G01165 | 0,96 | -0,31 | 2,14 | -0,15 | 8,84E-01 |
| AT5G01195 | 1,58 | -0,96 | 1,64 | -0,58 | 5,60E-01 |
| AT5G06230 | 0,49 | 2,10 | 3,92 | 0,54 | 5,92E-01 |
| AT5G06260 | 0,46 | 0,31 | 3,35 | 0,09 | 9,27E-01 |
| AT5G06330 | 0,08 | 0,00 | 4,41 | 0,00 | 1,00E+00 |
| AT5G06410 | 0,25 | 1,39 | 4,41 | 0,32 | 7,52E-01 |
| AT5G06430 | 0,51 | 2,52 | 4,35 | 0,58 | 5,63E-01 |
| AT5G06440 | 0,83 | 2,87 | 3,46 | 0,83 | 4,08E-01 |
| AT5G06520 | 0,44 | 0,00 | 4,41 | 0,00 | 1,00E+00 |
| AT5G06540 | 0,48 | 0,00 | 4,41 | 0,00 | 1,00E+00 |
| AT5G06560 | 0,78 | 1,81 | 2,27 | 0,80 | 4,24E-01 |
| AT5G06670 | 0,24 | -0,40 | 4,38 | -0,09 | 9,26E-01 |
| AT5G06690 | 0,35 | -0,44 | 3,61 | -0,12 | 9,03E-01 |
| AT5G06750 | 0,11 | 0,00 | 4,41 | 0,00 | 1,00E+00 |
| AT5G06770 | 0,36 | -0,54 | 4,41 | -0,12 | 9,02E-01 |
| AT5G06865 | 0,22 | -0,48 | 4,40 | -0,11 | 9,14E-01 |
| AT5G06970 | 0,87 | 1,26 | 2,11 | 0,60 | 5,50E-01 |
| AT5G01295 | 0,28 | 1,84 | 4,39 | 0,42 | 6,74E-01 |
| AT5G07120 | 0,92 | 2,67 | 2,47 | 1,08 | 2,79E-01 |
| AT5G07220 | 0,71 | -0,58 | 2,40 | -0,24 | 8,10E-01 |
| AT5G07250 | 0,62 | -1,72 | 2,98 | -0,58 | 5,65E-01 |
| AT5G07360 | 0,50 | -0,41 | 3,30 | -0,12 | 9,01E-01 |
| AT5G07590 | 0,15 | 0,00 | 4,41 | 0,00 | 1,00E+00 |

| | | | | | |
|---|---|---|---|---|---|
| AT5G07620 | 0,13 | -0,43 | 4,39 | -0,10 | 9,23E-01 |
| AT5G07675 | 0,20 | 0,00 | 4,41 | 0,00 | 1,00E+00 |
| AT5G07810 | 0,26 | 1,38 | 4,41 | 0,31 | 7,54E-01 |
| AT5G07950 | 0,39 | 1,79 | 4,37 | 0,41 | 6,81E-01 |
| AT5G07980 | 1,38 | 2,63 | 1,95 | 1,35 | 1,77E-01 |
| AT5G08040 | 0,14 | -0,78 | 4,37 | -0,18 | 8,58E-01 |
| AT5G08075 | 0,48 | 1,78 | 4,08 | 0,44 | 6,63E-01 |
| AT5G08110 | 0,94 | 3,25 | 2,33 | 1,40 | 1,63E-01 |
| AT5G08130 | 1,09 | 0,34 | 2,21 | 0,15 | 8,80E-01 |
| AT5G08160 | 1,05 | 2,44 | 2,06 | 1,19 | 2,36E-01 |
| AT5G08170 | 0,71 | 3,20 | 3,58 | 0,89 | 3,71E-01 |
| AT5G01495 | 0,69 | 1,21 | 2,10 | 0,58 | 5,64E-01 |
| AT5G01505 | 1,02 | 0,00 | 1,81 | 0,00 | 1,00E+00 |
| AT5G01515 | 0,11 | 0,00 | 4,41 | 0,00 | 1,00E+00 |
| AT5G08230 | 0,62 | 2,36 | 3,15 | 0,75 | 4,54E-01 |
| AT5G08280 | 1,65 | -3,45 | 1,73 | -2,00 | 4,55E-02 |
| AT5G08305 | 0,42 | 1,32 | 4,21 | 0,31 | 7,55E-01 |
| AT5G08315 | 0,44 | 3,08 | 4,32 | 0,71 | 4,75E-01 |
| AT5G08350 | 0,26 | 1,38 | 4,41 | 0,31 | 7,54E-01 |
| AT5G08450 | 0,66 | -0,94 | 2,89 | -0,33 | 7,45E-01 |
| AT5G08520 | 1,35 | -0,80 | 1,49 | -0,54 | 5,91E-01 |
| AT5G08560 | 0,55 | 0,54 | 2,84 | 0,19 | 8,49E-01 |
| AT5G08650 | 1,18 | 0,18 | 1,85 | 0,10 | 9,24E-01 |
| AT5G08770 | 1,04 | -1,26 | 2,14 | -0,59 | 5,56E-01 |
| AT5G08790 | 1,51 | 2,86 | 1,79 | 1,60 | 1,10E-01 |
| AT5G09330 | 1,10 | 2,36 | 1,79 | 1,32 | 1,88E-01 |
| AT5G09443 | 0,26 | 1,38 | 4,41 | 0,31 | 7,54E-01 |
| AT5G09460 | 0,93 | 1,90 | 1,99 | 0,96 | 3,39E-01 |
| AT5G09510 | 0,57 | 0,85 | 4,06 | 0,21 | 8,34E-01 |
| AT5G09620 | 0,88 | 0,43 | 2,53 | 0,17 | 8,66E-01 |
| AT5G09650 | 0,62 | 0,43 | 2,65 | 0,16 | 8,70E-01 |
| AT5G09820 | 0,63 | 3,06 | 3,11 | 0,99 | 3,24E-01 |
| AT5G09850 | 0,67 | 1,38 | 3,19 | 0,43 | 6,66E-01 |
| AT5G09880 | 0,62 | 0,00 | 3,06 | 0,00 | 1,00E+00 |
| AT5G09890 | 0,52 | 0,00 | 3,99 | 0,00 | 1,00E+00 |
| AT5G09930 | 0,08 | 0,00 | 4,41 | 0,00 | 1,00E+00 |
| AT5G10030 | 0,52 | 1,40 | 4,01 | 0,35 | 7,26E-01 |
| AT5G10235 | 0,28 | 0,50 | 4,40 | 0,11 | 9,10E-01 |
| AT5G10290 | 0,76 | -0,54 | 2,55 | -0,21 | 8,31E-01 |
| AT5G10330 | 0,57 | 0,48 | 2,79 | 0,17 | 8,64E-01 |
| AT5G10350 | 0,66 | 1,42 | 4,35 | 0,33 | 7,44E-01 |
| AT5G10360 | 1,44 | 0,98 | 1,55 | 0,63 | 5,26E-01 |
| AT5G10390 | 0,27 | 1,27 | 4,39 | 0,29 | 7,72E-01 |
| AT5G10400 | 0,23 | 0,00 | 4,41 | 0,00 | 1,00E+00 |
| AT5G10470 | 0,57 | 2,46 | 3,24 | 0,76 | 4,49E-01 |
| AT5G10490 | 0,10 | 0,00 | 4,41 | 0,00 | 1,00E+00 |
| AT5G10550 | 0,86 | -0,54 | 2,48 | -0,22 | 8,27E-01 |
| AT5G10605 | 0,65 | 2,85 | 3,12 | 0,91 | 3,62E-01 |
| AT5G10650 | 0,25 | 1,30 | 4,40 | 0,30 | 7,67E-01 |
| AT5G10690 | 0,38 | 1,38 | 4,40 | 0,31 | 7,54E-01 |
| AT5G01815 | 0,80 | -1,38 | 2,26 | -0,61 | 5,42E-01 |
| AT5G10700 | 0,56 | 2,21 | 3,25 | 0,68 | 4,96E-01 |
| AT5G10730 | 0,23 | 0,00 | 4,38 | 0,00 | 1,00E+00 |
| AT5G10770 | 0,94 | 1,35 | 2,45 | 0,55 | 5,82E-01 |
| AT5G10800 | 0,30 | 1,38 | 4,41 | 0,31 | 7,54E-01 |
| AT5G10820 | 0,63 | 0,00 | 4,41 | 0,00 | 1,00E+00 |

| | | | | | |
|---|---|---|---|---|---|
| AT5G10930 | 1,14 | 2,49 | 1,89 | 1,32 | 1,88E-01 |
| AT5G10940 | 0,77 | 2,72 | 3,60 | 0,75 | 4,51E-01 |
| AT5G10965 | 0,75 | 3,19 | 2,39 | 1,34 | 1,82E-01 |
| AT5G10980 | 0,55 | -0,25 | 3,85 | -0,07 | 9,48E-01 |
| AT5G11000 | 0,45 | 1,90 | 4,24 | 0,45 | 6,54E-01 |
| AT5G11070 | 1,00 | 2,36 | 2,22 | 1,06 | 2,88E-01 |
| AT5G11180 | 0,81 | -0,92 | 2,33 | -0,40 | 6,92E-01 |
| AT5G11210 | 1,06 | 3,83 | 3,13 | 1,23 | 2,20E-01 |
| AT5G11250 | 1,58 | 1,41 | 1,62 | 0,87 | 3,83E-01 |
| AT5G11270 | 0,22 | 1,39 | 4,41 | 0,32 | 7,52E-01 |
| AT5G11330 | 1,01 | 1,54 | 1,96 | 0,78 | 4,33E-01 |
| AT5G11480 | 0,46 | -1,86 | 3,22 | -0,58 | 5,62E-01 |
| AT5G11520 | 1,24 | -0,01 | 1,94 | -0,01 | 9,94E-01 |
| AT5G11550 | 0,28 | 1,34 | 4,40 | 0,30 | 7,61E-01 |
| AT5G11770 | 0,54 | -1,76 | 2,79 | -0,63 | 5,29E-01 |
| AT5G11850 | 0,25 | 0,00 | 4,41 | 0,00 | 1,00E+00 |
| AT5G11860 | 0,12 | 0,00 | 4,41 | 0,00 | 1,00E+00 |
| AT5G11910 | 0,27 | 1,34 | 4,40 | 0,30 | 7,61E-01 |
| AT5G11960 | 0,36 | 0,00 | 4,41 | 0,00 | 1,00E+00 |
| AT5G12010 | 0,59 | 1,33 | 3,12 | 0,43 | 6,71E-01 |
| AT5G12080 | 0,25 | 1,34 | 4,40 | 0,30 | 7,61E-01 |
| AT5G12140 | 0,22 | 0,00 | 4,41 | 0,00 | 1,00E+00 |
| AT5G12200 | 0,44 | -1,26 | 3,30 | -0,38 | 7,02E-01 |
| AT5G12310 | 0,42 | 1,29 | 3,42 | 0,38 | 7,05E-01 |
| AT5G12390 | 0,11 | 0,00 | 4,41 | 0,00 | 1,00E+00 |
| AT5G12430 | 1,28 | 4,27 | 2,28 | 1,87 | 6,15E-02 |
| AT5G12440 | 0,88 | 3,34 | 2,67 | 1,25 | 2,11E-01 |
| AT5G12480 | 0,34 | -0,54 | 4,41 | -0,12 | 9,02E-01 |
| AT5G12890 | 0,77 | 2,88 | 3,63 | 0,79 | 4,27E-01 |
| AT5G12900 | 0,23 | 0,00 | 4,41 | 0,00 | 1,00E+00 |
| AT5G13020 | 0,81 | 1,24 | 2,17 | 0,57 | 5,68E-01 |
| AT5G13050 | 0,42 | 1,46 | 4,22 | 0,35 | 7,28E-01 |
| AT5G13090 | 0,10 | 0,00 | 4,41 | 0,00 | 1,00E+00 |
| AT5G13120 | 0,17 | -0,82 | 4,37 | -0,19 | 8,51E-01 |
| AT5G13140 | 0,23 | -1,17 | 4,37 | -0,27 | 7,88E-01 |
| AT5G13170 | 0,48 | 1,38 | 4,41 | 0,31 | 7,54E-01 |
| AT5G13190 | 0,99 | 1,23 | 2,09 | 0,59 | 5,57E-01 |
| AT5G02055 | 0,48 | -1,63 | 2,79 | -0,58 | 5,59E-01 |
| AT5G13225 | 0,48 | -1,63 | 2,79 | -0,58 | 5,59E-01 |
| AT5G13270 | 0,40 | 2,13 | 4,20 | 0,51 | 6,13E-01 |
| AT5G13300 | 0,77 | 2,17 | 2,52 | 0,86 | 3,89E-01 |
| AT5G13400 | 0,43 | 0,00 | 4,41 | 0,00 | 1,00E+00 |
| AT5G13430 | 0,54 | 2,22 | 3,83 | 0,58 | 5,62E-01 |
| AT5G13440 | 0,29 | 0,00 | 4,41 | 0,00 | 1,00E+00 |
| AT5G13450 | 0,96 | -0,20 | 2,19 | -0,09 | 9,26E-01 |
| AT5G13550 | 0,46 | 1,50 | 4,05 | 0,37 | 7,12E-01 |
| AT5G13560 | 1,05 | 1,25 | 2,06 | 0,60 | 5,46E-01 |
| AT5G13590 | 1,32 | 2,62 | 2,00 | 1,31 | 1,91E-01 |
| AT5G13680 | 1,03 | 1,91 | 1,96 | 0,97 | 3,30E-01 |
| AT5G13710 | 0,59 | 2,44 | 3,11 | 0,79 | 4,32E-01 |
| AT5G13780 | 0,62 | 2,09 | 3,14 | 0,67 | 5,06E-01 |
| AT5G13800 | 0,74 | 0,36 | 2,47 | 0,14 | 8,85E-01 |
| AT5G13825 | 0,39 | -2,26 | 4,23 | -0,53 | 5,93E-01 |
| AT5G13950 | 1,63 | 1,46 | 1,64 | 0,89 | 3,73E-01 |
| AT5G13990 | 0,12 | 0,00 | 4,41 | 0,00 | 1,00E+00 |
| AT5G14100 | 0,11 | 0,00 | 4,41 | 0,00 | 1,00E+00 |

| | | | | | |
|---|---|---|---|---|---|
| AT5G14170 | 1,07 | 3,66 | 2,18 | 1,68 | 9,30E-02 |
| AT5G14210 | 0,50 | -0,46 | 3,38 | -0,14 | 8,92E-01 |
| AT5G14240 | 0,57 | 1,53 | 2,75 | 0,56 | 5,77E-01 |
| AT5G14320 | 0,97 | -0,13 | 1,82 | -0,07 | 9,41E-01 |
| AT5G14390 | 0,52 | 2,31 | 4,14 | 0,56 | 5,77E-01 |
| AT5G14420 | 0,18 | 0,00 | 4,41 | 0,00 | 1,00E+00 |
| AT5G14530 | 0,86 | 1,41 | 2,96 | 0,48 | 6,35E-01 |
| AT5G14540 | 0,54 | 2,71 | 3,93 | 0,69 | 4,91E-01 |
| AT5G02245 | 0,10 | 0,00 | 4,41 | 0,00 | 1,00E+00 |
| AT5G14580 | 1,26 | 3,52 | 2,67 | 1,32 | 1,88E-01 |
| AT5G14590 | 0,98 | -0,01 | 2,01 | 0,00 | 9,97E-01 |
| AT5G14700 | 0,24 | 0,00 | 4,41 | 0,00 | 1,00E+00 |
| AT5G14720 | 0,96 | 2,19 | 2,07 | 1,06 | 2,90E-01 |
| AT5G14770 | 0,20 | -0,44 | 4,39 | -0,10 | 9,20E-01 |
| AT5G14820 | 0,29 | 1,38 | 4,41 | 0,31 | 7,54E-01 |
| AT5G14940 | 1,18 | 1,40 | 2,46 | 0,57 | 5,69E-01 |
| AT5G14950 | 1,52 | 1,38 | 1,63 | 0,85 | 3,96E-01 |
| AT5G15050 | 0,10 | 0,00 | 4,41 | 0,00 | 1,00E+00 |
| AT5G15210 | 0,28 | 1,38 | 4,41 | 0,31 | 7,54E-01 |
| AT5G15250 | 0,05 | 0,00 | 4,41 | 0,00 | 1,00E+00 |
| AT5G15360 | 0,52 | -0,51 | 2,73 | -0,19 | 8,51E-01 |
| AT5G15410 | 1,41 | -0,18 | 1,88 | -0,10 | 9,22E-01 |
| AT5G15420 | 1,66 | -1,69 | 1,65 | -1,03 | 3,05E-01 |
| AT5G15450 | 1,23 | -0,33 | 2,05 | -0,16 | 8,74E-01 |
| AT5G15460 | 0,08 | 0,00 | 4,41 | 0,00 | 1,00E+00 |
| AT5G15470 | 0,21 | -0,54 | 4,41 | -0,12 | 9,02E-01 |
| AT5G15490 | 0,40 | 1,76 | 3,43 | 0,51 | 6,09E-01 |
| AT5G15537 | 0,21 | 1,38 | 4,41 | 0,31 | 7,54E-01 |
| AT5G15540 | 1,01 | 1,63 | 2,09 | 0,78 | 4,34E-01 |
| AT5G15580 | 0,70 | 0,71 | 2,45 | 0,29 | 7,73E-01 |
| AT5G15690 | 0,53 | 1,24 | 2,72 | 0,46 | 6,49E-01 |
| AT5G15700 | 0,61 | 0,43 | 2,46 | 0,17 | 8,62E-01 |
| AT5G15740 | 0,28 | 1,37 | 4,41 | 0,31 | 7,55E-01 |
| AT5G15780 | 0,22 | -0,48 | 4,40 | -0,11 | 9,14E-01 |
| AT5G15800 | 0,08 | 0,00 | 4,41 | 0,00 | 1,00E+00 |
| AT5G15802 | 0,08 | 0,00 | 4,41 | 0,00 | 1,00E+00 |
| AT5G15860 | 0,19 | 0,00 | 4,41 | 0,00 | 1,00E+00 |
| AT5G16050 | 0,47 | -1,35 | 2,83 | -0,48 | 6,33E-01 |
| AT5G16070 | 0,93 | 2,13 | 2,08 | 1,02 | 3,06E-01 |
| AT5G16110 | 1,09 | 2,51 | 1,88 | 1,33 | 1,83E-01 |
| AT5G16200 | 0,36 | 2,79 | 4,33 | 0,64 | 5,19E-01 |
| AT5G16220 | 0,18 | 0,00 | 4,41 | 0,00 | 1,00E+00 |
| AT5G16280 | 0,67 | 2,88 | 2,88 | 1,00 | 3,17E-01 |
| AT5G16360 | 0,08 | 0,00 | 4,41 | 0,00 | 1,00E+00 |
| AT5G16370 | 1,57 | 2,36 | 1,67 | 1,41 | 1,58E-01 |
| AT5G16380 | 0,25 | -0,54 | 4,41 | -0,12 | 9,02E-01 |
| AT5G16390 | 0,61 | 1,57 | 3,67 | 0,43 | 6,70E-01 |
| AT5G16480 | 0,56 | 2,13 | 4,34 | 0,49 | 6,23E-01 |
| AT5G16490 | 0,08 | 0,00 | 4,41 | 0,00 | 1,00E+00 |
| AT5G16520 | 0,47 | -0,42 | 4,22 | -0,10 | 9,21E-01 |
| AT5G16560 | 0,42 | -1,46 | 4,37 | -0,33 | 7,39E-01 |
| AT5G16610 | 0,08 | 0,00 | 4,41 | 0,00 | 1,00E+00 |
| AT5G16660 | 0,23 | 0,00 | 4,41 | 0,00 | 1,00E+00 |
| AT5G16680 | 0,78 | 2,19 | 2,33 | 0,94 | 3,49E-01 |
| AT5G16740 | 0,42 | 1,92 | 4,39 | 0,44 | 6,62E-01 |
| AT5G16750 | 0,89 | 1,55 | 2,17 | 0,71 | 4,75E-01 |

| | | | | | |
|---|---|---|---|---|---|
| AT5G16780 | 0,54 | 1,56 | 3,99 | 0,39 | 6,96E-01 |
| AT5G16800 | 0,21 | 1,25 | 4,39 | 0,29 | 7,75E-01 |
| AT5G16830 | 0,10 | 0,00 | 4,41 | 0,00 | 1,00E+00 |
| AT5G16980 | 0,08 | 0,00 | 4,41 | 0,00 | 1,00E+00 |
| AT5G16990 | 0,20 | 0,00 | 4,41 | 0,00 | 1,00E+00 |
| AT5G17000 | 0,20 | 1,40 | 4,41 | 0,32 | 7,51E-01 |
| AT5G17010 | 0,24 | 0,00 | 4,41 | 0,00 | 1,00E+00 |
| AT5G02655 | 0,55 | -2,12 | 2,61 | -0,81 | 4,17E-01 |
| AT5G02665 | 1,25 | 2,24 | 2,20 | 1,02 | 3,10E-01 |
| AT5G02675 | 0,37 | 1,23 | 4,33 | 0,28 | 7,76E-01 |
| AT5G17290 | 0,65 | 0,93 | 3,15 | 0,30 | 7,67E-01 |
| AT5G17300 | 0,69 | 2,59 | 2,99 | 0,87 | 3,87E-01 |
| AT5G17320 | 0,38 | 0,31 | 4,39 | 0,07 | 9,44E-01 |
| AT5G17380 | 0,80 | -0,08 | 2,65 | -0,03 | 9,75E-01 |
| AT5G17420 | 0,40 | 0,00 | 4,41 | 0,00 | 1,00E+00 |
| AT5G17460 | 0,52 | -2,95 | 4,30 | -0,69 | 4,93E-01 |
| AT5G17520 | 0,17 | -1,42 | 4,35 | -0,33 | 7,45E-01 |
| AT5G17550 | 0,08 | 0,00 | 4,41 | 0,00 | 1,00E+00 |
| AT5G17600 | 0,90 | 2,16 | 2,31 | 0,94 | 3,48E-01 |
| AT5G17620 | 0,42 | 1,92 | 4,39 | 0,44 | 6,62E-01 |
| AT5G17640 | 0,35 | 1,96 | 4,38 | 0,45 | 6,54E-01 |
| AT5G17690 | 0,20 | 0,00 | 4,41 | 0,00 | 1,00E+00 |
| AT5G17770 | 0,49 | 1,37 | 3,30 | 0,41 | 6,79E-01 |
| AT5G17860 | 0,63 | 0,00 | 2,98 | 0,00 | 1,00E+00 |
| AT5G17880 | 1,61 | 4,55 | 1,97 | 2,31 | 2,07E-02 |
| AT5G17930 | 0,48 | 0,00 | 4,41 | 0,00 | 1,00E+00 |
| AT5G18050 | 0,24 | 1,96 | 4,39 | 0,45 | 6,55E-01 |
| AT5G18110 | 0,39 | -0,54 | 4,41 | -0,12 | 9,02E-01 |
| AT5G18190 | 0,63 | 1,20 | 2,64 | 0,46 | 6,48E-01 |
| AT5G18230 | 0,99 | -0,46 | 2,94 | -0,16 | 8,77E-01 |
| AT5G18240 | 1,40 | 0,00 | 2,97 | 0,00 | 1,00E+00 |
| AT5G18260 | 0,49 | 2,51 | 4,36 | 0,57 | 5,66E-01 |
| AT5G18280 | 0,49 | 1,38 | 3,23 | 0,43 | 6,70E-01 |
| AT5G18290 | 0,36 | 2,37 | 4,36 | 0,54 | 5,86E-01 |
| AT5G18340 | 0,58 | 3,38 | 3,93 | 0,86 | 3,90E-01 |
| AT5G18400 | 0,32 | 1,38 | 4,41 | 0,31 | 7,54E-01 |
| AT5G02895 | 0,88 | 2,03 | 2,79 | 0,73 | 4,66E-01 |
| AT5G18475 | 0,21 | 0,00 | 4,41 | 0,00 | 1,00E+00 |
| AT5G18480 | 0,62 | 0,02 | 3,03 | 0,01 | 9,94E-01 |
| AT5G18570 | 0,43 | 0,00 | 4,10 | 0,00 | 1,00E+00 |
| AT5G18590 | 0,48 | 1,33 | 3,35 | 0,40 | 6,92E-01 |
| AT5G18620 | 0,19 | 0,00 | 4,41 | 0,00 | 1,00E+00 |
| AT5G18700 | 0,89 | 3,41 | 2,75 | 1,24 | 2,15E-01 |
| AT5G02935 | 0,05 | 0,00 | 4,41 | 0,00 | 1,00E+00 |
| AT5G19010 | 0,59 | 0,43 | 2,65 | 0,16 | 8,70E-01 |
| AT5G19130 | 0,62 | 1,44 | 2,68 | 0,54 | 5,91E-01 |
| AT5G19220 | 0,51 | 2,37 | 4,35 | 0,55 | 5,85E-01 |
| AT5G19360 | 0,72 | 3,54 | 4,30 | 0,82 | 4,11E-01 |
| AT5G19410 | 0,62 | 2,88 | 2,94 | 0,98 | 3,28E-01 |
| AT5G19420 | 1,09 | 3,27 | 1,93 | 1,69 | 9,08E-02 |
| AT5G19450 | 0,87 | 1,91 | 1,93 | 0,99 | 3,25E-01 |
| AT5G19485 | 0,20 | 1,38 | 4,41 | 0,31 | 7,54E-01 |
| AT5G19590 | 0,42 | 0,00 | 3,35 | 0,00 | 1,00E+00 |
| AT5G19660 | 0,36 | 1,30 | 4,40 | 0,30 | 7,68E-01 |
| AT5G19760 | 0,90 | 2,77 | 2,48 | 1,12 | 2,63E-01 |
| AT5G19850 | 0,75 | 0,00 | 3,74 | 0,00 | 1,00E+00 |

| | | | | | |
|---|---|---|---|---|---|
| AT5G19860 | 0,51 | 1,38 | 4,05 | 0,34 | 7,33E-01 |
| AT5G19900 | 0,73 | 3,18 | 3,04 | 1,05 | 2,96E-01 |
| AT5G19920 | 0,37 | 2,19 | 4,38 | 0,50 | 6,16E-01 |
| AT5G19930 | 0,12 | 0,00 | 4,41 | 0,00 | 1,00E+00 |
| AT5G19950 | 0,27 | 1,30 | 4,40 | 0,29 | 7,68E-01 |
| AT5G19980 | 0,33 | 1,16 | 4,37 | 0,27 | 7,91E-01 |
| AT5G20040 | 0,12 | 0,00 | 4,41 | 0,00 | 1,00E+00 |
| AT5G20160 | 0,08 | 0,00 | 4,41 | 0,00 | 1,00E+00 |
| AT5G20170 | 0,19 | 0,00 | 4,41 | 0,00 | 1,00E+00 |
| AT5G20225 | 0,17 | 0,00 | 4,41 | 0,00 | 1,00E+00 |
| AT5G20260 | 0,52 | 1,38 | 4,41 | 0,31 | 7,54E-01 |
| AT5G20270 | 0,14 | 0,00 | 4,41 | 0,00 | 1,00E+00 |
| AT5G20300 | 1,26 | 1,55 | 2,60 | 0,60 | 5,52E-01 |
| AT5G20320 | 1,21 | 2,19 | 2,59 | 0,85 | 3,97E-01 |
| AT5G20360 | 0,79 | -0,19 | 2,20 | -0,08 | 9,32E-01 |
| AT5G20380 | 0,45 | -1,65 | 4,35 | -0,38 | 7,04E-01 |
| AT5G20490 | 1,59 | -1,45 | 1,92 | -0,76 | 4,48E-01 |
| AT5G20520 | 1,55 | 0,69 | 1,63 | 0,42 | 6,75E-01 |
| AT5G20570 | 0,58 | 2,72 | 3,08 | 0,88 | 3,78E-01 |
| AT5G20600 | 1,15 | 2,86 | 2,19 | 1,31 | 1,91E-01 |
| AT5G20620 | 0,96 | 2,71 | 2,47 | 1,10 | 2,72E-01 |
| AT5G20630 | 1,03 | 1,74 | 1,80 | 0,96 | 3,35E-01 |
| AT5G20650 | 0,87 | 1,32 | 3,13 | 0,42 | 6,74E-01 |
| AT5G20820 | 0,08 | 0,00 | 4,41 | 0,00 | 1,00E+00 |
| AT5G20830 | 0,97 | 2,08 | 2,43 | 0,85 | 3,93E-01 |
| AT5G20840 | 1,18 | 1,89 | 1,84 | 1,03 | 3,03E-01 |
| AT5G20890 | 1,15 | 2,92 | 2,27 | 1,28 | 1,99E-01 |
| AT5G20920 | 0,64 | 0,77 | 3,25 | 0,24 | 8,13E-01 |
| AT5G20940 | 0,27 | 1,30 | 4,40 | 0,30 | 7,67E-01 |
| AT5G03265 | 0,23 | 0,00 | 4,41 | 0,00 | 1,00E+00 |
| AT5G20980 | 0,81 | 1,41 | 2,99 | 0,47 | 6,38E-01 |
| AT5G20990 | 1,65 | 3,82 | 1,91 | 2,00 | 4,57E-02 |
| AT5G21040 | 0,32 | 1,38 | 4,41 | 0,31 | 7,54E-01 |
| AT5G21160 | 0,41 | 0,00 | 4,41 | 0,00 | 1,00E+00 |
| AT5G21274 | 0,23 | 0,00 | 4,41 | 0,00 | 1,00E+00 |
| AT5G21920 | 0,36 | -0,58 | 4,40 | -0,13 | 8,94E-01 |
| AT5G21950 | 0,08 | 0,00 | 4,41 | 0,00 | 1,00E+00 |
| AT5G22010 | 0,59 | 1,35 | 3,11 | 0,44 | 6,64E-01 |
| AT5G22020 | 0,55 | 1,41 | 3,85 | 0,37 | 7,14E-01 |
| AT5G22030 | 0,76 | 2,89 | 3,39 | 0,85 | 3,94E-01 |
| AT5G22060 | 1,34 | 1,32 | 2,47 | 0,54 | 5,92E-01 |
| AT5G22080 | 0,35 | 0,00 | 3,60 | 0,00 | 1,00E+00 |
| AT5G22090 | 0,53 | 0,52 | 3,37 | 0,15 | 8,77E-01 |
| AT5G22250 | 1,15 | 1,36 | 2,31 | 0,59 | 5,57E-01 |
| AT5G22350 | 0,24 | 0,00 | 4,41 | 0,00 | 1,00E+00 |
| AT5G22360 | 0,12 | 0,00 | 4,41 | 0,00 | 1,00E+00 |
| AT5G22390 | 0,49 | -0,29 | 3,19 | -0,09 | 9,28E-01 |
| AT5G22440 | 0,73 | 3,30 | 2,95 | 1,12 | 2,63E-01 |
| AT5G22450 | 0,87 | 1,25 | 2,08 | 0,60 | 5,50E-01 |
| AT5G22505 | 1,58 | -3,46 | 1,88 | -1,84 | 6,63E-02 |
| AT5G22510 | 0,44 | 2,41 | 4,22 | 0,57 | 5,67E-01 |
| AT5G03425 | 0,62 | -1,43 | 3,58 | -0,40 | 6,89E-01 |
| AT5G22540 | 0,22 | 1,33 | 4,40 | 0,30 | 7,62E-01 |
| AT5G22570 | 0,26 | 1,29 | 4,39 | 0,29 | 7,69E-01 |
| AT5G22700 | 0,44 | 0,00 | 4,17 | 0,00 | 1,00E+00 |
| AT5G22740 | 0,58 | 2,68 | 3,87 | 0,69 | 4,89E-01 |

| | | | | | |
|---|---|---|---|---|---|
| AT5G22750 | 0,55 | 0,00 | 4,39 | 0,00 | 1,00E+00 |
| AT5G03475 | 1,21 | 0,25 | 1,85 | 0,14 | 8,92E-01 |
| AT5G22780 | 1,47 | 2,18 | 1,78 | 1,23 | 2,19E-01 |
| AT5G22850 | 0,23 | 0,00 | 4,41 | 0,00 | 1,00E+00 |
| AT5G22860 | 0,35 | -0,54 | 3,55 | -0,15 | 8,79E-01 |
| AT5G23040 | 0,63 | -1,33 | 2,60 | -0,51 | 6,10E-01 |
| AT5G23050 | 0,45 | 2,57 | 4,08 | 0,63 | 5,29E-01 |
| AT5G23060 | 1,11 | -1,12 | 1,88 | -0,59 | 5,53E-01 |
| AT5G23080 | 1,07 | 2,97 | 2,62 | 1,14 | 2,56E-01 |
| AT5G23240 | 0,89 | -0,75 | 2,30 | -0,32 | 7,45E-01 |
| AT5G23300 | 0,32 | 1,38 | 4,41 | 0,31 | 7,54E-01 |
| AT5G23390 | 0,47 | -0,54 | 4,03 | -0,13 | 8,93E-01 |
| AT5G23405 | 0,14 | 0,00 | 4,41 | 0,00 | 1,00E+00 |
| AT5G23430 | 0,90 | 2,11 | 2,31 | 0,91 | 3,61E-01 |
| AT5G23440 | 0,16 | -0,55 | 4,41 | -0,12 | 9,01E-01 |
| AT5G23450 | 0,83 | 2,19 | 2,96 | 0,74 | 4,60E-01 |
| AT5G23580 | 0,83 | 1,84 | 2,24 | 0,82 | 4,10E-01 |
| AT5G23590 | 0,61 | 1,36 | 3,75 | 0,36 | 7,18E-01 |
| AT5G23610 | 1,31 | 1,01 | 2,05 | 0,49 | 6,22E-01 |
| AT5G23660 | 0,79 | 0,00 | 3,75 | 0,00 | 1,00E+00 |
| AT5G23740 | 0,24 | 0,00 | 4,41 | 0,00 | 1,00E+00 |
| AT5G23900 | 0,57 | -0,85 | 2,78 | -0,31 | 7,60E-01 |
| AT5G23940 | 0,08 | 0,00 | 4,41 | 0,00 | 1,00E+00 |
| AT5G23990 | 0,29 | -0,45 | 4,39 | -0,10 | 9,18E-01 |
| AT5G24120 | 0,77 | 2,17 | 2,55 | 0,85 | 3,94E-01 |
| AT5G24140 | 0,08 | 0,00 | 4,41 | 0,00 | 1,00E+00 |
| AT5G24150 | 0,21 | 0,00 | 4,41 | 0,00 | 1,00E+00 |
| AT5G24160 | 0,65 | 1,69 | 2,98 | 0,57 | 5,71E-01 |
| AT5G24240 | 1,01 | -1,48 | 1,91 | -0,78 | 4,36E-01 |
| AT5G24260 | 1,06 | 3,77 | 2,13 | 1,77 | 7,68E-02 |
| AT5G24270 | 1,64 | 0,00 | 1,52 | 0,00 | 9,99E-01 |
| AT5G24320 | 0,20 | -0,44 | 4,39 | -0,10 | 9,19E-01 |
| AT5G24355 | 0,08 | 0,00 | 4,41 | 0,00 | 1,00E+00 |
| AT5G24430 | 0,56 | -1,11 | 4,00 | -0,28 | 7,81E-01 |
| AT5G24470 | 1,29 | 0,23 | 2,68 | 0,09 | 9,32E-01 |
| AT5G24480 | 0,23 | 0,00 | 4,41 | 0,00 | 1,00E+00 |
| AT5G24500 | 0,24 | 1,36 | 4,40 | 0,31 | 7,57E-01 |
| AT5G24520 | 0,95 | 0,52 | 2,29 | 0,23 | 8,22E-01 |
| AT5G24590 | 1,51 | 2,71 | 1,96 | 1,38 | 1,67E-01 |
| AT5G24655 | 0,37 | 0,00 | 4,41 | 0,00 | 1,00E+00 |
| AT5G24735 | 0,57 | 0,45 | 3,37 | 0,13 | 8,95E-01 |
| AT5G24740 | 1,40 | 0,05 | 1,84 | 0,03 | 9,77E-01 |
| AT5G24770 | 0,62 | -1,38 | 2,41 | -0,57 | 5,66E-01 |
| AT5G24780 | 0,08 | 0,00 | 4,41 | 0,00 | 1,00E+00 |
| AT5G24940 | 0,08 | 0,00 | 4,41 | 0,00 | 1,00E+00 |
| AT5G24960 | 0,36 | 0,00 | 4,41 | 0,00 | 1,00E+00 |
| AT5G25000 | 0,16 | 0,00 | 4,41 | 0,00 | 1,00E+00 |
| AT5G25070 | 1,26 | 3,62 | 1,82 | 1,99 | 4,68E-02 |
| AT5G25100 | 1,32 | -1,25 | 1,71 | -0,73 | 4,63E-01 |
| AT5G25200 | 0,60 | 0,45 | 2,73 | 0,17 | 8,68E-01 |
| AT5G25265 | 0,67 | 0,42 | 3,03 | 0,14 | 8,90E-01 |
| AT5G25350 | 1,15 | 1,26 | 2,14 | 0,59 | 5,55E-01 |
| AT5G25370 | 0,71 | -2,89 | 2,85 | -1,02 | 3,10E-01 |
| AT5G03985 | 0,37 | -1,68 | 4,36 | -0,38 | 7,01E-01 |
| AT5G25480 | 0,27 | 0,23 | 4,36 | 0,05 | 9,58E-01 |
| AT5G25510 | 0,49 | -1,60 | 3,28 | -0,49 | 6,25E-01 |

| | | | | | |
|---|---|---|---|---|---|
| AT5G25530 | 1,06 | 2,36 | 3,28 | 0,72 | 4,72E-01 |
| AT5G25756 | 0,35 | 0,00 | 4,41 | 0,00 | 1,00E+00 |
| AT5G25757 | 0,87 | -0,45 | 1,95 | -0,23 | 8,15E-01 |
| AT5G25770 | 0,28 | 2,07 | 4,38 | 0,47 | 6,37E-01 |
| AT5G04085 | 0,59 | 0,00 | 4,41 | 0,00 | 1,00E+00 |
| AT5G25780 | 1,13 | 0,79 | 1,65 | 0,48 | 6,33E-01 |
| AT5G25820 | 0,43 | 2,05 | 4,36 | 0,47 | 6,39E-01 |
| AT5G25830 | 0,05 | 0,00 | 4,41 | 0,00 | 1,00E+00 |
| AT5G25920 | 0,08 | 0,00 | 4,41 | 0,00 | 1,00E+00 |
| AT5G26170 | 0,17 | 0,00 | 4,41 | 0,00 | 1,00E+00 |
| AT5G26210 | 0,43 | 1,81 | 4,22 | 0,43 | 6,68E-01 |
| AT5G26230 | 0,08 | 0,00 | 4,41 | 0,00 | 1,00E+00 |
| AT5G26680 | 0,15 | 0,00 | 4,41 | 0,00 | 1,00E+00 |
| AT5G04245 | 1,57 | 0,00 | 2,45 | 0,00 | 1,00E+00 |
| AT5G26640 | 0,17 | 0,00 | 4,41 | 0,00 | 1,00E+00 |
| AT5G26622 | 0,46 | 1,86 | 4,38 | 0,43 | 6,70E-01 |
| AT5G04255 | 0,21 | 1,25 | 4,39 | 0,29 | 7,75E-01 |
| AT5G26760 | 0,42 | 2,18 | 4,18 | 0,52 | 6,03E-01 |
| AT5G26770 | 1,06 | -0,59 | 1,95 | -0,30 | 7,63E-01 |
| AT5G26880 | 0,63 | 1,63 | 3,05 | 0,53 | 5,94E-01 |
| AT5G26910 | 0,66 | 0,98 | 2,55 | 0,38 | 7,01E-01 |
| AT5G26960 | 0,27 | -0,42 | 4,39 | -0,10 | 9,24E-01 |
| AT5G26970 | 0,84 | -1,37 | 1,98 | -0,69 | 4,89E-01 |
| AT5G27030 | 1,03 | 3,06 | 2,06 | 1,49 | 1,36E-01 |
| AT5G27100 | 0,25 | 1,38 | 4,41 | 0,31 | 7,54E-01 |
| AT5G27238 | 0,05 | 0,00 | 4,41 | 0,00 | 1,00E+00 |
| AT5G04375 | 0,24 | 1,36 | 4,40 | 0,31 | 7,57E-01 |
| AT5G27330 | 0,05 | 0,00 | 4,41 | 0,00 | 1,00E+00 |
| AT5G27380 | 0,63 | -0,83 | 3,65 | -0,23 | 8,20E-01 |
| AT5G27550 | 0,34 | 1,46 | 4,40 | 0,33 | 7,39E-01 |
| AT5G27640 | 1,34 | 1,98 | 1,94 | 1,02 | 3,05E-01 |
| AT5G27700 | 0,49 | 2,09 | 3,29 | 0,64 | 5,25E-01 |
| AT5G27715 | 0,29 | 1,38 | 4,41 | 0,31 | 7,54E-01 |
| AT5G04435 | 0,39 | -0,79 | 4,27 | -0,19 | 8,53E-01 |
| AT5G27790 | 0,75 | 0,00 | 3,62 | 0,00 | 1,00E+00 |
| AT5G27840 | 0,72 | 1,38 | 2,94 | 0,47 | 6,38E-01 |
| AT5G27870 | 1,24 | 3,46 | 2,00 | 1,73 | 8,43E-02 |
| AT5G27920 | 0,08 | 0,00 | 4,41 | 0,00 | 1,00E+00 |
| AT5G27960 | 0,74 | 0,00 | 2,84 | 0,00 | 1,00E+00 |
| AT5G28020 | 0,34 | -1,00 | 3,59 | -0,28 | 7,81E-01 |
| AT5G28030 | 0,66 | 1,38 | 3,03 | 0,46 | 6,49E-01 |
| AT5G28050 | 0,28 | 1,34 | 4,40 | 0,30 | 7,61E-01 |
| AT5G04635 | 0,50 | -0,54 | 3,25 | -0,17 | 8,67E-01 |
| AT5G28060 | 0,65 | 0,44 | 2,32 | 0,19 | 8,50E-01 |
| AT5G04685 | 0,19 | 0,00 | 4,41 | 0,00 | 1,00E+00 |
| AT5G28190 | 0,36 | 0,00 | 4,41 | 0,00 | 1,00E+00 |
| AT5G28220 | 0,30 | -0,66 | 4,39 | -0,15 | 8,80E-01 |
| AT5G28235 | 0,34 | 0,00 | 4,41 | 0,00 | 1,00E+00 |
| AT5G28300 | 1,51 | -3,18 | 1,74 | -1,82 | 6,80E-02 |
| AT5G28310 | 0,08 | 0,00 | 4,41 | 0,00 | 1,00E+00 |
| AT5G28400 | 1,06 | 2,30 | 1,88 | 1,22 | 2,22E-01 |
| AT5G28442 | 0,36 | 0,00 | 3,49 | 0,00 | 1,00E+00 |
| AT5G28450 | 0,85 | -1,90 | 2,42 | -0,78 | 4,33E-01 |
| AT5G28530 | 0,22 | 0,00 | 4,41 | 0,00 | 1,00E+00 |
| AT5G28550 | 0,21 | 0,00 | 4,41 | 0,00 | 1,00E+00 |
| AT5G28640 | 0,08 | 0,00 | 4,41 | 0,00 | 1,00E+00 |

| | | | | | |
|---|---|---|---|---|---|
| AT5G28750 | 0,65 | 0,90 | 2,27 | 0,39 | 6,93E-01 |
| AT5G28770 | 0,50 | 0,00 | 3,99 | 0,00 | 1,00E+00 |
| AT5G28824 | 1,47 | 0,49 | 2,05 | 0,24 | 8,09E-01 |
| AT5G28850 | 0,48 | 1,30 | 3,31 | 0,39 | 6,95E-01 |
| AT5G28910 | 0,82 | 1,57 | 2,46 | 0,64 | 5,24E-01 |
| AT5G04925 | 0,18 | -0,42 | 4,39 | -0,10 | 9,23E-01 |
| AT5G32470 | 0,85 | 1,48 | 2,20 | 0,68 | 5,00E-01 |
| AT5G32590 | 0,68 | 0,36 | 2,37 | 0,15 | 8,78E-01 |
| AT5G33210 | 0,08 | 0,00 | 4,41 | 0,00 | 1,00E+00 |
| AT5G33439 | 0,11 | 0,00 | 4,41 | 0,00 | 1,00E+00 |
| AT5G34850 | 1,64 | 0,14 | 1,57 | 0,09 | 9,28E-01 |
| AT5G35100 | 0,30 | -0,53 | 3,72 | -0,14 | 8,87E-01 |
| AT5G35200 | 0,52 | 1,38 | 3,20 | 0,43 | 6,67E-01 |
| AT5G35220 | 0,25 | 1,34 | 4,40 | 0,30 | 7,61E-01 |
| AT5G35230 | 0,96 | 0,00 | 2,80 | 0,00 | 1,00E+00 |
| AT5G35370 | 0,70 | -0,58 | 2,14 | -0,27 | 7,88E-01 |
| AT5G35475 | 0,36 | -0,52 | 4,40 | -0,12 | 9,05E-01 |
| AT5G35480 | 0,65 | 1,38 | 3,18 | 0,43 | 6,65E-01 |
| AT5G35530 | 0,88 | 0,26 | 2,57 | 0,10 | 9,20E-01 |
| AT5G35560 | 0,64 | 0,44 | 2,34 | 0,19 | 8,52E-01 |
| AT5G35570 | 0,29 | 0,00 | 4,41 | 0,00 | 1,00E+00 |
| AT5G35590 | 0,12 | 0,00 | 4,41 | 0,00 | 1,00E+00 |
| AT5G35603 | 0,59 | -0,54 | 4,19 | -0,13 | 8,97E-01 |
| AT5G35604 | 0,45 | 0,00 | 4,03 | 0,00 | 1,00E+00 |
| AT5G35620 | 0,26 | -0,49 | 4,40 | -0,11 | 9,12E-01 |
| AT5G35690 | 0,37 | 1,38 | 4,38 | 0,32 | 7,53E-01 |
| AT5G35700 | 0,37 | 1,27 | 4,33 | 0,29 | 7,70E-01 |
| AT5G35750 | 1,20 | 1,11 | 2,05 | 0,54 | 5,88E-01 |
| AT5G35753 | 0,77 | 2,36 | 2,15 | 1,10 | 2,73E-01 |
| AT5G35790 | 0,34 | 0,10 | 4,35 | 0,02 | 9,82E-01 |
| AT5G35810 | 0,15 | 0,00 | 4,41 | 0,00 | 1,00E+00 |
| AT5G35890 | 0,19 | 0,00 | 4,41 | 0,00 | 1,00E+00 |
| AT5G35910 | 0,15 | 0,00 | 4,41 | 0,00 | 1,00E+00 |
| AT5G35913 | 0,08 | 0,00 | 4,41 | 0,00 | 1,00E+00 |
| AT5G35970 | 1,60 | -1,12 | 1,55 | -0,73 | 4,68E-01 |
| AT5G35995 | 0,29 | 0,00 | 4,41 | 0,00 | 1,00E+00 |
| AT5G36002 | 0,12 | 0,00 | 4,41 | 0,00 | 1,00E+00 |
| AT5G36120 | 0,05 | 0,00 | 4,41 | 0,00 | 1,00E+00 |
| AT5G36150 | 0,29 | -1,60 | 4,35 | -0,37 | 7,14E-01 |
| AT5G36160 | 0,29 | 1,26 | 4,39 | 0,29 | 7,74E-01 |
| AT5G36200 | 1,52 | -0,50 | 2,17 | -0,23 | 8,17E-01 |
| AT5G36210 | 1,11 | -1,68 | 2,00 | -0,84 | 4,01E-01 |
| AT5G36220 | 0,80 | 0,51 | 2,32 | 0,22 | 8,25E-01 |
| AT5G36700 | 0,58 | 1,52 | 2,64 | 0,58 | 5,64E-01 |
| AT5G36790 | 0,17 | -0,46 | 4,40 | -0,10 | 9,17E-01 |
| AT5G36880 | 0,93 | 3,42 | 2,24 | 1,53 | 1,26E-01 |
| AT5G36930 | 1,17 | -0,51 | 2,32 | -0,22 | 8,25E-01 |
| AT5G37260 | 0,78 | 2,47 | 3,37 | 0,73 | 4,64E-01 |
| AT5G37310 | 0,77 | 0,00 | 3,48 | 0,00 | 1,00E+00 |
| AT5G37370 | 0,59 | 0,52 | 3,77 | 0,14 | 8,90E-01 |
| AT5G37480 | 0,13 | 0,00 | 4,41 | 0,00 | 1,00E+00 |
| AT5G37590 | 0,66 | 2,03 | 3,63 | 0,56 | 5,76E-01 |
| AT5G37720 | 0,46 | 2,41 | 3,32 | 0,73 | 4,67E-01 |
| AT5G37770 | 1,11 | 0,00 | 2,86 | 0,00 | 1,00E+00 |
| AT5G37780 | 0,80 | 0,27 | 2,26 | 0,12 | 9,06E-01 |
| AT5G05545 | 0,31 | 0,00 | 4,41 | 0,00 | 1,00E+00 |

| | | | | | |
|---|---|---|---|---|---|
| AT5G37830 | 1,35 | -0,50 | 2,26 | -0,22 | 8,24E-01 |
| AT5G37890 | 0,36 | 0,00 | 4,41 | 0,00 | 1,00E+00 |
| AT5G05565 | 0,45 | 0,00 | 4,19 | 0,00 | 1,00E+00 |
| AT5G38020 | 0,42 | -1,17 | 4,32 | -0,27 | 7,86E-01 |
| AT5G38080 | 0,08 | 0,00 | 4,41 | 0,00 | 1,00E+00 |
| AT5G38140 | 0,10 | 0,00 | 4,41 | 0,00 | 1,00E+00 |
| AT5G38212 | 1,00 | 0,00 | 2,40 | 0,00 | 1,00E+00 |
| AT5G38240 | 0,41 | 2,61 | 4,35 | 0,60 | 5,49E-01 |
| AT5G38280 | 0,74 | -2,64 | 3,65 | -0,72 | 4,69E-01 |
| AT5G38450 | 0,46 | -0,40 | 4,18 | -0,09 | 9,25E-01 |
| AT5G05755 | 0,05 | 0,00 | 4,41 | 0,00 | 1,00E+00 |
| AT5G38510 | 0,76 | -0,66 | 2,54 | -0,26 | 7,94E-01 |
| AT5G38640 | 0,50 | 0,29 | 2,76 | 0,10 | 9,17E-01 |
| AT5G38660 | 1,59 | 1,88 | 1,71 | 1,10 | 2,72E-01 |
| AT5G38650 | 1,00 | 1,87 | 2,31 | 0,81 | 4,18E-01 |
| AT5G38830 | 0,49 | -1,34 | 4,37 | -0,31 | 7,60E-01 |
| AT5G39130 | 0,12 | 0,00 | 4,41 | 0,00 | 1,00E+00 |
| AT5G39160 | 0,34 | 0,00 | 4,41 | 0,00 | 1,00E+00 |
| AT5G05895 | 0,62 | -0,64 | 3,09 | -0,21 | 8,35E-01 |
| AT5G39480 | 0,08 | 0,00 | 4,41 | 0,00 | 1,00E+00 |
| AT5G39520 | 0,33 | 1,38 | 4,41 | 0,31 | 7,54E-01 |
| AT5G39530 | 0,77 | -1,11 | 2,88 | -0,39 | 6,99E-01 |
| AT5G39600 | 0,66 | -1,14 | 2,26 | -0,50 | 6,14E-01 |
| AT5G39610 | 0,52 | 2,47 | 3,92 | 0,63 | 5,28E-01 |
| AT5G05995 | 0,40 | 0,00 | 4,25 | 0,00 | 1,00E+00 |
| AT5G39680 | 0,42 | 1,38 | 4,41 | 0,31 | 7,54E-01 |
| AT5G39693 | 0,43 | -1,63 | 3,33 | -0,49 | 6,25E-01 |
| AT5G39920 | 0,25 | -1,39 | 4,37 | -0,32 | 7,50E-01 |
| AT5G39950 | 0,95 | 0,49 | 2,52 | 0,20 | 8,45E-01 |
| AT5G40170 | 0,68 | 1,10 | 2,31 | 0,48 | 6,34E-01 |
| AT5G40240 | 0,14 | 0,00 | 4,41 | 0,00 | 1,00E+00 |
| AT5G40370 | 0,37 | -0,44 | 3,47 | -0,13 | 8,99E-01 |
| AT5G40470 | 0,27 | 1,84 | 4,38 | 0,42 | 6,75E-01 |
| AT5G40580 | 0,60 | 2,17 | 4,36 | 0,50 | 6,18E-01 |
| AT5G40660 | 0,29 | 1,35 | 4,40 | 0,31 | 7,59E-01 |
| AT5G40740 | 0,85 | 1,38 | 2,54 | 0,54 | 5,87E-01 |
| AT5G40770 | 1,47 | 1,87 | 1,61 | 1,16 | 2,46E-01 |
| AT5G40820 | 0,95 | 3,22 | 2,34 | 1,37 | 1,69E-01 |
| AT5G40830 | 0,63 | 0,42 | 2,80 | 0,15 | 8,82E-01 |
| AT5G40940 | 0,56 | 1,38 | 3,88 | 0,36 | 7,22E-01 |
| AT5G41020 | 0,73 | 2,17 | 2,68 | 0,81 | 4,19E-01 |
| AT5G41130 | 0,21 | 0,00 | 4,41 | 0,00 | 1,00E+00 |
| AT5G41150 | 0,33 | 1,37 | 4,41 | 0,31 | 7,56E-01 |
| AT5G41190 | 0,83 | 3,58 | 2,88 | 1,24 | 2,14E-01 |
| AT5G41330 | 0,40 | 2,11 | 3,39 | 0,62 | 5,35E-01 |
| AT5G41400 | 0,65 | 2,16 | 2,74 | 0,79 | 4,30E-01 |
| AT5G41460 | 0,17 | -1,42 | 4,35 | -0,33 | 7,45E-01 |
| AT5G41520 | 0,58 | 2,32 | 2,79 | 0,83 | 4,05E-01 |
| AT5G41670 | 0,43 | 1,38 | 2,92 | 0,47 | 6,36E-01 |
| AT5G06235 | 0,77 | 0,00 | 4,41 | 0,00 | 1,00E+00 |
| AT5G41870 | 0,38 | -0,75 | 4,33 | -0,17 | 8,63E-01 |
| AT5G41990 | 1,56 | 2,36 | 1,76 | 1,34 | 1,82E-01 |
| AT5G42100 | 1,07 | -0,71 | 1,76 | -0,41 | 6,85E-01 |
| AT5G42110 | 0,47 | 1,38 | 3,36 | 0,41 | 6,81E-01 |
| AT5G42130 | 0,21 | 0,00 | 4,41 | 0,00 | 1,00E+00 |
| AT5G42140 | 0,12 | 0,00 | 4,41 | 0,00 | 1,00E+00 |

| | | | | | |
|---|---|---|---|---|---|
| AT5G42190 | 0,65 | 0,00 | 2,67 | 0,00 | 9,99E-01 |
| AT5G42280 | 0,14 | 0,00 | 4,41 | 0,00 | 1,00E+00 |
| AT5G42300 | 0,48 | 0,00 | 3,99 | 0,00 | 1,00E+00 |
| AT5G42360 | 1,47 | 3,64 | 1,77 | 2,05 | 4,03E-02 |
| AT5G42380 | 1,28 | 3,42 | 2,13 | 1,61 | 1,08E-01 |
| AT5G42400 | 0,21 | 0,00 | 4,41 | 0,00 | 1,00E+00 |
| AT5G06305 | 1,02 | -2,40 | 1,88 | -1,28 | 2,02E-01 |
| AT5G42490 | 0,56 | 1,36 | 3,93 | 0,35 | 7,29E-01 |
| AT5G42567 | 0,58 | 0,00 | 4,41 | 0,00 | 1,00E+00 |
| AT5G42570 | 0,15 | -0,43 | 4,39 | -0,10 | 9,23E-01 |
| AT5G42660 | 0,12 | 0,00 | 4,41 | 0,00 | 1,00E+00 |
| AT5G42750 | 1,69 | -0,52 | 1,88 | -0,28 | 7,82E-01 |
| AT5G42790 | 0,93 | -1,59 | 2,16 | -0,74 | 4,62E-01 |
| AT5G42800 | 0,38 | 1,38 | 4,41 | 0,31 | 7,54E-01 |
| AT5G42850 | 1,15 | 2,19 | 2,06 | 1,06 | 2,88E-01 |
| AT5G42870 | 0,81 | 1,34 | 2,48 | 0,54 | 5,90E-01 |
| AT5G42890 | 0,08 | 0,00 | 4,41 | 0,00 | 1,00E+00 |
| AT5G42900 | 0,25 | 1,26 | 4,39 | 0,29 | 7,74E-01 |
| AT5G42905 | 0,52 | 0,00 | 3,90 | 0,00 | 1,00E+00 |
| AT5G42940 | 0,81 | 1,93 | 2,51 | 0,77 | 4,42E-01 |
| AT5G42955 | 0,65 | -1,41 | 3,77 | -0,37 | 7,09E-01 |
| AT5G42957 | 0,83 | -0,77 | 2,46 | -0,31 | 7,54E-01 |
| AT5G42990 | 0,17 | 0,00 | 4,41 | 0,00 | 1,00E+00 |
| AT5G43010 | 0,13 | -0,43 | 4,39 | -0,10 | 9,23E-01 |
| AT5G43020 | 0,41 | 1,24 | 4,18 | 0,30 | 7,67E-01 |
| AT5G43130 | 1,30 | 3,20 | 2,19 | 1,46 | 1,44E-01 |
| AT5G43190 | 1,41 | 1,59 | 1,74 | 0,91 | 3,60E-01 |
| AT5G43260 | 0,52 | 0,36 | 2,82 | 0,13 | 8,99E-01 |
| AT5G43270 | 0,76 | 0,58 | 3,13 | 0,19 | 8,52E-01 |
| AT5G43285 | 0,35 | -1,96 | 4,33 | -0,45 | 6,51E-01 |
| AT5G43320 | 0,66 | 1,40 | 2,74 | 0,51 | 6,09E-01 |
| AT5G43440 | 0,40 | 0,42 | 4,41 | 0,09 | 9,25E-01 |
| AT5G43670 | 0,40 | 0,44 | 4,21 | 0,11 | 9,16E-01 |
| AT5G43710 | 0,21 | 0,00 | 4,41 | 0,00 | 1,00E+00 |
| AT5G43745 | 0,36 | 0,97 | 4,36 | 0,22 | 8,24E-01 |
| AT5G43750 | 0,47 | 2,40 | 4,36 | 0,55 | 5,82E-01 |
| AT5G43780 | 0,63 | 2,15 | 2,77 | 0,78 | 4,37E-01 |
| AT5G43790 | 0,45 | 1,11 | 4,27 | 0,26 | 7,95E-01 |
| AT5G43830 | 0,91 | 0,46 | 1,90 | 0,24 | 8,09E-01 |
| AT5G43850 | 0,29 | -1,22 | 3,68 | -0,33 | 7,40E-01 |
| AT5G43880 | 0,19 | -0,43 | 4,39 | -0,10 | 9,21E-01 |
| AT5G43900 | 1,55 | 2,27 | 1,76 | 1,29 | 1,96E-01 |
| AT5G43950 | 0,65 | 1,50 | 2,66 | 0,56 | 5,75E-01 |
| AT5G43960 | 0,54 | -0,43 | 3,17 | -0,14 | 8,91E-01 |
| AT5G44000 | 0,95 | 0,00 | 2,43 | 0,00 | 1,00E+00 |
| AT5G44005 | 0,16 | 0,00 | 4,41 | 0,00 | 1,00E+00 |
| AT5G44100 | 0,66 | 0,31 | 2,70 | 0,12 | 9,08E-01 |
| AT5G44250 | 0,12 | 0,00 | 4,41 | 0,00 | 1,00E+00 |
| AT5G44260 | 0,22 | -0,84 | 4,38 | -0,19 | 8,47E-01 |
| AT5G44283 | 0,91 | -0,86 | 2,58 | -0,33 | 7,41E-01 |
| AT5G44290 | 1,25 | 0,52 | 2,04 | 0,26 | 7,98E-01 |
| AT5G44306 | 0,93 | 1,38 | 3,03 | 0,46 | 6,49E-01 |
| AT5G44495 | 1,23 | -1,72 | 1,73 | -0,99 | 3,20E-01 |
| AT5G44510 | 0,61 | 0,00 | 4,41 | 0,00 | 1,00E+00 |
| AT5G44568 | 0,48 | -0,45 | 2,89 | -0,16 | 8,77E-01 |
| AT5G06595 | 1,38 | 1,77 | 1,63 | 1,09 | 2,76E-01 |

| | | | | | |
|---|---|---|---|---|---|
| AT5G44670 | 0,13 | 0,00 | 4,41 | 0,00 | 1,00E+00 |
| AT5G44680 | 0,48 | -2,53 | 4,31 | -0,59 | 5,56E-01 |
| AT5G44785 | 0,10 | 0,00 | 4,41 | 0,00 | 1,00E+00 |
| AT5G44790 | 1,35 | 2,28 | 1,81 | 1,26 | 2,07E-01 |
| AT5G44820 | 0,08 | 0,00 | 4,41 | 0,00 | 1,00E+00 |
| AT5G44840 | 0,14 | -0,78 | 4,37 | -0,18 | 8,58E-01 |
| AT5G45010 | 0,58 | 1,36 | 3,15 | 0,43 | 6,66E-01 |
| AT5G45040 | 0,08 | 0,00 | 4,41 | 0,00 | 1,00E+00 |
| AT5G45050 | 0,86 | 1,41 | 2,21 | 0,64 | 5,22E-01 |
| AT5G45170 | 0,11 | 0,00 | 4,41 | 0,00 | 1,00E+00 |
| AT5G45180 | 0,66 | -1,94 | 3,89 | -0,50 | 6,19E-01 |
| AT5G45230 | 0,97 | 0,09 | 1,99 | 0,05 | 9,63E-01 |
| AT5G45250 | 0,93 | 3,29 | 2,08 | 1,58 | 1,14E-01 |
| AT5G45275 | 0,48 | 0,00 | 4,40 | 0,00 | 1,00E+00 |
| AT5G45290 | 1,09 | 2,00 | 2,16 | 0,92 | 3,55E-01 |
| AT5G45340 | 0,20 | 0,00 | 4,41 | 0,00 | 1,00E+00 |
| AT5G45360 | 0,08 | 0,00 | 4,41 | 0,00 | 1,00E+00 |
| AT5G45410 | 0,46 | 0,85 | 4,07 | 0,21 | 8,34E-01 |
| AT5G45420 | 0,27 | -0,42 | 4,39 | -0,10 | 9,24E-01 |
| AT5G45440 | 0,74 | 1,38 | 3,53 | 0,39 | 6,96E-01 |
| AT5G45460 | 0,18 | 1,38 | 4,41 | 0,31 | 7,54E-01 |
| AT5G45573 | 0,61 | 0,00 | 3,03 | 0,00 | 1,00E+00 |
| AT5G45630 | 0,05 | 0,00 | 4,41 | 0,00 | 1,00E+00 |
| AT5G45650 | 0,61 | 1,85 | 3,29 | 0,56 | 5,73E-01 |
| AT5G45775 | 0,86 | 0,95 | 2,04 | 0,46 | 6,43E-01 |
| AT5G45800 | 0,52 | 0,41 | 2,50 | 0,16 | 8,69E-01 |
| AT5G45820 | 1,25 | -0,19 | 1,77 | -0,11 | 9,13E-01 |
| AT5G46020 | 1,24 | -1,35 | 2,45 | -0,55 | 5,83E-01 |
| AT5G46040 | 0,11 | 0,00 | 4,41 | 0,00 | 1,00E+00 |
| AT5G46050 | 0,37 | 1,76 | 4,38 | 0,40 | 6,87E-01 |
| AT5G46280 | 0,57 | -0,39 | 3,10 | -0,13 | 8,99E-01 |
| AT5G46290 | 0,63 | -0,56 | 3,12 | -0,18 | 8,58E-01 |
| AT5G46340 | 0,61 | -1,40 | 3,93 | -0,36 | 7,21E-01 |
| AT5G46395 | 0,11 | 0,00 | 4,41 | 0,00 | 1,00E+00 |
| AT5G46580 | 0,75 | -2,30 | 2,44 | -0,94 | 3,45E-01 |
| AT5G46620 | 0,64 | -2,51 | 3,05 | -0,82 | 4,12E-01 |
| AT5G46750 | 0,66 | 2,03 | 3,80 | 0,53 | 5,94E-01 |
| AT5G46790 | 0,53 | -0,54 | 3,15 | -0,17 | 8,63E-01 |
| AT5G46800 | 0,83 | -0,44 | 2,27 | -0,20 | 8,45E-01 |
| AT5G46860 | 0,46 | -1,76 | 4,09 | -0,43 | 6,68E-01 |
| AT5G46970 | 0,65 | -1,52 | 2,66 | -0,57 | 5,68E-01 |
| AT5G47040 | 1,34 | 2,92 | 1,84 | 1,59 | 1,12E-01 |
| AT5G47070 | 0,97 | 2,41 | 2,41 | 1,00 | 3,17E-01 |
| AT5G47080 | 0,13 | 0,00 | 4,41 | 0,00 | 1,00E+00 |
| AT5G47110 | 1,25 | 2,44 | 1,77 | 1,38 | 1,67E-01 |
| AT5G47190 | 0,93 | -0,74 | 2,12 | -0,35 | 7,26E-01 |
| AT5G47210 | 1,48 | 1,28 | 2,02 | 0,63 | 5,26E-01 |
| AT5G47220 | 0,78 | 3,15 | 2,55 | 1,23 | 2,17E-01 |
| AT5G47250 | 0,28 | -0,93 | 4,36 | -0,21 | 8,32E-01 |
| AT5G47370 | 0,28 | 1,84 | 4,39 | 0,42 | 6,74E-01 |
| AT5G47460 | 0,15 | 0,00 | 4,41 | 0,00 | 1,00E+00 |
| AT5G47570 | 0,30 | -0,53 | 4,41 | -0,12 | 9,04E-01 |
| AT5G47580 | 0,64 | 2,60 | 3,63 | 0,72 | 4,74E-01 |
| AT5G47650 | 0,20 | 1,38 | 4,41 | 0,31 | 7,54E-01 |
| AT5G47700 | 0,72 | -0,78 | 2,28 | -0,34 | 7,34E-01 |
| AT5G47730 | 0,16 | 0,00 | 4,41 | 0,00 | 1,00E+00 |

| | | | | | |
|---|---|---|---|---|---|
| AT5G47750 | 1,50 | 3,14 | 2,30 | 1,36 | 1,73E-01 |
| AT5G47760 | 0,12 | 0,00 | 4,41 | 0,00 | 1,00E+00 |
| AT5G47810 | 0,55 | -0,37 | 2,74 | -0,13 | 8,93E-01 |
| AT5G47830 | 0,54 | -0,85 | 4,38 | -0,19 | 8,46E-01 |
| AT5G47850 | 0,49 | 1,38 | 4,08 | 0,34 | 7,35E-01 |
| AT5G47880 | 0,55 | 1,38 | 3,26 | 0,42 | 6,72E-01 |
| AT5G47930 | 0,53 | 0,44 | 2,75 | 0,16 | 8,74E-01 |
| AT5G48000 | 0,12 | 0,00 | 4,41 | 0,00 | 1,00E+00 |
| AT5G07055 | 0,58 | -1,88 | 3,13 | -0,60 | 5,48E-01 |
| AT5G48120 | 1,18 | 2,48 | 1,88 | 1,32 | 1,87E-01 |
| AT5G48150 | 0,85 | 2,11 | 3,46 | 0,61 | 5,42E-01 |
| AT5G48180 | 0,98 | 0,44 | 2,25 | 0,20 | 8,44E-01 |
| AT5G48375 | 0,26 | 1,38 | 4,41 | 0,31 | 7,54E-01 |
| AT5G48412 | 0,64 | 0,70 | 2,66 | 0,26 | 7,91E-01 |
| AT5G48440 | 0,43 | 0,00 | 4,27 | 0,00 | 1,00E+00 |
| AT5G48540 | 0,08 | 0,00 | 4,41 | 0,00 | 1,00E+00 |
| AT5G48580 | 0,41 | 1,38 | 4,21 | 0,33 | 7,43E-01 |
| AT5G48640 | 0,19 | -0,54 | 4,41 | -0,12 | 9,02E-01 |
| AT5G48657 | 0,40 | 1,29 | 3,40 | 0,38 | 7,03E-01 |
| AT5G07265 | 0,12 | 0,00 | 4,41 | 0,00 | 1,00E+00 |
| AT5G48770 | 0,08 | 0,00 | 4,41 | 0,00 | 1,00E+00 |
| AT5G07275 | 0,71 | 1,38 | 4,41 | 0,31 | 7,54E-01 |
| AT5G48810 | 0,80 | 1,45 | 2,43 | 0,60 | 5,52E-01 |
| AT5G48830 | 0,28 | 1,39 | 4,41 | 0,32 | 7,52E-01 |
| AT5G48880 | 0,58 | 2,05 | 4,35 | 0,47 | 6,38E-01 |
| AT5G48910 | 0,76 | 0,45 | 2,38 | 0,19 | 8,49E-01 |
| AT5G48930 | 1,24 | 1,42 | 1,74 | 0,81 | 4,15E-01 |
| AT5G49100 | 1,11 | -0,02 | 2,07 | -0,01 | 9,92E-01 |
| AT5G49300 | 0,36 | 0,00 | 4,41 | 0,00 | 1,00E+00 |
| AT5G49350 | 0,08 | 0,00 | 4,41 | 0,00 | 1,00E+00 |
| AT5G49420 | 0,35 | -0,54 | 4,41 | -0,12 | 9,02E-01 |
| AT5G49430 | 1,30 | 2,38 | 1,79 | 1,33 | 1,82E-01 |
| AT5G49540 | 0,90 | 0,11 | 2,52 | 0,04 | 9,65E-01 |
| AT5G49550 | 0,27 | 1,40 | 4,40 | 0,32 | 7,51E-01 |
| AT5G49555 | 0,40 | 0,10 | 4,35 | 0,02 | 9,82E-01 |
| AT5G49570 | 0,11 | 0,00 | 4,41 | 0,00 | 1,00E+00 |
| AT5G49580 | 0,46 | -1,16 | 4,01 | -0,29 | 7,72E-01 |
| AT5G49630 | 0,19 | 0,00 | 4,41 | 0,00 | 1,00E+00 |
| AT5G49645 | 0,08 | 0,00 | 4,41 | 0,00 | 1,00E+00 |
| AT5G49660 | 0,65 | 0,29 | 2,65 | 0,11 | 9,12E-01 |
| AT5G49665 | 0,50 | 2,48 | 4,01 | 0,62 | 5,36E-01 |
| AT5G49830 | 1,27 | 1,72 | 1,92 | 0,89 | 3,71E-01 |
| AT5G49890 | 0,22 | -0,47 | 4,40 | -0,11 | 9,16E-01 |
| AT5G49900 | 0,41 | 1,51 | 4,21 | 0,36 | 7,21E-01 |
| AT5G49930 | 0,74 | -0,41 | 2,28 | -0,18 | 8,56E-01 |
| AT5G49950 | 0,25 | 0,50 | 4,40 | 0,11 | 9,09E-01 |
| AT5G50000 | 0,84 | -1,07 | 2,10 | -0,51 | 6,11E-01 |
| AT5G50150 | 0,10 | 0,00 | 4,41 | 0,00 | 1,00E+00 |
| AT5G50220 | 0,28 | 2,12 | 4,38 | 0,48 | 6,28E-01 |
| AT5G50320 | 0,77 | 2,70 | 2,99 | 0,90 | 3,67E-01 |
| AT5G50340 | 0,25 | 1,38 | 4,41 | 0,31 | 7,54E-01 |
| AT5G50350 | 0,29 | -0,54 | 4,41 | -0,12 | 9,02E-01 |
| AT5G50380 | 0,97 | 0,44 | 2,28 | 0,19 | 8,49E-01 |
| AT5G50390 | 0,17 | 0,00 | 4,41 | 0,00 | 1,00E+00 |
| AT5G50400 | 0,54 | 3,01 | 3,92 | 0,77 | 4,43E-01 |
| AT5G50460 | 0,36 | 1,14 | 4,37 | 0,26 | 7,94E-01 |

| | | | | | |
|---|---|---|---|---|---|
| AT5G50480 | 0,53 | -2,78 | 3,78 | -0,74 | 4,62E-01 |
| AT5G50570 | 0,31 | 0,44 | 4,39 | 0,10 | 9,20E-01 |
| AT5G50630 | 0,05 | 0,00 | 4,41 | 0,00 | 1,00E+00 |
| AT5G50640 | 0,24 | 1,38 | 4,41 | 0,31 | 7,54E-01 |
| AT5G50670 | 0,93 | 1,66 | 2,42 | 0,69 | 4,93E-01 |
| AT5G50860 | 0,49 | 2,10 | 3,93 | 0,53 | 5,94E-01 |
| AT5G50870 | 0,08 | 0,00 | 4,41 | 0,00 | 1,00E+00 |
| AT5G50900 | 0,83 | 0,88 | 2,54 | 0,35 | 7,27E-01 |
| AT5G51000 | 0,61 | -0,54 | 3,12 | -0,17 | 8,62E-01 |
| AT5G51020 | 0,37 | 1,27 | 4,33 | 0,29 | 7,70E-01 |
| AT5G51110 | 0,78 | -1,11 | 2,36 | -0,47 | 6,37E-01 |
| AT5G51170 | 0,12 | 0,00 | 4,41 | 0,00 | 1,00E+00 |
| AT5G51195 | 0,57 | -1,32 | 2,63 | -0,50 | 6,14E-01 |
| AT5G51220 | 0,23 | 0,00 | 4,41 | 0,00 | 1,00E+00 |
| AT5G51290 | 0,61 | 3,33 | 2,99 | 1,11 | 2,65E-01 |
| AT5G51390 | 0,29 | 1,74 | 4,38 | 0,40 | 6,90E-01 |
| AT5G51400 | 1,21 | 0,58 | 1,60 | 0,37 | 7,14E-01 |
| AT5G51550 | 1,02 | -0,06 | 2,11 | -0,03 | 9,78E-01 |
| AT5G51640 | 0,39 | 0,54 | 3,53 | 0,15 | 8,78E-01 |
| AT5G51650 | 0,32 | 0,00 | 3,61 | 0,00 | 1,00E+00 |
| AT5G51690 | 0,72 | 2,69 | 2,52 | 1,06 | 2,88E-01 |
| AT5G51710 | 0,10 | 0,00 | 4,41 | 0,00 | 1,00E+00 |
| AT5G51820 | 0,69 | 0,12 | 2,51 | 0,05 | 9,62E-01 |
| AT5G52060 | 0,54 | 0,43 | 2,76 | 0,16 | 8,76E-01 |
| AT5G52090 | 0,73 | 0,44 | 2,37 | 0,19 | 8,51E-01 |
| AT5G52120 | 0,47 | 0,40 | 4,35 | 0,09 | 9,26E-01 |
| AT5G52180 | 0,61 | 2,95 | 3,07 | 0,96 | 3,37E-01 |
| AT5G52330 | 0,15 | 0,00 | 4,39 | 0,00 | 1,00E+00 |
| AT5G52370 | 0,37 | 1,29 | 4,34 | 0,30 | 7,67E-01 |
| AT5G52450 | 0,44 | 2,09 | 3,28 | 0,64 | 5,23E-01 |
| AT5G52540 | 1,60 | 1,54 | 1,78 | 0,86 | 3,87E-01 |
| AT5G52550 | 0,25 | 0,00 | 4,41 | 0,00 | 1,00E+00 |
| AT5G52650 | 0,64 | 1,80 | 3,64 | 0,49 | 6,21E-01 |
| AT5G52780 | 0,46 | 0,11 | 3,29 | 0,03 | 9,74E-01 |
| AT5G52797 | 0,48 | 1,35 | 3,31 | 0,41 | 6,84E-01 |
| AT5G53000 | 0,29 | 1,38 | 4,41 | 0,31 | 7,54E-01 |
| AT5G53010 | 0,50 | 1,58 | 3,31 | 0,48 | 6,33E-01 |
| AT5G53050 | 0,49 | -1,27 | 3,18 | -0,40 | 6,90E-01 |
| AT5G53130 | 0,72 | 0,00 | 3,01 | 0,00 | 1,00E+00 |
| AT5G07955 | 0,08 | 0,00 | 4,41 | 0,00 | 1,00E+00 |
| AT5G53300 | 0,89 | 2,85 | 2,69 | 1,06 | 2,88E-01 |
| AT5G53320 | 0,58 | 2,88 | 2,99 | 0,96 | 3,37E-01 |
| AT5G07975 | 0,08 | 0,00 | 4,41 | 0,00 | 1,00E+00 |
| AT5G53490 | 0,65 | 1,40 | 2,31 | 0,61 | 5,44E-01 |
| AT5G53500 | 0,52 | -0,05 | 3,23 | -0,01 | 9,88E-01 |
| AT5G53530 | 0,37 | 0,59 | 4,36 | 0,14 | 8,92E-01 |
| AT5G53570 | 0,72 | 0,29 | 2,54 | 0,11 | 9,09E-01 |
| AT5G53580 | 0,36 | 0,52 | 3,62 | 0,14 | 8,87E-01 |
| AT5G53730 | 0,14 | 0,00 | 4,41 | 0,00 | 1,00E+00 |
| AT5G53850 | 0,47 | 0,49 | 4,34 | 0,11 | 9,10E-01 |
| AT5G53870 | 0,19 | 0,00 | 4,40 | 0,00 | 1,00E+00 |
| AT5G53900 | 0,79 | 1,66 | 2,30 | 0,72 | 4,70E-01 |
| AT5G54030 | 0,27 | 0,00 | 4,41 | 0,00 | 1,00E+00 |
| AT5G54040 | 0,27 | 0,00 | 4,41 | 0,00 | 1,00E+00 |
| AT5G54050 | 0,38 | 0,00 | 4,41 | 0,00 | 1,00E+00 |
| AT5G54060 | 0,12 | 0,00 | 4,41 | 0,00 | 1,00E+00 |

| | | | | | |
|---|---|---|---|---|---|
| AT5G54090 | 1,00 | 1,91 | 2,30 | 0,83 | 4,05E-01 |
| AT5G54100 | 0,65 | 1,29 | 3,03 | 0,43 | 6,70E-01 |
| AT5G54170 | 0,64 | 1,11 | 2,34 | 0,48 | 6,33E-01 |
| AT5G54280 | 1,29 | 2,11 | 1,80 | 1,17 | 2,41E-01 |
| AT5G54380 | 1,11 | 1,19 | 2,12 | 0,56 | 5,76E-01 |
| AT5G54410 | 0,82 | -1,38 | 2,26 | -0,61 | 5,42E-01 |
| AT5G54430 | 1,12 | 1,14 | 2,64 | 0,43 | 6,64E-01 |
| AT5G54460 | 1,09 | 0,00 | 3,40 | 0,00 | 1,00E+00 |
| AT5G54470 | 0,35 | 1,28 | 3,53 | 0,36 | 7,16E-01 |
| AT5G54510 | 0,08 | 0,00 | 4,41 | 0,00 | 1,00E+00 |
| AT5G54610 | 0,49 | 1,25 | 3,93 | 0,32 | 7,51E-01 |
| AT5G54620 | 0,12 | 0,00 | 4,41 | 0,00 | 1,00E+00 |
| AT5G54630 | 0,29 | -1,22 | 4,35 | -0,28 | 7,79E-01 |
| AT5G54650 | 0,47 | 1,94 | 4,39 | 0,44 | 6,59E-01 |
| AT5G54690 | 0,39 | 1,76 | 4,38 | 0,40 | 6,87E-01 |
| AT5G54700 | 0,89 | 1,79 | 2,31 | 0,77 | 4,39E-01 |
| AT5G54710 | 0,79 | 0,53 | 2,62 | 0,20 | 8,40E-01 |
| AT5G54720 | 0,73 | 0,52 | 2,48 | 0,21 | 8,35E-01 |
| AT5G54780 | 1,16 | 2,80 | 1,86 | 1,50 | 1,33E-01 |
| AT5G54930 | 0,40 | 0,46 | 4,23 | 0,11 | 9,13E-01 |
| AT5G54960 | 0,98 | -1,63 | 2,28 | -0,71 | 4,76E-01 |
| AT5G55040 | 0,74 | 0,38 | 2,34 | 0,16 | 8,70E-01 |
| AT5G55055 | 0,05 | 0,00 | 4,41 | 0,00 | 1,00E+00 |
| AT5G55100 | 1,12 | 3,86 | 2,41 | 1,60 | 1,10E-01 |
| AT5G55130 | 0,31 | 0,00 | 3,68 | 0,00 | 1,00E+00 |
| AT5G55150 | 1,21 | 4,09 | 2,42 | 1,69 | 9,13E-02 |
| AT5G55160 | 0,37 | 1,38 | 4,41 | 0,31 | 7,54E-01 |
| AT5G55210 | 0,08 | 0,00 | 4,41 | 0,00 | 1,00E+00 |
| AT5G55250 | 0,40 | -1,10 | 4,37 | -0,25 | 8,00E-01 |
| AT5G55280 | 1,13 | -3,11 | 2,39 | -1,30 | 1,93E-01 |
| AT5G55290 | 0,30 | 1,38 | 4,41 | 0,31 | 7,54E-01 |
| AT5G55300 | 0,70 | 0,02 | 2,24 | 0,01 | 9,91E-01 |
| AT5G55340 | 0,19 | 0,00 | 4,40 | 0,00 | 1,00E+00 |
| AT5G55390 | 0,36 | 0,00 | 4,41 | 0,00 | 1,00E+00 |
| AT5G08275 | 0,59 | 1,62 | 3,80 | 0,43 | 6,69E-01 |
| AT5G08285 | 0,30 | 0,42 | 4,36 | 0,10 | 9,23E-01 |
| AT5G55510 | 0,05 | 0,00 | 4,41 | 0,00 | 1,00E+00 |
| AT5G55540 | 1,63 | 1,30 | 1,66 | 0,78 | 4,33E-01 |
| AT5G55550 | 0,34 | 1,38 | 4,41 | 0,31 | 7,54E-01 |
| AT5G55570 | 1,30 | -1,71 | 2,89 | -0,59 | 5,54E-01 |
| AT5G55600 | 0,31 | -0,52 | 3,70 | -0,14 | 8,88E-01 |
| AT5G55670 | 0,30 | 0,00 | 4,41 | 0,00 | 1,00E+00 |
| AT5G55710 | 0,92 | 0,37 | 2,60 | 0,14 | 8,88E-01 |
| AT5G55740 | 0,27 | 0,00 | 4,41 | 0,00 | 1,00E+00 |
| AT5G55790 | 0,33 | 0,00 | 4,41 | 0,00 | 1,00E+00 |
| AT5G55835 | 0,61 | 0,39 | 2,45 | 0,16 | 8,75E-01 |
| AT5G55910 | 1,51 | 0,50 | 1,46 | 0,35 | 7,29E-01 |
| AT5G55920 | 0,81 | 1,97 | 2,55 | 0,77 | 4,39E-01 |
| AT5G55940 | 0,50 | 2,24 | 3,93 | 0,57 | 5,69E-01 |
| AT5G56000 | 1,26 | -0,94 | 1,95 | -0,48 | 6,28E-01 |
| AT5G56180 | 0,23 | 1,38 | 4,41 | 0,31 | 7,54E-01 |
| AT5G56240 | 1,23 | 3,04 | 2,30 | 1,32 | 1,86E-01 |
| AT5G56250 | 0,87 | 0,00 | 2,94 | 0,00 | 1,00E+00 |
| AT5G56260 | 0,25 | 0,50 | 4,40 | 0,11 | 9,09E-01 |
| AT5G56280 | 0,08 | 0,00 | 4,41 | 0,00 | 1,00E+00 |
| AT5G56350 | 0,73 | -0,33 | 1,98 | -0,17 | 8,69E-01 |

| | | | | | |
|---|---|---|---|---|---|
| AT5G56360 | 0,87 | -2,23 | 2,86 | -0,78 | 4,36E-01 |
| AT5G56410 | 0,30 | 1,61 | 4,39 | 0,37 | 7,14E-01 |
| AT5G56450 | 0,20 | 0,00 | 4,41 | 0,00 | 1,00E+00 |
| AT5G56710 | 0,79 | 3,09 | 3,38 | 0,92 | 3,60E-01 |
| AT5G56730 | 0,65 | 1,33 | 2,57 | 0,52 | 6,05E-01 |
| AT5G56760 | 0,72 | 1,20 | 2,79 | 0,43 | 6,68E-01 |
| AT5G56860 | 0,23 | 0,00 | 4,41 | 0,00 | 1,00E+00 |
| AT5G56890 | 0,16 | 0,00 | 4,41 | 0,00 | 1,00E+00 |
| AT5G56900 | 1,69 | 4,01 | 1,76 | 2,27 | 2,29E-02 |
| AT5G56980 | 0,28 | 1,38 | 4,41 | 0,31 | 7,54E-01 |
| AT5G57040 | 0,80 | 3,23 | 2,11 | 1,53 | 1,27E-01 |
| AT5G57160 | 0,81 | 1,78 | 2,46 | 0,72 | 4,69E-01 |
| AT5G57170 | 0,08 | 0,00 | 4,41 | 0,00 | 1,00E+00 |
| AT5G57250 | 0,54 | 1,34 | 3,96 | 0,34 | 7,35E-01 |
| AT5G57270 | 0,45 | 0,81 | 4,37 | 0,19 | 8,52E-01 |
| AT5G57290 | 0,69 | 0,38 | 2,56 | 0,15 | 8,83E-01 |
| AT5G57345 | 0,70 | 0,52 | 2,39 | 0,22 | 8,27E-01 |
| AT5G57580 | 0,90 | 0,04 | 2,51 | 0,02 | 9,87E-01 |
| AT5G57630 | 0,49 | 0,64 | 3,41 | 0,19 | 8,52E-01 |
| AT5G57670 | 0,16 | 0,00 | 4,41 | 0,00 | 1,00E+00 |
| AT5G57780 | 0,36 | 1,26 | 4,39 | 0,29 | 7,74E-01 |
| AT5G57785 | 0,56 | 0,00 | 2,77 | 0,00 | 1,00E+00 |
| AT5G57790 | 0,05 | 0,00 | 4,41 | 0,00 | 1,00E+00 |
| AT5G57850 | 0,51 | -0,14 | 4,33 | -0,03 | 9,73E-01 |
| AT5G57900 | 0,27 | -0,40 | 4,38 | -0,09 | 9,27E-01 |
| AT5G57960 | 0,12 | 0,00 | 4,41 | 0,00 | 1,00E+00 |
| AT5G58030 | 0,11 | 0,00 | 4,41 | 0,00 | 1,00E+00 |
| AT5G58050 | 0,17 | 0,00 | 4,41 | 0,00 | 1,00E+00 |
| AT5G58090 | 0,76 | 2,54 | 2,45 | 1,04 | 2,99E-01 |
| AT5G58130 | 0,84 | 1,38 | 2,82 | 0,49 | 6,24E-01 |
| AT5G58170 | 0,90 | 1,36 | 3,32 | 0,41 | 6,83E-01 |
| AT5G58220 | 0,12 | 0,00 | 4,41 | 0,00 | 1,00E+00 |
| AT5G58260 | 0,45 | -0,44 | 3,40 | -0,13 | 8,98E-01 |
| AT5G58380 | 0,93 | 1,18 | 2,33 | 0,51 | 6,11E-01 |
| AT5G58440 | 0,43 | -0,31 | 2,89 | -0,11 | 9,13E-01 |
| AT5G58570 | 0,61 | 0,00 | 3,98 | 0,00 | 1,00E+00 |
| AT5G08860 | 0,08 | 0,00 | 4,41 | 0,00 | 1,00E+00 |
| AT5G58600 | 0,29 | 1,85 | 4,39 | 0,42 | 6,73E-01 |
| AT5G58650 | 0,16 | 0,00 | 4,41 | 0,00 | 1,00E+00 |
| AT5G58670 | 0,69 | 0,72 | 2,20 | 0,33 | 7,45E-01 |
| AT5G58720 | 0,64 | 2,90 | 2,53 | 1,15 | 2,51E-01 |
| AT5G58730 | 0,50 | 2,01 | 4,02 | 0,50 | 6,17E-01 |
| AT5G58870 | 0,47 | -0,64 | 3,46 | -0,18 | 8,54E-01 |
| AT5G58900 | 0,28 | 1,84 | 4,39 | 0,42 | 6,74E-01 |
| AT5G58930 | 1,32 | 1,67 | 2,03 | 0,82 | 4,11E-01 |
| AT5G59030 | 0,49 | 2,11 | 3,93 | 0,54 | 5,90E-01 |
| AT5G59130 | 0,21 | 0,00 | 4,41 | 0,00 | 1,00E+00 |
| AT5G59220 | 0,62 | 3,27 | 3,72 | 0,88 | 3,79E-01 |
| AT5G59290 | 0,24 | 1,36 | 4,40 | 0,31 | 7,57E-01 |
| AT5G08920 | 0,49 | 2,01 | 4,05 | 0,50 | 6,19E-01 |
| AT5G59480 | 0,17 | 0,00 | 4,41 | 0,00 | 1,00E+00 |
| AT5G59550 | 1,55 | 2,51 | 1,90 | 1,32 | 1,87E-01 |
| AT5G59662 | 0,05 | 0,00 | 4,41 | 0,00 | 1,00E+00 |
| AT5G59700 | 0,33 | 2,08 | 4,37 | 0,48 | 6,34E-01 |
| AT5G59770 | 0,46 | 1,15 | 3,24 | 0,36 | 7,22E-01 |
| AT5G59780 | 1,19 | 2,89 | 2,04 | 1,42 | 1,57E-01 |

| | | | | | |
|---|---|---|---|---|---|
| AT5G59840 | 0,17 | -0,48 | 4,39 | -0,11 | 9,13E-01 |
| AT5G59850 | 0,69 | 0,92 | 2,50 | 0,37 | 7,12E-01 |
| AT5G59870 | 0,19 | 0,00 | 4,41 | 0,00 | 1,00E+00 |
| AT5G59900 | 0,16 | -0,55 | 4,41 | -0,12 | 9,01E-01 |
| AT5G59960 | 0,11 | 0,00 | 4,41 | 0,00 | 1,00E+00 |
| AT5G60050 | 0,15 | -0,42 | 4,39 | -0,10 | 9,23E-01 |
| AT5G60100 | 0,30 | -0,78 | 4,37 | -0,18 | 8,58E-01 |
| AT5G60160 | 0,49 | -0,54 | 4,14 | -0,13 | 8,95E-01 |
| AT5G60180 | 0,21 | 1,27 | 4,39 | 0,29 | 7,72E-01 |
| AT5G60210 | 0,29 | -0,57 | 4,40 | -0,13 | 8,97E-01 |
| AT5G60340 | 0,36 | 0,00 | 4,41 | 0,00 | 1,00E+00 |
| AT5G60400 | 0,12 | 0,00 | 4,41 | 0,00 | 1,00E+00 |
| AT5G60460 | 0,27 | -0,42 | 4,39 | -0,10 | 9,24E-01 |
| AT5G60670 | 0,64 | 2,04 | 2,26 | 0,90 | 3,67E-01 |
| AT5G60700 | 1,32 | 3,94 | 2,26 | 1,74 | 8,19E-02 |
| AT5G60710 | 1,48 | 2,07 | 1,56 | 1,33 | 1,85E-01 |
| AT5G60800 | 0,66 | 1,50 | 3,06 | 0,49 | 6,23E-01 |
| AT5G60840 | 0,12 | 0,00 | 4,41 | 0,00 | 1,00E+00 |
| AT5G60930 | 0,24 | 1,57 | 4,39 | 0,36 | 7,20E-01 |
| AT5G60960 | 0,29 | 1,47 | 4,40 | 0,33 | 7,39E-01 |
| AT5G60990 | 0,16 | 0,00 | 4,41 | 0,00 | 1,00E+00 |
| AT5G61040 | 0,88 | 1,59 | 4,33 | 0,37 | 7,14E-01 |
| AT5G61060 | 0,37 | 1,29 | 3,50 | 0,37 | 7,13E-01 |
| AT5G61130 | 0,17 | 0,00 | 4,41 | 0,00 | 1,00E+00 |
| AT5G61150 | 0,86 | 1,04 | 2,72 | 0,38 | 7,03E-01 |
| AT5G61250 | 0,23 | 1,38 | 4,41 | 0,31 | 7,54E-01 |
| AT5G61390 | 0,43 | 1,90 | 4,39 | 0,43 | 6,65E-01 |
| AT5G09005 | 1,51 | -2,88 | 1,66 | -1,73 | 8,36E-02 |
| AT5G61460 | 0,21 | 0,00 | 4,41 | 0,00 | 1,00E+00 |
| AT5G61480 | 0,83 | 1,88 | 2,21 | 0,85 | 3,94E-01 |
| AT5G61500 | 0,76 | 2,94 | 2,89 | 1,02 | 3,10E-01 |
| AT5G61510 | 0,56 | 1,64 | 3,27 | 0,50 | 6,15E-01 |
| AT5G61520 | 0,33 | 1,26 | 4,39 | 0,29 | 7,74E-01 |
| AT5G61590 | 1,28 | 0,43 | 1,68 | 0,25 | 7,99E-01 |
| AT5G61600 | 0,57 | -0,56 | 2,72 | -0,21 | 8,36E-01 |
| AT5G61800 | 0,12 | 0,00 | 4,41 | 0,00 | 1,00E+00 |
| AT5G61810 | 0,48 | 2,43 | 3,34 | 0,73 | 4,68E-01 |
| AT5G61820 | 1,07 | 2,86 | 2,39 | 1,20 | 2,31E-01 |
| AT5G61840 | 0,89 | 2,22 | 2,34 | 0,95 | 3,42E-01 |
| AT5G61910 | 1,60 | 1,27 | 1,88 | 0,67 | 5,01E-01 |
| AT5G61900 | 1,09 | 0,36 | 2,11 | 0,17 | 8,64E-01 |
| AT5G61960 | 0,94 | 1,16 | 2,58 | 0,45 | 6,54E-01 |
| AT5G61990 | 0,30 | 1,44 | 4,40 | 0,33 | 7,43E-01 |
| AT5G62090 | 0,32 | -1,23 | 4,36 | -0,28 | 7,77E-01 |
| AT5G62110 | 0,34 | 0,00 | 4,40 | 0,00 | 1,00E+00 |
| AT5G62130 | 0,45 | -0,46 | 3,29 | -0,14 | 8,89E-01 |
| AT5G62140 | 1,15 | -1,67 | 1,92 | -0,87 | 3,85E-01 |
| AT5G62180 | 0,08 | 0,00 | 4,41 | 0,00 | 1,00E+00 |
| AT5G62220 | 1,34 | 0,22 | 1,86 | 0,12 | 9,04E-01 |
| AT5G62300 | 1,07 | 0,87 | 1,77 | 0,49 | 6,23E-01 |
| AT5G62430 | 0,29 | 1,23 | 4,38 | 0,28 | 7,79E-01 |
| AT5G62470 | 0,30 | -1,22 | 4,36 | -0,28 | 7,79E-01 |
| AT5G62530 | 0,56 | -1,66 | 3,10 | -0,54 | 5,93E-01 |
| AT5G62600 | 0,50 | 2,01 | 4,02 | 0,50 | 6,17E-01 |
| AT5G62610 | 0,31 | 1,32 | 4,40 | 0,30 | 7,63E-01 |
| AT5G62630 | 0,37 | 0,02 | 4,37 | 0,00 | 9,96E-01 |

| | | | | | |
|---|---|---|---|---|---|
| AT5G62650 | 0,39 | 0,32 | 4,32 | 0,07 | 9,41E-01 |
| AT5G62680 | 0,33 | 1,32 | 4,40 | 0,30 | 7,64E-01 |
| AT5G62790 | 0,60 | -1,56 | 2,62 | -0,59 | 5,54E-01 |
| AT5G63080 | 0,23 | -0,40 | 4,38 | -0,09 | 9,28E-01 |
| AT5G63100 | 0,29 | 1,29 | 4,39 | 0,29 | 7,69E-01 |
| AT5G63110 | 0,58 | -1,06 | 3,15 | -0,34 | 7,37E-01 |
| AT5G63120 | 0,65 | 1,19 | 3,16 | 0,38 | 7,07E-01 |
| AT5G63160 | 0,51 | 0,00 | 3,23 | 0,00 | 1,00E+00 |
| AT5G63180 | 0,25 | -1,23 | 4,36 | -0,28 | 7,79E-01 |
| AT5G63290 | 0,56 | 2,06 | 4,38 | 0,47 | 6,38E-01 |
| AT5G63310 | 0,47 | 0,32 | 4,37 | 0,07 | 9,41E-01 |
| AT5G63320 | 0,52 | 0,00 | 3,97 | 0,00 | 1,00E+00 |
| AT5G63370 | 1,26 | 2,56 | 1,80 | 1,42 | 1,55E-01 |
| AT5G63400 | 0,30 | 0,01 | 4,37 | 0,00 | 9,98E-01 |
| AT5G63410 | 0,46 | 1,38 | 2,84 | 0,49 | 6,27E-01 |
| AT5G63470 | 0,44 | 1,37 | 4,11 | 0,33 | 7,38E-01 |
| AT5G63610 | 0,51 | 2,16 | 3,99 | 0,54 | 5,88E-01 |
| AT5G63625 | 1,28 | -2,93 | 1,98 | -1,48 | 1,38E-01 |
| AT5G63760 | 0,90 | 2,49 | 2,45 | 1,01 | 3,10E-01 |
| AT5G63790 | 0,54 | 1,75 | 4,38 | 0,40 | 6,89E-01 |
| AT5G63860 | 0,75 | 1,46 | 2,27 | 0,64 | 5,21E-01 |
| AT5G63870 | 0,08 | 0,00 | 4,41 | 0,00 | 1,00E+00 |
| AT5G63890 | 0,39 | -1,89 | 4,27 | -0,44 | 6,58E-01 |
| AT5G63910 | 0,66 | 1,73 | 2,99 | 0,58 | 5,64E-01 |
| AT5G64020 | 0,23 | 1,38 | 4,41 | 0,31 | 7,54E-01 |
| AT5G64050 | 1,12 | -0,21 | 1,80 | -0,12 | 9,06E-01 |
| AT5G64070 | 1,53 | 2,21 | 1,73 | 1,27 | 2,03E-01 |
| AT5G64130 | 0,35 | -1,23 | 3,62 | -0,34 | 7,33E-01 |
| AT5G64140 | 1,06 | 1,92 | 1,87 | 1,03 | 3,05E-01 |
| AT5G64170 | 0,64 | 1,38 | 3,27 | 0,42 | 6,74E-01 |
| AT5G64200 | 0,28 | 1,34 | 4,40 | 0,30 | 7,61E-01 |
| AT5G64280 | 1,39 | 4,07 | 1,74 | 2,34 | 1,95E-02 |
| AT5G64290 | 0,42 | -0,54 | 4,19 | -0,13 | 8,97E-01 |
| AT5G64340 | 1,19 | 1,05 | 1,90 | 0,55 | 5,79E-01 |
| AT5G64350 | 0,41 | -1,31 | 4,26 | -0,31 | 7,58E-01 |
| AT5G64360 | 0,62 | 3,30 | 3,64 | 0,91 | 3,64E-01 |
| AT5G64410 | 0,79 | -0,54 | 2,37 | -0,23 | 8,19E-01 |
| AT5G64430 | 0,96 | 0,22 | 2,33 | 0,10 | 9,23E-01 |
| AT5G64460 | 0,08 | 0,00 | 4,41 | 0,00 | 1,00E+00 |
| AT5G64550 | 0,25 | 1,38 | 4,41 | 0,31 | 7,54E-01 |
| AT5G64570 | 1,49 | 2,19 | 1,64 | 1,34 | 1,80E-01 |
| AT5G64580 | 0,77 | 2,30 | 2,63 | 0,87 | 3,82E-01 |
| AT5G64610 | 0,28 | 0,00 | 4,41 | 0,00 | 1,00E+00 |
| AT5G64660 | 0,37 | -0,53 | 3,51 | -0,15 | 8,79E-01 |
| AT5G64670 | 0,17 | -0,49 | 4,40 | -0,11 | 9,11E-01 |
| AT5G64760 | 0,83 | 1,41 | 3,10 | 0,45 | 6,49E-01 |
| AT5G64830 | 0,77 | 1,98 | 2,46 | 0,81 | 4,20E-01 |
| AT5G64860 | 0,28 | 0,00 | 4,41 | 0,00 | 1,00E+00 |
| AT5G64870 | 0,67 | 0,00 | 2,97 | 0,00 | 1,00E+00 |
| AT5G64920 | 1,21 | 3,50 | 1,92 | 1,82 | 6,87E-02 |
| AT5G64930 | 0,39 | 1,79 | 4,37 | 0,41 | 6,81E-01 |
| AT5G64980 | 0,12 | 0,00 | 4,41 | 0,00 | 1,00E+00 |
| AT5G65110 | 0,46 | -1,05 | 3,25 | -0,32 | 7,48E-01 |
| AT5G65200 | 0,16 | 0,00 | 4,41 | 0,00 | 1,00E+00 |
| AT5G65240 | 0,27 | 0,00 | 3,81 | 0,00 | 1,00E+00 |
| AT5G65310 | 1,18 | 1,70 | 2,01 | 0,85 | 3,98E-01 |

| | | | | | |
|---|---|---|---|---|---|
| AT5G65360 | 0,52 | 2,19 | 3,19 | 0,68 | 4,93E-01 |
| AT5G65430 | 0,62 | 2,86 | 2,65 | 1,08 | 2,81E-01 |
| AT5G65560 | 0,37 | 1,23 | 4,38 | 0,28 | 7,79E-01 |
| AT5G65685 | 0,97 | -0,39 | 2,21 | -0,18 | 8,60E-01 |
| AT5G65840 | 0,17 | 0,00 | 4,41 | 0,00 | 1,00E+00 |
| AT5G65910 | 1,14 | 1,73 | 3,26 | 0,53 | 5,95E-01 |
| AT5G65930 | 0,99 | 0,28 | 2,30 | 0,12 | 9,02E-01 |
| AT5G09535 | 0,24 | 1,37 | 4,41 | 0,31 | 7,56E-01 |
| AT5G65970 | 0,45 | 2,52 | 4,35 | 0,58 | 5,63E-01 |
| AT5G66055 | 0,63 | -0,47 | 2,60 | -0,18 | 8,58E-01 |
| AT5G66120 | 0,85 | -0,68 | 2,35 | -0,29 | 7,72E-01 |
| AT5G66210 | 1,05 | 1,77 | 2,28 | 0,77 | 4,39E-01 |
| AT5G66300 | 0,13 | 0,00 | 4,41 | 0,00 | 1,00E+00 |
| AT5G66310 | 0,12 | 0,00 | 4,41 | 0,00 | 1,00E+00 |
| AT5G66320 | 0,48 | 2,07 | 3,38 | 0,61 | 5,40E-01 |
| AT5G66330 | 0,11 | 0,00 | 4,41 | 0,00 | 1,00E+00 |
| AT5G66400 | 0,42 | -2,21 | 4,19 | -0,53 | 5,97E-01 |
| AT5G66440 | 0,33 | 0,00 | 4,41 | 0,00 | 1,00E+00 |
| AT5G66460 | 0,57 | 1,72 | 4,38 | 0,39 | 6,94E-01 |
| AT5G66500 | 0,15 | 0,00 | 4,41 | 0,00 | 1,00E+00 |
| AT5G66530 | 0,97 | 0,40 | 2,37 | 0,17 | 8,65E-01 |
| AT5G66540 | 0,21 | 0,00 | 4,41 | 0,00 | 1,00E+00 |
| AT5G66558 | 1,38 | 2,81 | 1,79 | 1,57 | 1,17E-01 |
| AT5G66595 | 0,74 | 0,50 | 2,52 | 0,20 | 8,42E-01 |
| AT5G66607 | 0,08 | 0,00 | 4,41 | 0,00 | 1,00E+00 |
| AT5G66650 | 0,26 | 1,38 | 4,41 | 0,31 | 7,54E-01 |
| AT5G66675 | 0,51 | 1,38 | 3,21 | 0,43 | 6,67E-01 |
| AT5G66680 | 0,76 | 2,48 | 3,72 | 0,67 | 5,05E-01 |
| AT5G66720 | 0,41 | 2,78 | 4,34 | 0,64 | 5,21E-01 |
| AT5G66730 | 0,31 | -0,41 | 4,38 | -0,09 | 9,26E-01 |
| AT5G66850 | 0,24 | -0,45 | 4,39 | -0,10 | 9,18E-01 |
| AT5G66890 | 0,75 | 1,67 | 3,53 | 0,47 | 6,36E-01 |
| AT5G66900 | 1,52 | 1,78 | 1,95 | 0,91 | 3,61E-01 |
| AT5G66910 | 0,38 | 1,38 | 3,49 | 0,39 | 6,94E-01 |
| AT5G67120 | 0,79 | 0,00 | 2,88 | 0,00 | 1,00E+00 |
| AT5G67150 | 0,35 | 1,32 | 4,40 | 0,30 | 7,63E-01 |
| AT5G67190 | 0,91 | 0,80 | 2,56 | 0,31 | 7,54E-01 |
| AT5G67220 | 0,60 | 0,38 | 2,60 | 0,14 | 8,85E-01 |
| AT5G67240 | 1,06 | -0,24 | 1,99 | -0,12 | 9,06E-01 |
| AT5G67245 | 0,45 | 1,38 | 4,41 | 0,31 | 7,54E-01 |
| AT5G67250 | 1,03 | 1,38 | 2,27 | 0,61 | 5,43E-01 |
| AT5G67280 | 0,58 | 0,33 | 2,70 | 0,12 | 9,02E-01 |
| AT5G67290 | 0,82 | 2,20 | 2,54 | 0,86 | 3,87E-01 |
| AT5G09855 | 0,69 | 2,30 | 2,52 | 0,91 | 3,61E-01 |
| AT5G67310 | 0,78 | 1,28 | 2,88 | 0,44 | 6,57E-01 |
| AT5G67350 | 0,75 | -0,70 | 2,69 | -0,26 | 7,95E-01 |
| AT5G67380 | 0,30 | 1,27 | 4,39 | 0,29 | 7,73E-01 |
| AT5G67460 | 0,10 | 0,00 | 4,41 | 0,00 | 1,00E+00 |
| AT5G67480 | 1,04 | 2,49 | 2,60 | 0,96 | 3,36E-01 |
| AT5G67490 | 1,16 | 0,32 | 2,02 | 0,16 | 8,73E-01 |
| AT5G67580 | 0,39 | -1,28 | 2,91 | -0,44 | 6,60E-01 |
| AT5G67610 | 0,37 | 2,72 | 4,34 | 0,63 | 5,31E-01 |

**vild type infected by TCV relative to uninfected wild type.**

| |
|---|
| type |
| protein_coding |
| miRNA_precursor |
| miRNA_precursor |
| protein_coding |
| miRNA_precursor |
| miRNA_precursor |
| protein_coding |
| protein_coding |
| protein_coding |
| ncRNA |
| protein_coding |
| miRNA_precursor |
| miRNA_precursor |
| protein_coding |
| miRNA_precursor |
| miRNA_precursor |
| protein_coding |
| protein_coding |
| ncRNA |
| ncRNA |
| miRNA_precursor |
| miRNA_precursor |
| protein_coding |
| miRNA_precursor |
| miRNA_precursor |
| protein_coding |
| protein_coding |
| miRNA_precursor |
| miRNA_precursor |
| protein_coding |
| protein_coding |
| ncRNA |
| miRNA_precursor |
| ncRNA |
| protein_coding |
| protein_coding |
| miRNA_precursor |
| protein_coding |
| protein_coding |
| protein_coding |
| miRNA_precursor |
| protein_coding |
| protein_coding |
| protein_coding |
| ribosomal_rna |
| protein_coding |
| ncRNA |
| ncRNA |
| miRNA_precursor |
| protein_coding |
| miRNA_precursor |
| protein_coding |
| protein_coding |

small_nucle(ol)ar_rna
miRNA_precursor
ncRNA
small_nucle(ol)ar_rna
protein_coding
protein_coding
protein_coding
protein_coding
small_nucle(ol)ar_rna
protein_coding
protein_coding
protein_coding
protein_coding
small_nucle(ol)ar_rna
ncRNA
ncRNA
protein_coding
protein_coding
miRNA_precursor
miRNA_precursor
miRNA_precursor
ncRNA
protein_coding
miRNA_precursor
protein_coding
protein_coding
ncRNA
protein_coding
protein_coding
protein_coding
small_nucle(ol)ar_rna
protein_coding
protein_coding
protein_coding
protein_coding
protein_coding
small_nucle(ol)ar_rna
protein_coding
protein_coding
protein_coding
miRNA_precursor
protein_coding
protein_coding
protein_coding
protein_coding
protein_coding
ncRNA
ncRNA
miRNA_precursor
protein_coding
protein_coding
protein_coding
miRNA_precursor
ncRNA
protein_coding
protein_coding

miRNA_precursor
protein_coding
protein_coding
protein_coding
protein_coding
protein_coding
protein_coding
small_nucle(ol)ar_rna
protein_coding
protein_coding
miRNA_precursor
protein_coding
protein_coding
protein_coding
protein_coding
miRNA_precursor
miRNA_precursor
miRNA_precursor
protein_coding
protein_coding
protein_coding
protein_coding
protein_coding
miRNA_precursor
protein_coding
protein_coding
protein_coding
protein_coding
ncRNA
protein_coding
protein_coding
miRNA_precursor
protein_coding
protein_coding
protein_coding
protein_coding
protein_coding
protein_coding
protein_coding
small_nucle(ol)ar_rna
protein_coding
protein_coding
protein_coding
protein_coding
protein_coding
protein_coding
protein_coding
protein_coding
protein_coding
protein_coding
protein_coding
protein_coding
protein_coding
protein_coding
ribosomal_rna
miRNA_precursor

protein_coding
ncRNA
protein_coding
protein_coding
protein_coding
protein_coding
miRNA_precursor
small_nucle(ol)ar_rna
ncRNA
protein_coding
ncRNA
protein_coding
protein_coding
protein_coding
protein_coding
protein_coding
protein_coding
protein_coding
protein_coding
protein_coding
protein_coding
protein_coding
protein_coding
protein_coding
protein_coding
protein_coding
protein_coding
protein_coding
miRNA_precursor
protein_coding
ncRNA
protein_coding
protein_coding
protein_coding
protein_coding
miRNA_precursor
miRNA_precursor
protein_coding
protein_coding
protein_coding
protein_coding
protein_coding
miRNA_precursor
protein_coding
protein_coding
protein_coding
protein_coding
miRNA_precursor
protein_coding
miRNA_precursor
protein_coding
protein_coding
protein_coding
protein_coding
protein_coding
protein_coding

protein_coding
protein_coding
protein_coding
miRNA_precursor
protein_coding
protein_coding
small_nucle(ol)ar_rna
protein_coding
miRNA_precursor
protein_coding
protein_coding
protein_coding
protein_coding
protein_coding
protein_coding
protein_coding
protein_coding
protein_coding
protein_coding
protein_coding
protein_coding
ncRNA
protein_coding
protein_coding
protein_coding
protein_coding
small_nucle(ol)ar_rna
protein_coding
protein_coding
protein_coding
miRNA_precursor
protein_coding
protein_coding
miRNA_precursor
protein_coding
protein_coding
protein_coding
protein_coding
protein_coding
ncRNA
protein_coding
protein_coding
protein_coding
protein_coding
protein_coding
protein_coding
protein_coding
protein_coding
protein_coding
protein_coding
protein_coding
protein_coding
protein_coding
protein_coding
protein_coding
protein_coding

protein_coding
protein_coding
protein_coding
protein_coding
miRNA_precursor
protein_coding
protein_coding
protein_coding
protein_coding
protein_coding
small_nucle(ol)ar_rna
small_nucle(ol)ar_rna
protein_coding
protein_coding
protein_coding
protein_coding
protein_coding
miRNA_precursor
protein_coding
protein_coding
miRNA_precursor
protein_coding
protein_coding
protein_coding
protein_coding
protein_coding
protein_coding
protein_coding
miRNA_precursor
protein_coding
protein_coding
protein_coding
small_nucle(ol)ar_rna
ncRNA
protein_coding
protein_coding
protein_coding
protein_coding
protein_coding
protein_coding
protein_coding
protein_coding
ncRNA
protein_coding
protein_coding
protein_coding
protein_coding
protein_coding
protein_coding
protein_coding
protein_coding
protein_coding
protein_coding
protein_coding
protein_coding
protein_coding

protein_coding
protein_coding
protein_coding
protein_coding
protein_coding
protein_coding
protein_coding
protein_coding
protein_coding
protein_coding
protein_coding
protein_coding
protein_coding
protein_coding
protein_coding
protein_coding
protein_coding
protein_coding
protein_coding
miRNA_precursor
protein_coding
protein_coding
protein_coding
protein_coding
protein_coding
protein_coding
protein_coding
miRNA_precursor
protein_coding
protein_coding
small_nucle(ol)ar_rna
miRNA_precursor
protein_coding
protein_coding
protein_coding
protein_coding
protein_coding
protein_coding
protein_coding
protein_coding
protein_coding
pre_trna
protein_coding
protein_coding
ncRNA
protein_coding
protein_coding
protein_coding
protein_coding
protein_coding
protein_coding
protein_coding
small_nucle(ol)ar_rna
protein_coding
protein_coding
protein_coding

protein_coding
protein_coding
protein_coding
protein_coding
protein_coding
protein_coding
protein_coding
protein_coding
protein_coding
protein_coding
protein_coding
protein_coding
protein_coding
protein_coding
protein_coding
protein_coding
protein_coding
protein_coding
protein_coding
protein_coding
protein_coding
protein_coding
protein_coding
protein_coding
protein_coding
miRNA_precursor
miRNA_precursor
protein_coding
small_nucle(ol)ar_rna
protein_coding
protein_coding
protein_coding
protein_coding
protein_coding
protein_coding
protein_coding
protein_coding
ncRNA
protein_coding
protein_coding
protein_coding
protein_coding
protein_coding
protein_coding
protein_coding
protein_coding
protein_coding
protein_coding
protein_coding
protein_coding
protein_coding
protein_coding
protein_coding
protein_coding
protein_coding
protein_coding

miRNA_precursor
ncRNA
protein_coding
protein_coding
protein_coding
protein_coding
protein_coding
miRNA_precursor
miRNA_precursor
protein_coding
protein_coding
protein_coding
protein_coding
protein_coding
protein_coding
protein_coding
protein_coding
small_nucle(ol)ar_rna
protein_coding
protein_coding
protein_coding
protein_coding
protein_coding
protein_coding
protein_coding
protein_coding
protein_coding
protein_coding
protein_coding
protein_coding
protein_coding
protein_coding
protein_coding
protein_coding
protein_coding
protein_coding
protein_coding
protein_coding
protein_coding
protein_coding
miRNA_precursor
protein_coding
protein_coding
protein_coding
protein_coding
protein_coding
protein_coding
protein_coding
protein_coding
protein_coding
protein_coding
protein_coding
miRNA_precursor
protein_coding
ncRNA

protein_coding
ncRNA
protein_coding
protein_coding
protein_coding
protein_coding
protein_coding
protein_coding
protein_coding
protein_coding
protein_coding
protein_coding
protein_coding
protein_coding
protein_coding
protein_coding
ncRNA
protein_coding
protein_coding
protein_coding
protein_coding
protein_coding
small_nucle(ol)ar_rna
protein_coding
protein_coding
protein_coding
protein_coding
protein_coding
protein_coding
ncRNA
miRNA_precursor
protein_coding
protein_coding
protein_coding
protein_coding
protein_coding
protein_coding
protein_coding
protein_coding
protein_coding
pre_trna
protein_coding
protein_coding
protein_coding
protein_coding
protein_coding
protein_coding
miRNA_precursor
protein_coding
protein_coding
protein_coding
protein_coding
protein_coding
protein_coding
protein_coding
protein_coding

protein_coding
protein_coding
protein_coding
protein_coding
ncRNA
protein_coding
ncRNA
protein_coding
protein_coding
protein_coding
protein_coding
protein_coding
small_nucle(ol)ar_rna
miRNA_precursor
protein_coding
miRNA_precursor
protein_coding
protein_coding
protein_coding
protein_coding
protein_coding
protein_coding
protein_coding
protein_coding
protein_coding
protein_coding
protein_coding
protein_coding
protein_coding
protein_coding
protein_coding
protein_coding
protein_coding
protein_coding
protein_coding
protein_coding
protein_coding
miRNA_precursor
miRNA_precursor
protein_coding
protein_coding
miRNA_precursor
protein_coding
protein_coding
protein_coding
protein_coding
protein_coding
protein_coding
ncRNA
protein_coding
protein_coding
protein_coding
protein_coding
miRNA_precursor
protein_coding
miRNA_precursor

small_nucle(ol)ar_rna
protein_coding
protein_coding
ncRNA
protein_coding
protein_coding
small_nucle(ol)ar_rna
small_nucle(ol)ar_rna
protein_coding
protein_coding
protein_coding
protein_coding
protein_coding
miRNA_precursor
protein_coding
protein_coding
protein_coding
miRNA_precursor
protein_coding
protein_coding
protein_coding
protein_coding
protein_coding
protein_coding
protein_coding
protein_coding
ncRNA
protein_coding
protein_coding
pre_trna
protein_coding
protein_coding
protein_coding
small_nucle(ol)ar_rna
protein_coding
protein_coding
protein_coding
protein_coding
protein_coding
protein_coding
ribosomal_rna
protein_coding
protein_coding
ncRNA
small_nucle(ol)ar_rna
protein_coding
miRNA_precursor
protein_coding
miRNA_precursor
protein_coding
protein_coding
protein_coding
small_nucle(ol)ar_rna
protein_coding
protein_coding
miRNA_precursor

ncRNA
protein_coding
protein_coding
protein_coding
ncRNA
protein_coding
protein_coding
protein_coding
protein_coding
protein_coding
protein_coding
protein_coding
miRNA_precursor
protein_coding
protein_coding
protein_coding
protein_coding
protein_coding
protein_coding
protein_coding
protein_coding
protein_coding
protein_coding
protein_coding
protein_coding
protein_coding
protein_coding
protein_coding
protein_coding
protein_coding
protein_coding
protein_coding
protein_coding
protein_coding
protein_coding
small_nucle(ol)ar_rna
protein_coding
protein_coding
protein_coding
protein_coding
protein_coding
protein_coding
protein_coding
protein_coding
protein_coding
protein_coding
protein_coding
protein_coding
protein_coding
protein_coding
protein_coding
protein_coding
protein_coding
protein_coding
protein_coding
protein_coding
protein_coding

protein_coding
protein_coding
protein_coding
protein_coding
protein_coding
protein_coding
miRNA_precursor
protein_coding
protein_coding
protein_coding
protein_coding
miRNA_precursor
protein_coding
protein_coding
protein_coding
protein_coding
small_nucle(ol)ar_rna
protein_coding
protein_coding
protein_coding
protein_coding
protein_coding
protein_coding
protein_coding
protein_coding
protein_coding
protein_coding
protein_coding
protein_coding
ncRNA
protein_coding
protein_coding
protein_coding
protein_coding
protein_coding
protein_coding
protein_coding
protein_coding
protein_coding
protein_coding
protein_coding
protein_coding
protein_coding
protein_coding
protein_coding
protein_coding
protein_coding
protein_coding
small_nucle(ol)ar_rna
protein_coding
protein_coding
protein_coding
protein_coding
protein_coding
protein_coding
protein_coding

protein_coding
protein_coding
protein_coding
protein_coding
protein_coding
protein_coding
protein_coding
protein_coding
protein_coding
protein_coding
protein_coding
miRNA_precursor
protein_coding
protein_coding
protein_coding
miRNA_precursor
protein_coding
protein_coding
protein_coding
protein_coding
protein_coding
protein_coding
protein_coding
ncRNA
protein_coding
protein_coding
protein_coding
ncRNA
protein_coding
small_nucle(ol)ar_rna
protein_coding
protein_coding
protein_coding
protein_coding
protein_coding
protein_coding
protein_coding
protein_coding
ncRNA
protein_coding
protein_coding
protein_coding
protein_coding
protein_coding
protein_coding
protein_coding
protein_coding
protein_coding
protein_coding
protein_coding
protein_coding
protein_coding
protein_coding
protein_coding
ncRNA
protein_coding

protein_coding
protein_coding
protein_coding
protein_coding
protein_coding
protein_coding
protein_coding
protein_coding
protein_coding
protein_coding
protein_coding
protein_coding
protein_coding
protein_coding
protein_coding
protein_coding
protein_coding
protein_coding
small_nucle(ol)ar_rna
small_nucle(ol)ar_rna
protein_coding
protein_coding
ncRNA
protein_coding
protein_coding
protein_coding
protein_coding
protein_coding
protein_coding
protein_coding
protein_coding
protein_coding
protein_coding
protein_coding
protein_coding
protein_coding
protein_coding
small_nucle(ol)ar_rna
ncRNA
protein_coding
protein_coding
small_nucle(ol)ar_rna
protein_coding
protein_coding
protein_coding
protein_coding
protein_coding
protein_coding
protein_coding
protein_coding
protein_coding
protein_coding
protein_coding
protein_coding
protein_coding
protein_coding

protein_coding
protein_coding
protein_coding
miRNA_precursor
ncRNA
miRNA_precursor
protein_coding
protein_coding
miRNA_precursor
protein_coding
protein_coding
protein_coding
ncRNA
protein_coding
protein_coding
protein_coding
protein_coding
small_nucle(ol)ar_rna
protein_coding
protein_coding
protein_coding
protein_coding
protein_coding
small_nucle(ol)ar_rna
ncRNA
protein_coding
protein_coding
miRNA_precursor
protein_coding
protein_coding
protein_coding
protein_coding
protein_coding
protein_coding
protein_coding
protein_coding
protein_coding
protein_coding
protein_coding
miRNA_precursor
protein_coding
small_nucle(ol)ar_rna
protein_coding
protein_coding
protein_coding
protein_coding
protein_coding
protein_coding
protein_coding
protein_coding
protein_coding
protein_coding
protein_coding
ncRNA
ncRNA
protein_coding

small_nucle(ol)ar_rna
protein_coding
protein_coding
ncRNA
protein_coding
protein_coding
protein_coding
protein_coding
protein_coding
protein_coding
protein_coding
protein_coding
protein_coding
protein_coding
protein_coding
protein_coding
protein_coding
protein_coding
small_nucle(ol)ar_rna
protein_coding
protein_coding
small_nucle(ol)ar_rna
protein_coding
protein_coding
protein_coding
protein_coding
protein_coding
protein_coding
miRNA_precursor
protein_coding
protein_coding
protein_coding
miRNA_precursor
protein_coding
protein_coding
ncRNA
protein_coding
protein_coding
protein_coding
miRNA_precursor
protein_coding
protein_coding
protein_coding
protein_coding
protein_coding
protein_coding
protein_coding
protein_coding
protein_coding
protein_coding
protein_coding
protein_coding
protein_coding
small_nucle(ol)ar_rna
ncRNA
protein_coding

small_nucle(ol)ar_rna
protein_coding
protein_coding
protein_coding
protein_coding
protein_coding
protein_coding
protein_coding
protein_coding
protein_coding
protein_coding
protein_coding
protein_coding
protein_coding
protein_coding
protein_coding
protein_coding
protein_coding
protein_coding
protein_coding
protein_coding
small_nucle(ol)ar_rna
protein_coding
miRNA_precursor
protein_coding
protein_coding
protein_coding
protein_coding
protein_coding
protein_coding
protein_coding
protein_coding
protein_coding
protein_coding
small_nucle(ol)ar_rna
protein_coding
miRNA_precursor
protein_coding
protein_coding
protein_coding
protein_coding
miRNA_precursor
protein_coding
protein_coding
small_nucle(ol)ar_rna
protein_coding
protein_coding
protein_coding
protein_coding
protein_coding
protein_coding
protein_coding
protein_coding
ncRNA
protein_coding
protein_coding

protein_coding
protein_coding
protein_coding
miRNA_precursor
protein_coding
protein_coding
protein_coding
protein_coding
protein_coding
ncRNA
protein_coding
protein_coding
protein_coding
protein_coding
protein_coding
protein_coding
protein_coding
protein_coding
small_nucle(ol)ar_rna
protein_coding
protein_coding
protein_coding
small_nucle(ol)ar_rna
protein_coding
protein_coding
protein_coding
protein_coding
protein_coding
ncRNA
protein_coding
protein_coding
protein_coding
protein_coding
protein_coding
protein_coding
protein_coding
ncRNA
protein_coding
protein_coding
protein_coding
small_nucle(ol)ar_rna
protein_coding
small_nucle(ol)ar_rna
protein_coding
protein_coding
protein_coding
miRNA_precursor
protein_coding
ncRNA
protein_coding
protein_coding
protein_coding
protein_coding
small_nucle(ol)ar_rna
protein_coding
ncRNA

protein_coding
protein_coding
protein_coding
protein_coding
protein_coding
protein_coding
small_nucle(ol)ar_rna
protein_coding
protein_coding
small_nucle(ol)ar_rna
protein_coding
protein_coding
small_nucle(ol)ar_rna
protein_coding
small_nucle(ol)ar_rna
protein_coding
ncRNA
protein_coding
protein_coding
protein_coding
protein_coding
protein_coding
protein_coding
protein_coding
protein_coding
protein_coding
protein_coding
protein_coding
protein_coding
small_nucle(ol)ar_rna
protein_coding
protein_coding
protein_coding
protein_coding
protein_coding
protein_coding
protein_coding
protein_coding
protein_coding
protein_coding
protein_coding
protein_coding
miRNA_precursor
protein_coding
protein_coding
protein_coding
protein_coding
protein_coding
protein_coding
ncRNA
protein_coding
protein_coding
protein_coding
protein_coding
protein_coding
protein_coding

protein_coding
protein_coding
protein_coding
protein_coding
protein_coding
protein_coding
protein_coding
protein_coding
protein_coding
protein_coding
protein_coding
protein_coding
protein_coding
protein_coding
protein_coding
protein_coding
protein_coding
protein_coding
protein_coding
protein_coding
protein_coding
protein_coding
protein_coding
protein_coding
protein_coding
protein_coding
miRNA_precursor
protein_coding
protein_coding
protein_coding
protein_coding
protein_coding
protein_coding
protein_coding
miRNA_precursor
miRNA_precursor
protein_coding
protein_coding
miRNA_precursor
protein_coding
protein_coding
protein_coding
protein_coding
protein_coding
protein_coding
protein_coding
protein_coding
protein_coding
protein_coding
protein_coding
protein_coding
protein_coding
protein_coding
miRNA_precursor
protein_coding
protein_coding

small_nucle(ol)ar_rna
protein_coding
protein_coding
protein_coding
protein_coding
protein_coding
protein_coding
protein_coding
small_nucle(ol)ar_rna
protein_coding
protein_coding
protein_coding
protein_coding
protein_coding
protein_coding
protein_coding
protein_coding
protein_coding
protein_coding
protein_coding
ncRNA
protein_coding
protein_coding
protein_coding
small_nucle(ol)ar_rna
protein_coding
protein_coding
protein_coding
protein_coding
protein_coding
small_nucle(ol)ar_rna
protein_coding
protein_coding
protein_coding
protein_coding
protein_coding
protein_coding
protein_coding
protein_coding
ncRNA
protein_coding
protein_coding
protein_coding
protein_coding
miRNA_precursor
protein_coding
protein_coding
miRNA_precursor
miRNA_precursor
protein_coding
protein_coding
protein_coding
small_nucle(ol)ar_rna
protein_coding
protein_coding
protein_coding

protein_coding
protein_coding
protein_coding
protein_coding
protein_coding
protein_coding
protein_coding
protein_coding
protein_coding
protein_coding
protein_coding
protein_coding
protein_coding
protein_coding
small_nucle(ol)ar_rna
protein_coding
protein_coding
protein_coding
small_nucle(ol)ar_rna
miRNA_precursor
protein_coding
protein_coding
protein_coding
protein_coding
protein_coding
protein_coding
protein_coding
protein_coding
protein_coding
protein_coding
protein_coding
protein_coding
protein_coding
protein_coding
protein_coding
protein_coding
protein_coding
protein_coding
protein_coding
protein_coding
protein_coding
protein_coding
protein_coding
ncRNA
protein_coding
protein_coding
protein_coding
protein_coding
protein_coding
small_nucle(ol)ar_rna
protein_coding
protein_coding
small_nucle(ol)ar_rna
protein_coding
protein_coding
protein_coding

protein_coding
protein_coding
protein_coding
protein_coding
protein_coding
protein_coding
protein_coding
protein_coding
protein_coding
protein_coding
protein_coding
protein_coding
protein_coding
protein_coding
protein_coding
ncRNA
protein_coding
protein_coding
miRNA_precursor
protein_coding
protein_coding
ncRNA
small_nucle(ol)ar_rna
ncRNA
protein_coding
protein_coding
protein_coding
small_nucle(ol)ar_rna
protein_coding
small_nucle(ol)ar_rna
protein_coding
protein_coding
protein_coding
protein_coding
protein_coding
protein_coding
protein_coding
protein_coding
protein_coding
small_nucle(ol)ar_rna
protein_coding
protein_coding
protein_coding
protein_coding
ncRNA
small_nucle(ol)ar_rna
protein_coding
protein_coding
protein_coding
protein_coding
protein_coding
protein_coding
protein_coding
protein_coding
protein_coding
protein_coding

protein_coding
protein_coding
protein_coding
protein_coding
protein_coding
protein_coding
protein_coding
ncRNA
small_nucle(ol)ar_rna
ncRNA
protein_coding
miRNA_precursor
protein_coding
miRNA_precursor
protein_coding
protein_coding
protein_coding
protein_coding
protein_coding
protein_coding
protein_coding
protein_coding
protein_coding
protein_coding
protein_coding
protein_coding
protein_coding
protein_coding
small_nucle(ol)ar_rna
protein_coding
protein_coding
protein_coding
protein_coding
protein_coding
small_nucle(ol)ar_rna
protein_coding
protein_coding
protein_coding
protein_coding
protein_coding
protein_coding
protein_coding
ncRNA
protein_coding
protein_coding
small_nucle(ol)ar_rna
protein_coding
protein_coding
protein_coding
protein_coding
miRNA_precursor
ncRNA
protein_coding
small_nucle(ol)ar_rna
protein_coding
protein_coding

ncRNA
protein_coding
protein_coding
protein_coding
protein_coding
miRNA_precursor
miRNA_precursor
protein_coding
protein_coding
protein_coding
protein_coding
protein_coding
protein_coding
protein_coding
protein_coding
protein_coding
protein_coding
protein_coding
protein_coding
protein_coding
protein_coding
protein_coding
protein_coding
protein_coding
protein_coding
protein_coding
ncRNA
small_nucle(ol)ar_rna
protein_coding
protein_coding
protein_coding
protein_coding
protein_coding
protein_coding
protein_coding
protein_coding
protein_coding
protein_coding
protein_coding
protein_coding
protein_coding
protein_coding
protein_coding
protein_coding
protein_coding
protein_coding
ncRNA
protein_coding
small_nucle(ol)ar_rna
protein_coding
pre_trna
small_nucle(ol)ar_rna
protein_coding
protein_coding
protein_coding
protein_coding

protein_coding
protein_coding
protein_coding
protein_coding
protein_coding
protein_coding
protein_coding
protein_coding
protein_coding
protein_coding
protein_coding
protein_coding
protein_coding
protein_coding
protein_coding
protein_coding
protein_coding
protein_coding
protein_coding
protein_coding
protein_coding
protein_coding
protein_coding
protein_coding
protein_coding
protein_coding
protein_coding
small_nucle(ol)ar_rna
protein_coding
protein_coding
protein_coding
protein_coding
protein_coding
small_nucle(ol)ar_rna
protein_coding
small_nucle(ol)ar_rna
protein_coding
protein_coding
protein_coding
protein_coding
protein_coding
miRNA_precursor
ncRNA
protein_coding
protein_coding
protein_coding
ncRNA
protein_coding
protein_coding
protein_coding
protein_coding
protein_coding
miRNA_precursor
ncRNA
protein_coding
protein_coding

small_nucle(ol)ar_rna
protein_coding
ncRNA
protein_coding
protein_coding
protein_coding
protein_coding
protein_coding
protein_coding
ncRNA
small_nucle(ol)ar_rna
small_nucle(ol)ar_rna
protein_coding
protein_coding
ncRNA
protein_coding
protein_coding
protein_coding
protein_coding
protein_coding
protein_coding
ncRNA
protein_coding
protein_coding
protein_coding
protein_coding
ncRNA
protein_coding
protein_coding
protein_coding
protein_coding
protein_coding
protein_coding
protein_coding
protein_coding
protein_coding
protein_coding
protein_coding
protein_coding
protein_coding
protein_coding
protein_coding
protein_coding
protein_coding
protein_coding
protein_coding
protein_coding
small_nucle(ol)ar_rna
protein_coding
small_nucle(ol)ar_rna
protein_coding
small_nucle(ol)ar_rna
protein_coding
protein_coding
protein_coding
protein_coding

protein_coding
protein_coding
protein_coding
protein_coding
ncRNA
protein_coding
protein_coding
protein_coding
protein_coding
protein_coding
protein_coding
protein_coding
protein_coding
protein_coding
small_nucle(ol)ar_rna
small_nucle(ol)ar_rna
protein_coding
protein_coding
protein_coding
ncRNA
protein_coding
protein_coding
protein_coding
protein_coding
protein_coding
miRNA_precursor
protein_coding
ncRNA
ncRNA
protein_coding
protein_coding
small_nucle(ol)ar_rna
protein_coding
protein_coding
protein_coding
protein_coding
protein_coding
protein_coding
protein_coding
protein_coding
protein_coding
protein_coding
protein_coding
protein_coding
ncRNA
protein_coding
protein_coding
protein_coding
protein_coding
protein_coding
ncRNA
protein_coding
protein_coding
protein_coding
protein_coding
protein_coding

protein_coding
small_nucle(ol)ar_rna
protein_coding
protein_coding
miRNA_precursor
protein_coding
protein_coding
small_nucle(ol)ar_rna
protein_coding
ncRNA
protein_coding
protein_coding
protein_coding
miRNA_precursor
ncRNA
protein_coding
protein_coding
protein_coding
ncRNA
protein_coding
protein_coding
protein_coding
ncRNA
protein_coding
miRNA_precursor
protein_coding
protein_coding
protein_coding
protein_coding
protein_coding
protein_coding
ncRNA
small_nucle(ol)ar_rna
protein_coding
protein_coding
protein_coding
protein_coding
protein_coding
small_nucle(ol)ar_rna
protein_coding
protein_coding
protein_coding
pre_trna
protein_coding
protein_coding
protein_coding
protein_coding
protein_coding
protein_coding
miRNA_precursor
protein_coding
protein_coding
protein_coding
ncRNA
protein_coding
protein_coding

protein_coding
protein_coding
protein_coding
ncRNA
protein_coding
protein_coding
protein_coding
protein_coding
protein_coding
protein_coding
protein_coding
protein_coding
protein_coding
protein_coding
protein_coding
protein_coding
protein_coding
protein_coding
protein_coding
protein_coding
protein_coding
protein_coding
protein_coding
protein_coding
miRNA_precursor
protein_coding
protein_coding
small_nucle(ol)ar_rna
miRNA_precursor
protein_coding
protein_coding
protein_coding
protein_coding
protein_coding
protein_coding
protein_coding
protein_coding
protein_coding
protein_coding
protein_coding
protein_coding
protein_coding
miRNA_precursor
protein_coding
protein_coding
protein_coding
protein_coding
protein_coding
protein_coding
protein_coding
protein_coding
protein_coding
protein_coding
protein_coding
protein_coding
protein_coding

protein_coding
protein_coding
protein_coding
protein_coding
protein_coding
protein_coding
protein_coding
protein_coding
protein_coding
protein_coding
protein_coding
protein_coding
protein_coding
protein_coding
protein_coding
protein_coding
protein_coding
protein_coding
ncRNA
protein_coding
miRNA_precursor
protein_coding
protein_coding
protein_coding
protein_coding
ncRNA
ncRNA
protein_coding
protein_coding
protein_coding
small_nucle(ol)ar_rna
protein_coding
protein_coding
protein_coding
protein_coding
protein_coding
protein_coding
ncRNA
protein_coding
protein_coding
protein_coding
miRNA_precursor
protein_coding
protein_coding
small_nucle(ol)ar_rna
protein_coding
protein_coding
small_nucle(ol)ar_rna
protein_coding
protein_coding
protein_coding
protein_coding
protein_coding
protein_coding
protein_coding
protein_coding

protein_coding
protein_coding
protein_coding
protein_coding
ncRNA
protein_coding
protein_coding
protein_coding
protein_coding
small_nucle(ol)ar_rna
protein_coding
protein_coding
protein_coding
protein_coding
protein_coding
protein_coding
protein_coding
protein_coding
protein_coding
protein_coding
small_nucle(ol)ar_rna
miRNA_precursor
protein_coding
ncRNA
protein_coding
protein_coding
protein_coding
protein_coding
ncRNA
ncRNA
protein_coding
protein_coding
protein_coding
protein_coding
protein_coding
ncRNA
protein_coding
protein_coding
protein_coding
miRNA_precursor
protein_coding
protein_coding
protein_coding
ncRNA
protein_coding
ncRNA
protein_coding
protein_coding
miRNA_precursor
protein_coding
protein_coding
protein_coding
protein_coding
protein_coding
small_nucle(ol)ar_rna
protein_coding

protein_coding
protein_coding
protein_coding
small_nucle(ol)ar_rna
protein_coding
protein_coding
protein_coding
protein_coding
protein_coding
miRNA_precursor
ncRNA
protein_coding
protein_coding
protein_coding
protein_coding
ncRNA
protein_coding
protein_coding
protein_coding
protein_coding
protein_coding
protein_coding
small_nucle(ol)ar_rna
protein_coding
protein_coding
small_nucle(ol)ar_rna
protein_coding
protein_coding
protein_coding
protein_coding
protein_coding
protein_coding
protein_coding
protein_coding
protein_coding
protein_coding
protein_coding
protein_coding
protein_coding
protein_coding
protein_coding
protein_coding
protein_coding
protein_coding
protein_coding
protein_coding
protein_coding
protein_coding
protein_coding
protein_coding
protein_coding
protein_coding
protein_coding
protein_coding
protein_coding
protein_coding
protein_coding

protein_coding
miRNA_precursor
protein_coding
protein_coding
protein_coding
protein_coding
small_nucle(ol)ar_rna
protein_coding
protein_coding
protein_coding
protein_coding
small_nucle(ol)ar_rna
protein_coding
protein_coding
protein_coding
protein_coding
protein_coding
small_nucle(ol)ar_rna
small_nucle(ol)ar_rna
protein_coding
protein_coding
protein_coding
protein_coding
protein_coding
small_nucle(ol)ar_rna
protein_coding
protein_coding
protein_coding
protein_coding
protein_coding
protein_coding
protein_coding
protein_coding
ncRNA
protein_coding
miRNA_precursor
protein_coding
protein_coding
miRNA_precursor
ncRNA
protein_coding
protein_coding
protein_coding
protein_coding
protein_coding
ncRNA
protein_coding
protein_coding
protein_coding
protein_coding
miRNA_precursor
ncRNA
protein_coding
ncRNA
protein_coding
small_nucle(ol)ar_rna

protein_coding
protein_coding
protein_coding
protein_coding
miRNA_precursor
protein_coding
protein_coding
protein_coding
small_nucle(ol)ar_rna
protein_coding
protein_coding
protein_coding
protein_coding
protein_coding
protein_coding
protein_coding
protein_coding
protein_coding
protein_coding
protein_coding
protein_coding
small_nucle(ol)ar_rna
protein_coding
protein_coding
protein_coding
protein_coding
protein_coding
protein_coding
miRNA_precursor
protein_coding
small_nucle(ol)ar_rna
ncRNA
protein_coding
protein_coding
protein_coding
protein_coding
protein_coding
protein_coding
protein_coding
protein_coding
protein_coding
miRNA_precursor
protein_coding
miRNA_precursor
protein_coding
protein_coding
small_nucle(ol)ar_rna
protein_coding
miRNA_precursor
ncRNA
protein_coding
protein_coding
protein_coding
protein_coding
protein_coding
protein_coding

protein_coding
protein_coding
protein_coding
protein_coding
protein_coding
protein_coding
protein_coding
protein_coding
protein_coding
small_nucle(ol)ar_rna
protein_coding
protein_coding
protein_coding
protein_coding
small_nucle(ol)ar_rna
protein_coding
protein_coding
small_nucle(ol)ar_rna
protein_coding
protein_coding
miRNA_precursor
protein_coding
protein_coding
protein_coding
protein_coding
miRNA_precursor
protein_coding
protein_coding
protein_coding
protein_coding
small_nucle(ol)ar_rna
protein_coding
protein_coding
protein_coding
protein_coding
small_nucle(ol)ar_rna
small_nucle(ol)ar_rna
protein_coding
protein_coding
protein_coding
protein_coding
protein_coding
protein_coding
protein_coding
protein_coding
protein_coding
protein_coding
protein_coding
protein_coding
protein_coding
small_nucle(ol)ar_rna
protein_coding
ribosomal_rna
miRNA_precursor
protein_coding
ncRNA

protein_coding
protein_coding
protein_coding
miRNA_precursor
protein_coding
protein_coding
protein_coding
protein_coding
protein_coding
protein_coding
protein_coding
small_nucle(ol)ar_rna
ncRNA
protein_coding
protein_coding
protein_coding
protein_coding
protein_coding
protein_coding
protein_coding
protein_coding
protein_coding
protein_coding
protein_coding
protein_coding
small_nucle(ol)ar_rna
protein_coding
protein_coding
protein_coding
protein_coding
protein_coding
protein_coding
protein_coding
ncRNA
protein_coding
protein_coding
protein_coding
protein_coding
ncRNA
protein_coding
protein_coding
protein_coding
small_nucle(ol)ar_rna
protein_coding
protein_coding
protein_coding
miRNA_precursor
protein_coding
protein_coding
protein_coding
protein_coding
protein_coding
protein_coding
protein_coding
protein_coding
protein_coding

protein_coding
protein_coding
protein_coding
protein_coding
protein_coding
miRNA_precursor
protein_coding
protein_coding
protein_coding
protein_coding
protein_coding
protein_coding
protein_coding
small_nucle(ol)ar_rna
protein_coding
miRNA_precursor
protein_coding
protein_coding
protein_coding
ncRNA
protein_coding
ncRNA
ncRNA
protein_coding
protein_coding
protein_coding
protein_coding
protein_coding
protein_coding
protein_coding
protein_coding
protein_coding
protein_coding
protein_coding
ncRNA
protein_coding
protein_coding
ncRNA
protein_coding
protein_coding
protein_coding
protein_coding
protein_coding
protein_coding
protein_coding
protein_coding
protein_coding
small_nucle(ol)ar_rna
protein_coding
miRNA_precursor
protein_coding
pre_trna
ncRNA
small_nucle(ol)ar_rna
protein_coding
pre_trna

protein_coding
protein_coding
protein_coding
ncRNA
small_nucle(ol)ar_rna
protein_coding
protein_coding
protein_coding
small_nucle(ol)ar_rna
protein_coding
protein_coding
small_nucle(ol)ar_rna
protein_coding
protein_coding
protein_coding
protein_coding
pre_trna
protein_coding
protein_coding
protein_coding
small_nucle(ol)ar_rna
protein_coding
ncRNA
protein_coding
protein_coding
protein_coding
protein_coding
protein_coding
protein_coding
protein_coding
protein_coding
protein_coding
protein_coding
protein_coding
miRNA_precursor
protein_coding
protein_coding
small_nucle(ol)ar_rna
protein_coding
protein_coding
ncRNA
protein_coding
protein_coding
protein_coding
protein_coding
protein_coding
miRNA_precursor
protein_coding
protein_coding
protein_coding
miRNA_precursor
small_nucle(ol)ar_rna
miRNA_precursor
protein_coding
protein_coding
protein_coding

protein_coding
ncRNA
protein_coding
protein_coding
ncRNA
ncRNA
protein_coding
protein_coding
miRNA_precursor
small_nucle(ol)ar_rna
protein_coding
protein_coding
protein_coding
protein_coding
protein_coding
miRNA_precursor
protein_coding
protein_coding
protein_coding
protein_coding
protein_coding
protein_coding
protein_coding
protein_coding
protein_coding
ncRNA
protein_coding
protein_coding
protein_coding
protein_coding
protein_coding
protein_coding
protein_coding
ncRNA
protein_coding
protein_coding
protein_coding
protein_coding
protein_coding
protein_coding
protein_coding
protein_coding
protein_coding
ncRNA
protein_coding
protein_coding
ncRNA
protein_coding
protein_coding
protein_coding
protein_coding
protein_coding
protein_coding
protein_coding
protein_coding
protein_coding

protein_coding
protein_coding
protein_coding
protein_coding
protein_coding
protein_coding
protein_coding
protein_coding
protein_coding
protein_coding
ncRNA
protein_coding
protein_coding
protein_coding
protein_coding
protein_coding
protein_coding
protein_coding
protein_coding
protein_coding
protein_coding
small_nucle(ol)ar_rna
small_nucle(ol)ar_rna
small_nucle(ol)ar_rna
small_nucle(ol)ar_rna
protein_coding
protein_coding
protein_coding
protein_coding
protein_coding
ncRNA
protein_coding
ncRNA
protein_coding
protein_coding
protein_coding
small_nucle(ol)ar_rna
small_nucle(ol)ar_rna
protein_coding
protein_coding
protein_coding
protein_coding
protein_coding
protein_coding
protein_coding
protein_coding
protein_coding
protein_coding
protein_coding
protein_coding
protein_coding
protein_coding
ncRNA
protein_coding
miRNA_precursor
protein_coding

protein_coding
protein_coding
protein_coding
protein_coding
pre_trna
ncRNA
protein_coding
protein_coding
protein_coding
protein_coding
protein_coding
protein_coding
protein_coding
protein_coding
protein_coding
protein_coding
protein_coding
protein_coding
protein_coding
protein_coding
protein_coding
ncRNA
protein_coding
protein_coding
protein_coding
protein_coding
protein_coding
protein_coding
protein_coding
protein_coding
protein_coding
protein_coding
protein_coding
ncRNA
protein_coding
protein_coding
protein_coding
protein_coding
protein_coding
protein_coding
protein_coding
protein_coding
protein_coding
protein_coding
protein_coding
protein_coding
protein_coding
protein_coding
protein_coding
protein_coding
protein_coding
protein_coding
protein_coding
protein_coding
protein_coding
protein_coding

protein_coding
protein_coding
protein_coding
protein_coding
protein_coding
protein_coding
protein_coding
protein_coding
protein_coding
protein_coding
protein_coding
protein_coding
protein_coding
protein_coding
protein_coding
protein_coding
ncRNA
protein_coding
protein_coding
protein_coding
protein_coding
protein_coding
protein_coding
protein_coding
protein_coding
protein_coding
protein_coding
protein_coding
protein_coding
protein_coding
protein_coding
protein_coding
protein_coding
protein_coding
protein_coding
protein_coding
protein_coding
protein_coding
protein_coding
protein_coding
protein_coding
protein_coding
protein_coding
protein_coding
protein_coding
protein_coding
protein_coding
protein_coding
protein_coding
protein_coding
protein_coding
protein_coding
protein_coding
protein_coding
protein_coding
protein_coding

protein_coding
protein_coding
protein_coding
protein_coding
protein_coding
protein_coding
protein_coding
protein_coding
protein_coding
protein_coding
protein_coding
protein_coding
protein_coding
protein_coding
protein_coding
protein_coding
protein_coding
small_nucle(ol)ar_rna
protein_coding
protein_coding
protein_coding
protein_coding
protein_coding
protein_coding
protein_coding
protein_coding
protein_coding
protein_coding
protein_coding
protein_coding
protein_coding
protein_coding
protein_coding
protein_coding
protein_coding
protein_coding
protein_coding
protein_coding
protein_coding
protein_coding
protein_coding
protein_coding
protein_coding
protein_coding
protein_coding
protein_coding
protein_coding
protein_coding
protein_coding
protein_coding
pre_trna
protein_coding
protein_coding
protein_coding
protein_coding
ncRNA

protein_coding
protein_coding
protein_coding
protein_coding
protein_coding
protein_coding
protein_coding
protein_coding
protein_coding
protein_coding
protein_coding
protein_coding
protein_coding
protein_coding
protein_coding
protein_coding
protein_coding
protein_coding
protein_coding
protein_coding
protein_coding
protein_coding
protein_coding
protein_coding
pre_trna
ncRNA
protein_coding
protein_coding
protein_coding
protein_coding
protein_coding
protein_coding
protein_coding
protein_coding
protein_coding
protein_coding
protein_coding
protein_coding
protein_coding
protein_coding
protein_coding
protein_coding
protein_coding
protein_coding
protein_coding
protein_coding
protein_coding
protein_coding
protein_coding
protein_coding
protein_coding
protein_coding
protein_coding
protein_coding
protein_coding
protein_coding

protein_coding
protein_coding
protein_coding
protein_coding
protein_coding
protein_coding
protein_coding
protein_coding
ncRNA
protein_coding
protein_coding
protein_coding
protein_coding
protein_coding
protein_coding
protein_coding
protein_coding
protein_coding
protein_coding
pre_trna
protein_coding
protein_coding
protein_coding
protein_coding
protein_coding
protein_coding
protein_coding
protein_coding
protein_coding
protein_coding
protein_coding
protein_coding
protein_coding
protein_coding
protein_coding
protein_coding
protein_coding
protein_coding
protein_coding
protein_coding
protein_coding
protein_coding
protein_coding
protein_coding
protein_coding
protein_coding
pre_trna
protein_coding
miRNA_precursor
protein_coding
protein_coding
protein_coding
protein_coding
protein_coding
protein_coding
protein_coding

protein_coding
protein_coding
protein_coding
protein_coding
small_nucle(ol)ar_rna
miRNA_precursor
protein_coding
ncRNA
protein_coding
protein_coding
protein_coding
protein_coding
protein_coding
protein_coding
protein_coding
protein_coding
ncRNA
protein_coding
protein_coding
protein_coding
protein_coding
protein_coding
miRNA_precursor
protein_coding
protein_coding
protein_coding
protein_coding
protein_coding
protein_coding
protein_coding
protein_coding
protein_coding
protein_coding
protein_coding
protein_coding
protein_coding
protein_coding
protein_coding
protein_coding
protein_coding
protein_coding
protein_coding
protein_coding
protein_coding
protein_coding
protein_coding
protein_coding
protein_coding
protein_coding
protein_coding
protein_coding
protein_coding
protein_coding
protein_coding
protein_coding
protein_coding

protein_coding
protein_coding
protein_coding
protein_coding
protein_coding
protein_coding
protein_coding
protein_coding
protein_coding
protein_coding
protein_coding
protein_coding
ncRNA
protein_coding
protein_coding
protein_coding
protein_coding
protein_coding
protein_coding
protein_coding
protein_coding
protein_coding
protein_coding
protein_coding
protein_coding
protein_coding
protein_coding
protein_coding
protein_coding
protein_coding
ncRNA
protein_coding
protein_coding
protein_coding
protein_coding
protein_coding
protein_coding
protein_coding
protein_coding
protein_coding
protein_coding
protein_coding
protein_coding
protein_coding
protein_coding
protein_coding
protein_coding
protein_coding
protein_coding
protein_coding
protein_coding
protein_coding
protein_coding
protein_coding
protein_coding

protein_coding
protein_coding
protein_coding
protein_coding
protein_coding
protein_coding
protein_coding
protein_coding
protein_coding
protein_coding
protein_coding
protein_coding
protein_coding
protein_coding
protein_coding
protein_coding
protein_coding
protein_coding
protein_coding
protein_coding
protein_coding
protein_coding
protein_coding
small_nucle(ol)ar_rna
protein_coding
protein_coding
protein_coding
protein_coding
ncRNA
protein_coding
protein_coding
protein_coding
protein_coding
protein_coding
protein_coding
protein_coding
protein_coding
protein_coding
protein_coding
protein_coding
protein_coding
protein_coding
protein_coding
ncRNA
protein_coding
protein_coding
protein_coding
pre_trna
protein_coding
protein_coding
protein_coding
protein_coding
protein_coding
protein_coding
protein_coding
protein_coding

small_nucle(ol)ar_rna
small_nucle(ol)ar_rna
small_nucle(ol)ar_rna
small_nucle(ol)ar_rna
protein_coding
protein_coding
protein_coding
protein_coding
protein_coding
protein_coding
protein_coding
protein_coding
protein_coding
protein_coding
protein_coding
protein_coding
pre_trna
protein_coding
protein_coding
protein_coding
protein_coding
protein_coding
protein_coding
protein_coding
protein_coding
protein_coding
protein_coding
protein_coding
ncRNA
protein_coding
protein_coding
protein_coding
protein_coding
protein_coding
protein_coding
protein_coding
protein_coding
ncRNA
protein_coding
protein_coding
protein_coding
protein_coding
protein_coding
protein_coding
protein_coding
protein_coding
protein_coding
protein_coding
protein_coding
protein_coding
protein_coding
small_nucle(ol)ar_rna
small_nucle(ol)ar_rna
protein_coding
protein_coding
protein_coding

miRNA_precursor
protein_coding
protein_coding
miRNA_precursor
protein_coding
protein_coding
protein_coding
protein_coding
protein_coding
protein_coding
protein_coding
protein_coding
protein_coding
protein_coding
protein_coding
protein_coding
protein_coding
protein_coding
protein_coding
protein_coding
protein_coding
protein_coding
protein_coding
protein_coding
protein_coding
protein_coding
protein_coding
protein_coding
ncRNA
protein_coding
protein_coding
protein_coding
protein_coding
protein_coding
protein_coding
protein_coding
protein_coding
protein_coding
ncRNA
protein_coding
protein_coding
protein_coding
protein_coding
protein_coding
protein_coding
protein_coding
protein_coding
ncRNA
protein_coding
protein_coding
protein_coding
protein_coding
ncRNA
protein_coding
protein_coding
protein_coding

protein_coding
protein_coding
protein_coding
protein_coding
protein_coding
protein_coding
protein_coding
protein_coding
protein_coding
protein_coding
protein_coding
protein_coding
protein_coding
ncRNA
protein_coding
protein_coding
ncRNA
protein_coding
protein_coding
ncRNA
protein_coding
protein_coding
protein_coding
protein_coding
protein_coding
ncRNA
protein_coding
protein_coding
protein_coding
miRNA_precursor
protein_coding
protein_coding
protein_coding
protein_coding
protein_coding
protein_coding
protein_coding
protein_coding
protein_coding
protein_coding
protein_coding
protein_coding
protein_coding
protein_coding
protein_coding
protein_coding
protein_coding
protein_coding
protein_coding
protein_coding
protein_coding
protein_coding
protein_coding
protein_coding
protein_coding
protein_coding

protein_coding
protein_coding
protein_coding
protein_coding
ncRNA
protein_coding
ncRNA
protein_coding
protein_coding
miRNA_precursor
protein_coding
protein_coding
protein_coding
protein_coding
protein_coding
protein_coding
protein_coding
protein_coding
protein_coding
protein_coding
protein_coding
protein_coding
protein_coding
protein_coding
protein_coding
protein_coding
ncRNA
protein_coding
protein_coding
small_nucle(ol)ar_rna
protein_coding
protein_coding
protein_coding
protein_coding
protein_coding
protein_coding
protein_coding
protein_coding
protein_coding
protein_coding
protein_coding
protein_coding
protein_coding
protein_coding
protein_coding
protein_coding
protein_coding
protein_coding
protein_coding
protein_coding
protein_coding
protein_coding
protein_coding
protein_coding
protein_coding
protein_coding

protein_coding
protein_coding
miRNA_precursor
protein_coding
protein_coding
protein_coding
protein_coding
protein_coding
protein_coding
protein_coding
protein_coding
protein_coding
protein_coding
protein_coding
protein_coding
protein_coding
protein_coding
protein_coding
protein_coding
protein_coding
protein_coding
protein_coding
protein_coding
protein_coding
ncRNA
protein_coding
protein_coding
protein_coding
protein_coding
protein_coding
ncRNA
protein_coding
protein_coding
protein_coding
protein_coding
protein_coding
protein_coding
protein_coding
protein_coding
protein_coding
protein_coding
protein_coding
protein_coding
protein_coding
protein_coding
protein_coding
protein_coding
protein_coding
protein_coding
protein_coding
protein_coding
protein_coding
protein_coding
protein_coding
protein_coding
protein_coding

protein_coding
protein_coding
protein_coding
protein_coding
ncRNA
protein_coding
protein_coding
protein_coding
protein_coding
protein_coding
protein_coding
protein_coding
ncRNA
protein_coding
ncRNA
small_nucle(ol)ar_rna
protein_coding
protein_coding
protein_coding
protein_coding
protein_coding
protein_coding
protein_coding
protein_coding
protein_coding
protein_coding
protein_coding
protein_coding
protein_coding
protein_coding
protein_coding
protein_coding
small_nucle(ol)ar_rna
protein_coding
protein_coding
protein_coding
protein_coding
protein_coding
protein_coding
protein_coding
protein_coding
protein_coding
protein_coding
protein_coding
protein_coding
protein_coding
protein_coding
protein_coding
protein_coding
protein_coding
protein_coding
protein_coding
protein_coding
protein_coding
protein_coding
protein_coding

protein_coding
protein_coding
protein_coding
protein_coding
protein_coding
protein_coding
protein_coding
ncRNA
protein_coding
protein_coding
protein_coding
ncRNA
protein_coding
protein_coding
protein_coding
protein_coding
protein_coding
protein_coding
protein_coding
protein_coding
protein_coding
protein_coding
protein_coding
protein_coding
protein_coding
protein_coding
protein_coding
protein_coding
protein_coding
protein_coding
protein_coding
protein_coding
protein_coding
protein_coding
protein_coding
protein_coding
protein_coding
protein_coding
protein_coding
protein_coding
protein_coding
protein_coding
ncRNA
protein_coding
protein_coding
protein_coding
protein_coding
protein_coding
protein_coding
protein_coding
protein_coding
protein_coding
protein_coding
protein_coding
miRNA_precursor
miRNA_precursor

protein_coding
protein_coding
protein_coding
protein_coding
ncRNA
protein_coding
protein_coding
protein_coding
small_nucle(ol)ar_rna
protein_coding
protein_coding
protein_coding
protein_coding
protein_coding
protein_coding
miRNA_precursor
protein_coding
protein_coding
protein_coding
protein_coding
protein_coding
protein_coding
protein_coding
protein_coding
protein_coding
protein_coding
protein_coding
protein_coding
protein_coding
ncRNA
protein_coding
ncRNA
protein_coding
protein_coding
protein_coding
protein_coding
protein_coding
protein_coding
protein_coding
protein_coding
protein_coding
protein_coding
protein_coding
protein_coding
protein_coding
protein_coding
protein_coding
protein_coding
protein_coding
protein_coding
protein_coding
protein_coding
protein_coding
protein_coding
protein_coding
protein_coding

small_nucle(ol)ar_rna
protein_coding
protein_coding
protein_coding
protein_coding
protein_coding
protein_coding
pre_trna
protein_coding
ncRNA
protein_coding
protein_coding
protein_coding
protein_coding
protein_coding
protein_coding
protein_coding
ncRNA
protein_coding
protein_coding
protein_coding
protein_coding
protein_coding
protein_coding
protein_coding
protein_coding
protein_coding
protein_coding
protein_coding
protein_coding
protein_coding
protein_coding
protein_coding
protein_coding
protein_coding
protein_coding
protein_coding
ncRNA
protein_coding
protein_coding
protein_coding
protein_coding
protein_coding
protein_coding
protein_coding
protein_coding
miRNA_precursor
protein_coding
protein_coding
protein_coding
protein_coding
protein_coding
protein_coding
protein_coding
protein_coding
protein_coding

protein_coding
protein_coding
protein_coding
protein_coding
protein_coding
ncRNA
protein_coding
ncRNA
protein_coding
small_nucle(ol)ar_rna
protein_coding
protein_coding
protein_coding
protein_coding
protein_coding
protein_coding
protein_coding
protein_coding
ncRNA
protein_coding
protein_coding
ncRNA
protein_coding
protein_coding
protein_coding
protein_coding
protein_coding
protein_coding
protein_coding
protein_coding
protein_coding
protein_coding
protein_coding
protein_coding
protein_coding
protein_coding
protein_coding
protein_coding
protein_coding
protein_coding
protein_coding
protein_coding
protein_coding
ncRNA
protein_coding
protein_coding
protein_coding
protein_coding
protein_coding
protein_coding
protein_coding
protein_coding
protein_coding
protein_coding
protein_coding

small_nucle(ol)ar_rna
protein_coding
protein_coding
protein_coding
protein_coding
protein_coding
protein_coding
protein_coding
protein_coding
protein_coding
protein_coding
protein_coding
protein_coding
protein_coding
protein_coding
protein_coding
protein_coding
protein_coding
protein_coding
protein_coding
protein_coding
protein_coding
miRNA_precursor
protein_coding
protein_coding
protein_coding
protein_coding
protein_coding
protein_coding
protein_coding
protein_coding
protein_coding
protein_coding
protein_coding
protein_coding
protein_coding
protein_coding
protein_coding
protein_coding
protein_coding
protein_coding
protein_coding
protein_coding
ncRNA
protein_coding
protein_coding
protein_coding
protein_coding
protein_coding
protein_coding
protein_coding
protein_coding
protein_coding
protein_coding
protein_coding
protein_coding

protein_coding
protein_coding
protein_coding
protein_coding
protein_coding
protein_coding
protein_coding
protein_coding
protein_coding
protein_coding
protein_coding
small_nucle(ol)ar_rna
protein_coding
protein_coding
protein_coding
protein_coding
protein_coding
ncRNA
protein_coding
protein_coding
protein_coding
protein_coding
protein_coding
protein_coding
protein_coding
protein_coding
protein_coding
protein_coding
protein_coding
protein_coding
protein_coding
protein_coding
protein_coding
protein_coding
protein_coding
protein_coding
protein_coding
protein_coding
protein_coding
protein_coding
protein_coding
protein_coding
protein_coding
protein_coding
protein_coding
protein_coding
protein_coding
protein_coding
protein_coding
protein_coding
protein_coding
protein_coding
protein_coding
protein_coding
protein_coding
protein_coding

protein_coding
protein_coding
protein_coding
protein_coding
protein_coding
small_nucle(ol)ar_rna
small_nucle(ol)ar_rna
small_nucle(ol)ar_rna
protein_coding
ncRNA
protein_coding
protein_coding
protein_coding
protein_coding
protein_coding
protein_coding
protein_coding
protein_coding
protein_coding
protein_coding
protein_coding
protein_coding
protein_coding
protein_coding
protein_coding
protein_coding
pre_trna
protein_coding
protein_coding
protein_coding
protein_coding
protein_coding
protein_coding
protein_coding
protein_coding
protein_coding
protein_coding
protein_coding
protein_coding
protein_coding
protein_coding
protein_coding
protein_coding
protein_coding
protein_coding
protein_coding
protein_coding
protein_coding
protein_coding
protein_coding
protein_coding
protein_coding
protein_coding
protein_coding
protein_coding
protein_coding

protein_coding
protein_coding
protein_coding
protein_coding
protein_coding
protein_coding
protein_coding
protein_coding
protein_coding
protein_coding
protein_coding
protein_coding
protein_coding
protein_coding
protein_coding
protein_coding
protein_coding
protein_coding
protein_coding
protein_coding
protein_coding
protein_coding
protein_coding
protein_coding
protein_coding
protein_coding
protein_coding
protein_coding
protein_coding
protein_coding
protein_coding
protein_coding
protein_coding
protein_coding
ncRNA
protein_coding
protein_coding
pre_trna
protein_coding
protein_coding
protein_coding
protein_coding
protein_coding
protein_coding
ncRNA
protein_coding
protein_coding
protein_coding
protein_coding
protein_coding
protein_coding
protein_coding
protein_coding
protein_coding
protein_coding
protein_coding

protein_coding
protein_coding
protein_coding
protein_coding
ncRNA
pre_trna
protein_coding
protein_coding
protein_coding
protein_coding
protein_coding
protein_coding
protein_coding
protein_coding
protein_coding
protein_coding
protein_coding
protein_coding
protein_coding
protein_coding
ncRNA
protein_coding
protein_coding
protein_coding
protein_coding
protein_coding
protein_coding
protein_coding
protein_coding
protein_coding
protein_coding
protein_coding
protein_coding
protein_coding
protein_coding
protein_coding
protein_coding
protein_coding
ncRNA
protein_coding
protein_coding
protein_coding
protein_coding
ncRNA
ncRNA
protein_coding
protein_coding
protein_coding
protein_coding
protein_coding
protein_coding
protein_coding
protein_coding
protein_coding
protein_coding
protein_coding

protein_coding
protein_coding
protein_coding
protein_coding
protein_coding
protein_coding
protein_coding
protein_coding
protein_coding
protein_coding
protein_coding
protein_coding
ncRNA
protein_coding
protein_coding
protein_coding
protein_coding
protein_coding
protein_coding
protein_coding
protein_coding
protein_coding
protein_coding
protein_coding
protein_coding
protein_coding
protein_coding
protein_coding
protein_coding
protein_coding
protein_coding
protein_coding
protein_coding
protein_coding
protein_coding
protein_coding
protein_coding
ncRNA
miRNA_precursor
protein_coding
ncRNA
protein_coding
ncRNA
protein_coding
protein_coding
protein_coding
ncRNA
protein_coding
protein_coding
protein_coding
protein_coding
protein_coding
protein_coding
protein_coding
protein_coding
protein_coding

protein_coding
protein_coding
protein_coding
pre_trna
protein_coding
protein_coding
protein_coding
protein_coding
protein_coding
protein_coding
ncRNA
protein_coding
protein_coding
protein_coding
protein_coding
protein_coding
protein_coding
protein_coding
protein_coding
protein_coding
protein_coding
protein_coding
protein_coding
pre_trna
protein_coding
protein_coding
protein_coding
ncRNA
protein_coding
protein_coding
protein_coding
protein_coding
protein_coding
protein_coding
protein_coding
protein_coding
protein_coding
ncRNA
protein_coding
protein_coding
ncRNA
protein_coding
protein_coding
protein_coding
ncRNA
protein_coding
protein_coding
protein_coding
protein_coding
ncRNA
protein_coding
protein_coding
protein_coding
protein_coding
protein_coding
protein_coding

protein_coding
protein_coding
protein_coding
protein_coding
protein_coding
ncRNA
protein_coding
ncRNA
protein_coding
protein_coding
protein_coding
protein_coding
protein_coding
protein_coding
protein_coding
protein_coding
protein_coding
protein_coding
protein_coding
protein_coding
protein_coding
ncRNA
protein_coding
protein_coding
protein_coding
protein_coding
protein_coding
protein_coding
protein_coding
protein_coding
protein_coding
protein_coding
protein_coding
protein_coding
protein_coding
protein_coding
protein_coding
protein_coding
protein_coding
protein_coding
protein_coding
protein_coding
protein_coding
ncRNA
protein_coding
protein_coding
protein_coding
protein_coding
protein_coding
protein_coding
protein_coding
protein_coding
protein_coding
protein_coding
protein_coding
protein_coding

protein_coding
protein_coding
protein_coding
protein_coding
protein_coding
protein_coding
protein_coding
protein_coding
protein_coding
protein_coding
protein_coding
protein_coding
protein_coding
protein_coding
protein_coding
protein_coding
protein_coding
protein_coding
protein_coding
protein_coding
protein_coding
ncRNA
protein_coding
protein_coding
protein_coding
protein_coding
ncRNA
ncRNA
protein_coding
protein_coding
protein_coding
protein_coding
protein_coding
protein_coding
protein_coding
protein_coding
protein_coding
protein_coding
miRNA_precursor
protein_coding
protein_coding
protein_coding
ncRNA
ncRNA
protein_coding
protein_coding
protein_coding
protein_coding
protein_coding
protein_coding
protein_coding
protein_coding
protein_coding
protein_coding
protein_coding
protein_coding

protein_coding
protein_coding
protein_coding
protein_coding
protein_coding
protein_coding
protein_coding
protein_coding
protein_coding
protein_coding
ncRNA
protein_coding
protein_coding
protein_coding
protein_coding
protein_coding
protein_coding
protein_coding
protein_coding
ncRNA
protein_coding
protein_coding
protein_coding
protein_coding
protein_coding
protein_coding
protein_coding
protein_coding
protein_coding
protein_coding
protein_coding
protein_coding
protein_coding
ncRNA
protein_coding
ncRNA
protein_coding
protein_coding
protein_coding
protein_coding
protein_coding
protein_coding
protein_coding
protein_coding
protein_coding
ncRNA
protein_coding
protein_coding
ncRNA
protein_coding
protein_coding
protein_coding
protein_coding
protein_coding
protein_coding
protein_coding

miRNA_precursor
protein_coding
protein_coding
protein_coding
miRNA_precursor
ncRNA
protein_coding
protein_coding
protein_coding
protein_coding
protein_coding
protein_coding
protein_coding
protein_coding
protein_coding
miRNA_precursor
protein_coding
protein_coding
pre_trna
protein_coding
ncRNA
ncRNA
protein_coding
protein_coding
ncRNA
protein_coding
protein_coding
protein_coding
protein_coding
protein_coding
protein_coding
protein_coding
protein_coding
protein_coding
protein_coding
protein_coding
protein_coding
protein_coding
protein_coding
protein_coding
protein_coding
protein_coding
protein_coding
protein_coding
protein_coding
protein_coding
protein_coding
protein_coding
protein_coding
protein_coding
protein_coding
protein_coding
protein_coding
protein_coding
protein_coding
protein_coding
protein_coding

ncRNA
protein_coding
protein_coding
protein_coding
protein_coding
protein_coding
protein_coding
protein_coding
protein_coding
protein_coding
protein_coding
protein_coding
protein_coding
protein_coding
protein_coding
protein_coding
protein_coding
protein_coding
protein_coding
protein_coding
protein_coding
protein_coding
protein_coding
protein_coding
protein_coding
protein_coding
protein_coding
protein_coding
protein_coding
protein_coding
protein_coding
protein_coding
protein_coding
protein_coding
protein_coding
protein_coding
protein_coding
protein_coding
protein_coding
protein_coding
protein_coding
protein_coding
pre_trna
protein_coding
protein_coding
protein_coding
protein_coding
protein_coding
protein_coding
protein_coding
protein_coding
protein_coding
protein_coding
protein_coding
protein_coding
protein_coding

protein_coding
protein_coding
protein_coding
protein_coding
protein_coding
protein_coding
protein_coding
protein_coding
protein_coding
protein_coding
pre_trna
small_nucle(ol)ar_rna
protein_coding
ncRNA
protein_coding
protein_coding
protein_coding
protein_coding
protein_coding
protein_coding
protein_coding
protein_coding
protein_coding
protein_coding
protein_coding
protein_coding
protein_coding
protein_coding
protein_coding
protein_coding
protein_coding
protein_coding
protein_coding
ncRNA
protein_coding
ncRNA
protein_coding
protein_coding
protein_coding
protein_coding
protein_coding
protein_coding
protein_coding
protein_coding
protein_coding
protein_coding
protein_coding
protein_coding
protein_coding
protein_coding
protein_coding
miRNA_precursor
protein_coding
protein_coding
protein_coding
protein_coding

protein_coding
protein_coding
protein_coding
protein_coding
protein_coding
protein_coding
protein_coding
small_nucle(ol)ar_rna
protein_coding
protein_coding
protein_coding
protein_coding
protein_coding
protein_coding
pre_trna
protein_coding
protein_coding
protein_coding
protein_coding
protein_coding
pre_trna
protein_coding
protein_coding
protein_coding
protein_coding
protein_coding
protein_coding
protein_coding
protein_coding
protein_coding
protein_coding
protein_coding
protein_coding
protein_coding
protein_coding
protein_coding
protein_coding
protein_coding
protein_coding
protein_coding
ncRNA
protein_coding
protein_coding
protein_coding
protein_coding
protein_coding
protein_coding
protein_coding
protein_coding
protein_coding
protein_coding
protein_coding
protein_coding
protein_coding
protein_coding
protein_coding

protein_coding
protein_coding
protein_coding
protein_coding
pre_trna
protein_coding
protein_coding
pre_trna
protein_coding
ncRNA
protein_coding
protein_coding
protein_coding
protein_coding
protein_coding
protein_coding
protein_coding
protein_coding
protein_coding
protein_coding
protein_coding
small_nucle(ol)ar_rna
protein_coding
protein_coding
ncRNA
protein_coding
protein_coding
protein_coding
ncRNA
protein_coding
protein_coding
protein_coding
protein_coding
protein_coding
protein_coding
protein_coding
protein_coding
protein_coding
protein_coding
protein_coding
protein_coding
protein_coding
protein_coding
protein_coding
protein_coding
protein_coding
protein_coding
protein_coding
protein_coding
protein_coding
protein_coding
protein_coding
protein_coding
protein_coding
protein_coding
protein_coding

protein_coding
protein_coding
protein_coding
protein_coding
protein_coding
protein_coding
protein_coding
protein_coding
protein_coding
protein_coding
protein_coding
protein_coding
protein_coding
protein_coding
protein_coding
protein_coding
protein_coding
protein_coding
protein_coding
protein_coding
protein_coding
protein_coding
protein_coding
protein_coding
protein_coding
miRNA_precursor
protein_coding
protein_coding
protein_coding
protein_coding
protein_coding
protein_coding
protein_coding
protein_coding
protein_coding
protein_coding
protein_coding
protein_coding
protein_coding
protein_coding
protein_coding
protein_coding
protein_coding
protein_coding
protein_coding
protein_coding
ncRNA
protein_coding
protein_coding
protein_coding
protein_coding
protein_coding
protein_coding
protein_coding
protein_coding
protein_coding

protein_coding
protein_coding
protein_coding
protein_coding
protein_coding
protein_coding
protein_coding
protein_coding
protein_coding
protein_coding
protein_coding
protein_coding
protein_coding
protein_coding
protein_coding
protein_coding
protein_coding
protein_coding
protein_coding
protein_coding
protein_coding
protein_coding
protein_coding
protein_coding
protein_coding
small_nucle(ol)ar_rna
small_nucle(ol)ar_rna
small_nucle(ol)ar_rna
protein_coding
protein_coding
protein_coding
protein_coding
ncRNA
protein_coding
protein_coding
protein_coding
protein_coding
protein_coding
protein_coding
protein_coding
protein_coding
protein_coding
protein_coding
protein_coding
protein_coding
protein_coding
protein_coding
protein_coding
protein_coding
protein_coding
protein_coding
protein_coding
protein_coding
protein_coding
protein_coding
protein_coding

protein_coding
protein_coding
protein_coding
protein_coding
protein_coding
protein_coding
protein_coding
protein_coding
protein_coding
protein_coding
protein_coding
protein_coding
protein_coding
protein_coding
protein_coding
pre_trna
protein_coding
protein_coding
protein_coding
protein_coding
protein_coding
protein_coding
pre_trna
protein_coding
protein_coding
protein_coding
ncRNA
protein_coding
protein_coding
protein_coding
protein_coding
protein_coding
protein_coding
protein_coding
protein_coding
protein_coding
protein_coding
protein_coding
protein_coding
protein_coding
small_nucle(ol)ar_rna
small_nucle(ol)ar_rna
protein_coding
protein_coding
protein_coding
protein_coding
protein_coding
protein_coding
protein_coding
protein_coding
protein_coding
protein_coding
protein_coding
protein_coding
protein_coding
protein_coding

protein_coding
protein_coding
protein_coding
protein_coding
protein_coding
protein_coding
protein_coding
protein_coding
protein_coding
protein_coding
protein_coding
small_nucle(ol)ar_rna
protein_coding
protein_coding
protein_coding
protein_coding
protein_coding
protein_coding
protein_coding
protein_coding
protein_coding
protein_coding
protein_coding
protein_coding
protein_coding
protein_coding
protein_coding
protein_coding
protein_coding
protein_coding
protein_coding
protein_coding
protein_coding
protein_coding
protein_coding
protein_coding
protein_coding
protein_coding
protein_coding
protein_coding
protein_coding
protein_coding
protein_coding
protein_coding
protein_coding
protein_coding
protein_coding
protein_coding
protein_coding
protein_coding
protein_coding
protein_coding
protein_coding
protein_coding
protein_coding
protein_coding

protein_coding
protein_coding
protein_coding
protein_coding
protein_coding
protein_coding
protein_coding
protein_coding
protein_coding
protein_coding
protein_coding
protein_coding
protein_coding
protein_coding
protein_coding
protein_coding
protein_coding
protein_coding
protein_coding
protein_coding
protein_coding
protein_coding
protein_coding
ncRNA
small_nucle(ol)ar_rna
protein_coding
protein_coding
protein_coding
protein_coding
protein_coding
protein_coding
protein_coding
protein_coding
protein_coding
protein_coding
protein_coding
protein_coding
protein_coding
protein_coding
protein_coding
protein_coding
protein_coding
protein_coding
protein_coding
protein_coding
protein_coding
protein_coding
protein_coding
protein_coding
protein_coding
protein_coding
protein_coding
protein_coding
protein_coding
protein_coding
protein_coding

ncRNA
ncRNA
protein_coding
protein_coding
protein_coding
protein_coding
protein_coding
protein_coding
protein_coding
protein_coding
protein_coding
protein_coding
protein_coding
protein_coding
protein_coding
protein_coding
protein_coding
protein_coding
protein_coding
protein_coding
protein_coding
protein_coding
ncRNA
ncRNA
protein_coding
protein_coding
protein_coding
protein_coding
protein_coding
protein_coding
protein_coding
protein_coding
protein_coding
protein_coding
protein_coding
protein_coding
protein_coding
protein_coding
protein_coding
protein_coding
protein_coding
protein_coding
protein_coding
protein_coding
protein_coding
protein_coding
protein_coding
protein_coding
protein_coding
protein_coding
protein_coding
protein_coding
protein_coding
protein_coding
protein_coding
protein_coding

protein_coding
protein_coding
protein_coding
protein_coding
protein_coding
protein_coding
protein_coding
protein_coding
protein_coding
protein_coding
protein_coding
protein_coding
protein_coding
protein_coding
protein_coding
protein_coding
protein_coding
protein_coding
protein_coding
protein_coding
protein_coding
protein_coding
protein_coding
protein_coding
protein_coding
protein_coding
ncRNA
protein_coding
protein_coding
protein_coding
protein_coding
protein_coding
protein_coding
protein_coding
protein_coding
protein_coding
protein_coding
protein_coding
protein_coding
protein_coding
protein_coding
protein_coding
protein_coding
protein_coding
protein_coding
protein_coding
protein_coding
protein_coding
protein_coding
protein_coding
protein_coding
protein_coding
protein_coding
protein_coding
protein_coding
protein_coding

protein_coding
protein_coding
protein_coding
protein_coding
protein_coding
protein_coding
protein_coding
protein_coding
protein_coding
protein_coding
protein_coding
protein_coding
protein_coding
protein_coding
protein_coding
protein_coding
protein_coding
protein_coding
protein_coding
protein_coding
protein_coding
protein_coding
protein_coding
ncRNA
protein_coding
protein_coding
protein_coding
protein_coding
protein_coding
protein_coding
protein_coding
protein_coding
protein_coding
protein_coding
protein_coding
protein_coding
protein_coding
protein_coding
protein_coding
protein_coding
protein_coding
ncRNA
protein_coding
protein_coding
protein_coding
protein_coding
protein_coding
protein_coding
protein_coding
protein_coding
protein_coding
protein_coding
protein_coding
protein_coding
protein_coding
protein_coding

protein_coding
protein_coding
protein_coding
protein_coding
protein_coding
small_nucle(ol)ar_rna
protein_coding
protein_coding
protein_coding
protein_coding
protein_coding
protein_coding
protein_coding
protein_coding
protein_coding
protein_coding
protein_coding
protein_coding
protein_coding
protein_coding
protein_coding
protein_coding
protein_coding
protein_coding
protein_coding
protein_coding
protein_coding
protein_coding
protein_coding
protein_coding
protein_coding
protein_coding
protein_coding
protein_coding
protein_coding
protein_coding
protein_coding
protein_coding
protein_coding
protein_coding
protein_coding
protein_coding
protein_coding
protein_coding
protein_coding
protein_coding
protein_coding
protein_coding
protein_coding
protein_coding
protein_coding
protein_coding
protein_coding
ncRNA
protein_coding
protein_coding

protein_coding
protein_coding
protein_coding
protein_coding
protein_coding
protein_coding
protein_coding
protein_coding
protein_coding
protein_coding
protein_coding
protein_coding
protein_coding
protein_coding
protein_coding
protein_coding
protein_coding
protein_coding
protein_coding
protein_coding
protein_coding
protein_coding
protein_coding
protein_coding
protein_coding
protein_coding
protein_coding
protein_coding
protein_coding
protein_coding
ncRNA
protein_coding
protein_coding
protein_coding
protein_coding
protein_coding
protein_coding
protein_coding
protein_coding
protein_coding
protein_coding
protein_coding
protein_coding
protein_coding
protein_coding
ncRNA
protein_coding
protein_coding
protein_coding
protein_coding
protein_coding
protein_coding
protein_coding
protein_coding
protein_coding
protein_coding

protein_coding
protein_coding
protein_coding
protein_coding
protein_coding
protein_coding
miRNA_precursor
protein_coding
protein_coding
protein_coding
protein_coding
protein_coding
protein_coding
protein_coding
protein_coding
protein_coding
protein_coding
protein_coding
miRNA_precursor
protein_coding
protein_coding
protein_coding
protein_coding
protein_coding
protein_coding
protein_coding
protein_coding
protein_coding
protein_coding
protein_coding
protein_coding
protein_coding
protein_coding
protein_coding
protein_coding
protein_coding
protein_coding
protein_coding
protein_coding
protein_coding
protein_coding
protein_coding
protein_coding
protein_coding
protein_coding
protein_coding
protein_coding
protein_coding
protein_coding
protein_coding
protein_coding
protein_coding
protein_coding
protein_coding
protein_coding
protein_coding

protein_coding
ncRNA
protein_coding
protein_coding
protein_coding
protein_coding
protein_coding
protein_coding
protein_coding
protein_coding
protein_coding
protein_coding
protein_coding
protein_coding
protein_coding
protein_coding
protein_coding
protein_coding
small_nucle(ol)ar_rna
protein_coding
ncRNA
protein_coding
protein_coding
protein_coding
protein_coding
ncRNA
protein_coding
protein_coding
protein_coding
protein_coding
protein_coding
protein_coding
protein_coding
protein_coding
protein_coding
protein_coding
protein_coding
protein_coding
protein_coding
protein_coding
protein_coding
protein_coding
ncRNA
protein_coding
protein_coding
protein_coding
protein_coding
protein_coding
protein_coding
protein_coding
protein_coding
protein_coding
protein_coding
protein_coding
protein_coding
protein_coding

protein_coding
protein_coding
protein_coding
protein_coding
protein_coding
protein_coding
protein_coding
protein_coding
protein_coding
protein_coding
protein_coding
protein_coding
protein_coding
protein_coding
protein_coding
protein_coding
protein_coding
protein_coding
protein_coding
protein_coding
protein_coding
protein_coding
protein_coding
protein_coding
protein_coding
protein_coding
protein_coding
protein_coding
protein_coding
protein_coding
protein_coding
protein_coding
protein_coding
ncRNA
protein_coding
protein_coding
protein_coding
protein_coding
protein_coding
protein_coding
protein_coding
protein_coding
protein_coding
protein_coding
protein_coding
protein_coding
protein_coding
ncRNA
small_nucle(ol)ar_rna
protein_coding
protein_coding
protein_coding
protein_coding
protein_coding
protein_coding
protein_coding

protein_coding
protein_coding
protein_coding
protein_coding
protein_coding
protein_coding
protein_coding
protein_coding
protein_coding
protein_coding
protein_coding
protein_coding
protein_coding
protein_coding
protein_coding
protein_coding
protein_coding
protein_coding
protein_coding
protein_coding
protein_coding
protein_coding
protein_coding
protein_coding
protein_coding
pre_trna
protein_coding
ncRNA
protein_coding
protein_coding
protein_coding
protein_coding
protein_coding
protein_coding
protein_coding
protein_coding
protein_coding
protein_coding
protein_coding
protein_coding
protein_coding
protein_coding
protein_coding
protein_coding
protein_coding
protein_coding
protein_coding
protein_coding
protein_coding
miRNA_precursor
protein_coding
protein_coding
protein_coding
protein_coding
protein_coding
protein_coding

protein_coding
protein_coding
protein_coding
protein_coding
ncRNA
protein_coding
protein_coding
protein_coding
protein_coding
protein_coding
protein_coding
protein_coding
small_nucle(ol)ar_rna
protein_coding
protein_coding
protein_coding
protein_coding
protein_coding
protein_coding
protein_coding
ncRNA
protein_coding
protein_coding
protein_coding
protein_coding
protein_coding
protein_coding
protein_coding
protein_coding
protein_coding
protein_coding
protein_coding
protein_coding
protein_coding
protein_coding
protein_coding
protein_coding
protein_coding
protein_coding
protein_coding
protein_coding
protein_coding
protein_coding
protein_coding
protein_coding
ncRNA
protein_coding
protein_coding
protein_coding
protein_coding
protein_coding
ncRNA
protein_coding
ncRNA
protein_coding
protein_coding
ncRNA

protein_coding
protein_coding
protein_coding
protein_coding
protein_coding
protein_coding
protein_coding
protein_coding
protein_coding
protein_coding
protein_coding
protein_coding
protein_coding
protein_coding
protein_coding
protein_coding
protein_coding
protein_coding
protein_coding
ncRNA
protein_coding
protein_coding
protein_coding
protein_coding
protein_coding
protein_coding
protein_coding
protein_coding
protein_coding
protein_coding
protein_coding
protein_coding
protein_coding
protein_coding
pre_trna
protein_coding
protein_coding
protein_coding
protein_coding
protein_coding
protein_coding
protein_coding
protein_coding
protein_coding
protein_coding
protein_coding
ncRNA
protein_coding
ncRNA
ncRNA
ncRNA
ncRNA
protein_coding
protein_coding
protein_coding
miRNA_precursor

protein_coding
protein_coding
small_nucle(ol)ar_rna
protein_coding
protein_coding
protein_coding
protein_coding
protein_coding
protein_coding
protein_coding
ncRNA
ncRNA
protein_coding
protein_coding
protein_coding
protein_coding
protein_coding
protein_coding
protein_coding
protein_coding
protein_coding
protein_coding
protein_coding
protein_coding
protein_coding
protein_coding
protein_coding
protein_coding
protein_coding
protein_coding
protein_coding
protein_coding
protein_coding
protein_coding
protein_coding
protein_coding
ncRNA
protein_coding
protein_coding
protein_coding
protein_coding
protein_coding
protein_coding
protein_coding
protein_coding
protein_coding
protein_coding
protein_coding
protein_coding
protein_coding
protein_coding
protein_coding
protein_coding
protein_coding
protein_coding
protein_coding

protein_coding
protein_coding
protein_coding
protein_coding
protein_coding
protein_coding
protein_coding
protein_coding
protein_coding
protein_coding
protein_coding
protein_coding
protein_coding
protein_coding
protein_coding
protein_coding
protein_coding
protein_coding
protein_coding
protein_coding
protein_coding
protein_coding
protein_coding
protein_coding
protein_coding
protein_coding
protein_coding
protein_coding
protein_coding
protein_coding
protein_coding
protein_coding
protein_coding
protein_coding
protein_coding
protein_coding
protein_coding
protein_coding
protein_coding
protein_coding
protein_coding
protein_coding
protein_coding
protein_coding
protein_coding
protein_coding
protein_coding
protein_coding
protein_coding
protein_coding
protein_coding
protein_coding
protein_coding
small_nucle(ol)ar_rna
protein_coding
protein_coding

protein_coding
miRNA_precursor
ncRNA
protein_coding
protein_coding
protein_coding
protein_coding
protein_coding
protein_coding
protein_coding
protein_coding
protein_coding
protein_coding
protein_coding
protein_coding
protein_coding
protein_coding
protein_coding
protein_coding
protein_coding
protein_coding
protein_coding
protein_coding
protein_coding
protein_coding
protein_coding
protein_coding
protein_coding
ncRNA
protein_coding
protein_coding
protein_coding
protein_coding
ncRNA
protein_coding
protein_coding
protein_coding
protein_coding
protein_coding
protein_coding
protein_coding
protein_coding
protein_coding
protein_coding
protein_coding
protein_coding
protein_coding
protein_coding
protein_coding
ncRNA
small_nucle(ol)ar_rna
protein_coding
protein_coding
protein_coding
protein_coding
protein_coding

protein_coding
protein_coding
protein_coding
protein_coding
protein_coding
protein_coding
protein_coding
protein_coding
protein_coding
protein_coding
protein_coding
ncRNA
protein_coding
protein_coding
protein_coding
protein_coding
protein_coding
protein_coding
protein_coding
protein_coding
protein_coding
protein_coding
ncRNA
protein_coding
protein_coding
protein_coding
protein_coding
protein_coding
protein_coding
ncRNA
protein_coding
protein_coding
protein_coding
protein_coding
protein_coding
protein_coding
protein_coding
protein_coding
protein_coding
ncRNA
protein_coding
protein_coding
protein_coding
protein_coding
protein_coding
protein_coding
protein_coding
protein_coding
protein_coding
protein_coding
protein_coding
protein_coding
protein_coding
protein_coding
protein_coding
protein_coding

protein_coding
ncRNA
protein_coding
protein_coding
protein_coding
protein_coding
protein_coding
protein_coding
protein_coding
protein_coding
protein_coding
protein_coding
protein_coding
protein_coding
protein_coding
protein_coding
protein_coding
protein_coding
protein_coding
protein_coding
protein_coding
protein_coding
protein_coding
protein_coding
protein_coding
protein_coding
protein_coding
protein_coding
protein_coding
protein_coding
protein_coding
protein_coding
protein_coding
protein_coding
protein_coding
protein_coding
protein_coding
protein_coding
protein_coding
miRNA_precursor
protein_coding
ncRNA
protein_coding
protein_coding
protein_coding
protein_coding
protein_coding
protein_coding
protein_coding
protein_coding
protein_coding
protein_coding
protein_coding
protein_coding
protein_coding
protein_coding

protein_coding
protein_coding
protein_coding
protein_coding
protein_coding
protein_coding
pre_trna
protein_coding
protein_coding
protein_coding
protein_coding
protein_coding
protein_coding
protein_coding
pre_trna
protein_coding
protein_coding
protein_coding
protein_coding
protein_coding
protein_coding
ncRNA
ncRNA
protein_coding
protein_coding
protein_coding
protein_coding
protein_coding
protein_coding
protein_coding
protein_coding
protein_coding
protein_coding
protein_coding
protein_coding
protein_coding
protein_coding
protein_coding
protein_coding
protein_coding
protein_coding
protein_coding
ncRNA
protein_coding
protein_coding
protein_coding
protein_coding
protein_coding
protein_coding
protein_coding
protein_coding
protein_coding
protein_coding
protein_coding
protein_coding
protein_coding

protein_coding
protein_coding
protein_coding
protein_coding
protein_coding
protein_coding
protein_coding
protein_coding
protein_coding
protein_coding
protein_coding
protein_coding
protein_coding
protein_coding
protein_coding
protein_coding
protein_coding
protein_coding
protein_coding
protein_coding
protein_coding
protein_coding
ncRNA
small_nucle(ol)ar_rna
protein_coding
protein_coding
protein_coding
protein_coding
protein_coding
protein_coding
protein_coding
protein_coding
protein_coding
protein_coding
protein_coding
protein_coding
protein_coding
protein_coding
protein_coding
protein_coding
protein_coding
protein_coding
protein_coding
protein_coding
protein_coding
ncRNA
protein_coding
ncRNA
protein_coding
protein_coding
protein_coding
protein_coding
protein_coding
protein_coding
protein_coding
protein_coding

protein_coding
protein_coding
protein_coding
protein_coding
protein_coding
ncRNA
protein_coding
protein_coding
protein_coding
protein_coding
protein_coding
protein_coding
pre_trna
protein_coding
protein_coding
protein_coding
protein_coding
protein_coding
protein_coding
protein_coding
protein_coding
protein_coding
protein_coding
ncRNA
ncRNA
protein_coding
protein_coding
protein_coding
protein_coding
protein_coding
protein_coding
protein_coding
protein_coding
protein_coding
protein_coding
protein_coding
protein_coding
protein_coding
protein_coding
protein_coding
protein_coding
miRNA_precursor
protein_coding
protein_coding
protein_coding
protein_coding
protein_coding
protein_coding
protein_coding
protein_coding
protein_coding
protein_coding
protein_coding
protein_coding
miRNA_precursor
protein_coding

protein_coding
protein_coding
protein_coding
protein_coding
protein_coding
protein_coding
miRNA_precursor
protein_coding
protein_coding
protein_coding
protein_coding
protein_coding
protein_coding
ncRNA
protein_coding
protein_coding
protein_coding
protein_coding
protein_coding
protein_coding
protein_coding
protein_coding
protein_coding
protein_coding
ncRNA
protein_coding
protein_coding
protein_coding
protein_coding
protein_coding
protein_coding
protein_coding
protein_coding
protein_coding
protein_coding
protein_coding
protein_coding
protein_coding
protein_coding
protein_coding
ncRNA
protein_coding
protein_coding
protein_coding
protein_coding
protein_coding
protein_coding
protein_coding
protein_coding
protein_coding
protein_coding
protein_coding
protein_coding
protein_coding
protein_coding
protein_coding

protein_coding
protein_coding
protein_coding
protein_coding
protein_coding
protein_coding
protein_coding
protein_coding
ncRNA
protein_coding
protein_coding
protein_coding
protein_coding
protein_coding
protein_coding
protein_coding
protein_coding
protein_coding
protein_coding
protein_coding
ncRNA
protein_coding
protein_coding
protein_coding
protein_coding
protein_coding
protein_coding
protein_coding
protein_coding
ncRNA
protein_coding
protein_coding
small_nucle(ol)ar_rna
small_nucle(ol)ar_rna
small_nucle(ol)ar_rna
protein_coding
protein_coding
protein_coding
protein_coding
protein_coding
protein_coding
protein_coding
protein_coding
protein_coding
protein_coding
protein_coding
protein_coding
protein_coding
protein_coding
protein_coding
protein_coding
protein_coding
protein_coding
protein_coding
protein_coding
protein_coding

protein_coding
ncRNA
protein_coding
protein_coding
protein_coding
protein_coding
small_nucle(ol)ar_rna
small_nucle(ol)ar_rna
miRNA_precursor
protein_coding
protein_coding
protein_coding
protein_coding
protein_coding
protein_coding
protein_coding
protein_coding
protein_coding
protein_coding
protein_coding
protein_coding
protein_coding
protein_coding
protein_coding
protein_coding
protein_coding
protein_coding
protein_coding
protein_coding
protein_coding
protein_coding
protein_coding
protein_coding
protein_coding
protein_coding
protein_coding
protein_coding
pre_trna
protein_coding
protein_coding
protein_coding
protein_coding
protein_coding
protein_coding
protein_coding
protein_coding
protein_coding
protein_coding
protein_coding
protein_coding
protein_coding
protein_coding
protein_coding
protein_coding
protein_coding
protein_coding

protein_coding
protein_coding
protein_coding
protein_coding
protein_coding
protein_coding
protein_coding
protein_coding
protein_coding
protein_coding
protein_coding
protein_coding
protein_coding
protein_coding
ncRNA
ncRNA
protein_coding
protein_coding
protein_coding
protein_coding
protein_coding
protein_coding
protein_coding
protein_coding
protein_coding
protein_coding
protein_coding
protein_coding
protein_coding
protein_coding
protein_coding
protein_coding
protein_coding
protein_coding
protein_coding
protein_coding
protein_coding
protein_coding
protein_coding
protein_coding
ncRNA
protein_coding
protein_coding
protein_coding
pre_trna
protein_coding
protein_coding
protein_coding
protein_coding
protein_coding
protein_coding
protein_coding
protein_coding
protein_coding
miRNA_precursor
protein_coding

protein_coding
protein_coding
protein_coding
protein_coding
protein_coding
protein_coding
protein_coding
protein_coding
protein_coding
protein_coding
protein_coding
protein_coding
protein_coding
protein_coding
protein_coding
protein_coding
protein_coding
protein_coding
ncRNA
protein_coding
protein_coding
protein_coding
protein_coding
protein_coding
protein_coding
protein_coding
protein_coding
protein_coding
protein_coding
protein_coding
protein_coding
protein_coding
protein_coding
protein_coding
protein_coding
protein_coding
protein_coding
protein_coding
protein_coding
protein_coding
protein_coding
protein_coding
protein_coding
protein_coding
protein_coding
protein_coding
protein_coding
ncRNA
protein_coding
protein_coding
protein_coding
ncRNA
protein_coding
protein_coding
small_nucle(ol)ar_rna
miRNA_precursor

protein_coding
protein_coding
protein_coding
protein_coding
protein_coding
protein_coding
protein_coding
protein_coding
protein_coding
protein_coding
protein_coding
protein_coding
protein_coding
protein_coding
protein_coding
protein_coding
protein_coding
protein_coding
protein_coding
protein_coding
protein_coding
protein_coding
protein_coding
protein_coding
ncRNA
protein_coding
protein_coding
protein_coding
protein_coding
protein_coding
protein_coding
protein_coding
protein_coding
protein_coding
protein_coding
protein_coding
protein_coding
protein_coding
protein_coding
protein_coding
protein_coding
protein_coding
protein_coding
protein_coding
protein_coding
protein_coding
protein_coding
protein_coding
protein_coding
ncRNA
protein_coding
protein_coding
protein_coding
protein_coding
protein_coding
protein_coding

protein_coding
protein_coding
protein_coding
protein_coding
protein_coding
protein_coding
protein_coding
protein_coding
protein_coding
protein_coding
small_nucle(ol)ar_rna
protein_coding
protein_coding
protein_coding
protein_coding
protein_coding
protein_coding
protein_coding
protein_coding
protein_coding
protein_coding
protein_coding
protein_coding
protein_coding
protein_coding
protein_coding
protein_coding
protein_coding
protein_coding
protein_coding
protein_coding
protein_coding
protein_coding
protein_coding
protein_coding
protein_coding
protein_coding
ncRNA
protein_coding
protein_coding
protein_coding
protein_coding
protein_coding
pre_trna
protein_coding
protein_coding
protein_coding
protein_coding
protein_coding
protein_coding
protein_coding
protein_coding
protein_coding
miRNA_precursor
protein_coding
protein_coding

protein_coding
protein_coding
protein_coding
protein_coding
protein_coding
protein_coding
protein_coding
protein_coding
pre_trna
protein_coding
protein_coding
protein_coding
protein_coding
protein_coding
protein_coding
protein_coding
protein_coding
protein_coding
protein_coding
protein_coding
protein_coding
protein_coding
protein_coding
protein_coding
protein_coding
protein_coding
protein_coding
protein_coding
protein_coding
protein_coding
protein_coding
protein_coding
protein_coding
protein_coding
protein_coding
protein_coding
protein_coding
protein_coding
ncRNA
ncRNA
protein_coding
protein_coding
protein_coding
protein_coding
protein_coding
protein_coding
protein_coding
protein_coding
ncRNA
protein_coding
protein_coding
protein_coding
protein_coding
protein_coding
protein_coding
protein_coding

protein_coding
small_nucle(ol)ar_rna
protein_coding
protein_coding
protein_coding
protein_coding
protein_coding
protein_coding
protein_coding
protein_coding
protein_coding
protein_coding
protein_coding
protein_coding
protein_coding
protein_coding
protein_coding
protein_coding
small_nucle(ol)ar_rna
small_nucle(ol)ar_rna
protein_coding
protein_coding
protein_coding
protein_coding
protein_coding
protein_coding
protein_coding
protein_coding
pre_trna
protein_coding
protein_coding
protein_coding
protein_coding
protein_coding
protein_coding
protein_coding
protein_coding
protein_coding
protein_coding
protein_coding
protein_coding
protein_coding
protein_coding
protein_coding
protein_coding
protein_coding
protein_coding
protein_coding
protein_coding
protein_coding
protein_coding
protein_coding
ncRNA
protein_coding
protein_coding
protein_coding

protein_coding
protein_coding
protein_coding
protein_coding
protein_coding
protein_coding
protein_coding
protein_coding
protein_coding
protein_coding
protein_coding
protein_coding
protein_coding
pre_trna
protein_coding
protein_coding
protein_coding
protein_coding
protein_coding
protein_coding
protein_coding
protein_coding
protein_coding
protein_coding
protein_coding
protein_coding
protein_coding
protein_coding
protein_coding
protein_coding
protein_coding
protein_coding
protein_coding
protein_coding
protein_coding
protein_coding
protein_coding
protein_coding
protein_coding
protein_coding
protein_coding
protein_coding
protein_coding
protein_coding
protein_coding
protein_coding
protein_coding
protein_coding
protein_coding
protein_coding
ncRNA
ncRNA
protein_coding
protein_coding
protein_coding

protein_coding
protein_coding
protein_coding
protein_coding
protein_coding
protein_coding
ncRNA
protein_coding
protein_coding
protein_coding
protein_coding
protein_coding
protein_coding
protein_coding
protein_coding
protein_coding
protein_coding
protein_coding
protein_coding
protein_coding
protein_coding
protein_coding
protein_coding
protein_coding
protein_coding
protein_coding
protein_coding
protein_coding
protein_coding
protein_coding
protein_coding
protein_coding
ncRNA
ncRNA
protein_coding
protein_coding
protein_coding
protein_coding
protein_coding
protein_coding
protein_coding
protein_coding
protein_coding
protein_coding
protein_coding
protein_coding
protein_coding
protein_coding
protein_coding
protein_coding
protein_coding
protein_coding
protein_coding
protein_coding
protein_coding
protein_coding

protein_coding
protein_coding
protein_coding
protein_coding
protein_coding
protein_coding
protein_coding
protein_coding
protein_coding
protein_coding
protein_coding
protein_coding
protein_coding
ncRNA
protein_coding
protein_coding
protein_coding
protein_coding
protein_coding
protein_coding
ncRNA
protein_coding
ncRNA
protein_coding
ncRNA
protein_coding
protein_coding
protein_coding
protein_coding
protein_coding
protein_coding
miRNA_precursor
protein_coding
protein_coding
protein_coding
ncRNA
protein_coding
protein_coding
protein_coding
protein_coding
protein_coding
protein_coding
protein_coding
protein_coding
protein_coding
protein_coding
protein_coding
protein_coding
protein_coding
protein_coding
protein_coding
protein_coding
protein_coding
protein_coding
protein_coding
protein_coding

protein_coding
protein_coding
protein_coding
protein_coding
protein_coding
protein_coding
protein_coding
protein_coding
protein_coding
protein_coding
protein_coding
protein_coding
protein_coding
protein_coding
protein_coding
protein_coding
protein_coding
protein_coding
protein_coding
protein_coding
protein_coding
ncRNA
protein_coding
protein_coding
protein_coding
protein_coding
protein_coding
protein_coding
protein_coding
ncRNA
protein_coding
protein_coding
protein_coding
protein_coding
protein_coding
protein_coding
protein_coding
protein_coding
protein_coding
protein_coding
protein_coding
protein_coding
protein_coding
protein_coding
protein_coding
protein_coding
protein_coding
protein_coding
protein_coding
protein_coding
protein_coding
pre_trna
protein_coding
protein_coding
protein_coding
protein_coding

protein_coding
protein_coding
protein_coding
protein_coding
protein_coding
ncRNA
protein_coding
protein_coding
protein_coding
protein_coding
protein_coding
protein_coding
protein_coding
protein_coding
protein_coding
protein_coding
protein_coding
protein_coding
protein_coding
protein_coding
protein_coding
protein_coding
protein_coding
ncRNA
protein_coding
protein_coding
protein_coding
protein_coding
protein_coding
protein_coding
protein_coding
protein_coding
protein_coding
protein_coding
protein_coding
protein_coding
protein_coding
protein_coding
protein_coding
protein_coding
protein_coding
protein_coding
protein_coding
protein_coding
protein_coding
protein_coding
protein_coding
protein_coding
ncRNA
protein_coding
protein_coding
protein_coding
protein_coding
protein_coding
protein_coding
protein_coding

protein_coding
protein_coding
protein_coding
protein_coding
ncRNA
protein_coding
protein_coding
protein_coding
protein_coding
protein_coding
protein_coding
protein_coding
protein_coding
protein_coding
protein_coding
protein_coding
protein_coding
protein_coding
protein_coding
protein_coding
protein_coding
protein_coding
protein_coding
protein_coding
protein_coding
protein_coding
protein_coding
protein_coding
protein_coding
protein_coding
ncRNA
protein_coding
protein_coding
protein_coding
protein_coding
protein_coding
protein_coding
protein_coding
protein_coding
protein_coding
protein_coding
protein_coding
protein_coding
protein_coding
protein_coding
protein_coding
protein_coding
protein_coding
protein_coding
protein_coding
protein_coding
protein_coding
protein_coding
protein_coding
protein_coding
protein_coding

protein_coding
protein_coding
protein_coding
protein_coding
protein_coding
protein_coding
protein_coding
miRNA_precursor
protein_coding
protein_coding
protein_coding
protein_coding
protein_coding
protein_coding
protein_coding
protein_coding
protein_coding
protein_coding
protein_coding
protein_coding
protein_coding
protein_coding
protein_coding
protein_coding
protein_coding
protein_coding
protein_coding
protein_coding
protein_coding
protein_coding
protein_coding
protein_coding
protein_coding
protein_coding
protein_coding
protein_coding
protein_coding
protein_coding
protein_coding
protein_coding
protein_coding
protein_coding
protein_coding
protein_coding
protein_coding
protein_coding
protein_coding
protein_coding
protein_coding
protein_coding
protein_coding
protein_coding
protein_coding
protein_coding
protein_coding
protein_coding

protein_coding
protein_coding
protein_coding
small_nucle(ol)ar_rna
protein_coding
protein_coding
protein_coding
protein_coding
protein_coding
protein_coding
protein_coding
protein_coding
protein_coding
protein_coding
protein_coding
protein_coding
protein_coding
protein_coding
protein_coding
protein_coding
protein_coding
protein_coding
protein_coding
protein_coding
protein_coding
protein_coding
protein_coding
protein_coding
protein_coding
protein_coding
protein_coding
protein_coding
protein_coding
protein_coding
ncRNA
protein_coding
protein_coding
protein_coding
protein_coding
protein_coding
protein_coding
protein_coding
protein_coding
protein_coding
protein_coding
protein_coding
protein_coding
protein_coding
protein_coding
protein_coding
protein_coding
protein_coding
protein_coding
protein_coding
protein_coding
protein_coding

protein_coding
protein_coding
protein_coding
protein_coding
protein_coding
protein_coding
protein_coding
protein_coding
protein_coding
pre_trna
protein_coding
protein_coding
protein_coding
protein_coding
protein_coding
ncRNA
ncRNA
protein_coding
protein_coding
protein_coding
protein_coding
protein_coding
protein_coding
protein_coding
protein_coding
protein_coding
protein_coding
protein_coding
protein_coding
protein_coding
protein_coding
protein_coding
protein_coding
protein_coding
protein_coding
protein_coding
protein_coding
protein_coding
protein_coding
ncRNA
protein_coding
protein_coding
protein_coding
protein_coding
protein_coding
protein_coding
protein_coding
protein_coding
protein_coding
protein_coding
protein_coding
protein_coding
protein_coding
protein_coding
protein_coding
protein_coding

protein_coding
protein_coding
protein_coding
protein_coding
protein_coding
protein_coding
protein_coding
protein_coding
protein_coding
protein_coding
protein_coding
protein_coding
protein_coding
protein_coding
ncRNA
protein_coding
protein_coding
protein_coding
protein_coding
protein_coding
protein_coding
protein_coding
protein_coding
protein_coding
protein_coding
protein_coding
protein_coding
protein_coding
pre_trna
protein_coding
protein_coding
protein_coding
protein_coding
protein_coding
protein_coding
protein_coding
protein_coding
protein_coding
protein_coding
protein_coding
protein_coding
protein_coding
protein_coding
ncRNA
protein_coding
protein_coding
protein_coding
protein_coding
protein_coding
protein_coding
protein_coding
protein_coding
protein_coding
protein_coding
protein_coding
protein_coding

protein_coding
protein_coding
protein_coding
protein_coding
protein_coding
protein_coding
protein_coding
protein_coding
protein_coding
protein_coding
protein_coding
protein_coding
protein_coding
protein_coding
protein_coding
protein_coding
protein_coding
protein_coding
protein_coding
ncRNA
protein_coding
protein_coding
protein_coding
protein_coding
protein_coding
protein_coding
protein_coding
protein_coding
protein_coding
protein_coding
protein_coding
protein_coding
protein_coding
protein_coding
protein_coding
protein_coding
miRNA_precursor
protein_coding
protein_coding
protein_coding
protein_coding
miRNA_precursor
ncRNA
miRNA_precursor
protein_coding
protein_coding
protein_coding
protein_coding
protein_coding
protein_coding
protein_coding
protein_coding
protein_coding
protein_coding
protein_coding
protein_coding

protein_coding
protein_coding
protein_coding
protein_coding
protein_coding
protein_coding
protein_coding
protein_coding
protein_coding
protein_coding
protein_coding
protein_coding
protein_coding
protein_coding
protein_coding
protein_coding
protein_coding
protein_coding
protein_coding
protein_coding
protein_coding
protein_coding
protein_coding
protein_coding
protein_coding
protein_coding
protein_coding
protein_coding
protein_coding
protein_coding
protein_coding
protein_coding
protein_coding
protein_coding
protein_coding
protein_coding
ncRNA
ncRNA
protein_coding
ncRNA
protein_coding
protein_coding
protein_coding
protein_coding
protein_coding
ncRNA
protein_coding
protein_coding
protein_coding
protein_coding
protein_coding
miRNA_precursor
protein_coding
protein_coding
protein_coding
protein_coding

protein_coding
protein_coding
ncRNA
protein_coding
protein_coding
protein_coding
protein_coding
protein_coding
protein_coding
protein_coding
protein_coding
protein_coding
protein_coding
ncRNA
protein_coding
protein_coding
protein_coding
protein_coding
protein_coding
protein_coding
protein_coding
protein_coding
protein_coding
protein_coding
protein_coding
protein_coding
protein_coding
protein_coding
protein_coding
protein_coding
protein_coding
protein_coding
protein_coding
protein_coding
protein_coding
protein_coding
protein_coding
protein_coding
protein_coding
pre_trna
protein_coding
protein_coding
protein_coding
protein_coding
protein_coding
protein_coding
protein_coding
protein_coding
protein_coding
protein_coding
protein_coding
protein_coding
protein_coding
protein_coding
protein_coding
protein_coding

protein_coding
protein_coding
protein_coding
protein_coding
protein_coding
protein_coding
protein_coding
protein_coding
protein_coding
protein_coding
protein_coding
protein_coding
protein_coding
protein_coding
protein_coding
protein_coding
protein_coding
protein_coding
protein_coding
protein_coding
protein_coding
protein_coding
protein_coding
protein_coding
protein_coding
protein_coding
small_nucle(ol)ar_rna
protein_coding
protein_coding
protein_coding
protein_coding
protein_coding
protein_coding
protein_coding
protein_coding
protein_coding
protein_coding
protein_coding
protein_coding
protein_coding
protein_coding
protein_coding
protein_coding
protein_coding
protein_coding
protein_coding
protein_coding
protein_coding
protein_coding
protein_coding
protein_coding
protein_coding
protein_coding
protein_coding
protein_coding
protein_coding

protein_coding
protein_coding
protein_coding
protein_coding
protein_coding
protein_coding
protein_coding
protein_coding
protein_coding
protein_coding
protein_coding
ncRNA
protein_coding
protein_coding
protein_coding
protein_coding
protein_coding
ncRNA
protein_coding
protein_coding
protein_coding
protein_coding
protein_coding
protein_coding
protein_coding
protein_coding
protein_coding
protein_coding
protein_coding
protein_coding
protein_coding
protein_coding
protein_coding
protein_coding
protein_coding
protein_coding
small_nucle(ol)ar_rna
small_nucle(ol)ar_rna
protein_coding
protein_coding
protein_coding
protein_coding
miRNA_precursor
protein_coding
protein_coding
protein_coding
protein_coding
protein_coding
protein_coding
protein_coding
ncRNA
protein_coding
protein_coding
protein_coding
protein_coding
protein_coding

protein_coding
protein_coding
protein_coding
protein_coding
ncRNA
protein_coding
protein_coding
protein_coding
protein_coding
protein_coding
protein_coding
protein_coding
protein_coding
protein_coding
protein_coding
protein_coding
small_nucle(ol)ar_rna
protein_coding
protein_coding
protein_coding
small_nucle(ol)ar_rna
protein_coding
protein_coding
protein_coding
protein_coding
protein_coding
protein_coding
protein_coding
protein_coding
protein_coding
protein_coding
protein_coding
protein_coding
protein_coding
protein_coding
protein_coding
protein_coding
protein_coding
protein_coding
protein_coding
protein_coding
protein_coding
protein_coding
protein_coding
protein_coding
protein_coding
protein_coding
protein_coding
protein_coding
protein_coding
protein_coding
protein_coding
protein_coding
protein_coding
protein_coding
protein_coding

protein_coding
protein_coding
protein_coding
protein_coding
protein_coding
protein_coding
protein_coding
protein_coding
small_nucle(ol)ar_rna
small_nucle(ol)ar_rna
protein_coding
protein_coding
protein_coding
ncRNA
protein_coding
protein_coding
protein_coding
protein_coding
protein_coding
protein_coding
protein_coding
protein_coding
protein_coding
protein_coding
protein_coding
protein_coding
protein_coding
protein_coding
protein_coding
protein_coding
protein_coding
protein_coding
protein_coding
protein_coding
protein_coding
protein_coding
protein_coding
protein_coding
protein_coding
protein_coding
protein_coding
ncRNA
protein_coding
protein_coding
protein_coding
miRNA_precursor
protein_coding
protein_coding
protein_coding
protein_coding
protein_coding
protein_coding
pre_trna
protein_coding
protein_coding
protein_coding

protein_coding
protein_coding
protein_coding
protein_coding
protein_coding
protein_coding
protein_coding
protein_coding
protein_coding
protein_coding
protein_coding
protein_coding
protein_coding
protein_coding
protein_coding
protein_coding
protein_coding
protein_coding
protein_coding
small_nucle(ol)ar_rna
protein_coding
protein_coding
protein_coding
protein_coding
protein_coding
protein_coding
protein_coding
protein_coding
ncRNA
protein_coding
protein_coding
protein_coding
ncRNA
protein_coding
protein_coding
protein_coding
protein_coding
protein_coding
protein_coding
protein_coding
protein_coding
protein_coding
protein_coding
protein_coding
protein_coding
protein_coding
protein_coding
protein_coding
protein_coding
protein_coding
protein_coding
protein_coding
protein_coding
protein_coding
protein_coding
protein_coding

protein_coding
protein_coding
protein_coding
protein_coding
protein_coding
protein_coding
protein_coding
protein_coding
protein_coding
protein_coding
ncRNA
protein_coding
protein_coding
protein_coding
protein_coding
protein_coding
protein_coding
protein_coding
protein_coding
protein_coding
protein_coding
protein_coding
protein_coding
protein_coding
protein_coding
protein_coding
protein_coding
protein_coding
protein_coding
protein_coding
protein_coding
protein_coding
protein_coding
protein_coding
protein_coding
protein_coding
protein_coding
protein_coding
protein_coding
protein_coding
protein_coding
protein_coding
protein_coding
protein_coding
protein_coding
protein_coding
protein_coding
protein_coding
protein_coding
small_nucle(ol)ar_rna
protein_coding
protein_coding
protein_coding
protein_coding
protein_coding
protein_coding

protein_coding
protein_coding
protein_coding
protein_coding
protein_coding
protein_coding
protein_coding
protein_coding
protein_coding
protein_coding
protein_coding
protein_coding
protein_coding
protein_coding
protein_coding
protein_coding
protein_coding
protein_coding
protein_coding
protein_coding
protein_coding
protein_coding
ncRNA
protein_coding
small_nucle(ol)ar_rna
protein_coding
protein_coding
protein_coding
protein_coding
protein_coding
protein_coding
protein_coding
protein_coding
protein_coding
ncRNA
protein_coding
protein_coding
protein_coding
protein_coding
protein_coding
protein_coding
protein_coding
protein_coding
protein_coding
protein_coding
protein_coding
protein_coding
pre_trna
protein_coding
protein_coding
ncRNA
protein_coding
protein_coding
protein_coding
protein_coding
protein_coding

protein_coding
protein_coding
protein_coding
protein_coding
protein_coding
protein_coding
protein_coding
protein_coding
protein_coding
protein_coding
protein_coding
protein_coding
protein_coding
protein_coding
protein_coding
protein_coding
protein_coding
protein_coding
protein_coding
protein_coding
protein_coding
protein_coding
protein_coding
protein_coding
protein_coding
protein_coding
protein_coding
protein_coding
protein_coding
protein_coding
protein_coding
protein_coding
protein_coding
protein_coding
protein_coding
protein_coding
protein_coding
protein_coding
pre_trna
protein_coding
protein_coding
protein_coding
protein_coding
protein_coding
protein_coding
protein_coding
protein_coding
protein_coding
protein_coding
protein_coding
protein_coding
protein_coding
protein_coding
protein_coding
protein_coding
protein_coding

protein_coding
protein_coding
protein_coding
protein_coding
protein_coding
protein_coding
protein_coding
protein_coding
protein_coding
protein_coding
protein_coding
protein_coding
protein_coding
protein_coding
protein_coding
protein_coding
protein_coding
protein_coding
protein_coding
ncRNA
protein_coding
protein_coding
protein_coding
protein_coding
protein_coding
ncRNA
protein_coding
protein_coding
protein_coding
protein_coding
protein_coding
protein_coding
protein_coding
protein_coding
protein_coding
protein_coding
protein_coding
protein_coding
protein_coding
protein_coding
protein_coding
protein_coding
protein_coding
protein_coding
protein_coding
protein_coding
protein_coding
protein_coding
protein_coding
protein_coding
protein_coding
protein_coding
pre_trna
ncRNA
protein_coding
protein_coding

protein_coding
ncRNA
protein_coding
protein_coding
protein_coding
pre_trna
protein_coding
protein_coding
protein_coding
protein_coding
protein_coding
protein_coding
protein_coding
protein_coding
protein_coding
protein_coding
protein_coding
protein_coding
protein_coding
protein_coding
protein_coding
protein_coding
protein_coding
protein_coding
protein_coding
protein_coding
protein_coding
protein_coding
protein_coding
protein_coding
protein_coding
protein_coding
protein_coding
protein_coding
protein_coding
protein_coding
protein_coding
small_nucle(ol)ar_rna
small_nucle(ol)ar_rna
pre_trna
protein_coding
protein_coding
protein_coding
protein_coding
protein_coding
protein_coding
protein_coding
protein_coding
protein_coding
protein_coding
protein_coding
protein_coding
protein_coding
protein_coding
pre_trna
protein_coding

ncRNA
protein_coding
protein_coding
protein_coding
protein_coding
protein_coding
protein_coding
ncRNA
protein_coding
protein_coding
protein_coding
protein_coding
protein_coding
protein_coding
protein_coding
protein_coding
protein_coding
protein_coding
protein_coding
protein_coding
protein_coding
protein_coding
protein_coding
protein_coding
miRNA_precursor
protein_coding
protein_coding
ncRNA
protein_coding
protein_coding
protein_coding
pre_trna
protein_coding
protein_coding
protein_coding
protein_coding
protein_coding
protein_coding
protein_coding
protein_coding
ncRNA
protein_coding
protein_coding
protein_coding
protein_coding
protein_coding
protein_coding
protein_coding
protein_coding
protein_coding
protein_coding
protein_coding
protein_coding
protein_coding
protein_coding
protein_coding

protein_coding
protein_coding
protein_coding
protein_coding
protein_coding
protein_coding
protein_coding
protein_coding
protein_coding
small_nucle(ol)ar_rna
protein_coding
protein_coding
protein_coding
protein_coding
ncRNA
protein_coding
protein_coding
protein_coding
protein_coding
protein_coding
protein_coding
protein_coding
protein_coding
protein_coding
protein_coding
protein_coding
protein_coding
protein_coding
protein_coding
protein_coding
protein_coding
protein_coding
protein_coding
protein_coding
small_nucle(ol)ar_rna
protein_coding
protein_coding
protein_coding
protein_coding
protein_coding
protein_coding
protein_coding
protein_coding
protein_coding
protein_coding
protein_coding
protein_coding
protein_coding
protein_coding
protein_coding
protein_coding
protein_coding
protein_coding
protein_coding
protein_coding
protein_coding

protein_coding
protein_coding
protein_coding
protein_coding
protein_coding
protein_coding
protein_coding
protein_coding
protein_coding
protein_coding
protein_coding
protein_coding
protein_coding
protein_coding
protein_coding
protein_coding
protein_coding
protein_coding
protein_coding
protein_coding
protein_coding
ncRNA
protein_coding
protein_coding
protein_coding
protein_coding
protein_coding
protein_coding
ncRNA
protein_coding
protein_coding
protein_coding
protein_coding
protein_coding
protein_coding
protein_coding
protein_coding
protein_coding
protein_coding
protein_coding
protein_coding
protein_coding
protein_coding
ncRNA
protein_coding
protein_coding
ncRNA
protein_coding
protein_coding
protein_coding
protein_coding
protein_coding
protein_coding
protein_coding
protein_coding
protein_coding

protein_coding
protein_coding
protein_coding
protein_coding
protein_coding
protein_coding
protein_coding
protein_coding
protein_coding
protein_coding
protein_coding
protein_coding
protein_coding
protein_coding
protein_coding
ncRNA
protein_coding
protein_coding
protein_coding
protein_coding
protein_coding
protein_coding
protein_coding
protein_coding
protein_coding
protein_coding
protein_coding
protein_coding
protein_coding
protein_coding
protein_coding
protein_coding
protein_coding
protein_coding
protein_coding
protein_coding
ncRNA
protein_coding
ncRNA
protein_coding
protein_coding
protein_coding
protein_coding
protein_coding
protein_coding
protein_coding
protein_coding
protein_coding
protein_coding
protein_coding
protein_coding
protein_coding
protein_coding
protein_coding
protein_coding
protein_coding

protein_coding
protein_coding
protein_coding
protein_coding
protein_coding
protein_coding
protein_coding
protein_coding
protein_coding
protein_coding
protein_coding
protein_coding
protein_coding
protein_coding
protein_coding
protein_coding
protein_coding
ncRNA
protein_coding
protein_coding
protein_coding
protein_coding
protein_coding
protein_coding
protein_coding
protein_coding
protein_coding
protein_coding
protein_coding
protein_coding
protein_coding
protein_coding
protein_coding
protein_coding
protein_coding
protein_coding
protein_coding
protein_coding
protein_coding
protein_coding
pre_trna
protein_coding
protein_coding
protein_coding
protein_coding
protein_coding
protein_coding
protein_coding
protein_coding
protein_coding
protein_coding
protein_coding
protein_coding
protein_coding
protein_coding
protein_coding

protein_coding
protein_coding
protein_coding
protein_coding
protein_coding
protein_coding
protein_coding
protein_coding
protein_coding
protein_coding
protein_coding
protein_coding
protein_coding
protein_coding
protein_coding
protein_coding
protein_coding
protein_coding
ncRNA
protein_coding
protein_coding
protein_coding
protein_coding
protein_coding
protein_coding
protein_coding
protein_coding
protein_coding
protein_coding
protein_coding
protein_coding
protein_coding
small_nucle(ol)ar_rna
protein_coding
protein_coding
protein_coding
protein_coding
ncRNA
protein_coding
protein_coding
protein_coding
protein_coding
protein_coding
protein_coding
protein_coding
protein_coding
protein_coding
protein_coding
ncRNA
protein_coding
protein_coding
protein_coding
ncRNA
protein_coding
protein_coding
protein_coding

protein_coding
protein_coding
protein_coding
protein_coding
protein_coding
protein_coding
protein_coding
protein_coding
protein_coding
protein_coding
protein_coding
protein_coding
protein_coding
protein_coding
protein_coding
protein_coding
protein_coding
protein_coding
protein_coding
ncRNA
protein_coding
protein_coding
protein_coding
protein_coding
ncRNA
protein_coding
protein_coding
protein_coding
protein_coding
protein_coding
pre_trna
protein_coding
protein_coding
protein_coding
protein_coding
ncRNA
protein_coding
protein_coding
ncRNA
protein_coding
protein_coding
protein_coding
protein_coding
protein_coding
protein_coding
protein_coding
protein_coding
protein_coding
protein_coding
protein_coding
protein_coding
protein_coding
protein_coding
ncRNA
protein_coding
protein_coding

protein_coding
small_nucle(ol)ar_rna
protein_coding
protein_coding
protein_coding
protein_coding
protein_coding
protein_coding
protein_coding
protein_coding
protein_coding
protein_coding
protein_coding
protein_coding
protein_coding
protein_coding
protein_coding
protein_coding
protein_coding
protein_coding
protein_coding
protein_coding
protein_coding
protein_coding
protein_coding
protein_coding
protein_coding
protein_coding
pre_trna
protein_coding
protein_coding
protein_coding
protein_coding
protein_coding
protein_coding
protein_coding
protein_coding
protein_coding
protein_coding
protein_coding
protein_coding
protein_coding
protein_coding
protein_coding
protein_coding
protein_coding
protein_coding
miRNA_precursor
protein_coding
protein_coding
protein_coding
protein_coding
protein_coding
protein_coding
protein_coding
protein_coding

protein_coding
protein_coding
protein_coding
protein_coding
protein_coding
protein_coding
protein_coding
protein_coding
protein_coding
protein_coding
protein_coding
protein_coding
protein_coding
protein_coding
protein_coding
protein_coding
protein_coding
protein_coding
ncRNA
protein_coding
ncRNA
protein_coding
protein_coding
protein_coding
protein_coding
protein_coding
protein_coding
protein_coding
protein_coding
protein_coding
protein_coding
protein_coding
protein_coding
protein_coding
protein_coding
protein_coding
protein_coding
protein_coding
protein_coding
protein_coding
protein_coding
protein_coding
protein_coding
small_nucle(ol)ar_rna
protein_coding
protein_coding
protein_coding
protein_coding
protein_coding
protein_coding
protein_coding
protein_coding
protein_coding
protein_coding
protein_coding
protein_coding

protein_coding
protein_coding
ncRNA
ncRNA
small_nucle(ol)ar_rna
protein_coding
protein_coding
protein_coding
protein_coding
protein_coding
protein_coding
protein_coding
protein_coding
protein_coding
small_nucle(ol)ar_rna
protein_coding
protein_coding
protein_coding
protein_coding
protein_coding
protein_coding
protein_coding
protein_coding
protein_coding
protein_coding
protein_coding
protein_coding
ncRNA
protein_coding
protein_coding
protein_coding
protein_coding
protein_coding
protein_coding
protein_coding
protein_coding
protein_coding
protein_coding
protein_coding
protein_coding
protein_coding
protein_coding
protein_coding
protein_coding
protein_coding
protein_coding
protein_coding
protein_coding
protein_coding
protein_coding
protein_coding
protein_coding
protein_coding
protein_coding
protein_coding
ncRNA

protein_coding
protein_coding
protein_coding
small_nucle(ol)ar_rna
protein_coding
protein_coding
protein_coding
protein_coding
protein_coding
protein_coding
protein_coding
protein_coding
protein_coding
protein_coding
protein_coding
protein_coding
protein_coding
protein_coding
protein_coding
protein_coding
protein_coding
protein_coding
protein_coding
protein_coding
protein_coding
ncRNA
protein_coding
protein_coding
protein_coding
protein_coding
protein_coding
protein_coding
protein_coding
protein_coding
ncRNA
protein_coding
protein_coding
protein_coding
protein_coding
protein_coding
protein_coding
protein_coding
protein_coding
protein_coding
protein_coding
protein_coding
protein_coding
protein_coding
protein_coding
protein_coding
protein_coding
protein_coding
protein_coding
protein_coding
small_nucle(ol)ar_rna
protein_coding

small_nucle(ol)ar_rna
protein_coding
protein_coding
ncRNA
protein_coding
protein_coding
protein_coding
protein_coding
protein_coding
protein_coding
ncRNA
protein_coding
protein_coding
ncRNA
protein_coding
protein_coding
protein_coding
protein_coding
ncRNA
miRNA_precursor
protein_coding
protein_coding
protein_coding
protein_coding
protein_coding
protein_coding
miRNA_precursor
protein_coding
protein_coding
small_nucle(ol)ar_rna
protein_coding
protein_coding
protein_coding
protein_coding
ncRNA
protein_coding
protein_coding
protein_coding
protein_coding
protein_coding
protein_coding
protein_coding
protein_coding
protein_coding
protein_coding
protein_coding
protein_coding
protein_coding
protein_coding
protein_coding
protein_coding
protein_coding
protein_coding
protein_coding
protein_coding
protein_coding

protein_coding
protein_coding
ncRNA
ncRNA
miRNA_precursor
protein_coding
ncRNA
protein_coding
protein_coding
protein_coding
protein_coding
protein_coding
protein_coding
protein_coding
protein_coding
protein_coding
protein_coding
protein_coding
protein_coding
protein_coding
protein_coding
ncRNA
protein_coding
protein_coding
protein_coding
protein_coding
protein_coding
protein_coding
protein_coding
protein_coding
protein_coding
protein_coding
protein_coding
protein_coding
protein_coding
protein_coding
protein_coding
protein_coding
protein_coding
protein_coding
protein_coding
protein_coding
protein_coding
protein_coding
ncRNA
protein_coding
protein_coding
protein_coding
protein_coding
protein_coding
protein_coding
protein_coding
protein_coding
protein_coding
protein_coding
protein_coding

protein_coding
ncRNA
protein_coding
protein_coding
ncRNA
protein_coding
protein_coding
ncRNA
protein_coding
protein_coding
ncRNA
small_nucle(ol)ar_rna
protein_coding
protein_coding
protein_coding
protein_coding
protein_coding
protein_coding
protein_coding
protein_coding
protein_coding
protein_coding
protein_coding
protein_coding
protein_coding
protein_coding
protein_coding
protein_coding
protein_coding
protein_coding
protein_coding
protein_coding
protein_coding
protein_coding
protein_coding
miRNA_precursor
protein_coding
protein_coding
protein_coding
protein_coding
protein_coding
protein_coding
protein_coding
protein_coding
protein_coding
protein_coding
protein_coding
protein_coding
protein_coding
protein_coding
protein_coding
protein_coding
protein_coding
protein_coding
protein_coding
ncRNA

protein_coding
protein_coding
protein_coding
protein_coding
protein_coding
protein_coding
ncRNA
protein_coding
protein_coding
protein_coding
protein_coding
ncRNA
protein_coding
protein_coding
protein_coding
ncRNA
protein_coding
protein_coding
protein_coding
protein_coding
protein_coding
protein_coding
ncRNA
protein_coding
protein_coding
protein_coding
protein_coding
protein_coding
protein_coding
protein_coding
protein_coding
ncRNA
protein_coding
protein_coding
protein_coding
protein_coding
protein_coding
protein_coding
protein_coding
protein_coding
protein_coding
protein_coding
miRNA_precursor
protein_coding
protein_coding
protein_coding
protein_coding
protein_coding
protein_coding
protein_coding
protein_coding
protein_coding
protein_coding
protein_coding
protein_coding
protein_coding

protein_coding
protein_coding
protein_coding
protein_coding
protein_coding
protein_coding
protein_coding
protein_coding
protein_coding
protein_coding
protein_coding
protein_coding
protein_coding
protein_coding
protein_coding
protein_coding
protein_coding
protein_coding
protein_coding
protein_coding
protein_coding
protein_coding
protein_coding
protein_coding
protein_coding
protein_coding
protein_coding
protein_coding
protein_coding
protein_coding
protein_coding
protein_coding
protein_coding
protein_coding
protein_coding
protein_coding
protein_coding
ncRNA
protein_coding
protein_coding
protein_coding
protein_coding
protein_coding
protein_coding
protein_coding
protein_coding
protein_coding
protein_coding
protein_coding
protein_coding
protein_coding
protein_coding
protein_coding
protein_coding
protein_coding
protein_coding

protein_coding
protein_coding
protein_coding
protein_coding
protein_coding
ncRNA
protein_coding
small_nucle(ol)ar_rna
small_nucle(ol)ar_rna
small_nucle(ol)ar_rna
protein_coding
protein_coding
protein_coding
protein_coding
protein_coding
protein_coding
protein_coding
protein_coding
protein_coding
protein_coding
protein_coding
protein_coding
protein_coding
protein_coding
protein_coding
protein_coding
protein_coding
protein_coding
protein_coding
protein_coding
protein_coding
protein_coding
protein_coding
protein_coding
protein_coding
protein_coding
protein_coding
protein_coding
protein_coding
protein_coding
protein_coding
protein_coding
protein_coding
protein_coding
protein_coding
protein_coding
protein_coding
protein_coding
protein_coding
protein_coding
protein_coding
protein_coding
protein_coding
protein_coding
protein_coding
protein_coding

ncRNA
protein_coding
protein_coding
protein_coding
protein_coding
protein_coding
protein_coding
protein_coding
protein_coding
protein_coding
protein_coding
protein_coding
protein_coding
protein_coding
protein_coding
protein_coding
protein_coding
protein_coding
protein_coding
protein_coding
protein_coding
protein_coding
protein_coding
protein_coding
protein_coding
protein_coding
protein_coding
protein_coding
protein_coding
protein_coding
protein_coding
ncRNA
protein_coding
protein_coding
protein_coding
protein_coding
protein_coding
protein_coding
protein_coding
protein_coding
protein_coding
protein_coding
protein_coding
protein_coding
protein_coding
protein_coding
protein_coding
protein_coding
protein_coding
protein_coding
protein_coding
protein_coding
protein_coding
protein_coding
protein_coding
protein_coding

protein_coding
protein_coding
protein_coding
protein_coding
ncRNA
protein_coding
protein_coding
protein_coding
protein_coding
protein_coding
protein_coding
protein_coding
protein_coding
protein_coding
protein_coding
protein_coding
protein_coding
protein_coding
protein_coding
protein_coding
protein_coding
protein_coding
protein_coding
protein_coding
protein_coding
protein_coding
protein_coding
protein_coding
protein_coding
protein_coding
ncRNA
protein_coding
protein_coding
protein_coding
protein_coding
protein_coding
protein_coding
protein_coding
protein_coding
protein_coding
protein_coding
protein_coding
ncRNA
protein_coding
protein_coding
protein_coding
protein_coding
protein_coding
protein_coding
protein_coding
protein_coding
protein_coding
protein_coding
protein_coding
protein_coding
protein_coding

protein_coding
protein_coding
protein_coding
protein_coding
protein_coding
protein_coding
protein_coding
protein_coding
protein_coding
protein_coding
protein_coding
protein_coding
protein_coding
protein_coding
protein_coding
protein_coding
protein_coding
protein_coding
protein_coding
protein_coding
miRNA_precursor
protein_coding
protein_coding
protein_coding
protein_coding
protein_coding
protein_coding
protein_coding
protein_coding
protein_coding
protein_coding
protein_coding
protein_coding
protein_coding
protein_coding
protein_coding
protein_coding
protein_coding
protein_coding
ncRNA
protein_coding
protein_coding
protein_coding
protein_coding
protein_coding
protein_coding
protein_coding
protein_coding
protein_coding
protein_coding
protein_coding
protein_coding
protein_coding
protein_coding
protein_coding
protein_coding

protein_coding
protein_coding
protein_coding
miRNA_precursor
protein_coding
protein_coding
protein_coding
protein_coding
protein_coding
protein_coding
protein_coding
protein_coding
protein_coding
protein_coding
protein_coding
protein_coding
protein_coding
protein_coding
protein_coding
protein_coding
protein_coding
protein_coding
protein_coding
protein_coding
protein_coding
protein_coding
protein_coding
protein_coding
protein_coding
protein_coding
protein_coding
protein_coding
protein_coding
protein_coding
protein_coding
protein_coding
protein_coding
protein_coding
protein_coding
protein_coding
ncRNA
protein_coding
protein_coding
protein_coding
protein_coding
protein_coding
protein_coding
protein_coding
protein_coding
protein_coding
protein_coding
protein_coding
protein_coding
protein_coding
protein_coding
protein_coding

protein_coding
protein_coding
ncRNA
protein_coding
protein_coding
protein_coding
protein_coding
protein_coding
protein_coding
protein_coding
ncRNA
protein_coding
protein_coding
protein_coding
protein_coding
protein_coding
protein_coding
protein_coding
protein_coding
protein_coding
protein_coding
protein_coding
protein_coding
protein_coding
protein_coding
protein_coding
protein_coding
protein_coding
protein_coding
protein_coding
protein_coding
protein_coding
protein_coding
protein_coding
protein_coding
protein_coding
protein_coding
protein_coding
protein_coding
protein_coding
protein_coding
protein_coding
small_nucle(ol)ar_rna
protein_coding
protein_coding
protein_coding
protein_coding
protein_coding
protein_coding
protein_coding
protein_coding
protein_coding
protein_coding
protein_coding
protein_coding
protein_coding

protein_coding
protein_coding
protein_coding
protein_coding
protein_coding
protein_coding
protein_coding
protein_coding
protein_coding
protein_coding
protein_coding
protein_coding
protein_coding
protein_coding
protein_coding
protein_coding
protein_coding
protein_coding
protein_coding
protein_coding
protein_coding
protein_coding
protein_coding
ncRNA
protein_coding
protein_coding
protein_coding
protein_coding
protein_coding
protein_coding
protein_coding
protein_coding
ncRNA
protein_coding
protein_coding
protein_coding
protein_coding
protein_coding
protein_coding
protein_coding
protein_coding
protein_coding
protein_coding
protein_coding
protein_coding
protein_coding
protein_coding
protein_coding
protein_coding
protein_coding
protein_coding
protein_coding
protein_coding
protein_coding
protein_coding
protein_coding

protein_coding
protein_coding
protein_coding
protein_coding
protein_coding
protein_coding
protein_coding
protein_coding
protein_coding
protein_coding
protein_coding
protein_coding
protein_coding
ncRNA
protein_coding
protein_coding
protein_coding
protein_coding
protein_coding
protein_coding
protein_coding
protein_coding
protein_coding
protein_coding
protein_coding
protein_coding
protein_coding
small_nucle(ol)ar_rna
small_nucle(ol)ar_rna
protein_coding
protein_coding
protein_coding
protein_coding
protein_coding
protein_coding
protein_coding
protein_coding
protein_coding
protein_coding
protein_coding
protein_coding
protein_coding
protein_coding
protein_coding
ncRNA
protein_coding
protein_coding
protein_coding
protein_coding
protein_coding
protein_coding
protein_coding
protein_coding
protein_coding
protein_coding
protein_coding

protein_coding
protein_coding
protein_coding
protein_coding
protein_coding
protein_coding
protein_coding
protein_coding
protein_coding
protein_coding
protein_coding
protein_coding
protein_coding
protein_coding
protein_coding
protein_coding
protein_coding
protein_coding
protein_coding
protein_coding
protein_coding
protein_coding
protein_coding
protein_coding
protein_coding
protein_coding
protein_coding
protein_coding
protein_coding
protein_coding
protein_coding
protein_coding
ncRNA
protein_coding
protein_coding
protein_coding
protein_coding
protein_coding
protein_coding
protein_coding
protein_coding
protein_coding
protein_coding
protein_coding
protein_coding
protein_coding
protein_coding
protein_coding
protein_coding
protein_coding
protein_coding
protein_coding
protein_coding
protein_coding
protein_coding
ncRNA

protein_coding
protein_coding
protein_coding
protein_coding
protein_coding
protein_coding
protein_coding
protein_coding
protein_coding
protein_coding
protein_coding
pre_trna
protein_coding
protein_coding
protein_coding
protein_coding
protein_coding
protein_coding
protein_coding
protein_coding
ncRNA
protein_coding
protein_coding
protein_coding
protein_coding
protein_coding
protein_coding
protein_coding
protein_coding
protein_coding
protein_coding
protein_coding
protein_coding
protein_coding
protein_coding
protein_coding
protein_coding
protein_coding
protein_coding
protein_coding
protein_coding
protein_coding
ncRNA
protein_coding
ncRNA
protein_coding
ncRNA
protein_coding
protein_coding
protein_coding
protein_coding
protein_coding
protein_coding
protein_coding
protein_coding
protein_coding

protein_coding
protein_coding
protein_coding
protein_coding
protein_coding
protein_coding
protein_coding
protein_coding
protein_coding
protein_coding
protein_coding
protein_coding
protein_coding
protein_coding
ncRNA
protein_coding
protein_coding
protein_coding
protein_coding
pre_trna
protein_coding
protein_coding
protein_coding
protein_coding
protein_coding
protein_coding
ncRNA
ncRNA
protein_coding
protein_coding
small_nucle(ol)ar_rna
protein_coding
protein_coding
protein_coding
protein_coding
ncRNA
protein_coding
protein_coding
protein_coding
ncRNA
protein_coding
ncRNA
protein_coding
protein_coding
protein_coding
protein_coding
protein_coding
protein_coding
protein_coding
protein_coding
protein_coding
protein_coding
protein_coding
protein_coding
protein_coding
protein_coding

protein_coding
protein_coding
protein_coding
protein_coding
protein_coding
protein_coding
ncRNA
protein_coding
protein_coding
ncRNA
protein_coding
protein_coding
ncRNA
protein_coding
protein_coding
protein_coding
protein_coding
protein_coding
ncRNA
protein_coding
protein_coding
ncRNA
protein_coding
protein_coding
protein_coding
protein_coding
protein_coding
protein_coding
protein_coding
protein_coding
protein_coding
protein_coding
protein_coding
protein_coding
protein_coding
protein_coding
protein_coding
protein_coding
protein_coding
protein_coding
protein_coding
protein_coding
protein_coding
protein_coding
protein_coding
protein_coding
protein_coding
ncRNA
protein_coding
protein_coding
protein_coding
protein_coding
protein_coding
protein_coding
protein_coding
protein_coding

protein_coding
protein_coding
protein_coding
protein_coding
protein_coding
protein_coding
protein_coding
pre_trna
protein_coding
protein_coding
protein_coding
protein_coding
protein_coding
protein_coding
protein_coding
protein_coding
ncRNA
ncRNA
protein_coding
protein_coding
protein_coding
protein_coding
protein_coding
miRNA_precursor
protein_coding
protein_coding
protein_coding
protein_coding
protein_coding
protein_coding
protein_coding
protein_coding
protein_coding
protein_coding
protein_coding
protein_coding
protein_coding
protein_coding
protein_coding
protein_coding
protein_coding
protein_coding
protein_coding
protein_coding
protein_coding
protein_coding
ncRNA
protein_coding
small_nucle(ol)ar_rna
protein_coding
protein_coding
protein_coding
protein_coding
protein_coding
protein_coding
protein_coding

protein_coding
protein_coding
protein_coding
protein_coding
protein_coding
ncRNA
protein_coding
protein_coding
protein_coding
protein_coding
protein_coding
protein_coding
protein_coding
protein_coding
protein_coding
protein_coding
protein_coding
protein_coding
protein_coding
protein_coding
protein_coding
protein_coding
protein_coding
protein_coding
protein_coding
ncRNA
protein_coding
protein_coding
protein_coding
protein_coding
protein_coding
protein_coding
protein_coding
protein_coding
protein_coding
protein_coding
protein_coding
protein_coding
protein_coding
protein_coding
protein_coding
protein_coding
protein_coding
miRNA_precursor
protein_coding
protein_coding
protein_coding
protein_coding
protein_coding
protein_coding
protein_coding
protein_coding
protein_coding
protein_coding
protein_coding
protein_coding

protein_coding
protein_coding
protein_coding
protein_coding
protein_coding
ncRNA
ncRNA
protein_coding
protein_coding
protein_coding
protein_coding
ncRNA
protein_coding
protein_coding
protein_coding
protein_coding
miRNA_precursor
protein_coding
protein_coding
ncRNA
protein_coding
protein_coding
protein_coding
protein_coding
protein_coding
protein_coding
protein_coding
protein_coding
protein_coding
protein_coding
protein_coding
protein_coding
protein_coding
protein_coding
protein_coding
protein_coding
protein_coding
protein_coding
protein_coding
protein_coding
protein_coding
protein_coding
protein_coding
protein_coding
protein_coding
ncRNA
protein_coding
protein_coding
protein_coding
protein_coding
protein_coding
protein_coding
protein_coding
protein_coding
protein_coding
protein_coding

protein_coding
protein_coding
protein_coding
protein_coding
protein_coding
protein_coding
protein_coding
protein_coding
protein_coding
protein_coding
protein_coding
protein_coding
protein_coding
protein_coding
protein_coding
protein_coding
protein_coding
protein_coding
protein_coding
protein_coding
protein_coding
protein_coding
protein_coding
protein_coding
protein_coding
miRNA_precursor
protein_coding
protein_coding
protein_coding
protein_coding
protein_coding
ncRNA
protein_coding
protein_coding
protein_coding
protein_coding
pre_trna
protein_coding
protein_coding
protein_coding
protein_coding
ncRNA
protein_coding
protein_coding
protein_coding
protein_coding
protein_coding
protein_coding
protein_coding
protein_coding
protein_coding
protein_coding
protein_coding
protein_coding
protein_coding
protein_coding

protein_coding
protein_coding
protein_coding
protein_coding
protein_coding
protein_coding
protein_coding
protein_coding
pre_trna
protein_coding
ncRNA
protein_coding
protein_coding
protein_coding
protein_coding
protein_coding
protein_coding
protein_coding
protein_coding
protein_coding
ncRNA
protein_coding
protein_coding
protein_coding
protein_coding
protein_coding
protein_coding
protein_coding
protein_coding
protein_coding
ncRNA
protein_coding
protein_coding
protein_coding
protein_coding
ncRNA
protein_coding
protein_coding
protein_coding
protein_coding
protein_coding
protein_coding
protein_coding
protein_coding
protein_coding
protein_coding
protein_coding
protein_coding
protein_coding
protein_coding
protein_coding
protein_coding
protein_coding
protein_coding
protein_coding
protein_coding

protein_coding
ncRNA
protein_coding
protein_coding
protein_coding
protein_coding
protein_coding
protein_coding
protein_coding
protein_coding
protein_coding
protein_coding
protein_coding
protein_coding
protein_coding
protein_coding
protein_coding
protein_coding
protein_coding
protein_coding
protein_coding
ncRNA
protein_coding
protein_coding
protein_coding
protein_coding
protein_coding
protein_coding
protein_coding
protein_coding
protein_coding
protein_coding
protein_coding
protein_coding
protein_coding
protein_coding
protein_coding
protein_coding
protein_coding
protein_coding
protein_coding
protein_coding
protein_coding
protein_coding
protein_coding
protein_coding
protein_coding
protein_coding
protein_coding
protein_coding
protein_coding
protein_coding
protein_coding
protein_coding
small_nucle(ol)ar_rna
small_nucle(ol)ar_rna

small_nucle(ol)ar_rna
ncRNA
protein_coding
protein_coding
protein_coding
protein_coding
protein_coding
protein_coding
protein_coding
protein_coding
protein_coding
protein_coding
protein_coding
protein_coding
protein_coding
protein_coding
protein_coding
protein_coding
protein_coding
protein_coding
protein_coding
protein_coding
protein_coding
protein_coding
protein_coding
protein_coding
protein_coding
protein_coding
protein_coding
protein_coding
protein_coding
protein_coding
protein_coding
protein_coding
protein_coding
protein_coding
protein_coding
protein_coding
protein_coding
protein_coding
protein_coding
protein_coding
protein_coding
protein_coding
protein_coding
protein_coding
protein_coding
protein_coding
ncRNA
small_nucle(ol)ar_rna
small_nucle(ol)ar_rna
ncRNA
protein_coding
protein_coding
protein_coding
protein_coding

protein_coding
protein_coding
protein_coding
protein_coding
protein_coding
protein_coding
protein_coding
protein_coding
protein_coding
protein_coding
protein_coding
protein_coding
protein_coding
protein_coding
protein_coding
protein_coding
protein_coding
ncRNA
protein_coding
protein_coding
protein_coding
protein_coding
ncRNA
protein_coding
protein_coding
protein_coding
protein_coding
protein_coding
protein_coding
miRNA_precursor
protein_coding
protein_coding
ncRNA
protein_coding
protein_coding
protein_coding
protein_coding
protein_coding
protein_coding
protein_coding
protein_coding
protein_coding
protein_coding
protein_coding
protein_coding
protein_coding
protein_coding
protein_coding
protein_coding
protein_coding
protein_coding
protein_coding
protein_coding
protein_coding
protein_coding
protein_coding

protein_coding
protein_coding
protein_coding
protein_coding
ncRNA
protein_coding
protein_coding
protein_coding
protein_coding
protein_coding
protein_coding
protein_coding
protein_coding
miRNA_precursor
protein_coding
protein_coding
protein_coding
protein_coding
protein_coding
protein_coding
protein_coding
protein_coding
protein_coding
protein_coding
protein_coding
protein_coding
pre_trna
protein_coding
protein_coding
protein_coding
ncRNA
protein_coding
protein_coding
protein_coding
protein_coding
protein_coding
protein_coding
protein_coding
protein_coding
protein_coding
protein_coding
protein_coding
protein_coding
protein_coding
protein_coding
protein_coding
protein_coding
protein_coding
protein_coding
protein_coding
protein_coding
protein_coding
protein_coding
protein_coding
protein_coding
protein_coding

protein_coding
protein_coding
protein_coding
protein_coding
protein_coding
protein_coding
ncRNA
protein_coding
protein_coding
protein_coding
protein_coding
ncRNA
protein_coding
protein_coding
protein_coding
ncRNA
protein_coding
protein_coding
protein_coding
ncRNA
protein_coding
protein_coding
protein_coding
protein_coding
protein_coding
protein_coding
protein_coding
protein_coding
protein_coding
protein_coding
protein_coding
protein_coding
protein_coding
protein_coding
protein_coding
protein_coding
protein_coding
protein_coding
protein_coding
ncRNA
protein_coding
protein_coding
protein_coding
protein_coding
protein_coding
protein_coding
protein_coding
protein_coding
protein_coding
protein_coding
protein_coding
protein_coding
protein_coding
protein_coding
protein_coding
protein_coding

protein_coding
protein_coding
protein_coding
protein_coding
protein_coding
protein_coding
protein_coding
protein_coding
protein_coding
protein_coding
protein_coding
protein_coding
protein_coding
protein_coding
protein_coding
protein_coding
protein_coding
protein_coding
protein_coding
protein_coding
protein_coding
protein_coding
protein_coding
protein_coding
protein_coding
protein_coding
protein_coding
protein_coding
protein_coding
protein_coding
protein_coding
protein_coding
protein_coding
protein_coding
protein_coding
protein_coding
protein_coding
ncRNA
protein_coding
protein_coding
protein_coding
protein_coding
protein_coding
protein_coding
protein_coding
protein_coding
protein_coding
protein_coding
protein_coding
protein_coding
protein_coding
protein_coding
protein_coding
protein_coding
protein_coding
protein_coding

protein_coding
protein_coding
protein_coding
protein_coding
protein_coding
protein_coding
protein_coding
protein_coding
protein_coding
protein_coding
protein_coding
protein_coding
protein_coding
protein_coding
protein_coding
protein_coding
protein_coding
protein_coding
protein_coding
protein_coding
protein_coding
protein_coding
protein_coding
protein_coding
ncRNA
protein_coding
protein_coding
protein_coding
protein_coding
protein_coding
protein_coding
protein_coding
protein_coding
protein_coding
protein_coding
protein_coding
protein_coding
protein_coding
protein_coding
protein_coding
protein_coding
protein_coding
protein_coding
protein_coding
ncRNA
protein_coding
protein_coding
protein_coding
protein_coding
protein_coding
protein_coding
protein_coding
protein_coding
protein_coding
protein_coding
protein_coding

protein_coding
protein_coding
protein_coding
protein_coding
protein_coding
protein_coding
protein_coding
protein_coding
protein_coding
protein_coding
pre_trna
protein_coding
protein_coding
protein_coding
protein_coding
protein_coding
protein_coding
protein_coding
small_nucle(ol)ar_rna
protein_coding
protein_coding
protein_coding
protein_coding
protein_coding
protein_coding
pre_trna
protein_coding
ncRNA
protein_coding
protein_coding
protein_coding
protein_coding
protein_coding
protein_coding
protein_coding
protein_coding
protein_coding
protein_coding
protein_coding
protein_coding
protein_coding
protein_coding
protein_coding
protein_coding
protein_coding
protein_coding
protein_coding
protein_coding
protein_coding
protein_coding
protein_coding
protein_coding
protein_coding
protein_coding
protein_coding
protein_coding

protein_coding
protein_coding
protein_coding
protein_coding
protein_coding
protein_coding
protein_coding
protein_coding
protein_coding
protein_coding
protein_coding
protein_coding
protein_coding
protein_coding
protein_coding
protein_coding
protein_coding
protein_coding
protein_coding
protein_coding
protein_coding
protein_coding
protein_coding
protein_coding
protein_coding
protein_coding
protein_coding
protein_coding
protein_coding
protein_coding
protein_coding
protein_coding
protein_coding
protein_coding
protein_coding
protein_coding
protein_coding
protein_coding
protein_coding
protein_coding
protein_coding
protein_coding
protein_coding
protein_coding
protein_coding
protein_coding
protein_coding
miRNA_precursor
protein_coding
protein_coding
small_nucle(ol)ar_rna
small_nucle(ol)ar_rna
ncRNA
protein_coding
protein_coding
protein_coding

protein_coding
protein_coding
protein_coding
protein_coding
protein_coding
protein_coding
protein_coding
protein_coding
protein_coding
protein_coding
protein_coding
protein_coding
protein_coding
protein_coding
protein_coding
protein_coding
protein_coding
protein_coding
protein_coding
protein_coding
protein_coding
protein_coding
protein_coding
protein_coding
protein_coding
protein_coding
protein_coding
protein_coding
protein_coding
protein_coding
protein_coding
protein_coding
protein_coding
ncRNA
protein_coding
protein_coding
protein_coding
protein_coding
protein_coding
ncRNA
protein_coding
protein_coding
protein_coding
protein_coding
protein_coding
protein_coding
protein_coding
protein_coding
protein_coding
protein_coding
protein_coding
protein_coding
protein_coding
protein_coding
protein_coding
protein_coding

protein_coding
protein_coding
protein_coding
protein_coding
protein_coding
protein_coding
protein_coding
protein_coding
protein_coding
protein_coding
protein_coding
protein_coding
protein_coding
protein_coding
protein_coding
protein_coding
protein_coding
protein_coding
protein_coding
protein_coding
protein_coding
protein_coding
protein_coding
protein_coding
protein_coding
protein_coding
protein_coding
protein_coding
protein_coding
protein_coding
ncRNA
protein_coding
protein_coding
protein_coding
protein_coding
protein_coding
protein_coding
protein_coding
protein_coding
protein_coding
protein_coding
protein_coding
protein_coding
protein_coding
protein_coding
miRNA_precursor
protein_coding
protein_coding
protein_coding
protein_coding
protein_coding
protein_coding
protein_coding
protein_coding
protein_coding
protein_coding

protein_coding
protein_coding
protein_coding
protein_coding
small_nucle(ol)ar_rna
small_nucle(ol)ar_rna
protein_coding
protein_coding
protein_coding
protein_coding
protein_coding
protein_coding
protein_coding
protein_coding
protein_coding
protein_coding
small_nucle(ol)ar_rna
protein_coding
protein_coding
protein_coding
protein_coding
protein_coding
protein_coding
protein_coding
protein_coding
protein_coding
protein_coding
protein_coding
protein_coding
protein_coding
protein_coding
protein_coding
protein_coding
protein_coding
protein_coding
protein_coding
protein_coding
ncRNA
protein_coding
protein_coding
protein_coding
protein_coding
protein_coding
protein_coding
protein_coding
protein_coding
protein_coding
protein_coding
protein_coding
protein_coding
protein_coding
protein_coding
protein_coding
protein_coding
protein_coding
protein_coding

protein_coding
protein_coding
protein_coding
protein_coding
protein_coding
protein_coding
protein_coding
protein_coding
protein_coding
protein_coding
protein_coding
protein_coding
protein_coding
pre_trna
protein_coding
ncRNA
protein_coding
protein_coding
protein_coding
protein_coding
protein_coding
protein_coding
protein_coding
protein_coding
protein_coding
protein_coding
protein_coding
protein_coding
protein_coding
protein_coding
protein_coding
protein_coding
protein_coding
protein_coding
protein_coding
protein_coding
protein_coding
protein_coding
protein_coding
protein_coding
protein_coding
protein_coding
protein_coding
protein_coding
protein_coding
protein_coding
protein_coding
protein_coding
protein_coding
protein_coding
protein_coding
protein_coding
protein_coding
protein_coding
protein_coding
protein_coding

protein_coding
protein_coding
protein_coding
protein_coding
protein_coding
protein_coding
protein_coding
protein_coding
protein_coding
protein_coding
protein_coding
protein_coding
ncRNA
protein_coding
protein_coding
protein_coding
protein_coding
protein_coding
protein_coding
protein_coding
protein_coding
protein_coding
protein_coding
protein_coding
protein_coding
protein_coding
protein_coding
protein_coding
protein_coding
protein_coding
protein_coding
protein_coding
ncRNA
small_nucle(ol)ar_rna
protein_coding
protein_coding
protein_coding
protein_coding
protein_coding
protein_coding
protein_coding
protein_coding
protein_coding
protein_coding
protein_coding
protein_coding
ncRNA
protein_coding
protein_coding
protein_coding
protein_coding
protein_coding
protein_coding
protein_coding
protein_coding
protein_coding

protein_coding
protein_coding
protein_coding
protein_coding
protein_coding
protein_coding
protein_coding
protein_coding
protein_coding
protein_coding
protein_coding
protein_coding
protein_coding
protein_coding
protein_coding
protein_coding
protein_coding
miRNA_precursor
miRNA_precursor
protein_coding
protein_coding
protein_coding
protein_coding
protein_coding
protein_coding
protein_coding
protein_coding
protein_coding
protein_coding
protein_coding
protein_coding
protein_coding
protein_coding
ncRNA
ncRNA
protein_coding
protein_coding
protein_coding
protein_coding
protein_coding
protein_coding
protein_coding
protein_coding
protein_coding
protein_coding
ncRNA
protein_coding
protein_coding
protein_coding
protein_coding
pre_trna
protein_coding
protein_coding
protein_coding
protein_coding
protein_coding

protein_coding
protein_coding
protein_coding
protein_coding
protein_coding
protein_coding
ncRNA
protein_coding
protein_coding
protein_coding
protein_coding
protein_coding
protein_coding
ncRNA
protein_coding
protein_coding
ncRNA
protein_coding
protein_coding
protein_coding
protein_coding
protein_coding
protein_coding
protein_coding
protein_coding
protein_coding
protein_coding
protein_coding
protein_coding
protein_coding
protein_coding
protein_coding
protein_coding
protein_coding
protein_coding
protein_coding
protein_coding
protein_coding
protein_coding
protein_coding
protein_coding
protein_coding
protein_coding
protein_coding
protein_coding
miRNA_precursor
protein_coding
protein_coding
protein_coding
protein_coding
protein_coding
ncRNA
ncRNA
protein_coding
protein_coding
protein_coding

protein_coding
protein_coding
protein_coding
protein_coding
protein_coding
protein_coding
protein_coding
protein_coding
protein_coding
protein_coding
protein_coding
protein_coding
protein_coding
protein_coding
protein_coding
protein_coding
protein_coding
protein_coding
protein_coding
protein_coding
protein_coding
ncRNA
protein_coding
protein_coding
miRNA_precursor
protein_coding
miRNA_precursor
protein_coding
protein_coding
protein_coding
protein_coding
protein_coding
protein_coding
protein_coding
protein_coding
protein_coding
protein_coding
protein_coding
protein_coding
protein_coding
protein_coding
protein_coding
protein_coding
protein_coding
ncRNA
protein_coding
protein_coding
protein_coding
protein_coding
protein_coding
protein_coding
protein_coding
protein_coding
protein_coding
protein_coding
protein_coding

protein_coding
protein_coding
protein_coding
protein_coding
protein_coding
protein_coding
protein_coding
protein_coding
protein_coding
protein_coding
protein_coding
protein_coding
protein_coding
protein_coding
protein_coding
protein_coding
protein_coding
protein_coding
protein_coding
protein_coding
protein_coding
protein_coding
protein_coding
protein_coding
protein_coding
protein_coding
protein_coding
protein_coding
protein_coding
protein_coding
protein_coding
protein_coding
protein_coding
protein_coding
protein_coding
protein_coding
protein_coding
protein_coding
protein_coding
protein_coding
protein_coding
ncRNA
protein_coding
protein_coding
protein_coding
protein_coding
protein_coding
protein_coding
protein_coding
protein_coding
protein_coding
protein_coding
protein_coding
protein_coding
protein_coding
protein_coding

protein_coding
protein_coding
protein_coding
protein_coding
protein_coding
protein_coding
protein_coding
protein_coding
protein_coding
protein_coding
protein_coding
protein_coding
protein_coding
protein_coding
protein_coding
protein_coding
protein_coding
protein_coding
protein_coding
protein_coding
protein_coding
protein_coding
small_nucle(ol)ar_rna
protein_coding
protein_coding
protein_coding
pre_trna
protein_coding
pre_trna
protein_coding
protein_coding
protein_coding
protein_coding
protein_coding
protein_coding
protein_coding
protein_coding
protein_coding
protein_coding
protein_coding
ncRNA
protein_coding
protein_coding
ncRNA
ncRNA
protein_coding
protein_coding
protein_coding
protein_coding
protein_coding
protein_coding
protein_coding
protein_coding
protein_coding
protein_coding
protein_coding

protein_coding
miRNA_precursor
miRNA_precursor
protein_coding
protein_coding
protein_coding
protein_coding
protein_coding
protein_coding
protein_coding
protein_coding
protein_coding
protein_coding
protein_coding
protein_coding
protein_coding
protein_coding
protein_coding
protein_coding
protein_coding
protein_coding
protein_coding
protein_coding
protein_coding
protein_coding
protein_coding
protein_coding
protein_coding
protein_coding
protein_coding
protein_coding
protein_coding
protein_coding
protein_coding
protein_coding
protein_coding
protein_coding
protein_coding
protein_coding
protein_coding
protein_coding
protein_coding
protein_coding
protein_coding
protein_coding
protein_coding
protein_coding
protein_coding
protein_coding
protein_coding
protein_coding
protein_coding
protein_coding
protein_coding
protein_coding
protein_coding

protein_coding
protein_coding
protein_coding
protein_coding
pre_trna
protein_coding
protein_coding
protein_coding
protein_coding
protein_coding
protein_coding
protein_coding
protein_coding
protein_coding
protein_coding
protein_coding
protein_coding
protein_coding
protein_coding
protein_coding
protein_coding
protein_coding
protein_coding
protein_coding
protein_coding
protein_coding
protein_coding
protein_coding
protein_coding
ncRNA
protein_coding
ncRNA
protein_coding
protein_coding
protein_coding
protein_coding
protein_coding
protein_coding
ncRNA
protein_coding
protein_coding
protein_coding
protein_coding
protein_coding
protein_coding
small_nucle(ol)ar_rna
protein_coding
protein_coding
protein_coding
protein_coding
protein_coding
protein_coding
ncRNA
protein_coding
protein_coding
protein_coding

protein_coding
protein_coding
protein_coding
protein_coding
protein_coding
protein_coding
protein_coding
protein_coding
protein_coding
protein_coding
protein_coding
protein_coding
protein_coding
protein_coding
protein_coding
protein_coding
protein_coding
protein_coding
protein_coding
protein_coding
protein_coding
protein_coding
protein_coding
protein_coding
protein_coding
protein_coding
protein_coding
protein_coding
protein_coding
protein_coding
ncRNA
protein_coding
protein_coding
protein_coding
ncRNA
protein_coding
protein_coding
protein_coding
protein_coding
protein_coding
protein_coding
protein_coding
protein_coding
protein_coding
protein_coding
protein_coding
ncRNA
protein_coding
protein_coding
protein_coding
protein_coding
protein_coding
protein_coding
protein_coding
protein_coding
protein_coding

protein_coding
protein_coding
protein_coding
protein_coding
protein_coding
protein_coding
protein_coding
protein_coding
protein_coding
protein_coding
protein_coding
protein_coding
protein_coding
protein_coding
protein_coding
protein_coding
protein_coding
ncRNA
protein_coding
protein_coding
ncRNA
ncRNA
protein_coding
protein_coding
protein_coding
protein_coding
protein_coding
protein_coding
protein_coding
protein_coding
protein_coding
protein_coding
protein_coding
protein_coding
protein_coding
protein_coding
protein_coding
protein_coding
protein_coding
protein_coding
protein_coding
protein_coding
protein_coding
protein_coding
protein_coding
protein_coding
protein_coding
protein_coding
protein_coding
protein_coding
protein_coding
protein_coding
ncRNA
protein_coding
protein_coding
protein_coding

protein_coding
protein_coding
protein_coding
protein_coding
protein_coding
protein_coding
protein_coding
protein_coding
protein_coding
pre_trna
protein_coding
protein_coding
protein_coding
protein_coding
protein_coding
protein_coding
protein_coding
protein_coding
protein_coding
protein_coding
protein_coding
protein_coding
protein_coding
protein_coding
ncRNA
protein_coding
protein_coding
protein_coding
protein_coding
protein_coding
protein_coding
protein_coding
protein_coding
protein_coding
protein_coding
protein_coding
protein_coding
protein_coding
protein_coding
protein_coding
small_nucle(ol)ar_rna
protein_coding
protein_coding
protein_coding
protein_coding
protein_coding
protein_coding
protein_coding
protein_coding
protein_coding
protein_coding
protein_coding
protein_coding
protein_coding
protein_coding
protein_coding

protein_coding
protein_coding
protein_coding
protein_coding
protein_coding
small_nucle(ol)ar_rna
small_nucle(ol)ar_rna
protein_coding
protein_coding
protein_coding
protein_coding
pre_trna
miRNA_precursor
protein_coding
protein_coding
protein_coding
protein_coding
protein_coding
protein_coding
protein_coding
protein_coding
ncRNA
protein_coding
ncRNA
ncRNA
protein_coding
protein_coding
protein_coding
protein_coding
protein_coding
protein_coding
protein_coding
protein_coding
protein_coding
protein_coding
protein_coding
protein_coding
protein_coding
protein_coding
protein_coding
protein_coding
protein_coding
ncRNA
protein_coding
protein_coding
protein_coding
protein_coding
protein_coding
protein_coding
protein_coding
protein_coding
protein_coding
protein_coding
protein_coding
protein_coding
ncRNA

protein_coding
protein_coding
protein_coding
miRNA_precursor
protein_coding
protein_coding
protein_coding
ncRNA
protein_coding
protein_coding
protein_coding
protein_coding
protein_coding
protein_coding
protein_coding
protein_coding
protein_coding
protein_coding
protein_coding
protein_coding
protein_coding
protein_coding
protein_coding
protein_coding
protein_coding
protein_coding
protein_coding
ncRNA
protein_coding
protein_coding
protein_coding
protein_coding
protein_coding
protein_coding
protein_coding
protein_coding
protein_coding
protein_coding
protein_coding
protein_coding
protein_coding
protein_coding
protein_coding
miRNA_precursor
protein_coding
protein_coding
protein_coding
protein_coding
protein_coding
protein_coding
protein_coding
protein_coding
protein_coding
protein_coding
protein_coding
protein_coding

miRNA_precursor
pre_trna
protein_coding
protein_coding
protein_coding
protein_coding
protein_coding
ncRNA
protein_coding
protein_coding
protein_coding
miRNA_precursor
protein_coding
protein_coding
protein_coding
protein_coding
protein_coding
miRNA_precursor
protein_coding
ncRNA
protein_coding
protein_coding
protein_coding
protein_coding
protein_coding
protein_coding
protein_coding
protein_coding
protein_coding
protein_coding
protein_coding
protein_coding
protein_coding
protein_coding
protein_coding
protein_coding
protein_coding
protein_coding
protein_coding
protein_coding
protein_coding
protein_coding
protein_coding
protein_coding
protein_coding
protein_coding
protein_coding
protein_coding
protein_coding
protein_coding
protein_coding
protein_coding
protein_coding
protein_coding
protein_coding
protein_coding

protein_coding
protein_coding
protein_coding
protein_coding
protein_coding
protein_coding
protein_coding
protein_coding
protein_coding
protein_coding
protein_coding
protein_coding
protein_coding
protein_coding
protein_coding
protein_coding
miRNA_precursor
protein_coding
protein_coding
protein_coding
protein_coding
protein_coding
protein_coding
protein_coding
protein_coding
protein_coding
protein_coding
ncRNA
small_nucle(ol)ar_rna
small_nucle(ol)ar_rna
small_nucle(ol)ar_rna
protein_coding
protein_coding
protein_coding
protein_coding
protein_coding
ncRNA
protein_coding
protein_coding
protein_coding
protein_coding
protein_coding
protein_coding
protein_coding
protein_coding
protein_coding
protein_coding
protein_coding
protein_coding
protein_coding
protein_coding
protein_coding
protein_coding
protein_coding
protein_coding
protein_coding

protein_coding
protein_coding
protein_coding
protein_coding
protein_coding
ncRNA
protein_coding
protein_coding
protein_coding
protein_coding
protein_coding
protein_coding
protein_coding
protein_coding
protein_coding
protein_coding
protein_coding
protein_coding
protein_coding
protein_coding
protein_coding
protein_coding
protein_coding
protein_coding
protein_coding
protein_coding
protein_coding
protein_coding
protein_coding
protein_coding
protein_coding
protein_coding
protein_coding
protein_coding
protein_coding
protein_coding
protein_coding
protein_coding
protein_coding
protein_coding
protein_coding
pre_trna
protein_coding
protein_coding
protein_coding
protein_coding
protein_coding
protein_coding
protein_coding
protein_coding
protein_coding
protein_coding
protein_coding
protein_coding
protein_coding
protein_coding

ncRNA
protein_coding
protein_coding
protein_coding
protein_coding
protein_coding
protein_coding
protein_coding
protein_coding
miRNA_precursor
protein_coding
protein_coding
protein_coding
protein_coding
protein_coding
protein_coding
protein_coding
protein_coding
protein_coding
protein_coding
ncRNA
protein_coding
protein_coding
protein_coding
protein_coding
protein_coding
protein_coding
protein_coding
protein_coding
protein_coding
protein_coding
protein_coding
ncRNA
protein_coding
protein_coding
protein_coding
protein_coding
protein_coding
protein_coding
protein_coding
protein_coding
protein_coding
protein_coding
protein_coding
protein_coding
protein_coding
protein_coding
protein_coding
protein_coding
protein_coding
protein_coding
protein_coding
protein_coding
protein_coding
pre_trna
protein_coding

protein_coding
protein_coding
protein_coding
protein_coding
protein_coding
protein_coding
protein_coding
protein_coding
protein_coding
protein_coding
protein_coding
protein_coding
protein_coding
protein_coding
protein_coding
protein_coding
protein_coding
protein_coding
ncRNA
protein_coding
protein_coding
protein_coding
protein_coding
protein_coding
protein_coding
protein_coding
protein_coding
protein_coding
protein_coding
ncRNA
protein_coding
protein_coding
ncRNA
small_nucle(ol)ar_rna
protein_coding
protein_coding
protein_coding
protein_coding
protein_coding
protein_coding
protein_coding
protein_coding
protein_coding
protein_coding
protein_coding
ncRNA
protein_coding
protein_coding
protein_coding
ncRNA
protein_coding
protein_coding
protein_coding
protein_coding
protein_coding
protein_coding

protein_coding
protein_coding
protein_coding
protein_coding
protein_coding
protein_coding
protein_coding
ncRNA
protein_coding
protein_coding
protein_coding
protein_coding
protein_coding
protein_coding
protein_coding
protein_coding
protein_coding
protein_coding
pre_trna
protein_coding
protein_coding
protein_coding
protein_coding
protein_coding
protein_coding
protein_coding
protein_coding
protein_coding
protein_coding
protein_coding
protein_coding
protein_coding
protein_coding
protein_coding
protein_coding
miRNA_precursor
small_nucle(ol)ar_rna
small_nucle(ol)ar_rna
protein_coding
pre_trna
protein_coding
protein_coding
protein_coding
protein_coding
protein_coding
protein_coding
protein_coding
ncRNA
protein_coding
protein_coding
protein_coding
protein_coding
protein_coding
ncRNA
protein_coding
protein_coding

protein_coding
protein_coding
protein_coding
protein_coding
protein_coding
protein_coding
protein_coding
protein_coding
protein_coding
protein_coding
protein_coding
protein_coding
protein_coding
pre_trna
protein_coding
protein_coding
protein_coding
protein_coding
protein_coding
protein_coding
protein_coding
protein_coding
protein_coding
protein_coding
protein_coding
protein_coding
protein_coding
protein_coding
protein_coding
protein_coding
protein_coding
protein_coding
protein_coding
protein_coding
protein_coding
protein_coding
protein_coding
protein_coding
small_nucle(ol)ar_rna
protein_coding
protein_coding
protein_coding
protein_coding
protein_coding
protein_coding
protein_coding
protein_coding
protein_coding
protein_coding
protein_coding
protein_coding
protein_coding
protein_coding
protein_coding
protein_coding
protein_coding

| |
|---|
| protein_coding |
| protein_coding |
| protein_coding |
| protein_coding |
| protein_coding |
| protein_coding |
| protein_coding |
| protein_coding |
| protein_coding |
| protein_coding |
| protein_coding |
| protein_coding |
| protein_coding |
| protein_coding |
| protein_coding |
| protein_coding |
| protein_coding |
| protein_coding |
| protein_coding |
| protein_coding |
| protein_coding |
| protein_coding |
| protein_coding |
| protein_coding |
| protein_coding |
| protein_coding |
| protein_coding |
| protein_coding |
| protein_coding |
| protein_coding |
| protein_coding |
| protein_coding |
| protein_coding |
| protein_coding |
| protein_coding |
| protein_coding |
| protein_coding |
| protein_coding |
| protein_coding |
| protein_coding |
| protein_coding |
| protein_coding |
| protein_coding |
| protein_coding |
| protein_coding |
| protein_coding |
| protein_coding |
| protein_coding |
| protein_coding |
| protein_coding |
| protein_coding |
| protein_coding |
| protein_coding |
| protein_coding |
| protein_coding |
| protein_coding |

protein_coding
protein_coding
protein_coding
protein_coding
protein_coding
protein_coding
protein_coding
protein_coding
protein_coding
protein_coding
protein_coding
protein_coding
protein_coding
protein_coding
protein_coding
protein_coding
protein_coding
protein_coding
protein_coding
protein_coding
protein_coding
protein_coding
protein_coding
protein_coding
protein_coding
protein_coding
protein_coding
protein_coding
protein_coding
protein_coding
protein_coding
ncRNA
protein_coding
protein_coding
ncRNA
protein_coding
protein_coding
protein_coding
protein_coding
protein_coding
protein_coding
protein_coding
protein_coding
protein_coding
protein_coding
protein_coding
protein_coding
protein_coding
protein_coding
protein_coding
protein_coding
protein_coding
protein_coding
protein_coding
protein_coding
protein_coding

protein_coding
protein_coding
protein_coding
protein_coding
protein_coding
protein_coding
protein_coding
protein_coding
protein_coding
protein_coding
protein_coding
protein_coding
protein_coding
protein_coding
protein_coding
protein_coding
protein_coding
protein_coding
protein_coding
protein_coding
protein_coding
protein_coding
protein_coding
protein_coding
ncRNA
protein_coding
protein_coding
protein_coding
protein_coding
protein_coding
protein_coding
protein_coding
protein_coding
ncRNA
protein_coding
protein_coding
protein_coding
ncRNA
protein_coding
protein_coding
protein_coding
protein_coding
protein_coding
protein_coding
protein_coding
protein_coding
protein_coding
protein_coding
protein_coding
protein_coding
protein_coding
protein_coding
protein_coding
protein_coding
ncRNA
protein_coding

protein_coding
protein_coding
protein_coding
ncRNA
protein_coding
protein_coding
protein_coding
protein_coding
protein_coding
protein_coding
protein_coding
protein_coding
protein_coding
protein_coding
protein_coding
protein_coding
protein_coding
protein_coding
protein_coding
protein_coding
ncRNA
protein_coding
protein_coding
protein_coding
protein_coding
protein_coding
protein_coding
pre_trna
protein_coding
protein_coding
protein_coding
protein_coding
protein_coding
protein_coding
protein_coding
protein_coding
protein_coding
protein_coding
protein_coding
protein_coding
protein_coding
protein_coding
protein_coding
protein_coding
ncRNA
pre_trna
protein_coding
protein_coding
protein_coding
protein_coding
protein_coding
protein_coding
protein_coding
ncRNA
protein_coding
protein_coding

protein_coding
ncRNA
protein_coding
protein_coding
pre_trna
protein_coding
protein_coding
protein_coding
protein_coding
protein_coding
protein_coding
protein_coding
protein_coding
protein_coding
protein_coding
protein_coding
protein_coding
protein_coding
protein_coding
protein_coding
protein_coding
protein_coding
protein_coding
protein_coding
protein_coding
protein_coding
protein_coding
protein_coding
protein_coding
protein_coding
protein_coding
protein_coding
protein_coding
protein_coding
protein_coding
protein_coding
protein_coding
protein_coding
protein_coding
protein_coding
protein_coding
protein_coding
protein_coding
protein_coding
protein_coding
protein_coding
protein_coding
protein_coding
protein_coding
protein_coding
protein_coding
protein_coding
protein_coding
protein_coding
ncRNA
protein_coding

protein_coding
protein_coding
protein_coding
protein_coding
protein_coding
protein_coding
protein_coding
protein_coding
protein_coding
protein_coding
protein_coding
protein_coding
protein_coding
protein_coding
protein_coding
protein_coding
protein_coding
protein_coding
protein_coding
ncRNA
protein_coding
ncRNA
protein_coding
protein_coding
protein_coding
protein_coding
ncRNA
protein_coding
protein_coding
protein_coding
protein_coding
protein_coding
protein_coding
small_nucle(ol)ar_rna
small_nucle(ol)ar_rna
protein_coding
protein_coding
protein_coding
protein_coding
protein_coding
protein_coding
protein_coding
protein_coding
protein_coding
protein_coding
protein_coding
protein_coding
protein_coding
ncRNA
protein_coding
ncRNA
protein_coding
protein_coding
protein_coding
protein_coding
protein_coding

protein_coding
pre_trna
protein_coding
protein_coding
protein_coding
protein_coding
pre_trna
protein_coding
protein_coding
protein_coding
protein_coding
small_nucle(ol)ar_rna
small_nucle(ol)ar_rna
small_nucle(ol)ar_rna
protein_coding
protein_coding
protein_coding
protein_coding
protein_coding
protein_coding
protein_coding
protein_coding
protein_coding
protein_coding
protein_coding
protein_coding
ncRNA
protein_coding
protein_coding
protein_coding
protein_coding
protein_coding
protein_coding
protein_coding
protein_coding
protein_coding
protein_coding
pre_trna
protein_coding
protein_coding
protein_coding
protein_coding
protein_coding
protein_coding
protein_coding
protein_coding
protein_coding
protein_coding
protein_coding
protein_coding
ncRNA
protein_coding
protein_coding
protein_coding
protein_coding
protein_coding

protein_coding
protein_coding
ncRNA
protein_coding
protein_coding
protein_coding
protein_coding
protein_coding
protein_coding
protein_coding
protein_coding
protein_coding
protein_coding
protein_coding
protein_coding
protein_coding
protein_coding
protein_coding
protein_coding
protein_coding
protein_coding
protein_coding
protein_coding
protein_coding
protein_coding
protein_coding
protein_coding
protein_coding
protein_coding
protein_coding
protein_coding
protein_coding
protein_coding
protein_coding
protein_coding
protein_coding
protein_coding
ncRNA
small_nucle(ol)ar_rna
protein_coding
protein_coding
protein_coding
protein_coding
protein_coding
protein_coding
protein_coding
protein_coding
protein_coding
protein_coding
protein_coding
protein_coding
protein_coding
protein_coding
protein_coding
protein_coding
protein_coding

protein_coding
protein_coding
protein_coding
protein_coding
protein_coding
protein_coding
protein_coding
protein_coding
small_nucle(ol)ar_rna
protein_coding
protein_coding
protein_coding
protein_coding
protein_coding
protein_coding
protein_coding
protein_coding
protein_coding
protein_coding
protein_coding
protein_coding
protein_coding
protein_coding
protein_coding
protein_coding
protein_coding
protein_coding
protein_coding
protein_coding
protein_coding
protein_coding
protein_coding
protein_coding
protein_coding
protein_coding
protein_coding
protein_coding
protein_coding
protein_coding
protein_coding
protein_coding
protein_coding
protein_coding
protein_coding
protein_coding
protein_coding
protein_coding
protein_coding
protein_coding
protein_coding
protein_coding
protein_coding
protein_coding
protein_coding
protein_coding
protein_coding

protein_coding
protein_coding
protein_coding
protein_coding
protein_coding
protein_coding
protein_coding
small_nucle(ol)ar_rna
small_nucle(ol)ar_rna
small_nucle(ol)ar_rna
protein_coding
protein_coding
protein_coding
protein_coding
protein_coding
protein_coding
protein_coding
protein_coding
protein_coding
protein_coding
protein_coding
protein_coding
protein_coding
protein_coding
protein_coding
protein_coding
protein_coding
protein_coding
protein_coding
protein_coding
protein_coding
protein_coding
protein_coding
protein_coding
protein_coding
protein_coding
ncRNA
protein_coding
protein_coding
protein_coding
protein_coding
protein_coding
protein_coding
ncRNA
protein_coding
protein_coding
protein_coding
protein_coding
protein_coding
protein_coding
protein_coding
protein_coding
protein_coding
protein_coding
protein_coding
protein_coding

protein_coding
protein_coding
protein_coding
protein_coding
protein_coding
protein_coding
protein_coding
protein_coding
protein_coding
ncRNA
protein_coding
protein_coding
protein_coding
protein_coding
protein_coding
protein_coding
protein_coding
protein_coding
protein_coding
protein_coding
protein_coding
protein_coding
protein_coding
protein_coding
protein_coding
protein_coding
protein_coding
protein_coding
protein_coding
miRNA_precursor
protein_coding
protein_coding
protein_coding
protein_coding
protein_coding
protein_coding
protein_coding
protein_coding
protein_coding
protein_coding
protein_coding
protein_coding
protein_coding
protein_coding
protein_coding
protein_coding
protein_coding
protein_coding
protein_coding
protein_coding
protein_coding
miRNA_precursor
protein_coding
protein_coding
protein_coding
protein_coding

protein_coding
miRNA_precursor
protein_coding
protein_coding
protein_coding
protein_coding
protein_coding
protein_coding
protein_coding
protein_coding
protein_coding
protein_coding
protein_coding
protein_coding
protein_coding
protein_coding
protein_coding
protein_coding
protein_coding
protein_coding
protein_coding
protein_coding
protein_coding
protein_coding
protein_coding
protein_coding
protein_coding
protein_coding
protein_coding
protein_coding
protein_coding
protein_coding
protein_coding
protein_coding
protein_coding
protein_coding
protein_coding
protein_coding
protein_coding
protein_coding
ncRNA
protein_coding
protein_coding
protein_coding
protein_coding
protein_coding
protein_coding
protein_coding
protein_coding
protein_coding
protein_coding
protein_coding
protein_coding
ncRNA
protein_coding
protein_coding

protein_coding
protein_coding
protein_coding
protein_coding
small_nucle(ol)ar_rna
protein_coding
protein_coding
protein_coding
protein_coding
protein_coding
protein_coding
protein_coding
protein_coding
ncRNA
protein_coding
protein_coding
miRNA_precursor
protein_coding
protein_coding
protein_coding
protein_coding
protein_coding
protein_coding
protein_coding
protein_coding
protein_coding
ncRNA
protein_coding
protein_coding
protein_coding
protein_coding
protein_coding
pre_trna
ncRNA
protein_coding
protein_coding
protein_coding
protein_coding
protein_coding
protein_coding
protein_coding
protein_coding
ncRNA
protein_coding
ncRNA
protein_coding
protein_coding
protein_coding
protein_coding
protein_coding
protein_coding
protein_coding
protein_coding
protein_coding
protein_coding
protein_coding

protein_coding
protein_coding
ncRNA
protein_coding
protein_coding
ncRNA
protein_coding
protein_coding
protein_coding
ncRNA
protein_coding
protein_coding
protein_coding
protein_coding
protein_coding
protein_coding
protein_coding
protein_coding
protein_coding
protein_coding
protein_coding
protein_coding
protein_coding
protein_coding
protein_coding
protein_coding
protein_coding
protein_coding
protein_coding
protein_coding
protein_coding
protein_coding
protein_coding
protein_coding
protein_coding
protein_coding
ncRNA
protein_coding
protein_coding
protein_coding
protein_coding
protein_coding
protein_coding
protein_coding
protein_coding
protein_coding
protein_coding
protein_coding
protein_coding
protein_coding
protein_coding
protein_coding
protein_coding
protein_coding
protein_coding
ncRNA

protein_coding
protein_coding
ncRNA
protein_coding
protein_coding
protein_coding
ncRNA
protein_coding
protein_coding
protein_coding
ncRNA
protein_coding
protein_coding
protein_coding
protein_coding
protein_coding
protein_coding
protein_coding
ncRNA
protein_coding
protein_coding
protein_coding
protein_coding
protein_coding
ncRNA
protein_coding
miRNA_precursor
protein_coding
protein_coding
protein_coding
protein_coding
protein_coding
protein_coding
protein_coding
protein_coding
protein_coding
protein_coding
protein_coding
protein_coding
protein_coding
protein_coding
protein_coding
protein_coding
protein_coding
protein_coding
protein_coding
protein_coding
protein_coding
protein_coding
ncRNA
protein_coding
protein_coding
protein_coding
protein_coding
protein_coding
protein_coding

protein_coding
protein_coding
protein_coding
protein_coding
protein_coding
protein_coding
small_nucle(ol)ar_rna
protein_coding
protein_coding
protein_coding
protein_coding
protein_coding
protein_coding
protein_coding
protein_coding
protein_coding
protein_coding
protein_coding
protein_coding
protein_coding
protein_coding
protein_coding
protein_coding
protein_coding
protein_coding
protein_coding
protein_coding
protein_coding
protein_coding
protein_coding
protein_coding
protein_coding
protein_coding
protein_coding
protein_coding
protein_coding
protein_coding
protein_coding
protein_coding
protein_coding
protein_coding
protein_coding
protein_coding
protein_coding
protein_coding
protein_coding
protein_coding
protein_coding
protein_coding
pre_trna
protein_coding
protein_coding
protein_coding
protein_coding
protein_coding
ncRNA

protein_coding
protein_coding
protein_coding
protein_coding
protein_coding
protein_coding
protein_coding
protein_coding
protein_coding
protein_coding
protein_coding
protein_coding
protein_coding
protein_coding
protein_coding
protein_coding
protein_coding
protein_coding
protein_coding
protein_coding
protein_coding
protein_coding
protein_coding
protein_coding
protein_coding
protein_coding
protein_coding
protein_coding
protein_coding
protein_coding
protein_coding
protein_coding
protein_coding
protein_coding
protein_coding
protein_coding
protein_coding
protein_coding
protein_coding
protein_coding
protein_coding
protein_coding
protein_coding
protein_coding
protein_coding
protein_coding
protein_coding
protein_coding
protein_coding
protein_coding
protein_coding
protein_coding
protein_coding
protein_coding
protein_coding
protein_coding

protein_coding
protein_coding
protein_coding
protein_coding
protein_coding
protein_coding
protein_coding
protein_coding
miRNA_precursor
protein_coding
protein_coding
protein_coding
protein_coding
ncRNA
protein_coding
protein_coding
protein_coding
protein_coding
protein_coding
ncRNA
protein_coding
ncRNA
protein_coding
protein_coding
protein_coding
protein_coding
protein_coding
protein_coding
protein_coding
protein_coding
protein_coding
protein_coding
protein_coding
protein_coding
protein_coding
protein_coding
protein_coding
protein_coding
protein_coding
protein_coding
protein_coding
protein_coding
protein_coding
protein_coding
protein_coding
protein_coding
protein_coding
protein_coding
protein_coding
protein_coding
protein_coding
protein_coding
protein_coding
protein_coding
protein_coding
protein_coding

protein_coding
protein_coding
protein_coding
protein_coding
protein_coding
protein_coding
protein_coding
protein_coding
protein_coding
protein_coding
protein_coding
protein_coding
protein_coding
protein_coding
protein_coding
protein_coding
protein_coding
protein_coding
protein_coding
protein_coding
protein_coding
protein_coding
protein_coding
protein_coding
protein_coding
protein_coding
protein_coding
protein_coding
protein_coding
protein_coding
protein_coding
protein_coding
protein_coding
protein_coding
miRNA_precursor
protein_coding
protein_coding
protein_coding
protein_coding
ncRNA
protein_coding
protein_coding
ncRNA
protein_coding
protein_coding
protein_coding
protein_coding
protein_coding
protein_coding
protein_coding
protein_coding
protein_coding
protein_coding
protein_coding
protein_coding
protein_coding

protein_coding
protein_coding
protein_coding
protein_coding
protein_coding
protein_coding
protein_coding
protein_coding
protein_coding
protein_coding
protein_coding
protein_coding
protein_coding
protein_coding
protein_coding
protein_coding
protein_coding
protein_coding
protein_coding
protein_coding
protein_coding
protein_coding
pre_trna
protein_coding
protein_coding
protein_coding
protein_coding
protein_coding
protein_coding
protein_coding
protein_coding
protein_coding
protein_coding
protein_coding
ncRNA
ncRNA
protein_coding
protein_coding
protein_coding
protein_coding
protein_coding
protein_coding
protein_coding
protein_coding
protein_coding
miRNA_precursor
protein_coding
protein_coding
protein_coding
protein_coding
protein_coding
protein_coding
protein_coding
protein_coding
protein_coding
protein_coding

protein_coding
protein_coding
protein_coding
protein_coding
protein_coding
protein_coding
protein_coding
protein_coding
protein_coding
protein_coding
protein_coding
protein_coding
protein_coding
protein_coding
protein_coding
protein_coding
protein_coding
protein_coding
protein_coding
protein_coding
protein_coding
protein_coding
protein_coding
protein_coding
protein_coding
protein_coding
protein_coding
protein_coding
protein_coding
protein_coding
protein_coding
protein_coding
protein_coding
protein_coding
protein_coding
protein_coding
small_nucle(ol)ar_rna
protein_coding
protein_coding
protein_coding
protein_coding
protein_coding
protein_coding
protein_coding
protein_coding
protein_coding
protein_coding
protein_coding
protein_coding
ncRNA
protein_coding
protein_coding
ncRNA
protein_coding
protein_coding
protein_coding

protein_coding
protein_coding
protein_coding
protein_coding
protein_coding
protein_coding
protein_coding
protein_coding
protein_coding
protein_coding
protein_coding
protein_coding
protein_coding
protein_coding
protein_coding
protein_coding
protein_coding
protein_coding
protein_coding
protein_coding
protein_coding
protein_coding
protein_coding
protein_coding
protein_coding
protein_coding
protein_coding
small_nucle(ol)ar_rna
protein_coding
protein_coding
protein_coding
protein_coding
protein_coding
protein_coding
protein_coding
protein_coding
protein_coding
protein_coding
protein_coding
protein_coding
protein_coding
protein_coding
protein_coding
protein_coding
protein_coding
protein_coding
protein_coding
protein_coding
protein_coding
protein_coding
protein_coding
protein_coding
protein_coding
protein_coding
protein_coding
protein_coding

protein_coding
protein_coding
protein_coding
protein_coding
protein_coding
protein_coding
protein_coding
protein_coding
protein_coding
protein_coding
protein_coding
protein_coding
protein_coding
protein_coding
protein_coding
protein_coding
protein_coding
protein_coding
protein_coding
protein_coding
protein_coding
protein_coding
protein_coding
protein_coding
protein_coding
protein_coding
protein_coding
protein_coding
protein_coding
protein_coding
protein_coding
protein_coding
protein_coding
protein_coding
protein_coding
protein_coding
protein_coding
protein_coding
protein_coding
protein_coding
protein_coding
protein_coding
protein_coding
protein_coding
protein_coding
protein_coding
protein_coding
protein_coding
protein_coding
protein_coding
protein_coding
protein_coding
protein_coding
protein_coding
protein_coding
protein_coding

protein_coding
protein_coding
protein_coding
protein_coding
protein_coding
protein_coding
protein_coding
ncRNA
protein_coding
protein_coding
protein_coding
protein_coding
protein_coding
protein_coding
protein_coding
protein_coding
protein_coding
protein_coding
protein_coding
protein_coding
protein_coding
protein_coding
ncRNA
protein_coding
protein_coding
protein_coding
protein_coding
protein_coding
protein_coding
protein_coding
protein_coding
protein_coding
protein_coding
protein_coding
protein_coding
protein_coding
protein_coding
protein_coding
protein_coding
protein_coding
protein_coding
protein_coding
protein_coding
ncRNA
protein_coding
protein_coding
protein_coding
protein_coding
protein_coding
protein_coding
protein_coding
protein_coding

**Dataset 1.3 Differential analysis of 21/22nt accumulation on Araport11 genes in c**

| ID | baseMean | log2FoldChange | lfcSE | stat | pvalue | padj |
|---|---|---|---|---|---|---|
| AT5G13630 | 269,68 | 5,77 | 0,41 | 14,24 | 4,87E-46 | 1,12E-42 |
| AT1G31280 | 162,06 | 6,88 | 0,60 | 11,42 | 3,42E-30 | 2,29E-27 |
| AT5G38850 | 479,48 | -2,45 | 0,21 | -11,40 | 3,97E-30 | 2,29E-27 |
| AT2G10606 | 1219,01 | 3,15 | 0,28 | 11,41 | 3,62E-30 | 2,29E-27 |
| AT2G26211 | 125,25 | 3,55 | 0,33 | 10,92 | 8,84E-28 | 4,08E-25 |
| AT4G06810 | 54,17 | 4,50 | 0,43 | 10,40 | 2,36E-25 | 9,07E-23 |
| AT2G18960 | 62,63 | 6,03 | 0,58 | 10,37 | 3,50E-25 | 1,01E-22 |
| AT4G06805 | 54,05 | 4,50 | 0,43 | 10,38 | 3,07E-25 | 1,01E-22 |
| AT1G29265 | 54,27 | 3,54 | 0,36 | 9,91 | 3,77E-23 | 9,68E-21 |
| AT4G19395 | 418,73 | 1,95 | 0,20 | 9,56 | 1,22E-21 | 2,82E-19 |
| AT1G62914 | 457,48 | -3,22 | 0,34 | -9,53 | 1,57E-21 | 3,30E-19 |
| AT3G23325 | 69,65 | 3,62 | 0,39 | 9,37 | 7,59E-21 | 1,46E-18 |
| AT1G37130 | 39,12 | 3,34 | 0,36 | 9,35 | 8,51E-21 | 1,51E-18 |
| AT3G23326 | 69,32 | 3,67 | 0,40 | 9,29 | 1,54E-20 | 2,54E-18 |
| AT3G17185 | 156,12 | -2,57 | 0,28 | -9,23 | 2,77E-20 | 4,27E-18 |
| AT1G56070 | 55,72 | 2,97 | 0,33 | 9,10 | 8,69E-20 | 1,25E-17 |
| AT1G63130 | 240,21 | -2,52 | 0,29 | -8,60 | 7,68E-18 | 1,04E-15 |
| AT1G63150 | 176,58 | -2,55 | 0,30 | -8,48 | 2,28E-17 | 2,92E-15 |
| AT4G13575 | 426,34 | -3,62 | 0,43 | -8,40 | 4,46E-17 | 5,42E-15 |
| AT1G15750 | 35,69 | 6,20 | 0,74 | 8,35 | 6,80E-17 | 7,84E-15 |
| AT3G25795 | 22,21 | 5,67 | 0,70 | 8,13 | 4,28E-16 | 4,71E-14 |
| AT3G51375 | 135,65 | 3,24 | 0,40 | 8,11 | 5,01E-16 | 5,25E-14 |
| AT4G06075 | 405,29 | -2,65 | 0,33 | -8,08 | 6,51E-16 | 6,53E-14 |
| AT1G62910 | 98,21 | -2,87 | 0,36 | -8,07 | 7,12E-16 | 6,85E-14 |
| AT4G06130 | 392,23 | -3,52 | 0,44 | -7,93 | 2,12E-15 | 1,96E-13 |
| AT4G13495 | 869,17 | -2,60 | 0,34 | -7,68 | 1,53E-14 | 1,36E-12 |
| AT4G04825 | 145,71 | -9,67 | 1,30 | -7,42 | 1,15E-13 | 9,80E-12 |
| AT1G12294 | 124,95 | 1,96 | 0,26 | 7,42 | 1,22E-13 | 1,00E-11 |
| AT5G05170 | 36,40 | 4,44 | 0,61 | 7,31 | 2,63E-13 | 2,09E-11 |
| AT1G18075 | 4243,99 | -1,27 | 0,18 | -7,18 | 6,81E-13 | 5,24E-11 |
| AT5G03553 | 58,75 | -2,70 | 0,38 | -7,17 | 7,59E-13 | 5,65E-11 |
| AT5G03552 | 58,24 | -2,68 | 0,38 | -7,15 | 8,54E-13 | 6,16E-11 |
| AT5G14565 | 6609,25 | -1,89 | 0,27 | -7,11 | 1,14E-12 | 7,97E-11 |
| AT3G01845 | 219,09 | -1,44 | 0,20 | -7,10 | 1,29E-12 | 8,75E-11 |
| AT1G09797 | 19,74 | 3,54 | 0,51 | 7,00 | 2,59E-12 | 1,62E-10 |
| AT1G62590 | 20,75 | -4,10 | 0,59 | -7,00 | 2,60E-12 | 1,62E-10 |
| AT1G09793 | 19,74 | 3,54 | 0,51 | 7,00 | 2,59E-12 | 1,62E-10 |
| AT1G63080 | 109,45 | -2,08 | 0,30 | -6,94 | 3,89E-12 | 2,37E-10 |
| AT5G02500 | 62,24 | 2,22 | 0,32 | 6,94 | 4,05E-12 | 2,40E-10 |
| AT2G15530 | 40,01 | 2,31 | 0,34 | 6,86 | 7,04E-12 | 4,06E-10 |
| AT4G34450 | 21,43 | 4,46 | 0,65 | 6,84 | 8,10E-12 | 4,56E-10 |
| AT3G02260 | 23,16 | 3,49 | 0,52 | 6,70 | 2,06E-11 | 1,13E-09 |
| AT3G01540 | 20,21 | 4,42 | 0,67 | 6,62 | 3,57E-11 | 1,92E-09 |
| AT2G28290 | 15,57 | 4,54 | 0,69 | 6,61 | 3,85E-11 | 2,02E-09 |
| AT1G12820 | 53,20 | -2,10 | 0,32 | -6,60 | 4,02E-11 | 2,06E-09 |
| AT2G47275 | 830,57 | 2,04 | 0,31 | 6,60 | 4,14E-11 | 2,08E-09 |
| AT1G62930 | 53,37 | -2,85 | 0,43 | -6,59 | 4,29E-11 | 2,10E-09 |
| AT1G70320 | 18,40 | 3,24 | 0,49 | 6,55 | 5,63E-11 | 2,71E-09 |
| AT1G09620 | 17,50 | 4,58 | 0,70 | 6,54 | 6,21E-11 | 2,92E-09 |
| AT1G12290 | 131,54 | 1,66 | 0,26 | 6,51 | 7,62E-11 | 3,52E-09 |
| AT3G09840 | 22,25 | 2,74 | 0,44 | 6,29 | 3,11E-10 | 1,41E-08 |
| AT1G04977 | 45,78 | -2,20 | 0,35 | -6,29 | 3,18E-10 | 1,41E-08 |
| AT3G13080 | 13,73 | 3,99 | 0,64 | 6,28 | 3,44E-10 | 1,50E-08 |

| | | | | | | |
|---|---|---|---|---|---|---|
| AT5G46315 | 17,01 | 7,98 | 1,27 | 6,26 | 3,94E-10 | 1,69E-08 |
| AT3G48201 | 30,66 | 3,24 | 0,52 | 6,23 | 4,57E-10 | 1,92E-08 |
| AT2G23350 | 29,35 | 4,31 | 0,69 | 6,22 | 5,13E-10 | 2,11E-08 |
| AT4G13493 | 160,77 | -1,98 | 0,32 | -6,18 | 6,22E-10 | 2,47E-08 |
| AT5G08185 | 1387,13 | 1,93 | 0,31 | 6,19 | 6,20E-10 | 2,47E-08 |
| AT2G21390 | 23,77 | 4,35 | 0,71 | 6,16 | 7,16E-10 | 2,80E-08 |
| AT5G43740 | 103,64 | -2,98 | 0,49 | -6,10 | 1,08E-09 | 4,16E-08 |
| AT2G47020 | 1248,75 | 1,89 | 0,31 | 6,09 | 1,12E-09 | 4,23E-08 |
| AT2G47015 | 1248,43 | 1,89 | 0,31 | 6,09 | 1,15E-09 | 4,28E-08 |
| AT5G64270 | 15,19 | 4,70 | 0,78 | 6,01 | 1,85E-09 | 6,79E-08 |
| AT2G41900 | 16,69 | 4,83 | 0,82 | 5,92 | 3,14E-09 | 1,13E-07 |
| AT3G59884 | 100,21 | 1,87 | 0,32 | 5,90 | 3,73E-09 | 1,33E-07 |
| AT4G28080 | 23,67 | 4,14 | 0,71 | 5,83 | 5,39E-09 | 1,88E-07 |
| AT5G23110 | 11,58 | 3,63 | 0,63 | 5,74 | 9,55E-09 | 3,29E-07 |
| AT1G59870 | 72,89 | 1,89 | 0,33 | 5,70 | 1,21E-08 | 4,05E-07 |
| AT3G11130 | 25,04 | 3,12 | 0,55 | 5,70 | 1,20E-08 | 4,05E-07 |
| AT1G70770 | 12,70 | 4,15 | 0,73 | 5,68 | 1,36E-08 | 4,49E-07 |
| AT1G08597 | 22,78 | -4,32 | 0,76 | -5,67 | 1,46E-08 | 4,68E-07 |
| AT2G43970 | 15,13 | 3,11 | 0,55 | 5,67 | 1,46E-08 | 4,68E-07 |
| AT2G37190 | 11,90 | 4,40 | 0,78 | 5,66 | 1,53E-08 | 4,84E-07 |
| AT2G46255 | 72,33 | -2,37 | 0,42 | -5,62 | 1,87E-08 | 5,83E-07 |
| AT1G50055 | 182,36 | -1,67 | 0,30 | -5,61 | 2,02E-08 | 6,20E-07 |
| AT5G04140 | 18,61 | 3,19 | 0,57 | 5,61 | 2,07E-08 | 6,27E-07 |
| AT5G51174 | 65,86 | -2,42 | 0,43 | -5,59 | 2,33E-08 | 6,99E-07 |
| AT3G48870 | 10,82 | 7,37 | 1,33 | 5,55 | 2,82E-08 | 8,36E-07 |
| AT5G42950 | 18,25 | 4,15 | 0,76 | 5,45 | 4,99E-08 | 1,46E-06 |
| AT4G13494 | 321,70 | -2,66 | 0,49 | -5,39 | 7,13E-08 | 2,06E-06 |
| AT2G22125 | 19,76 | 3,50 | 0,66 | 5,29 | 1,22E-07 | 3,48E-06 |
| AT5G35407 | 1076,14 | 1,94 | 0,37 | 5,28 | 1,29E-07 | 3,64E-06 |
| AT2G42270 | 6,29 | 7,27 | 1,38 | 5,26 | 1,44E-07 | 3,99E-06 |
| AT4G02480 | 22,30 | 4,17 | 0,79 | 5,25 | 1,52E-07 | 4,18E-06 |
| AT1G63400 | 45,16 | -2,32 | 0,44 | -5,24 | 1,64E-07 | 4,45E-06 |
| AT3G08943 | 11,39 | 5,38 | 1,04 | 5,15 | 2,60E-07 | 6,99E-06 |
| AT1G08537 | 8,63 | -6,13 | 1,21 | -5,05 | 4,32E-07 | 1,15E-05 |
| AT3G06435 | 378,12 | -1,27 | 0,25 | -5,03 | 4,97E-07 | 1,30E-05 |
| AT3G01835 | 72,10 | -1,44 | 0,29 | -5,01 | 5,40E-07 | 1,40E-05 |
| AT4G17140 | 11,35 | 3,81 | 0,76 | 5,01 | 5,47E-07 | 1,40E-05 |
| AT4G27000 | 10,07 | 3,80 | 0,76 | 4,99 | 5,93E-07 | 1,50E-05 |
| AT5G43725 | 9,10 | -3,61 | 0,73 | -4,96 | 7,18E-07 | 1,78E-05 |
| AT5G43730 | 9,10 | -3,61 | 0,73 | -4,96 | 7,18E-07 | 1,78E-05 |
| AT3G23690 | 28,16 | -2,49 | 0,51 | -4,94 | 7,93E-07 | 1,95E-05 |
| AT3G41768 | 91,21 | -1,48 | 0,30 | -4,93 | 8,37E-07 | 2,03E-05 |
| AT5G60790 | 11,59 | 3,54 | 0,72 | 4,90 | 9,68E-07 | 2,33E-05 |
| AT1G10290 | 9,22 | 4,61 | 0,94 | 4,89 | 9,93E-07 | 2,36E-05 |
| AT2G43900 | 9,76 | 6,33 | 1,30 | 4,87 | 1,13E-06 | 2,66E-05 |
| AT1G13980 | 8,14 | 5,01 | 1,04 | 4,82 | 1,45E-06 | 3,38E-05 |
| AT1G64720 | 13,91 | 3,20 | 0,67 | 4,80 | 1,56E-06 | 3,59E-05 |
| AT1G12775 | 21,87 | -2,41 | 0,50 | -4,77 | 1,82E-06 | 4,12E-05 |
| AT1G31173 | 488,28 | 1,15 | 0,24 | 4,77 | 1,81E-06 | 4,12E-05 |
| AT5G53460 | 17,48 | 2,23 | 0,47 | 4,76 | 1,96E-06 | 4,39E-05 |
| AT1G22610 | 11,17 | 2,97 | 0,62 | 4,75 | 2,05E-06 | 4,51E-05 |
| AT3G43300 | 11,08 | 3,44 | 0,72 | 4,75 | 2,05E-06 | 4,51E-05 |
| AT1G22620 | 12,70 | 2,92 | 0,62 | 4,72 | 2,35E-06 | 5,08E-05 |
| AT5G10946 | 91,61 | -1,29 | 0,27 | -4,72 | 2,36E-06 | 5,08E-05 |
| AT5G10945 | 91,45 | -1,29 | 0,28 | -4,70 | 2,59E-06 | 5,53E-05 |
| AT1G01320 | 16,50 | 2,45 | 0,52 | 4,69 | 2,71E-06 | 5,74E-05 |

| | | | | | | |
|---|---|---|---|---|---|---|
| AT3G19820 | 10,30 | 6,05 | 1,30 | 4,66 | 3,14E-06 | 6,58E-05 |
| AT4G01800 | 8,35 | 5,47 | 1,17 | 4,66 | 3,16E-06 | 6,58E-05 |
| AT2G02741 | 19,79 | -2,25 | 0,48 | -4,65 | 3,36E-06 | 6,92E-05 |
| AT2G02750 | 20,13 | -2,23 | 0,48 | -4,64 | 3,56E-06 | 7,27E-05 |
| AT1G55860 | 12,82 | 3,28 | 0,71 | 4,63 | 3,60E-06 | 7,30E-05 |
| AT4G31570 | 11,16 | 4,00 | 0,87 | 4,61 | 4,00E-06 | 7,96E-05 |
| AT5G07745 | 20,93 | 2,54 | 0,55 | 4,61 | 3,99E-06 | 7,96E-05 |
| AT2G43680 | 14,92 | 2,85 | 0,62 | 4,59 | 4,38E-06 | 8,65E-05 |
| AT2G42600 | 13,57 | 2,52 | 0,55 | 4,58 | 4,55E-06 | 8,90E-05 |
| AT1G62670 | 9,93 | -3,65 | 0,80 | -4,58 | 4,65E-06 | 9,02E-05 |
| AT1G50480 | 9,27 | 3,40 | 0,74 | 4,57 | 4,85E-06 | 9,31E-05 |
| AT1G20960 | 9,33 | 3,23 | 0,71 | 4,57 | 4,88E-06 | 9,31E-05 |
| AT1G30410 | 4,36 | 6,56 | 1,44 | 4,55 | 5,29E-06 | 1,00E-04 |
| AT4G00885 | 370,66 | -1,33 | 0,29 | -4,55 | 5,49E-06 | 1,03E-04 |
| AT5G15680 | 7,75 | 4,18 | 0,92 | 4,53 | 5,85E-06 | 1,09E-04 |
| AT1G78478 | 10,59 | -2,79 | 0,62 | -4,51 | 6,54E-06 | 1,21E-04 |
| AT4G36080 | 9,35 | 3,43 | 0,76 | 4,50 | 6,73E-06 | 1,23E-04 |
| AT1G10170 | 9,09 | 3,56 | 0,79 | 4,50 | 6,82E-06 | 1,24E-04 |
| AT3G18780 | 17,06 | 2,45 | 0,55 | 4,47 | 7,72E-06 | 1,39E-04 |
| AT1G03060 | 7,63 | 3,71 | 0,83 | 4,47 | 7,76E-06 | 1,39E-04 |
| AT1G20760 | 7,86 | 4,09 | 0,92 | 4,46 | 8,23E-06 | 1,46E-04 |
| AT1G09817 | 10,25 | -2,87 | 0,64 | -4,45 | 8,50E-06 | 1,49E-04 |
| AT1G79920 | 8,70 | 3,93 | 0,88 | 4,45 | 8,46E-06 | 1,49E-04 |
| AT2G26890 | 8,74 | 3,49 | 0,79 | 4,41 | 1,04E-05 | 1,80E-04 |
| AT5G65950 | 6,84 | 6,64 | 1,51 | 4,40 | 1,09E-05 | 1,88E-04 |
| AT1G62020 | 12,54 | 3,32 | 0,75 | 4,40 | 1,11E-05 | 1,89E-04 |
| AT4G23387 | 136,04 | -1,93 | 0,44 | -4,39 | 1,14E-05 | 1,93E-04 |
| AT1G29900 | 11,05 | 2,68 | 0,61 | 4,37 | 1,22E-05 | 2,06E-04 |
| AT4G39420 | 9,97 | 3,54 | 0,81 | 4,36 | 1,29E-05 | 2,15E-04 |
| AT4G13940 | 35,08 | 1,77 | 0,41 | 4,35 | 1,34E-05 | 2,23E-04 |
| AT1G79930 | 9,47 | 3,91 | 0,90 | 4,34 | 1,40E-05 | 2,31E-04 |
| AT3G50370 | 9,94 | 2,92 | 0,68 | 4,32 | 1,58E-05 | 2,59E-04 |
| AT2G39730 | 47,53 | 1,71 | 0,40 | 4,31 | 1,63E-05 | 2,65E-04 |
| AT4G05420 | 7,60 | 3,82 | 0,89 | 4,30 | 1,69E-05 | 2,73E-04 |
| AT2G30520 | 8,24 | 4,14 | 0,96 | 4,30 | 1,74E-05 | 2,79E-04 |
| AT1G74260 | 9,63 | 2,89 | 0,67 | 4,29 | 1,78E-05 | 2,83E-04 |
| AT2G27400 | 624,49 | -1,43 | 0,34 | -4,28 | 1,86E-05 | 2,94E-04 |
| AT5G50290 | 244,15 | -1,58 | 0,37 | -4,28 | 1,91E-05 | 2,99E-04 |
| AT1G26985 | 10,26 | 3,36 | 0,79 | 4,27 | 1,93E-05 | 3,01E-04 |
| AT1G36160 | 9,82 | 3,09 | 0,72 | 4,26 | 2,05E-05 | 3,17E-04 |
| AT4G07975 | 9,02 | 3,23 | 0,76 | 4,24 | 2,21E-05 | 3,41E-04 |
| AT5G20700 | 23,90 | 1,68 | 0,40 | 4,23 | 2,29E-05 | 3,50E-04 |
| AT5G46250 | 6,03 | 4,35 | 1,03 | 4,23 | 2,35E-05 | 3,57E-04 |
| AT1G78476 | 11,35 | -2,48 | 0,59 | -4,23 | 2,39E-05 | 3,60E-04 |
| AT2G26080 | 10,37 | 3,11 | 0,74 | 4,20 | 2,65E-05 | 3,97E-04 |
| AT5G60900 | 5,69 | 4,27 | 1,02 | 4,19 | 2,79E-05 | 4,15E-04 |
| AT1G02500 | 10,85 | 3,38 | 0,81 | 4,19 | 2,84E-05 | 4,20E-04 |
| AT4G38600 | 8,47 | 3,80 | 0,91 | 4,17 | 3,05E-05 | 4,48E-04 |
| AT1G49240 | 7,10 | 4,15 | 1,00 | 4,15 | 3,31E-05 | 4,83E-04 |
| AT1G55591 | 5,61 | 6,00 | 1,45 | 4,15 | 3,38E-05 | 4,90E-04 |
| AT5G65750 | 4,51 | 5,98 | 1,44 | 4,15 | 3,40E-05 | 4,90E-04 |
| AT1G07810 | 6,32 | 3,66 | 0,89 | 4,12 | 3,82E-05 | 5,47E-04 |
| AT4G16940 | 4,86 | 6,18 | 1,51 | 4,10 | 4,05E-05 | 5,77E-04 |
| AT2G40030 | 5,36 | 5,14 | 1,25 | 4,10 | 4,10E-05 | 5,80E-04 |
| AT5G07350 | 9,52 | 3,52 | 0,86 | 4,09 | 4,37E-05 | 6,16E-04 |
| AT2G01020 | 7030,11 | -1,45 | 0,35 | -4,08 | 4,47E-05 | 6,25E-04 |

| | | | | | | |
|---|---|---|---|---|---|---|
| AT2G20190 | 8,93 | 4,17 | 1,02 | 4,07 | 4,67E-05 | 6,50E-04 |
| AT1G06190 | 5,18 | 4,16 | 1,03 | 4,05 | 5,20E-05 | 7,18E-04 |
| AT5G39500 | 6,69 | 4,02 | 0,99 | 4,04 | 5,30E-05 | 7,24E-04 |
| AT1G06227 | 343,33 | -1,49 | 0,37 | -4,04 | 5,30E-05 | 7,24E-04 |
| AT1G67120 | 8,25 | 2,95 | 0,73 | 4,04 | 5,33E-05 | 7,24E-04 |
| AT2G45540 | 12,39 | 2,78 | 0,69 | 4,02 | 5,84E-05 | 7,88E-04 |
| AT3G57410 | 3,70 | 5,94 | 1,49 | 3,99 | 6,51E-05 | 8,73E-04 |
| AT3G23680 | 9,12 | 3,27 | 0,82 | 3,98 | 6,77E-05 | 9,04E-04 |
| AT3G23125 | 239,13 | -1,79 | 0,45 | -3,98 | 6,87E-05 | 9,12E-04 |
| AT1G06237 | 336,80 | -1,50 | 0,38 | -3,97 | 7,31E-05 | 9,64E-04 |
| AT5G57990 | 5,76 | 4,80 | 1,21 | 3,96 | 7,46E-05 | 9,79E-04 |
| AT5G09410 | 3,14 | 6,07 | 1,53 | 3,96 | 7,57E-05 | 9,87E-04 |
| AT5G58140 | 8,36 | 3,27 | 0,83 | 3,95 | 7,87E-05 | 1,02E-03 |
| AT1G47720 | 6,21 | 3,37 | 0,85 | 3,95 | 7,83E-05 | 1,02E-03 |
| AT2G02950 | 6,29 | -3,43 | 0,87 | -3,95 | 7,95E-05 | 1,02E-03 |
| AT5G20280 | 6,18 | 3,35 | 0,85 | 3,94 | 8,25E-05 | 1,05E-03 |
| AT3G13300 | 8,42 | 4,39 | 1,12 | 3,93 | 8,55E-05 | 1,08E-03 |
| AT4G07070 | 70,24 | -1,93 | 0,49 | -3,92 | 8,89E-05 | 1,12E-03 |
| AT1G15500 | 7,05 | 4,36 | 1,11 | 3,91 | 9,10E-05 | 1,14E-03 |
| AT2G43160 | 6,80 | 4,24 | 1,08 | 3,91 | 9,20E-05 | 1,15E-03 |
| AT2G18320 | 12,44 | -2,58 | 0,66 | -3,91 | 9,29E-05 | 1,15E-03 |
| AT3G43990 | 14,30 | 1,92 | 0,49 | 3,90 | 9,46E-05 | 1,17E-03 |
| AT5G25270 | 3,67 | 6,05 | 1,56 | 3,89 | 9,92E-05 | 1,22E-03 |
| AT1G49040 | 3,74 | 5,52 | 1,42 | 3,88 | 1,06E-04 | 1,30E-03 |
| AT1G48267 | 15177,13 | -1,30 | 0,34 | -3,87 | 1,07E-04 | 1,30E-03 |
| AT4G27640 | 8,75 | 4,36 | 1,13 | 3,87 | 1,07E-04 | 1,30E-03 |
| AT1G75660 | 6,24 | 4,14 | 1,07 | 3,87 | 1,10E-04 | 1,32E-03 |
| AT3G48720 | 18,61 | -1,99 | 0,52 | -3,86 | 1,14E-04 | 1,36E-03 |
| AT3G18290 | 9,13 | 3,15 | 0,82 | 3,85 | 1,19E-04 | 1,41E-03 |
| AT1G15290 | 5,85 | 3,62 | 0,94 | 3,85 | 1,19E-04 | 1,41E-03 |
| AT1G63770 | 7,56 | 2,83 | 0,74 | 3,85 | 1,20E-04 | 1,42E-03 |
| AT4G31480 | 6,09 | 3,70 | 0,96 | 3,84 | 1,21E-04 | 1,42E-03 |
| AT5G41663 | 49,62 | -1,72 | 0,45 | -3,84 | 1,23E-04 | 1,44E-03 |
| AT3G08530 | 12,53 | 2,21 | 0,58 | 3,83 | 1,27E-04 | 1,47E-03 |
| AT5G46510 | 5,48 | 3,10 | 0,81 | 3,83 | 1,28E-04 | 1,48E-03 |
| AT5G15833 | 13,93 | 2,56 | 0,67 | 3,82 | 1,34E-04 | 1,53E-03 |
| AT1G10760 | 16,93 | 2,25 | 0,59 | 3,82 | 1,33E-04 | 1,53E-03 |
| AT2G36910 | 8,87 | 3,03 | 0,79 | 3,82 | 1,36E-04 | 1,54E-03 |
| AT3G01460 | 5,55 | 4,27 | 1,12 | 3,80 | 1,42E-04 | 1,60E-03 |
| AT3G63500 | 4,53 | 3,88 | 1,02 | 3,80 | 1,43E-04 | 1,61E-03 |
| AT4G39080 | 4,27 | 5,37 | 1,42 | 3,79 | 1,48E-04 | 1,66E-03 |
| AT5G17920 | 34,31 | 1,53 | 0,40 | 3,79 | 1,49E-04 | 1,66E-03 |
| AT5G62162 | 10,16 | 2,54 | 0,67 | 3,79 | 1,54E-04 | 1,70E-03 |
| AT3G52140 | 9,89 | 2,68 | 0,71 | 3,78 | 1,59E-04 | 1,75E-03 |
| AT5G57350 | 4,58 | 4,53 | 1,20 | 3,77 | 1,64E-04 | 1,80E-03 |
| AT2G22330 | 4,57 | 4,62 | 1,23 | 3,76 | 1,68E-04 | 1,84E-03 |
| AT1G48090 | 11,18 | 2,03 | 0,54 | 3,75 | 1,77E-04 | 1,92E-03 |
| AT5G45510 | 8,42 | 2,78 | 0,74 | 3,75 | 1,80E-04 | 1,95E-03 |
| AT4G32410 | 11,90 | 2,74 | 0,73 | 3,74 | 1,83E-04 | 1,98E-03 |
| AT1G06220 | 3,15 | 5,69 | 1,52 | 3,74 | 1,86E-04 | 2,00E-03 |
| AT1G49760 | 5,88 | 4,02 | 1,08 | 3,73 | 1,94E-04 | 2,08E-03 |
| AT4G32020 | 12,23 | 2,11 | 0,57 | 3,70 | 2,12E-04 | 2,25E-03 |
| AT5G64840 | 5,36 | 3,27 | 0,89 | 3,69 | 2,24E-04 | 2,37E-03 |
| AT4G30190 | 7,61 | 3,23 | 0,88 | 3,68 | 2,32E-04 | 2,45E-03 |
| AT1G21630 | 4,05 | 5,27 | 1,43 | 3,67 | 2,38E-04 | 2,48E-03 |
| AT5G57870 | 8,84 | 2,67 | 0,73 | 3,68 | 2,38E-04 | 2,48E-03 |

| | | | | | | |
|---|---|---|---|---|---|---|
| AT1G15130 | 4,06 | 4,62 | 1,27 | 3,65 | 2,63E-04 | 2,74E-03 |
| AT4G36630 | 4,11 | 5,97 | 1,64 | 3,65 | 2,67E-04 | 2,76E-03 |
| AT3G46780 | 9,83 | 2,69 | 0,74 | 3,64 | 2,73E-04 | 2,81E-03 |
| AT3G26810 | 23,04 | -1,73 | 0,47 | -3,63 | 2,78E-04 | 2,86E-03 |
| AT5G15022 | 21,95 | -1,93 | 0,53 | -3,63 | 2,80E-04 | 2,86E-03 |
| AT2G28000 | 11,21 | 2,51 | 0,69 | 3,63 | 2,83E-04 | 2,88E-03 |
| AT4G00630 | 4,40 | 4,44 | 1,22 | 3,63 | 2,85E-04 | 2,89E-03 |
| AT5G53480 | 6,73 | 3,21 | 0,89 | 3,61 | 3,02E-04 | 3,05E-03 |
| AT3G23000 | 7,09 | 3,12 | 0,87 | 3,59 | 3,26E-04 | 3,27E-03 |
| AT2G16586 | 20,98 | -1,88 | 0,52 | -3,59 | 3,29E-04 | 3,29E-03 |
| AT1G48300 | 8,14 | 3,82 | 1,06 | 3,59 | 3,32E-04 | 3,30E-03 |
| AT5G51430 | 4,74 | 5,65 | 1,58 | 3,58 | 3,38E-04 | 3,34E-03 |
| AT5G06845 | 6,24 | 3,20 | 0,89 | 3,58 | 3,48E-04 | 3,43E-03 |
| AT3G17850 | 4,74 | 3,99 | 1,12 | 3,57 | 3,54E-04 | 3,48E-03 |
| AT3G62360 | 4,39 | 4,09 | 1,15 | 3,55 | 3,83E-04 | 3,75E-03 |
| AT1G50200 | 4,92 | 4,71 | 1,33 | 3,54 | 3,93E-04 | 3,83E-03 |
| AT1G31580 | 10,89 | -2,26 | 0,64 | -3,54 | 4,03E-04 | 3,87E-03 |
| AT3G26740 | 15,30 | -2,20 | 0,62 | -3,54 | 4,02E-04 | 3,87E-03 |
| AT2G41740 | 6,26 | 3,70 | 1,04 | 3,54 | 4,00E-04 | 3,87E-03 |
| AT3G59940 | 11,46 | -2,01 | 0,57 | -3,53 | 4,09E-04 | 3,90E-03 |
| AT4G14910 | 7,50 | 2,74 | 0,78 | 3,53 | 4,09E-04 | 3,90E-03 |
| AT2G46560 | 4,43 | 3,21 | 0,91 | 3,53 | 4,19E-04 | 3,97E-03 |
| AT2G45940 | 2,82 | 5,12 | 1,47 | 3,49 | 4,77E-04 | 4,51E-03 |
| AT3G08580 | 9,77 | 2,37 | 0,68 | 3,49 | 4,86E-04 | 4,58E-03 |
| AT1G04907 | 10,89 | 2,42 | 0,69 | 3,48 | 4,94E-04 | 4,64E-03 |
| AT3G09275 | 11,41 | -1,99 | 0,57 | -3,48 | 5,01E-04 | 4,68E-03 |
| AT3G01345 | 6,06 | 2,75 | 0,79 | 3,47 | 5,15E-04 | 4,79E-03 |
| AT5G01820 | 9,75 | 2,73 | 0,79 | 3,45 | 5,51E-04 | 5,11E-03 |
| AT2G39800 | 6,12 | 3,42 | 0,99 | 3,45 | 5,55E-04 | 5,13E-03 |
| AT2G47070 | 5,92 | 2,67 | 0,77 | 3,45 | 5,67E-04 | 5,22E-03 |
| AT3G59020 | 5,79 | 3,19 | 0,92 | 3,45 | 5,70E-04 | 5,22E-03 |
| AT1G29930 | 124,68 | 1,33 | 0,39 | 3,43 | 6,13E-04 | 5,59E-03 |
| AT2G01270 | 3,86 | 4,99 | 1,46 | 3,42 | 6,19E-04 | 5,60E-03 |
| AT3G49500 | 2,34 | 5,76 | 1,68 | 3,42 | 6,18E-04 | 5,60E-03 |
| AT1G79280 | 4,52 | 3,38 | 0,99 | 3,42 | 6,25E-04 | 5,63E-03 |
| AT4G24280 | 6,53 | 3,22 | 0,94 | 3,42 | 6,28E-04 | 5,64E-03 |
| AT4G20850 | 8,05 | 2,47 | 0,72 | 3,42 | 6,35E-04 | 5,68E-03 |
| AT4G16890 | 15,81 | 2,02 | 0,59 | 3,41 | 6,39E-04 | 5,69E-03 |
| AT1G58250 | 4,39 | 3,28 | 0,96 | 3,41 | 6,48E-04 | 5,74E-03 |
| AT1G77300 | 4,17 | 4,37 | 1,28 | 3,41 | 6,49E-04 | 5,74E-03 |
| AT3G08947 | 5,24 | 3,63 | 1,07 | 3,40 | 6,72E-04 | 5,92E-03 |
| AT1G19835 | 5,86 | 3,65 | 1,07 | 3,40 | 6,76E-04 | 5,94E-03 |
| AT1G30230 | 5,95 | 3,17 | 0,94 | 3,39 | 6,95E-04 | 6,07E-03 |
| AT5G01920 | 2,56 | 5,34 | 1,58 | 3,38 | 7,35E-04 | 6,40E-03 |
| AT2G25430 | 4,01 | 4,36 | 1,29 | 3,37 | 7,39E-04 | 6,42E-03 |
| AT5G18370 | 3,23 | 5,32 | 1,58 | 3,37 | 7,53E-04 | 6,51E-03 |
| AT2G09670 | 5,26 | 3,54 | 1,05 | 3,36 | 7,67E-04 | 6,58E-03 |
| AT1G80410 | 5,85 | 4,01 | 1,19 | 3,36 | 7,66E-04 | 6,58E-03 |
| AT1G53687 | 5,07 | -4,75 | 1,41 | -3,36 | 7,79E-04 | 6,66E-03 |
| AT5G58410 | 3,13 | 4,47 | 1,33 | 3,36 | 7,82E-04 | 6,66E-03 |
| AT5G07970 | 3,91 | 4,60 | 1,37 | 3,36 | 7,91E-04 | 6,71E-03 |
| AT2G43890 | 2,70 | 5,75 | 1,72 | 3,34 | 8,34E-04 | 7,05E-03 |
| AT3G06530 | 4,41 | 3,61 | 1,08 | 3,34 | 8,43E-04 | 7,10E-03 |
| AT2G31141 | 64,55 | 0,94 | 0,28 | 3,34 | 8,51E-04 | 7,14E-03 |
| AT1G29465 | 25,62 | -1,58 | 0,47 | -3,33 | 8,72E-04 | 7,29E-03 |
| AT5G06600 | 5,36 | 3,21 | 0,97 | 3,32 | 8,95E-04 | 7,46E-03 |

| | | | | | | |
|---|---|---|---|---|---|---|
| AT2G41440 | 2,60 | 5,05 | 1,52 | 3,32 | 9,10E-04 | 7,52E-03 |
| AT4G09135 | 26,82 | 1,49 | 0,45 | 3,32 | 9,10E-04 | 7,52E-03 |
| AT3G55320 | 2,94 | 4,59 | 1,38 | 3,32 | 9,12E-04 | 7,52E-03 |
| AT3G55220 | 4,51 | 3,78 | 1,14 | 3,31 | 9,20E-04 | 7,56E-03 |
| AT3G03710 | 4,42 | 4,37 | 1,32 | 3,31 | 9,25E-04 | 7,57E-03 |
| AT1G24460 | 4,84 | 4,06 | 1,23 | 3,31 | 9,34E-04 | 7,61E-03 |
| AT2G48160 | 6,20 | 4,46 | 1,35 | 3,31 | 9,36E-04 | 7,61E-03 |
| AT1G80070 | 7,13 | 2,40 | 0,73 | 3,31 | 9,46E-04 | 7,64E-03 |
| AT5G07940 | 5,66 | 3,06 | 0,92 | 3,31 | 9,47E-04 | 7,64E-03 |
| AT3G19190 | 3,92 | 4,58 | 1,39 | 3,30 | 9,72E-04 | 7,82E-03 |
| AT1G09877 | 23,86 | -1,79 | 0,54 | -3,30 | 9,77E-04 | 7,82E-03 |
| AT5G48160 | 6,17 | 2,99 | 0,91 | 3,30 | 9,80E-04 | 7,82E-03 |
| AT2G09610 | 9,53 | -2,45 | 0,74 | -3,29 | 9,86E-04 | 7,85E-03 |
| AT3G47347 | 15,63 | -1,56 | 0,47 | -3,29 | 1,01E-03 | 8,03E-03 |
| AT4G35790 | 4,32 | 4,22 | 1,29 | 3,29 | 1,02E-03 | 8,04E-03 |
| AT3G02020 | 19,73 | -1,58 | 0,48 | -3,28 | 1,05E-03 | 8,27E-03 |
| AT4G08755 | 25,91 | 1,33 | 0,41 | 3,27 | 1,06E-03 | 8,32E-03 |
| AT1G13350 | 2,82 | 5,60 | 1,71 | 3,27 | 1,07E-03 | 8,39E-03 |
| AT3G03380 | 4,30 | 3,40 | 1,04 | 3,27 | 1,09E-03 | 8,45E-03 |
| AT1G79520 | 27,20 | -1,79 | 0,55 | -3,27 | 1,09E-03 | 8,45E-03 |
| AT4G14504 | 4,83 | -3,31 | 1,02 | -3,25 | 1,14E-03 | 8,78E-03 |
| AT4G01860 | 3,28 | 5,28 | 1,62 | 3,25 | 1,14E-03 | 8,78E-03 |
| AT4G12770 | 19,67 | -1,40 | 0,43 | -3,25 | 1,17E-03 | 8,99E-03 |
| AT5G36180 | 3,47 | -3,82 | 1,18 | -3,24 | 1,19E-03 | 9,15E-03 |
| AT5G42390 | 8,64 | 2,58 | 0,80 | 3,24 | 1,21E-03 | 9,26E-03 |
| AT5G44340 | 14,19 | 2,09 | 0,65 | 3,22 | 1,27E-03 | 9,67E-03 |
| AT3G20760 | 5,52 | 3,39 | 1,05 | 3,22 | 1,28E-03 | 9,70E-03 |
| AT2G40805 | 280,87 | -1,28 | 0,40 | -3,22 | 1,29E-03 | 9,73E-03 |
| AT5G05980 | 2,96 | 5,45 | 1,70 | 3,22 | 1,29E-03 | 9,73E-03 |
| AT1G04430 | 7,26 | 2,74 | 0,85 | 3,21 | 1,32E-03 | 9,92E-03 |
| AT5G04275 | 20,42 | -1,36 | 0,42 | -3,21 | 1,35E-03 | 1,01E-02 |
| AT2G03820 | 4,25 | 3,57 | 1,12 | 3,20 | 1,36E-03 | 1,02E-02 |
| AT4G18600 | 3,50 | 4,52 | 1,41 | 3,20 | 1,36E-03 | 1,02E-02 |
| AT3G04340 | 3,02 | 4,63 | 1,45 | 3,20 | 1,37E-03 | 1,02E-02 |
| AT5G45307 | 438,83 | 1,00 | 0,31 | 3,19 | 1,42E-03 | 1,05E-02 |
| AT2G31660 | 3,59 | 4,58 | 1,44 | 3,18 | 1,45E-03 | 1,07E-02 |
| AT4G35800 | 5,60 | 2,76 | 0,87 | 3,18 | 1,47E-03 | 1,08E-02 |
| AT5G19200 | 4,59 | 3,32 | 1,04 | 3,18 | 1,48E-03 | 1,09E-02 |
| AT3G22380 | 8,79 | 2,22 | 0,70 | 3,17 | 1,52E-03 | 1,11E-02 |
| AT2G17930 | 7,05 | 2,25 | 0,71 | 3,16 | 1,57E-03 | 1,14E-02 |
| AT5G53740 | 20,74 | 1,57 | 0,50 | 3,16 | 1,59E-03 | 1,15E-02 |
| AT2G07360 | 3,99 | 3,94 | 1,25 | 3,16 | 1,59E-03 | 1,15E-02 |
| AT5G43603 | 44,77 | 1,21 | 0,38 | 3,16 | 1,60E-03 | 1,15E-02 |
| AT5G02325 | 15,37 | -2,15 | 0,68 | -3,15 | 1,61E-03 | 1,16E-02 |
| AT1G79750 | 4,46 | 4,19 | 1,33 | 3,15 | 1,63E-03 | 1,17E-02 |
| AT1G76630 | 2,14 | 5,38 | 1,71 | 3,14 | 1,69E-03 | 1,20E-02 |
| AT5G45430 | 3,88 | 2,87 | 0,91 | 3,14 | 1,71E-03 | 1,22E-02 |
| AT1G23310 | 12,45 | 1,73 | 0,55 | 3,13 | 1,77E-03 | 1,26E-02 |
| AT3G62980 | 20,74 | 1,63 | 0,52 | 3,12 | 1,83E-03 | 1,30E-02 |
| AT1G66100 | 8,07 | -2,48 | 0,80 | -3,11 | 1,88E-03 | 1,33E-02 |
| AT3G52250 | 4,06 | 3,10 | 1,00 | 3,09 | 2,01E-03 | 1,41E-02 |
| AT2G34208 | 2,28 | 4,89 | 1,59 | 3,09 | 2,03E-03 | 1,42E-02 |
| AT5G18350 | 2,51 | 5,47 | 1,77 | 3,09 | 2,02E-03 | 1,42E-02 |
| AT1G64430 | 2,85 | 5,03 | 1,64 | 3,07 | 2,11E-03 | 1,47E-02 |
| AT1G04810 | 3,41 | 4,98 | 1,62 | 3,07 | 2,15E-03 | 1,49E-02 |
| AT2G47800 | 2,70 | 4,11 | 1,34 | 3,07 | 2,17E-03 | 1,51E-02 |

| | | | | | | |
|---|---|---|---|---|---|---|
| AT4G37900 | 1,96 | 5,47 | 1,81 | 3,03 | 2,46E-03 | 1,70E-02 |
| AT3G47350 | 23,89 | 1,37 | 0,45 | 3,03 | 2,47E-03 | 1,70E-02 |
| AT5G54810 | 4,59 | 2,97 | 0,98 | 3,02 | 2,50E-03 | 1,72E-02 |
| AT5G06530 | 2,30 | 4,70 | 1,55 | 3,02 | 2,52E-03 | 1,72E-02 |
| AT1G05960 | 4,07 | 4,62 | 1,53 | 3,01 | 2,58E-03 | 1,76E-02 |
| AT1G62750 | 9,17 | 2,24 | 0,75 | 3,01 | 2,59E-03 | 1,76E-02 |
| AT1G13640 | 2,82 | 4,57 | 1,52 | 3,01 | 2,64E-03 | 1,79E-02 |
| AT4G16950 | 9,81 | 1,82 | 0,61 | 3,00 | 2,70E-03 | 1,83E-02 |
| AT4G37910 | 5,28 | 2,66 | 0,89 | 3,00 | 2,72E-03 | 1,83E-02 |
| AT3G07100 | 6,58 | 2,12 | 0,71 | 3,00 | 2,73E-03 | 1,84E-02 |
| AT2G46460 | 2,59 | -5,18 | 1,73 | -2,99 | 2,77E-03 | 1,86E-02 |
| AT3G10650 | 3,29 | 4,42 | 1,48 | 2,99 | 2,79E-03 | 1,87E-02 |
| AT5G64420 | 4,10 | 3,05 | 1,02 | 2,99 | 2,81E-03 | 1,87E-02 |
| AT1G35501 | 18,58 | -1,65 | 0,55 | -2,98 | 2,86E-03 | 1,90E-02 |
| AT1G10950 | 6,24 | 2,53 | 0,85 | 2,98 | 2,88E-03 | 1,91E-02 |
| AT1G59610 | 3,98 | 3,15 | 1,06 | 2,98 | 2,88E-03 | 1,91E-02 |
| AT5G63715 | 27,38 | -1,48 | 0,50 | -2,96 | 3,07E-03 | 2,02E-02 |
| AT5G13770 | 2,99 | 4,06 | 1,38 | 2,95 | 3,13E-03 | 2,05E-02 |
| AT1G48650 | 3,21 | 3,72 | 1,26 | 2,95 | 3,14E-03 | 2,05E-02 |
| AT3G55610 | 4,81 | 3,20 | 1,08 | 2,95 | 3,13E-03 | 2,05E-02 |
| AT1G07051 | 29,68 | 1,64 | 0,56 | 2,93 | 3,38E-03 | 2,20E-02 |
| AT2G35744 | 14,81 | -1,65 | 0,56 | -2,93 | 3,38E-03 | 2,20E-02 |
| AT3G23685 | 5,28 | 2,39 | 0,82 | 2,91 | 3,58E-03 | 2,32E-02 |
| AT1G69792 | 2,23 | 5,28 | 1,82 | 2,91 | 3,63E-03 | 2,35E-02 |
| AT1G58290 | 5,16 | 2,46 | 0,85 | 2,91 | 3,67E-03 | 2,37E-02 |
| AT5G58040 | 3,29 | 3,47 | 1,20 | 2,90 | 3,73E-03 | 2,40E-02 |
| AT1G27320 | 2,96 | 3,65 | 1,26 | 2,90 | 3,74E-03 | 2,40E-02 |
| AT1G08813 | 8,42 | -2,10 | 0,73 | -2,89 | 3,80E-03 | 2,42E-02 |
| AT4G36140 | 2,98 | 3,59 | 1,24 | 2,89 | 3,79E-03 | 2,42E-02 |
| AT3G25510 | 1,96 | 4,78 | 1,66 | 2,89 | 3,88E-03 | 2,47E-02 |
| AT1G13210 | 5,46 | 2,41 | 0,83 | 2,89 | 3,90E-03 | 2,47E-02 |
| AT2G44120 | 4,00 | 4,25 | 1,47 | 2,89 | 3,91E-03 | 2,47E-02 |
| AT3G59140 | 3,91 | 5,05 | 1,75 | 2,89 | 3,90E-03 | 2,47E-02 |
| AT2G20570 | 7,71 | 1,81 | 0,63 | 2,88 | 3,95E-03 | 2,49E-02 |
| AT1G55540 | 2,72 | 4,28 | 1,49 | 2,88 | 3,99E-03 | 2,50E-02 |
| AT4G28652 | 2,08 | 5,32 | 1,85 | 2,87 | 4,07E-03 | 2,54E-02 |
| AT4G28650 | 2,08 | 5,32 | 1,85 | 2,87 | 4,07E-03 | 2,54E-02 |
| AT4G01850 | 4,27 | 3,81 | 1,33 | 2,87 | 4,11E-03 | 2,56E-02 |
| AT4G27440 | 13,15 | -1,72 | 0,60 | -2,86 | 4,21E-03 | 2,61E-02 |
| AT3G13090 | 2,27 | 5,94 | 2,08 | 2,86 | 4,21E-03 | 2,61E-02 |
| AT4G24690 | 16,56 | 1,46 | 0,51 | 2,86 | 4,23E-03 | 2,61E-02 |
| AT4G29380 | 1,74 | 4,63 | 1,62 | 2,86 | 4,30E-03 | 2,65E-02 |
| AT5G61140 | 4,17 | 2,52 | 0,88 | 2,85 | 4,35E-03 | 2,67E-02 |
| AT5G66045 | 69,41 | 1,39 | 0,49 | 2,84 | 4,47E-03 | 2,74E-02 |
| AT1G06247 | 2,53 | 4,51 | 1,59 | 2,84 | 4,48E-03 | 2,74E-02 |
| AT2G33100 | 2,25 | 5,04 | 1,78 | 2,83 | 4,59E-03 | 2,79E-02 |
| AT5G18360 | 2,96 | 3,51 | 1,24 | 2,83 | 4,64E-03 | 2,82E-02 |
| AT2G38325 | 156,77 | 0,66 | 0,23 | 2,82 | 4,76E-03 | 2,88E-02 |
| AT5G41905 | 1173,00 | -0,72 | 0,25 | -2,82 | 4,85E-03 | 2,93E-02 |
| AT4G16720 | 3,07 | 4,27 | 1,52 | 2,81 | 4,91E-03 | 2,96E-02 |
| AT1G26975 | 8,83 | 1,98 | 0,70 | 2,81 | 4,95E-03 | 2,97E-02 |
| AT5G23065 | 1776,68 | 0,75 | 0,27 | 2,81 | 4,97E-03 | 2,98E-02 |
| AT1G63070 | 3,60 | 3,04 | 1,08 | 2,81 | 4,99E-03 | 2,98E-02 |
| AT2G47410 | 3,41 | 4,39 | 1,56 | 2,81 | 5,01E-03 | 2,99E-02 |
| AT1G73687 | 3456,65 | -0,73 | 0,26 | -2,80 | 5,04E-03 | 3,00E-02 |
| AT3G44480 | 4,70 | 2,34 | 0,84 | 2,80 | 5,06E-03 | 3,00E-02 |

| | | | | | | |
|---|---|---|---|---|---|---|
| AT5G11560 | 3,42 | 2,79 | 1,00 | 2,80 | 5,07E-03 | 3,00E-02 |
| AT2G26780 | 3,27 | 3,00 | 1,08 | 2,78 | 5,38E-03 | 3,16E-02 |
| AT5G04930 | 5,77 | 2,58 | 0,93 | 2,78 | 5,37E-03 | 3,16E-02 |
| AT2G47940 | 2,47 | 4,45 | 1,60 | 2,78 | 5,39E-03 | 3,16E-02 |
| AT1G63330 | 7,00 | -2,19 | 0,79 | -2,78 | 5,42E-03 | 3,17E-02 |
| AT1G06267 | 15,22 | -1,58 | 0,57 | -2,77 | 5,54E-03 | 3,24E-02 |
| AT4G38220 | 1,96 | 4,57 | 1,65 | 2,76 | 5,70E-03 | 3,31E-02 |
| AT4G03455 | 16,01 | 3,95 | 1,43 | 2,76 | 5,70E-03 | 3,31E-02 |
| AT1G32583 | 28,27 | -1,62 | 0,59 | -2,76 | 5,79E-03 | 3,36E-02 |
| AT4G36760 | 3,13 | 3,30 | 1,20 | 2,75 | 5,91E-03 | 3,42E-02 |
| AT2G31880 | 6,93 | 2,22 | 0,81 | 2,75 | 5,99E-03 | 3,45E-02 |
| AT3G14470 | 2,64 | 4,37 | 1,59 | 2,74 | 6,07E-03 | 3,49E-02 |
| AT5G02480 | 2,67 | 3,86 | 1,41 | 2,74 | 6,06E-03 | 3,49E-02 |
| AT4G03445 | 45,78 | -0,76 | 0,28 | -2,74 | 6,10E-03 | 3,50E-02 |
| AT4G31880 | 2,70 | 4,26 | 1,56 | 2,74 | 6,24E-03 | 3,56E-02 |
| AT1G32582 | 26,60 | -1,64 | 0,60 | -2,73 | 6,29E-03 | 3,59E-02 |
| AT5G39770 | 12,51 | -1,49 | 0,55 | -2,73 | 6,35E-03 | 3,61E-02 |
| AT1G36730 | 3,83 | 2,70 | 0,99 | 2,72 | 6,47E-03 | 3,67E-02 |
| AT4G34071 | 2,35 | -4,41 | 1,62 | -2,72 | 6,53E-03 | 3,69E-02 |
| AT1G72390 | 2,40 | 3,89 | 1,43 | 2,72 | 6,55E-03 | 3,70E-02 |
| AT3G51050 | 3,03 | 4,34 | 1,60 | 2,71 | 6,76E-03 | 3,80E-02 |
| AT4G16340 | 3,35 | 3,66 | 1,35 | 2,71 | 6,80E-03 | 3,82E-02 |
| AT3G21805 | 21,83 | -1,95 | 0,72 | -2,71 | 6,82E-03 | 3,82E-02 |
| AT1G64140 | 5,12 | 2,46 | 0,91 | 2,70 | 6,84E-03 | 3,83E-02 |
| AT5G24350 | 3,06 | 4,22 | 1,56 | 2,70 | 6,98E-03 | 3,89E-02 |
| AT3G22910 | 2,19 | 4,11 | 1,53 | 2,70 | 7,02E-03 | 3,89E-02 |
| AT5G03240 | 5,81 | 2,10 | 0,78 | 2,70 | 7,00E-03 | 3,89E-02 |
| AT3G63380 | 3,14 | 3,81 | 1,41 | 2,69 | 7,06E-03 | 3,91E-02 |
| AT2G45380 | 11,33 | -1,57 | 0,58 | -2,69 | 7,20E-03 | 3,97E-02 |
| AT3G02470 | 10,40 | 1,51 | 0,56 | 2,69 | 7,24E-03 | 3,98E-02 |
| AT3G04240 | 3,51 | 3,17 | 1,18 | 2,69 | 7,25E-03 | 3,98E-02 |
| AT1G15940 | 2,10 | 3,76 | 1,40 | 2,67 | 7,52E-03 | 4,09E-02 |
| AT1G70185 | 5,33 | 2,39 | 0,89 | 2,67 | 7,53E-03 | 4,09E-02 |
| AT1G29370 | 3,57 | 2,89 | 1,08 | 2,67 | 7,55E-03 | 4,09E-02 |
| AT1G62310 | 4,32 | -2,83 | 1,06 | -2,67 | 7,50E-03 | 4,09E-02 |
| AT3G57330 | 3,24 | 2,94 | 1,10 | 2,67 | 7,49E-03 | 4,09E-02 |
| AT5G16210 | 1,91 | 4,41 | 1,65 | 2,67 | 7,56E-03 | 4,09E-02 |
| AT1G35710 | 7,53 | 2,44 | 0,91 | 2,67 | 7,69E-03 | 4,15E-02 |
| AT5G12370 | 4,35 | 3,01 | 1,13 | 2,66 | 7,86E-03 | 4,24E-02 |
| AT5G54310 | 4,38 | 2,80 | 1,06 | 2,66 | 7,89E-03 | 4,24E-02 |
| AT3G04130 | 2,12 | 5,36 | 2,02 | 2,65 | 7,94E-03 | 4,26E-02 |
| AT5G47910 | 5,04 | 2,35 | 0,88 | 2,65 | 7,96E-03 | 4,26E-02 |
| AT5G23380 | 2,25 | 4,67 | 1,76 | 2,65 | 7,99E-03 | 4,27E-02 |
| AT5G58540 | 1,83 | 4,53 | 1,71 | 2,65 | 8,14E-03 | 4,34E-02 |
| AT3G44570 | 48,95 | -3,11 | 1,18 | -2,64 | 8,22E-03 | 4,37E-02 |
| AT1G30490 | 1,77 | 4,97 | 1,88 | 2,64 | 8,35E-03 | 4,43E-02 |
| AT1G28420 | 2,78 | 3,42 | 1,30 | 2,64 | 8,38E-03 | 4,44E-02 |
| AT1G64790 | 7,36 | 1,93 | 0,73 | 2,63 | 8,43E-03 | 4,45E-02 |
| AT4G01050 | 4,58 | 3,00 | 1,14 | 2,63 | 8,48E-03 | 4,46E-02 |
| AT4G30160 | 4,90 | 2,26 | 0,86 | 2,63 | 8,46E-03 | 4,46E-02 |
| AT2G35384 | 13,85 | -1,78 | 0,67 | -2,63 | 8,49E-03 | 4,46E-02 |
| AT4G34670 | 4,28 | 2,63 | 1,00 | 2,63 | 8,58E-03 | 4,49E-02 |
| AT1G75335 | 3,97 | 2,96 | 1,13 | 2,62 | 8,81E-03 | 4,60E-02 |
| AT1G18485 | 1,73 | 5,28 | 2,02 | 2,62 | 8,86E-03 | 4,61E-02 |
| AT4G18700 | 2,69 | 2,92 | 1,12 | 2,62 | 8,92E-03 | 4,64E-02 |
| AT3G58640 | 2,56 | 4,27 | 1,63 | 2,61 | 8,94E-03 | 4,64E-02 |

| | | | | | | |
|---|---|---|---|---|---|---|
| AT3G13857 | 124,31 | 0,96 | 0,37 | 2,61 | 9,00E-03 | 4,64E-02 |
| AT3G13855 | 124,31 | 0,96 | 0,37 | 2,61 | 9,00E-03 | 4,64E-02 |
| AT3G08850 | 2,35 | 3,49 | 1,34 | 2,61 | 9,05E-03 | 4,66E-02 |
| AT4G00730 | 3,19 | 3,25 | 1,25 | 2,60 | 9,32E-03 | 4,78E-02 |
| AT5G14800 | 5,34 | 3,14 | 1,21 | 2,60 | 9,32E-03 | 4,78E-02 |
| AT2G17440 | 1,80 | 4,45 | 1,72 | 2,60 | 9,46E-03 | 4,82E-02 |
| AT3G15730 | 15,00 | 1,37 | 0,53 | 2,60 | 9,44E-03 | 4,82E-02 |
| AT5G15400 | 2,82 | 2,78 | 1,07 | 2,59 | 9,47E-03 | 4,82E-02 |
| AT4G11420 | 3,20 | 3,64 | 1,41 | 2,59 | 9,59E-03 | 4,88E-02 |
| AT4G25880 | 1,93 | 4,19 | 1,62 | 2,59 | 9,67E-03 | 4,90E-02 |
| AT5G61020 | 2,23 | 4,15 | 1,61 | 2,58 | 9,91E-03 | 5,02E-02 |
| AT2G39175 | 43,76 | 0,86 | 0,33 | 2,58 | 9,95E-03 | 5,03E-02 |
| AT4G18596 | 1,78 | 4,48 | 1,74 | 2,57 | 1,01E-02 | 5,08E-02 |
| AT1G60870 | 2,59 | 3,46 | 1,35 | 2,57 | 1,01E-02 | 5,09E-02 |
| AT2G19850 | 5,66 | -2,75 | 1,07 | -2,57 | 1,02E-02 | 5,10E-02 |
| AT4G33010 | 11,10 | 1,84 | 0,72 | 2,57 | 1,02E-02 | 5,10E-02 |
| AT1G30470 | 3,87 | 3,06 | 1,19 | 2,57 | 1,02E-02 | 5,12E-02 |
| AT4G24680 | 3,35 | 2,65 | 1,03 | 2,56 | 1,04E-02 | 5,17E-02 |
| AT3G16850 | 3,96 | 2,84 | 1,11 | 2,56 | 1,04E-02 | 5,17E-02 |
| AT5G24990 | 2,50 | -3,84 | 1,50 | -2,56 | 1,05E-02 | 5,19E-02 |
| AT3G59770 | 3,31 | 2,77 | 1,08 | 2,56 | 1,05E-02 | 5,19E-02 |
| AT3G02170 | 3,18 | 3,78 | 1,48 | 2,56 | 1,05E-02 | 5,21E-02 |
| AT4G25080 | 4,26 | 2,44 | 0,95 | 2,56 | 1,06E-02 | 5,22E-02 |
| AT1G69360 | 2,19 | 4,37 | 1,71 | 2,56 | 1,06E-02 | 5,22E-02 |
| AT5G62000 | 10,38 | 1,39 | 0,54 | 2,55 | 1,08E-02 | 5,28E-02 |
| AT1G55670 | 5,12 | -2,41 | 0,95 | -2,55 | 1,08E-02 | 5,28E-02 |
| AT4G35830 | 9,01 | 2,05 | 0,81 | 2,55 | 1,08E-02 | 5,28E-02 |
| AT5G66030 | 2,48 | 4,03 | 1,58 | 2,55 | 1,08E-02 | 5,28E-02 |
| AT1G47497 | 2,25 | 4,71 | 1,85 | 2,55 | 1,09E-02 | 5,30E-02 |
| AT5G26830 | 2,03 | 3,59 | 1,41 | 2,54 | 1,11E-02 | 5,38E-02 |
| AT5G44800 | 4,77 | 2,10 | 0,83 | 2,54 | 1,11E-02 | 5,38E-02 |
| AT1G01960 | 2,74 | 2,64 | 1,04 | 2,53 | 1,13E-02 | 5,48E-02 |
| AT3G23780 | 4,59 | 2,41 | 0,95 | 2,53 | 1,14E-02 | 5,52E-02 |
| AT2G32730 | 3,18 | 3,00 | 1,19 | 2,53 | 1,15E-02 | 5,53E-02 |
| AT3G19180 | 1,86 | 4,64 | 1,84 | 2,52 | 1,18E-02 | 5,67E-02 |
| AT3G13070 | 1,98 | 4,51 | 1,79 | 2,52 | 1,18E-02 | 5,68E-02 |
| AT3G20010 | 2,80 | 4,35 | 1,73 | 2,51 | 1,21E-02 | 5,77E-02 |
| AT5G35210 | 2,88 | 3,56 | 1,42 | 2,50 | 1,23E-02 | 5,86E-02 |
| AT5G63020 | 8,08 | -1,84 | 0,73 | -2,50 | 1,24E-02 | 5,91E-02 |
| AT5G04810 | 3,05 | 3,35 | 1,34 | 2,50 | 1,24E-02 | 5,91E-02 |
| AT2G28910 | 4,89 | 2,96 | 1,19 | 2,50 | 1,25E-02 | 5,92E-02 |
| AT1G09970 | 4,22 | 2,46 | 0,99 | 2,50 | 1,26E-02 | 5,96E-02 |
| AT5G17165 | 3,90 | -2,58 | 1,03 | -2,49 | 1,27E-02 | 5,98E-02 |
| AT4G11830 | 1,70 | 4,77 | 1,91 | 2,49 | 1,27E-02 | 6,00E-02 |
| AT4G36360 | 3,63 | 3,64 | 1,46 | 2,49 | 1,27E-02 | 6,00E-02 |
| AT3G06650 | 3,17 | 3,90 | 1,57 | 2,49 | 1,28E-02 | 6,01E-02 |
| AT2G25800 | 4,68 | 2,35 | 0,95 | 2,49 | 1,28E-02 | 6,03E-02 |
| AT3G56680 | 2,66 | 3,56 | 1,43 | 2,49 | 1,29E-02 | 6,03E-02 |
| AT3G22790 | 2,45 | 3,96 | 1,60 | 2,48 | 1,31E-02 | 6,11E-02 |
| AT2G35382 | 3,86 | -2,56 | 1,03 | -2,48 | 1,31E-02 | 6,12E-02 |
| AT3G52240 | 2,27 | 4,01 | 1,62 | 2,48 | 1,32E-02 | 6,13E-02 |
| AT1G09770 | 3,11 | 2,85 | 1,15 | 2,48 | 1,32E-02 | 6,13E-02 |
| AT3G50590 | 4,47 | 3,12 | 1,26 | 2,48 | 1,32E-02 | 6,13E-02 |
| AT1G17210 | 2,32 | 3,85 | 1,56 | 2,48 | 1,33E-02 | 6,13E-02 |
| AT1G79000 | 2,98 | 4,70 | 1,90 | 2,48 | 1,33E-02 | 6,13E-02 |
| AT5G46210 | 3,09 | 3,17 | 1,28 | 2,48 | 1,33E-02 | 6,13E-02 |

| | | | | | | |
|---|---|---|---|---|---|---|
| AT1G72645 | 2,19 | -3,27 | 1,32 | -2,47 | 1,35E-02 | 6,18E-02 |
| AT3G23120 | 2,24 | 3,01 | 1,22 | 2,47 | 1,35E-02 | 6,18E-02 |
| AT5G50920 | 18,45 | 1,21 | 0,49 | 2,47 | 1,35E-02 | 6,18E-02 |
| AT5G14430 | 2,60 | 3,45 | 1,40 | 2,46 | 1,37E-02 | 6,26E-02 |
| AT5G44180 | 3,26 | 3,29 | 1,34 | 2,46 | 1,37E-02 | 6,26E-02 |
| AT5G10380 | 7,30 | 1,83 | 0,74 | 2,46 | 1,38E-02 | 6,27E-02 |
| AT2G16485 | 2,25 | 3,29 | 1,34 | 2,46 | 1,38E-02 | 6,27E-02 |
| AT4G00800 | 2,43 | 3,20 | 1,30 | 2,46 | 1,38E-02 | 6,27E-02 |
| AT1G05562 | 6,13 | -2,06 | 0,84 | -2,46 | 1,41E-02 | 6,36E-02 |
| AT1G05560 | 6,13 | -2,06 | 0,84 | -2,46 | 1,41E-02 | 6,36E-02 |
| AT1G80680 | 4,25 | 2,70 | 1,10 | 2,45 | 1,42E-02 | 6,39E-02 |
| AT4G34830 | 2,44 | 3,23 | 1,32 | 2,45 | 1,44E-02 | 6,46E-02 |
| AT5G13010 | 4,55 | 2,55 | 1,04 | 2,45 | 1,44E-02 | 6,47E-02 |
| AT5G46470 | 3,68 | 2,49 | 1,02 | 2,44 | 1,46E-02 | 6,51E-02 |
| AT1G04625 | 1,86 | 4,79 | 1,96 | 2,44 | 1,45E-02 | 6,51E-02 |
| AT1G48410 | 43,79 | 0,84 | 0,35 | 2,44 | 1,46E-02 | 6,51E-02 |
| AT5G52882 | 2,07 | 4,25 | 1,74 | 2,44 | 1,46E-02 | 6,51E-02 |
| AT2G38230 | 6,88 | -1,88 | 0,77 | -2,44 | 1,47E-02 | 6,54E-02 |
| AT2G46830 | 2,43 | 3,66 | 1,50 | 2,44 | 1,48E-02 | 6,55E-02 |
| AT4G08470 | 1,82 | 4,82 | 1,98 | 2,44 | 1,48E-02 | 6,55E-02 |
| AT3G48000 | 4,79 | 2,24 | 0,92 | 2,43 | 1,49E-02 | 6,60E-02 |
| AT2G01008 | 28,92 | -0,97 | 0,40 | -2,43 | 1,50E-02 | 6,64E-02 |
| AT4G00570 | 3,02 | 3,28 | 1,36 | 2,42 | 1,53E-02 | 6,76E-02 |
| AT1G34300 | 2,71 | 3,10 | 1,28 | 2,42 | 1,54E-02 | 6,78E-02 |
| AT5G54270 | 15,64 | -1,26 | 0,52 | -2,42 | 1,55E-02 | 6,81E-02 |
| AT3G22890 | 4,47 | 1,86 | 0,77 | 2,42 | 1,56E-02 | 6,84E-02 |
| AT1G58230 | 3,69 | 2,62 | 1,08 | 2,42 | 1,57E-02 | 6,87E-02 |
| AT5G16640 | 4,15 | -2,27 | 0,94 | -2,41 | 1,57E-02 | 6,87E-02 |
| AT1G49700 | 4,39 | 2,02 | 0,84 | 2,41 | 1,58E-02 | 6,88E-02 |
| AT1G12240 | 2,62 | 3,32 | 1,38 | 2,41 | 1,59E-02 | 6,89E-02 |
| AT1G63630 | 3,43 | -2,50 | 1,04 | -2,41 | 1,59E-02 | 6,89E-02 |
| AT3G03050 | 2,11 | 4,11 | 1,70 | 2,41 | 1,59E-02 | 6,89E-02 |
| AT1G63615 | 3,43 | -2,50 | 1,04 | -2,41 | 1,59E-02 | 6,89E-02 |
| AT5G12470 | 4,58 | 2,83 | 1,18 | 2,41 | 1,61E-02 | 6,94E-02 |
| AT5G58150 | 3,53 | 3,71 | 1,54 | 2,41 | 1,61E-02 | 6,94E-02 |
| AT4G31180 | 4,29 | 2,30 | 0,96 | 2,40 | 1,62E-02 | 6,96E-02 |
| AT1G45160 | 1,73 | 3,89 | 1,62 | 2,40 | 1,64E-02 | 7,03E-02 |
| AT1G43910 | 2,35 | 3,63 | 1,51 | 2,40 | 1,65E-02 | 7,06E-02 |
| AT5G60410 | 2,90 | 2,79 | 1,16 | 2,40 | 1,65E-02 | 7,06E-02 |
| AT5G59505 | 3,52 | -3,48 | 1,45 | -2,39 | 1,67E-02 | 7,13E-02 |
| AT5G44320 | 2,18 | 3,17 | 1,33 | 2,39 | 1,67E-02 | 7,13E-02 |
| AT5G02490 | 4,18 | 2,20 | 0,92 | 2,39 | 1,69E-02 | 7,17E-02 |
| AT5G11490 | 1,97 | 3,50 | 1,47 | 2,39 | 1,70E-02 | 7,23E-02 |
| AT2G20990 | 4,50 | 2,57 | 1,08 | 2,38 | 1,71E-02 | 7,26E-02 |
| AT5G43310 | 2,37 | 3,17 | 1,33 | 2,38 | 1,73E-02 | 7,33E-02 |
| AT5G08670 | 5,95 | 1,82 | 0,77 | 2,37 | 1,76E-02 | 7,43E-02 |
| AT3G18217 | 147,37 | 0,56 | 0,24 | 2,37 | 1,78E-02 | 7,49E-02 |
| AT3G02090 | 3,68 | 2,23 | 0,94 | 2,37 | 1,80E-02 | 7,55E-02 |
| AT1G61690 | 1,91 | 3,67 | 1,55 | 2,37 | 1,80E-02 | 7,55E-02 |
| AT3G11910 | 1,81 | 3,78 | 1,60 | 2,36 | 1,80E-02 | 7,55E-02 |
| AT1G74456 | 11,08 | -1,53 | 0,65 | -2,36 | 1,81E-02 | 7,56E-02 |
| AT5G58550 | 1,86 | 4,42 | 1,87 | 2,36 | 1,81E-02 | 7,56E-02 |
| AT1G63230 | 3,31 | -2,50 | 1,06 | -2,36 | 1,84E-02 | 7,67E-02 |
| AT3G03780 | 4,16 | 2,58 | 1,09 | 2,36 | 1,85E-02 | 7,67E-02 |
| AT2G03445 | 309,93 | -0,61 | 0,26 | -2,35 | 1,86E-02 | 7,68E-02 |
| AT2G38440 | 3,79 | 2,18 | 0,93 | 2,36 | 1,85E-02 | 7,68E-02 |

| | | | | | | |
|---|---|---|---|---|---|---|
| AT1G21580 | 3,07 | 2,38 | 1,01 | 2,35 | 1,85E-02 | 7,68E-02 |
| AT5G62350 | 2,26 | -3,44 | 1,46 | -2,35 | 1,87E-02 | 7,71E-02 |
| AT3G44330 | 1,72 | 4,69 | 2,00 | 2,35 | 1,89E-02 | 7,80E-02 |
| AT4G08691 | 15,19 | -1,30 | 0,55 | -2,35 | 1,90E-02 | 7,80E-02 |
| AT5G11040 | 2,15 | 3,21 | 1,37 | 2,34 | 1,92E-02 | 7,87E-02 |
| AT1G07000 | 3,04 | 2,95 | 1,26 | 2,34 | 1,93E-02 | 7,92E-02 |
| AT5G60450 | 7,78 | -1,59 | 0,68 | -2,34 | 1,93E-02 | 7,92E-02 |
| AT5G04560 | 3,61 | 2,47 | 1,06 | 2,33 | 1,96E-02 | 8,01E-02 |
| AT3G61897 | 2,76 | -5,99 | 2,57 | -2,32 | 2,01E-02 | 8,18E-02 |
| AT4G37460 | 2,98 | 2,67 | 1,15 | 2,33 | 2,01E-02 | 8,18E-02 |
| AT1G15690 | 11,49 | 1,44 | 0,62 | 2,32 | 2,03E-02 | 8,23E-02 |
| AT3G08730 | 2,48 | 3,91 | 1,68 | 2,32 | 2,03E-02 | 8,23E-02 |
| AT3G13100 | 2,83 | 2,97 | 1,28 | 2,32 | 2,03E-02 | 8,23E-02 |
| AT2G35750 | 26,17 | -1,17 | 0,50 | -2,32 | 2,04E-02 | 8,23E-02 |
| AT1G07590 | 3,38 | 2,09 | 0,90 | 2,32 | 2,04E-02 | 8,25E-02 |
| AT1G11735 | 8,53 | -1,71 | 0,74 | -2,32 | 2,06E-02 | 8,30E-02 |
| AT3G61580 | 5,37 | 1,81 | 0,78 | 2,31 | 2,07E-02 | 8,33E-02 |
| AT4G10710 | 3,86 | 2,74 | 1,18 | 2,31 | 2,07E-02 | 8,33E-02 |
| AT3G45780 | 10,09 | 1,31 | 0,57 | 2,31 | 2,09E-02 | 8,38E-02 |
| AT5G54390 | 4,62 | 2,71 | 1,18 | 2,30 | 2,12E-02 | 8,44E-02 |
| AT1G05163 | 491,36 | -0,62 | 0,27 | -2,30 | 2,12E-02 | 8,44E-02 |
| AT1G15405 | 491,36 | -0,62 | 0,27 | -2,30 | 2,12E-02 | 8,44E-02 |
| AT5G19820 | 4,02 | 2,85 | 1,23 | 2,31 | 2,11E-02 | 8,44E-02 |
| AT5G05200 | 1,99 | 4,22 | 1,83 | 2,30 | 2,13E-02 | 8,45E-02 |
| AT1G54090 | 3,59 | 3,20 | 1,39 | 2,30 | 2,14E-02 | 8,49E-02 |
| AT1G62035 | 5,54 | -1,71 | 0,74 | -2,30 | 2,17E-02 | 8,60E-02 |
| AT3G44444 | 1,95 | -2,92 | 1,27 | -2,29 | 2,18E-02 | 8,62E-02 |
| AT1G80180 | 13,61 | -1,24 | 0,54 | -2,29 | 2,19E-02 | 8,63E-02 |
| AT2G40930 | 1,74 | 4,32 | 1,88 | 2,29 | 2,20E-02 | 8,65E-02 |
| AT5G13100 | 3,43 | 2,90 | 1,27 | 2,29 | 2,22E-02 | 8,71E-02 |
| AT1G27430 | 3,11 | 2,72 | 1,19 | 2,29 | 2,22E-02 | 8,71E-02 |
| AT3G07160 | 3,87 | 2,32 | 1,02 | 2,29 | 2,22E-02 | 8,71E-02 |
| AT4G14500 | 7,04 | -1,68 | 0,73 | -2,28 | 2,25E-02 | 8,77E-02 |
| AT5G07875 | 4,76 | -2,15 | 0,94 | -2,28 | 2,25E-02 | 8,77E-02 |
| AT1G05917 | 8,49 | -1,78 | 0,78 | -2,28 | 2,29E-02 | 8,91E-02 |
| AT3G12120 | 5,50 | 2,39 | 1,05 | 2,28 | 2,29E-02 | 8,91E-02 |
| AT1G66795 | 14,00 | 1,20 | 0,53 | 2,27 | 2,30E-02 | 8,92E-02 |
| AT2G45685 | 1,79 | 3,83 | 1,68 | 2,27 | 2,30E-02 | 8,92E-02 |
| AT2G45680 | 1,79 | 3,83 | 1,68 | 2,27 | 2,30E-02 | 8,92E-02 |
| AT3G13870 | 4,03 | 3,46 | 1,53 | 2,27 | 2,32E-02 | 8,96E-02 |
| AT1G80000 | 1,90 | 4,53 | 2,00 | 2,27 | 2,34E-02 | 9,03E-02 |
| AT2G01010 | 3,38 | -2,18 | 0,96 | -2,26 | 2,38E-02 | 9,06E-02 |
| AT2G38040 | 4,80 | 2,26 | 1,00 | 2,26 | 2,38E-02 | 9,06E-02 |
| AT1G72440 | 1,79 | 3,62 | 1,60 | 2,26 | 2,38E-02 | 9,06E-02 |
| AT2G33730 | 2,24 | 2,89 | 1,28 | 2,26 | 2,36E-02 | 9,06E-02 |
| AT2G34100 | 7,49 | -1,64 | 0,72 | -2,26 | 2,36E-02 | 9,06E-02 |
| AT3G09200 | 3,98 | 2,35 | 1,04 | 2,26 | 2,36E-02 | 9,06E-02 |
| AT4G24520 | 3,90 | 3,48 | 1,54 | 2,26 | 2,37E-02 | 9,06E-02 |
| AT5G13000 | 3,30 | 2,12 | 0,94 | 2,26 | 2,38E-02 | 9,06E-02 |
| AT5G03305 | 5,94 | -1,68 | 0,75 | -2,26 | 2,39E-02 | 9,07E-02 |
| AT2G18280 | 2,10 | 3,91 | 1,74 | 2,25 | 2,43E-02 | 9,20E-02 |
| AT4G02380 | 7,09 | 1,45 | 0,65 | 2,25 | 2,42E-02 | 9,20E-02 |
| AT2G24790 | 3,54 | 2,73 | 1,21 | 2,25 | 2,43E-02 | 9,21E-02 |
| AT4G13560 | 52,08 | -0,77 | 0,34 | -2,25 | 2,46E-02 | 9,28E-02 |
| AT2G34410 | 2,89 | 2,66 | 1,19 | 2,24 | 2,48E-02 | 9,36E-02 |
| AT3G20770 | 2,68 | 2,86 | 1,28 | 2,24 | 2,49E-02 | 9,37E-02 |

| | | | | | | |
|---|---|---|---|---|---|---|
| AT1G16880 | 2,91 | 2,76 | 1,24 | 2,24 | 2,53E-02 | 9,49E-02 |
| AT4G12290 | 3,10 | 2,19 | 0,98 | 2,23 | 2,55E-02 | 9,55E-02 |
| AT1G32750 | 1,94 | 3,26 | 1,46 | 2,23 | 2,55E-02 | 9,55E-02 |
| AT1G21680 | 4,80 | 2,00 | 0,90 | 2,23 | 2,57E-02 | 9,55E-02 |
| AT1G77815 | 2,65 | -3,19 | 1,43 | -2,23 | 2,55E-02 | 9,55E-02 |
| AT4G06110 | 22,73 | -1,13 | 0,51 | -2,23 | 2,56E-02 | 9,55E-02 |
| AT5G42820 | 2,08 | 3,60 | 1,61 | 2,23 | 2,56E-02 | 9,55E-02 |
| AT3G62250 | 1,91 | 3,65 | 1,64 | 2,23 | 2,59E-02 | 9,61E-02 |
| AT4G11373 | 5,92 | -1,68 | 0,76 | -2,23 | 2,60E-02 | 9,62E-02 |
| AT1G79090 | 2,69 | 3,24 | 1,45 | 2,23 | 2,60E-02 | 9,62E-02 |
| AT4G23713 | 128,07 | -0,71 | 0,32 | -2,22 | 2,62E-02 | 9,64E-02 |
| AT4G31490 | 3,09 | 2,74 | 1,23 | 2,22 | 2,62E-02 | 9,64E-02 |
| AT5G36230 | 2,63 | 3,07 | 1,38 | 2,22 | 2,63E-02 | 9,64E-02 |
| AT5G37510 | 3,25 | 2,21 | 0,99 | 2,22 | 2,63E-02 | 9,64E-02 |
| AT5G61010 | 3,05 | 2,40 | 1,08 | 2,22 | 2,62E-02 | 9,64E-02 |
| AT1G08380 | 7,50 | -1,95 | 0,88 | -2,22 | 2,62E-02 | 9,64E-02 |
| AT2G08770 | 10,72 | -1,58 | 0,71 | -2,22 | 2,63E-02 | 9,64E-02 |
| AT2G23347 | 3,51 | -2,23 | 1,01 | -2,22 | 2,65E-02 | 9,69E-02 |
| AT3G20690 | 3,62 | -2,68 | 1,21 | -2,21 | 2,68E-02 | 9,79E-02 |
| AT5G12120 | 2,73 | 2,96 | 1,34 | 2,21 | 2,69E-02 | 9,82E-02 |
| AT5G15020 | 2,14 | 3,27 | 1,48 | 2,21 | 2,70E-02 | 9,84E-02 |
| AT3G07870 | 5,57 | 2,34 | 1,06 | 2,21 | 2,71E-02 | 9,86E-02 |
| AT5G58160 | 1,77 | 5,40 | 2,44 | 2,21 | 2,72E-02 | 9,86E-02 |
| AT4G19110 | 3,30 | 2,85 | 1,29 | 2,21 | 2,74E-02 | 9,90E-02 |
| AT5G03040 | 2,14 | 3,18 | 1,44 | 2,21 | 2,73E-02 | 9,90E-02 |
| AT4G02320 | 3,77 | 2,66 | 1,21 | 2,20 | 2,77E-02 | 1,00E-01 |
| AT1G32230 | 5,15 | 1,88 | 0,86 | 2,20 | 2,78E-02 | 1,00E-01 |
| AT2G23348 | 3,88 | -2,23 | 1,02 | -2,20 | 2,78E-02 | 1,00E-01 |
| AT5G66200 | 3,05 | 2,47 | 1,12 | 2,20 | 2,79E-02 | 1,00E-01 |
| AT1G52780 | 3,16 | 2,29 | 1,04 | 2,19 | 2,82E-02 | 1,01E-01 |
| AT5G61780 | 4,25 | 2,07 | 0,95 | 2,19 | 2,84E-02 | 1,02E-01 |
| AT3G22960 | 3,17 | 2,78 | 1,27 | 2,18 | 2,89E-02 | 1,04E-01 |
| AT3G13920 | 8,10 | 1,81 | 0,83 | 2,18 | 2,93E-02 | 1,05E-01 |
| AT3G07725 | 10,80 | -1,16 | 0,53 | -2,18 | 2,96E-02 | 1,06E-01 |
| AT4G03030 | 1,74 | 3,53 | 1,62 | 2,17 | 2,97E-02 | 1,06E-01 |
| AT2G35050 | 3,09 | 2,68 | 1,23 | 2,17 | 2,99E-02 | 1,06E-01 |
| AT2G45930 | 2,64 | 3,46 | 1,59 | 2,17 | 2,99E-02 | 1,06E-01 |
| AT3G28540 | 2,73 | 2,40 | 1,11 | 2,17 | 3,03E-02 | 1,08E-01 |
| AT3G60860 | 2,94 | 2,71 | 1,25 | 2,16 | 3,04E-02 | 1,08E-01 |
| AT5G19390 | 2,78 | 3,48 | 1,61 | 2,16 | 3,06E-02 | 1,08E-01 |
| AT4G17330 | 4,91 | 2,44 | 1,13 | 2,16 | 3,06E-02 | 1,08E-01 |
| AT3G13460 | 10,31 | 1,25 | 0,58 | 2,16 | 3,08E-02 | 1,09E-01 |
| AT1G61190 | 2,38 | 2,63 | 1,22 | 2,16 | 3,09E-02 | 1,09E-01 |
| AT1G16710 | 1,85 | 3,56 | 1,65 | 2,15 | 3,15E-02 | 1,11E-01 |
| AT4G04640 | 5,86 | -1,94 | 0,90 | -2,15 | 3,15E-02 | 1,11E-01 |
| AT1G55673 | 3,49 | -2,52 | 1,17 | -2,15 | 3,17E-02 | 1,11E-01 |
| AT1G79990 | 2,18 | 3,36 | 1,57 | 2,15 | 3,18E-02 | 1,11E-01 |
| AT5G24690 | 2,42 | 3,03 | 1,41 | 2,15 | 3,19E-02 | 1,11E-01 |
| AT1G29920 | 50,33 | 0,89 | 0,42 | 2,14 | 3,20E-02 | 1,11E-01 |
| AT1G48953 | 8,24 | -1,77 | 0,82 | -2,14 | 3,20E-02 | 1,11E-01 |
| AT1G79040 | 10,12 | -1,81 | 0,84 | -2,14 | 3,22E-02 | 1,12E-01 |
| AT2G01600 | 1,95 | 3,33 | 1,56 | 2,14 | 3,23E-02 | 1,12E-01 |
| AT5G05750 | 1,85 | 4,46 | 2,08 | 2,14 | 3,24E-02 | 1,12E-01 |
| AT3G23920 | 2,81 | 2,30 | 1,08 | 2,14 | 3,27E-02 | 1,13E-01 |
| AT4G00975 | 11,27 | 1,12 | 0,53 | 2,13 | 3,28E-02 | 1,13E-01 |
| AT1G78900 | 8,95 | 1,42 | 0,66 | 2,13 | 3,30E-02 | 1,14E-01 |

| | | | | | | |
|---|---|---|---|---|---|---|
| AT3G60415 | 1,73 | 3,81 | 1,79 | 2,13 | 3,30E-02 | 1,14E-01 |
| AT5G24280 | 2,00 | 6,39 | 3,00 | 2,13 | 3,30E-02 | 1,14E-01 |
| AT1G05913 | 9,06 | -1,68 | 0,79 | -2,13 | 3,32E-02 | 1,14E-01 |
| AT4G01810 | 2,76 | 2,84 | 1,34 | 2,13 | 3,32E-02 | 1,14E-01 |
| AT1G12620 | 2,40 | -2,81 | 1,32 | -2,13 | 3,33E-02 | 1,14E-01 |
| AT1G77680 | 1,90 | 4,16 | 1,96 | 2,12 | 3,36E-02 | 1,15E-01 |
| AT4G31877 | 9,33 | -1,49 | 0,70 | -2,12 | 3,37E-02 | 1,15E-01 |
| AT4G02660 | 2,17 | 3,08 | 1,45 | 2,12 | 3,38E-02 | 1,15E-01 |
| AT3G57520 | 3,92 | 2,74 | 1,29 | 2,12 | 3,39E-02 | 1,15E-01 |
| AT1G31330 | 6,03 | -1,65 | 0,78 | -2,12 | 3,40E-02 | 1,16E-01 |
| AT4G13570 | 36,83 | -0,79 | 0,37 | -2,11 | 3,46E-02 | 1,17E-01 |
| AT4G38200 | 2,17 | 3,44 | 1,63 | 2,11 | 3,47E-02 | 1,18E-01 |
| AT3G06720 | 1,89 | 3,97 | 1,88 | 2,11 | 3,48E-02 | 1,18E-01 |
| AT1G72930 | 3,01 | -3,22 | 1,53 | -2,11 | 3,51E-02 | 1,19E-01 |
| AT3G53460 | 6,35 | 1,71 | 0,81 | 2,11 | 3,53E-02 | 1,19E-01 |
| AT2G34357 | 1,72 | 3,27 | 1,56 | 2,10 | 3,58E-02 | 1,20E-01 |
| AT3G09105 | 30,37 | -1,05 | 0,50 | -2,10 | 3,58E-02 | 1,21E-01 |
| AT3G58196 | 21,50 | -1,00 | 0,48 | -2,10 | 3,60E-02 | 1,21E-01 |
| AT5G63190 | 2,74 | 2,73 | 1,30 | 2,09 | 3,62E-02 | 1,21E-01 |
| AT5G46260 | 1,90 | 2,74 | 1,31 | 2,09 | 3,62E-02 | 1,21E-01 |
| AT4G03550 | 3,00 | 2,81 | 1,34 | 2,09 | 3,66E-02 | 1,22E-01 |
| AT5G13650 | 5,78 | 1,71 | 0,82 | 2,09 | 3,67E-02 | 1,22E-01 |
| AT5G06175 | 3,94 | -1,88 | 0,90 | -2,09 | 3,67E-02 | 1,22E-01 |
| AT4G23375 | 8,41 | -1,34 | 0,64 | -2,08 | 3,71E-02 | 1,24E-01 |
| AT1G43170 | 3,66 | 1,86 | 0,89 | 2,08 | 3,74E-02 | 1,24E-01 |
| AT1G18160 | 2,26 | 2,78 | 1,34 | 2,08 | 3,79E-02 | 1,26E-01 |
| AT3G17240 | 1,76 | 3,70 | 1,78 | 2,08 | 3,79E-02 | 1,26E-01 |
| AT4G29410 | 17,21 | -1,12 | 0,54 | -2,08 | 3,79E-02 | 1,26E-01 |
| AT4G16490 | 2,72 | 3,34 | 1,61 | 2,07 | 3,83E-02 | 1,27E-01 |
| AT5G10960 | 3,18 | 2,18 | 1,06 | 2,07 | 3,85E-02 | 1,27E-01 |
| AT1G08450 | 3,29 | 2,52 | 1,22 | 2,07 | 3,85E-02 | 1,27E-01 |
| AT1G11640 | 3,15 | 2,70 | 1,31 | 2,07 | 3,85E-02 | 1,27E-01 |
| AT1G12300 | 2,68 | -2,45 | 1,18 | -2,07 | 3,86E-02 | 1,27E-01 |
| AT4G13577 | 2,60 | -3,25 | 1,57 | -2,07 | 3,85E-02 | 1,27E-01 |
| AT4G30100 | 3,56 | 2,17 | 1,05 | 2,07 | 3,88E-02 | 1,27E-01 |
| AT1G33680 | 2,02 | 3,80 | 1,84 | 2,06 | 3,89E-02 | 1,27E-01 |
| AT2G43710 | 3,74 | 2,34 | 1,13 | 2,06 | 3,90E-02 | 1,27E-01 |
| AT5G01960 | 1,82 | 3,97 | 1,92 | 2,06 | 3,91E-02 | 1,28E-01 |
| AT3G17510 | 2,73 | 3,13 | 1,52 | 2,06 | 3,91E-02 | 1,28E-01 |
| AT1G04210 | 3,48 | 2,06 | 1,00 | 2,06 | 3,92E-02 | 1,28E-01 |
| AT2G04030 | 4,28 | 1,94 | 0,94 | 2,06 | 3,97E-02 | 1,28E-01 |
| AT4G11380 | 1,75 | 3,17 | 1,54 | 2,06 | 3,96E-02 | 1,28E-01 |
| AT1G67960 | 1,96 | 3,44 | 1,68 | 2,06 | 3,98E-02 | 1,28E-01 |
| AT2G07645 | 4,48 | -1,75 | 0,85 | -2,06 | 3,97E-02 | 1,28E-01 |
| AT3G13330 | 3,62 | 2,07 | 1,01 | 2,06 | 3,96E-02 | 1,28E-01 |
| AT4G23650 | 3,54 | 2,21 | 1,08 | 2,06 | 3,97E-02 | 1,28E-01 |
| AT5G18525 | 1,70 | 3,73 | 1,81 | 2,06 | 3,96E-02 | 1,28E-01 |
| AT2G48060 | 4,01 | 2,03 | 0,99 | 2,06 | 3,98E-02 | 1,28E-01 |
| AT3G08165 | 3,95 | -2,36 | 1,15 | -2,05 | 4,00E-02 | 1,28E-01 |
| AT4G38510 | 2,20 | 2,95 | 1,44 | 2,05 | 4,00E-02 | 1,28E-01 |
| AT3G23450 | 2,19 | -2,81 | 1,37 | -2,05 | 4,03E-02 | 1,29E-01 |
| AT2G25660 | 3,83 | 2,40 | 1,17 | 2,05 | 4,04E-02 | 1,29E-01 |
| AT3G51770 | 2,18 | 2,94 | 1,43 | 2,05 | 4,04E-02 | 1,29E-01 |
| AT2G45160 | 10,70 | -1,46 | 0,71 | -2,05 | 4,06E-02 | 1,30E-01 |
| AT1G64710 | 2,31 | 2,93 | 1,43 | 2,05 | 4,07E-02 | 1,30E-01 |
| AT1G06277 | 5,95 | -1,54 | 0,75 | -2,04 | 4,12E-02 | 1,31E-01 |

| | | | | | | |
|---|---|---|---|---|---|---|
| AT3G20362 | 7,16 | -2,04 | 1,00 | -2,04 | 4,12E-02 | 1,31E-01 |
| AT1G05240 | 2,51 | -2,49 | 1,23 | -2,04 | 4,17E-02 | 1,32E-01 |
| AT3G03870 | 3,66 | 2,33 | 1,15 | 2,03 | 4,20E-02 | 1,33E-01 |
| AT4G25530 | 5,35 | -1,68 | 0,83 | -2,03 | 4,26E-02 | 1,35E-01 |
| AT2G21330 | 7,94 | 1,46 | 0,72 | 2,02 | 4,29E-02 | 1,36E-01 |
| AT5G49980 | 4,39 | 2,02 | 1,00 | 2,02 | 4,30E-02 | 1,36E-01 |
| AT4G04970 | 2,63 | 3,42 | 1,69 | 2,02 | 4,31E-02 | 1,36E-01 |
| AT4G38440 | 2,75 | 2,32 | 1,15 | 2,02 | 4,32E-02 | 1,36E-01 |
| AT4G16860 | 6,11 | 1,48 | 0,74 | 2,02 | 4,39E-02 | 1,38E-01 |
| AT4G13885 | 7,77 | -1,46 | 0,73 | -2,01 | 4,42E-02 | 1,39E-01 |
| AT2G47980 | 2,39 | 2,36 | 1,18 | 2,01 | 4,44E-02 | 1,39E-01 |
| AT3G28899 | 2,18 | -2,88 | 1,43 | -2,01 | 4,45E-02 | 1,39E-01 |
| AT5G46450 | 2,49 | 2,55 | 1,27 | 2,00 | 4,51E-02 | 1,41E-01 |
| AT2G44540 | 4,48 | -1,79 | 0,89 | -2,00 | 4,56E-02 | 1,42E-01 |
| AT5G53170 | 2,82 | 2,12 | 1,06 | 2,00 | 4,57E-02 | 1,42E-01 |
| AT2G39675 | 2246,76 | -0,68 | 0,34 | -2,00 | 4,58E-02 | 1,43E-01 |
| AT1G76550 | 2,61 | 2,38 | 1,19 | 2,00 | 4,60E-02 | 1,43E-01 |
| AT4G34890 | 2,69 | 2,41 | 1,21 | 1,99 | 4,61E-02 | 1,43E-01 |
| AT1G18745 | 15,99 | -1,10 | 0,55 | -1,99 | 4,64E-02 | 1,44E-01 |
| AT2G40840 | 4,12 | 1,79 | 0,90 | 1,99 | 4,66E-02 | 1,44E-01 |
| AT4G00231 | 1,99 | 2,47 | 1,24 | 1,99 | 4,66E-02 | 1,44E-01 |
| AT5G26742 | 9,59 | 1,42 | 0,71 | 1,99 | 4,67E-02 | 1,44E-01 |
| AT4G04565 | 64,98 | 0,59 | 0,30 | 1,99 | 4,71E-02 | 1,45E-01 |
| AT3G27865 | 3,38 | -2,14 | 1,08 | -1,98 | 4,73E-02 | 1,45E-01 |
| AT3G04470 | 2,70 | 2,20 | 1,11 | 1,98 | 4,73E-02 | 1,45E-01 |
| AT4G08165 | 5,10 | -2,14 | 1,08 | -1,98 | 4,73E-02 | 1,45E-01 |
| AT1G19680 | 2,21 | 3,40 | 1,71 | 1,98 | 4,75E-02 | 1,46E-01 |
| AT3G05355 | 1,75 | -3,06 | 1,55 | -1,98 | 4,79E-02 | 1,47E-01 |
| AT1G52230 | 2,95 | -2,38 | 1,20 | -1,98 | 4,82E-02 | 1,47E-01 |
| AT4G00450 | 3,39 | 2,19 | 1,11 | 1,97 | 4,83E-02 | 1,47E-01 |
| AT1G77180 | 2,45 | 2,38 | 1,21 | 1,97 | 4,83E-02 | 1,47E-01 |
| AT5G15845 | 2,91 | 2,21 | 1,12 | 1,97 | 4,90E-02 | 1,49E-01 |
| AT5G64220 | 2,50 | 3,28 | 1,67 | 1,97 | 4,91E-02 | 1,49E-01 |
| AT5G41765 | 5,08 | -1,60 | 0,81 | -1,97 | 4,92E-02 | 1,49E-01 |
| AT4G08175 | 7,31 | 1,76 | 0,90 | 1,96 | 4,94E-02 | 1,50E-01 |
| AT4G32640 | 2,41 | 2,86 | 1,46 | 1,96 | 4,96E-02 | 1,50E-01 |
| AT2G04280 | 2,22 | 2,89 | 1,47 | 1,96 | 4,96E-02 | 1,50E-01 |
| AT2G46020 | 3,28 | 2,02 | 1,03 | 1,96 | 5,00E-02 | 1,51E-01 |
| AT5G09590 | 2,74 | 3,18 | 1,62 | 1,96 | 4,99E-02 | 1,51E-01 |
| AT5G05190 | 2,00 | 2,78 | 1,42 | 1,96 | 5,03E-02 | 1,52E-01 |
| AT1G14610 | 2,42 | 2,59 | 1,33 | 1,96 | 5,05E-02 | 1,52E-01 |
| AT3G01313 | 12,85 | -1,12 | 0,57 | -1,96 | 5,05E-02 | 1,52E-01 |
| AT3G58050 | 1,93 | 2,85 | 1,46 | 1,95 | 5,07E-02 | 1,53E-01 |
| AT3G48860 | 1,96 | 4,52 | 2,31 | 1,95 | 5,09E-02 | 1,53E-01 |
| AT4G30960 | 5,13 | 1,42 | 0,73 | 1,95 | 5,09E-02 | 1,53E-01 |
| AT3G13530 | 2,08 | 2,76 | 1,42 | 1,95 | 5,14E-02 | 1,54E-01 |
| AT4G16980 | 2,88 | -2,13 | 1,09 | -1,94 | 5,18E-02 | 1,55E-01 |
| AT2G16500 | 6,48 | 1,39 | 0,72 | 1,94 | 5,23E-02 | 1,56E-01 |
| AT3G18490 | 8,11 | -1,23 | 0,64 | -1,94 | 5,22E-02 | 1,56E-01 |
| AT5G09730 | 35,07 | 0,85 | 0,44 | 1,94 | 5,22E-02 | 1,56E-01 |
| AT5G02880 | 3,00 | 2,07 | 1,06 | 1,94 | 5,23E-02 | 1,56E-01 |
| AT5G28740 | 1,92 | 2,93 | 1,51 | 1,94 | 5,26E-02 | 1,56E-01 |
| AT2G27150 | 1,72 | 3,82 | 1,97 | 1,94 | 5,27E-02 | 1,56E-01 |
| AT4G19520 | 3,67 | 1,64 | 0,85 | 1,94 | 5,29E-02 | 1,57E-01 |
| AT1G02860 | 4,18 | 2,41 | 1,24 | 1,93 | 5,31E-02 | 1,57E-01 |
| AT2G09690 | 14,22 | -1,49 | 0,77 | -1,93 | 5,32E-02 | 1,57E-01 |

| | | | | | | |
|---|---|---|---|---|---|---|
| AT2G25320 | 3,21 | 2,45 | 1,27 | 1,93 | 5,30E-02 | 1,57E-01 |
| AT5G19330 | 2,99 | 2,64 | 1,37 | 1,93 | 5,32E-02 | 1,57E-01 |
| AT1G23900 | 1,85 | 2,83 | 1,47 | 1,93 | 5,38E-02 | 1,58E-01 |
| AT4G13340 | 14,47 | 0,89 | 0,46 | 1,93 | 5,38E-02 | 1,58E-01 |
| AT5G04315 | 6,08 | 1,40 | 0,73 | 1,93 | 5,37E-02 | 1,58E-01 |
| AT5G04325 | 6,08 | 1,40 | 0,73 | 1,93 | 5,37E-02 | 1,58E-01 |
| AT5G46520 | 3,26 | 2,16 | 1,12 | 1,93 | 5,40E-02 | 1,58E-01 |
| AT5G51070 | 3,71 | 1,76 | 0,92 | 1,92 | 5,45E-02 | 1,59E-01 |
| AT1G59760 | 3,05 | 2,42 | 1,26 | 1,92 | 5,44E-02 | 1,59E-01 |
| AT3G10815 | 5,65 | 1,46 | 0,76 | 1,92 | 5,45E-02 | 1,59E-01 |
| AT4G13564 | 27,21 | -0,73 | 0,38 | -1,92 | 5,45E-02 | 1,59E-01 |
| AT1G06680 | 10,44 | -1,38 | 0,72 | -1,92 | 5,51E-02 | 1,60E-01 |
| AT1G65720 | 3,55 | 1,57 | 0,82 | 1,92 | 5,49E-02 | 1,60E-01 |
| AT4G00720 | 2,07 | 2,71 | 1,41 | 1,92 | 5,50E-02 | 1,60E-01 |
| AT1G21130 | 2,17 | -2,90 | 1,51 | -1,92 | 5,52E-02 | 1,60E-01 |
| AT1G47128 | 7,64 | 1,10 | 0,57 | 1,92 | 5,54E-02 | 1,60E-01 |
| AT5G15850 | 2,49 | 2,25 | 1,17 | 1,91 | 5,56E-02 | 1,61E-01 |
| AT2G22990 | 2,03 | -2,52 | 1,32 | -1,91 | 5,59E-02 | 1,61E-01 |
| AT2G44550 | 3,60 | -1,99 | 1,04 | -1,91 | 5,61E-02 | 1,62E-01 |
| AT2G04755 | 2,18 | 2,83 | 1,48 | 1,91 | 5,63E-02 | 1,62E-01 |
| AT3G13750 | 2,68 | 2,28 | 1,19 | 1,91 | 5,63E-02 | 1,62E-01 |
| AT1G63950 | 1,81 | -3,51 | 1,84 | -1,91 | 5,64E-02 | 1,62E-01 |
| AT1G66880 | 6,48 | 1,55 | 0,81 | 1,91 | 5,64E-02 | 1,62E-01 |
| AT1G74720 | 1,98 | 3,03 | 1,59 | 1,91 | 5,65E-02 | 1,62E-01 |
| AT3G57300 | 2,34 | 2,47 | 1,29 | 1,91 | 5,66E-02 | 1,62E-01 |
| AT5G52070 | 6,76 | -2,08 | 1,09 | -1,91 | 5,67E-02 | 1,62E-01 |
| AT3G14210 | 12,80 | -1,89 | 0,99 | -1,90 | 5,70E-02 | 1,63E-01 |
| AT3G17900 | 1,92 | 2,89 | 1,52 | 1,90 | 5,69E-02 | 1,63E-01 |
| AT5G24810 | 2,74 | 2,33 | 1,22 | 1,90 | 5,71E-02 | 1,63E-01 |
| AT1G77740 | 1,74 | 3,10 | 1,63 | 1,90 | 5,74E-02 | 1,63E-01 |
| AT1G14580 | 3,99 | -2,05 | 1,08 | -1,90 | 5,75E-02 | 1,63E-01 |
| AT3G44300 | 1,82 | 2,41 | 1,27 | 1,90 | 5,75E-02 | 1,63E-01 |
| AT2G43150 | 1,94 | 2,54 | 1,34 | 1,90 | 5,80E-02 | 1,64E-01 |
| AT5G09040 | 2,31 | -2,33 | 1,23 | -1,90 | 5,80E-02 | 1,64E-01 |
| AT3G14067 | 4,84 | 1,52 | 0,81 | 1,89 | 5,83E-02 | 1,65E-01 |
| AT1G29350 | 2,20 | 2,48 | 1,31 | 1,89 | 5,92E-02 | 1,67E-01 |
| AT1G05780 | 4,77 | -1,56 | 0,83 | -1,89 | 5,93E-02 | 1,67E-01 |
| AT5G07290 | 1,99 | 4,11 | 2,19 | 1,88 | 5,97E-02 | 1,68E-01 |
| AT4G16983 | 2,85 | -2,07 | 1,10 | -1,88 | 5,99E-02 | 1,68E-01 |
| AT4G38970 | 12,35 | 0,93 | 0,50 | 1,88 | 5,99E-02 | 1,68E-01 |
| AT4G34710 | 3,32 | 2,04 | 1,09 | 1,88 | 6,01E-02 | 1,69E-01 |
| AT5G27850 | 1,94 | 2,50 | 1,33 | 1,88 | 6,04E-02 | 1,69E-01 |
| AT1G42970 | 13,13 | 1,05 | 0,56 | 1,87 | 6,09E-02 | 1,70E-01 |
| AT4G16190 | 11,22 | 1,10 | 0,58 | 1,87 | 6,08E-02 | 1,70E-01 |
| AT5G49615 | 2,21 | -3,10 | 1,65 | -1,87 | 6,08E-02 | 1,70E-01 |
| AT3G47470 | 25,61 | -0,92 | 0,49 | -1,87 | 6,09E-02 | 1,70E-01 |
| AT5G43470 | 3,38 | 2,17 | 1,16 | 1,88 | 6,07E-02 | 1,70E-01 |
| AT1G24706 | 2,66 | 2,08 | 1,11 | 1,87 | 6,16E-02 | 1,71E-01 |
| AT4G08920 | 2,64 | 2,39 | 1,28 | 1,87 | 6,16E-02 | 1,71E-01 |
| AT3G54730 | 3,74 | -1,71 | 0,91 | -1,87 | 6,17E-02 | 1,71E-01 |
| AT3G09465 | 2,48 | -2,18 | 1,17 | -1,87 | 6,19E-02 | 1,72E-01 |
| AT4G12800 | 10,28 | -1,47 | 0,79 | -1,87 | 6,19E-02 | 1,72E-01 |
| AT1G07510 | 1,74 | 2,67 | 1,43 | 1,86 | 6,26E-02 | 1,73E-01 |
| AT1G05423 | 13,10 | -1,05 | 0,56 | -1,86 | 6,30E-02 | 1,74E-01 |
| AT1G28270 | 2,06 | -2,92 | 1,57 | -1,86 | 6,32E-02 | 1,74E-01 |
| AT4G12570 | 2,25 | 2,61 | 1,40 | 1,86 | 6,32E-02 | 1,74E-01 |

| | | | | | | |
|---|---|---|---|---|---|---|
| AT4G28760 | 2,13 | 2,65 | 1,43 | 1,86 | 6,33E-02 | 1,74E-01 |
| AT4G27870 | 2,09 | 2,58 | 1,39 | 1,85 | 6,36E-02 | 1,75E-01 |
| AT2G43570 | 2,25 | 2,47 | 1,33 | 1,86 | 6,35E-02 | 1,75E-01 |
| AT3G63460 | 4,10 | 1,73 | 0,93 | 1,86 | 6,36E-02 | 1,75E-01 |
| AT1G02890 | 1,79 | 2,55 | 1,38 | 1,85 | 6,39E-02 | 1,75E-01 |
| AT4G02570 | 1,99 | 2,46 | 1,33 | 1,85 | 6,40E-02 | 1,75E-01 |
| AT4G26530 | 4,43 | 1,42 | 0,77 | 1,85 | 6,41E-02 | 1,75E-01 |
| AT5G51630 | 1,97 | 2,76 | 1,49 | 1,85 | 6,44E-02 | 1,76E-01 |
| AT3G55734 | 132,05 | -0,55 | 0,29 | -1,85 | 6,46E-02 | 1,76E-01 |
| AT1G61226 | 20,72 | -0,93 | 0,51 | -1,85 | 6,47E-02 | 1,76E-01 |
| AT2G44410 | 2,51 | 3,15 | 1,71 | 1,85 | 6,48E-02 | 1,76E-01 |
| AT2G16030 | 25,88 | -0,87 | 0,47 | -1,85 | 6,50E-02 | 1,77E-01 |
| AT2G46340 | 1,90 | 3,02 | 1,64 | 1,84 | 6,51E-02 | 1,77E-01 |
| AT1G14320 | 3,20 | 2,03 | 1,10 | 1,84 | 6,58E-02 | 1,78E-01 |
| AT4G03260 | 2,29 | 2,25 | 1,23 | 1,83 | 6,68E-02 | 1,81E-01 |
| AT3G04455 | 2,46 | -2,93 | 1,60 | -1,83 | 6,68E-02 | 1,81E-01 |
| AT1G11700 | 23,46 | -0,74 | 0,40 | -1,83 | 6,69E-02 | 1,81E-01 |
| AT5G61455 | 11,96 | 1,02 | 0,56 | 1,83 | 6,72E-02 | 1,81E-01 |
| AT1G49050 | 1,80 | 3,16 | 1,73 | 1,83 | 6,74E-02 | 1,82E-01 |
| AT3G49120 | 5,10 | 1,39 | 0,76 | 1,83 | 6,76E-02 | 1,82E-01 |
| AT4G02510 | 15,59 | 1,01 | 0,55 | 1,83 | 6,76E-02 | 1,82E-01 |
| AT5G47480 | 1,95 | 2,58 | 1,41 | 1,83 | 6,73E-02 | 1,82E-01 |
| AT4G03580 | 13,67 | -1,12 | 0,61 | -1,83 | 6,77E-02 | 1,82E-01 |
| AT4G03020 | 2,29 | 2,02 | 1,11 | 1,83 | 6,77E-02 | 1,82E-01 |
| AT3G24180 | 1,99 | 2,38 | 1,31 | 1,82 | 6,81E-02 | 1,82E-01 |
| AT3G11540 | 2,22 | 2,94 | 1,62 | 1,82 | 6,88E-02 | 1,83E-01 |
| AT3G25690 | 2,72 | 1,92 | 1,05 | 1,82 | 6,86E-02 | 1,83E-01 |
| AT5G47490 | 2,21 | 2,16 | 1,19 | 1,82 | 6,88E-02 | 1,83E-01 |
| AT1G07200 | 1,82 | 2,65 | 1,45 | 1,82 | 6,86E-02 | 1,83E-01 |
| AT3G45190 | 2,88 | 2,04 | 1,12 | 1,82 | 6,86E-02 | 1,83E-01 |
| AT5G45500 | 2,29 | 3,23 | 1,77 | 1,82 | 6,87E-02 | 1,83E-01 |
| AT3G05420 | 2,70 | 2,33 | 1,28 | 1,82 | 6,94E-02 | 1,84E-01 |
| AT4G31210 | 1,90 | 2,97 | 1,63 | 1,81 | 6,96E-02 | 1,84E-01 |
| AT5G01980 | 1,71 | 2,99 | 1,64 | 1,81 | 6,95E-02 | 1,84E-01 |
| AT5G10180 | 2,74 | 2,33 | 1,28 | 1,82 | 6,95E-02 | 1,84E-01 |
| AT3G41762 | 25,36 | -0,80 | 0,44 | -1,81 | 7,00E-02 | 1,85E-01 |
| AT1G33560 | 2,19 | 3,16 | 1,75 | 1,81 | 7,07E-02 | 1,87E-01 |
| AT4G29200 | 3,91 | -1,56 | 0,86 | -1,81 | 7,08E-02 | 1,87E-01 |
| AT2G26530 | 1,88 | -2,89 | 1,60 | -1,81 | 7,10E-02 | 1,87E-01 |
| AT2G39885 | 9,28 | 1,38 | 0,77 | 1,80 | 7,12E-02 | 1,87E-01 |
| AT4G04205 | 2,87 | -1,80 | 1,00 | -1,80 | 7,14E-02 | 1,88E-01 |
| AT3G63375 | 13,10 | 0,97 | 0,54 | 1,80 | 7,15E-02 | 1,88E-01 |
| AT1G20970 | 4,69 | 2,08 | 1,16 | 1,80 | 7,20E-02 | 1,89E-01 |
| AT4G32285 | 2,40 | 2,10 | 1,16 | 1,80 | 7,21E-02 | 1,89E-01 |
| AT5G24710 | 4,46 | 1,92 | 1,07 | 1,80 | 7,24E-02 | 1,89E-01 |
| AT5G04720 | 2,75 | 2,67 | 1,49 | 1,79 | 7,30E-02 | 1,91E-01 |
| AT1G71220 | 2,96 | 2,16 | 1,21 | 1,79 | 7,35E-02 | 1,92E-01 |
| AT3G63490 | 1,82 | 2,92 | 1,64 | 1,79 | 7,40E-02 | 1,93E-01 |
| AT5G21940 | 4,72 | -1,37 | 0,76 | -1,79 | 7,40E-02 | 1,93E-01 |
| AT2G35940 | 16,85 | -0,92 | 0,52 | -1,79 | 7,42E-02 | 1,93E-01 |
| AT5G24300 | 2,35 | 2,31 | 1,29 | 1,78 | 7,44E-02 | 1,93E-01 |
| AT2G36850 | 3,05 | 2,28 | 1,28 | 1,78 | 7,44E-02 | 1,93E-01 |
| AT1G02080 | 3,21 | 2,38 | 1,34 | 1,78 | 7,47E-02 | 1,93E-01 |
| AT1G05630 | 1,84 | 2,68 | 1,50 | 1,78 | 7,49E-02 | 1,93E-01 |
| AT1G07920 | 7,76 | 1,37 | 0,77 | 1,78 | 7,49E-02 | 1,93E-01 |
| AT1G20100 | 1,77 | 2,78 | 1,56 | 1,78 | 7,47E-02 | 1,93E-01 |

| | | | | | | |
|---|---|---|---|---|---|---|
| AT4G00150 | 10,48 | -0,95 | 0,53 | -1,78 | 7,49E-02 | 1,93E-01 |
| AT1G54115 | 2,18 | 2,60 | 1,46 | 1,78 | 7,52E-02 | 1,94E-01 |
| AT5G14520 | 3,02 | 2,95 | 1,66 | 1,78 | 7,54E-02 | 1,94E-01 |
| AT3G61430 | 1,94 | 3,14 | 1,77 | 1,78 | 7,57E-02 | 1,95E-01 |
| AT5G42080 | 4,02 | 1,68 | 0,95 | 1,78 | 7,58E-02 | 1,95E-01 |
| AT1G09310 | 8,77 | -1,41 | 0,80 | -1,77 | 7,60E-02 | 1,95E-01 |
| AT4G09735 | 2,23 | -2,51 | 1,42 | -1,77 | 7,60E-02 | 1,95E-01 |
| AT4G35770 | 2,85 | -2,23 | 1,26 | -1,77 | 7,65E-02 | 1,96E-01 |
| AT3G25020 | 3,41 | -1,64 | 0,92 | -1,77 | 7,64E-02 | 1,96E-01 |
| AT3G04740 | 2,26 | 2,25 | 1,27 | 1,77 | 7,74E-02 | 1,97E-01 |
| AT4G19170 | 5,23 | -1,37 | 0,78 | -1,77 | 7,76E-02 | 1,97E-01 |
| AT1G20340 | 11,85 | -1,08 | 0,61 | -1,77 | 7,73E-02 | 1,97E-01 |
| AT2G43410 | 2,84 | 2,00 | 1,13 | 1,77 | 7,75E-02 | 1,97E-01 |
| AT5G12150 | 4,21 | 2,20 | 1,25 | 1,77 | 7,76E-02 | 1,97E-01 |
| AT5G18490 | 1,80 | 2,88 | 1,63 | 1,77 | 7,72E-02 | 1,97E-01 |
| AT2G20410 | 6,12 | 1,30 | 0,74 | 1,76 | 7,77E-02 | 1,97E-01 |
| AT5G21100 | 6,61 | -1,23 | 0,70 | -1,76 | 7,78E-02 | 1,97E-01 |
| AT2G38800 | 2,71 | 2,19 | 1,24 | 1,76 | 7,81E-02 | 1,98E-01 |
| AT2G27720 | 1,70 | 3,25 | 1,85 | 1,76 | 7,82E-02 | 1,98E-01 |
| AT2G18020 | 2,64 | 2,08 | 1,18 | 1,76 | 7,91E-02 | 2,00E-01 |
| AT2G35945 | 15,59 | -0,93 | 0,53 | -1,76 | 7,92E-02 | 2,00E-01 |
| AT2G05070 | 19,17 | -1,05 | 0,60 | -1,76 | 7,91E-02 | 2,00E-01 |
| AT5G65380 | 2,69 | 2,35 | 1,34 | 1,75 | 7,93E-02 | 2,00E-01 |
| AT5G02575 | 6,74 | -1,56 | 0,89 | -1,75 | 7,94E-02 | 2,00E-01 |
| AT5G54130 | 6,22 | -1,53 | 0,87 | -1,75 | 7,98E-02 | 2,00E-01 |
| AT1G09780 | 4,32 | 1,44 | 0,82 | 1,75 | 7,98E-02 | 2,00E-01 |
| AT1G31540 | 3,00 | 2,04 | 1,16 | 1,75 | 7,98E-02 | 2,00E-01 |
| AT1G64583 | 1,87 | -2,71 | 1,54 | -1,75 | 7,97E-02 | 2,00E-01 |
| AT5G08550 | 1,83 | 2,50 | 1,43 | 1,75 | 8,03E-02 | 2,01E-01 |
| AT2G46500 | 2,79 | 2,46 | 1,41 | 1,75 | 8,05E-02 | 2,01E-01 |
| AT2G08665 | 53,97 | 0,71 | 0,40 | 1,75 | 8,08E-02 | 2,01E-01 |
| AT4G03420 | 2,29 | 2,62 | 1,50 | 1,75 | 8,07E-02 | 2,01E-01 |
| AT2G34420 | 53,97 | 0,71 | 0,40 | 1,75 | 8,08E-02 | 2,01E-01 |
| AT2G31400 | 2,56 | 2,11 | 1,21 | 1,74 | 8,15E-02 | 2,03E-01 |
| AT3G03790 | 1,73 | 2,58 | 1,48 | 1,74 | 8,21E-02 | 2,04E-01 |
| AT5G03280 | 3,28 | 1,99 | 1,14 | 1,74 | 8,22E-02 | 2,04E-01 |
| AT5G42092 | 3,68 | 1,65 | 0,95 | 1,73 | 8,29E-02 | 2,06E-01 |
| AT4G05180 | 7,70 | -1,55 | 0,89 | -1,73 | 8,32E-02 | 2,06E-01 |
| AT4G14030 | 1,74 | 2,51 | 1,45 | 1,73 | 8,35E-02 | 2,07E-01 |
| AT2G25450 | 4,94 | 1,40 | 0,81 | 1,73 | 8,37E-02 | 2,07E-01 |
| AT3G54280 | 2,37 | 2,40 | 1,39 | 1,73 | 8,40E-02 | 2,07E-01 |
| AT5G38420 | 5,05 | 1,59 | 0,92 | 1,73 | 8,40E-02 | 2,07E-01 |
| AT5G41740 | 6,05 | 1,54 | 0,90 | 1,73 | 8,44E-02 | 2,08E-01 |
| AT3G54660 | 2,19 | 2,93 | 1,70 | 1,73 | 8,45E-02 | 2,08E-01 |
| AT1G19090 | 4,93 | -1,45 | 0,84 | -1,72 | 8,50E-02 | 2,09E-01 |
| AT1G05035 | 3,55 | -1,87 | 1,09 | -1,72 | 8,60E-02 | 2,11E-01 |
| AT5G09670 | 2,53 | 1,99 | 1,16 | 1,72 | 8,61E-02 | 2,11E-01 |
| AT1G27595 | 2,01 | 2,40 | 1,40 | 1,71 | 8,67E-02 | 2,12E-01 |
| AT5G19400 | 2,38 | 2,87 | 1,67 | 1,71 | 8,67E-02 | 2,12E-01 |
| AT1G18010 | 5,80 | -1,32 | 0,77 | -1,71 | 8,75E-02 | 2,14E-01 |
| AT2G41840 | 3,13 | 2,11 | 1,24 | 1,70 | 8,86E-02 | 2,17E-01 |
| AT1G47657 | 2,52 | -2,25 | 1,33 | -1,70 | 8,99E-02 | 2,19E-01 |
| AT3G62700 | 3,87 | 1,55 | 0,91 | 1,70 | 8,98E-02 | 2,19E-01 |
| AT5G22500 | 2,19 | -2,67 | 1,58 | -1,69 | 9,01E-02 | 2,20E-01 |
| AT2G06555 | 20,57 | -0,77 | 0,46 | -1,69 | 9,03E-02 | 2,20E-01 |
| AT2G02840 | 4,55 | -1,89 | 1,12 | -1,69 | 9,09E-02 | 2,21E-01 |

| | | | | | | |
|---|---|---|---|---|---|---|
| AT2G26355 | 2,39 | 1,90 | 1,12 | 1,69 | 9,10E-02 | 2,21E-01 |
| AT1G08420 | 1,72 | 2,61 | 1,54 | 1,69 | 9,11E-02 | 2,21E-01 |
| AT3G48050 | 3,14 | 1,50 | 0,89 | 1,69 | 9,18E-02 | 2,23E-01 |
| AT5G54440 | 2,06 | 2,18 | 1,30 | 1,68 | 9,21E-02 | 2,23E-01 |
| AT2G03250 | 2,57 | -2,20 | 1,31 | -1,68 | 9,20E-02 | 2,23E-01 |
| AT1G26558 | 2,72 | -2,11 | 1,26 | -1,68 | 9,30E-02 | 2,24E-01 |
| AT1G08115 | 40,91 | -0,70 | 0,42 | -1,68 | 9,30E-02 | 2,24E-01 |
| AT5G26360 | 2,70 | 1,91 | 1,14 | 1,68 | 9,29E-02 | 2,24E-01 |
| AT5G54075 | 15,55 | -0,86 | 0,51 | -1,68 | 9,29E-02 | 2,24E-01 |
| AT2G35387 | 52,44 | -0,60 | 0,36 | -1,67 | 9,42E-02 | 2,26E-01 |
| AT3G55020 | 1,94 | 2,42 | 1,44 | 1,67 | 9,41E-02 | 2,26E-01 |
| AT2G30950 | 3,17 | 1,66 | 1,00 | 1,67 | 9,48E-02 | 2,27E-01 |
| AT3G28956 | 2,37 | -1,94 | 1,16 | -1,67 | 9,48E-02 | 2,27E-01 |
| AT1G33950 | 1,88 | -2,57 | 1,54 | -1,67 | 9,49E-02 | 2,28E-01 |
| AT5G54250 | 1,88 | 2,97 | 1,78 | 1,67 | 9,51E-02 | 2,28E-01 |
| AT1G60072 | 5,42 | 1,33 | 0,80 | 1,67 | 9,54E-02 | 2,28E-01 |
| AT5G64040 | 3,44 | -1,86 | 1,12 | -1,66 | 9,61E-02 | 2,30E-01 |
| AT1G54410 | 19,25 | -1,13 | 0,68 | -1,66 | 9,67E-02 | 2,31E-01 |
| AT2G34750 | 2,36 | 2,21 | 1,33 | 1,66 | 9,68E-02 | 2,31E-01 |
| AT4G22410 | 2,93 | -1,78 | 1,07 | -1,66 | 9,66E-02 | 2,31E-01 |
| AT2G36160 | 1,81 | 2,65 | 1,60 | 1,66 | 9,73E-02 | 2,31E-01 |
| AT1G25440 | 6,25 | 1,53 | 0,93 | 1,66 | 9,79E-02 | 2,33E-01 |
| AT1G07600 | 1,87 | 2,05 | 1,24 | 1,65 | 9,83E-02 | 2,33E-01 |
| AT4G38770 | 7,08 | -1,12 | 0,68 | -1,65 | 9,82E-02 | 2,33E-01 |
| AT5G12860 | 2,32 | 2,87 | 1,74 | 1,65 | 9,83E-02 | 2,33E-01 |
| AT2G40140 | 4,76 | 1,55 | 0,94 | 1,65 | 9,86E-02 | 2,33E-01 |
| AT1G51570 | 1,72 | 2,50 | 1,51 | 1,65 | 9,86E-02 | 2,33E-01 |
| AT3G19380 | 2,79 | -1,82 | 1,11 | -1,65 | 9,87E-02 | 2,33E-01 |
| AT3G20820 | 3,82 | -1,83 | 1,11 | -1,65 | 9,91E-02 | 2,34E-01 |
| AT2G46192 | 21,12 | -0,99 | 0,60 | -1,65 | 9,93E-02 | 2,34E-01 |
| AT3G62290 | 5,55 | 1,36 | 0,83 | 1,65 | 9,93E-02 | 2,34E-01 |
| AT5G47260 | 4,17 | -1,39 | 0,85 | -1,65 | 9,95E-02 | 2,34E-01 |
| AT1G32630 | 4,04 | -1,44 | 0,87 | -1,64 | 1,00E-01 | 2,35E-01 |
| AT1G32640 | 4,04 | -1,44 | 0,87 | -1,64 | 1,00E-01 | 2,35E-01 |
| AT4G35310 | 1,84 | 2,04 | 1,24 | 1,64 | 1,00E-01 | 2,36E-01 |
| AT3G56910 | 1,94 | 2,53 | 1,54 | 1,64 | 1,01E-01 | 2,36E-01 |
| AT5G54070 | 15,72 | -0,85 | 0,52 | -1,64 | 1,01E-01 | 2,37E-01 |
| AT2G25730 | 2,88 | 1,74 | 1,06 | 1,64 | 1,01E-01 | 2,37E-01 |
| AT1G62540 | 2,48 | -2,54 | 1,56 | -1,64 | 1,02E-01 | 2,38E-01 |
| AT3G22886 | 587,34 | -0,41 | 0,25 | -1,63 | 1,02E-01 | 2,38E-01 |
| AT1G32010 | 1,81 | -2,47 | 1,51 | -1,63 | 1,02E-01 | 2,38E-01 |
| AT3G01715 | 19,56 | 1,02 | 0,62 | 1,63 | 1,02E-01 | 2,38E-01 |
| AT3G48430 | 1,70 | 3,06 | 1,88 | 1,63 | 1,02E-01 | 2,38E-01 |
| AT1G69550 | 1,87 | 2,36 | 1,44 | 1,63 | 1,03E-01 | 2,38E-01 |
| AT1G09640 | 2,66 | 2,53 | 1,55 | 1,63 | 1,03E-01 | 2,38E-01 |
| AT5G36228 | 1,95 | -2,79 | 1,71 | -1,63 | 1,03E-01 | 2,39E-01 |
| AT3G11010 | 3,23 | 1,96 | 1,20 | 1,63 | 1,04E-01 | 2,40E-01 |
| AT5G05010 | 2,08 | 2,36 | 1,45 | 1,63 | 1,04E-01 | 2,41E-01 |
| AT5G13390 | 2,05 | 2,32 | 1,43 | 1,62 | 1,05E-01 | 2,42E-01 |
| AT2G04965 | 3,81 | -1,43 | 0,88 | -1,62 | 1,05E-01 | 2,42E-01 |
| AT3G25500 | 1,85 | 2,03 | 1,25 | 1,62 | 1,05E-01 | 2,42E-01 |
| AT2G26250 | 1,96 | -2,07 | 1,28 | -1,62 | 1,06E-01 | 2,44E-01 |
| AT1G12090 | 8,60 | -1,24 | 0,77 | -1,62 | 1,06E-01 | 2,44E-01 |
| AT5G04590 | 4,30 | 1,53 | 0,95 | 1,62 | 1,06E-01 | 2,44E-01 |
| AT5G27606 | 14,40 | -0,85 | 0,53 | -1,61 | 1,07E-01 | 2,46E-01 |
| AT4G27060 | 1,71 | 2,39 | 1,48 | 1,61 | 1,07E-01 | 2,46E-01 |

| | | | | | | |
|---|---|---|---|---|---|---|
| AT1G31730 | 1,97 | 3,01 | 1,87 | 1,61 | 1,07E-01 | 2,47E-01 |
| AT3G01310 | 2,04 | 2,76 | 1,72 | 1,61 | 1,08E-01 | 2,47E-01 |
| AT1G08833 | 15,74 | -0,86 | 0,54 | -1,60 | 1,09E-01 | 2,48E-01 |
| AT5G62670 | 1,85 | 2,79 | 1,74 | 1,60 | 1,09E-01 | 2,48E-01 |
| AT4G06240 | 6,18 | -1,75 | 1,09 | -1,60 | 1,09E-01 | 2,49E-01 |
| AT1G33140 | 1,84 | 2,37 | 1,48 | 1,60 | 1,09E-01 | 2,49E-01 |
| AT4G00060 | 1,77 | 2,35 | 1,47 | 1,60 | 1,09E-01 | 2,49E-01 |
| AT2G01910 | 2,66 | -2,45 | 1,53 | -1,60 | 1,11E-01 | 2,52E-01 |
| AT5G23630 | 2,35 | 1,98 | 1,24 | 1,59 | 1,11E-01 | 2,53E-01 |
| AT2G18790 | 2,11 | 2,08 | 1,30 | 1,59 | 1,11E-01 | 2,53E-01 |
| AT2G33830 | 1,76 | -2,73 | 1,72 | -1,59 | 1,12E-01 | 2,53E-01 |
| AT3G25520 | 3,31 | 1,76 | 1,11 | 1,59 | 1,12E-01 | 2,53E-01 |
| AT1G29940 | 2,05 | 2,51 | 1,58 | 1,59 | 1,12E-01 | 2,54E-01 |
| AT1G27340 | 2,10 | 2,42 | 1,52 | 1,59 | 1,12E-01 | 2,54E-01 |
| AT3G27690 | 1,98 | -2,55 | 1,61 | -1,59 | 1,12E-01 | 2,55E-01 |
| AT1G17470 | 1,97 | 2,43 | 1,53 | 1,58 | 1,13E-01 | 2,55E-01 |
| AT2G46685 | 3232,16 | -0,42 | 0,26 | -1,58 | 1,13E-01 | 2,55E-01 |
| AT5G08610 | 1,71 | 2,53 | 1,60 | 1,58 | 1,14E-01 | 2,56E-01 |
| AT1G79570 | 2,57 | 1,76 | 1,12 | 1,58 | 1,14E-01 | 2,57E-01 |
| AT3G27400 | 1,80 | -2,21 | 1,40 | -1,58 | 1,15E-01 | 2,59E-01 |
| AT3G19420 | 2,24 | 2,50 | 1,59 | 1,58 | 1,15E-01 | 2,59E-01 |
| AT1G66970 | 4,60 | 1,23 | 0,78 | 1,58 | 1,15E-01 | 2,59E-01 |
| AT1G68520 | 2,00 | 1,77 | 1,13 | 1,57 | 1,15E-01 | 2,59E-01 |
| AT1G04425 | 1,91 | -2,11 | 1,34 | -1,57 | 1,16E-01 | 2,59E-01 |
| AT3G23820 | 9,86 | -0,88 | 0,56 | -1,57 | 1,16E-01 | 2,59E-01 |
| AT1G74453 | 1,99 | -1,92 | 1,22 | -1,57 | 1,16E-01 | 2,60E-01 |
| AT1G80250 | 2,27 | 2,63 | 1,68 | 1,57 | 1,17E-01 | 2,62E-01 |
| AT5G04540 | 1,96 | 2,50 | 1,59 | 1,57 | 1,17E-01 | 2,62E-01 |
| AT2G36350 | 2,13 | 2,39 | 1,53 | 1,56 | 1,18E-01 | 2,62E-01 |
| AT3G49230 | 1,94 | -2,40 | 1,54 | -1,56 | 1,18E-01 | 2,62E-01 |
| AT2G35747 | 15,41 | -1,00 | 0,64 | -1,56 | 1,18E-01 | 2,62E-01 |
| AT5G01400 | 1,70 | 2,98 | 1,91 | 1,56 | 1,18E-01 | 2,62E-01 |
| AT1G01183 | 317,62 | -0,56 | 0,36 | -1,56 | 1,18E-01 | 2,62E-01 |
| AT1G10522 | 24,13 | -0,68 | 0,43 | -1,56 | 1,18E-01 | 2,63E-01 |
| AT1G76135 | 2,70 | -2,02 | 1,30 | -1,56 | 1,19E-01 | 2,64E-01 |
| AT5G02310 | 4,95 | 1,18 | 0,76 | 1,56 | 1,19E-01 | 2,64E-01 |
| AT3G44630 | 4,39 | 1,38 | 0,89 | 1,56 | 1,19E-01 | 2,64E-01 |
| AT1G69797 | 12,98 | -0,84 | 0,54 | -1,56 | 1,19E-01 | 2,64E-01 |
| AT3G41761 | 2,51 | -2,77 | 1,78 | -1,56 | 1,20E-01 | 2,64E-01 |
| AT5G51300 | 1,97 | 3,25 | 2,09 | 1,56 | 1,20E-01 | 2,65E-01 |
| AT5G53370 | 2,35 | 1,91 | 1,23 | 1,55 | 1,20E-01 | 2,65E-01 |
| AT4G05320 | 17,67 | 0,78 | 0,50 | 1,55 | 1,21E-01 | 2,66E-01 |
| AT3G56410 | 2,56 | 1,75 | 1,13 | 1,55 | 1,22E-01 | 2,69E-01 |
| AT2G05100 | 10,97 | -0,85 | 0,55 | -1,54 | 1,22E-01 | 2,69E-01 |
| AT1G27535 | 2,34 | -1,81 | 1,17 | -1,54 | 1,23E-01 | 2,69E-01 |
| AT4G15258 | 2,27 | -1,89 | 1,22 | -1,54 | 1,23E-01 | 2,69E-01 |
| AT3G18830 | 2,46 | 1,68 | 1,09 | 1,54 | 1,23E-01 | 2,71E-01 |
| AT5G63195 | 1,93 | 2,70 | 1,75 | 1,54 | 1,23E-01 | 2,71E-01 |
| AT3G04930 | 1,93 | 2,24 | 1,45 | 1,54 | 1,24E-01 | 2,72E-01 |
| AT3G14010 | 2,74 | 2,11 | 1,37 | 1,54 | 1,24E-01 | 2,72E-01 |
| AT3G59780 | 2,03 | 2,14 | 1,39 | 1,53 | 1,25E-01 | 2,73E-01 |
| AT2G33150 | 2,47 | 1,79 | 1,16 | 1,53 | 1,25E-01 | 2,73E-01 |
| AT5G65610 | 2,72 | -2,02 | 1,32 | -1,53 | 1,25E-01 | 2,73E-01 |
| AT4G12320 | 1,93 | 2,57 | 1,68 | 1,53 | 1,25E-01 | 2,73E-01 |
| AT5G06640 | 3,20 | -1,69 | 1,10 | -1,53 | 1,26E-01 | 2,73E-01 |
| AT5G52170 | 2,49 | 2,02 | 1,32 | 1,53 | 1,26E-01 | 2,74E-01 |

| | | | | | | |
|---|---|---|---|---|---|---|
| AT3G55200 | 2,62 | 2,47 | 1,62 | 1,53 | 1,26E-01 | 2,75E-01 |
| AT3G13290 | 2,32 | 2,06 | 1,35 | 1,53 | 1,27E-01 | 2,76E-01 |
| AT1G21590 | 2,73 | 1,88 | 1,24 | 1,52 | 1,27E-01 | 2,76E-01 |
| AT1G14740 | 3,27 | 1,48 | 0,97 | 1,52 | 1,28E-01 | 2,77E-01 |
| AT2G47000 | 1,78 | 3,17 | 2,08 | 1,52 | 1,28E-01 | 2,77E-01 |
| AT3G05700 | 20,03 | 0,90 | 0,59 | 1,52 | 1,28E-01 | 2,78E-01 |
| AT1G65960 | 3,84 | 1,61 | 1,06 | 1,52 | 1,29E-01 | 2,78E-01 |
| AT1G53900 | 2,01 | -1,94 | 1,28 | -1,52 | 1,30E-01 | 2,80E-01 |
| AT4G16370 | 3,80 | 1,36 | 0,90 | 1,51 | 1,31E-01 | 2,81E-01 |
| AT5G17760 | 1,99 | 2,61 | 1,73 | 1,51 | 1,30E-01 | 2,81E-01 |
| AT4G21595 | 28,61 | 0,61 | 0,41 | 1,51 | 1,31E-01 | 2,82E-01 |
| AT3G54590 | 2,31 | -2,07 | 1,37 | -1,51 | 1,31E-01 | 2,82E-01 |
| AT3G51860 | 2,20 | 1,87 | 1,24 | 1,51 | 1,31E-01 | 2,82E-01 |
| AT3G63410 | 2,71 | 1,67 | 1,11 | 1,51 | 1,32E-01 | 2,84E-01 |
| AT1G17220 | 3,72 | 1,57 | 1,05 | 1,50 | 1,33E-01 | 2,84E-01 |
| AT3G22235 | 3,04 | 1,73 | 1,15 | 1,50 | 1,32E-01 | 2,84E-01 |
| AT1G19373 | 3,55 | -1,57 | 1,04 | -1,50 | 1,34E-01 | 2,87E-01 |
| AT3G57660 | 1,78 | 2,55 | 1,70 | 1,50 | 1,35E-01 | 2,87E-01 |
| AT4G24570 | 3,86 | -1,30 | 0,87 | -1,50 | 1,35E-01 | 2,87E-01 |
| AT2G33170 | 2,19 | 2,04 | 1,36 | 1,50 | 1,34E-01 | 2,87E-01 |
| AT5G20950 | 3,45 | 1,59 | 1,06 | 1,50 | 1,34E-01 | 2,87E-01 |
| AT3G59710 | 6,69 | -1,17 | 0,79 | -1,49 | 1,37E-01 | 2,92E-01 |
| AT3G09225 | 6,69 | -1,17 | 0,79 | -1,49 | 1,37E-01 | 2,92E-01 |
| AT1G26110 | 1,79 | 2,02 | 1,36 | 1,48 | 1,38E-01 | 2,94E-01 |
| AT1G33120 | 1,97 | 1,93 | 1,31 | 1,48 | 1,39E-01 | 2,96E-01 |
| AT3G06510 | 3,56 | 1,50 | 1,02 | 1,48 | 1,40E-01 | 2,97E-01 |
| AT4G16660 | 2,21 | 1,86 | 1,26 | 1,48 | 1,40E-01 | 2,97E-01 |
| AT2G46430 | 2,08 | 1,93 | 1,31 | 1,47 | 1,40E-01 | 2,97E-01 |
| AT3G58193 | 8,75 | -1,07 | 0,72 | -1,47 | 1,40E-01 | 2,97E-01 |
| AT3G07855 | 4,66 | -1,26 | 0,86 | -1,47 | 1,41E-01 | 2,98E-01 |
| AT1G12013 | 2,08 | -2,09 | 1,42 | -1,47 | 1,42E-01 | 2,99E-01 |
| AT1G63750 | 2,64 | 1,76 | 1,19 | 1,47 | 1,42E-01 | 2,99E-01 |
| AT3G56408 | 2,59 | 1,67 | 1,14 | 1,46 | 1,43E-01 | 3,01E-01 |
| AT5G05660 | 1,92 | 3,47 | 2,37 | 1,47 | 1,43E-01 | 3,01E-01 |
| AT5G08717 | 14,65 | -0,99 | 0,68 | -1,46 | 1,43E-01 | 3,02E-01 |
| AT1G63980 | 3,67 | 1,53 | 1,04 | 1,46 | 1,44E-01 | 3,02E-01 |
| AT3G13724 | 5,13 | -1,32 | 0,91 | -1,46 | 1,44E-01 | 3,02E-01 |
| AT2G22496 | 20,32 | 0,74 | 0,50 | 1,46 | 1,44E-01 | 3,02E-01 |
| AT2G46765 | 2,57 | -1,52 | 1,04 | -1,46 | 1,44E-01 | 3,03E-01 |
| AT1G51690 | 2,38 | 2,03 | 1,39 | 1,46 | 1,44E-01 | 3,03E-01 |
| AT2G03140 | 2,19 | 1,94 | 1,33 | 1,46 | 1,45E-01 | 3,04E-01 |
| AT3G06345 | 8,24 | -0,89 | 0,61 | -1,46 | 1,45E-01 | 3,04E-01 |
| AT4G27090 | 2,08 | 1,96 | 1,35 | 1,45 | 1,46E-01 | 3,05E-01 |
| AT1G07233 | 2,51 | -1,62 | 1,11 | -1,45 | 1,46E-01 | 3,06E-01 |
| AT5G42090 | 4,37 | 1,31 | 0,90 | 1,45 | 1,47E-01 | 3,07E-01 |
| AT1G18000 | 4,70 | -1,59 | 1,10 | -1,45 | 1,48E-01 | 3,07E-01 |
| AT1G31960 | 1,81 | -2,07 | 1,43 | -1,45 | 1,48E-01 | 3,07E-01 |
| AT1G09027 | 2,11 | -1,70 | 1,17 | -1,45 | 1,48E-01 | 3,07E-01 |
| AT3G15880 | 1,94 | 1,73 | 1,19 | 1,45 | 1,48E-01 | 3,07E-01 |
| AT3G52500 | 2,36 | -1,54 | 1,07 | -1,45 | 1,48E-01 | 3,08E-01 |
| AT5G67360 | 6,15 | 1,38 | 0,95 | 1,45 | 1,48E-01 | 3,08E-01 |
| AT3G45830 | 1,79 | 2,02 | 1,40 | 1,45 | 1,48E-01 | 3,08E-01 |
| AT3G29360 | 2,83 | 1,57 | 1,08 | 1,44 | 1,48E-01 | 3,08E-01 |
| AT4G18480 | 1,72 | 1,97 | 1,36 | 1,44 | 1,49E-01 | 3,08E-01 |
| AT2G47390 | 2,13 | 2,15 | 1,49 | 1,44 | 1,49E-01 | 3,08E-01 |
| AT4G27430 | 2,23 | 1,98 | 1,37 | 1,44 | 1,50E-01 | 3,09E-01 |

| | | | | | | |
|---|---|---|---|---|---|---|
| AT3G61690 | 2,25 | 1,90 | 1,32 | 1,44 | 1,50E-01 | 3,10E-01 |
| AT4G03295 | 3,55 | -1,39 | 0,97 | -1,44 | 1,50E-01 | 3,10E-01 |
| AT2G38170 | 3,76 | 1,44 | 1,00 | 1,43 | 1,51E-01 | 3,12E-01 |
| AT5G41510 | 1,76 | -2,15 | 1,50 | -1,43 | 1,52E-01 | 3,13E-01 |
| AT1G03680 | 1,73 | -2,15 | 1,50 | -1,43 | 1,52E-01 | 3,13E-01 |
| AT5G38190 | 2,86 | -1,44 | 1,01 | -1,43 | 1,53E-01 | 3,14E-01 |
| AT1G62390 | 2,10 | 2,48 | 1,73 | 1,43 | 1,53E-01 | 3,14E-01 |
| AT5G04550 | 2,33 | 1,79 | 1,26 | 1,43 | 1,53E-01 | 3,14E-01 |
| AT4G18197 | 1,87 | -1,82 | 1,27 | -1,43 | 1,53E-01 | 3,14E-01 |
| AT4G34110 | 4,15 | 1,14 | 0,80 | 1,43 | 1,54E-01 | 3,15E-01 |
| AT2G19360 | 2,70 | -1,71 | 1,20 | -1,42 | 1,55E-01 | 3,16E-01 |
| AT3G46740 | 3,41 | 1,54 | 1,08 | 1,42 | 1,55E-01 | 3,16E-01 |
| AT2G25900 | 2,31 | -1,82 | 1,28 | -1,42 | 1,55E-01 | 3,16E-01 |
| AT1G01510 | 2,79 | 2,06 | 1,45 | 1,42 | 1,56E-01 | 3,18E-01 |
| AT5G65480 | 2,00 | 1,80 | 1,27 | 1,42 | 1,56E-01 | 3,18E-01 |
| AT1G47660 | 2,01 | -1,86 | 1,31 | -1,42 | 1,56E-01 | 3,18E-01 |
| AT5G03940 | 2,83 | 1,59 | 1,12 | 1,42 | 1,56E-01 | 3,18E-01 |
| AT1G13320 | 2,17 | 1,86 | 1,32 | 1,42 | 1,57E-01 | 3,19E-01 |
| AT1G31910 | 2,79 | 1,47 | 1,04 | 1,41 | 1,57E-01 | 3,20E-01 |
| AT4G02970 | 72,42 | 0,44 | 0,31 | 1,41 | 1,58E-01 | 3,20E-01 |
| AT3G02200 | 1,75 | 2,42 | 1,71 | 1,41 | 1,58E-01 | 3,20E-01 |
| AT1G60950 | 5,66 | -1,29 | 0,91 | -1,41 | 1,58E-01 | 3,21E-01 |
| AT4G16180 | 1,78 | 3,04 | 2,16 | 1,41 | 1,59E-01 | 3,22E-01 |
| AT1G11650 | 3,30 | 1,36 | 0,97 | 1,41 | 1,59E-01 | 3,22E-01 |
| AT1G60140 | 2,14 | 1,92 | 1,36 | 1,41 | 1,60E-01 | 3,22E-01 |
| AT2G06520 | 10,63 | -1,17 | 0,83 | -1,41 | 1,60E-01 | 3,22E-01 |
| AT3G01770 | 1,96 | 1,92 | 1,37 | 1,41 | 1,60E-01 | 3,22E-01 |
| AT1G09010 | 3,02 | 1,73 | 1,23 | 1,40 | 1,60E-01 | 3,23E-01 |
| AT1G06230 | 2,14 | 1,96 | 1,40 | 1,40 | 1,60E-01 | 3,23E-01 |
| AT2G03150 | 2,08 | 1,91 | 1,36 | 1,40 | 1,60E-01 | 3,23E-01 |
| AT1G42470 | 1,71 | 2,64 | 1,88 | 1,40 | 1,61E-01 | 3,23E-01 |
| AT5G10540 | 2,27 | 1,78 | 1,27 | 1,40 | 1,61E-01 | 3,23E-01 |
| AT3G55512 | 2,56 | -2,07 | 1,48 | -1,40 | 1,62E-01 | 3,25E-01 |
| AT1G12280 | 1,78 | 1,66 | 1,18 | 1,40 | 1,62E-01 | 3,25E-01 |
| AT2G31820 | 4,59 | -1,08 | 0,77 | -1,40 | 1,62E-01 | 3,25E-01 |
| AT4G11570 | 2,16 | 1,91 | 1,37 | 1,40 | 1,62E-01 | 3,25E-01 |
| AT2G21100 | 3,61 | -1,50 | 1,08 | -1,40 | 1,62E-01 | 3,25E-01 |
| AT4G26110 | 2,20 | 1,79 | 1,28 | 1,40 | 1,63E-01 | 3,25E-01 |
| AT5G38990 | 2,97 | 1,43 | 1,02 | 1,39 | 1,63E-01 | 3,26E-01 |
| AT3G58030 | 2,17 | 1,94 | 1,39 | 1,39 | 1,64E-01 | 3,26E-01 |
| AT2G19480 | 1,78 | 1,95 | 1,40 | 1,39 | 1,64E-01 | 3,27E-01 |
| AT1G77560 | 2,14 | 1,75 | 1,26 | 1,39 | 1,64E-01 | 3,27E-01 |
| AT1G75800 | 1,71 | 2,61 | 1,88 | 1,39 | 1,65E-01 | 3,29E-01 |
| AT3G15353 | 5,91 | -1,24 | 0,89 | -1,39 | 1,65E-01 | 3,29E-01 |
| AT1G26973 | 11,41 | -0,90 | 0,65 | -1,38 | 1,66E-01 | 3,30E-01 |
| AT2G01422 | 2,93 | -2,03 | 1,47 | -1,38 | 1,67E-01 | 3,31E-01 |
| AT2G17510 | 1,71 | 1,77 | 1,28 | 1,38 | 1,67E-01 | 3,31E-01 |
| AT2G29080 | 1,97 | 1,81 | 1,31 | 1,38 | 1,67E-01 | 3,31E-01 |
| AT3G51950 | 4,09 | 1,16 | 0,84 | 1,38 | 1,67E-01 | 3,31E-01 |
| AT4G31160 | 2,16 | 2,21 | 1,60 | 1,38 | 1,67E-01 | 3,31E-01 |
| AT1G09690 | 1,78 | 1,77 | 1,29 | 1,38 | 1,67E-01 | 3,31E-01 |
| AT2G14080 | 4,62 | 1,42 | 1,03 | 1,38 | 1,68E-01 | 3,33E-01 |
| AT5G51090 | 2,18 | -1,70 | 1,23 | -1,38 | 1,69E-01 | 3,33E-01 |
| AT2G20780 | 1,93 | 1,89 | 1,37 | 1,37 | 1,69E-01 | 3,33E-01 |
| AT5G46110 | 7,87 | 0,99 | 0,72 | 1,37 | 1,70E-01 | 3,34E-01 |
| AT1G27770 | 5,67 | 1,28 | 0,93 | 1,37 | 1,70E-01 | 3,35E-01 |

| | | | | | | |
|---|---|---|---|---|---|---|
| AT2G37710 | 4,76 | 1,09 | 0,79 | 1,37 | 1,70E-01 | 3,35E-01 |
| AT3G05530 | 1,78 | 1,91 | 1,40 | 1,37 | 1,71E-01 | 3,35E-01 |
| AT4G16143 | 2,03 | 1,55 | 1,13 | 1,37 | 1,71E-01 | 3,35E-01 |
| AT2G22440 | 2,40 | -1,63 | 1,19 | -1,37 | 1,71E-01 | 3,36E-01 |
| AT4G32760 | 1,73 | 2,14 | 1,57 | 1,37 | 1,72E-01 | 3,37E-01 |
| AT5G27607 | 2,38 | -2,11 | 1,55 | -1,36 | 1,74E-01 | 3,40E-01 |
| AT1G08103 | 6,40 | -1,20 | 0,88 | -1,36 | 1,74E-01 | 3,40E-01 |
| AT5G14040 | 2,00 | 2,14 | 1,58 | 1,36 | 1,74E-01 | 3,40E-01 |
| AT4G21150 | 2,92 | 1,62 | 1,19 | 1,36 | 1,74E-01 | 3,40E-01 |
| AT5G40480 | 2,70 | 1,37 | 1,01 | 1,35 | 1,75E-01 | 3,42E-01 |
| AT4G00700 | 2,06 | 1,96 | 1,45 | 1,35 | 1,76E-01 | 3,44E-01 |
| AT1G79690 | 1,96 | 2,40 | 1,79 | 1,35 | 1,78E-01 | 3,48E-01 |
| AT1G07940 | 3,71 | 1,25 | 0,93 | 1,34 | 1,79E-01 | 3,49E-01 |
| AT1G47265 | 3,77 | -1,28 | 0,96 | -1,34 | 1,80E-01 | 3,49E-01 |
| AT2G07698 | 9,74 | -0,84 | 0,63 | -1,34 | 1,80E-01 | 3,49E-01 |
| AT2G36480 | 1,71 | 2,30 | 1,71 | 1,34 | 1,80E-01 | 3,49E-01 |
| AT5G05690 | 2,09 | 1,88 | 1,40 | 1,34 | 1,80E-01 | 3,49E-01 |
| AT5G48380 | 2,92 | 1,27 | 0,95 | 1,34 | 1,81E-01 | 3,50E-01 |
| AT5G49030 | 1,79 | 2,08 | 1,55 | 1,34 | 1,81E-01 | 3,50E-01 |
| AT1G68945 | 2,15 | -1,52 | 1,13 | -1,34 | 1,82E-01 | 3,51E-01 |
| AT4G06705 | 4,08 | -1,22 | 0,91 | -1,34 | 1,82E-01 | 3,51E-01 |
| AT4G26255 | 3,93 | -1,66 | 1,24 | -1,33 | 1,82E-01 | 3,52E-01 |
| AT2G33770 | 4,11 | 1,05 | 0,79 | 1,33 | 1,83E-01 | 3,53E-01 |
| AT5G20720 | 3,41 | 1,56 | 1,17 | 1,33 | 1,83E-01 | 3,53E-01 |
| AT4G26850 | 7,34 | 0,82 | 0,62 | 1,33 | 1,84E-01 | 3,54E-01 |
| AT2G41940 | 1,76 | -1,75 | 1,32 | -1,33 | 1,84E-01 | 3,54E-01 |
| AT5G14060 | 2,66 | -1,59 | 1,20 | -1,33 | 1,84E-01 | 3,54E-01 |
| AT2G41640 | 6,36 | -0,83 | 0,63 | -1,33 | 1,85E-01 | 3,55E-01 |
| AT1G20620 | 38,72 | -0,59 | 0,44 | -1,33 | 1,85E-01 | 3,55E-01 |
| AT3G28900 | 3,87 | -1,35 | 1,02 | -1,33 | 1,85E-01 | 3,55E-01 |
| AT4G04095 | 1,94 | 1,95 | 1,47 | 1,32 | 1,86E-01 | 3,56E-01 |
| AT3G07390 | 2,06 | -1,78 | 1,35 | -1,32 | 1,86E-01 | 3,56E-01 |
| AT1G28070 | 4,21 | 2,99 | 2,26 | 1,32 | 1,86E-01 | 3,56E-01 |
| AT3G55980 | 3,63 | -1,19 | 0,90 | -1,32 | 1,86E-01 | 3,56E-01 |
| AT1G06243 | 5,81 | -0,87 | 0,66 | -1,32 | 1,87E-01 | 3,57E-01 |
| AT4G06695 | 4,20 | -1,23 | 0,94 | -1,32 | 1,87E-01 | 3,57E-01 |
| AT5G62700 | 1,86 | -2,18 | 1,65 | -1,32 | 1,88E-01 | 3,58E-01 |
| AT5G46420 | 2,51 | 1,48 | 1,12 | 1,31 | 1,89E-01 | 3,59E-01 |
| AT1G66783 | 14,56 | 0,78 | 0,59 | 1,32 | 1,89E-01 | 3,59E-01 |
| AT5G47010 | 2,34 | 1,57 | 1,20 | 1,31 | 1,89E-01 | 3,59E-01 |
| AT3G48115 | 1,91 | 1,48 | 1,13 | 1,31 | 1,89E-01 | 3,59E-01 |
| AT3G63070 | 1,86 | 2,29 | 1,75 | 1,31 | 1,91E-01 | 3,63E-01 |
| AT1G30330 | 1,89 | -1,74 | 1,34 | -1,31 | 1,91E-01 | 3,63E-01 |
| AT1G12920 | 1,87 | 1,81 | 1,39 | 1,31 | 1,92E-01 | 3,63E-01 |
| AT4G30750 | 3,95 | 1,22 | 0,93 | 1,31 | 1,92E-01 | 3,63E-01 |
| AT1G70530 | 1,72 | 2,31 | 1,77 | 1,30 | 1,92E-01 | 3,64E-01 |
| AT1G12900 | 4,70 | -1,14 | 0,87 | -1,30 | 1,93E-01 | 3,64E-01 |
| AT1G30400 | 2,52 | 1,65 | 1,27 | 1,30 | 1,93E-01 | 3,65E-01 |
| AT1G20330 | 2,79 | 1,69 | 1,30 | 1,30 | 1,94E-01 | 3,67E-01 |
| AT2G36490 | 1,90 | 3,54 | 2,73 | 1,30 | 1,94E-01 | 3,67E-01 |
| AT1G68530 | 2,97 | -1,32 | 1,02 | -1,30 | 1,95E-01 | 3,67E-01 |
| AT4G20170 | 3,89 | -1,08 | 0,83 | -1,29 | 1,96E-01 | 3,68E-01 |
| AT5G49910 | 2,46 | 1,62 | 1,25 | 1,29 | 1,96E-01 | 3,68E-01 |
| AT2G14910 | 2,97 | 1,42 | 1,10 | 1,29 | 1,96E-01 | 3,68E-01 |
| AT2G22660 | 2,02 | 1,87 | 1,44 | 1,29 | 1,96E-01 | 3,69E-01 |
| AT4G15180 | 1,84 | 1,66 | 1,28 | 1,29 | 1,96E-01 | 3,69E-01 |

| | | | | | | |
|---|---|---|---|---|---|---|
| AT1G33420 | 2,44 | -1,29 | 1,00 | -1,29 | 1,96E-01 | 3,69E-01 |
| AT1G08117 | 4,98 | -1,45 | 1,12 | -1,29 | 1,97E-01 | 3,70E-01 |
| AT1G04393 | 5,00 | -1,07 | 0,83 | -1,29 | 1,98E-01 | 3,72E-01 |
| AT1G69572 | 2,44 | -1,59 | 1,24 | -1,28 | 1,99E-01 | 3,72E-01 |
| AT1G69570 | 2,44 | -1,59 | 1,24 | -1,28 | 1,99E-01 | 3,72E-01 |
| AT4G37640 | 1,71 | 1,71 | 1,33 | 1,28 | 1,99E-01 | 3,72E-01 |
| AT5G20290 | 5,09 | 1,10 | 0,86 | 1,28 | 2,00E-01 | 3,73E-01 |
| AT5G16270 | 1,71 | 1,74 | 1,36 | 1,28 | 2,00E-01 | 3,74E-01 |
| AT2G17840 | 2,90 | -1,66 | 1,29 | -1,28 | 2,00E-01 | 3,74E-01 |
| AT2G38220 | 2,26 | -1,45 | 1,13 | -1,28 | 2,01E-01 | 3,74E-01 |
| AT3G46620 | 4,33 | 1,12 | 0,88 | 1,28 | 2,02E-01 | 3,76E-01 |
| AT3G22940 | 1,72 | -1,93 | 1,51 | -1,27 | 2,02E-01 | 3,76E-01 |
| AT2G05765 | 3,87 | -1,30 | 1,02 | -1,27 | 2,03E-01 | 3,78E-01 |
| AT5G10760 | 4,94 | 0,99 | 0,78 | 1,27 | 2,04E-01 | 3,78E-01 |
| AT4G22710 | 1,76 | -2,22 | 1,75 | -1,27 | 2,05E-01 | 3,80E-01 |
| AT3G44703 | 4,72 | -1,30 | 1,03 | -1,27 | 2,05E-01 | 3,80E-01 |
| AT3G58610 | 2,68 | 1,36 | 1,07 | 1,27 | 2,05E-01 | 3,81E-01 |
| AT4G33300 | 3,93 | 1,32 | 1,04 | 1,27 | 2,06E-01 | 3,81E-01 |
| AT1G21270 | 2,66 | 1,86 | 1,47 | 1,26 | 2,06E-01 | 3,81E-01 |
| AT1G61100 | 1,97 | 1,83 | 1,45 | 1,26 | 2,06E-01 | 3,81E-01 |
| AT1G29670 | 1,73 | -2,04 | 1,62 | -1,26 | 2,07E-01 | 3,81E-01 |
| AT5G48620 | 2,46 | 1,70 | 1,34 | 1,26 | 2,07E-01 | 3,81E-01 |
| AT3G03105 | 2,11 | -1,67 | 1,32 | -1,26 | 2,08E-01 | 3,83E-01 |
| AT5G40910 | 2,64 | 1,24 | 0,98 | 1,26 | 2,08E-01 | 3,83E-01 |
| AT1G11860 | 2,12 | -1,77 | 1,41 | -1,26 | 2,09E-01 | 3,85E-01 |
| AT2G41650 | 5,84 | -0,84 | 0,67 | -1,26 | 2,09E-01 | 3,85E-01 |
| AT1G50440 | 6,47 | -0,89 | 0,71 | -1,25 | 2,10E-01 | 3,86E-01 |
| AT2G38540 | 3,59 | -1,42 | 1,13 | -1,25 | 2,10E-01 | 3,86E-01 |
| AT2G29630 | 5,51 | 0,85 | 0,68 | 1,25 | 2,11E-01 | 3,88E-01 |
| AT2G09555 | 2,08 | -1,53 | 1,23 | -1,25 | 2,12E-01 | 3,89E-01 |
| AT3G22970 | 2,77 | -1,29 | 1,03 | -1,25 | 2,13E-01 | 3,89E-01 |
| AT2G03855 | 12,69 | -0,70 | 0,56 | -1,25 | 2,13E-01 | 3,90E-01 |
| AT2G34250 | 2,94 | 1,27 | 1,02 | 1,24 | 2,13E-01 | 3,90E-01 |
| AT5G26340 | 2,20 | 1,60 | 1,29 | 1,24 | 2,15E-01 | 3,93E-01 |
| AT4G06225 | 6,57 | -1,28 | 1,03 | -1,24 | 2,16E-01 | 3,93E-01 |
| AT5G13530 | 2,40 | 1,36 | 1,10 | 1,24 | 2,16E-01 | 3,93E-01 |
| AT4G06800 | 2,19 | -1,35 | 1,09 | -1,24 | 2,15E-01 | 3,93E-01 |
| AT3G00980 | 2,10 | -2,17 | 1,75 | -1,24 | 2,16E-01 | 3,93E-01 |
| AT3G47348 | 6,89 | -1,32 | 1,07 | -1,24 | 2,17E-01 | 3,94E-01 |
| AT5G27950 | 2,87 | 1,40 | 1,14 | 1,24 | 2,17E-01 | 3,94E-01 |
| AT4G09650 | 3,00 | -1,76 | 1,43 | -1,23 | 2,18E-01 | 3,96E-01 |
| AT3G50820 | 7,18 | -1,10 | 0,89 | -1,23 | 2,18E-01 | 3,96E-01 |
| AT3G53740 | 2,01 | 2,33 | 1,89 | 1,23 | 2,18E-01 | 3,96E-01 |
| AT4G02770 | 8,37 | -0,81 | 0,66 | -1,23 | 2,18E-01 | 3,96E-01 |
| AT2G45960 | 8,71 | 0,83 | 0,67 | 1,23 | 2,19E-01 | 3,96E-01 |
| AT3G08740 | 1,74 | 1,63 | 1,32 | 1,23 | 2,19E-01 | 3,96E-01 |
| AT5G02645 | 2,33 | -1,41 | 1,14 | -1,23 | 2,19E-01 | 3,96E-01 |
| AT5G42930 | 2,68 | -1,38 | 1,13 | -1,23 | 2,19E-01 | 3,97E-01 |
| AT3G04775 | 14,57 | -0,57 | 0,47 | -1,23 | 2,20E-01 | 3,98E-01 |
| AT1G50030 | 2,19 | 1,64 | 1,34 | 1,22 | 2,21E-01 | 3,98E-01 |
| AT5G10450 | 3,14 | 1,25 | 1,02 | 1,22 | 2,21E-01 | 3,98E-01 |
| AT3G01500 | 4,00 | -1,41 | 1,15 | -1,22 | 2,21E-01 | 3,98E-01 |
| AT4G26480 | 2,39 | 1,62 | 1,33 | 1,22 | 2,21E-01 | 3,98E-01 |
| AT3G47910 | 2,73 | 1,31 | 1,07 | 1,22 | 2,22E-01 | 3,99E-01 |
| AT3G13525 | 2,38 | -1,33 | 1,09 | -1,22 | 2,22E-01 | 3,99E-01 |
| AT4G38760 | 3,63 | -1,19 | 0,98 | -1,22 | 2,22E-01 | 3,99E-01 |

| | | | | | | |
|---|---|---|---|---|---|---|
| AT5G20020 | 2,46 | 1,89 | 1,55 | 1,22 | 2,23E-01 | 4,01E-01 |
| AT5G08925 | 2,86 | -1,73 | 1,42 | -1,22 | 2,23E-01 | 4,01E-01 |
| AT4G30993 | 2,23 | 1,36 | 1,12 | 1,22 | 2,24E-01 | 4,01E-01 |
| AT1G08857 | 2,42 | 2,05 | 1,69 | 1,22 | 2,24E-01 | 4,01E-01 |
| AT3G01120 | 2,55 | 1,80 | 1,48 | 1,22 | 2,24E-01 | 4,01E-01 |
| AT4G39350 | 3,27 | 1,17 | 0,97 | 1,21 | 2,27E-01 | 4,05E-01 |
| AT5G08690 | 2,64 | 1,30 | 1,08 | 1,21 | 2,27E-01 | 4,05E-01 |
| AT5G07735 | 1,90 | -1,49 | 1,23 | -1,21 | 2,27E-01 | 4,05E-01 |
| AT2G33340 | 1,98 | 1,81 | 1,50 | 1,21 | 2,27E-01 | 4,05E-01 |
| AT1G05427 | 2,11 | -1,79 | 1,49 | -1,21 | 2,28E-01 | 4,06E-01 |
| AT2G38280 | 1,89 | 1,87 | 1,55 | 1,21 | 2,28E-01 | 4,06E-01 |
| AT3G26440 | 2,47 | -1,71 | 1,42 | -1,20 | 2,29E-01 | 4,07E-01 |
| AT5G62165 | 7,10 | -0,93 | 0,78 | -1,20 | 2,30E-01 | 4,08E-01 |
| AT5G64740 | 4,40 | 1,00 | 0,83 | 1,20 | 2,30E-01 | 4,08E-01 |
| AT1G65350 | 2,42 | -1,41 | 1,17 | -1,20 | 2,30E-01 | 4,08E-01 |
| AT5G17020 | 2,03 | 1,72 | 1,43 | 1,20 | 2,30E-01 | 4,08E-01 |
| AT4G23140 | 2,18 | 1,48 | 1,24 | 1,20 | 2,31E-01 | 4,09E-01 |
| AT4G34720 | 1,83 | 1,43 | 1,19 | 1,20 | 2,31E-01 | 4,09E-01 |
| AT5G60360 | 2,97 | 1,50 | 1,25 | 1,19 | 2,32E-01 | 4,10E-01 |
| AT1G29470 | 3,69 | 1,11 | 0,93 | 1,20 | 2,32E-01 | 4,10E-01 |
| AT1G22060 | 2,54 | 1,39 | 1,17 | 1,19 | 2,34E-01 | 4,13E-01 |
| AT1G03080 | 1,73 | 1,52 | 1,28 | 1,19 | 2,34E-01 | 4,13E-01 |
| AT4G16990 | 5,03 | 0,98 | 0,82 | 1,19 | 2,34E-01 | 4,13E-01 |
| AT1G65440 | 2,14 | 1,57 | 1,32 | 1,19 | 2,35E-01 | 4,14E-01 |
| AT3G50950 | 4,46 | 0,97 | 0,82 | 1,19 | 2,35E-01 | 4,14E-01 |
| AT1G28480 | 1,91 | 2,06 | 1,74 | 1,19 | 2,36E-01 | 4,15E-01 |
| AT2G13360 | 3,44 | -1,10 | 0,93 | -1,19 | 2,36E-01 | 4,15E-01 |
| AT2G03330 | 8,86 | -0,80 | 0,68 | -1,18 | 2,38E-01 | 4,17E-01 |
| AT4G00360 | 2,92 | -1,40 | 1,18 | -1,18 | 2,38E-01 | 4,17E-01 |
| AT3G10380 | 2,66 | 1,21 | 1,02 | 1,18 | 2,38E-01 | 4,17E-01 |
| AT5G42895 | 1,80 | -1,78 | 1,51 | -1,18 | 2,39E-01 | 4,20E-01 |
| AT2G02835 | 3,94 | -1,02 | 0,87 | -1,18 | 2,40E-01 | 4,20E-01 |
| AT1G05817 | 1,94 | -1,49 | 1,27 | -1,17 | 2,40E-01 | 4,21E-01 |
| AT3G00985 | 2,03 | -2,11 | 1,80 | -1,17 | 2,40E-01 | 4,21E-01 |
| AT2G44490 | 17,93 | -0,62 | 0,53 | -1,17 | 2,41E-01 | 4,21E-01 |
| AT1G72410 | 1,92 | 1,82 | 1,56 | 1,17 | 2,41E-01 | 4,21E-01 |
| AT4G08850 | 2,82 | 1,37 | 1,18 | 1,17 | 2,42E-01 | 4,23E-01 |
| AT1G18260 | 1,96 | 1,54 | 1,32 | 1,17 | 2,43E-01 | 4,24E-01 |
| AT3G55830 | 1,72 | 1,81 | 1,55 | 1,17 | 2,44E-01 | 4,25E-01 |
| AT3G53260 | 2,56 | 1,47 | 1,26 | 1,17 | 2,44E-01 | 4,25E-01 |
| AT4G18670 | 7,64 | -0,79 | 0,68 | -1,16 | 2,45E-01 | 4,27E-01 |
| AT5G02150 | 1,74 | 2,18 | 1,88 | 1,16 | 2,46E-01 | 4,27E-01 |
| AT3G13405 | 1,91 | -1,64 | 1,41 | -1,16 | 2,46E-01 | 4,28E-01 |
| AT5G20250 | 2,34 | 1,32 | 1,14 | 1,16 | 2,47E-01 | 4,28E-01 |
| AT5G04455 | 10,87 | 0,63 | 0,55 | 1,16 | 2,48E-01 | 4,29E-01 |
| AT5G04445 | 10,87 | 0,63 | 0,55 | 1,16 | 2,48E-01 | 4,29E-01 |
| AT3G26450 | 2,27 | -1,63 | 1,41 | -1,15 | 2,48E-01 | 4,30E-01 |
| AT2G26500 | 2,52 | 1,31 | 1,14 | 1,15 | 2,49E-01 | 4,31E-01 |
| AT5G02305 | 2,96 | -1,31 | 1,14 | -1,15 | 2,49E-01 | 4,32E-01 |
| AT5G07570 | 4,01 | -1,27 | 1,11 | -1,15 | 2,50E-01 | 4,32E-01 |
| AT3G10745 | 164,34 | -0,39 | 0,34 | -1,15 | 2,50E-01 | 4,32E-01 |
| AT4G10800 | 6,52 | 0,78 | 0,68 | 1,15 | 2,51E-01 | 4,33E-01 |
| AT2G28550 | 2,55 | -1,29 | 1,13 | -1,15 | 2,51E-01 | 4,33E-01 |
| AT5G19120 | 2,51 | -1,27 | 1,11 | -1,15 | 2,52E-01 | 4,34E-01 |
| AT5G01775 | 4,59 | -1,03 | 0,90 | -1,14 | 2,53E-01 | 4,34E-01 |
| AT1G07633 | 6,19 | -0,84 | 0,74 | -1,15 | 2,52E-01 | 4,34E-01 |

| | | | | | | |
|---|---|---|---|---|---|---|
| AT2G20010 | 1,89 | 1,56 | 1,36 | 1,14 | 2,52E-01 | 4,34E-01 |
| AT4G06235 | 7,69 | -1,14 | 1,00 | -1,14 | 2,53E-01 | 4,34E-01 |
| AT4G23100 | 1,83 | 1,36 | 1,19 | 1,14 | 2,53E-01 | 4,34E-01 |
| AT1G32720 | 2,47 | -1,28 | 1,12 | -1,14 | 2,54E-01 | 4,35E-01 |
| AT1G62630 | 2,10 | -1,40 | 1,22 | -1,14 | 2,54E-01 | 4,35E-01 |
| AT3G28180 | 8,56 | -1,27 | 1,11 | -1,14 | 2,54E-01 | 4,35E-01 |
| AT5G57660 | 4,42 | 0,86 | 0,75 | 1,14 | 2,55E-01 | 4,35E-01 |
| AT1G61740 | 2,13 | -1,42 | 1,25 | -1,14 | 2,55E-01 | 4,35E-01 |
| AT5G58330 | 1,92 | 1,65 | 1,44 | 1,14 | 2,54E-01 | 4,35E-01 |
| AT2G45180 | 2,18 | -1,50 | 1,32 | -1,14 | 2,56E-01 | 4,37E-01 |
| AT1G63360 | 4,56 | -1,00 | 0,88 | -1,14 | 2,56E-01 | 4,37E-01 |
| AT5G14570 | 2,87 | 1,10 | 0,97 | 1,13 | 2,56E-01 | 4,37E-01 |
| AT1G06430 | 1,82 | 1,63 | 1,44 | 1,13 | 2,57E-01 | 4,38E-01 |
| AT3G48200 | 2,11 | 1,90 | 1,67 | 1,13 | 2,57E-01 | 4,38E-01 |
| AT1G66725 | 1003,08 | -0,34 | 0,30 | -1,13 | 2,57E-01 | 4,38E-01 |
| AT3G20310 | 2,42 | -1,23 | 1,09 | -1,13 | 2,58E-01 | 4,39E-01 |
| AT4G08125 | 3,11 | -1,10 | 0,98 | -1,13 | 2,59E-01 | 4,40E-01 |
| AT2G18570 | 4,57 | -1,14 | 1,01 | -1,13 | 2,60E-01 | 4,40E-01 |
| AT1G77760 | 2,44 | 1,39 | 1,23 | 1,13 | 2,59E-01 | 4,40E-01 |
| AT3G44110 | 2,03 | 1,33 | 1,18 | 1,13 | 2,59E-01 | 4,40E-01 |
| AT5G41490 | 2,24 | -1,39 | 1,23 | -1,13 | 2,61E-01 | 4,42E-01 |
| AT2G38195 | 4,06 | -1,36 | 1,21 | -1,12 | 2,62E-01 | 4,43E-01 |
| AT5G27807 | 30,37 | 0,44 | 0,39 | 1,12 | 2,62E-01 | 4,44E-01 |
| AT1G48760 | 1,82 | 2,23 | 1,99 | 1,12 | 2,64E-01 | 4,46E-01 |
| AT3G01695 | 8,72 | 0,82 | 0,73 | 1,12 | 2,64E-01 | 4,47E-01 |
| AT5G56030 | 5,79 | 0,75 | 0,67 | 1,11 | 2,65E-01 | 4,48E-01 |
| AT1G67280 | 1,72 | 1,71 | 1,54 | 1,11 | 2,66E-01 | 4,48E-01 |
| AT5G09000 | 2,90 | -1,48 | 1,33 | -1,11 | 2,65E-01 | 4,48E-01 |
| AT4G02890 | 2,39 | 1,24 | 1,12 | 1,11 | 2,67E-01 | 4,49E-01 |
| AT5G19510 | 1,83 | 1,56 | 1,41 | 1,11 | 2,67E-01 | 4,49E-01 |
| AT3G49910 | 2,91 | 1,21 | 1,09 | 1,11 | 2,67E-01 | 4,50E-01 |
| AT1G05993 | 11,40 | -0,60 | 0,55 | -1,11 | 2,68E-01 | 4,50E-01 |
| AT2G17200 | 1,89 | 1,55 | 1,40 | 1,11 | 2,68E-01 | 4,50E-01 |
| AT5G17710 | 1,85 | 1,45 | 1,31 | 1,11 | 2,68E-01 | 4,50E-01 |
| AT5G63570 | 1,81 | 1,90 | 1,72 | 1,10 | 2,70E-01 | 4,51E-01 |
| AT1G76010 | 2,04 | 1,43 | 1,29 | 1,10 | 2,69E-01 | 4,51E-01 |
| AT3G05675 | 9,08 | 0,78 | 0,71 | 1,10 | 2,69E-01 | 4,51E-01 |
| AT5G11700 | 1,99 | 1,45 | 1,31 | 1,10 | 2,69E-01 | 4,51E-01 |
| AT3G26730 | 1,96 | 1,39 | 1,26 | 1,10 | 2,71E-01 | 4,53E-01 |
| AT3G09440 | 5,91 | 0,80 | 0,73 | 1,10 | 2,71E-01 | 4,53E-01 |
| AT4G04570 | 4,35 | -1,67 | 1,52 | -1,10 | 2,72E-01 | 4,53E-01 |
| AT1G11785 | 2,48 | -1,15 | 1,05 | -1,10 | 2,71E-01 | 4,53E-01 |
| AT2G05315 | 4,98 | -0,98 | 0,89 | -1,10 | 2,72E-01 | 4,54E-01 |
| AT3G24615 | 9,91 | -0,67 | 0,61 | -1,10 | 2,73E-01 | 4,54E-01 |
| AT4G21710 | 2,69 | 1,57 | 1,43 | 1,10 | 2,73E-01 | 4,54E-01 |
| AT5G20010 | 1,98 | 1,61 | 1,47 | 1,10 | 2,73E-01 | 4,54E-01 |
| AT1G09210 | 5,83 | 0,83 | 0,75 | 1,10 | 2,73E-01 | 4,54E-01 |
| AT1G59980 | 3,55 | -1,15 | 1,05 | -1,09 | 2,75E-01 | 4,57E-01 |
| AT4G36648 | 2,47 | 1,55 | 1,42 | 1,09 | 2,76E-01 | 4,58E-01 |
| AT2G16040 | 2,68 | -1,23 | 1,13 | -1,09 | 2,77E-01 | 4,60E-01 |
| AT5G02700 | 1,99 | -1,43 | 1,32 | -1,09 | 2,77E-01 | 4,60E-01 |
| AT3G07735 | 4,39 | -0,93 | 0,86 | -1,08 | 2,78E-01 | 4,60E-01 |
| AT4G17090 | 4,63 | 0,99 | 0,91 | 1,09 | 2,78E-01 | 4,60E-01 |
| AT5G09660 | 2,38 | -1,20 | 1,11 | -1,08 | 2,78E-01 | 4,60E-01 |
| AT1G77235 | 1,77 | -1,87 | 1,73 | -1,08 | 2,79E-01 | 4,61E-01 |
| AT4G05360 | 2,15 | -1,28 | 1,18 | -1,08 | 2,79E-01 | 4,61E-01 |

| | | | | | | |
|---|---|---|---|---|---|---|
| AT4G39090 | 7,03 | 0,79 | 0,73 | 1,08 | 2,80E-01 | 4,61E-01 |
| AT1G20015 | 7,84 | -0,83 | 0,77 | -1,08 | 2,80E-01 | 4,62E-01 |
| AT5G14270 | 2,74 | 1,16 | 1,08 | 1,07 | 2,83E-01 | 4,66E-01 |
| AT3G08210 | 3,75 | -0,98 | 0,91 | -1,07 | 2,83E-01 | 4,66E-01 |
| AT3G56150 | 2,36 | 1,34 | 1,25 | 1,07 | 2,83E-01 | 4,66E-01 |
| AT4G18030 | 2,74 | 1,40 | 1,30 | 1,07 | 2,83E-01 | 4,66E-01 |
| AT4G14400 | 8,78 | -0,92 | 0,86 | -1,07 | 2,84E-01 | 4,67E-01 |
| AT2G46220 | 1,83 | 1,61 | 1,50 | 1,07 | 2,85E-01 | 4,68E-01 |
| AT5G61420 | 2,01 | 1,64 | 1,54 | 1,07 | 2,85E-01 | 4,68E-01 |
| AT3G12980 | 1,82 | 1,90 | 1,78 | 1,07 | 2,86E-01 | 4,69E-01 |
| AT3G54925 | 1,76 | -1,77 | 1,66 | -1,07 | 2,87E-01 | 4,70E-01 |
| AT5G67030 | 1,94 | 1,30 | 1,22 | 1,06 | 2,87E-01 | 4,71E-01 |
| AT4G20260 | 2,30 | -1,27 | 1,20 | -1,06 | 2,88E-01 | 4,71E-01 |
| AT2G28056 | 34,24 | -0,41 | 0,39 | -1,06 | 2,88E-01 | 4,71E-01 |
| AT5G08530 | 3,76 | -1,16 | 1,09 | -1,06 | 2,88E-01 | 4,71E-01 |
| AT2G29980 | 3,37 | 1,36 | 1,28 | 1,06 | 2,88E-01 | 4,71E-01 |
| AT1G36180 | 2,20 | 1,43 | 1,36 | 1,06 | 2,91E-01 | 4,74E-01 |
| AT4G29060 | 2,00 | 1,44 | 1,36 | 1,06 | 2,91E-01 | 4,75E-01 |
| AT1G18080 | 2,60 | 1,27 | 1,20 | 1,05 | 2,92E-01 | 4,75E-01 |
| AT4G33650 | 1,75 | 1,66 | 1,57 | 1,05 | 2,92E-01 | 4,75E-01 |
| AT3G60630 | 5,65 | -0,80 | 0,76 | -1,05 | 2,92E-01 | 4,75E-01 |
| AT1G31163 | 6,82 | -0,70 | 0,66 | -1,05 | 2,94E-01 | 4,77E-01 |
| AT1G56340 | 4,49 | 0,84 | 0,80 | 1,05 | 2,94E-01 | 4,77E-01 |
| AT2G21920 | 4,05 | -0,93 | 0,88 | -1,05 | 2,94E-01 | 4,77E-01 |
| AT5G17890 | 1,77 | 1,80 | 1,72 | 1,05 | 2,95E-01 | 4,78E-01 |
| AT2G44560 | 4,56 | -0,80 | 0,77 | -1,05 | 2,95E-01 | 4,79E-01 |
| AT5G57710 | 4,28 | 0,80 | 0,77 | 1,04 | 2,97E-01 | 4,81E-01 |
| AT5G07325 | 8,05 | 0,63 | 0,61 | 1,04 | 2,97E-01 | 4,81E-01 |
| AT5G58950 | 2,42 | 1,23 | 1,18 | 1,04 | 2,97E-01 | 4,81E-01 |
| AT1G52360 | 2,02 | 1,38 | 1,32 | 1,04 | 2,99E-01 | 4,82E-01 |
| AT2G05710 | 3,35 | 1,03 | 0,99 | 1,04 | 2,98E-01 | 4,82E-01 |
| AT2G36390 | 2,07 | 1,47 | 1,41 | 1,04 | 2,98E-01 | 4,82E-01 |
| AT4G30972 | 11,83 | -0,52 | 0,51 | -1,04 | 2,99E-01 | 4,82E-01 |
| AT4G21960 | 15,77 | -0,50 | 0,48 | -1,04 | 2,99E-01 | 4,82E-01 |
| AT5G49740 | 2,42 | 1,18 | 1,14 | 1,04 | 2,99E-01 | 4,82E-01 |
| AT5G52190 | 1,80 | 1,78 | 1,71 | 1,04 | 2,99E-01 | 4,82E-01 |
| AT3G26650 | 12,31 | 0,63 | 0,61 | 1,04 | 3,00E-01 | 4,82E-01 |
| AT4G01010 | 2,44 | -1,37 | 1,32 | -1,03 | 3,01E-01 | 4,84E-01 |
| AT1G62180 | 2,66 | 1,07 | 1,03 | 1,03 | 3,01E-01 | 4,85E-01 |
| AT1G33415 | 2,72 | -0,99 | 0,96 | -1,03 | 3,02E-01 | 4,85E-01 |
| AT3G57710 | 4,92 | -0,75 | 0,73 | -1,03 | 3,02E-01 | 4,85E-01 |
| AT5G42650 | 3,24 | -1,25 | 1,21 | -1,03 | 3,02E-01 | 4,85E-01 |
| AT4G40040 | 3,01 | 1,04 | 1,01 | 1,03 | 3,03E-01 | 4,86E-01 |
| AT3G62200 | 2,36 | 1,38 | 1,35 | 1,03 | 3,04E-01 | 4,87E-01 |
| AT1G30380 | 5,72 | -1,12 | 1,10 | -1,03 | 3,05E-01 | 4,87E-01 |
| AT2G43375 | 6,65 | -0,68 | 0,67 | -1,03 | 3,05E-01 | 4,87E-01 |
| AT2G41410 | 4,84 | 0,96 | 0,94 | 1,03 | 3,05E-01 | 4,88E-01 |
| AT1G53210 | 1,83 | -1,54 | 1,50 | -1,02 | 3,06E-01 | 4,88E-01 |
| AT5G52310 | 5,67 | -1,47 | 1,43 | -1,02 | 3,05E-01 | 4,88E-01 |
| AT1G58602 | 11,97 | 0,55 | 0,54 | 1,02 | 3,07E-01 | 4,88E-01 |
| AT1G74470 | 19,25 | 0,51 | 0,50 | 1,02 | 3,06E-01 | 4,88E-01 |
| AT4G08335 | 4,06 | 0,93 | 0,90 | 1,02 | 3,06E-01 | 4,88E-01 |
| AT4G02520 | 4,48 | -0,89 | 0,87 | -1,02 | 3,06E-01 | 4,88E-01 |
| AT3G04120 | 5,88 | 0,72 | 0,70 | 1,02 | 3,07E-01 | 4,89E-01 |
| AT5G01785 | 2,53 | -1,50 | 1,47 | -1,02 | 3,08E-01 | 4,89E-01 |
| AT5G42050 | 3,62 | -0,96 | 0,95 | -1,02 | 3,08E-01 | 4,89E-01 |

| | | | | | | |
|---|---|---|---|---|---|---|
| AT5G63840 | 2,51 | 1,18 | 1,15 | 1,02 | 3,08E-01 | 4,89E-01 |
| AT3G04210 | 3,60 | 0,93 | 0,92 | 1,02 | 3,08E-01 | 4,89E-01 |
| AT5G26920 | 3,70 | 0,86 | 0,85 | 1,02 | 3,09E-01 | 4,90E-01 |
| AT2G04955 | 3,52 | -0,93 | 0,92 | -1,02 | 3,10E-01 | 4,91E-01 |
| AT2G16018 | 9,68 | -0,59 | 0,59 | -1,01 | 3,11E-01 | 4,93E-01 |
| AT5G62690 | 4,26 | 0,88 | 0,87 | 1,01 | 3,12E-01 | 4,93E-01 |
| AT2G07815 | 2,06 | -1,27 | 1,25 | -1,01 | 3,12E-01 | 4,93E-01 |
| AT1G21250 | 6,46 | 0,65 | 0,64 | 1,01 | 3,13E-01 | 4,94E-01 |
| AT5G09870 | 2,84 | 1,02 | 1,01 | 1,01 | 3,13E-01 | 4,94E-01 |
| AT3G30390 | 2,05 | 1,08 | 1,07 | 1,01 | 3,14E-01 | 4,96E-01 |
| AT3G52430 | 2,34 | 1,42 | 1,41 | 1,00 | 3,15E-01 | 4,97E-01 |
| AT2G08855 | 3,73 | -0,87 | 0,86 | -1,00 | 3,15E-01 | 4,97E-01 |
| AT5G61790 | 5,40 | 0,84 | 0,83 | 1,00 | 3,16E-01 | 4,97E-01 |
| AT3G57260 | 8,35 | 0,56 | 0,56 | 1,00 | 3,16E-01 | 4,98E-01 |
| AT4G30975 | 11,87 | -0,51 | 0,51 | -1,00 | 3,18E-01 | 5,00E-01 |
| AT2G34655 | 4,12 | -0,99 | 0,99 | -1,00 | 3,18E-01 | 5,00E-01 |
| AT1G15820 | 12,09 | -0,68 | 0,68 | -1,00 | 3,19E-01 | 5,01E-01 |
| AT4G28755 | 2,23 | -1,41 | 1,42 | -1,00 | 3,19E-01 | 5,01E-01 |
| AT3G29320 | 2,29 | 1,14 | 1,15 | 0,99 | 3,20E-01 | 5,02E-01 |
| AT5G39740 | 3,11 | 1,03 | 1,03 | 0,99 | 3,20E-01 | 5,02E-01 |
| AT1G16635 | 19,67 | -0,56 | 0,56 | -0,99 | 3,20E-01 | 5,02E-01 |
| AT3G07790 | 2,60 | 1,07 | 1,08 | 0,99 | 3,22E-01 | 5,04E-01 |
| AT1G19960 | 1,88 | 1,70 | 1,72 | 0,99 | 3,23E-01 | 5,05E-01 |
| AT5G46070 | 1,78 | 1,76 | 1,79 | 0,99 | 3,24E-01 | 5,06E-01 |
| AT2G21030 | 1,78 | -1,44 | 1,46 | -0,99 | 3,24E-01 | 5,06E-01 |
| AT2G30440 | 1,74 | -1,31 | 1,34 | -0,98 | 3,25E-01 | 5,07E-01 |
| AT1G29400 | 2,11 | 1,12 | 1,14 | 0,98 | 3,25E-01 | 5,07E-01 |
| AT2G07890 | 2,64 | -1,11 | 1,14 | -0,98 | 3,26E-01 | 5,09E-01 |
| AT1G61275 | 11,06 | -0,54 | 0,55 | -0,98 | 3,27E-01 | 5,09E-01 |
| AT3G11560 | 1,70 | 1,61 | 1,65 | 0,98 | 3,27E-01 | 5,09E-01 |
| AT5G25280 | 3,55 | 0,99 | 1,02 | 0,98 | 3,28E-01 | 5,10E-01 |
| AT5G47560 | 3,13 | -1,01 | 1,03 | -0,98 | 3,28E-01 | 5,10E-01 |
| AT1G61520 | 19,44 | -0,55 | 0,56 | -0,98 | 3,29E-01 | 5,11E-01 |
| AT2G25250 | 2,81 | -1,05 | 1,07 | -0,98 | 3,29E-01 | 5,11E-01 |
| AT5G27660 | 6,15 | 0,69 | 0,71 | 0,98 | 3,29E-01 | 5,11E-01 |
| AT3G44260 | 2,79 | -1,02 | 1,05 | -0,97 | 3,30E-01 | 5,11E-01 |
| AT1G05247 | 19,59 | -0,55 | 0,56 | -0,97 | 3,30E-01 | 5,11E-01 |
| AT1G49500 | 1,83 | -1,22 | 1,25 | -0,97 | 3,30E-01 | 5,11E-01 |
| AT1G21750 | 2,14 | 1,49 | 1,53 | 0,97 | 3,33E-01 | 5,14E-01 |
| AT1G06667 | 5,78 | -0,79 | 0,82 | -0,97 | 3,33E-01 | 5,14E-01 |
| AT2G20580 | 2,81 | 1,58 | 1,63 | 0,97 | 3,33E-01 | 5,14E-01 |
| AT3G14650 | 2,33 | 1,10 | 1,14 | 0,97 | 3,32E-01 | 5,14E-01 |
| AT5G58465 | 4,24 | -0,88 | 0,91 | -0,97 | 3,34E-01 | 5,15E-01 |
| AT5G65630 | 3,70 | 0,91 | 0,94 | 0,97 | 3,34E-01 | 5,15E-01 |
| AT4G10120 | 1,86 | 1,29 | 1,34 | 0,96 | 3,35E-01 | 5,15E-01 |
| AT2G18440 | 3,48 | 0,88 | 0,91 | 0,96 | 3,35E-01 | 5,15E-01 |
| AT3G21755 | 4,36 | -0,81 | 0,84 | -0,96 | 3,35E-01 | 5,15E-01 |
| AT5G46270 | 2,21 | 1,32 | 1,37 | 0,96 | 3,35E-01 | 5,15E-01 |
| AT1G35215 | 4,81 | -0,70 | 0,73 | -0,96 | 3,37E-01 | 5,18E-01 |
| AT4G25730 | 1,71 | 1,67 | 1,74 | 0,96 | 3,37E-01 | 5,18E-01 |
| AT4G39200 | 1,73 | 1,88 | 1,97 | 0,96 | 3,39E-01 | 5,20E-01 |
| AT5G38410 | 6,26 | -0,90 | 0,94 | -0,96 | 3,39E-01 | 5,20E-01 |
| AT3G24542 | 2,63 | -1,16 | 1,21 | -0,95 | 3,40E-01 | 5,21E-01 |
| AT2G32240 | 2,12 | 1,18 | 1,24 | 0,95 | 3,40E-01 | 5,21E-01 |
| AT3G44006 | 2,21 | -1,43 | 1,49 | -0,95 | 3,40E-01 | 5,21E-01 |
| AT5G52530 | 3,03 | 1,38 | 1,45 | 0,95 | 3,41E-01 | 5,21E-01 |

| | | | | | | |
|---|---|---|---|---|---|---|
| AT5G24890 | 2,39 | -1,06 | 1,11 | -0,95 | 3,41E-01 | 5,21E-01 |
| AT5G06165 | 4,41 | -0,84 | 0,89 | -0,95 | 3,41E-01 | 5,21E-01 |
| AT1G13930 | 7,00 | -0,73 | 0,76 | -0,95 | 3,42E-01 | 5,22E-01 |
| AT1G04733 | 20,09 | 0,43 | 0,45 | 0,95 | 3,42E-01 | 5,22E-01 |
| AT3G52850 | 1,76 | 1,30 | 1,37 | 0,95 | 3,42E-01 | 5,22E-01 |
| AT2G30920 | 2,18 | -1,16 | 1,22 | -0,95 | 3,42E-01 | 5,22E-01 |
| AT3G06325 | 6,42 | -0,70 | 0,74 | -0,95 | 3,44E-01 | 5,23E-01 |
| AT3G53400 | 1,89 | 1,49 | 1,57 | 0,95 | 3,44E-01 | 5,23E-01 |
| AT1G33240 | 1,91 | -1,42 | 1,50 | -0,95 | 3,44E-01 | 5,23E-01 |
| AT4G16920 | 1,96 | 1,21 | 1,28 | 0,94 | 3,45E-01 | 5,25E-01 |
| AT4G26090 | 1,90 | 1,17 | 1,24 | 0,94 | 3,46E-01 | 5,25E-01 |
| AT5G53440 | 2,08 | 1,17 | 1,24 | 0,94 | 3,46E-01 | 5,25E-01 |
| AT1G04080 | 1,89 | 1,26 | 1,34 | 0,94 | 3,47E-01 | 5,26E-01 |
| AT3G19170 | 4,99 | 0,91 | 0,97 | 0,94 | 3,47E-01 | 5,26E-01 |
| AT5G40395 | 2,01 | -1,08 | 1,15 | -0,94 | 3,49E-01 | 5,28E-01 |
| AT2G01021 | 3194,91 | -0,25 | 0,27 | -0,94 | 3,49E-01 | 5,29E-01 |
| AT5G43760 | 1,93 | 1,28 | 1,37 | 0,93 | 3,50E-01 | 5,29E-01 |
| AT1G44575 | 5,52 | -0,69 | 0,74 | -0,93 | 3,51E-01 | 5,30E-01 |
| AT1G09800 | 20,58 | 0,42 | 0,45 | 0,93 | 3,51E-01 | 5,30E-01 |
| AT2G01570 | 3,41 | 0,84 | 0,90 | 0,93 | 3,51E-01 | 5,30E-01 |
| AT5G42530 | 4,10 | -0,78 | 0,84 | -0,93 | 3,51E-01 | 5,30E-01 |
| AT2G09685 | 3,15 | -0,93 | 1,00 | -0,93 | 3,52E-01 | 5,31E-01 |
| AT1G64500 | 2,64 | -1,03 | 1,11 | -0,93 | 3,53E-01 | 5,32E-01 |
| AT5G18410 | 3,15 | -0,94 | 1,02 | -0,93 | 3,53E-01 | 5,32E-01 |
| AT5G05485 | 5,35 | -1,09 | 1,18 | -0,93 | 3,53E-01 | 5,32E-01 |
| AT2G05170 | 1,80 | 1,35 | 1,46 | 0,93 | 3,54E-01 | 5,33E-01 |
| AT1G02780 | 1,83 | -1,15 | 1,24 | -0,93 | 3,54E-01 | 5,33E-01 |
| AT1G50900 | 1,85 | 1,28 | 1,38 | 0,93 | 3,55E-01 | 5,33E-01 |
| AT1G23720 | 2,26 | -1,56 | 1,68 | -0,92 | 3,55E-01 | 5,33E-01 |
| AT5G46760 | 2,52 | 1,11 | 1,20 | 0,92 | 3,56E-01 | 5,33E-01 |
| AT1G64340 | 2,24 | -1,50 | 1,63 | -0,92 | 3,56E-01 | 5,34E-01 |
| AT3G21760 | 4,32 | -0,77 | 0,83 | -0,92 | 3,56E-01 | 5,34E-01 |
| AT3G04250 | 2,26 | -1,37 | 1,48 | -0,92 | 3,57E-01 | 5,34E-01 |
| AT3G09390 | 6,61 | 0,69 | 0,75 | 0,92 | 3,57E-01 | 5,34E-01 |
| AT1G08347 | 16,30 | -0,40 | 0,44 | -0,92 | 3,60E-01 | 5,38E-01 |
| AT1G03410 | 2,63 | -1,02 | 1,12 | -0,92 | 3,60E-01 | 5,38E-01 |
| AT3G25010 | 4,35 | -0,77 | 0,85 | -0,91 | 3,61E-01 | 5,39E-01 |
| AT1G18310 | 4,64 | 0,83 | 0,91 | 0,91 | 3,61E-01 | 5,39E-01 |
| AT4G04125 | 2,22 | 1,05 | 1,15 | 0,91 | 3,63E-01 | 5,41E-01 |
| AT5G14740 | 2,79 | 1,26 | 1,39 | 0,91 | 3,63E-01 | 5,41E-01 |
| AT1G75950 | 1,80 | 1,12 | 1,24 | 0,91 | 3,64E-01 | 5,41E-01 |
| AT4G26970 | 1,79 | 1,46 | 1,61 | 0,91 | 3,64E-01 | 5,41E-01 |
| AT3G47342 | 15,52 | -0,48 | 0,53 | -0,91 | 3,65E-01 | 5,42E-01 |
| AT3G03435 | 3,28 | 0,86 | 0,95 | 0,91 | 3,65E-01 | 5,42E-01 |
| AT5G24900 | 2,20 | -1,13 | 1,25 | -0,91 | 3,64E-01 | 5,42E-01 |
| AT2G37160 | 3,77 | -0,77 | 0,85 | -0,91 | 3,65E-01 | 5,42E-01 |
| AT4G09000 | 1,74 | -1,12 | 1,24 | -0,90 | 3,66E-01 | 5,43E-01 |
| AT3G52930 | 3,81 | 0,85 | 0,95 | 0,90 | 3,66E-01 | 5,43E-01 |
| AT4G34980 | 4,56 | -0,70 | 0,77 | -0,90 | 3,67E-01 | 5,44E-01 |
| AT1G01550 | 2,54 | -1,04 | 1,16 | -0,90 | 3,68E-01 | 5,45E-01 |
| AT3G56940 | 6,51 | 0,71 | 0,78 | 0,90 | 3,68E-01 | 5,45E-01 |
| AT5G41790 | 2,54 | 1,17 | 1,31 | 0,90 | 3,68E-01 | 5,45E-01 |
| AT5G00765 | 3,11 | 0,88 | 0,98 | 0,90 | 3,69E-01 | 5,45E-01 |
| AT5G09900 | 1,74 | 1,30 | 1,44 | 0,90 | 3,69E-01 | 5,45E-01 |
| AT2G07718 | 3,36 | -0,90 | 1,00 | -0,90 | 3,70E-01 | 5,47E-01 |
| AT4G37930 | 4,20 | 0,75 | 0,84 | 0,89 | 3,71E-01 | 5,47E-01 |

| | | | | | | |
|---|---|---|---|---|---|---|
| AT4G24413 | 2,10 | -1,12 | 1,26 | -0,89 | 3,71E-01 | 5,47E-01 |
| AT2G30860 | 6,26 | -0,67 | 0,75 | -0,89 | 3,72E-01 | 5,47E-01 |
| AT5G47430 | 3,26 | -0,86 | 0,96 | -0,89 | 3,72E-01 | 5,47E-01 |
| AT4G17340 | 2,05 | 1,23 | 1,38 | 0,89 | 3,73E-01 | 5,48E-01 |
| AT3G08170 | 3,83 | 0,81 | 0,91 | 0,89 | 3,73E-01 | 5,48E-01 |
| AT3G51800 | 4,28 | -0,76 | 0,86 | -0,89 | 3,73E-01 | 5,48E-01 |
| AT5G04105 | 18,20 | 0,44 | 0,50 | 0,89 | 3,73E-01 | 5,48E-01 |
| AT5G40450 | 4,44 | 0,80 | 0,90 | 0,89 | 3,73E-01 | 5,48E-01 |
| AT3G22120 | 4,08 | -0,90 | 1,01 | -0,89 | 3,74E-01 | 5,48E-01 |
| AT5G35630 | 4,18 | -0,92 | 1,04 | -0,89 | 3,75E-01 | 5,50E-01 |
| AT1G65930 | 2,39 | 1,17 | 1,32 | 0,89 | 3,76E-01 | 5,51E-01 |
| AT1G04183 | 2,59 | -0,99 | 1,12 | -0,88 | 3,76E-01 | 5,51E-01 |
| AT3G57645 | 1,71 | 1,35 | 1,53 | 0,88 | 3,77E-01 | 5,51E-01 |
| AT5G13850 | 2,26 | 1,07 | 1,22 | 0,88 | 3,78E-01 | 5,52E-01 |
| AT2G22500 | 4,35 | -0,83 | 0,95 | -0,88 | 3,79E-01 | 5,54E-01 |
| AT5G57560 | 1,92 | -1,07 | 1,22 | -0,88 | 3,80E-01 | 5,54E-01 |
| AT1G42550 | 2,32 | 1,02 | 1,16 | 0,87 | 3,82E-01 | 5,56E-01 |
| AT4G39330 | 1,70 | -1,16 | 1,33 | -0,88 | 3,81E-01 | 5,56E-01 |
| AT1G05030 | 1,78 | -1,31 | 1,50 | -0,88 | 3,81E-01 | 5,56E-01 |
| AT4G19410 | 1,78 | 1,20 | 1,37 | 0,87 | 3,82E-01 | 5,56E-01 |
| AT4G21640 | 1,99 | -1,32 | 1,51 | -0,88 | 3,81E-01 | 5,56E-01 |
| AT2G39681 | 1805,59 | -0,28 | 0,32 | -0,87 | 3,84E-01 | 5,58E-01 |
| AT3G59765 | 1,81 | 1,42 | 1,63 | 0,87 | 3,84E-01 | 5,58E-01 |
| AT3G59068 | 1,78 | -1,32 | 1,52 | -0,87 | 3,85E-01 | 5,59E-01 |
| AT3G59070 | 1,78 | -1,32 | 1,52 | -0,87 | 3,85E-01 | 5,59E-01 |
| AT2G04375 | 5,42 | 0,67 | 0,77 | 0,87 | 3,85E-01 | 5,59E-01 |
| AT3G60190 | 1,87 | 1,12 | 1,29 | 0,87 | 3,86E-01 | 5,60E-01 |
| AT1G30360 | 3,38 | 0,73 | 0,85 | 0,86 | 3,89E-01 | 5,63E-01 |
| AT5G54770 | 28,55 | -0,41 | 0,48 | -0,86 | 3,90E-01 | 5,64E-01 |
| AT3G28925 | 2,17 | -1,04 | 1,21 | -0,86 | 3,90E-01 | 5,65E-01 |
| AT3G09445 | 4,99 | -0,74 | 0,86 | -0,86 | 3,91E-01 | 5,65E-01 |
| AT4G12780 | 15,31 | -0,41 | 0,48 | -0,86 | 3,91E-01 | 5,65E-01 |
| AT5G25610 | 6,58 | -0,69 | 0,80 | -0,86 | 3,91E-01 | 5,65E-01 |
| AT4G14960 | 3,57 | 0,81 | 0,95 | 0,86 | 3,92E-01 | 5,65E-01 |
| AT5G26650 | 7,57 | -0,65 | 0,75 | -0,86 | 3,92E-01 | 5,65E-01 |
| AT4G02350 | 1,95 | 1,48 | 1,73 | 0,86 | 3,92E-01 | 5,66E-01 |
| AT3G61198 | 7,65 | -0,58 | 0,68 | -0,85 | 3,93E-01 | 5,66E-01 |
| AT4G05900 | 1,70 | -1,24 | 1,45 | -0,85 | 3,94E-01 | 5,67E-01 |
| AT4G27500 | 2,93 | 0,91 | 1,07 | 0,85 | 3,94E-01 | 5,67E-01 |
| AT3G06835 | 2,07 | -1,56 | 1,82 | -0,85 | 3,94E-01 | 5,67E-01 |
| AT4G07855 | 2,93 | 0,91 | 1,07 | 0,85 | 3,94E-01 | 5,67E-01 |
| AT3G22121 | 3,96 | -0,87 | 1,03 | -0,85 | 3,96E-01 | 5,68E-01 |
| AT2G03821 | 1,77 | -1,05 | 1,23 | -0,85 | 3,96E-01 | 5,68E-01 |
| AT4G10340 | 23,35 | 0,44 | 0,52 | 0,85 | 3,98E-01 | 5,71E-01 |
| AT3G16640 | 7,07 | 0,73 | 0,87 | 0,85 | 3,98E-01 | 5,71E-01 |
| AT5G56970 | 2,67 | -0,91 | 1,08 | -0,84 | 4,00E-01 | 5,73E-01 |
| AT3G05220 | 1,84 | -1,27 | 1,50 | -0,84 | 4,00E-01 | 5,73E-01 |
| AT1G26850 | 2,19 | 1,19 | 1,42 | 0,84 | 4,01E-01 | 5,73E-01 |
| AT5G61310 | 4,76 | -0,68 | 0,81 | -0,84 | 4,01E-01 | 5,74E-01 |
| AT5G17230 | 2,13 | 1,06 | 1,26 | 0,84 | 4,02E-01 | 5,74E-01 |
| AT1G76065 | 4,52 | -0,85 | 1,01 | -0,84 | 4,02E-01 | 5,75E-01 |
| AT5G45470 | 2,49 | 1,32 | 1,57 | 0,84 | 4,03E-01 | 5,75E-01 |
| AT2G47400 | 11,67 | 0,44 | 0,52 | 0,84 | 4,03E-01 | 5,75E-01 |
| AT5G07725 | 8,72 | -0,60 | 0,73 | -0,83 | 4,07E-01 | 5,80E-01 |
| AT5G11740 | 4,10 | 0,66 | 0,79 | 0,83 | 4,08E-01 | 5,81E-01 |
| AT1G50920 | 2,95 | 0,90 | 1,08 | 0,83 | 4,09E-01 | 5,81E-01 |

| | | | | | | |
|---|---|---|---|---|---|---|
| AT1G05010 | 2,06 | -1,30 | 1,57 | -0,83 | 4,08E-01 | 5,81E-01 |
| AT5G40890 | 1,74 | 1,19 | 1,44 | 0,83 | 4,09E-01 | 5,81E-01 |
| AT5G07740 | 1,77 | 1,04 | 1,26 | 0,82 | 4,10E-01 | 5,82E-01 |
| AT4G08593 | 1,89 | -1,24 | 1,51 | -0,82 | 4,12E-01 | 5,85E-01 |
| AT2G39940 | 1,84 | 1,53 | 1,87 | 0,82 | 4,13E-01 | 5,86E-01 |
| AT4G39364 | 2,19 | -1,13 | 1,38 | -0,82 | 4,13E-01 | 5,86E-01 |
| AT1G73170 | 1,76 | 1,35 | 1,65 | 0,82 | 4,13E-01 | 5,86E-01 |
| AT3G02540 | 1,81 | 1,20 | 1,47 | 0,82 | 4,14E-01 | 5,86E-01 |
| AT3G48350 | 1,84 | 1,60 | 1,96 | 0,81 | 4,15E-01 | 5,87E-01 |
| AT4G08630 | 1,97 | -1,11 | 1,36 | -0,82 | 4,15E-01 | 5,87E-01 |
| AT3G09162 | 2,15 | -1,46 | 1,79 | -0,81 | 4,16E-01 | 5,88E-01 |
| AT3G24929 | 2,38 | -0,97 | 1,19 | -0,81 | 4,17E-01 | 5,89E-01 |
| AT2G04885 | 2,17 | -1,00 | 1,23 | -0,81 | 4,18E-01 | 5,89E-01 |
| AT4G39850 | 2,19 | 1,12 | 1,38 | 0,81 | 4,17E-01 | 5,89E-01 |
| AT5G02565 | 2,69 | -1,01 | 1,25 | -0,81 | 4,19E-01 | 5,90E-01 |
| AT5G19940 | 2,37 | -1,06 | 1,31 | -0,81 | 4,19E-01 | 5,90E-01 |
| AT5G45100 | 1,72 | 1,45 | 1,79 | 0,81 | 4,18E-01 | 5,90E-01 |
| AT1G07140 | 1,81 | 1,24 | 1,53 | 0,81 | 4,19E-01 | 5,90E-01 |
| AT4G05050 | 3,49 | -0,76 | 0,94 | -0,81 | 4,19E-01 | 5,90E-01 |
| AT5G09515 | 9,09 | -0,55 | 0,69 | -0,81 | 4,20E-01 | 5,91E-01 |
| AT3G44860 | 2,37 | 0,93 | 1,16 | 0,80 | 4,21E-01 | 5,91E-01 |
| AT4G13918 | 2,19 | 0,96 | 1,20 | 0,81 | 4,21E-01 | 5,91E-01 |
| AT1G73870 | 2,16 | 1,09 | 1,36 | 0,80 | 4,22E-01 | 5,93E-01 |
| AT5G02810 | 1,73 | 1,36 | 1,70 | 0,80 | 4,22E-01 | 5,93E-01 |
| AT3G41979 | 6596,11 | -0,29 | 0,36 | -0,80 | 4,24E-01 | 5,94E-01 |
| AT3G04765 | 2,83 | -0,84 | 1,06 | -0,80 | 4,24E-01 | 5,94E-01 |
| AT3G06355 | 6711,54 | -0,28 | 0,36 | -0,80 | 4,26E-01 | 5,96E-01 |
| AT1G03130 | 1,99 | -0,96 | 1,21 | -0,79 | 4,27E-01 | 5,97E-01 |
| AT2G30570 | 3,97 | -0,71 | 0,89 | -0,80 | 4,26E-01 | 5,97E-01 |
| AT4G12650 | 2,23 | 0,91 | 1,15 | 0,80 | 4,27E-01 | 5,97E-01 |
| AT4G39400 | 3,54 | 0,74 | 0,93 | 0,79 | 4,28E-01 | 5,97E-01 |
| AT5G08180 | 2,30 | -0,80 | 1,01 | -0,79 | 4,28E-01 | 5,97E-01 |
| AT3G14840 | 1,81 | 1,07 | 1,35 | 0,79 | 4,29E-01 | 5,97E-01 |
| AT3G54050 | 3,78 | -0,87 | 1,10 | -0,79 | 4,28E-01 | 5,97E-01 |
| AT4G14160 | 2,71 | 1,03 | 1,31 | 0,79 | 4,28E-01 | 5,97E-01 |
| AT1G50250 | 2,35 | 0,85 | 1,07 | 0,79 | 4,29E-01 | 5,97E-01 |
| AT1G06950 | 2,18 | 1,48 | 1,88 | 0,79 | 4,30E-01 | 5,99E-01 |
| AT5G42270 | 4,20 | 0,71 | 0,90 | 0,79 | 4,31E-01 | 5,99E-01 |
| AT2G09465 | 4,22 | -0,68 | 0,86 | -0,79 | 4,31E-01 | 6,00E-01 |
| AT5G04415 | 4,71 | 0,59 | 0,75 | 0,78 | 4,33E-01 | 6,01E-01 |
| AT3G15210 | 1,70 | -0,97 | 1,24 | -0,78 | 4,33E-01 | 6,01E-01 |
| AT1G04480 | 2,74 | 0,91 | 1,17 | 0,78 | 4,35E-01 | 6,03E-01 |
| AT3G54890 | 13,73 | -0,50 | 0,64 | -0,78 | 4,35E-01 | 6,03E-01 |
| AT1G13110 | 2,32 | -0,93 | 1,19 | -0,78 | 4,37E-01 | 6,05E-01 |
| AT3G48057 | 47,97 | 0,42 | 0,53 | 0,78 | 4,38E-01 | 6,06E-01 |
| AT4G31980 | 2,53 | -1,00 | 1,29 | -0,78 | 4,38E-01 | 6,06E-01 |
| AT1G35720 | 3,50 | 0,73 | 0,94 | 0,77 | 4,39E-01 | 6,07E-01 |
| AT4G33240 | 2,23 | 1,10 | 1,42 | 0,77 | 4,39E-01 | 6,07E-01 |
| AT1G51805 | 2,63 | 0,87 | 1,13 | 0,77 | 4,40E-01 | 6,09E-01 |
| AT1G67450 | 1,86 | -0,92 | 1,19 | -0,77 | 4,41E-01 | 6,09E-01 |
| AT2G47470 | 1,77 | 1,20 | 1,56 | 0,77 | 4,42E-01 | 6,10E-01 |
| AT4G37260 | 2,49 | 0,87 | 1,13 | 0,77 | 4,43E-01 | 6,11E-01 |
| AT5G14930 | 1,80 | 1,62 | 2,11 | 0,77 | 4,43E-01 | 6,11E-01 |
| AT5G64940 | 2,50 | 1,09 | 1,42 | 0,76 | 4,45E-01 | 6,13E-01 |
| AT1G07637 | 2,62 | -0,82 | 1,08 | -0,76 | 4,46E-01 | 6,14E-01 |
| AT2G06420 | 4,36 | -0,71 | 0,94 | -0,76 | 4,46E-01 | 6,14E-01 |

| | | | | | | |
|---|---|---|---|---|---|---|
| AT1G53310 | 3,45 | 0,72 | 0,95 | 0,76 | 4,46E-01 | 6,14E-01 |
| AT2G02560 | 1,80 | 1,06 | 1,39 | 0,76 | 4,48E-01 | 6,15E-01 |
| AT2G34430 | 28,33 | 0,34 | 0,45 | 0,76 | 4,49E-01 | 6,16E-01 |
| AT4G21280 | 3,20 | -0,90 | 1,19 | -0,76 | 4,48E-01 | 6,16E-01 |
| AT4G35750 | 1,89 | 0,91 | 1,20 | 0,76 | 4,49E-01 | 6,16E-01 |
| AT5G67300 | 3,74 | 0,66 | 0,87 | 0,76 | 4,49E-01 | 6,16E-01 |
| AT4G35090 | 2,83 | -0,79 | 1,05 | -0,76 | 4,50E-01 | 6,16E-01 |
| AT5G01410 | 2,12 | 0,99 | 1,31 | 0,75 | 4,50E-01 | 6,17E-01 |
| AT3G15920 | 8,54 | 0,45 | 0,60 | 0,75 | 4,52E-01 | 6,19E-01 |
| AT3G12630 | 8,51 | 0,43 | 0,58 | 0,75 | 4,55E-01 | 6,22E-01 |
| AT5G67420 | 2,71 | -0,85 | 1,13 | -0,75 | 4,55E-01 | 6,22E-01 |
| AT5G13840 | 2,03 | 0,97 | 1,30 | 0,75 | 4,56E-01 | 6,23E-01 |
| AT5G19690 | 1,74 | 0,90 | 1,20 | 0,75 | 4,56E-01 | 6,23E-01 |
| AT1G07593 | 3,78 | 1,11 | 1,50 | 0,74 | 4,57E-01 | 6,23E-01 |
| AT5G45110 | 1,72 | 0,92 | 1,24 | 0,74 | 4,58E-01 | 6,25E-01 |
| AT5G60408 | 515,37 | 0,21 | 0,29 | 0,74 | 4,60E-01 | 6,26E-01 |
| AT4G11900 | 2,31 | -0,78 | 1,06 | -0,74 | 4,60E-01 | 6,26E-01 |
| AT5G60390 | 16,82 | 0,38 | 0,51 | 0,74 | 4,61E-01 | 6,27E-01 |
| AT2G27860 | 4,64 | 0,62 | 0,84 | 0,74 | 4,62E-01 | 6,28E-01 |
| AT1G22530 | 2,83 | 1,33 | 1,82 | 0,73 | 4,63E-01 | 6,30E-01 |
| AT1G44446 | 1,72 | 1,05 | 1,44 | 0,73 | 4,64E-01 | 6,30E-01 |
| AT4G29090 | 2,85 | -0,83 | 1,13 | -0,73 | 4,64E-01 | 6,30E-01 |
| AT5G10860 | 1,76 | 1,06 | 1,44 | 0,73 | 4,64E-01 | 6,30E-01 |
| AT1G61140 | 2,12 | 0,85 | 1,16 | 0,73 | 4,65E-01 | 6,31E-01 |
| AT1G78020 | 1,89 | -0,88 | 1,20 | -0,73 | 4,66E-01 | 6,32E-01 |
| AT3G08370 | 6,58 | -0,58 | 0,80 | -0,73 | 4,67E-01 | 6,33E-01 |
| AT5G15650 | 3,70 | 0,73 | 1,01 | 0,73 | 4,68E-01 | 6,34E-01 |
| AT4G26095 | 1,81 | 0,93 | 1,28 | 0,73 | 4,68E-01 | 6,34E-01 |
| AT5G44270 | 2,68 | -0,76 | 1,05 | -0,72 | 4,69E-01 | 6,34E-01 |
| AT2G04060 | 1,72 | -1,32 | 1,83 | -0,72 | 4,70E-01 | 6,35E-01 |
| AT5G65860 | 5,04 | 0,60 | 0,84 | 0,72 | 4,73E-01 | 6,39E-01 |
| AT4G22690 | 3,37 | -0,66 | 0,93 | -0,72 | 4,74E-01 | 6,40E-01 |
| AT4G02760 | 2,62 | -0,90 | 1,25 | -0,72 | 4,74E-01 | 6,40E-01 |
| AT5G66190 | 3,89 | -0,78 | 1,09 | -0,72 | 4,74E-01 | 6,40E-01 |
| AT4G39363 | 6,83 | -0,51 | 0,72 | -0,71 | 4,75E-01 | 6,41E-01 |
| AT4G34390 | 1,80 | 0,99 | 1,39 | 0,71 | 4,77E-01 | 6,43E-01 |
| AT2G07629 | 5,41 | -0,67 | 0,95 | -0,71 | 4,78E-01 | 6,43E-01 |
| AT3G62010 | 2,16 | 0,92 | 1,30 | 0,71 | 4,78E-01 | 6,43E-01 |
| AT3G29190 | 2,69 | 0,83 | 1,16 | 0,71 | 4,78E-01 | 6,43E-01 |
| AT4G29780 | 2,35 | -1,05 | 1,48 | -0,71 | 4,79E-01 | 6,44E-01 |
| AT3G11940 | 3,79 | 0,72 | 1,01 | 0,71 | 4,79E-01 | 6,44E-01 |
| AT4G32250 | 2,07 | 0,95 | 1,34 | 0,71 | 4,80E-01 | 6,44E-01 |
| AT4G15470 | 1,75 | -0,98 | 1,39 | -0,71 | 4,81E-01 | 6,45E-01 |
| AT2G20722 | 3,24 | -0,73 | 1,04 | -0,70 | 4,81E-01 | 6,45E-01 |
| AT2G42590 | 2,06 | 0,87 | 1,24 | 0,70 | 4,81E-01 | 6,45E-01 |
| AT3G23110 | 1,95 | 1,06 | 1,50 | 0,70 | 4,81E-01 | 6,45E-01 |
| AT2G41430 | 6,85 | -0,52 | 0,75 | -0,70 | 4,82E-01 | 6,45E-01 |
| AT4G28570 | 3,15 | -0,75 | 1,06 | -0,70 | 4,82E-01 | 6,46E-01 |
| AT3G22440 | 1,87 | 0,81 | 1,15 | 0,70 | 4,83E-01 | 6,46E-01 |
| AT2G41105 | 5,08 | 0,58 | 0,84 | 0,70 | 4,85E-01 | 6,47E-01 |
| AT5G06630 | 1,85 | -0,97 | 1,39 | -0,70 | 4,84E-01 | 6,47E-01 |
| AT1G26880 | 2,18 | -1,07 | 1,54 | -0,70 | 4,86E-01 | 6,48E-01 |
| AT2G29670 | 3,81 | 0,71 | 1,02 | 0,70 | 4,86E-01 | 6,48E-01 |
| AT3G24982 | 2,19 | -0,85 | 1,21 | -0,70 | 4,86E-01 | 6,48E-01 |
| AT4G20830 | 3,02 | 0,64 | 0,92 | 0,70 | 4,86E-01 | 6,48E-01 |
| AT4G04925 | 1,79 | -0,97 | 1,39 | -0,69 | 4,87E-01 | 6,49E-01 |

| | | | | | | |
|---|---|---|---|---|---|---|
| AT1G11910 | 5,12 | 0,67 | 0,97 | 0,69 | 4,88E-01 | 6,50E-01 |
| AT1G76062 | 4,01 | -0,76 | 1,10 | -0,69 | 4,88E-01 | 6,50E-01 |
| AT5G60600 | 2,43 | 0,68 | 0,98 | 0,69 | 4,89E-01 | 6,50E-01 |
| AT5G10840 | 2,05 | 0,95 | 1,38 | 0,69 | 4,90E-01 | 6,50E-01 |
| AT5G28840 | 2,50 | 0,71 | 1,03 | 0,69 | 4,90E-01 | 6,50E-01 |
| AT5G19150 | 6,35 | -0,46 | 0,66 | -0,69 | 4,91E-01 | 6,52E-01 |
| AT5G15970 | 2,85 | -1,07 | 1,55 | -0,69 | 4,92E-01 | 6,52E-01 |
| AT1G72600 | 18,04 | 0,31 | 0,45 | 0,69 | 4,92E-01 | 6,53E-01 |
| AT3G14420 | 4,42 | -0,66 | 0,97 | -0,68 | 4,94E-01 | 6,54E-01 |
| AT5G39570 | 4,04 | -0,69 | 1,00 | -0,68 | 4,94E-01 | 6,54E-01 |
| AT3G03425 | 4,95 | -0,49 | 0,72 | -0,68 | 4,94E-01 | 6,54E-01 |
| AT4G30440 | 3,67 | -0,68 | 0,99 | -0,68 | 4,94E-01 | 6,54E-01 |
| AT2G23810 | 3,17 | 0,74 | 1,09 | 0,68 | 4,95E-01 | 6,54E-01 |
| AT1G43850 | 1,82 | 0,95 | 1,39 | 0,68 | 4,95E-01 | 6,54E-01 |
| AT1G69940 | 3,40 | -0,61 | 0,90 | -0,68 | 4,98E-01 | 6,58E-01 |
| AT5G20730 | 2,22 | 0,80 | 1,19 | 0,67 | 5,00E-01 | 6,60E-01 |
| AT2G06510 | 1,80 | 0,86 | 1,27 | 0,67 | 5,00E-01 | 6,60E-01 |
| AT5G59732 | 3,71 | -0,71 | 1,07 | -0,67 | 5,03E-01 | 6,62E-01 |
| AT2G40000 | 5,19 | -0,50 | 0,75 | -0,67 | 5,03E-01 | 6,62E-01 |
| AT3G11670 | 1,77 | 1,13 | 1,68 | 0,67 | 5,03E-01 | 6,62E-01 |
| AT4G16900 | 2,10 | 0,81 | 1,20 | 0,67 | 5,03E-01 | 6,62E-01 |
| AT5G59730 | 3,71 | -0,71 | 1,07 | -0,67 | 5,03E-01 | 6,62E-01 |
| AT1G80440 | 3,83 | -0,72 | 1,08 | -0,67 | 5,04E-01 | 6,62E-01 |
| AT4G21060 | 1,98 | -0,87 | 1,30 | -0,67 | 5,04E-01 | 6,62E-01 |
| AT5G46845 | 66,07 | 0,19 | 0,29 | 0,67 | 5,04E-01 | 6,62E-01 |
| AT4G23170 | 6,38 | -0,56 | 0,85 | -0,66 | 5,07E-01 | 6,65E-01 |
| AT1G54040 | 2,21 | -0,81 | 1,24 | -0,66 | 5,11E-01 | 6,70E-01 |
| AT1G78570 | 2,32 | 0,79 | 1,20 | 0,66 | 5,11E-01 | 6,70E-01 |
| AT3G23100 | 2,14 | 0,91 | 1,39 | 0,66 | 5,12E-01 | 6,70E-01 |
| AT2G41310 | 487,34 | -0,18 | 0,28 | -0,65 | 5,15E-01 | 6,74E-01 |
| AT1G76090 | 1,72 | -0,94 | 1,46 | -0,65 | 5,19E-01 | 6,79E-01 |
| AT2G28840 | 1,76 | 0,99 | 1,55 | 0,64 | 5,20E-01 | 6,80E-01 |
| AT2G09250 | 487,12 | -0,18 | 0,28 | -0,64 | 5,21E-01 | 6,80E-01 |
| AT3G08510 | 2,33 | 0,97 | 1,51 | 0,64 | 5,21E-01 | 6,80E-01 |
| AT4G13920 | 2,03 | 0,85 | 1,33 | 0,64 | 5,22E-01 | 6,81E-01 |
| AT5G56070 | 3,26 | -0,80 | 1,25 | -0,64 | 5,22E-01 | 6,81E-01 |
| AT1G29910 | 27,17 | 0,33 | 0,51 | 0,64 | 5,22E-01 | 6,81E-01 |
| AT4G03560 | 2,24 | 0,79 | 1,24 | 0,64 | 5,23E-01 | 6,81E-01 |
| AT2G35110 | 1,99 | 0,90 | 1,41 | 0,64 | 5,24E-01 | 6,81E-01 |
| AT4G04365 | 93,18 | -0,27 | 0,43 | -0,64 | 5,24E-01 | 6,81E-01 |
| AT4G13810 | 2,84 | 0,79 | 1,23 | 0,64 | 5,24E-01 | 6,81E-01 |
| AT5G28920 | 2,84 | 0,67 | 1,06 | 0,64 | 5,24E-01 | 6,82E-01 |
| AT2G26975 | 1,93 | -0,77 | 1,20 | -0,64 | 5,25E-01 | 6,82E-01 |
| AT4G03039 | 93,29 | -0,27 | 0,43 | -0,63 | 5,26E-01 | 6,82E-01 |
| AT4G03380 | 7,25 | -0,42 | 0,66 | -0,63 | 5,26E-01 | 6,82E-01 |
| AT4G34150 | 1,90 | 0,77 | 1,22 | 0,63 | 5,28E-01 | 6,84E-01 |
| AT2G14610 | 3,13 | 0,59 | 0,93 | 0,63 | 5,27E-01 | 6,84E-01 |
| AT5G28040 | 2,94 | 0,71 | 1,12 | 0,63 | 5,27E-01 | 6,84E-01 |
| AT4G03038 | 93,33 | -0,27 | 0,43 | -0,63 | 5,28E-01 | 6,84E-01 |
| AT4G06320 | 7,79 | -0,41 | 0,65 | -0,63 | 5,29E-01 | 6,85E-01 |
| AT2G05520 | 18,30 | -0,24 | 0,39 | -0,63 | 5,30E-01 | 6,85E-01 |
| AT2G25520 | 1,73 | 1,08 | 1,72 | 0,63 | 5,30E-01 | 6,85E-01 |
| AT1G31835 | 2,35 | -0,73 | 1,17 | -0,63 | 5,32E-01 | 6,87E-01 |
| AT4G25100 | 2,11 | 0,79 | 1,26 | 0,63 | 5,32E-01 | 6,87E-01 |
| AT1G57720 | 1,72 | 0,92 | 1,47 | 0,62 | 5,32E-01 | 6,87E-01 |
| AT3G45140 | 15,78 | 0,31 | 0,50 | 0,62 | 5,33E-01 | 6,87E-01 |

| | | | | | | |
|---|---|---|---|---|---|---|
| AT2G36460 | 1,86 | 0,90 | 1,45 | 0,62 | 5,33E-01 | 6,87E-01 |
| AT4G39800 | 2,68 | -0,75 | 1,21 | -0,62 | 5,37E-01 | 6,92E-01 |
| AT1G22930 | 1,70 | 0,93 | 1,51 | 0,62 | 5,37E-01 | 6,92E-01 |
| AT3G10985 | 6,51 | -0,41 | 0,66 | -0,61 | 5,39E-01 | 6,94E-01 |
| AT2G21660 | 24,65 | 0,29 | 0,48 | 0,61 | 5,39E-01 | 6,94E-01 |
| AT4G14365 | 1,73 | 0,89 | 1,45 | 0,61 | 5,40E-01 | 6,94E-01 |
| AT5G18380 | 4,02 | -0,57 | 0,93 | -0,61 | 5,40E-01 | 6,94E-01 |
| AT5G01155 | 4,02 | -0,56 | 0,91 | -0,61 | 5,40E-01 | 6,94E-01 |
| AT5G09495 | 4,86 | 0,52 | 0,85 | 0,61 | 5,41E-01 | 6,94E-01 |
| AT1G05805 | 1,94 | -1,22 | 2,00 | -0,61 | 5,42E-01 | 6,95E-01 |
| AT1G43320 | 2,75 | 0,86 | 1,40 | 0,61 | 5,42E-01 | 6,95E-01 |
| AT2G47450 | 6,02 | 0,45 | 0,75 | 0,61 | 5,43E-01 | 6,95E-01 |
| AT5G49730 | 3,55 | 0,69 | 1,14 | 0,61 | 5,43E-01 | 6,95E-01 |
| AT1G63740 | 1,93 | -0,73 | 1,20 | -0,61 | 5,44E-01 | 6,97E-01 |
| AT3G46600 | 2,19 | 0,65 | 1,08 | 0,61 | 5,45E-01 | 6,97E-01 |
| AT5G47690 | 3,16 | 0,69 | 1,13 | 0,61 | 5,44E-01 | 6,97E-01 |
| AT1G72710 | 1,73 | 0,83 | 1,37 | 0,60 | 5,45E-01 | 6,97E-01 |
| AT2G20950 | 3,17 | -0,64 | 1,05 | -0,60 | 5,46E-01 | 6,98E-01 |
| AT4G20835 | 2,66 | 0,58 | 0,96 | 0,60 | 5,49E-01 | 7,01E-01 |
| AT2G44570 | 2,88 | -0,70 | 1,17 | -0,60 | 5,50E-01 | 7,01E-01 |
| AT5G01350 | 8,44 | 0,37 | 0,62 | 0,60 | 5,50E-01 | 7,02E-01 |
| AT3G01316 | 2,03 | -0,77 | 1,30 | -0,60 | 5,52E-01 | 7,03E-01 |
| AT5G49440 | 5,06 | -0,54 | 0,92 | -0,59 | 5,53E-01 | 7,04E-01 |
| AT2G39770 | 2,97 | 0,62 | 1,06 | 0,59 | 5,55E-01 | 7,06E-01 |
| AT3G44720 | 3,49 | 0,63 | 1,06 | 0,59 | 5,55E-01 | 7,06E-01 |
| AT4G29900 | 4,72 | 0,44 | 0,74 | 0,59 | 5,55E-01 | 7,06E-01 |
| AT1G72610 | 17,51 | 0,27 | 0,46 | 0,59 | 5,55E-01 | 7,06E-01 |
| AT2G15420 | 2,17 | -0,71 | 1,20 | -0,59 | 5,56E-01 | 7,07E-01 |
| AT3G52700 | 2,20 | -0,63 | 1,06 | -0,59 | 5,57E-01 | 7,07E-01 |
| AT4G31170 | 2,81 | 0,80 | 1,36 | 0,59 | 5,57E-01 | 7,07E-01 |
| AT3G26000 | 2,02 | 0,81 | 1,38 | 0,58 | 5,59E-01 | 7,09E-01 |
| AT5G20230 | 2,06 | 0,77 | 1,33 | 0,58 | 5,61E-01 | 7,11E-01 |
| AT4G18150 | 2,31 | -0,76 | 1,31 | -0,58 | 5,63E-01 | 7,12E-01 |
| AT4G13840 | 1,76 | -0,91 | 1,56 | -0,58 | 5,62E-01 | 7,12E-01 |
| AT5G04853 | 3,91 | -0,51 | 0,87 | -0,58 | 5,62E-01 | 7,12E-01 |
| AT5G03145 | 2,30 | 0,75 | 1,31 | 0,58 | 5,65E-01 | 7,14E-01 |
| AT1G33590 | 1,70 | -0,72 | 1,25 | -0,57 | 5,66E-01 | 7,16E-01 |
| AT2G36530 | 5,68 | 0,44 | 0,77 | 0,57 | 5,67E-01 | 7,17E-01 |
| AT5G25060 | 1,92 | 0,70 | 1,22 | 0,57 | 5,68E-01 | 7,17E-01 |
| AT4G05150 | 3,81 | 0,61 | 1,07 | 0,57 | 5,68E-01 | 7,17E-01 |
| AT3G09630 | 1,89 | 0,81 | 1,43 | 0,57 | 5,72E-01 | 7,20E-01 |
| AT3G16830 | 1,75 | 0,80 | 1,42 | 0,56 | 5,72E-01 | 7,20E-01 |
| AT3G61470 | 27,14 | -0,27 | 0,48 | -0,57 | 5,72E-01 | 7,20E-01 |
| AT3G52880 | 2,23 | 0,61 | 1,09 | 0,56 | 5,72E-01 | 7,20E-01 |
| AT5G12250 | 2,28 | 0,63 | 1,11 | 0,57 | 5,72E-01 | 7,20E-01 |
| AT4G01840 | 3,87 | -0,59 | 1,05 | -0,56 | 5,74E-01 | 7,22E-01 |
| AT3G16785 | 1,89 | 0,62 | 1,11 | 0,56 | 5,75E-01 | 7,23E-01 |
| AT3G17390 | 4,31 | -0,48 | 0,87 | -0,56 | 5,78E-01 | 7,26E-01 |
| AT2G44310 | 1,85 | -0,63 | 1,14 | -0,56 | 5,79E-01 | 7,27E-01 |
| AT5G62190 | 2,05 | 0,86 | 1,55 | 0,55 | 5,79E-01 | 7,27E-01 |
| AT3G19030 | 2,04 | 0,66 | 1,19 | 0,55 | 5,80E-01 | 7,27E-01 |
| AT5G41750 | 8,05 | 0,33 | 0,60 | 0,55 | 5,80E-01 | 7,27E-01 |
| AT1G27870 | 1,97 | -0,76 | 1,37 | -0,55 | 5,80E-01 | 7,27E-01 |
| AT1G75040 | 3,07 | 0,55 | 1,00 | 0,55 | 5,81E-01 | 7,28E-01 |
| AT3G60770 | 1,96 | 0,77 | 1,40 | 0,55 | 5,82E-01 | 7,28E-01 |
| AT5G02380 | 2,77 | 0,72 | 1,30 | 0,55 | 5,82E-01 | 7,28E-01 |

| | | | | | | |
|---|---|---|---|---|---|---|
| AT4G26542 | 2,99 | 0,55 | 1,00 | 0,55 | 5,83E-01 | 7,28E-01 |
| AT4G30760 | 1,76 | 0,70 | 1,28 | 0,55 | 5,82E-01 | 7,28E-01 |
| AT5G46430 | 2,53 | -0,66 | 1,21 | -0,55 | 5,85E-01 | 7,30E-01 |
| AT2G16600 | 11,35 | 0,33 | 0,60 | 0,55 | 5,85E-01 | 7,31E-01 |
| AT1G67840 | 1,96 | 0,74 | 1,36 | 0,54 | 5,87E-01 | 7,33E-01 |
| AT3G04155 | 3,42 | -0,53 | 0,97 | -0,54 | 5,88E-01 | 7,33E-01 |
| AT4G27600 | 12,18 | 0,32 | 0,59 | 0,54 | 5,89E-01 | 7,34E-01 |
| AT3G16370 | 1,93 | -0,74 | 1,38 | -0,54 | 5,91E-01 | 7,36E-01 |
| AT1G55490 | 3,72 | -0,60 | 1,12 | -0,54 | 5,92E-01 | 7,37E-01 |
| AT1G29418 | 9,45 | -0,28 | 0,53 | -0,53 | 5,94E-01 | 7,39E-01 |
| AT5G28919 | 2,64 | 0,56 | 1,06 | 0,53 | 5,97E-01 | 7,41E-01 |
| AT5G48385 | 2,37 | 0,76 | 1,43 | 0,53 | 5,96E-01 | 7,41E-01 |
| AT5G04905 | 2,64 | 0,56 | 1,06 | 0,53 | 5,97E-01 | 7,41E-01 |
| AT5G38540 | 8,36 | -0,65 | 1,24 | -0,53 | 5,96E-01 | 7,41E-01 |
| AT2G24270 | 3,15 | 0,55 | 1,04 | 0,53 | 5,97E-01 | 7,41E-01 |
| AT1G09240 | 1,84 | 1,03 | 1,96 | 0,53 | 5,98E-01 | 7,42E-01 |
| AT1G07930 | 3,92 | -0,51 | 0,96 | -0,53 | 5,99E-01 | 7,42E-01 |
| AT5G28560 | 3,51 | -0,47 | 0,90 | -0,52 | 6,00E-01 | 7,43E-01 |
| AT1G23190 | 1,73 | 0,92 | 1,76 | 0,52 | 6,02E-01 | 7,45E-01 |
| AT3G59970 | 1,71 | 0,63 | 1,21 | 0,52 | 6,03E-01 | 7,46E-01 |
| AT3G57480 | 3,07 | 0,73 | 1,40 | 0,52 | 6,03E-01 | 7,46E-01 |
| AT2G07605 | 7,80 | -0,35 | 0,68 | -0,52 | 6,04E-01 | 7,46E-01 |
| AT5G45472 | 4,93 | 0,51 | 0,99 | 0,52 | 6,04E-01 | 7,46E-01 |
| AT3G11410 | 4,48 | 0,56 | 1,09 | 0,52 | 6,04E-01 | 7,46E-01 |
| AT5G57110 | 2,43 | 0,70 | 1,36 | 0,52 | 6,04E-01 | 7,46E-01 |
| AT2G04605 | 3,46 | -0,54 | 1,05 | -0,52 | 6,06E-01 | 7,47E-01 |
| AT1G15125 | 1,84 | 0,74 | 1,44 | 0,51 | 6,07E-01 | 7,48E-01 |
| AT3G53990 | 1,83 | 0,76 | 1,48 | 0,51 | 6,07E-01 | 7,48E-01 |
| AT4G07120 | 22,20 | -0,27 | 0,52 | -0,51 | 6,08E-01 | 7,48E-01 |
| AT4G00430 | 2,16 | 0,62 | 1,21 | 0,51 | 6,08E-01 | 7,49E-01 |
| AT2G27310 | 2,22 | 0,55 | 1,08 | 0,51 | 6,10E-01 | 7,51E-01 |
| AT3G55800 | 2,45 | -0,64 | 1,27 | -0,51 | 6,11E-01 | 7,51E-01 |
| AT5G19140 | 6,43 | -0,33 | 0,65 | -0,51 | 6,11E-01 | 7,51E-01 |
| AT3G50930 | 3,98 | 0,42 | 0,83 | 0,51 | 6,12E-01 | 7,52E-01 |
| AT4G24415 | 10,68 | 0,28 | 0,56 | 0,51 | 6,13E-01 | 7,52E-01 |
| AT5G04410 | 1,77 | 1,05 | 2,08 | 0,51 | 6,13E-01 | 7,53E-01 |
| AT3G05325 | 9,46 | -0,31 | 0,63 | -0,50 | 6,15E-01 | 7,53E-01 |
| AT3G50480 | 1,71 | -0,65 | 1,30 | -0,50 | 6,15E-01 | 7,53E-01 |
| AT5G39050 | 4,52 | 0,49 | 0,97 | 0,50 | 6,14E-01 | 7,53E-01 |
| AT5G08525 | 4,34 | 0,51 | 1,01 | 0,50 | 6,16E-01 | 7,53E-01 |
| AT1G08053 | 2,32 | -0,63 | 1,25 | -0,50 | 6,16E-01 | 7,53E-01 |
| AT1G68560 | 3,34 | 0,51 | 1,02 | 0,50 | 6,16E-01 | 7,53E-01 |
| AT2G22430 | 4,62 | 0,43 | 0,87 | 0,50 | 6,17E-01 | 7,53E-01 |
| AT2G36880 | 2,29 | 0,57 | 1,13 | 0,50 | 6,16E-01 | 7,53E-01 |
| AT4G00140 | 2,38 | -0,62 | 1,24 | -0,50 | 6,15E-01 | 7,53E-01 |
| AT3G17400 | 6,94 | -0,38 | 0,77 | -0,50 | 6,17E-01 | 7,53E-01 |
| AT2G35260 | 3,89 | -0,48 | 0,96 | -0,50 | 6,19E-01 | 7,55E-01 |
| AT1G21200 | 5,07 | -0,44 | 0,88 | -0,50 | 6,19E-01 | 7,55E-01 |
| AT1G72180 | 2,71 | 0,50 | 1,02 | 0,50 | 6,20E-01 | 7,56E-01 |
| AT1G77230 | 3,05 | 0,53 | 1,07 | 0,49 | 6,21E-01 | 7,56E-01 |
| AT1G35560 | 2,43 | 0,57 | 1,15 | 0,50 | 6,20E-01 | 7,56E-01 |
| AT2G39010 | 4,44 | 0,53 | 1,07 | 0,49 | 6,22E-01 | 7,57E-01 |
| AT1G67090 | 31,55 | 0,22 | 0,45 | 0,49 | 6,24E-01 | 7,58E-01 |
| AT3G14050 | 1,74 | 0,59 | 1,21 | 0,49 | 6,25E-01 | 7,59E-01 |
| AT2G15970 | 3,79 | -0,58 | 1,19 | -0,49 | 6,26E-01 | 7,60E-01 |
| AT3G43160 | 4,62 | -0,37 | 0,76 | -0,49 | 6,26E-01 | 7,60E-01 |

| | | | | | | |
|---|---|---|---|---|---|---|
| AT1G44100 | 2,02 | -0,59 | 1,20 | -0,49 | 6,26E-01 | 7,60E-01 |
| AT5G49360 | 3,80 | 0,44 | 0,90 | 0,49 | 6,27E-01 | 7,61E-01 |
| AT3G13440 | 3,42 | 0,47 | 0,96 | 0,48 | 6,28E-01 | 7,61E-01 |
| AT4G34950 | 4,16 | -0,43 | 0,89 | -0,48 | 6,32E-01 | 7,65E-01 |
| AT1G05273 | 8,42 | 0,27 | 0,56 | 0,48 | 6,33E-01 | 7,66E-01 |
| AT1G16820 | 8,42 | 0,27 | 0,56 | 0,48 | 6,33E-01 | 7,66E-01 |
| AT1G03743 | 2,10 | -0,62 | 1,29 | -0,48 | 6,33E-01 | 7,66E-01 |
| AT1G09937 | 6,18 | -0,33 | 0,70 | -0,48 | 6,34E-01 | 7,66E-01 |
| AT2G26910 | 1,87 | -0,59 | 1,24 | -0,47 | 6,35E-01 | 7,68E-01 |
| AT2G31890 | 2,01 | 0,69 | 1,47 | 0,47 | 6,36E-01 | 7,68E-01 |
| AT3G29796 | 2,77 | -0,52 | 1,10 | -0,47 | 6,37E-01 | 7,69E-01 |
| AT5G42020 | 4,47 | 0,38 | 0,81 | 0,47 | 6,39E-01 | 7,71E-01 |
| AT4G13770 | 2,50 | 0,60 | 1,30 | 0,47 | 6,41E-01 | 7,72E-01 |
| AT4G32940 | 4,72 | 0,34 | 0,72 | 0,47 | 6,40E-01 | 7,72E-01 |
| AT2G43110 | 4,82 | -0,42 | 0,89 | -0,47 | 6,42E-01 | 7,73E-01 |
| AT3G18370 | 2,55 | 0,67 | 1,46 | 0,46 | 6,43E-01 | 7,74E-01 |
| AT3G46530 | 2,60 | 0,49 | 1,07 | 0,46 | 6,43E-01 | 7,74E-01 |
| AT1G40104 | 7,42 | 0,34 | 0,74 | 0,46 | 6,44E-01 | 7,75E-01 |
| AT1G06040 | 2,85 | 0,57 | 1,24 | 0,46 | 6,44E-01 | 7,75E-01 |
| AT2G04405 | 3,40 | 0,45 | 0,99 | 0,46 | 6,47E-01 | 7,77E-01 |
| AT5G14780 | 2,20 | 0,50 | 1,09 | 0,46 | 6,48E-01 | 7,78E-01 |
| AT4G03415 | 1,93 | 0,72 | 1,58 | 0,46 | 6,49E-01 | 7,79E-01 |
| AT1G56510 | 2,98 | -0,58 | 1,28 | -0,45 | 6,50E-01 | 7,80E-01 |
| AT2G31610 | 1,80 | 0,63 | 1,39 | 0,45 | 6,51E-01 | 7,80E-01 |
| AT1G56280 | 2,59 | 0,50 | 1,11 | 0,45 | 6,52E-01 | 7,81E-01 |
| AT4G31700 | 1,83 | -0,64 | 1,43 | -0,45 | 6,52E-01 | 7,81E-01 |
| AT1G29070 | 1,74 | 0,63 | 1,40 | 0,45 | 6,53E-01 | 7,81E-01 |
| AT1G33070 | 1,82 | -0,72 | 1,60 | -0,45 | 6,54E-01 | 7,81E-01 |
| AT3G04165 | 2,58 | -0,49 | 1,09 | -0,45 | 6,53E-01 | 7,81E-01 |
| AT4G17788 | 20,18 | -0,23 | 0,51 | -0,45 | 6,54E-01 | 7,81E-01 |
| AT5G54940 | 2,86 | -0,43 | 0,96 | -0,45 | 6,54E-01 | 7,81E-01 |
| AT3G60320 | 3,74 | -0,40 | 0,90 | -0,45 | 6,55E-01 | 7,81E-01 |
| AT4G06375 | 3,09 | 0,56 | 1,26 | 0,44 | 6,57E-01 | 7,84E-01 |
| AT1G07977 | 4,37 | -0,57 | 1,28 | -0,44 | 6,58E-01 | 7,85E-01 |
| AT1G21202 | 4,89 | -0,43 | 0,98 | -0,44 | 6,59E-01 | 7,85E-01 |
| AT1G05637 | 4,89 | -0,43 | 0,98 | -0,44 | 6,59E-01 | 7,85E-01 |
| AT1G15670 | 2,03 | -0,52 | 1,18 | -0,44 | 6,60E-01 | 7,86E-01 |
| AT4G31500 | 6,34 | 0,30 | 0,68 | 0,44 | 6,62E-01 | 7,88E-01 |
| AT4G33050 | 2,86 | 0,43 | 0,98 | 0,44 | 6,62E-01 | 7,88E-01 |
| AT4G04020 | 1,87 | -0,83 | 1,92 | -0,43 | 6,64E-01 | 7,90E-01 |
| AT3G60750 | 11,99 | 0,24 | 0,55 | 0,43 | 6,67E-01 | 7,93E-01 |
| AT3G05335 | 1,73 | -0,54 | 1,27 | -0,43 | 6,69E-01 | 7,93E-01 |
| AT5G61440 | 4,44 | -0,36 | 0,84 | -0,43 | 6,68E-01 | 7,93E-01 |
| AT3G29130 | 2,05 | -0,61 | 1,42 | -0,43 | 6,68E-01 | 7,93E-01 |
| AT5G49720 | 3,89 | -0,41 | 0,96 | -0,43 | 6,69E-01 | 7,93E-01 |
| AT3G07570 | 1,83 | -0,60 | 1,41 | -0,43 | 6,71E-01 | 7,95E-01 |
| AT5G58070 | 2,01 | 0,61 | 1,44 | 0,43 | 6,71E-01 | 7,95E-01 |
| AT5G56010 | 6,62 | 0,34 | 0,79 | 0,42 | 6,72E-01 | 7,95E-01 |
| AT3G08940 | 8,84 | -0,34 | 0,80 | -0,42 | 6,74E-01 | 7,97E-01 |
| AT1G07670 | 1,74 | 0,58 | 1,39 | 0,42 | 6,74E-01 | 7,98E-01 |
| AT1G65450 | 2,07 | -0,61 | 1,46 | -0,42 | 6,75E-01 | 7,98E-01 |
| AT1G20690 | 2,85 | -0,43 | 1,03 | -0,42 | 6,77E-01 | 7,99E-01 |
| AT1G07793 | 2,22 | -0,54 | 1,30 | -0,42 | 6,76E-01 | 7,99E-01 |
| AT1G20691 | 2,85 | -0,43 | 1,03 | -0,42 | 6,77E-01 | 7,99E-01 |
| AT1G07250 | 2,07 | -0,49 | 1,18 | -0,42 | 6,78E-01 | 7,99E-01 |
| AT1G55450 | 2,26 | -0,56 | 1,35 | -0,42 | 6,78E-01 | 7,99E-01 |

| | | | | | | |
|---|---|---|---|---|---|---|
| AT1G70490 | 1,85 | -0,56 | 1,34 | -0,42 | 6,78E-01 | 7,99E-01 |
| AT4G36040 | 3,49 | 0,38 | 0,93 | 0,41 | 6,80E-01 | 8,01E-01 |
| AT1G55920 | 3,12 | -0,45 | 1,11 | -0,41 | 6,82E-01 | 8,02E-01 |
| AT3G14270 | 2,35 | 0,47 | 1,16 | 0,41 | 6,83E-01 | 8,02E-01 |
| AT3G14415 | 1,72 | -0,57 | 1,39 | -0,41 | 6,81E-01 | 8,02E-01 |
| AT1G67740 | 3,89 | 0,37 | 0,90 | 0,41 | 6,83E-01 | 8,02E-01 |
| AT3G05655 | 1,87 | -0,57 | 1,40 | -0,41 | 6,82E-01 | 8,02E-01 |
| AT3G47500 | 2,57 | -0,46 | 1,12 | -0,41 | 6,83E-01 | 8,02E-01 |
| AT5G00760 | 4,31 | -0,34 | 0,82 | -0,41 | 6,82E-01 | 8,02E-01 |
| AT5G15200 | 4,33 | 0,34 | 0,84 | 0,41 | 6,83E-01 | 8,02E-01 |
| AT5G01530 | 20,05 | -0,24 | 0,60 | -0,41 | 6,84E-01 | 8,02E-01 |
| AT5G16620 | 2,01 | 0,46 | 1,14 | 0,41 | 6,85E-01 | 8,02E-01 |
| AT4G26540 | 3,42 | 0,41 | 1,00 | 0,40 | 6,86E-01 | 8,04E-01 |
| AT1G15010 | 2,66 | -0,41 | 1,03 | -0,40 | 6,87E-01 | 8,04E-01 |
| AT4G24715 | 2,06 | -0,50 | 1,24 | -0,40 | 6,87E-01 | 8,04E-01 |
| AT3G21870 | 1,78 | -0,65 | 1,65 | -0,40 | 6,92E-01 | 8,10E-01 |
| AT4G23630 | 1,89 | -0,47 | 1,19 | -0,39 | 6,94E-01 | 8,11E-01 |
| AT1G64385 | 1,99 | -0,69 | 1,76 | -0,39 | 6,95E-01 | 8,12E-01 |
| AT3G03475 | 2,58 | 0,43 | 1,10 | 0,39 | 6,96E-01 | 8,13E-01 |
| AT1G18880 | 1,72 | 0,48 | 1,23 | 0,39 | 6,97E-01 | 8,14E-01 |
| AT3G24480 | 4,54 | 0,34 | 0,88 | 0,39 | 6,98E-01 | 8,14E-01 |
| AT2G41530 | 2,42 | -0,51 | 1,31 | -0,39 | 6,98E-01 | 8,14E-01 |
| AT2G16020 | 3,14 | -0,40 | 1,05 | -0,39 | 6,99E-01 | 8,15E-01 |
| AT4G21366 | 2,02 | -0,46 | 1,21 | -0,38 | 7,01E-01 | 8,17E-01 |
| AT4G17390 | 6,32 | -0,30 | 0,79 | -0,38 | 7,01E-01 | 8,17E-01 |
| AT1G20440 | 3,88 | 0,33 | 0,85 | 0,38 | 7,03E-01 | 8,18E-01 |
| AT1G32060 | 5,29 | -0,38 | 1,00 | -0,38 | 7,05E-01 | 8,20E-01 |
| AT3G54920 | 2,20 | 0,42 | 1,11 | 0,38 | 7,05E-01 | 8,20E-01 |
| AT2G18480 | 2,00 | -0,56 | 1,47 | -0,38 | 7,05E-01 | 8,20E-01 |
| AT1G13440 | 5,54 | -0,29 | 0,77 | -0,37 | 7,08E-01 | 8,22E-01 |
| AT5G24660 | 2,17 | -0,48 | 1,29 | -0,37 | 7,08E-01 | 8,22E-01 |
| AT1G14920 | 1,77 | 0,55 | 1,47 | 0,37 | 7,10E-01 | 8,24E-01 |
| AT3G24612 | 3,02 | 0,35 | 0,95 | 0,37 | 7,10E-01 | 8,24E-01 |
| AT2G42540 | 2,58 | -0,72 | 1,95 | -0,37 | 7,12E-01 | 8,25E-01 |
| AT1G06460 | 1,79 | -0,56 | 1,52 | -0,37 | 7,12E-01 | 8,25E-01 |
| AT4G38775 | 2,66 | -0,38 | 1,04 | -0,37 | 7,13E-01 | 8,26E-01 |
| AT5G40340 | 2,24 | -0,43 | 1,17 | -0,37 | 7,14E-01 | 8,26E-01 |
| AT4G04465 | 2,27 | -0,41 | 1,13 | -0,37 | 7,14E-01 | 8,27E-01 |
| AT1G20510 | 2,74 | -0,45 | 1,23 | -0,36 | 7,17E-01 | 8,29E-01 |
| AT2G07981 | 2,13 | -0,58 | 1,61 | -0,36 | 7,18E-01 | 8,29E-01 |
| AT2G08986 | 10,75 | 0,19 | 0,53 | 0,36 | 7,18E-01 | 8,29E-01 |
| AT5G30510 | 1,83 | -0,56 | 1,54 | -0,36 | 7,18E-01 | 8,29E-01 |
| AT1G23390 | 2,44 | 0,41 | 1,15 | 0,36 | 7,19E-01 | 8,29E-01 |
| AT1G27090 | 1,85 | 0,43 | 1,19 | 0,36 | 7,20E-01 | 8,30E-01 |
| AT2G36470 | 2,06 | -0,55 | 1,54 | -0,36 | 7,20E-01 | 8,30E-01 |
| AT3G55850 | 2,66 | -0,46 | 1,27 | -0,36 | 7,20E-01 | 8,30E-01 |
| AT1G78206 | 14,29 | 0,18 | 0,50 | 0,36 | 7,21E-01 | 8,30E-01 |
| AT5G11790 | 2,58 | -0,37 | 1,03 | -0,36 | 7,21E-01 | 8,30E-01 |
| AT1G49750 | 3,00 | -0,41 | 1,15 | -0,36 | 7,21E-01 | 8,30E-01 |
| AT1G73805 | 1,81 | 0,51 | 1,44 | 0,36 | 7,22E-01 | 8,30E-01 |
| AT3G53420 | 2,26 | 0,45 | 1,26 | 0,35 | 7,24E-01 | 8,32E-01 |
| AT4G22240 | 1,83 | -0,47 | 1,34 | -0,35 | 7,24E-01 | 8,32E-01 |
| AT1G78830 | 2,12 | -0,48 | 1,38 | -0,35 | 7,25E-01 | 8,33E-01 |
| AT3G03175 | 7,33 | 0,22 | 0,63 | 0,35 | 7,27E-01 | 8,34E-01 |
| AT2G08725 | 34,05 | -0,20 | 0,58 | -0,35 | 7,27E-01 | 8,34E-01 |
| AT3G46970 | 7,07 | 0,23 | 0,67 | 0,35 | 7,28E-01 | 8,35E-01 |

| | | | | | | |
|---|---|---|---|---|---|---|
| AT4G20360 | 5,64 | -0,26 | 0,75 | -0,35 | 7,30E-01 | 8,36E-01 |
| AT4G20362 | 5,64 | -0,26 | 0,75 | -0,35 | 7,30E-01 | 8,36E-01 |
| AT5G26030 | 2,16 | -0,38 | 1,11 | -0,34 | 7,31E-01 | 8,36E-01 |
| AT2G29450 | 2,19 | -0,51 | 1,49 | -0,34 | 7,31E-01 | 8,36E-01 |
| AT5G55565 | 4,30 | -0,33 | 0,95 | -0,34 | 7,31E-01 | 8,36E-01 |
| AT4G01150 | 2,10 | 0,38 | 1,10 | 0,34 | 7,32E-01 | 8,37E-01 |
| AT2G41090 | 3,05 | -0,35 | 1,03 | -0,34 | 7,33E-01 | 8,37E-01 |
| AT4G15560 | 2,23 | -0,43 | 1,26 | -0,34 | 7,33E-01 | 8,37E-01 |
| AT3G07515 | 2,88 | -0,43 | 1,26 | -0,34 | 7,35E-01 | 8,39E-01 |
| AT1G48920 | 4,08 | 0,31 | 0,92 | 0,33 | 7,39E-01 | 8,41E-01 |
| AT1G66730 | 1,91 | 0,38 | 1,13 | 0,33 | 7,39E-01 | 8,41E-01 |
| AT1G46696 | 1,91 | -0,46 | 1,39 | -0,33 | 7,39E-01 | 8,41E-01 |
| AT1G76180 | 6,03 | -0,22 | 0,66 | -0,33 | 7,39E-01 | 8,41E-01 |
| AT3G47250 | 1,84 | 0,43 | 1,29 | 0,33 | 7,38E-01 | 8,41E-01 |
| AT2G15620 | 3,38 | 0,36 | 1,10 | 0,33 | 7,40E-01 | 8,42E-01 |
| AT1G04527 | 11,72 | 0,17 | 0,51 | 0,33 | 7,40E-01 | 8,42E-01 |
| AT1G50010 | 2,93 | 0,36 | 1,10 | 0,33 | 7,42E-01 | 8,44E-01 |
| AT1G08623 | 2,21 | -0,50 | 1,53 | -0,33 | 7,45E-01 | 8,46E-01 |
| AT5G10572 | 2,06 | -0,39 | 1,21 | -0,32 | 7,46E-01 | 8,47E-01 |
| AT2G34110 | 1,88 | 0,40 | 1,24 | 0,32 | 7,47E-01 | 8,48E-01 |
| AT3G43833 | 7,44 | -0,26 | 0,81 | -0,32 | 7,47E-01 | 8,48E-01 |
| AT2G45245 | 2,00 | -0,36 | 1,12 | -0,32 | 7,47E-01 | 8,48E-01 |
| AT2G08390 | 5,34 | -0,28 | 0,88 | -0,32 | 7,48E-01 | 8,48E-01 |
| AT5G58430 | 1,95 | 0,39 | 1,24 | 0,32 | 7,50E-01 | 8,50E-01 |
| AT2G47170 | 1,90 | -0,40 | 1,27 | -0,32 | 7,51E-01 | 8,50E-01 |
| AT1G15002 | 1,82 | -0,40 | 1,28 | -0,32 | 7,51E-01 | 8,50E-01 |
| AT3G07765 | 2,13 | 0,37 | 1,17 | 0,32 | 7,53E-01 | 8,51E-01 |
| AT2G16660 | 2,17 | -0,34 | 1,08 | -0,31 | 7,53E-01 | 8,51E-01 |
| AT4G01290 | 1,74 | 0,46 | 1,48 | 0,31 | 7,53E-01 | 8,51E-01 |
| AT5G06320 | 5,00 | 0,31 | 1,00 | 0,31 | 7,55E-01 | 8,52E-01 |
| AT1G28440 | 1,72 | -0,39 | 1,26 | -0,31 | 7,56E-01 | 8,53E-01 |
| AT1G79550 | 2,41 | 0,34 | 1,10 | 0,31 | 7,56E-01 | 8,53E-01 |
| AT4G04890 | 2,53 | -0,55 | 1,78 | -0,31 | 7,56E-01 | 8,53E-01 |
| AT1G13470 | 1,76 | -0,45 | 1,45 | -0,31 | 7,57E-01 | 8,54E-01 |
| AT1G19410 | 3,18 | -0,28 | 0,91 | -0,31 | 7,58E-01 | 8,54E-01 |
| AT2G37270 | 1,75 | 0,50 | 1,64 | 0,31 | 7,58E-01 | 8,54E-01 |
| AT2G32690 | 2,01 | -0,38 | 1,23 | -0,31 | 7,58E-01 | 8,54E-01 |
| AT3G59410 | 1,72 | -0,45 | 1,46 | -0,31 | 7,59E-01 | 8,54E-01 |
| AT4G35000 | 3,13 | -0,41 | 1,35 | -0,31 | 7,60E-01 | 8,55E-01 |
| AT5G04425 | 2,72 | -0,31 | 1,03 | -0,30 | 7,61E-01 | 8,55E-01 |
| AT1G69830 | 1,91 | 0,37 | 1,23 | 0,30 | 7,61E-01 | 8,55E-01 |
| AT2G42490 | 2,30 | 0,35 | 1,17 | 0,30 | 7,61E-01 | 8,55E-01 |
| AT2G25171 | 1,84 | -0,46 | 1,51 | -0,30 | 7,62E-01 | 8,55E-01 |
| AT4G02555 | 3,06 | 0,31 | 1,04 | 0,30 | 7,62E-01 | 8,55E-01 |
| AT3G16140 | 1,91 | -0,41 | 1,35 | -0,30 | 7,62E-01 | 8,55E-01 |
| AT2G23930 | 1,70 | 0,49 | 1,65 | 0,30 | 7,65E-01 | 8,58E-01 |
| AT4G03280 | 3,09 | -0,32 | 1,08 | -0,30 | 7,65E-01 | 8,58E-01 |
| AT1G70290 | 2,53 | 0,33 | 1,10 | 0,30 | 7,66E-01 | 8,58E-01 |
| AT3G47620 | 3,09 | 0,30 | 1,02 | 0,30 | 7,67E-01 | 8,58E-01 |
| AT1G52110 | 2,50 | -0,38 | 1,31 | -0,29 | 7,70E-01 | 8,61E-01 |
| AT3G01290 | 2,12 | -0,33 | 1,12 | -0,29 | 7,70E-01 | 8,61E-01 |
| AT4G32990 | 2,22 | 0,34 | 1,17 | 0,29 | 7,70E-01 | 8,61E-01 |
| AT2G41720 | 1,87 | -0,46 | 1,59 | -0,29 | 7,72E-01 | 8,62E-01 |
| AT3G17190 | 1,85 | -0,38 | 1,32 | -0,29 | 7,73E-01 | 8,62E-01 |
| AT5G24530 | 1,77 | -0,36 | 1,26 | -0,29 | 7,73E-01 | 8,62E-01 |
| AT5G39080 | 2,20 | -0,38 | 1,33 | -0,29 | 7,73E-01 | 8,62E-01 |

| | | | | | | |
|---|---|---|---|---|---|---|
| AT4G39260 | 6,03 | -0,20 | 0,71 | -0,29 | 7,74E-01 | 8,63E-01 |
| AT2G22300 | 2,59 | 0,38 | 1,34 | 0,28 | 7,76E-01 | 8,64E-01 |
| AT2G47240 | 1,70 | -0,38 | 1,35 | -0,28 | 7,76E-01 | 8,64E-01 |
| AT2G30110 | 3,88 | 0,27 | 0,95 | 0,28 | 7,76E-01 | 8,64E-01 |
| AT5G35735 | 2,19 | -0,31 | 1,09 | -0,28 | 7,76E-01 | 8,64E-01 |
| AT5G40384 | 2,72 | 0,37 | 1,30 | 0,28 | 7,77E-01 | 8,64E-01 |
| AT5G19770 | 1,75 | -0,49 | 1,74 | -0,28 | 7,77E-01 | 8,64E-01 |
| AT1G14670 | 2,24 | 0,33 | 1,17 | 0,28 | 7,79E-01 | 8,65E-01 |
| AT1G20450 | 3,27 | 0,30 | 1,08 | 0,28 | 7,78E-01 | 8,65E-01 |
| AT1G74100 | 5,98 | -0,23 | 0,82 | -0,28 | 7,80E-01 | 8,66E-01 |
| AT5G01915 | 2,27 | -0,35 | 1,26 | -0,28 | 7,81E-01 | 8,66E-01 |
| AT2G05455 | 4,14 | -0,24 | 0,87 | -0,28 | 7,83E-01 | 8,68E-01 |
| AT1G02470 | 5,92 | -0,25 | 0,91 | -0,27 | 7,84E-01 | 8,69E-01 |
| AT5G60640 | 2,58 | -0,34 | 1,25 | -0,27 | 7,84E-01 | 8,69E-01 |
| AT2G14560 | 27,90 | -0,13 | 0,48 | -0,27 | 7,85E-01 | 8,69E-01 |
| AT5G02870 | 2,41 | 0,36 | 1,32 | 0,27 | 7,87E-01 | 8,71E-01 |
| AT2G04455 | 3,39 | -0,25 | 0,96 | -0,26 | 7,92E-01 | 8,74E-01 |
| AT3G45640 | 1,79 | 0,32 | 1,21 | 0,26 | 7,91E-01 | 8,74E-01 |
| AT5G66570 | 20,06 | -0,16 | 0,62 | -0,26 | 7,92E-01 | 8,74E-01 |
| AT1G04897 | 2,68 | -0,29 | 1,11 | -0,26 | 7,92E-01 | 8,74E-01 |
| AT2G10940 | 5,77 | -0,27 | 1,01 | -0,27 | 7,91E-01 | 8,74E-01 |
| AT4G25520 | 4,54 | -0,24 | 0,89 | -0,26 | 7,92E-01 | 8,74E-01 |
| AT4G32260 | 3,86 | -0,26 | 1,00 | -0,26 | 7,91E-01 | 8,74E-01 |
| AT3G52590 | 1,74 | -0,36 | 1,39 | -0,26 | 7,94E-01 | 8,75E-01 |
| AT4G34100 | 3,45 | 0,28 | 1,07 | 0,26 | 7,95E-01 | 8,76E-01 |
| AT5G37600 | 1,93 | 0,38 | 1,46 | 0,26 | 7,95E-01 | 8,76E-01 |
| AT2G24980 | 3,78 | -0,26 | 1,03 | -0,26 | 7,97E-01 | 8,76E-01 |
| AT4G07875 | 11,03 | 0,16 | 0,63 | 0,26 | 7,96E-01 | 8,76E-01 |
| AT1G07493 | 1,97 | -0,42 | 1,65 | -0,26 | 7,97E-01 | 8,76E-01 |
| AT1G01040 | 4,73 | 0,20 | 0,78 | 0,25 | 8,00E-01 | 8,80E-01 |
| AT1G20010 | 2,01 | -0,31 | 1,26 | -0,25 | 8,04E-01 | 8,83E-01 |
| AT2G13560 | 2,19 | -0,39 | 1,59 | -0,24 | 8,07E-01 | 8,84E-01 |
| AT3G21750 | 1,77 | 0,29 | 1,21 | 0,24 | 8,07E-01 | 8,84E-01 |
| AT3G52800 | 1,93 | -0,33 | 1,34 | -0,25 | 8,04E-01 | 8,84E-01 |
| AT4G23915 | 12,58 | -0,13 | 0,54 | -0,24 | 8,07E-01 | 8,84E-01 |
| AT5G08712 | 123,44 | 0,08 | 0,33 | 0,24 | 8,08E-01 | 8,84E-01 |
| AT5G27720 | 3,01 | -0,25 | 1,01 | -0,25 | 8,06E-01 | 8,84E-01 |
| AT5G63145 | 2,83 | -0,26 | 1,08 | -0,24 | 8,07E-01 | 8,84E-01 |
| AT3G22231 | 4,08 | -0,21 | 0,84 | -0,25 | 8,06E-01 | 8,84E-01 |
| AT5G48860 | 3,76 | -0,24 | 0,95 | -0,25 | 8,05E-01 | 8,84E-01 |
| AT2G25170 | 3,04 | 0,28 | 1,16 | 0,24 | 8,09E-01 | 8,85E-01 |
| AT1G20860 | 3,03 | -0,24 | 1,01 | -0,24 | 8,11E-01 | 8,86E-01 |
| AT2G29925 | 2,27 | 0,40 | 1,65 | 0,24 | 8,10E-01 | 8,86E-01 |
| AT2G37220 | 3,40 | -0,25 | 1,04 | -0,24 | 8,11E-01 | 8,86E-01 |
| AT3G44310 | 10,52 | 0,15 | 0,66 | 0,24 | 8,14E-01 | 8,87E-01 |
| AT4G39361 | 1,80 | -0,37 | 1,56 | -0,24 | 8,12E-01 | 8,87E-01 |
| AT2G41560 | 2,99 | 0,24 | 1,02 | 0,24 | 8,13E-01 | 8,87E-01 |
| AT4G13505 | 4,72 | 0,18 | 0,77 | 0,24 | 8,14E-01 | 8,87E-01 |
| AT4G24770 | 3,23 | 0,25 | 1,05 | 0,24 | 8,14E-01 | 8,87E-01 |
| AT4G27520 | 5,05 | -0,22 | 0,95 | -0,24 | 8,13E-01 | 8,87E-01 |
| AT5G28823 | 1,86 | 0,36 | 1,56 | 0,23 | 8,16E-01 | 8,88E-01 |
| AT4G11600 | 2,32 | 0,26 | 1,13 | 0,23 | 8,16E-01 | 8,88E-01 |
| AT4G17410 | 2,55 | 0,34 | 1,47 | 0,23 | 8,18E-01 | 8,89E-01 |
| AT1G05563 | 2,75 | -0,25 | 1,08 | -0,23 | 8,18E-01 | 8,89E-01 |
| AT3G17800 | 1,86 | 0,33 | 1,41 | 0,23 | 8,17E-01 | 8,89E-01 |
| AT3G11830 | 1,90 | 0,32 | 1,38 | 0,23 | 8,19E-01 | 8,89E-01 |

| | | | | | | |
|---|---|---|---|---|---|---|
| AT4G04835 | 19,24 | -0,13 | 0,56 | -0,23 | 8,18E-01 | 8,89E-01 |
| AT3G61630 | 1,88 | 0,36 | 1,58 | 0,23 | 8,19E-01 | 8,89E-01 |
| AT4G29190 | 1,79 | -0,36 | 1,59 | -0,23 | 8,20E-01 | 8,90E-01 |
| AT5G44090 | 1,96 | 0,35 | 1,56 | 0,23 | 8,21E-01 | 8,90E-01 |
| AT5G52470 | 2,57 | -0,27 | 1,19 | -0,22 | 8,23E-01 | 8,91E-01 |
| AT5G04305 | 5,44 | 0,15 | 0,67 | 0,22 | 8,23E-01 | 8,91E-01 |
| AT5G38430 | 4,72 | 0,19 | 0,86 | 0,22 | 8,23E-01 | 8,91E-01 |
| AT2G07671 | 2,57 | -0,27 | 1,25 | -0,22 | 8,26E-01 | 8,94E-01 |
| AT1G05850 | 2,93 | 0,28 | 1,29 | 0,22 | 8,27E-01 | 8,94E-01 |
| AT4G24190 | 5,76 | 0,15 | 0,69 | 0,22 | 8,27E-01 | 8,94E-01 |
| AT2G08370 | 4,11 | 0,17 | 0,79 | 0,22 | 8,29E-01 | 8,95E-01 |
| AT3G56825 | 56,81 | 0,07 | 0,34 | 0,21 | 8,30E-01 | 8,97E-01 |
| AT3G44580 | 31,33 | -0,11 | 0,54 | -0,21 | 8,32E-01 | 8,98E-01 |
| AT5G22770 | 2,48 | 0,25 | 1,17 | 0,21 | 8,32E-01 | 8,98E-01 |
| AT3G55410 | 3,63 | 0,23 | 1,10 | 0,21 | 8,33E-01 | 8,98E-01 |
| AT2G46820 | 2,21 | -0,25 | 1,18 | -0,21 | 8,33E-01 | 8,98E-01 |
| AT4G23850 | 2,43 | 0,21 | 1,02 | 0,21 | 8,36E-01 | 9,01E-01 |
| AT5G65005 | 2,56 | -0,23 | 1,15 | -0,20 | 8,38E-01 | 9,03E-01 |
| AT1G63880 | 4,32 | -0,15 | 0,75 | -0,20 | 8,43E-01 | 9,07E-01 |
| AT4G04340 | 2,36 | 0,33 | 1,70 | 0,19 | 8,46E-01 | 9,10E-01 |
| AT1G69620 | 2,14 | -0,24 | 1,24 | -0,19 | 8,49E-01 | 9,12E-01 |
| AT2G29920 | 2,43 | 0,29 | 1,53 | 0,19 | 8,49E-01 | 9,12E-01 |
| AT2G16676 | 2,63 | 0,21 | 1,12 | 0,19 | 8,49E-01 | 9,12E-01 |
| AT3G53430 | 2,55 | -0,20 | 1,06 | -0,19 | 8,48E-01 | 9,12E-01 |
| AT1G53655 | 1,78 | -0,28 | 1,53 | -0,18 | 8,55E-01 | 9,17E-01 |
| AT3G12780 | 5,73 | 0,13 | 0,73 | 0,18 | 8,54E-01 | 9,17E-01 |
| AT3G05985 | 2,35 | 0,23 | 1,28 | 0,18 | 8,56E-01 | 9,18E-01 |
| AT3G44430 | 1,70 | 0,24 | 1,35 | 0,18 | 8,60E-01 | 9,21E-01 |
| AT1G08200 | 1,82 | -0,26 | 1,48 | -0,18 | 8,61E-01 | 9,22E-01 |
| AT1G07243 | 1,82 | -0,22 | 1,25 | -0,17 | 8,63E-01 | 9,24E-01 |
| AT2G04435 | 3,56 | -0,16 | 0,93 | -0,17 | 8,65E-01 | 9,26E-01 |
| AT3G56360 | 3,83 | 0,19 | 1,12 | 0,17 | 8,66E-01 | 9,27E-01 |
| AT5G13890 | 14,03 | 0,12 | 0,73 | 0,17 | 8,67E-01 | 9,27E-01 |
| AT1G05207 | 1,81 | 0,24 | 1,44 | 0,16 | 8,69E-01 | 9,28E-01 |
| AT4G39730 | 3,06 | 0,20 | 1,22 | 0,16 | 8,69E-01 | 9,28E-01 |
| AT5G08680 | 1,77 | 0,27 | 1,63 | 0,17 | 8,68E-01 | 9,28E-01 |
| AT1G55350 | 1,89 | 0,36 | 2,18 | 0,17 | 8,68E-01 | 9,28E-01 |
| AT1G04517 | 11,17 | 0,09 | 0,53 | 0,16 | 8,70E-01 | 9,28E-01 |
| AT5G41471 | 1,79 | -0,27 | 1,67 | -0,16 | 8,71E-01 | 9,29E-01 |
| AT1G08327 | 2,78 | 0,22 | 1,39 | 0,16 | 8,72E-01 | 9,29E-01 |
| AT5G13887 | 13,72 | 0,13 | 0,85 | 0,16 | 8,74E-01 | 9,31E-01 |
| AT1G68010 | 2,53 | 0,18 | 1,24 | 0,15 | 8,82E-01 | 9,34E-01 |
| AT3G06436 | 20,96 | 0,06 | 0,40 | 0,15 | 8,82E-01 | 9,34E-01 |
| AT3G07860 | 5,76 | 0,12 | 0,79 | 0,15 | 8,82E-01 | 9,34E-01 |
| AT4G13510 | 4,76 | 0,12 | 0,78 | 0,15 | 8,81E-01 | 9,34E-01 |
| AT5G11670 | 3,41 | 0,13 | 0,91 | 0,15 | 8,83E-01 | 9,34E-01 |
| AT5G49760 | 3,34 | 0,15 | 1,04 | 0,15 | 8,83E-01 | 9,34E-01 |
| AT2G25490 | 3,68 | 0,15 | 0,98 | 0,15 | 8,79E-01 | 9,34E-01 |
| AT3G14385 | 15,96 | -0,07 | 0,45 | -0,15 | 8,81E-01 | 9,34E-01 |
| AT3G46510 | 1,83 | 0,19 | 1,26 | 0,15 | 8,80E-01 | 9,34E-01 |
| AT3G48560 | 3,89 | -0,15 | 0,97 | -0,15 | 8,78E-01 | 9,34E-01 |
| AT3G51550 | 4,70 | 0,13 | 0,82 | 0,15 | 8,78E-01 | 9,34E-01 |
| AT3G54580 | 1,94 | -0,20 | 1,34 | -0,15 | 8,83E-01 | 9,34E-01 |
| AT3G62420 | 1,96 | 0,26 | 1,69 | 0,15 | 8,79E-01 | 9,34E-01 |
| AT3G63003 | 4,44 | 0,15 | 1,02 | 0,15 | 8,80E-01 | 9,34E-01 |
| AT4G18880 | 2,25 | 0,23 | 1,51 | 0,15 | 8,78E-01 | 9,34E-01 |

| | | | | | | |
|---|---|---|---|---|---|---|
| AT5G28540 | 7,23 | -0,10 | 0,68 | -0,15 | 8,82E-01 | 9,34E-01 |
| AT5G05505 | 4,00 | -0,17 | 1,18 | -0,15 | 8,82E-01 | 9,34E-01 |
| AT3G57400 | 1,77 | -0,20 | 1,36 | -0,14 | 8,85E-01 | 9,36E-01 |
| AT4G23420 | 23,63 | -0,07 | 0,47 | -0,14 | 8,86E-01 | 9,36E-01 |
| AT1G11840 | 1,70 | -0,18 | 1,27 | -0,14 | 8,87E-01 | 9,37E-01 |
| AT1G72370 | 2,39 | 0,16 | 1,14 | 0,14 | 8,88E-01 | 9,38E-01 |
| AT3G23810 | 2,25 | -0,19 | 1,35 | -0,14 | 8,90E-01 | 9,39E-01 |
| AT1G55325 | 2,49 | -0,14 | 1,04 | -0,14 | 8,91E-01 | 9,39E-01 |
| AT2G46400 | 1,95 | 0,18 | 1,35 | 0,14 | 8,91E-01 | 9,39E-01 |
| AT1G13360 | 23,38 | 0,05 | 0,39 | 0,14 | 8,92E-01 | 9,39E-01 |
| AT1G76030 | 2,84 | 0,16 | 1,23 | 0,13 | 8,93E-01 | 9,40E-01 |
| AT1G71002 | 2,81 | -0,15 | 1,13 | -0,13 | 8,93E-01 | 9,40E-01 |
| AT1G78040 | 2,48 | 0,16 | 1,18 | 0,13 | 8,95E-01 | 9,41E-01 |
| AT4G39160 | 1,99 | 0,16 | 1,25 | 0,13 | 8,97E-01 | 9,43E-01 |
| AT5G24930 | 2,10 | -0,18 | 1,40 | -0,13 | 8,97E-01 | 9,43E-01 |
| AT1G17290 | 2,38 | -0,15 | 1,14 | -0,13 | 8,98E-01 | 9,43E-01 |
| AT2G09700 | 2,16 | 0,17 | 1,34 | 0,13 | 8,99E-01 | 9,44E-01 |
| AT3G06055 | 2,49 | -0,13 | 1,05 | -0,13 | 8,99E-01 | 9,44E-01 |
| AT1G07890 | 4,08 | -0,10 | 0,85 | -0,12 | 9,02E-01 | 9,46E-01 |
| AT1G06148 | 2,19 | -0,15 | 1,25 | -0,12 | 9,03E-01 | 9,47E-01 |
| AT5G09505 | 92,76 | 0,04 | 0,30 | 0,12 | 9,03E-01 | 9,47E-01 |
| AT4G39366 | 2,55 | -0,13 | 1,12 | -0,12 | 9,04E-01 | 9,48E-01 |
| AT1G78080 | 3,84 | -0,12 | 1,02 | -0,12 | 9,05E-01 | 9,48E-01 |
| AT4G11270 | 3,79 | 0,12 | 1,00 | 0,12 | 9,06E-01 | 9,48E-01 |
| AT2G24103 | 3,17 | -0,12 | 1,04 | -0,12 | 9,08E-01 | 9,49E-01 |
| AT3G49470 | 2,41 | -0,16 | 1,34 | -0,12 | 9,08E-01 | 9,49E-01 |
| AT5G45480 | 2,31 | -0,16 | 1,35 | -0,12 | 9,07E-01 | 9,49E-01 |
| AT1G05570 | 10,12 | 0,06 | 0,56 | 0,11 | 9,11E-01 | 9,52E-01 |
| AT1G52400 | 1,99 | -0,18 | 1,61 | -0,11 | 9,12E-01 | 9,53E-01 |
| AT2G01850 | 1,74 | 0,14 | 1,31 | 0,11 | 9,13E-01 | 9,53E-01 |
| AT1G09340 | 1,73 | 0,15 | 1,36 | 0,11 | 9,14E-01 | 9,53E-01 |
| AT1G70940 | 2,12 | 0,14 | 1,36 | 0,10 | 9,17E-01 | 9,56E-01 |
| AT3G62410 | 1,88 | -0,13 | 1,26 | -0,10 | 9,18E-01 | 9,57E-01 |
| AT1G09070 | 2,87 | -0,11 | 1,09 | -0,10 | 9,20E-01 | 9,58E-01 |
| AT3G54810 | 2,87 | 0,11 | 1,08 | 0,10 | 9,19E-01 | 9,58E-01 |
| AT5G63800 | 1,87 | 0,12 | 1,20 | 0,10 | 9,20E-01 | 9,58E-01 |
| AT3G57730 | 5,17 | 0,07 | 0,75 | 0,10 | 9,21E-01 | 9,59E-01 |
| AT4G20735 | 3,19 | -0,13 | 1,31 | -0,10 | 9,22E-01 | 9,59E-01 |
| AT4G18210 | 1,90 | -0,15 | 1,56 | -0,10 | 9,24E-01 | 9,60E-01 |
| AT5G61410 | 2,49 | 0,09 | 1,01 | 0,09 | 9,26E-01 | 9,62E-01 |
| AT3G03990 | 2,37 | -0,12 | 1,27 | -0,09 | 9,26E-01 | 9,62E-01 |
| AT3G14930 | 2,04 | -0,12 | 1,31 | -0,09 | 9,27E-01 | 9,62E-01 |
| AT1G75380 | 2,78 | 0,09 | 1,00 | 0,09 | 9,27E-01 | 9,62E-01 |
| AT2G20721 | 2,89 | 0,09 | 0,94 | 0,09 | 9,28E-01 | 9,62E-01 |
| AT4G13430 | 3,55 | -0,08 | 0,91 | -0,09 | 9,28E-01 | 9,62E-01 |
| AT3G20950 | 1,75 | 0,11 | 1,28 | 0,09 | 9,30E-01 | 9,64E-01 |
| AT2G44060 | 2,21 | 0,09 | 1,15 | 0,08 | 9,37E-01 | 9,69E-01 |
| AT4G10210 | 2,08 | 0,10 | 1,28 | 0,08 | 9,38E-01 | 9,69E-01 |
| AT4G13880 | 2,00 | 0,11 | 1,42 | 0,08 | 9,37E-01 | 9,69E-01 |
| AT5G45095 | 2,04 | -0,10 | 1,27 | -0,08 | 9,37E-01 | 9,69E-01 |
| AT1G04820 | 2,61 | -0,11 | 1,36 | -0,08 | 9,38E-01 | 9,69E-01 |
| AT3G56070 | 2,75 | 0,08 | 1,01 | 0,08 | 9,38E-01 | 9,69E-01 |
| AT5G52320 | 1,77 | 0,11 | 1,35 | 0,08 | 9,36E-01 | 9,69E-01 |
| AT4G30210 | 1,86 | -0,12 | 1,57 | -0,08 | 9,39E-01 | 9,70E-01 |
| AT5G53902 | 6,85 | 0,06 | 0,78 | 0,08 | 9,40E-01 | 9,70E-01 |
| AT1G09570 | 2,05 | 0,09 | 1,26 | 0,07 | 9,40E-01 | 9,70E-01 |

| | | | | | | |
|---|---|---|---|---|---|---|
| AT5G53905 | 6,85 | 0,06 | 0,78 | 0,08 | 9,40E-01 | 9,70E-01 |
| AT1G32270 | 1,96 | 0,11 | 1,47 | 0,07 | 9,42E-01 | 9,70E-01 |
| AT2G47440 | 2,19 | 0,10 | 1,43 | 0,07 | 9,42E-01 | 9,70E-01 |
| AT5G03350 | 2,45 | -0,08 | 1,12 | -0,07 | 9,41E-01 | 9,70E-01 |
| AT1G71880 | 1,96 | -0,10 | 1,42 | -0,07 | 9,43E-01 | 9,71E-01 |
| AT1G18740 | 2,60 | 0,07 | 1,04 | 0,07 | 9,48E-01 | 9,71E-01 |
| AT1G67430 | 1,82 | 0,09 | 1,24 | 0,07 | 9,45E-01 | 9,71E-01 |
| AT1G72150 | 7,68 | 0,05 | 0,64 | 0,07 | 9,44E-01 | 9,71E-01 |
| AT3G05590 | 1,96 | -0,09 | 1,33 | -0,07 | 9,48E-01 | 9,71E-01 |
| AT5G01747 | 60,67 | 0,02 | 0,27 | 0,07 | 9,47E-01 | 9,71E-01 |
| AT5G64260 | 2,41 | 0,07 | 1,12 | 0,07 | 9,47E-01 | 9,71E-01 |
| AT1G07135 | 2,46 | 0,08 | 1,11 | 0,07 | 9,45E-01 | 9,71E-01 |
| AT1G51150 | 4,36 | -0,13 | 1,84 | -0,07 | 9,45E-01 | 9,71E-01 |
| AT3G28550 | 2,18 | -0,10 | 1,49 | -0,07 | 9,46E-01 | 9,71E-01 |
| AT4G31990 | 1,81 | 0,12 | 1,77 | 0,07 | 9,47E-01 | 9,71E-01 |
| AT5G26000 | 4,62 | 0,05 | 0,77 | 0,07 | 9,44E-01 | 9,71E-01 |
| AT5G60920 | 1,88 | 0,08 | 1,23 | 0,07 | 9,48E-01 | 9,71E-01 |
| AT4G40100 | 3,82 | -0,06 | 0,93 | -0,06 | 9,49E-01 | 9,71E-01 |
| AT5G09810 | 3,96 | 0,07 | 1,11 | 0,06 | 9,50E-01 | 9,71E-01 |
| AT5G09065 | 4,44 | -0,05 | 0,80 | -0,06 | 9,50E-01 | 9,71E-01 |
| AT4G02130 | 3,77 | -0,05 | 0,87 | -0,06 | 9,50E-01 | 9,71E-01 |
| AT3G56400 | 2,09 | 0,07 | 1,20 | 0,06 | 9,52E-01 | 9,72E-01 |
| AT5G14550 | 7872,48 | 0,01 | 0,21 | 0,06 | 9,52E-01 | 9,72E-01 |
| AT5G13180 | 1,85 | -0,08 | 1,37 | -0,06 | 9,53E-01 | 9,73E-01 |
| AT2G03890 | 2,03 | 0,07 | 1,15 | 0,06 | 9,55E-01 | 9,74E-01 |
| AT5G02960 | 2,13 | 0,08 | 1,38 | 0,06 | 9,55E-01 | 9,74E-01 |
| AT5G09585 | 3,20 | -0,05 | 0,96 | -0,06 | 9,56E-01 | 9,74E-01 |
| AT5G14545 | 7868,55 | 0,01 | 0,20 | 0,05 | 9,57E-01 | 9,75E-01 |
| AT2G01320 | 2,02 | 0,07 | 1,34 | 0,05 | 9,57E-01 | 9,75E-01 |
| AT1G74290 | 1,78 | -0,08 | 1,46 | -0,05 | 9,58E-01 | 9,75E-01 |
| AT4G19120 | 2,16 | 0,06 | 1,19 | 0,05 | 9,58E-01 | 9,75E-01 |
| AT3G02045 | 5,72 | 0,04 | 0,79 | 0,05 | 9,59E-01 | 9,76E-01 |
| AT2G07806 | 2,71 | -0,06 | 1,28 | -0,05 | 9,60E-01 | 9,76E-01 |
| AT1G54270 | 2,59 | 0,04 | 0,96 | 0,05 | 9,63E-01 | 9,79E-01 |
| AT2G07725 | 2,14 | -0,05 | 1,15 | -0,04 | 9,66E-01 | 9,80E-01 |
| AT1G04410 | 3,62 | 0,05 | 1,05 | 0,04 | 9,65E-01 | 9,80E-01 |
| AT1G63005 | 3,14 | 0,05 | 1,19 | 0,04 | 9,66E-01 | 9,80E-01 |
| AT3G26520 | 6,24 | -0,04 | 0,81 | -0,04 | 9,65E-01 | 9,80E-01 |
| AT4G14360 | 1,93 | 0,05 | 1,28 | 0,04 | 9,67E-01 | 9,81E-01 |
| AT1G11870 | 3,10 | -0,04 | 0,94 | -0,04 | 9,68E-01 | 9,81E-01 |
| AT3G04400 | 2,05 | -0,05 | 1,29 | -0,04 | 9,69E-01 | 9,81E-01 |
| AT3G56705 | 15,48 | 0,02 | 0,51 | 0,04 | 9,71E-01 | 9,83E-01 |
| AT2G25510 | 7,52 | -0,03 | 0,75 | -0,03 | 9,73E-01 | 9,85E-01 |
| AT4G26730 | 2,38 | -0,05 | 1,42 | -0,03 | 9,73E-01 | 9,85E-01 |
| AT1G24510 | 1,77 | -0,04 | 1,28 | -0,03 | 9,75E-01 | 9,86E-01 |
| AT1G76490 | 3,44 | -0,03 | 1,16 | -0,03 | 9,80E-01 | 9,88E-01 |
| AT2G07727 | 3,34 | 0,04 | 1,31 | 0,03 | 9,78E-01 | 9,88E-01 |
| AT2G05355 | 1,78 | 0,04 | 1,40 | 0,03 | 9,77E-01 | 9,88E-01 |
| AT2G41100 | 6,69 | -0,02 | 0,70 | -0,03 | 9,80E-01 | 9,88E-01 |
| AT4G38550 | 2,97 | 0,03 | 1,05 | 0,03 | 9,79E-01 | 9,88E-01 |
| AT2G05685 | 2,44 | -0,03 | 1,30 | -0,03 | 9,80E-01 | 9,88E-01 |
| AT4G38552 | 2,97 | 0,03 | 1,05 | 0,03 | 9,79E-01 | 9,88E-01 |
| AT1G20020 | 2,67 | 0,02 | 1,00 | 0,02 | 9,82E-01 | 9,89E-01 |
| AT1G74090 | 6,35 | -0,02 | 0,72 | -0,02 | 9,83E-01 | 9,89E-01 |
| AT3G01370 | 2,22 | -0,03 | 1,18 | -0,02 | 9,83E-01 | 9,89E-01 |
| AT3G57720 | 13,95 | -0,01 | 0,61 | -0,02 | 9,83E-01 | 9,89E-01 |

| | | | | | | |
|---|---|---|---|---|---|---|
| AT1G51980 | 1,85 | 0,03 | 1,38 | 0,02 | 9,82E-01 | 9,89E-01 |
| AT1G53640 | 2,14 | 0,03 | 1,46 | 0,02 | 9,83E-01 | 9,89E-01 |
| AT4G28750 | 4,56 | -0,02 | 1,08 | -0,02 | 9,84E-01 | 9,89E-01 |
| AT3G14310 | 2,07 | 0,02 | 1,64 | 0,01 | 9,90E-01 | 9,95E-01 |
| AT1G09687 | 2,04 | -0,01 | 1,14 | -0,01 | 9,91E-01 | 9,95E-01 |
| AT5G44870 | 1,81 | -0,01 | 1,24 | -0,01 | 9,93E-01 | 9,97E-01 |
| AT4G08620 | 2,91 | 0,01 | 1,21 | 0,01 | 9,94E-01 | 9,97E-01 |
| AT3G19010 | 1,81 | 0,01 | 1,49 | 0,01 | 9,94E-01 | 9,97E-01 |
| AT1G20260 | 2,71 | 0,00 | 1,20 | 0,00 | 9,98E-01 | 1,00E+00 |
| AT1G06273 | 2,17 | 0,00 | 2,08 | 0,00 | 1,00E+00 | 1,00E+00 |
| AT5G65700 | 1,88 | 0,00 | 1,27 | 0,00 | 1,00E+00 | 1,00E+00 |
| AT1G53683 | 15,44 | 0,00 | 1,74 | 0,00 | 1,00E+00 | 1,00E+00 |
| AT1G56630 | 1,84 | 0,00 | 2,38 | 0,00 | 1,00E+00 | 1,00E+00 |
| AT3G25855 | 1,70 | 0,00 | 2,19 | 0,00 | 1,00E+00 | 1,00E+00 |
| AT5G22800 | 1,71 | 0,00 | 1,43 | 0,00 | 9,99E-01 | 1,00E+00 |
| AT1G01046 | 1,63 | -0,65 | 1,32 | -0,49 | 6,23E-01 | |
| AT1G01160 | 0,43 | -1,49 | 3,18 | -0,47 | 6,40E-01 | |
| AT1G01180 | 0,85 | 1,73 | 2,15 | 0,81 | 4,20E-01 | |
| AT1G01220 | 0,11 | 1,33 | 4,36 | 0,31 | 7,59E-01 | |
| AT1G01225 | 0,31 | 0,83 | 4,35 | 0,19 | 8,49E-01 | |
| AT1G01230 | 0,37 | 0,09 | 4,38 | 0,02 | 9,84E-01 | |
| AT1G01240 | 0,19 | -0,39 | 4,29 | -0,09 | 9,27E-01 | |
| AT1G01260 | 0,53 | 0,87 | 2,64 | 0,33 | 7,42E-01 | |
| AT1G01300 | 1,08 | -0,41 | 1,68 | -0,24 | 8,09E-01 | |
| AT1G01350 | 0,29 | 1,86 | 4,33 | 0,43 | 6,67E-01 | |
| AT1G01448 | 0,59 | 3,63 | 4,32 | 0,84 | 4,00E-01 | |
| AT1G01480 | 0,12 | 2,87 | 4,34 | 0,66 | 5,08E-01 | |
| AT1G01520 | 0,12 | 2,87 | 4,34 | 0,66 | 5,08E-01 | |
| AT1G01540 | 0,22 | 0,81 | 4,27 | 0,19 | 8,50E-01 | |
| AT1G01560 | 1,60 | 2,97 | 1,78 | 1,67 | 9,46E-02 | |
| AT1G01630 | 0,58 | -0,07 | 3,84 | -0,02 | 9,85E-01 | |
| AT1G01710 | 0,82 | -0,33 | 3,10 | -0,11 | 9,14E-01 | |
| AT1G01720 | 1,37 | -0,08 | 1,35 | -0,06 | 9,54E-01 | |
| AT1G04047 | 0,70 | 0,23 | 2,33 | 0,10 | 9,21E-01 | |
| AT1G01730 | 0,40 | 1,64 | 4,38 | 0,37 | 7,08E-01 | |
| AT1G01790 | 0,83 | 0,73 | 1,78 | 0,41 | 6,80E-01 | |
| AT1G01810 | 0,12 | 2,81 | 4,34 | 0,65 | 5,18E-01 | |
| AT1G01950 | 0,18 | 0,26 | 4,30 | 0,06 | 9,51E-01 | |
| AT1G02010 | 0,10 | 0,39 | 4,35 | 0,09 | 9,29E-01 | |
| AT1G02100 | 0,49 | 1,65 | 3,06 | 0,54 | 5,89E-01 | |
| AT1G02120 | 0,36 | 1,51 | 3,39 | 0,44 | 6,57E-01 | |
| AT1G02150 | 1,38 | 0,03 | 1,62 | 0,02 | 9,85E-01 | |
| AT1G02205 | 1,06 | -0,27 | 1,82 | -0,15 | 8,82E-01 | |
| AT1G04093 | 0,08 | -0,56 | 4,38 | -0,13 | 8,99E-01 | |
| AT1G02300 | 0,34 | 1,12 | 4,32 | 0,26 | 7,96E-01 | |
| AT1G02305 | 0,80 | 2,42 | 2,44 | 0,99 | 3,21E-01 | |
| AT1G04103 | 0,08 | 2,33 | 4,36 | 0,53 | 5,93E-01 | |
| AT1G02390 | 0,19 | -0,01 | 4,32 | 0,00 | 9,98E-01 | |
| AT1G02420 | 0,52 | 3,95 | 3,86 | 1,02 | 3,06E-01 | |
| AT1G02560 | 1,57 | -0,46 | 1,52 | -0,30 | 7,62E-01 | |
| AT1G02690 | 0,11 | -1,28 | 4,35 | -0,29 | 7,69E-01 | |
| AT1G02750 | 0,51 | 1,71 | 4,40 | 0,39 | 6,98E-01 | |
| AT1G02816 | 0,38 | 1,00 | 3,27 | 0,31 | 7,59E-01 | |
| AT1G02850 | 0,62 | 3,35 | 2,87 | 1,17 | 2,42E-01 | |
| AT1G03160 | 0,95 | -0,24 | 2,16 | -0,11 | 9,10E-01 | |
| AT1G03190 | 0,39 | 1,35 | 3,37 | 0,40 | 6,89E-01 | |

| | | | | | |
|---|---|---|---|---|---|
| AT1G03220 | 1,11 | 0,86 | 1,91 | 0,45 | 6,51E-01 |
| AT1G03280 | 0,25 | 2,55 | 4,32 | 0,59 | 5,55E-01 |
| AT1G03290 | 0,75 | 1,29 | 2,21 | 0,58 | 5,59E-01 |
| AT1G03310 | 0,40 | 1,65 | 4,19 | 0,39 | 6,93E-01 |
| AT1G03360 | 0,35 | 0,65 | 3,40 | 0,19 | 8,48E-01 |
| AT1G03365 | 0,40 | 2,10 | 3,36 | 0,62 | 5,33E-01 |
| AT1G03370 | 0,51 | 0,63 | 3,85 | 0,16 | 8,70E-01 |
| AT1G03380 | 1,12 | 1,10 | 1,92 | 0,57 | 5,68E-01 |
| AT1G03430 | 0,37 | 0,18 | 4,35 | 0,04 | 9,67E-01 |
| AT1G03440 | 0,84 | -0,62 | 3,11 | -0,20 | 8,41E-01 |
| AT1G03545 | 0,70 | -0,51 | 2,29 | -0,22 | 8,25E-01 |
| AT1G03600 | 1,40 | -0,43 | 1,55 | -0,28 | 7,81E-01 |
| AT1G03610 | 0,72 | 0,88 | 3,10 | 0,28 | 7,76E-01 |
| AT1G03630 | 0,51 | -0,87 | 2,16 | -0,40 | 6,88E-01 |
| AT1G03687 | 0,11 | -1,28 | 4,35 | -0,29 | 7,69E-01 |
| AT1G03740 | 0,96 | -0,38 | 1,78 | -0,21 | 8,33E-01 |
| AT1G03810 | 0,77 | -1,32 | 2,04 | -0,65 | 5,19E-01 |
| AT1G03900 | 1,11 | 0,59 | 1,84 | 0,32 | 7,47E-01 |
| AT1G03905 | 0,24 | 0,00 | 4,41 | 0,00 | 1,00E+00 |
| AT1G03910 | 0,50 | -0,12 | 3,08 | -0,04 | 9,69E-01 |
| AT1G03930 | 0,53 | 1,40 | 2,63 | 0,53 | 5,94E-01 |
| AT1G04217 | 0,82 | 0,95 | 1,83 | 0,52 | 6,04E-01 |
| AT1G03935 | 0,16 | -0,80 | 4,37 | -0,18 | 8,54E-01 |
| AT1G04223 | 0,98 | -2,87 | 1,92 | -1,49 | 1,35E-01 |
| AT1G04227 | 0,32 | 0,86 | 3,60 | 0,24 | 8,12E-01 |
| AT1G03960 | 0,15 | 2,31 | 4,37 | 0,53 | 5,98E-01 |
| AT1G03970 | 0,36 | -0,33 | 4,23 | -0,08 | 9,39E-01 |
| AT1G04020 | 0,33 | 0,00 | 4,41 | 0,00 | 1,00E+00 |
| AT1G04130 | 0,50 | -0,89 | 2,51 | -0,35 | 7,24E-01 |
| AT1G04170 | 1,02 | 2,96 | 2,06 | 1,44 | 1,49E-01 |
| AT1G04247 | 0,30 | -1,30 | 4,35 | -0,30 | 7,65E-01 |
| AT1G04230 | 0,70 | 0,27 | 2,99 | 0,09 | 9,29E-01 |
| AT1G04295 | 0,43 | 0,66 | 2,62 | 0,25 | 8,01E-01 |
| AT1G04350 | 0,25 | -1,26 | 4,36 | -0,29 | 7,73E-01 |
| AT1G04420 | 1,20 | 1,39 | 1,66 | 0,84 | 4,02E-01 |
| AT1G04440 | 0,50 | -0,53 | 2,59 | -0,20 | 8,38E-01 |
| AT1G04273 | 0,37 | -0,46 | 3,44 | -0,14 | 8,93E-01 |
| AT1G04277 | 0,93 | -0,17 | 2,12 | -0,08 | 9,36E-01 |
| AT1G04510 | 0,67 | 2,08 | 2,98 | 0,70 | 4,85E-01 |
| AT1G04540 | 1,05 | 2,70 | 2,14 | 1,26 | 2,07E-01 |
| AT1G04690 | 0,77 | 0,75 | 2,20 | 0,34 | 7,35E-01 |
| AT1G04710 | 0,47 | 2,93 | 4,35 | 0,67 | 5,00E-01 |
| AT1G04760 | 0,36 | 0,31 | 4,35 | 0,07 | 9,42E-01 |
| AT1G04945 | 0,47 | 1,25 | 3,23 | 0,39 | 6,99E-01 |
| AT1G04970 | 0,34 | 2,55 | 4,35 | 0,59 | 5,58E-01 |
| AT1G05087 | 0,44 | 2,87 | 4,05 | 0,71 | 4,78E-01 |
| AT1G05090 | 0,12 | 2,87 | 4,34 | 0,66 | 5,08E-01 |
| AT1G05140 | 0,22 | -1,21 | 4,35 | -0,28 | 7,81E-01 |
| AT1G05200 | 1,12 | 1,04 | 1,90 | 0,55 | 5,82E-01 |
| AT1G05360 | 0,08 | 2,33 | 4,36 | 0,53 | 5,93E-01 |
| AT1G05380 | 1,42 | 2,92 | 1,93 | 1,51 | 1,30E-01 |
| AT1G05500 | 0,37 | -0,36 | 4,32 | -0,08 | 9,34E-01 |
| AT1G04397 | 0,08 | 2,33 | 4,36 | 0,53 | 5,93E-01 |
| AT1G05600 | 0,08 | -0,56 | 4,38 | -0,13 | 8,99E-01 |
| AT1G04423 | 1,18 | -1,80 | 1,75 | -1,02 | 3,05E-01 |
| AT1G05790 | 0,76 | 3,09 | 2,43 | 1,27 | 2,03E-01 |

| | | | | | |
|---|---|---|---|---|---|
| AT1G05830 | 0,53 | 0,49 | 3,23 | 0,15 | 8,80E-01 |
| AT1G05890 | 0,68 | 1,82 | 4,40 | 0,41 | 6,79E-01 |
| AT1G05900 | 0,21 | -0,24 | 4,35 | -0,06 | 9,56E-01 |
| AT1G05910 | 0,59 | 2,86 | 3,79 | 0,75 | 4,51E-01 |
| AT1G05980 | 0,23 | -1,37 | 4,35 | -0,32 | 7,52E-01 |
| AT1G06002 | 0,95 | -0,10 | 1,92 | -0,05 | 9,57E-01 |
| AT1G06030 | 0,96 | 2,92 | 3,37 | 0,87 | 3,85E-01 |
| AT1G06290 | 0,60 | 0,80 | 2,66 | 0,30 | 7,62E-01 |
| AT1G06380 | 0,45 | -1,48 | 4,17 | -0,36 | 7,22E-01 |
| AT1G06390 | 0,40 | 0,50 | 3,36 | 0,15 | 8,82E-01 |
| AT1G06410 | 1,61 | 0,53 | 1,42 | 0,37 | 7,09E-01 |
| AT1G06530 | 0,96 | 1,56 | 2,19 | 0,71 | 4,76E-01 |
| AT1G06640 | 1,12 | -0,54 | 1,57 | -0,35 | 7,29E-01 |
| AT1G06645 | 0,50 | 2,29 | 3,95 | 0,58 | 5,62E-01 |
| AT1G06650 | 0,69 | 1,50 | 2,90 | 0,52 | 6,05E-01 |
| AT1G06670 | 1,38 | 1,00 | 1,82 | 0,55 | 5,84E-01 |
| AT1G06750 | 0,24 | 2,45 | 4,35 | 0,56 | 5,73E-01 |
| AT1G06780 | 0,12 | 0,63 | 4,33 | 0,15 | 8,85E-01 |
| AT1G06910 | 0,23 | 1,68 | 4,33 | 0,39 | 6,98E-01 |
| AT1G06930 | 0,08 | 2,33 | 4,36 | 0,53 | 5,93E-01 |
| AT1G07010 | 0,86 | 0,92 | 2,81 | 0,33 | 7,43E-01 |
| AT1G04537 | 0,24 | 0,00 | 4,41 | 0,00 | 1,00E+00 |
| AT1G07080 | 0,86 | -0,03 | 2,00 | -0,01 | 9,90E-01 |
| AT1G07130 | 0,28 | 1,09 | 4,31 | 0,25 | 8,00E-01 |
| AT1G07170 | 0,78 | 2,25 | 2,03 | 1,11 | 2,69E-01 |
| AT1G07180 | 0,37 | -0,52 | 3,42 | -0,15 | 8,80E-01 |
| AT1G07210 | 0,17 | 3,04 | 4,33 | 0,70 | 4,83E-01 |
| AT1G07240 | 0,59 | 0,80 | 3,19 | 0,25 | 8,02E-01 |
| AT1G07280 | 1,33 | 3,58 | 1,96 | 1,82 | 6,87E-02 |
| AT1G07320 | 1,43 | 0,91 | 1,54 | 0,59 | 5,54E-01 |
| AT1G07470 | 0,61 | 2,83 | 3,73 | 0,76 | 4,49E-01 |
| AT1G07630 | 0,70 | 0,42 | 2,80 | 0,15 | 8,82E-01 |
| AT1G07705 | 0,12 | 1,65 | 4,34 | 0,38 | 7,04E-01 |
| AT1G07728 | 1,03 | 3,54 | 2,52 | 1,40 | 1,60E-01 |
| AT1G07790 | 0,25 | -0,13 | 4,36 | -0,03 | 9,77E-01 |
| AT1G07830 | 0,43 | 1,63 | 4,11 | 0,40 | 6,92E-01 |
| AT1G07990 | 0,78 | 1,52 | 2,91 | 0,52 | 6,03E-01 |
| AT1G08110 | 0,41 | 1,93 | 4,28 | 0,45 | 6,52E-01 |
| AT1G08190 | 0,34 | -0,46 | 3,48 | -0,13 | 8,95E-01 |
| AT1G08260 | 0,51 | 1,34 | 4,00 | 0,34 | 7,37E-01 |
| AT1G08360 | 1,45 | 2,86 | 1,59 | 1,79 | 7,28E-02 |
| AT1G08370 | 0,71 | 3,05 | 2,45 | 1,25 | 2,12E-01 |
| AT1G08410 | 0,87 | 0,90 | 2,49 | 0,36 | 7,19E-01 |
| AT1G08460 | 0,84 | 0,45 | 1,91 | 0,24 | 8,12E-01 |
| AT1G08465 | 0,35 | 1,15 | 4,31 | 0,27 | 7,90E-01 |
| AT1G08480 | 0,59 | 2,67 | 3,09 | 0,86 | 3,87E-01 |
| AT1G08520 | 1,05 | 1,19 | 1,69 | 0,70 | 4,82E-01 |
| AT1G08530 | 0,48 | 3,24 | 3,96 | 0,82 | 4,13E-01 |
| AT1G08550 | 0,85 | 2,31 | 2,24 | 1,03 | 3,04E-01 |
| AT1G08570 | 0,49 | 0,85 | 4,09 | 0,21 | 8,35E-01 |
| AT1G08590 | 0,74 | 3,49 | 2,84 | 1,23 | 2,20E-01 |
| AT1G08600 | 0,40 | 1,87 | 3,22 | 0,58 | 5,62E-01 |
| AT1G08620 | 1,06 | 1,69 | 1,76 | 0,96 | 3,37E-01 |
| AT1G08650 | 0,89 | 1,53 | 2,89 | 0,53 | 5,97E-01 |
| AT1G08680 | 0,17 | -1,97 | 4,31 | -0,46 | 6,47E-01 |
| AT1G08710 | 0,30 | 2,39 | 4,37 | 0,55 | 5,84E-01 |

| | | | | | |
|---|---|---|---|---|---|
| AT1G08720 | 1,46 | 4,09 | 2,04 | 2,01 | 4,43E-02 |
| AT1G08760 | 0,68 | 1,92 | 2,94 | 0,65 | 5,13E-01 |
| AT1G08770 | 0,33 | -1,15 | 3,46 | -0,33 | 7,38E-01 |
| AT1G08830 | 1,29 | 3,14 | 2,05 | 1,53 | 1,26E-01 |
| AT1G08900 | 0,34 | 0,78 | 3,39 | 0,23 | 8,18E-01 |
| AT1G08910 | 0,43 | -1,54 | 4,16 | -0,37 | 7,12E-01 |
| AT1G08920 | 0,59 | 2,82 | 3,16 | 0,89 | 3,73E-01 |
| AT1G08930 | 1,31 | 1,09 | 1,42 | 0,77 | 4,44E-01 |
| AT1G08970 | 0,82 | 1,82 | 2,52 | 0,72 | 4,70E-01 |
| AT1G08980 | 0,91 | 1,73 | 2,10 | 0,82 | 4,10E-01 |
| AT1G09040 | 0,45 | 1,89 | 3,28 | 0,57 | 5,65E-01 |
| AT1G09250 | 0,32 | 0,74 | 4,37 | 0,17 | 8,65E-01 |
| AT1G09260 | 0,62 | 0,00 | 4,41 | 0,00 | 1,00E+00 |
| AT1G09270 | 0,12 | 2,87 | 4,34 | 0,66 | 5,08E-01 |
| AT1G09300 | 0,36 | 1,44 | 4,33 | 0,33 | 7,40E-01 |
| AT1G09390 | 0,23 | 0,18 | 4,27 | 0,04 | 9,66E-01 |
| AT1G09421 | 0,17 | 1,41 | 4,35 | 0,32 | 7,46E-01 |
| AT1G09430 | 0,79 | 0,13 | 2,27 | 0,06 | 9,55E-01 |
| AT1G09450 | 0,24 | 0,06 | 4,31 | 0,01 | 9,89E-01 |
| AT1G09490 | 0,05 | -0,58 | 4,38 | -0,13 | 8,95E-01 |
| AT1G09500 | 0,08 | 2,33 | 4,36 | 0,53 | 5,93E-01 |
| AT1G09530 | 0,63 | 2,92 | 3,70 | 0,79 | 4,29E-01 |
| AT1G09560 | 0,29 | 3,11 | 4,33 | 0,72 | 4,73E-01 |
| AT1G09590 | 0,27 | -0,51 | 4,31 | -0,12 | 9,07E-01 |
| AT1G09600 | 0,94 | 4,30 | 2,79 | 1,54 | 1,24E-01 |
| AT1G09610 | 0,36 | 2,79 | 4,34 | 0,64 | 5,21E-01 |
| AT1G09645 | 0,33 | 2,37 | 4,35 | 0,55 | 5,85E-01 |
| AT1G09665 | 0,25 | 0,00 | 4,40 | 0,00 | 1,00E+00 |
| AT1G09710 | 0,91 | 1,52 | 2,86 | 0,53 | 5,95E-01 |
| AT1G09750 | 0,82 | -0,88 | 1,83 | -0,48 | 6,33E-01 |
| AT1G09795 | 0,58 | -2,11 | 3,94 | -0,53 | 5,93E-01 |
| AT1G09815 | 0,21 | -1,23 | 4,36 | -0,28 | 7,78E-01 |
| AT1G10050 | 0,78 | 1,88 | 2,26 | 0,83 | 4,06E-01 |
| AT1G10070 | 0,24 | 2,85 | 4,34 | 0,66 | 5,11E-01 |
| AT1G10130 | 1,07 | 0,75 | 1,73 | 0,43 | 6,65E-01 |
| AT1G10150 | 1,47 | 1,32 | 1,28 | 1,03 | 3,02E-01 |
| AT1G10270 | 0,58 | 0,85 | 2,60 | 0,33 | 7,44E-01 |
| AT1G10450 | 0,33 | 2,56 | 4,31 | 0,59 | 5,52E-01 |
| AT1G10490 | 0,65 | 2,88 | 2,44 | 1,18 | 2,39E-01 |
| AT1G10570 | 0,16 | 1,27 | 4,37 | 0,29 | 7,71E-01 |
| AT1G10600 | 1,27 | 3,03 | 1,85 | 1,64 | 1,01E-01 |
| AT1G10610 | 0,90 | 3,59 | 3,34 | 1,07 | 2,83E-01 |
| AT1G10660 | 0,54 | 0,71 | 4,04 | 0,18 | 8,60E-01 |
| AT1G10670 | 0,76 | -1,62 | 3,57 | -0,45 | 6,50E-01 |
| AT1G10700 | 0,45 | 0,91 | 4,34 | 0,21 | 8,34E-01 |
| AT1G10720 | 0,48 | 1,67 | 2,54 | 0,66 | 5,11E-01 |
| AT1G10745 | 1,57 | -3,41 | 1,78 | -1,92 | 5,51E-02 |
| AT1G10790 | 0,08 | 2,33 | 4,36 | 0,53 | 5,93E-01 |
| AT1G10850 | 0,85 | 3,53 | 3,49 | 1,01 | 3,12E-01 |
| AT1G10910 | 0,08 | 2,33 | 4,36 | 0,53 | 5,93E-01 |
| AT1G10960 | 0,86 | 3,09 | 2,86 | 1,08 | 2,80E-01 |
| AT1G11020 | 0,27 | 0,31 | 4,36 | 0,07 | 9,44E-01 |
| AT1G11050 | 0,22 | -1,04 | 4,30 | -0,24 | 8,08E-01 |
| AT1G11260 | 1,39 | 0,48 | 1,62 | 0,30 | 7,67E-01 |
| AT1G11300 | 0,32 | 2,54 | 4,33 | 0,59 | 5,57E-01 |
| AT1G11303 | 0,25 | 3,58 | 4,32 | 0,83 | 4,07E-01 |

| | | | | | |
|---|---|---|---|---|---|
| AT1G11320 | 0,08 | 2,08 | 4,38 | 0,47 | 6,35E-01 |
| AT1G11330 | 0,64 | 1,57 | 2,81 | 0,56 | 5,77E-01 |
| AT1G11400 | 0,26 | -0,08 | 4,30 | -0,02 | 9,85E-01 |
| AT1G11410 | 0,36 | 0,98 | 3,34 | 0,29 | 7,70E-01 |
| AT1G11430 | 0,35 | -0,01 | 3,56 | 0,00 | 9,98E-01 |
| AT1G11440 | 0,54 | 0,44 | 3,04 | 0,14 | 8,85E-01 |
| AT1G11580 | 0,31 | 2,37 | 4,31 | 0,55 | 5,82E-01 |
| AT1G11590 | 0,67 | -3,03 | 2,41 | -1,26 | 2,08E-01 |
| AT1G11593 | 0,40 | -1,40 | 4,09 | -0,34 | 7,32E-01 |
| AT1G11660 | 1,14 | 2,96 | 1,99 | 1,49 | 1,37E-01 |
| AT1G11680 | 0,83 | 3,17 | 2,71 | 1,17 | 2,42E-01 |
| AT1G11710 | 0,17 | 3,16 | 4,33 | 0,73 | 4,66E-01 |
| AT1G11750 | 0,56 | 0,32 | 3,20 | 0,10 | 9,20E-01 |
| AT1G11790 | 0,42 | 2,01 | 4,37 | 0,46 | 6,45E-01 |
| AT1G11890 | 1,50 | 0,54 | 1,68 | 0,32 | 7,49E-01 |
| AT1G11905 | 0,75 | 1,77 | 2,12 | 0,83 | 4,04E-01 |
| AT1G11930 | 0,68 | 1,92 | 2,21 | 0,87 | 3,85E-01 |
| AT1G12015 | 0,46 | -0,31 | 4,32 | -0,07 | 9,42E-01 |
| AT1G12050 | 1,22 | -1,47 | 1,55 | -0,95 | 3,44E-01 |
| AT1G12060 | 0,12 | 1,15 | 4,35 | 0,26 | 7,92E-01 |
| AT1G12140 | 0,12 | 2,87 | 4,34 | 0,66 | 5,08E-01 |
| AT1G12210 | 0,35 | 0,65 | 4,35 | 0,15 | 8,81E-01 |
| AT1G12220 | 1,65 | -1,44 | 1,30 | -1,11 | 2,67E-01 |
| AT1G12230 | 0,35 | 0,33 | 3,43 | 0,10 | 9,22E-01 |
| AT1G12250 | 0,19 | 0,03 | 4,35 | 0,01 | 9,95E-01 |
| AT1G12270 | 0,62 | 1,19 | 2,59 | 0,46 | 6,46E-01 |
| AT1G12320 | 0,82 | -0,83 | 2,61 | -0,32 | 7,50E-01 |
| AT1G12360 | 0,65 | 0,51 | 3,82 | 0,13 | 8,93E-01 |
| AT1G12410 | 0,67 | 1,73 | 3,81 | 0,45 | 6,50E-01 |
| AT1G12460 | 0,83 | -1,49 | 2,05 | -0,73 | 4,67E-01 |
| AT1G12580 | 0,43 | 0,46 | 2,67 | 0,17 | 8,64E-01 |
| AT1G12760 | 0,76 | 0,41 | 2,03 | 0,20 | 8,40E-01 |
| AT1G12810 | 0,42 | -0,42 | 2,33 | -0,18 | 8,57E-01 |
| AT1G12840 | 1,05 | 2,37 | 1,70 | 1,39 | 1,64E-01 |
| AT1G12860 | 0,23 | 0,82 | 4,35 | 0,19 | 8,50E-01 |
| AT1G12990 | 1,14 | 2,17 | 1,87 | 1,16 | 2,47E-01 |
| AT1G13040 | 0,24 | 0,39 | 4,34 | 0,09 | 9,28E-01 |
| AT1G13060 | 0,83 | 0,54 | 3,09 | 0,17 | 8,62E-01 |
| AT1G13090 | 1,29 | -0,61 | 1,75 | -0,35 | 7,25E-01 |
| AT1G13100 | 0,26 | -1,46 | 4,34 | -0,34 | 7,37E-01 |
| AT1G13180 | 0,14 | 0,87 | 4,31 | 0,20 | 8,40E-01 |
| AT1G13190 | 1,26 | 0,01 | 2,24 | 0,01 | 9,96E-01 |
| AT1G13220 | 0,56 | 2,33 | 3,73 | 0,62 | 5,33E-01 |
| AT1G13260 | 1,59 | 0,39 | 1,59 | 0,25 | 8,04E-01 |
| AT1G13270 | 0,60 | 2,47 | 3,07 | 0,80 | 4,22E-01 |
| AT1G13330 | 0,70 | 1,21 | 2,43 | 0,50 | 6,18E-01 |
| AT1G13400 | 0,23 | -0,64 | 4,39 | -0,15 | 8,84E-01 |
| AT1G13430 | 0,17 | 2,62 | 4,35 | 0,60 | 5,47E-01 |
| AT1G13460 | 0,37 | 2,51 | 4,33 | 0,58 | 5,62E-01 |
| AT1G13820 | 0,11 | 1,28 | 4,35 | 0,29 | 7,68E-01 |
| AT1G13850 | 0,13 | 2,37 | 4,36 | 0,54 | 5,86E-01 |
| AT1G13880 | 0,45 | 0,83 | 3,27 | 0,25 | 8,00E-01 |
| AT1G13990 | 0,47 | 1,47 | 3,22 | 0,46 | 6,48E-01 |
| AT1G14000 | 0,81 | 1,97 | 2,20 | 0,90 | 3,70E-01 |
| AT1G14020 | 0,41 | 1,06 | 4,21 | 0,25 | 8,01E-01 |
| AT1G05047 | 0,12 | 2,87 | 4,34 | 0,66 | 5,08E-01 |

| | | | | | |
|---|---|---|---|---|---|
| AT1G14110 | 0,52 | -0,45 | 3,70 | -0,12 | 9,03E-01 |
| AT1G14150 | 0,05 | -0,58 | 4,38 | -0,13 | 8,95E-01 |
| AT1G14185 | 0,05 | -0,58 | 4,38 | -0,13 | 8,95E-01 |
| AT1G14230 | 0,08 | -0,87 | 4,38 | -0,20 | 8,43E-01 |
| AT1G14250 | 0,32 | -0,56 | 4,38 | -0,13 | 8,99E-01 |
| AT1G14280 | 0,63 | -1,85 | 2,93 | -0,63 | 5,27E-01 |
| AT1G14310 | 0,25 | 1,98 | 4,32 | 0,46 | 6,47E-01 |
| AT1G14345 | 0,53 | 0,11 | 2,59 | 0,04 | 9,65E-01 |
| AT1G14350 | 0,37 | 1,15 | 4,34 | 0,27 | 7,90E-01 |
| AT1G14370 | 0,91 | 0,52 | 1,71 | 0,30 | 7,62E-01 |
| AT1G14470 | 0,37 | 0,00 | 4,40 | 0,00 | 1,00E+00 |
| AT1G14480 | 0,08 | -0,56 | 4,38 | -0,13 | 8,99E-01 |
| AT1G14560 | 0,21 | 1,20 | 4,36 | 0,28 | 7,83E-01 |
| AT1G14570 | 1,23 | 1,65 | 1,60 | 1,03 | 3,02E-01 |
| AT1G14650 | 1,32 | 3,60 | 2,11 | 1,71 | 8,77E-02 |
| AT1G14800 | 1,13 | -0,57 | 1,67 | -0,34 | 7,34E-01 |
| AT1G14860 | 0,08 | -0,56 | 4,38 | -0,13 | 8,99E-01 |
| AT1G15030 | 0,23 | 0,98 | 4,29 | 0,23 | 8,20E-01 |
| AT1G15120 | 0,30 | 1,38 | 4,29 | 0,32 | 7,48E-01 |
| AT1G15190 | 0,17 | 2,95 | 4,34 | 0,68 | 4,96E-01 |
| AT1G15240 | 0,58 | 1,46 | 2,52 | 0,58 | 5,62E-01 |
| AT1G15260 | 0,61 | 1,33 | 3,14 | 0,42 | 6,72E-01 |
| AT1G15390 | 0,48 | 3,50 | 4,33 | 0,81 | 4,19E-01 |
| AT1G15430 | 0,55 | 2,04 | 3,88 | 0,53 | 5,98E-01 |
| AT1G15650 | 0,36 | 0,00 | 4,41 | 0,00 | 1,00E+00 |
| AT1G05183 | 0,40 | 1,95 | 3,33 | 0,58 | 5,59E-01 |
| AT1G15720 | 0,33 | 1,28 | 4,29 | 0,30 | 7,65E-01 |
| AT1G15740 | 0,65 | -2,10 | 2,35 | -0,89 | 3,71E-01 |
| AT1G15810 | 0,77 | 0,32 | 1,84 | 0,18 | 8,61E-01 |
| AT1G15890 | 1,49 | -1,01 | 1,73 | -0,58 | 5,59E-01 |
| AT1G15910 | 0,69 | -0,14 | 3,64 | -0,04 | 9,70E-01 |
| AT1G15920 | 0,80 | 3,46 | 2,75 | 1,26 | 2,08E-01 |
| AT1G15930 | 1,08 | 2,82 | 2,00 | 1,41 | 1,58E-01 |
| AT1G15950 | 0,56 | 1,14 | 2,52 | 0,45 | 6,50E-01 |
| AT1G15980 | 1,06 | -1,06 | 1,79 | -0,59 | 5,53E-01 |
| AT1G16060 | 0,12 | 0,63 | 4,33 | 0,15 | 8,85E-01 |
| AT1G16080 | 0,31 | 2,98 | 4,33 | 0,69 | 4,92E-01 |
| AT1G16090 | 0,22 | 1,72 | 4,29 | 0,40 | 6,89E-01 |
| AT1G16110 | 0,61 | 3,95 | 3,61 | 1,09 | 2,74E-01 |
| AT1G16180 | 1,23 | 1,79 | 1,64 | 1,09 | 2,76E-01 |
| AT1G16250 | 0,81 | 0,63 | 2,10 | 0,30 | 7,63E-01 |
| AT1G16270 | 1,59 | 1,80 | 1,50 | 1,20 | 2,31E-01 |
| AT1G16350 | 0,39 | 0,71 | 4,27 | 0,17 | 8,68E-01 |
| AT1G16400 | 0,30 | -1,48 | 4,34 | -0,34 | 7,33E-01 |
| AT1G16410 | 1,19 | -2,34 | 1,60 | -1,46 | 1,45E-01 |
| AT1G16470 | 0,19 | -1,45 | 4,34 | -0,33 | 7,38E-01 |
| AT1G16500 | 0,25 | 0,42 | 4,34 | 0,10 | 9,23E-01 |
| AT1G16560 | 0,44 | 2,82 | 3,27 | 0,86 | 3,90E-01 |
| AT1G16670 | 0,35 | -1,24 | 4,35 | -0,29 | 7,75E-01 |
| AT1G16680 | 0,85 | 2,05 | 2,37 | 0,87 | 3,87E-01 |
| AT1G16690 | 0,12 | 1,15 | 4,35 | 0,26 | 7,92E-01 |
| AT1G16700 | 0,62 | 0,72 | 3,01 | 0,24 | 8,10E-01 |
| AT1G16720 | 1,64 | 3,61 | 1,69 | 2,14 | 3,25E-02 |
| AT1G16740 | 0,37 | -0,14 | 4,41 | -0,03 | 9,75E-01 |
| AT1G16810 | 0,12 | 2,87 | 4,34 | 0,66 | 5,08E-01 |
| AT1G16860 | 0,38 | 1,15 | 4,25 | 0,27 | 7,86E-01 |

| | | | | | |
|---|---|---|---|---|---|
| AT1G17050 | 0,20 | 2,39 | 4,37 | 0,55 | 5,84E-01 |
| AT1G17070 | 0,73 | 2,98 | 3,56 | 0,84 | 4,03E-01 |
| AT1G17100 | 0,86 | 0,62 | 1,69 | 0,37 | 7,14E-01 |
| AT1G17120 | 0,36 | -0,10 | 4,33 | -0,02 | 9,82E-01 |
| AT1G17150 | 0,40 | 0,16 | 3,30 | 0,05 | 9,61E-01 |
| AT1G17160 | 0,36 | 2,47 | 4,28 | 0,58 | 5,64E-01 |
| AT1G17200 | 0,19 | -1,34 | 4,35 | -0,31 | 7,58E-01 |
| AT1G17230 | 0,51 | -0,67 | 2,92 | -0,23 | 8,18E-01 |
| AT1G17255 | 1,02 | 3,36 | 2,56 | 1,31 | 1,89E-01 |
| AT1G17260 | 0,13 | 2,37 | 4,36 | 0,54 | 5,86E-01 |
| AT1G17300 | 0,61 | 3,95 | 3,61 | 1,09 | 2,74E-01 |
| AT1G17340 | 0,57 | -1,44 | 2,21 | -0,65 | 5,14E-01 |
| AT1G17360 | 1,17 | 3,26 | 2,10 | 1,56 | 1,20E-01 |
| AT1G17420 | 1,07 | -0,21 | 1,94 | -0,11 | 9,13E-01 |
| AT1G17460 | 0,31 | 0,77 | 4,34 | 0,18 | 8,58E-01 |
| AT1G17490 | 0,39 | -0,73 | 3,38 | -0,22 | 8,29E-01 |
| AT1G17500 | 0,69 | 2,02 | 2,14 | 0,94 | 3,45E-01 |
| AT1G17530 | 0,57 | 0,63 | 3,03 | 0,21 | 8,36E-01 |
| AT1G17550 | 1,05 | 1,25 | 1,66 | 0,75 | 4,52E-01 |
| AT1G17570 | 0,08 | -0,56 | 4,38 | -0,13 | 8,99E-01 |
| AT1G17580 | 1,15 | 2,13 | 2,06 | 1,03 | 3,02E-01 |
| AT1G17610 | 0,33 | 0,07 | 4,25 | 0,02 | 9,86E-01 |
| AT1G17620 | 1,43 | -0,55 | 1,54 | -0,36 | 7,22E-01 |
| AT1G17680 | 0,28 | 3,38 | 4,33 | 0,78 | 4,35E-01 |
| AT1G17745 | 0,96 | 0,13 | 1,75 | 0,07 | 9,42E-01 |
| AT1G17750 | 0,21 | 2,34 | 4,32 | 0,54 | 5,89E-01 |
| AT1G17840 | 1,51 | 0,76 | 1,51 | 0,50 | 6,14E-01 |
| AT1G17860 | 0,25 | 3,04 | 4,33 | 0,70 | 4,83E-01 |
| AT1G17970 | 0,47 | -1,39 | 4,09 | -0,34 | 7,33E-01 |
| AT1G18020 | 0,78 | -1,64 | 2,89 | -0,57 | 5,70E-01 |
| AT1G18090 | 0,88 | 4,26 | 3,50 | 1,22 | 2,24E-01 |
| AT1G18130 | 0,21 | 0,78 | 4,34 | 0,18 | 8,58E-01 |
| AT1G18170 | 0,31 | 2,79 | 4,34 | 0,64 | 5,20E-01 |
| AT1G18300 | 0,38 | -0,31 | 4,22 | -0,07 | 9,41E-01 |
| AT1G18390 | 0,88 | -0,11 | 1,85 | -0,06 | 9,52E-01 |
| AT1G18450 | 0,27 | 0,39 | 4,34 | 0,09 | 9,28E-01 |
| AT1G18460 | 0,79 | 1,50 | 1,98 | 0,76 | 4,50E-01 |
| AT1G18480 | 0,17 | -0,44 | 4,37 | -0,10 | 9,19E-01 |
| AT1G18490 | 0,64 | 4,08 | 4,31 | 0,95 | 3,44E-01 |
| AT1G18500 | 0,63 | 0,27 | 2,99 | 0,09 | 9,27E-01 |
| AT1G18570 | 0,88 | 1,03 | 1,83 | 0,56 | 5,74E-01 |
| AT1G18580 | 0,40 | 3,74 | 4,32 | 0,87 | 3,87E-01 |
| AT1G18600 | 0,74 | 1,83 | 2,40 | 0,76 | 4,45E-01 |
| AT1G18620 | 0,88 | 0,57 | 1,96 | 0,29 | 7,72E-01 |
| AT1G18700 | 1,03 | 2,22 | 2,29 | 0,97 | 3,32E-01 |
| AT1G18730 | 0,47 | -1,04 | 4,21 | -0,25 | 8,05E-01 |
| AT1G18820 | 0,52 | -1,03 | 4,34 | -0,24 | 8,12E-01 |
| AT1G18870 | 0,46 | 2,80 | 4,36 | 0,64 | 5,21E-01 |
| AT1G18879 | 1,60 | 0,54 | 1,36 | 0,40 | 6,92E-01 |
| AT1G18900 | 0,70 | 3,44 | 3,52 | 0,98 | 3,28E-01 |
| AT1G18950 | 0,83 | -0,11 | 1,94 | -0,06 | 9,55E-01 |
| AT1G19020 | 0,56 | 0,39 | 2,20 | 0,18 | 8,60E-01 |
| AT1G19025 | 0,65 | 1,44 | 3,05 | 0,47 | 6,36E-01 |
| AT1G19086 | 0,19 | -2,20 | 4,31 | -0,51 | 6,10E-01 |
| AT1G19110 | 0,80 | 2,03 | 2,23 | 0,91 | 3,64E-01 |
| AT1G19150 | 0,10 | -1,26 | 4,35 | -0,29 | 7,72E-01 |

| | | | | | |
|---|---|---|---|---|---|
| AT1G19180 | 1,37 | -2,09 | 1,71 | -1,23 | 2,20E-01 |
| AT1G19190 | 0,39 | 1,78 | 4,41 | 0,40 | 6,86E-01 |
| AT1G19290 | 0,57 | 3,63 | 3,73 | 0,97 | 3,30E-01 |
| AT1G19350 | 0,52 | -0,57 | 3,93 | -0,14 | 8,85E-01 |
| AT1G19376 | 0,56 | -0,47 | 2,86 | -0,16 | 8,70E-01 |
| AT1G19464 | 0,18 | -1,99 | 4,31 | -0,46 | 6,44E-01 |
| AT1G19580 | 0,28 | 2,59 | 4,35 | 0,59 | 5,53E-01 |
| AT1G05507 | 0,08 | -0,56 | 4,38 | -0,13 | 8,99E-01 |
| AT1G19710 | 0,37 | 2,79 | 4,28 | 0,65 | 5,14E-01 |
| AT1G19770 | 0,78 | -0,64 | 2,54 | -0,25 | 8,02E-01 |
| AT1G19800 | 0,29 | 1,93 | 4,36 | 0,44 | 6,58E-01 |
| AT1G19830 | 0,33 | -0,63 | 4,24 | -0,15 | 8,82E-01 |
| AT1G19850 | 0,68 | 1,93 | 2,75 | 0,70 | 4,83E-01 |
| AT1G19910 | 0,96 | 2,68 | 2,14 | 1,25 | 2,10E-01 |
| AT1G19990 | 0,28 | -1,11 | 4,29 | -0,26 | 7,96E-01 |
| AT1G20050 | 0,40 | 1,82 | 3,39 | 0,54 | 5,93E-01 |
| AT1G05523 | 0,59 | 1,69 | 2,35 | 0,72 | 4,71E-01 |
| AT1G20090 | 0,64 | -1,53 | 2,64 | -0,58 | 5,61E-01 |
| AT1G20110 | 0,46 | 1,42 | 2,66 | 0,53 | 5,93E-01 |
| AT1G20280 | 0,77 | -1,29 | 1,87 | -0,69 | 4,90E-01 |
| AT1G20300 | 0,15 | -0,67 | 4,39 | -0,15 | 8,78E-01 |
| AT1G20310 | 0,17 | 3,04 | 4,33 | 0,70 | 4,83E-01 |
| AT1G20375 | 1,50 | -0,01 | 1,52 | -0,01 | 9,95E-01 |
| AT1G20630 | 1,62 | -0,42 | 1,74 | -0,24 | 8,07E-01 |
| AT1G20640 | 1,05 | -0,75 | 1,95 | -0,38 | 7,02E-01 |
| AT1G20693 | 1,05 | -2,21 | 2,12 | -1,04 | 2,98E-01 |
| AT1G20696 | 0,69 | 0,55 | 2,18 | 0,25 | 8,01E-01 |
| AT1G20780 | 1,16 | 3,17 | 1,84 | 1,73 | 8,38E-02 |
| AT1G20823 | 0,81 | -1,39 | 2,01 | -0,69 | 4,90E-01 |
| AT1G20920 | 0,49 | 3,03 | 3,15 | 0,96 | 3,37E-01 |
| AT1G20980 | 1,44 | 2,17 | 1,71 | 1,27 | 2,04E-01 |
| AT1G21000 | 0,77 | -0,13 | 1,97 | -0,06 | 9,49E-01 |
| AT1G21050 | 0,58 | -2,03 | 2,87 | -0,71 | 4,80E-01 |
| AT1G21065 | 0,84 | 0,28 | 1,89 | 0,15 | 8,83E-01 |
| AT1G21140 | 0,08 | 2,33 | 4,36 | 0,53 | 5,93E-01 |
| AT1G21170 | 0,55 | -0,67 | 3,79 | -0,18 | 8,59E-01 |
| AT1G21210 | 0,63 | 3,36 | 2,87 | 1,17 | 2,41E-01 |
| AT1G21230 | 0,43 | -1,24 | 3,12 | -0,40 | 6,92E-01 |
| AT1G21240 | 0,38 | 2,29 | 4,23 | 0,54 | 5,89E-01 |
| AT1G21370 | 0,21 | 1,45 | 4,28 | 0,34 | 7,35E-01 |
| AT1G21430 | 0,53 | 3,29 | 4,33 | 0,76 | 4,47E-01 |
| AT1G21560 | 0,41 | 3,13 | 4,11 | 0,76 | 4,46E-01 |
| AT1G21640 | 0,89 | 0,90 | 1,70 | 0,53 | 5,96E-01 |
| AT1G21660 | 0,28 | 1,87 | 4,32 | 0,43 | 6,65E-01 |
| AT1G21670 | 0,86 | 2,13 | 3,36 | 0,63 | 5,27E-01 |
| AT1G21690 | 0,29 | 1,67 | 4,30 | 0,39 | 6,98E-01 |
| AT1G21720 | 0,34 | 1,12 | 4,32 | 0,26 | 7,96E-01 |
| AT1G21722 | 0,08 | 2,33 | 4,36 | 0,53 | 5,93E-01 |
| AT1G21780 | 0,75 | 0,16 | 2,37 | 0,07 | 9,45E-01 |
| AT1G21835 | 0,94 | -1,51 | 2,04 | -0,74 | 4,58E-01 |
| AT1G21840 | 0,12 | 2,87 | 4,34 | 0,66 | 5,08E-01 |
| AT1G21880 | 0,12 | 1,65 | 4,34 | 0,38 | 7,04E-01 |
| AT1G21910 | 0,33 | 0,00 | 4,41 | 0,00 | 1,00E+00 |
| AT1G21980 | 0,19 | 0,42 | 4,28 | 0,10 | 9,21E-01 |
| AT1G22040 | 0,05 | -0,58 | 4,38 | -0,13 | 8,95E-01 |
| AT1G22070 | 0,41 | 0,80 | 4,26 | 0,19 | 8,51E-01 |

| | | | | | |
|---|---|---|---|---|---|
| AT1G22190 | 1,08 | 1,07 | 1,66 | 0,65 | 5,17E-01 |
| AT1G22280 | 0,53 | -0,24 | 3,23 | -0,07 | 9,41E-01 |
| AT1G22330 | 0,11 | -1,28 | 4,35 | -0,29 | 7,69E-01 |
| AT1G22410 | 1,37 | -1,72 | 1,52 | -1,13 | 2,57E-01 |
| AT1G22500 | 0,40 | 3,80 | 4,32 | 0,88 | 3,79E-01 |
| AT1G22590 | 0,24 | -0,10 | 4,27 | -0,02 | 9,82E-01 |
| AT1G22640 | 0,56 | -1,63 | 3,69 | -0,44 | 6,59E-01 |
| AT1G22730 | 0,55 | 2,47 | 3,87 | 0,64 | 5,23E-01 |
| AT1G22750 | 0,35 | -0,57 | 3,41 | -0,17 | 8,67E-01 |
| AT1G22770 | 1,61 | -2,80 | 1,52 | -1,84 | 6,53E-02 |
| AT1G22780 | 0,54 | 0,23 | 3,18 | 0,07 | 9,41E-01 |
| AT1G22790 | 0,64 | 2,19 | 3,84 | 0,57 | 5,69E-01 |
| AT1G05767 | 0,23 | 1,96 | 4,31 | 0,45 | 6,50E-01 |
| AT1G22840 | 0,62 | 1,18 | 3,09 | 0,38 | 7,03E-01 |
| AT1G22860 | 0,86 | 1,91 | 1,97 | 0,97 | 3,33E-01 |
| AT1G22870 | 0,66 | 3,04 | 2,87 | 1,06 | 2,90E-01 |
| AT1G22880 | 1,11 | 1,77 | 2,92 | 0,61 | 5,43E-01 |
| AT1G22882 | 0,84 | 4,56 | 3,28 | 1,39 | 1,65E-01 |
| AT1G22940 | 0,55 | 1,97 | 2,69 | 0,73 | 4,64E-01 |
| AT1G23030 | 0,27 | 2,11 | 4,38 | 0,48 | 6,30E-01 |
| AT1G23040 | 0,34 | 1,06 | 4,28 | 0,25 | 8,05E-01 |
| AT1G23050 | 1,19 | -2,22 | 2,31 | -0,96 | 3,37E-01 |
| AT1G23100 | 0,08 | 2,33 | 4,36 | 0,53 | 5,93E-01 |
| AT1G23130 | 0,08 | 2,33 | 4,36 | 0,53 | 5,93E-01 |
| AT1G23230 | 1,20 | 0,63 | 1,51 | 0,41 | 6,79E-01 |
| AT1G23290 | 1,36 | 0,54 | 1,57 | 0,34 | 7,31E-01 |
| AT1G23440 | 0,65 | 4,07 | 2,96 | 1,38 | 1,69E-01 |
| AT1G23450 | 0,21 | 2,61 | 4,34 | 0,60 | 5,48E-01 |
| AT1G23490 | 1,31 | 0,02 | 1,36 | 0,02 | 9,86E-01 |
| AT1G23710 | 1,29 | -1,09 | 1,54 | -0,71 | 4,77E-01 |
| AT1G23820 | 1,10 | 2,23 | 2,88 | 0,78 | 4,38E-01 |
| AT1G05837 | 0,53 | 1,06 | 3,12 | 0,34 | 7,34E-01 |
| AT1G23870 | 0,51 | 1,96 | 3,74 | 0,52 | 6,01E-01 |
| AT1G23880 | 0,08 | 2,08 | 4,38 | 0,47 | 6,35E-01 |
| AT1G23915 | 1,01 | 3,12 | 2,36 | 1,32 | 1,86E-01 |
| AT1G24145 | 0,48 | 1,61 | 3,02 | 0,53 | 5,95E-01 |
| AT1G24147 | 0,16 | 1,27 | 4,37 | 0,29 | 7,71E-01 |
| AT1G24150 | 0,48 | 2,53 | 3,89 | 0,65 | 5,16E-01 |
| AT1G24280 | 0,44 | 3,47 | 4,33 | 0,80 | 4,23E-01 |
| AT1G24340 | 0,28 | 0,02 | 3,67 | 0,01 | 9,95E-01 |
| AT1G24360 | 0,34 | -0,79 | 4,32 | -0,18 | 8,54E-01 |
| AT1G24380 | 0,38 | -0,35 | 2,75 | -0,13 | 9,00E-01 |
| AT1G24490 | 0,45 | 1,07 | 3,14 | 0,34 | 7,33E-01 |
| AT1G24530 | 0,17 | 0,69 | 4,34 | 0,16 | 8,73E-01 |
| AT1G24560 | 0,68 | 0,30 | 1,96 | 0,15 | 8,77E-01 |
| AT1G24570 | 0,08 | 2,33 | 4,36 | 0,53 | 5,93E-01 |
| AT1G25275 | 1,34 | -1,10 | 1,63 | -0,67 | 5,02E-01 |
| AT1G25280 | 0,98 | 1,55 | 2,38 | 0,65 | 5,16E-01 |
| AT1G25480 | 0,19 | -2,00 | 4,31 | -0,46 | 6,43E-01 |
| AT1G25490 | 1,64 | 0,62 | 1,28 | 0,48 | 6,28E-01 |
| AT1G25520 | 0,81 | 2,29 | 3,53 | 0,65 | 5,17E-01 |
| AT1G25550 | 1,33 | -1,05 | 1,73 | -0,61 | 5,45E-01 |
| AT1G26130 | 0,54 | 3,11 | 3,81 | 0,81 | 4,15E-01 |
| AT1G26220 | 0,37 | -1,03 | 4,27 | -0,24 | 8,10E-01 |
| AT1G26250 | 0,44 | -1,55 | 3,98 | -0,39 | 6,96E-01 |
| AT1G26460 | 0,79 | 2,66 | 2,82 | 0,94 | 3,45E-01 |

| | | | | | |
|---|---|---|---|---|---|
| AT1G26470 | 0,34 | 3,20 | 4,33 | 0,74 | 4,60E-01 |
| AT1G26550 | 0,37 | -1,53 | 4,27 | -0,36 | 7,20E-01 |
| AT1G26560 | 0,24 | 0,80 | 4,32 | 0,18 | 8,54E-01 |
| AT1G26580 | 0,53 | 0,17 | 2,60 | 0,07 | 9,48E-01 |
| AT1G26630 | 1,29 | -0,52 | 1,59 | -0,33 | 7,42E-01 |
| AT1G26650 | 1,02 | 2,40 | 2,16 | 1,11 | 2,65E-01 |
| AT1G26665 | 0,17 | 3,16 | 4,33 | 0,73 | 4,66E-01 |
| AT1G26670 | 1,04 | 2,13 | 2,16 | 0,99 | 3,24E-01 |
| AT1G26770 | 0,19 | -2,00 | 4,31 | -0,46 | 6,43E-01 |
| AT1G26830 | 0,72 | 0,78 | 2,22 | 0,35 | 7,25E-01 |
| AT1G26920 | 0,73 | -1,29 | 2,11 | -0,61 | 5,42E-01 |
| AT1G26940 | 0,05 | -0,58 | 4,38 | -0,13 | 8,95E-01 |
| AT1G27000 | 0,38 | 1,39 | 4,36 | 0,32 | 7,50E-01 |
| AT1G27060 | 0,73 | 4,22 | 2,86 | 1,48 | 1,39E-01 |
| AT1G27130 | 0,51 | -0,89 | 3,07 | -0,29 | 7,72E-01 |
| AT1G27150 | 0,43 | 0,73 | 4,38 | 0,17 | 8,68E-01 |
| AT1G27170 | 0,35 | 1,45 | 3,36 | 0,43 | 6,66E-01 |
| AT1G27180 | 0,99 | 4,32 | 2,55 | 1,69 | 9,04E-02 |
| AT1G27290 | 1,54 | -0,86 | 1,25 | -0,68 | 4,93E-01 |
| AT1G27300 | 0,40 | 1,22 | 4,18 | 0,29 | 7,70E-01 |
| AT1G27330 | 0,50 | 2,19 | 4,38 | 0,50 | 6,17E-01 |
| AT1G27360 | 0,08 | 2,08 | 4,38 | 0,47 | 6,35E-01 |
| AT1G27400 | 0,67 | 0,91 | 1,97 | 0,46 | 6,44E-01 |
| AT1G06087 | 0,12 | 1,15 | 4,35 | 0,26 | 7,92E-01 |
| AT1G27450 | 0,37 | 1,75 | 4,30 | 0,41 | 6,83E-01 |
| AT1G27460 | 0,11 | 1,33 | 4,36 | 0,31 | 7,59E-01 |
| AT1G27480 | 0,41 | 2,51 | 4,18 | 0,60 | 5,49E-01 |
| AT1G27520 | 0,14 | 0,78 | 4,34 | 0,18 | 8,57E-01 |
| AT1G06097 | 0,36 | 0,27 | 4,37 | 0,06 | 9,52E-01 |
| AT1G27570 | 1,68 | -0,83 | 1,60 | -0,52 | 6,03E-01 |
| AT1G27630 | 0,58 | -1,17 | 3,92 | -0,30 | 7,64E-01 |
| AT1G27650 | 0,97 | 2,26 | 2,03 | 1,12 | 2,65E-01 |
| AT1G27750 | 0,63 | -0,24 | 2,49 | -0,10 | 9,22E-01 |
| AT1G27752 | 1,43 | 2,08 | 1,93 | 1,08 | 2,82E-01 |
| AT1G27760 | 0,65 | 1,97 | 2,44 | 0,81 | 4,19E-01 |
| AT1G27850 | 0,84 | 1,07 | 2,16 | 0,50 | 6,20E-01 |
| AT1G27950 | 0,55 | -1,57 | 3,74 | -0,42 | 6,75E-01 |
| AT1G28005 | 0,32 | -1,76 | 4,26 | -0,41 | 6,79E-01 |
| AT1G28060 | 0,32 | 1,89 | 4,31 | 0,44 | 6,61E-01 |
| AT1G28135 | 0,08 | -0,56 | 4,38 | -0,13 | 8,99E-01 |
| AT1G28340 | 0,87 | 3,70 | 2,66 | 1,39 | 1,63E-01 |
| AT1G28350 | 1,15 | -0,62 | 2,07 | -0,30 | 7,64E-01 |
| AT1G28380 | 1,21 | 1,57 | 1,96 | 0,80 | 4,25E-01 |
| AT1G06163 | 0,36 | -1,99 | 4,32 | -0,46 | 6,44E-01 |
| AT1G28490 | 0,17 | 2,04 | 4,33 | 0,47 | 6,37E-01 |
| AT1G28520 | 0,40 | 1,17 | 3,31 | 0,35 | 7,24E-01 |
| AT1G28680 | 0,55 | 0,63 | 3,13 | 0,20 | 8,40E-01 |
| AT1G28780 | 0,49 | 0,32 | 3,96 | 0,08 | 9,36E-01 |
| AT1G28760 | 0,10 | 0,39 | 4,35 | 0,09 | 9,29E-01 |
| AT1G29120 | 0,19 | 2,25 | 4,32 | 0,52 | 6,02E-01 |
| AT1G29150 | 0,99 | 0,24 | 2,31 | 0,10 | 9,19E-01 |
| AT1G29200 | 0,08 | 2,33 | 4,36 | 0,53 | 5,93E-01 |
| AT1G29260 | 0,92 | 0,93 | 1,62 | 0,58 | 5,64E-01 |
| AT1G29310 | 0,68 | 3,57 | 2,96 | 1,21 | 2,28E-01 |
| AT1G29340 | 0,94 | 0,13 | 1,94 | 0,07 | 9,48E-01 |
| AT1G29380 | 0,65 | 2,27 | 4,37 | 0,52 | 6,03E-01 |

| | | | | | |
|---|---|---|---|---|---|
| AT1G06233 | 0,68 | 0,06 | 2,48 | 0,02 | 9,81E-01 |
| AT1G06253 | 1,09 | 0,17 | 1,53 | 0,11 | 9,11E-01 |
| AT1G06263 | 0,52 | 0,88 | 4,08 | 0,21 | 8,30E-01 |
| AT1G06283 | 1,64 | -1,72 | 1,31 | -1,31 | 1,91E-01 |
| AT1G29620 | 0,47 | -0,72 | 3,15 | -0,23 | 8,18E-01 |
| AT1G29630 | 0,24 | 2,22 | 4,34 | 0,51 | 6,09E-01 |
| AT1G29660 | 1,40 | -2,34 | 1,60 | -1,46 | 1,44E-01 |
| AT1G29760 | 0,40 | 1,58 | 4,27 | 0,37 | 7,11E-01 |
| AT1G29770 | 0,71 | 3,87 | 4,31 | 0,90 | 3,68E-01 |
| AT1G29800 | 0,74 | 0,72 | 3,77 | 0,19 | 8,49E-01 |
| AT1G29830 | 1,13 | 0,86 | 1,77 | 0,49 | 6,27E-01 |
| AT1G29950 | 0,49 | 2,57 | 4,34 | 0,59 | 5,54E-01 |
| AT1G30010 | 0,08 | -0,56 | 4,38 | -0,13 | 8,99E-01 |
| AT1G30200 | 0,70 | 1,88 | 2,13 | 0,88 | 3,77E-01 |
| AT1G30260 | 1,16 | -1,01 | 2,25 | -0,45 | 6,52E-01 |
| AT1G30420 | 0,38 | 2,54 | 4,35 | 0,58 | 5,60E-01 |
| AT1G30430 | 0,16 | -0,80 | 4,37 | -0,18 | 8,54E-01 |
| AT1G30440 | 1,10 | -0,95 | 1,82 | -0,52 | 6,02E-01 |
| AT1G30450 | 0,86 | 2,36 | 2,06 | 1,14 | 2,52E-01 |
| AT1G30520 | 0,41 | 1,64 | 4,38 | 0,37 | 7,08E-01 |
| AT1G30530 | 0,46 | -0,76 | 3,24 | -0,23 | 8,14E-01 |
| AT1G30610 | 0,23 | 0,67 | 4,33 | 0,16 | 8,77E-01 |
| AT1G30620 | 0,56 | 0,76 | 3,15 | 0,24 | 8,09E-01 |
| AT1G30700 | 0,12 | 2,81 | 4,34 | 0,65 | 5,18E-01 |
| AT1G30720 | 0,53 | 3,29 | 4,33 | 0,76 | 4,47E-01 |
| AT1G30757 | 0,16 | -1,66 | 4,33 | -0,38 | 7,01E-01 |
| AT1G30800 | 0,20 | 2,11 | 4,37 | 0,48 | 6,29E-01 |
| AT1G30910 | 0,25 | 0,31 | 4,32 | 0,07 | 9,44E-01 |
| AT1G06393 | 0,32 | 0,36 | 4,35 | 0,08 | 9,34E-01 |
| AT1G31095 | 0,66 | -2,80 | 2,88 | -0,97 | 3,31E-01 |
| AT1G31170 | 0,22 | 1,27 | 4,31 | 0,29 | 7,69E-01 |
| AT1G31175 | 0,74 | 2,09 | 2,48 | 0,85 | 3,98E-01 |
| AT1G31190 | 0,86 | -0,15 | 2,22 | -0,07 | 9,47E-01 |
| AT1G31220 | 0,22 | -1,21 | 4,35 | -0,28 | 7,81E-01 |
| AT1G31290 | 0,15 | 0,40 | 4,33 | 0,09 | 9,27E-01 |
| AT1G31420 | 0,89 | 2,84 | 2,14 | 1,33 | 1,84E-01 |
| AT1G31480 | 0,40 | 3,80 | 4,32 | 0,88 | 3,79E-01 |
| AT1G06447 | 0,24 | 0,27 | 4,35 | 0,06 | 9,50E-01 |
| AT1G31710 | 0,82 | -1,44 | 3,34 | -0,43 | 6,66E-01 |
| AT1G31780 | 0,28 | 1,59 | 4,31 | 0,37 | 7,13E-01 |
| AT1G31800 | 0,33 | 1,51 | 3,46 | 0,44 | 6,62E-01 |
| AT1G31850 | 1,13 | 1,93 | 2,07 | 0,94 | 3,50E-01 |
| AT1G31930 | 0,12 | 2,81 | 4,34 | 0,65 | 5,18E-01 |
| AT1G31970 | 0,74 | 1,47 | 3,05 | 0,48 | 6,31E-01 |
| AT1G32050 | 0,65 | 0,99 | 2,20 | 0,45 | 6,53E-01 |
| AT1G32160 | 0,70 | 1,43 | 2,91 | 0,49 | 6,23E-01 |
| AT1G32220 | 0,21 | -1,82 | 4,32 | -0,42 | 6,73E-01 |
| AT1G32225 | 0,70 | -2,59 | 2,28 | -1,13 | 2,57E-01 |
| AT1G32320 | 0,23 | -0,93 | 4,38 | -0,21 | 8,32E-01 |
| AT1G32337 | 0,30 | 0,02 | 4,30 | 0,00 | 9,97E-01 |
| AT1G32360 | 0,41 | 1,85 | 3,36 | 0,55 | 5,83E-01 |
| AT1G06547 | 0,19 | -1,86 | 4,32 | -0,43 | 6,67E-01 |
| AT1G32385 | 0,41 | -0,80 | 2,77 | -0,29 | 7,74E-01 |
| AT1G32415 | 0,72 | 2,84 | 2,88 | 0,99 | 3,24E-01 |
| AT1G32460 | 0,41 | 0,11 | 4,24 | 0,03 | 9,79E-01 |
| AT1G32580 | 0,32 | 2,03 | 4,37 | 0,46 | 6,42E-01 |

| | | | | | |
|---|---|---|---|---|---|
| AT1G06597 | 1,64 | -0,34 | 1,46 | -0,23 | 8,17E-01 |
| AT1G32700 | 0,46 | 1,57 | 3,22 | 0,49 | 6,27E-01 |
| AT1G32810 | 0,36 | 2,21 | 3,43 | 0,64 | 5,20E-01 |
| AT1G06623 | 0,26 | -2,36 | 4,30 | -0,55 | 5,83E-01 |
| AT1G32870 | 0,59 | 2,48 | 3,80 | 0,65 | 5,14E-01 |
| AT1G32920 | 1,13 | -2,37 | 1,73 | -1,37 | 1,72E-01 |
| AT1G32928 | 0,50 | 0,50 | 4,04 | 0,12 | 9,01E-01 |
| AT1G32940 | 0,41 | 3,00 | 4,33 | 0,69 | 4,88E-01 |
| AT1G32950 | 0,62 | 3,84 | 3,64 | 1,06 | 2,91E-01 |
| AT1G32960 | 0,35 | 1,64 | 4,29 | 0,38 | 7,02E-01 |
| AT1G32990 | 0,48 | 0,11 | 4,38 | 0,03 | 9,79E-01 |
| AT1G33010 | 0,19 | -2,00 | 4,31 | -0,46 | 6,43E-01 |
| AT1G33040 | 0,24 | 1,12 | 4,32 | 0,26 | 7,95E-01 |
| AT1G33110 | 0,55 | 2,52 | 3,84 | 0,66 | 5,12E-01 |
| AT1G33170 | 0,44 | 2,88 | 3,28 | 0,88 | 3,80E-01 |
| AT1G33260 | 0,52 | 2,31 | 3,97 | 0,58 | 5,61E-01 |
| AT1G33360 | 0,47 | 0,35 | 3,99 | 0,09 | 9,30E-01 |
| AT1G33470 | 0,71 | 2,84 | 3,54 | 0,80 | 4,23E-01 |
| AT1G33600 | 1,39 | -1,06 | 1,46 | -0,72 | 4,70E-01 |
| AT1G33610 | 0,15 | 1,94 | 4,32 | 0,45 | 6,54E-01 |
| AT1G33612 | 0,17 | -0,14 | 4,41 | -0,03 | 9,75E-01 |
| AT1G33770 | 0,49 | 0,83 | 4,00 | 0,21 | 8,36E-01 |
| AT1G33811 | 0,18 | 0,41 | 4,34 | 0,10 | 9,24E-01 |
| AT1G33910 | 0,08 | -0,56 | 4,38 | -0,13 | 8,99E-01 |
| AT1G33930 | 0,36 | -1,76 | 4,32 | -0,41 | 6,83E-01 |
| AT1G33960 | 1,02 | 0,76 | 3,19 | 0,24 | 8,12E-01 |
| AT1G33980 | 0,92 | 1,83 | 4,28 | 0,43 | 6,69E-01 |
| AT1G33990 | 0,95 | 0,60 | 1,95 | 0,31 | 7,59E-01 |
| AT1G06723 | 0,59 | 0,79 | 2,69 | 0,29 | 7,68E-01 |
| AT1G34020 | 0,78 | 2,66 | 2,40 | 1,11 | 2,68E-01 |
| AT1G34070 | 0,46 | -2,12 | 3,98 | -0,53 | 5,94E-01 |
| AT1G34130 | 1,45 | 1,85 | 1,45 | 1,28 | 2,01E-01 |
| AT1G34150 | 1,09 | -1,66 | 2,43 | -0,68 | 4,95E-01 |
| AT1G34190 | 1,14 | 1,90 | 1,74 | 1,10 | 2,73E-01 |
| AT1G34200 | 0,21 | 2,29 | 4,32 | 0,53 | 5,96E-01 |
| AT1G34210 | 0,51 | 2,01 | 4,34 | 0,46 | 6,44E-01 |
| AT1G34260 | 1,42 | 2,56 | 1,79 | 1,43 | 1,53E-01 |
| AT1G34290 | 0,25 | -1,37 | 4,28 | -0,32 | 7,49E-01 |
| AT1G06747 | 0,08 | -0,56 | 4,38 | -0,13 | 8,99E-01 |
| AT1G34355 | 0,86 | 2,08 | 2,82 | 0,74 | 4,62E-01 |
| AT1G34370 | 1,49 | 4,22 | 1,93 | 2,18 | 2,91E-02 |
| AT1G34390 | 0,68 | -1,42 | 3,56 | -0,40 | 6,91E-01 |
| AT1G34410 | 0,91 | -3,15 | 3,16 | -1,00 | 3,19E-01 |
| AT1G34420 | 0,36 | 0,88 | 4,19 | 0,21 | 8,34E-01 |
| AT1G34460 | 0,31 | -1,73 | 4,27 | -0,41 | 6,85E-01 |
| AT1G34580 | 0,53 | 2,03 | 4,37 | 0,46 | 6,42E-01 |
| AT1G34770 | 0,12 | 2,87 | 4,34 | 0,66 | 5,08E-01 |
| AT1G34844 | 0,08 | 2,33 | 4,36 | 0,53 | 5,93E-01 |
| AT1G35030 | 0,42 | 1,01 | 3,24 | 0,31 | 7,56E-01 |
| AT1G35040 | 0,23 | -1,89 | 4,31 | -0,44 | 6,61E-01 |
| AT1G35140 | 0,39 | -2,38 | 4,18 | -0,57 | 5,69E-01 |
| AT1G35160 | 0,59 | 2,29 | 2,54 | 0,90 | 3,66E-01 |
| AT1G06827 | 0,56 | -0,37 | 2,61 | -0,14 | 8,87E-01 |
| AT1G35220 | 0,89 | 3,00 | 2,35 | 1,28 | 2,01E-01 |
| AT1G35230 | 1,07 | 1,43 | 2,09 | 0,68 | 4,95E-01 |
| AT1G35242 | 0,19 | 1,04 | 4,29 | 0,24 | 8,09E-01 |

| | | | | | |
|---|---|---|---|---|---|
| AT1G35420 | 0,41 | 0,26 | 3,33 | 0,08 | 9,37E-01 |
| AT1G35460 | 0,12 | 2,87 | 4,34 | 0,66 | 5,08E-01 |
| AT1G35500 | 0,14 | -1,69 | 4,33 | -0,39 | 6,96E-01 |
| AT1G35510 | 0,12 | 0,31 | 4,35 | 0,07 | 9,44E-01 |
| AT1G35515 | 0,52 | 0,62 | 2,19 | 0,28 | 7,79E-01 |
| AT1G35516 | 0,52 | 0,62 | 2,19 | 0,28 | 7,79E-01 |
| AT1G35530 | 0,08 | 2,33 | 4,36 | 0,53 | 5,93E-01 |
| AT1G35540 | 0,95 | -1,60 | 1,77 | -0,90 | 3,66E-01 |
| AT1G35614 | 1,12 | -0,63 | 2,01 | -0,31 | 7,55E-01 |
| AT1G35620 | 0,51 | 0,05 | 3,19 | 0,02 | 9,87E-01 |
| AT1G35660 | 0,28 | 2,49 | 4,33 | 0,57 | 5,66E-01 |
| AT1G35670 | 0,21 | 2,29 | 4,32 | 0,53 | 5,96E-01 |
| AT1G35680 | 0,81 | -1,13 | 2,31 | -0,49 | 6,24E-01 |
| AT1G06927 | 0,83 | -0,05 | 2,55 | -0,02 | 9,86E-01 |
| AT1G35850 | 0,23 | -2,25 | 4,30 | -0,52 | 6,00E-01 |
| AT1G36050 | 0,26 | 1,91 | 4,30 | 0,45 | 6,56E-01 |
| AT1G06963 | 0,05 | -0,58 | 4,38 | -0,13 | 8,95E-01 |
| AT1G36320 | 0,22 | -0,39 | 4,32 | -0,09 | 9,28E-01 |
| AT1G36510 | 0,10 | -1,26 | 4,35 | -0,29 | 7,72E-01 |
| AT1G06987 | 0,58 | 0,97 | 4,32 | 0,22 | 8,23E-01 |
| AT1G36675 | 0,81 | 1,04 | 2,34 | 0,44 | 6,57E-01 |
| AT1G36745 | 0,98 | 0,39 | 1,99 | 0,19 | 8,46E-01 |
| AT1G36756 | 0,94 | -1,25 | 2,24 | -0,56 | 5,76E-01 |
| AT1G36940 | 0,22 | -2,03 | 4,31 | -0,47 | 6,37E-01 |
| AT1G36980 | 0,52 | 2,82 | 3,13 | 0,90 | 3,68E-01 |
| AT1G36992 | 0,32 | 1,02 | 4,36 | 0,23 | 8,15E-01 |
| AT1G37113 | 0,08 | 2,08 | 4,38 | 0,47 | 6,35E-01 |
| AT1G42525 | 0,30 | -0,32 | 4,26 | -0,07 | 9,41E-01 |
| AT1G42540 | 1,07 | 2,68 | 2,25 | 1,19 | 2,33E-01 |
| AT1G07137 | 0,05 | -0,58 | 4,38 | -0,13 | 8,95E-01 |
| AT1G42960 | 0,66 | -0,13 | 4,41 | -0,03 | 9,77E-01 |
| AT1G42980 | 1,30 | 2,81 | 1,78 | 1,58 | 1,15E-01 |
| AT1G43040 | 0,16 | -1,66 | 4,33 | -0,38 | 7,01E-01 |
| AT1G43145 | 1,44 | 0,34 | 1,40 | 0,24 | 8,10E-01 |
| AT1G43171 | 0,72 | 0,00 | 4,41 | 0,00 | 1,00E+00 |
| AT1G43330 | 0,87 | -2,10 | 2,38 | -0,88 | 3,78E-01 |
| AT1G43670 | 0,63 | 0,40 | 2,57 | 0,15 | 8,77E-01 |
| AT1G43680 | 0,25 | 0,00 | 4,40 | 0,00 | 1,00E+00 |
| AT1G43700 | 0,72 | -1,45 | 1,96 | -0,74 | 4,59E-01 |
| AT1G43722 | 0,18 | -1,99 | 4,31 | -0,46 | 6,44E-01 |
| AT1G43770 | 1,22 | 1,33 | 1,61 | 0,82 | 4,10E-01 |
| AT1G43780 | 0,25 | 0,00 | 4,41 | 0,00 | 1,00E+00 |
| AT1G43815 | 1,28 | -3,65 | 1,94 | -1,88 | 6,08E-02 |
| AT1G43890 | 0,40 | 1,53 | 4,30 | 0,36 | 7,22E-01 |
| AT1G43970 | 0,43 | -3,40 | 4,27 | -0,80 | 4,26E-01 |
| AT1G44120 | 0,95 | 2,39 | 2,53 | 0,94 | 3,45E-01 |
| AT1G44750 | 0,33 | 3,89 | 4,32 | 0,90 | 3,68E-01 |
| AT1G44780 | 0,29 | 1,93 | 4,36 | 0,44 | 6,58E-01 |
| AT1G44810 | 0,26 | 1,11 | 3,70 | 0,30 | 7,63E-01 |
| AT1G44890 | 0,08 | -0,56 | 4,38 | -0,13 | 8,99E-01 |
| AT1G44900 | 0,12 | 2,87 | 4,34 | 0,66 | 5,08E-01 |
| AT1G44910 | 0,50 | -0,31 | 2,26 | -0,14 | 8,90E-01 |
| AT1G44960 | 0,10 | 0,39 | 4,35 | 0,09 | 9,29E-01 |
| AT1G45000 | 0,47 | 0,44 | 2,72 | 0,16 | 8,71E-01 |
| AT1G45165 | 0,70 | 1,85 | 2,35 | 0,79 | 4,31E-01 |
| AT1G45180 | 0,30 | 2,84 | 4,35 | 0,65 | 5,14E-01 |

| | | | | | |
|---|---|---|---|---|---|
| AT1G45474 | 0,41 | -1,96 | 3,30 | -0,59 | 5,53E-01 |
| AT1G45688 | 0,89 | -0,17 | 2,13 | -0,08 | 9,37E-01 |
| AT1G45976 | 0,41 | 1,08 | 4,23 | 0,26 | 7,98E-01 |
| AT1G46264 | 0,10 | 0,39 | 4,35 | 0,09 | 9,29E-01 |
| AT1G46554 | 0,28 | 1,20 | 4,33 | 0,28 | 7,82E-01 |
| AT1G47210 | 0,93 | 2,76 | 2,52 | 1,10 | 2,73E-01 |
| AT1G07327 | 1,09 | -1,71 | 2,11 | -0,81 | 4,18E-01 |
| AT1G47250 | 0,35 | 2,00 | 4,32 | 0,46 | 6,44E-01 |
| AT1G47270 | 0,13 | 2,37 | 4,36 | 0,54 | 5,86E-01 |
| AT1G07357 | 0,11 | -1,28 | 4,35 | -0,29 | 7,69E-01 |
| AT1G47480 | 1,07 | 0,71 | 1,84 | 0,38 | 7,00E-01 |
| AT1G47490 | 0,30 | 2,84 | 4,35 | 0,65 | 5,14E-01 |
| AT1G47500 | 0,37 | 2,86 | 4,28 | 0,67 | 5,04E-01 |
| AT1G47530 | 1,40 | 1,26 | 1,63 | 0,77 | 4,39E-01 |
| AT1G47550 | 0,50 | 0,37 | 4,07 | 0,09 | 9,28E-01 |
| AT1G47680 | 0,12 | -0,77 | 4,36 | -0,18 | 8,61E-01 |
| AT1G47690 | 0,26 | 1,88 | 4,28 | 0,44 | 6,61E-01 |
| AT1G47770 | 0,12 | -0,77 | 4,36 | -0,18 | 8,61E-01 |
| AT1G47890 | 1,24 | 2,05 | 1,95 | 1,05 | 2,92E-01 |
| AT1G47940 | 0,36 | -3,01 | 4,28 | -0,70 | 4,82E-01 |
| AT1G47970 | 1,63 | 1,13 | 1,40 | 0,81 | 4,21E-01 |
| AT1G48050 | 0,54 | 0,00 | 4,41 | 0,00 | 1,00E+00 |
| AT1G48100 | 0,12 | 1,65 | 4,34 | 0,38 | 7,04E-01 |
| AT1G48170 | 0,32 | 0,53 | 3,51 | 0,15 | 8,81E-01 |
| AT1G48210 | 0,24 | 2,12 | 4,34 | 0,49 | 6,26E-01 |
| AT1G48230 | 0,28 | 1,30 | 4,37 | 0,30 | 7,66E-01 |
| AT1G48315 | 0,17 | 3,14 | 4,33 | 0,73 | 4,68E-01 |
| AT1G48360 | 1,34 | 2,95 | 2,10 | 1,41 | 1,60E-01 |
| AT1G48460 | 0,51 | 0,19 | 2,95 | 0,06 | 9,50E-01 |
| AT1G07487 | 0,32 | 0,41 | 3,51 | 0,12 | 9,07E-01 |
| AT1G48480 | 0,38 | -1,30 | 4,31 | -0,30 | 7,63E-01 |
| AT1G48540 | 1,33 | 1,81 | 1,70 | 1,07 | 2,87E-01 |
| AT1G48560 | 0,14 | -1,69 | 4,33 | -0,39 | 6,96E-01 |
| AT1G48600 | 0,62 | 1,47 | 2,08 | 0,71 | 4,80E-01 |
| AT1G48640 | 0,26 | 2,30 | 4,37 | 0,53 | 5,98E-01 |
| AT1G48725 | 1,45 | -2,71 | 1,53 | -1,77 | 7,59E-02 |
| AT1G48820 | 0,24 | -0,14 | 4,41 | -0,03 | 9,75E-01 |
| AT1G49010 | 0,27 | 0,11 | 4,27 | 0,03 | 9,79E-01 |
| AT1G49032 | 0,71 | 0,13 | 2,03 | 0,06 | 9,50E-01 |
| AT1G49190 | 1,12 | -2,18 | 1,99 | -1,10 | 2,73E-01 |
| AT1G49200 | 0,08 | -0,56 | 4,38 | -0,13 | 8,99E-01 |
| AT1G49210 | 0,15 | -0,64 | 4,39 | -0,15 | 8,84E-01 |
| AT1G49230 | 0,38 | -0,33 | 3,38 | -0,10 | 9,21E-01 |
| AT1G49245 | 0,90 | -0,89 | 2,09 | -0,43 | 6,70E-01 |
| AT1G49300 | 0,19 | 0,37 | 4,28 | 0,09 | 9,31E-01 |
| AT1G49340 | 1,22 | -0,36 | 1,95 | -0,19 | 8,52E-01 |
| AT1G49390 | 0,08 | 2,08 | 4,38 | 0,47 | 6,35E-01 |
| AT1G49470 | 0,15 | -0,19 | 4,31 | -0,04 | 9,64E-01 |
| AT1G49510 | 0,25 | 0,00 | 4,41 | 0,00 | 1,00E+00 |
| AT1G49680 | 0,43 | -0,48 | 3,17 | -0,15 | 8,80E-01 |
| AT1G49720 | 0,23 | 2,14 | 4,38 | 0,49 | 6,25E-01 |
| AT1G49820 | 0,28 | 3,41 | 4,33 | 0,79 | 4,31E-01 |
| AT1G49890 | 0,23 | 1,20 | 4,34 | 0,28 | 7,82E-01 |
| AT1G49930 | 0,39 | 1,71 | 4,40 | 0,39 | 6,98E-01 |
| AT1G49975 | 1,21 | -0,80 | 2,23 | -0,36 | 7,20E-01 |
| AT1G49980 | 0,53 | 1,30 | 3,16 | 0,41 | 6,82E-01 |

| | | | | | |
|---|---|---|---|---|---|
| AT1G49990 | 0,80 | 4,79 | 4,25 | 1,13 | 2,59E-01 |
| AT1G50050 | 0,43 | -2,55 | 3,21 | -0,80 | 4,26E-01 |
| AT1G07613 | 1,47 | -2,33 | 1,52 | -1,53 | 1,25E-01 |
| AT1G50120 | 0,25 | 1,15 | 4,34 | 0,27 | 7,91E-01 |
| AT1G50170 | 0,08 | 2,08 | 4,38 | 0,47 | 6,35E-01 |
| AT1G50240 | 1,24 | 3,10 | 1,86 | 1,67 | 9,58E-02 |
| AT1G50270 | 0,45 | 0,18 | 3,30 | 0,05 | 9,57E-01 |
| AT1G50350 | 0,30 | -2,32 | 4,30 | -0,54 | 5,89E-01 |
| AT1G50360 | 0,23 | -0,46 | 4,30 | -0,11 | 9,15E-01 |
| AT1G50370 | 0,39 | 0,77 | 3,37 | 0,23 | 8,20E-01 |
| AT1G50380 | 0,57 | 1,53 | 3,04 | 0,50 | 6,16E-01 |
| AT1G50410 | 1,51 | 3,00 | 2,33 | 1,29 | 1,98E-01 |
| AT1G50560 | 0,41 | 1,78 | 4,26 | 0,42 | 6,76E-01 |
| AT1G50660 | 1,10 | 0,07 | 1,61 | 0,04 | 9,67E-01 |
| AT1G50730 | 0,12 | 2,87 | 4,34 | 0,66 | 5,08E-01 |
| AT1G50740 | 0,72 | 2,48 | 2,87 | 0,86 | 3,88E-01 |
| AT1G51060 | 0,42 | -1,45 | 4,25 | -0,34 | 7,33E-01 |
| AT1G51100 | 0,17 | -1,97 | 4,31 | -0,46 | 6,47E-01 |
| AT1G51110 | 0,52 | -0,04 | 3,19 | -0,01 | 9,89E-01 |
| AT1G51160 | 0,67 | 2,83 | 2,99 | 0,95 | 3,44E-01 |
| AT1G51172 | 1,67 | -1,55 | 1,42 | -1,09 | 2,76E-01 |
| AT1G51200 | 1,27 | -0,54 | 2,22 | -0,24 | 8,08E-01 |
| AT1G51350 | 0,44 | 1,51 | 3,21 | 0,47 | 6,38E-01 |
| AT1G51390 | 0,24 | -0,70 | 4,38 | -0,16 | 8,72E-01 |
| AT1G07737 | 0,83 | -2,24 | 2,26 | -0,99 | 3,21E-01 |
| AT1G51402 | 1,01 | -1,94 | 2,14 | -0,90 | 3,66E-01 |
| AT1G51440 | 0,14 | -0,45 | 4,33 | -0,10 | 9,17E-01 |
| AT1G51540 | 0,08 | -0,56 | 4,38 | -0,13 | 8,99E-01 |
| AT1G51580 | 0,35 | 0,20 | 4,33 | 0,05 | 9,62E-01 |
| AT1G51600 | 0,08 | 2,33 | 4,36 | 0,53 | 5,93E-01 |
| AT1G51645 | 0,08 | 2,33 | 4,36 | 0,53 | 5,93E-01 |
| AT1G51660 | 0,75 | 1,59 | 2,53 | 0,63 | 5,29E-01 |
| AT1G51670 | 0,25 | 2,97 | 4,33 | 0,69 | 4,93E-01 |
| AT1G51720 | 0,29 | 1,98 | 4,32 | 0,46 | 6,46E-01 |
| AT1G51745 | 0,21 | 2,82 | 4,35 | 0,65 | 5,16E-01 |
| AT1G51760 | 0,56 | -0,01 | 2,47 | -0,01 | 9,95E-01 |
| AT1G51950 | 0,22 | 0,08 | 4,27 | 0,02 | 9,85E-01 |
| AT1G52080 | 0,08 | 2,33 | 4,36 | 0,53 | 5,93E-01 |
| AT1G52190 | 0,40 | 1,94 | 3,32 | 0,58 | 5,59E-01 |
| AT1G52310 | 0,73 | 2,00 | 2,38 | 0,84 | 4,01E-01 |
| AT1G52320 | 0,99 | -0,05 | 1,61 | -0,03 | 9,76E-01 |
| AT1G52380 | 0,74 | 2,30 | 2,89 | 0,79 | 4,27E-01 |
| AT1G52410 | 0,68 | 1,55 | 2,59 | 0,60 | 5,50E-01 |
| AT1G52420 | 0,99 | 2,29 | 2,69 | 0,85 | 3,94E-01 |
| AT1G52500 | 0,39 | 2,36 | 4,35 | 0,54 | 5,88E-01 |
| AT1G52600 | 0,08 | 2,33 | 4,36 | 0,53 | 5,93E-01 |
| AT1G52618 | 0,68 | -2,06 | 2,70 | -0,76 | 4,45E-01 |
| AT1G52620 | 0,78 | 0,79 | 4,34 | 0,18 | 8,55E-01 |
| AT1G52720 | 0,19 | -2,00 | 4,31 | -0,46 | 6,43E-01 |
| AT1G52740 | 1,01 | -0,44 | 1,91 | -0,23 | 8,18E-01 |
| AT1G52760 | 0,33 | -0,43 | 3,55 | -0,12 | 9,03E-01 |
| AT1G52870 | 0,92 | -0,78 | 2,32 | -0,34 | 7,35E-01 |
| AT1G52905 | 0,68 | -0,29 | 2,63 | -0,11 | 9,11E-01 |
| AT1G52930 | 0,12 | 2,81 | 4,34 | 0,65 | 5,18E-01 |
| AT1G52940 | 0,76 | 0,02 | 2,29 | 0,01 | 9,92E-01 |
| AT1G52950 | 0,39 | -0,74 | 3,35 | -0,22 | 8,24E-01 |

| | | | | | |
|---|---|---|---|---|---|
| AT1G52980 | 1,44 | 2,68 | 1,70 | 1,57 | 1,16E-01 |
| AT1G53050 | 0,64 | 1,83 | 4,40 | 0,42 | 6,77E-01 |
| AT1G53090 | 0,57 | 1,85 | 4,35 | 0,43 | 6,70E-01 |
| AT1G53165 | 0,43 | 0,95 | 2,73 | 0,35 | 7,29E-01 |
| AT1G53233 | 0,99 | 1,75 | 2,01 | 0,87 | 3,86E-01 |
| AT1G53300 | 0,67 | -2,12 | 2,51 | -0,84 | 3,98E-01 |
| AT1G53330 | 0,35 | 0,00 | 4,41 | 0,00 | 1,00E+00 |
| AT1G53350 | 0,25 | 1,81 | 4,28 | 0,42 | 6,72E-01 |
| AT1G53390 | 0,38 | -1,39 | 4,33 | -0,32 | 7,48E-01 |
| AT1G53400 | 0,40 | 3,80 | 4,32 | 0,88 | 3,79E-01 |
| AT1G53430 | 1,51 | -0,09 | 1,68 | -0,05 | 9,59E-01 |
| AT1G53440 | 1,17 | -0,71 | 1,50 | -0,47 | 6,37E-01 |
| AT1G07887 | 0,84 | -1,33 | 2,94 | -0,45 | 6,52E-01 |
| AT1G53490 | 0,92 | -0,76 | 2,36 | -0,32 | 7,46E-01 |
| AT1G07893 | 0,39 | 1,59 | 4,31 | 0,37 | 7,13E-01 |
| AT1G07897 | 0,95 | 0,46 | 1,59 | 0,29 | 7,75E-01 |
| AT1G53570 | 0,96 | 0,81 | 2,40 | 0,34 | 7,36E-01 |
| AT1G53590 | 0,26 | -0,06 | 4,32 | -0,01 | 9,89E-01 |
| AT1G53670 | 0,12 | 2,81 | 4,34 | 0,65 | 5,18E-01 |
| AT1G53720 | 0,26 | 1,78 | 4,41 | 0,40 | 6,86E-01 |
| AT1G53730 | 1,50 | 0,11 | 2,10 | 0,05 | 9,58E-01 |
| AT1G53840 | 1,21 | -1,10 | 1,83 | -0,60 | 5,47E-01 |
| AT1G53870 | 0,49 | -0,34 | 3,80 | -0,09 | 9,28E-01 |
| AT1G53875 | 1,66 | -1,48 | 1,38 | -1,08 | 2,81E-01 |
| AT1G53885 | 0,24 | 1,12 | 4,32 | 0,26 | 7,95E-01 |
| AT1G53890 | 1,37 | -1,01 | 1,40 | -0,72 | 4,71E-01 |
| AT1G53910 | 0,62 | -0,97 | 2,30 | -0,42 | 6,72E-01 |
| AT1G54030 | 0,19 | -2,00 | 4,31 | -0,46 | 6,43E-01 |
| AT1G54130 | 1,27 | 2,86 | 1,92 | 1,49 | 1,37E-01 |
| AT1G54150 | 0,44 | 0,16 | 3,36 | 0,05 | 9,62E-01 |
| AT1G54200 | 0,80 | -1,02 | 2,30 | -0,44 | 6,58E-01 |
| AT1G54215 | 0,08 | 2,33 | 4,36 | 0,53 | 5,93E-01 |
| AT1G07957 | 0,34 | -0,31 | 3,33 | -0,09 | 9,26E-01 |
| AT1G54310 | 0,08 | 2,08 | 4,38 | 0,47 | 6,35E-01 |
| AT1G54340 | 0,28 | 0,88 | 4,30 | 0,20 | 8,38E-01 |
| AT1G54350 | 0,37 | 2,25 | 4,27 | 0,53 | 5,98E-01 |
| AT1G54360 | 0,57 | 1,65 | 4,39 | 0,38 | 7,06E-01 |
| AT1G54420 | 1,07 | -2,56 | 2,46 | -1,04 | 2,99E-01 |
| AT1G54440 | 0,25 | 2,55 | 4,32 | 0,59 | 5,55E-01 |
| AT1G54490 | 1,36 | 0,83 | 1,63 | 0,51 | 6,13E-01 |
| AT1G54550 | 0,56 | -0,46 | 2,75 | -0,17 | 8,68E-01 |
| AT1G54680 | 0,26 | 2,30 | 4,37 | 0,53 | 5,98E-01 |
| AT1G54720 | 0,66 | -1,49 | 2,91 | -0,51 | 6,09E-01 |
| AT1G54780 | 1,22 | -1,85 | 1,65 | -1,12 | 2,61E-01 |
| AT1G54820 | 0,40 | 0,48 | 4,28 | 0,11 | 9,10E-01 |
| AT1G54920 | 1,30 | 3,66 | 1,78 | 2,06 | 3,95E-02 |
| AT1G55000 | 0,70 | 2,15 | 2,50 | 0,86 | 3,91E-01 |
| AT1G55020 | 0,29 | 0,87 | 4,32 | 0,20 | 8,41E-01 |
| AT1G55040 | 0,41 | 2,34 | 4,16 | 0,56 | 5,73E-01 |
| AT1G55090 | 0,81 | 1,83 | 2,80 | 0,65 | 5,13E-01 |
| AT1G55120 | 1,23 | -2,54 | 1,86 | -1,37 | 1,71E-01 |
| AT1G55150 | 0,30 | 2,84 | 4,35 | 0,65 | 5,14E-01 |
| AT1G55160 | 0,99 | 0,52 | 2,18 | 0,24 | 8,10E-01 |
| AT1G55175 | 0,08 | 2,33 | 4,36 | 0,53 | 5,93E-01 |
| AT1G55320 | 0,25 | 2,25 | 4,37 | 0,52 | 6,06E-01 |
| AT1G55335 | 0,51 | 1,57 | 3,19 | 0,49 | 6,21E-01 |

| | | | | | |
|---|---|---|---|---|---|
| AT1G55370 | 0,24 | 0,00 | 4,41 | 0,00 | 1,00E+00 |
| AT1G55460 | 0,32 | 0,92 | 4,30 | 0,21 | 8,31E-01 |
| AT1G55480 | 0,48 | 0,58 | 4,06 | 0,14 | 8,86E-01 |
| AT1G55500 | 0,61 | 1,16 | 2,21 | 0,53 | 5,99E-01 |
| AT1G55510 | 0,20 | 2,83 | 4,35 | 0,65 | 5,14E-01 |
| AT1G55840 | 0,33 | 0,84 | 4,34 | 0,19 | 8,46E-01 |
| AT1G55850 | 0,50 | 3,80 | 4,32 | 0,88 | 3,78E-01 |
| AT1G08073 | 0,33 | 2,78 | 4,34 | 0,64 | 5,21E-01 |
| AT1G55890 | 0,12 | 1,15 | 4,35 | 0,26 | 7,92E-01 |
| AT1G55910 | 0,98 | 0,51 | 1,74 | 0,29 | 7,69E-01 |
| AT1G55915 | 0,81 | 0,79 | 2,84 | 0,28 | 7,80E-01 |
| AT1G08077 | 0,62 | -0,73 | 2,85 | -0,26 | 7,99E-01 |
| AT1G56045 | 0,75 | 0,82 | 2,64 | 0,31 | 7,55E-01 |
| AT1G56050 | 1,07 | 4,00 | 2,65 | 1,51 | 1,32E-01 |
| AT1G56190 | 1,11 | 1,26 | 1,96 | 0,64 | 5,22E-01 |
| AT1G56200 | 1,42 | 1,16 | 1,65 | 0,70 | 4,82E-01 |
| AT1G56210 | 0,05 | -0,58 | 4,38 | -0,13 | 8,95E-01 |
| AT1G56220 | 1,48 | -0,98 | 1,53 | -0,64 | 5,23E-01 |
| AT1G56450 | 0,83 | 1,48 | 2,03 | 0,73 | 4,66E-01 |
| AT1G56460 | 0,70 | 1,93 | 2,77 | 0,70 | 4,85E-01 |
| AT1G56500 | 0,95 | 1,34 | 1,80 | 0,74 | 4,58E-01 |
| AT1G56540 | 0,56 | 2,82 | 4,35 | 0,65 | 5,16E-01 |
| AT1G56560 | 0,52 | 3,84 | 3,86 | 0,99 | 3,20E-01 |
| AT1G56570 | 0,28 | 0,88 | 4,30 | 0,20 | 8,38E-01 |
| AT1G56600 | 0,19 | -1,00 | 4,37 | -0,23 | 8,19E-01 |
| AT1G56610 | 0,17 | 0,00 | 4,41 | 0,00 | 1,00E+00 |
| AT1G56660 | 1,02 | -0,10 | 1,53 | -0,06 | 9,49E-01 |
| AT1G56700 | 0,33 | 0,30 | 3,52 | 0,08 | 9,32E-01 |
| AT1G57620 | 0,88 | 2,80 | 3,00 | 0,93 | 3,50E-01 |
| AT1G57630 | 0,41 | 0,53 | 4,32 | 0,12 | 9,03E-01 |
| AT1G57650 | 0,46 | -0,49 | 2,70 | -0,18 | 8,57E-01 |
| AT1G57760 | 0,12 | 0,06 | 4,34 | 0,01 | 9,88E-01 |
| AT1G57765 | 0,15 | 0,69 | 4,31 | 0,16 | 8,73E-01 |
| AT1G58190 | 0,62 | -0,09 | 2,49 | -0,03 | 9,72E-01 |
| AT1G58215 | 0,57 | 3,32 | 4,33 | 0,77 | 4,43E-01 |
| AT1G58223 | 0,54 | 0,72 | 3,90 | 0,19 | 8,53E-01 |
| AT1G58260 | 0,12 | 0,63 | 4,33 | 0,15 | 8,85E-01 |
| AT1G58360 | 0,57 | -0,82 | 3,92 | -0,21 | 8,35E-01 |
| AT1G58525 | 0,11 | 1,33 | 4,36 | 0,31 | 7,59E-01 |
| AT1G58520 | 0,55 | 1,64 | 3,18 | 0,52 | 6,05E-01 |
| AT1G58643 | 0,42 | -0,03 | 4,39 | -0,01 | 9,94E-01 |
| AT1G58807 | 1,55 | 1,38 | 1,48 | 0,93 | 3,50E-01 |
| AT1G58808 | 1,22 | -0,13 | 2,07 | -0,06 | 9,50E-01 |
| AT1G58848 | 1,34 | 2,16 | 1,77 | 1,22 | 2,23E-01 |
| AT1G59124 | 1,27 | 1,84 | 1,52 | 1,21 | 2,26E-01 |
| AT1G59218 | 1,28 | 0,37 | 1,69 | 0,22 | 8,26E-01 |
| AT1G59590 | 0,59 | 0,64 | 3,10 | 0,20 | 8,38E-01 |
| AT1G59620 | 0,81 | 3,00 | 4,31 | 0,70 | 4,87E-01 |
| AT1G59650 | 0,54 | 1,15 | 3,86 | 0,30 | 7,66E-01 |
| AT1G59710 | 0,56 | 1,64 | 3,25 | 0,50 | 6,15E-01 |
| AT1G59750 | 1,67 | 5,26 | 2,40 | 2,19 | 2,84E-02 |
| AT1G59820 | 1,45 | 0,58 | 1,88 | 0,31 | 7,56E-01 |
| AT1G59890 | 0,93 | 1,33 | 2,28 | 0,58 | 5,60E-01 |
| AT1G59900 | 1,24 | 0,62 | 1,59 | 0,39 | 6,98E-01 |
| AT1G60025 | 0,05 | -0,58 | 4,38 | -0,13 | 8,95E-01 |
| AT1G60073 | 1,58 | 0,09 | 1,55 | 0,06 | 9,52E-01 |

| | | | | | |
|---|---|---|---|---|---|
| AT1G60170 | 1,29 | 1,37 | 2,16 | 0,64 | 5,25E-01 |
| AT1G60190 | 0,89 | 2,06 | 2,10 | 0,98 | 3,26E-01 |
| AT1G60220 | 0,08 | 2,08 | 4,38 | 0,47 | 6,35E-01 |
| AT1G60420 | 0,70 | 3,30 | 2,76 | 1,20 | 2,32E-01 |
| AT1G60545 | 1,07 | 4,15 | 2,98 | 1,39 | 1,64E-01 |
| AT1G60650 | 0,21 | 2,01 | 4,37 | 0,46 | 6,45E-01 |
| AT1G60720 | 1,62 | -2,30 | 1,66 | -1,38 | 1,66E-01 |
| AT1G60900 | 0,65 | 1,18 | 3,16 | 0,37 | 7,09E-01 |
| AT1G08353 | 0,98 | 0,89 | 2,00 | 0,44 | 6,57E-01 |
| AT1G61030 | 0,52 | 3,63 | 3,91 | 0,93 | 3,53E-01 |
| AT1G61040 | 0,54 | 1,15 | 3,86 | 0,30 | 7,66E-01 |
| AT1G61150 | 0,81 | 1,33 | 3,65 | 0,36 | 7,16E-01 |
| AT1G61170 | 0,33 | 1,90 | 4,27 | 0,44 | 6,56E-01 |
| AT1G61180 | 0,58 | 1,59 | 2,65 | 0,60 | 5,48E-01 |
| AT1G61210 | 1,07 | 1,34 | 1,73 | 0,77 | 4,40E-01 |
| AT1G61224 | 1,62 | 0,14 | 1,54 | 0,09 | 9,25E-01 |
| AT1G61250 | 0,18 | -0,57 | 4,30 | -0,13 | 8,94E-01 |
| AT1G61280 | 0,12 | 2,81 | 4,34 | 0,65 | 5,18E-01 |
| AT1G61350 | 0,76 | -1,88 | 2,18 | -0,86 | 3,88E-01 |
| AT1G61450 | 0,12 | 2,69 | 4,35 | 0,62 | 5,36E-01 |
| AT1G61470 | 0,18 | -1,39 | 4,34 | -0,32 | 7,49E-01 |
| AT1G61610 | 0,12 | 0,06 | 4,34 | 0,01 | 9,88E-01 |
| AT1G61620 | 0,37 | 1,64 | 4,38 | 0,37 | 7,08E-01 |
| AT1G61790 | 1,42 | 2,09 | 1,83 | 1,14 | 2,54E-01 |
| AT1G61800 | 0,56 | 3,81 | 4,32 | 0,88 | 3,78E-01 |
| AT1G61820 | 0,08 | 2,08 | 4,38 | 0,47 | 6,35E-01 |
| AT1G61850 | 0,80 | 2,37 | 2,25 | 1,05 | 2,92E-01 |
| AT1G61990 | 0,32 | 2,78 | 4,31 | 0,64 | 5,19E-01 |
| AT1G62170 | 0,36 | 0,31 | 4,35 | 0,07 | 9,42E-01 |
| AT1G08477 | 1,64 | 0,52 | 1,34 | 0,39 | 6,97E-01 |
| AT1G62270 | 0,29 | -2,65 | 4,29 | -0,62 | 5,38E-01 |
| AT1G08517 | 0,77 | -1,36 | 2,41 | -0,56 | 5,73E-01 |
| AT1G62380 | 1,16 | 0,67 | 1,53 | 0,44 | 6,64E-01 |
| AT1G62410 | 0,32 | -1,42 | 4,34 | -0,33 | 7,44E-01 |
| AT1G62480 | 0,12 | 0,06 | 4,34 | 0,01 | 9,88E-01 |
| AT1G62560 | 1,49 | -2,46 | 3,02 | -0,82 | 4,15E-01 |
| AT1G62570 | 0,39 | 0,80 | 4,25 | 0,19 | 8,51E-01 |
| AT1G62620 | 0,57 | 1,96 | 3,02 | 0,65 | 5,16E-01 |
| AT1G62660 | 1,32 | 2,61 | 1,76 | 1,48 | 1,38E-01 |
| AT1G62720 | 0,24 | -1,59 | 4,33 | -0,37 | 7,13E-01 |
| AT1G62790 | 0,51 | 1,71 | 4,40 | 0,39 | 6,98E-01 |
| AT1G62895 | 1,43 | 2,58 | 1,67 | 1,55 | 1,22E-01 |
| AT1G62920 | 0,97 | -0,08 | 2,79 | -0,03 | 9,77E-01 |
| AT1G62970 | 0,33 | 0,65 | 3,43 | 0,19 | 8,49E-01 |
| AT1G62975 | 0,08 | 2,08 | 4,38 | 0,47 | 6,35E-01 |
| AT1G63000 | 0,67 | -1,06 | 3,73 | -0,29 | 7,76E-01 |
| AT1G63110 | 0,32 | 2,20 | 4,32 | 0,51 | 6,10E-01 |
| AT1G63170 | 0,08 | 2,33 | 4,36 | 0,53 | 5,93E-01 |
| AT1G63430 | 0,74 | 2,51 | 3,80 | 0,66 | 5,09E-01 |
| AT1G63500 | 0,59 | 0,19 | 2,57 | 0,08 | 9,40E-01 |
| AT1G63540 | 0,37 | -2,99 | 4,28 | -0,70 | 4,84E-01 |
| AT1G63680 | 0,43 | 1,07 | 3,15 | 0,34 | 7,35E-01 |
| AT1G63690 | 1,03 | 0,00 | 1,67 | 0,00 | 1,00E+00 |
| AT1G63730 | 0,24 | -2,52 | 4,29 | -0,59 | 5,57E-01 |
| AT1G63830 | 0,49 | 0,32 | 3,96 | 0,08 | 9,36E-01 |
| AT1G63850 | 0,53 | 1,95 | 3,94 | 0,49 | 6,21E-01 |

| | | | | | |
|---|---|---|---|---|---|
| AT1G08643 | 0,79 | 0,32 | 2,77 | 0,12 | 9,07E-01 |
| AT1G63900 | 0,95 | 0,56 | 1,99 | 0,28 | 7,78E-01 |
| AT1G63940 | 0,81 | 0,62 | 1,90 | 0,33 | 7,43E-01 |
| AT1G64060 | 0,86 | 1,84 | 2,19 | 0,84 | 4,00E-01 |
| AT1G64065 | 0,29 | -0,42 | 4,33 | -0,10 | 9,22E-01 |
| AT1G64090 | 0,96 | 3,93 | 2,10 | 1,88 | 6,06E-02 |
| AT1G64100 | 0,61 | 3,12 | 3,70 | 0,84 | 3,99E-01 |
| AT1G64120 | 0,13 | -1,34 | 4,35 | -0,31 | 7,58E-01 |
| AT1G64130 | 0,12 | 2,81 | 4,34 | 0,65 | 5,18E-01 |
| AT1G08657 | 0,60 | 0,63 | 2,58 | 0,24 | 8,08E-01 |
| AT1G64230 | 1,39 | 3,17 | 1,78 | 1,79 | 7,37E-02 |
| AT1G64370 | 0,17 | -1,47 | 4,34 | -0,34 | 7,34E-01 |
| AT1G64390 | 1,48 | 0,72 | 2,13 | 0,34 | 7,36E-01 |
| AT1G64520 | 0,27 | -0,17 | 4,29 | -0,04 | 9,68E-01 |
| AT1G64530 | 0,54 | 3,20 | 3,08 | 1,04 | 2,99E-01 |
| AT1G64550 | 0,64 | 1,59 | 2,52 | 0,63 | 5,28E-01 |
| AT1G64570 | 0,71 | 3,02 | 2,43 | 1,24 | 2,13E-01 |
| AT1G64572 | 0,51 | 1,70 | 4,40 | 0,39 | 6,98E-01 |
| AT1G64620 | 0,45 | -1,20 | 3,15 | -0,38 | 7,02E-01 |
| AT1G64680 | 1,48 | 1,23 | 1,64 | 0,75 | 4,53E-01 |
| AT1G64740 | 0,25 | -1,37 | 4,28 | -0,32 | 7,49E-01 |
| AT1G64770 | 1,29 | -0,50 | 1,54 | -0,32 | 7,48E-01 |
| AT1G64795 | 0,74 | -0,52 | 1,80 | -0,29 | 7,71E-01 |
| AT1G64810 | 0,44 | 1,19 | 4,29 | 0,28 | 7,82E-01 |
| AT1G64820 | 0,08 | 2,33 | 4,36 | 0,53 | 5,93E-01 |
| AT1G64860 | 0,65 | 2,71 | 3,09 | 0,88 | 3,81E-01 |
| AT1G64900 | 1,03 | 2,34 | 1,89 | 1,24 | 2,17E-01 |
| AT1G64950 | 0,34 | 1,01 | 3,37 | 0,30 | 7,65E-01 |
| AT1G64960 | 0,60 | 2,78 | 3,90 | 0,71 | 4,75E-01 |
| AT1G65010 | 1,62 | 1,38 | 1,61 | 0,86 | 3,91E-01 |
| AT1G65220 | 0,59 | 1,04 | 2,59 | 0,40 | 6,87E-01 |
| AT1G65230 | 1,04 | -0,22 | 2,04 | -0,11 | 9,16E-01 |
| AT1G65260 | 0,21 | -2,30 | 4,30 | -0,54 | 5,92E-01 |
| AT1G65270 | 0,21 | -0,12 | 4,28 | -0,03 | 9,77E-01 |
| AT1G65280 | 0,39 | 0,65 | 4,24 | 0,15 | 8,79E-01 |
| AT1G65295 | 0,05 | -0,58 | 4,38 | -0,13 | 8,95E-01 |
| AT1G65360 | 0,90 | 1,68 | 2,45 | 0,69 | 4,93E-01 |
| AT1G08753 | 0,15 | 2,31 | 4,37 | 0,53 | 5,98E-01 |
| AT1G65490 | 0,49 | -0,94 | 4,34 | -0,22 | 8,28E-01 |
| AT1G65590 | 0,23 | 1,20 | 4,34 | 0,28 | 7,82E-01 |
| AT1G65681 | 0,20 | -2,22 | 4,30 | -0,52 | 6,06E-01 |
| AT1G65730 | 0,08 | 2,33 | 4,36 | 0,53 | 5,93E-01 |
| AT1G65790 | 0,93 | 1,42 | 1,72 | 0,82 | 4,10E-01 |
| AT1G65800 | 1,17 | -0,44 | 1,56 | -0,28 | 7,79E-01 |
| AT1G65820 | 0,38 | -0,35 | 2,74 | -0,13 | 8,99E-01 |
| AT1G65840 | 0,61 | 0,81 | 3,79 | 0,21 | 8,30E-01 |
| AT1G08793 | 0,15 | -0,19 | 4,31 | -0,04 | 9,64E-01 |
| AT1G66090 | 0,59 | 1,59 | 3,13 | 0,51 | 6,11E-01 |
| AT1G66150 | 1,55 | 0,63 | 1,26 | 0,50 | 6,19E-01 |
| AT1G66160 | 0,48 | 2,36 | 4,03 | 0,58 | 5,59E-01 |
| AT1G66180 | 0,53 | -1,29 | 3,06 | -0,42 | 6,72E-01 |
| AT1G66230 | 0,24 | 0,00 | 4,41 | 0,00 | 1,00E+00 |
| AT1G66340 | 0,60 | 2,96 | 2,91 | 1,02 | 3,10E-01 |
| AT1G66345 | 0,47 | 3,10 | 3,97 | 0,78 | 4,35E-01 |
| AT1G66430 | 0,50 | -0,14 | 4,41 | -0,03 | 9,75E-01 |
| AT1G66465 | 0,33 | 3,50 | 4,32 | 0,81 | 4,18E-01 |

| | | | | | |
|---|---|---|---|---|---|
| AT1G66580 | 1,55 | 1,70 | 1,45 | 1,17 | 2,43E-01 |
| AT1G66640 | 0,21 | -0,46 | 4,34 | -0,11 | 9,16E-01 |
| AT1G66750 | 0,31 | 2,98 | 4,33 | 0,69 | 4,92E-01 |
| AT1G66820 | 0,53 | 2,42 | 4,36 | 0,56 | 5,78E-01 |
| AT1G66900 | 0,20 | 1,57 | 4,34 | 0,36 | 7,17E-01 |
| AT1G67105 | 0,86 | 0,34 | 1,70 | 0,20 | 8,43E-01 |
| AT1G67190 | 0,34 | 0,88 | 4,26 | 0,21 | 8,36E-01 |
| AT1G08923 | 0,11 | -1,28 | 4,35 | -0,29 | 7,69E-01 |
| AT1G67420 | 0,13 | 2,37 | 4,36 | 0,54 | 5,86E-01 |
| AT1G08937 | 0,46 | -0,26 | 3,11 | -0,08 | 9,33E-01 |
| AT1G67470 | 0,43 | 1,18 | 3,27 | 0,36 | 7,19E-01 |
| AT1G67480 | 0,49 | 1,50 | 4,29 | 0,35 | 7,27E-01 |
| AT1G67530 | 0,54 | 0,79 | 3,95 | 0,20 | 8,42E-01 |
| AT1G67560 | 0,64 | 2,40 | 4,37 | 0,55 | 5,83E-01 |
| AT1G67620 | 0,15 | -0,41 | 4,32 | -0,09 | 9,25E-01 |
| AT1G67635 | 1,69 | -0,69 | 1,47 | -0,47 | 6,41E-01 |
| AT1G67700 | 0,46 | -2,63 | 3,91 | -0,67 | 5,01E-01 |
| AT1G67720 | 0,53 | 3,71 | 3,76 | 0,99 | 3,24E-01 |
| AT1G67792 | 0,38 | 2,54 | 4,35 | 0,58 | 5,60E-01 |
| AT1G67900 | 1,04 | 0,64 | 2,29 | 0,28 | 7,80E-01 |
| AT1G67940 | 0,66 | 2,82 | 2,94 | 0,96 | 3,37E-01 |
| AT1G08963 | 0,39 | 1,78 | 4,41 | 0,40 | 6,86E-01 |
| AT1G67980 | 0,18 | 1,77 | 4,41 | 0,40 | 6,88E-01 |
| AT1G67990 | 0,55 | 1,79 | 3,88 | 0,46 | 6,44E-01 |
| AT1G68050 | 0,24 | 2,00 | 4,34 | 0,46 | 6,45E-01 |
| AT1G68100 | 0,43 | 2,86 | 4,12 | 0,69 | 4,87E-01 |
| AT1G68110 | 0,27 | 2,11 | 4,38 | 0,48 | 6,30E-01 |
| AT1G68310 | 0,12 | 1,65 | 4,34 | 0,38 | 7,04E-01 |
| AT1G68440 | 1,32 | -1,32 | 1,52 | -0,87 | 3,83E-01 |
| AT1G68490 | 0,97 | 1,39 | 2,37 | 0,59 | 5,58E-01 |
| AT1G68570 | 0,58 | 0,34 | 2,41 | 0,14 | 8,87E-01 |
| AT1G68580 | 1,44 | 1,12 | 1,42 | 0,78 | 4,33E-01 |
| AT1G68585 | 0,27 | 0,31 | 4,36 | 0,07 | 9,44E-01 |
| AT1G68620 | 0,33 | 3,46 | 4,32 | 0,80 | 4,23E-01 |
| AT1G68650 | 0,32 | 2,47 | 4,35 | 0,57 | 5,69E-01 |
| AT1G68670 | 0,63 | 1,87 | 2,82 | 0,66 | 5,06E-01 |
| AT1G68720 | 1,50 | 2,90 | 2,03 | 1,43 | 1,54E-01 |
| AT1G68840 | 1,23 | -1,36 | 1,97 | -0,69 | 4,91E-01 |
| AT1G68890 | 1,09 | 3,42 | 1,96 | 1,74 | 8,11E-02 |
| AT1G68980 | 0,24 | 2,85 | 4,34 | 0,66 | 5,11E-01 |
| AT1G69085 | 0,08 | -0,87 | 4,38 | -0,20 | 8,43E-01 |
| AT1G69160 | 0,39 | -1,82 | 3,24 | -0,56 | 5,75E-01 |
| AT1G69170 | 0,40 | 3,80 | 4,32 | 0,88 | 3,79E-01 |
| AT1G69200 | 0,21 | -0,24 | 4,35 | -0,06 | 9,56E-01 |
| AT1G69252 | 0,90 | -2,54 | 2,96 | -0,86 | 3,92E-01 |
| AT1G69260 | 0,63 | -1,53 | 2,65 | -0,58 | 5,62E-01 |
| AT1G69350 | 0,78 | 2,83 | 2,34 | 1,21 | 2,27E-01 |
| AT1G69390 | 0,12 | 0,31 | 4,35 | 0,07 | 9,44E-01 |
| AT1G69410 | 0,27 | 1,58 | 4,35 | 0,36 | 7,16E-01 |
| AT1G69420 | 0,65 | 1,56 | 2,53 | 0,62 | 5,37E-01 |
| AT1G69510 | 0,10 | -1,26 | 4,35 | -0,29 | 7,72E-01 |
| AT1G69523 | 0,10 | -1,26 | 4,35 | -0,29 | 7,72E-01 |
| AT1G69526 | 0,08 | 2,33 | 4,36 | 0,53 | 5,93E-01 |
| AT1G69530 | 0,93 | -3,26 | 1,99 | -1,64 | 1,01E-01 |
| AT1G69580 | 0,26 | -0,20 | 4,35 | -0,05 | 9,64E-01 |
| AT1G69610 | 0,21 | 0,95 | 4,33 | 0,22 | 8,27E-01 |

| | | | | | |
|---|---|---|---|---|---|
| AT1G09087 | 0,37 | 1,12 | 3,43 | 0,33 | 7,44E-01 |
| AT1G69690 | 0,67 | -0,85 | 2,38 | -0,36 | 7,19E-01 |
| AT1G69740 | 0,99 | -0,64 | 1,69 | -0,38 | 7,05E-01 |
| AT1G69750 | 0,10 | -1,26 | 4,35 | -0,29 | 7,72E-01 |
| AT1G69870 | 0,38 | 2,12 | 4,31 | 0,49 | 6,23E-01 |
| AT1G70060 | 1,48 | 2,33 | 1,92 | 1,21 | 2,25E-01 |
| AT1G70090 | 0,99 | -0,87 | 1,92 | -0,45 | 6,50E-01 |
| AT1G70100 | 0,38 | 2,04 | 4,29 | 0,48 | 6,34E-01 |
| AT1G70160 | 1,19 | 1,72 | 1,51 | 1,14 | 2,55E-01 |
| AT1G70180 | 1,28 | 3,29 | 1,89 | 1,74 | 8,14E-02 |
| AT1G70190 | 0,40 | 2,19 | 4,34 | 0,51 | 6,13E-01 |
| AT1G70210 | 0,79 | -1,56 | 2,21 | -0,71 | 4,80E-01 |
| AT1G70230 | 0,59 | -1,17 | 2,61 | -0,45 | 6,53E-01 |
| AT1G70250 | 0,71 | 3,40 | 2,92 | 1,17 | 2,44E-01 |
| AT1G70330 | 0,33 | -0,39 | 3,46 | -0,11 | 9,11E-01 |
| AT1G70480 | 1,12 | 1,83 | 1,69 | 1,08 | 2,78E-01 |
| AT1G70510 | 0,12 | 2,69 | 4,35 | 0,62 | 5,36E-01 |
| AT1G70550 | 0,14 | -0,31 | 4,33 | -0,07 | 9,43E-01 |
| AT1G70570 | 0,36 | 1,73 | 4,40 | 0,39 | 6,94E-01 |
| AT1G70580 | 0,81 | 3,58 | 2,41 | 1,48 | 1,38E-01 |
| AT1G70620 | 1,32 | 3,04 | 2,03 | 1,50 | 1,33E-01 |
| AT1G70630 | 0,36 | 1,35 | 4,32 | 0,31 | 7,55E-01 |
| AT1G70645 | 0,05 | -0,58 | 4,38 | -0,13 | 8,95E-01 |
| AT1G70650 | 0,26 | 0,80 | 4,33 | 0,18 | 8,54E-01 |
| AT1G70660 | 0,20 | 2,11 | 4,37 | 0,48 | 6,29E-01 |
| AT1G70700 | 0,60 | -1,73 | 2,53 | -0,68 | 4,95E-01 |
| AT1G70760 | 0,35 | 1,44 | 3,42 | 0,42 | 6,73E-01 |
| AT1G70810 | 0,62 | 3,19 | 2,60 | 1,23 | 2,20E-01 |
| AT1G70820 | 0,43 | -0,43 | 2,76 | -0,16 | 8,75E-01 |
| AT1G70895 | 0,08 | -0,56 | 4,38 | -0,13 | 8,99E-01 |
| AT1G70900 | 1,03 | 1,59 | 1,91 | 0,84 | 4,03E-01 |
| AT1G70960 | 0,12 | 0,06 | 4,34 | 0,01 | 9,88E-01 |
| AT1G70980 | 0,40 | 0,22 | 4,36 | 0,05 | 9,61E-01 |
| AT1G71010 | 1,50 | 0,90 | 1,63 | 0,55 | 5,79E-01 |
| AT1G71090 | 0,35 | 1,12 | 3,47 | 0,32 | 7,47E-01 |
| AT1G71100 | 1,11 | 0,46 | 1,62 | 0,28 | 7,78E-01 |
| AT1G71110 | 0,15 | 0,40 | 4,33 | 0,09 | 9,27E-01 |
| AT1G71130 | 0,37 | 1,48 | 4,32 | 0,34 | 7,32E-01 |
| AT1G71180 | 0,42 | 1,87 | 4,21 | 0,44 | 6,57E-01 |
| AT1G71270 | 0,45 | 0,72 | 4,10 | 0,18 | 8,60E-01 |
| AT1G71300 | 0,38 | 2,05 | 4,37 | 0,47 | 6,40E-01 |
| AT1G71320 | 0,39 | -2,20 | 4,10 | -0,54 | 5,92E-01 |
| AT1G71480 | 0,80 | 2,84 | 2,81 | 1,01 | 3,11E-01 |
| AT1G09257 | 0,17 | 3,14 | 4,33 | 0,73 | 4,68E-01 |
| AT1G71500 | 0,95 | -1,11 | 1,97 | -0,57 | 5,72E-01 |
| AT1G71695 | 0,82 | 0,35 | 2,44 | 0,14 | 8,87E-01 |
| AT1G71720 | 0,53 | 1,71 | 3,88 | 0,44 | 6,59E-01 |
| AT1G71800 | 0,67 | 3,42 | 3,62 | 0,94 | 3,45E-01 |
| AT1G71820 | 1,53 | 0,50 | 1,30 | 0,39 | 7,00E-01 |
| AT1G71840 | 1,05 | -2,64 | 1,98 | -1,34 | 1,81E-01 |
| AT1G71900 | 0,51 | 1,70 | 4,40 | 0,39 | 6,98E-01 |
| AT1G71910 | 0,44 | -2,04 | 3,15 | -0,65 | 5,16E-01 |
| AT1G71920 | 0,73 | -1,07 | 2,22 | -0,48 | 6,30E-01 |
| AT1G71930 | 1,13 | -0,77 | 1,86 | -0,41 | 6,78E-01 |
| AT1G71970 | 0,14 | 0,87 | 4,31 | 0,20 | 8,40E-01 |
| AT1G71980 | 0,46 | 0,07 | 2,99 | 0,02 | 9,81E-01 |

| | | | | | |
|---|---|---|---|---|---|
| AT1G72010 | 0,42 | 0,78 | 3,20 | 0,24 | 8,07E-01 |
| AT1G72030 | 0,75 | 1,00 | 3,65 | 0,28 | 7,83E-01 |
| AT1G72050 | 0,75 | 3,56 | 3,62 | 0,98 | 3,26E-01 |
| AT1G72090 | 0,34 | 2,00 | 4,34 | 0,46 | 6,45E-01 |
| AT1G72240 | 0,60 | 1,94 | 3,83 | 0,51 | 6,13E-01 |
| AT1G72310 | 0,25 | 2,64 | 4,34 | 0,61 | 5,43E-01 |
| AT1G72330 | 0,67 | 1,73 | 2,82 | 0,61 | 5,39E-01 |
| AT1G72470 | 0,43 | 2,50 | 4,37 | 0,57 | 5,67E-01 |
| AT1G72510 | 0,57 | -0,34 | 3,14 | -0,11 | 9,14E-01 |
| AT1G72520 | 0,91 | 0,02 | 1,76 | 0,01 | 9,89E-01 |
| AT1G72530 | 0,05 | -0,58 | 4,38 | -0,13 | 8,95E-01 |
| AT1G09333 | 1,54 | -2,59 | 1,64 | -1,58 | 1,14E-01 |
| AT1G72650 | 0,79 | 2,33 | 2,19 | 1,06 | 2,88E-01 |
| AT1G72700 | 1,31 | 1,48 | 1,55 | 0,96 | 3,37E-01 |
| AT1G72750 | 0,63 | -0,31 | 2,04 | -0,15 | 8,80E-01 |
| AT1G72770 | 1,51 | 1,22 | 1,36 | 0,90 | 3,67E-01 |
| AT1G72790 | 0,71 | 0,80 | 2,84 | 0,28 | 7,77E-01 |
| AT1G72855 | 0,40 | 3,80 | 4,32 | 0,88 | 3,79E-01 |
| AT1G72900 | 0,32 | 1,45 | 3,47 | 0,42 | 6,76E-01 |
| AT1G72910 | 1,29 | 0,17 | 1,54 | 0,11 | 9,10E-01 |
| AT1G72920 | 0,44 | 0,32 | 3,35 | 0,10 | 9,24E-01 |
| AT1G72940 | 0,97 | 5,21 | 3,96 | 1,32 | 1,88E-01 |
| AT1G72970 | 0,16 | 1,18 | 4,32 | 0,27 | 7,85E-01 |
| AT1G73030 | 0,82 | -1,77 | 2,32 | -0,76 | 4,45E-01 |
| AT1G73060 | 0,24 | -1,17 | 4,35 | -0,27 | 7,89E-01 |
| AT1G73080 | 1,38 | 0,70 | 1,39 | 0,51 | 6,13E-01 |
| AT1G73100 | 0,70 | 0,96 | 2,50 | 0,39 | 7,00E-01 |
| AT1G73180 | 1,25 | 3,03 | 1,95 | 1,55 | 1,20E-01 |
| AT1G73430 | 1,05 | 1,64 | 2,11 | 0,77 | 4,38E-01 |
| AT1G73450 | 1,12 | 1,79 | 1,72 | 1,04 | 2,99E-01 |
| AT1G73460 | 0,79 | 1,45 | 2,37 | 0,61 | 5,41E-01 |
| AT1G73470 | 0,23 | 1,51 | 4,34 | 0,35 | 7,28E-01 |
| AT1G73480 | 1,37 | 1,55 | 2,15 | 0,72 | 4,69E-01 |
| AT1G73600 | 1,37 | -1,67 | 2,13 | -0,78 | 4,33E-01 |
| AT1G73630 | 0,10 | 0,34 | 4,35 | 0,08 | 9,37E-01 |
| AT1G73670 | 0,46 | 1,99 | 3,32 | 0,60 | 5,49E-01 |
| AT1G73710 | 0,22 | 1,72 | 4,29 | 0,40 | 6,89E-01 |
| AT1G73750 | 0,72 | 4,43 | 4,29 | 1,03 | 3,02E-01 |
| AT1G73770 | 0,08 | 2,33 | 4,36 | 0,53 | 5,93E-01 |
| AT1G73850 | 0,17 | -0,44 | 4,37 | -0,10 | 9,19E-01 |
| AT1G73875 | 0,39 | 1,70 | 4,18 | 0,41 | 6,84E-01 |
| AT1G73885 | 0,62 | 1,19 | 2,97 | 0,40 | 6,88E-01 |
| AT1G73920 | 0,62 | -0,98 | 2,50 | -0,39 | 6,94E-01 |
| AT1G73970 | 0,95 | -0,28 | 2,29 | -0,12 | 9,03E-01 |
| AT1G74160 | 1,08 | 1,87 | 1,81 | 1,03 | 3,02E-01 |
| AT1G74250 | 1,42 | 1,01 | 1,51 | 0,67 | 5,05E-01 |
| AT1G74360 | 1,21 | 3,78 | 2,39 | 1,58 | 1,14E-01 |
| AT1G74380 | 1,29 | -0,33 | 1,54 | -0,21 | 8,32E-01 |
| AT1G74430 | 0,38 | -1,32 | 4,35 | -0,30 | 7,61E-01 |
| AT1G74450 | 0,99 | -0,64 | 1,72 | -0,37 | 7,09E-01 |
| AT1G74580 | 0,25 | 1,05 | 4,33 | 0,24 | 8,08E-01 |
| AT1G74680 | 0,57 | 0,21 | 3,20 | 0,07 | 9,47E-01 |
| AT1G74710 | 0,60 | 0,98 | 2,48 | 0,40 | 6,91E-01 |
| AT1G74730 | 0,44 | 0,49 | 3,28 | 0,15 | 8,82E-01 |
| AT1G74750 | 0,50 | 2,63 | 4,35 | 0,60 | 5,46E-01 |
| AT1G74780 | 0,63 | 3,43 | 2,88 | 1,19 | 2,34E-01 |

| | | | | | |
|---|---|---|---|---|---|
| AT1G74790 | 0,51 | 0,19 | 3,90 | 0,05 | 9,60E-01 |
| AT1G74870 | 1,10 | 3,83 | 3,20 | 1,20 | 2,31E-01 |
| AT1G74930 | 0,81 | -3,10 | 2,62 | -1,18 | 2,36E-01 |
| AT1G74940 | 0,58 | 0,29 | 2,96 | 0,10 | 9,23E-01 |
| AT1G75080 | 0,21 | 2,34 | 4,32 | 0,54 | 5,89E-01 |
| AT1G09533 | 1,00 | -0,33 | 1,71 | -0,19 | 8,45E-01 |
| AT1G75163 | 0,94 | 0,37 | 1,81 | 0,20 | 8,38E-01 |
| AT1G75166 | 1,22 | -0,33 | 1,77 | -0,19 | 8,52E-01 |
| AT1G75170 | 0,12 | 1,65 | 4,34 | 0,38 | 7,04E-01 |
| AT1G09543 | 0,38 | -0,40 | 3,27 | -0,12 | 9,02E-01 |
| AT1G75190 | 0,22 | -1,61 | 4,34 | -0,37 | 7,10E-01 |
| AT1G75210 | 0,77 | -0,09 | 3,02 | -0,03 | 9,77E-01 |
| AT1G75280 | 0,22 | -2,03 | 4,31 | -0,47 | 6,37E-01 |
| AT1G75310 | 0,47 | 1,72 | 4,40 | 0,39 | 6,96E-01 |
| AT1G75350 | 0,83 | 2,03 | 1,95 | 1,04 | 2,98E-01 |
| AT1G75400 | 0,32 | 1,30 | 4,25 | 0,31 | 7,60E-01 |
| AT1G75420 | 0,31 | 0,77 | 4,34 | 0,18 | 8,58E-01 |
| AT1G75440 | 0,28 | 1,26 | 4,36 | 0,29 | 7,73E-01 |
| AT1G75460 | 0,84 | 2,22 | 2,24 | 0,99 | 3,20E-01 |
| AT1G75580 | 0,17 | 2,29 | 4,36 | 0,53 | 5,98E-01 |
| AT1G75710 | 0,31 | -0,80 | 4,29 | -0,19 | 8,52E-01 |
| AT1G75785 | 0,12 | 0,63 | 4,33 | 0,15 | 8,85E-01 |
| AT1G75810 | 0,60 | 1,52 | 2,65 | 0,57 | 5,66E-01 |
| AT1G75840 | 0,75 | 0,45 | 2,23 | 0,20 | 8,39E-01 |
| AT1G75900 | 0,32 | -0,50 | 4,30 | -0,12 | 9,07E-01 |
| AT1G75960 | 0,45 | 1,04 | 4,17 | 0,25 | 8,04E-01 |
| AT1G76000 | 0,58 | 2,15 | 2,87 | 0,75 | 4,53E-01 |
| AT1G76050 | 0,66 | 0,79 | 2,97 | 0,27 | 7,89E-01 |
| AT1G76140 | 0,48 | 0,37 | 3,95 | 0,09 | 9,26E-01 |
| AT1G76185 | 0,16 | 0,49 | 4,33 | 0,11 | 9,09E-01 |
| AT1G76190 | 0,11 | -1,28 | 4,35 | -0,29 | 7,69E-01 |
| AT1G76320 | 0,32 | 1,90 | 4,27 | 0,44 | 6,57E-01 |
| AT1G76350 | 0,12 | 1,15 | 4,35 | 0,26 | 7,92E-01 |
| AT1G76390 | 0,51 | 1,91 | 3,08 | 0,62 | 5,35E-01 |
| AT1G76450 | 0,08 | 2,08 | 4,38 | 0,47 | 6,35E-01 |
| AT1G76520 | 0,73 | -0,10 | 2,27 | -0,04 | 9,65E-01 |
| AT1G76580 | 0,80 | 0,39 | 3,50 | 0,11 | 9,12E-01 |
| AT1G76590 | 0,69 | -1,43 | 2,89 | -0,49 | 6,22E-01 |
| AT1G76600 | 0,49 | 0,97 | 3,09 | 0,31 | 7,54E-01 |
| AT1G76620 | 0,25 | 0,43 | 4,36 | 0,10 | 9,21E-01 |
| AT1G76680 | 0,14 | -0,45 | 4,38 | -0,10 | 9,18E-01 |
| AT1G76740 | 0,12 | 2,81 | 4,34 | 0,65 | 5,18E-01 |
| AT1G76780 | 1,47 | 1,56 | 1,69 | 0,92 | 3,56E-01 |
| AT1G76820 | 0,53 | 2,84 | 3,11 | 0,91 | 3,61E-01 |
| AT1G76850 | 1,15 | 0,71 | 1,84 | 0,39 | 6,99E-01 |
| AT1G76880 | 1,06 | -0,80 | 1,69 | -0,47 | 6,37E-01 |
| AT1G76900 | 0,91 | 2,15 | 2,72 | 0,79 | 4,31E-01 |
| AT1G76920 | 0,59 | 2,52 | 3,78 | 0,67 | 5,05E-01 |
| AT1G76950 | 0,83 | 0,83 | 2,83 | 0,29 | 7,69E-01 |
| AT1G76990 | 1,49 | -0,96 | 1,36 | -0,71 | 4,79E-01 |
| AT1G77080 | 0,41 | -1,09 | 3,27 | -0,33 | 7,39E-01 |
| AT1G77140 | 0,25 | -0,67 | 4,39 | -0,15 | 8,78E-01 |
| AT1G77320 | 0,26 | 2,13 | 4,38 | 0,49 | 6,27E-01 |
| AT1G77350 | 0,15 | -0,19 | 4,31 | -0,04 | 9,64E-01 |
| AT1G77420 | 0,12 | 2,87 | 4,34 | 0,66 | 5,08E-01 |
| AT1G77460 | 0,94 | 1,71 | 2,53 | 0,68 | 4,99E-01 |

| | | | | | |
|---|---|---|---|---|---|
| AT1G77490 | 0,64 | -2,10 | 2,96 | -0,71 | 4,77E-01 |
| AT1G77500 | 0,37 | 1,61 | 4,36 | 0,37 | 7,12E-01 |
| AT1G77510 | 1,26 | 1,96 | 1,67 | 1,18 | 2,38E-01 |
| AT1G77610 | 0,53 | 0,59 | 4,32 | 0,14 | 8,92E-01 |
| AT1G77710 | 0,32 | -1,10 | 4,30 | -0,26 | 7,98E-01 |
| AT1G77800 | 1,13 | 2,44 | 1,80 | 1,36 | 1,74E-01 |
| AT1G77840 | 0,31 | 1,18 | 3,52 | 0,33 | 7,38E-01 |
| AT1G77850 | 1,01 | 0,15 | 1,94 | 0,08 | 9,40E-01 |
| AT1G77890 | 0,98 | 1,54 | 2,01 | 0,76 | 4,45E-01 |
| AT1G77930 | 1,14 | 4,05 | 2,12 | 1,92 | 5,54E-02 |
| AT1G78010 | 0,90 | 1,09 | 2,11 | 0,52 | 6,06E-01 |
| AT1G78070 | 1,66 | -1,01 | 1,51 | -0,67 | 5,02E-01 |
| AT1G78100 | 0,64 | -0,38 | 2,05 | -0,19 | 8,53E-01 |
| AT1G78150 | 0,22 | 0,42 | 4,34 | 0,10 | 9,24E-01 |
| AT1G78200 | 0,54 | -0,36 | 2,61 | -0,14 | 8,89E-01 |
| AT1G78280 | 1,59 | 2,87 | 2,00 | 1,44 | 1,50E-01 |
| AT1G78300 | 0,54 | 0,89 | 3,04 | 0,29 | 7,71E-01 |
| AT1G78420 | 0,54 | 1,28 | 2,64 | 0,49 | 6,27E-01 |
| AT1G78490 | 0,05 | -0,58 | 4,38 | -0,13 | 8,95E-01 |
| AT1G78500 | 0,17 | 3,04 | 4,33 | 0,70 | 4,83E-01 |
| AT1G78600 | 0,20 | -0,80 | 4,37 | -0,18 | 8,55E-01 |
| AT1G78630 | 1,31 | -0,90 | 1,76 | -0,51 | 6,09E-01 |
| AT1G78670 | 0,52 | 2,11 | 3,04 | 0,69 | 4,88E-01 |
| AT1G78680 | 0,90 | -0,37 | 1,72 | -0,22 | 8,28E-01 |
| AT1G78700 | 0,37 | 2,29 | 4,31 | 0,53 | 5,94E-01 |
| AT1G78710 | 0,38 | 2,18 | 4,37 | 0,50 | 6,18E-01 |
| AT1G78770 | 0,21 | 2,34 | 4,32 | 0,54 | 5,89E-01 |
| AT1G78870 | 0,28 | 1,40 | 4,31 | 0,32 | 7,45E-01 |
| AT1G78890 | 0,36 | 1,64 | 4,38 | 0,37 | 7,08E-01 |
| AT1G78910 | 0,26 | 2,30 | 4,37 | 0,53 | 5,98E-01 |
| AT1G78920 | 1,61 | 2,53 | 1,51 | 1,67 | 9,47E-02 |
| AT1G79060 | 0,14 | -1,69 | 4,33 | -0,39 | 6,96E-01 |
| AT1G79103 | 0,05 | -0,58 | 4,38 | -0,13 | 8,95E-01 |
| AT1G79110 | 1,67 | 0,13 | 1,42 | 0,09 | 9,27E-01 |
| AT1G09847 | 0,44 | -2,16 | 3,22 | -0,67 | 5,02E-01 |
| AT1G79190 | 0,88 | 3,10 | 2,73 | 1,14 | 2,56E-01 |
| AT1G79230 | 0,71 | 2,17 | 2,90 | 0,75 | 4,54E-01 |
| AT1G79240 | 0,20 | 1,88 | 4,35 | 0,43 | 6,66E-01 |
| AT1G79270 | 1,18 | 2,13 | 1,58 | 1,35 | 1,77E-01 |
| AT1G79340 | 0,91 | -1,22 | 1,94 | -0,63 | 5,30E-01 |
| AT1G79380 | 0,91 | 2,80 | 2,74 | 1,02 | 3,08E-01 |
| AT1G79560 | 0,28 | 0,99 | 4,29 | 0,23 | 8,17E-01 |
| AT1G79590 | 0,18 | 1,51 | 4,31 | 0,35 | 7,25E-01 |
| AT1G79660 | 0,63 | 1,67 | 3,71 | 0,45 | 6,51E-01 |
| AT1G09903 | 0,08 | 2,33 | 4,36 | 0,53 | 5,93E-01 |
| AT1G79810 | 0,41 | 0,18 | 3,17 | 0,06 | 9,54E-01 |
| AT1G79840 | 0,77 | 2,91 | 2,91 | 1,00 | 3,17E-01 |
| AT1G79870 | 0,21 | 1,17 | 4,29 | 0,27 | 7,85E-01 |
| AT1G79890 | 0,21 | 2,01 | 4,37 | 0,46 | 6,45E-01 |
| AT1G79910 | 0,41 | 3,02 | 4,14 | 0,73 | 4,65E-01 |
| AT1G79940 | 1,20 | -0,02 | 1,62 | -0,01 | 9,90E-01 |
| AT1G79950 | 0,49 | 1,64 | 3,98 | 0,41 | 6,80E-01 |
| AT1G79970 | 0,27 | 0,14 | 4,29 | 0,03 | 9,75E-01 |
| AT1G79985 | 0,75 | 0,41 | 2,36 | 0,17 | 8,62E-01 |
| AT1G80010 | 1,24 | 2,93 | 1,61 | 1,82 | 6,88E-02 |
| AT1G80160 | 0,37 | 3,00 | 4,30 | 0,70 | 4,86E-01 |

| | | | | | |
|---|---|---|---|---|---|
| AT1G80260 | 0,59 | 0,84 | 3,03 | 0,28 | 7,81E-01 |
| AT1G80300 | 1,04 | 4,30 | 3,24 | 1,33 | 1,84E-01 |
| AT1G80310 | 0,64 | 0,87 | 2,53 | 0,35 | 7,30E-01 |
| AT1G80380 | 1,15 | 0,29 | 1,84 | 0,16 | 8,77E-01 |
| AT1G09963 | 0,96 | 2,37 | 2,50 | 0,95 | 3,44E-01 |
| AT1G80430 | 0,19 | 0,37 | 4,34 | 0,09 | 9,32E-01 |
| AT1G80510 | 0,61 | 0,81 | 3,79 | 0,21 | 8,30E-01 |
| AT1G80620 | 0,35 | 0,28 | 4,39 | 0,06 | 9,49E-01 |
| AT1G80770 | 0,54 | 0,22 | 3,12 | 0,07 | 9,44E-01 |
| AT1G80780 | 0,60 | 0,74 | 3,00 | 0,25 | 8,06E-01 |
| AT1G80790 | 0,69 | 3,95 | 2,82 | 1,40 | 1,62E-01 |
| AT1G80810 | 0,38 | 2,29 | 4,23 | 0,54 | 5,89E-01 |
| AT1G80840 | 1,09 | -1,42 | 2,20 | -0,65 | 5,17E-01 |
| AT1G80870 | 0,27 | 0,68 | 4,30 | 0,16 | 8,74E-01 |
| AT2G01070 | 0,81 | 2,43 | 2,47 | 0,98 | 3,27E-01 |
| AT2G01080 | 0,35 | 0,33 | 3,43 | 0,10 | 9,22E-01 |
| AT2G01100 | 0,12 | 0,06 | 4,34 | 0,01 | 9,88E-01 |
| AT2G01120 | 0,92 | 4,36 | 2,76 | 1,58 | 1,14E-01 |
| AT2G01190 | 0,50 | 0,47 | 2,98 | 0,16 | 8,75E-01 |
| AT2G01400 | 0,39 | -0,14 | 4,41 | -0,03 | 9,75E-01 |
| AT2G03975 | 0,34 | -0,46 | 3,51 | -0,13 | 8,96E-01 |
| AT2G01480 | 0,34 | 0,37 | 4,35 | 0,09 | 9,32E-01 |
| AT2G01620 | 0,60 | -0,02 | 2,43 | -0,01 | 9,95E-01 |
| AT2G01680 | 0,39 | 0,01 | 4,29 | 0,00 | 9,98E-01 |
| AT2G01750 | 0,24 | -0,94 | 4,31 | -0,22 | 8,28E-01 |
| AT2G01860 | 0,98 | 1,69 | 2,24 | 0,75 | 4,51E-01 |
| AT2G01900 | 1,22 | 0,38 | 1,51 | 0,25 | 8,01E-01 |
| AT2G01980 | 0,63 | 1,35 | 3,04 | 0,44 | 6,57E-01 |
| AT2G02040 | 0,10 | 0,34 | 4,35 | 0,08 | 9,37E-01 |
| AT2G02050 | 0,58 | -1,86 | 2,49 | -0,75 | 4,54E-01 |
| AT2G02070 | 0,84 | -0,30 | 1,83 | -0,16 | 8,71E-01 |
| AT2G02090 | 1,22 | 1,43 | 2,21 | 0,65 | 5,17E-01 |
| AT2G02100 | 0,54 | 1,17 | 3,94 | 0,30 | 7,67E-01 |
| AT2G02440 | 0,68 | -1,04 | 3,07 | -0,34 | 7,36E-01 |
| AT2G02450 | 0,66 | -1,13 | 2,44 | -0,46 | 6,45E-01 |
| AT2G02470 | 0,18 | -0,70 | 4,38 | -0,16 | 8,72E-01 |
| AT2G02550 | 1,01 | -1,34 | 1,84 | -0,73 | 4,68E-01 |
| AT2G02700 | 0,39 | 1,78 | 4,41 | 0,40 | 6,86E-01 |
| AT2G04295 | 0,12 | 0,31 | 4,35 | 0,07 | 9,44E-01 |
| AT2G02760 | 0,54 | -0,73 | 2,65 | -0,27 | 7,84E-01 |
| AT2G02790 | 0,39 | 1,15 | 4,23 | 0,27 | 7,86E-01 |
| AT2G02795 | 0,14 | -0,31 | 4,33 | -0,07 | 9,43E-01 |
| AT2G02810 | 0,62 | 2,34 | 2,94 | 0,80 | 4,26E-01 |
| AT2G04355 | 0,47 | 0,00 | 3,24 | 0,00 | 9,99E-01 |
| AT2G04365 | 0,97 | -0,84 | 2,16 | -0,39 | 6,96E-01 |
| AT2G02870 | 1,53 | 1,16 | 1,63 | 0,71 | 4,76E-01 |
| AT2G02955 | 0,36 | 0,85 | 4,32 | 0,20 | 8,44E-01 |
| AT2G03070 | 0,25 | 2,97 | 4,33 | 0,69 | 4,93E-01 |
| AT2G03120 | 0,64 | -0,10 | 2,67 | -0,04 | 9,69E-01 |
| AT2G03130 | 0,12 | 2,69 | 4,35 | 0,62 | 5,36E-01 |
| AT2G03270 | 0,39 | -0,30 | 2,72 | -0,11 | 9,13E-01 |
| AT2G03410 | 0,43 | 2,73 | 4,06 | 0,67 | 5,01E-01 |
| AT2G03450 | 0,32 | -0,22 | 4,40 | -0,05 | 9,61E-01 |
| AT2G03500 | 0,52 | -2,13 | 3,16 | -0,67 | 5,01E-01 |
| AT2G03520 | 0,08 | 2,33 | 4,36 | 0,53 | 5,93E-01 |
| AT2G03550 | 0,42 | 0,81 | 4,22 | 0,19 | 8,48E-01 |

| | | | | | |
|---|---|---|---|---|---|
| AT2G03567 | 0,47 | 0,35 | 3,24 | 0,11 | 9,13E-01 |
| AT2G03680 | 0,82 | -0,71 | 2,33 | -0,30 | 7,62E-01 |
| AT2G03710 | 0,17 | 3,16 | 4,33 | 0,73 | 4,66E-01 |
| AT2G03823 | 0,37 | -0,54 | 2,74 | -0,20 | 8,45E-01 |
| AT2G03870 | 0,41 | 3,13 | 4,11 | 0,76 | 4,46E-01 |
| AT2G03932 | 0,20 | -2,22 | 4,30 | -0,52 | 6,06E-01 |
| AT2G03931 | 0,05 | -0,58 | 4,38 | -0,13 | 8,95E-01 |
| AT2G03933 | 0,14 | -1,69 | 4,33 | -0,39 | 6,96E-01 |
| AT2G03936 | 0,18 | 0,08 | 4,29 | 0,02 | 9,85E-01 |
| AT2G03965 | 0,22 | -2,22 | 4,30 | -0,52 | 6,05E-01 |
| AT2G03980 | 0,82 | -0,54 | 2,57 | -0,21 | 8,35E-01 |
| AT2G04039 | 0,42 | 2,82 | 4,11 | 0,68 | 4,94E-01 |
| AT2G04735 | 0,15 | -0,97 | 4,37 | -0,22 | 8,24E-01 |
| AT2G04235 | 0,05 | -0,58 | 4,38 | -0,13 | 8,95E-01 |
| AT2G04270 | 1,17 | 1,64 | 2,11 | 0,78 | 4,36E-01 |
| AT2G04340 | 0,38 | 2,13 | 4,38 | 0,49 | 6,27E-01 |
| AT2G04350 | 1,50 | 2,31 | 1,57 | 1,47 | 1,42E-01 |
| AT2G04380 | 0,56 | -0,61 | 2,66 | -0,23 | 8,19E-01 |
| AT2G04390 | 0,85 | 1,78 | 1,98 | 0,90 | 3,69E-01 |
| AT2G04400 | 0,61 | 1,33 | 2,38 | 0,56 | 5,77E-01 |
| AT2G04450 | 0,21 | 2,34 | 4,32 | 0,54 | 5,89E-01 |
| AT2G04495 | 0,43 | 1,52 | 3,28 | 0,46 | 6,44E-01 |
| AT2G04550 | 0,15 | 2,31 | 4,37 | 0,53 | 5,98E-01 |
| AT2G04700 | 0,71 | 1,36 | 2,10 | 0,65 | 5,17E-01 |
| AT2G04740 | 0,38 | 2,54 | 4,35 | 0,58 | 5,60E-01 |
| AT2G04780 | 0,24 | 0,27 | 4,35 | 0,06 | 9,50E-01 |
| AT2G04795 | 0,15 | -0,19 | 4,31 | -0,04 | 9,64E-01 |
| AT2G04850 | 0,08 | 2,33 | 4,36 | 0,53 | 5,93E-01 |
| AT2G04865 | 0,25 | 3,51 | 4,32 | 0,81 | 4,16E-01 |
| AT2G04900 | 0,19 | 0,09 | 4,37 | 0,02 | 9,83E-01 |
| AT2G05060 | 0,21 | 2,01 | 4,37 | 0,46 | 6,45E-01 |
| AT2G05180 | 0,08 | 2,08 | 4,38 | 0,47 | 6,35E-01 |
| AT2G05210 | 0,12 | 0,63 | 4,33 | 0,15 | 8,85E-01 |
| AT2G05320 | 0,92 | 0,42 | 1,90 | 0,22 | 8,24E-01 |
| AT2G05350 | 0,23 | 0,91 | 4,30 | 0,21 | 8,32E-01 |
| AT2G05380 | 0,99 | 1,53 | 2,06 | 0,74 | 4,57E-01 |
| AT2G05540 | 0,30 | -2,05 | 4,31 | -0,48 | 6,35E-01 |
| AT2G04915 | 0,39 | -0,42 | 3,42 | -0,12 | 9,03E-01 |
| AT2G04935 | 1,06 | -0,98 | 1,61 | -0,61 | 5,44E-01 |
| AT2G05635 | 0,15 | -0,41 | 4,32 | -0,09 | 9,25E-01 |
| AT2G04985 | 0,95 | -0,94 | 1,98 | -0,48 | 6,35E-01 |
| AT2G05790 | 0,44 | 1,22 | 3,18 | 0,38 | 7,02E-01 |
| AT2G05812 | 1,46 | 0,05 | 1,42 | 0,03 | 9,73E-01 |
| AT2G05830 | 0,15 | 0,40 | 4,33 | 0,09 | 9,27E-01 |
| AT2G05840 | 0,17 | 2,29 | 4,36 | 0,53 | 5,98E-01 |
| AT2G05990 | 0,47 | 1,18 | 2,32 | 0,51 | 6,10E-01 |
| AT2G05995 | 0,63 | -1,44 | 3,08 | -0,47 | 6,40E-01 |
| AT2G06025 | 0,59 | -0,88 | 2,52 | -0,35 | 7,28E-01 |
| AT2G06095 | 0,70 | -0,12 | 3,59 | -0,03 | 9,74E-01 |
| AT2G06105 | 0,23 | -2,04 | 4,31 | -0,47 | 6,36E-01 |
| AT2G06845 | 0,18 | 0,41 | 4,28 | 0,10 | 9,23E-01 |
| AT2G06850 | 0,75 | 0,66 | 2,50 | 0,26 | 7,93E-01 |
| AT2G06904 | 0,38 | -1,13 | 4,23 | -0,27 | 7,90E-01 |
| AT2G06990 | 1,10 | -0,34 | 1,76 | -0,19 | 8,46E-01 |
| AT2G07050 | 0,64 | -0,04 | 2,79 | -0,01 | 9,88E-01 |
| AT2G07200 | 0,15 | -1,60 | 4,33 | -0,37 | 7,11E-01 |

| | | | | | |
|---|---|---|---|---|---|
| AT2G07565 | 0,91 | -3,29 | 2,36 | -1,40 | 1,62E-01 |
| AT2G07749 | 0,59 | -1,39 | 3,13 | -0,44 | 6,57E-01 |
| AT2G07672 | 0,25 | -1,37 | 4,28 | -0,32 | 7,49E-01 |
| AT2G07745 | 0,19 | -0,02 | 4,39 | 0,00 | 9,97E-01 |
| AT2G07674 | 0,81 | 1,32 | 3,56 | 0,37 | 7,12E-01 |
| AT2G07623 | 0,23 | 0,91 | 4,30 | 0,21 | 8,32E-01 |
| AT2G07678 | 0,32 | 0,00 | 4,40 | 0,00 | 1,00E+00 |
| AT2G07669 | 0,33 | -1,16 | 4,35 | -0,27 | 7,89E-01 |
| AT2G07681 | 0,75 | -1,00 | 1,98 | -0,50 | 6,15E-01 |
| AT2G07631 | 0,18 | -1,21 | 4,35 | -0,28 | 7,81E-01 |
| AT2G05215 | 0,68 | -0,04 | 2,41 | -0,01 | 9,88E-01 |
| AT2G07695 | 1,27 | 0,66 | 1,64 | 0,40 | 6,89E-01 |
| AT2G07798 | 1,27 | 0,23 | 1,54 | 0,15 | 8,84E-01 |
| AT2G07667 | 1,00 | 0,02 | 2,13 | 0,01 | 9,93E-01 |
| AT2G07638 | 1,46 | 2,24 | 1,72 | 1,30 | 1,93E-01 |
| AT2G07707 | 0,32 | 0,29 | 4,30 | 0,07 | 9,46E-01 |
| AT2G07708 | 0,57 | 0,54 | 4,37 | 0,12 | 9,01E-01 |
| AT2G07715 | 0,94 | -0,41 | 1,56 | -0,26 | 7,94E-01 |
| AT2G07728 | 0,11 | 1,28 | 4,35 | 0,29 | 7,68E-01 |
| AT2G07656 | 0,60 | -1,82 | 2,86 | -0,64 | 5,24E-01 |
| AT2G07827 | 0,36 | 0,50 | 3,44 | 0,15 | 8,84E-01 |
| AT2G07787 | 0,79 | 0,09 | 1,92 | 0,05 | 9,61E-01 |
| AT2G07659 | 0,28 | -1,22 | 4,29 | -0,28 | 7,77E-01 |
| AT2G07765 | 0,12 | 2,87 | 4,34 | 0,66 | 5,08E-01 |
| AT2G07741 | 0,27 | -1,91 | 4,32 | -0,44 | 6,59E-01 |
| AT2G07680 | 0,80 | -0,90 | 2,19 | -0,41 | 6,80E-01 |
| AT2G09990 | 0,53 | 1,69 | 3,03 | 0,56 | 5,77E-01 |
| AT2G05485 | 0,27 | -1,02 | 4,29 | -0,24 | 8,11E-01 |
| AT2G10545 | 0,08 | -0,56 | 4,38 | -0,13 | 8,99E-01 |
| AT2G10556 | 0,88 | 2,93 | 2,52 | 1,16 | 2,46E-01 |
| AT2G10965 | 0,86 | -1,20 | 1,86 | -0,64 | 5,20E-01 |
| AT2G11010 | 0,65 | -1,28 | 2,88 | -0,44 | 6,57E-01 |
| AT2G11035 | 0,48 | 0,45 | 4,02 | 0,11 | 9,11E-01 |
| AT2G11045 | 0,08 | -0,56 | 4,38 | -0,13 | 8,99E-01 |
| AT2G11225 | 0,91 | 0,00 | 1,63 | 0,00 | 9,99E-01 |
| AT2G11620 | 0,14 | -1,69 | 4,33 | -0,39 | 6,96E-01 |
| AT2G11851 | 0,57 | -1,35 | 2,80 | -0,48 | 6,30E-01 |
| AT2G05825 | 0,69 | -0,52 | 2,12 | -0,24 | 8,07E-01 |
| AT2G13100 | 0,26 | -0,18 | 4,35 | -0,04 | 9,67E-01 |
| AT2G13290 | 0,12 | 0,31 | 4,35 | 0,07 | 9,44E-01 |
| AT2G05885 | 0,14 | -1,69 | 4,33 | -0,39 | 6,96E-01 |
| AT2G13430 | 0,44 | 1,24 | 3,11 | 0,40 | 6,91E-01 |
| AT2G13500 | 0,11 | -1,28 | 4,35 | -0,29 | 7,69E-01 |
| AT2G13540 | 0,64 | 1,10 | 2,23 | 0,49 | 6,22E-01 |
| AT2G05925 | 0,12 | 0,06 | 4,34 | 0,01 | 9,88E-01 |
| AT2G13790 | 0,49 | 0,26 | 3,18 | 0,08 | 9,35E-01 |
| AT2G13845 | 0,67 | 1,33 | 2,10 | 0,63 | 5,25E-01 |
| AT2G13960 | 0,27 | -1,63 | 4,34 | -0,38 | 7,07E-01 |
| AT2G14070 | 0,19 | -1,34 | 4,35 | -0,31 | 7,58E-01 |
| AT2G14115 | 0,49 | -0,09 | 4,31 | -0,02 | 9,84E-01 |
| AT2G14395 | 0,63 | 0,88 | 4,34 | 0,20 | 8,39E-01 |
| AT2G14440 | 0,43 | -1,41 | 4,11 | -0,34 | 7,31E-01 |
| AT2G14530 | 0,21 | -1,82 | 4,32 | -0,42 | 6,73E-01 |
| AT2G14700 | 0,64 | -1,34 | 2,76 | -0,48 | 6,28E-01 |
| AT2G14740 | 1,25 | 3,84 | 2,35 | 1,64 | 1,02E-01 |
| AT2G14750 | 1,64 | 0,81 | 1,31 | 0,62 | 5,35E-01 |

| | | | | | |
|---|---|---|---|---|---|
| AT2G14765 | 0,40 | -1,36 | 4,25 | -0,32 | 7,48E-01 |
| AT2G14825 | 0,08 | 2,08 | 4,38 | 0,47 | 6,35E-01 |
| AT2G14835 | 0,23 | 2,47 | 4,31 | 0,57 | 5,67E-01 |
| AT2G14850 | 0,21 | 2,18 | 4,32 | 0,51 | 6,13E-01 |
| AT2G14870 | 0,17 | 0,00 | 4,41 | 0,00 | 1,00E+00 |
| AT2G14878 | 1,23 | -0,33 | 1,41 | -0,24 | 8,13E-01 |
| AT2G14890 | 0,77 | 0,27 | 1,97 | 0,14 | 8,90E-01 |
| AT2G06305 | 0,18 | -1,39 | 4,34 | -0,32 | 7,49E-01 |
| AT2G15042 | 0,11 | 1,28 | 4,35 | 0,29 | 7,68E-01 |
| AT2G15050 | 0,64 | -2,87 | 2,84 | -1,01 | 3,12E-01 |
| AT2G15080 | 0,92 | 0,71 | 1,91 | 0,37 | 7,10E-01 |
| AT2G15090 | 1,27 | 0,49 | 2,82 | 0,17 | 8,63E-01 |
| AT2G15300 | 0,31 | 1,65 | 4,39 | 0,38 | 7,06E-01 |
| AT2G15430 | 0,16 | 0,33 | 4,31 | 0,08 | 9,39E-01 |
| AT2G15450 | 0,29 | 0,46 | 4,32 | 0,11 | 9,15E-01 |
| AT2G15470 | 0,34 | -1,56 | 4,27 | -0,37 | 7,15E-01 |
| AT2G15580 | 0,29 | -0,09 | 3,59 | -0,03 | 9,79E-01 |
| AT2G15710 | 0,40 | -2,27 | 4,30 | -0,53 | 5,98E-01 |
| AT2G15730 | 0,08 | -0,56 | 4,38 | -0,13 | 8,99E-01 |
| AT2G15900 | 0,47 | 0,32 | 2,63 | 0,12 | 9,04E-01 |
| AT2G15960 | 1,09 | 2,06 | 1,82 | 1,14 | 2,56E-01 |
| AT2G06535 | 1,38 | 0,35 | 1,65 | 0,21 | 8,34E-01 |
| AT2G16019 | 1,05 | -1,55 | 1,91 | -0,81 | 4,15E-01 |
| AT2G16280 | 0,93 | -1,37 | 2,04 | -0,67 | 5,01E-01 |
| AT2G16365 | 0,96 | 1,53 | 2,12 | 0,72 | 4,71E-01 |
| AT2G16380 | 0,44 | -1,60 | 4,23 | -0,38 | 7,06E-01 |
| AT2G16390 | 0,40 | 3,73 | 4,32 | 0,86 | 3,87E-01 |
| AT2G16630 | 0,14 | 0,87 | 4,31 | 0,20 | 8,40E-01 |
| AT2G16700 | 0,28 | 2,34 | 4,33 | 0,54 | 5,89E-01 |
| AT2G16710 | 0,72 | 1,53 | 3,63 | 0,42 | 6,73E-01 |
| AT2G16800 | 0,26 | -1,21 | 4,36 | -0,28 | 7,81E-01 |
| AT2G16950 | 1,26 | 3,53 | 2,17 | 1,62 | 1,05E-01 |
| AT2G16970 | 1,25 | 0,79 | 2,02 | 0,39 | 6,96E-01 |
| AT2G17033 | 0,44 | 2,92 | 4,16 | 0,70 | 4,82E-01 |
| AT2G17040 | 1,57 | 0,61 | 1,52 | 0,40 | 6,91E-01 |
| AT2G17120 | 0,23 | 0,18 | 4,27 | 0,04 | 9,66E-01 |
| AT2G17140 | 0,42 | 1,21 | 3,33 | 0,36 | 7,17E-01 |
| AT2G17170 | 0,08 | 2,08 | 4,38 | 0,47 | 6,35E-01 |
| AT2G17240 | 0,52 | 0,56 | 3,18 | 0,18 | 8,60E-01 |
| AT2G17250 | 0,25 | 2,98 | 4,33 | 0,69 | 4,92E-01 |
| AT2G17265 | 1,56 | 1,91 | 1,41 | 1,35 | 1,76E-01 |
| AT2G17270 | 0,25 | 0,42 | 4,34 | 0,10 | 9,23E-01 |
| AT2G17290 | 0,78 | 1,44 | 2,24 | 0,64 | 5,21E-01 |
| AT2G06905 | 1,12 | 0,82 | 2,11 | 0,39 | 6,98E-01 |
| AT2G17360 | 0,51 | -0,52 | 3,18 | -0,16 | 8,71E-01 |
| AT2G17390 | 0,66 | 3,26 | 2,95 | 1,10 | 2,70E-01 |
| AT2G17410 | 1,45 | 1,33 | 1,65 | 0,81 | 4,20E-01 |
| AT2G17420 | 1,05 | 0,07 | 1,81 | 0,04 | 9,68E-01 |
| AT2G17520 | 1,17 | 1,02 | 1,61 | 0,63 | 5,26E-01 |
| AT2G17530 | 0,50 | 3,49 | 3,97 | 0,88 | 3,79E-01 |
| AT2G17570 | 0,42 | 2,98 | 4,15 | 0,72 | 4,73E-01 |
| AT2G17580 | 0,66 | 3,37 | 3,03 | 1,11 | 2,66E-01 |
| AT2G17705 | 0,42 | -1,18 | 3,18 | -0,37 | 7,09E-01 |
| AT2G17720 | 0,20 | 2,30 | 4,36 | 0,53 | 5,98E-01 |
| AT2G17760 | 0,65 | 1,48 | 2,92 | 0,51 | 6,11E-01 |
| AT2G17787 | 0,33 | 1,93 | 4,30 | 0,45 | 6,53E-01 |

| | | | | | |
|---|---|---|---|---|---|
| AT2G17790 | 0,80 | 1,02 | 2,37 | 0,43 | 6,67E-01 |
| AT2G17905 | 0,41 | -2,21 | 3,31 | -0,67 | 5,03E-01 |
| AT2G17920 | 0,15 | -0,14 | 4,34 | -0,03 | 9,74E-01 |
| AT2G17980 | 1,01 | 4,09 | 2,49 | 1,64 | 1,00E-01 |
| AT2G17990 | 0,69 | 1,78 | 2,43 | 0,73 | 4,63E-01 |
| AT2G18000 | 0,29 | 3,04 | 4,33 | 0,70 | 4,83E-01 |
| AT2G18030 | 0,68 | 3,31 | 2,92 | 1,13 | 2,58E-01 |
| AT2G18040 | 0,20 | -0,47 | 4,33 | -0,11 | 9,13E-01 |
| AT2G18090 | 1,01 | 1,56 | 1,66 | 0,94 | 3,49E-01 |
| AT2G18100 | 0,48 | 1,83 | 3,96 | 0,46 | 6,44E-01 |
| AT2G18110 | 0,62 | 0,02 | 3,06 | 0,01 | 9,96E-01 |
| AT2G18390 | 0,28 | 1,87 | 4,32 | 0,43 | 6,65E-01 |
| AT2G18420 | 0,81 | 3,22 | 3,46 | 0,93 | 3,53E-01 |
| AT2G18530 | 0,38 | 1,31 | 3,35 | 0,39 | 6,96E-01 |
| AT2G18560 | 1,54 | 0,89 | 2,30 | 0,39 | 7,00E-01 |
| AT2G18630 | 0,16 | 1,16 | 4,38 | 0,27 | 7,90E-01 |
| AT2G18670 | 1,13 | -0,37 | 1,52 | -0,24 | 8,10E-01 |
| AT2G18680 | 0,16 | 1,97 | 4,33 | 0,45 | 6,50E-01 |
| AT2G18690 | 1,29 | 0,09 | 1,59 | 0,05 | 9,57E-01 |
| AT2G18700 | 0,27 | 1,15 | 4,37 | 0,26 | 7,92E-01 |
| AT2G18730 | 0,58 | 0,81 | 2,40 | 0,34 | 7,35E-01 |
| AT2G18735 | 0,22 | 0,35 | 4,34 | 0,08 | 9,36E-01 |
| AT2G18750 | 0,67 | 0,18 | 1,91 | 0,10 | 9,24E-01 |
| AT2G18760 | 1,48 | 1,36 | 1,43 | 0,96 | 3,39E-01 |
| AT2G18850 | 0,52 | 0,00 | 4,41 | 0,00 | 1,00E+00 |
| AT2G18880 | 0,25 | 3,58 | 4,32 | 0,83 | 4,07E-01 |
| AT2G18917 | 0,08 | -0,56 | 4,38 | -0,13 | 8,99E-01 |
| AT2G07095 | 0,55 | 2,55 | 2,95 | 0,86 | 3,87E-01 |
| AT2G18938 | 0,72 | 0,29 | 2,42 | 0,12 | 9,05E-01 |
| AT2G18950 | 0,45 | 0,72 | 4,10 | 0,18 | 8,60E-01 |
| AT2G19110 | 0,15 | 1,64 | 4,33 | 0,38 | 7,06E-01 |
| AT2G19130 | 0,89 | -2,32 | 2,20 | -1,06 | 2,90E-01 |
| AT2G19160 | 0,39 | 2,54 | 4,33 | 0,59 | 5,58E-01 |
| AT2G19260 | 0,46 | 0,26 | 4,01 | 0,06 | 9,49E-01 |
| AT2G19280 | 0,36 | 3,22 | 4,34 | 0,74 | 4,59E-01 |
| AT2G19420 | 0,28 | -0,99 | 4,30 | -0,23 | 8,18E-01 |
| AT2G19430 | 0,24 | -0,80 | 4,38 | -0,18 | 8,55E-01 |
| AT2G19450 | 0,42 | 1,63 | 4,25 | 0,38 | 7,01E-01 |
| AT2G07205 | 0,75 | -1,52 | 3,53 | -0,43 | 6,68E-01 |
| AT2G19490 | 0,36 | 3,47 | 4,33 | 0,80 | 4,23E-01 |
| AT2G19520 | 0,55 | -0,65 | 3,97 | -0,16 | 8,70E-01 |
| AT2G19540 | 0,46 | 0,57 | 2,65 | 0,21 | 8,30E-01 |
| AT2G19572 | 0,52 | -1,06 | 2,88 | -0,37 | 7,14E-01 |
| AT2G19582 | 0,25 | -1,47 | 4,28 | -0,34 | 7,32E-01 |
| AT2G19600 | 0,64 | -0,13 | 3,81 | -0,03 | 9,73E-01 |
| AT2G19640 | 0,53 | 3,18 | 3,84 | 0,83 | 4,07E-01 |
| AT2G19710 | 1,44 | -0,31 | 1,91 | -0,17 | 8,69E-01 |
| AT2G19730 | 0,57 | -1,06 | 4,03 | -0,26 | 7,92E-01 |
| AT2G19740 | 0,67 | -0,48 | 2,12 | -0,23 | 8,19E-01 |
| AT2G19810 | 0,34 | -1,32 | 3,45 | -0,38 | 7,01E-01 |
| AT2G19880 | 0,75 | 2,58 | 2,43 | 1,06 | 2,90E-01 |
| AT2G20020 | 1,07 | 1,76 | 2,04 | 0,86 | 3,87E-01 |
| AT2G20100 | 0,33 | 2,03 | 4,34 | 0,47 | 6,39E-01 |
| AT2G20110 | 0,83 | 3,67 | 4,31 | 0,85 | 3,95E-01 |
| AT2G20142 | 0,28 | 1,78 | 4,33 | 0,41 | 6,80E-01 |
| AT2G20230 | 0,26 | -0,52 | 4,38 | -0,12 | 9,06E-01 |

| | | | | | |
|---|---|---|---|---|---|
| AT2G20260 | 1,52 | -0,21 | 1,67 | -0,13 | 9,00E-01 |
| AT2G20340 | 0,57 | 2,34 | 3,93 | 0,60 | 5,51E-01 |
| AT2G20360 | 0,22 | 1,27 | 4,31 | 0,29 | 7,69E-01 |
| AT2G20362 | 0,08 | 2,33 | 4,36 | 0,53 | 5,93E-01 |
| AT2G20370 | 1,01 | 0,36 | 2,37 | 0,15 | 8,78E-01 |
| AT2G20420 | 0,62 | 0,63 | 2,12 | 0,30 | 7,64E-01 |
| AT2G20450 | 0,40 | 3,40 | 4,14 | 0,82 | 4,11E-01 |
| AT2G20530 | 0,21 | 0,79 | 4,39 | 0,18 | 8,57E-01 |
| AT2G20690 | 1,05 | 1,70 | 2,33 | 0,73 | 4,65E-01 |
| AT2G20710 | 0,08 | 2,08 | 4,38 | 0,47 | 6,35E-01 |
| AT2G07335 | 0,46 | 2,47 | 4,17 | 0,59 | 5,53E-01 |
| AT2G20750 | 0,59 | 1,64 | 3,82 | 0,43 | 6,69E-01 |
| AT2G20770 | 0,17 | 3,14 | 4,33 | 0,73 | 4,68E-01 |
| AT2G20820 | 0,25 | 1,98 | 4,32 | 0,46 | 6,47E-01 |
| AT2G20860 | 1,36 | 2,19 | 1,55 | 1,41 | 1,58E-01 |
| AT2G20890 | 1,04 | 0,82 | 1,84 | 0,44 | 6,58E-01 |
| AT2G20920 | 0,65 | 2,49 | 2,46 | 1,01 | 3,12E-01 |
| AT2G20960 | 0,82 | 0,97 | 1,97 | 0,49 | 6,22E-01 |
| AT2G21160 | 0,91 | 0,88 | 3,01 | 0,29 | 7,69E-01 |
| AT2G21187 | 0,89 | -0,98 | 1,65 | -0,60 | 5,52E-01 |
| AT2G21190 | 0,23 | -0,91 | 4,37 | -0,21 | 8,36E-01 |
| AT2G21270 | 0,21 | 3,16 | 4,33 | 0,73 | 4,66E-01 |
| AT2G21380 | 1,04 | -1,00 | 1,98 | -0,51 | 6,12E-01 |
| AT2G21410 | 1,46 | 1,51 | 1,50 | 1,00 | 3,16E-01 |
| AT2G21520 | 0,84 | 1,70 | 1,91 | 0,89 | 3,76E-01 |
| AT2G21530 | 0,61 | -0,38 | 2,92 | -0,13 | 8,96E-01 |
| AT2G21550 | 0,95 | 0,29 | 2,56 | 0,11 | 9,09E-01 |
| AT2G21580 | 1,04 | 4,39 | 2,73 | 1,61 | 1,08E-01 |
| AT2G21620 | 1,16 | -1,65 | 2,20 | -0,75 | 4,53E-01 |
| AT2G21630 | 0,32 | 2,03 | 4,37 | 0,46 | 6,42E-01 |
| AT2G21770 | 0,18 | -0,70 | 4,38 | -0,16 | 8,72E-01 |
| AT2G21790 | 0,74 | 2,01 | 2,40 | 0,84 | 4,01E-01 |
| AT2G21850 | 1,02 | 2,19 | 2,58 | 0,85 | 3,95E-01 |
| AT2G07475 | 0,24 | 2,24 | 4,33 | 0,52 | 6,05E-01 |
| AT2G21960 | 0,80 | 1,44 | 2,54 | 0,57 | 5,70E-01 |
| AT2G07495 | 0,12 | 1,65 | 4,34 | 0,38 | 7,04E-01 |
| AT2G22080 | 0,65 | 2,95 | 2,54 | 1,16 | 2,45E-01 |
| AT2G22120 | 0,32 | 2,03 | 4,37 | 0,46 | 6,42E-01 |
| AT2G22425 | 0,10 | -1,26 | 4,35 | -0,29 | 7,72E-01 |
| AT2G22450 | 0,24 | 1,11 | 4,33 | 0,26 | 7,97E-01 |
| AT2G22480 | 0,23 | 1,20 | 4,34 | 0,28 | 7,82E-01 |
| AT2G22540 | 1,53 | -1,29 | 1,46 | -0,88 | 3,78E-01 |
| AT2G22670 | 0,80 | -2,17 | 2,12 | -1,03 | 3,04E-01 |
| AT2G22680 | 0,25 | -0,67 | 4,39 | -0,15 | 8,78E-01 |
| AT2G22720 | 0,99 | 3,22 | 2,69 | 1,20 | 2,32E-01 |
| AT2G07845 | 0,43 | 2,89 | 4,08 | 0,71 | 4,78E-01 |
| AT2G22790 | 0,08 | 2,33 | 4,36 | 0,53 | 5,93E-01 |
| AT2G22840 | 1,11 | 3,83 | 2,09 | 1,83 | 6,66E-02 |
| AT2G07850 | 0,72 | 0,10 | 1,87 | 0,05 | 9,57E-01 |
| AT2G22910 | 0,32 | 0,61 | 4,33 | 0,14 | 8,87E-01 |
| AT2G23080 | 0,38 | 1,13 | 4,27 | 0,27 | 7,91E-01 |
| AT2G23120 | 0,92 | 2,01 | 2,06 | 0,98 | 3,29E-01 |
| AT2G23130 | 0,41 | -0,37 | 2,68 | -0,14 | 8,89E-01 |
| AT2G23142 | 1,46 | 1,76 | 2,55 | 0,69 | 4,89E-01 |
| AT2G23200 | 1,64 | 1,69 | 1,54 | 1,09 | 2,74E-01 |
| AT2G23310 | 0,84 | 1,45 | 2,90 | 0,50 | 6,17E-01 |

| | | | | | |
|---|---|---|---|---|---|
| AT2G07895 | 0,15 | 1,93 | 4,33 | 0,45 | 6,55E-01 |
| AT2G23320 | 1,39 | 1,00 | 1,50 | 0,67 | 5,03E-01 |
| AT2G23420 | 0,47 | 2,40 | 4,37 | 0,55 | 5,83E-01 |
| AT2G23460 | 1,50 | 2,04 | 1,55 | 1,32 | 1,87E-01 |
| AT2G07915 | 1,63 | -4,39 | 2,30 | -1,91 | 5,65E-02 |
| AT2G23672 | 1,50 | 1,05 | 1,56 | 0,68 | 4,99E-01 |
| AT2G23680 | 0,31 | 2,25 | 4,37 | 0,51 | 6,07E-01 |
| AT2G23740 | 0,12 | 0,63 | 4,33 | 0,15 | 8,85E-01 |
| AT2G23890 | 0,37 | 2,03 | 3,44 | 0,59 | 5,55E-01 |
| AT2G23985 | 0,17 | 0,00 | 4,41 | 0,00 | 1,00E+00 |
| AT2G24090 | 0,67 | 1,00 | 2,53 | 0,39 | 6,93E-01 |
| AT2G24180 | 1,36 | 3,04 | 2,50 | 1,21 | 2,25E-01 |
| AT2G24240 | 0,31 | 0,28 | 4,30 | 0,06 | 9,48E-01 |
| AT2G24280 | 1,45 | 0,62 | 1,49 | 0,42 | 6,78E-01 |
| AT2G24300 | 0,15 | 0,40 | 4,33 | 0,09 | 9,27E-01 |
| AT2G08015 | 0,35 | 2,28 | 4,28 | 0,53 | 5,94E-01 |
| AT2G24350 | 0,50 | 2,46 | 3,99 | 0,62 | 5,37E-01 |
| AT2G24390 | 0,92 | 3,19 | 2,91 | 1,10 | 2,73E-01 |
| AT2G24380 | 0,65 | 2,07 | 3,82 | 0,54 | 5,88E-01 |
| AT2G24420 | 1,14 | 2,10 | 2,00 | 1,05 | 2,94E-01 |
| AT2G08045 | 0,23 | 0,83 | 4,27 | 0,20 | 8,45E-01 |
| AT2G08050 | 0,32 | 0,22 | 4,24 | 0,05 | 9,59E-01 |
| AT2G24540 | 0,88 | -0,68 | 1,65 | -0,41 | 6,80E-01 |
| AT2G24590 | 0,49 | 2,90 | 3,93 | 0,74 | 4,61E-01 |
| AT2G24592 | 0,83 | 1,66 | 2,01 | 0,83 | 4,08E-01 |
| AT2G24765 | 0,69 | 3,66 | 2,86 | 1,28 | 2,01E-01 |
| AT2G25000 | 0,52 | 3,38 | 3,80 | 0,89 | 3,73E-01 |
| AT2G25080 | 0,61 | -1,19 | 2,57 | -0,46 | 6,44E-01 |
| AT2G25110 | 0,19 | 1,14 | 4,29 | 0,26 | 7,91E-01 |
| AT2G25190 | 0,14 | -0,45 | 4,33 | -0,10 | 9,17E-01 |
| AT2G25370 | 1,47 | -4,18 | 2,03 | -2,06 | 3,98E-02 |
| AT2G25410 | 1,30 | -0,51 | 1,96 | -0,26 | 7,96E-01 |
| AT2G25480 | 0,05 | -0,58 | 4,38 | -0,13 | 8,95E-01 |
| AT2G25560 | 0,73 | 3,53 | 2,77 | 1,27 | 2,02E-01 |
| AT2G25580 | 0,08 | 2,08 | 4,38 | 0,47 | 6,35E-01 |
| AT2G25590 | 0,21 | 2,18 | 4,32 | 0,51 | 6,13E-01 |
| AT2G25625 | 0,15 | 1,94 | 4,32 | 0,45 | 6,54E-01 |
| AT2G25630 | 0,50 | -0,58 | 2,94 | -0,20 | 8,43E-01 |
| AT2G25650 | 0,70 | 3,80 | 2,87 | 1,32 | 1,86E-01 |
| AT2G25790 | 0,18 | -0,78 | 4,31 | -0,18 | 8,57E-01 |
| AT2G25840 | 0,59 | -0,87 | 3,88 | -0,22 | 8,23E-01 |
| AT2G25870 | 1,69 | 1,49 | 1,49 | 1,00 | 3,18E-01 |
| AT2G25930 | 1,26 | 1,71 | 1,50 | 1,14 | 2,54E-01 |
| AT2G26060 | 1,00 | 3,02 | 4,34 | 0,70 | 4,87E-01 |
| AT2G26110 | 0,61 | -2,16 | 2,84 | -0,76 | 4,46E-01 |
| AT2G26170 | 0,47 | 1,44 | 3,14 | 0,46 | 6,47E-01 |
| AT2G26270 | 0,21 | 2,01 | 4,37 | 0,46 | 6,45E-01 |
| AT2G26300 | 1,09 | 5,25 | 2,42 | 2,17 | 3,02E-02 |
| AT2G26340 | 0,57 | 1,82 | 3,08 | 0,59 | 5,55E-01 |
| AT2G26440 | 1,40 | 3,71 | 1,77 | 2,10 | 3,55E-02 |
| AT2G26510 | 0,24 | 1,11 | 4,33 | 0,26 | 7,97E-01 |
| AT2G26570 | 0,43 | 1,55 | 3,32 | 0,47 | 6,40E-01 |
| AT2G26600 | 0,45 | 1,69 | 3,14 | 0,54 | 5,90E-01 |
| AT2G26610 | 0,30 | 1,72 | 4,30 | 0,40 | 6,89E-01 |
| AT2G26640 | 0,71 | 3,02 | 3,61 | 0,83 | 4,04E-01 |
| AT2G26660 | 0,22 | 0,78 | 4,33 | 0,18 | 8,58E-01 |

| | | | | | |
|---|---|---|---|---|---|
| AT2G26692 | 0,08 | 2,33 | 4,36 | 0,53 | 5,93E-01 |
| AT2G26730 | 0,85 | -0,28 | 1,97 | -0,14 | 8,86E-01 |
| AT2G26920 | 0,60 | 2,81 | 3,04 | 0,93 | 3,55E-01 |
| AT2G27030 | 0,72 | -0,88 | 2,42 | -0,36 | 7,17E-01 |
| AT2G27050 | 0,99 | 0,96 | 1,62 | 0,59 | 5,53E-01 |
| AT2G27060 | 0,17 | 2,95 | 4,34 | 0,68 | 4,96E-01 |
| AT2G27080 | 0,39 | -1,32 | 4,11 | -0,32 | 7,48E-01 |
| AT2G27100 | 0,72 | 3,63 | 3,56 | 1,02 | 3,07E-01 |
| AT2G27130 | 0,30 | -2,05 | 4,31 | -0,48 | 6,35E-01 |
| AT2G27200 | 0,21 | 1,45 | 4,28 | 0,34 | 7,35E-01 |
| AT2G27210 | 0,89 | 1,88 | 2,12 | 0,89 | 3,75E-01 |
| AT2G27230 | 1,31 | 0,55 | 1,50 | 0,37 | 7,14E-01 |
| AT2G27260 | 0,40 | 1,64 | 3,24 | 0,51 | 6,12E-01 |
| AT2G27320 | 0,85 | -0,88 | 2,26 | -0,39 | 6,97E-01 |
| AT2G27380 | 0,13 | -1,34 | 4,35 | -0,31 | 7,58E-01 |
| AT2G27460 | 0,28 | 2,83 | 4,34 | 0,65 | 5,15E-01 |
| AT2G27540 | 0,95 | -3,02 | 2,98 | -1,01 | 3,12E-01 |
| AT2G27570 | 0,28 | 1,64 | 4,38 | 0,38 | 7,08E-01 |
| AT2G27600 | 1,39 | 1,08 | 1,66 | 0,65 | 5,15E-01 |
| AT2G27690 | 0,16 | 1,18 | 4,32 | 0,27 | 7,85E-01 |
| AT2G27710 | 0,57 | 0,40 | 3,15 | 0,13 | 8,98E-01 |
| AT2G27760 | 0,08 | 2,08 | 4,38 | 0,47 | 6,35E-01 |
| AT2G27810 | 0,54 | 0,19 | 2,58 | 0,07 | 9,41E-01 |
| AT2G27820 | 0,23 | 0,69 | 4,30 | 0,16 | 8,72E-01 |
| AT2G27900 | 1,32 | 1,14 | 1,39 | 0,82 | 4,13E-01 |
| AT2G28070 | 0,87 | 2,09 | 2,88 | 0,73 | 4,67E-01 |
| AT2G28120 | 0,29 | 3,20 | 4,33 | 0,74 | 4,60E-01 |
| AT2G28130 | 0,59 | 3,78 | 4,31 | 0,88 | 3,81E-01 |
| AT2G28190 | 0,42 | -0,57 | 4,25 | -0,13 | 8,93E-01 |
| AT2G28250 | 0,08 | 2,33 | 4,36 | 0,53 | 5,93E-01 |
| AT2G28310 | 0,30 | 0,27 | 4,35 | 0,06 | 9,50E-01 |
| AT2G28320 | 0,43 | 0,71 | 4,11 | 0,17 | 8,63E-01 |
| AT2G28350 | 0,96 | 2,58 | 1,91 | 1,35 | 1,78E-01 |
| AT2G28370 | 0,25 | 1,98 | 4,32 | 0,46 | 6,47E-01 |
| AT2G28460 | 0,18 | -1,50 | 4,34 | -0,35 | 7,30E-01 |
| AT2G28520 | 0,65 | 2,15 | 2,79 | 0,77 | 4,41E-01 |
| AT2G28540 | 0,87 | 3,37 | 2,34 | 1,44 | 1,49E-01 |
| AT2G28605 | 0,37 | -1,56 | 4,33 | -0,36 | 7,18E-01 |
| AT2G28660 | 0,10 | -1,26 | 4,35 | -0,29 | 7,72E-01 |
| AT2G28671 | 0,51 | 0,49 | 2,91 | 0,17 | 8,65E-01 |
| AT2G28710 | 0,24 | 0,93 | 4,29 | 0,22 | 8,29E-01 |
| AT2G28720 | 0,49 | -2,46 | 4,14 | -0,59 | 5,53E-01 |
| AT2G28730 | 1,02 | -0,72 | 1,55 | -0,46 | 6,45E-01 |
| AT2G28810 | 0,71 | 0,41 | 2,45 | 0,17 | 8,68E-01 |
| AT2G28930 | 0,12 | 1,15 | 4,35 | 0,26 | 7,92E-01 |
| AT2G29020 | 0,11 | -1,28 | 4,35 | -0,29 | 7,69E-01 |
| AT2G29065 | 0,30 | 2,68 | 4,35 | 0,61 | 5,39E-01 |
| AT2G29070 | 0,15 | 1,93 | 4,33 | 0,45 | 6,55E-01 |
| AT2G29140 | 0,94 | 2,31 | 2,34 | 0,99 | 3,24E-01 |
| AT2G29180 | 0,47 | 1,02 | 4,10 | 0,25 | 8,04E-01 |
| AT2G29210 | 1,16 | 2,35 | 2,01 | 1,17 | 2,41E-01 |
| AT2G29310 | 0,46 | -0,92 | 3,34 | -0,28 | 7,83E-01 |
| AT2G29350 | 0,65 | 2,87 | 3,77 | 0,76 | 4,46E-01 |
| AT2G29360 | 0,51 | 1,70 | 4,40 | 0,39 | 6,98E-01 |
| AT2G29510 | 0,82 | 4,56 | 3,29 | 1,38 | 1,66E-01 |
| AT2G29600 | 0,47 | -0,45 | 3,02 | -0,15 | 8,81E-01 |

| | | | | | |
|---|---|---|---|---|---|
| AT2G29640 | 0,37 | 2,79 | 4,28 | 0,65 | 5,14E-01 |
| AT2G29660 | 0,20 | 2,30 | 4,36 | 0,53 | 5,98E-01 |
| AT2G29690 | 0,14 | -0,45 | 4,38 | -0,10 | 9,18E-01 |
| AT2G29700 | 0,24 | -0,70 | 4,38 | -0,16 | 8,72E-01 |
| AT2G29970 | 1,50 | 0,24 | 1,29 | 0,19 | 8,51E-01 |
| AT2G30020 | 0,83 | -1,39 | 3,34 | -0,42 | 6,76E-01 |
| AT2G30040 | 0,38 | 1,13 | 3,25 | 0,35 | 7,28E-01 |
| AT2G30050 | 0,66 | -1,36 | 2,57 | -0,53 | 5,96E-01 |
| AT2G30140 | 0,54 | 3,14 | 3,82 | 0,82 | 4,11E-01 |
| AT2G30150 | 0,10 | 0,39 | 4,35 | 0,09 | 9,29E-01 |
| AT2G30180 | 1,06 | 0,23 | 1,65 | 0,14 | 8,87E-01 |
| AT2G08375 | 1,24 | -2,32 | 1,76 | -1,32 | 1,88E-01 |
| AT2G30230 | 0,50 | 0,85 | 3,16 | 0,27 | 7,88E-01 |
| AT2G30362 | 0,41 | -1,28 | 4,09 | -0,31 | 7,54E-01 |
| AT2G30410 | 0,72 | -1,43 | 2,48 | -0,58 | 5,64E-01 |
| AT2G30530 | 0,20 | 2,30 | 4,36 | 0,53 | 5,98E-01 |
| AT2G30550 | 1,44 | 1,42 | 1,86 | 0,76 | 4,46E-01 |
| AT2G30600 | 0,51 | -0,40 | 4,37 | -0,09 | 9,27E-01 |
| AT2G30620 | 0,85 | 0,42 | 2,19 | 0,19 | 8,47E-01 |
| AT2G30730 | 0,89 | -0,67 | 2,41 | -0,28 | 7,80E-01 |
| AT2G30780 | 0,62 | 2,57 | 3,64 | 0,71 | 4,81E-01 |
| AT2G30870 | 1,65 | 0,10 | 1,23 | 0,08 | 9,34E-01 |
| AT2G30930 | 0,49 | -0,55 | 2,63 | -0,21 | 8,33E-01 |
| AT2G30970 | 0,46 | 1,26 | 4,38 | 0,29 | 7,73E-01 |
| AT2G31010 | 0,95 | 3,33 | 2,34 | 1,42 | 1,54E-01 |
| AT2G31020 | 0,26 | 2,13 | 4,38 | 0,49 | 6,27E-01 |
| AT2G31200 | 0,16 | -0,97 | 4,35 | -0,22 | 8,24E-01 |
| AT2G31215 | 0,52 | -0,54 | 2,52 | -0,21 | 8,31E-01 |
| AT2G31280 | 1,47 | 3,59 | 2,00 | 1,79 | 7,27E-02 |
| AT2G31290 | 0,71 | 1,90 | 3,17 | 0,60 | 5,47E-01 |
| AT2G31350 | 0,38 | 2,34 | 4,28 | 0,55 | 5,85E-01 |
| AT2G31370 | 0,19 | 1,67 | 4,33 | 0,39 | 6,99E-01 |
| AT2G31380 | 1,06 | 2,87 | 1,91 | 1,51 | 1,32E-01 |
| AT2G08435 | 0,79 | 2,99 | 3,53 | 0,85 | 3,97E-01 |
| AT2G31560 | 0,67 | -1,49 | 2,36 | -0,63 | 5,28E-01 |
| AT2G31585 | 0,47 | 2,99 | 3,96 | 0,76 | 4,50E-01 |
| AT2G31750 | 0,75 | 1,89 | 2,90 | 0,65 | 5,15E-01 |
| AT2G31790 | 0,30 | -0,02 | 3,51 | -0,01 | 9,95E-01 |
| AT2G31810 | 1,10 | 0,59 | 2,05 | 0,29 | 7,72E-01 |
| AT2G31960 | 1,68 | 1,04 | 1,39 | 0,75 | 4,53E-01 |
| AT2G31970 | 0,18 | 1,51 | 4,31 | 0,35 | 7,25E-01 |
| AT2G32010 | 0,37 | 0,27 | 4,27 | 0,06 | 9,50E-01 |
| AT2G32040 | 0,08 | 2,08 | 4,38 | 0,47 | 6,35E-01 |
| AT2G32100 | 1,15 | -3,68 | 2,03 | -1,82 | 6,94E-02 |
| AT2G32150 | 1,00 | -0,57 | 2,19 | -0,26 | 7,94E-01 |
| AT2G32250 | 0,51 | 2,19 | 3,14 | 0,70 | 4,85E-01 |
| AT2G32260 | 0,44 | 2,02 | 4,05 | 0,50 | 6,18E-01 |
| AT2G32295 | 0,53 | 0,20 | 2,64 | 0,07 | 9,41E-01 |
| AT2G32320 | 0,15 | 0,40 | 4,33 | 0,09 | 9,27E-01 |
| AT2G32440 | 0,16 | 1,27 | 4,37 | 0,29 | 7,71E-01 |
| AT2G32450 | 0,56 | 1,20 | 2,43 | 0,49 | 6,21E-01 |
| AT2G32698 | 0,12 | 2,81 | 4,34 | 0,65 | 5,18E-01 |
| AT2G32700 | 1,61 | 2,08 | 1,29 | 1,61 | 1,08E-01 |
| AT2G32800 | 1,51 | 1,38 | 1,42 | 0,98 | 3,29E-01 |
| AT2G32900 | 0,29 | 2,24 | 4,31 | 0,52 | 6,03E-01 |
| AT2G32910 | 0,68 | 1,68 | 2,90 | 0,58 | 5,61E-01 |

| | | | | | |
|---|---|---|---|---|---|
| AT2G33080 | 0,74 | 4,26 | 3,44 | 1,24 | 2,16E-01 |
| AT2G33160 | 0,25 | 2,97 | 4,33 | 0,69 | 4,93E-01 |
| AT2G33220 | 0,54 | 2,33 | 3,13 | 0,75 | 4,56E-01 |
| AT2G33250 | 0,76 | 2,58 | 4,32 | 0,60 | 5,51E-01 |
| AT2G33290 | 0,29 | -0,09 | 3,59 | -0,03 | 9,79E-01 |
| AT2G33390 | 0,25 | -0,67 | 4,39 | -0,15 | 8,78E-01 |
| AT2G33420 | 0,37 | 2,79 | 4,28 | 0,65 | 5,14E-01 |
| AT2G08585 | 0,32 | -0,16 | 3,57 | -0,04 | 9,64E-01 |
| AT2G33435 | 1,31 | 0,49 | 1,68 | 0,29 | 7,73E-01 |
| AT2G33450 | 0,32 | 0,29 | 4,31 | 0,07 | 9,47E-01 |
| AT2G33490 | 0,35 | 2,51 | 4,27 | 0,59 | 5,57E-01 |
| AT2G33510 | 0,26 | 1,37 | 4,31 | 0,32 | 7,51E-01 |
| AT2G33570 | 0,17 | -0,71 | 4,32 | -0,16 | 8,69E-01 |
| AT2G33610 | 0,37 | 1,93 | 4,29 | 0,45 | 6,53E-01 |
| AT2G33650 | 0,29 | 0,00 | 4,41 | 0,00 | 1,00E+00 |
| AT2G33700 | 0,56 | -1,40 | 3,94 | -0,36 | 7,22E-01 |
| AT2G33760 | 0,64 | 2,87 | 4,35 | 0,66 | 5,09E-01 |
| AT2G33810 | 0,37 | -0,84 | 4,36 | -0,19 | 8,48E-01 |
| AT2G33840 | 0,52 | 0,42 | 3,18 | 0,13 | 8,95E-01 |
| AT2G33845 | 0,47 | 1,90 | 4,00 | 0,48 | 6,34E-01 |
| AT2G33950 | 0,39 | 0,63 | 4,29 | 0,15 | 8,83E-01 |
| AT2G34120 | 0,27 | -1,27 | 4,27 | -0,30 | 7,67E-01 |
| AT2G34240 | 0,12 | 2,69 | 4,35 | 0,62 | 5,36E-01 |
| AT2G34460 | 0,99 | -0,61 | 1,63 | -0,37 | 7,09E-01 |
| AT2G34660 | 0,69 | 1,03 | 2,48 | 0,42 | 6,77E-01 |
| AT2G34730 | 0,50 | 1,52 | 2,59 | 0,59 | 5,58E-01 |
| AT2G34770 | 0,22 | 0,78 | 4,33 | 0,18 | 8,58E-01 |
| AT2G34780 | 0,99 | 1,50 | 1,94 | 0,77 | 4,41E-01 |
| AT2G34810 | 0,16 | 1,18 | 4,32 | 0,27 | 7,85E-01 |
| AT2G34860 | 1,10 | 1,92 | 2,18 | 0,88 | 3,77E-01 |
| AT2G34970 | 0,22 | 0,42 | 4,34 | 0,10 | 9,24E-01 |
| AT2G35020 | 0,32 | 0,34 | 4,34 | 0,08 | 9,37E-01 |
| AT2G35170 | 0,10 | 0,39 | 4,35 | 0,09 | 9,29E-01 |
| AT2G35230 | 0,50 | 0,46 | 4,31 | 0,11 | 9,16E-01 |
| AT2G35330 | 0,35 | -1,38 | 4,34 | -0,32 | 7,51E-01 |
| AT2G35340 | 0,80 | 3,27 | 2,79 | 1,17 | 2,42E-01 |
| AT2G35370 | 0,43 | 0,86 | 4,26 | 0,20 | 8,39E-01 |
| AT2G35630 | 1,16 | 0,89 | 1,76 | 0,50 | 6,14E-01 |
| AT2G35635 | 0,64 | 0,00 | 4,41 | 0,00 | 1,00E+00 |
| AT2G35650 | 0,68 | 2,23 | 2,22 | 1,00 | 3,15E-01 |
| AT2G08750 | 0,08 | 2,08 | 4,38 | 0,47 | 6,35E-01 |
| AT2G35720 | 0,19 | 0,60 | 4,30 | 0,14 | 8,89E-01 |
| AT2G35760 | 0,17 | -0,44 | 4,37 | -0,10 | 9,19E-01 |
| AT2G35800 | 1,02 | 1,09 | 1,70 | 0,64 | 5,19E-01 |
| AT2G35820 | 0,71 | 3,95 | 4,31 | 0,92 | 3,59E-01 |
| AT2G35830 | 1,03 | 1,83 | 1,76 | 1,04 | 2,97E-01 |
| AT2G35840 | 0,63 | 0,40 | 2,36 | 0,17 | 8,65E-01 |
| AT2G35860 | 0,74 | -0,42 | 2,19 | -0,19 | 8,46E-01 |
| AT2G35920 | 0,77 | 2,77 | 2,35 | 1,18 | 2,39E-01 |
| AT2G35935 | 0,49 | -2,53 | 3,81 | -0,66 | 5,07E-01 |
| AT2G35960 | 0,20 | -0,87 | 4,29 | -0,20 | 8,40E-01 |
| AT2G36000 | 0,80 | 1,64 | 2,40 | 0,68 | 4,95E-01 |
| AT2G36010 | 0,59 | 0,74 | 3,13 | 0,24 | 8,14E-01 |
| AT2G36026 | 0,70 | 0,38 | 3,72 | 0,10 | 9,19E-01 |
| AT2G36090 | 0,69 | -2,56 | 3,43 | -0,75 | 4,55E-01 |
| AT2G36130 | 0,56 | 2,82 | 4,35 | 0,65 | 5,16E-01 |

| | | | | | |
|---|---|---|---|---|---|
| AT2G36320 | 1,39 | 0,62 | 1,54 | 0,40 | 6,87E-01 |
| AT2G36330 | 0,55 | 3,10 | 4,34 | 0,71 | 4,75E-01 |
| AT2G36430 | 1,44 | -1,09 | 1,69 | -0,65 | 5,18E-01 |
| AT2G36500 | 0,54 | 2,88 | 3,10 | 0,93 | 3,53E-01 |
| AT2G36510 | 0,37 | 2,86 | 4,28 | 0,67 | 5,04E-01 |
| AT2G36570 | 0,52 | -0,11 | 4,40 | -0,03 | 9,80E-01 |
| AT2G36580 | 1,14 | 0,77 | 1,43 | 0,54 | 5,90E-01 |
| AT2G36600 | 0,66 | 1,61 | 2,28 | 0,71 | 4,80E-01 |
| AT2G36720 | 1,25 | 3,23 | 2,07 | 1,56 | 1,18E-01 |
| AT2G36792 | 0,34 | -0,46 | 4,23 | -0,11 | 9,13E-01 |
| AT2G36830 | 0,86 | 1,13 | 2,12 | 0,54 | 5,92E-01 |
| AT2G36920 | 0,12 | 2,87 | 4,34 | 0,66 | 5,08E-01 |
| AT2G36950 | 0,34 | 3,18 | 4,34 | 0,73 | 4,63E-01 |
| AT2G36960 | 0,18 | 0,41 | 4,28 | 0,10 | 9,23E-01 |
| AT2G36970 | 0,19 | 1,14 | 4,29 | 0,26 | 7,91E-01 |
| AT2G36980 | 0,05 | -0,58 | 4,38 | -0,13 | 8,95E-01 |
| AT2G37025 | 0,19 | -0,02 | 4,39 | 0,00 | 9,97E-01 |
| AT2G37050 | 0,93 | 2,88 | 2,30 | 1,25 | 2,11E-01 |
| AT2G37110 | 1,16 | 0,11 | 2,06 | 0,05 | 9,58E-01 |
| AT2G37180 | 0,08 | 2,08 | 4,38 | 0,47 | 6,35E-01 |
| AT2G37250 | 0,74 | 1,43 | 2,10 | 0,68 | 4,97E-01 |
| AT2G08860 | 0,31 | 1,03 | 3,57 | 0,29 | 7,73E-01 |
| AT2G37320 | 0,08 | 2,33 | 4,36 | 0,53 | 5,93E-01 |
| AT2G37330 | 0,66 | 2,74 | 3,05 | 0,90 | 3,69E-01 |
| AT2G37362 | 0,21 | 3,40 | 4,32 | 0,79 | 4,32E-01 |
| AT2G37370 | 0,40 | -2,01 | 4,26 | -0,47 | 6,37E-01 |
| AT2G37470 | 0,26 | 2,02 | 4,29 | 0,47 | 6,37E-01 |
| AT2G37480 | 1,35 | 2,03 | 1,97 | 1,03 | 3,01E-01 |
| AT2G37555 | 0,61 | 2,69 | 2,97 | 0,91 | 3,65E-01 |
| AT2G37580 | 0,12 | 0,63 | 4,33 | 0,15 | 8,85E-01 |
| AT2G37650 | 1,11 | 1,07 | 1,88 | 0,57 | 5,70E-01 |
| AT2G37690 | 0,31 | 0,00 | 4,34 | 0,00 | 1,00E+00 |
| AT2G37770 | 0,28 | 2,67 | 4,31 | 0,62 | 5,35E-01 |
| AT2G37860 | 0,83 | -0,70 | 2,09 | -0,33 | 7,39E-01 |
| AT2G37940 | 0,73 | -2,17 | 2,97 | -0,73 | 4,66E-01 |
| AT2G37970 | 0,77 | 0,60 | 2,55 | 0,23 | 8,15E-01 |
| AT2G37990 | 0,11 | 1,28 | 4,35 | 0,29 | 7,68E-01 |
| AT2G38010 | 0,53 | 1,51 | 2,96 | 0,51 | 6,10E-01 |
| AT2G38090 | 0,20 | 2,11 | 4,37 | 0,48 | 6,29E-01 |
| AT2G38120 | 0,38 | 0,78 | 2,69 | 0,29 | 7,72E-01 |
| AT2G38140 | 0,26 | 0,46 | 3,69 | 0,13 | 9,00E-01 |
| AT2G38210 | 1,49 | -1,26 | 1,88 | -0,67 | 5,04E-01 |
| AT2G38310 | 0,71 | 0,84 | 2,95 | 0,28 | 7,77E-01 |
| AT2G38365 | 0,20 | 2,30 | 4,36 | 0,53 | 5,98E-01 |
| AT2G38420 | 0,26 | 2,13 | 4,38 | 0,49 | 6,27E-01 |
| AT2G38470 | 1,30 | 0,34 | 1,59 | 0,21 | 8,31E-01 |
| AT2G38560 | 1,56 | 2,81 | 1,79 | 1,57 | 1,16E-01 |
| AT2G38590 | 0,78 | -1,35 | 2,51 | -0,54 | 5,90E-01 |
| AT2G38670 | 0,86 | 2,02 | 2,17 | 0,93 | 3,52E-01 |
| AT2G38760 | 0,08 | 2,33 | 4,36 | 0,53 | 5,93E-01 |
| AT2G38780 | 0,82 | -0,24 | 1,69 | -0,14 | 8,86E-01 |
| AT2G38820 | 0,12 | 1,65 | 4,34 | 0,38 | 7,04E-01 |
| AT2G38840 | 0,33 | 1,76 | 4,33 | 0,41 | 6,85E-01 |
| AT2G38940 | 0,25 | 2,79 | 4,34 | 0,64 | 5,20E-01 |
| AT2G38950 | 0,65 | 1,88 | 2,63 | 0,72 | 4,75E-01 |
| AT2G39190 | 0,15 | 0,69 | 4,31 | 0,16 | 8,73E-01 |

| | | | | | |
|---|---|---|---|---|---|
| AT2G39230 | 0,19 | -1,45 | 4,34 | -0,33 | 7,38E-01 |
| AT2G39270 | 0,08 | 2,08 | 4,38 | 0,47 | 6,35E-01 |
| AT2G39340 | 0,19 | -1,45 | 4,34 | -0,33 | 7,38E-01 |
| AT2G39400 | 0,15 | 1,64 | 4,33 | 0,38 | 7,06E-01 |
| AT2G39420 | 0,56 | 0,24 | 3,18 | 0,08 | 9,40E-01 |
| AT2G39430 | 0,52 | 3,80 | 3,86 | 0,98 | 3,25E-01 |
| AT2G39460 | 0,89 | -0,26 | 1,75 | -0,15 | 8,81E-01 |
| AT2G39470 | 0,81 | 2,04 | 2,22 | 0,92 | 3,58E-01 |
| AT2G39570 | 1,60 | 0,63 | 2,78 | 0,23 | 8,21E-01 |
| AT2G39580 | 1,16 | 3,73 | 2,14 | 1,75 | 8,06E-02 |
| AT2G39670 | 1,03 | 3,34 | 2,09 | 1,60 | 1,10E-01 |
| AT2G39725 | 0,50 | 3,40 | 3,91 | 0,87 | 3,85E-01 |
| AT2G39740 | 0,63 | 0,74 | 2,99 | 0,25 | 8,04E-01 |
| AT2G39780 | 1,17 | 1,69 | 2,14 | 0,79 | 4,31E-01 |
| AT2G39810 | 1,48 | 2,07 | 2,32 | 0,89 | 3,71E-01 |
| AT2G39840 | 0,84 | 1,32 | 2,32 | 0,57 | 5,71E-01 |
| AT2G39850 | 0,54 | -1,54 | 4,31 | -0,36 | 7,21E-01 |
| AT2G39870 | 0,43 | -0,53 | 4,27 | -0,12 | 9,01E-01 |
| AT2G39930 | 1,27 | 0,93 | 1,69 | 0,55 | 5,82E-01 |
| AT2G40060 | 0,99 | 1,29 | 2,19 | 0,59 | 5,56E-01 |
| AT2G40095 | 0,64 | 2,77 | 3,07 | 0,90 | 3,67E-01 |
| AT2G40130 | 0,14 | 1,12 | 4,32 | 0,26 | 7,96E-01 |
| AT2G40150 | 0,08 | 2,08 | 4,38 | 0,47 | 6,35E-01 |
| AT2G40160 | 0,42 | 2,83 | 4,13 | 0,69 | 4,93E-01 |
| AT2G40200 | 0,38 | -0,39 | 4,28 | -0,09 | 9,28E-01 |
| AT2G09120 | 0,31 | 0,00 | 4,41 | 0,00 | 1,00E+00 |
| AT2G40310 | 0,83 | -0,36 | 1,98 | -0,18 | 8,56E-01 |
| AT2G40400 | 0,82 | 2,73 | 2,75 | 1,00 | 3,19E-01 |
| AT2G40410 | 0,54 | 0,09 | 4,05 | 0,02 | 9,83E-01 |
| AT2G40435 | 0,46 | 1,79 | 4,41 | 0,41 | 6,85E-01 |
| AT2G40460 | 0,63 | -0,10 | 2,43 | -0,04 | 9,66E-01 |
| AT2G40480 | 0,31 | 0,84 | 4,33 | 0,19 | 8,46E-01 |
| AT2G40490 | 1,23 | -0,60 | 1,90 | -0,31 | 7,54E-01 |
| AT2G40540 | 0,78 | 0,55 | 2,36 | 0,23 | 8,15E-01 |
| AT2G40550 | 0,20 | 2,30 | 4,36 | 0,53 | 5,98E-01 |
| AT2G40660 | 0,08 | -0,56 | 4,38 | -0,13 | 8,99E-01 |
| AT2G40670 | 0,10 | 0,39 | 4,35 | 0,09 | 9,29E-01 |
| AT2G40760 | 0,35 | 3,20 | 4,33 | 0,74 | 4,60E-01 |
| AT2G40765 | 0,08 | 2,08 | 4,38 | 0,47 | 6,35E-01 |
| AT2G40810 | 0,08 | 2,33 | 4,36 | 0,53 | 5,93E-01 |
| AT2G40830 | 0,33 | 3,37 | 4,32 | 0,78 | 4,36E-01 |
| AT2G40880 | 1,07 | 2,16 | 2,10 | 1,03 | 3,03E-01 |
| AT2G40955 | 1,38 | -1,28 | 1,57 | -0,81 | 4,16E-01 |
| AT2G40980 | 0,41 | 2,51 | 4,18 | 0,60 | 5,49E-01 |
| AT2G41080 | 0,59 | 0,04 | 2,48 | 0,02 | 9,87E-01 |
| AT2G41110 | 0,26 | 0,36 | 3,73 | 0,10 | 9,24E-01 |
| AT2G09230 | 0,12 | 2,87 | 4,34 | 0,66 | 5,08E-01 |
| AT2G41140 | 0,37 | 2,25 | 4,27 | 0,53 | 5,98E-01 |
| AT2G41200 | 0,84 | -1,00 | 2,17 | -0,46 | 6,45E-01 |
| AT2G41210 | 0,42 | 2,29 | 4,13 | 0,55 | 5,79E-01 |
| AT2G41220 | 0,64 | 0,65 | 2,80 | 0,23 | 8,17E-01 |
| AT2G41460 | 0,13 | 2,37 | 4,36 | 0,54 | 5,86E-01 |
| AT2G41510 | 0,08 | 2,08 | 4,38 | 0,47 | 6,35E-01 |
| AT2G41520 | 0,59 | 1,98 | 2,98 | 0,67 | 5,06E-01 |
| AT2G41590 | 1,66 | 0,46 | 1,42 | 0,32 | 7,46E-01 |
| AT2G41670 | 0,12 | 2,87 | 4,34 | 0,66 | 5,08E-01 |

| | | | | | |
|---|---|---|---|---|---|
| AT2G41730 | 0,99 | 2,28 | 1,95 | 1,17 | 2,43E-01 |
| AT2G41820 | 0,34 | 3,08 | 4,34 | 0,71 | 4,78E-01 |
| AT2G41880 | 0,26 | 0,36 | 3,73 | 0,10 | 9,24E-01 |
| AT2G41920 | 1,42 | -2,21 | 1,66 | -1,33 | 1,84E-01 |
| AT2G42130 | 0,48 | -0,42 | 3,29 | -0,13 | 8,99E-01 |
| AT2G42220 | 0,98 | 1,08 | 2,01 | 0,54 | 5,89E-01 |
| AT2G42310 | 0,61 | 1,27 | 2,40 | 0,53 | 5,97E-01 |
| AT2G42320 | 0,63 | 2,15 | 2,55 | 0,84 | 3,99E-01 |
| AT2G42430 | 0,21 | -2,30 | 4,30 | -0,54 | 5,92E-01 |
| AT2G42580 | 1,32 | 2,91 | 1,79 | 1,62 | 1,04E-01 |
| AT2G42610 | 0,29 | 1,50 | 4,34 | 0,35 | 7,29E-01 |
| AT2G42620 | 1,45 | 2,44 | 2,27 | 1,08 | 2,81E-01 |
| AT2G42700 | 0,72 | 2,88 | 2,99 | 0,96 | 3,35E-01 |
| AT2G42710 | 0,12 | 1,15 | 4,35 | 0,26 | 7,92E-01 |
| AT2G42750 | 0,57 | 3,93 | 3,74 | 1,05 | 2,93E-01 |
| AT2G42780 | 0,28 | -1,30 | 4,35 | -0,30 | 7,65E-01 |
| AT2G42890 | 0,32 | 2,51 | 4,30 | 0,58 | 5,60E-01 |
| AT2G42900 | 0,43 | -0,74 | 4,38 | -0,17 | 8,66E-01 |
| AT2G43010 | 1,28 | 0,60 | 1,82 | 0,33 | 7,43E-01 |
| AT2G43020 | 1,32 | -0,70 | 1,74 | -0,40 | 6,87E-01 |
| AT2G43030 | 0,58 | -1,36 | 3,10 | -0,44 | 6,61E-01 |
| AT2G43060 | 1,34 | -1,43 | 1,54 | -0,93 | 3,53E-01 |
| AT2G43080 | 0,63 | 0,97 | 2,98 | 0,32 | 7,46E-01 |
| AT2G43090 | 0,61 | -0,19 | 2,57 | -0,08 | 9,40E-01 |
| AT2G43100 | 0,56 | 0,23 | 3,22 | 0,07 | 9,43E-01 |
| AT2G43137 | 1,43 | 0,32 | 1,72 | 0,18 | 8,53E-01 |
| AT2G43138 | 1,21 | -1,35 | 1,79 | -0,76 | 4,49E-01 |
| AT2G43141 | 1,64 | -0,60 | 1,77 | -0,34 | 7,33E-01 |
| AT2G43210 | 0,63 | 3,94 | 2,93 | 1,34 | 1,79E-01 |
| AT2G43230 | 0,55 | 2,52 | 3,84 | 0,66 | 5,12E-01 |
| AT2G43250 | 0,31 | 2,17 | 4,31 | 0,50 | 6,14E-01 |
| AT2G43280 | 0,56 | 1,88 | 4,36 | 0,43 | 6,66E-01 |
| AT2G09440 | 0,08 | 2,33 | 4,36 | 0,53 | 5,93E-01 |
| AT2G43340 | 0,44 | -0,53 | 3,34 | -0,16 | 8,74E-01 |
| AT2G43400 | 0,37 | 2,26 | 4,27 | 0,53 | 5,97E-01 |
| AT2G43440 | 0,11 | -1,08 | 4,37 | -0,25 | 8,05E-01 |
| AT2G43540 | 1,53 | -0,69 | 1,56 | -0,44 | 6,57E-01 |
| AT2G43630 | 0,44 | 2,87 | 4,05 | 0,71 | 4,78E-01 |
| AT2G43650 | 0,83 | 1,83 | 2,36 | 0,78 | 4,38E-01 |
| AT2G43690 | 0,68 | 4,19 | 4,31 | 0,97 | 3,30E-01 |
| AT2G43790 | 0,39 | 1,42 | 2,75 | 0,52 | 6,04E-01 |
| AT2G43800 | 0,08 | -0,56 | 4,38 | -0,13 | 8,99E-01 |
| AT2G43820 | 0,45 | 2,11 | 3,21 | 0,66 | 5,10E-01 |
| AT2G44080 | 0,60 | -0,61 | 3,80 | -0,16 | 8,73E-01 |
| AT2G44100 | 0,65 | 1,68 | 2,46 | 0,68 | 4,95E-01 |
| AT2G44130 | 0,72 | -0,61 | 2,34 | -0,26 | 7,94E-01 |
| AT2G44160 | 0,80 | -0,22 | 2,26 | -0,10 | 9,22E-01 |
| AT2G44175 | 0,43 | -2,17 | 4,03 | -0,54 | 5,91E-01 |
| AT2G44200 | 0,72 | 1,69 | 2,39 | 0,71 | 4,79E-01 |
| AT2G44210 | 1,13 | -1,76 | 1,76 | -1,00 | 3,18E-01 |
| AT2G44230 | 0,76 | -1,02 | 2,71 | -0,38 | 7,06E-01 |
| AT2G44240 | 0,17 | -0,40 | 4,34 | -0,09 | 9,27E-01 |
| AT2G44260 | 0,20 | 1,94 | 4,34 | 0,45 | 6,55E-01 |
| AT2G44350 | 0,40 | -1,33 | 3,31 | -0,40 | 6,88E-01 |
| AT2G44420 | 0,20 | 2,30 | 4,36 | 0,53 | 5,98E-01 |
| AT2G44430 | 0,96 | -0,93 | 1,69 | -0,55 | 5,82E-01 |

| | | | | | |
|---|---|---|---|---|---|
| AT2G44500 | 0,91 | 1,06 | 2,66 | 0,40 | 6,90E-01 |
| AT2G44520 | 0,23 | 1,68 | 4,33 | 0,39 | 6,98E-01 |
| AT2G44530 | 0,99 | 2,66 | 2,13 | 1,25 | 2,12E-01 |
| AT2G44600 | 0,10 | 0,39 | 4,35 | 0,09 | 9,29E-01 |
| AT2G44640 | 1,50 | 2,49 | 1,65 | 1,51 | 1,30E-01 |
| AT2G44670 | 1,15 | -0,05 | 1,82 | -0,03 | 9,78E-01 |
| AT2G44710 | 1,40 | 1,44 | 1,55 | 0,93 | 3,53E-01 |
| AT2G44735 | 0,42 | 2,48 | 4,09 | 0,61 | 5,44E-01 |
| AT2G44770 | 0,15 | 1,94 | 4,32 | 0,45 | 6,54E-01 |
| AT2G44800 | 0,19 | 0,10 | 4,37 | 0,02 | 9,82E-01 |
| AT2G44870 | 0,51 | -0,72 | 2,66 | -0,27 | 7,87E-01 |
| AT2G44880 | 0,91 | 4,03 | 2,71 | 1,49 | 1,36E-01 |
| AT2G44925 | 0,92 | -1,12 | 1,66 | -0,67 | 5,00E-01 |
| AT2G44940 | 0,52 | 0,78 | 2,73 | 0,29 | 7,75E-01 |
| AT2G45000 | 0,38 | -2,09 | 3,38 | -0,62 | 5,36E-01 |
| AT2G45020 | 0,36 | 1,00 | 4,34 | 0,23 | 8,17E-01 |
| AT2G45030 | 0,91 | 2,28 | 2,67 | 0,85 | 3,94E-01 |
| AT2G45140 | 0,33 | 2,30 | 4,38 | 0,52 | 6,00E-01 |
| AT2G45240 | 0,80 | 2,59 | 3,00 | 0,86 | 3,89E-01 |
| AT2G45250 | 0,26 | 2,13 | 4,38 | 0,49 | 6,27E-01 |
| AT2G45300 | 0,10 | 0,34 | 4,35 | 0,08 | 9,37E-01 |
| AT2G45310 | 0,67 | 0,35 | 2,33 | 0,15 | 8,82E-01 |
| AT2G45390 | 0,17 | 0,00 | 4,41 | 0,00 | 1,00E+00 |
| AT2G45440 | 0,53 | 0,94 | 3,87 | 0,24 | 8,08E-01 |
| AT2G45510 | 0,84 | 0,42 | 1,95 | 0,22 | 8,29E-01 |
| AT2G45530 | 0,98 | 0,88 | 1,88 | 0,47 | 6,38E-01 |
| AT2G09650 | 0,36 | -0,28 | 4,28 | -0,07 | 9,48E-01 |
| AT2G45590 | 0,37 | 0,39 | 4,29 | 0,09 | 9,27E-01 |
| AT2G45620 | 1,35 | 2,70 | 1,90 | 1,42 | 1,56E-01 |
| AT2G45630 | 0,80 | 1,70 | 2,42 | 0,70 | 4,82E-01 |
| AT2G45710 | 0,45 | -0,88 | 4,37 | -0,20 | 8,40E-01 |
| AT2G45720 | 0,29 | 0,32 | 4,33 | 0,07 | 9,41E-01 |
| AT2G45740 | 0,35 | 0,74 | 3,54 | 0,21 | 8,34E-01 |
| AT2G45790 | 0,17 | -0,14 | 4,41 | -0,03 | 9,75E-01 |
| AT2G45810 | 0,90 | 1,71 | 2,20 | 0,78 | 4,37E-01 |
| AT2G45910 | 0,97 | 0,61 | 1,80 | 0,34 | 7,33E-01 |
| AT2G46100 | 0,37 | -1,10 | 2,80 | -0,39 | 6,94E-01 |
| AT2G46130 | 1,45 | -0,51 | 1,56 | -0,33 | 7,43E-01 |
| AT2G46170 | 0,69 | 1,18 | 3,70 | 0,32 | 7,50E-01 |
| AT2G46190 | 0,70 | -1,40 | 2,01 | -0,70 | 4,87E-01 |
| AT2G09695 | 1,56 | 0,10 | 1,47 | 0,07 | 9,45E-01 |
| AT2G09705 | 0,89 | -0,91 | 2,40 | -0,38 | 7,04E-01 |
| AT2G46225 | 0,27 | 0,48 | 4,31 | 0,11 | 9,11E-01 |
| AT2G46240 | 0,70 | 3,01 | 2,50 | 1,21 | 2,28E-01 |
| AT2G46260 | 0,08 | 2,08 | 4,38 | 0,47 | 6,35E-01 |
| AT2G46270 | 0,25 | 2,25 | 4,37 | 0,52 | 6,06E-01 |
| AT2G46370 | 0,44 | -0,63 | 3,36 | -0,19 | 8,51E-01 |
| AT2G46380 | 1,50 | 4,19 | 2,20 | 1,91 | 5,67E-02 |
| AT2G46420 | 0,84 | -1,42 | 2,91 | -0,49 | 6,26E-01 |
| AT2G46440 | 1,43 | 0,10 | 1,33 | 0,07 | 9,42E-01 |
| AT2G46450 | 1,09 | 2,46 | 2,04 | 1,21 | 2,28E-01 |
| AT2G46490 | 0,75 | 1,99 | 2,70 | 0,74 | 4,61E-01 |
| AT2G46520 | 1,16 | 1,39 | 1,52 | 0,91 | 3,60E-01 |
| AT2G46570 | 0,51 | 2,87 | 3,97 | 0,72 | 4,70E-01 |
| AT2G46590 | 0,52 | -0,26 | 2,85 | -0,09 | 9,26E-01 |
| AT2G46600 | 1,11 | 1,21 | 1,73 | 0,70 | 4,84E-01 |

| | | | | | |
|---|---|---|---|---|---|
| AT2G46630 | 0,28 | -0,37 | 4,28 | -0,09 | 9,31E-01 |
| AT2G46650 | 0,42 | 0,36 | 3,21 | 0,11 | 9,12E-01 |
| AT2G46690 | 0,43 | 0,71 | 4,11 | 0,17 | 8,63E-01 |
| AT2G46710 | 0,30 | -1,00 | 3,61 | -0,28 | 7,81E-01 |
| AT2G46790 | 0,54 | 1,55 | 2,50 | 0,62 | 5,36E-01 |
| AT2G46800 | 1,02 | 0,22 | 1,74 | 0,13 | 8,97E-01 |
| AT2G46810 | 0,59 | -1,06 | 3,10 | -0,34 | 7,33E-01 |
| AT2G46900 | 1,50 | 3,68 | 2,09 | 1,76 | 7,89E-02 |
| AT2G46910 | 0,44 | -0,86 | 4,17 | -0,21 | 8,36E-01 |
| AT2G46990 | 0,08 | 2,08 | 4,38 | 0,47 | 6,35E-01 |
| AT2G47110 | 0,66 | -1,12 | 2,07 | -0,54 | 5,88E-01 |
| AT2G09815 | 0,23 | -1,06 | 4,29 | -0,25 | 8,04E-01 |
| AT2G47190 | 0,40 | 3,74 | 4,32 | 0,87 | 3,87E-01 |
| AT2G47220 | 0,25 | 3,04 | 4,33 | 0,70 | 4,83E-01 |
| AT2G47320 | 0,28 | 0,21 | 4,39 | 0,05 | 9,62E-01 |
| AT2G47420 | 0,42 | 3,23 | 4,12 | 0,78 | 4,33E-01 |
| AT2G47510 | 1,59 | 2,76 | 1,68 | 1,64 | 1,01E-01 |
| AT2G47610 | 1,29 | 3,02 | 1,71 | 1,76 | 7,80E-02 |
| AT2G47620 | 0,26 | 0,36 | 3,73 | 0,10 | 9,24E-01 |
| AT2G47630 | 0,42 | 0,16 | 2,75 | 0,06 | 9,55E-01 |
| AT2G47640 | 0,28 | -1,16 | 4,35 | -0,27 | 7,89E-01 |
| AT2G47690 | 0,55 | 1,21 | 4,01 | 0,30 | 7,63E-01 |
| AT2G47730 | 0,80 | -1,97 | 2,16 | -0,92 | 3,60E-01 |
| AT2G47820 | 1,37 | 2,57 | 2,38 | 1,08 | 2,81E-01 |
| AT2G47850 | 0,53 | -0,41 | 3,08 | -0,13 | 8,95E-01 |
| AT2G47910 | 0,92 | -0,24 | 1,79 | -0,14 | 8,92E-01 |
| AT2G47935 | 0,72 | 2,17 | 2,75 | 0,79 | 4,30E-01 |
| AT2G47960 | 1,39 | 0,79 | 1,65 | 0,48 | 6,31E-01 |
| AT2G47970 | 0,70 | 3,88 | 3,46 | 1,12 | 2,63E-01 |
| AT2G48010 | 0,33 | 0,86 | 4,33 | 0,20 | 8,43E-01 |
| AT2G48020 | 0,56 | 0,97 | 3,15 | 0,31 | 7,58E-01 |
| AT2G48110 | 1,17 | 1,30 | 2,06 | 0,63 | 5,28E-01 |
| AT2G48120 | 0,32 | 2,63 | 4,35 | 0,60 | 5,46E-01 |
| AT3G01040 | 0,51 | 2,82 | 3,09 | 0,91 | 3,61E-01 |
| AT3G01050 | 0,18 | 0,77 | 4,33 | 0,18 | 8,59E-01 |
| AT3G01080 | 0,17 | 1,41 | 4,35 | 0,32 | 7,46E-01 |
| AT3G01085 | 1,09 | 0,22 | 1,63 | 0,14 | 8,93E-01 |
| AT3G01150 | 0,45 | -0,45 | 3,25 | -0,14 | 8,89E-01 |
| AT3G01170 | 0,42 | 2,29 | 4,13 | 0,55 | 5,79E-01 |
| AT3G01175 | 0,08 | 2,33 | 4,36 | 0,53 | 5,93E-01 |
| AT3G01210 | 0,10 | 0,34 | 4,35 | 0,08 | 9,37E-01 |
| AT3G01280 | 1,02 | -1,11 | 1,81 | -0,61 | 5,39E-01 |
| AT3G01320 | 1,14 | 1,46 | 1,62 | 0,90 | 3,67E-01 |
| AT3G01440 | 0,31 | -1,59 | 4,33 | -0,37 | 7,13E-01 |
| AT3G01475 | 1,05 | 2,08 | 1,84 | 1,13 | 2,58E-01 |
| AT3G01480 | 1,15 | -0,39 | 1,91 | -0,20 | 8,40E-01 |
| AT3G01560 | 0,68 | 1,54 | 2,44 | 0,63 | 5,26E-01 |
| AT3G01590 | 0,40 | 2,19 | 4,34 | 0,51 | 6,13E-01 |
| AT3G01610 | 0,25 | 1,14 | 4,33 | 0,26 | 7,92E-01 |
| AT3G01660 | 0,12 | 1,65 | 4,34 | 0,38 | 7,04E-01 |
| AT3G01690 | 1,08 | -1,64 | 1,85 | -0,88 | 3,77E-01 |
| AT3G01740 | 0,28 | 2,36 | 4,31 | 0,55 | 5,83E-01 |
| AT3G01750 | 0,72 | 3,22 | 3,00 | 1,07 | 2,83E-01 |
| AT3G01780 | 1,57 | 1,74 | 1,43 | 1,21 | 2,25E-01 |
| AT3G01810 | 0,37 | 1,90 | 4,39 | 0,43 | 6,65E-01 |
| AT3G01830 | 0,94 | 0,56 | 2,09 | 0,27 | 7,89E-01 |

| | | | | | |
|---|---|---|---|---|---|
| AT3G01920 | 0,37 | 0,38 | 4,38 | 0,09 | 9,31E-01 |
| AT3G02010 | 0,37 | 2,76 | 4,34 | 0,64 | 5,24E-01 |
| AT3G02030 | 0,12 | 0,63 | 4,33 | 0,15 | 8,85E-01 |
| AT3G02050 | 0,37 | 2,86 | 4,28 | 0,67 | 5,04E-01 |
| AT3G02060 | 1,23 | 0,99 | 1,89 | 0,52 | 6,00E-01 |
| AT3G02065 | 0,54 | 3,56 | 3,77 | 0,94 | 3,45E-01 |
| AT3G02070 | 0,35 | -0,23 | 3,46 | -0,07 | 9,47E-01 |
| AT3G02130 | 1,18 | 2,86 | 2,08 | 1,38 | 1,68E-01 |
| AT3G02280 | 0,25 | -0,67 | 4,39 | -0,15 | 8,78E-01 |
| AT3G02290 | 0,16 | 1,16 | 4,38 | 0,27 | 7,90E-01 |
| AT3G02300 | 0,16 | -1,42 | 4,34 | -0,33 | 7,43E-01 |
| AT3G02340 | 0,50 | 0,65 | 4,28 | 0,15 | 8,80E-01 |
| AT3G02350 | 0,84 | -0,95 | 2,05 | -0,46 | 6,45E-01 |
| AT3G02360 | 1,53 | 2,12 | 1,79 | 1,19 | 2,35E-01 |
| AT3G02370 | 0,47 | 1,48 | 3,31 | 0,45 | 6,54E-01 |
| AT3G02400 | 0,66 | 0,00 | 4,41 | 0,00 | 1,00E+00 |
| AT3G02490 | 0,33 | 1,66 | 4,35 | 0,38 | 7,02E-01 |
| AT3G02560 | 0,49 | 2,81 | 3,89 | 0,72 | 4,69E-01 |
| AT3G02610 | 0,81 | 0,16 | 2,81 | 0,06 | 9,56E-01 |
| AT3G02630 | 0,37 | 1,15 | 4,30 | 0,27 | 7,89E-01 |
| AT3G02690 | 0,66 | 1,12 | 2,52 | 0,45 | 6,56E-01 |
| AT3G02720 | 0,46 | 1,06 | 4,32 | 0,24 | 8,07E-01 |
| AT3G02830 | 1,22 | 1,32 | 1,99 | 0,66 | 5,08E-01 |
| AT3G02832 | 1,51 | -0,91 | 1,41 | -0,65 | 5,18E-01 |
| AT3G01145 | 0,57 | -0,90 | 2,37 | -0,38 | 7,06E-01 |
| AT3G02880 | 1,19 | 1,04 | 1,65 | 0,63 | 5,30E-01 |
| AT3G02900 | 0,28 | -2,55 | 4,29 | -0,60 | 5,52E-01 |
| AT3G02910 | 0,77 | 0,38 | 2,21 | 0,17 | 8,62E-01 |
| AT3G03020 | 0,43 | 1,63 | 4,11 | 0,40 | 6,92E-01 |
| AT3G03030 | 0,40 | 3,74 | 4,32 | 0,87 | 3,87E-01 |
| AT3G03040 | 0,24 | 0,06 | 4,31 | 0,01 | 9,89E-01 |
| AT3G03110 | 0,55 | 3,53 | 4,32 | 0,82 | 4,15E-01 |
| AT3G03160 | 0,47 | 2,09 | 3,93 | 0,53 | 5,95E-01 |
| AT3G03210 | 0,10 | -1,26 | 4,35 | -0,29 | 7,72E-01 |
| AT3G03320 | 0,15 | -0,64 | 4,39 | -0,15 | 8,84E-01 |
| AT3G03340 | 0,40 | 2,81 | 4,32 | 0,65 | 5,15E-01 |
| AT3G03350 | 0,24 | 2,80 | 4,34 | 0,65 | 5,18E-01 |
| AT3G03360 | 0,08 | 2,33 | 4,36 | 0,53 | 5,93E-01 |
| AT3G03610 | 0,27 | 1,73 | 4,40 | 0,39 | 6,94E-01 |
| AT3G03640 | 0,25 | -0,34 | 4,34 | -0,08 | 9,37E-01 |
| AT3G03680 | 1,60 | 0,26 | 1,51 | 0,17 | 8,64E-01 |
| AT3G03860 | 0,25 | -1,47 | 4,28 | -0,34 | 7,32E-01 |
| AT3G03880 | 0,28 | -0,01 | 4,28 | 0,00 | 9,97E-01 |
| AT3G03950 | 0,32 | 1,84 | 3,56 | 0,52 | 6,05E-01 |
| AT3G03960 | 1,45 | 2,19 | 1,47 | 1,49 | 1,37E-01 |
| AT3G04140 | 0,99 | 4,03 | 2,28 | 1,77 | 7,65E-02 |
| AT3G04490 | 0,26 | -1,00 | 4,37 | -0,23 | 8,18E-01 |
| AT3G04530 | 0,08 | 2,33 | 4,36 | 0,53 | 5,93E-01 |
| AT3G04550 | 0,64 | 0,73 | 2,10 | 0,35 | 7,28E-01 |
| AT3G04790 | 1,26 | 1,61 | 2,07 | 0,78 | 4,37E-01 |
| AT3G04810 | 0,59 | 3,63 | 4,32 | 0,84 | 4,00E-01 |
| AT3G04840 | 0,80 | 1,05 | 2,23 | 0,47 | 6,38E-01 |
| AT3G04870 | 0,49 | -1,78 | 3,14 | -0,57 | 5,71E-01 |
| AT3G04910 | 0,86 | -0,50 | 1,62 | -0,31 | 7,56E-01 |
| AT3G04920 | 0,51 | 0,72 | 4,08 | 0,18 | 8,60E-01 |
| AT3G04940 | 0,44 | 2,92 | 4,16 | 0,70 | 4,82E-01 |

| | | | | | |
|---|---|---|---|---|---|
| AT3G01555 | 0,12 | 1,15 | 4,35 | 0,26 | 7,92E-01 |
| AT3G01575 | 0,10 | -1,26 | 4,35 | -0,29 | 7,72E-01 |
| AT3G05070 | 0,17 | 0,31 | 4,36 | 0,07 | 9,43E-01 |
| AT3G05120 | 0,72 | 0,63 | 2,04 | 0,31 | 7,58E-01 |
| AT3G05200 | 0,41 | 0,22 | 4,19 | 0,05 | 9,59E-01 |
| AT3G05230 | 0,28 | 1,69 | 4,34 | 0,39 | 6,97E-01 |
| AT3G05380 | 0,92 | 2,99 | 2,22 | 1,34 | 1,79E-01 |
| AT3G05410 | 1,68 | -0,47 | 1,27 | -0,37 | 7,11E-01 |
| AT3G05430 | 0,21 | 3,22 | 4,33 | 0,74 | 4,58E-01 |
| AT3G05490 | 0,28 | -1,30 | 4,35 | -0,30 | 7,65E-01 |
| AT3G05500 | 0,55 | 0,80 | 2,57 | 0,31 | 7,55E-01 |
| AT3G05520 | 0,43 | -0,02 | 2,75 | -0,01 | 9,95E-01 |
| AT3G05545 | 0,82 | 0,45 | 2,06 | 0,22 | 8,29E-01 |
| AT3G05560 | 0,45 | 0,33 | 3,25 | 0,10 | 9,19E-01 |
| AT3G05630 | 0,21 | 3,40 | 4,32 | 0,79 | 4,32E-01 |
| AT3G05670 | 0,41 | 1,77 | 4,15 | 0,43 | 6,70E-01 |
| AT3G05710 | 0,95 | 0,89 | 2,01 | 0,44 | 6,57E-01 |
| AT3G05725 | 0,64 | -2,30 | 2,35 | -0,98 | 3,28E-01 |
| AT3G05750 | 0,61 | 3,35 | 3,63 | 0,92 | 3,56E-01 |
| AT3G05760 | 0,68 | 3,27 | 3,63 | 0,90 | 3,67E-01 |
| AT3G05830 | 0,23 | -0,93 | 4,38 | -0,21 | 8,32E-01 |
| AT3G05840 | 0,50 | 2,33 | 3,99 | 0,58 | 5,59E-01 |
| AT3G05905 | 0,51 | -0,21 | 3,23 | -0,07 | 9,48E-01 |
| AT3G06125 | 0,60 | 0,76 | 2,13 | 0,36 | 7,22E-01 |
| AT3G06170 | 0,51 | 0,51 | 2,93 | 0,17 | 8,62E-01 |
| AT3G06180 | 0,25 | 3,58 | 4,32 | 0,83 | 4,07E-01 |
| AT3G06200 | 0,41 | 0,04 | 4,27 | 0,01 | 9,93E-01 |
| AT3G06210 | 0,20 | 2,30 | 4,36 | 0,53 | 5,98E-01 |
| AT3G06240 | 0,61 | 1,83 | 2,29 | 0,80 | 4,25E-01 |
| AT3G06380 | 0,55 | 3,19 | 3,86 | 0,83 | 4,09E-01 |
| AT3G06400 | 0,95 | 0,62 | 2,38 | 0,26 | 7,93E-01 |
| AT3G06455 | 0,58 | 0,87 | 4,36 | 0,20 | 8,41E-01 |
| AT3G06470 | 0,36 | 2,05 | 4,31 | 0,48 | 6,34E-01 |
| AT3G06500 | 1,24 | 0,02 | 1,88 | 0,01 | 9,92E-01 |
| AT3G06520 | 0,50 | 3,27 | 3,90 | 0,84 | 4,02E-01 |
| AT3G06610 | 0,15 | -0,19 | 4,31 | -0,04 | 9,64E-01 |
| AT3G06660 | 0,08 | -0,56 | 4,38 | -0,13 | 8,99E-01 |
| AT3G06730 | 0,97 | 0,12 | 3,06 | 0,04 | 9,70E-01 |
| AT3G06740 | 0,50 | -0,64 | 4,06 | -0,16 | 8,76E-01 |
| AT3G06810 | 1,67 | 3,57 | 1,70 | 2,09 | 3,62E-02 |
| AT3G06830 | 0,08 | 2,33 | 4,36 | 0,53 | 5,93E-01 |
| AT3G06860 | 1,03 | 0,54 | 1,84 | 0,29 | 7,70E-01 |
| AT3G06868 | 0,29 | -0,80 | 4,37 | -0,18 | 8,55E-01 |
| AT3G06895 | 1,19 | -0,70 | 1,69 | -0,41 | 6,79E-01 |
| AT3G06920 | 0,42 | 1,49 | 4,26 | 0,35 | 7,26E-01 |
| AT3G06980 | 0,81 | 3,83 | 2,95 | 1,30 | 1,95E-01 |
| AT3G07110 | 1,39 | 2,03 | 1,68 | 1,21 | 2,27E-01 |
| AT3G07200 | 0,08 | 2,33 | 4,36 | 0,53 | 5,93E-01 |
| AT3G07230 | 0,29 | 0,93 | 4,39 | 0,21 | 8,32E-01 |
| AT3G07270 | 0,85 | 2,26 | 2,14 | 1,06 | 2,90E-01 |
| AT3G07350 | 0,42 | 1,41 | 3,27 | 0,43 | 6,65E-01 |
| AT3G07360 | 1,00 | 0,07 | 2,22 | 0,03 | 9,75E-01 |
| AT3G07400 | 0,87 | 1,67 | 2,09 | 0,80 | 4,23E-01 |
| AT3G07430 | 0,32 | -0,26 | 4,30 | -0,06 | 9,51E-01 |
| AT3G07440 | 0,08 | 2,33 | 4,36 | 0,53 | 5,93E-01 |
| AT3G07480 | 0,45 | 1,04 | 4,21 | 0,25 | 8,04E-01 |

| | | | | | |
|---|---|---|---|---|---|
| AT3G07565 | 0,19 | -0,01 | 4,32 | 0,00 | 9,98E-01 |
| AT3G07610 | 0,83 | -0,51 | 2,48 | -0,20 | 8,38E-01 |
| AT3G07640 | 0,17 | 3,16 | 4,33 | 0,73 | 4,66E-01 |
| AT3G07650 | 0,97 | 3,77 | 3,32 | 1,13 | 2,57E-01 |
| AT3G07680 | 0,59 | 0,63 | 3,15 | 0,20 | 8,42E-01 |
| AT3G07690 | 0,32 | 2,42 | 4,35 | 0,56 | 5,77E-01 |
| AT3G07770 | 1,44 | 0,82 | 1,57 | 0,52 | 6,01E-01 |
| AT3G07780 | 1,15 | 1,69 | 1,98 | 0,85 | 3,93E-01 |
| AT3G07810 | 0,90 | -0,48 | 1,83 | -0,26 | 7,95E-01 |
| AT3G07880 | 0,28 | 1,72 | 4,27 | 0,40 | 6,87E-01 |
| AT3G07950 | 0,43 | 3,68 | 4,32 | 0,85 | 3,94E-01 |
| AT3G08010 | 0,17 | -1,47 | 4,34 | -0,34 | 7,34E-01 |
| AT3G08030 | 1,37 | -1,23 | 1,53 | -0,81 | 4,20E-01 |
| AT3G08550 | 0,31 | -0,43 | 4,28 | -0,10 | 9,21E-01 |
| AT3G08560 | 1,47 | 3,75 | 1,94 | 1,93 | 5,32E-02 |
| AT3G08600 | 0,52 | 0,87 | 3,81 | 0,23 | 8,19E-01 |
| AT3G08640 | 0,20 | 2,39 | 4,37 | 0,55 | 5,84E-01 |
| AT3G08690 | 0,22 | -0,11 | 4,34 | -0,02 | 9,80E-01 |
| AT3G08920 | 0,32 | -0,58 | 4,31 | -0,13 | 8,93E-01 |
| AT3G08930 | 0,51 | 1,24 | 3,13 | 0,40 | 6,92E-01 |
| AT3G08950 | 0,44 | -1,78 | 4,18 | -0,43 | 6,70E-01 |
| AT3G08980 | 0,33 | 0,84 | 4,34 | 0,19 | 8,46E-01 |
| AT3G09020 | 0,57 | 2,67 | 3,94 | 0,68 | 4,99E-01 |
| AT3G09030 | 0,51 | 0,45 | 2,50 | 0,18 | 8,57E-01 |
| AT3G09035 | 0,19 | 1,20 | 4,29 | 0,28 | 7,79E-01 |
| AT3G09085 | 0,40 | 1,83 | 3,31 | 0,55 | 5,80E-01 |
| AT3G02105 | 0,40 | 3,73 | 4,32 | 0,86 | 3,87E-01 |
| AT3G09180 | 0,23 | 1,20 | 4,34 | 0,28 | 7,82E-01 |
| AT3G09300 | 1,07 | 1,80 | 2,53 | 0,71 | 4,77E-01 |
| AT3G09350 | 0,51 | 0,00 | 4,41 | 0,00 | 1,00E+00 |
| AT3G09370 | 0,29 | 3,11 | 4,33 | 0,72 | 4,73E-01 |
| AT3G09430 | 1,27 | 5,17 | 2,03 | 2,54 | 1,11E-02 |
| AT3G09500 | 0,81 | 1,78 | 2,16 | 0,82 | 4,12E-01 |
| AT3G09510 | 0,52 | -1,74 | 3,01 | -0,58 | 5,63E-01 |
| AT3G09570 | 0,70 | 2,76 | 2,49 | 1,11 | 2,68E-01 |
| AT3G09600 | 0,41 | 1,90 | 3,30 | 0,58 | 5,65E-01 |
| AT3G09650 | 0,58 | 4,18 | 4,31 | 0,97 | 3,33E-01 |
| AT3G09670 | 0,48 | 3,77 | 4,32 | 0,87 | 3,83E-01 |
| AT3G09690 | 0,55 | 2,14 | 4,38 | 0,49 | 6,25E-01 |
| AT3G09810 | 0,32 | -0,04 | 4,31 | -0,01 | 9,92E-01 |
| AT3G09820 | 0,77 | 1,46 | 2,31 | 0,63 | 5,26E-01 |
| AT3G09830 | 0,46 | -0,60 | 3,15 | -0,19 | 8,49E-01 |
| AT3G09880 | 0,75 | 2,38 | 2,41 | 0,99 | 3,24E-01 |
| AT3G09890 | 0,43 | 3,25 | 4,33 | 0,75 | 4,52E-01 |
| AT3G09900 | 0,19 | 2,19 | 4,32 | 0,51 | 6,13E-01 |
| AT3G10060 | 0,36 | 0,25 | 4,29 | 0,06 | 9,53E-01 |
| AT3G10120 | 0,12 | 2,69 | 4,35 | 0,62 | 5,36E-01 |
| AT3G10190 | 0,32 | 2,79 | 4,30 | 0,65 | 5,17E-01 |
| AT3G10200 | 0,08 | 2,33 | 4,36 | 0,53 | 5,93E-01 |
| AT3G10300 | 0,68 | -2,13 | 2,81 | -0,76 | 4,48E-01 |
| AT3G10310 | 0,16 | 1,27 | 4,37 | 0,29 | 7,71E-01 |
| AT3G10340 | 0,80 | -2,08 | 2,89 | -0,72 | 4,72E-01 |
| AT3G10370 | 0,28 | 0,97 | 4,27 | 0,23 | 8,20E-01 |
| AT3G10390 | 0,49 | 0,52 | 3,19 | 0,16 | 8,70E-01 |
| AT3G10420 | 1,49 | 2,06 | 1,50 | 1,37 | 1,70E-01 |
| AT3G10480 | 0,72 | 4,10 | 3,48 | 1,18 | 2,38E-01 |

| | | | | | |
|---|---|---|---|---|---|
| AT3G10490 | 0,18 | 0,77 | 4,33 | 0,18 | 8,59E-01 |
| AT3G10527 | 0,27 | 2,17 | 4,33 | 0,50 | 6,16E-01 |
| AT3G10540 | 0,56 | -0,17 | 2,57 | -0,07 | 9,47E-01 |
| AT3G10570 | 0,12 | 1,15 | 4,35 | 0,26 | 7,92E-01 |
| AT3G10760 | 0,48 | 0,03 | 3,25 | 0,01 | 9,92E-01 |
| AT3G10800 | 1,54 | 2,15 | 1,76 | 1,22 | 2,22E-01 |
| AT3G10820 | 0,24 | 0,00 | 4,41 | 0,00 | 1,00E+00 |
| AT3G10860 | 0,23 | 0,83 | 4,27 | 0,20 | 8,45E-01 |
| AT3G10920 | 1,12 | 0,72 | 1,92 | 0,37 | 7,09E-01 |
| AT3G10950 | 0,19 | -0,65 | 4,31 | -0,15 | 8,80E-01 |
| AT3G10980 | 0,79 | 1,26 | 3,04 | 0,41 | 6,78E-01 |
| AT3G11020 | 0,05 | -0,58 | 4,38 | -0,13 | 8,95E-01 |
| AT3G11080 | 0,18 | -0,57 | 4,30 | -0,13 | 8,94E-01 |
| AT3G11090 | 0,60 | 1,23 | 3,09 | 0,40 | 6,91E-01 |
| AT3G11120 | 0,39 | -2,18 | 3,33 | -0,66 | 5,12E-01 |
| AT3G11170 | 1,02 | -0,68 | 1,61 | -0,42 | 6,73E-01 |
| AT3G11230 | 0,53 | 0,00 | 4,41 | 0,00 | 1,00E+00 |
| AT3G11250 | 0,11 | -1,28 | 4,35 | -0,29 | 7,69E-01 |
| AT3G11270 | 0,10 | 0,34 | 4,35 | 0,08 | 9,37E-01 |
| AT3G11397 | 0,44 | -0,14 | 4,19 | -0,03 | 9,74E-01 |
| AT3G11400 | 0,87 | 1,07 | 2,27 | 0,47 | 6,39E-01 |
| AT3G11420 | 0,26 | -0,84 | 4,30 | -0,20 | 8,45E-01 |
| AT3G11440 | 0,65 | 3,76 | 2,96 | 1,27 | 2,04E-01 |
| AT3G02465 | 0,30 | 0,51 | 4,27 | 0,12 | 9,04E-01 |
| AT3G11630 | 0,96 | 0,46 | 1,57 | 0,29 | 7,70E-01 |
| AT3G11700 | 0,50 | 0,04 | 3,11 | 0,01 | 9,90E-01 |
| AT3G11760 | 0,23 | 1,68 | 4,33 | 0,39 | 6,98E-01 |
| AT3G11810 | 0,52 | -0,04 | 3,75 | -0,01 | 9,91E-01 |
| AT3G11900 | 0,13 | 2,37 | 4,36 | 0,54 | 5,86E-01 |
| AT3G11930 | 1,35 | 1,69 | 3,43 | 0,49 | 6,22E-01 |
| AT3G11960 | 1,04 | 3,46 | 1,97 | 1,75 | 7,96E-02 |
| AT3G12010 | 0,40 | -0,77 | 4,31 | -0,18 | 8,59E-01 |
| AT3G12020 | 0,79 | 2,44 | 2,38 | 1,03 | 3,04E-01 |
| AT3G12050 | 0,32 | -0,18 | 3,40 | -0,05 | 9,57E-01 |
| AT3G12080 | 1,08 | 2,97 | 2,44 | 1,21 | 2,25E-01 |
| AT3G12270 | 0,77 | 0,76 | 2,60 | 0,29 | 7,71E-01 |
| AT3G12290 | 0,81 | -1,12 | 2,36 | -0,48 | 6,33E-01 |
| AT3G12320 | 0,90 | 1,91 | 2,12 | 0,90 | 3,67E-01 |
| AT3G12360 | 0,71 | -0,21 | 2,42 | -0,09 | 9,32E-01 |
| AT3G12390 | 0,69 | 2,19 | 2,40 | 0,91 | 3,62E-01 |
| AT3G12480 | 0,26 | 2,19 | 4,32 | 0,51 | 6,13E-01 |
| AT3G12502 | 0,08 | 2,33 | 4,36 | 0,53 | 5,93E-01 |
| AT3G12570 | 1,24 | 3,95 | 2,65 | 1,49 | 1,37E-01 |
| AT3G12600 | 0,05 | -0,58 | 4,38 | -0,13 | 8,95E-01 |
| AT3G12640 | 0,60 | 1,62 | 2,16 | 0,75 | 4,55E-01 |
| AT3G12700 | 0,20 | -0,80 | 4,37 | -0,18 | 8,55E-01 |
| AT3G12740 | 0,52 | -0,40 | 3,17 | -0,13 | 9,00E-01 |
| AT3G12760 | 0,80 | 2,53 | 2,28 | 1,11 | 2,68E-01 |
| AT3G12810 | 1,56 | 2,33 | 1,57 | 1,48 | 1,39E-01 |
| AT3G12915 | 0,53 | 3,69 | 4,32 | 0,85 | 3,93E-01 |
| AT3G13060 | 0,87 | -0,36 | 2,05 | -0,17 | 8,61E-01 |
| AT3G13062 | 1,32 | 0,73 | 1,41 | 0,52 | 6,06E-01 |
| AT3G13065 | 0,88 | 4,37 | 3,37 | 1,30 | 1,94E-01 |
| AT3G13200 | 1,18 | 0,13 | 1,53 | 0,09 | 9,32E-01 |
| AT3G13210 | 0,13 | 2,37 | 4,36 | 0,54 | 5,86E-01 |
| AT3G13340 | 1,00 | -3,04 | 2,08 | -1,46 | 1,43E-01 |

| | | | | | |
|---|---|---|---|---|---|
| AT3G13370 | 0,63 | -3,81 | 4,27 | -0,89 | 3,72E-01 |
| AT3G13380 | 0,67 | 1,22 | 2,26 | 0,54 | 5,89E-01 |
| AT3G13437 | 0,08 | 2,08 | 4,38 | 0,47 | 6,35E-01 |
| AT3G13445 | 0,19 | 1,20 | 4,34 | 0,28 | 7,82E-01 |
| AT3G13470 | 0,61 | 1,34 | 3,80 | 0,35 | 7,24E-01 |
| AT3G02725 | 1,24 | -1,06 | 1,84 | -0,58 | 5,63E-01 |
| AT3G13662 | 0,34 | -0,96 | 4,29 | -0,22 | 8,23E-01 |
| AT3G13690 | 0,67 | 0,06 | 3,85 | 0,02 | 9,87E-01 |
| AT3G13760 | 0,08 | 2,08 | 4,38 | 0,47 | 6,35E-01 |
| AT3G13770 | 0,43 | 1,35 | 4,33 | 0,31 | 7,56E-01 |
| AT3G13810 | 0,42 | -1,26 | 4,13 | -0,31 | 7,60E-01 |
| AT3G13930 | 0,81 | -1,62 | 2,17 | -0,75 | 4,55E-01 |
| AT3G13950 | 0,55 | -0,41 | 3,88 | -0,10 | 9,17E-01 |
| AT3G14020 | 0,35 | 0,00 | 4,41 | 0,00 | 1,00E+00 |
| AT3G14100 | 0,69 | 0,52 | 2,38 | 0,22 | 8,27E-01 |
| AT3G14120 | 1,41 | 3,17 | 2,04 | 1,55 | 1,20E-01 |
| AT3G14160 | 0,19 | 1,14 | 4,29 | 0,26 | 7,91E-01 |
| AT3G14220 | 0,64 | -0,05 | 3,84 | -0,01 | 9,90E-01 |
| AT3G14290 | 0,42 | 3,19 | 4,12 | 0,77 | 4,39E-01 |
| AT3G14560 | 0,20 | 2,30 | 4,36 | 0,53 | 5,98E-01 |
| AT3G14595 | 1,01 | 1,52 | 1,99 | 0,76 | 4,44E-01 |
| AT3G14600 | 1,18 | 1,71 | 1,60 | 1,07 | 2,86E-01 |
| AT3G14610 | 0,28 | 1,64 | 4,38 | 0,38 | 7,08E-01 |
| AT3G14620 | 0,63 | -0,41 | 2,39 | -0,17 | 8,66E-01 |
| AT3G14640 | 0,33 | 2,56 | 4,35 | 0,59 | 5,57E-01 |
| AT3G14660 | 1,69 | 0,55 | 1,16 | 0,47 | 6,35E-01 |
| AT3G14670 | 1,07 | -1,06 | 1,74 | -0,61 | 5,43E-01 |
| AT3G14690 | 0,63 | 0,16 | 2,08 | 0,08 | 9,39E-01 |
| AT3G14720 | 0,35 | -0,87 | 4,30 | -0,20 | 8,40E-01 |
| AT3G14790 | 0,27 | 0,03 | 4,38 | 0,01 | 9,94E-01 |
| AT3G14830 | 0,30 | 1,72 | 4,30 | 0,40 | 6,89E-01 |
| AT3G14870 | 0,25 | 1,84 | 4,30 | 0,43 | 6,68E-01 |
| AT3G14920 | 0,30 | 2,91 | 4,34 | 0,67 | 5,04E-01 |
| AT3G14940 | 0,68 | 2,83 | 3,49 | 0,81 | 4,18E-01 |
| AT3G14990 | 0,39 | 0,58 | 2,72 | 0,21 | 8,32E-01 |
| AT3G15000 | 0,95 | 0,51 | 2,20 | 0,23 | 8,17E-01 |
| AT3G15010 | 0,55 | 2,48 | 3,03 | 0,82 | 4,13E-01 |
| AT3G15020 | 1,02 | 2,70 | 2,42 | 1,12 | 2,64E-01 |
| AT3G15030 | 0,69 | 0,65 | 3,08 | 0,21 | 8,33E-01 |
| AT3G15090 | 0,12 | 2,87 | 4,34 | 0,66 | 5,08E-01 |
| AT3G15095 | 1,44 | 0,77 | 1,39 | 0,56 | 5,77E-01 |
| AT3G15160 | 1,08 | 1,80 | 2,14 | 0,84 | 4,00E-01 |
| AT3G15180 | 0,21 | -0,24 | 4,35 | -0,06 | 9,56E-01 |
| AT3G15190 | 1,26 | 0,92 | 1,64 | 0,56 | 5,76E-01 |
| AT3G15260 | 0,39 | 2,59 | 4,25 | 0,61 | 5,43E-01 |
| AT3G15290 | 0,26 | 2,30 | 4,37 | 0,53 | 5,98E-01 |
| AT3G15350 | 1,19 | 2,52 | 2,31 | 1,09 | 2,75E-01 |
| AT3G15355 | 0,91 | 2,71 | 3,68 | 0,73 | 4,62E-01 |
| AT3G15360 | 1,20 | -0,55 | 1,69 | -0,33 | 7,43E-01 |
| AT3G15380 | 0,12 | 2,87 | 4,34 | 0,66 | 5,08E-01 |
| AT3G15450 | 0,77 | 1,90 | 2,18 | 0,87 | 3,84E-01 |
| AT3G15518 | 0,17 | 2,62 | 4,35 | 0,60 | 5,47E-01 |
| AT3G15550 | 0,17 | 2,11 | 4,37 | 0,48 | 6,28E-01 |
| AT3G03145 | 0,66 | -2,67 | 2,92 | -0,92 | 3,60E-01 |
| AT3G15635 | 0,25 | -0,31 | 4,33 | -0,07 | 9,42E-01 |
| AT3G15750 | 0,08 | 2,08 | 4,38 | 0,47 | 6,35E-01 |

| | | | | | |
|---|---|---|---|---|---|
| AT3G15760 | 0,84 | 1,23 | 2,45 | 0,50 | 6,15E-01 |
| AT3G15790 | 0,67 | 3,55 | 2,81 | 1,26 | 2,06E-01 |
| AT3G15820 | 0,32 | 1,90 | 4,27 | 0,44 | 6,57E-01 |
| AT3G15850 | 0,57 | -1,56 | 2,60 | -0,60 | 5,48E-01 |
| AT3G15930 | 0,12 | 2,81 | 4,34 | 0,65 | 5,18E-01 |
| AT3G16010 | 0,15 | -0,41 | 4,32 | -0,09 | 9,25E-01 |
| AT3G16060 | 0,47 | 0,63 | 4,07 | 0,16 | 8,76E-01 |
| AT3G16080 | 0,49 | 0,74 | 3,22 | 0,23 | 8,18E-01 |
| AT3G16130 | 0,87 | 0,03 | 1,88 | 0,02 | 9,87E-01 |
| AT3G16175 | 0,08 | 2,08 | 4,38 | 0,47 | 6,35E-01 |
| AT3G16240 | 1,39 | -3,48 | 1,73 | -2,02 | 4,37E-02 |
| AT3G16260 | 0,72 | 3,58 | 3,51 | 1,02 | 3,08E-01 |
| AT3G16270 | 0,71 | 1,68 | 2,35 | 0,71 | 4,75E-01 |
| AT3G16340 | 0,41 | 3,15 | 4,33 | 0,73 | 4,67E-01 |
| AT3G16420 | 0,28 | -0,24 | 4,40 | -0,05 | 9,57E-01 |
| AT3G16460 | 0,31 | -0,70 | 4,29 | -0,16 | 8,70E-01 |
| AT3G16525 | 0,84 | -2,13 | 2,08 | -1,03 | 3,05E-01 |
| AT3G16570 | 0,68 | 0,80 | 2,38 | 0,34 | 7,36E-01 |
| AT3G16710 | 0,43 | 1,26 | 4,21 | 0,30 | 7,65E-01 |
| AT3G16720 | 0,43 | 0,39 | 4,08 | 0,10 | 9,23E-01 |
| AT3G16740 | 0,38 | 1,31 | 3,35 | 0,39 | 6,96E-01 |
| AT3G16760 | 0,57 | 1,57 | 3,71 | 0,42 | 6,72E-01 |
| AT3G16770 | 1,33 | 1,03 | 2,30 | 0,45 | 6,53E-01 |
| AT3G16800 | 0,16 | -0,80 | 4,37 | -0,18 | 8,54E-01 |
| AT3G16857 | 0,72 | 2,66 | 3,78 | 0,70 | 4,81E-01 |
| AT3G16940 | 0,78 | 1,55 | 2,88 | 0,54 | 5,90E-01 |
| AT3G17100 | 1,17 | -0,59 | 1,91 | -0,31 | 7,56E-01 |
| AT3G17160 | 0,19 | 2,19 | 4,32 | 0,51 | 6,12E-01 |
| AT3G17205 | 0,35 | -1,02 | 4,30 | -0,24 | 8,13E-01 |
| AT3G17210 | 0,32 | 1,02 | 4,36 | 0,23 | 8,15E-01 |
| AT3G03465 | 0,80 | -0,02 | 2,10 | -0,01 | 9,92E-01 |
| AT3G17360 | 0,65 | 1,44 | 3,05 | 0,47 | 6,36E-01 |
| AT3G17380 | 0,28 | 1,87 | 4,37 | 0,43 | 6,69E-01 |
| AT3G17410 | 0,79 | 0,96 | 1,73 | 0,56 | 5,77E-01 |
| AT3G17470 | 0,76 | -0,60 | 3,79 | -0,16 | 8,74E-01 |
| AT3G17490 | 0,56 | -1,44 | 3,73 | -0,39 | 7,00E-01 |
| AT3G17626 | 0,20 | 2,30 | 4,36 | 0,53 | 5,98E-01 |
| AT3G17640 | 0,16 | -0,80 | 4,37 | -0,18 | 8,54E-01 |
| AT3G17650 | 0,88 | 2,21 | 1,87 | 1,18 | 2,38E-01 |
| AT3G17690 | 0,12 | 2,81 | 4,34 | 0,65 | 5,18E-01 |
| AT3G17700 | 0,35 | 0,51 | 4,28 | 0,12 | 9,05E-01 |
| AT3G17712 | 0,66 | 0,79 | 3,76 | 0,21 | 8,34E-01 |
| AT3G17750 | 1,26 | 1,65 | 1,64 | 1,01 | 3,14E-01 |
| AT3G17765 | 0,21 | 2,61 | 4,34 | 0,60 | 5,48E-01 |
| AT3G17790 | 0,83 | 2,79 | 2,44 | 1,14 | 2,53E-01 |
| AT3G03535 | 0,92 | -1,81 | 2,32 | -0,78 | 4,37E-01 |
| AT3G17810 | 0,53 | 0,60 | 2,65 | 0,23 | 8,19E-01 |
| AT3G17820 | 0,62 | 0,06 | 3,21 | 0,02 | 9,86E-01 |
| AT3G17860 | 0,36 | 0,00 | 4,25 | 0,00 | 1,00E+00 |
| AT3G17880 | 0,42 | 0,68 | 3,12 | 0,22 | 8,28E-01 |
| AT3G17890 | 0,10 | 0,34 | 4,35 | 0,08 | 9,37E-01 |
| AT3G17970 | 0,77 | 1,51 | 2,97 | 0,51 | 6,12E-01 |
| AT3G17980 | 0,35 | 0,00 | 4,41 | 0,00 | 1,00E+00 |
| AT3G18000 | 0,28 | 1,78 | 4,33 | 0,41 | 6,80E-01 |
| AT3G18020 | 0,41 | 1,47 | 4,23 | 0,35 | 7,28E-01 |
| AT3G18040 | 0,26 | 0,26 | 4,34 | 0,06 | 9,52E-01 |

| | | | | | |
|---|---|---|---|---|---|
| AT3G18050 | 1,52 | 1,75 | 1,52 | 1,15 | 2,51E-01 |
| AT3G18060 | 1,35 | 0,80 | 1,46 | 0,55 | 5,85E-01 |
| AT3G18070 | 0,11 | 1,33 | 4,36 | 0,31 | 7,59E-01 |
| AT3G18080 | 1,62 | 1,30 | 1,30 | 1,00 | 3,15E-01 |
| AT3G18090 | 1,26 | 5,12 | 2,93 | 1,75 | 8,08E-02 |
| AT3G18100 | 0,29 | 3,75 | 4,32 | 0,87 | 3,85E-01 |
| AT3G03625 | 0,40 | -1,33 | 3,31 | -0,40 | 6,88E-01 |
| AT3G18190 | 1,46 | 1,11 | 1,54 | 0,72 | 4,70E-01 |
| AT3G18240 | 0,43 | 1,39 | 4,18 | 0,33 | 7,39E-01 |
| AT3G18270 | 0,50 | 0,27 | 3,98 | 0,07 | 9,45E-01 |
| AT3G18340 | 0,19 | -2,00 | 4,31 | -0,46 | 6,43E-01 |
| AT3G18390 | 1,15 | 0,40 | 1,82 | 0,22 | 8,24E-01 |
| AT3G18440 | 0,08 | 2,33 | 4,36 | 0,53 | 5,93E-01 |
| AT3G18480 | 0,44 | 1,80 | 3,15 | 0,57 | 5,68E-01 |
| AT3G18740 | 0,73 | 2,02 | 2,15 | 0,94 | 3,48E-01 |
| AT3G18790 | 0,65 | 2,58 | 2,41 | 1,07 | 2,85E-01 |
| AT3G18800 | 0,33 | 2,56 | 4,35 | 0,59 | 5,57E-01 |
| AT3G18820 | 0,39 | -0,68 | 4,17 | -0,16 | 8,70E-01 |
| AT3G18827 | 0,57 | -0,86 | 3,15 | -0,27 | 7,86E-01 |
| AT3G18860 | 0,59 | 0,06 | 2,38 | 0,02 | 9,81E-01 |
| AT3G18890 | 0,86 | 0,65 | 2,21 | 0,29 | 7,68E-01 |
| AT3G18915 | 0,30 | -0,68 | 4,26 | -0,16 | 8,74E-01 |
| AT3G18960 | 0,08 | 2,33 | 4,36 | 0,53 | 5,93E-01 |
| AT3G18980 | 1,15 | 1,29 | 2,02 | 0,64 | 5,22E-01 |
| AT3G19050 | 0,62 | 2,45 | 3,01 | 0,81 | 4,16E-01 |
| AT3G19100 | 1,08 | 2,65 | 1,90 | 1,39 | 1,63E-01 |
| AT3G19240 | 1,33 | 0,35 | 1,34 | 0,26 | 7,96E-01 |
| AT3G19290 | 0,28 | -0,16 | 4,40 | -0,04 | 9,70E-01 |
| AT3G19390 | 0,35 | 0,00 | 4,41 | 0,00 | 1,00E+00 |
| AT3G19470 | 0,30 | -0,86 | 4,36 | -0,20 | 8,45E-01 |
| AT3G19480 | 0,52 | 1,32 | 3,10 | 0,43 | 6,70E-01 |
| AT3G19520 | 0,05 | -0,58 | 4,38 | -0,13 | 8,95E-01 |
| AT3G19540 | 1,20 | 0,41 | 1,88 | 0,22 | 8,26E-01 |
| AT3G19560 | 1,58 | -1,70 | 1,92 | -0,88 | 3,77E-01 |
| AT3G19640 | 0,86 | 2,38 | 2,42 | 0,98 | 3,26E-01 |
| AT3G19680 | 0,92 | 1,49 | 2,00 | 0,74 | 4,57E-01 |
| AT3G19760 | 1,20 | 0,25 | 1,66 | 0,15 | 8,80E-01 |
| AT3G19770 | 0,75 | 2,22 | 3,00 | 0,74 | 4,60E-01 |
| AT3G19780 | 1,34 | 3,62 | 2,05 | 1,77 | 7,69E-02 |
| AT3G19800 | 0,78 | 3,64 | 2,46 | 1,48 | 1,38E-01 |
| AT3G19860 | 0,48 | 2,67 | 4,02 | 0,66 | 5,07E-01 |
| AT3G19870 | 0,37 | 1,62 | 4,24 | 0,38 | 7,02E-01 |
| AT3G19910 | 1,61 | 1,15 | 1,32 | 0,87 | 3,87E-01 |
| AT3G19930 | 1,47 | -0,02 | 1,38 | -0,02 | 9,88E-01 |
| AT3G19960 | 1,65 | 0,80 | 1,44 | 0,56 | 5,77E-01 |
| AT3G19980 | 0,93 | 1,42 | 2,32 | 0,61 | 5,40E-01 |
| AT3G19990 | 0,32 | 2,47 | 4,35 | 0,57 | 5,69E-01 |
| AT3G20000 | 0,74 | 3,47 | 3,60 | 0,96 | 3,36E-01 |
| AT3G20030 | 0,79 | -0,84 | 2,19 | -0,38 | 7,02E-01 |
| AT3G20140 | 0,20 | 2,11 | 4,37 | 0,48 | 6,29E-01 |
| AT3G20150 | 0,54 | 0,00 | 4,41 | 0,00 | 1,00E+00 |
| AT3G20200 | 0,21 | 3,22 | 4,33 | 0,74 | 4,58E-01 |
| AT3G20410 | 0,37 | -1,56 | 4,33 | -0,36 | 7,18E-01 |
| AT3G20500 | 0,33 | 2,37 | 4,35 | 0,55 | 5,85E-01 |
| AT3G20520 | 0,48 | 3,77 | 4,32 | 0,87 | 3,83E-01 |
| AT3G20540 | 0,90 | 1,89 | 2,32 | 0,82 | 4,14E-01 |

| | | | | | |
|---|---|---|---|---|---|
| AT3G20600 | 0,25 | -0,67 | 4,39 | -0,15 | 8,78E-01 |
| AT3G03995 | 0,62 | -3,52 | 4,27 | -0,82 | 4,10E-01 |
| AT3G20720 | 0,42 | -0,13 | 3,30 | -0,04 | 9,69E-01 |
| AT3G20730 | 0,40 | 3,73 | 4,32 | 0,86 | 3,87E-01 |
| AT3G20790 | 0,23 | 2,49 | 4,37 | 0,57 | 5,68E-01 |
| AT3G20890 | 0,34 | 0,68 | 4,34 | 0,16 | 8,75E-01 |
| AT3G20970 | 0,63 | 2,09 | 3,00 | 0,70 | 4,86E-01 |
| AT3G21130 | 0,35 | -0,88 | 4,28 | -0,20 | 8,38E-01 |
| AT3G21170 | 1,15 | -1,80 | 2,12 | -0,85 | 3,95E-01 |
| AT3G21175 | 0,18 | -0,78 | 4,31 | -0,18 | 8,57E-01 |
| AT3G21220 | 0,51 | -2,02 | 4,02 | -0,50 | 6,15E-01 |
| AT3G21270 | 0,29 | 1,50 | 4,34 | 0,35 | 7,29E-01 |
| AT3G21295 | 0,41 | 0,59 | 3,37 | 0,18 | 8,60E-01 |
| AT3G21310 | 0,36 | 3,47 | 4,33 | 0,80 | 4,23E-01 |
| AT3G21465 | 0,14 | -0,45 | 4,38 | -0,10 | 9,18E-01 |
| AT3G21473 | 0,48 | -1,97 | 3,96 | -0,50 | 6,19E-01 |
| AT3G21560 | 0,31 | 0,77 | 4,34 | 0,18 | 8,58E-01 |
| AT3G21690 | 0,96 | 0,39 | 1,73 | 0,22 | 8,22E-01 |
| AT3G04175 | 0,99 | -0,93 | 1,87 | -0,50 | 6,17E-01 |
| AT3G21990 | 0,46 | -3,13 | 4,28 | -0,73 | 4,65E-01 |
| AT3G22072 | 0,17 | 0,00 | 4,41 | 0,00 | 1,00E+00 |
| AT3G22104 | 0,62 | 1,26 | 2,88 | 0,44 | 6,61E-01 |
| AT3G22125 | 1,40 | -0,35 | 1,51 | -0,23 | 8,19E-01 |
| AT3G22150 | 0,49 | 2,81 | 3,95 | 0,71 | 4,76E-01 |
| AT3G22200 | 0,76 | 1,23 | 2,07 | 0,59 | 5,54E-01 |
| AT3G04255 | 0,24 | -0,70 | 4,38 | -0,16 | 8,72E-01 |
| AT3G22270 | 0,79 | 0,14 | 2,54 | 0,06 | 9,56E-01 |
| AT3G22290 | 0,33 | 2,56 | 4,35 | 0,59 | 5,57E-01 |
| AT3G22330 | 0,85 | 0,14 | 2,10 | 0,07 | 9,48E-01 |
| AT3G22350 | 0,91 | -1,30 | 2,35 | -0,55 | 5,80E-01 |
| AT3G22370 | 0,72 | -0,44 | 2,46 | -0,18 | 8,59E-01 |
| AT3G22425 | 0,12 | 0,31 | 4,35 | 0,07 | 9,44E-01 |
| AT3G22430 | 0,19 | -0,79 | 4,32 | -0,18 | 8,54E-01 |
| AT3G22520 | 0,75 | 1,71 | 2,47 | 0,69 | 4,89E-01 |
| AT3G22760 | 0,51 | 1,11 | 3,19 | 0,35 | 7,27E-01 |
| AT3G22810 | 0,55 | 3,10 | 4,34 | 0,71 | 4,75E-01 |
| AT3G22845 | 0,51 | 1,71 | 4,40 | 0,39 | 6,98E-01 |
| AT3G22870 | 0,92 | -1,53 | 1,93 | -0,79 | 4,27E-01 |
| AT3G22900 | 0,24 | 2,85 | 4,34 | 0,66 | 5,11E-01 |
| AT3G22935 | 0,26 | -2,63 | 4,29 | -0,61 | 5,40E-01 |
| AT3G23010 | 0,77 | 1,06 | 3,58 | 0,29 | 7,68E-01 |
| AT3G23050 | 0,64 | -0,02 | 2,51 | -0,01 | 9,94E-01 |
| AT3G04395 | 0,16 | 1,18 | 4,32 | 0,27 | 7,85E-01 |
| AT3G23250 | 0,25 | 0,51 | 4,26 | 0,12 | 9,05E-01 |
| AT3G23280 | 0,45 | -0,31 | 3,99 | -0,08 | 9,39E-01 |
| AT3G23300 | 0,75 | 0,98 | 2,00 | 0,49 | 6,25E-01 |
| AT3G23310 | 0,63 | -0,33 | 2,16 | -0,15 | 8,79E-01 |
| AT3G23380 | 0,21 | -0,16 | 4,33 | -0,04 | 9,70E-01 |
| AT3G23390 | 0,15 | -0,41 | 4,32 | -0,09 | 9,25E-01 |
| AT3G23410 | 0,36 | 0,35 | 4,33 | 0,08 | 9,36E-01 |
| AT3G23530 | 1,09 | 0,96 | 1,97 | 0,49 | 6,27E-01 |
| AT3G23580 | 0,57 | 0,34 | 2,19 | 0,15 | 8,77E-01 |
| AT3G23590 | 1,64 | 3,14 | 1,70 | 1,85 | 6,44E-02 |
| AT3G23600 | 1,16 | 1,16 | 1,69 | 0,69 | 4,92E-01 |
| AT3G23640 | 0,83 | 0,19 | 2,58 | 0,07 | 9,41E-01 |
| AT3G23750 | 0,94 | -0,08 | 1,68 | -0,05 | 9,62E-01 |

| | | | | | |
|---|---|---|---|---|---|
| AT3G23790 | 0,38 | -0,18 | 4,33 | -0,04 | 9,66E-01 |
| AT3G23800 | 0,20 | 2,11 | 4,37 | 0,48 | 6,29E-01 |
| AT3G23890 | 0,56 | 1,79 | 3,94 | 0,45 | 6,50E-01 |
| AT3G23990 | 0,73 | 3,13 | 3,66 | 0,85 | 3,93E-01 |
| AT3G24000 | 0,34 | 1,89 | 4,34 | 0,44 | 6,63E-01 |
| AT3G24050 | 0,88 | 1,39 | 1,88 | 0,74 | 4,59E-01 |
| AT3G24070 | 0,38 | 3,26 | 4,34 | 0,75 | 4,52E-01 |
| AT3G24160 | 0,45 | -0,62 | 3,37 | -0,18 | 8,55E-01 |
| AT3G24190 | 0,59 | 2,20 | 3,07 | 0,72 | 4,73E-01 |
| AT3G24255 | 0,43 | 1,45 | 2,67 | 0,54 | 5,87E-01 |
| AT3G24490 | 0,64 | 4,37 | 3,54 | 1,23 | 2,17E-01 |
| AT3G24500 | 0,22 | -2,03 | 4,31 | -0,47 | 6,37E-01 |
| AT3G24514 | 0,29 | -2,62 | 4,29 | -0,61 | 5,41E-01 |
| AT3G24520 | 0,92 | -1,67 | 2,41 | -0,69 | 4,88E-01 |
| AT3G24550 | 0,60 | 1,27 | 2,78 | 0,46 | 6,48E-01 |
| AT3G24580 | 0,47 | 1,17 | 3,97 | 0,29 | 7,69E-01 |
| AT3G24590 | 0,21 | 2,01 | 4,37 | 0,46 | 6,45E-01 |
| AT3G24830 | 0,92 | 1,70 | 1,91 | 0,89 | 3,75E-01 |
| AT3G25012 | 0,95 | 0,34 | 1,75 | 0,19 | 8,46E-01 |
| AT3G25013 | 0,18 | -0,78 | 4,31 | -0,18 | 8,57E-01 |
| AT3G25030 | 0,21 | 1,07 | 4,33 | 0,25 | 8,06E-01 |
| AT3G25040 | 0,17 | 0,31 | 4,36 | 0,07 | 9,43E-01 |
| AT3G25070 | 0,31 | 0,53 | 3,49 | 0,15 | 8,79E-01 |
| AT3G25130 | 0,57 | 0,00 | 4,41 | 0,00 | 1,00E+00 |
| AT3G25140 | 1,51 | 3,64 | 2,05 | 1,78 | 7,54E-02 |
| AT3G25250 | 0,60 | 2,72 | 2,58 | 1,05 | 2,92E-01 |
| AT3G25290 | 0,29 | 1,51 | 4,34 | 0,35 | 7,27E-01 |
| AT3G25440 | 0,23 | -0,64 | 4,39 | -0,15 | 8,84E-01 |
| AT3G25470 | 0,67 | 2,39 | 3,75 | 0,64 | 5,24E-01 |
| AT3G25490 | 0,05 | -0,58 | 4,38 | -0,13 | 8,95E-01 |
| AT3G25585 | 0,71 | 1,72 | 2,63 | 0,65 | 5,13E-01 |
| AT3G25590 | 0,75 | 0,03 | 2,19 | 0,01 | 9,91E-01 |
| AT3G25700 | 0,66 | 0,75 | 2,17 | 0,35 | 7,28E-01 |
| AT3G04955 | 0,15 | 1,64 | 4,33 | 0,38 | 7,06E-01 |
| AT3G25720 | 0,26 | -1,25 | 4,29 | -0,29 | 7,71E-01 |
| AT3G25760 | 0,49 | 0,00 | 4,41 | 0,00 | 1,00E+00 |
| AT3G25770 | 1,09 | -0,78 | 1,84 | -0,42 | 6,71E-01 |
| AT3G25780 | 0,49 | -1,04 | 2,49 | -0,42 | 6,76E-01 |
| AT3G25860 | 0,63 | 0,54 | 3,14 | 0,17 | 8,64E-01 |
| AT3G25890 | 0,89 | 1,11 | 2,00 | 0,55 | 5,80E-01 |
| AT3G25910 | 0,36 | 2,98 | 4,30 | 0,69 | 4,89E-01 |
| AT3G26020 | 0,28 | -0,12 | 4,33 | -0,03 | 9,78E-01 |
| AT3G26030 | 1,16 | 2,47 | 1,79 | 1,38 | 1,66E-01 |
| AT3G26070 | 0,46 | -0,99 | 4,21 | -0,24 | 8,14E-01 |
| AT3G26080 | 0,26 | 2,30 | 4,37 | 0,53 | 5,98E-01 |
| AT3G26085 | 0,30 | 1,15 | 4,36 | 0,26 | 7,92E-01 |
| AT3G26090 | 0,21 | 0,95 | 4,33 | 0,22 | 8,27E-01 |
| AT3G04975 | 1,19 | -1,14 | 1,80 | -0,63 | 5,26E-01 |
| AT3G04985 | 0,58 | -0,70 | 2,46 | -0,28 | 7,77E-01 |
| AT3G26370 | 0,48 | 1,88 | 2,69 | 0,70 | 4,86E-01 |
| AT3G26395 | 0,16 | 1,97 | 4,33 | 0,45 | 6,50E-01 |
| AT3G26400 | 0,95 | -1,52 | 2,44 | -0,62 | 5,33E-01 |
| AT3G26430 | 1,60 | 1,95 | 1,39 | 1,41 | 1,59E-01 |
| AT3G26420 | 1,60 | 1,95 | 1,39 | 1,41 | 1,59E-01 |
| AT3G26512 | 0,38 | -0,52 | 3,21 | -0,16 | 8,72E-01 |
| AT3G26570 | 1,54 | 1,34 | 1,70 | 0,79 | 4,30E-01 |

| | | | | | |
|---|---|---|---|---|---|
| AT3G26580 | 0,73 | 0,76 | 2,11 | 0,36 | 7,20E-01 |
| AT3G26600 | 1,07 | 0,67 | 1,85 | 0,36 | 7,18E-01 |
| AT3G26640 | 0,37 | -1,37 | 3,32 | -0,41 | 6,79E-01 |
| AT3G26710 | 1,14 | 3,29 | 2,07 | 1,59 | 1,12E-01 |
| AT3G26720 | 0,82 | 1,68 | 2,44 | 0,69 | 4,91E-01 |
| AT3G26742 | 0,18 | 0,41 | 4,28 | 0,10 | 9,23E-01 |
| AT3G26780 | 1,24 | 2,06 | 1,82 | 1,13 | 2,57E-01 |
| AT3G26782 | 0,41 | 2,49 | 4,11 | 0,60 | 5,45E-01 |
| AT3G26800 | 0,34 | -0,42 | 3,47 | -0,12 | 9,04E-01 |
| AT3G26840 | 1,06 | 3,55 | 2,22 | 1,60 | 1,11E-01 |
| AT3G27090 | 1,24 | 2,22 | 1,78 | 1,25 | 2,13E-01 |
| AT3G27160 | 0,41 | -0,44 | 3,31 | -0,13 | 8,94E-01 |
| AT3G27170 | 0,68 | 4,15 | 2,82 | 1,47 | 1,41E-01 |
| AT3G27230 | 0,45 | 1,49 | 3,27 | 0,45 | 6,50E-01 |
| AT3G27250 | 0,85 | -0,89 | 2,51 | -0,36 | 7,22E-01 |
| AT3G27260 | 1,09 | 0,41 | 1,81 | 0,23 | 8,18E-01 |
| AT3G27270 | 0,62 | 0,83 | 3,86 | 0,22 | 8,30E-01 |
| AT3G27280 | 0,60 | 2,61 | 2,97 | 0,88 | 3,80E-01 |
| AT3G27325 | 0,56 | 0,27 | 2,59 | 0,10 | 9,18E-01 |
| AT3G27329 | 0,81 | 0,15 | 2,01 | 0,07 | 9,41E-01 |
| AT3G27331 | 0,96 | -2,44 | 3,03 | -0,81 | 4,20E-01 |
| AT3G27350 | 0,51 | 1,46 | 3,87 | 0,38 | 7,06E-01 |
| AT3G27430 | 0,35 | -0,53 | 3,52 | -0,15 | 8,81E-01 |
| AT3G27470 | 1,23 | 1,14 | 1,54 | 0,74 | 4,59E-01 |
| AT3G27540 | 0,29 | 0,25 | 3,58 | 0,07 | 9,45E-01 |
| AT3G27555 | 0,30 | -1,84 | 4,32 | -0,43 | 6,70E-01 |
| AT3G27670 | 1,19 | 1,55 | 2,10 | 0,74 | 4,61E-01 |
| AT3G05295 | 0,48 | -0,48 | 2,75 | -0,17 | 8,62E-01 |
| AT3G27820 | 1,03 | 1,42 | 1,65 | 0,86 | 3,91E-01 |
| AT3G27830 | 0,47 | -0,83 | 3,25 | -0,25 | 7,99E-01 |
| AT3G27850 | 1,55 | 0,06 | 1,66 | 0,04 | 9,70E-01 |
| AT3G27860 | 0,12 | 2,69 | 4,35 | 0,62 | 5,36E-01 |
| AT3G27870 | 0,75 | 1,39 | 2,48 | 0,56 | 5,74E-01 |
| AT3G27940 | 0,56 | -0,89 | 2,76 | -0,32 | 7,48E-01 |
| AT3G27980 | 0,34 | -2,46 | 4,29 | -0,57 | 5,66E-01 |
| AT3G27999 | 0,43 | -3,05 | 4,28 | -0,71 | 4,76E-01 |
| AT3G28040 | 1,16 | 0,69 | 2,11 | 0,33 | 7,45E-01 |
| AT3G28050 | 0,37 | 3,01 | 4,31 | 0,70 | 4,85E-01 |
| AT3G28080 | 0,70 | -0,82 | 2,13 | -0,39 | 7,00E-01 |
| AT3G28100 | 1,10 | -1,70 | 1,67 | -1,02 | 3,09E-01 |
| AT3G28170 | 0,75 | 3,53 | 3,52 | 1,00 | 3,16E-01 |
| AT3G28200 | 0,56 | 2,60 | 4,34 | 0,60 | 5,50E-01 |
| AT3G28220 | 0,69 | 0,29 | 3,02 | 0,10 | 9,22E-01 |
| AT3G28270 | 0,14 | -0,45 | 4,33 | -0,10 | 9,17E-01 |
| AT3G28290 | 1,10 | 0,77 | 2,62 | 0,29 | 7,69E-01 |
| AT3G28300 | 1,56 | 0,00 | 2,86 | 0,00 | 1,00E+00 |
| AT3G28450 | 0,78 | -0,55 | 2,38 | -0,23 | 8,17E-01 |
| AT3G28500 | 0,30 | 0,19 | 4,25 | 0,04 | 9,65E-01 |
| AT3G28510 | 0,99 | 2,38 | 2,44 | 0,97 | 3,30E-01 |
| AT3G05515 | 0,60 | -3,49 | 2,89 | -1,21 | 2,27E-01 |
| AT3G28580 | 0,77 | 2,21 | 3,31 | 0,67 | 5,05E-01 |
| AT3G28590 | 0,05 | -0,58 | 4,38 | -0,13 | 8,95E-01 |
| AT3G28650 | 0,20 | -0,80 | 4,37 | -0,18 | 8,55E-01 |
| AT3G28670 | 0,33 | 0,55 | 4,36 | 0,13 | 8,99E-01 |
| AT3G28690 | 0,39 | 1,63 | 4,17 | 0,39 | 6,97E-01 |
| AT3G28715 | 0,62 | 1,11 | 2,28 | 0,48 | 6,28E-01 |

| | | | | | |
|---|---|---|---|---|---|
| AT3G28720 | 0,91 | 3,04 | 3,13 | 0,97 | 3,32E-01 |
| AT3G28730 | 0,64 | 1,94 | 2,90 | 0,67 | 5,04E-01 |
| AT3G28870 | 0,74 | 1,54 | 3,66 | 0,42 | 6,73E-01 |
| AT3G28910 | 0,61 | 0,49 | 2,52 | 0,19 | 8,47E-01 |
| AT3G05595 | 0,35 | 1,42 | 4,31 | 0,33 | 7,42E-01 |
| AT3G28922 | 0,12 | 0,63 | 4,33 | 0,15 | 8,85E-01 |
| AT3G28958 | 0,58 | -0,59 | 2,94 | -0,20 | 8,41E-01 |
| AT3G29000 | 0,25 | 1,44 | 4,29 | 0,33 | 7,38E-01 |
| AT3G29010 | 0,38 | 2,05 | 4,37 | 0,47 | 6,40E-01 |
| AT3G29033 | 0,57 | 0,25 | 2,75 | 0,09 | 9,28E-01 |
| AT3G29080 | 1,28 | -0,41 | 1,39 | -0,30 | 7,66E-01 |
| AT3G29180 | 1,27 | 2,00 | 2,78 | 0,72 | 4,71E-01 |
| AT3G05715 | 0,40 | 0,04 | 4,23 | 0,01 | 9,93E-01 |
| AT3G29240 | 1,56 | -0,02 | 1,56 | -0,01 | 9,89E-01 |
| AT3G29270 | 0,39 | 1,71 | 4,40 | 0,39 | 6,98E-01 |
| AT3G29290 | 0,20 | 1,94 | 4,34 | 0,45 | 6,55E-01 |
| AT3G29370 | 0,05 | -0,58 | 4,38 | -0,13 | 8,95E-01 |
| AT3G29390 | 0,18 | 0,77 | 4,33 | 0,18 | 8,59E-01 |
| AT3G29450 | 0,14 | -1,69 | 4,33 | -0,39 | 6,96E-01 |
| AT3G29455 | 0,27 | -2,66 | 4,29 | -0,62 | 5,35E-01 |
| AT3G05805 | 0,31 | 0,36 | 4,25 | 0,08 | 9,33E-01 |
| AT3G29638 | 0,16 | 1,14 | 4,30 | 0,27 | 7,90E-01 |
| AT3G29670 | 0,94 | -3,51 | 2,20 | -1,60 | 1,10E-01 |
| AT3G30210 | 0,11 | 1,28 | 4,35 | 0,29 | 7,68E-01 |
| AT3G30320 | 0,40 | -1,22 | 4,26 | -0,29 | 7,75E-01 |
| AT3G30380 | 0,49 | 1,39 | 3,26 | 0,43 | 6,69E-01 |
| AT3G30383 | 0,12 | 2,69 | 4,35 | 0,62 | 5,36E-01 |
| AT3G30520 | 1,35 | -2,75 | 2,07 | -1,33 | 1,84E-01 |
| AT3G30525 | 1,14 | -1,10 | 2,47 | -0,45 | 6,56E-01 |
| AT3G30720 | 0,61 | 1,26 | 3,05 | 0,41 | 6,80E-01 |
| AT3G30770 | 0,13 | -1,34 | 4,35 | -0,31 | 7,58E-01 |
| AT3G30820 | 0,18 | -0,57 | 4,30 | -0,13 | 8,94E-01 |
| AT3G31950 | 0,59 | -0,27 | 2,17 | -0,13 | 9,00E-01 |
| AT3G32150 | 0,20 | -2,26 | 4,30 | -0,52 | 6,00E-01 |
| AT3G32280 | 0,39 | -1,80 | 4,24 | -0,43 | 6,70E-01 |
| AT3G32896 | 0,28 | -1,84 | 4,33 | -0,43 | 6,70E-01 |
| AT3G33530 | 1,04 | 0,00 | 1,69 | 0,00 | 9,98E-01 |
| AT3G42075 | 0,24 | 1,11 | 4,33 | 0,26 | 7,97E-01 |
| AT3G42170 | 0,48 | 2,47 | 3,11 | 0,79 | 4,28E-01 |
| AT3G42310 | 0,36 | -1,31 | 4,27 | -0,31 | 7,58E-01 |
| AT3G42560 | 0,10 | 0,34 | 4,35 | 0,08 | 9,37E-01 |
| AT3G42640 | 0,44 | -0,20 | 2,56 | -0,08 | 9,37E-01 |
| AT3G42870 | 1,35 | -2,28 | 1,84 | -1,24 | 2,14E-01 |
| AT3G42950 | 0,12 | 2,81 | 4,34 | 0,65 | 5,18E-01 |
| AT3G06495 | 0,51 | -0,27 | 3,02 | -0,09 | 9,30E-01 |
| AT3G43148 | 0,82 | -0,71 | 2,03 | -0,35 | 7,25E-01 |
| AT3G43150 | 0,30 | -0,65 | 3,50 | -0,19 | 8,52E-01 |
| AT3G43210 | 0,25 | 2,97 | 4,33 | 0,69 | 4,93E-01 |
| AT3G43230 | 1,49 | 3,47 | 1,60 | 2,17 | 3,00E-02 |
| AT3G43260 | 1,20 | -0,82 | 1,80 | -0,46 | 6,48E-01 |
| AT3G06565 | 0,05 | -0,58 | 4,38 | -0,13 | 8,95E-01 |
| AT3G43290 | 0,21 | 0,16 | 4,27 | 0,04 | 9,70E-01 |
| AT3G06625 | 0,13 | -0,63 | 4,39 | -0,14 | 8,85E-01 |
| AT3G43490 | 1,48 | -0,36 | 1,75 | -0,21 | 8,35E-01 |
| AT3G43503 | 0,30 | 1,11 | 3,61 | 0,31 | 7,59E-01 |
| AT3G06635 | 0,40 | -1,17 | 4,24 | -0,28 | 7,82E-01 |

| | | | | | |
|---|---|---|---|---|---|
| AT3G43670 | 0,53 | -0,25 | 2,63 | -0,09 | 9,25E-01 |
| AT3G43682 | 0,33 | 0,27 | 4,28 | 0,06 | 9,49E-01 |
| AT3G43740 | 0,28 | -0,70 | 4,31 | -0,16 | 8,70E-01 |
| AT3G43800 | 0,45 | 1,94 | 3,23 | 0,60 | 5,48E-01 |
| AT3G43837 | 0,47 | -1,04 | 3,97 | -0,26 | 7,94E-01 |
| AT3G43920 | 0,47 | -1,02 | 3,21 | -0,32 | 7,51E-01 |
| AT3G43950 | 0,37 | -0,36 | 4,32 | -0,08 | 9,34E-01 |
| AT3G44130 | 0,22 | -0,51 | 4,34 | -0,12 | 9,06E-01 |
| AT3G44150 | 0,18 | 1,77 | 4,41 | 0,40 | 6,88E-01 |
| AT3G44210 | 1,26 | -4,60 | 2,42 | -1,90 | 5,71E-02 |
| AT3G44230 | 0,25 | -0,07 | 4,37 | -0,02 | 9,87E-01 |
| AT3G44235 | 0,05 | -0,58 | 4,38 | -0,13 | 8,95E-01 |
| AT3G44320 | 0,76 | 3,35 | 2,85 | 1,18 | 2,40E-01 |
| AT3G44405 | 0,08 | 2,08 | 4,38 | 0,47 | 6,35E-01 |
| AT3G44550 | 0,99 | 1,18 | 2,20 | 0,54 | 5,91E-01 |
| AT3G44560 | 0,14 | -0,45 | 4,33 | -0,10 | 9,17E-01 |
| AT3G44620 | 0,30 | -1,12 | 3,56 | -0,32 | 7,53E-01 |
| AT3G44670 | 1,62 | 2,32 | 1,43 | 1,62 | 1,04E-01 |
| AT3G44730 | 0,52 | -0,27 | 2,24 | -0,12 | 9,02E-01 |
| AT3G44798 | 0,43 | 1,63 | 4,11 | 0,40 | 6,92E-01 |
| AT3G44880 | 0,54 | -1,77 | 4,00 | -0,44 | 6,59E-01 |
| AT3G44900 | 0,86 | 3,94 | 2,35 | 1,67 | 9,43E-02 |
| AT3G44935 | 1,28 | -1,06 | 1,78 | -0,60 | 5,51E-01 |
| AT3G44980 | 0,39 | 0,33 | 3,38 | 0,10 | 9,23E-01 |
| AT3G45010 | 0,31 | -0,66 | 4,31 | -0,15 | 8,78E-01 |
| AT3G45400 | 0,33 | 1,16 | 4,28 | 0,27 | 7,86E-01 |
| AT3G45620 | 1,31 | 2,24 | 2,32 | 0,97 | 3,34E-01 |
| AT3G45650 | 1,57 | 4,95 | 1,95 | 2,54 | 1,10E-02 |
| AT3G45660 | 1,01 | 2,84 | 2,59 | 1,09 | 2,74E-01 |
| AT3G45670 | 0,15 | -0,19 | 4,31 | -0,04 | 9,64E-01 |
| AT3G45730 | 0,15 | 1,93 | 4,33 | 0,45 | 6,55E-01 |
| AT3G45740 | 0,36 | 2,79 | 4,34 | 0,64 | 5,21E-01 |
| AT3G45870 | 0,08 | 2,33 | 4,36 | 0,53 | 5,93E-01 |
| AT3G45930 | 0,32 | 2,40 | 4,37 | 0,55 | 5,82E-01 |
| AT3G45965 | 0,21 | 1,85 | 4,38 | 0,42 | 6,73E-01 |
| AT3G45970 | 0,98 | 1,67 | 2,74 | 0,61 | 5,41E-01 |
| AT3G46040 | 0,85 | 2,36 | 2,15 | 1,10 | 2,73E-01 |
| AT3G46060 | 1,03 | -0,89 | 1,97 | -0,45 | 6,51E-01 |
| AT3G46130 | 0,12 | 2,81 | 4,34 | 0,65 | 5,18E-01 |
| AT3G46220 | 0,60 | 1,64 | 2,62 | 0,63 | 5,31E-01 |
| AT3G46290 | 1,13 | 2,33 | 1,82 | 1,28 | 2,02E-01 |
| AT3G46370 | 0,25 | -1,95 | 4,32 | -0,45 | 6,51E-01 |
| AT3G46490 | 0,32 | -1,93 | 4,32 | -0,45 | 6,56E-01 |
| AT3G46520 | 0,37 | 3,04 | 4,33 | 0,70 | 4,82E-01 |
| AT3G46560 | 0,33 | -1,81 | 4,27 | -0,43 | 6,71E-01 |
| AT3G46590 | 1,40 | 3,41 | 1,65 | 2,07 | 3,89E-02 |
| AT3G07505 | 1,14 | -2,22 | 1,90 | -1,16 | 2,44E-01 |
| AT3G46640 | 1,12 | 2,86 | 2,19 | 1,31 | 1,91E-01 |
| AT3G46658 | 0,59 | 1,25 | 2,50 | 0,50 | 6,17E-01 |
| AT3G46668 | 0,57 | 1,42 | 3,23 | 0,44 | 6,59E-01 |
| AT3G07555 | 1,02 | 0,71 | 1,92 | 0,37 | 7,12E-01 |
| AT3G07585 | 1,61 | -1,23 | 1,48 | -0,83 | 4,07E-01 |
| AT3G46930 | 0,21 | 3,22 | 4,33 | 0,74 | 4,58E-01 |
| AT3G47060 | 0,35 | 2,99 | 4,34 | 0,69 | 4,91E-01 |
| AT3G47140 | 0,11 | -1,28 | 4,35 | -0,29 | 7,69E-01 |
| AT3G07695 | 0,33 | 1,66 | 4,26 | 0,39 | 6,97E-01 |

| | | | | | |
|---|---|---|---|---|---|
| AT3G47290 | 0,08 | 2,33 | 4,36 | 0,53 | 5,93E-01 |
| AT3G47295 | 0,08 | -0,56 | 4,38 | -0,13 | 8,99E-01 |
| AT3G07715 | 0,48 | 0,19 | 2,63 | 0,07 | 9,43E-01 |
| AT3G47490 | 0,22 | -0,51 | 4,34 | -0,12 | 9,06E-01 |
| AT3G47520 | 0,29 | 2,11 | 4,32 | 0,49 | 6,26E-01 |
| AT3G47550 | 0,50 | 0,37 | 2,98 | 0,12 | 9,01E-01 |
| AT3G47570 | 0,50 | 2,50 | 3,21 | 0,78 | 4,35E-01 |
| AT3G47610 | 0,61 | -1,33 | 2,47 | -0,54 | 5,91E-01 |
| AT3G47650 | 0,65 | 3,76 | 2,96 | 1,27 | 2,04E-01 |
| AT3G47660 | 0,72 | 2,84 | 2,88 | 0,99 | 3,24E-01 |
| AT3G07795 | 0,21 | 2,34 | 4,32 | 0,54 | 5,89E-01 |
| AT3G07805 | 0,89 | 4,52 | 2,66 | 1,70 | 9,00E-02 |
| AT3G47770 | 0,17 | 3,14 | 4,33 | 0,73 | 4,68E-01 |
| AT3G47800 | 1,07 | 0,46 | 1,76 | 0,26 | 7,92E-01 |
| AT3G47820 | 1,04 | -0,50 | 1,75 | -0,29 | 7,74E-01 |
| AT3G47833 | 0,22 | -0,56 | 4,38 | -0,13 | 8,98E-01 |
| AT3G47920 | 1,60 | -1,48 | 1,69 | -0,88 | 3,79E-01 |
| AT3G47930 | 0,52 | 3,80 | 3,86 | 0,98 | 3,25E-01 |
| AT3G47950 | 0,34 | 0,44 | 4,33 | 0,10 | 9,19E-01 |
| AT3G48040 | 0,17 | -0,44 | 4,37 | -0,10 | 9,19E-01 |
| AT3G48070 | 0,40 | 2,00 | 4,30 | 0,46 | 6,43E-01 |
| AT3G48110 | 0,52 | 2,00 | 3,10 | 0,65 | 5,18E-01 |
| AT3G48140 | 0,97 | 1,00 | 1,95 | 0,51 | 6,09E-01 |
| AT3G48190 | 1,19 | 1,60 | 2,39 | 0,67 | 5,03E-01 |
| AT3G48310 | 0,40 | 2,21 | 3,34 | 0,66 | 5,08E-01 |
| AT3G48380 | 0,47 | 1,02 | 3,12 | 0,33 | 7,43E-01 |
| AT3G48420 | 0,42 | 1,67 | 4,21 | 0,40 | 6,93E-01 |
| AT3G48460 | 0,51 | -15,91 | 4,37 | -3,64 | 2,68E-04 |
| AT3G48500 | 0,57 | 1,72 | 3,92 | 0,44 | 6,60E-01 |
| AT3G48530 | 0,79 | -0,10 | 2,03 | -0,05 | 9,62E-01 |
| AT3G48620 | 0,08 | 2,08 | 4,38 | 0,47 | 6,35E-01 |
| AT3G48680 | 0,15 | 1,94 | 4,32 | 0,45 | 6,54E-01 |
| AT3G48710 | 0,27 | 1,29 | 4,29 | 0,30 | 7,64E-01 |
| AT3G48730 | 0,57 | 0,18 | 2,66 | 0,07 | 9,47E-01 |
| AT3G48750 | 0,54 | 1,69 | 2,65 | 0,64 | 5,23E-01 |
| AT3G48760 | 0,76 | 2,78 | 2,45 | 1,14 | 2,56E-01 |
| AT3G07995 | 0,83 | -1,69 | 2,08 | -0,81 | 4,16E-01 |
| AT3G48780 | 0,82 | 0,37 | 2,57 | 0,14 | 8,86E-01 |
| AT3G48810 | 0,58 | 3,94 | 3,64 | 1,08 | 2,79E-01 |
| AT3G48890 | 0,66 | 2,18 | 2,70 | 0,81 | 4,20E-01 |
| AT3G49000 | 0,27 | 1,09 | 4,29 | 0,26 | 7,99E-01 |
| AT3G49110 | 0,08 | 2,33 | 4,36 | 0,53 | 5,93E-01 |
| AT3G49130 | 1,37 | 4,53 | 2,37 | 1,91 | 5,56E-02 |
| AT3G49220 | 0,92 | 0,33 | 2,02 | 0,16 | 8,71E-01 |
| AT3G49240 | 0,54 | 0,00 | 4,15 | 0,00 | 9,99E-01 |
| AT3G49260 | 0,64 | 1,83 | 4,40 | 0,42 | 6,77E-01 |
| AT3G49430 | 0,93 | 2,25 | 2,09 | 1,07 | 2,83E-01 |
| AT3G49520 | 1,37 | -0,59 | 1,53 | -0,39 | 6,98E-01 |
| AT3G49590 | 1,44 | 0,86 | 1,60 | 0,54 | 5,91E-01 |
| AT3G49670 | 1,02 | 1,69 | 2,40 | 0,70 | 4,81E-01 |
| AT3G49680 | 0,45 | -0,70 | 3,23 | -0,22 | 8,29E-01 |
| AT3G49710 | 0,32 | 2,47 | 4,35 | 0,57 | 5,69E-01 |
| AT3G49780 | 0,25 | -1,00 | 4,30 | -0,23 | 8,16E-01 |
| AT3G50000 | 0,43 | 1,89 | 4,14 | 0,46 | 6,48E-01 |
| AT3G50080 | 0,21 | 1,60 | 4,34 | 0,37 | 7,13E-01 |
| AT3G50100 | 0,05 | -0,58 | 4,38 | -0,13 | 8,95E-01 |

| | | | | | |
|---|---|---|---|---|---|
| AT3G50270 | 0,60 | 3,10 | 3,06 | 1,01 | 3,12E-01 |
| AT3G50350 | 0,42 | -0,30 | 3,34 | -0,09 | 9,28E-01 |
| AT3G50360 | 0,31 | 0,44 | 4,33 | 0,10 | 9,18E-01 |
| AT3G50430 | 0,16 | -1,98 | 4,31 | -0,46 | 6,47E-01 |
| AT3G50530 | 0,25 | 1,74 | 4,31 | 0,40 | 6,87E-01 |
| AT3G50540 | 1,39 | 0,75 | 1,65 | 0,45 | 6,50E-01 |
| AT3G50550 | 0,51 | -0,11 | 3,21 | -0,03 | 9,73E-01 |
| AT3G50630 | 0,26 | 2,25 | 4,33 | 0,52 | 6,03E-01 |
| AT3G50700 | 0,53 | 0,24 | 2,51 | 0,10 | 9,24E-01 |
| AT3G50770 | 1,04 | 3,19 | 2,23 | 1,43 | 1,53E-01 |
| AT3G50890 | 0,25 | -1,33 | 4,28 | -0,31 | 7,56E-01 |
| AT3G50910 | 0,94 | 3,18 | 2,31 | 1,37 | 1,69E-01 |
| AT3G50970 | 0,39 | -0,14 | 4,41 | -0,03 | 9,75E-01 |
| AT3G51110 | 0,46 | 0,72 | 3,99 | 0,18 | 8,58E-01 |
| AT3G51140 | 0,97 | 0,02 | 2,04 | 0,01 | 9,94E-01 |
| AT3G51180 | 0,25 | -0,67 | 4,39 | -0,15 | 8,78E-01 |
| AT3G51260 | 0,46 | 1,94 | 3,28 | 0,59 | 5,55E-01 |
| AT3G51270 | 0,33 | 1,60 | 4,31 | 0,37 | 7,10E-01 |
| AT3G51325 | 0,36 | 0,57 | 4,32 | 0,13 | 8,95E-01 |
| AT3G51370 | 1,46 | 1,87 | 1,71 | 1,09 | 2,74E-01 |
| AT3G51420 | 0,66 | -1,44 | 2,26 | -0,64 | 5,24E-01 |
| AT3G51430 | 0,32 | 0,10 | 4,36 | 0,02 | 9,82E-01 |
| AT3G51440 | 0,80 | 2,00 | 2,11 | 0,95 | 3,43E-01 |
| AT3G51450 | 0,43 | -0,43 | 2,76 | -0,16 | 8,75E-01 |
| AT3G51460 | 0,27 | 0,29 | 4,37 | 0,07 | 9,46E-01 |
| AT3G51480 | 0,45 | 2,51 | 4,34 | 0,58 | 5,64E-01 |
| AT3G51500 | 0,44 | 4,02 | 4,31 | 0,93 | 3,51E-01 |
| AT3G51510 | 0,44 | 0,16 | 3,36 | 0,05 | 9,62E-01 |
| AT3G51540 | 0,25 | 2,97 | 4,33 | 0,69 | 4,93E-01 |
| AT3G51620 | 1,34 | 3,37 | 1,84 | 1,83 | 6,73E-02 |
| AT3G51630 | 0,87 | 4,29 | 3,21 | 1,34 | 1,81E-01 |
| AT3G51640 | 0,80 | 3,64 | 2,67 | 1,36 | 1,72E-01 |
| AT3G51650 | 1,02 | 3,02 | 2,14 | 1,41 | 1,58E-01 |
| AT3G51730 | 1,38 | 1,18 | 1,50 | 0,79 | 4,30E-01 |
| AT3G51830 | 0,25 | 2,28 | 4,32 | 0,53 | 5,97E-01 |
| AT3G51850 | 1,69 | 2,11 | 1,69 | 1,25 | 2,10E-01 |
| AT3G51870 | 1,46 | 2,94 | 2,05 | 1,43 | 1,52E-01 |
| AT3G51970 | 0,40 | 1,50 | 3,30 | 0,46 | 6,48E-01 |
| AT3G52080 | 0,33 | 2,10 | 4,31 | 0,49 | 6,26E-01 |
| AT3G52100 | 0,15 | 0,40 | 4,33 | 0,09 | 9,27E-01 |
| AT3G52105 | 0,19 | -0,39 | 4,29 | -0,09 | 9,27E-01 |
| AT3G52120 | 1,03 | -0,05 | 2,17 | -0,02 | 9,82E-01 |
| AT3G52150 | 0,83 | -3,17 | 2,78 | -1,14 | 2,54E-01 |
| AT3G52155 | 0,45 | 1,66 | 4,39 | 0,38 | 7,05E-01 |
| AT3G52200 | 0,34 | -0,85 | 4,29 | -0,20 | 8,42E-01 |
| AT3G52210 | 0,11 | 1,33 | 4,36 | 0,31 | 7,59E-01 |
| AT3G52230 | 0,83 | 0,96 | 2,97 | 0,32 | 7,47E-01 |
| AT3G52300 | 0,40 | 2,04 | 4,31 | 0,47 | 6,35E-01 |
| AT3G52340 | 0,38 | 0,82 | 4,30 | 0,19 | 8,48E-01 |
| AT3G52370 | 0,08 | 2,08 | 4,38 | 0,47 | 6,35E-01 |
| AT3G52380 | 1,54 | 2,42 | 1,94 | 1,25 | 2,12E-01 |
| AT3G52470 | 0,30 | -0,81 | 4,25 | -0,19 | 8,50E-01 |
| AT3G52570 | 0,29 | 0,74 | 4,25 | 0,17 | 8,62E-01 |
| AT3G08275 | 0,60 | -1,04 | 2,99 | -0,35 | 7,27E-01 |
| AT3G52610 | 0,30 | 2,91 | 4,34 | 0,67 | 5,04E-01 |
| AT3G52680 | 0,59 | -1,71 | 2,44 | -0,70 | 4,82E-01 |

| | | | | | |
|---|---|---|---|---|---|
| AT3G52710 | 0,88 | 0,87 | 1,78 | 0,49 | 6,24E-01 |
| AT3G08285 | 1,59 | -1,04 | 1,38 | -0,75 | 4,51E-01 |
| AT3G08295 | 0,73 | -2,95 | 3,32 | -0,89 | 3,74E-01 |
| AT3G52830 | 1,16 | -0,78 | 1,59 | -0,49 | 6,23E-01 |
| AT3G52840 | 0,82 | -0,02 | 2,70 | -0,01 | 9,94E-01 |
| AT3G52950 | 0,45 | 1,86 | 3,26 | 0,57 | 5,69E-01 |
| AT3G52960 | 0,80 | 0,41 | 1,73 | 0,23 | 8,14E-01 |
| AT3G52990 | 1,11 | -0,01 | 1,54 | 0,00 | 9,97E-01 |
| AT3G53000 | 0,31 | 1,06 | 4,33 | 0,25 | 8,06E-01 |
| AT3G53030 | 0,57 | 1,47 | 3,90 | 0,38 | 7,07E-01 |
| AT3G53050 | 0,05 | -0,58 | 4,38 | -0,13 | 8,95E-01 |
| AT3G53065 | 0,19 | -1,45 | 4,34 | -0,33 | 7,38E-01 |
| AT3G53090 | 0,36 | -0,51 | 4,38 | -0,12 | 9,08E-01 |
| AT3G53110 | 0,40 | -0,38 | 3,34 | -0,12 | 9,08E-01 |
| AT3G53180 | 0,20 | -0,66 | 4,32 | -0,15 | 8,78E-01 |
| AT3G53190 | 0,21 | -0,24 | 4,35 | -0,06 | 9,56E-01 |
| AT3G53230 | 1,14 | 2,81 | 2,74 | 1,03 | 3,04E-01 |
| AT3G53240 | 0,55 | 1,09 | 3,90 | 0,28 | 7,79E-01 |
| AT3G53305 | 0,72 | -0,29 | 2,00 | -0,14 | 8,87E-01 |
| AT3G53360 | 0,08 | 2,08 | 4,38 | 0,47 | 6,35E-01 |
| AT3G53390 | 0,27 | -0,47 | 4,30 | -0,11 | 9,14E-01 |
| AT3G53470 | 0,77 | -0,01 | 2,94 | 0,00 | 9,96E-01 |
| AT3G53520 | 0,83 | 2,08 | 2,53 | 0,82 | 4,12E-01 |
| AT3G53670 | 0,90 | 0,80 | 1,91 | 0,42 | 6,74E-01 |
| AT3G53700 | 0,94 | 1,03 | 1,75 | 0,59 | 5,57E-01 |
| AT3G53870 | 0,80 | 0,27 | 2,50 | 0,11 | 9,15E-01 |
| AT3G54220 | 0,50 | 2,23 | 3,15 | 0,71 | 4,79E-01 |
| AT3G54230 | 1,41 | 1,82 | 1,89 | 0,96 | 3,35E-01 |
| AT3G54366 | 1,11 | -0,05 | 1,83 | -0,03 | 9,77E-01 |
| AT3G54420 | 0,17 | 2,04 | 4,33 | 0,47 | 6,37E-01 |
| AT3G54540 | 1,42 | 3,06 | 1,76 | 1,74 | 8,24E-02 |
| AT3G54600 | 0,51 | -0,54 | 3,97 | -0,13 | 8,93E-01 |
| AT3G54610 | 0,47 | 3,71 | 3,96 | 0,94 | 3,48E-01 |
| AT3G54625 | 0,68 | 3,43 | 4,33 | 0,79 | 4,29E-01 |
| AT3G54680 | 0,75 | -0,14 | 2,85 | -0,05 | 9,62E-01 |
| AT3G54800 | 0,59 | -1,16 | 2,44 | -0,48 | 6,33E-01 |
| AT3G54820 | 0,15 | 0,69 | 4,31 | 0,16 | 8,73E-01 |
| AT3G54850 | 0,81 | 1,23 | 2,45 | 0,50 | 6,17E-01 |
| AT3G54980 | 0,41 | 1,06 | 4,21 | 0,25 | 8,01E-01 |
| AT3G55030 | 0,39 | 2,48 | 4,18 | 0,59 | 5,53E-01 |
| AT3G55250 | 0,63 | 0,32 | 2,23 | 0,14 | 8,86E-01 |
| AT3G55260 | 0,43 | -2,89 | 4,00 | -0,72 | 4,70E-01 |
| AT3G55270 | 1,69 | 3,75 | 1,80 | 2,08 | 3,73E-02 |
| AT3G55280 | 0,61 | -1,27 | 3,66 | -0,35 | 7,29E-01 |
| AT3G55370 | 0,12 | 0,63 | 4,33 | 0,15 | 8,85E-01 |
| AT3G55420 | 0,14 | -0,31 | 4,33 | -0,07 | 9,43E-01 |
| AT3G55440 | 1,26 | 0,44 | 1,55 | 0,29 | 7,74E-01 |
| AT3G55450 | 0,15 | 1,94 | 4,32 | 0,45 | 6,54E-01 |
| AT3G55460 | 1,02 | 1,74 | 1,80 | 0,97 | 3,32E-01 |
| AT3G55485 | 0,83 | -2,99 | 2,31 | -1,29 | 1,96E-01 |
| AT3G08485 | 0,83 | -2,99 | 2,31 | -1,29 | 1,96E-01 |
| AT3G55520 | 0,40 | 3,49 | 4,33 | 0,81 | 4,20E-01 |
| AT3G55620 | 0,54 | 0,78 | 2,52 | 0,31 | 7,56E-01 |
| AT3G55630 | 0,42 | 3,64 | 4,32 | 0,84 | 4,00E-01 |
| AT3G55677 | 0,24 | -1,17 | 4,35 | -0,27 | 7,89E-01 |
| AT3G55760 | 0,99 | 3,11 | 2,05 | 1,51 | 1,30E-01 |

| | | | | | |
|---|---|---|---|---|---|
| AT3G55770 | 0,98 | 1,00 | 1,86 | 0,54 | 5,89E-01 |
| AT3G55780 | 0,67 | 2,31 | 3,65 | 0,63 | 5,27E-01 |
| AT3G55880 | 0,30 | -0,02 | 3,51 | -0,01 | 9,95E-01 |
| AT3G55940 | 0,10 | 0,39 | 4,35 | 0,09 | 9,29E-01 |
| AT3G56130 | 0,33 | 0,67 | 4,28 | 0,16 | 8,75E-01 |
| AT3G56140 | 0,74 | 2,31 | 2,30 | 1,01 | 3,14E-01 |
| AT3G56180 | 0,21 | -0,24 | 4,35 | -0,06 | 9,56E-01 |
| AT3G56310 | 0,76 | 2,36 | 2,23 | 1,06 | 2,90E-01 |
| AT3G56330 | 0,52 | 3,79 | 3,84 | 0,99 | 3,24E-01 |
| AT3G56390 | 0,30 | 2,84 | 4,35 | 0,65 | 5,14E-01 |
| AT3G56450 | 0,39 | 2,04 | 4,37 | 0,47 | 6,41E-01 |
| AT3G08665 | 0,76 | 3,16 | 2,80 | 1,13 | 2,59E-01 |
| AT3G56470 | 0,12 | 2,81 | 4,34 | 0,65 | 5,18E-01 |
| AT3G56490 | 0,11 | 1,33 | 4,36 | 0,31 | 7,59E-01 |
| AT3G56590 | 0,57 | -1,13 | 2,95 | -0,38 | 7,01E-01 |
| AT3G56640 | 0,59 | 1,92 | 3,05 | 0,63 | 5,30E-01 |
| AT3G56650 | 0,53 | 0,83 | 4,00 | 0,21 | 8,36E-01 |
| AT3G56720 | 0,17 | 0,30 | 4,37 | 0,07 | 9,44E-01 |
| AT3G56740 | 0,49 | 1,60 | 3,94 | 0,41 | 6,85E-01 |
| AT3G56780 | 0,32 | 0,40 | 4,35 | 0,09 | 9,27E-01 |
| AT3G56840 | 0,64 | 1,87 | 2,38 | 0,78 | 4,33E-01 |
| AT3G56880 | 1,69 | 0,61 | 1,25 | 0,49 | 6,24E-01 |
| AT3G08785 | 1,51 | 2,48 | 1,83 | 1,35 | 1,77E-01 |
| AT3G56960 | 0,60 | 2,14 | 4,38 | 0,49 | 6,25E-01 |
| AT3G57040 | 1,37 | -1,32 | 1,60 | -0,83 | 4,09E-01 |
| AT3G57062 | 1,47 | 2,32 | 1,64 | 1,41 | 1,58E-01 |
| AT3G57070 | 0,24 | -2,23 | 4,30 | -0,52 | 6,04E-01 |
| AT3G57080 | 0,21 | 2,61 | 4,34 | 0,60 | 5,48E-01 |
| AT3G57090 | 0,59 | 0,26 | 2,08 | 0,13 | 8,99E-01 |
| AT3G08845 | 0,12 | 1,15 | 4,35 | 0,26 | 7,92E-01 |
| AT3G57230 | 0,38 | 0,25 | 2,79 | 0,09 | 9,29E-01 |
| AT3G57340 | 0,42 | 1,01 | 3,34 | 0,30 | 7,62E-01 |
| AT3G57390 | 0,11 | 1,33 | 4,36 | 0,31 | 7,59E-01 |
| AT3G57450 | 1,58 | 0,13 | 1,34 | 0,10 | 9,22E-01 |
| AT3G57550 | 0,50 | 1,24 | 3,07 | 0,40 | 6,86E-01 |
| AT3G57570 | 0,40 | 0,71 | 4,23 | 0,17 | 8,67E-01 |
| AT3G57587 | 0,73 | -0,70 | 2,05 | -0,34 | 7,33E-01 |
| AT3G57650 | 0,53 | 0,59 | 2,50 | 0,24 | 8,13E-01 |
| AT3G57700 | 1,50 | 0,80 | 1,24 | 0,65 | 5,18E-01 |
| AT3G08975 | 0,42 | 2,82 | 4,11 | 0,68 | 4,94E-01 |
| AT3G57760 | 0,08 | 2,33 | 4,36 | 0,53 | 5,93E-01 |
| AT3G57770 | 1,17 | -0,77 | 1,45 | -0,54 | 5,92E-01 |
| AT3G57810 | 0,21 | 1,07 | 4,33 | 0,25 | 8,06E-01 |
| AT3G57830 | 0,31 | 0,03 | 4,34 | 0,01 | 9,95E-01 |
| AT3G57890 | 1,04 | 0,95 | 1,86 | 0,51 | 6,09E-01 |
| AT3G57940 | 0,20 | -0,66 | 4,32 | -0,15 | 8,78E-01 |
| AT3G57970 | 0,17 | 3,16 | 4,33 | 0,73 | 4,66E-01 |
| AT3G58020 | 0,08 | 2,33 | 4,36 | 0,53 | 5,93E-01 |
| AT3G58110 | 1,07 | 4,76 | 3,17 | 1,50 | 1,33E-01 |
| AT3G58160 | 0,08 | 2,08 | 4,38 | 0,47 | 6,35E-01 |
| AT3G58490 | 0,30 | -0,88 | 4,27 | -0,21 | 8,37E-01 |
| AT3G58510 | 0,99 | 0,62 | 1,82 | 0,34 | 7,34E-01 |
| AT3G58530 | 0,46 | 0,40 | 3,99 | 0,10 | 9,20E-01 |
| AT3G58570 | 0,71 | 1,16 | 2,38 | 0,49 | 6,27E-01 |
| AT3G58680 | 0,61 | -0,80 | 2,55 | -0,31 | 7,54E-01 |
| AT3G58690 | 0,58 | 1,60 | 3,10 | 0,52 | 6,06E-01 |

| | | | | | |
|---|---|---|---|---|---|
| AT3G58700 | 0,20 | 2,11 | 4,37 | 0,48 | 6,29E-01 |
| AT3G58795 | 0,23 | -0,91 | 4,37 | -0,21 | 8,36E-01 |
| AT3G59030 | 0,64 | 3,81 | 3,65 | 1,04 | 2,96E-01 |
| AT3G59080 | 0,19 | 0,10 | 4,37 | 0,02 | 9,82E-01 |
| AT3G59090 | 0,80 | 1,32 | 2,46 | 0,54 | 5,91E-01 |
| AT3G59100 | 0,79 | 0,90 | 2,88 | 0,31 | 7,54E-01 |
| AT3G59110 | 0,41 | 0,01 | 4,19 | 0,00 | 9,98E-01 |
| AT3G59210 | 0,24 | 0,93 | 4,29 | 0,22 | 8,29E-01 |
| AT3G59290 | 1,04 | 0,37 | 1,76 | 0,21 | 8,34E-01 |
| AT3G59300 | 0,19 | 0,37 | 4,28 | 0,09 | 9,31E-01 |
| AT3G59350 | 0,86 | 2,65 | 2,40 | 1,10 | 2,70E-01 |
| AT3G59380 | 0,24 | 0,93 | 4,29 | 0,22 | 8,29E-01 |
| AT3G59500 | 0,47 | -0,52 | 3,39 | -0,15 | 8,78E-01 |
| AT3G59520 | 0,08 | 2,33 | 4,36 | 0,53 | 5,93E-01 |
| AT3G59530 | 0,53 | 1,65 | 4,38 | 0,38 | 7,07E-01 |
| AT3G59670 | 0,30 | -1,21 | 4,35 | -0,28 | 7,81E-01 |
| AT3G59700 | 0,10 | -1,26 | 4,35 | -0,29 | 7,72E-01 |
| AT3G59810 | 0,46 | 1,73 | 4,40 | 0,39 | 6,93E-01 |
| AT3G59920 | 0,37 | -0,71 | 4,27 | -0,17 | 8,67E-01 |
| AT3G60080 | 1,12 | -0,22 | 1,70 | -0,13 | 8,98E-01 |
| AT3G60110 | 0,48 | 0,61 | 4,36 | 0,14 | 8,88E-01 |
| AT3G60130 | 0,63 | 2,55 | 3,17 | 0,80 | 4,22E-01 |
| AT3G60140 | 0,58 | 2,69 | 3,81 | 0,71 | 4,80E-01 |
| AT3G60240 | 1,46 | 0,70 | 1,31 | 0,54 | 5,90E-01 |
| AT3G60245 | 0,58 | 0,43 | 2,25 | 0,19 | 8,48E-01 |
| AT3G60290 | 0,27 | -1,08 | 4,36 | -0,25 | 8,04E-01 |
| AT3G60300 | 0,58 | 0,92 | 2,61 | 0,35 | 7,25E-01 |
| AT3G60310 | 0,49 | 2,08 | 3,89 | 0,53 | 5,93E-01 |
| AT3G60410 | 0,24 | 2,80 | 4,34 | 0,65 | 5,18E-01 |
| AT3G60420 | 0,55 | 3,69 | 3,81 | 0,97 | 3,33E-01 |
| AT3G60440 | 0,83 | 1,96 | 3,47 | 0,57 | 5,71E-01 |
| AT3G60450 | 1,00 | 4,69 | 2,60 | 1,81 | 7,08E-02 |
| AT3G60500 | 0,28 | 1,78 | 4,33 | 0,41 | 6,80E-01 |
| AT3G60530 | 0,83 | -2,12 | 2,01 | -1,06 | 2,90E-01 |
| AT3G60600 | 1,58 | 3,62 | 1,65 | 2,20 | 2,77E-02 |
| AT3G60690 | 1,35 | 1,28 | 1,56 | 0,82 | 4,12E-01 |
| AT3G60910 | 0,08 | -0,87 | 4,38 | -0,20 | 8,43E-01 |
| AT3G60950 | 1,11 | -1,78 | 1,86 | -0,96 | 3,37E-01 |
| AT3G60960 | 1,54 | 0,24 | 1,44 | 0,17 | 8,68E-01 |
| AT3G09515 | 1,27 | 0,15 | 1,69 | 0,09 | 9,27E-01 |
| AT3G60961 | 1,33 | 4,44 | 2,04 | 2,18 | 2,93E-02 |
| AT3G09525 | 0,12 | 2,81 | 4,34 | 0,65 | 5,18E-01 |
| AT3G60970 | 0,12 | 2,81 | 4,34 | 0,65 | 5,18E-01 |
| AT3G61030 | 0,61 | -0,36 | 1,97 | -0,18 | 8,56E-01 |
| AT3G61050 | 1,51 | -0,13 | 1,59 | -0,08 | 9,35E-01 |
| AT3G61080 | 0,12 | 2,87 | 4,34 | 0,66 | 5,08E-01 |
| AT3G61110 | 1,40 | 1,14 | 1,71 | 0,67 | 5,03E-01 |
| AT3G61130 | 1,07 | -0,48 | 2,36 | -0,20 | 8,40E-01 |
| AT3G61140 | 0,73 | 0,60 | 1,93 | 0,31 | 7,55E-01 |
| AT3G61150 | 0,68 | 2,77 | 2,77 | 1,00 | 3,18E-01 |
| AT3G61180 | 0,94 | 1,99 | 2,38 | 0,83 | 4,04E-01 |
| AT3G61240 | 1,00 | 0,70 | 1,73 | 0,40 | 6,86E-01 |
| AT3G61280 | 0,41 | 1,43 | 3,29 | 0,43 | 6,64E-01 |
| AT3G61320 | 0,08 | -0,56 | 4,38 | -0,13 | 8,99E-01 |
| AT3G61440 | 1,04 | 1,69 | 2,24 | 0,76 | 4,50E-01 |
| AT3G61480 | 0,44 | 2,88 | 3,28 | 0,88 | 3,80E-01 |

| | | | | | |
|---|---|---|---|---|---|
| AT3G61490 | 0,55 | 2,14 | 4,38 | 0,49 | 6,25E-01 |
| AT3G61540 | 0,29 | 1,14 | 4,28 | 0,27 | 7,89E-01 |
| AT3G61550 | 0,37 | 0,97 | 3,40 | 0,29 | 7,75E-01 |
| AT3G61600 | 1,22 | 3,79 | 1,80 | 2,10 | 3,53E-02 |
| AT3G61670 | 0,29 | 1,74 | 4,31 | 0,40 | 6,87E-01 |
| AT3G61700 | 1,46 | -1,80 | 1,47 | -1,22 | 2,21E-01 |
| AT3G61755 | 0,16 | 0,33 | 4,31 | 0,08 | 9,39E-01 |
| AT3G61810 | 0,05 | -0,58 | 4,38 | -0,13 | 8,95E-01 |
| AT3G61850 | 0,15 | 0,69 | 4,31 | 0,16 | 8,73E-01 |
| AT3G61870 | 1,16 | -0,45 | 1,52 | -0,30 | 7,65E-01 |
| AT3G61950 | 0,11 | 1,33 | 4,36 | 0,31 | 7,59E-01 |
| AT3G62030 | 0,98 | -0,79 | 2,14 | -0,37 | 7,11E-01 |
| AT3G62130 | 0,39 | -0,61 | 4,29 | -0,14 | 8,86E-01 |
| AT3G62190 | 0,24 | 1,12 | 4,33 | 0,26 | 7,96E-01 |
| AT3G62245 | 0,42 | -1,40 | 4,01 | -0,35 | 7,26E-01 |
| AT3G62300 | 0,93 | 2,75 | 3,42 | 0,80 | 4,21E-01 |
| AT3G62370 | 0,90 | 2,25 | 2,04 | 1,10 | 2,70E-01 |
| AT3G62530 | 0,72 | 1,55 | 2,28 | 0,68 | 4,96E-01 |
| AT3G62550 | 0,70 | 1,49 | 3,74 | 0,40 | 6,89E-01 |
| AT3G62560 | 0,61 | -0,39 | 3,12 | -0,13 | 9,00E-01 |
| AT3G62580 | 0,64 | 0,40 | 2,05 | 0,19 | 8,46E-01 |
| AT3G09815 | 0,08 | 2,08 | 4,38 | 0,47 | 6,35E-01 |
| AT3G09835 | 1,22 | -2,43 | 1,77 | -1,38 | 1,69E-01 |
| AT3G62620 | 0,29 | -1,08 | 4,34 | -0,25 | 8,03E-01 |
| AT3G62650 | 1,34 | 1,30 | 1,63 | 0,79 | 4,27E-01 |
| AT3G62660 | 1,25 | -0,61 | 1,54 | -0,40 | 6,92E-01 |
| AT3G62720 | 0,40 | 0,59 | 4,01 | 0,15 | 8,83E-01 |
| AT3G62750 | 0,68 | -1,98 | 2,45 | -0,81 | 4,19E-01 |
| AT3G62820 | 0,26 | -0,55 | 4,31 | -0,13 | 8,98E-01 |
| AT3G62830 | 0,86 | 0,67 | 1,79 | 0,38 | 7,06E-01 |
| AT3G62890 | 0,20 | 0,36 | 4,30 | 0,08 | 9,34E-01 |
| AT3G63000 | 0,33 | 2,42 | 4,31 | 0,56 | 5,74E-01 |
| AT3G63010 | 0,38 | 0,18 | 3,38 | 0,05 | 9,59E-01 |
| AT3G63080 | 0,53 | 0,66 | 2,61 | 0,25 | 8,00E-01 |
| AT3G63130 | 0,43 | 0,54 | 4,09 | 0,13 | 8,94E-01 |
| AT3G63170 | 0,32 | -0,22 | 4,40 | -0,05 | 9,61E-01 |
| AT3G63215 | 0,08 | -0,56 | 4,38 | -0,13 | 8,99E-01 |
| AT3G63300 | 0,72 | 2,84 | 2,88 | 0,99 | 3,24E-01 |
| AT3G63370 | 0,57 | 2,08 | 3,10 | 0,67 | 5,02E-01 |
| AT3G63400 | 0,95 | 1,76 | 2,08 | 0,85 | 3,98E-01 |
| AT3G63420 | 0,46 | 3,13 | 4,33 | 0,72 | 4,70E-01 |
| AT3G63470 | 0,42 | 2,85 | 4,17 | 0,68 | 4,95E-01 |
| AT3G09995 | 0,45 | 3,59 | 4,03 | 0,89 | 3,72E-01 |
| AT3G63520 | 0,87 | 0,57 | 2,25 | 0,25 | 8,00E-01 |
| AT4G00030 | 0,38 | -0,11 | 4,34 | -0,02 | 9,81E-01 |
| AT4G00100 | 1,39 | 1,37 | 1,93 | 0,71 | 4,80E-01 |
| AT4G00163 | 0,65 | -1,73 | 2,64 | -0,66 | 5,11E-01 |
| AT4G00230 | 0,69 | 1,86 | 4,40 | 0,42 | 6,73E-01 |
| AT4G00300 | 1,08 | -2,38 | 1,76 | -1,35 | 1,78E-01 |
| AT4G00340 | 0,31 | -0,60 | 4,32 | -0,14 | 8,91E-01 |
| AT4G00440 | 0,33 | 1,91 | 4,34 | 0,44 | 6,59E-01 |
| AT4G00490 | 0,44 | 2,07 | 4,35 | 0,48 | 6,34E-01 |
| AT4G00580 | 0,48 | -1,20 | 3,94 | -0,30 | 7,61E-01 |
| AT4G00660 | 1,18 | 2,68 | 1,96 | 1,37 | 1,72E-01 |
| AT4G00710 | 0,48 | 2,96 | 4,34 | 0,68 | 4,95E-01 |
| AT4G00750 | 0,96 | 0,05 | 1,62 | 0,03 | 9,74E-01 |

| | | | | | |
|---|---|---|---|---|---|
| AT4G00820 | 0,05 | -0,58 | 4,38 | -0,13 | 8,95E-01 |
| AT4G00830 | 0,92 | 0,51 | 2,03 | 0,25 | 8,01E-01 |
| AT4G00889 | 0,23 | -2,41 | 4,30 | -0,56 | 5,75E-01 |
| AT4G00930 | 0,17 | 3,14 | 4,33 | 0,73 | 4,68E-01 |
| AT4G00953 | 0,67 | 0,38 | 2,52 | 0,15 | 8,79E-01 |
| AT4G00955 | 0,42 | -0,80 | 2,76 | -0,29 | 7,71E-01 |
| AT4G00960 | 1,30 | -2,89 | 1,91 | -1,51 | 1,31E-01 |
| AT4G00970 | 0,33 | -1,68 | 4,27 | -0,39 | 6,94E-01 |
| AT4G00990 | 0,61 | 2,23 | 3,92 | 0,57 | 5,69E-01 |
| AT4G01020 | 0,37 | 0,68 | 4,29 | 0,16 | 8,74E-01 |
| AT4G01026 | 0,56 | 1,27 | 2,51 | 0,50 | 6,14E-01 |
| AT4G01040 | 0,12 | 1,65 | 4,34 | 0,38 | 7,04E-01 |
| AT4G01100 | 0,48 | -0,21 | 3,07 | -0,07 | 9,44E-01 |
| AT4G01130 | 0,11 | -1,28 | 4,35 | -0,29 | 7,69E-01 |
| AT4G01280 | 0,59 | 0,87 | 3,07 | 0,28 | 7,77E-01 |
| AT4G01320 | 0,90 | 3,06 | 2,13 | 1,43 | 1,52E-01 |
| AT4G01330 | 0,80 | -0,32 | 2,30 | -0,14 | 8,90E-01 |
| AT4G01340 | 1,48 | 0,29 | 1,64 | 0,18 | 8,59E-01 |
| AT4G01360 | 0,89 | 0,27 | 2,75 | 0,10 | 9,21E-01 |
| AT4G01370 | 0,27 | 0,51 | 4,25 | 0,12 | 9,04E-01 |
| AT4G01460 | 0,10 | 0,34 | 4,35 | 0,08 | 9,37E-01 |
| AT4G01570 | 0,24 | 2,03 | 4,32 | 0,47 | 6,38E-01 |
| AT4G03905 | 0,55 | 0,49 | 3,79 | 0,13 | 8,97E-01 |
| AT4G03925 | 0,76 | -0,08 | 2,10 | -0,04 | 9,69E-01 |
| AT4G01610 | 0,62 | 0,11 | 2,28 | 0,05 | 9,61E-01 |
| AT4G01690 | 1,14 | 0,82 | 1,63 | 0,51 | 6,14E-01 |
| AT4G01720 | 0,44 | 0,72 | 4,13 | 0,17 | 8,61E-01 |
| AT4G01995 | 0,08 | 2,08 | 4,38 | 0,47 | 6,35E-01 |
| AT4G02000 | 0,69 | -0,45 | 2,02 | -0,22 | 8,25E-01 |
| AT4G02010 | 0,14 | 0,78 | 4,34 | 0,18 | 8,57E-01 |
| AT4G02020 | 0,55 | 0,63 | 3,13 | 0,20 | 8,40E-01 |
| AT4G02030 | 0,75 | 3,44 | 2,78 | 1,23 | 2,17E-01 |
| AT4G02070 | 0,12 | 2,69 | 4,35 | 0,62 | 5,36E-01 |
| AT4G02080 | 0,57 | 2,46 | 2,61 | 0,94 | 3,46E-01 |
| AT4G02100 | 0,08 | -0,56 | 4,38 | -0,13 | 8,99E-01 |
| AT4G02110 | 0,21 | 3,22 | 4,33 | 0,74 | 4,58E-01 |
| AT4G02200 | 0,55 | 1,10 | 3,76 | 0,29 | 7,70E-01 |
| AT4G02220 | 0,29 | 1,03 | 4,31 | 0,24 | 8,10E-01 |
| AT4G02260 | 0,56 | -0,41 | 2,96 | -0,14 | 8,89E-01 |
| AT4G02330 | 0,22 | -1,07 | 4,36 | -0,25 | 8,06E-01 |
| AT4G02340 | 1,02 | -0,27 | 2,17 | -0,12 | 9,01E-01 |
| AT4G02370 | 0,72 | 0,26 | 2,96 | 0,09 | 9,31E-01 |
| AT4G02390 | 0,10 | 0,34 | 4,35 | 0,08 | 9,37E-01 |
| AT4G02400 | 0,19 | 1,14 | 4,29 | 0,26 | 7,91E-01 |
| AT4G02440 | 0,90 | -1,00 | 1,74 | -0,58 | 5,64E-01 |
| AT4G04135 | 0,05 | -0,58 | 4,38 | -0,13 | 8,95E-01 |
| AT4G02500 | 1,37 | 3,50 | 2,73 | 1,28 | 2,01E-01 |
| AT4G04195 | 0,38 | 0,01 | 3,24 | 0,00 | 9,97E-01 |
| AT4G02540 | 0,29 | -0,28 | 3,66 | -0,08 | 9,39E-01 |
| AT4G02580 | 1,19 | 0,58 | 2,19 | 0,27 | 7,91E-01 |
| AT4G02600 | 0,66 | -0,15 | 2,21 | -0,07 | 9,44E-01 |
| AT4G02610 | 0,31 | 0,91 | 4,32 | 0,21 | 8,32E-01 |
| AT4G02650 | 0,24 | 2,80 | 4,34 | 0,65 | 5,18E-01 |
| AT4G02700 | 1,40 | -0,26 | 1,58 | -0,16 | 8,71E-01 |
| AT4G02725 | 0,69 | 1,18 | 3,01 | 0,39 | 6,96E-01 |
| AT4G02790 | 0,08 | 2,33 | 4,36 | 0,53 | 5,93E-01 |

| | | | | | |
|---|---|---|---|---|---|
| AT4G02880 | 0,95 | 0,92 | 2,00 | 0,46 | 6,47E-01 |
| AT4G02920 | 1,02 | 2,84 | 2,19 | 1,30 | 1,95E-01 |
| AT4G02940 | 0,12 | -0,77 | 4,36 | -0,18 | 8,61E-01 |
| AT4G02990 | 0,69 | -1,23 | 2,50 | -0,49 | 6,22E-01 |
| AT4G03000 | 0,64 | 2,26 | 2,94 | 0,77 | 4,42E-01 |
| AT4G04425 | 0,08 | 2,08 | 4,38 | 0,47 | 6,35E-01 |
| AT4G03100 | 0,38 | -0,41 | 4,37 | -0,09 | 9,24E-01 |
| AT4G03120 | 0,23 | 1,51 | 4,34 | 0,35 | 7,28E-01 |
| AT4G03180 | 0,27 | 1,57 | 4,36 | 0,36 | 7,19E-01 |
| AT4G03200 | 0,82 | 3,95 | 2,37 | 1,67 | 9,51E-02 |
| AT4G03205 | 0,48 | 1,61 | 3,18 | 0,51 | 6,13E-01 |
| AT4G03210 | 0,53 | 3,29 | 4,33 | 0,76 | 4,47E-01 |
| AT4G03285 | 0,16 | 1,18 | 4,32 | 0,27 | 7,85E-01 |
| AT4G03410 | 0,58 | -2,36 | 3,75 | -0,63 | 5,30E-01 |
| AT4G03500 | 0,47 | -0,19 | 4,35 | -0,04 | 9,65E-01 |
| AT4G03830 | 0,61 | -0,46 | 2,43 | -0,19 | 8,49E-01 |
| AT4G03930 | 0,65 | 2,83 | 2,98 | 0,95 | 3,42E-01 |
| AT4G03940 | 0,17 | 2,04 | 4,33 | 0,47 | 6,37E-01 |
| AT4G03945 | 0,92 | 0,27 | 2,08 | 0,13 | 8,98E-01 |
| AT4G04110 | 0,65 | 0,63 | 4,39 | 0,14 | 8,86E-01 |
| AT4G04180 | 0,08 | -0,56 | 4,38 | -0,13 | 8,99E-01 |
| AT4G04210 | 0,39 | 0,43 | 2,74 | 0,16 | 8,76E-01 |
| AT4G04220 | 0,89 | 1,23 | 2,01 | 0,61 | 5,40E-01 |
| AT4G04795 | 0,87 | -0,83 | 2,29 | -0,36 | 7,19E-01 |
| AT4G04315 | 0,10 | 0,34 | 4,35 | 0,08 | 9,37E-01 |
| AT4G04350 | 0,42 | -0,56 | 3,36 | -0,17 | 8,67E-01 |
| AT4G04450 | 0,17 | -0,14 | 4,41 | -0,03 | 9,75E-01 |
| AT4G04510 | 0,17 | -0,44 | 4,37 | -0,10 | 9,19E-01 |
| AT4G04540 | 0,26 | 0,26 | 4,34 | 0,06 | 9,52E-01 |
| AT4G04610 | 0,73 | 0,39 | 2,00 | 0,20 | 8,44E-01 |
| AT4G04670 | 0,48 | 3,50 | 4,33 | 0,81 | 4,19E-01 |
| AT4G04750 | 0,90 | -0,49 | 1,71 | -0,29 | 7,75E-01 |
| AT4G04760 | 1,18 | -1,92 | 1,93 | -0,99 | 3,21E-01 |
| AT4G04800 | 0,52 | 2,17 | 3,84 | 0,56 | 5,73E-01 |
| AT4G04840 | 1,00 | -1,98 | 1,99 | -1,00 | 3,19E-01 |
| AT4G04850 | 0,79 | 1,52 | 1,90 | 0,80 | 4,24E-01 |
| AT4G04860 | 0,28 | 1,11 | 3,65 | 0,30 | 7,62E-01 |
| AT4G04920 | 1,70 | 2,85 | 1,58 | 1,80 | 7,25E-02 |
| AT4G04960 | 0,39 | 0,31 | 2,73 | 0,11 | 9,09E-01 |
| AT4G05020 | 0,40 | 0,94 | 4,19 | 0,22 | 8,22E-01 |
| AT4G05080 | 0,23 | -2,25 | 4,30 | -0,52 | 6,00E-01 |
| AT4G05105 | 0,17 | 2,95 | 4,34 | 0,68 | 4,96E-01 |
| AT4G05160 | 0,46 | 2,51 | 4,03 | 0,62 | 5,33E-01 |
| AT4G05612 | 0,12 | 2,81 | 4,34 | 0,65 | 5,18E-01 |
| AT4G05632 | 0,08 | -0,56 | 4,38 | -0,13 | 8,99E-01 |
| AT4G06479 | 0,10 | -1,26 | 4,35 | -0,29 | 7,72E-01 |
| AT4G06598 | 0,74 | -0,14 | 3,57 | -0,04 | 9,70E-01 |
| AT4G06688 | 0,68 | 0,67 | 2,27 | 0,29 | 7,68E-01 |
| AT4G07390 | 0,27 | 0,31 | 3,64 | 0,08 | 9,33E-01 |
| AT4G07455 | 0,24 | 0,00 | 4,41 | 0,00 | 1,00E+00 |
| AT4G07825 | 1,05 | 1,40 | 2,58 | 0,54 | 5,88E-01 |
| AT4G07932 | 0,51 | -2,57 | 4,03 | -0,64 | 5,25E-01 |
| AT4G07965 | 0,15 | 1,94 | 4,32 | 0,45 | 6,54E-01 |
| AT4G08025 | 0,16 | 0,85 | 4,31 | 0,20 | 8,44E-01 |
| AT4G05305 | 0,12 | -1,57 | 4,34 | -0,36 | 7,18E-01 |
| AT4G08097 | 0,66 | -1,37 | 2,42 | -0,57 | 5,71E-01 |

| | | | | | |
|---|---|---|---|---|---|
| AT4G08180 | 1,01 | 2,99 | 2,11 | 1,41 | 1,57E-01 |
| AT4G08267 | 0,63 | 1,65 | 4,36 | 0,38 | 7,05E-01 |
| AT4G08270 | 0,17 | 1,41 | 4,35 | 0,32 | 7,46E-01 |
| AT4G08320 | 0,31 | -2,20 | 4,30 | -0,51 | 6,09E-01 |
| AT4G08350 | 0,65 | 0,73 | 2,40 | 0,30 | 7,62E-01 |
| AT4G08395 | 0,29 | -1,27 | 4,26 | -0,30 | 7,66E-01 |
| AT4G05455 | 0,68 | -1,14 | 2,45 | -0,47 | 6,40E-01 |
| AT4G08450 | 0,11 | -1,08 | 4,37 | -0,25 | 8,05E-01 |
| AT4G08455 | 0,63 | 2,83 | 4,31 | 0,66 | 5,11E-01 |
| AT4G08510 | 0,33 | 2,30 | 4,32 | 0,53 | 5,95E-01 |
| AT4G08545 | 1,36 | -0,59 | 1,46 | -0,41 | 6,83E-01 |
| AT4G08560 | 0,64 | 0,00 | 4,41 | 0,00 | 1,00E+00 |
| AT4G08780 | 0,43 | 1,02 | 4,15 | 0,25 | 8,06E-01 |
| AT4G05605 | 0,39 | -2,11 | 4,31 | -0,49 | 6,24E-01 |
| AT4G08930 | 0,71 | 3,52 | 2,37 | 1,48 | 1,38E-01 |
| AT4G08950 | 0,37 | -0,44 | 4,25 | -0,10 | 9,17E-01 |
| AT4G08980 | 0,65 | 1,26 | 2,77 | 0,45 | 6,50E-01 |
| AT4G08990 | 0,69 | -1,15 | 3,55 | -0,32 | 7,46E-01 |
| AT4G09020 | 1,04 | 0,58 | 1,76 | 0,33 | 7,41E-01 |
| AT4G09030 | 0,16 | 0,85 | 4,31 | 0,20 | 8,44E-01 |
| AT4G09150 | 0,62 | 2,27 | 2,91 | 0,78 | 4,34E-01 |
| AT4G09160 | 0,36 | 0,00 | 4,41 | 0,00 | 1,00E+00 |
| AT4G09250 | 0,12 | 1,65 | 4,34 | 0,38 | 7,04E-01 |
| AT4G09310 | 0,21 | 2,65 | 4,34 | 0,61 | 5,42E-01 |
| AT4G05705 | 0,10 | -1,26 | 4,35 | -0,29 | 7,72E-01 |
| AT4G09320 | 1,06 | -0,23 | 1,44 | -0,16 | 8,73E-01 |
| AT4G09360 | 0,71 | -0,57 | 2,21 | -0,26 | 7,94E-01 |
| AT4G09450 | 1,44 | 1,57 | 2,00 | 0,79 | 4,31E-01 |
| AT4G09460 | 0,37 | 1,99 | 4,30 | 0,46 | 6,43E-01 |
| AT4G09545 | 0,31 | -2,29 | 4,30 | -0,53 | 5,94E-01 |
| AT4G09570 | 0,54 | -1,12 | 2,27 | -0,49 | 6,23E-01 |
| AT4G09630 | 0,87 | -1,89 | 1,95 | -0,97 | 3,32E-01 |
| AT4G09660 | 0,30 | -1,67 | 4,27 | -0,39 | 6,96E-01 |
| AT4G09680 | 0,45 | 3,33 | 3,98 | 0,84 | 4,03E-01 |
| AT4G09690 | 1,50 | -1,00 | 1,60 | -0,63 | 5,31E-01 |
| AT4G09730 | 0,51 | 0,36 | 3,20 | 0,11 | 9,10E-01 |
| AT4G09800 | 0,41 | -0,83 | 4,26 | -0,19 | 8,46E-01 |
| AT4G09820 | 0,31 | -0,79 | 4,39 | -0,18 | 8,56E-01 |
| AT4G09830 | 0,41 | 2,07 | 3,36 | 0,62 | 5,38E-01 |
| AT4G09850 | 0,21 | 0,41 | 4,33 | 0,09 | 9,25E-01 |
| AT4G05745 | 0,45 | 0,52 | 3,04 | 0,17 | 8,63E-01 |
| AT4G09890 | 0,18 | 1,48 | 4,31 | 0,34 | 7,32E-01 |
| AT4G10060 | 0,79 | 1,15 | 2,04 | 0,56 | 5,74E-01 |
| AT4G10170 | 0,08 | 2,33 | 4,36 | 0,53 | 5,93E-01 |
| AT4G10180 | 0,21 | 2,34 | 4,32 | 0,54 | 5,89E-01 |
| AT4G10465 | 0,60 | -0,60 | 2,41 | -0,25 | 8,04E-01 |
| AT4G10610 | 1,30 | -0,18 | 2,10 | -0,09 | 9,30E-01 |
| AT4G10613 | 0,77 | -0,59 | 1,86 | -0,32 | 7,51E-01 |
| AT4G10620 | 0,24 | 0,00 | 4,41 | 0,00 | 1,00E+00 |
| AT4G10660 | 0,11 | -1,28 | 4,35 | -0,29 | 7,69E-01 |
| AT4G10840 | 1,51 | 2,50 | 1,88 | 1,33 | 1,85E-01 |
| AT4G11000 | 0,26 | 1,27 | 4,31 | 0,29 | 7,68E-01 |
| AT4G11070 | 0,37 | -0,36 | 4,37 | -0,08 | 9,34E-01 |
| AT4G11100 | 0,32 | 0,73 | 4,27 | 0,17 | 8,65E-01 |
| AT4G11110 | 0,51 | 1,04 | 2,59 | 0,40 | 6,88E-01 |
| AT4G11130 | 0,54 | 2,69 | 3,91 | 0,69 | 4,91E-01 |

| | | | | | |
|---|---|---|---|---|---|
| AT4G11150 | 1,03 | -0,94 | 1,56 | -0,60 | 5,48E-01 |
| AT4G11160 | 1,61 | 1,79 | 1,59 | 1,13 | 2,59E-01 |
| AT4G11170 | 0,50 | -1,42 | 3,14 | -0,45 | 6,51E-01 |
| AT4G11240 | 1,11 | 3,14 | 1,98 | 1,59 | 1,13E-01 |
| AT4G11280 | 0,67 | -0,74 | 2,99 | -0,25 | 8,05E-01 |
| AT4G11330 | 0,53 | 1,56 | 3,16 | 0,49 | 6,22E-01 |
| AT4G11350 | 0,19 | 1,04 | 4,29 | 0,24 | 8,09E-01 |
| AT4G11360 | 0,31 | 2,14 | 4,38 | 0,49 | 6,25E-01 |
| AT4G05885 | 0,80 | 0,80 | 2,78 | 0,29 | 7,73E-01 |
| AT4G11660 | 0,57 | 0,33 | 3,11 | 0,11 | 9,16E-01 |
| AT4G11720 | 0,60 | 0,45 | 3,00 | 0,15 | 8,81E-01 |
| AT4G11740 | 0,35 | -0,19 | 4,26 | -0,05 | 9,64E-01 |
| AT4G11880 | 0,28 | -1,30 | 4,35 | -0,30 | 7,65E-01 |
| AT4G11890 | 0,62 | 3,20 | 3,00 | 1,07 | 2,85E-01 |
| AT4G11910 | 0,12 | 2,87 | 4,34 | 0,66 | 5,08E-01 |
| AT4G12020 | 1,23 | 3,66 | 2,45 | 1,50 | 1,34E-01 |
| AT4G12030 | 0,68 | 3,82 | 3,45 | 1,11 | 2,68E-01 |
| AT4G12040 | 1,45 | 2,14 | 1,63 | 1,32 | 1,88E-01 |
| AT4G12070 | 0,19 | -1,34 | 4,35 | -0,31 | 7,58E-01 |
| AT4G12240 | 0,26 | 2,03 | 4,29 | 0,47 | 6,36E-01 |
| AT4G05985 | 0,10 | 0,34 | 4,35 | 0,08 | 9,37E-01 |
| AT4G12390 | 0,50 | 0,10 | 4,36 | 0,02 | 9,82E-01 |
| AT4G12600 | 0,92 | 3,17 | 2,94 | 1,08 | 2,80E-01 |
| AT4G12640 | 0,26 | 1,18 | 3,71 | 0,32 | 7,51E-01 |
| AT4G12680 | 0,21 | 2,82 | 4,35 | 0,65 | 5,16E-01 |
| AT4G12690 | 0,12 | 1,15 | 4,35 | 0,26 | 7,92E-01 |
| AT4G12720 | 1,54 | 0,39 | 1,42 | 0,27 | 7,85E-01 |
| AT4G12830 | 0,34 | -1,15 | 4,35 | -0,26 | 7,91E-01 |
| AT4G12900 | 0,05 | -0,58 | 4,38 | -0,13 | 8,95E-01 |
| AT4G05995 | 0,98 | -1,00 | 2,24 | -0,45 | 6,54E-01 |
| AT4G13010 | 0,63 | -1,28 | 3,64 | -0,35 | 7,26E-01 |
| AT4G13020 | 0,54 | 1,63 | 3,11 | 0,53 | 5,99E-01 |
| AT4G06010 | 0,28 | 1,09 | 4,31 | 0,25 | 8,00E-01 |
| AT4G06015 | 0,10 | 0,39 | 4,35 | 0,09 | 9,29E-01 |
| AT4G13245 | 0,48 | -1,99 | 3,19 | -0,62 | 5,33E-01 |
| AT4G13270 | 0,50 | 2,75 | 3,14 | 0,88 | 3,80E-01 |
| AT4G13390 | 1,53 | -1,77 | 1,61 | -1,10 | 2,72E-01 |
| AT4G13440 | 0,58 | 0,87 | 3,95 | 0,22 | 8,25E-01 |
| AT4G13530 | 0,95 | 1,95 | 1,91 | 1,02 | 3,07E-01 |
| AT4G13572 | 0,31 | 0,00 | 4,41 | 0,00 | 1,00E+00 |
| AT4G13610 | 0,37 | 0,04 | 2,72 | 0,02 | 9,88E-01 |
| AT4G13615 | 0,92 | 2,10 | 1,85 | 1,13 | 2,57E-01 |
| AT4G13670 | 0,92 | 4,83 | 3,14 | 1,54 | 1,24E-01 |
| AT4G13730 | 0,24 | 0,39 | 4,34 | 0,09 | 9,29E-01 |
| AT4G13750 | 0,39 | 0,04 | 3,27 | 0,01 | 9,90E-01 |
| AT4G13760 | 0,54 | 0,69 | 3,02 | 0,23 | 8,20E-01 |
| AT4G13830 | 0,70 | -0,88 | 2,81 | -0,31 | 7,53E-01 |
| AT4G13850 | 0,81 | 0,67 | 2,01 | 0,33 | 7,39E-01 |
| AT4G13860 | 0,43 | 2,91 | 4,10 | 0,71 | 4,78E-01 |
| AT4G13992 | 1,30 | -0,41 | 1,73 | -0,24 | 8,11E-01 |
| AT4G14000 | 0,12 | -1,57 | 4,34 | -0,36 | 7,18E-01 |
| AT4G14140 | 0,81 | -1,51 | 1,93 | -0,78 | 4,34E-01 |
| AT4G14220 | 0,46 | 0,05 | 4,38 | 0,01 | 9,90E-01 |
| AT4G14270 | 0,92 | -1,09 | 1,66 | -0,65 | 5,14E-01 |
| AT4G14300 | 0,77 | 2,72 | 3,56 | 0,77 | 4,44E-01 |
| AT4G14330 | 0,14 | 0,78 | 4,34 | 0,18 | 8,57E-01 |

| | | | | | |
|---|---|---|---|---|---|
| AT4G14540 | 0,71 | -1,85 | 2,41 | -0,77 | 4,44E-01 |
| AT4G14548 | 1,46 | -2,67 | 1,63 | -1,64 | 1,00E-01 |
| AT4G14560 | 0,12 | 0,08 | 4,33 | 0,02 | 9,84E-01 |
| AT4G14570 | 0,70 | 2,85 | 2,76 | 1,03 | 3,02E-01 |
| AT4G14630 | 0,43 | -3,05 | 3,98 | -0,77 | 4,44E-01 |
| AT4G14700 | 0,12 | 2,87 | 4,34 | 0,66 | 5,08E-01 |
| AT4G06245 | 1,52 | 3,57 | 2,16 | 1,65 | 9,83E-02 |
| AT4G06255 | 0,39 | 0,01 | 4,29 | 0,00 | 9,98E-01 |
| AT4G06260 | 0,42 | -0,70 | 3,24 | -0,22 | 8,29E-01 |
| AT4G14760 | 0,91 | 2,59 | 2,35 | 1,10 | 2,70E-01 |
| AT4G14800 | 0,34 | 2,03 | 4,34 | 0,47 | 6,39E-01 |
| AT4G14810 | 0,45 | -0,09 | 3,27 | -0,03 | 9,79E-01 |
| AT4G14850 | 0,15 | -0,41 | 4,32 | -0,09 | 9,25E-01 |
| AT4G14870 | 0,48 | -1,46 | 2,60 | -0,56 | 5,73E-01 |
| AT4G14890 | 0,73 | -0,07 | 2,15 | -0,03 | 9,74E-01 |
| AT4G14950 | 0,54 | 0,73 | 2,51 | 0,29 | 7,71E-01 |
| AT4G15000 | 1,11 | -0,32 | 1,85 | -0,17 | 8,62E-01 |
| AT4G15020 | 0,86 | 3,44 | 2,30 | 1,50 | 1,34E-01 |
| AT4G15030 | 0,61 | 0,46 | 2,26 | 0,20 | 8,39E-01 |
| AT4G15090 | 0,08 | 2,33 | 4,36 | 0,53 | 5,93E-01 |
| AT4G15096 | 1,15 | -3,60 | 2,08 | -1,73 | 8,42E-02 |
| AT4G15165 | 0,40 | -0,28 | 4,39 | -0,06 | 9,49E-01 |
| AT4G15233 | 0,90 | 2,71 | 2,37 | 1,14 | 2,52E-01 |
| AT4G15245 | 0,49 | -0,91 | 2,53 | -0,36 | 7,19E-01 |
| AT4G15260 | 0,57 | -0,81 | 2,95 | -0,28 | 7,83E-01 |
| AT4G15280 | 1,02 | 0,05 | 2,03 | 0,03 | 9,78E-01 |
| AT4G15415 | 0,24 | 0,58 | 4,32 | 0,13 | 8,93E-01 |
| AT4G15420 | 1,24 | 3,55 | 2,11 | 1,68 | 9,23E-02 |
| AT4G15440 | 0,99 | -2,79 | 2,35 | -1,19 | 2,34E-01 |
| AT4G15450 | 0,24 | -1,17 | 4,35 | -0,27 | 7,89E-01 |
| AT4G15475 | 0,96 | 0,00 | 4,14 | 0,00 | 1,00E+00 |
| AT4G15540 | 0,39 | -0,21 | 3,38 | -0,06 | 9,51E-01 |
| AT4G15545 | 1,17 | 1,09 | 1,47 | 0,74 | 4,60E-01 |
| AT4G15660 | 0,10 | 0,34 | 4,35 | 0,08 | 9,37E-01 |
| AT4G15700 | 0,08 | -0,56 | 4,38 | -0,13 | 8,99E-01 |
| AT4G15720 | 0,08 | 2,33 | 4,36 | 0,53 | 5,93E-01 |
| AT4G15770 | 0,18 | -0,70 | 4,38 | -0,16 | 8,72E-01 |
| AT4G15775 | 0,24 | -0,14 | 4,41 | -0,03 | 9,75E-01 |
| AT4G15800 | 0,76 | -0,06 | 2,03 | -0,03 | 9,77E-01 |
| AT4G15820 | 0,10 | 0,39 | 4,35 | 0,09 | 9,29E-01 |
| AT4G15840 | 0,73 | 1,10 | 2,01 | 0,55 | 5,84E-01 |
| AT4G15850 | 0,38 | 2,13 | 4,38 | 0,49 | 6,27E-01 |
| AT4G15880 | 0,12 | 0,63 | 4,33 | 0,15 | 8,85E-01 |
| AT4G15940 | 0,10 | -1,26 | 4,35 | -0,29 | 7,72E-01 |
| AT4G16008 | 0,08 | 2,33 | 4,36 | 0,53 | 5,93E-01 |
| AT4G16015 | 0,19 | 1,14 | 4,29 | 0,26 | 7,91E-01 |
| AT4G16130 | 1,05 | 2,59 | 2,03 | 1,27 | 2,02E-01 |
| AT4G16140 | 0,34 | -0,27 | 4,25 | -0,06 | 9,49E-01 |
| AT4G16150 | 0,50 | 1,74 | 4,04 | 0,43 | 6,67E-01 |
| AT4G16310 | 1,01 | 0,95 | 1,79 | 0,53 | 5,96E-01 |
| AT4G16360 | 0,61 | 1,54 | 2,64 | 0,58 | 5,61E-01 |
| AT4G16380 | 0,81 | -0,19 | 2,34 | -0,08 | 9,35E-01 |
| AT4G16390 | 0,67 | 1,29 | 2,17 | 0,59 | 5,53E-01 |
| AT4G16400 | 1,31 | 1,69 | 1,92 | 0,88 | 3,80E-01 |
| AT4G16430 | 0,28 | -0,16 | 4,40 | -0,04 | 9,70E-01 |
| AT4G16480 | 0,65 | 3,75 | 3,51 | 1,07 | 2,86E-01 |

| | | | | | |
|---|---|---|---|---|---|
| AT4G16510 | 0,53 | 2,80 | 4,01 | 0,70 | 4,85E-01 |
| AT4G16590 | 0,08 | 2,33 | 4,36 | 0,53 | 5,93E-01 |
| AT4G16650 | 0,41 | 2,25 | 4,24 | 0,53 | 5,95E-01 |
| AT4G16670 | 0,65 | 2,01 | 2,25 | 0,90 | 3,70E-01 |
| AT4G16710 | 0,63 | 2,17 | 2,43 | 0,89 | 3,73E-01 |
| AT4G16780 | 0,60 | 1,91 | 3,88 | 0,49 | 6,22E-01 |
| AT4G16830 | 0,78 | -1,52 | 3,13 | -0,49 | 6,27E-01 |
| AT4G16845 | 0,11 | -1,28 | 4,35 | -0,29 | 7,69E-01 |
| AT4G16850 | 0,29 | 3,25 | 4,33 | 0,75 | 4,53E-01 |
| AT4G16855 | 0,17 | 3,16 | 4,33 | 0,73 | 4,66E-01 |
| AT4G16955 | 0,78 | 1,52 | 3,00 | 0,51 | 6,13E-01 |
| AT4G17010 | 0,55 | 0,00 | 4,40 | 0,00 | 1,00E+00 |
| AT4G17080 | 0,57 | 3,10 | 4,32 | 0,72 | 4,74E-01 |
| AT4G17085 | 0,42 | 1,65 | 4,37 | 0,38 | 7,07E-01 |
| AT4G06665 | 1,22 | 1,25 | 1,74 | 0,72 | 4,70E-01 |
| AT4G17098 | 0,61 | -0,71 | 2,56 | -0,28 | 7,82E-01 |
| AT4G17245 | 0,66 | 0,86 | 2,46 | 0,35 | 7,26E-01 |
| AT4G17250 | 0,41 | 2,51 | 4,31 | 0,58 | 5,59E-01 |
| AT4G17270 | 0,64 | -2,24 | 2,98 | -0,75 | 4,52E-01 |
| AT4G17300 | 0,71 | -0,27 | 2,10 | -0,13 | 8,97E-01 |
| AT4G17380 | 0,44 | 1,19 | 2,67 | 0,45 | 6,56E-01 |
| AT4G17500 | 0,96 | -0,21 | 1,96 | -0,11 | 9,16E-01 |
| AT4G17520 | 1,02 | -1,02 | 2,66 | -0,38 | 7,02E-01 |
| AT4G17600 | 1,35 | 1,29 | 1,47 | 0,88 | 3,79E-01 |
| AT4G17616 | 0,78 | 3,78 | 3,41 | 1,11 | 2,67E-01 |
| AT4G17620 | 1,00 | 3,02 | 2,33 | 1,30 | 1,94E-01 |
| AT4G17650 | 0,32 | 2,96 | 4,34 | 0,68 | 4,95E-01 |
| AT4G17770 | 1,15 | 0,66 | 1,81 | 0,36 | 7,17E-01 |
| AT4G17810 | 0,14 | 0,00 | 4,41 | 0,00 | 1,00E+00 |
| AT4G17830 | 0,83 | 1,65 | 2,37 | 0,69 | 4,87E-01 |
| AT4G17870 | 1,03 | -0,32 | 2,07 | -0,16 | 8,76E-01 |
| AT4G17900 | 0,47 | 1,50 | 2,69 | 0,56 | 5,76E-01 |
| AT4G18070 | 1,02 | 0,02 | 2,15 | 0,01 | 9,92E-01 |
| AT4G18160 | 0,40 | -0,70 | 4,39 | -0,16 | 8,73E-01 |
| AT4G18195 | 0,22 | -1,30 | 4,35 | -0,30 | 7,65E-01 |
| AT4G18205 | 1,32 | -1,76 | 1,49 | -1,19 | 2,36E-01 |
| AT4G18220 | 1,16 | 0,35 | 1,52 | 0,23 | 8,17E-01 |
| AT4G18240 | 1,25 | 0,58 | 1,78 | 0,33 | 7,43E-01 |
| AT4G18370 | 0,87 | 2,17 | 2,30 | 0,94 | 3,47E-01 |
| AT4G18380 | 0,33 | 2,17 | 4,38 | 0,50 | 6,20E-01 |
| AT4G06795 | 0,56 | 2,17 | 2,99 | 0,73 | 4,67E-01 |
| AT4G18530 | 0,35 | 1,42 | 4,31 | 0,33 | 7,42E-01 |
| AT4G18570 | 0,37 | 0,38 | 2,81 | 0,14 | 8,91E-01 |
| AT4G18710 | 1,19 | 2,57 | 1,78 | 1,44 | 1,49E-01 |
| AT4G18725 | 0,19 | -1,45 | 4,34 | -0,33 | 7,38E-01 |
| AT4G18730 | 1,33 | 1,42 | 1,76 | 0,81 | 4,18E-01 |
| AT4G18750 | 0,50 | 1,41 | 4,02 | 0,35 | 7,25E-01 |
| AT4G18820 | 1,31 | 3,85 | 1,91 | 2,02 | 4,37E-02 |
| AT4G18830 | 0,19 | 0,42 | 4,28 | 0,10 | 9,21E-01 |
| AT4G18890 | 0,44 | -2,81 | 4,00 | -0,70 | 4,83E-01 |
| AT4G18905 | 0,35 | 1,15 | 4,31 | 0,27 | 7,90E-01 |
| AT4G18930 | 0,31 | 0,12 | 4,33 | 0,03 | 9,78E-01 |
| AT4G18950 | 1,00 | 0,60 | 1,83 | 0,33 | 7,44E-01 |
| AT4G19020 | 0,17 | 2,04 | 4,33 | 0,47 | 6,37E-01 |
| AT4G06865 | 0,10 | 0,34 | 4,35 | 0,08 | 9,37E-01 |
| AT4G19090 | 1,32 | 1,22 | 1,57 | 0,78 | 4,36E-01 |

| | | | | | |
|---|---|---|---|---|---|
| AT4G19100 | 1,38 | -3,03 | 2,04 | -1,49 | 1,37E-01 |
| AT4G19190 | 0,39 | 2,78 | 4,18 | 0,67 | 5,05E-01 |
| AT4G19200 | 0,86 | -0,15 | 2,11 | -0,07 | 9,44E-01 |
| AT4G19210 | 1,66 | 3,70 | 1,48 | 2,51 | 1,22E-02 |
| AT4G19450 | 0,65 | 2,66 | 4,28 | 0,62 | 5,35E-01 |
| AT4G19490 | 0,82 | 2,07 | 2,66 | 0,78 | 4,36E-01 |
| AT4G19500 | 0,89 | 2,15 | 2,15 | 1,00 | 3,17E-01 |
| AT4G19510 | 1,30 | 3,35 | 1,83 | 1,83 | 6,72E-02 |
| AT4G19530 | 1,16 | -0,52 | 1,78 | -0,29 | 7,72E-01 |
| AT4G19540 | 0,68 | 2,96 | 2,88 | 1,03 | 3,04E-01 |
| AT4G19580 | 0,44 | 2,87 | 4,05 | 0,71 | 4,78E-01 |
| AT4G19600 | 0,21 | 1,65 | 4,38 | 0,38 | 7,07E-01 |
| AT4G19610 | 0,37 | 1,39 | 3,40 | 0,41 | 6,82E-01 |
| AT4G19650 | 0,19 | 2,19 | 4,32 | 0,51 | 6,12E-01 |
| AT4G19710 | 1,47 | 2,62 | 1,59 | 1,65 | 9,89E-02 |
| AT4G19960 | 0,50 | 1,34 | 3,10 | 0,43 | 6,66E-01 |
| AT4G20110 | 0,83 | 1,22 | 2,09 | 0,58 | 5,60E-01 |
| AT4G20140 | 0,84 | 4,92 | 4,17 | 1,18 | 2,38E-01 |
| AT4G06950 | 0,12 | 0,31 | 4,35 | 0,07 | 9,44E-01 |
| AT4G20270 | 0,28 | 1,72 | 4,27 | 0,40 | 6,87E-01 |
| AT4G20300 | 0,68 | -0,98 | 2,41 | -0,41 | 6,85E-01 |
| AT4G20310 | 0,08 | 2,33 | 4,36 | 0,53 | 5,93E-01 |
| AT4G20400 | 1,23 | 2,84 | 2,16 | 1,32 | 1,87E-01 |
| AT4G20720 | 0,34 | 1,89 | 4,34 | 0,44 | 6,63E-01 |
| AT4G20840 | 1,22 | 4,02 | 2,46 | 1,64 | 1,02E-01 |
| AT4G20860 | 0,52 | 1,46 | 2,99 | 0,49 | 6,27E-01 |
| AT4G20890 | 0,68 | -0,57 | 2,56 | -0,22 | 8,24E-01 |
| AT4G20940 | 0,56 | 2,82 | 4,35 | 0,65 | 5,16E-01 |
| AT4G20960 | 0,48 | 0,10 | 2,57 | 0,04 | 9,70E-01 |
| AT4G20980 | 1,36 | 1,89 | 1,85 | 1,02 | 3,07E-01 |
| AT4G21105 | 0,54 | 1,03 | 4,32 | 0,24 | 8,11E-01 |
| AT4G21160 | 0,15 | -0,67 | 4,39 | -0,15 | 8,78E-01 |
| AT4G21180 | 0,08 | 2,08 | 4,38 | 0,47 | 6,35E-01 |
| AT4G21210 | 0,33 | 1,01 | 4,29 | 0,24 | 8,14E-01 |
| AT4G21300 | 0,05 | -0,58 | 4,38 | -0,13 | 8,95E-01 |
| AT4G21323 | 0,38 | 2,13 | 4,38 | 0,49 | 6,27E-01 |
| AT4G21450 | 0,15 | 1,69 | 4,34 | 0,39 | 6,98E-01 |
| AT4G21470 | 0,32 | 2,28 | 4,33 | 0,53 | 5,98E-01 |
| AT4G21515 | 0,05 | -0,58 | 4,38 | -0,13 | 8,95E-01 |
| AT4G21540 | 0,68 | 2,65 | 2,77 | 0,96 | 3,37E-01 |
| AT4G21550 | 0,37 | -0,14 | 4,41 | -0,03 | 9,75E-01 |
| AT4G21620 | 0,72 | 0,76 | 2,45 | 0,31 | 7,57E-01 |
| AT4G21670 | 1,62 | -0,42 | 1,31 | -0,32 | 7,50E-01 |
| AT4G21740 | 0,47 | 0,17 | 2,32 | 0,07 | 9,40E-01 |
| AT4G21750 | 0,81 | 3,95 | 2,75 | 1,44 | 1,51E-01 |
| AT4G21790 | 0,64 | 0,44 | 2,44 | 0,18 | 8,57E-01 |
| AT4G21800 | 0,53 | -1,20 | 3,86 | -0,31 | 7,56E-01 |
| AT4G07030 | 0,38 | 3,39 | 4,33 | 0,78 | 4,34E-01 |
| AT4G21940 | 0,15 | 1,93 | 4,33 | 0,45 | 6,55E-01 |
| AT4G22010 | 0,22 | 1,67 | 4,35 | 0,38 | 7,02E-01 |
| AT4G22050 | 0,20 | -0,87 | 4,29 | -0,20 | 8,40E-01 |
| AT4G07040 | 0,12 | -0,77 | 4,36 | -0,18 | 8,61E-01 |
| AT4G22130 | 0,44 | -0,33 | 2,65 | -0,12 | 9,01E-01 |
| AT4G22270 | 0,20 | 2,30 | 4,36 | 0,53 | 5,98E-01 |
| AT4G07075 | 1,23 | 0,36 | 1,47 | 0,25 | 8,04E-01 |
| AT4G07080 | 1,23 | 0,36 | 1,47 | 0,25 | 8,04E-01 |

| | | | | | |
|---|---|---|---|---|---|
| AT4G22330 | 0,17 | 3,16 | 4,33 | 0,73 | 4,66E-01 |
| AT4G22467 | 0,12 | 0,31 | 4,35 | 0,07 | 9,44E-01 |
| AT4G22475 | 0,49 | -2,96 | 3,85 | -0,77 | 4,42E-01 |
| AT4G22670 | 0,32 | -0,19 | 4,26 | -0,04 | 9,65E-01 |
| AT4G22720 | 0,28 | 1,78 | 4,33 | 0,41 | 6,80E-01 |
| AT4G22860 | 0,08 | 2,33 | 4,36 | 0,53 | 5,93E-01 |
| AT4G22890 | 1,50 | -1,30 | 1,71 | -0,76 | 4,49E-01 |
| AT4G22910 | 0,08 | 2,08 | 4,38 | 0,47 | 6,35E-01 |
| AT4G22960 | 0,12 | 0,31 | 4,35 | 0,07 | 9,44E-01 |
| AT4G23030 | 0,17 | -0,44 | 4,37 | -0,10 | 9,19E-01 |
| AT4G23050 | 0,25 | 2,79 | 4,34 | 0,64 | 5,20E-01 |
| AT4G23060 | 0,21 | 2,01 | 4,37 | 0,46 | 6,45E-01 |
| AT4G23150 | 0,15 | -0,97 | 4,37 | -0,22 | 8,24E-01 |
| AT4G23160 | 0,75 | -1,87 | 4,30 | -0,43 | 6,64E-01 |
| AT4G23180 | 1,57 | 0,20 | 1,27 | 0,16 | 8,75E-01 |
| AT4G23270 | 0,94 | 2,76 | 1,93 | 1,43 | 1,51E-01 |
| AT4G23400 | 0,56 | 0,42 | 2,12 | 0,20 | 8,42E-01 |
| AT4G23440 | 0,12 | 2,81 | 4,34 | 0,65 | 5,18E-01 |
| AT4G23470 | 0,84 | 0,82 | 1,81 | 0,45 | 6,50E-01 |
| AT4G23530 | 1,11 | 1,41 | 1,97 | 0,72 | 4,74E-01 |
| AT4G23570 | 0,44 | 0,42 | 3,34 | 0,13 | 8,99E-01 |
| AT4G23600 | 1,02 | 1,10 | 2,69 | 0,41 | 6,83E-01 |
| AT4G23740 | 0,40 | 1,98 | 4,31 | 0,46 | 6,46E-01 |
| AT4G23860 | 0,25 | -0,67 | 4,39 | -0,15 | 8,78E-01 |
| AT4G07190 | 0,45 | -0,10 | 3,41 | -0,03 | 9,77E-01 |
| AT4G23895 | 0,24 | 1,12 | 4,32 | 0,26 | 7,95E-01 |
| AT4G23940 | 1,06 | 3,58 | 2,18 | 1,64 | 1,02E-01 |
| AT4G23950 | 0,16 | 1,16 | 4,38 | 0,27 | 7,90E-01 |
| AT4G23980 | 1,47 | 1,39 | 1,45 | 0,96 | 3,37E-01 |
| AT4G24000 | 0,08 | 2,33 | 4,36 | 0,53 | 5,93E-01 |
| AT4G24010 | 0,33 | 2,17 | 4,38 | 0,50 | 6,20E-01 |
| AT4G24020 | 1,49 | 0,44 | 1,46 | 0,30 | 7,63E-01 |
| AT4G24060 | 0,12 | 2,81 | 4,34 | 0,65 | 5,18E-01 |
| AT4G24090 | 0,53 | 0,00 | 4,41 | 0,00 | 1,00E+00 |
| AT4G24120 | 1,68 | 4,94 | 2,10 | 2,36 | 1,85E-02 |
| AT4G24130 | 0,24 | 0,00 | 4,41 | 0,00 | 1,00E+00 |
| AT4G24160 | 0,15 | 0,40 | 4,33 | 0,09 | 9,27E-01 |
| AT4G24175 | 0,21 | 2,61 | 4,34 | 0,60 | 5,48E-01 |
| AT4G24220 | 1,52 | 1,04 | 1,41 | 0,74 | 4,58E-01 |
| AT4G24290 | 0,72 | 2,30 | 2,33 | 0,99 | 3,22E-01 |
| AT4G24350 | 0,25 | -1,00 | 4,30 | -0,23 | 8,16E-01 |
| AT4G24440 | 0,43 | 2,08 | 4,20 | 0,50 | 6,20E-01 |
| AT4G24460 | 0,18 | 1,77 | 4,41 | 0,40 | 6,88E-01 |
| AT4G24470 | 0,57 | 2,62 | 3,84 | 0,68 | 4,96E-01 |
| AT4G24550 | 0,38 | 2,18 | 3,38 | 0,65 | 5,18E-01 |
| AT4G24790 | 0,10 | 0,39 | 4,35 | 0,09 | 9,29E-01 |
| AT4G24800 | 0,98 | 2,76 | 2,56 | 1,08 | 2,81E-01 |
| AT4G24805 | 0,88 | -1,66 | 2,46 | -0,67 | 5,01E-01 |
| AT4G24880 | 0,08 | -0,56 | 4,38 | -0,13 | 8,99E-01 |
| AT4G07295 | 0,21 | 0,57 | 4,32 | 0,13 | 8,94E-01 |
| AT4G25030 | 0,40 | -0,30 | 2,70 | -0,11 | 9,12E-01 |
| AT4G25050 | 1,37 | 1,07 | 1,56 | 0,69 | 4,93E-01 |
| AT4G25070 | 0,53 | 3,05 | 4,35 | 0,70 | 4,84E-01 |
| AT4G25120 | 0,50 | 1,26 | 3,30 | 0,38 | 7,03E-01 |
| AT4G25170 | 1,05 | -1,18 | 2,00 | -0,59 | 5,53E-01 |
| AT4G25210 | 0,75 | -2,85 | 2,79 | -1,02 | 3,07E-01 |

| | | | | | |
|---|---|---|---|---|---|
| AT4G25230 | 0,77 | 1,04 | 2,03 | 0,51 | 6,07E-01 |
| AT4G25360 | 0,25 | 2,47 | 4,32 | 0,57 | 5,67E-01 |
| AT4G25370 | 0,59 | 3,11 | 3,85 | 0,81 | 4,19E-01 |
| AT4G25390 | 0,17 | -0,71 | 4,32 | -0,16 | 8,69E-01 |
| AT4G25400 | 0,31 | -2,62 | 4,29 | -0,61 | 5,41E-01 |
| AT4G25500 | 1,12 | 0,14 | 1,95 | 0,07 | 9,43E-01 |
| AT4G25515 | 0,32 | 1,58 | 4,35 | 0,36 | 7,17E-01 |
| AT4G25550 | 0,38 | 2,13 | 4,38 | 0,49 | 6,27E-01 |
| AT4G25610 | 0,54 | 2,22 | 3,15 | 0,71 | 4,81E-01 |
| AT4G25630 | 1,57 | -0,24 | 1,65 | -0,15 | 8,83E-01 |
| AT4G25631 | 1,33 | -0,31 | 1,89 | -0,16 | 8,71E-01 |
| AT4G25660 | 0,81 | 0,78 | 2,29 | 0,34 | 7,33E-01 |
| AT4G25810 | 0,12 | 2,69 | 4,35 | 0,62 | 5,36E-01 |
| AT4G25835 | 0,21 | 1,20 | 4,36 | 0,28 | 7,83E-01 |
| AT4G25900 | 0,36 | 0,48 | 4,30 | 0,11 | 9,11E-01 |
| AT4G25910 | 0,69 | 3,32 | 3,50 | 0,95 | 3,42E-01 |
| AT4G25960 | 1,53 | -2,25 | 1,65 | -1,36 | 1,73E-01 |
| AT4G25970 | 1,00 | 0,79 | 1,70 | 0,47 | 6,41E-01 |
| AT4G26000 | 0,96 | 2,31 | 2,34 | 0,99 | 3,24E-01 |
| AT4G26120 | 0,24 | 0,00 | 4,41 | 0,00 | 1,00E+00 |
| AT4G26170 | 0,08 | 2,08 | 4,38 | 0,47 | 6,35E-01 |
| AT4G26210 | 0,36 | 3,47 | 4,33 | 0,80 | 4,23E-01 |
| AT4G26230 | 0,82 | 1,62 | 2,53 | 0,64 | 5,23E-01 |
| AT4G26450 | 0,45 | 0,08 | 3,24 | 0,02 | 9,81E-01 |
| AT4G26455 | 0,54 | 3,68 | 3,76 | 0,98 | 3,27E-01 |
| AT4G26510 | 0,72 | 0,50 | 2,49 | 0,20 | 8,41E-01 |
| AT4G26520 | 1,37 | 2,60 | 2,05 | 1,27 | 2,03E-01 |
| AT4G26650 | 0,56 | 1,08 | 2,27 | 0,48 | 6,32E-01 |
| AT4G26670 | 0,33 | 0,78 | 4,34 | 0,18 | 8,57E-01 |
| AT4G26680 | 0,40 | 2,20 | 4,08 | 0,54 | 5,89E-01 |
| AT4G26700 | 0,28 | 0,85 | 4,32 | 0,20 | 8,44E-01 |
| AT4G26790 | 0,42 | -1,55 | 3,99 | -0,39 | 6,98E-01 |
| AT4G26800 | 0,12 | 0,06 | 4,34 | 0,01 | 9,88E-01 |
| AT4G26840 | 0,82 | 0,91 | 2,03 | 0,45 | 6,53E-01 |
| AT4G26870 | 0,92 | 1,76 | 2,07 | 0,85 | 3,94E-01 |
| AT4G27010 | 0,79 | 0,81 | 1,95 | 0,42 | 6,78E-01 |
| AT4G27030 | 0,63 | 2,66 | 2,53 | 1,05 | 2,94E-01 |
| AT4G07775 | 0,58 | -0,03 | 2,76 | -0,01 | 9,91E-01 |
| AT4G27080 | 1,09 | -0,52 | 2,19 | -0,24 | 8,11E-01 |
| AT4G27260 | 0,26 | 2,30 | 4,37 | 0,53 | 5,98E-01 |
| AT4G27290 | 0,45 | 2,02 | 4,37 | 0,46 | 6,44E-01 |
| AT4G27310 | 0,93 | -0,54 | 1,88 | -0,29 | 7,73E-01 |
| AT4G27380 | 0,21 | 1,85 | 4,38 | 0,42 | 6,73E-01 |
| AT4G27395 | 0,73 | 2,80 | 2,85 | 0,98 | 3,25E-01 |
| AT4G27652 | 0,57 | -2,03 | 3,60 | -0,56 | 5,73E-01 |
| AT4G27654 | 0,56 | 0,34 | 2,45 | 0,14 | 8,89E-01 |
| AT4G27657 | 0,12 | 0,31 | 4,35 | 0,07 | 9,44E-01 |
| AT4G27680 | 0,93 | 2,03 | 1,95 | 1,04 | 2,97E-01 |
| AT4G27710 | 0,52 | 1,56 | 3,19 | 0,49 | 6,25E-01 |
| AT4G27720 | 1,38 | 0,06 | 1,59 | 0,04 | 9,72E-01 |
| AT4G27740 | 0,52 | -2,50 | 3,72 | -0,67 | 5,02E-01 |
| AT4G27750 | 0,69 | 1,79 | 4,41 | 0,41 | 6,85E-01 |
| AT4G27760 | 0,28 | 1,64 | 4,38 | 0,38 | 7,08E-01 |
| AT4G27765 | 0,37 | 2,79 | 4,28 | 0,65 | 5,14E-01 |
| AT4G27780 | 0,50 | 2,54 | 3,95 | 0,64 | 5,20E-01 |
| AT4G27860 | 0,21 | 0,84 | 4,28 | 0,20 | 8,44E-01 |

| | | | | | |
|---|---|---|---|---|---|
| AT4G27880 | 0,08 | 2,08 | 4,38 | 0,47 | 6,35E-01 |
| AT4G27910 | 0,46 | 3,27 | 4,05 | 0,81 | 4,20E-01 |
| AT4G27940 | 1,14 | 2,49 | 2,21 | 1,12 | 2,61E-01 |
| AT4G28010 | 0,17 | 1,41 | 4,35 | 0,32 | 7,46E-01 |
| AT4G28210 | 0,08 | -0,56 | 4,38 | -0,13 | 8,99E-01 |
| AT4G28220 | 0,27 | 0,11 | 4,27 | 0,03 | 9,79E-01 |
| AT4G28260 | 0,80 | 3,29 | 2,86 | 1,15 | 2,51E-01 |
| AT4G28300 | 0,21 | 2,34 | 4,32 | 0,54 | 5,89E-01 |
| AT4G28362 | 0,21 | 2,01 | 4,37 | 0,46 | 6,45E-01 |
| AT4G28400 | 0,26 | -0,39 | 4,33 | -0,09 | 9,28E-01 |
| AT4G28440 | 0,16 | 0,03 | 4,34 | 0,01 | 9,94E-01 |
| AT4G28490 | 0,74 | 2,03 | 3,49 | 0,58 | 5,61E-01 |
| AT4G28600 | 1,03 | 2,69 | 2,05 | 1,31 | 1,90E-01 |
| AT4G28660 | 0,48 | -0,83 | 4,08 | -0,20 | 8,39E-01 |
| AT4G28706 | 0,39 | -1,54 | 4,30 | -0,36 | 7,20E-01 |
| AT4G28710 | 0,41 | 1,55 | 4,13 | 0,37 | 7,08E-01 |
| AT4G28740 | 0,90 | 0,38 | 2,39 | 0,16 | 8,73E-01 |
| AT4G28780 | 0,15 | -0,19 | 4,31 | -0,04 | 9,64E-01 |
| AT4G28860 | 0,76 | 1,79 | 3,09 | 0,58 | 5,63E-01 |
| AT4G28870 | 0,12 | -1,57 | 4,34 | -0,36 | 7,18E-01 |
| AT4G28880 | 1,20 | 0,11 | 1,59 | 0,07 | 9,44E-01 |
| AT4G28980 | 0,73 | 2,81 | 2,44 | 1,16 | 2,48E-01 |
| AT4G29020 | 0,17 | -0,44 | 4,37 | -0,10 | 9,19E-01 |
| AT4G29035 | 0,22 | -1,30 | 4,35 | -0,30 | 7,65E-01 |
| AT4G29040 | 1,10 | 0,53 | 1,78 | 0,30 | 7,67E-01 |
| AT4G29070 | 0,33 | -0,05 | 4,32 | -0,01 | 9,90E-01 |
| AT4G29110 | 0,63 | -1,67 | 2,78 | -0,60 | 5,49E-01 |
| AT4G29120 | 0,68 | 2,41 | 2,72 | 0,89 | 3,75E-01 |
| AT4G29160 | 0,87 | 1,38 | 1,78 | 0,77 | 4,39E-01 |
| AT4G29170 | 0,92 | 3,50 | 4,32 | 0,81 | 4,18E-01 |
| AT4G29180 | 0,96 | 3,19 | 2,64 | 1,21 | 2,26E-01 |
| AT4G29210 | 0,68 | 1,98 | 2,57 | 0,77 | 4,41E-01 |
| AT4G29350 | 0,47 | -0,12 | 3,98 | -0,03 | 9,77E-01 |
| AT4G29400 | 0,21 | -0,24 | 4,35 | -0,06 | 9,56E-01 |
| AT4G29510 | 0,71 | 1,37 | 2,18 | 0,63 | 5,31E-01 |
| AT4G29740 | 1,04 | -1,47 | 2,05 | -0,72 | 4,71E-01 |
| AT4G29750 | 0,66 | -0,36 | 2,61 | -0,14 | 8,91E-01 |
| AT4G29790 | 1,30 | 5,08 | 1,89 | 2,69 | 7,24E-03 |
| AT4G08245 | 0,54 | -0,54 | 2,99 | -0,18 | 8,57E-01 |
| AT4G08255 | 0,51 | -0,48 | 4,21 | -0,11 | 9,09E-01 |
| AT4G29950 | 0,57 | 0,47 | 2,48 | 0,19 | 8,51E-01 |
| AT4G30010 | 0,55 | 1,10 | 3,11 | 0,35 | 7,25E-01 |
| AT4G30020 | 1,12 | 1,79 | 2,05 | 0,87 | 3,82E-01 |
| AT4G30130 | 0,24 | 0,81 | 4,34 | 0,19 | 8,53E-01 |
| AT4G30150 | 1,08 | 2,82 | 1,99 | 1,42 | 1,56E-01 |
| AT4G30310 | 0,60 | 2,95 | 3,67 | 0,80 | 4,22E-01 |
| AT4G30350 | 0,36 | -0,04 | 4,25 | -0,01 | 9,92E-01 |
| AT4G30400 | 0,15 | -0,41 | 4,32 | -0,09 | 9,25E-01 |
| AT4G08285 | 0,08 | 2,08 | 4,38 | 0,47 | 6,35E-01 |
| AT4G30480 | 0,31 | 0,00 | 4,41 | 0,00 | 1,00E+00 |
| AT4G30530 | 1,06 | -1,23 | 1,65 | -0,74 | 4,58E-01 |
| AT4G30650 | 0,58 | -0,78 | 2,67 | -0,29 | 7,71E-01 |
| AT4G30660 | 0,47 | -0,15 | 2,72 | -0,05 | 9,57E-01 |
| AT4G30780 | 0,23 | -0,64 | 4,39 | -0,15 | 8,84E-01 |
| AT4G30800 | 0,17 | 0,00 | 4,41 | 0,00 | 1,00E+00 |
| AT4G30940 | 0,31 | -0,60 | 4,32 | -0,14 | 8,91E-01 |

| | | | | | |
|---|---|---|---|---|---|
| AT4G30990 | 1,34 | 3,35 | 1,84 | 1,82 | 6,90E-02 |
| AT4G08425 | 1,45 | 0,56 | 1,26 | 0,44 | 6,57E-01 |
| AT4G31080 | 0,70 | 2,22 | 2,79 | 0,79 | 4,27E-01 |
| AT4G31115 | 0,40 | 2,10 | 4,25 | 0,49 | 6,22E-01 |
| AT4G31300 | 0,37 | -2,19 | 4,30 | -0,51 | 6,11E-01 |
| AT4G31340 | 0,63 | 2,62 | 3,81 | 0,69 | 4,91E-01 |
| AT4G31390 | 0,47 | 0,44 | 2,72 | 0,16 | 8,71E-01 |
| AT4G31430 | 0,50 | 1,94 | 3,97 | 0,49 | 6,25E-01 |
| AT4G31530 | 0,77 | -0,67 | 2,42 | -0,28 | 7,81E-01 |
| AT4G31580 | 0,97 | 1,05 | 1,86 | 0,56 | 5,72E-01 |
| AT4G31710 | 0,50 | 2,54 | 3,95 | 0,64 | 5,20E-01 |
| AT4G31715 | 0,08 | 2,33 | 4,36 | 0,53 | 5,93E-01 |
| AT4G31780 | 1,36 | 1,30 | 2,71 | 0,48 | 6,32E-01 |
| AT4G31790 | 0,26 | 0,26 | 4,29 | 0,06 | 9,51E-01 |
| AT4G31800 | 1,69 | -0,51 | 1,50 | -0,34 | 7,32E-01 |
| AT4G31805 | 0,98 | 4,11 | 3,30 | 1,25 | 2,13E-01 |
| AT4G31850 | 0,54 | 0,35 | 3,19 | 0,11 | 9,14E-01 |
| AT4G31910 | 0,49 | -1,28 | 2,60 | -0,49 | 6,23E-01 |
| AT4G08615 | 0,91 | -1,17 | 2,15 | -0,54 | 5,86E-01 |
| AT4G08625 | 1,54 | -0,79 | 1,37 | -0,58 | 5,63E-01 |
| AT4G31985 | 0,37 | 2,89 | 4,33 | 0,67 | 5,05E-01 |
| AT4G32010 | 1,47 | 2,77 | 1,99 | 1,39 | 1,65E-01 |
| AT4G32030 | 0,47 | 0,63 | 4,18 | 0,15 | 8,80E-01 |
| AT4G32040 | 0,84 | 2,93 | 3,46 | 0,85 | 3,97E-01 |
| AT4G32060 | 1,61 | -0,34 | 1,29 | -0,26 | 7,95E-01 |
| AT4G32160 | 0,93 | 1,34 | 1,83 | 0,73 | 4,66E-01 |
| AT4G32190 | 0,86 | 3,75 | 2,65 | 1,42 | 1,57E-01 |
| AT4G32200 | 0,65 | 3,89 | 3,59 | 1,08 | 2,79E-01 |
| AT4G32265 | 1,11 | 0,70 | 1,73 | 0,40 | 6,86E-01 |
| AT4G32300 | 0,70 | 0,32 | 1,99 | 0,16 | 8,72E-01 |
| AT4G32330 | 0,80 | -0,46 | 1,96 | -0,24 | 8,14E-01 |
| AT4G32360 | 0,05 | -0,58 | 4,38 | -0,13 | 8,95E-01 |
| AT4G32390 | 1,34 | 2,37 | 1,68 | 1,41 | 1,58E-01 |
| AT4G32430 | 0,12 | 0,31 | 4,35 | 0,07 | 9,44E-01 |
| AT4G32551 | 0,49 | 0,61 | 2,61 | 0,23 | 8,16E-01 |
| AT4G32600 | 0,53 | 0,57 | 3,18 | 0,18 | 8,57E-01 |
| AT4G32620 | 1,61 | 3,64 | 2,00 | 1,82 | 6,85E-02 |
| AT4G32630 | 0,68 | 1,72 | 4,40 | 0,39 | 6,96E-01 |
| AT4G32700 | 0,67 | 4,00 | 3,62 | 1,11 | 2,68E-01 |
| AT4G32720 | 0,62 | 2,41 | 3,02 | 0,80 | 4,24E-01 |
| AT4G32730 | 0,45 | 3,37 | 3,98 | 0,85 | 3,98E-01 |
| AT4G32770 | 0,32 | 0,80 | 4,29 | 0,19 | 8,51E-01 |
| AT4G32800 | 0,31 | 0,23 | 4,33 | 0,05 | 9,57E-01 |
| AT4G32850 | 1,20 | 1,60 | 1,79 | 0,89 | 3,74E-01 |
| AT4G32910 | 0,98 | 4,26 | 2,05 | 2,07 | 3,80E-02 |
| AT4G32930 | 0,91 | 1,91 | 2,37 | 0,81 | 4,19E-01 |
| AT4G33000 | 0,14 | -1,69 | 4,33 | -0,39 | 6,96E-01 |
| AT4G33030 | 0,99 | 4,13 | 2,53 | 1,63 | 1,04E-01 |
| AT4G33060 | 0,15 | 1,94 | 4,32 | 0,45 | 6,54E-01 |
| AT4G33180 | 0,05 | -0,58 | 4,38 | -0,13 | 8,95E-01 |
| AT4G33220 | 0,80 | 0,05 | 3,10 | 0,02 | 9,88E-01 |
| AT4G33290 | 1,50 | -1,86 | 1,50 | -1,24 | 2,15E-01 |
| AT4G08785 | 0,69 | 0,78 | 3,69 | 0,21 | 8,32E-01 |
| AT4G33310 | 0,49 | -0,94 | 4,34 | -0,22 | 8,28E-01 |
| AT4G33380 | 0,54 | 2,04 | 3,21 | 0,64 | 5,24E-01 |
| AT4G33410 | 0,32 | 1,90 | 4,34 | 0,44 | 6,62E-01 |

| | | | | | |
|---|---|---|---|---|---|
| AT4G33510 | 1,23 | 0,56 | 1,37 | 0,41 | 6,80E-01 |
| AT4G33520 | 1,37 | 3,67 | 2,12 | 1,73 | 8,33E-02 |
| AT4G33540 | 0,41 | 0,84 | 4,18 | 0,20 | 8,41E-01 |
| AT4G33565 | 0,80 | -2,12 | 2,68 | -0,79 | 4,29E-01 |
| AT4G33580 | 1,02 | -1,32 | 3,10 | -0,43 | 6,69E-01 |
| AT4G33620 | 0,33 | 2,42 | 4,31 | 0,56 | 5,74E-01 |
| AT4G33630 | 1,15 | 0,90 | 1,98 | 0,45 | 6,50E-01 |
| AT4G33666 | 0,24 | 0,06 | 4,31 | 0,01 | 9,89E-01 |
| AT4G33690 | 0,48 | -2,31 | 3,98 | -0,58 | 5,62E-01 |
| AT4G33890 | 1,49 | 1,22 | 1,84 | 0,67 | 5,06E-01 |
| AT4G33920 | 0,65 | -1,57 | 3,48 | -0,45 | 6,52E-01 |
| AT4G33960 | 0,45 | 0,00 | 4,41 | 0,00 | 1,00E+00 |
| AT4G34030 | 0,40 | -0,38 | 3,36 | -0,11 | 9,10E-01 |
| AT4G34035 | 0,32 | 2,47 | 4,35 | 0,57 | 5,69E-01 |
| AT4G34050 | 1,48 | -1,57 | 1,49 | -1,06 | 2,91E-01 |
| AT4G34060 | 0,12 | 2,81 | 4,34 | 0,65 | 5,18E-01 |
| AT4G34120 | 1,07 | 0,02 | 1,80 | 0,01 | 9,93E-01 |
| AT4G34160 | 0,73 | 1,78 | 2,24 | 0,79 | 4,27E-01 |
| AT4G34240 | 0,30 | -1,48 | 4,34 | -0,34 | 7,33E-01 |
| AT4G34260 | 1,45 | 2,60 | 2,05 | 1,27 | 2,04E-01 |
| AT4G34280 | 0,96 | 4,06 | 2,74 | 1,48 | 1,38E-01 |
| AT4G34290 | 0,10 | 0,34 | 4,35 | 0,08 | 9,37E-01 |
| AT4G34310 | 0,63 | 1,26 | 4,37 | 0,29 | 7,72E-01 |
| AT4G34330 | 0,08 | 2,33 | 4,36 | 0,53 | 5,93E-01 |
| AT4G34370 | 0,50 | 2,54 | 3,95 | 0,64 | 5,20E-01 |
| AT4G34400 | 0,59 | 2,86 | 3,75 | 0,76 | 4,45E-01 |
| AT4G34410 | 0,77 | -1,63 | 2,61 | -0,63 | 5,32E-01 |
| AT4G34430 | 1,22 | 4,53 | 2,15 | 2,11 | 3,53E-02 |
| AT4G34440 | 0,20 | 2,30 | 4,36 | 0,53 | 5,98E-01 |
| AT4G34640 | 0,93 | 4,40 | 2,76 | 1,60 | 1,11E-01 |
| AT4G34680 | 0,14 | -0,45 | 4,33 | -0,10 | 9,17E-01 |
| AT4G34700 | 0,83 | -0,70 | 2,47 | -0,28 | 7,76E-01 |
| AT4G34870 | 1,35 | 0,18 | 1,43 | 0,12 | 9,02E-01 |
| AT4G34881 | 0,53 | -0,61 | 2,70 | -0,23 | 8,20E-01 |
| AT4G34910 | 0,61 | 3,45 | 3,00 | 1,15 | 2,50E-01 |
| AT4G34920 | 0,49 | 0,72 | 3,11 | 0,23 | 8,17E-01 |
| AT4G35040 | 0,31 | 2,37 | 4,31 | 0,55 | 5,82E-01 |
| AT4G35100 | 1,16 | 0,72 | 2,50 | 0,29 | 7,74E-01 |
| AT4G35370 | 0,13 | 2,37 | 4,36 | 0,54 | 5,86E-01 |
| AT4G35450 | 0,43 | -0,16 | 3,29 | -0,05 | 9,61E-01 |
| AT4G35460 | 0,66 | 2,82 | 2,94 | 0,96 | 3,37E-01 |
| AT4G35470 | 0,67 | 1,17 | 2,45 | 0,48 | 6,34E-01 |
| AT4G35500 | 0,79 | 3,55 | 2,75 | 1,29 | 1,97E-01 |
| AT4G35560 | 0,72 | 2,56 | 2,91 | 0,88 | 3,81E-01 |
| AT4G35600 | 1,15 | 2,10 | 1,81 | 1,16 | 2,45E-01 |
| AT4G35630 | 0,33 | 0,91 | 3,54 | 0,26 | 7,97E-01 |
| AT4G35850 | 0,77 | 2,34 | 2,18 | 1,07 | 2,84E-01 |
| AT4G35870 | 0,76 | 1,12 | 2,47 | 0,45 | 6,51E-01 |
| AT4G35880 | 0,17 | 2,62 | 4,35 | 0,60 | 5,47E-01 |
| AT4G35890 | 1,08 | 1,04 | 1,92 | 0,54 | 5,87E-01 |
| AT4G35940 | 1,33 | 0,55 | 1,63 | 0,34 | 7,36E-01 |
| AT4G36032 | 1,21 | -1,84 | 2,04 | -0,90 | 3,65E-01 |
| AT4G36052 | 0,12 | 2,87 | 4,34 | 0,66 | 5,08E-01 |
| AT4G36130 | 1,27 | 3,57 | 1,99 | 1,79 | 7,36E-02 |
| AT4G36150 | 1,48 | 5,20 | 2,51 | 2,07 | 3,81E-02 |
| AT4G36190 | 0,22 | -1,85 | 4,32 | -0,43 | 6,69E-01 |

| | | | | | |
|---|---|---|---|---|---|
| AT4G36195 | 0,86 | -0,48 | 2,06 | -0,23 | 8,15E-01 |
| AT4G36210 | 1,25 | 0,29 | 1,54 | 0,19 | 8,51E-01 |
| AT4G36250 | 0,69 | -1,25 | 2,39 | -0,52 | 6,01E-01 |
| AT4G36540 | 0,24 | 2,85 | 4,34 | 0,66 | 5,11E-01 |
| AT4G36710 | 1,27 | 2,34 | 2,41 | 0,97 | 3,30E-01 |
| AT4G36720 | 0,82 | 3,33 | 2,80 | 1,19 | 2,34E-01 |
| AT4G09375 | 0,14 | -0,45 | 4,38 | -0,10 | 9,18E-01 |
| AT4G36810 | 0,84 | 2,35 | 2,28 | 1,03 | 3,04E-01 |
| AT4G36870 | 0,25 | -1,95 | 4,32 | -0,45 | 6,51E-01 |
| AT4G36920 | 0,14 | -0,45 | 4,38 | -0,10 | 9,18E-01 |
| AT4G36925 | 1,06 | -0,37 | 1,72 | -0,22 | 8,28E-01 |
| AT4G36990 | 0,59 | 3,23 | 2,95 | 1,10 | 2,73E-01 |
| AT4G37000 | 0,23 | -0,64 | 4,39 | -0,15 | 8,84E-01 |
| AT4G37020 | 0,15 | 1,69 | 4,34 | 0,39 | 6,98E-01 |
| AT4G37080 | 0,19 | 1,62 | 4,35 | 0,37 | 7,09E-01 |
| AT4G37130 | 0,62 | 3,89 | 3,62 | 1,08 | 2,82E-01 |
| AT4G37180 | 0,47 | 0,37 | 3,99 | 0,09 | 9,25E-01 |
| AT4G37190 | 0,32 | 2,99 | 4,34 | 0,69 | 4,90E-01 |
| AT4G37210 | 0,15 | 2,31 | 4,37 | 0,53 | 5,98E-01 |
| AT4G37300 | 1,24 | 0,38 | 2,00 | 0,19 | 8,50E-01 |
| AT4G37430 | 0,86 | 3,26 | 3,34 | 0,97 | 3,30E-01 |
| AT4G37440 | 0,34 | -0,93 | 4,37 | -0,21 | 8,31E-01 |
| AT4G37550 | 0,88 | 0,26 | 1,97 | 0,13 | 8,95E-01 |
| AT4G37560 | 0,42 | 2,99 | 4,34 | 0,69 | 4,91E-01 |
| AT4G37650 | 1,47 | -2,21 | 1,70 | -1,29 | 1,95E-01 |
| AT4G37680 | 0,49 | 0,54 | 3,06 | 0,18 | 8,59E-01 |
| AT4G37730 | 0,75 | 0,03 | 1,89 | 0,02 | 9,88E-01 |
| AT4G37760 | 0,63 | -0,61 | 2,21 | -0,28 | 7,82E-01 |
| AT4G37790 | 0,15 | -0,64 | 4,39 | -0,15 | 8,84E-01 |
| AT4G37980 | 0,67 | -0,39 | 1,97 | -0,20 | 8,43E-01 |
| AT4G38040 | 0,20 | 2,30 | 4,36 | 0,53 | 5,98E-01 |
| AT4G38170 | 0,97 | 0,43 | 2,15 | 0,20 | 8,39E-01 |
| AT4G09685 | 0,46 | 3,03 | 3,99 | 0,76 | 4,48E-01 |
| AT4G38215 | 0,75 | 3,54 | 4,32 | 0,82 | 4,12E-01 |
| AT4G38225 | 0,78 | 0,24 | 2,34 | 0,10 | 9,20E-01 |
| AT4G38250 | 1,25 | 1,59 | 1,54 | 1,04 | 2,99E-01 |
| AT4G38270 | 0,51 | 3,06 | 3,07 | 1,00 | 3,19E-01 |
| AT4G38290 | 0,11 | 1,33 | 4,36 | 0,31 | 7,59E-01 |
| AT4G38320 | 0,19 | 1,89 | 4,32 | 0,44 | 6,61E-01 |
| AT4G38330 | 0,12 | 2,81 | 4,34 | 0,65 | 5,18E-01 |
| AT4G38360 | 0,15 | 1,94 | 4,32 | 0,45 | 6,54E-01 |
| AT4G38430 | 0,81 | 3,23 | 2,63 | 1,23 | 2,19E-01 |
| AT4G38460 | 1,41 | -0,74 | 1,56 | -0,47 | 6,35E-01 |
| AT4G38500 | 0,68 | 3,67 | 3,46 | 1,06 | 2,89E-01 |
| AT4G38540 | 0,10 | 0,39 | 4,35 | 0,09 | 9,29E-01 |
| AT4G38560 | 1,22 | 1,23 | 2,55 | 0,48 | 6,29E-01 |
| AT4G38580 | 0,50 | 0,00 | 4,41 | 0,00 | 1,00E+00 |
| AT4G38620 | 0,24 | 1,30 | 4,37 | 0,30 | 7,67E-01 |
| AT4G38790 | 0,69 | 3,19 | 2,98 | 1,07 | 2,85E-01 |
| AT4G38860 | 0,38 | -0,63 | 4,26 | -0,15 | 8,83E-01 |
| AT4G38880 | 0,39 | 1,71 | 4,40 | 0,39 | 6,98E-01 |
| AT4G38920 | 0,99 | 0,95 | 2,03 | 0,47 | 6,40E-01 |
| AT4G38930 | 0,41 | 0,79 | 4,39 | 0,18 | 8,58E-01 |
| AT4G39020 | 0,55 | 0,72 | 4,24 | 0,17 | 8,65E-01 |
| AT4G39030 | 0,29 | 1,64 | 4,38 | 0,37 | 7,08E-01 |
| AT4G39050 | 1,56 | 0,76 | 1,43 | 0,53 | 5,94E-01 |

| | | | | | |
|---|---|---|---|---|---|
| AT4G39210 | 1,07 | 3,52 | 2,21 | 1,59 | 1,11E-01 |
| AT4G39270 | 1,16 | 0,04 | 1,50 | 0,03 | 9,78E-01 |
| AT4G39620 | 0,18 | 1,51 | 4,31 | 0,35 | 7,25E-01 |
| AT4G39640 | 0,76 | 0,35 | 2,43 | 0,14 | 8,87E-01 |
| AT4G39670 | 0,55 | 0,88 | 3,05 | 0,29 | 7,72E-01 |
| AT4G39840 | 1,07 | -0,28 | 1,73 | -0,16 | 8,69E-01 |
| AT4G39860 | 0,51 | -0,32 | 2,26 | -0,14 | 8,86E-01 |
| AT4G39870 | 0,35 | 2,69 | 4,31 | 0,62 | 5,33E-01 |
| AT4G39940 | 0,36 | 1,44 | 4,33 | 0,33 | 7,40E-01 |
| AT4G39950 | 1,14 | 1,30 | 1,71 | 0,76 | 4,45E-01 |
| AT4G39952 | 0,28 | 1,64 | 4,38 | 0,38 | 7,08E-01 |
| AT4G39980 | 0,50 | 0,34 | 3,05 | 0,11 | 9,12E-01 |
| AT4G40063 | 0,73 | 3,91 | 2,93 | 1,33 | 1,82E-01 |
| AT5G00730 | 1,64 | -2,40 | 1,74 | -1,38 | 1,68E-01 |
| AT5G01170 | 0,10 | 0,34 | 4,35 | 0,08 | 9,37E-01 |
| AT5G01210 | 1,09 | -0,51 | 1,55 | -0,33 | 7,41E-01 |
| AT5G01240 | 0,22 | -0,13 | 4,30 | -0,03 | 9,76E-01 |
| AT5G01260 | 0,54 | 1,11 | 2,32 | 0,48 | 6,33E-01 |
| AT5G01290 | 0,28 | 2,54 | 4,30 | 0,59 | 5,55E-01 |
| AT5G01460 | 0,28 | -0,24 | 4,40 | -0,05 | 9,57E-01 |
| AT5G01542 | 0,46 | 1,39 | 4,03 | 0,34 | 7,30E-01 |
| AT5G01590 | 0,94 | 3,23 | 2,06 | 1,57 | 1,17E-01 |
| AT5G01595 | 0,81 | 1,79 | 2,29 | 0,78 | 4,35E-01 |
| AT5G01690 | 0,08 | 2,08 | 4,38 | 0,47 | 6,35E-01 |
| AT5G01732 | 0,29 | 3,20 | 4,33 | 0,74 | 4,60E-01 |
| AT5G01740 | 0,15 | -1,60 | 4,33 | -0,37 | 7,11E-01 |
| AT5G01750 | 1,69 | -0,55 | 1,67 | -0,33 | 7,44E-01 |
| AT5G01760 | 0,79 | 2,79 | 2,78 | 1,01 | 3,14E-01 |
| AT5G01800 | 0,74 | 2,81 | 2,41 | 1,17 | 2,44E-01 |
| AT5G01810 | 0,55 | 3,69 | 3,81 | 0,97 | 3,33E-01 |
| AT5G01850 | 0,24 | 0,67 | 4,34 | 0,15 | 8,78E-01 |
| AT5G02035 | 0,47 | 3,65 | 3,95 | 0,92 | 3,56E-01 |
| AT5G02040 | 0,25 | 2,79 | 4,34 | 0,64 | 5,20E-01 |
| AT5G02100 | 0,70 | 3,12 | 3,70 | 0,84 | 3,99E-01 |
| AT5G02120 | 0,39 | -0,73 | 4,35 | -0,17 | 8,67E-01 |
| AT5G00820 | 1,31 | 2,03 | 2,66 | 0,76 | 4,46E-01 |
| AT5G02160 | 1,30 | -0,36 | 1,64 | -0,22 | 8,24E-01 |
| AT5G02240 | 0,32 | 1,41 | 4,34 | 0,32 | 7,45E-01 |
| AT5G02250 | 1,33 | 2,38 | 2,24 | 1,06 | 2,88E-01 |
| AT5G02280 | 0,11 | 1,33 | 4,36 | 0,31 | 7,59E-01 |
| AT5G02420 | 0,12 | 1,65 | 4,34 | 0,38 | 7,04E-01 |
| AT5G02450 | 0,59 | 1,90 | 3,12 | 0,61 | 5,43E-01 |
| AT5G02620 | 0,32 | -0,75 | 4,35 | -0,17 | 8,63E-01 |
| AT5G02630 | 0,30 | 2,84 | 4,35 | 0,65 | 5,14E-01 |
| AT5G02650 | 0,40 | 3,80 | 4,32 | 0,88 | 3,79E-01 |
| AT5G02760 | 0,40 | 1,31 | 3,43 | 0,38 | 7,02E-01 |
| AT5G02790 | 0,39 | -1,62 | 3,38 | -0,48 | 6,31E-01 |
| AT5G02840 | 0,40 | 2,39 | 4,13 | 0,58 | 5,64E-01 |
| AT5G02850 | 0,54 | 2,33 | 3,07 | 0,76 | 4,47E-01 |
| AT5G02860 | 1,11 | 2,44 | 1,82 | 1,34 | 1,81E-01 |
| AT5G02950 | 0,61 | 3,59 | 3,79 | 0,95 | 3,43E-01 |
| AT5G02995 | 0,13 | -0,63 | 4,39 | -0,14 | 8,85E-01 |
| AT5G03020 | 0,28 | 1,64 | 4,38 | 0,38 | 7,08E-01 |
| AT5G03160 | 0,68 | 0,02 | 2,28 | 0,01 | 9,91E-01 |
| AT5G03190 | 0,92 | 0,79 | 1,89 | 0,42 | 6,76E-01 |
| AT5G03290 | 1,24 | 0,67 | 1,46 | 0,46 | 6,46E-01 |

| | | | | | |
|---|---|---|---|---|---|
| AT5G03300 | 0,81 | 1,06 | 2,28 | 0,46 | 6,43E-01 |
| AT5G03330 | 0,61 | 1,90 | 3,06 | 0,62 | 5,34E-01 |
| AT5G03355 | 0,46 | -1,69 | 3,32 | -0,51 | 6,12E-01 |
| AT5G03415 | 0,36 | 1,26 | 4,28 | 0,29 | 7,68E-01 |
| AT5G03420 | 0,63 | 1,86 | 2,44 | 0,76 | 4,46E-01 |
| AT5G03455 | 0,25 | 2,64 | 4,34 | 0,61 | 5,43E-01 |
| AT5G03460 | 0,54 | 1,16 | 3,12 | 0,37 | 7,11E-01 |
| AT5G03470 | 0,40 | 1,45 | 4,36 | 0,33 | 7,40E-01 |
| AT5G03490 | 0,11 | 1,28 | 4,35 | 0,29 | 7,68E-01 |
| AT5G03540 | 1,05 | 3,28 | 2,72 | 1,21 | 2,27E-01 |
| AT5G03650 | 0,91 | 0,08 | 1,66 | 0,05 | 9,61E-01 |
| AT5G03905 | 0,30 | 0,59 | 4,36 | 0,14 | 8,92E-01 |
| AT5G04040 | 1,27 | 3,07 | 1,92 | 1,60 | 1,10E-01 |
| AT5G04050 | 0,10 | 0,34 | 4,35 | 0,08 | 9,37E-01 |
| AT5G04060 | 0,37 | 1,48 | 4,32 | 0,34 | 7,32E-01 |
| AT5G04170 | 0,76 | 0,17 | 2,46 | 0,07 | 9,44E-01 |
| AT5G04190 | 0,13 | -1,34 | 4,35 | -0,31 | 7,58E-01 |
| AT5G04240 | 0,88 | 4,81 | 3,25 | 1,48 | 1,39E-01 |
| AT5G04280 | 1,25 | 2,14 | 1,72 | 1,24 | 2,14E-01 |
| AT5G04290 | 0,66 | -0,19 | 3,16 | -0,06 | 9,53E-01 |
| AT5G04360 | 0,45 | 2,56 | 4,15 | 0,62 | 5,38E-01 |
| AT5G00960 | 0,96 | 0,75 | 1,91 | 0,39 | 6,94E-01 |
| AT5G04460 | 0,56 | 2,88 | 3,08 | 0,94 | 3,49E-01 |
| AT5G04490 | 0,46 | -0,41 | 3,24 | -0,13 | 9,00E-01 |
| AT5G04510 | 0,58 | 1,99 | 2,66 | 0,75 | 4,54E-01 |
| AT5G04600 | 0,71 | 0,42 | 2,23 | 0,19 | 8,50E-01 |
| AT5G04620 | 0,43 | 2,13 | 4,38 | 0,49 | 6,26E-01 |
| AT5G04630 | 0,57 | 3,70 | 3,74 | 0,99 | 3,22E-01 |
| AT5G04660 | 0,24 | 1,12 | 4,33 | 0,26 | 7,96E-01 |
| AT5G00985 | 0,28 | 1,87 | 4,37 | 0,43 | 6,69E-01 |
| AT5G04800 | 0,67 | 0,01 | 2,06 | 0,01 | 9,95E-01 |
| AT5G04885 | 0,27 | 0,31 | 3,64 | 0,08 | 9,33E-01 |
| AT5G04910 | 0,42 | -0,01 | 4,08 | 0,00 | 9,98E-01 |
| AT5G05070 | 0,40 | 1,32 | 4,27 | 0,31 | 7,57E-01 |
| AT5G05090 | 0,35 | -0,54 | 3,48 | -0,15 | 8,77E-01 |
| AT5G05160 | 0,72 | 2,38 | 2,78 | 0,86 | 3,92E-01 |
| AT5G05210 | 0,28 | -0,21 | 4,31 | -0,05 | 9,62E-01 |
| AT5G05220 | 0,08 | 2,08 | 4,38 | 0,47 | 6,35E-01 |
| AT5G05230 | 0,08 | 2,08 | 4,38 | 0,47 | 6,35E-01 |
| AT5G05310 | 0,18 | -1,39 | 4,34 | -0,32 | 7,49E-01 |
| AT5G05440 | 0,16 | 1,16 | 4,38 | 0,27 | 7,90E-01 |
| AT5G05490 | 0,55 | 2,06 | 4,37 | 0,47 | 6,37E-01 |
| AT5G05560 | 1,14 | 0,75 | 1,73 | 0,43 | 6,66E-01 |
| AT5G05570 | 0,62 | 2,93 | 4,31 | 0,68 | 4,96E-01 |
| AT5G05600 | 0,31 | -0,67 | 4,39 | -0,15 | 8,79E-01 |
| AT5G05630 | 0,08 | -0,56 | 4,38 | -0,13 | 8,99E-01 |
| AT5G05680 | 1,56 | 3,24 | 1,84 | 1,75 | 7,94E-02 |
| AT5G05710 | 0,20 | 2,11 | 4,37 | 0,48 | 6,29E-01 |
| AT5G05780 | 0,96 | 3,65 | 2,46 | 1,49 | 1,37E-01 |
| AT5G05950 | 0,83 | 1,04 | 2,31 | 0,45 | 6,55E-01 |
| AT5G05970 | 0,30 | 1,05 | 4,33 | 0,24 | 8,09E-01 |
| AT5G05985 | 0,55 | 2,52 | 3,84 | 0,66 | 5,12E-01 |
| AT5G06000 | 0,23 | -1,63 | 4,33 | -0,38 | 7,06E-01 |
| AT5G06050 | 0,39 | 2,48 | 4,27 | 0,58 | 5,61E-01 |
| AT5G06100 | 0,21 | 2,34 | 4,32 | 0,54 | 5,89E-01 |
| AT5G06120 | 1,17 | 3,96 | 2,42 | 1,64 | 1,01E-01 |

| | | | | | |
|---|---|---|---|---|---|
| AT5G06170 | 0,31 | -2,37 | 4,30 | -0,55 | 5,81E-01 |
| AT5G06220 | 0,84 | 2,85 | 2,46 | 1,16 | 2,46E-01 |
| AT5G06265 | 0,26 | -0,69 | 4,29 | -0,16 | 8,72E-01 |
| AT5G06290 | 1,37 | 1,80 | 1,84 | 0,98 | 3,28E-01 |
| AT5G06310 | 0,33 | 0,43 | 4,24 | 0,10 | 9,20E-01 |
| AT5G01255 | 0,70 | 0,77 | 2,64 | 0,29 | 7,72E-01 |
| AT5G06350 | 0,36 | 0,38 | 3,38 | 0,11 | 9,10E-01 |
| AT5G06360 | 0,40 | 1,32 | 4,27 | 0,31 | 7,57E-01 |
| AT5G06380 | 0,08 | 2,08 | 4,38 | 0,47 | 6,35E-01 |
| AT5G06390 | 0,14 | -0,31 | 4,33 | -0,07 | 9,43E-01 |
| AT5G06460 | 1,60 | 3,11 | 1,79 | 1,73 | 8,34E-02 |
| AT5G06645 | 0,58 | -3,73 | 4,27 | -0,87 | 3,82E-01 |
| AT5G06660 | 0,96 | 2,22 | 3,44 | 0,64 | 5,19E-01 |
| AT5G06680 | 0,39 | -0,05 | 4,31 | -0,01 | 9,90E-01 |
| AT5G06700 | 0,91 | 0,14 | 1,71 | 0,08 | 9,35E-01 |
| AT5G06811 | 0,11 | 1,33 | 4,36 | 0,31 | 7,59E-01 |
| AT5G06820 | 0,12 | 2,81 | 4,34 | 0,65 | 5,18E-01 |
| AT5G06860 | 0,22 | -0,39 | 4,32 | -0,09 | 9,28E-01 |
| AT5G06870 | 0,67 | -1,59 | 2,71 | -0,59 | 5,56E-01 |
| AT5G06950 | 0,61 | 1,96 | 3,09 | 0,64 | 5,25E-01 |
| AT5G06980 | 0,16 | 1,97 | 4,33 | 0,45 | 6,50E-01 |
| AT5G07090 | 1,54 | -0,68 | 1,46 | -0,46 | 6,43E-01 |
| AT5G07110 | 0,85 | 3,60 | 3,35 | 1,07 | 2,83E-01 |
| AT5G01315 | 0,64 | 1,95 | 2,97 | 0,66 | 5,11E-01 |
| AT5G07130 | 0,33 | 2,78 | 4,34 | 0,64 | 5,21E-01 |
| AT5G07140 | 1,24 | 4,15 | 2,47 | 1,68 | 9,33E-02 |
| AT5G07210 | 1,09 | -0,88 | 1,85 | -0,48 | 6,35E-01 |
| AT5G07240 | 1,19 | 3,36 | 2,08 | 1,62 | 1,06E-01 |
| AT5G07270 | 0,15 | -0,41 | 4,32 | -0,09 | 9,25E-01 |
| AT5G07340 | 0,98 | 2,23 | 2,14 | 1,04 | 2,97E-01 |
| AT5G07410 | 1,62 | -1,51 | 1,55 | -0,97 | 3,30E-01 |
| AT5G07430 | 0,50 | -1,97 | 3,87 | -0,51 | 6,12E-01 |
| AT5G07440 | 0,36 | 2,51 | 4,35 | 0,58 | 5,64E-01 |
| AT5G07580 | 0,75 | -0,07 | 2,07 | -0,03 | 9,73E-01 |
| AT5G07610 | 0,48 | 0,17 | 3,93 | 0,04 | 9,65E-01 |
| AT5G07760 | 0,08 | 2,33 | 4,36 | 0,53 | 5,93E-01 |
| AT5G07770 | 0,16 | 1,27 | 4,37 | 0,29 | 7,71E-01 |
| AT5G07790 | 0,58 | 2,52 | 4,32 | 0,58 | 5,60E-01 |
| AT5G07830 | 0,86 | 0,06 | 2,51 | 0,03 | 9,79E-01 |
| AT5G07860 | 0,58 | 1,44 | 3,91 | 0,37 | 7,12E-01 |
| AT5G07870 | 0,49 | 2,86 | 4,33 | 0,66 | 5,09E-01 |
| AT5G07960 | 0,12 | 1,65 | 4,34 | 0,38 | 7,04E-01 |
| AT5G08050 | 0,48 | 1,60 | 4,12 | 0,39 | 6,98E-01 |
| AT5G08060 | 0,10 | 0,34 | 4,35 | 0,08 | 9,37E-01 |
| AT5G08070 | 0,35 | 0,00 | 4,41 | 0,00 | 1,00E+00 |
| AT5G08139 | 0,37 | -0,89 | 4,21 | -0,21 | 8,32E-01 |
| AT5G08190 | 0,29 | 3,27 | 4,33 | 0,76 | 4,49E-01 |
| AT5G08260 | 0,90 | 1,10 | 2,42 | 0,45 | 6,51E-01 |
| AT5G01545 | 0,37 | 0,00 | 4,40 | 0,00 | 1,00E+00 |
| AT5G08290 | 0,74 | 1,09 | 1,98 | 0,55 | 5,82E-01 |
| AT5G08300 | 0,94 | -0,11 | 2,22 | -0,05 | 9,61E-01 |
| AT5G08330 | 1,56 | -1,15 | 1,60 | -0,72 | 4,74E-01 |
| AT5G08335 | 0,18 | 0,41 | 4,28 | 0,10 | 9,23E-01 |
| AT5G08340 | 0,58 | 2,44 | 3,97 | 0,62 | 5,38E-01 |
| AT5G08390 | 0,26 | 1,88 | 4,28 | 0,44 | 6,61E-01 |
| AT5G08430 | 0,57 | 3,33 | 4,32 | 0,77 | 4,41E-01 |

| | | | | | |
|---|---|---|---|---|---|
| AT5G08440 | 0,59 | 1,82 | 3,89 | 0,47 | 6,40E-01 |
| AT5G08470 | 0,81 | 2,95 | 2,71 | 1,09 | 2,76E-01 |
| AT5G08510 | 0,51 | 0,89 | 3,29 | 0,27 | 7,86E-01 |
| AT5G08540 | 0,41 | 0,92 | 3,31 | 0,28 | 7,82E-01 |
| AT5G01555 | 0,36 | 0,28 | 4,34 | 0,06 | 9,48E-01 |
| AT5G08570 | 1,20 | 0,76 | 1,56 | 0,49 | 6,25E-01 |
| AT5G08590 | 0,63 | 0,84 | 2,61 | 0,32 | 7,48E-01 |
| AT5G08600 | 0,28 | 1,87 | 4,37 | 0,43 | 6,69E-01 |
| AT5G08620 | 1,12 | 1,44 | 1,74 | 0,83 | 4,08E-01 |
| AT5G08660 | 0,33 | -0,43 | 4,37 | -0,10 | 9,22E-01 |
| AT5G08695 | 0,20 | 2,11 | 4,37 | 0,48 | 6,29E-01 |
| AT5G08750 | 0,11 | 1,28 | 4,35 | 0,29 | 7,68E-01 |
| AT5G08760 | 0,55 | -1,23 | 3,92 | -0,31 | 7,53E-01 |
| AT5G08780 | 0,49 | 2,21 | 3,15 | 0,70 | 4,82E-01 |
| AT5G09220 | 0,82 | -0,90 | 2,95 | -0,31 | 7,60E-01 |
| AT5G09400 | 1,22 | 2,62 | 1,88 | 1,39 | 1,64E-01 |
| AT5G09440 | 0,77 | 0,38 | 2,18 | 0,17 | 8,63E-01 |
| AT5G09445 | 0,26 | 2,13 | 4,38 | 0,49 | 6,27E-01 |
| AT5G09450 | 1,25 | 0,95 | 2,15 | 0,44 | 6,59E-01 |
| AT5G09600 | 0,62 | 2,49 | 3,03 | 0,82 | 4,11E-01 |
| AT5G09760 | 0,14 | -0,31 | 4,33 | -0,07 | 9,43E-01 |
| AT5G10010 | 0,47 | -0,58 | 3,37 | -0,17 | 8,63E-01 |
| AT5G10020 | 0,41 | 3,16 | 4,33 | 0,73 | 4,66E-01 |
| AT5G10100 | 0,16 | 0,49 | 4,33 | 0,11 | 9,09E-01 |
| AT5G10190 | 0,41 | 1,90 | 3,30 | 0,58 | 5,65E-01 |
| AT5G10200 | 0,81 | 3,07 | 3,43 | 0,90 | 3,70E-01 |
| AT5G10340 | 1,40 | -2,29 | 2,16 | -1,06 | 2,89E-01 |
| AT5G10370 | 1,36 | 3,14 | 1,67 | 1,88 | 5,97E-02 |
| AT5G10410 | 0,05 | -0,58 | 4,38 | -0,13 | 8,95E-01 |
| AT5G10460 | 0,08 | 2,33 | 4,36 | 0,53 | 5,93E-01 |
| AT5G01765 | 1,15 | 1,75 | 1,73 | 1,01 | 3,12E-01 |
| AT5G10600 | 0,40 | 0,21 | 4,22 | 0,05 | 9,60E-01 |
| AT5G10630 | 0,28 | 1,13 | 4,28 | 0,27 | 7,91E-01 |
| AT5G10660 | 0,46 | 0,14 | 2,66 | 0,05 | 9,57E-01 |
| AT5G10695 | 0,85 | -1,20 | 1,89 | -0,63 | 5,26E-01 |
| AT5G10745 | 0,35 | 0,28 | 3,38 | 0,08 | 9,33E-01 |
| AT5G10750 | 1,12 | 0,99 | 2,12 | 0,46 | 6,42E-01 |
| AT5G10780 | 0,76 | 1,01 | 2,52 | 0,40 | 6,88E-01 |
| AT5G10790 | 0,93 | 1,50 | 1,94 | 0,77 | 4,39E-01 |
| AT5G10920 | 0,91 | 0,91 | 2,14 | 0,42 | 6,72E-01 |
| AT5G11060 | 1,02 | -0,57 | 1,46 | -0,39 | 6,99E-01 |
| AT5G11090 | 0,49 | 0,33 | 2,61 | 0,13 | 9,00E-01 |
| AT5G11110 | 0,50 | -0,05 | 3,95 | -0,01 | 9,90E-01 |
| AT5G11170 | 1,42 | 1,50 | 1,99 | 0,75 | 4,50E-01 |
| AT5G11200 | 0,33 | 2,56 | 4,31 | 0,59 | 5,52E-01 |
| AT5G11280 | 0,20 | 0,30 | 4,34 | 0,07 | 9,46E-01 |
| AT5G11340 | 0,28 | 1,96 | 4,32 | 0,45 | 6,51E-01 |
| AT5G11350 | 0,82 | 3,04 | 2,93 | 1,04 | 2,99E-01 |
| AT5G11390 | 0,30 | 2,91 | 4,34 | 0,67 | 5,04E-01 |
| AT5G11420 | 1,15 | -1,40 | 1,93 | -0,73 | 4,66E-01 |
| AT5G11430 | 0,70 | 0,65 | 2,16 | 0,30 | 7,62E-01 |
| AT5G11500 | 0,94 | 2,50 | 1,85 | 1,35 | 1,78E-01 |
| AT5G11510 | 0,81 | 3,19 | 2,35 | 1,36 | 1,75E-01 |
| AT5G11530 | 0,81 | 2,02 | 2,31 | 0,87 | 3,83E-01 |
| AT5G11580 | 0,12 | 1,15 | 4,35 | 0,26 | 7,92E-01 |
| AT5G11650 | 0,38 | 0,41 | 3,34 | 0,12 | 9,03E-01 |

| | | | | | |
|---|---|---|---|---|---|
| AT5G11680 | 0,19 | 1,14 | 4,29 | 0,26 | 7,91E-01 |
| AT5G11710 | 1,24 | 0,59 | 1,51 | 0,39 | 6,96E-01 |
| AT5G11720 | 1,68 | 3,40 | 2,06 | 1,65 | 9,87E-02 |
| AT5G11730 | 0,89 | -0,21 | 1,82 | -0,12 | 9,08E-01 |
| AT5G11760 | 0,22 | 0,37 | 4,35 | 0,08 | 9,32E-01 |
| AT5G11890 | 0,82 | -0,39 | 1,98 | -0,19 | 8,46E-01 |
| AT5G11920 | 0,12 | 1,15 | 4,35 | 0,26 | 7,92E-01 |
| AT5G11977 | 0,12 | 2,87 | 4,34 | 0,66 | 5,08E-01 |
| AT5G12090 | 0,79 | -2,84 | 2,27 | -1,25 | 2,11E-01 |
| AT5G12100 | 0,44 | 3,94 | 4,31 | 0,91 | 3,61E-01 |
| AT5G12110 | 1,52 | 2,37 | 1,47 | 1,61 | 1,08E-01 |
| AT5G12210 | 0,65 | 2,21 | 3,66 | 0,60 | 5,47E-01 |
| AT5G12260 | 0,32 | 1,41 | 4,34 | 0,32 | 7,45E-01 |
| AT5G12350 | 0,49 | 2,12 | 4,13 | 0,51 | 6,07E-01 |
| AT5G12400 | 0,93 | 2,82 | 2,75 | 1,03 | 3,04E-01 |
| AT5G12850 | 0,97 | -0,66 | 1,78 | -0,37 | 7,12E-01 |
| AT5G12950 | 0,53 | -0,40 | 2,35 | -0,17 | 8,66E-01 |
| AT5G12960 | 0,27 | 2,11 | 4,38 | 0,48 | 6,30E-01 |
| AT5G12970 | 0,24 | 0,00 | 4,41 | 0,00 | 1,00E+00 |
| AT5G13030 | 0,96 | -0,44 | 1,85 | -0,24 | 8,12E-01 |
| AT5G13200 | 1,26 | -0,78 | 1,52 | -0,51 | 6,07E-01 |
| AT5G13210 | 0,35 | 1,45 | 3,36 | 0,43 | 6,66E-01 |
| AT5G13260 | 0,92 | 1,49 | 2,16 | 0,69 | 4,90E-01 |
| AT5G13280 | 0,45 | 0,87 | 4,14 | 0,21 | 8,34E-01 |
| AT5G13310 | 0,31 | 2,33 | 4,31 | 0,54 | 5,88E-01 |
| AT5G13320 | 1,27 | 1,67 | 1,68 | 1,00 | 3,19E-01 |
| AT5G13330 | 0,37 | 3,54 | 4,32 | 0,82 | 4,12E-01 |
| AT5G13370 | 0,68 | 0,05 | 3,21 | 0,01 | 9,88E-01 |
| AT5G13380 | 0,54 | 3,73 | 4,32 | 0,86 | 3,88E-01 |
| AT5G13500 | 0,32 | -0,08 | 4,30 | -0,02 | 9,86E-01 |
| AT5G13510 | 0,48 | 1,92 | 3,30 | 0,58 | 5,62E-01 |
| AT5G13580 | 0,29 | 3,75 | 4,32 | 0,87 | 3,85E-01 |
| AT5G13640 | 0,84 | 1,55 | 3,00 | 0,52 | 6,05E-01 |
| AT5G13690 | 0,61 | 2,36 | 3,07 | 0,77 | 4,42E-01 |
| AT5G13720 | 0,21 | 0,78 | 4,40 | 0,18 | 8,60E-01 |
| AT5G13740 | 0,19 | 1,14 | 4,29 | 0,26 | 7,91E-01 |
| AT5G13750 | 0,26 | 2,30 | 4,37 | 0,53 | 5,98E-01 |
| AT5G13760 | 0,26 | -0,12 | 4,30 | -0,03 | 9,77E-01 |
| AT5G13810 | 0,93 | 0,04 | 1,67 | 0,02 | 9,81E-01 |
| AT5G13820 | 0,28 | 2,64 | 4,31 | 0,61 | 5,40E-01 |
| AT5G13930 | 1,07 | 4,26 | 2,22 | 1,92 | 5,49E-02 |
| AT5G13980 | 0,97 | 1,66 | 2,29 | 0,72 | 4,69E-01 |
| AT5G14030 | 0,98 | 2,74 | 2,36 | 1,16 | 2,46E-01 |
| AT5G14050 | 0,50 | 1,73 | 3,07 | 0,56 | 5,73E-01 |
| AT5G14120 | 1,26 | -1,42 | 1,54 | -0,92 | 3,55E-01 |
| AT5G14200 | 1,42 | 0,33 | 1,37 | 0,25 | 8,06E-01 |
| AT5G14230 | 0,39 | 0,06 | 3,28 | 0,02 | 9,86E-01 |
| AT5G14260 | 0,68 | 1,15 | 2,42 | 0,47 | 6,35E-01 |
| AT5G14280 | 0,43 | 2,30 | 4,38 | 0,52 | 6,00E-01 |
| AT5G14285 | 0,21 | 3,22 | 4,33 | 0,74 | 4,58E-01 |
| AT5G14480 | 0,25 | 1,74 | 4,31 | 0,40 | 6,87E-01 |
| AT5G14610 | 0,64 | 0,38 | 3,12 | 0,12 | 9,02E-01 |
| AT5G14670 | 0,08 | 2,33 | 4,36 | 0,53 | 5,93E-01 |
| AT5G14880 | 0,37 | 1,18 | 3,39 | 0,35 | 7,29E-01 |
| AT5G14890 | 0,99 | 4,83 | 4,01 | 1,20 | 2,28E-01 |
| AT5G14910 | 0,77 | 1,37 | 2,51 | 0,54 | 5,87E-01 |

| | | | | | |
|---|---|---|---|---|---|
| AT5G14970 | 0,18 | 1,48 | 4,31 | 0,34 | 7,32E-01 |
| AT5G14990 | 0,42 | 2,85 | 4,17 | 0,68 | 4,95E-01 |
| AT5G15010 | 0,38 | -0,07 | 4,40 | -0,02 | 9,87E-01 |
| AT5G02295 | 1,40 | -2,31 | 1,49 | -1,55 | 1,21E-01 |
| AT5G15080 | 0,92 | 2,91 | 2,92 | 1,00 | 3,19E-01 |
| AT5G15230 | 0,74 | 3,08 | 2,88 | 1,07 | 2,85E-01 |
| AT5G15240 | 0,05 | -0,58 | 4,38 | -0,13 | 8,95E-01 |
| AT5G15270 | 0,61 | 1,31 | 3,05 | 0,43 | 6,69E-01 |
| AT5G15280 | 0,54 | 2,81 | 3,84 | 0,73 | 4,64E-01 |
| AT5G15320 | 0,21 | 1,12 | 4,33 | 0,26 | 7,95E-01 |
| AT5G15330 | 0,11 | 1,33 | 4,36 | 0,31 | 7,59E-01 |
| AT5G15530 | 0,12 | 2,81 | 4,34 | 0,65 | 5,18E-01 |
| AT5G15570 | 0,44 | 4,02 | 4,31 | 0,93 | 3,51E-01 |
| AT5G15610 | 0,89 | 0,54 | 1,74 | 0,31 | 7,57E-01 |
| AT5G15685 | 0,72 | 0,05 | 1,85 | 0,03 | 9,77E-01 |
| AT5G15710 | 0,74 | 2,16 | 3,55 | 0,61 | 5,44E-01 |
| AT5G15730 | 0,59 | 2,18 | 2,50 | 0,87 | 3,83E-01 |
| AT5G15750 | 0,54 | -0,84 | 3,89 | -0,22 | 8,29E-01 |
| AT5G15820 | 0,22 | 0,78 | 4,33 | 0,18 | 8,58E-01 |
| AT5G15930 | 0,32 | 1,93 | 4,34 | 0,44 | 6,57E-01 |
| AT5G15950 | 1,23 | 4,84 | 2,07 | 2,33 | 1,96E-02 |
| AT5G15960 | 0,51 | 0,00 | 4,41 | 0,00 | 1,00E+00 |
| AT5G16010 | 0,36 | -0,10 | 4,33 | -0,02 | 9,82E-01 |
| AT5G16030 | 0,58 | -0,73 | 3,30 | -0,22 | 8,24E-01 |
| AT5G16120 | 0,30 | 1,62 | 4,35 | 0,37 | 7,10E-01 |
| AT5G16130 | 1,11 | 2,31 | 2,05 | 1,13 | 2,59E-01 |
| AT5G16150 | 1,33 | 0,34 | 1,48 | 0,23 | 8,16E-01 |
| AT5G16260 | 0,11 | -1,28 | 4,35 | -0,29 | 7,69E-01 |
| AT5G16290 | 0,43 | 1,19 | 4,26 | 0,28 | 7,80E-01 |
| AT5G16300 | 1,23 | 1,97 | 1,52 | 1,30 | 1,94E-01 |
| AT5G16440 | 0,67 | 0,67 | 2,54 | 0,26 | 7,91E-01 |
| AT5G16470 | 0,69 | -0,06 | 2,46 | -0,03 | 9,79E-01 |
| AT5G16540 | 0,32 | 2,42 | 4,35 | 0,56 | 5,77E-01 |
| AT5G16550 | 0,57 | 1,70 | 2,63 | 0,65 | 5,17E-01 |
| AT5G02555 | 0,05 | -0,58 | 4,38 | -0,13 | 8,95E-01 |
| AT5G16590 | 0,87 | 2,07 | 2,07 | 1,00 | 3,18E-01 |
| AT5G16630 | 0,92 | 3,42 | 2,61 | 1,31 | 1,90E-01 |
| AT5G16715 | 0,74 | 1,51 | 2,49 | 0,61 | 5,43E-01 |
| AT5G16730 | 1,05 | 0,55 | 1,78 | 0,31 | 7,56E-01 |
| AT5G16760 | 1,05 | 1,31 | 2,00 | 0,66 | 5,12E-01 |
| AT5G16820 | 0,94 | 2,88 | 2,29 | 1,26 | 2,07E-01 |
| AT5G16880 | 1,27 | 1,69 | 1,57 | 1,08 | 2,81E-01 |
| AT5G16900 | 0,33 | 3,34 | 4,32 | 0,77 | 4,40E-01 |
| AT5G16910 | 0,98 | 4,38 | 2,54 | 1,73 | 8,42E-02 |
| AT5G17167 | 0,20 | -0,66 | 4,32 | -0,15 | 8,78E-01 |
| AT5G17170 | 0,33 | 0,14 | 4,34 | 0,03 | 9,75E-01 |
| AT5G17190 | 0,51 | -2,35 | 3,10 | -0,76 | 4,48E-01 |
| AT5G17270 | 0,61 | 2,47 | 4,35 | 0,57 | 5,70E-01 |
| AT5G17280 | 0,29 | 1,74 | 4,31 | 0,40 | 6,87E-01 |
| AT5G17305 | 0,08 | 2,33 | 4,36 | 0,53 | 5,93E-01 |
| AT5G17440 | 0,24 | 1,18 | 4,36 | 0,27 | 7,87E-01 |
| AT5G17530 | 0,59 | 1,29 | 3,10 | 0,42 | 6,78E-01 |
| AT5G17580 | 0,17 | 3,04 | 4,33 | 0,70 | 4,83E-01 |
| AT5G17630 | 0,33 | 1,34 | 4,39 | 0,30 | 7,60E-01 |
| AT5G17670 | 0,39 | 1,78 | 4,41 | 0,40 | 6,86E-01 |
| AT5G17680 | 0,30 | 2,91 | 4,34 | 0,67 | 5,04E-01 |

| | | | | | |
|---|---|---|---|---|---|
| AT5G17850 | 0,18 | -2,02 | 4,31 | -0,47 | 6,40E-01 |
| AT5G17870 | 0,86 | -0,21 | 2,21 | -0,10 | 9,24E-01 |
| AT5G17900 | 0,30 | 0,09 | 4,30 | 0,02 | 9,84E-01 |
| AT5G17910 | 1,47 | 1,69 | 1,75 | 0,96 | 3,36E-01 |
| AT5G17980 | 0,38 | 2,20 | 4,19 | 0,53 | 5,99E-01 |
| AT5G17990 | 0,55 | 1,50 | 3,98 | 0,38 | 7,05E-01 |
| AT5G18040 | 0,71 | 2,17 | 2,34 | 0,93 | 3,54E-01 |
| AT5G18065 | 0,30 | 1,11 | 3,61 | 0,31 | 7,59E-01 |
| AT5G18070 | 0,62 | 0,83 | 2,56 | 0,32 | 7,46E-01 |
| AT5G18085 | 0,24 | 0,00 | 4,41 | 0,00 | 1,00E+00 |
| AT5G18100 | 0,33 | 1,01 | 4,29 | 0,24 | 8,14E-01 |
| AT5G18130 | 1,11 | 2,80 | 2,14 | 1,31 | 1,91E-01 |
| AT5G18140 | 1,06 | 2,35 | 2,50 | 0,94 | 3,48E-01 |
| AT5G18170 | 0,16 | 1,14 | 4,30 | 0,27 | 7,90E-01 |
| AT5G18200 | 0,08 | 2,08 | 4,38 | 0,47 | 6,35E-01 |
| AT5G18245 | 1,46 | -4,03 | 2,64 | -1,53 | 1,27E-01 |
| AT5G18255 | 1,25 | 1,62 | 1,58 | 1,03 | 3,04E-01 |
| AT5G18310 | 0,76 | -2,37 | 2,30 | -1,03 | 3,03E-01 |
| AT5G18390 | 0,33 | 1,52 | 4,34 | 0,35 | 7,26E-01 |
| AT5G18470 | 1,32 | 2,15 | 1,64 | 1,32 | 1,88E-01 |
| AT5G18500 | 0,37 | 0,68 | 4,29 | 0,16 | 8,74E-01 |
| AT5G18520 | 1,46 | 0,61 | 1,40 | 0,44 | 6,61E-01 |
| AT5G18580 | 0,90 | 1,04 | 2,03 | 0,51 | 6,09E-01 |
| AT5G18610 | 1,43 | 1,66 | 1,97 | 0,84 | 3,99E-01 |
| AT5G18630 | 0,28 | -1,71 | 4,32 | -0,39 | 6,93E-01 |
| AT5G18636 | 0,58 | -1,29 | 2,27 | -0,57 | 5,71E-01 |
| AT5G18650 | 0,68 | 0,24 | 3,04 | 0,08 | 9,36E-01 |
| AT5G18670 | 0,21 | 3,36 | 4,33 | 0,78 | 4,37E-01 |
| AT5G18750 | 0,77 | 3,17 | 3,30 | 0,96 | 3,37E-01 |
| AT5G18830 | 0,51 | 1,63 | 3,17 | 0,51 | 6,07E-01 |
| AT5G18890 | 0,98 | 1,79 | 4,33 | 0,41 | 6,80E-01 |
| AT5G18937 | 0,54 | -3,71 | 4,27 | -0,87 | 3,84E-01 |
| AT5G18940 | 1,00 | 2,42 | 2,72 | 0,89 | 3,75E-01 |
| AT5G18960 | 0,35 | 1,33 | 3,40 | 0,39 | 6,96E-01 |
| AT5G19000 | 0,44 | 0,51 | 3,35 | 0,15 | 8,79E-01 |
| AT5G19090 | 0,78 | -1,80 | 2,32 | -0,78 | 4,38E-01 |
| AT5G19180 | 0,77 | 0,82 | 2,07 | 0,40 | 6,91E-01 |
| AT5G19190 | 0,37 | 0,15 | 4,32 | 0,03 | 9,73E-01 |
| AT5G19210 | 0,49 | -1,30 | 3,26 | -0,40 | 6,90E-01 |
| AT5G19221 | 0,47 | 0,27 | 4,35 | 0,06 | 9,51E-01 |
| AT5G19240 | 0,65 | -0,16 | 1,96 | -0,08 | 9,35E-01 |
| AT5G19250 | 0,98 | -2,11 | 2,74 | -0,77 | 4,42E-01 |
| AT5G19290 | 0,41 | 0,62 | 3,28 | 0,19 | 8,49E-01 |
| AT5G19440 | 0,36 | 1,93 | 4,34 | 0,44 | 6,57E-01 |
| AT5G19500 | 1,08 | 2,42 | 2,27 | 1,06 | 2,87E-01 |
| AT5G19540 | 0,77 | 0,28 | 1,98 | 0,14 | 8,89E-01 |
| AT5G19550 | 0,90 | 1,19 | 1,86 | 0,64 | 5,23E-01 |
| AT5G19570 | 0,52 | 1,44 | 3,92 | 0,37 | 7,13E-01 |
| AT5G19620 | 0,50 | -0,54 | 3,97 | -0,14 | 8,91E-01 |
| AT5G19680 | 0,19 | 1,20 | 4,34 | 0,28 | 7,82E-01 |
| AT5G19750 | 0,55 | 4,02 | 4,30 | 0,93 | 3,50E-01 |
| AT5G19780 | 1,35 | 1,12 | 1,75 | 0,64 | 5,21E-01 |
| AT5G19990 | 0,77 | -0,07 | 2,28 | -0,03 | 9,77E-01 |
| AT5G20080 | 0,48 | -0,52 | 3,27 | -0,16 | 8,74E-01 |
| AT5G20120 | 0,55 | 3,10 | 4,34 | 0,71 | 4,75E-01 |
| AT5G20130 | 0,68 | 2,80 | 3,74 | 0,75 | 4,53E-01 |

| | | | | | |
|---|---|---|---|---|---|
| AT5G20150 | 0,56 | 1,69 | 2,51 | 0,67 | 5,00E-01 |
| AT5G20190 | 1,21 | 1,00 | 1,75 | 0,57 | 5,67E-01 |
| AT5G20200 | 1,58 | 4,28 | 1,79 | 2,39 | 1,67E-02 |
| AT5G20220 | 0,21 | 2,34 | 4,32 | 0,54 | 5,89E-01 |
| AT5G20350 | 1,36 | 2,24 | 1,56 | 1,44 | 1,50E-01 |
| AT5G20410 | 0,63 | 1,73 | 3,72 | 0,47 | 6,41E-01 |
| AT5G20480 | 0,15 | -0,41 | 4,32 | -0,09 | 9,25E-01 |
| AT5G20500 | 0,33 | 0,26 | 4,27 | 0,06 | 9,51E-01 |
| AT5G20590 | 0,85 | 1,77 | 2,14 | 0,83 | 4,09E-01 |
| AT5G20610 | 1,39 | 2,55 | 1,71 | 1,49 | 1,37E-01 |
| AT5G20660 | 1,27 | 2,20 | 1,75 | 1,26 | 2,09E-01 |
| AT5G20680 | 0,31 | 0,03 | 4,34 | 0,01 | 9,95E-01 |
| AT5G20710 | 0,12 | 2,81 | 4,34 | 0,65 | 5,18E-01 |
| AT5G20900 | 0,45 | -0,42 | 3,27 | -0,13 | 8,97E-01 |
| AT5G03255 | 0,31 | 2,17 | 4,31 | 0,50 | 6,14E-01 |
| AT5G20960 | 1,02 | 2,32 | 2,38 | 0,98 | 3,30E-01 |
| AT5G21020 | 1,50 | -0,29 | 1,33 | -0,22 | 8,29E-01 |
| AT5G21090 | 1,39 | -0,20 | 1,65 | -0,12 | 9,03E-01 |
| AT5G21105 | 1,68 | 0,68 | 1,44 | 0,47 | 6,35E-01 |
| AT5G21150 | 0,12 | 2,81 | 4,34 | 0,65 | 5,18E-01 |
| AT5G21222 | 1,01 | 2,34 | 2,66 | 0,88 | 3,80E-01 |
| AT5G21326 | 0,90 | 3,14 | 2,67 | 1,18 | 2,39E-01 |
| AT5G21482 | 0,24 | -0,10 | 4,27 | -0,02 | 9,82E-01 |
| AT5G21930 | 1,60 | 1,04 | 1,45 | 0,72 | 4,72E-01 |
| AT5G22000 | 1,27 | 1,56 | 1,71 | 0,92 | 3,59E-01 |
| AT5G22050 | 0,36 | -0,23 | 4,30 | -0,05 | 9,58E-01 |
| AT5G22070 | 0,57 | -1,39 | 2,93 | -0,47 | 6,36E-01 |
| AT5G22300 | 0,30 | -0,23 | 3,59 | -0,06 | 9,50E-01 |
| AT5G22315 | 0,20 | -0,80 | 4,37 | -0,18 | 8,55E-01 |
| AT5G22340 | 0,95 | 2,65 | 2,17 | 1,22 | 2,22E-01 |
| AT5G22420 | 0,42 | 2,98 | 4,15 | 0,72 | 4,73E-01 |
| AT5G22580 | 0,46 | 0,91 | 4,38 | 0,21 | 8,36E-01 |
| AT5G22590 | 1,03 | -3,36 | 2,49 | -1,35 | 1,77E-01 |
| AT5G22630 | 1,52 | 1,28 | 1,56 | 0,82 | 4,10E-01 |
| AT5G22640 | 1,19 | 2,76 | 1,65 | 1,67 | 9,50E-02 |
| AT5G22650 | 0,66 | -1,58 | 2,41 | -0,65 | 5,12E-01 |
| AT5G22690 | 0,12 | 2,81 | 4,34 | 0,65 | 5,18E-01 |
| AT5G22760 | 1,25 | 2,87 | 1,82 | 1,58 | 1,15E-01 |
| AT5G22794 | 0,08 | 2,33 | 4,36 | 0,53 | 5,93E-01 |
| AT5G22830 | 0,08 | -0,56 | 4,38 | -0,13 | 8,99E-01 |
| AT5G22950 | 0,12 | 0,31 | 4,35 | 0,07 | 9,44E-01 |
| AT5G23010 | 0,84 | -2,13 | 2,09 | -1,02 | 3,07E-01 |
| AT5G23070 | 0,19 | 0,28 | 4,33 | 0,07 | 9,48E-01 |
| AT5G03575 | 0,26 | 1,78 | 4,41 | 0,40 | 6,86E-01 |
| AT5G23120 | 0,65 | 1,08 | 2,06 | 0,52 | 6,01E-01 |
| AT5G23140 | 0,95 | 2,44 | 2,26 | 1,08 | 2,80E-01 |
| AT5G23150 | 1,41 | 2,46 | 1,99 | 1,24 | 2,16E-01 |
| AT5G23210 | 0,16 | 1,18 | 4,32 | 0,27 | 7,85E-01 |
| AT5G23280 | 1,01 | -0,18 | 2,43 | -0,08 | 9,40E-01 |
| AT5G23320 | 0,08 | 2,33 | 4,36 | 0,53 | 5,93E-01 |
| AT5G23330 | 0,43 | 2,13 | 4,38 | 0,49 | 6,26E-01 |
| AT5G23340 | 1,40 | 3,37 | 1,87 | 1,80 | 7,20E-02 |
| AT5G23490 | 0,49 | 0,48 | 2,55 | 0,19 | 8,50E-01 |
| AT5G23540 | 0,96 | -1,38 | 1,89 | -0,73 | 4,63E-01 |
| AT5G23570 | 0,58 | 0,16 | 2,29 | 0,07 | 9,45E-01 |
| AT5G23575 | 1,00 | -0,24 | 1,68 | -0,14 | 8,86E-01 |

| | | | | | |
|---|---|---|---|---|---|
| AT5G23600 | 0,08 | 2,33 | 4,36 | 0,53 | 5,93E-01 |
| AT5G23670 | 0,72 | 2,76 | 2,97 | 0,93 | 3,53E-01 |
| AT5G23720 | 0,51 | 0,89 | 3,29 | 0,27 | 7,86E-01 |
| AT5G23730 | 0,26 | 1,27 | 4,31 | 0,29 | 7,68E-01 |
| AT5G23890 | 0,99 | 2,90 | 2,68 | 1,08 | 2,80E-01 |
| AT5G23950 | 0,35 | -1,24 | 4,35 | -0,29 | 7,75E-01 |
| AT5G24020 | 0,16 | 1,18 | 4,32 | 0,27 | 7,85E-01 |
| AT5G24200 | 1,34 | 0,01 | 1,39 | 0,01 | 9,94E-01 |
| AT5G24210 | 0,96 | -0,44 | 2,13 | -0,20 | 8,38E-01 |
| AT5G24360 | 0,40 | 2,41 | 3,31 | 0,73 | 4,66E-01 |
| AT5G24380 | 0,41 | 2,07 | 3,36 | 0,62 | 5,38E-01 |
| AT5G24450 | 0,69 | 0,96 | 3,56 | 0,27 | 7,87E-01 |
| AT5G24460 | 0,88 | 1,87 | 1,91 | 0,98 | 3,27E-01 |
| AT5G24490 | 1,61 | 1,24 | 1,88 | 0,66 | 5,08E-01 |
| AT5G24610 | 0,55 | 1,46 | 2,62 | 0,56 | 5,77E-01 |
| AT5G24830 | 0,08 | 2,33 | 4,36 | 0,53 | 5,93E-01 |
| AT5G24980 | 0,43 | 1,13 | 3,32 | 0,34 | 7,33E-01 |
| AT5G25040 | 0,67 | -1,84 | 2,23 | -0,82 | 4,10E-01 |
| AT5G25120 | 0,51 | -1,35 | 3,07 | -0,44 | 6,59E-01 |
| AT5G03925 | 0,17 | -1,97 | 4,31 | -0,46 | 6,47E-01 |
| AT5G25130 | 1,42 | -1,96 | 1,46 | -1,34 | 1,80E-01 |
| AT5G25140 | 0,28 | 2,67 | 4,31 | 0,62 | 5,35E-01 |
| AT5G25150 | 0,27 | -0,36 | 4,31 | -0,08 | 9,34E-01 |
| AT5G25170 | 0,33 | 2,23 | 4,28 | 0,52 | 6,03E-01 |
| AT5G25210 | 0,42 | 1,01 | 3,24 | 0,31 | 7,56E-01 |
| AT5G25220 | 1,05 | 0,98 | 1,79 | 0,55 | 5,83E-01 |
| AT5G25230 | 1,66 | 3,00 | 1,64 | 1,83 | 6,74E-02 |
| AT5G25250 | 1,43 | -0,55 | 1,63 | -0,34 | 7,34E-01 |
| AT5G25260 | 1,02 | 4,60 | 2,67 | 1,72 | 8,49E-02 |
| AT5G25340 | 0,46 | 3,27 | 4,05 | 0,81 | 4,20E-01 |
| AT5G25460 | 1,53 | -1,31 | 1,98 | -0,66 | 5,08E-01 |
| AT5G25520 | 1,48 | 4,47 | 2,41 | 1,86 | 6,34E-02 |
| AT5G25560 | 0,53 | 1,50 | 3,18 | 0,47 | 6,39E-01 |
| AT5G25580 | 0,12 | 2,87 | 4,34 | 0,66 | 5,08E-01 |
| AT5G25630 | 0,47 | 2,34 | 3,12 | 0,75 | 4,55E-01 |
| AT5G25754 | 1,48 | 0,54 | 1,29 | 0,42 | 6,77E-01 |
| AT5G04145 | 0,89 | 0,49 | 2,30 | 0,21 | 8,32E-01 |
| AT5G25900 | 0,33 | 2,86 | 4,33 | 0,66 | 5,09E-01 |
| AT5G25910 | 0,60 | 4,12 | 3,67 | 1,12 | 2,62E-01 |
| AT5G25930 | 0,70 | 0,04 | 2,30 | 0,02 | 9,86E-01 |
| AT5G25980 | 1,32 | -0,51 | 1,52 | -0,34 | 7,35E-01 |
| AT5G26038 | 1,13 | -0,86 | 1,88 | -0,45 | 6,50E-01 |
| AT5G26146 | 0,89 | 0,70 | 2,08 | 0,33 | 7,39E-01 |
| AT5G26147 | 0,89 | 0,70 | 2,08 | 0,33 | 7,39E-01 |
| AT5G26200 | 0,12 | 2,81 | 4,34 | 0,65 | 5,18E-01 |
| AT5G26240 | 0,84 | 2,91 | 2,01 | 1,44 | 1,49E-01 |
| AT5G26270 | 0,26 | -0,28 | 4,32 | -0,06 | 9,49E-01 |
| AT5G26290 | 0,79 | -1,16 | 2,13 | -0,54 | 5,86E-01 |
| AT5G26570 | 1,51 | 2,64 | 1,75 | 1,51 | 1,32E-01 |
| AT5G26740 | 0,58 | 1,57 | 2,60 | 0,60 | 5,46E-01 |
| AT5G26731 | 0,38 | -0,25 | 4,31 | -0,06 | 9,53E-01 |
| AT5G26710 | 0,73 | 2,71 | 2,49 | 1,09 | 2,77E-01 |
| AT5G26670 | 0,26 | 0,00 | 4,41 | 0,00 | 1,00E+00 |
| AT5G26610 | 0,16 | -1,98 | 4,31 | -0,46 | 6,47E-01 |
| AT5G26600 | 0,82 | 1,53 | 2,49 | 0,62 | 5,37E-01 |
| AT5G26780 | 0,39 | 0,41 | 2,75 | 0,15 | 8,81E-01 |

| | | | | | |
|---|---|---|---|---|---|
| AT5G26820 | 0,42 | 2,45 | 4,36 | 0,56 | 5,74E-01 |
| AT5G26850 | 1,21 | 3,66 | 2,27 | 1,61 | 1,07E-01 |
| AT5G26860 | 1,16 | 2,23 | 1,66 | 1,35 | 1,78E-01 |
| AT5G26930 | 0,33 | 2,11 | 4,36 | 0,48 | 6,29E-01 |
| AT5G26940 | 0,66 | 2,87 | 3,72 | 0,77 | 4,40E-01 |
| AT5G26990 | 0,23 | -0,64 | 4,39 | -0,15 | 8,84E-01 |
| AT5G27120 | 0,91 | 3,52 | 3,47 | 1,01 | 3,11E-01 |
| AT5G27150 | 0,57 | 1,68 | 2,90 | 0,58 | 5,61E-01 |
| AT5G27170 | 0,25 | -1,14 | 4,30 | -0,26 | 7,92E-01 |
| AT5G27230 | 0,08 | -0,56 | 4,38 | -0,13 | 8,99E-01 |
| AT5G27240 | 0,05 | -0,58 | 4,38 | -0,13 | 8,95E-01 |
| AT5G27270 | 0,17 | 0,31 | 4,36 | 0,07 | 9,43E-01 |
| AT5G27280 | 0,34 | 0,34 | 3,55 | 0,10 | 9,24E-01 |
| AT5G27300 | 0,08 | -0,56 | 4,38 | -0,13 | 8,99E-01 |
| AT5G27320 | 0,08 | -0,56 | 4,38 | -0,13 | 8,99E-01 |
| AT5G27340 | 0,54 | -1,94 | 3,64 | -0,53 | 5,94E-01 |
| AT5G27350 | 0,64 | -0,69 | 2,41 | -0,29 | 7,75E-01 |
| AT5G27410 | 0,42 | -1,23 | 4,11 | -0,30 | 7,65E-01 |
| AT5G27420 | 1,44 | 2,33 | 1,52 | 1,53 | 1,26E-01 |
| AT5G27470 | 0,51 | 1,88 | 3,03 | 0,62 | 5,35E-01 |
| AT5G27530 | 0,32 | -1,71 | 4,33 | -0,39 | 6,94E-01 |
| AT5G27540 | 0,68 | 1,45 | 3,52 | 0,41 | 6,81E-01 |
| AT5G27600 | 0,18 | 0,26 | 4,30 | 0,06 | 9,51E-01 |
| AT5G27610 | 0,08 | -0,56 | 4,38 | -0,13 | 8,99E-01 |
| AT5G27630 | 0,17 | -0,71 | 4,32 | -0,16 | 8,69E-01 |
| AT5G27650 | 0,94 | 2,67 | 2,67 | 1,00 | 3,18E-01 |
| AT5G27710 | 0,67 | 0,18 | 2,36 | 0,08 | 9,38E-01 |
| AT5G27730 | 0,69 | -0,52 | 1,96 | -0,27 | 7,90E-01 |
| AT5G27760 | 0,27 | 0,29 | 4,37 | 0,07 | 9,46E-01 |
| AT5G27770 | 0,86 | 0,56 | 2,43 | 0,23 | 8,17E-01 |
| AT5G27800 | 0,16 | -0,65 | 4,31 | -0,15 | 8,80E-01 |
| AT5G27830 | 0,23 | 1,51 | 4,34 | 0,35 | 7,28E-01 |
| AT5G27890 | 0,40 | -0,81 | 4,24 | -0,19 | 8,48E-01 |
| AT5G27893 | 1,07 | -1,29 | 1,82 | -0,71 | 4,78E-01 |
| AT5G27970 | 1,51 | 2,27 | 2,04 | 1,11 | 2,67E-01 |
| AT5G28010 | 1,26 | 0,10 | 1,42 | 0,07 | 9,44E-01 |
| AT5G04625 | 0,10 | -1,26 | 4,35 | -0,29 | 7,72E-01 |
| AT5G04665 | 0,11 | -1,28 | 4,35 | -0,29 | 7,69E-01 |
| AT5G28090 | 0,29 | 0,10 | 4,26 | 0,02 | 9,82E-01 |
| AT5G04675 | 0,19 | -2,00 | 4,31 | -0,46 | 6,43E-01 |
| AT5G28320 | 0,89 | 4,71 | 2,62 | 1,80 | 7,27E-02 |
| AT5G28350 | 0,24 | 0,92 | 4,35 | 0,21 | 8,32E-01 |
| AT5G28410 | 0,36 | 0,84 | 4,33 | 0,19 | 8,46E-01 |
| AT5G28463 | 0,37 | -1,04 | 4,23 | -0,24 | 8,07E-01 |
| AT5G28500 | 0,70 | -1,14 | 2,51 | -0,45 | 6,50E-01 |
| AT5G04815 | 0,05 | -0,58 | 4,38 | -0,13 | 8,95E-01 |
| AT5G28810 | 0,66 | -2,70 | 2,72 | -0,99 | 3,22E-01 |
| AT5G28830 | 0,05 | -0,58 | 4,38 | -0,13 | 8,95E-01 |
| AT5G28900 | 0,40 | 1,72 | 4,28 | 0,40 | 6,88E-01 |
| AT5G29624 | 0,12 | 2,87 | 4,34 | 0,66 | 5,08E-01 |
| AT5G30341 | 0,12 | -0,77 | 4,36 | -0,18 | 8,61E-01 |
| AT5G04955 | 1,06 | -4,69 | 3,80 | -1,24 | 2,17E-01 |
| AT5G32440 | 1,33 | 0,50 | 1,62 | 0,31 | 7,55E-01 |
| AT5G32613 | 0,26 | -1,25 | 4,29 | -0,29 | 7,71E-01 |
| AT5G33290 | 0,62 | 0,94 | 3,91 | 0,24 | 8,09E-01 |
| AT5G33320 | 0,61 | 0,92 | 2,19 | 0,42 | 6,74E-01 |

| | | | | | |
|---|---|---|---|---|---|
| AT5G33390 | 0,38 | 2,91 | 4,34 | 0,67 | 5,03E-01 |
| AT5G34869 | 0,19 | 1,20 | 4,29 | 0,28 | 7,79E-01 |
| AT5G34871 | 0,12 | 1,65 | 4,34 | 0,38 | 7,04E-01 |
| AT5G34870 | 0,38 | 1,31 | 3,35 | 0,39 | 6,96E-01 |
| AT5G35160 | 1,68 | 0,62 | 1,33 | 0,46 | 6,42E-01 |
| AT5G35170 | 0,47 | 0,17 | 2,32 | 0,07 | 9,40E-01 |
| AT5G35180 | 0,52 | 0,79 | 2,54 | 0,31 | 7,56E-01 |
| AT5G35190 | 0,84 | 0,23 | 2,57 | 0,09 | 9,29E-01 |
| AT5G35300 | 0,23 | -1,13 | 4,29 | -0,26 | 7,92E-01 |
| AT5G35320 | 0,21 | -0,24 | 4,35 | -0,06 | 9,56E-01 |
| AT5G35360 | 1,57 | 2,57 | 1,52 | 1,70 | 8,95E-02 |
| AT5G35430 | 1,24 | 3,23 | 2,26 | 1,43 | 1,53E-01 |
| AT5G35450 | 0,40 | 3,49 | 4,33 | 0,81 | 4,20E-01 |
| AT5G05195 | 0,23 | 0,17 | 4,26 | 0,04 | 9,68E-01 |
| AT5G35490 | 1,40 | -1,11 | 2,04 | -0,54 | 5,87E-01 |
| AT5G35730 | 0,82 | 3,15 | 2,75 | 1,14 | 2,53E-01 |
| AT5G35840 | 1,35 | 2,63 | 2,05 | 1,28 | 1,99E-01 |
| AT5G35917 | 0,29 | -1,47 | 4,29 | -0,34 | 7,31E-01 |
| AT5G35920 | 1,48 | -0,50 | 1,42 | -0,35 | 7,27E-01 |
| AT5G35926 | 1,15 | -0,19 | 1,65 | -0,12 | 9,06E-01 |
| AT5G35980 | 1,15 | -0,88 | 1,76 | -0,50 | 6,14E-01 |
| AT5G36001 | 0,12 | 2,87 | 4,34 | 0,66 | 5,08E-01 |
| AT5G36080 | 1,49 | 0,12 | 1,51 | 0,08 | 9,34E-01 |
| AT5G36290 | 0,47 | -1,14 | 2,65 | -0,43 | 6,67E-01 |
| AT5G36300 | 0,87 | 0,30 | 2,13 | 0,14 | 8,88E-01 |
| AT5G36740 | 0,08 | 2,33 | 4,36 | 0,53 | 5,93E-01 |
| AT5G36870 | 1,07 | -1,72 | 1,97 | -0,87 | 3,82E-01 |
| AT5G36900 | 0,54 | 1,26 | 2,97 | 0,42 | 6,71E-01 |
| AT5G36940 | 0,51 | 3,24 | 3,85 | 0,84 | 3,99E-01 |
| AT5G37020 | 0,62 | 0,22 | 2,45 | 0,09 | 9,29E-01 |
| AT5G37070 | 0,43 | 0,00 | 4,41 | 0,00 | 1,00E+00 |
| AT5G37130 | 0,66 | 0,73 | 2,53 | 0,29 | 7,74E-01 |
| AT5G37150 | 0,98 | -3,09 | 2,11 | -1,46 | 1,43E-01 |
| AT5G37540 | 0,17 | 2,62 | 4,35 | 0,60 | 5,47E-01 |
| AT5G37630 | 0,94 | 3,44 | 2,22 | 1,55 | 1,21E-01 |
| AT5G37760 | 0,15 | 1,93 | 4,33 | 0,45 | 6,55E-01 |
| AT5G37793 | 0,51 | 0,13 | 2,56 | 0,05 | 9,58E-01 |
| AT5G37850 | 0,30 | 2,84 | 4,35 | 0,65 | 5,14E-01 |
| AT5G37950 | 0,11 | -1,28 | 4,35 | -0,29 | 7,69E-01 |
| AT5G38155 | 0,12 | 0,31 | 4,35 | 0,07 | 9,44E-01 |
| AT5G38210 | 1,00 | -0,42 | 1,92 | -0,22 | 8,25E-01 |
| AT5G38380 | 0,15 | -0,67 | 4,39 | -0,15 | 8,78E-01 |
| AT5G38470 | 0,73 | 1,47 | 3,66 | 0,40 | 6,89E-01 |
| AT5G38480 | 1,20 | 1,76 | 1,60 | 1,09 | 2,74E-01 |
| AT5G38560 | 0,34 | 0,05 | 3,40 | 0,01 | 9,88E-01 |
| AT5G38910 | 0,16 | -1,42 | 4,34 | -0,33 | 7,43E-01 |
| AT5G38980 | 0,57 | -0,34 | 2,53 | -0,14 | 8,92E-01 |
| AT5G39000 | 0,70 | -0,68 | 4,39 | -0,16 | 8,76E-01 |
| AT5G39030 | 0,81 | 3,34 | 2,78 | 1,20 | 2,29E-01 |
| AT5G39040 | 0,29 | 3,11 | 4,33 | 0,72 | 4,73E-01 |
| AT5G39250 | 0,34 | 0,00 | 4,34 | 0,00 | 9,99E-01 |
| AT5G39320 | 0,82 | 2,41 | 4,37 | 0,55 | 5,82E-01 |
| AT5G39350 | 0,21 | 0,06 | 4,31 | 0,01 | 9,89E-01 |
| AT5G39510 | 0,77 | -1,07 | 2,23 | -0,48 | 6,31E-01 |
| AT5G39590 | 0,82 | 0,36 | 2,23 | 0,16 | 8,71E-01 |
| AT5G39660 | 0,40 | 2,81 | 4,32 | 0,65 | 5,15E-01 |

| | | | | | |
|---|---|---|---|---|---|
| AT5G39670 | 0,40 | 1,01 | 4,07 | 0,25 | 8,05E-01 |
| AT5G39690 | 1,03 | -1,24 | 2,00 | -0,62 | 5,35E-01 |
| AT5G39710 | 0,25 | 2,97 | 4,33 | 0,69 | 4,93E-01 |
| AT5G39730 | 0,19 | -0,65 | 4,31 | -0,15 | 8,80E-01 |
| AT5G39775 | 0,53 | -1,01 | 4,10 | -0,25 | 8,05E-01 |
| AT5G39850 | 0,08 | 2,33 | 4,36 | 0,53 | 5,93E-01 |
| AT5G39960 | 0,21 | 2,61 | 4,34 | 0,60 | 5,48E-01 |
| AT5G39990 | 0,08 | 2,08 | 4,38 | 0,47 | 6,35E-01 |
| AT5G40060 | 1,33 | 3,06 | 2,46 | 1,24 | 2,13E-01 |
| AT5G40140 | 0,31 | 0,32 | 3,54 | 0,09 | 9,28E-01 |
| AT5G40200 | 0,62 | 0,41 | 2,65 | 0,16 | 8,76E-01 |
| AT5G40380 | 0,46 | -2,72 | 3,92 | -0,69 | 4,88E-01 |
| AT5G40400 | 1,55 | -1,33 | 1,29 | -1,03 | 3,04E-01 |
| AT5G40650 | 0,74 | 0,75 | 2,87 | 0,26 | 7,95E-01 |
| AT5G40670 | 1,01 | 1,81 | 2,18 | 0,83 | 4,08E-01 |
| AT5G40680 | 0,17 | 3,14 | 4,33 | 0,73 | 4,68E-01 |
| AT5G40690 | 0,62 | 3,02 | 3,62 | 0,83 | 4,04E-01 |
| AT5G40720 | 0,32 | -0,68 | 4,24 | -0,16 | 8,73E-01 |
| AT5G40760 | 1,16 | 2,16 | 1,93 | 1,12 | 2,64E-01 |
| AT5G40780 | 1,36 | 1,48 | 1,58 | 0,94 | 3,49E-01 |
| AT5G40810 | 1,32 | 1,37 | 1,70 | 0,81 | 4,20E-01 |
| AT5G40850 | 0,08 | -0,87 | 4,38 | -0,20 | 8,43E-01 |
| AT5G40870 | 0,14 | -0,31 | 4,33 | -0,07 | 9,43E-01 |
| AT5G40880 | 0,51 | 2,36 | 3,94 | 0,60 | 5,49E-01 |
| AT5G40900 | 0,15 | -0,64 | 4,39 | -0,15 | 8,84E-01 |
| AT5G41010 | 0,28 | -0,34 | 4,30 | -0,08 | 9,37E-01 |
| AT5G06115 | 0,20 | -0,47 | 4,33 | -0,11 | 9,13E-01 |
| AT5G41070 | 0,20 | -0,80 | 4,37 | -0,18 | 8,55E-01 |
| AT5G41100 | 0,28 | -1,53 | 4,33 | -0,35 | 7,24E-01 |
| AT5G41110 | 0,22 | -0,11 | 4,34 | -0,02 | 9,80E-01 |
| AT5G41140 | 0,59 | 0,13 | 3,82 | 0,03 | 9,72E-01 |
| AT5G41260 | 0,61 | 0,31 | 2,21 | 0,14 | 8,89E-01 |
| AT5G41360 | 0,66 | 0,76 | 3,75 | 0,20 | 8,40E-01 |
| AT5G41370 | 0,15 | -0,19 | 4,31 | -0,04 | 9,64E-01 |
| AT5G41410 | 1,13 | 0,46 | 1,70 | 0,27 | 7,85E-01 |
| AT5G41500 | 1,05 | -1,32 | 1,82 | -0,72 | 4,69E-01 |
| AT5G41660 | 0,71 | -1,78 | 2,80 | -0,63 | 5,26E-01 |
| AT5G41700 | 1,24 | 0,12 | 1,98 | 0,06 | 9,52E-01 |
| AT5G41761 | 0,83 | -1,94 | 2,15 | -0,90 | 3,67E-01 |
| AT5G41770 | 0,61 | 1,73 | 2,49 | 0,70 | 4,86E-01 |
| AT5G41860 | 0,77 | 3,27 | 4,26 | 0,77 | 4,43E-01 |
| AT5G41920 | 0,61 | -0,11 | 4,31 | -0,02 | 9,80E-01 |
| AT5G41940 | 0,98 | 2,67 | 1,99 | 1,35 | 1,78E-01 |
| AT5G41950 | 1,46 | 2,72 | 1,98 | 1,38 | 1,69E-01 |
| AT5G41960 | 0,15 | 2,31 | 4,37 | 0,53 | 5,98E-01 |
| AT5G41970 | 0,85 | 2,31 | 3,43 | 0,67 | 5,01E-01 |
| AT5G42010 | 0,60 | 2,25 | 3,05 | 0,74 | 4,61E-01 |
| AT5G42220 | 0,70 | 0,68 | 2,17 | 0,31 | 7,53E-01 |
| AT5G42250 | 0,33 | 3,34 | 4,32 | 0,77 | 4,40E-01 |
| AT5G42260 | 0,28 | -2,74 | 4,29 | -0,64 | 5,23E-01 |
| AT5G42290 | 0,08 | 2,33 | 4,36 | 0,53 | 5,93E-01 |
| AT5G42310 | 0,50 | -2,19 | 4,10 | -0,53 | 5,94E-01 |
| AT5G42350 | 0,37 | 1,62 | 4,32 | 0,38 | 7,08E-01 |
| AT5G42420 | 0,74 | 3,22 | 3,63 | 0,89 | 3,74E-01 |
| AT5G42460 | 0,33 | 2,76 | 4,34 | 0,64 | 5,25E-01 |
| AT5G42470 | 0,22 | -0,56 | 4,38 | -0,13 | 8,98E-01 |

| | | | | | |
|---|---|---|---|---|---|
| AT5G42480 | 0,45 | 3,33 | 4,32 | 0,77 | 4,41E-01 |
| AT5G42504 | 0,17 | -0,14 | 4,41 | -0,03 | 9,75E-01 |
| AT5G42520 | 0,71 | 0,81 | 2,29 | 0,35 | 7,23E-01 |
| AT5G42540 | 1,02 | 3,37 | 2,32 | 1,45 | 1,46E-01 |
| AT5G42620 | 0,80 | 1,18 | 2,31 | 0,51 | 6,09E-01 |
| AT5G42640 | 0,08 | -0,56 | 4,38 | -0,13 | 8,99E-01 |
| AT5G42670 | 0,16 | -1,66 | 4,33 | -0,38 | 7,01E-01 |
| AT5G42680 | 0,80 | -1,17 | 3,12 | -0,38 | 7,07E-01 |
| AT5G42720 | 0,19 | 0,10 | 4,37 | 0,02 | 9,82E-01 |
| AT5G42740 | 1,16 | 2,24 | 2,76 | 0,81 | 4,18E-01 |
| AT5G42760 | 0,12 | 0,63 | 4,33 | 0,15 | 8,85E-01 |
| AT5G06395 | 1,55 | 4,38 | 1,86 | 2,35 | 1,89E-02 |
| AT5G42825 | 0,29 | 2,37 | 4,35 | 0,55 | 5,85E-01 |
| AT5G42965 | 0,19 | 1,62 | 4,35 | 0,37 | 7,09E-01 |
| AT5G42980 | 0,59 | 0,37 | 2,59 | 0,14 | 8,87E-01 |
| AT5G43100 | 0,78 | 1,47 | 2,05 | 0,72 | 4,72E-01 |
| AT5G43150 | 0,61 | 2,25 | 3,00 | 0,75 | 4,52E-01 |
| AT5G06505 | 0,36 | -0,33 | 4,31 | -0,08 | 9,39E-01 |
| AT5G43460 | 1,01 | 2,71 | 2,01 | 1,35 | 1,78E-01 |
| AT5G43500 | 1,47 | 0,06 | 1,70 | 0,04 | 9,72E-01 |
| AT5G43525 | 0,69 | -0,93 | 2,60 | -0,36 | 7,20E-01 |
| AT5G43560 | 0,55 | -1,02 | 3,11 | -0,33 | 7,43E-01 |
| AT5G43610 | 1,12 | -0,74 | 2,11 | -0,35 | 7,27E-01 |
| AT5G43620 | 0,36 | 2,77 | 4,26 | 0,65 | 5,16E-01 |
| AT5G43700 | 0,52 | 2,80 | 3,12 | 0,90 | 3,70E-01 |
| AT5G43810 | 1,04 | 2,87 | 2,60 | 1,11 | 2,69E-01 |
| AT5G43930 | 0,73 | -0,25 | 2,56 | -0,10 | 9,23E-01 |
| AT5G43940 | 1,17 | -0,16 | 1,86 | -0,09 | 9,31E-01 |
| AT5G44020 | 1,03 | -0,94 | 1,97 | -0,48 | 6,32E-01 |
| AT5G44050 | 1,57 | -1,33 | 2,27 | -0,59 | 5,57E-01 |
| AT5G44060 | 0,47 | -0,50 | 2,63 | -0,19 | 8,50E-01 |
| AT5G44070 | 1,32 | -0,19 | 1,52 | -0,12 | 9,01E-01 |
| AT5G44130 | 0,80 | -0,14 | 2,97 | -0,05 | 9,63E-01 |
| AT5G44190 | 1,58 | -3,35 | 1,90 | -1,76 | 7,84E-02 |
| AT5G44230 | 0,12 | 2,87 | 4,34 | 0,66 | 5,08E-01 |
| AT5G44240 | 1,02 | -0,08 | 2,05 | -0,04 | 9,69E-01 |
| AT5G06565 | 0,63 | -0,44 | 2,56 | -0,17 | 8,63E-01 |
| AT5G44286 | 0,85 | -1,37 | 2,08 | -0,66 | 5,12E-01 |
| AT5G44530 | 1,17 | 4,52 | 2,59 | 1,74 | 8,16E-02 |
| AT5G44574 | 0,91 | -3,06 | 3,12 | -0,98 | 3,27E-01 |
| AT5G44580 | 0,52 | -0,83 | 3,99 | -0,21 | 8,35E-01 |
| AT5G44585 | 1,38 | -1,42 | 1,76 | -0,81 | 4,18E-01 |
| AT5G06605 | 1,31 | -0,89 | 1,81 | -0,49 | 6,24E-01 |
| AT5G44640 | 1,59 | -1,74 | 1,56 | -1,11 | 2,66E-01 |
| AT5G44650 | 0,46 | 2,90 | 4,04 | 0,72 | 4,72E-01 |
| AT5G44720 | 0,16 | 0,85 | 4,31 | 0,20 | 8,44E-01 |
| AT5G44750 | 0,74 | 2,72 | 2,21 | 1,23 | 2,18E-01 |
| AT5G44760 | 0,36 | 3,22 | 4,34 | 0,74 | 4,59E-01 |
| AT5G44770 | 0,22 | -0,30 | 4,29 | -0,07 | 9,45E-01 |
| AT5G45000 | 0,38 | 0,82 | 4,32 | 0,19 | 8,49E-01 |
| AT5G06695 | 0,59 | 3,40 | 3,04 | 1,12 | 2,63E-01 |
| AT5G45030 | 0,59 | 3,40 | 3,04 | 1,12 | 2,63E-01 |
| AT5G45060 | 1,56 | 1,67 | 1,63 | 1,03 | 3,05E-01 |
| AT5G45130 | 0,78 | 3,29 | 2,94 | 1,12 | 2,64E-01 |
| AT5G45150 | 0,08 | 2,08 | 4,38 | 0,47 | 6,35E-01 |
| AT5G45240 | 0,12 | 2,87 | 4,34 | 0,66 | 5,08E-01 |

| | | | | | |
|---|---|---|---|---|---|
| AT5G45260 | 0,75 | 2,69 | 2,87 | 0,94 | 3,49E-01 |
| AT5G45390 | 0,62 | -1,17 | 2,23 | -0,52 | 6,00E-01 |
| AT5G45400 | 0,76 | 2,17 | 2,50 | 0,87 | 3,87E-01 |
| AT5G45450 | 0,08 | 2,08 | 4,38 | 0,47 | 6,35E-01 |
| AT5G45475 | 0,74 | -0,20 | 2,14 | -0,09 | 9,27E-01 |
| AT5G45490 | 1,25 | -0,47 | 1,60 | -0,30 | 7,68E-01 |
| AT5G45570 | 0,68 | -1,22 | 2,25 | -0,54 | 5,90E-01 |
| AT5G45620 | 0,59 | 1,17 | 2,97 | 0,39 | 6,93E-01 |
| AT5G45670 | 0,10 | -1,26 | 4,35 | -0,29 | 7,72E-01 |
| AT5G45680 | 0,26 | -1,00 | 4,37 | -0,23 | 8,18E-01 |
| AT5G45710 | 0,55 | 0,76 | 3,84 | 0,20 | 8,43E-01 |
| AT5G45750 | 0,19 | -2,00 | 4,31 | -0,46 | 6,43E-01 |
| AT5G45770 | 0,12 | 2,81 | 4,34 | 0,65 | 5,18E-01 |
| AT5G45920 | 1,21 | 0,62 | 1,99 | 0,31 | 7,57E-01 |
| AT5G46330 | 0,47 | 2,09 | 3,93 | 0,53 | 5,95E-01 |
| AT5G46410 | 0,44 | 0,11 | 4,38 | 0,02 | 9,81E-01 |
| AT5G06825 | 0,67 | -2,50 | 2,91 | -0,86 | 3,90E-01 |
| AT5G46460 | 0,90 | 4,87 | 3,23 | 1,51 | 1,32E-01 |
| AT5G46490 | 1,46 | 1,05 | 1,42 | 0,73 | 4,62E-01 |
| AT5G46500 | 1,11 | 4,39 | 2,37 | 1,86 | 6,34E-02 |
| AT5G06835 | 1,13 | 1,75 | 2,63 | 0,67 | 5,05E-01 |
| AT5G46630 | 1,04 | 2,14 | 1,99 | 1,07 | 2,83E-01 |
| AT5G46640 | 1,47 | 2,84 | 2,08 | 1,36 | 1,74E-01 |
| AT5G46660 | 1,02 | -4,48 | 3,97 | -1,13 | 2,60E-01 |
| AT5G46670 | 0,74 | -3,10 | 2,64 | -1,18 | 2,39E-01 |
| AT5G46730 | 0,08 | -0,56 | 4,38 | -0,13 | 8,99E-01 |
| AT5G46780 | 0,35 | 0,55 | 4,28 | 0,13 | 8,97E-01 |
| AT5G46825 | 1,18 | -1,34 | 1,46 | -0,92 | 3,58E-01 |
| AT5G46840 | 0,62 | 0,78 | 3,09 | 0,25 | 8,00E-01 |
| AT5G46850 | 0,29 | 1,98 | 4,32 | 0,46 | 6,46E-01 |
| AT5G47020 | 1,57 | 0,80 | 1,61 | 0,49 | 6,22E-01 |
| AT5G47030 | 0,63 | 1,60 | 3,79 | 0,42 | 6,73E-01 |
| AT5G47120 | 1,06 | 0,10 | 1,77 | 0,05 | 9,57E-01 |
| AT5G47130 | 0,25 | 1,15 | 4,30 | 0,27 | 7,89E-01 |
| AT5G47140 | 0,26 | -0,06 | 4,32 | -0,01 | 9,89E-01 |
| AT5G47200 | 0,34 | 1,77 | 4,32 | 0,41 | 6,82E-01 |
| AT5G47230 | 0,27 | -1,33 | 4,27 | -0,31 | 7,56E-01 |
| AT5G47240 | 0,14 | -1,69 | 4,33 | -0,39 | 6,96E-01 |
| AT5G47280 | 1,15 | -1,40 | 1,68 | -0,83 | 4,06E-01 |
| AT5G47300 | 1,47 | -1,68 | 1,54 | -1,09 | 2,75E-01 |
| AT5G47320 | 0,19 | -0,39 | 4,29 | -0,09 | 9,27E-01 |
| AT5G47390 | 0,64 | 0,63 | 1,89 | 0,33 | 7,39E-01 |
| AT5G47420 | 0,50 | 2,46 | 3,99 | 0,62 | 5,37E-01 |
| AT5G47435 | 1,48 | 3,15 | 1,97 | 1,60 | 1,10E-01 |
| AT5G47640 | 0,70 | 0,11 | 2,22 | 0,05 | 9,59E-01 |
| AT5G47660 | 0,08 | -0,56 | 4,38 | -0,13 | 8,99E-01 |
| AT5G47720 | 0,16 | 1,97 | 4,33 | 0,45 | 6,50E-01 |
| AT5G47770 | 0,39 | -0,57 | 4,19 | -0,14 | 8,91E-01 |
| AT5G47780 | 0,52 | 0,86 | 2,57 | 0,34 | 7,37E-01 |
| AT5G47790 | 0,28 | 3,25 | 4,33 | 0,75 | 4,53E-01 |
| AT5G47800 | 0,24 | 2,80 | 4,34 | 0,65 | 5,18E-01 |
| AT5G47820 | 0,70 | 1,38 | 3,68 | 0,38 | 7,07E-01 |
| AT5G47840 | 0,45 | -1,53 | 2,62 | -0,58 | 5,59E-01 |
| AT5G47860 | 0,53 | 2,68 | 3,82 | 0,70 | 4,83E-01 |
| AT5G47890 | 1,07 | 4,50 | 2,00 | 2,25 | 2,45E-02 |
| AT5G48190 | 1,46 | -0,59 | 1,71 | -0,34 | 7,32E-01 |

| | | | | | |
|---|---|---|---|---|---|
| AT5G48220 | 0,16 | 1,16 | 4,38 | 0,27 | 7,90E-01 |
| AT5G48290 | 0,36 | 0,00 | 4,41 | 0,00 | 1,00E+00 |
| AT5G48300 | 1,23 | -0,54 | 1,88 | -0,29 | 7,74E-01 |
| AT5G48340 | 0,30 | 2,84 | 4,35 | 0,65 | 5,14E-01 |
| AT5G48360 | 0,08 | -0,56 | 4,38 | -0,13 | 8,99E-01 |
| AT5G48390 | 0,69 | 3,32 | 3,50 | 0,95 | 3,42E-01 |
| AT5G48400 | 0,56 | 4,04 | 3,76 | 1,07 | 2,83E-01 |
| AT5G48410 | 0,12 | 2,81 | 4,34 | 0,65 | 5,18E-01 |
| AT5G07095 | 1,59 | 0,40 | 1,33 | 0,30 | 7,65E-01 |
| AT5G07105 | 0,86 | -2,02 | 2,02 | -1,00 | 3,17E-01 |
| AT5G48545 | 0,30 | -1,00 | 3,61 | -0,28 | 7,81E-01 |
| AT5G48720 | 0,16 | 1,27 | 4,37 | 0,29 | 7,71E-01 |
| AT5G48730 | 0,12 | 2,81 | 4,34 | 0,65 | 5,18E-01 |
| AT5G48775 | 0,08 | -0,56 | 4,38 | -0,13 | 8,99E-01 |
| AT5G48800 | 0,90 | 3,52 | 2,83 | 1,24 | 2,15E-01 |
| AT5G48900 | 0,13 | -0,63 | 4,39 | -0,14 | 8,85E-01 |
| AT5G49010 | 1,17 | -2,76 | 2,02 | -1,37 | 1,72E-01 |
| AT5G49020 | 0,22 | -0,13 | 4,30 | -0,03 | 9,76E-01 |
| AT5G49060 | 0,17 | 2,04 | 4,33 | 0,47 | 6,37E-01 |
| AT5G49160 | 0,93 | 1,37 | 2,43 | 0,56 | 5,73E-01 |
| AT5G49215 | 0,08 | -0,56 | 4,38 | -0,13 | 8,99E-01 |
| AT5G49250 | 0,66 | -0,82 | 2,58 | -0,32 | 7,50E-01 |
| AT5G49260 | 0,08 | -0,56 | 4,38 | -0,13 | 8,99E-01 |
| AT5G49450 | 0,19 | 2,25 | 4,32 | 0,52 | 6,02E-01 |
| AT5G49460 | 0,62 | 1,95 | 2,99 | 0,65 | 5,13E-01 |
| AT5G49470 | 0,29 | 1,98 | 4,32 | 0,46 | 6,46E-01 |
| AT5G49480 | 1,15 | 1,48 | 2,22 | 0,67 | 5,04E-01 |
| AT5G49510 | 0,19 | 0,73 | 4,31 | 0,17 | 8,66E-01 |
| AT5G49520 | 0,12 | 0,63 | 4,33 | 0,15 | 8,85E-01 |
| AT5G49530 | 0,46 | 2,51 | 4,03 | 0,62 | 5,33E-01 |
| AT5G49590 | 0,86 | -2,82 | 2,76 | -1,02 | 3,06E-01 |
| AT5G49640 | 0,08 | 2,08 | 4,38 | 0,47 | 6,35E-01 |
| AT5G49650 | 0,61 | 1,01 | 2,20 | 0,46 | 6,47E-01 |
| AT5G49810 | 1,47 | 2,54 | 2,04 | 1,24 | 2,13E-01 |
| AT5G49840 | 0,10 | 0,39 | 4,35 | 0,09 | 9,29E-01 |
| AT5G49945 | 0,51 | 3,95 | 3,81 | 1,04 | 3,00E-01 |
| AT5G49970 | 0,29 | 1,68 | 4,27 | 0,39 | 6,94E-01 |
| AT5G50020 | 0,47 | 0,00 | 4,41 | 0,00 | 1,00E+00 |
| AT5G50160 | 1,06 | 1,30 | 2,06 | 0,63 | 5,29E-01 |
| AT5G50170 | 0,08 | 2,08 | 4,38 | 0,47 | 6,35E-01 |
| AT5G50200 | 0,08 | -0,56 | 4,38 | -0,13 | 8,99E-01 |
| AT5G07605 | 0,25 | -0,88 | 4,31 | -0,21 | 8,38E-01 |
| AT5G50310 | 0,21 | 2,34 | 4,32 | 0,54 | 5,89E-01 |
| AT5G50580 | 0,65 | 3,83 | 3,62 | 1,06 | 2,90E-01 |
| AT5G50680 | 1,09 | 1,69 | 2,11 | 0,80 | 4,23E-01 |
| AT5G50717 | 0,11 | -1,28 | 4,35 | -0,29 | 7,69E-01 |
| AT5G50780 | 0,59 | 2,79 | 4,33 | 0,64 | 5,19E-01 |
| AT5G50850 | 0,82 | 2,80 | 2,22 | 1,26 | 2,06E-01 |
| AT5G50950 | 1,52 | -0,80 | 1,28 | -0,62 | 5,34E-01 |
| AT5G50960 | 0,50 | 2,54 | 3,95 | 0,64 | 5,20E-01 |
| AT5G51010 | 0,08 | 2,33 | 4,36 | 0,53 | 5,93E-01 |
| AT5G51050 | 0,20 | 1,88 | 4,35 | 0,43 | 6,66E-01 |
| AT5G51055 | 0,34 | -2,09 | 3,49 | -0,60 | 5,50E-01 |
| AT5G51120 | 0,08 | 2,33 | 4,36 | 0,53 | 5,93E-01 |
| AT5G51150 | 0,28 | 1,26 | 4,36 | 0,29 | 7,73E-01 |
| AT5G51180 | 0,38 | 0,25 | 2,79 | 0,09 | 9,29E-01 |

| | | | | | |
|---|---|---|---|---|---|
| AT5G51200 | 1,04 | 2,36 | 2,05 | 1,15 | 2,49E-01 |
| AT5G51230 | 0,21 | 0,57 | 4,32 | 0,13 | 8,94E-01 |
| AT5G51280 | 0,48 | 0,05 | 2,54 | 0,02 | 9,85E-01 |
| AT5G51340 | 0,39 | 1,21 | 3,38 | 0,36 | 7,20E-01 |
| AT5G51450 | 0,53 | 2,91 | 3,95 | 0,74 | 4,62E-01 |
| AT5G51560 | 0,28 | -0,24 | 4,40 | -0,05 | 9,57E-01 |
| AT5G51570 | 0,40 | -0,92 | 3,32 | -0,28 | 7,81E-01 |
| AT5G51660 | 1,44 | 1,11 | 1,62 | 0,69 | 4,92E-01 |
| AT5G51670 | 0,12 | 1,65 | 4,34 | 0,38 | 7,04E-01 |
| AT5G51680 | 0,64 | 1,23 | 2,55 | 0,48 | 6,30E-01 |
| AT5G51750 | 0,75 | 0,41 | 3,60 | 0,11 | 9,10E-01 |
| AT5G51760 | 1,35 | 0,58 | 1,57 | 0,37 | 7,12E-01 |
| AT5G51830 | 0,27 | 0,51 | 4,25 | 0,12 | 9,04E-01 |
| AT5G51960 | 0,11 | -1,28 | 4,35 | -0,29 | 7,69E-01 |
| AT5G51970 | 1,04 | -0,19 | 1,59 | -0,12 | 9,03E-01 |
| AT5G52050 | 0,61 | -0,70 | 2,33 | -0,30 | 7,65E-01 |
| AT5G52115 | 1,12 | -3,97 | 2,54 | -1,56 | 1,19E-01 |
| AT5G52240 | 1,16 | 1,87 | 1,96 | 0,95 | 3,40E-01 |
| AT5G52280 | 0,34 | 0,46 | 4,33 | 0,11 | 9,16E-01 |
| AT5G52471 | 1,40 | -2,30 | 1,84 | -1,25 | 2,11E-01 |
| AT5G52510 | 0,69 | 2,47 | 3,00 | 0,82 | 4,11E-01 |
| AT5G52520 | 0,23 | -1,11 | 4,34 | -0,25 | 7,99E-01 |
| AT5G52580 | 0,50 | 0,17 | 3,13 | 0,05 | 9,58E-01 |
| AT5G52630 | 0,08 | 2,08 | 4,38 | 0,47 | 6,35E-01 |
| AT5G52700 | 0,14 | -1,69 | 4,33 | -0,39 | 6,96E-01 |
| AT5G52750 | 0,33 | 0,53 | 4,24 | 0,13 | 9,00E-01 |
| AT5G52760 | 0,52 | -0,60 | 2,42 | -0,25 | 8,04E-01 |
| AT5G52920 | 1,32 | 2,25 | 1,98 | 1,14 | 2,55E-01 |
| AT5G07945 | 0,52 | 0,08 | 3,83 | 0,02 | 9,83E-01 |
| AT5G52940 | 0,11 | 1,28 | 4,35 | 0,29 | 7,68E-01 |
| AT5G53020 | 0,27 | 0,48 | 4,31 | 0,11 | 9,11E-01 |
| AT5G53030 | 0,24 | -0,70 | 4,38 | -0,16 | 8,72E-01 |
| AT5G53048 | 0,25 | 1,15 | 4,31 | 0,27 | 7,90E-01 |
| AT5G53060 | 0,48 | 2,17 | 3,91 | 0,56 | 5,79E-01 |
| AT5G53080 | 0,12 | 2,81 | 4,34 | 0,65 | 5,18E-01 |
| AT5G53110 | 0,31 | 2,14 | 4,38 | 0,49 | 6,25E-01 |
| AT5G53120 | 0,82 | 1,36 | 2,40 | 0,57 | 5,70E-01 |
| AT5G53140 | 1,45 | 1,57 | 1,62 | 0,97 | 3,30E-01 |
| AT5G53150 | 0,95 | 0,95 | 2,18 | 0,43 | 6,64E-01 |
| AT5G53160 | 1,14 | 0,12 | 1,57 | 0,08 | 9,38E-01 |
| AT5G53220 | 0,48 | 1,48 | 3,01 | 0,49 | 6,24E-01 |
| AT5G53280 | 0,44 | 0,92 | 4,14 | 0,22 | 8,24E-01 |
| AT5G53350 | 0,54 | 0,30 | 3,22 | 0,09 | 9,26E-01 |
| AT5G53360 | 0,08 | 2,08 | 4,38 | 0,47 | 6,35E-01 |
| AT5G53410 | 1,13 | 1,13 | 1,61 | 0,70 | 4,85E-01 |
| AT5G53420 | 0,48 | 0,56 | 3,27 | 0,17 | 8,64E-01 |
| AT5G53430 | 1,00 | 1,09 | 1,81 | 0,60 | 5,46E-01 |
| AT5G53450 | 0,71 | 0,22 | 2,58 | 0,09 | 9,31E-01 |
| AT5G53550 | 0,99 | 2,75 | 2,15 | 1,28 | 2,01E-01 |
| AT5G53560 | 1,33 | 0,54 | 1,50 | 0,36 | 7,21E-01 |
| AT5G53590 | 0,48 | 3,00 | 4,01 | 0,75 | 4,54E-01 |
| AT5G53620 | 1,09 | 1,77 | 1,73 | 1,03 | 3,05E-01 |
| AT5G53860 | 0,25 | -0,36 | 3,69 | -0,10 | 9,23E-01 |
| AT5G53890 | 0,71 | 1,89 | 2,32 | 0,81 | 4,15E-01 |
| AT5G54035 | 0,15 | -1,81 | 4,32 | -0,42 | 6,75E-01 |
| AT5G54043 | 0,22 | -1,04 | 4,30 | -0,24 | 8,08E-01 |

| | | | | | |
|---|---|---|---|---|---|
| AT5G54080 | 0,99 | 2,72 | 2,07 | 1,32 | 1,87E-01 |
| AT5G54110 | 0,33 | 3,45 | 4,32 | 0,80 | 4,24E-01 |
| AT5G54140 | 0,24 | 2,85 | 4,34 | 0,66 | 5,11E-01 |
| AT5G54145 | 0,12 | -1,57 | 4,34 | -0,36 | 7,18E-01 |
| AT5G54160 | 0,46 | 1,42 | 4,09 | 0,35 | 7,29E-01 |
| AT5G54180 | 0,12 | 0,08 | 4,33 | 0,02 | 9,84E-01 |
| AT5G54290 | 0,66 | -0,85 | 2,61 | -0,33 | 7,45E-01 |
| AT5G54500 | 0,31 | 0,30 | 4,33 | 0,07 | 9,44E-01 |
| AT5G54540 | 0,58 | 1,68 | 2,53 | 0,66 | 5,07E-01 |
| AT5G54600 | 0,87 | 1,72 | 1,94 | 0,89 | 3,76E-01 |
| AT5G08225 | 0,36 | 0,80 | 4,35 | 0,18 | 8,55E-01 |
| AT5G54640 | 0,05 | -0,58 | 4,38 | -0,13 | 8,95E-01 |
| AT5G54680 | 0,72 | -1,14 | 2,53 | -0,45 | 6,52E-01 |
| AT5G54750 | 0,55 | 1,59 | 3,86 | 0,41 | 6,80E-01 |
| AT5G54760 | 0,49 | -1,15 | 2,67 | -0,43 | 6,68E-01 |
| AT5G54800 | 0,08 | -0,87 | 4,38 | -0,20 | 8,43E-01 |
| AT5G54830 | 0,76 | 3,82 | 2,70 | 1,41 | 1,57E-01 |
| AT5G54855 | 0,18 | 1,77 | 4,41 | 0,40 | 6,88E-01 |
| AT5G54860 | 0,26 | 2,30 | 4,37 | 0,53 | 5,98E-01 |
| AT5G54900 | 0,82 | 2,81 | 2,39 | 1,18 | 2,39E-01 |
| AT5G54920 | 0,83 | 3,67 | 4,31 | 0,85 | 3,95E-01 |
| AT5G55060 | 0,14 | -0,31 | 4,33 | -0,07 | 9,43E-01 |
| AT5G55070 | 1,01 | 2,01 | 1,76 | 1,14 | 2,55E-01 |
| AT5G55190 | 0,80 | 2,35 | 2,25 | 1,04 | 2,96E-01 |
| AT5G55220 | 0,88 | -1,68 | 2,62 | -0,64 | 5,23E-01 |
| AT5G55230 | 0,95 | -0,23 | 1,85 | -0,12 | 9,02E-01 |
| AT5G55450 | 0,62 | 0,92 | 3,05 | 0,30 | 7,62E-01 |
| AT5G55460 | 0,37 | -0,59 | 3,38 | -0,18 | 8,61E-01 |
| AT5G55500 | 0,57 | 3,19 | 3,06 | 1,04 | 2,96E-01 |
| AT5G55530 | 0,30 | 1,73 | 4,30 | 0,40 | 6,88E-01 |
| AT5G55580 | 0,70 | 2,00 | 3,86 | 0,52 | 6,05E-01 |
| AT5G55620 | 0,43 | -1,47 | 4,18 | -0,35 | 7,26E-01 |
| AT5G55630 | 0,12 | 1,65 | 4,34 | 0,38 | 7,04E-01 |
| AT5G55660 | 0,90 | 0,22 | 2,19 | 0,10 | 9,21E-01 |
| AT5G55840 | 0,22 | 0,75 | 4,27 | 0,18 | 8,60E-01 |
| AT5G55850 | 0,24 | -0,77 | 4,31 | -0,18 | 8,58E-01 |
| AT5G55900 | 0,10 | -1,26 | 4,35 | -0,29 | 7,72E-01 |
| AT5G55930 | 0,70 | 2,78 | 2,86 | 0,97 | 3,31E-01 |
| AT5G55960 | 0,59 | -0,87 | 3,03 | -0,29 | 7,74E-01 |
| AT5G55990 | 0,33 | 1,89 | 4,30 | 0,44 | 6,59E-01 |
| AT5G56020 | 0,32 | 2,42 | 4,35 | 0,56 | 5,77E-01 |
| AT5G56040 | 0,73 | 1,99 | 2,27 | 0,88 | 3,81E-01 |
| AT5G56060 | 0,67 | 1,15 | 4,32 | 0,27 | 7,91E-01 |
| AT5G56140 | 0,98 | 0,48 | 1,84 | 0,26 | 7,95E-01 |
| AT5G56160 | 0,49 | 0,00 | 4,41 | 0,00 | 1,00E+00 |
| AT5G56170 | 0,53 | -0,21 | 4,34 | -0,05 | 9,61E-01 |
| AT5G56270 | 0,76 | 3,08 | 2,83 | 1,09 | 2,76E-01 |
| AT5G56290 | 0,52 | 0,38 | 2,26 | 0,17 | 8,66E-01 |
| AT5G56340 | 0,44 | 2,88 | 3,28 | 0,88 | 3,80E-01 |
| AT5G08495 | 0,43 | -1,91 | 4,16 | -0,46 | 6,46E-01 |
| AT5G56460 | 0,74 | 0,56 | 3,93 | 0,14 | 8,87E-01 |
| AT5G56500 | 0,32 | -0,22 | 4,40 | -0,05 | 9,61E-01 |
| AT5G56590 | 0,24 | 0,27 | 4,35 | 0,06 | 9,50E-01 |
| AT5G56630 | 0,71 | 1,10 | 2,54 | 0,43 | 6,65E-01 |
| AT5G56650 | 0,64 | -0,02 | 2,22 | -0,01 | 9,92E-01 |
| AT5G56680 | 1,22 | 2,13 | 1,80 | 1,18 | 2,37E-01 |

| | | | | | |
|---|---|---|---|---|---|
| AT5G56720 | 0,12 | 1,15 | 4,35 | 0,26 | 7,92E-01 |
| AT5G56870 | 0,19 | 0,42 | 4,28 | 0,10 | 9,21E-01 |
| AT5G56910 | 0,15 | 1,94 | 4,32 | 0,45 | 6,54E-01 |
| AT5G56920 | 0,65 | -1,22 | 2,88 | -0,42 | 6,72E-01 |
| AT5G56940 | 0,08 | -0,87 | 4,38 | -0,20 | 8,43E-01 |
| AT5G56950 | 0,44 | 3,02 | 3,25 | 0,93 | 3,53E-01 |
| AT5G57015 | 0,32 | -0,44 | 4,28 | -0,10 | 9,19E-01 |
| AT5G57020 | 1,25 | 1,57 | 1,54 | 1,02 | 3,09E-01 |
| AT5G57030 | 0,86 | 4,18 | 2,31 | 1,81 | 7,02E-02 |
| AT5G57035 | 1,04 | 1,93 | 2,57 | 0,75 | 4,54E-01 |
| AT5G57070 | 0,42 | 1,29 | 4,22 | 0,30 | 7,61E-01 |
| AT5G57150 | 0,25 | 2,98 | 4,33 | 0,69 | 4,92E-01 |
| AT5G57180 | 1,05 | -1,02 | 1,87 | -0,55 | 5,86E-01 |
| AT5G57320 | 0,08 | 2,08 | 4,38 | 0,47 | 6,35E-01 |
| AT5G57330 | 0,49 | 0,07 | 2,77 | 0,02 | 9,81E-01 |
| AT5G57340 | 0,20 | -0,87 | 4,29 | -0,20 | 8,40E-01 |
| AT5G57360 | 0,96 | 1,55 | 1,68 | 0,92 | 3,55E-01 |
| AT5G57370 | 0,31 | 1,06 | 4,33 | 0,25 | 8,06E-01 |
| AT5G57460 | 1,07 | 1,68 | 1,66 | 1,01 | 3,12E-01 |
| AT5G57520 | 0,31 | -1,89 | 4,27 | -0,44 | 6,58E-01 |
| AT5G57610 | 0,89 | 1,36 | 1,73 | 0,78 | 4,33E-01 |
| AT5G57655 | 1,67 | 0,53 | 1,46 | 0,36 | 7,16E-01 |
| AT5G57735 | 0,19 | -1,70 | 4,33 | -0,39 | 6,96E-01 |
| AT5G57800 | 0,61 | -2,22 | 2,56 | -0,86 | 3,87E-01 |
| AT5G08805 | 1,03 | 0,32 | 2,21 | 0,14 | 8,86E-01 |
| AT5G57880 | 0,48 | 1,69 | 4,12 | 0,41 | 6,81E-01 |
| AT5G57930 | 1,66 | 4,76 | 1,89 | 2,51 | 1,19E-02 |
| AT5G57940 | 0,35 | 1,35 | 3,47 | 0,39 | 6,97E-01 |
| AT5G58000 | 0,08 | 2,33 | 4,36 | 0,53 | 5,93E-01 |
| AT5G58020 | 0,39 | 1,67 | 4,39 | 0,38 | 7,04E-01 |
| AT5G58060 | 0,59 | 0,79 | 2,44 | 0,32 | 7,47E-01 |
| AT5G58100 | 0,97 | -0,35 | 1,72 | -0,20 | 8,39E-01 |
| AT5G58110 | 0,27 | 0,05 | 4,28 | 0,01 | 9,90E-01 |
| AT5G58120 | 1,53 | 1,11 | 1,22 | 0,91 | 3,61E-01 |
| AT5G08820 | 0,19 | 1,89 | 4,32 | 0,44 | 6,61E-01 |
| AT5G58200 | 0,40 | -0,11 | 4,40 | -0,02 | 9,81E-01 |
| AT5G58230 | 0,77 | 2,61 | 2,37 | 1,10 | 2,71E-01 |
| AT5G58250 | 0,32 | 0,20 | 4,29 | 0,05 | 9,63E-01 |
| AT5G58270 | 0,43 | 1,64 | 4,18 | 0,39 | 6,95E-01 |
| AT5G58290 | 1,43 | 0,59 | 1,54 | 0,39 | 7,00E-01 |
| AT5G58300 | 0,99 | 2,57 | 2,24 | 1,15 | 2,52E-01 |
| AT5G58320 | 0,24 | 2,45 | 4,35 | 0,56 | 5,73E-01 |
| AT5G58350 | 1,12 | 2,71 | 1,78 | 1,53 | 1,27E-01 |
| AT5G58370 | 0,29 | 0,22 | 4,36 | 0,05 | 9,59E-01 |
| AT5G58375 | 0,48 | 0,77 | 4,35 | 0,18 | 8,59E-01 |
| AT5G58420 | 1,59 | -0,33 | 1,85 | -0,18 | 8,57E-01 |
| AT5G58450 | 0,68 | 2,18 | 2,91 | 0,75 | 4,54E-01 |
| AT5G58495 | 0,12 | 2,81 | 4,34 | 0,65 | 5,18E-01 |
| AT5G58520 | 0,39 | -0,08 | 4,32 | -0,02 | 9,84E-01 |
| AT5G58560 | 0,47 | 1,13 | 3,12 | 0,36 | 7,18E-01 |
| AT5G08855 | 0,43 | -2,29 | 4,17 | -0,55 | 5,83E-01 |
| AT5G58620 | 0,34 | 2,58 | 4,37 | 0,59 | 5,54E-01 |
| AT5G58640 | 0,71 | 1,62 | 3,01 | 0,54 | 5,91E-01 |
| AT5G58710 | 0,66 | 0,89 | 3,02 | 0,29 | 7,69E-01 |
| AT5G58990 | 0,29 | -1,08 | 4,34 | -0,25 | 8,03E-01 |
| AT5G59160 | 0,32 | 2,58 | 4,30 | 0,60 | 5,49E-01 |

| | | | | | |
|---|---|---|---|---|---|
| AT5G59170 | 0,26 | 2,30 | 4,37 | 0,53 | 5,98E-01 |
| AT5G59250 | 0,50 | 1,16 | 3,21 | 0,36 | 7,19E-01 |
| AT5G59350 | 0,35 | 1,71 | 4,32 | 0,40 | 6,92E-01 |
| AT5G59420 | 1,43 | 1,76 | 1,68 | 1,05 | 2,95E-01 |
| AT5G59430 | 0,64 | 1,02 | 2,41 | 0,42 | 6,72E-01 |
| AT5G59450 | 0,61 | 1,78 | 3,77 | 0,47 | 6,36E-01 |
| AT5G59610 | 0,51 | 2,64 | 3,96 | 0,67 | 5,04E-01 |
| AT5G59660 | 0,05 | -0,58 | 4,38 | -0,13 | 8,95E-01 |
| AT5G59670 | 0,28 | -1,06 | 3,63 | -0,29 | 7,70E-01 |
| AT5G59680 | 0,64 | 1,99 | 4,39 | 0,45 | 6,50E-01 |
| AT5G59690 | 0,64 | 1,54 | 2,80 | 0,55 | 5,81E-01 |
| AT5G59820 | 0,42 | 0,78 | 3,20 | 0,24 | 8,07E-01 |
| AT5G59830 | 0,85 | 2,95 | 2,92 | 1,01 | 3,12E-01 |
| AT5G59860 | 0,43 | 2,13 | 4,38 | 0,49 | 6,26E-01 |
| AT5G59880 | 0,87 | 1,09 | 2,50 | 0,44 | 6,62E-01 |
| AT5G59910 | 0,91 | -1,69 | 1,99 | -0,85 | 3,96E-01 |
| AT5G59920 | 0,58 | 3,76 | 3,70 | 1,02 | 3,09E-01 |
| AT5G59950 | 1,16 | 1,32 | 1,77 | 0,74 | 4,57E-01 |
| AT5G59980 | 0,39 | 1,78 | 4,41 | 0,40 | 6,86E-01 |
| AT5G60040 | 0,79 | 2,44 | 2,38 | 1,03 | 3,04E-01 |
| AT5G60110 | 0,25 | 2,97 | 4,33 | 0,69 | 4,93E-01 |
| AT5G60120 | 0,67 | 2,61 | 2,76 | 0,95 | 3,43E-01 |
| AT5G60170 | 1,68 | 4,12 | 1,74 | 2,36 | 1,81E-02 |
| AT5G60270 | 0,48 | 4,17 | 4,30 | 0,97 | 3,33E-01 |
| AT5G60280 | 0,76 | 3,04 | 3,41 | 0,89 | 3,74E-01 |
| AT5G60300 | 0,52 | 2,98 | 3,85 | 0,78 | 4,38E-01 |
| AT5G60320 | 0,25 | 2,98 | 4,33 | 0,69 | 4,92E-01 |
| AT5G60550 | 0,49 | 2,29 | 3,13 | 0,73 | 4,65E-01 |
| AT5G60560 | 0,60 | -0,63 | 2,91 | -0,22 | 8,28E-01 |
| AT5G60570 | 0,88 | -1,73 | 2,06 | -0,84 | 4,00E-01 |
| AT5G60610 | 0,72 | 2,48 | 3,60 | 0,69 | 4,91E-01 |
| AT5G60620 | 0,40 | 1,16 | 3,41 | 0,34 | 7,34E-01 |
| AT5G60680 | 1,59 | 1,45 | 1,55 | 0,94 | 3,50E-01 |
| AT5G60780 | 1,02 | 2,45 | 2,17 | 1,13 | 2,61E-01 |
| AT5G60820 | 0,45 | 2,03 | 4,09 | 0,50 | 6,19E-01 |
| AT5G60850 | 1,63 | 1,24 | 1,81 | 0,68 | 4,95E-01 |
| AT5G60890 | 0,24 | 0,00 | 4,41 | 0,00 | 1,00E+00 |
| AT5G60945 | 1,24 | -1,68 | 1,57 | -1,07 | 2,84E-01 |
| AT5G60963 | 0,50 | 1,71 | 2,58 | 0,66 | 5,08E-01 |
| AT5G60964 | 0,93 | -2,82 | 3,29 | -0,86 | 3,92E-01 |
| AT5G60980 | 1,13 | 0,97 | 2,10 | 0,46 | 6,44E-01 |
| AT5G61190 | 0,65 | 3,55 | 2,46 | 1,44 | 1,50E-01 |
| AT5G61200 | 0,25 | 0,00 | 4,40 | 0,00 | 1,00E+00 |
| AT5G61210 | 1,18 | 1,60 | 1,59 | 1,00 | 3,16E-01 |
| AT5G61230 | 1,00 | 2,25 | 2,01 | 1,12 | 2,63E-01 |
| AT5G61240 | 0,29 | 0,87 | 4,32 | 0,20 | 8,41E-01 |
| AT5G61380 | 0,78 | 3,17 | 2,38 | 1,33 | 1,82E-01 |
| AT5G61400 | 0,78 | 2,67 | 3,50 | 0,76 | 4,45E-01 |
| AT5G61470 | 0,39 | -1,21 | 3,19 | -0,38 | 7,05E-01 |
| AT5G61490 | 0,17 | 3,16 | 4,33 | 0,73 | 4,66E-01 |
| AT5G61550 | 0,85 | 4,65 | 3,29 | 1,41 | 1,57E-01 |
| AT5G61570 | 0,19 | -1,34 | 4,35 | -0,31 | 7,58E-01 |
| AT5G61580 | 0,26 | 1,27 | 4,31 | 0,29 | 7,68E-01 |
| AT5G61950 | 0,32 | 2,03 | 4,37 | 0,46 | 6,42E-01 |
| AT5G61970 | 0,49 | 0,07 | 3,32 | 0,02 | 9,83E-01 |
| AT5G61980 | 0,55 | 2,06 | 4,37 | 0,47 | 6,37E-01 |

| | | | | | |
|---|---|---|---|---|---|
| AT5G62030 | 0,41 | 0,12 | 4,11 | 0,03 | 9,76E-01 |
| AT5G62270 | 0,20 | 0,36 | 4,30 | 0,08 | 9,34E-01 |
| AT5G62290 | 0,13 | -1,34 | 4,35 | -0,31 | 7,58E-01 |
| AT5G62360 | 0,27 | -0,36 | 4,31 | -0,08 | 9,34E-01 |
| AT5G62460 | 0,44 | 2,26 | 4,14 | 0,55 | 5,84E-01 |
| AT5G62500 | 0,05 | -0,58 | 4,38 | -0,13 | 8,95E-01 |
| AT5G62520 | 0,28 | 3,41 | 4,33 | 0,79 | 4,31E-01 |
| AT5G62560 | 0,37 | -0,52 | 3,42 | -0,15 | 8,80E-01 |
| AT5G62570 | 0,48 | 0,19 | 2,52 | 0,07 | 9,41E-01 |
| AT5G62575 | 0,21 | 3,35 | 4,33 | 0,77 | 4,39E-01 |
| AT5G62620 | 0,35 | -0,12 | 4,28 | -0,03 | 9,77E-01 |
| AT5G62710 | 0,23 | 1,20 | 4,34 | 0,28 | 7,82E-01 |
| AT5G62720 | 0,31 | 1,49 | 4,30 | 0,35 | 7,28E-01 |
| AT5G62740 | 0,43 | 2,73 | 4,07 | 0,67 | 5,03E-01 |
| AT5G62770 | 0,31 | 1,70 | 4,32 | 0,39 | 6,94E-01 |
| AT5G62890 | 1,11 | 1,58 | 1,86 | 0,85 | 3,96E-01 |
| AT5G62910 | 0,36 | 2,31 | 4,33 | 0,54 | 5,93E-01 |
| AT5G62970 | 0,35 | -0,05 | 3,46 | -0,01 | 9,89E-01 |
| AT5G63010 | 0,41 | 1,48 | 3,25 | 0,46 | 6,48E-01 |
| AT5G09120 | 1,28 | -2,08 | 1,76 | -1,18 | 2,37E-01 |
| AT5G63050 | 0,55 | 1,15 | 4,02 | 0,29 | 7,74E-01 |
| AT5G63130 | 0,64 | 2,80 | 2,97 | 0,94 | 3,47E-01 |
| AT5G63200 | 0,75 | -0,92 | 3,58 | -0,26 | 7,97E-01 |
| AT5G63260 | 0,08 | 2,33 | 4,36 | 0,53 | 5,93E-01 |
| AT5G63280 | 0,12 | 2,81 | 4,34 | 0,65 | 5,18E-01 |
| AT5G09150 | 0,24 | 0,00 | 4,41 | 0,00 | 1,00E+00 |
| AT5G63300 | 0,47 | 2,86 | 4,35 | 0,66 | 5,11E-01 |
| AT5G63340 | 0,26 | -1,25 | 4,29 | -0,29 | 7,71E-01 |
| AT5G63420 | 0,69 | 1,16 | 3,14 | 0,37 | 7,11E-01 |
| AT5G63480 | 0,12 | -1,57 | 4,34 | -0,36 | 7,18E-01 |
| AT5G63510 | 0,37 | 2,25 | 4,28 | 0,53 | 5,98E-01 |
| AT5G63640 | 0,36 | 2,48 | 4,36 | 0,57 | 5,69E-01 |
| AT5G63680 | 0,63 | 1,46 | 3,08 | 0,47 | 6,35E-01 |
| AT5G63770 | 0,64 | 1,01 | 2,96 | 0,34 | 7,32E-01 |
| AT5G63780 | 0,69 | -1,05 | 2,20 | -0,48 | 6,32E-01 |
| AT5G63810 | 0,68 | 1,05 | 3,09 | 0,34 | 7,33E-01 |
| AT5G63940 | 0,43 | -1,44 | 3,24 | -0,44 | 6,57E-01 |
| AT5G63960 | 0,45 | -0,14 | 4,41 | -0,03 | 9,75E-01 |
| AT5G63980 | 0,08 | 2,33 | 4,36 | 0,53 | 5,93E-01 |
| AT5G64030 | 1,58 | 2,25 | 1,50 | 1,50 | 1,34E-01 |
| AT5G64090 | 0,72 | 2,75 | 2,42 | 1,14 | 2,55E-01 |
| AT5G64160 | 0,20 | 1,15 | 4,36 | 0,26 | 7,92E-01 |
| AT5G64190 | 0,08 | 2,33 | 4,36 | 0,53 | 5,93E-01 |
| AT5G64240 | 0,28 | 1,09 | 4,31 | 0,25 | 8,00E-01 |
| AT5G64300 | 1,43 | 0,68 | 1,48 | 0,46 | 6,49E-01 |
| AT5G64310 | 0,80 | 0,04 | 2,55 | 0,02 | 9,87E-01 |
| AT5G64330 | 1,11 | 0,13 | 1,96 | 0,07 | 9,46E-01 |
| AT5G64370 | 0,51 | 2,37 | 3,90 | 0,61 | 5,44E-01 |
| AT5G64380 | 0,10 | 0,34 | 4,35 | 0,08 | 9,37E-01 |
| AT5G64400 | 0,52 | 1,80 | 4,11 | 0,44 | 6,61E-01 |
| AT5G64440 | 0,42 | -1,61 | 3,25 | -0,50 | 6,19E-01 |
| AT5G64500 | 0,82 | 1,19 | 2,23 | 0,53 | 5,94E-01 |
| AT5G64505 | 0,26 | 2,30 | 4,37 | 0,53 | 5,98E-01 |
| AT5G64572 | 1,22 | 0,70 | 1,66 | 0,42 | 6,75E-01 |
| AT5G64690 | 0,11 | -1,28 | 4,35 | -0,29 | 7,69E-01 |
| AT5G64750 | 0,12 | 2,87 | 4,34 | 0,66 | 5,08E-01 |

| | | | | | |
|---|---|---|---|---|---|
| AT5G64813 | 0,60 | 1,51 | 2,47 | 0,61 | 5,41E-01 |
| AT5G09285 | 0,45 | 3,33 | 3,98 | 0,84 | 4,03E-01 |
| AT5G64950 | 0,48 | 3,83 | 4,32 | 0,89 | 3,75E-01 |
| AT5G64960 | 0,59 | 2,88 | 3,12 | 0,92 | 3,55E-01 |
| AT5G65010 | 1,00 | 1,03 | 1,76 | 0,58 | 5,60E-01 |
| AT5G65015 | 0,39 | 2,44 | 4,32 | 0,56 | 5,72E-01 |
| AT5G65210 | 0,26 | 2,30 | 4,37 | 0,53 | 5,98E-01 |
| AT5G65220 | 1,17 | -0,42 | 2,08 | -0,20 | 8,42E-01 |
| AT5G65230 | 0,08 | 2,08 | 4,38 | 0,47 | 6,35E-01 |
| AT5G65270 | 0,10 | 0,34 | 4,35 | 0,08 | 9,37E-01 |
| AT5G65290 | 0,47 | 0,00 | 4,41 | 0,00 | 1,00E+00 |
| AT5G65400 | 0,17 | 2,29 | 4,36 | 0,53 | 5,98E-01 |
| AT5G65440 | 0,58 | -0,08 | 3,03 | -0,03 | 9,78E-01 |
| AT5G65470 | 0,30 | 0,75 | 4,25 | 0,18 | 8,59E-01 |
| AT5G65510 | 0,12 | 0,31 | 4,35 | 0,07 | 9,44E-01 |
| AT5G65570 | 0,25 | 2,79 | 4,34 | 0,64 | 5,20E-01 |
| AT5G65620 | 1,40 | -0,93 | 1,59 | -0,59 | 5,58E-01 |
| AT5G65640 | 0,08 | 2,33 | 4,36 | 0,53 | 5,93E-01 |
| AT5G65670 | 0,91 | 0,50 | 2,02 | 0,25 | 8,03E-01 |
| AT5G65710 | 0,77 | 0,47 | 1,93 | 0,24 | 8,08E-01 |
| AT5G65720 | 1,65 | 3,28 | 1,45 | 2,26 | 2,41E-02 |
| AT5G65760 | 0,55 | 0,41 | 2,44 | 0,17 | 8,65E-01 |
| AT5G65770 | 1,30 | -0,15 | 1,85 | -0,08 | 9,34E-01 |
| AT5G65780 | 1,41 | 0,13 | 1,80 | 0,07 | 9,41E-01 |
| AT5G65890 | 0,05 | -0,58 | 4,38 | -0,13 | 8,95E-01 |
| AT5G65960 | 1,37 | 4,43 | 1,90 | 2,34 | 1,95E-02 |
| AT5G65990 | 0,70 | 1,16 | 1,90 | 0,61 | 5,40E-01 |
| AT5G66052 | 0,53 | 1,45 | 4,02 | 0,36 | 7,19E-01 |
| AT5G66070 | 0,10 | 0,34 | 4,35 | 0,08 | 9,37E-01 |
| AT5G66100 | 0,24 | 0,93 | 4,29 | 0,22 | 8,29E-01 |
| AT5G66110 | 0,30 | 2,68 | 4,35 | 0,61 | 5,39E-01 |
| AT5G66160 | 0,31 | 0,02 | 4,37 | 0,00 | 9,97E-01 |
| AT5G66170 | 0,33 | 2,76 | 4,34 | 0,64 | 5,25E-01 |
| AT5G66240 | 1,02 | 2,20 | 2,15 | 1,02 | 3,06E-01 |
| AT5G66360 | 0,60 | 3,56 | 3,78 | 0,94 | 3,46E-01 |
| AT5G66450 | 0,53 | -0,95 | 3,11 | -0,30 | 7,60E-01 |
| AT5G66560 | 1,38 | 1,94 | 1,59 | 1,22 | 2,21E-01 |
| AT5G66562 | 1,20 | -0,03 | 1,42 | -0,02 | 9,86E-01 |
| AT5G66564 | 1,37 | -1,13 | 1,42 | -0,80 | 4,25E-01 |
| AT5G66568 | 0,20 | -0,66 | 4,32 | -0,15 | 8,78E-01 |
| AT5G66760 | 1,08 | 2,53 | 1,93 | 1,31 | 1,90E-01 |
| AT5G66770 | 0,43 | 2,33 | 4,28 | 0,54 | 5,86E-01 |
| AT5G66790 | 0,74 | 1,94 | 2,18 | 0,89 | 3,74E-01 |
| AT5G66810 | 0,67 | 2,54 | 2,48 | 1,02 | 3,07E-01 |
| AT5G66930 | 0,12 | 1,15 | 4,35 | 0,26 | 7,92E-01 |
| AT5G66950 | 1,10 | -0,11 | 1,55 | -0,07 | 9,43E-01 |
| AT5G67070 | 0,51 | -1,98 | 3,89 | -0,51 | 6,11E-01 |
| AT5G67080 | 0,30 | 0,59 | 4,36 | 0,14 | 8,92E-01 |
| AT5G67170 | 0,08 | 2,08 | 4,38 | 0,47 | 6,35E-01 |
| AT5G67260 | 0,65 | -0,32 | 3,83 | -0,08 | 9,33E-01 |
| AT5G67320 | 0,96 | 3,65 | 2,56 | 1,42 | 1,55E-01 |
| AT5G67330 | 1,26 | 2,06 | 1,65 | 1,25 | 2,12E-01 |
| AT5G67340 | 0,24 | 2,22 | 4,34 | 0,51 | 6,09E-01 |
| AT5G67385 | 0,80 | -0,25 | 2,06 | -0,12 | 9,05E-01 |
| AT5G67455 | 0,10 | 0,34 | 4,35 | 0,08 | 9,37E-01 |
| AT5G67470 | 1,09 | 1,36 | 1,85 | 0,73 | 4,63E-01 |

| | | | | | |
|---|---|---|---|---|---|
| AT5G67488 | 1,20 | 0,78 | 1,65 | 0,47 | 6,36E-01 |
| AT5G67500 | 0,21 | 2,12 | 4,32 | 0,49 | 6,25E-01 |
| AT5G67530 | 0,08 | 2,08 | 4,38 | 0,47 | 6,35E-01 |
| AT5G67560 | 0,70 | 1,78 | 3,62 | 0,49 | 6,24E-01 |
| AT5G67570 | 0,48 | 0,19 | 4,13 | 0,05 | 9,63E-01 |
| AT5G67590 | 0,32 | -1,22 | 4,35 | -0,28 | 7,78E-01 |
| AT1G01020 | 0,15 | -0,41 | 4,32 | -0,09 | 9,25E-01 |
| AT1G03993 | 0,11 | 1,33 | 4,36 | 0,31 | 7,59E-01 |
| AT1G01050 | 0,66 | 2,82 | 2,94 | 0,96 | 3,37E-01 |
| AT1G01060 | 0,69 | 2,68 | 2,41 | 1,11 | 2,66E-01 |
| AT1G01090 | 1,61 | 1,14 | 1,44 | 0,79 | 4,29E-01 |
| AT1G01100 | 1,47 | 1,17 | 1,46 | 0,80 | 4,23E-01 |
| AT1G01120 | 1,43 | -0,70 | 1,36 | -0,51 | 6,09E-01 |
| AT1G01130 | 0,29 | -0,58 | 4,33 | -0,13 | 8,94E-01 |
| AT1G01140 | 0,54 | -0,76 | 2,39 | -0,32 | 7,51E-01 |
| AT1G01170 | 0,58 | -0,59 | 2,94 | -0,20 | 8,41E-01 |
| AT1G01440 | 0,35 | 0,89 | 4,30 | 0,21 | 8,36E-01 |
| AT1G01450 | 0,59 | 3,63 | 4,32 | 0,84 | 4,00E-01 |
| AT1G01470 | 1,29 | -0,61 | 1,41 | -0,43 | 6,65E-01 |
| AT1G01490 | 0,15 | -0,67 | 4,39 | -0,15 | 8,78E-01 |
| AT1G01530 | 0,08 | -0,87 | 4,38 | -0,20 | 8,43E-01 |
| AT1G01610 | 0,73 | -0,98 | 2,44 | -0,40 | 6,88E-01 |
| AT1G01620 | 1,23 | -1,50 | 1,97 | -0,76 | 4,45E-01 |
| AT1G01650 | 0,59 | 0,89 | 2,74 | 0,33 | 7,44E-01 |
| AT1G04027 | 0,46 | 0,47 | 2,97 | 0,16 | 8,75E-01 |
| AT1G01680 | 0,72 | 0,29 | 2,46 | 0,12 | 9,05E-01 |
| AT1G01725 | 1,09 | 1,04 | 1,70 | 0,61 | 5,43E-01 |
| AT1G04063 | 0,34 | -1,57 | 4,33 | -0,36 | 7,16E-01 |
| AT1G01820 | 0,86 | 0,16 | 2,27 | 0,07 | 9,42E-01 |
| AT1G01830 | 1,22 | 2,16 | 1,79 | 1,21 | 2,27E-01 |
| AT1G01880 | 1,08 | 1,49 | 2,36 | 0,63 | 5,28E-01 |
| AT1G01890 | 1,04 | 0,66 | 2,76 | 0,24 | 8,10E-01 |
| AT1G01930 | 0,16 | 1,16 | 4,38 | 0,27 | 7,90E-01 |
| AT1G01990 | 0,21 | 2,29 | 4,32 | 0,53 | 5,96E-01 |
| AT1G02020 | 0,45 | 2,45 | 4,07 | 0,60 | 5,47E-01 |
| AT1G02090 | 0,84 | 3,15 | 2,71 | 1,16 | 2,46E-01 |
| AT1G02140 | 0,21 | 1,12 | 4,33 | 0,26 | 7,95E-01 |
| AT1G02260 | 0,54 | 1,61 | 3,21 | 0,50 | 6,17E-01 |
| AT1G02280 | 0,61 | 2,88 | 3,01 | 0,95 | 3,40E-01 |
| AT1G02310 | 0,21 | 2,61 | 4,34 | 0,60 | 5,48E-01 |
| AT1G04117 | 0,17 | 2,04 | 4,33 | 0,47 | 6,37E-01 |
| AT1G02430 | 0,17 | 1,90 | 4,33 | 0,44 | 6,61E-01 |
| AT1G02450 | 0,20 | 2,90 | 4,34 | 0,67 | 5,04E-01 |
| AT1G02610 | 0,33 | -0,51 | 4,38 | -0,12 | 9,07E-01 |
| AT1G02660 | 0,36 | 0,18 | 4,34 | 0,04 | 9,67E-01 |
| AT1G02770 | 0,41 | 0,00 | 4,41 | 0,00 | 1,00E+00 |
| AT1G02870 | 0,35 | 1,66 | 4,39 | 0,38 | 7,06E-01 |
| AT1G02910 | 0,70 | 2,44 | 2,51 | 0,97 | 3,31E-01 |
| AT1G02920 | 0,41 | -2,09 | 3,26 | -0,64 | 5,23E-01 |
| AT1G02930 | 0,49 | 0,03 | 3,25 | 0,01 | 9,93E-01 |
| AT1G02960 | 0,08 | 2,33 | 4,36 | 0,53 | 5,93E-01 |
| AT1G03000 | 0,70 | 1,85 | 2,54 | 0,73 | 4,67E-01 |
| AT1G03040 | 0,62 | 0,30 | 2,60 | 0,12 | 9,07E-01 |
| AT1G03140 | 0,90 | 0,39 | 1,63 | 0,24 | 8,10E-01 |
| AT1G03240 | 1,51 | -3,97 | 2,17 | -1,83 | 6,76E-02 |
| AT1G03300 | 0,12 | 2,87 | 4,34 | 0,66 | 5,08E-01 |

| | | | | | |
|---|---|---|---|---|---|
| AT1G03400 | 0,49 | 1,14 | 2,68 | 0,42 | 6,71E-01 |
| AT1G03445 | 0,28 | 2,49 | 4,33 | 0,57 | 5,66E-01 |
| AT1G03475 | 0,69 | -0,79 | 2,21 | -0,36 | 7,22E-01 |
| AT1G03520 | 0,48 | 2,00 | 3,97 | 0,50 | 6,15E-01 |
| AT1G03530 | 0,41 | 2,69 | 4,34 | 0,62 | 5,36E-01 |
| AT1G03550 | 0,70 | -0,51 | 2,29 | -0,22 | 8,25E-01 |
| AT1G03590 | 0,41 | 1,06 | 4,21 | 0,25 | 8,01E-01 |
| AT1G03660 | 0,52 | 1,21 | 2,64 | 0,46 | 6,47E-01 |
| AT1G03730 | 1,35 | 1,31 | 2,34 | 0,56 | 5,75E-01 |
| AT1G03746 | 0,46 | -1,57 | 4,08 | -0,39 | 7,00E-01 |
| AT1G03760 | 0,08 | -0,56 | 4,38 | -0,13 | 8,99E-01 |
| AT1G03830 | 0,65 | 2,45 | 2,48 | 0,99 | 3,23E-01 |
| AT1G03870 | 0,58 | 1,22 | 3,16 | 0,38 | 7,00E-01 |
| AT1G03940 | 1,20 | 1,41 | 1,71 | 0,83 | 4,07E-01 |
| AT1G04233 | 0,08 | 2,08 | 4,38 | 0,47 | 6,35E-01 |
| AT1G03980 | 0,11 | -1,28 | 4,35 | -0,29 | 7,69E-01 |
| AT1G04000 | 0,55 | -1,42 | 3,07 | -0,46 | 6,44E-01 |
| AT1G04010 | 0,05 | -0,58 | 4,38 | -0,13 | 8,95E-01 |
| AT1G04050 | 0,08 | -0,56 | 4,38 | -0,13 | 8,99E-01 |
| AT1G04070 | 0,16 | 1,27 | 4,37 | 0,29 | 7,71E-01 |
| AT1G04110 | 0,35 | 0,00 | 4,41 | 0,00 | 1,00E+00 |
| AT1G04120 | 1,63 | 2,34 | 1,39 | 1,68 | 9,27E-02 |
| AT1G04140 | 1,54 | 2,91 | 1,62 | 1,79 | 7,27E-02 |
| AT1G04190 | 0,13 | -0,63 | 4,39 | -0,14 | 8,85E-01 |
| AT1G04220 | 0,35 | -0,36 | 4,34 | -0,08 | 9,34E-01 |
| AT1G04240 | 0,91 | -1,30 | 2,35 | -0,55 | 5,82E-01 |
| AT1G04260 | 0,47 | 1,66 | 2,62 | 0,64 | 5,25E-01 |
| AT1G04270 | 1,04 | -0,14 | 2,05 | -0,07 | 9,44E-01 |
| AT1G04280 | 0,49 | -0,20 | 3,24 | -0,06 | 9,52E-01 |
| AT1G04290 | 0,43 | 0,66 | 2,62 | 0,25 | 8,01E-01 |
| AT1G04300 | 0,83 | 1,70 | 2,95 | 0,58 | 5,64E-01 |
| AT1G04380 | 0,20 | 2,11 | 4,37 | 0,48 | 6,29E-01 |
| AT1G04390 | 0,19 | 1,14 | 4,29 | 0,26 | 7,91E-01 |
| AT1G04400 | 0,89 | -1,17 | 2,38 | -0,49 | 6,23E-01 |
| AT1G04263 | 1,35 | 4,37 | 2,36 | 1,85 | 6,41E-02 |
| AT1G04530 | 0,42 | 0,90 | 3,22 | 0,28 | 7,79E-01 |
| AT1G04570 | 0,11 | 1,28 | 4,35 | 0,29 | 7,68E-01 |
| AT1G04630 | 0,66 | 3,57 | 3,71 | 0,96 | 3,36E-01 |
| AT1G04680 | 0,74 | -0,53 | 3,00 | -0,18 | 8,60E-01 |
| AT1G04730 | 0,74 | 2,32 | 2,48 | 0,93 | 3,50E-01 |
| AT1G04750 | 0,67 | -0,71 | 2,24 | -0,32 | 7,50E-01 |
| AT1G04770 | 0,89 | -0,13 | 2,09 | -0,06 | 9,50E-01 |
| AT1G04780 | 0,86 | 0,32 | 1,89 | 0,17 | 8,64E-01 |
| AT1G04840 | 0,63 | 3,98 | 3,69 | 1,08 | 2,81E-01 |
| AT1G04850 | 0,86 | 1,34 | 2,66 | 0,50 | 6,14E-01 |
| AT1G04860 | 0,57 | 2,03 | 3,92 | 0,52 | 6,04E-01 |
| AT1G04870 | 0,84 | 1,63 | 3,04 | 0,54 | 5,92E-01 |
| AT1G04880 | 0,21 | -1,23 | 4,36 | -0,28 | 7,78E-01 |
| AT1G04910 | 0,78 | 2,64 | 2,74 | 0,96 | 3,35E-01 |
| AT1G04950 | 0,50 | 2,10 | 3,23 | 0,65 | 5,14E-01 |
| AT1G04960 | 0,36 | -0,08 | 4,40 | -0,02 | 9,86E-01 |
| AT1G04980 | 0,36 | 0,00 | 4,41 | 0,00 | 1,00E+00 |
| AT1G04985 | 0,26 | -0,08 | 4,30 | -0,02 | 9,85E-01 |
| AT1G04990 | 0,84 | 2,89 | 2,38 | 1,21 | 2,25E-01 |
| AT1G05120 | 0,53 | 2,97 | 4,33 | 0,69 | 4,93E-01 |
| AT1G05150 | 0,85 | 1,25 | 2,07 | 0,60 | 5,47E-01 |

| | | | | | |
|---|---|---|---|---|---|
| AT1G05160 | 0,59 | 3,63 | 4,32 | 0,84 | 4,00E-01 |
| AT1G05170 | 0,47 | -0,14 | 4,41 | -0,03 | 9,75E-01 |
| AT1G05190 | 0,78 | -0,99 | 2,31 | -0,43 | 6,68E-01 |
| AT1G05205 | 0,08 | 2,33 | 4,36 | 0,53 | 5,93E-01 |
| AT1G05230 | 0,45 | 2,45 | 4,07 | 0,60 | 5,47E-01 |
| AT1G05300 | 0,18 | 0,08 | 4,29 | 0,02 | 9,85E-01 |
| AT1G05350 | 0,28 | 1,78 | 4,33 | 0,41 | 6,80E-01 |
| AT1G05520 | 0,61 | 1,59 | 2,97 | 0,53 | 5,93E-01 |
| AT1G05575 | 0,16 | -0,65 | 4,31 | -0,15 | 8,80E-01 |
| AT1G05640 | 0,20 | -0,80 | 4,37 | -0,18 | 8,55E-01 |
| AT1G05670 | 0,15 | 1,69 | 4,34 | 0,39 | 6,98E-01 |
| AT1G05720 | 0,13 | -0,63 | 4,39 | -0,14 | 8,85E-01 |
| AT1G05783 | 0,96 | 1,28 | 1,71 | 0,75 | 4,54E-01 |
| AT1G05840 | 0,75 | -0,14 | 2,01 | -0,07 | 9,44E-01 |
| AT1G05870 | 0,42 | -0,85 | 4,13 | -0,21 | 8,36E-01 |
| AT1G05950 | 0,12 | 2,87 | 4,34 | 0,66 | 5,08E-01 |
| AT1G05970 | 0,08 | 2,08 | 4,38 | 0,47 | 6,35E-01 |
| AT1G06000 | 0,95 | -0,10 | 1,92 | -0,05 | 9,57E-01 |
| AT1G06130 | 0,62 | 2,48 | 3,06 | 0,81 | 4,19E-01 |
| AT1G06150 | 1,38 | 2,58 | 1,70 | 1,51 | 1,30E-01 |
| AT1G06250 | 0,72 | 0,00 | 4,41 | 0,00 | 1,00E+00 |
| AT1G06265 | 0,08 | 2,33 | 4,36 | 0,53 | 5,93E-01 |
| AT1G06360 | 0,33 | -1,05 | 4,36 | -0,24 | 8,10E-01 |
| AT1G06400 | 0,21 | 0,16 | 4,27 | 0,04 | 9,70E-01 |
| AT1G06440 | 0,58 | 1,07 | 2,59 | 0,41 | 6,78E-01 |
| AT1G06540 | 1,04 | 2,40 | 2,12 | 1,13 | 2,58E-01 |
| AT1G06570 | 0,97 | 3,51 | 2,48 | 1,42 | 1,57E-01 |
| AT1G06580 | 1,59 | 0,19 | 1,37 | 0,14 | 8,89E-01 |
| AT1G04487 | 0,39 | 1,78 | 4,16 | 0,43 | 6,68E-01 |
| AT1G06660 | 0,50 | 2,46 | 3,99 | 0,62 | 5,37E-01 |
| AT1G06690 | 0,36 | 2,48 | 4,36 | 0,57 | 5,69E-01 |
| AT1G06700 | 1,69 | 2,20 | 1,56 | 1,41 | 1,57E-01 |
| AT1G06710 | 0,26 | 2,13 | 4,38 | 0,49 | 6,27E-01 |
| AT1G06720 | 0,55 | 0,17 | 4,00 | 0,04 | 9,66E-01 |
| AT1G06760 | 1,45 | -1,16 | 1,52 | -0,77 | 4,42E-01 |
| AT1G06810 | 0,08 | 2,08 | 4,38 | 0,47 | 6,35E-01 |
| AT1G06840 | 0,17 | 0,30 | 4,37 | 0,07 | 9,44E-01 |
| AT1G06890 | 0,40 | -0,02 | 2,82 | -0,01 | 9,95E-01 |
| AT1G06900 | 1,35 | 2,44 | 1,55 | 1,57 | 1,16E-01 |
| AT1G07020 | 0,55 | 0,00 | 4,41 | 0,00 | 1,00E+00 |
| AT1G07050 | 0,42 | 2,88 | 4,11 | 0,70 | 4,83E-01 |
| AT1G07090 | 0,43 | 0,83 | 4,26 | 0,19 | 8,46E-01 |
| AT1G07110 | 1,24 | 3,87 | 2,35 | 1,65 | 9,94E-02 |
| AT1G07128 | 0,28 | 1,09 | 4,31 | 0,25 | 8,00E-01 |
| AT1G07220 | 0,25 | 0,00 | 4,41 | 0,00 | 1,00E+00 |
| AT1G07260 | 0,48 | 0,87 | 4,31 | 0,20 | 8,40E-01 |
| AT1G04573 | 0,66 | 1,88 | 4,37 | 0,43 | 6,67E-01 |
| AT1G07310 | 0,69 | 2,03 | 2,57 | 0,79 | 4,29E-01 |
| AT1G07350 | 0,45 | 1,86 | 3,27 | 0,57 | 5,70E-01 |
| AT1G07360 | 0,57 | -0,84 | 3,13 | -0,27 | 7,89E-01 |
| AT1G07430 | 0,32 | 2,96 | 4,34 | 0,68 | 4,95E-01 |
| AT1G07440 | 0,75 | 1,58 | 2,40 | 0,66 | 5,11E-01 |
| AT1G07480 | 0,80 | 0,81 | 2,19 | 0,37 | 7,10E-01 |
| AT1G07500 | 0,08 | -0,56 | 4,38 | -0,13 | 8,99E-01 |
| AT1G07520 | 1,20 | 4,47 | 2,07 | 2,16 | 3,09E-02 |
| AT1G07530 | 1,00 | 2,09 | 1,74 | 1,20 | 2,30E-01 |

| | | | | | |
|---|---|---|---|---|---|
| AT1G07640 | 0,24 | -0,04 | 4,34 | -0,01 | 9,93E-01 |
| AT1G07720 | 1,26 | -0,64 | 1,46 | -0,44 | 6,61E-01 |
| AT1G07725 | 1,03 | 3,54 | 2,52 | 1,40 | 1,60E-01 |
| AT1G07740 | 0,29 | 3,14 | 4,33 | 0,72 | 4,69E-01 |
| AT1G07730 | 0,29 | 3,14 | 4,33 | 0,72 | 4,69E-01 |
| AT1G07750 | 0,25 | 0,00 | 4,41 | 0,00 | 1,00E+00 |
| AT1G07770 | 0,87 | 0,00 | 1,62 | 0,00 | 1,00E+00 |
| AT1G07870 | 0,12 | 1,65 | 4,34 | 0,38 | 7,04E-01 |
| AT1G07970 | 0,62 | 3,59 | 3,62 | 0,99 | 3,22E-01 |
| AT1G08040 | 0,40 | 1,92 | 4,28 | 0,45 | 6,54E-01 |
| AT1G08050 | 0,33 | 2,56 | 4,31 | 0,59 | 5,52E-01 |
| AT1G08060 | 1,18 | 2,88 | 2,00 | 1,43 | 1,51E-01 |
| AT1G08125 | 0,32 | 0,40 | 4,35 | 0,09 | 9,27E-01 |
| AT1G08510 | 1,46 | 0,63 | 1,41 | 0,45 | 6,56E-01 |
| AT1G08580 | 0,24 | 0,00 | 4,41 | 0,00 | 1,00E+00 |
| AT1G08592 | 0,74 | 3,49 | 2,84 | 1,23 | 2,20E-01 |
| AT1G08610 | 0,28 | 0,84 | 4,35 | 0,19 | 8,46E-01 |
| AT1G08640 | 0,47 | 1,06 | 3,96 | 0,27 | 7,89E-01 |
| AT1G08645 | 0,64 | 0,74 | 3,76 | 0,20 | 8,43E-01 |
| AT1G08660 | 0,10 | -1,26 | 4,35 | -0,29 | 7,72E-01 |
| AT1G08700 | 1,22 | 4,53 | 2,08 | 2,18 | 2,91E-02 |
| AT1G08800 | 0,74 | 3,13 | 2,88 | 1,09 | 2,77E-01 |
| AT1G08820 | 0,89 | -2,57 | 2,60 | -0,99 | 3,23E-01 |
| AT1G08880 | 0,12 | 2,81 | 4,34 | 0,65 | 5,18E-01 |
| AT1G09020 | 0,78 | 1,62 | 2,09 | 0,78 | 4,38E-01 |
| AT1G09060 | 1,04 | 2,79 | 2,22 | 1,26 | 2,09E-01 |
| AT1G09080 | 0,40 | 0,55 | 4,26 | 0,13 | 8,98E-01 |
| AT1G09130 | 0,46 | -0,63 | 3,20 | -0,20 | 8,43E-01 |
| AT1G09160 | 0,40 | -0,01 | 3,39 | 0,00 | 9,98E-01 |
| AT1G09190 | 0,17 | 3,16 | 4,33 | 0,73 | 4,66E-01 |
| AT1G09195 | 0,70 | 2,12 | 2,88 | 0,74 | 4,61E-01 |
| AT1G09200 | 0,34 | -1,70 | 4,26 | -0,40 | 6,90E-01 |
| AT1G09230 | 0,23 | -0,64 | 4,39 | -0,15 | 8,84E-01 |
| AT1G09280 | 1,11 | 0,85 | 1,71 | 0,49 | 6,22E-01 |
| AT1G09420 | 0,88 | 2,41 | 3,28 | 0,74 | 4,62E-01 |
| AT1G09440 | 0,21 | 1,65 | 4,38 | 0,38 | 7,07E-01 |
| AT1G04707 | 0,36 | 2,79 | 4,34 | 0,64 | 5,21E-01 |
| AT1G09630 | 0,08 | 2,08 | 4,38 | 0,47 | 6,35E-01 |
| AT1G04717 | 0,54 | 3,60 | 3,79 | 0,95 | 3,42E-01 |
| AT1G09680 | 1,19 | 4,17 | 2,48 | 1,69 | 9,19E-02 |
| AT1G09730 | 0,83 | 3,89 | 2,63 | 1,48 | 1,40E-01 |
| AT1G09760 | 0,17 | 1,90 | 4,33 | 0,44 | 6,61E-01 |
| AT1G09810 | 0,42 | 0,78 | 2,69 | 0,29 | 7,71E-01 |
| AT1G09880 | 0,33 | 2,11 | 4,36 | 0,48 | 6,29E-01 |
| AT1G09910 | 0,78 | 2,45 | 2,35 | 1,04 | 2,98E-01 |
| AT1G10020 | 0,10 | 0,34 | 4,35 | 0,08 | 9,37E-01 |
| AT1G10090 | 0,57 | 1,62 | 2,58 | 0,63 | 5,29E-01 |
| AT1G10155 | 1,47 | 1,32 | 1,28 | 1,03 | 3,02E-01 |
| AT1G10180 | 0,61 | 0,19 | 2,61 | 0,07 | 9,42E-01 |
| AT1G10190 | 0,94 | -3,73 | 2,66 | -1,40 | 1,61E-01 |
| AT1G10200 | 0,57 | 2,12 | 3,79 | 0,56 | 5,76E-01 |
| AT1G10310 | 0,42 | 2,45 | 4,36 | 0,56 | 5,74E-01 |
| AT1G10320 | 0,47 | 3,13 | 3,94 | 0,79 | 4,27E-01 |
| AT1G10340 | 1,00 | 0,25 | 2,19 | 0,11 | 9,10E-01 |
| AT1G10350 | 0,52 | 3,80 | 3,86 | 0,98 | 3,25E-01 |
| AT1G10390 | 1,30 | 2,24 | 1,68 | 1,33 | 1,82E-01 |

| | | | | | |
|---|---|---|---|---|---|
| AT1G10400 | 0,08 | 2,33 | 4,36 | 0,53 | 5,93E-01 |
| AT1G10410 | 0,96 | 0,55 | 1,77 | 0,31 | 7,58E-01 |
| AT1G10430 | 0,87 | 0,28 | 1,95 | 0,15 | 8,84E-01 |
| AT1G10470 | 1,21 | -1,06 | 1,76 | -0,61 | 5,45E-01 |
| AT1G10500 | 1,39 | 4,00 | 2,38 | 1,68 | 9,31E-02 |
| AT1G10580 | 0,57 | 3,77 | 3,84 | 0,98 | 3,27E-01 |
| AT1G10586 | 1,03 | -1,25 | 1,89 | -0,66 | 5,09E-01 |
| AT1G10590 | 0,69 | 0,63 | 2,93 | 0,22 | 8,28E-01 |
| AT1G10630 | 0,65 | 1,96 | 2,65 | 0,74 | 4,59E-01 |
| AT1G10640 | 0,05 | -0,58 | 4,38 | -0,13 | 8,95E-01 |
| AT1G10680 | 0,08 | 2,08 | 4,38 | 0,47 | 6,35E-01 |
| AT1G10840 | 0,88 | 0,45 | 1,82 | 0,25 | 8,05E-01 |
| AT1G10870 | 1,11 | 0,88 | 1,94 | 0,45 | 6,50E-01 |
| AT1G10900 | 1,03 | 1,09 | 2,92 | 0,37 | 7,09E-01 |
| AT1G10920 | 0,82 | 4,19 | 4,31 | 0,97 | 3,31E-01 |
| AT1G10930 | 0,74 | 2,67 | 2,88 | 0,93 | 3,54E-01 |
| AT1G10940 | 0,48 | 0,33 | 3,25 | 0,10 | 9,20E-01 |
| AT1G04833 | 0,22 | -1,04 | 4,30 | -0,24 | 8,08E-01 |
| AT1G11100 | 0,45 | 1,64 | 3,29 | 0,50 | 6,19E-01 |
| AT1G11210 | 0,56 | -0,98 | 2,34 | -0,42 | 6,76E-01 |
| AT1G11240 | 0,05 | -0,58 | 4,38 | -0,13 | 8,95E-01 |
| AT1G11270 | 0,83 | -2,24 | 2,08 | -1,08 | 2,82E-01 |
| AT1G11280 | 0,45 | -0,46 | 2,73 | -0,17 | 8,67E-01 |
| AT1G11290 | 0,63 | 3,97 | 3,63 | 1,09 | 2,74E-01 |
| AT1G11310 | 1,46 | 1,64 | 1,43 | 1,15 | 2,51E-01 |
| AT1G11340 | 0,08 | 2,33 | 4,36 | 0,53 | 5,93E-01 |
| AT1G11350 | 0,52 | 1,91 | 3,15 | 0,61 | 5,43E-01 |
| AT1G11360 | 0,33 | -0,38 | 4,31 | -0,09 | 9,29E-01 |
| AT1G11480 | 0,70 | 0,59 | 2,08 | 0,28 | 7,77E-01 |
| AT1G11630 | 0,35 | 0,00 | 4,41 | 0,00 | 1,00E+00 |
| AT1G11820 | 0,25 | 1,45 | 4,32 | 0,34 | 7,38E-01 |
| AT1G12000 | 0,52 | 1,05 | 2,92 | 0,36 | 7,20E-01 |
| AT1G12020 | 0,63 | -1,02 | 2,09 | -0,49 | 6,26E-01 |
| AT1G12120 | 0,70 | -0,60 | 2,55 | -0,23 | 8,14E-01 |
| AT1G12211 | 0,63 | 1,71 | 3,13 | 0,55 | 5,83E-01 |
| AT1G12310 | 0,63 | 0,30 | 3,00 | 0,10 | 9,20E-01 |
| AT1G12380 | 0,49 | 0,32 | 3,96 | 0,08 | 9,36E-01 |
| AT1G12440 | 1,21 | -0,18 | 1,63 | -0,11 | 9,12E-01 |
| AT1G12500 | 0,25 | 1,81 | 4,28 | 0,42 | 6,72E-01 |
| AT1G12520 | 0,51 | 2,02 | 4,37 | 0,46 | 6,45E-01 |
| AT1G12590 | 0,78 | -0,88 | 1,70 | -0,52 | 6,04E-01 |
| AT1G12610 | 0,19 | 0,42 | 4,28 | 0,10 | 9,21E-01 |
| AT1G12615 | 1,59 | -2,17 | 1,79 | -1,21 | 2,27E-01 |
| AT1G12640 | 0,58 | 3,84 | 4,32 | 0,89 | 3,74E-01 |
| AT1G12680 | 1,43 | -1,59 | 1,62 | -0,98 | 3,29E-01 |
| AT1G12700 | 0,22 | 0,81 | 4,27 | 0,19 | 8,50E-01 |
| AT1G12780 | 0,71 | 0,99 | 2,09 | 0,47 | 6,37E-01 |
| AT1G12800 | 0,97 | 0,55 | 2,55 | 0,22 | 8,29E-01 |
| AT1G12910 | 1,07 | 3,53 | 2,00 | 1,76 | 7,77E-02 |
| AT1G13020 | 0,59 | 1,87 | 2,59 | 0,72 | 4,72E-01 |
| AT1G13120 | 0,23 | 2,38 | 4,31 | 0,55 | 5,80E-01 |
| AT1G13170 | 0,46 | 1,95 | 3,21 | 0,61 | 5,43E-01 |
| AT1G13200 | 0,66 | 2,60 | 3,64 | 0,71 | 4,76E-01 |
| AT1G13245 | 0,44 | -0,14 | 4,14 | -0,03 | 9,74E-01 |
| AT1G13310 | 0,35 | 0,00 | 4,41 | 0,00 | 1,00E+00 |
| AT1G13340 | 0,95 | 0,94 | 2,02 | 0,47 | 6,42E-01 |

| | | | | | |
|---|---|---|---|---|---|
| AT1G13390 | 0,17 | 3,14 | 4,33 | 0,73 | 4,68E-01 |
| AT1G13450 | 0,76 | 1,81 | 2,15 | 0,84 | 4,02E-01 |
| AT1G13520 | 0,54 | 1,38 | 4,35 | 0,32 | 7,51E-01 |
| AT1G13560 | 0,43 | 1,35 | 4,33 | 0,31 | 7,56E-01 |
| AT1G13825 | 0,76 | 2,76 | 3,49 | 0,79 | 4,28E-01 |
| AT1G13860 | 0,10 | 0,39 | 4,35 | 0,09 | 9,29E-01 |
| AT1G13870 | 0,12 | -0,77 | 4,36 | -0,18 | 8,61E-01 |
| AT1G13900 | 0,21 | 2,12 | 4,32 | 0,49 | 6,25E-01 |
| AT1G13940 | 0,86 | 1,94 | 2,25 | 0,86 | 3,89E-01 |
| AT1G14010 | 0,36 | 0,25 | 4,29 | 0,06 | 9,53E-01 |
| AT1G14030 | 0,62 | 2,82 | 3,08 | 0,92 | 3,60E-01 |
| AT1G14290 | 0,26 | 2,13 | 4,38 | 0,49 | 6,27E-01 |
| AT1G14300 | 0,22 | -0,07 | 4,40 | -0,02 | 9,88E-01 |
| AT1G14330 | 0,59 | 1,88 | 2,56 | 0,73 | 4,64E-01 |
| AT1G14360 | 0,18 | -0,70 | 4,38 | -0,16 | 8,72E-01 |
| AT1G14400 | 0,66 | 2,36 | 3,04 | 0,77 | 4,38E-01 |
| AT1G14450 | 0,11 | -1,08 | 4,37 | -0,25 | 8,05E-01 |
| AT1G14455 | 0,14 | 0,00 | 4,41 | 0,00 | 1,00E+00 |
| AT1G14518 | 0,84 | -1,61 | 2,13 | -0,75 | 4,51E-01 |
| AT1G14590 | 0,42 | 2,85 | 4,17 | 0,68 | 4,95E-01 |
| AT1G14690 | 0,78 | 1,51 | 2,33 | 0,65 | 5,16E-01 |
| AT1G14710 | 0,51 | 3,85 | 3,84 | 1,00 | 3,16E-01 |
| AT1G14720 | 0,26 | 0,00 | 4,41 | 0,00 | 1,00E+00 |
| AT1G14790 | 1,42 | 2,96 | 1,99 | 1,49 | 1,37E-01 |
| AT1G14810 | 1,02 | 2,11 | 2,09 | 1,01 | 3,13E-01 |
| AT1G14830 | 0,90 | 1,91 | 2,65 | 0,72 | 4,70E-01 |
| AT1G14840 | 0,51 | 0,31 | 2,25 | 0,14 | 8,90E-01 |
| AT1G14850 | 1,26 | 1,59 | 1,70 | 0,93 | 3,51E-01 |
| AT1G14880 | 1,19 | -0,32 | 1,61 | -0,20 | 8,43E-01 |
| AT1G14900 | 0,38 | 1,68 | 4,39 | 0,38 | 7,02E-01 |
| AT1G14910 | 0,80 | 2,29 | 2,44 | 0,94 | 3,49E-01 |
| AT1G14980 | 0,57 | 3,34 | 3,05 | 1,09 | 2,74E-01 |
| AT1G05137 | 0,28 | 1,59 | 4,31 | 0,37 | 7,13E-01 |
| AT1G15020 | 0,98 | 4,24 | 2,56 | 1,65 | 9,82E-02 |
| AT1G15040 | 0,21 | 1,45 | 4,28 | 0,34 | 7,35E-01 |
| AT1G15100 | 1,26 | -0,76 | 2,44 | -0,31 | 7,56E-01 |
| AT1G15140 | 0,76 | 1,87 | 2,09 | 0,90 | 3,71E-01 |
| AT1G15175 | 0,27 | 1,46 | 4,33 | 0,34 | 7,36E-01 |
| AT1G15210 | 0,17 | 3,16 | 4,33 | 0,73 | 4,66E-01 |
| AT1G15220 | 0,11 | 1,28 | 4,35 | 0,29 | 7,68E-01 |
| AT1G15340 | 0,21 | -0,58 | 4,32 | -0,13 | 8,94E-01 |
| AT1G15350 | 1,25 | 0,98 | 2,02 | 0,49 | 6,25E-01 |
| AT1G15400 | 0,43 | 0,16 | 2,64 | 0,06 | 9,51E-01 |
| AT1G15440 | 0,87 | 1,74 | 2,16 | 0,80 | 4,22E-01 |
| AT1G15520 | 0,46 | 0,43 | 3,16 | 0,14 | 8,92E-01 |
| AT1G15530 | 1,02 | 3,30 | 2,32 | 1,42 | 1,56E-01 |
| AT1G15550 | 0,24 | 0,00 | 4,41 | 0,00 | 1,00E+00 |
| AT1G15700 | 0,40 | 1,95 | 3,33 | 0,58 | 5,59E-01 |
| AT1G05187 | 0,24 | 2,85 | 4,34 | 0,66 | 5,11E-01 |
| AT1G15730 | 0,44 | -2,54 | 4,06 | -0,63 | 5,32E-01 |
| AT1G15780 | 1,01 | 0,91 | 1,85 | 0,49 | 6,24E-01 |
| AT1G15790 | 1,05 | 0,33 | 2,75 | 0,12 | 9,05E-01 |
| AT1G05203 | 0,26 | 2,30 | 4,37 | 0,53 | 5,98E-01 |
| AT1G15880 | 0,20 | -0,80 | 4,37 | -0,18 | 8,55E-01 |
| AT1G15970 | 0,11 | -1,28 | 4,35 | -0,29 | 7,69E-01 |
| AT1G16010 | 0,99 | 4,04 | 2,23 | 1,81 | 7,08E-02 |

| | | | | | |
|---|---|---|---|---|---|
| AT1G16070 | 0,40 | 1,98 | 4,31 | 0,46 | 6,46E-01 |
| AT1G16210 | 0,60 | 0,71 | 3,12 | 0,23 | 8,21E-01 |
| AT1G16240 | 0,80 | 2,69 | 2,35 | 1,14 | 2,52E-01 |
| AT1G16260 | 1,63 | 1,76 | 1,58 | 1,12 | 2,64E-01 |
| AT1G16460 | 0,57 | 0,31 | 2,20 | 0,14 | 8,87E-01 |
| AT1G16570 | 0,28 | 1,09 | 4,31 | 0,25 | 8,00E-01 |
| AT1G16610 | 1,02 | 1,79 | 2,99 | 0,60 | 5,48E-01 |
| AT1G16750 | 0,49 | 2,46 | 3,10 | 0,79 | 4,28E-01 |
| AT1G16780 | 0,12 | 2,87 | 4,34 | 0,66 | 5,08E-01 |
| AT1G16800 | 1,44 | 3,05 | 2,22 | 1,38 | 1,68E-01 |
| AT1G16870 | 0,95 | 3,03 | 3,35 | 0,91 | 3,65E-01 |
| AT1G16890 | 1,30 | -0,73 | 1,61 | -0,45 | 6,51E-01 |
| AT1G16900 | 0,65 | 4,05 | 3,67 | 1,10 | 2,71E-01 |
| AT1G16920 | 0,22 | 0,62 | 4,28 | 0,14 | 8,85E-01 |
| AT1G16970 | 0,37 | 1,49 | 3,43 | 0,43 | 6,64E-01 |
| AT1G17080 | 0,60 | 0,47 | 3,19 | 0,15 | 8,83E-01 |
| AT1G17110 | 0,57 | 2,15 | 2,59 | 0,83 | 4,06E-01 |
| AT1G17140 | 0,95 | 2,80 | 3,44 | 0,81 | 4,16E-01 |
| AT1G17147 | 0,12 | -1,57 | 4,34 | -0,36 | 7,18E-01 |
| AT1G17232 | 0,51 | -0,67 | 2,92 | -0,23 | 8,18E-01 |
| AT1G17240 | 0,05 | -0,58 | 4,38 | -0,13 | 8,95E-01 |
| AT1G17250 | 0,99 | 3,95 | 2,53 | 1,56 | 1,19E-01 |
| AT1G17370 | 0,81 | 1,00 | 2,55 | 0,39 | 6,94E-01 |
| AT1G17380 | 0,50 | -0,95 | 2,49 | -0,38 | 7,02E-01 |
| AT1G05313 | 0,32 | -0,43 | 4,25 | -0,10 | 9,20E-01 |
| AT1G17440 | 0,46 | -0,17 | 3,18 | -0,05 | 9,58E-01 |
| AT1G17450 | 1,22 | 2,86 | 1,72 | 1,67 | 9,58E-02 |
| AT1G17590 | 0,46 | 1,57 | 3,22 | 0,49 | 6,27E-01 |
| AT1G17600 | 1,00 | -0,29 | 1,92 | -0,15 | 8,82E-01 |
| AT1G17665 | 0,20 | 2,68 | 4,35 | 0,61 | 5,39E-01 |
| AT1G17670 | 0,37 | 1,22 | 4,35 | 0,28 | 7,80E-01 |
| AT1G17690 | 0,25 | 2,79 | 4,34 | 0,64 | 5,20E-01 |
| AT1G17720 | 0,66 | 0,66 | 2,09 | 0,32 | 7,52E-01 |
| AT1G17744 | 0,16 | 1,16 | 4,38 | 0,27 | 7,90E-01 |
| AT1G17790 | 0,63 | 0,72 | 3,18 | 0,23 | 8,20E-01 |
| AT1G05333 | 0,71 | 2,84 | 3,54 | 0,80 | 4,23E-01 |
| AT1G17820 | 0,71 | 2,84 | 3,54 | 0,80 | 4,23E-01 |
| AT1G17850 | 0,36 | 1,28 | 4,34 | 0,30 | 7,68E-01 |
| AT1G05347 | 0,21 | 2,80 | 4,34 | 0,65 | 5,18E-01 |
| AT1G17870 | 0,39 | 2,17 | 4,23 | 0,51 | 6,09E-01 |
| AT1G17880 | 0,59 | 2,18 | 3,92 | 0,56 | 5,78E-01 |
| AT1G17890 | 0,42 | 0,79 | 4,40 | 0,18 | 8,58E-01 |
| AT1G17930 | 0,20 | 2,30 | 4,36 | 0,53 | 5,98E-01 |
| AT1G17980 | 1,06 | -0,42 | 2,12 | -0,20 | 8,43E-01 |
| AT1G17990 | 0,53 | -0,41 | 3,08 | -0,13 | 8,95E-01 |
| AT1G18050 | 0,60 | 1,28 | 2,36 | 0,54 | 5,89E-01 |
| AT1G18060 | 0,60 | 0,72 | 2,19 | 0,33 | 7,44E-01 |
| AT1G18070 | 0,60 | 2,35 | 2,50 | 0,94 | 3,46E-01 |
| AT1G18100 | 0,36 | 3,47 | 4,33 | 0,80 | 4,23E-01 |
| AT1G18150 | 1,19 | 2,55 | 2,27 | 1,12 | 2,62E-01 |
| AT1G18210 | 0,61 | -1,69 | 2,18 | -0,78 | 4,36E-01 |
| AT1G18330 | 0,22 | 0,78 | 4,33 | 0,18 | 8,58E-01 |
| AT1G18360 | 1,25 | -0,10 | 1,67 | -0,06 | 9,53E-01 |
| AT1G18382 | 0,44 | -0,13 | 2,60 | -0,05 | 9,60E-01 |
| AT1G18470 | 0,48 | -0,22 | 2,53 | -0,09 | 9,32E-01 |
| AT1G18540 | 1,15 | 0,78 | 1,92 | 0,40 | 6,87E-01 |

| | | | | | |
|---|---|---|---|---|---|
| AT1G18720 | 0,23 | 0,83 | 4,27 | 0,20 | 8,45E-01 |
| AT1G05433 | 0,53 | -1,71 | 2,91 | -0,59 | 5,58E-01 |
| AT1G18840 | 0,43 | 1,19 | 3,28 | 0,36 | 7,16E-01 |
| AT1G18890 | 0,28 | 1,26 | 4,36 | 0,29 | 7,73E-01 |
| AT1G18990 | 0,45 | 2,07 | 4,15 | 0,50 | 6,18E-01 |
| AT1G19000 | 0,86 | -0,18 | 1,66 | -0,11 | 9,13E-01 |
| AT1G19130 | 0,49 | 0,87 | 2,71 | 0,32 | 7,49E-01 |
| AT1G19140 | 0,44 | 1,05 | 4,35 | 0,24 | 8,10E-01 |
| AT1G19210 | 0,10 | -1,26 | 4,35 | -0,29 | 7,72E-01 |
| AT1G19260 | 0,97 | -0,77 | 2,34 | -0,33 | 7,41E-01 |
| AT1G19300 | 0,88 | 1,59 | 2,26 | 0,70 | 4,81E-01 |
| AT1G19360 | 0,50 | 1,94 | 3,97 | 0,49 | 6,25E-01 |
| AT1G19370 | 0,41 | 1,98 | 4,26 | 0,46 | 6,43E-01 |
| AT1G19400 | 1,69 | 0,71 | 1,14 | 0,62 | 5,32E-01 |
| AT1G19430 | 0,39 | 0,79 | 4,24 | 0,19 | 8,52E-01 |
| AT1G19450 | 0,33 | -0,88 | 4,37 | -0,20 | 8,40E-01 |
| AT1G19570 | 0,89 | -0,69 | 1,91 | -0,36 | 7,20E-01 |
| AT1G19640 | 0,14 | 0,00 | 4,41 | 0,00 | 1,00E+00 |
| AT1G19660 | 0,55 | -1,07 | 2,56 | -0,42 | 6,74E-01 |
| AT1G19670 | 1,59 | -1,33 | 1,78 | -0,74 | 4,57E-01 |
| AT1G19700 | 0,25 | 0,64 | 4,35 | 0,15 | 8,82E-01 |
| AT1G19715 | 1,55 | 4,02 | 2,04 | 1,97 | 4,90E-02 |
| AT1G19720 | 1,66 | 1,86 | 1,46 | 1,27 | 2,05E-01 |
| AT1G19780 | 0,50 | -0,98 | 3,98 | -0,25 | 8,05E-01 |
| AT1G19860 | 0,15 | 0,40 | 4,33 | 0,09 | 9,27E-01 |
| AT1G19870 | 0,80 | 2,17 | 2,43 | 0,89 | 3,72E-01 |
| AT1G19880 | 0,41 | -1,28 | 3,31 | -0,39 | 6,99E-01 |
| AT1G19920 | 1,00 | 2,53 | 2,29 | 1,11 | 2,69E-01 |
| AT1G20040 | 0,34 | 0,91 | 4,35 | 0,21 | 8,35E-01 |
| AT1G20070 | 0,59 | 1,69 | 2,35 | 0,72 | 4,71E-01 |
| AT1G20160 | 0,33 | 3,34 | 4,32 | 0,77 | 4,40E-01 |
| AT1G20190 | 0,64 | 0,51 | 3,24 | 0,16 | 8,74E-01 |
| AT1G20200 | 0,33 | 0,95 | 3,45 | 0,28 | 7,82E-01 |
| AT1G20320 | 0,05 | -0,58 | 4,38 | -0,13 | 8,95E-01 |
| AT1G20400 | 1,62 | -2,17 | 1,40 | -1,55 | 1,21E-01 |
| AT1G20430 | 0,89 | 3,83 | 2,82 | 1,36 | 1,75E-01 |
| AT1G20490 | 0,48 | 2,96 | 4,34 | 0,68 | 4,95E-01 |
| AT1G20560 | 0,25 | 1,66 | 4,31 | 0,39 | 6,99E-01 |
| AT1G20570 | 0,08 | 2,33 | 4,36 | 0,53 | 5,93E-01 |
| AT1G20650 | 0,41 | -1,14 | 4,36 | -0,26 | 7,94E-01 |
| AT1G20670 | 0,31 | 1,26 | 4,34 | 0,29 | 7,72E-01 |
| AT1G20770 | 0,16 | 1,16 | 4,38 | 0,27 | 7,90E-01 |
| AT1G20840 | 1,09 | -0,28 | 1,75 | -0,16 | 8,72E-01 |
| AT1G20880 | 0,25 | 2,05 | 4,37 | 0,47 | 6,40E-01 |
| AT1G20890 | 0,48 | 2,74 | 3,96 | 0,69 | 4,90E-01 |
| AT1G20950 | 0,69 | 2,51 | 2,43 | 1,03 | 3,02E-01 |
| AT1G21080 | 0,61 | -1,60 | 2,48 | -0,65 | 5,19E-01 |
| AT1G05627 | 0,05 | -0,58 | 4,38 | -0,13 | 8,95E-01 |
| AT1G21160 | 0,26 | 0,60 | 4,28 | 0,14 | 8,89E-01 |
| AT1G21190 | 0,45 | -0,04 | 4,10 | -0,01 | 9,92E-01 |
| AT1G21310 | 0,44 | 1,09 | 2,72 | 0,40 | 6,90E-01 |
| AT1G21380 | 1,01 | 0,79 | 2,09 | 0,38 | 7,04E-01 |
| AT1G21390 | 0,08 | 2,33 | 4,36 | 0,53 | 5,93E-01 |
| AT1G21440 | 1,10 | -1,91 | 2,66 | -0,72 | 4,73E-01 |
| AT1G21500 | 0,51 | -2,42 | 3,08 | -0,78 | 4,33E-01 |
| AT1G21520 | 0,17 | 1,90 | 4,33 | 0,44 | 6,61E-01 |

| | | | | | |
|---|---|---|---|---|---|
| AT1G21550 | 0,17 | 2,04 | 4,33 | 0,47 | 6,37E-01 |
| AT1G21600 | 0,77 | 0,02 | 1,90 | 0,01 | 9,92E-01 |
| AT1G21650 | 0,46 | 1,89 | 3,21 | 0,59 | 5,55E-01 |
| AT1G21770 | 0,66 | -0,69 | 2,52 | -0,27 | 7,84E-01 |
| AT1G21890 | 0,36 | -1,88 | 4,26 | -0,44 | 6,59E-01 |
| AT1G21900 | 0,26 | 0,26 | 4,29 | 0,06 | 9,51E-01 |
| AT1G21920 | 0,41 | 1,94 | 3,27 | 0,59 | 5,53E-01 |
| AT1G21928 | 0,29 | 0,55 | 4,28 | 0,13 | 8,98E-01 |
| AT1G22180 | 0,30 | -0,65 | 3,50 | -0,19 | 8,52E-01 |
| AT1G22200 | 0,54 | 2,92 | 3,87 | 0,76 | 4,50E-01 |
| AT1G22300 | 0,58 | -0,71 | 2,48 | -0,29 | 7,73E-01 |
| AT1G22360 | 0,63 | 0,98 | 2,68 | 0,36 | 7,16E-01 |
| AT1G22400 | 0,23 | 0,97 | 4,30 | 0,23 | 8,21E-01 |
| AT1G22550 | 0,19 | -0,39 | 4,29 | -0,09 | 9,27E-01 |
| AT1G22570 | 0,46 | 2,66 | 3,98 | 0,67 | 5,03E-01 |
| AT1G22710 | 0,50 | -0,38 | 2,48 | -0,15 | 8,80E-01 |
| AT1G22850 | 1,38 | 4,54 | 1,84 | 2,46 | 1,37E-02 |
| AT1G23020 | 0,20 | 2,11 | 4,37 | 0,48 | 6,29E-01 |
| AT1G23052 | 1,19 | -2,22 | 2,31 | -0,96 | 3,37E-01 |
| AT1G23080 | 0,61 | -0,63 | 3,19 | -0,20 | 8,44E-01 |
| AT1G05793 | 0,08 | -0,56 | 4,38 | -0,13 | 8,99E-01 |
| AT1G23205 | 0,35 | -1,24 | 4,36 | -0,28 | 7,76E-01 |
| AT1G23260 | 0,22 | 1,81 | 4,30 | 0,42 | 6,74E-01 |
| AT1G23300 | 1,57 | 0,19 | 1,48 | 0,13 | 8,96E-01 |
| AT1G23330 | 0,10 | -1,26 | 4,35 | -0,29 | 7,72E-01 |
| AT1G23650 | 0,08 | 2,08 | 4,38 | 0,47 | 6,35E-01 |
| AT1G23670 | 0,22 | -2,38 | 4,30 | -0,55 | 5,80E-01 |
| AT1G23740 | 1,33 | 0,49 | 1,83 | 0,27 | 7,88E-01 |
| AT1G23780 | 0,31 | 2,37 | 4,31 | 0,55 | 5,82E-01 |
| AT1G23830 | 0,53 | 1,06 | 3,12 | 0,34 | 7,34E-01 |
| AT1G23840 | 0,48 | 2,47 | 3,10 | 0,80 | 4,26E-01 |
| AT1G23850 | 0,35 | 1,47 | 4,32 | 0,34 | 7,34E-01 |
| AT1G23960 | 0,23 | -0,26 | 4,31 | -0,06 | 9,52E-01 |
| AT1G24040 | 0,10 | 0,39 | 4,35 | 0,09 | 9,29E-01 |
| AT1G24100 | 1,00 | 0,51 | 1,71 | 0,30 | 7,66E-01 |
| AT1G24120 | 0,29 | -0,57 | 4,33 | -0,13 | 8,96E-01 |
| AT1G24140 | 0,74 | 1,58 | 3,00 | 0,53 | 5,99E-01 |
| AT1G24160 | 0,84 | 4,56 | 3,28 | 1,39 | 1,65E-01 |
| AT1G24190 | 0,84 | 1,77 | 2,60 | 0,68 | 4,95E-01 |
| AT1G24267 | 0,48 | 2,58 | 3,15 | 0,82 | 4,13E-01 |
| AT1G24290 | 0,17 | 1,90 | 4,33 | 0,44 | 6,61E-01 |
| AT1G24300 | 1,68 | 1,51 | 1,55 | 0,97 | 3,30E-01 |
| AT1G24350 | 0,14 | -0,45 | 4,38 | -0,10 | 9,18E-01 |
| AT1G05907 | 0,77 | -0,89 | 2,30 | -0,39 | 7,00E-01 |
| AT1G24764 | 0,53 | 1,04 | 3,21 | 0,32 | 7,46E-01 |
| AT1G24822 | 0,94 | 0,00 | 4,41 | 0,00 | 1,00E+00 |
| AT1G25220 | 0,36 | 3,47 | 4,33 | 0,80 | 4,23E-01 |
| AT1G25350 | 1,21 | 1,23 | 1,74 | 0,71 | 4,80E-01 |
| AT1G25370 | 0,12 | 2,81 | 4,34 | 0,65 | 5,18E-01 |
| AT1G25390 | 0,51 | 1,60 | 2,96 | 0,54 | 5,89E-01 |
| AT1G25400 | 0,63 | 1,57 | 2,74 | 0,57 | 5,67E-01 |
| AT1G25500 | 0,25 | 1,24 | 4,28 | 0,29 | 7,72E-01 |
| AT1G25540 | 0,94 | 1,90 | 2,10 | 0,91 | 3,65E-01 |
| AT1G25560 | 0,90 | -0,52 | 1,81 | -0,29 | 7,73E-01 |
| AT1G25570 | 0,49 | 0,00 | 4,41 | 0,00 | 1,00E+00 |
| AT1G26150 | 0,88 | 0,87 | 2,15 | 0,40 | 6,87E-01 |

| | | | | | |
|---|---|---|---|---|---|
| AT1G26160 | 0,24 | 0,06 | 4,31 | 0,01 | 9,89E-01 |
| AT1G26170 | 0,54 | 1,25 | 3,06 | 0,41 | 6,83E-01 |
| AT1G26208 | 0,43 | -0,35 | 4,28 | -0,08 | 9,36E-01 |
| AT1G26218 | 0,37 | -1,03 | 4,27 | -0,24 | 8,10E-01 |
| AT1G26233 | 0,08 | -0,56 | 4,38 | -0,13 | 8,99E-01 |
| AT1G26240 | 0,88 | -1,73 | 2,40 | -0,72 | 4,71E-01 |
| AT1G26255 | 0,44 | -1,55 | 3,98 | -0,39 | 6,96E-01 |
| AT1G26270 | 1,28 | 0,73 | 2,10 | 0,35 | 7,28E-01 |
| AT1G26380 | 0,46 | -0,90 | 4,37 | -0,21 | 8,36E-01 |
| AT1G26390 | 0,25 | 0,42 | 4,34 | 0,10 | 9,23E-01 |
| AT1G26420 | 0,40 | 0,04 | 4,23 | 0,01 | 9,93E-01 |
| AT1G26690 | 0,33 | 2,56 | 4,31 | 0,59 | 5,52E-01 |
| AT1G26761 | 0,85 | 3,10 | 4,30 | 0,72 | 4,70E-01 |
| AT1G26800 | 0,42 | -0,81 | 2,76 | -0,29 | 7,68E-01 |
| AT1G06033 | 1,64 | -1,70 | 1,76 | -0,97 | 3,33E-01 |
| AT1G26930 | 0,36 | 1,52 | 4,24 | 0,36 | 7,20E-01 |
| AT1G27008 | 0,08 | 2,33 | 4,36 | 0,53 | 5,93E-01 |
| AT1G27100 | 1,46 | -1,15 | 1,45 | -0,79 | 4,28E-01 |
| AT1G27190 | 0,77 | 1,26 | 2,02 | 0,62 | 5,33E-01 |
| AT1G27200 | 0,57 | 0,38 | 2,61 | 0,15 | 8,83E-01 |
| AT1G27210 | 0,26 | -0,17 | 4,26 | -0,04 | 9,68E-01 |
| AT1G27350 | 0,20 | -0,65 | 4,36 | -0,15 | 8,82E-01 |
| AT1G27370 | 0,51 | 0,83 | 4,04 | 0,21 | 8,37E-01 |
| AT1G27385 | 0,43 | 0,00 | 4,38 | 0,00 | 1,00E+00 |
| AT1G27440 | 0,24 | 1,18 | 4,36 | 0,27 | 7,87E-01 |
| AT1G27500 | 0,29 | 3,27 | 4,33 | 0,76 | 4,49E-01 |
| AT1G27510 | 0,44 | 1,47 | 3,13 | 0,47 | 6,39E-01 |
| AT1G06093 | 0,29 | 0,27 | 4,37 | 0,06 | 9,50E-01 |
| AT1G27600 | 1,12 | 2,73 | 2,14 | 1,27 | 2,03E-01 |
| AT1G27700 | 0,50 | 2,63 | 4,35 | 0,60 | 5,46E-01 |
| AT1G27720 | 0,50 | 3,40 | 3,91 | 0,87 | 3,85E-01 |
| AT1G27730 | 1,60 | -0,76 | 1,24 | -0,61 | 5,39E-01 |
| AT1G27900 | 0,40 | -0,58 | 3,17 | -0,18 | 8,55E-01 |
| AT1G27930 | 0,53 | 2,69 | 3,04 | 0,89 | 3,75E-01 |
| AT1G28000 | 0,35 | -1,93 | 4,26 | -0,45 | 6,50E-01 |
| AT1G28120 | 0,73 | 4,11 | 2,85 | 1,44 | 1,49E-01 |
| AT1G28140 | 1,15 | 3,49 | 1,86 | 1,88 | 6,00E-02 |
| AT1G28150 | 0,60 | 2,71 | 2,98 | 0,91 | 3,63E-01 |
| AT1G28190 | 0,15 | 1,64 | 4,33 | 0,38 | 7,06E-01 |
| AT1G28200 | 0,24 | 1,23 | 4,34 | 0,28 | 7,77E-01 |
| AT1G28230 | 0,78 | 0,43 | 2,02 | 0,21 | 8,30E-01 |
| AT1G28240 | 0,33 | -0,37 | 4,32 | -0,09 | 9,31E-01 |
| AT1G28260 | 1,27 | 3,06 | 1,85 | 1,65 | 9,79E-02 |
| AT1G28280 | 1,03 | 0,05 | 1,74 | 0,03 | 9,76E-01 |
| AT1G28290 | 0,27 | -1,63 | 4,34 | -0,38 | 7,07E-01 |
| AT1G28310 | 0,66 | -1,06 | 1,98 | -0,54 | 5,92E-01 |
| AT1G28330 | 0,60 | -0,14 | 3,17 | -0,05 | 9,64E-01 |
| AT1G28370 | 0,92 | -0,88 | 2,12 | -0,41 | 6,79E-01 |
| AT1G28390 | 0,25 | 0,00 | 4,40 | 0,00 | 1,00E+00 |
| AT1G28400 | 0,35 | -0,61 | 3,43 | -0,18 | 8,59E-01 |
| AT1G28570 | 0,12 | 1,65 | 4,34 | 0,38 | 7,04E-01 |
| AT1G28580 | 0,48 | 2,90 | 3,87 | 0,75 | 4,53E-01 |
| AT1G28600 | 1,10 | 0,71 | 1,70 | 0,42 | 6,76E-01 |
| AT1G28630 | 0,79 | -2,61 | 2,77 | -0,94 | 3,46E-01 |
| AT1G28660 | 0,08 | 2,33 | 4,36 | 0,53 | 5,93E-01 |
| AT1G28685 | 0,55 | 0,63 | 3,13 | 0,20 | 8,40E-01 |

| | | | | | |
|---|---|---|---|---|---|
| AT1G28765 | 0,17 | 3,14 | 4,33 | 0,73 | 4,68E-01 |
| AT1G28960 | 0,56 | 2,82 | 3,89 | 0,72 | 4,69E-01 |
| AT1G29050 | 0,14 | -1,69 | 4,33 | -0,39 | 6,96E-01 |
| AT1G29071 | 1,34 | -1,01 | 1,45 | -0,70 | 4,87E-01 |
| AT1G29170 | 0,29 | 1,00 | 4,27 | 0,23 | 8,15E-01 |
| AT1G29240 | 0,32 | 2,47 | 4,35 | 0,57 | 5,69E-01 |
| AT1G29300 | 0,08 | -0,87 | 4,38 | -0,20 | 8,43E-01 |
| AT1G29390 | 0,82 | 2,61 | 2,10 | 1,24 | 2,13E-01 |
| AT1G29395 | 0,39 | 2,12 | 3,36 | 0,63 | 5,29E-01 |
| AT1G29530 | 0,36 | 1,29 | 3,45 | 0,37 | 7,09E-01 |
| AT1G29690 | 0,27 | -0,66 | 4,31 | -0,15 | 8,78E-01 |
| AT1G29700 | 0,34 | 0,00 | 4,41 | 0,00 | 1,00E+00 |
| AT1G29720 | 0,71 | 0,87 | 2,56 | 0,34 | 7,33E-01 |
| AT1G29750 | 0,26 | 0,92 | 4,30 | 0,21 | 8,31E-01 |
| AT1G29880 | 1,19 | 0,96 | 1,72 | 0,56 | 5,77E-01 |
| AT1G29970 | 0,83 | 1,29 | 2,47 | 0,52 | 6,02E-01 |
| AT1G30000 | 0,84 | 1,27 | 2,55 | 0,50 | 6,17E-01 |
| AT1G30060 | 0,17 | -1,97 | 4,31 | -0,46 | 6,47E-01 |
| AT1G30070 | 0,15 | -0,64 | 4,39 | -0,15 | 8,84E-01 |
| AT1G30090 | 0,16 | 1,18 | 4,32 | 0,27 | 7,85E-01 |
| AT1G30120 | 1,11 | 0,94 | 1,74 | 0,54 | 5,89E-01 |
| AT1G30210 | 0,29 | 0,68 | 4,34 | 0,16 | 8,75E-01 |
| AT1G30250 | 0,60 | -1,45 | 2,42 | -0,60 | 5,47E-01 |
| AT1G30370 | 0,55 | 2,89 | 3,07 | 0,94 | 3,47E-01 |
| AT1G30500 | 0,64 | 1,93 | 3,81 | 0,51 | 6,13E-01 |
| AT1G06347 | 0,94 | -0,49 | 2,03 | -0,24 | 8,09E-01 |
| AT1G30540 | 0,42 | -0,59 | 4,19 | -0,14 | 8,88E-01 |
| AT1G30580 | 1,07 | 1,31 | 2,00 | 0,66 | 5,11E-01 |
| AT1G30590 | 0,66 | -2,73 | 2,75 | -0,99 | 3,20E-01 |
| AT1G30755 | 0,62 | -0,94 | 2,60 | -0,36 | 7,19E-01 |
| AT1G30780 | 0,83 | -1,15 | 2,08 | -0,55 | 5,81E-01 |
| AT1G30810 | 0,83 | 1,44 | 1,93 | 0,74 | 4,57E-01 |
| AT1G30880 | 0,08 | 2,08 | 4,38 | 0,47 | 6,35E-01 |
| AT1G30960 | 0,37 | 2,73 | 4,34 | 0,63 | 5,29E-01 |
| AT1G06407 | 0,34 | -0,06 | 3,50 | -0,02 | 9,85E-01 |
| AT1G31130 | 1,47 | 1,18 | 1,41 | 0,84 | 4,01E-01 |
| AT1G31160 | 0,42 | 2,83 | 4,13 | 0,69 | 4,93E-01 |
| AT1G31180 | 0,17 | 1,90 | 4,33 | 0,44 | 6,61E-01 |
| AT1G31230 | 1,31 | -1,00 | 1,70 | -0,59 | 5,57E-01 |
| AT1G31240 | 0,22 | -1,85 | 4,32 | -0,43 | 6,69E-01 |
| AT1G31350 | 0,45 | 2,11 | 3,21 | 0,66 | 5,10E-01 |
| AT1G31410 | 0,08 | 2,08 | 4,38 | 0,47 | 6,35E-01 |
| AT1G31470 | 0,15 | 1,93 | 4,33 | 0,45 | 6,55E-01 |
| AT1G31550 | 0,81 | 2,78 | 2,55 | 1,09 | 2,76E-01 |
| AT1G31600 | 0,81 | 1,03 | 2,87 | 0,36 | 7,21E-01 |
| AT1G31640 | 0,23 | 0,18 | 4,27 | 0,04 | 9,66E-01 |
| AT1G31650 | 0,25 | -0,44 | 4,34 | -0,10 | 9,20E-01 |
| AT1G31660 | 0,51 | 2,15 | 4,36 | 0,49 | 6,21E-01 |
| AT1G31740 | 0,22 | 0,81 | 4,27 | 0,19 | 8,50E-01 |
| AT1G31810 | 0,57 | 1,05 | 2,25 | 0,46 | 6,43E-01 |
| AT1G31812 | 0,29 | -1,08 | 4,34 | -0,25 | 8,03E-01 |
| AT1G31814 | 0,37 | 3,05 | 4,33 | 0,70 | 4,81E-01 |
| AT1G31820 | 0,94 | 3,41 | 2,67 | 1,28 | 2,01E-01 |
| AT1G31830 | 0,66 | 2,15 | 2,54 | 0,85 | 3,97E-01 |
| AT1G06453 | 1,48 | -1,03 | 1,73 | -0,59 | 5,52E-01 |
| AT1G31860 | 1,08 | -0,50 | 1,57 | -0,32 | 7,50E-01 |

| | | | | | |
|---|---|---|---|---|---|
| AT1G06463 | 0,27 | 0,50 | 4,27 | 0,12 | 9,08E-01 |
| AT1G31920 | 0,52 | 1,12 | 2,60 | 0,43 | 6,68E-01 |
| AT1G31940 | 0,35 | -0,68 | 4,39 | -0,15 | 8,77E-01 |
| AT1G06473 | 0,38 | 0,98 | 4,34 | 0,23 | 8,21E-01 |
| AT1G32000 | 0,35 | -1,02 | 4,30 | -0,24 | 8,13E-01 |
| AT1G32070 | 0,34 | 2,94 | 4,34 | 0,68 | 4,98E-01 |
| AT1G32080 | 0,85 | 0,21 | 1,82 | 0,12 | 9,07E-01 |
| AT1G32130 | 1,27 | 1,81 | 1,94 | 0,93 | 3,51E-01 |
| AT1G32200 | 0,75 | -2,02 | 2,17 | -0,93 | 3,50E-01 |
| AT1G32340 | 0,12 | 1,15 | 4,35 | 0,26 | 7,92E-01 |
| AT1G06537 | 1,54 | 2,93 | 1,73 | 1,70 | 8,92E-02 |
| AT1G32400 | 0,47 | -0,33 | 2,56 | -0,13 | 8,97E-01 |
| AT1G32450 | 0,27 | 0,42 | 4,31 | 0,10 | 9,21E-01 |
| AT1G06573 | 0,41 | 0,11 | 4,24 | 0,03 | 9,79E-01 |
| AT1G32470 | 0,50 | -0,29 | 2,52 | -0,11 | 9,10E-01 |
| AT1G32490 | 0,70 | 1,57 | 2,06 | 0,76 | 4,46E-01 |
| AT1G32500 | 0,52 | -0,39 | 2,71 | -0,14 | 8,87E-01 |
| AT1G32530 | 0,53 | 1,40 | 2,63 | 0,53 | 5,94E-01 |
| AT1G06577 | 0,16 | -0,80 | 4,37 | -0,18 | 8,54E-01 |
| AT1G32713 | 1,03 | -0,66 | 2,06 | -0,32 | 7,47E-01 |
| AT1G32790 | 1,57 | 3,71 | 1,84 | 2,02 | 4,37E-02 |
| AT1G32860 | 0,05 | -0,58 | 4,38 | -0,13 | 8,95E-01 |
| AT1G32880 | 0,26 | 2,30 | 4,37 | 0,53 | 5,98E-01 |
| AT1G32900 | 0,61 | -0,99 | 2,99 | -0,33 | 7,39E-01 |
| AT1G32970 | 0,17 | 3,16 | 4,33 | 0,73 | 4,66E-01 |
| AT1G33050 | 0,95 | 1,82 | 1,89 | 0,96 | 3,38E-01 |
| AT1G06643 | 0,36 | 2,51 | 4,35 | 0,58 | 5,64E-01 |
| AT1G33250 | 0,90 | 2,30 | 2,08 | 1,11 | 2,68E-01 |
| AT1G33270 | 0,24 | 2,80 | 4,34 | 0,65 | 5,18E-01 |
| AT1G06673 | 0,47 | -0,69 | 3,18 | -0,22 | 8,28E-01 |
| AT1G33390 | 1,05 | 1,71 | 2,39 | 0,72 | 4,73E-01 |
| AT1G33410 | 1,37 | 1,29 | 1,46 | 0,88 | 3,77E-01 |
| AT1G33430 | 0,08 | 2,08 | 4,38 | 0,47 | 6,35E-01 |
| AT1G33520 | 0,47 | 2,80 | 3,15 | 0,89 | 3,74E-01 |
| AT1G33615 | 0,17 | -0,14 | 4,41 | -0,03 | 9,75E-01 |
| AT1G33700 | 0,88 | 2,07 | 2,03 | 1,02 | 3,08E-01 |
| AT1G33720 | 1,30 | 0,55 | 1,74 | 0,32 | 7,50E-01 |
| AT1G33780 | 0,08 | -0,87 | 4,38 | -0,20 | 8,43E-01 |
| AT1G33870 | 0,60 | 1,52 | 2,45 | 0,62 | 5,36E-01 |
| AT1G33970 | 0,67 | 0,21 | 2,51 | 0,08 | 9,34E-01 |
| AT1G34030 | 0,44 | 1,11 | 2,79 | 0,40 | 6,91E-01 |
| AT1G34042 | 0,29 | -0,46 | 4,25 | -0,11 | 9,14E-01 |
| AT1G34110 | 0,91 | 0,52 | 1,71 | 0,30 | 7,62E-01 |
| AT1G34170 | 0,20 | 1,39 | 4,37 | 0,32 | 7,51E-01 |
| AT1G34220 | 1,01 | 3,60 | 2,80 | 1,28 | 1,99E-01 |
| AT1G34270 | 0,12 | 2,69 | 4,35 | 0,62 | 5,36E-01 |
| AT1G34310 | 0,97 | -2,73 | 2,49 | -1,10 | 2,73E-01 |
| AT1G34315 | 0,23 | 2,48 | 4,31 | 0,57 | 5,66E-01 |
| AT1G34380 | 0,08 | 2,08 | 4,38 | 0,47 | 6,35E-01 |
| AT1G34440 | 1,54 | -1,63 | 1,49 | -1,09 | 2,74E-01 |
| AT1G34550 | 0,63 | 0,04 | 2,29 | 0,02 | 9,87E-01 |
| AT1G34750 | 0,23 | -0,22 | 4,28 | -0,05 | 9,60E-01 |
| AT1G34780 | 0,68 | 3,19 | 2,82 | 1,13 | 2,58E-01 |
| AT1G35150 | 0,16 | -0,58 | 4,32 | -0,14 | 8,92E-01 |
| AT1G35240 | 1,26 | 0,68 | 1,56 | 0,44 | 6,63E-01 |
| AT1G35250 | 0,19 | -1,86 | 4,32 | -0,43 | 6,67E-01 |

| | | | | | |
|---|---|---|---|---|---|
| AT1G35320 | 0,59 | -0,93 | 2,76 | -0,34 | 7,36E-01 |
| AT1G35350 | 0,96 | -0,37 | 1,69 | -0,22 | 8,28E-01 |
| AT1G06883 | 0,19 | -2,20 | 4,31 | -0,51 | 6,10E-01 |
| AT1G06897 | 0,91 | -2,55 | 3,41 | -0,75 | 4,54E-01 |
| AT1G06903 | 0,10 | -1,26 | 4,35 | -0,29 | 7,72E-01 |
| AT1G35520 | 1,01 | -1,11 | 2,25 | -0,49 | 6,21E-01 |
| AT1G35545 | 0,38 | -1,62 | 3,29 | -0,49 | 6,21E-01 |
| AT1G35580 | 1,07 | 0,43 | 1,53 | 0,28 | 7,77E-01 |
| AT1G35610 | 0,22 | 0,35 | 4,34 | 0,08 | 9,36E-01 |
| AT1G35625 | 0,08 | 2,08 | 4,38 | 0,47 | 6,35E-01 |
| AT1G35730 | 0,17 | 3,14 | 4,33 | 0,73 | 4,68E-01 |
| AT1G35820 | 1,40 | -3,31 | 2,69 | -1,23 | 2,18E-01 |
| AT1G35895 | 0,55 | 2,21 | 3,11 | 0,71 | 4,77E-01 |
| AT1G36070 | 0,25 | 2,97 | 4,33 | 0,69 | 4,93E-01 |
| AT1G36085 | 0,68 | -4,04 | 4,26 | -0,95 | 3,43E-01 |
| AT1G36230 | 0,12 | -1,57 | 4,34 | -0,36 | 7,18E-01 |
| AT1G36325 | 1,41 | -1,05 | 1,53 | -0,68 | 4,93E-01 |
| AT1G36370 | 0,71 | 1,29 | 2,26 | 0,57 | 5,66E-01 |
| AT1G36390 | 0,17 | 2,29 | 4,36 | 0,53 | 5,98E-01 |
| AT1G36622 | 0,40 | 0,48 | 3,41 | 0,14 | 8,89E-01 |
| AT1G36990 | 1,02 | 1,30 | 1,85 | 0,70 | 4,83E-01 |
| AT1G07013 | 0,15 | -0,64 | 4,39 | -0,15 | 8,84E-01 |
| AT1G40087 | 0,19 | -2,00 | 4,31 | -0,46 | 6,43E-01 |
| AT1G41830 | 0,28 | -0,26 | 4,25 | -0,06 | 9,51E-01 |
| AT1G41920 | 0,51 | 0,85 | 3,14 | 0,27 | 7,86E-01 |
| AT1G42440 | 0,46 | -0,18 | 3,28 | -0,05 | 9,57E-01 |
| AT1G42560 | 0,05 | -0,58 | 4,38 | -0,13 | 8,95E-01 |
| AT1G42990 | 1,44 | -0,11 | 1,39 | -0,08 | 9,35E-01 |
| AT1G43190 | 0,38 | 0,92 | 4,32 | 0,21 | 8,32E-01 |
| AT1G43560 | 0,22 | -0,13 | 4,30 | -0,03 | 9,76E-01 |
| AT1G43580 | 0,21 | 1,12 | 4,33 | 0,26 | 7,95E-01 |
| AT1G43620 | 0,44 | 0,88 | 4,15 | 0,21 | 8,32E-01 |
| AT1G43665 | 0,88 | 0,40 | 3,04 | 0,13 | 8,95E-01 |
| AT1G43710 | 0,50 | 1,70 | 3,14 | 0,54 | 5,89E-01 |
| AT1G43730 | 0,40 | -0,75 | 4,18 | -0,18 | 8,58E-01 |
| AT1G43760 | 0,66 | 0,39 | 2,40 | 0,16 | 8,70E-01 |
| AT1G43825 | 0,81 | -3,04 | 2,69 | -1,13 | 2,58E-01 |
| AT1G43860 | 1,69 | 1,40 | 1,63 | 0,86 | 3,90E-01 |
| AT1G43950 | 0,46 | -0,98 | 3,95 | -0,25 | 8,05E-01 |
| AT1G44000 | 0,08 | 2,08 | 4,38 | 0,47 | 6,35E-01 |
| AT1G44030 | 0,25 | -0,89 | 4,30 | -0,21 | 8,36E-01 |
| AT1G44080 | 1,35 | -1,68 | 2,08 | -0,81 | 4,19E-01 |
| AT1G44085 | 0,65 | -1,88 | 2,72 | -0,69 | 4,90E-01 |
| AT1G44090 | 0,05 | -0,58 | 4,38 | -0,13 | 8,95E-01 |
| AT1G44125 | 0,91 | 2,69 | 2,54 | 1,06 | 2,90E-01 |
| AT1G44170 | 0,30 | -1,67 | 4,27 | -0,39 | 6,96E-01 |
| AT1G44350 | 0,41 | 1,08 | 4,23 | 0,26 | 7,98E-01 |
| AT1G44542 | 0,40 | -1,83 | 4,32 | -0,42 | 6,72E-01 |
| AT1G44770 | 0,41 | 0,56 | 3,22 | 0,17 | 8,61E-01 |
| AT1G44790 | 0,21 | -1,08 | 4,30 | -0,25 | 8,02E-01 |
| AT1G44920 | 0,44 | -0,20 | 3,25 | -0,06 | 9,50E-01 |
| AT1G44980 | 0,59 | -0,23 | 2,59 | -0,09 | 9,29E-01 |
| AT1G45063 | 0,43 | 1,66 | 4,39 | 0,38 | 7,05E-01 |
| AT1G45145 | 0,72 | 1,70 | 3,00 | 0,57 | 5,70E-01 |
| AT1G45150 | 0,25 | -0,44 | 4,34 | -0,10 | 9,20E-01 |
| AT1G45163 | 0,70 | 1,85 | 2,35 | 0,79 | 4,31E-01 |

| | | | | | |
|---|---|---|---|---|---|
| AT1G45207 | 0,37 | 0,00 | 4,41 | 0,00 | 1,00E+00 |
| AT1G07323 | 0,55 | -0,11 | 2,56 | -0,04 | 9,66E-01 |
| AT1G47200 | 0,16 | -1,42 | 4,34 | -0,33 | 7,43E-01 |
| AT1G47240 | 0,43 | 2,89 | 4,08 | 0,71 | 4,78E-01 |
| AT1G07333 | 0,81 | -3,90 | 3,23 | -1,21 | 2,27E-01 |
| AT1G47260 | 1,01 | -0,09 | 1,51 | -0,06 | 9,54E-01 |
| AT1G47280 | 1,31 | -3,55 | 1,99 | -1,79 | 7,40E-02 |
| AT1G07347 | 0,18 | -1,99 | 4,31 | -0,46 | 6,44E-01 |
| AT1G47380 | 0,18 | 0,77 | 4,33 | 0,18 | 8,59E-01 |
| AT1G47420 | 0,71 | 1,34 | 2,04 | 0,66 | 5,10E-01 |
| AT1G07383 | 0,55 | -1,39 | 2,58 | -0,54 | 5,89E-01 |
| AT1G07393 | 1,67 | -0,78 | 1,60 | -0,48 | 6,28E-01 |
| AT1G47640 | 0,35 | 0,33 | 3,43 | 0,10 | 9,22E-01 |
| AT1G47670 | 0,05 | -0,58 | 4,38 | -0,13 | 8,95E-01 |
| AT1G47765 | 0,55 | -1,26 | 2,28 | -0,55 | 5,81E-01 |
| AT1G47900 | 0,45 | 1,92 | 4,14 | 0,46 | 6,43E-01 |
| AT1G47960 | 1,43 | -0,01 | 1,38 | 0,00 | 9,96E-01 |
| AT1G48000 | 0,50 | 0,24 | 4,37 | 0,05 | 9,57E-01 |
| AT1G48030 | 1,26 | 2,56 | 1,93 | 1,32 | 1,85E-01 |
| AT1G48040 | 0,63 | 4,08 | 4,30 | 0,95 | 3,42E-01 |
| AT1G48110 | 0,42 | 2,32 | 4,31 | 0,54 | 5,90E-01 |
| AT1G48120 | 0,82 | 2,95 | 2,26 | 1,30 | 1,92E-01 |
| AT1G48160 | 0,15 | 0,40 | 4,33 | 0,09 | 9,27E-01 |
| AT1G48285 | 0,80 | 0,38 | 2,10 | 0,18 | 8,56E-01 |
| AT1G48320 | 0,17 | 3,14 | 4,33 | 0,73 | 4,68E-01 |
| AT1G48380 | 0,10 | 0,34 | 4,35 | 0,08 | 9,37E-01 |
| AT1G48405 | 0,82 | 4,60 | 3,28 | 1,40 | 1,60E-01 |
| AT1G48430 | 0,15 | 1,94 | 4,32 | 0,45 | 6,54E-01 |
| AT1G48450 | 1,45 | 0,84 | 1,78 | 0,47 | 6,35E-01 |
| AT1G48550 | 1,08 | 2,80 | 2,62 | 1,07 | 2,86E-01 |
| AT1G48570 | 0,26 | 1,71 | 4,30 | 0,40 | 6,90E-01 |
| AT1G48580 | 0,08 | 2,33 | 4,36 | 0,53 | 5,93E-01 |
| AT1G48620 | 0,46 | -2,30 | 3,17 | -0,72 | 4,69E-01 |
| AT1G48630 | 1,25 | 3,84 | 1,80 | 2,13 | 3,28E-02 |
| AT1G48730 | 0,75 | -2,84 | 2,22 | -1,28 | 2,01E-01 |
| AT1G48742 | 0,46 | 1,97 | 3,16 | 0,62 | 5,33E-01 |
| AT1G48790 | 0,44 | 1,87 | 3,14 | 0,59 | 5,52E-01 |
| AT1G48850 | 0,39 | 1,01 | 4,31 | 0,23 | 8,15E-01 |
| AT1G48900 | 0,23 | 0,71 | 4,38 | 0,16 | 8,71E-01 |
| AT1G48930 | 0,71 | 3,68 | 2,83 | 1,30 | 1,92E-01 |
| AT1G48970 | 1,37 | 2,02 | 1,75 | 1,15 | 2,49E-01 |
| AT1G49005 | 0,38 | -0,21 | 4,26 | -0,05 | 9,61E-01 |
| AT1G49140 | 0,12 | 0,31 | 4,35 | 0,07 | 9,44E-01 |
| AT1G49180 | 0,08 | 2,08 | 4,38 | 0,47 | 6,35E-01 |
| AT1G49205 | 0,08 | -0,56 | 4,38 | -0,13 | 8,99E-01 |
| AT1G49225 | 0,48 | -1,03 | 3,13 | -0,33 | 7,41E-01 |
| AT1G49270 | 0,10 | -1,26 | 4,35 | -0,29 | 7,72E-01 |
| AT1G49490 | 0,32 | 0,74 | 4,37 | 0,17 | 8,65E-01 |
| AT1G49540 | 0,83 | 2,88 | 2,39 | 1,20 | 2,28E-01 |
| AT1G49600 | 0,82 | 2,21 | 1,99 | 1,11 | 2,66E-01 |
| AT1G49670 | 0,26 | -1,42 | 4,27 | -0,33 | 7,39E-01 |
| AT1G49710 | 0,34 | 0,30 | 3,37 | 0,09 | 9,28E-01 |
| AT1G49780 | 1,22 | 0,77 | 1,48 | 0,52 | 6,04E-01 |
| AT1G49850 | 0,39 | 2,44 | 4,32 | 0,56 | 5,72E-01 |
| AT1G49970 | 0,66 | 0,07 | 2,02 | 0,03 | 9,74E-01 |
| AT1G07603 | 0,61 | 1,28 | 3,15 | 0,41 | 6,83E-01 |

| | | | | | |
|---|---|---|---|---|---|
| AT1G50020 | 0,58 | -0,52 | 4,02 | -0,13 | 8,97E-01 |
| AT1G50140 | 0,20 | 2,68 | 4,35 | 0,61 | 5,39E-01 |
| AT1G50190 | 0,32 | 1,93 | 4,34 | 0,44 | 6,57E-01 |
| AT1G50260 | 0,32 | 1,34 | 4,33 | 0,31 | 7,56E-01 |
| AT1G50280 | 0,61 | -1,15 | 2,26 | -0,51 | 6,11E-01 |
| AT1G50320 | 0,25 | 1,45 | 4,32 | 0,34 | 7,38E-01 |
| AT1G07627 | 0,18 | -1,39 | 4,34 | -0,32 | 7,49E-01 |
| AT1G50420 | 0,49 | 2,57 | 4,34 | 0,59 | 5,54E-01 |
| AT1G50430 | 1,02 | 1,31 | 1,68 | 0,78 | 4,37E-01 |
| AT1G50450 | 0,63 | 1,16 | 2,96 | 0,39 | 6,96E-01 |
| AT1G50500 | 0,57 | 1,99 | 3,13 | 0,64 | 5,24E-01 |
| AT1G07647 | 0,08 | 2,08 | 4,38 | 0,47 | 6,35E-01 |
| AT1G50570 | 0,65 | -0,26 | 2,60 | -0,10 | 9,19E-01 |
| AT1G50600 | 0,88 | 1,29 | 1,86 | 0,69 | 4,87E-01 |
| AT1G50640 | 1,08 | 1,12 | 1,78 | 0,63 | 5,29E-01 |
| AT1G50659 | 0,43 | -0,44 | 3,09 | -0,14 | 8,86E-01 |
| AT1G50750 | 0,08 | -0,56 | 4,38 | -0,13 | 8,99E-01 |
| AT1G50790 | 0,12 | 2,81 | 4,34 | 0,65 | 5,18E-01 |
| AT1G50820 | 0,50 | 2,33 | 3,99 | 0,58 | 5,59E-01 |
| AT1G50910 | 1,39 | -0,71 | 1,56 | -0,45 | 6,50E-01 |
| AT1G50980 | 0,39 | -0,14 | 4,41 | -0,03 | 9,75E-01 |
| AT1G51080 | 0,73 | 0,16 | 2,48 | 0,06 | 9,50E-01 |
| AT1G07733 | 0,24 | -0,70 | 4,38 | -0,16 | 8,72E-01 |
| AT1G51400 | 0,93 | -2,36 | 2,18 | -1,08 | 2,79E-01 |
| AT1G51480 | 0,33 | 2,11 | 4,36 | 0,48 | 6,29E-01 |
| AT1G51500 | 1,21 | -0,52 | 1,88 | -0,28 | 7,83E-01 |
| AT1G51510 | 0,91 | -1,00 | 2,07 | -0,49 | 6,27E-01 |
| AT1G51560 | 0,45 | 1,93 | 3,29 | 0,59 | 5,58E-01 |
| AT1G51590 | 0,26 | -1,25 | 4,29 | -0,29 | 7,71E-01 |
| AT1G51630 | 0,52 | -1,13 | 3,10 | -0,36 | 7,15E-01 |
| AT1G51680 | 0,80 | 0,16 | 1,92 | 0,08 | 9,34E-01 |
| AT1G07763 | 0,56 | 0,00 | 4,41 | 0,00 | 1,00E+00 |
| AT1G51710 | 0,45 | 2,53 | 4,11 | 0,61 | 5,39E-01 |
| AT1G51730 | 0,47 | 1,52 | 3,19 | 0,48 | 6,33E-01 |
| AT1G51790 | 0,25 | -0,67 | 4,39 | -0,15 | 8,78E-01 |
| AT1G51890 | 0,23 | 2,47 | 4,31 | 0,57 | 5,67E-01 |
| AT1G51940 | 1,08 | 1,47 | 2,46 | 0,60 | 5,51E-01 |
| AT1G52000 | 0,46 | -0,13 | 4,11 | -0,03 | 9,74E-01 |
| AT1G52040 | 0,54 | 0,00 | 3,98 | 0,00 | 1,00E+00 |
| AT1G52150 | 1,30 | 3,60 | 2,28 | 1,58 | 1,14E-01 |
| AT1G52160 | 0,61 | -0,54 | 3,03 | -0,18 | 8,59E-01 |
| AT1G52180 | 1,00 | -0,70 | 1,93 | -0,36 | 7,17E-01 |
| AT1G52185 | 0,32 | 0,56 | 4,37 | 0,13 | 8,98E-01 |
| AT1G52200 | 0,25 | 3,58 | 4,32 | 0,83 | 4,08E-01 |
| AT1G52220 | 0,76 | 0,98 | 2,09 | 0,47 | 6,40E-01 |
| AT1G52290 | 0,41 | 2,21 | 4,19 | 0,53 | 5,97E-01 |
| AT1G52300 | 0,35 | 1,03 | 4,26 | 0,24 | 8,09E-01 |
| AT1G52340 | 0,08 | -0,56 | 4,38 | -0,13 | 8,99E-01 |
| AT1G52342 | 0,45 | 3,85 | 4,32 | 0,89 | 3,72E-01 |
| AT1G52510 | 0,47 | 1,33 | 4,16 | 0,32 | 7,49E-01 |
| AT1G52570 | 1,59 | 3,98 | 2,27 | 1,75 | 7,93E-02 |
| AT1G52750 | 0,31 | 2,33 | 4,31 | 0,54 | 5,88E-01 |
| AT1G52880 | 0,40 | 2,53 | 3,37 | 0,75 | 4,53E-01 |
| AT1G52890 | 0,42 | 2,35 | 4,14 | 0,57 | 5,70E-01 |
| AT1G52970 | 1,13 | -1,37 | 2,11 | -0,65 | 5,17E-01 |
| AT1G52990 | 0,41 | -3,24 | 4,28 | -0,76 | 4,49E-01 |

| | | | | | |
|---|---|---|---|---|---|
| AT1G53000 | 0,49 | 1,21 | 4,02 | 0,30 | 7,64E-01 |
| AT1G53025 | 0,43 | -0,27 | 3,33 | -0,08 | 9,35E-01 |
| AT1G53035 | 0,12 | 2,81 | 4,34 | 0,65 | 5,18E-01 |
| AT1G53170 | 0,65 | -0,81 | 2,46 | -0,33 | 7,42E-01 |
| AT1G53200 | 0,24 | 2,80 | 4,34 | 0,65 | 5,18E-01 |
| AT1G53230 | 1,22 | 1,76 | 1,92 | 0,91 | 3,60E-01 |
| AT1G53240 | 1,16 | -0,07 | 1,69 | -0,04 | 9,68E-01 |
| AT1G53265 | 0,53 | 1,26 | 4,03 | 0,31 | 7,54E-01 |
| AT1G53320 | 0,35 | 1,15 | 4,31 | 0,27 | 7,90E-01 |
| AT1G53450 | 0,86 | -2,38 | 1,97 | -1,21 | 2,26E-01 |
| AT1G53480 | 0,84 | -1,33 | 2,94 | -0,45 | 6,52E-01 |
| AT1G53500 | 0,50 | 0,52 | 3,25 | 0,16 | 8,73E-01 |
| AT1G53510 | 0,76 | -0,37 | 2,41 | -0,15 | 8,79E-01 |
| AT1G53541 | 0,64 | 1,02 | 2,29 | 0,44 | 6,58E-01 |
| AT1G53580 | 0,12 | 0,63 | 4,33 | 0,15 | 8,85E-01 |
| AT1G53710 | 0,67 | 2,87 | 2,93 | 0,98 | 3,26E-01 |
| AT1G53750 | 1,03 | 0,38 | 1,87 | 0,20 | 8,39E-01 |
| AT1G53760 | 0,08 | 2,33 | 4,36 | 0,53 | 5,93E-01 |
| AT1G53770 | 0,33 | 2,56 | 4,35 | 0,59 | 5,57E-01 |
| AT1G53780 | 1,40 | 3,51 | 1,84 | 1,91 | 5,65E-02 |
| AT1G54050 | 0,12 | 0,63 | 4,33 | 0,15 | 8,85E-01 |
| AT1G54080 | 1,16 | 3,52 | 2,02 | 1,74 | 8,13E-02 |
| AT1G54100 | 1,08 | 0,16 | 1,88 | 0,08 | 9,33E-01 |
| AT1G54140 | 0,15 | 0,40 | 4,33 | 0,09 | 9,27E-01 |
| AT1G54170 | 0,82 | 2,19 | 2,17 | 1,01 | 3,12E-01 |
| AT1G54320 | 0,67 | 1,24 | 2,31 | 0,54 | 5,93E-01 |
| AT1G54370 | 0,38 | 2,34 | 4,28 | 0,55 | 5,85E-01 |
| AT1G54500 | 0,85 | -0,80 | 1,85 | -0,43 | 6,67E-01 |
| AT1G54590 | 0,10 | 0,34 | 4,35 | 0,08 | 9,37E-01 |
| AT1G54630 | 0,35 | 0,65 | 4,35 | 0,15 | 8,81E-01 |
| AT1G54690 | 0,48 | 0,30 | 2,67 | 0,11 | 9,09E-01 |
| AT1G54775 | 0,57 | -1,66 | 3,79 | -0,44 | 6,61E-01 |
| AT1G54840 | 0,49 | -0,14 | 4,41 | -0,03 | 9,75E-01 |
| AT1G54950 | 0,44 | -2,18 | 4,01 | -0,54 | 5,87E-01 |
| AT1G55152 | 0,78 | 0,41 | 2,28 | 0,18 | 8,56E-01 |
| AT1G55180 | 0,78 | 3,18 | 2,75 | 1,15 | 2,49E-01 |
| AT1G55200 | 0,08 | 2,33 | 4,36 | 0,53 | 5,93E-01 |
| AT1G08043 | 0,30 | 2,91 | 4,34 | 0,67 | 5,04E-01 |
| AT1G55270 | 1,00 | 3,53 | 3,17 | 1,11 | 2,65E-01 |
| AT1G55300 | 0,10 | 0,39 | 4,35 | 0,09 | 9,29E-01 |
| AT1G55330 | 0,51 | 1,57 | 3,19 | 0,49 | 6,21E-01 |
| AT1G55360 | 0,31 | 0,19 | 4,29 | 0,04 | 9,65E-01 |
| AT1G55530 | 0,83 | 1,94 | 2,11 | 0,92 | 3,58E-01 |
| AT1G55535 | 0,40 | 2,56 | 4,23 | 0,61 | 5,45E-01 |
| AT1G55590 | 0,61 | 1,02 | 2,58 | 0,40 | 6,92E-01 |
| AT1G55610 | 0,12 | 0,63 | 4,33 | 0,15 | 8,85E-01 |
| AT1G55620 | 0,57 | 0,35 | 3,03 | 0,12 | 9,07E-01 |
| AT1G55675 | 0,68 | -1,41 | 3,74 | -0,38 | 7,07E-01 |
| AT1G55680 | 1,44 | 4,69 | 1,97 | 2,38 | 1,72E-02 |
| AT1G55690 | 0,12 | 2,81 | 4,34 | 0,65 | 5,18E-01 |
| AT1G55830 | 0,23 | 0,75 | 4,27 | 0,17 | 8,61E-01 |
| AT1G55880 | 0,36 | 3,22 | 4,34 | 0,74 | 4,59E-01 |
| AT1G55950 | 0,93 | 0,10 | 1,90 | 0,05 | 9,58E-01 |
| AT1G55960 | 0,25 | -0,51 | 4,38 | -0,12 | 9,08E-01 |
| AT1G55970 | 0,17 | 3,16 | 4,33 | 0,73 | 4,66E-01 |
| AT1G56010 | 0,42 | 2,55 | 4,09 | 0,62 | 5,34E-01 |

| | | | | | |
|---|---|---|---|---|---|
| AT1G56100 | 0,86 | -0,31 | 2,88 | -0,11 | 9,15E-01 |
| AT1G56110 | 1,35 | 0,28 | 1,87 | 0,15 | 8,81E-01 |
| AT1G56120 | 0,47 | 0,56 | 4,03 | 0,14 | 8,90E-01 |
| AT1G56140 | 0,44 | 0,70 | 3,17 | 0,22 | 8,24E-01 |
| AT1G56145 | 0,80 | 3,42 | 2,74 | 1,25 | 2,12E-01 |
| AT1G56165 | 0,20 | 2,11 | 4,37 | 0,48 | 6,29E-01 |
| AT1G56230 | 0,19 | 0,58 | 4,36 | 0,13 | 8,94E-01 |
| AT1G08107 | 0,31 | -1,45 | 3,54 | -0,41 | 6,83E-01 |
| AT1G08113 | 0,55 | -0,22 | 3,19 | -0,07 | 9,46E-01 |
| AT1G08123 | 0,56 | -0,08 | 2,31 | -0,03 | 9,73E-01 |
| AT1G56300 | 0,85 | -0,79 | 2,77 | -0,29 | 7,74E-01 |
| AT1G56330 | 0,78 | -2,56 | 2,45 | -1,04 | 2,96E-01 |
| AT1G56345 | 0,19 | -1,45 | 4,34 | -0,33 | 7,38E-01 |
| AT1G56440 | 0,27 | 0,42 | 4,31 | 0,10 | 9,21E-01 |
| AT1G56520 | 1,21 | 1,18 | 2,07 | 0,57 | 5,68E-01 |
| AT1G56550 | 0,17 | 2,29 | 4,36 | 0,53 | 5,98E-01 |
| AT1G56580 | 0,24 | 0,06 | 4,31 | 0,01 | 9,89E-01 |
| AT1G56650 | 1,01 | 5,29 | 3,90 | 1,36 | 1,74E-01 |
| AT1G57680 | 1,45 | 2,32 | 1,77 | 1,31 | 1,90E-01 |
| AT1G57775 | 0,38 | -1,62 | 3,29 | -0,49 | 6,21E-01 |
| AT1G57777 | 0,31 | 0,61 | 3,51 | 0,17 | 8,62E-01 |
| AT1G57860 | 0,30 | 1,74 | 4,40 | 0,40 | 6,93E-01 |
| AT1G57870 | 0,40 | 0,31 | 4,18 | 0,07 | 9,41E-01 |
| AT1G57943 | 0,08 | 2,08 | 4,38 | 0,47 | 6,35E-01 |
| AT1G57990 | 0,77 | 1,04 | 2,37 | 0,44 | 6,61E-01 |
| AT1G58030 | 0,73 | 1,47 | 2,41 | 0,61 | 5,41E-01 |
| AT1G58060 | 0,60 | 1,57 | 3,00 | 0,52 | 6,01E-01 |
| AT1G58080 | 0,29 | 2,35 | 4,31 | 0,54 | 5,86E-01 |
| AT1G58100 | 1,20 | 1,32 | 1,93 | 0,68 | 4,94E-01 |
| AT1G58110 | 0,53 | 2,47 | 3,91 | 0,63 | 5,28E-01 |
| AT1G58200 | 0,16 | -1,42 | 4,34 | -0,33 | 7,43E-01 |
| AT1G58220 | 0,54 | 0,72 | 3,90 | 0,19 | 8,53E-01 |
| AT1G58300 | 1,33 | -1,46 | 1,78 | -0,82 | 4,13E-01 |
| AT1G58350 | 0,80 | 0,02 | 2,45 | 0,01 | 9,93E-01 |
| AT1G58390 | 0,70 | -1,88 | 2,92 | -0,64 | 5,19E-01 |
| AT1G58440 | 0,58 | -1,99 | 2,98 | -0,67 | 5,04E-01 |
| AT1G58684 | 0,49 | -1,59 | 4,33 | -0,37 | 7,14E-01 |
| AT1G58725 | 0,33 | -0,98 | 4,28 | -0,23 | 8,19E-01 |
| AT1G58766 | 0,16 | 1,18 | 4,32 | 0,27 | 7,85E-01 |
| AT1G59030 | 0,44 | 0,27 | 4,19 | 0,06 | 9,48E-01 |
| AT1G59077 | 0,16 | -0,80 | 4,37 | -0,18 | 8,54E-01 |
| AT1G59406 | 0,14 | -0,45 | 4,33 | -0,10 | 9,17E-01 |
| AT1G59453 | 0,25 | -0,54 | 4,27 | -0,13 | 8,99E-01 |
| AT1G59720 | 0,61 | -0,06 | 2,62 | -0,02 | 9,82E-01 |
| AT1G59780 | 0,52 | -1,20 | 3,11 | -0,39 | 7,00E-01 |
| AT1G59835 | 1,47 | -0,66 | 2,04 | -0,32 | 7,46E-01 |
| AT1G59910 | 1,37 | 0,32 | 1,80 | 0,18 | 8,60E-01 |
| AT1G59920 | 0,42 | -1,82 | 3,19 | -0,57 | 5,68E-01 |
| AT1G59930 | 1,11 | -0,20 | 2,13 | -0,10 | 9,24E-01 |
| AT1G59990 | 1,13 | 1,74 | 2,09 | 0,83 | 4,04E-01 |
| AT1G60000 | 0,38 | -0,08 | 3,45 | -0,02 | 9,80E-01 |
| AT1G60070 | 1,35 | 4,13 | 2,01 | 2,05 | 4,00E-02 |
| AT1G60160 | 0,43 | 0,78 | 3,10 | 0,25 | 8,01E-01 |
| AT1G60200 | 0,68 | -1,45 | 3,14 | -0,46 | 6,44E-01 |
| AT1G60230 | 0,48 | -0,91 | 4,05 | -0,22 | 8,23E-01 |
| AT1G60260 | 0,77 | -0,99 | 2,43 | -0,41 | 6,83E-01 |

| | | | | | |
|---|---|---|---|---|---|
| AT1G08317 | 0,08 | 2,08 | 4,38 | 0,47 | 6,35E-01 |
| AT1G60430 | 0,45 | 3,33 | 3,98 | 0,84 | 4,03E-01 |
| AT1G60490 | 0,25 | 1,45 | 4,32 | 0,34 | 7,38E-01 |
| AT1G60550 | 0,38 | 2,13 | 4,38 | 0,49 | 6,27E-01 |
| AT1G60560 | 0,17 | 3,04 | 4,33 | 0,70 | 4,83E-01 |
| AT1G60590 | 0,17 | 0,00 | 4,41 | 0,00 | 1,00E+00 |
| AT1G60630 | 0,08 | 2,08 | 4,38 | 0,47 | 6,35E-01 |
| AT1G60640 | 0,12 | 1,65 | 4,34 | 0,38 | 7,04E-01 |
| AT1G60700 | 0,48 | 3,77 | 4,32 | 0,87 | 3,83E-01 |
| AT1G60710 | 0,41 | 1,68 | 4,39 | 0,38 | 7,02E-01 |
| AT1G60770 | 1,20 | 4,16 | 2,11 | 1,97 | 4,86E-02 |
| AT1G60780 | 0,33 | 0,06 | 4,24 | 0,01 | 9,90E-01 |
| AT1G60800 | 0,56 | 2,53 | 3,09 | 0,82 | 4,13E-01 |
| AT1G60810 | 0,12 | -1,57 | 4,34 | -0,36 | 7,18E-01 |
| AT1G60850 | 0,11 | 1,33 | 4,36 | 0,31 | 7,59E-01 |
| AT1G60890 | 0,24 | 3,20 | 4,33 | 0,74 | 4,60E-01 |
| AT1G60930 | 0,24 | 2,45 | 4,35 | 0,56 | 5,73E-01 |
| AT1G60940 | 0,93 | 0,47 | 2,09 | 0,23 | 8,20E-01 |
| AT1G60960 | 1,57 | 0,91 | 1,67 | 0,55 | 5,83E-01 |
| AT1G60990 | 0,24 | 2,80 | 4,34 | 0,65 | 5,18E-01 |
| AT1G60995 | 0,35 | -1,02 | 4,30 | -0,24 | 8,13E-01 |
| AT1G61010 | 0,89 | 0,06 | 2,35 | 0,03 | 9,80E-01 |
| AT1G61200 | 1,13 | -0,75 | 1,52 | -0,50 | 6,19E-01 |
| AT1G61260 | 0,47 | -0,80 | 3,13 | -0,25 | 7,99E-01 |
| AT1G61300 | 1,06 | 1,11 | 2,17 | 0,51 | 6,08E-01 |
| AT1G61310 | 0,33 | -0,49 | 4,25 | -0,12 | 9,08E-01 |
| AT1G61360 | 0,15 | 0,40 | 4,33 | 0,09 | 9,27E-01 |
| AT1G61370 | 0,17 | 0,69 | 4,34 | 0,16 | 8,73E-01 |
| AT1G61380 | 0,34 | 0,82 | 4,25 | 0,19 | 8,47E-01 |
| AT1G61420 | 0,12 | 2,69 | 4,35 | 0,62 | 5,36E-01 |
| AT1G61560 | 0,37 | 1,72 | 4,31 | 0,40 | 6,90E-01 |
| AT1G08417 | 0,75 | -2,62 | 2,77 | -0,94 | 3,45E-01 |
| AT1G61660 | 0,65 | 0,29 | 2,80 | 0,10 | 9,17E-01 |
| AT1G61730 | 1,18 | 3,86 | 2,18 | 1,77 | 7,72E-02 |
| AT1G61890 | 1,56 | 0,08 | 1,49 | 0,05 | 9,59E-01 |
| AT1G61900 | 0,26 | -1,30 | 4,35 | -0,30 | 7,66E-01 |
| AT1G62030 | 1,53 | -2,36 | 1,56 | -1,51 | 1,31E-01 |
| AT1G62085 | 0,60 | 1,28 | 3,10 | 0,41 | 6,80E-01 |
| AT1G62300 | 0,27 | -1,50 | 4,34 | -0,35 | 7,29E-01 |
| AT1G62422 | 0,51 | 1,44 | 2,52 | 0,57 | 5,68E-01 |
| AT1G62610 | 0,76 | 1,69 | 2,32 | 0,73 | 4,66E-01 |
| AT1G62680 | 1,24 | 1,43 | 1,82 | 0,78 | 4,34E-01 |
| AT1G62760 | 0,29 | 0,44 | 4,26 | 0,10 | 9,18E-01 |
| AT1G62780 | 0,49 | -0,68 | 3,15 | -0,22 | 8,29E-01 |
| AT1G62810 | 0,97 | 2,64 | 1,99 | 1,33 | 1,84E-01 |
| AT1G62915 | 0,64 | -1,25 | 2,86 | -0,44 | 6,63E-01 |
| AT1G62960 | 0,36 | -0,51 | 4,38 | -0,12 | 9,08E-01 |
| AT1G63010 | 0,16 | -0,80 | 4,37 | -0,18 | 8,54E-01 |
| AT1G63020 | 0,44 | 4,02 | 4,31 | 0,93 | 3,51E-01 |
| AT1G63050 | 0,34 | 0,68 | 4,34 | 0,16 | 8,75E-01 |
| AT1G63090 | 1,34 | 0,62 | 1,44 | 0,43 | 6,67E-01 |
| AT1G63250 | 0,13 | 2,37 | 4,36 | 0,54 | 5,86E-01 |
| AT1G63290 | 0,12 | 1,65 | 4,34 | 0,38 | 7,04E-01 |
| AT1G63320 | 0,20 | -2,26 | 4,30 | -0,52 | 6,00E-01 |
| AT1G63410 | 0,62 | 0,69 | 2,50 | 0,28 | 7,82E-01 |
| AT1G63490 | 0,53 | 1,08 | 3,12 | 0,35 | 7,29E-01 |

| | | | | | |
|---|---|---|---|---|---|
| AT1G63660 | 0,92 | 0,73 | 2,07 | 0,35 | 7,25E-01 |
| AT1G63670 | 0,18 | -0,70 | 4,38 | -0,16 | 8,72E-01 |
| AT1G63700 | 1,36 | 2,03 | 1,36 | 1,49 | 1,35E-01 |
| AT1G63720 | 0,19 | 0,28 | 4,33 | 0,07 | 9,48E-01 |
| AT1G08627 | 1,18 | 0,78 | 2,10 | 0,37 | 7,09E-01 |
| AT1G63800 | 0,47 | -0,96 | 4,36 | -0,22 | 8,26E-01 |
| AT1G63810 | 0,31 | 1,92 | 4,39 | 0,44 | 6,62E-01 |
| AT1G63840 | 0,40 | 2,19 | 4,34 | 0,51 | 6,13E-01 |
| AT1G63860 | 1,09 | 0,91 | 1,77 | 0,51 | 6,09E-01 |
| AT1G63870 | 0,58 | 1,91 | 3,00 | 0,64 | 5,24E-01 |
| AT1G63970 | 0,51 | 0,89 | 3,20 | 0,28 | 7,81E-01 |
| AT1G64080 | 0,15 | -0,41 | 4,32 | -0,09 | 9,25E-01 |
| AT1G64190 | 1,11 | 1,14 | 1,62 | 0,70 | 4,83E-01 |
| AT1G64200 | 0,78 | 3,57 | 4,32 | 0,83 | 4,08E-01 |
| AT1G64280 | 0,86 | 2,61 | 2,42 | 1,08 | 2,81E-01 |
| AT1G64380 | 0,37 | -2,19 | 4,30 | -0,51 | 6,11E-01 |
| AT1G64460 | 0,72 | 2,75 | 2,42 | 1,14 | 2,55E-01 |
| AT1G64480 | 0,18 | -1,39 | 4,34 | -0,32 | 7,49E-01 |
| AT1G64510 | 0,32 | -0,56 | 4,31 | -0,13 | 8,96E-01 |
| AT1G64580 | 0,24 | 1,30 | 4,37 | 0,30 | 7,67E-01 |
| AT1G64610 | 0,62 | -0,37 | 2,63 | -0,14 | 8,89E-01 |
| AT1G64640 | 0,11 | 1,28 | 4,35 | 0,29 | 7,68E-01 |
| AT1G64650 | 0,59 | -0,52 | 2,26 | -0,23 | 8,19E-01 |
| AT1G64670 | 0,10 | 0,39 | 4,35 | 0,09 | 9,29E-01 |
| AT1G64750 | 0,36 | 1,79 | 4,41 | 0,41 | 6,85E-01 |
| AT1G64970 | 0,63 | 1,40 | 2,51 | 0,56 | 5,78E-01 |
| AT1G64980 | 1,26 | -1,80 | 1,72 | -1,05 | 2,95E-01 |
| AT1G65040 | 0,28 | -1,17 | 3,63 | -0,32 | 7,47E-01 |
| AT1G65150 | 0,20 | 2,90 | 4,34 | 0,67 | 5,04E-01 |
| AT1G65190 | 0,43 | 0,71 | 4,10 | 0,17 | 8,63E-01 |
| AT1G08747 | 0,81 | 1,12 | 2,89 | 0,39 | 6,98E-01 |
| AT1G65290 | 0,77 | 1,25 | 2,08 | 0,60 | 5,48E-01 |
| AT1G65320 | 0,39 | 2,16 | 4,29 | 0,50 | 6,14E-01 |
| AT1G65346 | 0,34 | -0,49 | 4,25 | -0,11 | 9,09E-01 |
| AT1G65420 | 0,15 | 2,31 | 4,37 | 0,53 | 5,98E-01 |
| AT1G65430 | 0,52 | 2,19 | 3,06 | 0,71 | 4,75E-01 |
| AT1G65520 | 0,29 | -0,21 | 4,31 | -0,05 | 9,62E-01 |
| AT1G65540 | 1,25 | 3,65 | 1,97 | 1,85 | 6,38E-02 |
| AT1G65550 | 0,21 | 2,85 | 4,34 | 0,66 | 5,11E-01 |
| AT1G65580 | 0,46 | -0,58 | 2,74 | -0,21 | 8,32E-01 |
| AT1G65690 | 0,12 | 2,87 | 4,34 | 0,66 | 5,08E-01 |
| AT1G65950 | 0,28 | 1,64 | 4,38 | 0,38 | 7,08E-01 |
| AT1G08797 | 0,63 | 3,13 | 4,30 | 0,73 | 4,67E-01 |
| AT1G65980 | 1,27 | 2,13 | 2,24 | 0,95 | 3,42E-01 |
| AT1G65985 | 0,33 | 0,89 | 4,31 | 0,21 | 8,36E-01 |
| AT1G65990 | 1,24 | -0,84 | 1,74 | -0,48 | 6,29E-01 |
| AT1G66200 | 1,38 | -1,61 | 1,52 | -1,06 | 2,91E-01 |
| AT1G66235 | 0,12 | 0,31 | 4,35 | 0,07 | 9,44E-01 |
| AT1G66240 | 0,12 | 1,65 | 4,34 | 0,38 | 7,04E-01 |
| AT1G66260 | 1,28 | 1,67 | 1,45 | 1,15 | 2,49E-01 |
| AT1G66290 | 0,49 | 0,86 | 3,27 | 0,26 | 7,92E-01 |
| AT1G66330 | 0,52 | 0,85 | 2,67 | 0,32 | 7,50E-01 |
| AT1G66410 | 1,68 | -1,19 | 1,55 | -0,76 | 4,45E-01 |
| AT1G66460 | 0,25 | 3,14 | 4,33 | 0,72 | 4,69E-01 |
| AT1G66490 | 0,90 | 0,04 | 2,52 | 0,01 | 9,89E-01 |
| AT1G66500 | 0,25 | 0,00 | 4,40 | 0,00 | 1,00E+00 |

| | | | | | |
|---|---|---|---|---|---|
| AT1G66570 | 1,67 | -0,35 | 1,37 | -0,26 | 7,98E-01 |
| AT1G66660 | 0,44 | 1,01 | 4,11 | 0,25 | 8,06E-01 |
| AT1G66670 | 1,37 | -0,12 | 1,58 | -0,08 | 9,39E-01 |
| AT1G66680 | 0,59 | 3,24 | 4,33 | 0,75 | 4,54E-01 |
| AT1G66700 | 0,46 | 2,47 | 4,29 | 0,58 | 5,64E-01 |
| AT1G66830 | 0,68 | 1,05 | 3,76 | 0,28 | 7,80E-01 |
| AT1G66910 | 0,65 | 1,36 | 2,17 | 0,63 | 5,31E-01 |
| AT1G66940 | 0,34 | -2,02 | 4,31 | -0,47 | 6,39E-01 |
| AT1G66980 | 1,51 | 0,20 | 1,49 | 0,14 | 8,91E-01 |
| AT1G67000 | 0,91 | -2,04 | 1,88 | -1,09 | 2,78E-01 |
| AT1G67050 | 0,28 | -1,06 | 3,63 | -0,29 | 7,70E-01 |
| AT1G67080 | 0,36 | 1,61 | 4,33 | 0,37 | 7,10E-01 |
| AT1G67140 | 0,95 | 2,40 | 2,44 | 0,98 | 3,26E-01 |
| AT1G67230 | 0,62 | 0,22 | 2,45 | 0,09 | 9,29E-01 |
| AT1G67235 | 0,29 | -2,72 | 4,29 | -0,63 | 5,26E-01 |
| AT1G67250 | 0,34 | -0,81 | 4,39 | -0,18 | 8,54E-01 |
| AT1G67300 | 0,37 | 2,19 | 4,29 | 0,51 | 6,10E-01 |
| AT1G67310 | 0,33 | 2,56 | 4,31 | 0,59 | 5,52E-01 |
| AT1G67325 | 0,61 | 3,31 | 2,94 | 1,13 | 2,60E-01 |
| AT1G67360 | 0,73 | -1,89 | 2,88 | -0,66 | 5,12E-01 |
| AT1G67481 | 1,30 | 0,47 | 1,71 | 0,28 | 7,81E-01 |
| AT1G67490 | 0,57 | -0,45 | 3,12 | -0,14 | 8,85E-01 |
| AT1G67500 | 0,52 | 1,92 | 3,99 | 0,48 | 6,31E-01 |
| AT1G67510 | 0,38 | -0,05 | 4,22 | -0,01 | 9,91E-01 |
| AT1G67580 | 1,21 | 1,18 | 1,81 | 0,65 | 5,13E-01 |
| AT1G67590 | 0,17 | -0,44 | 4,37 | -0,10 | 9,19E-01 |
| AT1G67680 | 0,43 | 2,45 | 4,30 | 0,57 | 5,69E-01 |
| AT1G67785 | 0,15 | -0,67 | 4,39 | -0,15 | 8,78E-01 |
| AT1G67790 | 0,26 | 2,30 | 4,37 | 0,53 | 5,98E-01 |
| AT1G67860 | 0,58 | 1,33 | 3,02 | 0,44 | 6,59E-01 |
| AT1G67865 | 0,24 | 2,03 | 4,34 | 0,47 | 6,39E-01 |
| AT1G67870 | 0,89 | -1,76 | 1,83 | -0,97 | 3,34E-01 |
| AT1G67930 | 1,11 | 1,37 | 1,55 | 0,88 | 3,76E-01 |
| AT1G67950 | 0,39 | 1,78 | 4,41 | 0,40 | 6,86E-01 |
| AT1G68060 | 0,56 | 1,38 | 3,13 | 0,44 | 6,60E-01 |
| AT1G68140 | 0,87 | 1,44 | 1,89 | 0,76 | 4,46E-01 |
| AT1G68325 | 0,05 | -0,58 | 4,38 | -0,13 | 8,95E-01 |
| AT1G68370 | 0,32 | 2,99 | 4,34 | 0,69 | 4,90E-01 |
| AT1G68550 | 0,87 | 0,44 | 1,78 | 0,25 | 8,04E-01 |
| AT1G08997 | 0,58 | 1,40 | 3,74 | 0,37 | 7,09E-01 |
| AT1G68590 | 1,41 | 0,88 | 1,58 | 0,56 | 5,76E-01 |
| AT1G68660 | 0,71 | 0,80 | 2,22 | 0,36 | 7,18E-01 |
| AT1G68710 | 0,46 | 3,38 | 4,33 | 0,78 | 4,35E-01 |
| AT1G68830 | 0,57 | 1,48 | 2,58 | 0,57 | 5,66E-01 |
| AT1G68910 | 0,08 | 2,33 | 4,36 | 0,53 | 5,93E-01 |
| AT1G69020 | 0,30 | 2,91 | 4,34 | 0,67 | 5,04E-01 |
| AT1G69070 | 0,46 | 1,86 | 3,30 | 0,56 | 5,73E-01 |
| AT1G69220 | 1,28 | 3,36 | 1,99 | 1,68 | 9,23E-02 |
| AT1G69270 | 0,78 | 2,02 | 3,05 | 0,66 | 5,07E-01 |
| AT1G69280 | 0,21 | 2,85 | 4,34 | 0,66 | 5,11E-01 |
| AT1G69290 | 0,15 | 1,93 | 4,33 | 0,45 | 6,55E-01 |
| AT1G69310 | 0,25 | 2,55 | 4,32 | 0,59 | 5,55E-01 |
| AT1G69450 | 0,38 | -1,74 | 4,30 | -0,41 | 6,85E-01 |
| AT1G69480 | 0,55 | 3,10 | 4,34 | 0,71 | 4,75E-01 |
| AT1G69630 | 0,59 | 0,00 | 4,41 | 0,00 | 1,00E+00 |
| AT1G69640 | 0,90 | 1,20 | 2,25 | 0,53 | 5,95E-01 |

| | | | | | |
|---|---|---|---|---|---|
| AT1G69730 | 0,73 | -0,39 | 2,12 | -0,18 | 8,54E-01 |
| AT1G69760 | 0,47 | -2,61 | 3,11 | -0,84 | 4,00E-01 |
| AT1G69770 | 1,05 | 2,28 | 2,31 | 0,99 | 3,23E-01 |
| AT1G69795 | 1,39 | 5,07 | 2,33 | 2,18 | 2,95E-02 |
| AT1G69840 | 0,65 | -0,28 | 2,90 | -0,10 | 9,23E-01 |
| AT1G69850 | 0,22 | 0,08 | 4,27 | 0,02 | 9,85E-01 |
| AT1G69890 | 1,49 | 0,75 | 1,50 | 0,50 | 6,16E-01 |
| AT1G69960 | 0,51 | 2,32 | 3,12 | 0,74 | 4,58E-01 |
| AT1G70000 | 0,98 | 3,32 | 2,01 | 1,66 | 9,77E-02 |
| AT1G70070 | 1,07 | 0,86 | 2,11 | 0,41 | 6,83E-01 |
| AT1G70200 | 0,37 | -1,27 | 4,35 | -0,29 | 7,70E-01 |
| AT1G70280 | 0,17 | 3,04 | 4,33 | 0,70 | 4,83E-01 |
| AT1G70300 | 0,38 | 2,12 | 4,26 | 0,50 | 6,19E-01 |
| AT1G70310 | 0,89 | 2,16 | 3,06 | 0,71 | 4,79E-01 |
| AT1G70340 | 0,34 | 0,60 | 3,52 | 0,17 | 8,65E-01 |
| AT1G70370 | 0,57 | -0,76 | 2,55 | -0,30 | 7,66E-01 |
| AT1G70410 | 0,70 | -1,74 | 2,61 | -0,67 | 5,05E-01 |
| AT1G70420 | 0,44 | -0,86 | 4,08 | -0,21 | 8,34E-01 |
| AT1G70520 | 0,53 | 3,38 | 4,32 | 0,78 | 4,34E-01 |
| AT1G70600 | 0,97 | 3,13 | 2,87 | 1,09 | 2,75E-01 |
| AT1G70730 | 0,91 | 3,19 | 2,36 | 1,35 | 1,76E-01 |
| AT1G70740 | 0,10 | -1,26 | 4,35 | -0,29 | 7,72E-01 |
| AT1G70750 | 0,44 | 3,68 | 4,32 | 0,85 | 3,95E-01 |
| AT1G70780 | 0,73 | 0,94 | 2,62 | 0,36 | 7,18E-01 |
| AT1G70830 | 0,27 | 1,09 | 4,29 | 0,26 | 7,99E-01 |
| AT1G70890 | 1,35 | -0,87 | 1,70 | -0,51 | 6,10E-01 |
| AT1G71020 | 0,94 | 3,68 | 2,32 | 1,59 | 1,13E-01 |
| AT1G71030 | 0,60 | -0,23 | 2,46 | -0,10 | 9,24E-01 |
| AT1G71040 | 0,83 | 0,39 | 1,88 | 0,21 | 8,36E-01 |
| AT1G71060 | 0,73 | 4,59 | 3,39 | 1,35 | 1,76E-01 |
| AT1G71070 | 0,49 | 2,86 | 3,17 | 0,90 | 3,66E-01 |
| AT1G71080 | 0,90 | 2,91 | 2,85 | 1,02 | 3,07E-01 |
| AT1G71160 | 0,69 | 4,16 | 3,53 | 1,18 | 2,38E-01 |
| AT1G71190 | 0,19 | 1,04 | 4,29 | 0,24 | 8,09E-01 |
| AT1G71210 | 0,73 | 2,51 | 2,82 | 0,89 | 3,73E-01 |
| AT1G71310 | 0,27 | 2,92 | 4,35 | 0,67 | 5,02E-01 |
| AT1G71330 | 0,30 | 2,68 | 4,35 | 0,61 | 5,39E-01 |
| AT1G71340 | 0,92 | -0,98 | 2,28 | -0,43 | 6,66E-01 |
| AT1G71350 | 0,21 | 3,40 | 4,32 | 0,79 | 4,32E-01 |
| AT1G71410 | 1,45 | 0,76 | 1,87 | 0,40 | 6,86E-01 |
| AT1G09253 | 0,32 | 1,70 | 4,33 | 0,39 | 6,95E-01 |
| AT1G71490 | 0,42 | 2,56 | 4,16 | 0,62 | 5,38E-01 |
| AT1G71710 | 0,44 | 0,60 | 2,79 | 0,22 | 8,30E-01 |
| AT1G71770 | 0,40 | 1,19 | 4,05 | 0,29 | 7,69E-01 |
| AT1G71870 | 0,88 | 0,08 | 2,64 | 0,03 | 9,76E-01 |
| AT1G71960 | 0,26 | 1,18 | 3,71 | 0,32 | 7,51E-01 |
| AT1G72020 | 0,56 | -0,26 | 2,54 | -0,10 | 9,19E-01 |
| AT1G72040 | 0,83 | 1,27 | 1,98 | 0,64 | 5,23E-01 |
| AT1G72160 | 0,64 | -1,36 | 2,18 | -0,62 | 5,33E-01 |
| AT1G72190 | 0,52 | 4,03 | 3,84 | 1,05 | 2,94E-01 |
| AT1G72270 | 0,12 | 1,15 | 4,35 | 0,26 | 7,92E-01 |
| AT1G72300 | 0,54 | 2,33 | 3,07 | 0,76 | 4,47E-01 |
| AT1G72320 | 0,39 | 0,77 | 3,37 | 0,23 | 8,20E-01 |
| AT1G72450 | 1,12 | -0,57 | 1,57 | -0,36 | 7,16E-01 |
| AT1G72480 | 0,35 | 2,99 | 4,34 | 0,69 | 4,91E-01 |
| AT1G72500 | 0,46 | 0,05 | 4,38 | 0,01 | 9,90E-01 |

| | | | | | |
|---|---|---|---|---|---|
| AT1G72550 | 0,79 | 1,96 | 2,52 | 0,78 | 4,37E-01 |
| AT1G72560 | 1,09 | -0,42 | 2,37 | -0,18 | 8,59E-01 |
| AT1G09327 | 0,20 | 2,39 | 4,37 | 0,55 | 5,84E-01 |
| AT1G72670 | 0,20 | 2,11 | 4,37 | 0,48 | 6,29E-01 |
| AT1G72720 | 0,35 | 0,00 | 4,41 | 0,00 | 1,00E+00 |
| AT1G72730 | 0,32 | 2,47 | 4,35 | 0,57 | 5,69E-01 |
| AT1G72760 | 0,65 | 4,71 | 4,29 | 1,10 | 2,72E-01 |
| AT1G72840 | 0,41 | 2,87 | 4,32 | 0,67 | 5,06E-01 |
| AT1G72860 | 0,44 | 4,02 | 4,31 | 0,93 | 3,51E-01 |
| AT1G73010 | 0,40 | 3,80 | 4,32 | 0,88 | 3,79E-01 |
| AT1G09363 | 0,41 | -1,58 | 3,24 | -0,49 | 6,25E-01 |
| AT1G73110 | 0,63 | 3,23 | 3,81 | 0,85 | 3,97E-01 |
| AT1G73200 | 0,73 | 1,13 | 2,22 | 0,51 | 6,11E-01 |
| AT1G73230 | 0,17 | 3,16 | 4,33 | 0,73 | 4,66E-01 |
| AT1G73260 | 0,63 | 3,24 | 2,97 | 1,09 | 2,76E-01 |
| AT1G73370 | 0,52 | 4,34 | 4,30 | 1,01 | 3,13E-01 |
| AT1G73390 | 0,25 | 1,06 | 4,31 | 0,25 | 8,05E-01 |
| AT1G73490 | 0,37 | 2,14 | 4,31 | 0,50 | 6,20E-01 |
| AT1G73500 | 0,92 | 0,19 | 1,93 | 0,10 | 9,22E-01 |
| AT1G73510 | 0,45 | 2,45 | 4,07 | 0,60 | 5,47E-01 |
| AT1G73530 | 0,46 | -1,45 | 4,18 | -0,35 | 7,28E-01 |
| AT1G73540 | 0,81 | -0,35 | 1,95 | -0,18 | 8,56E-01 |
| AT1G73650 | 0,96 | 3,83 | 2,40 | 1,60 | 1,10E-01 |
| AT1G73655 | 0,74 | 2,19 | 3,69 | 0,59 | 5,53E-01 |
| AT1G73660 | 1,58 | 0,86 | 1,77 | 0,48 | 6,28E-01 |
| AT1G73680 | 0,85 | 1,24 | 1,93 | 0,64 | 5,21E-01 |
| AT1G73730 | 0,17 | 3,04 | 4,33 | 0,70 | 4,83E-01 |
| AT1G73760 | 0,39 | -0,35 | 4,20 | -0,08 | 9,34E-01 |
| AT1G73930 | 0,25 | 2,64 | 4,34 | 0,61 | 5,43E-01 |
| AT1G73950 | 0,45 | 2,76 | 4,36 | 0,63 | 5,27E-01 |
| AT1G73960 | 0,29 | 2,24 | 4,31 | 0,52 | 6,03E-01 |
| AT1G73980 | 0,67 | 0,58 | 2,42 | 0,24 | 8,11E-01 |
| AT1G74050 | 0,43 | 1,61 | 3,20 | 0,50 | 6,14E-01 |
| AT1G74060 | 0,72 | -1,51 | 4,34 | -0,35 | 7,28E-01 |
| AT1G74070 | 0,42 | -1,72 | 4,22 | -0,41 | 6,83E-01 |
| AT1G74120 | 0,82 | 0,38 | 2,83 | 0,14 | 8,92E-01 |
| AT1G74170 | 0,18 | 1,48 | 4,31 | 0,34 | 7,32E-01 |
| AT1G74340 | 0,31 | 2,98 | 4,33 | 0,69 | 4,92E-01 |
| AT1G74370 | 0,53 | 0,02 | 3,89 | 0,01 | 9,96E-01 |
| AT1G74390 | 0,95 | 3,96 | 2,67 | 1,48 | 1,39E-01 |
| AT1G74420 | 0,45 | 2,02 | 4,37 | 0,46 | 6,44E-01 |
| AT1G74440 | 0,80 | -0,75 | 2,89 | -0,26 | 7,95E-01 |
| AT1G09467 | 0,50 | 2,23 | 4,11 | 0,54 | 5,88E-01 |
| AT1G74560 | 0,37 | -1,56 | 3,37 | -0,46 | 6,44E-01 |
| AT1G74690 | 0,95 | 3,42 | 2,07 | 1,65 | 9,82E-02 |
| AT1G74770 | 0,42 | 2,85 | 4,17 | 0,68 | 4,95E-01 |
| AT1G74800 | 0,56 | 1,22 | 3,02 | 0,40 | 6,86E-01 |
| AT1G74830 | 0,12 | 2,81 | 4,34 | 0,65 | 5,18E-01 |
| AT1G74850 | 0,94 | 1,41 | 2,02 | 0,70 | 4,83E-01 |
| AT1G74880 | 0,11 | 1,33 | 4,36 | 0,31 | 7,59E-01 |
| AT1G74920 | 0,86 | -0,73 | 2,56 | -0,28 | 7,77E-01 |
| AT1G74950 | 0,28 | -0,66 | 4,30 | -0,15 | 8,78E-01 |
| AT1G74960 | 1,51 | 3,57 | 2,66 | 1,34 | 1,80E-01 |
| AT1G74970 | 0,46 | 0,07 | 4,33 | 0,02 | 9,87E-01 |
| AT1G75010 | 0,20 | -2,26 | 4,30 | -0,52 | 6,00E-01 |
| AT1G75100 | 0,60 | -0,04 | 2,07 | -0,02 | 9,83E-01 |

| | | | | | |
|---|---|---|---|---|---|
| AT1G75130 | 0,12 | 1,65 | 4,34 | 0,38 | 7,04E-01 |
| AT1G75180 | 0,38 | -0,40 | 3,27 | -0,12 | 9,02E-01 |
| AT1G75200 | 0,15 | 1,69 | 4,34 | 0,39 | 6,98E-01 |
| AT1G75220 | 0,55 | 1,52 | 3,04 | 0,50 | 6,16E-01 |
| AT1G75370 | 1,12 | 0,06 | 1,55 | 0,04 | 9,71E-01 |
| AT1G75410 | 0,90 | 2,28 | 2,40 | 0,95 | 3,43E-01 |
| AT1G75450 | 0,11 | 1,28 | 4,35 | 0,29 | 7,68E-01 |
| AT1G75500 | 0,97 | 0,02 | 1,73 | 0,01 | 9,90E-01 |
| AT1G75680 | 0,99 | -1,16 | 1,65 | -0,71 | 4,80E-01 |
| AT1G75690 | 0,84 | -0,89 | 1,98 | -0,45 | 6,55E-01 |
| AT1G75750 | 1,13 | 3,91 | 2,26 | 1,73 | 8,43E-02 |
| AT1G75780 | 0,25 | 0,84 | 4,30 | 0,20 | 8,44E-01 |
| AT1G75820 | 1,17 | 2,56 | 2,20 | 1,16 | 2,45E-01 |
| AT1G75860 | 0,67 | 1,09 | 2,22 | 0,49 | 6,22E-01 |
| AT1G75945 | 0,28 | 1,62 | 4,35 | 0,37 | 7,10E-01 |
| AT1G76060 | 0,20 | -0,80 | 4,37 | -0,18 | 8,55E-01 |
| AT1G76080 | 1,03 | -0,20 | 1,54 | -0,13 | 8,97E-01 |
| AT1G76100 | 1,22 | -0,90 | 1,88 | -0,48 | 6,31E-01 |
| AT1G76120 | 0,15 | 1,64 | 4,33 | 0,38 | 7,06E-01 |
| AT1G76150 | 0,72 | 2,22 | 2,74 | 0,81 | 4,19E-01 |
| AT1G76160 | 0,88 | 2,10 | 1,86 | 1,13 | 2,59E-01 |
| AT1G76270 | 0,08 | 2,08 | 4,38 | 0,47 | 6,35E-01 |
| AT1G76280 | 0,12 | 1,65 | 4,34 | 0,38 | 7,04E-01 |
| AT1G09627 | 0,45 | 2,59 | 4,01 | 0,65 | 5,17E-01 |
| AT1G76400 | 0,57 | 1,65 | 2,94 | 0,56 | 5,74E-01 |
| AT1G76460 | 0,53 | 0,41 | 3,27 | 0,13 | 9,00E-01 |
| AT1G76640 | 0,08 | 2,08 | 4,38 | 0,47 | 6,35E-01 |
| AT1G76650 | 0,42 | -1,98 | 3,26 | -0,61 | 5,43E-01 |
| AT1G76660 | 0,13 | -0,63 | 4,39 | -0,14 | 8,85E-01 |
| AT1G76670 | 1,15 | 1,61 | 2,10 | 0,77 | 4,42E-01 |
| AT1G76700 | 0,39 | 2,04 | 4,37 | 0,47 | 6,41E-01 |
| AT1G76810 | 1,64 | -1,56 | 1,36 | -1,15 | 2,50E-01 |
| AT1G76878 | 1,06 | -0,80 | 1,69 | -0,47 | 6,37E-01 |
| AT1G76890 | 0,74 | -0,67 | 1,79 | -0,37 | 7,08E-01 |
| AT1G76955 | 0,30 | 2,68 | 4,35 | 0,61 | 5,39E-01 |
| AT1G76960 | 0,40 | -0,21 | 3,36 | -0,06 | 9,50E-01 |
| AT1G76965 | 0,25 | 2,25 | 4,37 | 0,52 | 6,06E-01 |
| AT1G76970 | 0,43 | 0,48 | 3,26 | 0,15 | 8,84E-01 |
| AT1G76980 | 0,73 | 1,75 | 2,96 | 0,59 | 5,55E-01 |
| AT1G77030 | 0,32 | -1,10 | 4,30 | -0,26 | 7,98E-01 |
| AT1G77060 | 0,12 | 0,63 | 4,33 | 0,15 | 8,85E-01 |
| AT1G77100 | 0,82 | 0,62 | 3,51 | 0,18 | 8,59E-01 |
| AT1G77145 | 0,08 | 2,33 | 4,36 | 0,53 | 5,93E-01 |
| AT1G77170 | 0,63 | 3,04 | 2,94 | 1,04 | 3,00E-01 |
| AT1G77240 | 1,26 | 3,86 | 1,90 | 2,03 | 4,25E-02 |
| AT1G77250 | 0,55 | 3,55 | 3,76 | 0,94 | 3,45E-01 |
| AT1G77260 | 0,57 | 2,28 | 3,78 | 0,60 | 5,47E-01 |
| AT1G77480 | 0,33 | 0,00 | 4,41 | 0,00 | 1,00E+00 |
| AT1G77580 | 0,18 | 0,26 | 4,30 | 0,06 | 9,51E-01 |
| AT1G77600 | 0,35 | 1,58 | 4,28 | 0,37 | 7,12E-01 |
| AT1G77660 | 0,33 | 2,42 | 4,31 | 0,56 | 5,74E-01 |
| AT1G77810 | 0,58 | -0,43 | 3,81 | -0,11 | 9,09E-01 |
| AT1G77932 | 0,47 | 2,93 | 4,35 | 0,67 | 5,00E-01 |
| AT1G77940 | 0,55 | 1,41 | 2,45 | 0,58 | 5,65E-01 |
| AT1G77960 | 1,37 | 0,12 | 1,52 | 0,08 | 9,35E-01 |
| AT1G77980 | 0,77 | -2,33 | 2,00 | -1,17 | 2,43E-01 |

| | | | | | |
|---|---|---|---|---|---|
| AT1G77990 | 0,37 | 0,71 | 4,24 | 0,17 | 8,67E-01 |
| AT1G78060 | 0,43 | -0,74 | 4,35 | -0,17 | 8,64E-01 |
| AT1G78210 | 0,32 | -0,41 | 4,32 | -0,10 | 9,24E-01 |
| AT1G78240 | 0,51 | 1,56 | 2,72 | 0,57 | 5,68E-01 |
| AT1G78370 | 1,22 | 0,44 | 1,98 | 0,22 | 8,26E-01 |
| AT1G78380 | 1,48 | -0,46 | 1,56 | -0,30 | 7,67E-01 |
| AT1G78460 | 0,60 | 0,68 | 2,65 | 0,26 | 7,97E-01 |
| AT1G78480 | 0,26 | 1,37 | 4,31 | 0,32 | 7,51E-01 |
| AT1G78510 | 0,69 | 1,18 | 3,76 | 0,31 | 7,54E-01 |
| AT1G78580 | 0,70 | 2,24 | 2,22 | 1,01 | 3,13E-01 |
| AT1G78800 | 0,21 | 2,29 | 4,32 | 0,53 | 5,96E-01 |
| AT1G78820 | 0,54 | 0,35 | 2,23 | 0,16 | 8,76E-01 |
| AT1G78850 | 0,94 | -0,49 | 2,04 | -0,24 | 8,10E-01 |
| AT1G78865 | 0,13 | -1,34 | 4,35 | -0,31 | 7,58E-01 |
| AT1G78880 | 0,88 | 3,02 | 2,72 | 1,11 | 2,68E-01 |
| AT1G78915 | 0,18 | -1,50 | 4,34 | -0,35 | 7,30E-01 |
| AT1G78990 | 0,38 | 2,18 | 4,37 | 0,50 | 6,18E-01 |
| AT1G78995 | 0,24 | 1,12 | 4,33 | 0,26 | 7,96E-01 |
| AT1G79020 | 0,25 | 0,00 | 4,40 | 0,00 | 1,00E+00 |
| AT1G79030 | 0,53 | -1,19 | 3,96 | -0,30 | 7,63E-01 |
| AT1G79050 | 0,58 | 0,40 | 3,14 | 0,13 | 8,99E-01 |
| AT1G79080 | 1,23 | 4,88 | 2,16 | 2,26 | 2,41E-02 |
| AT1G79120 | 0,10 | -1,26 | 4,35 | -0,29 | 7,72E-01 |
| AT1G79150 | 0,35 | 2,35 | 4,33 | 0,54 | 5,87E-01 |
| AT1G79350 | 0,49 | 1,11 | 2,66 | 0,42 | 6,76E-01 |
| AT1G79430 | 0,17 | -0,44 | 4,37 | -0,10 | 9,19E-01 |
| AT1G79440 | 0,39 | 1,29 | 3,45 | 0,37 | 7,09E-01 |
| AT1G09867 | 1,21 | -1,29 | 2,11 | -0,61 | 5,41E-01 |
| AT1G09883 | 0,79 | -2,97 | 2,84 | -1,05 | 2,96E-01 |
| AT1G79540 | 0,53 | 2,48 | 3,10 | 0,80 | 4,24E-01 |
| AT1G79600 | 1,43 | 2,21 | 1,67 | 1,32 | 1,87E-01 |
| AT1G79640 | 0,40 | 2,19 | 4,34 | 0,51 | 6,13E-01 |
| AT1G79670 | 0,90 | 3,27 | 2,47 | 1,33 | 1,85E-01 |
| AT1G79700 | 0,08 | 2,33 | 4,36 | 0,53 | 5,93E-01 |
| AT1G79720 | 0,47 | 2,32 | 3,26 | 0,71 | 4,77E-01 |
| AT1G79730 | 0,63 | 0,12 | 3,79 | 0,03 | 9,75E-01 |
| AT1G79740 | 0,25 | 0,00 | 4,40 | 0,00 | 1,00E+00 |
| AT1G79790 | 0,05 | -0,58 | 4,38 | -0,13 | 8,95E-01 |
| AT1G79830 | 1,06 | 0,01 | 1,64 | 0,01 | 9,96E-01 |
| AT1G79880 | 0,11 | 1,33 | 4,36 | 0,31 | 7,59E-01 |
| AT1G79900 | 0,42 | 2,85 | 4,17 | 0,68 | 4,95E-01 |
| AT1G80030 | 1,07 | 3,02 | 2,18 | 1,39 | 1,66E-01 |
| AT1G80040 | 0,61 | 1,23 | 2,59 | 0,48 | 6,35E-01 |
| AT1G80080 | 0,50 | 1,22 | 3,09 | 0,39 | 6,93E-01 |
| AT1G80325 | 1,08 | 2,08 | 1,89 | 1,10 | 2,72E-01 |
| AT1G80350 | 0,38 | 2,18 | 4,37 | 0,50 | 6,18E-01 |
| AT1G80420 | 0,34 | 0,19 | 4,32 | 0,05 | 9,64E-01 |
| AT1G80460 | 0,24 | 1,11 | 4,33 | 0,26 | 7,97E-01 |
| AT1G80480 | 0,56 | 1,25 | 3,02 | 0,41 | 6,79E-01 |
| AT1G80490 | 1,23 | 1,66 | 1,64 | 1,01 | 3,12E-01 |
| AT1G80530 | 0,68 | 2,95 | 2,94 | 1,00 | 3,16E-01 |
| AT1G80600 | 1,40 | 0,48 | 1,97 | 0,24 | 8,07E-01 |
| AT1G80610 | 0,35 | -0,25 | 3,32 | -0,07 | 9,40E-01 |
| AT1G80630 | 0,28 | 1,26 | 4,36 | 0,29 | 7,73E-01 |
| AT1G80660 | 1,14 | 2,94 | 2,04 | 1,44 | 1,49E-01 |
| AT1G80670 | 0,39 | 1,78 | 4,41 | 0,40 | 6,86E-01 |

| | | | | | |
|---|---|---|---|---|---|
| AT1G80760 | 0,62 | 1,37 | 3,74 | 0,37 | 7,14E-01 |
| AT1G80830 | 0,27 | -0,47 | 4,30 | -0,11 | 9,14E-01 |
| AT1G80850 | 0,20 | -2,22 | 4,30 | -0,52 | 6,06E-01 |
| AT1G80920 | 0,74 | 0,21 | 1,94 | 0,11 | 9,13E-01 |
| AT1G80930 | 0,69 | -0,72 | 2,27 | -0,32 | 7,51E-01 |
| AT1G80950 | 0,35 | -0,09 | 4,30 | -0,02 | 9,84E-01 |
| AT2G01023 | 0,35 | 0,26 | 2,84 | 0,09 | 9,26E-01 |
| AT2G01050 | 0,45 | -1,05 | 4,28 | -0,24 | 8,07E-01 |
| AT2G01060 | 0,57 | 0,26 | 2,41 | 0,11 | 9,14E-01 |
| AT2G01110 | 0,90 | 2,91 | 2,35 | 1,24 | 2,15E-01 |
| AT2G01130 | 1,10 | 3,38 | 2,01 | 1,68 | 9,34E-02 |
| AT2G01140 | 1,55 | 1,50 | 1,57 | 0,95 | 3,41E-01 |
| AT2G01180 | 0,86 | -0,97 | 1,71 | -0,57 | 5,71E-01 |
| AT2G01250 | 0,52 | 3,31 | 3,90 | 0,85 | 3,97E-01 |
| AT2G01275 | 0,19 | -0,65 | 4,31 | -0,15 | 8,80E-01 |
| AT2G01290 | 0,18 | 1,51 | 4,31 | 0,35 | 7,25E-01 |
| AT2G01300 | 0,08 | 2,08 | 4,38 | 0,47 | 6,35E-01 |
| AT2G01310 | 0,05 | -0,58 | 4,38 | -0,13 | 8,95E-01 |
| AT2G01350 | 0,12 | -1,57 | 4,34 | -0,36 | 7,18E-01 |
| AT2G01410 | 0,34 | -0,46 | 3,51 | -0,13 | 8,96E-01 |
| AT2G01420 | 1,24 | -2,32 | 1,71 | -1,36 | 1,75E-01 |
| AT2G01440 | 1,25 | 3,37 | 2,35 | 1,43 | 1,53E-01 |
| AT2G01450 | 1,15 | -0,07 | 1,48 | -0,04 | 9,65E-01 |
| AT2G01505 | 0,30 | 0,40 | 4,27 | 0,09 | 9,25E-01 |
| AT2G01630 | 0,87 | 2,69 | 2,78 | 0,97 | 3,34E-01 |
| AT2G01650 | 1,03 | 1,29 | 1,98 | 0,65 | 5,15E-01 |
| AT2G01670 | 0,39 | 0,72 | 3,20 | 0,22 | 8,22E-01 |
| AT2G01690 | 0,55 | -0,84 | 3,75 | -0,22 | 8,24E-01 |
| AT2G01720 | 1,57 | 3,82 | 2,14 | 1,79 | 7,36E-02 |
| AT2G01755 | 0,14 | -0,45 | 4,33 | -0,10 | 9,17E-01 |
| AT2G01820 | 1,26 | 4,40 | 2,34 | 1,88 | 6,06E-02 |
| AT2G01830 | 0,91 | 0,80 | 1,83 | 0,44 | 6,64E-01 |
| AT2G04125 | 0,12 | 0,31 | 4,35 | 0,07 | 9,44E-01 |
| AT2G01870 | 0,33 | 1,34 | 4,39 | 0,30 | 7,60E-01 |
| AT2G01918 | 1,01 | -2,09 | 2,04 | -1,02 | 3,06E-01 |
| AT2G01930 | 0,16 | 1,16 | 4,38 | 0,27 | 7,90E-01 |
| AT2G01970 | 1,49 | 0,47 | 1,55 | 0,30 | 7,63E-01 |
| AT2G01990 | 0,25 | -0,49 | 4,27 | -0,11 | 9,10E-01 |
| AT2G02080 | 1,03 | -0,28 | 2,49 | -0,11 | 9,11E-01 |
| AT2G02103 | 0,70 | -0,87 | 2,33 | -0,37 | 7,09E-01 |
| AT2G02148 | 0,42 | 2,82 | 4,11 | 0,68 | 4,94E-01 |
| AT2G02150 | 0,39 | 2,17 | 4,23 | 0,51 | 6,09E-01 |
| AT2G02160 | 0,60 | 1,52 | 2,38 | 0,64 | 5,22E-01 |
| AT2G02220 | 0,80 | 1,35 | 2,43 | 0,56 | 5,77E-01 |
| AT2G02230 | 0,17 | 1,90 | 4,33 | 0,44 | 6,61E-01 |
| AT2G02340 | 1,23 | -0,90 | 2,08 | -0,43 | 6,66E-01 |
| AT2G02360 | 0,16 | 1,18 | 4,32 | 0,27 | 7,85E-01 |
| AT2G02400 | 0,84 | -0,59 | 2,02 | -0,29 | 7,70E-01 |
| AT2G02520 | 0,08 | -0,87 | 4,38 | -0,20 | 8,43E-01 |
| AT2G02570 | 0,25 | -0,54 | 4,27 | -0,13 | 8,99E-01 |
| AT2G02635 | 0,17 | -0,65 | 4,36 | -0,15 | 8,81E-01 |
| AT2G02695 | 0,58 | -0,91 | 2,81 | -0,32 | 7,46E-01 |
| AT2G02710 | 0,66 | 0,84 | 2,37 | 0,35 | 7,23E-01 |
| AT2G02780 | 0,28 | -1,65 | 4,33 | -0,38 | 7,02E-01 |
| AT2G02800 | 0,78 | -0,12 | 2,41 | -0,05 | 9,60E-01 |
| AT2G04345 | 0,62 | -1,18 | 2,46 | -0,48 | 6,31E-01 |

| | | | | | |
|---|---|---|---|---|---|
| AT2G02930 | 0,49 | 0,97 | 3,09 | 0,31 | 7,54E-01 |
| AT2G03050 | 0,25 | 2,54 | 4,32 | 0,59 | 5,56E-01 |
| AT2G03090 | 0,08 | 2,33 | 4,36 | 0,53 | 5,93E-01 |
| AT2G03240 | 0,24 | 2,85 | 4,34 | 0,66 | 5,11E-01 |
| AT2G03340 | 0,45 | 0,27 | 3,27 | 0,08 | 9,35E-01 |
| AT2G03390 | 0,61 | 1,41 | 2,26 | 0,63 | 5,31E-01 |
| AT2G03420 | 0,43 | 2,73 | 4,06 | 0,67 | 5,01E-01 |
| AT2G03430 | 0,52 | 2,21 | 4,37 | 0,51 | 6,14E-01 |
| AT2G03440 | 0,35 | -1,10 | 4,29 | -0,26 | 7,98E-01 |
| AT2G03460 | 0,08 | 2,33 | 4,36 | 0,53 | 5,93E-01 |
| AT2G03470 | 1,22 | 1,23 | 2,25 | 0,54 | 5,86E-01 |
| AT2G03490 | 0,08 | 2,33 | 4,36 | 0,53 | 5,93E-01 |
| AT2G03620 | 0,33 | 2,76 | 4,34 | 0,64 | 5,25E-01 |
| AT2G03640 | 0,13 | 2,37 | 4,36 | 0,54 | 5,86E-01 |
| AT2G03690 | 0,23 | -0,64 | 4,39 | -0,15 | 8,84E-01 |
| AT2G03730 | 0,29 | 0,31 | 4,29 | 0,07 | 9,43E-01 |
| AT2G03750 | 1,29 | -1,45 | 1,50 | -0,97 | 3,33E-01 |
| AT2G03760 | 0,93 | -0,38 | 3,36 | -0,11 | 9,11E-01 |
| AT2G03770 | 0,15 | 1,94 | 4,32 | 0,45 | 6,54E-01 |
| AT2G03930 | 1,60 | -1,30 | 1,31 | -1,00 | 3,20E-01 |
| AT2G04655 | 0,14 | -1,69 | 4,33 | -0,39 | 6,96E-01 |
| AT2G04031 | 0,15 | -1,60 | 4,33 | -0,37 | 7,11E-01 |
| AT2G04037 | 0,16 | -0,97 | 4,35 | -0,22 | 8,24E-01 |
| AT2G04050 | 0,12 | 2,87 | 4,34 | 0,66 | 5,08E-01 |
| AT2G04062 | 0,70 | -1,44 | 2,72 | -0,53 | 5,98E-01 |
| AT2G04063 | 0,69 | -0,92 | 2,54 | -0,36 | 7,18E-01 |
| AT2G04066 | 0,42 | 0,67 | 4,28 | 0,16 | 8,75E-01 |
| AT2G04070 | 0,33 | 2,11 | 4,36 | 0,48 | 6,29E-01 |
| AT2G04080 | 0,61 | 0,89 | 2,42 | 0,37 | 7,13E-01 |
| AT2G04160 | 0,43 | -1,07 | 3,97 | -0,27 | 7,89E-01 |
| AT2G04230 | 1,09 | 3,20 | 2,08 | 1,54 | 1,24E-01 |
| AT2G04360 | 0,70 | -0,06 | 2,52 | -0,02 | 9,81E-01 |
| AT2G04378 | 0,56 | -0,61 | 2,66 | -0,23 | 8,19E-01 |
| AT2G04520 | 0,26 | -1,46 | 4,34 | -0,34 | 7,37E-01 |
| AT2G04620 | 0,17 | 3,16 | 4,33 | 0,73 | 4,66E-01 |
| AT2G04630 | 0,19 | 0,60 | 4,30 | 0,14 | 8,89E-01 |
| AT2G04842 | 0,88 | 2,22 | 2,94 | 0,76 | 4,50E-01 |
| AT2G04870 | 0,55 | -1,37 | 2,62 | -0,52 | 6,00E-01 |
| AT2G04890 | 0,11 | 1,28 | 4,35 | 0,29 | 7,68E-01 |
| AT2G05120 | 0,91 | 2,91 | 2,06 | 1,41 | 1,58E-01 |
| AT2G05220 | 0,38 | 1,63 | 4,31 | 0,38 | 7,06E-01 |
| AT2G05250 | 0,41 | 3,15 | 4,33 | 0,73 | 4,67E-01 |
| AT2G04895 | 0,17 | -0,71 | 4,32 | -0,16 | 8,69E-01 |
| AT2G05752 | 0,90 | -0,61 | 2,10 | -0,29 | 7,74E-01 |
| AT2G04995 | 1,03 | -1,12 | 1,94 | -0,58 | 5,63E-01 |
| AT2G05810 | 1,10 | -0,23 | 1,67 | -0,14 | 8,92E-01 |
| AT2G05914 | 0,51 | -0,56 | 4,31 | -0,13 | 8,97E-01 |
| AT2G05920 | 1,42 | 1,21 | 1,36 | 0,89 | 3,74E-01 |
| AT2G05940 | 0,97 | -0,31 | 1,70 | -0,18 | 8,55E-01 |
| AT2G06000 | 0,52 | 1,70 | 2,50 | 0,68 | 4,95E-01 |
| AT2G06040 | 0,24 | 1,89 | 4,34 | 0,43 | 6,64E-01 |
| AT2G06050 | 0,61 | -1,94 | 2,92 | -0,66 | 5,07E-01 |
| AT2G06530 | 0,61 | 0,51 | 3,18 | 0,16 | 8,73E-01 |
| AT2G06541 | 0,49 | 0,16 | 4,00 | 0,04 | 9,68E-01 |
| AT2G06645 | 0,15 | -0,97 | 4,37 | -0,22 | 8,24E-01 |
| AT2G06675 | 0,19 | -2,00 | 4,31 | -0,46 | 6,43E-01 |

| | | | | | |
|---|---|---|---|---|---|
| AT2G06906 | 0,12 | 1,65 | 4,34 | 0,38 | 7,04E-01 |
| AT2G06925 | 0,35 | -0,61 | 3,48 | -0,18 | 8,61E-01 |
| AT2G06960 | 0,68 | -2,82 | 2,74 | -1,03 | 3,05E-01 |
| AT2G07020 | 0,35 | -2,79 | 4,28 | -0,65 | 5,15E-01 |
| AT2G07120 | 0,12 | 0,06 | 4,34 | 0,01 | 9,88E-01 |
| AT2G07180 | 1,49 | 0,57 | 1,89 | 0,30 | 7,61E-01 |
| AT2G07240 | 1,04 | -0,63 | 1,72 | -0,37 | 7,14E-01 |
| AT2G07560 | 0,12 | 1,15 | 4,35 | 0,26 | 7,92E-01 |
| AT2G07771 | 0,58 | -0,59 | 2,57 | -0,23 | 8,18E-01 |
| AT2G07777 | 0,89 | 1,15 | 1,96 | 0,59 | 5,58E-01 |
| AT2G07779 | 0,28 | -0,06 | 4,33 | -0,01 | 9,90E-01 |
| AT2G07621 | 0,12 | 0,31 | 4,35 | 0,07 | 9,44E-01 |
| AT2G07774 | 0,25 | -1,62 | 4,33 | -0,37 | 7,09E-01 |
| AT2G07687 | 1,11 | 1,33 | 1,88 | 0,71 | 4,78E-01 |
| AT2G05205 | 0,68 | -0,04 | 2,41 | -0,01 | 9,88E-01 |
| AT2G07599 | 0,90 | -0,15 | 2,02 | -0,07 | 9,42E-01 |
| AT2G07706 | 1,10 | -2,00 | 2,29 | -0,88 | 3,81E-01 |
| AT2G07641 | 0,45 | 0,25 | 2,69 | 0,09 | 9,27E-01 |
| AT2G07719 | 0,62 | 1,77 | 2,54 | 0,69 | 4,88E-01 |
| AT2G07756 | 1,02 | -0,97 | 1,74 | -0,56 | 5,77E-01 |
| AT2G07734 | 0,71 | -1,63 | 2,90 | -0,56 | 5,73E-01 |
| AT2G07658 | 0,65 | 0,71 | 2,35 | 0,30 | 7,64E-01 |
| AT2G07662 | 1,06 | 0,85 | 2,04 | 0,42 | 6,76E-01 |
| AT2G07835 | 0,78 | 1,76 | 2,87 | 0,61 | 5,40E-01 |
| AT2G07690 | 0,71 | 1,97 | 2,40 | 0,82 | 4,13E-01 |
| AT2G07760 | 0,23 | 1,34 | 4,38 | 0,30 | 7,61E-01 |
| AT2G09795 | 0,34 | 1,89 | 4,34 | 0,44 | 6,63E-01 |
| AT2G05465 | 0,27 | -1,02 | 4,29 | -0,24 | 8,11E-01 |
| AT2G10557 | 1,21 | 5,62 | 3,63 | 1,55 | 1,21E-01 |
| AT2G10608 | 0,36 | -0,45 | 4,38 | -0,10 | 9,19E-01 |
| AT2G05565 | 0,43 | -2,89 | 4,00 | -0,72 | 4,70E-01 |
| AT2G10625 | 0,25 | -2,34 | 4,30 | -0,54 | 5,87E-01 |
| AT2G10950 | 0,47 | -1,90 | 3,86 | -0,49 | 6,23E-01 |
| AT2G10955 | 0,26 | -1,73 | 4,32 | -0,40 | 6,89E-01 |
| AT2G11000 | 0,08 | -0,87 | 4,38 | -0,20 | 8,43E-01 |
| AT2G11852 | 0,26 | -0,14 | 4,26 | -0,03 | 9,74E-01 |
| AT2G11910 | 1,43 | 2,16 | 1,52 | 1,42 | 1,54E-01 |
| AT2G12400 | 0,28 | -0,12 | 4,33 | -0,03 | 9,78E-01 |
| AT2G12405 | 0,18 | -1,50 | 4,34 | -0,35 | 7,30E-01 |
| AT2G05705 | 0,21 | -1,08 | 4,36 | -0,25 | 8,04E-01 |
| AT2G12461 | 0,12 | 2,87 | 4,34 | 0,66 | 5,08E-01 |
| AT2G05835 | 0,29 | -0,37 | 4,34 | -0,09 | 9,32E-01 |
| AT2G12945 | 0,05 | -0,58 | 4,38 | -0,13 | 8,95E-01 |
| AT2G13370 | 1,20 | 1,22 | 2,22 | 0,55 | 5,83E-01 |
| AT2G13450 | 1,51 | -1,07 | 1,68 | -0,64 | 5,24E-01 |
| AT2G13550 | 0,22 | 0,41 | 4,28 | 0,10 | 9,24E-01 |
| AT2G13640 | 0,11 | 1,33 | 4,36 | 0,31 | 7,59E-01 |
| AT2G13650 | 0,22 | -1,21 | 4,35 | -0,28 | 7,81E-01 |
| AT2G13820 | 1,49 | -4,00 | 2,67 | -1,50 | 1,34E-01 |
| AT2G14100 | 0,15 | -0,19 | 4,31 | -0,04 | 9,64E-01 |
| AT2G14120 | 0,98 | 1,20 | 1,83 | 0,65 | 5,13E-01 |
| AT2G14170 | 1,22 | 1,48 | 1,87 | 0,79 | 4,28E-01 |
| AT2G14260 | 0,26 | 2,02 | 4,29 | 0,47 | 6,37E-01 |
| AT2G14265 | 1,52 | -1,51 | 1,59 | -0,95 | 3,43E-01 |
| AT2G14288 | 0,20 | -1,75 | 4,32 | -0,41 | 6,85E-01 |
| AT2G14510 | 0,15 | -0,19 | 4,31 | -0,04 | 9,64E-01 |

| | | | | | |
|---|---|---|---|---|---|
| AT2G14540 | 0,12 | 1,65 | 4,34 | 0,38 | 7,04E-01 |
| AT2G14670 | 1,21 | -4,29 | 2,41 | -1,78 | 7,57E-02 |
| AT2G14720 | 1,05 | 1,00 | 2,15 | 0,47 | 6,40E-01 |
| AT2G14755 | 0,32 | -2,82 | 4,28 | -0,66 | 5,11E-01 |
| AT2G15000 | 0,23 | -0,64 | 4,39 | -0,15 | 8,84E-01 |
| AT2G15020 | 0,51 | 1,25 | 3,23 | 0,39 | 7,00E-01 |
| AT2G06355 | 0,23 | -0,46 | 4,30 | -0,11 | 9,15E-01 |
| AT2G15110 | 0,63 | -0,52 | 2,81 | -0,19 | 8,53E-01 |
| AT2G15290 | 0,58 | 3,41 | 3,74 | 0,91 | 3,63E-01 |
| AT2G15292 | 1,13 | 1,76 | 1,52 | 1,15 | 2,49E-01 |
| AT2G15320 | 1,25 | -0,58 | 1,40 | -0,42 | 6,78E-01 |
| AT2G15390 | 0,37 | 1,32 | 4,27 | 0,31 | 7,58E-01 |
| AT2G15555 | 1,03 | 1,02 | 1,59 | 0,64 | 5,22E-01 |
| AT2G15560 | 0,05 | -0,58 | 4,38 | -0,13 | 8,95E-01 |
| AT2G15790 | 0,41 | 0,59 | 4,14 | 0,14 | 8,86E-01 |
| AT2G15860 | 0,36 | 0,13 | 4,25 | 0,03 | 9,76E-01 |
| AT2G15880 | 1,49 | 2,29 | 1,76 | 1,30 | 1,94E-01 |
| AT2G15890 | 1,14 | 1,20 | 1,44 | 0,83 | 4,05E-01 |
| AT2G06515 | 0,69 | -0,04 | 3,75 | -0,01 | 9,92E-01 |
| AT2G15980 | 0,29 | 1,20 | 3,64 | 0,33 | 7,42E-01 |
| AT2G16016 | 1,06 | -0,36 | 1,88 | -0,19 | 8,49E-01 |
| AT2G06575 | 0,75 | -1,87 | 1,91 | -0,98 | 3,28E-01 |
| AT2G16070 | 0,64 | 1,09 | 3,79 | 0,29 | 7,74E-01 |
| AT2G16090 | 0,55 | 0,72 | 2,39 | 0,30 | 7,62E-01 |
| AT2G16370 | 0,11 | 1,33 | 4,36 | 0,31 | 7,59E-01 |
| AT2G16400 | 1,00 | 1,09 | 1,75 | 0,62 | 5,33E-01 |
| AT2G16405 | 0,20 | 2,90 | 4,34 | 0,67 | 5,04E-01 |
| AT2G16440 | 1,04 | 2,13 | 3,50 | 0,61 | 5,43E-01 |
| AT2G16450 | 0,33 | 2,76 | 4,34 | 0,64 | 5,25E-01 |
| AT2G16490 | 1,37 | -3,01 | 2,36 | -1,27 | 2,04E-01 |
| AT2G16505 | 0,85 | -0,16 | 2,03 | -0,08 | 9,37E-01 |
| AT2G16510 | 1,63 | 2,41 | 1,53 | 1,58 | 1,15E-01 |
| AT2G16620 | 0,18 | -0,57 | 4,30 | -0,13 | 8,94E-01 |
| AT2G16640 | 1,05 | 2,27 | 1,96 | 1,16 | 2,45E-01 |
| AT2G16650 | 0,37 | 4,01 | 4,31 | 0,93 | 3,52E-01 |
| AT2G16870 | 1,12 | 2,09 | 1,91 | 1,09 | 2,74E-01 |
| AT2G16880 | 0,51 | 3,24 | 4,34 | 0,75 | 4,55E-01 |
| AT2G16920 | 0,25 | 2,90 | 4,34 | 0,67 | 5,03E-01 |
| AT2G16940 | 0,88 | 1,38 | 2,00 | 0,69 | 4,90E-01 |
| AT2G17020 | 0,78 | 1,90 | 1,95 | 0,97 | 3,30E-01 |
| AT2G17110 | 0,49 | 0,16 | 2,61 | 0,06 | 9,50E-01 |
| AT2G17130 | 0,45 | 1,73 | 4,18 | 0,42 | 6,78E-01 |
| AT2G17190 | 0,94 | 0,78 | 2,24 | 0,35 | 7,28E-01 |
| AT2G17210 | 0,23 | 1,68 | 4,33 | 0,39 | 6,98E-01 |
| AT2G17220 | 0,39 | -0,04 | 4,39 | -0,01 | 9,92E-01 |
| AT2G17230 | 0,92 | 1,06 | 2,26 | 0,47 | 6,41E-01 |
| AT2G17300 | 0,17 | 1,90 | 4,33 | 0,44 | 6,61E-01 |
| AT2G06935 | 0,05 | -0,58 | 4,38 | -0,13 | 8,95E-01 |
| AT2G17340 | 0,86 | 2,27 | 2,17 | 1,05 | 2,96E-01 |
| AT2G17350 | 0,34 | -0,65 | 4,39 | -0,15 | 8,81E-01 |
| AT2G17450 | 0,63 | 0,29 | 2,16 | 0,13 | 8,95E-01 |
| AT2G17470 | 0,10 | 0,39 | 4,35 | 0,09 | 9,29E-01 |
| AT2G17550 | 0,71 | 2,66 | 2,32 | 1,15 | 2,51E-01 |
| AT2G17560 | 0,46 | 1,26 | 4,36 | 0,29 | 7,73E-01 |
| AT2G17700 | 0,43 | 1,16 | 4,08 | 0,28 | 7,76E-01 |
| AT2G17820 | 0,96 | 0,99 | 2,04 | 0,49 | 6,26E-01 |

| | | | | | |
|---|---|---|---|---|---|
| AT2G17870 | 0,45 | -0,13 | 4,10 | -0,03 | 9,74E-01 |
| AT2G18130 | 0,22 | -1,04 | 4,30 | -0,24 | 8,08E-01 |
| AT2G18160 | 1,66 | -0,19 | 1,46 | -0,13 | 8,94E-01 |
| AT2G18170 | 0,50 | 2,10 | 3,23 | 0,65 | 5,14E-01 |
| AT2G18193 | 1,29 | 3,20 | 2,13 | 1,50 | 1,33E-01 |
| AT2G18290 | 0,19 | 1,20 | 4,29 | 0,28 | 7,79E-01 |
| AT2G18300 | 1,33 | -0,91 | 2,08 | -0,44 | 6,62E-01 |
| AT2G18310 | 0,43 | 0,49 | 4,38 | 0,11 | 9,12E-01 |
| AT2G18520 | 0,25 | 0,00 | 4,40 | 0,00 | 1,00E+00 |
| AT2G18600 | 0,24 | 0,00 | 4,41 | 0,00 | 1,00E+00 |
| AT2G18660 | 0,89 | 0,25 | 1,92 | 0,13 | 8,95E-01 |
| AT2G18685 | 0,08 | 2,33 | 4,36 | 0,53 | 5,93E-01 |
| AT2G18900 | 1,01 | 1,43 | 1,83 | 0,78 | 4,35E-01 |
| AT2G18915 | 0,29 | 1,38 | 4,32 | 0,32 | 7,49E-01 |
| AT2G18940 | 0,31 | 1,12 | 3,54 | 0,32 | 7,52E-01 |
| AT2G19120 | 0,71 | 1,65 | 3,61 | 0,46 | 6,48E-01 |
| AT2G07135 | 0,16 | -1,98 | 4,31 | -0,46 | 6,47E-01 |
| AT2G07165 | 0,21 | 0,81 | 4,32 | 0,19 | 8,52E-01 |
| AT2G19290 | 0,05 | -0,58 | 4,38 | -0,13 | 8,95E-01 |
| AT2G19310 | 0,12 | 0,06 | 4,34 | 0,01 | 9,88E-01 |
| AT2G19340 | 0,43 | 2,32 | 4,33 | 0,54 | 5,92E-01 |
| AT2G19365 | 0,81 | -0,43 | 1,96 | -0,22 | 8,27E-01 |
| AT2G19390 | 0,76 | 0,67 | 2,14 | 0,31 | 7,56E-01 |
| AT2G07195 | 0,51 | -0,22 | 2,47 | -0,09 | 9,29E-01 |
| AT2G19400 | 0,15 | -0,64 | 4,39 | -0,15 | 8,84E-01 |
| AT2G19470 | 0,28 | 0,53 | 4,29 | 0,12 | 9,02E-01 |
| AT2G19560 | 0,20 | 1,94 | 4,34 | 0,45 | 6,55E-01 |
| AT2G19570 | 0,52 | -1,06 | 2,88 | -0,37 | 7,14E-01 |
| AT2G19580 | 0,25 | -1,47 | 4,28 | -0,34 | 7,32E-01 |
| AT2G19760 | 0,25 | -1,09 | 4,27 | -0,26 | 7,98E-01 |
| AT2G19780 | 0,19 | -1,34 | 4,35 | -0,31 | 7,58E-01 |
| AT2G19910 | 0,11 | 1,33 | 4,36 | 0,31 | 7,59E-01 |
| AT2G20000 | 0,44 | 2,74 | 3,26 | 0,84 | 4,01E-01 |
| AT2G20050 | 0,77 | 2,06 | 2,20 | 0,94 | 3,49E-01 |
| AT2G20120 | 0,38 | -0,41 | 4,37 | -0,09 | 9,24E-01 |
| AT2G20180 | 0,53 | 0,47 | 2,51 | 0,19 | 8,51E-01 |
| AT2G20240 | 1,31 | 3,41 | 2,45 | 1,39 | 1,64E-01 |
| AT2G20270 | 0,73 | 0,97 | 3,04 | 0,32 | 7,49E-01 |
| AT2G20280 | 0,90 | 1,79 | 2,73 | 0,66 | 5,12E-01 |
| AT2G20300 | 0,57 | 0,80 | 2,54 | 0,32 | 7,51E-01 |
| AT2G20310 | 0,15 | 1,93 | 4,33 | 0,45 | 6,55E-01 |
| AT2G20380 | 0,08 | 2,08 | 4,38 | 0,47 | 6,35E-01 |
| AT2G20562 | 0,11 | -1,28 | 4,35 | -0,29 | 7,69E-01 |
| AT2G20610 | 0,62 | -0,69 | 2,44 | -0,28 | 7,76E-01 |
| AT2G20630 | 0,57 | -0,83 | 2,60 | -0,32 | 7,50E-01 |
| AT2G20670 | 0,99 | 2,91 | 2,12 | 1,37 | 1,70E-01 |
| AT2G07325 | 0,08 | 2,08 | 4,38 | 0,47 | 6,35E-01 |
| AT2G20720 | 0,50 | 2,93 | 3,21 | 0,91 | 3,61E-01 |
| AT2G20723 | 1,21 | -0,99 | 1,57 | -0,63 | 5,27E-01 |
| AT2G20725 | 0,17 | 2,62 | 4,35 | 0,60 | 5,47E-01 |
| AT2G20760 | 1,00 | 0,31 | 2,12 | 0,15 | 8,82E-01 |
| AT2G20790 | 0,65 | 0,07 | 3,77 | 0,02 | 9,84E-01 |
| AT2G20825 | 0,24 | 2,45 | 4,35 | 0,56 | 5,73E-01 |
| AT2G20830 | 0,42 | -1,69 | 3,18 | -0,53 | 5,95E-01 |
| AT2G20850 | 0,80 | 0,81 | 3,45 | 0,24 | 8,14E-01 |
| AT2G20900 | 0,70 | 1,75 | 2,99 | 0,59 | 5,57E-01 |

| | | | | | |
|---|---|---|---|---|---|
| AT2G21060 | 0,37 | -0,90 | 4,31 | -0,21 | 8,35E-01 |
| AT2G21090 | 0,12 | 2,81 | 4,34 | 0,65 | 5,18E-01 |
| AT2G21120 | 1,14 | 0,07 | 1,61 | 0,04 | 9,64E-01 |
| AT2G21170 | 1,20 | 1,39 | 1,55 | 0,90 | 3,71E-01 |
| AT2G21185 | 0,86 | -0,88 | 1,66 | -0,53 | 5,97E-01 |
| AT2G21188 | 0,47 | 1,93 | 4,21 | 0,46 | 6,46E-01 |
| AT2G21210 | 1,12 | 0,89 | 1,96 | 0,46 | 6,48E-01 |
| AT2G21240 | 0,62 | 0,81 | 2,18 | 0,37 | 7,10E-01 |
| AT2G21280 | 0,38 | 2,18 | 4,37 | 0,50 | 6,18E-01 |
| AT2G21300 | 0,99 | 2,39 | 2,17 | 1,10 | 2,71E-01 |
| AT2G21350 | 0,05 | -0,58 | 4,38 | -0,13 | 8,95E-01 |
| AT2G21370 | 0,82 | -0,35 | 1,86 | -0,19 | 8,51E-01 |
| AT2G21440 | 0,17 | 3,14 | 4,33 | 0,73 | 4,68E-01 |
| AT2G21480 | 0,25 | 2,97 | 4,33 | 0,69 | 4,93E-01 |
| AT2G21500 | 0,51 | 0,44 | 2,94 | 0,15 | 8,80E-01 |
| AT2G21560 | 0,27 | -2,69 | 4,29 | -0,63 | 5,30E-01 |
| AT2G21870 | 1,17 | 0,15 | 2,23 | 0,07 | 9,48E-01 |
| AT2G21950 | 0,24 | 2,24 | 4,33 | 0,52 | 6,05E-01 |
| AT2G21970 | 0,61 | 2,78 | 3,87 | 0,72 | 4,73E-01 |
| AT2G22010 | 0,65 | 0,30 | 2,34 | 0,13 | 8,98E-01 |
| AT2G22090 | 0,34 | -0,81 | 4,37 | -0,19 | 8,52E-01 |
| AT2G22140 | 0,05 | -0,58 | 4,38 | -0,13 | 8,95E-01 |
| AT2G22160 | 0,35 | -0,88 | 4,23 | -0,21 | 8,36E-01 |
| AT2G22240 | 0,21 | 0,06 | 4,31 | 0,01 | 9,89E-01 |
| AT2G22250 | 0,26 | 2,13 | 4,38 | 0,49 | 6,27E-01 |
| AT2G07555 | 0,37 | -2,81 | 4,23 | -0,66 | 5,06E-01 |
| AT2G22400 | 0,55 | 0,41 | 3,78 | 0,11 | 9,14E-01 |
| AT2G22470 | 0,19 | 1,89 | 4,32 | 0,44 | 6,61E-01 |
| AT2G22490 | 1,17 | 3,40 | 2,39 | 1,42 | 1,55E-01 |
| AT2G22740 | 0,43 | 2,89 | 4,08 | 0,71 | 4,78E-01 |
| AT2G22850 | 0,76 | -0,15 | 1,84 | -0,08 | 9,34E-01 |
| AT2G22890 | 0,53 | -2,68 | 3,01 | -0,89 | 3,72E-01 |
| AT2G22940 | 0,87 | -0,97 | 2,41 | -0,40 | 6,87E-01 |
| AT2G23070 | 0,53 | 0,07 | 2,06 | 0,04 | 9,72E-01 |
| AT2G23090 | 0,40 | 1,53 | 4,30 | 0,36 | 7,22E-01 |
| AT2G23100 | 0,16 | 1,16 | 4,38 | 0,27 | 7,90E-01 |
| AT2G23140 | 0,38 | 1,12 | 4,15 | 0,27 | 7,87E-01 |
| AT2G23210 | 0,71 | 3,30 | 2,80 | 1,18 | 2,39E-01 |
| AT2G23290 | 0,64 | -0,01 | 2,50 | 0,00 | 9,97E-01 |
| AT2G23390 | 0,32 | 1,89 | 4,31 | 0,44 | 6,61E-01 |
| AT2G23430 | 0,47 | -0,37 | 3,24 | -0,11 | 9,10E-01 |
| AT2G23450 | 0,62 | 0,21 | 2,13 | 0,10 | 9,22E-01 |
| AT2G23520 | 0,43 | -0,34 | 3,34 | -0,10 | 9,19E-01 |
| AT2G07910 | 1,63 | -4,39 | 2,30 | -1,91 | 5,65E-02 |
| AT2G23550 | 1,59 | -4,33 | 2,32 | -1,87 | 6,17E-02 |
| AT2G23600 | 1,45 | -0,13 | 1,78 | -0,07 | 9,42E-01 |
| AT2G23610 | 0,63 | -1,16 | 2,98 | -0,39 | 6,98E-01 |
| AT2G23670 | 1,50 | 1,05 | 1,56 | 0,68 | 4,99E-01 |
| AT2G23760 | 0,34 | -1,55 | 4,34 | -0,36 | 7,20E-01 |
| AT2G23790 | 0,79 | 2,48 | 2,89 | 0,86 | 3,91E-01 |
| AT2G23840 | 0,70 | 0,19 | 3,07 | 0,06 | 9,51E-01 |
| AT2G23987 | 0,17 | 0,00 | 4,41 | 0,00 | 1,00E+00 |
| AT2G24120 | 0,81 | -1,55 | 2,09 | -0,74 | 4,58E-01 |
| AT2G24150 | 0,79 | 1,44 | 2,08 | 0,69 | 4,90E-01 |
| AT2G24170 | 0,17 | 2,04 | 4,33 | 0,47 | 6,37E-01 |
| AT2G24200 | 1,48 | 1,00 | 1,42 | 0,71 | 4,79E-01 |

| | | | | | |
|---|---|---|---|---|---|
| AT2G24230 | 0,59 | -1,70 | 2,50 | -0,68 | 4,97E-01 |
| AT2G24330 | 0,33 | -0,88 | 4,37 | -0,20 | 8,40E-01 |
| AT2G24340 | 1,46 | -0,94 | 1,44 | -0,65 | 5,16E-01 |
| AT2G24360 | 1,19 | 0,63 | 1,73 | 0,37 | 7,14E-01 |
| AT2G24500 | 1,13 | 0,95 | 2,10 | 0,45 | 6,51E-01 |
| AT2G08055 | 0,33 | 0,82 | 4,27 | 0,19 | 8,47E-01 |
| AT2G24545 | 1,16 | 0,04 | 1,50 | 0,03 | 9,78E-01 |
| AT2G24550 | 1,50 | 0,59 | 1,43 | 0,42 | 6,77E-01 |
| AT2G24600 | 1,31 | 2,35 | 2,04 | 1,15 | 2,51E-01 |
| AT2G24640 | 0,24 | 2,80 | 4,34 | 0,65 | 5,18E-01 |
| AT2G24710 | 0,19 | 2,25 | 4,32 | 0,52 | 6,02E-01 |
| AT2G24755 | 0,71 | 1,23 | 2,46 | 0,50 | 6,16E-01 |
| AT2G24780 | 0,92 | -0,18 | 2,33 | -0,08 | 9,40E-01 |
| AT2G24850 | 1,45 | 0,35 | 1,56 | 0,22 | 8,22E-01 |
| AT2G24870 | 1,09 | -3,37 | 2,40 | -1,40 | 1,60E-01 |
| AT2G24880 | 1,30 | 0,14 | 1,54 | 0,09 | 9,26E-01 |
| AT2G25011 | 1,24 | -0,12 | 1,64 | -0,07 | 9,41E-01 |
| AT2G25050 | 0,47 | 0,79 | 3,22 | 0,25 | 8,06E-01 |
| AT2G25070 | 0,45 | -0,33 | 2,66 | -0,12 | 9,01E-01 |
| AT2G08105 | 0,74 | 1,94 | 2,41 | 0,80 | 4,22E-01 |
| AT2G25180 | 0,76 | 2,23 | 3,01 | 0,74 | 4,59E-01 |
| AT2G25200 | 0,69 | -0,71 | 3,81 | -0,19 | 8,52E-01 |
| AT2G25300 | 0,15 | 2,31 | 4,37 | 0,53 | 5,98E-01 |
| AT2G25310 | 0,20 | -0,80 | 4,37 | -0,18 | 8,55E-01 |
| AT2G25570 | 0,45 | 0,59 | 4,35 | 0,14 | 8,92E-01 |
| AT2G25610 | 0,53 | -0,13 | 3,11 | -0,04 | 9,68E-01 |
| AT2G25670 | 0,93 | 1,69 | 2,15 | 0,78 | 4,33E-01 |
| AT2G25760 | 0,93 | 2,50 | 2,82 | 0,89 | 3,75E-01 |
| AT2G25850 | 0,16 | 1,18 | 4,32 | 0,27 | 7,85E-01 |
| AT2G25910 | 0,91 | -2,87 | 2,42 | -1,18 | 2,37E-01 |
| AT2G25970 | 1,54 | 1,42 | 1,67 | 0,85 | 3,95E-01 |
| AT2G26030 | 0,31 | -0,01 | 3,56 | 0,00 | 9,97E-01 |
| AT2G26070 | 0,39 | 1,71 | 4,40 | 0,39 | 6,98E-01 |
| AT2G26140 | 1,61 | 1,87 | 1,57 | 1,20 | 2,32E-01 |
| AT2G26190 | 1,38 | -1,28 | 1,67 | -0,76 | 4,44E-01 |
| AT2G26280 | 1,11 | -0,06 | 2,29 | -0,03 | 9,78E-01 |
| AT2G26430 | 0,38 | 2,12 | 4,26 | 0,50 | 6,19E-01 |
| AT2G26540 | 0,42 | 0,25 | 4,31 | 0,06 | 9,54E-01 |
| AT2G26560 | 1,67 | 0,06 | 1,34 | 0,05 | 9,61E-01 |
| AT2G26580 | 0,26 | -0,20 | 4,35 | -0,05 | 9,64E-01 |
| AT2G26590 | 0,25 | 0,16 | 4,31 | 0,04 | 9,70E-01 |
| AT2G26620 | 0,18 | -0,78 | 4,31 | -0,18 | 8,57E-01 |
| AT2G26680 | 0,33 | 2,76 | 4,34 | 0,64 | 5,25E-01 |
| AT2G26690 | 0,68 | 2,56 | 2,44 | 1,05 | 2,93E-01 |
| AT2G26740 | 0,33 | 1,16 | 4,28 | 0,27 | 7,86E-01 |
| AT2G08175 | 1,14 | -1,77 | 1,76 | -1,01 | 3,14E-01 |
| AT2G26790 | 0,21 | 3,35 | 4,33 | 0,77 | 4,39E-01 |
| AT2G26800 | 0,41 | 1,88 | 3,37 | 0,56 | 5,77E-01 |
| AT2G26840 | 0,08 | 2,33 | 4,36 | 0,53 | 5,93E-01 |
| AT2G26900 | 1,34 | 0,52 | 1,63 | 0,32 | 7,51E-01 |
| AT2G26930 | 0,69 | 1,18 | 3,01 | 0,39 | 6,96E-01 |
| AT2G26980 | 0,61 | 0,19 | 2,25 | 0,09 | 9,32E-01 |
| AT2G26990 | 0,56 | 2,57 | 3,09 | 0,83 | 4,04E-01 |
| AT2G27020 | 0,74 | 1,85 | 2,48 | 0,75 | 4,55E-01 |
| AT2G27040 | 0,44 | -0,36 | 3,27 | -0,11 | 9,12E-01 |
| AT2G27070 | 0,56 | 0,83 | 3,13 | 0,27 | 7,90E-01 |

| | | | | | |
|---|---|---|---|---|---|
| AT2G27090 | 0,77 | 3,25 | 2,95 | 1,10 | 2,70E-01 |
| AT2G27170 | 0,61 | 0,42 | 2,49 | 0,17 | 8,67E-01 |
| AT2G27190 | 0,39 | -1,54 | 4,30 | -0,36 | 7,20E-01 |
| AT2G27290 | 0,57 | 0,67 | 2,61 | 0,26 | 7,96E-01 |
| AT2G27330 | 0,10 | -1,26 | 4,35 | -0,29 | 7,72E-01 |
| AT2G27350 | 0,23 | 0,75 | 4,27 | 0,17 | 8,61E-01 |
| AT2G27385 | 1,03 | -1,58 | 2,02 | -0,78 | 4,33E-01 |
| AT2G27500 | 0,69 | -2,05 | 2,37 | -0,87 | 3,87E-01 |
| AT2G27510 | 0,44 | 1,00 | 4,13 | 0,24 | 8,08E-01 |
| AT2G27530 | 0,92 | 0,39 | 1,74 | 0,23 | 8,21E-01 |
| AT2G27580 | 0,43 | 1,89 | 4,14 | 0,46 | 6,48E-01 |
| AT2G27660 | 0,41 | 0,03 | 4,39 | 0,01 | 9,95E-01 |
| AT2G27680 | 0,72 | 0,91 | 1,91 | 0,48 | 6,32E-01 |
| AT2G27770 | 0,38 | -0,66 | 4,26 | -0,16 | 8,77E-01 |
| AT2G27950 | 1,06 | 1,97 | 1,86 | 1,06 | 2,90E-01 |
| AT2G28080 | 0,87 | -2,42 | 2,65 | -0,92 | 3,60E-01 |
| AT2G28170 | 0,05 | -0,58 | 4,38 | -0,13 | 8,95E-01 |
| AT2G28360 | 1,42 | -0,74 | 1,50 | -0,49 | 6,23E-01 |
| AT2G28380 | 0,74 | 2,18 | 2,83 | 0,77 | 4,40E-01 |
| AT2G28390 | 0,41 | 1,92 | 4,37 | 0,44 | 6,61E-01 |
| AT2G28400 | 0,81 | 2,91 | 2,42 | 1,20 | 2,29E-01 |
| AT2G28450 | 1,48 | 2,27 | 1,86 | 1,22 | 2,21E-01 |
| AT2G28470 | 0,87 | -2,41 | 2,59 | -0,93 | 3,51E-01 |
| AT2G28480 | 0,12 | 1,15 | 4,35 | 0,26 | 7,92E-01 |
| AT2G28620 | 1,12 | 0,37 | 1,86 | 0,20 | 8,41E-01 |
| AT2G08275 | 1,12 | 0,37 | 1,86 | 0,20 | 8,41E-01 |
| AT2G28630 | 0,37 | -0,57 | 4,24 | -0,13 | 8,93E-01 |
| AT2G28640 | 0,62 | -2,16 | 2,94 | -0,74 | 4,62E-01 |
| AT2G28670 | 0,51 | 0,49 | 2,91 | 0,17 | 8,65E-01 |
| AT2G28740 | 0,44 | 0,58 | 4,08 | 0,14 | 8,87E-01 |
| AT2G28800 | 0,62 | -1,70 | 2,88 | -0,59 | 5,54E-01 |
| AT2G08300 | 0,20 | -0,80 | 4,37 | -0,18 | 8,55E-01 |
| AT2G28880 | 0,33 | 3,22 | 4,33 | 0,74 | 4,56E-01 |
| AT2G28890 | 0,77 | 0,60 | 2,00 | 0,30 | 7,66E-01 |
| AT2G28900 | 0,81 | -2,71 | 2,54 | -1,07 | 2,86E-01 |
| AT2G28950 | 0,45 | 0,06 | 4,10 | 0,02 | 9,88E-01 |
| AT2G29030 | 0,32 | -1,88 | 4,32 | -0,44 | 6,63E-01 |
| AT2G29040 | 0,08 | -0,56 | 4,38 | -0,13 | 8,99E-01 |
| AT2G29110 | 0,41 | 2,03 | 4,10 | 0,49 | 6,21E-01 |
| AT2G29120 | 1,16 | 4,05 | 2,10 | 1,92 | 5,43E-02 |
| AT2G29170 | 0,18 | 0,08 | 4,29 | 0,02 | 9,85E-01 |
| AT2G08330 | 0,26 | 2,13 | 4,38 | 0,49 | 6,27E-01 |
| AT2G29190 | 0,61 | 1,62 | 2,57 | 0,63 | 5,28E-01 |
| AT2G29220 | 0,39 | 1,71 | 4,40 | 0,39 | 6,98E-01 |
| AT2G29390 | 0,71 | 2,28 | 2,44 | 0,94 | 3,49E-01 |
| AT2G29400 | 0,98 | 1,85 | 1,71 | 1,08 | 2,78E-01 |
| AT2G29550 | 0,75 | -0,06 | 2,11 | -0,03 | 9,76E-01 |
| AT2G29570 | 0,14 | -0,31 | 4,33 | -0,07 | 9,43E-01 |
| AT2G29605 | 0,65 | -2,65 | 2,76 | -0,96 | 3,36E-01 |
| AT2G29650 | 0,77 | 1,94 | 2,51 | 0,77 | 4,40E-01 |
| AT2G30250 | 0,71 | 1,36 | 2,24 | 0,61 | 5,43E-01 |
| AT2G30260 | 0,44 | 2,22 | 3,26 | 0,68 | 4,96E-01 |
| AT2G30270 | 0,18 | 1,17 | 4,30 | 0,27 | 7,86E-01 |
| AT2G30360 | 0,37 | -0,78 | 4,28 | -0,18 | 8,55E-01 |
| AT2G30390 | 0,49 | 1,23 | 3,30 | 0,37 | 7,10E-01 |
| AT2G30470 | 0,73 | 2,21 | 2,18 | 1,01 | 3,11E-01 |

| | | | | | |
|---|---|---|---|---|---|
| AT2G30490 | 0,91 | 0,74 | 3,00 | 0,25 | 8,05E-01 |
| AT2G30500 | 1,56 | 4,49 | 1,79 | 2,51 | 1,20E-02 |
| AT2G30505 | 0,35 | 2,20 | 4,32 | 0,51 | 6,11E-01 |
| AT2G30700 | 0,36 | 2,71 | 4,27 | 0,63 | 5,26E-01 |
| AT2G30710 | 0,35 | 1,71 | 4,32 | 0,40 | 6,92E-01 |
| AT2G30720 | 0,49 | 1,39 | 3,20 | 0,43 | 6,64E-01 |
| AT2G30790 | 0,21 | -0,24 | 4,35 | -0,06 | 9,56E-01 |
| AT2G30800 | 1,27 | 0,68 | 1,97 | 0,35 | 7,30E-01 |
| AT2G30850 | 0,41 | 0,00 | 4,39 | 0,00 | 1,00E+00 |
| AT2G30880 | 0,34 | 1,57 | 4,30 | 0,37 | 7,14E-01 |
| AT2G08420 | 0,54 | 2,77 | 4,00 | 0,69 | 4,89E-01 |
| AT2G30980 | 0,41 | 1,62 | 4,33 | 0,37 | 7,09E-01 |
| AT2G31040 | 0,19 | 0,58 | 4,36 | 0,13 | 8,94E-01 |
| AT2G31060 | 0,49 | 3,35 | 4,30 | 0,78 | 4,36E-01 |
| AT2G31070 | 0,79 | 0,15 | 2,04 | 0,07 | 9,41E-01 |
| AT2G31130 | 0,17 | -0,40 | 4,34 | -0,09 | 9,27E-01 |
| AT2G31170 | 0,45 | 1,59 | 4,34 | 0,37 | 7,14E-01 |
| AT2G31240 | 0,25 | 2,97 | 4,33 | 0,69 | 4,93E-01 |
| AT2G31250 | 0,39 | 1,78 | 4,41 | 0,40 | 6,86E-01 |
| AT2G31260 | 0,53 | -1,36 | 2,47 | -0,55 | 5,83E-01 |
| AT2G08430 | 0,08 | 2,33 | 4,36 | 0,53 | 5,93E-01 |
| AT2G31360 | 1,30 | 1,30 | 1,86 | 0,70 | 4,83E-01 |
| AT2G31390 | 0,86 | 2,33 | 2,88 | 0,81 | 4,18E-01 |
| AT2G31410 | 0,37 | 0,00 | 4,41 | 0,00 | 1,00E+00 |
| AT2G31440 | 0,44 | 3,92 | 4,31 | 0,91 | 3,64E-01 |
| AT2G31510 | 0,63 | -0,85 | 3,15 | -0,27 | 7,86E-01 |
| AT2G31570 | 0,14 | 0,78 | 4,34 | 0,18 | 8,57E-01 |
| AT2G31670 | 0,73 | 3,04 | 2,47 | 1,23 | 2,18E-01 |
| AT2G31730 | 0,53 | 0,23 | 4,36 | 0,05 | 9,57E-01 |
| AT2G31740 | 0,52 | 2,46 | 3,21 | 0,77 | 4,44E-01 |
| AT2G31751 | 0,97 | 2,29 | 2,31 | 0,99 | 3,21E-01 |
| AT2G31800 | 0,85 | 0,34 | 1,89 | 0,18 | 8,58E-01 |
| AT2G31830 | 0,34 | 2,03 | 4,34 | 0,47 | 6,39E-01 |
| AT2G31865 | 0,37 | 3,61 | 4,32 | 0,84 | 4,04E-01 |
| AT2G31870 | 0,27 | 0,36 | 3,65 | 0,10 | 9,22E-01 |
| AT2G08475 | 0,81 | 1,25 | 2,27 | 0,55 | 5,81E-01 |
| AT2G31990 | 0,31 | 0,75 | 4,30 | 0,17 | 8,62E-01 |
| AT2G32000 | 0,55 | -0,36 | 2,60 | -0,14 | 8,90E-01 |
| AT2G32030 | 0,22 | 0,81 | 4,27 | 0,19 | 8,50E-01 |
| AT2G32050 | 0,08 | 2,08 | 4,38 | 0,47 | 6,35E-01 |
| AT2G32060 | 0,34 | 0,04 | 4,25 | 0,01 | 9,92E-01 |
| AT2G32070 | 0,64 | 2,49 | 2,44 | 1,02 | 3,08E-01 |
| AT2G32080 | 0,57 | 1,96 | 2,43 | 0,81 | 4,20E-01 |
| AT2G32130 | 0,28 | 2,67 | 4,31 | 0,62 | 5,35E-01 |
| AT2G32230 | 0,70 | 3,11 | 2,86 | 1,09 | 2,77E-01 |
| AT2G32235 | 0,12 | 2,81 | 4,34 | 0,65 | 5,18E-01 |
| AT2G32390 | 0,36 | 3,22 | 4,34 | 0,74 | 4,59E-01 |
| AT2G32400 | 0,29 | 3,75 | 4,32 | 0,87 | 3,85E-01 |
| AT2G32415 | 0,42 | 2,52 | 4,36 | 0,58 | 5,64E-01 |
| AT2G32480 | 0,87 | 1,27 | 2,22 | 0,57 | 5,68E-01 |
| AT2G32520 | 0,35 | 2,20 | 4,29 | 0,51 | 6,08E-01 |
| AT2G32600 | 0,08 | 2,33 | 4,36 | 0,53 | 5,93E-01 |
| AT2G32680 | 1,55 | 0,80 | 1,45 | 0,55 | 5,80E-01 |
| AT2G32810 | 0,70 | 0,13 | 2,32 | 0,06 | 9,55E-01 |
| AT2G32920 | 1,12 | 0,33 | 1,68 | 0,19 | 8,46E-01 |
| AT2G32940 | 0,12 | 2,81 | 4,34 | 0,65 | 5,18E-01 |

| | | | | | |
|---|---|---|---|---|---|
| AT2G33040 | 0,74 | -0,02 | 2,15 | -0,01 | 9,92E-01 |
| AT2G33051 | 0,64 | 3,25 | 3,86 | 0,84 | 4,00E-01 |
| AT2G33070 | 0,08 | 2,08 | 4,38 | 0,47 | 6,35E-01 |
| AT2G33110 | 0,40 | 3,80 | 4,32 | 0,88 | 3,79E-01 |
| AT2G33120 | 0,77 | -0,09 | 1,92 | -0,04 | 9,64E-01 |
| AT2G33175 | 0,67 | -1,52 | 2,34 | -0,65 | 5,16E-01 |
| AT2G33210 | 0,70 | 0,88 | 2,33 | 0,38 | 7,06E-01 |
| AT2G33260 | 0,24 | 2,45 | 4,35 | 0,56 | 5,73E-01 |
| AT2G33320 | 0,54 | 0,70 | 2,40 | 0,29 | 7,70E-01 |
| AT2G33370 | 0,46 | -0,29 | 2,66 | -0,11 | 9,12E-01 |
| AT2G33385 | 0,08 | -0,56 | 4,38 | -0,13 | 8,99E-01 |
| AT2G33460 | 0,28 | 1,96 | 4,32 | 0,45 | 6,51E-01 |
| AT2G33470 | 0,39 | 1,06 | 3,24 | 0,33 | 7,44E-01 |
| AT2G33500 | 0,51 | 0,00 | 4,41 | 0,00 | 1,00E+00 |
| AT2G33530 | 0,99 | 1,18 | 1,80 | 0,66 | 5,11E-01 |
| AT2G33540 | 0,96 | 0,21 | 1,59 | 0,13 | 8,95E-01 |
| AT2G33550 | 0,30 | -1,84 | 4,32 | -0,43 | 6,70E-01 |
| AT2G33580 | 0,75 | -0,40 | 2,02 | -0,20 | 8,43E-01 |
| AT2G33630 | 0,50 | 0,09 | 3,24 | 0,03 | 9,77E-01 |
| AT2G33655 | 0,29 | 0,00 | 4,41 | 0,00 | 1,00E+00 |
| AT2G33800 | 0,66 | 1,14 | 2,56 | 0,45 | 6,56E-01 |
| AT2G33815 | 0,37 | -0,84 | 4,36 | -0,19 | 8,48E-01 |
| AT2G33860 | 0,43 | 0,71 | 4,11 | 0,17 | 8,63E-01 |
| AT2G33980 | 0,47 | 1,62 | 3,21 | 0,51 | 6,13E-01 |
| AT2G34040 | 1,26 | 2,29 | 1,47 | 1,56 | 1,18E-01 |
| AT2G34070 | 0,35 | 0,63 | 3,47 | 0,18 | 8,55E-01 |
| AT2G34090 | 0,11 | -1,28 | 4,35 | -0,29 | 7,69E-01 |
| AT2G34290 | 0,45 | -1,13 | 3,91 | -0,29 | 7,73E-01 |
| AT2G34300 | 0,72 | 2,87 | 4,31 | 0,67 | 5,05E-01 |
| AT2G34450 | 0,60 | 3,56 | 3,78 | 0,94 | 3,46E-01 |
| AT2G34470 | 0,95 | 2,56 | 2,01 | 1,27 | 2,03E-01 |
| AT2G34480 | 1,66 | 0,16 | 1,25 | 0,13 | 8,98E-01 |
| AT2G34490 | 0,49 | 0,63 | 3,96 | 0,16 | 8,73E-01 |
| AT2G34510 | 0,33 | 1,70 | 4,38 | 0,39 | 6,97E-01 |
| AT2G34520 | 0,36 | 0,00 | 4,41 | 0,00 | 1,00E+00 |
| AT2G34585 | 1,42 | 0,90 | 1,36 | 0,66 | 5,08E-01 |
| AT2G34680 | 1,27 | 0,89 | 1,99 | 0,45 | 6,54E-01 |
| AT2G34710 | 1,10 | 2,93 | 2,42 | 1,21 | 2,26E-01 |
| AT2G34790 | 0,43 | -1,84 | 4,09 | -0,45 | 6,53E-01 |
| AT2G34920 | 0,05 | -0,58 | 4,38 | -0,13 | 8,95E-01 |
| AT2G34930 | 0,44 | -1,70 | 3,38 | -0,50 | 6,14E-01 |
| AT2G35030 | 0,48 | 3,18 | 3,98 | 0,80 | 4,23E-01 |
| AT2G35040 | 0,75 | 2,19 | 2,95 | 0,74 | 4,59E-01 |
| AT2G35060 | 1,08 | -0,60 | 1,65 | -0,36 | 7,17E-01 |
| AT2G35120 | 0,10 | -1,26 | 4,35 | -0,29 | 7,72E-01 |
| AT2G35250 | 1,05 | 1,11 | 1,75 | 0,63 | 5,26E-01 |
| AT2G35360 | 0,45 | 0,87 | 4,14 | 0,21 | 8,34E-01 |
| AT2G35390 | 0,25 | 2,25 | 4,37 | 0,52 | 6,06E-01 |
| AT2G35420 | 0,17 | 1,90 | 4,33 | 0,44 | 6,61E-01 |
| AT2G35490 | 0,67 | 0,24 | 2,00 | 0,12 | 9,05E-01 |
| AT2G35510 | 0,57 | 0,60 | 3,19 | 0,19 | 8,51E-01 |
| AT2G35610 | 0,17 | 2,04 | 4,33 | 0,47 | 6,37E-01 |
| AT2G35680 | 0,73 | 1,27 | 2,21 | 0,58 | 5,65E-01 |
| AT2G35730 | 0,35 | 0,00 | 4,41 | 0,00 | 1,00E+00 |
| AT2G35742 | 0,34 | -1,49 | 3,44 | -0,43 | 6,65E-01 |
| AT2G08755 | 0,29 | 1,50 | 4,28 | 0,35 | 7,27E-01 |

| | | | | | |
|---|---|---|---|---|---|
| AT2G08765 | 0,51 | -1,66 | 3,18 | -0,52 | 6,02E-01 |
| AT2G35859 | 0,71 | 0,29 | 2,63 | 0,11 | 9,11E-01 |
| AT2G35880 | 0,24 | 2,85 | 4,34 | 0,66 | 5,11E-01 |
| AT2G35930 | 0,76 | -1,38 | 2,66 | -0,52 | 6,03E-01 |
| AT2G36030 | 0,62 | -0,72 | 4,35 | -0,17 | 8,68E-01 |
| AT2G36050 | 0,69 | -1,65 | 2,83 | -0,58 | 5,62E-01 |
| AT2G36060 | 0,48 | 0,52 | 2,70 | 0,19 | 8,48E-01 |
| AT2G36070 | 0,34 | 0,68 | 4,34 | 0,16 | 8,75E-01 |
| AT2G36080 | 0,43 | -0,10 | 3,27 | -0,03 | 9,77E-01 |
| AT2G36180 | 0,76 | -2,52 | 3,30 | -0,77 | 4,44E-01 |
| AT2G36200 | 0,33 | 0,20 | 4,33 | 0,05 | 9,63E-01 |
| AT2G36230 | 0,20 | -0,53 | 4,30 | -0,12 | 9,03E-01 |
| AT2G36250 | 0,55 | -2,04 | 3,16 | -0,64 | 5,19E-01 |
| AT2G36300 | 0,44 | -0,96 | 3,25 | -0,30 | 7,67E-01 |
| AT2G36360 | 0,43 | -2,05 | 3,28 | -0,63 | 5,31E-01 |
| AT2G36400 | 0,83 | 1,76 | 2,92 | 0,60 | 5,46E-01 |
| AT2G36620 | 0,25 | 1,05 | 4,33 | 0,24 | 8,08E-01 |
| AT2G36680 | 0,35 | 0,23 | 4,36 | 0,05 | 9,57E-01 |
| AT2G36710 | 0,27 | -1,91 | 4,32 | -0,44 | 6,59E-01 |
| AT2G36750 | 0,21 | 3,40 | 4,32 | 0,79 | 4,32E-01 |
| AT2G36790 | 0,34 | -0,46 | 4,23 | -0,11 | 9,13E-01 |
| AT2G36800 | 0,85 | 3,30 | 2,28 | 1,45 | 1,48E-01 |
| AT2G36810 | 1,10 | 1,99 | 2,00 | 1,00 | 3,19E-01 |
| AT2G36835 | 0,30 | -0,34 | 3,57 | -0,10 | 9,24E-01 |
| AT2G36895 | 0,35 | 0,00 | 4,41 | 0,00 | 1,00E+00 |
| AT2G36900 | 0,55 | 2,16 | 3,74 | 0,58 | 5,63E-01 |
| AT2G36930 | 0,54 | 1,15 | 3,86 | 0,30 | 7,66E-01 |
| AT2G36990 | 1,19 | 1,50 | 2,58 | 0,58 | 5,62E-01 |
| AT2G37035 | 0,78 | 2,01 | 2,27 | 0,88 | 3,77E-01 |
| AT2G37040 | 1,58 | 2,38 | 1,49 | 1,59 | 1,11E-01 |
| AT2G37080 | 0,12 | 2,69 | 4,35 | 0,62 | 5,36E-01 |
| AT2G37130 | 0,68 | -0,16 | 2,94 | -0,05 | 9,57E-01 |
| AT2G37150 | 0,75 | 3,90 | 3,41 | 1,14 | 2,53E-01 |
| AT2G37170 | 0,90 | 3,47 | 2,27 | 1,52 | 1,27E-01 |
| AT2G37200 | 0,31 | 1,18 | 3,52 | 0,33 | 7,38E-01 |
| AT2G37230 | 0,56 | 0,80 | 3,08 | 0,26 | 7,96E-01 |
| AT2G37310 | 0,77 | 4,45 | 3,47 | 1,28 | 2,00E-01 |
| AT2G37340 | 1,12 | 3,43 | 2,24 | 1,53 | 1,26E-01 |
| AT2G37360 | 0,21 | 3,40 | 4,32 | 0,79 | 4,32E-01 |
| AT2G37400 | 0,25 | 2,79 | 4,34 | 0,64 | 5,20E-01 |
| AT2G37410 | 0,19 | 2,25 | 4,32 | 0,52 | 6,02E-01 |
| AT2G37520 | 1,03 | 0,89 | 1,70 | 0,53 | 5,99E-01 |
| AT2G37540 | 0,69 | 4,04 | 2,86 | 1,41 | 1,58E-01 |
| AT2G37550 | 1,38 | 2,98 | 1,77 | 1,68 | 9,28E-02 |
| AT2G37570 | 0,98 | 1,29 | 1,72 | 0,75 | 4,55E-01 |
| AT2G37630 | 0,98 | 1,91 | 1,88 | 1,02 | 3,09E-01 |
| AT2G37660 | 1,35 | -1,21 | 1,63 | -0,75 | 4,56E-01 |
| AT2G37678 | 0,57 | 0,97 | 4,38 | 0,22 | 8,25E-01 |
| AT2G08895 | 1,00 | 0,24 | 2,02 | 0,12 | 9,05E-01 |
| AT2G08900 | 0,86 | -0,13 | 1,82 | -0,07 | 9,44E-01 |
| AT2G08910 | 1,52 | -3,24 | 1,58 | -2,05 | 4,06E-02 |
| AT2G08915 | 0,73 | 0,49 | 2,60 | 0,19 | 8,49E-01 |
| AT2G38000 | 0,32 | -0,62 | 4,32 | -0,14 | 8,86E-01 |
| AT2G38020 | 1,29 | 2,52 | 2,22 | 1,14 | 2,55E-01 |
| AT2G38160 | 0,87 | 1,69 | 3,35 | 0,51 | 6,13E-01 |
| AT2G38270 | 0,22 | -1,85 | 4,32 | -0,43 | 6,69E-01 |

| | | | | | |
|---|---|---|---|---|---|
| AT2G38290 | 0,66 | 2,92 | 2,84 | 1,03 | 3,05E-01 |
| AT2G38340 | 0,17 | 0,69 | 4,34 | 0,16 | 8,73E-01 |
| AT2G38360 | 0,39 | -0,80 | 3,26 | -0,25 | 8,05E-01 |
| AT2G38410 | 0,35 | 1,98 | 3,41 | 0,58 | 5,61E-01 |
| AT2G38480 | 0,17 | 2,04 | 4,33 | 0,47 | 6,37E-01 |
| AT2G38610 | 0,83 | 3,82 | 2,38 | 1,60 | 1,09E-01 |
| AT2G38650 | 1,16 | 1,74 | 1,93 | 0,90 | 3,66E-01 |
| AT2G38680 | 0,08 | -0,56 | 4,38 | -0,13 | 8,99E-01 |
| AT2G38750 | 0,05 | -0,58 | 4,38 | -0,13 | 8,95E-01 |
| AT2G38770 | 1,53 | 2,09 | 1,54 | 1,35 | 1,77E-01 |
| AT2G38790 | 0,55 | -0,27 | 2,20 | -0,12 | 9,01E-01 |
| AT2G38860 | 0,41 | 1,14 | 3,33 | 0,34 | 7,32E-01 |
| AT2G39000 | 1,06 | 4,29 | 2,87 | 1,50 | 1,35E-01 |
| AT2G39080 | 0,20 | 2,11 | 4,37 | 0,48 | 6,29E-01 |
| AT2G39130 | 0,08 | -0,56 | 4,38 | -0,13 | 8,99E-01 |
| AT2G39160 | 0,24 | 2,08 | 4,31 | 0,48 | 6,29E-01 |
| AT2G39170 | 0,19 | 0,09 | 4,37 | 0,02 | 9,83E-01 |
| AT2G09010 | 0,08 | -0,56 | 4,38 | -0,13 | 8,99E-01 |
| AT2G39210 | 0,91 | 3,89 | 2,72 | 1,43 | 1,52E-01 |
| AT2G39260 | 1,36 | 1,25 | 1,42 | 0,88 | 3,77E-01 |
| AT2G39350 | 0,49 | 1,74 | 3,13 | 0,56 | 5,78E-01 |
| AT2G39360 | 0,80 | 2,81 | 3,50 | 0,80 | 4,22E-01 |
| AT2G09045 | 0,52 | 3,80 | 3,86 | 0,98 | 3,25E-01 |
| AT2G39450 | 0,29 | 2,99 | 4,34 | 0,69 | 4,90E-01 |
| AT2G39480 | 1,05 | 0,16 | 1,90 | 0,08 | 9,35E-01 |
| AT2G39650 | 0,49 | 0,33 | 3,91 | 0,08 | 9,33E-01 |
| AT2G39710 | 0,60 | 1,32 | 3,13 | 0,42 | 6,74E-01 |
| AT2G39720 | 1,30 | 2,83 | 2,33 | 1,21 | 2,25E-01 |
| AT2G39750 | 1,24 | 0,69 | 1,93 | 0,36 | 7,21E-01 |
| AT2G09080 | 0,50 | 2,50 | 3,21 | 0,78 | 4,35E-01 |
| AT2G39805 | 0,30 | 2,91 | 4,34 | 0,67 | 5,04E-01 |
| AT2G39920 | 0,18 | 1,26 | 4,36 | 0,29 | 7,73E-01 |
| AT2G09090 | 0,82 | 2,64 | 2,89 | 0,91 | 3,61E-01 |
| AT2G39950 | 0,75 | 2,51 | 2,56 | 0,98 | 3,27E-01 |
| AT2G39960 | 0,25 | 2,44 | 4,35 | 0,56 | 5,74E-01 |
| AT2G39990 | 1,46 | 2,13 | 2,55 | 0,84 | 4,03E-01 |
| AT2G40070 | 0,52 | 3,95 | 3,86 | 1,02 | 3,06E-01 |
| AT2G40080 | 0,23 | 0,75 | 4,27 | 0,17 | 8,61E-01 |
| AT2G40165 | 0,38 | 2,13 | 4,38 | 0,49 | 6,27E-01 |
| AT2G40205 | 0,27 | -1,50 | 4,34 | -0,35 | 7,29E-01 |
| AT2G40270 | 1,34 | 1,20 | 1,39 | 0,86 | 3,89E-01 |
| AT2G40280 | 0,63 | 2,77 | 2,99 | 0,93 | 3,54E-01 |
| AT2G40350 | 0,12 | 2,87 | 4,34 | 0,66 | 5,08E-01 |
| AT2G40360 | 0,28 | -0,99 | 4,27 | -0,23 | 8,17E-01 |
| AT2G40430 | 0,45 | -1,16 | 4,17 | -0,28 | 7,80E-01 |
| AT2G40475 | 0,89 | -0,64 | 2,02 | -0,32 | 7,51E-01 |
| AT2G40590 | 0,50 | 0,36 | 2,70 | 0,13 | 8,94E-01 |
| AT2G40600 | 0,11 | 1,28 | 4,35 | 0,29 | 7,68E-01 |
| AT2G40610 | 0,37 | -0,10 | 4,40 | -0,02 | 9,82E-01 |
| AT2G40650 | 0,77 | -0,90 | 2,04 | -0,44 | 6,60E-01 |
| AT2G40730 | 0,34 | -0,68 | 3,40 | -0,20 | 8,41E-01 |
| AT2G40750 | 0,46 | 2,24 | 4,16 | 0,54 | 5,91E-01 |
| AT2G40770 | 0,51 | -0,51 | 4,02 | -0,13 | 9,00E-01 |
| AT2G40815 | 0,24 | 2,85 | 4,34 | 0,66 | 5,11E-01 |
| AT2G40860 | 0,30 | 2,91 | 4,34 | 0,67 | 5,04E-01 |
| AT2G40890 | 0,53 | 2,36 | 3,82 | 0,62 | 5,37E-01 |

| | | | | | |
|---|---|---|---|---|---|
| AT2G40920 | 0,35 | 2,27 | 4,31 | 0,53 | 5,99E-01 |
| AT2G40925 | 0,65 | 3,20 | 3,66 | 0,87 | 3,82E-01 |
| AT2G40950 | 0,74 | 3,32 | 3,46 | 0,96 | 3,37E-01 |
| AT2G40970 | 0,42 | 0,00 | 4,41 | 0,00 | 1,00E+00 |
| AT2G09225 | 0,24 | 1,12 | 4,33 | 0,26 | 7,96E-01 |
| AT2G41120 | 0,21 | 3,40 | 4,32 | 0,79 | 4,32E-01 |
| AT2G41160 | 0,39 | 2,44 | 4,32 | 0,56 | 5,72E-01 |
| AT2G41250 | 0,87 | 0,30 | 2,18 | 0,14 | 8,92E-01 |
| AT2G41420 | 0,11 | -1,28 | 4,35 | -0,29 | 7,69E-01 |
| AT2G41475 | 0,71 | 0,03 | 2,47 | 0,01 | 9,91E-01 |
| AT2G41500 | 0,96 | 2,47 | 2,11 | 1,17 | 2,43E-01 |
| AT2G41600 | 0,58 | 0,87 | 2,54 | 0,34 | 7,33E-01 |
| AT2G41620 | 1,68 | 3,84 | 1,81 | 2,11 | 3,45E-02 |
| AT2G41616 | 0,33 | 3,34 | 4,32 | 0,77 | 4,40E-01 |
| AT2G41630 | 0,92 | 0,35 | 2,80 | 0,12 | 9,02E-01 |
| AT2G41680 | 0,32 | 1,34 | 4,25 | 0,31 | 7,53E-01 |
| AT2G41700 | 1,62 | 1,83 | 1,81 | 1,01 | 3,13E-01 |
| AT2G41770 | 0,40 | 0,80 | 4,24 | 0,19 | 8,50E-01 |
| AT2G41790 | 0,58 | 0,88 | 2,50 | 0,35 | 7,26E-01 |
| AT2G41830 | 0,40 | 3,41 | 4,15 | 0,82 | 4,11E-01 |
| AT2G41835 | 0,70 | 2,46 | 3,64 | 0,68 | 4,99E-01 |
| AT2G41870 | 0,29 | 1,86 | 4,33 | 0,43 | 6,67E-01 |
| AT2G41890 | 0,43 | -0,22 | 4,11 | -0,05 | 9,58E-01 |
| AT2G41945 | 0,60 | 2,67 | 3,87 | 0,69 | 4,91E-01 |
| AT2G41960 | 0,92 | 2,89 | 1,92 | 1,50 | 1,33E-01 |
| AT2G42010 | 0,72 | -0,05 | 1,90 | -0,02 | 9,80E-01 |
| AT2G42020 | 0,48 | -0,80 | 4,36 | -0,18 | 8,54E-01 |
| AT2G42070 | 0,23 | -0,22 | 4,28 | -0,05 | 9,60E-01 |
| AT2G42230 | 0,42 | 2,55 | 4,09 | 0,62 | 5,34E-01 |
| AT2G42280 | 0,33 | 2,56 | 4,35 | 0,59 | 5,57E-01 |
| AT2G42365 | 0,35 | 0,00 | 4,41 | 0,00 | 1,00E+00 |
| AT2G42400 | 1,01 | 3,01 | 2,07 | 1,45 | 1,47E-01 |
| AT2G42500 | 0,82 | 0,13 | 2,16 | 0,06 | 9,50E-01 |
| AT2G42530 | 0,98 | -0,28 | 2,11 | -0,13 | 8,94E-01 |
| AT2G42690 | 1,34 | 0,25 | 1,36 | 0,18 | 8,54E-01 |
| AT2G42760 | 0,47 | -1,22 | 2,50 | -0,49 | 6,25E-01 |
| AT2G42790 | 0,53 | 1,97 | 2,67 | 0,74 | 4,60E-01 |
| AT2G42800 | 0,18 | -1,99 | 4,31 | -0,46 | 6,44E-01 |
| AT2G42810 | 0,84 | 1,61 | 1,93 | 0,83 | 4,06E-01 |
| AT2G42840 | 0,22 | -2,03 | 4,31 | -0,47 | 6,37E-01 |
| AT2G42865 | 0,08 | 2,33 | 4,36 | 0,53 | 5,93E-01 |
| AT2G42870 | 0,10 | 0,39 | 4,35 | 0,09 | 9,29E-01 |
| AT2G42880 | 0,41 | 0,75 | 3,36 | 0,22 | 8,23E-01 |
| AT2G43180 | 0,70 | 3,08 | 2,90 | 1,06 | 2,89E-01 |
| AT2G43260 | 0,20 | 2,30 | 4,36 | 0,53 | 5,98E-01 |
| AT2G43290 | 0,50 | 1,70 | 3,08 | 0,55 | 5,82E-01 |
| AT2G43320 | 0,22 | 1,81 | 4,30 | 0,42 | 6,74E-01 |
| AT2G43360 | 0,33 | 2,93 | 4,33 | 0,68 | 4,98E-01 |
| AT2G43420 | 0,11 | 1,28 | 4,35 | 0,29 | 7,68E-01 |
| AT2G43430 | 0,59 | -1,31 | 2,63 | -0,50 | 6,18E-01 |
| AT2G43460 | 0,41 | -0,48 | 3,36 | -0,14 | 8,87E-01 |
| AT2G43490 | 0,44 | 2,87 | 4,05 | 0,71 | 4,78E-01 |
| AT2G43560 | 0,25 | 1,15 | 4,34 | 0,27 | 7,91E-01 |
| AT2G43750 | 1,46 | 0,75 | 1,44 | 0,52 | 6,04E-01 |
| AT2G43865 | 0,15 | -0,19 | 4,31 | -0,04 | 9,64E-01 |
| AT2G43910 | 0,26 | 1,11 | 3,70 | 0,30 | 7,63E-01 |

| | | | | | |
|---|---|---|---|---|---|
| AT2G43940 | 0,08 | 2,08 | 4,38 | 0,47 | 6,35E-01 |
| AT2G43945 | 0,47 | 1,77 | 4,40 | 0,40 | 6,88E-01 |
| AT2G43950 | 0,24 | 2,80 | 4,34 | 0,65 | 5,18E-01 |
| AT2G44090 | 0,69 | 2,13 | 2,61 | 0,81 | 4,15E-01 |
| AT2G44140 | 0,91 | -1,14 | 2,12 | -0,54 | 5,91E-01 |
| AT2G44180 | 0,46 | -0,23 | 3,18 | -0,07 | 9,42E-01 |
| AT2G44255 | 0,20 | 1,94 | 4,34 | 0,45 | 6,55E-01 |
| AT2G44360 | 0,63 | 0,96 | 4,27 | 0,22 | 8,22E-01 |
| AT2G44610 | 0,48 | 1,60 | 4,12 | 0,39 | 6,98E-01 |
| AT2G44660 | 0,35 | 1,44 | 3,41 | 0,42 | 6,72E-01 |
| AT2G44760 | 0,32 | 1,31 | 3,51 | 0,37 | 7,10E-01 |
| AT2G44798 | 0,36 | 0,04 | 4,32 | 0,01 | 9,93E-01 |
| AT2G44950 | 0,77 | 2,37 | 2,23 | 1,06 | 2,87E-01 |
| AT2G45070 | 0,80 | -0,51 | 1,97 | -0,26 | 7,94E-01 |
| AT2G45220 | 0,54 | 3,99 | 4,32 | 0,92 | 3,55E-01 |
| AT2G45315 | 0,67 | 0,35 | 2,33 | 0,15 | 8,82E-01 |
| AT2G45460 | 0,20 | -1,01 | 4,30 | -0,23 | 8,15E-01 |
| AT2G45470 | 1,65 | -2,06 | 1,99 | -1,04 | 3,00E-01 |
| AT2G45520 | 0,22 | -0,39 | 4,32 | -0,09 | 9,28E-01 |
| AT2G09640 | 1,68 | 1,39 | 1,96 | 0,71 | 4,79E-01 |
| AT2G45560 | 0,47 | 0,00 | 4,41 | 0,00 | 1,00E+00 |
| AT2G45660 | 0,55 | 0,14 | 2,13 | 0,06 | 9,49E-01 |
| AT2G45670 | 0,95 | 4,47 | 2,27 | 1,97 | 4,89E-02 |
| AT2G45700 | 0,38 | 2,04 | 4,29 | 0,48 | 6,34E-01 |
| AT2G45820 | 0,98 | -0,63 | 1,75 | -0,36 | 7,17E-01 |
| AT2G45970 | 0,56 | 1,27 | 2,51 | 0,50 | 6,14E-01 |
| AT2G45980 | 0,45 | 1,91 | 4,03 | 0,47 | 6,35E-01 |
| AT2G46030 | 0,57 | 3,85 | 4,31 | 0,89 | 3,72E-01 |
| AT2G46040 | 0,20 | 2,11 | 4,37 | 0,48 | 6,29E-01 |
| AT2G46060 | 0,49 | 0,00 | 4,41 | 0,00 | 1,00E+00 |
| AT2G46080 | 0,51 | 0,10 | 3,19 | 0,03 | 9,76E-01 |
| AT2G46180 | 0,35 | 0,15 | 4,31 | 0,03 | 9,73E-01 |
| AT2G46280 | 1,16 | 2,46 | 1,94 | 1,27 | 2,03E-01 |
| AT2G46330 | 0,93 | -0,04 | 1,94 | -0,02 | 9,82E-01 |
| AT2G46390 | 0,46 | 3,10 | 4,07 | 0,76 | 4,46E-01 |
| AT2G46505 | 0,89 | 0,06 | 2,35 | 0,03 | 9,80E-01 |
| AT2G46510 | 1,07 | -1,92 | 1,92 | -1,00 | 3,17E-01 |
| AT2G46540 | 0,12 | -0,77 | 4,36 | -0,18 | 8,61E-01 |
| AT2G46550 | 0,57 | 0,94 | 2,71 | 0,35 | 7,28E-01 |
| AT2G46572 | 0,51 | 2,87 | 3,97 | 0,72 | 4,70E-01 |
| AT2G46610 | 0,85 | -1,10 | 1,95 | -0,56 | 5,73E-01 |
| AT2G46620 | 1,45 | 1,41 | 1,60 | 0,88 | 3,78E-01 |
| AT2G46680 | 0,94 | 1,17 | 2,19 | 0,53 | 5,94E-01 |
| AT2G46840 | 0,08 | 2,33 | 4,36 | 0,53 | 5,93E-01 |
| AT2G46920 | 0,44 | 2,92 | 3,99 | 0,73 | 4,64E-01 |
| AT2G46980 | 0,24 | 0,00 | 4,41 | 0,00 | 1,00E+00 |
| AT2G47090 | 0,25 | 3,58 | 4,32 | 0,83 | 4,07E-01 |
| AT2G47180 | 1,41 | 0,41 | 1,64 | 0,25 | 8,03E-01 |
| AT2G47250 | 1,15 | 4,61 | 2,27 | 2,03 | 4,20E-02 |
| AT2G47330 | 0,60 | 1,05 | 3,03 | 0,35 | 7,29E-01 |
| AT2G47600 | 0,61 | 1,95 | 2,44 | 0,80 | 4,23E-01 |
| AT2G47650 | 1,46 | 2,63 | 1,65 | 1,59 | 1,11E-01 |
| AT2G47680 | 0,89 | 1,30 | 1,96 | 0,66 | 5,08E-01 |
| AT2G47700 | 0,93 | 0,93 | 2,26 | 0,41 | 6,82E-01 |
| AT2G47840 | 0,52 | -0,40 | 3,29 | -0,12 | 9,03E-01 |
| AT2G47990 | 0,88 | 3,14 | 2,69 | 1,17 | 2,43E-01 |

| | | | | | |
|---|---|---|---|---|---|
| AT2G48000 | 1,00 | 4,84 | 3,32 | 1,46 | 1,45E-01 |
| AT2G48030 | 0,73 | 2,60 | 2,31 | 1,12 | 2,62E-01 |
| AT2G48130 | 0,91 | -1,19 | 1,97 | -0,61 | 5,45E-01 |
| AT3G01090 | 0,63 | -0,19 | 1,98 | -0,10 | 9,22E-01 |
| AT3G01100 | 0,90 | 0,82 | 2,12 | 0,39 | 6,97E-01 |
| AT3G01130 | 0,21 | -0,24 | 4,35 | -0,06 | 9,56E-01 |
| AT3G01160 | 0,24 | 2,08 | 4,31 | 0,48 | 6,29E-01 |
| AT3G01180 | 0,84 | 1,19 | 1,77 | 0,67 | 5,01E-01 |
| AT3G01300 | 0,08 | 2,33 | 4,36 | 0,53 | 5,93E-01 |
| AT3G01340 | 0,50 | 2,28 | 3,15 | 0,72 | 4,69E-01 |
| AT3G01380 | 1,09 | 2,67 | 2,00 | 1,34 | 1,81E-01 |
| AT3G01390 | 0,38 | 0,98 | 4,34 | 0,23 | 8,21E-01 |
| AT3G01415 | 0,40 | -1,85 | 4,30 | -0,43 | 6,68E-01 |
| AT3G01420 | 0,31 | 2,17 | 4,31 | 0,50 | 6,14E-01 |
| AT3G01430 | 0,79 | 1,02 | 2,89 | 0,35 | 7,25E-01 |
| AT3G01470 | 1,49 | 1,94 | 1,49 | 1,30 | 1,93E-01 |
| AT3G01550 | 0,10 | 0,34 | 4,35 | 0,08 | 9,37E-01 |
| AT3G01720 | 0,80 | 2,40 | 2,76 | 0,87 | 3,85E-01 |
| AT3G01820 | 0,81 | 3,31 | 3,58 | 0,92 | 3,56E-01 |
| AT3G01840 | 0,88 | 1,44 | 2,14 | 0,68 | 4,99E-01 |
| AT3G01910 | 0,46 | 2,63 | 4,04 | 0,65 | 5,15E-01 |
| AT3G01930 | 0,62 | 2,55 | 3,02 | 0,84 | 3,98E-01 |
| AT3G01980 | 0,16 | 1,97 | 4,33 | 0,45 | 6,50E-01 |
| AT3G02040 | 0,68 | 0,38 | 2,36 | 0,16 | 8,72E-01 |
| AT3G02080 | 1,20 | 1,22 | 2,04 | 0,60 | 5,50E-01 |
| AT3G02190 | 0,44 | 0,58 | 3,31 | 0,17 | 8,61E-01 |
| AT3G02210 | 0,52 | 1,87 | 3,96 | 0,47 | 6,36E-01 |
| AT3G02250 | 0,34 | -0,38 | 4,36 | -0,09 | 9,31E-01 |
| AT3G02380 | 0,91 | 1,52 | 2,22 | 0,68 | 4,95E-01 |
| AT3G02410 | 0,66 | 0,00 | 4,41 | 0,00 | 1,00E+00 |
| AT3G02450 | 0,68 | 0,23 | 2,23 | 0,10 | 9,20E-01 |
| AT3G02520 | 1,11 | 1,09 | 1,57 | 0,69 | 4,88E-01 |
| AT3G02530 | 0,67 | 1,18 | 2,54 | 0,46 | 6,43E-01 |
| AT3G02700 | 0,39 | 0,84 | 4,21 | 0,20 | 8,42E-01 |
| AT3G02730 | 0,92 | -0,21 | 2,91 | -0,07 | 9,43E-01 |
| AT3G02750 | 1,16 | 1,25 | 1,83 | 0,68 | 4,95E-01 |
| AT3G02760 | 0,94 | 3,10 | 2,69 | 1,15 | 2,50E-01 |
| AT3G01205 | 0,80 | 0,17 | 2,16 | 0,08 | 9,37E-01 |
| AT3G03090 | 0,29 | 0,00 | 4,41 | 0,00 | 1,00E+00 |
| AT3G03150 | 0,22 | -1,61 | 4,34 | -0,37 | 7,10E-01 |
| AT3G03220 | 0,16 | -1,98 | 4,31 | -0,46 | 6,47E-01 |
| AT3G03250 | 1,00 | 0,44 | 1,58 | 0,28 | 7,82E-01 |
| AT3G03300 | 0,33 | 3,85 | 4,32 | 0,89 | 3,72E-01 |
| AT3G03305 | 0,32 | 1,93 | 4,34 | 0,44 | 6,57E-01 |
| AT3G03310 | 0,29 | 3,04 | 4,33 | 0,70 | 4,83E-01 |
| AT3G03330 | 0,30 | -0,29 | 4,36 | -0,07 | 9,48E-01 |
| AT3G03405 | 0,16 | -1,98 | 4,31 | -0,46 | 6,47E-01 |
| AT3G03460 | 0,17 | 2,62 | 4,35 | 0,60 | 5,47E-01 |
| AT3G03470 | 1,68 | 1,75 | 1,50 | 1,17 | 2,44E-01 |
| AT3G03480 | 0,08 | 2,33 | 4,36 | 0,53 | 5,93E-01 |
| AT3G03550 | 0,10 | 0,34 | 4,35 | 0,08 | 9,37E-01 |
| AT3G03560 | 0,61 | 3,08 | 2,94 | 1,05 | 2,94E-01 |
| AT3G03580 | 0,26 | 2,30 | 4,37 | 0,53 | 5,98E-01 |
| AT3G03590 | 0,08 | 2,33 | 4,36 | 0,53 | 5,93E-01 |
| AT3G03600 | 0,30 | -1,36 | 4,35 | -0,31 | 7,54E-01 |
| AT3G03720 | 0,26 | 0,99 | 4,26 | 0,23 | 8,17E-01 |

| | | | | | |
|---|---|---|---|---|---|
| AT3G03770 | 0,99 | 4,29 | 3,01 | 1,42 | 1,54E-01 |
| AT3G03890 | 0,79 | 1,31 | 2,33 | 0,56 | 5,75E-01 |
| AT3G03920 | 0,42 | -0,90 | 3,35 | -0,27 | 7,89E-01 |
| AT3G03940 | 0,41 | -0,79 | 2,68 | -0,30 | 7,68E-01 |
| AT3G04030 | 0,25 | -2,53 | 4,29 | -0,59 | 5,55E-01 |
| AT3G04080 | 1,01 | 3,00 | 2,01 | 1,49 | 1,37E-01 |
| AT3G04150 | 0,26 | 2,30 | 4,37 | 0,53 | 5,98E-01 |
| AT3G04200 | 0,62 | 3,54 | 4,32 | 0,82 | 4,13E-01 |
| AT3G04220 | 0,33 | 2,17 | 4,29 | 0,51 | 6,13E-01 |
| AT3G04260 | 0,61 | -0,04 | 4,39 | -0,01 | 9,94E-01 |
| AT3G04290 | 0,84 | 2,30 | 3,03 | 0,76 | 4,48E-01 |
| AT3G04350 | 0,35 | -0,88 | 4,23 | -0,21 | 8,36E-01 |
| AT3G04480 | 0,84 | 1,87 | 2,18 | 0,86 | 3,90E-01 |
| AT3G04500 | 0,20 | -0,80 | 4,37 | -0,18 | 8,55E-01 |
| AT3G04580 | 1,03 | 3,25 | 2,03 | 1,60 | 1,09E-01 |
| AT3G04640 | 0,85 | -1,41 | 1,95 | -0,72 | 4,72E-01 |
| AT3G04690 | 0,22 | -0,76 | 4,38 | -0,17 | 8,63E-01 |
| AT3G04720 | 0,50 | 0,21 | 2,92 | 0,07 | 9,42E-01 |
| AT3G04730 | 1,02 | 0,34 | 1,56 | 0,22 | 8,29E-01 |
| AT3G04750 | 0,57 | 0,79 | 2,59 | 0,30 | 7,61E-01 |
| AT3G04760 | 0,89 | 1,38 | 1,79 | 0,77 | 4,41E-01 |
| AT3G04860 | 0,12 | 2,69 | 4,35 | 0,62 | 5,36E-01 |
| AT3G04980 | 0,46 | 3,53 | 4,33 | 0,81 | 4,15E-01 |
| AT3G05030 | 0,28 | 0,97 | 4,27 | 0,23 | 8,20E-01 |
| AT3G01585 | 0,10 | -1,26 | 4,35 | -0,29 | 7,72E-01 |
| AT3G05040 | 1,58 | 1,47 | 1,51 | 0,97 | 3,32E-01 |
| AT3G05050 | 0,38 | 2,27 | 4,33 | 0,53 | 6,00E-01 |
| AT3G05060 | 0,60 | 2,55 | 3,66 | 0,70 | 4,86E-01 |
| AT3G05090 | 0,52 | -0,01 | 2,45 | 0,00 | 9,98E-01 |
| AT3G05165 | 0,83 | 0,38 | 1,85 | 0,21 | 8,37E-01 |
| AT3G05240 | 0,13 | 2,37 | 4,36 | 0,54 | 5,86E-01 |
| AT3G05270 | 0,58 | 1,88 | 2,58 | 0,73 | 4,67E-01 |
| AT3G05280 | 0,59 | 1,64 | 3,82 | 0,43 | 6,69E-01 |
| AT3G05540 | 1,04 | -0,80 | 1,75 | -0,46 | 6,47E-01 |
| AT3G05650 | 0,98 | 1,88 | 2,09 | 0,90 | 3,68E-01 |
| AT3G05660 | 0,79 | 3,13 | 2,65 | 1,18 | 2,38E-01 |
| AT3G05680 | 1,68 | 2,50 | 1,42 | 1,76 | 7,88E-02 |
| AT3G05690 | 0,11 | -1,28 | 4,35 | -0,29 | 7,69E-01 |
| AT3G05770 | 0,46 | -0,17 | 3,18 | -0,05 | 9,58E-01 |
| AT3G05810 | 0,21 | 1,12 | 4,33 | 0,26 | 7,95E-01 |
| AT3G05850 | 0,90 | 0,69 | 2,43 | 0,29 | 7,76E-01 |
| AT3G05880 | 0,55 | 0,90 | 3,20 | 0,28 | 7,78E-01 |
| AT3G05900 | 0,51 | -0,21 | 3,23 | -0,07 | 9,48E-01 |
| AT3G05910 | 0,68 | 0,09 | 2,66 | 0,03 | 9,74E-01 |
| AT3G05932 | 0,27 | 2,08 | 4,31 | 0,48 | 6,29E-01 |
| AT3G05970 | 0,71 | -1,14 | 2,19 | -0,52 | 6,04E-01 |
| AT3G06030 | 0,29 | 3,75 | 4,32 | 0,87 | 3,85E-01 |
| AT3G06050 | 0,18 | -1,50 | 4,34 | -0,35 | 7,30E-01 |
| AT3G06060 | 0,17 | -0,44 | 4,37 | -0,10 | 9,19E-01 |
| AT3G06080 | 1,38 | -1,87 | 1,54 | -1,22 | 2,24E-01 |
| AT3G06150 | 0,98 | 0,36 | 1,58 | 0,23 | 8,21E-01 |
| AT3G06190 | 0,71 | -0,34 | 3,04 | -0,11 | 9,10E-01 |
| AT3G06250 | 0,74 | 2,48 | 2,80 | 0,89 | 3,75E-01 |
| AT3G06290 | 0,72 | 1,78 | 2,30 | 0,77 | 4,41E-01 |
| AT3G06310 | 0,08 | 2,08 | 4,38 | 0,47 | 6,35E-01 |
| AT3G06350 | 0,64 | 3,83 | 2,96 | 1,29 | 1,97E-01 |

| | | | | | |
|---|---|---|---|---|---|
| AT3G06430 | 0,12 | 2,87 | 4,34 | 0,66 | 5,08E-01 |
| AT3G06440 | 0,12 | 2,69 | 4,35 | 0,62 | 5,36E-01 |
| AT3G06450 | 0,61 | 1,42 | 2,54 | 0,56 | 5,75E-01 |
| AT3G06480 | 1,05 | 1,35 | 1,83 | 0,74 | 4,61E-01 |
| AT3G06483 | 0,45 | -1,23 | 2,70 | -0,46 | 6,49E-01 |
| AT3G06540 | 0,08 | 2,08 | 4,38 | 0,47 | 6,35E-01 |
| AT3G06550 | 0,32 | -0,08 | 4,30 | -0,02 | 9,86E-01 |
| AT3G06580 | 0,15 | 1,93 | 4,33 | 0,45 | 6,55E-01 |
| AT3G06590 | 0,40 | -0,77 | 4,37 | -0,18 | 8,60E-01 |
| AT3G06620 | 0,12 | 2,87 | 4,34 | 0,66 | 5,08E-01 |
| AT3G06665 | 0,25 | -2,53 | 4,29 | -0,59 | 5,55E-01 |
| AT3G06670 | 1,42 | 2,68 | 1,63 | 1,65 | 9,93E-02 |
| AT3G06680 | 0,28 | -1,48 | 4,35 | -0,34 | 7,33E-01 |
| AT3G06690 | 0,20 | -0,80 | 4,37 | -0,18 | 8,55E-01 |
| AT3G06700 | 0,57 | -0,29 | 2,65 | -0,11 | 9,14E-01 |
| AT3G06790 | 0,16 | 1,16 | 4,38 | 0,27 | 7,90E-01 |
| AT3G06820 | 0,08 | 2,33 | 4,36 | 0,53 | 5,93E-01 |
| AT3G06870 | 0,21 | -1,08 | 4,36 | -0,25 | 8,04E-01 |
| AT3G06900 | 1,19 | -0,70 | 1,69 | -0,41 | 6,79E-01 |
| AT3G06930 | 0,40 | 1,22 | 4,18 | 0,29 | 7,70E-01 |
| AT3G06960 | 0,89 | -1,40 | 2,47 | -0,57 | 5,71E-01 |
| AT3G07010 | 0,41 | 0,17 | 4,24 | 0,04 | 9,68E-01 |
| AT3G07020 | 0,54 | 3,68 | 3,76 | 0,98 | 3,27E-01 |
| AT3G07040 | 1,04 | 3,84 | 2,14 | 1,80 | 7,26E-02 |
| AT3G07050 | 0,53 | -0,25 | 3,16 | -0,08 | 9,37E-01 |
| AT3G07115 | 0,08 | -0,56 | 4,38 | -0,13 | 8,99E-01 |
| AT3G07180 | 0,12 | 2,87 | 4,34 | 0,66 | 5,08E-01 |
| AT3G07215 | 0,55 | -2,17 | 2,48 | -0,88 | 3,81E-01 |
| AT3G07273 | 0,08 | 2,33 | 4,36 | 0,53 | 5,93E-01 |
| AT3G07290 | 0,69 | 4,41 | 3,46 | 1,27 | 2,03E-01 |
| AT3G07300 | 0,29 | 2,14 | 4,36 | 0,49 | 6,23E-01 |
| AT3G07330 | 0,98 | 1,06 | 2,30 | 0,46 | 6,46E-01 |
| AT3G07380 | 1,22 | -1,61 | 1,46 | -1,10 | 2,71E-01 |
| AT3G07460 | 0,63 | -1,28 | 2,55 | -0,50 | 6,15E-01 |
| AT3G07470 | 0,58 | 0,51 | 3,20 | 0,16 | 8,74E-01 |
| AT3G07520 | 0,51 | 0,81 | 3,10 | 0,26 | 7,93E-01 |
| AT3G07540 | 0,08 | 2,33 | 4,36 | 0,53 | 5,93E-01 |
| AT3G07660 | 1,40 | 2,06 | 1,74 | 1,18 | 2,38E-01 |
| AT3G07700 | 0,94 | 1,14 | 1,63 | 0,70 | 4,85E-01 |
| AT3G07800 | 0,53 | 1,55 | 2,63 | 0,59 | 5,55E-01 |
| AT3G07890 | 0,34 | 0,88 | 4,26 | 0,21 | 8,36E-01 |
| AT3G08570 | 1,10 | 4,41 | 2,14 | 2,06 | 3,94E-02 |
| AT3G08590 | 1,52 | 1,42 | 1,54 | 0,92 | 3,55E-01 |
| AT3G08610 | 0,59 | 1,17 | 3,19 | 0,37 | 7,14E-01 |
| AT3G08650 | 0,24 | 0,81 | 4,34 | 0,19 | 8,53E-01 |
| AT3G08720 | 1,37 | 2,37 | 1,74 | 1,36 | 1,73E-01 |
| AT3G08760 | 0,70 | 0,94 | 2,12 | 0,44 | 6,58E-01 |
| AT3G08840 | 0,67 | 2,11 | 3,02 | 0,70 | 4,84E-01 |
| AT3G08870 | 0,75 | 4,10 | 3,42 | 1,20 | 2,30E-01 |
| AT3G08960 | 0,38 | -0,45 | 4,25 | -0,11 | 9,16E-01 |
| AT3G09040 | 0,17 | 3,14 | 4,33 | 0,73 | 4,68E-01 |
| AT3G09070 | 1,31 | -0,01 | 1,89 | -0,01 | 9,96E-01 |
| AT3G09090 | 1,29 | 3,63 | 1,88 | 1,93 | 5,39E-02 |
| AT3G09190 | 0,55 | 0,79 | 3,74 | 0,21 | 8,32E-01 |
| AT3G09210 | 0,27 | 0,00 | 4,41 | 0,00 | 1,00E+00 |
| AT3G09270 | 0,53 | 0,85 | 2,57 | 0,33 | 7,40E-01 |

| | | | | | |
|---|---|---|---|---|---|
| AT3G09320 | 0,85 | 2,57 | 2,22 | 1,16 | 2,47E-01 |
| AT3G09410 | 0,82 | 3,65 | 2,39 | 1,53 | 1,27E-01 |
| AT3G09470 | 0,82 | 3,41 | 3,41 | 1,00 | 3,17E-01 |
| AT3G09560 | 0,97 | 2,12 | 2,15 | 0,99 | 3,24E-01 |
| AT3G09580 | 0,48 | 0,46 | 3,35 | 0,14 | 8,91E-01 |
| AT3G09770 | 0,38 | 1,63 | 4,31 | 0,38 | 7,06E-01 |
| AT3G09780 | 0,23 | -0,64 | 4,39 | -0,15 | 8,84E-01 |
| AT3G09850 | 1,65 | 3,25 | 1,64 | 1,98 | 4,76E-02 |
| AT3G09870 | 0,08 | 2,33 | 4,36 | 0,53 | 5,93E-01 |
| AT3G09920 | 0,16 | 1,27 | 4,37 | 0,29 | 7,71E-01 |
| AT3G10010 | 0,35 | -0,68 | 4,39 | -0,15 | 8,77E-01 |
| AT3G10020 | 0,27 | 1,61 | 4,36 | 0,37 | 7,11E-01 |
| AT3G10030 | 0,88 | 2,67 | 2,89 | 0,92 | 3,56E-01 |
| AT3G10050 | 0,25 | 1,85 | 4,33 | 0,43 | 6,69E-01 |
| AT3G10070 | 0,46 | 0,74 | 4,22 | 0,18 | 8,60E-01 |
| AT3G10130 | 0,21 | 2,34 | 4,32 | 0,54 | 5,89E-01 |
| AT3G10160 | 0,36 | 0,78 | 3,34 | 0,23 | 8,15E-01 |
| AT3G10230 | 1,13 | 2,79 | 2,04 | 1,37 | 1,71E-01 |
| AT3G10250 | 0,08 | 2,33 | 4,36 | 0,53 | 5,93E-01 |
| AT3G10270 | 0,56 | 0,00 | 3,16 | 0,00 | 9,99E-01 |
| AT3G10320 | 0,12 | 2,87 | 4,34 | 0,66 | 5,08E-01 |
| AT3G10360 | 0,63 | 1,53 | 3,78 | 0,41 | 6,85E-01 |
| AT3G10405 | 0,68 | -0,14 | 4,41 | -0,03 | 9,75E-01 |
| AT3G10410 | 1,05 | 0,96 | 1,76 | 0,55 | 5,86E-01 |
| AT3G10520 | 0,15 | -0,64 | 4,39 | -0,15 | 8,84E-01 |
| AT3G10525 | 0,39 | 2,36 | 3,38 | 0,70 | 4,86E-01 |
| AT3G10630 | 0,99 | 2,56 | 1,82 | 1,41 | 1,59E-01 |
| AT3G10640 | 0,33 | -0,37 | 4,32 | -0,09 | 9,31E-01 |
| AT3G10660 | 0,27 | 1,46 | 4,33 | 0,34 | 7,36E-01 |
| AT3G10690 | 0,77 | 3,13 | 3,46 | 0,90 | 3,66E-01 |
| AT3G10720 | 0,36 | -1,90 | 4,32 | -0,44 | 6,60E-01 |
| AT3G10730 | 0,58 | 0,83 | 3,07 | 0,27 | 7,87E-01 |
| AT3G10770 | 0,74 | 2,99 | 2,44 | 1,22 | 2,21E-01 |
| AT3G10810 | 0,46 | 2,27 | 4,18 | 0,54 | 5,87E-01 |
| AT3G10840 | 0,41 | 0,00 | 4,41 | 0,00 | 1,00E+00 |
| AT3G10850 | 0,36 | -0,02 | 4,26 | 0,00 | 9,96E-01 |
| AT3G10900 | 0,31 | -1,49 | 4,26 | -0,35 | 7,26E-01 |
| AT3G10915 | 0,24 | 0,39 | 4,34 | 0,09 | 9,28E-01 |
| AT3G10930 | 0,61 | -0,59 | 2,30 | -0,26 | 7,98E-01 |
| AT3G10960 | 1,23 | 0,02 | 1,43 | 0,02 | 9,87E-01 |
| AT3G11030 | 0,56 | 0,35 | 3,80 | 0,09 | 9,27E-01 |
| AT3G11100 | 0,47 | 2,95 | 3,96 | 0,74 | 4,57E-01 |
| AT3G11280 | 0,31 | 0,21 | 4,36 | 0,05 | 9,62E-01 |
| AT3G11290 | 0,30 | 1,56 | 4,30 | 0,36 | 7,17E-01 |
| AT3G11310 | 0,31 | -1,15 | 4,25 | -0,27 | 7,86E-01 |
| AT3G11320 | 0,17 | 1,90 | 4,33 | 0,44 | 6,61E-01 |
| AT3G11330 | 0,97 | 4,56 | 2,57 | 1,77 | 7,62E-02 |
| AT3G11435 | 0,23 | 0,00 | 4,41 | 0,00 | 1,00E+00 |
| AT3G11450 | 1,01 | 0,56 | 1,79 | 0,32 | 7,53E-01 |
| AT3G11510 | 1,34 | -0,66 | 1,86 | -0,36 | 7,21E-01 |
| AT3G02435 | 1,34 | -3,24 | 1,89 | -1,72 | 8,62E-02 |
| AT3G02445 | 0,88 | -3,68 | 3,39 | -1,09 | 2,78E-01 |
| AT3G02455 | 0,99 | 1,87 | 2,78 | 0,67 | 5,02E-01 |
| AT3G11570 | 0,77 | 1,09 | 2,98 | 0,36 | 7,16E-01 |
| AT3G11580 | 0,26 | 0,00 | 4,27 | 0,00 | 1,00E+00 |
| AT3G11650 | 0,48 | -1,02 | 3,08 | -0,33 | 7,39E-01 |

| | | | | | |
|---|---|---|---|---|---|
| AT3G11660 | 1,33 | 0,36 | 3,18 | 0,11 | 9,11E-01 |
| AT3G11680 | 0,22 | 0,08 | 4,27 | 0,02 | 9,85E-01 |
| AT3G11710 | 1,08 | -0,06 | 1,97 | -0,03 | 9,76E-01 |
| AT3G11730 | 0,25 | 2,97 | 4,33 | 0,69 | 4,93E-01 |
| AT3G11770 | 0,28 | 0,84 | 4,35 | 0,19 | 8,46E-01 |
| AT3G11780 | 0,21 | 1,46 | 4,31 | 0,34 | 7,35E-01 |
| AT3G11820 | 1,51 | 1,42 | 1,60 | 0,89 | 3,72E-01 |
| AT3G11840 | 0,47 | 2,19 | 4,04 | 0,54 | 5,88E-01 |
| AT3G11964 | 0,92 | 0,82 | 1,82 | 0,45 | 6,52E-01 |
| AT3G12030 | 0,12 | 2,87 | 4,34 | 0,66 | 5,08E-01 |
| AT3G12130 | 0,80 | 1,48 | 2,51 | 0,59 | 5,55E-01 |
| AT3G12140 | 0,25 | -0,25 | 4,29 | -0,06 | 9,53E-01 |
| AT3G12145 | 0,15 | -0,19 | 4,31 | -0,04 | 9,64E-01 |
| AT3G12200 | 0,53 | -0,52 | 3,89 | -0,13 | 8,93E-01 |
| AT3G12260 | 0,32 | -1,53 | 4,33 | -0,35 | 7,23E-01 |
| AT3G12280 | 1,55 | 2,45 | 1,43 | 1,71 | 8,65E-02 |
| AT3G12400 | 0,71 | 0,47 | 1,82 | 0,26 | 7,96E-01 |
| AT3G12490 | 0,47 | 0,16 | 2,28 | 0,07 | 9,45E-01 |
| AT3G12500 | 0,08 | 2,33 | 4,36 | 0,53 | 5,93E-01 |
| AT3G12580 | 0,43 | 2,50 | 4,37 | 0,57 | 5,67E-01 |
| AT3G12587 | 0,28 | -0,24 | 4,40 | -0,05 | 9,57E-01 |
| AT3G12590 | 0,82 | 2,39 | 2,44 | 0,98 | 3,26E-01 |
| AT3G12610 | 0,59 | -0,15 | 2,29 | -0,07 | 9,48E-01 |
| AT3G12670 | 1,09 | 2,72 | 2,45 | 1,11 | 2,68E-01 |
| AT3G12680 | 0,32 | 0,10 | 4,36 | 0,02 | 9,82E-01 |
| AT3G12690 | 0,57 | 4,41 | 4,30 | 1,03 | 3,05E-01 |
| AT3G12750 | 0,45 | 3,37 | 3,98 | 0,85 | 3,98E-01 |
| AT3G12770 | 0,67 | 1,16 | 2,91 | 0,40 | 6,91E-01 |
| AT3G12800 | 0,96 | -0,45 | 1,66 | -0,27 | 7,85E-01 |
| AT3G12820 | 0,21 | 2,61 | 4,34 | 0,60 | 5,48E-01 |
| AT3G12920 | 1,14 | -0,98 | 2,57 | -0,38 | 7,02E-01 |
| AT3G12940 | 0,28 | 0,85 | 4,27 | 0,20 | 8,42E-01 |
| AT3G12950 | 0,48 | 3,77 | 4,32 | 0,87 | 3,83E-01 |
| AT3G12965 | 0,28 | -1,48 | 4,35 | -0,34 | 7,33E-01 |
| AT3G13000 | 0,24 | 0,00 | 4,41 | 0,00 | 1,00E+00 |
| AT3G13010 | 0,29 | 3,20 | 4,33 | 0,74 | 4,60E-01 |
| AT3G13020 | 0,33 | 1,99 | 4,37 | 0,46 | 6,49E-01 |
| AT3G13030 | 0,97 | 2,86 | 2,33 | 1,22 | 2,21E-01 |
| AT3G13040 | 0,58 | -1,21 | 2,59 | -0,46 | 6,42E-01 |
| AT3G13061 | 0,23 | -0,84 | 4,31 | -0,19 | 8,46E-01 |
| AT3G13110 | 1,06 | 3,41 | 1,99 | 1,71 | 8,65E-02 |
| AT3G13120 | 0,60 | 0,83 | 3,05 | 0,27 | 7,84E-01 |
| AT3G13150 | 0,37 | 2,80 | 4,34 | 0,65 | 5,18E-01 |
| AT3G13160 | 0,08 | 2,08 | 4,38 | 0,47 | 6,35E-01 |
| AT3G13220 | 0,37 | 3,50 | 4,32 | 0,81 | 4,18E-01 |
| AT3G13222 | 0,48 | 4,19 | 4,30 | 0,97 | 3,30E-01 |
| AT3G13225 | 0,32 | 2,83 | 4,30 | 0,66 | 5,10E-01 |
| AT3G13226 | 0,08 | 2,33 | 4,36 | 0,53 | 5,93E-01 |
| AT3G13235 | 0,44 | -0,10 | 2,70 | -0,04 | 9,69E-01 |
| AT3G13310 | 0,25 | 1,03 | 4,31 | 0,24 | 8,11E-01 |
| AT3G13390 | 0,05 | -0,58 | 4,38 | -0,13 | 8,95E-01 |
| AT3G13404 | 0,46 | -2,18 | 4,06 | -0,54 | 5,92E-01 |
| AT3G13410 | 1,64 | -0,63 | 2,03 | -0,31 | 7,59E-01 |
| AT3G13450 | 0,75 | 3,44 | 2,78 | 1,23 | 2,17E-01 |
| AT3G13480 | 0,20 | 0,30 | 4,34 | 0,07 | 9,46E-01 |
| AT3G13490 | 0,57 | 2,41 | 3,04 | 0,79 | 4,27E-01 |

| | | | | | |
|---|---|---|---|---|---|
| AT3G13570 | 0,44 | 1,93 | 4,20 | 0,46 | 6,47E-01 |
| AT3G13720 | 0,96 | -1,67 | 1,59 | -1,05 | 2,93E-01 |
| AT3G13772 | 0,72 | 3,37 | 2,76 | 1,22 | 2,22E-01 |
| AT3G13860 | 0,89 | 2,31 | 1,92 | 1,20 | 2,29E-01 |
| AT3G13990 | 1,05 | 3,75 | 2,22 | 1,69 | 9,12E-02 |
| AT3G14070 | 0,23 | 2,10 | 4,31 | 0,49 | 6,27E-01 |
| AT3G14075 | 0,26 | 2,19 | 4,32 | 0,51 | 6,13E-01 |
| AT3G14090 | 0,68 | -0,13 | 3,12 | -0,04 | 9,67E-01 |
| AT3G14110 | 0,61 | 1,71 | 2,56 | 0,67 | 5,05E-01 |
| AT3G14172 | 1,38 | 2,81 | 1,88 | 1,50 | 1,34E-01 |
| AT3G14180 | 0,40 | 3,49 | 4,33 | 0,81 | 4,20E-01 |
| AT3G14205 | 0,51 | 0,28 | 2,68 | 0,11 | 9,15E-01 |
| AT3G14230 | 0,40 | 1,22 | 4,18 | 0,29 | 7,70E-01 |
| AT3G14240 | 1,34 | -0,03 | 1,82 | -0,02 | 9,87E-01 |
| AT3G14330 | 0,56 | 3,61 | 2,99 | 1,21 | 2,27E-01 |
| AT3G14350 | 1,44 | -1,26 | 1,47 | -0,86 | 3,90E-01 |
| AT3G14400 | 1,24 | 4,85 | 3,16 | 1,53 | 1,25E-01 |
| AT3G14430 | 0,25 | 2,90 | 4,34 | 0,67 | 5,03E-01 |
| AT3G14440 | 0,33 | 1,07 | 4,25 | 0,25 | 8,01E-01 |
| AT3G14460 | 0,50 | 1,55 | 3,91 | 0,40 | 6,93E-01 |
| AT3G14590 | 0,14 | -0,31 | 4,33 | -0,07 | 9,43E-01 |
| AT3G14810 | 0,79 | 3,16 | 2,32 | 1,36 | 1,74E-01 |
| AT3G14900 | 0,17 | 2,95 | 4,34 | 0,68 | 4,96E-01 |
| AT3G15110 | 0,15 | -0,64 | 4,39 | -0,15 | 8,84E-01 |
| AT3G15120 | 0,82 | 1,46 | 2,34 | 0,63 | 5,31E-01 |
| AT3G15150 | 0,08 | -0,56 | 4,38 | -0,13 | 8,99E-01 |
| AT3G15220 | 0,77 | 1,40 | 2,45 | 0,57 | 5,68E-01 |
| AT3G15354 | 0,91 | 2,48 | 2,06 | 1,21 | 2,27E-01 |
| AT3G15356 | 0,36 | 0,85 | 4,39 | 0,19 | 8,46E-01 |
| AT3G15470 | 0,41 | -0,07 | 2,70 | -0,02 | 9,80E-01 |
| AT3G03125 | 0,17 | 2,62 | 4,35 | 0,60 | 5,47E-01 |
| AT3G15530 | 1,09 | 0,51 | 2,20 | 0,23 | 8,17E-01 |
| AT3G15570 | 0,83 | -3,39 | 2,69 | -1,26 | 2,07E-01 |
| AT3G15590 | 0,41 | 1,60 | 4,22 | 0,38 | 7,04E-01 |
| AT3G15630 | 0,46 | -1,27 | 3,08 | -0,41 | 6,79E-01 |
| AT3G15720 | 0,20 | -0,53 | 4,30 | -0,12 | 9,03E-01 |
| AT3G15810 | 0,58 | -2,11 | 2,41 | -0,87 | 3,82E-01 |
| AT3G15840 | 1,45 | -0,54 | 1,33 | -0,41 | 6,83E-01 |
| AT3G15900 | 0,29 | -1,08 | 4,34 | -0,25 | 8,03E-01 |
| AT3G15940 | 1,43 | 3,29 | 1,92 | 1,71 | 8,73E-02 |
| AT3G15970 | 0,63 | 1,71 | 2,48 | 0,69 | 4,90E-01 |
| AT3G15980 | 1,59 | -0,39 | 1,34 | -0,29 | 7,71E-01 |
| AT3G16000 | 0,51 | -0,86 | 3,13 | -0,28 | 7,83E-01 |
| AT3G16110 | 0,21 | 2,82 | 4,35 | 0,65 | 5,16E-01 |
| AT3G16190 | 0,21 | 2,01 | 4,37 | 0,46 | 6,45E-01 |
| AT3G03235 | 0,34 | 3,08 | 4,34 | 0,71 | 4,78E-01 |
| AT3G16290 | 0,84 | 0,16 | 1,82 | 0,09 | 9,28E-01 |
| AT3G16310 | 0,55 | 2,98 | 3,76 | 0,79 | 4,28E-01 |
| AT3G16500 | 0,17 | 0,69 | 4,34 | 0,16 | 8,73E-01 |
| AT3G16510 | 0,65 | 0,15 | 2,93 | 0,05 | 9,59E-01 |
| AT3G16520 | 0,76 | -0,09 | 1,97 | -0,05 | 9,64E-01 |
| AT3G16530 | 0,27 | -0,47 | 4,30 | -0,11 | 9,14E-01 |
| AT3G16560 | 0,66 | 2,15 | 2,41 | 0,89 | 3,72E-01 |
| AT3G16630 | 1,01 | 0,20 | 1,86 | 0,11 | 9,14E-01 |
| AT3G16820 | 1,66 | -2,17 | 1,27 | -1,71 | 8,69E-02 |
| AT3G16840 | 0,19 | 0,42 | 4,28 | 0,10 | 9,21E-01 |

| | | | | | |
|---|---|---|---|---|---|
| AT3G16860 | 0,47 | -2,19 | 3,99 | -0,55 | 5,84E-01 |
| AT3G16870 | 0,39 | 1,78 | 4,41 | 0,40 | 6,86E-01 |
| AT3G16890 | 0,19 | 0,25 | 4,35 | 0,06 | 9,54E-01 |
| AT3G17020 | 1,21 | 0,23 | 2,03 | 0,11 | 9,10E-01 |
| AT3G03415 | 1,34 | 0,03 | 1,60 | 0,02 | 9,86E-01 |
| AT3G03445 | 0,77 | 1,86 | 2,50 | 0,74 | 4,57E-01 |
| AT3G17170 | 0,31 | 0,70 | 4,39 | 0,16 | 8,73E-01 |
| AT3G17320 | 0,12 | 0,08 | 4,33 | 0,02 | 9,84E-01 |
| AT3G17330 | 0,20 | 1,94 | 4,34 | 0,45 | 6,55E-01 |
| AT3G17420 | 0,17 | 2,11 | 4,37 | 0,48 | 6,28E-01 |
| AT3G17440 | 0,08 | 2,33 | 4,36 | 0,53 | 5,93E-01 |
| AT3G17450 | 0,60 | 2,46 | 2,92 | 0,84 | 3,99E-01 |
| AT3G17655 | 0,19 | 2,19 | 4,32 | 0,51 | 6,13E-01 |
| AT3G17740 | 0,37 | 2,19 | 4,29 | 0,51 | 6,10E-01 |
| AT3G17760 | 0,21 | 2,61 | 4,34 | 0,60 | 5,48E-01 |
| AT3G17770 | 1,51 | 1,47 | 1,35 | 1,09 | 2,77E-01 |
| AT3G03585 | 0,05 | -0,58 | 4,38 | -0,13 | 8,95E-01 |
| AT3G17940 | 0,81 | 1,26 | 2,52 | 0,50 | 6,18E-01 |
| AT3G18030 | 0,52 | 1,87 | 3,96 | 0,47 | 6,36E-01 |
| AT3G18035 | 1,23 | -0,50 | 1,51 | -0,33 | 7,39E-01 |
| AT3G18110 | 1,16 | 4,90 | 2,44 | 2,01 | 4,47E-02 |
| AT3G18130 | 0,88 | 1,66 | 2,32 | 0,71 | 4,75E-01 |
| AT3G18140 | 0,30 | -1,73 | 4,32 | -0,40 | 6,88E-01 |
| AT3G18250 | 0,55 | -0,69 | 4,39 | -0,16 | 8,76E-01 |
| AT3G18380 | 0,25 | 2,97 | 4,33 | 0,69 | 4,93E-01 |
| AT3G18420 | 0,20 | 0,36 | 4,30 | 0,08 | 9,34E-01 |
| AT3G18535 | 0,67 | -0,45 | 2,24 | -0,20 | 8,41E-01 |
| AT3G18600 | 0,25 | 0,00 | 4,40 | 0,00 | 1,00E+00 |
| AT3G18610 | 0,55 | 3,10 | 4,34 | 0,71 | 4,75E-01 |
| AT3G18630 | 0,20 | 2,11 | 4,37 | 0,48 | 6,29E-01 |
| AT3G18700 | 1,19 | 1,24 | 1,79 | 0,69 | 4,89E-01 |
| AT3G18750 | 0,17 | 3,04 | 4,33 | 0,70 | 4,83E-01 |
| AT3G18930 | 0,90 | -2,59 | 2,13 | -1,22 | 2,24E-01 |
| AT3G18970 | 0,08 | 2,33 | 4,36 | 0,53 | 5,93E-01 |
| AT3G19000 | 0,24 | 0,95 | 4,31 | 0,22 | 8,26E-01 |
| AT3G19020 | 0,54 | 0,31 | 3,08 | 0,10 | 9,19E-01 |
| AT3G19150 | 0,16 | 0,03 | 4,34 | 0,01 | 9,94E-01 |
| AT3G03815 | 0,25 | 0,00 | 4,40 | 0,00 | 1,00E+00 |
| AT3G19300 | 0,12 | 1,65 | 4,34 | 0,38 | 7,04E-01 |
| AT3G19340 | 0,12 | 0,31 | 4,35 | 0,07 | 9,44E-01 |
| AT3G19430 | 0,25 | 0,00 | 4,40 | 0,00 | 1,00E+00 |
| AT3G19490 | 0,72 | 2,37 | 2,39 | 1,00 | 3,20E-01 |
| AT3G19510 | 0,12 | 0,63 | 4,33 | 0,15 | 8,85E-01 |
| AT3G19515 | 0,59 | 2,45 | 3,11 | 0,79 | 4,31E-01 |
| AT3G19553 | 0,42 | -1,79 | 4,10 | -0,44 | 6,62E-01 |
| AT3G19580 | 0,72 | 0,72 | 2,26 | 0,32 | 7,49E-01 |
| AT3G19620 | 0,32 | 2,42 | 4,35 | 0,56 | 5,77E-01 |
| AT3G19630 | 0,39 | 1,78 | 4,41 | 0,40 | 6,86E-01 |
| AT3G19710 | 0,69 | 0,90 | 2,31 | 0,39 | 6,98E-01 |
| AT3G19720 | 1,17 | 3,05 | 2,28 | 1,34 | 1,80E-01 |
| AT3G19740 | 0,59 | 1,33 | 3,05 | 0,44 | 6,62E-01 |
| AT3G19790 | 0,71 | 0,09 | 1,86 | 0,05 | 9,61E-01 |
| AT3G19810 | 0,70 | 1,79 | 4,41 | 0,41 | 6,85E-01 |
| AT3G19830 | 0,67 | -0,32 | 2,80 | -0,11 | 9,09E-01 |
| AT3G19850 | 0,50 | -0,27 | 3,10 | -0,09 | 9,31E-01 |
| AT3G19880 | 0,35 | 0,85 | 3,33 | 0,26 | 7,99E-01 |

| | | | | | |
|---|---|---|---|---|---|
| AT3G19895 | 0,23 | -1,06 | 4,29 | -0,25 | 8,04E-01 |
| AT3G19900 | 0,17 | 3,14 | 4,33 | 0,73 | 4,68E-01 |
| AT3G20015 | 0,62 | -1,77 | 3,06 | -0,58 | 5,62E-01 |
| AT3G20050 | 0,74 | 1,46 | 2,41 | 0,61 | 5,44E-01 |
| AT3G20060 | 0,33 | 0,16 | 4,29 | 0,04 | 9,69E-01 |
| AT3G20230 | 0,35 | 1,56 | 4,27 | 0,37 | 7,14E-01 |
| AT3G20250 | 1,61 | 2,27 | 1,69 | 1,35 | 1,78E-01 |
| AT3G20290 | 0,63 | 1,01 | 2,90 | 0,35 | 7,27E-01 |
| AT3G20320 | 0,51 | 0,30 | 2,66 | 0,11 | 9,10E-01 |
| AT3G20390 | 0,62 | -0,64 | 2,12 | -0,30 | 7,62E-01 |
| AT3G20470 | 0,38 | -0,01 | 4,34 | 0,00 | 9,98E-01 |
| AT3G20620 | 0,68 | 3,98 | 3,55 | 1,12 | 2,63E-01 |
| AT3G20630 | 1,20 | 1,45 | 1,57 | 0,92 | 3,58E-01 |
| AT3G20655 | 0,30 | 2,68 | 4,35 | 0,61 | 5,39E-01 |
| AT3G20700 | 1,05 | -2,40 | 2,24 | -1,07 | 2,83E-01 |
| AT3G03985 | 0,54 | -3,60 | 4,27 | -0,84 | 4,00E-01 |
| AT3G20830 | 0,60 | 0,63 | 3,24 | 0,19 | 8,46E-01 |
| AT3G21055 | 1,34 | 0,37 | 1,67 | 0,22 | 8,26E-01 |
| AT3G21070 | 0,23 | 0,75 | 4,27 | 0,17 | 8,61E-01 |
| AT3G21140 | 0,25 | 1,77 | 4,34 | 0,41 | 6,83E-01 |
| AT3G21160 | 0,11 | 1,33 | 4,36 | 0,31 | 7,59E-01 |
| AT3G21215 | 0,11 | 1,28 | 4,35 | 0,29 | 7,68E-01 |
| AT3G21300 | 0,80 | 0,32 | 2,18 | 0,15 | 8,82E-01 |
| AT3G21430 | 0,61 | 0,00 | 3,05 | 0,00 | 9,99E-01 |
| AT3G21540 | 0,20 | 2,39 | 4,37 | 0,55 | 5,84E-01 |
| AT3G21570 | 0,87 | -2,92 | 2,87 | -1,02 | 3,09E-01 |
| AT3G21630 | 0,12 | 0,31 | 4,35 | 0,07 | 9,44E-01 |
| AT3G21650 | 0,33 | 3,34 | 4,32 | 0,77 | 4,40E-01 |
| AT3G21670 | 1,42 | 3,91 | 2,30 | 1,70 | 8,83E-02 |
| AT3G21740 | 0,24 | 1,18 | 4,36 | 0,27 | 7,87E-01 |
| AT3G21790 | 0,96 | 1,15 | 1,91 | 0,60 | 5,48E-01 |
| AT3G21865 | 0,48 | 0,60 | 2,58 | 0,23 | 8,16E-01 |
| AT3G22070 | 0,17 | 0,00 | 4,41 | 0,00 | 1,00E+00 |
| AT3G22110 | 0,73 | -0,45 | 2,38 | -0,19 | 8,50E-01 |
| AT3G22142 | 1,40 | -0,35 | 1,51 | -0,23 | 8,19E-01 |
| AT3G22170 | 1,10 | 2,28 | 1,85 | 1,23 | 2,19E-01 |
| AT3G22190 | 0,78 | 0,63 | 2,95 | 0,21 | 8,31E-01 |
| AT3G22210 | 0,48 | -0,34 | 3,32 | -0,10 | 9,18E-01 |
| AT3G22220 | 0,36 | 1,85 | 3,46 | 0,53 | 5,94E-01 |
| AT3G22230 | 0,95 | 0,99 | 1,76 | 0,56 | 5,76E-01 |
| AT3G22240 | 1,25 | 0,13 | 1,55 | 0,08 | 9,34E-01 |
| AT3G22320 | 0,41 | 1,60 | 4,22 | 0,38 | 7,04E-01 |
| AT3G22421 | 0,96 | 1,04 | 2,47 | 0,42 | 6,75E-01 |
| AT3G22470 | 0,59 | 1,58 | 2,52 | 0,63 | 5,30E-01 |
| AT3G22530 | 0,38 | 2,42 | 4,37 | 0,55 | 5,80E-01 |
| AT3G22550 | 0,60 | 2,25 | 3,00 | 0,75 | 4,52E-01 |
| AT3G22580 | 0,15 | -0,64 | 4,39 | -0,15 | 8,84E-01 |
| AT3G22600 | 0,17 | 3,16 | 4,33 | 0,73 | 4,66E-01 |
| AT3G22630 | 0,68 | 0,49 | 2,08 | 0,24 | 8,14E-01 |
| AT3G22660 | 0,44 | -0,27 | 4,39 | -0,06 | 9,50E-01 |
| AT3G22690 | 0,29 | 1,79 | 4,31 | 0,41 | 6,78E-01 |
| AT3G22710 | 1,14 | -0,28 | 1,77 | -0,16 | 8,76E-01 |
| AT3G22720 | 0,08 | -0,56 | 4,38 | -0,13 | 8,99E-01 |
| AT3G22730 | 1,61 | -2,46 | 1,46 | -1,69 | 9,19E-02 |
| AT3G22770 | 0,68 | 0,77 | 3,16 | 0,24 | 8,07E-01 |
| AT3G22950 | 0,15 | -0,67 | 4,39 | -0,15 | 8,78E-01 |

| | | | | | |
|---|---|---|---|---|---|
| AT3G22980 | 0,74 | 2,05 | 2,44 | 0,84 | 4,02E-01 |
| AT3G23020 | 0,53 | 3,07 | 3,81 | 0,81 | 4,20E-01 |
| AT3G23030 | 0,69 | 0,35 | 1,98 | 0,18 | 8,60E-01 |
| AT3G23080 | 0,08 | 2,33 | 4,36 | 0,53 | 5,93E-01 |
| AT3G23170 | 0,50 | 0,39 | 2,50 | 0,16 | 8,75E-01 |
| AT3G23172 | 0,24 | 0,64 | 4,28 | 0,15 | 8,80E-01 |
| AT3G23400 | 0,58 | 0,14 | 3,13 | 0,05 | 9,64E-01 |
| AT3G23420 | 0,32 | -0,60 | 4,25 | -0,14 | 8,88E-01 |
| AT3G23490 | 0,26 | -0,79 | 4,27 | -0,18 | 8,54E-01 |
| AT3G23570 | 0,32 | 2,20 | 4,32 | 0,51 | 6,10E-01 |
| AT3G23633 | 0,12 | -1,57 | 4,34 | -0,36 | 7,18E-01 |
| AT3G23650 | 0,55 | -0,82 | 2,76 | -0,30 | 7,66E-01 |
| AT3G04585 | 0,72 | -0,99 | 2,63 | -0,38 | 7,07E-01 |
| AT3G23660 | 0,75 | 2,40 | 2,87 | 0,84 | 4,03E-01 |
| AT3G23700 | 0,24 | -0,36 | 4,34 | -0,08 | 9,34E-01 |
| AT3G23740 | 0,08 | 2,08 | 4,38 | 0,47 | 6,35E-01 |
| AT3G23830 | 0,27 | 2,01 | 4,37 | 0,46 | 6,46E-01 |
| AT3G23900 | 0,77 | 2,78 | 2,66 | 1,05 | 2,95E-01 |
| AT3G23970 | 0,08 | 2,08 | 4,38 | 0,47 | 6,35E-01 |
| AT3G23980 | 0,76 | -0,66 | 2,48 | -0,27 | 7,90E-01 |
| AT3G24080 | 0,22 | 1,72 | 4,29 | 0,40 | 6,89E-01 |
| AT3G24120 | 0,83 | 2,81 | 3,45 | 0,81 | 4,16E-01 |
| AT3G24140 | 0,26 | -1,01 | 4,30 | -0,24 | 8,14E-01 |
| AT3G24170 | 1,33 | 2,31 | 1,69 | 1,37 | 1,71E-01 |
| AT3G24200 | 0,11 | 1,33 | 4,36 | 0,31 | 7,59E-01 |
| AT3G24260 | 0,15 | -1,60 | 4,33 | -0,37 | 7,11E-01 |
| AT3G24270 | 0,91 | 0,88 | 2,08 | 0,42 | 6,73E-01 |
| AT3G24320 | 0,63 | 2,55 | 3,02 | 0,84 | 3,99E-01 |
| AT3G24430 | 0,54 | -0,31 | 2,62 | -0,12 | 9,05E-01 |
| AT3G24440 | 0,50 | 2,22 | 3,18 | 0,70 | 4,86E-01 |
| AT3G24465 | 0,32 | 0,24 | 3,56 | 0,07 | 9,46E-01 |
| AT3G24503 | 1,13 | -1,29 | 1,70 | -0,76 | 4,50E-01 |
| AT3G24518 | 0,96 | -1,80 | 2,39 | -0,76 | 4,49E-01 |
| AT3G24614 | 0,96 | -0,69 | 1,70 | -0,41 | 6,85E-01 |
| AT3G24670 | 0,20 | 1,88 | 4,35 | 0,43 | 6,66E-01 |
| AT3G24870 | 1,16 | -0,03 | 1,96 | -0,01 | 9,89E-01 |
| AT3G24880 | 0,67 | -1,23 | 3,03 | -0,41 | 6,85E-01 |
| AT3G24900 | 0,51 | 1,28 | 3,03 | 0,42 | 6,72E-01 |
| AT3G25090 | 0,87 | -2,17 | 1,85 | -1,18 | 2,39E-01 |
| AT3G25150 | 1,03 | 3,18 | 2,48 | 1,28 | 1,99E-01 |
| AT3G25180 | 0,15 | -0,64 | 4,39 | -0,15 | 8,84E-01 |
| AT3G25460 | 1,48 | -0,79 | 1,52 | -0,52 | 6,01E-01 |
| AT3G25530 | 0,19 | -0,79 | 4,32 | -0,18 | 8,54E-01 |
| AT3G25560 | 0,41 | 0,79 | 4,39 | 0,18 | 8,58E-01 |
| AT3G25610 | 1,05 | 0,72 | 2,45 | 0,30 | 7,68E-01 |
| AT3G25660 | 0,47 | 3,65 | 3,95 | 0,92 | 3,56E-01 |
| AT3G04935 | 0,15 | 1,64 | 4,33 | 0,38 | 7,06E-01 |
| AT3G25800 | 1,14 | 3,03 | 2,00 | 1,51 | 1,31E-01 |
| AT3G25840 | 1,32 | 0,73 | 1,60 | 0,46 | 6,47E-01 |
| AT3G25870 | 0,65 | 0,55 | 2,28 | 0,24 | 8,11E-01 |
| AT3G25880 | 1,38 | 3,77 | 2,55 | 1,48 | 1,40E-01 |
| AT3G25882 | 0,50 | 1,80 | 3,98 | 0,45 | 6,51E-01 |
| AT3G25920 | 0,58 | 0,59 | 2,19 | 0,27 | 7,87E-01 |
| AT3G25990 | 0,75 | 1,68 | 3,52 | 0,48 | 6,32E-01 |
| AT3G26170 | 1,29 | 0,50 | 1,54 | 0,33 | 7,45E-01 |
| AT3G26180 | 0,36 | 0,07 | 4,26 | 0,02 | 9,88E-01 |

| | | | | | |
|---|---|---|---|---|---|
| AT3G26210 | 1,30 | 0,56 | 1,80 | 0,31 | 7,57E-01 |
| AT3G26220 | 0,49 | -0,47 | 3,94 | -0,12 | 9,04E-01 |
| AT3G26230 | 0,79 | -1,56 | 2,44 | -0,64 | 5,25E-01 |
| AT3G26280 | 1,16 | -1,54 | 1,71 | -0,90 | 3,69E-01 |
| AT3G26290 | 0,78 | -0,27 | 1,80 | -0,15 | 8,83E-01 |
| AT3G26300 | 0,36 | 1,64 | 4,38 | 0,37 | 7,08E-01 |
| AT3G26340 | 0,62 | -0,14 | 3,85 | -0,04 | 9,72E-01 |
| AT3G26410 | 0,31 | 0,00 | 4,41 | 0,00 | 1,00E+00 |
| AT3G26500 | 1,32 | -0,12 | 1,62 | -0,07 | 9,42E-01 |
| AT3G26510 | 0,66 | -0,87 | 1,96 | -0,45 | 6,56E-01 |
| AT3G26550 | 0,20 | 2,30 | 4,36 | 0,53 | 5,98E-01 |
| AT3G26560 | 0,40 | -0,28 | 3,23 | -0,09 | 9,32E-01 |
| AT3G26590 | 0,41 | -0,92 | 4,33 | -0,21 | 8,31E-01 |
| AT3G26616 | 0,23 | -0,22 | 4,28 | -0,05 | 9,60E-01 |
| AT3G26630 | 0,69 | 1,93 | 2,44 | 0,79 | 4,29E-01 |
| AT3G26670 | 0,54 | 1,01 | 3,23 | 0,31 | 7,56E-01 |
| AT3G26744 | 0,39 | 2,48 | 4,27 | 0,58 | 5,61E-01 |
| AT3G26812 | 0,17 | -0,71 | 4,32 | -0,16 | 8,69E-01 |
| AT3G26819 | 0,05 | -0,58 | 4,38 | -0,13 | 8,95E-01 |
| AT3G26860 | 0,30 | 0,00 | 4,41 | 0,00 | 1,00E+00 |
| AT3G26870 | 1,55 | -0,86 | 2,06 | -0,42 | 6,76E-01 |
| AT3G26890 | 0,29 | -0,39 | 4,29 | -0,09 | 9,28E-01 |
| AT3G26932 | 0,90 | -0,24 | 2,14 | -0,11 | 9,10E-01 |
| AT3G26935 | 0,27 | 0,42 | 4,31 | 0,10 | 9,21E-01 |
| AT3G26980 | 0,14 | 1,12 | 4,32 | 0,26 | 7,96E-01 |
| AT3G27010 | 0,67 | 2,97 | 3,65 | 0,81 | 4,16E-01 |
| AT3G27020 | 0,51 | 0,32 | 2,23 | 0,14 | 8,86E-01 |
| AT3G27040 | 1,43 | -2,06 | 1,67 | -1,24 | 2,17E-01 |
| AT3G27060 | 0,90 | 0,06 | 2,20 | 0,03 | 9,77E-01 |
| AT3G27100 | 0,74 | 1,79 | 4,41 | 0,41 | 6,85E-01 |
| AT3G27240 | 0,37 | -1,37 | 3,32 | -0,41 | 6,79E-01 |
| AT3G27300 | 0,49 | 1,71 | 3,29 | 0,52 | 6,02E-01 |
| AT3G27380 | 0,45 | 1,59 | 4,34 | 0,37 | 7,14E-01 |
| AT3G05235 | 0,99 | -2,85 | 2,42 | -1,18 | 2,39E-01 |
| AT3G05245 | 0,33 | -1,83 | 4,33 | -0,42 | 6,71E-01 |
| AT3G27530 | 0,76 | 1,81 | 2,56 | 0,71 | 4,79E-01 |
| AT3G27600 | 1,14 | -1,57 | 2,07 | -0,76 | 4,47E-01 |
| AT3G27610 | 0,33 | 0,55 | 4,36 | 0,13 | 8,99E-01 |
| AT3G27680 | 0,86 | -3,38 | 2,25 | -1,50 | 1,34E-01 |
| AT3G27700 | 1,39 | 1,79 | 1,74 | 1,03 | 3,03E-01 |
| AT3G27740 | 0,52 | 3,17 | 4,00 | 0,79 | 4,27E-01 |
| AT3G27770 | 0,43 | 1,32 | 2,74 | 0,48 | 6,29E-01 |
| AT3G27906 | 1,35 | -1,21 | 1,57 | -0,77 | 4,43E-01 |
| AT3G27925 | 0,65 | 0,95 | 3,00 | 0,32 | 7,52E-01 |
| AT3G27930 | 0,20 | -0,53 | 4,30 | -0,12 | 9,03E-01 |
| AT3G27990 | 0,64 | -2,86 | 2,43 | -1,18 | 2,39E-01 |
| AT3G28340 | 0,22 | -1,01 | 4,28 | -0,24 | 8,13E-01 |
| AT3G28380 | 0,08 | 2,08 | 4,38 | 0,47 | 6,35E-01 |
| AT3G28410 | 0,14 | -0,45 | 4,33 | -0,10 | 9,17E-01 |
| AT3G28674 | 0,28 | -0,34 | 4,30 | -0,08 | 9,37E-01 |
| AT3G28695 | 0,43 | -1,50 | 4,11 | -0,37 | 7,14E-01 |
| AT3G28710 | 0,99 | 0,13 | 1,54 | 0,08 | 9,34E-01 |
| AT3G28780 | 0,61 | -0,16 | 2,99 | -0,05 | 9,58E-01 |
| AT3G28860 | 1,33 | 2,21 | 2,30 | 0,96 | 3,37E-01 |
| AT3G28890 | 0,43 | 2,70 | 4,34 | 0,62 | 5,34E-01 |
| AT3G28920 | 1,35 | 0,31 | 1,70 | 0,18 | 8,57E-01 |

| | | | | | |
|---|---|---|---|---|---|
| AT3G28930 | 0,77 | 0,28 | 2,81 | 0,10 | 9,20E-01 |
| AT3G28940 | 1,30 | -0,11 | 1,50 | -0,07 | 9,41E-01 |
| AT3G28950 | 0,97 | -0,85 | 1,91 | -0,45 | 6,56E-01 |
| AT3G28960 | 0,30 | -1,14 | 4,30 | -0,27 | 7,90E-01 |
| AT3G28970 | 0,52 | 3,13 | 3,96 | 0,79 | 4,30E-01 |
| AT3G29030 | 0,33 | -2,33 | 4,30 | -0,54 | 5,88E-01 |
| AT3G05665 | 0,80 | -3,13 | 2,55 | -1,23 | 2,19E-01 |
| AT3G29110 | 0,53 | -1,97 | 2,61 | -0,75 | 4,51E-01 |
| AT3G29160 | 0,59 | -15,39 | 4,37 | -3,52 | 4,24E-04 |
| AT3G29200 | 0,59 | 2,91 | 3,03 | 0,96 | 3,38E-01 |
| AT3G29350 | 0,17 | -0,44 | 4,37 | -0,10 | 9,19E-01 |
| AT3G29400 | 0,67 | -0,20 | 2,41 | -0,08 | 9,35E-01 |
| AT3G29570 | 0,38 | -0,25 | 4,31 | -0,06 | 9,53E-01 |
| AT3G05745 | 0,14 | 0,87 | 4,31 | 0,20 | 8,40E-01 |
| AT3G29633 | 0,14 | 0,00 | 4,41 | 0,00 | 1,00E+00 |
| AT3G29639 | 0,16 | -0,65 | 4,31 | -0,15 | 8,80E-01 |
| AT3G05895 | 0,08 | -0,56 | 4,38 | -0,13 | 8,99E-01 |
| AT3G29680 | 0,25 | -1,58 | 4,34 | -0,37 | 7,15E-01 |
| AT3G29720 | 0,40 | 0,22 | 4,36 | 0,05 | 9,61E-01 |
| AT3G29760 | 0,67 | 2,11 | 3,02 | 0,70 | 4,84E-01 |
| AT3G29785 | 0,22 | -0,79 | 4,28 | -0,18 | 8,54E-01 |
| AT3G30160 | 0,58 | -2,45 | 2,86 | -0,86 | 3,91E-01 |
| AT3G30200 | 0,12 | 0,63 | 4,33 | 0,15 | 8,85E-01 |
| AT3G30220 | 0,08 | -0,56 | 4,38 | -0,13 | 8,99E-01 |
| AT3G30230 | 0,54 | 0,45 | 3,80 | 0,12 | 9,05E-01 |
| AT3G30300 | 0,53 | 1,07 | 4,32 | 0,25 | 8,05E-01 |
| AT3G30840 | 0,21 | 2,29 | 4,32 | 0,53 | 5,96E-01 |
| AT3G32190 | 0,34 | 0,78 | 3,39 | 0,23 | 8,18E-01 |
| AT3G32420 | 0,10 | 0,34 | 4,35 | 0,08 | 9,37E-01 |
| AT3G32904 | 0,65 | -1,39 | 3,05 | -0,45 | 6,50E-01 |
| AT3G33528 | 0,61 | 1,47 | 2,37 | 0,62 | 5,36E-01 |
| AT3G42050 | 1,44 | 1,37 | 1,86 | 0,74 | 4,62E-01 |
| AT3G42060 | 0,32 | -0,82 | 4,27 | -0,19 | 8,47E-01 |
| AT3G42140 | 0,14 | -0,45 | 4,33 | -0,10 | 9,17E-01 |
| AT3G42155 | 0,34 | -1,89 | 4,30 | -0,44 | 6,60E-01 |
| AT3G42550 | 0,12 | -0,77 | 4,36 | -0,18 | 8,61E-01 |
| AT3G42723 | 0,12 | 0,31 | 4,35 | 0,07 | 9,44E-01 |
| AT3G42790 | 0,59 | 0,33 | 3,04 | 0,11 | 9,15E-01 |
| AT3G43153 | 0,99 | 2,04 | 2,33 | 0,87 | 3,82E-01 |
| AT3G43190 | 0,26 | -1,00 | 4,37 | -0,23 | 8,18E-01 |
| AT3G43220 | 0,79 | 2,69 | 2,87 | 0,94 | 3,48E-01 |
| AT3G43240 | 0,86 | 2,80 | 2,68 | 1,04 | 2,97E-01 |
| AT3G43270 | 0,68 | -2,02 | 2,45 | -0,83 | 4,08E-01 |
| AT3G43280 | 0,11 | -1,28 | 4,35 | -0,29 | 7,69E-01 |
| AT3G43291 | 0,65 | -2,00 | 2,42 | -0,83 | 4,08E-01 |
| AT3G06595 | 0,10 | 0,34 | 4,35 | 0,08 | 9,37E-01 |
| AT3G43432 | 0,16 | -0,58 | 4,32 | -0,14 | 8,92E-01 |
| AT3G43470 | 0,45 | -1,73 | 3,86 | -0,45 | 6,54E-01 |
| AT3G43500 | 0,45 | -2,38 | 3,90 | -0,61 | 5,42E-01 |
| AT3G43600 | 0,98 | 1,85 | 2,30 | 0,80 | 4,22E-01 |
| AT3G43610 | 0,74 | 1,40 | 2,46 | 0,57 | 5,69E-01 |
| AT3G06645 | 0,23 | -0,93 | 4,38 | -0,21 | 8,32E-01 |
| AT3G06655 | 1,06 | -0,93 | 1,84 | -0,50 | 6,15E-01 |
| AT3G43720 | 0,87 | 0,00 | 2,86 | 0,00 | 1,00E+00 |
| AT3G43810 | 0,51 | -2,08 | 3,14 | -0,66 | 5,06E-01 |
| AT3G43880 | 0,10 | 0,39 | 4,35 | 0,09 | 9,29E-01 |

| | | | | | |
|---|---|---|---|---|---|
| AT3G43930 | 0,08 | 2,08 | 4,38 | 0,47 | 6,35E-01 |
| AT3G43940 | 0,53 | -1,51 | 3,81 | -0,40 | 6,92E-01 |
| AT3G44010 | 1,00 | -0,67 | 1,82 | -0,37 | 7,12E-01 |
| AT3G44070 | 0,37 | -0,44 | 3,38 | -0,13 | 8,97E-01 |
| AT3G44160 | 0,43 | 0,54 | 4,09 | 0,13 | 8,94E-01 |
| AT3G44265 | 0,40 | -0,66 | 4,20 | -0,16 | 8,75E-01 |
| AT3G44340 | 1,56 | 1,27 | 1,44 | 0,88 | 3,77E-01 |
| AT3G44400 | 0,82 | 1,12 | 2,56 | 0,44 | 6,63E-01 |
| AT3G44435 | 0,70 | 0,71 | 3,65 | 0,19 | 8,47E-01 |
| AT3G44450 | 0,56 | -1,47 | 3,00 | -0,49 | 6,25E-01 |
| AT3G44460 | 0,11 | -1,28 | 4,35 | -0,29 | 7,69E-01 |
| AT3G44635 | 1,31 | 4,43 | 2,90 | 1,53 | 1,26E-01 |
| AT3G44680 | 0,12 | 1,65 | 4,34 | 0,38 | 7,04E-01 |
| AT3G44753 | 0,14 | -0,31 | 4,33 | -0,07 | 9,43E-01 |
| AT3G44780 | 0,10 | -1,26 | 4,35 | -0,29 | 7,72E-01 |
| AT3G44820 | 0,38 | 1,61 | 4,35 | 0,37 | 7,11E-01 |
| AT3G44990 | 0,60 | -0,08 | 3,74 | -0,02 | 9,83E-01 |
| AT3G45030 | 0,91 | 1,32 | 2,39 | 0,55 | 5,81E-01 |
| AT3G45040 | 0,29 | -1,27 | 4,26 | -0,30 | 7,66E-01 |
| AT3G45060 | 0,08 | 2,33 | 4,36 | 0,53 | 5,93E-01 |
| AT3G45090 | 1,49 | 3,00 | 1,96 | 1,53 | 1,25E-01 |
| AT3G06955 | 0,21 | -2,30 | 4,30 | -0,54 | 5,92E-01 |
| AT3G45130 | 0,82 | -2,88 | 2,28 | -1,26 | 2,07E-01 |
| AT3G45180 | 1,00 | -0,47 | 1,80 | -0,26 | 7,92E-01 |
| AT3G07065 | 0,05 | -0,58 | 4,38 | -0,13 | 8,95E-01 |
| AT3G45240 | 0,25 | 2,79 | 4,34 | 0,64 | 5,20E-01 |
| AT3G07095 | 0,20 | 2,90 | 4,34 | 0,67 | 5,04E-01 |
| AT3G45260 | 1,01 | 0,95 | 1,78 | 0,54 | 5,92E-01 |
| AT3G45300 | 0,17 | -0,14 | 4,41 | -0,03 | 9,75E-01 |
| AT3G45430 | 0,20 | 2,30 | 4,36 | 0,53 | 5,98E-01 |
| AT3G45443 | 0,08 | -0,56 | 4,38 | -0,13 | 8,99E-01 |
| AT3G45600 | 0,92 | 1,68 | 1,90 | 0,89 | 3,75E-01 |
| AT3G45630 | 0,80 | 1,08 | 2,11 | 0,51 | 6,10E-01 |
| AT3G45770 | 0,14 | -0,45 | 4,33 | -0,10 | 9,17E-01 |
| AT3G45820 | 0,59 | -2,41 | 2,96 | -0,81 | 4,16E-01 |
| AT3G45850 | 1,07 | 1,66 | 2,34 | 0,71 | 4,78E-01 |
| AT3G45860 | 0,95 | -0,73 | 2,09 | -0,35 | 7,26E-01 |
| AT3G45910 | 1,00 | 0,47 | 2,27 | 0,21 | 8,34E-01 |
| AT3G45990 | 0,46 | 0,89 | 3,30 | 0,27 | 7,88E-01 |
| AT3G46000 | 0,30 | 1,11 | 3,61 | 0,31 | 7,59E-01 |
| AT3G46030 | 0,35 | -1,12 | 4,30 | -0,26 | 7,94E-01 |
| AT3G46090 | 0,72 | -1,12 | 2,19 | -0,51 | 6,09E-01 |
| AT3G46100 | 0,51 | -1,23 | 4,00 | -0,31 | 7,59E-01 |
| AT3G46210 | 0,56 | -0,49 | 3,77 | -0,13 | 8,97E-01 |
| AT3G07375 | 0,21 | 2,65 | 4,34 | 0,61 | 5,42E-01 |
| AT3G46320 | 0,61 | 1,24 | 4,28 | 0,29 | 7,73E-01 |
| AT3G46360 | 0,82 | -1,39 | 2,88 | -0,48 | 6,30E-01 |
| AT3G46390 | 0,25 | 1,38 | 4,27 | 0,32 | 7,47E-01 |
| AT3G46440 | 0,45 | 2,51 | 4,34 | 0,58 | 5,64E-01 |
| AT3G46450 | 0,53 | 2,24 | 3,81 | 0,59 | 5,57E-01 |
| AT3G46470 | 0,38 | 0,39 | 4,16 | 0,09 | 9,26E-01 |
| AT3G46540 | 0,31 | 0,44 | 4,33 | 0,10 | 9,18E-01 |
| AT3G46550 | 1,58 | -0,46 | 1,23 | -0,37 | 7,09E-01 |
| AT3G46610 | 1,41 | 0,60 | 1,49 | 0,40 | 6,87E-01 |
| AT3G46630 | 0,81 | 1,24 | 2,18 | 0,57 | 5,69E-01 |
| AT3G46650 | 0,58 | 3,56 | 3,11 | 1,15 | 2,51E-01 |

| | | | | | |
|---|---|---|---|---|---|
| AT3G46660 | 0,59 | 1,25 | 2,50 | 0,50 | 6,17E-01 |
| AT3G46670 | 0,57 | 1,42 | 3,23 | 0,44 | 6,59E-01 |
| AT3G46690 | 0,12 | 2,87 | 4,34 | 0,66 | 5,08E-01 |
| AT3G46730 | 0,12 | 2,81 | 4,34 | 0,65 | 5,18E-01 |
| AT3G46790 | 0,53 | 3,29 | 4,33 | 0,76 | 4,47E-01 |
| AT3G46845 | 1,29 | 0,37 | 1,61 | 0,23 | 8,17E-01 |
| AT3G46920 | 0,08 | 2,08 | 4,38 | 0,47 | 6,35E-01 |
| AT3G46960 | 0,92 | 1,57 | 2,14 | 0,73 | 4,64E-01 |
| AT3G47000 | 0,33 | -0,43 | 4,37 | -0,10 | 9,22E-01 |
| AT3G47070 | 0,67 | -0,96 | 2,56 | -0,37 | 7,08E-01 |
| AT3G47090 | 0,54 | 2,72 | 3,73 | 0,73 | 4,66E-01 |
| AT3G47160 | 0,74 | -0,87 | 2,38 | -0,37 | 7,15E-01 |
| AT3G47370 | 0,35 | -0,25 | 4,32 | -0,06 | 9,54E-01 |
| AT3G47420 | 0,51 | 0,45 | 2,50 | 0,18 | 8,57E-01 |
| AT3G47480 | 0,70 | -0,93 | 2,81 | -0,33 | 7,39E-01 |
| AT3G47670 | 0,97 | 1,82 | 1,73 | 1,05 | 2,94E-01 |
| AT3G47675 | 0,21 | 2,01 | 4,37 | 0,46 | 6,45E-01 |
| AT3G47680 | 0,08 | 2,08 | 4,38 | 0,47 | 6,35E-01 |
| AT3G47700 | 0,43 | 0,66 | 2,62 | 0,25 | 8,01E-01 |
| AT3G47730 | 0,93 | 4,65 | 2,64 | 1,76 | 7,85E-02 |
| AT3G47810 | 0,27 | 1,46 | 4,33 | 0,34 | 7,36E-01 |
| AT3G47860 | 0,23 | -1,34 | 4,35 | -0,31 | 7,58E-01 |
| AT3G47890 | 1,54 | 2,85 | 1,56 | 1,83 | 6,72E-02 |
| AT3G47940 | 0,35 | 1,07 | 4,34 | 0,25 | 8,06E-01 |
| AT3G47960 | 1,01 | -0,31 | 1,71 | -0,18 | 8,58E-01 |
| AT3G47990 | 0,32 | 1,58 | 4,35 | 0,36 | 7,17E-01 |
| AT3G48060 | 1,66 | 1,25 | 1,78 | 0,70 | 4,82E-01 |
| AT3G48080 | 1,04 | 2,08 | 2,00 | 1,04 | 2,99E-01 |
| AT3G48090 | 1,08 | 2,85 | 2,63 | 1,08 | 2,79E-01 |
| AT3G48100 | 0,85 | -0,04 | 4,36 | -0,01 | 9,94E-01 |
| AT3G48131 | 0,31 | -0,87 | 4,26 | -0,20 | 8,39E-01 |
| AT3G48150 | 0,55 | 2,52 | 3,84 | 0,66 | 5,12E-01 |
| AT3G48170 | 0,90 | 1,38 | 2,55 | 0,54 | 5,89E-01 |
| AT3G48195 | 0,48 | 1,61 | 3,02 | 0,53 | 5,95E-01 |
| AT3G48360 | 1,03 | 3,27 | 2,85 | 1,15 | 2,50E-01 |
| AT3G48520 | 0,38 | 3,15 | 4,34 | 0,73 | 4,67E-01 |
| AT3G48570 | 0,28 | -1,53 | 4,33 | -0,35 | 7,24E-01 |
| AT3G48640 | 0,25 | 1,05 | 4,33 | 0,24 | 8,08E-01 |
| AT3G48690 | 0,79 | 2,55 | 2,46 | 1,04 | 2,99E-01 |
| AT3G48740 | 0,68 | 0,81 | 1,96 | 0,41 | 6,81E-01 |
| AT3G48770 | 1,49 | -1,16 | 1,43 | -0,81 | 4,17E-01 |
| AT3G08005 | 0,05 | -0,58 | 4,38 | -0,13 | 8,95E-01 |
| AT3G48840 | 0,12 | 1,15 | 4,35 | 0,26 | 7,92E-01 |
| AT3G48930 | 0,53 | 1,36 | 2,59 | 0,53 | 5,99E-01 |
| AT3G48990 | 0,58 | 3,52 | 2,98 | 1,18 | 2,38E-01 |
| AT3G49010 | 1,04 | 0,83 | 1,64 | 0,51 | 6,12E-01 |
| AT3G49055 | 0,42 | 2,30 | 4,37 | 0,53 | 5,99E-01 |
| AT3G49060 | 0,44 | -2,37 | 3,23 | -0,73 | 4,63E-01 |
| AT3G49070 | 1,57 | 0,00 | 3,15 | 0,00 | 1,00E+00 |
| AT3G49140 | 0,93 | 1,05 | 2,16 | 0,49 | 6,27E-01 |
| AT3G49160 | 0,11 | 1,33 | 4,36 | 0,31 | 7,59E-01 |
| AT3G49250 | 0,54 | 0,00 | 4,15 | 0,00 | 9,99E-01 |
| AT3G49350 | 0,36 | 2,32 | 4,30 | 0,54 | 5,90E-01 |
| AT3G49490 | 1,14 | 2,68 | 2,05 | 1,31 | 1,91E-01 |
| AT3G49530 | 1,18 | 0,42 | 1,94 | 0,22 | 8,28E-01 |
| AT3G49600 | 1,57 | 3,39 | 1,90 | 1,79 | 7,34E-02 |

| | | | | | |
|---|---|---|---|---|---|
| AT3G49601 | 0,27 | -0,51 | 4,25 | -0,12 | 9,04E-01 |
| AT3G49650 | 0,08 | 2,08 | 4,38 | 0,47 | 6,35E-01 |
| AT3G49720 | 1,09 | -2,91 | 2,16 | -1,35 | 1,78E-01 |
| AT3G49790 | 0,38 | -0,52 | 3,21 | -0,16 | 8,72E-01 |
| AT3G49800 | 1,37 | 1,00 | 1,37 | 0,73 | 4,66E-01 |
| AT3G49870 | 0,08 | 2,08 | 4,38 | 0,47 | 6,35E-01 |
| AT3G49980 | 0,21 | 0,06 | 4,31 | 0,01 | 9,89E-01 |
| AT3G49990 | 0,57 | 2,70 | 3,69 | 0,73 | 4,64E-01 |
| AT3G50060 | 0,43 | -1,17 | 2,69 | -0,43 | 6,65E-01 |
| AT3G50110 | 0,18 | 0,41 | 4,28 | 0,10 | 9,23E-01 |
| AT3G50120 | 0,78 | 0,59 | 2,22 | 0,27 | 7,91E-01 |
| AT3G50240 | 0,71 | 3,87 | 4,31 | 0,90 | 3,68E-01 |
| AT3G50320 | 1,17 | -2,03 | 1,90 | -1,07 | 2,86E-01 |
| AT3G50380 | 1,07 | 0,77 | 2,39 | 0,32 | 7,46E-01 |
| AT3G50500 | 1,35 | 0,60 | 1,88 | 0,32 | 7,48E-01 |
| AT3G50670 | 0,33 | 1,55 | 4,30 | 0,36 | 7,19E-01 |
| AT3G50690 | 0,85 | -0,28 | 1,76 | -0,16 | 8,76E-01 |
| AT3G50740 | 0,53 | -0,49 | 3,18 | -0,16 | 8,77E-01 |
| AT3G50750 | 1,00 | -1,41 | 1,95 | -0,72 | 4,70E-01 |
| AT3G50780 | 0,08 | 2,33 | 4,36 | 0,53 | 5,93E-01 |
| AT3G50790 | 0,29 | 0,17 | 4,33 | 0,04 | 9,68E-01 |
| AT3G50808 | 0,10 | -1,26 | 4,35 | -0,29 | 7,72E-01 |
| AT3G50825 | 0,41 | -3,13 | 4,28 | -0,73 | 4,64E-01 |
| AT3G50920 | 0,48 | 1,70 | 4,09 | 0,42 | 6,78E-01 |
| AT3G51000 | 0,52 | 0,56 | 3,08 | 0,18 | 8,56E-01 |
| AT3G51120 | 1,56 | 4,13 | 1,80 | 2,30 | 2,14E-02 |
| AT3G51135 | 0,51 | -0,06 | 2,60 | -0,02 | 9,82E-01 |
| AT3G51160 | 0,58 | -0,92 | 3,15 | -0,29 | 7,71E-01 |
| AT3G51265 | 0,13 | -1,34 | 4,35 | -0,31 | 7,58E-01 |
| AT3G51310 | 0,51 | 1,85 | 3,17 | 0,59 | 5,58E-01 |
| AT3G51580 | 0,25 | 2,55 | 4,32 | 0,59 | 5,55E-01 |
| AT3G51600 | 1,48 | -1,59 | 1,65 | -0,96 | 3,36E-01 |
| AT3G51660 | 0,43 | -1,44 | 4,15 | -0,35 | 7,29E-01 |
| AT3G51790 | 0,60 | 4,23 | 3,63 | 1,16 | 2,45E-01 |
| AT3G51820 | 0,59 | 3,17 | 3,81 | 0,83 | 4,05E-01 |
| AT3G51840 | 0,33 | 1,16 | 4,31 | 0,27 | 7,88E-01 |
| AT3G51880 | 0,48 | 1,61 | 3,18 | 0,51 | 6,13E-01 |
| AT3G51890 | 1,41 | 2,04 | 2,67 | 0,77 | 4,44E-01 |
| AT3G51895 | 0,82 | 4,48 | 2,70 | 1,66 | 9,75E-02 |
| AT3G51920 | 0,46 | 0,30 | 2,58 | 0,12 | 9,08E-01 |
| AT3G08215 | 0,08 | 2,33 | 4,36 | 0,53 | 5,93E-01 |
| AT3G51980 | 0,42 | -0,90 | 3,35 | -0,27 | 7,89E-01 |
| AT3G52060 | 0,46 | 1,07 | 2,74 | 0,39 | 6,95E-01 |
| AT3G52072 | 0,22 | 0,08 | 4,27 | 0,02 | 9,85E-01 |
| AT3G08230 | 0,12 | 0,63 | 4,33 | 0,15 | 8,85E-01 |
| AT3G52110 | 0,12 | 1,65 | 4,34 | 0,38 | 7,04E-01 |
| AT3G52180 | 0,66 | 0,62 | 2,53 | 0,25 | 8,06E-01 |
| AT3G52190 | 0,13 | 2,37 | 4,36 | 0,54 | 5,86E-01 |
| AT3G52220 | 0,11 | 1,33 | 4,36 | 0,31 | 7,59E-01 |
| AT3G52260 | 0,64 | 4,37 | 3,54 | 1,23 | 2,17E-01 |
| AT3G52400 | 1,57 | -0,13 | 1,29 | -0,10 | 9,20E-01 |
| AT3G52480 | 0,53 | -0,63 | 2,85 | -0,22 | 8,24E-01 |
| AT3G52560 | 0,47 | 1,16 | 3,29 | 0,35 | 7,25E-01 |
| AT3G52640 | 0,21 | 3,36 | 4,33 | 0,78 | 4,37E-01 |
| AT3G52730 | 0,69 | 1,18 | 3,76 | 0,31 | 7,54E-01 |
| AT3G52740 | 0,17 | -0,44 | 4,37 | -0,10 | 9,19E-01 |

| | | | | | |
|---|---|---|---|---|---|
| AT3G52870 | 0,19 | -2,00 | 4,31 | -0,46 | 6,43E-01 |
| AT3G53015 | 1,03 | 3,71 | 2,20 | 1,68 | 9,23E-02 |
| AT3G53016 | 0,52 | -1,61 | 4,03 | -0,40 | 6,90E-01 |
| AT3G53100 | 0,69 | -1,72 | 3,07 | -0,56 | 5,74E-01 |
| AT3G53380 | 0,40 | -2,32 | 4,16 | -0,56 | 5,77E-01 |
| AT3G53410 | 0,47 | 2,86 | 4,35 | 0,66 | 5,11E-01 |
| AT3G53500 | 0,57 | -0,33 | 2,72 | -0,12 | 9,04E-01 |
| AT3G53510 | 0,25 | 3,58 | 4,32 | 0,83 | 4,07E-01 |
| AT3G53540 | 1,08 | 4,07 | 2,23 | 1,83 | 6,79E-02 |
| AT3G53560 | 0,23 | -1,11 | 4,34 | -0,25 | 7,99E-01 |
| AT3G53570 | 0,21 | 2,01 | 4,37 | 0,46 | 6,45E-01 |
| AT3G53610 | 0,53 | -0,20 | 3,03 | -0,07 | 9,47E-01 |
| AT3G53710 | 0,37 | 1,24 | 3,35 | 0,37 | 7,10E-01 |
| AT3G53730 | 0,67 | 0,37 | 2,25 | 0,17 | 8,69E-01 |
| AT3G53810 | 0,44 | 0,35 | 4,18 | 0,08 | 9,33E-01 |
| AT3G53890 | 0,52 | 2,46 | 3,21 | 0,77 | 4,44E-01 |
| AT3G53910 | 1,01 | -0,59 | 1,90 | -0,31 | 7,56E-01 |
| AT3G53920 | 0,24 | 2,45 | 4,35 | 0,56 | 5,73E-01 |
| AT3G53960 | 0,98 | 3,02 | 2,46 | 1,23 | 2,20E-01 |
| AT3G54100 | 0,47 | 2,16 | 3,16 | 0,68 | 4,95E-01 |
| AT3G54120 | 0,20 | 2,11 | 4,37 | 0,48 | 6,29E-01 |
| AT3G54140 | 0,70 | 0,85 | 2,47 | 0,35 | 7,29E-01 |
| AT3G54210 | 1,01 | -0,66 | 1,55 | -0,42 | 6,73E-01 |
| AT3G54290 | 0,42 | 0,26 | 4,25 | 0,06 | 9,50E-01 |
| AT3G54350 | 1,02 | 2,10 | 1,81 | 1,16 | 2,47E-01 |
| AT3G54360 | 1,14 | 3,20 | 1,92 | 1,67 | 9,49E-02 |
| AT3G54400 | 0,32 | 2,82 | 4,34 | 0,65 | 5,16E-01 |
| AT3G54440 | 0,61 | -2,31 | 2,22 | -1,04 | 2,98E-01 |
| AT3G54460 | 0,97 | 3,57 | 2,69 | 1,33 | 1,83E-01 |
| AT3G54470 | 0,44 | 2,88 | 3,28 | 0,88 | 3,80E-01 |
| AT3G54500 | 1,37 | 3,31 | 2,06 | 1,60 | 1,09E-01 |
| AT3G54620 | 0,24 | 2,85 | 4,34 | 0,66 | 5,11E-01 |
| AT3G54630 | 0,68 | 3,43 | 4,33 | 0,79 | 4,29E-01 |
| AT3G54640 | 0,22 | -1,21 | 4,35 | -0,28 | 7,81E-01 |
| AT3G54650 | 0,62 | 0,70 | 3,11 | 0,22 | 8,23E-01 |
| AT3G54750 | 0,48 | 3,10 | 4,35 | 0,71 | 4,76E-01 |
| AT3G54760 | 1,04 | 1,05 | 1,80 | 0,58 | 5,60E-01 |
| AT3G54790 | 1,31 | 0,72 | 1,82 | 0,40 | 6,92E-01 |
| AT3G54860 | 0,55 | 1,19 | 2,88 | 0,41 | 6,78E-01 |
| AT3G54900 | 0,95 | 2,47 | 1,86 | 1,33 | 1,84E-01 |
| AT3G54960 | 0,90 | 0,05 | 1,79 | 0,03 | 9,78E-01 |
| AT3G55040 | 0,21 | 0,00 | 4,41 | 0,00 | 1,00E+00 |
| AT3G55050 | 0,45 | 1,41 | 3,19 | 0,44 | 6,60E-01 |
| AT3G55060 | 0,14 | -0,45 | 4,33 | -0,10 | 9,17E-01 |
| AT3G55130 | 0,51 | -2,00 | 3,12 | -0,64 | 5,21E-01 |
| AT3G55160 | 0,95 | 2,11 | 1,80 | 1,17 | 2,43E-01 |
| AT3G55252 | 0,66 | -1,24 | 2,18 | -0,57 | 5,69E-01 |
| AT3G55254 | 0,80 | 0,56 | 2,06 | 0,27 | 7,85E-01 |
| AT3G55330 | 0,44 | 1,09 | 2,72 | 0,40 | 6,90E-01 |
| AT3G55360 | 0,60 | 0,38 | 2,36 | 0,16 | 8,73E-01 |
| AT3G55400 | 0,18 | 1,70 | 4,39 | 0,39 | 6,99E-01 |
| AT3G55430 | 0,77 | 0,00 | 1,99 | 0,00 | 1,00E+00 |
| AT3G55480 | 1,64 | 3,23 | 1,67 | 1,93 | 5,30E-02 |
| AT3G55840 | 1,41 | 2,13 | 1,68 | 1,27 | 2,03E-01 |
| AT3G55860 | 0,83 | -0,91 | 1,93 | -0,47 | 6,36E-01 |
| AT3G55935 | 0,46 | 0,05 | 2,58 | 0,02 | 9,83E-01 |

| | | | | | |
|---|---|---|---|---|---|
| AT3G55950 | 0,63 | -1,32 | 2,33 | -0,57 | 5,70E-01 |
| AT3G55960 | 0,64 | 0,98 | 2,51 | 0,39 | 6,98E-01 |
| AT3G56040 | 0,77 | 1,34 | 2,05 | 0,65 | 5,13E-01 |
| AT3G56050 | 0,97 | 1,08 | 1,86 | 0,58 | 5,62E-01 |
| AT3G56085 | 0,12 | 0,31 | 4,35 | 0,07 | 9,44E-01 |
| AT3G56090 | 0,34 | -0,09 | 3,54 | -0,02 | 9,80E-01 |
| AT3G56110 | 0,41 | -0,56 | 3,27 | -0,17 | 8,64E-01 |
| AT3G56190 | 1,16 | 0,41 | 1,62 | 0,25 | 8,01E-01 |
| AT3G56210 | 0,45 | -0,69 | 4,36 | -0,16 | 8,74E-01 |
| AT3G56240 | 0,35 | -0,58 | 2,77 | -0,21 | 8,35E-01 |
| AT3G56290 | 1,19 | 2,28 | 2,20 | 1,04 | 3,00E-01 |
| AT3G56320 | 0,58 | 1,33 | 3,86 | 0,35 | 7,30E-01 |
| AT3G56340 | 0,44 | -0,36 | 3,27 | -0,11 | 9,12E-01 |
| AT3G56370 | 0,84 | 2,24 | 2,24 | 1,00 | 3,19E-01 |
| AT3G56460 | 1,07 | 2,08 | 1,83 | 1,14 | 2,55E-01 |
| AT3G56480 | 0,39 | 0,58 | 2,72 | 0,21 | 8,32E-01 |
| AT3G56670 | 0,38 | 0,15 | 4,24 | 0,03 | 9,73E-01 |
| AT3G56690 | 0,43 | 0,42 | 4,13 | 0,10 | 9,18E-01 |
| AT3G56790 | 0,33 | 0,55 | 4,36 | 0,13 | 8,99E-01 |
| AT3G56800 | 1,39 | -1,11 | 1,39 | -0,80 | 4,25E-01 |
| AT3G56820 | 0,25 | 0,00 | 4,40 | 0,00 | 1,00E+00 |
| AT3G56830 | 0,17 | 3,04 | 4,33 | 0,70 | 4,83E-01 |
| AT3G56850 | 0,48 | 0,33 | 3,25 | 0,10 | 9,20E-01 |
| AT3G56860 | 0,49 | -0,49 | 2,57 | -0,19 | 8,50E-01 |
| AT3G56950 | 0,79 | 0,80 | 2,16 | 0,37 | 7,12E-01 |
| AT3G56970 | 0,12 | 2,81 | 4,34 | 0,65 | 5,18E-01 |
| AT3G56990 | 0,46 | 1,16 | 3,25 | 0,36 | 7,21E-01 |
| AT3G57020 | 0,42 | -0,03 | 4,39 | -0,01 | 9,94E-01 |
| AT3G57030 | 0,05 | -0,58 | 4,38 | -0,13 | 8,95E-01 |
| AT3G08825 | 0,72 | -0,45 | 2,92 | -0,15 | 8,78E-01 |
| AT3G57050 | 0,52 | 1,10 | 2,50 | 0,44 | 6,61E-01 |
| AT3G08855 | 0,12 | 1,15 | 4,35 | 0,26 | 7,92E-01 |
| AT3G57140 | 0,48 | 2,13 | 3,04 | 0,70 | 4,84E-01 |
| AT3G57240 | 0,24 | 0,00 | 4,41 | 0,00 | 1,00E+00 |
| AT3G57290 | 1,20 | 0,16 | 2,12 | 0,08 | 9,39E-01 |
| AT3G57350 | 0,44 | 2,20 | 3,26 | 0,67 | 5,01E-01 |
| AT3G57420 | 0,74 | -0,24 | 2,15 | -0,11 | 9,12E-01 |
| AT3G57460 | 0,24 | -1,45 | 4,28 | -0,34 | 7,35E-01 |
| AT3G08935 | 1,61 | -1,21 | 2,06 | -0,59 | 5,56E-01 |
| AT3G57470 | 0,21 | 0,78 | 4,40 | 0,18 | 8,60E-01 |
| AT3G57530 | 1,08 | -0,72 | 1,82 | -0,40 | 6,92E-01 |
| AT3G57540 | 0,08 | -0,56 | 4,38 | -0,13 | 8,99E-01 |
| AT3G57580 | 0,45 | -1,52 | 3,25 | -0,47 | 6,41E-01 |
| AT3G57610 | 1,05 | 1,94 | 4,32 | 0,45 | 6,53E-01 |
| AT3G57640 | 0,11 | -1,28 | 4,35 | -0,29 | 7,69E-01 |
| AT3G57765 | 1,26 | -0,36 | 1,37 | -0,26 | 7,95E-01 |
| AT3G57780 | 0,43 | 0,00 | 4,38 | 0,00 | 1,00E+00 |
| AT3G57790 | 0,44 | 2,48 | 4,30 | 0,58 | 5,63E-01 |
| AT3G57800 | 0,37 | -1,53 | 4,25 | -0,36 | 7,18E-01 |
| AT3G57870 | 0,53 | 1,65 | 4,38 | 0,38 | 7,07E-01 |
| AT3G57880 | 0,23 | 0,83 | 4,27 | 0,20 | 8,45E-01 |
| AT3G57930 | 0,30 | 0,02 | 4,35 | 0,00 | 9,96E-01 |
| AT3G57965 | 0,28 | 3,41 | 4,33 | 0,79 | 4,31E-01 |
| AT3G57980 | 0,74 | 0,87 | 2,63 | 0,33 | 7,39E-01 |
| AT3G57990 | 0,40 | -0,36 | 3,23 | -0,11 | 9,12E-01 |
| AT3G58100 | 1,03 | 4,57 | 3,28 | 1,39 | 1,63E-01 |

| | | | | | |
|---|---|---|---|---|---|
| AT3G58120 | 0,15 | 2,31 | 4,37 | 0,53 | 5,98E-01 |
| AT3G58140 | 0,42 | -1,20 | 3,31 | -0,36 | 7,16E-01 |
| AT3G58170 | 0,42 | 0,61 | 4,30 | 0,14 | 8,88E-01 |
| AT3G58210 | 0,64 | 1,83 | 4,40 | 0,42 | 6,77E-01 |
| AT3G58270 | 0,99 | 2,25 | 2,44 | 0,92 | 3,55E-01 |
| AT3G58500 | 1,34 | 1,73 | 1,40 | 1,24 | 2,16E-01 |
| AT3G58560 | 0,23 | -0,23 | 4,29 | -0,05 | 9,58E-01 |
| AT3G58580 | 0,38 | 2,25 | 4,27 | 0,53 | 5,99E-01 |
| AT3G58670 | 0,12 | 1,15 | 4,35 | 0,26 | 7,92E-01 |
| AT3G58730 | 1,29 | -0,12 | 1,61 | -0,08 | 9,40E-01 |
| AT3G58750 | 0,61 | 2,55 | 3,85 | 0,66 | 5,07E-01 |
| AT3G58790 | 0,34 | -0,01 | 4,31 | 0,00 | 9,99E-01 |
| AT3G58970 | 1,53 | 5,04 | 2,34 | 2,16 | 3,10E-02 |
| AT3G58990 | 0,12 | 0,63 | 4,33 | 0,15 | 8,85E-01 |
| AT3G59040 | 1,65 | 3,62 | 1,82 | 1,99 | 4,64E-02 |
| AT3G59050 | 0,52 | 1,12 | 3,00 | 0,37 | 7,10E-01 |
| AT3G59060 | 1,47 | -0,35 | 1,38 | -0,25 | 8,02E-01 |
| AT3G59120 | 0,10 | 0,34 | 4,35 | 0,08 | 9,37E-01 |
| AT3G59280 | 0,32 | 0,61 | 4,33 | 0,14 | 8,87E-01 |
| AT3G59360 | 0,61 | 1,96 | 2,82 | 0,69 | 4,87E-01 |
| AT3G59400 | 1,33 | -0,36 | 1,46 | -0,25 | 8,03E-01 |
| AT3G59540 | 1,10 | 1,76 | 1,96 | 0,90 | 3,68E-01 |
| AT3G59570 | 0,57 | 1,23 | 2,90 | 0,42 | 6,71E-01 |
| AT3G59660 | 1,15 | 1,64 | 1,71 | 0,96 | 3,38E-01 |
| AT3G59760 | 0,20 | -0,70 | 4,38 | -0,16 | 8,72E-01 |
| AT3G59840 | 0,15 | -0,41 | 4,32 | -0,09 | 9,25E-01 |
| AT3G59910 | 0,48 | 1,07 | 4,08 | 0,26 | 7,94E-01 |
| AT3G59950 | 1,22 | -1,43 | 2,33 | -0,61 | 5,40E-01 |
| AT3G59980 | 0,35 | 1,33 | 3,40 | 0,39 | 6,96E-01 |
| AT3G60030 | 1,14 | 3,11 | 2,11 | 1,48 | 1,40E-01 |
| AT3G09335 | 0,23 | 0,69 | 4,30 | 0,16 | 8,72E-01 |
| AT3G60050 | 0,12 | 2,69 | 4,35 | 0,62 | 5,36E-01 |
| AT3G60070 | 0,35 | 1,66 | 4,39 | 0,38 | 7,06E-01 |
| AT3G60160 | 1,27 | 2,09 | 1,75 | 1,20 | 2,31E-01 |
| AT3G60176 | 0,85 | -0,18 | 2,52 | -0,07 | 9,44E-01 |
| AT3G60220 | 0,15 | -1,60 | 4,33 | -0,37 | 7,11E-01 |
| AT3G60250 | 1,02 | 1,88 | 2,07 | 0,91 | 3,64E-01 |
| AT3G60350 | 1,30 | 3,87 | 1,98 | 1,95 | 5,09E-02 |
| AT3G60380 | 0,49 | 0,39 | 3,92 | 0,10 | 9,21E-01 |
| AT3G60540 | 0,08 | -0,56 | 4,38 | -0,13 | 8,99E-01 |
| AT3G60670 | 0,12 | 0,63 | 4,33 | 0,15 | 8,85E-01 |
| AT3G60740 | 0,66 | -0,81 | 2,27 | -0,35 | 7,23E-01 |
| AT3G60760 | 0,12 | 2,87 | 4,34 | 0,66 | 5,08E-01 |
| AT3G60820 | 0,15 | -0,64 | 4,39 | -0,15 | 8,84E-01 |
| AT3G60990 | 0,73 | 0,55 | 2,58 | 0,21 | 8,31E-01 |
| AT3G61070 | 0,50 | 1,41 | 4,02 | 0,35 | 7,25E-01 |
| AT3G09565 | 0,50 | -0,97 | 3,91 | -0,25 | 8,03E-01 |
| AT3G61190 | 1,32 | 0,37 | 1,57 | 0,24 | 8,12E-01 |
| AT3G61200 | 0,69 | 2,17 | 2,38 | 0,91 | 3,62E-01 |
| AT3G61260 | 0,83 | -1,24 | 2,52 | -0,49 | 6,23E-01 |
| AT3G61310 | 0,38 | 2,13 | 4,38 | 0,49 | 6,27E-01 |
| AT3G61380 | 0,18 | 1,13 | 4,32 | 0,26 | 7,94E-01 |
| AT3G61420 | 0,37 | -0,14 | 4,41 | -0,03 | 9,75E-01 |
| AT3G61460 | 0,83 | -0,61 | 1,75 | -0,35 | 7,28E-01 |
| AT3G61530 | 0,25 | 3,04 | 4,33 | 0,70 | 4,83E-01 |
| AT3G61640 | 0,38 | -0,08 | 3,21 | -0,02 | 9,80E-01 |

| | | | | | |
|---|---|---|---|---|---|
| AT3G61740 | 0,78 | 3,00 | 2,39 | 1,25 | 2,10E-01 |
| AT3G61750 | 0,24 | 0,00 | 4,41 | 0,00 | 1,00E+00 |
| AT3G61780 | 1,45 | 2,66 | 1,56 | 1,70 | 8,83E-02 |
| AT3G61800 | 0,23 | -1,21 | 4,29 | -0,28 | 7,77E-01 |
| AT3G61820 | 1,36 | -0,07 | 1,57 | -0,04 | 9,65E-01 |
| AT3G61860 | 0,39 | 0,79 | 4,29 | 0,18 | 8,55E-01 |
| AT3G61890 | 0,44 | -0,34 | 3,45 | -0,10 | 9,21E-01 |
| AT3G61960 | 1,02 | -0,21 | 1,93 | -0,11 | 9,13E-01 |
| AT3G61990 | 0,15 | 0,69 | 4,31 | 0,16 | 8,73E-01 |
| AT3G62070 | 0,12 | -1,57 | 4,34 | -0,36 | 7,18E-01 |
| AT3G62120 | 1,00 | 2,06 | 2,01 | 1,03 | 3,05E-01 |
| AT3G62240 | 0,93 | 2,26 | 2,73 | 0,83 | 4,07E-01 |
| AT3G62260 | 0,41 | 1,52 | 4,09 | 0,37 | 7,11E-01 |
| AT3G62310 | 0,33 | 2,48 | 4,35 | 0,57 | 5,68E-01 |
| AT3G62330 | 0,73 | 2,66 | 2,90 | 0,92 | 3,59E-01 |
| AT3G62470 | 0,24 | 2,80 | 4,34 | 0,65 | 5,18E-01 |
| AT3G62540 | 0,28 | 1,64 | 4,38 | 0,38 | 7,08E-01 |
| AT3G09795 | 0,21 | -1,08 | 4,30 | -0,25 | 8,01E-01 |
| AT3G62590 | 0,64 | 1,54 | 2,85 | 0,54 | 5,88E-01 |
| AT3G62600 | 0,95 | 0,20 | 1,80 | 0,11 | 9,11E-01 |
| AT3G09825 | 1,22 | -2,43 | 1,77 | -1,38 | 1,69E-01 |
| AT3G09845 | 0,08 | -0,56 | 4,38 | -0,13 | 8,99E-01 |
| AT3G62630 | 0,91 | -0,75 | 2,19 | -0,34 | 7,31E-01 |
| AT3G62800 | 0,37 | 3,54 | 4,32 | 0,82 | 4,12E-01 |
| AT3G62840 | 0,66 | 1,85 | 3,07 | 0,60 | 5,46E-01 |
| AT3G62860 | 0,81 | 2,76 | 3,52 | 0,78 | 4,34E-01 |
| AT3G62870 | 1,22 | 3,25 | 1,88 | 1,73 | 8,40E-02 |
| AT3G62900 | 0,94 | 2,22 | 2,14 | 1,04 | 3,01E-01 |
| AT3G62910 | 0,49 | -0,98 | 4,06 | -0,24 | 8,08E-01 |
| AT3G62940 | 0,31 | -1,44 | 4,34 | -0,33 | 7,41E-01 |
| AT3G63052 | 0,42 | -1,20 | 3,31 | -0,36 | 7,16E-01 |
| AT3G63110 | 0,56 | -1,49 | 2,56 | -0,58 | 5,61E-01 |
| AT3G63140 | 0,93 | -1,39 | 1,97 | -0,71 | 4,80E-01 |
| AT3G63150 | 0,90 | 1,00 | 2,33 | 0,43 | 6,68E-01 |
| AT3G63160 | 1,15 | -1,42 | 1,85 | -0,77 | 4,43E-01 |
| AT3G63180 | 1,24 | 2,23 | 1,83 | 1,22 | 2,22E-01 |
| AT3G63190 | 0,25 | -0,89 | 4,30 | -0,21 | 8,36E-01 |
| AT3G63210 | 0,47 | -3,19 | 4,27 | -0,75 | 4,55E-01 |
| AT3G63310 | 1,42 | 2,49 | 1,92 | 1,30 | 1,95E-01 |
| AT3G63340 | 1,13 | 3,66 | 2,47 | 1,48 | 1,38E-01 |
| AT3G63390 | 0,24 | 0,16 | 4,28 | 0,04 | 9,69E-01 |
| AT3G63540 | 1,40 | -1,03 | 1,59 | -0,65 | 5,16E-01 |
| AT4G00005 | 0,19 | 0,58 | 4,36 | 0,13 | 8,94E-01 |
| AT4G00020 | 0,79 | -0,11 | 2,13 | -0,05 | 9,57E-01 |
| AT4G00070 | 0,43 | 1,39 | 4,18 | 0,33 | 7,39E-01 |
| AT4G00110 | 1,14 | -0,48 | 2,46 | -0,20 | 8,45E-01 |
| AT4G00120 | 1,32 | -3,33 | 1,73 | -1,92 | 5,49E-02 |
| AT4G00165 | 0,65 | -1,73 | 2,64 | -0,66 | 5,11E-01 |
| AT4G00180 | 0,45 | 1,59 | 4,34 | 0,37 | 7,14E-01 |
| AT4G00280 | 0,25 | 0,00 | 4,40 | 0,00 | 1,00E+00 |
| AT4G00290 | 0,50 | -0,32 | 2,74 | -0,12 | 9,06E-01 |
| AT4G00330 | 0,80 | 0,70 | 1,94 | 0,36 | 7,17E-01 |
| AT4G03585 | 1,51 | -1,60 | 1,59 | -1,01 | 3,13E-01 |
| AT4G00370 | 0,63 | 2,94 | 2,59 | 1,14 | 2,55E-01 |
| AT4G00400 | 0,42 | -0,60 | 2,69 | -0,22 | 8,24E-01 |
| AT4G00500 | 0,24 | 1,23 | 4,34 | 0,28 | 7,77E-01 |

| | | | | | |
|---|---|---|---|---|---|
| AT4G00540 | 0,54 | -2,27 | 2,50 | -0,91 | 3,64E-01 |
| AT4G00550 | 0,15 | 1,94 | 4,32 | 0,45 | 6,54E-01 |
| AT4G00560 | 0,40 | 3,10 | 4,30 | 0,72 | 4,71E-01 |
| AT4G00620 | 0,40 | -1,21 | 4,19 | -0,29 | 7,72E-01 |
| AT4G00740 | 0,31 | -0,48 | 4,34 | -0,11 | 9,13E-01 |
| AT4G00810 | 0,25 | 2,79 | 4,34 | 0,64 | 5,20E-01 |
| AT4G00890 | 0,23 | -2,41 | 4,30 | -0,56 | 5,75E-01 |
| AT4G00900 | 1,09 | 3,51 | 2,00 | 1,75 | 7,99E-02 |
| AT4G00980 | 0,33 | 1,52 | 4,34 | 0,35 | 7,26E-01 |
| AT4G00985 | 0,26 | 0,00 | 4,41 | 0,00 | 1,00E+00 |
| AT4G01000 | 0,50 | 2,70 | 3,85 | 0,70 | 4,82E-01 |
| AT4G01030 | 0,51 | 1,07 | 2,94 | 0,36 | 7,17E-01 |
| AT4G01037 | 0,21 | -0,52 | 4,38 | -0,12 | 9,05E-01 |
| AT4G01070 | 1,43 | -0,07 | 1,58 | -0,05 | 9,64E-01 |
| AT4G01090 | 0,44 | 1,87 | 3,17 | 0,59 | 5,55E-01 |
| AT4G01110 | 0,08 | 2,08 | 4,38 | 0,47 | 6,35E-01 |
| AT4G01120 | 0,18 | 1,77 | 4,41 | 0,40 | 6,88E-01 |
| AT4G01180 | 0,08 | 2,08 | 4,38 | 0,47 | 6,35E-01 |
| AT4G01190 | 0,20 | 2,11 | 4,37 | 0,48 | 6,29E-01 |
| AT4G01200 | 0,46 | 2,03 | 4,38 | 0,46 | 6,43E-01 |
| AT4G01210 | 1,50 | 1,81 | 1,66 | 1,09 | 2,78E-01 |
| AT4G01240 | 0,94 | -1,38 | 1,92 | -0,72 | 4,71E-01 |
| AT4G01310 | 0,66 | -2,33 | 3,04 | -0,77 | 4,44E-01 |
| AT4G01395 | 0,71 | -1,28 | 2,17 | -0,59 | 5,57E-01 |
| AT4G01533 | 1,62 | -2,00 | 1,72 | -1,16 | 2,45E-01 |
| AT4G01540 | 0,20 | 0,30 | 4,34 | 0,07 | 9,46E-01 |
| AT4G01550 | 0,48 | 2,75 | 4,36 | 0,63 | 5,28E-01 |
| AT4G01700 | 0,60 | 1,29 | 3,12 | 0,41 | 6,79E-01 |
| AT4G01870 | 0,45 | 2,56 | 4,15 | 0,62 | 5,38E-01 |
| AT4G01950 | 0,49 | -1,81 | 2,62 | -0,69 | 4,89E-01 |
| AT4G01985 | 0,41 | -0,46 | 4,37 | -0,10 | 9,17E-01 |
| AT4G02195 | 0,71 | 0,10 | 2,56 | 0,04 | 9,70E-01 |
| AT4G02210 | 0,65 | 0,08 | 2,74 | 0,03 | 9,76E-01 |
| AT4G02230 | 0,28 | 1,26 | 4,36 | 0,29 | 7,73E-01 |
| AT4G02405 | 0,05 | -0,58 | 4,38 | -0,13 | 8,95E-01 |
| AT4G02410 | 0,84 | 2,01 | 2,80 | 0,72 | 4,74E-01 |
| AT4G02420 | 1,44 | -0,70 | 1,54 | -0,46 | 6,49E-01 |
| AT4G02450 | 0,79 | -0,41 | 2,26 | -0,18 | 8,57E-01 |
| AT4G02460 | 0,19 | 1,89 | 4,32 | 0,44 | 6,61E-01 |
| AT4G02530 | 0,43 | -1,17 | 3,28 | -0,36 | 7,22E-01 |
| AT4G04185 | 0,38 | 0,01 | 3,24 | 0,00 | 9,97E-01 |
| AT4G02550 | 0,12 | 0,63 | 4,33 | 0,15 | 8,85E-01 |
| AT4G02560 | 0,57 | 2,13 | 2,66 | 0,80 | 4,24E-01 |
| AT4G02590 | 0,27 | 2,11 | 4,38 | 0,48 | 6,30E-01 |
| AT4G02620 | 0,23 | 2,14 | 4,38 | 0,49 | 6,25E-01 |
| AT4G02680 | 0,66 | 2,29 | 3,78 | 0,61 | 5,45E-01 |
| AT4G02710 | 0,71 | 3,09 | 2,87 | 1,08 | 2,82E-01 |
| AT4G02715 | 1,63 | 3,74 | 1,88 | 1,99 | 4,61E-02 |
| AT4G02735 | 0,41 | 0,00 | 4,41 | 0,00 | 1,00E+00 |
| AT4G02740 | 0,60 | 1,26 | 2,94 | 0,43 | 6,67E-01 |
| AT4G02750 | 0,98 | 4,45 | 2,19 | 2,03 | 4,26E-02 |
| AT4G02910 | 0,98 | 0,36 | 2,19 | 0,16 | 8,70E-01 |
| AT4G02930 | 0,46 | 2,71 | 4,35 | 0,62 | 5,34E-01 |
| AT4G03080 | 1,69 | 2,23 | 1,29 | 1,73 | 8,28E-02 |
| AT4G03090 | 0,91 | -0,02 | 1,72 | -0,01 | 9,92E-01 |
| AT4G03110 | 0,81 | 2,00 | 2,90 | 0,69 | 4,90E-01 |

| | | | | | |
|---|---|---|---|---|---|
| AT4G03130 | 0,08 | 2,08 | 4,38 | 0,47 | 6,35E-01 |
| AT4G03150 | 0,26 | 2,13 | 4,38 | 0,49 | 6,27E-01 |
| AT4G03190 | 0,84 | 1,56 | 2,30 | 0,68 | 4,96E-01 |
| AT4G03230 | 0,47 | -0,58 | 4,03 | -0,14 | 8,85E-01 |
| AT4G03270 | 0,81 | 3,67 | 2,77 | 1,32 | 1,86E-01 |
| AT4G04475 | 0,42 | -1,18 | 3,27 | -0,36 | 7,19E-01 |
| AT4G04485 | 0,66 | 1,96 | 2,29 | 0,85 | 3,93E-01 |
| AT4G03340 | 0,12 | 2,69 | 4,35 | 0,62 | 5,36E-01 |
| AT4G03390 | 0,56 | 1,45 | 2,62 | 0,56 | 5,79E-01 |
| AT4G03400 | 0,95 | 0,60 | 1,91 | 0,32 | 7,52E-01 |
| AT4G03430 | 1,44 | 0,83 | 1,33 | 0,63 | 5,32E-01 |
| AT4G03435 | 0,08 | -0,56 | 4,38 | -0,13 | 8,99E-01 |
| AT4G03438 | 0,26 | -1,57 | 4,34 | -0,36 | 7,17E-01 |
| AT4G03450 | 0,60 | -0,72 | 2,64 | -0,27 | 7,85E-01 |
| AT4G03490 | 0,42 | -0,40 | 4,28 | -0,09 | 9,25E-01 |
| AT4G03505 | 0,25 | 0,00 | 4,40 | 0,00 | 1,00E+00 |
| AT4G03520 | 1,13 | 2,04 | 1,81 | 1,13 | 2,60E-01 |
| AT4G03570 | 0,98 | -1,55 | 1,60 | -0,97 | 3,33E-01 |
| AT4G03590 | 1,16 | -2,12 | 2,05 | -1,03 | 3,01E-01 |
| AT4G03728 | 0,69 | -1,30 | 3,59 | -0,36 | 7,18E-01 |
| AT4G03740 | 1,37 | -1,24 | 1,67 | -0,74 | 4,58E-01 |
| AT4G04675 | 1,07 | 0,15 | 2,00 | 0,07 | 9,42E-01 |
| AT4G04030 | 0,82 | -1,76 | 2,43 | -0,72 | 4,69E-01 |
| AT4G04221 | 1,00 | 0,32 | 1,94 | 0,17 | 8,69E-01 |
| AT4G04805 | 1,38 | -1,45 | 1,98 | -0,74 | 4,62E-01 |
| AT4G04370 | 1,03 | 2,77 | 2,08 | 1,33 | 1,83E-01 |
| AT4G04470 | 0,08 | -0,56 | 4,38 | -0,13 | 8,99E-01 |
| AT4G04580 | 0,11 | 1,28 | 4,35 | 0,29 | 7,68E-01 |
| AT4G04614 | 0,51 | 1,66 | 4,39 | 0,38 | 7,06E-01 |
| AT4G04632 | 0,44 | 0,76 | 3,99 | 0,19 | 8,50E-01 |
| AT4G04650 | 0,82 | -2,94 | 2,23 | -1,32 | 1,87E-01 |
| AT4G04710 | 0,47 | 2,16 | 3,16 | 0,68 | 4,95E-01 |
| AT4G04720 | 1,28 | 2,95 | 2,07 | 1,42 | 1,55E-01 |
| AT4G04740 | 0,87 | 1,72 | 1,77 | 0,97 | 3,33E-01 |
| AT4G04745 | 1,61 | -2,97 | 1,62 | -1,84 | 6,55E-02 |
| AT4G04770 | 1,41 | 1,31 | 1,71 | 0,77 | 4,43E-01 |
| AT4G04910 | 0,75 | -0,86 | 2,18 | -0,40 | 6,93E-01 |
| AT4G04950 | 0,83 | 1,79 | 3,71 | 0,48 | 6,30E-01 |
| AT4G04957 | 0,91 | -1,28 | 2,39 | -0,54 | 5,92E-01 |
| AT4G04980 | 0,11 | -1,28 | 4,35 | -0,29 | 7,69E-01 |
| AT4G05070 | 1,30 | 0,43 | 1,57 | 0,27 | 7,85E-01 |
| AT4G05110 | 0,17 | 2,95 | 4,34 | 0,68 | 4,96E-01 |
| AT4G05085 | 0,05 | -0,58 | 4,38 | -0,13 | 8,95E-01 |
| AT4G05330 | 0,44 | 1,81 | 4,05 | 0,45 | 6,55E-01 |
| AT4G05410 | 0,51 | 1,05 | 4,07 | 0,26 | 7,95E-01 |
| AT4G05470 | 0,12 | 1,65 | 4,34 | 0,38 | 7,04E-01 |
| AT4G05631 | 0,41 | 0,36 | 3,39 | 0,11 | 9,15E-01 |
| AT4G06526 | 0,56 | -1,90 | 3,91 | -0,49 | 6,26E-01 |
| AT4G06639 | 0,21 | -0,16 | 4,33 | -0,04 | 9,70E-01 |
| AT4G06599 | 0,16 | -0,97 | 4,35 | -0,22 | 8,24E-01 |
| AT4G07350 | 0,08 | -0,56 | 4,38 | -0,13 | 8,99E-01 |
| AT4G07400 | 0,28 | 1,04 | 4,29 | 0,24 | 8,09E-01 |
| AT4G07410 | 1,28 | 1,02 | 2,04 | 0,50 | 6,18E-01 |
| AT4G07524 | 0,19 | 1,20 | 4,29 | 0,28 | 7,79E-01 |
| AT4G07526 | 0,83 | 0,80 | 2,48 | 0,32 | 7,48E-01 |
| AT4G05225 | 0,74 | -2,50 | 3,01 | -0,83 | 4,06E-01 |

| | | | | | |
|---|---|---|---|---|---|
| AT4G07990 | 0,20 | -0,80 | 4,37 | -0,18 | 8,55E-01 |
| AT4G08230 | 0,30 | -1,62 | 4,28 | -0,38 | 7,05E-01 |
| AT4G08310 | 0,85 | -1,30 | 1,95 | -0,66 | 5,07E-01 |
| AT4G05415 | 0,95 | 4,14 | 2,61 | 1,59 | 1,13E-01 |
| AT4G08370 | 0,46 | 0,34 | 2,53 | 0,13 | 8,93E-01 |
| AT4G08400 | 0,54 | -1,74 | 2,84 | -0,61 | 5,41E-01 |
| AT4G08410 | 1,29 | -1,30 | 2,02 | -0,64 | 5,21E-01 |
| AT4G05535 | 0,95 | 0,76 | 2,40 | 0,32 | 7,51E-01 |
| AT4G08460 | 0,62 | 1,54 | 2,54 | 0,61 | 5,44E-01 |
| AT4G08480 | 0,67 | -1,08 | 3,03 | -0,36 | 7,21E-01 |
| AT4G08485 | 0,63 | -0,14 | 3,80 | -0,04 | 9,71E-01 |
| AT4G08590 | 0,56 | 3,19 | 4,33 | 0,74 | 4,61E-01 |
| AT4G08760 | 1,19 | -2,65 | 2,09 | -1,27 | 2,04E-01 |
| AT4G08810 | 0,77 | 2,67 | 2,56 | 1,05 | 2,96E-01 |
| AT4G08870 | 1,44 | 2,32 | 1,79 | 1,30 | 1,95E-01 |
| AT4G08874 | 0,12 | 0,31 | 4,35 | 0,07 | 9,44E-01 |
| AT4G08940 | 0,46 | 1,37 | 3,26 | 0,42 | 6,75E-01 |
| AT4G09010 | 1,11 | 0,75 | 1,78 | 0,42 | 6,72E-01 |
| AT4G09012 | 0,55 | 1,70 | 3,15 | 0,54 | 5,90E-01 |
| AT4G09035 | 0,16 | 0,85 | 4,31 | 0,20 | 8,44E-01 |
| AT4G09040 | 0,47 | -0,01 | 4,35 | 0,00 | 9,99E-01 |
| AT4G09340 | 0,44 | 1,95 | 3,23 | 0,60 | 5,46E-01 |
| AT4G09350 | 0,26 | -0,20 | 4,35 | -0,05 | 9,64E-01 |
| AT4G09490 | 0,39 | 0,00 | 4,41 | 0,00 | 1,00E+00 |
| AT4G09510 | 0,38 | 1,19 | 3,38 | 0,35 | 7,25E-01 |
| AT4G09620 | 0,40 | 3,80 | 4,32 | 0,88 | 3,79E-01 |
| AT4G09640 | 0,98 | -0,73 | 1,74 | -0,42 | 6,73E-01 |
| AT4G05740 | 1,41 | -1,20 | 1,47 | -0,82 | 4,14E-01 |
| AT4G09900 | 0,13 | -0,63 | 4,39 | -0,14 | 8,85E-01 |
| AT4G09960 | 0,46 | 0,91 | 4,38 | 0,21 | 8,36E-01 |
| AT4G09980 | 0,57 | 1,55 | 2,54 | 0,61 | 5,41E-01 |
| AT4G10070 | 1,18 | 2,72 | 2,19 | 1,24 | 2,16E-01 |
| AT4G10080 | 1,04 | -0,49 | 2,04 | -0,24 | 8,10E-01 |
| AT4G10130 | 0,55 | 0,82 | 3,83 | 0,21 | 8,31E-01 |
| AT4G10320 | 1,43 | 0,21 | 1,27 | 0,17 | 8,66E-01 |
| AT4G10400 | 0,39 | 1,78 | 4,41 | 0,40 | 6,86E-01 |
| AT4G10420 | 1,07 | 3,21 | 2,68 | 1,20 | 2,30E-01 |
| AT4G10440 | 0,08 | 2,08 | 4,38 | 0,47 | 6,35E-01 |
| AT4G10480 | 0,41 | 0,85 | 4,35 | 0,20 | 8,44E-01 |
| AT4G10540 | 0,12 | 2,87 | 4,34 | 0,66 | 5,08E-01 |
| AT4G10550 | 0,52 | 3,84 | 3,86 | 0,99 | 3,20E-01 |
| AT4G10570 | 1,14 | 0,45 | 1,61 | 0,28 | 7,78E-01 |
| AT4G10590 | 1,40 | 3,28 | 1,94 | 1,70 | 8,99E-02 |
| AT4G05800 | 1,37 | -0,26 | 1,63 | -0,16 | 8,72E-01 |
| AT4G10700 | 0,54 | -2,69 | 3,66 | -0,74 | 4,62E-01 |
| AT4G10730 | 0,28 | 1,64 | 4,38 | 0,38 | 7,08E-01 |
| AT4G10740 | 1,27 | -2,27 | 2,03 | -1,12 | 2,64E-01 |
| AT4G10760 | 0,19 | -2,00 | 4,31 | -0,46 | 6,43E-01 |
| AT4G10770 | 0,10 | 0,34 | 4,35 | 0,08 | 9,37E-01 |
| AT4G10790 | 1,30 | 2,15 | 1,48 | 1,46 | 1,45E-01 |
| AT4G10920 | 0,27 | 0,36 | 3,65 | 0,10 | 9,22E-01 |
| AT4G10960 | 0,05 | -0,58 | 4,38 | -0,13 | 8,95E-01 |
| AT4G10970 | 0,10 | 0,34 | 4,35 | 0,08 | 9,37E-01 |
| AT4G11010 | 0,17 | 1,90 | 4,33 | 0,44 | 6,61E-01 |
| AT4G11050 | 0,37 | -0,59 | 3,38 | -0,18 | 8,61E-01 |
| AT4G11080 | 0,05 | -0,58 | 4,38 | -0,13 | 8,95E-01 |

| | | | | | |
|---|---|---|---|---|---|
| AT4G05855 | 1,27 | 1,86 | 2,18 | 0,86 | 3,92E-01 |
| AT4G11216 | 0,26 | 2,13 | 4,38 | 0,49 | 6,27E-01 |
| AT4G11220 | 0,29 | -1,43 | 4,34 | -0,33 | 7,41E-01 |
| AT4G11260 | 0,66 | 0,75 | 4,30 | 0,18 | 8,61E-01 |
| AT4G11300 | 0,45 | 1,78 | 4,09 | 0,43 | 6,64E-01 |
| AT4G11340 | 1,18 | 0,34 | 1,49 | 0,23 | 8,22E-01 |
| AT4G11410 | 0,24 | 2,80 | 4,34 | 0,65 | 5,18E-01 |
| AT4G05880 | 0,80 | 0,80 | 2,78 | 0,29 | 7,73E-01 |
| AT4G11650 | 0,43 | 2,13 | 4,38 | 0,49 | 6,26E-01 |
| AT4G11670 | 0,44 | 1,19 | 2,67 | 0,45 | 6,56E-01 |
| AT4G11800 | 1,18 | 0,57 | 1,52 | 0,38 | 7,06E-01 |
| AT4G05925 | 1,28 | -2,91 | 1,94 | -1,50 | 1,34E-01 |
| AT4G11820 | 0,25 | 0,00 | 4,41 | 0,00 | 1,00E+00 |
| AT4G11840 | 1,34 | 1,75 | 1,49 | 1,17 | 2,41E-01 |
| AT4G11850 | 1,56 | 0,71 | 1,49 | 0,47 | 6,35E-01 |
| AT4G11860 | 0,25 | 2,98 | 4,33 | 0,69 | 4,92E-01 |
| AT4G11920 | 0,40 | 0,16 | 4,04 | 0,04 | 9,69E-01 |
| AT4G05930 | 0,37 | 0,32 | 4,13 | 0,08 | 9,37E-01 |
| AT4G12010 | 0,59 | 3,56 | 3,63 | 0,98 | 3,27E-01 |
| AT4G05980 | 0,40 | 1,98 | 4,31 | 0,46 | 6,46E-01 |
| AT4G12300 | 1,00 | 0,87 | 2,01 | 0,44 | 6,63E-01 |
| AT4G12310 | 0,40 | 0,23 | 2,75 | 0,08 | 9,34E-01 |
| AT4G12330 | 0,81 | -0,90 | 2,77 | -0,32 | 7,47E-01 |
| AT4G12420 | 1,53 | -0,06 | 1,85 | -0,03 | 9,76E-01 |
| AT4G12590 | 0,64 | -2,09 | 2,52 | -0,83 | 4,06E-01 |
| AT4G12710 | 0,37 | 2,89 | 4,33 | 0,67 | 5,05E-01 |
| AT4G12730 | 0,13 | -0,63 | 4,39 | -0,14 | 8,85E-01 |
| AT4G12750 | 1,05 | 2,72 | 2,04 | 1,34 | 1,81E-01 |
| AT4G12790 | 0,53 | 0,18 | 3,17 | 0,06 | 9,55E-01 |
| AT4G12820 | 0,35 | 0,00 | 4,41 | 0,00 | 1,00E+00 |
| AT4G12880 | 0,35 | 0,00 | 4,41 | 0,00 | 1,00E+00 |
| AT4G12980 | 0,57 | -0,02 | 2,56 | -0,01 | 9,95E-01 |
| AT4G13150 | 0,55 | 2,15 | 4,38 | 0,49 | 6,23E-01 |
| AT4G13250 | 1,54 | 1,04 | 1,86 | 0,56 | 5,74E-01 |
| AT4G13350 | 0,30 | -0,22 | 4,27 | -0,05 | 9,59E-01 |
| AT4G13395 | 0,15 | -1,60 | 4,33 | -0,37 | 7,11E-01 |
| AT4G13420 | 0,12 | 2,81 | 4,34 | 0,65 | 5,18E-01 |
| AT4G13550 | 0,15 | 1,93 | 4,33 | 0,45 | 6,55E-01 |
| AT4G13590 | 0,56 | -1,21 | 2,55 | -0,48 | 6,34E-01 |
| AT4G13630 | 0,80 | 4,36 | 3,40 | 1,28 | 2,00E-01 |
| AT4G13640 | 0,21 | 3,40 | 4,32 | 0,79 | 4,32E-01 |
| AT4G13780 | 1,18 | 2,43 | 1,82 | 1,34 | 1,81E-01 |
| AT4G13800 | 0,45 | 2,28 | 3,20 | 0,71 | 4,77E-01 |
| AT4G13820 | 0,22 | 0,75 | 4,27 | 0,18 | 8,60E-01 |
| AT4G13870 | 0,08 | 2,08 | 4,38 | 0,47 | 6,35E-01 |
| AT4G13930 | 1,56 | 0,63 | 1,45 | 0,44 | 6,61E-01 |
| AT4G13950 | 0,08 | 2,33 | 4,36 | 0,53 | 5,93E-01 |
| AT4G13970 | 0,50 | 2,31 | 3,84 | 0,60 | 5,48E-01 |
| AT4G14010 | 0,10 | -1,26 | 4,35 | -0,29 | 7,72E-01 |
| AT4G14040 | 0,70 | 0,02 | 2,58 | 0,01 | 9,94E-01 |
| AT4G14050 | 0,14 | 1,12 | 4,32 | 0,26 | 7,96E-01 |
| AT4G14070 | 0,95 | 2,59 | 2,61 | 0,99 | 3,22E-01 |
| AT4G14090 | 0,47 | -0,13 | 2,68 | -0,05 | 9,61E-01 |
| AT4G14110 | 0,26 | 2,30 | 4,37 | 0,53 | 5,98E-01 |
| AT4G14200 | 0,21 | 1,20 | 4,30 | 0,28 | 7,81E-01 |
| AT4G14210 | 0,70 | -0,37 | 1,94 | -0,19 | 8,50E-01 |

| | | | | | |
|---|---|---|---|---|---|
| AT4G14230 | 0,73 | 2,46 | 2,96 | 0,83 | 4,06E-01 |
| AT4G14240 | 0,36 | 0,82 | 4,35 | 0,19 | 8,51E-01 |
| AT4G14340 | 0,26 | 2,30 | 4,37 | 0,53 | 5,98E-01 |
| AT4G14342 | 0,72 | 4,50 | 4,29 | 1,05 | 2,94E-01 |
| AT4G14350 | 0,16 | 0,03 | 4,34 | 0,01 | 9,94E-01 |
| AT4G14370 | 0,47 | -0,34 | 2,69 | -0,13 | 8,99E-01 |
| AT4G14385 | 0,08 | 2,33 | 4,36 | 0,53 | 5,93E-01 |
| AT4G14420 | 0,43 | 2,13 | 4,38 | 0,49 | 6,26E-01 |
| AT4G14430 | 1,12 | 2,38 | 1,64 | 1,45 | 1,47E-01 |
| AT4G14440 | 0,19 | -1,86 | 4,32 | -0,43 | 6,67E-01 |
| AT4G14510 | 1,36 | 2,25 | 1,79 | 1,26 | 2,08E-01 |
| AT4G14680 | 0,20 | 2,11 | 4,37 | 0,48 | 6,29E-01 |
| AT4G14710 | 0,27 | -0,62 | 4,33 | -0,14 | 8,86E-01 |
| AT4G14740 | 0,35 | 0,28 | 4,39 | 0,06 | 9,49E-01 |
| AT4G14746 | 0,16 | 0,33 | 4,31 | 0,08 | 9,39E-01 |
| AT4G14790 | 0,23 | -0,64 | 4,39 | -0,15 | 8,84E-01 |
| AT4G14811 | 0,45 | -0,09 | 3,27 | -0,03 | 9,79E-01 |
| AT4G14880 | 1,32 | -1,46 | 1,50 | -0,97 | 3,31E-01 |
| AT4G14900 | 1,30 | 0,57 | 1,60 | 0,36 | 7,22E-01 |
| AT4G14905 | 0,68 | 0,06 | 2,57 | 0,03 | 9,80E-01 |
| AT4G14920 | 0,44 | 2,26 | 4,14 | 0,55 | 5,84E-01 |
| AT4G14990 | 0,87 | 1,67 | 1,80 | 0,93 | 3,52E-01 |
| AT4G15050 | 0,52 | -0,02 | 3,68 | -0,01 | 9,95E-01 |
| AT4G15053 | 0,24 | -2,00 | 4,31 | -0,46 | 6,42E-01 |
| AT4G15080 | 0,43 | 0,59 | 4,09 | 0,14 | 8,86E-01 |
| AT4G15200 | 0,36 | 3,22 | 4,34 | 0,74 | 4,59E-01 |
| AT4G15210 | 0,12 | 2,81 | 4,34 | 0,65 | 5,18E-01 |
| AT4G15242 | 0,72 | -1,61 | 1,96 | -0,82 | 4,13E-01 |
| AT4G06305 | 0,44 | -0,86 | 4,08 | -0,21 | 8,34E-01 |
| AT4G06310 | 0,35 | -0,53 | 4,38 | -0,12 | 9,04E-01 |
| AT4G06315 | 1,00 | -4,45 | 2,99 | -1,49 | 1,36E-01 |
| AT4G15270 | 0,91 | 0,89 | 2,18 | 0,41 | 6,84E-01 |
| AT4G15490 | 0,92 | 1,89 | 1,82 | 1,04 | 2,99E-01 |
| AT4G15530 | 1,47 | 1,49 | 1,63 | 0,92 | 3,60E-01 |
| AT4G15550 | 0,31 | 0,91 | 4,32 | 0,21 | 8,32E-01 |
| AT4G15810 | 0,14 | -0,45 | 4,33 | -0,10 | 9,17E-01 |
| AT4G06340 | 0,10 | 0,39 | 4,35 | 0,09 | 9,29E-01 |
| AT4G15920 | 0,36 | -0,81 | 4,27 | -0,19 | 8,50E-01 |
| AT4G15955 | 0,12 | 2,87 | 4,34 | 0,66 | 5,08E-01 |
| AT4G16070 | 0,52 | 1,39 | 2,60 | 0,54 | 5,91E-01 |
| AT4G16141 | 0,51 | -0,13 | 4,08 | -0,03 | 9,75E-01 |
| AT4G16144 | 1,36 | 0,44 | 2,08 | 0,21 | 8,32E-01 |
| AT4G16155 | 0,30 | 0,27 | 4,35 | 0,06 | 9,50E-01 |
| AT4G16215 | 0,40 | -1,21 | 3,17 | -0,38 | 7,02E-01 |
| AT4G16250 | 0,73 | 2,11 | 3,02 | 0,70 | 4,83E-01 |
| AT4G16280 | 0,45 | 2,11 | 3,21 | 0,66 | 5,10E-01 |
| AT4G16330 | 0,88 | 2,26 | 2,70 | 0,84 | 4,02E-01 |
| AT4G16410 | 1,15 | 1,58 | 1,98 | 0,80 | 4,25E-01 |
| AT4G16470 | 0,11 | 1,33 | 4,36 | 0,31 | 7,59E-01 |
| AT4G16500 | 1,23 | -0,96 | 1,51 | -0,63 | 5,26E-01 |
| AT4G16515 | 0,18 | 1,77 | 4,41 | 0,40 | 6,88E-01 |
| AT4G16520 | 0,32 | -0,22 | 4,30 | -0,05 | 9,59E-01 |
| AT4G16540 | 0,20 | -0,80 | 4,37 | -0,18 | 8,55E-01 |
| AT4G16580 | 0,33 | -0,43 | 3,55 | -0,12 | 9,03E-01 |
| AT4G16630 | 0,66 | -0,32 | 3,79 | -0,09 | 9,32E-01 |
| AT4G16680 | 0,40 | 3,73 | 4,32 | 0,86 | 3,87E-01 |

| | | | | | |
|---|---|---|---|---|---|
| AT4G16690 | 0,88 | -0,29 | 1,93 | -0,15 | 8,82E-01 |
| AT4G16750 | 0,15 | -0,67 | 4,39 | -0,15 | 8,78E-01 |
| AT4G16760 | 1,69 | 2,19 | 1,46 | 1,50 | 1,33E-01 |
| AT4G16790 | 0,35 | 1,03 | 3,44 | 0,30 | 7,66E-01 |
| AT4G16880 | 1,01 | -0,52 | 1,93 | -0,27 | 7,86E-01 |
| AT4G16892 | 1,06 | 2,17 | 2,25 | 0,97 | 3,34E-01 |
| AT4G16957 | 0,81 | 4,56 | 3,37 | 1,35 | 1,77E-01 |
| AT4G16960 | 0,92 | 5,12 | 4,02 | 1,27 | 2,03E-01 |
| AT4G17040 | 0,44 | 1,93 | 4,20 | 0,46 | 6,47E-01 |
| AT4G17060 | 0,64 | 3,24 | 3,65 | 0,89 | 3,74E-01 |
| AT4G17070 | 0,40 | 1,64 | 3,24 | 0,51 | 6,12E-01 |
| AT4G17100 | 0,77 | -0,68 | 2,37 | -0,29 | 7,76E-01 |
| AT4G17190 | 0,27 | 0,11 | 4,27 | 0,03 | 9,79E-01 |
| AT4G17230 | 1,13 | -0,87 | 1,57 | -0,55 | 5,82E-01 |
| AT4G17243 | 0,66 | 0,86 | 2,46 | 0,35 | 7,26E-01 |
| AT4G17280 | 0,46 | 2,18 | 3,26 | 0,67 | 5,03E-01 |
| AT4G17430 | 0,16 | -1,66 | 4,33 | -0,38 | 7,01E-01 |
| AT4G17486 | 0,51 | 1,71 | 4,40 | 0,39 | 6,98E-01 |
| AT4G17490 | 0,05 | -0,58 | 4,38 | -0,13 | 8,95E-01 |
| AT4G17510 | 0,15 | 0,69 | 4,31 | 0,16 | 8,73E-01 |
| AT4G17550 | 0,46 | 1,17 | 4,13 | 0,28 | 7,78E-01 |
| AT4G17570 | 0,79 | -1,35 | 2,34 | -0,58 | 5,64E-01 |
| AT4G17590 | 0,76 | 1,20 | 2,59 | 0,46 | 6,43E-01 |
| AT4G17610 | 0,22 | 0,81 | 4,27 | 0,19 | 8,50E-01 |
| AT4G17670 | 0,33 | 3,22 | 4,33 | 0,74 | 4,56E-01 |
| AT4G17720 | 0,25 | -0,51 | 4,38 | -0,12 | 9,08E-01 |
| AT4G17740 | 0,24 | 1,12 | 4,32 | 0,26 | 7,95E-01 |
| AT4G17840 | 0,82 | 1,99 | 2,12 | 0,94 | 3,49E-01 |
| AT4G17880 | 0,51 | 2,87 | 3,12 | 0,92 | 3,59E-01 |
| AT4G17950 | 0,50 | 2,00 | 4,11 | 0,49 | 6,26E-01 |
| AT4G18010 | 1,46 | 1,30 | 1,62 | 0,80 | 4,21E-01 |
| AT4G18020 | 0,33 | 1,91 | 4,34 | 0,44 | 6,59E-01 |
| AT4G18040 | 0,22 | -0,13 | 4,30 | -0,03 | 9,76E-01 |
| AT4G18100 | 0,24 | -0,94 | 4,31 | -0,22 | 8,28E-01 |
| AT4G18110 | 0,16 | 1,16 | 4,38 | 0,27 | 7,90E-01 |
| AT4G18130 | 0,68 | 2,86 | 2,78 | 1,03 | 3,05E-01 |
| AT4G18140 | 0,37 | 0,35 | 4,38 | 0,08 | 9,37E-01 |
| AT4G18215 | 1,61 | 0,76 | 1,41 | 0,54 | 5,91E-01 |
| AT4G18230 | 0,15 | 1,94 | 4,32 | 0,45 | 6,54E-01 |
| AT4G18250 | 0,30 | 2,91 | 4,34 | 0,67 | 5,04E-01 |
| AT4G18253 | 0,68 | 1,99 | 2,39 | 0,83 | 4,06E-01 |
| AT4G18255 | 0,35 | 0,00 | 4,41 | 0,00 | 1,00E+00 |
| AT4G18390 | 1,03 | 0,91 | 2,12 | 0,43 | 6,68E-01 |
| AT4G18440 | 0,69 | -1,82 | 3,48 | -0,52 | 6,02E-01 |
| AT4G18520 | 0,18 | 1,51 | 4,31 | 0,35 | 7,25E-01 |
| AT4G18580 | 0,08 | 2,08 | 4,38 | 0,47 | 6,35E-01 |
| AT4G18790 | 0,08 | 2,33 | 4,36 | 0,53 | 5,93E-01 |
| AT4G18810 | 0,53 | -0,66 | 2,79 | -0,24 | 8,13E-01 |
| AT4G18840 | 0,38 | 0,53 | 3,35 | 0,16 | 8,73E-01 |
| AT4G19006 | 0,25 | 1,15 | 4,34 | 0,27 | 7,91E-01 |
| AT4G19040 | 0,87 | 2,81 | 3,23 | 0,87 | 3,83E-01 |
| AT4G19045 | 0,12 | 2,69 | 4,35 | 0,62 | 5,36E-01 |
| AT4G19050 | 0,78 | 3,82 | 2,71 | 1,41 | 1,58E-01 |
| AT4G19060 | 0,46 | 1,72 | 3,97 | 0,43 | 6,64E-01 |
| AT4G19080 | 1,21 | 2,69 | 1,78 | 1,51 | 1,31E-01 |
| AT4G19095 | 0,58 | -1,38 | 2,55 | -0,54 | 5,87E-01 |

| | | | | | |
|---|---|---|---|---|---|
| AT4G06870 | 0,16 | -0,65 | 4,31 | -0,15 | 8,80E-01 |
| AT4G19180 | 0,15 | 1,93 | 4,33 | 0,45 | 6,55E-01 |
| AT4G19220 | 0,37 | 2,86 | 4,28 | 0,67 | 5,04E-01 |
| AT4G19240 | 0,33 | 1,60 | 4,31 | 0,37 | 7,10E-01 |
| AT4G19420 | 0,68 | 0,05 | 2,28 | 0,02 | 9,82E-01 |
| AT4G19440 | 0,20 | 1,94 | 4,34 | 0,45 | 6,55E-01 |
| AT4G19640 | 0,35 | -1,16 | 3,39 | -0,34 | 7,33E-01 |
| AT4G19660 | 1,11 | 0,78 | 1,79 | 0,43 | 6,64E-01 |
| AT4G19670 | 0,77 | 3,50 | 3,52 | 0,99 | 3,20E-01 |
| AT4G06915 | 0,34 | 1,51 | 4,25 | 0,35 | 7,23E-01 |
| AT4G19700 | 1,20 | -0,09 | 1,80 | -0,05 | 9,59E-01 |
| AT4G19720 | 0,28 | -1,14 | 4,36 | -0,26 | 7,93E-01 |
| AT4G19860 | 1,15 | 1,57 | 1,66 | 0,94 | 3,46E-01 |
| AT4G19870 | 0,10 | 0,39 | 4,35 | 0,09 | 9,29E-01 |
| AT4G19880 | 0,52 | 1,27 | 2,45 | 0,52 | 6,05E-01 |
| AT4G19890 | 0,56 | 2,21 | 3,04 | 0,73 | 4,66E-01 |
| AT4G19900 | 0,17 | 2,62 | 4,35 | 0,60 | 5,47E-01 |
| AT4G19985 | 0,13 | 2,37 | 4,36 | 0,54 | 5,86E-01 |
| AT4G20095 | 0,45 | 0,00 | 4,41 | 0,00 | 1,00E+00 |
| AT4G20150 | 0,58 | 0,30 | 2,62 | 0,11 | 9,10E-01 |
| AT4G06945 | 1,00 | 2,54 | 2,71 | 0,94 | 3,47E-01 |
| AT4G20210 | 0,49 | -1,84 | 2,59 | -0,71 | 4,77E-01 |
| AT4G20330 | 0,48 | 2,03 | 2,66 | 0,76 | 4,45E-01 |
| AT4G20410 | 1,25 | 0,35 | 1,94 | 0,18 | 8,56E-01 |
| AT4G20440 | 0,64 | 0,58 | 2,08 | 0,28 | 7,81E-01 |
| AT4G20690 | 0,21 | 3,40 | 4,32 | 0,79 | 4,32E-01 |
| AT4G20880 | 0,74 | -1,34 | 1,79 | -0,75 | 4,54E-01 |
| AT4G20910 | 0,27 | 0,03 | 4,38 | 0,01 | 9,94E-01 |
| AT4G20953 | 0,51 | -0,07 | 3,08 | -0,02 | 9,82E-01 |
| AT4G21100 | 1,08 | 0,62 | 1,76 | 0,35 | 7,24E-01 |
| AT4G21140 | 0,20 | 2,11 | 4,37 | 0,48 | 6,29E-01 |
| AT4G21170 | 0,28 | -2,16 | 4,31 | -0,50 | 6,17E-01 |
| AT4G06985 | 0,08 | 2,08 | 4,38 | 0,47 | 6,35E-01 |
| AT4G21190 | 0,08 | 2,33 | 4,36 | 0,53 | 5,93E-01 |
| AT4G21350 | 0,08 | 2,33 | 4,36 | 0,53 | 5,93E-01 |
| AT4G21380 | 1,64 | 2,24 | 1,40 | 1,60 | 1,10E-01 |
| AT4G21400 | 0,16 | 1,27 | 4,37 | 0,29 | 7,71E-01 |
| AT4G21430 | 0,22 | -1,61 | 4,34 | -0,37 | 7,10E-01 |
| AT4G21510 | 0,05 | -0,58 | 4,38 | -0,13 | 8,95E-01 |
| AT4G21520 | 0,22 | 0,78 | 4,33 | 0,18 | 8,58E-01 |
| AT4G21560 | 0,38 | -0,81 | 4,37 | -0,19 | 8,53E-01 |
| AT4G21570 | 1,42 | 0,88 | 1,80 | 0,49 | 6,25E-01 |
| AT4G21660 | 1,13 | 1,22 | 2,04 | 0,60 | 5,50E-01 |
| AT4G21810 | 1,02 | 1,41 | 1,92 | 0,74 | 4,62E-01 |
| AT4G21860 | 0,50 | 1,49 | 3,33 | 0,45 | 6,55E-01 |
| AT4G21890 | 0,12 | 2,87 | 4,34 | 0,66 | 5,08E-01 |
| AT4G21910 | 0,54 | 1,31 | 4,28 | 0,31 | 7,60E-01 |
| AT4G21990 | 0,81 | 0,76 | 2,42 | 0,31 | 7,54E-01 |
| AT4G22120 | 0,12 | 2,81 | 4,34 | 0,65 | 5,18E-01 |
| AT4G22150 | 0,22 | -0,07 | 4,40 | -0,02 | 9,88E-01 |
| AT4G22190 | 0,98 | 1,71 | 1,82 | 0,93 | 3,50E-01 |
| AT4G22200 | 0,21 | 0,81 | 4,32 | 0,19 | 8,52E-01 |
| AT4G22250 | 0,16 | 1,18 | 4,32 | 0,27 | 7,85E-01 |
| AT4G22260 | 0,18 | -1,21 | 4,35 | -0,28 | 7,81E-01 |
| AT4G22265 | 0,20 | 2,30 | 4,36 | 0,53 | 5,98E-01 |
| AT4G22285 | 0,20 | 1,15 | 4,36 | 0,26 | 7,92E-01 |

| | | | | | |
|---|---|---|---|---|---|
| AT4G22290 | 0,23 | -1,06 | 4,29 | -0,25 | 8,04E-01 |
| AT4G22350 | 0,69 | -0,47 | 3,68 | -0,13 | 8,99E-01 |
| AT4G22380 | 0,51 | -0,51 | 2,48 | -0,20 | 8,39E-01 |
| AT4G22470 | 0,12 | 0,31 | 4,35 | 0,07 | 9,44E-01 |
| AT4G22485 | 0,49 | -2,96 | 3,85 | -0,77 | 4,42E-01 |
| AT4G22550 | 0,26 | 1,18 | 3,71 | 0,32 | 7,51E-01 |
| AT4G22570 | 0,24 | 1,18 | 4,36 | 0,27 | 7,87E-01 |
| AT4G22745 | 0,42 | 2,83 | 4,13 | 0,69 | 4,93E-01 |
| AT4G22750 | 1,03 | 1,78 | 2,22 | 0,80 | 4,24E-01 |
| AT4G22755 | 0,47 | 2,86 | 4,35 | 0,66 | 5,11E-01 |
| AT4G22770 | 1,29 | 1,35 | 1,36 | 0,99 | 3,22E-01 |
| AT4G22780 | 0,24 | 0,16 | 4,33 | 0,04 | 9,70E-01 |
| AT4G22970 | 0,52 | 2,59 | 3,79 | 0,68 | 4,95E-01 |
| AT4G22980 | 0,47 | 1,10 | 3,84 | 0,29 | 7,74E-01 |
| AT4G23010 | 0,83 | 1,86 | 2,35 | 0,79 | 4,27E-01 |
| AT4G23040 | 0,28 | 1,15 | 4,37 | 0,26 | 7,92E-01 |
| AT4G23130 | 0,55 | 3,17 | 3,89 | 0,82 | 4,15E-01 |
| AT4G23190 | 1,18 | 0,39 | 1,72 | 0,23 | 8,21E-01 |
| AT4G23205 | 0,08 | 2,33 | 4,36 | 0,53 | 5,93E-01 |
| AT4G23200 | 0,43 | 2,30 | 4,38 | 0,52 | 6,00E-01 |
| AT4G23210 | 0,21 | 2,29 | 4,32 | 0,53 | 5,96E-01 |
| AT4G23220 | 0,76 | 1,89 | 2,24 | 0,85 | 3,98E-01 |
| AT4G23240 | 0,37 | 3,54 | 4,32 | 0,82 | 4,12E-01 |
| AT4G23250 | 0,45 | 1,08 | 3,83 | 0,28 | 7,77E-01 |
| AT4G23260 | 0,19 | 1,89 | 4,32 | 0,44 | 6,61E-01 |
| AT4G23290 | 0,10 | -1,26 | 4,35 | -0,29 | 7,72E-01 |
| AT4G23320 | 0,08 | 2,33 | 4,36 | 0,53 | 5,93E-01 |
| AT4G23460 | 1,64 | 1,36 | 1,57 | 0,86 | 3,88E-01 |
| AT4G23490 | 0,15 | -0,67 | 4,39 | -0,15 | 8,78E-01 |
| AT4G07130 | 0,46 | 1,58 | 4,12 | 0,38 | 7,01E-01 |
| AT4G23540 | 0,67 | 1,25 | 3,16 | 0,39 | 6,94E-01 |
| AT4G23560 | 0,10 | -1,26 | 4,35 | -0,29 | 7,72E-01 |
| AT4G07135 | 1,25 | 1,98 | 1,88 | 1,06 | 2,91E-01 |
| AT4G07145 | 0,96 | 4,13 | 2,63 | 1,57 | 1,16E-01 |
| AT4G23620 | 0,31 | 0,86 | 4,30 | 0,20 | 8,41E-01 |
| AT4G23640 | 0,77 | 0,97 | 2,20 | 0,44 | 6,58E-01 |
| AT4G23670 | 1,50 | -3,12 | 1,79 | -1,74 | 8,10E-02 |
| AT4G23700 | 0,29 | 2,14 | 4,36 | 0,49 | 6,23E-01 |
| AT4G23770 | 0,22 | -0,29 | 4,32 | -0,07 | 9,47E-01 |
| AT4G23810 | 0,35 | 1,15 | 4,31 | 0,27 | 7,90E-01 |
| AT4G23820 | 0,63 | -1,58 | 2,18 | -0,73 | 4,67E-01 |
| AT4G23880 | 0,47 | -2,81 | 3,12 | -0,90 | 3,68E-01 |
| AT4G23890 | 0,66 | 0,16 | 2,71 | 0,06 | 9,53E-01 |
| AT4G24180 | 0,37 | 2,80 | 4,34 | 0,65 | 5,18E-01 |
| AT4G24200 | 0,70 | 1,45 | 2,87 | 0,50 | 6,15E-01 |
| AT4G07250 | 0,17 | 0,00 | 4,41 | 0,00 | 1,00E+00 |
| AT4G24300 | 0,45 | -2,06 | 4,24 | -0,48 | 6,28E-01 |
| AT4G24320 | 0,33 | 3,55 | 4,32 | 0,82 | 4,11E-01 |
| AT4G24330 | 0,84 | -0,37 | 1,74 | -0,21 | 8,33E-01 |
| AT4G07255 | 0,10 | -1,26 | 4,35 | -0,29 | 7,72E-01 |
| AT4G24390 | 0,29 | 1,16 | 4,30 | 0,27 | 7,87E-01 |
| AT4G24490 | 0,72 | 1,71 | 2,41 | 0,71 | 4,77E-01 |
| AT4G24530 | 0,08 | 2,33 | 4,36 | 0,53 | 5,93E-01 |
| AT4G24560 | 1,21 | 2,96 | 1,97 | 1,50 | 1,33E-01 |
| AT4G24580 | 0,46 | 0,48 | 4,10 | 0,12 | 9,08E-01 |
| AT4G24610 | 0,67 | 3,39 | 2,93 | 1,16 | 2,47E-01 |

| | | | | | |
|---|---|---|---|---|---|
| AT4G24620 | 0,42 | -0,87 | 4,16 | -0,21 | 8,34E-01 |
| AT4G24660 | 1,04 | -0,99 | 2,06 | -0,48 | 6,31E-01 |
| AT4G24670 | 0,08 | -0,56 | 4,38 | -0,13 | 8,99E-01 |
| AT4G24730 | 0,45 | 2,45 | 4,07 | 0,60 | 5,47E-01 |
| AT4G24740 | 0,63 | 0,76 | 3,03 | 0,25 | 8,02E-01 |
| AT4G24750 | 0,28 | 1,64 | 4,38 | 0,38 | 7,08E-01 |
| AT4G24780 | 1,27 | -1,99 | 1,98 | -1,00 | 3,15E-01 |
| AT4G07285 | 0,27 | -0,89 | 4,37 | -0,20 | 8,39E-01 |
| AT4G24820 | 1,43 | 1,18 | 1,53 | 0,77 | 4,42E-01 |
| AT4G24830 | 0,25 | -0,89 | 4,30 | -0,21 | 8,36E-01 |
| AT4G24840 | 0,53 | -0,81 | 3,03 | -0,27 | 7,90E-01 |
| AT4G24920 | 0,86 | 3,97 | 2,72 | 1,46 | 1,45E-01 |
| AT4G24990 | 0,31 | -0,60 | 4,32 | -0,14 | 8,91E-01 |
| AT4G25090 | 0,29 | 3,75 | 4,32 | 0,87 | 3,85E-01 |
| AT4G25270 | 0,45 | 2,45 | 4,07 | 0,60 | 5,47E-01 |
| AT4G25280 | 0,71 | 3,87 | 4,31 | 0,90 | 3,68E-01 |
| AT4G25290 | 0,52 | 2,17 | 3,84 | 0,56 | 5,73E-01 |
| AT4G25340 | 0,19 | 1,67 | 4,33 | 0,39 | 6,99E-01 |
| AT4G25435 | 0,88 | 0,00 | 2,83 | 0,00 | 1,00E+00 |
| AT4G25540 | 0,13 | 2,37 | 4,36 | 0,54 | 5,86E-01 |
| AT4G25570 | 0,43 | 0,08 | 4,16 | 0,02 | 9,85E-01 |
| AT4G25640 | 1,48 | 1,85 | 1,52 | 1,22 | 2,24E-01 |
| AT4G25650 | 1,11 | 1,70 | 1,60 | 1,06 | 2,88E-01 |
| AT4G25670 | 0,79 | 2,77 | 2,82 | 0,98 | 3,26E-01 |
| AT4G25690 | 0,56 | 0,23 | 3,25 | 0,07 | 9,44E-01 |
| AT4G25740 | 0,38 | 0,92 | 4,32 | 0,21 | 8,32E-01 |
| AT4G25870 | 0,26 | 2,13 | 4,38 | 0,49 | 6,27E-01 |
| AT4G25940 | 0,39 | 2,88 | 4,34 | 0,66 | 5,06E-01 |
| AT4G25990 | 0,26 | -1,61 | 4,34 | -0,37 | 7,10E-01 |
| AT4G26080 | 0,87 | 1,87 | 2,46 | 0,76 | 4,48E-01 |
| AT4G26100 | 0,81 | 0,78 | 1,90 | 0,41 | 6,81E-01 |
| AT4G26130 | 1,45 | 2,08 | 1,65 | 1,26 | 2,08E-01 |
| AT4G26140 | 0,21 | -0,24 | 4,35 | -0,06 | 9,56E-01 |
| AT4G26190 | 0,25 | 0,51 | 4,26 | 0,12 | 9,05E-01 |
| AT4G26300 | 0,69 | 2,82 | 2,56 | 1,10 | 2,69E-01 |
| AT4G07565 | 0,79 | -1,84 | 1,91 | -0,96 | 3,35E-01 |
| AT4G07625 | 0,79 | 1,21 | 2,54 | 0,48 | 6,35E-01 |
| AT4G07635 | 0,12 | 0,31 | 4,35 | 0,07 | 9,44E-01 |
| AT4G26370 | 0,48 | -1,01 | 4,06 | -0,25 | 8,04E-01 |
| AT4G26375 | 0,40 | 3,80 | 4,32 | 0,88 | 3,79E-01 |
| AT4G26400 | 0,60 | 2,10 | 3,92 | 0,54 | 5,91E-01 |
| AT4G26620 | 0,28 | 2,17 | 4,31 | 0,50 | 6,14E-01 |
| AT4G26630 | 0,72 | 1,08 | 2,26 | 0,48 | 6,32E-01 |
| AT4G26640 | 1,00 | 2,85 | 2,24 | 1,28 | 2,02E-01 |
| AT4G26690 | 1,51 | 0,26 | 1,47 | 0,18 | 8,58E-01 |
| AT4G26750 | 0,43 | 2,34 | 4,16 | 0,56 | 5,73E-01 |
| AT4G26760 | 0,64 | 2,78 | 3,03 | 0,92 | 3,60E-01 |
| AT4G26795 | 0,42 | -1,55 | 3,99 | -0,39 | 6,98E-01 |
| AT4G26860 | 0,37 | 0,76 | 4,27 | 0,18 | 8,59E-01 |
| AT4G26910 | 1,33 | 2,94 | 1,77 | 1,66 | 9,72E-02 |
| AT4G26940 | 0,43 | 1,16 | 4,08 | 0,28 | 7,76E-01 |
| AT4G26980 | 0,78 | 3,00 | 2,39 | 1,25 | 2,10E-01 |
| AT4G27020 | 0,48 | 1,00 | 3,28 | 0,30 | 7,61E-01 |
| AT4G07765 | 0,66 | -0,37 | 2,62 | -0,14 | 8,88E-01 |
| AT4G27130 | 0,28 | -0,59 | 3,58 | -0,16 | 8,70E-01 |
| AT4G27270 | 0,66 | 3,20 | 3,79 | 0,84 | 3,99E-01 |

| | | | | | |
|---|---|---|---|---|---|
| AT4G27280 | 0,84 | -1,84 | 2,62 | -0,70 | 4,84E-01 |
| AT4G27300 | 0,83 | 0,48 | 2,31 | 0,21 | 8,34E-01 |
| AT4G27320 | 1,19 | 1,71 | 2,13 | 0,80 | 4,21E-01 |
| AT4G27340 | 0,21 | 1,46 | 4,31 | 0,34 | 7,35E-01 |
| AT4G27410 | 0,98 | 3,89 | 2,32 | 1,68 | 9,34E-02 |
| AT4G27450 | 0,72 | 0,21 | 2,27 | 0,09 | 9,27E-01 |
| AT4G27480 | 0,65 | 4,43 | 3,54 | 1,25 | 2,11E-01 |
| AT4G27650 | 0,39 | 1,51 | 3,42 | 0,44 | 6,59E-01 |
| AT4G27700 | 0,47 | -0,45 | 3,27 | -0,14 | 8,90E-01 |
| AT4G27730 | 0,37 | 3,04 | 4,33 | 0,70 | 4,82E-01 |
| AT4G27800 | 1,09 | 1,52 | 1,98 | 0,77 | 4,43E-01 |
| AT4G27830 | 0,84 | -0,44 | 1,76 | -0,25 | 8,01E-01 |
| AT4G27840 | 0,43 | -0,90 | 4,36 | -0,21 | 8,36E-01 |
| AT4G27885 | 0,16 | -1,66 | 4,33 | -0,38 | 7,01E-01 |
| AT4G27960 | 0,38 | -1,81 | 4,21 | -0,43 | 6,67E-01 |
| AT4G28030 | 0,45 | 1,09 | 2,69 | 0,41 | 6,85E-01 |
| AT4G28050 | 0,43 | 3,32 | 4,33 | 0,77 | 4,43E-01 |
| AT4G28060 | 0,25 | 0,00 | 4,40 | 0,00 | 1,00E+00 |
| AT4G28180 | 0,69 | -0,22 | 2,37 | -0,09 | 9,26E-01 |
| AT4G28200 | 0,59 | 2,00 | 2,66 | 0,75 | 4,53E-01 |
| AT4G28230 | 0,33 | 3,31 | 4,32 | 0,77 | 4,44E-01 |
| AT4G28240 | 0,66 | -0,40 | 2,03 | -0,20 | 8,45E-01 |
| AT4G28320 | 0,20 | 2,11 | 4,37 | 0,48 | 6,29E-01 |
| AT4G28390 | 0,33 | 3,34 | 4,32 | 0,77 | 4,40E-01 |
| AT4G28450 | 0,61 | 0,00 | 4,41 | 0,00 | 1,00E+00 |
| AT4G28470 | 0,62 | 2,01 | 3,04 | 0,66 | 5,09E-01 |
| AT4G28610 | 0,93 | 1,63 | 2,48 | 0,66 | 5,10E-01 |
| AT4G28630 | 0,12 | 2,81 | 4,34 | 0,65 | 5,18E-01 |
| AT4G28770 | 0,20 | -0,47 | 4,33 | -0,11 | 9,13E-01 |
| AT4G28910 | 1,06 | 3,49 | 2,40 | 1,45 | 1,46E-01 |
| AT4G29010 | 1,63 | 0,54 | 1,51 | 0,36 | 7,22E-01 |
| AT4G29050 | 0,13 | -0,63 | 4,39 | -0,14 | 8,85E-01 |
| AT4G29080 | 0,28 | 0,53 | 4,29 | 0,12 | 9,02E-01 |
| AT4G29130 | 1,04 | 1,72 | 1,91 | 0,90 | 3,70E-01 |
| AT4G29220 | 0,63 | 1,24 | 3,07 | 0,40 | 6,86E-01 |
| AT4G29260 | 0,48 | -1,67 | 3,88 | -0,43 | 6,66E-01 |
| AT4G29300 | 1,28 | -1,36 | 1,58 | -0,86 | 3,88E-01 |
| AT4G29305 | 0,76 | -2,38 | 2,24 | -1,07 | 2,87E-01 |
| AT4G08155 | 1,40 | -1,73 | 1,40 | -1,24 | 2,17E-01 |
| AT4G29440 | 0,66 | 0,52 | 2,84 | 0,18 | 8,55E-01 |
| AT4G29520 | 0,77 | 0,77 | 2,50 | 0,31 | 7,58E-01 |
| AT4G29654 | 1,07 | -0,32 | 2,39 | -0,14 | 8,92E-01 |
| AT4G29670 | 0,65 | -0,19 | 2,17 | -0,09 | 9,29E-01 |
| AT4G29680 | 0,85 | 0,85 | 3,02 | 0,28 | 7,78E-01 |
| AT4G29720 | 0,17 | 2,11 | 4,37 | 0,48 | 6,28E-01 |
| AT4G29735 | 0,17 | 2,29 | 4,36 | 0,53 | 5,98E-01 |
| AT4G29810 | 0,27 | 0,92 | 4,32 | 0,21 | 8,32E-01 |
| AT4G29820 | 0,24 | 0,06 | 4,31 | 0,01 | 9,89E-01 |
| AT4G29840 | 0,58 | 0,31 | 2,24 | 0,14 | 8,89E-01 |
| AT4G29880 | 0,21 | 0,86 | 4,30 | 0,20 | 8,42E-01 |
| AT4G29905 | 1,17 | -1,37 | 1,65 | -0,83 | 4,08E-01 |
| AT4G29910 | 0,08 | -0,56 | 4,38 | -0,13 | 8,99E-01 |
| AT4G29920 | 0,05 | -0,58 | 4,38 | -0,13 | 8,95E-01 |
| AT4G29940 | 0,28 | 1,64 | 4,38 | 0,38 | 7,08E-01 |
| AT4G29960 | 0,50 | 3,40 | 3,91 | 0,87 | 3,85E-01 |
| AT4G29990 | 0,08 | 2,08 | 4,38 | 0,47 | 6,35E-01 |

| | | | | | |
|---|---|---|---|---|---|
| AT4G30000 | 0,38 | 2,18 | 4,37 | 0,50 | 6,18E-01 |
| AT4G30030 | 0,45 | 1,75 | 4,25 | 0,41 | 6,81E-01 |
| AT4G30060 | 0,08 | 2,33 | 4,36 | 0,53 | 5,93E-01 |
| AT4G30080 | 0,31 | 0,19 | 4,29 | 0,04 | 9,65E-01 |
| AT4G30110 | 1,05 | 4,48 | 3,12 | 1,43 | 1,52E-01 |
| AT4G30200 | 1,47 | 2,14 | 1,54 | 1,39 | 1,63E-01 |
| AT4G30240 | 0,30 | 1,56 | 4,30 | 0,36 | 7,17E-01 |
| AT4G30260 | 0,47 | 2,78 | 3,14 | 0,89 | 3,76E-01 |
| AT4G30270 | 1,68 | 1,49 | 1,27 | 1,18 | 2,39E-01 |
| AT4G30280 | 0,12 | -0,77 | 4,36 | -0,18 | 8,61E-01 |
| AT4G30360 | 0,21 | -2,30 | 4,30 | -0,54 | 5,92E-01 |
| AT4G30410 | 0,12 | 0,63 | 4,33 | 0,15 | 8,85E-01 |
| AT4G30430 | 0,08 | 2,08 | 4,38 | 0,47 | 6,35E-01 |
| AT4G30560 | 0,74 | 3,37 | 3,49 | 0,96 | 3,35E-01 |
| AT4G30580 | 0,92 | 4,41 | 3,36 | 1,31 | 1,90E-01 |
| AT4G30600 | 1,60 | -0,11 | 1,43 | -0,08 | 9,37E-01 |
| AT4G30620 | 0,23 | -0,64 | 4,39 | -0,15 | 8,84E-01 |
| AT4G30700 | 0,08 | 2,33 | 4,36 | 0,53 | 5,93E-01 |
| AT4G30710 | 0,12 | 2,81 | 4,34 | 0,65 | 5,18E-01 |
| AT4G30720 | 1,29 | 2,48 | 1,60 | 1,55 | 1,20E-01 |
| AT4G30740 | 1,03 | -1,65 | 2,17 | -0,76 | 4,47E-01 |
| AT4G30790 | 0,99 | 1,36 | 1,99 | 0,68 | 4,95E-01 |
| AT4G30890 | 0,60 | 0,33 | 2,98 | 0,11 | 9,11E-01 |
| AT4G30920 | 1,19 | 1,22 | 1,82 | 0,67 | 5,04E-01 |
| AT4G30950 | 1,64 | 2,30 | 1,49 | 1,55 | 1,21E-01 |
| AT4G31020 | 0,29 | 1,64 | 4,38 | 0,37 | 7,08E-01 |
| AT4G31120 | 0,87 | 2,25 | 2,43 | 0,93 | 3,55E-01 |
| AT4G31130 | 0,59 | 0,87 | 3,07 | 0,28 | 7,78E-01 |
| AT4G31420 | 0,28 | 0,84 | 4,35 | 0,19 | 8,46E-01 |
| AT4G31450 | 0,46 | 2,24 | 4,16 | 0,54 | 5,91E-01 |
| AT4G31540 | 0,43 | -0,97 | 4,09 | -0,24 | 8,12E-01 |
| AT4G31550 | 1,02 | 0,27 | 1,74 | 0,15 | 8,77E-01 |
| AT4G31590 | 0,63 | 1,02 | 2,49 | 0,41 | 6,81E-01 |
| AT4G31810 | 0,08 | 2,33 | 4,36 | 0,53 | 5,93E-01 |
| AT4G31820 | 0,43 | 1,59 | 2,71 | 0,58 | 5,59E-01 |
| AT4G32070 | 0,58 | 1,44 | 3,91 | 0,37 | 7,12E-01 |
| AT4G32120 | 0,15 | 0,40 | 4,33 | 0,09 | 9,27E-01 |
| AT4G32130 | 0,18 | -0,57 | 4,30 | -0,13 | 8,94E-01 |
| AT4G32150 | 1,17 | 0,09 | 1,53 | 0,06 | 9,55E-01 |
| AT4G32180 | 1,00 | 4,06 | 2,51 | 1,62 | 1,05E-01 |
| AT4G32208 | 0,37 | 2,86 | 4,28 | 0,67 | 5,04E-01 |
| AT4G32340 | 1,02 | -0,78 | 1,61 | -0,48 | 6,28E-01 |
| AT4G32420 | 0,78 | 2,74 | 2,47 | 1,11 | 2,66E-01 |
| AT4G32460 | 0,31 | 1,10 | 4,33 | 0,26 | 7,99E-01 |
| AT4G32470 | 0,29 | -0,86 | 4,30 | -0,20 | 8,42E-01 |
| AT4G32510 | 0,56 | 4,04 | 3,73 | 1,08 | 2,78E-01 |
| AT4G32520 | 1,00 | 1,58 | 1,81 | 0,87 | 3,84E-01 |
| AT4G32530 | 0,89 | 1,97 | 2,44 | 0,81 | 4,18E-01 |
| AT4G32670 | 0,15 | 1,64 | 4,33 | 0,38 | 7,06E-01 |
| AT4G32750 | 0,18 | 1,30 | 4,37 | 0,30 | 7,65E-01 |
| AT4G32820 | 0,83 | 3,60 | 2,62 | 1,38 | 1,69E-01 |
| AT4G32880 | 1,09 | 0,40 | 1,99 | 0,20 | 8,42E-01 |
| AT4G32900 | 0,12 | 1,65 | 4,34 | 0,38 | 7,04E-01 |
| AT4G32920 | 0,70 | 2,08 | 2,71 | 0,77 | 4,43E-01 |
| AT4G33040 | 0,37 | 1,70 | 4,31 | 0,39 | 6,94E-01 |
| AT4G33090 | 0,64 | 0,25 | 2,91 | 0,09 | 9,32E-01 |

| | | | | | |
|---|---|---|---|---|---|
| AT4G33110 | 0,17 | 0,57 | 4,29 | 0,13 | 8,95E-01 |
| AT4G33150 | 0,65 | 4,53 | 3,60 | 1,26 | 2,09E-01 |
| AT4G33200 | 0,29 | 3,04 | 4,33 | 0,70 | 4,83E-01 |
| AT4G33210 | 1,11 | 3,14 | 1,98 | 1,59 | 1,13E-01 |
| AT4G33250 | 0,89 | 1,04 | 1,89 | 0,55 | 5,84E-01 |
| AT4G33260 | 0,12 | 2,81 | 4,34 | 0,65 | 5,18E-01 |
| AT4G33270 | 0,12 | -0,77 | 4,36 | -0,18 | 8,61E-01 |
| AT4G33400 | 0,33 | 2,78 | 4,34 | 0,64 | 5,21E-01 |
| AT4G33430 | 1,19 | 0,78 | 1,76 | 0,45 | 6,56E-01 |
| AT4G33470 | 0,67 | 2,94 | 2,86 | 1,03 | 3,04E-01 |
| AT4G33495 | 0,29 | 3,04 | 4,33 | 0,70 | 4,83E-01 |
| AT4G33500 | 0,96 | 1,78 | 2,51 | 0,71 | 4,79E-01 |
| AT4G33530 | 1,44 | 2,41 | 1,57 | 1,53 | 1,26E-01 |
| AT4G33585 | 1,20 | 1,26 | 2,99 | 0,42 | 6,72E-01 |
| AT4G33625 | 0,16 | 1,14 | 4,30 | 0,27 | 7,90E-01 |
| AT4G33670 | 0,76 | 0,72 | 2,03 | 0,35 | 7,23E-01 |
| AT4G33680 | 0,20 | 1,57 | 4,34 | 0,36 | 7,17E-01 |
| AT4G33700 | 0,28 | 0,24 | 4,32 | 0,06 | 9,55E-01 |
| AT4G33790 | 0,16 | 1,27 | 4,37 | 0,29 | 7,71E-01 |
| AT4G33860 | 0,68 | 3,51 | 3,61 | 0,97 | 3,32E-01 |
| AT4G33865 | 0,81 | 1,85 | 1,99 | 0,93 | 3,53E-01 |
| AT4G33910 | 0,26 | 2,13 | 4,38 | 0,49 | 6,27E-01 |
| AT4G34020 | 0,41 | 2,16 | 4,37 | 0,49 | 6,21E-01 |
| AT4G34040 | 0,82 | 2,55 | 2,15 | 1,19 | 2,36E-01 |
| AT4G34070 | 0,72 | 1,52 | 2,69 | 0,57 | 5,72E-01 |
| AT4G34090 | 0,71 | 3,88 | 4,30 | 0,90 | 3,67E-01 |
| AT4G34131 | 0,52 | 3,84 | 3,86 | 0,99 | 3,20E-01 |
| AT4G34180 | 0,68 | 2,66 | 2,83 | 0,94 | 3,46E-01 |
| AT4G34190 | 1,52 | 0,63 | 1,57 | 0,40 | 6,89E-01 |
| AT4G34200 | 1,37 | 2,21 | 1,59 | 1,39 | 1,65E-01 |
| AT4G34220 | 1,59 | -0,65 | 1,43 | -0,45 | 6,51E-01 |
| AT4G34332 | 0,08 | 2,33 | 4,36 | 0,53 | 5,93E-01 |
| AT4G34340 | 0,12 | 2,81 | 4,34 | 0,65 | 5,18E-01 |
| AT4G34350 | 1,55 | 2,25 | 1,41 | 1,59 | 1,11E-01 |
| AT4G08985 | 0,45 | -1,34 | 3,08 | -0,43 | 6,64E-01 |
| AT4G34460 | 0,42 | 0,24 | 3,16 | 0,08 | 9,38E-01 |
| AT4G34480 | 1,19 | 1,52 | 1,66 | 0,92 | 3,58E-01 |
| AT4G34490 | 1,39 | 2,18 | 1,49 | 1,47 | 1,42E-01 |
| AT4G34610 | 0,37 | -0,72 | 4,34 | -0,17 | 8,69E-01 |
| AT4G34620 | 1,51 | 0,33 | 1,88 | 0,17 | 8,63E-01 |
| AT4G34630 | 0,16 | -0,97 | 4,35 | -0,22 | 8,24E-01 |
| AT4G34690 | 1,55 | -0,66 | 1,59 | -0,42 | 6,75E-01 |
| AT4G34730 | 0,55 | -0,77 | 2,57 | -0,30 | 7,64E-01 |
| AT4G34740 | 0,69 | 0,78 | 2,93 | 0,27 | 7,89E-01 |
| AT4G34860 | 0,08 | 2,08 | 4,38 | 0,47 | 6,35E-01 |
| AT4G34900 | 0,65 | 3,12 | 3,59 | 0,87 | 3,85E-01 |
| AT4G35060 | 0,12 | 2,81 | 4,34 | 0,65 | 5,18E-01 |
| AT4G35140 | 0,36 | 0,27 | 4,37 | 0,06 | 9,52E-01 |
| AT4G35180 | 0,66 | 1,26 | 2,95 | 0,43 | 6,68E-01 |
| AT4G35230 | 1,11 | 0,00 | 1,76 | 0,00 | 1,00E+00 |
| AT4G35240 | 1,02 | 0,76 | 2,16 | 0,35 | 7,24E-01 |
| AT4G35260 | 0,53 | -1,77 | 3,12 | -0,57 | 5,69E-01 |
| AT4G35300 | 1,43 | -0,23 | 2,09 | -0,11 | 9,12E-01 |
| AT4G35320 | 0,51 | -1,81 | 3,02 | -0,60 | 5,50E-01 |
| AT4G35360 | 0,54 | -0,13 | 4,07 | -0,03 | 9,74E-01 |
| AT4G35440 | 0,24 | 0,16 | 4,28 | 0,04 | 9,69E-01 |

| | | | | | |
|---|---|---|---|---|---|
| AT4G35570 | 0,29 | 1,74 | 4,31 | 0,40 | 6,87E-01 |
| AT4G35580 | 0,56 | 0,50 | 3,04 | 0,17 | 8,68E-01 |
| AT4G35780 | 0,15 | 1,93 | 4,33 | 0,45 | 6,55E-01 |
| AT4G35785 | 0,84 | 3,50 | 2,77 | 1,27 | 2,06E-01 |
| AT4G35920 | 0,70 | 1,78 | 4,39 | 0,41 | 6,85E-01 |
| AT4G36010 | 0,74 | 0,13 | 2,64 | 0,05 | 9,61E-01 |
| AT4G36030 | 1,21 | -1,84 | 2,04 | -0,90 | 3,65E-01 |
| AT4G36050 | 0,52 | 4,37 | 4,30 | 1,02 | 3,10E-01 |
| AT4G36070 | 0,08 | 2,08 | 4,38 | 0,47 | 6,35E-01 |
| AT4G36220 | 1,38 | 0,38 | 1,38 | 0,27 | 7,85E-01 |
| AT4G36270 | 0,14 | 0,87 | 4,31 | 0,20 | 8,40E-01 |
| AT4G36280 | 0,08 | 2,08 | 4,38 | 0,47 | 6,35E-01 |
| AT4G36380 | 0,24 | -0,14 | 4,41 | -0,03 | 9,75E-01 |
| AT4G36390 | 0,60 | 0,67 | 2,42 | 0,28 | 7,81E-01 |
| AT4G36460 | 0,50 | -0,94 | 3,03 | -0,31 | 7,58E-01 |
| AT4G36500 | 0,97 | -0,31 | 2,13 | -0,15 | 8,83E-01 |
| AT4G36520 | 1,04 | 1,00 | 1,53 | 0,65 | 5,16E-01 |
| AT4G36550 | 0,53 | 1,52 | 3,05 | 0,50 | 6,18E-01 |
| AT4G36640 | 1,54 | 4,72 | 2,02 | 2,33 | 1,97E-02 |
| AT4G36670 | 0,44 | -0,33 | 3,22 | -0,10 | 9,18E-01 |
| AT4G36690 | 0,56 | -0,15 | 3,18 | -0,05 | 9,62E-01 |
| AT4G36730 | 0,51 | -1,40 | 3,16 | -0,44 | 6,57E-01 |
| AT4G36770 | 0,41 | 1,48 | 4,13 | 0,36 | 7,20E-01 |
| AT4G36800 | 0,60 | 1,89 | 3,00 | 0,63 | 5,29E-01 |
| AT4G36808 | 0,88 | 2,03 | 2,26 | 0,90 | 3,67E-01 |
| AT4G36860 | 0,41 | 0,81 | 4,30 | 0,19 | 8,50E-01 |
| AT4G36890 | 0,60 | 1,24 | 3,89 | 0,32 | 7,49E-01 |
| AT4G36970 | 0,18 | -0,70 | 4,38 | -0,16 | 8,72E-01 |
| AT4G36980 | 0,30 | 2,84 | 4,35 | 0,65 | 5,14E-01 |
| AT4G37100 | 0,62 | 0,43 | 3,04 | 0,14 | 8,87E-01 |
| AT4G37270 | 0,72 | 2,07 | 2,43 | 0,85 | 3,94E-01 |
| AT4G37280 | 0,63 | 1,67 | 3,71 | 0,45 | 6,51E-01 |
| AT4G37370 | 0,86 | 2,13 | 2,52 | 0,85 | 3,97E-01 |
| AT4G37432 | 0,86 | 3,26 | 3,34 | 0,97 | 3,30E-01 |
| AT4G37470 | 1,39 | -0,89 | 1,53 | -0,58 | 5,62E-01 |
| AT4G37510 | 0,72 | 1,52 | 2,29 | 0,66 | 5,07E-01 |
| AT4G37540 | 0,41 | -1,98 | 4,33 | -0,46 | 6,48E-01 |
| AT4G37553 | 0,42 | 2,99 | 4,34 | 0,69 | 4,91E-01 |
| AT4G37590 | 0,76 | 0,89 | 2,31 | 0,38 | 7,01E-01 |
| AT4G37610 | 0,33 | 0,91 | 3,54 | 0,26 | 7,97E-01 |
| AT4G37740 | 0,08 | -0,56 | 4,38 | -0,13 | 8,99E-01 |
| AT4G37800 | 0,23 | -1,06 | 4,29 | -0,25 | 8,04E-01 |
| AT4G37830 | 0,60 | -0,48 | 2,55 | -0,19 | 8,51E-01 |
| AT4G37850 | 0,24 | 0,00 | 4,41 | 0,00 | 1,00E+00 |
| AT4G37870 | 0,70 | 0,43 | 2,03 | 0,21 | 8,31E-01 |
| AT4G37920 | 0,32 | 2,25 | 4,32 | 0,52 | 6,03E-01 |
| AT4G38010 | 0,08 | 2,08 | 4,38 | 0,47 | 6,35E-01 |
| AT4G38050 | 0,35 | 3,20 | 4,33 | 0,74 | 4,60E-01 |
| AT4G38120 | 0,89 | 2,03 | 2,14 | 0,95 | 3,43E-01 |
| AT4G38130 | 0,71 | 0,77 | 1,81 | 0,43 | 6,70E-01 |
| AT4G38180 | 0,60 | 1,33 | 2,22 | 0,60 | 5,51E-01 |
| AT4G38210 | 0,75 | 3,54 | 4,32 | 0,82 | 4,12E-01 |
| AT4G38230 | 0,40 | 0,50 | 3,36 | 0,15 | 8,82E-01 |
| AT4G38240 | 0,62 | 3,84 | 3,64 | 1,06 | 2,91E-01 |
| AT4G09695 | 1,25 | 1,59 | 1,54 | 1,04 | 2,99E-01 |
| AT4G38300 | 0,68 | 3,98 | 3,55 | 1,12 | 2,63E-01 |

| | | | | | |
|---|---|---|---|---|---|
| AT4G38350 | 1,14 | 2,29 | 1,85 | 1,24 | 2,15E-01 |
| AT4G38380 | 0,43 | 2,86 | 4,12 | 0,69 | 4,87E-01 |
| AT4G38520 | 0,55 | -0,30 | 2,20 | -0,14 | 8,90E-01 |
| AT4G38545 | 0,17 | 0,57 | 4,29 | 0,13 | 8,95E-01 |
| AT4G38570 | 0,81 | 3,34 | 2,78 | 1,20 | 2,29E-01 |
| AT4G38630 | 0,74 | 1,06 | 1,99 | 0,53 | 5,95E-01 |
| AT4G38640 | 0,43 | 0,74 | 3,30 | 0,22 | 8,23E-01 |
| AT4G38660 | 0,37 | -1,23 | 4,36 | -0,28 | 7,78E-01 |
| AT4G38670 | 0,49 | 2,88 | 3,17 | 0,91 | 3,64E-01 |
| AT4G38680 | 1,18 | -0,56 | 1,48 | -0,38 | 7,03E-01 |
| AT4G38700 | 0,34 | -2,02 | 4,31 | -0,47 | 6,39E-01 |
| AT4G38710 | 1,00 | -1,13 | 1,54 | -0,73 | 4,63E-01 |
| AT4G38730 | 0,38 | 2,13 | 4,38 | 0,49 | 6,27E-01 |
| AT4G38740 | 1,15 | -2,34 | 1,77 | -1,32 | 1,87E-01 |
| AT4G38780 | 1,10 | 3,51 | 2,40 | 1,47 | 1,43E-01 |
| AT4G38800 | 0,65 | -1,39 | 2,51 | -0,55 | 5,79E-01 |
| AT4G38810 | 0,68 | -0,03 | 2,58 | -0,01 | 9,91E-01 |
| AT4G38870 | 0,21 | 2,29 | 4,32 | 0,53 | 5,96E-01 |
| AT4G38890 | 0,48 | -0,92 | 3,27 | -0,28 | 7,79E-01 |
| AT4G38900 | 0,33 | -0,09 | 4,34 | -0,02 | 9,84E-01 |
| AT4G38932 | 0,58 | 1,02 | 2,65 | 0,39 | 6,99E-01 |
| AT4G38950 | 1,10 | 2,74 | 2,05 | 1,34 | 1,81E-01 |
| AT4G39100 | 0,33 | 3,36 | 4,33 | 0,78 | 4,37E-01 |
| AT4G39120 | 0,19 | 1,90 | 4,33 | 0,44 | 6,60E-01 |
| AT4G39140 | 0,65 | 0,84 | 3,79 | 0,22 | 8,25E-01 |
| AT4G39180 | 0,48 | 1,97 | 4,31 | 0,46 | 6,49E-01 |
| AT4G39190 | 0,69 | -1,96 | 2,25 | -0,87 | 3,85E-01 |
| AT4G39195 | 0,79 | 3,29 | 2,87 | 1,14 | 2,52E-01 |
| AT4G39240 | 1,33 | 3,97 | 2,17 | 1,83 | 6,71E-02 |
| AT4G39280 | 1,06 | -1,21 | 1,58 | -0,76 | 4,45E-01 |
| AT4G39460 | 0,85 | 3,37 | 2,43 | 1,39 | 1,65E-01 |
| AT4G39500 | 0,20 | 2,11 | 4,37 | 0,48 | 6,29E-01 |
| AT4G39510 | 0,43 | 1,79 | 4,41 | 0,41 | 6,85E-01 |
| AT4G39520 | 1,02 | 0,79 | 1,51 | 0,52 | 6,00E-01 |
| AT4G39550 | 0,17 | 3,16 | 4,33 | 0,73 | 4,66E-01 |
| AT4G39560 | 0,59 | 2,86 | 3,79 | 0,75 | 4,51E-01 |
| AT4G39570 | 0,55 | 4,17 | 3,71 | 1,12 | 2,61E-01 |
| AT4G39580 | 0,15 | 1,64 | 4,33 | 0,38 | 7,06E-01 |
| AT4G39590 | 0,08 | 2,08 | 4,38 | 0,47 | 6,35E-01 |
| AT4G39630 | 0,05 | -0,58 | 4,38 | -0,13 | 8,95E-01 |
| AT4G39680 | 0,30 | -0,81 | 4,25 | -0,19 | 8,50E-01 |
| AT4G39710 | 0,56 | -0,76 | 3,88 | -0,20 | 8,44E-01 |
| AT4G39780 | 0,21 | 1,12 | 4,33 | 0,26 | 7,95E-01 |
| AT4G39820 | 0,46 | 2,42 | 4,04 | 0,60 | 5,49E-01 |
| AT4G39838 | 1,07 | -0,28 | 1,73 | -0,16 | 8,69E-01 |
| AT4G39865 | 0,25 | 0,00 | 4,40 | 0,00 | 1,00E+00 |
| AT4G39910 | 0,28 | 0,37 | 4,35 | 0,08 | 9,33E-01 |
| AT4G40000 | 0,43 | 2,79 | 4,12 | 0,68 | 4,99E-01 |
| AT4G40020 | 0,75 | 3,39 | 2,77 | 1,22 | 2,21E-01 |
| AT4G40030 | 1,32 | 0,43 | 1,66 | 0,26 | 7,95E-01 |
| AT4G40045 | 0,12 | 2,81 | 4,34 | 0,65 | 5,18E-01 |
| AT4G40050 | 1,65 | 1,90 | 1,49 | 1,28 | 2,02E-01 |
| AT4G40060 | 1,22 | -1,20 | 1,58 | -0,76 | 4,50E-01 |
| AT4G40065 | 1,04 | 2,52 | 1,76 | 1,43 | 1,52E-01 |
| AT5G01010 | 1,03 | 3,57 | 2,30 | 1,55 | 1,20E-01 |
| AT5G01020 | 1,61 | 1,91 | 1,45 | 1,31 | 1,89E-01 |

| | | | | | |
|---|---|---|---|---|---|
| AT5G01100 | 0,27 | -0,47 | 4,30 | -0,11 | 9,14E-01 |
| AT5G01215 | 1,09 | -0,51 | 1,55 | -0,33 | 7,41E-01 |
| AT5G01220 | 0,12 | 2,87 | 4,34 | 0,66 | 5,08E-01 |
| AT5G01270 | 0,34 | 2,00 | 4,34 | 0,46 | 6,45E-01 |
| AT5G01365 | 0,66 | 3,35 | 4,33 | 0,78 | 4,38E-01 |
| AT5G01540 | 0,46 | 1,39 | 4,03 | 0,34 | 7,30E-01 |
| AT5G01560 | 1,57 | 2,51 | 1,61 | 1,56 | 1,20E-01 |
| AT5G01600 | 1,52 | 0,25 | 1,42 | 0,18 | 8,60E-01 |
| AT5G01630 | 1,03 | 3,09 | 2,55 | 1,21 | 2,27E-01 |
| AT5G01700 | 0,62 | 2,94 | 4,34 | 0,68 | 4,98E-01 |
| AT5G01710 | 1,65 | -0,24 | 1,51 | -0,16 | 8,75E-01 |
| AT5G01720 | 0,49 | 1,11 | 3,89 | 0,29 | 7,75E-01 |
| AT5G01730 | 0,29 | 3,20 | 4,33 | 0,74 | 4,60E-01 |
| AT5G01770 | 0,54 | 3,15 | 3,78 | 0,83 | 4,05E-01 |
| AT5G01780 | 0,19 | 0,37 | 4,28 | 0,09 | 9,31E-01 |
| AT5G01890 | 0,64 | 1,03 | 2,49 | 0,41 | 6,78E-01 |
| AT5G01930 | 1,04 | 2,28 | 3,15 | 0,72 | 4,69E-01 |
| AT5G01950 | 0,59 | 0,03 | 2,63 | 0,01 | 9,92E-01 |
| AT5G01970 | 1,20 | 4,06 | 2,03 | 2,00 | 4,52E-02 |
| AT5G01990 | 1,12 | 2,62 | 2,17 | 1,21 | 2,27E-01 |
| AT5G02130 | 0,08 | -0,56 | 4,38 | -0,13 | 8,99E-01 |
| AT5G02290 | 0,48 | 0,30 | 2,67 | 0,11 | 9,09E-01 |
| AT5G02502 | 0,33 | 0,16 | 4,29 | 0,04 | 9,69E-01 |
| AT5G02530 | 0,37 | 1,60 | 4,35 | 0,37 | 7,12E-01 |
| AT5G02670 | 0,44 | 1,51 | 3,21 | 0,47 | 6,38E-01 |
| AT5G02770 | 0,75 | -0,41 | 2,18 | -0,19 | 8,50E-01 |
| AT5G02800 | 0,65 | 1,22 | 3,00 | 0,41 | 6,85E-01 |
| AT5G02830 | 0,85 | 0,39 | 1,75 | 0,22 | 8,22E-01 |
| AT5G02890 | 0,18 | -1,21 | 4,35 | -0,28 | 7,81E-01 |
| AT5G02940 | 0,88 | 0,67 | 2,22 | 0,30 | 7,64E-01 |
| AT5G03140 | 0,48 | 0,90 | 2,55 | 0,35 | 7,24E-01 |
| AT5G03170 | 0,32 | 0,00 | 4,40 | 0,00 | 1,00E+00 |
| AT5G03204 | 0,29 | -1,08 | 4,34 | -0,25 | 8,03E-01 |
| AT5G03210 | 0,71 | -0,17 | 2,29 | -0,08 | 9,40E-01 |
| AT5G03340 | 1,55 | 2,26 | 1,56 | 1,45 | 1,47E-01 |
| AT5G03380 | 0,84 | 1,27 | 2,12 | 0,60 | 5,50E-01 |
| AT5G03400 | 1,62 | 0,11 | 1,35 | 0,08 | 9,37E-01 |
| AT5G03430 | 0,65 | 0,42 | 2,98 | 0,14 | 8,88E-01 |
| AT5G03450 | 0,39 | 0,95 | 4,38 | 0,22 | 8,29E-01 |
| AT5G03555 | 1,08 | 2,42 | 1,99 | 1,22 | 2,24E-01 |
| AT5G03560 | 0,43 | 0,39 | 4,08 | 0,10 | 9,23E-01 |
| AT5G03630 | 0,55 | 0,29 | 3,08 | 0,10 | 9,24E-01 |
| AT5G03700 | 0,11 | 1,33 | 4,36 | 0,31 | 7,59E-01 |
| AT5G03730 | 0,62 | 2,25 | 2,55 | 0,88 | 3,78E-01 |
| AT5G03760 | 0,44 | -0,81 | 4,13 | -0,20 | 8,44E-01 |
| AT5G03800 | 0,30 | 2,91 | 4,34 | 0,67 | 5,04E-01 |
| AT5G03850 | 0,83 | -1,65 | 3,01 | -0,55 | 5,84E-01 |
| AT5G03880 | 0,57 | 0,69 | 3,06 | 0,23 | 8,22E-01 |
| AT5G03910 | 0,83 | 2,59 | 2,34 | 1,11 | 2,69E-01 |
| AT5G03970 | 0,19 | 2,19 | 4,32 | 0,51 | 6,12E-01 |
| AT5G04020 | 0,78 | 1,07 | 1,82 | 0,58 | 5,59E-01 |
| AT5G04130 | 0,05 | -0,58 | 4,38 | -0,13 | 8,95E-01 |
| AT5G04267 | 0,16 | -1,50 | 4,35 | -0,35 | 7,30E-01 |
| AT5G04340 | 1,00 | 0,14 | 1,90 | 0,07 | 9,41E-01 |
| AT5G00955 | 1,17 | 0,56 | 1,58 | 0,36 | 7,22E-01 |
| AT5G04420 | 0,49 | 2,81 | 3,95 | 0,71 | 4,76E-01 |

| | | | | | |
|---|---|---|---|---|---|
| AT5G04430 | 1,54 | 2,56 | 1,60 | 1,59 | 1,11E-01 |
| AT5G04470 | 0,16 | 1,18 | 4,32 | 0,27 | 7,85E-01 |
| AT5G04480 | 0,41 | 0,30 | 3,17 | 0,09 | 9,25E-01 |
| AT5G04670 | 1,32 | 3,22 | 1,82 | 1,77 | 7,71E-02 |
| AT5G04700 | 0,30 | 1,62 | 4,35 | 0,37 | 7,10E-01 |
| AT5G04710 | 0,12 | 2,81 | 4,34 | 0,65 | 5,18E-01 |
| AT5G04730 | 1,06 | 3,50 | 2,11 | 1,66 | 9,63E-02 |
| AT5G04760 | 0,48 | 3,32 | 4,33 | 0,77 | 4,43E-01 |
| AT5G04830 | 0,60 | 1,53 | 3,06 | 0,50 | 6,16E-01 |
| AT5G04850 | 0,17 | 1,90 | 4,33 | 0,44 | 6,61E-01 |
| AT5G04860 | 0,46 | -0,01 | 4,18 | 0,00 | 9,99E-01 |
| AT5G04870 | 0,78 | 1,11 | 2,08 | 0,53 | 5,95E-01 |
| AT5G04940 | 0,64 | 0,54 | 2,92 | 0,19 | 8,53E-01 |
| AT5G04950 | 0,11 | -1,28 | 4,35 | -0,29 | 7,69E-01 |
| AT5G04990 | 0,74 | -0,09 | 3,53 | -0,03 | 9,79E-01 |
| AT5G05060 | 0,37 | 0,00 | 4,40 | 0,00 | 1,00E+00 |
| AT5G05080 | 0,77 | 0,54 | 4,36 | 0,12 | 9,02E-01 |
| AT5G05085 | 0,35 | -0,54 | 3,48 | -0,15 | 8,77E-01 |
| AT5G05100 | 0,72 | 3,20 | 2,94 | 1,09 | 2,77E-01 |
| AT5G01005 | 0,75 | 2,78 | 2,89 | 0,96 | 3,37E-01 |
| AT5G05260 | 0,23 | 0,72 | 4,27 | 0,17 | 8,67E-01 |
| AT5G01055 | 0,10 | -1,26 | 4,35 | -0,29 | 7,72E-01 |
| AT5G05480 | 0,31 | 1,18 | 3,52 | 0,33 | 7,38E-01 |
| AT5G05670 | 0,17 | 1,41 | 4,35 | 0,32 | 7,46E-01 |
| AT5G05700 | 0,69 | 3,30 | 2,90 | 1,14 | 2,56E-01 |
| AT5G05730 | 0,58 | 0,75 | 3,24 | 0,23 | 8,16E-01 |
| AT5G01095 | 0,95 | 4,51 | 4,29 | 1,05 | 2,93E-01 |
| AT5G05800 | 0,36 | 3,22 | 4,34 | 0,74 | 4,59E-01 |
| AT5G06043 | 0,95 | -0,85 | 2,62 | -0,33 | 7,45E-01 |
| AT5G06060 | 0,53 | -0,42 | 3,19 | -0,13 | 8,94E-01 |
| AT5G06110 | 0,69 | 2,08 | 2,88 | 0,72 | 4,69E-01 |
| AT5G06140 | 0,36 | 2,29 | 4,35 | 0,52 | 6,00E-01 |
| AT5G06160 | 0,43 | 0,72 | 2,66 | 0,27 | 7,86E-01 |
| AT5G01165 | 0,96 | -0,53 | 1,78 | -0,30 | 7,66E-01 |
| AT5G01195 | 1,58 | -1,30 | 1,35 | -0,96 | 3,36E-01 |
| AT5G06230 | 0,49 | 3,54 | 3,86 | 0,92 | 3,59E-01 |
| AT5G06260 | 0,46 | 1,74 | 3,20 | 0,54 | 5,86E-01 |
| AT5G06330 | 0,08 | -0,56 | 4,38 | -0,13 | 8,99E-01 |
| AT5G06410 | 0,25 | 2,64 | 4,34 | 0,61 | 5,43E-01 |
| AT5G06430 | 0,51 | 1,70 | 4,40 | 0,39 | 6,98E-01 |
| AT5G06440 | 0,83 | 1,79 | 3,64 | 0,49 | 6,23E-01 |
| AT5G06520 | 0,44 | 4,01 | 4,31 | 0,93 | 3,52E-01 |
| AT5G06540 | 0,48 | 4,17 | 4,30 | 0,97 | 3,33E-01 |
| AT5G06560 | 0,78 | 1,23 | 2,16 | 0,57 | 5,69E-01 |
| AT5G06670 | 0,24 | -0,36 | 4,34 | -0,08 | 9,34E-01 |
| AT5G06690 | 0,35 | -0,05 | 3,46 | -0,01 | 9,89E-01 |
| AT5G06750 | 0,11 | 1,28 | 4,35 | 0,29 | 7,68E-01 |
| AT5G06770 | 0,36 | -0,23 | 4,30 | -0,05 | 9,58E-01 |
| AT5G06865 | 0,22 | -0,39 | 4,32 | -0,09 | 9,28E-01 |
| AT5G06970 | 0,87 | 1,58 | 1,83 | 0,86 | 3,88E-01 |
| AT5G01295 | 0,28 | 1,64 | 4,38 | 0,38 | 7,08E-01 |
| AT5G07120 | 0,92 | 1,35 | 2,26 | 0,60 | 5,50E-01 |
| AT5G07220 | 0,71 | 0,58 | 2,03 | 0,29 | 7,74E-01 |
| AT5G07250 | 0,62 | -0,13 | 3,12 | -0,04 | 9,66E-01 |
| AT5G07360 | 0,50 | 0,24 | 3,17 | 0,07 | 9,40E-01 |
| AT5G07590 | 0,15 | 1,94 | 4,32 | 0,45 | 6,54E-01 |

| | | | | | |
|---|---|---|---|---|---|
| AT5G07620 | 0,13 | -0,63 | 4,39 | -0,14 | 8,85E-01 |
| AT5G07675 | 0,20 | -1,75 | 4,32 | -0,41 | 6,85E-01 |
| AT5G07810 | 0,26 | 2,13 | 4,38 | 0,49 | 6,27E-01 |
| AT5G07950 | 0,39 | 1,71 | 4,40 | 0,39 | 6,98E-01 |
| AT5G07980 | 1,38 | 2,57 | 1,84 | 1,40 | 1,63E-01 |
| AT5G08040 | 0,14 | 0,00 | 4,41 | 0,00 | 1,00E+00 |
| AT5G08075 | 0,48 | -0,70 | 4,01 | -0,17 | 8,62E-01 |
| AT5G08110 | 0,94 | 3,75 | 2,34 | 1,60 | 1,09E-01 |
| AT5G08130 | 1,09 | -2,30 | 1,94 | -1,18 | 2,36E-01 |
| AT5G08160 | 1,05 | -0,61 | 2,14 | -0,29 | 7,75E-01 |
| AT5G08170 | 0,71 | 1,25 | 3,62 | 0,34 | 7,31E-01 |
| AT5G01495 | 0,69 | -0,53 | 1,79 | -0,30 | 7,66E-01 |
| AT5G01505 | 1,02 | -1,81 | 1,56 | -1,16 | 2,45E-01 |
| AT5G01515 | 0,11 | 1,33 | 4,36 | 0,31 | 7,59E-01 |
| AT5G08230 | 0,62 | 0,89 | 3,04 | 0,29 | 7,70E-01 |
| AT5G08280 | 1,65 | 1,82 | 1,58 | 1,15 | 2,48E-01 |
| AT5G08305 | 0,42 | 2,35 | 4,14 | 0,57 | 5,70E-01 |
| AT5G08315 | 0,44 | -0,14 | 4,41 | -0,03 | 9,75E-01 |
| AT5G08350 | 0,26 | 2,30 | 4,37 | 0,53 | 5,98E-01 |
| AT5G08450 | 0,66 | 1,24 | 2,64 | 0,47 | 6,38E-01 |
| AT5G08520 | 1,35 | -1,17 | 1,69 | -0,69 | 4,92E-01 |
| AT5G08560 | 0,55 | -0,16 | 2,62 | -0,06 | 9,53E-01 |
| AT5G08650 | 1,18 | 0,32 | 1,93 | 0,17 | 8,68E-01 |
| AT5G08770 | 1,04 | -0,13 | 1,87 | -0,07 | 9,43E-01 |
| AT5G08790 | 1,51 | 1,62 | 1,35 | 1,20 | 2,29E-01 |
| AT5G09330 | 1,10 | -0,26 | 1,58 | -0,16 | 8,70E-01 |
| AT5G09443 | 0,26 | 2,13 | 4,38 | 0,49 | 6,27E-01 |
| AT5G09460 | 0,93 | 2,44 | 1,82 | 1,34 | 1,80E-01 |
| AT5G09510 | 0,57 | 0,28 | 3,97 | 0,07 | 9,44E-01 |
| AT5G09620 | 0,88 | -1,77 | 2,25 | -0,79 | 4,31E-01 |
| AT5G09650 | 0,62 | 0,69 | 2,50 | 0,28 | 7,82E-01 |
| AT5G09820 | 0,63 | -0,15 | 3,04 | -0,05 | 9,61E-01 |
| AT5G09850 | 0,67 | 0,58 | 2,94 | 0,20 | 8,43E-01 |
| AT5G09880 | 0,62 | 1,80 | 2,80 | 0,64 | 5,20E-01 |
| AT5G09890 | 0,52 | 2,17 | 3,82 | 0,57 | 5,71E-01 |
| AT5G09930 | 0,08 | 2,33 | 4,36 | 0,53 | 5,93E-01 |
| AT5G10030 | 0,52 | 2,37 | 3,93 | 0,60 | 5,46E-01 |
| AT5G10235 | 0,28 | 2,59 | 4,35 | 0,59 | 5,53E-01 |
| AT5G10290 | 0,76 | 1,99 | 2,24 | 0,89 | 3,73E-01 |
| AT5G10330 | 0,57 | 1,03 | 2,55 | 0,41 | 6,84E-01 |
| AT5G10350 | 0,66 | 2,72 | 4,37 | 0,62 | 5,33E-01 |
| AT5G10360 | 1,44 | 0,10 | 1,42 | 0,07 | 9,46E-01 |
| AT5G10390 | 0,27 | -1,28 | 4,33 | -0,29 | 7,68E-01 |
| AT5G10400 | 0,23 | -2,41 | 4,30 | -0,56 | 5,75E-01 |
| AT5G10470 | 0,57 | 1,15 | 3,18 | 0,36 | 7,18E-01 |
| AT5G10490 | 0,10 | 0,34 | 4,35 | 0,08 | 9,37E-01 |
| AT5G10550 | 0,86 | 1,99 | 2,15 | 0,93 | 3,54E-01 |
| AT5G10605 | 0,65 | -0,36 | 3,04 | -0,12 | 9,06E-01 |
| AT5G10650 | 0,25 | 2,79 | 4,34 | 0,64 | 5,20E-01 |
| AT5G10690 | 0,38 | 0,38 | 4,27 | 0,09 | 9,29E-01 |
| AT5G01815 | 0,80 | -0,97 | 1,89 | -0,52 | 6,06E-01 |
| AT5G10700 | 0,56 | -0,97 | 3,10 | -0,31 | 7,53E-01 |
| AT5G10730 | 0,23 | -0,93 | 4,38 | -0,21 | 8,32E-01 |
| AT5G10770 | 0,94 | 0,78 | 2,03 | 0,39 | 7,00E-01 |
| AT5G10800 | 0,30 | 2,84 | 4,35 | 0,65 | 5,14E-01 |
| AT5G10820 | 0,63 | 3,92 | 4,31 | 0,91 | 3,63E-01 |

| | | | | | |
|---|---|---|---|---|---|
| AT5G10930 | 1,14 | 1,43 | 1,66 | 0,86 | 3,90E-01 |
| AT5G10940 | 0,77 | 2,62 | 3,59 | 0,73 | 4,65E-01 |
| AT5G10965 | 0,75 | 2,30 | 2,38 | 0,97 | 3,33E-01 |
| AT5G10980 | 0,55 | 1,50 | 3,76 | 0,40 | 6,91E-01 |
| AT5G11000 | 0,45 | 0,40 | 4,13 | 0,10 | 9,22E-01 |
| AT5G11070 | 1,00 | -0,04 | 1,82 | -0,02 | 9,83E-01 |
| AT5G11180 | 0,81 | 1,36 | 2,15 | 0,63 | 5,26E-01 |
| AT5G11210 | 1,06 | 2,40 | 3,23 | 0,74 | 4,57E-01 |
| AT5G11250 | 1,58 | 1,26 | 1,34 | 0,94 | 3,47E-01 |
| AT5G11270 | 0,22 | 0,37 | 4,35 | 0,08 | 9,32E-01 |
| AT5G11330 | 1,01 | 0,51 | 1,61 | 0,32 | 7,52E-01 |
| AT5G11480 | 0,46 | -0,45 | 3,21 | -0,14 | 8,89E-01 |
| AT5G11520 | 1,24 | 0,72 | 1,83 | 0,40 | 6,93E-01 |
| AT5G11550 | 0,28 | 1,40 | 4,31 | 0,32 | 7,45E-01 |
| AT5G11770 | 0,54 | 0,01 | 2,61 | 0,00 | 9,97E-01 |
| AT5G11850 | 0,25 | 1,81 | 4,28 | 0,42 | 6,72E-01 |
| AT5G11860 | 0,12 | 2,81 | 4,34 | 0,65 | 5,18E-01 |
| AT5G11910 | 0,27 | 2,08 | 4,31 | 0,48 | 6,29E-01 |
| AT5G11960 | 0,36 | 3,47 | 4,33 | 0,80 | 4,23E-01 |
| AT5G12010 | 0,59 | 1,95 | 3,03 | 0,64 | 5,20E-01 |
| AT5G12080 | 0,25 | 2,97 | 4,33 | 0,69 | 4,93E-01 |
| AT5G12140 | 0,22 | 0,62 | 4,28 | 0,14 | 8,85E-01 |
| AT5G12200 | 0,44 | 2,46 | 3,25 | 0,76 | 4,49E-01 |
| AT5G12310 | 0,42 | -0,46 | 3,25 | -0,14 | 8,88E-01 |
| AT5G12390 | 0,11 | 1,33 | 4,36 | 0,31 | 7,59E-01 |
| AT5G12430 | 1,28 | 3,40 | 2,36 | 1,44 | 1,49E-01 |
| AT5G12440 | 0,88 | 3,81 | 2,68 | 1,42 | 1,56E-01 |
| AT5G12480 | 0,34 | 0,48 | 4,26 | 0,11 | 9,11E-01 |
| AT5G12890 | 0,77 | 3,45 | 3,63 | 0,95 | 3,43E-01 |
| AT5G12900 | 0,23 | 0,91 | 4,30 | 0,21 | 8,32E-01 |
| AT5G13020 | 0,81 | 1,75 | 2,21 | 0,79 | 4,29E-01 |
| AT5G13050 | 0,42 | 0,65 | 4,15 | 0,16 | 8,77E-01 |
| AT5G13090 | 0,10 | 0,39 | 4,35 | 0,09 | 9,29E-01 |
| AT5G13120 | 0,17 | -0,14 | 4,41 | -0,03 | 9,75E-01 |
| AT5G13140 | 0,23 | -0,91 | 4,37 | -0,21 | 8,36E-01 |
| AT5G13170 | 0,48 | 3,32 | 4,33 | 0,77 | 4,43E-01 |
| AT5G13190 | 0,99 | 0,41 | 1,63 | 0,25 | 8,03E-01 |
| AT5G02055 | 0,48 | -1,05 | 2,70 | -0,39 | 6,97E-01 |
| AT5G13225 | 0,48 | -1,05 | 2,70 | -0,39 | 6,97E-01 |
| AT5G13270 | 0,40 | 1,22 | 4,18 | 0,29 | 7,70E-01 |
| AT5G13300 | 0,77 | 2,46 | 2,36 | 1,04 | 2,97E-01 |
| AT5G13400 | 0,43 | 2,75 | 4,31 | 0,64 | 5,24E-01 |
| AT5G13430 | 0,54 | 3,56 | 3,77 | 0,94 | 3,45E-01 |
| AT5G13440 | 0,29 | -0,87 | 4,27 | -0,20 | 8,39E-01 |
| AT5G13450 | 0,96 | -0,16 | 1,96 | -0,08 | 9,35E-01 |
| AT5G13550 | 0,46 | 2,63 | 3,97 | 0,66 | 5,07E-01 |
| AT5G13560 | 1,05 | 2,45 | 2,16 | 1,13 | 2,57E-01 |
| AT5G13590 | 1,32 | 2,56 | 1,93 | 1,33 | 1,84E-01 |
| AT5G13680 | 1,03 | 0,89 | 1,66 | 0,54 | 5,92E-01 |
| AT5G13710 | 0,59 | 0,35 | 2,99 | 0,12 | 9,07E-01 |
| AT5G13780 | 0,62 | 2,25 | 2,99 | 0,75 | 4,51E-01 |
| AT5G13800 | 0,74 | -2,49 | 2,47 | -1,01 | 3,15E-01 |
| AT5G13825 | 0,39 | -0,59 | 4,28 | -0,14 | 8,91E-01 |
| AT5G13950 | 1,63 | 1,68 | 1,45 | 1,16 | 2,47E-01 |
| AT5G13990 | 0,12 | 0,31 | 4,35 | 0,07 | 9,44E-01 |
| AT5G14100 | 0,11 | 1,33 | 4,36 | 0,31 | 7,59E-01 |

| | | | | | |
|---|---|---|---|---|---|
| AT5G14170 | 1,07 | 3,06 | 2,18 | 1,40 | 1,60E-01 |
| AT5G14210 | 0,50 | -0,75 | 3,31 | -0,23 | 8,21E-01 |
| AT5G14240 | 0,57 | -0,75 | 2,66 | -0,28 | 7,79E-01 |
| AT5G14320 | 0,97 | -0,04 | 1,60 | -0,02 | 9,80E-01 |
| AT5G14390 | 0,52 | 0,86 | 4,04 | 0,21 | 8,32E-01 |
| AT5G14420 | 0,18 | 1,51 | 4,31 | 0,35 | 7,25E-01 |
| AT5G14530 | 0,86 | 2,05 | 2,77 | 0,74 | 4,60E-01 |
| AT5G14540 | 0,54 | 1,11 | 3,93 | 0,28 | 7,78E-01 |
| AT5G02245 | 0,10 | -1,26 | 4,35 | -0,29 | 7,72E-01 |
| AT5G14580 | 1,26 | 2,08 | 2,66 | 0,78 | 4,34E-01 |
| AT5G14590 | 0,98 | 1,83 | 1,83 | 1,00 | 3,18E-01 |
| AT5G14700 | 0,24 | -1,45 | 4,28 | -0,34 | 7,35E-01 |
| AT5G14720 | 0,96 | 1,82 | 1,70 | 1,07 | 2,85E-01 |
| AT5G14770 | 0,20 | 2,83 | 4,35 | 0,65 | 5,14E-01 |
| AT5G14820 | 0,29 | 1,74 | 4,31 | 0,40 | 6,87E-01 |
| AT5G14940 | 1,18 | 2,65 | 2,18 | 1,21 | 2,24E-01 |
| AT5G14950 | 1,52 | 1,46 | 1,33 | 1,09 | 2,74E-01 |
| AT5G15050 | 0,10 | 0,34 | 4,35 | 0,08 | 9,37E-01 |
| AT5G15210 | 0,28 | 1,62 | 4,35 | 0,37 | 7,10E-01 |
| AT5G15250 | 0,05 | -0,58 | 4,38 | -0,13 | 8,95E-01 |
| AT5G15360 | 0,52 | -0,91 | 2,40 | -0,38 | 7,06E-01 |
| AT5G15410 | 1,41 | 3,44 | 1,78 | 1,94 | 5,27E-02 |
| AT5G15420 | 1,66 | 0,49 | 1,42 | 0,34 | 7,31E-01 |
| AT5G15450 | 1,23 | 2,24 | 1,66 | 1,35 | 1,78E-01 |
| AT5G15460 | 0,08 | 2,33 | 4,36 | 0,53 | 5,93E-01 |
| AT5G15470 | 0,21 | 1,20 | 4,36 | 0,28 | 7,83E-01 |
| AT5G15490 | 0,40 | 0,86 | 3,32 | 0,26 | 7,95E-01 |
| AT5G15537 | 0,21 | 2,61 | 4,34 | 0,60 | 5,48E-01 |
| AT5G15540 | 1,01 | 2,52 | 1,88 | 1,34 | 1,82E-01 |
| AT5G15580 | 0,70 | 0,85 | 2,51 | 0,34 | 7,33E-01 |
| AT5G15690 | 0,53 | -0,86 | 2,66 | -0,33 | 7,45E-01 |
| AT5G15700 | 0,61 | -0,01 | 2,13 | -0,01 | 9,96E-01 |
| AT5G15740 | 0,28 | -1,84 | 4,33 | -0,43 | 6,70E-01 |
| AT5G15780 | 0,22 | -0,39 | 4,32 | -0,09 | 9,28E-01 |
| AT5G15800 | 0,08 | -0,87 | 4,38 | -0,20 | 8,43E-01 |
| AT5G15802 | 0,08 | -0,56 | 4,38 | -0,13 | 8,99E-01 |
| AT5G15860 | 0,19 | 2,19 | 4,32 | 0,51 | 6,12E-01 |
| AT5G16050 | 0,47 | 0,11 | 2,62 | 0,04 | 9,66E-01 |
| AT5G16070 | 0,93 | 1,56 | 2,24 | 0,70 | 4,86E-01 |
| AT5G16110 | 1,09 | 1,42 | 1,69 | 0,84 | 4,00E-01 |
| AT5G16200 | 0,36 | 0,00 | 4,41 | 0,00 | 1,00E+00 |
| AT5G16220 | 0,18 | -1,99 | 4,31 | -0,46 | 6,44E-01 |
| AT5G16280 | 0,67 | 3,61 | 2,86 | 1,26 | 2,07E-01 |
| AT5G16360 | 0,08 | 2,33 | 4,36 | 0,53 | 5,93E-01 |
| AT5G16370 | 1,57 | 2,92 | 1,55 | 1,89 | 5,92E-02 |
| AT5G16380 | 0,25 | 1,85 | 4,33 | 0,43 | 6,69E-01 |
| AT5G16390 | 0,61 | 2,40 | 3,74 | 0,64 | 5,21E-01 |
| AT5G16480 | 0,56 | -0,14 | 4,41 | -0,03 | 9,75E-01 |
| AT5G16490 | 0,08 | -0,56 | 4,38 | -0,13 | 8,99E-01 |
| AT5G16520 | 0,47 | 1,78 | 4,15 | 0,43 | 6,68E-01 |
| AT5G16560 | 0,42 | 0,01 | 4,30 | 0,00 | 9,98E-01 |
| AT5G16610 | 0,08 | 2,33 | 4,36 | 0,53 | 5,93E-01 |
| AT5G16660 | 0,23 | 1,37 | 4,28 | 0,32 | 7,50E-01 |
| AT5G16680 | 0,78 | -0,02 | 2,09 | -0,01 | 9,92E-01 |
| AT5G16740 | 0,42 | 0,79 | 4,40 | 0,18 | 8,58E-01 |
| AT5G16750 | 0,89 | 1,26 | 1,93 | 0,65 | 5,13E-01 |

| | | | | | |
|---|---|---|---|---|---|
| AT5G16780 | 0,54 | -1,00 | 3,84 | -0,26 | 7,94E-01 |
| AT5G16800 | 0,21 | -0,24 | 4,35 | -0,06 | 9,56E-01 |
| AT5G16830 | 0,10 | 0,34 | 4,35 | 0,08 | 9,37E-01 |
| AT5G16980 | 0,08 | 2,08 | 4,38 | 0,47 | 6,35E-01 |
| AT5G16990 | 0,20 | 1,57 | 4,34 | 0,36 | 7,17E-01 |
| AT5G17000 | 0,20 | -0,80 | 4,37 | -0,18 | 8,55E-01 |
| AT5G17010 | 0,24 | 0,54 | 4,33 | 0,12 | 9,01E-01 |
| AT5G02655 | 0,55 | -2,30 | 2,56 | -0,90 | 3,68E-01 |
| AT5G02665 | 1,25 | -0,42 | 2,03 | -0,21 | 8,37E-01 |
| AT5G02675 | 0,37 | 1,36 | 4,25 | 0,32 | 7,49E-01 |
| AT5G17290 | 0,65 | 1,35 | 2,96 | 0,45 | 6,49E-01 |
| AT5G17300 | 0,69 | 2,72 | 2,91 | 0,93 | 3,50E-01 |
| AT5G17320 | 0,38 | 1,20 | 4,35 | 0,28 | 7,83E-01 |
| AT5G17380 | 0,80 | 2,24 | 2,38 | 0,94 | 3,47E-01 |
| AT5G17420 | 0,40 | 2,81 | 4,32 | 0,65 | 5,15E-01 |
| AT5G17460 | 0,52 | 0,00 | 4,41 | 0,00 | 1,00E+00 |
| AT5G17520 | 0,17 | 0,00 | 4,41 | 0,00 | 1,00E+00 |
| AT5G17550 | 0,08 | 2,33 | 4,36 | 0,53 | 5,93E-01 |
| AT5G17600 | 0,90 | -0,09 | 1,97 | -0,04 | 9,65E-01 |
| AT5G17620 | 0,42 | 0,79 | 4,40 | 0,18 | 8,58E-01 |
| AT5G17640 | 0,35 | -1,26 | 4,35 | -0,29 | 7,72E-01 |
| AT5G17690 | 0,20 | 1,88 | 4,35 | 0,43 | 6,66E-01 |
| AT5G17770 | 0,49 | 0,34 | 3,07 | 0,11 | 9,11E-01 |
| AT5G17860 | 0,63 | 0,64 | 2,66 | 0,24 | 8,09E-01 |
| AT5G17880 | 1,61 | 1,86 | 1,93 | 0,96 | 3,36E-01 |
| AT5G17930 | 0,48 | 4,11 | 4,30 | 0,95 | 3,40E-01 |
| AT5G18050 | 0,24 | 0,00 | 4,41 | 0,00 | 1,00E+00 |
| AT5G18110 | 0,39 | 0,24 | 4,27 | 0,06 | 9,56E-01 |
| AT5G18190 | 0,63 | 1,97 | 2,56 | 0,77 | 4,43E-01 |
| AT5G18230 | 0,99 | 3,09 | 2,71 | 1,14 | 2,54E-01 |
| AT5G18240 | 1,40 | -3,97 | 2,69 | -1,47 | 1,40E-01 |
| AT5G18260 | 0,49 | 2,35 | 4,36 | 0,54 | 5,90E-01 |
| AT5G18280 | 0,49 | 1,92 | 3,05 | 0,63 | 5,29E-01 |
| AT5G18290 | 0,36 | 0,00 | 4,41 | 0,00 | 1,00E+00 |
| AT5G18340 | 0,58 | 1,47 | 3,97 | 0,37 | 7,12E-01 |
| AT5G18400 | 0,32 | -0,62 | 4,28 | -0,14 | 8,85E-01 |
| AT5G02895 | 0,88 | 1,88 | 2,57 | 0,73 | 4,65E-01 |
| AT5G18475 | 0,21 | 3,35 | 4,33 | 0,77 | 4,39E-01 |
| AT5G18480 | 0,62 | 2,83 | 2,95 | 0,96 | 3,38E-01 |
| AT5G18570 | 0,43 | 1,31 | 3,92 | 0,33 | 7,38E-01 |
| AT5G18590 | 0,48 | 0,60 | 3,19 | 0,19 | 8,51E-01 |
| AT5G18620 | 0,19 | 0,37 | 4,28 | 0,09 | 9,31E-01 |
| AT5G18700 | 0,89 | 1,80 | 2,73 | 0,66 | 5,10E-01 |
| AT5G02935 | 0,05 | -0,58 | 4,38 | -0,13 | 8,95E-01 |
| AT5G19010 | 0,59 | 0,14 | 2,37 | 0,06 | 9,55E-01 |
| AT5G19130 | 0,62 | 1,29 | 2,56 | 0,50 | 6,15E-01 |
| AT5G19220 | 0,51 | 0,68 | 4,34 | 0,16 | 8,75E-01 |
| AT5G19360 | 0,72 | 0,00 | 4,41 | 0,00 | 1,00E+00 |
| AT5G19410 | 0,62 | 3,42 | 2,93 | 1,17 | 2,43E-01 |
| AT5G19420 | 1,09 | 1,34 | 1,79 | 0,75 | 4,52E-01 |
| AT5G19450 | 0,87 | 0,35 | 1,76 | 0,20 | 8,44E-01 |
| AT5G19485 | 0,20 | 2,30 | 4,36 | 0,53 | 5,98E-01 |
| AT5G19590 | 0,42 | -0,28 | 3,10 | -0,09 | 9,27E-01 |
| AT5G19660 | 0,36 | 1,09 | 4,26 | 0,25 | 7,99E-01 |
| AT5G19760 | 0,90 | 0,50 | 2,47 | 0,20 | 8,40E-01 |
| AT5G19850 | 0,75 | 2,34 | 3,57 | 0,66 | 5,12E-01 |

| | | | | | |
|---|---|---|---|---|---|
| AT5G19860 | 0,51 | 2,64 | 3,96 | 0,67 | 5,04E-01 |
| AT5G19900 | 0,73 | -0,72 | 2,98 | -0,24 | 8,09E-01 |
| AT5G19920 | 0,37 | 2,44 | 4,35 | 0,56 | 5,74E-01 |
| AT5G19930 | 0,12 | 0,63 | 4,33 | 0,15 | 8,85E-01 |
| AT5G19950 | 0,27 | 0,14 | 4,29 | 0,03 | 9,75E-01 |
| AT5G19980 | 0,33 | 0,79 | 4,32 | 0,18 | 8,55E-01 |
| AT5G20040 | 0,12 | 2,87 | 4,34 | 0,66 | 5,08E-01 |
| AT5G20160 | 0,08 | -0,56 | 4,38 | -0,13 | 8,99E-01 |
| AT5G20170 | 0,19 | -0,01 | 4,32 | 0,00 | 9,98E-01 |
| AT5G20225 | 0,17 | 3,14 | 4,33 | 0,73 | 4,68E-01 |
| AT5G20260 | 0,52 | 3,61 | 4,33 | 0,83 | 4,04E-01 |
| AT5G20270 | 0,14 | -0,31 | 4,33 | -0,07 | 9,43E-01 |
| AT5G20300 | 1,26 | 1,80 | 2,38 | 0,76 | 4,49E-01 |
| AT5G20320 | 1,21 | 5,22 | 2,42 | 2,16 | 3,08E-02 |
| AT5G20360 | 0,79 | 1,18 | 2,09 | 0,57 | 5,72E-01 |
| AT5G20380 | 0,45 | 2,77 | 4,34 | 0,64 | 5,24E-01 |
| AT5G20490 | 1,59 | 0,42 | 1,71 | 0,25 | 8,04E-01 |
| AT5G20520 | 1,55 | 1,26 | 1,51 | 0,83 | 4,06E-01 |
| AT5G20570 | 0,58 | 1,59 | 3,10 | 0,51 | 6,08E-01 |
| AT5G20600 | 1,15 | 4,92 | 2,09 | 2,35 | 1,86E-02 |
| AT5G20620 | 0,96 | 2,31 | 2,34 | 0,99 | 3,24E-01 |
| AT5G20630 | 1,03 | -0,44 | 1,84 | -0,24 | 8,12E-01 |
| AT5G20650 | 0,87 | 2,56 | 2,90 | 0,88 | 3,77E-01 |
| AT5G20820 | 0,08 | -0,56 | 4,38 | -0,13 | 8,99E-01 |
| AT5G20830 | 0,97 | 1,29 | 2,36 | 0,55 | 5,85E-01 |
| AT5G20840 | 1,18 | -0,23 | 1,41 | -0,16 | 8,70E-01 |
| AT5G20890 | 1,15 | 2,04 | 2,04 | 1,00 | 3,18E-01 |
| AT5G20920 | 0,64 | 0,68 | 3,18 | 0,21 | 8,32E-01 |
| AT5G20940 | 0,27 | 1,96 | 4,31 | 0,45 | 6,50E-01 |
| AT5G03265 | 0,23 | 1,05 | 4,31 | 0,24 | 8,08E-01 |
| AT5G20980 | 0,81 | 3,01 | 2,76 | 1,09 | 2,74E-01 |
| AT5G20990 | 1,65 | 1,37 | 1,64 | 0,84 | 4,02E-01 |
| AT5G21040 | 0,32 | 1,90 | 4,34 | 0,44 | 6,62E-01 |
| AT5G21160 | 0,41 | -0,80 | 4,29 | -0,19 | 8,52E-01 |
| AT5G21274 | 0,23 | 1,37 | 4,28 | 0,32 | 7,50E-01 |
| AT5G21920 | 0,36 | 1,39 | 4,29 | 0,32 | 7,47E-01 |
| AT5G21950 | 0,08 | 2,08 | 4,38 | 0,47 | 6,35E-01 |
| AT5G22010 | 0,59 | 3,23 | 2,95 | 1,10 | 2,73E-01 |
| AT5G22020 | 0,55 | -0,11 | 3,92 | -0,03 | 9,79E-01 |
| AT5G22030 | 0,76 | 0,71 | 3,39 | 0,21 | 8,34E-01 |
| AT5G22060 | 1,34 | 2,15 | 2,05 | 1,05 | 2,94E-01 |
| AT5G22080 | 0,35 | -0,20 | 3,38 | -0,06 | 9,54E-01 |
| AT5G22090 | 0,53 | 1,18 | 3,26 | 0,36 | 7,16E-01 |
| AT5G22250 | 1,15 | -0,21 | 1,79 | -0,12 | 9,06E-01 |
| AT5G22350 | 0,24 | -0,04 | 4,27 | -0,01 | 9,93E-01 |
| AT5G22360 | 0,12 | 2,87 | 4,34 | 0,66 | 5,08E-01 |
| AT5G22390 | 0,49 | 3,14 | 3,17 | 0,99 | 3,22E-01 |
| AT5G22440 | 0,73 | -0,04 | 3,03 | -0,01 | 9,91E-01 |
| AT5G22450 | 0,87 | -0,23 | 1,69 | -0,14 | 8,90E-01 |
| AT5G22505 | 1,58 | -2,66 | 1,81 | -1,47 | 1,43E-01 |
| AT5G22510 | 0,44 | -1,78 | 4,18 | -0,43 | 6,70E-01 |
| AT5G03425 | 0,62 | 1,64 | 3,67 | 0,45 | 6,54E-01 |
| AT5G22540 | 0,22 | -0,11 | 4,34 | -0,02 | 9,80E-01 |
| AT5G22570 | 0,26 | 1,27 | 4,31 | 0,29 | 7,68E-01 |
| AT5G22700 | 0,44 | 3,01 | 4,07 | 0,74 | 4,59E-01 |
| AT5G22740 | 0,58 | 1,40 | 3,85 | 0,36 | 7,16E-01 |

| | | | | | |
|---|---|---|---|---|---|
| AT5G22750 | 0,55 | 3,10 | 4,34 | 0,71 | 4,75E-01 |
| AT5G03475 | 1,21 | -0,61 | 1,66 | -0,37 | 7,12E-01 |
| AT5G22780 | 1,47 | -0,64 | 1,41 | -0,45 | 6,50E-01 |
| AT5G22850 | 0,23 | 2,48 | 4,31 | 0,57 | 5,66E-01 |
| AT5G22860 | 0,35 | 1,07 | 3,37 | 0,32 | 7,51E-01 |
| AT5G23040 | 0,63 | -1,75 | 2,43 | -0,72 | 4,72E-01 |
| AT5G23050 | 0,45 | 2,45 | 4,07 | 0,60 | 5,47E-01 |
| AT5G23060 | 1,11 | 1,35 | 1,56 | 0,87 | 3,84E-01 |
| AT5G23080 | 1,07 | 3,09 | 2,55 | 1,22 | 2,24E-01 |
| AT5G23240 | 0,89 | 0,02 | 2,03 | 0,01 | 9,94E-01 |
| AT5G23300 | 0,32 | 1,70 | 4,33 | 0,39 | 6,95E-01 |
| AT5G23390 | 0,47 | 2,85 | 3,91 | 0,73 | 4,66E-01 |
| AT5G23405 | 0,14 | 1,12 | 4,32 | 0,26 | 7,96E-01 |
| AT5G23430 | 0,90 | 0,84 | 1,94 | 0,43 | 6,67E-01 |
| AT5G23440 | 0,16 | -0,80 | 4,37 | -0,18 | 8,54E-01 |
| AT5G23450 | 0,83 | 2,70 | 2,80 | 0,96 | 3,35E-01 |
| AT5G23580 | 0,83 | 0,97 | 2,21 | 0,44 | 6,62E-01 |
| AT5G23590 | 0,61 | 4,32 | 3,63 | 1,19 | 2,34E-01 |
| AT5G23610 | 1,31 | 3,54 | 1,95 | 1,81 | 6,95E-02 |
| AT5G23660 | 0,79 | 0,11 | 3,49 | 0,03 | 9,74E-01 |
| AT5G23740 | 0,24 | -1,45 | 4,28 | -0,34 | 7,35E-01 |
| AT5G23900 | 0,57 | 0,91 | 2,70 | 0,34 | 7,36E-01 |
| AT5G23940 | 0,08 | -0,56 | 4,38 | -0,13 | 8,99E-01 |
| AT5G23990 | 0,29 | -0,95 | 4,30 | -0,22 | 8,25E-01 |
| AT5G24120 | 0,77 | 2,61 | 2,37 | 1,10 | 2,71E-01 |
| AT5G24140 | 0,08 | -0,56 | 4,38 | -0,13 | 8,99E-01 |
| AT5G24150 | 0,21 | 3,22 | 4,33 | 0,74 | 4,58E-01 |
| AT5G24160 | 0,65 | 1,45 | 2,78 | 0,52 | 6,02E-01 |
| AT5G24240 | 1,01 | -0,04 | 1,66 | -0,02 | 9,81E-01 |
| AT5G24260 | 1,06 | 0,57 | 2,04 | 0,28 | 7,82E-01 |
| AT5G24270 | 1,64 | -0,15 | 1,30 | -0,12 | 9,06E-01 |
| AT5G24320 | 0,20 | 1,15 | 4,36 | 0,26 | 7,92E-01 |
| AT5G24355 | 0,08 | 2,33 | 4,36 | 0,53 | 5,93E-01 |
| AT5G24430 | 0,56 | 0,19 | 3,88 | 0,05 | 9,62E-01 |
| AT5G24470 | 1,29 | 0,74 | 2,54 | 0,29 | 7,70E-01 |
| AT5G24480 | 0,23 | 1,49 | 4,28 | 0,35 | 7,29E-01 |
| AT5G24500 | 0,24 | 1,11 | 4,33 | 0,26 | 7,97E-01 |
| AT5G24520 | 0,95 | 1,18 | 1,90 | 0,62 | 5,34E-01 |
| AT5G24590 | 1,51 | -1,14 | 1,50 | -0,76 | 4,50E-01 |
| AT5G24655 | 0,37 | -0,45 | 4,25 | -0,11 | 9,15E-01 |
| AT5G24735 | 0,57 | -0,78 | 3,16 | -0,25 | 8,04E-01 |
| AT5G24740 | 1,40 | 2,05 | 1,58 | 1,30 | 1,94E-01 |
| AT5G24770 | 0,62 | -0,12 | 2,11 | -0,06 | 9,56E-01 |
| AT5G24780 | 0,08 | -0,56 | 4,38 | -0,13 | 8,99E-01 |
| AT5G24940 | 0,08 | 2,08 | 4,38 | 0,47 | 6,35E-01 |
| AT5G24960 | 0,36 | -1,19 | 4,23 | -0,28 | 7,79E-01 |
| AT5G25000 | 0,16 | 0,85 | 4,31 | 0,20 | 8,44E-01 |
| AT5G25070 | 1,26 | 2,32 | 1,73 | 1,34 | 1,82E-01 |
| AT5G25100 | 1,32 | 1,66 | 1,56 | 1,06 | 2,89E-01 |
| AT5G25200 | 0,60 | -1,86 | 2,47 | -0,75 | 4,52E-01 |
| AT5G25265 | 0,67 | 2,76 | 2,88 | 0,96 | 3,37E-01 |
| AT5G25350 | 1,15 | 3,97 | 1,95 | 2,04 | 4,15E-02 |
| AT5G25370 | 0,71 | -1,89 | 2,88 | -0,66 | 5,10E-01 |
| AT5G03985 | 0,37 | 0,00 | 4,40 | 0,00 | 1,00E+00 |
| AT5G25480 | 0,27 | 0,00 | 4,41 | 0,00 | 1,00E+00 |
| AT5G25510 | 0,49 | 1,80 | 3,19 | 0,56 | 5,73E-01 |

| | | | | | |
|---|---|---|---|---|---|
| AT5G25530 | 1,06 | 3,42 | 3,27 | 1,05 | 2,95E-01 |
| AT5G25756 | 0,35 | -3,07 | 4,28 | -0,72 | 4,73E-01 |
| AT5G25757 | 0,87 | 0,61 | 2,04 | 0,30 | 7,65E-01 |
| AT5G25770 | 0,28 | 1,87 | 4,37 | 0,43 | 6,69E-01 |
| AT5G04085 | 0,59 | -3,60 | 4,27 | -0,84 | 3,99E-01 |
| AT5G25780 | 1,13 | 2,04 | 1,91 | 1,07 | 2,85E-01 |
| AT5G25820 | 0,43 | -0,74 | 4,38 | -0,17 | 8,66E-01 |
| AT5G25830 | 0,05 | -0,58 | 4,38 | -0,13 | 8,95E-01 |
| AT5G25920 | 0,08 | 2,08 | 4,38 | 0,47 | 6,35E-01 |
| AT5G26170 | 0,17 | 2,04 | 4,33 | 0,47 | 6,37E-01 |
| AT5G26210 | 0,43 | 2,46 | 4,13 | 0,60 | 5,51E-01 |
| AT5G26230 | 0,08 | -0,87 | 4,38 | -0,20 | 8,43E-01 |
| AT5G26680 | 0,15 | -0,41 | 4,32 | -0,09 | 9,25E-01 |
| AT5G04245 | 1,57 | -1,03 | 1,93 | -0,54 | 5,92E-01 |
| AT5G26640 | 0,17 | 2,62 | 4,35 | 0,60 | 5,47E-01 |
| AT5G26622 | 0,46 | -1,68 | 4,33 | -0,39 | 6,98E-01 |
| AT5G04255 | 0,21 | -0,24 | 4,35 | -0,06 | 9,56E-01 |
| AT5G26760 | 0,42 | 1,22 | 4,12 | 0,30 | 7,67E-01 |
| AT5G26770 | 1,06 | -1,29 | 1,84 | -0,70 | 4,83E-01 |
| AT5G26880 | 0,63 | 0,59 | 3,14 | 0,19 | 8,50E-01 |
| AT5G26910 | 0,66 | 1,37 | 2,53 | 0,54 | 5,87E-01 |
| AT5G26960 | 0,27 | -0,97 | 4,31 | -0,23 | 8,21E-01 |
| AT5G26970 | 0,84 | -1,46 | 1,89 | -0,77 | 4,41E-01 |
| AT5G27030 | 1,03 | 0,49 | 1,82 | 0,27 | 7,86E-01 |
| AT5G27100 | 0,25 | 2,98 | 4,33 | 0,69 | 4,92E-01 |
| AT5G27238 | 0,05 | -0,58 | 4,38 | -0,13 | 8,95E-01 |
| AT5G04375 | 0,24 | 1,11 | 4,33 | 0,26 | 7,97E-01 |
| AT5G27330 | 0,05 | -0,58 | 4,38 | -0,13 | 8,95E-01 |
| AT5G27380 | 0,63 | 4,17 | 3,56 | 1,17 | 2,41E-01 |
| AT5G27550 | 0,34 | 1,89 | 4,34 | 0,44 | 6,63E-01 |
| AT5G27640 | 1,34 | 1,97 | 1,66 | 1,18 | 2,37E-01 |
| AT5G27700 | 0,49 | 2,45 | 3,20 | 0,77 | 4,44E-01 |
| AT5G27715 | 0,29 | -1,95 | 4,32 | -0,45 | 6,51E-01 |
| AT5G04435 | 0,39 | -0,98 | 4,21 | -0,23 | 8,16E-01 |
| AT5G27790 | 0,75 | -1,99 | 3,36 | -0,59 | 5,54E-01 |
| AT5G27840 | 0,72 | 3,35 | 2,76 | 1,21 | 2,25E-01 |
| AT5G27870 | 1,24 | 2,92 | 1,91 | 1,53 | 1,26E-01 |
| AT5G27920 | 0,08 | 2,33 | 4,36 | 0,53 | 5,93E-01 |
| AT5G27960 | 0,74 | -1,06 | 2,47 | -0,43 | 6,68E-01 |
| AT5G28020 | 0,34 | 1,17 | 3,50 | 0,33 | 7,38E-01 |
| AT5G28030 | 0,66 | 3,04 | 2,87 | 1,06 | 2,90E-01 |
| AT5G28050 | 0,28 | 0,99 | 4,29 | 0,23 | 8,17E-01 |
| AT5G04635 | 0,50 | 1,48 | 3,02 | 0,49 | 6,24E-01 |
| AT5G28060 | 0,65 | 0,33 | 2,15 | 0,15 | 8,78E-01 |
| AT5G04685 | 0,19 | -2,00 | 4,31 | -0,46 | 6,43E-01 |
| AT5G28190 | 0,36 | -0,28 | 4,26 | -0,07 | 9,48E-01 |
| AT5G28220 | 0,30 | -2,32 | 4,30 | -0,54 | 5,89E-01 |
| AT5G28235 | 0,34 | -2,98 | 4,28 | -0,70 | 4,87E-01 |
| AT5G28300 | 1,51 | -0,51 | 1,45 | -0,35 | 7,23E-01 |
| AT5G28310 | 0,08 | 2,08 | 4,38 | 0,47 | 6,35E-01 |
| AT5G28400 | 1,06 | 2,21 | 1,75 | 1,26 | 2,08E-01 |
| AT5G28442 | 0,36 | -0,52 | 3,25 | -0,16 | 8,74E-01 |
| AT5G28450 | 0,85 | -2,22 | 2,23 | -1,00 | 3,19E-01 |
| AT5G28530 | 0,22 | -0,30 | 4,29 | -0,07 | 9,45E-01 |
| AT5G28550 | 0,21 | -0,12 | 4,28 | -0,03 | 9,77E-01 |
| AT5G28640 | 0,08 | -0,56 | 4,38 | -0,13 | 8,99E-01 |

| | | | | | |
|---|---|---|---|---|---|
| AT5G28750 | 0,65 | -1,06 | 2,23 | -0,48 | 6,34E-01 |
| AT5G28770 | 0,50 | 2,70 | 3,85 | 0,70 | 4,82E-01 |
| AT5G28824 | 1,47 | 1,65 | 1,54 | 1,07 | 2,84E-01 |
| AT5G28850 | 0,48 | 0,72 | 3,17 | 0,23 | 8,20E-01 |
| AT5G28910 | 0,82 | -1,00 | 2,42 | -0,41 | 6,80E-01 |
| AT5G04925 | 0,18 | 0,41 | 4,34 | 0,10 | 9,24E-01 |
| AT5G32470 | 0,85 | 0,62 | 1,93 | 0,32 | 7,47E-01 |
| AT5G32590 | 0,68 | -0,90 | 2,05 | -0,44 | 6,62E-01 |
| AT5G33210 | 0,08 | -0,56 | 4,38 | -0,13 | 8,99E-01 |
| AT5G33439 | 0,11 | -1,28 | 4,35 | -0,29 | 7,69E-01 |
| AT5G34850 | 1,64 | 1,31 | 1,32 | 1,00 | 3,19E-01 |
| AT5G35100 | 0,30 | 0,78 | 3,56 | 0,22 | 8,26E-01 |
| AT5G35200 | 0,52 | 2,18 | 2,99 | 0,73 | 4,67E-01 |
| AT5G35220 | 0,25 | 2,97 | 4,33 | 0,69 | 4,93E-01 |
| AT5G35230 | 0,96 | -1,84 | 2,46 | -0,75 | 4,53E-01 |
| AT5G35370 | 0,70 | 0,48 | 1,91 | 0,25 | 8,01E-01 |
| AT5G35475 | 0,36 | -1,04 | 4,27 | -0,24 | 8,07E-01 |
| AT5G35480 | 0,65 | -1,30 | 2,94 | -0,44 | 6,58E-01 |
| AT5G35530 | 0,88 | 2,40 | 2,33 | 1,03 | 3,03E-01 |
| AT5G35560 | 0,64 | 0,70 | 2,28 | 0,31 | 7,60E-01 |
| AT5G35570 | 0,29 | 3,73 | 4,32 | 0,86 | 3,88E-01 |
| AT5G35590 | 0,12 | -0,77 | 4,36 | -0,18 | 8,61E-01 |
| AT5G35603 | 0,59 | -0,62 | 3,98 | -0,16 | 8,76E-01 |
| AT5G35604 | 0,45 | 0,73 | 3,83 | 0,19 | 8,48E-01 |
| AT5G35620 | 0,26 | 0,66 | 4,33 | 0,15 | 8,78E-01 |
| AT5G35690 | 0,37 | 2,18 | 4,30 | 0,51 | 6,13E-01 |
| AT5G35700 | 0,37 | 2,71 | 4,24 | 0,64 | 5,23E-01 |
| AT5G35750 | 1,20 | 3,65 | 1,93 | 1,89 | 5,90E-02 |
| AT5G35753 | 0,77 | 1,94 | 2,21 | 0,88 | 3,79E-01 |
| AT5G35790 | 0,34 | -0,19 | 4,40 | -0,04 | 9,66E-01 |
| AT5G35810 | 0,15 | 0,69 | 4,31 | 0,16 | 8,73E-01 |
| AT5G35890 | 0,19 | -0,39 | 4,29 | -0,09 | 9,27E-01 |
| AT5G35910 | 0,15 | 1,69 | 4,34 | 0,39 | 6,98E-01 |
| AT5G35913 | 0,08 | 2,33 | 4,36 | 0,53 | 5,93E-01 |
| AT5G35970 | 1,60 | 0,45 | 1,25 | 0,36 | 7,21E-01 |
| AT5G35995 | 0,29 | 2,61 | 4,31 | 0,61 | 5,45E-01 |
| AT5G36002 | 0,12 | 2,87 | 4,34 | 0,66 | 5,08E-01 |
| AT5G36120 | 0,05 | -0,58 | 4,38 | -0,13 | 8,95E-01 |
| AT5G36150 | 0,29 | -0,96 | 4,38 | -0,22 | 8,26E-01 |
| AT5G36160 | 0,29 | 1,67 | 4,30 | 0,39 | 6,98E-01 |
| AT5G36200 | 1,52 | -0,40 | 1,60 | -0,25 | 8,03E-01 |
| AT5G36210 | 1,11 | 3,14 | 1,94 | 1,62 | 1,05E-01 |
| AT5G36220 | 0,80 | 1,44 | 2,08 | 0,69 | 4,87E-01 |
| AT5G36700 | 0,58 | 1,48 | 2,59 | 0,57 | 5,67E-01 |
| AT5G36790 | 0,17 | 0,30 | 4,37 | 0,07 | 9,44E-01 |
| AT5G36880 | 0,93 | 2,68 | 2,20 | 1,22 | 2,24E-01 |
| AT5G36930 | 1,17 | 0,79 | 1,98 | 0,40 | 6,90E-01 |
| AT5G37260 | 0,78 | 4,09 | 3,32 | 1,23 | 2,17E-01 |
| AT5G37310 | 0,77 | 4,06 | 3,32 | 1,22 | 2,21E-01 |
| AT5G37370 | 0,59 | 3,68 | 3,67 | 1,00 | 3,16E-01 |
| AT5G37480 | 0,13 | -1,34 | 4,35 | -0,31 | 7,58E-01 |
| AT5G37590 | 0,66 | 2,98 | 3,64 | 0,82 | 4,12E-01 |
| AT5G37720 | 0,46 | 1,94 | 3,28 | 0,59 | 5,55E-01 |
| AT5G37770 | 1,11 | 0,87 | 2,45 | 0,36 | 7,21E-01 |
| AT5G37780 | 0,80 | 1,09 | 1,92 | 0,57 | 5,71E-01 |
| AT5G05545 | 0,31 | -2,62 | 4,29 | -0,61 | 5,41E-01 |

| | | | | | |
|---|---|---|---|---|---|
| AT5G37830 | 1,35 | 3,29 | 1,84 | 1,79 | 7,39E-02 |
| AT5G37890 | 0,36 | 3,47 | 4,33 | 0,80 | 4,23E-01 |
| AT5G05565 | 0,45 | 0,32 | 4,03 | 0,08 | 9,36E-01 |
| AT5G38020 | 0,42 | 0,00 | 4,41 | 0,00 | 1,00E+00 |
| AT5G38080 | 0,08 | 2,08 | 4,38 | 0,47 | 6,35E-01 |
| AT5G38140 | 0,10 | -1,26 | 4,35 | -0,29 | 7,72E-01 |
| AT5G38212 | 1,00 | -0,42 | 1,92 | -0,22 | 8,25E-01 |
| AT5G38240 | 0,41 | 1,71 | 4,38 | 0,39 | 6,97E-01 |
| AT5G38280 | 0,74 | -1,69 | 3,68 | -0,46 | 6,46E-01 |
| AT5G38450 | 0,46 | -0,66 | 4,11 | -0,16 | 8,72E-01 |
| AT5G05755 | 0,05 | -0,58 | 4,38 | -0,13 | 8,95E-01 |
| AT5G38510 | 0,76 | 0,85 | 2,47 | 0,34 | 7,31E-01 |
| AT5G38640 | 0,50 | 1,08 | 2,59 | 0,42 | 6,78E-01 |
| AT5G38660 | 1,59 | 1,37 | 1,57 | 0,88 | 3,81E-01 |
| AT5G38650 | 1,00 | 1,98 | 2,14 | 0,93 | 3,55E-01 |
| AT5G38830 | 0,49 | 2,82 | 4,34 | 0,65 | 5,17E-01 |
| AT5G39130 | 0,12 | 0,63 | 4,33 | 0,15 | 8,85E-01 |
| AT5G39160 | 0,34 | -2,77 | 4,28 | -0,65 | 5,18E-01 |
| AT5G05895 | 0,62 | -2,03 | 2,88 | -0,70 | 4,82E-01 |
| AT5G39480 | 0,08 | -0,56 | 4,38 | -0,13 | 8,99E-01 |
| AT5G39520 | 0,33 | 2,03 | 4,34 | 0,47 | 6,39E-01 |
| AT5G39530 | 0,77 | 3,35 | 2,89 | 1,16 | 2,47E-01 |
| AT5G39600 | 0,66 | 1,83 | 2,28 | 0,80 | 4,21E-01 |
| AT5G39610 | 0,52 | 1,63 | 3,87 | 0,42 | 6,74E-01 |
| AT5G05995 | 0,40 | 1,01 | 4,07 | 0,25 | 8,05E-01 |
| AT5G39680 | 0,42 | 3,55 | 4,33 | 0,82 | 4,11E-01 |
| AT5G39693 | 0,43 | -1,28 | 3,26 | -0,39 | 6,96E-01 |
| AT5G39920 | 0,25 | 0,73 | 4,38 | 0,17 | 8,67E-01 |
| AT5G39950 | 0,95 | 1,68 | 2,34 | 0,72 | 4,74E-01 |
| AT5G40170 | 0,68 | -0,91 | 2,20 | -0,41 | 6,78E-01 |
| AT5G40240 | 0,14 | 1,12 | 4,32 | 0,26 | 7,96E-01 |
| AT5G40370 | 0,37 | 1,02 | 3,34 | 0,30 | 7,61E-01 |
| AT5G40470 | 0,27 | 0,03 | 4,38 | 0,01 | 9,94E-01 |
| AT5G40580 | 0,60 | 2,52 | 4,37 | 0,58 | 5,65E-01 |
| AT5G40660 | 0,29 | 3,27 | 4,33 | 0,76 | 4,49E-01 |
| AT5G40740 | 0,85 | 2,57 | 2,22 | 1,16 | 2,47E-01 |
| AT5G40770 | 1,47 | 3,24 | 1,73 | 1,87 | 6,12E-02 |
| AT5G40820 | 0,95 | 2,17 | 2,21 | 0,98 | 3,25E-01 |
| AT5G40830 | 0,63 | 0,30 | 2,51 | 0,12 | 9,04E-01 |
| AT5G40940 | 0,56 | 4,04 | 3,76 | 1,07 | 2,83E-01 |
| AT5G41020 | 0,73 | 0,38 | 2,47 | 0,15 | 8,79E-01 |
| AT5G41130 | 0,21 | 3,16 | 4,33 | 0,73 | 4,66E-01 |
| AT5G41150 | 0,33 | 3,34 | 4,32 | 0,77 | 4,40E-01 |
| AT5G41190 | 0,83 | 0,18 | 2,88 | 0,06 | 9,51E-01 |
| AT5G41330 | 0,40 | 1,94 | 3,32 | 0,58 | 5,59E-01 |
| AT5G41400 | 0,65 | -1,12 | 2,51 | -0,44 | 6,56E-01 |
| AT5G41460 | 0,17 | 0,00 | 4,41 | 0,00 | 1,00E+00 |
| AT5G41520 | 0,58 | 0,36 | 2,61 | 0,14 | 8,91E-01 |
| AT5G41670 | 0,43 | -0,15 | 2,64 | -0,06 | 9,53E-01 |
| AT5G06235 | 0,77 | 3,27 | 4,26 | 0,77 | 4,43E-01 |
| AT5G41870 | 0,38 | 0,20 | 4,27 | 0,05 | 9,62E-01 |
| AT5G41990 | 1,56 | 4,31 | 1,71 | 2,51 | 1,20E-02 |
| AT5G42100 | 1,07 | -1,34 | 1,61 | -0,83 | 4,06E-01 |
| AT5G42110 | 0,47 | -0,85 | 3,15 | -0,27 | 7,88E-01 |
| AT5G42130 | 0,21 | 2,29 | 4,32 | 0,53 | 5,96E-01 |
| AT5G42140 | 0,12 | 0,31 | 4,35 | 0,07 | 9,44E-01 |

| | | | | | |
|---|---|---|---|---|---|
| AT5G42190 | 0,65 | 0,83 | 2,42 | 0,34 | 7,30E-01 |
| AT5G42280 | 0,14 | 1,12 | 4,32 | 0,26 | 7,96E-01 |
| AT5G42300 | 0,48 | 0,22 | 3,79 | 0,06 | 9,54E-01 |
| AT5G42360 | 1,47 | 3,03 | 1,63 | 1,86 | 6,27E-02 |
| AT5G42380 | 1,28 | -0,12 | 1,91 | -0,06 | 9,52E-01 |
| AT5G42400 | 0,21 | 1,57 | 4,30 | 0,37 | 7,14E-01 |
| AT5G06305 | 1,02 | -0,15 | 1,55 | -0,09 | 9,25E-01 |
| AT5G42490 | 0,56 | 2,36 | 3,85 | 0,61 | 5,39E-01 |
| AT5G42567 | 0,58 | -3,80 | 4,27 | -0,89 | 3,73E-01 |
| AT5G42570 | 0,15 | 2,31 | 4,37 | 0,53 | 5,98E-01 |
| AT5G42660 | 0,12 | 0,31 | 4,35 | 0,07 | 9,44E-01 |
| AT5G42750 | 1,69 | -1,53 | 1,30 | -1,18 | 2,40E-01 |
| AT5G42790 | 0,93 | 0,45 | 1,80 | 0,25 | 8,02E-01 |
| AT5G42800 | 0,38 | 3,45 | 4,33 | 0,80 | 4,26E-01 |
| AT5G42850 | 1,15 | 2,58 | 2,07 | 1,25 | 2,13E-01 |
| AT5G42870 | 0,81 | 1,63 | 2,13 | 0,77 | 4,44E-01 |
| AT5G42890 | 0,08 | -0,56 | 4,38 | -0,13 | 8,99E-01 |
| AT5G42900 | 0,25 | 2,25 | 4,37 | 0,52 | 6,06E-01 |
| AT5G42905 | 0,52 | 0,42 | 3,68 | 0,11 | 9,10E-01 |
| AT5G42940 | 0,81 | 2,75 | 2,52 | 1,09 | 2,75E-01 |
| AT5G42955 | 0,65 | 1,95 | 3,85 | 0,51 | 6,12E-01 |
| AT5G42957 | 0,83 | -3,30 | 2,33 | -1,42 | 1,56E-01 |
| AT5G42990 | 0,17 | 3,16 | 4,33 | 0,73 | 4,66E-01 |
| AT5G43010 | 0,13 | -0,63 | 4,39 | -0,14 | 8,85E-01 |
| AT5G43020 | 0,41 | 0,12 | 4,11 | 0,03 | 9,76E-01 |
| AT5G43130 | 1,30 | 3,88 | 2,12 | 1,83 | 6,67E-02 |
| AT5G43190 | 1,41 | -0,17 | 1,55 | -0,11 | 9,13E-01 |
| AT5G43260 | 0,52 | -0,85 | 2,60 | -0,33 | 7,44E-01 |
| AT5G43270 | 0,76 | -1,31 | 3,01 | -0,43 | 6,64E-01 |
| AT5G43285 | 0,35 | -0,03 | 4,39 | -0,01 | 9,95E-01 |
| AT5G43320 | 0,66 | -0,16 | 2,61 | -0,06 | 9,52E-01 |
| AT5G43440 | 0,40 | 0,16 | 4,28 | 0,04 | 9,71E-01 |
| AT5G43670 | 0,40 | -1,38 | 4,20 | -0,33 | 7,42E-01 |
| AT5G43710 | 0,21 | 3,22 | 4,33 | 0,74 | 4,58E-01 |
| AT5G43745 | 0,36 | 1,64 | 4,38 | 0,37 | 7,08E-01 |
| AT5G43750 | 0,47 | -1,39 | 4,34 | -0,32 | 7,49E-01 |
| AT5G43780 | 0,63 | -0,73 | 2,52 | -0,29 | 7,73E-01 |
| AT5G43790 | 0,45 | 1,34 | 4,22 | 0,32 | 7,51E-01 |
| AT5G43830 | 0,91 | 0,87 | 1,87 | 0,46 | 6,43E-01 |
| AT5G43850 | 0,29 | 1,20 | 3,64 | 0,33 | 7,42E-01 |
| AT5G43880 | 0,19 | -1,81 | 4,32 | -0,42 | 6,75E-01 |
| AT5G43900 | 1,55 | 3,12 | 1,53 | 2,04 | 4,15E-02 |
| AT5G43950 | 0,65 | 1,83 | 2,55 | 0,72 | 4,73E-01 |
| AT5G43960 | 0,54 | 2,98 | 3,06 | 0,97 | 3,30E-01 |
| AT5G44000 | 0,95 | 2,23 | 2,03 | 1,09 | 2,74E-01 |
| AT5G44005 | 0,16 | 1,27 | 4,37 | 0,29 | 7,71E-01 |
| AT5G44100 | 0,66 | 1,84 | 2,47 | 0,74 | 4,57E-01 |
| AT5G44250 | 0,12 | 2,81 | 4,34 | 0,65 | 5,18E-01 |
| AT5G44260 | 0,22 | -0,76 | 4,38 | -0,17 | 8,63E-01 |
| AT5G44283 | 0,91 | 0,12 | 2,40 | 0,05 | 9,61E-01 |
| AT5G44290 | 1,25 | 1,31 | 1,75 | 0,75 | 4,54E-01 |
| AT5G44306 | 0,93 | 0,17 | 2,73 | 0,06 | 9,50E-01 |
| AT5G44495 | 1,23 | -1,51 | 1,42 | -1,06 | 2,89E-01 |
| AT5G44510 | 0,61 | 4,50 | 4,29 | 1,05 | 2,94E-01 |
| AT5G44568 | 0,48 | -0,83 | 2,78 | -0,30 | 7,65E-01 |
| AT5G06595 | 1,38 | -1,42 | 1,76 | -0,81 | 4,18E-01 |

| | | | | | |
|---|---|---|---|---|---|
| AT5G44670 | 0,13 | 2,37 | 4,36 | 0,54 | 5,86E-01 |
| AT5G44680 | 0,48 | -0,13 | 4,41 | -0,03 | 9,77E-01 |
| AT5G44785 | 0,10 | 0,34 | 4,35 | 0,08 | 9,37E-01 |
| AT5G44790 | 1,35 | 2,25 | 1,54 | 1,46 | 1,45E-01 |
| AT5G44820 | 0,08 | 2,08 | 4,38 | 0,47 | 6,35E-01 |
| AT5G44840 | 0,14 | 0,00 | 4,41 | 0,00 | 1,00E+00 |
| AT5G45010 | 0,58 | -0,10 | 3,18 | -0,03 | 9,76E-01 |
| AT5G45040 | 0,08 | 2,33 | 4,36 | 0,53 | 5,93E-01 |
| AT5G45050 | 0,86 | 1,85 | 1,96 | 0,94 | 3,46E-01 |
| AT5G45170 | 0,11 | -1,28 | 4,35 | -0,29 | 7,69E-01 |
| AT5G45180 | 0,66 | 0,75 | 4,00 | 0,19 | 8,52E-01 |
| AT5G45230 | 0,97 | -1,53 | 1,69 | -0,90 | 3,66E-01 |
| AT5G45250 | 0,93 | 1,21 | 1,99 | 0,61 | 5,43E-01 |
| AT5G45275 | 0,48 | 2,35 | 4,33 | 0,54 | 5,87E-01 |
| AT5G45290 | 1,09 | 2,50 | 2,04 | 1,22 | 2,22E-01 |
| AT5G45340 | 0,20 | -0,87 | 4,29 | -0,20 | 8,40E-01 |
| AT5G45360 | 0,08 | 2,33 | 4,36 | 0,53 | 5,93E-01 |
| AT5G45410 | 0,46 | 2,78 | 4,02 | 0,69 | 4,89E-01 |
| AT5G45420 | 0,27 | 0,48 | 4,31 | 0,11 | 9,11E-01 |
| AT5G45440 | 0,74 | 4,47 | 3,38 | 1,32 | 1,85E-01 |
| AT5G45460 | 0,18 | -0,70 | 4,38 | -0,16 | 8,72E-01 |
| AT5G45573 | 0,61 | -0,77 | 2,71 | -0,28 | 7,77E-01 |
| AT5G45630 | 0,05 | -0,58 | 4,38 | -0,13 | 8,95E-01 |
| AT5G45650 | 0,61 | 0,26 | 3,08 | 0,08 | 9,34E-01 |
| AT5G45775 | 0,86 | 1,25 | 1,91 | 0,65 | 5,13E-01 |
| AT5G45800 | 0,52 | 0,62 | 2,19 | 0,28 | 7,79E-01 |
| AT5G45820 | 1,25 | -0,76 | 1,52 | -0,50 | 6,16E-01 |
| AT5G46020 | 1,24 | 0,07 | 2,44 | 0,03 | 9,77E-01 |
| AT5G46040 | 0,11 | 1,28 | 4,35 | 0,29 | 7,68E-01 |
| AT5G46050 | 0,37 | -0,14 | 4,41 | -0,03 | 9,75E-01 |
| AT5G46280 | 0,57 | -0,99 | 3,03 | -0,33 | 7,44E-01 |
| AT5G46290 | 0,63 | 2,01 | 3,12 | 0,64 | 5,20E-01 |
| AT5G46340 | 0,61 | 0,84 | 4,01 | 0,21 | 8,34E-01 |
| AT5G46395 | 0,11 | -1,28 | 4,35 | -0,29 | 7,69E-01 |
| AT5G46580 | 0,75 | 0,04 | 2,31 | 0,02 | 9,85E-01 |
| AT5G46620 | 0,64 | 0,23 | 3,06 | 0,07 | 9,40E-01 |
| AT5G46750 | 0,66 | 1,19 | 3,75 | 0,32 | 7,51E-01 |
| AT5G46790 | 0,53 | 0,94 | 2,90 | 0,32 | 7,46E-01 |
| AT5G46800 | 0,83 | -0,55 | 2,30 | -0,24 | 8,10E-01 |
| AT5G46860 | 0,46 | -1,89 | 4,05 | -0,47 | 6,40E-01 |
| AT5G46970 | 0,65 | -0,42 | 2,41 | -0,17 | 8,61E-01 |
| AT5G47040 | 1,34 | 0,81 | 1,51 | 0,54 | 5,90E-01 |
| AT5G47070 | 0,97 | -0,36 | 2,10 | -0,17 | 8,64E-01 |
| AT5G47080 | 0,13 | 2,37 | 4,36 | 0,54 | 5,86E-01 |
| AT5G47110 | 1,25 | -0,99 | 1,48 | -0,67 | 5,04E-01 |
| AT5G47190 | 0,93 | 0,87 | 2,10 | 0,41 | 6,79E-01 |
| AT5G47210 | 1,48 | -0,50 | 1,88 | -0,27 | 7,90E-01 |
| AT5G47220 | 0,78 | 0,71 | 2,43 | 0,29 | 7,69E-01 |
| AT5G47250 | 0,28 | -0,12 | 4,35 | -0,03 | 9,77E-01 |
| AT5G47370 | 0,28 | 1,64 | 4,38 | 0,38 | 7,08E-01 |
| AT5G47460 | 0,15 | 1,94 | 4,32 | 0,45 | 6,54E-01 |
| AT5G47570 | 0,30 | 0,65 | 4,28 | 0,15 | 8,79E-01 |
| AT5G47580 | 0,64 | 3,06 | 3,63 | 0,84 | 4,00E-01 |
| AT5G47650 | 0,20 | 2,30 | 4,36 | 0,53 | 5,98E-01 |
| AT5G47700 | 0,72 | 1,48 | 2,07 | 0,71 | 4,75E-01 |
| AT5G47730 | 0,16 | 1,18 | 4,32 | 0,27 | 7,85E-01 |

| | | | | | |
|---|---|---|---|---|---|
| AT5G47750 | 1,50 | 3,76 | 2,17 | 1,73 | 8,29E-02 |
| AT5G47760 | 0,12 | 0,31 | 4,35 | 0,07 | 9,44E-01 |
| AT5G47810 | 0,55 | 1,77 | 2,63 | 0,67 | 5,02E-01 |
| AT5G47830 | 0,54 | 3,04 | 4,35 | 0,70 | 4,84E-01 |
| AT5G47850 | 0,49 | 0,10 | 3,98 | 0,03 | 9,79E-01 |
| AT5G47880 | 0,55 | 1,79 | 3,09 | 0,58 | 5,62E-01 |
| AT5G47930 | 0,53 | -1,64 | 2,50 | -0,65 | 5,13E-01 |
| AT5G48000 | 0,12 | -0,77 | 4,36 | -0,18 | 8,61E-01 |
| AT5G07055 | 0,58 | -0,84 | 3,02 | -0,28 | 7,81E-01 |
| AT5G48120 | 1,18 | 1,90 | 1,69 | 1,13 | 2,60E-01 |
| AT5G48150 | 0,85 | 1,97 | 3,59 | 0,55 | 5,83E-01 |
| AT5G48180 | 0,98 | 1,36 | 1,78 | 0,76 | 4,46E-01 |
| AT5G48375 | 0,26 | 2,30 | 4,37 | 0,53 | 5,98E-01 |
| AT5G48412 | 0,64 | -1,05 | 2,59 | -0,41 | 6,85E-01 |
| AT5G48440 | 0,43 | 1,89 | 4,14 | 0,46 | 6,48E-01 |
| AT5G48540 | 0,08 | 2,08 | 4,38 | 0,47 | 6,35E-01 |
| AT5G48580 | 0,41 | 2,49 | 4,11 | 0,60 | 5,45E-01 |
| AT5G48640 | 0,19 | -1,34 | 4,35 | -0,31 | 7,58E-01 |
| AT5G48657 | 0,40 | -0,94 | 3,24 | -0,29 | 7,73E-01 |
| AT5G07265 | 0,12 | 2,81 | 4,34 | 0,65 | 5,18E-01 |
| AT5G48770 | 0,08 | -0,56 | 4,38 | -0,13 | 8,99E-01 |
| AT5G07275 | 0,71 | 3,95 | 4,31 | 0,92 | 3,59E-01 |
| AT5G48810 | 0,80 | 1,17 | 2,19 | 0,53 | 5,95E-01 |
| AT5G48830 | 0,28 | 1,26 | 4,36 | 0,29 | 7,73E-01 |
| AT5G48880 | 0,58 | 0,51 | 4,33 | 0,12 | 9,06E-01 |
| AT5G48910 | 0,76 | 0,62 | 2,01 | 0,31 | 7,57E-01 |
| AT5G48930 | 1,24 | 1,56 | 1,67 | 0,94 | 3,49E-01 |
| AT5G49100 | 1,11 | -2,99 | 1,84 | -1,62 | 1,05E-01 |
| AT5G49300 | 0,36 | 3,22 | 4,34 | 0,74 | 4,59E-01 |
| AT5G49350 | 0,08 | 2,33 | 4,36 | 0,53 | 5,93E-01 |
| AT5G49420 | 0,35 | -0,87 | 4,30 | -0,20 | 8,39E-01 |
| AT5G49430 | 1,30 | 0,68 | 1,55 | 0,44 | 6,62E-01 |
| AT5G49540 | 0,90 | 2,49 | 2,56 | 0,97 | 3,30E-01 |
| AT5G49550 | 0,27 | 0,31 | 4,36 | 0,07 | 9,44E-01 |
| AT5G49555 | 0,40 | -0,98 | 4,36 | -0,22 | 8,22E-01 |
| AT5G49570 | 0,11 | 1,33 | 4,36 | 0,31 | 7,59E-01 |
| AT5G49580 | 0,46 | 0,96 | 3,98 | 0,24 | 8,09E-01 |
| AT5G49630 | 0,19 | -2,00 | 4,31 | -0,46 | 6,43E-01 |
| AT5G49645 | 0,08 | 2,08 | 4,38 | 0,47 | 6,35E-01 |
| AT5G49660 | 0,65 | 0,92 | 2,39 | 0,39 | 7,00E-01 |
| AT5G49665 | 0,50 | 1,39 | 4,01 | 0,35 | 7,29E-01 |
| AT5G49830 | 1,27 | 2,71 | 1,61 | 1,68 | 9,24E-02 |
| AT5G49890 | 0,22 | -0,52 | 4,33 | -0,12 | 9,05E-01 |
| AT5G49900 | 0,41 | 2,55 | 4,12 | 0,62 | 5,36E-01 |
| AT5G49930 | 0,74 | 1,99 | 2,08 | 0,96 | 3,38E-01 |
| AT5G49950 | 0,25 | -0,71 | 4,39 | -0,16 | 8,72E-01 |
| AT5G50000 | 0,84 | -0,77 | 1,92 | -0,40 | 6,87E-01 |
| AT5G50150 | 0,10 | 0,39 | 4,35 | 0,09 | 9,29E-01 |
| AT5G50220 | 0,28 | -0,24 | 4,40 | -0,05 | 9,57E-01 |
| AT5G50320 | 0,77 | 1,26 | 2,94 | 0,43 | 6,68E-01 |
| AT5G50340 | 0,25 | 2,98 | 4,33 | 0,69 | 4,92E-01 |
| AT5G50350 | 0,29 | -1,08 | 4,34 | -0,25 | 8,03E-01 |
| AT5G50380 | 0,97 | 0,64 | 1,85 | 0,35 | 7,29E-01 |
| AT5G50390 | 0,17 | 3,16 | 4,33 | 0,73 | 4,66E-01 |
| AT5G50400 | 0,54 | -0,76 | 3,95 | -0,19 | 8,47E-01 |
| AT5G50460 | 0,36 | -0,08 | 4,40 | -0,02 | 9,86E-01 |

| | | | | | |
|---|---|---|---|---|---|
| AT5G50480 | 0,53 | -0,14 | 3,90 | -0,03 | 9,72E-01 |
| AT5G50570 | 0,31 | -1,44 | 4,34 | -0,33 | 7,41E-01 |
| AT5G50630 | 0,05 | -0,58 | 4,38 | -0,13 | 8,95E-01 |
| AT5G50640 | 0,24 | 2,85 | 4,34 | 0,66 | 5,11E-01 |
| AT5G50670 | 0,93 | -0,74 | 2,42 | -0,31 | 7,59E-01 |
| AT5G50860 | 0,49 | 3,55 | 3,87 | 0,92 | 3,60E-01 |
| AT5G50870 | 0,08 | 2,08 | 4,38 | 0,47 | 6,35E-01 |
| AT5G50900 | 0,83 | 1,01 | 2,39 | 0,42 | 6,73E-01 |
| AT5G51000 | 0,61 | -0,71 | 2,82 | -0,25 | 8,00E-01 |
| AT5G51020 | 0,37 | 2,71 | 4,24 | 0,64 | 5,23E-01 |
| AT5G51110 | 0,78 | 0,88 | 2,22 | 0,40 | 6,92E-01 |
| AT5G51170 | 0,12 | 2,87 | 4,34 | 0,66 | 5,08E-01 |
| AT5G51195 | 0,57 | 0,14 | 2,38 | 0,06 | 9,53E-01 |
| AT5G51220 | 0,23 | -0,84 | 4,31 | -0,19 | 8,46E-01 |
| AT5G51290 | 0,61 | 0,80 | 3,08 | 0,26 | 7,95E-01 |
| AT5G51390 | 0,29 | 1,64 | 4,38 | 0,37 | 7,08E-01 |
| AT5G51400 | 1,21 | 0,30 | 1,56 | 0,19 | 8,47E-01 |
| AT5G51550 | 1,02 | -0,38 | 1,67 | -0,23 | 8,18E-01 |
| AT5G51640 | 0,39 | -0,71 | 3,45 | -0,21 | 8,36E-01 |
| AT5G51650 | 0,32 | -0,18 | 3,40 | -0,05 | 9,57E-01 |
| AT5G51690 | 0,72 | 1,78 | 2,43 | 0,73 | 4,62E-01 |
| AT5G51710 | 0,10 | -1,26 | 4,35 | -0,29 | 7,72E-01 |
| AT5G51820 | 0,69 | 1,16 | 2,57 | 0,45 | 6,53E-01 |
| AT5G52060 | 0,54 | 0,86 | 2,49 | 0,34 | 7,30E-01 |
| AT5G52090 | 0,73 | -0,80 | 1,96 | -0,41 | 6,82E-01 |
| AT5G52120 | 0,47 | -0,14 | 4,41 | -0,03 | 9,75E-01 |
| AT5G52180 | 0,61 | 1,90 | 3,06 | 0,62 | 5,34E-01 |
| AT5G52330 | 0,15 | -0,64 | 4,39 | -0,15 | 8,84E-01 |
| AT5G52370 | 0,37 | 1,99 | 4,30 | 0,46 | 6,43E-01 |
| AT5G52450 | 0,44 | 2,25 | 3,20 | 0,70 | 4,81E-01 |
| AT5G52540 | 1,60 | 1,35 | 1,46 | 0,92 | 3,57E-01 |
| AT5G52550 | 0,25 | 1,77 | 4,34 | 0,41 | 6,83E-01 |
| AT5G52650 | 0,64 | 0,14 | 3,70 | 0,04 | 9,70E-01 |
| AT5G52780 | 0,46 | -0,57 | 3,22 | -0,18 | 8,60E-01 |
| AT5G52797 | 0,48 | -0,74 | 3,08 | -0,24 | 8,11E-01 |
| AT5G53000 | 0,29 | 1,74 | 4,31 | 0,40 | 6,87E-01 |
| AT5G53010 | 0,50 | 1,26 | 3,31 | 0,38 | 7,05E-01 |
| AT5G53050 | 0,49 | 0,18 | 3,10 | 0,06 | 9,54E-01 |
| AT5G53130 | 0,72 | 2,57 | 2,74 | 0,94 | 3,49E-01 |
| AT5G07955 | 0,08 | 2,08 | 4,38 | 0,47 | 6,35E-01 |
| AT5G53300 | 0,89 | 2,93 | 2,60 | 1,12 | 2,61E-01 |
| AT5G53320 | 0,58 | 3,18 | 2,99 | 1,06 | 2,89E-01 |
| AT5G07975 | 0,08 | 2,08 | 4,38 | 0,47 | 6,35E-01 |
| AT5G53490 | 0,65 | -1,65 | 2,18 | -0,76 | 4,50E-01 |
| AT5G53500 | 0,52 | -2,41 | 3,14 | -0,77 | 4,44E-01 |
| AT5G53530 | 0,37 | 1,22 | 4,35 | 0,28 | 7,80E-01 |
| AT5G53570 | 0,72 | 1,94 | 2,35 | 0,83 | 4,08E-01 |
| AT5G53580 | 0,36 | 0,75 | 3,49 | 0,21 | 8,31E-01 |
| AT5G53730 | 0,14 | -0,31 | 4,33 | -0,07 | 9,43E-01 |
| AT5G53850 | 0,47 | 1,72 | 4,40 | 0,39 | 6,96E-01 |
| AT5G53870 | 0,19 | -1,45 | 4,34 | -0,33 | 7,38E-01 |
| AT5G53900 | 0,79 | -0,43 | 2,15 | -0,20 | 8,43E-01 |
| AT5G54030 | 0,27 | -2,66 | 4,29 | -0,62 | 5,35E-01 |
| AT5G54040 | 0,27 | -1,55 | 4,28 | -0,36 | 7,16E-01 |
| AT5G54050 | 0,38 | 0,00 | 4,25 | 0,00 | 1,00E+00 |
| AT5G54060 | 0,12 | 2,81 | 4,34 | 0,65 | 5,18E-01 |

| | | | | | |
|---|---|---|---|---|---|
| AT5G54090 | 1,00 | 4,41 | 2,25 | 1,96 | 4,95E-02 |
| AT5G54100 | 0,65 | 3,54 | 2,88 | 1,23 | 2,18E-01 |
| AT5G54170 | 0,64 | 0,14 | 2,09 | 0,06 | 9,48E-01 |
| AT5G54280 | 1,29 | -0,19 | 1,45 | -0,13 | 8,97E-01 |
| AT5G54380 | 1,11 | 1,10 | 1,76 | 0,62 | 5,33E-01 |
| AT5G54410 | 0,82 | -0,96 | 1,87 | -0,51 | 6,07E-01 |
| AT5G54430 | 1,12 | 1,47 | 2,39 | 0,62 | 5,38E-01 |
| AT5G54460 | 1,09 | -1,30 | 3,12 | -0,42 | 6,76E-01 |
| AT5G54470 | 0,35 | -1,16 | 3,39 | -0,34 | 7,33E-01 |
| AT5G54510 | 0,08 | 2,33 | 4,36 | 0,53 | 5,93E-01 |
| AT5G54610 | 0,49 | 2,11 | 3,81 | 0,55 | 5,79E-01 |
| AT5G54620 | 0,12 | 2,69 | 4,35 | 0,62 | 5,36E-01 |
| AT5G54630 | 0,29 | 0,79 | 4,34 | 0,18 | 8,56E-01 |
| AT5G54650 | 0,47 | 2,93 | 4,35 | 0,67 | 5,00E-01 |
| AT5G54690 | 0,39 | 1,78 | 4,41 | 0,40 | 6,86E-01 |
| AT5G54700 | 0,89 | -3,12 | 2,24 | -1,39 | 1,65E-01 |
| AT5G54710 | 0,79 | -2,40 | 2,41 | -0,99 | 3,20E-01 |
| AT5G54720 | 0,73 | -2,45 | 2,30 | -1,06 | 2,87E-01 |
| AT5G54780 | 1,16 | 2,05 | 1,77 | 1,15 | 2,48E-01 |
| AT5G54930 | 0,40 | 3,38 | 4,15 | 0,82 | 4,15E-01 |
| AT5G54960 | 0,98 | 0,44 | 2,09 | 0,21 | 8,32E-01 |
| AT5G55040 | 0,74 | 0,69 | 1,97 | 0,35 | 7,24E-01 |
| AT5G55055 | 0,05 | -0,58 | 4,38 | -0,13 | 8,95E-01 |
| AT5G55100 | 1,12 | 3,80 | 2,45 | 1,55 | 1,22E-01 |
| AT5G55130 | 0,31 | 0,53 | 3,49 | 0,15 | 8,79E-01 |
| AT5G55150 | 1,21 | 3,58 | 2,48 | 1,44 | 1,50E-01 |
| AT5G55160 | 0,37 | 2,34 | 4,33 | 0,54 | 5,89E-01 |
| AT5G55210 | 0,08 | -0,56 | 4,38 | -0,13 | 8,99E-01 |
| AT5G55250 | 0,40 | -0,01 | 4,26 | 0,00 | 9,99E-01 |
| AT5G55280 | 1,13 | -0,94 | 2,27 | -0,41 | 6,79E-01 |
| AT5G55290 | 0,30 | -1,84 | 4,32 | -0,43 | 6,70E-01 |
| AT5G55300 | 0,70 | 1,94 | 2,30 | 0,84 | 3,99E-01 |
| AT5G55340 | 0,19 | 0,09 | 4,37 | 0,02 | 9,83E-01 |
| AT5G55390 | 0,36 | -0,67 | 4,23 | -0,16 | 8,74E-01 |
| AT5G08275 | 0,59 | 1,94 | 3,81 | 0,51 | 6,11E-01 |
| AT5G08285 | 0,30 | 0,49 | 4,33 | 0,11 | 9,10E-01 |
| AT5G55510 | 0,05 | -0,58 | 4,38 | -0,13 | 8,95E-01 |
| AT5G55540 | 1,63 | 3,23 | 1,59 | 2,03 | 4,20E-02 |
| AT5G55550 | 0,34 | 3,18 | 4,34 | 0,73 | 4,63E-01 |
| AT5G55570 | 1,30 | 0,00 | 3,15 | 0,00 | 1,00E+00 |
| AT5G55600 | 0,31 | 1,46 | 3,57 | 0,41 | 6,84E-01 |
| AT5G55670 | 0,30 | -0,32 | 4,26 | -0,07 | 9,41E-01 |
| AT5G55710 | 0,92 | 3,39 | 2,39 | 1,42 | 1,56E-01 |
| AT5G55740 | 0,27 | 1,09 | 4,29 | 0,26 | 7,99E-01 |
| AT5G55790 | 0,33 | 1,14 | 4,29 | 0,26 | 7,91E-01 |
| AT5G55835 | 0,61 | 0,81 | 2,18 | 0,37 | 7,09E-01 |
| AT5G55910 | 1,51 | 1,67 | 1,51 | 1,11 | 2,68E-01 |
| AT5G55920 | 0,81 | 1,71 | 2,36 | 0,73 | 4,68E-01 |
| AT5G55940 | 0,50 | 3,39 | 3,88 | 0,87 | 3,82E-01 |
| AT5G56000 | 1,26 | -3,30 | 1,97 | -1,67 | 9,42E-02 |
| AT5G56180 | 0,23 | -0,64 | 4,39 | -0,15 | 8,84E-01 |
| AT5G56240 | 1,23 | 2,47 | 2,12 | 1,17 | 2,43E-01 |
| AT5G56250 | 0,87 | 2,64 | 2,66 | 0,99 | 3,20E-01 |
| AT5G56260 | 0,25 | -0,71 | 4,39 | -0,16 | 8,72E-01 |
| AT5G56280 | 0,08 | -0,56 | 4,38 | -0,13 | 8,99E-01 |
| AT5G56350 | 0,73 | 1,87 | 2,04 | 0,92 | 3,58E-01 |

| | | | | | |
|---|---|---|---|---|---|
| AT5G56360 | 0,87 | 2,81 | 2,83 | 0,99 | 3,21E-01 |
| AT5G56410 | 0,30 | 0,27 | 4,35 | 0,06 | 9,50E-01 |
| AT5G56450 | 0,20 | 1,57 | 4,34 | 0,36 | 7,17E-01 |
| AT5G56710 | 0,79 | 3,27 | 3,39 | 0,96 | 3,35E-01 |
| AT5G56730 | 0,65 | 2,82 | 2,51 | 1,12 | 2,61E-01 |
| AT5G56760 | 0,72 | 0,81 | 2,81 | 0,29 | 7,74E-01 |
| AT5G56860 | 0,23 | -1,06 | 4,29 | -0,25 | 8,04E-01 |
| AT5G56890 | 0,16 | 1,97 | 4,33 | 0,45 | 6,50E-01 |
| AT5G56900 | 1,69 | 1,36 | 1,53 | 0,89 | 3,74E-01 |
| AT5G56980 | 0,28 | 1,62 | 4,35 | 0,37 | 7,10E-01 |
| AT5G57040 | 0,80 | 1,05 | 1,97 | 0,53 | 5,94E-01 |
| AT5G57160 | 0,81 | 1,82 | 2,48 | 0,73 | 4,64E-01 |
| AT5G57170 | 0,08 | -0,56 | 4,38 | -0,13 | 8,99E-01 |
| AT5G57250 | 0,54 | 3,10 | 3,85 | 0,81 | 4,20E-01 |
| AT5G57270 | 0,45 | -0,02 | 4,39 | -0,01 | 9,96E-01 |
| AT5G57290 | 0,69 | -0,64 | 2,53 | -0,25 | 8,00E-01 |
| AT5G57345 | 0,70 | 0,80 | 2,16 | 0,37 | 7,11E-01 |
| AT5G57580 | 0,90 | 3,30 | 2,38 | 1,38 | 1,67E-01 |
| AT5G57630 | 0,49 | 0,43 | 3,31 | 0,13 | 8,96E-01 |
| AT5G57670 | 0,16 | -0,65 | 4,31 | -0,15 | 8,80E-01 |
| AT5G57780 | 0,36 | -0,17 | 4,28 | -0,04 | 9,69E-01 |
| AT5G57785 | 0,56 | 0,34 | 2,45 | 0,14 | 8,89E-01 |
| AT5G57790 | 0,05 | -0,58 | 4,38 | -0,13 | 8,95E-01 |
| AT5G57850 | 0,51 | -0,04 | 4,39 | -0,01 | 9,94E-01 |
| AT5G57900 | 0,27 | 1,98 | 4,35 | 0,46 | 6,48E-01 |
| AT5G57960 | 0,12 | 0,08 | 4,33 | 0,02 | 9,84E-01 |
| AT5G58030 | 0,11 | 1,33 | 4,36 | 0,31 | 7,59E-01 |
| AT5G58050 | 0,17 | 3,04 | 4,33 | 0,70 | 4,83E-01 |
| AT5G58090 | 0,76 | 1,88 | 2,28 | 0,82 | 4,11E-01 |
| AT5G58130 | 0,84 | 3,90 | 2,62 | 1,49 | 1,36E-01 |
| AT5G58170 | 0,90 | 5,11 | 3,16 | 1,62 | 1,06E-01 |
| AT5G58220 | 0,12 | 0,06 | 4,34 | 0,01 | 9,88E-01 |
| AT5G58260 | 0,45 | 0,10 | 3,39 | 0,03 | 9,76E-01 |
| AT5G58380 | 0,93 | 1,58 | 2,01 | 0,79 | 4,30E-01 |
| AT5G58440 | 0,43 | 0,69 | 2,79 | 0,25 | 8,06E-01 |
| AT5G58570 | 0,61 | 1,77 | 3,79 | 0,47 | 6,41E-01 |
| AT5G08860 | 0,08 | -0,56 | 4,38 | -0,13 | 8,99E-01 |
| AT5G58600 | 0,29 | 2,14 | 4,36 | 0,49 | 6,23E-01 |
| AT5G58650 | 0,16 | 0,03 | 4,34 | 0,01 | 9,94E-01 |
| AT5G58670 | 0,69 | 1,84 | 2,21 | 0,83 | 4,05E-01 |
| AT5G58720 | 0,64 | 1,76 | 2,46 | 0,72 | 4,74E-01 |
| AT5G58730 | 0,50 | 1,94 | 3,97 | 0,49 | 6,25E-01 |
| AT5G58870 | 0,47 | 0,74 | 3,26 | 0,23 | 8,20E-01 |
| AT5G58900 | 0,28 | 1,64 | 4,38 | 0,38 | 7,08E-01 |
| AT5G58930 | 1,32 | 3,76 | 1,87 | 2,01 | 4,41E-02 |
| AT5G59030 | 0,49 | 3,48 | 3,87 | 0,90 | 3,68E-01 |
| AT5G59130 | 0,21 | 3,36 | 4,33 | 0,78 | 4,37E-01 |
| AT5G59220 | 0,62 | 2,43 | 3,77 | 0,65 | 5,18E-01 |
| AT5G59290 | 0,24 | 1,11 | 4,33 | 0,26 | 7,97E-01 |
| AT5G08920 | 0,49 | 2,15 | 3,96 | 0,54 | 5,88E-01 |
| AT5G59480 | 0,17 | 3,14 | 4,33 | 0,73 | 4,68E-01 |
| AT5G59550 | 1,55 | 1,48 | 1,46 | 1,01 | 3,12E-01 |
| AT5G59662 | 0,05 | -0,58 | 4,38 | -0,13 | 8,95E-01 |
| AT5G59700 | 0,33 | -0,14 | 4,41 | -0,03 | 9,75E-01 |
| AT5G59770 | 0,46 | 0,33 | 3,27 | 0,10 | 9,19E-01 |
| AT5G59780 | 1,19 | 3,26 | 1,87 | 1,74 | 8,15E-02 |

| | | | | | |
|---|---|---|---|---|---|
| AT5G59840 | 0,17 | 0,31 | 4,36 | 0,07 | 9,43E-01 |
| AT5G59850 | 0,69 | -1,03 | 2,62 | -0,39 | 6,94E-01 |
| AT5G59870 | 0,19 | 0,12 | 4,32 | 0,03 | 9,77E-01 |
| AT5G59900 | 0,16 | -0,80 | 4,37 | -0,18 | 8,54E-01 |
| AT5G59960 | 0,11 | 1,33 | 4,36 | 0,31 | 7,59E-01 |
| AT5G60050 | 0,15 | 2,12 | 4,38 | 0,48 | 6,28E-01 |
| AT5G60100 | 0,30 | -0,01 | 4,35 | 0,00 | 9,99E-01 |
| AT5G60160 | 0,49 | 0,48 | 3,96 | 0,12 | 9,04E-01 |
| AT5G60180 | 0,21 | 2,01 | 4,37 | 0,46 | 6,45E-01 |
| AT5G60210 | 0,29 | 1,50 | 4,34 | 0,35 | 7,29E-01 |
| AT5G60340 | 0,36 | 0,31 | 4,35 | 0,07 | 9,42E-01 |
| AT5G60400 | 0,12 | 2,87 | 4,34 | 0,66 | 5,08E-01 |
| AT5G60460 | 0,27 | -0,51 | 4,31 | -0,12 | 9,07E-01 |
| AT5G60670 | 0,64 | 1,12 | 2,24 | 0,50 | 6,18E-01 |
| AT5G60700 | 1,32 | 4,16 | 2,29 | 1,82 | 6,93E-02 |
| AT5G60710 | 1,48 | 2,17 | 1,50 | 1,45 | 1,48E-01 |
| AT5G60800 | 0,66 | 1,61 | 3,02 | 0,53 | 5,95E-01 |
| AT5G60840 | 0,12 | -1,57 | 4,34 | -0,36 | 7,18E-01 |
| AT5G60930 | 0,24 | 1,18 | 4,36 | 0,27 | 7,87E-01 |
| AT5G60960 | 0,29 | 1,86 | 4,33 | 0,43 | 6,67E-01 |
| AT5G60990 | 0,16 | 1,18 | 4,32 | 0,27 | 7,85E-01 |
| AT5G61040 | 0,88 | -0,06 | 4,34 | -0,01 | 9,88E-01 |
| AT5G61060 | 0,37 | 1,06 | 3,40 | 0,31 | 7,55E-01 |
| AT5G61130 | 0,17 | 3,14 | 4,33 | 0,73 | 4,68E-01 |
| AT5G61150 | 0,86 | 0,67 | 2,46 | 0,27 | 7,85E-01 |
| AT5G61250 | 0,23 | 1,51 | 4,34 | 0,35 | 7,28E-01 |
| AT5G61390 | 0,43 | 2,50 | 4,37 | 0,57 | 5,67E-01 |
| AT5G09005 | 1,51 | 0,50 | 1,26 | 0,40 | 6,90E-01 |
| AT5G61460 | 0,21 | 3,36 | 4,33 | 0,78 | 4,37E-01 |
| AT5G61480 | 0,83 | -0,24 | 2,09 | -0,12 | 9,08E-01 |
| AT5G61500 | 0,76 | 0,48 | 2,83 | 0,17 | 8,65E-01 |
| AT5G61510 | 0,56 | 1,38 | 3,26 | 0,42 | 6,73E-01 |
| AT5G61520 | 0,33 | 1,31 | 4,30 | 0,30 | 7,61E-01 |
| AT5G61590 | 1,28 | -2,57 | 1,87 | -1,38 | 1,69E-01 |
| AT5G61600 | 0,57 | -1,04 | 2,40 | -0,43 | 6,65E-01 |
| AT5G61800 | 0,12 | 2,69 | 4,35 | 0,62 | 5,36E-01 |
| AT5G61810 | 0,48 | 0,33 | 3,25 | 0,10 | 9,20E-01 |
| AT5G61820 | 1,07 | 2,44 | 2,22 | 1,10 | 2,71E-01 |
| AT5G61840 | 0,89 | 1,28 | 2,03 | 0,63 | 5,28E-01 |
| AT5G61910 | 1,60 | 1,36 | 1,32 | 1,03 | 3,04E-01 |
| AT5G61900 | 1,09 | 1,08 | 1,59 | 0,68 | 4,98E-01 |
| AT5G61960 | 0,94 | 0,60 | 2,51 | 0,24 | 8,12E-01 |
| AT5G61990 | 0,30 | 2,39 | 4,37 | 0,55 | 5,84E-01 |
| AT5G62090 | 0,32 | 1,54 | 4,32 | 0,36 | 7,23E-01 |
| AT5G62110 | 0,34 | 0,54 | 4,24 | 0,13 | 8,99E-01 |
| AT5G62130 | 0,45 | -2,01 | 3,11 | -0,65 | 5,18E-01 |
| AT5G62140 | 1,15 | 0,23 | 1,46 | 0,16 | 8,72E-01 |
| AT5G62180 | 0,08 | 2,33 | 4,36 | 0,53 | 5,93E-01 |
| AT5G62220 | 1,34 | -2,50 | 1,60 | -1,56 | 1,18E-01 |
| AT5G62300 | 1,07 | 1,42 | 1,97 | 0,72 | 4,73E-01 |
| AT5G62430 | 0,29 | 0,17 | 4,33 | 0,04 | 9,68E-01 |
| AT5G62470 | 0,30 | -1,84 | 4,32 | -0,43 | 6,70E-01 |
| AT5G62530 | 0,56 | 3,00 | 3,02 | 0,99 | 3,21E-01 |
| AT5G62600 | 0,50 | 1,94 | 3,97 | 0,49 | 6,25E-01 |
| AT5G62610 | 0,31 | 0,05 | 4,28 | 0,01 | 9,92E-01 |
| AT5G62630 | 0,37 | 2,24 | 4,35 | 0,52 | 6,06E-01 |

| | | | | | |
|---|---|---|---|---|---|
| AT5G62650 | 0,39 | 0,00 | 4,41 | 0,00 | 1,00E+00 |
| AT5G62680 | 0,33 | 1,61 | 4,30 | 0,37 | 7,08E-01 |
| AT5G62790 | 0,60 | -1,81 | 2,46 | -0,74 | 4,61E-01 |
| AT5G63080 | 0,23 | 1,34 | 4,38 | 0,30 | 7,61E-01 |
| AT5G63100 | 0,29 | 2,35 | 4,31 | 0,54 | 5,86E-01 |
| AT5G63110 | 0,58 | 2,35 | 3,05 | 0,77 | 4,41E-01 |
| AT5G63120 | 0,65 | 1,61 | 3,03 | 0,53 | 5,96E-01 |
| AT5G63160 | 0,51 | -1,44 | 2,98 | -0,48 | 6,29E-01 |
| AT5G63180 | 0,25 | 0,64 | 4,35 | 0,15 | 8,82E-01 |
| AT5G63290 | 0,56 | 3,19 | 4,33 | 0,74 | 4,61E-01 |
| AT5G63310 | 0,47 | 2,24 | 4,35 | 0,52 | 6,06E-01 |
| AT5G63320 | 0,52 | -1,80 | 3,77 | -0,48 | 6,33E-01 |
| AT5G63370 | 1,26 | 2,68 | 1,65 | 1,63 | 1,04E-01 |
| AT5G63400 | 0,30 | 1,69 | 4,39 | 0,38 | 7,01E-01 |
| AT5G63410 | 0,46 | -0,94 | 2,55 | -0,37 | 7,12E-01 |
| AT5G63470 | 0,44 | -2,86 | 3,98 | -0,72 | 4,72E-01 |
| AT5G63610 | 0,51 | 2,08 | 3,95 | 0,53 | 5,99E-01 |
| AT5G63625 | 1,28 | -0,13 | 1,68 | -0,07 | 9,40E-01 |
| AT5G63760 | 0,90 | 1,23 | 2,20 | 0,56 | 5,76E-01 |
| AT5G63790 | 0,54 | 3,75 | 4,32 | 0,87 | 3,86E-01 |
| AT5G63860 | 0,75 | 1,74 | 2,09 | 0,83 | 4,06E-01 |
| AT5G63870 | 0,08 | 2,08 | 4,38 | 0,47 | 6,35E-01 |
| AT5G63890 | 0,39 | 1,44 | 4,29 | 0,34 | 7,38E-01 |
| AT5G63910 | 0,66 | -1,07 | 2,97 | -0,36 | 7,19E-01 |
| AT5G64020 | 0,23 | 1,51 | 4,34 | 0,35 | 7,28E-01 |
| AT5G64050 | 1,12 | 1,30 | 1,61 | 0,81 | 4,20E-01 |
| AT5G64070 | 1,53 | 1,07 | 1,35 | 0,79 | 4,31E-01 |
| AT5G64130 | 0,35 | -0,01 | 3,56 | 0,00 | 9,98E-01 |
| AT5G64140 | 1,06 | -1,26 | 1,56 | -0,81 | 4,18E-01 |
| AT5G64170 | 0,64 | 0,24 | 3,05 | 0,08 | 9,37E-01 |
| AT5G64200 | 0,28 | 1,40 | 4,31 | 0,32 | 7,45E-01 |
| AT5G64280 | 1,39 | 1,46 | 1,58 | 0,92 | 3,56E-01 |
| AT5G64290 | 0,42 | 2,33 | 4,09 | 0,57 | 5,68E-01 |
| AT5G64340 | 1,19 | 0,37 | 1,63 | 0,23 | 8,21E-01 |
| AT5G64350 | 0,41 | 1,24 | 4,23 | 0,29 | 7,69E-01 |
| AT5G64360 | 0,62 | -0,14 | 3,76 | -0,04 | 9,71E-01 |
| AT5G64410 | 0,79 | -0,15 | 2,08 | -0,07 | 9,43E-01 |
| AT5G64430 | 0,96 | 1,97 | 2,14 | 0,92 | 3,57E-01 |
| AT5G64460 | 0,08 | -0,56 | 4,38 | -0,13 | 8,99E-01 |
| AT5G64550 | 0,25 | 2,98 | 4,33 | 0,69 | 4,92E-01 |
| AT5G64570 | 1,49 | 0,71 | 1,45 | 0,49 | 6,26E-01 |
| AT5G64580 | 0,77 | 1,14 | 2,58 | 0,44 | 6,59E-01 |
| AT5G64610 | 0,28 | 0,51 | 4,28 | 0,12 | 9,06E-01 |
| AT5G64660 | 0,37 | -0,01 | 3,30 | 0,00 | 9,98E-01 |
| AT5G64670 | 0,17 | 2,11 | 4,37 | 0,48 | 6,28E-01 |
| AT5G64760 | 0,83 | 1,76 | 2,92 | 0,60 | 5,46E-01 |
| AT5G64830 | 0,77 | 2,70 | 2,45 | 1,10 | 2,70E-01 |
| AT5G64860 | 0,28 | 1,54 | 4,28 | 0,36 | 7,19E-01 |
| AT5G64870 | 0,67 | -0,46 | 2,62 | -0,17 | 8,61E-01 |
| AT5G64920 | 1,21 | 2,14 | 1,71 | 1,26 | 2,09E-01 |
| AT5G64930 | 0,39 | 1,71 | 4,40 | 0,39 | 6,98E-01 |
| AT5G64980 | 0,12 | 2,81 | 4,34 | 0,65 | 5,18E-01 |
| AT5G65110 | 0,46 | 1,87 | 3,12 | 0,60 | 5,50E-01 |
| AT5G65200 | 0,16 | 1,18 | 4,32 | 0,27 | 7,85E-01 |
| AT5G65240 | 0,27 | 0,31 | 3,64 | 0,08 | 9,33E-01 |
| AT5G65310 | 1,18 | -1,15 | 2,16 | -0,53 | 5,95E-01 |

| | | | | | |
|---|---|---|---|---|---|
| AT5G65360 | 0,52 | -0,58 | 3,07 | -0,19 | 8,51E-01 |
| AT5G65430 | 0,62 | -0,94 | 2,60 | -0,36 | 7,19E-01 |
| AT5G65560 | 0,37 | 1,00 | 4,30 | 0,23 | 8,16E-01 |
| AT5G65685 | 0,97 | -0,32 | 2,03 | -0,16 | 8,75E-01 |
| AT5G65840 | 0,17 | 0,69 | 4,34 | 0,16 | 8,73E-01 |
| AT5G65910 | 1,14 | 3,90 | 3,23 | 1,21 | 2,27E-01 |
| AT5G65930 | 0,99 | 2,09 | 1,99 | 1,05 | 2,92E-01 |
| AT5G09535 | 0,24 | 2,80 | 4,34 | 0,65 | 5,18E-01 |
| AT5G65970 | 0,45 | 1,66 | 4,39 | 0,38 | 7,05E-01 |
| AT5G66055 | 0,63 | 2,83 | 2,51 | 1,13 | 2,58E-01 |
| AT5G66120 | 0,85 | 1,12 | 1,98 | 0,57 | 5,71E-01 |
| AT5G66210 | 1,05 | -0,76 | 1,99 | -0,38 | 7,03E-01 |
| AT5G66300 | 0,13 | -1,34 | 4,35 | -0,31 | 7,58E-01 |
| AT5G66310 | 0,12 | 2,81 | 4,34 | 0,65 | 5,18E-01 |
| AT5G66320 | 0,48 | 0,74 | 3,24 | 0,23 | 8,20E-01 |
| AT5G66330 | 0,11 | -1,08 | 4,37 | -0,25 | 8,05E-01 |
| AT5G66400 | 0,42 | 2,36 | 4,22 | 0,56 | 5,76E-01 |
| AT5G66440 | 0,33 | 0,55 | 4,36 | 0,13 | 8,99E-01 |
| AT5G66460 | 0,57 | 1,61 | 4,34 | 0,37 | 7,11E-01 |
| AT5G66500 | 0,15 | 1,93 | 4,33 | 0,45 | 6,55E-01 |
| AT5G66530 | 0,97 | 0,72 | 2,62 | 0,27 | 7,83E-01 |
| AT5G66540 | 0,21 | 1,45 | 4,28 | 0,34 | 7,35E-01 |
| AT5G66558 | 1,38 | 1,94 | 1,59 | 1,22 | 2,21E-01 |
| AT5G66595 | 0,74 | -2,34 | 2,26 | -1,04 | 3,00E-01 |
| AT5G66607 | 0,08 | -0,56 | 4,38 | -0,13 | 8,99E-01 |
| AT5G66650 | 0,26 | 2,30 | 4,37 | 0,53 | 5,98E-01 |
| AT5G66675 | 0,51 | -1,94 | 2,99 | -0,65 | 5,16E-01 |
| AT5G66680 | 0,76 | 0,19 | 3,58 | 0,05 | 9,58E-01 |
| AT5G66720 | 0,41 | -0,13 | 4,41 | -0,03 | 9,77E-01 |
| AT5G66730 | 0,31 | 0,65 | 4,33 | 0,15 | 8,81E-01 |
| AT5G66850 | 0,24 | 3,20 | 4,33 | 0,74 | 4,60E-01 |
| AT5G66890 | 0,75 | 3,61 | 3,46 | 1,04 | 2,97E-01 |
| AT5G66900 | 1,52 | 2,01 | 1,56 | 1,29 | 1,97E-01 |
| AT5G66910 | 0,38 | -0,21 | 3,30 | -0,06 | 9,48E-01 |
| AT5G67120 | 0,79 | -0,01 | 2,51 | 0,00 | 9,97E-01 |
| AT5G67150 | 0,35 | -0,36 | 4,34 | -0,08 | 9,34E-01 |
| AT5G67190 | 0,91 | 0,54 | 2,30 | 0,23 | 8,14E-01 |
| AT5G67220 | 0,60 | 1,25 | 2,43 | 0,51 | 6,07E-01 |
| AT5G67240 | 1,06 | 1,17 | 1,88 | 0,62 | 5,35E-01 |
| AT5G67245 | 0,45 | 0,59 | 4,35 | 0,14 | 8,92E-01 |
| AT5G67250 | 1,03 | -0,09 | 1,75 | -0,05 | 9,57E-01 |
| AT5G67280 | 0,58 | -1,14 | 2,57 | -0,44 | 6,57E-01 |
| AT5G67290 | 0,82 | 1,06 | 2,27 | 0,47 | 6,41E-01 |
| AT5G09855 | 0,69 | 0,48 | 2,27 | 0,21 | 8,33E-01 |
| AT5G67310 | 0,78 | 4,34 | 2,78 | 1,56 | 1,18E-01 |
| AT5G67350 | 0,75 | -0,06 | 2,38 | -0,03 | 9,79E-01 |
| AT5G67380 | 0,30 | -0,80 | 4,30 | -0,19 | 8,52E-01 |
| AT5G67460 | 0,10 | 0,34 | 4,35 | 0,08 | 9,37E-01 |
| AT5G67480 | 1,04 | 2,66 | 2,43 | 1,09 | 2,74E-01 |
| AT5G67490 | 1,16 | 0,65 | 1,66 | 0,39 | 6,94E-01 |
| AT5G67580 | 0,39 | 0,63 | 2,75 | 0,23 | 8,19E-01 |
| AT5G67610 | 0,37 | 0,00 | 4,41 | 0,00 | 1,00E+00 |

**chr11 chr17 infected by TCV relative to uninfected *chr11 chr17*.**

| |
|---|
| type |
| protein_coding |
| protein_coding |
| protein_coding |
| miRNA_precursor |
| miRNA_precursor |
| ncRNA |
| protein_coding |
| ncRNA |
| miRNA_precursor |
| miRNA_precursor |
| protein_coding |
| protein_coding |
| protein_coding |
| miRNA_precursor |
| ncRNA |
| protein_coding |
| protein_coding |
| protein_coding |
| protein_coding |
| protein_coding |
| ncRNA |
| miRNA_precursor |
| miRNA_precursor |
| protein_coding |
| miRNA_precursor |
| ncRNA |
| miRNA_precursor |
| miRNA_precursor |
| protein_coding |
| miRNA_precursor |
| protein_coding |
| miRNA_precursor |
| miRNA_precursor |
| ncRNA |
| ncRNA |
| protein_coding |
| ncRNA |
| protein_coding |
| protein_coding |
| protein_coding |
| protein_coding |
| protein_coding |
| protein_coding |
| protein_coding |
| protein_coding |
| miRNA_precursor |
| protein_coding |
| protein_coding |
| protein_coding |
| protein_coding |
| protein_coding |
| ncRNA |
| protein_coding |

small_nucle(ol)ar_rna
miRNA_precursor
protein_coding
miRNA_precursor
miRNA_precursor
protein_coding
protein_coding
protein_coding
miRNA_precursor
protein_coding
protein_coding
miRNA_precursor
protein_coding
protein_coding
protein_coding
protein_coding
protein_coding
miRNA_precursor
protein_coding
protein_coding
miRNA_precursor
ncRNA
protein_coding
small_nucle(ol)ar_rna
protein_coding
protein_coding
miRNA_precursor
protein_coding
miRNA_precursor
protein_coding
protein_coding
protein_coding
protein_coding
ncRNA
protein_coding
ncRNA
protein_coding
protein_coding
ncRNA
protein_coding
protein_coding
ribosomal_rna
protein_coding
protein_coding
protein_coding
protein_coding
protein_coding
protein_coding
miRNA_precursor
protein_coding
protein_coding
protein_coding
protein_coding
protein_coding
miRNA_precursor
protein_coding

protein_coding
protein_coding
miRNA_precursor
protein_coding
protein_coding
protein_coding
ncRNA
protein_coding
protein_coding
protein_coding
protein_coding
protein_coding
protein_coding
miRNA_precursor
protein_coding
miRNA_precursor
protein_coding
protein_coding
protein_coding
protein_coding
protein_coding
miRNA_precursor
protein_coding
protein_coding
protein_coding
protein_coding
miRNA_precursor
protein_coding
protein_coding
protein_coding
protein_coding
protein_coding
protein_coding
protein_coding
protein_coding
protein_coding
ncRNA
protein_coding
miRNA_precursor
protein_coding
ncRNA
protein_coding
protein_coding
protein_coding
protein_coding
protein_coding
protein_coding
protein_coding
protein_coding
miRNA_precursor
protein_coding
protein_coding
protein_coding
protein_coding
protein_coding
ribosomal_rna

protein_coding
protein_coding
protein_coding
ncRNA
protein_coding
protein_coding
protein_coding
protein_coding
miRNA_precursor
small_nucle(ol)ar_rna
protein_coding
protein_coding
protein_coding
protein_coding
protein_coding
protein_coding
protein_coding
ncRNA
protein_coding
protein_coding
protein_coding
protein_coding
protein_coding
protein_coding
miRNA_precursor
protein_coding
protein_coding
protein_coding
protein_coding
protein_coding
protein_coding
protein_coding
miRNA_precursor
protein_coding
protein_coding
miRNA_precursor
protein_coding
protein_coding
protein_coding
protein_coding
protein_coding
protein_coding
miRNA_precursor
protein_coding
protein_coding
protein_coding
protein_coding
protein_coding
protein_coding
protein_coding
protein_coding
protein_coding
protein_coding
protein_coding
protein_coding
protein_coding

protein_coding
protein_coding
protein_coding
protein_coding
ncRNA
protein_coding
protein_coding
protein_coding
protein_coding
protein_coding
protein_coding
protein_coding
ncRNA
protein_coding
protein_coding
protein_coding
protein_coding
protein_coding
protein_coding
protein_coding
protein_coding
protein_coding
protein_coding
protein_coding
small_nucle(ol)ar_rna
ncRNA
protein_coding
protein_coding
protein_coding
protein_coding
protein_coding
protein_coding
protein_coding
protein_coding
protein_coding
protein_coding
protein_coding
protein_coding
protein_coding
protein_coding
protein_coding
protein_coding
protein_coding
protein_coding
protein_coding
protein_coding
miRNA_precursor
protein_coding
miRNA_precursor
protein_coding
protein_coding
protein_coding
protein_coding
protein_coding
protein_coding
protein_coding

protein_coding
small_nucle(ol)ar_rna
protein_coding
protein_coding
protein_coding
protein_coding
protein_coding
protein_coding
protein_coding
protein_coding
small_nucle(ol)ar_rna
protein_coding
ncRNA
small_nucle(ol)ar_rna
protein_coding
protein_coding
small_nucle(ol)ar_rna
protein_coding
protein_coding
protein_coding
miRNA_precursor
protein_coding
protein_coding
protein_coding
protein_coding
protein_coding
protein_coding
miRNA_precursor
protein_coding
protein_coding
miRNA_precursor
protein_coding
protein_coding
protein_coding
miRNA_precursor
protein_coding
protein_coding
protein_coding
protein_coding
protein_coding
protein_coding
protein_coding
miRNA_precursor
small_nucle(ol)ar_rna
protein_coding
protein_coding
protein_coding
protein_coding
protein_coding
protein_coding
protein_coding
miRNA_precursor
protein_coding
protein_coding
protein_coding
protein_coding

protein_coding
protein_coding
protein_coding
protein_coding
protein_coding
protein_coding
protein_coding
protein_coding
protein_coding
protein_coding
protein_coding
protein_coding
protein_coding
miRNA_precursor
protein_coding
protein_coding
miRNA_precursor
protein_coding
protein_coding
protein_coding
miRNA_precursor
small_nucle(ol)ar_rna
protein_coding
miRNA_precursor
protein_coding
protein_coding
protein_coding
miRNA_precursor
protein_coding
protein_coding
protein_coding
protein_coding
protein_coding
protein_coding
protein_coding
ncRNA
protein_coding
protein_coding
protein_coding
protein_coding
protein_coding
protein_coding
protein_coding
miRNA_precursor
small_nucle(ol)ar_rna
protein_coding
protein_coding
miRNA_precursor
miRNA_precursor
protein_coding
miRNA_precursor
miRNA_precursor
protein_coding
protein_coding
miRNA_precursor
protein_coding

protein_coding
protein_coding
protein_coding
protein_coding
protein_coding
small_nucle(ol)ar_rna
protein_coding
miRNA_precursor
protein_coding
protein_coding
protein_coding
protein_coding
protein_coding
miRNA_precursor
protein_coding
miRNA_precursor
protein_coding
protein_coding
ncRNA
protein_coding
protein_coding
protein_coding
small_nucle(ol)ar_rna
protein_coding
protein_coding
protein_coding
protein_coding
protein_coding
protein_coding
protein_coding
protein_coding
protein_coding
ncRNA
protein_coding
protein_coding
protein_coding
protein_coding
protein_coding
protein_coding
protein_coding
protein_coding
protein_coding
protein_coding
protein_coding
protein_coding
protein_coding
protein_coding
protein_coding
protein_coding
protein_coding
small_nucle(ol)ar_rna
protein_coding
protein_coding
protein_coding
protein_coding
protein_coding

protein_coding
small_nucle(ol)ar_rna
protein_coding
protein_coding
protein_coding
protein_coding
protein_coding
protein_coding
protein_coding
protein_coding
protein_coding
miRNA_precursor
protein_coding
protein_coding
protein_coding
protein_coding
protein_coding
protein_coding
protein_coding
protein_coding
protein_coding
protein_coding
protein_coding
protein_coding
protein_coding
protein_coding
protein_coding
protein_coding
protein_coding
protein_coding
protein_coding
protein_coding
protein_coding
protein_coding
protein_coding
protein_coding
protein_coding
protein_coding
protein_coding
protein_coding
protein_coding
protein_coding
protein_coding
protein_coding
protein_coding
protein_coding
protein_coding
protein_coding
protein_coding
small_nucle(ol)ar_rna
protein_coding
protein_coding
protein_coding
protein_coding
protein_coding
protein_coding

protein_coding
protein_coding
protein_coding
protein_coding
protein_coding
protein_coding
protein_coding
protein_coding
ncRNA
protein_coding
protein_coding
protein_coding
protein_coding
protein_coding
protein_coding
protein_coding
protein_coding
protein_coding
protein_coding
protein_coding
protein_coding
protein_coding
protein_coding
protein_coding
protein_coding
protein_coding
protein_coding
protein_coding
protein_coding
protein_coding
protein_coding
protein_coding
protein_coding
protein_coding
protein_coding
protein_coding
protein_coding
protein_coding
protein_coding
miRNA_precursor
protein_coding
protein_coding
protein_coding
protein_coding
protein_coding
protein_coding
miRNA_precursor
protein_coding
protein_coding
protein_coding
small_nucle(ol)ar_rna
protein_coding
protein_coding
protein_coding
miRNA_precursor
protein_coding

protein_coding
protein_coding
protein_coding
protein_coding
protein_coding
protein_coding
protein_coding
protein_coding
miRNA_precursor
protein_coding
protein_coding
protein_coding
protein_coding
protein_coding
protein_coding
miRNA_precursor
protein_coding
protein_coding
protein_coding
protein_coding
ncRNA
ncRNA
protein_coding
protein_coding
protein_coding
miRNA_precursor
miRNA_precursor
protein_coding
protein_coding
protein_coding
protein_coding
protein_coding
protein_coding
miRNA_precursor
small_nucle(ol)ar_rna
protein_coding
miRNA_precursor
ncRNA
protein_coding
protein_coding
protein_coding
ribosomal_rna
protein_coding
protein_coding
protein_coding
protein_coding
protein_coding
protein_coding
protein_coding
miRNA_precursor
protein_coding
protein_coding
protein_coding
protein_coding
protein_coding
protein_coding

protein_coding
protein_coding
protein_coding
protein_coding
protein_coding
miRNA_precursor
protein_coding
protein_coding
protein_coding
protein_coding
miRNA_precursor
protein_coding
protein_coding
protein_coding
protein_coding
protein_coding
small_nucle(ol)ar_rna
miRNA_precursor
protein_coding
protein_coding
protein_coding
protein_coding
protein_coding
protein_coding
protein_coding
protein_coding
protein_coding
protein_coding
protein_coding
protein_coding
protein_coding
protein_coding
protein_coding
small_nucle(ol)ar_rna
protein_coding
protein_coding
protein_coding
protein_coding
protein_coding
protein_coding
protein_coding
protein_coding
protein_coding
protein_coding
protein_coding
protein_coding
protein_coding
protein_coding
protein_coding
protein_coding
protein_coding
protein_coding
protein_coding
protein_coding
ncRNA
protein_coding

protein_coding
protein_coding
ncRNA
protein_coding
protein_coding
protein_coding
miRNA_precursor
protein_coding
protein_coding
protein_coding
protein_coding
protein_coding
protein_coding
protein_coding
protein_coding
protein_coding
ncRNA
small_nucle(ol)ar_rna
protein_coding
protein_coding
protein_coding
protein_coding
miRNA_precursor
miRNA_precursor
protein_coding
protein_coding
protein_coding
protein_coding
protein_coding
protein_coding
protein_coding
pre_trna
protein_coding
protein_coding
protein_coding
protein_coding
protein_coding
protein_coding
protein_coding
protein_coding
protein_coding
protein_coding
protein_coding
ncRNA
protein_coding
protein_coding
protein_coding
protein_coding
small_nucle(ol)ar_rna
protein_coding
protein_coding
protein_coding
protein_coding
protein_coding
protein_coding
small_nucle(ol)ar_rna

protein_coding
protein_coding
protein_coding
protein_coding
protein_coding
protein_coding
protein_coding
protein_coding
protein_coding
protein_coding
protein_coding
protein_coding
protein_coding
protein_coding
protein_coding
ncRNA
protein_coding
protein_coding
ncRNA
protein_coding
protein_coding
protein_coding
ncRNA
small_nucle(ol)ar_rna
protein_coding
small_nucle(ol)ar_rna
protein_coding
small_nucle(ol)ar_rna
protein_coding
protein_coding
protein_coding
ncRNA
protein_coding
protein_coding
small_nucle(ol)ar_rna
protein_coding
protein_coding
protein_coding
protein_coding
protein_coding
protein_coding
small_nucle(ol)ar_rna
protein_coding
protein_coding
protein_coding
protein_coding
protein_coding
protein_coding
protein_coding
protein_coding
protein_coding
protein_coding
protein_coding
protein_coding
protein_coding
small_nucle(ol)ar_rna

protein_coding
protein_coding
protein_coding
protein_coding
small_nucle(ol)ar_rna
ncRNA
protein_coding
protein_coding
protein_coding
protein_coding
miRNA_precursor
protein_coding
protein_coding
protein_coding
protein_coding
protein_coding
protein_coding
protein_coding
protein_coding
ncRNA
protein_coding
protein_coding
protein_coding
protein_coding
protein_coding
protein_coding
protein_coding
protein_coding
protein_coding
protein_coding
protein_coding
protein_coding
protein_coding
ncRNA
protein_coding
protein_coding
protein_coding
protein_coding
protein_coding
protein_coding
protein_coding
protein_coding
protein_coding
protein_coding
ncRNA
protein_coding
protein_coding
protein_coding
protein_coding
protein_coding
ncRNA
protein_coding
protein_coding
small_nucle(ol)ar_rna
protein_coding
protein_coding

protein_coding
protein_coding
protein_coding
protein_coding
protein_coding
protein_coding
protein_coding
protein_coding
miRNA_precursor
miRNA_precursor
protein_coding
protein_coding
protein_coding
protein_coding
protein_coding
ncRNA
protein_coding
small_nucle(ol)ar_rna
protein_coding
protein_coding
protein_coding
protein_coding
protein_coding
protein_coding
protein_coding
protein_coding
protein_coding
protein_coding
protein_coding
protein_coding
protein_coding
protein_coding
protein_coding
protein_coding
protein_coding
protein_coding
protein_coding
protein_coding
protein_coding
miRNA_precursor
small_nucle(ol)ar_rna
miRNA_precursor
protein_coding
protein_coding
protein_coding
protein_coding
protein_coding
protein_coding
protein_coding
protein_coding
protein_coding
protein_coding
protein_coding
protein_coding
protein_coding
protein_coding

protein_coding
protein_coding
protein_coding
protein_coding
protein_coding
protein_coding
miRNA_precursor
protein_coding
protein_coding
protein_coding
protein_coding
protein_coding
protein_coding
protein_coding
protein_coding
protein_coding
protein_coding
protein_coding
protein_coding
protein_coding
ncRNA
protein_coding
protein_coding
small_nucle(ol)ar_rna
protein_coding
protein_coding
protein_coding
protein_coding
protein_coding
protein_coding
ncRNA
protein_coding
protein_coding
protein_coding
protein_coding
protein_coding
ncRNA
protein_coding
protein_coding
protein_coding
protein_coding
protein_coding
protein_coding
protein_coding
protein_coding
protein_coding
protein_coding
protein_coding
protein_coding
protein_coding
protein_coding
protein_coding
protein_coding
protein_coding
protein_coding
protein_coding

ncRNA
protein_coding
protein_coding
protein_coding
protein_coding
ncRNA
small_nucle(ol)ar_rna
protein_coding
small_nucle(ol)ar_rna
small_nucle(ol)ar_rna
protein_coding
protein_coding
protein_coding
protein_coding
protein_coding
miRNA_precursor
protein_coding
protein_coding
protein_coding
protein_coding
protein_coding
protein_coding
protein_coding
protein_coding
protein_coding
protein_coding
protein_coding
protein_coding
protein_coding
ncRNA
protein_coding
protein_coding
protein_coding
protein_coding
protein_coding
protein_coding
protein_coding
protein_coding
protein_coding
miRNA_precursor
protein_coding
small_nucle(ol)ar_rna
protein_coding
protein_coding
protein_coding
protein_coding
protein_coding
protein_coding
protein_coding
small_nucle(ol)ar_rna
protein_coding
protein_coding
protein_coding
protein_coding
protein_coding
protein_coding

protein_coding
protein_coding
ncRNA
protein_coding
small_nucle(ol)ar_rna
protein_coding
protein_coding
protein_coding
protein_coding
protein_coding
protein_coding
protein_coding
protein_coding
protein_coding
protein_coding
protein_coding
miRNA_precursor
protein_coding
protein_coding
protein_coding
protein_coding
protein_coding
protein_coding
ncRNA
protein_coding
small_nucle(ol)ar_rna
pre_trna
protein_coding
protein_coding
protein_coding
small_nucle(ol)ar_rna
protein_coding
miRNA_precursor
protein_coding
miRNA_precursor
protein_coding
protein_coding
miRNA_precursor
ncRNA
protein_coding
protein_coding
protein_coding
protein_coding
protein_coding
protein_coding
small_nucle(ol)ar_rna
protein_coding
ncRNA
protein_coding
protein_coding
protein_coding
protein_coding
protein_coding
protein_coding
protein_coding
protein_coding

protein_coding
protein_coding
protein_coding
protein_coding
protein_coding
protein_coding
protein_coding
protein_coding
protein_coding
protein_coding
miRNA_precursor
protein_coding
protein_coding
protein_coding
protein_coding
protein_coding
small_nucle(ol)ar_rna
protein_coding
protein_coding
protein_coding
protein_coding
protein_coding
small_nucle(ol)ar_rna
protein_coding
protein_coding
protein_coding
protein_coding
protein_coding
small_nucle(ol)ar_rna
ncRNA
small_nucle(ol)ar_rna
protein_coding
ncRNA
protein_coding
miRNA_precursor
protein_coding
miRNA_precursor
miRNA_precursor
protein_coding
protein_coding
protein_coding
miRNA_precursor
protein_coding
ncRNA
protein_coding
protein_coding
protein_coding
small_nucle(ol)ar_rna
protein_coding
protein_coding
protein_coding
protein_coding
protein_coding
protein_coding
protein_coding
protein_coding

protein_coding
small_nucle(ol)ar_rna
protein_coding
protein_coding
protein_coding
protein_coding
protein_coding
protein_coding
protein_coding
protein_coding
protein_coding
protein_coding
protein_coding
protein_coding
protein_coding
protein_coding
protein_coding
protein_coding
protein_coding
protein_coding
protein_coding
protein_coding
protein_coding
protein_coding
protein_coding
protein_coding
protein_coding
protein_coding
protein_coding
protein_coding
protein_coding
protein_coding
miRNA_precursor
protein_coding
protein_coding
protein_coding
protein_coding
protein_coding
protein_coding
protein_coding
protein_coding
pre_trna
protein_coding
protein_coding
miRNA_precursor
ncRNA
protein_coding
protein_coding
protein_coding
protein_coding
protein_coding
protein_coding
protein_coding
protein_coding
protein_coding
protein_coding

protein_coding
protein_coding
protein_coding
protein_coding
protein_coding
protein_coding
ncRNA
protein_coding
protein_coding
protein_coding
protein_coding
protein_coding
protein_coding
protein_coding
protein_coding
protein_coding
protein_coding
protein_coding
protein_coding
protein_coding
small_nucle(ol)ar_rna
ncRNA
protein_coding
protein_coding
protein_coding
protein_coding
protein_coding
protein_coding
protein_coding
protein_coding
miRNA_precursor
protein_coding
protein_coding
protein_coding
small_nucle(ol)ar_rna
ncRNA
protein_coding
protein_coding
miRNA_precursor
protein_coding
ncRNA
protein_coding
protein_coding
protein_coding
protein_coding
protein_coding
protein_coding
protein_coding
protein_coding
protein_coding
protein_coding
protein_coding
protein_coding
protein_coding
protein_coding
protein_coding

protein_coding
small_nucle(ol)ar_rna
miRNA_precursor
ncRNA
protein_coding
protein_coding
protein_coding
protein_coding
protein_coding
protein_coding
protein_coding
protein_coding
small_nucle(ol)ar_rna
protein_coding
protein_coding
protein_coding
protein_coding
protein_coding
protein_coding
protein_coding
protein_coding
protein_coding
ncRNA
protein_coding
protein_coding
protein_coding
protein_coding
protein_coding
protein_coding
ncRNA
protein_coding
miRNA_precursor
protein_coding
protein_coding
ncRNA
protein_coding
ncRNA
ncRNA
small_nucle(ol)ar_rna
protein_coding
protein_coding
protein_coding
protein_coding
protein_coding
protein_coding
protein_coding
ncRNA
protein_coding
ncRNA
protein_coding
protein_coding
protein_coding
protein_coding
protein_coding
small_nucle(ol)ar_rna
protein_coding

protein_coding
miRNA_precursor
protein_coding
ncRNA
protein_coding
protein_coding
protein_coding
small_nucle(ol)ar_rna
protein_coding
small_nucle(ol)ar_rna
protein_coding
protein_coding
protein_coding
protein_coding
protein_coding
protein_coding
protein_coding
protein_coding
protein_coding
protein_coding
protein_coding
protein_coding
protein_coding
protein_coding
protein_coding
protein_coding
protein_coding
protein_coding
protein_coding
protein_coding
protein_coding
protein_coding
ncRNA
small_nucle(ol)ar_rna
protein_coding
protein_coding
protein_coding
protein_coding
protein_coding
protein_coding
protein_coding
protein_coding
miRNA_precursor
protein_coding
small_nucle(ol)ar_rna
ncRNA
protein_coding
protein_coding
small_nucle(ol)ar_rna
protein_coding
miRNA_precursor
protein_coding
protein_coding
protein_coding
ncRNA
miRNA_precursor

protein_coding
ncRNA
protein_coding
protein_coding
protein_coding
protein_coding
protein_coding
protein_coding
protein_coding
protein_coding
protein_coding
protein_coding
protein_coding
protein_coding
miRNA_precursor
protein_coding
small_nucle(ol)ar_rna
protein_coding
protein_coding
protein_coding
protein_coding
protein_coding
miRNA_precursor
protein_coding
small_nucle(ol)ar_rna
protein_coding
protein_coding
small_nucle(ol)ar_rna
protein_coding
protein_coding
protein_coding
small_nucle(ol)ar_rna
protein_coding
protein_coding
protein_coding
protein_coding
protein_coding
protein_coding
protein_coding
protein_coding
protein_coding
protein_coding
pre_trna
small_nucle(ol)ar_rna
protein_coding
protein_coding
protein_coding
protein_coding
ncRNA
protein_coding
protein_coding
small_nucle(ol)ar_rna
protein_coding
protein_coding
miRNA_precursor
protein_coding

protein_coding
small_nucle(ol)ar_rna
protein_coding
small_nucle(ol)ar_rna
protein_coding
protein_coding
protein_coding
protein_coding
protein_coding
protein_coding
protein_coding
protein_coding
protein_coding
miRNA_precursor
protein_coding
protein_coding
protein_coding
protein_coding
protein_coding
protein_coding
protein_coding
protein_coding
protein_coding
protein_coding
protein_coding
protein_coding
protein_coding
ncRNA
protein_coding
protein_coding
protein_coding
protein_coding
miRNA_precursor
protein_coding
protein_coding
protein_coding
protein_coding
protein_coding
protein_coding
ncRNA
protein_coding
protein_coding
protein_coding
protein_coding
protein_coding
ncRNA
protein_coding
protein_coding
protein_coding
protein_coding
protein_coding
miRNA_precursor
protein_coding
protein_coding
small_nucle(ol)ar_rna
protein_coding

protein_coding
protein_coding
protein_coding
small_nucle(ol)ar_rna
protein_coding
protein_coding
protein_coding
protein_coding
protein_coding
protein_coding
protein_coding
miRNA_precursor
protein_coding
protein_coding
ncRNA
protein_coding
protein_coding
protein_coding
protein_coding
protein_coding
ncRNA
protein_coding
protein_coding
protein_coding
protein_coding
protein_coding
protein_coding
ncRNA
small_nucle(ol)ar_rna
protein_coding
protein_coding
protein_coding
protein_coding
protein_coding
protein_coding
protein_coding
small_nucle(ol)ar_rna
protein_coding
protein_coding
small_nucle(ol)ar_rna
protein_coding
protein_coding
miRNA_precursor
protein_coding
protein_coding
ncRNA
ncRNA
protein_coding
protein_coding
protein_coding
protein_coding
protein_coding
protein_coding
protein_coding
protein_coding
protein_coding

protein_coding
ncRNA
protein_coding
small_nucle(ol)ar_rna
protein_coding
protein_coding
ncRNA
protein_coding
protein_coding
protein_coding
protein_coding
protein_coding
protein_coding
protein_coding
small_nucle(ol)ar_rna
protein_coding
protein_coding
protein_coding
protein_coding
protein_coding
protein_coding
small_nucle(ol)ar_rna
protein_coding
protein_coding
ncRNA
protein_coding
protein_coding
protein_coding
protein_coding
protein_coding
protein_coding
protein_coding
protein_coding
protein_coding
small_nucle(ol)ar_rna
protein_coding
protein_coding
protein_coding
miRNA_precursor
protein_coding
protein_coding
protein_coding
small_nucle(ol)ar_rna
ncRNA
protein_coding
protein_coding
protein_coding
protein_coding
protein_coding
protein_coding
protein_coding
protein_coding
small_nucle(ol)ar_rna
protein_coding
protein_coding
protein_coding

protein_coding
protein_coding
protein_coding
protein_coding
ncRNA
protein_coding
small_nucle(ol)ar_rna
protein_coding
protein_coding
protein_coding
protein_coding
ncRNA
small_nucle(ol)ar_rna
protein_coding
protein_coding
protein_coding
protein_coding
protein_coding
protein_coding
protein_coding
protein_coding
ncRNA
ncRNA
ncRNA
protein_coding
small_nucle(ol)ar_rna
protein_coding
protein_coding
protein_coding
protein_coding
ncRNA
protein_coding
protein_coding
protein_coding
protein_coding
protein_coding
ncRNA
ncRNA
protein_coding
ncRNA
ncRNA
ncRNA
protein_coding
protein_coding
protein_coding
protein_coding
protein_coding
protein_coding
protein_coding
protein_coding
protein_coding
protein_coding
protein_coding
small_nucle(ol)ar_rna
protein_coding
protein_coding

protein_coding
protein_coding
protein_coding
protein_coding
protein_coding
small_nucle(ol)ar_rna
protein_coding
protein_coding
protein_coding
protein_coding
protein_coding
protein_coding
ncRNA
protein_coding
small_nucle(ol)ar_rna
protein_coding
protein_coding
protein_coding
protein_coding
small_nucle(ol)ar_rna
protein_coding
ncRNA
protein_coding
protein_coding
ribosomal_rna
miRNA_precursor
ncRNA
protein_coding
protein_coding
protein_coding
protein_coding
protein_coding
protein_coding
protein_coding
protein_coding
protein_coding
protein_coding
protein_coding
small_nucle(ol)ar_rna
miRNA_precursor
protein_coding
protein_coding
protein_coding
protein_coding
miRNA_precursor
protein_coding
protein_coding
protein_coding
protein_coding
protein_coding
protein_coding
protein_coding
protein_coding
protein_coding
ncRNA
protein_coding

protein_coding
protein_coding
protein_coding
protein_coding
protein_coding
protein_coding
protein_coding
protein_coding
protein_coding
protein_coding
protein_coding
protein_coding
protein_coding
small_nucle(ol)ar_rna
protein_coding
miRNA_precursor
protein_coding
protein_coding
protein_coding
protein_coding
protein_coding
protein_coding
protein_coding
protein_coding
protein_coding
ncRNA
protein_coding
ncRNA
protein_coding
protein_coding
protein_coding
protein_coding
protein_coding
protein_coding
small_nucle(ol)ar_rna
protein_coding
protein_coding
protein_coding
protein_coding
protein_coding
protein_coding
protein_coding
protein_coding
small_nucle(ol)ar_rna
protein_coding
protein_coding
protein_coding
protein_coding
protein_coding
protein_coding
protein_coding
protein_coding
protein_coding
protein_coding
protein_coding
protein_coding

protein_coding
miRNA_precursor
protein_coding
protein_coding
protein_coding
protein_coding
protein_coding
protein_coding
protein_coding
protein_coding
small_nucle(ol)ar_rna
protein_coding
protein_coding
protein_coding
protein_coding
protein_coding
protein_coding
ncRNA
protein_coding
protein_coding
protein_coding
protein_coding
protein_coding
protein_coding
miRNA_precursor
protein_coding
protein_coding
protein_coding
protein_coding
protein_coding
protein_coding
protein_coding
ncRNA
protein_coding
protein_coding
protein_coding
protein_coding
protein_coding
protein_coding
miRNA_precursor
protein_coding
protein_coding
protein_coding
miRNA_precursor
protein_coding
protein_coding
protein_coding
protein_coding
ncRNA
small_nucle(ol)ar_rna
protein_coding
protein_coding
protein_coding
protein_coding
protein_coding
protein_coding

protein_coding
protein_coding
protein_coding
protein_coding
protein_coding
protein_coding
protein_coding
ncRNA
small_nucle(ol)ar_rna
protein_coding
protein_coding
protein_coding
protein_coding
protein_coding
protein_coding
protein_coding
protein_coding
protein_coding
protein_coding
protein_coding
protein_coding
small_nucle(ol)ar_rna
protein_coding
protein_coding
protein_coding
protein_coding
protein_coding
protein_coding
protein_coding
protein_coding
protein_coding
protein_coding
protein_coding
protein_coding
protein_coding
miRNA_precursor
protein_coding
protein_coding
protein_coding
protein_coding
protein_coding
protein_coding
protein_coding
protein_coding
protein_coding
protein_coding
protein_coding
protein_coding
protein_coding
protein_coding
protein_coding
protein_coding
protein_coding
protein_coding
protein_coding
protein_coding

ncRNA
protein_coding
protein_coding
protein_coding
protein_coding
small_nucle(ol)ar_rna
protein_coding
protein_coding
protein_coding
protein_coding
protein_coding
protein_coding
ncRNA
protein_coding
protein_coding
protein_coding
protein_coding
protein_coding
protein_coding
protein_coding
protein_coding
small_nucle(ol)ar_rna
ncRNA
protein_coding
protein_coding
ncRNA
protein_coding
protein_coding
small_nucle(ol)ar_rna
protein_coding
protein_coding
protein_coding
protein_coding
protein_coding
miRNA_precursor
protein_coding
small_nucle(ol)ar_rna
protein_coding
protein_coding
small_nucle(ol)ar_rna
miRNA_precursor
protein_coding
protein_coding
protein_coding
protein_coding
protein_coding
protein_coding
protein_coding
protein_coding
protein_coding
protein_coding
protein_coding
protein_coding
protein_coding
protein_coding
protein_coding

protein_coding
protein_coding
protein_coding
protein_coding
small_nucle(ol)ar_rna
protein_coding
small_nucle(ol)ar_rna
ncRNA
protein_coding
protein_coding
protein_coding
protein_coding
protein_coding
protein_coding
protein_coding
protein_coding
protein_coding
protein_coding
protein_coding
ncRNA
protein_coding
protein_coding
protein_coding
protein_coding
protein_coding
protein_coding
protein_coding
protein_coding
small_nucle(ol)ar_rna
miRNA_precursor
protein_coding
protein_coding
small_nucle(ol)ar_rna
ncRNA
miRNA_precursor
miRNA_precursor
protein_coding
protein_coding
protein_coding
protein_coding
protein_coding
small_nucle(ol)ar_rna
protein_coding
protein_coding
protein_coding
protein_coding
protein_coding
protein_coding
protein_coding
protein_coding
protein_coding
protein_coding
ncRNA
ncRNA
protein_coding
protein_coding

protein_coding
protein_coding
protein_coding
protein_coding
protein_coding
protein_coding
ncRNA
protein_coding
small_nucle(ol)ar_rna
protein_coding
protein_coding
protein_coding
protein_coding
protein_coding
protein_coding
protein_coding
protein_coding
protein_coding
ncRNA
protein_coding
protein_coding
protein_coding
protein_coding
protein_coding
protein_coding
protein_coding
protein_coding
protein_coding
protein_coding
protein_coding
protein_coding
protein_coding
small_nucle(ol)ar_rna
protein_coding
protein_coding
protein_coding
protein_coding
small_nucle(ol)ar_rna
protein_coding
protein_coding
protein_coding
protein_coding
protein_coding
protein_coding
protein_coding
protein_coding
miRNA_precursor
protein_coding
protein_coding
protein_coding
protein_coding
protein_coding
protein_coding
small_nucle(ol)ar_rna
small_nucle(ol)ar_rna
protein_coding

protein_coding
ncRNA
protein_coding
protein_coding
protein_coding
protein_coding
protein_coding
protein_coding
miRNA_precursor
protein_coding
protein_coding
protein_coding
protein_coding
protein_coding
protein_coding
small_nucle(ol)ar_rna
protein_coding
ncRNA
small_nucle(ol)ar_rna
protein_coding
protein_coding
ncRNA
small_nucle(ol)ar_rna
protein_coding
protein_coding
ncRNA
ncRNA
protein_coding
protein_coding
protein_coding
protein_coding
protein_coding
protein_coding
protein_coding
protein_coding
protein_coding
protein_coding
protein_coding
protein_coding
small_nucle(ol)ar_rna
protein_coding
protein_coding
miRNA_precursor
small_nucle(ol)ar_rna
protein_coding
protein_coding
protein_coding
protein_coding
protein_coding
protein_coding
protein_coding
protein_coding
protein_coding
protein_coding
protein_coding
protein_coding

protein_coding
protein_coding
protein_coding
protein_coding
protein_coding
miRNA_precursor
protein_coding
protein_coding
protein_coding
protein_coding
miRNA_precursor
miRNA_precursor
protein_coding
protein_coding
protein_coding
protein_coding
small_nucle(ol)ar_rna
protein_coding
protein_coding
small_nucle(ol)ar_rna
protein_coding
protein_coding
protein_coding
protein_coding
protein_coding
protein_coding
protein_coding
small_nucle(ol)ar_rna
ncRNA
protein_coding
protein_coding
protein_coding
protein_coding
protein_coding
pre_trna
miRNA_precursor
protein_coding
pre_trna
protein_coding
protein_coding
protein_coding
protein_coding
protein_coding
protein_coding
protein_coding
small_nucle(ol)ar_rna
protein_coding
ncRNA
protein_coding
protein_coding
protein_coding
protein_coding
protein_coding
small_nucle(ol)ar_rna
protein_coding
protein_coding

miRNA_precursor
protein_coding
protein_coding
protein_coding
protein_coding
small_nucle(ol)ar_rna
protein_coding
protein_coding
protein_coding
protein_coding
small_nucle(ol)ar_rna
small_nucle(ol)ar_rna
protein_coding
protein_coding
protein_coding
protein_coding
protein_coding
protein_coding
protein_coding
protein_coding
protein_coding
protein_coding
protein_coding
protein_coding
protein_coding
protein_coding
miRNA_precursor
protein_coding
protein_coding
miRNA_precursor
ncRNA
protein_coding
protein_coding
ncRNA
protein_coding
protein_coding
protein_coding
small_nucle(ol)ar_rna
small_nucle(ol)ar_rna
small_nucle(ol)ar_rna
miRNA_precursor
protein_coding
protein_coding
protein_coding
protein_coding
protein_coding
protein_coding
protein_coding
miRNA_precursor
protein_coding
protein_coding
protein_coding
protein_coding
protein_coding
pre_trna
protein_coding

protein_coding
miRNA_precursor
protein_coding
protein_coding
protein_coding
protein_coding
protein_coding
protein_coding
protein_coding
protein_coding
protein_coding
miRNA_precursor
protein_coding
protein_coding
protein_coding
protein_coding
small_nucle(ol)ar_rna
ncRNA
protein_coding
protein_coding
ncRNA
small_nucle(ol)ar_rna
protein_coding
protein_coding
miRNA_precursor
protein_coding
protein_coding
protein_coding
protein_coding
protein_coding
protein_coding
protein_coding
protein_coding
protein_coding
protein_coding
protein_coding
protein_coding
protein_coding
protein_coding
protein_coding
protein_coding
protein_coding
protein_coding
small_nucle(ol)ar_rna
protein_coding
protein_coding
protein_coding
protein_coding
protein_coding
protein_coding
protein_coding
protein_coding
protein_coding
protein_coding
small_nucle(ol)ar_rna
protein_coding

protein_coding
protein_coding
protein_coding
protein_coding
protein_coding
protein_coding
protein_coding
protein_coding
protein_coding
miRNA_precursor
protein_coding
protein_coding
protein_coding
protein_coding
protein_coding
protein_coding
protein_coding
protein_coding
protein_coding
ncRNA
protein_coding
protein_coding
protein_coding
protein_coding
protein_coding
protein_coding
small_nucle(ol)ar_rna
miRNA_precursor
protein_coding
protein_coding
protein_coding
small_nucle(ol)ar_rna
protein_coding
protein_coding
protein_coding
protein_coding
miRNA_precursor
protein_coding
protein_coding
protein_coding
protein_coding
small_nucle(ol)ar_rna
protein_coding
protein_coding
protein_coding
protein_coding
protein_coding
pre_trna
protein_coding
protein_coding
ncRNA
ncRNA
protein_coding
protein_coding
protein_coding
protein_coding

protein_coding
protein_coding
protein_coding
protein_coding
miRNA_precursor
protein_coding
protein_coding
protein_coding
protein_coding
small_nucle(ol)ar_rna
protein_coding
miRNA_precursor
protein_coding
protein_coding
protein_coding
miRNA_precursor
protein_coding
protein_coding
protein_coding
protein_coding
protein_coding
protein_coding
protein_coding
protein_coding
protein_coding
ncRNA
protein_coding
protein_coding
protein_coding
protein_coding
protein_coding
protein_coding
protein_coding
ncRNA
protein_coding
protein_coding
protein_coding
protein_coding
protein_coding
protein_coding
protein_coding
protein_coding
protein_coding
ncRNA
protein_coding
protein_coding
ncRNA
protein_coding
protein_coding
protein_coding
protein_coding
protein_coding
protein_coding
protein_coding
protein_coding
protein_coding

protein_coding
protein_coding
protein_coding
protein_coding
protein_coding
protein_coding
protein_coding
protein_coding
protein_coding
protein_coding
ncRNA
protein_coding
protein_coding
protein_coding
protein_coding
protein_coding
protein_coding
protein_coding
protein_coding
protein_coding
protein_coding
small_nucle(ol)ar_rna
small_nucle(ol)ar_rna
small_nucle(ol)ar_rna
small_nucle(ol)ar_rna
protein_coding
protein_coding
protein_coding
protein_coding
protein_coding
ncRNA
protein_coding
ncRNA
protein_coding
protein_coding
protein_coding
small_nucle(ol)ar_rna
small_nucle(ol)ar_rna
protein_coding
protein_coding
protein_coding
protein_coding
protein_coding
protein_coding
protein_coding
protein_coding
protein_coding
protein_coding
protein_coding
protein_coding
protein_coding
protein_coding
ncRNA
protein_coding
miRNA_precursor
protein_coding

protein_coding
protein_coding
protein_coding
protein_coding
pre_trna
ncRNA
protein_coding
protein_coding
protein_coding
protein_coding
protein_coding
protein_coding
protein_coding
protein_coding
protein_coding
protein_coding
protein_coding
protein_coding
protein_coding
protein_coding
protein_coding
ncRNA
protein_coding
protein_coding
protein_coding
protein_coding
protein_coding
protein_coding
protein_coding
protein_coding
protein_coding
protein_coding
protein_coding
ncRNA
protein_coding
protein_coding
protein_coding
protein_coding
protein_coding
protein_coding
protein_coding
protein_coding
protein_coding
protein_coding
protein_coding
protein_coding
protein_coding
protein_coding
protein_coding
protein_coding
protein_coding
protein_coding
protein_coding
protein_coding
protein_coding
protein_coding

protein_coding
protein_coding
protein_coding
protein_coding
protein_coding
protein_coding
protein_coding
protein_coding
protein_coding
protein_coding
protein_coding
protein_coding
protein_coding
protein_coding
protein_coding
protein_coding
ncRNA
protein_coding
protein_coding
protein_coding
protein_coding
protein_coding
protein_coding
protein_coding
protein_coding
protein_coding
protein_coding
protein_coding
protein_coding
protein_coding
protein_coding
protein_coding
protein_coding
protein_coding
protein_coding
protein_coding
protein_coding
protein_coding
protein_coding
protein_coding
protein_coding
protein_coding
protein_coding
protein_coding
protein_coding
protein_coding
protein_coding
protein_coding
protein_coding
protein_coding
protein_coding
protein_coding
protein_coding
protein_coding
protein_coding
protein_coding

protein_coding
protein_coding
protein_coding
protein_coding
protein_coding
protein_coding
protein_coding
protein_coding
protein_coding
protein_coding
protein_coding
protein_coding
protein_coding
protein_coding
protein_coding
protein_coding
protein_coding
small_nucle(ol)ar_rna
protein_coding
protein_coding
protein_coding
protein_coding
protein_coding
protein_coding
protein_coding
protein_coding
protein_coding
protein_coding
protein_coding
protein_coding
protein_coding
protein_coding
protein_coding
protein_coding
protein_coding
protein_coding
protein_coding
protein_coding
protein_coding
protein_coding
protein_coding
protein_coding
protein_coding
protein_coding
protein_coding
protein_coding
protein_coding
protein_coding
protein_coding
protein_coding
pre_trna
protein_coding
protein_coding
protein_coding
protein_coding
ncRNA

protein_coding
protein_coding
protein_coding
protein_coding
protein_coding
protein_coding
protein_coding
protein_coding
protein_coding
protein_coding
protein_coding
protein_coding
protein_coding
protein_coding
protein_coding
protein_coding
protein_coding
protein_coding
protein_coding
protein_coding
protein_coding
protein_coding
protein_coding
protein_coding
pre_trna
ncRNA
protein_coding
protein_coding
protein_coding
protein_coding
protein_coding
protein_coding
protein_coding
protein_coding
protein_coding
protein_coding
protein_coding
protein_coding
protein_coding
protein_coding
protein_coding
protein_coding
protein_coding
protein_coding
protein_coding
protein_coding
protein_coding
protein_coding
protein_coding
protein_coding
protein_coding
protein_coding
protein_coding
protein_coding
protein_coding
protein_coding

protein_coding
protein_coding
protein_coding
protein_coding
protein_coding
protein_coding
protein_coding
protein_coding
ncRNA
protein_coding
protein_coding
protein_coding
protein_coding
protein_coding
protein_coding
protein_coding
protein_coding
protein_coding
protein_coding
pre_trna
protein_coding
protein_coding
protein_coding
protein_coding
protein_coding
protein_coding
protein_coding
protein_coding
protein_coding
protein_coding
protein_coding
protein_coding
protein_coding
protein_coding
protein_coding
protein_coding
protein_coding
protein_coding
protein_coding
protein_coding
protein_coding
protein_coding
protein_coding
protein_coding
protein_coding
protein_coding
pre_trna
protein_coding
miRNA_precursor
protein_coding
protein_coding
protein_coding
protein_coding
protein_coding
protein_coding
protein_coding

protein_coding
protein_coding
protein_coding
protein_coding
small_nucle(ol)ar_rna
miRNA_precursor
protein_coding
ncRNA
protein_coding
protein_coding
protein_coding
protein_coding
protein_coding
protein_coding
protein_coding
protein_coding
ncRNA
protein_coding
protein_coding
protein_coding
protein_coding
protein_coding
miRNA_precursor
protein_coding
protein_coding
protein_coding
protein_coding
protein_coding
protein_coding
protein_coding
protein_coding
protein_coding
protein_coding
protein_coding
protein_coding
protein_coding
protein_coding
protein_coding
protein_coding
protein_coding
protein_coding
protein_coding
protein_coding
protein_coding
protein_coding
protein_coding
protein_coding
protein_coding
protein_coding
protein_coding
protein_coding
protein_coding
protein_coding
protein_coding
protein_coding
protein_coding

protein_coding
protein_coding
protein_coding
protein_coding
protein_coding
protein_coding
protein_coding
protein_coding
protein_coding
protein_coding
protein_coding
protein_coding
ncRNA
protein_coding
protein_coding
protein_coding
protein_coding
protein_coding
protein_coding
protein_coding
protein_coding
protein_coding
protein_coding
protein_coding
protein_coding
protein_coding
protein_coding
protein_coding
protein_coding
protein_coding
protein_coding
ncRNA
protein_coding
protein_coding
protein_coding
protein_coding
protein_coding
protein_coding
protein_coding
protein_coding
protein_coding
protein_coding
protein_coding
protein_coding
protein_coding
protein_coding
protein_coding
protein_coding
protein_coding
protein_coding
protein_coding
protein_coding
protein_coding
protein_coding
protein_coding
protein_coding

protein_coding
protein_coding
protein_coding
protein_coding
protein_coding
protein_coding
protein_coding
protein_coding
protein_coding
protein_coding
protein_coding
protein_coding
protein_coding
protein_coding
protein_coding
protein_coding
protein_coding
protein_coding
protein_coding
protein_coding
protein_coding
protein_coding
protein_coding
small_nucle(ol)ar_rna
protein_coding
protein_coding
protein_coding
protein_coding
ncRNA
protein_coding
protein_coding
protein_coding
protein_coding
protein_coding
protein_coding
protein_coding
protein_coding
protein_coding
protein_coding
protein_coding
protein_coding
protein_coding
protein_coding
ncRNA
protein_coding
protein_coding
protein_coding
pre_trna
protein_coding
protein_coding
protein_coding
protein_coding
protein_coding
protein_coding
protein_coding
protein_coding

small_nucle(ol)ar_rna
small_nucle(ol)ar_rna
small_nucle(ol)ar_rna
small_nucle(ol)ar_rna
protein_coding
protein_coding
protein_coding
protein_coding
protein_coding
protein_coding
protein_coding
protein_coding
protein_coding
protein_coding
protein_coding
protein_coding
pre_trna
protein_coding
protein_coding
protein_coding
protein_coding
protein_coding
protein_coding
protein_coding
protein_coding
protein_coding
protein_coding
protein_coding
ncRNA
protein_coding
protein_coding
protein_coding
protein_coding
protein_coding
protein_coding
protein_coding
protein_coding
ncRNA
protein_coding
protein_coding
protein_coding
protein_coding
protein_coding
protein_coding
protein_coding
protein_coding
protein_coding
protein_coding
protein_coding
protein_coding
protein_coding
small_nucle(ol)ar_rna
small_nucle(ol)ar_rna
protein_coding
protein_coding
protein_coding

miRNA_precursor
protein_coding
protein_coding
miRNA_precursor
protein_coding
protein_coding
protein_coding
protein_coding
protein_coding
protein_coding
protein_coding
protein_coding
protein_coding
protein_coding
protein_coding
protein_coding
protein_coding
protein_coding
protein_coding
protein_coding
protein_coding
protein_coding
protein_coding
protein_coding
protein_coding
protein_coding
protein_coding
protein_coding
ncRNA
protein_coding
protein_coding
protein_coding
protein_coding
protein_coding
protein_coding
protein_coding
protein_coding
protein_coding
ncRNA
protein_coding
protein_coding
protein_coding
protein_coding
protein_coding
protein_coding
protein_coding
protein_coding
ncRNA
protein_coding
protein_coding
protein_coding
protein_coding
ncRNA
protein_coding
protein_coding
protein_coding

protein_coding
protein_coding
protein_coding
protein_coding
protein_coding
protein_coding
protein_coding
protein_coding
protein_coding
protein_coding
protein_coding
protein_coding
protein_coding
ncRNA
protein_coding
protein_coding
ncRNA
protein_coding
protein_coding
ncRNA
protein_coding
protein_coding
protein_coding
protein_coding
protein_coding
ncRNA
protein_coding
protein_coding
protein_coding
miRNA_precursor
protein_coding
protein_coding
protein_coding
protein_coding
protein_coding
protein_coding
protein_coding
protein_coding
protein_coding
protein_coding
protein_coding
protein_coding
protein_coding
protein_coding
protein_coding
protein_coding
protein_coding
protein_coding
protein_coding
protein_coding
protein_coding
protein_coding
protein_coding
protein_coding
protein_coding
protein_coding

protein_coding
protein_coding
protein_coding
protein_coding
ncRNA
protein_coding
ncRNA
protein_coding
protein_coding
miRNA_precursor
protein_coding
protein_coding
protein_coding
protein_coding
protein_coding
protein_coding
protein_coding
protein_coding
protein_coding
protein_coding
protein_coding
protein_coding
protein_coding
protein_coding
protein_coding
protein_coding
ncRNA
protein_coding
protein_coding
small_nucle(ol)ar_rna
protein_coding
protein_coding
protein_coding
protein_coding
protein_coding
protein_coding
protein_coding
protein_coding
protein_coding
protein_coding
protein_coding
protein_coding
protein_coding
protein_coding
protein_coding
protein_coding
protein_coding
protein_coding
protein_coding
protein_coding
protein_coding
protein_coding
protein_coding
protein_coding
protein_coding
protein_coding

protein_coding
protein_coding
miRNA_precursor
protein_coding
protein_coding
protein_coding
protein_coding
protein_coding
protein_coding
protein_coding
protein_coding
protein_coding
protein_coding
protein_coding
protein_coding
protein_coding
protein_coding
protein_coding
protein_coding
protein_coding
protein_coding
protein_coding
protein_coding
protein_coding
ncRNA
protein_coding
protein_coding
protein_coding
protein_coding
protein_coding
ncRNA
protein_coding
protein_coding
protein_coding
protein_coding
protein_coding
protein_coding
protein_coding
protein_coding
protein_coding
protein_coding
protein_coding
protein_coding
protein_coding
protein_coding
protein_coding
protein_coding
protein_coding
protein_coding
protein_coding
protein_coding
protein_coding
protein_coding
protein_coding
protein_coding
protein_coding

protein_coding
protein_coding
protein_coding
protein_coding
ncRNA
protein_coding
protein_coding
protein_coding
protein_coding
protein_coding
protein_coding
protein_coding
ncRNA
protein_coding
ncRNA
small_nucle(ol)ar_rna
protein_coding
protein_coding
protein_coding
protein_coding
protein_coding
protein_coding
protein_coding
protein_coding
protein_coding
protein_coding
protein_coding
protein_coding
protein_coding
protein_coding
protein_coding
protein_coding
small_nucle(ol)ar_rna
protein_coding
protein_coding
protein_coding
protein_coding
protein_coding
protein_coding
protein_coding
protein_coding
protein_coding
protein_coding
protein_coding
protein_coding
protein_coding
protein_coding
protein_coding
protein_coding
protein_coding
protein_coding
protein_coding
protein_coding
protein_coding
protein_coding
protein_coding

protein_coding
protein_coding
protein_coding
protein_coding
protein_coding
protein_coding
protein_coding
ncRNA
protein_coding
protein_coding
protein_coding
ncRNA
protein_coding
protein_coding
protein_coding
protein_coding
protein_coding
protein_coding
protein_coding
protein_coding
protein_coding
protein_coding
protein_coding
protein_coding
protein_coding
protein_coding
protein_coding
protein_coding
protein_coding
protein_coding
protein_coding
protein_coding
protein_coding
protein_coding
protein_coding
protein_coding
protein_coding
protein_coding
protein_coding
protein_coding
protein_coding
protein_coding
ncRNA
protein_coding
protein_coding
protein_coding
protein_coding
protein_coding
protein_coding
protein_coding
protein_coding
protein_coding
protein_coding
protein_coding
miRNA_precursor
miRNA_precursor

protein_coding
protein_coding
protein_coding
protein_coding
ncRNA
protein_coding
protein_coding
protein_coding
small_nucle(ol)ar_rna
protein_coding
protein_coding
protein_coding
protein_coding
protein_coding
protein_coding
miRNA_precursor
protein_coding
protein_coding
protein_coding
protein_coding
protein_coding
protein_coding
protein_coding
protein_coding
protein_coding
protein_coding
protein_coding
protein_coding
protein_coding
ncRNA
protein_coding
ncRNA
protein_coding
protein_coding
protein_coding
protein_coding
protein_coding
protein_coding
protein_coding
protein_coding
protein_coding
protein_coding
protein_coding
protein_coding
protein_coding
protein_coding
protein_coding
protein_coding
protein_coding
protein_coding
protein_coding
protein_coding
protein_coding
protein_coding
protein_coding
protein_coding

small_nucle(ol)ar_rna
protein_coding
protein_coding
protein_coding
protein_coding
protein_coding
protein_coding
pre_trna
protein_coding
ncRNA
protein_coding
protein_coding
protein_coding
protein_coding
protein_coding
protein_coding
protein_coding
ncRNA
protein_coding
protein_coding
protein_coding
protein_coding
protein_coding
protein_coding
protein_coding
protein_coding
protein_coding
protein_coding
protein_coding
protein_coding
protein_coding
protein_coding
protein_coding
protein_coding
protein_coding
protein_coding
protein_coding
ncRNA
protein_coding
protein_coding
protein_coding
protein_coding
protein_coding
protein_coding
protein_coding
protein_coding
miRNA_precursor
protein_coding
protein_coding
protein_coding
protein_coding
protein_coding
protein_coding
protein_coding
protein_coding
protein_coding

protein_coding
protein_coding
protein_coding
protein_coding
protein_coding
ncRNA
protein_coding
ncRNA
protein_coding
small_nucle(ol)ar_rna
protein_coding
protein_coding
protein_coding
protein_coding
protein_coding
protein_coding
protein_coding
protein_coding
ncRNA
protein_coding
protein_coding
ncRNA
protein_coding
protein_coding
protein_coding
protein_coding
protein_coding
protein_coding
protein_coding
protein_coding
protein_coding
protein_coding
protein_coding
protein_coding
protein_coding
protein_coding
protein_coding
protein_coding
protein_coding
protein_coding
protein_coding
protein_coding
protein_coding
protein_coding
ncRNA
protein_coding
protein_coding
protein_coding
protein_coding
protein_coding
protein_coding
protein_coding
protein_coding
protein_coding
protein_coding
protein_coding

small_nucle(ol)ar_rna
protein_coding
protein_coding
protein_coding
protein_coding
protein_coding
protein_coding
protein_coding
protein_coding
protein_coding
protein_coding
protein_coding
protein_coding
protein_coding
protein_coding
protein_coding
protein_coding
protein_coding
protein_coding
protein_coding
protein_coding
protein_coding
miRNA_precursor
protein_coding
protein_coding
protein_coding
protein_coding
protein_coding
protein_coding
protein_coding
protein_coding
protein_coding
protein_coding
protein_coding
protein_coding
protein_coding
protein_coding
protein_coding
protein_coding
protein_coding
protein_coding
protein_coding
protein_coding
ncRNA
protein_coding
protein_coding
protein_coding
protein_coding
protein_coding
protein_coding
protein_coding
protein_coding
protein_coding
protein_coding
protein_coding
protein_coding

protein_coding
protein_coding
protein_coding
protein_coding
protein_coding
protein_coding
protein_coding
protein_coding
protein_coding
protein_coding
protein_coding
small_nucle(ol)ar_rna
protein_coding
protein_coding
protein_coding
protein_coding
protein_coding
ncRNA
protein_coding
protein_coding
protein_coding
protein_coding
protein_coding
protein_coding
protein_coding
protein_coding
protein_coding
protein_coding
protein_coding
protein_coding
protein_coding
protein_coding
protein_coding
protein_coding
protein_coding
protein_coding
protein_coding
protein_coding
protein_coding
protein_coding
protein_coding
protein_coding
protein_coding
protein_coding
protein_coding
protein_coding
protein_coding
protein_coding
protein_coding
protein_coding
protein_coding
protein_coding
protein_coding
protein_coding
protein_coding
protein_coding

protein_coding
protein_coding
protein_coding
protein_coding
protein_coding
small_nucle(ol)ar_rna
small_nucle(ol)ar_rna
small_nucle(ol)ar_rna
protein_coding
ncRNA
protein_coding
protein_coding
protein_coding
protein_coding
protein_coding
protein_coding
protein_coding
protein_coding
protein_coding
protein_coding
protein_coding
protein_coding
protein_coding
protein_coding
protein_coding
protein_coding
pre_trna
protein_coding
protein_coding
protein_coding
protein_coding
protein_coding
protein_coding
protein_coding
protein_coding
protein_coding
protein_coding
protein_coding
protein_coding
protein_coding
protein_coding
protein_coding
protein_coding
protein_coding
protein_coding
protein_coding
protein_coding
protein_coding
protein_coding
protein_coding
protein_coding
protein_coding
protein_coding
protein_coding
protein_coding
protein_coding

protein_coding
protein_coding
protein_coding
protein_coding
protein_coding
protein_coding
protein_coding
protein_coding
protein_coding
protein_coding
protein_coding
protein_coding
protein_coding
protein_coding
protein_coding
protein_coding
protein_coding
protein_coding
protein_coding
protein_coding
protein_coding
protein_coding
protein_coding
protein_coding
protein_coding
protein_coding
protein_coding
protein_coding
protein_coding
protein_coding
protein_coding
protein_coding
protein_coding
protein_coding
ncRNA
protein_coding
protein_coding
pre_trna
protein_coding
protein_coding
protein_coding
protein_coding
protein_coding
protein_coding
ncRNA
protein_coding
protein_coding
protein_coding
protein_coding
protein_coding
protein_coding
protein_coding
protein_coding
protein_coding
protein_coding
protein_coding

protein_coding
protein_coding
protein_coding
protein_coding
ncRNA
pre_trna
protein_coding
protein_coding
protein_coding
protein_coding
protein_coding
protein_coding
protein_coding
protein_coding
protein_coding
protein_coding
protein_coding
protein_coding
protein_coding
protein_coding
ncRNA
protein_coding
protein_coding
protein_coding
protein_coding
protein_coding
protein_coding
protein_coding
protein_coding
protein_coding
protein_coding
protein_coding
protein_coding
protein_coding
protein_coding
protein_coding
protein_coding
protein_coding
ncRNA
protein_coding
protein_coding
protein_coding
protein_coding
ncRNA
ncRNA
protein_coding
protein_coding
protein_coding
protein_coding
protein_coding
protein_coding
protein_coding
protein_coding
protein_coding
protein_coding
protein_coding

protein_coding
protein_coding
protein_coding
protein_coding
protein_coding
protein_coding
protein_coding
protein_coding
protein_coding
protein_coding
protein_coding
protein_coding
ncRNA
protein_coding
protein_coding
protein_coding
protein_coding
protein_coding
protein_coding
protein_coding
protein_coding
protein_coding
protein_coding
protein_coding
protein_coding
protein_coding
protein_coding
protein_coding
protein_coding
protein_coding
protein_coding
protein_coding
protein_coding
protein_coding
protein_coding
protein_coding
protein_coding
ncRNA
miRNA_precursor
protein_coding
ncRNA
protein_coding
ncRNA
protein_coding
protein_coding
protein_coding
ncRNA
protein_coding
protein_coding
protein_coding
protein_coding
protein_coding
protein_coding
protein_coding
protein_coding
protein_coding

protein_coding
protein_coding
protein_coding
pre_trna
protein_coding
protein_coding
protein_coding
protein_coding
protein_coding
protein_coding
ncRNA
protein_coding
protein_coding
protein_coding
protein_coding
protein_coding
protein_coding
protein_coding
protein_coding
protein_coding
protein_coding
protein_coding
protein_coding
pre_trna
protein_coding
protein_coding
protein_coding
ncRNA
protein_coding
protein_coding
protein_coding
protein_coding
protein_coding
protein_coding
protein_coding
protein_coding
protein_coding
ncRNA
protein_coding
protein_coding
ncRNA
protein_coding
protein_coding
protein_coding
ncRNA
protein_coding
protein_coding
protein_coding
protein_coding
ncRNA
protein_coding
protein_coding
protein_coding
protein_coding
protein_coding
protein_coding

protein_coding
protein_coding
protein_coding
protein_coding
protein_coding
ncRNA
protein_coding
ncRNA
protein_coding
protein_coding
protein_coding
protein_coding
protein_coding
protein_coding
protein_coding
protein_coding
protein_coding
protein_coding
protein_coding
protein_coding
protein_coding
ncRNA
protein_coding
protein_coding
protein_coding
protein_coding
protein_coding
protein_coding
protein_coding
protein_coding
protein_coding
protein_coding
protein_coding
protein_coding
protein_coding
protein_coding
protein_coding
protein_coding
protein_coding
protein_coding
protein_coding
protein_coding
protein_coding
ncRNA
protein_coding
protein_coding
protein_coding
protein_coding
protein_coding
protein_coding
protein_coding
protein_coding
protein_coding
protein_coding
protein_coding
protein_coding

protein_coding
protein_coding
protein_coding
protein_coding
protein_coding
protein_coding
protein_coding
protein_coding
protein_coding
protein_coding
protein_coding
protein_coding
protein_coding
protein_coding
protein_coding
protein_coding
protein_coding
protein_coding
protein_coding
protein_coding
protein_coding
ncRNA
protein_coding
protein_coding
protein_coding
protein_coding
ncRNA
ncRNA
protein_coding
protein_coding
protein_coding
protein_coding
protein_coding
protein_coding
protein_coding
protein_coding
protein_coding
protein_coding
miRNA_precursor
protein_coding
protein_coding
protein_coding
ncRNA
ncRNA
protein_coding
protein_coding
protein_coding
protein_coding
protein_coding
protein_coding
protein_coding
protein_coding
protein_coding
protein_coding
protein_coding
protein_coding

protein_coding
protein_coding
protein_coding
protein_coding
protein_coding
protein_coding
protein_coding
protein_coding
protein_coding
protein_coding
ncRNA
protein_coding
protein_coding
protein_coding
protein_coding
protein_coding
protein_coding
protein_coding
protein_coding
ncRNA
protein_coding
protein_coding
protein_coding
protein_coding
protein_coding
protein_coding
protein_coding
protein_coding
protein_coding
protein_coding
protein_coding
protein_coding
protein_coding
ncRNA
protein_coding
ncRNA
protein_coding
protein_coding
protein_coding
protein_coding
protein_coding
protein_coding
protein_coding
protein_coding
protein_coding
ncRNA
protein_coding
protein_coding
ncRNA
protein_coding
protein_coding
protein_coding
protein_coding
protein_coding
protein_coding
protein_coding

miRNA_precursor
protein_coding
protein_coding
protein_coding
miRNA_precursor
ncRNA
protein_coding
protein_coding
protein_coding
protein_coding
protein_coding
protein_coding
protein_coding
protein_coding
protein_coding
miRNA_precursor
protein_coding
protein_coding
pre_trna
protein_coding
ncRNA
ncRNA
protein_coding
protein_coding
ncRNA
protein_coding
protein_coding
protein_coding
protein_coding
protein_coding
protein_coding
protein_coding
protein_coding
protein_coding
protein_coding
protein_coding
protein_coding
protein_coding
protein_coding
protein_coding
protein_coding
protein_coding
protein_coding
protein_coding
protein_coding
protein_coding
protein_coding
protein_coding
protein_coding
protein_coding
protein_coding
protein_coding
protein_coding
protein_coding
protein_coding
protein_coding
protein_coding
protein_coding

ncRNA
protein_coding
protein_coding
protein_coding
protein_coding
protein_coding
protein_coding
protein_coding
protein_coding
protein_coding
protein_coding
protein_coding
protein_coding
protein_coding
protein_coding
protein_coding
protein_coding
protein_coding
protein_coding
protein_coding
protein_coding
protein_coding
protein_coding
protein_coding
protein_coding
protein_coding
protein_coding
protein_coding
protein_coding
protein_coding
protein_coding
protein_coding
protein_coding
protein_coding
protein_coding
protein_coding
protein_coding
protein_coding
protein_coding
protein_coding
protein_coding
protein_coding
pre_trna
protein_coding
protein_coding
protein_coding
protein_coding
protein_coding
protein_coding
protein_coding
protein_coding
protein_coding
protein_coding
protein_coding
protein_coding
protein_coding

protein_coding
protein_coding
protein_coding
protein_coding
protein_coding
protein_coding
protein_coding
protein_coding
protein_coding
protein_coding
pre_trna
small_nucle(ol)ar_rna
protein_coding
ncRNA
protein_coding
protein_coding
protein_coding
protein_coding
protein_coding
protein_coding
protein_coding
protein_coding
protein_coding
protein_coding
protein_coding
protein_coding
protein_coding
protein_coding
protein_coding
protein_coding
protein_coding
protein_coding
protein_coding
ncRNA
protein_coding
ncRNA
protein_coding
protein_coding
protein_coding
protein_coding
protein_coding
protein_coding
protein_coding
protein_coding
protein_coding
protein_coding
protein_coding
protein_coding
protein_coding
protein_coding
protein_coding
miRNA_precursor
protein_coding
protein_coding
protein_coding
protein_coding

protein_coding
protein_coding
protein_coding
protein_coding
protein_coding
protein_coding
protein_coding
small_nucle(ol)ar_rna
protein_coding
protein_coding
protein_coding
protein_coding
protein_coding
protein_coding
pre_trna
protein_coding
protein_coding
protein_coding
protein_coding
protein_coding
pre_trna
protein_coding
protein_coding
protein_coding
protein_coding
protein_coding
protein_coding
protein_coding
protein_coding
protein_coding
protein_coding
protein_coding
protein_coding
protein_coding
protein_coding
protein_coding
protein_coding
protein_coding
protein_coding
protein_coding
ncRNA
protein_coding
protein_coding
protein_coding
protein_coding
protein_coding
protein_coding
protein_coding
protein_coding
protein_coding
protein_coding
protein_coding
protein_coding
protein_coding
protein_coding
protein_coding

protein_coding
protein_coding
protein_coding
protein_coding
pre_trna
protein_coding
protein_coding
pre_trna
protein_coding
ncRNA
protein_coding
protein_coding
protein_coding
protein_coding
protein_coding
protein_coding
protein_coding
protein_coding
protein_coding
protein_coding
protein_coding
small_nucle(ol)ar_rna
protein_coding
protein_coding
ncRNA
protein_coding
protein_coding
protein_coding
ncRNA
protein_coding
protein_coding
protein_coding
protein_coding
protein_coding
protein_coding
protein_coding
protein_coding
protein_coding
protein_coding
protein_coding
protein_coding
protein_coding
protein_coding
protein_coding
protein_coding
protein_coding
protein_coding
protein_coding
protein_coding
protein_coding
protein_coding
protein_coding
protein_coding
protein_coding
protein_coding
protein_coding
protein_coding

protein_coding
protein_coding
protein_coding
protein_coding
protein_coding
protein_coding
protein_coding
protein_coding
protein_coding
protein_coding
protein_coding
protein_coding
protein_coding
protein_coding
protein_coding
protein_coding
protein_coding
protein_coding
protein_coding
protein_coding
protein_coding
protein_coding
protein_coding
protein_coding
protein_coding
miRNA_precursor
protein_coding
protein_coding
protein_coding
protein_coding
protein_coding
protein_coding
protein_coding
protein_coding
protein_coding
protein_coding
protein_coding
protein_coding
protein_coding
protein_coding
protein_coding
protein_coding
protein_coding
protein_coding
protein_coding
protein_coding
ncRNA
protein_coding
protein_coding
protein_coding
protein_coding
protein_coding
protein_coding
protein_coding
protein_coding
protein_coding

protein_coding
protein_coding
protein_coding
protein_coding
protein_coding
protein_coding
protein_coding
protein_coding
protein_coding
protein_coding
protein_coding
protein_coding
protein_coding
protein_coding
protein_coding
protein_coding
protein_coding
protein_coding
protein_coding
protein_coding
protein_coding
protein_coding
protein_coding
protein_coding
protein_coding
small_nucle(ol)ar_rna
small_nucle(ol)ar_rna
small_nucle(ol)ar_rna
protein_coding
protein_coding
protein_coding
protein_coding
ncRNA
protein_coding
protein_coding
protein_coding
protein_coding
protein_coding
protein_coding
protein_coding
protein_coding
protein_coding
protein_coding
protein_coding
protein_coding
protein_coding
protein_coding
protein_coding
protein_coding
protein_coding
protein_coding
protein_coding
protein_coding
protein_coding
protein_coding
protein_coding

protein_coding
protein_coding
protein_coding
protein_coding
protein_coding
protein_coding
protein_coding
protein_coding
protein_coding
protein_coding
protein_coding
protein_coding
protein_coding
protein_coding
protein_coding
pre_trna
protein_coding
protein_coding
protein_coding
protein_coding
protein_coding
protein_coding
pre_trna
protein_coding
protein_coding
protein_coding
ncRNA
protein_coding
protein_coding
protein_coding
protein_coding
protein_coding
protein_coding
protein_coding
protein_coding
protein_coding
protein_coding
protein_coding
protein_coding
protein_coding
small_nucle(ol)ar_rna
small_nucle(ol)ar_rna
protein_coding
protein_coding
protein_coding
protein_coding
protein_coding
protein_coding
protein_coding
protein_coding
protein_coding
protein_coding
protein_coding
protein_coding
protein_coding
protein_coding

protein_coding
protein_coding
protein_coding
protein_coding
protein_coding
protein_coding
protein_coding
protein_coding
protein_coding
protein_coding
protein_coding
small_nucle(ol)ar_rna
protein_coding
protein_coding
protein_coding
protein_coding
protein_coding
protein_coding
protein_coding
protein_coding
protein_coding
protein_coding
protein_coding
protein_coding
protein_coding
protein_coding
protein_coding
protein_coding
protein_coding
protein_coding
protein_coding
protein_coding
protein_coding
protein_coding
protein_coding
protein_coding
protein_coding
protein_coding
protein_coding
protein_coding
protein_coding
protein_coding
protein_coding
protein_coding
protein_coding
protein_coding
protein_coding
protein_coding
protein_coding
protein_coding
protein_coding
protein_coding
protein_coding
protein_coding
protein_coding
protein_coding

protein_coding
protein_coding
protein_coding
protein_coding
protein_coding
protein_coding
protein_coding
protein_coding
protein_coding
protein_coding
protein_coding
protein_coding
protein_coding
protein_coding
protein_coding
protein_coding
protein_coding
protein_coding
protein_coding
protein_coding
protein_coding
protein_coding
protein_coding
ncRNA
small_nucle(ol)ar_rna
protein_coding
protein_coding
protein_coding
protein_coding
protein_coding
protein_coding
protein_coding
protein_coding
protein_coding
protein_coding
protein_coding
protein_coding
protein_coding
protein_coding
protein_coding
protein_coding
protein_coding
protein_coding
protein_coding
protein_coding
protein_coding
protein_coding
protein_coding
protein_coding
protein_coding
protein_coding
protein_coding
protein_coding
protein_coding
protein_coding
protein_coding

ncRNA
ncRNA
protein_coding
protein_coding
protein_coding
protein_coding
protein_coding
protein_coding
protein_coding
protein_coding
protein_coding
protein_coding
protein_coding
protein_coding
protein_coding
protein_coding
protein_coding
protein_coding
protein_coding
protein_coding
protein_coding
protein_coding
ncRNA
ncRNA
protein_coding
protein_coding
protein_coding
protein_coding
protein_coding
protein_coding
protein_coding
protein_coding
protein_coding
protein_coding
protein_coding
protein_coding
protein_coding
protein_coding
protein_coding
protein_coding
protein_coding
protein_coding
protein_coding
protein_coding
protein_coding
protein_coding
protein_coding
protein_coding
protein_coding
protein_coding
protein_coding
protein_coding
protein_coding
protein_coding
protein_coding
protein_coding

protein_coding
protein_coding
protein_coding
protein_coding
protein_coding
protein_coding
protein_coding
protein_coding
protein_coding
protein_coding
protein_coding
protein_coding
protein_coding
protein_coding
protein_coding
protein_coding
protein_coding
protein_coding
protein_coding
protein_coding
protein_coding
protein_coding
protein_coding
protein_coding
protein_coding
protein_coding
ncRNA
protein_coding
protein_coding
protein_coding
protein_coding
protein_coding
protein_coding
protein_coding
protein_coding
protein_coding
protein_coding
protein_coding
protein_coding
protein_coding
protein_coding
protein_coding
protein_coding
protein_coding
protein_coding
protein_coding
protein_coding
protein_coding
protein_coding
protein_coding
protein_coding
protein_coding
protein_coding
protein_coding
protein_coding
protein_coding

protein_coding
protein_coding
protein_coding
protein_coding
protein_coding
protein_coding
protein_coding
protein_coding
protein_coding
protein_coding
protein_coding
protein_coding
protein_coding
protein_coding
protein_coding
protein_coding
protein_coding
protein_coding
protein_coding
protein_coding
protein_coding
protein_coding
protein_coding
ncRNA
protein_coding
protein_coding
protein_coding
protein_coding
protein_coding
protein_coding
protein_coding
protein_coding
protein_coding
protein_coding
protein_coding
protein_coding
protein_coding
protein_coding
protein_coding
protein_coding
protein_coding
ncRNA
protein_coding
protein_coding
protein_coding
protein_coding
protein_coding
protein_coding
protein_coding
protein_coding
protein_coding
protein_coding
protein_coding
protein_coding
protein_coding
protein_coding

protein_coding
protein_coding
protein_coding
protein_coding
protein_coding
small_nucle(ol)ar_rna
protein_coding
protein_coding
protein_coding
protein_coding
protein_coding
protein_coding
protein_coding
protein_coding
protein_coding
protein_coding
protein_coding
protein_coding
protein_coding
protein_coding
protein_coding
protein_coding
protein_coding
protein_coding
protein_coding
protein_coding
protein_coding
protein_coding
protein_coding
protein_coding
protein_coding
protein_coding
protein_coding
protein_coding
protein_coding
protein_coding
protein_coding
protein_coding
protein_coding
protein_coding
protein_coding
protein_coding
protein_coding
protein_coding
protein_coding
protein_coding
protein_coding
protein_coding
protein_coding
protein_coding
protein_coding
protein_coding
protein_coding
ncRNA
protein_coding
protein_coding

protein_coding
protein_coding
protein_coding
protein_coding
protein_coding
protein_coding
protein_coding
protein_coding
protein_coding
protein_coding
protein_coding
protein_coding
protein_coding
protein_coding
protein_coding
protein_coding
protein_coding
protein_coding
protein_coding
protein_coding
protein_coding
protein_coding
protein_coding
protein_coding
protein_coding
protein_coding
protein_coding
protein_coding
protein_coding
protein_coding
ncRNA
protein_coding
protein_coding
protein_coding
protein_coding
protein_coding
protein_coding
protein_coding
protein_coding
protein_coding
protein_coding
protein_coding
protein_coding
protein_coding
protein_coding
ncRNA
protein_coding
protein_coding
protein_coding
protein_coding
protein_coding
protein_coding
protein_coding
protein_coding
protein_coding
protein_coding

protein_coding
protein_coding
protein_coding
protein_coding
protein_coding
protein_coding
miRNA_precursor
protein_coding
protein_coding
protein_coding
protein_coding
protein_coding
protein_coding
protein_coding
protein_coding
protein_coding
protein_coding
protein_coding
miRNA_precursor
protein_coding
protein_coding
protein_coding
protein_coding
protein_coding
protein_coding
protein_coding
protein_coding
protein_coding
protein_coding
protein_coding
protein_coding
protein_coding
protein_coding
protein_coding
protein_coding
protein_coding
protein_coding
protein_coding
protein_coding
protein_coding
protein_coding
protein_coding
protein_coding
protein_coding
protein_coding
protein_coding
protein_coding
protein_coding
protein_coding
protein_coding
protein_coding
protein_coding
protein_coding
protein_coding
protein_coding
protein_coding

protein_coding
ncRNA
protein_coding
protein_coding
protein_coding
protein_coding
protein_coding
protein_coding
protein_coding
protein_coding
protein_coding
protein_coding
protein_coding
protein_coding
protein_coding
protein_coding
protein_coding
protein_coding
small_nucle(ol)ar_rna
protein_coding
ncRNA
protein_coding
protein_coding
protein_coding
protein_coding
ncRNA
protein_coding
protein_coding
protein_coding
protein_coding
protein_coding
protein_coding
protein_coding
protein_coding
protein_coding
protein_coding
protein_coding
protein_coding
protein_coding
protein_coding
protein_coding
protein_coding
ncRNA
protein_coding
protein_coding
protein_coding
protein_coding
protein_coding
protein_coding
protein_coding
protein_coding
protein_coding
protein_coding
protein_coding
protein_coding
protein_coding

protein_coding
protein_coding
protein_coding
protein_coding
protein_coding
protein_coding
protein_coding
protein_coding
protein_coding
protein_coding
protein_coding
protein_coding
protein_coding
protein_coding
protein_coding
protein_coding
protein_coding
protein_coding
protein_coding
protein_coding
protein_coding
protein_coding
protein_coding
protein_coding
protein_coding
protein_coding
protein_coding
protein_coding
protein_coding
protein_coding
protein_coding
protein_coding
protein_coding
ncRNA
protein_coding
protein_coding
protein_coding
protein_coding
protein_coding
protein_coding
protein_coding
protein_coding
protein_coding
protein_coding
protein_coding
protein_coding
protein_coding
ncRNA
small_nucle(ol)ar_rna
protein_coding
protein_coding
protein_coding
protein_coding
protein_coding
protein_coding
protein_coding

protein_coding
protein_coding
protein_coding
protein_coding
protein_coding
protein_coding
protein_coding
protein_coding
protein_coding
protein_coding
protein_coding
protein_coding
protein_coding
protein_coding
protein_coding
protein_coding
protein_coding
protein_coding
protein_coding
protein_coding
protein_coding
protein_coding
protein_coding
protein_coding
protein_coding
pre_trna
protein_coding
ncRNA
protein_coding
protein_coding
protein_coding
protein_coding
protein_coding
protein_coding
protein_coding
protein_coding
protein_coding
protein_coding
protein_coding
protein_coding
protein_coding
protein_coding
protein_coding
protein_coding
protein_coding
protein_coding
protein_coding
protein_coding
protein_coding
miRNA_precursor
protein_coding
protein_coding
protein_coding
protein_coding
protein_coding
protein_coding

protein_coding
protein_coding
protein_coding
protein_coding
ncRNA
protein_coding
protein_coding
protein_coding
protein_coding
protein_coding
protein_coding
protein_coding
small_nucle(ol)ar_rna
protein_coding
protein_coding
protein_coding
protein_coding
protein_coding
protein_coding
protein_coding
ncRNA
protein_coding
protein_coding
protein_coding
protein_coding
protein_coding
protein_coding
protein_coding
protein_coding
protein_coding
protein_coding
protein_coding
protein_coding
protein_coding
protein_coding
protein_coding
protein_coding
protein_coding
protein_coding
protein_coding
protein_coding
protein_coding
protein_coding
protein_coding
ncRNA
protein_coding
protein_coding
protein_coding
protein_coding
protein_coding
ncRNA
protein_coding
ncRNA
protein_coding
protein_coding
ncRNA

protein_coding
protein_coding
protein_coding
protein_coding
protein_coding
protein_coding
protein_coding
protein_coding
protein_coding
protein_coding
protein_coding
protein_coding
protein_coding
protein_coding
protein_coding
protein_coding
protein_coding
protein_coding
protein_coding
ncRNA
protein_coding
protein_coding
protein_coding
protein_coding
protein_coding
protein_coding
protein_coding
protein_coding
protein_coding
protein_coding
protein_coding
protein_coding
protein_coding
protein_coding
pre_trna
protein_coding
protein_coding
protein_coding
protein_coding
protein_coding
protein_coding
protein_coding
protein_coding
protein_coding
protein_coding
protein_coding
ncRNA
protein_coding
ncRNA
ncRNA
ncRNA
ncRNA
protein_coding
protein_coding
protein_coding
miRNA_precursor

protein_coding
protein_coding
small_nucle(ol)ar_rna
protein_coding
protein_coding
protein_coding
protein_coding
protein_coding
protein_coding
protein_coding
ncRNA
ncRNA
protein_coding
protein_coding
protein_coding
protein_coding
protein_coding
protein_coding
protein_coding
protein_coding
protein_coding
protein_coding
protein_coding
protein_coding
protein_coding
protein_coding
protein_coding
protein_coding
protein_coding
protein_coding
protein_coding
protein_coding
protein_coding
protein_coding
protein_coding
protein_coding
ncRNA
protein_coding
protein_coding
protein_coding
protein_coding
protein_coding
protein_coding
protein_coding
protein_coding
protein_coding
protein_coding
protein_coding
protein_coding
protein_coding
protein_coding
protein_coding
protein_coding
protein_coding
protein_coding
protein_coding

protein_coding
protein_coding
protein_coding
protein_coding
protein_coding
protein_coding
protein_coding
protein_coding
protein_coding
protein_coding
protein_coding
protein_coding
protein_coding
protein_coding
protein_coding
protein_coding
protein_coding
protein_coding
protein_coding
protein_coding
protein_coding
protein_coding
protein_coding
protein_coding
protein_coding
protein_coding
protein_coding
protein_coding
protein_coding
protein_coding
protein_coding
protein_coding
protein_coding
protein_coding
protein_coding
protein_coding
protein_coding
protein_coding
protein_coding
protein_coding
protein_coding
protein_coding
protein_coding
protein_coding
protein_coding
protein_coding
protein_coding
protein_coding
protein_coding
protein_coding
protein_coding
protein_coding
protein_coding
small_nucle(ol)ar_rna
protein_coding
protein_coding

protein_coding
miRNA_precursor
ncRNA
protein_coding
protein_coding
protein_coding
protein_coding
protein_coding
protein_coding
protein_coding
protein_coding
protein_coding
protein_coding
protein_coding
protein_coding
protein_coding
protein_coding
protein_coding
protein_coding
protein_coding
protein_coding
protein_coding
protein_coding
protein_coding
protein_coding
protein_coding
protein_coding
protein_coding
protein_coding
ncRNA
protein_coding
protein_coding
protein_coding
protein_coding
ncRNA
protein_coding
protein_coding
protein_coding
protein_coding
protein_coding
protein_coding
protein_coding
protein_coding
protein_coding
protein_coding
protein_coding
protein_coding
protein_coding
protein_coding
protein_coding
ncRNA
small_nucle(ol)ar_rna
protein_coding
protein_coding
protein_coding
protein_coding
protein_coding

protein_coding
protein_coding
protein_coding
protein_coding
protein_coding
protein_coding
protein_coding
protein_coding
protein_coding
protein_coding
protein_coding
ncRNA
protein_coding
protein_coding
protein_coding
protein_coding
protein_coding
protein_coding
protein_coding
protein_coding
protein_coding
protein_coding
ncRNA
protein_coding
protein_coding
protein_coding
protein_coding
protein_coding
protein_coding
ncRNA
protein_coding
protein_coding
protein_coding
protein_coding
protein_coding
protein_coding
protein_coding
protein_coding
protein_coding
ncRNA
protein_coding
protein_coding
protein_coding
protein_coding
protein_coding
protein_coding
protein_coding
protein_coding
protein_coding
protein_coding
protein_coding
protein_coding
protein_coding
protein_coding
protein_coding
protein_coding

protein_coding
ncRNA
protein_coding
protein_coding
protein_coding
protein_coding
protein_coding
protein_coding
protein_coding
protein_coding
protein_coding
protein_coding
protein_coding
protein_coding
protein_coding
protein_coding
protein_coding
protein_coding
protein_coding
protein_coding
protein_coding
protein_coding
protein_coding
protein_coding
protein_coding
protein_coding
protein_coding
protein_coding
protein_coding
protein_coding
protein_coding
protein_coding
protein_coding
protein_coding
protein_coding
protein_coding
protein_coding
protein_coding
protein_coding
miRNA_precursor
protein_coding
ncRNA
protein_coding
protein_coding
protein_coding
protein_coding
protein_coding
protein_coding
protein_coding
protein_coding
protein_coding
protein_coding
protein_coding
protein_coding
protein_coding
protein_coding

protein_coding
protein_coding
protein_coding
protein_coding
protein_coding
protein_coding
pre_trna
protein_coding
protein_coding
protein_coding
protein_coding
protein_coding
protein_coding
protein_coding
pre_trna
protein_coding
protein_coding
protein_coding
protein_coding
protein_coding
protein_coding
ncRNA
ncRNA
protein_coding
protein_coding
protein_coding
protein_coding
protein_coding
protein_coding
protein_coding
protein_coding
protein_coding
protein_coding
protein_coding
protein_coding
protein_coding
protein_coding
protein_coding
protein_coding
protein_coding
protein_coding
protein_coding
ncRNA
protein_coding
protein_coding
protein_coding
protein_coding
protein_coding
protein_coding
protein_coding
protein_coding
protein_coding
protein_coding
protein_coding
protein_coding
protein_coding

protein_coding
protein_coding
protein_coding
protein_coding
protein_coding
protein_coding
protein_coding
protein_coding
protein_coding
protein_coding
protein_coding
protein_coding
protein_coding
protein_coding
protein_coding
protein_coding
protein_coding
protein_coding
protein_coding
protein_coding
protein_coding
protein_coding
ncRNA
small_nucle(ol)ar_rna
protein_coding
protein_coding
protein_coding
protein_coding
protein_coding
protein_coding
protein_coding
protein_coding
protein_coding
protein_coding
protein_coding
protein_coding
protein_coding
protein_coding
protein_coding
protein_coding
protein_coding
protein_coding
protein_coding
protein_coding
protein_coding
ncRNA
protein_coding
ncRNA
protein_coding
protein_coding
protein_coding
protein_coding
protein_coding
protein_coding
protein_coding
protein_coding

protein_coding
protein_coding
protein_coding
protein_coding
protein_coding
ncRNA
protein_coding
protein_coding
protein_coding
protein_coding
protein_coding
protein_coding
pre_trna
protein_coding
protein_coding
protein_coding
protein_coding
protein_coding
protein_coding
protein_coding
protein_coding
protein_coding
protein_coding
ncRNA
ncRNA
protein_coding
protein_coding
protein_coding
protein_coding
protein_coding
protein_coding
protein_coding
protein_coding
protein_coding
protein_coding
protein_coding
protein_coding
protein_coding
protein_coding
protein_coding
protein_coding
miRNA_precursor
protein_coding
protein_coding
protein_coding
protein_coding
protein_coding
protein_coding
protein_coding
protein_coding
protein_coding
protein_coding
protein_coding
protein_coding
miRNA_precursor
protein_coding

protein_coding
protein_coding
protein_coding
protein_coding
protein_coding
protein_coding
miRNA_precursor
protein_coding
protein_coding
protein_coding
protein_coding
protein_coding
protein_coding
ncRNA
protein_coding
protein_coding
protein_coding
protein_coding
protein_coding
protein_coding
protein_coding
protein_coding
protein_coding
protein_coding
ncRNA
protein_coding
protein_coding
protein_coding
protein_coding
protein_coding
protein_coding
protein_coding
protein_coding
protein_coding
protein_coding
protein_coding
protein_coding
protein_coding
protein_coding
protein_coding
ncRNA
protein_coding
protein_coding
protein_coding
protein_coding
protein_coding
protein_coding
protein_coding
protein_coding
protein_coding
protein_coding
protein_coding
protein_coding
protein_coding
protein_coding
protein_coding

protein_coding
protein_coding
protein_coding
protein_coding
protein_coding
protein_coding
protein_coding
protein_coding
ncRNA
protein_coding
protein_coding
protein_coding
protein_coding
protein_coding
protein_coding
protein_coding
protein_coding
protein_coding
protein_coding
protein_coding
ncRNA
protein_coding
protein_coding
protein_coding
protein_coding
protein_coding
protein_coding
protein_coding
protein_coding
ncRNA
protein_coding
protein_coding
small_nucle(ol)ar_rna
small_nucle(ol)ar_rna
small_nucle(ol)ar_rna
protein_coding
protein_coding
protein_coding
protein_coding
protein_coding
protein_coding
protein_coding
protein_coding
protein_coding
protein_coding
protein_coding
protein_coding
protein_coding
protein_coding
protein_coding
protein_coding
protein_coding
protein_coding
protein_coding
protein_coding
protein_coding

protein_coding
ncRNA
protein_coding
protein_coding
protein_coding
protein_coding
small_nucle(ol)ar_rna
small_nucle(ol)ar_rna
miRNA_precursor
protein_coding
protein_coding
protein_coding
protein_coding
protein_coding
protein_coding
protein_coding
protein_coding
protein_coding
protein_coding
protein_coding
protein_coding
protein_coding
protein_coding
protein_coding
protein_coding
protein_coding
protein_coding
protein_coding
protein_coding
protein_coding
protein_coding
protein_coding
protein_coding
protein_coding
protein_coding
protein_coding
protein_coding
pre_trna
protein_coding
protein_coding
protein_coding
protein_coding
protein_coding
protein_coding
protein_coding
protein_coding
protein_coding
protein_coding
protein_coding
protein_coding
protein_coding
protein_coding
protein_coding
protein_coding
protein_coding
protein_coding

protein_coding
protein_coding
protein_coding
protein_coding
protein_coding
protein_coding
protein_coding
protein_coding
protein_coding
protein_coding
protein_coding
protein_coding
protein_coding
protein_coding
ncRNA
ncRNA
protein_coding
protein_coding
protein_coding
protein_coding
protein_coding
protein_coding
protein_coding
protein_coding
protein_coding
protein_coding
protein_coding
protein_coding
protein_coding
protein_coding
protein_coding
protein_coding
protein_coding
protein_coding
protein_coding
protein_coding
protein_coding
protein_coding
protein_coding
protein_coding
ncRNA
protein_coding
protein_coding
protein_coding
pre_trna
protein_coding
protein_coding
protein_coding
protein_coding
protein_coding
protein_coding
protein_coding
protein_coding
protein_coding
miRNA_precursor
protein_coding

protein_coding
protein_coding
protein_coding
protein_coding
protein_coding
protein_coding
protein_coding
protein_coding
protein_coding
protein_coding
protein_coding
protein_coding
protein_coding
protein_coding
protein_coding
protein_coding
protein_coding
protein_coding
ncRNA
protein_coding
protein_coding
protein_coding
protein_coding
protein_coding
protein_coding
protein_coding
protein_coding
protein_coding
protein_coding
protein_coding
protein_coding
protein_coding
protein_coding
protein_coding
protein_coding
protein_coding
protein_coding
protein_coding
protein_coding
protein_coding
protein_coding
protein_coding
protein_coding
protein_coding
protein_coding
protein_coding
protein_coding
ncRNA
protein_coding
protein_coding
protein_coding
ncRNA
protein_coding
protein_coding
small_nucle(ol)ar_rna
miRNA_precursor

protein_coding
protein_coding
protein_coding
protein_coding
protein_coding
protein_coding
protein_coding
protein_coding
protein_coding
protein_coding
protein_coding
protein_coding
protein_coding
protein_coding
protein_coding
protein_coding
protein_coding
protein_coding
protein_coding
protein_coding
protein_coding
protein_coding
protein_coding
protein_coding
ncRNA
protein_coding
protein_coding
protein_coding
protein_coding
protein_coding
protein_coding
protein_coding
protein_coding
protein_coding
protein_coding
protein_coding
protein_coding
protein_coding
protein_coding
protein_coding
protein_coding
protein_coding
protein_coding
protein_coding
protein_coding
protein_coding
protein_coding
protein_coding
protein_coding
ncRNA
protein_coding
protein_coding
protein_coding
protein_coding
protein_coding
protein_coding

protein_coding
protein_coding
protein_coding
protein_coding
protein_coding
protein_coding
protein_coding
protein_coding
protein_coding
protein_coding
small_nucle(ol)ar_rna
protein_coding
protein_coding
protein_coding
protein_coding
protein_coding
protein_coding
protein_coding
protein_coding
protein_coding
protein_coding
protein_coding
protein_coding
protein_coding
protein_coding
protein_coding
protein_coding
protein_coding
protein_coding
protein_coding
protein_coding
protein_coding
protein_coding
protein_coding
protein_coding
protein_coding
protein_coding
ncRNA
protein_coding
protein_coding
protein_coding
protein_coding
protein_coding
pre_trna
protein_coding
protein_coding
protein_coding
protein_coding
protein_coding
protein_coding
protein_coding
protein_coding
protein_coding
miRNA_precursor
protein_coding
protein_coding

protein_coding
protein_coding
protein_coding
protein_coding
protein_coding
protein_coding
protein_coding
protein_coding
pre_trna
protein_coding
protein_coding
protein_coding
protein_coding
protein_coding
protein_coding
protein_coding
protein_coding
protein_coding
protein_coding
protein_coding
protein_coding
protein_coding
protein_coding
protein_coding
protein_coding
protein_coding
protein_coding
protein_coding
protein_coding
protein_coding
protein_coding
protein_coding
protein_coding
protein_coding
protein_coding
protein_coding
protein_coding
protein_coding
ncRNA
ncRNA
protein_coding
protein_coding
protein_coding
protein_coding
protein_coding
protein_coding
protein_coding
protein_coding
ncRNA
protein_coding
protein_coding
protein_coding
protein_coding
protein_coding
protein_coding
protein_coding

protein_coding
small_nucle(ol)ar_rna
protein_coding
protein_coding
protein_coding
protein_coding
protein_coding
protein_coding
protein_coding
protein_coding
protein_coding
protein_coding
protein_coding
protein_coding
protein_coding
protein_coding
protein_coding
protein_coding
small_nucle(ol)ar_rna
small_nucle(ol)ar_rna
protein_coding
protein_coding
protein_coding
protein_coding
protein_coding
protein_coding
protein_coding
protein_coding
pre_trna
protein_coding
protein_coding
protein_coding
protein_coding
protein_coding
protein_coding
protein_coding
protein_coding
protein_coding
protein_coding
protein_coding
protein_coding
protein_coding
protein_coding
protein_coding
protein_coding
protein_coding
protein_coding
protein_coding
protein_coding
protein_coding
protein_coding
protein_coding
ncRNA
protein_coding
protein_coding
protein_coding

protein_coding
protein_coding
protein_coding
protein_coding
protein_coding
protein_coding
protein_coding
protein_coding
protein_coding
protein_coding
protein_coding
protein_coding
protein_coding
pre_trna
protein_coding
protein_coding
protein_coding
protein_coding
protein_coding
protein_coding
protein_coding
protein_coding
protein_coding
protein_coding
protein_coding
protein_coding
protein_coding
protein_coding
protein_coding
protein_coding
protein_coding
protein_coding
protein_coding
protein_coding
protein_coding
protein_coding
protein_coding
protein_coding
protein_coding
protein_coding
protein_coding
protein_coding
protein_coding
protein_coding
protein_coding
protein_coding
protein_coding
protein_coding
protein_coding
protein_coding
protein_coding
ncRNA
ncRNA
protein_coding
protein_coding
protein_coding

protein_coding
protein_coding
protein_coding
protein_coding
protein_coding
protein_coding
ncRNA
protein_coding
protein_coding
protein_coding
protein_coding
protein_coding
protein_coding
protein_coding
protein_coding
protein_coding
protein_coding
protein_coding
protein_coding
protein_coding
protein_coding
protein_coding
protein_coding
protein_coding
protein_coding
protein_coding
protein_coding
protein_coding
protein_coding
protein_coding
protein_coding
protein_coding
ncRNA
ncRNA
protein_coding
protein_coding
protein_coding
protein_coding
protein_coding
protein_coding
protein_coding
protein_coding
protein_coding
protein_coding
protein_coding
protein_coding
protein_coding
protein_coding
protein_coding
protein_coding
protein_coding
protein_coding
protein_coding
protein_coding
protein_coding
protein_coding

protein_coding
protein_coding
protein_coding
protein_coding
protein_coding
protein_coding
protein_coding
protein_coding
protein_coding
protein_coding
protein_coding
protein_coding
protein_coding
ncRNA
protein_coding
protein_coding
protein_coding
protein_coding
protein_coding
protein_coding
ncRNA
protein_coding
ncRNA
protein_coding
ncRNA
protein_coding
protein_coding
protein_coding
protein_coding
protein_coding
protein_coding
miRNA_precursor
protein_coding
protein_coding
protein_coding
ncRNA
protein_coding
protein_coding
protein_coding
protein_coding
protein_coding
protein_coding
protein_coding
protein_coding
protein_coding
protein_coding
protein_coding
protein_coding
protein_coding
protein_coding
protein_coding
protein_coding
protein_coding
protein_coding
protein_coding
protein_coding

protein_coding
protein_coding
protein_coding
protein_coding
protein_coding
protein_coding
protein_coding
protein_coding
protein_coding
protein_coding
protein_coding
protein_coding
protein_coding
protein_coding
protein_coding
protein_coding
protein_coding
protein_coding
protein_coding
protein_coding
protein_coding
ncRNA
protein_coding
protein_coding
protein_coding
protein_coding
protein_coding
protein_coding
protein_coding
ncRNA
protein_coding
protein_coding
protein_coding
protein_coding
protein_coding
protein_coding
protein_coding
protein_coding
protein_coding
protein_coding
protein_coding
protein_coding
protein_coding
protein_coding
protein_coding
protein_coding
protein_coding
protein_coding
protein_coding
protein_coding
protein_coding
pre_trna
protein_coding
protein_coding
protein_coding
protein_coding

protein_coding
protein_coding
protein_coding
protein_coding
protein_coding
ncRNA
protein_coding
protein_coding
protein_coding
protein_coding
protein_coding
protein_coding
protein_coding
protein_coding
protein_coding
protein_coding
protein_coding
protein_coding
protein_coding
protein_coding
protein_coding
protein_coding
protein_coding
ncRNA
protein_coding
protein_coding
protein_coding
protein_coding
protein_coding
protein_coding
protein_coding
protein_coding
protein_coding
protein_coding
protein_coding
protein_coding
protein_coding
protein_coding
protein_coding
protein_coding
protein_coding
protein_coding
protein_coding
protein_coding
protein_coding
protein_coding
protein_coding
protein_coding
ncRNA
protein_coding
protein_coding
protein_coding
protein_coding
protein_coding
protein_coding
protein_coding

protein_coding
protein_coding
protein_coding
protein_coding
ncRNA
protein_coding
protein_coding
protein_coding
protein_coding
protein_coding
protein_coding
protein_coding
protein_coding
protein_coding
protein_coding
protein_coding
protein_coding
protein_coding
protein_coding
protein_coding
protein_coding
protein_coding
protein_coding
protein_coding
protein_coding
protein_coding
protein_coding
protein_coding
protein_coding
protein_coding
ncRNA
protein_coding
protein_coding
protein_coding
protein_coding
protein_coding
protein_coding
protein_coding
protein_coding
protein_coding
protein_coding
protein_coding
protein_coding
protein_coding
protein_coding
protein_coding
protein_coding
protein_coding
protein_coding
protein_coding
protein_coding
protein_coding
protein_coding
protein_coding
protein_coding
protein_coding

protein_coding
protein_coding
protein_coding
protein_coding
protein_coding
protein_coding
protein_coding
miRNA_precursor
protein_coding
protein_coding
protein_coding
protein_coding
protein_coding
protein_coding
protein_coding
protein_coding
protein_coding
protein_coding
protein_coding
protein_coding
protein_coding
protein_coding
protein_coding
protein_coding
protein_coding
protein_coding
protein_coding
protein_coding
protein_coding
protein_coding
protein_coding
protein_coding
protein_coding
protein_coding
protein_coding
protein_coding
protein_coding
protein_coding
protein_coding
protein_coding
protein_coding
protein_coding
protein_coding
protein_coding
protein_coding
protein_coding
protein_coding
protein_coding
protein_coding
protein_coding
protein_coding
protein_coding
protein_coding
protein_coding
protein_coding
protein_coding

protein_coding
protein_coding
protein_coding
small_nucle(ol)ar_rna
protein_coding
protein_coding
protein_coding
protein_coding
protein_coding
protein_coding
protein_coding
protein_coding
protein_coding
protein_coding
protein_coding
protein_coding
protein_coding
protein_coding
protein_coding
protein_coding
protein_coding
protein_coding
protein_coding
protein_coding
protein_coding
protein_coding
protein_coding
protein_coding
protein_coding
protein_coding
protein_coding
protein_coding
protein_coding
protein_coding
ncRNA
protein_coding
protein_coding
protein_coding
protein_coding
protein_coding
protein_coding
protein_coding
protein_coding
protein_coding
protein_coding
protein_coding
protein_coding
protein_coding
protein_coding
protein_coding
protein_coding
protein_coding
protein_coding
protein_coding
protein_coding
protein_coding

protein_coding
protein_coding
protein_coding
protein_coding
protein_coding
protein_coding
protein_coding
protein_coding
protein_coding
pre_trna
protein_coding
protein_coding
protein_coding
protein_coding
protein_coding
ncRNA
ncRNA
protein_coding
protein_coding
protein_coding
protein_coding
protein_coding
protein_coding
protein_coding
protein_coding
protein_coding
protein_coding
protein_coding
protein_coding
protein_coding
protein_coding
protein_coding
protein_coding
protein_coding
protein_coding
protein_coding
protein_coding
protein_coding
protein_coding
ncRNA
protein_coding
protein_coding
protein_coding
protein_coding
protein_coding
protein_coding
protein_coding
protein_coding
protein_coding
protein_coding
protein_coding
protein_coding
protein_coding
protein_coding
protein_coding
protein_coding

protein_coding
protein_coding
protein_coding
protein_coding
protein_coding
protein_coding
protein_coding
protein_coding
protein_coding
protein_coding
protein_coding
protein_coding
protein_coding
protein_coding
ncRNA
protein_coding
protein_coding
protein_coding
protein_coding
protein_coding
protein_coding
protein_coding
protein_coding
protein_coding
protein_coding
protein_coding
protein_coding
protein_coding
pre_trna
protein_coding
protein_coding
protein_coding
protein_coding
protein_coding
protein_coding
protein_coding
protein_coding
protein_coding
protein_coding
protein_coding
protein_coding
protein_coding
protein_coding
ncRNA
protein_coding
protein_coding
protein_coding
protein_coding
protein_coding
protein_coding
protein_coding
protein_coding
protein_coding
protein_coding
protein_coding
protein_coding

protein_coding
protein_coding
protein_coding
protein_coding
protein_coding
protein_coding
protein_coding
protein_coding
protein_coding
protein_coding
protein_coding
protein_coding
protein_coding
protein_coding
protein_coding
protein_coding
protein_coding
protein_coding
protein_coding
ncRNA
protein_coding
protein_coding
protein_coding
protein_coding
protein_coding
protein_coding
protein_coding
protein_coding
protein_coding
protein_coding
protein_coding
protein_coding
protein_coding
protein_coding
protein_coding
protein_coding
miRNA_precursor
protein_coding
protein_coding
protein_coding
protein_coding
miRNA_precursor
ncRNA
miRNA_precursor
protein_coding
protein_coding
protein_coding
protein_coding
protein_coding
protein_coding
protein_coding
protein_coding
protein_coding
protein_coding
protein_coding
protein_coding

protein_coding
protein_coding
protein_coding
protein_coding
protein_coding
protein_coding
protein_coding
protein_coding
protein_coding
protein_coding
protein_coding
protein_coding
protein_coding
protein_coding
protein_coding
protein_coding
protein_coding
protein_coding
protein_coding
protein_coding
protein_coding
protein_coding
protein_coding
protein_coding
protein_coding
protein_coding
protein_coding
protein_coding
protein_coding
protein_coding
protein_coding
protein_coding
protein_coding
protein_coding
protein_coding
protein_coding
ncRNA
ncRNA
protein_coding
ncRNA
protein_coding
protein_coding
protein_coding
protein_coding
protein_coding
ncRNA
protein_coding
protein_coding
protein_coding
protein_coding
protein_coding
miRNA_precursor
protein_coding
protein_coding
protein_coding
protein_coding

protein_coding
protein_coding
ncRNA
protein_coding
protein_coding
protein_coding
protein_coding
protein_coding
protein_coding
protein_coding
protein_coding
protein_coding
protein_coding
ncRNA
protein_coding
protein_coding
protein_coding
protein_coding
protein_coding
protein_coding
protein_coding
protein_coding
protein_coding
protein_coding
protein_coding
protein_coding
protein_coding
protein_coding
protein_coding
protein_coding
protein_coding
protein_coding
protein_coding
protein_coding
protein_coding
protein_coding
protein_coding
protein_coding
protein_coding
pre_trna
protein_coding
protein_coding
protein_coding
protein_coding
protein_coding
protein_coding
protein_coding
protein_coding
protein_coding
protein_coding
protein_coding
protein_coding
protein_coding
protein_coding
protein_coding
protein_coding

protein_coding
protein_coding
protein_coding
protein_coding
protein_coding
protein_coding
protein_coding
protein_coding
protein_coding
protein_coding
protein_coding
protein_coding
protein_coding
protein_coding
protein_coding
protein_coding
protein_coding
protein_coding
protein_coding
protein_coding
protein_coding
protein_coding
protein_coding
protein_coding
protein_coding
protein_coding
small_nucle(ol)ar_rna
protein_coding
protein_coding
protein_coding
protein_coding
protein_coding
protein_coding
protein_coding
protein_coding
protein_coding
protein_coding
protein_coding
protein_coding
protein_coding
protein_coding
protein_coding
protein_coding
protein_coding
protein_coding
protein_coding
protein_coding
protein_coding
protein_coding
protein_coding
protein_coding
protein_coding
protein_coding
protein_coding
protein_coding
protein_coding

protein_coding
protein_coding
protein_coding
protein_coding
protein_coding
protein_coding
protein_coding
protein_coding
protein_coding
protein_coding
protein_coding
ncRNA
protein_coding
protein_coding
protein_coding
protein_coding
protein_coding
ncRNA
protein_coding
protein_coding
protein_coding
protein_coding
protein_coding
protein_coding
protein_coding
protein_coding
protein_coding
protein_coding
protein_coding
protein_coding
protein_coding
protein_coding
protein_coding
protein_coding
protein_coding
protein_coding
small_nucle(ol)ar_rna
small_nucle(ol)ar_rna
protein_coding
protein_coding
protein_coding
protein_coding
miRNA_precursor
protein_coding
protein_coding
protein_coding
protein_coding
protein_coding
protein_coding
protein_coding
ncRNA
protein_coding
protein_coding
protein_coding
protein_coding
protein_coding

protein_coding
protein_coding
protein_coding
protein_coding
ncRNA
protein_coding
protein_coding
protein_coding
protein_coding
protein_coding
protein_coding
protein_coding
protein_coding
protein_coding
protein_coding
protein_coding
small_nucle(ol)ar_rna
protein_coding
protein_coding
protein_coding
small_nucle(ol)ar_rna
protein_coding
protein_coding
protein_coding
protein_coding
protein_coding
protein_coding
protein_coding
protein_coding
protein_coding
protein_coding
protein_coding
protein_coding
protein_coding
protein_coding
protein_coding
protein_coding
protein_coding
protein_coding
protein_coding
protein_coding
protein_coding
protein_coding
protein_coding
protein_coding
protein_coding
protein_coding
protein_coding
protein_coding
protein_coding
protein_coding
protein_coding
protein_coding
protein_coding
protein_coding
protein_coding

protein_coding
protein_coding
protein_coding
protein_coding
protein_coding
protein_coding
protein_coding
protein_coding
small_nucle(ol)ar_rna
small_nucle(ol)ar_rna
protein_coding
protein_coding
protein_coding
ncRNA
protein_coding
protein_coding
protein_coding
protein_coding
protein_coding
protein_coding
protein_coding
protein_coding
protein_coding
protein_coding
protein_coding
protein_coding
protein_coding
protein_coding
protein_coding
protein_coding
protein_coding
protein_coding
protein_coding
protein_coding
protein_coding
protein_coding
protein_coding
protein_coding
protein_coding
protein_coding
protein_coding
ncRNA
protein_coding
protein_coding
protein_coding
miRNA_precursor
protein_coding
protein_coding
protein_coding
protein_coding
protein_coding
protein_coding
pre_trna
protein_coding
protein_coding
protein_coding

protein_coding
protein_coding
protein_coding
protein_coding
protein_coding
protein_coding
protein_coding
protein_coding
protein_coding
protein_coding
protein_coding
protein_coding
protein_coding
protein_coding
protein_coding
protein_coding
protein_coding
protein_coding
protein_coding
small_nucle(ol)ar_rna
protein_coding
protein_coding
protein_coding
protein_coding
protein_coding
protein_coding
protein_coding
protein_coding
ncRNA
protein_coding
protein_coding
protein_coding
ncRNA
protein_coding
protein_coding
protein_coding
protein_coding
protein_coding
protein_coding
protein_coding
protein_coding
protein_coding
protein_coding
protein_coding
protein_coding
protein_coding
protein_coding
protein_coding
protein_coding
protein_coding
protein_coding
protein_coding
protein_coding
protein_coding
protein_coding
protein_coding

protein_coding
protein_coding
protein_coding
protein_coding
protein_coding
protein_coding
protein_coding
protein_coding
protein_coding
protein_coding
ncRNA
protein_coding
protein_coding
protein_coding
protein_coding
protein_coding
protein_coding
protein_coding
protein_coding
protein_coding
protein_coding
protein_coding
protein_coding
protein_coding
protein_coding
protein_coding
protein_coding
protein_coding
protein_coding
protein_coding
protein_coding
protein_coding
protein_coding
protein_coding
protein_coding
protein_coding
protein_coding
protein_coding
protein_coding
protein_coding
protein_coding
protein_coding
protein_coding
protein_coding
protein_coding
protein_coding
protein_coding
protein_coding
protein_coding
small_nucle(ol)ar_rna
protein_coding
protein_coding
protein_coding
protein_coding
protein_coding
protein_coding

protein_coding
protein_coding
protein_coding
protein_coding
protein_coding
protein_coding
protein_coding
protein_coding
protein_coding
protein_coding
protein_coding
protein_coding
protein_coding
protein_coding
protein_coding
protein_coding
protein_coding
protein_coding
protein_coding
protein_coding
protein_coding
protein_coding
ncRNA
protein_coding
small_nucle(ol)ar_rna
protein_coding
protein_coding
protein_coding
protein_coding
protein_coding
protein_coding
protein_coding
protein_coding
protein_coding
ncRNA
protein_coding
protein_coding
protein_coding
protein_coding
protein_coding
protein_coding
protein_coding
protein_coding
protein_coding
protein_coding
protein_coding
protein_coding
pre_trna
protein_coding
protein_coding
ncRNA
protein_coding
protein_coding
protein_coding
protein_coding
protein_coding

protein_coding
protein_coding
protein_coding
protein_coding
protein_coding
protein_coding
protein_coding
protein_coding
protein_coding
protein_coding
protein_coding
protein_coding
protein_coding
protein_coding
protein_coding
protein_coding
protein_coding
protein_coding
protein_coding
protein_coding
protein_coding
protein_coding
protein_coding
protein_coding
protein_coding
protein_coding
protein_coding
protein_coding
protein_coding
protein_coding
protein_coding
protein_coding
protein_coding
protein_coding
protein_coding
protein_coding
protein_coding
protein_coding
pre_trna
protein_coding
protein_coding
protein_coding
protein_coding
protein_coding
protein_coding
protein_coding
protein_coding
protein_coding
protein_coding
protein_coding
protein_coding
protein_coding
protein_coding
protein_coding
protein_coding
protein_coding

protein_coding
protein_coding
protein_coding
protein_coding
protein_coding
protein_coding
protein_coding
protein_coding
protein_coding
protein_coding
protein_coding
protein_coding
protein_coding
protein_coding
protein_coding
protein_coding
protein_coding
protein_coding
protein_coding
ncRNA
protein_coding
protein_coding
protein_coding
protein_coding
protein_coding
ncRNA
protein_coding
protein_coding
protein_coding
protein_coding
protein_coding
protein_coding
protein_coding
protein_coding
protein_coding
protein_coding
protein_coding
protein_coding
protein_coding
protein_coding
protein_coding
protein_coding
protein_coding
protein_coding
protein_coding
protein_coding
protein_coding
protein_coding
protein_coding
protein_coding
protein_coding
protein_coding
pre_trna
ncRNA
protein_coding
protein_coding

protein_coding
ncRNA
protein_coding
protein_coding
protein_coding
pre_trna
protein_coding
protein_coding
protein_coding
protein_coding
protein_coding
protein_coding
protein_coding
protein_coding
protein_coding
protein_coding
protein_coding
protein_coding
protein_coding
protein_coding
protein_coding
protein_coding
protein_coding
protein_coding
protein_coding
protein_coding
protein_coding
protein_coding
protein_coding
protein_coding
protein_coding
protein_coding
protein_coding
protein_coding
protein_coding
protein_coding
protein_coding
small_nucle(ol)ar_rna
small_nucle(ol)ar_rna
pre_trna
protein_coding
protein_coding
protein_coding
protein_coding
protein_coding
protein_coding
protein_coding
protein_coding
protein_coding
protein_coding
protein_coding
protein_coding
protein_coding
protein_coding
pre_trna
protein_coding

ncRNA
protein_coding
protein_coding
protein_coding
protein_coding
protein_coding
protein_coding
ncRNA
protein_coding
protein_coding
protein_coding
protein_coding
protein_coding
protein_coding
protein_coding
protein_coding
protein_coding
protein_coding
protein_coding
protein_coding
protein_coding
protein_coding
protein_coding
protein_coding
miRNA_precursor
protein_coding
protein_coding
ncRNA
protein_coding
protein_coding
protein_coding
pre_trna
protein_coding
protein_coding
protein_coding
protein_coding
protein_coding
protein_coding
protein_coding
protein_coding
ncRNA
protein_coding
protein_coding
protein_coding
protein_coding
protein_coding
protein_coding
protein_coding
protein_coding
protein_coding
protein_coding
protein_coding
protein_coding
protein_coding
protein_coding
protein_coding

protein_coding
protein_coding
protein_coding
protein_coding
protein_coding
protein_coding
protein_coding
protein_coding
protein_coding
small_nucle(ol)ar_rna
protein_coding
protein_coding
protein_coding
protein_coding
ncRNA
protein_coding
protein_coding
protein_coding
protein_coding
protein_coding
protein_coding
protein_coding
protein_coding
protein_coding
protein_coding
protein_coding
protein_coding
protein_coding
protein_coding
protein_coding
protein_coding
protein_coding
protein_coding
protein_coding
small_nucle(ol)ar_rna
protein_coding
protein_coding
protein_coding
protein_coding
protein_coding
protein_coding
protein_coding
protein_coding
protein_coding
protein_coding
protein_coding
protein_coding
protein_coding
protein_coding
protein_coding
protein_coding
protein_coding
protein_coding
protein_coding
protein_coding
protein_coding

protein_coding
protein_coding
protein_coding
protein_coding
protein_coding
protein_coding
protein_coding
protein_coding
protein_coding
protein_coding
protein_coding
protein_coding
protein_coding
protein_coding
protein_coding
protein_coding
protein_coding
protein_coding
protein_coding
protein_coding
protein_coding
ncRNA
protein_coding
protein_coding
protein_coding
protein_coding
protein_coding
protein_coding
ncRNA
protein_coding
protein_coding
protein_coding
protein_coding
protein_coding
protein_coding
protein_coding
protein_coding
protein_coding
protein_coding
protein_coding
protein_coding
protein_coding
protein_coding
ncRNA
protein_coding
protein_coding
ncRNA
protein_coding
protein_coding
protein_coding
protein_coding
protein_coding
protein_coding
protein_coding
protein_coding
protein_coding

protein_coding
protein_coding
protein_coding
protein_coding
protein_coding
protein_coding
protein_coding
protein_coding
protein_coding
protein_coding
protein_coding
protein_coding
protein_coding
protein_coding
protein_coding
ncRNA
protein_coding
protein_coding
protein_coding
protein_coding
protein_coding
protein_coding
protein_coding
protein_coding
protein_coding
protein_coding
protein_coding
protein_coding
protein_coding
protein_coding
protein_coding
protein_coding
protein_coding
protein_coding
protein_coding
protein_coding
ncRNA
protein_coding
ncRNA
protein_coding
protein_coding
protein_coding
protein_coding
protein_coding
protein_coding
protein_coding
protein_coding
protein_coding
protein_coding
protein_coding
protein_coding
protein_coding
protein_coding
protein_coding
protein_coding
protein_coding

protein_coding
protein_coding
protein_coding
protein_coding
protein_coding
protein_coding
protein_coding
protein_coding
protein_coding
protein_coding
protein_coding
protein_coding
protein_coding
protein_coding
protein_coding
protein_coding
protein_coding
ncRNA
protein_coding
protein_coding
protein_coding
protein_coding
protein_coding
protein_coding
protein_coding
protein_coding
protein_coding
protein_coding
protein_coding
protein_coding
protein_coding
protein_coding
protein_coding
protein_coding
protein_coding
protein_coding
protein_coding
protein_coding
protein_coding
protein_coding
pre_trna
protein_coding
protein_coding
protein_coding
protein_coding
protein_coding
protein_coding
protein_coding
protein_coding
protein_coding
protein_coding
protein_coding
protein_coding
protein_coding
protein_coding
protein_coding

protein_coding
protein_coding
protein_coding
protein_coding
protein_coding
protein_coding
protein_coding
protein_coding
protein_coding
protein_coding
protein_coding
protein_coding
protein_coding
protein_coding
protein_coding
protein_coding
protein_coding
protein_coding
ncRNA
protein_coding
protein_coding
protein_coding
protein_coding
protein_coding
protein_coding
protein_coding
protein_coding
protein_coding
protein_coding
protein_coding
protein_coding
protein_coding
small_nucle(ol)ar_rna
protein_coding
protein_coding
protein_coding
protein_coding
ncRNA
protein_coding
protein_coding
protein_coding
protein_coding
protein_coding
protein_coding
protein_coding
protein_coding
protein_coding
protein_coding
ncRNA
protein_coding
protein_coding
protein_coding
ncRNA
protein_coding
protein_coding
protein_coding

protein_coding
protein_coding
protein_coding
protein_coding
protein_coding
protein_coding
protein_coding
protein_coding
protein_coding
protein_coding
protein_coding
protein_coding
protein_coding
protein_coding
protein_coding
protein_coding
protein_coding
protein_coding
protein_coding
ncRNA
protein_coding
protein_coding
protein_coding
protein_coding
ncRNA
protein_coding
protein_coding
protein_coding
protein_coding
protein_coding
pre_trna
protein_coding
protein_coding
protein_coding
protein_coding
ncRNA
protein_coding
protein_coding
ncRNA
protein_coding
protein_coding
protein_coding
protein_coding
protein_coding
protein_coding
protein_coding
protein_coding
protein_coding
protein_coding
protein_coding
protein_coding
protein_coding
protein_coding
ncRNA
protein_coding
protein_coding

protein_coding
small_nucle(ol)ar_rna
protein_coding
protein_coding
protein_coding
protein_coding
protein_coding
protein_coding
protein_coding
protein_coding
protein_coding
protein_coding
protein_coding
protein_coding
protein_coding
protein_coding
protein_coding
protein_coding
protein_coding
protein_coding
protein_coding
protein_coding
protein_coding
protein_coding
protein_coding
protein_coding
protein_coding
protein_coding
pre_trna
protein_coding
protein_coding
protein_coding
protein_coding
protein_coding
protein_coding
protein_coding
protein_coding
protein_coding
protein_coding
protein_coding
protein_coding
protein_coding
protein_coding
protein_coding
protein_coding
protein_coding
protein_coding
miRNA_precursor
protein_coding
protein_coding
protein_coding
protein_coding
protein_coding
protein_coding
protein_coding
protein_coding

protein_coding
protein_coding
protein_coding
protein_coding
protein_coding
protein_coding
protein_coding
protein_coding
protein_coding
protein_coding
protein_coding
protein_coding
protein_coding
protein_coding
protein_coding
protein_coding
protein_coding
protein_coding
ncRNA
protein_coding
ncRNA
protein_coding
protein_coding
protein_coding
protein_coding
protein_coding
protein_coding
protein_coding
protein_coding
protein_coding
protein_coding
protein_coding
protein_coding
protein_coding
protein_coding
protein_coding
protein_coding
protein_coding
protein_coding
protein_coding
protein_coding
protein_coding
protein_coding
small_nucle(ol)ar_rna
protein_coding
protein_coding
protein_coding
protein_coding
protein_coding
protein_coding
protein_coding
protein_coding
protein_coding
protein_coding
protein_coding
protein_coding

protein_coding
protein_coding
ncRNA
ncRNA
small_nucle(ol)ar_rna
protein_coding
protein_coding
protein_coding
protein_coding
protein_coding
protein_coding
protein_coding
protein_coding
protein_coding
small_nucle(ol)ar_rna
protein_coding
protein_coding
protein_coding
protein_coding
protein_coding
protein_coding
protein_coding
protein_coding
protein_coding
protein_coding
protein_coding
protein_coding
ncRNA
protein_coding
protein_coding
protein_coding
protein_coding
protein_coding
protein_coding
protein_coding
protein_coding
protein_coding
protein_coding
protein_coding
protein_coding
protein_coding
protein_coding
protein_coding
protein_coding
protein_coding
protein_coding
protein_coding
protein_coding
protein_coding
protein_coding
protein_coding
protein_coding
protein_coding
protein_coding
protein_coding
ncRNA

protein_coding
protein_coding
protein_coding
small_nucle(ol)ar_rna
protein_coding
protein_coding
protein_coding
protein_coding
protein_coding
protein_coding
protein_coding
protein_coding
protein_coding
protein_coding
protein_coding
protein_coding
protein_coding
protein_coding
protein_coding
protein_coding
protein_coding
protein_coding
protein_coding
protein_coding
protein_coding
ncRNA
protein_coding
protein_coding
protein_coding
protein_coding
protein_coding
protein_coding
protein_coding
protein_coding
ncRNA
protein_coding
protein_coding
protein_coding
protein_coding
protein_coding
protein_coding
protein_coding
protein_coding
protein_coding
protein_coding
protein_coding
protein_coding
protein_coding
protein_coding
protein_coding
protein_coding
protein_coding
protein_coding
protein_coding
small_nucle(ol)ar_rna
protein_coding

small_nucle(ol)ar_rna
protein_coding
protein_coding
ncRNA
protein_coding
protein_coding
protein_coding
protein_coding
protein_coding
protein_coding
ncRNA
protein_coding
protein_coding
ncRNA
protein_coding
protein_coding
protein_coding
protein_coding
ncRNA
miRNA_precursor
protein_coding
protein_coding
protein_coding
protein_coding
protein_coding
protein_coding
miRNA_precursor
protein_coding
protein_coding
small_nucle(ol)ar_rna
protein_coding
protein_coding
protein_coding
protein_coding
ncRNA
protein_coding
protein_coding
protein_coding
protein_coding
protein_coding
protein_coding
protein_coding
protein_coding
protein_coding
protein_coding
protein_coding
protein_coding
protein_coding
protein_coding
protein_coding
protein_coding
protein_coding
protein_coding
protein_coding
protein_coding
protein_coding

protein_coding
protein_coding
ncRNA
ncRNA
miRNA_precursor
protein_coding
ncRNA
protein_coding
protein_coding
protein_coding
protein_coding
protein_coding
protein_coding
protein_coding
protein_coding
protein_coding
protein_coding
protein_coding
protein_coding
protein_coding
protein_coding
ncRNA
protein_coding
protein_coding
protein_coding
protein_coding
protein_coding
protein_coding
protein_coding
protein_coding
protein_coding
protein_coding
protein_coding
protein_coding
protein_coding
protein_coding
protein_coding
protein_coding
protein_coding
protein_coding
protein_coding
protein_coding
protein_coding
protein_coding
ncRNA
protein_coding
protein_coding
protein_coding
protein_coding
protein_coding
protein_coding
protein_coding
protein_coding
protein_coding
protein_coding
protein_coding

protein_coding
ncRNA
protein_coding
protein_coding
ncRNA
protein_coding
protein_coding
ncRNA
protein_coding
protein_coding
ncRNA
small_nucle(ol)ar_rna
protein_coding
protein_coding
protein_coding
protein_coding
protein_coding
protein_coding
protein_coding
protein_coding
protein_coding
protein_coding
protein_coding
protein_coding
protein_coding
protein_coding
protein_coding
protein_coding
protein_coding
protein_coding
protein_coding
protein_coding
protein_coding
protein_coding
protein_coding
miRNA_precursor
protein_coding
protein_coding
protein_coding
protein_coding
protein_coding
protein_coding
protein_coding
protein_coding
protein_coding
protein_coding
protein_coding
protein_coding
protein_coding
protein_coding
protein_coding
protein_coding
protein_coding
protein_coding
protein_coding
ncRNA

protein_coding
protein_coding
protein_coding
protein_coding
protein_coding
protein_coding
ncRNA
protein_coding
protein_coding
protein_coding
protein_coding
ncRNA
protein_coding
protein_coding
protein_coding
ncRNA
protein_coding
protein_coding
protein_coding
protein_coding
protein_coding
protein_coding
ncRNA
protein_coding
protein_coding
protein_coding
protein_coding
protein_coding
protein_coding
protein_coding
protein_coding
ncRNA
protein_coding
protein_coding
protein_coding
protein_coding
protein_coding
protein_coding
protein_coding
protein_coding
protein_coding
protein_coding
miRNA_precursor
protein_coding
protein_coding
protein_coding
protein_coding
protein_coding
protein_coding
protein_coding
protein_coding
protein_coding
protein_coding
protein_coding
protein_coding
protein_coding

protein_coding
protein_coding
protein_coding
protein_coding
protein_coding
protein_coding
protein_coding
protein_coding
protein_coding
protein_coding
protein_coding
protein_coding
protein_coding
protein_coding
protein_coding
protein_coding
protein_coding
protein_coding
protein_coding
protein_coding
protein_coding
protein_coding
protein_coding
protein_coding
protein_coding
protein_coding
protein_coding
protein_coding
protein_coding
protein_coding
protein_coding
protein_coding
protein_coding
protein_coding
protein_coding
protein_coding
protein_coding
ncRNA
protein_coding
protein_coding
protein_coding
protein_coding
protein_coding
protein_coding
protein_coding
protein_coding
protein_coding
protein_coding
protein_coding
protein_coding
protein_coding
protein_coding
protein_coding
protein_coding
protein_coding
protein_coding

protein_coding
protein_coding
protein_coding
protein_coding
protein_coding
ncRNA
protein_coding
small_nucle(ol)ar_rna
small_nucle(ol)ar_rna
small_nucle(ol)ar_rna
protein_coding
protein_coding
protein_coding
protein_coding
protein_coding
protein_coding
protein_coding
protein_coding
protein_coding
protein_coding
protein_coding
protein_coding
protein_coding
protein_coding
protein_coding
protein_coding
protein_coding
protein_coding
protein_coding
protein_coding
protein_coding
protein_coding
protein_coding
protein_coding
protein_coding
protein_coding
protein_coding
protein_coding
protein_coding
protein_coding
protein_coding
protein_coding
protein_coding
protein_coding
protein_coding
protein_coding
protein_coding
protein_coding
protein_coding
protein_coding
protein_coding
protein_coding
protein_coding
protein_coding
protein_coding
protein_coding

ncRNA
protein_coding
protein_coding
protein_coding
protein_coding
protein_coding
protein_coding
protein_coding
protein_coding
protein_coding
protein_coding
protein_coding
protein_coding
protein_coding
protein_coding
protein_coding
protein_coding
protein_coding
protein_coding
protein_coding
protein_coding
protein_coding
protein_coding
protein_coding
protein_coding
protein_coding
protein_coding
protein_coding
protein_coding
protein_coding
protein_coding
ncRNA
protein_coding
protein_coding
protein_coding
protein_coding
protein_coding
protein_coding
protein_coding
protein_coding
protein_coding
protein_coding
protein_coding
protein_coding
protein_coding
protein_coding
protein_coding
protein_coding
protein_coding
protein_coding
protein_coding
protein_coding
protein_coding
protein_coding
protein_coding
protein_coding

protein_coding
protein_coding
protein_coding
protein_coding
ncRNA
protein_coding
protein_coding
protein_coding
protein_coding
protein_coding
protein_coding
protein_coding
protein_coding
protein_coding
protein_coding
protein_coding
protein_coding
protein_coding
protein_coding
protein_coding
protein_coding
protein_coding
protein_coding
protein_coding
protein_coding
protein_coding
protein_coding
protein_coding
protein_coding
protein_coding
ncRNA
protein_coding
protein_coding
protein_coding
protein_coding
protein_coding
protein_coding
protein_coding
protein_coding
protein_coding
protein_coding
protein_coding
ncRNA
protein_coding
protein_coding
protein_coding
protein_coding
protein_coding
protein_coding
protein_coding
protein_coding
protein_coding
protein_coding
protein_coding
protein_coding
protein_coding

protein_coding
protein_coding
protein_coding
protein_coding
protein_coding
protein_coding
protein_coding
protein_coding
protein_coding
protein_coding
protein_coding
protein_coding
protein_coding
protein_coding
protein_coding
protein_coding
protein_coding
protein_coding
protein_coding
protein_coding
miRNA_precursor
protein_coding
protein_coding
protein_coding
protein_coding
protein_coding
protein_coding
protein_coding
protein_coding
protein_coding
protein_coding
protein_coding
protein_coding
protein_coding
protein_coding
protein_coding
protein_coding
protein_coding
protein_coding
ncRNA
protein_coding
protein_coding
protein_coding
protein_coding
protein_coding
protein_coding
protein_coding
protein_coding
protein_coding
protein_coding
protein_coding
protein_coding
protein_coding
protein_coding
protein_coding
protein_coding

protein_coding
protein_coding
protein_coding
miRNA_precursor
protein_coding
protein_coding
protein_coding
protein_coding
protein_coding
protein_coding
protein_coding
protein_coding
protein_coding
protein_coding
protein_coding
protein_coding
protein_coding
protein_coding
protein_coding
protein_coding
protein_coding
protein_coding
protein_coding
protein_coding
protein_coding
protein_coding
protein_coding
protein_coding
protein_coding
protein_coding
protein_coding
protein_coding
protein_coding
protein_coding
protein_coding
protein_coding
protein_coding
protein_coding
protein_coding
protein_coding
ncRNA
protein_coding
protein_coding
protein_coding
protein_coding
protein_coding
protein_coding
protein_coding
protein_coding
protein_coding
protein_coding
protein_coding
protein_coding
protein_coding
protein_coding
protein_coding

protein_coding
protein_coding
ncRNA
protein_coding
protein_coding
protein_coding
protein_coding
protein_coding
protein_coding
protein_coding
ncRNA
protein_coding
protein_coding
protein_coding
protein_coding
protein_coding
protein_coding
protein_coding
protein_coding
protein_coding
protein_coding
protein_coding
protein_coding
protein_coding
protein_coding
protein_coding
protein_coding
protein_coding
protein_coding
protein_coding
protein_coding
protein_coding
protein_coding
protein_coding
protein_coding
protein_coding
protein_coding
protein_coding
protein_coding
protein_coding
protein_coding
protein_coding
small_nucle(ol)ar_rna
protein_coding
protein_coding
protein_coding
protein_coding
protein_coding
protein_coding
protein_coding
protein_coding
protein_coding
protein_coding
protein_coding
protein_coding
protein_coding

protein_coding
protein_coding
protein_coding
protein_coding
protein_coding
protein_coding
protein_coding
protein_coding
protein_coding
protein_coding
protein_coding
protein_coding
protein_coding
protein_coding
protein_coding
protein_coding
protein_coding
protein_coding
protein_coding
protein_coding
protein_coding
protein_coding
protein_coding
ncRNA
protein_coding
protein_coding
protein_coding
protein_coding
protein_coding
protein_coding
protein_coding
protein_coding
ncRNA
protein_coding
protein_coding
protein_coding
protein_coding
protein_coding
protein_coding
protein_coding
protein_coding
protein_coding
protein_coding
protein_coding
protein_coding
protein_coding
protein_coding
protein_coding
protein_coding
protein_coding
protein_coding
protein_coding
protein_coding
protein_coding
protein_coding
protein_coding

protein_coding
protein_coding
protein_coding
protein_coding
protein_coding
protein_coding
protein_coding
protein_coding
protein_coding
protein_coding
protein_coding
protein_coding
protein_coding
ncRNA
protein_coding
protein_coding
protein_coding
protein_coding
protein_coding
protein_coding
protein_coding
protein_coding
protein_coding
protein_coding
protein_coding
protein_coding
protein_coding
small_nucle(ol)ar_rna
small_nucle(ol)ar_rna
protein_coding
protein_coding
protein_coding
protein_coding
protein_coding
protein_coding
protein_coding
protein_coding
protein_coding
protein_coding
protein_coding
protein_coding
protein_coding
protein_coding
protein_coding
ncRNA
protein_coding
protein_coding
protein_coding
protein_coding
protein_coding
protein_coding
protein_coding
protein_coding
protein_coding
protein_coding
protein_coding

protein_coding
protein_coding
protein_coding
protein_coding
protein_coding
protein_coding
protein_coding
protein_coding
protein_coding
protein_coding
protein_coding
protein_coding
protein_coding
protein_coding
protein_coding
protein_coding
protein_coding
protein_coding
protein_coding
protein_coding
protein_coding
protein_coding
protein_coding
protein_coding
protein_coding
protein_coding
protein_coding
protein_coding
protein_coding
protein_coding
protein_coding
protein_coding
ncRNA
protein_coding
protein_coding
protein_coding
protein_coding
protein_coding
protein_coding
protein_coding
protein_coding
protein_coding
protein_coding
protein_coding
protein_coding
protein_coding
protein_coding
protein_coding
protein_coding
protein_coding
protein_coding
protein_coding
protein_coding
protein_coding
protein_coding
ncRNA

protein_coding
protein_coding
protein_coding
protein_coding
protein_coding
protein_coding
protein_coding
protein_coding
protein_coding
protein_coding
protein_coding
pre_trna
protein_coding
protein_coding
protein_coding
protein_coding
protein_coding
protein_coding
protein_coding
protein_coding
ncRNA
protein_coding
protein_coding
protein_coding
protein_coding
protein_coding
protein_coding
protein_coding
protein_coding
protein_coding
protein_coding
protein_coding
protein_coding
protein_coding
protein_coding
protein_coding
protein_coding
protein_coding
protein_coding
protein_coding
protein_coding
protein_coding
ncRNA
protein_coding
ncRNA
protein_coding
ncRNA
protein_coding
protein_coding
protein_coding
protein_coding
protein_coding
protein_coding
protein_coding
protein_coding
protein_coding

protein_coding
protein_coding
protein_coding
protein_coding
protein_coding
protein_coding
protein_coding
protein_coding
protein_coding
protein_coding
protein_coding
protein_coding
protein_coding
protein_coding
ncRNA
protein_coding
protein_coding
protein_coding
protein_coding
pre_trna
protein_coding
protein_coding
protein_coding
protein_coding
protein_coding
protein_coding
ncRNA
ncRNA
protein_coding
protein_coding
small_nucle(ol)ar_rna
protein_coding
protein_coding
protein_coding
protein_coding
ncRNA
protein_coding
protein_coding
protein_coding
ncRNA
protein_coding
ncRNA
protein_coding
protein_coding
protein_coding
protein_coding
protein_coding
protein_coding
protein_coding
protein_coding
protein_coding
protein_coding
protein_coding
protein_coding
protein_coding
protein_coding

protein_coding
protein_coding
protein_coding
protein_coding
protein_coding
protein_coding
ncRNA
protein_coding
protein_coding
ncRNA
protein_coding
protein_coding
ncRNA
protein_coding
protein_coding
protein_coding
protein_coding
protein_coding
ncRNA
protein_coding
protein_coding
ncRNA
protein_coding
protein_coding
protein_coding
protein_coding
protein_coding
protein_coding
protein_coding
protein_coding
protein_coding
protein_coding
protein_coding
protein_coding
protein_coding
protein_coding
protein_coding
protein_coding
protein_coding
protein_coding
protein_coding
protein_coding
protein_coding
protein_coding
protein_coding
protein_coding
protein_coding
ncRNA
protein_coding
protein_coding
protein_coding
protein_coding
protein_coding
protein_coding
protein_coding
protein_coding

protein_coding
protein_coding
protein_coding
protein_coding
protein_coding
protein_coding
protein_coding
pre_trna
protein_coding
protein_coding
protein_coding
protein_coding
protein_coding
protein_coding
protein_coding
protein_coding
ncRNA
ncRNA
protein_coding
protein_coding
protein_coding
protein_coding
protein_coding
miRNA_precursor
protein_coding
protein_coding
protein_coding
protein_coding
protein_coding
protein_coding
protein_coding
protein_coding
protein_coding
protein_coding
protein_coding
protein_coding
protein_coding
protein_coding
protein_coding
protein_coding
protein_coding
protein_coding
protein_coding
protein_coding
protein_coding
protein_coding
ncRNA
protein_coding
small_nucle(ol)ar_rna
protein_coding
protein_coding
protein_coding
protein_coding
protein_coding
protein_coding
protein_coding

protein_coding
protein_coding
protein_coding
protein_coding
protein_coding
ncRNA
protein_coding
protein_coding
protein_coding
protein_coding
protein_coding
protein_coding
protein_coding
protein_coding
protein_coding
protein_coding
protein_coding
protein_coding
protein_coding
protein_coding
protein_coding
protein_coding
protein_coding
protein_coding
protein_coding
ncRNA
protein_coding
protein_coding
protein_coding
protein_coding
protein_coding
protein_coding
protein_coding
protein_coding
protein_coding
protein_coding
protein_coding
protein_coding
protein_coding
protein_coding
protein_coding
protein_coding
protein_coding
miRNA_precursor
protein_coding
protein_coding
protein_coding
protein_coding
protein_coding
protein_coding
protein_coding
protein_coding
protein_coding
protein_coding
protein_coding
protein_coding

protein_coding
protein_coding
protein_coding
protein_coding
protein_coding
ncRNA
ncRNA
protein_coding
protein_coding
protein_coding
protein_coding
ncRNA
protein_coding
protein_coding
protein_coding
protein_coding
miRNA_precursor
protein_coding
protein_coding
ncRNA
protein_coding
protein_coding
protein_coding
protein_coding
protein_coding
protein_coding
protein_coding
protein_coding
protein_coding
protein_coding
protein_coding
protein_coding
protein_coding
protein_coding
protein_coding
protein_coding
protein_coding
protein_coding
protein_coding
protein_coding
protein_coding
protein_coding
protein_coding
protein_coding
protein_coding
ncRNA
protein_coding
protein_coding
protein_coding
protein_coding
protein_coding
protein_coding
protein_coding
protein_coding
protein_coding
protein_coding

protein_coding
protein_coding
protein_coding
protein_coding
protein_coding
protein_coding
protein_coding
protein_coding
protein_coding
protein_coding
protein_coding
protein_coding
protein_coding
protein_coding
protein_coding
protein_coding
protein_coding
protein_coding
protein_coding
protein_coding
protein_coding
protein_coding
protein_coding
protein_coding
protein_coding
miRNA_precursor
protein_coding
protein_coding
protein_coding
protein_coding
protein_coding
ncRNA
protein_coding
protein_coding
protein_coding
protein_coding
pre_trna
protein_coding
protein_coding
protein_coding
protein_coding
ncRNA
protein_coding
protein_coding
protein_coding
protein_coding
protein_coding
protein_coding
protein_coding
protein_coding
protein_coding
protein_coding
protein_coding
protein_coding
protein_coding
protein_coding

protein_coding
protein_coding
protein_coding
protein_coding
protein_coding
protein_coding
protein_coding
protein_coding
pre_trna
protein_coding
ncRNA
protein_coding
protein_coding
protein_coding
protein_coding
protein_coding
protein_coding
protein_coding
protein_coding
protein_coding
ncRNA
protein_coding
protein_coding
protein_coding
protein_coding
protein_coding
protein_coding
protein_coding
protein_coding
protein_coding
ncRNA
protein_coding
protein_coding
protein_coding
protein_coding
ncRNA
protein_coding
protein_coding
protein_coding
protein_coding
protein_coding
protein_coding
protein_coding
protein_coding
protein_coding
protein_coding
protein_coding
protein_coding
protein_coding
protein_coding
protein_coding
protein_coding
protein_coding
protein_coding
protein_coding
protein_coding

protein_coding
ncRNA
protein_coding
protein_coding
protein_coding
protein_coding
protein_coding
protein_coding
protein_coding
protein_coding
protein_coding
protein_coding
protein_coding
protein_coding
protein_coding
protein_coding
protein_coding
protein_coding
protein_coding
protein_coding
protein_coding
ncRNA
protein_coding
protein_coding
protein_coding
protein_coding
protein_coding
protein_coding
protein_coding
protein_coding
protein_coding
protein_coding
protein_coding
protein_coding
protein_coding
protein_coding
protein_coding
protein_coding
protein_coding
protein_coding
protein_coding
protein_coding
protein_coding
protein_coding
protein_coding
protein_coding
protein_coding
protein_coding
protein_coding
protein_coding
protein_coding
protein_coding
protein_coding
protein_coding
small_nucle(ol)ar_rna
small_nucle(ol)ar_rna

small_nucle(ol)ar_rna
ncRNA
protein_coding
protein_coding
protein_coding
protein_coding
protein_coding
protein_coding
protein_coding
protein_coding
protein_coding
protein_coding
protein_coding
protein_coding
protein_coding
protein_coding
protein_coding
protein_coding
protein_coding
protein_coding
protein_coding
protein_coding
protein_coding
protein_coding
protein_coding
protein_coding
protein_coding
protein_coding
protein_coding
protein_coding
protein_coding
protein_coding
protein_coding
protein_coding
protein_coding
protein_coding
protein_coding
protein_coding
protein_coding
protein_coding
protein_coding
protein_coding
protein_coding
protein_coding
protein_coding
protein_coding
protein_coding
protein_coding
ncRNA
small_nucle(ol)ar_rna
small_nucle(ol)ar_rna
ncRNA
protein_coding
protein_coding
protein_coding
protein_coding

protein_coding
protein_coding
protein_coding
protein_coding
protein_coding
protein_coding
protein_coding
protein_coding
protein_coding
protein_coding
protein_coding
protein_coding
protein_coding
protein_coding
protein_coding
protein_coding
protein_coding
ncRNA
protein_coding
protein_coding
protein_coding
protein_coding
ncRNA
protein_coding
protein_coding
protein_coding
protein_coding
protein_coding
protein_coding
miRNA_precursor
protein_coding
protein_coding
ncRNA
protein_coding
protein_coding
protein_coding
protein_coding
protein_coding
protein_coding
protein_coding
protein_coding
protein_coding
protein_coding
protein_coding
protein_coding
protein_coding
protein_coding
protein_coding
protein_coding
protein_coding
protein_coding
protein_coding
protein_coding
protein_coding
protein_coding
protein_coding

protein_coding
protein_coding
protein_coding
protein_coding
ncRNA
protein_coding
protein_coding
protein_coding
protein_coding
protein_coding
protein_coding
protein_coding
protein_coding
miRNA_precursor
protein_coding
protein_coding
protein_coding
protein_coding
protein_coding
protein_coding
protein_coding
protein_coding
protein_coding
protein_coding
protein_coding
protein_coding
pre_trna
protein_coding
protein_coding
protein_coding
ncRNA
protein_coding
protein_coding
protein_coding
protein_coding
protein_coding
protein_coding
protein_coding
protein_coding
protein_coding
protein_coding
protein_coding
protein_coding
protein_coding
protein_coding
protein_coding
protein_coding
protein_coding
protein_coding
protein_coding
protein_coding
protein_coding
protein_coding
protein_coding
protein_coding
protein_coding

protein_coding
protein_coding
protein_coding
protein_coding
protein_coding
protein_coding
ncRNA
protein_coding
protein_coding
protein_coding
protein_coding
ncRNA
protein_coding
protein_coding
protein_coding
ncRNA
protein_coding
protein_coding
protein_coding
ncRNA
protein_coding
protein_coding
protein_coding
protein_coding
protein_coding
protein_coding
protein_coding
protein_coding
protein_coding
protein_coding
protein_coding
protein_coding
protein_coding
protein_coding
protein_coding
protein_coding
protein_coding
protein_coding
protein_coding
ncRNA
protein_coding
protein_coding
protein_coding
protein_coding
protein_coding
protein_coding
protein_coding
protein_coding
protein_coding
protein_coding
protein_coding
protein_coding
protein_coding
protein_coding
protein_coding
protein_coding

protein_coding
protein_coding
protein_coding
protein_coding
protein_coding
protein_coding
protein_coding
protein_coding
protein_coding
protein_coding
protein_coding
protein_coding
protein_coding
protein_coding
protein_coding
protein_coding
protein_coding
protein_coding
protein_coding
protein_coding
protein_coding
protein_coding
protein_coding
protein_coding
protein_coding
protein_coding
protein_coding
protein_coding
protein_coding
protein_coding
protein_coding
protein_coding
protein_coding
protein_coding
protein_coding
protein_coding
protein_coding
ncRNA
protein_coding
protein_coding
protein_coding
protein_coding
protein_coding
protein_coding
protein_coding
protein_coding
protein_coding
protein_coding
protein_coding
protein_coding
protein_coding
protein_coding
protein_coding
protein_coding
protein_coding
protein_coding

protein_coding
protein_coding
protein_coding
protein_coding
protein_coding
protein_coding
protein_coding
protein_coding
protein_coding
protein_coding
protein_coding
protein_coding
protein_coding
protein_coding
protein_coding
protein_coding
protein_coding
protein_coding
protein_coding
protein_coding
protein_coding
protein_coding
protein_coding
protein_coding
ncRNA
protein_coding
protein_coding
protein_coding
protein_coding
protein_coding
protein_coding
protein_coding
protein_coding
protein_coding
protein_coding
protein_coding
protein_coding
protein_coding
protein_coding
protein_coding
protein_coding
protein_coding
protein_coding
protein_coding
ncRNA
protein_coding
protein_coding
protein_coding
protein_coding
protein_coding
protein_coding
protein_coding
protein_coding
protein_coding
protein_coding
protein_coding

protein_coding
protein_coding
protein_coding
protein_coding
protein_coding
protein_coding
protein_coding
protein_coding
protein_coding
protein_coding
pre_trna
protein_coding
protein_coding
protein_coding
protein_coding
protein_coding
protein_coding
protein_coding
small_nucle(ol)ar_rna
protein_coding
protein_coding
protein_coding
protein_coding
protein_coding
protein_coding
pre_trna
protein_coding
ncRNA
protein_coding
protein_coding
protein_coding
protein_coding
protein_coding
protein_coding
protein_coding
protein_coding
protein_coding
protein_coding
protein_coding
protein_coding
protein_coding
protein_coding
protein_coding
protein_coding
protein_coding
protein_coding
protein_coding
protein_coding
protein_coding
protein_coding
protein_coding
protein_coding
protein_coding
protein_coding
protein_coding
protein_coding

protein_coding
protein_coding
protein_coding
protein_coding
protein_coding
protein_coding
protein_coding
protein_coding
protein_coding
protein_coding
protein_coding
protein_coding
protein_coding
protein_coding
protein_coding
protein_coding
protein_coding
protein_coding
protein_coding
protein_coding
protein_coding
protein_coding
protein_coding
protein_coding
protein_coding
protein_coding
protein_coding
protein_coding
protein_coding
protein_coding
protein_coding
protein_coding
protein_coding
protein_coding
protein_coding
protein_coding
protein_coding
protein_coding
protein_coding
protein_coding
protein_coding
protein_coding
protein_coding
protein_coding
protein_coding
protein_coding
protein_coding
miRNA_precursor
protein_coding
protein_coding
small_nucle(ol)ar_rna
small_nucle(ol)ar_rna
ncRNA
protein_coding
protein_coding
protein_coding

protein_coding
protein_coding
protein_coding
protein_coding
protein_coding
protein_coding
protein_coding
protein_coding
protein_coding
protein_coding
protein_coding
protein_coding
protein_coding
protein_coding
protein_coding
protein_coding
protein_coding
protein_coding
protein_coding
protein_coding
protein_coding
protein_coding
protein_coding
protein_coding
protein_coding
protein_coding
protein_coding
protein_coding
protein_coding
protein_coding
protein_coding
protein_coding
protein_coding
ncRNA
protein_coding
protein_coding
protein_coding
protein_coding
protein_coding
ncRNA
protein_coding
protein_coding
protein_coding
protein_coding
protein_coding
protein_coding
protein_coding
protein_coding
protein_coding
protein_coding
protein_coding
protein_coding
protein_coding
protein_coding
protein_coding
protein_coding

protein_coding
protein_coding
protein_coding
protein_coding
protein_coding
protein_coding
protein_coding
protein_coding
protein_coding
protein_coding
protein_coding
protein_coding
protein_coding
protein_coding
protein_coding
protein_coding
protein_coding
protein_coding
protein_coding
protein_coding
protein_coding
protein_coding
protein_coding
protein_coding
protein_coding
protein_coding
protein_coding
protein_coding
protein_coding
protein_coding
ncRNA
protein_coding
protein_coding
protein_coding
protein_coding
protein_coding
protein_coding
protein_coding
protein_coding
protein_coding
protein_coding
protein_coding
protein_coding
protein_coding
protein_coding
miRNA_precursor
protein_coding
protein_coding
protein_coding
protein_coding
protein_coding
protein_coding
protein_coding
protein_coding
protein_coding
protein_coding

protein_coding
protein_coding
protein_coding
protein_coding
small_nucle(ol)ar_rna
small_nucle(ol)ar_rna
protein_coding
protein_coding
protein_coding
protein_coding
protein_coding
protein_coding
protein_coding
protein_coding
protein_coding
protein_coding
small_nucle(ol)ar_rna
protein_coding
protein_coding
protein_coding
protein_coding
protein_coding
protein_coding
protein_coding
protein_coding
protein_coding
protein_coding
protein_coding
protein_coding
protein_coding
protein_coding
protein_coding
protein_coding
protein_coding
protein_coding
protein_coding
protein_coding
ncRNA
protein_coding
protein_coding
protein_coding
protein_coding
protein_coding
protein_coding
protein_coding
protein_coding
protein_coding
protein_coding
protein_coding
protein_coding
protein_coding
protein_coding
protein_coding
protein_coding
protein_coding
protein_coding

protein_coding
protein_coding
protein_coding
protein_coding
protein_coding
protein_coding
protein_coding
protein_coding
protein_coding
protein_coding
protein_coding
protein_coding
protein_coding
pre_trna
protein_coding
ncRNA
protein_coding
protein_coding
protein_coding
protein_coding
protein_coding
protein_coding
protein_coding
protein_coding
protein_coding
protein_coding
protein_coding
protein_coding
protein_coding
protein_coding
protein_coding
protein_coding
protein_coding
protein_coding
protein_coding
protein_coding
protein_coding
protein_coding
protein_coding
protein_coding
protein_coding
protein_coding
protein_coding
protein_coding
protein_coding
protein_coding
protein_coding
protein_coding
protein_coding
protein_coding
protein_coding
protein_coding
protein_coding
protein_coding
protein_coding
protein_coding

protein_coding
protein_coding
protein_coding
protein_coding
protein_coding
protein_coding
protein_coding
protein_coding
protein_coding
protein_coding
protein_coding
protein_coding
ncRNA
protein_coding
protein_coding
protein_coding
protein_coding
protein_coding
protein_coding
protein_coding
protein_coding
protein_coding
protein_coding
protein_coding
protein_coding
protein_coding
protein_coding
protein_coding
protein_coding
protein_coding
protein_coding
protein_coding
ncRNA
small_nucle(ol)ar_rna
protein_coding
protein_coding
protein_coding
protein_coding
protein_coding
protein_coding
protein_coding
protein_coding
protein_coding
protein_coding
protein_coding
protein_coding
ncRNA
protein_coding
protein_coding
protein_coding
protein_coding
protein_coding
protein_coding
protein_coding
protein_coding
protein_coding

| |
|---|
| protein_coding |
| protein_coding |
| protein_coding |
| protein_coding |
| protein_coding |
| protein_coding |
| protein_coding |
| protein_coding |
| protein_coding |
| protein_coding |
| protein_coding |
| protein_coding |
| protein_coding |
| protein_coding |
| protein_coding |
| protein_coding |
| protein_coding |
| miRNA_precursor |
| miRNA_precursor |
| protein_coding |
| protein_coding |
| protein_coding |
| protein_coding |
| protein_coding |
| protein_coding |
| protein_coding |
| protein_coding |
| protein_coding |
| protein_coding |
| protein_coding |
| protein_coding |
| protein_coding |
| protein_coding |
| ncRNA |
| ncRNA |
| protein_coding |
| protein_coding |
| protein_coding |
| protein_coding |
| protein_coding |
| protein_coding |
| protein_coding |
| protein_coding |
| protein_coding |
| protein_coding |
| ncRNA |
| protein_coding |
| protein_coding |
| protein_coding |
| protein_coding |
| pre_trna |
| protein_coding |
| protein_coding |
| protein_coding |
| protein_coding |
| protein_coding |

protein_coding
protein_coding
protein_coding
protein_coding
protein_coding
protein_coding
ncRNA
protein_coding
protein_coding
protein_coding
protein_coding
protein_coding
protein_coding
ncRNA
protein_coding
protein_coding
ncRNA
protein_coding
protein_coding
protein_coding
protein_coding
protein_coding
protein_coding
protein_coding
protein_coding
protein_coding
protein_coding
protein_coding
protein_coding
protein_coding
protein_coding
protein_coding
protein_coding
protein_coding
protein_coding
protein_coding
protein_coding
protein_coding
protein_coding
protein_coding
protein_coding
protein_coding
protein_coding
protein_coding
protein_coding
miRNA_precursor
protein_coding
protein_coding
protein_coding
protein_coding
protein_coding
ncRNA
ncRNA
protein_coding
protein_coding
protein_coding

protein_coding
protein_coding
protein_coding
protein_coding
protein_coding
protein_coding
protein_coding
protein_coding
protein_coding
protein_coding
protein_coding
protein_coding
protein_coding
protein_coding
protein_coding
protein_coding
protein_coding
protein_coding
protein_coding
protein_coding
protein_coding
ncRNA
protein_coding
protein_coding
miRNA_precursor
protein_coding
miRNA_precursor
protein_coding
protein_coding
protein_coding
protein_coding
protein_coding
protein_coding
protein_coding
protein_coding
protein_coding
protein_coding
protein_coding
protein_coding
protein_coding
protein_coding
protein_coding
protein_coding
protein_coding
ncRNA
protein_coding
protein_coding
protein_coding
protein_coding
protein_coding
protein_coding
protein_coding
protein_coding
protein_coding
protein_coding
protein_coding

protein_coding
protein_coding
protein_coding
protein_coding
protein_coding
protein_coding
protein_coding
protein_coding
protein_coding
protein_coding
protein_coding
protein_coding
protein_coding
protein_coding
protein_coding
protein_coding
protein_coding
protein_coding
protein_coding
protein_coding
protein_coding
protein_coding
protein_coding
protein_coding
protein_coding
protein_coding
protein_coding
protein_coding
protein_coding
protein_coding
protein_coding
protein_coding
protein_coding
protein_coding
protein_coding
protein_coding
protein_coding
protein_coding
protein_coding
protein_coding
protein_coding
ncRNA
protein_coding
protein_coding
protein_coding
protein_coding
protein_coding
protein_coding
protein_coding
protein_coding
protein_coding
protein_coding
protein_coding
protein_coding
protein_coding
protein_coding

protein_coding
protein_coding
protein_coding
protein_coding
protein_coding
protein_coding
protein_coding
protein_coding
protein_coding
protein_coding
protein_coding
protein_coding
protein_coding
protein_coding
protein_coding
protein_coding
protein_coding
protein_coding
protein_coding
protein_coding
protein_coding
protein_coding
small_nucle(ol)ar_rna
protein_coding
protein_coding
protein_coding
pre_trna
protein_coding
pre_trna
protein_coding
protein_coding
protein_coding
protein_coding
protein_coding
protein_coding
protein_coding
protein_coding
protein_coding
protein_coding
protein_coding
ncRNA
protein_coding
protein_coding
ncRNA
ncRNA
protein_coding
protein_coding
protein_coding
protein_coding
protein_coding
protein_coding
protein_coding
protein_coding
protein_coding
protein_coding
protein_coding

protein_coding
miRNA_precursor
miRNA_precursor
protein_coding
protein_coding
protein_coding
protein_coding
protein_coding
protein_coding
protein_coding
protein_coding
protein_coding
protein_coding
protein_coding
protein_coding
protein_coding
protein_coding
protein_coding
protein_coding
protein_coding
protein_coding
protein_coding
protein_coding
protein_coding
protein_coding
protein_coding
protein_coding
protein_coding
protein_coding
protein_coding
protein_coding
protein_coding
protein_coding
protein_coding
protein_coding
protein_coding
protein_coding
protein_coding
protein_coding
protein_coding
protein_coding
protein_coding
protein_coding
protein_coding
protein_coding
protein_coding
protein_coding
protein_coding
protein_coding
protein_coding
protein_coding
protein_coding
protein_coding
protein_coding
protein_coding
protein_coding

protein_coding
protein_coding
protein_coding
protein_coding
pre_trna
protein_coding
protein_coding
protein_coding
protein_coding
protein_coding
protein_coding
protein_coding
protein_coding
protein_coding
protein_coding
protein_coding
protein_coding
protein_coding
protein_coding
protein_coding
protein_coding
protein_coding
protein_coding
protein_coding
protein_coding
protein_coding
protein_coding
protein_coding
protein_coding
ncRNA
protein_coding
ncRNA
protein_coding
protein_coding
protein_coding
protein_coding
protein_coding
protein_coding
ncRNA
protein_coding
protein_coding
protein_coding
protein_coding
protein_coding
protein_coding
small_nucle(ol)ar_rna
protein_coding
protein_coding
protein_coding
protein_coding
protein_coding
protein_coding
ncRNA
protein_coding
protein_coding
protein_coding

protein_coding
protein_coding
protein_coding
protein_coding
protein_coding
protein_coding
protein_coding
protein_coding
protein_coding
protein_coding
protein_coding
protein_coding
protein_coding
protein_coding
protein_coding
protein_coding
protein_coding
protein_coding
protein_coding
protein_coding
protein_coding
protein_coding
protein_coding
protein_coding
protein_coding
protein_coding
protein_coding
protein_coding
protein_coding
protein_coding
ncRNA
protein_coding
protein_coding
protein_coding
ncRNA
protein_coding
protein_coding
protein_coding
protein_coding
protein_coding
protein_coding
protein_coding
protein_coding
protein_coding
protein_coding
protein_coding
ncRNA
protein_coding
protein_coding
protein_coding
protein_coding
protein_coding
protein_coding
protein_coding
protein_coding
protein_coding

protein_coding
protein_coding
protein_coding
protein_coding
protein_coding
protein_coding
protein_coding
protein_coding
protein_coding
protein_coding
protein_coding
protein_coding
protein_coding
protein_coding
protein_coding
protein_coding
protein_coding
ncRNA
protein_coding
protein_coding
ncRNA
ncRNA
protein_coding
protein_coding
protein_coding
protein_coding
protein_coding
protein_coding
protein_coding
protein_coding
protein_coding
protein_coding
protein_coding
protein_coding
protein_coding
protein_coding
protein_coding
protein_coding
protein_coding
protein_coding
protein_coding
protein_coding
protein_coding
protein_coding
protein_coding
protein_coding
protein_coding
protein_coding
protein_coding
protein_coding
protein_coding
protein_coding
ncRNA
protein_coding
protein_coding
protein_coding

protein_coding
protein_coding
protein_coding
protein_coding
protein_coding
protein_coding
protein_coding
protein_coding
protein_coding
pre_trna
protein_coding
protein_coding
protein_coding
protein_coding
protein_coding
protein_coding
protein_coding
protein_coding
protein_coding
protein_coding
protein_coding
protein_coding
protein_coding
protein_coding
ncRNA
protein_coding
protein_coding
protein_coding
protein_coding
protein_coding
protein_coding
protein_coding
protein_coding
protein_coding
protein_coding
protein_coding
protein_coding
protein_coding
protein_coding
protein_coding
small_nucle(ol)ar_rna
protein_coding
protein_coding
protein_coding
protein_coding
protein_coding
protein_coding
protein_coding
protein_coding
protein_coding
protein_coding
protein_coding
protein_coding
protein_coding
protein_coding
protein_coding

protein_coding
protein_coding
protein_coding
protein_coding
protein_coding
small_nucle(ol)ar_rna
small_nucle(ol)ar_rna
protein_coding
protein_coding
protein_coding
protein_coding
pre_trna
miRNA_precursor
protein_coding
protein_coding
protein_coding
protein_coding
protein_coding
protein_coding
protein_coding
protein_coding
ncRNA
protein_coding
ncRNA
ncRNA
protein_coding
protein_coding
protein_coding
protein_coding
protein_coding
protein_coding
protein_coding
protein_coding
protein_coding
protein_coding
protein_coding
protein_coding
protein_coding
protein_coding
protein_coding
protein_coding
protein_coding
ncRNA
protein_coding
protein_coding
protein_coding
protein_coding
protein_coding
protein_coding
protein_coding
protein_coding
protein_coding
protein_coding
protein_coding
protein_coding
ncRNA

protein_coding
protein_coding
protein_coding
miRNA_precursor
protein_coding
protein_coding
protein_coding
ncRNA
protein_coding
protein_coding
protein_coding
protein_coding
protein_coding
protein_coding
protein_coding
protein_coding
protein_coding
protein_coding
protein_coding
protein_coding
protein_coding
protein_coding
protein_coding
protein_coding
protein_coding
protein_coding
protein_coding
ncRNA
protein_coding
protein_coding
protein_coding
protein_coding
protein_coding
protein_coding
protein_coding
protein_coding
protein_coding
protein_coding
protein_coding
protein_coding
protein_coding
protein_coding
protein_coding
miRNA_precursor
protein_coding
protein_coding
protein_coding
protein_coding
protein_coding
protein_coding
protein_coding
protein_coding
protein_coding
protein_coding
protein_coding
protein_coding

miRNA_precursor
pre_trna
protein_coding
protein_coding
protein_coding
protein_coding
protein_coding
ncRNA
protein_coding
protein_coding
protein_coding
miRNA_precursor
protein_coding
protein_coding
protein_coding
protein_coding
protein_coding
miRNA_precursor
protein_coding
ncRNA
protein_coding
protein_coding
protein_coding
protein_coding
protein_coding
protein_coding
protein_coding
protein_coding
protein_coding
protein_coding
protein_coding
protein_coding
protein_coding
protein_coding
protein_coding
protein_coding
protein_coding
protein_coding
protein_coding
protein_coding
protein_coding
protein_coding
protein_coding
protein_coding
protein_coding
protein_coding
protein_coding
protein_coding
protein_coding
protein_coding
protein_coding
protein_coding
protein_coding
protein_coding
protein_coding
protein_coding

protein_coding
protein_coding
protein_coding
protein_coding
protein_coding
protein_coding
protein_coding
protein_coding
protein_coding
protein_coding
protein_coding
protein_coding
protein_coding
protein_coding
protein_coding
protein_coding
miRNA_precursor
protein_coding
protein_coding
protein_coding
protein_coding
protein_coding
protein_coding
protein_coding
protein_coding
protein_coding
protein_coding
ncRNA
small_nucle(ol)ar_rna
small_nucle(ol)ar_rna
small_nucle(ol)ar_rna
protein_coding
protein_coding
protein_coding
protein_coding
protein_coding
ncRNA
protein_coding
protein_coding
protein_coding
protein_coding
protein_coding
protein_coding
protein_coding
protein_coding
protein_coding
protein_coding
protein_coding
protein_coding
protein_coding
protein_coding
protein_coding
protein_coding
protein_coding
protein_coding
protein_coding

protein_coding
protein_coding
protein_coding
protein_coding
protein_coding
ncRNA
protein_coding
protein_coding
protein_coding
protein_coding
protein_coding
protein_coding
protein_coding
protein_coding
protein_coding
protein_coding
protein_coding
protein_coding
protein_coding
protein_coding
protein_coding
protein_coding
protein_coding
protein_coding
protein_coding
protein_coding
protein_coding
protein_coding
protein_coding
protein_coding
protein_coding
protein_coding
protein_coding
protein_coding
protein_coding
protein_coding
protein_coding
protein_coding
protein_coding
protein_coding
protein_coding
pre_trna
protein_coding
protein_coding
protein_coding
protein_coding
protein_coding
protein_coding
protein_coding
protein_coding
protein_coding
protein_coding
protein_coding
protein_coding
protein_coding
protein_coding

ncRNA
protein_coding
protein_coding
protein_coding
protein_coding
protein_coding
protein_coding
protein_coding
protein_coding
miRNA_precursor
protein_coding
protein_coding
protein_coding
protein_coding
protein_coding
protein_coding
protein_coding
protein_coding
protein_coding
protein_coding
ncRNA
protein_coding
protein_coding
protein_coding
protein_coding
protein_coding
protein_coding
protein_coding
protein_coding
protein_coding
protein_coding
protein_coding
ncRNA
protein_coding
protein_coding
protein_coding
protein_coding
protein_coding
protein_coding
protein_coding
protein_coding
protein_coding
protein_coding
protein_coding
protein_coding
protein_coding
protein_coding
protein_coding
protein_coding
protein_coding
protein_coding
protein_coding
protein_coding
protein_coding
pre_trna
protein_coding

protein_coding
protein_coding
protein_coding
protein_coding
protein_coding
protein_coding
protein_coding
protein_coding
protein_coding
protein_coding
protein_coding
protein_coding
protein_coding
protein_coding
protein_coding
protein_coding
protein_coding
protein_coding
ncRNA
protein_coding
protein_coding
protein_coding
protein_coding
protein_coding
protein_coding
protein_coding
protein_coding
protein_coding
protein_coding
ncRNA
protein_coding
protein_coding
ncRNA
small_nucle(ol)ar_rna
protein_coding
protein_coding
protein_coding
protein_coding
protein_coding
protein_coding
protein_coding
protein_coding
protein_coding
protein_coding
protein_coding
ncRNA
protein_coding
protein_coding
protein_coding
ncRNA
protein_coding
protein_coding
protein_coding
protein_coding
protein_coding
protein_coding

protein_coding
protein_coding
protein_coding
protein_coding
protein_coding
protein_coding
protein_coding
ncRNA
protein_coding
protein_coding
protein_coding
protein_coding
protein_coding
protein_coding
protein_coding
protein_coding
protein_coding
protein_coding
pre_trna
protein_coding
protein_coding
protein_coding
protein_coding
protein_coding
protein_coding
protein_coding
protein_coding
protein_coding
protein_coding
protein_coding
protein_coding
protein_coding
protein_coding
protein_coding
protein_coding
miRNA_precursor
small_nucle(ol)ar_rna
small_nucle(ol)ar_rna
protein_coding
pre_trna
protein_coding
protein_coding
protein_coding
protein_coding
protein_coding
protein_coding
protein_coding
ncRNA
protein_coding
protein_coding
protein_coding
protein_coding
protein_coding
ncRNA
protein_coding
protein_coding

protein_coding
protein_coding
protein_coding
protein_coding
protein_coding
protein_coding
protein_coding
protein_coding
protein_coding
protein_coding
protein_coding
protein_coding
protein_coding
pre_trna
protein_coding
protein_coding
protein_coding
protein_coding
protein_coding
protein_coding
protein_coding
protein_coding
protein_coding
protein_coding
protein_coding
protein_coding
protein_coding
protein_coding
protein_coding
protein_coding
protein_coding
protein_coding
protein_coding
protein_coding
protein_coding
protein_coding
protein_coding
protein_coding
small_nucle(ol)ar_rna
protein_coding
protein_coding
protein_coding
protein_coding
protein_coding
protein_coding
protein_coding
protein_coding
protein_coding
protein_coding
protein_coding
protein_coding
protein_coding
protein_coding
protein_coding
protein_coding
protein_coding

protein_coding
protein_coding
protein_coding
protein_coding
protein_coding
protein_coding
protein_coding
protein_coding
protein_coding
protein_coding
protein_coding
protein_coding
protein_coding
protein_coding
protein_coding
protein_coding
protein_coding
protein_coding
protein_coding
protein_coding
protein_coding
protein_coding
protein_coding
protein_coding
protein_coding
protein_coding
protein_coding
protein_coding
protein_coding
protein_coding
protein_coding
protein_coding
protein_coding
protein_coding
protein_coding
protein_coding
protein_coding
protein_coding
protein_coding
protein_coding
protein_coding
protein_coding
protein_coding
protein_coding
protein_coding
protein_coding
protein_coding
protein_coding
protein_coding
protein_coding
protein_coding
protein_coding
protein_coding
protein_coding
protein_coding
protein_coding

protein_coding
protein_coding
protein_coding
protein_coding
protein_coding
protein_coding
protein_coding
protein_coding
protein_coding
protein_coding
protein_coding
protein_coding
protein_coding
protein_coding
protein_coding
protein_coding
protein_coding
protein_coding
protein_coding
protein_coding
protein_coding
protein_coding
protein_coding
protein_coding
protein_coding
protein_coding
protein_coding
protein_coding
protein_coding
protein_coding
protein_coding
ncRNA
protein_coding
protein_coding
ncRNA
protein_coding
protein_coding
protein_coding
protein_coding
protein_coding
protein_coding
protein_coding
protein_coding
protein_coding
protein_coding
protein_coding
protein_coding
protein_coding
protein_coding
protein_coding
protein_coding
protein_coding
protein_coding
protein_coding
protein_coding
protein_coding

protein_coding
protein_coding
protein_coding
protein_coding
protein_coding
protein_coding
protein_coding
protein_coding
protein_coding
protein_coding
protein_coding
protein_coding
protein_coding
protein_coding
protein_coding
protein_coding
protein_coding
protein_coding
protein_coding
protein_coding
protein_coding
protein_coding
protein_coding
protein_coding
ncRNA
protein_coding
protein_coding
protein_coding
protein_coding
protein_coding
protein_coding
protein_coding
protein_coding
ncRNA
protein_coding
protein_coding
protein_coding
ncRNA
protein_coding
protein_coding
protein_coding
protein_coding
protein_coding
protein_coding
protein_coding
protein_coding
protein_coding
protein_coding
protein_coding
protein_coding
protein_coding
protein_coding
protein_coding
protein_coding
ncRNA
protein_coding

protein_coding
protein_coding
protein_coding
ncRNA
protein_coding
protein_coding
protein_coding
protein_coding
protein_coding
protein_coding
protein_coding
protein_coding
protein_coding
protein_coding
protein_coding
protein_coding
protein_coding
protein_coding
protein_coding
protein_coding
ncRNA
protein_coding
protein_coding
protein_coding
protein_coding
protein_coding
protein_coding
pre_trna
protein_coding
protein_coding
protein_coding
protein_coding
protein_coding
protein_coding
protein_coding
protein_coding
protein_coding
protein_coding
protein_coding
protein_coding
protein_coding
protein_coding
protein_coding
protein_coding
ncRNA
pre_trna
protein_coding
protein_coding
protein_coding
protein_coding
protein_coding
protein_coding
protein_coding
ncRNA
protein_coding
protein_coding

protein_coding
ncRNA
protein_coding
protein_coding
pre_trna
protein_coding
protein_coding
protein_coding
protein_coding
protein_coding
protein_coding
protein_coding
protein_coding
protein_coding
protein_coding
protein_coding
protein_coding
protein_coding
protein_coding
protein_coding
protein_coding
protein_coding
protein_coding
protein_coding
protein_coding
protein_coding
protein_coding
protein_coding
protein_coding
protein_coding
protein_coding
protein_coding
protein_coding
protein_coding
protein_coding
protein_coding
protein_coding
protein_coding
protein_coding
protein_coding
protein_coding
protein_coding
protein_coding
protein_coding
protein_coding
protein_coding
protein_coding
protein_coding
protein_coding
protein_coding
protein_coding
protein_coding
protein_coding
protein_coding
ncRNA
protein_coding

protein_coding
protein_coding
protein_coding
protein_coding
protein_coding
protein_coding
protein_coding
protein_coding
protein_coding
protein_coding
protein_coding
protein_coding
protein_coding
protein_coding
protein_coding
protein_coding
protein_coding
protein_coding
protein_coding
ncRNA
protein_coding
ncRNA
protein_coding
protein_coding
protein_coding
protein_coding
ncRNA
protein_coding
protein_coding
protein_coding
protein_coding
protein_coding
protein_coding
small_nucle(ol)ar_rna
small_nucle(ol)ar_rna
protein_coding
protein_coding
protein_coding
protein_coding
protein_coding
protein_coding
protein_coding
protein_coding
protein_coding
protein_coding
protein_coding
protein_coding
protein_coding
ncRNA
protein_coding
ncRNA
protein_coding
protein_coding
protein_coding
protein_coding
protein_coding

protein_coding
pre_trna
protein_coding
protein_coding
protein_coding
protein_coding
pre_trna
protein_coding
protein_coding
protein_coding
protein_coding
small_nucle(ol)ar_rna
small_nucle(ol)ar_rna
small_nucle(ol)ar_rna
protein_coding
protein_coding
protein_coding
protein_coding
protein_coding
protein_coding
protein_coding
protein_coding
protein_coding
protein_coding
protein_coding
protein_coding
ncRNA
protein_coding
protein_coding
protein_coding
protein_coding
protein_coding
protein_coding
protein_coding
protein_coding
protein_coding
protein_coding
pre_trna
protein_coding
protein_coding
protein_coding
protein_coding
protein_coding
protein_coding
protein_coding
protein_coding
protein_coding
protein_coding
protein_coding
protein_coding
ncRNA
protein_coding
protein_coding
protein_coding
protein_coding
protein_coding

protein_coding
protein_coding
ncRNA
protein_coding
protein_coding
protein_coding
protein_coding
protein_coding
protein_coding
protein_coding
protein_coding
protein_coding
protein_coding
protein_coding
protein_coding
protein_coding
protein_coding
protein_coding
protein_coding
protein_coding
protein_coding
protein_coding
protein_coding
protein_coding
protein_coding
protein_coding
protein_coding
protein_coding
protein_coding
protein_coding
protein_coding
protein_coding
protein_coding
protein_coding
protein_coding
protein_coding
protein_coding
ncRNA
small_nucle(ol)ar_rna
protein_coding
protein_coding
protein_coding
protein_coding
protein_coding
protein_coding
protein_coding
protein_coding
protein_coding
protein_coding
protein_coding
protein_coding
protein_coding
protein_coding
protein_coding
protein_coding
protein_coding

protein_coding
protein_coding
protein_coding
protein_coding
protein_coding
protein_coding
protein_coding
protein_coding
small_nucle(ol)ar_rna
protein_coding
protein_coding
protein_coding
protein_coding
protein_coding
protein_coding
protein_coding
protein_coding
protein_coding
protein_coding
protein_coding
protein_coding
protein_coding
protein_coding
protein_coding
protein_coding
protein_coding
protein_coding
protein_coding
protein_coding
protein_coding
protein_coding
protein_coding
protein_coding
protein_coding
protein_coding
protein_coding
protein_coding
protein_coding
protein_coding
protein_coding
protein_coding
protein_coding
protein_coding
protein_coding
protein_coding
protein_coding
protein_coding
protein_coding
protein_coding
protein_coding
protein_coding
protein_coding
protein_coding
protein_coding
protein_coding
protein_coding

protein_coding
protein_coding
protein_coding
protein_coding
protein_coding
protein_coding
protein_coding
small_nucle(ol)ar_rna
small_nucle(ol)ar_rna
small_nucle(ol)ar_rna
protein_coding
protein_coding
protein_coding
protein_coding
protein_coding
protein_coding
protein_coding
protein_coding
protein_coding
protein_coding
protein_coding
protein_coding
protein_coding
protein_coding
protein_coding
protein_coding
protein_coding
protein_coding
protein_coding
protein_coding
protein_coding
protein_coding
protein_coding
protein_coding
protein_coding
protein_coding
ncRNA
protein_coding
protein_coding
protein_coding
protein_coding
protein_coding
protein_coding
ncRNA
protein_coding
protein_coding
protein_coding
protein_coding
protein_coding
protein_coding
protein_coding
protein_coding
protein_coding
protein_coding
protein_coding
protein_coding

protein_coding
protein_coding
protein_coding
protein_coding
protein_coding
protein_coding
protein_coding
protein_coding
protein_coding
ncRNA
protein_coding
protein_coding
protein_coding
protein_coding
protein_coding
protein_coding
protein_coding
protein_coding
protein_coding
protein_coding
protein_coding
protein_coding
protein_coding
protein_coding
protein_coding
protein_coding
protein_coding
protein_coding
protein_coding
miRNA_precursor
protein_coding
protein_coding
protein_coding
protein_coding
protein_coding
protein_coding
protein_coding
protein_coding
protein_coding
protein_coding
protein_coding
protein_coding
protein_coding
protein_coding
protein_coding
protein_coding
protein_coding
protein_coding
protein_coding
protein_coding
protein_coding
miRNA_precursor
protein_coding
protein_coding
protein_coding
protein_coding

protein_coding
miRNA_precursor
protein_coding
protein_coding
protein_coding
protein_coding
protein_coding
protein_coding
protein_coding
protein_coding
protein_coding
protein_coding
protein_coding
protein_coding
protein_coding
protein_coding
protein_coding
protein_coding
protein_coding
protein_coding
protein_coding
protein_coding
protein_coding
protein_coding
protein_coding
protein_coding
protein_coding
protein_coding
protein_coding
protein_coding
protein_coding
protein_coding
protein_coding
protein_coding
protein_coding
protein_coding
protein_coding
protein_coding
protein_coding
protein_coding
ncRNA
protein_coding
protein_coding
protein_coding
protein_coding
protein_coding
protein_coding
protein_coding
protein_coding
protein_coding
protein_coding
protein_coding
protein_coding
ncRNA
protein_coding
protein_coding

protein_coding
protein_coding
protein_coding
protein_coding
small_nucle(ol)ar_rna
protein_coding
protein_coding
protein_coding
protein_coding
protein_coding
protein_coding
protein_coding
protein_coding
ncRNA
protein_coding
protein_coding
miRNA_precursor
protein_coding
protein_coding
protein_coding
protein_coding
protein_coding
protein_coding
protein_coding
protein_coding
protein_coding
ncRNA
protein_coding
protein_coding
protein_coding
protein_coding
protein_coding
pre_trna
ncRNA
protein_coding
protein_coding
protein_coding
protein_coding
protein_coding
protein_coding
protein_coding
protein_coding
ncRNA
protein_coding
ncRNA
protein_coding
protein_coding
protein_coding
protein_coding
protein_coding
protein_coding
protein_coding
protein_coding
protein_coding
protein_coding
protein_coding

protein_coding
protein_coding
ncRNA
protein_coding
protein_coding
ncRNA
protein_coding
protein_coding
protein_coding
ncRNA
protein_coding
protein_coding
protein_coding
protein_coding
protein_coding
protein_coding
protein_coding
protein_coding
protein_coding
protein_coding
protein_coding
protein_coding
protein_coding
protein_coding
protein_coding
protein_coding
protein_coding
protein_coding
protein_coding
protein_coding
protein_coding
protein_coding
protein_coding
protein_coding
protein_coding
protein_coding
ncRNA
protein_coding
protein_coding
protein_coding
protein_coding
protein_coding
protein_coding
protein_coding
protein_coding
protein_coding
protein_coding
protein_coding
protein_coding
protein_coding
protein_coding
protein_coding
protein_coding
protein_coding
protein_coding
ncRNA

protein_coding
protein_coding
ncRNA
protein_coding
protein_coding
protein_coding
ncRNA
protein_coding
protein_coding
protein_coding
ncRNA
protein_coding
protein_coding
protein_coding
protein_coding
protein_coding
protein_coding
protein_coding
ncRNA
protein_coding
protein_coding
protein_coding
protein_coding
protein_coding
ncRNA
protein_coding
miRNA_precursor
protein_coding
protein_coding
protein_coding
protein_coding
protein_coding
protein_coding
protein_coding
protein_coding
protein_coding
protein_coding
protein_coding
protein_coding
protein_coding
protein_coding
protein_coding
protein_coding
protein_coding
protein_coding
protein_coding
protein_coding
protein_coding
protein_coding
ncRNA
protein_coding
protein_coding
protein_coding
protein_coding
protein_coding
protein_coding

protein_coding
protein_coding
protein_coding
protein_coding
protein_coding
protein_coding
small_nucle(ol)ar_rna
protein_coding
protein_coding
protein_coding
protein_coding
protein_coding
protein_coding
protein_coding
protein_coding
protein_coding
protein_coding
protein_coding
protein_coding
protein_coding
protein_coding
protein_coding
protein_coding
protein_coding
protein_coding
protein_coding
protein_coding
protein_coding
protein_coding
protein_coding
protein_coding
protein_coding
protein_coding
protein_coding
protein_coding
protein_coding
protein_coding
protein_coding
protein_coding
protein_coding
protein_coding
protein_coding
protein_coding
protein_coding
protein_coding
protein_coding
protein_coding
protein_coding
protein_coding
pre_trna
protein_coding
protein_coding
protein_coding
protein_coding
protein_coding
ncRNA

protein_coding
protein_coding
protein_coding
protein_coding
protein_coding
protein_coding
protein_coding
protein_coding
protein_coding
protein_coding
protein_coding
protein_coding
protein_coding
protein_coding
protein_coding
protein_coding
protein_coding
protein_coding
protein_coding
protein_coding
protein_coding
protein_coding
protein_coding
protein_coding
protein_coding
protein_coding
protein_coding
protein_coding
protein_coding
protein_coding
protein_coding
protein_coding
protein_coding
protein_coding
protein_coding
protein_coding
protein_coding
protein_coding
protein_coding
protein_coding
protein_coding
protein_coding
protein_coding
protein_coding
protein_coding
protein_coding
protein_coding
protein_coding
protein_coding
protein_coding
protein_coding
protein_coding
protein_coding
protein_coding
protein_coding
protein_coding

protein_coding
protein_coding
protein_coding
protein_coding
protein_coding
protein_coding
protein_coding
protein_coding
miRNA_precursor
protein_coding
protein_coding
protein_coding
protein_coding
ncRNA
protein_coding
protein_coding
protein_coding
protein_coding
protein_coding
ncRNA
protein_coding
ncRNA
protein_coding
protein_coding
protein_coding
protein_coding
protein_coding
protein_coding
protein_coding
protein_coding
protein_coding
protein_coding
protein_coding
protein_coding
protein_coding
protein_coding
protein_coding
protein_coding
protein_coding
protein_coding
protein_coding
protein_coding
protein_coding
protein_coding
protein_coding
protein_coding
protein_coding
protein_coding
protein_coding
protein_coding
protein_coding
protein_coding
protein_coding
protein_coding
protein_coding
protein_coding

protein_coding
protein_coding
protein_coding
protein_coding
protein_coding
protein_coding
protein_coding
protein_coding
protein_coding
protein_coding
protein_coding
protein_coding
protein_coding
protein_coding
protein_coding
protein_coding
protein_coding
protein_coding
protein_coding
protein_coding
protein_coding
protein_coding
protein_coding
protein_coding
protein_coding
protein_coding
protein_coding
protein_coding
protein_coding
protein_coding
protein_coding
protein_coding
protein_coding
protein_coding
miRNA_precursor
protein_coding
protein_coding
protein_coding
protein_coding
ncRNA
protein_coding
protein_coding
ncRNA
protein_coding
protein_coding
protein_coding
protein_coding
protein_coding
protein_coding
protein_coding
protein_coding
protein_coding
protein_coding
protein_coding
protein_coding
protein_coding

protein_coding
protein_coding
protein_coding
protein_coding
protein_coding
protein_coding
protein_coding
protein_coding
protein_coding
protein_coding
protein_coding
protein_coding
protein_coding
protein_coding
protein_coding
protein_coding
protein_coding
protein_coding
protein_coding
protein_coding
protein_coding
protein_coding
pre_trna
protein_coding
protein_coding
protein_coding
protein_coding
protein_coding
protein_coding
protein_coding
protein_coding
protein_coding
protein_coding
protein_coding
ncRNA
ncRNA
protein_coding
protein_coding
protein_coding
protein_coding
protein_coding
protein_coding
protein_coding
protein_coding
protein_coding
miRNA_precursor
protein_coding
protein_coding
protein_coding
protein_coding
protein_coding
protein_coding
protein_coding
protein_coding
protein_coding
protein_coding

protein_coding
protein_coding
protein_coding
protein_coding
protein_coding
protein_coding
protein_coding
protein_coding
protein_coding
protein_coding
protein_coding
protein_coding
protein_coding
protein_coding
protein_coding
protein_coding
protein_coding
protein_coding
protein_coding
protein_coding
protein_coding
protein_coding
protein_coding
protein_coding
protein_coding
protein_coding
protein_coding
protein_coding
protein_coding
protein_coding
protein_coding
protein_coding
protein_coding
protein_coding
protein_coding
protein_coding
small_nucle(ol)ar_rna
protein_coding
protein_coding
protein_coding
protein_coding
protein_coding
protein_coding
protein_coding
protein_coding
protein_coding
protein_coding
protein_coding
protein_coding
ncRNA
protein_coding
protein_coding
ncRNA
protein_coding
protein_coding
protein_coding

protein_coding
protein_coding
protein_coding
protein_coding
protein_coding
protein_coding
protein_coding
protein_coding
protein_coding
protein_coding
protein_coding
protein_coding
protein_coding
protein_coding
protein_coding
protein_coding
protein_coding
protein_coding
protein_coding
protein_coding
protein_coding
protein_coding
protein_coding
protein_coding
protein_coding
protein_coding
protein_coding
small_nucle(ol)ar_rna
protein_coding
protein_coding
protein_coding
protein_coding
protein_coding
protein_coding
protein_coding
protein_coding
protein_coding
protein_coding
protein_coding
protein_coding
protein_coding
protein_coding
protein_coding
protein_coding
protein_coding
protein_coding
protein_coding
protein_coding
protein_coding
protein_coding
protein_coding
protein_coding
protein_coding
protein_coding
protein_coding
protein_coding

protein_coding
protein_coding
protein_coding
protein_coding
protein_coding
protein_coding
protein_coding
protein_coding
protein_coding
protein_coding
protein_coding
protein_coding
protein_coding
protein_coding
protein_coding
protein_coding
protein_coding
protein_coding
protein_coding
protein_coding
protein_coding
protein_coding
protein_coding
protein_coding
protein_coding
protein_coding
protein_coding
protein_coding
protein_coding
protein_coding
protein_coding
protein_coding
protein_coding
protein_coding
protein_coding
protein_coding
protein_coding
protein_coding
protein_coding
protein_coding
protein_coding
protein_coding
protein_coding
protein_coding
protein_coding
protein_coding
protein_coding
protein_coding
protein_coding
protein_coding
protein_coding
protein_coding
protein_coding
protein_coding
protein_coding
protein_coding

protein_coding
protein_coding
protein_coding
protein_coding
protein_coding
protein_coding
protein_coding
ncRNA
protein_coding
protein_coding
protein_coding
protein_coding
protein_coding
protein_coding
protein_coding
protein_coding
protein_coding
protein_coding
protein_coding
protein_coding
protein_coding
protein_coding
ncRNA
protein_coding
protein_coding
protein_coding
protein_coding
protein_coding
protein_coding
protein_coding
protein_coding
protein_coding
protein_coding
protein_coding
protein_coding
protein_coding
protein_coding
protein_coding
protein_coding
protein_coding
protein_coding
protein_coding
protein_coding
ncRNA
protein_coding
protein_coding
protein_coding
protein_coding
protein_coding
protein_coding
protein_coding
protein_coding

**Dataset 1.4 Differential analysis of 21/22nt accumulation on Araport11 genes in** [illegible]

| ID | baseMean | log2FoldChange | lfcSE | stat | pvalue | padj |
|---|---|---|---|---|---|---|
| AT5G23065 | 1776,68 | -3,27 | 0,26 | -12,35 | 4,74E-35 | 2,95E-32 |
| AT4G23713 | 128,07 | 6,05 | 0,72 | 8,44 | 3,12E-17 | 9,70E-15 |
| AT2G26211 | 125,25 | -2,00 | 0,26 | -7,81 | 5,66E-15 | 1,17E-12 |
| AT4G03039 | 93,29 | 6,06 | 0,79 | 7,65 | 2,00E-14 | 2,22E-12 |
| AT4G03038 | 93,33 | 6,07 | 0,79 | 7,65 | 1,98E-14 | 2,22E-12 |
| AT4G04365 | 93,18 | 6,06 | 0,79 | 7,64 | 2,14E-14 | 2,22E-12 |
| AT5G60408 | 515,37 | 2,25 | 0,30 | 7,56 | 4,04E-14 | 3,59E-12 |
| AT4G13493 | 160,77 | -2,40 | 0,32 | -7,40 | 1,36E-13 | 1,05E-11 |
| AT3G44570 | 48,95 | -7,85 | 1,13 | -6,96 | 3,45E-12 | 2,38E-10 |
| AT3G44580 | 31,33 | -3,18 | 0,49 | -6,47 | 9,74E-11 | 6,06E-09 |
| AT1G66725 | 1003,08 | 1,95 | 0,31 | 6,34 | 2,26E-10 | 1,28E-08 |
| AT3G25795 | 22,21 | 8,16 | 1,29 | 6,33 | 2,48E-10 | 1,28E-08 |
| AT4G21595 | 28,61 | 7,80 | 1,24 | 6,28 | 3,48E-10 | 1,66E-08 |
| AT3G10745 | 164,34 | -2,12 | 0,34 | -6,25 | 4,06E-10 | 1,81E-08 |
| AT1G53683 | 15,44 | -7,97 | 1,29 | -6,20 | 5,54E-10 | 2,30E-08 |
| AT4G04825 | 145,71 | -7,98 | 1,31 | -6,08 | 1,17E-09 | 4,55E-08 |
| AT4G23387 | 136,04 | 3,00 | 0,53 | 5,66 | 1,49E-08 | 5,44E-07 |
| AT5G08185 | 1387,13 | -1,76 | 0,31 | -5,65 | 1,60E-08 | 5,52E-07 |
| AT3G01845 | 219,09 | 1,28 | 0,24 | 5,36 | 8,43E-08 | 2,76E-06 |
| AT4G03455 | 16,01 | -4,41 | 0,83 | -5,33 | 9,90E-08 | 2,93E-06 |
| AT4G13560 | 52,08 | -1,85 | 0,35 | -5,34 | 9,43E-08 | 2,93E-06 |
| AT5G41905 | 1173,00 | -1,34 | 0,26 | -5,25 | 1,48E-07 | 4,19E-06 |
| AT3G23325 | 69,65 | -1,54 | 0,30 | -5,12 | 3,09E-07 | 8,06E-06 |
| AT4G19395 | 418,73 | 1,06 | 0,21 | 5,12 | 3,11E-07 | 8,06E-06 |
| AT3G23326 | 69,32 | -1,55 | 0,31 | -5,07 | 4,00E-07 | 9,95E-06 |
| AT1G31173 | 488,28 | 1,22 | 0,24 | 4,98 | 6,36E-07 | 1,52E-05 |
| AT4G13494 | 321,70 | -2,37 | 0,50 | -4,78 | 1,80E-06 | 4,14E-05 |
| AT5G27606 | 14,40 | 6,06 | 1,29 | 4,71 | 2,49E-06 | 5,54E-05 |
| AT4G13575 | 426,34 | -2,05 | 0,44 | -4,65 | 3,26E-06 | 6,99E-05 |
| AT3G14210 | 12,80 | -4,43 | 0,96 | -4,61 | 4,12E-06 | 8,54E-05 |
| AT4G13570 | 36,83 | -1,75 | 0,38 | -4,59 | 4,46E-06 | 8,67E-05 |
| AT2G16030 | 25,88 | 5,03 | 1,09 | 4,59 | 4,34E-06 | 8,67E-05 |
| AT1G63400 | 45,16 | 5,73 | 1,26 | 4,56 | 5,10E-06 | 9,44E-05 |
| AT2G40805 | 280,87 | 1,90 | 0,42 | 4,56 | 5,16E-06 | 9,44E-05 |
| AT4G04835 | 19,24 | -2,45 | 0,54 | -4,50 | 6,67E-06 | 1,19E-04 |
| AT4G08691 | 15,19 | 5,82 | 1,30 | 4,48 | 7,44E-06 | 1,29E-04 |
| AT3G01345 | 6,06 | 6,06 | 1,38 | 4,40 | 1,06E-05 | 1,78E-04 |
| AT4G13564 | 27,21 | -1,71 | 0,39 | -4,39 | 1,11E-05 | 1,82E-04 |
| AT2G05070 | 19,17 | -2,63 | 0,60 | -4,37 | 1,22E-05 | 1,95E-04 |
| AT4G13495 | 869,17 | -1,47 | 0,34 | -4,33 | 1,52E-05 | 2,30E-04 |
| AT2G06555 | 20,57 | 3,43 | 0,79 | 4,33 | 1,49E-05 | 2,30E-04 |
| AT3G43990 | 14,30 | 4,71 | 1,12 | 4,22 | 2,49E-05 | 3,68E-04 |
| AT1G26973 | 11,41 | -2,69 | 0,65 | -4,15 | 3,28E-05 | 4,75E-04 |
| AT2G31141 | 64,55 | 1,27 | 0,31 | 4,03 | 5,46E-05 | 7,72E-04 |
| AT4G06110 | 22,73 | -2,07 | 0,51 | -4,02 | 5,82E-05 | 8,05E-04 |
| AT4G06130 | 392,23 | -1,82 | 0,45 | -4,01 | 6,04E-05 | 8,17E-04 |
| AT5G53902 | 6,85 | 5,54 | 1,40 | 3,94 | 8,02E-05 | 1,02E-03 |
| AT3G05700 | 20,03 | -2,22 | 0,56 | -3,94 | 8,03E-05 | 1,02E-03 |
| AT5G53905 | 6,85 | 5,54 | 1,40 | 3,94 | 8,02E-05 | 1,02E-03 |

| | | | | | | |
|---|---|---|---|---|---|---|
| AT1G01183 | 317,62 | -1,41 | 0,36 | -3,93 | 8,48E-05 | 1,05E-03 |
| AT3G01715 | 19,56 | -2,29 | 0,59 | -3,91 | 9,33E-05 | 1,14E-03 |
| AT1G11700 | 23,46 | 2,34 | 0,60 | 3,89 | 1,01E-04 | 1,20E-03 |
| AT2G44490 | 17,93 | -2,06 | 0,53 | -3,87 | 1,11E-04 | 1,30E-03 |
| AT2G39175 | 43,76 | 1,43 | 0,38 | 3,79 | 1,52E-04 | 1,75E-03 |
| AT2G01020 | 7030,11 | -1,34 | 0,35 | -3,77 | 1,66E-04 | 1,87E-03 |
| AT5G02575 | 6,74 | -3,28 | 0,89 | -3,69 | 2,25E-04 | 2,49E-03 |
| AT4G12770 | 19,67 | 2,64 | 0,73 | 3,64 | 2,74E-04 | 2,99E-03 |
| AT3G08370 | 6,58 | 5,08 | 1,41 | 3,59 | 3,27E-04 | 3,51E-03 |
| AT5G14545 | 7868,55 | 0,73 | 0,20 | 3,57 | 3,54E-04 | 3,73E-03 |
| AT5G63715 | 27,38 | 4,52 | 1,27 | 3,55 | 3,88E-04 | 4,03E-03 |
| AT3G47470 | 25,61 | -1,77 | 0,50 | -3,52 | 4,38E-04 | 4,46E-03 |
| AT2G23350 | 29,35 | -1,73 | 0,49 | -3,51 | 4,46E-04 | 4,47E-03 |
| AT4G17788 | 20,18 | -1,78 | 0,51 | -3,47 | 5,14E-04 | 5,05E-03 |
| AT5G14550 | 7872,48 | 0,73 | 0,21 | 3,47 | 5,20E-04 | 5,05E-03 |
| AT3G51375 | 135,65 | -1,27 | 0,37 | -3,44 | 5,74E-04 | 5,49E-03 |
| AT1G11735 | 8,53 | -2,58 | 0,76 | -3,41 | 6,54E-04 | 6,17E-03 |
| AT2G47275 | 830,57 | -1,03 | 0,31 | -3,36 | 7,69E-04 | 7,11E-03 |
| AT1G29920 | 50,33 | -1,37 | 0,41 | -3,36 | 7,78E-04 | 7,11E-03 |
| AT2G46685 | 3232,16 | -0,89 | 0,26 | -3,35 | 8,15E-04 | 7,35E-03 |
| AT1G07051 | 29,68 | -1,71 | 0,52 | -3,31 | 9,32E-04 | 8,29E-03 |
| AT5G38850 | 479,48 | 0,80 | 0,24 | 3,30 | 9,63E-04 | 8,43E-03 |
| AT1G05163 | 491,36 | 0,88 | 0,28 | 3,20 | 1,37E-03 | 1,17E-02 |
| AT1G15405 | 491,36 | 0,88 | 0,28 | 3,20 | 1,37E-03 | 1,17E-02 |
| AT1G63330 | 7,00 | -2,60 | 0,83 | -3,15 | 1,64E-03 | 1,37E-02 |
| AT2G03445 | 309,93 | 0,85 | 0,27 | 3,15 | 1,65E-03 | 1,37E-02 |
| AT3G43833 | 7,44 | 4,46 | 1,42 | 3,14 | 1,69E-03 | 1,38E-02 |
| AT4G03580 | 13,67 | 3,00 | 0,96 | 3,13 | 1,75E-03 | 1,40E-02 |
| AT5G27807 | 30,37 | 1,44 | 0,46 | 3,13 | 1,77E-03 | 1,40E-02 |
| AT5G45307 | 438,83 | -0,98 | 0,31 | -3,12 | 1,78E-03 | 1,40E-02 |
| AT2G35945 | 15,59 | 2,32 | 0,75 | 3,09 | 2,03E-03 | 1,58E-02 |
| AT2G16018 | 9,68 | 3,11 | 1,01 | 3,08 | 2,07E-03 | 1,59E-02 |
| AT5G26650 | 7,57 | 4,27 | 1,39 | 3,07 | 2,17E-03 | 1,65E-02 |
| AT2G47015 | 1248,43 | 0,95 | 0,31 | 3,05 | 2,28E-03 | 1,69E-02 |
| AT2G47020 | 1248,75 | 0,95 | 0,31 | 3,05 | 2,28E-03 | 1,69E-02 |
| AT1G29930 | 124,68 | -1,16 | 0,38 | -3,04 | 2,33E-03 | 1,70E-02 |
| AT3G56825 | 56,81 | 1,16 | 0,38 | 3,03 | 2,43E-03 | 1,76E-02 |
| AT2G18320 | 12,44 | -2,17 | 0,73 | -2,97 | 2,93E-03 | 2,10E-02 |
| AT2G35940 | 16,85 | 2,07 | 0,70 | 2,96 | 3,07E-03 | 2,17E-02 |
| AT5G38540 | 8,36 | -3,38 | 1,15 | -2,95 | 3,22E-03 | 2,25E-02 |
| AT5G53740 | 20,74 | 1,79 | 0,61 | 2,94 | 3,32E-03 | 2,27E-02 |
| AT5G62165 | 7,10 | 4,11 | 1,40 | 2,94 | 3,31E-03 | 2,27E-02 |
| AT4G23375 | 8,41 | 3,91 | 1,34 | 2,92 | 3,48E-03 | 2,35E-02 |
| AT5G41663 | 49,62 | 1,56 | 0,54 | 2,90 | 3,73E-03 | 2,50E-02 |
| AT5G54130 | 6,22 | -2,54 | 0,89 | -2,85 | 4,44E-03 | 2,94E-02 |
| AT3G13080 | 13,73 | 1,66 | 0,59 | 2,80 | 5,11E-03 | 3,34E-02 |
| AT3G23125 | 239,13 | -1,26 | 0,45 | -2,78 | 5,49E-03 | 3,56E-02 |
| AT1G63080 | 109,45 | 1,03 | 0,37 | 2,75 | 6,03E-03 | 3,86E-02 |
| AT3G06436 | 20,96 | 1,35 | 0,50 | 2,72 | 6,46E-03 | 4,10E-02 |
| AT5G43603 | 44,77 | 1,10 | 0,41 | 2,70 | 6,90E-03 | 4,29E-02 |
| AT5G66045 | 69,41 | -1,30 | 0,48 | -2,70 | 6,86E-03 | 4,29E-02 |
| AT1G13360 | 23,38 | 1,28 | 0,47 | 2,70 | 7,01E-03 | 4,32E-02 |

| | | | | | | |
|---|---|---|---|---|---|---|
| AT2G22125 | 19,76 | -1,38 | 0,51 | -2,68 | 7,39E-03 | 4,50E-02 |
| AT5G45510 | 8,42 | 2,17 | 0,82 | 2,66 | 7,89E-03 | 4,76E-02 |
| AT1G32583 | 28,27 | -1,60 | 0,60 | -2,64 | 8,35E-03 | 4,95E-02 |
| AT1G62910 | 98,21 | 1,21 | 0,46 | 2,64 | 8,29E-03 | 4,95E-02 |
| AT2G27400 | 624,49 | -0,89 | 0,34 | -2,62 | 8,67E-03 | 5,04E-02 |
| AT3G01313 | 12,85 | 2,25 | 0,86 | 2,63 | 8,61E-03 | 5,04E-02 |
| AT2G22496 | 20,32 | -1,30 | 0,50 | -2,61 | 8,93E-03 | 5,14E-02 |
| AT1G32582 | 26,60 | -1,61 | 0,62 | -2,60 | 9,40E-03 | 5,36E-02 |
| AT2G08986 | 10,75 | 1,81 | 0,70 | 2,58 | 9,97E-03 | 5,64E-02 |
| AT2G09610 | 9,53 | -2,07 | 0,81 | -2,57 | 1,02E-02 | 5,72E-02 |
| AT5G54270 | 15,64 | -1,43 | 0,56 | -2,56 | 1,04E-02 | 5,76E-02 |
| AT5G01820 | 9,75 | 2,16 | 0,85 | 2,55 | 1,09E-02 | 6,00E-02 |
| AT1G01320 | 16,50 | -1,12 | 0,44 | -2,53 | 1,13E-02 | 6,18E-02 |
| AT1G80180 | 13,61 | 2,01 | 0,80 | 2,52 | 1,16E-02 | 6,29E-02 |
| AT4G12780 | 15,31 | 1,54 | 0,61 | 2,52 | 1,18E-02 | 6,33E-02 |
| AT5G43740 | 103,64 | -1,32 | 0,53 | -2,51 | 1,22E-02 | 6,47E-02 |
| AT1G06227 | 343,33 | 0,95 | 0,38 | 2,49 | 1,26E-02 | 6,66E-02 |
| AT1G40104 | 7,42 | 2,51 | 1,01 | 2,49 | 1,29E-02 | 6,72E-02 |
| AT3G20362 | 7,16 | -2,59 | 1,04 | -2,48 | 1,31E-02 | 6,78E-02 |
| AT3G12630 | 8,51 | 1,87 | 0,76 | 2,48 | 1,32E-02 | 6,80E-02 |
| AT1G63150 | 176,58 | 0,89 | 0,36 | 2,47 | 1,35E-02 | 6,88E-02 |
| AT1G06237 | 336,80 | 0,96 | 0,39 | 2,44 | 1,46E-02 | 7,37E-02 |
| AT2G34430 | 28,33 | -1,10 | 0,45 | -2,43 | 1,51E-02 | 7,39E-02 |
| AT3G26740 | 15,30 | -1,67 | 0,68 | -2,44 | 1,48E-02 | 7,39E-02 |
| AT1G54410 | 19,25 | -1,72 | 0,71 | -2,43 | 1,49E-02 | 7,39E-02 |
| AT1G09877 | 23,86 | -1,40 | 0,58 | -2,43 | 1,50E-02 | 7,39E-02 |
| AT3G48201 | 30,66 | -1,06 | 0,44 | -2,42 | 1,56E-02 | 7,59E-02 |
| AT2G41650 | 5,84 | 3,24 | 1,34 | 2,41 | 1,58E-02 | 7,60E-02 |
| AT3G41979 | 6596,11 | -0,86 | 0,36 | -2,40 | 1,64E-02 | 7,83E-02 |
| AT1G79520 | 27,20 | -1,39 | 0,58 | -2,40 | 1,65E-02 | 7,83E-02 |
| AT2G41100 | 6,69 | 2,57 | 1,08 | 2,39 | 1,70E-02 | 7,89E-02 |
| AT3G61470 | 27,14 | -1,17 | 0,49 | -2,38 | 1,71E-02 | 7,89E-02 |
| AT4G14400 | 8,78 | -2,09 | 0,88 | -2,39 | 1,71E-02 | 7,89E-02 |
| AT1G62590 | 20,75 | 3,13 | 1,31 | 2,38 | 1,71E-02 | 7,89E-02 |
| AT3G06355 | 6711,54 | -0,84 | 0,36 | -2,37 | 1,77E-02 | 8,08E-02 |
| AT1G29910 | 27,17 | -1,21 | 0,51 | -2,36 | 1,81E-02 | 8,21E-02 |
| AT5G44340 | 14,19 | -1,37 | 0,58 | -2,35 | 1,90E-02 | 8,54E-02 |
| AT1G66100 | 8,07 | -2,01 | 0,86 | -2,33 | 1,95E-02 | 8,75E-02 |
| AT1G06680 | 10,44 | -1,77 | 0,76 | -2,31 | 2,06E-02 | 9,17E-02 |
| AT1G56070 | 55,72 | -0,65 | 0,28 | -2,31 | 2,10E-02 | 9,27E-02 |
| AT1G61226 | 20,72 | -1,22 | 0,53 | -2,28 | 2,24E-02 | 9,77E-02 |
| AT4G06810 | 54,17 | -0,70 | 0,31 | -2,28 | 2,25E-02 | 9,77E-02 |
| AT1G66795 | 14,00 | 1,33 | 0,59 | 2,25 | 2,47E-02 | 1,04E-01 |
| AT5G17920 | 34,31 | -0,88 | 0,39 | -2,25 | 2,47E-02 | 1,04E-01 |
| AT2G45380 | 11,33 | -1,43 | 0,64 | -2,25 | 2,45E-02 | 1,04E-01 |
| AT4G03445 | 45,78 | 0,75 | 0,34 | 2,25 | 2,46E-02 | 1,04E-01 |
| AT4G06805 | 54,05 | -0,69 | 0,31 | -2,25 | 2,45E-02 | 1,04E-01 |
| AT4G10800 | 6,52 | 2,14 | 0,96 | 2,23 | 2,60E-02 | 1,09E-01 |
| AT5G04105 | 18,20 | -1,10 | 0,50 | -2,21 | 2,72E-02 | 1,13E-01 |
| AT1G37130 | 39,12 | 0,72 | 0,33 | 2,20 | 2,79E-02 | 1,14E-01 |
| AT4G11373 | 5,92 | 3,06 | 1,39 | 2,20 | 2,79E-02 | 1,14E-01 |
| AT5G09730 | 35,07 | 1,04 | 0,47 | 2,20 | 2,78E-02 | 1,14E-01 |

| | | | | | | |
|---|---|---|---|---|---|---|
| AT1G12090 | 8,60 | -1,77 | 0,81 | -2,20 | 2,81E-02 | 1,14E-01 |
| AT5G05170 | 36,40 | -0,99 | 0,45 | -2,19 | 2,85E-02 | 1,14E-01 |
| AT1G66783 | 14,56 | 1,44 | 0,66 | 2,19 | 2,88E-02 | 1,15E-01 |
| AT5G07745 | 20,93 | -1,06 | 0,49 | -2,18 | 2,93E-02 | 1,16E-01 |
| AT1G08813 | 8,42 | -1,69 | 0,78 | -2,16 | 3,07E-02 | 1,21E-01 |
| AT1G12820 | 53,20 | 1,00 | 0,47 | 2,15 | 3,13E-02 | 1,22E-01 |
| AT5G04455 | 10,87 | 1,38 | 0,64 | 2,14 | 3,22E-02 | 1,24E-01 |
| AT5G04445 | 10,87 | 1,38 | 0,64 | 2,14 | 3,22E-02 | 1,24E-01 |
| AT4G23420 | 23,63 | -1,04 | 0,48 | -2,14 | 3,24E-02 | 1,24E-01 |
| AT2G41640 | 6,36 | 2,52 | 1,18 | 2,14 | 3,26E-02 | 1,24E-01 |
| AT2G28000 | 11,21 | -1,26 | 0,59 | -2,13 | 3,33E-02 | 1,26E-01 |
| AT4G07120 | 22,20 | -1,12 | 0,53 | -2,10 | 3,56E-02 | 1,34E-01 |
| AT1G74470 | 19,25 | -1,05 | 0,51 | -2,09 | 3,68E-02 | 1,38E-01 |
| AT1G61275 | 11,06 | 1,56 | 0,76 | 2,05 | 4,05E-02 | 1,51E-01 |
| AT1G23310 | 12,45 | -1,03 | 0,51 | -2,03 | 4,19E-02 | 1,55E-01 |
| AT1G20620 | 38,72 | -0,93 | 0,46 | -2,03 | 4,24E-02 | 1,56E-01 |
| AT3G22886 | 587,34 | 0,51 | 0,25 | 2,01 | 4,48E-02 | 1,64E-01 |
| AT4G23915 | 12,58 | 1,33 | 0,67 | 2,00 | 4,52E-02 | 1,64E-01 |
| AT2G38230 | 6,88 | -1,69 | 0,85 | -1,99 | 4,71E-02 | 1,70E-01 |
| AT2G01008 | 28,92 | 1,03 | 0,52 | 1,98 | 4,74E-02 | 1,70E-01 |
| AT4G00150 | 10,48 | 1,44 | 0,73 | 1,98 | 4,77E-02 | 1,70E-01 |
| AT5G15833 | 13,93 | -1,16 | 0,59 | -1,97 | 4,84E-02 | 1,71E-01 |
| AT3G14385 | 15,96 | -0,92 | 0,47 | -1,98 | 4,82E-02 | 1,71E-01 |
| AT3G23690 | 28,16 | 1,90 | 0,97 | 1,96 | 4,95E-02 | 1,74E-01 |
| AT1G08115 | 40,91 | -0,85 | 0,43 | -1,96 | 5,06E-02 | 1,77E-01 |
| AT1G50440 | 6,47 | 1,94 | 1,00 | 1,94 | 5,22E-02 | 1,80E-01 |
| AT3G52140 | 9,89 | -1,16 | 0,60 | -1,94 | 5,22E-02 | 1,80E-01 |
| AT1G18010 | 5,80 | 2,63 | 1,37 | 1,92 | 5,44E-02 | 1,85E-01 |
| AT5G51174 | 65,86 | -0,89 | 0,46 | -1,93 | 5,38E-02 | 1,85E-01 |
| AT1G10760 | 16,93 | -1,02 | 0,53 | -1,93 | 5,42E-02 | 1,85E-01 |
| AT1G20340 | 11,85 | -1,28 | 0,67 | -1,92 | 5,46E-02 | 1,85E-01 |
| AT2G35384 | 13,85 | -1,37 | 0,72 | -1,92 | 5,49E-02 | 1,85E-01 |
| AT5G21100 | 6,61 | 2,04 | 1,07 | 1,91 | 5,55E-02 | 1,86E-01 |
| AT2G21390 | 23,77 | -1,03 | 0,54 | -1,91 | 5,61E-02 | 1,87E-01 |
| AT3G19820 | 10,30 | -1,10 | 0,58 | -1,89 | 5,90E-02 | 1,95E-01 |
| AT1G07633 | 6,19 | 1,87 | 1,01 | 1,85 | 6,47E-02 | 2,12E-01 |
| AT3G54890 | 13,73 | -1,23 | 0,66 | -1,85 | 6,46E-02 | 2,12E-01 |
| AT3G05675 | 9,08 | -1,27 | 0,69 | -1,84 | 6,62E-02 | 2,16E-01 |
| AT3G13920 | 8,10 | -1,40 | 0,76 | -1,83 | 6,71E-02 | 2,17E-01 |
| AT4G00885 | 370,66 | 0,57 | 0,31 | 1,82 | 6,90E-02 | 2,22E-01 |
| AT1G04977 | 45,78 | 0,93 | 0,51 | 1,81 | 6,99E-02 | 2,23E-01 |
| AT5G35407 | 1076,14 | 0,67 | 0,37 | 1,81 | 6,99E-02 | 2,23E-01 |
| AT1G05993 | 11,40 | 1,28 | 0,71 | 1,79 | 7,43E-02 | 2,36E-01 |
| AT1G48410 | 43,79 | 0,65 | 0,37 | 1,78 | 7,55E-02 | 2,37E-01 |
| AT3G11130 | 25,04 | -0,84 | 0,47 | -1,78 | 7,53E-02 | 2,37E-01 |
| AT5G07350 | 9,52 | -1,17 | 0,66 | -1,77 | 7,61E-02 | 2,38E-01 |
| AT1G04733 | 20,09 | 0,89 | 0,51 | 1,74 | 8,22E-02 | 2,56E-01 |
| AT1G09310 | 8,77 | -1,51 | 0,87 | -1,73 | 8,28E-02 | 2,56E-01 |
| AT4G04640 | 5,86 | -1,70 | 0,98 | -1,73 | 8,33E-02 | 2,56E-01 |
| AT1G74456 | 11,08 | -1,21 | 0,70 | -1,73 | 8,42E-02 | 2,58E-01 |
| AT3G62980 | 20,74 | -0,86 | 0,50 | -1,71 | 8,67E-02 | 2,64E-01 |
| AT3G26650 | 12,31 | -1,04 | 0,61 | -1,71 | 8,76E-02 | 2,66E-01 |

| | | | | | | |
|---|---|---|---|---|---|---|
| AT3G01695 | 8,72 | -1,22 | 0,72 | -1,70 | 8,83E-02 | 2,67E-01 |
| AT3G23820 | 9,86 | -1,04 | 0,61 | -1,70 | 8,88E-02 | 2,67E-01 |
| AT2G45160 | 10,70 | -1,36 | 0,80 | -1,69 | 9,02E-02 | 2,70E-01 |
| AT4G30975 | 11,87 | 1,09 | 0,64 | 1,69 | 9,10E-02 | 2,70E-01 |
| AT5G01530 | 20,05 | -1,04 | 0,62 | -1,69 | 9,13E-02 | 2,70E-01 |
| AT4G30972 | 11,83 | 1,07 | 0,64 | 1,67 | 9,58E-02 | 2,82E-01 |
| AT4G27640 | 8,75 | -1,29 | 0,79 | -1,64 | 1,01E-01 | 2,96E-01 |
| AT1G62930 | 53,37 | -0,83 | 0,51 | -1,63 | 1,03E-01 | 2,97E-01 |
| AT2G38325 | 156,77 | 0,40 | 0,24 | 1,64 | 1,02E-01 | 2,97E-01 |
| AT3G28180 | 8,56 | -1,87 | 1,15 | -1,63 | 1,03E-01 | 2,97E-01 |
| AT4G05180 | 7,70 | -1,58 | 0,97 | -1,63 | 1,03E-01 | 2,97E-01 |
| AT3G41768 | 91,21 | -0,53 | 0,33 | -1,62 | 1,05E-01 | 3,01E-01 |
| AT5G46315 | 17,01 | -0,83 | 0,51 | -1,61 | 1,07E-01 | 3,04E-01 |
| AT1G02470 | 5,92 | -1,48 | 0,92 | -1,61 | 1,07E-01 | 3,04E-01 |
| AT1G15820 | 12,09 | -1,15 | 0,71 | -1,61 | 1,07E-01 | 3,04E-01 |
| AT2G20190 | 8,93 | -1,15 | 0,72 | -1,59 | 1,11E-01 | 3,13E-01 |
| AT4G10340 | 23,35 | -0,82 | 0,52 | -1,57 | 1,15E-01 | 3,23E-01 |
| AT1G61520 | 19,44 | -0,93 | 0,60 | -1,56 | 1,18E-01 | 3,26E-01 |
| AT3G47350 | 23,89 | 0,74 | 0,47 | 1,56 | 1,18E-01 | 3,26E-01 |
| AT3G58196 | 21,50 | 0,89 | 0,57 | 1,56 | 1,18E-01 | 3,26E-01 |
| AT2G21330 | 7,94 | -1,06 | 0,68 | -1,56 | 1,19E-01 | 3,26E-01 |
| AT3G01540 | 20,21 | -0,72 | 0,46 | -1,56 | 1,19E-01 | 3,26E-01 |
| AT1G09800 | 20,58 | 0,79 | 0,51 | 1,55 | 1,21E-01 | 3,29E-01 |
| AT4G32410 | 11,90 | -0,97 | 0,63 | -1,54 | 1,25E-01 | 3,39E-01 |
| AT3G10985 | 6,51 | 1,33 | 0,87 | 1,53 | 1,26E-01 | 3,40E-01 |
| AT2G46255 | 72,33 | 0,79 | 0,52 | 1,52 | 1,28E-01 | 3,45E-01 |
| AT3G63375 | 13,10 | -0,81 | 0,53 | -1,52 | 1,29E-01 | 3,47E-01 |
| AT2G16600 | 11,35 | 1,03 | 0,69 | 1,51 | 1,32E-01 | 3,52E-01 |
| AT3G50370 | 9,94 | -0,83 | 0,56 | -1,50 | 1,33E-01 | 3,54E-01 |
| AT3G09105 | 30,37 | 0,85 | 0,57 | 1,49 | 1,35E-01 | 3,58E-01 |
| AT2G20410 | 6,12 | 1,31 | 0,88 | 1,49 | 1,38E-01 | 3,61E-01 |
| AT4G31877 | 9,33 | -1,16 | 0,78 | -1,48 | 1,38E-01 | 3,61E-01 |
| AT4G13940 | 35,08 | -0,58 | 0,39 | -1,48 | 1,40E-01 | 3,63E-01 |
| AT2G39675 | 2246,76 | -0,50 | 0,34 | -1,48 | 1,39E-01 | 3,63E-01 |
| AT5G03552 | 58,24 | -0,66 | 0,45 | -1,47 | 1,43E-01 | 3,70E-01 |
| AT5G03553 | 58,75 | -0,65 | 0,45 | -1,46 | 1,44E-01 | 3,72E-01 |
| AT1G63130 | 240,21 | 0,48 | 0,33 | 1,45 | 1,46E-01 | 3,75E-01 |
| AT1G49760 | 5,88 | -1,03 | 0,72 | -1,43 | 1,52E-01 | 3,90E-01 |
| AT3G06325 | 6,42 | 1,39 | 0,97 | 1,43 | 1,54E-01 | 3,92E-01 |
| AT1G31580 | 10,89 | -1,04 | 0,73 | -1,42 | 1,56E-01 | 3,96E-01 |
| AT5G20700 | 23,90 | 0,58 | 0,41 | 1,42 | 1,57E-01 | 3,97E-01 |
| AT3G47342 | 15,52 | 0,89 | 0,63 | 1,41 | 1,58E-01 | 3,99E-01 |
| AT3G05325 | 9,46 | 1,10 | 0,79 | 1,40 | 1,62E-01 | 4,05E-01 |
| AT5G03305 | 5,94 | 1,52 | 1,09 | 1,40 | 1,62E-01 | 4,05E-01 |
| AT4G07975 | 9,02 | -0,83 | 0,60 | -1,39 | 1,63E-01 | 4,07E-01 |
| AT5G14565 | 6609,25 | 0,37 | 0,27 | 1,39 | 1,64E-01 | 4,07E-01 |
| AT1G18075 | 4243,99 | 0,25 | 0,18 | 1,39 | 1,65E-01 | 4,08E-01 |
| AT1G59870 | 72,89 | 0,46 | 0,33 | 1,38 | 1,67E-01 | 4,08E-01 |
| AT5G66570 | 20,06 | -0,89 | 0,65 | -1,38 | 1,66E-01 | 4,08E-01 |
| AT5G46845 | 66,07 | 0,44 | 0,32 | 1,38 | 1,67E-01 | 4,08E-01 |
| AT1G12775 | 21,87 | 0,98 | 0,71 | 1,37 | 1,69E-01 | 4,11E-01 |
| AT4G16860 | 6,11 | 1,11 | 0,81 | 1,37 | 1,72E-01 | 4,17E-01 |

| | | | | | | |
|---|---|---|---|---|---|---|
| AT1G48267 | 15177,13 | -0,45 | 0,34 | -1,35 | 1,78E-01 | 4,19E-01 |
| AT2G18960 | 62,63 | 0,58 | 0,43 | 1,35 | 1,77E-01 | 4,19E-01 |
| AT2G41310 | 487,34 | -0,38 | 0,28 | -1,36 | 1,75E-01 | 4,19E-01 |
| AT3G13857 | 124,31 | -0,50 | 0,37 | -1,35 | 1,77E-01 | 4,19E-01 |
| AT3G13855 | 124,31 | -0,50 | 0,37 | -1,35 | 1,77E-01 | 4,19E-01 |
| AT3G56705 | 15,48 | 0,79 | 0,59 | 1,35 | 1,78E-01 | 4,19E-01 |
| AT4G12800 | 10,28 | -1,16 | 0,86 | -1,35 | 1,76E-01 | 4,19E-01 |
| AT2G09250 | 487,12 | -0,38 | 0,28 | -1,35 | 1,79E-01 | 4,19E-01 |
| AT5G07725 | 8,72 | 1,19 | 0,89 | 1,34 | 1,80E-01 | 4,21E-01 |
| AT5G52070 | 6,76 | -1,53 | 1,15 | -1,34 | 1,81E-01 | 4,21E-01 |
| AT5G62162 | 10,16 | 0,90 | 0,67 | 1,34 | 1,81E-01 | 4,21E-01 |
| AT1G06267 | 15,22 | 0,98 | 0,76 | 1,30 | 1,93E-01 | 4,43E-01 |
| AT3G08940 | 8,84 | -1,08 | 0,83 | -1,30 | 1,93E-01 | 4,43E-01 |
| AT4G04565 | 64,98 | 0,41 | 0,31 | 1,30 | 1,92E-01 | 4,43E-01 |
| AT5G10946 | 91,61 | 0,43 | 0,33 | 1,30 | 1,94E-01 | 4,43E-01 |
| AT1G09620 | 17,50 | -0,59 | 0,46 | -1,29 | 1,99E-01 | 4,51E-01 |
| AT1G67090 | 31,55 | -0,59 | 0,46 | -1,29 | 1,99E-01 | 4,51E-01 |
| AT2G06520 | 10,63 | -1,14 | 0,89 | -1,28 | 2,01E-01 | 4,56E-01 |
| AT1G26975 | 8,83 | -0,81 | 0,64 | -1,27 | 2,06E-01 | 4,60E-01 |
| AT2G35744 | 14,81 | -0,80 | 0,63 | -1,27 | 2,05E-01 | 4,60E-01 |
| AT5G10945 | 91,45 | 0,42 | 0,33 | 1,27 | 2,05E-01 | 4,60E-01 |
| AT3G02020 | 19,73 | 0,75 | 0,59 | 1,26 | 2,07E-01 | 4,61E-01 |
| AT1G78206 | 14,29 | -0,65 | 0,52 | -1,24 | 2,14E-01 | 4,72E-01 |
| AT5G57870 | 8,84 | -0,77 | 0,62 | -1,24 | 2,14E-01 | 4,72E-01 |
| AT4G21960 | 15,77 | -0,66 | 0,53 | -1,24 | 2,14E-01 | 4,72E-01 |
| AT3G47348 | 6,89 | 1,69 | 1,36 | 1,24 | 2,16E-01 | 4,74E-01 |
| AT3G48057 | 47,97 | 0,68 | 0,55 | 1,22 | 2,21E-01 | 4,83E-01 |
| AT1G47128 | 7,64 | 0,80 | 0,65 | 1,22 | 2,21E-01 | 4,83E-01 |
| AT2G05520 | 18,30 | 0,60 | 0,49 | 1,22 | 2,23E-01 | 4,84E-01 |
| AT3G18290 | 9,13 | -0,81 | 0,67 | -1,21 | 2,28E-01 | 4,92E-01 |
| AT4G08175 | 7,31 | -0,99 | 0,82 | -1,21 | 2,27E-01 | 4,92E-01 |
| AT5G46250 | 6,03 | 1,04 | 0,86 | 1,20 | 2,29E-01 | 4,92E-01 |
| AT4G03380 | 7,25 | 1,00 | 0,83 | 1,20 | 2,30E-01 | 4,92E-01 |
| AT3G18780 | 17,06 | -0,59 | 0,50 | -1,19 | 2,34E-01 | 5,00E-01 |
| AT5G02500 | 62,24 | -0,36 | 0,30 | -1,18 | 2,39E-01 | 5,09E-01 |
| AT1G07920 | 7,76 | -0,85 | 0,74 | -1,16 | 2,47E-01 | 5,22E-01 |
| AT2G20570 | 7,71 | 0,77 | 0,67 | 1,16 | 2,47E-01 | 5,22E-01 |
| AT3G56940 | 6,51 | -0,91 | 0,79 | -1,16 | 2,46E-01 | 5,22E-01 |
| AT1G66880 | 6,48 | 1,01 | 0,87 | 1,15 | 2,49E-01 | 5,22E-01 |
| AT4G13885 | 7,77 | -0,96 | 0,83 | -1,15 | 2,49E-01 | 5,22E-01 |
| AT1G08833 | 15,74 | -0,69 | 0,60 | -1,14 | 2,53E-01 | 5,22E-01 |
| AT1G75660 | 6,24 | -0,83 | 0,73 | -1,14 | 2,52E-01 | 5,22E-01 |
| AT4G07070 | 70,24 | 0,65 | 0,57 | 1,15 | 2,52E-01 | 5,22E-01 |
| AT3G16640 | 7,07 | -0,99 | 0,86 | -1,15 | 2,51E-01 | 5,22E-01 |
| AT3G48870 | 10,82 | -0,73 | 0,64 | -1,13 | 2,56E-01 | 5,28E-01 |
| AT4G34450 | 21,43 | -0,53 | 0,47 | -1,13 | 2,59E-01 | 5,32E-01 |
| AT4G33010 | 11,10 | -0,76 | 0,68 | -1,13 | 2,60E-01 | 5,32E-01 |
| AT2G34100 | 7,49 | 1,07 | 0,96 | 1,12 | 2,62E-01 | 5,35E-01 |
| AT2G16586 | 20,98 | 0,69 | 0,62 | 1,11 | 2,68E-01 | 5,39E-01 |
| AT5G43725 | 9,10 | 1,51 | 1,36 | 1,11 | 2,68E-01 | 5,39E-01 |
| AT5G43730 | 9,10 | 1,51 | 1,36 | 1,11 | 2,68E-01 | 5,39E-01 |
| AT2G35747 | 15,41 | 0,85 | 0,76 | 1,11 | 2,66E-01 | 5,39E-01 |

| | | | | | | |
|---|---|---|---|---|---|---|
| AT2G15530 | 40,01 | -0,34 | 0,31 | -1,09 | 2,74E-01 | 5,48E-01 |
| AT5G64270 | 15,19 | -0,56 | 0,52 | -1,09 | 2,74E-01 | 5,48E-01 |
| AT4G36080 | 9,35 | -0,63 | 0,58 | -1,09 | 2,77E-01 | 5,52E-01 |
| AT5G03240 | 5,81 | 0,88 | 0,81 | 1,08 | 2,81E-01 | 5,58E-01 |
| AT1G15750 | 35,69 | -0,41 | 0,39 | -1,07 | 2,83E-01 | 5,59E-01 |
| AT3G57260 | 8,35 | -0,62 | 0,58 | -1,07 | 2,83E-01 | 5,59E-01 |
| AT3G22380 | 8,79 | 0,75 | 0,71 | 1,06 | 2,90E-01 | 5,70E-01 |
| AT1G02500 | 10,85 | -0,71 | 0,67 | -1,05 | 2,92E-01 | 5,72E-01 |
| AT3G58193 | 8,75 | 0,91 | 0,88 | 1,04 | 2,99E-01 | 5,85E-01 |
| AT5G15022 | 21,95 | 0,68 | 0,65 | 1,04 | 3,00E-01 | 5,85E-01 |
| AT1G73687 | 3456,65 | 0,27 | 0,26 | 1,03 | 3,01E-01 | 5,85E-01 |
| AT1G79920 | 8,70 | -0,65 | 0,63 | -1,03 | 3,05E-01 | 5,91E-01 |
| AT1G04430 | 7,26 | -0,74 | 0,73 | -1,01 | 3,12E-01 | 6,01E-01 |
| AT3G43300 | 11,08 | -0,59 | 0,58 | -1,01 | 3,11E-01 | 6,01E-01 |
| AT3G09840 | 22,25 | 0,41 | 0,41 | 1,00 | 3,15E-01 | 6,02E-01 |
| AT3G21805 | 21,83 | 0,85 | 0,85 | 1,01 | 3,14E-01 | 6,02E-01 |
| AT3G15920 | 8,54 | 0,69 | 0,69 | 1,00 | 3,15E-01 | 6,02E-01 |
| AT1G55860 | 12,82 | 0,66 | 0,66 | 1,00 | 3,16E-01 | 6,02E-01 |
| AT3G46780 | 9,83 | -0,65 | 0,65 | -1,00 | 3,19E-01 | 6,04E-01 |
| AT5G61455 | 11,96 | -0,56 | 0,56 | -1,00 | 3,20E-01 | 6,04E-01 |
| AT1G06243 | 5,81 | 0,88 | 0,90 | 0,98 | 3,26E-01 | 6,11E-01 |
| AT1G19835 | 5,86 | -0,81 | 0,82 | -0,98 | 3,26E-01 | 6,11E-01 |
| AT3G15353 | 5,91 | 1,12 | 1,13 | 0,98 | 3,25E-01 | 6,11E-01 |
| AT1G48300 | 8,14 | -0,83 | 0,86 | -0,97 | 3,31E-01 | 6,12E-01 |
| AT1G04907 | 10,89 | 0,67 | 0,69 | 0,97 | 3,31E-01 | 6,12E-01 |
| AT1G09937 | 6,18 | -0,74 | 0,75 | -0,98 | 3,29E-01 | 6,12E-01 |
| AT2G43900 | 9,76 | -0,63 | 0,65 | -0,97 | 3,31E-01 | 6,12E-01 |
| AT4G02480 | 22,30 | -0,66 | 0,68 | -0,97 | 3,33E-01 | 6,15E-01 |
| AT3G45780 | 10,09 | -0,54 | 0,56 | -0,96 | 3,36E-01 | 6,16E-01 |
| AT4G07875 | 11,03 | -0,63 | 0,65 | -0,96 | 3,37E-01 | 6,16E-01 |
| AT1G25440 | 6,25 | -0,86 | 0,89 | -0,96 | 3,37E-01 | 6,16E-01 |
| AT1G31280 | 162,06 | -0,44 | 0,46 | -0,95 | 3,42E-01 | 6,25E-01 |
| AT1G31330 | 6,03 | -0,86 | 0,92 | -0,94 | 3,48E-01 | 6,32E-01 |
| AT4G28080 | 23,67 | -0,55 | 0,59 | -0,93 | 3,50E-01 | 6,35E-01 |
| AT2G39885 | 9,28 | -0,69 | 0,75 | -0,92 | 3,56E-01 | 6,39E-01 |
| AT2G43160 | 6,80 | -0,70 | 0,76 | -0,92 | 3,55E-01 | 6,39E-01 |
| AT3G17400 | 6,94 | 0,86 | 0,93 | 0,93 | 3,54E-01 | 6,39E-01 |
| AT5G65950 | 6,84 | -0,90 | 0,98 | -0,92 | 3,56E-01 | 6,39E-01 |
| AT2G39681 | 1805,59 | 0,30 | 0,32 | 0,92 | 3,60E-01 | 6,43E-01 |
| AT1G62670 | 9,93 | 1,29 | 1,42 | 0,91 | 3,61E-01 | 6,44E-01 |
| AT1G10522 | 24,13 | 0,44 | 0,49 | 0,90 | 3,67E-01 | 6,48E-01 |
| AT1G72150 | 7,68 | 0,70 | 0,77 | 0,91 | 3,65E-01 | 6,48E-01 |
| AT2G08770 | 10,72 | -0,73 | 0,81 | -0,90 | 3,66E-01 | 6,48E-01 |
| AT1G15500 | 7,05 | -0,70 | 0,78 | -0,90 | 3,69E-01 | 6,49E-01 |
| AT2G14560 | 27,90 | -0,44 | 0,50 | -0,89 | 3,73E-01 | 6,49E-01 |
| AT3G02470 | 10,40 | 0,53 | 0,59 | 0,89 | 3,73E-01 | 6,49E-01 |
| AT1G16635 | 19,67 | 0,57 | 0,64 | 0,90 | 3,71E-01 | 6,49E-01 |
| AT1G05247 | 19,59 | 0,57 | 0,64 | 0,89 | 3,72E-01 | 6,49E-01 |
| AT2G21660 | 24,65 | -0,44 | 0,50 | -0,89 | 3,71E-01 | 6,49E-01 |
| AT1G29265 | 54,27 | 0,28 | 0,32 | 0,88 | 3,80E-01 | 6,53E-01 |
| AT3G01835 | 72,10 | 0,29 | 0,33 | 0,88 | 3,78E-01 | 6,53E-01 |
| AT3G26520 | 6,24 | 0,86 | 0,97 | 0,88 | 3,78E-01 | 6,53E-01 |

| | | | | | | |
|---|---|---|---|---|---|---|
| AT3G61198 | 7,65 | 0,70 | 0,80 | 0,88 | 3,80E-01 | 6,53E-01 |
| AT5G25610 | 6,58 | -0,79 | 0,90 | -0,88 | 3,81E-01 | 6,53E-01 |
| AT1G35710 | 7,53 | 0,79 | 0,91 | 0,87 | 3,85E-01 | 6,57E-01 |
| AT4G05320 | 17,67 | 0,47 | 0,54 | 0,87 | 3,85E-01 | 6,57E-01 |
| AT5G63020 | 8,08 | 0,94 | 1,09 | 0,86 | 3,88E-01 | 6,60E-01 |
| AT4G13340 | 14,47 | -0,41 | 0,47 | -0,86 | 3,89E-01 | 6,60E-01 |
| AT2G28056 | 34,24 | 0,38 | 0,45 | 0,85 | 3,97E-01 | 6,69E-01 |
| AT5G60790 | 11,59 | -0,49 | 0,58 | -0,85 | 3,96E-01 | 6,69E-01 |
| AT5G09505 | 92,76 | 0,27 | 0,32 | 0,84 | 4,00E-01 | 6,73E-01 |
| AT1G62750 | 9,17 | -0,58 | 0,69 | -0,84 | 4,03E-01 | 6,76E-01 |
| AT4G08755 | 25,91 | 0,35 | 0,42 | 0,83 | 4,08E-01 | 6,83E-01 |
| AT1G10290 | 9,22 | -0,51 | 0,63 | -0,81 | 4,16E-01 | 6,93E-01 |
| AT1G08347 | 16,30 | 0,42 | 0,53 | 0,80 | 4,24E-01 | 7,05E-01 |
| AT4G39090 | 7,03 | 0,65 | 0,82 | 0,79 | 4,27E-01 | 7,08E-01 |
| AT2G30860 | 6,26 | -0,64 | 0,82 | -0,78 | 4,34E-01 | 7,14E-01 |
| AT5G23110 | 11,58 | 0,42 | 0,53 | 0,78 | 4,33E-01 | 7,14E-01 |
| AT3G18490 | 8,11 | 0,66 | 0,84 | 0,78 | 4,33E-01 | 7,14E-01 |
| AT1G10950 | 6,24 | -0,59 | 0,76 | -0,77 | 4,41E-01 | 7,17E-01 |
| AT3G04120 | 5,88 | -0,56 | 0,72 | -0,77 | 4,42E-01 | 7,17E-01 |
| AT4G24415 | 10,68 | 0,52 | 0,67 | 0,77 | 4,43E-01 | 7,17E-01 |
| AT1G08380 | 7,50 | -0,79 | 1,03 | -0,77 | 4,41E-01 | 7,17E-01 |
| AT3G07100 | 6,58 | 0,55 | 0,71 | 0,77 | 4,40E-01 | 7,17E-01 |
| AT5G54075 | 15,55 | 0,45 | 0,58 | 0,77 | 4,40E-01 | 7,17E-01 |
| AT5G53480 | 6,73 | 0,62 | 0,82 | 0,76 | 4,49E-01 | 7,26E-01 |
| AT5G26742 | 9,59 | -0,53 | 0,70 | -0,75 | 4,53E-01 | 7,30E-01 |
| AT2G08725 | 34,05 | 0,46 | 0,61 | 0,75 | 4,55E-01 | 7,31E-01 |
| AT3G23680 | 9,12 | -0,52 | 0,69 | -0,75 | 4,56E-01 | 7,31E-01 |
| AT2G30520 | 8,24 | -0,51 | 0,69 | -0,74 | 4,58E-01 | 7,32E-01 |
| AT3G08943 | 11,39 | -0,47 | 0,63 | -0,74 | 4,60E-01 | 7,34E-01 |
| AT1G62914 | 457,48 | 0,26 | 0,36 | 0,73 | 4,64E-01 | 7,37E-01 |
| AT3G13460 | 10,31 | -0,43 | 0,58 | -0,73 | 4,64E-01 | 7,37E-01 |
| AT1G20015 | 7,84 | -0,60 | 0,85 | -0,71 | 4,76E-01 | 7,38E-01 |
| AT2G01021 | 3194,91 | -0,19 | 0,27 | -0,72 | 4,74E-01 | 7,38E-01 |
| AT3G48720 | 18,61 | 0,45 | 0,63 | 0,72 | 4,73E-01 | 7,38E-01 |
| AT5G60390 | 16,82 | -0,38 | 0,53 | -0,71 | 4,76E-01 | 7,38E-01 |
| AT1G03060 | 7,63 | 0,51 | 0,71 | 0,73 | 4,68E-01 | 7,38E-01 |
| AT1G04517 | 11,17 | 0,43 | 0,61 | 0,71 | 4,75E-01 | 7,38E-01 |
| AT2G02950 | 6,29 | -0,80 | 1,12 | -0,71 | 4,76E-01 | 7,38E-01 |
| AT4G27600 | 12,18 | -0,44 | 0,61 | -0,72 | 4,71E-01 | 7,38E-01 |
| AT5G01350 | 8,44 | -0,47 | 0,66 | -0,72 | 4,74E-01 | 7,38E-01 |
| AT5G41740 | 6,05 | 0,65 | 0,93 | 0,70 | 4,81E-01 | 7,42E-01 |
| AT5G46110 | 7,87 | -0,51 | 0,73 | -0,71 | 4,81E-01 | 7,42E-01 |
| AT1G15290 | 5,85 | -0,50 | 0,72 | -0,70 | 4,83E-01 | 7,44E-01 |
| AT5G02325 | 15,37 | 0,56 | 0,81 | 0,69 | 4,89E-01 | 7,51E-01 |
| AT1G05273 | 8,42 | 0,44 | 0,65 | 0,68 | 4,95E-01 | 7,51E-01 |
| AT2G42270 | 6,29 | 0,54 | 0,80 | 0,68 | 4,93E-01 | 7,51E-01 |
| AT4G32020 | 12,23 | -0,37 | 0,54 | -0,68 | 4,95E-01 | 7,51E-01 |
| AT1G16820 | 8,42 | 0,44 | 0,65 | 0,68 | 4,95E-01 | 7,51E-01 |
| AT2G26080 | 10,37 | 0,47 | 0,68 | 0,68 | 4,94E-01 | 7,51E-01 |
| AT2G05100 | 10,97 | 0,48 | 0,71 | 0,68 | 4,97E-01 | 7,53E-01 |
| AT4G18670 | 7,64 | 0,59 | 0,88 | 0,68 | 4,99E-01 | 7,54E-01 |
| AT5G54070 | 15,72 | 0,40 | 0,59 | 0,67 | 5,01E-01 | 7,55E-01 |

| | | | | | | |
|---|---|---|---|---|---|---|
| AT1G47720 | 6,21 | 0,50 | 0,75 | 0,67 | 5,04E-01 | 7,57E-01 |
| AT1G64720 | 13,91 | -0,39 | 0,58 | -0,67 | 5,05E-01 | 7,58E-01 |
| AT3G17185 | 156,12 | 0,22 | 0,33 | 0,66 | 5,08E-01 | 7,60E-01 |
| AT3G15730 | 15,00 | -0,34 | 0,53 | -0,65 | 5,13E-01 | 7,66E-01 |
| AT4G14500 | 7,04 | 0,63 | 0,98 | 0,65 | 5,16E-01 | 7,68E-01 |
| AT4G38770 | 7,08 | 0,63 | 0,97 | 0,65 | 5,18E-01 | 7,68E-01 |
| AT1G72600 | 18,04 | -0,31 | 0,49 | -0,64 | 5,21E-01 | 7,71E-01 |
| AT3G03175 | 7,33 | 0,44 | 0,71 | 0,62 | 5,32E-01 | 7,79E-01 |
| AT4G31480 | 6,09 | 0,52 | 0,84 | 0,63 | 5,32E-01 | 7,79E-01 |
| AT3G06435 | 378,12 | 0,16 | 0,26 | 0,63 | 5,31E-01 | 7,79E-01 |
| AT4G16950 | 9,81 | 0,39 | 0,62 | 0,63 | 5,30E-01 | 7,79E-01 |
| AT5G04275 | 20,42 | 0,31 | 0,50 | 0,63 | 5,29E-01 | 7,79E-01 |
| AT1G79930 | 9,47 | -0,44 | 0,71 | -0,62 | 5,37E-01 | 7,84E-01 |
| AT1G15690 | 11,49 | 0,40 | 0,65 | 0,61 | 5,39E-01 | 7,85E-01 |
| AT1G29418 | 9,45 | 0,39 | 0,65 | 0,61 | 5,45E-01 | 7,91E-01 |
| AT1G78900 | 8,95 | -0,39 | 0,66 | -0,60 | 5,50E-01 | 7,96E-01 |
| AT1G80410 | 5,85 | -0,55 | 0,92 | -0,60 | 5,50E-01 | 7,96E-01 |
| AT2G03855 | 12,69 | 0,41 | 0,69 | 0,59 | 5,58E-01 | 8,00E-01 |
| AT3G55734 | 132,05 | -0,18 | 0,30 | -0,58 | 5,60E-01 | 8,00E-01 |
| AT4G06225 | 6,57 | 0,76 | 1,30 | 0,58 | 5,60E-01 | 8,00E-01 |
| AT4G39420 | 9,97 | -0,39 | 0,67 | -0,58 | 5,59E-01 | 8,00E-01 |
| AT3G02260 | 23,16 | 0,27 | 0,46 | 0,59 | 5,57E-01 | 8,00E-01 |
| AT5G08670 | 5,95 | -0,43 | 0,73 | -0,59 | 5,57E-01 | 8,00E-01 |
| AT1G50055 | 182,36 | 0,18 | 0,32 | 0,58 | 5,63E-01 | 8,02E-01 |
| AT1G79040 | 10,12 | -0,54 | 0,95 | -0,57 | 5,72E-01 | 8,04E-01 |
| AT2G07605 | 7,80 | 0,45 | 0,81 | 0,55 | 5,80E-01 | 8,04E-01 |
| AT2G09690 | 14,22 | -0,45 | 0,83 | -0,55 | 5,86E-01 | 8,04E-01 |
| AT3G59940 | 11,46 | 0,40 | 0,73 | 0,55 | 5,82E-01 | 8,04E-01 |
| AT4G00975 | 11,27 | -0,30 | 0,54 | -0,56 | 5,75E-01 | 8,04E-01 |
| AT4G27440 | 13,15 | -0,40 | 0,72 | -0,55 | 5,83E-01 | 8,04E-01 |
| AT5G50290 | 244,15 | 0,21 | 0,38 | 0,55 | 5,82E-01 | 8,04E-01 |
| AT5G62000 | 10,38 | 0,32 | 0,57 | 0,56 | 5,75E-01 | 8,04E-01 |
| AT1G09210 | 5,83 | -0,44 | 0,77 | -0,57 | 5,69E-01 | 8,04E-01 |
| AT1G70770 | 12,70 | -0,29 | 0,53 | -0,55 | 5,85E-01 | 8,04E-01 |
| AT1G72610 | 17,51 | -0,28 | 0,50 | -0,56 | 5,77E-01 | 8,04E-01 |
| AT2G02741 | 19,79 | 0,36 | 0,64 | 0,56 | 5,74E-01 | 8,04E-01 |
| AT2G45540 | 12,39 | 0,36 | 0,66 | 0,55 | 5,80E-01 | 8,04E-01 |
| AT3G09275 | 11,41 | 0,41 | 0,74 | 0,55 | 5,81E-01 | 8,04E-01 |
| AT4G01800 | 8,35 | -0,36 | 0,65 | -0,55 | 5,84E-01 | 8,04E-01 |
| AT5G15680 | 7,75 | -0,38 | 0,66 | -0,57 | 5,71E-01 | 8,04E-01 |
| AT5G53460 | 17,48 | -0,24 | 0,44 | -0,54 | 5,89E-01 | 8,05E-01 |
| AT1G10170 | 9,09 | -0,34 | 0,64 | -0,54 | 5,89E-01 | 8,05E-01 |
| AT5G56010 | 6,62 | -0,44 | 0,83 | -0,53 | 5,99E-01 | 8,17E-01 |
| AT1G13930 | 7,00 | -0,45 | 0,86 | -0,52 | 6,02E-01 | 8,19E-01 |
| AT3G23000 | 7,09 | -0,38 | 0,74 | -0,52 | 6,06E-01 | 8,21E-01 |
| AT5G39500 | 6,69 | -0,38 | 0,74 | -0,52 | 6,06E-01 | 8,21E-01 |
| AT2G17930 | 7,05 | -0,33 | 0,66 | -0,51 | 6,09E-01 | 8,24E-01 |
| AT2G36910 | 8,87 | -0,35 | 0,70 | -0,51 | 6,13E-01 | 8,25E-01 |
| AT2G41430 | 6,85 | 0,44 | 0,86 | 0,51 | 6,12E-01 | 8,25E-01 |
| AT4G02970 | 72,42 | 0,16 | 0,32 | 0,50 | 6,16E-01 | 8,25E-01 |
| AT5G13630 | 269,68 | -0,18 | 0,35 | -0,50 | 6,15E-01 | 8,25E-01 |
| AT2G43680 | 14,92 | -0,28 | 0,56 | -0,50 | 6,20E-01 | 8,30E-01 |

| | | | | | | |
|---|---|---|---|---|---|---|
| AT5G42950 | 18,25 | -0,31 | 0,64 | -0,49 | 6,22E-01 | 8,30E-01 |
| AT5G67360 | 6,15 | -0,46 | 0,94 | -0,49 | 6,23E-01 | 8,30E-01 |
| AT5G28540 | 7,23 | -0,36 | 0,74 | -0,49 | 6,26E-01 | 8,32E-01 |
| AT1G05570 | 10,12 | -0,29 | 0,60 | -0,48 | 6,28E-01 | 8,33E-01 |
| AT2G07698 | 9,74 | -0,34 | 0,71 | -0,48 | 6,33E-01 | 8,37E-01 |
| AT1G20760 | 7,86 | 0,34 | 0,75 | 0,46 | 6,46E-01 | 8,46E-01 |
| AT1G50480 | 9,27 | 0,29 | 0,64 | 0,45 | 6,49E-01 | 8,46E-01 |
| AT1G69797 | 12,98 | 0,28 | 0,62 | 0,46 | 6,49E-01 | 8,46E-01 |
| AT3G13300 | 8,42 | 0,43 | 0,95 | 0,45 | 6,50E-01 | 8,46E-01 |
| AT3G26810 | 23,04 | -0,26 | 0,57 | -0,46 | 6,44E-01 | 8,46E-01 |
| AT3G41762 | 25,36 | 0,24 | 0,51 | 0,47 | 6,42E-01 | 8,46E-01 |
| AT3G53460 | 6,35 | -0,36 | 0,79 | -0,46 | 6,48E-01 | 8,46E-01 |
| AT4G17140 | 11,35 | -0,28 | 0,60 | -0,46 | 6,48E-01 | 8,46E-01 |
| AT5G01747 | 60,67 | 0,13 | 0,30 | 0,45 | 6,54E-01 | 8,47E-01 |
| AT3G09390 | 6,61 | 0,36 | 0,81 | 0,45 | 6,55E-01 | 8,47E-01 |
| AT5G09515 | 9,09 | 0,37 | 0,82 | 0,45 | 6,52E-01 | 8,47E-01 |
| AT4G24690 | 16,56 | 0,23 | 0,52 | 0,44 | 6,57E-01 | 8,48E-01 |
| AT5G06845 | 6,24 | -0,33 | 0,75 | -0,44 | 6,60E-01 | 8,48E-01 |
| AT5G58140 | 8,36 | 0,33 | 0,75 | 0,44 | 6,60E-01 | 8,48E-01 |
| AT2G45960 | 8,71 | 0,31 | 0,72 | 0,43 | 6,65E-01 | 8,48E-01 |
| AT2G43970 | 15,13 | 0,21 | 0,49 | 0,43 | 6,66E-01 | 8,48E-01 |
| AT2G48160 | 6,20 | -0,47 | 1,07 | -0,44 | 6,63E-01 | 8,48E-01 |
| AT4G27000 | 10,07 | -0,25 | 0,58 | -0,44 | 6,63E-01 | 8,48E-01 |
| AT1G22610 | 11,17 | -0,23 | 0,54 | -0,43 | 6,70E-01 | 8,49E-01 |
| AT4G09135 | 26,82 | -0,19 | 0,45 | -0,42 | 6,74E-01 | 8,49E-01 |
| AT1G70320 | 18,40 | -0,17 | 0,41 | -0,42 | 6,78E-01 | 8,49E-01 |
| AT2G02750 | 20,13 | 0,26 | 0,62 | 0,42 | 6,77E-01 | 8,49E-01 |
| AT2G10606 | 1219,01 | -0,11 | 0,27 | -0,41 | 6,80E-01 | 8,49E-01 |
| AT2G35387 | 52,44 | -0,16 | 0,39 | -0,42 | 6,76E-01 | 8,49E-01 |
| AT2G35750 | 26,17 | 0,25 | 0,58 | 0,43 | 6,70E-01 | 8,49E-01 |
| AT2G37190 | 11,90 | 0,25 | 0,59 | 0,42 | 6,76E-01 | 8,49E-01 |
| AT3G46970 | 7,07 | -0,30 | 0,73 | -0,41 | 6,79E-01 | 8,49E-01 |
| AT4G14910 | 7,50 | -0,29 | 0,70 | -0,42 | 6,75E-01 | 8,49E-01 |
| AT1G74090 | 6,35 | -0,32 | 0,78 | -0,41 | 6,85E-01 | 8,52E-01 |
| AT3G07725 | 10,80 | 0,26 | 0,65 | 0,40 | 6,89E-01 | 8,52E-01 |
| AT3G47347 | 15,63 | 0,23 | 0,59 | 0,40 | 6,92E-01 | 8,52E-01 |
| AT5G41750 | 8,05 | 0,26 | 0,66 | 0,40 | 6,87E-01 | 8,52E-01 |
| AT4G06235 | 7,69 | 0,48 | 1,21 | 0,40 | 6,92E-01 | 8,52E-01 |
| AT5G38410 | 6,26 | -0,41 | 1,03 | -0,40 | 6,88E-01 | 8,52E-01 |
| AT5G60450 | 7,78 | -0,32 | 0,82 | -0,40 | 6,91E-01 | 8,52E-01 |
| AT4G35830 | 9,01 | -0,30 | 0,77 | -0,39 | 6,95E-01 | 8,53E-01 |
| AT5G07325 | 8,05 | 0,26 | 0,66 | 0,39 | 6,95E-01 | 8,53E-01 |
| AT1G05917 | 8,49 | -0,34 | 0,89 | -0,38 | 7,04E-01 | 8,62E-01 |
| AT4G06320 | 7,79 | 0,29 | 0,78 | 0,38 | 7,06E-01 | 8,63E-01 |
| AT3G45140 | 15,78 | 0,20 | 0,55 | 0,37 | 7,12E-01 | 8,63E-01 |
| AT5G10380 | 7,30 | -0,27 | 0,72 | -0,37 | 7,08E-01 | 8,63E-01 |
| AT1G48090 | 11,18 | 0,20 | 0,53 | 0,37 | 7,09E-01 | 8,63E-01 |
| AT2G39800 | 6,12 | -0,30 | 0,81 | -0,37 | 7,11E-01 | 8,63E-01 |
| AT3G60750 | 11,99 | -0,21 | 0,60 | -0,36 | 7,21E-01 | 8,72E-01 |
| AT1G05913 | 9,06 | -0,31 | 0,89 | -0,35 | 7,26E-01 | 8,77E-01 |
| AT1G29465 | 25,62 | 0,19 | 0,56 | 0,34 | 7,33E-01 | 8,80E-01 |
| AT1G80070 | 7,13 | -0,23 | 0,67 | -0,34 | 7,35E-01 | 8,80E-01 |

| | | | | | | |
|---|---|---|---|---|---|---|
| AT3G24615 | 9,91 | -0,23 | 0,69 | -0,34 | 7,36E-01 | 8,80E-01 |
| AT4G17390 | 6,32 | -0,29 | 0,86 | -0,34 | 7,35E-01 | 8,80E-01 |
| AT5G50920 | 18,45 | 0,17 | 0,51 | 0,34 | 7,33E-01 | 8,80E-01 |
| AT5G19150 | 6,35 | -0,25 | 0,76 | -0,33 | 7,41E-01 | 8,84E-01 |
| AT2G26890 | 8,74 | -0,20 | 0,64 | -0,32 | 7,50E-01 | 8,93E-01 |
| AT1G62020 | 12,54 | -0,21 | 0,67 | -0,31 | 7,56E-01 | 8,99E-01 |
| AT1G20960 | 9,33 | 0,19 | 0,61 | 0,30 | 7,62E-01 | 9,03E-01 |
| AT1G76180 | 6,03 | 0,24 | 0,78 | 0,30 | 7,61E-01 | 9,03E-01 |
| AT4G16190 | 11,22 | 0,18 | 0,61 | 0,30 | 7,65E-01 | 9,04E-01 |
| AT4G02510 | 15,59 | 0,16 | 0,57 | 0,28 | 7,76E-01 | 9,15E-01 |
| AT4G05420 | 7,60 | -0,19 | 0,68 | -0,28 | 7,78E-01 | 9,15E-01 |
| AT3G09440 | 5,91 | 0,22 | 0,79 | 0,28 | 7,78E-01 | 9,15E-01 |
| AT2G46192 | 21,12 | -0,18 | 0,65 | -0,28 | 7,82E-01 | 9,16E-01 |
| AT2G41740 | 6,26 | -0,24 | 0,85 | -0,28 | 7,80E-01 | 9,16E-01 |
| AT2G47070 | 5,92 | 0,19 | 0,71 | 0,26 | 7,92E-01 | 9,21E-01 |
| AT1G67120 | 8,25 | -0,17 | 0,63 | -0,27 | 7,89E-01 | 9,21E-01 |
| AT3G18217 | 147,37 | -0,06 | 0,24 | -0,26 | 7,91E-01 | 9,21E-01 |
| AT4G26850 | 7,34 | 0,18 | 0,67 | 0,26 | 7,92E-01 | 9,21E-01 |
| AT4G29410 | 17,21 | -0,16 | 0,61 | -0,26 | 7,93E-01 | 9,21E-01 |
| AT2G47400 | 11,67 | -0,14 | 0,56 | -0,26 | 7,97E-01 | 9,22E-01 |
| AT5G13887 | 13,72 | 0,23 | 0,88 | 0,26 | 7,97E-01 | 9,22E-01 |
| AT1G26985 | 10,26 | 0,16 | 0,69 | 0,23 | 8,18E-01 | 9,24E-01 |
| AT1G35501 | 18,58 | -0,16 | 0,65 | -0,25 | 8,01E-01 | 9,24E-01 |
| AT2G03330 | 8,86 | 0,19 | 0,80 | 0,24 | 8,11E-01 | 9,24E-01 |
| AT2G28290 | 15,57 | 0,12 | 0,50 | 0,24 | 8,11E-01 | 9,24E-01 |
| AT2G08665 | 53,97 | 0,10 | 0,42 | 0,23 | 8,16E-01 | 9,24E-01 |
| AT2G41900 | 16,69 | -0,15 | 0,61 | -0,25 | 8,03E-01 | 9,24E-01 |
| AT4G31570 | 11,16 | -0,18 | 0,72 | -0,25 | 8,06E-01 | 9,24E-01 |
| AT4G39363 | 6,83 | -0,18 | 0,80 | -0,23 | 8,20E-01 | 9,24E-01 |
| AT2G16500 | 6,48 | 0,17 | 0,74 | 0,23 | 8,20E-01 | 9,24E-01 |
| AT2G34420 | 53,97 | 0,10 | 0,42 | 0,23 | 8,16E-01 | 9,24E-01 |
| AT2G42600 | 13,57 | 0,12 | 0,52 | 0,23 | 8,18E-01 | 9,24E-01 |
| AT4G02770 | 8,37 | -0,18 | 0,78 | -0,23 | 8,20E-01 | 9,24E-01 |
| AT4G20850 | 8,05 | -0,17 | 0,67 | -0,25 | 8,04E-01 | 9,24E-01 |
| AT5G13890 | 14,03 | 0,18 | 0,77 | 0,23 | 8,20E-01 | 9,24E-01 |
| AT1G21250 | 6,46 | 0,15 | 0,70 | 0,22 | 8,30E-01 | 9,30E-01 |
| AT1G78478 | 10,59 | -0,17 | 0,79 | -0,22 | 8,29E-01 | 9,30E-01 |
| AT2G43375 | 6,65 | -0,17 | 0,77 | -0,22 | 8,28E-01 | 9,30E-01 |
| AT1G36160 | 9,82 | -0,13 | 0,63 | -0,21 | 8,36E-01 | 9,36E-01 |
| AT4G06240 | 6,18 | 0,27 | 1,35 | 0,20 | 8,39E-01 | 9,37E-01 |
| AT1G12290 | 131,54 | 0,05 | 0,26 | 0,20 | 8,44E-01 | 9,40E-01 |
| AT1G12294 | 124,95 | 0,05 | 0,26 | 0,20 | 8,45E-01 | 9,40E-01 |
| AT4G31500 | 6,34 | 0,14 | 0,74 | 0,19 | 8,46E-01 | 9,40E-01 |
| AT1G31163 | 6,82 | -0,15 | 0,79 | -0,19 | 8,49E-01 | 9,41E-01 |
| AT1G08103 | 6,40 | -0,18 | 1,00 | -0,18 | 8,53E-01 | 9,44E-01 |
| AT5G04140 | 18,61 | 0,09 | 0,51 | 0,18 | 8,55E-01 | 9,45E-01 |
| AT1G42970 | 13,13 | -0,10 | 0,58 | -0,18 | 8,57E-01 | 9,45E-01 |
| AT4G24280 | 6,53 | 0,15 | 0,84 | 0,17 | 8,62E-01 | 9,49E-01 |
| AT1G07810 | 6,32 | 0,12 | 0,73 | 0,17 | 8,66E-01 | 9,50E-01 |
| AT5G19140 | 6,43 | -0,13 | 0,75 | -0,17 | 8,65E-01 | 9,50E-01 |
| AT1G29900 | 11,05 | -0,09 | 0,56 | -0,16 | 8,75E-01 | 9,52E-01 |
| AT1G09797 | 19,74 | 0,06 | 0,42 | 0,15 | 8,77E-01 | 9,52E-01 |

| | | | | | | |
|---|---|---|---|---|---|---|
| AT1G78476 | 11,35 | -0,11 | 0,73 | -0,16 | 8,76E-01 | 9,52E-01 |
| AT3G44310 | 10,52 | 0,11 | 0,72 | 0,15 | 8,80E-01 | 9,52E-01 |
| AT5G08712 | 123,44 | -0,05 | 0,34 | -0,14 | 8,88E-01 | 9,52E-01 |
| AT1G22620 | 12,70 | -0,09 | 0,55 | -0,16 | 8,72E-01 | 9,52E-01 |
| AT1G08597 | 22,78 | 0,15 | 1,03 | 0,14 | 8,87E-01 | 9,52E-01 |
| AT1G74100 | 5,98 | 0,13 | 0,91 | 0,15 | 8,82E-01 | 9,52E-01 |
| AT1G09793 | 19,74 | 0,06 | 0,42 | 0,15 | 8,77E-01 | 9,52E-01 |
| AT3G08530 | 12,53 | -0,08 | 0,55 | -0,14 | 8,86E-01 | 9,52E-01 |
| AT4G02380 | 7,09 | 0,10 | 0,66 | 0,16 | 8,76E-01 | 9,52E-01 |
| AT5G04315 | 6,08 | -0,11 | 0,73 | -0,15 | 8,84E-01 | 9,52E-01 |
| AT5G04325 | 6,08 | -0,11 | 0,73 | -0,15 | 8,84E-01 | 9,52E-01 |
| AT1G05562 | 6,13 | -0,15 | 1,09 | -0,13 | 8,94E-01 | 9,55E-01 |
| AT1G05560 | 6,13 | -0,15 | 1,09 | -0,13 | 8,94E-01 | 9,55E-01 |
| AT1G30230 | 5,95 | -0,11 | 0,80 | -0,13 | 8,96E-01 | 9,56E-01 |
| AT4G38600 | 8,47 | -0,09 | 0,76 | -0,12 | 9,08E-01 | 9,66E-01 |
| AT4G39260 | 6,03 | -0,09 | 0,81 | -0,12 | 9,08E-01 | 9,66E-01 |
| AT5G54770 | 28,55 | 0,06 | 0,53 | 0,11 | 9,13E-01 | 9,67E-01 |
| AT1G04527 | 11,72 | -0,06 | 0,57 | -0,11 | 9,13E-01 | 9,67E-01 |
| AT1G06277 | 5,95 | 0,08 | 0,93 | 0,09 | 9,31E-01 | 9,70E-01 |
| AT1G49240 | 7,10 | 0,07 | 0,77 | 0,10 | 9,23E-01 | 9,70E-01 |
| AT5G42390 | 8,64 | -0,07 | 0,74 | -0,09 | 9,29E-01 | 9,70E-01 |
| AT1G48953 | 8,24 | 0,09 | 1,01 | 0,09 | 9,31E-01 | 9,70E-01 |
| AT1G64790 | 7,36 | 0,07 | 0,72 | 0,10 | 9,20E-01 | 9,70E-01 |
| AT1G74260 | 9,63 | -0,06 | 0,60 | -0,10 | 9,20E-01 | 9,70E-01 |
| AT2G39730 | 47,53 | 0,04 | 0,40 | 0,09 | 9,29E-01 | 9,70E-01 |
| AT3G04775 | 14,57 | -0,05 | 0,54 | -0,09 | 9,30E-01 | 9,70E-01 |
| AT4G30190 | 7,61 | -0,07 | 0,76 | -0,09 | 9,28E-01 | 9,70E-01 |
| AT4G38970 | 12,35 | -0,05 | 0,53 | -0,09 | 9,31E-01 | 9,70E-01 |
| AT1G05423 | 13,10 | -0,05 | 0,68 | -0,08 | 9,37E-01 | 9,72E-01 |
| AT4G16890 | 15,81 | -0,05 | 0,58 | -0,08 | 9,37E-01 | 9,72E-01 |
| AT5G48160 | 6,17 | 0,06 | 0,82 | 0,08 | 9,37E-01 | 9,72E-01 |
| AT1G13980 | 8,14 | 0,05 | 0,67 | 0,07 | 9,41E-01 | 9,73E-01 |
| AT5G39770 | 12,51 | 0,05 | 0,65 | 0,07 | 9,41E-01 | 9,73E-01 |
| AT1G63770 | 7,56 | 0,04 | 0,66 | 0,06 | 9,51E-01 | 9,76E-01 |
| AT1G09817 | 10,25 | -0,05 | 0,84 | -0,06 | 9,51E-01 | 9,76E-01 |
| AT3G08580 | 9,77 | 0,04 | 0,64 | 0,06 | 9,51E-01 | 9,76E-01 |
| AT3G59884 | 100,21 | 0,02 | 0,32 | 0,06 | 9,51E-01 | 9,76E-01 |
| AT1G08537 | 8,63 | 0,08 | 1,69 | 0,05 | 9,61E-01 | 9,78E-01 |
| AT2G31880 | 6,93 | 0,04 | 0,78 | 0,05 | 9,59E-01 | 9,78E-01 |
| AT5G08717 | 14,65 | 0,03 | 0,72 | 0,05 | 9,62E-01 | 9,78E-01 |
| AT3G59710 | 6,69 | 0,05 | 0,93 | 0,05 | 9,60E-01 | 9,78E-01 |
| AT3G09225 | 6,69 | 0,05 | 0,93 | 0,05 | 9,60E-01 | 9,78E-01 |
| AT5G20280 | 6,18 | 0,04 | 0,71 | 0,05 | 9,58E-01 | 9,78E-01 |
| AT2G25510 | 7,52 | -0,03 | 0,81 | -0,04 | 9,72E-01 | 9,83E-01 |
| AT4G06075 | 405,29 | 0,01 | 0,34 | 0,03 | 9,73E-01 | 9,83E-01 |
| AT3G50820 | 7,18 | -0,04 | 1,04 | -0,04 | 9,70E-01 | 9,83E-01 |
| AT5G27660 | 6,15 | 0,03 | 0,76 | 0,04 | 9,70E-01 | 9,83E-01 |
| AT3G57720 | 13,95 | -0,02 | 0,65 | -0,03 | 9,76E-01 | 9,84E-01 |
| AT1G58602 | 11,97 | 0,01 | 0,57 | 0,02 | 9,81E-01 | 9,86E-01 |
| AT1G18745 | 15,99 | 0,02 | 0,65 | 0,02 | 9,80E-01 | 9,86E-01 |
| AT3G06345 | 8,24 | 0,01 | 0,70 | 0,01 | 9,90E-01 | 9,93E-01 |
| AT4G23170 | 6,38 | -0,01 | 0,93 | -0,01 | 9,93E-01 | 9,94E-01 |

| | | | | | | |
|---|---|---|---|---|---|---|
| AT2G47450 | 6,02 | 0,00 | 0,80 | 0,00 | 9,99E-01 | 9,99E-01 |
| AT1G01040 | 4,73 | 0,92 | 0,94 | 0,99 | 3,24E-01 | |
| AT1G01046 | 1,63 | 0,23 | 1,54 | 0,15 | 8,82E-01 | |
| AT1G01160 | 0,43 | -0,85 | 3,29 | -0,26 | 7,96E-01 | |
| AT1G01180 | 0,85 | 2,86 | 2,32 | 1,23 | 2,18E-01 | |
| AT1G01220 | 0,11 | 0,65 | 4,36 | 0,15 | 8,81E-01 | |
| AT1G01225 | 0,31 | -1,33 | 4,35 | -0,31 | 7,60E-01 | |
| AT1G01230 | 0,37 | -2,04 | 4,37 | -0,47 | 6,40E-01 | |
| AT1G01240 | 0,19 | 0,36 | 4,36 | 0,08 | 9,34E-01 | |
| AT1G01260 | 0,53 | -0,13 | 2,71 | -0,05 | 9,62E-01 | |
| AT1G01300 | 1,08 | 2,24 | 2,01 | 1,12 | 2,64E-01 | |
| AT1G01350 | 0,29 | 0,10 | 4,32 | 0,02 | 9,82E-01 | |
| AT1G01448 | 0,59 | 1,99 | 4,32 | 0,46 | 6,44E-01 | |
| AT1G01480 | 0,12 | 1,23 | 4,34 | 0,28 | 7,76E-01 | |
| AT1G01510 | 2,79 | -1,75 | 1,26 | -1,39 | 1,66E-01 | |
| AT1G01520 | 0,12 | 1,23 | 4,34 | 0,28 | 7,76E-01 | |
| AT1G01540 | 0,22 | 1,19 | 4,33 | 0,27 | 7,84E-01 | |
| AT1G01550 | 2,54 | -2,12 | 1,19 | -1,78 | 7,49E-02 | |
| AT1G01560 | 1,60 | 0,43 | 1,66 | 0,26 | 7,97E-01 | |
| AT1G01630 | 0,58 | -2,96 | 3,78 | -0,78 | 4,34E-01 | |
| AT1G01710 | 0,82 | -1,49 | 3,15 | -0,47 | 6,35E-01 | |
| AT1G01720 | 1,37 | -0,16 | 1,52 | -0,10 | 9,17E-01 | |
| AT1G04047 | 0,70 | 2,09 | 2,54 | 0,82 | 4,11E-01 | |
| AT1G01730 | 0,40 | -1,72 | 4,35 | -0,40 | 6,92E-01 | |
| AT1G01790 | 0,83 | -0,26 | 1,85 | -0,14 | 8,89E-01 | |
| AT1G01810 | 0,12 | 1,16 | 4,34 | 0,27 | 7,89E-01 | |
| AT1G01950 | 0,18 | 0,65 | 4,35 | 0,15 | 8,82E-01 | |
| AT1G02010 | 0,10 | 0,21 | 4,38 | 0,05 | 9,62E-01 | |
| AT1G02080 | 3,21 | -0,34 | 1,26 | -0,27 | 7,85E-01 | |
| AT1G02100 | 0,49 | 2,14 | 3,14 | 0,68 | 4,96E-01 | |
| AT1G02120 | 0,36 | 1,57 | 3,46 | 0,46 | 6,49E-01 | |
| AT1G02150 | 1,38 | -0,71 | 1,71 | -0,41 | 6,80E-01 | |
| AT1G02205 | 1,06 | 1,65 | 2,09 | 0,79 | 4,30E-01 | |
| AT1G04093 | 0,08 | 0,00 | 4,40 | 0,00 | 1,00E+00 | |
| AT1G02300 | 0,34 | 0,12 | 4,35 | 0,03 | 9,78E-01 | |
| AT1G02305 | 0,80 | 0,46 | 2,42 | 0,19 | 8,49E-01 | |
| AT1G04103 | 0,08 | 0,69 | 4,36 | 0,16 | 8,75E-01 | |
| AT1G02390 | 0,19 | 0,38 | 4,37 | 0,09 | 9,31E-01 | |
| AT1G02420 | 0,52 | 1,36 | 3,86 | 0,35 | 7,24E-01 | |
| AT1G02560 | 1,57 | -1,55 | 1,55 | -1,00 | 3,18E-01 | |
| AT1G02690 | 0,11 | 0,00 | 4,41 | 0,00 | 1,00E+00 | |
| AT1G02750 | 0,51 | -2,36 | 4,32 | -0,55 | 5,84E-01 | |
| AT1G02816 | 0,38 | 0,48 | 3,34 | 0,14 | 8,85E-01 | |
| AT1G02850 | 0,62 | 2,64 | 2,88 | 0,92 | 3,59E-01 | |
| AT1G02860 | 4,18 | 1,87 | 1,32 | 1,42 | 1,57E-01 | |
| AT1G03160 | 0,95 | -1,05 | 2,25 | -0,47 | 6,41E-01 | |
| AT1G03190 | 0,39 | 0,61 | 3,43 | 0,18 | 8,58E-01 | |
| AT1G03220 | 1,11 | 0,53 | 2,04 | 0,26 | 7,93E-01 | |
| AT1G03280 | 0,25 | 1,82 | 4,32 | 0,42 | 6,73E-01 | |
| AT1G03290 | 0,75 | 2,75 | 2,42 | 1,14 | 2,54E-01 | |
| AT1G03310 | 0,40 | -0,23 | 4,19 | -0,06 | 9,56E-01 | |
| AT1G03360 | 0,35 | 0,23 | 3,48 | 0,07 | 9,46E-01 | |

| | | | | | |
|---|---|---|---|---|---|
| AT1G03365 | 0,40 | 0,41 | 3,36 | 0,12 | 9,03E-01 |
| AT1G03370 | 0,51 | 1,97 | 3,95 | 0,50 | 6,18E-01 |
| AT1G03380 | 1,12 | 0,42 | 1,99 | 0,21 | 8,33E-01 |
| AT1G04183 | 2,59 | 0,74 | 1,38 | 0,54 | 5,89E-01 |
| AT1G03430 | 0,37 | -1,03 | 4,36 | -0,24 | 8,14E-01 |
| AT1G03440 | 0,84 | -1,79 | 3,14 | -0,57 | 5,68E-01 |
| AT1G03545 | 0,70 | -2,13 | 2,29 | -0,93 | 3,52E-01 |
| AT1G03600 | 1,40 | 1,04 | 1,79 | 0,58 | 5,64E-01 |
| AT1G03610 | 0,72 | 1,65 | 3,21 | 0,52 | 6,06E-01 |
| AT1G03630 | 0,51 | -0,15 | 2,38 | -0,06 | 9,51E-01 |
| AT1G03687 | 0,11 | 0,00 | 4,41 | 0,00 | 1,00E+00 |
| AT1G03740 | 0,96 | 1,03 | 2,01 | 0,52 | 6,06E-01 |
| AT1G03810 | 0,77 | -1,87 | 2,13 | -0,88 | 3,80E-01 |
| AT1G03900 | 1,11 | 0,84 | 2,04 | 0,41 | 6,79E-01 |
| AT1G03905 | 0,24 | -2,36 | 4,39 | -0,54 | 5,91E-01 |
| AT1G03910 | 0,50 | -0,91 | 3,11 | -0,29 | 7,68E-01 |
| AT1G03930 | 0,53 | -0,61 | 2,62 | -0,23 | 8,16E-01 |
| AT1G04217 | 0,82 | 0,27 | 1,95 | 0,14 | 8,92E-01 |
| AT1G03935 | 0,16 | 0,00 | 4,41 | 0,00 | 1,00E+00 |
| AT1G04223 | 0,98 | -1,80 | 2,14 | -0,84 | 4,00E-01 |
| AT1G04227 | 0,32 | 0,83 | 3,64 | 0,23 | 8,19E-01 |
| AT1G03960 | 0,15 | 0,67 | 4,37 | 0,15 | 8,79E-01 |
| AT1G03970 | 0,36 | 1,22 | 4,34 | 0,28 | 7,80E-01 |
| AT1G04020 | 0,33 | -2,31 | 4,39 | -0,53 | 5,98E-01 |
| AT1G04080 | 1,89 | -1,29 | 1,26 | -1,03 | 3,04E-01 |
| AT1G04130 | 0,50 | -0,13 | 2,70 | -0,05 | 9,61E-01 |
| AT1G04170 | 1,02 | -0,62 | 1,83 | -0,34 | 7,36E-01 |
| AT1G04210 | 3,48 | -0,93 | 0,89 | -1,05 | 2,94E-01 |
| AT1G04247 | 0,30 | -1,71 | 4,40 | -0,39 | 6,98E-01 |
| AT1G04230 | 0,70 | 0,16 | 3,04 | 0,05 | 9,59E-01 |
| AT1G04295 | 0,43 | 0,64 | 2,74 | 0,24 | 8,14E-01 |
| AT1G04350 | 0,25 | 0,00 | 4,41 | 0,00 | 1,00E+00 |
| AT1G04420 | 1,20 | -0,25 | 1,67 | -0,15 | 8,79E-01 |
| AT1G04425 | 1,91 | -0,49 | 1,65 | -0,30 | 7,67E-01 |
| AT1G04440 | 0,50 | -1,09 | 2,68 | -0,41 | 6,84E-01 |
| AT1G04480 | 2,74 | -0,79 | 1,17 | -0,68 | 4,99E-01 |
| AT1G04273 | 0,37 | -0,87 | 3,52 | -0,25 | 8,04E-01 |
| AT1G04277 | 0,93 | -1,39 | 2,16 | -0,64 | 5,20E-01 |
| AT1G04510 | 0,67 | -0,74 | 2,90 | -0,25 | 7,99E-01 |
| AT1G04540 | 1,05 | -0,25 | 2,04 | -0,12 | 9,04E-01 |
| AT1G04690 | 0,77 | 0,77 | 2,34 | 0,33 | 7,42E-01 |
| AT1G04710 | 0,47 | -0,23 | 4,33 | -0,05 | 9,58E-01 |
| AT1G04760 | 0,36 | 0,42 | 4,38 | 0,10 | 9,23E-01 |
| AT1G04810 | 3,41 | -0,19 | 1,22 | -0,16 | 8,74E-01 |
| AT1G04945 | 0,47 | -1,21 | 3,20 | -0,38 | 7,04E-01 |
| AT1G04970 | 0,34 | 0,91 | 4,35 | 0,21 | 8,35E-01 |
| AT1G05035 | 3,55 | -0,39 | 1,29 | -0,31 | 7,59E-01 |
| AT1G05087 | 0,44 | 2,16 | 4,05 | 0,53 | 5,95E-01 |
| AT1G05090 | 0,12 | 1,23 | 4,34 | 0,28 | 7,76E-01 |
| AT1G05140 | 0,22 | -1,77 | 4,41 | -0,40 | 6,88E-01 |
| AT1G05200 | 1,12 | -0,64 | 1,91 | -0,33 | 7,39E-01 |
| AT1G05240 | 2,51 | 0,30 | 1,63 | 0,19 | 8,53E-01 |

| | | | | | |
|---|---|---|---|---|---|
| AT1G05360 | 0,08 | 0,69 | 4,36 | 0,16 | 8,75E-01 |
| AT1G05380 | 1,42 | 0,89 | 1,88 | 0,47 | 6,35E-01 |
| AT1G05500 | 0,37 | 0,31 | 4,39 | 0,07 | 9,44E-01 |
| AT1G04397 | 0,08 | 0,69 | 4,36 | 0,16 | 8,75E-01 |
| AT1G05600 | 0,08 | 0,00 | 4,40 | 0,00 | 1,00E+00 |
| AT1G05630 | 1,84 | 0,18 | 1,40 | 0,13 | 8,98E-01 |
| AT1G05780 | 4,77 | 0,72 | 1,05 | 0,69 | 4,92E-01 |
| AT1G04423 | 1,18 | -0,62 | 1,94 | -0,32 | 7,50E-01 |
| AT1G05790 | 0,76 | 0,63 | 2,39 | 0,26 | 7,93E-01 |
| AT1G05805 | 1,94 | 2,58 | 2,30 | 1,12 | 2,63E-01 |
| AT1G05830 | 0,53 | 0,15 | 3,30 | 0,04 | 9,65E-01 |
| AT1G05890 | 0,68 | -2,39 | 4,32 | -0,55 | 5,80E-01 |
| AT1G05900 | 0,21 | -1,41 | 4,36 | -0,32 | 7,46E-01 |
| AT1G05910 | 0,59 | -1,22 | 3,70 | -0,33 | 7,41E-01 |
| AT1G05980 | 0,23 | 0,00 | 4,41 | 0,00 | 1,00E+00 |
| AT1G06002 | 0,95 | -1,31 | 1,96 | -0,67 | 5,05E-01 |
| AT1G06030 | 0,96 | -1,03 | 3,28 | -0,31 | 7,53E-01 |
| AT1G06220 | 3,15 | -0,51 | 1,05 | -0,49 | 6,26E-01 |
| AT1G06230 | 2,14 | -0,62 | 1,31 | -0,47 | 6,37E-01 |
| AT1G06290 | 0,60 | -0,41 | 2,71 | -0,15 | 8,81E-01 |
| AT1G06380 | 0,45 | -1,76 | 4,24 | -0,41 | 6,78E-01 |
| AT1G06390 | 0,40 | -1,21 | 3,37 | -0,36 | 7,19E-01 |
| AT1G06410 | 1,61 | 0,43 | 1,56 | 0,27 | 7,85E-01 |
| AT1G06530 | 0,96 | -2,21 | 1,97 | -1,12 | 2,62E-01 |
| AT1G06640 | 1,12 | 0,12 | 1,82 | 0,06 | 9,49E-01 |
| AT1G06645 | 0,50 | 1,07 | 3,97 | 0,27 | 7,87E-01 |
| AT1G06650 | 0,69 | -0,04 | 2,90 | -0,01 | 9,88E-01 |
| AT1G06670 | 1,38 | -1,03 | 1,79 | -0,57 | 5,66E-01 |
| AT1G06750 | 0,24 | -0,08 | 4,34 | -0,02 | 9,85E-01 |
| AT1G06780 | 0,12 | 0,49 | 4,36 | 0,11 | 9,11E-01 |
| AT1G06910 | 0,23 | -0,05 | 4,34 | -0,01 | 9,91E-01 |
| AT1G06930 | 0,08 | 0,69 | 4,36 | 0,16 | 8,75E-01 |
| AT1G07010 | 0,86 | -0,77 | 2,82 | -0,27 | 7,84E-01 |
| AT1G04537 | 0,24 | -2,36 | 4,39 | -0,54 | 5,91E-01 |
| AT1G07080 | 0,86 | -0,15 | 2,15 | -0,07 | 9,45E-01 |
| AT1G07130 | 0,28 | 1,27 | 4,35 | 0,29 | 7,69E-01 |
| AT1G07170 | 0,78 | 1,52 | 2,12 | 0,72 | 4,74E-01 |
| AT1G07180 | 0,37 | -0,62 | 3,52 | -0,18 | 8,61E-01 |
| AT1G07210 | 0,17 | 1,40 | 4,33 | 0,32 | 7,47E-01 |
| AT1G07240 | 0,59 | -0,72 | 3,21 | -0,22 | 8,22E-01 |
| AT1G07250 | 2,07 | -1,44 | 1,24 | -1,16 | 2,44E-01 |
| AT1G07280 | 1,33 | -0,89 | 1,74 | -0,51 | 6,11E-01 |
| AT1G07320 | 1,43 | -1,44 | 1,46 | -0,99 | 3,23E-01 |
| AT1G07470 | 0,61 | -1,33 | 3,64 | -0,37 | 7,15E-01 |
| AT1G07510 | 1,74 | 1,94 | 1,56 | 1,24 | 2,15E-01 |
| AT1G07630 | 0,70 | -0,51 | 2,84 | -0,18 | 8,59E-01 |
| AT1G07705 | 0,12 | 0,97 | 4,34 | 0,22 | 8,23E-01 |
| AT1G07728 | 1,03 | -1,48 | 2,31 | -0,64 | 5,21E-01 |
| AT1G07790 | 0,25 | 0,04 | 4,39 | 0,01 | 9,93E-01 |
| AT1G07830 | 0,43 | 1,70 | 4,15 | 0,41 | 6,82E-01 |
| AT1G07890 | 4,08 | -0,47 | 0,91 | -0,51 | 6,07E-01 |
| AT1G07930 | 3,92 | -0,87 | 1,02 | -0,86 | 3,91E-01 |

| | | | | | |
|---|---|---|---|---|---|
| AT1G07990 | 0,78 | 0,95 | 2,96 | 0,32 | 7,48E-01 |
| AT1G08110 | 0,41 | -0,36 | 4,26 | -0,08 | 9,33E-01 |
| AT1G08190 | 0,34 | 0,73 | 3,59 | 0,20 | 8,38E-01 |
| AT1G08260 | 0,51 | 2,16 | 4,08 | 0,53 | 5,97E-01 |
| AT1G08360 | 1,45 | 1,08 | 1,57 | 0,68 | 4,94E-01 |
| AT1G08370 | 0,71 | 0,60 | 2,39 | 0,25 | 8,03E-01 |
| AT1G08410 | 0,87 | -1,31 | 2,47 | -0,53 | 5,94E-01 |
| AT1G08420 | 1,72 | -0,23 | 1,40 | -0,16 | 8,70E-01 |
| AT1G08460 | 0,84 | 0,71 | 2,13 | 0,34 | 7,37E-01 |
| AT1G08465 | 0,35 | 0,16 | 4,34 | 0,04 | 9,71E-01 |
| AT1G08480 | 0,59 | 2,56 | 3,12 | 0,82 | 4,11E-01 |
| AT1G08520 | 1,05 | -0,94 | 1,65 | -0,57 | 5,69E-01 |
| AT1G08530 | 0,48 | 2,56 | 3,96 | 0,65 | 5,18E-01 |
| AT1G08550 | 0,85 | 1,36 | 2,30 | 0,59 | 5,55E-01 |
| AT1G08570 | 0,49 | 0,14 | 4,10 | 0,03 | 9,73E-01 |
| AT1G08590 | 0,74 | 1,84 | 2,84 | 0,65 | 5,16E-01 |
| AT1G08600 | 0,40 | 1,87 | 3,30 | 0,57 | 5,70E-01 |
| AT1G08620 | 1,06 | -0,38 | 1,71 | -0,22 | 8,24E-01 |
| AT1G08650 | 0,89 | 0,44 | 2,93 | 0,15 | 8,80E-01 |
| AT1G08680 | 0,17 | 0,00 | 4,41 | 0,00 | 1,00E+00 |
| AT1G08710 | 0,30 | -0,27 | 4,36 | -0,06 | 9,50E-01 |
| AT1G08720 | 1,46 | 0,81 | 1,90 | 0,43 | 6,69E-01 |
| AT1G08760 | 0,68 | 0,91 | 2,97 | 0,31 | 7,59E-01 |
| AT1G08770 | 0,33 | -0,91 | 3,56 | -0,26 | 7,98E-01 |
| AT1G08830 | 1,29 | 1,30 | 2,03 | 0,64 | 5,22E-01 |
| AT1G08900 | 0,34 | 0,24 | 3,46 | 0,07 | 9,44E-01 |
| AT1G08910 | 0,43 | -2,56 | 4,21 | -0,61 | 5,43E-01 |
| AT1G08920 | 0,59 | -0,86 | 3,09 | -0,28 | 7,80E-01 |
| AT1G08930 | 1,31 | 0,41 | 1,54 | 0,27 | 7,91E-01 |
| AT1G08970 | 0,82 | 1,21 | 2,60 | 0,47 | 6,42E-01 |
| AT1G08980 | 0,91 | -1,23 | 1,95 | -0,63 | 5,27E-01 |
| AT1G09040 | 0,45 | -0,55 | 3,24 | -0,17 | 8,66E-01 |
| AT1G09070 | 2,87 | 2,23 | 1,47 | 1,52 | 1,29E-01 |
| AT1G09240 | 1,84 | 1,48 | 2,14 | 0,69 | 4,89E-01 |
| AT1G09250 | 0,32 | -1,93 | 4,33 | -0,45 | 6,55E-01 |
| AT1G09260 | 0,62 | 0,00 | 4,41 | 0,00 | 1,00E+00 |
| AT1G09270 | 0,12 | 1,23 | 4,34 | 0,28 | 7,76E-01 |
| AT1G09300 | 0,36 | -0,56 | 4,32 | -0,13 | 8,97E-01 |
| AT1G09340 | 1,73 | -1,39 | 1,37 | -1,02 | 3,10E-01 |
| AT1G09390 | 0,23 | 0,86 | 4,34 | 0,20 | 8,43E-01 |
| AT1G09421 | 0,17 | 0,61 | 4,37 | 0,14 | 8,88E-01 |
| AT1G09430 | 0,79 | -0,12 | 2,38 | -0,05 | 9,61E-01 |
| AT1G09450 | 0,24 | -0,73 | 4,37 | -0,17 | 8,68E-01 |
| AT1G09490 | 0,05 | 0,00 | 4,41 | 0,00 | 1,00E+00 |
| AT1G09500 | 0,08 | 0,69 | 4,36 | 0,16 | 8,75E-01 |
| AT1G09530 | 0,63 | 0,21 | 3,67 | 0,06 | 9,54E-01 |
| AT1G09560 | 0,29 | 0,51 | 4,33 | 0,12 | 9,07E-01 |
| AT1G09590 | 0,27 | 0,29 | 4,39 | 0,07 | 9,48E-01 |
| AT1G09600 | 0,94 | 0,18 | 2,66 | 0,07 | 9,46E-01 |
| AT1G09610 | 0,36 | -0,29 | 4,32 | -0,07 | 9,46E-01 |
| AT1G09640 | 2,66 | -1,35 | 1,36 | -0,99 | 3,20E-01 |
| AT1G09645 | 0,33 | -0,97 | 4,32 | -0,23 | 8,21E-01 |

| | | | | | |
|---|---|---|---|---|---|
| AT1G09665 | 0,25 | -2,19 | 4,38 | -0,50 | 6,17E-01 |
| AT1G09710 | 0,91 | -1,36 | 2,77 | -0,49 | 6,23E-01 |
| AT1G09750 | 0,82 | -0,13 | 2,09 | -0,06 | 9,51E-01 |
| AT1G09770 | 3,11 | 0,37 | 1,06 | 0,35 | 7,30E-01 |
| AT1G09795 | 0,58 | -2,38 | 4,01 | -0,59 | 5,53E-01 |
| AT1G09815 | 0,21 | 0,00 | 4,41 | 0,00 | 1,00E+00 |
| AT1G09970 | 4,22 | 1,15 | 1,01 | 1,14 | 2,53E-01 |
| AT1G10050 | 0,78 | -0,42 | 2,24 | -0,19 | 8,52E-01 |
| AT1G10070 | 0,24 | 0,24 | 4,34 | 0,06 | 9,55E-01 |
| AT1G10130 | 1,07 | 0,19 | 1,82 | 0,10 | 9,17E-01 |
| AT1G10150 | 1,47 | 0,66 | 1,41 | 0,47 | 6,42E-01 |
| AT1G10270 | 0,58 | 0,04 | 2,66 | 0,01 | 9,88E-01 |
| AT1G10450 | 0,33 | 0,92 | 4,31 | 0,21 | 8,30E-01 |
| AT1G10490 | 0,65 | 0,42 | 2,39 | 0,18 | 8,61E-01 |
| AT1G10570 | 0,16 | 0,65 | 4,37 | 0,15 | 8,82E-01 |
| AT1G10600 | 1,27 | 0,78 | 1,76 | 0,44 | 6,60E-01 |
| AT1G10610 | 0,90 | -1,01 | 3,20 | -0,31 | 7,54E-01 |
| AT1G10660 | 0,54 | -2,15 | 3,97 | -0,54 | 5,87E-01 |
| AT1G10670 | 0,76 | 0,00 | 3,66 | 0,00 | 1,00E+00 |
| AT1G10700 | 0,45 | -0,34 | 4,37 | -0,08 | 9,38E-01 |
| AT1G10720 | 0,48 | 0,75 | 2,62 | 0,29 | 7,74E-01 |
| AT1G10745 | 1,57 | -2,22 | 1,95 | -1,14 | 2,55E-01 |
| AT1G10790 | 0,08 | 0,69 | 4,36 | 0,16 | 8,75E-01 |
| AT1G10850 | 0,85 | -0,91 | 3,37 | -0,27 | 7,87E-01 |
| AT1G10910 | 0,08 | 0,69 | 4,36 | 0,16 | 8,75E-01 |
| AT1G10960 | 0,86 | -0,20 | 2,79 | -0,07 | 9,44E-01 |
| AT1G11020 | 0,27 | -0,96 | 4,39 | -0,22 | 8,27E-01 |
| AT1G11050 | 0,22 | 0,00 | 4,38 | 0,00 | 1,00E+00 |
| AT1G11260 | 1,39 | -0,15 | 1,76 | -0,09 | 9,31E-01 |
| AT1G11300 | 0,32 | 1,86 | 4,33 | 0,43 | 6,67E-01 |
| AT1G11303 | 0,25 | 1,94 | 4,32 | 0,45 | 6,54E-01 |
| AT1G11320 | 0,08 | 0,44 | 4,38 | 0,10 | 9,21E-01 |
| AT1G11330 | 0,64 | 0,69 | 2,88 | 0,24 | 8,09E-01 |
| AT1G11400 | 0,26 | -0,61 | 4,36 | -0,14 | 8,89E-01 |
| AT1G11410 | 0,36 | 1,40 | 3,43 | 0,41 | 6,82E-01 |
| AT1G11430 | 0,35 | 0,58 | 3,64 | 0,16 | 8,74E-01 |
| AT1G11440 | 0,54 | 0,10 | 3,07 | 0,03 | 9,75E-01 |
| AT1G11580 | 0,31 | 0,65 | 4,32 | 0,15 | 8,81E-01 |
| AT1G11590 | 0,67 | 0,00 | 2,63 | 0,00 | 1,00E+00 |
| AT1G11593 | 0,40 | 0,64 | 4,22 | 0,15 | 8,80E-01 |
| AT1G11650 | 3,30 | 0,00 | 0,99 | 0,00 | 1,00E+00 |
| AT1G11660 | 1,14 | -0,89 | 1,73 | -0,52 | 6,06E-01 |
| AT1G11680 | 0,83 | 0,67 | 2,70 | 0,25 | 8,05E-01 |
| AT1G11710 | 0,17 | 1,52 | 4,33 | 0,35 | 7,26E-01 |
| AT1G11750 | 0,56 | 0,33 | 3,26 | 0,10 | 9,20E-01 |
| AT1G11790 | 0,42 | -0,97 | 4,34 | -0,22 | 8,23E-01 |
| AT1G11840 | 1,70 | 0,38 | 1,49 | 0,26 | 7,98E-01 |
| AT1G11860 | 2,12 | -1,68 | 1,57 | -1,07 | 2,84E-01 |
| AT1G11870 | 3,10 | 1,16 | 1,17 | 0,99 | 3,24E-01 |
| AT1G11890 | 1,50 | -0,11 | 1,78 | -0,06 | 9,49E-01 |
| AT1G11905 | 0,75 | -0,47 | 2,09 | -0,22 | 8,22E-01 |
| AT1G11930 | 0,68 | -1,03 | 2,11 | -0,49 | 6,27E-01 |

| | | | | | |
|---|---|---|---|---|---|
| AT1G12013 | 2,08 | -1,70 | 1,59 | -1,07 | 2,86E-01 |
| AT1G12015 | 0,46 | 0,00 | 4,38 | 0,00 | 1,00E+00 |
| AT1G12050 | 1,22 | -2,35 | 1,61 | -1,46 | 1,43E-01 |
| AT1G12060 | 0,12 | 0,37 | 4,37 | 0,08 | 9,33E-01 |
| AT1G12140 | 0,12 | 1,23 | 4,34 | 0,28 | 7,76E-01 |
| AT1G12210 | 0,35 | -0,57 | 4,38 | -0,13 | 8,96E-01 |
| AT1G12220 | 1,65 | -0,01 | 1,56 | 0,00 | 9,97E-01 |
| AT1G12230 | 0,35 | -0,15 | 3,50 | -0,04 | 9,66E-01 |
| AT1G12240 | 2,62 | 0,77 | 1,26 | 0,61 | 5,43E-01 |
| AT1G12250 | 0,19 | 0,01 | 4,38 | 0,00 | 9,99E-01 |
| AT1G12270 | 0,62 | 0,28 | 2,66 | 0,11 | 9,16E-01 |
| AT1G12320 | 0,82 | 1,81 | 2,83 | 0,64 | 5,22E-01 |
| AT1G12360 | 0,65 | -1,65 | 3,77 | -0,44 | 6,62E-01 |
| AT1G12410 | 0,67 | -0,87 | 3,81 | -0,23 | 8,19E-01 |
| AT1G12460 | 0,83 | -0,98 | 2,24 | -0,44 | 6,61E-01 |
| AT1G12580 | 0,43 | 0,48 | 2,79 | 0,17 | 8,64E-01 |
| AT1G12760 | 0,76 | -0,99 | 2,05 | -0,48 | 6,29E-01 |
| AT1G12810 | 0,42 | 0,82 | 2,50 | 0,33 | 7,44E-01 |
| AT1G12840 | 1,05 | 0,68 | 1,73 | 0,39 | 6,95E-01 |
| AT1G12860 | 0,23 | 0,00 | 4,38 | 0,00 | 9,99E-01 |
| AT1G12990 | 1,14 | 1,28 | 1,99 | 0,64 | 5,20E-01 |
| AT1G13040 | 0,24 | 0,59 | 4,38 | 0,13 | 8,93E-01 |
| AT1G13060 | 0,83 | -1,20 | 3,10 | -0,39 | 6,97E-01 |
| AT1G13090 | 1,29 | -2,11 | 1,76 | -1,20 | 2,30E-01 |
| AT1G13100 | 0,26 | 0,00 | 4,41 | 0,00 | 1,00E+00 |
| AT1G13110 | 2,32 | 1,06 | 1,55 | 0,68 | 4,96E-01 |
| AT1G13180 | 0,14 | 0,76 | 4,35 | 0,18 | 8,61E-01 |
| AT1G13190 | 1,26 | 2,70 | 2,49 | 1,08 | 2,78E-01 |
| AT1G13220 | 0,56 | 2,36 | 3,78 | 0,62 | 5,33E-01 |
| AT1G13260 | 1,59 | 0,19 | 1,72 | 0,11 | 9,13E-01 |
| AT1G13270 | 0,60 | 0,83 | 3,07 | 0,27 | 7,88E-01 |
| AT1G13330 | 0,70 | -0,03 | 2,48 | -0,01 | 9,90E-01 |
| AT1G13400 | 0,23 | -1,78 | 4,41 | -0,40 | 6,87E-01 |
| AT1G13430 | 0,17 | 0,98 | 4,35 | 0,23 | 8,21E-01 |
| AT1G13460 | 0,37 | 0,02 | 4,32 | 0,01 | 9,96E-01 |
| AT1G13820 | 0,11 | 0,48 | 4,37 | 0,11 | 9,12E-01 |
| AT1G13850 | 0,13 | 0,73 | 4,36 | 0,17 | 8,67E-01 |
| AT1G13880 | 0,45 | 1,83 | 3,38 | 0,54 | 5,88E-01 |
| AT1G13990 | 0,47 | -0,14 | 3,23 | -0,04 | 9,66E-01 |
| AT1G14000 | 0,81 | 1,13 | 2,26 | 0,50 | 6,18E-01 |
| AT1G14020 | 0,41 | -1,28 | 4,19 | -0,31 | 7,60E-01 |
| AT1G05047 | 0,12 | 1,23 | 4,34 | 0,28 | 7,76E-01 |
| AT1G14110 | 0,52 | 1,47 | 3,83 | 0,38 | 7,01E-01 |
| AT1G14150 | 0,05 | 0,00 | 4,41 | 0,00 | 1,00E+00 |
| AT1G14185 | 0,05 | 0,00 | 4,41 | 0,00 | 1,00E+00 |
| AT1G14230 | 0,08 | 0,00 | 4,41 | 0,00 | 1,00E+00 |
| AT1G14250 | 0,32 | 0,00 | 4,41 | 0,00 | 1,00E+00 |
| AT1G14280 | 0,63 | -1,76 | 3,04 | -0,58 | 5,62E-01 |
| AT1G14310 | 0,25 | 0,34 | 4,32 | 0,08 | 9,37E-01 |
| AT1G14320 | 3,20 | -0,46 | 1,03 | -0,44 | 6,57E-01 |
| AT1G14345 | 0,53 | -0,51 | 2,67 | -0,19 | 8,48E-01 |
| AT1G14350 | 0,37 | -0,59 | 4,33 | -0,14 | 8,92E-01 |

| | | | | | |
|---|---|---|---|---|---|
| AT1G14370 | 0,91 | 1,16 | 1,93 | 0,60 | 5,49E-01 |
| AT1G14470 | 0,37 | 0,00 | 4,40 | 0,00 | 1,00E+00 |
| AT1G14480 | 0,08 | 0,00 | 4,40 | 0,00 | 1,00E+00 |
| AT1G14560 | 0,21 | 0,45 | 4,37 | 0,10 | 9,17E-01 |
| AT1G14570 | 1,23 | -0,11 | 1,59 | -0,07 | 9,43E-01 |
| AT1G14580 | 3,99 | -0,86 | 1,23 | -0,70 | 4,83E-01 |
| AT1G14650 | 1,32 | 1,86 | 2,12 | 0,88 | 3,81E-01 |
| AT1G14670 | 2,24 | -0,57 | 1,23 | -0,47 | 6,41E-01 |
| AT1G14800 | 1,13 | -0,86 | 1,77 | -0,49 | 6,27E-01 |
| AT1G14860 | 0,08 | 0,00 | 4,40 | 0,00 | 1,00E+00 |
| AT1G14920 | 1,77 | -0,70 | 1,52 | -0,46 | 6,45E-01 |
| AT1G15010 | 2,66 | 2,72 | 1,54 | 1,77 | 7,71E-02 |
| AT1G15030 | 0,23 | 1,14 | 4,34 | 0,26 | 7,92E-01 |
| AT1G15120 | 0,30 | 0,23 | 4,33 | 0,05 | 9,57E-01 |
| AT1G15125 | 1,84 | -0,71 | 1,46 | -0,49 | 6,26E-01 |
| AT1G15190 | 0,17 | 1,31 | 4,34 | 0,30 | 7,63E-01 |
| AT1G15240 | 0,58 | 2,14 | 2,64 | 0,81 | 4,17E-01 |
| AT1G15260 | 0,61 | -0,36 | 3,15 | -0,11 | 9,10E-01 |
| AT1G15390 | 0,48 | 0,90 | 4,33 | 0,21 | 8,36E-01 |
| AT1G15430 | 0,55 | -0,01 | 3,85 | 0,00 | 9,97E-01 |
| AT1G15650 | 0,36 | -2,77 | 4,36 | -0,64 | 5,25E-01 |
| AT1G15670 | 2,03 | -0,84 | 1,27 | -0,66 | 5,12E-01 |
| AT1G05183 | 0,40 | -0,49 | 3,30 | -0,15 | 8,81E-01 |
| AT1G15720 | 0,33 | 0,00 | 4,31 | 0,00 | 1,00E+00 |
| AT1G15740 | 0,65 | -1,60 | 2,50 | -0,64 | 5,21E-01 |
| AT1G15810 | 0,77 | -0,19 | 1,99 | -0,10 | 9,22E-01 |
| AT1G05207 | 1,81 | -0,80 | 1,53 | -0,53 | 5,99E-01 |
| AT1G15890 | 1,49 | -2,09 | 1,76 | -1,19 | 2,34E-01 |
| AT1G15910 | 0,69 | -3,94 | 3,51 | -1,12 | 2,61E-01 |
| AT1G15920 | 0,80 | -1,07 | 2,58 | -0,42 | 6,77E-01 |
| AT1G15930 | 1,08 | 0,28 | 1,89 | 0,15 | 8,83E-01 |
| AT1G15940 | 2,10 | 0,48 | 1,22 | 0,39 | 6,93E-01 |
| AT1G15950 | 0,56 | -1,01 | 2,50 | -0,40 | 6,86E-01 |
| AT1G15980 | 1,06 | 0,04 | 2,06 | 0,02 | 9,86E-01 |
| AT1G16060 | 0,12 | 0,49 | 4,36 | 0,11 | 9,11E-01 |
| AT1G16080 | 0,31 | 1,34 | 4,33 | 0,31 | 7,57E-01 |
| AT1G16090 | 0,22 | 1,51 | 4,33 | 0,35 | 7,27E-01 |
| AT1G16110 | 0,61 | 1,36 | 3,61 | 0,38 | 7,05E-01 |
| AT1G16180 | 1,23 | 0,98 | 1,76 | 0,56 | 5,77E-01 |
| AT1G16250 | 0,81 | 0,48 | 2,26 | 0,21 | 8,31E-01 |
| AT1G16270 | 1,59 | -0,71 | 1,39 | -0,51 | 6,11E-01 |
| AT1G16350 | 0,39 | 0,03 | 4,27 | 0,01 | 9,94E-01 |
| AT1G16400 | 0,30 | -1,78 | 4,41 | -0,40 | 6,87E-01 |
| AT1G16410 | 1,19 | -1,70 | 1,85 | -0,92 | 3,58E-01 |
| AT1G16470 | 0,19 | 0,00 | 4,41 | 0,00 | 1,00E+00 |
| AT1G16500 | 0,25 | -0,79 | 4,34 | -0,18 | 8,55E-01 |
| AT1G16560 | 0,44 | -0,63 | 3,23 | -0,19 | 8,46E-01 |
| AT1G16670 | 0,35 | -2,53 | 4,37 | -0,58 | 5,63E-01 |
| AT1G16680 | 0,85 | 0,91 | 2,42 | 0,38 | 7,06E-01 |
| AT1G16690 | 0,12 | 0,37 | 4,37 | 0,08 | 9,33E-01 |
| AT1G16700 | 0,62 | -0,32 | 3,05 | -0,11 | 9,16E-01 |
| AT1G16710 | 1,85 | 0,81 | 1,57 | 0,52 | 6,06E-01 |

| | | | | | |
|---|---|---|---|---|---|
| AT1G16720 | 1,64 | 0,14 | 1,42 | 0,10 | 9,22E-01 |
| AT1G16740 | 0,37 | -2,24 | 4,39 | -0,51 | 6,09E-01 |
| AT1G16810 | 0,12 | 1,23 | 4,34 | 0,28 | 7,76E-01 |
| AT1G16860 | 0,38 | 0,35 | 4,27 | 0,08 | 9,35E-01 |
| AT1G16880 | 2,91 | 0,61 | 1,18 | 0,52 | 6,03E-01 |
| AT1G17050 | 0,20 | 0,75 | 4,37 | 0,17 | 8,64E-01 |
| AT1G17070 | 0,73 | 2,30 | 3,56 | 0,65 | 5,19E-01 |
| AT1G17100 | 0,86 | 0,02 | 1,82 | 0,01 | 9,93E-01 |
| AT1G17120 | 0,36 | -1,28 | 4,34 | -0,29 | 7,68E-01 |
| AT1G17150 | 0,40 | 1,21 | 3,41 | 0,35 | 7,23E-01 |
| AT1G17160 | 0,36 | 0,78 | 4,28 | 0,18 | 8,56E-01 |
| AT1G17200 | 0,19 | 0,00 | 4,41 | 0,00 | 1,00E+00 |
| AT1G17220 | 3,72 | -0,90 | 0,98 | -0,92 | 3,56E-01 |
| AT1G17230 | 0,51 | 1,49 | 3,10 | 0,48 | 6,30E-01 |
| AT1G17255 | 1,02 | 2,39 | 2,62 | 0,91 | 3,61E-01 |
| AT1G17260 | 0,13 | 0,73 | 4,36 | 0,17 | 8,67E-01 |
| AT1G17290 | 2,38 | 0,01 | 1,28 | 0,01 | 9,94E-01 |
| AT1G17300 | 0,61 | 1,36 | 3,61 | 0,38 | 7,05E-01 |
| AT1G17340 | 0,57 | -1,81 | 2,34 | -0,77 | 4,40E-01 |
| AT1G17360 | 1,17 | 0,85 | 2,04 | 0,42 | 6,78E-01 |
| AT1G17420 | 1,07 | 0,25 | 2,13 | 0,12 | 9,07E-01 |
| AT1G17460 | 0,31 | -0,64 | 4,36 | -0,15 | 8,83E-01 |
| AT1G17470 | 1,97 | -0,79 | 1,41 | -0,56 | 5,74E-01 |
| AT1G17490 | 0,39 | 0,63 | 3,52 | 0,18 | 8,57E-01 |
| AT1G17500 | 0,69 | 0,29 | 2,15 | 0,13 | 8,94E-01 |
| AT1G17530 | 0,57 | 0,31 | 3,13 | 0,10 | 9,22E-01 |
| AT1G17550 | 1,05 | -0,79 | 1,65 | -0,48 | 6,32E-01 |
| AT1G17570 | 0,08 | 0,00 | 4,40 | 0,00 | 1,00E+00 |
| AT1G17580 | 1,15 | -0,38 | 1,96 | -0,19 | 8,45E-01 |
| AT1G17610 | 0,33 | 1,15 | 4,33 | 0,27 | 7,91E-01 |
| AT1G17620 | 1,43 | 0,89 | 1,79 | 0,50 | 6,20E-01 |
| AT1G17680 | 0,28 | 1,74 | 4,33 | 0,40 | 6,88E-01 |
| AT1G17745 | 0,96 | -0,19 | 1,89 | -0,10 | 9,19E-01 |
| AT1G17750 | 0,21 | 1,63 | 4,33 | 0,38 | 7,06E-01 |
| AT1G17840 | 1,51 | 1,76 | 1,79 | 0,98 | 3,27E-01 |
| AT1G17860 | 0,25 | 1,40 | 4,33 | 0,32 | 7,47E-01 |
| AT1G17970 | 0,47 | -1,84 | 4,14 | -0,44 | 6,57E-01 |
| AT1G18020 | 0,78 | -1,16 | 3,03 | -0,38 | 7,02E-01 |
| AT1G18080 | 2,60 | -0,27 | 1,21 | -0,22 | 8,23E-01 |
| AT1G18090 | 0,88 | 0,69 | 3,45 | 0,20 | 8,42E-01 |
| AT1G18130 | 0,21 | 0,43 | 4,37 | 0,10 | 9,23E-01 |
| AT1G18160 | 2,26 | 1,97 | 1,46 | 1,35 | 1,77E-01 |
| AT1G18170 | 0,31 | 1,15 | 4,34 | 0,27 | 7,91E-01 |
| AT1G18300 | 0,38 | -0,48 | 4,29 | -0,11 | 9,11E-01 |
| AT1G18390 | 0,88 | -1,08 | 1,91 | -0,56 | 5,73E-01 |
| AT1G18450 | 0,27 | 0,44 | 4,38 | 0,10 | 9,20E-01 |
| AT1G18460 | 0,79 | 0,73 | 2,07 | 0,36 | 7,22E-01 |
| AT1G18480 | 0,17 | -1,70 | 4,40 | -0,39 | 7,00E-01 |
| AT1G18485 | 1,73 | 0,18 | 1,74 | 0,10 | 9,19E-01 |
| AT1G18490 | 0,64 | 1,51 | 4,31 | 0,35 | 7,26E-01 |
| AT1G18500 | 0,63 | -0,61 | 3,03 | -0,20 | 8,41E-01 |
| AT1G18570 | 0,88 | 0,56 | 1,97 | 0,28 | 7,76E-01 |

| | | | | | |
|---|---|---|---|---|---|
| AT1G18580 | 0,40 | 2,10 | 4,32 | 0,49 | 6,27E-01 |
| AT1G18600 | 0,74 | -0,30 | 2,40 | -0,12 | 9,01E-01 |
| AT1G18620 | 0,88 | 0,98 | 2,11 | 0,46 | 6,44E-01 |
| AT1G18700 | 1,03 | -2,04 | 2,07 | -0,98 | 3,26E-01 |
| AT1G18730 | 0,47 | 0,00 | 4,25 | 0,00 | 1,00E+00 |
| AT1G18740 | 2,60 | 0,55 | 1,18 | 0,46 | 6,43E-01 |
| AT1G18820 | 0,52 | 0,00 | 4,39 | 0,00 | 1,00E+00 |
| AT1G18870 | 0,46 | 1,16 | 4,36 | 0,27 | 7,91E-01 |
| AT1G18880 | 1,72 | 2,41 | 1,66 | 1,45 | 1,46E-01 |
| AT1G18879 | 1,60 | 3,41 | 1,75 | 1,95 | 5,16E-02 |
| AT1G18900 | 0,70 | 2,73 | 3,52 | 0,77 | 4,39E-01 |
| AT1G18950 | 0,83 | 0,84 | 2,18 | 0,38 | 7,00E-01 |
| AT1G19020 | 0,56 | -0,55 | 2,29 | -0,24 | 8,11E-01 |
| AT1G19025 | 0,65 | -2,12 | 2,91 | -0,73 | 4,67E-01 |
| AT1G19086 | 0,19 | 0,00 | 4,41 | 0,00 | 1,00E+00 |
| AT1G19090 | 4,93 | 0,92 | 1,10 | 0,83 | 4,05E-01 |
| AT1G19110 | 0,80 | -1,10 | 2,07 | -0,53 | 5,94E-01 |
| AT1G19150 | 0,10 | 0,00 | 4,41 | 0,00 | 1,00E+00 |
| AT1G19180 | 1,37 | -1,26 | 1,87 | -0,67 | 5,03E-01 |
| AT1G19190 | 0,39 | -1,28 | 4,38 | -0,29 | 7,70E-01 |
| AT1G19290 | 0,57 | 1,07 | 3,73 | 0,29 | 7,73E-01 |
| AT1G19350 | 0,52 | -1,28 | 3,98 | -0,32 | 7,48E-01 |
| AT1G19373 | 3,55 | 1,18 | 1,45 | 0,81 | 4,17E-01 |
| AT1G19376 | 0,56 | 1,70 | 3,04 | 0,56 | 5,76E-01 |
| AT1G19464 | 0,18 | 0,00 | 4,41 | 0,00 | 1,00E+00 |
| AT1G19580 | 0,28 | -0,02 | 4,35 | 0,00 | 9,97E-01 |
| AT1G19680 | 2,21 | 1,54 | 1,69 | 0,91 | 3,63E-01 |
| AT1G05507 | 0,08 | 0,00 | 4,40 | 0,00 | 1,00E+00 |
| AT1G19710 | 0,37 | -0,56 | 4,24 | -0,13 | 8,95E-01 |
| AT1G19770 | 0,78 | 1,93 | 2,76 | 0,70 | 4,85E-01 |
| AT1G19800 | 0,29 | -0,48 | 4,35 | -0,11 | 9,12E-01 |
| AT1G19830 | 0,33 | 0,84 | 4,34 | 0,19 | 8,47E-01 |
| AT1G19850 | 0,68 | 2,85 | 2,87 | 0,99 | 3,21E-01 |
| AT1G19910 | 0,96 | 2,50 | 2,26 | 1,10 | 2,69E-01 |
| AT1G19960 | 1,88 | 1,65 | 1,86 | 0,89 | 3,74E-01 |
| AT1G19990 | 0,28 | 0,24 | 4,39 | 0,05 | 9,57E-01 |
| AT1G20020 | 2,67 | -0,81 | 1,07 | -0,76 | 4,48E-01 |
| AT1G20050 | 0,40 | 0,68 | 3,41 | 0,20 | 8,42E-01 |
| AT1G05523 | 0,59 | 1,17 | 2,44 | 0,48 | 6,31E-01 |
| AT1G20090 | 0,64 | -2,56 | 2,70 | -0,95 | 3,42E-01 |
| AT1G20110 | 0,46 | 1,48 | 2,73 | 0,54 | 5,87E-01 |
| AT1G20260 | 2,71 | -0,35 | 1,31 | -0,26 | 7,92E-01 |
| AT1G20280 | 0,77 | -0,16 | 2,15 | -0,07 | 9,42E-01 |
| AT1G20300 | 0,15 | 0,00 | 4,41 | 0,00 | 1,00E+00 |
| AT1G20310 | 0,17 | 1,40 | 4,33 | 0,32 | 7,47E-01 |
| AT1G20375 | 1,50 | -1,75 | 1,52 | -1,15 | 2,49E-01 |
| AT1G20630 | 1,62 | -1,74 | 1,75 | -0,99 | 3,20E-01 |
| AT1G20640 | 1,05 | -2,09 | 1,97 | -1,06 | 2,90E-01 |
| AT1G20690 | 2,85 | 0,81 | 1,29 | 0,63 | 5,31E-01 |
| AT1G20693 | 1,05 | -0,70 | 2,37 | -0,30 | 7,67E-01 |
| AT1G20696 | 0,69 | -0,11 | 2,28 | -0,05 | 9,61E-01 |
| AT1G20780 | 1,16 | 1,53 | 1,83 | 0,83 | 4,04E-01 |

| | | | | | |
|---|---|---|---|---|---|
| AT1G20823 | 0,81 | 1,57 | 2,30 | 0,68 | 4,95E-01 |
| AT1G20920 | 0,49 | -0,30 | 3,09 | -0,10 | 9,22E-01 |
| AT1G20970 | 4,69 | -1,05 | 1,08 | -0,97 | 3,31E-01 |
| AT1G20980 | 1,44 | 1,15 | 1,78 | 0,64 | 5,19E-01 |
| AT1G21000 | 0,77 | 0,72 | 2,22 | 0,33 | 7,44E-01 |
| AT1G21050 | 0,58 | 0,47 | 3,06 | 0,15 | 8,79E-01 |
| AT1G21065 | 0,84 | 1,96 | 2,14 | 0,92 | 3,59E-01 |
| AT1G21140 | 0,08 | 0,69 | 4,36 | 0,16 | 8,75E-01 |
| AT1G21170 | 0,55 | 0,15 | 3,87 | 0,04 | 9,70E-01 |
| AT1G21200 | 5,07 | 0,48 | 0,99 | 0,48 | 6,31E-01 |
| AT1G21202 | 4,89 | 0,47 | 1,09 | 0,44 | 6,62E-01 |
| AT1G21210 | 0,63 | 1,65 | 2,89 | 0,57 | 5,68E-01 |
| AT1G21230 | 0,43 | 0,82 | 3,29 | 0,25 | 8,03E-01 |
| AT1G21240 | 0,38 | 0,60 | 4,23 | 0,14 | 8,88E-01 |
| AT1G21270 | 2,66 | 0,18 | 1,46 | 0,12 | 9,03E-01 |
| AT1G21370 | 0,21 | 1,42 | 4,33 | 0,33 | 7,43E-01 |
| AT1G21430 | 0,53 | 0,75 | 4,32 | 0,17 | 8,61E-01 |
| AT1G21560 | 0,41 | -0,20 | 4,07 | -0,05 | 9,61E-01 |
| AT1G21630 | 4,05 | -0,29 | 0,97 | -0,30 | 7,66E-01 |
| AT1G21640 | 0,89 | 0,37 | 1,82 | 0,20 | 8,39E-01 |
| AT1G21660 | 0,28 | 0,13 | 4,33 | 0,03 | 9,77E-01 |
| AT1G21670 | 0,86 | -2,20 | 3,25 | -0,68 | 4,99E-01 |
| AT1G21680 | 4,80 | -0,49 | 0,83 | -0,59 | 5,56E-01 |
| AT1G21690 | 0,29 | 0,50 | 4,31 | 0,12 | 9,07E-01 |
| AT1G21720 | 0,34 | 0,12 | 4,35 | 0,03 | 9,78E-01 |
| AT1G21722 | 0,08 | 0,69 | 4,36 | 0,16 | 8,75E-01 |
| AT1G21750 | 2,14 | 0,78 | 1,60 | 0,49 | 6,25E-01 |
| AT1G21780 | 0,75 | 0,02 | 2,53 | 0,01 | 9,94E-01 |
| AT1G21835 | 0,94 | -0,93 | 2,23 | -0,42 | 6,77E-01 |
| AT1G21840 | 0,12 | 1,23 | 4,34 | 0,28 | 7,76E-01 |
| AT1G21880 | 0,12 | 0,97 | 4,34 | 0,22 | 8,23E-01 |
| AT1G21910 | 0,33 | 0,00 | 4,41 | 0,00 | 1,00E+00 |
| AT1G21980 | 0,19 | 0,84 | 4,35 | 0,19 | 8,46E-01 |
| AT1G22040 | 0,05 | 0,00 | 4,41 | 0,00 | 1,00E+00 |
| AT1G22070 | 0,41 | 0,24 | 4,31 | 0,06 | 9,55E-01 |
| AT1G22190 | 1,08 | -0,61 | 1,68 | -0,36 | 7,15E-01 |
| AT1G22280 | 0,53 | -1,20 | 3,28 | -0,37 | 7,14E-01 |
| AT1G22330 | 0,11 | 0,00 | 4,41 | 0,00 | 1,00E+00 |
| AT1G22410 | 1,37 | -1,75 | 1,66 | -1,06 | 2,91E-01 |
| AT1G22500 | 0,40 | 2,16 | 4,32 | 0,50 | 6,17E-01 |
| AT1G22590 | 0,24 | 0,82 | 4,35 | 0,19 | 8,50E-01 |
| AT1G22640 | 0,56 | 0,19 | 3,82 | 0,05 | 9,61E-01 |
| AT1G22730 | 0,55 | -1,50 | 3,79 | -0,40 | 6,93E-01 |
| AT1G22750 | 0,35 | -0,48 | 3,49 | -0,14 | 8,90E-01 |
| AT1G22770 | 1,61 | -3,12 | 1,60 | -1,95 | 5,06E-02 |
| AT1G22780 | 0,54 | -0,93 | 3,22 | -0,29 | 7,74E-01 |
| AT1G22790 | 0,64 | -0,88 | 3,76 | -0,23 | 8,16E-01 |
| AT1G05767 | 0,23 | 1,14 | 4,34 | 0,26 | 7,92E-01 |
| AT1G22840 | 0,62 | 2,25 | 3,22 | 0,70 | 4,84E-01 |
| AT1G22860 | 0,86 | -0,90 | 1,87 | -0,48 | 6,29E-01 |
| AT1G22870 | 0,66 | 1,40 | 2,87 | 0,49 | 6,27E-01 |
| AT1G22880 | 1,11 | -2,93 | 2,73 | -1,07 | 2,83E-01 |

| | | | | | |
|---|---|---|---|---|---|
| AT1G22882 | 0,84 | 1,96 | 3,28 | 0,60 | 5,51E-01 |
| AT1G22930 | 1,70 | 0,45 | 1,61 | 0,28 | 7,82E-01 |
| AT1G22940 | 0,55 | -0,57 | 2,62 | -0,22 | 8,28E-01 |
| AT1G23030 | 0,27 | -0,47 | 4,37 | -0,11 | 9,14E-01 |
| AT1G23040 | 0,34 | 1,59 | 4,33 | 0,37 | 7,13E-01 |
| AT1G23050 | 1,19 | -0,61 | 2,55 | -0,24 | 8,12E-01 |
| AT1G23100 | 0,08 | 0,69 | 4,36 | 0,16 | 8,75E-01 |
| AT1G23130 | 0,08 | 0,69 | 4,36 | 0,16 | 8,75E-01 |
| AT1G23190 | 1,73 | -1,97 | 1,65 | -1,19 | 2,33E-01 |
| AT1G23230 | 1,20 | 0,63 | 1,74 | 0,36 | 7,19E-01 |
| AT1G23290 | 1,36 | 1,26 | 1,86 | 0,68 | 5,00E-01 |
| AT1G23440 | 0,65 | 1,47 | 2,96 | 0,50 | 6,20E-01 |
| AT1G23450 | 0,21 | 0,01 | 4,34 | 0,00 | 9,98E-01 |
| AT1G23490 | 1,31 | 0,87 | 1,63 | 0,54 | 5,91E-01 |
| AT1G23710 | 1,29 | 1,13 | 1,90 | 0,60 | 5,51E-01 |
| AT1G23720 | 2,26 | 2,88 | 2,04 | 1,41 | 1,58E-01 |
| AT1G23820 | 1,10 | -0,47 | 2,78 | -0,17 | 8,67E-01 |
| AT1G05837 | 0,53 | 0,49 | 3,18 | 0,15 | 8,79E-01 |
| AT1G23870 | 0,51 | 2,36 | 3,82 | 0,62 | 5,37E-01 |
| AT1G23880 | 0,08 | 0,44 | 4,38 | 0,10 | 9,21E-01 |
| AT1G23900 | 1,85 | 0,65 | 1,40 | 0,47 | 6,41E-01 |
| AT1G23915 | 1,01 | 3,57 | 2,48 | 1,44 | 1,50E-01 |
| AT1G24145 | 0,48 | 1,11 | 3,09 | 0,36 | 7,19E-01 |
| AT1G24147 | 0,16 | 0,65 | 4,37 | 0,15 | 8,82E-01 |
| AT1G24150 | 0,48 | 0,37 | 3,89 | 0,09 | 9,25E-01 |
| AT1G24280 | 0,44 | 1,83 | 4,33 | 0,42 | 6,73E-01 |
| AT1G24340 | 0,28 | 0,81 | 3,75 | 0,22 | 8,29E-01 |
| AT1G24360 | 0,34 | -0,92 | 4,39 | -0,21 | 8,35E-01 |
| AT1G24380 | 0,38 | 0,86 | 2,91 | 0,30 | 7,66E-01 |
| AT1G24460 | 4,84 | -0,01 | 0,94 | -0,02 | 9,88E-01 |
| AT1G24490 | 0,45 | 0,56 | 3,21 | 0,18 | 8,61E-01 |
| AT1G24530 | 0,17 | 0,35 | 4,37 | 0,08 | 9,37E-01 |
| AT1G24560 | 0,68 | 0,83 | 2,19 | 0,38 | 7,03E-01 |
| AT1G24570 | 0,08 | 0,69 | 4,36 | 0,16 | 8,75E-01 |
| AT1G24706 | 2,66 | -0,57 | 1,02 | -0,56 | 5,77E-01 |
| AT1G25275 | 1,34 | -0,63 | 1,86 | -0,34 | 7,34E-01 |
| AT1G25280 | 0,98 | -1,37 | 2,24 | -0,61 | 5,40E-01 |
| AT1G25480 | 0,19 | 0,00 | 4,41 | 0,00 | 1,00E+00 |
| AT1G25490 | 1,64 | -0,03 | 1,39 | -0,02 | 9,86E-01 |
| AT1G25520 | 0,81 | 0,81 | 3,52 | 0,23 | 8,19E-01 |
| AT1G25550 | 1,33 | 1,09 | 2,05 | 0,53 | 5,96E-01 |
| AT1G26130 | 0,54 | 1,49 | 3,81 | 0,39 | 6,95E-01 |
| AT1G26220 | 0,37 | 0,43 | 4,38 | 0,10 | 9,21E-01 |
| AT1G26250 | 0,44 | 0,69 | 4,11 | 0,17 | 8,66E-01 |
| AT1G26460 | 0,79 | 0,40 | 2,76 | 0,15 | 8,84E-01 |
| AT1G26470 | 0,34 | 0,60 | 4,34 | 0,14 | 8,90E-01 |
| AT1G26550 | 0,37 | -0,91 | 4,38 | -0,21 | 8,35E-01 |
| AT1G26558 | 2,72 | -1,85 | 1,36 | -1,36 | 1,74E-01 |
| AT1G26560 | 0,24 | 0,85 | 4,35 | 0,19 | 8,45E-01 |
| AT1G26580 | 0,53 | -0,88 | 2,65 | -0,33 | 7,39E-01 |
| AT1G26630 | 1,29 | -0,11 | 1,79 | -0,06 | 9,52E-01 |
| AT1G26650 | 1,02 | -0,56 | 2,05 | -0,28 | 7,83E-01 |

| | | | | | |
|---|---|---|---|---|---|
| AT1G26665 | 0,17 | 1,52 | 4,33 | 0,35 | 7,26E-01 |
| AT1G26670 | 1,04 | -0,25 | 2,11 | -0,12 | 9,06E-01 |
| AT1G26770 | 0,19 | 0,00 | 4,41 | 0,00 | 1,00E+00 |
| AT1G26830 | 0,72 | 1,47 | 2,39 | 0,62 | 5,37E-01 |
| AT1G26920 | 0,73 | -0,37 | 2,35 | -0,16 | 8,75E-01 |
| AT1G26940 | 0,05 | 0,00 | 4,41 | 0,00 | 1,00E+00 |
| AT1G27000 | 0,38 | -0,30 | 4,36 | -0,07 | 9,45E-01 |
| AT1G27060 | 0,73 | 0,83 | 2,81 | 0,30 | 7,66E-01 |
| AT1G27130 | 0,51 | 0,29 | 3,20 | 0,09 | 9,28E-01 |
| AT1G27150 | 0,43 | -1,72 | 4,33 | -0,40 | 6,91E-01 |
| AT1G27170 | 0,35 | 0,54 | 3,42 | 0,16 | 8,75E-01 |
| AT1G27180 | 0,99 | 0,41 | 2,44 | 0,17 | 8,68E-01 |
| AT1G27290 | 1,54 | 0,04 | 1,45 | 0,03 | 9,75E-01 |
| AT1G27300 | 0,40 | -1,30 | 4,16 | -0,31 | 7,55E-01 |
| AT1G27320 | 2,96 | 0,48 | 1,05 | 0,45 | 6,51E-01 |
| AT1G27330 | 0,50 | -0,85 | 4,35 | -0,20 | 8,44E-01 |
| AT1G27340 | 2,10 | -1,22 | 1,33 | -0,92 | 3,56E-01 |
| AT1G27360 | 0,08 | 0,44 | 4,38 | 0,10 | 9,21E-01 |
| AT1G27400 | 0,67 | -0,02 | 2,04 | -0,01 | 9,92E-01 |
| AT1G06087 | 0,12 | 0,37 | 4,37 | 0,08 | 9,33E-01 |
| AT1G27450 | 0,37 | 1,40 | 4,33 | 0,32 | 7,46E-01 |
| AT1G27460 | 0,11 | 0,65 | 4,36 | 0,15 | 8,81E-01 |
| AT1G27480 | 0,41 | -0,47 | 4,16 | -0,11 | 9,10E-01 |
| AT1G27520 | 0,14 | 0,40 | 4,37 | 0,09 | 9,27E-01 |
| AT1G27535 | 2,34 | 0,01 | 1,53 | 0,01 | 9,93E-01 |
| AT1G06097 | 0,36 | -0,91 | 4,39 | -0,21 | 8,35E-01 |
| AT1G27570 | 1,68 | 2,89 | 1,96 | 1,48 | 1,40E-01 |
| AT1G27595 | 2,01 | 0,40 | 1,36 | 0,29 | 7,71E-01 |
| AT1G27630 | 0,58 | -1,83 | 3,97 | -0,46 | 6,45E-01 |
| AT1G27650 | 0,97 | 0,13 | 1,97 | 0,07 | 9,47E-01 |
| AT1G27750 | 0,63 | -0,66 | 2,60 | -0,25 | 7,99E-01 |
| AT1G27752 | 1,43 | 2,41 | 2,15 | 1,12 | 2,63E-01 |
| AT1G27760 | 0,65 | 0,35 | 2,44 | 0,14 | 8,87E-01 |
| AT1G27850 | 0,84 | -1,05 | 2,10 | -0,50 | 6,17E-01 |
| AT1G27950 | 0,55 | 0,00 | 3,85 | 0,00 | 1,00E+00 |
| AT1G28005 | 0,32 | 0,14 | 4,38 | 0,03 | 9,74E-01 |
| AT1G28060 | 0,32 | 1,07 | 4,35 | 0,25 | 8,06E-01 |
| AT1G28070 | 4,21 | -3,97 | 1,96 | -2,03 | 4,22E-02 |
| AT1G28135 | 0,08 | 0,00 | 4,40 | 0,00 | 1,00E+00 |
| AT1G28270 | 2,06 | -2,40 | 1,72 | -1,39 | 1,64E-01 |
| AT1G28340 | 0,87 | 3,02 | 2,66 | 1,14 | 2,55E-01 |
| AT1G28350 | 1,15 | -2,82 | 2,03 | -1,39 | 1,65E-01 |
| AT1G28380 | 1,21 | 0,81 | 2,06 | 0,39 | 6,93E-01 |
| AT1G28420 | 2,78 | 2,34 | 1,39 | 1,68 | 9,23E-02 |
| AT1G28440 | 1,72 | -1,18 | 1,33 | -0,89 | 3,76E-01 |
| AT1G06163 | 0,36 | -1,78 | 4,41 | -0,40 | 6,87E-01 |
| AT1G28490 | 0,17 | 1,33 | 4,33 | 0,31 | 7,59E-01 |
| AT1G28520 | 0,40 | 0,29 | 3,36 | 0,09 | 9,32E-01 |
| AT1G28680 | 0,55 | 0,05 | 3,20 | 0,01 | 9,89E-01 |
| AT1G28780 | 0,49 | 1,58 | 4,07 | 0,39 | 6,98E-01 |
| AT1G28760 | 0,10 | 0,21 | 4,38 | 0,05 | 9,62E-01 |
| AT1G29070 | 1,74 | -1,09 | 1,40 | -0,78 | 4,35E-01 |

| | | | | | |
|---|---|---|---|---|---|
| AT1G29120 | 0,19 | 1,50 | 4,33 | 0,35 | 7,30E-01 |
| AT1G29150 | 0,99 | -2,03 | 2,27 | -0,90 | 3,70E-01 |
| AT1G29200 | 0,08 | 0,69 | 4,36 | 0,16 | 8,75E-01 |
| AT1G29260 | 0,92 | -0,30 | 1,68 | -0,18 | 8,60E-01 |
| AT1G29310 | 0,68 | 0,49 | 2,92 | 0,17 | 8,66E-01 |
| AT1G29340 | 0,94 | 1,23 | 2,12 | 0,58 | 5,60E-01 |
| AT1G29380 | 0,65 | -1,27 | 4,31 | -0,29 | 7,68E-01 |
| AT1G06233 | 0,68 | -0,40 | 2,58 | -0,15 | 8,77E-01 |
| AT1G06247 | 2,53 | -0,39 | 1,22 | -0,32 | 7,46E-01 |
| AT1G06253 | 1,09 | 0,87 | 1,79 | 0,48 | 6,29E-01 |
| AT1G06263 | 0,52 | 0,11 | 4,09 | 0,03 | 9,78E-01 |
| AT1G06273 | 2,17 | -4,45 | 1,81 | -2,46 | 1,39E-02 |
| AT1G06283 | 1,64 | 0,92 | 1,74 | 0,53 | 5,96E-01 |
| AT1G29620 | 0,47 | 0,23 | 3,26 | 0,07 | 9,45E-01 |
| AT1G29630 | 0,24 | 1,54 | 4,34 | 0,35 | 7,23E-01 |
| AT1G29660 | 1,40 | -1,74 | 1,84 | -0,95 | 3,45E-01 |
| AT1G29670 | 1,73 | -1,84 | 1,77 | -1,03 | 3,01E-01 |
| AT1G29760 | 0,40 | -0,14 | 4,28 | -0,03 | 9,75E-01 |
| AT1G29770 | 0,71 | 1,27 | 4,31 | 0,30 | 7,68E-01 |
| AT1G29800 | 0,74 | -3,45 | 3,64 | -0,95 | 3,43E-01 |
| AT1G29830 | 1,13 | -1,08 | 1,76 | -0,61 | 5,42E-01 |
| AT1G29950 | 0,49 | -1,37 | 4,28 | -0,32 | 7,49E-01 |
| AT1G30010 | 0,08 | 0,00 | 4,40 | 0,00 | 1,00E+00 |
| AT1G30200 | 0,70 | -0,07 | 2,12 | -0,03 | 9,73E-01 |
| AT1G30260 | 1,16 | -0,83 | 2,40 | -0,35 | 7,28E-01 |
| AT1G30360 | 3,38 | 0,09 | 0,92 | 0,10 | 9,20E-01 |
| AT1G30380 | 5,72 | -0,59 | 1,21 | -0,49 | 6,23E-01 |
| AT1G30400 | 2,52 | 0,33 | 1,30 | 0,26 | 7,99E-01 |
| AT1G30410 | 4,36 | 0,35 | 0,92 | 0,38 | 7,01E-01 |
| AT1G30420 | 0,38 | -0,91 | 4,32 | -0,21 | 8,34E-01 |
| AT1G30430 | 0,16 | 0,00 | 4,41 | 0,00 | 1,00E+00 |
| AT1G30440 | 1,10 | -1,47 | 1,93 | -0,76 | 4,45E-01 |
| AT1G30450 | 0,86 | 1,23 | 2,09 | 0,59 | 5,57E-01 |
| AT1G30470 | 3,87 | -0,76 | 1,00 | -0,76 | 4,46E-01 |
| AT1G30520 | 0,41 | -2,17 | 4,32 | -0,50 | 6,15E-01 |
| AT1G30530 | 0,46 | 0,17 | 3,35 | 0,05 | 9,60E-01 |
| AT1G30610 | 0,23 | 0,81 | 4,37 | 0,19 | 8,53E-01 |
| AT1G30620 | 0,56 | -0,12 | 3,19 | -0,04 | 9,69E-01 |
| AT1G30700 | 0,12 | 1,16 | 4,34 | 0,27 | 7,89E-01 |
| AT1G30720 | 0,53 | 0,75 | 4,32 | 0,17 | 8,61E-01 |
| AT1G30757 | 0,16 | 0,00 | 4,41 | 0,00 | 1,00E+00 |
| AT1G30800 | 0,20 | -0,44 | 4,36 | -0,10 | 9,19E-01 |
| AT1G30910 | 0,25 | 0,73 | 4,37 | 0,17 | 8,68E-01 |
| AT1G06393 | 0,32 | 0,00 | 4,38 | 0,00 | 1,00E+00 |
| AT1G31095 | 0,66 | -2,17 | 3,02 | -0,72 | 4,73E-01 |
| AT1G31170 | 0,22 | 0,88 | 4,35 | 0,20 | 8,39E-01 |
| AT1G31175 | 0,74 | -0,69 | 2,38 | -0,29 | 7,72E-01 |
| AT1G31190 | 0,86 | 1,09 | 2,46 | 0,44 | 6,57E-01 |
| AT1G31220 | 0,22 | -1,77 | 4,41 | -0,40 | 6,88E-01 |
| AT1G31290 | 0,15 | 0,45 | 4,37 | 0,10 | 9,19E-01 |
| AT1G31420 | 0,89 | 0,36 | 2,09 | 0,17 | 8,62E-01 |
| AT1G31480 | 0,40 | 2,16 | 4,32 | 0,50 | 6,17E-01 |

| | | | | | |
|---|---|---|---|---|---|
| AT1G06447 | 0,24 | -1,19 | 4,35 | -0,27 | 7,84E-01 |
| AT1G31710 | 0,82 | 1,61 | 3,52 | 0,46 | 6,48E-01 |
| AT1G31730 | 1,97 | -2,19 | 1,54 | -1,42 | 1,55E-01 |
| AT1G31780 | 0,28 | 1,42 | 4,34 | 0,33 | 7,43E-01 |
| AT1G31800 | 0,33 | 1,16 | 3,50 | 0,33 | 7,41E-01 |
| AT1G31850 | 1,13 | -1,37 | 1,95 | -0,70 | 4,82E-01 |
| AT1G31910 | 2,79 | -0,38 | 1,03 | -0,37 | 7,09E-01 |
| AT1G31930 | 0,12 | 1,16 | 4,34 | 0,27 | 7,89E-01 |
| AT1G31970 | 0,74 | -0,51 | 3,04 | -0,17 | 8,67E-01 |
| AT1G32050 | 0,65 | 0,52 | 2,34 | 0,22 | 8,25E-01 |
| AT1G32060 | 5,29 | 0,48 | 1,16 | 0,41 | 6,78E-01 |
| AT1G32160 | 0,70 | 1,48 | 2,98 | 0,50 | 6,20E-01 |
| AT1G32220 | 0,21 | 0,00 | 4,41 | 0,00 | 1,00E+00 |
| AT1G32225 | 0,70 | -0,88 | 2,48 | -0,35 | 7,23E-01 |
| AT1G32230 | 5,15 | -0,68 | 0,80 | -0,84 | 3,99E-01 |
| AT1G32270 | 1,96 | 0,63 | 1,66 | 0,38 | 7,05E-01 |
| AT1G32320 | 0,23 | 0,00 | 4,41 | 0,00 | 1,00E+00 |
| AT1G32337 | 0,30 | 0,37 | 4,36 | 0,09 | 9,32E-01 |
| AT1G32360 | 0,41 | 1,17 | 3,36 | 0,35 | 7,28E-01 |
| AT1G06547 | 0,19 | 0,00 | 4,41 | 0,00 | 1,00E+00 |
| AT1G32385 | 0,41 | 0,07 | 2,89 | 0,02 | 9,80E-01 |
| AT1G32415 | 0,72 | -1,68 | 2,72 | -0,62 | 5,37E-01 |
| AT1G32460 | 0,41 | 1,23 | 4,33 | 0,28 | 7,77E-01 |
| AT1G32580 | 0,32 | -0,98 | 4,35 | -0,23 | 8,22E-01 |
| AT1G06597 | 1,64 | -0,55 | 1,62 | -0,34 | 7,36E-01 |
| AT1G32630 | 4,04 | 0,48 | 1,11 | 0,43 | 6,64E-01 |
| AT1G32700 | 0,46 | 0,29 | 3,24 | 0,09 | 9,29E-01 |
| AT1G32720 | 2,47 | 2,93 | 1,61 | 1,83 | 6,78E-02 |
| AT1G32810 | 0,36 | 0,52 | 3,45 | 0,15 | 8,80E-01 |
| AT1G06623 | 0,26 | 0,00 | 4,41 | 0,00 | 1,00E+00 |
| AT1G32870 | 0,59 | 0,42 | 3,77 | 0,11 | 9,11E-01 |
| AT1G32920 | 1,13 | -0,64 | 1,97 | -0,33 | 7,44E-01 |
| AT1G32928 | 0,50 | -1,01 | 4,04 | -0,25 | 8,03E-01 |
| AT1G32940 | 0,41 | -0,04 | 4,31 | -0,01 | 9,92E-01 |
| AT1G32950 | 0,62 | 1,27 | 3,63 | 0,35 | 7,28E-01 |
| AT1G32960 | 0,35 | 1,78 | 4,33 | 0,41 | 6,81E-01 |
| AT1G32990 | 0,48 | 0,00 | 4,39 | 0,00 | 1,00E+00 |
| AT1G33010 | 0,19 | 0,00 | 4,41 | 0,00 | 1,00E+00 |
| AT1G33040 | 0,24 | 1,08 | 4,35 | 0,25 | 8,03E-01 |
| AT1G33070 | 1,82 | 0,22 | 1,85 | 0,12 | 9,06E-01 |
| AT1G33110 | 0,55 | -1,30 | 3,78 | -0,34 | 7,31E-01 |
| AT1G33120 | 1,97 | 0,83 | 1,40 | 0,59 | 5,53E-01 |
| AT1G33140 | 1,84 | -0,31 | 1,37 | -0,23 | 8,19E-01 |
| AT1G33170 | 0,44 | -0,54 | 3,23 | -0,17 | 8,67E-01 |
| AT1G33260 | 0,52 | 1,14 | 3,98 | 0,29 | 7,74E-01 |
| AT1G06667 | 5,78 | 0,63 | 1,01 | 0,63 | 5,29E-01 |
| AT1G33360 | 0,47 | 1,19 | 4,07 | 0,29 | 7,70E-01 |
| AT1G33415 | 2,72 | 0,52 | 1,23 | 0,43 | 6,69E-01 |
| AT1G33470 | 0,71 | -1,53 | 3,43 | -0,45 | 6,56E-01 |
| AT1G33560 | 2,19 | -0,58 | 1,54 | -0,38 | 7,07E-01 |
| AT1G33590 | 1,70 | 1,20 | 1,68 | 0,72 | 4,73E-01 |
| AT1G33600 | 1,39 | -0,41 | 1,72 | -0,24 | 8,10E-01 |

| | | | | | |
|---|---|---|---|---|---|
| AT1G33610 | 0,15 | 1,16 | 4,34 | 0,27 | 7,88E-01 |
| AT1G33612 | 0,17 | 0,00 | 4,41 | 0,00 | 1,00E+00 |
| AT1G33680 | 2,02 | -1,14 | 1,51 | -0,75 | 4,52E-01 |
| AT1G33770 | 0,49 | -0,42 | 4,03 | -0,10 | 9,16E-01 |
| AT1G33811 | 0,18 | 0,00 | 4,38 | 0,00 | 1,00E+00 |
| AT1G33910 | 0,08 | 0,00 | 4,40 | 0,00 | 1,00E+00 |
| AT1G33930 | 0,36 | 0,00 | 4,41 | 0,00 | 1,00E+00 |
| AT1G33960 | 1,02 | -2,92 | 3,09 | -0,95 | 3,44E-01 |
| AT1G33980 | 0,92 | -2,03 | 4,22 | -0,48 | 6,30E-01 |
| AT1G33990 | 0,95 | -0,90 | 1,99 | -0,45 | 6,51E-01 |
| AT1G06723 | 0,59 | -1,91 | 2,62 | -0,73 | 4,65E-01 |
| AT1G34020 | 0,78 | 0,99 | 2,40 | 0,41 | 6,78E-01 |
| AT1G34070 | 0,46 | 0,21 | 4,12 | 0,05 | 9,59E-01 |
| AT1G34130 | 1,45 | 1,19 | 1,62 | 0,73 | 4,63E-01 |
| AT1G34150 | 1,09 | -2,02 | 2,50 | -0,81 | 4,19E-01 |
| AT1G34190 | 1,14 | 0,40 | 1,76 | 0,22 | 8,22E-01 |
| AT1G34200 | 0,21 | 1,55 | 4,33 | 0,36 | 7,20E-01 |
| AT1G34210 | 0,51 | -0,30 | 4,32 | -0,07 | 9,46E-01 |
| AT1G34260 | 1,42 | 1,69 | 1,91 | 0,88 | 3,78E-01 |
| AT1G34290 | 0,25 | 0,13 | 4,39 | 0,03 | 9,77E-01 |
| AT1G34300 | 2,71 | -0,09 | 1,12 | -0,08 | 9,37E-01 |
| AT1G06747 | 0,08 | 0,00 | 4,40 | 0,00 | 1,00E+00 |
| AT1G34355 | 0,86 | -1,74 | 2,68 | -0,65 | 5,15E-01 |
| AT1G34370 | 1,49 | 0,63 | 1,75 | 0,36 | 7,18E-01 |
| AT1G34390 | 0,68 | 0,97 | 3,71 | 0,26 | 7,94E-01 |
| AT1G34410 | 0,91 | 0,28 | 3,35 | 0,08 | 9,34E-01 |
| AT1G34420 | 0,36 | 1,68 | 4,27 | 0,39 | 6,94E-01 |
| AT1G34460 | 0,31 | 0,00 | 4,38 | 0,00 | 1,00E+00 |
| AT1G34580 | 0,53 | 0,39 | 4,37 | 0,09 | 9,29E-01 |
| AT1G34770 | 0,12 | 1,23 | 4,34 | 0,28 | 7,76E-01 |
| AT1G34844 | 0,08 | 0,69 | 4,36 | 0,16 | 8,75E-01 |
| AT1G35030 | 0,42 | 1,37 | 3,33 | 0,41 | 6,81E-01 |
| AT1G35040 | 0,23 | 0,00 | 4,41 | 0,00 | 1,00E+00 |
| AT1G35140 | 0,39 | -1,75 | 4,29 | -0,41 | 6,83E-01 |
| AT1G35160 | 0,59 | 1,53 | 2,57 | 0,60 | 5,51E-01 |
| AT1G06827 | 0,56 | -0,44 | 2,71 | -0,16 | 8,72E-01 |
| AT1G35220 | 0,89 | 0,02 | 2,21 | 0,01 | 9,94E-01 |
| AT1G35230 | 1,07 | -0,96 | 1,99 | -0,48 | 6,28E-01 |
| AT1G35242 | 0,19 | 1,09 | 4,34 | 0,25 | 8,01E-01 |
| AT1G35420 | 0,41 | 0,38 | 3,38 | 0,11 | 9,10E-01 |
| AT1G35460 | 0,12 | 1,23 | 4,34 | 0,28 | 7,76E-01 |
| AT1G35500 | 0,14 | 0,00 | 4,41 | 0,00 | 1,00E+00 |
| AT1G35510 | 0,12 | 0,00 | 4,38 | 0,00 | 1,00E+00 |
| AT1G35515 | 0,52 | 0,54 | 2,35 | 0,23 | 8,17E-01 |
| AT1G35516 | 0,52 | 0,54 | 2,35 | 0,23 | 8,17E-01 |
| AT1G35530 | 0,08 | 0,69 | 4,36 | 0,16 | 8,75E-01 |
| AT1G35540 | 0,95 | -0,15 | 2,05 | -0,07 | 9,42E-01 |
| AT1G35614 | 1,12 | 0,26 | 2,22 | 0,12 | 9,08E-01 |
| AT1G35620 | 0,51 | 1,21 | 3,32 | 0,37 | 7,15E-01 |
| AT1G35660 | 0,28 | 1,81 | 4,33 | 0,42 | 6,76E-01 |
| AT1G35670 | 0,21 | 1,55 | 4,33 | 0,36 | 7,20E-01 |
| AT1G35680 | 0,81 | -3,72 | 2,21 | -1,68 | 9,21E-02 |

| | | | | | |
|---|---|---|---|---|---|
| AT1G35720 | 3,50 | -0,68 | 0,96 | -0,70 | 4,82E-01 |
| AT1G06927 | 0,83 | 2,44 | 2,77 | 0,88 | 3,78E-01 |
| AT1G35850 | 0,23 | 0,00 | 4,41 | 0,00 | 1,00E+00 |
| AT1G36050 | 0,26 | 1,54 | 4,33 | 0,36 | 7,21E-01 |
| AT1G06963 | 0,05 | 0,00 | 4,41 | 0,00 | 1,00E+00 |
| AT1G36180 | 2,20 | -1,24 | 1,28 | -0,97 | 3,31E-01 |
| AT1G36320 | 0,22 | 0,24 | 4,38 | 0,05 | 9,57E-01 |
| AT1G36510 | 0,10 | 0,00 | 4,41 | 0,00 | 1,00E+00 |
| AT1G06987 | 0,58 | 0,90 | 4,36 | 0,21 | 8,37E-01 |
| AT1G36675 | 0,81 | 1,19 | 2,47 | 0,48 | 6,29E-01 |
| AT1G36745 | 0,98 | 2,73 | 2,26 | 1,21 | 2,27E-01 |
| AT1G36756 | 0,94 | 1,92 | 2,50 | 0,77 | 4,42E-01 |
| AT1G36940 | 0,22 | 0,00 | 4,41 | 0,00 | 1,00E+00 |
| AT1G36980 | 0,52 | 2,14 | 3,13 | 0,68 | 4,95E-01 |
| AT1G36992 | 0,32 | -1,28 | 4,34 | -0,29 | 7,68E-01 |
| AT1G37113 | 0,08 | 0,44 | 4,38 | 0,10 | 9,21E-01 |
| AT1G42470 | 1,71 | 0,02 | 1,81 | 0,01 | 9,91E-01 |
| AT1G42525 | 0,30 | 0,85 | 4,34 | 0,20 | 8,45E-01 |
| AT1G42540 | 1,07 | -0,52 | 2,11 | -0,25 | 8,05E-01 |
| AT1G42550 | 2,32 | 0,22 | 1,24 | 0,18 | 8,57E-01 |
| AT1G07137 | 0,05 | 0,00 | 4,41 | 0,00 | 1,00E+00 |
| AT1G42960 | 0,66 | 0,00 | 4,41 | 0,00 | 1,00E+00 |
| AT1G42980 | 1,30 | 0,74 | 1,71 | 0,44 | 6,64E-01 |
| AT1G43040 | 0,16 | 0,00 | 4,41 | 0,00 | 1,00E+00 |
| AT1G43145 | 1,44 | 3,08 | 1,78 | 1,73 | 8,38E-02 |
| AT1G43170 | 3,66 | 0,43 | 0,91 | 0,47 | 6,35E-01 |
| AT1G43171 | 0,72 | -3,93 | 4,30 | -0,91 | 3,60E-01 |
| AT1G43320 | 2,75 | 2,79 | 1,73 | 1,61 | 1,07E-01 |
| AT1G43330 | 0,87 | 1,20 | 2,63 | 0,46 | 6,48E-01 |
| AT1G43670 | 0,63 | -0,58 | 2,60 | -0,22 | 8,23E-01 |
| AT1G43680 | 0,25 | -2,19 | 4,38 | -0,50 | 6,17E-01 |
| AT1G43700 | 0,72 | -1,76 | 2,11 | -0,84 | 4,03E-01 |
| AT1G43722 | 0,18 | 0,00 | 4,41 | 0,00 | 1,00E+00 |
| AT1G43770 | 1,22 | 0,02 | 1,68 | 0,01 | 9,90E-01 |
| AT1G43780 | 0,25 | 0,00 | 4,41 | 0,00 | 1,00E+00 |
| AT1G43815 | 1,28 | -0,84 | 2,25 | -0,37 | 7,10E-01 |
| AT1G43850 | 1,82 | 0,22 | 1,46 | 0,15 | 8,81E-01 |
| AT1G07233 | 2,51 | -1,58 | 1,22 | -1,30 | 1,94E-01 |
| AT1G43890 | 0,40 | -1,07 | 4,26 | -0,25 | 8,03E-01 |
| AT1G43970 | 0,43 | 0,00 | 4,41 | 0,00 | 1,00E+00 |
| AT1G44120 | 0,95 | 3,22 | 2,66 | 1,21 | 2,27E-01 |
| AT1G44575 | 5,52 | -0,57 | 0,85 | -0,68 | 4,98E-01 |
| AT1G44750 | 0,33 | 2,24 | 4,32 | 0,52 | 6,03E-01 |
| AT1G44780 | 0,29 | -0,48 | 4,35 | -0,11 | 9,12E-01 |
| AT1G44810 | 0,26 | 1,17 | 3,74 | 0,31 | 7,55E-01 |
| AT1G44890 | 0,08 | 0,00 | 4,40 | 0,00 | 1,00E+00 |
| AT1G44900 | 0,12 | 1,23 | 4,34 | 0,28 | 7,76E-01 |
| AT1G44910 | 0,50 | -0,06 | 2,43 | -0,02 | 9,81E-01 |
| AT1G44960 | 0,10 | 0,21 | 4,38 | 0,05 | 9,62E-01 |
| AT1G45000 | 0,47 | -1,27 | 2,74 | -0,46 | 6,42E-01 |
| AT1G45165 | 0,70 | 1,50 | 2,46 | 0,61 | 5,41E-01 |
| AT1G45180 | 0,30 | 0,24 | 4,35 | 0,05 | 9,57E-01 |

| | | | | | |
|---|---|---|---|---|---|
| AT1G45474 | 0,41 | 0,00 | 3,42 | 0,00 | 1,00E+00 |
| AT1G45688 | 0,89 | 1,13 | 2,36 | 0,48 | 6,30E-01 |
| AT1G45976 | 0,41 | 0,06 | 4,23 | 0,01 | 9,89E-01 |
| AT1G46264 | 0,10 | 0,21 | 4,38 | 0,05 | 9,62E-01 |
| AT1G46554 | 0,28 | -0,12 | 4,34 | -0,03 | 9,78E-01 |
| AT1G47210 | 0,93 | -0,72 | 2,42 | -0,30 | 7,67E-01 |
| AT1G07327 | 1,09 | 1,43 | 2,38 | 0,60 | 5,49E-01 |
| AT1G47250 | 0,35 | 1,19 | 4,34 | 0,27 | 7,84E-01 |
| AT1G47265 | 3,77 | -0,89 | 1,07 | -0,83 | 4,04E-01 |
| AT1G47270 | 0,13 | 0,73 | 4,36 | 0,17 | 8,67E-01 |
| AT1G07357 | 0,11 | 0,00 | 4,41 | 0,00 | 1,00E+00 |
| AT1G47480 | 1,07 | 2,27 | 2,08 | 1,09 | 2,75E-01 |
| AT1G47490 | 0,30 | 0,24 | 4,35 | 0,05 | 9,57E-01 |
| AT1G47500 | 0,37 | -0,47 | 4,25 | -0,11 | 9,11E-01 |
| AT1G47530 | 1,40 | -1,38 | 1,51 | -0,91 | 3,62E-01 |
| AT1G47550 | 0,50 | -0,41 | 4,12 | -0,10 | 9,21E-01 |
| AT1G47657 | 2,52 | 2,21 | 1,76 | 1,26 | 2,09E-01 |
| AT1G47680 | 0,12 | 0,00 | 4,39 | 0,00 | 1,00E+00 |
| AT1G47690 | 0,26 | 1,71 | 4,32 | 0,39 | 6,93E-01 |
| AT1G47770 | 0,12 | 0,00 | 4,39 | 0,00 | 1,00E+00 |
| AT1G47890 | 1,24 | -1,41 | 1,75 | -0,81 | 4,20E-01 |
| AT1G47940 | 0,36 | 0,00 | 4,41 | 0,00 | 1,00E+00 |
| AT1G47970 | 1,63 | -0,50 | 1,40 | -0,35 | 7,23E-01 |
| AT1G48050 | 0,54 | -1,73 | 4,40 | -0,39 | 6,94E-01 |
| AT1G48100 | 0,12 | 0,97 | 4,34 | 0,22 | 8,23E-01 |
| AT1G48170 | 0,32 | 0,04 | 3,56 | 0,01 | 9,91E-01 |
| AT1G48210 | 0,24 | 1,44 | 4,34 | 0,33 | 7,41E-01 |
| AT1G48230 | 0,28 | -0,34 | 4,37 | -0,08 | 9,38E-01 |
| AT1G48315 | 0,17 | 1,50 | 4,33 | 0,35 | 7,29E-01 |
| AT1G48360 | 1,34 | -1,36 | 1,89 | -0,72 | 4,71E-01 |
| AT1G48460 | 0,51 | 1,75 | 3,11 | 0,56 | 5,74E-01 |
| AT1G07487 | 0,32 | 1,20 | 3,60 | 0,33 | 7,39E-01 |
| AT1G48480 | 0,38 | -1,68 | 4,35 | -0,39 | 7,00E-01 |
| AT1G48540 | 1,33 | 0,70 | 1,76 | 0,40 | 6,90E-01 |
| AT1G48560 | 0,14 | 0,00 | 4,41 | 0,00 | 1,00E+00 |
| AT1G48600 | 0,62 | 0,97 | 2,18 | 0,45 | 6,55E-01 |
| AT1G48640 | 0,26 | -0,30 | 4,37 | -0,07 | 9,46E-01 |
| AT1G48725 | 1,45 | 1,22 | 1,91 | 0,64 | 5,22E-01 |
| AT1G48820 | 0,24 | 0,00 | 4,41 | 0,00 | 1,00E+00 |
| AT1G48920 | 4,08 | -1,99 | 0,89 | -2,25 | 2,47E-02 |
| AT1G49010 | 0,27 | 1,03 | 4,35 | 0,24 | 8,13E-01 |
| AT1G49032 | 0,71 | -0,75 | 2,10 | -0,36 | 7,19E-01 |
| AT1G49050 | 1,80 | 1,45 | 1,72 | 0,84 | 3,99E-01 |
| AT1G49190 | 1,12 | 1,14 | 2,28 | 0,50 | 6,17E-01 |
| AT1G49200 | 0,08 | 0,00 | 4,40 | 0,00 | 1,00E+00 |
| AT1G49210 | 0,15 | 0,00 | 4,41 | 0,00 | 1,00E+00 |
| AT1G49230 | 0,38 | -1,59 | 3,41 | -0,47 | 6,41E-01 |
| AT1G49245 | 0,90 | -1,14 | 2,21 | -0,52 | 6,06E-01 |
| AT1G49300 | 0,19 | 0,86 | 4,34 | 0,20 | 8,42E-01 |
| AT1G49340 | 1,22 | -2,05 | 1,95 | -1,05 | 2,92E-01 |
| AT1G49390 | 0,08 | 0,44 | 4,38 | 0,10 | 9,21E-01 |
| AT1G49470 | 0,15 | 0,29 | 4,37 | 0,07 | 9,47E-01 |

| | | | | | |
|---|---|---|---|---|---|
| AT1G49510 | 0,25 | 0,00 | 4,41 | 0,00 | 1,00E+00 |
| AT1G49680 | 0,43 | 1,17 | 3,32 | 0,35 | 7,25E-01 |
| AT1G49720 | 0,23 | 0,50 | 4,38 | 0,11 | 9,09E-01 |
| AT1G49820 | 0,28 | 1,76 | 4,33 | 0,41 | 6,84E-01 |
| AT1G49890 | 0,23 | -0,48 | 4,35 | -0,11 | 9,12E-01 |
| AT1G49930 | 0,39 | -1,40 | 4,37 | -0,32 | 7,49E-01 |
| AT1G49975 | 1,21 | 0,47 | 2,45 | 0,19 | 8,49E-01 |
| AT1G49980 | 0,53 | -2,00 | 3,08 | -0,65 | 5,16E-01 |
| AT1G49990 | 0,80 | 3,15 | 4,25 | 0,74 | 4,58E-01 |
| AT1G50010 | 2,93 | -1,36 | 1,10 | -1,24 | 2,16E-01 |
| AT1G50050 | 0,43 | -1,78 | 3,36 | -0,53 | 5,97E-01 |
| AT1G07613 | 1,47 | 0,73 | 1,88 | 0,39 | 6,97E-01 |
| AT1G50120 | 0,25 | -0,49 | 4,34 | -0,11 | 9,11E-01 |
| AT1G50170 | 0,08 | 0,44 | 4,38 | 0,10 | 9,21E-01 |
| AT1G50240 | 1,24 | -0,09 | 1,70 | -0,06 | 9,56E-01 |
| AT1G50270 | 0,45 | -1,73 | 3,30 | -0,52 | 6,01E-01 |
| AT1G50350 | 0,30 | 0,00 | 4,41 | 0,00 | 1,00E+00 |
| AT1G50360 | 0,23 | 0,13 | 4,37 | 0,03 | 9,77E-01 |
| AT1G50370 | 0,39 | -0,66 | 3,40 | -0,20 | 8,45E-01 |
| AT1G50380 | 0,57 | 1,45 | 3,13 | 0,46 | 6,44E-01 |
| AT1G50410 | 1,51 | 0,06 | 2,17 | 0,03 | 9,79E-01 |
| AT1G50560 | 0,41 | -0,62 | 4,24 | -0,15 | 8,84E-01 |
| AT1G50660 | 1,10 | 0,73 | 1,85 | 0,40 | 6,93E-01 |
| AT1G50730 | 0,12 | 1,23 | 4,34 | 0,28 | 7,76E-01 |
| AT1G50740 | 0,72 | 0,02 | 2,83 | 0,01 | 9,95E-01 |
| AT1G50900 | 1,85 | -0,42 | 1,38 | -0,30 | 7,62E-01 |
| AT1G50920 | 2,95 | -0,52 | 1,11 | -0,47 | 6,38E-01 |
| AT1G51060 | 0,42 | 0,00 | 4,32 | 0,00 | 1,00E+00 |
| AT1G51100 | 0,17 | 0,00 | 4,41 | 0,00 | 1,00E+00 |
| AT1G51110 | 0,52 | -1,03 | 3,23 | -0,32 | 7,51E-01 |
| AT1G51160 | 0,67 | -1,07 | 2,90 | -0,37 | 7,11E-01 |
| AT1G51172 | 1,67 | 1,34 | 1,82 | 0,74 | 4,62E-01 |
| AT1G51200 | 1,27 | -0,44 | 2,35 | -0,19 | 8,50E-01 |
| AT1G51350 | 0,44 | 0,65 | 3,24 | 0,20 | 8,42E-01 |
| AT1G51390 | 0,24 | 0,00 | 4,41 | 0,00 | 1,00E+00 |
| AT1G07737 | 0,83 | 0,13 | 2,50 | 0,05 | 9,60E-01 |
| AT1G51402 | 1,01 | 0,24 | 2,39 | 0,10 | 9,19E-01 |
| AT1G51440 | 0,14 | 0,03 | 4,39 | 0,01 | 9,95E-01 |
| AT1G51540 | 0,08 | 0,00 | 4,40 | 0,00 | 1,00E+00 |
| AT1G51580 | 0,35 | 0,39 | 4,37 | 0,09 | 9,30E-01 |
| AT1G51600 | 0,08 | 0,69 | 4,36 | 0,16 | 8,75E-01 |
| AT1G51645 | 0,08 | 0,69 | 4,36 | 0,16 | 8,75E-01 |
| AT1G51660 | 0,75 | -0,61 | 2,51 | -0,24 | 8,07E-01 |
| AT1G51670 | 0,25 | 0,41 | 4,33 | 0,09 | 9,25E-01 |
| AT1G51690 | 2,38 | -0,76 | 1,25 | -0,61 | 5,43E-01 |
| AT1G51720 | 0,29 | 0,35 | 4,32 | 0,08 | 9,36E-01 |
| AT1G51745 | 0,21 | 1,18 | 4,35 | 0,27 | 7,86E-01 |
| AT1G51760 | 0,56 | 0,21 | 2,63 | 0,08 | 9,35E-01 |
| AT1G51950 | 0,22 | 0,83 | 4,34 | 0,19 | 8,49E-01 |
| AT1G52080 | 0,08 | 0,69 | 4,36 | 0,16 | 8,75E-01 |
| AT1G07793 | 2,22 | -0,39 | 1,56 | -0,25 | 8,04E-01 |
| AT1G52190 | 0,40 | -0,54 | 3,29 | -0,16 | 8,71E-01 |

| | | | | | |
|---|---|---|---|---|---|
| AT1G52230 | 2,95 | -2,38 | 1,32 | -1,81 | 7,09E-02 |
| AT1G52310 | 0,73 | 0,31 | 2,40 | 0,13 | 8,99E-01 |
| AT1G52320 | 0,99 | 1,12 | 1,90 | 0,59 | 5,57E-01 |
| AT1G52360 | 2,02 | -1,08 | 1,25 | -0,86 | 3,90E-01 |
| AT1G52380 | 0,74 | -0,36 | 2,85 | -0,13 | 8,98E-01 |
| AT1G52400 | 1,99 | 0,09 | 1,84 | 0,05 | 9,61E-01 |
| AT1G52410 | 0,68 | -0,09 | 2,59 | -0,04 | 9,72E-01 |
| AT1G52420 | 0,99 | 1,26 | 2,74 | 0,46 | 6,45E-01 |
| AT1G52500 | 0,39 | -0,25 | 4,35 | -0,06 | 9,55E-01 |
| AT1G52600 | 0,08 | 0,69 | 4,36 | 0,16 | 8,75E-01 |
| AT1G52618 | 0,68 | 0,80 | 2,91 | 0,28 | 7,82E-01 |
| AT1G52620 | 0,78 | 0,26 | 4,35 | 0,06 | 9,52E-01 |
| AT1G52720 | 0,19 | 0,00 | 4,41 | 0,00 | 1,00E+00 |
| AT1G52740 | 1,01 | 0,56 | 2,17 | 0,26 | 7,97E-01 |
| AT1G52760 | 0,33 | -0,91 | 3,60 | -0,25 | 8,01E-01 |
| AT1G52780 | 3,16 | 2,51 | 1,28 | 1,97 | 4,93E-02 |
| AT1G52870 | 0,92 | 2,15 | 2,57 | 0,84 | 4,03E-01 |
| AT1G52905 | 0,68 | 2,07 | 2,84 | 0,73 | 4,66E-01 |
| AT1G52930 | 0,12 | 1,16 | 4,34 | 0,27 | 7,89E-01 |
| AT1G52940 | 0,76 | -0,03 | 2,44 | -0,01 | 9,90E-01 |
| AT1G52950 | 0,39 | 0,83 | 3,49 | 0,24 | 8,12E-01 |
| AT1G52980 | 1,44 | -0,52 | 1,53 | -0,34 | 7,34E-01 |
| AT1G53050 | 0,64 | -2,46 | 4,31 | -0,57 | 5,68E-01 |
| AT1G53090 | 0,57 | -0,48 | 4,33 | -0,11 | 9,11E-01 |
| AT1G53165 | 0,43 | 0,26 | 2,80 | 0,09 | 9,25E-01 |
| AT1G53210 | 1,83 | 0,95 | 1,86 | 0,51 | 6,09E-01 |
| AT1G53233 | 0,99 | 2,06 | 2,21 | 0,93 | 3,51E-01 |
| AT1G53300 | 0,67 | -1,72 | 2,66 | -0,65 | 5,17E-01 |
| AT1G53330 | 0,35 | -2,17 | 4,38 | -0,49 | 6,21E-01 |
| AT1G53350 | 0,25 | 1,68 | 4,32 | 0,39 | 6,97E-01 |
| AT1G53390 | 0,38 | -1,86 | 4,37 | -0,43 | 6,70E-01 |
| AT1G53400 | 0,40 | 2,16 | 4,32 | 0,50 | 6,17E-01 |
| AT1G53430 | 1,51 | -1,39 | 1,72 | -0,80 | 4,21E-01 |
| AT1G53440 | 1,17 | -0,49 | 1,74 | -0,28 | 7,78E-01 |
| AT1G07887 | 0,84 | 0,05 | 3,09 | 0,02 | 9,87E-01 |
| AT1G53490 | 0,92 | 0,67 | 2,57 | 0,26 | 7,93E-01 |
| AT1G07893 | 0,39 | 1,04 | 4,32 | 0,24 | 8,09E-01 |
| AT1G07897 | 0,95 | 0,34 | 1,74 | 0,20 | 8,43E-01 |
| AT1G53570 | 0,96 | -3,10 | 2,22 | -1,40 | 1,61E-01 |
| AT1G53590 | 0,26 | 0,41 | 4,37 | 0,09 | 9,25E-01 |
| AT1G53655 | 1,78 | 2,24 | 1,88 | 1,19 | 2,32E-01 |
| AT1G53670 | 0,12 | 1,16 | 4,34 | 0,27 | 7,89E-01 |
| AT1G53687 | 5,07 | -3,91 | 1,53 | -2,55 | 1,08E-02 |
| AT1G53720 | 0,26 | 0,14 | 4,41 | 0,03 | 9,74E-01 |
| AT1G53730 | 1,50 | -2,11 | 2,05 | -1,02 | 3,05E-01 |
| AT1G53840 | 1,21 | 0,78 | 2,13 | 0,37 | 7,15E-01 |
| AT1G53870 | 0,49 | 1,46 | 3,93 | 0,37 | 7,11E-01 |
| AT1G53875 | 1,66 | 0,41 | 1,77 | 0,23 | 8,17E-01 |
| AT1G53885 | 0,24 | 1,08 | 4,35 | 0,25 | 8,03E-01 |
| AT1G53890 | 1,37 | 2,21 | 1,79 | 1,23 | 2,17E-01 |
| AT1G53900 | 2,01 | -0,77 | 1,51 | -0,51 | 6,08E-01 |
| AT1G53910 | 0,62 | 1,15 | 2,52 | 0,45 | 6,50E-01 |

| | | | | | |
|---|---|---|---|---|---|
| AT1G54030 | 0,19 | 0,00 | 4,41 | 0,00 | 1,00E+00 |
| AT1G54090 | 3,59 | -0,93 | 1,17 | -0,79 | 4,27E-01 |
| AT1G54115 | 2,18 | 0,31 | 1,39 | 0,22 | 8,22E-01 |
| AT1G54130 | 1,27 | -0,16 | 1,80 | -0,09 | 9,28E-01 |
| AT1G54150 | 0,44 | -0,75 | 3,42 | -0,22 | 8,26E-01 |
| AT1G54200 | 0,80 | -0,79 | 2,43 | -0,32 | 7,46E-01 |
| AT1G54215 | 0,08 | 0,69 | 4,36 | 0,16 | 8,75E-01 |
| AT1G54270 | 2,59 | 1,52 | 1,36 | 1,12 | 2,63E-01 |
| AT1G07957 | 0,34 | 1,16 | 3,47 | 0,33 | 7,38E-01 |
| AT1G54310 | 0,08 | 0,44 | 4,38 | 0,10 | 9,21E-01 |
| AT1G54340 | 0,28 | -0,22 | 4,33 | -0,05 | 9,59E-01 |
| AT1G54350 | 0,37 | 0,57 | 4,27 | 0,13 | 8,93E-01 |
| AT1G54360 | 0,57 | -2,42 | 4,31 | -0,56 | 5,75E-01 |
| AT1G54420 | 1,07 | 0,88 | 2,71 | 0,33 | 7,45E-01 |
| AT1G54440 | 0,25 | 1,82 | 4,32 | 0,42 | 6,73E-01 |
| AT1G54490 | 1,36 | -1,26 | 1,60 | -0,79 | 4,31E-01 |
| AT1G54550 | 0,56 | 1,77 | 2,94 | 0,60 | 5,47E-01 |
| AT1G07977 | 4,37 | -1,01 | 1,42 | -0,71 | 4,78E-01 |
| AT1G54680 | 0,26 | -0,30 | 4,37 | -0,07 | 9,46E-01 |
| AT1G54720 | 0,66 | 1,20 | 3,11 | 0,39 | 6,98E-01 |
| AT1G54780 | 1,22 | -0,32 | 1,96 | -0,16 | 8,71E-01 |
| AT1G54820 | 0,40 | -0,69 | 4,32 | -0,16 | 8,73E-01 |
| AT1G54920 | 1,30 | 0,17 | 1,52 | 0,11 | 9,13E-01 |
| AT1G55000 | 0,70 | 2,26 | 2,56 | 0,88 | 3,77E-01 |
| AT1G55020 | 0,29 | -0,02 | 4,33 | -0,01 | 9,96E-01 |
| AT1G55040 | 0,41 | 0,25 | 4,14 | 0,06 | 9,51E-01 |
| AT1G55090 | 0,81 | 0,53 | 2,84 | 0,19 | 8,52E-01 |
| AT1G55120 | 1,23 | -2,66 | 2,03 | -1,31 | 1,90E-01 |
| AT1G55150 | 0,30 | 0,24 | 4,35 | 0,05 | 9,57E-01 |
| AT1G55160 | 0,99 | 0,66 | 2,36 | 0,28 | 7,81E-01 |
| AT1G55175 | 0,08 | 0,69 | 4,36 | 0,16 | 8,75E-01 |
| AT1G55320 | 0,25 | -0,23 | 4,35 | -0,05 | 9,58E-01 |
| AT1G55325 | 2,49 | -1,00 | 1,09 | -0,91 | 3,61E-01 |
| AT1G55335 | 0,51 | -0,40 | 3,19 | -0,13 | 8,99E-01 |
| AT1G55370 | 0,24 | -2,36 | 4,39 | -0,54 | 5,91E-01 |
| AT1G55460 | 0,32 | 0,82 | 4,34 | 0,19 | 8,50E-01 |
| AT1G55480 | 0,48 | -1,04 | 4,08 | -0,25 | 7,99E-01 |
| AT1G55500 | 0,61 | -1,26 | 2,14 | -0,59 | 5,57E-01 |
| AT1G55510 | 0,20 | 1,19 | 4,35 | 0,27 | 7,84E-01 |
| AT1G55591 | 5,61 | -1,32 | 0,88 | -1,49 | 1,36E-01 |
| AT1G55673 | 3,49 | -2,92 | 1,24 | -2,35 | 1,89E-02 |
| AT1G55840 | 0,33 | -1,22 | 4,34 | -0,28 | 7,79E-01 |
| AT1G55850 | 0,50 | 2,16 | 4,32 | 0,50 | 6,17E-01 |
| AT1G08073 | 0,33 | 0,26 | 4,33 | 0,06 | 9,52E-01 |
| AT1G55890 | 0,12 | 0,37 | 4,37 | 0,08 | 9,33E-01 |
| AT1G55910 | 0,98 | -1,18 | 1,74 | -0,68 | 4,99E-01 |
| AT1G55915 | 0,81 | 0,10 | 2,89 | 0,04 | 9,72E-01 |
| AT1G55920 | 3,12 | 0,56 | 1,28 | 0,43 | 6,64E-01 |
| AT1G08077 | 0,62 | 0,97 | 3,02 | 0,32 | 7,48E-01 |
| AT1G56045 | 0,75 | -0,15 | 2,69 | -0,05 | 9,57E-01 |
| AT1G56050 | 1,07 | -0,46 | 2,49 | -0,18 | 8,54E-01 |
| AT1G56190 | 1,11 | 0,63 | 2,10 | 0,30 | 7,63E-01 |

| | | | | | |
|---|---|---|---|---|---|
| AT1G56200 | 1,42 | -0,22 | 1,69 | -0,13 | 8,99E-01 |
| AT1G56210 | 0,05 | 0,00 | 4,41 | 0,00 | 1,00E+00 |
| AT1G56220 | 1,48 | -0,79 | 1,69 | -0,47 | 6,40E-01 |
| AT1G56450 | 0,83 | 1,32 | 2,19 | 0,60 | 5,47E-01 |
| AT1G56460 | 0,70 | 2,54 | 2,86 | 0,89 | 3,76E-01 |
| AT1G56500 | 0,95 | 0,54 | 1,90 | 0,29 | 7,76E-01 |
| AT1G56510 | 2,98 | -1,01 | 1,33 | -0,76 | 4,47E-01 |
| AT1G56540 | 0,56 | -0,38 | 4,32 | -0,09 | 9,29E-01 |
| AT1G56560 | 0,52 | 1,24 | 3,86 | 0,32 | 7,49E-01 |
| AT1G56570 | 0,28 | -0,22 | 4,33 | -0,05 | 9,59E-01 |
| AT1G56600 | 0,19 | 0,00 | 4,40 | 0,00 | 1,00E+00 |
| AT1G56610 | 0,17 | 0,00 | 4,41 | 0,00 | 1,00E+00 |
| AT1G56660 | 1,02 | 1,29 | 1,87 | 0,69 | 4,90E-01 |
| AT1G56700 | 0,33 | 0,74 | 3,59 | 0,21 | 8,36E-01 |
| AT1G57620 | 0,88 | 1,47 | 3,01 | 0,49 | 6,25E-01 |
| AT1G57630 | 0,41 | -0,29 | 4,33 | -0,07 | 9,46E-01 |
| AT1G57650 | 0,46 | 0,47 | 2,82 | 0,17 | 8,67E-01 |
| AT1G57720 | 1,72 | -0,98 | 1,45 | -0,68 | 4,98E-01 |
| AT1G57760 | 0,12 | 0,00 | 4,38 | 0,00 | 1,00E+00 |
| AT1G57765 | 0,15 | 0,74 | 4,35 | 0,17 | 8,65E-01 |
| AT1G58190 | 0,62 | -1,93 | 2,47 | -0,78 | 4,35E-01 |
| AT1G58215 | 0,57 | 0,78 | 4,32 | 0,18 | 8,56E-01 |
| AT1G58223 | 0,54 | -2,15 | 3,83 | -0,56 | 5,75E-01 |
| AT1G58230 | 3,69 | 0,82 | 1,07 | 0,77 | 4,42E-01 |
| AT1G58260 | 0,12 | 0,49 | 4,36 | 0,11 | 9,11E-01 |
| AT1G58360 | 0,57 | -2,22 | 3,95 | -0,56 | 5,74E-01 |
| AT1G58525 | 0,11 | 0,65 | 4,36 | 0,15 | 8,81E-01 |
| AT1G58520 | 0,55 | 1,08 | 3,26 | 0,33 | 7,40E-01 |
| AT1G58643 | 0,42 | -2,05 | 4,37 | -0,47 | 6,39E-01 |
| AT1G58807 | 1,55 | -0,68 | 1,44 | -0,47 | 6,36E-01 |
| AT1G58808 | 1,22 | -2,98 | 1,93 | -1,54 | 1,23E-01 |
| AT1G58848 | 1,34 | 0,22 | 1,74 | 0,13 | 9,00E-01 |
| AT1G59124 | 1,27 | -0,43 | 1,43 | -0,30 | 7,66E-01 |
| AT1G59218 | 1,28 | 0,12 | 1,82 | 0,07 | 9,46E-01 |
| AT1G59590 | 0,59 | -0,90 | 3,11 | -0,29 | 7,73E-01 |
| AT1G59610 | 3,98 | 0,22 | 0,92 | 0,24 | 8,10E-01 |
| AT1G59620 | 0,81 | -0,59 | 4,28 | -0,14 | 8,90E-01 |
| AT1G59650 | 0,54 | 0,09 | 3,89 | 0,02 | 9,81E-01 |
| AT1G59710 | 0,56 | -0,01 | 3,25 | 0,00 | 9,99E-01 |
| AT1G59750 | 1,67 | 0,44 | 2,19 | 0,20 | 8,40E-01 |
| AT1G59820 | 1,45 | -1,10 | 1,87 | -0,59 | 5,54E-01 |
| AT1G59890 | 0,93 | 1,51 | 2,45 | 0,62 | 5,37E-01 |
| AT1G59900 | 1,24 | 2,71 | 1,91 | 1,42 | 1,55E-01 |
| AT1G59980 | 3,55 | -1,38 | 1,12 | -1,23 | 2,20E-01 |
| AT1G60025 | 0,05 | 0,00 | 4,41 | 0,00 | 1,00E+00 |
| AT1G60072 | 5,42 | 0,92 | 0,86 | 1,06 | 2,87E-01 |
| AT1G60073 | 1,58 | 3,40 | 1,91 | 1,78 | 7,56E-02 |
| AT1G60170 | 1,29 | 0,66 | 2,25 | 0,29 | 7,70E-01 |
| AT1G60190 | 0,89 | -0,08 | 2,04 | -0,04 | 9,71E-01 |
| AT1G60220 | 0,08 | 0,44 | 4,38 | 0,10 | 9,21E-01 |
| AT1G60420 | 0,70 | 2,97 | 2,82 | 1,05 | 2,92E-01 |
| AT1G60545 | 1,07 | -0,17 | 2,86 | -0,06 | 9,51E-01 |

| | | | | | |
|---|---|---|---|---|---|
| AT1G60650 | 0,21 | -0,49 | 4,35 | -0,11 | 9,11E-01 |
| AT1G60720 | 1,62 | 0,74 | 2,01 | 0,37 | 7,11E-01 |
| AT1G60870 | 2,59 | 1,09 | 1,24 | 0,88 | 3,78E-01 |
| AT1G60900 | 0,65 | 0,42 | 3,21 | 0,13 | 8,95E-01 |
| AT1G08353 | 0,98 | -1,39 | 1,92 | -0,72 | 4,70E-01 |
| AT1G60950 | 5,66 | -0,67 | 1,03 | -0,65 | 5,15E-01 |
| AT1G61030 | 0,52 | 1,08 | 3,90 | 0,28 | 7,82E-01 |
| AT1G61040 | 0,54 | 0,09 | 3,89 | 0,02 | 9,81E-01 |
| AT1G61150 | 0,81 | -1,44 | 3,59 | -0,40 | 6,89E-01 |
| AT1G61170 | 0,33 | 1,92 | 4,32 | 0,44 | 6,57E-01 |
| AT1G61180 | 0,58 | 0,09 | 2,67 | 0,03 | 9,74E-01 |
| AT1G61190 | 2,38 | 1,15 | 1,25 | 0,92 | 3,55E-01 |
| AT1G61210 | 1,07 | -1,14 | 1,68 | -0,68 | 4,95E-01 |
| AT1G61224 | 1,62 | -0,31 | 1,63 | -0,19 | 8,51E-01 |
| AT1G61250 | 0,18 | 0,21 | 4,37 | 0,05 | 9,62E-01 |
| AT1G61280 | 0,12 | 1,16 | 4,34 | 0,27 | 7,89E-01 |
| AT1G61350 | 0,76 | -0,12 | 2,44 | -0,05 | 9,60E-01 |
| AT1G61450 | 0,12 | 1,05 | 4,35 | 0,24 | 8,10E-01 |
| AT1G61470 | 0,18 | 0,00 | 4,41 | 0,00 | 1,00E+00 |
| AT1G61610 | 0,12 | 0,00 | 4,38 | 0,00 | 1,00E+00 |
| AT1G61620 | 0,37 | -1,39 | 4,35 | -0,32 | 7,50E-01 |
| AT1G61790 | 1,42 | -2,25 | 1,54 | -1,46 | 1,45E-01 |
| AT1G61800 | 0,56 | 1,22 | 4,32 | 0,28 | 7,78E-01 |
| AT1G61820 | 0,08 | 0,44 | 4,38 | 0,10 | 9,21E-01 |
| AT1G61850 | 0,80 | 0,23 | 2,23 | 0,10 | 9,17E-01 |
| AT1G61990 | 0,32 | 2,04 | 4,32 | 0,47 | 6,36E-01 |
| AT1G62170 | 0,36 | 0,42 | 4,38 | 0,10 | 9,23E-01 |
| AT1G08477 | 1,64 | 1,05 | 1,62 | 0,65 | 5,17E-01 |
| AT1G62270 | 0,29 | 0,00 | 4,41 | 0,00 | 1,00E+00 |
| AT1G08517 | 0,77 | -0,94 | 2,58 | -0,37 | 7,15E-01 |
| AT1G62380 | 1,16 | 2,41 | 1,84 | 1,31 | 1,90E-01 |
| AT1G62410 | 0,32 | 0,00 | 4,41 | 0,00 | 1,00E+00 |
| AT1G62480 | 0,12 | 0,00 | 4,38 | 0,00 | 1,00E+00 |
| AT1G62540 | 2,48 | -0,83 | 1,79 | -0,46 | 6,45E-01 |
| AT1G62560 | 1,49 | -1,79 | 3,11 | -0,58 | 5,64E-01 |
| AT1G62570 | 0,39 | 1,16 | 4,32 | 0,27 | 7,88E-01 |
| AT1G62620 | 0,57 | -0,99 | 2,97 | -0,33 | 7,40E-01 |
| AT1G62630 | 2,10 | 0,02 | 1,51 | 0,01 | 9,90E-01 |
| AT1G62660 | 1,32 | -0,25 | 1,61 | -0,16 | 8,76E-01 |
| AT1G62720 | 0,24 | 0,00 | 4,41 | 0,00 | 1,00E+00 |
| AT1G62790 | 0,51 | -2,36 | 4,32 | -0,55 | 5,84E-01 |
| AT1G62895 | 1,43 | 1,04 | 1,67 | 0,63 | 5,31E-01 |
| AT1G62920 | 0,97 | 1,99 | 2,98 | 0,67 | 5,04E-01 |
| AT1G62970 | 0,33 | 1,25 | 3,52 | 0,35 | 7,24E-01 |
| AT1G62975 | 0,08 | 0,44 | 4,38 | 0,10 | 9,21E-01 |
| AT1G63000 | 0,67 | -0,36 | 3,86 | -0,09 | 9,25E-01 |
| AT1G63110 | 0,32 | 0,55 | 4,32 | 0,13 | 8,98E-01 |
| AT1G63170 | 0,08 | 0,69 | 4,36 | 0,16 | 8,75E-01 |
| AT1G63230 | 3,31 | -0,87 | 1,32 | -0,66 | 5,12E-01 |
| AT1G63430 | 0,74 | 1,72 | 3,84 | 0,45 | 6,54E-01 |
| AT1G63500 | 0,59 | 0,40 | 2,72 | 0,15 | 8,82E-01 |
| AT1G63540 | 0,37 | 0,00 | 4,41 | 0,00 | 1,00E+00 |

| | | | | | |
|---|---|---|---|---|---|
| AT1G63630 | 3,43 | -0,56 | 1,30 | -0,43 | 6,64E-01 |
| AT1G63680 | 0,43 | 0,56 | 3,23 | 0,17 | 8,63E-01 |
| AT1G63690 | 1,03 | -1,79 | 1,65 | -1,08 | 2,79E-01 |
| AT1G08623 | 2,21 | -1,53 | 1,60 | -0,95 | 3,41E-01 |
| AT1G63730 | 0,24 | 0,00 | 4,41 | 0,00 | 1,00E+00 |
| AT1G63740 | 1,93 | 1,14 | 1,56 | 0,73 | 4,64E-01 |
| AT1G63750 | 2,64 | -0,08 | 1,18 | -0,07 | 9,45E-01 |
| AT1G63830 | 0,49 | 1,58 | 4,07 | 0,39 | 6,98E-01 |
| AT1G63850 | 0,53 | -0,21 | 3,94 | -0,05 | 9,58E-01 |
| AT1G08643 | 0,79 | 2,39 | 2,96 | 0,80 | 4,21E-01 |
| AT1G63900 | 0,95 | -1,02 | 2,00 | -0,51 | 6,10E-01 |
| AT1G63940 | 0,81 | -0,81 | 1,94 | -0,42 | 6,77E-01 |
| AT1G63950 | 1,81 | -0,85 | 2,16 | -0,39 | 6,93E-01 |
| AT1G63980 | 3,67 | -0,69 | 1,01 | -0,68 | 4,98E-01 |
| AT1G64060 | 0,86 | 1,83 | 2,35 | 0,78 | 4,36E-01 |
| AT1G64065 | 0,29 | -0,89 | 4,37 | -0,20 | 8,39E-01 |
| AT1G64090 | 0,96 | -0,07 | 1,93 | -0,04 | 9,71E-01 |
| AT1G64100 | 0,61 | -0,05 | 3,67 | -0,01 | 9,90E-01 |
| AT1G64120 | 0,13 | 0,00 | 4,41 | 0,00 | 1,00E+00 |
| AT1G64130 | 0,12 | 1,16 | 4,34 | 0,27 | 7,89E-01 |
| AT1G08657 | 0,60 | 0,12 | 2,67 | 0,04 | 9,64E-01 |
| AT1G64230 | 1,39 | 0,16 | 1,60 | 0,10 | 9,21E-01 |
| AT1G64340 | 2,24 | 3,00 | 2,00 | 1,50 | 1,34E-01 |
| AT1G64370 | 0,17 | 0,00 | 4,41 | 0,00 | 1,00E+00 |
| AT1G64385 | 1,99 | -1,81 | 1,78 | -1,02 | 3,09E-01 |
| AT1G64390 | 1,48 | -1,39 | 2,08 | -0,67 | 5,04E-01 |
| AT1G64430 | 2,85 | -1,28 | 1,20 | -1,06 | 2,88E-01 |
| AT1G64500 | 2,64 | -0,56 | 1,24 | -0,45 | 6,50E-01 |
| AT1G64520 | 0,27 | 0,72 | 4,37 | 0,17 | 8,69E-01 |
| AT1G64530 | 0,54 | -0,23 | 3,04 | -0,07 | 9,41E-01 |
| AT1G64550 | 0,64 | 0,58 | 2,58 | 0,22 | 8,24E-01 |
| AT1G64570 | 0,71 | 1,30 | 2,40 | 0,54 | 5,88E-01 |
| AT1G64572 | 0,51 | -2,04 | 4,34 | -0,47 | 6,38E-01 |
| AT1G64620 | 0,45 | -0,46 | 3,28 | -0,14 | 8,88E-01 |
| AT1G64680 | 1,48 | 0,71 | 1,79 | 0,40 | 6,92E-01 |
| AT1G64710 | 2,31 | -1,40 | 1,19 | -1,18 | 2,39E-01 |
| AT1G64740 | 0,25 | 0,13 | 4,39 | 0,03 | 9,77E-01 |
| AT1G64770 | 1,29 | -1,11 | 1,62 | -0,69 | 4,92E-01 |
| AT1G64795 | 0,74 | 0,35 | 2,07 | 0,17 | 8,67E-01 |
| AT1G64810 | 0,44 | 0,62 | 4,32 | 0,14 | 8,86E-01 |
| AT1G64820 | 0,08 | 0,69 | 4,36 | 0,16 | 8,75E-01 |
| AT1G64860 | 0,65 | 0,99 | 3,11 | 0,32 | 7,50E-01 |
| AT1G64900 | 1,03 | 0,47 | 1,87 | 0,25 | 8,02E-01 |
| AT1G64950 | 0,34 | 1,36 | 3,45 | 0,40 | 6,93E-01 |
| AT1G64960 | 0,60 | 0,75 | 3,89 | 0,19 | 8,48E-01 |
| AT1G65010 | 1,62 | -0,09 | 1,64 | -0,06 | 9,56E-01 |
| AT1G65220 | 0,59 | 0,77 | 2,65 | 0,29 | 7,72E-01 |
| AT1G65230 | 1,04 | -2,39 | 1,99 | -1,20 | 2,28E-01 |
| AT1G65260 | 0,21 | 0,00 | 4,41 | 0,00 | 1,00E+00 |
| AT1G65270 | 0,21 | 0,64 | 4,35 | 0,15 | 8,84E-01 |
| AT1G65280 | 0,39 | 0,00 | 4,27 | 0,00 | 9,99E-01 |
| AT1G65295 | 0,05 | 0,00 | 4,41 | 0,00 | 1,00E+00 |

| | | | | | |
|---|---|---|---|---|---|
| AT1G65360 | 0,90 | -2,21 | 2,32 | -0,95 | 3,41E-01 |
| AT1G08753 | 0,15 | 0,67 | 4,37 | 0,15 | 8,79E-01 |
| AT1G65450 | 2,07 | -2,41 | 1,45 | -1,66 | 9,69E-02 |
| AT1G65490 | 0,49 | -3,04 | 4,33 | -0,70 | 4,83E-01 |
| AT1G65590 | 0,23 | -0,48 | 4,35 | -0,11 | 9,12E-01 |
| AT1G65681 | 0,20 | 0,00 | 4,41 | 0,00 | 1,00E+00 |
| AT1G65730 | 0,08 | 0,69 | 4,36 | 0,16 | 8,75E-01 |
| AT1G65790 | 0,93 | 0,04 | 1,76 | 0,02 | 9,84E-01 |
| AT1G65800 | 1,17 | 0,38 | 1,82 | 0,21 | 8,33E-01 |
| AT1G65820 | 0,38 | -0,15 | 2,88 | -0,05 | 9,58E-01 |
| AT1G65840 | 0,61 | -0,03 | 3,82 | -0,01 | 9,94E-01 |
| AT1G65930 | 2,39 | 0,10 | 1,37 | 0,07 | 9,44E-01 |
| AT1G65960 | 3,84 | 1,59 | 1,21 | 1,31 | 1,90E-01 |
| AT1G08793 | 0,15 | 0,29 | 4,37 | 0,07 | 9,47E-01 |
| AT1G66090 | 0,59 | 0,46 | 3,17 | 0,14 | 8,85E-01 |
| AT1G66150 | 1,55 | -0,06 | 1,39 | -0,04 | 9,65E-01 |
| AT1G66160 | 0,48 | 1,68 | 4,03 | 0,42 | 6,78E-01 |
| AT1G66180 | 0,53 | -1,04 | 3,18 | -0,33 | 7,43E-01 |
| AT1G66230 | 0,24 | -2,36 | 4,39 | -0,54 | 5,91E-01 |
| AT1G66340 | 0,60 | 1,32 | 2,91 | 0,45 | 6,51E-01 |
| AT1G66345 | 0,47 | 2,40 | 3,98 | 0,60 | 5,45E-01 |
| AT1G66430 | 0,50 | 0,00 | 4,41 | 0,00 | 1,00E+00 |
| AT1G66465 | 0,33 | 1,86 | 4,32 | 0,43 | 6,67E-01 |
| AT1G66580 | 1,55 | 0,40 | 1,51 | 0,27 | 7,90E-01 |
| AT1G66640 | 0,21 | 0,05 | 4,39 | 0,01 | 9,90E-01 |
| AT1G66730 | 1,91 | 1,03 | 1,46 | 0,71 | 4,78E-01 |
| AT1G66750 | 0,31 | 1,34 | 4,33 | 0,31 | 7,58E-01 |
| AT1G66820 | 0,53 | -0,05 | 4,33 | -0,01 | 9,91E-01 |
| AT1G66900 | 0,20 | 1,00 | 4,36 | 0,23 | 8,18E-01 |
| AT1G67105 | 0,86 | 1,89 | 1,97 | 0,96 | 3,35E-01 |
| AT1G67190 | 0,34 | 0,31 | 4,33 | 0,07 | 9,42E-01 |
| AT1G08923 | 0,11 | 0,00 | 4,41 | 0,00 | 1,00E+00 |
| AT1G67420 | 0,13 | 0,73 | 4,36 | 0,17 | 8,67E-01 |
| AT1G67430 | 1,82 | 0,49 | 1,43 | 0,34 | 7,34E-01 |
| AT1G08937 | 0,46 | 1,44 | 3,27 | 0,44 | 6,59E-01 |
| AT1G67450 | 1,86 | 1,61 | 1,64 | 0,98 | 3,27E-01 |
| AT1G67470 | 0,43 | -0,89 | 3,27 | -0,27 | 7,87E-01 |
| AT1G67480 | 0,49 | -0,04 | 4,30 | -0,01 | 9,93E-01 |
| AT1G67530 | 0,54 | -2,42 | 3,87 | -0,62 | 5,32E-01 |
| AT1G67560 | 0,64 | -1,29 | 4,31 | -0,30 | 7,64E-01 |
| AT1G67620 | 0,15 | 0,00 | 4,38 | 0,00 | 1,00E+00 |
| AT1G67635 | 1,69 | 3,06 | 1,86 | 1,64 | 1,01E-01 |
| AT1G67700 | 0,46 | -1,72 | 4,04 | -0,43 | 6,69E-01 |
| AT1G67720 | 0,53 | 0,39 | 3,71 | 0,11 | 9,16E-01 |
| AT1G67792 | 0,38 | -0,91 | 4,32 | -0,21 | 8,34E-01 |
| AT1G67840 | 1,96 | 1,29 | 1,63 | 0,79 | 4,28E-01 |
| AT1G67900 | 1,04 | -0,16 | 2,31 | -0,07 | 9,45E-01 |
| AT1G67940 | 0,66 | -1,69 | 2,79 | -0,61 | 5,43E-01 |
| AT1G08963 | 0,39 | -1,28 | 4,38 | -0,29 | 7,70E-01 |
| AT1G67980 | 0,18 | 0,13 | 4,41 | 0,03 | 9,76E-01 |
| AT1G67990 | 0,55 | -1,61 | 3,84 | -0,42 | 6,74E-01 |
| AT1G68010 | 2,53 | 0,30 | 1,39 | 0,22 | 8,27E-01 |

| | | | | | |
|---|---|---|---|---|---|
| AT1G68050 | 0,24 | 1,32 | 4,34 | 0,30 | 7,61E-01 |
| AT1G68100 | 0,43 | 0,26 | 4,12 | 0,06 | 9,50E-01 |
| AT1G68110 | 0,27 | -0,47 | 4,37 | -0,11 | 9,14E-01 |
| AT1G68310 | 0,12 | 0,97 | 4,34 | 0,22 | 8,23E-01 |
| AT1G68440 | 1,32 | -1,45 | 1,62 | -0,89 | 3,72E-01 |
| AT1G68490 | 0,97 | -0,37 | 2,36 | -0,16 | 8,74E-01 |
| AT1G68570 | 0,58 | 1,51 | 2,57 | 0,59 | 5,57E-01 |
| AT1G68580 | 1,44 | -0,33 | 1,45 | -0,23 | 8,21E-01 |
| AT1G68585 | 0,27 | -0,96 | 4,39 | -0,22 | 8,27E-01 |
| AT1G68620 | 0,33 | 0,91 | 4,32 | 0,21 | 8,33E-01 |
| AT1G68650 | 0,32 | -0,65 | 4,33 | -0,15 | 8,81E-01 |
| AT1G68670 | 0,63 | 2,61 | 2,93 | 0,89 | 3,73E-01 |
| AT1G68720 | 1,50 | 2,28 | 2,16 | 1,05 | 2,92E-01 |
| AT1G68840 | 1,23 | -3,66 | 1,90 | -1,93 | 5,38E-02 |
| AT1G68890 | 1,09 | 0,53 | 1,82 | 0,29 | 7,69E-01 |
| AT1G68945 | 2,15 | 1,32 | 1,61 | 0,82 | 4,12E-01 |
| AT1G68980 | 0,24 | 0,24 | 4,34 | 0,06 | 9,55E-01 |
| AT1G69085 | 0,08 | 0,00 | 4,41 | 0,00 | 1,00E+00 |
| AT1G69160 | 0,39 | 0,01 | 3,39 | 0,00 | 9,98E-01 |
| AT1G69170 | 0,40 | 2,16 | 4,32 | 0,50 | 6,17E-01 |
| AT1G69200 | 0,21 | -1,41 | 4,36 | -0,32 | 7,46E-01 |
| AT1G69252 | 0,90 | -2,61 | 3,07 | -0,85 | 3,95E-01 |
| AT1G69260 | 0,63 | -0,82 | 2,83 | -0,29 | 7,72E-01 |
| AT1G69350 | 0,78 | 0,62 | 2,32 | 0,27 | 7,89E-01 |
| AT1G69390 | 0,12 | 0,00 | 4,38 | 0,00 | 1,00E+00 |
| AT1G69410 | 0,27 | 0,86 | 4,35 | 0,20 | 8,43E-01 |
| AT1G69420 | 0,65 | -1,99 | 2,38 | -0,84 | 4,03E-01 |
| AT1G69510 | 0,10 | 0,00 | 4,41 | 0,00 | 1,00E+00 |
| AT1G69523 | 0,10 | 0,00 | 4,41 | 0,00 | 1,00E+00 |
| AT1G69526 | 0,08 | 0,69 | 4,36 | 0,16 | 8,75E-01 |
| AT1G69530 | 0,93 | -3,14 | 2,11 | -1,49 | 1,37E-01 |
| AT1G69572 | 2,44 | 0,13 | 1,49 | 0,09 | 9,28E-01 |
| AT1G69580 | 0,26 | 0,00 | 4,38 | 0,00 | 1,00E+00 |
| AT1G69610 | 0,21 | 0,58 | 4,36 | 0,13 | 8,94E-01 |
| AT1G69620 | 2,14 | 0,26 | 1,40 | 0,19 | 8,51E-01 |
| AT1G09087 | 0,37 | 0,03 | 3,46 | 0,01 | 9,92E-01 |
| AT1G69690 | 0,67 | 0,86 | 2,57 | 0,34 | 7,37E-01 |
| AT1G69740 | 0,99 | -0,17 | 1,93 | -0,09 | 9,31E-01 |
| AT1G69750 | 0,10 | 0,00 | 4,41 | 0,00 | 1,00E+00 |
| AT1G69870 | 0,38 | 1,33 | 4,33 | 0,31 | 7,58E-01 |
| AT1G70060 | 1,48 | 0,31 | 1,88 | 0,16 | 8,69E-01 |
| AT1G70090 | 0,99 | -0,43 | 2,11 | -0,20 | 8,40E-01 |
| AT1G70100 | 0,38 | 1,36 | 4,29 | 0,32 | 7,51E-01 |
| AT1G70160 | 1,19 | 0,79 | 1,60 | 0,49 | 6,23E-01 |
| AT1G70180 | 1,28 | -0,54 | 1,64 | -0,33 | 7,43E-01 |
| AT1G70185 | 5,33 | -0,59 | 0,79 | -0,74 | 4,60E-01 |
| AT1G70190 | 0,40 | 1,45 | 4,35 | 0,33 | 7,38E-01 |
| AT1G70210 | 0,79 | 1,03 | 2,47 | 0,42 | 6,78E-01 |
| AT1G70230 | 0,59 | -0,92 | 2,77 | -0,33 | 7,40E-01 |
| AT1G70250 | 0,71 | -0,68 | 2,80 | -0,24 | 8,07E-01 |
| AT1G70330 | 0,33 | 0,83 | 3,58 | 0,23 | 8,17E-01 |
| AT1G70480 | 1,12 | -0,41 | 1,62 | -0,25 | 8,00E-01 |

| | | | | | |
|---|---|---|---|---|---|
| AT1G70510 | 0,12 | 1,05 | 4,35 | 0,24 | 8,10E-01 |
| AT1G70550 | 0,14 | 0,09 | 4,38 | 0,02 | 9,84E-01 |
| AT1G70570 | 0,36 | -0,77 | 4,38 | -0,18 | 8,61E-01 |
| AT1G70580 | 0,81 | 0,05 | 2,33 | 0,02 | 9,83E-01 |
| AT1G70620 | 1,32 | 0,58 | 1,91 | 0,30 | 7,61E-01 |
| AT1G70630 | 0,36 | -0,86 | 4,30 | -0,20 | 8,42E-01 |
| AT1G70645 | 0,05 | 0,00 | 4,41 | 0,00 | 1,00E+00 |
| AT1G70650 | 0,26 | -0,48 | 4,35 | -0,11 | 9,12E-01 |
| AT1G70660 | 0,20 | -0,44 | 4,36 | -0,10 | 9,19E-01 |
| AT1G70700 | 0,60 | 0,14 | 2,73 | 0,05 | 9,58E-01 |
| AT1G70760 | 0,35 | 1,63 | 3,47 | 0,47 | 6,38E-01 |
| AT1G70810 | 0,62 | -0,34 | 2,53 | -0,14 | 8,92E-01 |
| AT1G70820 | 0,43 | -0,89 | 2,83 | -0,32 | 7,52E-01 |
| AT1G70895 | 0,08 | 0,00 | 4,40 | 0,00 | 1,00E+00 |
| AT1G70900 | 1,03 | -0,43 | 1,86 | -0,23 | 8,17E-01 |
| AT1G70940 | 2,12 | -1,97 | 1,33 | -1,48 | 1,40E-01 |
| AT1G70960 | 0,12 | 0,00 | 4,38 | 0,00 | 1,00E+00 |
| AT1G70980 | 0,40 | 0,23 | 4,40 | 0,05 | 9,59E-01 |
| AT1G71010 | 1,50 | 0,28 | 1,71 | 0,16 | 8,70E-01 |
| AT1G71090 | 0,35 | 0,33 | 3,52 | 0,09 | 9,25E-01 |
| AT1G71100 | 1,11 | 0,89 | 1,86 | 0,48 | 6,33E-01 |
| AT1G71110 | 0,15 | 0,45 | 4,37 | 0,10 | 9,19E-01 |
| AT1G71130 | 0,37 | 1,21 | 4,35 | 0,28 | 7,81E-01 |
| AT1G71180 | 0,42 | 0,71 | 4,23 | 0,17 | 8,67E-01 |
| AT1G71220 | 2,96 | 0,26 | 1,19 | 0,22 | 8,24E-01 |
| AT1G71270 | 0,45 | 1,33 | 4,17 | 0,32 | 7,50E-01 |
| AT1G71300 | 0,38 | -1,27 | 4,33 | -0,29 | 7,68E-01 |
| AT1G71320 | 0,39 | 0,01 | 4,24 | 0,00 | 9,98E-01 |
| AT1G71480 | 0,80 | 0,75 | 2,79 | 0,27 | 7,88E-01 |
| AT1G09257 | 0,17 | 1,50 | 4,33 | 0,35 | 7,29E-01 |
| AT1G71500 | 0,95 | -2,45 | 2,00 | -1,22 | 2,22E-01 |
| AT1G71695 | 0,82 | -1,43 | 2,44 | -0,59 | 5,58E-01 |
| AT1G71720 | 0,53 | 0,07 | 3,88 | 0,02 | 9,86E-01 |
| AT1G71800 | 0,67 | -0,69 | 3,52 | -0,20 | 8,44E-01 |
| AT1G71820 | 1,53 | 0,89 | 1,58 | 0,56 | 5,73E-01 |
| AT1G71840 | 1,05 | -0,15 | 2,28 | -0,07 | 9,48E-01 |
| AT1G71880 | 1,96 | 0,71 | 1,59 | 0,45 | 6,53E-01 |
| AT1G71900 | 0,51 | -2,04 | 4,34 | -0,47 | 6,38E-01 |
| AT1G71910 | 0,44 | 0,02 | 3,31 | 0,01 | 9,95E-01 |
| AT1G71920 | 0,73 | 1,47 | 2,47 | 0,60 | 5,51E-01 |
| AT1G71930 | 1,13 | 1,79 | 2,13 | 0,84 | 4,03E-01 |
| AT1G71970 | 0,14 | 0,76 | 4,35 | 0,18 | 8,61E-01 |
| AT1G71980 | 0,46 | 1,77 | 3,15 | 0,56 | 5,75E-01 |
| AT1G72010 | 0,42 | 1,14 | 3,29 | 0,35 | 7,28E-01 |
| AT1G72030 | 0,75 | -0,55 | 3,64 | -0,15 | 8,81E-01 |
| AT1G72050 | 0,75 | -0,71 | 3,51 | -0,20 | 8,39E-01 |
| AT1G72090 | 0,34 | 0,36 | 4,34 | 0,08 | 9,34E-01 |
| AT1G72180 | 2,71 | -0,26 | 1,10 | -0,24 | 8,12E-01 |
| AT1G72240 | 0,60 | -0,16 | 3,83 | -0,04 | 9,66E-01 |
| AT1G72310 | 0,25 | 0,03 | 4,34 | 0,01 | 9,95E-01 |
| AT1G72330 | 0,67 | -0,68 | 2,80 | -0,24 | 8,08E-01 |
| AT1G72390 | 2,40 | 0,67 | 1,18 | 0,56 | 5,72E-01 |

| | | | | | |
|---|---|---|---|---|---|
| AT1G72470 | 0,43 | -0,62 | 4,35 | -0,14 | 8,86E-01 |
| AT1G72510 | 0,57 | 0,77 | 3,26 | 0,24 | 8,13E-01 |
| AT1G72520 | 0,91 | 1,09 | 2,05 | 0,53 | 5,96E-01 |
| AT1G72530 | 0,05 | 0,00 | 4,41 | 0,00 | 1,00E+00 |
| AT1G72645 | 2,19 | -1,07 | 1,64 | -0,65 | 5,14E-01 |
| AT1G09333 | 1,54 | 0,77 | 1,99 | 0,39 | 6,97E-01 |
| AT1G72650 | 0,79 | 1,91 | 2,32 | 0,83 | 4,09E-01 |
| AT1G72700 | 1,31 | 2,30 | 1,85 | 1,24 | 2,15E-01 |
| AT1G72710 | 1,73 | 0,10 | 1,46 | 0,07 | 9,44E-01 |
| AT1G72750 | 0,63 | -0,57 | 2,19 | -0,26 | 7,94E-01 |
| AT1G72770 | 1,51 | 0,59 | 1,48 | 0,40 | 6,91E-01 |
| AT1G72790 | 0,71 | 2,12 | 2,99 | 0,71 | 4,78E-01 |
| AT1G72855 | 0,40 | 2,16 | 4,32 | 0,50 | 6,17E-01 |
| AT1G72900 | 0,32 | 1,47 | 3,53 | 0,42 | 6,76E-01 |
| AT1G72910 | 1,29 | 1,22 | 1,81 | 0,68 | 4,98E-01 |
| AT1G72920 | 0,44 | -1,27 | 3,36 | -0,38 | 7,07E-01 |
| AT1G72930 | 3,01 | -1,43 | 1,70 | -0,85 | 3,98E-01 |
| AT1G72940 | 0,97 | 3,57 | 3,96 | 0,90 | 3,67E-01 |
| AT1G72970 | 0,16 | 0,92 | 4,35 | 0,21 | 8,33E-01 |
| AT1G73030 | 0,82 | -2,57 | 2,37 | -1,08 | 2,79E-01 |
| AT1G73060 | 0,24 | -1,85 | 4,40 | -0,42 | 6,74E-01 |
| AT1G73080 | 1,38 | -0,61 | 1,43 | -0,43 | 6,69E-01 |
| AT1G73100 | 0,70 | -0,55 | 2,51 | -0,22 | 8,25E-01 |
| AT1G73180 | 1,25 | -0,98 | 1,73 | -0,57 | 5,70E-01 |
| AT1G73430 | 1,05 | 0,41 | 2,17 | 0,19 | 8,50E-01 |
| AT1G73450 | 1,12 | 0,28 | 1,74 | 0,16 | 8,73E-01 |
| AT1G73460 | 0,79 | -0,05 | 2,38 | -0,02 | 9,83E-01 |
| AT1G73470 | 0,23 | -0,21 | 4,35 | -0,05 | 9,61E-01 |
| AT1G73480 | 1,37 | -1,51 | 2,03 | -0,75 | 4,56E-01 |
| AT1G73600 | 1,37 | -1,35 | 2,31 | -0,58 | 5,60E-01 |
| AT1G73630 | 0,10 | 0,01 | 4,38 | 0,00 | 9,99E-01 |
| AT1G73670 | 0,46 | 0,37 | 3,31 | 0,11 | 9,10E-01 |
| AT1G73710 | 0,22 | 1,51 | 4,33 | 0,35 | 7,27E-01 |
| AT1G73750 | 0,72 | 2,79 | 4,29 | 0,65 | 5,16E-01 |
| AT1G73770 | 0,08 | 0,69 | 4,36 | 0,16 | 8,75E-01 |
| AT1G73850 | 0,17 | -1,70 | 4,40 | -0,39 | 7,00E-01 |
| AT1G73870 | 2,16 | -1,60 | 1,27 | -1,26 | 2,07E-01 |
| AT1G73875 | 0,39 | 0,38 | 4,20 | 0,09 | 9,28E-01 |
| AT1G73885 | 0,62 | 2,07 | 3,08 | 0,67 | 5,02E-01 |
| AT1G73920 | 0,62 | -2,36 | 2,52 | -0,94 | 3,49E-01 |
| AT1G73970 | 0,95 | 1,47 | 2,49 | 0,59 | 5,55E-01 |
| AT1G74160 | 1,08 | -1,60 | 1,65 | -0,97 | 3,32E-01 |
| AT1G74250 | 1,42 | 1,40 | 1,76 | 0,80 | 4,26E-01 |
| AT1G74290 | 1,78 | 1,13 | 1,70 | 0,67 | 5,06E-01 |
| AT1G74360 | 1,21 | 1,58 | 2,34 | 0,68 | 5,00E-01 |
| AT1G74380 | 1,29 | -1,10 | 1,61 | -0,68 | 4,95E-01 |
| AT1G74430 | 0,38 | -1,78 | 4,41 | -0,40 | 6,87E-01 |
| AT1G74450 | 0,99 | 1,00 | 2,04 | 0,49 | 6,22E-01 |
| AT1G74580 | 0,25 | -0,12 | 4,34 | -0,03 | 9,78E-01 |
| AT1G74680 | 0,57 | -2,19 | 3,14 | -0,70 | 4,86E-01 |
| AT1G74710 | 0,60 | 0,82 | 2,61 | 0,31 | 7,54E-01 |
| AT1G74720 | 1,98 | -0,43 | 1,41 | -0,31 | 7,58E-01 |

| | | | | | |
|---|---|---|---|---|---|
| AT1G74730 | 0,44 | -0,80 | 3,30 | -0,24 | 8,08E-01 |
| AT1G74750 | 0,50 | -1,32 | 4,28 | -0,31 | 7,58E-01 |
| AT1G74780 | 0,63 | 1,71 | 2,90 | 0,59 | 5,55E-01 |
| AT1G74790 | 0,51 | -1,22 | 3,91 | -0,31 | 7,55E-01 |
| AT1G74870 | 1,10 | 2,11 | 3,21 | 0,66 | 5,11E-01 |
| AT1G74930 | 0,81 | 0,03 | 2,85 | 0,01 | 9,91E-01 |
| AT1G74940 | 0,58 | 1,48 | 3,10 | 0,48 | 6,32E-01 |
| AT1G75040 | 3,07 | -0,99 | 1,01 | -0,98 | 3,29E-01 |
| AT1G75080 | 0,21 | 1,63 | 4,33 | 0,38 | 7,06E-01 |
| AT1G09533 | 1,00 | -0,12 | 1,92 | -0,06 | 9,48E-01 |
| AT1G75163 | 0,94 | 2,30 | 2,08 | 1,11 | 2,69E-01 |
| AT1G75166 | 1,22 | 0,87 | 2,04 | 0,43 | 6,68E-01 |
| AT1G75170 | 0,12 | 0,97 | 4,34 | 0,22 | 8,23E-01 |
| AT1G09543 | 0,38 | 0,83 | 3,38 | 0,25 | 8,06E-01 |
| AT1G75190 | 0,22 | 0,00 | 4,41 | 0,00 | 1,00E+00 |
| AT1G75210 | 0,77 | -1,66 | 3,01 | -0,55 | 5,80E-01 |
| AT1G75280 | 0,22 | 0,00 | 4,41 | 0,00 | 1,00E+00 |
| AT1G75310 | 0,47 | -1,39 | 4,37 | -0,32 | 7,51E-01 |
| AT1G75335 | 3,97 | -0,74 | 0,96 | -0,77 | 4,39E-01 |
| AT1G75350 | 0,83 | 1,69 | 2,10 | 0,81 | 4,21E-01 |
| AT1G75400 | 0,32 | 1,76 | 4,32 | 0,41 | 6,84E-01 |
| AT1G75420 | 0,31 | -0,64 | 4,36 | -0,15 | 8,83E-01 |
| AT1G75440 | 0,28 | -0,49 | 4,38 | -0,11 | 9,11E-01 |
| AT1G75460 | 0,84 | -0,94 | 2,08 | -0,45 | 6,51E-01 |
| AT1G75580 | 0,17 | 0,65 | 4,36 | 0,15 | 8,81E-01 |
| AT1G75710 | 0,31 | 0,34 | 4,39 | 0,08 | 9,38E-01 |
| AT1G75785 | 0,12 | 0,49 | 4,36 | 0,11 | 9,11E-01 |
| AT1G75800 | 1,71 | -1,97 | 1,63 | -1,21 | 2,26E-01 |
| AT1G75810 | 0,60 | -1,46 | 2,53 | -0,58 | 5,65E-01 |
| AT1G75840 | 0,75 | -1,43 | 2,20 | -0,65 | 5,16E-01 |
| AT1G75900 | 0,32 | -0,59 | 4,37 | -0,14 | 8,92E-01 |
| AT1G75950 | 1,80 | 0,17 | 1,34 | 0,12 | 9,01E-01 |
| AT1G75960 | 0,45 | -0,45 | 4,19 | -0,11 | 9,15E-01 |
| AT1G76000 | 0,58 | 2,60 | 2,96 | 0,88 | 3,79E-01 |
| AT1G76030 | 2,84 | -0,94 | 1,27 | -0,74 | 4,58E-01 |
| AT1G76050 | 0,66 | -1,65 | 2,94 | -0,56 | 5,75E-01 |
| AT1G76135 | 2,70 | 1,97 | 1,73 | 1,14 | 2,54E-01 |
| AT1G76140 | 0,48 | 1,04 | 4,02 | 0,26 | 7,96E-01 |
| AT1G76185 | 0,16 | 0,38 | 4,37 | 0,09 | 9,30E-01 |
| AT1G76190 | 0,11 | 0,00 | 4,41 | 0,00 | 1,00E+00 |
| AT1G76320 | 0,32 | 1,91 | 4,32 | 0,44 | 6,59E-01 |
| AT1G76350 | 0,12 | 0,37 | 4,37 | 0,08 | 9,33E-01 |
| AT1G76390 | 0,51 | 1,20 | 3,15 | 0,38 | 7,03E-01 |
| AT1G76450 | 0,08 | 0,44 | 4,38 | 0,10 | 9,21E-01 |
| AT1G76490 | 3,44 | -2,40 | 1,11 | -2,15 | 3,12E-02 |
| AT1G76520 | 0,73 | -2,16 | 2,23 | -0,97 | 3,33E-01 |
| AT1G76580 | 0,80 | -1,41 | 3,49 | -0,40 | 6,86E-01 |
| AT1G76590 | 0,69 | -1,45 | 2,97 | -0,49 | 6,26E-01 |
| AT1G76600 | 0,49 | 1,88 | 3,20 | 0,59 | 5,56E-01 |
| AT1G76620 | 0,25 | 0,03 | 4,38 | 0,01 | 9,95E-01 |
| AT1G76630 | 2,14 | 0,54 | 1,40 | 0,38 | 7,01E-01 |
| AT1G76680 | 0,14 | 0,00 | 4,41 | 0,00 | 1,00E+00 |

| | | | | | |
|---|---|---|---|---|---|
| AT1G76740 | 0,12 | 1,16 | 4,34 | 0,27 | 7,89E-01 |
| AT1G76780 | 1,47 | 0,77 | 1,82 | 0,42 | 6,71E-01 |
| AT1G76820 | 0,53 | 1,27 | 3,09 | 0,41 | 6,80E-01 |
| AT1G76850 | 1,15 | -1,79 | 1,75 | -1,02 | 3,07E-01 |
| AT1G76880 | 1,06 | -0,48 | 1,90 | -0,25 | 7,99E-01 |
| AT1G76900 | 0,91 | 0,53 | 2,71 | 0,19 | 8,46E-01 |
| AT1G09687 | 2,04 | 0,57 | 1,35 | 0,42 | 6,74E-01 |
| AT1G76920 | 0,59 | -0,06 | 3,73 | -0,02 | 9,87E-01 |
| AT1G76950 | 0,83 | -2,16 | 2,74 | -0,79 | 4,30E-01 |
| AT1G76990 | 1,49 | -0,22 | 1,59 | -0,14 | 8,90E-01 |
| AT1G77080 | 0,41 | 0,08 | 3,39 | 0,02 | 9,82E-01 |
| AT1G77140 | 0,25 | -1,82 | 4,40 | -0,41 | 6,79E-01 |
| AT1G77230 | 3,05 | -0,42 | 1,11 | -0,37 | 7,08E-01 |
| AT1G77235 | 1,77 | -1,66 | 1,85 | -0,89 | 3,71E-01 |
| AT1G77320 | 0,26 | -0,48 | 4,38 | -0,11 | 9,13E-01 |
| AT1G77350 | 0,15 | 0,29 | 4,37 | 0,07 | 9,47E-01 |
| AT1G77420 | 0,12 | 1,23 | 4,34 | 0,28 | 7,76E-01 |
| AT1G77460 | 0,94 | -0,83 | 2,49 | -0,33 | 7,40E-01 |
| AT1G77490 | 0,64 | -2,53 | 3,05 | -0,83 | 4,07E-01 |
| AT1G77500 | 0,37 | -0,03 | 4,36 | -0,01 | 9,95E-01 |
| AT1G77510 | 1,26 | -0,67 | 1,55 | -0,43 | 6,65E-01 |
| AT1G77610 | 0,53 | -1,14 | 4,32 | -0,26 | 7,91E-01 |
| AT1G77710 | 0,32 | -0,91 | 4,39 | -0,21 | 8,36E-01 |
| AT1G77740 | 1,74 | 0,16 | 1,49 | 0,11 | 9,13E-01 |
| AT1G77800 | 1,13 | -0,27 | 1,66 | -0,16 | 8,71E-01 |
| AT1G77840 | 0,31 | 0,23 | 3,59 | 0,06 | 9,49E-01 |
| AT1G77850 | 1,01 | 1,49 | 2,17 | 0,68 | 4,94E-01 |
| AT1G77890 | 0,98 | -2,40 | 1,78 | -1,35 | 1,77E-01 |
| AT1G77930 | 1,14 | 1,96 | 2,06 | 0,95 | 3,41E-01 |
| AT1G78010 | 0,90 | -0,74 | 2,10 | -0,35 | 7,25E-01 |
| AT1G78020 | 1,89 | -0,67 | 1,33 | -0,50 | 6,15E-01 |
| AT1G78040 | 2,48 | 0,60 | 1,36 | 0,44 | 6,63E-01 |
| AT1G78070 | 1,66 | -0,83 | 1,70 | -0,49 | 6,24E-01 |
| AT1G78080 | 3,84 | -1,30 | 1,04 | -1,24 | 2,14E-01 |
| AT1G78100 | 0,64 | 0,22 | 2,26 | 0,10 | 9,21E-01 |
| AT1G78150 | 0,22 | -0,86 | 4,36 | -0,20 | 8,44E-01 |
| AT1G78200 | 0,54 | -1,62 | 2,64 | -0,62 | 5,38E-01 |
| AT1G78280 | 1,59 | 2,01 | 2,08 | 0,96 | 3,35E-01 |
| AT1G78300 | 0,54 | 1,97 | 3,17 | 0,62 | 5,34E-01 |
| AT1G78420 | 0,54 | -0,26 | 2,63 | -0,10 | 9,22E-01 |
| AT1G78490 | 0,05 | 0,00 | 4,41 | 0,00 | 1,00E+00 |
| AT1G78500 | 0,17 | 1,40 | 4,33 | 0,32 | 7,47E-01 |
| AT1G78570 | 2,32 | -0,17 | 1,25 | -0,14 | 8,92E-01 |
| AT1G78600 | 0,20 | -1,80 | 4,41 | -0,41 | 6,83E-01 |
| AT1G78630 | 1,31 | -2,13 | 1,79 | -1,19 | 2,34E-01 |
| AT1G78670 | 0,52 | 0,64 | 3,07 | 0,21 | 8,35E-01 |
| AT1G78680 | 0,90 | 0,69 | 2,00 | 0,34 | 7,31E-01 |
| AT1G78700 | 0,37 | 0,40 | 4,30 | 0,09 | 9,26E-01 |
| AT1G78710 | 0,38 | -1,52 | 4,32 | -0,35 | 7,24E-01 |
| AT1G78770 | 0,21 | 1,63 | 4,33 | 0,38 | 7,06E-01 |
| AT1G78870 | 0,28 | 0,09 | 4,33 | 0,02 | 9,83E-01 |
| AT1G78890 | 0,36 | -1,42 | 4,35 | -0,33 | 7,45E-01 |

| | | | | | |
|---|---|---|---|---|---|
| AT1G78910 | 0,26 | -0,30 | 4,37 | -0,07 | 9,46E-01 |
| AT1G78920 | 1,61 | 1,11 | 1,55 | 0,72 | 4,74E-01 |
| AT1G79060 | 0,14 | 0,00 | 4,41 | 0,00 | 1,00E+00 |
| AT1G79103 | 0,05 | 0,00 | 4,41 | 0,00 | 1,00E+00 |
| AT1G79110 | 1,67 | 0,19 | 1,58 | 0,12 | 9,06E-01 |
| AT1G09847 | 0,44 | 0,00 | 3,35 | 0,00 | 1,00E+00 |
| AT1G79190 | 0,88 | -1,81 | 2,54 | -0,71 | 4,75E-01 |
| AT1G79230 | 0,71 | -0,61 | 2,81 | -0,22 | 8,29E-01 |
| AT1G79240 | 0,20 | 1,20 | 4,35 | 0,28 | 7,83E-01 |
| AT1G79270 | 1,18 | 0,28 | 1,57 | 0,18 | 8,59E-01 |
| AT1G79340 | 0,91 | 0,83 | 2,20 | 0,38 | 7,05E-01 |
| AT1G79380 | 0,91 | 0,95 | 2,70 | 0,35 | 7,26E-01 |
| AT1G79560 | 0,28 | 0,08 | 4,33 | 0,02 | 9,86E-01 |
| AT1G79590 | 0,18 | 1,24 | 4,33 | 0,29 | 7,74E-01 |
| AT1G79660 | 0,63 | -2,82 | 3,59 | -0,79 | 4,32E-01 |
| AT1G09903 | 0,08 | 0,69 | 4,36 | 0,16 | 8,75E-01 |
| AT1G79690 | 1,96 | -1,18 | 1,58 | -0,74 | 4,58E-01 |
| AT1G79810 | 0,41 | 0,25 | 3,27 | 0,08 | 9,40E-01 |
| AT1G79840 | 0,77 | 0,49 | 2,87 | 0,17 | 8,66E-01 |
| AT1G79870 | 0,21 | 1,26 | 4,33 | 0,29 | 7,71E-01 |
| AT1G79890 | 0,21 | -0,49 | 4,35 | -0,11 | 9,11E-01 |
| AT1G79910 | 0,41 | -0,33 | 4,10 | -0,08 | 9,35E-01 |
| AT1G79940 | 1,20 | -1,22 | 1,68 | -0,72 | 4,69E-01 |
| AT1G79950 | 0,49 | -2,07 | 3,92 | -0,53 | 5,97E-01 |
| AT1G79970 | 0,27 | -0,36 | 4,34 | -0,08 | 9,34E-01 |
| AT1G79985 | 0,75 | 0,69 | 2,50 | 0,28 | 7,82E-01 |
| AT1G79990 | 2,18 | -0,36 | 1,34 | -0,27 | 7,91E-01 |
| AT1G80000 | 1,90 | 0,74 | 1,81 | 0,41 | 6,81E-01 |
| AT1G80010 | 1,24 | 0,71 | 1,56 | 0,46 | 6,47E-01 |
| AT1G80160 | 0,37 | 2,32 | 4,31 | 0,54 | 5,91E-01 |
| AT1G80260 | 0,59 | -1,22 | 3,02 | -0,40 | 6,87E-01 |
| AT1G80300 | 1,04 | -0,33 | 3,09 | -0,11 | 9,15E-01 |
| AT1G80310 | 0,64 | 0,13 | 2,58 | 0,05 | 9,60E-01 |
| AT1G80380 | 1,15 | -1,28 | 1,86 | -0,69 | 4,90E-01 |
| AT1G09963 | 0,96 | -1,16 | 2,33 | -0,50 | 6,17E-01 |
| AT1G80430 | 0,19 | 0,49 | 4,38 | 0,11 | 9,11E-01 |
| AT1G80440 | 3,83 | 1,98 | 1,48 | 1,34 | 1,81E-01 |
| AT1G80510 | 0,61 | -0,03 | 3,82 | -0,01 | 9,94E-01 |
| AT1G80620 | 0,35 | 0,15 | 4,41 | 0,03 | 9,74E-01 |
| AT1G80680 | 4,25 | -0,24 | 1,00 | -0,24 | 8,13E-01 |
| AT1G80770 | 0,54 | -0,48 | 3,17 | -0,15 | 8,79E-01 |
| AT1G80780 | 0,60 | 0,82 | 3,12 | 0,26 | 7,92E-01 |
| AT1G80790 | 0,69 | 1,42 | 2,80 | 0,51 | 6,12E-01 |
| AT1G80810 | 0,38 | 0,60 | 4,23 | 0,14 | 8,88E-01 |
| AT1G80840 | 1,09 | 1,99 | 2,47 | 0,80 | 4,22E-01 |
| AT1G80870 | 0,27 | -0,32 | 4,34 | -0,07 | 9,41E-01 |
| AT2G01010 | 3,38 | 0,88 | 1,39 | 0,63 | 5,28E-01 |
| AT2G01070 | 0,81 | -0,54 | 2,35 | -0,23 | 8,19E-01 |
| AT2G01080 | 0,35 | -0,15 | 3,50 | -0,04 | 9,66E-01 |
| AT2G01100 | 0,12 | 0,00 | 4,38 | 0,00 | 1,00E+00 |
| AT2G01120 | 0,92 | 0,80 | 2,69 | 0,30 | 7,66E-01 |
| AT2G01190 | 0,50 | 1,62 | 3,11 | 0,52 | 6,02E-01 |

| | | | | | |
|---|---|---|---|---|---|
| AT2G01270 | 3,86 | -0,54 | 0,94 | -0,58 | 5,64E-01 |
| AT2G01400 | 0,39 | -2,30 | 4,38 | -0,53 | 5,99E-01 |
| AT2G03975 | 0,34 | -0,93 | 3,58 | -0,26 | 7,94E-01 |
| AT2G01422 | 2,93 | 0,04 | 1,72 | 0,02 | 9,84E-01 |
| AT2G01480 | 0,34 | -1,75 | 4,35 | -0,40 | 6,88E-01 |
| AT2G01600 | 1,95 | 0,64 | 1,41 | 0,45 | 6,51E-01 |
| AT2G01620 | 0,60 | -1,28 | 2,46 | -0,52 | 6,03E-01 |
| AT2G01680 | 0,39 | -0,94 | 4,33 | -0,22 | 8,28E-01 |
| AT2G01750 | 0,24 | 0,05 | 4,39 | 0,01 | 9,91E-01 |
| AT2G01850 | 1,74 | 1,36 | 1,64 | 0,83 | 4,06E-01 |
| AT2G01860 | 0,98 | 2,07 | 2,43 | 0,85 | 3,95E-01 |
| AT2G01900 | 1,22 | 2,50 | 1,84 | 1,36 | 1,73E-01 |
| AT2G01910 | 2,66 | 0,06 | 1,79 | 0,03 | 9,75E-01 |
| AT2G01980 | 0,63 | -0,07 | 3,07 | -0,02 | 9,82E-01 |
| AT2G02040 | 0,10 | 0,01 | 4,38 | 0,00 | 9,99E-01 |
| AT2G02050 | 0,58 | -0,89 | 2,69 | -0,33 | 7,42E-01 |
| AT2G02070 | 0,84 | -0,63 | 1,99 | -0,32 | 7,52E-01 |
| AT2G02090 | 1,22 | -0,58 | 2,18 | -0,27 | 7,88E-01 |
| AT2G02100 | 0,54 | 0,41 | 3,95 | 0,10 | 9,18E-01 |
| AT2G02440 | 0,68 | -1,79 | 3,12 | -0,57 | 5,66E-01 |
| AT2G02450 | 0,66 | 0,92 | 2,66 | 0,35 | 7,28E-01 |
| AT2G02470 | 0,18 | -1,78 | 4,41 | -0,40 | 6,87E-01 |
| AT2G02550 | 1,01 | 1,69 | 2,15 | 0,78 | 4,33E-01 |
| AT2G02560 | 1,80 | -1,28 | 1,32 | -0,97 | 3,33E-01 |
| AT2G02700 | 0,39 | -1,28 | 4,38 | -0,29 | 7,70E-01 |
| AT2G04295 | 0,12 | 0,00 | 4,38 | 0,00 | 1,00E+00 |
| AT2G02760 | 0,54 | 0,03 | 2,77 | 0,01 | 9,92E-01 |
| AT2G02790 | 0,39 | -0,49 | 4,23 | -0,11 | 9,09E-01 |
| AT2G02795 | 0,14 | 0,09 | 4,38 | 0,02 | 9,84E-01 |
| AT2G02810 | 0,62 | 1,59 | 3,00 | 0,53 | 5,96E-01 |
| AT2G04355 | 0,47 | -0,84 | 3,29 | -0,26 | 7,97E-01 |
| AT2G02840 | 4,55 | -1,48 | 1,17 | -1,26 | 2,06E-01 |
| AT2G04365 | 0,97 | 1,91 | 2,42 | 0,79 | 4,30E-01 |
| AT2G02870 | 1,53 | 1,23 | 1,80 | 0,68 | 4,94E-01 |
| AT2G04375 | 5,42 | 0,07 | 0,83 | 0,09 | 9,29E-01 |
| AT2G04405 | 3,40 | 0,49 | 1,13 | 0,43 | 6,67E-01 |
| AT2G04435 | 3,56 | 0,78 | 1,11 | 0,70 | 4,84E-01 |
| AT2G04455 | 3,39 | 0,97 | 1,18 | 0,83 | 4,09E-01 |
| AT2G02955 | 0,36 | -0,85 | 4,33 | -0,20 | 8,44E-01 |
| AT2G03070 | 0,25 | 0,41 | 4,33 | 0,09 | 9,25E-01 |
| AT2G03120 | 0,64 | -1,50 | 2,69 | -0,56 | 5,76E-01 |
| AT2G03130 | 0,12 | 1,05 | 4,35 | 0,24 | 8,10E-01 |
| AT2G03140 | 2,19 | -0,24 | 1,28 | -0,19 | 8,51E-01 |
| AT2G03150 | 2,08 | -1,08 | 1,25 | -0,87 | 3,86E-01 |
| AT2G03270 | 0,39 | -0,03 | 2,85 | -0,01 | 9,91E-01 |
| AT2G03410 | 0,43 | 1,94 | 4,08 | 0,48 | 6,35E-01 |
| AT2G03450 | 0,32 | -2,61 | 4,37 | -0,60 | 5,51E-01 |
| AT2G03500 | 0,52 | -1,78 | 3,30 | -0,54 | 5,90E-01 |
| AT2G03520 | 0,08 | 0,69 | 4,36 | 0,16 | 8,75E-01 |
| AT2G03550 | 0,42 | 0,01 | 4,24 | 0,00 | 9,99E-01 |
| AT2G03567 | 0,47 | 1,42 | 3,36 | 0,42 | 6,71E-01 |
| AT2G03680 | 0,82 | -2,21 | 2,35 | -0,94 | 3,46E-01 |

| | | | | | |
|---|---|---|---|---|---|
| AT2G03710 | 0,17 | 1,52 | 4,33 | 0,35 | 7,26E-01 |
| AT2G03823 | 0,37 | 0,81 | 2,89 | 0,28 | 7,78E-01 |
| AT2G03870 | 0,41 | -0,20 | 4,07 | -0,05 | 9,61E-01 |
| AT2G03890 | 2,03 | 0,46 | 1,33 | 0,34 | 7,31E-01 |
| AT2G03932 | 0,20 | 0,00 | 4,41 | 0,00 | 1,00E+00 |
| AT2G03931 | 0,05 | 0,00 | 4,41 | 0,00 | 1,00E+00 |
| AT2G03933 | 0,14 | 0,00 | 4,41 | 0,00 | 1,00E+00 |
| AT2G03936 | 0,18 | 0,56 | 4,36 | 0,13 | 8,97E-01 |
| AT2G03965 | 0,22 | 0,00 | 4,41 | 0,00 | 1,00E+00 |
| AT2G03980 | 0,82 | -1,80 | 2,60 | -0,69 | 4,89E-01 |
| AT2G04030 | 4,28 | -0,43 | 0,89 | -0,48 | 6,30E-01 |
| AT2G04039 | 0,42 | -0,54 | 4,08 | -0,13 | 8,95E-01 |
| AT2G04735 | 0,15 | 0,00 | 4,41 | 0,00 | 1,00E+00 |
| AT2G04235 | 0,05 | 0,00 | 4,41 | 0,00 | 1,00E+00 |
| AT2G04270 | 1,17 | 1,36 | 2,26 | 0,60 | 5,46E-01 |
| AT2G04755 | 2,18 | 1,39 | 1,50 | 0,92 | 3,55E-01 |
| AT2G04340 | 0,38 | -1,21 | 4,34 | -0,28 | 7,81E-01 |
| AT2G04350 | 1,50 | 0,19 | 1,48 | 0,13 | 8,95E-01 |
| AT2G04380 | 0,56 | -1,34 | 2,75 | -0,49 | 6,25E-01 |
| AT2G04390 | 0,85 | 0,60 | 2,04 | 0,29 | 7,69E-01 |
| AT2G04400 | 0,61 | 0,94 | 2,48 | 0,38 | 7,04E-01 |
| AT2G04450 | 0,21 | 1,63 | 4,33 | 0,38 | 7,06E-01 |
| AT2G04495 | 0,43 | 0,83 | 3,35 | 0,25 | 8,05E-01 |
| AT2G04550 | 0,15 | 0,67 | 4,37 | 0,15 | 8,79E-01 |
| AT2G04700 | 0,71 | 1,07 | 2,23 | 0,48 | 6,32E-01 |
| AT2G04740 | 0,38 | -0,91 | 4,32 | -0,21 | 8,34E-01 |
| AT2G04780 | 0,24 | -1,19 | 4,35 | -0,27 | 7,84E-01 |
| AT2G04795 | 0,15 | 0,29 | 4,37 | 0,07 | 9,47E-01 |
| AT2G04850 | 0,08 | 0,69 | 4,36 | 0,16 | 8,75E-01 |
| AT2G04865 | 0,25 | 1,87 | 4,32 | 0,43 | 6,65E-01 |
| AT2G04900 | 0,19 | 0,13 | 4,41 | 0,03 | 9,76E-01 |
| AT2G05060 | 0,21 | -0,49 | 4,35 | -0,11 | 9,11E-01 |
| AT2G05170 | 1,80 | -0,74 | 1,42 | -0,52 | 6,00E-01 |
| AT2G05180 | 0,08 | 0,44 | 4,38 | 0,10 | 9,21E-01 |
| AT2G05210 | 0,12 | 0,49 | 4,36 | 0,11 | 9,11E-01 |
| AT2G04885 | 2,17 | 3,20 | 1,68 | 1,90 | 5,73E-02 |
| AT2G05320 | 0,92 | 2,22 | 2,15 | 1,03 | 3,03E-01 |
| AT2G05350 | 0,23 | 1,21 | 4,34 | 0,28 | 7,81E-01 |
| AT2G05380 | 0,99 | -1,50 | 1,90 | -0,79 | 4,29E-01 |
| AT2G05540 | 0,30 | -1,78 | 4,41 | -0,40 | 6,87E-01 |
| AT2G04915 | 0,39 | 0,08 | 3,50 | 0,02 | 9,82E-01 |
| AT2G04935 | 1,06 | 0,18 | 1,83 | 0,10 | 9,22E-01 |
| AT2G05635 | 0,15 | 0,00 | 4,38 | 0,00 | 1,00E+00 |
| AT2G05710 | 3,35 | 0,26 | 1,05 | 0,24 | 8,07E-01 |
| AT2G04985 | 0,95 | 1,28 | 2,24 | 0,57 | 5,69E-01 |
| AT2G05790 | 0,44 | 0,38 | 3,21 | 0,12 | 9,07E-01 |
| AT2G05812 | 1,46 | 0,35 | 1,60 | 0,22 | 8,28E-01 |
| AT2G05830 | 0,15 | 0,45 | 4,37 | 0,10 | 9,19E-01 |
| AT2G05840 | 0,17 | 0,65 | 4,36 | 0,15 | 8,81E-01 |
| AT2G05990 | 0,47 | 0,29 | 2,41 | 0,12 | 9,03E-01 |
| AT2G05995 | 0,63 | -1,78 | 3,18 | -0,56 | 5,76E-01 |
| AT2G06025 | 0,59 | -0,96 | 2,64 | -0,36 | 7,16E-01 |

| | | | | | |
|---|---|---|---|---|---|
| AT2G06095 | 0,70 | 0,00 | 3,59 | 0,00 | 1,00E+00 |
| AT2G06105 | 0,23 | 0,00 | 4,40 | 0,00 | 1,00E+00 |
| AT2G06420 | 4,36 | 4,42 | 1,49 | 2,96 | 3,10E-03 |
| AT2G06510 | 1,80 | 1,11 | 1,45 | 0,77 | 4,44E-01 |
| AT2G06845 | 0,18 | 0,81 | 4,34 | 0,19 | 8,52E-01 |
| AT2G06850 | 0,75 | 0,17 | 2,60 | 0,06 | 9,49E-01 |
| AT2G06904 | 0,38 | 0,82 | 4,35 | 0,19 | 8,51E-01 |
| AT2G06990 | 1,10 | 1,16 | 2,04 | 0,57 | 5,69E-01 |
| AT2G07050 | 0,64 | 1,98 | 2,97 | 0,67 | 5,05E-01 |
| AT2G07200 | 0,15 | 0,00 | 4,41 | 0,00 | 1,00E+00 |
| AT2G07565 | 0,91 | -1,78 | 2,59 | -0,69 | 4,93E-01 |
| AT2G07749 | 0,59 | 0,41 | 3,29 | 0,13 | 9,00E-01 |
| AT2G07672 | 0,25 | 0,13 | 4,39 | 0,03 | 9,77E-01 |
| AT2G07745 | 0,19 | 0,00 | 4,41 | 0,00 | 1,00E+00 |
| AT2G07674 | 0,81 | 1,41 | 3,67 | 0,39 | 7,00E-01 |
| AT2G07623 | 0,23 | 1,21 | 4,34 | 0,28 | 7,81E-01 |
| AT2G07678 | 0,32 | -2,18 | 4,38 | -0,50 | 6,18E-01 |
| AT2G07669 | 0,33 | -1,63 | 4,39 | -0,37 | 7,10E-01 |
| AT2G07681 | 0,75 | -0,61 | 2,18 | -0,28 | 7,79E-01 |
| AT2G07631 | 0,18 | 0,00 | 4,41 | 0,00 | 1,00E+00 |
| AT2G05215 | 0,68 | 1,40 | 2,60 | 0,54 | 5,90E-01 |
| AT2G07695 | 1,27 | 2,85 | 1,95 | 1,46 | 1,45E-01 |
| AT2G07798 | 1,27 | 1,53 | 1,86 | 0,82 | 4,10E-01 |
| AT2G07667 | 1,00 | 1,53 | 2,34 | 0,65 | 5,14E-01 |
| AT2G07638 | 1,46 | -0,26 | 1,64 | -0,16 | 8,74E-01 |
| AT2G07707 | 0,32 | 0,49 | 4,36 | 0,11 | 9,10E-01 |
| AT2G07708 | 0,57 | -0,14 | 4,38 | -0,03 | 9,75E-01 |
| AT2G07715 | 0,94 | -0,22 | 1,76 | -0,12 | 9,01E-01 |
| AT2G07718 | 3,36 | -0,62 | 1,10 | -0,56 | 5,76E-01 |
| AT2G05315 | 4,98 | 0,97 | 1,09 | 0,89 | 3,72E-01 |
| AT2G07725 | 2,14 | -0,26 | 1,28 | -0,20 | 8,40E-01 |
| AT2G07727 | 3,34 | -1,25 | 1,33 | -0,94 | 3,47E-01 |
| AT2G07728 | 0,11 | 0,48 | 4,37 | 0,11 | 9,12E-01 |
| AT2G07656 | 0,60 | 0,64 | 3,05 | 0,21 | 8,33E-01 |
| AT2G07827 | 0,36 | -0,36 | 3,48 | -0,10 | 9,18E-01 |
| AT2G07787 | 0,79 | 0,82 | 2,17 | 0,38 | 7,05E-01 |
| AT2G07659 | 0,28 | 0,07 | 4,39 | 0,02 | 9,87E-01 |
| AT2G07806 | 2,71 | -1,30 | 1,31 | -0,99 | 3,20E-01 |
| AT2G05355 | 1,78 | -1,13 | 1,46 | -0,77 | 4,39E-01 |
| AT2G07765 | 0,12 | 1,23 | 4,34 | 0,28 | 7,76E-01 |
| AT2G07741 | 0,27 | 0,00 | 4,41 | 0,00 | 1,00E+00 |
| AT2G07680 | 0,80 | 1,85 | 2,45 | 0,76 | 4,50E-01 |
| AT2G07981 | 2,13 | 3,48 | 1,97 | 1,76 | 7,79E-02 |
| AT2G05455 | 4,14 | 2,67 | 1,44 | 1,86 | 6,36E-02 |
| AT2G09990 | 0,53 | 2,14 | 3,11 | 0,69 | 4,91E-01 |
| AT2G05485 | 0,27 | 0,00 | 4,38 | 0,00 | 1,00E+00 |
| AT2G10545 | 0,08 | 0,00 | 4,40 | 0,00 | 1,00E+00 |
| AT2G10556 | 0,88 | 3,41 | 2,63 | 1,29 | 1,96E-01 |
| AT2G10965 | 0,86 | 1,22 | 2,15 | 0,57 | 5,70E-01 |
| AT2G11010 | 0,65 | -0,49 | 3,04 | -0,16 | 8,73E-01 |
| AT2G11035 | 0,48 | -0,93 | 4,03 | -0,23 | 8,17E-01 |
| AT2G11045 | 0,08 | 0,00 | 4,40 | 0,00 | 1,00E+00 |

| | | | | | |
|---|---|---|---|---|---|
| AT2G11225 | 0,91 | -0,61 | 1,73 | -0,35 | 7,25E-01 |
| AT2G11620 | 0,14 | 0,00 | 4,41 | 0,00 | 1,00E+00 |
| AT2G11851 | 0,57 | 1,15 | 3,00 | 0,38 | 7,02E-01 |
| AT2G05825 | 0,69 | -0,85 | 2,26 | -0,38 | 7,07E-01 |
| AT2G13100 | 0,26 | -1,45 | 4,37 | -0,33 | 7,39E-01 |
| AT2G13290 | 0,12 | 0,00 | 4,38 | 0,00 | 1,00E+00 |
| AT2G05885 | 0,14 | 0,00 | 4,41 | 0,00 | 1,00E+00 |
| AT2G13430 | 0,44 | 1,94 | 3,20 | 0,60 | 5,45E-01 |
| AT2G13500 | 0,11 | 0,00 | 4,41 | 0,00 | 1,00E+00 |
| AT2G13540 | 0,64 | 0,58 | 2,35 | 0,25 | 8,06E-01 |
| AT2G13560 | 2,19 | -0,52 | 1,70 | -0,31 | 7,60E-01 |
| AT2G05925 | 0,12 | 0,00 | 4,38 | 0,00 | 1,00E+00 |
| AT2G13790 | 0,49 | 0,33 | 3,30 | 0,10 | 9,20E-01 |
| AT2G13845 | 0,67 | 1,18 | 2,26 | 0,52 | 6,00E-01 |
| AT2G13960 | 0,27 | -1,78 | 4,41 | -0,40 | 6,87E-01 |
| AT2G14070 | 0,19 | 0,00 | 4,41 | 0,00 | 1,00E+00 |
| AT2G14080 | 4,62 | 1,24 | 1,11 | 1,12 | 2,63E-01 |
| AT2G14115 | 0,49 | -1,02 | 4,33 | -0,23 | 8,14E-01 |
| AT2G14395 | 0,63 | -0,07 | 4,35 | -0,02 | 9,87E-01 |
| AT2G14440 | 0,43 | 0,00 | 4,18 | 0,00 | 1,00E+00 |
| AT2G14530 | 0,21 | 0,00 | 4,41 | 0,00 | 1,00E+00 |
| AT2G14700 | 0,64 | 1,26 | 2,97 | 0,42 | 6,72E-01 |
| AT2G14740 | 1,25 | 2,63 | 2,39 | 1,10 | 2,72E-01 |
| AT2G14750 | 1,64 | -0,40 | 1,36 | -0,29 | 7,70E-01 |
| AT2G14765 | 0,40 | 0,46 | 4,36 | 0,11 | 9,16E-01 |
| AT2G14825 | 0,08 | 0,44 | 4,38 | 0,10 | 9,21E-01 |
| AT2G14835 | 0,23 | 1,72 | 4,32 | 0,40 | 6,90E-01 |
| AT2G14850 | 0,21 | 1,58 | 4,33 | 0,37 | 7,15E-01 |
| AT2G14870 | 0,17 | 0,00 | 4,41 | 0,00 | 1,00E+00 |
| AT2G14878 | 1,23 | 0,24 | 1,67 | 0,14 | 8,86E-01 |
| AT2G14890 | 0,77 | -0,04 | 2,13 | -0,02 | 9,85E-01 |
| AT2G06305 | 0,18 | 0,00 | 4,41 | 0,00 | 1,00E+00 |
| AT2G15042 | 0,11 | 0,48 | 4,37 | 0,11 | 9,12E-01 |
| AT2G15050 | 0,64 | 0,00 | 3,03 | 0,00 | 1,00E+00 |
| AT2G15080 | 0,92 | 0,83 | 2,05 | 0,41 | 6,85E-01 |
| AT2G15090 | 1,27 | 2,81 | 3,02 | 0,93 | 3,52E-01 |
| AT2G15300 | 0,31 | -0,92 | 4,38 | -0,21 | 8,33E-01 |
| AT2G15430 | 0,16 | 0,54 | 4,36 | 0,12 | 9,01E-01 |
| AT2G15450 | 0,29 | 0,46 | 4,35 | 0,11 | 9,15E-01 |
| AT2G15470 | 0,34 | 0,00 | 4,38 | 0,00 | 9,99E-01 |
| AT2G15580 | 0,29 | 0,82 | 3,69 | 0,22 | 8,24E-01 |
| AT2G15620 | 3,38 | -2,22 | 1,05 | -2,12 | 3,38E-02 |
| AT2G15710 | 0,40 | 0,00 | 4,41 | 0,00 | 1,00E+00 |
| AT2G15730 | 0,08 | 0,00 | 4,40 | 0,00 | 1,00E+00 |
| AT2G15900 | 0,47 | 0,24 | 2,73 | 0,09 | 9,29E-01 |
| AT2G15960 | 1,09 | -0,61 | 1,67 | -0,37 | 7,15E-01 |
| AT2G15970 | 3,79 | -1,14 | 1,29 | -0,88 | 3,77E-01 |
| AT2G06535 | 1,38 | 3,29 | 1,99 | 1,66 | 9,73E-02 |
| AT2G16019 | 1,05 | 1,79 | 2,21 | 0,81 | 4,19E-01 |
| AT2G16020 | 3,14 | 3,05 | 1,56 | 1,96 | 5,03E-02 |
| AT2G16040 | 2,68 | 3,33 | 1,62 | 2,06 | 3,97E-02 |
| AT2G16280 | 0,93 | 1,65 | 2,33 | 0,71 | 4,77E-01 |

| | | | | | |
|---|---|---|---|---|---|
| AT2G16365 | 0,96 | 1,50 | 2,28 | 0,66 | 5,12E-01 |
| AT2G16380 | 0,44 | -2,26 | 4,28 | -0,53 | 5,97E-01 |
| AT2G16390 | 0,40 | 2,09 | 4,32 | 0,48 | 6,28E-01 |
| AT2G16630 | 0,14 | 0,76 | 4,35 | 0,18 | 8,61E-01 |
| AT2G16700 | 0,28 | 1,66 | 4,33 | 0,38 | 7,02E-01 |
| AT2G16710 | 0,72 | 2,06 | 3,70 | 0,56 | 5,77E-01 |
| AT2G16800 | 0,26 | 0,00 | 4,41 | 0,00 | 1,00E+00 |
| AT2G16950 | 1,26 | 1,68 | 2,18 | 0,77 | 4,41E-01 |
| AT2G16970 | 1,25 | -2,34 | 1,85 | -1,26 | 2,06E-01 |
| AT2G17033 | 0,44 | -0,30 | 4,14 | -0,07 | 9,41E-01 |
| AT2G17040 | 1,57 | -0,67 | 1,55 | -0,43 | 6,65E-01 |
| AT2G17120 | 0,23 | 0,86 | 4,34 | 0,20 | 8,43E-01 |
| AT2G17140 | 0,42 | 1,44 | 3,39 | 0,43 | 6,70E-01 |
| AT2G17170 | 0,08 | 0,44 | 4,38 | 0,10 | 9,21E-01 |
| AT2G17240 | 0,52 | -0,90 | 3,19 | -0,28 | 7,79E-01 |
| AT2G17250 | 0,25 | 0,38 | 4,33 | 0,09 | 9,31E-01 |
| AT2G17265 | 1,56 | 0,90 | 1,50 | 0,60 | 5,50E-01 |
| AT2G17270 | 0,25 | -0,79 | 4,34 | -0,18 | 8,55E-01 |
| AT2G17290 | 0,78 | -0,11 | 2,24 | -0,05 | 9,60E-01 |
| AT2G06905 | 1,12 | -3,28 | 1,91 | -1,72 | 8,63E-02 |
| AT2G17360 | 0,51 | -0,24 | 3,31 | -0,07 | 9,41E-01 |
| AT2G17390 | 0,66 | -0,54 | 2,85 | -0,19 | 8,49E-01 |
| AT2G17410 | 1,45 | -1,77 | 1,49 | -1,19 | 2,34E-01 |
| AT2G17420 | 1,05 | 0,09 | 1,95 | 0,04 | 9,65E-01 |
| AT2G17440 | 1,80 | 0,73 | 1,46 | 0,50 | 6,16E-01 |
| AT2G17520 | 1,17 | 1,58 | 1,86 | 0,85 | 3,94E-01 |
| AT2G17530 | 0,50 | 0,01 | 3,93 | 0,00 | 9,98E-01 |
| AT2G17570 | 0,42 | -0,40 | 4,12 | -0,10 | 9,23E-01 |
| AT2G17580 | 0,66 | -0,52 | 2,93 | -0,18 | 8,59E-01 |
| AT2G17705 | 0,42 | -0,48 | 3,30 | -0,15 | 8,84E-01 |
| AT2G17720 | 0,20 | -0,31 | 4,36 | -0,07 | 9,44E-01 |
| AT2G17760 | 0,65 | -1,13 | 2,87 | -0,39 | 6,94E-01 |
| AT2G17787 | 0,33 | 0,50 | 4,33 | 0,12 | 9,08E-01 |
| AT2G17790 | 0,80 | 0,91 | 2,50 | 0,36 | 7,17E-01 |
| AT2G17905 | 0,41 | -1,78 | 3,44 | -0,52 | 6,05E-01 |
| AT2G17920 | 0,15 | 0,00 | 4,38 | 0,00 | 9,99E-01 |
| AT2G17980 | 1,01 | 2,50 | 2,50 | 1,00 | 3,18E-01 |
| AT2G17990 | 0,69 | 0,45 | 2,44 | 0,18 | 8,54E-01 |
| AT2G18000 | 0,29 | 0,47 | 4,33 | 0,11 | 9,14E-01 |
| AT2G18020 | 2,64 | 1,82 | 1,36 | 1,34 | 1,80E-01 |
| AT2G18030 | 0,68 | 1,76 | 2,94 | 0,60 | 5,49E-01 |
| AT2G18040 | 0,20 | 0,00 | 4,38 | 0,00 | 1,00E+00 |
| AT2G18090 | 1,01 | 0,98 | 1,82 | 0,54 | 5,89E-01 |
| AT2G18100 | 0,48 | 2,28 | 4,02 | 0,57 | 5,71E-01 |
| AT2G18110 | 0,62 | 0,15 | 3,10 | 0,05 | 9,62E-01 |
| AT2G18280 | 2,10 | -0,66 | 1,38 | -0,48 | 6,30E-01 |
| AT2G18390 | 0,28 | 0,13 | 4,33 | 0,03 | 9,77E-01 |
| AT2G18420 | 0,81 | -1,19 | 3,34 | -0,35 | 7,23E-01 |
| AT2G18530 | 0,38 | 1,32 | 3,42 | 0,39 | 7,00E-01 |
| AT2G18560 | 1,54 | 1,19 | 2,45 | 0,49 | 6,26E-01 |
| AT2G18570 | 4,57 | 1,84 | 1,34 | 1,38 | 1,69E-01 |
| AT2G18630 | 0,16 | 0,49 | 4,38 | 0,11 | 9,12E-01 |

| | | | | | |
|---|---|---|---|---|---|
| AT2G18670 | 1,13 | 1,23 | 1,86 | 0,66 | 5,08E-01 |
| AT2G18680 | 0,16 | 1,33 | 4,33 | 0,31 | 7,59E-01 |
| AT2G18690 | 1,29 | 1,02 | 1,85 | 0,55 | 5,80E-01 |
| AT2G18700 | 0,27 | 0,41 | 4,37 | 0,09 | 9,25E-01 |
| AT2G18730 | 0,58 | 1,47 | 2,55 | 0,58 | 5,63E-01 |
| AT2G18735 | 0,22 | 0,44 | 4,38 | 0,10 | 9,19E-01 |
| AT2G18750 | 0,67 | -0,50 | 2,04 | -0,25 | 8,06E-01 |
| AT2G18760 | 1,48 | 0,23 | 1,48 | 0,15 | 8,78E-01 |
| AT2G18790 | 2,11 | 0,23 | 1,29 | 0,18 | 8,58E-01 |
| AT2G18850 | 0,52 | -2,44 | 4,37 | -0,56 | 5,77E-01 |
| AT2G18880 | 0,25 | 1,94 | 4,32 | 0,45 | 6,54E-01 |
| AT2G18917 | 0,08 | 0,00 | 4,40 | 0,00 | 1,00E+00 |
| AT2G07095 | 0,55 | 2,61 | 3,01 | 0,87 | 3,87E-01 |
| AT2G18938 | 0,72 | -1,19 | 2,44 | -0,49 | 6,28E-01 |
| AT2G18950 | 0,45 | 1,33 | 4,17 | 0,32 | 7,50E-01 |
| AT2G19110 | 0,15 | 0,83 | 4,35 | 0,19 | 8,49E-01 |
| AT2G19130 | 0,89 | -3,10 | 2,27 | -1,36 | 1,73E-01 |
| AT2G19160 | 0,39 | 1,86 | 4,33 | 0,43 | 6,68E-01 |
| AT2G19260 | 0,46 | -0,49 | 4,04 | -0,12 | 9,03E-01 |
| AT2G19280 | 0,36 | 1,57 | 4,34 | 0,36 | 7,17E-01 |
| AT2G19360 | 2,70 | -0,74 | 1,37 | -0,54 | 5,88E-01 |
| AT2G19420 | 0,28 | 0,01 | 4,38 | 0,00 | 9,99E-01 |
| AT2G19430 | 0,24 | 0,00 | 4,41 | 0,00 | 1,00E+00 |
| AT2G19450 | 0,42 | 0,99 | 4,26 | 0,23 | 8,17E-01 |
| AT2G07205 | 0,75 | -0,32 | 3,67 | -0,09 | 9,30E-01 |
| AT2G19480 | 1,78 | 1,63 | 1,60 | 1,02 | 3,09E-01 |
| AT2G19490 | 0,36 | 1,83 | 4,33 | 0,42 | 6,73E-01 |
| AT2G19520 | 0,55 | -0,77 | 4,02 | -0,19 | 8,48E-01 |
| AT2G19540 | 0,46 | 0,28 | 2,77 | 0,10 | 9,19E-01 |
| AT2G19572 | 0,52 | 1,23 | 3,07 | 0,40 | 6,89E-01 |
| AT2G19582 | 0,25 | 0,02 | 4,39 | 0,00 | 9,97E-01 |
| AT2G19600 | 0,64 | -0,09 | 3,90 | -0,02 | 9,81E-01 |
| AT2G19640 | 0,53 | -0,54 | 3,77 | -0,14 | 8,85E-01 |
| AT2G19710 | 1,44 | -1,93 | 1,91 | -1,01 | 3,11E-01 |
| AT2G19730 | 0,57 | 0,77 | 4,15 | 0,19 | 8,53E-01 |
| AT2G19740 | 0,67 | 0,27 | 2,34 | 0,12 | 9,07E-01 |
| AT2G19810 | 0,34 | -0,89 | 3,56 | -0,25 | 8,02E-01 |
| AT2G19850 | 5,66 | -0,78 | 1,23 | -0,63 | 5,28E-01 |
| AT2G19880 | 0,75 | 0,00 | 2,42 | 0,00 | 9,99E-01 |
| AT2G20020 | 1,07 | 0,34 | 2,07 | 0,17 | 8,68E-01 |
| AT2G20100 | 0,33 | 0,39 | 4,34 | 0,09 | 9,28E-01 |
| AT2G20110 | 0,83 | 2,03 | 4,31 | 0,47 | 6,39E-01 |
| AT2G20142 | 0,28 | 0,23 | 4,33 | 0,05 | 9,58E-01 |
| AT2G20230 | 0,26 | -1,82 | 4,40 | -0,41 | 6,80E-01 |
| AT2G20260 | 1,52 | -2,48 | 1,61 | -1,54 | 1,23E-01 |
| AT2G20340 | 0,57 | 1,81 | 3,95 | 0,46 | 6,48E-01 |
| AT2G20360 | 0,22 | 0,88 | 4,35 | 0,20 | 8,39E-01 |
| AT2G20362 | 0,08 | 0,69 | 4,36 | 0,16 | 8,75E-01 |
| AT2G20370 | 1,01 | -1,46 | 2,35 | -0,62 | 5,35E-01 |
| AT2G20420 | 0,62 | 0,65 | 2,25 | 0,29 | 7,71E-01 |
| AT2G20450 | 0,40 | 0,84 | 4,13 | 0,20 | 8,39E-01 |
| AT2G20530 | 0,21 | 0,05 | 4,39 | 0,01 | 9,91E-01 |

| | | | | | |
|---|---|---|---|---|---|
| AT2G20580 | 2,81 | -1,40 | 1,55 | -0,90 | 3,68E-01 |
| AT2G20690 | 1,05 | 2,77 | 2,49 | 1,11 | 2,67E-01 |
| AT2G20710 | 0,08 | 0,44 | 4,38 | 0,10 | 9,21E-01 |
| AT2G07335 | 0,46 | -1,04 | 4,12 | -0,25 | 8,01E-01 |
| AT2G20750 | 0,59 | -2,41 | 3,74 | -0,64 | 5,20E-01 |
| AT2G20770 | 0,17 | 1,50 | 4,33 | 0,35 | 7,29E-01 |
| AT2G20820 | 0,25 | 0,34 | 4,32 | 0,08 | 9,37E-01 |
| AT2G20860 | 1,36 | 0,23 | 1,53 | 0,15 | 8,81E-01 |
| AT2G20890 | 1,04 | -0,25 | 1,93 | -0,13 | 8,95E-01 |
| AT2G20920 | 0,65 | -0,71 | 2,33 | -0,31 | 7,60E-01 |
| AT2G20950 | 3,17 | 0,30 | 1,30 | 0,23 | 8,15E-01 |
| AT2G20960 | 0,82 | -0,50 | 2,00 | -0,25 | 8,02E-01 |
| AT2G20990 | 4,50 | -0,24 | 0,99 | -0,25 | 8,04E-01 |
| AT2G21160 | 0,91 | 0,30 | 3,10 | 0,10 | 9,24E-01 |
| AT2G21187 | 0,89 | -0,31 | 1,88 | -0,17 | 8,68E-01 |
| AT2G21190 | 0,23 | 0,00 | 4,41 | 0,00 | 1,00E+00 |
| AT2G21270 | 0,21 | 1,51 | 4,33 | 0,35 | 7,27E-01 |
| AT2G21380 | 1,04 | -1,97 | 2,06 | -0,96 | 3,40E-01 |
| AT2G21410 | 1,46 | -1,65 | 1,35 | -1,23 | 2,20E-01 |
| AT2G21520 | 0,84 | 1,54 | 2,06 | 0,75 | 4,56E-01 |
| AT2G21530 | 0,61 | 0,56 | 3,05 | 0,18 | 8,53E-01 |
| AT2G21550 | 0,95 | 1,17 | 2,70 | 0,43 | 6,66E-01 |
| AT2G21580 | 1,04 | 0,29 | 2,60 | 0,11 | 9,10E-01 |
| AT2G21620 | 1,16 | -2,61 | 2,28 | -1,14 | 2,53E-01 |
| AT2G21630 | 0,32 | -0,98 | 4,35 | -0,23 | 8,22E-01 |
| AT2G21770 | 0,18 | -1,78 | 4,41 | -0,40 | 6,87E-01 |
| AT2G21790 | 0,74 | -0,32 | 2,36 | -0,13 | 8,93E-01 |
| AT2G21850 | 1,02 | -1,69 | 2,44 | -0,69 | 4,89E-01 |
| AT2G07475 | 0,24 | 1,56 | 4,34 | 0,36 | 7,18E-01 |
| AT2G21960 | 0,80 | -1,84 | 2,44 | -0,75 | 4,51E-01 |
| AT2G07495 | 0,12 | 0,97 | 4,34 | 0,22 | 8,23E-01 |
| AT2G22080 | 0,65 | 1,31 | 2,53 | 0,52 | 6,05E-01 |
| AT2G22120 | 0,32 | -0,98 | 4,35 | -0,23 | 8,22E-01 |
| AT2G22300 | 2,59 | -1,48 | 1,33 | -1,11 | 2,66E-01 |
| AT2G22330 | 4,57 | -0,24 | 0,87 | -0,28 | 7,80E-01 |
| AT2G22425 | 0,10 | 0,00 | 4,41 | 0,00 | 1,00E+00 |
| AT2G22440 | 2,40 | -0,88 | 1,41 | -0,62 | 5,32E-01 |
| AT2G22450 | 0,24 | -0,27 | 4,35 | -0,06 | 9,51E-01 |
| AT2G22480 | 0,23 | -0,48 | 4,35 | -0,11 | 9,12E-01 |
| AT2G22540 | 1,53 | -2,32 | 1,50 | -1,55 | 1,20E-01 |
| AT2G22660 | 2,02 | -0,33 | 1,38 | -0,24 | 8,10E-01 |
| AT2G22670 | 0,80 | -0,14 | 2,39 | -0,06 | 9,54E-01 |
| AT2G22680 | 0,25 | -1,82 | 4,40 | -0,41 | 6,79E-01 |
| AT2G22720 | 0,99 | -0,45 | 2,55 | -0,18 | 8,59E-01 |
| AT2G07845 | 0,43 | 2,16 | 4,09 | 0,53 | 5,97E-01 |
| AT2G22790 | 0,08 | 0,69 | 4,36 | 0,16 | 8,75E-01 |
| AT2G22840 | 1,11 | 2,24 | 2,07 | 1,08 | 2,80E-01 |
| AT2G07850 | 0,72 | 1,30 | 2,09 | 0,62 | 5,34E-01 |
| AT2G22910 | 0,32 | -0,25 | 4,36 | -0,06 | 9,55E-01 |
| AT2G22990 | 2,03 | -2,37 | 1,42 | -1,66 | 9,62E-02 |
| AT2G23080 | 0,38 | 0,58 | 4,31 | 0,13 | 8,93E-01 |
| AT2G23120 | 0,92 | 1,00 | 2,14 | 0,47 | 6,40E-01 |

| | | | | | |
|---|---|---|---|---|---|
| AT2G23130 | 0,41 | 0,79 | 2,82 | 0,28 | 7,78E-01 |
| AT2G23142 | 1,46 | -0,84 | 2,55 | -0,33 | 7,42E-01 |
| AT2G07890 | 2,64 | -0,56 | 1,29 | -0,43 | 6,66E-01 |
| AT2G23200 | 1,64 | -0,46 | 1,49 | -0,31 | 7,59E-01 |
| AT2G23310 | 0,84 | -1,83 | 2,79 | -0,66 | 5,12E-01 |
| AT2G07895 | 0,15 | 1,20 | 4,34 | 0,28 | 7,81E-01 |
| AT2G23320 | 1,39 | 0,46 | 1,62 | 0,28 | 7,78E-01 |
| AT2G23420 | 0,47 | -0,92 | 4,32 | -0,21 | 8,31E-01 |
| AT2G23460 | 1,50 | -0,56 | 1,49 | -0,38 | 7,08E-01 |
| AT2G07915 | 1,63 | 0,00 | 2,56 | 0,00 | 1,00E+00 |
| AT2G23672 | 1,50 | 0,74 | 1,74 | 0,42 | 6,72E-01 |
| AT2G23680 | 0,31 | 0,60 | 4,37 | 0,14 | 8,90E-01 |
| AT2G23740 | 0,12 | 0,49 | 4,36 | 0,11 | 9,11E-01 |
| AT2G23890 | 0,37 | 0,41 | 3,44 | 0,12 | 9,06E-01 |
| AT2G23930 | 1,70 | -1,57 | 1,61 | -0,98 | 3,30E-01 |
| AT2G23985 | 0,17 | 0,00 | 4,41 | 0,00 | 1,00E+00 |
| AT2G24090 | 0,67 | 0,25 | 2,59 | 0,10 | 9,22E-01 |
| AT2G24103 | 3,17 | -0,99 | 1,07 | -0,92 | 3,59E-01 |
| AT2G24180 | 1,36 | 2,45 | 2,54 | 0,96 | 3,35E-01 |
| AT2G24240 | 0,31 | 0,87 | 4,37 | 0,20 | 8,43E-01 |
| AT2G24280 | 1,45 | 0,10 | 1,65 | 0,06 | 9,49E-01 |
| AT2G24300 | 0,15 | 0,45 | 4,37 | 0,10 | 9,19E-01 |
| AT2G08015 | 0,35 | 2,07 | 4,32 | 0,48 | 6,32E-01 |
| AT2G24350 | 0,50 | -1,51 | 3,91 | -0,39 | 7,00E-01 |
| AT2G24390 | 0,92 | -0,15 | 2,85 | -0,05 | 9,58E-01 |
| AT2G24380 | 0,65 | -1,00 | 3,79 | -0,26 | 7,92E-01 |
| AT2G24420 | 1,14 | 1,40 | 2,11 | 0,66 | 5,07E-01 |
| AT2G08045 | 0,23 | 1,27 | 4,33 | 0,29 | 7,70E-01 |
| AT2G08050 | 0,32 | 1,34 | 4,34 | 0,31 | 7,57E-01 |
| AT2G24540 | 0,88 | -1,63 | 1,73 | -0,94 | 3,46E-01 |
| AT2G24590 | 0,49 | -0,82 | 3,87 | -0,21 | 8,32E-01 |
| AT2G24592 | 0,83 | 0,44 | 2,07 | 0,21 | 8,32E-01 |
| AT2G24765 | 0,69 | 0,29 | 2,81 | 0,10 | 9,18E-01 |
| AT2G24790 | 3,54 | 0,19 | 1,14 | 0,17 | 8,66E-01 |
| AT2G24980 | 3,78 | 4,46 | 1,55 | 2,88 | 3,95E-03 |
| AT2G25000 | 0,52 | 2,68 | 3,80 | 0,71 | 4,80E-01 |
| AT2G25080 | 0,61 | -0,88 | 2,72 | -0,32 | 7,45E-01 |
| AT2G25110 | 0,19 | 1,15 | 4,34 | 0,27 | 7,91E-01 |
| AT2G25170 | 3,04 | 0,49 | 1,29 | 0,38 | 7,06E-01 |
| AT2G25171 | 1,84 | 1,34 | 1,78 | 0,75 | 4,53E-01 |
| AT2G25190 | 0,14 | 0,03 | 4,39 | 0,01 | 9,95E-01 |
| AT2G25250 | 2,81 | 0,60 | 1,34 | 0,45 | 6,56E-01 |
| AT2G25370 | 1,47 | -1,78 | 2,32 | -0,76 | 4,44E-01 |
| AT2G25410 | 1,30 | 2,81 | 2,26 | 1,24 | 2,13E-01 |
| AT2G25430 | 4,01 | -0,49 | 0,89 | -0,56 | 5,78E-01 |
| AT2G25480 | 0,05 | 0,00 | 4,41 | 0,00 | 1,00E+00 |
| AT2G25520 | 1,73 | 0,46 | 1,79 | 0,26 | 7,96E-01 |
| AT2G25560 | 0,73 | 1,89 | 2,77 | 0,68 | 4,95E-01 |
| AT2G25580 | 0,08 | 0,44 | 4,38 | 0,10 | 9,21E-01 |
| AT2G25590 | 0,21 | 1,58 | 4,33 | 0,37 | 7,15E-01 |
| AT2G25625 | 0,15 | 1,16 | 4,34 | 0,27 | 7,88E-01 |
| AT2G25630 | 0,50 | 1,52 | 3,12 | 0,49 | 6,25E-01 |

| | | | | | |
|---|---|---|---|---|---|
| AT2G25650 | 0,70 | 0,45 | 2,82 | 0,16 | 8,74E-01 |
| AT2G25790 | 0,18 | 0,00 | 4,38 | 0,00 | 9,99E-01 |
| AT2G25840 | 0,59 | -1,21 | 3,96 | -0,31 | 7,59E-01 |
| AT2G25870 | 1,69 | -0,15 | 1,49 | -0,10 | 9,18E-01 |
| AT2G25900 | 2,31 | 0,60 | 1,71 | 0,35 | 7,24E-01 |
| AT2G25930 | 1,26 | 0,48 | 1,55 | 0,31 | 7,59E-01 |
| AT2G26060 | 1,00 | 0,57 | 4,31 | 0,13 | 8,95E-01 |
| AT2G26110 | 0,61 | -0,87 | 3,03 | -0,29 | 7,73E-01 |
| AT2G26170 | 0,47 | -0,02 | 3,16 | -0,01 | 9,95E-01 |
| AT2G26270 | 0,21 | -0,49 | 4,35 | -0,11 | 9,11E-01 |
| AT2G26300 | 1,09 | 2,67 | 2,42 | 1,10 | 2,70E-01 |
| AT2G26340 | 0,57 | -1,12 | 2,98 | -0,38 | 7,07E-01 |
| AT2G26355 | 2,39 | 1,54 | 1,26 | 1,23 | 2,20E-01 |
| AT2G26440 | 1,40 | 3,57 | 1,85 | 1,93 | 5,38E-02 |
| AT2G26500 | 2,52 | -0,13 | 1,15 | -0,11 | 9,11E-01 |
| AT2G26510 | 0,24 | -0,27 | 4,35 | -0,06 | 9,51E-01 |
| AT2G26570 | 0,43 | -0,05 | 3,32 | -0,01 | 9,89E-01 |
| AT2G26600 | 0,45 | 0,78 | 3,20 | 0,24 | 8,07E-01 |
| AT2G26610 | 0,30 | 0,46 | 4,32 | 0,11 | 9,16E-01 |
| AT2G26640 | 0,71 | 1,37 | 3,61 | 0,38 | 7,04E-01 |
| AT2G26660 | 0,22 | -0,47 | 4,35 | -0,11 | 9,13E-01 |
| AT2G26692 | 0,08 | 0,69 | 4,36 | 0,16 | 8,75E-01 |
| AT2G26730 | 0,85 | 1,39 | 2,20 | 0,63 | 5,27E-01 |
| AT2G26780 | 3,27 | 1,23 | 1,06 | 1,16 | 2,48E-01 |
| AT2G26910 | 1,87 | -1,51 | 1,30 | -1,17 | 2,43E-01 |
| AT2G26920 | 0,60 | 0,99 | 3,03 | 0,33 | 7,45E-01 |
| AT2G26975 | 1,93 | -0,33 | 1,35 | -0,25 | 8,04E-01 |
| AT2G27030 | 0,72 | -0,81 | 2,56 | -0,32 | 7,52E-01 |
| AT2G27050 | 0,99 | 0,61 | 1,80 | 0,34 | 7,35E-01 |
| AT2G27060 | 0,17 | 1,31 | 4,34 | 0,30 | 7,63E-01 |
| AT2G27080 | 0,39 | 0,41 | 4,23 | 0,10 | 9,23E-01 |
| AT2G27100 | 0,72 | -0,46 | 3,46 | -0,13 | 8,94E-01 |
| AT2G27130 | 0,30 | -1,78 | 4,41 | -0,40 | 6,87E-01 |
| AT2G27150 | 1,72 | 0,41 | 1,84 | 0,22 | 8,24E-01 |
| AT2G27200 | 0,21 | 1,42 | 4,33 | 0,33 | 7,43E-01 |
| AT2G27210 | 0,89 | 2,98 | 2,31 | 1,29 | 1,97E-01 |
| AT2G27230 | 1,31 | -1,47 | 1,46 | -1,01 | 3,15E-01 |
| AT2G27260 | 0,40 | 0,76 | 3,31 | 0,23 | 8,18E-01 |
| AT2G27320 | 0,85 | 0,08 | 2,42 | 0,03 | 9,75E-01 |
| AT2G27380 | 0,13 | 0,00 | 4,41 | 0,00 | 1,00E+00 |
| AT2G27460 | 0,28 | 1,19 | 4,34 | 0,27 | 7,85E-01 |
| AT2G27540 | 0,95 | 0,42 | 3,19 | 0,13 | 8,95E-01 |
| AT2G27570 | 0,28 | -1,42 | 4,36 | -0,33 | 7,44E-01 |
| AT2G27600 | 1,39 | 0,97 | 1,85 | 0,53 | 5,99E-01 |
| AT2G27690 | 0,16 | 0,92 | 4,35 | 0,21 | 8,33E-01 |
| AT2G27710 | 0,57 | -1,16 | 3,17 | -0,37 | 7,14E-01 |
| AT2G27720 | 1,70 | -2,57 | 1,49 | -1,72 | 8,50E-02 |
| AT2G27760 | 0,08 | 0,44 | 4,38 | 0,10 | 9,21E-01 |
| AT2G27810 | 0,54 | -0,33 | 2,67 | -0,12 | 9,01E-01 |
| AT2G27820 | 0,23 | 0,70 | 4,35 | 0,16 | 8,73E-01 |
| AT2G27900 | 1,32 | -0,54 | 1,39 | -0,39 | 6,96E-01 |
| AT2G28070 | 0,87 | -1,14 | 2,76 | -0,41 | 6,79E-01 |

| | | | | | |
|---|---|---|---|---|---|
| AT2G28120 | 0,29 | 0,65 | 4,32 | 0,15 | 8,80E-01 |
| AT2G28130 | 0,59 | 2,14 | 4,31 | 0,50 | 6,20E-01 |
| AT2G28190 | 0,42 | -1,21 | 4,31 | -0,28 | 7,80E-01 |
| AT2G28250 | 0,08 | 0,69 | 4,36 | 0,16 | 8,75E-01 |
| AT2G28310 | 0,30 | -1,19 | 4,35 | -0,27 | 7,84E-01 |
| AT2G28320 | 0,43 | 1,23 | 4,17 | 0,29 | 7,68E-01 |
| AT2G28350 | 0,96 | 0,41 | 1,88 | 0,22 | 8,29E-01 |
| AT2G28370 | 0,25 | 0,34 | 4,32 | 0,08 | 9,37E-01 |
| AT2G28460 | 0,18 | 0,00 | 4,41 | 0,00 | 1,00E+00 |
| AT2G28520 | 0,65 | 0,86 | 2,82 | 0,30 | 7,62E-01 |
| AT2G28540 | 0,87 | -1,04 | 2,15 | -0,48 | 6,30E-01 |
| AT2G28605 | 0,37 | -1,78 | 4,41 | -0,40 | 6,87E-01 |
| AT2G28660 | 0,10 | 0,00 | 4,41 | 0,00 | 1,00E+00 |
| AT2G28671 | 0,51 | 1,71 | 3,05 | 0,56 | 5,74E-01 |
| AT2G28710 | 0,24 | 1,30 | 4,34 | 0,30 | 7,64E-01 |
| AT2G28720 | 0,49 | -1,74 | 4,25 | -0,41 | 6,82E-01 |
| AT2G28730 | 1,02 | -1,59 | 1,62 | -0,98 | 3,27E-01 |
| AT2G28810 | 0,71 | -0,33 | 2,53 | -0,13 | 8,95E-01 |
| AT2G28840 | 1,76 | -0,63 | 1,55 | -0,41 | 6,85E-01 |
| AT2G28930 | 0,12 | 0,37 | 4,37 | 0,08 | 9,33E-01 |
| AT2G29020 | 0,11 | 0,00 | 4,41 | 0,00 | 1,00E+00 |
| AT2G29065 | 0,30 | 0,07 | 4,35 | 0,02 | 9,87E-01 |
| AT2G29070 | 0,15 | 1,20 | 4,34 | 0,28 | 7,81E-01 |
| AT2G29140 | 0,94 | 2,14 | 2,44 | 0,88 | 3,80E-01 |
| AT2G29180 | 0,47 | -0,64 | 4,10 | -0,16 | 8,76E-01 |
| AT2G29210 | 1,16 | -0,75 | 1,89 | -0,40 | 6,91E-01 |
| AT2G29310 | 0,46 | -2,75 | 3,32 | -0,83 | 4,07E-01 |
| AT2G29350 | 0,65 | 3,06 | 3,82 | 0,80 | 4,23E-01 |
| AT2G29360 | 0,51 | -2,04 | 4,34 | -0,47 | 6,38E-01 |
| AT2G29510 | 0,82 | 1,53 | 3,26 | 0,47 | 6,40E-01 |
| AT2G29600 | 0,47 | 1,50 | 3,18 | 0,47 | 6,37E-01 |
| AT2G29630 | 5,51 | 0,58 | 0,77 | 0,76 | 4,49E-01 |
| AT2G29640 | 0,37 | -0,56 | 4,24 | -0,13 | 8,95E-01 |
| AT2G29660 | 0,20 | -0,31 | 4,36 | -0,07 | 9,44E-01 |
| AT2G29690 | 0,14 | 0,00 | 4,41 | 0,00 | 1,00E+00 |
| AT2G29700 | 0,24 | 0,00 | 4,41 | 0,00 | 1,00E+00 |
| AT2G29920 | 2,43 | 3,99 | 1,90 | 2,09 | 3,62E-02 |
| AT2G29970 | 1,50 | 2,60 | 1,68 | 1,55 | 1,22E-01 |
| AT2G30020 | 0,83 | 1,56 | 3,51 | 0,44 | 6,56E-01 |
| AT2G30040 | 0,38 | 0,57 | 3,32 | 0,17 | 8,63E-01 |
| AT2G30050 | 0,66 | -2,45 | 2,60 | -0,94 | 3,46E-01 |
| AT2G30140 | 0,54 | 1,52 | 3,81 | 0,40 | 6,89E-01 |
| AT2G30150 | 0,10 | 0,21 | 4,38 | 0,05 | 9,62E-01 |
| AT2G30180 | 1,06 | -0,50 | 1,73 | -0,29 | 7,74E-01 |
| AT2G08370 | 4,11 | 0,12 | 0,91 | 0,14 | 8,91E-01 |
| AT2G08375 | 1,24 | -0,63 | 1,99 | -0,32 | 7,51E-01 |
| AT2G30230 | 0,50 | 1,29 | 3,23 | 0,40 | 6,90E-01 |
| AT2G30362 | 0,41 | 0,58 | 4,21 | 0,14 | 8,91E-01 |
| AT2G30410 | 0,72 | -2,47 | 2,53 | -0,98 | 3,28E-01 |
| AT2G30440 | 1,74 | -0,67 | 1,49 | -0,45 | 6,51E-01 |
| AT2G30530 | 0,20 | -0,31 | 4,36 | -0,07 | 9,44E-01 |
| AT2G30550 | 1,44 | -1,62 | 1,75 | -0,92 | 3,56E-01 |

| | | | | | |
|---|---|---|---|---|---|
| AT2G30600 | 0,51 | -2,24 | 4,37 | -0,51 | 6,08E-01 |
| AT2G30620 | 0,85 | 1,68 | 2,37 | 0,71 | 4,78E-01 |
| AT2G30730 | 0,89 | -0,41 | 2,56 | -0,16 | 8,72E-01 |
| AT2G30780 | 0,62 | 2,27 | 3,69 | 0,61 | 5,39E-01 |
| AT2G30870 | 1,65 | 2,01 | 1,65 | 1,22 | 2,23E-01 |
| AT2G30930 | 0,49 | -1,69 | 2,67 | -0,63 | 5,26E-01 |
| AT2G30950 | 3,17 | -0,27 | 0,97 | -0,28 | 7,77E-01 |
| AT2G30970 | 0,46 | -1,03 | 4,35 | -0,24 | 8,13E-01 |
| AT2G31010 | 0,95 | -1,09 | 2,16 | -0,51 | 6,12E-01 |
| AT2G31020 | 0,26 | -0,48 | 4,38 | -0,11 | 9,13E-01 |
| AT2G31200 | 0,16 | 0,00 | 4,39 | 0,00 | 1,00E+00 |
| AT2G31215 | 0,52 | 0,90 | 2,70 | 0,33 | 7,39E-01 |
| AT2G31280 | 1,47 | 0,70 | 1,87 | 0,37 | 7,09E-01 |
| AT2G31290 | 0,71 | -1,02 | 3,10 | -0,33 | 7,43E-01 |
| AT2G31350 | 0,38 | 0,70 | 4,28 | 0,16 | 8,71E-01 |
| AT2G31370 | 0,19 | 0,90 | 4,34 | 0,21 | 8,35E-01 |
| AT2G31380 | 1,06 | -1,03 | 1,71 | -0,60 | 5,45E-01 |
| AT2G08435 | 0,79 | -0,72 | 3,47 | -0,21 | 8,36E-01 |
| AT2G31560 | 0,67 | -0,48 | 2,56 | -0,19 | 8,53E-01 |
| AT2G31585 | 0,47 | 0,53 | 3,93 | 0,13 | 8,94E-01 |
| AT2G31610 | 1,80 | -0,57 | 1,42 | -0,40 | 6,90E-01 |
| AT2G31660 | 3,59 | -1,03 | 0,98 | -1,05 | 2,94E-01 |
| AT2G31750 | 0,75 | 2,31 | 2,98 | 0,78 | 4,38E-01 |
| AT2G31790 | 0,30 | 1,17 | 3,63 | 0,32 | 7,46E-01 |
| AT2G31810 | 1,10 | -1,60 | 1,99 | -0,81 | 4,20E-01 |
| AT2G31820 | 4,59 | 0,38 | 0,97 | 0,39 | 6,96E-01 |
| AT2G31960 | 1,68 | -0,26 | 1,42 | -0,18 | 8,56E-01 |
| AT2G31970 | 0,18 | 1,24 | 4,33 | 0,29 | 7,74E-01 |
| AT2G32010 | 0,37 | 1,10 | 4,34 | 0,25 | 8,00E-01 |
| AT2G32040 | 0,08 | 0,44 | 4,38 | 0,10 | 9,21E-01 |
| AT2G32100 | 1,15 | 0,05 | 2,32 | 0,02 | 9,83E-01 |
| AT2G32150 | 1,00 | 0,36 | 2,38 | 0,15 | 8,79E-01 |
| AT2G32250 | 0,51 | 0,03 | 3,11 | 0,01 | 9,92E-01 |
| AT2G32260 | 0,44 | 0,83 | 4,08 | 0,20 | 8,39E-01 |
| AT2G32295 | 0,53 | -0,17 | 2,75 | -0,06 | 9,52E-01 |
| AT2G32320 | 0,15 | 0,45 | 4,37 | 0,10 | 9,19E-01 |
| AT2G32440 | 0,16 | 0,65 | 4,37 | 0,15 | 8,82E-01 |
| AT2G32450 | 0,56 | 1,16 | 2,57 | 0,45 | 6,51E-01 |
| AT2G32698 | 0,12 | 1,16 | 4,34 | 0,27 | 7,89E-01 |
| AT2G32700 | 1,61 | 0,86 | 1,37 | 0,63 | 5,31E-01 |
| AT2G32730 | 3,18 | 0,25 | 1,08 | 0,23 | 8,15E-01 |
| AT2G32800 | 1,51 | -0,50 | 1,39 | -0,36 | 7,17E-01 |
| AT2G32900 | 0,29 | 0,60 | 4,31 | 0,14 | 8,89E-01 |
| AT2G32910 | 0,68 | 0,83 | 2,95 | 0,28 | 7,79E-01 |
| AT2G33080 | 0,74 | 1,66 | 3,44 | 0,48 | 6,29E-01 |
| AT2G33160 | 0,25 | 0,41 | 4,33 | 0,09 | 9,25E-01 |
| AT2G33220 | 0,54 | -0,20 | 3,08 | -0,06 | 9,48E-01 |
| AT2G33250 | 0,76 | 1,90 | 4,32 | 0,44 | 6,61E-01 |
| AT2G33290 | 0,29 | 0,82 | 3,69 | 0,22 | 8,24E-01 |
| AT2G33390 | 0,25 | -1,82 | 4,40 | -0,41 | 6,79E-01 |
| AT2G33420 | 0,37 | -0,56 | 4,24 | -0,13 | 8,95E-01 |
| AT2G08585 | 0,32 | 0,81 | 3,67 | 0,22 | 8,25E-01 |

| | | | | | |
|---|---|---|---|---|---|
| AT2G33435 | 1,31 | -1,63 | 1,63 | -1,00 | 3,19E-01 |
| AT2G33450 | 0,32 | 0,46 | 4,36 | 0,11 | 9,16E-01 |
| AT2G33490 | 0,35 | 0,81 | 4,28 | 0,19 | 8,50E-01 |
| AT2G33510 | 0,26 | 0,12 | 4,33 | 0,03 | 9,79E-01 |
| AT2G33570 | 0,17 | 0,07 | 4,40 | 0,02 | 9,88E-01 |
| AT2G33610 | 0,37 | 0,34 | 4,28 | 0,08 | 9,37E-01 |
| AT2G33650 | 0,29 | 0,00 | 4,41 | 0,00 | 1,00E+00 |
| AT2G33700 | 0,56 | -1,40 | 4,05 | -0,35 | 7,29E-01 |
| AT2G33760 | 0,64 | -0,72 | 4,31 | -0,17 | 8,67E-01 |
| AT2G33810 | 0,37 | -1,96 | 4,38 | -0,45 | 6,54E-01 |
| AT2G33840 | 0,52 | -1,52 | 3,17 | -0,48 | 6,32E-01 |
| AT2G33845 | 0,47 | 1,97 | 4,04 | 0,49 | 6,26E-01 |
| AT2G33950 | 0,39 | -1,37 | 4,28 | -0,32 | 7,49E-01 |
| AT2G34110 | 1,88 | 0,53 | 1,43 | 0,37 | 7,10E-01 |
| AT2G34120 | 0,27 | 0,25 | 4,38 | 0,06 | 9,55E-01 |
| AT2G34208 | 2,28 | 0,96 | 1,28 | 0,75 | 4,50E-01 |
| AT2G34240 | 0,12 | 1,05 | 4,35 | 0,24 | 8,10E-01 |
| AT2G34250 | 2,94 | 0,41 | 1,09 | 0,38 | 7,05E-01 |
| AT2G34357 | 1,72 | 1,70 | 1,55 | 1,09 | 2,74E-01 |
| AT2G34410 | 2,89 | 1,27 | 1,22 | 1,04 | 2,96E-01 |
| AT2G34460 | 0,99 | -0,31 | 1,85 | -0,17 | 8,65E-01 |
| AT2G34655 | 4,12 | 1,42 | 1,38 | 1,03 | 3,03E-01 |
| AT2G34660 | 0,69 | 0,89 | 2,62 | 0,34 | 7,33E-01 |
| AT2G34730 | 0,50 | 0,65 | 2,65 | 0,24 | 8,07E-01 |
| AT2G34770 | 0,22 | -0,47 | 4,35 | -0,11 | 9,13E-01 |
| AT2G34780 | 0,99 | 0,12 | 1,97 | 0,06 | 9,50E-01 |
| AT2G34810 | 0,16 | 0,92 | 4,35 | 0,21 | 8,33E-01 |
| AT2G34860 | 1,10 | -1,81 | 2,00 | -0,90 | 3,66E-01 |
| AT2G34970 | 0,22 | -0,86 | 4,36 | -0,20 | 8,44E-01 |
| AT2G35020 | 0,32 | -0,59 | 4,37 | -0,14 | 8,92E-01 |
| AT2G35050 | 3,09 | 0,02 | 1,13 | 0,02 | 9,84E-01 |
| AT2G35170 | 0,10 | 0,21 | 4,38 | 0,05 | 9,62E-01 |
| AT2G35230 | 0,50 | -0,59 | 4,33 | -0,14 | 8,92E-01 |
| AT2G35260 | 3,89 | -0,57 | 1,07 | -0,53 | 5,93E-01 |
| AT2G35330 | 0,35 | -2,42 | 4,38 | -0,55 | 5,81E-01 |
| AT2G35340 | 0,80 | -1,29 | 2,63 | -0,49 | 6,24E-01 |
| AT2G35370 | 0,43 | -0,63 | 4,27 | -0,15 | 8,82E-01 |
| AT2G35630 | 1,16 | -1,13 | 1,73 | -0,65 | 5,14E-01 |
| AT2G35635 | 0,64 | -3,09 | 4,34 | -0,71 | 4,76E-01 |
| AT2G35650 | 0,68 | -0,96 | 2,09 | -0,46 | 6,45E-01 |
| AT2G08750 | 0,08 | 0,44 | 4,38 | 0,10 | 9,21E-01 |
| AT2G35720 | 0,19 | 0,85 | 4,35 | 0,20 | 8,45E-01 |
| AT2G35760 | 0,17 | -1,70 | 4,40 | -0,39 | 7,00E-01 |
| AT2G35800 | 1,02 | -0,66 | 1,69 | -0,39 | 6,96E-01 |
| AT2G35820 | 0,71 | 1,35 | 4,31 | 0,31 | 7,54E-01 |
| AT2G35830 | 1,03 | 1,21 | 1,87 | 0,65 | 5,18E-01 |
| AT2G35840 | 0,63 | -1,01 | 2,38 | -0,42 | 6,72E-01 |
| AT2G35860 | 0,74 | 0,09 | 2,40 | 0,04 | 9,70E-01 |
| AT2G35920 | 0,77 | 0,21 | 2,31 | 0,09 | 9,27E-01 |
| AT2G35935 | 0,49 | 0,04 | 3,96 | 0,01 | 9,92E-01 |
| AT2G35960 | 0,20 | 0,15 | 4,37 | 0,04 | 9,72E-01 |
| AT2G36000 | 0,80 | 1,51 | 2,54 | 0,59 | 5,52E-01 |

| | | | | | |
|---|---|---|---|---|---|
| AT2G36010 | 0,59 | -1,90 | 3,05 | -0,62 | 5,33E-01 |
| AT2G36026 | 0,70 | 0,73 | 3,80 | 0,19 | 8,48E-01 |
| AT2G36090 | 0,69 | 0,52 | 3,61 | 0,15 | 8,84E-01 |
| AT2G36130 | 0,56 | -0,38 | 4,32 | -0,09 | 9,29E-01 |
| AT2G36160 | 1,81 | -1,15 | 1,37 | -0,84 | 4,02E-01 |
| AT2G36320 | 1,39 | 1,36 | 1,84 | 0,74 | 4,61E-01 |
| AT2G36330 | 0,55 | 1,46 | 4,34 | 0,34 | 7,37E-01 |
| AT2G36350 | 2,13 | 0,10 | 1,47 | 0,07 | 9,45E-01 |
| AT2G36390 | 2,07 | 0,62 | 1,49 | 0,42 | 6,76E-01 |
| AT2G36430 | 1,44 | 0,61 | 1,95 | 0,31 | 7,56E-01 |
| AT2G36500 | 0,54 | -1,21 | 2,98 | -0,41 | 6,84E-01 |
| AT2G36510 | 0,37 | -0,47 | 4,25 | -0,11 | 9,11E-01 |
| AT2G36570 | 0,52 | -3,40 | 4,32 | -0,79 | 4,31E-01 |
| AT2G36580 | 1,14 | 0,20 | 1,55 | 0,13 | 8,98E-01 |
| AT2G36600 | 0,66 | -0,03 | 2,28 | -0,01 | 9,90E-01 |
| AT2G36720 | 1,25 | -0,89 | 1,84 | -0,48 | 6,28E-01 |
| AT2G36792 | 0,34 | 1,06 | 4,34 | 0,24 | 8,07E-01 |
| AT2G36830 | 0,86 | -1,31 | 2,04 | -0,64 | 5,22E-01 |
| AT2G36920 | 0,12 | 1,23 | 4,34 | 0,28 | 7,76E-01 |
| AT2G36950 | 0,34 | 0,58 | 4,34 | 0,13 | 8,94E-01 |
| AT2G36960 | 0,18 | 0,81 | 4,34 | 0,19 | 8,52E-01 |
| AT2G36970 | 0,19 | 1,15 | 4,34 | 0,27 | 7,91E-01 |
| AT2G36980 | 0,05 | 0,00 | 4,41 | 0,00 | 1,00E+00 |
| AT2G37025 | 0,19 | 0,00 | 4,41 | 0,00 | 1,00E+00 |
| AT2G37050 | 0,93 | -0,55 | 2,15 | -0,25 | 7,99E-01 |
| AT2G37110 | 1,16 | 1,57 | 2,30 | 0,68 | 4,95E-01 |
| AT2G37160 | 3,77 | 0,64 | 1,08 | 0,59 | 5,53E-01 |
| AT2G08855 | 3,73 | 0,54 | 1,10 | 0,49 | 6,21E-01 |
| AT2G37180 | 0,08 | 0,44 | 4,38 | 0,10 | 9,21E-01 |
| AT2G37250 | 0,74 | 1,04 | 2,23 | 0,47 | 6,42E-01 |
| AT2G08860 | 0,31 | 1,16 | 3,61 | 0,32 | 7,47E-01 |
| AT2G37320 | 0,08 | 0,69 | 4,36 | 0,16 | 8,75E-01 |
| AT2G37330 | 0,66 | 0,07 | 2,98 | 0,02 | 9,81E-01 |
| AT2G37362 | 0,21 | 1,76 | 4,32 | 0,41 | 6,85E-01 |
| AT2G37370 | 0,40 | 0,00 | 4,38 | 0,00 | 1,00E+00 |
| AT2G37470 | 0,26 | 1,69 | 4,32 | 0,39 | 6,96E-01 |
| AT2G37480 | 1,35 | 1,06 | 2,03 | 0,52 | 6,02E-01 |
| AT2G37555 | 0,61 | 1,05 | 2,97 | 0,35 | 7,24E-01 |
| AT2G37580 | 0,12 | 0,49 | 4,36 | 0,11 | 9,11E-01 |
| AT2G37650 | 1,11 | -0,56 | 1,88 | -0,30 | 7,66E-01 |
| AT2G37690 | 0,31 | -0,76 | 4,38 | -0,17 | 8,62E-01 |
| AT2G37770 | 0,28 | 1,91 | 4,32 | 0,44 | 6,58E-01 |
| AT2G37860 | 0,83 | -1,80 | 2,13 | -0,85 | 3,98E-01 |
| AT2G37940 | 0,73 | -3,20 | 3,01 | -1,06 | 2,88E-01 |
| AT2G37970 | 0,77 | 1,86 | 2,69 | 0,69 | 4,89E-01 |
| AT2G37990 | 0,11 | 0,48 | 4,37 | 0,11 | 9,12E-01 |
| AT2G38010 | 0,53 | 0,98 | 3,04 | 0,32 | 7,47E-01 |
| AT2G38040 | 4,80 | -0,49 | 0,92 | -0,53 | 5,94E-01 |
| AT2G38090 | 0,20 | -0,44 | 4,36 | -0,10 | 9,19E-01 |
| AT2G38120 | 0,38 | 1,59 | 2,82 | 0,57 | 5,72E-01 |
| AT2G38140 | 0,26 | 0,82 | 3,76 | 0,22 | 8,27E-01 |
| AT2G38195 | 4,06 | 0,73 | 1,47 | 0,50 | 6,18E-01 |

| | | | | | |
|---|---|---|---|---|---|
| AT2G38210 | 1,49 | 0,04 | 2,13 | 0,02 | 9,87E-01 |
| AT2G38220 | 2,26 | 0,19 | 1,38 | 0,14 | 8,92E-01 |
| AT2G38310 | 0,71 | -0,85 | 2,95 | -0,29 | 7,73E-01 |
| AT2G38365 | 0,20 | -0,31 | 4,36 | -0,07 | 9,44E-01 |
| AT2G38420 | 0,26 | -0,48 | 4,38 | -0,11 | 9,13E-01 |
| AT2G38440 | 3,79 | -0,27 | 0,87 | -0,32 | 7,52E-01 |
| AT2G38470 | 1,30 | -0,24 | 1,68 | -0,14 | 8,87E-01 |
| AT2G38540 | 3,59 | -2,22 | 1,19 | -1,87 | 6,12E-02 |
| AT2G38560 | 1,56 | 0,56 | 1,73 | 0,32 | 7,46E-01 |
| AT2G38590 | 0,78 | 1,60 | 2,74 | 0,59 | 5,58E-01 |
| AT2G38670 | 0,86 | -0,42 | 2,11 | -0,20 | 8,42E-01 |
| AT2G38760 | 0,08 | 0,69 | 4,36 | 0,16 | 8,75E-01 |
| AT2G38780 | 0,82 | -0,48 | 1,88 | -0,25 | 7,99E-01 |
| AT2G38800 | 2,71 | 1,99 | 1,39 | 1,43 | 1,54E-01 |
| AT2G38820 | 0,12 | 0,97 | 4,34 | 0,22 | 8,23E-01 |
| AT2G38840 | 0,33 | -0,27 | 4,32 | -0,06 | 9,50E-01 |
| AT2G38940 | 0,25 | 0,26 | 4,33 | 0,06 | 9,51E-01 |
| AT2G38950 | 0,65 | -1,40 | 2,48 | -0,56 | 5,73E-01 |
| AT2G39010 | 4,44 | -0,72 | 1,09 | -0,66 | 5,10E-01 |
| AT2G39190 | 0,15 | 0,74 | 4,35 | 0,17 | 8,65E-01 |
| AT2G39230 | 0,19 | 0,00 | 4,41 | 0,00 | 1,00E+00 |
| AT2G39270 | 0,08 | 0,44 | 4,38 | 0,10 | 9,21E-01 |
| AT2G39340 | 0,19 | 0,00 | 4,41 | 0,00 | 1,00E+00 |
| AT2G39400 | 0,15 | 0,83 | 4,35 | 0,19 | 8,49E-01 |
| AT2G39420 | 0,56 | -1,61 | 3,18 | -0,51 | 6,11E-01 |
| AT2G39430 | 0,52 | 1,20 | 3,86 | 0,31 | 7,57E-01 |
| AT2G39460 | 0,89 | 0,75 | 2,01 | 0,37 | 7,08E-01 |
| AT2G39470 | 0,81 | 0,36 | 2,23 | 0,16 | 8,73E-01 |
| AT2G39570 | 1,60 | -1,07 | 2,78 | -0,39 | 7,00E-01 |
| AT2G39580 | 1,16 | -1,35 | 1,88 | -0,72 | 4,71E-01 |
| AT2G39670 | 1,03 | 3,42 | 2,18 | 1,57 | 1,16E-01 |
| AT2G39725 | 0,50 | -0,16 | 3,86 | -0,04 | 9,66E-01 |
| AT2G39740 | 0,63 | -0,43 | 3,01 | -0,14 | 8,86E-01 |
| AT2G39770 | 2,97 | -0,35 | 1,10 | -0,32 | 7,51E-01 |
| AT2G39780 | 1,17 | -0,67 | 2,07 | -0,33 | 7,45E-01 |
| AT2G39810 | 1,48 | -0,98 | 2,20 | -0,44 | 6,57E-01 |
| AT2G39840 | 0,84 | 0,81 | 2,44 | 0,33 | 7,41E-01 |
| AT2G39850 | 0,54 | 0,00 | 4,38 | 0,00 | 1,00E+00 |
| AT2G39870 | 0,43 | -0,75 | 4,34 | -0,17 | 8,64E-01 |
| AT2G39930 | 1,27 | -1,63 | 1,58 | -1,03 | 3,01E-01 |
| AT2G40030 | 5,36 | 0,43 | 0,82 | 0,52 | 6,01E-01 |
| AT2G40060 | 0,99 | 1,57 | 2,38 | 0,66 | 5,08E-01 |
| AT2G40095 | 0,64 | 0,33 | 3,03 | 0,11 | 9,14E-01 |
| AT2G40130 | 0,14 | 0,83 | 4,35 | 0,19 | 8,49E-01 |
| AT2G40140 | 4,76 | -0,41 | 0,92 | -0,44 | 6,59E-01 |
| AT2G40150 | 0,08 | 0,44 | 4,38 | 0,10 | 9,21E-01 |
| AT2G40160 | 0,42 | -0,56 | 4,10 | -0,14 | 8,91E-01 |
| AT2G40200 | 0,38 | -0,37 | 4,36 | -0,08 | 9,33E-01 |
| AT2G09120 | 0,31 | 0,00 | 4,41 | 0,00 | 1,00E+00 |
| AT2G40310 | 0,83 | 2,24 | 2,26 | 0,99 | 3,23E-01 |
| AT2G40400 | 0,82 | 0,68 | 2,71 | 0,25 | 8,02E-01 |
| AT2G40410 | 0,54 | -2,28 | 4,01 | -0,57 | 5,70E-01 |

| | | | | | |
|---|---|---|---|---|---|
| AT2G40435 | 0,46 | -1,19 | 4,38 | -0,27 | 7,86E-01 |
| AT2G40460 | 0,63 | 1,23 | 2,60 | 0,47 | 6,35E-01 |
| AT2G40480 | 0,31 | -0,19 | 4,36 | -0,04 | 9,65E-01 |
| AT2G40490 | 1,23 | -0,78 | 2,00 | -0,39 | 6,98E-01 |
| AT2G40540 | 0,78 | 0,67 | 2,46 | 0,27 | 7,85E-01 |
| AT2G40550 | 0,20 | -0,31 | 4,36 | -0,07 | 9,44E-01 |
| AT2G40660 | 0,08 | 0,00 | 4,40 | 0,00 | 1,00E+00 |
| AT2G40670 | 0,10 | 0,21 | 4,38 | 0,05 | 9,62E-01 |
| AT2G40760 | 0,35 | 1,56 | 4,33 | 0,36 | 7,19E-01 |
| AT2G40765 | 0,08 | 0,44 | 4,38 | 0,10 | 9,21E-01 |
| AT2G40810 | 0,08 | 0,69 | 4,36 | 0,16 | 8,75E-01 |
| AT2G40830 | 0,33 | 0,76 | 4,32 | 0,18 | 8,60E-01 |
| AT2G40840 | 4,12 | -0,40 | 0,86 | -0,47 | 6,40E-01 |
| AT2G40880 | 1,07 | 0,39 | 2,11 | 0,19 | 8,52E-01 |
| AT2G40955 | 1,38 | 1,21 | 1,94 | 0,63 | 5,32E-01 |
| AT2G40980 | 0,41 | -0,47 | 4,16 | -0,11 | 9,10E-01 |
| AT2G41080 | 0,59 | -0,79 | 2,55 | -0,31 | 7,58E-01 |
| AT2G41090 | 3,05 | -0,06 | 1,17 | -0,05 | 9,60E-01 |
| AT2G41110 | 0,26 | 0,84 | 3,79 | 0,22 | 8,25E-01 |
| AT2G09230 | 0,12 | 1,23 | 4,34 | 0,28 | 7,76E-01 |
| AT2G41140 | 0,37 | 0,57 | 4,27 | 0,13 | 8,93E-01 |
| AT2G41200 | 0,84 | -1,51 | 2,29 | -0,66 | 5,09E-01 |
| AT2G41210 | 0,42 | 0,40 | 4,12 | 0,10 | 9,23E-01 |
| AT2G41220 | 0,64 | 1,02 | 2,95 | 0,35 | 7,29E-01 |
| AT2G41440 | 2,60 | 0,91 | 1,16 | 0,79 | 4,30E-01 |
| AT2G41460 | 0,13 | 0,73 | 4,36 | 0,17 | 8,67E-01 |
| AT2G41510 | 0,08 | 0,44 | 4,38 | 0,10 | 9,21E-01 |
| AT2G41520 | 0,59 | -0,87 | 2,93 | -0,30 | 7,67E-01 |
| AT2G41590 | 1,66 | 0,18 | 1,57 | 0,12 | 9,08E-01 |
| AT2G41670 | 0,12 | 1,23 | 4,34 | 0,28 | 7,76E-01 |
| AT2G41730 | 0,99 | -0,65 | 1,82 | -0,35 | 7,24E-01 |
| AT2G41820 | 0,34 | 0,47 | 4,34 | 0,11 | 9,13E-01 |
| AT2G41880 | 0,26 | 0,84 | 3,79 | 0,22 | 8,25E-01 |
| AT2G41920 | 1,42 | 0,79 | 2,01 | 0,39 | 6,95E-01 |
| AT2G41940 | 1,76 | 0,03 | 1,62 | 0,02 | 9,87E-01 |
| AT2G42130 | 0,48 | -0,82 | 3,36 | -0,25 | 8,06E-01 |
| AT2G42220 | 0,98 | 0,05 | 2,09 | 0,02 | 9,82E-01 |
| AT2G42310 | 0,61 | 0,62 | 2,50 | 0,25 | 8,05E-01 |
| AT2G42320 | 0,63 | 0,71 | 2,59 | 0,28 | 7,83E-01 |
| AT2G42430 | 0,21 | 0,00 | 4,41 | 0,00 | 1,00E+00 |
| AT2G42580 | 1,32 | 0,78 | 1,75 | 0,45 | 6,55E-01 |
| AT2G42610 | 0,29 | 0,74 | 4,35 | 0,17 | 8,65E-01 |
| AT2G42620 | 1,45 | 2,09 | 2,37 | 0,88 | 3,77E-01 |
| AT2G42700 | 0,72 | 0,43 | 2,95 | 0,15 | 8,85E-01 |
| AT2G42710 | 0,12 | 0,37 | 4,37 | 0,08 | 9,33E-01 |
| AT2G42750 | 0,57 | 1,38 | 3,73 | 0,37 | 7,12E-01 |
| AT2G42780 | 0,28 | 0,00 | 4,41 | 0,00 | 1,00E+00 |
| AT2G42890 | 0,32 | 1,69 | 4,32 | 0,39 | 6,95E-01 |
| AT2G42900 | 0,43 | -2,45 | 4,36 | -0,56 | 5,75E-01 |
| AT2G43010 | 1,28 | -1,78 | 1,72 | -1,03 | 3,01E-01 |
| AT2G43020 | 1,32 | 1,59 | 2,07 | 0,77 | 4,43E-01 |
| AT2G43030 | 0,58 | -0,85 | 3,21 | -0,27 | 7,91E-01 |

| | | | | | |
|---|---|---|---|---|---|
| AT2G43060 | 1,34 | -0,14 | 1,84 | -0,08 | 9,39E-01 |
| AT2G43080 | 0,63 | 0,71 | 3,08 | 0,23 | 8,19E-01 |
| AT2G43090 | 0,61 | 0,05 | 2,73 | 0,02 | 9,86E-01 |
| AT2G43100 | 0,56 | 0,43 | 3,35 | 0,13 | 8,98E-01 |
| AT2G43110 | 4,82 | -0,34 | 0,97 | -0,35 | 7,24E-01 |
| AT2G43137 | 1,43 | 0,51 | 1,93 | 0,26 | 7,93E-01 |
| AT2G43138 | 1,21 | 0,71 | 2,11 | 0,33 | 7,38E-01 |
| AT2G43141 | 1,64 | -0,78 | 1,87 | -0,42 | 6,75E-01 |
| AT2G43150 | 1,94 | 0,84 | 1,33 | 0,63 | 5,29E-01 |
| AT2G43210 | 0,63 | 1,33 | 2,93 | 0,46 | 6,49E-01 |
| AT2G43230 | 0,55 | -1,30 | 3,78 | -0,34 | 7,31E-01 |
| AT2G43250 | 0,31 | 0,48 | 4,32 | 0,11 | 9,11E-01 |
| AT2G43280 | 0,56 | -1,50 | 4,33 | -0,35 | 7,29E-01 |
| AT2G09440 | 0,08 | 0,69 | 4,36 | 0,16 | 8,75E-01 |
| AT2G43340 | 0,44 | -1,76 | 3,37 | -0,52 | 6,00E-01 |
| AT2G43400 | 0,37 | 0,75 | 4,30 | 0,17 | 8,62E-01 |
| AT2G43440 | 0,11 | 0,00 | 4,41 | 0,00 | 1,00E+00 |
| AT2G43540 | 1,53 | 1,06 | 1,86 | 0,57 | 5,67E-01 |
| AT2G43630 | 0,44 | 2,16 | 4,05 | 0,53 | 5,95E-01 |
| AT2G43650 | 0,83 | 1,39 | 2,48 | 0,56 | 5,75E-01 |
| AT2G43690 | 0,68 | 1,62 | 4,30 | 0,38 | 7,06E-01 |
| AT2G43710 | 3,74 | -0,90 | 1,01 | -0,89 | 3,74E-01 |
| AT2G43790 | 0,39 | 1,46 | 2,82 | 0,52 | 6,04E-01 |
| AT2G43800 | 0,08 | 0,00 | 4,40 | 0,00 | 1,00E+00 |
| AT2G43820 | 0,45 | -0,42 | 3,18 | -0,13 | 8,96E-01 |
| AT2G43890 | 2,70 | 0,24 | 1,36 | 0,18 | 8,57E-01 |
| AT2G44060 | 2,21 | -0,34 | 1,27 | -0,27 | 7,86E-01 |
| AT2G44080 | 0,60 | -3,70 | 3,71 | -1,00 | 3,19E-01 |
| AT2G44100 | 0,65 | 2,10 | 2,56 | 0,82 | 4,11E-01 |
| AT2G44130 | 0,72 | 0,79 | 2,52 | 0,31 | 7,54E-01 |
| AT2G44160 | 0,80 | -0,82 | 2,34 | -0,35 | 7,25E-01 |
| AT2G44175 | 0,43 | 0,19 | 4,16 | 0,05 | 9,63E-01 |
| AT2G44200 | 0,72 | 1,42 | 2,52 | 0,56 | 5,73E-01 |
| AT2G44210 | 1,13 | -2,19 | 1,88 | -1,17 | 2,43E-01 |
| AT2G44230 | 0,76 | -0,01 | 2,84 | -0,01 | 9,96E-01 |
| AT2G44240 | 0,17 | 0,03 | 4,39 | 0,01 | 9,95E-01 |
| AT2G44260 | 0,20 | 1,26 | 4,34 | 0,29 | 7,71E-01 |
| AT2G44310 | 1,85 | 0,84 | 1,50 | 0,56 | 5,77E-01 |
| AT2G44350 | 0,40 | 0,00 | 3,44 | 0,00 | 9,99E-01 |
| AT2G44410 | 2,51 | 0,40 | 1,60 | 0,25 | 8,03E-01 |
| AT2G44420 | 0,20 | -0,31 | 4,36 | -0,07 | 9,44E-01 |
| AT2G44430 | 0,96 | 0,82 | 1,96 | 0,42 | 6,74E-01 |
| AT2G44500 | 0,91 | 2,84 | 2,85 | 1,00 | 3,19E-01 |
| AT2G44520 | 0,23 | -0,05 | 4,34 | -0,01 | 9,91E-01 |
| AT2G44530 | 0,99 | 1,95 | 2,22 | 0,88 | 3,78E-01 |
| AT2G44600 | 0,10 | 0,21 | 4,38 | 0,05 | 9,62E-01 |
| AT2G44640 | 1,50 | -1,01 | 1,45 | -0,70 | 4,84E-01 |
| AT2G44670 | 1,15 | -0,06 | 1,95 | -0,03 | 9,75E-01 |
| AT2G44710 | 1,40 | -1,01 | 1,46 | -0,69 | 4,90E-01 |
| AT2G44735 | 0,42 | 0,72 | 4,08 | 0,18 | 8,59E-01 |
| AT2G44770 | 0,15 | 1,16 | 4,34 | 0,27 | 7,88E-01 |
| AT2G44800 | 0,19 | 0,07 | 4,40 | 0,02 | 9,88E-01 |

| | | | | | |
|---|---|---|---|---|---|
| AT2G44870 | 0,51 | -0,88 | 2,73 | -0,32 | 7,46E-01 |
| AT2G44880 | 0,91 | 0,21 | 2,61 | 0,08 | 9,37E-01 |
| AT2G44925 | 0,92 | -0,92 | 1,87 | -0,49 | 6,22E-01 |
| AT2G44940 | 0,52 | -0,94 | 2,72 | -0,34 | 7,31E-01 |
| AT2G45000 | 0,38 | -1,78 | 3,51 | -0,51 | 6,12E-01 |
| AT2G45020 | 0,36 | 0,56 | 4,36 | 0,13 | 8,97E-01 |
| AT2G45030 | 0,91 | 0,81 | 2,68 | 0,30 | 7,64E-01 |
| AT2G45140 | 0,33 | -0,25 | 4,38 | -0,06 | 9,55E-01 |
| AT2G45180 | 2,18 | 0,76 | 1,73 | 0,44 | 6,60E-01 |
| AT2G45240 | 0,80 | 1,20 | 3,01 | 0,40 | 6,90E-01 |
| AT2G45250 | 0,26 | -0,48 | 4,38 | -0,11 | 9,13E-01 |
| AT2G45300 | 0,10 | 0,01 | 4,38 | 0,00 | 9,99E-01 |
| AT2G45310 | 0,67 | 0,66 | 2,50 | 0,26 | 7,93E-01 |
| AT2G45390 | 0,17 | 0,00 | 4,41 | 0,00 | 1,00E+00 |
| AT2G45440 | 0,53 | -1,66 | 3,83 | -0,43 | 6,64E-01 |
| AT2G45510 | 0,84 | -0,55 | 2,02 | -0,27 | 7,85E-01 |
| AT2G45530 | 0,98 | 0,04 | 1,99 | 0,02 | 9,85E-01 |
| AT2G09650 | 0,36 | 0,45 | 4,36 | 0,10 | 9,18E-01 |
| AT2G45590 | 0,37 | -0,54 | 4,31 | -0,12 | 9,01E-01 |
| AT2G45620 | 1,35 | -0,83 | 1,67 | -0,50 | 6,19E-01 |
| AT2G45630 | 0,80 | 0,63 | 2,49 | 0,25 | 7,99E-01 |
| AT2G45685 | 1,79 | 0,90 | 1,52 | 0,59 | 5,54E-01 |
| AT2G45710 | 0,45 | -2,11 | 4,38 | -0,48 | 6,29E-01 |
| AT2G45720 | 0,29 | 0,36 | 4,37 | 0,08 | 9,35E-01 |
| AT2G45740 | 0,35 | 0,88 | 3,58 | 0,24 | 8,07E-01 |
| AT2G45790 | 0,17 | 0,00 | 4,41 | 0,00 | 1,00E+00 |
| AT2G45810 | 0,90 | 2,81 | 2,38 | 1,18 | 2,39E-01 |
| AT2G45910 | 0,97 | -0,01 | 1,90 | 0,00 | 9,97E-01 |
| AT2G09670 | 5,26 | -0,83 | 0,80 | -1,05 | 2,94E-01 |
| AT2G45940 | 2,82 | 0,58 | 1,07 | 0,54 | 5,89E-01 |
| AT2G46020 | 3,28 | -0,59 | 0,96 | -0,62 | 5,39E-01 |
| AT2G46100 | 0,37 | 0,04 | 2,94 | 0,01 | 9,90E-01 |
| AT2G46130 | 1,45 | -0,78 | 1,70 | -0,46 | 6,48E-01 |
| AT2G46170 | 0,69 | -1,77 | 3,64 | -0,49 | 6,27E-01 |
| AT2G46190 | 0,70 | 0,76 | 2,26 | 0,33 | 7,38E-01 |
| AT2G09685 | 3,15 | 0,26 | 1,26 | 0,21 | 8,37E-01 |
| AT2G09695 | 1,56 | -1,00 | 1,53 | -0,65 | 5,14E-01 |
| AT2G09700 | 2,16 | 1,20 | 1,62 | 0,74 | 4,58E-01 |
| AT2G09705 | 0,89 | 1,18 | 2,61 | 0,45 | 6,53E-01 |
| AT2G46220 | 1,83 | -0,17 | 1,48 | -0,11 | 9,11E-01 |
| AT2G46225 | 0,27 | 0,84 | 4,36 | 0,19 | 8,48E-01 |
| AT2G46240 | 0,70 | 2,29 | 2,51 | 0,91 | 3,61E-01 |
| AT2G46260 | 0,08 | 0,44 | 4,38 | 0,10 | 9,21E-01 |
| AT2G46270 | 0,25 | -0,23 | 4,35 | -0,05 | 9,58E-01 |
| AT2G46370 | 0,44 | -1,44 | 3,42 | -0,42 | 6,73E-01 |
| AT2G46380 | 1,50 | -1,24 | 1,93 | -0,64 | 5,20E-01 |
| AT2G46420 | 0,84 | -1,20 | 3,03 | -0,40 | 6,91E-01 |
| AT2G46430 | 2,08 | -0,10 | 1,27 | -0,08 | 9,37E-01 |
| AT2G46440 | 1,43 | -0,84 | 1,39 | -0,60 | 5,48E-01 |
| AT2G46450 | 1,09 | 0,19 | 2,00 | 0,09 | 9,25E-01 |
| AT2G46460 | 2,59 | 0,00 | 2,08 | 0,00 | 1,00E+00 |
| AT2G46490 | 0,75 | 2,79 | 2,83 | 0,99 | 3,24E-01 |

| | | | | | |
|---|---|---|---|---|---|
| AT2G46520 | 1,16 | 1,14 | 1,73 | 0,66 | 5,10E-01 |
| AT2G46570 | 0,51 | 1,14 | 3,99 | 0,29 | 7,74E-01 |
| AT2G46590 | 0,52 | 1,74 | 3,03 | 0,57 | 5,66E-01 |
| AT2G46600 | 1,11 | -1,53 | 1,63 | -0,94 | 3,50E-01 |
| AT2G46630 | 0,28 | 0,60 | 4,36 | 0,14 | 8,91E-01 |
| AT2G46650 | 0,42 | 0,40 | 3,30 | 0,12 | 9,04E-01 |
| AT2G46690 | 0,43 | 1,23 | 4,17 | 0,29 | 7,68E-01 |
| AT2G46710 | 0,30 | 0,16 | 3,72 | 0,04 | 9,65E-01 |
| AT2G46765 | 2,57 | -0,35 | 1,23 | -0,29 | 7,75E-01 |
| AT2G46790 | 0,54 | 0,74 | 2,56 | 0,29 | 7,73E-01 |
| AT2G46800 | 1,02 | -0,58 | 1,81 | -0,32 | 7,47E-01 |
| AT2G46810 | 0,59 | 0,38 | 3,24 | 0,12 | 9,06E-01 |
| AT2G46820 | 2,21 | -0,20 | 1,37 | -0,14 | 8,87E-01 |
| AT2G46830 | 2,43 | -0,57 | 1,21 | -0,47 | 6,39E-01 |
| AT2G46900 | 1,50 | -0,81 | 1,88 | -0,43 | 6,67E-01 |
| AT2G46910 | 0,44 | 0,84 | 4,29 | 0,20 | 8,45E-01 |
| AT2G46990 | 0,08 | 0,44 | 4,38 | 0,10 | 9,21E-01 |
| AT2G47110 | 0,66 | -0,02 | 2,30 | -0,01 | 9,92E-01 |
| AT2G47170 | 1,90 | 0,63 | 1,56 | 0,40 | 6,88E-01 |
| AT2G09815 | 0,23 | 0,19 | 4,38 | 0,04 | 9,66E-01 |
| AT2G47190 | 0,40 | 2,10 | 4,32 | 0,49 | 6,27E-01 |
| AT2G47220 | 0,25 | 1,40 | 4,33 | 0,32 | 7,47E-01 |
| AT2G47240 | 1,70 | 0,23 | 1,59 | 0,14 | 8,85E-01 |
| AT2G47320 | 0,28 | 0,13 | 4,41 | 0,03 | 9,76E-01 |
| AT2G47410 | 3,41 | 0,74 | 1,30 | 0,57 | 5,72E-01 |
| AT2G47420 | 0,42 | 1,59 | 4,12 | 0,39 | 7,00E-01 |
| AT2G47440 | 2,19 | -0,73 | 1,49 | -0,49 | 6,24E-01 |
| AT2G47470 | 1,77 | 1,18 | 1,76 | 0,67 | 5,01E-01 |
| AT2G47510 | 1,59 | -0,45 | 1,49 | -0,30 | 7,65E-01 |
| AT2G47610 | 1,29 | 1,44 | 1,69 | 0,85 | 3,95E-01 |
| AT2G47620 | 0,26 | 0,84 | 3,79 | 0,22 | 8,25E-01 |
| AT2G47630 | 0,42 | -0,12 | 2,86 | -0,04 | 9,67E-01 |
| AT2G47640 | 0,28 | -1,60 | 4,38 | -0,36 | 7,16E-01 |
| AT2G47690 | 0,55 | -1,06 | 3,99 | -0,27 | 7,89E-01 |
| AT2G47730 | 0,80 | -0,50 | 2,34 | -0,21 | 8,32E-01 |
| AT2G47800 | 2,70 | 0,32 | 1,09 | 0,30 | 7,66E-01 |
| AT2G47820 | 1,37 | -0,26 | 2,28 | -0,11 | 9,11E-01 |
| AT2G47850 | 0,53 | -1,87 | 3,10 | -0,60 | 5,47E-01 |
| AT2G47910 | 0,92 | 1,04 | 2,08 | 0,50 | 6,17E-01 |
| AT2G47935 | 0,72 | -0,16 | 2,74 | -0,06 | 9,54E-01 |
| AT2G47960 | 1,39 | -1,05 | 1,63 | -0,64 | 5,20E-01 |
| AT2G47970 | 0,70 | 0,17 | 3,39 | 0,05 | 9,59E-01 |
| AT2G47980 | 2,39 | 0,90 | 1,20 | 0,75 | 4,55E-01 |
| AT2G48010 | 0,33 | 1,00 | 4,36 | 0,23 | 8,19E-01 |
| AT2G48020 | 0,56 | -0,30 | 3,18 | -0,09 | 9,24E-01 |
| AT2G48110 | 1,17 | -1,75 | 1,92 | -0,91 | 3,61E-01 |
| AT2G48120 | 0,32 | 0,02 | 4,35 | 0,01 | 9,96E-01 |
| AT3G01040 | 0,51 | 1,12 | 3,11 | 0,36 | 7,19E-01 |
| AT3G01050 | 0,18 | 0,43 | 4,36 | 0,10 | 9,22E-01 |
| AT3G01080 | 0,17 | 0,61 | 4,37 | 0,14 | 8,88E-01 |
| AT3G01085 | 1,09 | 1,37 | 1,94 | 0,70 | 4,81E-01 |
| AT3G01150 | 0,45 | -1,75 | 3,26 | -0,54 | 5,92E-01 |

| | | | | | |
|---|---|---|---|---|---|
| AT3G01170 | 0,42 | 0,40 | 4,12 | 0,10 | 9,23E-01 |
| AT3G01175 | 0,08 | 0,69 | 4,36 | 0,16 | 8,75E-01 |
| AT3G01210 | 0,10 | 0,01 | 4,38 | 0,00 | 9,99E-01 |
| AT3G01280 | 1,02 | -2,96 | 1,79 | -1,65 | 9,89E-02 |
| AT3G01320 | 1,14 | -0,30 | 1,61 | -0,19 | 8,51E-01 |
| AT3G01370 | 2,22 | 0,29 | 1,38 | 0,21 | 8,34E-01 |
| AT3G01440 | 0,31 | 0,00 | 4,41 | 0,00 | 1,00E+00 |
| AT3G01460 | 5,55 | -0,22 | 0,82 | -0,27 | 7,85E-01 |
| AT3G01475 | 1,05 | -0,70 | 1,71 | -0,41 | 6,81E-01 |
| AT3G01480 | 1,15 | -1,89 | 1,92 | -0,98 | 3,25E-01 |
| AT3G01560 | 0,68 | -0,16 | 2,45 | -0,06 | 9,49E-01 |
| AT3G01590 | 0,40 | 1,45 | 4,35 | 0,33 | 7,38E-01 |
| AT3G01610 | 0,25 | 0,74 | 4,36 | 0,17 | 8,66E-01 |
| AT3G01660 | 0,12 | 0,97 | 4,34 | 0,22 | 8,23E-01 |
| AT3G01690 | 1,08 | -1,07 | 2,01 | -0,53 | 5,96E-01 |
| AT3G01740 | 0,28 | 1,57 | 4,32 | 0,36 | 7,16E-01 |
| AT3G01750 | 0,72 | -0,87 | 2,88 | -0,30 | 7,63E-01 |
| AT3G01780 | 1,57 | -0,55 | 1,37 | -0,40 | 6,87E-01 |
| AT3G01810 | 0,37 | 0,26 | 4,39 | 0,06 | 9,53E-01 |
| AT3G01830 | 0,94 | -0,68 | 2,13 | -0,32 | 7,49E-01 |
| AT3G01920 | 0,37 | -0,69 | 4,39 | -0,16 | 8,76E-01 |
| AT3G02010 | 0,37 | -0,64 | 4,30 | -0,15 | 8,81E-01 |
| AT3G02030 | 0,12 | 0,49 | 4,36 | 0,11 | 9,11E-01 |
| AT3G02050 | 0,37 | -0,47 | 4,25 | -0,11 | 9,11E-01 |
| AT3G02060 | 1,23 | 0,58 | 1,98 | 0,29 | 7,69E-01 |
| AT3G02065 | 0,54 | 0,12 | 3,73 | 0,03 | 9,75E-01 |
| AT3G02070 | 0,35 | -0,31 | 3,56 | -0,09 | 9,30E-01 |
| AT3G02090 | 3,68 | -0,06 | 0,89 | -0,07 | 9,46E-01 |
| AT3G02130 | 1,18 | 0,61 | 2,05 | 0,30 | 7,67E-01 |
| AT3G02200 | 1,75 | -0,05 | 1,67 | -0,03 | 9,78E-01 |
| AT3G02280 | 0,25 | -1,82 | 4,40 | -0,41 | 6,79E-01 |
| AT3G02290 | 0,16 | 0,49 | 4,38 | 0,11 | 9,12E-01 |
| AT3G02300 | 0,16 | 0,00 | 4,41 | 0,00 | 1,00E+00 |
| AT3G02340 | 0,50 | 0,87 | 4,34 | 0,20 | 8,41E-01 |
| AT3G02350 | 0,84 | -0,81 | 2,24 | -0,36 | 7,19E-01 |
| AT3G02360 | 1,53 | -1,32 | 1,63 | -0,81 | 4,16E-01 |
| AT3G02370 | 0,47 | 0,55 | 3,34 | 0,17 | 8,68E-01 |
| AT3G02400 | 0,66 | -4,03 | 4,30 | -0,94 | 3,48E-01 |
| AT3G02490 | 0,33 | 0,02 | 4,35 | 0,01 | 9,96E-01 |
| AT3G02560 | 0,49 | 1,09 | 3,90 | 0,28 | 7,80E-01 |
| AT3G02610 | 0,81 | 0,73 | 2,91 | 0,25 | 8,02E-01 |
| AT3G02630 | 0,37 | -1,44 | 4,28 | -0,34 | 7,36E-01 |
| AT3G02690 | 0,66 | -1,27 | 2,44 | -0,52 | 6,02E-01 |
| AT3G02720 | 0,46 | -1,01 | 4,32 | -0,23 | 8,15E-01 |
| AT3G02830 | 1,22 | 0,50 | 2,07 | 0,24 | 8,11E-01 |
| AT3G02832 | 1,51 | 0,35 | 1,71 | 0,20 | 8,39E-01 |
| AT3G01145 | 0,57 | 1,31 | 2,60 | 0,51 | 6,13E-01 |
| AT3G02880 | 1,19 | -1,61 | 1,55 | -1,04 | 2,98E-01 |
| AT3G02900 | 0,28 | 0,00 | 4,41 | 0,00 | 1,00E+00 |
| AT3G02910 | 0,77 | 0,43 | 2,34 | 0,18 | 8,54E-01 |
| AT3G03020 | 0,43 | 1,70 | 4,15 | 0,41 | 6,82E-01 |
| AT3G03030 | 0,40 | 2,10 | 4,32 | 0,49 | 6,27E-01 |

| | | | | | |
|---|---|---|---|---|---|
| AT3G03040 | 0,24 | -0,73 | 4,37 | -0,17 | 8,68E-01 |
| AT3G03050 | 2,11 | 0,64 | 1,49 | 0,43 | 6,69E-01 |
| AT3G03110 | 0,55 | 0,33 | 4,31 | 0,08 | 9,38E-01 |
| AT3G03160 | 0,47 | 1,11 | 3,98 | 0,28 | 7,80E-01 |
| AT3G03210 | 0,10 | 0,00 | 4,41 | 0,00 | 1,00E+00 |
| AT3G03320 | 0,15 | 0,00 | 4,41 | 0,00 | 1,00E+00 |
| AT3G03340 | 0,40 | 2,13 | 4,32 | 0,49 | 6,22E-01 |
| AT3G03350 | 0,24 | 0,21 | 4,34 | 0,05 | 9,62E-01 |
| AT3G03360 | 0,08 | 0,69 | 4,36 | 0,16 | 8,75E-01 |
| AT3G03380 | 4,30 | 0,04 | 0,87 | 0,05 | 9,63E-01 |
| AT3G03610 | 0,27 | 0,09 | 4,40 | 0,02 | 9,84E-01 |
| AT3G03640 | 0,25 | 0,00 | 4,38 | 0,00 | 1,00E+00 |
| AT3G03680 | 1,60 | 0,08 | 1,62 | 0,05 | 9,58E-01 |
| AT3G03710 | 4,42 | -0,63 | 0,91 | -0,69 | 4,88E-01 |
| AT3G03780 | 4,16 | -0,89 | 0,93 | -0,96 | 3,39E-01 |
| AT3G03790 | 1,73 | 0,10 | 1,37 | 0,08 | 9,40E-01 |
| AT3G03860 | 0,25 | 0,02 | 4,39 | 0,00 | 9,97E-01 |
| AT3G03870 | 3,66 | -1,33 | 0,96 | -1,39 | 1,65E-01 |
| AT3G03880 | 0,28 | 0,86 | 4,36 | 0,20 | 8,44E-01 |
| AT3G03950 | 0,32 | 1,16 | 3,56 | 0,33 | 7,44E-01 |
| AT3G03960 | 1,45 | 1,27 | 1,58 | 0,80 | 4,22E-01 |
| AT3G03990 | 2,37 | -1,01 | 1,32 | -0,77 | 4,42E-01 |
| AT3G04130 | 2,12 | -0,13 | 1,72 | -0,07 | 9,42E-01 |
| AT3G04140 | 0,99 | 0,20 | 2,14 | 0,09 | 9,27E-01 |
| AT3G04250 | 2,26 | 2,98 | 1,88 | 1,59 | 1,13E-01 |
| AT3G04400 | 2,05 | -1,44 | 1,32 | -1,09 | 2,77E-01 |
| AT3G04490 | 0,26 | -1,78 | 4,41 | -0,40 | 6,87E-01 |
| AT3G04530 | 0,08 | 0,69 | 4,36 | 0,16 | 8,75E-01 |
| AT3G04550 | 0,64 | 1,14 | 2,24 | 0,51 | 6,12E-01 |
| AT3G04740 | 2,26 | -0,67 | 1,16 | -0,58 | 5,62E-01 |
| AT3G04790 | 1,26 | -0,65 | 2,02 | -0,32 | 7,50E-01 |
| AT3G04810 | 0,59 | 1,99 | 4,32 | 0,46 | 6,44E-01 |
| AT3G04840 | 0,80 | -0,77 | 2,20 | -0,35 | 7,26E-01 |
| AT3G04870 | 0,49 | -1,72 | 3,24 | -0,53 | 5,94E-01 |
| AT3G04910 | 0,86 | 0,84 | 1,94 | 0,43 | 6,66E-01 |
| AT3G04920 | 0,51 | -1,61 | 4,04 | -0,40 | 6,91E-01 |
| AT3G04930 | 1,93 | -0,24 | 1,37 | -0,18 | 8,60E-01 |
| AT3G04940 | 0,44 | -0,30 | 4,14 | -0,07 | 9,41E-01 |
| AT3G01555 | 0,12 | 0,37 | 4,37 | 0,08 | 9,33E-01 |
| AT3G01575 | 0,10 | 0,00 | 4,41 | 0,00 | 1,00E+00 |
| AT3G05070 | 0,17 | 0,02 | 4,39 | 0,00 | 9,97E-01 |
| AT3G05120 | 0,72 | 0,88 | 2,22 | 0,39 | 6,93E-01 |
| AT3G05200 | 0,41 | -0,54 | 4,21 | -0,13 | 8,98E-01 |
| AT3G05220 | 1,84 | 1,53 | 1,81 | 0,85 | 3,98E-01 |
| AT3G05230 | 0,28 | 1,14 | 4,36 | 0,26 | 7,94E-01 |
| AT3G05380 | 0,92 | -0,31 | 2,08 | -0,15 | 8,81E-01 |
| AT3G05410 | 1,68 | 0,12 | 1,45 | 0,08 | 9,33E-01 |
| AT3G05420 | 2,70 | -1,19 | 1,09 | -1,09 | 2,75E-01 |
| AT3G05430 | 0,21 | 1,57 | 4,33 | 0,36 | 7,16E-01 |
| AT3G05490 | 0,28 | 0,00 | 4,41 | 0,00 | 1,00E+00 |
| AT3G05500 | 0,55 | -0,39 | 2,62 | -0,15 | 8,81E-01 |
| AT3G05520 | 0,43 | -0,13 | 2,87 | -0,04 | 9,64E-01 |

| | | | | | |
|---|---|---|---|---|---|
| AT3G05530 | 1,78 | -1,29 | 1,28 | -1,01 | 3,15E-01 |
| AT3G05545 | 0,82 | 1,27 | 2,23 | 0,57 | 5,69E-01 |
| AT3G05560 | 0,45 | 0,79 | 3,31 | 0,24 | 8,11E-01 |
| AT3G05590 | 1,96 | -1,87 | 1,31 | -1,43 | 1,54E-01 |
| AT3G05630 | 0,21 | 1,76 | 4,32 | 0,41 | 6,85E-01 |
| AT3G05670 | 0,41 | 0,54 | 4,18 | 0,13 | 8,97E-01 |
| AT3G05710 | 0,95 | -1,30 | 1,98 | -0,65 | 5,13E-01 |
| AT3G05725 | 0,64 | 0,01 | 2,58 | 0,00 | 9,97E-01 |
| AT3G05750 | 0,61 | 2,60 | 3,64 | 0,71 | 4,75E-01 |
| AT3G05760 | 0,68 | 2,59 | 3,63 | 0,71 | 4,75E-01 |
| AT3G05830 | 0,23 | 0,00 | 4,41 | 0,00 | 1,00E+00 |
| AT3G05840 | 0,50 | -1,77 | 3,90 | -0,45 | 6,51E-01 |
| AT3G05905 | 0,51 | -0,84 | 3,26 | -0,26 | 7,98E-01 |
| AT3G06125 | 0,60 | -0,26 | 2,21 | -0,12 | 9,08E-01 |
| AT3G06170 | 0,51 | 1,49 | 3,06 | 0,49 | 6,25E-01 |
| AT3G06180 | 0,25 | 1,94 | 4,32 | 0,45 | 6,54E-01 |
| AT3G06200 | 0,41 | -0,04 | 4,36 | -0,01 | 9,93E-01 |
| AT3G06210 | 0,20 | -0,31 | 4,36 | -0,07 | 9,44E-01 |
| AT3G06240 | 0,61 | -0,57 | 2,26 | -0,25 | 8,00E-01 |
| AT3G06380 | 0,55 | 0,55 | 3,86 | 0,14 | 8,87E-01 |
| AT3G06400 | 0,95 | 1,56 | 2,53 | 0,61 | 5,39E-01 |
| AT3G06455 | 0,58 | -0,65 | 4,35 | -0,15 | 8,81E-01 |
| AT3G06470 | 0,36 | 1,22 | 4,34 | 0,28 | 7,78E-01 |
| AT3G06500 | 1,24 | 0,29 | 2,05 | 0,14 | 8,87E-01 |
| AT3G06510 | 3,56 | 1,16 | 1,12 | 1,04 | 2,98E-01 |
| AT3G06520 | 0,50 | -0,11 | 3,87 | -0,03 | 9,78E-01 |
| AT3G06530 | 4,41 | 0,25 | 0,90 | 0,27 | 7,83E-01 |
| AT3G06610 | 0,15 | 0,29 | 4,37 | 0,07 | 9,47E-01 |
| AT3G06660 | 0,08 | 0,00 | 4,40 | 0,00 | 1,00E+00 |
| AT3G06720 | 1,89 | -0,57 | 1,55 | -0,36 | 7,15E-01 |
| AT3G06730 | 0,97 | -2,05 | 3,02 | -0,68 | 4,97E-01 |
| AT3G06740 | 0,50 | -1,36 | 4,13 | -0,33 | 7,41E-01 |
| AT3G06810 | 1,67 | -0,81 | 1,35 | -0,60 | 5,50E-01 |
| AT3G06830 | 0,08 | 0,69 | 4,36 | 0,16 | 8,75E-01 |
| AT3G06860 | 1,03 | -0,66 | 1,89 | -0,35 | 7,29E-01 |
| AT3G06868 | 0,29 | -1,87 | 4,40 | -0,43 | 6,70E-01 |
| AT3G06895 | 1,19 | -0,38 | 1,84 | -0,20 | 8,38E-01 |
| AT3G06920 | 0,42 | 0,04 | 4,28 | 0,01 | 9,92E-01 |
| AT3G06980 | 0,81 | 0,78 | 2,91 | 0,27 | 7,89E-01 |
| AT3G07110 | 1,39 | 0,22 | 1,66 | 0,13 | 8,93E-01 |
| AT3G07200 | 0,08 | 0,69 | 4,36 | 0,16 | 8,75E-01 |
| AT3G07230 | 0,29 | 0,23 | 4,39 | 0,05 | 9,58E-01 |
| AT3G07270 | 0,85 | -1,31 | 1,95 | -0,67 | 5,01E-01 |
| AT3G07350 | 0,42 | 0,25 | 3,30 | 0,08 | 9,40E-01 |
| AT3G07360 | 1,00 | 0,49 | 2,41 | 0,20 | 8,39E-01 |
| AT3G07390 | 2,06 | 0,05 | 1,66 | 0,03 | 9,76E-01 |
| AT3G07400 | 0,87 | -1,22 | 1,96 | -0,62 | 5,33E-01 |
| AT3G07430 | 0,32 | 0,22 | 4,36 | 0,05 | 9,60E-01 |
| AT3G07440 | 0,08 | 0,69 | 4,36 | 0,16 | 8,75E-01 |
| AT3G07480 | 0,45 | -0,10 | 4,23 | -0,02 | 9,81E-01 |
| AT3G07565 | 0,19 | 0,38 | 4,37 | 0,09 | 9,31E-01 |
| AT3G07610 | 0,83 | -1,87 | 2,50 | -0,75 | 4,54E-01 |

| | | | | | |
|---|---|---|---|---|---|
| AT3G07640 | 0,17 | 1,52 | 4,33 | 0,35 | 7,26E-01 |
| AT3G07650 | 0,97 | 1,38 | 3,29 | 0,42 | 6,75E-01 |
| AT3G07680 | 0,59 | -1,28 | 3,14 | -0,41 | 6,84E-01 |
| AT3G07690 | 0,32 | -0,94 | 4,31 | -0,22 | 8,28E-01 |
| AT3G07770 | 1,44 | -1,52 | 1,50 | -1,01 | 3,13E-01 |
| AT3G07780 | 1,15 | 0,89 | 2,10 | 0,42 | 6,71E-01 |
| AT3G07790 | 2,60 | -0,27 | 1,11 | -0,25 | 8,06E-01 |
| AT3G07810 | 0,90 | -1,74 | 1,87 | -0,93 | 3,53E-01 |
| AT3G07860 | 5,76 | 1,90 | 1,14 | 1,67 | 9,52E-02 |
| AT3G02045 | 5,72 | 1,83 | 1,14 | 1,61 | 1,08E-01 |
| AT3G07870 | 5,57 | -1,05 | 0,96 | -1,09 | 2,74E-01 |
| AT3G07880 | 0,28 | 1,73 | 4,32 | 0,40 | 6,89E-01 |
| AT3G07950 | 0,43 | 2,04 | 4,32 | 0,47 | 6,37E-01 |
| AT3G08010 | 0,17 | 0,00 | 4,41 | 0,00 | 1,00E+00 |
| AT3G08030 | 1,37 | -1,73 | 1,62 | -1,07 | 2,86E-01 |
| AT3G08550 | 0,31 | -0,55 | 4,35 | -0,13 | 8,99E-01 |
| AT3G08560 | 1,47 | -0,62 | 1,72 | -0,36 | 7,19E-01 |
| AT3G08600 | 0,52 | 1,29 | 3,88 | 0,33 | 7,39E-01 |
| AT3G08640 | 0,20 | 0,75 | 4,37 | 0,17 | 8,64E-01 |
| AT3G08690 | 0,22 | -0,91 | 4,38 | -0,21 | 8,35E-01 |
| AT3G08920 | 0,32 | -0,82 | 4,35 | -0,19 | 8,51E-01 |
| AT3G08930 | 0,51 | 0,73 | 3,21 | 0,23 | 8,20E-01 |
| AT3G08947 | 5,24 | -0,14 | 0,89 | -0,16 | 8,72E-01 |
| AT3G08950 | 0,44 | -2,81 | 4,22 | -0,67 | 5,05E-01 |
| AT3G08980 | 0,33 | -1,22 | 4,34 | -0,28 | 7,79E-01 |
| AT3G09020 | 0,57 | 2,61 | 3,98 | 0,66 | 5,12E-01 |
| AT3G09030 | 0,51 | -0,48 | 2,55 | -0,19 | 8,51E-01 |
| AT3G09035 | 0,19 | 1,22 | 4,33 | 0,28 | 7,78E-01 |
| AT3G09085 | 0,40 | 0,92 | 3,37 | 0,27 | 7,84E-01 |
| AT3G02105 | 0,40 | 2,09 | 4,32 | 0,48 | 6,28E-01 |
| AT3G09180 | 0,23 | -0,48 | 4,35 | -0,11 | 9,12E-01 |
| AT3G09300 | 1,07 | -1,74 | 2,37 | -0,73 | 4,63E-01 |
| AT3G09350 | 0,51 | -2,33 | 4,39 | -0,53 | 5,95E-01 |
| AT3G09370 | 0,29 | 0,51 | 4,33 | 0,12 | 9,07E-01 |
| AT3G09430 | 1,27 | 1,08 | 1,86 | 0,58 | 5,60E-01 |
| AT3G09500 | 0,81 | -1,39 | 2,02 | -0,69 | 4,90E-01 |
| AT3G09510 | 0,52 | 0,24 | 3,17 | 0,07 | 9,41E-01 |
| AT3G09570 | 0,70 | 0,36 | 2,45 | 0,15 | 8,82E-01 |
| AT3G09600 | 0,41 | -0,61 | 3,27 | -0,19 | 8,52E-01 |
| AT3G09630 | 1,89 | -0,38 | 1,49 | -0,26 | 7,97E-01 |
| AT3G09650 | 0,58 | 1,59 | 4,31 | 0,37 | 7,11E-01 |
| AT3G09670 | 0,48 | 1,17 | 4,32 | 0,27 | 7,86E-01 |
| AT3G09690 | 0,55 | -2,06 | 4,30 | -0,48 | 6,31E-01 |
| AT3G09810 | 0,32 | -0,49 | 4,34 | -0,11 | 9,11E-01 |
| AT3G09820 | 0,77 | 1,45 | 2,42 | 0,60 | 5,51E-01 |
| AT3G09830 | 0,46 | 0,81 | 3,29 | 0,25 | 8,06E-01 |
| AT3G09880 | 0,75 | 0,87 | 2,42 | 0,36 | 7,21E-01 |
| AT3G09890 | 0,43 | 0,75 | 4,31 | 0,17 | 8,63E-01 |
| AT3G09900 | 0,19 | 1,46 | 4,33 | 0,34 | 7,35E-01 |
| AT3G10060 | 0,36 | -0,12 | 4,35 | -0,03 | 9,78E-01 |
| AT3G10120 | 0,12 | 1,05 | 4,35 | 0,24 | 8,10E-01 |
| AT3G10190 | 0,32 | 2,00 | 4,32 | 0,46 | 6,44E-01 |

| | | | | | |
|---|---|---|---|---|---|
| AT3G10200 | 0,08 | 0,69 | 4,36 | 0,16 | 8,75E-01 |
| AT3G10300 | 0,68 | 0,02 | 3,00 | 0,01 | 9,93E-01 |
| AT3G10310 | 0,16 | 0,65 | 4,37 | 0,15 | 8,82E-01 |
| AT3G10340 | 0,80 | 0,36 | 3,08 | 0,12 | 9,06E-01 |
| AT3G10370 | 0,28 | 1,35 | 4,33 | 0,31 | 7,55E-01 |
| AT3G10390 | 0,49 | -1,15 | 3,19 | -0,36 | 7,19E-01 |
| AT3G10420 | 1,49 | 0,62 | 1,53 | 0,41 | 6,84E-01 |
| AT3G10480 | 0,72 | 0,99 | 3,45 | 0,29 | 7,74E-01 |
| AT3G10490 | 0,18 | 0,43 | 4,36 | 0,10 | 9,22E-01 |
| AT3G10527 | 0,27 | 1,49 | 4,33 | 0,34 | 7,30E-01 |
| AT3G10540 | 0,56 | -1,27 | 2,60 | -0,49 | 6,25E-01 |
| AT3G10570 | 0,12 | 0,37 | 4,37 | 0,08 | 9,33E-01 |
| AT3G10760 | 0,48 | -1,21 | 3,29 | -0,37 | 7,13E-01 |
| AT3G10800 | 1,54 | 0,24 | 1,73 | 0,14 | 8,90E-01 |
| AT3G10820 | 0,24 | -2,36 | 4,39 | -0,54 | 5,91E-01 |
| AT3G10860 | 0,23 | 1,27 | 4,33 | 0,29 | 7,70E-01 |
| AT3G10920 | 1,12 | -0,56 | 1,98 | -0,29 | 7,76E-01 |
| AT3G10950 | 0,19 | 0,00 | 4,38 | 0,00 | 9,99E-01 |
| AT3G10980 | 0,79 | -0,87 | 3,00 | -0,29 | 7,72E-01 |
| AT3G11020 | 0,05 | 0,00 | 4,41 | 0,00 | 1,00E+00 |
| AT3G11080 | 0,18 | 0,21 | 4,37 | 0,05 | 9,62E-01 |
| AT3G11090 | 0,60 | 0,73 | 3,17 | 0,23 | 8,17E-01 |
| AT3G11120 | 0,39 | -1,78 | 3,46 | -0,51 | 6,07E-01 |
| AT3G11170 | 1,02 | -0,58 | 1,82 | -0,32 | 7,50E-01 |
| AT3G11230 | 0,53 | -2,17 | 4,38 | -0,50 | 6,20E-01 |
| AT3G11250 | 0,11 | 0,00 | 4,41 | 0,00 | 1,00E+00 |
| AT3G11270 | 0,10 | 0,01 | 4,38 | 0,00 | 9,99E-01 |
| AT3G11397 | 0,44 | -1,85 | 4,19 | -0,44 | 6,58E-01 |
| AT3G11400 | 0,87 | -2,03 | 2,16 | -0,94 | 3,47E-01 |
| AT3G11420 | 0,26 | 0,00 | 4,38 | 0,00 | 9,99E-01 |
| AT3G11440 | 0,65 | 0,31 | 2,91 | 0,11 | 9,16E-01 |
| AT3G11540 | 2,22 | -0,47 | 1,47 | -0,32 | 7,49E-01 |
| AT3G11560 | 1,70 | 1,67 | 1,85 | 0,90 | 3,67E-01 |
| AT3G02465 | 0,30 | -0,04 | 4,34 | -0,01 | 9,93E-01 |
| AT3G11630 | 0,96 | 2,27 | 1,87 | 1,21 | 2,25E-01 |
| AT3G11700 | 0,50 | 0,95 | 3,22 | 0,30 | 7,68E-01 |
| AT3G11760 | 0,23 | -0,05 | 4,34 | -0,01 | 9,91E-01 |
| AT3G11810 | 0,52 | -0,66 | 3,78 | -0,18 | 8,61E-01 |
| AT3G11830 | 1,90 | -1,21 | 1,39 | -0,87 | 3,82E-01 |
| AT3G11900 | 0,13 | 0,73 | 4,36 | 0,17 | 8,67E-01 |
| AT3G11930 | 1,35 | -0,48 | 3,42 | -0,14 | 8,88E-01 |
| AT3G11960 | 1,04 | 0,50 | 1,86 | 0,27 | 7,88E-01 |
| AT3G12010 | 0,40 | -1,45 | 4,35 | -0,33 | 7,40E-01 |
| AT3G12020 | 0,79 | -1,10 | 2,21 | -0,50 | 6,19E-01 |
| AT3G12050 | 0,32 | 1,19 | 3,53 | 0,34 | 7,36E-01 |
| AT3G12080 | 1,08 | -0,54 | 2,35 | -0,23 | 8,20E-01 |
| AT3G12270 | 0,77 | 0,19 | 2,68 | 0,07 | 9,42E-01 |
| AT3G12290 | 0,81 | -1,73 | 2,42 | -0,71 | 4,75E-01 |
| AT3G12320 | 0,90 | -1,94 | 1,91 | -1,02 | 3,09E-01 |
| AT3G12360 | 0,71 | -0,79 | 2,49 | -0,32 | 7,50E-01 |
| AT3G12390 | 0,69 | -0,74 | 2,37 | -0,31 | 7,54E-01 |
| AT3G12480 | 0,26 | 1,43 | 4,34 | 0,33 | 7,41E-01 |

| | | | | | |
|---|---|---|---|---|---|
| AT3G12502 | 0,08 | 0,69 | 4,36 | 0,16 | 8,75E-01 |
| AT3G12570 | 1,24 | 3,27 | 2,65 | 1,23 | 2,18E-01 |
| AT3G12600 | 0,05 | 0,00 | 4,41 | 0,00 | 1,00E+00 |
| AT3G12640 | 0,60 | -0,06 | 2,18 | -0,03 | 9,76E-01 |
| AT3G12700 | 0,20 | -1,80 | 4,41 | -0,41 | 6,83E-01 |
| AT3G12740 | 0,52 | 0,25 | 3,32 | 0,08 | 9,39E-01 |
| AT3G12760 | 0,80 | -0,32 | 2,22 | -0,15 | 8,85E-01 |
| AT3G12810 | 1,56 | -0,34 | 1,46 | -0,23 | 8,18E-01 |
| AT3G12915 | 0,53 | 0,62 | 4,29 | 0,14 | 8,86E-01 |
| AT3G12980 | 1,82 | -1,94 | 1,62 | -1,20 | 2,31E-01 |
| AT3G13060 | 0,87 | -0,69 | 2,11 | -0,33 | 7,45E-01 |
| AT3G13062 | 1,32 | -0,08 | 1,51 | -0,05 | 9,59E-01 |
| AT3G13065 | 0,88 | 0,33 | 3,27 | 0,10 | 9,19E-01 |
| AT3G13200 | 1,18 | 0,90 | 1,78 | 0,50 | 6,15E-01 |
| AT3G13210 | 0,13 | 0,73 | 4,36 | 0,17 | 8,67E-01 |
| AT3G13290 | 2,32 | 0,96 | 1,41 | 0,68 | 4,97E-01 |
| AT3G13340 | 1,00 | -2,62 | 2,26 | -1,16 | 2,46E-01 |
| AT3G13370 | 0,63 | 0,00 | 4,41 | 0,00 | 1,00E+00 |
| AT3G13380 | 0,67 | 1,49 | 2,43 | 0,61 | 5,40E-01 |
| AT3G13437 | 0,08 | 0,44 | 4,38 | 0,10 | 9,21E-01 |
| AT3G13440 | 3,42 | 0,88 | 1,11 | 0,80 | 4,26E-01 |
| AT3G13445 | 0,19 | 0,37 | 4,37 | 0,09 | 9,32E-01 |
| AT3G13470 | 0,61 | 0,21 | 3,83 | 0,05 | 9,56E-01 |
| AT3G13525 | 2,38 | 1,22 | 1,57 | 0,78 | 4,38E-01 |
| AT3G02725 | 1,24 | -2,11 | 1,89 | -1,12 | 2,64E-01 |
| AT3G13662 | 0,34 | -0,78 | 4,36 | -0,18 | 8,58E-01 |
| AT3G13690 | 0,67 | -1,45 | 3,85 | -0,38 | 7,06E-01 |
| AT3G13750 | 2,68 | -0,08 | 1,13 | -0,07 | 9,41E-01 |
| AT3G13760 | 0,08 | 0,44 | 4,38 | 0,10 | 9,21E-01 |
| AT3G13770 | 0,43 | -1,19 | 4,32 | -0,27 | 7,84E-01 |
| AT3G13810 | 0,42 | -1,07 | 4,20 | -0,26 | 7,98E-01 |
| AT3G13930 | 0,81 | -3,59 | 2,15 | -1,67 | 9,49E-02 |
| AT3G13950 | 0,55 | -0,38 | 3,98 | -0,10 | 9,24E-01 |
| AT3G14010 | 2,74 | -0,25 | 1,32 | -0,19 | 8,52E-01 |
| AT3G14020 | 0,35 | -2,17 | 4,38 | -0,49 | 6,21E-01 |
| AT3G14100 | 0,69 | 2,16 | 2,58 | 0,84 | 4,03E-01 |
| AT3G14120 | 1,41 | -1,28 | 1,76 | -0,73 | 4,67E-01 |
| AT3G14160 | 0,19 | 1,15 | 4,34 | 0,27 | 7,91E-01 |
| AT3G14220 | 0,64 | -3,62 | 3,74 | -0,97 | 3,34E-01 |
| AT3G14270 | 2,35 | -0,27 | 1,22 | -0,22 | 8,24E-01 |
| AT3G14290 | 0,42 | 0,58 | 4,12 | 0,14 | 8,88E-01 |
| AT3G14415 | 1,72 | -0,18 | 1,56 | -0,12 | 9,07E-01 |
| AT3G14420 | 4,42 | -0,36 | 1,10 | -0,32 | 7,46E-01 |
| AT3G14470 | 2,64 | -0,62 | 1,16 | -0,54 | 5,92E-01 |
| AT3G14560 | 0,20 | -0,31 | 4,36 | -0,07 | 9,44E-01 |
| AT3G14595 | 1,01 | 0,62 | 2,04 | 0,30 | 7,62E-01 |
| AT3G14600 | 1,18 | 0,20 | 1,63 | 0,12 | 9,02E-01 |
| AT3G14610 | 0,28 | -1,42 | 4,36 | -0,33 | 7,44E-01 |
| AT3G14620 | 0,63 | 0,07 | 2,55 | 0,03 | 9,79E-01 |
| AT3G14640 | 0,33 | -0,45 | 4,32 | -0,10 | 9,16E-01 |
| AT3G14650 | 2,33 | -0,14 | 1,17 | -0,12 | 9,02E-01 |
| AT3G14660 | 1,69 | 1,29 | 1,47 | 0,88 | 3,80E-01 |

| | | | | | |
|---|---|---|---|---|---|
| AT3G14670 | 1,07 | 0,66 | 2,00 | 0,33 | 7,42E-01 |
| AT3G14690 | 0,63 | 1,62 | 2,30 | 0,71 | 4,80E-01 |
| AT3G14720 | 0,35 | -1,17 | 4,36 | -0,27 | 7,89E-01 |
| AT3G14790 | 0,27 | -2,20 | 4,38 | -0,50 | 6,16E-01 |
| AT3G14830 | 0,30 | 0,46 | 4,32 | 0,11 | 9,16E-01 |
| AT3G14840 | 1,81 | -0,59 | 1,35 | -0,44 | 6,61E-01 |
| AT3G14870 | 0,25 | 1,59 | 4,33 | 0,37 | 7,14E-01 |
| AT3G14920 | 0,30 | 0,30 | 4,34 | 0,07 | 9,45E-01 |
| AT3G14930 | 2,04 | -2,26 | 1,28 | -1,77 | 7,67E-02 |
| AT3G14940 | 0,68 | 1,47 | 3,52 | 0,42 | 6,75E-01 |
| AT3G14990 | 0,39 | 1,48 | 2,84 | 0,52 | 6,03E-01 |
| AT3G15000 | 0,95 | -0,16 | 2,31 | -0,07 | 9,45E-01 |
| AT3G15010 | 0,55 | 2,65 | 3,08 | 0,86 | 3,90E-01 |
| AT3G15020 | 1,02 | -0,92 | 2,23 | -0,41 | 6,79E-01 |
| AT3G15030 | 0,69 | -2,59 | 2,97 | -0,87 | 3,84E-01 |
| AT3G15090 | 0,12 | 1,23 | 4,34 | 0,28 | 7,76E-01 |
| AT3G15095 | 1,44 | -0,13 | 1,50 | -0,09 | 9,31E-01 |
| AT3G15160 | 1,08 | -0,45 | 2,06 | -0,22 | 8,25E-01 |
| AT3G15180 | 0,21 | -1,41 | 4,36 | -0,32 | 7,46E-01 |
| AT3G15190 | 1,26 | -1,37 | 1,56 | -0,88 | 3,79E-01 |
| AT3G15210 | 1,70 | 1,41 | 1,67 | 0,85 | 3,98E-01 |
| AT3G15260 | 0,39 | 1,80 | 4,27 | 0,42 | 6,73E-01 |
| AT3G15290 | 0,26 | -0,30 | 4,37 | -0,07 | 9,46E-01 |
| AT3G15350 | 1,19 | -1,82 | 2,09 | -0,87 | 3,84E-01 |
| AT3G03105 | 2,11 | -1,32 | 1,46 | -0,90 | 3,66E-01 |
| AT3G15355 | 0,91 | 1,01 | 3,69 | 0,27 | 7,84E-01 |
| AT3G15360 | 1,20 | -0,85 | 1,84 | -0,46 | 6,45E-01 |
| AT3G15380 | 0,12 | 1,23 | 4,34 | 0,28 | 7,76E-01 |
| AT3G15450 | 0,77 | -1,42 | 2,06 | -0,69 | 4,91E-01 |
| AT3G15518 | 0,17 | 0,98 | 4,35 | 0,23 | 8,21E-01 |
| AT3G15550 | 0,17 | 0,47 | 4,37 | 0,11 | 9,14E-01 |
| AT3G03145 | 0,66 | -2,04 | 3,04 | -0,67 | 5,03E-01 |
| AT3G15635 | 0,25 | 0,01 | 4,37 | 0,00 | 9,99E-01 |
| AT3G15750 | 0,08 | 0,44 | 4,38 | 0,10 | 9,21E-01 |
| AT3G15760 | 0,84 | 1,05 | 2,59 | 0,41 | 6,84E-01 |
| AT3G15790 | 0,67 | 1,84 | 2,83 | 0,65 | 5,16E-01 |
| AT3G15820 | 0,32 | 1,91 | 4,32 | 0,44 | 6,59E-01 |
| AT3G15850 | 0,57 | 0,02 | 2,78 | 0,01 | 9,95E-01 |
| AT3G15930 | 0,12 | 1,16 | 4,34 | 0,27 | 7,89E-01 |
| AT3G16010 | 0,15 | 0,00 | 4,38 | 0,00 | 1,00E+00 |
| AT3G16060 | 0,47 | -0,49 | 4,08 | -0,12 | 9,05E-01 |
| AT3G16080 | 0,49 | -0,89 | 3,23 | -0,28 | 7,82E-01 |
| AT3G16130 | 0,87 | -0,35 | 2,04 | -0,17 | 8,65E-01 |
| AT3G16175 | 0,08 | 0,44 | 4,38 | 0,10 | 9,21E-01 |
| AT3G16240 | 1,39 | -3,26 | 1,87 | -1,75 | 8,10E-02 |
| AT3G16260 | 0,72 | -0,32 | 3,43 | -0,09 | 9,26E-01 |
| AT3G16270 | 0,71 | 2,14 | 2,47 | 0,87 | 3,87E-01 |
| AT3G16340 | 0,41 | 0,04 | 4,30 | 0,01 | 9,93E-01 |
| AT3G16370 | 1,93 | -1,77 | 1,43 | -1,23 | 2,17E-01 |
| AT3G16420 | 0,28 | -2,52 | 4,38 | -0,58 | 5,65E-01 |
| AT3G16460 | 0,31 | -0,76 | 4,36 | -0,17 | 8,62E-01 |
| AT3G16525 | 0,84 | -0,85 | 2,33 | -0,36 | 7,16E-01 |

| | | | | | |
|---|---|---|---|---|---|
| AT3G16570 | 0,68 | 0,59 | 2,46 | 0,24 | 8,11E-01 |
| AT3G16710 | 0,43 | -0,13 | 4,24 | -0,03 | 9,75E-01 |
| AT3G16720 | 0,43 | -0,24 | 4,14 | -0,06 | 9,55E-01 |
| AT3G16740 | 0,38 | 1,32 | 3,42 | 0,39 | 7,00E-01 |
| AT3G16760 | 0,57 | 1,97 | 3,77 | 0,52 | 6,01E-01 |
| AT3G16770 | 1,33 | 2,28 | 2,48 | 0,92 | 3,58E-01 |
| AT3G16785 | 1,89 | -0,15 | 1,19 | -0,13 | 8,99E-01 |
| AT3G16800 | 0,16 | 0,00 | 4,41 | 0,00 | 1,00E+00 |
| AT3G16830 | 1,75 | -0,24 | 1,48 | -0,17 | 8,69E-01 |
| AT3G16850 | 3,96 | -0,59 | 0,93 | -0,63 | 5,26E-01 |
| AT3G16857 | 0,72 | -0,50 | 3,68 | -0,13 | 8,93E-01 |
| AT3G16940 | 0,78 | -0,20 | 2,87 | -0,07 | 9,44E-01 |
| AT3G17100 | 1,17 | -2,20 | 1,91 | -1,15 | 2,49E-01 |
| AT3G17160 | 0,19 | 1,40 | 4,33 | 0,32 | 7,46E-01 |
| AT3G17190 | 1,85 | 3,03 | 1,74 | 1,74 | 8,12E-02 |
| AT3G17205 | 0,35 | -0,94 | 4,39 | -0,21 | 8,31E-01 |
| AT3G17210 | 0,32 | -1,28 | 4,34 | -0,29 | 7,68E-01 |
| AT3G03465 | 0,80 | 2,23 | 2,35 | 0,95 | 3,43E-01 |
| AT3G17360 | 0,65 | -2,12 | 2,91 | -0,73 | 4,67E-01 |
| AT3G17380 | 0,28 | -1,42 | 4,35 | -0,33 | 7,44E-01 |
| AT3G03475 | 2,58 | 0,87 | 1,32 | 0,66 | 5,08E-01 |
| AT3G17410 | 0,79 | 0,37 | 1,84 | 0,20 | 8,41E-01 |
| AT3G17470 | 0,76 | -3,67 | 3,72 | -0,99 | 3,24E-01 |
| AT3G17490 | 0,56 | 1,13 | 3,88 | 0,29 | 7,71E-01 |
| AT3G17626 | 0,20 | -0,31 | 4,36 | -0,07 | 9,44E-01 |
| AT3G17640 | 0,16 | 0,00 | 4,41 | 0,00 | 1,00E+00 |
| AT3G17650 | 0,88 | 3,04 | 2,05 | 1,49 | 1,37E-01 |
| AT3G17690 | 0,12 | 1,16 | 4,34 | 0,27 | 7,89E-01 |
| AT3G17700 | 0,35 | 1,23 | 4,35 | 0,28 | 7,78E-01 |
| AT3G17712 | 0,66 | -2,88 | 3,65 | -0,79 | 4,29E-01 |
| AT3G17750 | 1,26 | 0,21 | 1,67 | 0,13 | 8,99E-01 |
| AT3G17765 | 0,21 | 0,01 | 4,34 | 0,00 | 9,98E-01 |
| AT3G17790 | 0,83 | 0,22 | 2,37 | 0,09 | 9,26E-01 |
| AT3G03535 | 0,92 | -3,37 | 2,33 | -1,45 | 1,48E-01 |
| AT3G17810 | 0,53 | 0,63 | 2,78 | 0,23 | 8,21E-01 |
| AT3G17820 | 0,62 | -2,61 | 3,15 | -0,83 | 4,08E-01 |
| AT3G17860 | 0,36 | 1,17 | 4,35 | 0,27 | 7,88E-01 |
| AT3G17880 | 0,42 | 1,90 | 3,25 | 0,58 | 5,60E-01 |
| AT3G17890 | 0,10 | 0,01 | 4,38 | 0,00 | 9,99E-01 |
| AT3G17900 | 1,92 | -1,03 | 1,26 | -0,81 | 4,18E-01 |
| AT3G17970 | 0,77 | 0,48 | 2,99 | 0,16 | 8,72E-01 |
| AT3G17980 | 0,35 | -2,17 | 4,38 | -0,49 | 6,21E-01 |
| AT3G18000 | 0,28 | 0,23 | 4,33 | 0,05 | 9,58E-01 |
| AT3G18020 | 0,41 | -0,57 | 4,23 | -0,13 | 8,93E-01 |
| AT3G18040 | 0,26 | 0,56 | 4,38 | 0,13 | 8,99E-01 |
| AT3G18050 | 1,52 | -0,36 | 1,50 | -0,24 | 8,11E-01 |
| AT3G18060 | 1,35 | -1,07 | 1,44 | -0,75 | 4,56E-01 |
| AT3G18070 | 0,11 | 0,65 | 4,36 | 0,15 | 8,81E-01 |
| AT3G18080 | 1,62 | 0,46 | 1,42 | 0,33 | 7,44E-01 |
| AT3G18090 | 1,26 | 1,75 | 2,87 | 0,61 | 5,43E-01 |
| AT3G18100 | 0,29 | 2,11 | 4,32 | 0,49 | 6,26E-01 |
| AT3G03625 | 0,40 | 0,00 | 3,44 | 0,00 | 9,99E-01 |

| | | | | | |
|---|---|---|---|---|---|
| AT3G18190 | 1,46 | -1,12 | 1,51 | -0,74 | 4,61E-01 |
| AT3G18240 | 0,43 | 0,15 | 4,18 | 0,04 | 9,72E-01 |
| AT3G18270 | 0,50 | -1,22 | 3,98 | -0,31 | 7,60E-01 |
| AT3G18340 | 0,19 | 0,00 | 4,41 | 0,00 | 1,00E+00 |
| AT3G18370 | 2,55 | -1,06 | 1,45 | -0,73 | 4,66E-01 |
| AT3G18390 | 1,15 | -0,64 | 1,89 | -0,34 | 7,35E-01 |
| AT3G18440 | 0,08 | 0,69 | 4,36 | 0,16 | 8,75E-01 |
| AT3G18480 | 0,44 | 0,93 | 3,21 | 0,29 | 7,72E-01 |
| AT3G18740 | 0,73 | 2,07 | 2,25 | 0,92 | 3,57E-01 |
| AT3G18790 | 0,65 | 0,08 | 2,37 | 0,03 | 9,74E-01 |
| AT3G18800 | 0,33 | -0,45 | 4,32 | -0,10 | 9,16E-01 |
| AT3G18820 | 0,39 | 0,08 | 4,24 | 0,02 | 9,85E-01 |
| AT3G18827 | 0,57 | -2,35 | 3,17 | -0,74 | 4,59E-01 |
| AT3G18860 | 0,59 | 0,60 | 2,57 | 0,23 | 8,14E-01 |
| AT3G18890 | 0,86 | -1,79 | 2,11 | -0,85 | 3,96E-01 |
| AT3G18915 | 0,30 | 0,63 | 4,35 | 0,14 | 8,85E-01 |
| AT3G18960 | 0,08 | 0,69 | 4,36 | 0,16 | 8,75E-01 |
| AT3G18980 | 1,15 | -1,79 | 1,91 | -0,94 | 3,48E-01 |
| AT3G19050 | 0,62 | -2,04 | 2,86 | -0,71 | 4,77E-01 |
| AT3G19100 | 1,08 | -0,33 | 1,77 | -0,19 | 8,51E-01 |
| AT3G19180 | 1,86 | 1,64 | 1,67 | 0,98 | 3,25E-01 |
| AT3G19190 | 3,92 | 0,40 | 1,06 | 0,38 | 7,05E-01 |
| AT3G19240 | 1,33 | -0,28 | 1,43 | -0,19 | 8,47E-01 |
| AT3G19290 | 0,28 | -2,31 | 4,39 | -0,53 | 5,99E-01 |
| AT3G19390 | 0,35 | -2,17 | 4,38 | -0,49 | 6,21E-01 |
| AT3G19420 | 2,24 | -1,43 | 1,36 | -1,05 | 2,93E-01 |
| AT3G19470 | 0,30 | -2,32 | 4,39 | -0,53 | 5,96E-01 |
| AT3G19480 | 0,52 | 1,01 | 3,18 | 0,32 | 7,50E-01 |
| AT3G19520 | 0,05 | 0,00 | 4,41 | 0,00 | 1,00E+00 |
| AT3G19540 | 1,20 | -1,30 | 1,87 | -0,70 | 4,86E-01 |
| AT3G19560 | 1,58 | -1,60 | 2,02 | -0,79 | 4,28E-01 |
| AT3G19640 | 0,86 | -0,54 | 2,30 | -0,23 | 8,15E-01 |
| AT3G19680 | 0,92 | -0,25 | 2,00 | -0,13 | 8,99E-01 |
| AT3G19760 | 1,20 | -0,30 | 1,75 | -0,17 | 8,62E-01 |
| AT3G19770 | 0,75 | -0,26 | 2,96 | -0,09 | 9,29E-01 |
| AT3G19780 | 1,34 | 1,11 | 2,00 | 0,56 | 5,78E-01 |
| AT3G19800 | 0,78 | 0,29 | 2,38 | 0,12 | 9,03E-01 |
| AT3G19860 | 0,48 | -0,82 | 3,97 | -0,21 | 8,36E-01 |
| AT3G19870 | 0,37 | 1,89 | 4,28 | 0,44 | 6,58E-01 |
| AT3G19910 | 1,61 | 1,40 | 1,57 | 0,89 | 3,74E-01 |
| AT3G19930 | 1,47 | -1,16 | 1,43 | -0,81 | 4,16E-01 |
| AT3G19960 | 1,65 | 0,01 | 1,52 | 0,01 | 9,93E-01 |
| AT3G19980 | 0,93 | -0,97 | 2,24 | -0,43 | 6,66E-01 |
| AT3G19990 | 0,32 | -0,65 | 4,33 | -0,15 | 8,81E-01 |
| AT3G20000 | 0,74 | 0,01 | 3,55 | 0,00 | 9,97E-01 |
| AT3G20010 | 2,80 | -0,58 | 1,35 | -0,43 | 6,66E-01 |
| AT3G20030 | 0,79 | 1,74 | 2,44 | 0,71 | 4,75E-01 |
| AT3G20140 | 0,20 | -0,44 | 4,36 | -0,10 | 9,19E-01 |
| AT3G20150 | 0,54 | -3,64 | 4,31 | -0,85 | 3,98E-01 |
| AT3G20200 | 0,21 | 1,57 | 4,33 | 0,36 | 7,16E-01 |
| AT3G20410 | 0,37 | -1,78 | 4,41 | -0,40 | 6,87E-01 |
| AT3G20500 | 0,33 | -0,97 | 4,32 | -0,23 | 8,21E-01 |

| | | | | | |
|---|---|---|---|---|---|
| AT3G20520 | 0,48 | 1,17 | 4,32 | 0,27 | 7,86E-01 |
| AT3G20540 | 0,90 | -1,10 | 2,21 | -0,50 | 6,18E-01 |
| AT3G20600 | 0,25 | -1,82 | 4,40 | -0,41 | 6,79E-01 |
| AT3G03995 | 0,62 | 0,00 | 4,41 | 0,00 | 1,00E+00 |
| AT3G20720 | 0,42 | 0,12 | 3,42 | 0,04 | 9,71E-01 |
| AT3G20730 | 0,40 | 2,09 | 4,32 | 0,48 | 6,28E-01 |
| AT3G20790 | 0,23 | 0,85 | 4,37 | 0,20 | 8,45E-01 |
| AT3G20820 | 3,82 | 0,27 | 1,35 | 0,20 | 8,42E-01 |
| AT3G20890 | 0,34 | -0,92 | 4,34 | -0,21 | 8,32E-01 |
| AT3G20950 | 1,75 | 2,34 | 1,69 | 1,38 | 1,67E-01 |
| AT3G20970 | 0,63 | 0,53 | 2,99 | 0,18 | 8,58E-01 |
| AT3G21130 | 0,35 | 0,39 | 4,37 | 0,09 | 9,29E-01 |
| AT3G21170 | 1,15 | 1,79 | 2,41 | 0,75 | 4,56E-01 |
| AT3G21175 | 0,18 | 0,00 | 4,38 | 0,00 | 9,99E-01 |
| AT3G21220 | 0,51 | -2,22 | 4,08 | -0,54 | 5,86E-01 |
| AT3G21270 | 0,29 | 0,74 | 4,35 | 0,17 | 8,65E-01 |
| AT3G21295 | 0,41 | -0,32 | 3,42 | -0,09 | 9,26E-01 |
| AT3G21310 | 0,36 | 1,83 | 4,33 | 0,42 | 6,73E-01 |
| AT3G21465 | 0,14 | 0,00 | 4,41 | 0,00 | 1,00E+00 |
| AT3G21473 | 0,48 | 0,00 | 4,05 | 0,00 | 1,00E+00 |
| AT3G21560 | 0,31 | -0,64 | 4,36 | -0,15 | 8,83E-01 |
| AT3G21690 | 0,96 | 2,61 | 2,03 | 1,29 | 1,98E-01 |
| AT3G21750 | 1,77 | 1,05 | 1,46 | 0,72 | 4,71E-01 |
| AT3G21760 | 4,32 | 0,62 | 1,07 | 0,58 | 5,65E-01 |
| AT3G04155 | 3,42 | -0,21 | 1,12 | -0,19 | 8,51E-01 |
| AT3G04165 | 2,58 | 0,58 | 1,29 | 0,45 | 6,53E-01 |
| AT3G04175 | 0,99 | -0,37 | 2,04 | -0,18 | 8,54E-01 |
| AT3G21990 | 0,46 | 0,00 | 4,41 | 0,00 | 1,00E+00 |
| AT3G22072 | 0,17 | 0,00 | 4,41 | 0,00 | 1,00E+00 |
| AT3G22104 | 0,62 | -0,93 | 2,87 | -0,32 | 7,47E-01 |
| AT3G22121 | 3,96 | -1,69 | 1,07 | -1,59 | 1,12E-01 |
| AT3G22125 | 1,40 | 2,67 | 1,87 | 1,42 | 1,54E-01 |
| AT3G22150 | 0,49 | -0,99 | 3,88 | -0,26 | 7,98E-01 |
| AT3G22200 | 0,76 | 0,73 | 2,21 | 0,33 | 7,42E-01 |
| AT3G04255 | 0,24 | 0,00 | 4,41 | 0,00 | 1,00E+00 |
| AT3G22270 | 0,79 | -1,38 | 2,54 | -0,54 | 5,87E-01 |
| AT3G22290 | 0,33 | -0,45 | 4,32 | -0,10 | 9,16E-01 |
| AT3G22330 | 0,85 | -0,50 | 2,19 | -0,23 | 8,21E-01 |
| AT3G22350 | 0,91 | 1,78 | 2,60 | 0,68 | 4,93E-01 |
| AT3G22370 | 0,72 | -1,67 | 2,50 | -0,67 | 5,05E-01 |
| AT3G22425 | 0,12 | 0,00 | 4,38 | 0,00 | 1,00E+00 |
| AT3G22430 | 0,19 | 0,01 | 4,38 | 0,00 | 9,98E-01 |
| AT3G22440 | 1,87 | 1,58 | 1,48 | 1,07 | 2,86E-01 |
| AT3G22520 | 0,75 | 0,19 | 2,50 | 0,08 | 9,38E-01 |
| AT3G22760 | 0,51 | 0,23 | 3,24 | 0,07 | 9,43E-01 |
| AT3G22810 | 0,55 | 1,46 | 4,34 | 0,34 | 7,37E-01 |
| AT3G22845 | 0,51 | -2,36 | 4,32 | -0,55 | 5,84E-01 |
| AT3G22870 | 0,92 | 0,55 | 2,21 | 0,25 | 8,04E-01 |
| AT3G22890 | 4,47 | 0,83 | 0,82 | 1,01 | 3,10E-01 |
| AT3G22900 | 0,24 | 0,24 | 4,34 | 0,06 | 9,55E-01 |
| AT3G22935 | 0,26 | 0,00 | 4,41 | 0,00 | 1,00E+00 |
| AT3G22940 | 1,72 | 2,06 | 1,89 | 1,09 | 2,77E-01 |

| | | | | | |
|---|---|---|---|---|---|
| AT3G22960 | 3,17 | -0,48 | 1,08 | -0,45 | 6,55E-01 |
| AT3G22970 | 2,77 | 0,27 | 1,27 | 0,22 | 8,29E-01 |
| AT3G23010 | 0,77 | -2,74 | 3,46 | -0,79 | 4,28E-01 |
| AT3G23050 | 0,64 | 0,19 | 2,67 | 0,07 | 9,44E-01 |
| AT3G23100 | 2,14 | -0,94 | 1,37 | -0,68 | 4,95E-01 |
| AT3G04395 | 0,16 | 0,92 | 4,35 | 0,21 | 8,33E-01 |
| AT3G23250 | 0,25 | 1,18 | 4,33 | 0,27 | 7,85E-01 |
| AT3G23280 | 0,45 | 0,85 | 4,08 | 0,21 | 8,35E-01 |
| AT3G23300 | 0,75 | -0,77 | 2,00 | -0,38 | 7,01E-01 |
| AT3G23310 | 0,63 | -0,07 | 2,30 | -0,03 | 9,74E-01 |
| AT3G23380 | 0,21 | 0,01 | 4,37 | 0,00 | 9,98E-01 |
| AT3G23390 | 0,15 | 0,00 | 4,38 | 0,00 | 1,00E+00 |
| AT3G23410 | 0,36 | 0,20 | 4,37 | 0,05 | 9,64E-01 |
| AT3G23450 | 2,19 | 0,33 | 1,79 | 0,19 | 8,53E-01 |
| AT3G23530 | 1,09 | -1,85 | 1,84 | -1,00 | 3,16E-01 |
| AT3G23580 | 0,57 | -0,12 | 2,31 | -0,05 | 9,58E-01 |
| AT3G23590 | 1,64 | 1,98 | 1,74 | 1,13 | 2,57E-01 |
| AT3G23600 | 1,16 | 0,51 | 1,79 | 0,28 | 7,77E-01 |
| AT3G23640 | 0,83 | -2,50 | 2,50 | -1,00 | 3,17E-01 |
| AT3G23685 | 5,28 | 0,13 | 0,78 | 0,16 | 8,69E-01 |
| AT3G23750 | 0,94 | -0,99 | 1,76 | -0,56 | 5,74E-01 |
| AT3G23790 | 0,38 | -1,32 | 4,34 | -0,30 | 7,62E-01 |
| AT3G23800 | 0,20 | -0,44 | 4,36 | -0,10 | 9,19E-01 |
| AT3G23890 | 0,56 | -0,66 | 3,91 | -0,17 | 8,66E-01 |
| AT3G23920 | 2,81 | 1,44 | 1,18 | 1,22 | 2,22E-01 |
| AT3G23990 | 0,73 | -0,90 | 3,57 | -0,25 | 8,00E-01 |
| AT3G24000 | 0,34 | 0,13 | 4,34 | 0,03 | 9,76E-01 |
| AT3G24050 | 0,88 | 1,50 | 2,07 | 0,72 | 4,69E-01 |
| AT3G24070 | 0,38 | 0,66 | 4,34 | 0,15 | 8,80E-01 |
| AT3G24160 | 0,45 | 0,80 | 3,50 | 0,23 | 8,18E-01 |
| AT3G24190 | 0,59 | 0,56 | 3,07 | 0,18 | 8,55E-01 |
| AT3G24255 | 0,43 | 0,55 | 2,75 | 0,20 | 8,41E-01 |
| AT3G24490 | 0,64 | 1,77 | 3,54 | 0,50 | 6,18E-01 |
| AT3G24500 | 0,22 | 0,00 | 4,41 | 0,00 | 1,00E+00 |
| AT3G24514 | 0,29 | 0,00 | 4,41 | 0,00 | 1,00E+00 |
| AT3G24520 | 0,92 | -1,45 | 2,53 | -0,57 | 5,68E-01 |
| AT3G24542 | 2,63 | -1,41 | 1,31 | -1,07 | 2,83E-01 |
| AT3G24550 | 0,60 | 2,48 | 2,93 | 0,85 | 3,96E-01 |
| AT3G24580 | 0,47 | 1,63 | 4,03 | 0,40 | 6,86E-01 |
| AT3G24590 | 0,21 | -0,49 | 4,35 | -0,11 | 9,11E-01 |
| AT3G24830 | 0,92 | 2,29 | 2,06 | 1,11 | 2,66E-01 |
| AT3G24929 | 2,38 | 2,92 | 1,65 | 1,77 | 7,73E-02 |
| AT3G25012 | 0,95 | -1,25 | 1,75 | -0,71 | 4,77E-01 |
| AT3G25013 | 0,18 | 0,00 | 4,38 | 0,00 | 9,99E-01 |
| AT3G25030 | 0,21 | 0,83 | 4,36 | 0,19 | 8,48E-01 |
| AT3G25040 | 0,17 | 0,02 | 4,39 | 0,00 | 9,97E-01 |
| AT3G25070 | 0,31 | 1,40 | 3,59 | 0,39 | 6,95E-01 |
| AT3G25130 | 0,57 | -1,79 | 4,41 | -0,41 | 6,85E-01 |
| AT3G25140 | 1,51 | 1,16 | 2,01 | 0,58 | 5,62E-01 |
| AT3G25250 | 0,60 | 2,04 | 2,58 | 0,79 | 4,29E-01 |
| AT3G25290 | 0,29 | -0,47 | 4,33 | -0,11 | 9,14E-01 |
| AT3G25440 | 0,23 | -1,78 | 4,41 | -0,40 | 6,87E-01 |

| | | | | | |
|---|---|---|---|---|---|
| AT3G25470 | 0,67 | -0,90 | 3,71 | -0,24 | 8,07E-01 |
| AT3G25490 | 0,05 | 0,00 | 4,41 | 0,00 | 1,00E+00 |
| AT3G25585 | 0,71 | -0,86 | 2,57 | -0,33 | 7,39E-01 |
| AT3G25590 | 0,75 | 1,20 | 2,39 | 0,50 | 6,16E-01 |
| AT3G25690 | 2,72 | 0,67 | 1,10 | 0,61 | 5,43E-01 |
| AT3G25700 | 0,66 | 0,36 | 2,23 | 0,16 | 8,70E-01 |
| AT3G04955 | 0,15 | 0,83 | 4,35 | 0,19 | 8,49E-01 |
| AT3G25720 | 0,26 | 0,00 | 4,38 | 0,00 | 9,99E-01 |
| AT3G25760 | 0,49 | -2,96 | 4,35 | -0,68 | 4,95E-01 |
| AT3G25770 | 1,09 | -0,19 | 2,08 | -0,09 | 9,27E-01 |
| AT3G25780 | 0,49 | -0,08 | 2,67 | -0,03 | 9,76E-01 |
| AT3G25860 | 0,63 | 0,97 | 3,23 | 0,30 | 7,63E-01 |
| AT3G25890 | 0,89 | 0,64 | 2,14 | 0,30 | 7,65E-01 |
| AT3G25910 | 0,36 | 2,21 | 4,32 | 0,51 | 6,09E-01 |
| AT3G26020 | 0,28 | 0,00 | 4,38 | 0,00 | 9,99E-01 |
| AT3G26030 | 1,16 | 0,00 | 1,74 | 0,00 | 9,98E-01 |
| AT3G26070 | 0,46 | -2,76 | 4,21 | -0,66 | 5,12E-01 |
| AT3G26080 | 0,26 | -0,30 | 4,37 | -0,07 | 9,46E-01 |
| AT3G26085 | 0,30 | -0,77 | 4,35 | -0,18 | 8,59E-01 |
| AT3G26090 | 0,21 | 0,58 | 4,36 | 0,13 | 8,94E-01 |
| AT3G04975 | 1,19 | 1,76 | 2,11 | 0,83 | 4,04E-01 |
| AT3G04985 | 0,58 | 1,25 | 2,66 | 0,47 | 6,38E-01 |
| AT3G26370 | 0,48 | -0,62 | 2,65 | -0,23 | 8,15E-01 |
| AT3G26395 | 0,16 | 1,33 | 4,33 | 0,31 | 7,59E-01 |
| AT3G26400 | 0,95 | -3,60 | 2,40 | -1,50 | 1,34E-01 |
| AT3G26430 | 1,60 | -0,36 | 1,31 | -0,27 | 7,85E-01 |
| AT3G26420 | 1,60 | -0,36 | 1,31 | -0,27 | 7,85E-01 |
| AT3G26440 | 2,47 | -1,75 | 1,54 | -1,14 | 2,55E-01 |
| AT3G26512 | 0,38 | 1,15 | 3,36 | 0,34 | 7,32E-01 |
| AT3G26570 | 1,54 | 0,41 | 1,78 | 0,23 | 8,19E-01 |
| AT3G26580 | 0,73 | 1,56 | 2,27 | 0,69 | 4,91E-01 |
| AT3G26600 | 1,07 | 0,42 | 2,01 | 0,21 | 8,35E-01 |
| AT3G26640 | 0,37 | -0,86 | 3,45 | -0,25 | 8,03E-01 |
| AT3G26710 | 1,14 | -1,53 | 1,83 | -0,84 | 4,03E-01 |
| AT3G26720 | 0,82 | 0,44 | 2,46 | 0,18 | 8,58E-01 |
| AT3G26730 | 1,96 | 1,56 | 1,53 | 1,02 | 3,06E-01 |
| AT3G26742 | 0,18 | 0,81 | 4,34 | 0,19 | 8,52E-01 |
| AT3G26780 | 1,24 | 0,79 | 1,87 | 0,42 | 6,74E-01 |
| AT3G26782 | 0,41 | 0,79 | 4,12 | 0,19 | 8,49E-01 |
| AT3G26800 | 0,34 | -0,85 | 3,51 | -0,24 | 8,10E-01 |
| AT3G26840 | 1,06 | 0,51 | 2,08 | 0,25 | 8,06E-01 |
| AT3G27090 | 1,24 | 0,17 | 1,77 | 0,10 | 9,24E-01 |
| AT3G27160 | 0,41 | -0,01 | 3,38 | 0,00 | 9,98E-01 |
| AT3G27170 | 0,68 | 1,55 | 2,82 | 0,55 | 5,83E-01 |
| AT3G27230 | 0,45 | -0,37 | 3,28 | -0,11 | 9,10E-01 |
| AT3G27250 | 0,85 | -2,02 | 2,53 | -0,80 | 4,24E-01 |
| AT3G27260 | 1,09 | 0,65 | 1,95 | 0,34 | 7,37E-01 |
| AT3G27270 | 0,62 | -1,35 | 3,85 | -0,35 | 7,26E-01 |
| AT3G27280 | 0,60 | 0,63 | 2,95 | 0,21 | 8,32E-01 |
| AT3G27325 | 0,56 | -0,83 | 2,64 | -0,31 | 7,53E-01 |
| AT3G27329 | 0,81 | 1,29 | 2,21 | 0,59 | 5,58E-01 |
| AT3G27331 | 0,96 | 0,93 | 3,23 | 0,29 | 7,73E-01 |

| | | | | | |
|---|---|---|---|---|---|
| AT3G27350 | 0,51 | 1,89 | 3,92 | 0,48 | 6,29E-01 |
| AT3G27400 | 1,80 | -1,10 | 1,68 | -0,66 | 5,11E-01 |
| AT3G27430 | 0,35 | 0,32 | 3,62 | 0,09 | 9,30E-01 |
| AT3G27470 | 1,23 | 2,28 | 1,86 | 1,22 | 2,22E-01 |
| AT3G27540 | 0,29 | 0,83 | 3,66 | 0,23 | 8,20E-01 |
| AT3G27555 | 0,30 | -1,78 | 4,41 | -0,40 | 6,87E-01 |
| AT3G27670 | 1,19 | -0,31 | 2,08 | -0,15 | 8,81E-01 |
| AT3G27690 | 1,98 | -2,46 | 1,74 | -1,41 | 1,58E-01 |
| AT3G05295 | 0,48 | -0,11 | 2,90 | -0,04 | 9,69E-01 |
| AT3G27820 | 1,03 | 3,12 | 1,94 | 1,61 | 1,08E-01 |
| AT3G27830 | 0,47 | 0,02 | 3,38 | 0,01 | 9,94E-01 |
| AT3G27850 | 1,55 | -0,78 | 1,75 | -0,45 | 6,53E-01 |
| AT3G27860 | 0,12 | 1,05 | 4,35 | 0,24 | 8,10E-01 |
| AT3G27865 | 3,38 | 1,37 | 1,58 | 0,87 | 3,85E-01 |
| AT3G05335 | 1,73 | 1,37 | 1,68 | 0,81 | 4,17E-01 |
| AT3G05355 | 1,75 | -0,90 | 1,86 | -0,49 | 6,27E-01 |
| AT3G27870 | 0,75 | -0,77 | 2,45 | -0,31 | 7,54E-01 |
| AT3G27940 | 0,56 | 1,51 | 2,96 | 0,51 | 6,09E-01 |
| AT3G27980 | 0,34 | 0,00 | 4,41 | 0,00 | 1,00E+00 |
| AT3G27999 | 0,43 | 0,00 | 4,41 | 0,00 | 1,00E+00 |
| AT3G28040 | 1,16 | -1,63 | 2,06 | -0,79 | 4,28E-01 |
| AT3G28050 | 0,37 | 2,26 | 4,32 | 0,52 | 6,00E-01 |
| AT3G28080 | 0,70 | -0,82 | 2,30 | -0,36 | 7,20E-01 |
| AT3G28100 | 1,10 | 0,41 | 1,93 | 0,21 | 8,32E-01 |
| AT3G28170 | 0,75 | -0,64 | 3,42 | -0,19 | 8,51E-01 |
| AT3G28200 | 0,56 | 0,96 | 4,34 | 0,22 | 8,26E-01 |
| AT3G28220 | 0,69 | 0,02 | 3,06 | 0,01 | 9,95E-01 |
| AT3G28270 | 0,14 | 0,03 | 4,39 | 0,01 | 9,95E-01 |
| AT3G28290 | 1,10 | 0,09 | 2,62 | 0,03 | 9,73E-01 |
| AT3G28300 | 1,56 | -2,12 | 2,81 | -0,75 | 4,50E-01 |
| AT3G28450 | 0,78 | 1,40 | 2,59 | 0,54 | 5,90E-01 |
| AT3G28500 | 0,30 | 1,14 | 4,33 | 0,26 | 7,92E-01 |
| AT3G28510 | 0,99 | 0,59 | 2,45 | 0,24 | 8,09E-01 |
| AT3G28540 | 2,73 | 0,46 | 1,08 | 0,42 | 6,72E-01 |
| AT3G05515 | 0,60 | 0,00 | 3,09 | 0,00 | 1,00E+00 |
| AT3G28580 | 0,77 | 2,92 | 3,41 | 0,85 | 3,93E-01 |
| AT3G28590 | 0,05 | 0,00 | 4,41 | 0,00 | 1,00E+00 |
| AT3G28650 | 0,20 | -1,80 | 4,41 | -0,41 | 6,83E-01 |
| AT3G28670 | 0,33 | 0,35 | 4,39 | 0,08 | 9,36E-01 |
| AT3G28690 | 0,39 | 1,18 | 4,20 | 0,28 | 7,79E-01 |
| AT3G28715 | 0,62 | -0,66 | 2,27 | -0,29 | 7,70E-01 |
| AT3G28720 | 0,91 | 1,02 | 3,14 | 0,33 | 7,45E-01 |
| AT3G28730 | 0,64 | -1,27 | 2,81 | -0,45 | 6,51E-01 |
| AT3G28870 | 0,74 | 1,56 | 3,69 | 0,42 | 6,72E-01 |
| AT3G28899 | 2,18 | -2,09 | 1,57 | -1,33 | 1,84E-01 |
| AT3G28910 | 0,61 | -0,35 | 2,59 | -0,14 | 8,92E-01 |
| AT3G05595 | 0,35 | 1,03 | 4,35 | 0,24 | 8,12E-01 |
| AT3G28922 | 0,12 | 0,49 | 4,36 | 0,11 | 9,11E-01 |
| AT3G28925 | 2,17 | 1,00 | 1,55 | 0,65 | 5,17E-01 |
| AT3G28958 | 0,58 | 1,00 | 3,10 | 0,32 | 7,47E-01 |
| AT3G29000 | 0,25 | 1,36 | 4,33 | 0,31 | 7,54E-01 |
| AT3G29010 | 0,38 | -1,27 | 4,33 | -0,29 | 7,68E-01 |

| | | | | | |
|---|---|---|---|---|---|
| AT3G29033 | 0,57 | 2,11 | 2,92 | 0,72 | 4,71E-01 |
| AT3G29080 | 1,28 | 1,43 | 1,78 | 0,81 | 4,20E-01 |
| AT3G29180 | 1,27 | -1,35 | 2,62 | -0,51 | 6,07E-01 |
| AT3G05715 | 0,40 | 0,68 | 4,29 | 0,16 | 8,73E-01 |
| AT3G29240 | 1,56 | -0,83 | 1,61 | -0,51 | 6,08E-01 |
| AT3G29270 | 0,39 | -1,40 | 4,37 | -0,32 | 7,49E-01 |
| AT3G29290 | 0,20 | 1,26 | 4,34 | 0,29 | 7,71E-01 |
| AT3G29320 | 2,29 | 1,23 | 1,40 | 0,88 | 3,78E-01 |
| AT3G29370 | 0,05 | 0,00 | 4,41 | 0,00 | 1,00E+00 |
| AT3G29390 | 0,18 | 0,43 | 4,36 | 0,10 | 9,22E-01 |
| AT3G29450 | 0,14 | 0,00 | 4,41 | 0,00 | 1,00E+00 |
| AT3G29455 | 0,27 | 0,00 | 4,41 | 0,00 | 1,00E+00 |
| AT3G05805 | 0,31 | 1,35 | 4,33 | 0,31 | 7,56E-01 |
| AT3G29638 | 0,16 | 1,07 | 4,34 | 0,25 | 8,05E-01 |
| AT3G29670 | 0,94 | 0,00 | 2,45 | 0,00 | 1,00E+00 |
| AT3G29796 | 2,77 | 3,82 | 1,59 | 2,40 | 1,65E-02 |
| AT3G30210 | 0,11 | 0,48 | 4,37 | 0,11 | 9,12E-01 |
| AT3G30320 | 0,40 | 0,29 | 4,36 | 0,07 | 9,47E-01 |
| AT3G30380 | 0,49 | -1,18 | 3,21 | -0,37 | 7,12E-01 |
| AT3G30383 | 0,12 | 1,05 | 4,35 | 0,24 | 8,10E-01 |
| AT3G30520 | 1,35 | 1,25 | 2,36 | 0,53 | 5,98E-01 |
| AT3G30525 | 1,14 | 2,19 | 2,71 | 0,81 | 4,20E-01 |
| AT3G30720 | 0,61 | 0,93 | 3,10 | 0,30 | 7,64E-01 |
| AT3G30770 | 0,13 | 0,00 | 4,41 | 0,00 | 1,00E+00 |
| AT3G30820 | 0,18 | 0,21 | 4,37 | 0,05 | 9,62E-01 |
| AT3G31950 | 0,59 | 0,87 | 2,36 | 0,37 | 7,11E-01 |
| AT3G32150 | 0,20 | 0,00 | 4,41 | 0,00 | 1,00E+00 |
| AT3G32280 | 0,39 | 0,00 | 4,32 | 0,00 | 1,00E+00 |
| AT3G32896 | 0,28 | -1,77 | 4,41 | -0,40 | 6,88E-01 |
| AT3G33530 | 1,04 | -0,58 | 1,78 | -0,32 | 7,46E-01 |
| AT3G41761 | 2,51 | -1,65 | 2,01 | -0,82 | 4,12E-01 |
| AT3G42075 | 0,24 | -0,27 | 4,35 | -0,06 | 9,51E-01 |
| AT3G42170 | 0,48 | 0,03 | 3,07 | 0,01 | 9,93E-01 |
| AT3G42310 | 0,36 | 0,30 | 4,38 | 0,07 | 9,45E-01 |
| AT3G42560 | 0,10 | 0,01 | 4,38 | 0,00 | 9,99E-01 |
| AT3G42640 | 0,44 | 0,23 | 2,71 | 0,08 | 9,34E-01 |
| AT3G42870 | 1,35 | 0,36 | 2,13 | 0,17 | 8,67E-01 |
| AT3G42950 | 0,12 | 1,16 | 4,34 | 0,27 | 7,89E-01 |
| AT3G06495 | 0,51 | 1,26 | 3,16 | 0,40 | 6,89E-01 |
| AT3G43148 | 0,82 | 1,93 | 2,31 | 0,84 | 4,03E-01 |
| AT3G43150 | 0,30 | 0,82 | 3,63 | 0,22 | 8,22E-01 |
| AT3G43160 | 4,62 | 3,54 | 1,39 | 2,55 | 1,09E-02 |
| AT3G43210 | 0,25 | 0,41 | 4,33 | 0,09 | 9,25E-01 |
| AT3G43230 | 1,49 | 1,14 | 1,49 | 0,76 | 4,45E-01 |
| AT3G43260 | 1,20 | 2,42 | 2,12 | 1,14 | 2,54E-01 |
| AT3G06565 | 0,05 | 0,00 | 4,41 | 0,00 | 1,00E+00 |
| AT3G43290 | 0,21 | 0,81 | 4,34 | 0,19 | 8,51E-01 |
| AT3G06625 | 0,13 | 0,00 | 4,41 | 0,00 | 1,00E+00 |
| AT3G43490 | 1,48 | 1,47 | 2,07 | 0,71 | 4,77E-01 |
| AT3G43503 | 0,30 | 1,26 | 3,65 | 0,35 | 7,30E-01 |
| AT3G06635 | 0,40 | 0,64 | 4,35 | 0,15 | 8,82E-01 |
| AT3G43670 | 0,53 | -1,16 | 2,69 | -0,43 | 6,67E-01 |

| | | | | | |
|---|---|---|---|---|---|
| AT3G43682 | 0,33 | 0,98 | 4,35 | 0,23 | 8,21E-01 |
| AT3G43740 | 0,28 | 0,02 | 4,39 | 0,00 | 9,97E-01 |
| AT3G43800 | 0,45 | 0,23 | 3,25 | 0,07 | 9,43E-01 |
| AT3G43837 | 0,47 | -0,73 | 4,06 | -0,18 | 8,57E-01 |
| AT3G43920 | 0,47 | -1,77 | 3,28 | -0,54 | 5,90E-01 |
| AT3G43950 | 0,37 | 0,31 | 4,39 | 0,07 | 9,44E-01 |
| AT3G44006 | 2,21 | -2,39 | 1,55 | -1,55 | 1,22E-01 |
| AT3G44130 | 0,22 | 0,07 | 4,40 | 0,02 | 9,88E-01 |
| AT3G44150 | 0,18 | 0,13 | 4,41 | 0,03 | 9,76E-01 |
| AT3G44210 | 1,26 | 0,00 | 2,67 | 0,00 | 1,00E+00 |
| AT3G44230 | 0,25 | 0,10 | 4,40 | 0,02 | 9,81E-01 |
| AT3G44235 | 0,05 | 0,00 | 4,41 | 0,00 | 1,00E+00 |
| AT3G44300 | 1,82 | 1,42 | 1,37 | 1,04 | 3,01E-01 |
| AT3G44320 | 0,76 | 0,03 | 2,79 | 0,01 | 9,93E-01 |
| AT3G44330 | 1,72 | 2,16 | 1,91 | 1,13 | 2,58E-01 |
| AT3G44405 | 0,08 | 0,44 | 4,38 | 0,10 | 9,21E-01 |
| AT3G44430 | 1,70 | -0,87 | 1,38 | -0,63 | 5,30E-01 |
| AT3G44550 | 0,99 | 2,89 | 2,42 | 1,19 | 2,32E-01 |
| AT3G44560 | 0,14 | 0,03 | 4,39 | 0,01 | 9,95E-01 |
| AT3G44620 | 0,30 | 0,03 | 3,68 | 0,01 | 9,94E-01 |
| AT3G44630 | 4,39 | -0,20 | 0,89 | -0,22 | 8,23E-01 |
| AT3G44670 | 1,62 | 0,65 | 1,43 | 0,45 | 6,51E-01 |
| AT3G44703 | 4,72 | -2,56 | 1,05 | -2,45 | 1,45E-02 |
| AT3G44720 | 3,49 | -0,69 | 1,08 | -0,64 | 5,21E-01 |
| AT3G44730 | 0,52 | 0,87 | 2,41 | 0,36 | 7,20E-01 |
| AT3G44798 | 0,43 | 1,70 | 4,15 | 0,41 | 6,82E-01 |
| AT3G44860 | 2,37 | 2,15 | 1,50 | 1,44 | 1,51E-01 |
| AT3G44880 | 0,54 | -2,30 | 4,07 | -0,57 | 5,72E-01 |
| AT3G44900 | 0,86 | -0,15 | 2,20 | -0,07 | 9,47E-01 |
| AT3G44935 | 1,28 | 2,15 | 2,10 | 1,02 | 3,06E-01 |
| AT3G44980 | 0,39 | -0,41 | 3,44 | -0,12 | 9,05E-01 |
| AT3G45010 | 0,31 | -1,10 | 4,36 | -0,25 | 8,02E-01 |
| AT3G45400 | 0,33 | 0,22 | 4,32 | 0,05 | 9,59E-01 |
| AT3G45620 | 1,31 | 1,41 | 2,42 | 0,58 | 5,60E-01 |
| AT3G45640 | 1,79 | 0,65 | 1,38 | 0,47 | 6,37E-01 |
| AT3G45650 | 1,57 | -0,07 | 1,67 | -0,04 | 9,69E-01 |
| AT3G45660 | 1,01 | -2,20 | 2,39 | -0,92 | 3,58E-01 |
| AT3G45670 | 0,15 | 0,29 | 4,37 | 0,07 | 9,47E-01 |
| AT3G45730 | 0,15 | 1,20 | 4,34 | 0,28 | 7,81E-01 |
| AT3G45740 | 0,36 | -0,29 | 4,32 | -0,07 | 9,46E-01 |
| AT3G45830 | 1,79 | 0,76 | 1,44 | 0,53 | 5,98E-01 |
| AT3G45870 | 0,08 | 0,69 | 4,36 | 0,16 | 8,75E-01 |
| AT3G45930 | 0,32 | -0,15 | 4,36 | -0,03 | 9,72E-01 |
| AT3G45965 | 0,21 | 0,21 | 4,38 | 0,05 | 9,62E-01 |
| AT3G45970 | 0,98 | -1,39 | 2,64 | -0,53 | 5,98E-01 |
| AT3G46040 | 0,85 | 0,59 | 2,11 | 0,28 | 7,78E-01 |
| AT3G46060 | 1,03 | -1,67 | 2,04 | -0,82 | 4,12E-01 |
| AT3G46130 | 0,12 | 1,16 | 4,34 | 0,27 | 7,89E-01 |
| AT3G46220 | 0,60 | -1,28 | 2,54 | -0,50 | 6,14E-01 |
| AT3G46290 | 1,13 | 0,42 | 1,82 | 0,23 | 8,15E-01 |
| AT3G46370 | 0,25 | 0,00 | 4,41 | 0,00 | 1,00E+00 |
| AT3G46490 | 0,32 | 0,00 | 4,41 | 0,00 | 1,00E+00 |

| | | | | | |
|---|---|---|---|---|---|
| AT3G46520 | 0,37 | 0,01 | 4,31 | 0,00 | 9,97E-01 |
| AT3G46560 | 0,33 | 0,02 | 4,39 | 0,01 | 9,96E-01 |
| AT3G46590 | 1,40 | 1,95 | 1,69 | 1,15 | 2,50E-01 |
| AT3G46600 | 2,19 | 0,10 | 1,18 | 0,09 | 9,31E-01 |
| AT3G07505 | 1,14 | -1,57 | 2,11 | -0,74 | 4,57E-01 |
| AT3G07515 | 2,88 | 2,22 | 1,63 | 1,36 | 1,75E-01 |
| AT3G46640 | 1,12 | -0,47 | 2,01 | -0,23 | 8,16E-01 |
| AT3G46658 | 0,59 | 0,84 | 2,62 | 0,32 | 7,47E-01 |
| AT3G46668 | 0,57 | -0,15 | 3,24 | -0,05 | 9,63E-01 |
| AT3G07555 | 1,02 | 1,66 | 2,17 | 0,76 | 4,45E-01 |
| AT3G07585 | 1,61 | 0,11 | 1,74 | 0,06 | 9,48E-01 |
| AT3G46930 | 0,21 | 1,57 | 4,33 | 0,36 | 7,16E-01 |
| AT3G47060 | 0,35 | 1,35 | 4,34 | 0,31 | 7,56E-01 |
| AT3G47140 | 0,11 | 0,00 | 4,41 | 0,00 | 1,00E+00 |
| AT3G07695 | 0,33 | 1,93 | 4,32 | 0,45 | 6,54E-01 |
| AT3G47290 | 0,08 | 0,69 | 4,36 | 0,16 | 8,75E-01 |
| AT3G47295 | 0,08 | 0,00 | 4,40 | 0,00 | 1,00E+00 |
| AT3G07715 | 0,48 | -0,05 | 2,73 | -0,02 | 9,84E-01 |
| AT3G07735 | 4,39 | -0,33 | 1,00 | -0,33 | 7,41E-01 |
| AT3G07765 | 2,13 | 1,48 | 1,58 | 0,94 | 3,49E-01 |
| AT3G47490 | 0,22 | 0,07 | 4,40 | 0,02 | 9,88E-01 |
| AT3G47520 | 0,29 | 1,38 | 4,33 | 0,32 | 7,50E-01 |
| AT3G47550 | 0,50 | 1,34 | 3,10 | 0,43 | 6,65E-01 |
| AT3G47570 | 0,50 | 0,01 | 3,17 | 0,00 | 9,98E-01 |
| AT3G47610 | 0,61 | -2,15 | 2,54 | -0,85 | 3,97E-01 |
| AT3G47620 | 3,09 | -0,07 | 1,10 | -0,06 | 9,49E-01 |
| AT3G47650 | 0,65 | 0,31 | 2,91 | 0,11 | 9,16E-01 |
| AT3G47660 | 0,72 | -1,68 | 2,72 | -0,62 | 5,37E-01 |
| AT3G07795 | 0,21 | 1,63 | 4,33 | 0,38 | 7,06E-01 |
| AT3G07805 | 0,89 | 2,87 | 2,66 | 1,08 | 2,81E-01 |
| AT3G47770 | 0,17 | 1,50 | 4,33 | 0,35 | 7,29E-01 |
| AT3G47800 | 1,07 | -0,17 | 1,85 | -0,09 | 9,27E-01 |
| AT3G47820 | 1,04 | -0,40 | 1,89 | -0,21 | 8,33E-01 |
| AT3G47833 | 0,22 | 0,00 | 4,41 | 0,00 | 1,00E+00 |
| AT3G47920 | 1,60 | -0,88 | 1,95 | -0,45 | 6,52E-01 |
| AT3G47930 | 0,52 | 1,20 | 3,86 | 0,31 | 7,57E-01 |
| AT3G47950 | 0,34 | -1,44 | 4,33 | -0,33 | 7,40E-01 |
| AT3G07855 | 4,66 | 0,46 | 1,17 | 0,39 | 6,95E-01 |
| AT3G48040 | 0,17 | -1,70 | 4,40 | -0,39 | 7,00E-01 |
| AT3G48070 | 0,40 | 0,36 | 4,30 | 0,08 | 9,34E-01 |
| AT3G48110 | 0,52 | -0,87 | 3,03 | -0,29 | 7,74E-01 |
| AT3G48140 | 0,97 | 0,81 | 2,12 | 0,38 | 7,02E-01 |
| AT3G48190 | 1,19 | -1,30 | 2,28 | -0,57 | 5,69E-01 |
| AT3G48310 | 0,40 | 0,57 | 3,34 | 0,17 | 8,65E-01 |
| AT3G48350 | 1,84 | -2,92 | 1,73 | -1,69 | 9,14E-02 |
| AT3G48380 | 0,47 | 0,63 | 3,21 | 0,20 | 8,43E-01 |
| AT3G48420 | 0,42 | 0,14 | 4,22 | 0,03 | 9,74E-01 |
| AT3G48430 | 1,70 | -1,08 | 1,66 | -0,65 | 5,16E-01 |
| AT3G48460 | 0,51 | -18,09 | 4,35 | -4,16 | 3,21E-05 |
| AT3G48500 | 0,57 | 1,21 | 3,94 | 0,31 | 7,59E-01 |
| AT3G48530 | 0,79 | -0,35 | 2,14 | -0,16 | 8,70E-01 |
| AT3G48620 | 0,08 | 0,44 | 4,38 | 0,10 | 9,21E-01 |

| | | | | | |
|---|---|---|---|---|---|
| AT3G48680 | 0,15 | 1,16 | 4,34 | 0,27 | 7,88E-01 |
| AT3G48710 | 0,27 | 1,53 | 4,33 | 0,35 | 7,24E-01 |
| AT3G48730 | 0,57 | -1,29 | 2,69 | -0,48 | 6,31E-01 |
| AT3G48750 | 0,54 | -0,70 | 2,62 | -0,27 | 7,88E-01 |
| AT3G48760 | 0,76 | 0,44 | 2,40 | 0,18 | 8,56E-01 |
| AT3G07995 | 0,83 | 0,21 | 2,32 | 0,09 | 9,30E-01 |
| AT3G48780 | 0,82 | 1,65 | 2,72 | 0,61 | 5,43E-01 |
| AT3G48810 | 0,58 | 2,30 | 3,64 | 0,63 | 5,28E-01 |
| AT3G48860 | 1,96 | -0,75 | 2,07 | -0,36 | 7,16E-01 |
| AT3G48890 | 0,66 | -0,27 | 2,65 | -0,10 | 9,19E-01 |
| AT3G49000 | 0,27 | 1,46 | 4,33 | 0,34 | 7,37E-01 |
| AT3G49110 | 0,08 | 0,69 | 4,36 | 0,16 | 8,75E-01 |
| AT3G49120 | 5,10 | 1,81 | 0,92 | 1,97 | 4,85E-02 |
| AT3G49130 | 1,37 | -0,59 | 2,14 | -0,28 | 7,81E-01 |
| AT3G49220 | 0,92 | -2,07 | 1,96 | -1,06 | 2,90E-01 |
| AT3G49230 | 1,94 | -0,32 | 1,81 | -0,18 | 8,60E-01 |
| AT3G49240 | 0,54 | 0,11 | 4,19 | 0,03 | 9,80E-01 |
| AT3G49260 | 0,64 | -2,46 | 4,31 | -0,57 | 5,68E-01 |
| AT3G49430 | 0,93 | 0,11 | 2,09 | 0,05 | 9,57E-01 |
| AT3G49470 | 2,41 | 0,91 | 1,61 | 0,56 | 5,74E-01 |
| AT3G49520 | 1,37 | 0,14 | 1,70 | 0,09 | 9,32E-01 |
| AT3G49590 | 1,44 | -1,72 | 1,53 | -1,13 | 2,60E-01 |
| AT3G49670 | 1,02 | -2,19 | 2,24 | -0,98 | 3,29E-01 |
| AT3G49680 | 0,45 | -0,01 | 3,31 | 0,00 | 9,98E-01 |
| AT3G49710 | 0,32 | -0,65 | 4,33 | -0,15 | 8,81E-01 |
| AT3G49780 | 0,25 | 0,02 | 4,39 | 0,01 | 9,96E-01 |
| AT3G49910 | 2,91 | -0,67 | 1,07 | -0,63 | 5,29E-01 |
| AT3G50000 | 0,43 | 1,96 | 4,18 | 0,47 | 6,40E-01 |
| AT3G50080 | 0,21 | 0,79 | 4,36 | 0,18 | 8,55E-01 |
| AT3G50100 | 0,05 | 0,00 | 4,41 | 0,00 | 1,00E+00 |
| AT3G50270 | 0,60 | -0,30 | 3,01 | -0,10 | 9,20E-01 |
| AT3G50350 | 0,42 | -0,41 | 3,45 | -0,12 | 9,06E-01 |
| AT3G50360 | 0,31 | -0,03 | 4,36 | -0,01 | 9,95E-01 |
| AT3G50430 | 0,16 | 0,00 | 4,41 | 0,00 | 1,00E+00 |
| AT3G50480 | 1,71 | 1,06 | 1,58 | 0,67 | 5,03E-01 |
| AT3G50530 | 0,25 | 1,20 | 4,33 | 0,28 | 7,81E-01 |
| AT3G50540 | 1,39 | 0,01 | 1,77 | 0,01 | 9,96E-01 |
| AT3G50550 | 0,51 | -2,30 | 3,17 | -0,73 | 4,68E-01 |
| AT3G50590 | 4,47 | -1,57 | 0,99 | -1,58 | 1,15E-01 |
| AT3G50630 | 0,26 | 1,53 | 4,33 | 0,35 | 7,24E-01 |
| AT3G50700 | 0,53 | 0,49 | 2,66 | 0,18 | 8,55E-01 |
| AT3G50770 | 1,04 | 1,73 | 2,26 | 0,76 | 4,44E-01 |
| AT3G50890 | 0,25 | 0,00 | 4,38 | 0,00 | 1,00E+00 |
| AT3G50910 | 0,94 | 0,06 | 2,16 | 0,03 | 9,77E-01 |
| AT3G50930 | 3,98 | 0,71 | 0,95 | 0,75 | 4,54E-01 |
| AT3G50950 | 4,46 | 0,21 | 0,87 | 0,24 | 8,12E-01 |
| AT3G50970 | 0,39 | -2,30 | 4,38 | -0,53 | 5,99E-01 |
| AT3G51050 | 3,03 | -0,84 | 1,26 | -0,67 | 5,06E-01 |
| AT3G51110 | 0,46 | -1,42 | 3,98 | -0,36 | 7,21E-01 |
| AT3G51140 | 0,97 | -1,85 | 2,01 | -0,92 | 3,57E-01 |
| AT3G51180 | 0,25 | -1,82 | 4,40 | -0,41 | 6,79E-01 |
| AT3G51260 | 0,46 | -0,73 | 3,22 | -0,23 | 8,20E-01 |

| | | | | | |
|---|---|---|---|---|---|
| AT3G51270 | 0,33 | 0,39 | 4,32 | 0,09 | 9,28E-01 |
| AT3G51325 | 0,36 | -0,05 | 4,36 | -0,01 | 9,91E-01 |
| AT3G51370 | 1,46 | -0,98 | 1,62 | -0,61 | 5,45E-01 |
| AT3G51420 | 0,66 | -1,49 | 2,38 | -0,63 | 5,30E-01 |
| AT3G51430 | 0,32 | -0,74 | 4,39 | -0,17 | 8,66E-01 |
| AT3G51440 | 0,80 | 0,31 | 2,09 | 0,15 | 8,82E-01 |
| AT3G51450 | 0,43 | -0,89 | 2,83 | -0,32 | 7,52E-01 |
| AT3G51460 | 0,27 | -0,79 | 4,39 | -0,18 | 8,58E-01 |
| AT3G51480 | 0,45 | -1,26 | 4,29 | -0,29 | 7,70E-01 |
| AT3G51500 | 0,44 | 2,38 | 4,31 | 0,55 | 5,81E-01 |
| AT3G51510 | 0,44 | -0,75 | 3,42 | -0,22 | 8,26E-01 |
| AT3G51540 | 0,25 | 0,41 | 4,33 | 0,09 | 9,25E-01 |
| AT3G51620 | 1,34 | 0,73 | 1,78 | 0,41 | 6,80E-01 |
| AT3G51630 | 0,87 | 0,60 | 3,12 | 0,19 | 8,49E-01 |
| AT3G51640 | 0,80 | 2,00 | 2,67 | 0,75 | 4,53E-01 |
| AT3G51650 | 1,02 | -0,78 | 1,97 | -0,40 | 6,92E-01 |
| AT3G51730 | 1,38 | 0,44 | 1,62 | 0,27 | 7,87E-01 |
| AT3G51830 | 0,25 | 1,51 | 4,33 | 0,35 | 7,28E-01 |
| AT3G51850 | 1,69 | -1,41 | 1,51 | -0,93 | 3,50E-01 |
| AT3G51860 | 2,20 | 0,57 | 1,27 | 0,45 | 6,53E-01 |
| AT3G51870 | 1,46 | -1,64 | 1,76 | -0,93 | 3,52E-01 |
| AT3G51970 | 0,40 | 0,68 | 3,35 | 0,20 | 8,39E-01 |
| AT3G52080 | 0,33 | 0,49 | 4,32 | 0,11 | 9,09E-01 |
| AT3G52100 | 0,15 | 0,45 | 4,37 | 0,10 | 9,19E-01 |
| AT3G52105 | 0,19 | 0,36 | 4,36 | 0,08 | 9,34E-01 |
| AT3G52120 | 1,03 | -2,94 | 2,06 | -1,43 | 1,53E-01 |
| AT3G52150 | 0,83 | 0,00 | 2,98 | 0,00 | 1,00E+00 |
| AT3G52155 | 0,45 | -2,08 | 4,33 | -0,48 | 6,32E-01 |
| AT3G52200 | 0,34 | -0,92 | 4,36 | -0,21 | 8,32E-01 |
| AT3G52210 | 0,11 | 0,65 | 4,36 | 0,15 | 8,81E-01 |
| AT3G52230 | 0,83 | -1,87 | 2,90 | -0,65 | 5,19E-01 |
| AT3G52250 | 4,06 | 0,32 | 0,90 | 0,36 | 7,22E-01 |
| AT3G52300 | 0,40 | 0,39 | 4,32 | 0,09 | 9,28E-01 |
| AT3G52340 | 0,38 | 1,14 | 4,34 | 0,26 | 7,93E-01 |
| AT3G52370 | 0,08 | 0,44 | 4,38 | 0,10 | 9,21E-01 |
| AT3G52380 | 1,54 | -0,48 | 1,80 | -0,26 | 7,91E-01 |
| AT3G52430 | 2,34 | -0,59 | 1,39 | -0,43 | 6,70E-01 |
| AT3G52470 | 0,30 | 0,71 | 4,36 | 0,16 | 8,71E-01 |
| AT3G52570 | 0,29 | 1,43 | 4,33 | 0,33 | 7,41E-01 |
| AT3G52590 | 1,74 | -0,88 | 1,46 | -0,60 | 5,46E-01 |
| AT3G08275 | 0,60 | -0,64 | 3,13 | -0,20 | 8,38E-01 |
| AT3G52610 | 0,30 | 0,30 | 4,34 | 0,07 | 9,45E-01 |
| AT3G52680 | 0,59 | -1,70 | 2,56 | -0,66 | 5,06E-01 |
| AT3G52710 | 0,88 | 1,16 | 2,00 | 0,58 | 5,60E-01 |
| AT3G08285 | 1,59 | -0,42 | 1,51 | -0,28 | 7,82E-01 |
| AT3G08295 | 0,73 | 0,04 | 3,50 | 0,01 | 9,91E-01 |
| AT3G52800 | 1,93 | 1,05 | 1,64 | 0,64 | 5,21E-01 |
| AT3G52830 | 1,16 | 1,99 | 1,93 | 1,03 | 3,02E-01 |
| AT3G52840 | 0,82 | -0,78 | 2,76 | -0,28 | 7,76E-01 |
| AT3G52850 | 1,76 | 2,10 | 1,68 | 1,25 | 2,10E-01 |
| AT3G52950 | 0,45 | -0,64 | 3,21 | -0,20 | 8,43E-01 |
| AT3G52960 | 0,80 | -0,22 | 1,88 | -0,12 | 9,05E-01 |

| | | | | | |
|---|---|---|---|---|---|
| AT3G52990 | 1,11 | 0,60 | 1,83 | 0,33 | 7,43E-01 |
| AT3G53000 | 0,31 | -0,13 | 4,36 | -0,03 | 9,77E-01 |
| AT3G53030 | 0,57 | 0,28 | 3,93 | 0,07 | 9,43E-01 |
| AT3G53050 | 0,05 | 0,00 | 4,41 | 0,00 | 1,00E+00 |
| AT3G53065 | 0,19 | 0,00 | 4,41 | 0,00 | 1,00E+00 |
| AT3G53090 | 0,36 | -1,75 | 4,40 | -0,40 | 6,91E-01 |
| AT3G53110 | 0,40 | -1,75 | 3,37 | -0,52 | 6,04E-01 |
| AT3G53180 | 0,20 | 0,02 | 4,39 | 0,00 | 9,97E-01 |
| AT3G53190 | 0,21 | -1,41 | 4,36 | -0,32 | 7,46E-01 |
| AT3G53230 | 1,14 | -1,03 | 2,57 | -0,40 | 6,88E-01 |
| AT3G53240 | 0,55 | -2,11 | 3,84 | -0,55 | 5,82E-01 |
| AT3G53305 | 0,72 | 1,71 | 2,25 | 0,76 | 4,49E-01 |
| AT3G53360 | 0,08 | 0,44 | 4,38 | 0,10 | 9,21E-01 |
| AT3G53390 | 0,27 | -0,81 | 4,37 | -0,19 | 8,53E-01 |
| AT3G53400 | 1,89 | -0,28 | 1,56 | -0,18 | 8,58E-01 |
| AT3G53470 | 0,77 | -2,84 | 2,85 | -1,00 | 3,18E-01 |
| AT3G53520 | 0,83 | -0,86 | 2,43 | -0,35 | 7,24E-01 |
| AT3G53670 | 0,90 | -0,31 | 1,97 | -0,16 | 8,74E-01 |
| AT3G53700 | 0,94 | 1,08 | 1,93 | 0,56 | 5,78E-01 |
| AT3G53870 | 0,80 | 0,21 | 2,64 | 0,08 | 9,38E-01 |
| AT3G54050 | 3,78 | -2,79 | 1,08 | -2,58 | 9,96E-03 |
| AT3G54220 | 0,50 | 0,74 | 3,17 | 0,23 | 8,15E-01 |
| AT3G54230 | 1,41 | 0,02 | 1,89 | 0,01 | 9,90E-01 |
| AT3G54280 | 2,37 | -0,11 | 1,28 | -0,08 | 9,33E-01 |
| AT3G54366 | 1,11 | 2,85 | 2,14 | 1,33 | 1,83E-01 |
| AT3G54420 | 0,17 | 1,33 | 4,33 | 0,31 | 7,59E-01 |
| AT3G54540 | 1,42 | 0,93 | 1,69 | 0,55 | 5,81E-01 |
| AT3G54590 | 2,31 | 1,49 | 1,79 | 0,83 | 4,05E-01 |
| AT3G54600 | 0,51 | 0,50 | 4,07 | 0,12 | 9,01E-01 |
| AT3G54610 | 0,47 | 2,07 | 3,96 | 0,52 | 6,01E-01 |
| AT3G54625 | 0,68 | 0,74 | 4,32 | 0,17 | 8,64E-01 |
| AT3G54680 | 0,75 | -1,70 | 2,85 | -0,60 | 5,51E-01 |
| AT3G54730 | 3,74 | 2,64 | 1,47 | 1,79 | 7,29E-02 |
| AT3G54800 | 0,59 | 1,19 | 2,66 | 0,45 | 6,53E-01 |
| AT3G54810 | 2,87 | 0,34 | 1,23 | 0,28 | 7,83E-01 |
| AT3G54820 | 0,15 | 0,74 | 4,35 | 0,17 | 8,65E-01 |
| AT3G54850 | 0,81 | -1,82 | 2,37 | -0,77 | 4,42E-01 |
| AT3G54920 | 2,20 | -0,77 | 1,14 | -0,68 | 4,98E-01 |
| AT3G54980 | 0,41 | -1,28 | 4,19 | -0,31 | 7,60E-01 |
| AT3G55030 | 0,39 | 1,79 | 4,18 | 0,43 | 6,69E-01 |
| AT3G55200 | 2,62 | -0,40 | 1,55 | -0,26 | 7,94E-01 |
| AT3G55250 | 0,63 | -0,21 | 2,34 | -0,09 | 9,30E-01 |
| AT3G55260 | 0,43 | -1,75 | 4,13 | -0,42 | 6,72E-01 |
| AT3G55270 | 1,69 | -0,63 | 1,46 | -0,43 | 6,66E-01 |
| AT3G55280 | 0,61 | -0,01 | 3,76 | 0,00 | 9,97E-01 |
| AT3G55370 | 0,12 | 0,49 | 4,36 | 0,11 | 9,11E-01 |
| AT3G55410 | 3,63 | -2,54 | 1,05 | -2,41 | 1,59E-02 |
| AT3G55420 | 0,14 | 0,09 | 4,38 | 0,02 | 9,84E-01 |
| AT3G55440 | 1,26 | 1,62 | 1,83 | 0,89 | 3,75E-01 |
| AT3G55450 | 0,15 | 1,16 | 4,34 | 0,27 | 7,88E-01 |
| AT3G55460 | 1,02 | 1,54 | 1,96 | 0,79 | 4,32E-01 |
| AT3G55485 | 0,83 | -1,74 | 2,53 | -0,69 | 4,92E-01 |

| | | | | | |
|---|---|---|---|---|---|
| AT3G08485 | 0,83 | -1,74 | 2,53 | -0,69 | 4,92E-01 |
| AT3G55512 | 2,56 | 2,74 | 1,88 | 1,45 | 1,46E-01 |
| AT3G55520 | 0,40 | 1,85 | 4,33 | 0,43 | 6,69E-01 |
| AT3G55620 | 0,54 | -0,33 | 2,57 | -0,13 | 8,98E-01 |
| AT3G55630 | 0,42 | 1,03 | 4,32 | 0,24 | 8,11E-01 |
| AT3G55677 | 0,24 | -1,85 | 4,40 | -0,42 | 6,74E-01 |
| AT3G55760 | 0,99 | -1,10 | 1,86 | -0,59 | 5,55E-01 |
| AT3G55770 | 0,98 | -0,89 | 1,83 | -0,49 | 6,26E-01 |
| AT3G55780 | 0,67 | -2,18 | 3,53 | -0,62 | 5,38E-01 |
| AT3G55800 | 2,45 | -1,29 | 1,37 | -0,94 | 3,45E-01 |
| AT3G55830 | 1,72 | -0,71 | 1,48 | -0,48 | 6,33E-01 |
| AT3G55880 | 0,30 | 1,17 | 3,63 | 0,32 | 7,46E-01 |
| AT3G55940 | 0,10 | 0,21 | 4,38 | 0,05 | 9,62E-01 |
| AT3G55980 | 3,63 | 0,64 | 1,13 | 0,56 | 5,73E-01 |
| AT3G56130 | 0,33 | 0,83 | 4,34 | 0,19 | 8,49E-01 |
| AT3G56140 | 0,74 | 1,32 | 2,39 | 0,55 | 5,79E-01 |
| AT3G56180 | 0,21 | -1,41 | 4,36 | -0,32 | 7,46E-01 |
| AT3G56310 | 0,76 | 1,91 | 2,33 | 0,82 | 4,14E-01 |
| AT3G56330 | 0,52 | 1,23 | 3,84 | 0,32 | 7,49E-01 |
| AT3G56360 | 3,83 | -1,99 | 1,10 | -1,81 | 7,03E-02 |
| AT3G56390 | 0,30 | 0,24 | 4,35 | 0,05 | 9,57E-01 |
| AT3G56408 | 2,59 | 2,54 | 1,48 | 1,72 | 8,56E-02 |
| AT3G56450 | 0,39 | -0,79 | 4,35 | -0,18 | 8,57E-01 |
| AT3G08665 | 0,76 | 1,66 | 2,83 | 0,59 | 5,57E-01 |
| AT3G56470 | 0,12 | 1,16 | 4,34 | 0,27 | 7,89E-01 |
| AT3G56490 | 0,11 | 0,65 | 4,36 | 0,15 | 8,81E-01 |
| AT3G56590 | 0,57 | -0,79 | 3,03 | -0,26 | 7,94E-01 |
| AT3G56640 | 0,59 | 0,82 | 3,08 | 0,26 | 7,91E-01 |
| AT3G56650 | 0,53 | -1,27 | 4,00 | -0,32 | 7,50E-01 |
| AT3G56680 | 2,66 | -0,60 | 1,11 | -0,54 | 5,88E-01 |
| AT3G56720 | 0,17 | 0,18 | 4,39 | 0,04 | 9,67E-01 |
| AT3G56740 | 0,49 | 0,59 | 3,98 | 0,15 | 8,83E-01 |
| AT3G56780 | 0,32 | -1,43 | 4,36 | -0,33 | 7,44E-01 |
| AT3G56840 | 0,64 | 0,39 | 2,42 | 0,16 | 8,71E-01 |
| AT3G56880 | 1,69 | 0,30 | 1,42 | 0,21 | 8,34E-01 |
| AT3G08785 | 1,51 | 0,42 | 1,81 | 0,23 | 8,18E-01 |
| AT3G56960 | 0,60 | -1,46 | 4,32 | -0,34 | 7,35E-01 |
| AT3G57040 | 1,37 | -1,01 | 1,80 | -0,56 | 5,75E-01 |
| AT3G57062 | 1,47 | 0,01 | 1,57 | 0,00 | 9,96E-01 |
| AT3G57070 | 0,24 | 0,00 | 4,41 | 0,00 | 1,00E+00 |
| AT3G57080 | 0,21 | 0,01 | 4,34 | 0,00 | 9,98E-01 |
| AT3G57090 | 0,59 | 0,72 | 2,27 | 0,32 | 7,51E-01 |
| AT3G08845 | 0,12 | 0,37 | 4,37 | 0,08 | 9,33E-01 |
| AT3G57230 | 0,38 | -0,16 | 2,89 | -0,06 | 9,55E-01 |
| AT3G57300 | 2,34 | -0,96 | 1,09 | -0,87 | 3,83E-01 |
| AT3G57340 | 0,42 | 0,46 | 3,41 | 0,13 | 8,93E-01 |
| AT3G57390 | 0,11 | 0,65 | 4,36 | 0,15 | 8,81E-01 |
| AT3G57400 | 1,77 | 0,69 | 1,56 | 0,44 | 6,61E-01 |
| AT3G57450 | 1,58 | -0,75 | 1,40 | -0,53 | 5,94E-01 |
| AT3G57550 | 0,50 | 0,90 | 3,17 | 0,28 | 7,76E-01 |
| AT3G57570 | 0,40 | -0,86 | 4,22 | -0,20 | 8,38E-01 |
| AT3G57587 | 0,73 | -0,19 | 2,26 | -0,09 | 9,32E-01 |

| | | | | | |
|---|---|---|---|---|---|
| AT3G57645 | 1,71 | 4,02 | 1,88 | 2,14 | 3,24E-02 |
| AT3G57650 | 0,53 | 1,75 | 2,66 | 0,66 | 5,10E-01 |
| AT3G57660 | 1,78 | -1,51 | 1,47 | -1,03 | 3,05E-01 |
| AT3G57700 | 1,50 | 1,22 | 1,52 | 0,81 | 4,20E-01 |
| AT3G08975 | 0,42 | -0,54 | 4,08 | -0,13 | 8,95E-01 |
| AT3G57760 | 0,08 | 0,69 | 4,36 | 0,16 | 8,75E-01 |
| AT3G57770 | 1,17 | 0,04 | 1,67 | 0,02 | 9,80E-01 |
| AT3G57810 | 0,21 | 0,83 | 4,36 | 0,19 | 8,48E-01 |
| AT3G57830 | 0,31 | 0,03 | 4,38 | 0,01 | 9,94E-01 |
| AT3G57890 | 1,04 | -0,83 | 1,87 | -0,44 | 6,58E-01 |
| AT3G57940 | 0,20 | 0,02 | 4,39 | 0,00 | 9,97E-01 |
| AT3G57970 | 0,17 | 1,52 | 4,33 | 0,35 | 7,26E-01 |
| AT3G58020 | 0,08 | 0,69 | 4,36 | 0,16 | 8,75E-01 |
| AT3G58030 | 2,17 | 1,02 | 1,47 | 0,70 | 4,87E-01 |
| AT3G58050 | 1,93 | -0,81 | 1,24 | -0,65 | 5,15E-01 |
| AT3G58110 | 1,07 | 2,16 | 3,17 | 0,68 | 4,96E-01 |
| AT3G58160 | 0,08 | 0,44 | 4,38 | 0,10 | 9,21E-01 |
| AT3G58490 | 0,30 | 0,45 | 4,38 | 0,10 | 9,18E-01 |
| AT3G58510 | 0,99 | 0,00 | 1,95 | 0,00 | 9,99E-01 |
| AT3G58530 | 0,46 | -2,07 | 3,97 | -0,52 | 6,03E-01 |
| AT3G58570 | 0,71 | 0,90 | 2,50 | 0,36 | 7,20E-01 |
| AT3G58610 | 2,68 | -0,61 | 1,05 | -0,58 | 5,60E-01 |
| AT3G58680 | 0,61 | -1,44 | 2,63 | -0,55 | 5,85E-01 |
| AT3G58690 | 0,58 | -1,22 | 3,01 | -0,40 | 6,87E-01 |
| AT3G58700 | 0,20 | -0,44 | 4,36 | -0,10 | 9,19E-01 |
| AT3G58795 | 0,23 | 0,00 | 4,41 | 0,00 | 1,00E+00 |
| AT3G59030 | 0,64 | 0,65 | 3,62 | 0,18 | 8,57E-01 |
| AT3G59068 | 1,78 | -0,31 | 1,72 | -0,18 | 8,55E-01 |
| AT3G59080 | 0,19 | 0,07 | 4,40 | 0,02 | 9,88E-01 |
| AT3G59090 | 0,80 | 0,78 | 2,56 | 0,31 | 7,60E-01 |
| AT3G59100 | 0,79 | -3,00 | 2,70 | -1,11 | 2,67E-01 |
| AT3G59110 | 0,41 | 1,01 | 4,27 | 0,24 | 8,12E-01 |
| AT3G59210 | 0,24 | 1,30 | 4,34 | 0,30 | 7,64E-01 |
| AT3G59290 | 1,04 | -0,17 | 1,89 | -0,09 | 9,28E-01 |
| AT3G59300 | 0,19 | 0,86 | 4,34 | 0,20 | 8,42E-01 |
| AT3G59350 | 0,86 | -0,90 | 2,28 | -0,39 | 6,95E-01 |
| AT3G59380 | 0,24 | 1,30 | 4,34 | 0,30 | 7,64E-01 |
| AT3G59410 | 1,72 | -0,96 | 1,54 | -0,62 | 5,34E-01 |
| AT3G59500 | 0,47 | 0,33 | 3,49 | 0,09 | 9,25E-01 |
| AT3G59520 | 0,08 | 0,69 | 4,36 | 0,16 | 8,75E-01 |
| AT3G59530 | 0,53 | -2,71 | 4,29 | -0,63 | 5,28E-01 |
| AT3G59670 | 0,30 | -1,78 | 4,41 | -0,40 | 6,87E-01 |
| AT3G59700 | 0,10 | 0,00 | 4,41 | 0,00 | 1,00E+00 |
| AT3G59780 | 2,03 | 1,17 | 1,50 | 0,78 | 4,36E-01 |
| AT3G59810 | 0,46 | -1,24 | 4,37 | -0,28 | 7,77E-01 |
| AT3G59920 | 0,37 | -0,44 | 4,34 | -0,10 | 9,19E-01 |
| AT3G59970 | 1,71 | 2,48 | 1,63 | 1,52 | 1,27E-01 |
| AT3G60080 | 1,12 | -0,19 | 1,83 | -0,10 | 9,19E-01 |
| AT3G60110 | 0,48 | -0,47 | 4,37 | -0,11 | 9,14E-01 |
| AT3G60130 | 0,63 | 0,91 | 3,17 | 0,29 | 7,75E-01 |
| AT3G60140 | 0,58 | 2,02 | 3,81 | 0,53 | 5,97E-01 |
| AT3G60240 | 1,46 | 1,09 | 1,53 | 0,72 | 4,75E-01 |

| | | | | | |
|---|---|---|---|---|---|
| AT3G60245 | 0,58 | -0,40 | 2,36 | -0,17 | 8,65E-01 |
| AT3G60290 | 0,27 | 0,00 | 4,41 | 0,00 | 1,00E+00 |
| AT3G60300 | 0,58 | 0,26 | 2,69 | 0,10 | 9,22E-01 |
| AT3G60310 | 0,49 | 1,87 | 3,94 | 0,47 | 6,35E-01 |
| AT3G60410 | 0,24 | 0,21 | 4,34 | 0,05 | 9,62E-01 |
| AT3G60415 | 1,73 | -0,50 | 1,46 | -0,34 | 7,34E-01 |
| AT3G60420 | 0,55 | 0,25 | 3,78 | 0,07 | 9,47E-01 |
| AT3G60440 | 0,83 | -2,52 | 3,35 | -0,75 | 4,51E-01 |
| AT3G60450 | 1,00 | 1,19 | 2,53 | 0,47 | 6,37E-01 |
| AT3G60500 | 0,28 | 0,23 | 4,33 | 0,05 | 9,58E-01 |
| AT3G09445 | 4,99 | -0,92 | 0,94 | -0,98 | 3,26E-01 |
| AT3G60530 | 0,83 | 0,03 | 2,27 | 0,02 | 9,88E-01 |
| AT3G09465 | 2,48 | -0,58 | 1,51 | -0,38 | 7,01E-01 |
| AT3G60600 | 1,58 | 0,70 | 1,50 | 0,46 | 6,44E-01 |
| AT3G60690 | 1,35 | 1,51 | 1,74 | 0,87 | 3,84E-01 |
| AT3G60860 | 2,94 | 0,25 | 1,17 | 0,22 | 8,29E-01 |
| AT3G60910 | 0,08 | 0,00 | 4,41 | 0,00 | 1,00E+00 |
| AT3G60950 | 1,11 | 1,07 | 2,15 | 0,50 | 6,20E-01 |
| AT3G60960 | 1,54 | -0,15 | 1,58 | -0,10 | 9,24E-01 |
| AT3G09515 | 1,27 | -0,45 | 1,79 | -0,25 | 8,00E-01 |
| AT3G60961 | 1,33 | 2,77 | 2,05 | 1,36 | 1,75E-01 |
| AT3G09525 | 0,12 | 1,16 | 4,34 | 0,27 | 7,89E-01 |
| AT3G60970 | 0,12 | 1,16 | 4,34 | 0,27 | 7,89E-01 |
| AT3G61030 | 0,61 | 0,81 | 2,18 | 0,37 | 7,09E-01 |
| AT3G61050 | 1,51 | -0,37 | 1,76 | -0,21 | 8,33E-01 |
| AT3G61080 | 0,12 | 1,23 | 4,34 | 0,28 | 7,76E-01 |
| AT3G61110 | 1,40 | 0,49 | 1,79 | 0,27 | 7,87E-01 |
| AT3G61130 | 1,07 | -3,21 | 2,28 | -1,41 | 1,58E-01 |
| AT3G61140 | 0,73 | -1,33 | 1,91 | -0,69 | 4,88E-01 |
| AT3G61150 | 0,68 | -0,10 | 2,71 | -0,04 | 9,72E-01 |
| AT3G61180 | 0,94 | 1,06 | 2,45 | 0,43 | 6,65E-01 |
| AT3G61240 | 1,00 | -0,17 | 1,81 | -0,09 | 9,26E-01 |
| AT3G61280 | 0,41 | 0,75 | 3,34 | 0,23 | 8,21E-01 |
| AT3G61320 | 0,08 | 0,00 | 4,40 | 0,00 | 1,00E+00 |
| AT3G61430 | 1,94 | 0,23 | 1,66 | 0,14 | 8,89E-01 |
| AT3G61440 | 1,04 | 0,22 | 2,26 | 0,10 | 9,21E-01 |
| AT3G61480 | 0,44 | -0,54 | 3,23 | -0,17 | 8,67E-01 |
| AT3G61490 | 0,55 | -2,06 | 4,30 | -0,48 | 6,31E-01 |
| AT3G61540 | 0,29 | 1,23 | 4,33 | 0,28 | 7,76E-01 |
| AT3G61550 | 0,37 | 1,51 | 3,48 | 0,43 | 6,64E-01 |
| AT3G61580 | 5,37 | -0,30 | 0,75 | -0,40 | 6,85E-01 |
| AT3G61600 | 1,22 | 0,66 | 1,60 | 0,41 | 6,79E-01 |
| AT3G61630 | 1,88 | -1,42 | 1,57 | -0,90 | 3,65E-01 |
| AT3G61670 | 0,29 | 0,24 | 4,33 | 0,06 | 9,55E-01 |
| AT3G61690 | 2,25 | -1,05 | 1,25 | -0,84 | 3,98E-01 |
| AT3G61700 | 1,46 | -0,96 | 1,65 | -0,58 | 5,60E-01 |
| AT3G61755 | 0,16 | 0,54 | 4,36 | 0,12 | 9,01E-01 |
| AT3G61810 | 0,05 | 0,00 | 4,41 | 0,00 | 1,00E+00 |
| AT3G61850 | 0,15 | 0,74 | 4,35 | 0,17 | 8,65E-01 |
| AT3G61870 | 1,16 | -0,65 | 1,68 | -0,38 | 7,00E-01 |
| AT3G61897 | 2,76 | 0,00 | 2,83 | 0,00 | 1,00E+00 |
| AT3G61950 | 0,11 | 0,65 | 4,36 | 0,15 | 8,81E-01 |

| | | | | | |
|---|---|---|---|---|---|
| AT3G62010 | 2,16 | -1,01 | 1,28 | -0,79 | 4,27E-01 |
| AT3G62030 | 0,98 | -1,54 | 2,24 | -0,69 | 4,93E-01 |
| AT3G62130 | 0,39 | -1,35 | 4,33 | -0,31 | 7,56E-01 |
| AT3G62190 | 0,24 | 0,92 | 4,36 | 0,21 | 8,32E-01 |
| AT3G62245 | 0,42 | 0,44 | 4,13 | 0,11 | 9,16E-01 |
| AT3G62250 | 1,91 | -0,46 | 1,30 | -0,36 | 7,21E-01 |
| AT3G62290 | 5,55 | -0,52 | 0,82 | -0,64 | 5,25E-01 |
| AT3G62300 | 0,93 | -0,09 | 3,38 | -0,03 | 9,79E-01 |
| AT3G62370 | 0,90 | 2,74 | 2,20 | 1,25 | 2,11E-01 |
| AT3G62410 | 1,88 | -0,20 | 1,42 | -0,14 | 8,85E-01 |
| AT3G62530 | 0,72 | 2,38 | 2,42 | 0,98 | 3,25E-01 |
| AT3G62550 | 0,70 | 0,71 | 3,76 | 0,19 | 8,50E-01 |
| AT3G62560 | 0,61 | -2,37 | 3,09 | -0,77 | 4,43E-01 |
| AT3G62580 | 0,64 | -0,21 | 2,18 | -0,10 | 9,23E-01 |
| AT3G09815 | 0,08 | 0,44 | 4,38 | 0,10 | 9,21E-01 |
| AT3G09835 | 1,22 | 0,02 | 2,07 | 0,01 | 9,92E-01 |
| AT3G62620 | 0,29 | 0,00 | 4,39 | 0,00 | 1,00E+00 |
| AT3G62650 | 1,34 | -0,59 | 1,60 | -0,37 | 7,13E-01 |
| AT3G62660 | 1,25 | 0,71 | 1,81 | 0,39 | 6,96E-01 |
| AT3G62720 | 0,40 | 1,69 | 4,10 | 0,41 | 6,80E-01 |
| AT3G62750 | 0,68 | -2,95 | 2,51 | -1,18 | 2,39E-01 |
| AT3G62820 | 0,26 | -0,87 | 4,36 | -0,20 | 8,42E-01 |
| AT3G62830 | 0,86 | 0,53 | 1,97 | 0,27 | 7,87E-01 |
| AT3G62890 | 0,20 | 0,81 | 4,36 | 0,19 | 8,52E-01 |
| AT3G63000 | 0,33 | 0,79 | 4,31 | 0,18 | 8,55E-01 |
| AT3G63010 | 0,38 | 0,22 | 3,48 | 0,06 | 9,50E-01 |
| AT3G63070 | 1,86 | -0,11 | 1,68 | -0,07 | 9,48E-01 |
| AT3G63080 | 0,53 | -0,78 | 2,63 | -0,30 | 7,67E-01 |
| AT3G63130 | 0,43 | -0,73 | 4,09 | -0,18 | 8,58E-01 |
| AT3G63170 | 0,32 | -2,61 | 4,37 | -0,60 | 5,51E-01 |
| AT3G63215 | 0,08 | 0,00 | 4,40 | 0,00 | 1,00E+00 |
| AT3G63300 | 0,72 | -1,68 | 2,72 | -0,62 | 5,37E-01 |
| AT3G63370 | 0,57 | -0,98 | 3,03 | -0,32 | 7,46E-01 |
| AT3G63400 | 0,95 | 0,43 | 2,13 | 0,20 | 8,40E-01 |
| AT3G63420 | 0,46 | 0,19 | 4,30 | 0,04 | 9,65E-01 |
| AT3G63470 | 0,42 | -0,70 | 4,13 | -0,17 | 8,66E-01 |
| AT3G63490 | 1,82 | 0,82 | 1,57 | 0,52 | 6,04E-01 |
| AT3G09995 | 0,45 | 1,00 | 4,02 | 0,25 | 8,03E-01 |
| AT3G63520 | 0,87 | -1,25 | 2,23 | -0,56 | 5,75E-01 |
| AT4G00030 | 0,38 | 0,01 | 4,39 | 0,00 | 9,98E-01 |
| AT4G00060 | 1,77 | 1,69 | 1,58 | 1,07 | 2,85E-01 |
| AT4G00100 | 1,39 | -0,55 | 1,90 | -0,29 | 7,70E-01 |
| AT4G00163 | 0,65 | -2,37 | 2,73 | -0,87 | 3,85E-01 |
| AT4G00230 | 0,69 | -2,06 | 4,34 | -0,48 | 6,34E-01 |
| AT4G00231 | 1,99 | 1,46 | 1,34 | 1,09 | 2,76E-01 |
| AT4G00300 | 1,08 | -1,59 | 2,00 | -0,80 | 4,25E-01 |
| AT4G00340 | 0,31 | -0,92 | 4,39 | -0,21 | 8,35E-01 |
| AT4G00360 | 2,92 | -1,80 | 1,27 | -1,41 | 1,58E-01 |
| AT4G00440 | 0,33 | 0,27 | 4,34 | 0,06 | 9,51E-01 |
| AT4G00450 | 3,39 | 0,23 | 1,09 | 0,21 | 8,34E-01 |
| AT4G00490 | 0,44 | 1,39 | 4,35 | 0,32 | 7,50E-01 |
| AT4G00580 | 0,48 | -0,87 | 4,04 | -0,22 | 8,29E-01 |

| | | | | | |
|---|---|---|---|---|---|
| AT4G00660 | 1,18 | 0,26 | 1,91 | 0,14 | 8,90E-01 |
| AT4G00700 | 2,06 | 0,55 | 1,47 | 0,37 | 7,11E-01 |
| AT4G00710 | 0,48 | 0,36 | 4,34 | 0,08 | 9,34E-01 |
| AT4G00750 | 0,96 | -0,22 | 1,80 | -0,12 | 9,02E-01 |
| AT4G00820 | 0,05 | 0,00 | 4,41 | 0,00 | 1,00E+00 |
| AT4G00830 | 0,92 | 0,01 | 2,12 | 0,01 | 9,95E-01 |
| AT4G00889 | 0,23 | 0,00 | 4,41 | 0,00 | 1,00E+00 |
| AT4G00930 | 0,17 | 1,50 | 4,33 | 0,35 | 7,29E-01 |
| AT4G00953 | 0,67 | 0,26 | 2,64 | 0,10 | 9,23E-01 |
| AT4G00955 | 0,42 | 0,59 | 2,92 | 0,20 | 8,39E-01 |
| AT4G00960 | 1,30 | -0,35 | 2,18 | -0,16 | 8,73E-01 |
| AT4G00970 | 0,33 | 0,00 | 4,38 | 0,00 | 1,00E+00 |
| AT4G00990 | 0,61 | -0,97 | 3,84 | -0,25 | 8,00E-01 |
| AT4G01020 | 0,37 | -0,40 | 4,33 | -0,09 | 9,26E-01 |
| AT4G01026 | 0,56 | 0,06 | 2,56 | 0,02 | 9,82E-01 |
| AT4G01040 | 0,12 | 0,97 | 4,34 | 0,22 | 8,23E-01 |
| AT4G01050 | 4,58 | -0,80 | 0,96 | -0,84 | 4,04E-01 |
| AT4G01100 | 0,48 | 0,90 | 3,19 | 0,28 | 7,78E-01 |
| AT4G01130 | 0,11 | 0,00 | 4,41 | 0,00 | 1,00E+00 |
| AT4G01150 | 2,10 | 0,19 | 1,24 | 0,15 | 8,78E-01 |
| AT4G01280 | 0,59 | 0,74 | 3,16 | 0,23 | 8,16E-01 |
| AT4G01320 | 0,90 | -0,07 | 1,96 | -0,04 | 9,70E-01 |
| AT4G01330 | 0,80 | 0,82 | 2,50 | 0,33 | 7,43E-01 |
| AT4G01340 | 1,48 | 0,19 | 1,85 | 0,10 | 9,20E-01 |
| AT4G01360 | 0,89 | 0,88 | 2,92 | 0,30 | 7,63E-01 |
| AT4G01370 | 0,27 | 1,29 | 4,33 | 0,30 | 7,65E-01 |
| AT4G01460 | 0,10 | 0,01 | 4,38 | 0,00 | 9,99E-01 |
| AT4G01570 | 0,24 | 1,26 | 4,33 | 0,29 | 7,71E-01 |
| AT4G03905 | 0,55 | 1,88 | 3,91 | 0,48 | 6,31E-01 |
| AT4G03925 | 0,76 | 0,23 | 2,30 | 0,10 | 9,21E-01 |
| AT4G01610 | 0,62 | 0,94 | 2,48 | 0,38 | 7,04E-01 |
| AT4G01690 | 1,14 | -0,82 | 1,64 | -0,50 | 6,17E-01 |
| AT4G01720 | 0,44 | -2,01 | 4,09 | -0,49 | 6,22E-01 |
| AT4G01840 | 3,87 | 3,39 | 1,56 | 2,18 | 2,95E-02 |
| AT4G01995 | 0,08 | 0,44 | 4,38 | 0,10 | 9,21E-01 |
| AT4G02000 | 0,69 | 1,24 | 2,26 | 0,55 | 5,82E-01 |
| AT4G02010 | 0,14 | 0,40 | 4,37 | 0,09 | 9,27E-01 |
| AT4G02020 | 0,55 | 0,05 | 3,20 | 0,01 | 9,89E-01 |
| AT4G02030 | 0,75 | -1,00 | 2,63 | -0,38 | 7,03E-01 |
| AT4G02070 | 0,12 | 1,05 | 4,35 | 0,24 | 8,10E-01 |
| AT4G02080 | 0,57 | 0,87 | 2,59 | 0,34 | 7,37E-01 |
| AT4G02100 | 0,08 | 0,00 | 4,40 | 0,00 | 1,00E+00 |
| AT4G02110 | 0,21 | 1,57 | 4,33 | 0,36 | 7,16E-01 |
| AT4G02200 | 0,55 | 1,84 | 3,84 | 0,48 | 6,32E-01 |
| AT4G02220 | 0,29 | 0,05 | 4,33 | 0,01 | 9,90E-01 |
| AT4G02260 | 0,56 | -0,90 | 3,00 | -0,30 | 7,64E-01 |
| AT4G04095 | 1,94 | -0,90 | 1,34 | -0,67 | 5,03E-01 |
| AT4G02330 | 0,22 | 0,00 | 4,41 | 0,00 | 1,00E+00 |
| AT4G02340 | 1,02 | -1,19 | 2,24 | -0,53 | 5,94E-01 |
| AT4G02350 | 1,95 | -2,63 | 1,55 | -1,69 | 9,10E-02 |
| AT4G02370 | 0,72 | 1,11 | 3,08 | 0,36 | 7,19E-01 |
| AT4G02390 | 0,10 | 0,01 | 4,38 | 0,00 | 9,99E-01 |

| | | | | | |
|---|---|---|---|---|---|
| AT4G02400 | 0,19 | 1,15 | 4,34 | 0,27 | 7,91E-01 |
| AT4G02440 | 0,90 | -0,63 | 1,94 | -0,32 | 7,48E-01 |
| AT4G04135 | 0,05 | 0,00 | 4,41 | 0,00 | 1,00E+00 |
| AT4G02500 | 1,37 | 2,59 | 2,74 | 0,95 | 3,44E-01 |
| AT4G04195 | 0,38 | 1,50 | 3,38 | 0,44 | 6,57E-01 |
| AT4G02540 | 0,29 | 0,16 | 3,72 | 0,04 | 9,65E-01 |
| AT4G02555 | 3,06 | 0,48 | 1,17 | 0,41 | 6,84E-01 |
| AT4G04205 | 2,87 | 0,14 | 1,28 | 0,11 | 9,15E-01 |
| AT4G02570 | 1,99 | 0,60 | 1,30 | 0,46 | 6,47E-01 |
| AT4G02580 | 1,19 | -1,41 | 2,15 | -0,66 | 5,12E-01 |
| AT4G02600 | 0,66 | -1,39 | 2,26 | -0,62 | 5,37E-01 |
| AT4G02610 | 0,31 | 1,01 | 4,35 | 0,23 | 8,16E-01 |
| AT4G02650 | 0,24 | 0,21 | 4,34 | 0,05 | 9,62E-01 |
| AT4G02700 | 1,40 | 0,39 | 1,79 | 0,22 | 8,27E-01 |
| AT4G02725 | 0,69 | -0,89 | 3,00 | -0,30 | 7,68E-01 |
| AT4G02790 | 0,08 | 0,69 | 4,36 | 0,16 | 8,75E-01 |
| AT4G02880 | 0,95 | -1,05 | 1,99 | -0,53 | 6,00E-01 |
| AT4G02920 | 1,02 | 2,17 | 2,31 | 0,94 | 3,46E-01 |
| AT4G02940 | 0,12 | 0,00 | 4,39 | 0,00 | 1,00E+00 |
| AT4G02990 | 0,69 | -2,17 | 2,56 | -0,85 | 3,98E-01 |
| AT4G03000 | 0,64 | 0,62 | 2,94 | 0,21 | 8,32E-01 |
| AT4G04425 | 0,08 | 0,44 | 4,38 | 0,10 | 9,21E-01 |
| AT4G03100 | 0,38 | -2,37 | 4,35 | -0,55 | 5,85E-01 |
| AT4G03120 | 0,23 | -0,21 | 4,35 | -0,05 | 9,61E-01 |
| AT4G03180 | 0,27 | 0,89 | 4,36 | 0,20 | 8,38E-01 |
| AT4G03200 | 0,82 | 1,35 | 2,37 | 0,57 | 5,69E-01 |
| AT4G03205 | 0,48 | -1,27 | 3,13 | -0,41 | 6,85E-01 |
| AT4G03210 | 0,53 | 0,75 | 4,32 | 0,17 | 8,61E-01 |
| AT4G03260 | 2,29 | 0,01 | 1,16 | 0,01 | 9,93E-01 |
| AT4G03280 | 3,09 | -0,29 | 1,19 | -0,25 | 8,04E-01 |
| AT4G03285 | 0,16 | 0,92 | 4,35 | 0,21 | 8,33E-01 |
| AT4G03410 | 0,58 | -1,85 | 3,86 | -0,48 | 6,31E-01 |
| AT4G03420 | 2,29 | 1,03 | 1,51 | 0,68 | 4,95E-01 |
| AT4G03500 | 0,47 | -1,20 | 4,38 | -0,27 | 7,85E-01 |
| AT4G03550 | 3,00 | 0,29 | 1,25 | 0,23 | 8,16E-01 |
| AT4G03560 | 2,24 | 0,75 | 1,47 | 0,51 | 6,13E-01 |
| AT4G03830 | 0,61 | -0,57 | 2,53 | -0,23 | 8,21E-01 |
| AT4G03930 | 0,65 | 2,72 | 3,01 | 0,90 | 3,66E-01 |
| AT4G03940 | 0,17 | 1,33 | 4,33 | 0,31 | 7,59E-01 |
| AT4G03945 | 0,92 | 2,62 | 2,34 | 1,12 | 2,62E-01 |
| AT4G04020 | 1,87 | 0,00 | 2,09 | 0,00 | 9,98E-01 |
| AT4G04110 | 0,65 | -1,11 | 4,38 | -0,25 | 7,99E-01 |
| AT4G04180 | 0,08 | 0,00 | 4,40 | 0,00 | 1,00E+00 |
| AT4G04210 | 0,39 | 0,38 | 2,86 | 0,13 | 8,95E-01 |
| AT4G04220 | 0,89 | -1,30 | 1,92 | -0,68 | 4,98E-01 |
| AT4G04795 | 0,87 | 2,14 | 2,55 | 0,84 | 4,02E-01 |
| AT4G04315 | 0,10 | 0,01 | 4,38 | 0,00 | 9,99E-01 |
| AT4G04340 | 2,36 | -1,76 | 1,66 | -1,06 | 2,89E-01 |
| AT4G04350 | 0,42 | -0,74 | 3,44 | -0,22 | 8,29E-01 |
| AT4G04450 | 0,17 | 0,00 | 4,41 | 0,00 | 1,00E+00 |
| AT4G04510 | 0,17 | -1,70 | 4,40 | -0,39 | 7,00E-01 |
| AT4G04540 | 0,26 | 0,56 | 4,38 | 0,13 | 8,99E-01 |

| | | | | | |
|---|---|---|---|---|---|
| AT4G04570 | 4,35 | -3,59 | 1,50 | -2,39 | 1,69E-02 |
| AT4G04610 | 0,73 | 0,16 | 2,15 | 0,07 | 9,41E-01 |
| AT4G04670 | 0,48 | 0,90 | 4,33 | 0,21 | 8,36E-01 |
| AT4G04750 | 0,90 | 1,62 | 2,01 | 0,81 | 4,19E-01 |
| AT4G04760 | 1,18 | -0,62 | 2,16 | -0,29 | 7,74E-01 |
| AT4G04800 | 0,52 | 0,53 | 3,84 | 0,14 | 8,90E-01 |
| AT4G04840 | 1,00 | -1,14 | 2,20 | -0,52 | 6,03E-01 |
| AT4G04850 | 0,79 | -0,56 | 1,87 | -0,30 | 7,67E-01 |
| AT4G04860 | 0,28 | 1,16 | 3,71 | 0,31 | 7,55E-01 |
| AT4G04920 | 1,70 | 0,32 | 1,47 | 0,22 | 8,29E-01 |
| AT4G04925 | 1,79 | 0,82 | 1,68 | 0,49 | 6,28E-01 |
| AT4G04960 | 0,39 | 0,25 | 2,86 | 0,09 | 9,31E-01 |
| AT4G04970 | 2,63 | 3,95 | 1,89 | 2,09 | 3,67E-02 |
| AT4G05020 | 0,40 | 0,52 | 4,26 | 0,12 | 9,04E-01 |
| AT4G05080 | 0,23 | 0,00 | 4,41 | 0,00 | 1,00E+00 |
| AT4G05105 | 0,17 | 1,31 | 4,34 | 0,30 | 7,63E-01 |
| AT4G05150 | 3,81 | 1,10 | 1,22 | 0,90 | 3,69E-01 |
| AT4G05160 | 0,46 | 2,28 | 4,05 | 0,56 | 5,73E-01 |
| AT4G05360 | 2,15 | 1,87 | 1,65 | 1,13 | 2,57E-01 |
| AT4G05612 | 0,12 | 1,16 | 4,34 | 0,27 | 7,89E-01 |
| AT4G05632 | 0,08 | 0,00 | 4,40 | 0,00 | 1,00E+00 |
| AT4G06479 | 0,10 | 0,00 | 4,41 | 0,00 | 1,00E+00 |
| AT4G06598 | 0,74 | -4,16 | 3,43 | -1,21 | 2,25E-01 |
| AT4G06688 | 0,68 | -0,06 | 2,36 | -0,03 | 9,79E-01 |
| AT4G07390 | 0,27 | 1,17 | 3,74 | 0,31 | 7,55E-01 |
| AT4G07455 | 0,24 | -2,36 | 4,39 | -0,54 | 5,91E-01 |
| AT4G07825 | 1,05 | -0,65 | 2,54 | -0,26 | 7,98E-01 |
| AT4G07932 | 0,51 | 0,00 | 4,16 | 0,00 | 1,00E+00 |
| AT4G07965 | 0,15 | 1,16 | 4,34 | 0,27 | 7,88E-01 |
| AT4G08025 | 0,16 | 0,84 | 4,35 | 0,19 | 8,46E-01 |
| AT4G05305 | 0,12 | 0,00 | 4,41 | 0,00 | 1,00E+00 |
| AT4G08097 | 0,66 | 0,70 | 2,64 | 0,26 | 7,92E-01 |
| AT4G08180 | 1,01 | 0,45 | 2,04 | 0,22 | 8,25E-01 |
| AT4G08267 | 0,63 | 0,71 | 4,37 | 0,16 | 8,71E-01 |
| AT4G08270 | 0,17 | 0,61 | 4,37 | 0,14 | 8,88E-01 |
| AT4G08320 | 0,31 | 0,00 | 4,41 | 0,00 | 1,00E+00 |
| AT4G08350 | 0,65 | 2,13 | 2,59 | 0,82 | 4,10E-01 |
| AT4G08395 | 0,29 | 0,26 | 4,37 | 0,06 | 9,52E-01 |
| AT4G05455 | 0,68 | 0,22 | 2,62 | 0,08 | 9,34E-01 |
| AT4G08450 | 0,11 | 0,00 | 4,41 | 0,00 | 1,00E+00 |
| AT4G08455 | 0,63 | 2,14 | 4,32 | 0,50 | 6,20E-01 |
| AT4G08510 | 0,33 | 1,55 | 4,33 | 0,36 | 7,19E-01 |
| AT4G08545 | 1,36 | 1,64 | 1,84 | 0,89 | 3,72E-01 |
| AT4G08560 | 0,64 | -3,82 | 4,31 | -0,89 | 3,75E-01 |
| AT4G08593 | 1,89 | 2,80 | 1,89 | 1,48 | 1,39E-01 |
| AT4G08620 | 2,91 | 3,19 | 1,66 | 1,92 | 5,53E-02 |
| AT4G08780 | 0,43 | -1,86 | 4,11 | -0,45 | 6,51E-01 |
| AT4G05605 | 0,39 | -1,78 | 4,41 | -0,40 | 6,87E-01 |
| AT4G08920 | 2,64 | -0,06 | 1,21 | -0,05 | 9,64E-01 |
| AT4G08930 | 0,71 | 1,82 | 2,40 | 0,76 | 4,48E-01 |
| AT4G08950 | 0,37 | -0,32 | 4,31 | -0,07 | 9,41E-01 |
| AT4G08980 | 0,65 | 2,18 | 2,91 | 0,75 | 4,52E-01 |

| | | | | | |
|---|---|---|---|---|---|
| AT4G08990 | 0,69 | 1,51 | 3,71 | 0,41 | 6,83E-01 |
| AT4G09000 | 1,74 | -0,53 | 1,41 | -0,38 | 7,04E-01 |
| AT4G09020 | 1,04 | 2,22 | 2,02 | 1,10 | 2,71E-01 |
| AT4G09030 | 0,16 | 0,84 | 4,35 | 0,19 | 8,46E-01 |
| AT4G09150 | 0,62 | 0,34 | 2,91 | 0,12 | 9,06E-01 |
| AT4G09160 | 0,36 | -3,19 | 4,33 | -0,74 | 4,61E-01 |
| AT4G09250 | 0,12 | 0,97 | 4,34 | 0,22 | 8,23E-01 |
| AT4G09310 | 0,21 | 1,01 | 4,34 | 0,23 | 8,17E-01 |
| AT4G05705 | 0,10 | 0,00 | 4,41 | 0,00 | 1,00E+00 |
| AT4G09320 | 1,06 | 0,91 | 1,77 | 0,52 | 6,06E-01 |
| AT4G09360 | 0,71 | 1,73 | 2,44 | 0,71 | 4,79E-01 |
| AT4G09450 | 1,44 | -0,94 | 1,91 | -0,49 | 6,23E-01 |
| AT4G09460 | 0,37 | 0,40 | 4,29 | 0,09 | 9,26E-01 |
| AT4G09545 | 0,31 | 0,00 | 4,41 | 0,00 | 1,00E+00 |
| AT4G09570 | 0,54 | -1,83 | 2,37 | -0,77 | 4,39E-01 |
| AT4G09630 | 0,87 | -1,25 | 2,11 | -0,59 | 5,54E-01 |
| AT4G09650 | 3,00 | 1,74 | 1,82 | 0,95 | 3,41E-01 |
| AT4G09660 | 0,30 | 0,01 | 4,38 | 0,00 | 9,99E-01 |
| AT4G09680 | 0,45 | -0,01 | 3,94 | 0,00 | 9,99E-01 |
| AT4G09690 | 1,50 | 0,31 | 1,92 | 0,16 | 8,72E-01 |
| AT4G09730 | 0,51 | -1,12 | 3,22 | -0,35 | 7,27E-01 |
| AT4G09800 | 0,41 | -2,51 | 4,27 | -0,59 | 5,57E-01 |
| AT4G09820 | 0,31 | 0,00 | 4,41 | 0,00 | 1,00E+00 |
| AT4G09830 | 0,41 | 0,43 | 3,36 | 0,13 | 8,99E-01 |
| AT4G09850 | 0,21 | 0,34 | 4,37 | 0,08 | 9,38E-01 |
| AT4G05745 | 0,45 | 0,56 | 3,15 | 0,18 | 8,59E-01 |
| AT4G09890 | 0,18 | 1,15 | 4,34 | 0,26 | 7,91E-01 |
| AT4G10060 | 0,79 | 0,50 | 2,14 | 0,23 | 8,17E-01 |
| AT4G10120 | 1,86 | -0,95 | 1,29 | -0,74 | 4,61E-01 |
| AT4G10170 | 0,08 | 0,69 | 4,36 | 0,16 | 8,75E-01 |
| AT4G10180 | 0,21 | 1,63 | 4,33 | 0,38 | 7,06E-01 |
| AT4G10210 | 2,08 | 0,03 | 1,42 | 0,02 | 9,85E-01 |
| AT4G10465 | 0,60 | 1,55 | 2,63 | 0,59 | 5,55E-01 |
| AT4G10610 | 1,30 | -2,04 | 2,08 | -0,98 | 3,28E-01 |
| AT4G10613 | 0,77 | 1,77 | 2,15 | 0,83 | 4,08E-01 |
| AT4G10620 | 0,24 | -2,36 | 4,39 | -0,54 | 5,91E-01 |
| AT4G10660 | 0,11 | 0,00 | 4,41 | 0,00 | 1,00E+00 |
| AT4G10840 | 1,51 | -0,54 | 1,77 | -0,31 | 7,59E-01 |
| AT4G11000 | 0,26 | 0,01 | 4,33 | 0,00 | 9,98E-01 |
| AT4G11070 | 0,37 | -2,13 | 4,37 | -0,49 | 6,27E-01 |
| AT4G11100 | 0,32 | 0,23 | 4,32 | 0,05 | 9,58E-01 |
| AT4G11110 | 0,51 | 1,14 | 2,67 | 0,43 | 6,70E-01 |
| AT4G11130 | 0,54 | -1,42 | 3,82 | -0,37 | 7,11E-01 |
| AT4G11150 | 1,03 | 1,53 | 1,90 | 0,80 | 4,21E-01 |
| AT4G11160 | 1,61 | -0,09 | 1,56 | -0,06 | 9,55E-01 |
| AT4G11170 | 0,50 | -1,72 | 3,22 | -0,53 | 5,94E-01 |
| AT4G11240 | 1,11 | -0,39 | 1,75 | -0,22 | 8,22E-01 |
| AT4G11270 | 3,79 | -0,12 | 1,10 | -0,11 | 9,12E-01 |
| AT4G11280 | 0,67 | -1,16 | 3,06 | -0,38 | 7,04E-01 |
| AT4G11330 | 0,53 | -0,58 | 3,13 | -0,19 | 8,52E-01 |
| AT4G11350 | 0,19 | 1,09 | 4,34 | 0,25 | 8,01E-01 |
| AT4G11360 | 0,31 | 0,50 | 4,38 | 0,11 | 9,09E-01 |

| | | | | | |
|---|---|---|---|---|---|
| AT4G11380 | 1,75 | 2,30 | 1,67 | 1,38 | 1,68E-01 |
| AT4G05885 | 0,80 | 0,01 | 2,81 | 0,00 | 9,96E-01 |
| AT4G11570 | 2,16 | -0,56 | 1,28 | -0,43 | 6,64E-01 |
| AT4G05900 | 1,70 | 1,09 | 1,79 | 0,61 | 5,43E-01 |
| AT4G11660 | 0,57 | -0,03 | 3,19 | -0,01 | 9,91E-01 |
| AT4G11720 | 0,60 | -1,28 | 3,01 | -0,43 | 6,70E-01 |
| AT4G11740 | 0,35 | -0,19 | 4,34 | -0,04 | 9,64E-01 |
| AT4G11880 | 0,28 | 0,00 | 4,41 | 0,00 | 1,00E+00 |
| AT4G11890 | 0,62 | 2,44 | 3,02 | 0,81 | 4,19E-01 |
| AT4G11910 | 0,12 | 1,23 | 4,34 | 0,28 | 7,76E-01 |
| AT4G12020 | 1,23 | 2,00 | 2,47 | 0,81 | 4,18E-01 |
| AT4G12030 | 0,68 | 3,07 | 3,46 | 0,89 | 3,74E-01 |
| AT4G12040 | 1,45 | 1,78 | 1,77 | 1,00 | 3,17E-01 |
| AT4G12070 | 0,19 | 0,00 | 4,41 | 0,00 | 1,00E+00 |
| AT4G12240 | 0,26 | 1,74 | 4,32 | 0,40 | 6,86E-01 |
| AT4G12290 | 3,10 | 2,22 | 1,20 | 1,86 | 6,31E-02 |
| AT4G05985 | 0,10 | 0,01 | 4,38 | 0,00 | 9,99E-01 |
| AT4G12390 | 0,50 | -0,67 | 4,39 | -0,15 | 8,78E-01 |
| AT4G12570 | 2,25 | 0,35 | 1,35 | 0,26 | 7,94E-01 |
| AT4G12600 | 0,92 | 1,51 | 2,94 | 0,51 | 6,07E-01 |
| AT4G12640 | 0,26 | 1,24 | 3,76 | 0,33 | 7,41E-01 |
| AT4G12680 | 0,21 | 1,18 | 4,35 | 0,27 | 7,86E-01 |
| AT4G12690 | 0,12 | 0,37 | 4,37 | 0,08 | 9,33E-01 |
| AT4G12720 | 1,54 | 0,52 | 1,57 | 0,33 | 7,43E-01 |
| AT4G12830 | 0,34 | 0,00 | 4,40 | 0,00 | 1,00E+00 |
| AT4G12900 | 0,05 | 0,00 | 4,41 | 0,00 | 1,00E+00 |
| AT4G05995 | 0,98 | 2,17 | 2,50 | 0,87 | 3,85E-01 |
| AT4G13010 | 0,63 | 0,00 | 3,71 | 0,00 | 1,00E+00 |
| AT4G13020 | 0,54 | 1,82 | 3,16 | 0,58 | 5,65E-01 |
| AT4G06010 | 0,28 | 1,27 | 4,35 | 0,29 | 7,69E-01 |
| AT4G06015 | 0,10 | 0,21 | 4,38 | 0,05 | 9,62E-01 |
| AT4G13245 | 0,48 | -1,78 | 3,31 | -0,54 | 5,91E-01 |
| AT4G13270 | 0,50 | 2,07 | 3,14 | 0,66 | 5,09E-01 |
| AT4G13390 | 1,53 | 1,15 | 1,97 | 0,58 | 5,61E-01 |
| AT4G13440 | 0,58 | -0,01 | 3,99 | 0,00 | 9,98E-01 |
| AT4G13510 | 4,76 | 0,46 | 0,90 | 0,50 | 6,14E-01 |
| AT4G13530 | 0,95 | 1,31 | 2,02 | 0,65 | 5,19E-01 |
| AT4G13572 | 0,31 | -2,59 | 4,37 | -0,59 | 5,53E-01 |
| AT4G13610 | 0,37 | 1,22 | 2,88 | 0,43 | 6,70E-01 |
| AT4G13615 | 0,92 | 0,54 | 1,86 | 0,29 | 7,73E-01 |
| AT4G13670 | 0,92 | 2,23 | 3,14 | 0,71 | 4,78E-01 |
| AT4G13730 | 0,24 | 0,50 | 4,38 | 0,11 | 9,10E-01 |
| AT4G13750 | 0,39 | 0,82 | 3,38 | 0,24 | 8,07E-01 |
| AT4G13760 | 0,54 | 1,89 | 3,16 | 0,60 | 5,48E-01 |
| AT4G13830 | 0,70 | 0,16 | 2,92 | 0,05 | 9,57E-01 |
| AT4G13850 | 0,81 | 0,87 | 2,20 | 0,39 | 6,93E-01 |
| AT4G13860 | 0,43 | 2,20 | 4,10 | 0,54 | 5,91E-01 |
| AT4G13880 | 2,00 | -0,04 | 1,54 | -0,03 | 9,80E-01 |
| AT4G13920 | 2,03 | 2,01 | 1,64 | 1,22 | 2,21E-01 |
| AT4G13992 | 1,30 | 1,89 | 2,03 | 0,93 | 3,52E-01 |
| AT4G14000 | 0,12 | 0,00 | 4,41 | 0,00 | 1,00E+00 |
| AT4G14140 | 0,81 | -0,14 | 2,20 | -0,06 | 9,49E-01 |

| | | | | | |
|---|---|---|---|---|---|
| AT4G14160 | 2,71 | -0,05 | 1,33 | -0,04 | 9,69E-01 |
| AT4G14220 | 0,46 | -2,73 | 4,34 | -0,63 | 5,29E-01 |
| AT4G14270 | 0,92 | -1,05 | 1,81 | -0,58 | 5,64E-01 |
| AT4G14300 | 0,77 | -0,97 | 3,48 | -0,28 | 7,82E-01 |
| AT4G14330 | 0,14 | 0,40 | 4,37 | 0,09 | 9,27E-01 |
| AT4G14504 | 4,83 | 0,84 | 1,55 | 0,54 | 5,86E-01 |
| AT4G14540 | 0,71 | 0,14 | 2,62 | 0,06 | 9,56E-01 |
| AT4G14548 | 1,46 | 0,22 | 1,99 | 0,11 | 9,10E-01 |
| AT4G14560 | 0,12 | 0,26 | 4,37 | 0,06 | 9,52E-01 |
| AT4G14570 | 0,70 | 0,03 | 2,70 | 0,01 | 9,90E-01 |
| AT4G14630 | 0,43 | 0,00 | 4,12 | 0,00 | 1,00E+00 |
| AT4G14700 | 0,12 | 1,23 | 4,34 | 0,28 | 7,76E-01 |
| AT4G06245 | 1,52 | 0,99 | 2,15 | 0,46 | 6,45E-01 |
| AT4G06255 | 0,39 | -0,94 | 4,33 | -0,22 | 8,28E-01 |
| AT4G06260 | 0,42 | -1,66 | 3,28 | -0,51 | 6,13E-01 |
| AT4G14760 | 0,91 | -0,93 | 2,22 | -0,42 | 6,75E-01 |
| AT4G14800 | 0,34 | 0,39 | 4,34 | 0,09 | 9,28E-01 |
| AT4G14810 | 0,45 | -0,71 | 3,35 | -0,21 | 8,33E-01 |
| AT4G14850 | 0,15 | 0,00 | 4,38 | 0,00 | 1,00E+00 |
| AT4G14870 | 0,48 | -0,92 | 2,77 | -0,33 | 7,41E-01 |
| AT4G14890 | 0,73 | 0,03 | 2,30 | 0,01 | 9,91E-01 |
| AT4G14950 | 0,54 | 0,29 | 2,59 | 0,11 | 9,11E-01 |
| AT4G15000 | 1,11 | -2,09 | 1,86 | -1,12 | 2,63E-01 |
| AT4G15020 | 0,86 | 1,89 | 2,30 | 0,82 | 4,11E-01 |
| AT4G15030 | 0,61 | 0,34 | 2,41 | 0,14 | 8,89E-01 |
| AT4G15090 | 0,08 | 0,69 | 4,36 | 0,16 | 8,75E-01 |
| AT4G15096 | 1,15 | 0,05 | 2,37 | 0,02 | 9,85E-01 |
| AT4G15165 | 0,40 | -2,96 | 4,35 | -0,68 | 4,97E-01 |
| AT4G15180 | 1,84 | 0,12 | 1,30 | 0,09 | 9,26E-01 |
| AT4G15233 | 0,90 | -0,29 | 2,24 | -0,13 | 8,98E-01 |
| AT4G15245 | 0,49 | 0,88 | 2,73 | 0,32 | 7,48E-01 |
| AT4G15260 | 0,57 | -1,92 | 2,98 | -0,64 | 5,20E-01 |
| AT4G15280 | 1,02 | -0,21 | 2,17 | -0,10 | 9,21E-01 |
| AT4G15415 | 0,24 | -0,38 | 4,35 | -0,09 | 9,31E-01 |
| AT4G15420 | 1,24 | -0,44 | 1,89 | -0,23 | 8,15E-01 |
| AT4G15440 | 0,99 | -3,19 | 2,43 | -1,31 | 1,89E-01 |
| AT4G15450 | 0,24 | -1,85 | 4,40 | -0,42 | 6,74E-01 |
| AT4G15470 | 1,75 | -1,72 | 1,45 | -1,19 | 2,35E-01 |
| AT4G15475 | 0,96 | -4,34 | 4,01 | -1,08 | 2,79E-01 |
| AT4G15540 | 0,39 | -0,11 | 3,50 | -0,03 | 9,75E-01 |
| AT4G15545 | 1,17 | 0,81 | 1,67 | 0,48 | 6,29E-01 |
| AT4G15560 | 2,23 | -1,38 | 1,32 | -1,05 | 2,96E-01 |
| AT4G15660 | 0,10 | 0,01 | 4,38 | 0,00 | 9,99E-01 |
| AT4G15700 | 0,08 | 0,00 | 4,40 | 0,00 | 1,00E+00 |
| AT4G15720 | 0,08 | 0,69 | 4,36 | 0,16 | 8,75E-01 |
| AT4G15770 | 0,18 | -1,78 | 4,41 | -0,40 | 6,87E-01 |
| AT4G15775 | 0,24 | 0,00 | 4,41 | 0,00 | 1,00E+00 |
| AT4G15800 | 0,76 | -2,02 | 2,00 | -1,01 | 3,14E-01 |
| AT4G15820 | 0,10 | 0,21 | 4,38 | 0,05 | 9,62E-01 |
| AT4G15840 | 0,73 | -1,27 | 1,92 | -0,66 | 5,10E-01 |
| AT4G15850 | 0,38 | -1,21 | 4,34 | -0,28 | 7,81E-01 |
| AT4G15880 | 0,12 | 0,49 | 4,36 | 0,11 | 9,11E-01 |

| | | | | | |
|---|---|---|---|---|---|
| AT4G15940 | 0,10 | 0,00 | 4,41 | 0,00 | 1,00E+00 |
| AT4G16008 | 0,08 | 0,69 | 4,36 | 0,16 | 8,75E-01 |
| AT4G16015 | 0,19 | 1,15 | 4,34 | 0,27 | 7,91E-01 |
| AT4G06375 | 3,09 | -0,13 | 1,31 | -0,10 | 9,19E-01 |
| AT4G16130 | 1,05 | 0,29 | 1,99 | 0,14 | 8,85E-01 |
| AT4G16140 | 0,34 | 0,96 | 4,35 | 0,22 | 8,26E-01 |
| AT4G16150 | 0,50 | 0,93 | 4,08 | 0,23 | 8,19E-01 |
| AT4G16310 | 1,01 | 1,52 | 2,02 | 0,75 | 4,53E-01 |
| AT4G16360 | 0,61 | -0,10 | 2,64 | -0,04 | 9,69E-01 |
| AT4G16380 | 0,81 | -0,75 | 2,43 | -0,31 | 7,58E-01 |
| AT4G16390 | 0,67 | -0,13 | 2,20 | -0,06 | 9,54E-01 |
| AT4G16400 | 1,31 | -1,18 | 1,80 | -0,66 | 5,11E-01 |
| AT4G16430 | 0,28 | -2,31 | 4,39 | -0,53 | 5,99E-01 |
| AT4G16480 | 0,65 | 3,04 | 3,52 | 0,86 | 3,88E-01 |
| AT4G16510 | 0,53 | 0,21 | 4,01 | 0,05 | 9,57E-01 |
| AT4G16590 | 0,08 | 0,69 | 4,36 | 0,16 | 8,75E-01 |
| AT4G16650 | 0,41 | -0,33 | 4,23 | -0,08 | 9,37E-01 |
| AT4G16660 | 2,21 | 0,83 | 1,32 | 0,63 | 5,31E-01 |
| AT4G16670 | 0,65 | -1,23 | 2,12 | -0,58 | 5,62E-01 |
| AT4G16710 | 0,63 | 1,34 | 2,49 | 0,54 | 5,90E-01 |
| AT4G16780 | 0,60 | 0,94 | 3,90 | 0,24 | 8,09E-01 |
| AT4G16830 | 0,78 | -2,58 | 3,18 | -0,81 | 4,17E-01 |
| AT4G16845 | 0,11 | 0,00 | 4,41 | 0,00 | 1,00E+00 |
| AT4G16850 | 0,29 | 0,64 | 4,33 | 0,15 | 8,82E-01 |
| AT4G16855 | 0,17 | 1,52 | 4,33 | 0,35 | 7,26E-01 |
| AT4G16955 | 0,78 | -1,83 | 2,90 | -0,63 | 5,27E-01 |
| AT4G16980 | 2,88 | -1,37 | 1,26 | -1,08 | 2,78E-01 |
| AT4G16990 | 5,03 | -0,04 | 0,86 | -0,04 | 9,65E-01 |
| AT4G17010 | 0,55 | 0,00 | 4,40 | 0,00 | 1,00E+00 |
| AT4G17080 | 0,57 | -0,16 | 4,29 | -0,04 | 9,70E-01 |
| AT4G17085 | 0,42 | -1,60 | 4,34 | -0,37 | 7,12E-01 |
| AT4G06665 | 1,22 | 0,58 | 1,86 | 0,31 | 7,55E-01 |
| AT4G17098 | 0,61 | 0,04 | 2,68 | 0,01 | 9,89E-01 |
| AT4G17245 | 0,66 | -0,43 | 2,51 | -0,17 | 8,65E-01 |
| AT4G17250 | 0,41 | 0,41 | 4,29 | 0,10 | 9,24E-01 |
| AT4G17270 | 0,64 | -3,29 | 3,01 | -1,09 | 2,74E-01 |
| AT4G17300 | 0,71 | -0,04 | 2,25 | -0,02 | 9,84E-01 |
| AT4G17340 | 2,05 | 1,76 | 1,65 | 1,07 | 2,86E-01 |
| AT4G17380 | 0,44 | -0,48 | 2,68 | -0,18 | 8,57E-01 |
| AT4G06695 | 4,20 | 0,08 | 1,13 | 0,07 | 9,41E-01 |
| AT4G17410 | 2,55 | -1,02 | 1,49 | -0,69 | 4,91E-01 |
| AT4G17500 | 0,96 | 1,38 | 2,24 | 0,61 | 5,39E-01 |
| AT4G17520 | 1,02 | -2,13 | 2,71 | -0,79 | 4,32E-01 |
| AT4G17600 | 1,35 | 1,52 | 1,74 | 0,87 | 3,83E-01 |
| AT4G17616 | 0,78 | -0,21 | 3,31 | -0,06 | 9,50E-01 |
| AT4G17620 | 1,00 | 1,06 | 2,31 | 0,46 | 6,47E-01 |
| AT4G17650 | 0,32 | 0,36 | 4,34 | 0,08 | 9,34E-01 |
| AT4G17770 | 1,15 | -0,22 | 1,91 | -0,12 | 9,06E-01 |
| AT4G17810 | 0,14 | 0,00 | 4,41 | 0,00 | 1,00E+00 |
| AT4G17830 | 0,83 | 1,37 | 2,49 | 0,55 | 5,80E-01 |
| AT4G17870 | 1,03 | 2,27 | 2,33 | 0,97 | 3,30E-01 |
| AT4G17900 | 0,47 | 0,68 | 2,75 | 0,25 | 8,04E-01 |

| | | | | | |
|---|---|---|---|---|---|
| AT4G18070 | 1,02 | 0,32 | 2,35 | 0,14 | 8,91E-01 |
| AT4G18150 | 2,31 | 0,46 | 1,54 | 0,30 | 7,66E-01 |
| AT4G18160 | 0,40 | -3,12 | 4,34 | -0,72 | 4,73E-01 |
| AT4G18195 | 0,22 | -1,75 | 4,40 | -0,40 | 6,91E-01 |
| AT4G18197 | 1,87 | 0,13 | 1,61 | 0,08 | 9,35E-01 |
| AT4G18205 | 1,32 | 0,21 | 1,86 | 0,12 | 9,08E-01 |
| AT4G18210 | 1,90 | 1,37 | 1,85 | 0,74 | 4,59E-01 |
| AT4G18220 | 1,16 | 0,14 | 1,67 | 0,09 | 9,32E-01 |
| AT4G18240 | 1,25 | 0,02 | 1,90 | 0,01 | 9,93E-01 |
| AT4G18370 | 0,87 | -0,58 | 2,24 | -0,26 | 7,97E-01 |
| AT4G18380 | 0,33 | -0,37 | 4,37 | -0,09 | 9,32E-01 |
| AT4G06795 | 0,56 | -0,70 | 2,94 | -0,24 | 8,13E-01 |
| AT4G18530 | 0,35 | 1,03 | 4,35 | 0,24 | 8,12E-01 |
| AT4G18570 | 0,37 | -0,15 | 2,90 | -0,05 | 9,60E-01 |
| AT4G18596 | 1,78 | 1,35 | 1,53 | 0,88 | 3,78E-01 |
| AT4G18710 | 1,19 | 1,00 | 1,80 | 0,56 | 5,78E-01 |
| AT4G18725 | 0,19 | 0,00 | 4,41 | 0,00 | 1,00E+00 |
| AT4G18730 | 1,33 | -2,25 | 1,56 | -1,44 | 1,50E-01 |
| AT4G18750 | 0,50 | 0,46 | 4,06 | 0,11 | 9,09E-01 |
| AT4G18820 | 1,31 | 1,08 | 1,78 | 0,61 | 5,43E-01 |
| AT4G18830 | 0,19 | 0,84 | 4,35 | 0,19 | 8,46E-01 |
| AT4G18890 | 0,44 | -1,77 | 4,13 | -0,43 | 6,69E-01 |
| AT4G18905 | 0,35 | 0,16 | 4,34 | 0,04 | 9,71E-01 |
| AT4G18930 | 0,31 | 0,48 | 4,38 | 0,11 | 9,12E-01 |
| AT4G18950 | 1,00 | 1,37 | 2,09 | 0,65 | 5,13E-01 |
| AT4G19020 | 0,17 | 1,33 | 4,33 | 0,31 | 7,59E-01 |
| AT4G06865 | 0,10 | 0,01 | 4,38 | 0,00 | 9,99E-01 |
| AT4G19090 | 1,32 | -0,14 | 1,60 | -0,09 | 9,32E-01 |
| AT4G19100 | 1,38 | 0,85 | 2,33 | 0,36 | 7,17E-01 |
| AT4G19170 | 5,23 | -0,04 | 0,95 | -0,04 | 9,64E-01 |
| AT4G19190 | 0,39 | 2,09 | 4,18 | 0,50 | 6,16E-01 |
| AT4G19200 | 0,86 | -0,64 | 2,21 | -0,29 | 7,72E-01 |
| AT4G19210 | 1,66 | 1,38 | 1,35 | 1,02 | 3,06E-01 |
| AT4G19450 | 0,65 | 2,47 | 4,30 | 0,57 | 5,67E-01 |
| AT4G19490 | 0,82 | -0,54 | 2,62 | -0,21 | 8,37E-01 |
| AT4G19500 | 0,89 | -0,29 | 2,11 | -0,14 | 8,92E-01 |
| AT4G19510 | 1,30 | 0,97 | 1,75 | 0,56 | 5,78E-01 |
| AT4G19530 | 1,16 | -1,31 | 1,86 | -0,70 | 4,82E-01 |
| AT4G19540 | 0,68 | 1,42 | 2,91 | 0,49 | 6,25E-01 |
| AT4G19580 | 0,44 | 2,16 | 4,05 | 0,53 | 5,95E-01 |
| AT4G19600 | 0,21 | 0,01 | 4,38 | 0,00 | 9,98E-01 |
| AT4G19610 | 0,37 | 0,23 | 3,43 | 0,07 | 9,46E-01 |
| AT4G19650 | 0,19 | 1,40 | 4,33 | 0,32 | 7,46E-01 |
| AT4G19710 | 1,47 | -0,43 | 1,43 | -0,30 | 7,61E-01 |
| AT4G19960 | 0,50 | 1,08 | 3,19 | 0,34 | 7,36E-01 |
| AT4G20110 | 0,83 | 0,03 | 2,16 | 0,02 | 9,87E-01 |
| AT4G20140 | 0,84 | 3,28 | 4,17 | 0,79 | 4,31E-01 |
| AT4G20170 | 3,89 | 1,38 | 1,17 | 1,18 | 2,38E-01 |
| AT4G06950 | 0,12 | 0,00 | 4,38 | 0,00 | 1,00E+00 |
| AT4G20260 | 2,30 | 0,53 | 1,49 | 0,35 | 7,24E-01 |
| AT4G20270 | 0,28 | 1,73 | 4,32 | 0,40 | 6,89E-01 |
| AT4G20300 | 0,68 | -1,32 | 2,48 | -0,53 | 5,94E-01 |

| | | | | | |
|---|---|---|---|---|---|
| AT4G20310 | 0,08 | 0,69 | 4,36 | 0,16 | 8,75E-01 |
| AT4G20360 | 5,64 | 0,14 | 0,89 | 0,16 | 8,76E-01 |
| AT4G20400 | 1,23 | -0,70 | 2,00 | -0,35 | 7,25E-01 |
| AT4G20720 | 0,34 | 0,13 | 4,34 | 0,03 | 9,76E-01 |
| AT4G20735 | 3,19 | 3,13 | 1,74 | 1,79 | 7,31E-02 |
| AT4G20830 | 3,02 | 1,75 | 1,19 | 1,47 | 1,41E-01 |
| AT4G20840 | 1,22 | -1,09 | 2,23 | -0,49 | 6,25E-01 |
| AT4G20860 | 0,52 | 1,23 | 3,08 | 0,40 | 6,91E-01 |
| AT4G20890 | 0,68 | -0,88 | 2,65 | -0,33 | 7,40E-01 |
| AT4G20940 | 0,56 | -0,38 | 4,32 | -0,09 | 9,29E-01 |
| AT4G20960 | 0,48 | 0,35 | 2,74 | 0,13 | 8,98E-01 |
| AT4G20980 | 1,36 | 1,20 | 1,93 | 0,62 | 5,33E-01 |
| AT4G21060 | 1,98 | -1,12 | 1,46 | -0,77 | 4,44E-01 |
| AT4G21105 | 0,54 | -0,93 | 4,32 | -0,22 | 8,29E-01 |
| AT4G21150 | 2,92 | -0,05 | 1,19 | -0,04 | 9,68E-01 |
| AT4G21160 | 0,15 | 0,00 | 4,41 | 0,00 | 1,00E+00 |
| AT4G21180 | 0,08 | 0,44 | 4,38 | 0,10 | 9,21E-01 |
| AT4G21210 | 0,33 | 1,01 | 4,34 | 0,23 | 8,16E-01 |
| AT4G21280 | 3,20 | -1,78 | 1,24 | -1,44 | 1,51E-01 |
| AT4G21300 | 0,05 | 0,00 | 4,41 | 0,00 | 1,00E+00 |
| AT4G21323 | 0,38 | -1,21 | 4,34 | -0,28 | 7,81E-01 |
| AT4G21366 | 2,02 | 0,73 | 1,43 | 0,51 | 6,09E-01 |
| AT4G21450 | 0,15 | 1,01 | 4,34 | 0,23 | 8,17E-01 |
| AT4G21470 | 0,32 | 1,54 | 4,34 | 0,35 | 7,23E-01 |
| AT4G21515 | 0,05 | 0,00 | 4,41 | 0,00 | 1,00E+00 |
| AT4G21540 | 0,68 | 2,84 | 2,82 | 1,01 | 3,15E-01 |
| AT4G21550 | 0,37 | -2,24 | 4,39 | -0,51 | 6,09E-01 |
| AT4G21620 | 0,72 | -0,30 | 2,50 | -0,12 | 9,05E-01 |
| AT4G21670 | 1,62 | -0,12 | 1,49 | -0,08 | 9,34E-01 |
| AT4G21740 | 0,47 | -0,17 | 2,45 | -0,07 | 9,45E-01 |
| AT4G21750 | 0,81 | 0,28 | 2,66 | 0,10 | 9,17E-01 |
| AT4G21790 | 0,64 | -0,03 | 2,53 | -0,01 | 9,90E-01 |
| AT4G21800 | 0,53 | -2,81 | 3,86 | -0,73 | 4,66E-01 |
| AT4G07030 | 0,38 | 0,78 | 4,33 | 0,18 | 8,56E-01 |
| AT4G21940 | 0,15 | 1,20 | 4,34 | 0,28 | 7,81E-01 |
| AT4G22010 | 0,22 | 0,99 | 4,35 | 0,23 | 8,21E-01 |
| AT4G22050 | 0,20 | 0,15 | 4,37 | 0,04 | 9,72E-01 |
| AT4G07040 | 0,12 | 0,00 | 4,39 | 0,00 | 1,00E+00 |
| AT4G22130 | 0,44 | -0,11 | 2,81 | -0,04 | 9,70E-01 |
| AT4G22270 | 0,20 | -0,31 | 4,36 | -0,07 | 9,44E-01 |
| AT4G07075 | 1,23 | 1,67 | 1,82 | 0,92 | 3,57E-01 |
| AT4G07080 | 1,23 | 1,67 | 1,82 | 0,92 | 3,57E-01 |
| AT4G22330 | 0,17 | 1,52 | 4,33 | 0,35 | 7,26E-01 |
| AT4G22467 | 0,12 | 0,00 | 4,38 | 0,00 | 1,00E+00 |
| AT4G22475 | 0,49 | 0,00 | 3,99 | 0,00 | 1,00E+00 |
| AT4G22670 | 0,32 | 0,73 | 4,34 | 0,17 | 8,67E-01 |
| AT4G22690 | 3,37 | -1,36 | 0,98 | -1,38 | 1,66E-01 |
| AT4G22710 | 1,76 | -0,92 | 1,91 | -0,48 | 6,30E-01 |
| AT4G22720 | 0,28 | 0,23 | 4,33 | 0,05 | 9,58E-01 |
| AT4G22860 | 0,08 | 0,69 | 4,36 | 0,16 | 8,75E-01 |
| AT4G22890 | 1,50 | -1,72 | 1,83 | -0,94 | 3,46E-01 |
| AT4G22910 | 0,08 | 0,44 | 4,38 | 0,10 | 9,21E-01 |

| | | | | | |
|---|---|---|---|---|---|
| AT4G22960 | 0,12 | 0,00 | 4,38 | 0,00 | 1,00E+00 |
| AT4G23030 | 0,17 | -1,70 | 4,40 | -0,39 | 7,00E-01 |
| AT4G23050 | 0,25 | 0,26 | 4,33 | 0,06 | 9,51E-01 |
| AT4G23060 | 0,21 | -0,49 | 4,35 | -0,11 | 9,11E-01 |
| AT4G23140 | 2,18 | 0,50 | 1,32 | 0,38 | 7,03E-01 |
| AT4G23150 | 0,15 | 0,00 | 4,41 | 0,00 | 1,00E+00 |
| AT4G23160 | 0,75 | -3,51 | 4,31 | -0,82 | 4,14E-01 |
| AT4G23180 | 1,57 | -0,31 | 1,38 | -0,23 | 8,22E-01 |
| AT4G23270 | 0,94 | 1,82 | 2,04 | 0,89 | 3,74E-01 |
| AT4G23400 | 0,56 | 1,58 | 2,31 | 0,68 | 4,94E-01 |
| AT4G23440 | 0,12 | 1,16 | 4,34 | 0,27 | 7,89E-01 |
| AT4G23470 | 0,84 | 1,53 | 2,07 | 0,74 | 4,60E-01 |
| AT4G23530 | 1,11 | -1,45 | 1,85 | -0,78 | 4,33E-01 |
| AT4G23570 | 0,44 | -0,50 | 3,38 | -0,15 | 8,82E-01 |
| AT4G23600 | 1,02 | 0,25 | 2,74 | 0,09 | 9,28E-01 |
| AT4G23630 | 1,89 | -0,61 | 1,31 | -0,47 | 6,41E-01 |
| AT4G23740 | 0,40 | 1,78 | 4,33 | 0,41 | 6,81E-01 |
| AT4G23860 | 0,25 | -1,82 | 4,40 | -0,41 | 6,79E-01 |
| AT4G07190 | 0,45 | -0,77 | 3,47 | -0,22 | 8,24E-01 |
| AT4G23895 | 0,24 | 1,08 | 4,35 | 0,25 | 8,03E-01 |
| AT4G23940 | 1,06 | 0,56 | 2,05 | 0,27 | 7,84E-01 |
| AT4G23950 | 0,16 | 0,49 | 4,38 | 0,11 | 9,12E-01 |
| AT4G23980 | 1,47 | -0,26 | 1,45 | -0,18 | 8,59E-01 |
| AT4G24000 | 0,08 | 0,69 | 4,36 | 0,16 | 8,75E-01 |
| AT4G24010 | 0,33 | -0,37 | 4,37 | -0,09 | 9,32E-01 |
| AT4G24020 | 1,49 | -1,07 | 1,47 | -0,73 | 4,67E-01 |
| AT4G24060 | 0,12 | 1,16 | 4,34 | 0,27 | 7,89E-01 |
| AT4G24090 | 0,53 | 0,00 | 4,41 | 0,00 | 1,00E+00 |
| AT4G24120 | 1,68 | 3,30 | 2,10 | 1,57 | 1,16E-01 |
| AT4G24130 | 0,24 | -2,36 | 4,39 | -0,54 | 5,91E-01 |
| AT4G24160 | 0,15 | 0,45 | 4,37 | 0,10 | 9,19E-01 |
| AT4G24175 | 0,21 | 0,01 | 4,34 | 0,00 | 9,98E-01 |
| AT4G24220 | 1,52 | 0,08 | 1,50 | 0,05 | 9,60E-01 |
| AT4G24290 | 0,72 | 0,90 | 2,36 | 0,38 | 7,02E-01 |
| AT4G24350 | 0,25 | 0,02 | 4,39 | 0,01 | 9,96E-01 |
| AT4G24440 | 0,43 | -0,05 | 4,20 | -0,01 | 9,90E-01 |
| AT4G24460 | 0,18 | 0,13 | 4,41 | 0,03 | 9,76E-01 |
| AT4G24470 | 0,57 | 0,98 | 3,84 | 0,25 | 8,00E-01 |
| AT4G24550 | 0,38 | 0,48 | 3,40 | 0,14 | 8,88E-01 |
| AT4G24570 | 3,86 | 2,38 | 1,45 | 1,64 | 1,01E-01 |
| AT4G24680 | 3,35 | -0,09 | 0,93 | -0,09 | 9,25E-01 |
| AT4G24790 | 0,10 | 0,21 | 4,38 | 0,05 | 9,62E-01 |
| AT4G24800 | 0,98 | -0,49 | 2,47 | -0,20 | 8,43E-01 |
| AT4G24805 | 0,88 | -1,47 | 2,58 | -0,57 | 5,70E-01 |
| AT4G24880 | 0,08 | 0,00 | 4,40 | 0,00 | 1,00E+00 |
| AT4G07295 | 0,21 | 0,50 | 4,36 | 0,12 | 9,08E-01 |
| AT4G25030 | 0,40 | 0,80 | 2,84 | 0,28 | 7,78E-01 |
| AT4G25050 | 1,37 | -0,51 | 1,57 | -0,33 | 7,43E-01 |
| AT4G25070 | 0,53 | 0,52 | 4,34 | 0,12 | 9,04E-01 |
| AT4G25080 | 4,26 | -0,21 | 0,87 | -0,24 | 8,08E-01 |
| AT4G25120 | 0,50 | -1,40 | 3,27 | -0,43 | 6,67E-01 |
| AT4G25170 | 1,05 | -2,38 | 2,03 | -1,17 | 2,42E-01 |

| | | | | | |
|---|---|---|---|---|---|
| AT4G25210 | 0,75 | -2,96 | 2,89 | -1,02 | 3,06E-01 |
| AT4G25230 | 0,77 | 2,19 | 2,23 | 0,98 | 3,26E-01 |
| AT4G25360 | 0,25 | 1,72 | 4,33 | 0,40 | 6,91E-01 |
| AT4G25370 | 0,59 | -0,21 | 3,81 | -0,06 | 9,55E-01 |
| AT4G25390 | 0,17 | 0,07 | 4,40 | 0,02 | 9,88E-01 |
| AT4G25400 | 0,31 | 0,00 | 4,41 | 0,00 | 1,00E+00 |
| AT4G25500 | 1,12 | -0,32 | 2,07 | -0,16 | 8,76E-01 |
| AT4G25515 | 0,32 | -0,25 | 4,36 | -0,06 | 9,55E-01 |
| AT4G25530 | 5,35 | 2,50 | 1,39 | 1,80 | 7,19E-02 |
| AT4G25550 | 0,38 | -1,21 | 4,34 | -0,28 | 7,81E-01 |
| AT4G25610 | 0,54 | -0,34 | 3,09 | -0,11 | 9,13E-01 |
| AT4G25630 | 1,57 | 0,18 | 1,79 | 0,10 | 9,19E-01 |
| AT4G25631 | 1,33 | -0,18 | 2,01 | -0,09 | 9,30E-01 |
| AT4G25660 | 0,81 | -1,37 | 2,27 | -0,60 | 5,46E-01 |
| AT4G25810 | 0,12 | 1,05 | 4,35 | 0,24 | 8,10E-01 |
| AT4G25835 | 0,21 | 0,45 | 4,37 | 0,10 | 9,17E-01 |
| AT4G25880 | 1,93 | 0,40 | 1,30 | 0,31 | 7,57E-01 |
| AT4G25900 | 0,36 | -0,20 | 4,35 | -0,05 | 9,63E-01 |
| AT4G25910 | 0,69 | -0,70 | 3,41 | -0,20 | 8,38E-01 |
| AT4G25960 | 1,53 | -1,97 | 1,84 | -1,07 | 2,85E-01 |
| AT4G25970 | 1,00 | 0,07 | 1,81 | 0,04 | 9,70E-01 |
| AT4G26000 | 0,96 | 0,13 | 2,28 | 0,06 | 9,53E-01 |
| AT4G26090 | 1,90 | -0,30 | 1,25 | -0,24 | 8,12E-01 |
| AT4G26110 | 2,20 | -0,03 | 1,27 | -0,02 | 9,84E-01 |
| AT4G26120 | 0,24 | -2,36 | 4,39 | -0,54 | 5,91E-01 |
| AT4G26170 | 0,08 | 0,44 | 4,38 | 0,10 | 9,21E-01 |
| AT4G26210 | 0,36 | 1,83 | 4,33 | 0,42 | 6,73E-01 |
| AT4G26230 | 0,82 | -0,98 | 2,43 | -0,40 | 6,87E-01 |
| AT4G26255 | 3,93 | -2,83 | 1,26 | -2,25 | 2,42E-02 |
| AT4G26450 | 0,45 | -0,49 | 3,31 | -0,15 | 8,83E-01 |
| AT4G26455 | 0,54 | 0,36 | 3,71 | 0,10 | 9,23E-01 |
| AT4G26510 | 0,72 | -1,07 | 2,47 | -0,43 | 6,64E-01 |
| AT4G26520 | 1,37 | -1,87 | 1,78 | -1,05 | 2,93E-01 |
| AT4G26530 | 4,43 | 0,92 | 0,86 | 1,07 | 2,84E-01 |
| AT4G26542 | 2,99 | 0,50 | 1,13 | 0,44 | 6,57E-01 |
| AT4G26650 | 0,56 | -0,54 | 2,26 | -0,24 | 8,12E-01 |
| AT4G26670 | 0,33 | 0,43 | 4,38 | 0,10 | 9,22E-01 |
| AT4G26680 | 0,40 | 2,21 | 4,13 | 0,53 | 5,93E-01 |
| AT4G26700 | 0,28 | 0,95 | 4,36 | 0,22 | 8,27E-01 |
| AT4G26730 | 2,38 | 2,75 | 1,78 | 1,55 | 1,22E-01 |
| AT4G26790 | 0,42 | 0,39 | 4,12 | 0,09 | 9,25E-01 |
| AT4G26800 | 0,12 | 0,00 | 4,38 | 0,00 | 1,00E+00 |
| AT4G26840 | 0,82 | -0,25 | 2,08 | -0,12 | 9,05E-01 |
| AT4G26870 | 0,92 | -0,33 | 2,05 | -0,16 | 8,72E-01 |
| AT4G26970 | 1,79 | 0,99 | 1,73 | 0,57 | 5,69E-01 |
| AT4G27010 | 0,79 | 1,06 | 2,13 | 0,50 | 6,18E-01 |
| AT4G27030 | 0,63 | 1,98 | 2,53 | 0,78 | 4,35E-01 |
| AT4G07775 | 0,58 | 1,95 | 2,94 | 0,66 | 5,07E-01 |
| AT4G27080 | 1,09 | -2,39 | 2,17 | -1,10 | 2,69E-01 |
| AT4G27260 | 0,26 | -0,30 | 4,37 | -0,07 | 9,46E-01 |
| AT4G27290 | 0,45 | -1,75 | 4,32 | -0,40 | 6,86E-01 |
| AT4G27310 | 0,93 | 0,61 | 2,14 | 0,29 | 7,74E-01 |

| | | | | | |
|---|---|---|---|---|---|
| AT4G27380 | 0,21 | 0,21 | 4,38 | 0,05 | 9,62E-01 |
| AT4G27395 | 0,73 | 1,16 | 2,85 | 0,41 | 6,84E-01 |
| AT4G27430 | 2,23 | -1,01 | 1,24 | -0,81 | 4,16E-01 |
| AT4G27500 | 2,93 | -0,08 | 1,11 | -0,07 | 9,45E-01 |
| AT4G27652 | 0,57 | 0,38 | 3,75 | 0,10 | 9,19E-01 |
| AT4G27654 | 0,56 | 1,75 | 2,64 | 0,66 | 5,06E-01 |
| AT4G27657 | 0,12 | 0,00 | 4,38 | 0,00 | 1,00E+00 |
| AT4G27680 | 0,93 | 0,64 | 1,99 | 0,32 | 7,49E-01 |
| AT4G27710 | 0,52 | 1,61 | 3,25 | 0,50 | 6,20E-01 |
| AT4G27720 | 1,38 | 0,46 | 1,80 | 0,26 | 7,96E-01 |
| AT4G27740 | 0,52 | 0,05 | 3,87 | 0,01 | 9,90E-01 |
| AT4G27750 | 0,69 | -1,05 | 4,37 | -0,24 | 8,11E-01 |
| AT4G27760 | 0,28 | -1,42 | 4,36 | -0,33 | 7,44E-01 |
| AT4G27765 | 0,37 | -0,56 | 4,24 | -0,13 | 8,95E-01 |
| AT4G27780 | 0,50 | -1,20 | 3,89 | -0,31 | 7,59E-01 |
| AT4G27860 | 0,21 | 1,13 | 4,34 | 0,26 | 7,95E-01 |
| AT4G27870 | 2,09 | -0,46 | 1,22 | -0,38 | 7,06E-01 |
| AT4G27880 | 0,08 | 0,44 | 4,38 | 0,10 | 9,21E-01 |
| AT4G27910 | 0,46 | -0,16 | 4,02 | -0,04 | 9,69E-01 |
| AT4G27940 | 1,14 | -1,38 | 2,03 | -0,68 | 4,98E-01 |
| AT4G28010 | 0,17 | 0,61 | 4,37 | 0,14 | 8,88E-01 |
| AT4G28210 | 0,08 | 0,00 | 4,40 | 0,00 | 1,00E+00 |
| AT4G28220 | 0,27 | 1,03 | 4,35 | 0,24 | 8,13E-01 |
| AT4G28260 | 0,80 | 1,65 | 2,86 | 0,57 | 5,66E-01 |
| AT4G28300 | 0,21 | 1,63 | 4,33 | 0,38 | 7,06E-01 |
| AT4G28362 | 0,21 | -0,49 | 4,35 | -0,11 | 9,11E-01 |
| AT4G28400 | 0,26 | -0,89 | 4,37 | -0,20 | 8,39E-01 |
| AT4G28440 | 0,16 | 0,00 | 4,37 | 0,00 | 9,99E-01 |
| AT4G28490 | 0,74 | 2,99 | 3,59 | 0,83 | 4,06E-01 |
| AT4G28570 | 3,15 | -0,42 | 1,23 | -0,34 | 7,31E-01 |
| AT4G28600 | 1,03 | 1,02 | 2,07 | 0,49 | 6,22E-01 |
| AT4G28652 | 2,08 | -0,12 | 1,52 | -0,08 | 9,39E-01 |
| AT4G28660 | 0,48 | -1,76 | 4,11 | -0,43 | 6,68E-01 |
| AT4G28706 | 0,39 | 0,00 | 4,38 | 0,00 | 1,00E+00 |
| AT4G28710 | 0,41 | 0,51 | 4,17 | 0,12 | 9,02E-01 |
| AT4G28740 | 0,90 | 0,75 | 2,56 | 0,29 | 7,70E-01 |
| AT4G28755 | 2,23 | -2,64 | 1,45 | -1,83 | 6,75E-02 |
| AT4G28760 | 2,13 | -0,81 | 1,22 | -0,66 | 5,08E-01 |
| AT4G28780 | 0,15 | 0,29 | 4,37 | 0,07 | 9,47E-01 |
| AT4G28860 | 0,76 | -1,52 | 2,97 | -0,51 | 6,08E-01 |
| AT4G28870 | 0,12 | 0,00 | 4,41 | 0,00 | 1,00E+00 |
| AT4G28880 | 1,20 | -2,16 | 1,53 | -1,41 | 1,59E-01 |
| AT4G28980 | 0,73 | 1,10 | 2,46 | 0,45 | 6,55E-01 |
| AT4G29020 | 0,17 | -1,70 | 4,40 | -0,39 | 7,00E-01 |
| AT4G29035 | 0,22 | -1,75 | 4,40 | -0,40 | 6,91E-01 |
| AT4G29040 | 1,10 | 1,59 | 2,06 | 0,77 | 4,41E-01 |
| AT4G29060 | 2,00 | 3,63 | 1,73 | 2,10 | 3,58E-02 |
| AT4G29070 | 0,33 | -0,48 | 4,36 | -0,11 | 9,12E-01 |
| AT4G29090 | 2,85 | 2,30 | 1,60 | 1,44 | 1,49E-01 |
| AT4G29110 | 0,63 | -0,54 | 2,96 | -0,18 | 8,54E-01 |
| AT4G29120 | 0,68 | 2,87 | 2,82 | 1,02 | 3,08E-01 |
| AT4G29160 | 0,87 | -0,12 | 1,80 | -0,07 | 9,46E-01 |

| | | | | | |
|---|---|---|---|---|---|
| AT4G29170 | 0,92 | -0,78 | 4,24 | -0,18 | 8,54E-01 |
| AT4G29180 | 0,96 | -1,65 | 2,45 | -0,68 | 4,99E-01 |
| AT4G29210 | 0,68 | -0,53 | 2,51 | -0,21 | 8,33E-01 |
| AT4G29350 | 0,47 | 1,03 | 4,07 | 0,25 | 8,00E-01 |
| AT4G29380 | 1,74 | 1,10 | 1,39 | 0,79 | 4,31E-01 |
| AT4G29400 | 0,21 | -1,41 | 4,36 | -0,32 | 7,46E-01 |
| AT4G29510 | 0,71 | -0,07 | 2,22 | -0,03 | 9,75E-01 |
| AT4G29740 | 1,04 | -2,07 | 2,14 | -0,97 | 3,34E-01 |
| AT4G29750 | 0,66 | -1,10 | 2,67 | -0,41 | 6,81E-01 |
| AT4G29780 | 2,35 | 3,22 | 1,87 | 1,72 | 8,50E-02 |
| AT4G29790 | 1,30 | 1,00 | 1,70 | 0,59 | 5,58E-01 |
| AT4G08245 | 0,54 | 0,24 | 3,16 | 0,08 | 9,40E-01 |
| AT4G08255 | 0,51 | 1,40 | 4,33 | 0,32 | 7,47E-01 |
| AT4G29950 | 0,57 | -0,22 | 2,56 | -0,08 | 9,33E-01 |
| AT4G30010 | 0,55 | 2,20 | 3,23 | 0,68 | 4,97E-01 |
| AT4G30020 | 1,12 | -1,34 | 1,88 | -0,71 | 4,77E-01 |
| AT4G30100 | 3,56 | 0,04 | 1,01 | 0,04 | 9,66E-01 |
| AT4G30130 | 0,24 | 0,86 | 4,37 | 0,20 | 8,43E-01 |
| AT4G30150 | 1,08 | 0,46 | 1,92 | 0,24 | 8,11E-01 |
| AT4G30160 | 4,90 | 1,09 | 0,90 | 1,22 | 2,24E-01 |
| AT4G30210 | 1,86 | 0,15 | 1,68 | 0,09 | 9,30E-01 |
| AT4G30310 | 0,60 | 0,14 | 3,64 | 0,04 | 9,70E-01 |
| AT4G30350 | 0,36 | 1,18 | 4,35 | 0,27 | 7,87E-01 |
| AT4G30400 | 0,15 | 0,00 | 4,38 | 0,00 | 1,00E+00 |
| AT4G08285 | 0,08 | 0,44 | 4,38 | 0,10 | 9,21E-01 |
| AT4G30480 | 0,31 | -2,59 | 4,37 | -0,59 | 5,53E-01 |
| AT4G30530 | 1,06 | 1,22 | 1,98 | 0,62 | 5,37E-01 |
| AT4G30650 | 0,58 | 0,34 | 2,82 | 0,12 | 9,04E-01 |
| AT4G30660 | 0,47 | -0,88 | 2,77 | -0,32 | 7,51E-01 |
| AT4G08335 | 4,06 | 4,02 | 1,45 | 2,78 | 5,46E-03 |
| AT4G30760 | 1,76 | 2,78 | 1,67 | 1,66 | 9,73E-02 |
| AT4G30780 | 0,23 | -1,78 | 4,41 | -0,40 | 6,87E-01 |
| AT4G30800 | 0,17 | 0,00 | 4,41 | 0,00 | 1,00E+00 |
| AT4G30940 | 0,31 | -0,92 | 4,39 | -0,21 | 8,35E-01 |
| AT4G30960 | 5,13 | 2,14 | 0,93 | 2,30 | 2,13E-02 |
| AT4G30990 | 1,34 | -0,41 | 1,59 | -0,26 | 7,96E-01 |
| AT4G30993 | 2,23 | 1,91 | 1,40 | 1,36 | 1,73E-01 |
| AT4G08425 | 1,45 | 1,76 | 1,65 | 1,07 | 2,87E-01 |
| AT4G31080 | 0,70 | 0,66 | 2,81 | 0,23 | 8,15E-01 |
| AT4G31115 | 0,40 | -1,46 | 4,21 | -0,35 | 7,28E-01 |
| AT4G31160 | 2,16 | -0,40 | 1,52 | -0,26 | 7,93E-01 |
| AT4G31180 | 4,29 | -0,61 | 0,86 | -0,71 | 4,78E-01 |
| AT4G31210 | 1,90 | -1,01 | 1,41 | -0,71 | 4,75E-01 |
| AT4G31300 | 0,37 | 0,00 | 4,41 | 0,00 | 1,00E+00 |
| AT4G31340 | 0,63 | -0,50 | 3,77 | -0,13 | 8,94E-01 |
| AT4G31390 | 0,47 | -1,27 | 2,74 | -0,46 | 6,42E-01 |
| AT4G31430 | 0,50 | -0,34 | 3,95 | -0,09 | 9,32E-01 |
| AT4G31490 | 3,09 | -0,26 | 1,09 | -0,24 | 8,09E-01 |
| AT4G31530 | 0,77 | -1,30 | 2,49 | -0,52 | 6,02E-01 |
| AT4G31580 | 0,97 | 0,65 | 2,01 | 0,32 | 7,48E-01 |
| AT4G31710 | 0,50 | -1,20 | 3,89 | -0,31 | 7,59E-01 |
| AT4G31715 | 0,08 | 0,69 | 4,36 | 0,16 | 8,75E-01 |

| | | | | | |
|---|---|---|---|---|---|
| AT4G31780 | 1,36 | -1,43 | 2,66 | -0,54 | 5,90E-01 |
| AT4G31790 | 0,26 | 0,84 | 4,35 | 0,19 | 8,46E-01 |
| AT4G31800 | 1,69 | -0,12 | 1,63 | -0,07 | 9,41E-01 |
| AT4G31805 | 0,98 | -0,25 | 3,17 | -0,08 | 9,36E-01 |
| AT4G31850 | 0,54 | -0,98 | 3,22 | -0,30 | 7,61E-01 |
| AT4G31910 | 0,49 | 0,60 | 2,80 | 0,21 | 8,31E-01 |
| AT4G31980 | 2,53 | 0,24 | 1,53 | 0,16 | 8,74E-01 |
| AT4G08615 | 0,91 | -0,13 | 2,39 | -0,05 | 9,57E-01 |
| AT4G08625 | 1,54 | 0,15 | 1,69 | 0,09 | 9,30E-01 |
| AT4G31985 | 0,37 | -0,30 | 4,31 | -0,07 | 9,44E-01 |
| AT4G32010 | 1,47 | -1,77 | 1,73 | -1,02 | 3,06E-01 |
| AT4G32030 | 0,47 | -0,63 | 4,20 | -0,15 | 8,80E-01 |
| AT4G32040 | 0,84 | 1,35 | 3,45 | 0,39 | 6,96E-01 |
| AT4G32060 | 1,61 | 1,54 | 1,68 | 0,92 | 3,59E-01 |
| AT4G32160 | 0,93 | 2,40 | 2,05 | 1,17 | 2,40E-01 |
| AT4G32190 | 0,86 | 2,03 | 2,67 | 0,76 | 4,47E-01 |
| AT4G32200 | 0,65 | 1,29 | 3,59 | 0,36 | 7,20E-01 |
| AT4G32265 | 1,11 | 0,14 | 1,87 | 0,08 | 9,38E-01 |
| AT4G32285 | 2,40 | 0,58 | 1,18 | 0,49 | 6,25E-01 |
| AT4G32300 | 0,70 | -0,03 | 2,14 | -0,01 | 9,88E-01 |
| AT4G32330 | 0,80 | -0,35 | 2,13 | -0,16 | 8,71E-01 |
| AT4G32360 | 0,05 | 0,00 | 4,41 | 0,00 | 1,00E+00 |
| AT4G32390 | 1,34 | 2,36 | 1,81 | 1,31 | 1,91E-01 |
| AT4G32430 | 0,12 | 0,00 | 4,38 | 0,00 | 1,00E+00 |
| AT4G32551 | 0,49 | 0,23 | 2,72 | 0,08 | 9,34E-01 |
| AT4G32600 | 0,53 | 0,86 | 3,30 | 0,26 | 7,93E-01 |
| AT4G32620 | 1,61 | 0,88 | 1,87 | 0,47 | 6,38E-01 |
| AT4G32630 | 0,68 | -2,36 | 4,32 | -0,54 | 5,86E-01 |
| AT4G32640 | 2,41 | -0,72 | 1,23 | -0,59 | 5,58E-01 |
| AT4G32700 | 0,67 | 0,36 | 3,56 | 0,10 | 9,19E-01 |
| AT4G32720 | 0,62 | 0,14 | 2,97 | 0,05 | 9,62E-01 |
| AT4G32730 | 0,45 | 0,04 | 3,94 | 0,01 | 9,91E-01 |
| AT4G32760 | 1,73 | -0,90 | 1,42 | -0,64 | 5,25E-01 |
| AT4G32770 | 0,32 | 0,19 | 4,34 | 0,04 | 9,65E-01 |
| AT4G32800 | 0,31 | -0,48 | 4,36 | -0,11 | 9,13E-01 |
| AT4G32850 | 1,20 | -0,60 | 1,77 | -0,34 | 7,35E-01 |
| AT4G32910 | 0,98 | 1,65 | 2,05 | 0,81 | 4,21E-01 |
| AT4G32930 | 0,91 | -1,24 | 2,26 | -0,55 | 5,81E-01 |
| AT4G32990 | 2,22 | 1,91 | 1,55 | 1,23 | 2,18E-01 |
| AT4G33000 | 0,14 | 0,00 | 4,41 | 0,00 | 1,00E+00 |
| AT4G33030 | 0,99 | 0,52 | 2,44 | 0,22 | 8,30E-01 |
| AT4G33060 | 0,15 | 1,16 | 4,34 | 0,27 | 7,88E-01 |
| AT4G33180 | 0,05 | 0,00 | 4,41 | 0,00 | 1,00E+00 |
| AT4G33220 | 0,80 | 1,88 | 3,26 | 0,58 | 5,64E-01 |
| AT4G33290 | 1,50 | -1,07 | 1,68 | -0,64 | 5,24E-01 |
| AT4G08785 | 0,69 | -1,34 | 3,69 | -0,36 | 7,16E-01 |
| AT4G33310 | 0,49 | -3,04 | 4,33 | -0,70 | 4,83E-01 |
| AT4G33380 | 0,54 | 1,13 | 3,27 | 0,35 | 7,30E-01 |
| AT4G33410 | 0,32 | 0,27 | 4,34 | 0,06 | 9,51E-01 |
| AT4G33510 | 1,23 | 0,51 | 1,58 | 0,32 | 7,49E-01 |
| AT4G33520 | 1,37 | 1,36 | 2,07 | 0,66 | 5,12E-01 |
| AT4G33540 | 0,41 | 1,39 | 4,25 | 0,33 | 7,44E-01 |

| | | | | | |
|---|---|---|---|---|---|
| AT4G33565 | 0,80 | -0,83 | 2,87 | -0,29 | 7,72E-01 |
| AT4G33580 | 1,02 | -2,18 | 3,13 | -0,70 | 4,87E-01 |
| AT4G33620 | 0,33 | 0,79 | 4,31 | 0,18 | 8,55E-01 |
| AT4G33630 | 1,15 | -1,95 | 1,87 | -1,04 | 2,99E-01 |
| AT4G33650 | 1,75 | -0,27 | 1,54 | -0,18 | 8,59E-01 |
| AT4G33666 | 0,24 | -0,73 | 4,37 | -0,17 | 8,68E-01 |
| AT4G33690 | 0,48 | -2,58 | 4,06 | -0,64 | 5,25E-01 |
| AT4G33890 | 1,49 | -1,53 | 1,74 | -0,88 | 3,79E-01 |
| AT4G33920 | 0,65 | 0,93 | 3,64 | 0,26 | 7,98E-01 |
| AT4G33960 | 0,45 | 0,00 | 4,41 | 0,00 | 1,00E+00 |
| AT4G34030 | 0,40 | 0,07 | 3,44 | 0,02 | 9,84E-01 |
| AT4G34035 | 0,32 | -0,65 | 4,33 | -0,15 | 8,81E-01 |
| AT4G34050 | 1,48 | 2,20 | 1,87 | 1,18 | 2,39E-01 |
| AT4G34060 | 0,12 | 1,16 | 4,34 | 0,27 | 7,89E-01 |
| AT4G34071 | 2,35 | -1,78 | 1,98 | -0,90 | 3,69E-01 |
| AT4G34100 | 3,45 | -0,56 | 1,11 | -0,51 | 6,12E-01 |
| AT4G34110 | 4,15 | -0,28 | 0,82 | -0,34 | 7,36E-01 |
| AT4G34120 | 1,07 | -1,60 | 1,80 | -0,89 | 3,74E-01 |
| AT4G34150 | 1,90 | 1,59 | 1,53 | 1,04 | 2,98E-01 |
| AT4G34160 | 0,73 | 1,61 | 2,39 | 0,67 | 5,01E-01 |
| AT4G34240 | 0,30 | -1,78 | 4,41 | -0,40 | 6,87E-01 |
| AT4G34260 | 1,45 | -2,00 | 1,82 | -1,10 | 2,71E-01 |
| AT4G34280 | 0,96 | -0,45 | 2,57 | -0,18 | 8,61E-01 |
| AT4G34290 | 0,10 | 0,01 | 4,38 | 0,00 | 9,99E-01 |
| AT4G34310 | 0,63 | -1,59 | 4,32 | -0,37 | 7,12E-01 |
| AT4G34330 | 0,08 | 0,69 | 4,36 | 0,16 | 8,75E-01 |
| AT4G34370 | 0,50 | -0,49 | 3,92 | -0,12 | 9,01E-01 |
| AT4G34390 | 1,80 | 0,43 | 1,47 | 0,29 | 7,72E-01 |
| AT4G34400 | 0,59 | -1,08 | 3,66 | -0,30 | 7,68E-01 |
| AT4G34410 | 0,77 | 1,29 | 2,83 | 0,45 | 6,49E-01 |
| AT4G34430 | 1,22 | 0,68 | 2,01 | 0,34 | 7,36E-01 |
| AT4G34440 | 0,20 | -0,31 | 4,36 | -0,07 | 9,44E-01 |
| AT4G34640 | 0,93 | 0,41 | 2,64 | 0,16 | 8,75E-01 |
| AT4G34670 | 4,28 | -1,00 | 0,83 | -1,21 | 2,28E-01 |
| AT4G34680 | 0,14 | 0,03 | 4,39 | 0,01 | 9,95E-01 |
| AT4G34700 | 0,83 | -1,11 | 2,55 | -0,44 | 6,63E-01 |
| AT4G34870 | 1,35 | -0,08 | 1,57 | -0,05 | 9,61E-01 |
| AT4G34881 | 0,53 | -0,70 | 2,82 | -0,25 | 8,04E-01 |
| AT4G34910 | 0,61 | 1,81 | 2,99 | 0,60 | 5,45E-01 |
| AT4G34920 | 0,49 | -0,13 | 3,17 | -0,04 | 9,67E-01 |
| AT4G35040 | 0,31 | 0,65 | 4,32 | 0,15 | 8,81E-01 |
| AT4G35100 | 1,16 | -1,26 | 2,50 | -0,50 | 6,15E-01 |
| AT4G35310 | 1,84 | 2,83 | 1,61 | 1,76 | 7,86E-02 |
| AT4G35370 | 0,13 | 0,73 | 4,36 | 0,17 | 8,67E-01 |
| AT4G35450 | 0,43 | -0,72 | 3,37 | -0,21 | 8,30E-01 |
| AT4G35460 | 0,66 | -1,69 | 2,79 | -0,61 | 5,43E-01 |
| AT4G35470 | 0,67 | 1,21 | 2,61 | 0,46 | 6,42E-01 |
| AT4G35500 | 0,79 | -0,97 | 2,59 | -0,37 | 7,09E-01 |
| AT4G35560 | 0,72 | 0,10 | 2,89 | 0,03 | 9,73E-01 |
| AT4G35600 | 1,15 | -1,20 | 1,67 | -0,72 | 4,73E-01 |
| AT4G35630 | 0,33 | -0,15 | 3,58 | -0,04 | 9,66E-01 |
| AT4G35770 | 2,85 | -1,34 | 1,41 | -0,95 | 3,40E-01 |

| | | | | | |
|---|---|---|---|---|---|
| AT4G35850 | 0,77 | -0,16 | 2,13 | -0,08 | 9,39E-01 |
| AT4G35870 | 0,76 | -0,27 | 2,47 | -0,11 | 9,13E-01 |
| AT4G35880 | 0,17 | 0,98 | 4,35 | 0,23 | 8,21E-01 |
| AT4G35890 | 1,08 | -0,65 | 1,92 | -0,34 | 7,36E-01 |
| AT4G35940 | 1,33 | -2,13 | 1,54 | -1,38 | 1,67E-01 |
| AT4G36032 | 1,21 | -1,22 | 2,17 | -0,56 | 5,73E-01 |
| AT4G36052 | 0,12 | 1,23 | 4,34 | 0,28 | 7,76E-01 |
| AT4G36130 | 1,27 | -1,46 | 1,72 | -0,85 | 3,96E-01 |
| AT4G36150 | 1,48 | 1,20 | 2,38 | 0,50 | 6,14E-01 |
| AT4G36190 | 0,22 | 0,00 | 4,41 | 0,00 | 1,00E+00 |
| AT4G36195 | 0,86 | 0,53 | 2,28 | 0,23 | 8,16E-01 |
| AT4G36210 | 1,25 | -0,94 | 1,57 | -0,60 | 5,50E-01 |
| AT4G36250 | 0,69 | -0,77 | 2,53 | -0,30 | 7,62E-01 |
| AT4G36540 | 0,24 | 0,24 | 4,34 | 0,06 | 9,55E-01 |
| AT4G36710 | 1,27 | 1,62 | 2,48 | 0,65 | 5,13E-01 |
| AT4G36720 | 0,82 | -0,48 | 2,71 | -0,18 | 8,58E-01 |
| AT4G09375 | 0,14 | 0,00 | 4,41 | 0,00 | 1,00E+00 |
| AT4G36760 | 3,13 | -0,01 | 1,00 | -0,01 | 9,88E-01 |
| AT4G36810 | 0,84 | 2,02 | 2,39 | 0,84 | 3,99E-01 |
| AT4G36870 | 0,25 | 0,00 | 4,41 | 0,00 | 1,00E+00 |
| AT4G36920 | 0,14 | 0,00 | 4,41 | 0,00 | 1,00E+00 |
| AT4G36925 | 1,06 | -1,05 | 1,82 | -0,58 | 5,64E-01 |
| AT4G36990 | 0,59 | 1,51 | 2,97 | 0,51 | 6,10E-01 |
| AT4G37000 | 0,23 | -1,78 | 4,41 | -0,40 | 6,87E-01 |
| AT4G37020 | 0,15 | 1,01 | 4,34 | 0,23 | 8,17E-01 |
| AT4G37080 | 0,19 | 0,94 | 4,35 | 0,22 | 8,28E-01 |
| AT4G37130 | 0,62 | 0,40 | 3,57 | 0,11 | 9,10E-01 |
| AT4G37180 | 0,47 | 1,30 | 4,08 | 0,32 | 7,50E-01 |
| AT4G37190 | 0,32 | 0,39 | 4,34 | 0,09 | 9,28E-01 |
| AT4G37210 | 0,15 | 0,67 | 4,37 | 0,15 | 8,79E-01 |
| AT4G37260 | 2,49 | 0,83 | 1,28 | 0,65 | 5,18E-01 |
| AT4G37300 | 1,24 | -1,88 | 1,95 | -0,97 | 3,33E-01 |
| AT4G37430 | 0,86 | 2,53 | 3,35 | 0,76 | 4,50E-01 |
| AT4G37440 | 0,34 | -2,39 | 4,38 | -0,55 | 5,85E-01 |
| AT4G37550 | 0,88 | -0,47 | 2,03 | -0,23 | 8,17E-01 |
| AT4G37560 | 0,42 | 0,03 | 4,31 | 0,01 | 9,94E-01 |
| AT4G37650 | 1,47 | -0,88 | 1,92 | -0,46 | 6,46E-01 |
| AT4G37680 | 0,49 | 0,55 | 3,18 | 0,17 | 8,62E-01 |
| AT4G37730 | 0,75 | 0,90 | 2,15 | 0,42 | 6,75E-01 |
| AT4G37760 | 0,63 | -0,27 | 2,40 | -0,11 | 9,11E-01 |
| AT4G37790 | 0,15 | 0,00 | 4,41 | 0,00 | 1,00E+00 |
| AT4G37980 | 0,67 | 1,52 | 2,22 | 0,68 | 4,95E-01 |
| AT4G38040 | 0,20 | -0,31 | 4,36 | -0,07 | 9,44E-01 |
| AT4G38170 | 0,97 | -1,12 | 2,16 | -0,52 | 6,03E-01 |
| AT4G09685 | 0,46 | -0,33 | 3,95 | -0,08 | 9,33E-01 |
| AT4G38200 | 2,17 | -1,31 | 1,24 | -1,05 | 2,92E-01 |
| AT4G38215 | 0,75 | 0,53 | 4,29 | 0,12 | 9,01E-01 |
| AT4G38220 | 1,96 | 0,47 | 1,30 | 0,36 | 7,19E-01 |
| AT4G38225 | 0,78 | -0,19 | 2,45 | -0,08 | 9,39E-01 |
| AT4G38250 | 1,25 | -0,20 | 1,54 | -0,13 | 8,99E-01 |
| AT4G38270 | 0,51 | 2,31 | 3,09 | 0,75 | 4,55E-01 |
| AT4G38290 | 0,11 | 0,65 | 4,36 | 0,15 | 8,81E-01 |

| | | | | | |
|---|---|---|---|---|---|
| AT4G38320 | 0,19 | 1,08 | 4,34 | 0,25 | 8,04E-01 |
| AT4G38330 | 0,12 | 1,16 | 4,34 | 0,27 | 7,89E-01 |
| AT4G38360 | 0,15 | 1,16 | 4,34 | 0,27 | 7,88E-01 |
| AT4G38430 | 0,81 | 0,37 | 2,56 | 0,15 | 8,84E-01 |
| AT4G38440 | 2,75 | -0,83 | 0,99 | -0,84 | 4,01E-01 |
| AT4G38460 | 1,41 | -0,82 | 1,66 | -0,49 | 6,23E-01 |
| AT4G38500 | 0,68 | 2,99 | 3,46 | 0,86 | 3,88E-01 |
| AT4G38540 | 0,10 | 0,21 | 4,38 | 0,05 | 9,62E-01 |
| AT4G38550 | 2,97 | 0,49 | 1,19 | 0,41 | 6,78E-01 |
| AT4G38560 | 1,22 | -2,92 | 2,34 | -1,25 | 2,12E-01 |
| AT4G38580 | 0,50 | -3,11 | 4,34 | -0,71 | 4,75E-01 |
| AT4G38620 | 0,24 | 0,62 | 4,37 | 0,14 | 8,88E-01 |
| AT4G09735 | 2,23 | 1,05 | 1,78 | 0,59 | 5,54E-01 |
| AT4G38775 | 2,66 | 1,65 | 1,53 | 1,08 | 2,79E-01 |
| AT4G38790 | 0,69 | 0,67 | 2,96 | 0,23 | 8,21E-01 |
| AT4G38860 | 0,38 | 0,67 | 4,36 | 0,15 | 8,78E-01 |
| AT4G38880 | 0,39 | -1,40 | 4,37 | -0,32 | 7,49E-01 |
| AT4G38920 | 0,99 | -0,02 | 2,10 | -0,01 | 9,92E-01 |
| AT4G38930 | 0,41 | -1,53 | 4,36 | -0,35 | 7,26E-01 |
| AT4G39020 | 0,55 | 1,82 | 4,32 | 0,42 | 6,73E-01 |
| AT4G39030 | 0,29 | -1,33 | 4,35 | -0,31 | 7,60E-01 |
| AT4G39050 | 1,56 | 1,10 | 1,68 | 0,65 | 5,13E-01 |
| AT4G39080 | 4,27 | -0,15 | 0,90 | -0,17 | 8,67E-01 |
| AT4G39160 | 1,99 | -0,49 | 1,32 | -0,37 | 7,11E-01 |
| AT4G39200 | 1,73 | -2,99 | 1,64 | -1,82 | 6,88E-02 |
| AT4G39210 | 1,07 | -0,14 | 2,07 | -0,07 | 9,45E-01 |
| AT4G39270 | 1,16 | -0,87 | 1,59 | -0,55 | 5,84E-01 |
| AT4G39330 | 1,70 | -1,82 | 1,39 | -1,31 | 1,91E-01 |
| AT4G39350 | 3,27 | -0,78 | 0,95 | -0,82 | 4,12E-01 |
| AT4G39361 | 1,80 | 2,23 | 1,91 | 1,16 | 2,44E-01 |
| AT4G39364 | 2,19 | 0,42 | 1,64 | 0,26 | 7,97E-01 |
| AT4G39366 | 2,55 | 1,12 | 1,44 | 0,78 | 4,37E-01 |
| AT4G39400 | 3,54 | 0,55 | 1,03 | 0,53 | 5,94E-01 |
| AT4G39620 | 0,18 | 1,24 | 4,33 | 0,29 | 7,74E-01 |
| AT4G39640 | 0,76 | -0,02 | 2,51 | -0,01 | 9,95E-01 |
| AT4G39670 | 0,55 | 1,94 | 3,18 | 0,61 | 5,41E-01 |
| AT4G39730 | 3,06 | 0,24 | 1,37 | 0,17 | 8,62E-01 |
| AT4G39840 | 1,07 | -0,45 | 1,86 | -0,24 | 8,11E-01 |
| AT4G39850 | 2,19 | -1,47 | 1,32 | -1,11 | 2,66E-01 |
| AT4G39860 | 0,51 | -0,05 | 2,43 | -0,02 | 9,84E-01 |
| AT4G39870 | 0,35 | 1,96 | 4,32 | 0,45 | 6,50E-01 |
| AT4G39940 | 0,36 | -0,56 | 4,32 | -0,13 | 8,97E-01 |
| AT4G39950 | 1,14 | 0,31 | 1,81 | 0,17 | 8,63E-01 |
| AT4G39952 | 0,28 | -1,42 | 4,36 | -0,33 | 7,44E-01 |
| AT4G39980 | 0,50 | -0,31 | 3,11 | -0,10 | 9,20E-01 |
| AT4G40063 | 0,73 | 0,73 | 2,90 | 0,25 | 8,01E-01 |
| AT4G40100 | 3,82 | 0,53 | 1,14 | 0,46 | 6,43E-01 |
| AT5G00730 | 1,64 | -0,81 | 2,04 | -0,40 | 6,91E-01 |
| AT5G01170 | 0,10 | 0,01 | 4,38 | 0,00 | 9,99E-01 |
| AT5G01210 | 1,09 | 1,00 | 1,87 | 0,53 | 5,93E-01 |
| AT5G01240 | 0,22 | 0,25 | 4,36 | 0,06 | 9,54E-01 |
| AT5G01260 | 0,54 | -0,65 | 2,30 | -0,28 | 7,78E-01 |

| | | | | | |
|---|---|---|---|---|---|
| AT5G01290 | 0,28 | 1,74 | 4,32 | 0,40 | 6,87E-01 |
| AT5G01460 | 0,28 | -2,52 | 4,38 | -0,58 | 5,65E-01 |
| AT5G01542 | 0,46 | 0,10 | 4,03 | 0,02 | 9,80E-01 |
| AT5G01590 | 0,94 | 1,65 | 2,07 | 0,80 | 4,25E-01 |
| AT5G01595 | 0,81 | 1,77 | 2,43 | 0,73 | 4,65E-01 |
| AT5G01690 | 0,08 | 0,44 | 4,38 | 0,10 | 9,21E-01 |
| AT5G01732 | 0,29 | 0,65 | 4,32 | 0,15 | 8,80E-01 |
| AT5G01740 | 0,15 | 0,00 | 4,41 | 0,00 | 1,00E+00 |
| AT5G01750 | 1,69 | 2,58 | 2,01 | 1,29 | 1,99E-01 |
| AT5G01760 | 0,79 | -1,85 | 2,61 | -0,71 | 4,78E-01 |
| AT5G01800 | 0,74 | -0,39 | 2,27 | -0,17 | 8,65E-01 |
| AT5G01810 | 0,55 | 0,25 | 3,78 | 0,07 | 9,47E-01 |
| AT5G01850 | 0,24 | 0,37 | 4,37 | 0,08 | 9,33E-01 |
| AT5G01920 | 2,56 | -0,30 | 1,16 | -0,26 | 7,98E-01 |
| AT5G01960 | 1,82 | 1,48 | 1,85 | 0,80 | 4,24E-01 |
| AT5G01980 | 1,71 | -0,03 | 1,47 | -0,02 | 9,83E-01 |
| AT5G02035 | 0,47 | 2,00 | 3,95 | 0,51 | 6,12E-01 |
| AT5G02040 | 0,25 | 0,26 | 4,33 | 0,06 | 9,51E-01 |
| AT5G02100 | 0,70 | 2,44 | 3,70 | 0,66 | 5,09E-01 |
| AT5G02120 | 0,39 | 0,00 | 4,39 | 0,00 | 1,00E+00 |
| AT5G00820 | 1,31 | -1,29 | 2,61 | -0,49 | 6,22E-01 |
| AT5G02160 | 1,30 | -0,25 | 1,86 | -0,13 | 8,94E-01 |
| AT5G02240 | 0,32 | -0,24 | 4,34 | -0,06 | 9,56E-01 |
| AT5G02250 | 1,33 | -0,77 | 2,12 | -0,36 | 7,17E-01 |
| AT5G02280 | 0,11 | 0,65 | 4,36 | 0,15 | 8,81E-01 |
| AT5G02310 | 4,95 | -0,10 | 0,78 | -0,12 | 9,02E-01 |
| AT5G02420 | 0,12 | 0,97 | 4,34 | 0,22 | 8,23E-01 |
| AT5G02450 | 0,59 | -0,11 | 3,13 | -0,03 | 9,73E-01 |
| AT5G02480 | 2,67 | 0,74 | 1,24 | 0,60 | 5,51E-01 |
| AT5G02620 | 0,32 | -2,05 | 4,37 | -0,47 | 6,38E-01 |
| AT5G02630 | 0,30 | 0,24 | 4,35 | 0,05 | 9,57E-01 |
| AT5G02650 | 0,40 | 2,16 | 4,32 | 0,50 | 6,17E-01 |
| AT5G02760 | 0,40 | -0,33 | 3,43 | -0,10 | 9,24E-01 |
| AT5G02790 | 0,39 | -0,90 | 3,52 | -0,26 | 7,97E-01 |
| AT5G02840 | 0,40 | 0,81 | 4,14 | 0,20 | 8,45E-01 |
| AT5G02850 | 0,54 | -0,05 | 3,02 | -0,02 | 9,86E-01 |
| AT5G02860 | 1,11 | 1,17 | 1,89 | 0,62 | 5,35E-01 |
| AT5G02870 | 2,41 | -0,90 | 1,35 | -0,67 | 5,04E-01 |
| AT5G02880 | 3,00 | 0,72 | 1,10 | 0,66 | 5,12E-01 |
| AT5G02950 | 0,61 | 1,95 | 3,79 | 0,52 | 6,06E-01 |
| AT5G02995 | 0,13 | 0,00 | 4,41 | 0,00 | 1,00E+00 |
| AT5G03020 | 0,28 | -1,42 | 4,36 | -0,33 | 7,44E-01 |
| AT5G03160 | 0,68 | -0,95 | 2,32 | -0,41 | 6,84E-01 |
| AT5G03190 | 0,92 | -0,85 | 1,89 | -0,45 | 6,54E-01 |
| AT5G03280 | 3,28 | 0,35 | 1,14 | 0,31 | 7,56E-01 |
| AT5G03290 | 1,24 | -0,43 | 1,52 | -0,28 | 7,80E-01 |
| AT5G03300 | 0,81 | 2,16 | 2,45 | 0,88 | 3,78E-01 |
| AT5G03330 | 0,61 | -1,44 | 2,98 | -0,48 | 6,28E-01 |
| AT5G03355 | 0,46 | -2,58 | 3,38 | -0,76 | 4,45E-01 |
| AT5G03415 | 0,36 | 0,17 | 4,32 | 0,04 | 9,68E-01 |
| AT5G03420 | 0,63 | 1,40 | 2,53 | 0,55 | 5,79E-01 |
| AT5G03455 | 0,25 | 0,03 | 4,34 | 0,01 | 9,95E-01 |

| | | | | | |
|---|---|---|---|---|---|
| AT5G03460 | 0,54 | -2,21 | 3,03 | -0,73 | 4,66E-01 |
| AT5G03470 | 0,40 | -1,14 | 4,31 | -0,26 | 7,92E-01 |
| AT5G03490 | 0,11 | 0,48 | 4,37 | 0,11 | 9,12E-01 |
| AT5G03540 | 1,05 | 0,38 | 2,63 | 0,14 | 8,86E-01 |
| AT5G03650 | 0,91 | -0,32 | 1,83 | -0,17 | 8,61E-01 |
| AT5G03905 | 0,30 | -0,50 | 4,38 | -0,11 | 9,09E-01 |
| AT5G04040 | 1,27 | 2,32 | 1,98 | 1,17 | 2,42E-01 |
| AT5G04050 | 0,10 | 0,01 | 4,38 | 0,00 | 9,99E-01 |
| AT5G04060 | 0,37 | 1,21 | 4,35 | 0,28 | 7,81E-01 |
| AT5G04170 | 0,76 | -1,55 | 2,48 | -0,63 | 5,32E-01 |
| AT5G04190 | 0,13 | 0,00 | 4,41 | 0,00 | 1,00E+00 |
| AT5G04240 | 0,88 | 3,17 | 3,25 | 0,98 | 3,29E-01 |
| AT5G04280 | 1,25 | 2,26 | 1,93 | 1,17 | 2,42E-01 |
| AT5G04290 | 0,66 | -1,10 | 3,22 | -0,34 | 7,33E-01 |
| AT5G04360 | 0,45 | -0,83 | 4,12 | -0,20 | 8,40E-01 |
| AT5G00960 | 0,96 | 2,95 | 2,19 | 1,35 | 1,77E-01 |
| AT5G04410 | 1,77 | 0,82 | 2,18 | 0,38 | 7,07E-01 |
| AT5G04460 | 0,56 | 1,24 | 3,08 | 0,40 | 6,86E-01 |
| AT5G04490 | 0,46 | -0,11 | 3,37 | -0,03 | 9,75E-01 |
| AT5G04510 | 0,58 | -0,38 | 2,62 | -0,15 | 8,84E-01 |
| AT5G04560 | 3,61 | -0,64 | 0,94 | -0,68 | 4,99E-01 |
| AT5G04590 | 4,30 | 0,52 | 0,99 | 0,53 | 5,98E-01 |
| AT5G04600 | 0,71 | -1,21 | 2,23 | -0,54 | 5,89E-01 |
| AT5G04620 | 0,43 | -1,11 | 4,34 | -0,25 | 7,99E-01 |
| AT5G04630 | 0,57 | 1,13 | 3,73 | 0,30 | 7,61E-01 |
| AT5G04660 | 0,24 | 0,92 | 4,36 | 0,21 | 8,32E-01 |
| AT5G00985 | 0,28 | -1,42 | 4,35 | -0,33 | 7,44E-01 |
| AT5G04720 | 2,75 | 1,47 | 1,52 | 0,96 | 3,35E-01 |
| AT5G04800 | 0,67 | -1,35 | 2,09 | -0,65 | 5,19E-01 |
| AT5G04810 | 3,05 | -0,73 | 0,99 | -0,74 | 4,59E-01 |
| AT5G04885 | 0,27 | 1,17 | 3,74 | 0,31 | 7,55E-01 |
| AT5G04910 | 0,42 | -0,90 | 4,11 | -0,22 | 8,26E-01 |
| AT5G04930 | 5,77 | 0,61 | 0,91 | 0,68 | 4,99E-01 |
| AT5G05010 | 2,08 | -0,27 | 1,36 | -0,20 | 8,45E-01 |
| AT5G05070 | 0,40 | -0,67 | 4,27 | -0,16 | 8,74E-01 |
| AT5G05090 | 0,35 | -0,45 | 3,57 | -0,13 | 9,00E-01 |
| AT5G05160 | 0,72 | -0,88 | 2,70 | -0,33 | 7,44E-01 |
| AT5G05190 | 2,00 | 0,27 | 1,31 | 0,20 | 8,39E-01 |
| AT5G05210 | 0,28 | 0,43 | 4,37 | 0,10 | 9,22E-01 |
| AT5G05220 | 0,08 | 0,44 | 4,38 | 0,10 | 9,21E-01 |
| AT5G05230 | 0,08 | 0,44 | 4,38 | 0,10 | 9,21E-01 |
| AT5G05310 | 0,18 | 0,00 | 4,41 | 0,00 | 1,00E+00 |
| AT5G05440 | 0,16 | 0,49 | 4,38 | 0,11 | 9,12E-01 |
| AT5G05490 | 0,55 | -1,70 | 4,31 | -0,40 | 6,93E-01 |
| AT5G05560 | 1,14 | -0,09 | 1,82 | -0,05 | 9,63E-01 |
| AT5G05570 | 0,62 | 2,23 | 4,31 | 0,52 | 6,05E-01 |
| AT5G05600 | 0,31 | 0,00 | 4,41 | 0,00 | 1,00E+00 |
| AT5G05630 | 0,08 | 0,00 | 4,40 | 0,00 | 1,00E+00 |
| AT5G05680 | 1,56 | -1,18 | 1,56 | -0,75 | 4,52E-01 |
| AT5G05710 | 0,20 | -0,44 | 4,36 | -0,10 | 9,19E-01 |
| AT5G05750 | 1,85 | 2,29 | 2,02 | 1,13 | 2,57E-01 |
| AT5G05780 | 0,96 | 2,90 | 2,48 | 1,17 | 2,42E-01 |

| | | | | | |
|---|---|---|---|---|---|
| AT5G05950 | 0,83 | -0,69 | 2,31 | -0,30 | 7,64E-01 |
| AT5G05970 | 0,30 | -0,24 | 4,34 | -0,06 | 9,56E-01 |
| AT5G05985 | 0,55 | -1,30 | 3,78 | -0,34 | 7,31E-01 |
| AT5G06000 | 0,23 | 0,00 | 4,41 | 0,00 | 1,00E+00 |
| AT5G06050 | 0,39 | -0,72 | 4,25 | -0,17 | 8,65E-01 |
| AT5G06100 | 0,21 | 1,63 | 4,33 | 0,38 | 7,06E-01 |
| AT5G06120 | 1,17 | -0,40 | 2,25 | -0,18 | 8,59E-01 |
| AT5G01155 | 4,02 | 0,33 | 1,09 | 0,31 | 7,58E-01 |
| AT5G06170 | 0,31 | 0,00 | 4,41 | 0,00 | 1,00E+00 |
| AT5G06220 | 0,84 | -0,23 | 2,34 | -0,10 | 9,23E-01 |
| AT5G06265 | 0,26 | 0,30 | 4,38 | 0,07 | 9,46E-01 |
| AT5G06290 | 1,37 | -0,88 | 1,73 | -0,51 | 6,10E-01 |
| AT5G06310 | 0,33 | 1,39 | 4,33 | 0,32 | 7,47E-01 |
| AT5G01255 | 0,70 | 1,24 | 2,78 | 0,45 | 6,55E-01 |
| AT5G06350 | 0,36 | 0,84 | 3,46 | 0,24 | 8,09E-01 |
| AT5G06360 | 0,40 | -0,67 | 4,27 | -0,16 | 8,74E-01 |
| AT5G06380 | 0,08 | 0,44 | 4,38 | 0,10 | 9,21E-01 |
| AT5G06390 | 0,14 | 0,09 | 4,38 | 0,02 | 9,84E-01 |
| AT5G06460 | 1,60 | -0,34 | 1,60 | -0,21 | 8,31E-01 |
| AT5G06645 | 0,58 | 0,00 | 4,41 | 0,00 | 1,00E+00 |
| AT5G06660 | 0,96 | 2,88 | 3,53 | 0,82 | 4,14E-01 |
| AT5G06680 | 0,39 | -1,73 | 4,32 | -0,40 | 6,88E-01 |
| AT5G06700 | 0,91 | -0,57 | 1,85 | -0,31 | 7,59E-01 |
| AT5G06811 | 0,11 | 0,65 | 4,36 | 0,15 | 8,81E-01 |
| AT5G06820 | 0,12 | 1,16 | 4,34 | 0,27 | 7,89E-01 |
| AT5G06860 | 0,22 | 0,24 | 4,38 | 0,05 | 9,57E-01 |
| AT5G06870 | 0,67 | -0,93 | 2,83 | -0,33 | 7,41E-01 |
| AT5G06950 | 0,61 | -1,12 | 2,98 | -0,38 | 7,06E-01 |
| AT5G06980 | 0,16 | 1,33 | 4,33 | 0,31 | 7,59E-01 |
| AT5G07090 | 1,54 | -2,33 | 1,46 | -1,60 | 1,09E-01 |
| AT5G07110 | 0,85 | -0,14 | 3,27 | -0,04 | 9,65E-01 |
| AT5G01315 | 0,64 | -1,04 | 2,89 | -0,36 | 7,18E-01 |
| AT5G07130 | 0,33 | 0,26 | 4,33 | 0,06 | 9,52E-01 |
| AT5G07140 | 1,24 | -0,53 | 2,28 | -0,23 | 8,16E-01 |
| AT5G07210 | 1,09 | 0,21 | 2,08 | 0,10 | 9,21E-01 |
| AT5G07240 | 1,19 | 0,73 | 1,97 | 0,37 | 7,10E-01 |
| AT5G07270 | 0,15 | 0,00 | 4,38 | 0,00 | 1,00E+00 |
| AT5G07290 | 1,99 | 0,06 | 1,94 | 0,03 | 9,76E-01 |
| AT5G07340 | 0,98 | 2,91 | 2,28 | 1,28 | 2,02E-01 |
| AT5G07410 | 1,62 | 2,40 | 1,93 | 1,25 | 2,13E-01 |
| AT5G07430 | 0,50 | -1,46 | 3,95 | -0,37 | 7,12E-01 |
| AT5G07440 | 0,36 | -0,09 | 4,35 | -0,02 | 9,83E-01 |
| AT5G07570 | 4,01 | 1,99 | 1,50 | 1,32 | 1,87E-01 |
| AT5G07580 | 0,75 | -0,33 | 2,21 | -0,15 | 8,83E-01 |
| AT5G07610 | 0,48 | 1,83 | 4,05 | 0,45 | 6,51E-01 |
| AT5G07760 | 0,08 | 0,69 | 4,36 | 0,16 | 8,75E-01 |
| AT5G07770 | 0,16 | 0,65 | 4,37 | 0,15 | 8,82E-01 |
| AT5G07790 | 0,58 | 1,84 | 4,32 | 0,43 | 6,71E-01 |
| AT5G07830 | 0,86 | -0,35 | 2,63 | -0,13 | 8,93E-01 |
| AT5G07860 | 0,58 | -1,69 | 3,82 | -0,44 | 6,57E-01 |
| AT5G07870 | 0,49 | -0,90 | 4,27 | -0,21 | 8,33E-01 |
| AT5G07940 | 5,66 | -0,07 | 0,81 | -0,08 | 9,36E-01 |

| | | | | | |
|---|---|---|---|---|---|
| AT5G07960 | 0,12 | 0,97 | 4,34 | 0,22 | 8,23E-01 |
| AT5G08050 | 0,48 | 1,64 | 4,16 | 0,39 | 6,94E-01 |
| AT5G08060 | 0,10 | 0,01 | 4,38 | 0,00 | 9,99E-01 |
| AT5G08070 | 0,35 | -2,17 | 4,38 | -0,49 | 6,21E-01 |
| AT5G08139 | 0,37 | 0,82 | 4,32 | 0,19 | 8,49E-01 |
| AT5G08190 | 0,29 | 0,70 | 4,32 | 0,16 | 8,72E-01 |
| AT5G08260 | 0,90 | -2,82 | 2,26 | -1,25 | 2,12E-01 |
| AT5G01545 | 0,37 | 0,00 | 4,40 | 0,00 | 1,00E+00 |
| AT5G08290 | 0,74 | 1,32 | 2,17 | 0,61 | 5,44E-01 |
| AT5G08300 | 0,94 | -0,56 | 2,33 | -0,24 | 8,10E-01 |
| AT5G08330 | 1,56 | -2,81 | 1,60 | -1,75 | 7,96E-02 |
| AT5G08335 | 0,18 | 0,81 | 4,34 | 0,19 | 8,52E-01 |
| AT5G08340 | 0,58 | -0,09 | 3,92 | -0,02 | 9,81E-01 |
| AT5G08390 | 0,26 | 1,71 | 4,32 | 0,39 | 6,93E-01 |
| AT5G08430 | 0,57 | -0,10 | 4,28 | -0,02 | 9,82E-01 |
| AT5G08440 | 0,59 | -1,17 | 3,81 | -0,31 | 7,58E-01 |
| AT5G08470 | 0,81 | 0,20 | 2,63 | 0,08 | 9,38E-01 |
| AT5G08510 | 0,51 | -2,16 | 3,20 | -0,68 | 4,99E-01 |
| AT5G08540 | 0,41 | 1,27 | 3,38 | 0,38 | 7,07E-01 |
| AT5G01555 | 0,36 | -0,38 | 4,38 | -0,09 | 9,31E-01 |
| AT5G08570 | 1,20 | -0,55 | 1,61 | -0,34 | 7,30E-01 |
| AT5G08590 | 0,63 | 0,27 | 2,69 | 0,10 | 9,20E-01 |
| AT5G08600 | 0,28 | -1,42 | 4,35 | -0,33 | 7,44E-01 |
| AT5G08610 | 1,71 | -0,64 | 1,41 | -0,45 | 6,50E-01 |
| AT5G08620 | 1,12 | -0,83 | 1,71 | -0,48 | 6,29E-01 |
| AT5G08660 | 0,33 | -2,47 | 4,36 | -0,57 | 5,71E-01 |
| AT5G08680 | 1,77 | -1,39 | 1,63 | -0,85 | 3,94E-01 |
| AT5G08690 | 2,64 | 0,51 | 1,15 | 0,45 | 6,55E-01 |
| AT5G08695 | 0,20 | -0,44 | 4,36 | -0,10 | 9,19E-01 |
| AT5G08750 | 0,11 | 0,48 | 4,37 | 0,11 | 9,12E-01 |
| AT5G08760 | 0,55 | -1,66 | 3,96 | -0,42 | 6,75E-01 |
| AT5G08780 | 0,49 | -0,23 | 3,11 | -0,07 | 9,42E-01 |
| AT5G09220 | 0,82 | 0,36 | 3,10 | 0,12 | 9,08E-01 |
| AT5G09400 | 1,22 | 0,71 | 1,84 | 0,38 | 7,02E-01 |
| AT5G09410 | 3,14 | 0,24 | 1,08 | 0,22 | 8,25E-01 |
| AT5G09440 | 0,77 | 0,43 | 2,29 | 0,19 | 8,50E-01 |
| AT5G09445 | 0,26 | -0,48 | 4,38 | -0,11 | 9,13E-01 |
| AT5G09450 | 1,25 | -2,19 | 1,98 | -1,10 | 2,70E-01 |
| AT5G09585 | 3,20 | 4,22 | 1,50 | 2,82 | 4,87E-03 |
| AT5G09590 | 2,74 | -0,71 | 1,34 | -0,53 | 5,96E-01 |
| AT5G09600 | 0,62 | 0,85 | 3,03 | 0,28 | 7,79E-01 |
| AT5G09760 | 0,14 | 0,09 | 4,38 | 0,02 | 9,84E-01 |
| AT5G09810 | 3,96 | -0,81 | 1,14 | -0,71 | 4,81E-01 |
| AT5G09870 | 2,84 | -0,79 | 1,00 | -0,79 | 4,29E-01 |
| AT5G10010 | 0,47 | -1,61 | 3,40 | -0,47 | 6,37E-01 |
| AT5G10020 | 0,41 | -0,10 | 4,29 | -0,02 | 9,82E-01 |
| AT5G10100 | 0,16 | 0,38 | 4,37 | 0,09 | 9,30E-01 |
| AT5G10180 | 2,74 | 0,37 | 1,25 | 0,30 | 7,67E-01 |
| AT5G10190 | 0,41 | -0,61 | 3,27 | -0,19 | 8,52E-01 |
| AT5G10200 | 0,81 | -1,11 | 3,33 | -0,33 | 7,39E-01 |
| AT5G10340 | 1,40 | -2,48 | 2,25 | -1,10 | 2,70E-01 |
| AT5G10370 | 1,36 | 2,32 | 1,75 | 1,32 | 1,86E-01 |

| | | | | | |
|---|---|---|---|---|---|
| AT5G10410 | 0,05 | 0,00 | 4,41 | 0,00 | 1,00E+00 |
| AT5G10460 | 0,08 | 0,69 | 4,36 | 0,16 | 8,75E-01 |
| AT5G10540 | 2,27 | -0,37 | 1,23 | -0,30 | 7,62E-01 |
| AT5G01765 | 1,15 | 0,47 | 1,79 | 0,26 | 7,93E-01 |
| AT5G01775 | 4,59 | 0,55 | 1,09 | 0,51 | 6,12E-01 |
| AT5G01785 | 2,53 | 0,80 | 1,75 | 0,45 | 6,49E-01 |
| AT5G10572 | 2,06 | 0,33 | 1,43 | 0,23 | 8,17E-01 |
| AT5G10600 | 0,40 | -0,40 | 4,26 | -0,09 | 9,26E-01 |
| AT5G10630 | 0,28 | 1,58 | 4,33 | 0,37 | 7,15E-01 |
| AT5G10660 | 0,46 | -0,19 | 2,76 | -0,07 | 9,46E-01 |
| AT5G10695 | 0,85 | 1,39 | 2,18 | 0,64 | 5,23E-01 |
| AT5G10745 | 0,35 | 1,54 | 3,50 | 0,44 | 6,60E-01 |
| AT5G10750 | 1,12 | -0,50 | 2,14 | -0,23 | 8,15E-01 |
| AT5G10780 | 0,76 | -0,32 | 2,54 | -0,13 | 8,99E-01 |
| AT5G10790 | 0,93 | -1,03 | 1,85 | -0,56 | 5,77E-01 |
| AT5G10920 | 0,91 | -0,83 | 2,12 | -0,39 | 6,94E-01 |
| AT5G10960 | 3,18 | -0,10 | 1,00 | -0,10 | 9,20E-01 |
| AT5G11040 | 2,15 | -0,02 | 1,21 | -0,02 | 9,86E-01 |
| AT5G11060 | 1,02 | 0,01 | 1,71 | 0,01 | 9,94E-01 |
| AT5G11090 | 0,49 | 0,01 | 2,70 | 0,00 | 9,96E-01 |
| AT5G11110 | 0,50 | -0,39 | 4,02 | -0,10 | 9,22E-01 |
| AT5G11170 | 1,42 | -1,50 | 1,89 | -0,79 | 4,28E-01 |
| AT5G11200 | 0,33 | 0,92 | 4,31 | 0,21 | 8,30E-01 |
| AT5G11280 | 0,20 | 0,41 | 4,37 | 0,09 | 9,26E-01 |
| AT5G11340 | 0,28 | 0,26 | 4,33 | 0,06 | 9,52E-01 |
| AT5G11350 | 0,82 | -1,31 | 2,79 | -0,47 | 6,38E-01 |
| AT5G11390 | 0,30 | 0,30 | 4,34 | 0,07 | 9,45E-01 |
| AT5G11420 | 1,15 | -1,23 | 2,08 | -0,59 | 5,56E-01 |
| AT5G11430 | 0,70 | -0,77 | 2,20 | -0,35 | 7,27E-01 |
| AT5G11490 | 1,97 | 0,55 | 1,24 | 0,44 | 6,57E-01 |
| AT5G11500 | 0,94 | 1,64 | 1,95 | 0,84 | 3,98E-01 |
| AT5G11510 | 0,81 | 0,68 | 2,28 | 0,30 | 7,66E-01 |
| AT5G11530 | 0,81 | -0,15 | 2,30 | -0,07 | 9,47E-01 |
| AT5G11580 | 0,12 | 0,37 | 4,37 | 0,08 | 9,33E-01 |
| AT5G11650 | 0,38 | 0,28 | 3,44 | 0,08 | 9,36E-01 |
| AT5G11670 | 3,41 | 0,13 | 1,03 | 0,12 | 9,03E-01 |
| AT5G11680 | 0,19 | 1,15 | 4,34 | 0,27 | 7,91E-01 |
| AT5G11710 | 1,24 | -0,52 | 1,58 | -0,33 | 7,41E-01 |
| AT5G11720 | 1,68 | -1,21 | 1,78 | -0,68 | 4,98E-01 |
| AT5G11730 | 0,89 | -1,18 | 1,89 | -0,62 | 5,33E-01 |
| AT5G11740 | 4,10 | 0,30 | 0,88 | 0,34 | 7,31E-01 |
| AT5G11760 | 0,22 | -0,88 | 4,38 | -0,20 | 8,40E-01 |
| AT5G11790 | 2,58 | 1,35 | 1,41 | 0,96 | 3,38E-01 |
| AT5G01915 | 2,27 | 2,14 | 1,69 | 1,26 | 2,06E-01 |
| AT5G11890 | 0,82 | -1,03 | 2,08 | -0,50 | 6,20E-01 |
| AT5G11920 | 0,12 | 0,37 | 4,37 | 0,08 | 9,33E-01 |
| AT5G11977 | 0,12 | 1,23 | 4,34 | 0,28 | 7,76E-01 |
| AT5G12090 | 0,79 | 0,02 | 2,52 | 0,01 | 9,93E-01 |
| AT5G12100 | 0,44 | 2,30 | 4,31 | 0,53 | 5,94E-01 |
| AT5G12110 | 1,52 | -0,02 | 1,40 | -0,02 | 9,87E-01 |
| AT5G12210 | 0,65 | 0,03 | 3,64 | 0,01 | 9,94E-01 |
| AT5G12260 | 0,32 | -0,24 | 4,34 | -0,06 | 9,56E-01 |

| | | | | | |
|---|---|---|---|---|---|
| AT5G12350 | 0,49 | -0,56 | 4,08 | -0,14 | 8,90E-01 |
| AT5G12400 | 0,93 | 1,37 | 2,77 | 0,49 | 6,21E-01 |
| AT5G12850 | 0,97 | -1,22 | 1,87 | -0,65 | 5,14E-01 |
| AT5G12950 | 0,53 | 1,48 | 2,56 | 0,58 | 5,62E-01 |
| AT5G12960 | 0,27 | -0,47 | 4,37 | -0,11 | 9,14E-01 |
| AT5G12970 | 0,24 | -2,36 | 4,39 | -0,54 | 5,91E-01 |
| AT5G13010 | 4,55 | -0,85 | 0,92 | -0,93 | 3,53E-01 |
| AT5G13030 | 0,96 | 1,10 | 2,09 | 0,52 | 6,00E-01 |
| AT5G13100 | 3,43 | -1,52 | 1,04 | -1,47 | 1,43E-01 |
| AT5G13180 | 1,85 | -0,33 | 1,48 | -0,22 | 8,22E-01 |
| AT5G13200 | 1,26 | 0,14 | 1,75 | 0,08 | 9,38E-01 |
| AT5G13210 | 0,35 | 0,54 | 3,42 | 0,16 | 8,75E-01 |
| AT5G13260 | 0,92 | -1,37 | 2,06 | -0,67 | 5,06E-01 |
| AT5G13280 | 0,45 | -1,86 | 4,10 | -0,45 | 6,50E-01 |
| AT5G13310 | 0,31 | 0,62 | 4,32 | 0,14 | 8,85E-01 |
| AT5G13320 | 1,27 | 1,49 | 1,85 | 0,80 | 4,21E-01 |
| AT5G13330 | 0,37 | 0,94 | 4,32 | 0,22 | 8,28E-01 |
| AT5G13370 | 0,68 | -0,78 | 3,26 | -0,24 | 8,11E-01 |
| AT5G13380 | 0,54 | 2,09 | 4,32 | 0,48 | 6,29E-01 |
| AT5G13390 | 2,05 | 0,08 | 1,40 | 0,06 | 9,52E-01 |
| AT5G13500 | 0,32 | -0,31 | 4,36 | -0,07 | 9,44E-01 |
| AT5G13510 | 0,48 | 0,27 | 3,30 | 0,08 | 9,34E-01 |
| AT5G13530 | 2,40 | 0,48 | 1,18 | 0,41 | 6,82E-01 |
| AT5G13580 | 0,29 | 2,11 | 4,32 | 0,49 | 6,26E-01 |
| AT5G13640 | 0,84 | -1,82 | 2,89 | -0,63 | 5,28E-01 |
| AT5G13650 | 5,78 | -0,02 | 0,81 | -0,02 | 9,82E-01 |
| AT5G13690 | 0,61 | -0,75 | 2,99 | -0,25 | 8,02E-01 |
| AT5G13720 | 0,21 | 0,10 | 4,40 | 0,02 | 9,82E-01 |
| AT5G13740 | 0,19 | 1,15 | 4,34 | 0,27 | 7,91E-01 |
| AT5G13750 | 0,26 | -0,30 | 4,37 | -0,07 | 9,46E-01 |
| AT5G13760 | 0,26 | -0,71 | 4,36 | -0,16 | 8,71E-01 |
| AT5G13770 | 2,99 | 1,42 | 1,24 | 1,15 | 2,51E-01 |
| AT5G13810 | 0,93 | -0,63 | 1,77 | -0,35 | 7,23E-01 |
| AT5G13820 | 0,28 | 1,90 | 4,32 | 0,44 | 6,60E-01 |
| AT5G13850 | 2,26 | -0,20 | 1,26 | -0,16 | 8,75E-01 |
| AT5G13930 | 1,07 | 0,43 | 2,10 | 0,20 | 8,39E-01 |
| AT5G13980 | 0,97 | -0,73 | 2,25 | -0,32 | 7,46E-01 |
| AT5G14030 | 0,98 | 0,63 | 2,33 | 0,27 | 7,88E-01 |
| AT5G14050 | 0,50 | 2,19 | 3,15 | 0,69 | 4,88E-01 |
| AT5G14060 | 2,66 | -0,51 | 1,38 | -0,37 | 7,11E-01 |
| AT5G14120 | 1,26 | -2,37 | 1,59 | -1,49 | 1,36E-01 |
| AT5G14200 | 1,42 | 0,88 | 1,61 | 0,55 | 5,85E-01 |
| AT5G14230 | 0,39 | 1,56 | 3,42 | 0,46 | 6,47E-01 |
| AT5G14260 | 0,68 | -0,32 | 2,44 | -0,13 | 8,95E-01 |
| AT5G14270 | 2,74 | -1,03 | 1,05 | -0,99 | 3,24E-01 |
| AT5G14280 | 0,43 | -0,82 | 4,36 | -0,19 | 8,50E-01 |
| AT5G14285 | 0,21 | 1,57 | 4,33 | 0,36 | 7,16E-01 |
| AT5G14430 | 2,60 | 0,26 | 1,22 | 0,21 | 8,34E-01 |
| AT5G14480 | 0,25 | 1,20 | 4,33 | 0,28 | 7,81E-01 |
| AT5G14610 | 0,64 | -0,37 | 3,19 | -0,12 | 9,08E-01 |
| AT5G14670 | 0,08 | 0,69 | 4,36 | 0,16 | 8,75E-01 |
| AT5G14740 | 2,79 | -2,65 | 1,18 | -2,25 | 2,43E-02 |

| | | | | | |
|---|---|---|---|---|---|
| AT5G14780 | 2,20 | 1,26 | 1,43 | 0,88 | 3,81E-01 |
| AT5G14880 | 0,37 | 0,28 | 3,44 | 0,08 | 9,35E-01 |
| AT5G14890 | 0,99 | 3,19 | 4,01 | 0,80 | 4,27E-01 |
| AT5G14910 | 0,77 | 0,66 | 2,59 | 0,26 | 7,98E-01 |
| AT5G14930 | 1,80 | -2,05 | 1,97 | -1,04 | 2,99E-01 |
| AT5G14970 | 0,18 | 1,15 | 4,34 | 0,26 | 7,91E-01 |
| AT5G14990 | 0,42 | -0,70 | 4,13 | -0,17 | 8,66E-01 |
| AT5G15010 | 0,38 | 0,00 | 4,41 | 0,00 | 1,00E+00 |
| AT5G02295 | 1,40 | 0,22 | 1,85 | 0,12 | 9,03E-01 |
| AT5G02305 | 2,96 | 0,44 | 1,44 | 0,30 | 7,62E-01 |
| AT5G15080 | 0,92 | -1,60 | 2,77 | -0,58 | 5,64E-01 |
| AT5G15230 | 0,74 | -0,67 | 2,80 | -0,24 | 8,12E-01 |
| AT5G15240 | 0,05 | 0,00 | 4,41 | 0,00 | 1,00E+00 |
| AT5G15270 | 0,61 | 1,73 | 3,13 | 0,55 | 5,80E-01 |
| AT5G15280 | 0,54 | -1,08 | 3,77 | -0,29 | 7,74E-01 |
| AT5G15320 | 0,21 | 0,68 | 4,35 | 0,16 | 8,76E-01 |
| AT5G15330 | 0,11 | 0,65 | 4,36 | 0,15 | 8,81E-01 |
| AT5G15530 | 0,12 | 1,16 | 4,34 | 0,27 | 7,89E-01 |
| AT5G15570 | 0,44 | 2,38 | 4,31 | 0,55 | 5,81E-01 |
| AT5G15610 | 0,89 | -0,94 | 1,76 | -0,53 | 5,93E-01 |
| AT5G15650 | 3,70 | -1,23 | 0,99 | -1,24 | 2,15E-01 |
| AT5G15685 | 0,72 | 0,70 | 2,10 | 0,33 | 7,38E-01 |
| AT5G15710 | 0,74 | 2,25 | 3,59 | 0,63 | 5,30E-01 |
| AT5G15730 | 0,59 | 1,31 | 2,57 | 0,51 | 6,10E-01 |
| AT5G15750 | 0,54 | -0,72 | 3,97 | -0,18 | 8,55E-01 |
| AT5G15820 | 0,22 | -0,47 | 4,35 | -0,11 | 9,13E-01 |
| AT5G15845 | 2,91 | -0,44 | 1,03 | -0,43 | 6,68E-01 |
| AT5G15930 | 0,32 | 0,23 | 4,34 | 0,05 | 9,58E-01 |
| AT5G15950 | 1,23 | 0,39 | 1,86 | 0,21 | 8,35E-01 |
| AT5G15960 | 0,51 | 0,00 | 4,41 | 0,00 | 1,00E+00 |
| AT5G15970 | 2,85 | -1,72 | 1,66 | -1,04 | 2,99E-01 |
| AT5G16010 | 0,36 | -1,28 | 4,34 | -0,29 | 7,68E-01 |
| AT5G16030 | 0,58 | 0,04 | 3,40 | 0,01 | 9,90E-01 |
| AT5G16120 | 0,30 | -0,02 | 4,35 | 0,00 | 9,96E-01 |
| AT5G16130 | 1,11 | -0,19 | 1,97 | -0,10 | 9,24E-01 |
| AT5G16150 | 1,33 | 0,25 | 1,66 | 0,15 | 8,79E-01 |
| AT5G16260 | 0,11 | 0,00 | 4,41 | 0,00 | 1,00E+00 |
| AT5G16270 | 1,71 | -0,26 | 1,33 | -0,20 | 8,45E-01 |
| AT5G16290 | 0,43 | 1,09 | 4,31 | 0,25 | 8,01E-01 |
| AT5G16300 | 1,23 | 1,41 | 1,68 | 0,84 | 4,00E-01 |
| AT5G16440 | 0,67 | -1,77 | 2,46 | -0,72 | 4,73E-01 |
| AT5G16470 | 0,69 | -1,69 | 2,47 | -0,68 | 4,94E-01 |
| AT5G16540 | 0,32 | -0,94 | 4,31 | -0,22 | 8,28E-01 |
| AT5G16550 | 0,57 | -0,18 | 2,61 | -0,07 | 9,46E-01 |
| AT5G02555 | 0,05 | 0,00 | 4,41 | 0,00 | 1,00E+00 |
| AT5G16590 | 0,87 | 1,07 | 2,14 | 0,50 | 6,18E-01 |
| AT5G02565 | 2,69 | -0,66 | 1,51 | -0,44 | 6,63E-01 |
| AT5G16620 | 2,01 | -0,31 | 1,25 | -0,25 | 8,06E-01 |
| AT5G16630 | 0,92 | 1,80 | 2,60 | 0,69 | 4,90E-01 |
| AT5G16640 | 4,15 | 0,60 | 1,37 | 0,44 | 6,61E-01 |
| AT5G16715 | 0,74 | -1,52 | 2,44 | -0,62 | 5,32E-01 |
| AT5G16730 | 1,05 | 0,96 | 2,03 | 0,47 | 6,37E-01 |

| | | | | | |
|---|---|---|---|---|---|
| AT5G16760 | 1,05 | 0,75 | 2,12 | 0,36 | 7,22E-01 |
| AT5G16820 | 0,94 | -0,02 | 2,16 | -0,01 | 9,91E-01 |
| AT5G16880 | 1,27 | -0,05 | 1,58 | -0,03 | 9,75E-01 |
| AT5G16900 | 0,33 | 0,75 | 4,32 | 0,17 | 8,62E-01 |
| AT5G16910 | 0,98 | 0,52 | 2,43 | 0,22 | 8,29E-01 |
| AT5G17020 | 2,03 | -1,62 | 1,25 | -1,30 | 1,94E-01 |
| AT5G17167 | 0,20 | 0,02 | 4,39 | 0,00 | 9,97E-01 |
| AT5G17170 | 0,33 | -1,24 | 4,35 | -0,29 | 7,75E-01 |
| AT5G17190 | 0,51 | -2,59 | 3,21 | -0,81 | 4,20E-01 |
| AT5G17270 | 0,61 | -1,82 | 4,26 | -0,43 | 6,70E-01 |
| AT5G17280 | 0,29 | 1,55 | 4,33 | 0,36 | 7,21E-01 |
| AT5G17305 | 0,08 | 0,69 | 4,36 | 0,16 | 8,75E-01 |
| AT5G17440 | 0,24 | -0,74 | 4,35 | -0,17 | 8,65E-01 |
| AT5G17530 | 0,59 | -1,26 | 3,09 | -0,41 | 6,84E-01 |
| AT5G17580 | 0,17 | 1,40 | 4,33 | 0,32 | 7,47E-01 |
| AT5G17630 | 0,33 | -0,23 | 4,38 | -0,05 | 9,59E-01 |
| AT5G17670 | 0,39 | -1,28 | 4,38 | -0,29 | 7,70E-01 |
| AT5G17680 | 0,30 | 0,30 | 4,34 | 0,07 | 9,45E-01 |
| AT5G17850 | 0,18 | 0,00 | 4,41 | 0,00 | 1,00E+00 |
| AT5G17870 | 0,86 | -0,28 | 2,33 | -0,12 | 9,05E-01 |
| AT5G17890 | 1,77 | -1,54 | 1,55 | -0,99 | 3,20E-01 |
| AT5G17900 | 0,30 | -0,39 | 4,34 | -0,09 | 9,29E-01 |
| AT5G17910 | 1,47 | -0,14 | 1,75 | -0,08 | 9,37E-01 |
| AT5G17980 | 0,38 | 0,94 | 4,21 | 0,22 | 8,23E-01 |
| AT5G17990 | 0,55 | -0,83 | 3,95 | -0,21 | 8,35E-01 |
| AT5G18040 | 0,71 | 1,53 | 2,43 | 0,63 | 5,30E-01 |
| AT5G18065 | 0,30 | 1,26 | 3,65 | 0,35 | 7,30E-01 |
| AT5G18070 | 0,62 | -1,14 | 2,51 | -0,45 | 6,49E-01 |
| AT5G18085 | 0,24 | -2,36 | 4,39 | -0,54 | 5,91E-01 |
| AT5G18100 | 0,33 | 1,01 | 4,34 | 0,23 | 8,16E-01 |
| AT5G18130 | 1,11 | 1,25 | 2,16 | 0,58 | 5,64E-01 |
| AT5G18140 | 1,06 | -0,34 | 2,46 | -0,14 | 8,89E-01 |
| AT5G18170 | 0,16 | 1,07 | 4,34 | 0,25 | 8,05E-01 |
| AT5G18200 | 0,08 | 0,44 | 4,38 | 0,10 | 9,21E-01 |
| AT5G18245 | 1,46 | 0,18 | 2,88 | 0,06 | 9,49E-01 |
| AT5G18255 | 1,25 | -0,50 | 1,55 | -0,32 | 7,49E-01 |
| AT5G18310 | 0,76 | 0,00 | 2,54 | 0,00 | 9,99E-01 |
| AT5G18390 | 0,33 | -0,58 | 4,33 | -0,13 | 8,93E-01 |
| AT5G18470 | 1,32 | 0,99 | 1,69 | 0,59 | 5,58E-01 |
| AT5G18500 | 0,37 | -0,40 | 4,33 | -0,09 | 9,26E-01 |
| AT5G18520 | 1,46 | 0,61 | 1,64 | 0,37 | 7,11E-01 |
| AT5G18580 | 0,90 | 0,17 | 2,14 | 0,08 | 9,36E-01 |
| AT5G18610 | 1,43 | -2,51 | 1,76 | -1,43 | 1,53E-01 |
| AT5G18630 | 0,28 | -1,78 | 4,41 | -0,40 | 6,87E-01 |
| AT5G18636 | 0,58 | -1,60 | 2,41 | -0,67 | 5,06E-01 |
| AT5G18650 | 0,68 | -1,08 | 3,07 | -0,35 | 7,25E-01 |
| AT5G18670 | 0,21 | 1,72 | 4,33 | 0,40 | 6,91E-01 |
| AT5G18750 | 0,77 | 3,17 | 3,35 | 0,95 | 3,44E-01 |
| AT5G18830 | 0,51 | 1,74 | 3,22 | 0,54 | 5,89E-01 |
| AT5G18890 | 0,98 | -1,33 | 4,29 | -0,31 | 7,56E-01 |
| AT5G18937 | 0,54 | 0,00 | 4,41 | 0,00 | 1,00E+00 |
| AT5G18940 | 1,00 | -1,36 | 2,58 | -0,53 | 5,99E-01 |

| | | | | | |
|---|---|---|---|---|---|
| AT5G18960 | 0,35 | 0,49 | 3,45 | 0,14 | 8,88E-01 |
| AT5G19000 | 0,44 | -1,55 | 3,33 | -0,47 | 6,40E-01 |
| AT5G19090 | 0,78 | -1,35 | 2,47 | -0,55 | 5,84E-01 |
| AT5G19120 | 2,51 | -1,45 | 1,19 | -1,21 | 2,24E-01 |
| AT5G19180 | 0,77 | 1,07 | 2,26 | 0,48 | 6,35E-01 |
| AT5G19190 | 0,37 | -1,05 | 4,33 | -0,24 | 8,08E-01 |
| AT5G19200 | 4,59 | 0,33 | 0,90 | 0,36 | 7,17E-01 |
| AT5G19210 | 0,49 | -1,75 | 3,35 | -0,52 | 6,01E-01 |
| AT5G19221 | 0,47 | -1,86 | 4,32 | -0,43 | 6,66E-01 |
| AT5G19240 | 0,65 | 1,58 | 2,21 | 0,72 | 4,74E-01 |
| AT5G19250 | 0,98 | -1,82 | 2,86 | -0,64 | 5,25E-01 |
| AT5G19290 | 0,41 | 0,34 | 3,33 | 0,10 | 9,18E-01 |
| AT5G19390 | 2,78 | -0,74 | 1,32 | -0,56 | 5,74E-01 |
| AT5G19400 | 2,38 | -0,93 | 1,49 | -0,62 | 5,34E-01 |
| AT5G19440 | 0,36 | 0,29 | 4,34 | 0,07 | 9,48E-01 |
| AT5G19500 | 1,08 | -0,54 | 2,15 | -0,25 | 8,03E-01 |
| AT5G19540 | 0,77 | -0,92 | 2,04 | -0,45 | 6,51E-01 |
| AT5G19550 | 0,90 | 1,73 | 2,08 | 0,83 | 4,05E-01 |
| AT5G19570 | 0,52 | 0,18 | 3,93 | 0,05 | 9,63E-01 |
| AT5G19620 | 0,50 | 0,50 | 4,07 | 0,12 | 9,02E-01 |
| AT5G19680 | 0,19 | 0,37 | 4,37 | 0,09 | 9,32E-01 |
| AT5G19690 | 1,74 | 0,93 | 1,42 | 0,66 | 5,10E-01 |
| AT5G19750 | 0,55 | 2,38 | 4,30 | 0,55 | 5,80E-01 |
| AT5G19780 | 1,35 | -0,33 | 1,78 | -0,18 | 8,54E-01 |
| AT5G19940 | 2,37 | -1,70 | 1,39 | -1,22 | 2,22E-01 |
| AT5G19990 | 0,77 | -2,94 | 2,15 | -1,37 | 1,72E-01 |
| AT5G20010 | 1,98 | -0,71 | 1,40 | -0,51 | 6,13E-01 |
| AT5G20020 | 2,46 | 0,95 | 1,60 | 0,59 | 5,55E-01 |
| AT5G20080 | 0,48 | -1,46 | 3,32 | -0,44 | 6,59E-01 |
| AT5G20120 | 0,55 | 1,46 | 4,34 | 0,34 | 7,37E-01 |
| AT5G20130 | 0,68 | -0,96 | 3,67 | -0,26 | 7,94E-01 |
| AT5G20150 | 0,56 | -0,01 | 2,52 | 0,00 | 9,97E-01 |
| AT5G20190 | 1,21 | 0,87 | 1,88 | 0,46 | 6,42E-01 |
| AT5G20200 | 1,58 | 0,74 | 1,54 | 0,48 | 6,29E-01 |
| AT5G20220 | 0,21 | 1,63 | 4,33 | 0,38 | 7,06E-01 |
| AT5G20230 | 2,06 | 0,62 | 1,46 | 0,43 | 6,69E-01 |
| AT5G20250 | 2,34 | 1,50 | 1,34 | 1,12 | 2,64E-01 |
| AT5G20350 | 1,36 | 1,16 | 1,66 | 0,69 | 4,87E-01 |
| AT5G20410 | 0,63 | -2,72 | 3,60 | -0,76 | 4,50E-01 |
| AT5G20480 | 0,15 | 0,00 | 4,38 | 0,00 | 1,00E+00 |
| AT5G03145 | 2,30 | 0,98 | 1,55 | 0,63 | 5,26E-01 |
| AT5G20500 | 0,33 | -0,06 | 4,34 | -0,01 | 9,89E-01 |
| AT5G20590 | 0,85 | 0,01 | 2,13 | 0,00 | 9,97E-01 |
| AT5G20610 | 1,39 | -0,48 | 1,53 | -0,31 | 7,55E-01 |
| AT5G20660 | 1,27 | -0,60 | 1,64 | -0,36 | 7,17E-01 |
| AT5G20680 | 0,31 | 0,03 | 4,38 | 0,01 | 9,94E-01 |
| AT5G20710 | 0,12 | 1,16 | 4,34 | 0,27 | 7,89E-01 |
| AT5G20720 | 3,41 | -1,09 | 1,10 | -0,99 | 3,23E-01 |
| AT5G20900 | 0,45 | -0,41 | 3,38 | -0,12 | 9,04E-01 |
| AT5G03255 | 0,31 | 0,48 | 4,32 | 0,11 | 9,11E-01 |
| AT5G20960 | 1,02 | 3,48 | 2,55 | 1,36 | 1,73E-01 |
| AT5G21020 | 1,50 | 1,31 | 1,72 | 0,76 | 4,45E-01 |

| | | | | | |
|---|---|---|---|---|---|
| AT5G21090 | 1,39 | -0,09 | 1,76 | -0,05 | 9,57E-01 |
| AT5G21105 | 1,68 | 1,54 | 1,74 | 0,88 | 3,77E-01 |
| AT5G21150 | 0,12 | 1,16 | 4,34 | 0,27 | 7,89E-01 |
| AT5G21222 | 1,01 | -0,84 | 2,56 | -0,33 | 7,43E-01 |
| AT5G21326 | 0,90 | 2,04 | 2,71 | 0,75 | 4,52E-01 |
| AT5G21482 | 0,24 | 0,82 | 4,35 | 0,19 | 8,50E-01 |
| AT5G21930 | 1,60 | 0,27 | 1,53 | 0,18 | 8,58E-01 |
| AT5G22000 | 1,27 | -1,26 | 1,58 | -0,80 | 4,23E-01 |
| AT5G22050 | 0,36 | 0,38 | 4,37 | 0,09 | 9,31E-01 |
| AT5G22070 | 0,57 | 0,36 | 3,09 | 0,12 | 9,07E-01 |
| AT5G22300 | 0,30 | 0,81 | 3,69 | 0,22 | 8,26E-01 |
| AT5G22315 | 0,20 | -1,80 | 4,41 | -0,41 | 6,83E-01 |
| AT5G22340 | 0,95 | 0,14 | 2,08 | 0,07 | 9,46E-01 |
| AT5G22420 | 0,42 | -0,40 | 4,12 | -0,10 | 9,23E-01 |
| AT5G22500 | 2,19 | -0,78 | 1,93 | -0,40 | 6,87E-01 |
| AT5G22580 | 0,46 | 0,29 | 4,39 | 0,07 | 9,48E-01 |
| AT5G22590 | 1,03 | -0,79 | 2,73 | -0,29 | 7,73E-01 |
| AT5G22630 | 1,52 | -1,29 | 1,46 | -0,88 | 3,77E-01 |
| AT5G22640 | 1,19 | 1,12 | 1,66 | 0,68 | 4,97E-01 |
| AT5G22650 | 0,66 | 0,22 | 2,62 | 0,09 | 9,32E-01 |
| AT5G22690 | 0,12 | 1,16 | 4,34 | 0,27 | 7,89E-01 |
| AT5G22760 | 1,25 | -0,38 | 1,65 | -0,23 | 8,17E-01 |
| AT5G22794 | 0,08 | 0,69 | 4,36 | 0,16 | 8,75E-01 |
| AT5G22830 | 0,08 | 0,00 | 4,40 | 0,00 | 1,00E+00 |
| AT5G22950 | 0,12 | 0,00 | 4,38 | 0,00 | 1,00E+00 |
| AT5G23010 | 0,84 | -1,42 | 2,30 | -0,62 | 5,36E-01 |
| AT5G23070 | 0,19 | 0,37 | 4,37 | 0,08 | 9,33E-01 |
| AT5G03575 | 0,26 | 0,14 | 4,41 | 0,03 | 9,74E-01 |
| AT5G23120 | 0,65 | 0,51 | 2,19 | 0,23 | 8,18E-01 |
| AT5G23140 | 0,95 | -1,53 | 2,08 | -0,74 | 4,61E-01 |
| AT5G23150 | 1,41 | 0,28 | 1,95 | 0,14 | 8,85E-01 |
| AT5G23210 | 0,16 | 0,92 | 4,35 | 0,21 | 8,33E-01 |
| AT5G23280 | 1,01 | -2,44 | 2,39 | -1,02 | 3,06E-01 |
| AT5G23320 | 0,08 | 0,69 | 4,36 | 0,16 | 8,75E-01 |
| AT5G23330 | 0,43 | -1,11 | 4,34 | -0,25 | 7,99E-01 |
| AT5G23340 | 1,40 | -0,76 | 1,57 | -0,49 | 6,28E-01 |
| AT5G23380 | 2,25 | -1,22 | 1,39 | -0,88 | 3,79E-01 |
| AT5G23490 | 0,49 | -0,51 | 2,60 | -0,20 | 8,44E-01 |
| AT5G23540 | 0,96 | -1,65 | 2,01 | -0,82 | 4,13E-01 |
| AT5G23570 | 0,58 | 0,85 | 2,42 | 0,35 | 7,25E-01 |
| AT5G23575 | 1,00 | -1,91 | 1,68 | -1,14 | 2,54E-01 |
| AT5G23600 | 0,08 | 0,69 | 4,36 | 0,16 | 8,75E-01 |
| AT5G23670 | 0,72 | 0,35 | 2,93 | 0,12 | 9,06E-01 |
| AT5G23720 | 0,51 | -2,16 | 3,20 | -0,68 | 4,99E-01 |
| AT5G23730 | 0,26 | 0,01 | 4,33 | 0,00 | 9,98E-01 |
| AT5G23890 | 0,99 | -0,37 | 2,60 | -0,14 | 8,88E-01 |
| AT5G23950 | 0,35 | -2,53 | 4,37 | -0,58 | 5,63E-01 |
| AT5G24020 | 0,16 | 0,92 | 4,35 | 0,21 | 8,33E-01 |
| AT5G24200 | 1,34 | -0,66 | 1,48 | -0,44 | 6,57E-01 |
| AT5G24210 | 0,96 | -2,51 | 2,11 | -1,19 | 2,35E-01 |
| AT5G24300 | 2,35 | 1,35 | 1,39 | 0,97 | 3,30E-01 |
| AT5G24350 | 3,06 | 0,04 | 1,27 | 0,03 | 9,75E-01 |

| | | | | | |
|---|---|---|---|---|---|
| AT5G24360 | 0,40 | 0,78 | 3,31 | 0,23 | 8,14E-01 |
| AT5G24380 | 0,41 | 0,43 | 3,36 | 0,13 | 8,99E-01 |
| AT5G24450 | 0,69 | 2,38 | 3,68 | 0,65 | 5,18E-01 |
| AT5G24460 | 0,88 | 0,28 | 1,89 | 0,15 | 8,80E-01 |
| AT5G24490 | 1,61 | 1,71 | 2,12 | 0,81 | 4,18E-01 |
| AT5G24530 | 1,77 | -0,52 | 1,38 | -0,37 | 7,08E-01 |
| AT5G24610 | 0,55 | 0,23 | 2,66 | 0,09 | 9,30E-01 |
| AT5G24830 | 0,08 | 0,69 | 4,36 | 0,16 | 8,75E-01 |
| AT5G24930 | 2,10 | -1,85 | 1,40 | -1,32 | 1,88E-01 |
| AT5G24980 | 0,43 | 0,89 | 3,37 | 0,26 | 7,92E-01 |
| AT5G24990 | 2,50 | 0,13 | 1,89 | 0,07 | 9,46E-01 |
| AT5G25040 | 0,67 | -0,18 | 2,48 | -0,07 | 9,44E-01 |
| AT5G25120 | 0,51 | 0,83 | 3,24 | 0,25 | 7,99E-01 |
| AT5G03925 | 0,17 | 0,00 | 4,41 | 0,00 | 1,00E+00 |
| AT5G25130 | 1,42 | -1,08 | 1,67 | -0,65 | 5,16E-01 |
| AT5G25140 | 0,28 | 1,91 | 4,32 | 0,44 | 6,58E-01 |
| AT5G25150 | 0,27 | -0,98 | 4,37 | -0,22 | 8,23E-01 |
| AT5G25170 | 0,33 | 1,99 | 4,32 | 0,46 | 6,44E-01 |
| AT5G25210 | 0,42 | 1,37 | 3,33 | 0,41 | 6,81E-01 |
| AT5G25220 | 1,05 | 3,18 | 2,08 | 1,53 | 1,27E-01 |
| AT5G25230 | 1,66 | -0,06 | 1,43 | -0,05 | 9,64E-01 |
| AT5G25250 | 1,43 | 1,87 | 1,98 | 0,95 | 3,44E-01 |
| AT5G25260 | 1,02 | 1,99 | 2,67 | 0,75 | 4,55E-01 |
| AT5G25280 | 3,55 | -1,46 | 0,97 | -1,50 | 1,34E-01 |
| AT5G25340 | 0,46 | -0,16 | 4,02 | -0,04 | 9,69E-01 |
| AT5G25460 | 1,53 | -0,60 | 2,18 | -0,28 | 7,83E-01 |
| AT5G25520 | 1,48 | -0,01 | 2,23 | 0,00 | 9,96E-01 |
| AT5G25560 | 0,53 | -1,52 | 3,08 | -0,49 | 6,21E-01 |
| AT5G25580 | 0,12 | 1,23 | 4,34 | 0,28 | 7,76E-01 |
| AT5G25630 | 0,47 | 1,06 | 3,15 | 0,34 | 7,36E-01 |
| AT5G25754 | 1,48 | 0,64 | 1,48 | 0,43 | 6,67E-01 |
| AT5G04145 | 0,89 | -2,39 | 2,23 | -1,07 | 2,85E-01 |
| AT5G25900 | 0,33 | 0,26 | 4,33 | 0,06 | 9,53E-01 |
| AT5G25910 | 0,60 | 1,51 | 3,67 | 0,41 | 6,80E-01 |
| AT5G25930 | 0,70 | 0,85 | 2,51 | 0,34 | 7,34E-01 |
| AT5G25980 | 1,32 | 2,44 | 1,88 | 1,30 | 1,95E-01 |
| AT5G26030 | 2,16 | 1,41 | 1,46 | 0,97 | 3,34E-01 |
| AT5G26038 | 1,13 | 0,85 | 2,11 | 0,40 | 6,87E-01 |
| AT5G26146 | 0,89 | 1,82 | 2,29 | 0,80 | 4,25E-01 |
| AT5G26147 | 0,89 | 1,82 | 2,29 | 0,80 | 4,25E-01 |
| AT5G26200 | 0,12 | 1,16 | 4,34 | 0,27 | 7,89E-01 |
| AT5G26240 | 0,84 | -0,01 | 1,94 | -0,01 | 9,96E-01 |
| AT5G26270 | 0,26 | 0,31 | 4,38 | 0,07 | 9,44E-01 |
| AT5G26290 | 0,79 | -1,76 | 2,23 | -0,79 | 4,29E-01 |
| AT5G26570 | 1,51 | -0,06 | 1,58 | -0,04 | 9,68E-01 |
| AT5G26740 | 0,58 | 1,68 | 2,66 | 0,63 | 5,28E-01 |
| AT5G26731 | 0,38 | 0,40 | 4,38 | 0,09 | 9,26E-01 |
| AT5G26710 | 0,73 | 0,06 | 2,40 | 0,02 | 9,81E-01 |
| AT5G26670 | 0,26 | -1,86 | 4,40 | -0,42 | 6,72E-01 |
| AT5G26610 | 0,16 | 0,00 | 4,41 | 0,00 | 1,00E+00 |
| AT5G26600 | 0,82 | 0,49 | 2,54 | 0,19 | 8,48E-01 |
| AT5G26780 | 0,39 | -0,15 | 2,86 | -0,05 | 9,58E-01 |

| | | | | | |
|---|---|---|---|---|---|
| AT5G26820 | 0,42 | -0,89 | 4,33 | -0,20 | 8,38E-01 |
| AT5G26830 | 2,03 | 0,12 | 1,21 | 0,10 | 9,24E-01 |
| AT5G26850 | 1,21 | -1,10 | 2,05 | -0,54 | 5,91E-01 |
| AT5G26860 | 1,16 | 1,30 | 1,77 | 0,74 | 4,61E-01 |
| AT5G26920 | 3,70 | -0,27 | 0,88 | -0,30 | 7,61E-01 |
| AT5G26930 | 0,33 | -1,30 | 4,33 | -0,30 | 7,64E-01 |
| AT5G26940 | 0,66 | -1,55 | 3,61 | -0,43 | 6,67E-01 |
| AT5G26990 | 0,23 | -1,78 | 4,41 | -0,40 | 6,87E-01 |
| AT5G27120 | 0,91 | -0,31 | 3,39 | -0,09 | 9,27E-01 |
| AT5G27150 | 0,57 | 1,18 | 2,99 | 0,40 | 6,92E-01 |
| AT5G27170 | 0,25 | 0,00 | 4,38 | 0,00 | 1,00E+00 |
| AT5G27230 | 0,08 | 0,00 | 4,40 | 0,00 | 1,00E+00 |
| AT5G27240 | 0,05 | 0,00 | 4,41 | 0,00 | 1,00E+00 |
| AT5G27270 | 0,17 | 0,02 | 4,39 | 0,00 | 9,97E-01 |
| AT5G27280 | 0,34 | -0,43 | 3,59 | -0,12 | 9,05E-01 |
| AT5G27300 | 0,08 | 0,00 | 4,40 | 0,00 | 1,00E+00 |
| AT5G27320 | 0,08 | 0,00 | 4,40 | 0,00 | 1,00E+00 |
| AT5G27340 | 0,54 | 0,40 | 3,80 | 0,10 | 9,17E-01 |
| AT5G27350 | 0,64 | 1,23 | 2,61 | 0,47 | 6,39E-01 |
| AT5G27410 | 0,42 | -2,49 | 4,13 | -0,60 | 5,46E-01 |
| AT5G27420 | 1,44 | 1,05 | 1,58 | 0,67 | 5,05E-01 |
| AT5G27470 | 0,51 | 1,12 | 3,08 | 0,36 | 7,17E-01 |
| AT5G27530 | 0,32 | 0,00 | 4,41 | 0,00 | 1,00E+00 |
| AT5G27540 | 0,68 | -1,64 | 3,48 | -0,47 | 6,38E-01 |
| AT5G27600 | 0,18 | 0,65 | 4,35 | 0,15 | 8,82E-01 |
| AT5G27607 | 2,38 | 2,51 | 1,94 | 1,30 | 1,95E-01 |
| AT5G27610 | 0,08 | 0,00 | 4,40 | 0,00 | 1,00E+00 |
| AT5G27630 | 0,17 | 0,07 | 4,40 | 0,02 | 9,88E-01 |
| AT5G27650 | 0,94 | -0,26 | 2,59 | -0,10 | 9,19E-01 |
| AT5G04415 | 4,71 | -0,33 | 0,79 | -0,42 | 6,73E-01 |
| AT5G27710 | 0,67 | 0,43 | 2,50 | 0,17 | 8,64E-01 |
| AT5G27720 | 3,01 | 0,63 | 1,20 | 0,52 | 6,01E-01 |
| AT5G04425 | 2,72 | 0,89 | 1,29 | 0,69 | 4,88E-01 |
| AT5G27730 | 0,69 | 1,57 | 2,22 | 0,70 | 4,81E-01 |
| AT5G27760 | 0,27 | -0,79 | 4,39 | -0,18 | 8,58E-01 |
| AT5G27770 | 0,86 | -2,58 | 2,34 | -1,10 | 2,70E-01 |
| AT5G27800 | 0,16 | 0,09 | 4,38 | 0,02 | 9,83E-01 |
| AT5G27830 | 0,23 | -0,21 | 4,35 | -0,05 | 9,61E-01 |
| AT5G27850 | 1,94 | 0,77 | 1,32 | 0,58 | 5,59E-01 |
| AT5G27890 | 0,40 | 0,83 | 4,34 | 0,19 | 8,49E-01 |
| AT5G27893 | 1,07 | -1,64 | 1,98 | -0,83 | 4,08E-01 |
| AT5G27950 | 2,87 | 1,73 | 1,40 | 1,23 | 2,17E-01 |
| AT5G27970 | 1,51 | 1,88 | 2,18 | 0,86 | 3,88E-01 |
| AT5G28010 | 1,26 | 0,46 | 1,63 | 0,28 | 7,79E-01 |
| AT5G04625 | 0,10 | 0,00 | 4,41 | 0,00 | 1,00E+00 |
| AT5G04665 | 0,11 | 0,00 | 4,41 | 0,00 | 1,00E+00 |
| AT5G28090 | 0,29 | 1,06 | 4,34 | 0,24 | 8,07E-01 |
| AT5G04675 | 0,19 | 0,00 | 4,41 | 0,00 | 1,00E+00 |
| AT5G28320 | 0,89 | 1,33 | 2,56 | 0,52 | 6,03E-01 |
| AT5G28350 | 0,24 | -0,46 | 4,36 | -0,10 | 9,17E-01 |
| AT5G28410 | 0,36 | -0,80 | 4,34 | -0,18 | 8,54E-01 |
| AT5G28463 | 0,37 | 0,41 | 4,34 | 0,09 | 9,25E-01 |

| | | | | | |
|---|---|---|---|---|---|
| AT5G28500 | 0,70 | -0,70 | 2,68 | -0,26 | 7,95E-01 |
| AT5G28560 | 3,51 | 4,08 | 1,46 | 2,79 | 5,31E-03 |
| AT5G28740 | 1,92 | -0,42 | 1,26 | -0,33 | 7,39E-01 |
| AT5G04815 | 0,05 | 0,00 | 4,41 | 0,00 | 1,00E+00 |
| AT5G28810 | 0,66 | 0,08 | 2,94 | 0,03 | 9,78E-01 |
| AT5G28830 | 0,05 | 0,00 | 4,41 | 0,00 | 1,00E+00 |
| AT5G28900 | 0,40 | -0,19 | 4,27 | -0,05 | 9,64E-01 |
| AT5G28919 | 2,64 | 0,74 | 1,21 | 0,61 | 5,41E-01 |
| AT5G29624 | 0,12 | 1,23 | 4,34 | 0,28 | 7,76E-01 |
| AT5G30341 | 0,12 | 0,00 | 4,39 | 0,00 | 1,00E+00 |
| AT5G04955 | 1,06 | 0,00 | 3,96 | 0,00 | 1,00E+00 |
| AT5G32440 | 1,33 | -0,30 | 1,69 | -0,18 | 8,60E-01 |
| AT5G32613 | 0,26 | 0,03 | 4,39 | 0,01 | 9,94E-01 |
| AT5G33290 | 0,62 | -2,50 | 3,84 | -0,65 | 5,15E-01 |
| AT5G33320 | 0,61 | -0,37 | 2,23 | -0,16 | 8,69E-01 |
| AT5G33390 | 0,38 | 1,27 | 4,34 | 0,29 | 7,70E-01 |
| AT5G34869 | 0,19 | 1,22 | 4,33 | 0,28 | 7,78E-01 |
| AT5G34871 | 0,12 | 0,97 | 4,34 | 0,22 | 8,23E-01 |
| AT5G34870 | 0,38 | 1,32 | 3,42 | 0,39 | 7,00E-01 |
| AT5G35160 | 1,68 | -1,19 | 1,32 | -0,90 | 3,69E-01 |
| AT5G35170 | 0,47 | -0,17 | 2,45 | -0,07 | 9,45E-01 |
| AT5G35180 | 0,52 | 0,22 | 2,65 | 0,08 | 9,34E-01 |
| AT5G35190 | 0,84 | 2,44 | 2,77 | 0,88 | 3,80E-01 |
| AT5G35300 | 0,23 | 0,01 | 4,38 | 0,00 | 9,98E-01 |
| AT5G35320 | 0,21 | -1,41 | 4,36 | -0,32 | 7,46E-01 |
| AT5G35360 | 1,57 | -0,28 | 1,39 | -0,20 | 8,41E-01 |
| AT5G35430 | 1,24 | -1,04 | 2,06 | -0,50 | 6,15E-01 |
| AT5G35450 | 0,40 | 1,85 | 4,33 | 0,43 | 6,69E-01 |
| AT5G05195 | 0,23 | 0,94 | 4,34 | 0,22 | 8,29E-01 |
| AT5G35490 | 1,40 | 2,36 | 2,33 | 1,01 | 3,10E-01 |
| AT5G35630 | 4,18 | -1,49 | 1,10 | -1,35 | 1,78E-01 |
| AT5G35730 | 0,82 | -1,08 | 2,62 | -0,41 | 6,80E-01 |
| AT5G35840 | 1,35 | 2,39 | 2,20 | 1,09 | 2,77E-01 |
| AT5G35917 | 0,29 | 0,07 | 4,40 | 0,02 | 9,88E-01 |
| AT5G35920 | 1,48 | 2,85 | 1,81 | 1,57 | 1,16E-01 |
| AT5G35926 | 1,15 | 0,16 | 1,87 | 0,09 | 9,31E-01 |
| AT5G35980 | 1,15 | -0,65 | 1,93 | -0,34 | 7,36E-01 |
| AT5G36001 | 0,12 | 1,23 | 4,34 | 0,28 | 7,76E-01 |
| AT5G36080 | 1,49 | 3,32 | 1,88 | 1,77 | 7,67E-02 |
| AT5G36180 | 3,47 | -1,58 | 1,42 | -1,11 | 2,66E-01 |
| AT5G36228 | 1,95 | -3,06 | 1,84 | -1,66 | 9,71E-02 |
| AT5G36230 | 2,63 | 0,28 | 1,24 | 0,22 | 8,23E-01 |
| AT5G36290 | 0,47 | -0,85 | 2,79 | -0,30 | 7,62E-01 |
| AT5G36300 | 0,87 | 2,50 | 2,38 | 1,05 | 2,93E-01 |
| AT5G36740 | 0,08 | 0,69 | 4,36 | 0,16 | 8,75E-01 |
| AT5G36870 | 1,07 | -0,85 | 2,21 | -0,38 | 7,02E-01 |
| AT5G36900 | 0,54 | 2,15 | 3,09 | 0,70 | 4,87E-01 |
| AT5G36940 | 0,51 | 2,48 | 3,86 | 0,64 | 5,20E-01 |
| AT5G37020 | 0,62 | 1,29 | 2,61 | 0,49 | 6,22E-01 |
| AT5G37070 | 0,43 | -2,17 | 4,38 | -0,50 | 6,20E-01 |
| AT5G37130 | 0,66 | -0,09 | 2,62 | -0,03 | 9,73E-01 |
| AT5G37150 | 0,98 | -0,80 | 2,39 | -0,33 | 7,39E-01 |

| | | | | | |
|---|---|---|---|---|---|
| AT5G37510 | 3,25 | 0,82 | 1,02 | 0,80 | 4,24E-01 |
| AT5G37540 | 0,17 | 0,98 | 4,35 | 0,23 | 8,21E-01 |
| AT5G37630 | 0,94 | 1,07 | 2,13 | 0,50 | 6,15E-01 |
| AT5G37760 | 0,15 | 1,20 | 4,34 | 0,28 | 7,81E-01 |
| AT5G37793 | 0,51 | 0,83 | 2,67 | 0,31 | 7,57E-01 |
| AT5G37850 | 0,30 | 0,24 | 4,35 | 0,05 | 9,57E-01 |
| AT5G37950 | 0,11 | 0,00 | 4,41 | 0,00 | 1,00E+00 |
| AT5G38155 | 0,12 | 0,00 | 4,38 | 0,00 | 1,00E+00 |
| AT5G38190 | 2,86 | 1,22 | 1,42 | 0,86 | 3,90E-01 |
| AT5G38210 | 1,00 | 2,33 | 2,21 | 1,05 | 2,91E-01 |
| AT5G38380 | 0,15 | 0,00 | 4,41 | 0,00 | 1,00E+00 |
| AT5G38470 | 0,73 | -1,12 | 3,61 | -0,31 | 7,57E-01 |
| AT5G38480 | 1,20 | -0,42 | 1,54 | -0,27 | 7,87E-01 |
| AT5G38560 | 0,34 | -0,13 | 3,49 | -0,04 | 9,71E-01 |
| AT5G38910 | 0,16 | 0,00 | 4,41 | 0,00 | 1,00E+00 |
| AT5G38980 | 0,57 | 0,40 | 2,72 | 0,15 | 8,84E-01 |
| AT5G38990 | 2,97 | 0,85 | 1,13 | 0,76 | 4,50E-01 |
| AT5G39000 | 0,70 | -3,55 | 4,31 | -0,82 | 4,11E-01 |
| AT5G39030 | 0,81 | 1,06 | 2,74 | 0,39 | 6,98E-01 |
| AT5G39040 | 0,29 | 1,46 | 4,33 | 0,34 | 7,35E-01 |
| AT5G39050 | 4,52 | 1,76 | 1,24 | 1,42 | 1,55E-01 |
| AT5G39080 | 2,20 | 0,79 | 1,59 | 0,49 | 6,22E-01 |
| AT5G39250 | 0,34 | -1,26 | 4,35 | -0,29 | 7,71E-01 |
| AT5G39320 | 0,82 | -1,55 | 4,30 | -0,36 | 7,18E-01 |
| AT5G39350 | 0,21 | 0,23 | 4,37 | 0,05 | 9,57E-01 |
| AT5G39510 | 0,77 | 0,01 | 2,44 | 0,01 | 9,95E-01 |
| AT5G39570 | 4,04 | -0,32 | 1,17 | -0,27 | 7,84E-01 |
| AT5G39590 | 0,82 | 0,92 | 2,41 | 0,38 | 7,03E-01 |
| AT5G39660 | 0,40 | 2,13 | 4,32 | 0,49 | 6,22E-01 |
| AT5G39670 | 0,40 | 1,87 | 4,16 | 0,45 | 6,53E-01 |
| AT5G39690 | 1,03 | -0,79 | 2,14 | -0,37 | 7,12E-01 |
| AT5G39710 | 0,25 | 0,41 | 4,33 | 0,09 | 9,25E-01 |
| AT5G39730 | 0,19 | 0,00 | 4,38 | 0,00 | 9,99E-01 |
| AT5G39740 | 3,11 | -1,43 | 0,98 | -1,46 | 1,43E-01 |
| AT5G39775 | 0,53 | -0,53 | 4,21 | -0,13 | 9,00E-01 |
| AT5G39850 | 0,08 | 0,69 | 4,36 | 0,16 | 8,75E-01 |
| AT5G39960 | 0,21 | 0,01 | 4,34 | 0,00 | 9,98E-01 |
| AT5G39990 | 0,08 | 0,44 | 4,38 | 0,10 | 9,21E-01 |
| AT5G40060 | 1,33 | 1,25 | 2,46 | 0,51 | 6,11E-01 |
| AT5G40140 | 0,31 | -0,15 | 3,62 | -0,04 | 9,67E-01 |
| AT5G40200 | 0,62 | 0,37 | 2,78 | 0,13 | 8,95E-01 |
| AT5G40380 | 0,46 | 0,00 | 4,05 | 0,00 | 1,00E+00 |
| AT5G40400 | 1,55 | -0,66 | 1,51 | -0,44 | 6,62E-01 |
| AT5G40480 | 2,70 | 0,58 | 1,09 | 0,53 | 5,97E-01 |
| AT5G40650 | 0,74 | -3,25 | 2,72 | -1,19 | 2,33E-01 |
| AT5G40670 | 1,01 | -0,14 | 2,16 | -0,06 | 9,48E-01 |
| AT5G40680 | 0,17 | 1,50 | 4,33 | 0,35 | 7,29E-01 |
| AT5G40690 | 0,62 | 1,57 | 3,66 | 0,43 | 6,68E-01 |
| AT5G40720 | 0,32 | 0,88 | 4,35 | 0,20 | 8,41E-01 |
| AT5G40760 | 1,16 | -0,58 | 1,81 | -0,32 | 7,48E-01 |
| AT5G40780 | 1,36 | -0,75 | 1,51 | -0,50 | 6,18E-01 |
| AT5G40810 | 1,32 | -0,02 | 1,76 | -0,01 | 9,90E-01 |

| | | | | | |
|---|---|---|---|---|---|
| AT5G40850 | 0,08 | 0,00 | 4,41 | 0,00 | 1,00E+00 |
| AT5G40870 | 0,14 | 0,09 | 4,38 | 0,02 | 9,84E-01 |
| AT5G40880 | 0,51 | 0,74 | 3,94 | 0,19 | 8,51E-01 |
| AT5G40900 | 0,15 | 0,00 | 4,41 | 0,00 | 1,00E+00 |
| AT5G40910 | 2,64 | 0,29 | 1,04 | 0,28 | 7,81E-01 |
| AT5G41010 | 0,28 | 0,36 | 4,37 | 0,08 | 9,34E-01 |
| AT5G06115 | 0,20 | 0,00 | 4,38 | 0,00 | 1,00E+00 |
| AT5G41070 | 0,20 | -1,80 | 4,41 | -0,41 | 6,83E-01 |
| AT5G41100 | 0,28 | -1,78 | 4,41 | -0,40 | 6,87E-01 |
| AT5G41110 | 0,22 | -0,91 | 4,38 | -0,21 | 8,35E-01 |
| AT5G41140 | 0,59 | -3,07 | 3,76 | -0,82 | 4,14E-01 |
| AT5G41260 | 0,61 | -1,69 | 2,18 | -0,77 | 4,39E-01 |
| AT5G41360 | 0,66 | -1,66 | 3,70 | -0,45 | 6,54E-01 |
| AT5G41370 | 0,15 | 0,29 | 4,37 | 0,07 | 9,47E-01 |
| AT5G41410 | 1,13 | 0,09 | 1,83 | 0,05 | 9,61E-01 |
| AT5G41471 | 1,79 | -0,10 | 1,87 | -0,05 | 9,58E-01 |
| AT5G41490 | 2,24 | 2,99 | 1,69 | 1,77 | 7,63E-02 |
| AT5G41500 | 1,05 | 1,94 | 2,13 | 0,91 | 3,64E-01 |
| AT5G41510 | 1,76 | 2,02 | 1,89 | 1,07 | 2,86E-01 |
| AT5G06175 | 3,94 | 0,40 | 1,21 | 0,33 | 7,40E-01 |
| AT5G41660 | 0,71 | 1,07 | 3,01 | 0,36 | 7,22E-01 |
| AT5G41700 | 1,24 | -1,85 | 1,95 | -0,95 | 3,43E-01 |
| AT5G41761 | 0,83 | -0,05 | 2,37 | -0,02 | 9,84E-01 |
| AT5G41770 | 0,61 | 2,05 | 2,57 | 0,80 | 4,26E-01 |
| AT5G41790 | 2,54 | 0,33 | 1,37 | 0,24 | 8,10E-01 |
| AT5G41860 | 0,77 | 3,08 | 4,29 | 0,72 | 4,72E-01 |
| AT5G41920 | 0,61 | -0,38 | 4,36 | -0,09 | 9,31E-01 |
| AT5G41940 | 0,98 | 0,18 | 1,91 | 0,09 | 9,26E-01 |
| AT5G41950 | 1,46 | -0,29 | 1,85 | -0,16 | 8,76E-01 |
| AT5G41960 | 0,15 | 0,67 | 4,37 | 0,15 | 8,79E-01 |
| AT5G41970 | 0,85 | -2,45 | 3,29 | -0,75 | 4,56E-01 |
| AT5G42010 | 0,60 | 1,30 | 3,11 | 0,42 | 6,75E-01 |
| AT5G42050 | 3,62 | -0,49 | 1,05 | -0,46 | 6,43E-01 |
| AT5G42090 | 4,37 | 0,76 | 0,99 | 0,77 | 4,43E-01 |
| AT5G42220 | 0,70 | 0,20 | 2,25 | 0,09 | 9,31E-01 |
| AT5G42250 | 0,33 | 0,75 | 4,32 | 0,17 | 8,62E-01 |
| AT5G42260 | 0,28 | 0,00 | 4,41 | 0,00 | 1,00E+00 |
| AT5G42270 | 4,20 | 2,66 | 1,28 | 2,07 | 3,82E-02 |
| AT5G42290 | 0,08 | 0,69 | 4,36 | 0,16 | 8,75E-01 |
| AT5G42310 | 0,50 | -2,56 | 4,17 | -0,61 | 5,39E-01 |
| AT5G42350 | 0,37 | -0,72 | 4,30 | -0,17 | 8,68E-01 |
| AT5G42420 | 0,74 | -0,56 | 3,56 | -0,16 | 8,76E-01 |
| AT5G42460 | 0,33 | -0,32 | 4,32 | -0,07 | 9,41E-01 |
| AT5G42470 | 0,22 | 0,00 | 4,41 | 0,00 | 1,00E+00 |
| AT5G42480 | 0,45 | 0,26 | 4,30 | 0,06 | 9,52E-01 |
| AT5G42504 | 0,17 | 0,00 | 4,41 | 0,00 | 1,00E+00 |
| AT5G42520 | 0,71 | -1,08 | 2,29 | -0,47 | 6,38E-01 |
| AT5G42540 | 1,02 | 0,34 | 2,18 | 0,16 | 8,75E-01 |
| AT5G42620 | 0,80 | -0,24 | 2,33 | -0,10 | 9,19E-01 |
| AT5G42640 | 0,08 | 0,00 | 4,40 | 0,00 | 1,00E+00 |
| AT5G42670 | 0,16 | 0,00 | 4,41 | 0,00 | 1,00E+00 |
| AT5G42680 | 0,80 | -0,28 | 3,26 | -0,09 | 9,31E-01 |

| | | | | | |
|---|---|---|---|---|---|
| AT5G42720 | 0,19 | 0,07 | 4,40 | 0,02 | 9,88E-01 |
| AT5G42740 | 1,16 | 2,34 | 2,81 | 0,83 | 4,05E-01 |
| AT5G42760 | 0,12 | 0,49 | 4,36 | 0,11 | 9,11E-01 |
| AT5G06395 | 1,55 | -0,96 | 1,54 | -0,62 | 5,35E-01 |
| AT5G42825 | 0,29 | -0,23 | 4,35 | -0,05 | 9,58E-01 |
| AT5G42895 | 1,80 | -1,74 | 1,60 | -1,09 | 2,76E-01 |
| AT5G42965 | 0,19 | 0,94 | 4,35 | 0,22 | 8,28E-01 |
| AT5G42980 | 0,59 | 0,28 | 2,72 | 0,10 | 9,17E-01 |
| AT5G43100 | 0,78 | 0,14 | 2,10 | 0,07 | 9,46E-01 |
| AT5G43150 | 0,61 | -0,40 | 2,94 | -0,14 | 8,91E-01 |
| AT5G06505 | 0,36 | -0,71 | 4,38 | -0,16 | 8,71E-01 |
| AT5G43460 | 1,01 | -1,12 | 1,77 | -0,63 | 5,26E-01 |
| AT5G43500 | 1,47 | 0,32 | 1,92 | 0,17 | 8,68E-01 |
| AT5G43525 | 0,69 | 1,74 | 2,82 | 0,62 | 5,37E-01 |
| AT5G43560 | 0,55 | -1,17 | 3,22 | -0,36 | 7,16E-01 |
| AT5G43610 | 1,12 | -0,71 | 2,22 | -0,32 | 7,49E-01 |
| AT5G43620 | 0,36 | 1,97 | 4,28 | 0,46 | 6,46E-01 |
| AT5G43700 | 0,52 | -0,26 | 3,08 | -0,08 | 9,34E-01 |
| AT5G43760 | 1,93 | 0,62 | 1,49 | 0,42 | 6,77E-01 |
| AT5G43810 | 1,04 | 1,15 | 2,61 | 0,44 | 6,61E-01 |
| AT5G43930 | 0,73 | -1,56 | 2,59 | -0,60 | 5,45E-01 |
| AT5G43940 | 1,17 | 0,33 | 2,02 | 0,16 | 8,71E-01 |
| AT5G44020 | 1,03 | -0,54 | 2,12 | -0,25 | 7,99E-01 |
| AT5G44050 | 1,57 | 0,00 | 2,38 | 0,00 | 1,00E+00 |
| AT5G44060 | 0,47 | -0,16 | 2,79 | -0,06 | 9,53E-01 |
| AT5G44070 | 1,32 | 0,34 | 1,75 | 0,19 | 8,48E-01 |
| AT5G44130 | 0,80 | -1,78 | 2,97 | -0,60 | 5,49E-01 |
| AT5G44180 | 3,26 | -0,64 | 1,08 | -0,59 | 5,53E-01 |
| AT5G44190 | 1,58 | -1,79 | 2,18 | -0,82 | 4,13E-01 |
| AT5G44230 | 0,12 | 1,23 | 4,34 | 0,28 | 7,76E-01 |
| AT5G44240 | 1,02 | -3,06 | 1,90 | -1,61 | 1,08E-01 |
| AT5G44270 | 2,68 | 0,57 | 1,29 | 0,44 | 6,60E-01 |
| AT5G06565 | 0,63 | 1,02 | 2,74 | 0,37 | 7,09E-01 |
| AT5G44286 | 0,85 | -0,82 | 2,24 | -0,36 | 7,16E-01 |
| AT5G44530 | 1,17 | 0,98 | 2,52 | 0,39 | 6,97E-01 |
| AT5G44574 | 0,91 | -2,55 | 3,26 | -0,78 | 4,33E-01 |
| AT5G44580 | 0,52 | -3,11 | 3,95 | -0,79 | 4,31E-01 |
| AT5G44585 | 1,38 | -3,49 | 1,72 | -2,03 | 4,28E-02 |
| AT5G06605 | 1,31 | -3,51 | 1,73 | -2,03 | 4,23E-02 |
| AT5G44640 | 1,59 | 2,03 | 1,93 | 1,05 | 2,93E-01 |
| AT5G44650 | 0,46 | 0,30 | 4,04 | 0,07 | 9,41E-01 |
| AT5G44720 | 0,16 | 0,84 | 4,35 | 0,19 | 8,46E-01 |
| AT5G44750 | 0,74 | 0,12 | 2,14 | 0,06 | 9,54E-01 |
| AT5G44760 | 0,36 | 1,57 | 4,34 | 0,36 | 7,17E-01 |
| AT5G44770 | 0,22 | 0,48 | 4,36 | 0,11 | 9,13E-01 |
| AT5G44870 | 1,81 | 0,45 | 1,44 | 0,31 | 7,53E-01 |
| AT5G45000 | 0,38 | -1,35 | 4,32 | -0,31 | 7,55E-01 |
| AT5G06695 | 0,59 | -0,21 | 2,97 | -0,07 | 9,45E-01 |
| AT5G45030 | 0,59 | -0,21 | 2,97 | -0,07 | 9,45E-01 |
| AT5G45060 | 1,56 | -0,01 | 1,64 | 0,00 | 9,97E-01 |
| AT5G45095 | 2,04 | 3,28 | 1,70 | 1,93 | 5,39E-02 |
| AT5G45100 | 1,72 | 0,99 | 1,89 | 0,52 | 6,00E-01 |

| | | | | | |
|---|---|---|---|---|---|
| AT5G45110 | 1,72 | 1,02 | 1,42 | 0,72 | 4,74E-01 |
| AT5G45130 | 0,78 | 1,72 | 2,94 | 0,58 | 5,59E-01 |
| AT5G45150 | 0,08 | 0,44 | 4,38 | 0,10 | 9,21E-01 |
| AT5G45240 | 0,12 | 1,23 | 4,34 | 0,28 | 7,76E-01 |
| AT5G45260 | 0,75 | -0,54 | 2,74 | -0,20 | 8,44E-01 |
| AT5G45390 | 0,62 | -0,89 | 2,37 | -0,38 | 7,07E-01 |
| AT5G45400 | 0,76 | -1,03 | 2,36 | -0,44 | 6,62E-01 |
| AT5G45430 | 3,88 | 1,05 | 0,90 | 1,17 | 2,43E-01 |
| AT5G45450 | 0,08 | 0,44 | 4,38 | 0,10 | 9,21E-01 |
| AT5G45472 | 4,93 | 1,37 | 1,15 | 1,19 | 2,35E-01 |
| AT5G45475 | 0,74 | 0,23 | 2,31 | 0,10 | 9,21E-01 |
| AT5G45490 | 1,25 | 0,35 | 1,83 | 0,19 | 8,48E-01 |
| AT5G45570 | 0,68 | 1,34 | 2,50 | 0,53 | 5,93E-01 |
| AT5G45620 | 0,59 | 2,00 | 3,09 | 0,65 | 5,17E-01 |
| AT5G45670 | 0,10 | 0,00 | 4,41 | 0,00 | 1,00E+00 |
| AT5G45680 | 0,26 | -1,78 | 4,41 | -0,40 | 6,87E-01 |
| AT5G45710 | 0,55 | -2,17 | 3,77 | -0,58 | 5,65E-01 |
| AT5G45750 | 0,19 | 0,00 | 4,41 | 0,00 | 1,00E+00 |
| AT5G45770 | 0,12 | 1,16 | 4,34 | 0,27 | 7,89E-01 |
| AT5G45920 | 1,21 | 0,84 | 2,09 | 0,40 | 6,89E-01 |
| AT5G46070 | 1,78 | -2,29 | 1,60 | -1,43 | 1,54E-01 |
| AT5G46330 | 0,47 | 1,11 | 3,98 | 0,28 | 7,80E-01 |
| AT5G46410 | 0,44 | -1,22 | 4,39 | -0,28 | 7,81E-01 |
| AT5G46420 | 2,51 | -0,77 | 1,07 | -0,72 | 4,75E-01 |
| AT5G46430 | 2,53 | 1,51 | 1,59 | 0,95 | 3,43E-01 |
| AT5G06825 | 0,67 | -0,47 | 3,09 | -0,15 | 8,79E-01 |
| AT5G46450 | 2,49 | -0,38 | 1,14 | -0,33 | 7,38E-01 |
| AT5G46460 | 0,90 | 1,82 | 3,19 | 0,57 | 5,69E-01 |
| AT5G46470 | 3,68 | -0,80 | 0,87 | -0,92 | 3,58E-01 |
| AT5G46490 | 1,46 | -0,16 | 1,46 | -0,11 | 9,12E-01 |
| AT5G46500 | 1,11 | 2,68 | 2,39 | 1,12 | 2,63E-01 |
| AT5G46510 | 5,48 | 1,07 | 0,77 | 1,38 | 1,67E-01 |
| AT5G06835 | 1,13 | -0,85 | 2,63 | -0,32 | 7,46E-01 |
| AT5G46520 | 3,26 | 0,46 | 1,11 | 0,41 | 6,82E-01 |
| AT5G46630 | 1,04 | -0,56 | 1,91 | -0,29 | 7,71E-01 |
| AT5G46640 | 1,47 | -2,82 | 1,78 | -1,58 | 1,14E-01 |
| AT5G46660 | 1,02 | 0,00 | 4,13 | 0,00 | 1,00E+00 |
| AT5G46670 | 0,74 | 0,04 | 2,86 | 0,01 | 9,88E-01 |
| AT5G46730 | 0,08 | 0,00 | 4,40 | 0,00 | 1,00E+00 |
| AT5G46760 | 2,52 | 0,29 | 1,27 | 0,23 | 8,17E-01 |
| AT5G46780 | 0,35 | 0,82 | 4,34 | 0,19 | 8,50E-01 |
| AT5G46825 | 1,18 | 0,27 | 1,82 | 0,15 | 8,82E-01 |
| AT5G46840 | 0,62 | -1,16 | 3,08 | -0,38 | 7,07E-01 |
| AT5G46850 | 0,29 | 0,35 | 4,32 | 0,08 | 9,36E-01 |
| AT5G47010 | 2,34 | -0,37 | 1,17 | -0,32 | 7,50E-01 |
| AT5G47020 | 1,57 | -0,75 | 1,62 | -0,46 | 6,45E-01 |
| AT5G47030 | 0,63 | -1,01 | 3,78 | -0,27 | 7,89E-01 |
| AT5G47120 | 1,06 | 2,79 | 2,09 | 1,34 | 1,81E-01 |
| AT5G47130 | 0,25 | 1,09 | 4,34 | 0,25 | 8,01E-01 |
| AT5G47140 | 0,26 | 0,41 | 4,37 | 0,09 | 9,25E-01 |
| AT5G47200 | 0,34 | 1,15 | 4,34 | 0,26 | 7,92E-01 |
| AT5G47230 | 0,27 | 0,18 | 4,37 | 0,04 | 9,68E-01 |

| | | | | | |
|---|---|---|---|---|---|
| AT5G47240 | 0,14 | 0,00 | 4,41 | 0,00 | 1,00E+00 |
| AT5G47260 | 4,17 | 2,24 | 1,43 | 1,56 | 1,19E-01 |
| AT5G47280 | 1,15 | 1,65 | 2,02 | 0,82 | 4,14E-01 |
| AT5G47300 | 1,47 | 0,98 | 1,91 | 0,51 | 6,10E-01 |
| AT5G47320 | 0,19 | 0,36 | 4,36 | 0,08 | 9,34E-01 |
| AT5G47390 | 0,64 | -0,24 | 1,98 | -0,12 | 9,04E-01 |
| AT5G47420 | 0,50 | -1,51 | 3,91 | -0,39 | 7,00E-01 |
| AT5G47435 | 1,48 | -1,46 | 1,71 | -0,86 | 3,91E-01 |
| AT5G47480 | 1,95 | -0,28 | 1,26 | -0,22 | 8,27E-01 |
| AT5G47490 | 2,21 | 0,04 | 1,14 | 0,04 | 9,71E-01 |
| AT5G47640 | 0,70 | -1,58 | 2,23 | -0,71 | 4,81E-01 |
| AT5G47660 | 0,08 | 0,00 | 4,40 | 0,00 | 1,00E+00 |
| AT5G47690 | 3,16 | -0,90 | 1,14 | -0,80 | 4,26E-01 |
| AT5G47720 | 0,16 | 1,33 | 4,33 | 0,31 | 7,59E-01 |
| AT5G47770 | 0,39 | 0,39 | 4,28 | 0,09 | 9,27E-01 |
| AT5G47780 | 0,52 | 0,66 | 2,67 | 0,25 | 8,06E-01 |
| AT5G47790 | 0,28 | 1,61 | 4,33 | 0,37 | 7,10E-01 |
| AT5G47800 | 0,24 | 0,21 | 4,34 | 0,05 | 9,62E-01 |
| AT5G47820 | 0,70 | -1,91 | 3,61 | -0,53 | 5,97E-01 |
| AT5G47840 | 0,45 | -0,91 | 2,79 | -0,33 | 7,44E-01 |
| AT5G47860 | 0,53 | 2,44 | 3,85 | 0,64 | 5,25E-01 |
| AT5G47890 | 1,07 | 0,40 | 1,82 | 0,22 | 8,24E-01 |
| AT5G47910 | 5,04 | 0,59 | 0,88 | 0,68 | 4,98E-01 |
| AT5G48190 | 1,46 | 0,26 | 1,99 | 0,13 | 8,95E-01 |
| AT5G48220 | 0,16 | 0,49 | 4,38 | 0,11 | 9,12E-01 |
| AT5G48290 | 0,36 | -2,77 | 4,36 | -0,64 | 5,25E-01 |
| AT5G48300 | 1,23 | -0,11 | 2,03 | -0,06 | 9,56E-01 |
| AT5G48340 | 0,30 | 0,24 | 4,35 | 0,05 | 9,57E-01 |
| AT5G48360 | 0,08 | 0,00 | 4,40 | 0,00 | 1,00E+00 |
| AT5G48385 | 2,37 | -0,87 | 1,43 | -0,61 | 5,44E-01 |
| AT5G48390 | 0,69 | -0,70 | 3,41 | -0,20 | 8,38E-01 |
| AT5G48400 | 0,56 | 1,44 | 3,76 | 0,38 | 7,02E-01 |
| AT5G48410 | 0,12 | 1,16 | 4,34 | 0,27 | 7,89E-01 |
| AT5G07095 | 1,59 | -0,60 | 1,40 | -0,43 | 6,69E-01 |
| AT5G07105 | 0,86 | -2,21 | 2,14 | -1,03 | 3,02E-01 |
| AT5G48545 | 0,30 | 0,16 | 3,72 | 0,04 | 9,65E-01 |
| AT5G48620 | 2,46 | 0,91 | 1,42 | 0,64 | 5,19E-01 |
| AT5G48720 | 0,16 | 0,65 | 4,37 | 0,15 | 8,82E-01 |
| AT5G48730 | 0,12 | 1,16 | 4,34 | 0,27 | 7,89E-01 |
| AT5G48775 | 0,08 | 0,00 | 4,40 | 0,00 | 1,00E+00 |
| AT5G48800 | 0,90 | 1,25 | 2,83 | 0,44 | 6,58E-01 |
| AT5G48900 | 0,13 | 0,00 | 4,41 | 0,00 | 1,00E+00 |
| AT5G49010 | 1,17 | -3,26 | 2,12 | -1,54 | 1,23E-01 |
| AT5G49020 | 0,22 | 0,25 | 4,36 | 0,06 | 9,54E-01 |
| AT5G49060 | 0,17 | 1,33 | 4,33 | 0,31 | 7,59E-01 |
| AT5G49160 | 0,93 | -0,34 | 2,42 | -0,14 | 8,87E-01 |
| AT5G49215 | 0,08 | 0,00 | 4,40 | 0,00 | 1,00E+00 |
| AT5G49250 | 0,66 | 1,76 | 2,80 | 0,63 | 5,30E-01 |
| AT5G49260 | 0,08 | 0,00 | 4,40 | 0,00 | 1,00E+00 |
| AT5G49440 | 5,06 | -1,26 | 0,97 | -1,29 | 1,96E-01 |
| AT5G49450 | 0,19 | 1,50 | 4,33 | 0,35 | 7,30E-01 |
| AT5G49460 | 0,62 | -1,11 | 2,91 | -0,38 | 7,03E-01 |

| | | | | | |
|---|---|---|---|---|---|
| AT5G49470 | 0,29 | 0,35 | 4,32 | 0,08 | 9,36E-01 |
| AT5G49480 | 1,15 | -2,02 | 2,09 | -0,97 | 3,34E-01 |
| AT5G49510 | 0,19 | 0,78 | 4,35 | 0,18 | 8,58E-01 |
| AT5G49520 | 0,12 | 0,49 | 4,36 | 0,11 | 9,11E-01 |
| AT5G49530 | 0,46 | 2,28 | 4,05 | 0,56 | 5,73E-01 |
| AT5G49590 | 0,86 | -1,77 | 2,91 | -0,61 | 5,43E-01 |
| AT5G49615 | 2,21 | 1,16 | 2,01 | 0,58 | 5,65E-01 |
| AT5G49640 | 0,08 | 0,44 | 4,38 | 0,10 | 9,21E-01 |
| AT5G49650 | 0,61 | 0,55 | 2,29 | 0,24 | 8,11E-01 |
| AT5G49760 | 3,34 | -1,41 | 1,05 | -1,35 | 1,77E-01 |
| AT5G49810 | 1,47 | -0,56 | 1,91 | -0,29 | 7,69E-01 |
| AT5G49840 | 0,10 | 0,21 | 4,38 | 0,05 | 9,62E-01 |
| AT5G49910 | 2,46 | -0,76 | 1,19 | -0,64 | 5,25E-01 |
| AT5G49945 | 0,51 | 2,31 | 3,81 | 0,61 | 5,45E-01 |
| AT5G49970 | 0,29 | 1,75 | 4,32 | 0,41 | 6,85E-01 |
| AT5G50020 | 0,47 | -2,98 | 4,35 | -0,69 | 4,93E-01 |
| AT5G50160 | 1,06 | 2,04 | 2,29 | 0,89 | 3,73E-01 |
| AT5G50170 | 0,08 | 0,44 | 4,38 | 0,10 | 9,21E-01 |
| AT5G50200 | 0,08 | 0,00 | 4,40 | 0,00 | 1,00E+00 |
| AT5G07605 | 0,25 | 0,05 | 4,39 | 0,01 | 9,92E-01 |
| AT5G50310 | 0,21 | 1,63 | 4,33 | 0,38 | 7,06E-01 |
| AT5G50580 | 0,65 | 0,79 | 3,59 | 0,22 | 8,25E-01 |
| AT5G50680 | 1,09 | 2,43 | 2,31 | 1,05 | 2,94E-01 |
| AT5G50717 | 0,11 | 0,00 | 4,41 | 0,00 | 1,00E+00 |
| AT5G50780 | 0,59 | -0,22 | 4,31 | -0,05 | 9,59E-01 |
| AT5G50850 | 0,82 | 2,85 | 2,31 | 1,23 | 2,18E-01 |
| AT5G50950 | 1,52 | 0,00 | 1,58 | 0,00 | 9,98E-01 |
| AT5G50960 | 0,50 | -0,49 | 3,92 | -0,12 | 9,01E-01 |
| AT5G51010 | 0,08 | 0,69 | 4,36 | 0,16 | 8,75E-01 |
| AT5G51050 | 0,20 | 1,20 | 4,35 | 0,28 | 7,83E-01 |
| AT5G51055 | 0,34 | 0,00 | 3,61 | 0,00 | 1,00E+00 |
| AT5G51070 | 3,71 | -0,20 | 0,89 | -0,23 | 8,20E-01 |
| AT5G51120 | 0,08 | 0,69 | 4,36 | 0,16 | 8,75E-01 |
| AT5G51150 | 0,28 | -0,49 | 4,38 | -0,11 | 9,11E-01 |
| AT5G07735 | 1,90 | 2,16 | 1,68 | 1,29 | 1,99E-01 |
| AT5G51180 | 0,38 | -0,16 | 2,89 | -0,06 | 9,55E-01 |
| AT5G51200 | 1,04 | 1,91 | 2,12 | 0,90 | 3,69E-01 |
| AT5G51230 | 0,21 | 0,50 | 4,36 | 0,12 | 9,08E-01 |
| AT5G51280 | 0,48 | 0,25 | 2,70 | 0,09 | 9,27E-01 |
| AT5G51340 | 0,39 | -0,18 | 3,41 | -0,05 | 9,57E-01 |
| AT5G51450 | 0,53 | 0,25 | 3,94 | 0,06 | 9,49E-01 |
| AT5G51560 | 0,28 | -2,52 | 4,38 | -0,58 | 5,65E-01 |
| AT5G51570 | 0,40 | 0,26 | 3,45 | 0,08 | 9,39E-01 |
| AT5G51630 | 1,97 | 0,84 | 1,45 | 0,57 | 5,65E-01 |
| AT5G51660 | 1,44 | -1,55 | 1,56 | -0,99 | 3,21E-01 |
| AT5G51670 | 0,12 | 0,97 | 4,34 | 0,22 | 8,23E-01 |
| AT5G51680 | 0,64 | -0,91 | 2,49 | -0,36 | 7,15E-01 |
| AT5G51750 | 0,75 | 0,96 | 3,67 | 0,26 | 7,94E-01 |
| AT5G51760 | 1,35 | -0,44 | 1,64 | -0,27 | 7,88E-01 |
| AT5G51830 | 0,27 | 1,29 | 4,33 | 0,30 | 7,65E-01 |
| AT5G51960 | 0,11 | 0,00 | 4,41 | 0,00 | 1,00E+00 |
| AT5G51970 | 1,04 | 0,19 | 1,82 | 0,10 | 9,18E-01 |

| | | | | | |
|---|---|---|---|---|---|
| AT5G52050 | 0,61 | 0,48 | 2,54 | 0,19 | 8,50E-01 |
| AT5G07875 | 4,76 | -1,97 | 1,00 | -1,97 | 4,94E-02 |
| AT5G52115 | 1,12 | 0,00 | 2,78 | 0,00 | 1,00E+00 |
| AT5G52170 | 2,49 | 0,36 | 1,32 | 0,27 | 7,85E-01 |
| AT5G52240 | 1,16 | 0,84 | 2,03 | 0,42 | 6,78E-01 |
| AT5G52280 | 0,34 | -1,28 | 4,34 | -0,29 | 7,69E-01 |
| AT5G52310 | 5,67 | -2,30 | 1,53 | -1,50 | 1,34E-01 |
| AT5G52470 | 2,57 | -0,89 | 1,26 | -0,71 | 4,77E-01 |
| AT5G52471 | 1,40 | -1,47 | 2,04 | -0,72 | 4,71E-01 |
| AT5G52510 | 0,69 | 1,79 | 3,00 | 0,60 | 5,51E-01 |
| AT5G52520 | 0,23 | -1,58 | 4,38 | -0,36 | 7,18E-01 |
| AT5G52530 | 3,03 | -0,73 | 1,42 | -0,51 | 6,08E-01 |
| AT5G52580 | 0,50 | 1,38 | 3,26 | 0,42 | 6,72E-01 |
| AT5G52630 | 0,08 | 0,44 | 4,38 | 0,10 | 9,21E-01 |
| AT5G52700 | 0,14 | 0,00 | 4,41 | 0,00 | 1,00E+00 |
| AT5G52750 | 0,33 | 1,48 | 4,32 | 0,34 | 7,32E-01 |
| AT5G52760 | 0,52 | 0,30 | 2,62 | 0,12 | 9,08E-01 |
| AT5G52920 | 1,32 | 1,05 | 2,03 | 0,52 | 6,04E-01 |
| AT5G07945 | 0,52 | 1,77 | 3,96 | 0,45 | 6,55E-01 |
| AT5G52940 | 0,11 | 0,48 | 4,37 | 0,11 | 9,12E-01 |
| AT5G53020 | 0,27 | 0,84 | 4,36 | 0,19 | 8,48E-01 |
| AT5G53030 | 0,24 | 0,00 | 4,41 | 0,00 | 1,00E+00 |
| AT5G53048 | 0,25 | 1,12 | 4,34 | 0,26 | 7,97E-01 |
| AT5G53060 | 0,48 | 2,19 | 3,95 | 0,55 | 5,80E-01 |
| AT5G53080 | 0,12 | 1,16 | 4,34 | 0,27 | 7,89E-01 |
| AT5G53110 | 0,31 | 0,50 | 4,38 | 0,11 | 9,09E-01 |
| AT5G53120 | 0,82 | 0,54 | 2,49 | 0,22 | 8,28E-01 |
| AT5G53140 | 1,45 | -0,38 | 1,59 | -0,24 | 8,09E-01 |
| AT5G53150 | 0,95 | -1,82 | 2,04 | -0,89 | 3,72E-01 |
| AT5G53160 | 1,14 | 1,65 | 1,91 | 0,86 | 3,88E-01 |
| AT5G53220 | 0,48 | 0,99 | 3,07 | 0,32 | 7,48E-01 |
| AT5G53280 | 0,44 | -2,04 | 4,10 | -0,50 | 6,18E-01 |
| AT5G53350 | 0,54 | -0,98 | 3,25 | -0,30 | 7,62E-01 |
| AT5G53360 | 0,08 | 0,44 | 4,38 | 0,10 | 9,21E-01 |
| AT5G53410 | 1,13 | 2,38 | 1,87 | 1,27 | 2,03E-01 |
| AT5G53420 | 0,48 | -1,28 | 3,26 | -0,39 | 6,93E-01 |
| AT5G53430 | 1,00 | -0,57 | 1,83 | -0,31 | 7,53E-01 |
| AT5G53450 | 0,71 | -2,15 | 2,52 | -0,85 | 3,93E-01 |
| AT5G53550 | 0,99 | 1,60 | 2,22 | 0,72 | 4,70E-01 |
| AT5G53560 | 1,33 | 1,30 | 1,76 | 0,74 | 4,59E-01 |
| AT5G53590 | 0,48 | 0,40 | 4,01 | 0,10 | 9,21E-01 |
| AT5G53620 | 1,09 | -0,94 | 1,60 | -0,59 | 5,56E-01 |
| AT5G53860 | 0,25 | 0,81 | 3,80 | 0,21 | 8,31E-01 |
| AT5G53890 | 0,71 | 2,47 | 2,44 | 1,01 | 3,11E-01 |
| AT5G54035 | 0,15 | 0,00 | 4,41 | 0,00 | 1,00E+00 |
| AT5G54043 | 0,22 | 0,00 | 4,38 | 0,00 | 1,00E+00 |
| AT5G54080 | 0,99 | 1,01 | 2,08 | 0,49 | 6,26E-01 |
| AT5G54110 | 0,33 | 0,85 | 4,32 | 0,20 | 8,44E-01 |
| AT5G54140 | 0,24 | 0,24 | 4,34 | 0,06 | 9,55E-01 |
| AT5G54145 | 0,12 | 0,00 | 4,41 | 0,00 | 1,00E+00 |
| AT5G54160 | 0,46 | 1,39 | 4,13 | 0,34 | 7,37E-01 |
| AT5G54180 | 0,12 | 0,26 | 4,37 | 0,06 | 9,52E-01 |

| | | | | | |
|---|---|---|---|---|---|
| AT5G54290 | 0,66 | -1,88 | 2,65 | -0,71 | 4,79E-01 |
| AT5G54390 | 4,62 | 1,62 | 1,21 | 1,33 | 1,82E-01 |
| AT5G54440 | 2,06 | 0,07 | 1,26 | 0,06 | 9,55E-01 |
| AT5G54500 | 0,31 | 0,51 | 4,37 | 0,12 | 9,07E-01 |
| AT5G54540 | 0,58 | 1,13 | 2,64 | 0,43 | 6,68E-01 |
| AT5G54600 | 0,87 | 0,05 | 1,94 | 0,03 | 9,78E-01 |
| AT5G08225 | 0,36 | 0,60 | 4,37 | 0,14 | 8,91E-01 |
| AT5G54640 | 0,05 | 0,00 | 4,41 | 0,00 | 1,00E+00 |
| AT5G54680 | 0,72 | -1,99 | 2,58 | -0,77 | 4,40E-01 |
| AT5G54750 | 0,55 | 1,41 | 3,88 | 0,36 | 7,17E-01 |
| AT5G54760 | 0,49 | 0,01 | 2,82 | 0,00 | 9,99E-01 |
| AT5G54800 | 0,08 | 0,00 | 4,41 | 0,00 | 1,00E+00 |
| AT5G54830 | 0,76 | 2,11 | 2,73 | 0,77 | 4,39E-01 |
| AT5G54855 | 0,18 | 0,13 | 4,41 | 0,03 | 9,76E-01 |
| AT5G54860 | 0,26 | -0,30 | 4,37 | -0,07 | 9,46E-01 |
| AT5G54900 | 0,82 | 0,02 | 2,29 | 0,01 | 9,94E-01 |
| AT5G54920 | 0,83 | 2,03 | 4,31 | 0,47 | 6,39E-01 |
| AT5G55060 | 0,14 | 0,09 | 4,38 | 0,02 | 9,84E-01 |
| AT5G55070 | 1,01 | 0,70 | 1,83 | 0,38 | 7,01E-01 |
| AT5G55190 | 0,80 | 1,10 | 2,29 | 0,48 | 6,29E-01 |
| AT5G55220 | 0,88 | -2,55 | 2,70 | -0,94 | 3,46E-01 |
| AT5G55230 | 0,95 | -1,30 | 1,90 | -0,68 | 4,96E-01 |
| AT5G55450 | 0,62 | -1,95 | 2,99 | -0,65 | 5,14E-01 |
| AT5G55460 | 0,37 | -0,83 | 3,43 | -0,24 | 8,08E-01 |
| AT5G55500 | 0,57 | 1,56 | 3,06 | 0,51 | 6,10E-01 |
| AT5G55530 | 0,30 | 0,48 | 4,32 | 0,11 | 9,12E-01 |
| AT5G55580 | 0,70 | 0,22 | 3,85 | 0,06 | 9,54E-01 |
| AT5G55620 | 0,43 | 0,04 | 4,28 | 0,01 | 9,92E-01 |
| AT5G55630 | 0,12 | 0,97 | 4,34 | 0,22 | 8,23E-01 |
| AT5G55660 | 0,90 | -1,46 | 2,19 | -0,66 | 5,06E-01 |
| AT5G55840 | 0,22 | 1,17 | 4,33 | 0,27 | 7,88E-01 |
| AT5G55850 | 0,24 | 0,00 | 4,38 | 0,00 | 9,99E-01 |
| AT5G55900 | 0,10 | 0,00 | 4,41 | 0,00 | 1,00E+00 |
| AT5G55930 | 0,70 | 1,01 | 2,87 | 0,35 | 7,24E-01 |
| AT5G55960 | 0,59 | -0,01 | 3,14 | 0,00 | 9,98E-01 |
| AT5G55990 | 0,33 | 0,71 | 4,31 | 0,16 | 8,69E-01 |
| AT5G56020 | 0,32 | -0,94 | 4,31 | -0,22 | 8,28E-01 |
| AT5G56030 | 5,79 | 0,40 | 0,74 | 0,54 | 5,87E-01 |
| AT5G56040 | 0,73 | 1,44 | 2,39 | 0,60 | 5,48E-01 |
| AT5G56060 | 0,67 | 1,22 | 4,36 | 0,28 | 7,80E-01 |
| AT5G56070 | 3,26 | 3,82 | 1,70 | 2,24 | 2,48E-02 |
| AT5G56140 | 0,98 | -0,72 | 1,90 | -0,38 | 7,06E-01 |
| AT5G56160 | 0,49 | -1,78 | 4,41 | -0,40 | 6,87E-01 |
| AT5G56170 | 0,53 | -0,62 | 4,38 | -0,14 | 8,87E-01 |
| AT5G56270 | 0,76 | -1,46 | 2,67 | -0,55 | 5,85E-01 |
| AT5G56290 | 0,52 | 0,27 | 2,41 | 0,11 | 9,12E-01 |
| AT5G56340 | 0,44 | -0,54 | 3,23 | -0,17 | 8,67E-01 |
| AT5G08495 | 0,43 | -1,83 | 4,25 | -0,43 | 6,67E-01 |
| AT5G08525 | 4,34 | 0,55 | 1,11 | 0,49 | 6,22E-01 |
| AT5G56460 | 0,74 | 0,88 | 3,98 | 0,22 | 8,24E-01 |
| AT5G56500 | 0,32 | -2,61 | 4,37 | -0,60 | 5,51E-01 |
| AT5G56590 | 0,24 | -1,19 | 4,35 | -0,27 | 7,84E-01 |

| | | | | | |
|---|---|---|---|---|---|
| AT5G56630 | 0,71 | -1,22 | 2,47 | -0,49 | 6,21E-01 |
| AT5G56650 | 0,64 | -1,52 | 2,23 | -0,68 | 4,97E-01 |
| AT5G56680 | 1,22 | -0,58 | 1,73 | -0,34 | 7,35E-01 |
| AT5G56720 | 0,12 | 0,37 | 4,37 | 0,08 | 9,33E-01 |
| AT5G56870 | 0,19 | 0,84 | 4,35 | 0,19 | 8,46E-01 |
| AT5G56910 | 0,15 | 1,16 | 4,34 | 0,27 | 7,88E-01 |
| AT5G56920 | 0,65 | 0,13 | 3,07 | 0,04 | 9,67E-01 |
| AT5G56940 | 0,08 | 0,00 | 4,41 | 0,00 | 1,00E+00 |
| AT5G56950 | 0,44 | 0,48 | 3,24 | 0,15 | 8,83E-01 |
| AT5G57015 | 0,32 | -0,68 | 4,35 | -0,16 | 8,77E-01 |
| AT5G57020 | 1,25 | 0,89 | 1,70 | 0,52 | 6,02E-01 |
| AT5G57030 | 0,86 | 1,64 | 2,29 | 0,72 | 4,73E-01 |
| AT5G57035 | 1,04 | -1,46 | 2,46 | -0,59 | 5,53E-01 |
| AT5G57070 | 0,42 | -1,11 | 4,20 | -0,26 | 7,91E-01 |
| AT5G57150 | 0,25 | 0,38 | 4,33 | 0,09 | 9,31E-01 |
| AT5G57180 | 1,05 | -1,76 | 1,95 | -0,90 | 3,67E-01 |
| AT5G57320 | 0,08 | 0,44 | 4,38 | 0,10 | 9,21E-01 |
| AT5G57330 | 0,49 | -0,06 | 2,86 | -0,02 | 9,84E-01 |
| AT5G57340 | 0,20 | 0,15 | 4,37 | 0,04 | 9,72E-01 |
| AT5G57360 | 0,96 | 0,86 | 1,81 | 0,47 | 6,36E-01 |
| AT5G57370 | 0,31 | -0,13 | 4,36 | -0,03 | 9,77E-01 |
| AT5G57460 | 1,07 | -0,07 | 1,65 | -0,04 | 9,68E-01 |
| AT5G57520 | 0,31 | 0,02 | 4,39 | 0,01 | 9,96E-01 |
| AT5G57610 | 0,89 | 0,49 | 1,82 | 0,27 | 7,88E-01 |
| AT5G57655 | 1,67 | 1,12 | 1,71 | 0,65 | 5,15E-01 |
| AT5G57660 | 4,42 | -0,06 | 0,81 | -0,08 | 9,38E-01 |
| AT5G57710 | 4,28 | 0,80 | 0,89 | 0,90 | 3,69E-01 |
| AT5G57735 | 0,19 | 0,00 | 4,41 | 0,00 | 1,00E+00 |
| AT5G57800 | 0,61 | -2,55 | 2,68 | -0,95 | 3,41E-01 |
| AT5G08805 | 1,03 | 1,12 | 2,40 | 0,47 | 6,40E-01 |
| AT5G57880 | 0,48 | -2,56 | 4,02 | -0,64 | 5,25E-01 |
| AT5G57930 | 1,66 | -0,16 | 1,61 | -0,10 | 9,22E-01 |
| AT5G57940 | 0,35 | 1,58 | 3,53 | 0,45 | 6,55E-01 |
| AT5G58000 | 0,08 | 0,69 | 4,36 | 0,16 | 8,75E-01 |
| AT5G58020 | 0,39 | 0,03 | 4,39 | 0,01 | 9,95E-01 |
| AT5G58040 | 3,29 | 0,09 | 0,98 | 0,09 | 9,27E-01 |
| AT5G58060 | 0,59 | -0,95 | 2,45 | -0,39 | 6,97E-01 |
| AT5G58100 | 0,97 | -1,08 | 1,82 | -0,59 | 5,54E-01 |
| AT5G58110 | 0,27 | 0,66 | 4,35 | 0,15 | 8,79E-01 |
| AT5G58120 | 1,53 | 1,14 | 1,48 | 0,77 | 4,40E-01 |
| AT5G08820 | 0,19 | 1,08 | 4,34 | 0,25 | 8,04E-01 |
| AT5G58160 | 1,77 | 0,67 | 2,24 | 0,30 | 7,65E-01 |
| AT5G58200 | 0,40 | 0,00 | 4,41 | 0,00 | 1,00E+00 |
| AT5G58230 | 0,77 | 0,67 | 2,37 | 0,28 | 7,76E-01 |
| AT5G58250 | 0,32 | 0,87 | 4,35 | 0,20 | 8,41E-01 |
| AT5G58270 | 0,43 | 0,29 | 4,19 | 0,07 | 9,46E-01 |
| AT5G58290 | 1,43 | 1,87 | 1,87 | 1,00 | 3,18E-01 |
| AT5G58300 | 0,99 | -1,42 | 2,06 | -0,69 | 4,91E-01 |
| AT5G58320 | 0,24 | -0,08 | 4,34 | -0,02 | 9,85E-01 |
| AT5G58350 | 1,12 | 1,39 | 1,84 | 0,76 | 4,50E-01 |
| AT5G58370 | 0,29 | -1,88 | 4,35 | -0,43 | 6,67E-01 |
| AT5G58375 | 0,48 | -1,37 | 4,33 | -0,32 | 7,51E-01 |

| | | | | | |
|---|---|---|---|---|---|
| AT5G58410 | 3,13 | 1,17 | 1,04 | 1,12 | 2,61E-01 |
| AT5G58420 | 1,59 | -1,72 | 1,87 | -0,92 | 3,57E-01 |
| AT5G58450 | 0,68 | -0,27 | 2,87 | -0,09 | 9,26E-01 |
| AT5G58465 | 4,24 | 0,02 | 1,08 | 0,02 | 9,82E-01 |
| AT5G58495 | 0,12 | 1,16 | 4,34 | 0,27 | 7,89E-01 |
| AT5G58520 | 0,39 | -1,16 | 4,36 | -0,27 | 7,91E-01 |
| AT5G58540 | 1,83 | 0,52 | 1,46 | 0,36 | 7,21E-01 |
| AT5G58560 | 0,47 | 0,80 | 3,19 | 0,25 | 8,03E-01 |
| AT5G08855 | 0,43 | -1,78 | 4,28 | -0,41 | 6,78E-01 |
| AT5G58620 | 0,34 | -0,13 | 4,35 | -0,03 | 9,76E-01 |
| AT5G58640 | 0,71 | -0,96 | 2,93 | -0,33 | 7,43E-01 |
| AT5G58710 | 0,66 | 1,16 | 3,12 | 0,37 | 7,09E-01 |
| AT5G58990 | 0,29 | 0,00 | 4,39 | 0,00 | 1,00E+00 |
| AT5G59160 | 0,32 | 1,79 | 4,32 | 0,41 | 6,79E-01 |
| AT5G59170 | 0,26 | -0,30 | 4,37 | -0,07 | 9,46E-01 |
| AT5G59250 | 0,50 | -0,85 | 3,20 | -0,26 | 7,91E-01 |
| AT5G59350 | 0,35 | -0,48 | 4,32 | -0,11 | 9,12E-01 |
| AT5G59420 | 1,43 | 1,25 | 1,84 | 0,68 | 4,97E-01 |
| AT5G59430 | 0,64 | 1,73 | 2,56 | 0,68 | 4,99E-01 |
| AT5G59450 | 0,61 | 0,42 | 3,79 | 0,11 | 9,12E-01 |
| AT5G59505 | 3,52 | -1,38 | 1,62 | -0,85 | 3,95E-01 |
| AT5G59610 | 0,51 | 0,98 | 3,96 | 0,25 | 8,04E-01 |
| AT5G59660 | 0,05 | 0,00 | 4,41 | 0,00 | 1,00E+00 |
| AT5G59670 | 0,28 | 0,09 | 3,74 | 0,02 | 9,82E-01 |
| AT5G59680 | 0,64 | -2,20 | 4,30 | -0,51 | 6,10E-01 |
| AT5G59690 | 0,64 | 1,12 | 2,89 | 0,39 | 6,98E-01 |
| AT5G59732 | 3,71 | -0,47 | 1,16 | -0,40 | 6,87E-01 |
| AT5G59820 | 0,42 | 1,14 | 3,29 | 0,35 | 7,28E-01 |
| AT5G59830 | 0,85 | -1,43 | 2,77 | -0,52 | 6,06E-01 |
| AT5G59860 | 0,43 | -1,11 | 4,34 | -0,25 | 7,99E-01 |
| AT5G59880 | 0,87 | 0,05 | 2,54 | 0,02 | 9,83E-01 |
| AT5G59910 | 0,91 | -0,29 | 2,20 | -0,13 | 8,96E-01 |
| AT5G59920 | 0,58 | 0,26 | 3,65 | 0,07 | 9,44E-01 |
| AT5G59950 | 1,16 | -0,93 | 1,75 | -0,53 | 5,96E-01 |
| AT5G59980 | 0,39 | -1,28 | 4,38 | -0,29 | 7,70E-01 |
| AT5G60040 | 0,79 | -1,10 | 2,21 | -0,50 | 6,19E-01 |
| AT5G60110 | 0,25 | 0,41 | 4,33 | 0,09 | 9,25E-01 |
| AT5G60120 | 0,67 | 0,89 | 2,77 | 0,32 | 7,47E-01 |
| AT5G60170 | 1,68 | 0,91 | 1,59 | 0,58 | 5,64E-01 |
| AT5G60270 | 0,48 | 2,53 | 4,30 | 0,59 | 5,57E-01 |
| AT5G60280 | 0,76 | 3,03 | 3,44 | 0,88 | 3,78E-01 |
| AT5G60300 | 0,52 | 2,30 | 3,85 | 0,60 | 5,49E-01 |
| AT5G60320 | 0,25 | 0,38 | 4,33 | 0,09 | 9,31E-01 |
| AT5G60360 | 2,97 | 0,22 | 1,29 | 0,17 | 8,66E-01 |
| AT5G60410 | 2,90 | -0,33 | 1,00 | -0,33 | 7,38E-01 |
| AT5G60550 | 0,49 | 0,92 | 3,16 | 0,29 | 7,72E-01 |
| AT5G60560 | 0,60 | 1,33 | 3,09 | 0,43 | 6,66E-01 |
| AT5G60570 | 0,88 | -2,93 | 2,09 | -1,40 | 1,61E-01 |
| AT5G60600 | 2,43 | 0,93 | 1,21 | 0,77 | 4,43E-01 |
| AT5G60610 | 0,72 | -1,83 | 3,49 | -0,52 | 6,01E-01 |
| AT5G60620 | 0,40 | 0,29 | 3,47 | 0,08 | 9,33E-01 |
| AT5G60680 | 1,59 | 0,67 | 1,65 | 0,41 | 6,83E-01 |

| | | | | | |
|---|---|---|---|---|---|
| AT5G60780 | 1,02 | -1,61 | 1,95 | -0,82 | 4,11E-01 |
| AT5G60820 | 0,45 | -0,02 | 4,08 | 0,00 | 9,96E-01 |
| AT5G60850 | 1,63 | -2,17 | 1,63 | -1,33 | 1,84E-01 |
| AT5G60890 | 0,24 | -2,36 | 4,39 | -0,54 | 5,91E-01 |
| AT5G60945 | 1,24 | -1,55 | 1,73 | -0,89 | 3,72E-01 |
| AT5G60963 | 0,50 | 0,86 | 2,64 | 0,33 | 7,44E-01 |
| AT5G60964 | 0,93 | -1,26 | 3,41 | -0,37 | 7,11E-01 |
| AT5G60980 | 1,13 | -1,21 | 2,05 | -0,59 | 5,56E-01 |
| AT5G61010 | 3,05 | -0,55 | 0,96 | -0,57 | 5,69E-01 |
| AT5G61140 | 4,17 | 0,27 | 0,83 | 0,32 | 7,46E-01 |
| AT5G61190 | 0,65 | 0,18 | 2,40 | 0,08 | 9,40E-01 |
| AT5G61200 | 0,25 | -2,19 | 4,38 | -0,50 | 6,17E-01 |
| AT5G61210 | 1,18 | 1,18 | 1,77 | 0,67 | 5,05E-01 |
| AT5G61230 | 1,00 | 2,30 | 2,12 | 1,08 | 2,79E-01 |
| AT5G61240 | 0,29 | 0,55 | 4,35 | 0,13 | 8,99E-01 |
| AT5G61380 | 0,78 | 0,86 | 2,36 | 0,36 | 7,17E-01 |
| AT5G61400 | 0,78 | -1,02 | 3,43 | -0,30 | 7,65E-01 |
| AT5G61440 | 4,44 | -0,69 | 0,95 | -0,73 | 4,62E-01 |
| AT5G61470 | 0,39 | 0,81 | 3,36 | 0,24 | 8,10E-01 |
| AT5G61490 | 0,17 | 1,52 | 4,33 | 0,35 | 7,26E-01 |
| AT5G61550 | 0,85 | 1,49 | 3,26 | 0,46 | 6,47E-01 |
| AT5G61570 | 0,19 | 0,00 | 4,41 | 0,00 | 1,00E+00 |
| AT5G61580 | 0,26 | 0,01 | 4,33 | 0,00 | 9,98E-01 |
| AT5G61780 | 4,25 | -0,51 | 0,88 | -0,58 | 5,65E-01 |
| AT5G61950 | 0,32 | -0,98 | 4,35 | -0,23 | 8,22E-01 |
| AT5G61970 | 0,49 | 0,04 | 3,42 | 0,01 | 9,91E-01 |
| AT5G61980 | 0,55 | -1,70 | 4,31 | -0,40 | 6,93E-01 |
| AT5G62030 | 0,41 | -0,25 | 4,14 | -0,06 | 9,51E-01 |
| AT5G62270 | 0,20 | 0,81 | 4,36 | 0,19 | 8,52E-01 |
| AT5G62290 | 0,13 | 0,00 | 4,41 | 0,00 | 1,00E+00 |
| AT5G62350 | 2,26 | -1,72 | 1,75 | -0,98 | 3,27E-01 |
| AT5G62360 | 0,27 | -0,98 | 4,37 | -0,22 | 8,23E-01 |
| AT5G09065 | 4,44 | 0,11 | 0,89 | 0,13 | 8,99E-01 |
| AT5G62460 | 0,44 | -0,01 | 4,11 | 0,00 | 9,99E-01 |
| AT5G62500 | 0,05 | 0,00 | 4,41 | 0,00 | 1,00E+00 |
| AT5G62520 | 0,28 | 1,76 | 4,33 | 0,41 | 6,84E-01 |
| AT5G62560 | 0,37 | -0,62 | 3,52 | -0,18 | 8,61E-01 |
| AT5G62570 | 0,48 | 0,48 | 2,65 | 0,18 | 8,55E-01 |
| AT5G62575 | 0,21 | 1,71 | 4,33 | 0,39 | 6,93E-01 |
| AT5G62620 | 0,35 | -0,67 | 4,33 | -0,15 | 8,78E-01 |
| AT5G62670 | 1,85 | -0,52 | 1,59 | -0,33 | 7,42E-01 |
| AT5G62690 | 4,26 | 0,15 | 0,93 | 0,16 | 8,69E-01 |
| AT5G62700 | 1,86 | -3,73 | 1,66 | -2,25 | 2,42E-02 |
| AT5G62710 | 0,23 | -0,48 | 4,35 | -0,11 | 9,12E-01 |
| AT5G62720 | 0,31 | 1,63 | 4,33 | 0,38 | 7,07E-01 |
| AT5G62740 | 0,43 | 2,05 | 4,07 | 0,50 | 6,15E-01 |
| AT5G62770 | 0,31 | 0,04 | 4,33 | 0,01 | 9,93E-01 |
| AT5G62890 | 1,11 | 1,62 | 2,01 | 0,80 | 4,22E-01 |
| AT5G62910 | 0,36 | 0,68 | 4,32 | 0,16 | 8,75E-01 |
| AT5G62970 | 0,35 | 0,91 | 3,57 | 0,26 | 7,98E-01 |
| AT5G63010 | 0,41 | 0,57 | 3,30 | 0,17 | 8,62E-01 |
| AT5G09120 | 1,28 | 0,81 | 2,08 | 0,39 | 6,95E-01 |

| | | | | | |
|---|---|---|---|---|---|
| AT5G63050 | 0,55 | -1,99 | 3,98 | -0,50 | 6,18E-01 |
| AT5G63130 | 0,64 | 1,19 | 2,97 | 0,40 | 6,88E-01 |
| AT5G63145 | 2,83 | 0,01 | 1,24 | 0,01 | 9,94E-01 |
| AT5G63190 | 2,74 | -0,25 | 1,18 | -0,21 | 8,32E-01 |
| AT5G63200 | 0,75 | -3,19 | 3,54 | -0,90 | 3,68E-01 |
| AT5G63260 | 0,08 | 0,69 | 4,36 | 0,16 | 8,75E-01 |
| AT5G63280 | 0,12 | 1,16 | 4,34 | 0,27 | 7,89E-01 |
| AT5G09150 | 0,24 | -2,36 | 4,39 | -0,54 | 5,91E-01 |
| AT5G63300 | 0,47 | -0,35 | 4,33 | -0,08 | 9,36E-01 |
| AT5G63340 | 0,26 | 0,03 | 4,39 | 0,01 | 9,94E-01 |
| AT5G63420 | 0,69 | -2,03 | 3,05 | -0,67 | 5,05E-01 |
| AT5G63480 | 0,12 | 0,00 | 4,41 | 0,00 | 1,00E+00 |
| AT5G63510 | 0,37 | 0,65 | 4,28 | 0,15 | 8,80E-01 |
| AT5G63570 | 1,81 | -1,05 | 1,58 | -0,67 | 5,04E-01 |
| AT5G63640 | 0,36 | -0,01 | 4,35 | 0,00 | 9,98E-01 |
| AT5G63680 | 0,63 | -0,04 | 3,10 | -0,01 | 9,90E-01 |
| AT5G63770 | 0,64 | 0,48 | 2,99 | 0,16 | 8,72E-01 |
| AT5G63780 | 0,69 | -1,71 | 2,31 | -0,74 | 4,59E-01 |
| AT5G63800 | 1,87 | 1,54 | 1,62 | 0,95 | 3,40E-01 |
| AT5G63810 | 0,68 | -1,23 | 3,03 | -0,41 | 6,84E-01 |
| AT5G63840 | 2,51 | -0,86 | 1,13 | -0,76 | 4,47E-01 |
| AT5G63940 | 0,43 | -0,85 | 3,38 | -0,25 | 8,01E-01 |
| AT5G63960 | 0,45 | -3,38 | 4,32 | -0,78 | 4,34E-01 |
| AT5G63980 | 0,08 | 0,69 | 4,36 | 0,16 | 8,75E-01 |
| AT5G64030 | 1,58 | -0,48 | 1,34 | -0,36 | 7,19E-01 |
| AT5G64090 | 0,72 | 0,29 | 2,37 | 0,12 | 9,02E-01 |
| AT5G64160 | 0,20 | 0,40 | 4,37 | 0,09 | 9,28E-01 |
| AT5G64190 | 0,08 | 0,69 | 4,36 | 0,16 | 8,75E-01 |
| AT5G64220 | 2,50 | -2,09 | 1,30 | -1,61 | 1,07E-01 |
| AT5G64240 | 0,28 | 1,27 | 4,35 | 0,29 | 7,69E-01 |
| AT5G64260 | 2,41 | 2,83 | 1,60 | 1,77 | 7,63E-02 |
| AT5G64300 | 1,43 | 1,36 | 1,75 | 0,78 | 4,35E-01 |
| AT5G64310 | 0,80 | 1,12 | 2,70 | 0,42 | 6,78E-01 |
| AT5G64330 | 1,11 | -1,94 | 1,93 | -1,01 | 3,14E-01 |
| AT5G64370 | 0,51 | 0,66 | 3,91 | 0,17 | 8,66E-01 |
| AT5G64380 | 0,10 | 0,01 | 4,38 | 0,00 | 9,99E-01 |
| AT5G64400 | 0,52 | 1,91 | 4,15 | 0,46 | 6,45E-01 |
| AT5G64420 | 4,10 | -0,22 | 0,87 | -0,25 | 8,04E-01 |
| AT5G64440 | 0,42 | 0,00 | 3,39 | 0,00 | 1,00E+00 |
| AT5G64500 | 0,82 | -2,17 | 2,08 | -1,04 | 2,98E-01 |
| AT5G64505 | 0,26 | -0,30 | 4,37 | -0,07 | 9,46E-01 |
| AT5G64572 | 1,22 | -1,07 | 1,65 | -0,65 | 5,15E-01 |
| AT5G64690 | 0,11 | 0,00 | 4,41 | 0,00 | 1,00E+00 |
| AT5G64740 | 4,40 | -1,00 | 0,81 | -1,23 | 2,19E-01 |
| AT5G64750 | 0,12 | 1,23 | 4,34 | 0,28 | 7,76E-01 |
| AT5G64813 | 0,60 | -0,58 | 2,46 | -0,24 | 8,13E-01 |
| AT5G09285 | 0,45 | -0,01 | 3,94 | 0,00 | 9,99E-01 |
| AT5G64940 | 2,50 | -1,67 | 1,35 | -1,24 | 2,16E-01 |
| AT5G64950 | 0,48 | 2,19 | 4,32 | 0,51 | 6,12E-01 |
| AT5G64960 | 0,59 | 1,25 | 3,12 | 0,40 | 6,89E-01 |
| AT5G65005 | 2,56 | -1,29 | 1,21 | -1,06 | 2,87E-01 |
| AT5G65010 | 1,00 | -1,23 | 1,67 | -0,74 | 4,60E-01 |

| | | | | | |
|---|---|---|---|---|---|
| AT5G65015 | 0,39 | 1,69 | 4,33 | 0,39 | 6,96E-01 |
| AT5G65210 | 0,26 | -0,30 | 4,37 | -0,07 | 9,46E-01 |
| AT5G65220 | 1,17 | -0,10 | 2,26 | -0,04 | 9,64E-01 |
| AT5G65230 | 0,08 | 0,44 | 4,38 | 0,10 | 9,21E-01 |
| AT5G65270 | 0,10 | 0,01 | 4,38 | 0,00 | 9,99E-01 |
| AT5G65290 | 0,47 | -2,88 | 4,35 | -0,66 | 5,09E-01 |
| AT5G65400 | 0,17 | 0,65 | 4,36 | 0,15 | 8,81E-01 |
| AT5G65440 | 0,58 | -2,02 | 3,00 | -0,68 | 5,00E-01 |
| AT5G65470 | 0,30 | 1,49 | 4,32 | 0,34 | 7,30E-01 |
| AT5G65480 | 2,00 | 0,45 | 1,30 | 0,34 | 7,31E-01 |
| AT5G65510 | 0,12 | 0,00 | 4,38 | 0,00 | 1,00E+00 |
| AT5G65570 | 0,25 | 0,26 | 4,33 | 0,06 | 9,51E-01 |
| AT5G65610 | 2,72 | 0,87 | 1,75 | 0,50 | 6,17E-01 |
| AT5G65620 | 1,40 | -2,39 | 1,60 | -1,49 | 1,35E-01 |
| AT5G65640 | 0,08 | 0,69 | 4,36 | 0,16 | 8,75E-01 |
| AT5G65670 | 0,91 | 0,39 | 2,14 | 0,18 | 8,56E-01 |
| AT5G65700 | 1,88 | -1,11 | 1,33 | -0,83 | 4,06E-01 |
| AT5G65710 | 0,77 | 1,02 | 2,16 | 0,47 | 6,36E-01 |
| AT5G65720 | 1,65 | 1,58 | 1,48 | 1,07 | 2,85E-01 |
| AT5G65750 | 4,51 | -0,74 | 0,90 | -0,83 | 4,09E-01 |
| AT5G65760 | 0,55 | 0,92 | 2,62 | 0,35 | 7,26E-01 |
| AT5G65770 | 1,30 | -1,33 | 1,89 | -0,71 | 4,81E-01 |
| AT5G65780 | 1,41 | -1,14 | 1,84 | -0,62 | 5,33E-01 |
| AT5G65890 | 0,05 | 0,00 | 4,41 | 0,00 | 1,00E+00 |
| AT5G65960 | 1,37 | 2,00 | 1,82 | 1,10 | 2,72E-01 |
| AT5G65990 | 0,70 | 1,98 | 2,07 | 0,96 | 3,39E-01 |
| AT5G66052 | 0,53 | -1,88 | 3,94 | -0,48 | 6,33E-01 |
| AT5G66070 | 0,10 | 0,01 | 4,38 | 0,00 | 9,99E-01 |
| AT5G66100 | 0,24 | 1,30 | 4,34 | 0,30 | 7,64E-01 |
| AT5G66110 | 0,30 | 0,07 | 4,35 | 0,02 | 9,87E-01 |
| AT5G66160 | 0,31 | -0,89 | 4,40 | -0,20 | 8,39E-01 |
| AT5G66170 | 0,33 | -0,32 | 4,32 | -0,07 | 9,41E-01 |
| AT5G66240 | 1,02 | 0,14 | 2,13 | 0,07 | 9,48E-01 |
| AT5G66360 | 0,60 | 0,37 | 3,76 | 0,10 | 9,21E-01 |
| AT5G66450 | 0,53 | -1,76 | 3,16 | -0,56 | 5,77E-01 |
| AT5G66560 | 1,38 | -0,67 | 1,51 | -0,44 | 6,58E-01 |
| AT5G66562 | 1,20 | 0,25 | 1,62 | 0,15 | 8,78E-01 |
| AT5G66564 | 1,37 | 1,00 | 1,81 | 0,56 | 5,79E-01 |
| AT5G66568 | 0,20 | 0,02 | 4,39 | 0,00 | 9,97E-01 |
| AT5G66760 | 1,08 | -1,04 | 1,68 | -0,62 | 5,37E-01 |
| AT5G66770 | 0,43 | 2,01 | 4,32 | 0,47 | 6,42E-01 |
| AT5G66790 | 0,74 | -1,06 | 2,11 | -0,50 | 6,16E-01 |
| AT5G66810 | 0,67 | -0,66 | 2,34 | -0,28 | 7,79E-01 |
| AT5G66930 | 0,12 | 0,37 | 4,37 | 0,08 | 9,33E-01 |
| AT5G66950 | 1,10 | 0,62 | 1,85 | 0,34 | 7,36E-01 |
| AT5G67070 | 0,51 | -1,86 | 3,97 | -0,47 | 6,41E-01 |
| AT5G67080 | 0,30 | -0,50 | 4,38 | -0,11 | 9,09E-01 |
| AT5G67170 | 0,08 | 0,44 | 4,38 | 0,10 | 9,21E-01 |
| AT5G67260 | 0,65 | -0,68 | 3,91 | -0,18 | 8,61E-01 |
| AT5G67300 | 3,74 | 1,08 | 1,01 | 1,06 | 2,88E-01 |
| AT5G67320 | 0,96 | 2,95 | 2,57 | 1,15 | 2,51E-01 |
| AT5G67330 | 1,26 | -0,22 | 1,62 | -0,14 | 8,90E-01 |

| | | | | | |
|---|---|---|---|---|---|
| AT5G67340 | 0,24 | 1,54 | 4,34 | 0,35 | 7,23E-01 |
| AT5G67385 | 0,80 | 0,79 | 2,31 | 0,34 | 7,31E-01 |
| AT5G67455 | 0,10 | 0,01 | 4,38 | 0,00 | 9,99E-01 |
| AT5G67470 | 1,09 | 0,23 | 1,90 | 0,12 | 9,04E-01 |
| AT5G67488 | 1,20 | 1,06 | 1,83 | 0,58 | 5,62E-01 |
| AT5G67500 | 0,21 | 1,35 | 4,34 | 0,31 | 7,56E-01 |
| AT5G67530 | 0,08 | 0,44 | 4,38 | 0,10 | 9,21E-01 |
| AT5G67560 | 0,70 | 1,82 | 3,67 | 0,49 | 6,21E-01 |
| AT5G67570 | 0,48 | -1,60 | 4,11 | -0,39 | 6,97E-01 |
| AT5G67590 | 0,32 | -1,72 | 4,40 | -0,39 | 6,95E-01 |
| AT1G01020 | 0,15 | 0,00 | 4,38 | 0,00 | 1,00E+00 |
| AT1G03993 | 0,11 | 0,65 | 4,36 | 0,15 | 8,81E-01 |
| AT1G01050 | 0,66 | -1,69 | 2,79 | -0,61 | 5,43E-01 |
| AT1G01060 | 0,69 | -0,57 | 2,29 | -0,25 | 8,03E-01 |
| AT1G01090 | 1,61 | -1,16 | 1,37 | -0,85 | 3,98E-01 |
| AT1G01100 | 1,47 | -1,29 | 1,38 | -0,93 | 3,52E-01 |
| AT1G01120 | 1,43 | 1,05 | 1,71 | 0,62 | 5,37E-01 |
| AT1G01130 | 0,29 | -1,27 | 4,35 | -0,29 | 7,70E-01 |
| AT1G01140 | 0,54 | 0,25 | 2,58 | 0,10 | 9,24E-01 |
| AT1G01170 | 0,58 | 1,00 | 3,10 | 0,32 | 7,47E-01 |
| AT1G01440 | 0,35 | 0,15 | 4,34 | 0,04 | 9,72E-01 |
| AT1G01450 | 0,59 | 1,99 | 4,32 | 0,46 | 6,44E-01 |
| AT1G01470 | 1,29 | -0,07 | 1,67 | -0,04 | 9,67E-01 |
| AT1G01490 | 0,15 | 0,00 | 4,41 | 0,00 | 1,00E+00 |
| AT1G01530 | 0,08 | 0,00 | 4,41 | 0,00 | 1,00E+00 |
| AT1G01610 | 0,73 | 0,03 | 2,59 | 0,01 | 9,90E-01 |
| AT1G01620 | 1,23 | -1,02 | 2,15 | -0,47 | 6,36E-01 |
| AT1G01650 | 0,59 | 2,37 | 2,90 | 0,82 | 4,13E-01 |
| AT1G04027 | 0,46 | 1,94 | 3,12 | 0,62 | 5,34E-01 |
| AT1G01680 | 0,72 | -0,74 | 2,49 | -0,29 | 7,68E-01 |
| AT1G01725 | 1,09 | 2,93 | 1,99 | 1,47 | 1,41E-01 |
| AT1G04063 | 0,34 | 0,00 | 4,39 | 0,00 | 1,00E+00 |
| AT1G01820 | 0,86 | -2,10 | 2,18 | -0,96 | 3,37E-01 |
| AT1G01830 | 1,22 | 2,22 | 1,98 | 1,12 | 2,63E-01 |
| AT1G01880 | 1,08 | 0,78 | 2,37 | 0,33 | 7,43E-01 |
| AT1G01890 | 1,04 | 0,02 | 2,77 | 0,01 | 9,95E-01 |
| AT1G01930 | 0,16 | 0,49 | 4,38 | 0,11 | 9,12E-01 |
| AT1G01960 | 2,74 | 1,02 | 1,05 | 0,97 | 3,32E-01 |
| AT1G01990 | 0,21 | 1,55 | 4,33 | 0,36 | 7,20E-01 |
| AT1G02020 | 0,45 | -1,34 | 4,01 | -0,33 | 7,38E-01 |
| AT1G02090 | 0,84 | 1,02 | 2,70 | 0,38 | 7,07E-01 |
| AT1G02140 | 0,21 | 0,68 | 4,35 | 0,16 | 8,76E-01 |
| AT1G02260 | 0,54 | -1,54 | 3,10 | -0,50 | 6,20E-01 |
| AT1G02280 | 0,61 | 2,20 | 3,01 | 0,73 | 4,66E-01 |
| AT1G02310 | 0,21 | 0,01 | 4,34 | 0,00 | 9,98E-01 |
| AT1G04117 | 0,17 | 1,33 | 4,33 | 0,31 | 7,59E-01 |
| AT1G02430 | 0,17 | 1,15 | 4,34 | 0,27 | 7,91E-01 |
| AT1G02450 | 0,20 | 1,26 | 4,34 | 0,29 | 7,72E-01 |
| AT1G02610 | 0,33 | -1,63 | 4,38 | -0,37 | 7,09E-01 |
| AT1G02660 | 0,36 | -2,06 | 4,33 | -0,48 | 6,33E-01 |
| AT1G02770 | 0,41 | -3,10 | 4,34 | -0,72 | 4,75E-01 |
| AT1G02780 | 1,83 | -1,34 | 1,37 | -0,98 | 3,26E-01 |

| | | | | | |
|---|---|---|---|---|---|
| AT1G02870 | 0,35 | -1,51 | 4,36 | -0,35 | 7,28E-01 |
| AT1G02890 | 1,79 | 0,80 | 1,36 | 0,59 | 5,56E-01 |
| AT1G02910 | 0,70 | -1,99 | 2,34 | -0,85 | 3,95E-01 |
| AT1G02920 | 0,41 | 0,06 | 3,43 | 0,02 | 9,85E-01 |
| AT1G02930 | 0,49 | -1,77 | 3,24 | -0,55 | 5,85E-01 |
| AT1G02960 | 0,08 | 0,69 | 4,36 | 0,16 | 8,75E-01 |
| AT1G03000 | 0,70 | 1,43 | 2,64 | 0,54 | 5,88E-01 |
| AT1G03040 | 0,62 | -0,85 | 2,65 | -0,32 | 7,48E-01 |
| AT1G03080 | 1,73 | 1,19 | 1,42 | 0,84 | 4,01E-01 |
| AT1G03130 | 1,99 | -0,57 | 1,40 | -0,41 | 6,84E-01 |
| AT1G03140 | 0,90 | 0,39 | 1,84 | 0,21 | 8,31E-01 |
| AT1G03240 | 1,51 | -2,38 | 2,38 | -1,00 | 3,17E-01 |
| AT1G03300 | 0,12 | 1,23 | 4,34 | 0,28 | 7,76E-01 |
| AT1G03400 | 0,49 | -0,81 | 2,67 | -0,30 | 7,61E-01 |
| AT1G03410 | 2,63 | 0,75 | 1,38 | 0,54 | 5,87E-01 |
| AT1G03445 | 0,28 | 1,81 | 4,33 | 0,42 | 6,76E-01 |
| AT1G03475 | 0,69 | -0,87 | 2,34 | -0,37 | 7,11E-01 |
| AT1G03520 | 0,48 | 0,05 | 3,98 | 0,01 | 9,91E-01 |
| AT1G03530 | 0,41 | -0,73 | 4,31 | -0,17 | 8,65E-01 |
| AT1G03550 | 0,70 | -2,13 | 2,29 | -0,93 | 3,52E-01 |
| AT1G03590 | 0,41 | -1,28 | 4,19 | -0,31 | 7,60E-01 |
| AT1G03660 | 0,52 | -0,41 | 2,64 | -0,16 | 8,76E-01 |
| AT1G03680 | 1,73 | -1,18 | 1,76 | -0,67 | 5,05E-01 |
| AT1G03730 | 1,35 | 2,45 | 2,51 | 0,98 | 3,29E-01 |
| AT1G03743 | 2,10 | 0,04 | 1,50 | 0,02 | 9,80E-01 |
| AT1G03746 | 0,46 | -1,67 | 4,14 | -0,40 | 6,86E-01 |
| AT1G03760 | 0,08 | 0,00 | 4,40 | 0,00 | 1,00E+00 |
| AT1G03830 | 0,65 | -0,13 | 2,46 | -0,05 | 9,57E-01 |
| AT1G03870 | 0,58 | -1,03 | 3,15 | -0,33 | 7,45E-01 |
| AT1G03940 | 1,20 | 1,09 | 1,88 | 0,58 | 5,62E-01 |
| AT1G04233 | 0,08 | 0,44 | 4,38 | 0,10 | 9,21E-01 |
| AT1G03980 | 0,11 | 0,00 | 4,41 | 0,00 | 1,00E+00 |
| AT1G04000 | 0,55 | 0,15 | 3,22 | 0,05 | 9,64E-01 |
| AT1G04010 | 0,05 | 0,00 | 4,41 | 0,00 | 1,00E+00 |
| AT1G04050 | 0,08 | 0,00 | 4,40 | 0,00 | 1,00E+00 |
| AT1G04070 | 0,16 | 0,65 | 4,37 | 0,15 | 8,82E-01 |
| AT1G04110 | 0,35 | -2,17 | 4,38 | -0,49 | 6,21E-01 |
| AT1G04120 | 1,63 | 1,27 | 1,47 | 0,86 | 3,88E-01 |
| AT1G04140 | 1,54 | -0,36 | 1,40 | -0,26 | 7,96E-01 |
| AT1G04190 | 0,13 | 0,00 | 4,41 | 0,00 | 1,00E+00 |
| AT1G04220 | 0,35 | -0,88 | 4,39 | -0,20 | 8,42E-01 |
| AT1G04240 | 0,91 | -1,75 | 2,47 | -0,71 | 4,77E-01 |
| AT1G04260 | 0,47 | 0,77 | 2,69 | 0,29 | 7,75E-01 |
| AT1G04270 | 1,04 | 0,10 | 2,24 | 0,05 | 9,63E-01 |
| AT1G04280 | 0,49 | 0,64 | 3,34 | 0,19 | 8,48E-01 |
| AT1G04290 | 0,43 | 0,64 | 2,74 | 0,24 | 8,14E-01 |
| AT1G04300 | 0,83 | -1,29 | 2,88 | -0,45 | 6,54E-01 |
| AT1G04380 | 0,20 | -0,44 | 4,36 | -0,10 | 9,19E-01 |
| AT1G04390 | 0,19 | 1,15 | 4,34 | 0,27 | 7,91E-01 |
| AT1G04400 | 0,89 | -3,20 | 2,35 | -1,36 | 1,73E-01 |
| AT1G04410 | 3,62 | 0,24 | 1,20 | 0,20 | 8,44E-01 |
| AT1G04263 | 1,35 | 1,76 | 2,36 | 0,75 | 4,55E-01 |

| | | | | | |
|---|---|---|---|---|---|
| AT1G04530 | 0,42 | 0,48 | 3,27 | 0,15 | 8,82E-01 |
| AT1G04570 | 0,11 | 0,48 | 4,37 | 0,11 | 9,12E-01 |
| AT1G04625 | 1,86 | -0,47 | 1,66 | -0,28 | 7,77E-01 |
| AT1G04630 | 0,66 | 0,21 | 3,67 | 0,06 | 9,54E-01 |
| AT1G04680 | 0,74 | 0,79 | 3,13 | 0,25 | 8,00E-01 |
| AT1G04730 | 0,74 | -0,80 | 2,34 | -0,34 | 7,34E-01 |
| AT1G04750 | 0,67 | -1,64 | 2,31 | -0,71 | 4,77E-01 |
| AT1G04770 | 0,89 | -0,15 | 2,24 | -0,07 | 9,47E-01 |
| AT1G04780 | 0,86 | 0,23 | 2,03 | 0,11 | 9,11E-01 |
| AT1G04820 | 2,61 | -2,78 | 1,27 | -2,19 | 2,87E-02 |
| AT1G04840 | 0,63 | 0,47 | 3,65 | 0,13 | 8,97E-01 |
| AT1G04850 | 0,86 | 0,08 | 2,69 | 0,03 | 9,77E-01 |
| AT1G04860 | 0,57 | -0,80 | 3,87 | -0,21 | 8,36E-01 |
| AT1G04870 | 0,84 | -1,12 | 2,98 | -0,38 | 7,06E-01 |
| AT1G04880 | 0,21 | 0,00 | 4,41 | 0,00 | 1,00E+00 |
| AT1G04910 | 0,78 | 1,31 | 2,75 | 0,47 | 6,35E-01 |
| AT1G04950 | 0,50 | -0,81 | 3,15 | -0,26 | 7,98E-01 |
| AT1G04960 | 0,36 | -2,50 | 4,37 | -0,57 | 5,67E-01 |
| AT1G04980 | 0,36 | -2,77 | 4,36 | -0,64 | 5,25E-01 |
| AT1G04985 | 0,26 | -0,61 | 4,36 | -0,14 | 8,89E-01 |
| AT1G04990 | 0,84 | 0,32 | 2,31 | 0,14 | 8,88E-01 |
| AT1G05010 | 2,06 | -1,05 | 1,70 | -0,62 | 5,38E-01 |
| AT1G05030 | 1,78 | 1,13 | 1,87 | 0,60 | 5,45E-01 |
| AT1G05120 | 0,53 | -0,35 | 4,30 | -0,08 | 9,36E-01 |
| AT1G05150 | 0,85 | -1,17 | 2,01 | -0,58 | 5,61E-01 |
| AT1G05160 | 0,59 | 1,99 | 4,32 | 0,46 | 6,44E-01 |
| AT1G05170 | 0,47 | -2,25 | 4,39 | -0,51 | 6,08E-01 |
| AT1G05190 | 0,78 | -1,43 | 2,39 | -0,60 | 5,49E-01 |
| AT1G05205 | 0,08 | 0,69 | 4,36 | 0,16 | 8,75E-01 |
| AT1G05230 | 0,45 | -1,34 | 4,01 | -0,33 | 7,38E-01 |
| AT1G05300 | 0,18 | 0,56 | 4,36 | 0,13 | 8,97E-01 |
| AT1G05350 | 0,28 | 0,23 | 4,33 | 0,05 | 9,58E-01 |
| AT1G05520 | 0,61 | -0,71 | 2,95 | -0,24 | 8,08E-01 |
| AT1G04393 | 5,00 | 0,73 | 1,02 | 0,72 | 4,73E-01 |
| AT1G05575 | 0,16 | 0,09 | 4,38 | 0,02 | 9,83E-01 |
| AT1G05640 | 0,20 | -1,80 | 4,41 | -0,41 | 6,83E-01 |
| AT1G05670 | 0,15 | 1,01 | 4,34 | 0,23 | 8,17E-01 |
| AT1G05720 | 0,13 | 0,00 | 4,41 | 0,00 | 1,00E+00 |
| AT1G05783 | 0,96 | 1,50 | 1,94 | 0,77 | 4,42E-01 |
| AT1G05840 | 0,75 | -1,30 | 2,05 | -0,64 | 5,25E-01 |
| AT1G05850 | 2,93 | 0,56 | 1,44 | 0,39 | 6,97E-01 |
| AT1G05870 | 0,42 | -1,35 | 4,18 | -0,32 | 7,46E-01 |
| AT1G05950 | 0,12 | 1,23 | 4,34 | 0,28 | 7,76E-01 |
| AT1G05960 | 4,07 | -1,25 | 1,03 | -1,22 | 2,24E-01 |
| AT1G05970 | 0,08 | 0,44 | 4,38 | 0,10 | 9,21E-01 |
| AT1G06000 | 0,95 | -1,31 | 1,96 | -0,67 | 5,05E-01 |
| AT1G06040 | 2,85 | -0,25 | 1,31 | -0,19 | 8,50E-01 |
| AT1G06130 | 0,62 | -1,69 | 2,95 | -0,57 | 5,67E-01 |
| AT1G06150 | 1,38 | -0,11 | 1,63 | -0,07 | 9,47E-01 |
| AT1G06148 | 2,19 | 1,89 | 1,66 | 1,13 | 2,57E-01 |
| AT1G06190 | 5,18 | 0,67 | 0,83 | 0,81 | 4,18E-01 |
| AT1G06250 | 0,72 | -3,93 | 4,30 | -0,91 | 3,60E-01 |

| | | | | | |
|---|---|---|---|---|---|
| AT1G06265 | 0,08 | 0,69 | 4,36 | 0,16 | 8,75E-01 |
| AT1G06360 | 0,33 | 0,00 | 4,41 | 0,00 | 1,00E+00 |
| AT1G06400 | 0,21 | 0,81 | 4,34 | 0,19 | 8,51E-01 |
| AT1G06430 | 1,82 | 0,90 | 1,57 | 0,57 | 5,68E-01 |
| AT1G06440 | 0,58 | -0,25 | 2,63 | -0,10 | 9,24E-01 |
| AT1G06460 | 1,79 | -2,29 | 1,51 | -1,51 | 1,31E-01 |
| AT1G06540 | 1,04 | -1,40 | 1,88 | -0,74 | 4,58E-01 |
| AT1G06570 | 0,97 | 0,12 | 2,41 | 0,05 | 9,62E-01 |
| AT1G06580 | 1,59 | 1,18 | 1,72 | 0,69 | 4,93E-01 |
| AT1G04487 | 0,39 | 0,58 | 4,18 | 0,14 | 8,90E-01 |
| AT1G06660 | 0,50 | -1,51 | 3,91 | -0,39 | 7,00E-01 |
| AT1G06690 | 0,36 | -0,01 | 4,35 | 0,00 | 9,98E-01 |
| AT1G06700 | 1,69 | -0,44 | 1,47 | -0,30 | 7,65E-01 |
| AT1G06710 | 0,26 | -0,48 | 4,38 | -0,11 | 9,13E-01 |
| AT1G06720 | 0,55 | -1,80 | 3,98 | -0,45 | 6,51E-01 |
| AT1G06760 | 1,45 | -2,15 | 1,57 | -1,37 | 1,70E-01 |
| AT1G06810 | 0,08 | 0,44 | 4,38 | 0,10 | 9,21E-01 |
| AT1G06840 | 0,17 | 0,18 | 4,39 | 0,04 | 9,67E-01 |
| AT1G06890 | 0,40 | 0,83 | 2,94 | 0,28 | 7,77E-01 |
| AT1G06900 | 1,35 | 0,73 | 1,56 | 0,47 | 6,37E-01 |
| AT1G06950 | 2,18 | -0,87 | 1,82 | -0,48 | 6,33E-01 |
| AT1G07000 | 3,04 | 0,06 | 1,13 | 0,05 | 9,57E-01 |
| AT1G07020 | 0,55 | -3,18 | 4,34 | -0,73 | 4,63E-01 |
| AT1G07050 | 0,42 | 1,24 | 4,11 | 0,30 | 7,62E-01 |
| AT1G07090 | 0,43 | -0,51 | 4,28 | -0,12 | 9,05E-01 |
| AT1G07110 | 1,24 | -1,24 | 2,11 | -0,59 | 5,57E-01 |
| AT1G07128 | 0,28 | 1,27 | 4,35 | 0,29 | 7,69E-01 |
| AT1G07135 | 2,46 | 0,60 | 1,29 | 0,46 | 6,42E-01 |
| AT1G07140 | 1,81 | -0,76 | 1,50 | -0,50 | 6,16E-01 |
| AT1G07200 | 1,82 | 0,83 | 1,43 | 0,58 | 5,63E-01 |
| AT1G07220 | 0,25 | 0,00 | 4,41 | 0,00 | 1,00E+00 |
| AT1G07260 | 0,48 | -0,90 | 4,32 | -0,21 | 8,34E-01 |
| AT1G04573 | 0,66 | -2,69 | 4,27 | -0,63 | 5,28E-01 |
| AT1G07310 | 0,69 | 0,02 | 2,56 | 0,01 | 9,93E-01 |
| AT1G07350 | 0,45 | -0,63 | 3,22 | -0,20 | 8,44E-01 |
| AT1G07360 | 0,57 | 0,06 | 3,25 | 0,02 | 9,85E-01 |
| AT1G07430 | 0,32 | 0,36 | 4,34 | 0,08 | 9,34E-01 |
| AT1G07440 | 0,75 | 0,44 | 2,46 | 0,18 | 8,59E-01 |
| AT1G07480 | 0,80 | -0,13 | 2,25 | -0,06 | 9,55E-01 |
| AT1G07500 | 0,08 | 0,00 | 4,40 | 0,00 | 1,00E+00 |
| AT1G07520 | 1,20 | 1,13 | 1,97 | 0,57 | 5,65E-01 |
| AT1G07530 | 1,00 | -0,20 | 1,63 | -0,12 | 9,03E-01 |
| AT1G07590 | 3,38 | 3,93 | 1,41 | 2,78 | 5,42E-03 |
| AT1G07600 | 1,87 | 4,18 | 1,63 | 2,57 | 1,01E-02 |
| AT1G07640 | 0,24 | 0,00 | 4,38 | 0,00 | 1,00E+00 |
| AT1G07670 | 1,74 | -1,46 | 1,36 | -1,07 | 2,84E-01 |
| AT1G07720 | 1,26 | 1,92 | 1,82 | 1,06 | 2,90E-01 |
| AT1G07725 | 1,03 | -1,48 | 2,31 | -0,64 | 5,21E-01 |
| AT1G07740 | 0,29 | 0,53 | 4,33 | 0,12 | 9,02E-01 |
| AT1G07730 | 0,29 | 0,53 | 4,33 | 0,12 | 9,02E-01 |
| AT1G07750 | 0,25 | 0,00 | 4,41 | 0,00 | 1,00E+00 |
| AT1G07770 | 0,87 | -0,10 | 1,81 | -0,06 | 9,54E-01 |

| | | | | | |
|---|---|---|---|---|---|
| AT1G07870 | 0,12 | 0,97 | 4,34 | 0,22 | 8,23E-01 |
| AT1G07940 | 3,71 | 0,15 | 0,98 | 0,15 | 8,81E-01 |
| AT1G07970 | 0,62 | -0,13 | 3,55 | -0,04 | 9,71E-01 |
| AT1G08040 | 0,40 | 0,37 | 4,29 | 0,09 | 9,32E-01 |
| AT1G08050 | 0,33 | 0,92 | 4,31 | 0,21 | 8,30E-01 |
| AT1G08060 | 1,18 | 0,43 | 1,96 | 0,22 | 8,28E-01 |
| AT1G08125 | 0,32 | -1,43 | 4,36 | -0,33 | 7,44E-01 |
| AT1G08200 | 1,82 | -0,81 | 1,56 | -0,52 | 6,01E-01 |
| AT1G08450 | 3,29 | 0,36 | 1,18 | 0,31 | 7,58E-01 |
| AT1G08510 | 1,46 | 1,51 | 1,74 | 0,87 | 3,85E-01 |
| AT1G08580 | 0,24 | -2,36 | 4,39 | -0,54 | 5,91E-01 |
| AT1G08592 | 0,74 | 1,84 | 2,84 | 0,65 | 5,16E-01 |
| AT1G08610 | 0,28 | -0,47 | 4,37 | -0,11 | 9,14E-01 |
| AT1G08640 | 0,47 | -1,11 | 3,95 | -0,28 | 7,78E-01 |
| AT1G08645 | 0,64 | -1,08 | 3,74 | -0,29 | 7,73E-01 |
| AT1G08660 | 0,10 | 0,00 | 4,41 | 0,00 | 1,00E+00 |
| AT1G08700 | 1,22 | 2,89 | 2,08 | 1,39 | 1,64E-01 |
| AT1G08800 | 0,74 | 0,92 | 2,85 | 0,32 | 7,48E-01 |
| AT1G08820 | 0,89 | -3,09 | 2,68 | -1,15 | 2,48E-01 |
| AT1G08880 | 0,12 | 1,16 | 4,34 | 0,27 | 7,89E-01 |
| AT1G09010 | 3,02 | -1,99 | 1,06 | -1,88 | 6,07E-02 |
| AT1G09020 | 0,78 | 0,34 | 2,14 | 0,16 | 8,74E-01 |
| AT1G09060 | 1,04 | 0,80 | 2,21 | 0,36 | 7,18E-01 |
| AT1G09080 | 0,40 | -0,07 | 4,31 | -0,02 | 9,87E-01 |
| AT1G09130 | 0,46 | -0,01 | 3,30 | 0,00 | 9,98E-01 |
| AT1G09160 | 0,40 | 0,44 | 3,47 | 0,13 | 9,00E-01 |
| AT1G09190 | 0,17 | 1,52 | 4,33 | 0,35 | 7,26E-01 |
| AT1G09195 | 0,70 | -0,48 | 2,83 | -0,17 | 8,65E-01 |
| AT1G09200 | 0,34 | 0,23 | 4,38 | 0,05 | 9,58E-01 |
| AT1G09230 | 0,23 | -1,78 | 4,41 | -0,40 | 6,87E-01 |
| AT1G09280 | 1,11 | -0,28 | 1,80 | -0,16 | 8,76E-01 |
| AT1G09420 | 0,88 | 2,46 | 3,33 | 0,74 | 4,61E-01 |
| AT1G09440 | 0,21 | 0,01 | 4,38 | 0,00 | 9,98E-01 |
| AT1G09570 | 2,05 | -2,41 | 1,20 | -2,01 | 4,47E-02 |
| AT1G04707 | 0,36 | -0,29 | 4,32 | -0,07 | 9,46E-01 |
| AT1G09630 | 0,08 | 0,44 | 4,38 | 0,10 | 9,21E-01 |
| AT1G04717 | 0,54 | 0,08 | 3,75 | 0,02 | 9,84E-01 |
| AT1G09680 | 1,19 | 1,86 | 2,44 | 0,76 | 4,46E-01 |
| AT1G09690 | 1,78 | 0,41 | 1,32 | 0,31 | 7,54E-01 |
| AT1G09730 | 0,83 | 3,11 | 2,66 | 1,17 | 2,42E-01 |
| AT1G09760 | 0,17 | 1,15 | 4,34 | 0,27 | 7,91E-01 |
| AT1G09780 | 4,32 | 0,28 | 0,85 | 0,33 | 7,43E-01 |
| AT1G09810 | 0,42 | 0,29 | 2,79 | 0,11 | 9,16E-01 |
| AT1G09880 | 0,33 | -1,30 | 4,33 | -0,30 | 7,64E-01 |
| AT1G09910 | 0,78 | -1,08 | 2,18 | -0,50 | 6,19E-01 |
| AT1G10020 | 0,10 | 0,01 | 4,38 | 0,00 | 9,99E-01 |
| AT1G10090 | 0,57 | 0,38 | 2,61 | 0,15 | 8,84E-01 |
| AT1G10155 | 1,47 | 0,66 | 1,41 | 0,47 | 6,42E-01 |
| AT1G10180 | 0,61 | -2,57 | 2,53 | -1,02 | 3,09E-01 |
| AT1G10190 | 0,94 | 0,00 | 2,88 | 0,00 | 1,00E+00 |
| AT1G10200 | 0,57 | 0,47 | 3,80 | 0,12 | 9,02E-01 |
| AT1G10310 | 0,42 | -0,89 | 4,33 | -0,20 | 8,38E-01 |

| | | | | | |
|---|---|---|---|---|---|
| AT1G10320 | 0,47 | 2,41 | 3,94 | 0,61 | 5,42E-01 |
| AT1G10340 | 1,00 | -2,45 | 2,10 | -1,17 | 2,43E-01 |
| AT1G10350 | 0,52 | 1,20 | 3,86 | 0,31 | 7,57E-01 |
| AT1G10390 | 1,30 | 1,73 | 1,85 | 0,94 | 3,49E-01 |
| AT1G10400 | 0,08 | 0,69 | 4,36 | 0,16 | 8,75E-01 |
| AT1G10410 | 0,96 | 0,81 | 1,99 | 0,41 | 6,83E-01 |
| AT1G10430 | 0,87 | 1,03 | 2,20 | 0,47 | 6,40E-01 |
| AT1G10470 | 1,21 | -1,48 | 1,85 | -0,80 | 4,24E-01 |
| AT1G10500 | 1,39 | 1,40 | 2,30 | 0,61 | 5,43E-01 |
| AT1G10580 | 0,57 | 2,13 | 3,84 | 0,55 | 5,80E-01 |
| AT1G10586 | 1,03 | 1,63 | 2,19 | 0,75 | 4,56E-01 |
| AT1G10590 | 0,69 | -1,29 | 2,93 | -0,44 | 6,60E-01 |
| AT1G10630 | 0,65 | -1,16 | 2,51 | -0,46 | 6,43E-01 |
| AT1G10640 | 0,05 | 0,00 | 4,41 | 0,00 | 1,00E+00 |
| AT1G10680 | 0,08 | 0,44 | 4,38 | 0,10 | 9,21E-01 |
| AT1G10840 | 0,88 | -1,07 | 1,84 | -0,58 | 5,63E-01 |
| AT1G10870 | 1,11 | -1,97 | 1,81 | -1,09 | 2,76E-01 |
| AT1G10900 | 1,03 | -2,85 | 2,74 | -1,04 | 2,98E-01 |
| AT1G10920 | 0,82 | 0,65 | 4,26 | 0,15 | 8,78E-01 |
| AT1G10930 | 0,74 | -0,44 | 2,81 | -0,16 | 8,75E-01 |
| AT1G10940 | 0,48 | -1,26 | 3,26 | -0,39 | 6,99E-01 |
| AT1G04833 | 0,22 | 0,00 | 4,38 | 0,00 | 1,00E+00 |
| AT1G11100 | 0,45 | 1,05 | 3,35 | 0,31 | 7,53E-01 |
| AT1G11210 | 0,56 | 1,17 | 2,56 | 0,46 | 6,48E-01 |
| AT1G11240 | 0,05 | 0,00 | 4,41 | 0,00 | 1,00E+00 |
| AT1G11270 | 0,83 | -0,16 | 2,35 | -0,07 | 9,45E-01 |
| AT1G11280 | 0,45 | -1,70 | 2,78 | -0,61 | 5,42E-01 |
| AT1G11290 | 0,63 | 0,56 | 3,59 | 0,16 | 8,75E-01 |
| AT1G11310 | 1,46 | 0,23 | 1,46 | 0,15 | 8,77E-01 |
| AT1G11340 | 0,08 | 0,69 | 4,36 | 0,16 | 8,75E-01 |
| AT1G11350 | 0,52 | 0,52 | 3,18 | 0,16 | 8,71E-01 |
| AT1G11360 | 0,33 | -0,82 | 4,36 | -0,19 | 8,51E-01 |
| AT1G11480 | 0,70 | -0,58 | 2,13 | -0,27 | 7,87E-01 |
| AT1G11630 | 0,35 | -2,17 | 4,38 | -0,49 | 6,21E-01 |
| AT1G11640 | 3,15 | -1,45 | 1,06 | -1,37 | 1,70E-01 |
| AT1G11785 | 2,48 | -0,12 | 1,30 | -0,10 | 9,24E-01 |
| AT1G11820 | 0,25 | 1,16 | 4,34 | 0,27 | 7,89E-01 |
| AT1G04897 | 2,68 | 0,83 | 1,31 | 0,63 | 5,27E-01 |
| AT1G11910 | 5,12 | 0,42 | 1,05 | 0,40 | 6,92E-01 |
| AT1G12000 | 0,52 | 2,27 | 3,06 | 0,74 | 4,58E-01 |
| AT1G12020 | 0,63 | -0,04 | 2,32 | -0,02 | 9,85E-01 |
| AT1G12120 | 0,70 | -2,17 | 2,56 | -0,85 | 3,97E-01 |
| AT1G12211 | 0,63 | 0,54 | 3,16 | 0,17 | 8,63E-01 |
| AT1G12280 | 1,78 | 0,28 | 1,23 | 0,22 | 8,22E-01 |
| AT1G12300 | 2,68 | 0,25 | 1,64 | 0,15 | 8,79E-01 |
| AT1G12310 | 0,63 | 2,00 | 3,16 | 0,63 | 5,26E-01 |
| AT1G12380 | 0,49 | 1,58 | 4,07 | 0,39 | 6,98E-01 |
| AT1G12440 | 1,21 | -1,49 | 1,66 | -0,90 | 3,69E-01 |
| AT1G12500 | 0,25 | 1,68 | 4,32 | 0,39 | 6,97E-01 |
| AT1G12520 | 0,51 | -1,36 | 4,33 | -0,31 | 7,53E-01 |
| AT1G12590 | 0,78 | -0,20 | 1,97 | -0,10 | 9,20E-01 |
| AT1G12610 | 0,19 | 0,84 | 4,35 | 0,19 | 8,46E-01 |

| | | | | | |
|---|---|---|---|---|---|
| AT1G12615 | 1,59 | 1,97 | 2,13 | 0,92 | 3,55E-01 |
| AT1G12620 | 2,40 | 0,54 | 1,75 | 0,31 | 7,56E-01 |
| AT1G12640 | 0,58 | 0,88 | 4,29 | 0,20 | 8,38E-01 |
| AT1G12680 | 1,43 | 0,67 | 1,97 | 0,34 | 7,34E-01 |
| AT1G12700 | 0,22 | 1,19 | 4,33 | 0,27 | 7,84E-01 |
| AT1G12780 | 0,71 | 1,35 | 2,30 | 0,59 | 5,58E-01 |
| AT1G12800 | 0,97 | -3,01 | 2,40 | -1,25 | 2,11E-01 |
| AT1G12900 | 4,70 | -1,31 | 0,96 | -1,37 | 1,70E-01 |
| AT1G12910 | 1,07 | 1,06 | 1,92 | 0,55 | 5,82E-01 |
| AT1G12920 | 1,87 | -0,82 | 1,28 | -0,64 | 5,20E-01 |
| AT1G13020 | 0,59 | -1,05 | 2,48 | -0,42 | 6,73E-01 |
| AT1G13120 | 0,23 | 1,59 | 4,33 | 0,37 | 7,13E-01 |
| AT1G13170 | 0,46 | -0,55 | 3,18 | -0,17 | 8,62E-01 |
| AT1G13200 | 0,66 | -1,82 | 3,53 | -0,52 | 6,05E-01 |
| AT1G13210 | 5,46 | 0,93 | 0,85 | 1,10 | 2,71E-01 |
| AT1G13245 | 0,44 | -1,71 | 4,13 | -0,41 | 6,79E-01 |
| AT1G13310 | 0,35 | -2,17 | 4,38 | -0,49 | 6,21E-01 |
| AT1G13320 | 2,17 | -0,85 | 1,21 | -0,70 | 4,83E-01 |
| AT1G13340 | 0,95 | 0,26 | 2,14 | 0,12 | 9,05E-01 |
| AT1G13350 | 2,82 | -0,50 | 1,31 | -0,38 | 7,04E-01 |
| AT1G13390 | 0,17 | 1,50 | 4,33 | 0,35 | 7,29E-01 |
| AT1G13440 | 5,54 | -0,55 | 0,84 | -0,65 | 5,17E-01 |
| AT1G13450 | 0,76 | 0,10 | 2,14 | 0,05 | 9,62E-01 |
| AT1G13470 | 1,76 | 0,11 | 1,61 | 0,07 | 9,44E-01 |
| AT1G13520 | 0,54 | -0,87 | 4,35 | -0,20 | 8,40E-01 |
| AT1G13560 | 0,43 | -1,19 | 4,32 | -0,27 | 7,84E-01 |
| AT1G13640 | 2,82 | -0,40 | 1,06 | -0,38 | 7,03E-01 |
| AT1G13825 | 0,76 | 2,82 | 3,53 | 0,80 | 4,24E-01 |
| AT1G13860 | 0,10 | 0,21 | 4,38 | 0,05 | 9,62E-01 |
| AT1G13870 | 0,12 | 0,00 | 4,39 | 0,00 | 1,00E+00 |
| AT1G13900 | 0,21 | 1,35 | 4,34 | 0,31 | 7,56E-01 |
| AT1G13940 | 0,86 | 1,10 | 2,36 | 0,46 | 6,42E-01 |
| AT1G14010 | 0,36 | -0,12 | 4,35 | -0,03 | 9,78E-01 |
| AT1G14030 | 0,62 | 1,09 | 3,06 | 0,36 | 7,21E-01 |
| AT1G14290 | 0,26 | -0,48 | 4,38 | -0,11 | 9,13E-01 |
| AT1G14300 | 0,22 | 0,00 | 4,41 | 0,00 | 1,00E+00 |
| AT1G14330 | 0,59 | 1,10 | 2,62 | 0,42 | 6,73E-01 |
| AT1G14360 | 0,18 | -1,78 | 4,41 | -0,40 | 6,87E-01 |
| AT1G14400 | 0,66 | 1,47 | 3,07 | 0,48 | 6,33E-01 |
| AT1G14450 | 0,11 | 0,00 | 4,41 | 0,00 | 1,00E+00 |
| AT1G14455 | 0,14 | 0,00 | 4,41 | 0,00 | 1,00E+00 |
| AT1G14518 | 0,84 | 0,09 | 2,37 | 0,04 | 9,71E-01 |
| AT1G14590 | 0,42 | -0,70 | 4,13 | -0,17 | 8,66E-01 |
| AT1G14610 | 2,42 | -0,93 | 1,13 | -0,82 | 4,10E-01 |
| AT1G14690 | 0,78 | -1,68 | 2,17 | -0,77 | 4,39E-01 |
| AT1G14710 | 0,51 | 2,21 | 3,84 | 0,57 | 5,65E-01 |
| AT1G14720 | 0,26 | -1,86 | 4,40 | -0,42 | 6,72E-01 |
| AT1G14740 | 3,27 | -0,63 | 0,94 | -0,67 | 5,03E-01 |
| AT1G14790 | 1,42 | 2,41 | 2,10 | 1,14 | 2,53E-01 |
| AT1G14810 | 1,02 | -0,81 | 1,95 | -0,41 | 6,78E-01 |
| AT1G14830 | 0,90 | 1,40 | 2,73 | 0,51 | 6,07E-01 |
| AT1G14840 | 0,51 | 1,19 | 2,40 | 0,50 | 6,20E-01 |

| | | | | | |
|---|---|---|---|---|---|
| AT1G14850 | 1,26 | -0,99 | 1,60 | -0,62 | 5,33E-01 |
| AT1G14880 | 1,19 | 1,69 | 1,95 | 0,86 | 3,88E-01 |
| AT1G14900 | 0,38 | -0,93 | 4,39 | -0,21 | 8,33E-01 |
| AT1G14910 | 0,80 | 1,74 | 2,54 | 0,69 | 4,92E-01 |
| AT1G14980 | 0,57 | -0,01 | 3,00 | 0,00 | 9,96E-01 |
| AT1G15002 | 1,82 | -0,20 | 1,47 | -0,13 | 8,93E-01 |
| AT1G05137 | 0,28 | 1,42 | 4,34 | 0,33 | 7,43E-01 |
| AT1G15020 | 0,98 | 0,46 | 2,46 | 0,19 | 8,51E-01 |
| AT1G15040 | 0,21 | 1,42 | 4,33 | 0,33 | 7,43E-01 |
| AT1G15100 | 1,26 | -1,57 | 2,49 | -0,63 | 5,30E-01 |
| AT1G15130 | 4,06 | 0,81 | 0,90 | 0,90 | 3,66E-01 |
| AT1G15140 | 0,76 | 1,09 | 2,13 | 0,51 | 6,09E-01 |
| AT1G15175 | 0,27 | -0,21 | 4,34 | -0,05 | 9,62E-01 |
| AT1G15210 | 0,17 | 1,52 | 4,33 | 0,35 | 7,26E-01 |
| AT1G15220 | 0,11 | 0,48 | 4,37 | 0,11 | 9,12E-01 |
| AT1G15340 | 0,21 | 0,04 | 4,39 | 0,01 | 9,93E-01 |
| AT1G15350 | 1,25 | -1,65 | 1,95 | -0,85 | 3,98E-01 |
| AT1G15400 | 0,43 | 0,44 | 2,78 | 0,16 | 8,76E-01 |
| AT1G15440 | 0,87 | -1,66 | 2,00 | -0,83 | 4,08E-01 |
| AT1G15520 | 0,46 | 0,25 | 3,26 | 0,08 | 9,40E-01 |
| AT1G15530 | 1,02 | -0,98 | 2,16 | -0,45 | 6,51E-01 |
| AT1G15550 | 0,24 | -2,36 | 4,39 | -0,54 | 5,91E-01 |
| AT1G15700 | 0,40 | -0,49 | 3,30 | -0,15 | 8,81E-01 |
| AT1G05187 | 0,24 | 0,24 | 4,34 | 0,06 | 9,55E-01 |
| AT1G15730 | 0,44 | -1,76 | 4,18 | -0,42 | 6,73E-01 |
| AT1G15780 | 1,01 | 1,20 | 2,07 | 0,58 | 5,62E-01 |
| AT1G15790 | 1,05 | -1,86 | 2,71 | -0,68 | 4,93E-01 |
| AT1G05203 | 0,26 | -0,30 | 4,37 | -0,07 | 9,46E-01 |
| AT1G15880 | 0,20 | -1,80 | 4,41 | -0,41 | 6,83E-01 |
| AT1G15970 | 0,11 | 0,00 | 4,41 | 0,00 | 1,00E+00 |
| AT1G16010 | 0,99 | 2,33 | 2,26 | 1,03 | 3,02E-01 |
| AT1G16070 | 0,40 | 1,78 | 4,33 | 0,41 | 6,81E-01 |
| AT1G16210 | 0,60 | -2,14 | 3,05 | -0,70 | 4,83E-01 |
| AT1G16240 | 0,80 | 2,41 | 2,41 | 1,00 | 3,18E-01 |
| AT1G16260 | 1,63 | -0,97 | 1,47 | -0,66 | 5,08E-01 |
| AT1G16460 | 0,57 | 0,24 | 2,35 | 0,10 | 9,19E-01 |
| AT1G16570 | 0,28 | 1,27 | 4,35 | 0,29 | 7,69E-01 |
| AT1G16610 | 1,02 | 0,00 | 2,97 | 0,00 | 9,99E-01 |
| AT1G16750 | 0,49 | -0,03 | 3,06 | -0,01 | 9,92E-01 |
| AT1G16780 | 0,12 | 1,23 | 4,34 | 0,28 | 7,76E-01 |
| AT1G16800 | 1,44 | 0,84 | 2,19 | 0,38 | 7,03E-01 |
| AT1G16870 | 0,95 | -1,99 | 3,19 | -0,63 | 5,32E-01 |
| AT1G16890 | 1,30 | -1,13 | 1,73 | -0,65 | 5,16E-01 |
| AT1G16900 | 0,65 | 0,98 | 3,65 | 0,27 | 7,88E-01 |
| AT1G16920 | 0,22 | 0,99 | 4,34 | 0,23 | 8,19E-01 |
| AT1G16970 | 0,37 | 0,25 | 3,46 | 0,07 | 9,43E-01 |
| AT1G17080 | 0,60 | -0,17 | 3,24 | -0,05 | 9,57E-01 |
| AT1G17110 | 0,57 | 0,63 | 2,59 | 0,24 | 8,07E-01 |
| AT1G17140 | 0,95 | -1,56 | 3,32 | -0,47 | 6,39E-01 |
| AT1G17147 | 0,12 | 0,00 | 4,41 | 0,00 | 1,00E+00 |
| AT1G17210 | 2,32 | -0,01 | 1,28 | -0,01 | 9,91E-01 |
| AT1G17232 | 0,51 | 1,49 | 3,10 | 0,48 | 6,30E-01 |

| | | | | | |
|---|---|---|---|---|---|
| AT1G17240 | 0,05 | 0,00 | 4,41 | 0,00 | 1,00E+00 |
| AT1G17250 | 0,99 | 2,31 | 2,53 | 0,91 | 3,62E-01 |
| AT1G17370 | 0,81 | 0,88 | 2,68 | 0,33 | 7,42E-01 |
| AT1G17380 | 0,50 | -0,03 | 2,66 | -0,01 | 9,90E-01 |
| AT1G05313 | 0,32 | 0,91 | 4,35 | 0,21 | 8,35E-01 |
| AT1G17440 | 0,46 | 0,36 | 3,32 | 0,11 | 9,14E-01 |
| AT1G17450 | 1,22 | 1,48 | 1,77 | 0,84 | 4,02E-01 |
| AT1G17590 | 0,46 | 0,29 | 3,24 | 0,09 | 9,29E-01 |
| AT1G17600 | 1,00 | 0,04 | 2,05 | 0,02 | 9,86E-01 |
| AT1G17665 | 0,20 | 1,03 | 4,35 | 0,24 | 8,12E-01 |
| AT1G17670 | 0,37 | -0,75 | 4,35 | -0,17 | 8,64E-01 |
| AT1G17690 | 0,25 | 0,26 | 4,33 | 0,06 | 9,51E-01 |
| AT1G17720 | 0,66 | 1,00 | 2,29 | 0,44 | 6,63E-01 |
| AT1G17744 | 0,16 | 0,49 | 4,38 | 0,11 | 9,12E-01 |
| AT1G17790 | 0,63 | -2,51 | 3,06 | -0,82 | 4,13E-01 |
| AT1G05333 | 0,71 | -1,53 | 3,43 | -0,45 | 6,56E-01 |
| AT1G17820 | 0,71 | -1,53 | 3,43 | -0,45 | 6,56E-01 |
| AT1G17850 | 0,36 | -1,49 | 4,30 | -0,35 | 7,30E-01 |
| AT1G05347 | 0,21 | 1,16 | 4,34 | 0,27 | 7,89E-01 |
| AT1G17870 | 0,39 | -0,03 | 4,23 | -0,01 | 9,94E-01 |
| AT1G17880 | 0,59 | -2,21 | 3,81 | -0,58 | 5,62E-01 |
| AT1G17890 | 0,42 | -1,40 | 4,38 | -0,32 | 7,50E-01 |
| AT1G17930 | 0,20 | -0,31 | 4,36 | -0,07 | 9,44E-01 |
| AT1G17980 | 1,06 | 0,34 | 2,35 | 0,14 | 8,85E-01 |
| AT1G17990 | 0,53 | -1,87 | 3,10 | -0,60 | 5,47E-01 |
| AT1G18000 | 4,70 | -1,04 | 1,19 | -0,87 | 3,82E-01 |
| AT1G18050 | 0,60 | 2,13 | 2,51 | 0,85 | 3,95E-01 |
| AT1G18060 | 0,60 | -0,77 | 2,23 | -0,35 | 7,30E-01 |
| AT1G18070 | 0,60 | 0,71 | 2,49 | 0,29 | 7,74E-01 |
| AT1G18100 | 0,36 | 1,83 | 4,33 | 0,42 | 6,73E-01 |
| AT1G18150 | 1,19 | -1,56 | 2,11 | -0,74 | 4,60E-01 |
| AT1G18210 | 0,61 | -0,75 | 2,41 | -0,31 | 7,56E-01 |
| AT1G18260 | 1,96 | 0,10 | 1,34 | 0,07 | 9,43E-01 |
| AT1G18310 | 4,64 | 4,93 | 1,47 | 3,37 | 7,65E-04 |
| AT1G18330 | 0,22 | -0,47 | 4,35 | -0,11 | 9,13E-01 |
| AT1G18360 | 1,25 | -1,77 | 1,66 | -1,06 | 2,87E-01 |
| AT1G18382 | 0,44 | 0,28 | 2,76 | 0,10 | 9,18E-01 |
| AT1G18470 | 0,48 | 1,16 | 2,70 | 0,43 | 6,66E-01 |
| AT1G18540 | 1,15 | 0,27 | 2,02 | 0,13 | 8,95E-01 |
| AT1G18720 | 0,23 | 1,27 | 4,33 | 0,29 | 7,70E-01 |
| AT1G05427 | 2,11 | -0,80 | 1,69 | -0,47 | 6,37E-01 |
| AT1G05433 | 0,53 | 0,80 | 3,10 | 0,26 | 7,95E-01 |
| AT1G18840 | 0,43 | -0,35 | 3,30 | -0,11 | 9,16E-01 |
| AT1G18890 | 0,28 | -0,49 | 4,38 | -0,11 | 9,11E-01 |
| AT1G18990 | 0,45 | -0,45 | 4,12 | -0,11 | 9,13E-01 |
| AT1G19000 | 0,86 | -0,48 | 1,84 | -0,26 | 7,93E-01 |
| AT1G19130 | 0,49 | 1,19 | 2,80 | 0,43 | 6,70E-01 |
| AT1G19140 | 0,44 | -1,56 | 4,31 | -0,36 | 7,17E-01 |
| AT1G19210 | 0,10 | 0,00 | 4,41 | 0,00 | 1,00E+00 |
| AT1G19260 | 0,97 | 2,17 | 2,58 | 0,84 | 4,01E-01 |
| AT1G19300 | 0,88 | 2,92 | 2,44 | 1,19 | 2,33E-01 |
| AT1G19360 | 0,50 | -0,34 | 3,95 | -0,09 | 9,32E-01 |

| | | | | | |
|---|---|---|---|---|---|
| AT1G19370 | 0,41 | 0,30 | 4,26 | 0,07 | 9,45E-01 |
| AT1G19400 | 1,69 | 1,54 | 1,47 | 1,04 | 2,97E-01 |
| AT1G19410 | 3,18 | 0,47 | 1,12 | 0,42 | 6,76E-01 |
| AT1G19430 | 0,39 | -0,88 | 4,25 | -0,21 | 8,36E-01 |
| AT1G19450 | 0,33 | -1,71 | 4,40 | -0,39 | 6,98E-01 |
| AT1G19570 | 0,89 | -1,08 | 2,06 | -0,52 | 6,00E-01 |
| AT1G19640 | 0,14 | 0,00 | 4,41 | 0,00 | 1,00E+00 |
| AT1G19660 | 0,55 | -0,45 | 2,73 | -0,16 | 8,69E-01 |
| AT1G19670 | 1,59 | 0,05 | 2,04 | 0,03 | 9,79E-01 |
| AT1G19700 | 0,25 | 0,37 | 4,37 | 0,09 | 9,32E-01 |
| AT1G19715 | 1,55 | 0,46 | 1,86 | 0,25 | 8,05E-01 |
| AT1G19720 | 1,66 | -0,33 | 1,44 | -0,23 | 8,21E-01 |
| AT1G19780 | 0,50 | -1,60 | 4,02 | -0,40 | 6,90E-01 |
| AT1G19860 | 0,15 | 0,45 | 4,37 | 0,10 | 9,19E-01 |
| AT1G19870 | 0,80 | 1,30 | 2,48 | 0,53 | 5,99E-01 |
| AT1G19880 | 0,41 | -0,48 | 3,43 | -0,14 | 8,89E-01 |
| AT1G19920 | 1,00 | -1,41 | 2,07 | -0,68 | 4,96E-01 |
| AT1G20010 | 2,01 | -2,08 | 1,25 | -1,66 | 9,61E-02 |
| AT1G20040 | 0,34 | 0,39 | 4,37 | 0,09 | 9,28E-01 |
| AT1G20070 | 0,59 | 1,17 | 2,44 | 0,48 | 6,31E-01 |
| AT1G20100 | 1,77 | 1,38 | 1,60 | 0,86 | 3,90E-01 |
| AT1G20160 | 0,33 | 0,75 | 4,32 | 0,17 | 8,62E-01 |
| AT1G20190 | 0,64 | 0,66 | 3,29 | 0,20 | 8,40E-01 |
| AT1G20200 | 0,33 | 1,44 | 3,52 | 0,41 | 6,83E-01 |
| AT1G20320 | 0,05 | 0,00 | 4,41 | 0,00 | 1,00E+00 |
| AT1G20330 | 2,79 | 0,69 | 1,35 | 0,51 | 6,08E-01 |
| AT1G20400 | 1,62 | -0,31 | 1,64 | -0,19 | 8,51E-01 |
| AT1G20430 | 0,89 | 1,31 | 2,80 | 0,47 | 6,40E-01 |
| AT1G20440 | 3,88 | -0,24 | 0,94 | -0,26 | 7,97E-01 |
| AT1G20450 | 3,27 | -0,10 | 1,22 | -0,09 | 9,32E-01 |
| AT1G20490 | 0,48 | 0,36 | 4,34 | 0,08 | 9,34E-01 |
| AT1G20510 | 2,74 | 0,78 | 1,41 | 0,56 | 5,78E-01 |
| AT1G20560 | 0,25 | 1,34 | 4,34 | 0,31 | 7,57E-01 |
| AT1G20570 | 0,08 | 0,69 | 4,36 | 0,16 | 8,75E-01 |
| AT1G20650 | 0,41 | -2,27 | 4,39 | -0,52 | 6,05E-01 |
| AT1G20670 | 0,31 | -0,98 | 4,34 | -0,23 | 8,22E-01 |
| AT1G20691 | 2,85 | 0,81 | 1,29 | 0,63 | 5,31E-01 |
| AT1G05563 | 2,75 | 0,81 | 1,32 | 0,61 | 5,41E-01 |
| AT1G20770 | 0,16 | 0,49 | 4,38 | 0,11 | 9,12E-01 |
| AT1G20840 | 1,09 | -0,17 | 1,97 | -0,09 | 9,29E-01 |
| AT1G20860 | 3,03 | 0,48 | 1,19 | 0,40 | 6,87E-01 |
| AT1G20880 | 0,25 | 0,41 | 4,37 | 0,09 | 9,26E-01 |
| AT1G20890 | 0,48 | 2,06 | 3,96 | 0,52 | 6,03E-01 |
| AT1G20950 | 0,69 | 0,30 | 2,42 | 0,12 | 9,02E-01 |
| AT1G21080 | 0,61 | -1,70 | 2,62 | -0,65 | 5,16E-01 |
| AT1G21130 | 2,17 | -2,59 | 1,63 | -1,59 | 1,13E-01 |
| AT1G05627 | 0,05 | 0,00 | 4,41 | 0,00 | 1,00E+00 |
| AT1G21160 | 0,26 | 1,17 | 4,34 | 0,27 | 7,87E-01 |
| AT1G21190 | 0,45 | 0,48 | 4,16 | 0,12 | 9,07E-01 |
| AT1G05637 | 4,89 | 0,47 | 1,09 | 0,44 | 6,62E-01 |
| AT1G21310 | 0,44 | 0,27 | 2,77 | 0,10 | 9,24E-01 |
| AT1G21380 | 1,01 | -0,42 | 2,13 | -0,20 | 8,43E-01 |

| | | | | | |
|---|---|---|---|---|---|
| AT1G21390 | 0,08 | 0,69 | 4,36 | 0,16 | 8,75E-01 |
| AT1G21440 | 1,10 | -3,22 | 2,68 | -1,20 | 2,30E-01 |
| AT1G21500 | 0,51 | -1,67 | 3,21 | -0,52 | 6,03E-01 |
| AT1G21520 | 0,17 | 1,15 | 4,34 | 0,27 | 7,91E-01 |
| AT1G21550 | 0,17 | 1,33 | 4,33 | 0,31 | 7,59E-01 |
| AT1G21580 | 3,07 | 0,65 | 1,00 | 0,65 | 5,17E-01 |
| AT1G21590 | 2,73 | -0,37 | 1,18 | -0,32 | 7,53E-01 |
| AT1G21600 | 0,77 | 0,20 | 2,08 | 0,09 | 9,25E-01 |
| AT1G21650 | 0,46 | 0,45 | 3,22 | 0,14 | 8,89E-01 |
| AT1G21770 | 0,66 | -0,67 | 2,68 | -0,25 | 8,02E-01 |
| AT1G21890 | 0,36 | 0,00 | 4,38 | 0,00 | 1,00E+00 |
| AT1G21900 | 0,26 | 0,84 | 4,35 | 0,19 | 8,46E-01 |
| AT1G21920 | 0,41 | 0,69 | 3,30 | 0,21 | 8,33E-01 |
| AT1G21928 | 0,29 | -0,14 | 4,34 | -0,03 | 9,74E-01 |
| AT1G22060 | 2,54 | -0,79 | 1,14 | -0,70 | 4,85E-01 |
| AT1G22180 | 0,30 | 0,82 | 3,63 | 0,22 | 8,22E-01 |
| AT1G22200 | 0,54 | -0,13 | 3,84 | -0,03 | 9,73E-01 |
| AT1G22300 | 0,58 | 0,83 | 2,66 | 0,31 | 7,55E-01 |
| AT1G22360 | 0,63 | 0,18 | 2,73 | 0,07 | 9,47E-01 |
| AT1G22400 | 0,23 | 1,02 | 4,34 | 0,24 | 8,14E-01 |
| AT1G22530 | 2,83 | -4,04 | 1,44 | -2,81 | 4,94E-03 |
| AT1G22550 | 0,19 | 0,36 | 4,36 | 0,08 | 9,34E-01 |
| AT1G22570 | 0,46 | 1,91 | 3,99 | 0,48 | 6,32E-01 |
| AT1G22710 | 0,50 | 1,42 | 2,67 | 0,53 | 5,96E-01 |
| AT1G22850 | 1,38 | 1,14 | 1,75 | 0,65 | 5,15E-01 |
| AT1G23020 | 0,20 | -0,44 | 4,36 | -0,10 | 9,19E-01 |
| AT1G23052 | 1,19 | -0,61 | 2,55 | -0,24 | 8,12E-01 |
| AT1G23080 | 0,61 | -1,19 | 3,26 | -0,37 | 7,14E-01 |
| AT1G05793 | 0,08 | 0,00 | 4,40 | 0,00 | 1,00E+00 |
| AT1G23205 | 0,35 | -1,66 | 4,39 | -0,38 | 7,06E-01 |
| AT1G23260 | 0,22 | 1,50 | 4,33 | 0,35 | 7,28E-01 |
| AT1G23300 | 1,57 | 0,40 | 1,72 | 0,23 | 8,17E-01 |
| AT1G23330 | 0,10 | 0,00 | 4,41 | 0,00 | 1,00E+00 |
| AT1G23390 | 2,44 | 0,38 | 1,30 | 0,29 | 7,72E-01 |
| AT1G05817 | 1,94 | -1,65 | 1,38 | -1,19 | 2,32E-01 |
| AT1G23650 | 0,08 | 0,44 | 4,38 | 0,10 | 9,21E-01 |
| AT1G23670 | 0,22 | 0,00 | 4,41 | 0,00 | 1,00E+00 |
| AT1G23740 | 1,33 | -0,66 | 1,90 | -0,35 | 7,28E-01 |
| AT1G23780 | 0,31 | 0,65 | 4,32 | 0,15 | 8,81E-01 |
| AT1G23830 | 0,53 | 0,49 | 3,18 | 0,15 | 8,79E-01 |
| AT1G23840 | 0,48 | 0,00 | 3,07 | 0,00 | 9,99E-01 |
| AT1G23850 | 0,35 | 0,30 | 4,34 | 0,07 | 9,44E-01 |
| AT1G23960 | 0,23 | 0,36 | 4,37 | 0,08 | 9,34E-01 |
| AT1G24040 | 0,10 | 0,21 | 4,38 | 0,05 | 9,62E-01 |
| AT1G24100 | 1,00 | 2,73 | 2,01 | 1,36 | 1,75E-01 |
| AT1G24120 | 0,29 | 0,11 | 4,40 | 0,02 | 9,81E-01 |
| AT1G24140 | 0,74 | -2,17 | 2,85 | -0,76 | 4,46E-01 |
| AT1G24160 | 0,84 | 1,96 | 3,28 | 0,60 | 5,51E-01 |
| AT1G24190 | 0,84 | 0,88 | 2,63 | 0,33 | 7,39E-01 |
| AT1G24267 | 0,48 | 0,91 | 3,15 | 0,29 | 7,73E-01 |
| AT1G24290 | 0,17 | 1,15 | 4,34 | 0,27 | 7,91E-01 |
| AT1G24300 | 1,68 | -0,03 | 1,57 | -0,02 | 9,83E-01 |

| | | | | | |
|---|---|---|---|---|---|
| AT1G24350 | 0,14 | 0,00 | 4,41 | 0,00 | 1,00E+00 |
| AT1G24510 | 1,77 | 0,47 | 1,48 | 0,32 | 7,52E-01 |
| AT1G05907 | 0,77 | 0,45 | 2,54 | 0,18 | 8,59E-01 |
| AT1G24764 | 0,53 | 1,48 | 3,28 | 0,45 | 6,52E-01 |
| AT1G24822 | 0,94 | -3,73 | 4,31 | -0,87 | 3,86E-01 |
| AT1G25220 | 0,36 | 1,83 | 4,33 | 0,42 | 6,73E-01 |
| AT1G25350 | 1,21 | -0,94 | 1,68 | -0,56 | 5,77E-01 |
| AT1G25370 | 0,12 | 1,16 | 4,34 | 0,27 | 7,89E-01 |
| AT1G25390 | 0,51 | 1,07 | 3,04 | 0,35 | 7,24E-01 |
| AT1G25400 | 0,63 | 1,34 | 2,85 | 0,47 | 6,37E-01 |
| AT1G25500 | 0,25 | 1,42 | 4,33 | 0,33 | 7,44E-01 |
| AT1G25540 | 0,94 | 2,06 | 2,29 | 0,90 | 3,67E-01 |
| AT1G25560 | 0,90 | 1,00 | 2,11 | 0,48 | 6,34E-01 |
| AT1G25570 | 0,49 | -3,61 | 4,31 | -0,84 | 4,03E-01 |
| AT1G26110 | 1,79 | 0,29 | 1,35 | 0,22 | 8,29E-01 |
| AT1G26150 | 0,88 | 0,25 | 2,25 | 0,11 | 9,12E-01 |
| AT1G26160 | 0,24 | -0,73 | 4,37 | -0,17 | 8,68E-01 |
| AT1G26170 | 0,54 | 1,68 | 3,13 | 0,54 | 5,93E-01 |
| AT1G26208 | 0,43 | 0,83 | 4,36 | 0,19 | 8,50E-01 |
| AT1G26218 | 0,37 | 0,43 | 4,38 | 0,10 | 9,21E-01 |
| AT1G26233 | 0,08 | 0,00 | 4,40 | 0,00 | 1,00E+00 |
| AT1G26240 | 0,88 | 1,47 | 2,64 | 0,56 | 5,79E-01 |
| AT1G26255 | 0,44 | 0,69 | 4,11 | 0,17 | 8,66E-01 |
| AT1G26270 | 1,28 | -2,99 | 1,93 | -1,55 | 1,21E-01 |
| AT1G26380 | 0,46 | -2,12 | 4,38 | -0,48 | 6,29E-01 |
| AT1G26390 | 0,25 | -0,79 | 4,34 | -0,18 | 8,55E-01 |
| AT1G26420 | 0,40 | 0,68 | 4,29 | 0,16 | 8,73E-01 |
| AT1G26690 | 0,33 | 0,92 | 4,31 | 0,21 | 8,30E-01 |
| AT1G26761 | 0,85 | 2,42 | 4,30 | 0,56 | 5,73E-01 |
| AT1G26800 | 0,42 | -0,61 | 2,91 | -0,21 | 8,34E-01 |
| AT1G26850 | 2,19 | -0,90 | 1,38 | -0,66 | 5,12E-01 |
| AT1G26880 | 2,18 | -1,74 | 1,62 | -1,07 | 2,83E-01 |
| AT1G06033 | 1,64 | -1,94 | 1,91 | -1,01 | 3,12E-01 |
| AT1G26930 | 0,36 | 1,95 | 4,31 | 0,45 | 6,52E-01 |
| AT1G27008 | 0,08 | 0,69 | 4,36 | 0,16 | 8,75E-01 |
| AT1G27090 | 1,85 | -0,18 | 1,31 | -0,14 | 8,90E-01 |
| AT1G27100 | 1,46 | 2,41 | 1,84 | 1,31 | 1,91E-01 |
| AT1G27190 | 0,77 | 0,29 | 2,10 | 0,14 | 8,89E-01 |
| AT1G27200 | 0,57 | 0,51 | 2,76 | 0,19 | 8,53E-01 |
| AT1G27210 | 0,26 | 0,82 | 4,34 | 0,19 | 8,50E-01 |
| AT1G27350 | 0,20 | -1,61 | 4,39 | -0,37 | 7,14E-01 |
| AT1G27370 | 0,51 | -1,22 | 4,04 | -0,30 | 7,62E-01 |
| AT1G27385 | 0,43 | -1,83 | 4,35 | -0,42 | 6,74E-01 |
| AT1G27430 | 3,11 | 0,29 | 1,11 | 0,26 | 7,93E-01 |
| AT1G27440 | 0,24 | -0,74 | 4,35 | -0,17 | 8,65E-01 |
| AT1G27500 | 0,29 | 0,70 | 4,32 | 0,16 | 8,72E-01 |
| AT1G27510 | 0,44 | 0,97 | 3,20 | 0,30 | 7,61E-01 |
| AT1G06093 | 0,29 | -0,92 | 4,39 | -0,21 | 8,34E-01 |
| AT1G27600 | 1,12 | -2,05 | 1,91 | -1,07 | 2,83E-01 |
| AT1G27700 | 0,50 | -1,32 | 4,28 | -0,31 | 7,58E-01 |
| AT1G27720 | 0,50 | -0,16 | 3,86 | -0,04 | 9,66E-01 |
| AT1G27730 | 1,60 | 1,29 | 1,63 | 0,79 | 4,29E-01 |

| | | | | | |
|---|---|---|---|---|---|
| AT1G27770 | 5,67 | 0,46 | 0,97 | 0,47 | 6,36E-01 |
| AT1G27870 | 1,97 | 1,49 | 1,77 | 0,84 | 3,99E-01 |
| AT1G27900 | 0,40 | 1,21 | 3,32 | 0,36 | 7,17E-01 |
| AT1G27930 | 0,53 | 0,83 | 3,05 | 0,27 | 7,86E-01 |
| AT1G28000 | 0,35 | 0,11 | 4,38 | 0,03 | 9,80E-01 |
| AT1G28120 | 0,73 | 0,69 | 2,80 | 0,25 | 8,06E-01 |
| AT1G28140 | 1,15 | 0,85 | 1,74 | 0,49 | 6,23E-01 |
| AT1G28150 | 0,60 | 1,06 | 2,98 | 0,35 | 7,23E-01 |
| AT1G28190 | 0,15 | 0,83 | 4,35 | 0,19 | 8,49E-01 |
| AT1G28200 | 0,24 | 0,43 | 4,36 | 0,10 | 9,22E-01 |
| AT1G28230 | 0,78 | 1,98 | 2,25 | 0,88 | 3,78E-01 |
| AT1G28240 | 0,33 | 0,01 | 4,37 | 0,00 | 9,99E-01 |
| AT1G28260 | 1,27 | 2,24 | 1,93 | 1,16 | 2,46E-01 |
| AT1G28280 | 1,03 | 0,74 | 1,98 | 0,37 | 7,08E-01 |
| AT1G28290 | 0,27 | -1,78 | 4,41 | -0,40 | 6,87E-01 |
| AT1G28310 | 0,66 | 1,16 | 2,24 | 0,52 | 6,06E-01 |
| AT1G28330 | 0,60 | -0,88 | 3,22 | -0,27 | 7,85E-01 |
| AT1G28370 | 0,92 | 1,66 | 2,38 | 0,70 | 4,85E-01 |
| AT1G28390 | 0,25 | -2,19 | 4,38 | -0,50 | 6,17E-01 |
| AT1G28400 | 0,35 | 0,00 | 3,51 | 0,00 | 9,99E-01 |
| AT1G28480 | 1,91 | 2,35 | 1,89 | 1,24 | 2,14E-01 |
| AT1G28570 | 0,12 | 0,97 | 4,34 | 0,22 | 8,23E-01 |
| AT1G28580 | 0,48 | 1,28 | 3,88 | 0,33 | 7,42E-01 |
| AT1G28600 | 1,10 | -1,65 | 1,66 | -0,99 | 3,22E-01 |
| AT1G28630 | 0,79 | 0,00 | 2,95 | 0,00 | 1,00E+00 |
| AT1G28660 | 0,08 | 0,69 | 4,36 | 0,16 | 8,75E-01 |
| AT1G28685 | 0,55 | 0,05 | 3,20 | 0,01 | 9,89E-01 |
| AT1G28765 | 0,17 | 1,50 | 4,33 | 0,35 | 7,29E-01 |
| AT1G28960 | 0,56 | 1,26 | 3,90 | 0,32 | 7,47E-01 |
| AT1G29050 | 0,14 | 0,00 | 4,41 | 0,00 | 1,00E+00 |
| AT1G29071 | 1,34 | 1,14 | 1,79 | 0,64 | 5,23E-01 |
| AT1G29170 | 0,29 | 1,44 | 4,33 | 0,33 | 7,40E-01 |
| AT1G29240 | 0,32 | -0,65 | 4,33 | -0,15 | 8,81E-01 |
| AT1G29300 | 0,08 | 0,00 | 4,41 | 0,00 | 1,00E+00 |
| AT1G29350 | 2,20 | -0,08 | 1,21 | -0,07 | 9,48E-01 |
| AT1G29370 | 3,57 | 0,17 | 0,97 | 0,18 | 8,57E-01 |
| AT1G29390 | 0,82 | 1,05 | 2,09 | 0,50 | 6,16E-01 |
| AT1G29395 | 0,39 | 1,33 | 3,38 | 0,39 | 6,94E-01 |
| AT1G29400 | 2,11 | 0,18 | 1,21 | 0,15 | 8,81E-01 |
| AT1G29470 | 3,69 | -0,56 | 0,93 | -0,60 | 5,47E-01 |
| AT1G29530 | 0,36 | 0,14 | 3,47 | 0,04 | 9,67E-01 |
| AT1G29690 | 0,27 | 0,04 | 4,38 | 0,01 | 9,93E-01 |
| AT1G29700 | 0,34 | -1,73 | 4,40 | -0,39 | 6,94E-01 |
| AT1G29720 | 0,71 | -1,87 | 2,49 | -0,75 | 4,52E-01 |
| AT1G29750 | 0,26 | -0,14 | 4,34 | -0,03 | 9,74E-01 |
| AT1G29880 | 1,19 | 1,03 | 1,90 | 0,54 | 5,89E-01 |
| AT1G29940 | 2,05 | -1,21 | 1,45 | -0,83 | 4,05E-01 |
| AT1G29970 | 0,83 | -0,90 | 2,43 | -0,37 | 7,11E-01 |
| AT1G30000 | 0,84 | -0,47 | 2,53 | -0,19 | 8,52E-01 |
| AT1G30060 | 0,17 | 0,00 | 4,41 | 0,00 | 1,00E+00 |
| AT1G30070 | 0,15 | 0,00 | 4,41 | 0,00 | 1,00E+00 |
| AT1G30090 | 0,16 | 0,92 | 4,35 | 0,21 | 8,33E-01 |

| | | | | | |
|---|---|---|---|---|---|
| AT1G30120 | 1,11 | 0,43 | 1,91 | 0,23 | 8,21E-01 |
| AT1G30210 | 0,29 | -0,92 | 4,34 | -0,21 | 8,32E-01 |
| AT1G30250 | 0,60 | -0,36 | 2,63 | -0,13 | 8,93E-01 |
| AT1G30330 | 1,89 | -0,21 | 1,61 | -0,13 | 8,95E-01 |
| AT1G30370 | 0,55 | 1,25 | 3,07 | 0,41 | 6,85E-01 |
| AT1G30490 | 1,77 | -0,19 | 1,57 | -0,12 | 9,03E-01 |
| AT1G30500 | 0,64 | 0,69 | 3,81 | 0,18 | 8,56E-01 |
| AT1G06347 | 0,94 | -0,52 | 2,17 | -0,24 | 8,10E-01 |
| AT1G30540 | 0,42 | -2,49 | 4,17 | -0,60 | 5,52E-01 |
| AT1G30580 | 1,07 | -0,33 | 2,02 | -0,17 | 8,68E-01 |
| AT1G30590 | 0,66 | -0,91 | 2,94 | -0,31 | 7,58E-01 |
| AT1G30755 | 0,62 | -2,31 | 2,62 | -0,88 | 3,77E-01 |
| AT1G30780 | 0,83 | -0,26 | 2,32 | -0,11 | 9,10E-01 |
| AT1G30810 | 0,83 | -1,35 | 1,85 | -0,73 | 4,66E-01 |
| AT1G30880 | 0,08 | 0,44 | 4,38 | 0,10 | 9,21E-01 |
| AT1G30960 | 0,37 | -0,62 | 4,30 | -0,14 | 8,86E-01 |
| AT1G06407 | 0,34 | -0,14 | 3,59 | -0,04 | 9,69E-01 |
| AT1G31130 | 1,47 | 2,56 | 1,77 | 1,45 | 1,48E-01 |
| AT1G31160 | 0,42 | -0,56 | 4,10 | -0,14 | 8,91E-01 |
| AT1G31180 | 0,17 | 1,15 | 4,34 | 0,27 | 7,91E-01 |
| AT1G31230 | 1,31 | -0,33 | 1,91 | -0,17 | 8,62E-01 |
| AT1G31240 | 0,22 | 0,00 | 4,41 | 0,00 | 1,00E+00 |
| AT1G31350 | 0,45 | -0,42 | 3,18 | -0,13 | 8,96E-01 |
| AT1G31410 | 0,08 | 0,44 | 4,38 | 0,10 | 9,21E-01 |
| AT1G31470 | 0,15 | 1,20 | 4,34 | 0,28 | 7,81E-01 |
| AT1G31540 | 3,00 | 0,86 | 1,20 | 0,72 | 4,74E-01 |
| AT1G31550 | 0,81 | 1,15 | 2,55 | 0,45 | 6,53E-01 |
| AT1G31600 | 0,81 | -0,13 | 2,89 | -0,04 | 9,65E-01 |
| AT1G31640 | 0,23 | 0,86 | 4,34 | 0,20 | 8,43E-01 |
| AT1G31650 | 0,25 | -1,33 | 4,36 | -0,31 | 7,60E-01 |
| AT1G31660 | 0,51 | -1,53 | 4,30 | -0,36 | 7,22E-01 |
| AT1G31740 | 0,22 | 1,19 | 4,33 | 0,27 | 7,84E-01 |
| AT1G31810 | 0,57 | -0,66 | 2,27 | -0,29 | 7,71E-01 |
| AT1G31812 | 0,29 | 0,00 | 4,39 | 0,00 | 1,00E+00 |
| AT1G31814 | 0,37 | 0,56 | 4,32 | 0,13 | 8,98E-01 |
| AT1G31820 | 0,94 | 1,31 | 2,67 | 0,49 | 6,23E-01 |
| AT1G31830 | 0,66 | 1,28 | 2,60 | 0,49 | 6,22E-01 |
| AT1G31835 | 2,35 | 0,55 | 1,49 | 0,37 | 7,10E-01 |
| AT1G06453 | 1,48 | 1,48 | 2,03 | 0,73 | 4,68E-01 |
| AT1G31860 | 1,08 | -0,10 | 1,79 | -0,06 | 9,54E-01 |
| AT1G06463 | 0,27 | 1,14 | 4,34 | 0,26 | 7,92E-01 |
| AT1G31920 | 0,52 | -0,49 | 2,60 | -0,19 | 8,50E-01 |
| AT1G31940 | 0,35 | -2,70 | 4,37 | -0,62 | 5,36E-01 |
| AT1G06473 | 0,38 | 0,70 | 4,37 | 0,16 | 8,72E-01 |
| AT1G31960 | 1,81 | 0,09 | 1,82 | 0,05 | 9,59E-01 |
| AT1G32000 | 0,35 | -0,94 | 4,39 | -0,21 | 8,31E-01 |
| AT1G32010 | 1,81 | 1,62 | 1,90 | 0,85 | 3,93E-01 |
| AT1G32070 | 0,34 | 0,34 | 4,34 | 0,08 | 9,38E-01 |
| AT1G32080 | 0,85 | -0,14 | 1,96 | -0,07 | 9,44E-01 |
| AT1G32130 | 1,27 | -0,95 | 1,86 | -0,51 | 6,08E-01 |
| AT1G32200 | 0,75 | -1,75 | 2,35 | -0,74 | 4,56E-01 |
| AT1G32340 | 0,12 | 0,37 | 4,37 | 0,08 | 9,33E-01 |

| | | | | | |
|---|---|---|---|---|---|
| AT1G06537 | 1,54 | 0,50 | 1,60 | 0,31 | 7,57E-01 |
| AT1G32400 | 0,47 | 0,19 | 2,73 | 0,07 | 9,43E-01 |
| AT1G32450 | 0,27 | -0,34 | 4,34 | -0,08 | 9,37E-01 |
| AT1G06573 | 0,41 | 1,23 | 4,33 | 0,28 | 7,77E-01 |
| AT1G32470 | 0,50 | 1,23 | 2,69 | 0,46 | 6,49E-01 |
| AT1G32490 | 0,70 | 1,05 | 2,21 | 0,48 | 6,34E-01 |
| AT1G32500 | 0,52 | -2,06 | 2,71 | -0,76 | 4,47E-01 |
| AT1G32530 | 0,53 | -0,61 | 2,62 | -0,23 | 8,16E-01 |
| AT1G06577 | 0,16 | 0,00 | 4,41 | 0,00 | 1,00E+00 |
| AT1G32640 | 4,04 | 0,48 | 1,11 | 0,43 | 6,64E-01 |
| AT1G32713 | 1,03 | 2,03 | 2,33 | 0,87 | 3,83E-01 |
| AT1G32750 | 1,94 | 0,67 | 1,38 | 0,48 | 6,28E-01 |
| AT1G32790 | 1,57 | 2,92 | 1,92 | 1,52 | 1,29E-01 |
| AT1G32860 | 0,05 | 0,00 | 4,41 | 0,00 | 1,00E+00 |
| AT1G32880 | 0,26 | -0,30 | 4,37 | -0,07 | 9,46E-01 |
| AT1G32900 | 0,61 | -0,45 | 3,09 | -0,15 | 8,83E-01 |
| AT1G32970 | 0,17 | 1,52 | 4,33 | 0,35 | 7,26E-01 |
| AT1G33050 | 0,95 | -0,49 | 1,82 | -0,27 | 7,89E-01 |
| AT1G06643 | 0,36 | -0,09 | 4,35 | -0,02 | 9,83E-01 |
| AT1G33240 | 1,91 | -1,79 | 1,57 | -1,14 | 2,54E-01 |
| AT1G33250 | 0,90 | 2,13 | 2,24 | 0,95 | 3,41E-01 |
| AT1G33270 | 0,24 | 0,21 | 4,34 | 0,05 | 9,62E-01 |
| AT1G06673 | 0,47 | 0,91 | 3,33 | 0,27 | 7,84E-01 |
| AT1G33390 | 1,05 | -1,27 | 2,29 | -0,55 | 5,79E-01 |
| AT1G33410 | 1,37 | -0,71 | 1,42 | -0,50 | 6,19E-01 |
| AT1G33420 | 2,44 | 0,70 | 1,35 | 0,52 | 6,05E-01 |
| AT1G33430 | 0,08 | 0,44 | 4,38 | 0,10 | 9,21E-01 |
| AT1G33520 | 0,47 | 1,07 | 3,17 | 0,34 | 7,35E-01 |
| AT1G33615 | 0,17 | 0,00 | 4,41 | 0,00 | 1,00E+00 |
| AT1G33700 | 0,88 | 0,63 | 2,04 | 0,31 | 7,56E-01 |
| AT1G33720 | 1,30 | 0,00 | 1,84 | 0,00 | 9,99E-01 |
| AT1G33780 | 0,08 | 0,00 | 4,41 | 0,00 | 1,00E+00 |
| AT1G33870 | 0,60 | 1,95 | 2,57 | 0,76 | 4,49E-01 |
| AT1G33950 | 1,88 | -0,71 | 1,80 | -0,40 | 6,92E-01 |
| AT1G33970 | 0,67 | -1,13 | 2,55 | -0,45 | 6,56E-01 |
| AT1G34030 | 0,44 | 0,00 | 2,81 | 0,00 | 9,99E-01 |
| AT1G34042 | 0,29 | 0,79 | 4,34 | 0,18 | 8,56E-01 |
| AT1G34110 | 0,91 | 1,16 | 1,93 | 0,60 | 5,49E-01 |
| AT1G34170 | 0,20 | 0,75 | 4,37 | 0,17 | 8,63E-01 |
| AT1G34220 | 1,01 | 2,76 | 2,82 | 0,98 | 3,28E-01 |
| AT1G34270 | 0,12 | 1,05 | 4,35 | 0,24 | 8,10E-01 |
| AT1G34310 | 0,97 | 0,41 | 2,73 | 0,15 | 8,81E-01 |
| AT1G34315 | 0,23 | 1,70 | 4,32 | 0,39 | 6,94E-01 |
| AT1G34380 | 0,08 | 0,44 | 4,38 | 0,10 | 9,21E-01 |
| AT1G34440 | 1,54 | 1,88 | 1,87 | 1,00 | 3,15E-01 |
| AT1G34550 | 0,63 | 0,45 | 2,43 | 0,18 | 8,53E-01 |
| AT1G34750 | 0,23 | 0,67 | 4,36 | 0,15 | 8,77E-01 |
| AT1G34780 | 0,68 | 1,55 | 2,82 | 0,55 | 5,82E-01 |
| AT1G35150 | 0,16 | 0,04 | 4,38 | 0,01 | 9,92E-01 |
| AT1G35215 | 4,81 | 2,49 | 1,24 | 2,00 | 4,54E-02 |
| AT1G35240 | 1,26 | 3,18 | 1,90 | 1,67 | 9,40E-02 |
| AT1G35250 | 0,19 | 0,00 | 4,41 | 0,00 | 1,00E+00 |

| | | | | | |
|---|---|---|---|---|---|
| AT1G35320 | 0,59 | 1,49 | 2,96 | 0,50 | 6,15E-01 |
| AT1G35350 | 0,96 | 1,31 | 2,01 | 0,65 | 5,15E-01 |
| AT1G06883 | 0,19 | 0,00 | 4,41 | 0,00 | 1,00E+00 |
| AT1G06897 | 0,91 | 0,00 | 3,54 | 0,00 | 1,00E+00 |
| AT1G06903 | 0,10 | 0,00 | 4,41 | 0,00 | 1,00E+00 |
| AT1G35520 | 1,01 | 2,13 | 2,51 | 0,85 | 3,96E-01 |
| AT1G35545 | 0,38 | 0,06 | 3,43 | 0,02 | 9,85E-01 |
| AT1G35560 | 2,43 | -1,07 | 1,16 | -0,92 | 3,56E-01 |
| AT1G35580 | 1,07 | 0,40 | 1,75 | 0,23 | 8,20E-01 |
| AT1G35610 | 0,22 | 0,44 | 4,38 | 0,10 | 9,19E-01 |
| AT1G35625 | 0,08 | 0,44 | 4,38 | 0,10 | 9,21E-01 |
| AT1G35730 | 0,17 | 1,50 | 4,33 | 0,35 | 7,29E-01 |
| AT1G35820 | 1,40 | 0,85 | 2,92 | 0,29 | 7,72E-01 |
| AT1G35895 | 0,55 | 2,47 | 3,17 | 0,78 | 4,36E-01 |
| AT1G36070 | 0,25 | 0,41 | 4,33 | 0,09 | 9,25E-01 |
| AT1G36085 | 0,68 | 0,00 | 4,41 | 0,00 | 1,00E+00 |
| AT1G36230 | 0,12 | 0,00 | 4,41 | 0,00 | 1,00E+00 |
| AT1G36325 | 1,41 | -1,66 | 1,61 | -1,03 | 3,02E-01 |
| AT1G36370 | 0,71 | 1,04 | 2,34 | 0,44 | 6,56E-01 |
| AT1G36390 | 0,17 | 0,65 | 4,36 | 0,15 | 8,81E-01 |
| AT1G36622 | 0,40 | -0,25 | 3,46 | -0,07 | 9,42E-01 |
| AT1G36730 | 3,83 | -0,15 | 0,89 | -0,17 | 8,64E-01 |
| AT1G36990 | 1,02 | -0,41 | 1,84 | -0,22 | 8,23E-01 |
| AT1G07013 | 0,15 | 0,00 | 4,41 | 0,00 | 1,00E+00 |
| AT1G40087 | 0,19 | 0,00 | 4,41 | 0,00 | 1,00E+00 |
| AT1G41830 | 0,28 | 0,98 | 4,35 | 0,23 | 8,21E-01 |
| AT1G41920 | 0,51 | 1,37 | 3,23 | 0,42 | 6,72E-01 |
| AT1G42440 | 0,46 | -0,61 | 3,36 | -0,18 | 8,56E-01 |
| AT1G42560 | 0,05 | 0,00 | 4,41 | 0,00 | 1,00E+00 |
| AT1G42990 | 1,44 | 0,26 | 1,58 | 0,16 | 8,71E-01 |
| AT1G43190 | 0,38 | -0,20 | 4,33 | -0,05 | 9,63E-01 |
| AT1G43560 | 0,22 | 0,25 | 4,36 | 0,06 | 9,54E-01 |
| AT1G43580 | 0,21 | 0,68 | 4,35 | 0,16 | 8,76E-01 |
| AT1G43620 | 0,44 | -0,09 | 4,19 | -0,02 | 9,83E-01 |
| AT1G43665 | 0,88 | 2,46 | 3,20 | 0,77 | 4,42E-01 |
| AT1G43710 | 0,50 | 0,97 | 3,20 | 0,30 | 7,61E-01 |
| AT1G43730 | 0,40 | 0,78 | 4,29 | 0,18 | 8,55E-01 |
| AT1G43760 | 0,66 | 1,52 | 2,56 | 0,59 | 5,53E-01 |
| AT1G43825 | 0,81 | -3,08 | 2,78 | -1,11 | 2,69E-01 |
| AT1G43860 | 1,69 | -0,11 | 1,64 | -0,07 | 9,45E-01 |
| AT1G43910 | 2,35 | 0,14 | 1,27 | 0,11 | 9,13E-01 |
| AT1G43950 | 0,46 | -0,31 | 4,04 | -0,08 | 9,39E-01 |
| AT1G44000 | 0,08 | 0,44 | 4,38 | 0,10 | 9,21E-01 |
| AT1G44030 | 0,25 | 0,13 | 4,39 | 0,03 | 9,76E-01 |
| AT1G44080 | 1,35 | -2,55 | 2,17 | -1,17 | 2,40E-01 |
| AT1G44085 | 0,65 | 0,84 | 2,94 | 0,29 | 7,75E-01 |
| AT1G44090 | 0,05 | 0,00 | 4,41 | 0,00 | 1,00E+00 |
| AT1G44100 | 2,02 | 2,17 | 1,66 | 1,31 | 1,90E-01 |
| AT1G07243 | 1,82 | 2,20 | 1,67 | 1,32 | 1,87E-01 |
| AT1G44125 | 0,91 | 3,20 | 2,64 | 1,21 | 2,26E-01 |
| AT1G44170 | 0,30 | 0,01 | 4,38 | 0,00 | 9,99E-01 |
| AT1G44350 | 0,41 | 0,06 | 4,23 | 0,01 | 9,89E-01 |

| | | | | | |
|---|---|---|---|---|---|
| AT1G44446 | 1,72 | 0,26 | 1,54 | 0,17 | 8,64E-01 |
| AT1G44542 | 0,40 | -2,20 | 4,38 | -0,50 | 6,15E-01 |
| AT1G44770 | 0,41 | 0,48 | 3,31 | 0,15 | 8,85E-01 |
| AT1G44790 | 0,21 | 0,01 | 4,38 | 0,00 | 9,98E-01 |
| AT1G44920 | 0,44 | -0,08 | 3,36 | -0,02 | 9,80E-01 |
| AT1G44980 | 0,59 | 1,30 | 2,77 | 0,47 | 6,39E-01 |
| AT1G45063 | 0,43 | 0,02 | 4,39 | 0,00 | 9,96E-01 |
| AT1G45145 | 0,72 | 1,60 | 3,11 | 0,51 | 6,07E-01 |
| AT1G45150 | 0,25 | -1,33 | 4,36 | -0,31 | 7,60E-01 |
| AT1G45160 | 1,73 | 1,28 | 1,53 | 0,83 | 4,05E-01 |
| AT1G45163 | 0,70 | 1,50 | 2,46 | 0,61 | 5,41E-01 |
| AT1G45207 | 0,37 | -3,12 | 4,34 | -0,72 | 4,72E-01 |
| AT1G46696 | 1,91 | 1,85 | 1,76 | 1,05 | 2,94E-01 |
| AT1G07323 | 0,55 | -0,46 | 2,66 | -0,17 | 8,62E-01 |
| AT1G47200 | 0,16 | 0,00 | 4,41 | 0,00 | 1,00E+00 |
| AT1G47240 | 0,43 | 2,16 | 4,09 | 0,53 | 5,97E-01 |
| AT1G07333 | 0,81 | 0,00 | 3,42 | 0,00 | 1,00E+00 |
| AT1G47260 | 1,01 | 0,09 | 1,74 | 0,05 | 9,60E-01 |
| AT1G47280 | 1,31 | -4,15 | 2,04 | -2,04 | 4,13E-02 |
| AT1G07347 | 0,18 | 0,00 | 4,41 | 0,00 | 1,00E+00 |
| AT1G47380 | 0,18 | 0,43 | 4,36 | 0,10 | 9,22E-01 |
| AT1G47420 | 0,71 | 1,24 | 2,20 | 0,57 | 5,71E-01 |
| AT1G47497 | 2,25 | 0,93 | 1,70 | 0,55 | 5,83E-01 |
| AT1G07383 | 0,55 | -0,87 | 2,75 | -0,32 | 7,52E-01 |
| AT1G07393 | 1,67 | -0,82 | 1,76 | -0,46 | 6,43E-01 |
| AT1G47640 | 0,35 | -0,15 | 3,50 | -0,04 | 9,66E-01 |
| AT1G47660 | 2,01 | 1,36 | 1,70 | 0,80 | 4,25E-01 |
| AT1G47670 | 0,05 | 0,00 | 4,41 | 0,00 | 1,00E+00 |
| AT1G47765 | 0,55 | -1,65 | 2,39 | -0,69 | 4,91E-01 |
| AT1G47900 | 0,45 | -1,41 | 4,10 | -0,34 | 7,31E-01 |
| AT1G47960 | 1,43 | 0,16 | 1,59 | 0,10 | 9,19E-01 |
| AT1G48000 | 0,50 | -0,05 | 4,39 | -0,01 | 9,91E-01 |
| AT1G48030 | 1,26 | 2,71 | 2,13 | 1,27 | 2,04E-01 |
| AT1G48040 | 0,63 | 2,44 | 4,30 | 0,57 | 5,70E-01 |
| AT1G48110 | 0,42 | 1,53 | 4,33 | 0,35 | 7,24E-01 |
| AT1G48120 | 0,82 | 0,99 | 2,25 | 0,44 | 6,61E-01 |
| AT1G48160 | 0,15 | 0,45 | 4,37 | 0,10 | 9,19E-01 |
| AT1G48285 | 0,80 | -0,16 | 2,21 | -0,07 | 9,42E-01 |
| AT1G48320 | 0,17 | 1,50 | 4,33 | 0,35 | 7,29E-01 |
| AT1G48380 | 0,10 | 0,01 | 4,38 | 0,00 | 9,99E-01 |
| AT1G48405 | 0,82 | 2,00 | 3,28 | 0,61 | 5,42E-01 |
| AT1G48430 | 0,15 | 1,16 | 4,34 | 0,27 | 7,88E-01 |
| AT1G48450 | 1,45 | -1,90 | 1,70 | -1,12 | 2,63E-01 |
| AT1G48550 | 1,08 | 0,97 | 2,62 | 0,37 | 7,10E-01 |
| AT1G48570 | 0,26 | 1,34 | 4,33 | 0,31 | 7,57E-01 |
| AT1G48580 | 0,08 | 0,69 | 4,36 | 0,16 | 8,75E-01 |
| AT1G07493 | 1,97 | -1,57 | 1,71 | -0,92 | 3,59E-01 |
| AT1G48620 | 0,46 | -1,80 | 3,31 | -0,54 | 5,86E-01 |
| AT1G48630 | 1,25 | 0,72 | 1,60 | 0,45 | 6,54E-01 |
| AT1G48650 | 3,21 | 0,17 | 1,03 | 0,17 | 8,68E-01 |
| AT1G48730 | 0,75 | 0,05 | 2,48 | 0,02 | 9,83E-01 |
| AT1G48742 | 0,46 | 1,17 | 3,19 | 0,37 | 7,14E-01 |

| | | | | | |
|---|---|---|---|---|---|
| AT1G48760 | 1,82 | -2,68 | 1,78 | -1,50 | 1,32E-01 |
| AT1G48790 | 0,44 | 0,95 | 3,20 | 0,30 | 7,66E-01 |
| AT1G48850 | 0,39 | -1,00 | 4,30 | -0,23 | 8,16E-01 |
| AT1G48900 | 0,23 | 0,01 | 4,39 | 0,00 | 9,98E-01 |
| AT1G48930 | 0,71 | -0,44 | 2,70 | -0,16 | 8,70E-01 |
| AT1G48970 | 1,37 | 2,07 | 1,92 | 1,08 | 2,82E-01 |
| AT1G49005 | 0,38 | 0,90 | 4,35 | 0,21 | 8,37E-01 |
| AT1G49040 | 3,74 | 0,47 | 1,01 | 0,47 | 6,42E-01 |
| AT1G49140 | 0,12 | 0,00 | 4,38 | 0,00 | 1,00E+00 |
| AT1G49180 | 0,08 | 0,44 | 4,38 | 0,10 | 9,21E-01 |
| AT1G49205 | 0,08 | 0,00 | 4,40 | 0,00 | 1,00E+00 |
| AT1G49225 | 0,48 | -1,54 | 3,19 | -0,48 | 6,30E-01 |
| AT1G49270 | 0,10 | 0,00 | 4,41 | 0,00 | 1,00E+00 |
| AT1G49490 | 0,32 | -1,93 | 4,33 | -0,45 | 6,55E-01 |
| AT1G49500 | 1,83 | -0,61 | 1,45 | -0,42 | 6,73E-01 |
| AT1G49540 | 0,83 | 2,71 | 2,46 | 1,10 | 2,70E-01 |
| AT1G49600 | 0,82 | -0,25 | 1,93 | -0,13 | 8,98E-01 |
| AT1G49670 | 0,26 | 0,09 | 4,38 | 0,02 | 9,84E-01 |
| AT1G49700 | 4,39 | -0,09 | 0,80 | -0,11 | 9,12E-01 |
| AT1G49710 | 0,34 | 1,47 | 3,49 | 0,42 | 6,74E-01 |
| AT1G49750 | 3,00 | 0,21 | 1,35 | 0,16 | 8,76E-01 |
| AT1G07593 | 3,78 | -2,21 | 1,39 | -1,59 | 1,12E-01 |
| AT1G49780 | 1,22 | 1,73 | 1,80 | 0,96 | 3,36E-01 |
| AT1G49850 | 0,39 | 1,69 | 4,33 | 0,39 | 6,96E-01 |
| AT1G49970 | 0,66 | 0,51 | 2,22 | 0,23 | 8,17E-01 |
| AT1G07603 | 0,61 | -1,11 | 3,11 | -0,36 | 7,21E-01 |
| AT1G50020 | 0,58 | -3,03 | 3,98 | -0,76 | 4,46E-01 |
| AT1G50030 | 2,19 | 0,39 | 1,37 | 0,29 | 7,74E-01 |
| AT1G50140 | 0,20 | 1,03 | 4,35 | 0,24 | 8,12E-01 |
| AT1G50190 | 0,32 | 0,23 | 4,34 | 0,05 | 9,58E-01 |
| AT1G50200 | 4,92 | -0,32 | 0,91 | -0,35 | 7,29E-01 |
| AT1G50250 | 2,35 | 0,57 | 1,22 | 0,47 | 6,40E-01 |
| AT1G50260 | 0,32 | -0,62 | 4,34 | -0,14 | 8,87E-01 |
| AT1G50280 | 0,61 | -0,95 | 2,43 | -0,39 | 6,95E-01 |
| AT1G50320 | 0,25 | 1,16 | 4,34 | 0,27 | 7,89E-01 |
| AT1G07627 | 0,18 | 0,00 | 4,41 | 0,00 | 1,00E+00 |
| AT1G50420 | 0,49 | -1,37 | 4,28 | -0,32 | 7,49E-01 |
| AT1G50430 | 1,02 | 0,12 | 1,74 | 0,07 | 9,47E-01 |
| AT1G07637 | 2,62 | 2,26 | 1,53 | 1,48 | 1,39E-01 |
| AT1G50450 | 0,63 | -1,69 | 2,91 | -0,58 | 5,60E-01 |
| AT1G50500 | 0,57 | 0,03 | 3,14 | 0,01 | 9,93E-01 |
| AT1G07647 | 0,08 | 0,44 | 4,38 | 0,10 | 9,21E-01 |
| AT1G50570 | 0,65 | -0,70 | 2,70 | -0,26 | 7,95E-01 |
| AT1G50600 | 0,88 | 1,65 | 2,08 | 0,79 | 4,27E-01 |
| AT1G50640 | 1,08 | -0,96 | 1,76 | -0,55 | 5,85E-01 |
| AT1G50659 | 0,43 | 1,42 | 3,25 | 0,44 | 6,63E-01 |
| AT1G50750 | 0,08 | 0,00 | 4,40 | 0,00 | 1,00E+00 |
| AT1G50790 | 0,12 | 1,16 | 4,34 | 0,27 | 7,89E-01 |
| AT1G50820 | 0,50 | -1,77 | 3,90 | -0,45 | 6,51E-01 |
| AT1G50910 | 1,39 | -1,94 | 1,59 | -1,22 | 2,24E-01 |
| AT1G50980 | 0,39 | -2,30 | 4,38 | -0,53 | 5,99E-01 |
| AT1G51080 | 0,73 | -1,06 | 2,52 | -0,42 | 6,74E-01 |

| | | | | | |
|---|---|---|---|---|---|
| AT1G51150 | 4,36 | -4,21 | 1,55 | -2,71 | 6,66E-03 |
| AT1G07733 | 0,24 | 0,00 | 4,41 | 0,00 | 1,00E+00 |
| AT1G51400 | 0,93 | 0,14 | 2,43 | 0,06 | 9,55E-01 |
| AT1G51480 | 0,33 | -1,30 | 4,33 | -0,30 | 7,64E-01 |
| AT1G51500 | 1,21 | 0,47 | 2,13 | 0,22 | 8,26E-01 |
| AT1G51510 | 0,91 | -0,96 | 2,22 | -0,43 | 6,65E-01 |
| AT1G51560 | 0,45 | -0,73 | 3,24 | -0,22 | 8,23E-01 |
| AT1G51570 | 1,72 | -0,77 | 1,38 | -0,56 | 5,76E-01 |
| AT1G51590 | 0,26 | 0,00 | 4,38 | 0,00 | 9,99E-01 |
| AT1G51630 | 0,52 | 0,15 | 3,24 | 0,05 | 9,64E-01 |
| AT1G51680 | 0,80 | 0,24 | 2,07 | 0,11 | 9,09E-01 |
| AT1G07763 | 0,56 | -1,78 | 4,41 | -0,40 | 6,87E-01 |
| AT1G51710 | 0,45 | -0,85 | 4,07 | -0,21 | 8,35E-01 |
| AT1G51730 | 0,47 | 0,38 | 3,21 | 0,12 | 9,06E-01 |
| AT1G51790 | 0,25 | -1,82 | 4,40 | -0,41 | 6,79E-01 |
| AT1G51805 | 2,63 | 0,96 | 1,32 | 0,73 | 4,64E-01 |
| AT1G51890 | 0,23 | 1,72 | 4,32 | 0,40 | 6,90E-01 |
| AT1G51940 | 1,08 | 1,39 | 2,60 | 0,53 | 5,94E-01 |
| AT1G51980 | 1,85 | 0,18 | 1,55 | 0,12 | 9,07E-01 |
| AT1G52000 | 0,46 | -1,77 | 4,11 | -0,43 | 6,67E-01 |
| AT1G52040 | 0,54 | -2,60 | 3,95 | -0,66 | 5,11E-01 |
| AT1G52110 | 2,50 | -0,71 | 1,46 | -0,49 | 6,23E-01 |
| AT1G52150 | 1,30 | 0,81 | 2,19 | 0,37 | 7,12E-01 |
| AT1G52160 | 0,61 | -0,75 | 3,15 | -0,24 | 8,12E-01 |
| AT1G52180 | 1,00 | -0,17 | 2,08 | -0,08 | 9,36E-01 |
| AT1G52185 | 0,32 | 0,04 | 4,39 | 0,01 | 9,93E-01 |
| AT1G52200 | 0,25 | 1,94 | 4,32 | 0,45 | 6,54E-01 |
| AT1G52220 | 0,76 | 0,65 | 2,23 | 0,29 | 7,72E-01 |
| AT1G52290 | 0,41 | 0,88 | 4,22 | 0,21 | 8,34E-01 |
| AT1G52300 | 0,35 | 1,40 | 4,33 | 0,32 | 7,45E-01 |
| AT1G52340 | 0,08 | 0,00 | 4,40 | 0,00 | 1,00E+00 |
| AT1G52342 | 0,45 | 1,25 | 4,31 | 0,29 | 7,72E-01 |
| AT1G52510 | 0,47 | -0,06 | 4,18 | -0,01 | 9,89E-01 |
| AT1G52570 | 1,59 | -1,72 | 1,99 | -0,86 | 3,89E-01 |
| AT1G52750 | 0,31 | 0,62 | 4,32 | 0,14 | 8,85E-01 |
| AT1G52880 | 0,40 | 0,90 | 3,37 | 0,27 | 7,90E-01 |
| AT1G52890 | 0,42 | 0,75 | 4,13 | 0,18 | 8,56E-01 |
| AT1G52970 | 1,13 | -2,50 | 2,14 | -1,17 | 2,42E-01 |
| AT1G52990 | 0,41 | 0,00 | 4,41 | 0,00 | 1,00E+00 |
| AT1G53000 | 0,49 | -1,99 | 3,97 | -0,50 | 6,16E-01 |
| AT1G53025 | 0,43 | -1,74 | 3,35 | -0,52 | 6,02E-01 |
| AT1G53035 | 0,12 | 1,16 | 4,34 | 0,27 | 7,89E-01 |
| AT1G53170 | 0,65 | 0,83 | 2,65 | 0,31 | 7,53E-01 |
| AT1G53200 | 0,24 | 0,21 | 4,34 | 0,05 | 9,62E-01 |
| AT1G53230 | 1,22 | 1,60 | 2,04 | 0,78 | 4,34E-01 |
| AT1G53240 | 1,16 | 1,92 | 1,98 | 0,97 | 3,33E-01 |
| AT1G53265 | 0,53 | 0,52 | 4,04 | 0,13 | 8,99E-01 |
| AT1G53310 | 3,45 | -1,20 | 0,93 | -1,29 | 1,98E-01 |
| AT1G53320 | 0,35 | 0,16 | 4,34 | 0,04 | 9,71E-01 |
| AT1G53450 | 0,86 | -2,57 | 2,13 | -1,21 | 2,27E-01 |
| AT1G53480 | 0,84 | 0,05 | 3,09 | 0,02 | 9,87E-01 |
| AT1G53500 | 0,50 | 1,26 | 3,34 | 0,38 | 7,07E-01 |

| | | | | | |
|---|---|---|---|---|---|
| AT1G53510 | 0,76 | -1,79 | 2,44 | -0,73 | 4,63E-01 |
| AT1G53541 | 0,64 | 0,15 | 2,35 | 0,06 | 9,49E-01 |
| AT1G53580 | 0,12 | 0,49 | 4,36 | 0,11 | 9,11E-01 |
| AT1G53640 | 2,14 | -1,82 | 1,45 | -1,26 | 2,07E-01 |
| AT1G53710 | 0,67 | -0,94 | 2,83 | -0,33 | 7,39E-01 |
| AT1G53750 | 1,03 | -1,33 | 1,87 | -0,71 | 4,77E-01 |
| AT1G53760 | 0,08 | 0,69 | 4,36 | 0,16 | 8,75E-01 |
| AT1G53770 | 0,33 | -0,45 | 4,32 | -0,10 | 9,16E-01 |
| AT1G53780 | 1,40 | 1,45 | 1,77 | 0,82 | 4,12E-01 |
| AT1G54040 | 2,21 | 3,09 | 1,68 | 1,83 | 6,70E-02 |
| AT1G54050 | 0,12 | 0,49 | 4,36 | 0,11 | 9,11E-01 |
| AT1G54080 | 1,16 | 0,88 | 1,95 | 0,45 | 6,50E-01 |
| AT1G54100 | 1,08 | 0,11 | 2,00 | 0,05 | 9,57E-01 |
| AT1G54140 | 0,15 | 0,45 | 4,37 | 0,10 | 9,19E-01 |
| AT1G54170 | 0,82 | 0,01 | 2,13 | 0,01 | 9,96E-01 |
| AT1G54320 | 0,67 | 1,64 | 2,46 | 0,67 | 5,06E-01 |
| AT1G54370 | 0,38 | 0,70 | 4,28 | 0,16 | 8,71E-01 |
| AT1G54500 | 0,85 | 0,00 | 2,08 | 0,00 | 1,00E+00 |
| AT1G54590 | 0,10 | 0,01 | 4,38 | 0,00 | 9,99E-01 |
| AT1G54630 | 0,35 | -0,57 | 4,38 | -0,13 | 8,96E-01 |
| AT1G54690 | 0,48 | -1,05 | 2,70 | -0,39 | 6,96E-01 |
| AT1G54775 | 0,57 | 0,00 | 3,88 | 0,00 | 1,00E+00 |
| AT1G54840 | 0,49 | -3,12 | 4,34 | -0,72 | 4,72E-01 |
| AT1G54950 | 0,44 | 0,00 | 4,12 | 0,00 | 1,00E+00 |
| AT1G55152 | 0,78 | 1,13 | 2,49 | 0,45 | 6,50E-01 |
| AT1G55180 | 0,78 | 0,41 | 2,69 | 0,15 | 8,78E-01 |
| AT1G55200 | 0,08 | 0,69 | 4,36 | 0,16 | 8,75E-01 |
| AT1G08043 | 0,30 | 0,30 | 4,34 | 0,07 | 9,45E-01 |
| AT1G55270 | 1,00 | -1,02 | 3,03 | -0,34 | 7,36E-01 |
| AT1G08053 | 2,32 | 0,24 | 1,56 | 0,16 | 8,77E-01 |
| AT1G55300 | 0,10 | 0,21 | 4,38 | 0,05 | 9,62E-01 |
| AT1G55330 | 0,51 | -0,40 | 3,19 | -0,13 | 8,99E-01 |
| AT1G55350 | 1,89 | -2,25 | 2,11 | -1,07 | 2,86E-01 |
| AT1G55360 | 0,31 | -0,54 | 4,33 | -0,12 | 9,02E-01 |
| AT1G55450 | 2,26 | 1,01 | 1,64 | 0,61 | 5,39E-01 |
| AT1G55490 | 3,72 | -2,50 | 1,10 | -2,27 | 2,32E-02 |
| AT1G55530 | 0,83 | 0,22 | 2,12 | 0,10 | 9,18E-01 |
| AT1G55535 | 0,40 | -0,51 | 4,21 | -0,12 | 9,03E-01 |
| AT1G55540 | 2,72 | -0,86 | 1,09 | -0,79 | 4,32E-01 |
| AT1G55590 | 0,61 | -1,07 | 2,56 | -0,42 | 6,76E-01 |
| AT1G55610 | 0,12 | 0,49 | 4,36 | 0,11 | 9,11E-01 |
| AT1G55620 | 0,57 | 1,55 | 3,17 | 0,49 | 6,26E-01 |
| AT1G55670 | 5,12 | -2,12 | 1,05 | -2,02 | 4,39E-02 |
| AT1G55675 | 0,68 | -3,75 | 3,69 | -1,02 | 3,10E-01 |
| AT1G55680 | 1,44 | 2,57 | 1,94 | 1,32 | 1,86E-01 |
| AT1G55690 | 0,12 | 1,16 | 4,34 | 0,27 | 7,89E-01 |
| AT1G55830 | 0,23 | 1,14 | 4,34 | 0,26 | 7,92E-01 |
| AT1G55880 | 0,36 | 1,57 | 4,34 | 0,36 | 7,17E-01 |
| AT1G55950 | 0,93 | 2,47 | 2,18 | 1,13 | 2,57E-01 |
| AT1G55960 | 0,25 | -1,75 | 4,40 | -0,40 | 6,91E-01 |
| AT1G55970 | 0,17 | 1,52 | 4,33 | 0,35 | 7,26E-01 |
| AT1G56010 | 0,42 | 0,90 | 4,09 | 0,22 | 8,26E-01 |

| | | | | | |
|---|---|---|---|---|---|
| AT1G56100 | 0,86 | -0,95 | 2,94 | -0,32 | 7,47E-01 |
| AT1G56110 | 1,35 | -1,13 | 1,88 | -0,60 | 5,47E-01 |
| AT1G56120 | 0,47 | -0,46 | 4,05 | -0,11 | 9,10E-01 |
| AT1G56140 | 0,44 | 0,44 | 3,27 | 0,13 | 8,94E-01 |
| AT1G56145 | 0,80 | -1,12 | 2,58 | -0,43 | 6,65E-01 |
| AT1G56165 | 0,20 | -0,44 | 4,36 | -0,10 | 9,19E-01 |
| AT1G56230 | 0,19 | 0,47 | 4,38 | 0,11 | 9,14E-01 |
| AT1G08107 | 0,31 | 0,07 | 3,68 | 0,02 | 9,85E-01 |
| AT1G08113 | 0,55 | 0,34 | 3,28 | 0,10 | 9,16E-01 |
| AT1G08117 | 4,98 | -0,93 | 1,21 | -0,77 | 4,44E-01 |
| AT1G08123 | 0,56 | 1,82 | 2,53 | 0,72 | 4,72E-01 |
| AT1G56280 | 2,59 | -0,71 | 1,15 | -0,62 | 5,35E-01 |
| AT1G56300 | 0,85 | -1,45 | 2,81 | -0,52 | 6,06E-01 |
| AT1G56330 | 0,78 | -3,29 | 2,52 | -1,31 | 1,91E-01 |
| AT1G56340 | 4,49 | 0,51 | 0,88 | 0,58 | 5,60E-01 |
| AT1G56345 | 0,19 | 0,00 | 4,41 | 0,00 | 1,00E+00 |
| AT1G56440 | 0,27 | -0,34 | 4,34 | -0,08 | 9,37E-01 |
| AT1G56520 | 1,21 | 0,01 | 2,11 | 0,01 | 9,96E-01 |
| AT1G56550 | 0,17 | 0,65 | 4,36 | 0,15 | 8,81E-01 |
| AT1G56580 | 0,24 | -0,73 | 4,37 | -0,17 | 8,68E-01 |
| AT1G56630 | 1,84 | -2,54 | 2,30 | -1,10 | 2,70E-01 |
| AT1G56650 | 1,01 | 3,65 | 3,90 | 0,94 | 3,49E-01 |
| AT1G57680 | 1,45 | 2,36 | 1,93 | 1,22 | 2,22E-01 |
| AT1G57775 | 0,38 | 0,06 | 3,43 | 0,02 | 9,85E-01 |
| AT1G57777 | 0,31 | 1,51 | 3,61 | 0,42 | 6,76E-01 |
| AT1G57860 | 0,30 | -0,86 | 4,40 | -0,20 | 8,44E-01 |
| AT1G57870 | 0,40 | 1,02 | 4,26 | 0,24 | 8,10E-01 |
| AT1G57943 | 0,08 | 0,44 | 4,38 | 0,10 | 9,21E-01 |
| AT1G57990 | 0,77 | 0,07 | 2,42 | 0,03 | 9,76E-01 |
| AT1G58030 | 0,73 | 0,10 | 2,45 | 0,04 | 9,68E-01 |
| AT1G58060 | 0,60 | 0,56 | 3,06 | 0,18 | 8,53E-01 |
| AT1G58080 | 0,29 | 0,71 | 4,31 | 0,17 | 8,68E-01 |
| AT1G58100 | 1,20 | -1,94 | 1,78 | -1,09 | 2,76E-01 |
| AT1G58110 | 0,53 | -0,72 | 3,88 | -0,18 | 8,53E-01 |
| AT1G58200 | 0,16 | 0,00 | 4,41 | 0,00 | 1,00E+00 |
| AT1G58220 | 0,54 | -2,15 | 3,83 | -0,56 | 5,75E-01 |
| AT1G58250 | 4,39 | 0,36 | 0,83 | 0,43 | 6,65E-01 |
| AT1G58290 | 5,16 | -0,47 | 0,75 | -0,62 | 5,33E-01 |
| AT1G58300 | 1,33 | -1,75 | 1,95 | -0,90 | 3,68E-01 |
| AT1G58350 | 0,80 | -1,32 | 2,48 | -0,53 | 5,95E-01 |
| AT1G58390 | 0,70 | 0,20 | 3,10 | 0,06 | 9,50E-01 |
| AT1G58440 | 0,58 | 0,00 | 3,12 | 0,00 | 1,00E+00 |
| AT1G58684 | 0,49 | 0,00 | 4,40 | 0,00 | 1,00E+00 |
| AT1G58725 | 0,33 | 0,20 | 4,36 | 0,05 | 9,64E-01 |
| AT1G58766 | 0,16 | 0,92 | 4,35 | 0,21 | 8,33E-01 |
| AT1G59030 | 0,44 | 1,31 | 4,28 | 0,31 | 7,59E-01 |
| AT1G59077 | 0,16 | 0,00 | 4,41 | 0,00 | 1,00E+00 |
| AT1G59406 | 0,14 | 0,03 | 4,39 | 0,01 | 9,95E-01 |
| AT1G59453 | 0,25 | 0,60 | 4,36 | 0,14 | 8,90E-01 |
| AT1G59720 | 0,61 | -1,18 | 2,66 | -0,44 | 6,58E-01 |
| AT1G59760 | 3,05 | -1,33 | 1,03 | -1,28 | 1,99E-01 |
| AT1G59780 | 0,52 | -1,41 | 3,20 | -0,44 | 6,59E-01 |

| | | | | | |
|---|---|---|---|---|---|
| AT1G59835 | 1,47 | -3,26 | 1,94 | -1,68 | 9,29E-02 |
| AT1G59910 | 1,37 | 0,38 | 1,96 | 0,19 | 8,46E-01 |
| AT1G59920 | 0,42 | 0,04 | 3,35 | 0,01 | 9,91E-01 |
| AT1G59930 | 1,11 | 2,62 | 2,39 | 1,10 | 2,72E-01 |
| AT1G59990 | 1,13 | -0,27 | 2,04 | -0,13 | 8,95E-01 |
| AT1G60000 | 0,38 | -0,33 | 3,53 | -0,09 | 9,24E-01 |
| AT1G60070 | 1,35 | 0,87 | 1,85 | 0,47 | 6,40E-01 |
| AT1G60140 | 2,14 | -1,35 | 1,19 | -1,14 | 2,55E-01 |
| AT1G60160 | 0,43 | 0,65 | 3,19 | 0,20 | 8,39E-01 |
| AT1G60200 | 0,68 | -2,37 | 3,18 | -0,74 | 4,56E-01 |
| AT1G60230 | 0,48 | -3,17 | 4,01 | -0,79 | 4,29E-01 |
| AT1G60260 | 0,77 | -1,62 | 2,49 | -0,65 | 5,17E-01 |
| AT1G08317 | 0,08 | 0,44 | 4,38 | 0,10 | 9,21E-01 |
| AT1G60430 | 0,45 | -0,01 | 3,94 | 0,00 | 9,99E-01 |
| AT1G60490 | 0,25 | 1,16 | 4,34 | 0,27 | 7,89E-01 |
| AT1G60550 | 0,38 | -1,21 | 4,34 | -0,28 | 7,81E-01 |
| AT1G60560 | 0,17 | 1,40 | 4,33 | 0,32 | 7,47E-01 |
| AT1G08327 | 2,78 | -2,26 | 1,34 | -1,68 | 9,21E-02 |
| AT1G60590 | 0,17 | 0,00 | 4,41 | 0,00 | 1,00E+00 |
| AT1G60630 | 0,08 | 0,44 | 4,38 | 0,10 | 9,21E-01 |
| AT1G60640 | 0,12 | 0,97 | 4,34 | 0,22 | 8,23E-01 |
| AT1G60700 | 0,48 | 1,17 | 4,32 | 0,27 | 7,86E-01 |
| AT1G60710 | 0,41 | -2,33 | 4,31 | -0,54 | 5,90E-01 |
| AT1G60770 | 1,20 | -0,62 | 1,87 | -0,33 | 7,42E-01 |
| AT1G60780 | 0,33 | 1,23 | 4,33 | 0,28 | 7,77E-01 |
| AT1G60800 | 0,56 | -0,51 | 3,05 | -0,17 | 8,68E-01 |
| AT1G60810 | 0,12 | 0,00 | 4,41 | 0,00 | 1,00E+00 |
| AT1G60850 | 0,11 | 0,65 | 4,36 | 0,15 | 8,81E-01 |
| AT1G60890 | 0,24 | 1,56 | 4,33 | 0,36 | 7,19E-01 |
| AT1G60930 | 0,24 | -0,08 | 4,34 | -0,02 | 9,85E-01 |
| AT1G60940 | 0,93 | -1,23 | 2,11 | -0,58 | 5,59E-01 |
| AT1G60960 | 1,57 | -1,64 | 1,57 | -1,05 | 2,96E-01 |
| AT1G60990 | 0,24 | 0,21 | 4,34 | 0,05 | 9,62E-01 |
| AT1G60995 | 0,35 | -0,94 | 4,39 | -0,21 | 8,31E-01 |
| AT1G61010 | 0,89 | -3,22 | 2,22 | -1,45 | 1,46E-01 |
| AT1G61100 | 1,97 | -0,10 | 1,44 | -0,07 | 9,46E-01 |
| AT1G61140 | 2,12 | -0,28 | 1,21 | -0,23 | 8,19E-01 |
| AT1G61200 | 1,13 | 0,56 | 1,86 | 0,30 | 7,63E-01 |
| AT1G61260 | 0,47 | -0,15 | 3,28 | -0,05 | 9,63E-01 |
| AT1G61300 | 1,06 | -0,05 | 2,23 | -0,02 | 9,82E-01 |
| AT1G61310 | 0,33 | 0,80 | 4,35 | 0,18 | 8,53E-01 |
| AT1G61360 | 0,15 | 0,45 | 4,37 | 0,10 | 9,19E-01 |
| AT1G61370 | 0,17 | 0,35 | 4,37 | 0,08 | 9,37E-01 |
| AT1G61380 | 0,34 | 1,55 | 4,32 | 0,36 | 7,20E-01 |
| AT1G61420 | 0,12 | 1,05 | 4,35 | 0,24 | 8,10E-01 |
| AT1G61560 | 0,37 | 1,53 | 4,33 | 0,35 | 7,25E-01 |
| AT1G08417 | 0,75 | -3,38 | 2,82 | -1,20 | 2,31E-01 |
| AT1G61660 | 0,65 | 1,74 | 2,94 | 0,59 | 5,56E-01 |
| AT1G61690 | 1,91 | 0,76 | 1,42 | 0,54 | 5,90E-01 |
| AT1G61730 | 1,18 | -0,26 | 2,02 | -0,13 | 8,96E-01 |
| AT1G61740 | 2,13 | -0,59 | 1,41 | -0,42 | 6,77E-01 |
| AT1G61890 | 1,56 | 1,54 | 1,75 | 0,88 | 3,81E-01 |

| | | | | | |
|---|---|---|---|---|---|
| AT1G61900 | 0,26 | 0,00 | 4,41 | 0,00 | 1,00E+00 |
| AT1G62030 | 1,53 | 1,48 | 1,93 | 0,76 | 4,45E-01 |
| AT1G62035 | 5,54 | -0,92 | 0,85 | -1,08 | 2,81E-01 |
| AT1G62085 | 0,60 | 0,85 | 3,16 | 0,27 | 7,87E-01 |
| AT1G62180 | 2,66 | 1,10 | 1,20 | 0,92 | 3,60E-01 |
| AT1G62300 | 0,27 | -1,78 | 4,41 | -0,40 | 6,87E-01 |
| AT1G62310 | 4,32 | -2,62 | 1,14 | -2,30 | 2,16E-02 |
| AT1G62390 | 2,10 | 0,50 | 1,70 | 0,30 | 7,68E-01 |
| AT1G62422 | 0,51 | 0,97 | 2,61 | 0,37 | 7,11E-01 |
| AT1G62610 | 0,76 | -1,41 | 2,21 | -0,64 | 5,24E-01 |
| AT1G62680 | 1,24 | 1,68 | 1,95 | 0,86 | 3,90E-01 |
| AT1G62760 | 0,29 | 1,22 | 4,33 | 0,28 | 7,79E-01 |
| AT1G62780 | 0,49 | -0,47 | 3,25 | -0,15 | 8,84E-01 |
| AT1G62810 | 0,97 | 0,55 | 1,94 | 0,28 | 7,79E-01 |
| AT1G62915 | 0,64 | 0,10 | 3,00 | 0,03 | 9,74E-01 |
| AT1G62960 | 0,36 | -1,75 | 4,40 | -0,40 | 6,91E-01 |
| AT1G63005 | 3,14 | 3,25 | 1,64 | 1,98 | 4,72E-02 |
| AT1G63010 | 0,16 | 0,00 | 4,41 | 0,00 | 1,00E+00 |
| AT1G63020 | 0,44 | 2,38 | 4,31 | 0,55 | 5,81E-01 |
| AT1G63050 | 0,34 | -0,92 | 4,34 | -0,21 | 8,32E-01 |
| AT1G63070 | 3,60 | 2,64 | 1,34 | 1,98 | 4,81E-02 |
| AT1G63090 | 1,34 | 0,97 | 1,69 | 0,58 | 5,65E-01 |
| AT1G63250 | 0,13 | 0,73 | 4,36 | 0,17 | 8,67E-01 |
| AT1G63290 | 0,12 | 0,97 | 4,34 | 0,22 | 8,23E-01 |
| AT1G63320 | 0,20 | 0,00 | 4,41 | 0,00 | 1,00E+00 |
| AT1G63360 | 4,56 | -0,51 | 1,02 | -0,50 | 6,15E-01 |
| AT1G63410 | 0,62 | 0,25 | 2,57 | 0,10 | 9,22E-01 |
| AT1G63490 | 0,53 | 0,55 | 3,20 | 0,17 | 8,63E-01 |
| AT1G63615 | 3,43 | -0,56 | 1,30 | -0,43 | 6,64E-01 |
| AT1G63660 | 0,92 | -1,13 | 2,05 | -0,55 | 5,81E-01 |
| AT1G63670 | 0,18 | -1,78 | 4,41 | -0,40 | 6,87E-01 |
| AT1G63700 | 1,36 | 0,80 | 1,42 | 0,56 | 5,76E-01 |
| AT1G63720 | 0,19 | 0,37 | 4,37 | 0,08 | 9,33E-01 |
| AT1G08627 | 1,18 | -0,97 | 2,11 | -0,46 | 6,45E-01 |
| AT1G63800 | 0,47 | -2,14 | 4,38 | -0,49 | 6,24E-01 |
| AT1G63810 | 0,31 | -0,68 | 4,39 | -0,16 | 8,77E-01 |
| AT1G63840 | 0,40 | 1,45 | 4,35 | 0,33 | 7,38E-01 |
| AT1G63860 | 1,09 | -1,30 | 1,72 | -0,76 | 4,50E-01 |
| AT1G63870 | 0,58 | 1,09 | 3,05 | 0,36 | 7,20E-01 |
| AT1G63880 | 4,32 | 1,31 | 0,97 | 1,36 | 1,75E-01 |
| AT1G63970 | 0,51 | -0,03 | 3,25 | -0,01 | 9,94E-01 |
| AT1G64080 | 0,15 | 0,00 | 4,38 | 0,00 | 1,00E+00 |
| AT1G64140 | 5,12 | -0,76 | 0,80 | -0,95 | 3,42E-01 |
| AT1G64190 | 1,11 | -0,19 | 1,66 | -0,11 | 9,10E-01 |
| AT1G64200 | 0,78 | 1,93 | 4,32 | 0,45 | 6,55E-01 |
| AT1G64280 | 0,86 | 0,97 | 2,44 | 0,40 | 6,89E-01 |
| AT1G64380 | 0,37 | 0,00 | 4,41 | 0,00 | 1,00E+00 |
| AT1G64460 | 0,72 | 0,29 | 2,37 | 0,12 | 9,02E-01 |
| AT1G64480 | 0,18 | 0,00 | 4,41 | 0,00 | 1,00E+00 |
| AT1G64510 | 0,32 | 0,00 | 4,38 | 0,00 | 9,99E-01 |
| AT1G64580 | 0,24 | 0,62 | 4,37 | 0,14 | 8,88E-01 |
| AT1G64583 | 1,87 | -0,90 | 1,89 | -0,48 | 6,34E-01 |

| | | | | | |
|---|---|---|---|---|---|
| AT1G64610 | 0,62 | -2,21 | 2,60 | -0,85 | 3,96E-01 |
| AT1G64640 | 0,11 | 0,48 | 4,37 | 0,11 | 9,12E-01 |
| AT1G64650 | 0,59 | 0,54 | 2,48 | 0,22 | 8,29E-01 |
| AT1G64670 | 0,10 | 0,21 | 4,38 | 0,05 | 9,62E-01 |
| AT1G64750 | 0,36 | -0,71 | 4,39 | -0,16 | 8,71E-01 |
| AT1G64970 | 0,63 | 0,63 | 2,57 | 0,25 | 8,05E-01 |
| AT1G64980 | 1,26 | -2,40 | 1,81 | -1,32 | 1,86E-01 |
| AT1G65040 | 0,28 | 0,06 | 3,75 | 0,02 | 9,87E-01 |
| AT1G65150 | 0,20 | 1,26 | 4,34 | 0,29 | 7,72E-01 |
| AT1G65190 | 0,43 | 0,34 | 4,14 | 0,08 | 9,35E-01 |
| AT1G08747 | 0,81 | -1,65 | 2,83 | -0,58 | 5,61E-01 |
| AT1G65290 | 0,77 | 0,70 | 2,20 | 0,32 | 7,49E-01 |
| AT1G65320 | 0,39 | 1,77 | 4,32 | 0,41 | 6,82E-01 |
| AT1G65350 | 2,42 | 0,50 | 1,48 | 0,34 | 7,36E-01 |
| AT1G65346 | 0,34 | 0,86 | 4,34 | 0,20 | 8,43E-01 |
| AT1G65420 | 0,15 | 0,67 | 4,37 | 0,15 | 8,79E-01 |
| AT1G65430 | 0,52 | 0,81 | 3,09 | 0,26 | 7,94E-01 |
| AT1G65440 | 2,14 | -0,36 | 1,29 | -0,28 | 7,82E-01 |
| AT1G65520 | 0,29 | 0,13 | 4,37 | 0,03 | 9,76E-01 |
| AT1G65540 | 1,25 | 2,03 | 1,96 | 1,04 | 3,00E-01 |
| AT1G65550 | 0,21 | 1,21 | 4,34 | 0,28 | 7,81E-01 |
| AT1G65580 | 0,46 | -0,92 | 2,84 | -0,32 | 7,47E-01 |
| AT1G65690 | 0,12 | 1,23 | 4,34 | 0,28 | 7,76E-01 |
| AT1G65720 | 3,55 | 1,42 | 0,97 | 1,47 | 1,42E-01 |
| AT1G65950 | 0,28 | -1,42 | 4,36 | -0,33 | 7,44E-01 |
| AT1G08797 | 0,63 | 2,45 | 4,30 | 0,57 | 5,69E-01 |
| AT1G65980 | 1,27 | -2,10 | 1,99 | -1,05 | 2,92E-01 |
| AT1G65985 | 0,33 | -0,12 | 4,33 | -0,03 | 9,77E-01 |
| AT1G65990 | 1,24 | 2,49 | 2,07 | 1,20 | 2,30E-01 |
| AT1G66200 | 1,38 | -2,05 | 1,61 | -1,27 | 2,03E-01 |
| AT1G66235 | 0,12 | 0,00 | 4,38 | 0,00 | 1,00E+00 |
| AT1G66240 | 0,12 | 0,97 | 4,34 | 0,22 | 8,23E-01 |
| AT1G66260 | 1,28 | 2,17 | 1,74 | 1,25 | 2,11E-01 |
| AT1G66290 | 0,49 | 0,15 | 3,27 | 0,05 | 9,63E-01 |
| AT1G66330 | 0,52 | 0,08 | 2,70 | 0,03 | 9,77E-01 |
| AT1G66410 | 1,68 | -1,54 | 1,64 | -0,94 | 3,48E-01 |
| AT1G66460 | 0,25 | 1,50 | 4,33 | 0,35 | 7,30E-01 |
| AT1G66490 | 0,90 | -3,29 | 2,41 | -1,37 | 1,72E-01 |
| AT1G66500 | 0,25 | -2,19 | 4,38 | -0,50 | 6,17E-01 |
| AT1G66570 | 1,67 | -0,43 | 1,49 | -0,29 | 7,74E-01 |
| AT1G66660 | 0,44 | 0,10 | 4,15 | 0,02 | 9,80E-01 |
| AT1G66670 | 1,37 | 1,48 | 1,90 | 0,78 | 4,36E-01 |
| AT1G66680 | 0,59 | -0,79 | 4,25 | -0,19 | 8,52E-01 |
| AT1G66700 | 0,46 | 2,12 | 4,32 | 0,49 | 6,23E-01 |
| AT1G66830 | 0,68 | -0,45 | 3,76 | -0,12 | 9,04E-01 |
| AT1G08857 | 2,42 | 2,05 | 1,83 | 1,12 | 2,61E-01 |
| AT1G66910 | 0,65 | -1,09 | 2,10 | -0,52 | 6,03E-01 |
| AT1G66940 | 0,34 | -1,74 | 4,40 | -0,40 | 6,93E-01 |
| AT1G66970 | 4,60 | 1,03 | 0,88 | 1,17 | 2,40E-01 |
| AT1G66980 | 1,51 | -0,98 | 1,53 | -0,64 | 5,21E-01 |
| AT1G67000 | 0,91 | 0,63 | 2,17 | 0,29 | 7,73E-01 |
| AT1G67050 | 0,28 | 0,09 | 3,74 | 0,02 | 9,82E-01 |

| | | | | | |
|---|---|---|---|---|---|
| AT1G67080 | 0,36 | -0,78 | 4,30 | -0,18 | 8,57E-01 |
| AT1G67140 | 0,95 | -0,24 | 2,35 | -0,10 | 9,18E-01 |
| AT1G67230 | 0,62 | 1,29 | 2,61 | 0,49 | 6,22E-01 |
| AT1G67235 | 0,29 | 0,00 | 4,41 | 0,00 | 1,00E+00 |
| AT1G67250 | 0,34 | -1,78 | 4,41 | -0,40 | 6,87E-01 |
| AT1G67280 | 1,72 | -1,02 | 1,47 | -0,69 | 4,90E-01 |
| AT1G67300 | 0,37 | 0,47 | 4,30 | 0,11 | 9,13E-01 |
| AT1G67310 | 0,33 | 0,92 | 4,31 | 0,21 | 8,30E-01 |
| AT1G67325 | 0,61 | 1,67 | 2,94 | 0,57 | 5,69E-01 |
| AT1G67360 | 0,73 | -1,72 | 2,99 | -0,57 | 5,67E-01 |
| AT1G67481 | 1,30 | -0,80 | 1,74 | -0,46 | 6,44E-01 |
| AT1G67490 | 0,57 | -2,08 | 3,11 | -0,67 | 5,04E-01 |
| AT1G67500 | 0,52 | -0,96 | 3,93 | -0,24 | 8,07E-01 |
| AT1G67510 | 0,38 | 0,32 | 4,27 | 0,08 | 9,40E-01 |
| AT1G67580 | 1,21 | -0,38 | 1,82 | -0,21 | 8,37E-01 |
| AT1G67590 | 0,17 | -1,70 | 4,40 | -0,39 | 7,00E-01 |
| AT1G67680 | 0,43 | 0,16 | 4,29 | 0,04 | 9,71E-01 |
| AT1G67740 | 3,89 | 0,21 | 1,07 | 0,19 | 8,46E-01 |
| AT1G67785 | 0,15 | 0,00 | 4,41 | 0,00 | 1,00E+00 |
| AT1G67790 | 0,26 | -0,30 | 4,37 | -0,07 | 9,46E-01 |
| AT1G67860 | 0,58 | 0,46 | 3,08 | 0,15 | 8,80E-01 |
| AT1G67865 | 0,24 | 1,35 | 4,34 | 0,31 | 7,55E-01 |
| AT1G67870 | 0,89 | 0,85 | 2,13 | 0,40 | 6,90E-01 |
| AT1G67930 | 1,11 | 0,01 | 1,60 | 0,01 | 9,94E-01 |
| AT1G67950 | 0,39 | -1,28 | 4,38 | -0,29 | 7,70E-01 |
| AT1G67960 | 1,96 | -0,02 | 1,46 | -0,01 | 9,89E-01 |
| AT1G68060 | 0,56 | 1,40 | 3,21 | 0,43 | 6,64E-01 |
| AT1G68140 | 0,87 | 1,90 | 2,10 | 0,90 | 3,66E-01 |
| AT1G68325 | 0,05 | 0,00 | 4,41 | 0,00 | 1,00E+00 |
| AT1G68370 | 0,32 | 0,39 | 4,34 | 0,09 | 9,28E-01 |
| AT1G68520 | 2,00 | 0,30 | 1,15 | 0,26 | 7,94E-01 |
| AT1G68530 | 2,97 | -0,60 | 1,18 | -0,51 | 6,10E-01 |
| AT1G68550 | 0,87 | 2,44 | 2,06 | 1,19 | 2,36E-01 |
| AT1G68560 | 3,34 | 1,19 | 1,26 | 0,95 | 3,44E-01 |
| AT1G08997 | 0,58 | 1,29 | 3,79 | 0,34 | 7,34E-01 |
| AT1G68590 | 1,41 | -0,86 | 1,57 | -0,55 | 5,85E-01 |
| AT1G68660 | 0,71 | 0,02 | 2,28 | 0,01 | 9,94E-01 |
| AT1G68710 | 0,46 | 0,40 | 4,30 | 0,09 | 9,25E-01 |
| AT1G68830 | 0,57 | 0,56 | 2,65 | 0,21 | 8,32E-01 |
| AT1G68910 | 0,08 | 0,69 | 4,36 | 0,16 | 8,75E-01 |
| AT1G09027 | 2,11 | 1,13 | 1,64 | 0,69 | 4,90E-01 |
| AT1G69020 | 0,30 | 0,30 | 4,34 | 0,07 | 9,45E-01 |
| AT1G69070 | 0,46 | -0,97 | 3,23 | -0,30 | 7,63E-01 |
| AT1G69220 | 1,28 | -0,64 | 1,73 | -0,37 | 7,09E-01 |
| AT1G69270 | 0,78 | -0,93 | 2,98 | -0,31 | 7,54E-01 |
| AT1G69280 | 0,21 | 1,21 | 4,34 | 0,28 | 7,81E-01 |
| AT1G69290 | 0,15 | 1,20 | 4,34 | 0,28 | 7,81E-01 |
| AT1G69310 | 0,25 | 1,82 | 4,32 | 0,42 | 6,73E-01 |
| AT1G69360 | 2,19 | 0,19 | 1,37 | 0,14 | 8,87E-01 |
| AT1G69450 | 0,38 | -2,35 | 4,36 | -0,54 | 5,89E-01 |
| AT1G69480 | 0,55 | 1,46 | 4,34 | 0,34 | 7,37E-01 |
| AT1G69550 | 1,87 | 0,29 | 1,40 | 0,21 | 8,37E-01 |

| | | | | | |
|---|---|---|---|---|---|
| AT1G69570 | 2,44 | 0,13 | 1,49 | 0,09 | 9,28E-01 |
| AT1G69630 | 0,59 | -3,49 | 4,32 | -0,81 | 4,18E-01 |
| AT1G69640 | 0,90 | 2,06 | 2,40 | 0,86 | 3,90E-01 |
| AT1G69730 | 0,73 | 0,96 | 2,36 | 0,41 | 6,83E-01 |
| AT1G69760 | 0,47 | -1,69 | 3,25 | -0,52 | 6,03E-01 |
| AT1G69770 | 1,05 | -1,66 | 2,12 | -0,79 | 4,32E-01 |
| AT1G69792 | 2,23 | 3,64 | 1,82 | 2,00 | 4,50E-02 |
| AT1G69795 | 1,39 | 3,43 | 2,33 | 1,47 | 1,41E-01 |
| AT1G69830 | 1,91 | -0,86 | 1,27 | -0,68 | 4,96E-01 |
| AT1G69840 | 0,65 | 0,32 | 3,04 | 0,11 | 9,15E-01 |
| AT1G69850 | 0,22 | 0,83 | 4,34 | 0,19 | 8,49E-01 |
| AT1G69890 | 1,49 | 3,43 | 1,86 | 1,85 | 6,43E-02 |
| AT1G69940 | 3,40 | 0,51 | 1,13 | 0,45 | 6,51E-01 |
| AT1G69960 | 0,51 | 0,35 | 3,11 | 0,11 | 9,10E-01 |
| AT1G70000 | 0,98 | 0,89 | 1,93 | 0,46 | 6,43E-01 |
| AT1G70070 | 1,07 | 0,03 | 2,20 | 0,01 | 9,90E-01 |
| AT1G70200 | 0,37 | -1,71 | 4,40 | -0,39 | 6,98E-01 |
| AT1G70280 | 0,17 | 1,40 | 4,33 | 0,32 | 7,47E-01 |
| AT1G70290 | 2,53 | -0,26 | 1,18 | -0,22 | 8,24E-01 |
| AT1G70300 | 0,38 | 0,28 | 4,25 | 0,07 | 9,47E-01 |
| AT1G70310 | 0,89 | -0,42 | 3,04 | -0,14 | 8,90E-01 |
| AT1G70340 | 0,34 | -0,32 | 3,57 | -0,09 | 9,30E-01 |
| AT1G70370 | 0,57 | 0,93 | 2,74 | 0,34 | 7,34E-01 |
| AT1G70410 | 0,70 | -2,90 | 2,66 | -1,09 | 2,75E-01 |
| AT1G70420 | 0,44 | 0,00 | 4,12 | 0,00 | 1,00E+00 |
| AT1G70490 | 1,85 | -1,42 | 1,40 | -1,01 | 3,12E-01 |
| AT1G70520 | 0,53 | 1,74 | 4,32 | 0,40 | 6,88E-01 |
| AT1G70530 | 1,72 | 0,60 | 1,76 | 0,34 | 7,31E-01 |
| AT1G70600 | 0,97 | 0,36 | 2,81 | 0,13 | 8,99E-01 |
| AT1G70730 | 0,91 | 0,83 | 2,27 | 0,36 | 7,15E-01 |
| AT1G70740 | 0,10 | 0,00 | 4,41 | 0,00 | 1,00E+00 |
| AT1G70750 | 0,44 | 2,04 | 4,32 | 0,47 | 6,38E-01 |
| AT1G70780 | 0,73 | 2,17 | 2,77 | 0,78 | 4,34E-01 |
| AT1G70830 | 0,27 | 1,46 | 4,33 | 0,34 | 7,37E-01 |
| AT1G70890 | 1,35 | -0,54 | 1,91 | -0,28 | 7,77E-01 |
| AT1G71002 | 2,81 | -1,08 | 1,18 | -0,91 | 3,60E-01 |
| AT1G71020 | 0,94 | -0,81 | 2,13 | -0,38 | 7,02E-01 |
| AT1G71030 | 0,60 | -0,48 | 2,53 | -0,19 | 8,49E-01 |
| AT1G71040 | 0,83 | -0,81 | 1,92 | -0,42 | 6,73E-01 |
| AT1G71060 | 0,73 | 1,99 | 3,39 | 0,58 | 5,59E-01 |
| AT1G71070 | 0,49 | -0,47 | 3,11 | -0,15 | 8,81E-01 |
| AT1G71080 | 0,90 | -0,07 | 2,76 | -0,03 | 9,79E-01 |
| AT1G71160 | 0,69 | 1,11 | 3,50 | 0,32 | 7,50E-01 |
| AT1G71190 | 0,19 | 1,09 | 4,34 | 0,25 | 8,01E-01 |
| AT1G71210 | 0,73 | 0,37 | 2,80 | 0,13 | 8,96E-01 |
| AT1G71310 | 0,27 | 1,28 | 4,35 | 0,29 | 7,69E-01 |
| AT1G71330 | 0,30 | 0,07 | 4,35 | 0,02 | 9,87E-01 |
| AT1G71340 | 0,92 | -0,46 | 2,45 | -0,19 | 8,51E-01 |
| AT1G71350 | 0,21 | 1,76 | 4,32 | 0,41 | 6,85E-01 |
| AT1G71410 | 1,45 | -2,34 | 1,76 | -1,33 | 1,82E-01 |
| AT1G09253 | 0,32 | 0,22 | 4,34 | 0,05 | 9,61E-01 |
| AT1G71490 | 0,42 | 0,92 | 4,16 | 0,22 | 8,25E-01 |

| | | | | | |
|---|---|---|---|---|---|
| AT1G71710 | 0,44 | -0,17 | 2,87 | -0,06 | 9,53E-01 |
| AT1G71770 | 0,40 | 1,82 | 4,13 | 0,44 | 6,59E-01 |
| AT1G71870 | 0,88 | 1,90 | 2,83 | 0,67 | 5,02E-01 |
| AT1G71960 | 0,26 | 1,24 | 3,76 | 0,33 | 7,41E-01 |
| AT1G72020 | 0,56 | -0,68 | 2,63 | -0,26 | 7,97E-01 |
| AT1G72040 | 0,83 | 1,03 | 2,15 | 0,48 | 6,32E-01 |
| AT1G72160 | 0,64 | -1,04 | 2,35 | -0,44 | 6,59E-01 |
| AT1G72190 | 0,52 | 2,39 | 3,84 | 0,62 | 5,34E-01 |
| AT1G72270 | 0,12 | 0,37 | 4,37 | 0,08 | 9,33E-01 |
| AT1G72300 | 0,54 | -0,05 | 3,02 | -0,02 | 9,86E-01 |
| AT1G72320 | 0,39 | -0,66 | 3,40 | -0,20 | 8,45E-01 |
| AT1G72370 | 2,39 | -0,85 | 1,18 | -0,71 | 4,76E-01 |
| AT1G72410 | 1,92 | -2,04 | 1,38 | -1,48 | 1,39E-01 |
| AT1G72440 | 1,79 | 1,89 | 1,59 | 1,19 | 2,35E-01 |
| AT1G72450 | 1,12 | -0,33 | 1,79 | -0,18 | 8,54E-01 |
| AT1G72480 | 0,35 | 1,35 | 4,34 | 0,31 | 7,56E-01 |
| AT1G72500 | 0,46 | -2,73 | 4,34 | -0,63 | 5,29E-01 |
| AT1G72550 | 0,79 | -1,09 | 2,39 | -0,46 | 6,47E-01 |
| AT1G72560 | 1,09 | -1,14 | 2,41 | -0,47 | 6,36E-01 |
| AT1G09327 | 0,20 | 0,75 | 4,37 | 0,17 | 8,64E-01 |
| AT1G72670 | 0,20 | -0,44 | 4,36 | -0,10 | 9,19E-01 |
| AT1G72720 | 0,35 | -2,17 | 4,38 | -0,49 | 6,21E-01 |
| AT1G72730 | 0,32 | -0,65 | 4,33 | -0,15 | 8,81E-01 |
| AT1G72760 | 0,65 | 3,07 | 4,29 | 0,72 | 4,74E-01 |
| AT1G72840 | 0,41 | 2,19 | 4,32 | 0,51 | 6,12E-01 |
| AT1G72860 | 0,44 | 2,38 | 4,31 | 0,55 | 5,81E-01 |
| AT1G73010 | 0,40 | 2,16 | 4,32 | 0,50 | 6,17E-01 |
| AT1G09363 | 0,41 | -0,92 | 3,37 | -0,27 | 7,86E-01 |
| AT1G73110 | 0,63 | 1,59 | 3,81 | 0,42 | 6,77E-01 |
| AT1G73170 | 1,76 | -0,05 | 1,67 | -0,03 | 9,78E-01 |
| AT1G73200 | 0,73 | -1,15 | 2,14 | -0,54 | 5,91E-01 |
| AT1G73230 | 0,17 | 1,52 | 4,33 | 0,35 | 7,26E-01 |
| AT1G73260 | 0,63 | -0,89 | 2,85 | -0,31 | 7,55E-01 |
| AT1G73370 | 0,52 | 2,69 | 4,30 | 0,63 | 5,31E-01 |
| AT1G73390 | 0,25 | 0,91 | 4,35 | 0,21 | 8,34E-01 |
| AT1G73490 | 0,37 | 0,04 | 4,30 | 0,01 | 9,93E-01 |
| AT1G73500 | 0,92 | -1,06 | 1,98 | -0,53 | 5,93E-01 |
| AT1G73510 | 0,45 | -1,34 | 4,01 | -0,33 | 7,38E-01 |
| AT1G73530 | 0,46 | 0,37 | 4,30 | 0,09 | 9,32E-01 |
| AT1G73540 | 0,81 | 0,99 | 2,21 | 0,45 | 6,55E-01 |
| AT1G73650 | 0,96 | 2,15 | 2,42 | 0,89 | 3,74E-01 |
| AT1G73655 | 0,74 | 1,35 | 3,73 | 0,36 | 7,18E-01 |
| AT1G73660 | 1,58 | -1,02 | 1,76 | -0,58 | 5,63E-01 |
| AT1G73680 | 0,85 | 2,57 | 2,15 | 1,20 | 2,32E-01 |
| AT1G73730 | 0,17 | 1,40 | 4,33 | 0,32 | 7,47E-01 |
| AT1G73760 | 0,39 | -0,76 | 4,25 | -0,18 | 8,58E-01 |
| AT1G73805 | 1,81 | 0,29 | 1,55 | 0,19 | 8,51E-01 |
| AT1G73930 | 0,25 | 0,03 | 4,34 | 0,01 | 9,95E-01 |
| AT1G73950 | 0,45 | 0,28 | 4,34 | 0,06 | 9,49E-01 |
| AT1G73960 | 0,29 | 0,60 | 4,31 | 0,14 | 8,89E-01 |
| AT1G73980 | 0,67 | 0,79 | 2,59 | 0,30 | 7,61E-01 |
| AT1G74050 | 0,43 | 0,82 | 3,24 | 0,25 | 8,01E-01 |

| | | | | | |
|---|---|---|---|---|---|
| AT1G74060 | 0,72 | -2,70 | 4,36 | -0,62 | 5,36E-01 |
| AT1G74070 | 0,42 | 0,00 | 4,30 | 0,00 | 1,00E+00 |
| AT1G74120 | 0,82 | -0,44 | 2,87 | -0,15 | 8,78E-01 |
| AT1G74170 | 0,18 | 1,15 | 4,34 | 0,26 | 7,91E-01 |
| AT1G74340 | 0,31 | 1,34 | 4,33 | 0,31 | 7,58E-01 |
| AT1G74370 | 0,53 | -0,58 | 3,95 | -0,15 | 8,82E-01 |
| AT1G74390 | 0,95 | -0,39 | 2,52 | -0,16 | 8,76E-01 |
| AT1G74420 | 0,45 | -1,75 | 4,32 | -0,40 | 6,86E-01 |
| AT1G74440 | 0,80 | -0,15 | 3,01 | -0,05 | 9,59E-01 |
| AT1G74453 | 1,99 | 0,13 | 1,54 | 0,08 | 9,33E-01 |
| AT1G09467 | 0,50 | 1,55 | 4,11 | 0,38 | 7,07E-01 |
| AT1G74560 | 0,37 | 0,02 | 3,51 | 0,01 | 9,95E-01 |
| AT1G74690 | 0,95 | 0,85 | 1,98 | 0,43 | 6,68E-01 |
| AT1G74770 | 0,42 | -0,70 | 4,13 | -0,17 | 8,66E-01 |
| AT1G74800 | 0,56 | 2,14 | 3,14 | 0,68 | 4,95E-01 |
| AT1G74830 | 0,12 | 1,16 | 4,34 | 0,27 | 7,89E-01 |
| AT1G74850 | 0,94 | -1,17 | 1,95 | -0,60 | 5,49E-01 |
| AT1G74880 | 0,11 | 0,65 | 4,36 | 0,15 | 8,81E-01 |
| AT1G74920 | 0,86 | -1,47 | 2,63 | -0,56 | 5,76E-01 |
| AT1G74950 | 0,28 | -0,83 | 4,37 | -0,19 | 8,49E-01 |
| AT1G74960 | 1,51 | -0,62 | 2,48 | -0,25 | 8,04E-01 |
| AT1G74970 | 0,46 | -1,63 | 4,34 | -0,38 | 7,07E-01 |
| AT1G75010 | 0,20 | 0,00 | 4,41 | 0,00 | 1,00E+00 |
| AT1G75100 | 0,60 | 0,46 | 2,24 | 0,20 | 8,39E-01 |
| AT1G75130 | 0,12 | 0,97 | 4,34 | 0,22 | 8,23E-01 |
| AT1G75180 | 0,38 | 0,83 | 3,38 | 0,25 | 8,06E-01 |
| AT1G75200 | 0,15 | 1,01 | 4,34 | 0,23 | 8,17E-01 |
| AT1G75220 | 0,55 | 2,32 | 3,15 | 0,74 | 4,62E-01 |
| AT1G75370 | 1,12 | -1,49 | 1,57 | -0,95 | 3,41E-01 |
| AT1G75380 | 2,78 | -0,02 | 1,13 | -0,02 | 9,83E-01 |
| AT1G75410 | 0,90 | 1,21 | 2,46 | 0,49 | 6,24E-01 |
| AT1G75450 | 0,11 | 0,48 | 4,37 | 0,11 | 9,12E-01 |
| AT1G75500 | 0,97 | -0,23 | 1,85 | -0,12 | 9,03E-01 |
| AT1G75680 | 0,99 | -0,14 | 1,95 | -0,07 | 9,44E-01 |
| AT1G75690 | 0,84 | -2,11 | 2,01 | -1,05 | 2,94E-01 |
| AT1G75750 | 1,13 | -0,44 | 2,08 | -0,21 | 8,32E-01 |
| AT1G75780 | 0,25 | 0,83 | 4,34 | 0,19 | 8,48E-01 |
| AT1G75820 | 1,17 | -0,10 | 2,10 | -0,05 | 9,64E-01 |
| AT1G75860 | 0,67 | -0,99 | 2,20 | -0,45 | 6,52E-01 |
| AT1G75945 | 0,28 | -0,02 | 4,35 | 0,00 | 9,96E-01 |
| AT1G76010 | 2,04 | -0,18 | 1,30 | -0,14 | 8,89E-01 |
| AT1G76060 | 0,20 | -1,80 | 4,41 | -0,41 | 6,83E-01 |
| AT1G76065 | 4,52 | 0,84 | 1,19 | 0,70 | 4,81E-01 |
| AT1G76062 | 4,01 | 1,02 | 1,31 | 0,78 | 4,35E-01 |
| AT1G76080 | 1,03 | 1,94 | 1,86 | 1,04 | 2,97E-01 |
| AT1G76090 | 1,72 | -1,15 | 1,56 | -0,73 | 4,63E-01 |
| AT1G76100 | 1,22 | -1,43 | 1,95 | -0,73 | 4,63E-01 |
| AT1G76120 | 0,15 | 0,83 | 4,35 | 0,19 | 8,49E-01 |
| AT1G76150 | 0,72 | 2,93 | 2,86 | 1,02 | 3,06E-01 |
| AT1G76160 | 0,88 | 0,55 | 1,86 | 0,30 | 7,68E-01 |
| AT1G76270 | 0,08 | 0,44 | 4,38 | 0,10 | 9,21E-01 |
| AT1G76280 | 0,12 | 0,97 | 4,34 | 0,22 | 8,23E-01 |

| | | | | | |
|---|---|---|---|---|---|
| AT1G09627 | 0,45 | 0,99 | 4,01 | 0,25 | 8,05E-01 |
| AT1G76400 | 0,57 | 1,20 | 3,03 | 0,40 | 6,92E-01 |
| AT1G76460 | 0,53 | -1,82 | 3,25 | -0,56 | 5,76E-01 |
| AT1G76550 | 2,61 | -0,33 | 1,10 | -0,30 | 7,63E-01 |
| AT1G76640 | 0,08 | 0,44 | 4,38 | 0,10 | 9,21E-01 |
| AT1G76650 | 0,42 | 0,06 | 3,41 | 0,02 | 9,85E-01 |
| AT1G76660 | 0,13 | 0,00 | 4,41 | 0,00 | 1,00E+00 |
| AT1G76670 | 1,15 | -1,26 | 1,96 | -0,64 | 5,20E-01 |
| AT1G76700 | 0,39 | -0,79 | 4,35 | -0,18 | 8,57E-01 |
| AT1G76810 | 1,64 | 0,47 | 1,70 | 0,27 | 7,83E-01 |
| AT1G76878 | 1,06 | -0,48 | 1,90 | -0,25 | 7,99E-01 |
| AT1G76890 | 0,74 | -0,60 | 1,98 | -0,30 | 7,62E-01 |
| AT1G76955 | 0,30 | 0,07 | 4,35 | 0,02 | 9,87E-01 |
| AT1G76960 | 0,40 | -0,34 | 3,43 | -0,10 | 9,22E-01 |
| AT1G76965 | 0,25 | -0,23 | 4,35 | -0,05 | 9,58E-01 |
| AT1G76970 | 0,43 | 1,22 | 3,35 | 0,36 | 7,16E-01 |
| AT1G76980 | 0,73 | 1,41 | 2,99 | 0,47 | 6,38E-01 |
| AT1G77030 | 0,32 | -0,91 | 4,39 | -0,21 | 8,36E-01 |
| AT1G77060 | 0,12 | 0,49 | 4,36 | 0,11 | 9,11E-01 |
| AT1G77100 | 0,82 | 2,48 | 3,65 | 0,68 | 4,97E-01 |
| AT1G77145 | 0,08 | 0,69 | 4,36 | 0,16 | 8,75E-01 |
| AT1G77170 | 0,63 | 0,53 | 2,90 | 0,18 | 8,56E-01 |
| AT1G77180 | 2,45 | 0,40 | 1,17 | 0,35 | 7,29E-01 |
| AT1G77240 | 1,26 | 0,32 | 1,68 | 0,19 | 8,48E-01 |
| AT1G77250 | 0,55 | 0,17 | 3,72 | 0,04 | 9,64E-01 |
| AT1G77260 | 0,57 | 0,81 | 3,80 | 0,21 | 8,30E-01 |
| AT1G77300 | 4,17 | -0,47 | 0,88 | -0,53 | 5,98E-01 |
| AT1G77480 | 0,33 | 0,00 | 4,41 | 0,00 | 1,00E+00 |
| AT1G77560 | 2,14 | 0,68 | 1,33 | 0,51 | 6,11E-01 |
| AT1G77580 | 0,18 | 0,65 | 4,35 | 0,15 | 8,82E-01 |
| AT1G77600 | 0,35 | 0,54 | 4,32 | 0,12 | 9,01E-01 |
| AT1G77660 | 0,33 | 0,79 | 4,31 | 0,18 | 8,55E-01 |
| AT1G77680 | 1,90 | -1,72 | 1,63 | -1,05 | 2,92E-01 |
| AT1G77760 | 2,44 | 0,49 | 1,29 | 0,38 | 7,01E-01 |
| AT1G77810 | 0,58 | -0,68 | 3,86 | -0,18 | 8,60E-01 |
| AT1G77815 | 2,65 | -0,27 | 1,67 | -0,16 | 8,71E-01 |
| AT1G77932 | 0,47 | -0,23 | 4,33 | -0,05 | 9,58E-01 |
| AT1G77940 | 0,55 | 0,97 | 2,57 | 0,38 | 7,06E-01 |
| AT1G77960 | 1,37 | -0,44 | 1,64 | -0,27 | 7,89E-01 |
| AT1G77980 | 0,77 | -0,90 | 2,22 | -0,40 | 6,86E-01 |
| AT1G77990 | 0,37 | 1,75 | 4,32 | 0,40 | 6,86E-01 |
| AT1G78060 | 0,43 | 0,00 | 4,39 | 0,00 | 1,00E+00 |
| AT1G78210 | 0,32 | -0,73 | 4,39 | -0,17 | 8,68E-01 |
| AT1G78240 | 0,51 | -0,90 | 2,68 | -0,34 | 7,36E-01 |
| AT1G78370 | 1,22 | -0,39 | 2,06 | -0,19 | 8,49E-01 |
| AT1G78380 | 1,48 | -0,34 | 1,78 | -0,19 | 8,48E-01 |
| AT1G78460 | 0,60 | -1,65 | 2,61 | -0,63 | 5,28E-01 |
| AT1G78480 | 0,26 | 0,12 | 4,33 | 0,03 | 9,79E-01 |
| AT1G78510 | 0,69 | -0,99 | 3,71 | -0,27 | 7,89E-01 |
| AT1G78580 | 0,70 | -0,08 | 2,20 | -0,04 | 9,71E-01 |
| AT1G78800 | 0,21 | 1,55 | 4,33 | 0,36 | 7,20E-01 |
| AT1G78820 | 0,54 | -0,19 | 2,36 | -0,08 | 9,35E-01 |

| | | | | | |
|---|---|---|---|---|---|
| AT1G78830 | 2,12 | -1,93 | 1,39 | -1,38 | 1,66E-01 |
| AT1G78850 | 0,94 | -0,90 | 2,12 | -0,42 | 6,72E-01 |
| AT1G78865 | 0,13 | 0,00 | 4,41 | 0,00 | 1,00E+00 |
| AT1G78880 | 0,88 | 1,54 | 2,75 | 0,56 | 5,76E-01 |
| AT1G78915 | 0,18 | 0,00 | 4,41 | 0,00 | 1,00E+00 |
| AT1G78990 | 0,38 | -1,52 | 4,32 | -0,35 | 7,24E-01 |
| AT1G78995 | 0,24 | 0,92 | 4,36 | 0,21 | 8,32E-01 |
| AT1G79000 | 2,98 | -0,45 | 1,57 | -0,28 | 7,76E-01 |
| AT1G79020 | 0,25 | -2,19 | 4,38 | -0,50 | 6,17E-01 |
| AT1G79030 | 0,53 | -2,64 | 3,98 | -0,66 | 5,06E-01 |
| AT1G79050 | 0,58 | -2,17 | 3,10 | -0,70 | 4,85E-01 |
| AT1G79080 | 1,23 | 1,45 | 2,09 | 0,69 | 4,89E-01 |
| AT1G79090 | 2,69 | -0,56 | 1,24 | -0,45 | 6,49E-01 |
| AT1G79120 | 0,10 | 0,00 | 4,41 | 0,00 | 1,00E+00 |
| AT1G79150 | 0,35 | 1,67 | 4,33 | 0,39 | 6,99E-01 |
| AT1G79280 | 4,52 | 0,36 | 0,85 | 0,43 | 6,68E-01 |
| AT1G79350 | 0,49 | -0,70 | 2,66 | -0,27 | 7,91E-01 |
| AT1G79430 | 0,17 | -1,70 | 4,40 | -0,39 | 7,00E-01 |
| AT1G79440 | 0,39 | 0,36 | 3,50 | 0,10 | 9,18E-01 |
| AT1G09867 | 1,21 | 1,78 | 2,39 | 0,74 | 4,57E-01 |
| AT1G09883 | 0,79 | -2,63 | 2,96 | -0,89 | 3,75E-01 |
| AT1G79540 | 0,53 | -0,20 | 3,04 | -0,06 | 9,48E-01 |
| AT1G79550 | 2,41 | -0,39 | 1,17 | -0,33 | 7,41E-01 |
| AT1G79570 | 2,57 | 0,13 | 1,12 | 0,12 | 9,07E-01 |
| AT1G79600 | 1,43 | -0,10 | 1,60 | -0,07 | 9,48E-01 |
| AT1G79640 | 0,40 | 1,45 | 4,35 | 0,33 | 7,38E-01 |
| AT1G79670 | 0,90 | 2,30 | 2,53 | 0,91 | 3,64E-01 |
| AT1G79700 | 0,08 | 0,69 | 4,36 | 0,16 | 8,75E-01 |
| AT1G79720 | 0,47 | 1,61 | 3,28 | 0,49 | 6,24E-01 |
| AT1G79730 | 0,63 | -0,86 | 3,83 | -0,22 | 8,22E-01 |
| AT1G79740 | 0,25 | -2,19 | 4,38 | -0,50 | 6,17E-01 |
| AT1G79750 | 4,46 | -0,76 | 0,94 | -0,81 | 4,20E-01 |
| AT1G79790 | 0,05 | 0,00 | 4,41 | 0,00 | 1,00E+00 |
| AT1G79830 | 1,06 | 0,38 | 1,82 | 0,21 | 8,36E-01 |
| AT1G79880 | 0,11 | 0,65 | 4,36 | 0,15 | 8,81E-01 |
| AT1G79900 | 0,42 | -0,70 | 4,13 | -0,17 | 8,66E-01 |
| AT1G80030 | 1,07 | 0,17 | 2,07 | 0,08 | 9,36E-01 |
| AT1G80040 | 0,61 | -2,15 | 2,50 | -0,86 | 3,90E-01 |
| AT1G80080 | 0,50 | -0,61 | 3,10 | -0,20 | 8,45E-01 |
| AT1G80250 | 2,27 | -2,61 | 1,32 | -1,97 | 4,88E-02 |
| AT1G80325 | 1,08 | 0,38 | 1,92 | 0,20 | 8,42E-01 |
| AT1G80350 | 0,38 | -1,52 | 4,32 | -0,35 | 7,24E-01 |
| AT1G80420 | 0,34 | 0,63 | 4,37 | 0,15 | 8,84E-01 |
| AT1G80460 | 0,24 | -0,27 | 4,35 | -0,06 | 9,51E-01 |
| AT1G80480 | 0,56 | 1,01 | 3,12 | 0,32 | 7,47E-01 |
| AT1G80490 | 1,23 | -1,02 | 1,56 | -0,65 | 5,13E-01 |
| AT1G80530 | 0,68 | 2,14 | 2,98 | 0,72 | 4,74E-01 |
| AT1G80600 | 1,40 | -1,75 | 1,90 | -0,92 | 3,57E-01 |
| AT1G80610 | 0,35 | 1,15 | 3,46 | 0,33 | 7,39E-01 |
| AT1G80630 | 0,28 | -0,49 | 4,38 | -0,11 | 9,11E-01 |
| AT1G80660 | 1,14 | -0,94 | 1,78 | -0,53 | 5,99E-01 |
| AT1G80670 | 0,39 | -1,28 | 4,38 | -0,29 | 7,70E-01 |

| | | | | | |
|---|---|---|---|---|---|
| AT1G80760 | 0,62 | -1,45 | 3,68 | -0,39 | 6,93E-01 |
| AT1G80830 | 0,27 | -0,81 | 4,37 | -0,19 | 8,53E-01 |
| AT1G80850 | 0,20 | 0,00 | 4,41 | 0,00 | 1,00E+00 |
| AT1G80920 | 0,74 | 0,19 | 2,10 | 0,09 | 9,28E-01 |
| AT1G80930 | 0,69 | 0,19 | 2,44 | 0,08 | 9,37E-01 |
| AT1G80950 | 0,35 | -0,44 | 4,37 | -0,10 | 9,20E-01 |
| AT2G01023 | 0,35 | 0,80 | 2,94 | 0,27 | 7,86E-01 |
| AT2G01050 | 0,45 | 0,37 | 4,39 | 0,08 | 9,32E-01 |
| AT2G01060 | 0,57 | 1,76 | 2,60 | 0,68 | 4,99E-01 |
| AT2G01110 | 0,90 | 0,52 | 2,30 | 0,23 | 8,21E-01 |
| AT2G01130 | 1,10 | -0,09 | 1,79 | -0,05 | 9,58E-01 |
| AT2G01140 | 1,55 | 1,60 | 1,76 | 0,91 | 3,64E-01 |
| AT2G01180 | 0,86 | 0,27 | 2,01 | 0,14 | 8,92E-01 |
| AT2G01250 | 0,52 | 0,70 | 3,90 | 0,18 | 8,57E-01 |
| AT2G01275 | 0,19 | 0,00 | 4,38 | 0,00 | 9,99E-01 |
| AT2G01290 | 0,18 | 1,24 | 4,33 | 0,29 | 7,74E-01 |
| AT2G01300 | 0,08 | 0,44 | 4,38 | 0,10 | 9,21E-01 |
| AT2G01310 | 0,05 | 0,00 | 4,41 | 0,00 | 1,00E+00 |
| AT2G01320 | 2,02 | -1,10 | 1,37 | -0,80 | 4,22E-01 |
| AT2G01350 | 0,12 | 0,00 | 4,41 | 0,00 | 1,00E+00 |
| AT2G01410 | 0,34 | -0,93 | 3,58 | -0,26 | 7,94E-01 |
| AT2G01420 | 1,24 | -0,75 | 2,02 | -0,37 | 7,10E-01 |
| AT2G01440 | 1,25 | 0,74 | 2,30 | 0,32 | 7,46E-01 |
| AT2G01450 | 1,15 | -0,19 | 1,63 | -0,12 | 9,05E-01 |
| AT2G01505 | 0,30 | 1,22 | 4,35 | 0,28 | 7,79E-01 |
| AT2G01570 | 3,41 | -0,56 | 0,92 | -0,61 | 5,43E-01 |
| AT2G01630 | 0,87 | -1,03 | 2,63 | -0,39 | 6,96E-01 |
| AT2G01650 | 1,03 | -0,17 | 2,00 | -0,09 | 9,31E-01 |
| AT2G01670 | 0,39 | 1,46 | 3,31 | 0,44 | 6,58E-01 |
| AT2G01690 | 0,55 | -1,54 | 3,79 | -0,40 | 6,86E-01 |
| AT2G01720 | 1,57 | 1,80 | 2,11 | 0,85 | 3,94E-01 |
| AT2G01755 | 0,14 | 0,03 | 4,39 | 0,01 | 9,95E-01 |
| AT2G01820 | 1,26 | 0,52 | 2,20 | 0,23 | 8,15E-01 |
| AT2G01830 | 0,91 | 0,98 | 2,04 | 0,48 | 6,32E-01 |
| AT2G04125 | 0,12 | 0,00 | 4,38 | 0,00 | 1,00E+00 |
| AT2G01870 | 0,33 | -0,23 | 4,38 | -0,05 | 9,59E-01 |
| AT2G01918 | 1,01 | -0,75 | 2,30 | -0,33 | 7,45E-01 |
| AT2G01930 | 0,16 | 0,49 | 4,38 | 0,11 | 9,12E-01 |
| AT2G01970 | 1,49 | -0,34 | 1,64 | -0,21 | 8,34E-01 |
| AT2G01990 | 0,25 | 0,55 | 4,35 | 0,13 | 8,99E-01 |
| AT2G02080 | 1,03 | -0,01 | 2,62 | 0,00 | 9,97E-01 |
| AT2G02103 | 0,70 | 1,21 | 2,55 | 0,48 | 6,34E-01 |
| AT2G02148 | 0,42 | -0,54 | 4,08 | -0,13 | 8,95E-01 |
| AT2G02150 | 0,39 | -0,03 | 4,23 | -0,01 | 9,94E-01 |
| AT2G02160 | 0,60 | 1,13 | 2,50 | 0,45 | 6,50E-01 |
| AT2G02220 | 0,80 | 1,01 | 2,53 | 0,40 | 6,89E-01 |
| AT2G02230 | 0,17 | 1,15 | 4,34 | 0,27 | 7,91E-01 |
| AT2G02340 | 1,23 | -1,80 | 2,14 | -0,84 | 4,00E-01 |
| AT2G02360 | 0,16 | 0,92 | 4,35 | 0,21 | 8,33E-01 |
| AT2G02400 | 0,84 | 1,03 | 2,27 | 0,45 | 6,50E-01 |
| AT2G02520 | 0,08 | 0,00 | 4,41 | 0,00 | 1,00E+00 |
| AT2G02570 | 0,25 | 0,60 | 4,36 | 0,14 | 8,90E-01 |

| | | | | | |
|---|---|---|---|---|---|
| AT2G02635 | 0,17 | 0,00 | 4,40 | 0,00 | 1,00E+00 |
| AT2G02695 | 0,58 | 1,49 | 3,00 | 0,50 | 6,19E-01 |
| AT2G02710 | 0,66 | -0,25 | 2,41 | -0,10 | 9,18E-01 |
| AT2G02780 | 0,28 | 0,00 | 4,41 | 0,00 | 1,00E+00 |
| AT2G02800 | 0,78 | 0,43 | 2,53 | 0,17 | 8,64E-01 |
| AT2G04345 | 0,62 | -0,83 | 2,60 | -0,32 | 7,51E-01 |
| AT2G02835 | 3,94 | -0,81 | 0,96 | -0,85 | 3,98E-01 |
| AT2G02930 | 0,49 | 1,88 | 3,20 | 0,59 | 5,56E-01 |
| AT2G03050 | 0,25 | 1,85 | 4,32 | 0,43 | 6,68E-01 |
| AT2G03090 | 0,08 | 0,69 | 4,36 | 0,16 | 8,75E-01 |
| AT2G03240 | 0,24 | 0,24 | 4,34 | 0,06 | 9,55E-01 |
| AT2G03250 | 2,57 | -0,53 | 1,63 | -0,33 | 7,44E-01 |
| AT2G03340 | 0,45 | -1,43 | 3,26 | -0,44 | 6,61E-01 |
| AT2G03390 | 0,61 | 0,49 | 2,35 | 0,21 | 8,34E-01 |
| AT2G03420 | 0,43 | 1,94 | 4,08 | 0,48 | 6,35E-01 |
| AT2G03430 | 0,52 | -0,98 | 4,33 | -0,23 | 8,22E-01 |
| AT2G03440 | 0,35 | -0,91 | 4,38 | -0,21 | 8,35E-01 |
| AT2G03460 | 0,08 | 0,69 | 4,36 | 0,16 | 8,75E-01 |
| AT2G03470 | 1,22 | -0,59 | 2,24 | -0,26 | 7,92E-01 |
| AT2G03490 | 0,08 | 0,69 | 4,36 | 0,16 | 8,75E-01 |
| AT2G03620 | 0,33 | -0,32 | 4,32 | -0,07 | 9,41E-01 |
| AT2G03640 | 0,13 | 0,73 | 4,36 | 0,17 | 8,67E-01 |
| AT2G03690 | 0,23 | -1,78 | 4,41 | -0,40 | 6,87E-01 |
| AT2G03730 | 0,29 | 0,89 | 4,35 | 0,20 | 8,39E-01 |
| AT2G03750 | 1,29 | 0,31 | 1,86 | 0,17 | 8,67E-01 |
| AT2G03760 | 0,93 | -0,63 | 3,41 | -0,19 | 8,52E-01 |
| AT2G03770 | 0,15 | 1,16 | 4,34 | 0,27 | 7,88E-01 |
| AT2G03820 | 4,25 | -0,35 | 0,90 | -0,39 | 6,94E-01 |
| AT2G03821 | 1,77 | 2,53 | 1,68 | 1,51 | 1,31E-01 |
| AT2G03930 | 1,60 | 2,04 | 1,73 | 1,18 | 2,38E-01 |
| AT2G04605 | 3,46 | 4,19 | 1,56 | 2,68 | 7,31E-03 |
| AT2G04655 | 0,14 | 0,00 | 4,41 | 0,00 | 1,00E+00 |
| AT2G04031 | 0,15 | 0,00 | 4,41 | 0,00 | 1,00E+00 |
| AT2G04037 | 0,16 | 0,00 | 4,39 | 0,00 | 1,00E+00 |
| AT2G04050 | 0,12 | 1,23 | 4,34 | 0,28 | 7,76E-01 |
| AT2G04060 | 1,72 | 2,72 | 2,16 | 1,26 | 2,08E-01 |
| AT2G04062 | 0,70 | 1,30 | 2,93 | 0,44 | 6,56E-01 |
| AT2G04063 | 0,69 | 1,75 | 2,76 | 0,63 | 5,27E-01 |
| AT2G04066 | 0,42 | 0,13 | 4,33 | 0,03 | 9,75E-01 |
| AT2G04070 | 0,33 | -1,30 | 4,33 | -0,30 | 7,64E-01 |
| AT2G04080 | 0,61 | -0,17 | 2,48 | -0,07 | 9,46E-01 |
| AT2G04160 | 0,43 | 0,67 | 4,10 | 0,16 | 8,70E-01 |
| AT2G04230 | 1,09 | 0,78 | 2,02 | 0,39 | 6,98E-01 |
| AT2G04280 | 2,22 | 1,48 | 1,49 | 0,99 | 3,22E-01 |
| AT2G04360 | 0,70 | -2,88 | 2,41 | -1,20 | 2,31E-01 |
| AT2G04378 | 0,56 | -1,34 | 2,75 | -0,49 | 6,25E-01 |
| AT2G04520 | 0,26 | 0,00 | 4,41 | 0,00 | 1,00E+00 |
| AT2G04620 | 0,17 | 1,52 | 4,33 | 0,35 | 7,26E-01 |
| AT2G04630 | 0,19 | 0,85 | 4,35 | 0,20 | 8,45E-01 |
| AT2G04842 | 0,88 | 2,71 | 3,02 | 0,90 | 3,70E-01 |
| AT2G04870 | 0,55 | -0,82 | 2,79 | -0,30 | 7,68E-01 |
| AT2G04890 | 0,11 | 0,48 | 4,37 | 0,11 | 9,12E-01 |

| | | | | | |
|---|---|---|---|---|---|
| AT2G05120 | 0,91 | 0,40 | 2,01 | 0,20 | 8,43E-01 |
| AT2G05220 | 0,38 | 0,38 | 4,32 | 0,09 | 9,31E-01 |
| AT2G05250 | 0,41 | 0,04 | 4,30 | 0,01 | 9,93E-01 |
| AT2G04895 | 0,17 | 0,07 | 4,40 | 0,02 | 9,88E-01 |
| AT2G05752 | 0,90 | 0,81 | 2,36 | 0,34 | 7,33E-01 |
| AT2G05765 | 3,87 | 0,20 | 1,20 | 0,16 | 8,70E-01 |
| AT2G04955 | 3,52 | -0,49 | 1,03 | -0,47 | 6,37E-01 |
| AT2G04965 | 3,81 | -0,75 | 1,03 | -0,73 | 4,67E-01 |
| AT2G04995 | 1,03 | 1,31 | 2,22 | 0,59 | 5,54E-01 |
| AT2G05810 | 1,10 | -0,14 | 1,82 | -0,08 | 9,40E-01 |
| AT2G05914 | 0,51 | 0,34 | 4,39 | 0,08 | 9,39E-01 |
| AT2G05920 | 1,42 | 3,30 | 1,72 | 1,91 | 5,56E-02 |
| AT2G05940 | 0,97 | 1,33 | 2,02 | 0,66 | 5,10E-01 |
| AT2G06000 | 0,52 | 0,03 | 2,51 | 0,01 | 9,90E-01 |
| AT2G06040 | 0,24 | 1,17 | 4,35 | 0,27 | 7,87E-01 |
| AT2G06050 | 0,61 | 0,75 | 3,12 | 0,24 | 8,09E-01 |
| AT2G06530 | 0,61 | 1,58 | 3,30 | 0,48 | 6,31E-01 |
| AT2G06541 | 0,49 | 1,50 | 4,11 | 0,36 | 7,16E-01 |
| AT2G06645 | 0,15 | 0,00 | 4,41 | 0,00 | 1,00E+00 |
| AT2G06675 | 0,19 | 0,00 | 4,41 | 0,00 | 1,00E+00 |
| AT2G06906 | 0,12 | 0,97 | 4,34 | 0,22 | 8,23E-01 |
| AT2G06925 | 0,35 | 0,03 | 3,57 | 0,01 | 9,93E-01 |
| AT2G06960 | 0,68 | 0,11 | 2,96 | 0,04 | 9,70E-01 |
| AT2G07020 | 0,35 | 0,00 | 4,41 | 0,00 | 1,00E+00 |
| AT2G07120 | 0,12 | 0,00 | 4,38 | 0,00 | 1,00E+00 |
| AT2G07180 | 1,49 | -1,15 | 1,88 | -0,61 | 5,43E-01 |
| AT2G07240 | 1,04 | 1,98 | 2,04 | 0,97 | 3,30E-01 |
| AT2G07360 | 3,99 | -1,06 | 0,85 | -1,25 | 2,12E-01 |
| AT2G07560 | 0,12 | 0,37 | 4,37 | 0,08 | 9,33E-01 |
| AT2G07771 | 0,58 | -1,15 | 2,67 | -0,43 | 6,67E-01 |
| AT2G07777 | 0,89 | 0,41 | 2,06 | 0,20 | 8,41E-01 |
| AT2G07671 | 2,57 | -0,16 | 1,47 | -0,11 | 9,13E-01 |
| AT2G07779 | 0,28 | -1,25 | 4,35 | -0,29 | 7,73E-01 |
| AT2G07621 | 0,12 | 0,00 | 4,38 | 0,00 | 1,00E+00 |
| AT2G07774 | 0,25 | 0,00 | 4,41 | 0,00 | 1,00E+00 |
| AT2G07687 | 1,11 | 0,99 | 2,00 | 0,50 | 6,20E-01 |
| AT2G07629 | 5,41 | -2,08 | 0,96 | -2,17 | 3,01E-02 |
| AT2G05205 | 0,68 | 1,40 | 2,60 | 0,54 | 5,90E-01 |
| AT2G07599 | 0,90 | 0,64 | 2,26 | 0,28 | 7,76E-01 |
| AT2G07706 | 1,10 | 0,18 | 2,52 | 0,07 | 9,44E-01 |
| AT2G07641 | 0,45 | -0,16 | 2,79 | -0,06 | 9,53E-01 |
| AT2G07719 | 0,62 | 0,89 | 2,61 | 0,34 | 7,32E-01 |
| AT2G07815 | 2,06 | 0,20 | 1,54 | 0,13 | 8,98E-01 |
| AT2G07756 | 1,02 | 0,11 | 2,00 | 0,05 | 9,57E-01 |
| AT2G07734 | 0,71 | -2,46 | 2,93 | -0,84 | 4,01E-01 |
| AT2G07658 | 0,65 | 0,52 | 2,49 | 0,21 | 8,34E-01 |
| AT2G07662 | 1,06 | 2,53 | 2,27 | 1,11 | 2,65E-01 |
| AT2G07835 | 0,78 | 1,87 | 2,92 | 0,64 | 5,22E-01 |
| AT2G07690 | 0,71 | 0,55 | 2,44 | 0,22 | 8,23E-01 |
| AT2G07760 | 0,23 | 0,66 | 4,38 | 0,15 | 8,81E-01 |
| AT2G09795 | 0,34 | 0,13 | 4,34 | 0,03 | 9,76E-01 |
| AT2G05465 | 0,27 | 0,00 | 4,38 | 0,00 | 1,00E+00 |

| | | | | | |
|---|---|---|---|---|---|
| AT2G10557 | 1,21 | 3,98 | 3,63 | 1,10 | 2,73E-01 |
| AT2G10608 | 0,36 | -2,23 | 4,38 | -0,51 | 6,10E-01 |
| AT2G05565 | 0,43 | -1,75 | 4,13 | -0,42 | 6,72E-01 |
| AT2G10625 | 0,25 | 0,00 | 4,41 | 0,00 | 1,00E+00 |
| AT2G10940 | 5,77 | -0,07 | 1,13 | -0,06 | 9,51E-01 |
| AT2G10950 | 0,47 | 0,42 | 4,00 | 0,10 | 9,17E-01 |
| AT2G10955 | 0,26 | 0,00 | 4,41 | 0,00 | 1,00E+00 |
| AT2G11000 | 0,08 | 0,00 | 4,41 | 0,00 | 1,00E+00 |
| AT2G11852 | 0,26 | 0,84 | 4,34 | 0,19 | 8,47E-01 |
| AT2G11910 | 1,43 | 1,47 | 1,68 | 0,87 | 3,83E-01 |
| AT2G05685 | 2,44 | 2,27 | 1,68 | 1,35 | 1,76E-01 |
| AT2G12400 | 0,28 | 0,00 | 4,38 | 0,00 | 9,99E-01 |
| AT2G12405 | 0,18 | 0,00 | 4,41 | 0,00 | 1,00E+00 |
| AT2G05705 | 0,21 | -1,78 | 4,41 | -0,40 | 6,87E-01 |
| AT2G12461 | 0,12 | 1,23 | 4,34 | 0,28 | 7,76E-01 |
| AT2G05835 | 0,29 | 0,10 | 4,40 | 0,02 | 9,83E-01 |
| AT2G12945 | 0,05 | 0,00 | 4,41 | 0,00 | 1,00E+00 |
| AT2G13360 | 3,44 | -1,08 | 1,03 | -1,05 | 2,94E-01 |
| AT2G13370 | 1,20 | -0,51 | 2,24 | -0,23 | 8,20E-01 |
| AT2G13450 | 1,51 | -1,71 | 1,79 | -0,95 | 3,40E-01 |
| AT2G13550 | 0,22 | 0,90 | 4,34 | 0,21 | 8,36E-01 |
| AT2G13640 | 0,11 | 0,65 | 4,36 | 0,15 | 8,81E-01 |
| AT2G13650 | 0,22 | -1,77 | 4,41 | -0,40 | 6,88E-01 |
| AT2G13820 | 1,49 | -0,01 | 2,90 | 0,00 | 9,99E-01 |
| AT2G14100 | 0,15 | 0,29 | 4,37 | 0,07 | 9,47E-01 |
| AT2G14120 | 0,98 | 1,00 | 2,01 | 0,50 | 6,19E-01 |
| AT2G14170 | 1,22 | 2,90 | 2,10 | 1,38 | 1,67E-01 |
| AT2G14260 | 0,26 | 1,69 | 4,32 | 0,39 | 6,96E-01 |
| AT2G14265 | 1,52 | 0,69 | 1,94 | 0,35 | 7,23E-01 |
| AT2G14288 | 0,20 | 0,00 | 4,41 | 0,00 | 1,00E+00 |
| AT2G14510 | 0,15 | 0,29 | 4,37 | 0,07 | 9,47E-01 |
| AT2G14540 | 0,12 | 0,97 | 4,34 | 0,22 | 8,23E-01 |
| AT2G14610 | 3,13 | 1,47 | 1,15 | 1,27 | 2,03E-01 |
| AT2G14670 | 1,21 | 0,00 | 2,67 | 0,00 | 1,00E+00 |
| AT2G14720 | 1,05 | -0,06 | 2,18 | -0,03 | 9,78E-01 |
| AT2G14755 | 0,32 | 0,00 | 4,41 | 0,00 | 1,00E+00 |
| AT2G14910 | 2,97 | 0,30 | 1,14 | 0,26 | 7,94E-01 |
| AT2G15000 | 0,23 | -1,78 | 4,41 | -0,40 | 6,87E-01 |
| AT2G15020 | 0,51 | -0,72 | 3,22 | -0,22 | 8,23E-01 |
| AT2G06355 | 0,23 | 0,13 | 4,37 | 0,03 | 9,77E-01 |
| AT2G15110 | 0,63 | 1,25 | 2,97 | 0,42 | 6,73E-01 |
| AT2G15290 | 0,58 | 0,35 | 3,72 | 0,09 | 9,26E-01 |
| AT2G15292 | 1,13 | 1,24 | 1,70 | 0,73 | 4,66E-01 |
| AT2G15320 | 1,25 | 1,22 | 1,78 | 0,69 | 4,93E-01 |
| AT2G15390 | 0,37 | 0,37 | 4,32 | 0,09 | 9,31E-01 |
| AT2G15420 | 2,17 | 2,21 | 1,66 | 1,33 | 1,83E-01 |
| AT2G15555 | 1,03 | 0,43 | 1,76 | 0,25 | 8,06E-01 |
| AT2G15560 | 0,05 | 0,00 | 4,41 | 0,00 | 1,00E+00 |
| AT2G15790 | 0,41 | -0,06 | 4,16 | -0,01 | 9,89E-01 |
| AT2G15860 | 0,36 | 1,06 | 4,33 | 0,24 | 8,07E-01 |
| AT2G15880 | 1,49 | -0,93 | 1,59 | -0,58 | 5,59E-01 |
| AT2G15890 | 1,14 | 0,93 | 1,63 | 0,57 | 5,70E-01 |

| | | | | | |
|---|---|---|---|---|---|
| AT2G06515 | 0,69 | -1,78 | 3,76 | -0,47 | 6,36E-01 |
| AT2G15980 | 0,29 | 0,87 | 3,67 | 0,24 | 8,13E-01 |
| AT2G16016 | 1,06 | 0,39 | 2,06 | 0,19 | 8,49E-01 |
| AT2G06575 | 0,75 | -0,15 | 2,20 | -0,07 | 9,46E-01 |
| AT2G16070 | 0,64 | -0,78 | 3,79 | -0,21 | 8,37E-01 |
| AT2G16090 | 0,55 | -0,22 | 2,45 | -0,09 | 9,30E-01 |
| AT2G16370 | 0,11 | 0,65 | 4,36 | 0,15 | 8,81E-01 |
| AT2G16400 | 1,00 | 1,46 | 1,99 | 0,73 | 4,64E-01 |
| AT2G16405 | 0,20 | 1,26 | 4,34 | 0,29 | 7,72E-01 |
| AT2G16440 | 1,04 | -0,79 | 3,41 | -0,23 | 8,17E-01 |
| AT2G16450 | 0,33 | -0,32 | 4,32 | -0,07 | 9,41E-01 |
| AT2G16485 | 2,25 | 0,76 | 1,26 | 0,60 | 5,47E-01 |
| AT2G16490 | 1,37 | -2,57 | 2,52 | -1,02 | 3,09E-01 |
| AT2G16505 | 0,85 | 1,95 | 2,29 | 0,85 | 3,95E-01 |
| AT2G16510 | 1,63 | 2,21 | 1,74 | 1,27 | 2,03E-01 |
| AT2G16620 | 0,18 | 0,21 | 4,37 | 0,05 | 9,62E-01 |
| AT2G16640 | 1,05 | 1,38 | 2,06 | 0,67 | 5,03E-01 |
| AT2G16650 | 0,37 | 2,37 | 4,31 | 0,55 | 5,83E-01 |
| AT2G16660 | 2,17 | -0,40 | 1,21 | -0,33 | 7,40E-01 |
| AT2G16676 | 2,63 | 0,20 | 1,28 | 0,16 | 8,73E-01 |
| AT2G16870 | 1,12 | -0,75 | 1,80 | -0,42 | 6,76E-01 |
| AT2G16880 | 0,51 | 0,04 | 4,32 | 0,01 | 9,93E-01 |
| AT2G16920 | 0,25 | 1,26 | 4,34 | 0,29 | 7,71E-01 |
| AT2G16940 | 0,88 | 0,27 | 2,08 | 0,13 | 8,96E-01 |
| AT2G17020 | 0,78 | 0,17 | 1,95 | 0,09 | 9,29E-01 |
| AT2G17110 | 0,49 | 0,25 | 2,71 | 0,09 | 9,26E-01 |
| AT2G17130 | 0,45 | -1,15 | 4,14 | -0,28 | 7,81E-01 |
| AT2G17190 | 0,94 | 1,52 | 2,45 | 0,62 | 5,34E-01 |
| AT2G17200 | 1,89 | 1,25 | 1,57 | 0,79 | 4,28E-01 |
| AT2G17210 | 0,23 | -0,05 | 4,34 | -0,01 | 9,91E-01 |
| AT2G17220 | 0,39 | -3,00 | 4,35 | -0,69 | 4,90E-01 |
| AT2G17230 | 0,92 | 2,56 | 2,46 | 1,04 | 2,98E-01 |
| AT2G17300 | 0,17 | 1,15 | 4,34 | 0,27 | 7,91E-01 |
| AT2G06935 | 0,05 | 0,00 | 4,41 | 0,00 | 1,00E+00 |
| AT2G17340 | 0,86 | 0,46 | 2,16 | 0,21 | 8,33E-01 |
| AT2G17350 | 0,34 | 0,00 | 4,41 | 0,00 | 1,00E+00 |
| AT2G17450 | 0,63 | -1,08 | 2,19 | -0,49 | 6,21E-01 |
| AT2G17470 | 0,10 | 0,21 | 4,38 | 0,05 | 9,62E-01 |
| AT2G17510 | 1,71 | 1,73 | 1,49 | 1,17 | 2,43E-01 |
| AT2G17550 | 0,71 | 1,75 | 2,40 | 0,73 | 4,68E-01 |
| AT2G17560 | 0,46 | -1,21 | 4,34 | -0,28 | 7,81E-01 |
| AT2G17700 | 0,43 | -0,48 | 4,07 | -0,12 | 9,05E-01 |
| AT2G17820 | 0,96 | 0,01 | 2,13 | 0,00 | 9,98E-01 |
| AT2G17840 | 2,90 | 0,85 | 1,68 | 0,50 | 6,14E-01 |
| AT2G17870 | 0,45 | -1,80 | 4,10 | -0,44 | 6,60E-01 |
| AT2G18130 | 0,22 | 0,00 | 4,38 | 0,00 | 1,00E+00 |
| AT2G18160 | 1,66 | -0,62 | 1,54 | -0,40 | 6,87E-01 |
| AT2G18170 | 0,50 | -0,81 | 3,15 | -0,26 | 7,98E-01 |
| AT2G18193 | 1,29 | -1,05 | 1,87 | -0,56 | 5,76E-01 |
| AT2G18290 | 0,19 | 1,22 | 4,33 | 0,28 | 7,78E-01 |
| AT2G18300 | 1,33 | -2,40 | 2,10 | -1,14 | 2,52E-01 |
| AT2G18310 | 0,43 | 0,04 | 4,40 | 0,01 | 9,92E-01 |

| | | | | | |
|---|---|---|---|---|---|
| AT2G18440 | 3,48 | 0,10 | 0,99 | 0,10 | 9,20E-01 |
| AT2G18480 | 2,00 | 0,08 | 1,64 | 0,05 | 9,61E-01 |
| AT2G18520 | 0,25 | -2,19 | 4,38 | -0,50 | 6,17E-01 |
| AT2G18600 | 0,24 | -2,36 | 4,39 | -0,54 | 5,91E-01 |
| AT2G18660 | 0,89 | 1,07 | 2,18 | 0,49 | 6,23E-01 |
| AT2G18685 | 0,08 | 0,69 | 4,36 | 0,16 | 8,75E-01 |
| AT2G18900 | 1,01 | -1,23 | 1,75 | -0,70 | 4,83E-01 |
| AT2G18915 | 0,29 | 0,19 | 4,32 | 0,04 | 9,65E-01 |
| AT2G18940 | 0,31 | 1,22 | 3,58 | 0,34 | 7,34E-01 |
| AT2G19120 | 0,71 | -2,43 | 3,52 | -0,69 | 4,91E-01 |
| AT2G07135 | 0,16 | 0,00 | 4,41 | 0,00 | 1,00E+00 |
| AT2G07165 | 0,21 | 0,67 | 4,36 | 0,15 | 8,78E-01 |
| AT2G19290 | 0,05 | 0,00 | 4,41 | 0,00 | 1,00E+00 |
| AT2G19310 | 0,12 | 0,00 | 4,38 | 0,00 | 1,00E+00 |
| AT2G19340 | 0,43 | 1,64 | 4,33 | 0,38 | 7,05E-01 |
| AT2G19365 | 0,81 | 0,33 | 2,17 | 0,15 | 8,78E-01 |
| AT2G19390 | 0,76 | 1,42 | 2,37 | 0,60 | 5,49E-01 |
| AT2G07195 | 0,51 | 0,31 | 2,66 | 0,12 | 9,07E-01 |
| AT2G19400 | 0,15 | 0,00 | 4,41 | 0,00 | 1,00E+00 |
| AT2G19470 | 0,28 | -0,18 | 4,34 | -0,04 | 9,66E-01 |
| AT2G19560 | 0,20 | 1,26 | 4,34 | 0,29 | 7,71E-01 |
| AT2G19570 | 0,52 | 1,23 | 3,07 | 0,40 | 6,89E-01 |
| AT2G19580 | 0,25 | 0,02 | 4,39 | 0,00 | 9,97E-01 |
| AT2G19760 | 0,25 | 0,22 | 4,37 | 0,05 | 9,60E-01 |
| AT2G19780 | 0,19 | 0,00 | 4,41 | 0,00 | 1,00E+00 |
| AT2G19910 | 0,11 | 0,65 | 4,36 | 0,15 | 8,81E-01 |
| AT2G20000 | 0,44 | 1,05 | 3,28 | 0,32 | 7,48E-01 |
| AT2G20010 | 1,89 | -1,16 | 1,27 | -0,91 | 3,63E-01 |
| AT2G20050 | 0,77 | 1,80 | 2,33 | 0,77 | 4,39E-01 |
| AT2G20120 | 0,38 | -2,37 | 4,35 | -0,55 | 5,85E-01 |
| AT2G20180 | 0,53 | 1,15 | 2,63 | 0,44 | 6,63E-01 |
| AT2G20240 | 1,31 | 1,92 | 2,48 | 0,77 | 4,39E-01 |
| AT2G20270 | 0,73 | 0,20 | 3,09 | 0,07 | 9,48E-01 |
| AT2G20280 | 0,90 | -1,26 | 2,64 | -0,48 | 6,34E-01 |
| AT2G20300 | 0,57 | 1,63 | 2,67 | 0,61 | 5,41E-01 |
| AT2G20310 | 0,15 | 1,20 | 4,34 | 0,28 | 7,81E-01 |
| AT2G20380 | 0,08 | 0,44 | 4,38 | 0,10 | 9,21E-01 |
| AT2G20562 | 0,11 | 0,00 | 4,41 | 0,00 | 1,00E+00 |
| AT2G20610 | 0,62 | -0,68 | 2,52 | -0,27 | 7,87E-01 |
| AT2G20630 | 0,57 | -1,02 | 2,70 | -0,38 | 7,06E-01 |
| AT2G20670 | 0,99 | -1,94 | 1,87 | -1,03 | 3,02E-01 |
| AT2G07325 | 0,08 | 0,44 | 4,38 | 0,10 | 9,21E-01 |
| AT2G20720 | 0,50 | -0,52 | 3,16 | -0,16 | 8,71E-01 |
| AT2G20721 | 2,89 | 2,82 | 1,48 | 1,91 | 5,62E-02 |
| AT2G20722 | 3,24 | 1,88 | 1,44 | 1,30 | 1,92E-01 |
| AT2G20723 | 1,21 | 0,74 | 1,91 | 0,39 | 7,00E-01 |
| AT2G20725 | 0,17 | 0,98 | 4,35 | 0,23 | 8,21E-01 |
| AT2G20760 | 1,00 | -0,16 | 2,20 | -0,07 | 9,40E-01 |
| AT2G20780 | 1,93 | 0,86 | 1,44 | 0,60 | 5,48E-01 |
| AT2G20790 | 0,65 | -1,79 | 3,77 | -0,47 | 6,35E-01 |
| AT2G20825 | 0,24 | -0,08 | 4,34 | -0,02 | 9,85E-01 |
| AT2G20830 | 0,42 | -0,87 | 3,30 | -0,26 | 7,92E-01 |

| | | | | | |
|---|---|---|---|---|---|
| AT2G20850 | 0,80 | -1,40 | 3,45 | -0,41 | 6,84E-01 |
| AT2G20900 | 0,70 | -1,61 | 2,87 | -0,56 | 5,75E-01 |
| AT2G21030 | 1,78 | 0,57 | 1,78 | 0,32 | 7,48E-01 |
| AT2G21060 | 0,37 | 0,11 | 4,40 | 0,02 | 9,80E-01 |
| AT2G21090 | 0,12 | 1,16 | 4,34 | 0,27 | 7,89E-01 |
| AT2G21100 | 3,61 | -0,59 | 1,31 | -0,45 | 6,52E-01 |
| AT2G21120 | 1,14 | -1,25 | 1,64 | -0,76 | 4,45E-01 |
| AT2G21170 | 1,20 | -0,16 | 1,57 | -0,10 | 9,19E-01 |
| AT2G21185 | 0,86 | -0,32 | 1,88 | -0,17 | 8,65E-01 |
| AT2G21188 | 0,47 | -0,53 | 4,18 | -0,13 | 8,99E-01 |
| AT2G21210 | 1,12 | -0,75 | 1,96 | -0,38 | 7,00E-01 |
| AT2G21240 | 0,62 | 1,71 | 2,34 | 0,73 | 4,64E-01 |
| AT2G21280 | 0,38 | -1,52 | 4,32 | -0,35 | 7,24E-01 |
| AT2G21300 | 0,99 | 0,69 | 2,19 | 0,32 | 7,52E-01 |
| AT2G21350 | 0,05 | 0,00 | 4,41 | 0,00 | 1,00E+00 |
| AT2G21370 | 0,82 | -1,32 | 1,93 | -0,69 | 4,93E-01 |
| AT2G21440 | 0,17 | 1,50 | 4,33 | 0,35 | 7,29E-01 |
| AT2G21480 | 0,25 | 0,41 | 4,33 | 0,09 | 9,25E-01 |
| AT2G21500 | 0,51 | 0,70 | 3,03 | 0,23 | 8,18E-01 |
| AT2G21560 | 0,27 | 0,00 | 4,41 | 0,00 | 1,00E+00 |
| AT2G21870 | 1,17 | -2,61 | 2,13 | -1,23 | 2,20E-01 |
| AT2G21920 | 4,05 | -1,25 | 0,96 | -1,31 | 1,92E-01 |
| AT2G21950 | 0,24 | 1,56 | 4,34 | 0,36 | 7,18E-01 |
| AT2G21970 | 0,61 | 0,23 | 3,87 | 0,06 | 9,53E-01 |
| AT2G22010 | 0,65 | 0,59 | 2,48 | 0,24 | 8,12E-01 |
| AT2G22090 | 0,34 | -1,85 | 4,40 | -0,42 | 6,74E-01 |
| AT2G22140 | 0,05 | 0,00 | 4,41 | 0,00 | 1,00E+00 |
| AT2G22160 | 0,35 | 0,88 | 4,35 | 0,20 | 8,40E-01 |
| AT2G22240 | 0,21 | 0,23 | 4,37 | 0,05 | 9,57E-01 |
| AT2G22250 | 0,26 | -0,48 | 4,38 | -0,11 | 9,13E-01 |
| AT2G07555 | 0,37 | 0,00 | 4,36 | 0,00 | 1,00E+00 |
| AT2G22400 | 0,55 | -1,22 | 3,79 | -0,32 | 7,48E-01 |
| AT2G22430 | 4,62 | -0,88 | 0,89 | -1,00 | 3,18E-01 |
| AT2G22470 | 0,19 | 1,08 | 4,34 | 0,25 | 8,04E-01 |
| AT2G22490 | 1,17 | -1,88 | 2,16 | -0,87 | 3,83E-01 |
| AT2G22500 | 4,35 | -0,34 | 1,04 | -0,33 | 7,40E-01 |
| AT2G07645 | 4,48 | 1,23 | 1,30 | 0,95 | 3,44E-01 |
| AT2G22740 | 0,43 | 2,16 | 4,09 | 0,53 | 5,97E-01 |
| AT2G22850 | 0,76 | 1,31 | 2,08 | 0,63 | 5,30E-01 |
| AT2G22890 | 0,53 | -2,55 | 3,13 | -0,81 | 4,15E-01 |
| AT2G22940 | 0,87 | 1,99 | 2,65 | 0,75 | 4,54E-01 |
| AT2G23070 | 0,53 | 1,55 | 2,28 | 0,68 | 4,96E-01 |
| AT2G23090 | 0,40 | -1,07 | 4,26 | -0,25 | 8,03E-01 |
| AT2G23100 | 0,16 | 0,49 | 4,38 | 0,11 | 9,12E-01 |
| AT2G23140 | 0,38 | 1,86 | 4,23 | 0,44 | 6,60E-01 |
| AT2G23210 | 0,71 | 1,05 | 2,80 | 0,38 | 7,06E-01 |
| AT2G23290 | 0,64 | 0,29 | 2,63 | 0,11 | 9,11E-01 |
| AT2G23348 | 3,88 | -2,44 | 1,07 | -2,29 | 2,22E-02 |
| AT2G23347 | 3,51 | -2,16 | 1,07 | -2,01 | 4,45E-02 |
| AT2G23390 | 0,32 | 1,07 | 4,35 | 0,25 | 8,06E-01 |
| AT2G23430 | 0,47 | 0,24 | 3,36 | 0,07 | 9,44E-01 |
| AT2G23450 | 0,62 | -0,90 | 2,18 | -0,41 | 6,79E-01 |

| | | | | | |
|---|---|---|---|---|---|
| AT2G23520 | 0,43 | -1,83 | 3,35 | -0,55 | 5,85E-01 |
| AT2G07910 | 1,63 | 0,00 | 2,56 | 0,00 | 1,00E+00 |
| AT2G23550 | 1,59 | 0,00 | 2,57 | 0,00 | 1,00E+00 |
| AT2G23600 | 1,45 | -1,98 | 1,76 | -1,12 | 2,62E-01 |
| AT2G23610 | 0,63 | -0,68 | 3,10 | -0,22 | 8,26E-01 |
| AT2G23670 | 1,50 | 0,74 | 1,74 | 0,42 | 6,72E-01 |
| AT2G23760 | 0,34 | -1,75 | 4,40 | -0,40 | 6,91E-01 |
| AT2G23790 | 0,79 | 2,43 | 2,94 | 0,83 | 4,08E-01 |
| AT2G23810 | 3,17 | 1,69 | 1,33 | 1,27 | 2,04E-01 |
| AT2G23840 | 0,70 | -1,50 | 3,07 | -0,49 | 6,25E-01 |
| AT2G23987 | 0,17 | 0,00 | 4,41 | 0,00 | 1,00E+00 |
| AT2G24120 | 0,81 | -0,47 | 2,29 | -0,20 | 8,39E-01 |
| AT2G24150 | 0,79 | -1,10 | 2,01 | -0,55 | 5,83E-01 |
| AT2G24170 | 0,17 | 1,33 | 4,33 | 0,31 | 7,59E-01 |
| AT2G24200 | 1,48 | 2,49 | 1,78 | 1,40 | 1,62E-01 |
| AT2G24230 | 0,59 | -0,86 | 2,70 | -0,32 | 7,50E-01 |
| AT2G24270 | 3,15 | -0,43 | 1,08 | -0,39 | 6,94E-01 |
| AT2G24330 | 0,33 | -1,71 | 4,40 | -0,39 | 6,98E-01 |
| AT2G24340 | 1,46 | 1,62 | 1,82 | 0,89 | 3,75E-01 |
| AT2G24360 | 1,19 | -0,16 | 1,81 | -0,09 | 9,32E-01 |
| AT2G24500 | 1,13 | -0,26 | 2,14 | -0,12 | 9,05E-01 |
| AT2G08055 | 0,33 | 0,17 | 4,33 | 0,04 | 9,68E-01 |
| AT2G24545 | 1,16 | -0,87 | 1,59 | -0,55 | 5,84E-01 |
| AT2G24550 | 1,50 | 2,00 | 1,77 | 1,13 | 2,58E-01 |
| AT2G24600 | 1,31 | 2,58 | 2,24 | 1,15 | 2,49E-01 |
| AT2G24640 | 0,24 | 0,21 | 4,34 | 0,05 | 9,62E-01 |
| AT2G24710 | 0,19 | 1,50 | 4,33 | 0,35 | 7,30E-01 |
| AT2G24755 | 0,71 | 0,18 | 2,52 | 0,07 | 9,44E-01 |
| AT2G24780 | 0,92 | -1,04 | 2,38 | -0,44 | 6,63E-01 |
| AT2G24850 | 1,45 | -0,86 | 1,60 | -0,53 | 5,93E-01 |
| AT2G24870 | 1,09 | -2,42 | 2,59 | -0,93 | 3,52E-01 |
| AT2G24880 | 1,30 | -0,31 | 1,71 | -0,18 | 8,54E-01 |
| AT2G25011 | 1,24 | 1,33 | 1,91 | 0,69 | 4,87E-01 |
| AT2G25050 | 0,47 | 0,43 | 3,27 | 0,13 | 8,95E-01 |
| AT2G25070 | 0,45 | -0,14 | 2,82 | -0,05 | 9,59E-01 |
| AT2G08105 | 0,74 | 0,18 | 2,43 | 0,07 | 9,41E-01 |
| AT2G25180 | 0,76 | 1,99 | 3,05 | 0,65 | 5,13E-01 |
| AT2G25200 | 0,69 | 0,00 | 3,88 | 0,00 | 9,99E-01 |
| AT2G25300 | 0,15 | 0,67 | 4,37 | 0,15 | 8,79E-01 |
| AT2G25310 | 0,20 | -1,80 | 4,41 | -0,41 | 6,83E-01 |
| AT2G25320 | 3,21 | -0,06 | 1,20 | -0,05 | 9,57E-01 |
| AT2G25450 | 4,94 | 0,94 | 0,89 | 1,06 | 2,89E-01 |
| AT2G25490 | 3,68 | 1,27 | 1,18 | 1,07 | 2,83E-01 |
| AT2G25570 | 0,45 | -0,48 | 4,37 | -0,11 | 9,13E-01 |
| AT2G25610 | 0,53 | -2,08 | 3,09 | -0,67 | 5,00E-01 |
| AT2G25660 | 3,83 | -1,53 | 1,00 | -1,54 | 1,24E-01 |
| AT2G25670 | 0,93 | 1,79 | 2,33 | 0,77 | 4,43E-01 |
| AT2G25730 | 2,88 | -0,65 | 1,00 | -0,65 | 5,17E-01 |
| AT2G25760 | 0,93 | 0,15 | 2,75 | 0,06 | 9,56E-01 |
| AT2G25800 | 4,68 | -0,78 | 0,84 | -0,92 | 3,57E-01 |
| AT2G25850 | 0,16 | 0,92 | 4,35 | 0,21 | 8,33E-01 |
| AT2G25910 | 0,91 | -1,78 | 2,64 | -0,67 | 5,01E-01 |

| | | | | | |
|---|---|---|---|---|---|
| AT2G25970 | 1,54 | -0,50 | 1,65 | -0,30 | 7,61E-01 |
| AT2G26030 | 0,31 | 0,85 | 3,65 | 0,23 | 8,16E-01 |
| AT2G26070 | 0,39 | -1,40 | 4,37 | -0,32 | 7,49E-01 |
| AT2G26140 | 1,61 | -0,45 | 1,49 | -0,30 | 7,65E-01 |
| AT2G26190 | 1,38 | -1,84 | 1,75 | -1,05 | 2,94E-01 |
| AT2G26250 | 1,96 | -0,65 | 1,59 | -0,41 | 6,84E-01 |
| AT2G26280 | 1,11 | -1,94 | 2,27 | -0,85 | 3,94E-01 |
| AT2G26430 | 0,38 | 0,28 | 4,25 | 0,07 | 9,47E-01 |
| AT2G26530 | 1,88 | 1,52 | 1,97 | 0,77 | 4,42E-01 |
| AT2G26540 | 0,42 | -1,72 | 4,30 | -0,40 | 6,90E-01 |
| AT2G26560 | 1,67 | 1,14 | 1,63 | 0,70 | 4,85E-01 |
| AT2G26580 | 0,26 | 0,00 | 4,38 | 0,00 | 1,00E+00 |
| AT2G26590 | 0,25 | 0,61 | 4,36 | 0,14 | 8,89E-01 |
| AT2G26620 | 0,18 | 0,00 | 4,38 | 0,00 | 9,99E-01 |
| AT2G26680 | 0,33 | -0,32 | 4,32 | -0,07 | 9,41E-01 |
| AT2G26690 | 0,68 | 1,63 | 2,52 | 0,65 | 5,19E-01 |
| AT2G26740 | 0,33 | 0,22 | 4,32 | 0,05 | 9,59E-01 |
| AT2G08175 | 1,14 | -1,08 | 1,97 | -0,55 | 5,84E-01 |
| AT2G26790 | 0,21 | 1,71 | 4,33 | 0,39 | 6,93E-01 |
| AT2G26800 | 0,41 | 0,27 | 3,38 | 0,08 | 9,36E-01 |
| AT2G26840 | 0,08 | 0,69 | 4,36 | 0,16 | 8,75E-01 |
| AT2G26900 | 1,34 | 0,47 | 1,78 | 0,26 | 7,93E-01 |
| AT2G26930 | 0,69 | -0,89 | 3,00 | -0,30 | 7,68E-01 |
| AT2G26980 | 0,61 | -0,13 | 2,38 | -0,05 | 9,56E-01 |
| AT2G26990 | 0,56 | 0,73 | 3,08 | 0,24 | 8,13E-01 |
| AT2G27020 | 0,74 | 2,06 | 2,55 | 0,81 | 4,19E-01 |
| AT2G27040 | 0,44 | -1,01 | 3,34 | -0,30 | 7,63E-01 |
| AT2G27070 | 0,56 | 0,14 | 3,13 | 0,04 | 9,64E-01 |
| AT2G27090 | 0,77 | 2,41 | 2,98 | 0,81 | 4,18E-01 |
| AT2G27170 | 0,61 | 0,78 | 2,61 | 0,30 | 7,65E-01 |
| AT2G27190 | 0,39 | 0,00 | 4,38 | 0,00 | 1,00E+00 |
| AT2G27290 | 0,57 | 1,29 | 2,72 | 0,47 | 6,36E-01 |
| AT2G27310 | 2,22 | 0,68 | 1,24 | 0,55 | 5,82E-01 |
| AT2G27330 | 0,10 | 0,00 | 4,41 | 0,00 | 1,00E+00 |
| AT2G27350 | 0,23 | 1,14 | 4,34 | 0,26 | 7,92E-01 |
| AT2G27385 | 1,03 | -0,90 | 2,23 | -0,41 | 6,85E-01 |
| AT2G27500 | 0,69 | -1,46 | 2,52 | -0,58 | 5,63E-01 |
| AT2G27510 | 0,44 | 0,08 | 4,18 | 0,02 | 9,85E-01 |
| AT2G27530 | 0,92 | -0,64 | 1,79 | -0,36 | 7,22E-01 |
| AT2G27580 | 0,43 | 1,96 | 4,18 | 0,47 | 6,40E-01 |
| AT2G27660 | 0,41 | -2,88 | 4,34 | -0,66 | 5,07E-01 |
| AT2G27680 | 0,72 | -0,34 | 1,97 | -0,17 | 8,62E-01 |
| AT2G27770 | 0,38 | 0,77 | 4,37 | 0,18 | 8,61E-01 |
| AT2G27860 | 4,64 | -1,31 | 0,83 | -1,58 | 1,14E-01 |
| AT2G27950 | 1,06 | -0,56 | 1,78 | -0,31 | 7,53E-01 |
| AT2G28080 | 0,87 | -0,57 | 2,87 | -0,20 | 8,42E-01 |
| AT2G28170 | 0,05 | 0,00 | 4,41 | 0,00 | 1,00E+00 |
| AT2G28360 | 1,42 | -2,83 | 1,47 | -1,93 | 5,34E-02 |
| AT2G28380 | 0,74 | 2,26 | 2,88 | 0,78 | 4,33E-01 |
| AT2G28390 | 0,41 | -1,90 | 4,31 | -0,44 | 6,58E-01 |
| AT2G28400 | 0,81 | 0,59 | 2,39 | 0,25 | 8,05E-01 |
| AT2G28450 | 1,48 | -1,47 | 1,64 | -0,90 | 3,69E-01 |

| | | | | | |
|---|---|---|---|---|---|
| AT2G28470 | 0,87 | 0,90 | 2,82 | 0,32 | 7,50E-01 |
| AT2G28480 | 0,12 | 0,37 | 4,37 | 0,08 | 9,33E-01 |
| AT2G28550 | 2,55 | -0,55 | 1,35 | -0,41 | 6,83E-01 |
| AT2G28620 | 1,12 | 2,78 | 2,15 | 1,29 | 1,95E-01 |
| AT2G08275 | 1,12 | 2,78 | 2,15 | 1,29 | 1,95E-01 |
| AT2G28630 | 0,37 | 0,02 | 4,30 | 0,00 | 9,97E-01 |
| AT2G28640 | 0,62 | -1,74 | 3,06 | -0,57 | 5,69E-01 |
| AT2G28670 | 0,51 | 1,71 | 3,05 | 0,56 | 5,74E-01 |
| AT2G28740 | 0,44 | 1,39 | 4,16 | 0,33 | 7,39E-01 |
| AT2G28800 | 0,62 | 0,76 | 3,07 | 0,25 | 8,06E-01 |
| AT2G08300 | 0,20 | -1,80 | 4,41 | -0,41 | 6,83E-01 |
| AT2G28880 | 0,33 | 0,66 | 4,32 | 0,15 | 8,78E-01 |
| AT2G28890 | 0,77 | -0,03 | 2,11 | -0,01 | 9,88E-01 |
| AT2G28900 | 0,81 | -1,78 | 2,74 | -0,65 | 5,16E-01 |
| AT2G28910 | 4,89 | 0,63 | 1,14 | 0,55 | 5,80E-01 |
| AT2G28950 | 0,45 | -1,10 | 4,11 | -0,27 | 7,88E-01 |
| AT2G29030 | 0,32 | -1,78 | 4,41 | -0,40 | 6,87E-01 |
| AT2G29040 | 0,08 | 0,00 | 4,40 | 0,00 | 1,00E+00 |
| AT2G29080 | 1,97 | 0,42 | 1,35 | 0,31 | 7,55E-01 |
| AT2G29110 | 0,41 | -0,04 | 4,09 | -0,01 | 9,92E-01 |
| AT2G29120 | 1,16 | 2,40 | 2,10 | 1,14 | 2,53E-01 |
| AT2G29170 | 0,18 | 0,56 | 4,36 | 0,13 | 8,97E-01 |
| AT2G08330 | 0,26 | -0,48 | 4,38 | -0,11 | 9,13E-01 |
| AT2G29190 | 0,61 | -0,13 | 2,55 | -0,05 | 9,58E-01 |
| AT2G29220 | 0,39 | -1,40 | 4,37 | -0,32 | 7,49E-01 |
| AT2G29390 | 0,71 | 1,01 | 2,48 | 0,41 | 6,84E-01 |
| AT2G29400 | 0,98 | 0,22 | 1,71 | 0,13 | 8,97E-01 |
| AT2G29450 | 2,19 | 0,55 | 1,68 | 0,33 | 7,43E-01 |
| AT2G29550 | 0,75 | 2,03 | 2,36 | 0,86 | 3,89E-01 |
| AT2G29570 | 0,14 | 0,09 | 4,38 | 0,02 | 9,84E-01 |
| AT2G29605 | 0,65 | 0,12 | 2,97 | 0,04 | 9,67E-01 |
| AT2G29650 | 0,77 | -0,63 | 2,50 | -0,25 | 8,00E-01 |
| AT2G29670 | 3,81 | -0,48 | 1,05 | -0,46 | 6,48E-01 |
| AT2G29925 | 2,27 | 3,90 | 2,00 | 1,95 | 5,10E-02 |
| AT2G29980 | 3,37 | -2,30 | 1,12 | -2,05 | 4,00E-02 |
| AT2G30110 | 3,88 | -1,38 | 0,95 | -1,45 | 1,46E-01 |
| AT2G30250 | 0,71 | -0,84 | 2,17 | -0,39 | 7,00E-01 |
| AT2G30260 | 0,44 | 0,53 | 3,26 | 0,16 | 8,70E-01 |
| AT2G30270 | 0,18 | 1,10 | 4,34 | 0,25 | 8,00E-01 |
| AT2G30360 | 0,37 | 0,50 | 4,38 | 0,11 | 9,09E-01 |
| AT2G30390 | 0,49 | -1,84 | 3,25 | -0,57 | 5,71E-01 |
| AT2G08390 | 5,34 | -0,12 | 0,98 | -0,12 | 9,05E-01 |
| AT2G30470 | 0,73 | 1,11 | 2,23 | 0,50 | 6,18E-01 |
| AT2G30490 | 0,91 | -0,03 | 3,04 | -0,01 | 9,93E-01 |
| AT2G30500 | 1,56 | 2,04 | 1,70 | 1,20 | 2,31E-01 |
| AT2G30505 | 0,35 | 1,38 | 4,34 | 0,32 | 7,50E-01 |
| AT2G30570 | 3,97 | -0,11 | 1,06 | -0,10 | 9,16E-01 |
| AT2G30700 | 0,36 | 2,02 | 4,28 | 0,47 | 6,38E-01 |
| AT2G30710 | 0,35 | -0,48 | 4,32 | -0,11 | 9,12E-01 |
| AT2G30720 | 0,49 | -0,23 | 3,21 | -0,07 | 9,43E-01 |
| AT2G30790 | 0,21 | -1,41 | 4,36 | -0,32 | 7,46E-01 |
| AT2G30800 | 1,27 | -2,50 | 1,84 | -1,36 | 1,74E-01 |

| | | | | | |
|---|---|---|---|---|---|
| AT2G30850 | 0,41 | -2,04 | 4,37 | -0,47 | 6,40E-01 |
| AT2G30880 | 0,34 | 0,09 | 4,32 | 0,02 | 9,83E-01 |
| AT2G30920 | 2,18 | 0,35 | 1,55 | 0,23 | 8,19E-01 |
| AT2G08420 | 0,54 | 0,17 | 4,00 | 0,04 | 9,67E-01 |
| AT2G30980 | 0,41 | -1,21 | 4,29 | -0,28 | 7,78E-01 |
| AT2G31040 | 0,19 | 0,47 | 4,38 | 0,11 | 9,14E-01 |
| AT2G31060 | 0,49 | 2,67 | 4,30 | 0,62 | 5,34E-01 |
| AT2G31070 | 0,79 | 1,34 | 2,28 | 0,59 | 5,55E-01 |
| AT2G31130 | 0,17 | 0,03 | 4,39 | 0,01 | 9,95E-01 |
| AT2G31170 | 0,45 | 0,78 | 4,36 | 0,18 | 8,58E-01 |
| AT2G31240 | 0,25 | 0,41 | 4,33 | 0,09 | 9,25E-01 |
| AT2G31250 | 0,39 | -1,28 | 4,38 | -0,29 | 7,70E-01 |
| AT2G31260 | 0,53 | 0,82 | 2,69 | 0,30 | 7,61E-01 |
| AT2G08430 | 0,08 | 0,69 | 4,36 | 0,16 | 8,75E-01 |
| AT2G31360 | 1,30 | 0,46 | 1,92 | 0,24 | 8,12E-01 |
| AT2G31390 | 0,86 | -0,70 | 2,77 | -0,25 | 7,99E-01 |
| AT2G31400 | 2,56 | 0,21 | 1,19 | 0,18 | 8,57E-01 |
| AT2G31410 | 0,37 | -3,12 | 4,34 | -0,72 | 4,72E-01 |
| AT2G31440 | 0,44 | 2,28 | 4,31 | 0,53 | 5,98E-01 |
| AT2G31510 | 0,63 | -1,78 | 3,20 | -0,56 | 5,78E-01 |
| AT2G31570 | 0,14 | 0,40 | 4,37 | 0,09 | 9,27E-01 |
| AT2G31670 | 0,73 | 2,29 | 2,49 | 0,92 | 3,58E-01 |
| AT2G31730 | 0,53 | -1,87 | 4,35 | -0,43 | 6,68E-01 |
| AT2G31740 | 0,52 | -1,58 | 3,10 | -0,51 | 6,11E-01 |
| AT2G31751 | 0,97 | 1,73 | 2,36 | 0,73 | 4,65E-01 |
| AT2G31800 | 0,85 | 0,12 | 2,05 | 0,06 | 9,55E-01 |
| AT2G31830 | 0,34 | 0,39 | 4,34 | 0,09 | 9,28E-01 |
| AT2G31865 | 0,37 | 1,07 | 4,31 | 0,25 | 8,04E-01 |
| AT2G31870 | 0,27 | 1,23 | 3,75 | 0,33 | 7,42E-01 |
| AT2G08475 | 0,81 | 0,64 | 2,34 | 0,28 | 7,83E-01 |
| AT2G31890 | 2,01 | -0,55 | 1,49 | -0,37 | 7,11E-01 |
| AT2G31990 | 0,31 | 0,00 | 4,33 | 0,00 | 1,00E+00 |
| AT2G32000 | 0,55 | -2,39 | 2,57 | -0,93 | 3,52E-01 |
| AT2G32030 | 0,22 | 1,19 | 4,33 | 0,27 | 7,84E-01 |
| AT2G32050 | 0,08 | 0,44 | 4,38 | 0,10 | 9,21E-01 |
| AT2G32060 | 0,34 | 1,21 | 4,34 | 0,28 | 7,80E-01 |
| AT2G32070 | 0,64 | 2,53 | 2,52 | 1,01 | 3,15E-01 |
| AT2G32080 | 0,57 | 1,06 | 2,51 | 0,42 | 6,73E-01 |
| AT2G32130 | 0,28 | 1,91 | 4,32 | 0,44 | 6,58E-01 |
| AT2G32230 | 0,70 | -1,33 | 2,70 | -0,49 | 6,22E-01 |
| AT2G32235 | 0,12 | 1,16 | 4,34 | 0,27 | 7,89E-01 |
| AT2G32240 | 2,12 | 2,09 | 1,54 | 1,35 | 1,76E-01 |
| AT2G32390 | 0,36 | 1,57 | 4,34 | 0,36 | 7,17E-01 |
| AT2G32400 | 0,29 | 2,11 | 4,32 | 0,49 | 6,26E-01 |
| AT2G32415 | 0,42 | -1,24 | 4,30 | -0,29 | 7,72E-01 |
| AT2G32480 | 0,87 | -0,69 | 2,20 | -0,31 | 7,53E-01 |
| AT2G32520 | 0,35 | 1,81 | 4,32 | 0,42 | 6,75E-01 |
| AT2G32600 | 0,08 | 0,69 | 4,36 | 0,16 | 8,75E-01 |
| AT2G32680 | 1,55 | 1,66 | 1,73 | 0,96 | 3,37E-01 |
| AT2G32690 | 2,01 | 0,44 | 1,50 | 0,29 | 7,70E-01 |
| AT2G32810 | 0,70 | 1,45 | 2,49 | 0,58 | 5,59E-01 |
| AT2G32920 | 1,12 | -0,80 | 1,75 | -0,46 | 6,48E-01 |

| | | | | | |
|---|---|---|---|---|---|
| AT2G32940 | 0,12 | 1,16 | 4,34 | 0,27 | 7,89E-01 |
| AT2G33040 | 0,74 | 1,98 | 2,38 | 0,83 | 4,06E-01 |
| AT2G33051 | 0,64 | -0,10 | 3,81 | -0,03 | 9,79E-01 |
| AT2G33070 | 0,08 | 0,44 | 4,38 | 0,10 | 9,21E-01 |
| AT2G33100 | 2,25 | -0,77 | 1,41 | -0,55 | 5,83E-01 |
| AT2G33110 | 0,40 | 2,16 | 4,32 | 0,50 | 6,17E-01 |
| AT2G33120 | 0,77 | -0,07 | 2,10 | -0,03 | 9,73E-01 |
| AT2G33150 | 2,47 | 0,87 | 1,26 | 0,69 | 4,89E-01 |
| AT2G33170 | 2,19 | 1,07 | 1,43 | 0,75 | 4,54E-01 |
| AT2G33175 | 0,67 | -0,28 | 2,55 | -0,11 | 9,14E-01 |
| AT2G33210 | 0,70 | 0,01 | 2,39 | 0,00 | 9,98E-01 |
| AT2G33260 | 0,24 | -0,08 | 4,34 | -0,02 | 9,85E-01 |
| AT2G33320 | 0,54 | 0,96 | 2,55 | 0,38 | 7,06E-01 |
| AT2G33340 | 1,98 | -1,68 | 1,36 | -1,23 | 2,17E-01 |
| AT2G33370 | 0,46 | -0,11 | 2,81 | -0,04 | 9,68E-01 |
| AT2G33385 | 0,08 | 0,00 | 4,40 | 0,00 | 1,00E+00 |
| AT2G33460 | 0,28 | 0,26 | 4,33 | 0,06 | 9,52E-01 |
| AT2G33470 | 0,39 | 0,60 | 3,32 | 0,18 | 8,55E-01 |
| AT2G33500 | 0,51 | -3,08 | 4,34 | -0,71 | 4,77E-01 |
| AT2G33530 | 0,99 | -1,26 | 1,73 | -0,73 | 4,64E-01 |
| AT2G33540 | 0,96 | 0,41 | 1,81 | 0,23 | 8,22E-01 |
| AT2G33550 | 0,30 | -1,78 | 4,41 | -0,40 | 6,87E-01 |
| AT2G33580 | 0,75 | 0,06 | 2,21 | 0,03 | 9,77E-01 |
| AT2G33630 | 0,50 | -2,34 | 3,21 | -0,73 | 4,67E-01 |
| AT2G33655 | 0,29 | 0,00 | 4,41 | 0,00 | 1,00E+00 |
| AT2G33730 | 2,24 | 0,09 | 1,11 | 0,08 | 9,35E-01 |
| AT2G33770 | 4,11 | 1,08 | 0,91 | 1,18 | 2,36E-01 |
| AT2G33800 | 0,66 | -0,12 | 2,58 | -0,05 | 9,63E-01 |
| AT2G33815 | 0,37 | -1,96 | 4,38 | -0,45 | 6,54E-01 |
| AT2G33830 | 1,76 | -3,21 | 1,83 | -1,76 | 7,89E-02 |
| AT2G33860 | 0,43 | 1,23 | 4,17 | 0,29 | 7,68E-01 |
| AT2G33980 | 0,47 | -0,45 | 3,21 | -0,14 | 8,88E-01 |
| AT2G34040 | 1,26 | 2,12 | 1,69 | 1,25 | 2,11E-01 |
| AT2G34070 | 0,35 | 1,38 | 3,57 | 0,39 | 6,99E-01 |
| AT2G34090 | 0,11 | 0,00 | 4,41 | 0,00 | 1,00E+00 |
| AT2G34290 | 0,45 | 0,84 | 4,04 | 0,21 | 8,35E-01 |
| AT2G34300 | 0,72 | 1,24 | 4,31 | 0,29 | 7,74E-01 |
| AT2G34450 | 0,60 | 0,37 | 3,76 | 0,10 | 9,21E-01 |
| AT2G34470 | 0,95 | -0,12 | 1,91 | -0,06 | 9,49E-01 |
| AT2G34480 | 1,66 | 0,16 | 1,40 | 0,12 | 9,08E-01 |
| AT2G34490 | 0,49 | 2,04 | 4,07 | 0,50 | 6,16E-01 |
| AT2G34510 | 0,33 | -0,91 | 4,38 | -0,21 | 8,35E-01 |
| AT2G34520 | 0,36 | -3,19 | 4,33 | -0,74 | 4,61E-01 |
| AT2G34585 | 1,42 | 1,46 | 1,64 | 0,89 | 3,72E-01 |
| AT2G34680 | 1,27 | 0,80 | 2,15 | 0,37 | 7,10E-01 |
| AT2G34710 | 1,10 | -0,32 | 2,30 | -0,14 | 8,88E-01 |
| AT2G34750 | 2,36 | -0,52 | 1,28 | -0,40 | 6,86E-01 |
| AT2G34790 | 0,43 | -2,54 | 4,16 | -0,61 | 5,42E-01 |
| AT2G34920 | 0,05 | 0,00 | 4,41 | 0,00 | 1,00E+00 |
| AT2G34930 | 0,44 | -0,89 | 3,50 | -0,26 | 7,99E-01 |
| AT2G35030 | 0,48 | 0,01 | 3,96 | 0,00 | 9,98E-01 |
| AT2G35040 | 0,75 | 0,71 | 2,98 | 0,24 | 8,11E-01 |

| | | | | | |
|---|---|---|---|---|---|
| AT2G35060 | 1,08 | 0,48 | 1,89 | 0,25 | 8,01E-01 |
| AT2G35110 | 1,99 | -1,82 | 1,33 | -1,37 | 1,72E-01 |
| AT2G35120 | 0,10 | 0,00 | 4,41 | 0,00 | 1,00E+00 |
| AT2G35250 | 1,05 | 2,24 | 1,98 | 1,13 | 2,58E-01 |
| AT2G35360 | 0,45 | -1,86 | 4,10 | -0,45 | 6,50E-01 |
| AT2G35382 | 3,86 | -1,36 | 1,20 | -1,13 | 2,57E-01 |
| AT2G35390 | 0,25 | -0,23 | 4,35 | -0,05 | 9,58E-01 |
| AT2G35420 | 0,17 | 1,15 | 4,34 | 0,27 | 7,91E-01 |
| AT2G35490 | 0,67 | 0,82 | 2,22 | 0,37 | 7,11E-01 |
| AT2G35510 | 0,57 | -2,00 | 3,13 | -0,64 | 5,24E-01 |
| AT2G35610 | 0,17 | 1,33 | 4,33 | 0,31 | 7,59E-01 |
| AT2G35680 | 0,73 | 1,26 | 2,34 | 0,54 | 5,89E-01 |
| AT2G35730 | 0,35 | -2,17 | 4,38 | -0,49 | 6,21E-01 |
| AT2G35742 | 0,34 | 0,06 | 3,57 | 0,02 | 9,87E-01 |
| AT2G08755 | 0,29 | 1,60 | 4,33 | 0,37 | 7,11E-01 |
| AT2G08765 | 0,51 | -1,78 | 3,28 | -0,54 | 5,88E-01 |
| AT2G35859 | 0,71 | 0,30 | 2,77 | 0,11 | 9,13E-01 |
| AT2G35880 | 0,24 | 0,24 | 4,34 | 0,06 | 9,55E-01 |
| AT2G35930 | 0,76 | 1,14 | 2,87 | 0,40 | 6,91E-01 |
| AT2G36030 | 0,62 | 0,00 | 4,39 | 0,00 | 1,00E+00 |
| AT2G36050 | 0,69 | -0,97 | 2,95 | -0,33 | 7,43E-01 |
| AT2G36060 | 0,48 | -1,11 | 2,71 | -0,41 | 6,82E-01 |
| AT2G36070 | 0,34 | -0,92 | 4,34 | -0,21 | 8,32E-01 |
| AT2G36080 | 0,43 | -0,56 | 3,34 | -0,17 | 8,67E-01 |
| AT2G36180 | 0,76 | 0,15 | 3,48 | 0,04 | 9,65E-01 |
| AT2G36200 | 0,33 | -0,49 | 4,35 | -0,11 | 9,11E-01 |
| AT2G36230 | 0,20 | 0,31 | 4,38 | 0,07 | 9,43E-01 |
| AT2G36250 | 0,55 | -1,77 | 3,30 | -0,54 | 5,91E-01 |
| AT2G36300 | 0,44 | -1,35 | 3,31 | -0,41 | 6,84E-01 |
| AT2G36360 | 0,43 | 0,00 | 3,41 | 0,00 | 1,00E+00 |
| AT2G36400 | 0,83 | 1,03 | 2,97 | 0,35 | 7,28E-01 |
| AT2G36460 | 1,86 | -0,36 | 1,48 | -0,24 | 8,06E-01 |
| AT2G36470 | 2,06 | 1,69 | 1,88 | 0,90 | 3,70E-01 |
| AT2G36480 | 1,71 | -1,48 | 1,49 | -0,99 | 3,21E-01 |
| AT2G36490 | 1,90 | -2,42 | 2,50 | -0,97 | 3,33E-01 |
| AT2G36530 | 5,68 | -1,35 | 0,77 | -1,76 | 7,76E-02 |
| AT2G36620 | 0,25 | -0,12 | 4,34 | -0,03 | 9,78E-01 |
| AT2G36680 | 0,35 | -1,89 | 4,36 | -0,43 | 6,64E-01 |
| AT2G36710 | 0,27 | 0,00 | 4,41 | 0,00 | 1,00E+00 |
| AT2G36750 | 0,21 | 1,76 | 4,32 | 0,41 | 6,85E-01 |
| AT2G36790 | 0,34 | 1,06 | 4,34 | 0,24 | 8,07E-01 |
| AT2G36800 | 0,85 | 0,86 | 2,22 | 0,39 | 6,98E-01 |
| AT2G36810 | 1,10 | 0,10 | 1,97 | 0,05 | 9,61E-01 |
| AT2G36835 | 0,30 | 0,49 | 3,67 | 0,13 | 8,94E-01 |
| AT2G36850 | 3,05 | -0,59 | 1,19 | -0,50 | 6,19E-01 |
| AT2G36880 | 2,29 | -0,55 | 1,17 | -0,47 | 6,38E-01 |
| AT2G36895 | 0,35 | -2,17 | 4,38 | -0,49 | 6,21E-01 |
| AT2G36900 | 0,55 | -0,26 | 3,73 | -0,07 | 9,44E-01 |
| AT2G36930 | 0,54 | 0,09 | 3,89 | 0,02 | 9,81E-01 |
| AT2G36990 | 1,19 | -3,22 | 2,40 | -1,35 | 1,78E-01 |
| AT2G37035 | 0,78 | 1,71 | 2,41 | 0,71 | 4,76E-01 |
| AT2G37040 | 1,58 | 0,45 | 1,46 | 0,31 | 7,56E-01 |

| | | | | | |
|---|---|---|---|---|---|
| AT2G37080 | 0,12 | 1,05 | 4,35 | 0,24 | 8,10E-01 |
| AT2G37130 | 0,68 | 0,08 | 3,07 | 0,03 | 9,78E-01 |
| AT2G37150 | 0,75 | 0,11 | 3,34 | 0,03 | 9,74E-01 |
| AT2G37170 | 0,90 | 1,05 | 2,20 | 0,48 | 6,33E-01 |
| AT2G37200 | 0,31 | 0,23 | 3,59 | 0,06 | 9,49E-01 |
| AT2G37220 | 3,40 | -0,13 | 1,18 | -0,11 | 9,12E-01 |
| AT2G37230 | 0,56 | 0,22 | 3,16 | 0,07 | 9,45E-01 |
| AT2G37270 | 1,75 | -1,49 | 1,60 | -0,93 | 3,53E-01 |
| AT2G37310 | 0,77 | 1,26 | 3,45 | 0,36 | 7,15E-01 |
| AT2G37340 | 1,12 | -1,61 | 2,00 | -0,80 | 4,21E-01 |
| AT2G37360 | 0,21 | 1,76 | 4,32 | 0,41 | 6,85E-01 |
| AT2G37400 | 0,25 | 0,26 | 4,33 | 0,06 | 9,51E-01 |
| AT2G37410 | 0,19 | 1,50 | 4,33 | 0,35 | 7,30E-01 |
| AT2G37520 | 1,03 | 1,15 | 1,88 | 0,61 | 5,39E-01 |
| AT2G37540 | 0,69 | 0,63 | 2,81 | 0,22 | 8,23E-01 |
| AT2G37550 | 1,38 | 0,91 | 1,70 | 0,54 | 5,91E-01 |
| AT2G37570 | 0,98 | 0,18 | 1,79 | 0,10 | 9,21E-01 |
| AT2G37630 | 0,98 | -1,18 | 1,75 | -0,67 | 5,02E-01 |
| AT2G37660 | 1,35 | -0,14 | 1,92 | -0,07 | 9,43E-01 |
| AT2G37678 | 0,57 | -0,06 | 4,39 | -0,01 | 9,89E-01 |
| AT2G37710 | 4,76 | 1,42 | 0,97 | 1,46 | 1,43E-01 |
| AT2G08895 | 1,00 | 2,24 | 2,27 | 0,99 | 3,24E-01 |
| AT2G08900 | 0,86 | 1,65 | 2,08 | 0,79 | 4,27E-01 |
| AT2G08910 | 1,52 | -2,30 | 1,76 | -1,31 | 1,91E-01 |
| AT2G08915 | 0,73 | 1,78 | 2,74 | 0,65 | 5,17E-01 |
| AT2G38000 | 0,32 | -0,87 | 4,37 | -0,20 | 8,41E-01 |
| AT2G38020 | 1,29 | -1,17 | 2,02 | -0,58 | 5,63E-01 |
| AT2G38160 | 0,87 | -3,30 | 3,20 | -1,03 | 3,01E-01 |
| AT2G38170 | 3,76 | 0,50 | 1,06 | 0,47 | 6,35E-01 |
| AT2G38270 | 0,22 | 0,00 | 4,41 | 0,00 | 1,00E+00 |
| AT2G38280 | 1,89 | -0,97 | 1,44 | -0,67 | 5,04E-01 |
| AT2G38290 | 0,66 | 0,39 | 2,81 | 0,14 | 8,91E-01 |
| AT2G38340 | 0,17 | 0,35 | 4,37 | 0,08 | 9,37E-01 |
| AT2G38360 | 0,39 | -0,48 | 3,35 | -0,14 | 8,85E-01 |
| AT2G38410 | 0,35 | 0,31 | 3,42 | 0,09 | 9,28E-01 |
| AT2G38480 | 0,17 | 1,33 | 4,33 | 0,31 | 7,59E-01 |
| AT2G38610 | 0,83 | 2,19 | 2,38 | 0,92 | 3,59E-01 |
| AT2G38650 | 1,16 | 2,76 | 2,12 | 1,30 | 1,93E-01 |
| AT2G38680 | 0,08 | 0,00 | 4,40 | 0,00 | 1,00E+00 |
| AT2G38750 | 0,05 | 0,00 | 4,41 | 0,00 | 1,00E+00 |
| AT2G38770 | 1,53 | -0,11 | 1,48 | -0,07 | 9,42E-01 |
| AT2G38790 | 0,55 | 0,21 | 2,40 | 0,09 | 9,29E-01 |
| AT2G38860 | 0,41 | 0,40 | 3,39 | 0,12 | 9,05E-01 |
| AT2G39000 | 1,06 | 0,80 | 2,81 | 0,29 | 7,75E-01 |
| AT2G39080 | 0,20 | -0,44 | 4,36 | -0,10 | 9,19E-01 |
| AT2G39130 | 0,08 | 0,00 | 4,40 | 0,00 | 1,00E+00 |
| AT2G39160 | 0,24 | 1,28 | 4,33 | 0,30 | 7,67E-01 |
| AT2G39170 | 0,19 | 0,13 | 4,41 | 0,03 | 9,76E-01 |
| AT2G09010 | 0,08 | 0,00 | 4,40 | 0,00 | 1,00E+00 |
| AT2G39210 | 0,91 | -0,07 | 2,60 | -0,03 | 9,79E-01 |
| AT2G39260 | 1,36 | 0,91 | 1,57 | 0,58 | 5,62E-01 |
| AT2G39350 | 0,49 | -0,18 | 3,13 | -0,06 | 9,54E-01 |

| | | | | | |
|---|---|---|---|---|---|
| AT2G39360 | 0,80 | -1,81 | 3,38 | -0,54 | 5,92E-01 |
| AT2G09045 | 0,52 | 1,20 | 3,86 | 0,31 | 7,57E-01 |
| AT2G39450 | 0,29 | 0,48 | 4,32 | 0,11 | 9,12E-01 |
| AT2G39480 | 1,05 | -0,51 | 1,99 | -0,26 | 7,96E-01 |
| AT2G39650 | 0,49 | 1,75 | 4,02 | 0,43 | 6,64E-01 |
| AT2G39710 | 0,60 | 1,07 | 3,23 | 0,33 | 7,40E-01 |
| AT2G39720 | 1,30 | -0,69 | 2,21 | -0,31 | 7,55E-01 |
| AT2G39750 | 1,24 | -1,81 | 1,83 | -0,99 | 3,24E-01 |
| AT2G09080 | 0,50 | 0,01 | 3,17 | 0,00 | 9,98E-01 |
| AT2G39805 | 0,30 | 0,30 | 4,34 | 0,07 | 9,45E-01 |
| AT2G39920 | 0,18 | 0,48 | 4,38 | 0,11 | 9,13E-01 |
| AT2G09090 | 0,82 | -1,16 | 2,80 | -0,41 | 6,79E-01 |
| AT2G39940 | 1,84 | -1,67 | 1,76 | -0,95 | 3,42E-01 |
| AT2G39950 | 0,75 | 0,18 | 2,52 | 0,07 | 9,43E-01 |
| AT2G39960 | 0,25 | 0,80 | 4,35 | 0,18 | 8,54E-01 |
| AT2G39990 | 1,46 | 0,89 | 2,58 | 0,35 | 7,29E-01 |
| AT2G40000 | 5,19 | 1,49 | 0,99 | 1,50 | 1,33E-01 |
| AT2G40070 | 0,52 | 1,36 | 3,86 | 0,35 | 7,24E-01 |
| AT2G40080 | 0,23 | 1,14 | 4,34 | 0,26 | 7,92E-01 |
| AT2G40165 | 0,38 | -1,21 | 4,34 | -0,28 | 7,81E-01 |
| AT2G40205 | 0,27 | -1,78 | 4,41 | -0,40 | 6,87E-01 |
| AT2G40270 | 1,34 | 1,34 | 1,63 | 0,82 | 4,11E-01 |
| AT2G40280 | 0,63 | -0,32 | 2,94 | -0,11 | 9,14E-01 |
| AT2G40350 | 0,12 | 1,23 | 4,34 | 0,28 | 7,76E-01 |
| AT2G40360 | 0,28 | 0,40 | 4,37 | 0,09 | 9,27E-01 |
| AT2G40430 | 0,45 | -2,47 | 4,19 | -0,59 | 5,56E-01 |
| AT2G40475 | 0,89 | 2,08 | 2,30 | 0,91 | 3,65E-01 |
| AT2G40590 | 0,50 | -1,77 | 2,70 | -0,66 | 5,12E-01 |
| AT2G40600 | 0,11 | 0,48 | 4,37 | 0,11 | 9,12E-01 |
| AT2G40610 | 0,37 | 0,00 | 4,41 | 0,00 | 1,00E+00 |
| AT2G40650 | 0,77 | 1,72 | 2,31 | 0,74 | 4,58E-01 |
| AT2G40730 | 0,34 | 0,46 | 3,52 | 0,13 | 8,96E-01 |
| AT2G40750 | 0,46 | -0,36 | 4,16 | -0,09 | 9,31E-01 |
| AT2G40770 | 0,51 | -1,80 | 4,03 | -0,45 | 6,54E-01 |
| AT2G40815 | 0,24 | 0,24 | 4,34 | 0,06 | 9,55E-01 |
| AT2G40860 | 0,30 | 0,30 | 4,34 | 0,07 | 9,45E-01 |
| AT2G40890 | 0,53 | 2,42 | 3,87 | 0,63 | 5,32E-01 |
| AT2G40920 | 0,35 | 1,48 | 4,33 | 0,34 | 7,33E-01 |
| AT2G40925 | 0,65 | -0,13 | 3,61 | -0,04 | 9,70E-01 |
| AT2G40930 | 1,74 | -1,29 | 1,55 | -0,84 | 4,04E-01 |
| AT2G40950 | 0,74 | 1,89 | 3,49 | 0,54 | 5,88E-01 |
| AT2G40970 | 0,42 | -2,91 | 4,35 | -0,67 | 5,03E-01 |
| AT2G41105 | 5,08 | 2,96 | 1,27 | 2,34 | 1,95E-02 |
| AT2G09225 | 0,24 | 0,92 | 4,36 | 0,21 | 8,32E-01 |
| AT2G41120 | 0,21 | 1,76 | 4,32 | 0,41 | 6,85E-01 |
| AT2G41160 | 0,39 | 1,69 | 4,33 | 0,39 | 6,96E-01 |
| AT2G41250 | 0,87 | 1,33 | 2,42 | 0,55 | 5,83E-01 |
| AT2G41410 | 4,84 | 0,80 | 1,02 | 0,78 | 4,33E-01 |
| AT2G41420 | 0,11 | 0,00 | 4,41 | 0,00 | 1,00E+00 |
| AT2G41475 | 0,71 | -1,16 | 2,52 | -0,46 | 6,46E-01 |
| AT2G41500 | 0,96 | -1,10 | 1,93 | -0,57 | 5,67E-01 |
| AT2G41530 | 2,42 | -1,85 | 1,33 | -1,40 | 1,63E-01 |

| | | | | | |
|---|---|---|---|---|---|
| AT2G41560 | 2,99 | -1,66 | 1,00 | -1,65 | 9,83E-02 |
| AT2G41600 | 0,58 | -0,34 | 2,57 | -0,13 | 8,95E-01 |
| AT2G41620 | 1,68 | -0,46 | 1,51 | -0,31 | 7,59E-01 |
| AT2G41616 | 0,33 | 0,75 | 4,32 | 0,17 | 8,62E-01 |
| AT2G41630 | 0,92 | -2,55 | 2,70 | -0,94 | 3,45E-01 |
| AT2G41680 | 0,32 | 1,76 | 4,32 | 0,41 | 6,84E-01 |
| AT2G41700 | 1,62 | 3,89 | 2,09 | 1,86 | 6,24E-02 |
| AT2G41720 | 1,87 | -0,37 | 1,72 | -0,21 | 8,31E-01 |
| AT2G41770 | 0,40 | -0,12 | 4,26 | -0,03 | 9,78E-01 |
| AT2G41790 | 0,58 | 0,65 | 2,62 | 0,25 | 8,05E-01 |
| AT2G41830 | 0,40 | 0,81 | 4,15 | 0,19 | 8,46E-01 |
| AT2G41835 | 0,70 | 2,55 | 3,67 | 0,70 | 4,87E-01 |
| AT2G41840 | 3,13 | -1,81 | 1,05 | -1,73 | 8,40E-02 |
| AT2G41870 | 0,29 | 0,10 | 4,32 | 0,02 | 9,82E-01 |
| AT2G41890 | 0,43 | -0,36 | 4,18 | -0,09 | 9,32E-01 |
| AT2G41945 | 0,60 | 1,72 | 3,90 | 0,44 | 6,59E-01 |
| AT2G41960 | 0,92 | 2,00 | 1,99 | 1,01 | 3,15E-01 |
| AT2G42010 | 0,72 | 0,31 | 2,13 | 0,15 | 8,83E-01 |
| AT2G42020 | 0,48 | 0,00 | 4,40 | 0,00 | 1,00E+00 |
| AT2G42070 | 0,23 | 0,67 | 4,36 | 0,15 | 8,77E-01 |
| AT2G42230 | 0,42 | 0,90 | 4,09 | 0,22 | 8,26E-01 |
| AT2G42280 | 0,33 | -0,45 | 4,32 | -0,10 | 9,16E-01 |
| AT2G42365 | 0,35 | -2,17 | 4,38 | -0,49 | 6,21E-01 |
| AT2G42400 | 1,01 | 0,12 | 1,94 | 0,06 | 9,49E-01 |
| AT2G42490 | 2,30 | -0,69 | 1,21 | -0,57 | 5,70E-01 |
| AT2G42500 | 0,82 | -0,46 | 2,29 | -0,20 | 8,40E-01 |
| AT2G42530 | 0,98 | -0,69 | 2,23 | -0,31 | 7,56E-01 |
| AT2G42540 | 2,58 | -2,54 | 1,95 | -1,30 | 1,94E-01 |
| AT2G42590 | 2,06 | 2,42 | 1,65 | 1,46 | 1,44E-01 |
| AT2G42690 | 1,34 | 1,33 | 1,72 | 0,77 | 4,39E-01 |
| AT2G42760 | 0,47 | 0,79 | 2,70 | 0,29 | 7,69E-01 |
| AT2G42790 | 0,53 | -0,57 | 2,60 | -0,22 | 8,27E-01 |
| AT2G42800 | 0,18 | 0,00 | 4,41 | 0,00 | 1,00E+00 |
| AT2G42810 | 0,84 | -0,73 | 1,84 | -0,40 | 6,89E-01 |
| AT2G42840 | 0,22 | 0,00 | 4,41 | 0,00 | 1,00E+00 |
| AT2G42865 | 0,08 | 0,69 | 4,36 | 0,16 | 8,75E-01 |
| AT2G42870 | 0,10 | 0,21 | 4,38 | 0,05 | 9,62E-01 |
| AT2G42880 | 0,41 | 0,58 | 3,43 | 0,17 | 8,65E-01 |
| AT2G43180 | 0,70 | -1,47 | 2,74 | -0,53 | 5,93E-01 |
| AT2G43260 | 0,20 | -0,31 | 4,36 | -0,07 | 9,44E-01 |
| AT2G43290 | 0,50 | 2,17 | 3,17 | 0,69 | 4,93E-01 |
| AT2G43320 | 0,22 | 1,50 | 4,33 | 0,35 | 7,28E-01 |
| AT2G43360 | 0,33 | 0,38 | 4,33 | 0,09 | 9,30E-01 |
| AT2G09465 | 4,22 | 0,03 | 1,00 | 0,03 | 9,76E-01 |
| AT2G43410 | 2,84 | 0,94 | 1,19 | 0,79 | 4,27E-01 |
| AT2G43420 | 0,11 | 0,48 | 4,37 | 0,11 | 9,12E-01 |
| AT2G43430 | 0,59 | -2,55 | 2,68 | -0,95 | 3,41E-01 |
| AT2G43460 | 0,41 | -0,93 | 3,43 | -0,27 | 7,86E-01 |
| AT2G43490 | 0,44 | 2,16 | 4,05 | 0,53 | 5,95E-01 |
| AT2G43560 | 0,25 | -0,49 | 4,34 | -0,11 | 9,11E-01 |
| AT2G43570 | 2,25 | 2,44 | 1,54 | 1,59 | 1,13E-01 |
| AT2G43750 | 1,46 | -0,95 | 1,44 | -0,66 | 5,09E-01 |

| | | | | | |
|---|---|---|---|---|---|
| AT2G43865 | 0,15 | 0,29 | 4,37 | 0,07 | 9,47E-01 |
| AT2G43910 | 0,26 | 1,17 | 3,74 | 0,31 | 7,55E-01 |
| AT2G43940 | 0,08 | 0,44 | 4,38 | 0,10 | 9,21E-01 |
| AT2G43945 | 0,47 | -1,31 | 4,38 | -0,30 | 7,65E-01 |
| AT2G43950 | 0,24 | 0,21 | 4,34 | 0,05 | 9,62E-01 |
| AT2G44090 | 0,69 | -1,22 | 2,46 | -0,49 | 6,21E-01 |
| AT2G44120 | 4,00 | -1,57 | 0,94 | -1,66 | 9,72E-02 |
| AT2G44140 | 0,91 | 0,75 | 2,34 | 0,32 | 7,47E-01 |
| AT2G44180 | 0,46 | 0,13 | 3,27 | 0,04 | 9,69E-01 |
| AT2G44255 | 0,20 | 1,26 | 4,34 | 0,29 | 7,71E-01 |
| AT2G44360 | 0,63 | 16,73 | 4,34 | 3,85 | 1,17E-04 |
| AT2G44540 | 4,48 | 3,03 | 1,47 | 2,07 | 3,85E-02 |
| AT2G44550 | 3,60 | -0,17 | 1,31 | -0,13 | 8,97E-01 |
| AT2G44560 | 4,56 | 0,19 | 0,97 | 0,19 | 8,46E-01 |
| AT2G44570 | 2,88 | 2,74 | 1,63 | 1,68 | 9,22E-02 |
| AT2G44610 | 0,48 | 1,64 | 4,16 | 0,39 | 6,94E-01 |
| AT2G44660 | 0,35 | 0,47 | 3,47 | 0,13 | 8,93E-01 |
| AT2G44760 | 0,32 | 0,42 | 3,56 | 0,12 | 9,06E-01 |
| AT2G44798 | 0,36 | -0,48 | 4,36 | -0,11 | 9,11E-01 |
| AT2G09555 | 2,08 | -0,35 | 1,45 | -0,24 | 8,07E-01 |
| AT2G44950 | 0,77 | -0,75 | 2,07 | -0,36 | 7,15E-01 |
| AT2G45070 | 0,80 | 1,67 | 2,23 | 0,75 | 4,55E-01 |
| AT2G45220 | 0,54 | 1,49 | 4,30 | 0,35 | 7,30E-01 |
| AT2G45245 | 2,00 | -0,62 | 1,23 | -0,51 | 6,12E-01 |
| AT2G45315 | 0,67 | 0,66 | 2,50 | 0,26 | 7,93E-01 |
| AT2G45460 | 0,20 | 0,09 | 4,39 | 0,02 | 9,84E-01 |
| AT2G45470 | 1,65 | -2,36 | 2,05 | -1,15 | 2,51E-01 |
| AT2G45520 | 0,22 | 0,24 | 4,38 | 0,05 | 9,57E-01 |
| AT2G09640 | 1,68 | 0,83 | 2,07 | 0,40 | 6,87E-01 |
| AT2G45560 | 0,47 | -2,98 | 4,35 | -0,69 | 4,93E-01 |
| AT2G45660 | 0,55 | 0,57 | 2,31 | 0,25 | 8,06E-01 |
| AT2G45670 | 0,95 | 0,99 | 2,19 | 0,45 | 6,53E-01 |
| AT2G45680 | 1,79 | 0,90 | 1,52 | 0,59 | 5,54E-01 |
| AT2G45700 | 0,38 | 1,36 | 4,29 | 0,32 | 7,51E-01 |
| AT2G45820 | 0,98 | -0,14 | 2,00 | -0,07 | 9,44E-01 |
| AT2G45930 | 2,64 | -1,78 | 1,18 | -1,51 | 1,30E-01 |
| AT2G45970 | 0,56 | 0,06 | 2,56 | 0,02 | 9,82E-01 |
| AT2G45980 | 0,45 | 0,80 | 4,08 | 0,20 | 8,45E-01 |
| AT2G46030 | 0,57 | 0,73 | 4,29 | 0,17 | 8,65E-01 |
| AT2G46040 | 0,20 | -0,44 | 4,36 | -0,10 | 9,19E-01 |
| AT2G46060 | 0,49 | -1,78 | 4,41 | -0,40 | 6,87E-01 |
| AT2G46080 | 0,51 | -1,49 | 3,19 | -0,47 | 6,42E-01 |
| AT2G46180 | 0,35 | -0,36 | 4,36 | -0,08 | 9,35E-01 |
| AT2G46280 | 1,16 | -0,66 | 1,75 | -0,38 | 7,05E-01 |
| AT2G46330 | 0,93 | 1,33 | 2,16 | 0,62 | 5,37E-01 |
| AT2G46340 | 1,90 | 2,51 | 1,77 | 1,42 | 1,55E-01 |
| AT2G46390 | 0,46 | -0,51 | 4,02 | -0,13 | 8,98E-01 |
| AT2G46400 | 1,95 | -2,14 | 1,31 | -1,64 | 1,01E-01 |
| AT2G46500 | 2,79 | -1,06 | 1,24 | -0,85 | 3,94E-01 |
| AT2G46505 | 0,89 | -3,22 | 2,22 | -1,45 | 1,46E-01 |
| AT2G46510 | 1,07 | -1,91 | 2,02 | -0,94 | 3,45E-01 |
| AT2G46540 | 0,12 | 0,00 | 4,39 | 0,00 | 1,00E+00 |

| | | | | | |
|---|---|---|---|---|---|
| AT2G46550 | 0,57 | -0,62 | 2,71 | -0,23 | 8,20E-01 |
| AT2G46560 | 4,43 | 0,86 | 0,84 | 1,03 | 3,05E-01 |
| AT2G46572 | 0,51 | 1,14 | 3,99 | 0,29 | 7,74E-01 |
| AT2G46610 | 0,85 | -2,76 | 1,95 | -1,42 | 1,57E-01 |
| AT2G46620 | 1,45 | -1,05 | 1,52 | -0,69 | 4,87E-01 |
| AT2G46680 | 0,94 | 1,00 | 2,24 | 0,44 | 6,56E-01 |
| AT2G46840 | 0,08 | 0,69 | 4,36 | 0,16 | 8,75E-01 |
| AT2G46920 | 0,44 | 1,21 | 4,00 | 0,30 | 7,62E-01 |
| AT2G46980 | 0,24 | -2,36 | 4,39 | -0,54 | 5,91E-01 |
| AT2G47000 | 1,78 | -1,14 | 1,86 | -0,61 | 5,40E-01 |
| AT2G47090 | 0,25 | 1,94 | 4,32 | 0,45 | 6,54E-01 |
| AT2G47180 | 1,41 | -1,44 | 1,62 | -0,89 | 3,76E-01 |
| AT2G47250 | 1,15 | 1,15 | 2,20 | 0,52 | 6,00E-01 |
| AT2G47330 | 0,60 | 1,55 | 3,11 | 0,50 | 6,18E-01 |
| AT2G47390 | 2,13 | 0,44 | 1,49 | 0,30 | 7,67E-01 |
| AT2G47600 | 0,61 | 0,16 | 2,45 | 0,07 | 9,47E-01 |
| AT2G47650 | 1,46 | -0,29 | 1,47 | -0,20 | 8,43E-01 |
| AT2G47680 | 0,89 | -1,21 | 1,87 | -0,64 | 5,19E-01 |
| AT2G47700 | 0,93 | -0,58 | 2,27 | -0,25 | 7,99E-01 |
| AT2G47840 | 0,52 | -0,86 | 3,37 | -0,26 | 7,99E-01 |
| AT2G47940 | 2,47 | 0,28 | 1,25 | 0,22 | 8,24E-01 |
| AT2G47990 | 0,88 | -1,68 | 2,50 | -0,67 | 5,02E-01 |
| AT2G48000 | 1,00 | 1,60 | 3,30 | 0,48 | 6,28E-01 |
| AT2G48030 | 0,73 | 2,64 | 2,40 | 1,10 | 2,71E-01 |
| AT2G48060 | 4,01 | -0,41 | 0,94 | -0,44 | 6,59E-01 |
| AT2G48130 | 0,91 | -0,21 | 2,20 | -0,10 | 9,24E-01 |
| AT3G01090 | 0,63 | 0,55 | 2,20 | 0,25 | 8,04E-01 |
| AT3G01100 | 0,90 | -1,90 | 2,00 | -0,95 | 3,43E-01 |
| AT3G01120 | 2,55 | -0,50 | 1,43 | -0,35 | 7,27E-01 |
| AT3G01130 | 0,21 | -1,41 | 4,36 | -0,32 | 7,46E-01 |
| AT3G01160 | 0,24 | 1,28 | 4,33 | 0,30 | 7,67E-01 |
| AT3G01180 | 0,84 | 0,09 | 1,86 | 0,05 | 9,63E-01 |
| AT3G01290 | 2,12 | 0,35 | 1,29 | 0,27 | 7,85E-01 |
| AT3G01300 | 0,08 | 0,69 | 4,36 | 0,16 | 8,75E-01 |
| AT3G01310 | 2,04 | 0,46 | 1,65 | 0,28 | 7,79E-01 |
| AT3G01316 | 2,03 | 2,12 | 1,71 | 1,24 | 2,15E-01 |
| AT3G00980 | 2,10 | 1,03 | 2,10 | 0,49 | 6,24E-01 |
| AT3G00985 | 2,03 | 1,01 | 2,13 | 0,47 | 6,37E-01 |
| AT3G01340 | 0,50 | 1,94 | 3,19 | 0,61 | 5,43E-01 |
| AT3G01380 | 1,09 | -0,58 | 1,85 | -0,31 | 7,55E-01 |
| AT3G01390 | 0,38 | 0,70 | 4,37 | 0,16 | 8,72E-01 |
| AT3G01415 | 0,40 | 0,00 | 4,39 | 0,00 | 1,00E+00 |
| AT3G01420 | 0,31 | 0,48 | 4,32 | 0,11 | 9,11E-01 |
| AT3G01430 | 0,79 | -0,83 | 2,87 | -0,29 | 7,72E-01 |
| AT3G01470 | 1,49 | -0,28 | 1,44 | -0,19 | 8,46E-01 |
| AT3G01500 | 4,00 | -1,00 | 1,29 | -0,77 | 4,40E-01 |
| AT3G01550 | 0,10 | 0,01 | 4,38 | 0,00 | 9,99E-01 |
| AT3G01720 | 0,80 | 1,66 | 2,83 | 0,59 | 5,58E-01 |
| AT3G01770 | 1,96 | 0,41 | 1,38 | 0,30 | 7,64E-01 |
| AT3G01820 | 0,81 | -0,24 | 3,51 | -0,07 | 9,45E-01 |
| AT3G01840 | 0,88 | -0,58 | 2,09 | -0,28 | 7,82E-01 |
| AT3G01910 | 0,46 | 1,85 | 4,06 | 0,46 | 6,48E-01 |

| | | | | | |
|---|---|---|---|---|---|
| AT3G01930 | 0,62 | -1,19 | 2,94 | -0,41 | 6,85E-01 |
| AT3G01980 | 0,16 | 1,33 | 4,33 | 0,31 | 7,59E-01 |
| AT3G02040 | 0,68 | -1,68 | 2,35 | -0,71 | 4,76E-01 |
| AT3G02080 | 1,20 | -1,37 | 1,99 | -0,68 | 4,94E-01 |
| AT3G02170 | 3,18 | 1,18 | 1,37 | 0,86 | 3,90E-01 |
| AT3G02190 | 0,44 | -0,20 | 3,36 | -0,06 | 9,53E-01 |
| AT3G02210 | 0,52 | 0,93 | 3,99 | 0,23 | 8,17E-01 |
| AT3G02250 | 0,34 | 0,00 | 4,39 | 0,00 | 1,00E+00 |
| AT3G02380 | 0,91 | -1,46 | 2,12 | -0,69 | 4,90E-01 |
| AT3G02410 | 0,66 | -4,03 | 4,30 | -0,94 | 3,48E-01 |
| AT3G02450 | 0,68 | -1,45 | 2,23 | -0,65 | 5,16E-01 |
| AT3G02520 | 1,11 | 0,44 | 1,69 | 0,26 | 7,94E-01 |
| AT3G02530 | 0,67 | 1,23 | 2,68 | 0,46 | 6,46E-01 |
| AT3G02540 | 1,81 | -1,51 | 1,37 | -1,10 | 2,72E-01 |
| AT3G02700 | 0,39 | -1,22 | 4,22 | -0,29 | 7,72E-01 |
| AT3G02730 | 0,92 | -3,98 | 2,75 | -1,45 | 1,48E-01 |
| AT3G02750 | 1,16 | -1,31 | 1,73 | -0,75 | 4,50E-01 |
| AT3G02760 | 0,94 | -0,04 | 2,57 | -0,02 | 9,87E-01 |
| AT3G01205 | 0,80 | 0,46 | 2,30 | 0,20 | 8,41E-01 |
| AT3G03090 | 0,29 | -1,78 | 4,41 | -0,40 | 6,87E-01 |
| AT3G03150 | 0,22 | 0,00 | 4,41 | 0,00 | 1,00E+00 |
| AT3G03220 | 0,16 | 0,00 | 4,41 | 0,00 | 1,00E+00 |
| AT3G03250 | 1,00 | 1,57 | 1,90 | 0,83 | 4,08E-01 |
| AT3G03300 | 0,33 | 2,21 | 4,32 | 0,51 | 6,08E-01 |
| AT3G03305 | 0,32 | 0,23 | 4,34 | 0,05 | 9,58E-01 |
| AT3G03310 | 0,29 | 0,47 | 4,33 | 0,11 | 9,14E-01 |
| AT3G03330 | 0,30 | -2,04 | 4,37 | -0,47 | 6,40E-01 |
| AT3G03405 | 0,16 | 0,00 | 4,41 | 0,00 | 1,00E+00 |
| AT3G03460 | 0,17 | 0,98 | 4,35 | 0,23 | 8,21E-01 |
| AT3G03470 | 1,68 | 2,05 | 1,75 | 1,17 | 2,42E-01 |
| AT3G03480 | 0,08 | 0,69 | 4,36 | 0,16 | 8,75E-01 |
| AT3G03550 | 0,10 | 0,01 | 4,38 | 0,00 | 9,99E-01 |
| AT3G03560 | 0,61 | 1,44 | 2,93 | 0,49 | 6,22E-01 |
| AT3G03580 | 0,26 | -0,30 | 4,37 | -0,07 | 9,46E-01 |
| AT3G03590 | 0,08 | 0,69 | 4,36 | 0,16 | 8,75E-01 |
| AT3G03600 | 0,30 | 0,00 | 4,41 | 0,00 | 1,00E+00 |
| AT3G03720 | 0,26 | 1,41 | 4,33 | 0,32 | 7,45E-01 |
| AT3G03770 | 0,99 | 3,56 | 3,02 | 1,18 | 2,39E-01 |
| AT3G03890 | 0,79 | -0,69 | 2,30 | -0,30 | 7,64E-01 |
| AT3G03920 | 0,42 | -1,78 | 3,39 | -0,52 | 6,00E-01 |
| AT3G03940 | 0,41 | 0,82 | 2,85 | 0,29 | 7,74E-01 |
| AT3G04030 | 0,25 | 0,00 | 4,41 | 0,00 | 1,00E+00 |
| AT3G04080 | 1,01 | 0,15 | 1,89 | 0,08 | 9,37E-01 |
| AT3G04150 | 0,26 | -0,30 | 4,37 | -0,07 | 9,46E-01 |
| AT3G04200 | 0,62 | 0,98 | 4,32 | 0,23 | 8,20E-01 |
| AT3G04210 | 3,60 | 1,32 | 1,08 | 1,23 | 2,20E-01 |
| AT3G04220 | 0,33 | 1,86 | 4,32 | 0,43 | 6,67E-01 |
| AT3G04240 | 3,51 | 0,46 | 1,07 | 0,43 | 6,70E-01 |
| AT3G04260 | 0,61 | -2,06 | 4,37 | -0,47 | 6,38E-01 |
| AT3G04290 | 0,84 | 0,96 | 3,04 | 0,31 | 7,53E-01 |
| AT3G04340 | 3,02 | 0,42 | 1,15 | 0,37 | 7,12E-01 |
| AT3G04350 | 0,35 | 0,88 | 4,35 | 0,20 | 8,40E-01 |

| | | | | | |
|---|---|---|---|---|---|
| AT3G04470 | 2,70 | -0,11 | 1,05 | -0,10 | 9,19E-01 |
| AT3G04480 | 0,84 | 1,69 | 2,27 | 0,75 | 4,55E-01 |
| AT3G04500 | 0,20 | -1,80 | 4,41 | -0,41 | 6,83E-01 |
| AT3G04580 | 1,03 | 0,77 | 1,95 | 0,40 | 6,91E-01 |
| AT3G04640 | 0,85 | 0,32 | 2,24 | 0,14 | 8,85E-01 |
| AT3G04690 | 0,22 | 0,00 | 4,41 | 0,00 | 1,00E+00 |
| AT3G04720 | 0,50 | 1,88 | 3,08 | 0,61 | 5,41E-01 |
| AT3G04730 | 1,02 | -0,40 | 1,67 | -0,24 | 8,11E-01 |
| AT3G04750 | 0,57 | -1,37 | 2,55 | -0,54 | 5,92E-01 |
| AT3G04760 | 0,89 | 1,85 | 2,04 | 0,91 | 3,63E-01 |
| AT3G04765 | 2,83 | -0,01 | 1,29 | -0,01 | 9,92E-01 |
| AT3G04860 | 0,12 | 1,05 | 4,35 | 0,24 | 8,10E-01 |
| AT3G04980 | 0,46 | 0,88 | 4,32 | 0,20 | 8,38E-01 |
| AT3G05030 | 0,28 | 1,35 | 4,33 | 0,31 | 7,55E-01 |
| AT3G01585 | 0,10 | 0,00 | 4,41 | 0,00 | 1,00E+00 |
| AT3G05040 | 1,58 | -1,73 | 1,38 | -1,25 | 2,12E-01 |
| AT3G05050 | 0,38 | 0,63 | 4,33 | 0,15 | 8,84E-01 |
| AT3G05060 | 0,60 | 0,09 | 3,66 | 0,03 | 9,80E-01 |
| AT3G05090 | 0,52 | 0,49 | 2,59 | 0,19 | 8,50E-01 |
| AT3G05165 | 0,83 | -0,35 | 1,97 | -0,18 | 8,58E-01 |
| AT3G05240 | 0,13 | 0,73 | 4,36 | 0,17 | 8,67E-01 |
| AT3G05270 | 0,58 | -1,36 | 2,46 | -0,55 | 5,79E-01 |
| AT3G05280 | 0,59 | -2,41 | 3,74 | -0,64 | 5,20E-01 |
| AT3G05540 | 1,04 | -0,27 | 1,98 | -0,14 | 8,92E-01 |
| AT3G05650 | 0,98 | 1,17 | 2,18 | 0,54 | 5,90E-01 |
| AT3G05660 | 0,79 | 2,02 | 2,71 | 0,74 | 4,57E-01 |
| AT3G05680 | 1,68 | 1,21 | 1,49 | 0,81 | 4,16E-01 |
| AT3G05690 | 0,11 | 0,00 | 4,41 | 0,00 | 1,00E+00 |
| AT3G05770 | 0,46 | 0,36 | 3,32 | 0,11 | 9,14E-01 |
| AT3G05810 | 0,21 | 0,68 | 4,35 | 0,16 | 8,76E-01 |
| AT3G05850 | 0,90 | -1,97 | 2,33 | -0,85 | 3,97E-01 |
| AT3G05880 | 0,55 | -0,70 | 3,21 | -0,22 | 8,28E-01 |
| AT3G05900 | 0,51 | -0,84 | 3,26 | -0,26 | 7,98E-01 |
| AT3G05910 | 0,68 | 0,33 | 2,72 | 0,12 | 9,05E-01 |
| AT3G05932 | 0,27 | 0,37 | 4,33 | 0,09 | 9,32E-01 |
| AT3G05970 | 0,71 | -0,72 | 2,37 | -0,30 | 7,63E-01 |
| AT3G06030 | 0,29 | 2,11 | 4,32 | 0,49 | 6,26E-01 |
| AT3G06050 | 0,18 | 0,00 | 4,41 | 0,00 | 1,00E+00 |
| AT3G06060 | 0,17 | -1,70 | 4,40 | -0,39 | 7,00E-01 |
| AT3G06080 | 1,38 | -1,69 | 1,69 | -1,00 | 3,16E-01 |
| AT3G06150 | 0,98 | -0,27 | 1,71 | -0,16 | 8,76E-01 |
| AT3G06190 | 0,71 | -1,56 | 3,07 | -0,51 | 6,12E-01 |
| AT3G06250 | 0,74 | 0,87 | 2,81 | 0,31 | 7,57E-01 |
| AT3G06290 | 0,72 | 1,60 | 2,41 | 0,66 | 5,07E-01 |
| AT3G06310 | 0,08 | 0,44 | 4,38 | 0,10 | 9,21E-01 |
| AT3G06350 | 0,64 | 0,39 | 2,91 | 0,13 | 8,94E-01 |
| AT3G06430 | 0,12 | 1,23 | 4,34 | 0,28 | 7,76E-01 |
| AT3G06440 | 0,12 | 1,05 | 4,35 | 0,24 | 8,10E-01 |
| AT3G06450 | 0,61 | 0,01 | 2,57 | 0,00 | 9,96E-01 |
| AT3G06480 | 1,05 | -1,18 | 1,74 | -0,67 | 5,00E-01 |
| AT3G06483 | 0,45 | -0,90 | 2,86 | -0,31 | 7,53E-01 |
| AT3G06540 | 0,08 | 0,44 | 4,38 | 0,10 | 9,21E-01 |

| | | | | | |
|---|---|---|---|---|---|
| AT3G06550 | 0,32 | 0,34 | 4,36 | 0,08 | 9,38E-01 |
| AT3G06580 | 0,15 | 1,20 | 4,34 | 0,28 | 7,81E-01 |
| AT3G06590 | 0,40 | -2,27 | 4,39 | -0,52 | 6,05E-01 |
| AT3G06620 | 0,12 | 1,23 | 4,34 | 0,28 | 7,76E-01 |
| AT3G06650 | 3,17 | -1,71 | 1,20 | -1,43 | 1,54E-01 |
| AT3G06665 | 0,25 | 0,00 | 4,41 | 0,00 | 1,00E+00 |
| AT3G06670 | 1,42 | -0,35 | 1,41 | -0,24 | 8,07E-01 |
| AT3G06680 | 0,28 | -1,75 | 4,40 | -0,40 | 6,91E-01 |
| AT3G06690 | 0,20 | -1,80 | 4,41 | -0,41 | 6,83E-01 |
| AT3G06700 | 0,57 | -1,28 | 2,70 | -0,47 | 6,37E-01 |
| AT3G06790 | 0,16 | 0,49 | 4,38 | 0,11 | 9,12E-01 |
| AT3G06820 | 0,08 | 0,69 | 4,36 | 0,16 | 8,75E-01 |
| AT3G06870 | 0,21 | -1,78 | 4,41 | -0,40 | 6,87E-01 |
| AT3G06900 | 1,19 | -0,38 | 1,84 | -0,20 | 8,38E-01 |
| AT3G06930 | 0,40 | -1,30 | 4,16 | -0,31 | 7,55E-01 |
| AT3G06960 | 0,89 | 0,30 | 2,66 | 0,11 | 9,09E-01 |
| AT3G07010 | 0,41 | -1,32 | 4,25 | -0,31 | 7,56E-01 |
| AT3G07020 | 0,54 | 0,36 | 3,71 | 0,10 | 9,23E-01 |
| AT3G07040 | 1,04 | 0,91 | 2,04 | 0,45 | 6,55E-01 |
| AT3G07050 | 0,53 | -0,27 | 3,27 | -0,08 | 9,33E-01 |
| AT3G07115 | 0,08 | 0,00 | 4,40 | 0,00 | 1,00E+00 |
| AT3G07160 | 3,87 | 0,49 | 1,00 | 0,48 | 6,28E-01 |
| AT3G07180 | 0,12 | 1,23 | 4,34 | 0,28 | 7,76E-01 |
| AT3G07215 | 0,55 | -0,91 | 2,68 | -0,34 | 7,34E-01 |
| AT3G07273 | 0,08 | 0,69 | 4,36 | 0,16 | 8,75E-01 |
| AT3G07290 | 0,69 | 1,86 | 3,45 | 0,54 | 5,89E-01 |
| AT3G07300 | 0,29 | -0,93 | 4,34 | -0,21 | 8,31E-01 |
| AT3G07330 | 0,98 | 1,65 | 2,50 | 0,66 | 5,09E-01 |
| AT3G07380 | 1,22 | -0,44 | 1,73 | -0,26 | 7,98E-01 |
| AT3G07460 | 0,63 | -0,88 | 2,72 | -0,32 | 7,48E-01 |
| AT3G07470 | 0,58 | -1,20 | 3,20 | -0,38 | 7,07E-01 |
| AT3G07520 | 0,51 | 0,87 | 3,22 | 0,27 | 7,88E-01 |
| AT3G07540 | 0,08 | 0,69 | 4,36 | 0,16 | 8,75E-01 |
| AT3G07570 | 1,83 | -1,93 | 1,44 | -1,34 | 1,80E-01 |
| AT3G07660 | 1,40 | -0,48 | 1,67 | -0,29 | 7,73E-01 |
| AT3G07700 | 0,94 | 0,07 | 1,73 | 0,04 | 9,69E-01 |
| AT3G07800 | 0,53 | -0,31 | 2,63 | -0,12 | 9,06E-01 |
| AT3G07890 | 0,34 | 0,31 | 4,33 | 0,07 | 9,42E-01 |
| AT3G08510 | 2,33 | -0,27 | 1,54 | -0,18 | 8,59E-01 |
| AT3G08570 | 1,10 | -0,05 | 1,93 | -0,03 | 9,79E-01 |
| AT3G08590 | 1,52 | -0,58 | 1,52 | -0,38 | 7,04E-01 |
| AT3G08610 | 0,59 | -1,60 | 3,12 | -0,51 | 6,08E-01 |
| AT3G08650 | 0,24 | 0,86 | 4,37 | 0,20 | 8,43E-01 |
| AT3G08720 | 1,37 | 1,70 | 1,88 | 0,90 | 3,67E-01 |
| AT3G08730 | 2,48 | -1,15 | 1,31 | -0,88 | 3,81E-01 |
| AT3G08740 | 1,74 | -0,10 | 1,32 | -0,08 | 9,37E-01 |
| AT3G08760 | 0,70 | 0,90 | 2,28 | 0,40 | 6,91E-01 |
| AT3G08840 | 0,67 | -1,33 | 2,89 | -0,46 | 6,47E-01 |
| AT3G08850 | 2,35 | 1,09 | 1,22 | 0,89 | 3,74E-01 |
| AT3G08870 | 0,75 | 2,46 | 3,42 | 0,72 | 4,71E-01 |
| AT3G08960 | 0,38 | -0,23 | 4,32 | -0,05 | 9,57E-01 |
| AT3G09040 | 0,17 | 1,50 | 4,33 | 0,35 | 7,29E-01 |

| | | | | | |
|---|---|---|---|---|---|
| AT3G09070 | 1,31 | 2,16 | 2,17 | 1,00 | 3,19E-01 |
| AT3G09090 | 1,29 | 1,34 | 1,84 | 0,73 | 4,68E-01 |
| AT3G09162 | 2,15 | -4,39 | 1,71 | -2,57 | 1,00E-02 |
| AT3G09190 | 0,55 | -1,67 | 3,72 | -0,45 | 6,53E-01 |
| AT3G09200 | 3,98 | -0,80 | 0,92 | -0,87 | 3,82E-01 |
| AT3G09210 | 0,27 | -1,86 | 4,40 | -0,42 | 6,72E-01 |
| AT3G09270 | 0,53 | 0,49 | 2,69 | 0,18 | 8,55E-01 |
| AT3G09320 | 0,85 | 1,99 | 2,32 | 0,86 | 3,92E-01 |
| AT3G09410 | 0,82 | -0,44 | 2,25 | -0,20 | 8,43E-01 |
| AT3G09470 | 0,82 | 1,70 | 3,43 | 0,50 | 6,20E-01 |
| AT3G09560 | 0,97 | 0,32 | 2,15 | 0,15 | 8,82E-01 |
| AT3G09580 | 0,48 | -1,13 | 3,35 | -0,34 | 7,36E-01 |
| AT3G09770 | 0,38 | 0,38 | 4,32 | 0,09 | 9,31E-01 |
| AT3G09780 | 0,23 | -1,78 | 4,41 | -0,40 | 6,87E-01 |
| AT3G09850 | 1,65 | -0,25 | 1,45 | -0,17 | 8,66E-01 |
| AT3G09870 | 0,08 | 0,69 | 4,36 | 0,16 | 8,75E-01 |
| AT3G09920 | 0,16 | 0,65 | 4,37 | 0,15 | 8,82E-01 |
| AT3G10010 | 0,35 | -2,70 | 4,37 | -0,62 | 5,36E-01 |
| AT3G10020 | 0,27 | 0,94 | 4,36 | 0,21 | 8,30E-01 |
| AT3G10030 | 0,88 | 0,05 | 2,88 | 0,02 | 9,86E-01 |
| AT3G10050 | 0,25 | 1,15 | 4,34 | 0,27 | 7,91E-01 |
| AT3G10070 | 0,46 | -1,13 | 4,22 | -0,27 | 7,89E-01 |
| AT3G10130 | 0,21 | 1,63 | 4,33 | 0,38 | 7,06E-01 |
| AT3G10160 | 0,36 | 0,30 | 3,40 | 0,09 | 9,30E-01 |
| AT3G10230 | 1,13 | 2,56 | 2,18 | 1,17 | 2,40E-01 |
| AT3G10250 | 0,08 | 0,69 | 4,36 | 0,16 | 8,75E-01 |
| AT3G10270 | 0,56 | -1,06 | 3,20 | -0,33 | 7,41E-01 |
| AT3G10320 | 0,12 | 1,23 | 4,34 | 0,28 | 7,76E-01 |
| AT3G10360 | 0,63 | -1,16 | 3,73 | -0,31 | 7,55E-01 |
| AT3G10380 | 2,66 | -0,46 | 1,02 | -0,45 | 6,52E-01 |
| AT3G10405 | 0,68 | -3,58 | 4,32 | -0,83 | 4,06E-01 |
| AT3G10410 | 1,05 | -0,10 | 1,84 | -0,05 | 9,59E-01 |
| AT3G10520 | 0,15 | 0,00 | 4,41 | 0,00 | 1,00E+00 |
| AT3G10525 | 0,39 | 0,64 | 3,40 | 0,19 | 8,50E-01 |
| AT3G10630 | 0,99 | 1,29 | 1,88 | 0,69 | 4,92E-01 |
| AT3G10640 | 0,33 | 0,01 | 4,37 | 0,00 | 9,99E-01 |
| AT3G10650 | 3,29 | -1,06 | 1,02 | -1,03 | 3,01E-01 |
| AT3G10660 | 0,27 | -0,21 | 4,34 | -0,05 | 9,62E-01 |
| AT3G10690 | 0,77 | -0,64 | 3,38 | -0,19 | 8,49E-01 |
| AT3G10720 | 0,36 | 0,00 | 4,41 | 0,00 | 1,00E+00 |
| AT3G10730 | 0,58 | -1,59 | 3,06 | -0,52 | 6,04E-01 |
| AT3G10770 | 0,74 | 1,34 | 2,44 | 0,55 | 5,82E-01 |
| AT3G10810 | 0,46 | -0,20 | 4,16 | -0,05 | 9,61E-01 |
| AT3G10815 | 5,65 | 2,71 | 1,01 | 2,68 | 7,40E-03 |
| AT3G10840 | 0,41 | -3,10 | 4,34 | -0,72 | 4,75E-01 |
| AT3G10850 | 0,36 | -0,01 | 4,35 | 0,00 | 9,97E-01 |
| AT3G10900 | 0,31 | 0,22 | 4,37 | 0,05 | 9,60E-01 |
| AT3G10915 | 0,24 | 0,59 | 4,38 | 0,13 | 8,93E-01 |
| AT3G10930 | 0,61 | 0,43 | 2,51 | 0,17 | 8,64E-01 |
| AT3G10960 | 1,23 | 0,76 | 1,71 | 0,44 | 6,57E-01 |
| AT3G11010 | 3,23 | 2,14 | 1,39 | 1,53 | 1,25E-01 |
| AT3G11030 | 0,56 | -2,18 | 3,76 | -0,58 | 5,62E-01 |

| | | | | | |
|---|---|---|---|---|---|
| AT3G11100 | 0,47 | 2,16 | 3,98 | 0,54 | 5,88E-01 |
| AT3G11280 | 0,31 | -1,08 | 4,37 | -0,25 | 8,04E-01 |
| AT3G11290 | 0,30 | 0,29 | 4,32 | 0,07 | 9,47E-01 |
| AT3G11310 | 0,31 | 0,58 | 4,36 | 0,13 | 8,95E-01 |
| AT3G11320 | 0,17 | 1,15 | 4,34 | 0,27 | 7,91E-01 |
| AT3G11330 | 0,97 | 0,71 | 2,46 | 0,29 | 7,74E-01 |
| AT3G11410 | 4,48 | -1,51 | 1,07 | -1,41 | 1,58E-01 |
| AT3G11435 | 0,23 | 0,00 | 4,41 | 0,00 | 1,00E+00 |
| AT3G11450 | 1,01 | -1,01 | 1,79 | -0,56 | 5,72E-01 |
| AT3G11510 | 1,34 | -1,64 | 1,91 | -0,86 | 3,90E-01 |
| AT3G02435 | 1,34 | -0,73 | 2,20 | -0,33 | 7,39E-01 |
| AT3G02445 | 0,88 | 0,00 | 3,56 | 0,00 | 1,00E+00 |
| AT3G02455 | 0,99 | 3,03 | 2,93 | 1,04 | 3,00E-01 |
| AT3G11570 | 0,77 | -1,25 | 2,92 | -0,43 | 6,70E-01 |
| AT3G11580 | 0,26 | 0,94 | 4,35 | 0,22 | 8,30E-01 |
| AT3G11650 | 0,48 | -0,15 | 3,24 | -0,05 | 9,63E-01 |
| AT3G11660 | 1,33 | 3,10 | 3,36 | 0,92 | 3,57E-01 |
| AT3G11670 | 1,77 | -1,77 | 1,58 | -1,12 | 2,63E-01 |
| AT3G11680 | 0,22 | 0,83 | 4,34 | 0,19 | 8,49E-01 |
| AT3G11710 | 1,08 | -0,54 | 2,07 | -0,26 | 7,96E-01 |
| AT3G11730 | 0,25 | 0,41 | 4,33 | 0,09 | 9,25E-01 |
| AT3G11770 | 0,28 | -0,47 | 4,37 | -0,11 | 9,14E-01 |
| AT3G11780 | 0,21 | 1,16 | 4,34 | 0,27 | 7,89E-01 |
| AT3G11820 | 1,51 | 0,25 | 1,65 | 0,15 | 8,82E-01 |
| AT3G11840 | 0,47 | 2,25 | 4,08 | 0,55 | 5,80E-01 |
| AT3G11910 | 1,81 | 0,29 | 1,31 | 0,22 | 8,25E-01 |
| AT3G11940 | 3,79 | -1,62 | 0,97 | -1,66 | 9,69E-02 |
| AT3G11964 | 0,92 | -0,50 | 1,85 | -0,27 | 7,85E-01 |
| AT3G12030 | 0,12 | 1,23 | 4,34 | 0,28 | 7,76E-01 |
| AT3G12120 | 5,50 | -1,52 | 0,83 | -1,83 | 6,66E-02 |
| AT3G12130 | 0,80 | -1,58 | 2,42 | -0,65 | 5,13E-01 |
| AT3G12140 | 0,25 | 0,29 | 4,36 | 0,07 | 9,47E-01 |
| AT3G12145 | 0,15 | 0,29 | 4,37 | 0,07 | 9,47E-01 |
| AT3G12200 | 0,53 | -0,75 | 3,97 | -0,19 | 8,50E-01 |
| AT3G12260 | 0,32 | 0,00 | 4,40 | 0,00 | 1,00E+00 |
| AT3G12280 | 1,55 | 1,08 | 1,47 | 0,74 | 4,62E-01 |
| AT3G12400 | 0,71 | -0,10 | 1,98 | -0,05 | 9,59E-01 |
| AT3G12490 | 0,47 | 0,23 | 2,41 | 0,10 | 9,23E-01 |
| AT3G12500 | 0,08 | 0,69 | 4,36 | 0,16 | 8,75E-01 |
| AT3G12580 | 0,43 | -0,62 | 4,35 | -0,14 | 8,86E-01 |
| AT3G12587 | 0,28 | -2,52 | 4,38 | -0,58 | 5,65E-01 |
| AT3G12590 | 0,82 | -1,22 | 2,25 | -0,54 | 5,86E-01 |
| AT3G12610 | 0,59 | -1,25 | 2,35 | -0,53 | 5,94E-01 |
| AT3G12670 | 1,09 | 1,87 | 2,48 | 0,75 | 4,50E-01 |
| AT3G12680 | 0,32 | -0,74 | 4,39 | -0,17 | 8,66E-01 |
| AT3G12690 | 0,57 | 2,77 | 4,30 | 0,64 | 5,20E-01 |
| AT3G12750 | 0,45 | 0,04 | 3,94 | 0,01 | 9,91E-01 |
| AT3G12770 | 0,67 | -2,44 | 2,80 | -0,87 | 3,84E-01 |
| AT3G12780 | 5,73 | -0,06 | 0,83 | -0,07 | 9,42E-01 |
| AT3G12800 | 0,96 | 0,90 | 1,97 | 0,46 | 6,48E-01 |
| AT3G12820 | 0,21 | 0,01 | 4,34 | 0,00 | 9,98E-01 |
| AT3G12920 | 1,14 | -0,08 | 2,72 | -0,03 | 9,76E-01 |

| | | | | | |
|---|---|---|---|---|---|
| AT3G12940 | 0,28 | 1,42 | 4,33 | 0,33 | 7,43E-01 |
| AT3G12950 | 0,48 | 1,17 | 4,32 | 0,27 | 7,86E-01 |
| AT3G12965 | 0,28 | -1,75 | 4,40 | -0,40 | 6,91E-01 |
| AT3G13000 | 0,24 | -2,36 | 4,39 | -0,54 | 5,91E-01 |
| AT3G13010 | 0,29 | 0,65 | 4,32 | 0,15 | 8,80E-01 |
| AT3G13020 | 0,33 | -0,94 | 4,34 | -0,22 | 8,28E-01 |
| AT3G13030 | 0,97 | -1,93 | 2,12 | -0,91 | 3,63E-01 |
| AT3G13040 | 0,58 | -0,62 | 2,75 | -0,23 | 8,22E-01 |
| AT3G13061 | 0,23 | 0,00 | 4,38 | 0,00 | 1,00E+00 |
| AT3G13070 | 1,98 | 0,59 | 1,49 | 0,40 | 6,92E-01 |
| AT3G13090 | 2,27 | 1,02 | 1,82 | 0,56 | 5,76E-01 |
| AT3G13100 | 2,83 | 2,69 | 1,49 | 1,80 | 7,12E-02 |
| AT3G13110 | 1,06 | 0,65 | 1,90 | 0,34 | 7,33E-01 |
| AT3G13120 | 0,60 | -1,23 | 3,03 | -0,41 | 6,85E-01 |
| AT3G13150 | 0,37 | -0,17 | 4,31 | -0,04 | 9,68E-01 |
| AT3G13160 | 0,08 | 0,44 | 4,38 | 0,10 | 9,21E-01 |
| AT3G13220 | 0,37 | 0,92 | 4,32 | 0,21 | 8,30E-01 |
| AT3G13222 | 0,48 | 2,55 | 4,30 | 0,59 | 5,53E-01 |
| AT3G13225 | 0,32 | 2,06 | 4,32 | 0,48 | 6,34E-01 |
| AT3G13226 | 0,08 | 0,69 | 4,36 | 0,16 | 8,75E-01 |
| AT3G13235 | 0,44 | -0,11 | 2,82 | -0,04 | 9,68E-01 |
| AT3G13310 | 0,25 | 0,91 | 4,34 | 0,21 | 8,35E-01 |
| AT3G13330 | 3,62 | -0,30 | 0,95 | -0,31 | 7,56E-01 |
| AT3G13390 | 0,05 | 0,00 | 4,41 | 0,00 | 1,00E+00 |
| AT3G13404 | 0,46 | -1,68 | 4,15 | -0,41 | 6,85E-01 |
| AT3G13405 | 1,91 | -1,16 | 1,60 | -0,72 | 4,69E-01 |
| AT3G13410 | 1,64 | -0,37 | 2,16 | -0,17 | 8,65E-01 |
| AT3G13450 | 0,75 | -1,00 | 2,63 | -0,38 | 7,03E-01 |
| AT3G13480 | 0,20 | 0,41 | 4,37 | 0,09 | 9,26E-01 |
| AT3G13490 | 0,57 | 0,07 | 3,00 | 0,02 | 9,82E-01 |
| AT3G13530 | 2,08 | 0,60 | 1,35 | 0,44 | 6,59E-01 |
| AT3G13570 | 0,44 | -1,32 | 4,18 | -0,32 | 7,51E-01 |
| AT3G13720 | 0,96 | 0,26 | 1,93 | 0,13 | 8,94E-01 |
| AT3G13724 | 5,13 | -0,67 | 1,05 | -0,64 | 5,23E-01 |
| AT3G13772 | 0,72 | 1,72 | 2,76 | 0,62 | 5,33E-01 |
| AT3G13860 | 0,89 | 0,20 | 1,90 | 0,11 | 9,15E-01 |
| AT3G13870 | 4,03 | -1,31 | 1,28 | -1,03 | 3,04E-01 |
| AT3G13990 | 1,05 | -1,09 | 1,99 | -0,55 | 5,85E-01 |
| AT3G14050 | 1,74 | 2,36 | 1,63 | 1,44 | 1,49E-01 |
| AT3G14067 | 4,84 | -0,33 | 0,79 | -0,41 | 6,80E-01 |
| AT3G14070 | 0,23 | 1,28 | 4,34 | 0,29 | 7,68E-01 |
| AT3G14075 | 0,26 | 1,43 | 4,34 | 0,33 | 7,41E-01 |
| AT3G14090 | 0,68 | -1,43 | 3,15 | -0,45 | 6,50E-01 |
| AT3G14110 | 0,61 | -1,22 | 2,52 | -0,48 | 6,28E-01 |
| AT3G14172 | 1,38 | -1,24 | 1,62 | -0,77 | 4,43E-01 |
| AT3G14180 | 0,40 | 1,85 | 4,33 | 0,43 | 6,69E-01 |
| AT3G14205 | 0,51 | -0,89 | 2,73 | -0,33 | 7,43E-01 |
| AT3G14230 | 0,40 | -1,30 | 4,16 | -0,31 | 7,55E-01 |
| AT3G14240 | 1,34 | -2,70 | 1,72 | -1,57 | 1,16E-01 |
| AT3G14310 | 2,07 | 0,24 | 1,85 | 0,13 | 8,95E-01 |
| AT3G14330 | 0,56 | 0,23 | 2,94 | 0,08 | 9,37E-01 |
| AT3G14350 | 1,44 | -2,57 | 1,49 | -1,72 | 8,46E-02 |

| | | | | | |
|---|---|---|---|---|---|
| AT3G14400 | 1,24 | 0,86 | 3,06 | 0,28 | 7,79E-01 |
| AT3G14430 | 0,25 | 1,26 | 4,34 | 0,29 | 7,71E-01 |
| AT3G14440 | 0,33 | 1,69 | 4,32 | 0,39 | 6,96E-01 |
| AT3G14460 | 0,50 | 0,20 | 3,92 | 0,05 | 9,59E-01 |
| AT3G14590 | 0,14 | 0,09 | 4,38 | 0,02 | 9,84E-01 |
| AT3G14810 | 0,79 | 1,52 | 2,32 | 0,65 | 5,14E-01 |
| AT3G14900 | 0,17 | 1,31 | 4,34 | 0,30 | 7,63E-01 |
| AT3G15110 | 0,15 | 0,00 | 4,41 | 0,00 | 1,00E+00 |
| AT3G15120 | 0,82 | -0,72 | 2,29 | -0,31 | 7,53E-01 |
| AT3G15150 | 0,08 | 0,00 | 4,40 | 0,00 | 1,00E+00 |
| AT3G15220 | 0,77 | -2,10 | 2,36 | -0,89 | 3,74E-01 |
| AT3G15354 | 0,91 | 1,62 | 2,14 | 0,76 | 4,47E-01 |
| AT3G15356 | 0,36 | -1,71 | 4,35 | -0,39 | 6,95E-01 |
| AT3G15470 | 0,41 | -0,16 | 2,83 | -0,06 | 9,53E-01 |
| AT3G03125 | 0,17 | 0,98 | 4,35 | 0,23 | 8,21E-01 |
| AT3G15530 | 1,09 | 1,81 | 2,44 | 0,74 | 4,58E-01 |
| AT3G15570 | 0,83 | -2,06 | 2,85 | -0,72 | 4,70E-01 |
| AT3G15590 | 0,41 | -0,39 | 4,21 | -0,09 | 9,26E-01 |
| AT3G15630 | 0,46 | 0,84 | 3,25 | 0,26 | 7,96E-01 |
| AT3G15720 | 0,20 | 0,31 | 4,38 | 0,07 | 9,43E-01 |
| AT3G15810 | 0,58 | -0,92 | 2,63 | -0,35 | 7,26E-01 |
| AT3G15840 | 1,45 | -1,20 | 1,43 | -0,84 | 3,99E-01 |
| AT3G15880 | 1,94 | 0,16 | 1,21 | 0,13 | 8,94E-01 |
| AT3G15900 | 0,29 | 0,00 | 4,39 | 0,00 | 1,00E+00 |
| AT3G15940 | 1,43 | 0,75 | 1,86 | 0,40 | 6,86E-01 |
| AT3G15970 | 0,63 | 1,06 | 2,57 | 0,41 | 6,78E-01 |
| AT3G15980 | 1,59 | -0,73 | 1,45 | -0,50 | 6,15E-01 |
| AT3G16000 | 0,51 | 0,61 | 3,28 | 0,19 | 8,52E-01 |
| AT3G16110 | 0,21 | 1,18 | 4,35 | 0,27 | 7,86E-01 |
| AT3G16140 | 1,91 | -1,24 | 1,42 | -0,87 | 3,83E-01 |
| AT3G16190 | 0,21 | -0,49 | 4,35 | -0,11 | 9,11E-01 |
| AT3G03235 | 0,34 | 0,47 | 4,34 | 0,11 | 9,13E-01 |
| AT3G16290 | 0,84 | 1,64 | 2,06 | 0,79 | 4,28E-01 |
| AT3G16310 | 0,55 | 1,18 | 3,76 | 0,31 | 7,53E-01 |
| AT3G16500 | 0,17 | 0,35 | 4,37 | 0,08 | 9,37E-01 |
| AT3G16510 | 0,65 | 0,28 | 3,06 | 0,09 | 9,27E-01 |
| AT3G16520 | 0,76 | 0,79 | 2,20 | 0,36 | 7,21E-01 |
| AT3G16530 | 0,27 | -0,81 | 4,37 | -0,19 | 8,53E-01 |
| AT3G16560 | 0,66 | 0,32 | 2,41 | 0,13 | 8,94E-01 |
| AT3G16630 | 1,01 | 0,27 | 2,07 | 0,13 | 8,98E-01 |
| AT3G16820 | 1,66 | -0,79 | 1,50 | -0,52 | 6,01E-01 |
| AT3G16840 | 0,19 | 0,84 | 4,35 | 0,19 | 8,46E-01 |
| AT3G16860 | 0,47 | -1,67 | 4,08 | -0,41 | 6,82E-01 |
| AT3G16870 | 0,39 | -1,28 | 4,38 | -0,29 | 7,70E-01 |
| AT3G16890 | 0,19 | 0,00 | 4,38 | 0,00 | 9,99E-01 |
| AT3G17020 | 1,21 | -0,94 | 2,06 | -0,45 | 6,49E-01 |
| AT3G03415 | 1,34 | -0,08 | 1,73 | -0,05 | 9,64E-01 |
| AT3G03425 | 4,95 | 2,38 | 1,24 | 1,92 | 5,43E-02 |
| AT3G03435 | 3,28 | 1,14 | 1,14 | 1,01 | 3,13E-01 |
| AT3G03445 | 0,77 | 0,27 | 2,49 | 0,11 | 9,13E-01 |
| AT3G17170 | 0,31 | 0,02 | 4,39 | 0,01 | 9,96E-01 |
| AT3G17240 | 1,76 | -0,25 | 1,48 | -0,17 | 8,68E-01 |

| | | | | | |
|---|---|---|---|---|---|
| AT3G17320 | 0,12 | 0,26 | 4,37 | 0,06 | 9,52E-01 |
| AT3G17330 | 0,20 | 1,26 | 4,34 | 0,29 | 7,71E-01 |
| AT3G17390 | 4,31 | 0,29 | 1,04 | 0,28 | 7,77E-01 |
| AT3G17420 | 0,17 | 0,47 | 4,37 | 0,11 | 9,14E-01 |
| AT3G17440 | 0,08 | 0,69 | 4,36 | 0,16 | 8,75E-01 |
| AT3G17450 | 0,60 | -0,43 | 2,84 | -0,15 | 8,80E-01 |
| AT3G17510 | 2,73 | -0,81 | 1,31 | -0,62 | 5,37E-01 |
| AT3G17655 | 0,19 | 1,46 | 4,33 | 0,34 | 7,35E-01 |
| AT3G17740 | 0,37 | 0,47 | 4,30 | 0,11 | 9,13E-01 |
| AT3G17760 | 0,21 | 0,01 | 4,34 | 0,00 | 9,98E-01 |
| AT3G17770 | 1,51 | 0,31 | 1,42 | 0,22 | 8,29E-01 |
| AT3G17800 | 1,86 | -0,69 | 1,46 | -0,47 | 6,39E-01 |
| AT3G17850 | 4,74 | 0,14 | 0,88 | 0,16 | 8,70E-01 |
| AT3G03585 | 0,05 | 0,00 | 4,41 | 0,00 | 1,00E+00 |
| AT3G17940 | 0,81 | -1,03 | 2,45 | -0,42 | 6,75E-01 |
| AT3G18030 | 0,52 | 0,93 | 3,99 | 0,23 | 8,17E-01 |
| AT3G18035 | 1,23 | -0,12 | 1,70 | -0,07 | 9,43E-01 |
| AT3G18110 | 1,16 | 1,03 | 2,32 | 0,44 | 6,57E-01 |
| AT3G18130 | 0,88 | -0,39 | 2,29 | -0,17 | 8,64E-01 |
| AT3G18140 | 0,30 | -1,59 | 4,38 | -0,36 | 7,17E-01 |
| AT3G18250 | 0,55 | -3,44 | 4,32 | -0,79 | 4,27E-01 |
| AT3G18380 | 0,25 | 0,41 | 4,33 | 0,09 | 9,25E-01 |
| AT3G18420 | 0,20 | 0,81 | 4,36 | 0,19 | 8,52E-01 |
| AT3G18535 | 0,67 | 0,64 | 2,47 | 0,26 | 7,95E-01 |
| AT3G18600 | 0,25 | -2,19 | 4,38 | -0,50 | 6,17E-01 |
| AT3G18610 | 0,55 | 1,46 | 4,34 | 0,34 | 7,37E-01 |
| AT3G18630 | 0,20 | -0,44 | 4,36 | -0,10 | 9,19E-01 |
| AT3G18700 | 1,19 | 3,33 | 2,07 | 1,61 | 1,08E-01 |
| AT3G18750 | 0,17 | 1,40 | 4,33 | 0,32 | 7,47E-01 |
| AT3G18830 | 2,46 | 1,67 | 1,29 | 1,29 | 1,96E-01 |
| AT3G18930 | 0,90 | -0,40 | 2,39 | -0,17 | 8,66E-01 |
| AT3G18970 | 0,08 | 0,69 | 4,36 | 0,16 | 8,75E-01 |
| AT3G19000 | 0,24 | 1,17 | 4,35 | 0,27 | 7,87E-01 |
| AT3G19010 | 1,81 | -0,64 | 1,57 | -0,41 | 6,84E-01 |
| AT3G19020 | 0,54 | -1,44 | 3,08 | -0,47 | 6,41E-01 |
| AT3G19030 | 2,04 | -0,16 | 1,26 | -0,13 | 8,97E-01 |
| AT3G19150 | 0,16 | 0,00 | 4,37 | 0,00 | 9,99E-01 |
| AT3G19170 | 4,99 | 2,85 | 1,40 | 2,04 | 4,17E-02 |
| AT3G03815 | 0,25 | -2,19 | 4,38 | -0,50 | 6,17E-01 |
| AT3G19300 | 0,12 | 0,97 | 4,34 | 0,22 | 8,23E-01 |
| AT3G19340 | 0,12 | 0,00 | 4,38 | 0,00 | 1,00E+00 |
| AT3G19380 | 2,79 | -0,20 | 1,29 | -0,15 | 8,78E-01 |
| AT3G19430 | 0,25 | -2,19 | 4,38 | -0,50 | 6,17E-01 |
| AT3G19490 | 0,72 | 1,53 | 2,44 | 0,63 | 5,30E-01 |
| AT3G19510 | 0,12 | 0,49 | 4,36 | 0,11 | 9,11E-01 |
| AT3G19515 | 0,59 | 0,81 | 3,11 | 0,26 | 7,95E-01 |
| AT3G19553 | 0,42 | -2,51 | 4,14 | -0,60 | 5,45E-01 |
| AT3G19580 | 0,72 | 1,39 | 2,46 | 0,57 | 5,70E-01 |
| AT3G19620 | 0,32 | -0,94 | 4,31 | -0,22 | 8,28E-01 |
| AT3G19630 | 0,39 | -1,28 | 4,38 | -0,29 | 7,70E-01 |
| AT3G19710 | 0,69 | 1,15 | 2,47 | 0,47 | 6,42E-01 |
| AT3G19720 | 1,17 | 3,79 | 2,42 | 1,57 | 1,17E-01 |

| | | | | | |
|---|---|---|---|---|---|
| AT3G19740 | 0,59 | 0,78 | 3,13 | 0,25 | 8,04E-01 |
| AT3G19790 | 0,71 | -1,22 | 1,89 | -0,64 | 5,20E-01 |
| AT3G19810 | 0,70 | -1,08 | 4,37 | -0,25 | 8,04E-01 |
| AT3G19830 | 0,67 | 1,78 | 2,99 | 0,59 | 5,53E-01 |
| AT3G19850 | 0,50 | -1,97 | 3,10 | -0,64 | 5,24E-01 |
| AT3G19880 | 0,35 | 1,73 | 3,43 | 0,50 | 6,14E-01 |
| AT3G19895 | 0,23 | 0,19 | 4,38 | 0,04 | 9,66E-01 |
| AT3G19900 | 0,17 | 1,50 | 4,33 | 0,35 | 7,29E-01 |
| AT3G20015 | 0,62 | -3,33 | 3,07 | -1,09 | 2,77E-01 |
| AT3G20050 | 0,74 | -0,06 | 2,44 | -0,02 | 9,82E-01 |
| AT3G20060 | 0,33 | 0,54 | 4,36 | 0,12 | 9,01E-01 |
| AT3G20230 | 0,35 | 1,92 | 4,32 | 0,44 | 6,57E-01 |
| AT3G20250 | 1,61 | -0,88 | 1,59 | -0,55 | 5,81E-01 |
| AT3G20290 | 0,63 | 2,17 | 3,04 | 0,71 | 4,76E-01 |
| AT3G20310 | 2,42 | 0,25 | 1,34 | 0,19 | 8,53E-01 |
| AT3G20320 | 0,51 | 0,29 | 2,76 | 0,11 | 9,15E-01 |
| AT3G20390 | 0,62 | 0,85 | 2,33 | 0,36 | 7,16E-01 |
| AT3G20470 | 0,38 | -0,76 | 4,38 | -0,17 | 8,62E-01 |
| AT3G20620 | 0,68 | 0,84 | 3,53 | 0,24 | 8,11E-01 |
| AT3G20630 | 1,20 | -0,78 | 1,52 | -0,52 | 6,07E-01 |
| AT3G20655 | 0,30 | 0,07 | 4,35 | 0,02 | 9,87E-01 |
| AT3G20690 | 3,62 | -0,50 | 1,56 | -0,32 | 7,47E-01 |
| AT3G20700 | 1,05 | 1,14 | 2,51 | 0,46 | 6,49E-01 |
| AT3G03985 | 0,54 | 0,00 | 4,41 | 0,00 | 1,00E+00 |
| AT3G20760 | 5,52 | -1,37 | 0,77 | -1,79 | 7,34E-02 |
| AT3G20770 | 2,68 | -0,56 | 1,09 | -0,51 | 6,07E-01 |
| AT3G20830 | 0,60 | 1,12 | 3,31 | 0,34 | 7,34E-01 |
| AT3G21055 | 1,34 | -0,91 | 1,72 | -0,53 | 5,94E-01 |
| AT3G21070 | 0,23 | 1,14 | 4,34 | 0,26 | 7,92E-01 |
| AT3G21140 | 0,25 | 0,96 | 4,36 | 0,22 | 8,25E-01 |
| AT3G21160 | 0,11 | 0,65 | 4,36 | 0,15 | 8,81E-01 |
| AT3G21215 | 0,11 | 0,48 | 4,37 | 0,11 | 9,12E-01 |
| AT3G21300 | 0,80 | -0,95 | 2,21 | -0,43 | 6,68E-01 |
| AT3G21430 | 0,61 | 0,24 | 3,19 | 0,08 | 9,39E-01 |
| AT3G21540 | 0,20 | 0,75 | 4,37 | 0,17 | 8,64E-01 |
| AT3G21570 | 0,87 | -1,75 | 3,04 | -0,57 | 5,66E-01 |
| AT3G21630 | 0,12 | 0,00 | 4,38 | 0,00 | 1,00E+00 |
| AT3G21650 | 0,33 | 0,75 | 4,32 | 0,17 | 8,62E-01 |
| AT3G21670 | 1,42 | 1,29 | 2,24 | 0,58 | 5,65E-01 |
| AT3G21740 | 0,24 | -0,74 | 4,35 | -0,17 | 8,65E-01 |
| AT3G21755 | 4,36 | 0,62 | 1,08 | 0,58 | 5,63E-01 |
| AT3G21790 | 0,96 | 1,70 | 2,15 | 0,79 | 4,27E-01 |
| AT3G21865 | 0,48 | 0,66 | 2,69 | 0,24 | 8,07E-01 |
| AT3G21870 | 1,78 | 0,43 | 1,95 | 0,22 | 8,25E-01 |
| AT3G22070 | 0,17 | 0,00 | 4,41 | 0,00 | 1,00E+00 |
| AT3G22110 | 0,73 | -1,32 | 2,44 | -0,54 | 5,89E-01 |
| AT3G22120 | 4,08 | -1,84 | 1,04 | -1,77 | 7,71E-02 |
| AT3G22142 | 1,40 | 2,67 | 1,87 | 1,42 | 1,54E-01 |
| AT3G22170 | 1,10 | 1,20 | 1,94 | 0,62 | 5,35E-01 |
| AT3G22190 | 0,78 | -2,07 | 2,87 | -0,72 | 4,70E-01 |
| AT3G22210 | 0,48 | -0,88 | 3,37 | -0,26 | 7,93E-01 |
| AT3G22220 | 0,36 | 0,21 | 3,46 | 0,06 | 9,53E-01 |

| | | | | | |
|---|---|---|---|---|---|
| AT3G22230 | 0,95 | -0,47 | 1,79 | -0,26 | 7,92E-01 |
| AT3G22231 | 4,08 | 0,91 | 1,08 | 0,84 | 4,01E-01 |
| AT3G22235 | 3,04 | 0,12 | 1,16 | 0,11 | 9,15E-01 |
| AT3G22240 | 1,25 | 1,78 | 1,89 | 0,94 | 3,48E-01 |
| AT3G22320 | 0,41 | -0,39 | 4,21 | -0,09 | 9,26E-01 |
| AT3G22421 | 0,96 | 3,06 | 2,67 | 1,15 | 2,52E-01 |
| AT3G22470 | 0,59 | 0,64 | 2,59 | 0,25 | 8,04E-01 |
| AT3G22530 | 0,38 | -0,24 | 4,36 | -0,05 | 9,56E-01 |
| AT3G22550 | 0,60 | 1,11 | 3,03 | 0,37 | 7,14E-01 |
| AT3G22580 | 0,15 | 0,00 | 4,41 | 0,00 | 1,00E+00 |
| AT3G22600 | 0,17 | 1,52 | 4,33 | 0,35 | 7,26E-01 |
| AT3G22630 | 0,68 | -0,76 | 2,13 | -0,36 | 7,19E-01 |
| AT3G22660 | 0,44 | -2,98 | 4,35 | -0,69 | 4,93E-01 |
| AT3G22690 | 0,29 | 1,54 | 4,33 | 0,35 | 7,23E-01 |
| AT3G22710 | 1,14 | -1,54 | 1,80 | -0,85 | 3,93E-01 |
| AT3G22720 | 0,08 | 0,00 | 4,40 | 0,00 | 1,00E+00 |
| AT3G22730 | 1,61 | 1,52 | 1,86 | 0,82 | 4,12E-01 |
| AT3G22770 | 0,68 | 1,74 | 3,27 | 0,53 | 5,96E-01 |
| AT3G22790 | 2,45 | -0,98 | 1,17 | -0,84 | 4,01E-01 |
| AT3G22910 | 2,19 | 1,00 | 1,29 | 0,77 | 4,39E-01 |
| AT3G22950 | 0,15 | 0,00 | 4,41 | 0,00 | 1,00E+00 |
| AT3G22980 | 0,74 | 0,83 | 2,46 | 0,34 | 7,37E-01 |
| AT3G23020 | 0,53 | -0,66 | 3,74 | -0,18 | 8,60E-01 |
| AT3G23030 | 0,69 | -0,95 | 2,03 | -0,46 | 6,42E-01 |
| AT3G23080 | 0,08 | 0,69 | 4,36 | 0,16 | 8,75E-01 |
| AT3G23110 | 1,95 | -0,92 | 1,48 | -0,62 | 5,33E-01 |
| AT3G23120 | 2,24 | 2,03 | 1,33 | 1,53 | 1,26E-01 |
| AT3G23170 | 0,50 | 0,67 | 2,64 | 0,25 | 7,99E-01 |
| AT3G23172 | 0,24 | 1,17 | 4,34 | 0,27 | 7,88E-01 |
| AT3G23400 | 0,58 | -0,12 | 3,22 | -0,04 | 9,71E-01 |
| AT3G23420 | 0,32 | 0,82 | 4,36 | 0,19 | 8,50E-01 |
| AT3G23490 | 0,26 | 0,48 | 4,37 | 0,11 | 9,13E-01 |
| AT3G04455 | 2,46 | -0,70 | 1,94 | -0,36 | 7,19E-01 |
| AT3G23570 | 0,32 | 0,55 | 4,32 | 0,13 | 8,98E-01 |
| AT3G23633 | 0,12 | 0,00 | 4,41 | 0,00 | 1,00E+00 |
| AT3G23650 | 0,55 | 1,52 | 2,96 | 0,51 | 6,08E-01 |
| AT3G04585 | 0,72 | 1,68 | 2,84 | 0,59 | 5,54E-01 |
| AT3G23660 | 0,75 | -0,62 | 2,80 | -0,22 | 8,24E-01 |
| AT3G23700 | 0,24 | 0,05 | 4,39 | 0,01 | 9,91E-01 |
| AT3G23740 | 0,08 | 0,44 | 4,38 | 0,10 | 9,21E-01 |
| AT3G23780 | 4,59 | -0,84 | 0,83 | -1,02 | 3,10E-01 |
| AT3G23810 | 2,25 | -0,87 | 1,44 | -0,60 | 5,47E-01 |
| AT3G23830 | 0,27 | 0,37 | 4,37 | 0,08 | 9,33E-01 |
| AT3G23900 | 0,77 | 1,79 | 2,74 | 0,65 | 5,13E-01 |
| AT3G23970 | 0,08 | 0,44 | 4,38 | 0,10 | 9,21E-01 |
| AT3G23980 | 0,76 | -1,60 | 2,54 | -0,63 | 5,28E-01 |
| AT3G24080 | 0,22 | 1,51 | 4,33 | 0,35 | 7,27E-01 |
| AT3G24120 | 0,83 | 1,23 | 3,44 | 0,36 | 7,21E-01 |
| AT3G24140 | 0,26 | 0,03 | 4,39 | 0,01 | 9,94E-01 |
| AT3G24170 | 1,33 | 0,47 | 1,66 | 0,28 | 7,77E-01 |
| AT3G24180 | 1,99 | -0,07 | 1,20 | -0,06 | 9,55E-01 |
| AT3G24200 | 0,11 | 0,65 | 4,36 | 0,15 | 8,81E-01 |

| | | | | | |
|---|---|---|---|---|---|
| AT3G24260 | 0,15 | 0,00 | 4,41 | 0,00 | 1,00E+00 |
| AT3G24270 | 0,91 | 2,90 | 2,33 | 1,25 | 2,12E-01 |
| AT3G24320 | 0,63 | 0,39 | 2,99 | 0,13 | 8,97E-01 |
| AT3G24430 | 0,54 | -1,01 | 2,71 | -0,37 | 7,10E-01 |
| AT3G24440 | 0,50 | -0,54 | 3,11 | -0,17 | 8,63E-01 |
| AT3G24465 | 0,32 | 0,82 | 3,64 | 0,23 | 8,21E-01 |
| AT3G24480 | 4,54 | -1,91 | 0,85 | -2,24 | 2,52E-02 |
| AT3G24503 | 1,13 | -1,32 | 1,83 | -0,72 | 4,71E-01 |
| AT3G24518 | 0,96 | -1,45 | 2,51 | -0,58 | 5,62E-01 |
| AT3G24612 | 3,02 | 0,72 | 1,11 | 0,65 | 5,15E-01 |
| AT3G24614 | 0,96 | -0,78 | 1,83 | -0,42 | 6,72E-01 |
| AT3G24670 | 0,20 | 1,20 | 4,35 | 0,28 | 7,83E-01 |
| AT3G24870 | 1,16 | 0,25 | 2,15 | 0,12 | 9,07E-01 |
| AT3G24880 | 0,67 | -2,51 | 3,05 | -0,82 | 4,12E-01 |
| AT3G24900 | 0,51 | 0,85 | 3,11 | 0,27 | 7,84E-01 |
| AT3G24982 | 2,19 | 2,72 | 1,66 | 1,64 | 1,02E-01 |
| AT3G25010 | 4,35 | -1,54 | 0,88 | -1,74 | 8,16E-02 |
| AT3G25020 | 3,41 | 1,36 | 1,36 | 1,00 | 3,18E-01 |
| AT3G25090 | 0,87 | -0,14 | 2,15 | -0,06 | 9,48E-01 |
| AT3G25150 | 1,03 | -0,08 | 2,38 | -0,03 | 9,74E-01 |
| AT3G25180 | 0,15 | 0,00 | 4,41 | 0,00 | 1,00E+00 |
| AT3G25460 | 1,48 | 2,03 | 1,87 | 1,08 | 2,80E-01 |
| AT3G25500 | 1,85 | 0,10 | 1,22 | 0,08 | 9,37E-01 |
| AT3G25510 | 1,96 | -0,38 | 1,30 | -0,29 | 7,72E-01 |
| AT3G25520 | 3,31 | -1,24 | 0,99 | -1,24 | 2,14E-01 |
| AT3G25530 | 0,19 | 0,01 | 4,38 | 0,00 | 9,98E-01 |
| AT3G25560 | 0,41 | -1,53 | 4,36 | -0,35 | 7,26E-01 |
| AT3G25610 | 1,05 | -2,65 | 2,29 | -1,16 | 2,47E-01 |
| AT3G25660 | 0,47 | 2,00 | 3,95 | 0,51 | 6,12E-01 |
| AT3G04935 | 0,15 | 0,83 | 4,35 | 0,19 | 8,49E-01 |
| AT3G25800 | 1,14 | 0,59 | 1,93 | 0,30 | 7,61E-01 |
| AT3G25840 | 1,32 | -1,31 | 1,56 | -0,84 | 4,01E-01 |
| AT3G25855 | 1,70 | -4,15 | 1,95 | -2,12 | 3,38E-02 |
| AT3G25870 | 0,65 | 1,49 | 2,45 | 0,61 | 5,45E-01 |
| AT3G25880 | 1,38 | -1,59 | 2,33 | -0,68 | 4,95E-01 |
| AT3G25882 | 0,50 | 0,45 | 3,99 | 0,11 | 9,10E-01 |
| AT3G25920 | 0,58 | -0,62 | 2,24 | -0,28 | 7,81E-01 |
| AT3G25990 | 0,75 | -3,23 | 3,38 | -0,96 | 3,39E-01 |
| AT3G26000 | 2,02 | -1,47 | 1,34 | -1,10 | 2,73E-01 |
| AT3G26170 | 1,29 | 1,53 | 1,86 | 0,82 | 4,12E-01 |
| AT3G26180 | 0,36 | 0,04 | 4,33 | 0,01 | 9,92E-01 |
| AT3G26210 | 1,30 | 1,15 | 2,01 | 0,57 | 5,69E-01 |
| AT3G26220 | 0,49 | 0,12 | 4,02 | 0,03 | 9,76E-01 |
| AT3G26230 | 0,79 | -2,07 | 2,52 | -0,82 | 4,11E-01 |
| AT3G26280 | 1,16 | -2,50 | 1,78 | -1,40 | 1,60E-01 |
| AT3G26290 | 0,78 | 0,24 | 2,05 | 0,12 | 9,08E-01 |
| AT3G26300 | 0,36 | -1,42 | 4,35 | -0,33 | 7,45E-01 |
| AT3G26340 | 0,62 | -3,53 | 3,75 | -0,94 | 3,46E-01 |
| AT3G26410 | 0,31 | -2,59 | 4,37 | -0,59 | 5,53E-01 |
| AT3G26450 | 2,27 | -1,48 | 1,55 | -0,95 | 3,42E-01 |
| AT3G26500 | 1,32 | 3,03 | 1,97 | 1,54 | 1,24E-01 |
| AT3G26510 | 0,66 | 0,31 | 2,19 | 0,14 | 8,86E-01 |

| | | | | | |
|---|---|---|---|---|---|
| AT3G26550 | 0,20 | -0,31 | 4,36 | -0,07 | 9,44E-01 |
| AT3G26560 | 0,40 | 1,16 | 3,37 | 0,35 | 7,30E-01 |
| AT3G26590 | 0,41 | -2,38 | 4,35 | -0,55 | 5,83E-01 |
| AT3G26616 | 0,23 | 0,67 | 4,36 | 0,15 | 8,77E-01 |
| AT3G26630 | 0,69 | -1,63 | 2,30 | -0,71 | 4,78E-01 |
| AT3G26670 | 0,54 | -0,86 | 3,23 | -0,26 | 7,91E-01 |
| AT3G26744 | 0,39 | -0,72 | 4,25 | -0,17 | 8,65E-01 |
| AT3G26812 | 0,17 | 0,07 | 4,40 | 0,02 | 9,88E-01 |
| AT3G26819 | 0,05 | 0,00 | 4,41 | 0,00 | 1,00E+00 |
| AT3G26860 | 0,30 | 0,00 | 4,41 | 0,00 | 1,00E+00 |
| AT3G26870 | 1,55 | 0,29 | 2,25 | 0,13 | 8,98E-01 |
| AT3G26890 | 0,29 | -0,67 | 4,36 | -0,15 | 8,78E-01 |
| AT3G26932 | 0,90 | 2,15 | 2,39 | 0,90 | 3,69E-01 |
| AT3G26935 | 0,27 | -0,34 | 4,34 | -0,08 | 9,37E-01 |
| AT3G26980 | 0,14 | 0,83 | 4,35 | 0,19 | 8,49E-01 |
| AT3G27010 | 0,67 | 0,48 | 3,62 | 0,13 | 8,94E-01 |
| AT3G27020 | 0,51 | -0,52 | 2,32 | -0,23 | 8,21E-01 |
| AT3G27040 | 1,43 | 1,54 | 2,02 | 0,76 | 4,46E-01 |
| AT3G27060 | 0,90 | 2,05 | 2,43 | 0,84 | 3,99E-01 |
| AT3G27100 | 0,74 | -1,93 | 4,36 | -0,44 | 6,58E-01 |
| AT3G27240 | 0,37 | -0,86 | 3,45 | -0,25 | 8,03E-01 |
| AT3G27300 | 0,49 | -0,53 | 3,28 | -0,16 | 8,71E-01 |
| AT3G27380 | 0,45 | 0,78 | 4,36 | 0,18 | 8,58E-01 |
| AT3G05235 | 0,99 | -0,83 | 2,65 | -0,31 | 7,53E-01 |
| AT3G05245 | 0,33 | 0,00 | 4,41 | 0,00 | 1,00E+00 |
| AT3G27530 | 0,76 | 0,29 | 2,57 | 0,11 | 9,10E-01 |
| AT3G27600 | 1,14 | -0,60 | 2,22 | -0,27 | 7,86E-01 |
| AT3G27610 | 0,33 | 0,35 | 4,39 | 0,08 | 9,36E-01 |
| AT3G27680 | 0,86 | -1,77 | 2,49 | -0,71 | 4,78E-01 |
| AT3G27700 | 1,39 | 2,32 | 2,00 | 1,16 | 2,46E-01 |
| AT3G27740 | 0,52 | -0,23 | 3,97 | -0,06 | 9,54E-01 |
| AT3G27770 | 0,43 | 0,48 | 2,80 | 0,17 | 8,65E-01 |
| AT3G27906 | 1,35 | -0,59 | 1,77 | -0,33 | 7,38E-01 |
| AT3G27925 | 0,65 | 0,95 | 3,07 | 0,31 | 7,58E-01 |
| AT3G27930 | 0,20 | 0,31 | 4,38 | 0,07 | 9,43E-01 |
| AT3G27990 | 0,64 | -1,78 | 2,64 | -0,67 | 5,01E-01 |
| AT3G28340 | 0,22 | 0,17 | 4,37 | 0,04 | 9,69E-01 |
| AT3G28380 | 0,08 | 0,44 | 4,38 | 0,10 | 9,21E-01 |
| AT3G28410 | 0,14 | 0,03 | 4,39 | 0,01 | 9,95E-01 |
| AT3G28550 | 2,18 | 2,49 | 1,83 | 1,36 | 1,75E-01 |
| AT3G28674 | 0,28 | 0,36 | 4,37 | 0,08 | 9,34E-01 |
| AT3G28695 | 0,43 | -1,03 | 4,21 | -0,24 | 8,07E-01 |
| AT3G28710 | 0,99 | 0,20 | 1,74 | 0,11 | 9,09E-01 |
| AT3G28780 | 0,61 | 1,68 | 3,16 | 0,53 | 5,95E-01 |
| AT3G28860 | 1,33 | -0,29 | 2,20 | -0,13 | 8,96E-01 |
| AT3G28890 | 0,43 | 0,12 | 4,33 | 0,03 | 9,78E-01 |
| AT3G28900 | 3,87 | -1,70 | 1,07 | -1,59 | 1,12E-01 |
| AT3G28920 | 1,35 | 0,46 | 1,88 | 0,24 | 8,08E-01 |
| AT3G28930 | 0,77 | 1,64 | 2,97 | 0,55 | 5,79E-01 |
| AT3G28940 | 1,30 | 0,63 | 1,76 | 0,36 | 7,19E-01 |
| AT3G28950 | 0,97 | -1,21 | 2,06 | -0,59 | 5,56E-01 |
| AT3G28956 | 2,37 | 1,79 | 1,63 | 1,10 | 2,72E-01 |

| | | | | | |
|---|---|---|---|---|---|
| AT3G28960 | 0,30 | 0,07 | 4,40 | 0,02 | 9,88E-01 |
| AT3G28970 | 0,52 | 1,49 | 3,96 | 0,38 | 7,06E-01 |
| AT3G29030 | 0,33 | 0,00 | 4,41 | 0,00 | 1,00E+00 |
| AT3G05655 | 1,87 | 0,70 | 1,64 | 0,43 | 6,68E-01 |
| AT3G05665 | 0,80 | -0,90 | 2,77 | -0,33 | 7,44E-01 |
| AT3G29110 | 0,53 | -1,74 | 2,75 | -0,63 | 5,26E-01 |
| AT3G29130 | 2,05 | 0,83 | 1,71 | 0,48 | 6,28E-01 |
| AT3G29160 | 0,59 | -16,61 | 4,39 | -3,78 | 1,54E-04 |
| AT3G29190 | 2,69 | 1,80 | 1,48 | 1,21 | 2,25E-01 |
| AT3G29200 | 0,59 | 0,40 | 3,00 | 0,13 | 8,93E-01 |
| AT3G29350 | 0,17 | -1,70 | 4,40 | -0,39 | 7,00E-01 |
| AT3G29360 | 2,83 | 0,22 | 1,11 | 0,19 | 8,46E-01 |
| AT3G29400 | 0,67 | -0,35 | 2,56 | -0,14 | 8,90E-01 |
| AT3G29570 | 0,38 | 0,40 | 4,38 | 0,09 | 9,26E-01 |
| AT3G05745 | 0,14 | 0,76 | 4,35 | 0,18 | 8,61E-01 |
| AT3G29633 | 0,14 | 0,00 | 4,41 | 0,00 | 1,00E+00 |
| AT3G29639 | 0,16 | 0,09 | 4,38 | 0,02 | 9,83E-01 |
| AT3G05895 | 0,08 | 0,00 | 4,40 | 0,00 | 1,00E+00 |
| AT3G29680 | 0,25 | -1,76 | 4,41 | -0,40 | 6,89E-01 |
| AT3G29720 | 0,40 | 0,23 | 4,40 | 0,05 | 9,59E-01 |
| AT3G29760 | 0,67 | -1,33 | 2,89 | -0,46 | 6,47E-01 |
| AT3G29785 | 0,22 | 0,28 | 4,37 | 0,06 | 9,49E-01 |
| AT3G05985 | 2,35 | 0,09 | 1,37 | 0,07 | 9,47E-01 |
| AT3G30160 | 0,58 | 0,12 | 3,06 | 0,04 | 9,68E-01 |
| AT3G30200 | 0,12 | 0,49 | 4,36 | 0,11 | 9,11E-01 |
| AT3G30220 | 0,08 | 0,00 | 4,40 | 0,00 | 1,00E+00 |
| AT3G30230 | 0,54 | 1,67 | 3,91 | 0,43 | 6,69E-01 |
| AT3G30300 | 0,53 | -0,63 | 4,32 | -0,15 | 8,84E-01 |
| AT3G30390 | 2,05 | 0,46 | 1,19 | 0,39 | 7,00E-01 |
| AT3G06055 | 2,49 | 1,79 | 1,41 | 1,27 | 2,04E-01 |
| AT3G30840 | 0,21 | 1,55 | 4,33 | 0,36 | 7,20E-01 |
| AT3G32190 | 0,34 | 0,24 | 3,46 | 0,07 | 9,44E-01 |
| AT3G32420 | 0,10 | 0,01 | 4,38 | 0,00 | 9,99E-01 |
| AT3G32904 | 0,65 | -0,89 | 3,18 | -0,28 | 7,79E-01 |
| AT3G33528 | 0,61 | 1,88 | 2,49 | 0,75 | 4,50E-01 |
| AT3G42050 | 1,44 | -1,72 | 1,74 | -0,99 | 3,23E-01 |
| AT3G42060 | 0,32 | 0,50 | 4,37 | 0,11 | 9,10E-01 |
| AT3G42140 | 0,14 | 0,03 | 4,39 | 0,01 | 9,95E-01 |
| AT3G42155 | 0,34 | -1,54 | 4,38 | -0,35 | 7,25E-01 |
| AT3G42550 | 0,12 | 0,00 | 4,39 | 0,00 | 1,00E+00 |
| AT3G42723 | 0,12 | 0,00 | 4,38 | 0,00 | 1,00E+00 |
| AT3G42790 | 0,59 | 0,15 | 3,15 | 0,05 | 9,62E-01 |
| AT3G43153 | 0,99 | 3,29 | 2,50 | 1,31 | 1,89E-01 |
| AT3G43190 | 0,26 | -1,78 | 4,41 | -0,40 | 6,87E-01 |
| AT3G43220 | 0,79 | 0,59 | 2,85 | 0,21 | 8,36E-01 |
| AT3G43240 | 0,86 | -0,38 | 2,60 | -0,15 | 8,83E-01 |
| AT3G43270 | 0,68 | 0,00 | 2,62 | 0,00 | 1,00E+00 |
| AT3G43280 | 0,11 | 0,00 | 4,41 | 0,00 | 1,00E+00 |
| AT3G43291 | 0,65 | -0,84 | 2,61 | -0,32 | 7,48E-01 |
| AT3G06595 | 0,10 | 0,01 | 4,38 | 0,00 | 9,99E-01 |
| AT3G43432 | 0,16 | 0,04 | 4,38 | 0,01 | 9,92E-01 |
| AT3G43470 | 0,45 | 0,41 | 4,00 | 0,10 | 9,18E-01 |

| | | | | | |
|---|---|---|---|---|---|
| AT3G43500 | 0,45 | 0,06 | 4,04 | 0,02 | 9,87E-01 |
| AT3G43600 | 0,98 | -2,24 | 2,14 | -1,04 | 2,96E-01 |
| AT3G43610 | 0,74 | -0,71 | 2,41 | -0,29 | 7,70E-01 |
| AT3G06645 | 0,23 | 0,00 | 4,41 | 0,00 | 1,00E+00 |
| AT3G06655 | 1,06 | 2,26 | 2,16 | 1,05 | 2,94E-01 |
| AT3G43720 | 0,87 | -2,55 | 2,81 | -0,91 | 3,63E-01 |
| AT3G43810 | 0,51 | -1,78 | 3,27 | -0,54 | 5,87E-01 |
| AT3G43880 | 0,10 | 0,21 | 4,38 | 0,05 | 9,62E-01 |
| AT3G43930 | 0,08 | 0,44 | 4,38 | 0,10 | 9,21E-01 |
| AT3G43940 | 0,53 | 0,29 | 3,95 | 0,07 | 9,41E-01 |
| AT3G44010 | 1,00 | -0,66 | 2,00 | -0,33 | 7,42E-01 |
| AT3G44070 | 0,37 | 0,00 | 3,45 | 0,00 | 9,99E-01 |
| AT3G44110 | 2,03 | -0,21 | 1,19 | -0,18 | 8,58E-01 |
| AT3G44160 | 0,43 | -0,73 | 4,09 | -0,18 | 8,58E-01 |
| AT3G44260 | 2,79 | 3,37 | 1,56 | 2,16 | 3,04E-02 |
| AT3G44265 | 0,40 | 0,91 | 4,31 | 0,21 | 8,33E-01 |
| AT3G44340 | 1,56 | 0,97 | 1,59 | 0,61 | 5,40E-01 |
| AT3G44400 | 0,82 | 0,71 | 2,63 | 0,27 | 7,87E-01 |
| AT3G44435 | 0,70 | 0,59 | 3,72 | 0,16 | 8,74E-01 |
| AT3G06835 | 2,07 | -4,51 | 1,70 | -2,65 | 8,08E-03 |
| AT3G44444 | 1,95 | -0,66 | 1,57 | -0,42 | 6,74E-01 |
| AT3G44450 | 0,56 | -0,84 | 3,10 | -0,27 | 7,87E-01 |
| AT3G44460 | 0,11 | 0,00 | 4,41 | 0,00 | 1,00E+00 |
| AT3G44480 | 4,70 | 0,12 | 0,79 | 0,15 | 8,81E-01 |
| AT3G44635 | 1,31 | -0,03 | 2,76 | -0,01 | 9,92E-01 |
| AT3G44680 | 0,12 | 0,97 | 4,34 | 0,22 | 8,23E-01 |
| AT3G44753 | 0,14 | 0,09 | 4,38 | 0,02 | 9,84E-01 |
| AT3G44780 | 0,10 | 0,00 | 4,41 | 0,00 | 1,00E+00 |
| AT3G44820 | 0,38 | -0,46 | 4,33 | -0,11 | 9,16E-01 |
| AT3G44990 | 0,60 | 0,00 | 3,75 | 0,00 | 1,00E+00 |
| AT3G45030 | 0,91 | 0,17 | 2,45 | 0,07 | 9,45E-01 |
| AT3G45040 | 0,29 | 0,26 | 4,37 | 0,06 | 9,52E-01 |
| AT3G45060 | 0,08 | 0,69 | 4,36 | 0,16 | 8,75E-01 |
| AT3G45090 | 1,49 | -1,15 | 1,68 | -0,69 | 4,92E-01 |
| AT3G06955 | 0,21 | 0,00 | 4,41 | 0,00 | 1,00E+00 |
| AT3G45130 | 0,82 | 0,07 | 2,54 | 0,03 | 9,79E-01 |
| AT3G45180 | 1,00 | -1,09 | 1,89 | -0,58 | 5,65E-01 |
| AT3G07065 | 0,05 | 0,00 | 4,41 | 0,00 | 1,00E+00 |
| AT3G45190 | 2,88 | -0,55 | 1,04 | -0,53 | 5,99E-01 |
| AT3G45240 | 0,25 | 0,26 | 4,33 | 0,06 | 9,51E-01 |
| AT3G07095 | 0,20 | 1,26 | 4,34 | 0,29 | 7,72E-01 |
| AT3G45260 | 1,01 | -1,03 | 1,74 | -0,59 | 5,55E-01 |
| AT3G45300 | 0,17 | 0,00 | 4,41 | 0,00 | 1,00E+00 |
| AT3G45430 | 0,20 | -0,31 | 4,36 | -0,07 | 9,44E-01 |
| AT3G45443 | 0,08 | 0,00 | 4,40 | 0,00 | 1,00E+00 |
| AT3G45600 | 0,92 | 0,05 | 1,92 | 0,03 | 9,79E-01 |
| AT3G45630 | 0,80 | -0,55 | 2,11 | -0,26 | 7,95E-01 |
| AT3G45770 | 0,14 | 0,03 | 4,39 | 0,01 | 9,95E-01 |
| AT3G45820 | 0,59 | 0,00 | 3,12 | 0,00 | 1,00E+00 |
| AT3G45850 | 1,07 | -0,79 | 2,30 | -0,34 | 7,32E-01 |
| AT3G45860 | 0,95 | -3,02 | 2,04 | -1,48 | 1,38E-01 |
| AT3G45910 | 1,00 | 2,53 | 2,50 | 1,01 | 3,11E-01 |

| | | | | | |
|---|---|---|---|---|---|
| AT3G45990 | 0,46 | 1,19 | 3,38 | 0,35 | 7,25E-01 |
| AT3G46000 | 0,30 | 1,26 | 3,65 | 0,35 | 7,30E-01 |
| AT3G46030 | 0,35 | -0,86 | 4,36 | -0,20 | 8,43E-01 |
| AT3G46090 | 0,72 | 0,30 | 2,44 | 0,12 | 9,02E-01 |
| AT3G46100 | 0,51 | 0,00 | 4,06 | 0,00 | 1,00E+00 |
| AT3G46210 | 0,56 | 1,27 | 3,90 | 0,33 | 7,44E-01 |
| AT3G07375 | 0,21 | 1,01 | 4,34 | 0,23 | 8,17E-01 |
| AT3G46320 | 0,61 | 15,90 | 4,34 | 3,66 | 2,52E-04 |
| AT3G46360 | 0,82 | 0,00 | 2,98 | 0,00 | 1,00E+00 |
| AT3G46390 | 0,25 | 1,53 | 4,33 | 0,35 | 7,24E-01 |
| AT3G46440 | 0,45 | -1,26 | 4,29 | -0,29 | 7,70E-01 |
| AT3G46450 | 0,53 | 0,81 | 3,83 | 0,21 | 8,33E-01 |
| AT3G46470 | 0,38 | 1,56 | 4,25 | 0,37 | 7,14E-01 |
| AT3G46510 | 1,83 | 0,29 | 1,44 | 0,20 | 8,38E-01 |
| AT3G46530 | 2,60 | -0,36 | 1,13 | -0,32 | 7,50E-01 |
| AT3G46540 | 0,31 | -0,03 | 4,36 | -0,01 | 9,95E-01 |
| AT3G46550 | 1,58 | 0,88 | 1,57 | 0,56 | 5,72E-01 |
| AT3G46610 | 1,41 | 0,27 | 1,63 | 0,16 | 8,69E-01 |
| AT3G46620 | 4,33 | 0,06 | 0,91 | 0,07 | 9,46E-01 |
| AT3G46630 | 0,81 | 1,32 | 2,26 | 0,59 | 5,58E-01 |
| AT3G46650 | 0,58 | 0,16 | 3,06 | 0,05 | 9,58E-01 |
| AT3G46660 | 0,59 | 0,84 | 2,62 | 0,32 | 7,47E-01 |
| AT3G46670 | 0,57 | -0,15 | 3,24 | -0,05 | 9,63E-01 |
| AT3G46690 | 0,12 | 1,23 | 4,34 | 0,28 | 7,76E-01 |
| AT3G46730 | 0,12 | 1,16 | 4,34 | 0,27 | 7,89E-01 |
| AT3G46740 | 3,41 | -1,42 | 0,99 | -1,43 | 1,52E-01 |
| AT3G46790 | 0,53 | 0,75 | 4,32 | 0,17 | 8,61E-01 |
| AT3G46845 | 1,29 | 0,82 | 1,84 | 0,45 | 6,55E-01 |
| AT3G46920 | 0,08 | 0,44 | 4,38 | 0,10 | 9,21E-01 |
| AT3G46960 | 0,92 | -0,16 | 2,15 | -0,07 | 9,41E-01 |
| AT3G47000 | 0,33 | -2,47 | 4,36 | -0,57 | 5,71E-01 |
| AT3G47070 | 0,67 | -1,15 | 2,65 | -0,43 | 6,66E-01 |
| AT3G47090 | 0,54 | 0,65 | 3,72 | 0,18 | 8,61E-01 |
| AT3G47160 | 0,74 | 1,19 | 2,60 | 0,46 | 6,47E-01 |
| AT3G47250 | 1,84 | -0,47 | 1,34 | -0,35 | 7,25E-01 |
| AT3G47370 | 0,35 | -1,37 | 4,35 | -0,31 | 7,53E-01 |
| AT3G47420 | 0,51 | -0,48 | 2,55 | -0,19 | 8,51E-01 |
| AT3G47480 | 0,70 | 0,05 | 2,97 | 0,02 | 9,86E-01 |
| AT3G47500 | 2,57 | -0,27 | 1,28 | -0,21 | 8,33E-01 |
| AT3G47670 | 0,97 | 1,67 | 1,93 | 0,87 | 3,86E-01 |
| AT3G47675 | 0,21 | -0,49 | 4,35 | -0,11 | 9,11E-01 |
| AT3G47680 | 0,08 | 0,44 | 4,38 | 0,10 | 9,21E-01 |
| AT3G47700 | 0,43 | 0,64 | 2,74 | 0,24 | 8,14E-01 |
| AT3G47730 | 0,93 | 3,01 | 2,64 | 1,14 | 2,55E-01 |
| AT3G47810 | 0,27 | -0,21 | 4,34 | -0,05 | 9,62E-01 |
| AT3G47860 | 0,23 | -1,78 | 4,41 | -0,40 | 6,87E-01 |
| AT3G47890 | 1,54 | 0,05 | 1,44 | 0,04 | 9,71E-01 |
| AT3G47910 | 2,73 | -0,90 | 1,04 | -0,87 | 3,85E-01 |
| AT3G47940 | 0,35 | -0,44 | 4,35 | -0,10 | 9,19E-01 |
| AT3G47960 | 1,01 | 0,03 | 1,90 | 0,02 | 9,87E-01 |
| AT3G47990 | 0,32 | -0,25 | 4,36 | -0,06 | 9,55E-01 |
| AT3G48000 | 4,79 | -0,43 | 0,85 | -0,51 | 6,12E-01 |

| | | | | | |
|---|---|---|---|---|---|
| AT3G48050 | 3,14 | 0,56 | 0,95 | 0,58 | 5,61E-01 |
| AT3G48060 | 1,66 | -0,53 | 1,76 | -0,30 | 7,65E-01 |
| AT3G48080 | 1,04 | -0,43 | 1,90 | -0,22 | 8,22E-01 |
| AT3G48090 | 1,08 | 1,12 | 2,60 | 0,43 | 6,66E-01 |
| AT3G48100 | 0,85 | -3,87 | 4,27 | -0,91 | 3,65E-01 |
| AT3G48115 | 1,91 | 1,83 | 1,43 | 1,27 | 2,03E-01 |
| AT3G48131 | 0,31 | 0,47 | 4,36 | 0,11 | 9,13E-01 |
| AT3G48150 | 0,55 | -1,30 | 3,78 | -0,34 | 7,31E-01 |
| AT3G48170 | 0,90 | -0,72 | 2,51 | -0,28 | 7,76E-01 |
| AT3G48195 | 0,48 | 1,11 | 3,09 | 0,36 | 7,19E-01 |
| AT3G48200 | 2,11 | -1,78 | 1,44 | -1,24 | 2,15E-01 |
| AT3G48360 | 1,03 | -1,31 | 2,69 | -0,49 | 6,27E-01 |
| AT3G48520 | 0,38 | 0,55 | 4,34 | 0,13 | 8,99E-01 |
| AT3G48560 | 3,89 | -1,32 | 1,01 | -1,31 | 1,90E-01 |
| AT3G48570 | 0,28 | -1,78 | 4,41 | -0,40 | 6,87E-01 |
| AT3G48640 | 0,25 | -0,12 | 4,34 | -0,03 | 9,78E-01 |
| AT3G48690 | 0,79 | -0,84 | 2,31 | -0,36 | 7,17E-01 |
| AT3G48740 | 0,68 | 1,24 | 2,18 | 0,57 | 5,71E-01 |
| AT3G48770 | 1,49 | -0,26 | 1,62 | -0,16 | 8,70E-01 |
| AT3G08005 | 0,05 | 0,00 | 4,41 | 0,00 | 1,00E+00 |
| AT3G48840 | 0,12 | 0,37 | 4,37 | 0,08 | 9,33E-01 |
| AT3G48930 | 0,53 | -0,42 | 2,59 | -0,16 | 8,71E-01 |
| AT3G48990 | 0,58 | 0,21 | 2,91 | 0,07 | 9,43E-01 |
| AT3G49010 | 1,04 | 0,32 | 1,81 | 0,18 | 8,59E-01 |
| AT3G49055 | 0,42 | -1,03 | 4,32 | -0,24 | 8,12E-01 |
| AT3G49060 | 0,44 | -1,69 | 3,35 | -0,50 | 6,14E-01 |
| AT3G49070 | 1,57 | -3,17 | 3,05 | -1,04 | 2,98E-01 |
| AT3G49140 | 0,93 | -1,09 | 2,12 | -0,51 | 6,08E-01 |
| AT3G49160 | 0,11 | 0,65 | 4,36 | 0,15 | 8,81E-01 |
| AT3G49250 | 0,54 | 0,11 | 4,19 | 0,03 | 9,80E-01 |
| AT3G49350 | 0,36 | 0,62 | 4,31 | 0,14 | 8,85E-01 |
| AT3G49490 | 1,14 | -0,11 | 1,94 | -0,06 | 9,53E-01 |
| AT3G49500 | 2,34 | 0,48 | 1,32 | 0,37 | 7,14E-01 |
| AT3G49530 | 1,18 | -1,01 | 1,96 | -0,51 | 6,07E-01 |
| AT3G49600 | 1,57 | -1,03 | 1,64 | -0,63 | 5,30E-01 |
| AT3G49601 | 0,27 | 0,77 | 4,35 | 0,18 | 8,59E-01 |
| AT3G49650 | 0,08 | 0,44 | 4,38 | 0,10 | 9,21E-01 |
| AT3G49720 | 1,09 | -0,21 | 2,38 | -0,09 | 9,28E-01 |
| AT3G49790 | 0,38 | 1,15 | 3,36 | 0,34 | 7,32E-01 |
| AT3G49800 | 1,37 | 0,14 | 1,48 | 0,10 | 9,24E-01 |
| AT3G49870 | 0,08 | 0,44 | 4,38 | 0,10 | 9,21E-01 |
| AT3G49980 | 0,21 | 0,23 | 4,37 | 0,05 | 9,57E-01 |
| AT3G49990 | 0,57 | 2,43 | 3,73 | 0,65 | 5,15E-01 |
| AT3G50060 | 0,43 | -0,90 | 2,82 | -0,32 | 7,49E-01 |
| AT3G50110 | 0,18 | 0,81 | 4,34 | 0,19 | 8,52E-01 |
| AT3G50120 | 0,78 | 0,84 | 2,30 | 0,36 | 7,16E-01 |
| AT3G50240 | 0,71 | 1,27 | 4,31 | 0,30 | 7,68E-01 |
| AT3G50320 | 1,17 | 1,59 | 2,21 | 0,72 | 4,71E-01 |
| AT3G50380 | 1,07 | -1,52 | 2,34 | -0,65 | 5,17E-01 |
| AT3G50500 | 1,35 | 1,44 | 2,13 | 0,68 | 4,97E-01 |
| AT3G50670 | 0,33 | 1,48 | 4,33 | 0,34 | 7,32E-01 |
| AT3G50690 | 0,85 | 0,23 | 2,01 | 0,12 | 9,07E-01 |

| | | | | | |
|---|---|---|---|---|---|
| AT3G50740 | 0,53 | 0,24 | 3,28 | 0,07 | 9,42E-01 |
| AT3G50750 | 1,00 | -0,91 | 2,12 | -0,43 | 6,67E-01 |
| AT3G50780 | 0,08 | 0,69 | 4,36 | 0,16 | 8,75E-01 |
| AT3G50790 | 0,29 | -0,77 | 4,34 | -0,18 | 8,58E-01 |
| AT3G50808 | 0,10 | 0,00 | 4,41 | 0,00 | 1,00E+00 |
| AT3G08165 | 3,95 | 1,25 | 1,63 | 0,77 | 4,42E-01 |
| AT3G50825 | 0,41 | 0,00 | 4,41 | 0,00 | 1,00E+00 |
| AT3G50920 | 0,48 | -2,37 | 4,00 | -0,59 | 5,53E-01 |
| AT3G08170 | 3,83 | 0,24 | 0,98 | 0,25 | 8,04E-01 |
| AT3G51000 | 0,52 | 0,70 | 3,21 | 0,22 | 8,27E-01 |
| AT3G51120 | 1,56 | 0,62 | 1,54 | 0,40 | 6,86E-01 |
| AT3G51135 | 0,51 | -1,18 | 2,65 | -0,44 | 6,57E-01 |
| AT3G51160 | 0,58 | -3,25 | 3,09 | -1,05 | 2,93E-01 |
| AT3G51265 | 0,13 | 0,00 | 4,41 | 0,00 | 1,00E+00 |
| AT3G51310 | 0,51 | 0,21 | 3,17 | 0,07 | 9,47E-01 |
| AT3G51550 | 4,70 | -1,16 | 0,84 | -1,38 | 1,67E-01 |
| AT3G51580 | 0,25 | 1,82 | 4,32 | 0,42 | 6,73E-01 |
| AT3G51600 | 1,48 | -2,15 | 1,76 | -1,22 | 2,23E-01 |
| AT3G51660 | 0,43 | -1,78 | 4,22 | -0,42 | 6,73E-01 |
| AT3G51770 | 2,18 | 0,42 | 1,32 | 0,32 | 7,51E-01 |
| AT3G51790 | 0,60 | 1,62 | 3,63 | 0,45 | 6,55E-01 |
| AT3G51800 | 4,28 | 1,23 | 1,20 | 1,02 | 3,06E-01 |
| AT3G08210 | 3,75 | 0,88 | 1,25 | 0,71 | 4,80E-01 |
| AT3G51820 | 0,59 | 1,49 | 3,82 | 0,39 | 6,96E-01 |
| AT3G51840 | 0,33 | 0,24 | 4,32 | 0,06 | 9,56E-01 |
| AT3G51880 | 0,48 | -1,27 | 3,13 | -0,41 | 6,85E-01 |
| AT3G51890 | 1,41 | -1,24 | 2,53 | -0,49 | 6,24E-01 |
| AT3G51895 | 0,82 | 1,88 | 2,70 | 0,69 | 4,87E-01 |
| AT3G51920 | 0,46 | 0,24 | 2,69 | 0,09 | 9,29E-01 |
| AT3G51950 | 4,09 | -0,01 | 0,88 | -0,01 | 9,90E-01 |
| AT3G08215 | 0,08 | 0,69 | 4,36 | 0,16 | 8,75E-01 |
| AT3G51980 | 0,42 | -1,78 | 3,39 | -0,52 | 6,00E-01 |
| AT3G52060 | 0,46 | 0,22 | 2,80 | 0,08 | 9,37E-01 |
| AT3G52072 | 0,22 | 0,83 | 4,34 | 0,19 | 8,49E-01 |
| AT3G08230 | 0,12 | 0,49 | 4,36 | 0,11 | 9,11E-01 |
| AT3G52110 | 0,12 | 0,97 | 4,34 | 0,22 | 8,23E-01 |
| AT3G52180 | 0,66 | 1,00 | 2,70 | 0,37 | 7,11E-01 |
| AT3G52190 | 0,13 | 0,73 | 4,36 | 0,17 | 8,67E-01 |
| AT3G52220 | 0,11 | 0,65 | 4,36 | 0,15 | 8,81E-01 |
| AT3G52240 | 2,27 | -0,31 | 1,24 | -0,25 | 8,00E-01 |
| AT3G52260 | 0,64 | 1,77 | 3,54 | 0,50 | 6,18E-01 |
| AT3G52400 | 1,57 | 0,26 | 1,49 | 0,18 | 8,59E-01 |
| AT3G52480 | 0,53 | 1,51 | 3,03 | 0,50 | 6,18E-01 |
| AT3G52500 | 2,36 | 0,61 | 1,44 | 0,42 | 6,71E-01 |
| AT3G52560 | 0,47 | 0,31 | 3,35 | 0,09 | 9,26E-01 |
| AT3G52640 | 0,21 | 1,72 | 4,33 | 0,40 | 6,91E-01 |
| AT3G52700 | 2,20 | 0,12 | 1,31 | 0,09 | 9,29E-01 |
| AT3G52730 | 0,69 | -0,99 | 3,71 | -0,27 | 7,89E-01 |
| AT3G52740 | 0,17 | -1,70 | 4,40 | -0,39 | 7,00E-01 |
| AT3G52870 | 0,19 | 0,00 | 4,41 | 0,00 | 1,00E+00 |
| AT3G52880 | 2,23 | 1,59 | 1,45 | 1,09 | 2,74E-01 |
| AT3G52930 | 3,81 | -0,13 | 0,99 | -0,13 | 8,97E-01 |

| | | | | | |
|---|---|---|---|---|---|
| AT3G53015 | 1,03 | 1,96 | 2,22 | 0,89 | 3,76E-01 |
| AT3G53016 | 0,52 | -2,10 | 4,08 | -0,51 | 6,07E-01 |
| AT3G53100 | 0,69 | -3,03 | 3,09 | -0,98 | 3,27E-01 |
| AT3G53260 | 2,56 | 0,21 | 1,29 | 0,16 | 8,73E-01 |
| AT3G53380 | 0,40 | -1,75 | 4,27 | -0,41 | 6,82E-01 |
| AT3G53410 | 0,47 | -0,35 | 4,33 | -0,08 | 9,36E-01 |
| AT3G53420 | 2,26 | 1,46 | 1,51 | 0,97 | 3,34E-01 |
| AT3G53430 | 2,55 | -0,96 | 1,13 | -0,85 | 3,98E-01 |
| AT3G53500 | 0,57 | -1,59 | 2,77 | -0,57 | 5,66E-01 |
| AT3G53510 | 0,25 | 1,94 | 4,32 | 0,45 | 6,54E-01 |
| AT3G53540 | 1,08 | 0,29 | 2,11 | 0,14 | 8,92E-01 |
| AT3G53560 | 0,23 | -1,58 | 4,38 | -0,36 | 7,18E-01 |
| AT3G53570 | 0,21 | -0,49 | 4,35 | -0,11 | 9,11E-01 |
| AT3G53610 | 0,53 | 1,32 | 3,18 | 0,41 | 6,79E-01 |
| AT3G53710 | 0,37 | 1,66 | 3,42 | 0,49 | 6,27E-01 |
| AT3G53730 | 0,67 | 1,18 | 2,47 | 0,48 | 6,32E-01 |
| AT3G53740 | 2,01 | -2,02 | 1,70 | -1,19 | 2,35E-01 |
| AT3G53810 | 0,44 | 0,41 | 4,26 | 0,10 | 9,23E-01 |
| AT3G53890 | 0,52 | -1,58 | 3,10 | -0,51 | 6,11E-01 |
| AT3G53910 | 1,01 | 2,22 | 2,20 | 1,01 | 3,13E-01 |
| AT3G53920 | 0,24 | -0,08 | 4,34 | -0,02 | 9,85E-01 |
| AT3G53960 | 0,98 | 0,31 | 2,37 | 0,13 | 8,95E-01 |
| AT3G53990 | 1,83 | -1,59 | 1,40 | -1,14 | 2,56E-01 |
| AT3G54100 | 0,47 | 2,22 | 3,21 | 0,69 | 4,89E-01 |
| AT3G54120 | 0,20 | -0,44 | 4,36 | -0,10 | 9,19E-01 |
| AT3G54140 | 0,70 | 0,40 | 2,55 | 0,16 | 8,74E-01 |
| AT3G54210 | 1,01 | 1,52 | 1,88 | 0,81 | 4,19E-01 |
| AT3G54290 | 0,42 | -2,34 | 4,21 | -0,56 | 5,78E-01 |
| AT3G54350 | 1,02 | 1,96 | 2,00 | 0,98 | 3,27E-01 |
| AT3G54360 | 1,14 | 1,08 | 1,92 | 0,56 | 5,74E-01 |
| AT3G54400 | 0,32 | 0,22 | 4,34 | 0,05 | 9,59E-01 |
| AT3G54440 | 0,61 | -2,63 | 2,35 | -1,12 | 2,63E-01 |
| AT3G54460 | 0,97 | 0,43 | 2,58 | 0,16 | 8,69E-01 |
| AT3G54470 | 0,44 | -0,54 | 3,23 | -0,17 | 8,67E-01 |
| AT3G54500 | 1,37 | -0,75 | 1,82 | -0,41 | 6,79E-01 |
| AT3G54580 | 1,94 | 3,42 | 1,76 | 1,94 | 5,18E-02 |
| AT3G54620 | 0,24 | 0,24 | 4,34 | 0,06 | 9,55E-01 |
| AT3G54630 | 0,68 | 0,74 | 4,32 | 0,17 | 8,64E-01 |
| AT3G54640 | 0,22 | -1,77 | 4,41 | -0,40 | 6,88E-01 |
| AT3G54650 | 0,62 | -0,33 | 3,15 | -0,10 | 9,17E-01 |
| AT3G54660 | 2,19 | -1,68 | 1,32 | -1,27 | 2,04E-01 |
| AT3G54750 | 0,48 | 0,56 | 4,34 | 0,13 | 8,98E-01 |
| AT3G54760 | 1,04 | 0,18 | 1,89 | 0,10 | 9,23E-01 |
| AT3G54790 | 1,31 | -2,17 | 1,66 | -1,31 | 1,91E-01 |
| AT3G54860 | 0,55 | 1,07 | 2,98 | 0,36 | 7,19E-01 |
| AT3G54900 | 0,95 | 1,58 | 1,97 | 0,80 | 4,22E-01 |
| AT3G54925 | 1,76 | -0,65 | 1,89 | -0,34 | 7,32E-01 |
| AT3G54960 | 0,90 | 0,90 | 2,06 | 0,44 | 6,62E-01 |
| AT3G55020 | 1,94 | -0,67 | 1,27 | -0,53 | 5,98E-01 |
| AT3G55040 | 0,21 | 0,00 | 4,41 | 0,00 | 1,00E+00 |
| AT3G55050 | 0,45 | 1,88 | 3,27 | 0,57 | 5,66E-01 |
| AT3G55060 | 0,14 | 0,03 | 4,39 | 0,01 | 9,95E-01 |

| | | | | | |
|---|---|---|---|---|---|
| AT3G55130 | 0,51 | 0,00 | 3,25 | 0,00 | 1,00E+00 |
| AT3G55160 | 0,95 | -0,07 | 1,77 | -0,04 | 9,68E-01 |
| AT3G55220 | 4,51 | -0,36 | 0,85 | -0,43 | 6,70E-01 |
| AT3G55252 | 0,66 | -1,44 | 2,33 | -0,62 | 5,36E-01 |
| AT3G55254 | 0,80 | 1,71 | 2,25 | 0,76 | 4,46E-01 |
| AT3G55320 | 2,94 | 2,44 | 1,27 | 1,92 | 5,49E-02 |
| AT3G55330 | 0,44 | 0,27 | 2,77 | 0,10 | 9,24E-01 |
| AT3G55360 | 0,60 | 1,35 | 2,53 | 0,53 | 5,94E-01 |
| AT3G55400 | 0,18 | 0,06 | 4,40 | 0,01 | 9,89E-01 |
| AT3G55430 | 0,77 | -0,78 | 2,09 | -0,37 | 7,08E-01 |
| AT3G55480 | 1,64 | -0,13 | 1,43 | -0,09 | 9,29E-01 |
| AT3G55610 | 4,81 | -1,10 | 0,85 | -1,30 | 1,94E-01 |
| AT3G55840 | 1,41 | -0,20 | 1,61 | -0,12 | 9,02E-01 |
| AT3G55850 | 2,66 | -0,90 | 1,32 | -0,68 | 4,96E-01 |
| AT3G55860 | 0,83 | 1,16 | 2,19 | 0,53 | 5,96E-01 |
| AT3G55935 | 0,46 | -0,88 | 2,63 | -0,33 | 7,38E-01 |
| AT3G55950 | 0,63 | -0,10 | 2,56 | -0,04 | 9,67E-01 |
| AT3G55960 | 0,64 | 0,72 | 2,60 | 0,28 | 7,82E-01 |
| AT3G56040 | 0,77 | 0,10 | 2,10 | 0,05 | 9,61E-01 |
| AT3G56050 | 0,97 | 0,71 | 1,99 | 0,35 | 7,23E-01 |
| AT3G56070 | 2,75 | 1,30 | 1,39 | 0,93 | 3,51E-01 |
| AT3G56085 | 0,12 | 0,00 | 4,38 | 0,00 | 1,00E+00 |
| AT3G56090 | 0,34 | 0,04 | 3,58 | 0,01 | 9,90E-01 |
| AT3G56110 | 0,41 | -0,49 | 3,36 | -0,15 | 8,83E-01 |
| AT3G56150 | 2,36 | 0,11 | 1,28 | 0,08 | 9,34E-01 |
| AT3G56190 | 1,16 | 1,49 | 1,91 | 0,78 | 4,36E-01 |
| AT3G56210 | 0,45 | -2,82 | 4,35 | -0,65 | 5,16E-01 |
| AT3G56240 | 0,35 | 0,83 | 2,93 | 0,28 | 7,77E-01 |
| AT3G56290 | 1,19 | -1,85 | 1,99 | -0,93 | 3,51E-01 |
| AT3G56320 | 0,58 | -1,73 | 3,81 | -0,45 | 6,50E-01 |
| AT3G56340 | 0,44 | -1,01 | 3,34 | -0,30 | 7,63E-01 |
| AT3G56370 | 0,84 | -0,83 | 2,13 | -0,39 | 6,95E-01 |
| AT3G56400 | 2,09 | 0,64 | 1,40 | 0,46 | 6,44E-01 |
| AT3G56410 | 2,56 | 2,53 | 1,47 | 1,73 | 8,42E-02 |
| AT3G56460 | 1,07 | 1,45 | 1,95 | 0,74 | 4,58E-01 |
| AT3G56480 | 0,39 | 1,48 | 2,84 | 0,52 | 6,03E-01 |
| AT3G56670 | 0,38 | 0,20 | 4,31 | 0,05 | 9,63E-01 |
| AT3G56690 | 0,43 | 0,58 | 4,17 | 0,14 | 8,88E-01 |
| AT3G56790 | 0,33 | 0,35 | 4,39 | 0,08 | 9,36E-01 |
| AT3G56800 | 1,39 | 0,24 | 1,66 | 0,15 | 8,84E-01 |
| AT3G56820 | 0,25 | -2,19 | 4,38 | -0,50 | 6,17E-01 |
| AT3G56830 | 0,17 | 1,40 | 4,33 | 0,32 | 7,47E-01 |
| AT3G56850 | 0,48 | -1,26 | 3,26 | -0,39 | 6,99E-01 |
| AT3G56860 | 0,49 | 0,88 | 2,73 | 0,32 | 7,48E-01 |
| AT3G56910 | 1,94 | 0,78 | 1,55 | 0,50 | 6,17E-01 |
| AT3G56950 | 0,79 | 0,19 | 2,27 | 0,08 | 9,34E-01 |
| AT3G56970 | 0,12 | 1,16 | 4,34 | 0,27 | 7,89E-01 |
| AT3G56990 | 0,46 | 0,38 | 3,27 | 0,12 | 9,06E-01 |
| AT3G57020 | 0,42 | -2,05 | 4,37 | -0,47 | 6,39E-01 |
| AT3G57030 | 0,05 | 0,00 | 4,41 | 0,00 | 1,00E+00 |
| AT3G08825 | 0,72 | 0,55 | 3,05 | 0,18 | 8,56E-01 |
| AT3G57050 | 0,52 | 0,53 | 2,60 | 0,21 | 8,37E-01 |

| | | | | | |
|---|---|---|---|---|---|
| AT3G08855 | 0,12 | 0,37 | 4,37 | 0,08 | 9,33E-01 |
| AT3G57140 | 0,48 | 1,25 | 3,10 | 0,40 | 6,88E-01 |
| AT3G57240 | 0,24 | -2,36 | 4,39 | -0,54 | 5,91E-01 |
| AT3G57290 | 1,20 | 0,99 | 2,32 | 0,43 | 6,69E-01 |
| AT3G57330 | 3,24 | 0,26 | 0,99 | 0,26 | 7,91E-01 |
| AT3G57350 | 0,44 | -0,23 | 3,22 | -0,07 | 9,43E-01 |
| AT3G57410 | 3,70 | -0,14 | 1,01 | -0,14 | 8,86E-01 |
| AT3G57420 | 0,74 | -0,87 | 2,27 | -0,39 | 7,00E-01 |
| AT3G57460 | 0,24 | 0,03 | 4,39 | 0,01 | 9,94E-01 |
| AT3G08935 | 1,61 | -3,50 | 1,99 | -1,76 | 7,83E-02 |
| AT3G57470 | 0,21 | 0,10 | 4,40 | 0,02 | 9,82E-01 |
| AT3G57480 | 3,07 | -2,59 | 1,23 | -2,11 | 3,51E-02 |
| AT3G57520 | 3,92 | -2,29 | 0,97 | -2,37 | 1,80E-02 |
| AT3G57530 | 1,08 | -0,09 | 2,03 | -0,04 | 9,65E-01 |
| AT3G57540 | 0,08 | 0,00 | 4,40 | 0,00 | 1,00E+00 |
| AT3G57580 | 0,45 | -0,88 | 3,40 | -0,26 | 7,96E-01 |
| AT3G57610 | 1,05 | -2,44 | 4,21 | -0,58 | 5,63E-01 |
| AT3G57640 | 0,11 | 0,00 | 4,41 | 0,00 | 1,00E+00 |
| AT3G57710 | 4,92 | -0,03 | 0,85 | -0,04 | 9,70E-01 |
| AT3G57730 | 5,17 | -0,33 | 0,81 | -0,41 | 6,81E-01 |
| AT3G57765 | 1,26 | 0,44 | 1,60 | 0,27 | 7,83E-01 |
| AT3G57780 | 0,43 | -1,83 | 4,35 | -0,42 | 6,74E-01 |
| AT3G57790 | 0,44 | 2,29 | 4,32 | 0,53 | 5,96E-01 |
| AT3G57800 | 0,37 | 0,33 | 4,36 | 0,08 | 9,40E-01 |
| AT3G57870 | 0,53 | -2,71 | 4,29 | -0,63 | 5,28E-01 |
| AT3G57880 | 0,23 | 1,27 | 4,33 | 0,29 | 7,70E-01 |
| AT3G57930 | 0,30 | -1,04 | 4,37 | -0,24 | 8,12E-01 |
| AT3G57965 | 0,28 | 1,76 | 4,33 | 0,41 | 6,84E-01 |
| AT3G57980 | 0,74 | 2,61 | 2,81 | 0,93 | 3,53E-01 |
| AT3G57990 | 0,40 | -0,06 | 3,34 | -0,02 | 9,86E-01 |
| AT3G58100 | 1,03 | 1,97 | 3,28 | 0,60 | 5,48E-01 |
| AT3G58120 | 0,15 | 0,67 | 4,37 | 0,15 | 8,79E-01 |
| AT3G58140 | 0,42 | -0,89 | 3,44 | -0,26 | 7,95E-01 |
| AT3G58170 | 0,42 | -0,91 | 4,31 | -0,21 | 8,32E-01 |
| AT3G58210 | 0,64 | -2,46 | 4,31 | -0,57 | 5,68E-01 |
| AT3G58270 | 0,99 | 2,10 | 2,55 | 0,82 | 4,10E-01 |
| AT3G58500 | 1,34 | 2,31 | 1,71 | 1,35 | 1,77E-01 |
| AT3G58560 | 0,23 | 0,55 | 4,36 | 0,13 | 9,00E-01 |
| AT3G58580 | 0,38 | 1,49 | 4,28 | 0,35 | 7,27E-01 |
| AT3G58640 | 2,56 | -0,59 | 1,22 | -0,48 | 6,30E-01 |
| AT3G58670 | 0,12 | 0,37 | 4,37 | 0,08 | 9,33E-01 |
| AT3G58730 | 1,29 | -0,05 | 1,84 | -0,02 | 9,81E-01 |
| AT3G58750 | 0,61 | -0,72 | 3,81 | -0,19 | 8,50E-01 |
| AT3G58790 | 0,34 | 0,40 | 4,36 | 0,09 | 9,26E-01 |
| AT3G58970 | 1,53 | 0,29 | 2,13 | 0,14 | 8,91E-01 |
| AT3G58990 | 0,12 | 0,49 | 4,36 | 0,11 | 9,11E-01 |
| AT3G59020 | 5,79 | -0,04 | 0,81 | -0,05 | 9,61E-01 |
| AT3G59040 | 1,65 | -0,77 | 1,48 | -0,52 | 6,03E-01 |
| AT3G59050 | 0,52 | 0,71 | 3,09 | 0,23 | 8,19E-01 |
| AT3G59060 | 1,47 | -1,09 | 1,44 | -0,76 | 4,49E-01 |
| AT3G59070 | 1,78 | -0,31 | 1,72 | -0,18 | 8,55E-01 |
| AT3G59120 | 0,10 | 0,01 | 4,38 | 0,00 | 9,99E-01 |

| | | | | | |
|---|---|---|---|---|---|
| AT3G59140 | 3,91 | 2,73 | 1,65 | 1,65 | 9,87E-02 |
| AT3G59280 | 0,32 | -0,25 | 4,36 | -0,06 | 9,55E-01 |
| AT3G59360 | 0,61 | 2,50 | 2,91 | 0,86 | 3,91E-01 |
| AT3G59400 | 1,33 | -0,74 | 1,57 | -0,47 | 6,37E-01 |
| AT3G59540 | 1,10 | -1,02 | 1,87 | -0,54 | 5,86E-01 |
| AT3G59570 | 0,57 | 2,14 | 3,02 | 0,71 | 4,80E-01 |
| AT3G59660 | 1,15 | 0,98 | 1,86 | 0,53 | 5,99E-01 |
| AT3G59760 | 0,20 | 0,00 | 4,41 | 0,00 | 1,00E+00 |
| AT3G59765 | 1,81 | -1,35 | 1,48 | -0,91 | 3,64E-01 |
| AT3G59770 | 3,31 | 1,66 | 1,15 | 1,43 | 1,52E-01 |
| AT3G59840 | 0,15 | 0,00 | 4,38 | 0,00 | 1,00E+00 |
| AT3G59910 | 0,48 | -1,98 | 4,02 | -0,49 | 6,23E-01 |
| AT3G59950 | 1,22 | -1,05 | 2,45 | -0,43 | 6,66E-01 |
| AT3G59980 | 0,35 | 0,49 | 3,45 | 0,14 | 8,88E-01 |
| AT3G60030 | 1,14 | -0,53 | 1,91 | -0,28 | 7,82E-01 |
| AT3G09335 | 0,23 | 0,70 | 4,35 | 0,16 | 8,73E-01 |
| AT3G60050 | 0,12 | 1,05 | 4,35 | 0,24 | 8,10E-01 |
| AT3G60070 | 0,35 | -1,51 | 4,36 | -0,35 | 7,28E-01 |
| AT3G60160 | 1,27 | 1,40 | 1,88 | 0,74 | 4,57E-01 |
| AT3G60176 | 0,85 | 2,23 | 2,73 | 0,81 | 4,15E-01 |
| AT3G60190 | 1,87 | 0,07 | 1,37 | 0,05 | 9,62E-01 |
| AT3G60220 | 0,15 | 0,00 | 4,41 | 0,00 | 1,00E+00 |
| AT3G60250 | 1,02 | -2,21 | 1,83 | -1,21 | 2,26E-01 |
| AT3G60320 | 3,74 | -1,52 | 0,93 | -1,64 | 1,00E-01 |
| AT3G60350 | 1,30 | -1,26 | 1,70 | -0,74 | 4,59E-01 |
| AT3G60380 | 0,49 | -2,09 | 3,91 | -0,54 | 5,92E-01 |
| AT3G60540 | 0,08 | 0,00 | 4,40 | 0,00 | 1,00E+00 |
| AT3G60630 | 5,65 | -0,69 | 0,83 | -0,84 | 4,03E-01 |
| AT3G60670 | 0,12 | 0,49 | 4,36 | 0,11 | 9,11E-01 |
| AT3G60740 | 0,66 | -0,26 | 2,44 | -0,11 | 9,14E-01 |
| AT3G60760 | 0,12 | 1,23 | 4,34 | 0,28 | 7,76E-01 |
| AT3G60770 | 1,96 | -1,05 | 1,39 | -0,75 | 4,50E-01 |
| AT3G60820 | 0,15 | 0,00 | 4,41 | 0,00 | 1,00E+00 |
| AT3G60990 | 0,73 | 0,33 | 2,70 | 0,12 | 9,02E-01 |
| AT3G61070 | 0,50 | 0,46 | 4,06 | 0,11 | 9,09E-01 |
| AT3G09565 | 0,50 | -1,81 | 3,94 | -0,46 | 6,46E-01 |
| AT3G61190 | 1,32 | 0,04 | 1,72 | 0,02 | 9,83E-01 |
| AT3G61200 | 0,69 | 2,37 | 2,45 | 0,97 | 3,34E-01 |
| AT3G61260 | 0,83 | -0,88 | 2,65 | -0,33 | 7,39E-01 |
| AT3G61310 | 0,38 | -1,21 | 4,34 | -0,28 | 7,81E-01 |
| AT3G61380 | 0,18 | 0,73 | 4,36 | 0,17 | 8,67E-01 |
| AT3G61420 | 0,37 | -2,24 | 4,39 | -0,51 | 6,09E-01 |
| AT3G61460 | 0,83 | 1,28 | 2,03 | 0,63 | 5,29E-01 |
| AT3G61530 | 0,25 | 1,40 | 4,33 | 0,32 | 7,47E-01 |
| AT3G61640 | 0,38 | 1,39 | 3,35 | 0,42 | 6,78E-01 |
| AT3G61740 | 0,78 | 0,39 | 2,31 | 0,17 | 8,66E-01 |
| AT3G61750 | 0,24 | -2,36 | 4,39 | -0,54 | 5,91E-01 |
| AT3G61780 | 1,45 | 0,51 | 1,49 | 0,34 | 7,33E-01 |
| AT3G61800 | 0,23 | 0,12 | 4,39 | 0,03 | 9,77E-01 |
| AT3G61820 | 1,36 | -0,52 | 1,70 | -0,31 | 7,58E-01 |
| AT3G61860 | 0,39 | -0,91 | 4,30 | -0,21 | 8,33E-01 |
| AT3G61890 | 0,44 | 0,83 | 3,57 | 0,23 | 8,16E-01 |

| | | | | | |
|---|---|---|---|---|---|
| AT3G61960 | 1,02 | -1,84 | 1,93 | -0,95 | 3,42E-01 |
| AT3G61990 | 0,15 | 0,74 | 4,35 | 0,17 | 8,65E-01 |
| AT3G62070 | 0,12 | 0,00 | 4,41 | 0,00 | 1,00E+00 |
| AT3G62120 | 1,00 | 1,32 | 2,05 | 0,64 | 5,22E-01 |
| AT3G62200 | 2,36 | -0,98 | 1,29 | -0,76 | 4,48E-01 |
| AT3G62240 | 0,93 | 1,37 | 2,81 | 0,49 | 6,28E-01 |
| AT3G62260 | 0,41 | 0,56 | 4,10 | 0,14 | 8,91E-01 |
| AT3G62310 | 0,33 | 0,84 | 4,35 | 0,19 | 8,46E-01 |
| AT3G62330 | 0,73 | 0,45 | 2,85 | 0,16 | 8,74E-01 |
| AT3G62360 | 4,39 | -0,12 | 0,85 | -0,14 | 8,89E-01 |
| AT3G62420 | 1,96 | -1,24 | 1,70 | -0,73 | 4,65E-01 |
| AT3G62470 | 0,24 | 0,21 | 4,34 | 0,05 | 9,62E-01 |
| AT3G62540 | 0,28 | -1,42 | 4,36 | -0,33 | 7,44E-01 |
| AT3G09795 | 0,21 | 0,05 | 4,39 | 0,01 | 9,90E-01 |
| AT3G62590 | 0,64 | 2,17 | 2,95 | 0,73 | 4,62E-01 |
| AT3G62600 | 0,95 | -0,42 | 1,90 | -0,22 | 8,25E-01 |
| AT3G09825 | 1,22 | 0,02 | 2,07 | 0,01 | 9,92E-01 |
| AT3G09845 | 0,08 | 0,00 | 4,40 | 0,00 | 1,00E+00 |
| AT3G62630 | 0,91 | -2,04 | 2,21 | -0,92 | 3,56E-01 |
| AT3G62700 | 3,87 | 0,55 | 0,98 | 0,57 | 5,70E-01 |
| AT3G62800 | 0,37 | 0,94 | 4,32 | 0,22 | 8,28E-01 |
| AT3G62840 | 0,66 | -1,18 | 2,98 | -0,40 | 6,91E-01 |
| AT3G62860 | 0,81 | -1,59 | 3,41 | -0,47 | 6,40E-01 |
| AT3G62870 | 1,22 | -0,36 | 1,64 | -0,22 | 8,27E-01 |
| AT3G62900 | 0,94 | 2,93 | 2,29 | 1,28 | 2,01E-01 |
| AT3G62910 | 0,49 | -3,25 | 4,02 | -0,81 | 4,19E-01 |
| AT3G62940 | 0,31 | -1,74 | 4,40 | -0,39 | 6,93E-01 |
| AT3G63003 | 4,44 | 1,56 | 1,40 | 1,12 | 2,63E-01 |
| AT3G63052 | 0,42 | -0,89 | 3,44 | -0,26 | 7,95E-01 |
| AT3G63110 | 0,56 | 0,22 | 2,74 | 0,08 | 9,37E-01 |
| AT3G63140 | 0,93 | -0,39 | 2,21 | -0,18 | 8,60E-01 |
| AT3G63150 | 0,90 | -0,23 | 2,37 | -0,10 | 9,22E-01 |
| AT3G63160 | 1,15 | 1,06 | 2,15 | 0,49 | 6,22E-01 |
| AT3G63180 | 1,24 | -1,77 | 1,61 | -1,09 | 2,74E-01 |
| AT3G63190 | 0,25 | 0,13 | 4,39 | 0,03 | 9,76E-01 |
| AT3G63210 | 0,47 | 0,00 | 4,41 | 0,00 | 1,00E+00 |
| AT3G63310 | 1,42 | -0,47 | 1,75 | -0,27 | 7,87E-01 |
| AT3G63340 | 1,13 | 0,46 | 2,38 | 0,19 | 8,47E-01 |
| AT3G63380 | 3,14 | -0,06 | 1,17 | -0,05 | 9,62E-01 |
| AT3G63390 | 0,24 | 0,88 | 4,36 | 0,20 | 8,41E-01 |
| AT3G63410 | 2,71 | -0,53 | 1,07 | -0,50 | 6,19E-01 |
| AT3G63460 | 4,10 | -1,22 | 0,85 | -1,44 | 1,50E-01 |
| AT3G63500 | 4,53 | 0,67 | 0,83 | 0,81 | 4,19E-01 |
| AT3G63540 | 1,40 | -0,95 | 1,77 | -0,54 | 5,89E-01 |
| AT4G00005 | 0,19 | 0,47 | 4,38 | 0,11 | 9,14E-01 |
| AT4G00020 | 0,79 | 0,27 | 2,33 | 0,11 | 9,09E-01 |
| AT4G00070 | 0,43 | 0,15 | 4,18 | 0,04 | 9,72E-01 |
| AT4G00110 | 1,14 | -3,60 | 2,34 | -1,54 | 1,25E-01 |
| AT4G00120 | 1,32 | -0,78 | 2,07 | -0,38 | 7,06E-01 |
| AT4G00140 | 2,38 | 3,62 | 1,69 | 2,14 | 3,25E-02 |
| AT4G00165 | 0,65 | -2,37 | 2,73 | -0,87 | 3,85E-01 |
| AT4G00180 | 0,45 | 0,78 | 4,36 | 0,18 | 8,58E-01 |

| | | | | | |
|---|---|---|---|---|---|
| AT4G00280 | 0,25 | -2,19 | 4,38 | -0,50 | 6,17E-01 |
| AT4G00290 | 0,50 | -0,72 | 2,82 | -0,26 | 7,98E-01 |
| AT4G00330 | 0,80 | 1,19 | 2,16 | 0,55 | 5,84E-01 |
| AT4G03585 | 1,51 | -1,51 | 1,75 | -0,86 | 3,89E-01 |
| AT4G00370 | 0,63 | 0,34 | 2,59 | 0,13 | 8,96E-01 |
| AT4G00400 | 0,42 | 0,56 | 2,84 | 0,20 | 8,43E-01 |
| AT4G00430 | 2,16 | 0,13 | 1,33 | 0,10 | 9,21E-01 |
| AT4G00500 | 0,24 | 0,43 | 4,36 | 0,10 | 9,22E-01 |
| AT4G00540 | 0,54 | 0,05 | 2,72 | 0,02 | 9,86E-01 |
| AT4G00550 | 0,15 | 1,16 | 4,34 | 0,27 | 7,88E-01 |
| AT4G00560 | 0,40 | 2,32 | 4,31 | 0,54 | 5,90E-01 |
| AT4G00570 | 3,02 | -0,51 | 1,11 | -0,46 | 6,45E-01 |
| AT4G00620 | 0,40 | -1,47 | 4,24 | -0,35 | 7,29E-01 |
| AT4G00630 | 4,40 | -0,19 | 0,80 | -0,24 | 8,08E-01 |
| AT4G00720 | 2,07 | 0,23 | 1,32 | 0,18 | 8,60E-01 |
| AT4G00730 | 3,19 | 0,03 | 1,09 | 0,02 | 9,81E-01 |
| AT4G00740 | 0,31 | -0,90 | 4,40 | -0,21 | 8,37E-01 |
| AT4G00800 | 2,43 | 0,16 | 1,13 | 0,15 | 8,85E-01 |
| AT4G00810 | 0,25 | 0,26 | 4,33 | 0,06 | 9,51E-01 |
| AT4G00890 | 0,23 | 0,00 | 4,41 | 0,00 | 1,00E+00 |
| AT4G00900 | 1,09 | 1,87 | 2,00 | 0,93 | 3,51E-01 |
| AT4G00980 | 0,33 | -0,58 | 4,33 | -0,13 | 8,93E-01 |
| AT4G00985 | 0,26 | -1,86 | 4,40 | -0,42 | 6,72E-01 |
| AT4G01000 | 0,50 | 2,45 | 3,88 | 0,63 | 5,28E-01 |
| AT4G01010 | 2,44 | -1,32 | 1,45 | -0,91 | 3,62E-01 |
| AT4G01030 | 0,51 | 0,93 | 3,05 | 0,31 | 7,60E-01 |
| AT4G01037 | 0,21 | 0,00 | 4,41 | 0,00 | 1,00E+00 |
| AT4G01070 | 1,43 | 0,30 | 1,82 | 0,16 | 8,69E-01 |
| AT4G01090 | 0,44 | 0,91 | 3,23 | 0,28 | 7,80E-01 |
| AT4G01110 | 0,08 | 0,44 | 4,38 | 0,10 | 9,21E-01 |
| AT4G01120 | 0,18 | 0,13 | 4,41 | 0,03 | 9,76E-01 |
| AT4G01180 | 0,08 | 0,44 | 4,38 | 0,10 | 9,21E-01 |
| AT4G01190 | 0,20 | -0,44 | 4,36 | -0,10 | 9,19E-01 |
| AT4G01200 | 0,46 | -1,83 | 4,32 | -0,42 | 6,72E-01 |
| AT4G01210 | 1,50 | -0,10 | 1,64 | -0,06 | 9,53E-01 |
| AT4G01240 | 0,94 | -1,53 | 2,03 | -0,76 | 4,50E-01 |
| AT4G01290 | 1,74 | -0,07 | 1,58 | -0,04 | 9,67E-01 |
| AT4G01310 | 0,66 | -3,23 | 3,09 | -1,05 | 2,96E-01 |
| AT4G01395 | 0,71 | -2,44 | 2,22 | -1,10 | 2,70E-01 |
| AT4G01533 | 1,62 | -0,63 | 1,92 | -0,33 | 7,41E-01 |
| AT4G01540 | 0,20 | 0,41 | 4,37 | 0,09 | 9,26E-01 |
| AT4G01550 | 0,48 | 1,11 | 4,36 | 0,25 | 8,00E-01 |
| AT4G01700 | 0,60 | 0,11 | 3,13 | 0,04 | 9,71E-01 |
| AT4G01810 | 2,76 | -0,61 | 1,16 | -0,53 | 5,97E-01 |
| AT4G01850 | 4,27 | -1,02 | 0,94 | -1,09 | 2,76E-01 |
| AT4G01860 | 3,28 | -1,27 | 1,17 | -1,08 | 2,79E-01 |
| AT4G01870 | 0,45 | -0,83 | 4,12 | -0,20 | 8,40E-01 |
| AT4G01950 | 0,49 | -0,91 | 2,81 | -0,32 | 7,47E-01 |
| AT4G01985 | 0,41 | 0,00 | 4,40 | 0,00 | 1,00E+00 |
| AT4G02130 | 3,77 | -0,59 | 0,93 | -0,63 | 5,28E-01 |
| AT4G02195 | 0,71 | -2,29 | 2,50 | -0,92 | 3,59E-01 |
| AT4G02210 | 0,65 | 0,78 | 2,85 | 0,27 | 7,84E-01 |

| | | | | | |
|---|---|---|---|---|---|
| AT4G02230 | 0,28 | -0,49 | 4,38 | -0,11 | 9,11E-01 |
| AT4G02320 | 3,77 | -1,43 | 0,95 | -1,50 | 1,35E-01 |
| AT4G04125 | 2,22 | 0,65 | 1,28 | 0,50 | 6,15E-01 |
| AT4G02405 | 0,05 | 0,00 | 4,41 | 0,00 | 1,00E+00 |
| AT4G02410 | 0,84 | -0,64 | 2,73 | -0,24 | 8,13E-01 |
| AT4G02420 | 1,44 | 1,51 | 1,91 | 0,79 | 4,29E-01 |
| AT4G02450 | 0,79 | 1,74 | 2,50 | 0,70 | 4,86E-01 |
| AT4G02460 | 0,19 | 1,08 | 4,34 | 0,25 | 8,04E-01 |
| AT4G02520 | 4,48 | -0,45 | 0,96 | -0,47 | 6,38E-01 |
| AT4G02530 | 0,43 | 0,10 | 3,41 | 0,03 | 9,76E-01 |
| AT4G04185 | 0,38 | 1,50 | 3,38 | 0,44 | 6,57E-01 |
| AT4G02550 | 0,12 | 0,49 | 4,36 | 0,11 | 9,11E-01 |
| AT4G02560 | 0,57 | -0,61 | 2,57 | -0,24 | 8,11E-01 |
| AT4G02590 | 0,27 | -0,47 | 4,37 | -0,11 | 9,14E-01 |
| AT4G02620 | 0,23 | 0,50 | 4,38 | 0,11 | 9,09E-01 |
| AT4G02660 | 2,17 | 0,41 | 1,32 | 0,31 | 7,58E-01 |
| AT4G02680 | 0,66 | -0,76 | 3,72 | -0,20 | 8,39E-01 |
| AT4G02710 | 0,71 | 0,34 | 2,79 | 0,12 | 9,02E-01 |
| AT4G02715 | 1,63 | 2,73 | 1,93 | 1,41 | 1,57E-01 |
| AT4G02735 | 0,41 | -3,10 | 4,34 | -0,72 | 4,75E-01 |
| AT4G02740 | 0,60 | 0,48 | 3,00 | 0,16 | 8,73E-01 |
| AT4G02750 | 0,98 | 0,36 | 2,03 | 0,18 | 8,60E-01 |
| AT4G02760 | 2,62 | -2,29 | 1,27 | -1,81 | 7,01E-02 |
| AT4G02890 | 2,39 | 0,59 | 1,24 | 0,48 | 6,33E-01 |
| AT4G02910 | 0,98 | 0,69 | 2,34 | 0,29 | 7,69E-01 |
| AT4G02930 | 0,46 | -0,64 | 4,32 | -0,15 | 8,83E-01 |
| AT4G03020 | 2,29 | 0,70 | 1,16 | 0,61 | 5,42E-01 |
| AT4G03030 | 1,74 | 0,49 | 1,50 | 0,33 | 7,44E-01 |
| AT4G03080 | 1,69 | 1,04 | 1,36 | 0,77 | 4,43E-01 |
| AT4G03090 | 0,91 | 1,62 | 1,99 | 0,81 | 4,17E-01 |
| AT4G03110 | 0,81 | -0,81 | 2,82 | -0,29 | 7,74E-01 |
| AT4G03130 | 0,08 | 0,44 | 4,38 | 0,10 | 9,21E-01 |
| AT4G03150 | 0,26 | -0,48 | 4,38 | -0,11 | 9,13E-01 |
| AT4G03190 | 0,84 | -1,69 | 2,13 | -0,79 | 4,27E-01 |
| AT4G03230 | 0,47 | -3,28 | 3,97 | -0,83 | 4,09E-01 |
| AT4G03270 | 0,81 | -0,20 | 2,67 | -0,08 | 9,40E-01 |
| AT4G04465 | 2,27 | 1,11 | 1,49 | 0,75 | 4,54E-01 |
| AT4G03295 | 3,55 | 0,02 | 1,24 | 0,01 | 9,89E-01 |
| AT4G04475 | 0,42 | -0,82 | 3,36 | -0,25 | 8,06E-01 |
| AT4G04485 | 0,66 | 1,48 | 2,41 | 0,61 | 5,39E-01 |
| AT4G03340 | 0,12 | 1,05 | 4,35 | 0,24 | 8,10E-01 |
| AT4G03390 | 0,56 | -0,18 | 2,62 | -0,07 | 9,47E-01 |
| AT4G03400 | 0,95 | 1,75 | 2,16 | 0,81 | 4,17E-01 |
| AT4G03415 | 1,93 | -1,07 | 1,57 | -0,68 | 4,94E-01 |
| AT4G03430 | 1,44 | -0,01 | 1,41 | 0,00 | 9,97E-01 |
| AT4G03435 | 0,08 | 0,00 | 4,40 | 0,00 | 1,00E+00 |
| AT4G03438 | 0,26 | -1,78 | 4,41 | -0,40 | 6,87E-01 |
| AT4G03450 | 0,60 | -2,39 | 2,64 | -0,91 | 3,64E-01 |
| AT4G03490 | 0,42 | -1,78 | 4,30 | -0,41 | 6,78E-01 |
| AT4G03505 | 0,25 | -2,19 | 4,38 | -0,50 | 6,17E-01 |
| AT4G03520 | 1,13 | 1,54 | 1,95 | 0,79 | 4,30E-01 |
| AT4G03570 | 0,98 | -0,19 | 1,92 | -0,10 | 9,20E-01 |

| | | | | | |
|---|---|---|---|---|---|
| AT4G03590 | 1,16 | 1,20 | 2,34 | 0,51 | 6,07E-01 |
| AT4G03728 | 0,69 | 0,54 | 3,73 | 0,14 | 8,86E-01 |
| AT4G03740 | 1,37 | 2,39 | 2,02 | 1,18 | 2,37E-01 |
| AT4G04675 | 1,07 | -0,50 | 2,11 | -0,24 | 8,12E-01 |
| AT4G04030 | 0,82 | 0,10 | 2,64 | 0,04 | 9,69E-01 |
| AT4G04221 | 1,00 | -1,30 | 1,94 | -0,67 | 5,00E-01 |
| AT4G04805 | 1,38 | 2,34 | 2,28 | 1,02 | 3,07E-01 |
| AT4G04370 | 1,03 | -0,74 | 1,86 | -0,40 | 6,90E-01 |
| AT4G04470 | 0,08 | 0,00 | 4,40 | 0,00 | 1,00E+00 |
| AT4G04580 | 0,11 | 0,48 | 4,37 | 0,11 | 9,12E-01 |
| AT4G04614 | 0,51 | -2,08 | 4,33 | -0,48 | 6,31E-01 |
| AT4G04632 | 0,44 | 1,81 | 4,08 | 0,44 | 6,57E-01 |
| AT4G04650 | 0,82 | -0,89 | 2,49 | -0,36 | 7,21E-01 |
| AT4G04710 | 0,47 | 2,22 | 3,21 | 0,69 | 4,89E-01 |
| AT4G04720 | 1,28 | 0,40 | 2,05 | 0,20 | 8,44E-01 |
| AT4G04740 | 0,87 | 1,64 | 1,95 | 0,84 | 3,99E-01 |
| AT4G04745 | 1,61 | -1,74 | 1,86 | -0,93 | 3,51E-01 |
| AT4G04770 | 1,41 | 1,28 | 1,84 | 0,70 | 4,87E-01 |
| AT4G04890 | 2,53 | -4,70 | 1,59 | -2,95 | 3,18E-03 |
| AT4G04910 | 0,75 | -1,76 | 2,25 | -0,78 | 4,34E-01 |
| AT4G04950 | 0,83 | -1,60 | 3,67 | -0,44 | 6,63E-01 |
| AT4G04957 | 0,91 | 1,88 | 2,64 | 0,71 | 4,77E-01 |
| AT4G04980 | 0,11 | 0,00 | 4,41 | 0,00 | 1,00E+00 |
| AT4G05050 | 3,49 | 1,09 | 1,17 | 0,93 | 3,51E-01 |
| AT4G05070 | 1,30 | -0,66 | 1,64 | -0,40 | 6,87E-01 |
| AT4G05110 | 0,17 | 1,31 | 4,34 | 0,30 | 7,63E-01 |
| AT4G05085 | 0,05 | 0,00 | 4,41 | 0,00 | 1,00E+00 |
| AT4G05330 | 0,44 | 0,00 | 4,05 | 0,00 | 9,99E-01 |
| AT4G05410 | 0,51 | -1,45 | 4,03 | -0,36 | 7,19E-01 |
| AT4G05470 | 0,12 | 0,97 | 4,34 | 0,22 | 8,23E-01 |
| AT4G05631 | 0,41 | -0,54 | 3,43 | -0,16 | 8,75E-01 |
| AT4G06526 | 0,56 | -1,79 | 4,00 | -0,45 | 6,54E-01 |
| AT4G06639 | 0,21 | 0,01 | 4,37 | 0,00 | 9,98E-01 |
| AT4G06599 | 0,16 | 0,00 | 4,39 | 0,00 | 1,00E+00 |
| AT4G07350 | 0,08 | 0,00 | 4,40 | 0,00 | 1,00E+00 |
| AT4G07400 | 0,28 | 1,04 | 4,34 | 0,24 | 8,11E-01 |
| AT4G07410 | 1,28 | -1,49 | 1,97 | -0,76 | 4,47E-01 |
| AT4G07524 | 0,19 | 1,22 | 4,33 | 0,28 | 7,78E-01 |
| AT4G07526 | 0,83 | -2,01 | 2,41 | -0,83 | 4,06E-01 |
| AT4G05225 | 0,74 | 0,00 | 3,18 | 0,00 | 1,00E+00 |
| AT4G07990 | 0,20 | -1,80 | 4,41 | -0,41 | 6,83E-01 |
| AT4G08230 | 0,30 | 0,02 | 4,39 | 0,01 | 9,96E-01 |
| AT4G08310 | 0,85 | -2,80 | 1,96 | -1,43 | 1,54E-01 |
| AT4G05415 | 0,95 | 0,21 | 2,49 | 0,08 | 9,34E-01 |
| AT4G08370 | 0,46 | 0,56 | 2,68 | 0,21 | 8,34E-01 |
| AT4G08400 | 0,54 | 0,82 | 3,04 | 0,27 | 7,88E-01 |
| AT4G08410 | 1,29 | 2,30 | 2,32 | 0,99 | 3,21E-01 |
| AT4G05535 | 0,95 | -1,43 | 2,38 | -0,60 | 5,48E-01 |
| AT4G08460 | 0,62 | -0,86 | 2,50 | -0,35 | 7,30E-01 |
| AT4G08470 | 1,82 | 1,56 | 1,82 | 0,86 | 3,91E-01 |
| AT4G08480 | 0,67 | -1,99 | 3,08 | -0,65 | 5,18E-01 |
| AT4G08485 | 0,63 | 0,00 | 3,80 | 0,00 | 1,00E+00 |

| | | | | | |
|---|---|---|---|---|---|
| AT4G08590 | 0,56 | -0,09 | 4,31 | -0,02 | 9,83E-01 |
| AT4G08630 | 1,97 | 3,01 | 1,78 | 1,69 | 9,03E-02 |
| AT4G08760 | 1,19 | 1,07 | 2,38 | 0,45 | 6,53E-01 |
| AT4G08810 | 0,77 | -0,58 | 2,42 | -0,24 | 8,11E-01 |
| AT4G08850 | 2,82 | 1,01 | 1,27 | 0,79 | 4,27E-01 |
| AT4G08870 | 1,44 | -0,74 | 1,66 | -0,45 | 6,53E-01 |
| AT4G08874 | 0,12 | 0,00 | 4,38 | 0,00 | 1,00E+00 |
| AT4G08940 | 0,46 | 0,70 | 3,33 | 0,21 | 8,34E-01 |
| AT4G09010 | 1,11 | -0,24 | 1,85 | -0,13 | 8,98E-01 |
| AT4G09012 | 0,55 | 1,92 | 3,21 | 0,60 | 5,49E-01 |
| AT4G09035 | 0,16 | 0,84 | 4,35 | 0,19 | 8,46E-01 |
| AT4G09040 | 0,47 | -1,76 | 4,34 | -0,41 | 6,85E-01 |
| AT4G09340 | 0,44 | -0,58 | 3,21 | -0,18 | 8,56E-01 |
| AT4G09350 | 0,26 | 0,00 | 4,38 | 0,00 | 1,00E+00 |
| AT4G09490 | 0,39 | 0,00 | 4,41 | 0,00 | 1,00E+00 |
| AT4G09510 | 0,38 | -0,48 | 3,40 | -0,14 | 8,88E-01 |
| AT4G09620 | 0,40 | 2,16 | 4,32 | 0,50 | 6,17E-01 |
| AT4G09640 | 0,98 | -0,81 | 1,88 | -0,43 | 6,68E-01 |
| AT4G05740 | 1,41 | -0,68 | 1,71 | -0,40 | 6,91E-01 |
| AT4G09900 | 0,13 | 0,00 | 4,41 | 0,00 | 1,00E+00 |
| AT4G09960 | 0,46 | 0,29 | 4,39 | 0,07 | 9,48E-01 |
| AT4G09980 | 0,57 | -0,20 | 2,54 | -0,08 | 9,36E-01 |
| AT4G10070 | 1,18 | -1,16 | 2,02 | -0,58 | 5,64E-01 |
| AT4G10080 | 1,04 | -1,71 | 2,08 | -0,82 | 4,10E-01 |
| AT4G10130 | 0,55 | -2,37 | 3,78 | -0,63 | 5,32E-01 |
| AT4G10320 | 1,43 | 0,90 | 1,56 | 0,57 | 5,67E-01 |
| AT4G10400 | 0,39 | -1,28 | 4,38 | -0,29 | 7,70E-01 |
| AT4G10420 | 1,07 | -1,69 | 2,49 | -0,68 | 4,98E-01 |
| AT4G10440 | 0,08 | 0,44 | 4,38 | 0,10 | 9,21E-01 |
| AT4G10480 | 0,41 | 0,53 | 4,37 | 0,12 | 9,03E-01 |
| AT4G10540 | 0,12 | 1,23 | 4,34 | 0,28 | 7,76E-01 |
| AT4G10550 | 0,52 | 1,24 | 3,86 | 0,32 | 7,49E-01 |
| AT4G10570 | 1,14 | 0,09 | 1,71 | 0,05 | 9,57E-01 |
| AT4G10590 | 1,40 | 1,20 | 1,88 | 0,64 | 5,24E-01 |
| AT4G05800 | 1,37 | 1,23 | 1,87 | 0,66 | 5,12E-01 |
| AT4G10700 | 0,54 | 0,02 | 3,82 | 0,00 | 9,97E-01 |
| AT4G10710 | 3,86 | -0,19 | 1,09 | -0,18 | 8,59E-01 |
| AT4G10730 | 0,28 | -1,42 | 4,36 | -0,33 | 7,44E-01 |
| AT4G10740 | 1,27 | 1,51 | 2,33 | 0,65 | 5,17E-01 |
| AT4G10760 | 0,19 | 0,00 | 4,41 | 0,00 | 1,00E+00 |
| AT4G10770 | 0,10 | 0,01 | 4,38 | 0,00 | 9,99E-01 |
| AT4G10790 | 1,30 | 2,95 | 1,70 | 1,74 | 8,19E-02 |
| AT4G10920 | 0,27 | 1,23 | 3,75 | 0,33 | 7,42E-01 |
| AT4G10960 | 0,05 | 0,00 | 4,41 | 0,00 | 1,00E+00 |
| AT4G10970 | 0,10 | 0,01 | 4,38 | 0,00 | 9,99E-01 |
| AT4G11010 | 0,17 | 1,15 | 4,34 | 0,27 | 7,91E-01 |
| AT4G11050 | 0,37 | -0,83 | 3,43 | -0,24 | 8,08E-01 |
| AT4G11080 | 0,05 | 0,00 | 4,41 | 0,00 | 1,00E+00 |
| AT4G05855 | 1,27 | -1,78 | 1,99 | -0,89 | 3,73E-01 |
| AT4G11216 | 0,26 | -0,48 | 4,38 | -0,11 | 9,13E-01 |
| AT4G11220 | 0,29 | -1,78 | 4,41 | -0,40 | 6,87E-01 |
| AT4G11260 | 0,66 | -0,86 | 4,31 | -0,20 | 8,42E-01 |

| | | | | | |
|---|---|---|---|---|---|
| AT4G11300 | 0,45 | -0,09 | 4,08 | -0,02 | 9,82E-01 |
| AT4G11340 | 1,18 | 0,99 | 1,77 | 0,56 | 5,75E-01 |
| AT4G11410 | 0,24 | 0,21 | 4,34 | 0,05 | 9,62E-01 |
| AT4G11420 | 3,20 | 0,67 | 1,25 | 0,53 | 5,93E-01 |
| AT4G05880 | 0,80 | 0,01 | 2,81 | 0,00 | 9,96E-01 |
| AT4G11600 | 2,32 | 0,69 | 1,44 | 0,48 | 6,33E-01 |
| AT4G11650 | 0,43 | -1,11 | 4,34 | -0,25 | 7,99E-01 |
| AT4G11670 | 0,44 | -0,48 | 2,68 | -0,18 | 8,57E-01 |
| AT4G11800 | 1,18 | 1,84 | 1,85 | 0,99 | 3,21E-01 |
| AT4G05925 | 1,28 | -2,19 | 2,11 | -1,04 | 2,99E-01 |
| AT4G11820 | 0,25 | 0,00 | 4,41 | 0,00 | 1,00E+00 |
| AT4G11830 | 1,70 | -0,24 | 1,63 | -0,14 | 8,85E-01 |
| AT4G11840 | 1,34 | 1,22 | 1,66 | 0,74 | 4,61E-01 |
| AT4G11850 | 1,56 | 0,37 | 1,62 | 0,23 | 8,19E-01 |
| AT4G11860 | 0,25 | 0,38 | 4,33 | 0,09 | 9,31E-01 |
| AT4G11900 | 2,31 | 0,09 | 1,26 | 0,07 | 9,40E-01 |
| AT4G11920 | 0,40 | 1,48 | 4,13 | 0,36 | 7,21E-01 |
| AT4G05930 | 0,37 | 1,48 | 4,22 | 0,35 | 7,25E-01 |
| AT4G12010 | 0,59 | 2,85 | 3,64 | 0,78 | 4,33E-01 |
| AT4G05980 | 0,40 | 1,78 | 4,33 | 0,41 | 6,81E-01 |
| AT4G12300 | 1,00 | 1,43 | 2,24 | 0,64 | 5,21E-01 |
| AT4G12310 | 0,40 | -0,10 | 2,85 | -0,03 | 9,73E-01 |
| AT4G12320 | 1,93 | -0,69 | 1,44 | -0,48 | 6,31E-01 |
| AT4G12330 | 0,81 | 0,00 | 2,83 | 0,00 | 1,00E+00 |
| AT4G12420 | 1,53 | -2,71 | 1,78 | -1,52 | 1,29E-01 |
| AT4G12590 | 0,64 | -0,89 | 2,73 | -0,33 | 7,43E-01 |
| AT4G12650 | 2,23 | -0,18 | 1,21 | -0,15 | 8,81E-01 |
| AT4G12710 | 0,37 | -0,30 | 4,31 | -0,07 | 9,44E-01 |
| AT4G12730 | 0,13 | 0,00 | 4,41 | 0,00 | 1,00E+00 |
| AT4G12750 | 1,05 | -0,79 | 1,82 | -0,43 | 6,66E-01 |
| AT4G12790 | 0,53 | 0,67 | 3,28 | 0,20 | 8,39E-01 |
| AT4G12820 | 0,35 | 0,00 | 4,41 | 0,00 | 1,00E+00 |
| AT4G12880 | 0,35 | 0,00 | 4,41 | 0,00 | 1,00E+00 |
| AT4G12980 | 0,57 | 1,26 | 2,72 | 0,46 | 6,44E-01 |
| AT4G13150 | 0,55 | -1,61 | 4,32 | -0,37 | 7,09E-01 |
| AT4G13250 | 1,54 | -0,82 | 1,84 | -0,44 | 6,57E-01 |
| AT4G13350 | 0,30 | 0,75 | 4,35 | 0,17 | 8,63E-01 |
| AT4G13395 | 0,15 | 0,00 | 4,41 | 0,00 | 1,00E+00 |
| AT4G13420 | 0,12 | 1,16 | 4,34 | 0,27 | 7,89E-01 |
| AT4G13430 | 3,55 | -0,46 | 0,99 | -0,46 | 6,44E-01 |
| AT4G13505 | 4,72 | 0,46 | 0,90 | 0,51 | 6,07E-01 |
| AT4G13550 | 0,15 | 1,20 | 4,34 | 0,28 | 7,81E-01 |
| AT4G13577 | 2,60 | -4,29 | 1,61 | -2,67 | 7,65E-03 |
| AT4G13590 | 0,56 | -1,55 | 2,66 | -0,58 | 5,61E-01 |
| AT4G13630 | 0,80 | 1,78 | 3,40 | 0,52 | 6,01E-01 |
| AT4G13640 | 0,21 | 1,76 | 4,32 | 0,41 | 6,85E-01 |
| AT4G13770 | 2,50 | 2,60 | 1,69 | 1,54 | 1,24E-01 |
| AT4G13780 | 1,18 | -0,22 | 1,72 | -0,13 | 8,96E-01 |
| AT4G13800 | 0,45 | -0,19 | 3,16 | -0,06 | 9,52E-01 |
| AT4G13810 | 2,84 | 3,16 | 1,67 | 1,89 | 5,84E-02 |
| AT4G13820 | 0,22 | 1,17 | 4,33 | 0,27 | 7,88E-01 |
| AT4G13840 | 1,76 | -0,81 | 1,72 | -0,47 | 6,38E-01 |

| | | | | | |
|---|---|---|---|---|---|
| AT4G13870 | 0,08 | 0,44 | 4,38 | 0,10 | 9,21E-01 |
| AT4G13918 | 2,19 | 1,42 | 1,40 | 1,01 | 3,11E-01 |
| AT4G13930 | 1,56 | -0,99 | 1,45 | -0,68 | 4,95E-01 |
| AT4G13950 | 0,08 | 0,69 | 4,36 | 0,16 | 8,75E-01 |
| AT4G13970 | 0,50 | 0,26 | 3,84 | 0,07 | 9,47E-01 |
| AT4G14010 | 0,10 | 0,00 | 4,41 | 0,00 | 1,00E+00 |
| AT4G14030 | 1,74 | 0,57 | 1,40 | 0,40 | 6,86E-01 |
| AT4G14040 | 0,70 | -2,53 | 2,49 | -1,02 | 3,09E-01 |
| AT4G14050 | 0,14 | 0,83 | 4,35 | 0,19 | 8,49E-01 |
| AT4G14070 | 0,95 | -0,20 | 2,54 | -0,08 | 9,37E-01 |
| AT4G14090 | 0,47 | -0,71 | 2,78 | -0,25 | 7,99E-01 |
| AT4G14110 | 0,26 | -0,30 | 4,37 | -0,07 | 9,46E-01 |
| AT4G14200 | 0,21 | 1,14 | 4,34 | 0,26 | 7,92E-01 |
| AT4G14210 | 0,70 | -1,52 | 1,99 | -0,76 | 4,45E-01 |
| AT4G14230 | 0,73 | 0,03 | 2,91 | 0,01 | 9,93E-01 |
| AT4G14240 | 0,36 | -1,19 | 4,36 | -0,27 | 7,85E-01 |
| AT4G14340 | 0,26 | -0,30 | 4,37 | -0,07 | 9,46E-01 |
| AT4G14342 | 0,72 | 2,86 | 4,29 | 0,67 | 5,04E-01 |
| AT4G14350 | 0,16 | 0,00 | 4,37 | 0,00 | 9,99E-01 |
| AT4G14360 | 1,93 | -1,43 | 1,29 | -1,11 | 2,69E-01 |
| AT4G14365 | 1,73 | 1,00 | 1,62 | 0,62 | 5,37E-01 |
| AT4G14370 | 0,47 | -0,16 | 2,84 | -0,06 | 9,55E-01 |
| AT4G14385 | 0,08 | 0,69 | 4,36 | 0,16 | 8,75E-01 |
| AT4G14420 | 0,43 | -1,11 | 4,34 | -0,25 | 7,99E-01 |
| AT4G14430 | 1,12 | 2,87 | 1,83 | 1,57 | 1,16E-01 |
| AT4G14440 | 0,19 | 0,00 | 4,41 | 0,00 | 1,00E+00 |
| AT4G14510 | 1,36 | 2,02 | 1,91 | 1,06 | 2,91E-01 |
| AT4G14680 | 0,20 | -0,44 | 4,36 | -0,10 | 9,19E-01 |
| AT4G14710 | 0,27 | 0,03 | 4,39 | 0,01 | 9,94E-01 |
| AT4G14740 | 0,35 | 0,15 | 4,41 | 0,03 | 9,74E-01 |
| AT4G14746 | 0,16 | 0,54 | 4,36 | 0,12 | 9,01E-01 |
| AT4G14790 | 0,23 | -1,78 | 4,41 | -0,40 | 6,87E-01 |
| AT4G14811 | 0,45 | -0,71 | 3,35 | -0,21 | 8,33E-01 |
| AT4G14880 | 1,32 | -1,42 | 1,64 | -0,87 | 3,84E-01 |
| AT4G14900 | 1,30 | -0,85 | 1,63 | -0,52 | 6,01E-01 |
| AT4G14905 | 0,68 | -1,98 | 2,54 | -0,78 | 4,35E-01 |
| AT4G14920 | 0,44 | -0,01 | 4,11 | 0,00 | 9,99E-01 |
| AT4G14960 | 3,57 | -0,02 | 1,01 | -0,02 | 9,86E-01 |
| AT4G14990 | 0,87 | -0,30 | 1,76 | -0,17 | 8,65E-01 |
| AT4G15050 | 0,52 | 1,68 | 3,80 | 0,44 | 6,58E-01 |
| AT4G15053 | 0,24 | 0,00 | 4,41 | 0,00 | 1,00E+00 |
| AT4G15080 | 0,43 | -0,54 | 4,11 | -0,13 | 8,95E-01 |
| AT4G15200 | 0,36 | 1,57 | 4,34 | 0,36 | 7,17E-01 |
| AT4G15210 | 0,12 | 1,16 | 4,34 | 0,27 | 7,89E-01 |
| AT4G15242 | 0,72 | -0,12 | 2,24 | -0,06 | 9,56E-01 |
| AT4G06305 | 0,44 | 0,00 | 4,12 | 0,00 | 1,00E+00 |
| AT4G06310 | 0,35 | -2,51 | 4,37 | -0,57 | 5,66E-01 |
| AT4G06315 | 1,00 | -1,78 | 3,20 | -0,56 | 5,78E-01 |
| AT4G15258 | 2,27 | -1,48 | 1,34 | -1,10 | 2,71E-01 |
| AT4G15270 | 0,91 | -1,53 | 2,09 | -0,73 | 4,64E-01 |
| AT4G15490 | 0,92 | 2,10 | 2,03 | 1,04 | 3,00E-01 |
| AT4G15530 | 1,47 | 0,42 | 1,69 | 0,25 | 8,04E-01 |

| | | | | | |
|---|---|---|---|---|---|
| AT4G15550 | 0,31 | 1,01 | 4,35 | 0,23 | 8,16E-01 |
| AT4G15810 | 0,14 | 0,03 | 4,39 | 0,01 | 9,95E-01 |
| AT4G06340 | 0,10 | 0,21 | 4,38 | 0,05 | 9,62E-01 |
| AT4G15920 | 0,36 | 0,00 | 4,34 | 0,00 | 1,00E+00 |
| AT4G15955 | 0,12 | 1,23 | 4,34 | 0,28 | 7,76E-01 |
| AT4G16070 | 0,52 | 0,86 | 2,69 | 0,32 | 7,49E-01 |
| AT4G16141 | 0,51 | -3,72 | 3,97 | -0,94 | 3,49E-01 |
| AT4G16143 | 2,03 | 0,68 | 1,22 | 0,56 | 5,78E-01 |
| AT4G16144 | 1,36 | 1,99 | 2,30 | 0,87 | 3,87E-01 |
| AT4G16155 | 0,30 | -1,19 | 4,35 | -0,27 | 7,84E-01 |
| AT4G16180 | 1,78 | 0,85 | 2,12 | 0,40 | 6,88E-01 |
| AT4G16215 | 0,40 | 0,82 | 3,33 | 0,25 | 8,06E-01 |
| AT4G16250 | 0,73 | -1,26 | 2,96 | -0,42 | 6,72E-01 |
| AT4G16280 | 0,45 | -0,42 | 3,18 | -0,13 | 8,96E-01 |
| AT4G16330 | 0,88 | 3,13 | 2,83 | 1,11 | 2,68E-01 |
| AT4G16340 | 3,35 | -1,19 | 0,99 | -1,20 | 2,30E-01 |
| AT4G16370 | 3,80 | -0,49 | 0,88 | -0,55 | 5,81E-01 |
| AT4G16410 | 1,15 | -1,08 | 1,87 | -0,58 | 5,63E-01 |
| AT4G16470 | 0,11 | 0,65 | 4,36 | 0,15 | 8,81E-01 |
| AT4G16490 | 2,72 | 0,72 | 1,51 | 0,48 | 6,33E-01 |
| AT4G16500 | 1,23 | -1,46 | 1,61 | -0,91 | 3,65E-01 |
| AT4G16515 | 0,18 | 0,13 | 4,41 | 0,03 | 9,76E-01 |
| AT4G16520 | 0,32 | -0,42 | 4,36 | -0,10 | 9,24E-01 |
| AT4G16540 | 0,20 | -1,80 | 4,41 | -0,41 | 6,83E-01 |
| AT4G16580 | 0,33 | -0,91 | 3,60 | -0,25 | 8,01E-01 |
| AT4G16630 | 0,66 | -3,45 | 3,71 | -0,93 | 3,53E-01 |
| AT4G16680 | 0,40 | 2,09 | 4,32 | 0,48 | 6,28E-01 |
| AT4G16690 | 0,88 | 0,70 | 2,18 | 0,32 | 7,49E-01 |
| AT4G16720 | 3,07 | -0,77 | 1,05 | -0,73 | 4,68E-01 |
| AT4G16750 | 0,15 | 0,00 | 4,41 | 0,00 | 1,00E+00 |
| AT4G16760 | 1,69 | 0,01 | 1,40 | 0,01 | 9,93E-01 |
| AT4G16790 | 0,35 | 0,24 | 3,48 | 0,07 | 9,46E-01 |
| AT4G16880 | 1,01 | -0,33 | 2,08 | -0,16 | 8,75E-01 |
| AT4G16892 | 1,06 | 1,63 | 2,35 | 0,69 | 4,88E-01 |
| AT4G16900 | 2,10 | -1,26 | 1,17 | -1,07 | 2,83E-01 |
| AT4G16920 | 1,96 | -0,21 | 1,30 | -0,16 | 8,73E-01 |
| AT4G16940 | 4,86 | -0,76 | 0,99 | -0,77 | 4,41E-01 |
| AT4G16957 | 0,81 | 1,39 | 3,35 | 0,41 | 6,79E-01 |
| AT4G16960 | 0,92 | 3,48 | 4,02 | 0,87 | 3,87E-01 |
| AT4G16983 | 2,85 | -1,38 | 1,27 | -1,08 | 2,79E-01 |
| AT4G17040 | 0,44 | -1,32 | 4,18 | -0,32 | 7,51E-01 |
| AT4G17060 | 0,64 | -0,67 | 3,57 | -0,19 | 8,51E-01 |
| AT4G17070 | 0,40 | 0,76 | 3,31 | 0,23 | 8,18E-01 |
| AT4G17090 | 4,63 | -0,04 | 0,96 | -0,04 | 9,69E-01 |
| AT4G17100 | 0,77 | 0,01 | 2,49 | 0,01 | 9,96E-01 |
| AT4G17190 | 0,27 | 1,03 | 4,35 | 0,24 | 8,13E-01 |
| AT4G17230 | 1,13 | 1,21 | 1,93 | 0,63 | 5,29E-01 |
| AT4G17243 | 0,66 | -0,43 | 2,51 | -0,17 | 8,65E-01 |
| AT4G17280 | 0,46 | -0,31 | 3,22 | -0,10 | 9,24E-01 |
| AT4G17330 | 4,91 | -0,21 | 1,07 | -0,19 | 8,45E-01 |
| AT4G06705 | 4,08 | 0,38 | 1,15 | 0,33 | 7,45E-01 |
| AT4G17430 | 0,16 | 0,00 | 4,41 | 0,00 | 1,00E+00 |

| | | | | | |
|---|---|---|---|---|---|
| AT4G17486 | 0,51 | -2,36 | 4,32 | -0,55 | 5,84E-01 |
| AT4G17490 | 0,05 | 0,00 | 4,41 | 0,00 | 1,00E+00 |
| AT4G17510 | 0,15 | 0,74 | 4,35 | 0,17 | 8,65E-01 |
| AT4G17550 | 0,46 | -0,33 | 4,15 | -0,08 | 9,37E-01 |
| AT4G17570 | 0,79 | -2,13 | 2,40 | -0,89 | 3,73E-01 |
| AT4G17590 | 0,76 | 2,76 | 2,77 | 1,00 | 3,18E-01 |
| AT4G17610 | 0,22 | 1,19 | 4,33 | 0,27 | 7,84E-01 |
| AT4G17670 | 0,33 | 0,66 | 4,32 | 0,15 | 8,78E-01 |
| AT4G17720 | 0,25 | -1,75 | 4,40 | -0,40 | 6,91E-01 |
| AT4G17740 | 0,24 | 1,08 | 4,35 | 0,25 | 8,03E-01 |
| AT4G17840 | 0,82 | 1,87 | 2,28 | 0,82 | 4,13E-01 |
| AT4G17880 | 0,51 | 2,19 | 3,12 | 0,70 | 4,84E-01 |
| AT4G17950 | 0,50 | -1,37 | 4,07 | -0,34 | 7,38E-01 |
| AT4G18010 | 1,46 | 3,46 | 1,93 | 1,79 | 7,31E-02 |
| AT4G18020 | 0,33 | 0,27 | 4,34 | 0,06 | 9,51E-01 |
| AT4G18030 | 2,74 | -0,62 | 1,27 | -0,49 | 6,23E-01 |
| AT4G18040 | 0,22 | 0,25 | 4,36 | 0,06 | 9,54E-01 |
| AT4G18100 | 0,24 | 0,05 | 4,39 | 0,01 | 9,91E-01 |
| AT4G18110 | 0,16 | 0,49 | 4,38 | 0,11 | 9,12E-01 |
| AT4G18130 | 0,68 | -0,01 | 2,72 | 0,00 | 9,98E-01 |
| AT4G18140 | 0,37 | -0,72 | 4,38 | -0,16 | 8,69E-01 |
| AT4G18215 | 1,61 | 1,09 | 1,58 | 0,69 | 4,92E-01 |
| AT4G18230 | 0,15 | 1,16 | 4,34 | 0,27 | 7,88E-01 |
| AT4G18250 | 0,30 | 0,30 | 4,34 | 0,07 | 9,45E-01 |
| AT4G18253 | 0,68 | -0,78 | 2,31 | -0,34 | 7,36E-01 |
| AT4G18255 | 0,35 | -2,17 | 4,38 | -0,49 | 6,21E-01 |
| AT4G18390 | 1,03 | 0,41 | 2,24 | 0,18 | 8,54E-01 |
| AT4G06800 | 2,19 | -0,10 | 1,35 | -0,07 | 9,43E-01 |
| AT4G18440 | 0,69 | 0,32 | 3,62 | 0,09 | 9,30E-01 |
| AT4G18480 | 1,72 | 1,10 | 1,48 | 0,75 | 4,56E-01 |
| AT4G18520 | 0,18 | 1,24 | 4,33 | 0,29 | 7,74E-01 |
| AT4G18580 | 0,08 | 0,44 | 4,38 | 0,10 | 9,21E-01 |
| AT4G18600 | 3,50 | 0,65 | 1,08 | 0,61 | 5,45E-01 |
| AT4G18700 | 2,69 | 0,64 | 1,03 | 0,62 | 5,37E-01 |
| AT4G18790 | 0,08 | 0,69 | 4,36 | 0,16 | 8,75E-01 |
| AT4G18810 | 0,53 | -0,81 | 2,91 | -0,28 | 7,82E-01 |
| AT4G18840 | 0,38 | 0,90 | 3,43 | 0,26 | 7,93E-01 |
| AT4G18880 | 2,25 | 0,24 | 1,59 | 0,15 | 8,78E-01 |
| AT4G19006 | 0,25 | -0,49 | 4,34 | -0,11 | 9,11E-01 |
| AT4G19040 | 0,87 | 2,65 | 3,26 | 0,81 | 4,15E-01 |
| AT4G19045 | 0,12 | 1,05 | 4,35 | 0,24 | 8,10E-01 |
| AT4G19050 | 0,78 | 0,08 | 2,59 | 0,03 | 9,75E-01 |
| AT4G19060 | 0,46 | 0,70 | 3,99 | 0,18 | 8,61E-01 |
| AT4G19080 | 1,21 | 1,08 | 1,81 | 0,59 | 5,52E-01 |
| AT4G19095 | 0,58 | -1,66 | 2,66 | -0,63 | 5,31E-01 |
| AT4G06870 | 0,16 | 0,09 | 4,38 | 0,02 | 9,83E-01 |
| AT4G19110 | 3,30 | 0,53 | 1,23 | 0,43 | 6,70E-01 |
| AT4G19120 | 2,16 | -1,32 | 1,22 | -1,08 | 2,79E-01 |
| AT4G19180 | 0,15 | 1,20 | 4,34 | 0,28 | 7,81E-01 |
| AT4G19220 | 0,37 | -0,47 | 4,25 | -0,11 | 9,11E-01 |
| AT4G19240 | 0,33 | 0,39 | 4,32 | 0,09 | 9,28E-01 |
| AT4G19410 | 1,78 | -0,97 | 1,32 | -0,73 | 4,64E-01 |

| | | | | | |
|---|---|---|---|---|---|
| AT4G19420 | 0,68 | 0,87 | 2,50 | 0,35 | 7,26E-01 |
| AT4G19440 | 0,20 | 1,26 | 4,34 | 0,29 | 7,71E-01 |
| AT4G19520 | 3,67 | 0,59 | 0,90 | 0,66 | 5,11E-01 |
| AT4G19640 | 0,35 | -0,87 | 3,49 | -0,25 | 8,03E-01 |
| AT4G19660 | 1,11 | -0,68 | 1,81 | -0,37 | 7,08E-01 |
| AT4G19670 | 0,77 | -0,46 | 3,43 | -0,13 | 8,94E-01 |
| AT4G06915 | 0,34 | 1,87 | 4,32 | 0,43 | 6,65E-01 |
| AT4G19700 | 1,20 | 1,35 | 2,00 | 0,68 | 4,99E-01 |
| AT4G19720 | 0,28 | 0,00 | 4,41 | 0,00 | 1,00E+00 |
| AT4G19860 | 1,15 | -0,01 | 1,68 | -0,01 | 9,95E-01 |
| AT4G19870 | 0,10 | 0,21 | 4,38 | 0,05 | 9,62E-01 |
| AT4G19880 | 0,52 | 0,77 | 2,55 | 0,30 | 7,63E-01 |
| AT4G19890 | 0,56 | -0,72 | 2,97 | -0,24 | 8,09E-01 |
| AT4G19900 | 0,17 | 0,98 | 4,35 | 0,23 | 8,21E-01 |
| AT4G19985 | 0,13 | 0,73 | 4,36 | 0,17 | 8,67E-01 |
| AT4G20095 | 0,45 | 0,00 | 4,41 | 0,00 | 1,00E+00 |
| AT4G20150 | 0,58 | -0,63 | 2,70 | -0,23 | 8,15E-01 |
| AT4G06945 | 1,00 | 0,75 | 2,71 | 0,28 | 7,81E-01 |
| AT4G20210 | 0,49 | 0,16 | 2,79 | 0,06 | 9,53E-01 |
| AT4G20330 | 0,48 | -0,45 | 2,62 | -0,17 | 8,64E-01 |
| AT4G20362 | 5,64 | 0,14 | 0,89 | 0,16 | 8,76E-01 |
| AT4G20410 | 1,25 | -2,45 | 1,83 | -1,34 | 1,80E-01 |
| AT4G20440 | 0,64 | 0,77 | 2,27 | 0,34 | 7,34E-01 |
| AT4G20690 | 0,21 | 1,76 | 4,32 | 0,41 | 6,85E-01 |
| AT4G20835 | 2,66 | 2,16 | 1,35 | 1,60 | 1,10E-01 |
| AT4G20880 | 0,74 | -1,01 | 2,00 | -0,51 | 6,13E-01 |
| AT4G20910 | 0,27 | -2,20 | 4,38 | -0,50 | 6,16E-01 |
| AT4G20953 | 0,51 | 1,82 | 3,24 | 0,56 | 5,75E-01 |
| AT4G21100 | 1,08 | -1,67 | 1,72 | -0,97 | 3,31E-01 |
| AT4G21140 | 0,20 | -0,44 | 4,36 | -0,10 | 9,19E-01 |
| AT4G21170 | 0,28 | 0,00 | 4,41 | 0,00 | 1,00E+00 |
| AT4G06985 | 0,08 | 0,44 | 4,38 | 0,10 | 9,21E-01 |
| AT4G21190 | 0,08 | 0,69 | 4,36 | 0,16 | 8,75E-01 |
| AT4G21350 | 0,08 | 0,69 | 4,36 | 0,16 | 8,75E-01 |
| AT4G21380 | 1,64 | 1,98 | 1,62 | 1,22 | 2,21E-01 |
| AT4G21400 | 0,16 | 0,65 | 4,37 | 0,15 | 8,82E-01 |
| AT4G21430 | 0,22 | 0,00 | 4,41 | 0,00 | 1,00E+00 |
| AT4G21510 | 0,05 | 0,00 | 4,41 | 0,00 | 1,00E+00 |
| AT4G21520 | 0,22 | -0,47 | 4,35 | -0,11 | 9,13E-01 |
| AT4G21560 | 0,38 | -2,51 | 4,37 | -0,57 | 5,65E-01 |
| AT4G21570 | 1,42 | 1,88 | 2,05 | 0,92 | 3,60E-01 |
| AT4G21640 | 1,99 | -0,36 | 1,80 | -0,20 | 8,39E-01 |
| AT4G21660 | 1,13 | -0,50 | 2,03 | -0,25 | 8,04E-01 |
| AT4G21710 | 2,69 | -1,94 | 1,29 | -1,51 | 1,32E-01 |
| AT4G21810 | 1,02 | -0,10 | 1,94 | -0,05 | 9,59E-01 |
| AT4G21860 | 0,50 | -0,09 | 3,34 | -0,03 | 9,78E-01 |
| AT4G21890 | 0,12 | 1,23 | 4,34 | 0,28 | 7,76E-01 |
| AT4G21910 | 0,54 | 1,74 | 4,33 | 0,40 | 6,88E-01 |
| AT4G21990 | 0,81 | -1,46 | 2,38 | -0,61 | 5,39E-01 |
| AT4G22120 | 0,12 | 1,16 | 4,34 | 0,27 | 7,89E-01 |
| AT4G22150 | 0,22 | 0,00 | 4,41 | 0,00 | 1,00E+00 |
| AT4G22190 | 0,98 | -0,60 | 1,72 | -0,35 | 7,29E-01 |

| | | | | | |
|---|---|---|---|---|---|
| AT4G22200 | 0,21 | 0,67 | 4,36 | 0,15 | 8,78E-01 |
| AT4G22240 | 1,83 | -1,91 | 1,35 | -1,41 | 1,59E-01 |
| AT4G22250 | 0,16 | 0,92 | 4,35 | 0,21 | 8,33E-01 |
| AT4G22260 | 0,18 | 0,00 | 4,41 | 0,00 | 1,00E+00 |
| AT4G22265 | 0,20 | -0,31 | 4,36 | -0,07 | 9,44E-01 |
| AT4G22285 | 0,20 | 0,40 | 4,37 | 0,09 | 9,28E-01 |
| AT4G22290 | 0,23 | 0,19 | 4,38 | 0,04 | 9,66E-01 |
| AT4G22350 | 0,69 | -1,14 | 3,72 | -0,31 | 7,60E-01 |
| AT4G22380 | 0,51 | 0,24 | 2,67 | 0,09 | 9,28E-01 |
| AT4G22410 | 2,93 | 2,04 | 1,58 | 1,29 | 1,97E-01 |
| AT4G22470 | 0,12 | 0,00 | 4,38 | 0,00 | 1,00E+00 |
| AT4G22485 | 0,49 | 0,00 | 3,99 | 0,00 | 1,00E+00 |
| AT4G22550 | 0,26 | 1,24 | 3,76 | 0,33 | 7,41E-01 |
| AT4G22570 | 0,24 | -0,74 | 4,35 | -0,17 | 8,65E-01 |
| AT4G22745 | 0,42 | -0,56 | 4,10 | -0,14 | 8,91E-01 |
| AT4G22750 | 1,03 | -0,06 | 2,23 | -0,03 | 9,77E-01 |
| AT4G22755 | 0,47 | -0,35 | 4,33 | -0,08 | 9,36E-01 |
| AT4G22770 | 1,29 | 1,92 | 1,68 | 1,14 | 2,53E-01 |
| AT4G22780 | 0,24 | -0,84 | 4,35 | -0,19 | 8,47E-01 |
| AT4G22970 | 0,52 | 2,33 | 3,82 | 0,61 | 5,43E-01 |
| AT4G22980 | 0,47 | 2,08 | 3,94 | 0,53 | 5,97E-01 |
| AT4G23010 | 0,83 | -2,04 | 2,16 | -0,94 | 3,45E-01 |
| AT4G23040 | 0,28 | 0,43 | 4,38 | 0,10 | 9,22E-01 |
| AT4G23100 | 1,83 | 0,37 | 1,27 | 0,29 | 7,73E-01 |
| AT4G23130 | 0,55 | 1,44 | 3,90 | 0,37 | 7,12E-01 |
| AT4G23190 | 1,18 | -0,21 | 1,82 | -0,12 | 9,08E-01 |
| AT4G23205 | 0,08 | 0,69 | 4,36 | 0,16 | 8,75E-01 |
| AT4G23200 | 0,43 | -0,82 | 4,36 | -0,19 | 8,50E-01 |
| AT4G23210 | 0,21 | 1,55 | 4,33 | 0,36 | 7,20E-01 |
| AT4G23220 | 0,76 | 1,71 | 2,34 | 0,73 | 4,65E-01 |
| AT4G23240 | 0,37 | 0,94 | 4,32 | 0,22 | 8,28E-01 |
| AT4G23250 | 0,45 | 2,02 | 3,92 | 0,52 | 6,06E-01 |
| AT4G23260 | 0,19 | 1,08 | 4,34 | 0,25 | 8,04E-01 |
| AT4G23290 | 0,10 | 0,00 | 4,41 | 0,00 | 1,00E+00 |
| AT4G23320 | 0,08 | 0,69 | 4,36 | 0,16 | 8,75E-01 |
| AT4G23460 | 1,64 | -1,59 | 1,45 | -1,10 | 2,73E-01 |
| AT4G23490 | 0,15 | 0,00 | 4,41 | 0,00 | 1,00E+00 |
| AT4G07130 | 0,46 | -0,76 | 4,10 | -0,19 | 8,53E-01 |
| AT4G23540 | 0,67 | -2,06 | 3,05 | -0,67 | 5,00E-01 |
| AT4G23560 | 0,10 | 0,00 | 4,41 | 0,00 | 1,00E+00 |
| AT4G07135 | 1,25 | -0,85 | 1,77 | -0,48 | 6,31E-01 |
| AT4G07145 | 0,96 | -0,44 | 2,45 | -0,18 | 8,59E-01 |
| AT4G23620 | 0,31 | 0,09 | 4,33 | 0,02 | 9,84E-01 |
| AT4G23640 | 0,77 | 0,71 | 2,32 | 0,31 | 7,60E-01 |
| AT4G23650 | 3,54 | -0,97 | 0,97 | -1,00 | 3,17E-01 |
| AT4G23670 | 1,50 | -1,76 | 2,07 | -0,85 | 3,94E-01 |
| AT4G23700 | 0,29 | -0,93 | 4,34 | -0,21 | 8,31E-01 |
| AT4G23770 | 0,22 | 0,26 | 4,38 | 0,06 | 9,52E-01 |
| AT4G23810 | 0,35 | 0,16 | 4,34 | 0,04 | 9,71E-01 |
| AT4G23820 | 0,63 | -1,72 | 2,31 | -0,74 | 4,57E-01 |
| AT4G23850 | 2,43 | 0,20 | 1,17 | 0,17 | 8,62E-01 |
| AT4G23880 | 0,47 | -1,78 | 3,29 | -0,54 | 5,89E-01 |

| | | | | | |
|---|---|---|---|---|---|
| AT4G23890 | 0,66 | -1,56 | 2,71 | -0,57 | 5,66E-01 |
| AT4G24180 | 0,37 | -0,17 | 4,31 | -0,04 | 9,68E-01 |
| AT4G24190 | 5,76 | 0,54 | 0,81 | 0,67 | 5,05E-01 |
| AT4G24200 | 0,70 | -0,58 | 2,87 | -0,20 | 8,39E-01 |
| AT4G07250 | 0,17 | 0,00 | 4,41 | 0,00 | 1,00E+00 |
| AT4G24300 | 0,45 | 0,22 | 4,37 | 0,05 | 9,59E-01 |
| AT4G24320 | 0,33 | 1,91 | 4,32 | 0,44 | 6,58E-01 |
| AT4G24330 | 0,84 | 0,71 | 2,03 | 0,35 | 7,27E-01 |
| AT4G07255 | 0,10 | 0,00 | 4,41 | 0,00 | 1,00E+00 |
| AT4G24390 | 0,29 | 0,21 | 4,32 | 0,05 | 9,60E-01 |
| AT4G24413 | 2,10 | -0,19 | 1,56 | -0,12 | 9,02E-01 |
| AT4G24490 | 0,72 | 0,01 | 2,41 | 0,00 | 9,97E-01 |
| AT4G24520 | 3,90 | -2,14 | 1,21 | -1,77 | 7,62E-02 |
| AT4G24530 | 0,08 | 0,69 | 4,36 | 0,16 | 8,75E-01 |
| AT4G24560 | 1,21 | 1,12 | 1,92 | 0,58 | 5,60E-01 |
| AT4G24580 | 0,46 | -1,85 | 4,08 | -0,45 | 6,50E-01 |
| AT4G24610 | 0,67 | -0,75 | 2,81 | -0,27 | 7,89E-01 |
| AT4G24620 | 0,42 | -1,31 | 4,19 | -0,31 | 7,55E-01 |
| AT4G24660 | 1,04 | 0,43 | 2,29 | 0,19 | 8,52E-01 |
| AT4G24670 | 0,08 | 0,00 | 4,40 | 0,00 | 1,00E+00 |
| AT4G24715 | 2,06 | -1,85 | 1,27 | -1,46 | 1,44E-01 |
| AT4G24730 | 0,45 | -1,34 | 4,01 | -0,33 | 7,38E-01 |
| AT4G24740 | 0,63 | -1,30 | 3,02 | -0,43 | 6,66E-01 |
| AT4G24750 | 0,28 | -1,42 | 4,36 | -0,33 | 7,44E-01 |
| AT4G24770 | 3,23 | 0,08 | 1,14 | 0,07 | 9,42E-01 |
| AT4G24780 | 1,27 | 0,22 | 2,21 | 0,10 | 9,22E-01 |
| AT4G07285 | 0,27 | 0,00 | 4,41 | 0,00 | 1,00E+00 |
| AT4G24820 | 1,43 | 0,29 | 1,60 | 0,18 | 8,55E-01 |
| AT4G24830 | 0,25 | 0,13 | 4,39 | 0,03 | 9,76E-01 |
| AT4G24840 | 0,53 | -0,43 | 3,13 | -0,14 | 8,90E-01 |
| AT4G24920 | 0,86 | 0,02 | 2,61 | 0,01 | 9,93E-01 |
| AT4G24990 | 0,31 | -0,92 | 4,39 | -0,21 | 8,35E-01 |
| AT4G25090 | 0,29 | 2,11 | 4,32 | 0,49 | 6,26E-01 |
| AT4G25100 | 2,11 | 3,65 | 1,68 | 2,18 | 2,95E-02 |
| AT4G25270 | 0,45 | -1,34 | 4,01 | -0,33 | 7,38E-01 |
| AT4G25280 | 0,71 | 1,27 | 4,31 | 0,30 | 7,68E-01 |
| AT4G25290 | 0,52 | 0,53 | 3,84 | 0,14 | 8,90E-01 |
| AT4G25340 | 0,19 | 0,90 | 4,34 | 0,21 | 8,35E-01 |
| AT4G25435 | 0,88 | -2,90 | 2,73 | -1,06 | 2,89E-01 |
| AT4G25520 | 4,54 | -0,72 | 0,97 | -0,75 | 4,54E-01 |
| AT4G25540 | 0,13 | 0,73 | 4,36 | 0,17 | 8,67E-01 |
| AT4G25570 | 0,43 | -0,81 | 4,17 | -0,19 | 8,46E-01 |
| AT4G25640 | 1,48 | -0,39 | 1,45 | -0,27 | 7,90E-01 |
| AT4G25650 | 1,11 | 0,47 | 1,68 | 0,28 | 7,81E-01 |
| AT4G25670 | 0,79 | -0,38 | 2,69 | -0,14 | 8,89E-01 |
| AT4G25690 | 0,56 | -2,04 | 3,21 | -0,63 | 5,26E-01 |
| AT4G25730 | 1,71 | -0,50 | 1,69 | -0,30 | 7,68E-01 |
| AT4G25740 | 0,38 | -0,20 | 4,33 | -0,05 | 9,63E-01 |
| AT4G25870 | 0,26 | -0,48 | 4,38 | -0,11 | 9,13E-01 |
| AT4G25940 | 0,39 | 0,37 | 4,32 | 0,09 | 9,31E-01 |
| AT4G25990 | 0,26 | -1,78 | 4,41 | -0,40 | 6,87E-01 |
| AT4G26080 | 0,87 | 0,82 | 2,53 | 0,32 | 7,46E-01 |

| | | | | | |
|---|---|---|---|---|---|
| AT4G26095 | 1,81 | -0,56 | 1,30 | -0,43 | 6,66E-01 |
| AT4G26100 | 0,81 | 0,06 | 1,99 | 0,03 | 9,77E-01 |
| AT4G26130 | 1,45 | 2,93 | 1,88 | 1,56 | 1,18E-01 |
| AT4G26140 | 0,21 | -1,41 | 4,36 | -0,32 | 7,46E-01 |
| AT4G26190 | 0,25 | 1,18 | 4,33 | 0,27 | 7,85E-01 |
| AT4G26300 | 0,69 | 0,42 | 2,49 | 0,17 | 8,67E-01 |
| AT4G07565 | 0,79 | 0,80 | 2,20 | 0,36 | 7,17E-01 |
| AT4G07625 | 0,79 | 0,69 | 2,64 | 0,26 | 7,93E-01 |
| AT4G07635 | 0,12 | 0,00 | 4,38 | 0,00 | 1,00E+00 |
| AT4G26370 | 0,48 | -1,73 | 4,10 | -0,42 | 6,74E-01 |
| AT4G26375 | 0,40 | 2,16 | 4,32 | 0,50 | 6,17E-01 |
| AT4G26400 | 0,60 | -0,50 | 3,87 | -0,13 | 8,97E-01 |
| AT4G26480 | 2,39 | 0,34 | 1,37 | 0,25 | 8,03E-01 |
| AT4G26540 | 3,42 | -0,16 | 1,07 | -0,15 | 8,82E-01 |
| AT4G26620 | 0,28 | 1,35 | 4,33 | 0,31 | 7,55E-01 |
| AT4G26630 | 0,72 | 1,51 | 2,43 | 0,62 | 5,35E-01 |
| AT4G26640 | 1,00 | 2,90 | 2,33 | 1,24 | 2,13E-01 |
| AT4G26690 | 1,51 | 1,17 | 1,75 | 0,67 | 5,04E-01 |
| AT4G26750 | 0,43 | 1,66 | 4,16 | 0,40 | 6,89E-01 |
| AT4G26760 | 0,64 | 1,20 | 3,05 | 0,39 | 6,94E-01 |
| AT4G26795 | 0,42 | 0,39 | 4,12 | 0,09 | 9,25E-01 |
| AT4G26860 | 0,37 | 1,13 | 4,33 | 0,26 | 7,94E-01 |
| AT4G26910 | 1,33 | -0,98 | 1,47 | -0,67 | 5,05E-01 |
| AT4G26940 | 0,43 | -0,48 | 4,07 | -0,12 | 9,05E-01 |
| AT4G26980 | 0,78 | 0,39 | 2,31 | 0,17 | 8,66E-01 |
| AT4G27020 | 0,48 | 0,58 | 3,37 | 0,17 | 8,63E-01 |
| AT4G07765 | 0,66 | 1,98 | 2,83 | 0,70 | 4,84E-01 |
| AT4G27060 | 1,71 | -0,01 | 1,40 | -0,01 | 9,95E-01 |
| AT4G27090 | 2,08 | 1,64 | 1,51 | 1,09 | 2,78E-01 |
| AT4G27130 | 0,28 | 0,81 | 3,70 | 0,22 | 8,27E-01 |
| AT4G27270 | 0,66 | 1,96 | 3,82 | 0,51 | 6,08E-01 |
| AT4G27280 | 0,84 | -0,10 | 2,82 | -0,04 | 9,71E-01 |
| AT4G27300 | 0,83 | 0,22 | 2,39 | 0,09 | 9,27E-01 |
| AT4G27320 | 1,19 | -0,84 | 2,06 | -0,41 | 6,85E-01 |
| AT4G27340 | 0,21 | 1,16 | 4,34 | 0,27 | 7,89E-01 |
| AT4G27410 | 0,98 | -0,65 | 2,12 | -0,31 | 7,58E-01 |
| AT4G27450 | 0,72 | -1,24 | 2,29 | -0,54 | 5,88E-01 |
| AT4G27480 | 0,65 | 1,86 | 3,53 | 0,53 | 5,99E-01 |
| AT4G07855 | 2,93 | -0,08 | 1,11 | -0,07 | 9,45E-01 |
| AT4G27520 | 5,05 | 0,23 | 1,07 | 0,22 | 8,30E-01 |
| AT4G27650 | 0,39 | -0,13 | 3,42 | -0,04 | 9,69E-01 |
| AT4G27700 | 0,47 | -1,87 | 3,29 | -0,57 | 5,71E-01 |
| AT4G27730 | 0,37 | 0,01 | 4,31 | 0,00 | 9,97E-01 |
| AT4G27800 | 1,09 | -0,71 | 1,89 | -0,38 | 7,07E-01 |
| AT4G27830 | 0,84 | -0,69 | 1,92 | -0,36 | 7,19E-01 |
| AT4G27840 | 0,43 | -2,67 | 4,37 | -0,61 | 5,40E-01 |
| AT4G27885 | 0,16 | 0,00 | 4,41 | 0,00 | 1,00E+00 |
| AT4G27960 | 0,38 | 0,12 | 4,34 | 0,03 | 9,78E-01 |
| AT4G28030 | 0,45 | 0,21 | 2,76 | 0,08 | 9,39E-01 |
| AT4G28050 | 0,43 | 1,68 | 4,33 | 0,39 | 6,98E-01 |
| AT4G28060 | 0,25 | -2,19 | 4,38 | -0,50 | 6,17E-01 |
| AT4G28180 | 0,69 | 0,52 | 2,57 | 0,20 | 8,39E-01 |

| | | | | | |
|---|---|---|---|---|---|
| AT4G28200 | 0,59 | 0,38 | 2,66 | 0,14 | 8,86E-01 |
| AT4G28230 | 0,33 | 0,71 | 4,32 | 0,16 | 8,70E-01 |
| AT4G28240 | 0,66 | 0,30 | 2,24 | 0,14 | 8,92E-01 |
| AT4G28320 | 0,20 | -0,44 | 4,36 | -0,10 | 9,19E-01 |
| AT4G28390 | 0,33 | 0,75 | 4,32 | 0,17 | 8,62E-01 |
| AT4G28450 | 0,61 | -3,93 | 4,30 | -0,91 | 3,60E-01 |
| AT4G28470 | 0,62 | -0,12 | 3,03 | -0,04 | 9,67E-01 |
| AT4G28610 | 0,93 | 0,10 | 2,49 | 0,04 | 9,67E-01 |
| AT4G28630 | 0,12 | 1,16 | 4,34 | 0,27 | 7,89E-01 |
| AT4G28650 | 2,08 | -0,12 | 1,52 | -0,08 | 9,39E-01 |
| AT4G28750 | 4,56 | -1,19 | 1,11 | -1,08 | 2,82E-01 |
| AT4G28770 | 0,20 | 0,00 | 4,38 | 0,00 | 1,00E+00 |
| AT4G28910 | 1,06 | -0,74 | 2,24 | -0,33 | 7,42E-01 |
| AT4G29010 | 1,63 | 2,01 | 1,79 | 1,12 | 2,61E-01 |
| AT4G29050 | 0,13 | 0,00 | 4,41 | 0,00 | 1,00E+00 |
| AT4G29080 | 0,28 | -0,18 | 4,34 | -0,04 | 9,66E-01 |
| AT4G29130 | 1,04 | -1,04 | 1,80 | -0,58 | 5,62E-01 |
| AT4G29190 | 1,79 | -2,89 | 1,51 | -1,91 | 5,57E-02 |
| AT4G29200 | 3,91 | 1,32 | 1,32 | 1,00 | 3,19E-01 |
| AT4G29220 | 0,63 | -0,01 | 3,11 | 0,00 | 9,99E-01 |
| AT4G29260 | 0,48 | 0,74 | 4,02 | 0,18 | 8,55E-01 |
| AT4G29300 | 1,28 | -0,11 | 1,85 | -0,06 | 9,52E-01 |
| AT4G29305 | 0,76 | -0,45 | 2,47 | -0,18 | 8,56E-01 |
| AT4G08125 | 3,11 | 0,66 | 1,38 | 0,48 | 6,35E-01 |
| AT4G08155 | 1,40 | -1,04 | 1,57 | -0,66 | 5,06E-01 |
| AT4G08165 | 5,10 | 3,60 | 1,59 | 2,26 | 2,37E-02 |
| AT4G29440 | 0,66 | -0,43 | 2,89 | -0,15 | 8,81E-01 |
| AT4G29520 | 0,77 | -0,69 | 2,51 | -0,28 | 7,83E-01 |
| AT4G29654 | 1,07 | 2,60 | 2,63 | 0,99 | 3,24E-01 |
| AT4G29670 | 0,65 | -1,00 | 2,25 | -0,44 | 6,58E-01 |
| AT4G29680 | 0,85 | -1,43 | 2,96 | -0,48 | 6,29E-01 |
| AT4G29720 | 0,17 | 0,47 | 4,37 | 0,11 | 9,14E-01 |
| AT4G29735 | 0,17 | 0,65 | 4,36 | 0,15 | 8,81E-01 |
| AT4G29810 | 0,27 | 1,06 | 4,35 | 0,24 | 8,07E-01 |
| AT4G29820 | 0,24 | -0,73 | 4,37 | -0,17 | 8,68E-01 |
| AT4G29840 | 0,58 | -0,82 | 2,30 | -0,35 | 7,23E-01 |
| AT4G29880 | 0,21 | 0,94 | 4,35 | 0,22 | 8,29E-01 |
| AT4G29900 | 4,72 | 0,49 | 0,84 | 0,58 | 5,63E-01 |
| AT4G29905 | 1,17 | 0,68 | 1,91 | 0,36 | 7,22E-01 |
| AT4G29910 | 0,08 | 0,00 | 4,40 | 0,00 | 1,00E+00 |
| AT4G29920 | 0,05 | 0,00 | 4,41 | 0,00 | 1,00E+00 |
| AT4G29940 | 0,28 | -1,42 | 4,36 | -0,33 | 7,44E-01 |
| AT4G29960 | 0,50 | -0,16 | 3,86 | -0,04 | 9,66E-01 |
| AT4G29990 | 0,08 | 0,44 | 4,38 | 0,10 | 9,21E-01 |
| AT4G30000 | 0,38 | -1,52 | 4,32 | -0,35 | 7,24E-01 |
| AT4G30030 | 0,45 | 1,23 | 4,29 | 0,29 | 7,74E-01 |
| AT4G30060 | 0,08 | 0,69 | 4,36 | 0,16 | 8,75E-01 |
| AT4G30080 | 0,31 | -0,54 | 4,33 | -0,12 | 9,02E-01 |
| AT4G30110 | 1,05 | 1,15 | 3,08 | 0,37 | 7,10E-01 |
| AT4G30200 | 1,47 | 0,92 | 1,60 | 0,58 | 5,65E-01 |
| AT4G30240 | 0,30 | 0,29 | 4,32 | 0,07 | 9,47E-01 |
| AT4G30260 | 0,47 | 1,07 | 3,16 | 0,34 | 7,36E-01 |

| | | | | | |
|---|---|---|---|---|---|
| AT4G30270 | 1,68 | 2,65 | 1,60 | 1,65 | 9,90E-02 |
| AT4G30280 | 0,12 | 0,00 | 4,39 | 0,00 | 1,00E+00 |
| AT4G30360 | 0,21 | 0,00 | 4,41 | 0,00 | 1,00E+00 |
| AT4G30410 | 0,12 | 0,49 | 4,36 | 0,11 | 9,11E-01 |
| AT4G30430 | 0,08 | 0,44 | 4,38 | 0,10 | 9,21E-01 |
| AT4G30440 | 3,67 | 1,34 | 1,26 | 1,06 | 2,89E-01 |
| AT4G30560 | 0,74 | -0,91 | 3,38 | -0,27 | 7,87E-01 |
| AT4G30580 | 0,92 | 1,16 | 3,32 | 0,35 | 7,28E-01 |
| AT4G30600 | 1,60 | -0,32 | 1,53 | -0,21 | 8,32E-01 |
| AT4G30620 | 0,23 | -1,78 | 4,41 | -0,40 | 6,87E-01 |
| AT4G30700 | 0,08 | 0,69 | 4,36 | 0,16 | 8,75E-01 |
| AT4G30710 | 0,12 | 1,16 | 4,34 | 0,27 | 7,89E-01 |
| AT4G30720 | 1,29 | 0,18 | 1,52 | 0,12 | 9,04E-01 |
| AT4G30740 | 1,03 | 1,77 | 2,45 | 0,72 | 4,70E-01 |
| AT4G30750 | 3,95 | 4,06 | 1,46 | 2,78 | 5,51E-03 |
| AT4G30790 | 0,99 | 1,95 | 2,22 | 0,88 | 3,81E-01 |
| AT4G30890 | 0,60 | -1,62 | 2,96 | -0,55 | 5,85E-01 |
| AT4G30920 | 1,19 | -0,06 | 1,86 | -0,03 | 9,73E-01 |
| AT4G30950 | 1,64 | 0,63 | 1,51 | 0,42 | 6,75E-01 |
| AT4G31020 | 0,29 | -1,33 | 4,35 | -0,31 | 7,60E-01 |
| AT4G31120 | 0,87 | -1,03 | 2,30 | -0,45 | 6,54E-01 |
| AT4G31130 | 0,59 | 1,32 | 3,18 | 0,42 | 6,78E-01 |
| AT4G31170 | 2,81 | -0,57 | 1,37 | -0,42 | 6,77E-01 |
| AT4G31420 | 0,28 | -0,47 | 4,37 | -0,11 | 9,14E-01 |
| AT4G31450 | 0,46 | -0,36 | 4,16 | -0,09 | 9,31E-01 |
| AT4G31540 | 0,43 | 0,23 | 4,18 | 0,06 | 9,55E-01 |
| AT4G31550 | 1,02 | -1,29 | 1,75 | -0,74 | 4,61E-01 |
| AT4G31590 | 0,63 | 0,78 | 2,62 | 0,30 | 7,66E-01 |
| AT4G31700 | 1,83 | 0,59 | 1,66 | 0,36 | 7,22E-01 |
| AT4G31810 | 0,08 | 0,69 | 4,36 | 0,16 | 8,75E-01 |
| AT4G31820 | 0,43 | 0,67 | 2,78 | 0,24 | 8,09E-01 |
| AT4G31880 | 2,70 | -0,86 | 1,14 | -0,76 | 4,48E-01 |
| AT4G31990 | 1,81 | -2,41 | 1,71 | -1,41 | 1,58E-01 |
| AT4G32070 | 0,58 | -1,69 | 3,82 | -0,44 | 6,57E-01 |
| AT4G32120 | 0,15 | 0,45 | 4,37 | 0,10 | 9,19E-01 |
| AT4G32130 | 0,18 | 0,21 | 4,37 | 0,05 | 9,62E-01 |
| AT4G32150 | 1,17 | -0,92 | 1,58 | -0,59 | 5,58E-01 |
| AT4G32180 | 1,00 | 0,63 | 2,45 | 0,26 | 7,96E-01 |
| AT4G32208 | 0,37 | -0,47 | 4,25 | -0,11 | 9,11E-01 |
| AT4G32250 | 2,07 | 0,82 | 1,47 | 0,56 | 5,74E-01 |
| AT4G32260 | 3,86 | 0,42 | 1,20 | 0,35 | 7,23E-01 |
| AT4G32340 | 1,02 | -0,45 | 1,80 | -0,25 | 8,04E-01 |
| AT4G32420 | 0,78 | 1,10 | 2,47 | 0,45 | 6,55E-01 |
| AT4G32460 | 0,31 | 0,76 | 4,36 | 0,17 | 8,62E-01 |
| AT4G32470 | 0,29 | -0,88 | 4,37 | -0,20 | 8,40E-01 |
| AT4G32510 | 0,56 | 1,44 | 3,73 | 0,39 | 6,99E-01 |
| AT4G32520 | 1,00 | 0,40 | 1,89 | 0,21 | 8,34E-01 |
| AT4G32530 | 0,89 | -0,54 | 2,35 | -0,23 | 8,17E-01 |
| AT4G32670 | 0,15 | 0,83 | 4,35 | 0,19 | 8,49E-01 |
| AT4G32750 | 0,18 | 0,62 | 4,37 | 0,14 | 8,86E-01 |
| AT4G32820 | 0,83 | 1,38 | 2,61 | 0,53 | 5,98E-01 |
| AT4G32880 | 1,09 | -2,07 | 1,90 | -1,09 | 2,75E-01 |

| | | | | | |
|---|---|---|---|---|---|
| AT4G32900 | 0,12 | 0,97 | 4,34 | 0,22 | 8,23E-01 |
| AT4G32920 | 0,70 | 1,53 | 2,80 | 0,55 | 5,85E-01 |
| AT4G32940 | 4,72 | 0,59 | 0,83 | 0,71 | 4,77E-01 |
| AT4G33040 | 0,37 | 0,04 | 4,32 | 0,01 | 9,93E-01 |
| AT4G33050 | 2,86 | 0,38 | 1,09 | 0,34 | 7,31E-01 |
| AT4G33090 | 0,64 | 1,78 | 3,07 | 0,58 | 5,62E-01 |
| AT4G33110 | 0,17 | 0,82 | 4,35 | 0,19 | 8,51E-01 |
| AT4G33150 | 0,65 | 1,93 | 3,60 | 0,54 | 5,92E-01 |
| AT4G33200 | 0,29 | 0,47 | 4,33 | 0,11 | 9,14E-01 |
| AT4G33210 | 1,11 | -0,39 | 1,75 | -0,22 | 8,22E-01 |
| AT4G33240 | 2,23 | -0,78 | 1,40 | -0,56 | 5,78E-01 |
| AT4G33250 | 0,89 | -0,72 | 1,88 | -0,39 | 7,00E-01 |
| AT4G33260 | 0,12 | 1,16 | 4,34 | 0,27 | 7,89E-01 |
| AT4G33270 | 0,12 | 0,00 | 4,39 | 0,00 | 1,00E+00 |
| AT4G33300 | 3,93 | -0,22 | 1,05 | -0,21 | 8,35E-01 |
| AT4G33400 | 0,33 | 0,26 | 4,33 | 0,06 | 9,52E-01 |
| AT4G33430 | 1,19 | 0,79 | 1,94 | 0,41 | 6,85E-01 |
| AT4G33470 | 0,67 | 0,71 | 2,86 | 0,25 | 8,04E-01 |
| AT4G33495 | 0,29 | 0,47 | 4,33 | 0,11 | 9,14E-01 |
| AT4G33500 | 0,96 | 1,84 | 2,58 | 0,71 | 4,75E-01 |
| AT4G33530 | 1,44 | 1,31 | 1,68 | 0,78 | 4,36E-01 |
| AT4G33585 | 1,20 | -2,81 | 2,83 | -0,99 | 3,21E-01 |
| AT4G33625 | 0,16 | 1,07 | 4,34 | 0,25 | 8,05E-01 |
| AT4G33670 | 0,76 | -1,66 | 1,97 | -0,84 | 4,01E-01 |
| AT4G33680 | 0,20 | 1,00 | 4,36 | 0,23 | 8,18E-01 |
| AT4G33700 | 0,28 | 0,28 | 4,36 | 0,06 | 9,49E-01 |
| AT4G33790 | 0,16 | 0,65 | 4,37 | 0,15 | 8,82E-01 |
| AT4G33860 | 0,68 | -0,48 | 3,52 | -0,14 | 8,92E-01 |
| AT4G33865 | 0,81 | 1,51 | 2,14 | 0,71 | 4,81E-01 |
| AT4G33910 | 0,26 | -0,48 | 4,38 | -0,11 | 9,13E-01 |
| AT4G34020 | 0,41 | 0,52 | 4,37 | 0,12 | 9,05E-01 |
| AT4G34040 | 0,82 | 2,03 | 2,28 | 0,89 | 3,73E-01 |
| AT4G34070 | 0,72 | -0,88 | 2,65 | -0,33 | 7,40E-01 |
| AT4G34090 | 0,71 | 2,24 | 4,30 | 0,52 | 6,03E-01 |
| AT4G34131 | 0,52 | 1,24 | 3,86 | 0,32 | 7,49E-01 |
| AT4G34180 | 0,68 | 2,49 | 2,86 | 0,87 | 3,85E-01 |
| AT4G34190 | 1,52 | -0,33 | 1,66 | -0,20 | 8,44E-01 |
| AT4G34200 | 1,37 | 0,61 | 1,61 | 0,38 | 7,06E-01 |
| AT4G34220 | 1,59 | -0,07 | 1,63 | -0,04 | 9,65E-01 |
| AT4G34332 | 0,08 | 0,69 | 4,36 | 0,16 | 8,75E-01 |
| AT4G34340 | 0,12 | 1,16 | 4,34 | 0,27 | 7,89E-01 |
| AT4G34350 | 1,55 | 1,77 | 1,62 | 1,09 | 2,75E-01 |
| AT4G08985 | 0,45 | 0,82 | 3,25 | 0,25 | 8,00E-01 |
| AT4G34460 | 0,42 | 0,27 | 3,24 | 0,08 | 9,34E-01 |
| AT4G34480 | 1,19 | 0,34 | 1,71 | 0,20 | 8,43E-01 |
| AT4G34490 | 1,39 | 0,71 | 1,52 | 0,47 | 6,40E-01 |
| AT4G34610 | 0,37 | -2,42 | 4,35 | -0,56 | 5,78E-01 |
| AT4G34620 | 1,51 | -0,53 | 1,93 | -0,28 | 7,83E-01 |
| AT4G34630 | 0,16 | 0,00 | 4,39 | 0,00 | 1,00E+00 |
| AT4G34690 | 1,55 | 2,96 | 1,95 | 1,52 | 1,29E-01 |
| AT4G34710 | 3,32 | -0,94 | 0,99 | -0,95 | 3,41E-01 |
| AT4G34720 | 1,83 | 0,46 | 1,30 | 0,36 | 7,22E-01 |

| | | | | | |
|---|---|---|---|---|---|
| AT4G34730 | 0,55 | -0,96 | 2,64 | -0,36 | 7,17E-01 |
| AT4G34740 | 0,69 | -1,62 | 2,89 | -0,56 | 5,75E-01 |
| AT4G34830 | 2,44 | 0,07 | 1,13 | 0,06 | 9,51E-01 |
| AT4G34860 | 0,08 | 0,44 | 4,38 | 0,10 | 9,21E-01 |
| AT4G34890 | 2,69 | 0,95 | 1,23 | 0,77 | 4,40E-01 |
| AT4G34900 | 0,65 | -0,99 | 3,50 | -0,28 | 7,77E-01 |
| AT4G34950 | 4,16 | -1,13 | 0,94 | -1,20 | 2,30E-01 |
| AT4G34980 | 4,56 | -0,85 | 0,84 | -1,01 | 3,11E-01 |
| AT4G35000 | 3,13 | -1,11 | 1,43 | -0,78 | 4,38E-01 |
| AT4G35060 | 0,12 | 1,16 | 4,34 | 0,27 | 7,89E-01 |
| AT4G35090 | 2,83 | -0,74 | 1,17 | -0,63 | 5,31E-01 |
| AT4G35140 | 0,36 | -0,91 | 4,39 | -0,21 | 8,35E-01 |
| AT4G35180 | 0,66 | -0,70 | 2,93 | -0,24 | 8,11E-01 |
| AT4G35230 | 1,11 | -0,37 | 1,86 | -0,20 | 8,42E-01 |
| AT4G35240 | 1,02 | -2,22 | 2,10 | -1,06 | 2,89E-01 |
| AT4G35260 | 0,53 | -2,24 | 3,19 | -0,70 | 4,82E-01 |
| AT4G35300 | 1,43 | -0,78 | 2,16 | -0,36 | 7,17E-01 |
| AT4G35320 | 0,51 | -0,59 | 3,19 | -0,18 | 8,54E-01 |
| AT4G35360 | 0,54 | -2,92 | 4,01 | -0,73 | 4,66E-01 |
| AT4G35440 | 0,24 | 0,88 | 4,36 | 0,20 | 8,41E-01 |
| AT4G35570 | 0,29 | 0,24 | 4,33 | 0,06 | 9,55E-01 |
| AT4G35580 | 0,56 | -0,73 | 3,05 | -0,24 | 8,12E-01 |
| AT4G35750 | 1,89 | 0,20 | 1,28 | 0,15 | 8,77E-01 |
| AT4G35780 | 0,15 | 1,20 | 4,34 | 0,28 | 7,81E-01 |
| AT4G35785 | 0,84 | 1,86 | 2,77 | 0,67 | 5,02E-01 |
| AT4G35790 | 4,32 | 0,58 | 1,05 | 0,55 | 5,79E-01 |
| AT4G35800 | 5,60 | -0,33 | 0,76 | -0,43 | 6,65E-01 |
| AT4G35920 | 0,70 | -3,05 | 4,27 | -0,71 | 4,76E-01 |
| AT4G36010 | 0,74 | 0,27 | 2,78 | 0,10 | 9,23E-01 |
| AT4G36030 | 1,21 | -1,22 | 2,17 | -0,56 | 5,73E-01 |
| AT4G36040 | 3,49 | -0,54 | 0,97 | -0,55 | 5,79E-01 |
| AT4G36050 | 0,52 | 2,72 | 4,30 | 0,63 | 5,26E-01 |
| AT4G36070 | 0,08 | 0,44 | 4,38 | 0,10 | 9,21E-01 |
| AT4G36140 | 2,98 | 0,66 | 1,07 | 0,62 | 5,37E-01 |
| AT4G36220 | 1,38 | 0,17 | 1,54 | 0,11 | 9,12E-01 |
| AT4G36270 | 0,14 | 0,76 | 4,35 | 0,18 | 8,61E-01 |
| AT4G36280 | 0,08 | 0,44 | 4,38 | 0,10 | 9,21E-01 |
| AT4G36360 | 3,63 | -0,78 | 1,15 | -0,68 | 4,99E-01 |
| AT4G36380 | 0,24 | 0,00 | 4,41 | 0,00 | 1,00E+00 |
| AT4G36390 | 0,60 | -0,49 | 2,46 | -0,20 | 8,43E-01 |
| AT4G36460 | 0,50 | 0,88 | 3,19 | 0,27 | 7,84E-01 |
| AT4G36500 | 0,97 | -0,99 | 2,22 | -0,44 | 6,57E-01 |
| AT4G36520 | 1,04 | 1,80 | 1,81 | 1,00 | 3,19E-01 |
| AT4G36550 | 0,53 | 1,14 | 3,14 | 0,36 | 7,17E-01 |
| AT4G36630 | 4,11 | -0,79 | 1,18 | -0,67 | 5,06E-01 |
| AT4G36640 | 1,54 | -0,34 | 1,75 | -0,19 | 8,48E-01 |
| AT4G36648 | 2,47 | 0,91 | 1,50 | 0,61 | 5,44E-01 |
| AT4G36670 | 0,44 | 0,86 | 3,35 | 0,26 | 7,98E-01 |
| AT4G36690 | 0,56 | -0,17 | 3,27 | -0,05 | 9,58E-01 |
| AT4G36730 | 0,51 | -1,70 | 3,24 | -0,53 | 5,99E-01 |
| AT4G36770 | 0,41 | -0,16 | 4,13 | -0,04 | 9,70E-01 |
| AT4G36800 | 0,60 | 1,06 | 3,03 | 0,35 | 7,27E-01 |

| | | | | | |
|---|---|---|---|---|---|
| AT4G36808 | 0,88 | 2,02 | 2,39 | 0,85 | 3,97E-01 |
| AT4G36860 | 0,41 | -1,12 | 4,30 | -0,26 | 7,94E-01 |
| AT4G36890 | 0,60 | -1,86 | 3,82 | -0,49 | 6,26E-01 |
| AT4G36970 | 0,18 | -1,78 | 4,41 | -0,40 | 6,87E-01 |
| AT4G36980 | 0,30 | 0,24 | 4,35 | 0,05 | 9,57E-01 |
| AT4G37100 | 0,62 | -1,78 | 2,99 | -0,60 | 5,51E-01 |
| AT4G37270 | 0,72 | 1,20 | 2,47 | 0,49 | 6,26E-01 |
| AT4G37280 | 0,63 | -2,82 | 3,59 | -0,79 | 4,32E-01 |
| AT4G37370 | 0,86 | -0,43 | 2,43 | -0,18 | 8,61E-01 |
| AT4G37432 | 0,86 | 2,53 | 3,35 | 0,76 | 4,50E-01 |
| AT4G37460 | 2,98 | 0,24 | 1,08 | 0,22 | 8,26E-01 |
| AT4G37470 | 1,39 | 0,35 | 1,78 | 0,20 | 8,44E-01 |
| AT4G37510 | 0,72 | 0,46 | 2,35 | 0,19 | 8,46E-01 |
| AT4G37540 | 0,41 | -1,78 | 4,41 | -0,40 | 6,87E-01 |
| AT4G37553 | 0,42 | 0,03 | 4,31 | 0,01 | 9,94E-01 |
| AT4G37590 | 0,76 | 2,12 | 2,49 | 0,85 | 3,95E-01 |
| AT4G37610 | 0,33 | -0,15 | 3,58 | -0,04 | 9,66E-01 |
| AT4G37640 | 1,71 | 0,82 | 1,43 | 0,57 | 5,66E-01 |
| AT4G37740 | 0,08 | 0,00 | 4,40 | 0,00 | 1,00E+00 |
| AT4G37800 | 0,23 | 0,19 | 4,38 | 0,04 | 9,66E-01 |
| AT4G37830 | 0,60 | -1,95 | 2,57 | -0,76 | 4,50E-01 |
| AT4G37850 | 0,24 | -2,36 | 4,39 | -0,54 | 5,91E-01 |
| AT4G37870 | 0,70 | 0,48 | 2,21 | 0,22 | 8,28E-01 |
| AT4G37900 | 1,96 | 0,45 | 1,50 | 0,30 | 7,65E-01 |
| AT4G37910 | 5,28 | -0,44 | 0,77 | -0,57 | 5,67E-01 |
| AT4G37920 | 0,32 | 0,53 | 4,33 | 0,12 | 9,02E-01 |
| AT4G37930 | 4,20 | 0,46 | 0,96 | 0,48 | 6,30E-01 |
| AT4G38010 | 0,08 | 0,44 | 4,38 | 0,10 | 9,21E-01 |
| AT4G38050 | 0,35 | 1,56 | 4,33 | 0,36 | 7,19E-01 |
| AT4G38120 | 0,89 | -0,27 | 2,11 | -0,13 | 8,98E-01 |
| AT4G38130 | 0,71 | 0,94 | 2,03 | 0,46 | 6,43E-01 |
| AT4G38180 | 0,60 | -0,29 | 2,21 | -0,13 | 8,95E-01 |
| AT4G38210 | 0,75 | 0,53 | 4,29 | 0,12 | 9,01E-01 |
| AT4G38230 | 0,40 | -1,21 | 3,37 | -0,36 | 7,19E-01 |
| AT4G38240 | 0,62 | 1,27 | 3,63 | 0,35 | 7,28E-01 |
| AT4G09695 | 1,25 | -0,20 | 1,54 | -0,13 | 8,99E-01 |
| AT4G38300 | 0,68 | 0,84 | 3,53 | 0,24 | 8,11E-01 |
| AT4G38350 | 1,14 | 0,79 | 1,88 | 0,42 | 6,74E-01 |
| AT4G38380 | 0,43 | 0,26 | 4,12 | 0,06 | 9,50E-01 |
| AT4G38510 | 2,20 | -0,44 | 1,28 | -0,35 | 7,29E-01 |
| AT4G38520 | 0,55 | -0,96 | 2,31 | -0,41 | 6,79E-01 |
| AT4G38545 | 0,17 | 0,82 | 4,35 | 0,19 | 8,51E-01 |
| AT4G38552 | 2,97 | 0,49 | 1,19 | 0,41 | 6,78E-01 |
| AT4G38570 | 0,81 | 1,06 | 2,74 | 0,39 | 6,98E-01 |
| AT4G38630 | 0,74 | -0,57 | 1,99 | -0,29 | 7,74E-01 |
| AT4G38640 | 0,43 | -0,58 | 3,32 | -0,17 | 8,62E-01 |
| AT4G38660 | 0,37 | -2,28 | 4,38 | -0,52 | 6,03E-01 |
| AT4G38670 | 0,49 | -0,48 | 3,13 | -0,16 | 8,77E-01 |
| AT4G38680 | 1,18 | 1,02 | 1,83 | 0,56 | 5,78E-01 |
| AT4G38700 | 0,34 | -1,74 | 4,40 | -0,40 | 6,93E-01 |
| AT4G38710 | 1,00 | 0,22 | 1,83 | 0,12 | 9,05E-01 |
| AT4G38730 | 0,38 | -1,21 | 4,34 | -0,28 | 7,81E-01 |

| | | | | | |
|---|---|---|---|---|---|
| AT4G38740 | 1,15 | -1,66 | 1,93 | -0,86 | 3,88E-01 |
| AT4G38760 | 3,63 | 0,82 | 1,21 | 0,68 | 4,99E-01 |
| AT4G38780 | 1,10 | 3,75 | 2,47 | 1,52 | 1,29E-01 |
| AT4G38800 | 0,65 | -1,05 | 2,66 | -0,40 | 6,92E-01 |
| AT4G38810 | 0,68 | -0,67 | 2,64 | -0,25 | 7,99E-01 |
| AT4G38870 | 0,21 | 1,55 | 4,33 | 0,36 | 7,20E-01 |
| AT4G38890 | 0,48 | -0,66 | 3,38 | -0,19 | 8,46E-01 |
| AT4G38900 | 0,33 | -1,72 | 4,34 | -0,40 | 6,92E-01 |
| AT4G38932 | 0,58 | -0,81 | 2,62 | -0,31 | 7,56E-01 |
| AT4G38950 | 1,10 | -1,10 | 1,80 | -0,61 | 5,40E-01 |
| AT4G39100 | 0,33 | 0,83 | 4,31 | 0,19 | 8,47E-01 |
| AT4G39120 | 0,19 | 1,25 | 4,34 | 0,29 | 7,72E-01 |
| AT4G39140 | 0,65 | -0,58 | 3,79 | -0,15 | 8,78E-01 |
| AT4G39180 | 0,48 | 1,66 | 4,34 | 0,38 | 7,02E-01 |
| AT4G39190 | 0,69 | -0,17 | 2,50 | -0,07 | 9,46E-01 |
| AT4G39195 | 0,79 | 1,65 | 2,87 | 0,57 | 5,67E-01 |
| AT4G39240 | 1,33 | 1,60 | 2,07 | 0,77 | 4,41E-01 |
| AT4G39280 | 1,06 | 0,67 | 1,93 | 0,35 | 7,28E-01 |
| AT4G39460 | 0,85 | -0,45 | 2,31 | -0,20 | 8,44E-01 |
| AT4G39500 | 0,20 | -0,44 | 4,36 | -0,10 | 9,19E-01 |
| AT4G39510 | 0,43 | -0,70 | 4,39 | -0,16 | 8,73E-01 |
| AT4G39520 | 1,02 | -0,53 | 1,56 | -0,34 | 7,34E-01 |
| AT4G39550 | 0,17 | 1,52 | 4,33 | 0,35 | 7,26E-01 |
| AT4G39560 | 0,59 | -1,22 | 3,70 | -0,33 | 7,41E-01 |
| AT4G39570 | 0,55 | 2,53 | 3,71 | 0,68 | 4,96E-01 |
| AT4G39580 | 0,15 | 0,83 | 4,35 | 0,19 | 8,49E-01 |
| AT4G39590 | 0,08 | 0,44 | 4,38 | 0,10 | 9,21E-01 |
| AT4G39630 | 0,05 | 0,00 | 4,41 | 0,00 | 1,00E+00 |
| AT4G39680 | 0,30 | 0,71 | 4,36 | 0,16 | 8,71E-01 |
| AT4G39710 | 0,56 | -0,65 | 3,97 | -0,16 | 8,71E-01 |
| AT4G39780 | 0,21 | 0,68 | 4,35 | 0,16 | 8,76E-01 |
| AT4G39800 | 2,68 | -2,03 | 1,24 | -1,64 | 1,01E-01 |
| AT4G39820 | 0,46 | 0,82 | 4,04 | 0,20 | 8,40E-01 |
| AT4G39838 | 1,07 | -0,45 | 1,86 | -0,24 | 8,11E-01 |
| AT4G39865 | 0,25 | -2,19 | 4,38 | -0,50 | 6,17E-01 |
| AT4G39910 | 0,28 | 0,08 | 4,38 | 0,02 | 9,85E-01 |
| AT4G40000 | 0,43 | 0,09 | 4,11 | 0,02 | 9,83E-01 |
| AT4G40020 | 0,75 | -1,07 | 2,61 | -0,41 | 6,83E-01 |
| AT4G40030 | 1,32 | -0,22 | 1,74 | -0,13 | 9,00E-01 |
| AT4G40040 | 3,01 | 3,36 | 1,50 | 2,24 | 2,53E-02 |
| AT4G40045 | 0,12 | 1,16 | 4,34 | 0,27 | 7,89E-01 |
| AT4G40050 | 1,65 | -0,14 | 1,44 | -0,10 | 9,22E-01 |
| AT4G40060 | 1,22 | 0,94 | 1,94 | 0,49 | 6,27E-01 |
| AT4G40065 | 1,04 | 0,33 | 1,72 | 0,19 | 8,47E-01 |
| AT5G01010 | 1,03 | 0,91 | 2,21 | 0,41 | 6,82E-01 |
| AT5G01020 | 1,61 | 0,09 | 1,43 | 0,06 | 9,50E-01 |
| AT5G01100 | 0,27 | -0,81 | 4,37 | -0,19 | 8,53E-01 |
| AT5G01215 | 1,09 | 1,00 | 1,87 | 0,53 | 5,93E-01 |
| AT5G01220 | 0,12 | 1,23 | 4,34 | 0,28 | 7,76E-01 |
| AT5G01270 | 0,34 | 0,36 | 4,34 | 0,08 | 9,34E-01 |
| AT5G00760 | 4,31 | -0,76 | 0,90 | -0,84 | 3,98E-01 |
| AT5G00765 | 3,11 | -0,02 | 1,05 | -0,02 | 9,85E-01 |

| | | | | | |
|---|---|---|---|---|---|
| AT5G01365 | 0,66 | 1,71 | 4,33 | 0,40 | 6,92E-01 |
| AT5G01400 | 1,70 | -0,26 | 1,77 | -0,15 | 8,82E-01 |
| AT5G01410 | 2,12 | 1,40 | 1,53 | 0,92 | 3,58E-01 |
| AT5G01540 | 0,46 | 0,10 | 4,03 | 0,02 | 9,80E-01 |
| AT5G01560 | 1,57 | -0,02 | 1,50 | -0,02 | 9,87E-01 |
| AT5G01600 | 1,52 | -0,22 | 1,52 | -0,14 | 8,87E-01 |
| AT5G01630 | 1,03 | 0,92 | 2,54 | 0,36 | 7,16E-01 |
| AT5G01700 | 0,62 | -0,78 | 4,29 | -0,18 | 8,56E-01 |
| AT5G01710 | 1,65 | -1,76 | 1,52 | -1,16 | 2,47E-01 |
| AT5G01720 | 0,49 | 1,52 | 3,95 | 0,38 | 7,00E-01 |
| AT5G01730 | 0,29 | 0,65 | 4,32 | 0,15 | 8,80E-01 |
| AT5G01770 | 0,54 | -0,56 | 3,71 | -0,15 | 8,80E-01 |
| AT5G01780 | 0,19 | 0,86 | 4,34 | 0,20 | 8,42E-01 |
| AT5G01890 | 0,64 | 0,80 | 2,62 | 0,31 | 7,59E-01 |
| AT5G01930 | 1,04 | -3,00 | 2,98 | -1,01 | 3,14E-01 |
| AT5G01950 | 0,59 | -1,69 | 2,62 | -0,65 | 5,18E-01 |
| AT5G01970 | 1,20 | 1,59 | 1,97 | 0,81 | 4,20E-01 |
| AT5G01990 | 1,12 | -1,49 | 1,92 | -0,77 | 4,39E-01 |
| AT5G02130 | 0,08 | 0,00 | 4,40 | 0,00 | 1,00E+00 |
| AT5G02150 | 1,74 | -1,10 | 1,74 | -0,63 | 5,27E-01 |
| AT5G02290 | 0,48 | -1,05 | 2,70 | -0,39 | 6,96E-01 |
| AT5G02380 | 2,77 | 1,03 | 1,47 | 0,70 | 4,85E-01 |
| AT5G02490 | 4,18 | 0,41 | 0,91 | 0,45 | 6,51E-01 |
| AT5G02502 | 0,33 | 0,54 | 4,36 | 0,12 | 9,01E-01 |
| AT5G02530 | 0,37 | -0,08 | 4,35 | -0,02 | 9,86E-01 |
| AT5G02670 | 0,44 | 0,65 | 3,24 | 0,20 | 8,42E-01 |
| AT5G02700 | 1,99 | -0,01 | 1,53 | -0,01 | 9,93E-01 |
| AT5G02770 | 0,75 | -0,95 | 2,25 | -0,42 | 6,72E-01 |
| AT5G02800 | 0,65 | -2,21 | 2,87 | -0,77 | 4,40E-01 |
| AT5G02810 | 1,73 | -1,50 | 1,58 | -0,95 | 3,41E-01 |
| AT5G02830 | 0,85 | 1,02 | 2,00 | 0,51 | 6,10E-01 |
| AT5G02890 | 0,18 | 0,00 | 4,41 | 0,00 | 1,00E+00 |
| AT5G02940 | 0,88 | 1,50 | 2,44 | 0,61 | 5,40E-01 |
| AT5G02960 | 2,13 | -1,27 | 1,41 | -0,90 | 3,68E-01 |
| AT5G03040 | 2,14 | -0,67 | 1,15 | -0,58 | 5,62E-01 |
| AT5G03140 | 0,48 | 0,56 | 2,66 | 0,21 | 8,34E-01 |
| AT5G03170 | 0,32 | -2,18 | 4,38 | -0,50 | 6,18E-01 |
| AT5G03204 | 0,29 | 0,00 | 4,39 | 0,00 | 1,00E+00 |
| AT5G03210 | 0,71 | 1,50 | 2,49 | 0,60 | 5,47E-01 |
| AT5G03340 | 1,55 | 2,17 | 1,79 | 1,21 | 2,26E-01 |
| AT5G03350 | 2,45 | 0,37 | 1,33 | 0,27 | 7,83E-01 |
| AT5G03380 | 0,84 | 2,51 | 2,32 | 1,08 | 2,78E-01 |
| AT5G03400 | 1,62 | 3,16 | 1,75 | 1,81 | 7,09E-02 |
| AT5G03430 | 0,65 | -0,48 | 3,04 | -0,16 | 8,74E-01 |
| AT5G03450 | 0,39 | -1,37 | 4,36 | -0,31 | 7,53E-01 |
| AT5G03555 | 1,08 | 0,20 | 1,90 | 0,11 | 9,14E-01 |
| AT5G03560 | 0,43 | -0,24 | 4,14 | -0,06 | 9,55E-01 |
| AT5G03630 | 0,55 | 0,42 | 3,13 | 0,14 | 8,93E-01 |
| AT5G03700 | 0,11 | 0,65 | 4,36 | 0,15 | 8,81E-01 |
| AT5G03730 | 0,62 | 0,61 | 2,55 | 0,24 | 8,10E-01 |
| AT5G03760 | 0,44 | -1,86 | 4,16 | -0,45 | 6,54E-01 |
| AT5G03800 | 0,30 | 0,30 | 4,34 | 0,07 | 9,45E-01 |

| | | | | | |
|---|---|---|---|---|---|
| AT5G03850 | 0,83 | -3,54 | 2,99 | -1,18 | 2,37E-01 |
| AT5G03880 | 0,57 | 1,81 | 3,19 | 0,57 | 5,71E-01 |
| AT5G03910 | 0,83 | 1,09 | 2,32 | 0,47 | 6,39E-01 |
| AT5G03940 | 2,83 | -0,86 | 1,04 | -0,82 | 4,10E-01 |
| AT5G03970 | 0,19 | 1,40 | 4,33 | 0,32 | 7,46E-01 |
| AT5G04020 | 0,78 | 0,02 | 1,90 | 0,01 | 9,90E-01 |
| AT5G04130 | 0,05 | 0,00 | 4,41 | 0,00 | 1,00E+00 |
| AT5G04267 | 0,16 | 0,00 | 4,41 | 0,00 | 1,00E+00 |
| AT5G04340 | 1,00 | -0,80 | 1,98 | -0,40 | 6,86E-01 |
| AT5G00955 | 1,17 | 2,02 | 1,90 | 1,06 | 2,89E-01 |
| AT5G04420 | 0,49 | -0,99 | 3,88 | -0,26 | 7,98E-01 |
| AT5G04430 | 1,54 | 1,38 | 1,69 | 0,81 | 4,15E-01 |
| AT5G04470 | 0,16 | 0,92 | 4,35 | 0,21 | 8,33E-01 |
| AT5G04480 | 0,41 | 0,37 | 3,27 | 0,11 | 9,09E-01 |
| AT5G04540 | 1,96 | -0,97 | 1,34 | -0,72 | 4,69E-01 |
| AT5G04550 | 2,33 | -0,31 | 1,24 | -0,25 | 8,02E-01 |
| AT5G04670 | 1,32 | 2,31 | 1,86 | 1,24 | 2,14E-01 |
| AT5G04700 | 0,30 | -0,02 | 4,35 | 0,00 | 9,96E-01 |
| AT5G04710 | 0,12 | 1,16 | 4,34 | 0,27 | 7,89E-01 |
| AT5G04730 | 1,06 | -0,07 | 1,95 | -0,04 | 9,72E-01 |
| AT5G04760 | 0,48 | 0,72 | 4,33 | 0,17 | 8,68E-01 |
| AT5G04830 | 0,60 | 0,79 | 3,11 | 0,25 | 7,99E-01 |
| AT5G04850 | 0,17 | 1,15 | 4,34 | 0,27 | 7,91E-01 |
| AT5G04853 | 3,91 | 3,40 | 1,45 | 2,34 | 1,93E-02 |
| AT5G04860 | 0,46 | -2,42 | 4,15 | -0,58 | 5,59E-01 |
| AT5G04870 | 0,78 | 0,21 | 2,20 | 0,10 | 9,23E-01 |
| AT5G04940 | 0,64 | -1,94 | 2,89 | -0,67 | 5,02E-01 |
| AT5G04950 | 0,11 | 0,00 | 4,41 | 0,00 | 1,00E+00 |
| AT5G04990 | 0,74 | -3,95 | 3,40 | -1,16 | 2,46E-01 |
| AT5G05060 | 0,37 | 0,00 | 4,40 | 0,00 | 1,00E+00 |
| AT5G05080 | 0,77 | -1,06 | 4,36 | -0,24 | 8,07E-01 |
| AT5G05085 | 0,35 | -0,45 | 3,57 | -0,13 | 9,00E-01 |
| AT5G05100 | 0,72 | -0,91 | 2,82 | -0,32 | 7,47E-01 |
| AT5G01005 | 0,75 | 0,21 | 2,88 | 0,07 | 9,41E-01 |
| AT5G05200 | 1,99 | 1,11 | 1,68 | 0,66 | 5,10E-01 |
| AT5G05260 | 0,23 | 1,22 | 4,33 | 0,28 | 7,79E-01 |
| AT5G01055 | 0,10 | 0,00 | 4,41 | 0,00 | 1,00E+00 |
| AT5G05480 | 0,31 | 0,23 | 3,59 | 0,06 | 9,49E-01 |
| AT5G05660 | 1,92 | -2,62 | 2,09 | -1,25 | 2,11E-01 |
| AT5G05670 | 0,17 | 0,61 | 4,37 | 0,14 | 8,88E-01 |
| AT5G05690 | 2,09 | -1,32 | 1,28 | -1,03 | 3,03E-01 |
| AT5G05700 | 0,69 | 1,58 | 2,89 | 0,54 | 5,86E-01 |
| AT5G05730 | 0,58 | -2,43 | 3,13 | -0,78 | 4,38E-01 |
| AT5G01095 | 0,95 | 2,86 | 4,29 | 0,67 | 5,04E-01 |
| AT5G05800 | 0,36 | 1,57 | 4,34 | 0,36 | 7,17E-01 |
| AT5G05980 | 2,96 | -0,61 | 1,29 | -0,47 | 6,37E-01 |
| AT5G06043 | 0,95 | -0,91 | 2,73 | -0,33 | 7,40E-01 |
| AT5G06060 | 0,53 | -0,85 | 3,27 | -0,26 | 7,94E-01 |
| AT5G06110 | 0,69 | -1,03 | 2,80 | -0,37 | 7,14E-01 |
| AT5G06140 | 0,36 | -0,65 | 4,33 | -0,15 | 8,81E-01 |
| AT5G06160 | 0,43 | 0,27 | 2,73 | 0,10 | 9,21E-01 |
| AT5G06165 | 4,41 | 0,32 | 1,07 | 0,30 | 7,62E-01 |

| | | | | | |
|---|---|---|---|---|---|
| AT5G01165 | 0,96 | 0,69 | 2,07 | 0,33 | 7,39E-01 |
| AT5G01195 | 1,58 | 0,09 | 1,66 | 0,05 | 9,59E-01 |
| AT5G06230 | 0,49 | 0,22 | 3,82 | 0,06 | 9,54E-01 |
| AT5G06260 | 0,46 | 0,69 | 3,23 | 0,21 | 8,32E-01 |
| AT5G06320 | 5,00 | 0,85 | 1,09 | 0,78 | 4,33E-01 |
| AT5G06330 | 0,08 | 0,00 | 4,40 | 0,00 | 1,00E+00 |
| AT5G06410 | 0,25 | 0,03 | 4,34 | 0,01 | 9,95E-01 |
| AT5G06430 | 0,51 | -2,04 | 4,34 | -0,47 | 6,38E-01 |
| AT5G06440 | 0,83 | -3,11 | 3,50 | -0,89 | 3,74E-01 |
| AT5G06520 | 0,44 | 2,37 | 4,31 | 0,55 | 5,83E-01 |
| AT5G06530 | 2,30 | 1,18 | 1,25 | 0,94 | 3,45E-01 |
| AT5G06540 | 0,48 | 2,53 | 4,30 | 0,59 | 5,57E-01 |
| AT5G06560 | 0,78 | -0,50 | 2,14 | -0,23 | 8,15E-01 |
| AT5G06600 | 5,36 | -0,24 | 0,82 | -0,30 | 7,66E-01 |
| AT5G06630 | 1,85 | 3,02 | 1,80 | 1,68 | 9,33E-02 |
| AT5G06640 | 3,20 | 3,28 | 1,60 | 2,05 | 4,03E-02 |
| AT5G06670 | 0,24 | 0,05 | 4,39 | 0,01 | 9,91E-01 |
| AT5G06690 | 0,35 | 0,91 | 3,57 | 0,26 | 7,98E-01 |
| AT5G06750 | 0,11 | 0,48 | 4,37 | 0,11 | 9,12E-01 |
| AT5G06770 | 0,36 | 0,38 | 4,37 | 0,09 | 9,31E-01 |
| AT5G06865 | 0,22 | 0,24 | 4,38 | 0,05 | 9,57E-01 |
| AT5G06970 | 0,87 | 0,34 | 1,90 | 0,18 | 8,58E-01 |
| AT5G01295 | 0,28 | -1,42 | 4,36 | -0,33 | 7,44E-01 |
| AT5G07120 | 0,92 | -0,01 | 2,30 | 0,00 | 9,97E-01 |
| AT5G07220 | 0,71 | 2,11 | 2,25 | 0,94 | 3,48E-01 |
| AT5G07250 | 0,62 | -1,78 | 3,12 | -0,57 | 5,69E-01 |
| AT5G07360 | 0,50 | 1,00 | 3,27 | 0,31 | 7,59E-01 |
| AT5G07590 | 0,15 | 1,16 | 4,34 | 0,27 | 7,88E-01 |
| AT5G07620 | 0,13 | 0,00 | 4,41 | 0,00 | 1,00E+00 |
| AT5G07675 | 0,20 | 0,00 | 4,41 | 0,00 | 1,00E+00 |
| AT5G07740 | 1,77 | -0,11 | 1,31 | -0,08 | 9,33E-01 |
| AT5G07810 | 0,26 | -0,48 | 4,38 | -0,11 | 9,13E-01 |
| AT5G07950 | 0,39 | -1,40 | 4,37 | -0,32 | 7,49E-01 |
| AT5G07970 | 3,91 | 0,89 | 1,04 | 0,86 | 3,89E-01 |
| AT5G07980 | 1,38 | -0,36 | 1,69 | -0,21 | 8,32E-01 |
| AT5G08040 | 0,14 | 0,00 | 4,41 | 0,00 | 1,00E+00 |
| AT5G08075 | 0,48 | -1,36 | 4,04 | -0,34 | 7,36E-01 |
| AT5G08110 | 0,94 | -0,73 | 2,15 | -0,34 | 7,35E-01 |
| AT5G08130 | 1,09 | -0,83 | 2,18 | -0,38 | 7,03E-01 |
| AT5G08160 | 1,05 | -3,07 | 2,07 | -1,49 | 1,37E-01 |
| AT5G08170 | 0,71 | -2,11 | 3,52 | -0,60 | 5,50E-01 |
| AT5G08180 | 2,30 | -0,59 | 1,15 | -0,52 | 6,04E-01 |
| AT5G01495 | 0,69 | -0,60 | 1,94 | -0,31 | 7,57E-01 |
| AT5G01505 | 1,02 | -1,00 | 1,82 | -0,55 | 5,80E-01 |
| AT5G01515 | 0,11 | 0,65 | 4,36 | 0,15 | 8,81E-01 |
| AT5G08230 | 0,62 | -0,54 | 3,05 | -0,18 | 8,59E-01 |
| AT5G08280 | 1,65 | 2,29 | 1,78 | 1,29 | 1,98E-01 |
| AT5G08305 | 0,42 | 0,75 | 4,13 | 0,18 | 8,56E-01 |
| AT5G08315 | 0,44 | -3,48 | 4,32 | -0,81 | 4,20E-01 |
| AT5G08350 | 0,26 | -0,30 | 4,37 | -0,07 | 9,46E-01 |
| AT5G08450 | 0,66 | 2,17 | 2,78 | 0,78 | 4,36E-01 |
| AT5G08520 | 1,35 | -2,38 | 1,74 | -1,37 | 1,71E-01 |

| | | | | | |
|---|---|---|---|---|---|
| AT5G08530 | 3,76 | 0,30 | 1,37 | 0,22 | 8,28E-01 |
| AT5G08550 | 1,83 | 0,35 | 1,36 | 0,26 | 7,98E-01 |
| AT5G08560 | 0,55 | 0,25 | 2,78 | 0,09 | 9,30E-01 |
| AT5G08650 | 1,18 | -1,20 | 1,92 | -0,63 | 5,32E-01 |
| AT5G08770 | 1,04 | 0,74 | 2,13 | 0,35 | 7,28E-01 |
| AT5G08790 | 1,51 | 0,67 | 1,45 | 0,47 | 6,41E-01 |
| AT5G09330 | 1,10 | -1,23 | 1,65 | -0,75 | 4,56E-01 |
| AT5G09443 | 0,26 | -0,48 | 4,38 | -0,11 | 9,13E-01 |
| AT5G09460 | 0,93 | 0,06 | 1,74 | 0,04 | 9,70E-01 |
| AT5G09510 | 0,57 | -0,26 | 4,01 | -0,07 | 9,48E-01 |
| AT5G09620 | 0,88 | 0,32 | 2,45 | 0,13 | 8,97E-01 |
| AT5G09650 | 0,62 | 0,25 | 2,57 | 0,10 | 9,22E-01 |
| AT5G09660 | 2,38 | 0,51 | 1,48 | 0,35 | 7,29E-01 |
| AT5G09670 | 2,53 | 0,05 | 1,13 | 0,05 | 9,62E-01 |
| AT5G09820 | 0,63 | -2,02 | 3,02 | -0,67 | 5,04E-01 |
| AT5G09850 | 0,67 | 0,82 | 3,08 | 0,26 | 7,91E-01 |
| AT5G09880 | 0,62 | 2,64 | 2,92 | 0,90 | 3,66E-01 |
| AT5G09890 | 0,52 | 2,51 | 3,88 | 0,65 | 5,17E-01 |
| AT5G09900 | 1,74 | -1,36 | 1,36 | -1,00 | 3,17E-01 |
| AT5G09930 | 0,08 | 0,69 | 4,36 | 0,16 | 8,75E-01 |
| AT5G10030 | 0,52 | 0,70 | 3,92 | 0,18 | 8,58E-01 |
| AT5G10235 | 0,28 | -0,02 | 4,35 | 0,00 | 9,97E-01 |
| AT5G10290 | 0,76 | 2,52 | 2,37 | 1,06 | 2,89E-01 |
| AT5G10330 | 0,57 | 0,67 | 2,66 | 0,25 | 8,02E-01 |
| AT5G10350 | 0,66 | -0,21 | 4,34 | -0,05 | 9,61E-01 |
| AT5G10360 | 1,44 | -0,99 | 1,48 | -0,67 | 5,01E-01 |
| AT5G10390 | 0,27 | -1,50 | 4,37 | -0,34 | 7,31E-01 |
| AT5G10400 | 0,23 | 0,00 | 4,41 | 0,00 | 1,00E+00 |
| AT5G10450 | 3,14 | -0,64 | 1,01 | -0,64 | 5,25E-01 |
| AT5G10470 | 0,57 | -0,76 | 3,16 | -0,24 | 8,09E-01 |
| AT5G10490 | 0,10 | 0,01 | 4,38 | 0,00 | 9,99E-01 |
| AT5G10550 | 0,86 | 2,66 | 2,30 | 1,16 | 2,46E-01 |
| AT5G10605 | 0,65 | -1,81 | 3,05 | -0,59 | 5,53E-01 |
| AT5G10650 | 0,25 | 0,26 | 4,33 | 0,06 | 9,51E-01 |
| AT5G10690 | 0,38 | 0,10 | 4,33 | 0,02 | 9,82E-01 |
| AT5G01815 | 0,80 | 1,38 | 2,17 | 0,64 | 5,24E-01 |
| AT5G10700 | 0,56 | -1,39 | 3,20 | -0,43 | 6,64E-01 |
| AT5G10730 | 0,23 | 0,00 | 4,41 | 0,00 | 1,00E+00 |
| AT5G10760 | 4,94 | 0,38 | 0,85 | 0,45 | 6,54E-01 |
| AT5G10770 | 0,94 | 1,71 | 2,26 | 0,76 | 4,50E-01 |
| AT5G10800 | 0,30 | 0,24 | 4,35 | 0,05 | 9,57E-01 |
| AT5G10820 | 0,63 | 2,28 | 4,31 | 0,53 | 5,97E-01 |
| AT5G10840 | 2,05 | -1,42 | 1,31 | -1,09 | 2,77E-01 |
| AT5G10860 | 1,76 | -0,15 | 1,48 | -0,10 | 9,18E-01 |
| AT5G10930 | 1,14 | -0,76 | 1,61 | -0,47 | 6,36E-01 |
| AT5G10940 | 0,77 | -0,36 | 3,50 | -0,10 | 9,19E-01 |
| AT5G10965 | 0,75 | -1,21 | 2,22 | -0,54 | 5,87E-01 |
| AT5G10980 | 0,55 | 1,31 | 3,79 | 0,35 | 7,30E-01 |
| AT5G11000 | 0,45 | -0,51 | 4,16 | -0,12 | 9,03E-01 |
| AT5G11070 | 1,00 | 0,32 | 2,02 | 0,16 | 8,75E-01 |
| AT5G11180 | 0,81 | 1,01 | 2,29 | 0,44 | 6,60E-01 |
| AT5G11210 | 1,06 | -2,65 | 3,07 | -0,86 | 3,88E-01 |

| | | | | | |
|---|---|---|---|---|---|
| AT5G11250 | 1,58 | -0,09 | 1,38 | -0,07 | 9,46E-01 |
| AT5G11270 | 0,22 | -0,88 | 4,38 | -0,20 | 8,40E-01 |
| AT5G11330 | 1,01 | -0,17 | 1,70 | -0,10 | 9,21E-01 |
| AT5G11480 | 0,46 | 0,00 | 3,28 | 0,00 | 9,99E-01 |
| AT5G11520 | 1,24 | -0,05 | 1,90 | -0,03 | 9,77E-01 |
| AT5G11550 | 0,28 | 0,09 | 4,33 | 0,02 | 9,83E-01 |
| AT5G11560 | 3,42 | 0,56 | 0,93 | 0,60 | 5,46E-01 |
| AT5G11700 | 1,99 | -0,04 | 1,33 | -0,03 | 9,79E-01 |
| AT5G11770 | 0,54 | 1,25 | 2,77 | 0,45 | 6,52E-01 |
| AT5G11850 | 0,25 | 1,68 | 4,32 | 0,39 | 6,97E-01 |
| AT5G11860 | 0,12 | 1,16 | 4,34 | 0,27 | 7,89E-01 |
| AT5G11910 | 0,27 | 0,37 | 4,33 | 0,09 | 9,32E-01 |
| AT5G11960 | 0,36 | 1,83 | 4,33 | 0,42 | 6,73E-01 |
| AT5G12010 | 0,59 | -0,13 | 3,01 | -0,04 | 9,66E-01 |
| AT5G12080 | 0,25 | 0,41 | 4,33 | 0,09 | 9,25E-01 |
| AT5G12120 | 2,73 | -0,64 | 1,14 | -0,56 | 5,73E-01 |
| AT5G12140 | 0,22 | 0,99 | 4,34 | 0,23 | 8,19E-01 |
| AT5G12150 | 4,21 | -0,51 | 1,19 | -0,43 | 6,70E-01 |
| AT5G12200 | 0,44 | 1,82 | 3,26 | 0,56 | 5,76E-01 |
| AT5G12250 | 2,28 | 0,64 | 1,27 | 0,51 | 6,11E-01 |
| AT5G12310 | 0,42 | -0,12 | 3,35 | -0,04 | 9,70E-01 |
| AT5G12370 | 4,35 | -0,88 | 0,94 | -0,94 | 3,48E-01 |
| AT5G12390 | 0,11 | 0,65 | 4,36 | 0,15 | 8,81E-01 |
| AT5G12430 | 1,28 | -2,09 | 2,11 | -0,99 | 3,20E-01 |
| AT5G12440 | 0,88 | -0,75 | 2,51 | -0,30 | 7,65E-01 |
| AT5G12470 | 4,58 | -1,63 | 0,96 | -1,70 | 8,94E-02 |
| AT5G12480 | 0,34 | 1,25 | 4,34 | 0,29 | 7,74E-01 |
| AT5G12860 | 2,32 | 0,60 | 1,66 | 0,36 | 7,18E-01 |
| AT5G12890 | 0,77 | -0,66 | 3,54 | -0,19 | 8,52E-01 |
| AT5G12900 | 0,23 | 1,21 | 4,34 | 0,28 | 7,81E-01 |
| AT5G13000 | 3,30 | 0,14 | 0,91 | 0,16 | 8,76E-01 |
| AT5G13020 | 0,81 | -1,41 | 2,08 | -0,68 | 4,98E-01 |
| AT5G13050 | 0,42 | -0,35 | 4,17 | -0,08 | 9,33E-01 |
| AT5G13090 | 0,10 | 0,21 | 4,38 | 0,05 | 9,62E-01 |
| AT5G13120 | 0,17 | 0,00 | 4,41 | 0,00 | 1,00E+00 |
| AT5G13140 | 0,23 | 0,00 | 4,41 | 0,00 | 1,00E+00 |
| AT5G13170 | 0,48 | 0,72 | 4,33 | 0,17 | 8,68E-01 |
| AT5G13190 | 0,99 | 0,65 | 1,85 | 0,35 | 7,27E-01 |
| AT5G02055 | 0,48 | 0,10 | 2,85 | 0,03 | 9,73E-01 |
| AT5G13225 | 0,48 | 0,10 | 2,85 | 0,03 | 9,73E-01 |
| AT5G13270 | 0,40 | -1,30 | 4,16 | -0,31 | 7,55E-01 |
| AT5G13300 | 0,77 | 0,47 | 2,34 | 0,20 | 8,42E-01 |
| AT5G13400 | 0,43 | 1,99 | 4,33 | 0,46 | 6,46E-01 |
| AT5G13430 | 0,54 | 0,12 | 3,73 | 0,03 | 9,75E-01 |
| AT5G13440 | 0,29 | 0,53 | 4,38 | 0,12 | 9,04E-01 |
| AT5G13450 | 0,96 | 0,71 | 2,15 | 0,33 | 7,41E-01 |
| AT5G13550 | 0,46 | 0,82 | 3,96 | 0,21 | 8,35E-01 |
| AT5G13560 | 1,05 | -0,73 | 2,03 | -0,36 | 7,19E-01 |
| AT5G13590 | 1,32 | -0,46 | 1,76 | -0,26 | 7,93E-01 |
| AT5G13680 | 1,03 | -0,38 | 1,71 | -0,22 | 8,24E-01 |
| AT5G13710 | 0,59 | -1,19 | 3,01 | -0,39 | 6,93E-01 |
| AT5G13780 | 0,62 | 0,50 | 3,00 | 0,17 | 8,68E-01 |

| | | | | | |
|---|---|---|---|---|---|
| AT5G13800 | 0,74 | -2,50 | 2,58 | -0,97 | 3,34E-01 |
| AT5G13825 | 0,39 | 0,00 | 4,31 | 0,00 | 1,00E+00 |
| AT5G13840 | 2,03 | -0,14 | 1,36 | -0,10 | 9,17E-01 |
| AT5G13950 | 1,63 | -0,26 | 1,44 | -0,18 | 8,58E-01 |
| AT5G13990 | 0,12 | 0,00 | 4,38 | 0,00 | 1,00E+00 |
| AT5G14040 | 2,00 | -1,33 | 1,38 | -0,96 | 3,37E-01 |
| AT5G14100 | 0,11 | 0,65 | 4,36 | 0,15 | 8,81E-01 |
| AT5G14170 | 1,07 | -0,91 | 1,96 | -0,46 | 6,42E-01 |
| AT5G14210 | 0,50 | -0,89 | 3,38 | -0,26 | 7,93E-01 |
| AT5G14240 | 0,57 | -2,01 | 2,70 | -0,74 | 4,57E-01 |
| AT5G14320 | 0,97 | -0,31 | 1,74 | -0,18 | 8,58E-01 |
| AT5G14390 | 0,52 | -0,66 | 4,05 | -0,16 | 8,71E-01 |
| AT5G14420 | 0,18 | 1,24 | 4,33 | 0,29 | 7,74E-01 |
| AT5G14520 | 3,02 | -2,81 | 1,24 | -2,27 | 2,33E-02 |
| AT5G14530 | 0,86 | 1,05 | 2,83 | 0,37 | 7,11E-01 |
| AT5G14540 | 0,54 | -1,66 | 3,88 | -0,43 | 6,68E-01 |
| AT5G02245 | 0,10 | 0,00 | 4,41 | 0,00 | 1,00E+00 |
| AT5G14570 | 2,87 | 1,63 | 1,21 | 1,35 | 1,78E-01 |
| AT5G14580 | 1,26 | -1,24 | 2,54 | -0,49 | 6,24E-01 |
| AT5G14590 | 0,98 | 0,43 | 1,87 | 0,23 | 8,18E-01 |
| AT5G14700 | 0,24 | 0,03 | 4,39 | 0,01 | 9,94E-01 |
| AT5G14720 | 0,96 | 0,86 | 1,81 | 0,47 | 6,36E-01 |
| AT5G14770 | 0,20 | 1,19 | 4,35 | 0,27 | 7,84E-01 |
| AT5G14800 | 5,34 | -1,61 | 0,97 | -1,65 | 9,85E-02 |
| AT5G14820 | 0,29 | 0,24 | 4,33 | 0,06 | 9,55E-01 |
| AT5G14940 | 1,18 | 1,85 | 2,24 | 0,83 | 4,08E-01 |
| AT5G14950 | 1,52 | 0,21 | 1,38 | 0,15 | 8,78E-01 |
| AT5G15020 | 2,14 | 1,12 | 1,40 | 0,80 | 4,23E-01 |
| AT5G15050 | 0,10 | 0,01 | 4,38 | 0,00 | 9,99E-01 |
| AT5G15200 | 4,33 | -0,92 | 0,86 | -1,07 | 2,84E-01 |
| AT5G15210 | 0,28 | -0,02 | 4,35 | 0,00 | 9,96E-01 |
| AT5G15250 | 0,05 | 0,00 | 4,41 | 0,00 | 1,00E+00 |
| AT5G15360 | 0,52 | 1,16 | 2,62 | 0,44 | 6,57E-01 |
| AT5G15400 | 2,82 | 0,56 | 1,01 | 0,55 | 5,81E-01 |
| AT5G15410 | 1,41 | 1,51 | 1,76 | 0,86 | 3,91E-01 |
| AT5G15420 | 1,66 | 0,94 | 1,68 | 0,56 | 5,76E-01 |
| AT5G15450 | 1,23 | 2,38 | 1,87 | 1,27 | 2,04E-01 |
| AT5G15460 | 0,08 | 0,69 | 4,36 | 0,16 | 8,75E-01 |
| AT5G15470 | 0,21 | 0,45 | 4,37 | 0,10 | 9,17E-01 |
| AT5G15490 | 0,40 | -0,44 | 3,34 | -0,13 | 8,94E-01 |
| AT5G15537 | 0,21 | 0,01 | 4,34 | 0,00 | 9,98E-01 |
| AT5G15540 | 1,01 | 0,55 | 1,87 | 0,29 | 7,69E-01 |
| AT5G15580 | 0,70 | -2,02 | 2,45 | -0,83 | 4,09E-01 |
| AT5G15690 | 0,53 | -1,67 | 2,72 | -0,61 | 5,39E-01 |
| AT5G15700 | 0,61 | 0,24 | 2,32 | 0,11 | 9,16E-01 |
| AT5G15740 | 0,28 | -1,77 | 4,41 | -0,40 | 6,88E-01 |
| AT5G15780 | 0,22 | 0,24 | 4,38 | 0,05 | 9,57E-01 |
| AT5G15800 | 0,08 | 0,00 | 4,41 | 0,00 | 1,00E+00 |
| AT5G15802 | 0,08 | 0,00 | 4,40 | 0,00 | 1,00E+00 |
| AT5G15850 | 2,49 | 0,08 | 1,12 | 0,07 | 9,42E-01 |
| AT5G15860 | 0,19 | 1,40 | 4,33 | 0,32 | 7,46E-01 |
| AT5G16050 | 0,47 | 1,14 | 2,76 | 0,41 | 6,80E-01 |

| | | | | | |
|---|---|---|---|---|---|
| AT5G16070 | 0,93 | -1,94 | 2,04 | -0,95 | 3,41E-01 |
| AT5G16110 | 1,09 | -1,20 | 1,62 | -0,74 | 4,57E-01 |
| AT5G16200 | 0,36 | -3,19 | 4,33 | -0,74 | 4,61E-01 |
| AT5G16210 | 1,91 | 0,15 | 1,37 | 0,11 | 9,14E-01 |
| AT5G16220 | 0,18 | 0,00 | 4,41 | 0,00 | 1,00E+00 |
| AT5G16280 | 0,67 | -0,49 | 2,74 | -0,18 | 8,57E-01 |
| AT5G16360 | 0,08 | 0,69 | 4,36 | 0,16 | 8,75E-01 |
| AT5G16370 | 1,57 | 0,34 | 1,38 | 0,25 | 8,05E-01 |
| AT5G16380 | 0,25 | 1,15 | 4,34 | 0,27 | 7,91E-01 |
| AT5G16390 | 0,61 | -1,31 | 3,67 | -0,36 | 7,22E-01 |
| AT5G16480 | 0,56 | -2,66 | 4,36 | -0,61 | 5,42E-01 |
| AT5G16490 | 0,08 | 0,00 | 4,40 | 0,00 | 1,00E+00 |
| AT5G16520 | 0,47 | 1,68 | 4,17 | 0,40 | 6,88E-01 |
| AT5G16560 | 0,42 | 0,57 | 4,36 | 0,13 | 8,97E-01 |
| AT5G16610 | 0,08 | 0,69 | 4,36 | 0,16 | 8,75E-01 |
| AT5G16660 | 0,23 | 1,38 | 4,33 | 0,32 | 7,50E-01 |
| AT5G16680 | 0,78 | -0,61 | 2,23 | -0,28 | 7,83E-01 |
| AT5G16740 | 0,42 | -1,40 | 4,38 | -0,32 | 7,50E-01 |
| AT5G16750 | 0,89 | -0,05 | 1,97 | -0,03 | 9,79E-01 |
| AT5G16780 | 0,54 | -0,44 | 3,94 | -0,11 | 9,12E-01 |
| AT5G16800 | 0,21 | -1,41 | 4,36 | -0,32 | 7,46E-01 |
| AT5G16830 | 0,10 | 0,01 | 4,38 | 0,00 | 9,99E-01 |
| AT5G16980 | 0,08 | 0,44 | 4,38 | 0,10 | 9,21E-01 |
| AT5G16990 | 0,20 | 1,00 | 4,36 | 0,23 | 8,18E-01 |
| AT5G17000 | 0,20 | -1,80 | 4,41 | -0,41 | 6,83E-01 |
| AT5G17010 | 0,24 | 0,66 | 4,37 | 0,15 | 8,80E-01 |
| AT5G17165 | 3,90 | 2,47 | 1,56 | 1,59 | 1,12E-01 |
| AT5G17230 | 2,13 | -0,86 | 1,24 | -0,69 | 4,89E-01 |
| AT5G02645 | 2,33 | -1,60 | 1,24 | -1,29 | 1,97E-01 |
| AT5G02655 | 0,55 | 0,00 | 2,74 | 0,00 | 1,00E+00 |
| AT5G02665 | 1,25 | -1,98 | 2,04 | -0,97 | 3,33E-01 |
| AT5G02675 | 0,37 | 0,42 | 4,27 | 0,10 | 9,21E-01 |
| AT5G17290 | 0,65 | 0,37 | 3,02 | 0,12 | 9,02E-01 |
| AT5G17300 | 0,69 | -0,25 | 2,85 | -0,09 | 9,29E-01 |
| AT5G17320 | 0,38 | -0,31 | 4,36 | -0,07 | 9,43E-01 |
| AT5G17380 | 0,80 | 1,95 | 2,50 | 0,78 | 4,35E-01 |
| AT5G17420 | 0,40 | 2,13 | 4,32 | 0,49 | 6,22E-01 |
| AT5G17460 | 0,52 | 0,00 | 4,41 | 0,00 | 1,00E+00 |
| AT5G17520 | 0,17 | 0,00 | 4,41 | 0,00 | 1,00E+00 |
| AT5G17550 | 0,08 | 0,69 | 4,36 | 0,16 | 8,75E-01 |
| AT5G17600 | 0,90 | -0,03 | 2,14 | -0,01 | 9,89E-01 |
| AT5G17620 | 0,42 | -1,40 | 4,38 | -0,32 | 7,50E-01 |
| AT5G17640 | 0,35 | -2,36 | 4,38 | -0,54 | 5,90E-01 |
| AT5G17690 | 0,20 | 1,20 | 4,35 | 0,28 | 7,83E-01 |
| AT5G17710 | 1,85 | -1,21 | 1,21 | -1,00 | 3,19E-01 |
| AT5G17760 | 1,99 | -0,38 | 1,59 | -0,24 | 8,13E-01 |
| AT5G17770 | 0,49 | 0,62 | 3,20 | 0,19 | 8,46E-01 |
| AT5G17860 | 0,63 | 2,31 | 2,84 | 0,81 | 4,16E-01 |
| AT5G17880 | 1,61 | -2,34 | 1,74 | -1,35 | 1,79E-01 |
| AT5G17930 | 0,48 | 2,46 | 4,30 | 0,57 | 5,67E-01 |
| AT5G18050 | 0,24 | -2,36 | 4,39 | -0,54 | 5,91E-01 |
| AT5G18110 | 0,39 | 0,85 | 4,34 | 0,20 | 8,44E-01 |

| | | | | | |
|---|---|---|---|---|---|
| AT5G18190 | 0,63 | -0,45 | 2,51 | -0,18 | 8,58E-01 |
| AT5G18230 | 0,99 | 3,35 | 2,77 | 1,21 | 2,26E-01 |
| AT5G18240 | 1,40 | 0,18 | 2,92 | 0,06 | 9,50E-01 |
| AT5G18260 | 0,49 | -1,38 | 4,31 | -0,32 | 7,49E-01 |
| AT5G18280 | 0,49 | 1,01 | 3,11 | 0,32 | 7,46E-01 |
| AT5G18290 | 0,36 | -2,77 | 4,36 | -0,64 | 5,25E-01 |
| AT5G18340 | 0,58 | -2,27 | 3,88 | -0,58 | 5,59E-01 |
| AT5G18350 | 2,51 | -0,30 | 1,40 | -0,21 | 8,33E-01 |
| AT5G18360 | 2,96 | 0,52 | 1,05 | 0,50 | 6,20E-01 |
| AT5G18370 | 3,23 | -0,94 | 1,13 | -0,84 | 4,01E-01 |
| AT5G18380 | 4,02 | -2,61 | 0,91 | -2,88 | 4,02E-03 |
| AT5G18400 | 0,32 | -0,70 | 4,36 | -0,16 | 8,72E-01 |
| AT5G18410 | 3,15 | -0,99 | 1,10 | -0,90 | 3,68E-01 |
| AT5G02895 | 0,88 | 0,96 | 2,62 | 0,37 | 7,15E-01 |
| AT5G18475 | 0,21 | 1,71 | 4,33 | 0,39 | 6,93E-01 |
| AT5G18480 | 0,62 | 0,94 | 2,96 | 0,32 | 7,50E-01 |
| AT5G18490 | 1,80 | 2,73 | 1,73 | 1,57 | 1,15E-01 |
| AT5G18525 | 1,70 | -0,54 | 1,49 | -0,36 | 7,16E-01 |
| AT5G18570 | 0,43 | 2,03 | 4,00 | 0,51 | 6,13E-01 |
| AT5G18590 | 0,48 | 0,36 | 3,28 | 0,11 | 9,13E-01 |
| AT5G18620 | 0,19 | 0,86 | 4,34 | 0,20 | 8,42E-01 |
| AT5G18700 | 0,89 | -1,80 | 2,61 | -0,69 | 4,92E-01 |
| AT5G02935 | 0,05 | 0,00 | 4,41 | 0,00 | 1,00E+00 |
| AT5G19010 | 0,59 | 0,54 | 2,52 | 0,21 | 8,31E-01 |
| AT5G19130 | 0,62 | -0,39 | 2,56 | -0,15 | 8,80E-01 |
| AT5G19220 | 0,51 | -1,60 | 4,31 | -0,37 | 7,11E-01 |
| AT5G19330 | 2,99 | -1,07 | 1,15 | -0,93 | 3,54E-01 |
| AT5G19360 | 0,72 | -3,93 | 4,30 | -0,91 | 3,60E-01 |
| AT5G19410 | 0,62 | -0,68 | 2,81 | -0,24 | 8,09E-01 |
| AT5G19420 | 1,09 | -0,89 | 1,71 | -0,52 | 6,05E-01 |
| AT5G19450 | 0,87 | -1,30 | 1,76 | -0,74 | 4,60E-01 |
| AT5G19485 | 0,20 | -0,31 | 4,36 | -0,07 | 9,44E-01 |
| AT5G19510 | 1,83 | -0,63 | 1,35 | -0,47 | 6,41E-01 |
| AT5G19590 | 0,42 | 1,38 | 3,25 | 0,42 | 6,71E-01 |
| AT5G19660 | 0,36 | 0,59 | 4,31 | 0,14 | 8,92E-01 |
| AT5G19760 | 0,90 | -2,34 | 2,38 | -0,98 | 3,26E-01 |
| AT5G19770 | 1,75 | -3,13 | 1,67 | -1,87 | 6,09E-02 |
| AT5G19820 | 4,02 | -0,39 | 1,11 | -0,35 | 7,26E-01 |
| AT5G19850 | 0,75 | 2,47 | 3,61 | 0,68 | 4,94E-01 |
| AT5G19860 | 0,51 | 0,98 | 3,96 | 0,25 | 8,04E-01 |
| AT5G19900 | 0,73 | -2,30 | 2,98 | -0,77 | 4,40E-01 |
| AT5G19920 | 0,37 | -0,97 | 4,32 | -0,23 | 8,22E-01 |
| AT5G19930 | 0,12 | 0,49 | 4,36 | 0,11 | 9,11E-01 |
| AT5G19950 | 0,27 | -0,36 | 4,34 | -0,08 | 9,34E-01 |
| AT5G19980 | 0,33 | -0,25 | 4,34 | -0,06 | 9,54E-01 |
| AT5G20040 | 0,12 | 1,23 | 4,34 | 0,28 | 7,76E-01 |
| AT5G20160 | 0,08 | 0,00 | 4,40 | 0,00 | 1,00E+00 |
| AT5G20170 | 0,19 | 0,38 | 4,37 | 0,09 | 9,31E-01 |
| AT5G20225 | 0,17 | 1,50 | 4,33 | 0,35 | 7,29E-01 |
| AT5G20260 | 0,52 | 1,01 | 4,33 | 0,23 | 8,16E-01 |
| AT5G20270 | 0,14 | 0,09 | 4,38 | 0,02 | 9,84E-01 |
| AT5G20290 | 5,09 | -0,68 | 0,85 | -0,80 | 4,23E-01 |

| | | | | | |
|---|---|---|---|---|---|
| AT5G20300 | 1,26 | 0,68 | 2,42 | 0,28 | 7,78E-01 |
| AT5G20320 | 1,21 | 1,80 | 2,36 | 0,76 | 4,44E-01 |
| AT5G20360 | 0,79 | 0,56 | 2,17 | 0,26 | 7,97E-01 |
| AT5G20380 | 0,45 | 1,12 | 4,34 | 0,26 | 7,95E-01 |
| AT5G20490 | 1,59 | 1,01 | 1,90 | 0,53 | 5,96E-01 |
| AT5G20520 | 1,55 | -0,34 | 1,53 | -0,22 | 8,24E-01 |
| AT5G20570 | 0,58 | -1,45 | 3,01 | -0,48 | 6,29E-01 |
| AT5G20600 | 1,15 | 0,83 | 1,92 | 0,44 | 6,63E-01 |
| AT5G20620 | 0,96 | 0,13 | 2,28 | 0,06 | 9,53E-01 |
| AT5G20630 | 1,03 | -2,46 | 1,80 | -1,37 | 1,70E-01 |
| AT5G20650 | 0,87 | 2,11 | 2,97 | 0,71 | 4,77E-01 |
| AT5G20730 | 2,22 | -1,47 | 1,15 | -1,28 | 2,02E-01 |
| AT5G20820 | 0,08 | 0,00 | 4,40 | 0,00 | 1,00E+00 |
| AT5G20830 | 0,97 | -1,41 | 2,29 | -0,62 | 5,36E-01 |
| AT5G20840 | 1,18 | -0,24 | 1,57 | -0,15 | 8,80E-01 |
| AT5G20890 | 1,15 | 0,18 | 2,05 | 0,09 | 9,30E-01 |
| AT5G20920 | 0,64 | -0,59 | 3,22 | -0,18 | 8,55E-01 |
| AT5G20940 | 0,27 | 0,26 | 4,33 | 0,06 | 9,52E-01 |
| AT5G20950 | 3,45 | -0,16 | 1,06 | -0,15 | 8,78E-01 |
| AT5G03265 | 0,23 | 1,19 | 4,34 | 0,27 | 7,83E-01 |
| AT5G20980 | 0,81 | 2,06 | 2,83 | 0,73 | 4,66E-01 |
| AT5G20990 | 1,65 | -0,73 | 1,61 | -0,45 | 6,53E-01 |
| AT5G21040 | 0,32 | 0,27 | 4,34 | 0,06 | 9,51E-01 |
| AT5G21160 | 0,41 | 0,33 | 4,39 | 0,07 | 9,41E-01 |
| AT5G21274 | 0,23 | 1,38 | 4,33 | 0,32 | 7,50E-01 |
| AT5G21920 | 0,36 | 1,19 | 4,33 | 0,28 | 7,83E-01 |
| AT5G21940 | 4,72 | 0,06 | 0,91 | 0,06 | 9,49E-01 |
| AT5G21950 | 0,08 | 0,44 | 4,38 | 0,10 | 9,21E-01 |
| AT5G22010 | 0,59 | 1,51 | 2,97 | 0,51 | 6,10E-01 |
| AT5G22020 | 0,55 | -2,91 | 3,87 | -0,75 | 4,52E-01 |
| AT5G22030 | 0,76 | -2,14 | 3,33 | -0,64 | 5,21E-01 |
| AT5G22060 | 1,34 | 2,52 | 2,23 | 1,13 | 2,59E-01 |
| AT5G22080 | 0,35 | 1,28 | 3,51 | 0,36 | 7,16E-01 |
| AT5G22090 | 0,53 | 0,64 | 3,31 | 0,19 | 8,46E-01 |
| AT5G22250 | 1,15 | 1,79 | 2,09 | 0,86 | 3,92E-01 |
| AT5G22350 | 0,24 | 0,90 | 4,35 | 0,21 | 8,37E-01 |
| AT5G22360 | 0,12 | 1,23 | 4,34 | 0,28 | 7,76E-01 |
| AT5G22390 | 0,49 | 0,54 | 3,17 | 0,17 | 8,65E-01 |
| AT5G22440 | 0,73 | -2,82 | 2,95 | -0,96 | 3,38E-01 |
| AT5G22450 | 0,87 | -0,16 | 1,88 | -0,08 | 9,33E-01 |
| AT5G22505 | 1,58 | 0,21 | 2,12 | 0,10 | 9,22E-01 |
| AT5G22510 | 0,44 | -2,81 | 4,22 | -0,67 | 5,05E-01 |
| AT5G03425 | 0,62 | -0,89 | 3,65 | -0,24 | 8,07E-01 |
| AT5G22540 | 0,22 | -0,91 | 4,38 | -0,21 | 8,35E-01 |
| AT5G22570 | 0,26 | 0,01 | 4,33 | 0,00 | 9,98E-01 |
| AT5G22700 | 0,44 | 2,32 | 4,07 | 0,57 | 5,68E-01 |
| AT5G22740 | 0,58 | -1,30 | 3,80 | -0,34 | 7,32E-01 |
| AT5G22750 | 0,55 | 1,46 | 4,34 | 0,34 | 7,37E-01 |
| AT5G22770 | 2,48 | -1,23 | 1,18 | -1,04 | 2,97E-01 |
| AT5G03475 | 1,21 | -0,64 | 1,79 | -0,36 | 7,20E-01 |
| AT5G22780 | 1,47 | -1,02 | 1,49 | -0,68 | 4,94E-01 |
| AT5G22800 | 1,71 | -0,83 | 1,52 | -0,55 | 5,85E-01 |

| | | | | | |
|---|---|---|---|---|---|
| AT5G22850 | 0,23 | 1,70 | 4,32 | 0,39 | 6,94E-01 |
| AT5G22860 | 0,35 | 1,54 | 3,45 | 0,45 | 6,55E-01 |
| AT5G23040 | 0,63 | 0,01 | 2,63 | 0,00 | 9,97E-01 |
| AT5G23050 | 0,45 | -1,34 | 4,01 | -0,33 | 7,38E-01 |
| AT5G23060 | 1,11 | 1,64 | 1,80 | 0,91 | 3,63E-01 |
| AT5G23080 | 1,07 | -0,01 | 2,44 | 0,00 | 9,98E-01 |
| AT5G23240 | 0,89 | 0,90 | 2,22 | 0,41 | 6,84E-01 |
| AT5G23300 | 0,32 | 0,22 | 4,34 | 0,05 | 9,61E-01 |
| AT5G23390 | 0,47 | 2,05 | 3,93 | 0,52 | 6,03E-01 |
| AT5G23405 | 0,14 | 0,83 | 4,35 | 0,19 | 8,49E-01 |
| AT5G23430 | 0,90 | 0,63 | 2,11 | 0,30 | 7,66E-01 |
| AT5G23440 | 0,16 | 0,00 | 4,41 | 0,00 | 1,00E+00 |
| AT5G23450 | 0,83 | 1,01 | 2,81 | 0,36 | 7,20E-01 |
| AT5G23580 | 0,83 | -1,36 | 2,13 | -0,64 | 5,23E-01 |
| AT5G23590 | 0,61 | 1,74 | 3,63 | 0,48 | 6,31E-01 |
| AT5G23610 | 1,31 | 1,10 | 1,88 | 0,59 | 5,57E-01 |
| AT5G23630 | 2,35 | -0,63 | 1,16 | -0,54 | 5,89E-01 |
| AT5G23660 | 0,79 | 2,38 | 3,64 | 0,65 | 5,14E-01 |
| AT5G23740 | 0,24 | 0,03 | 4,39 | 0,01 | 9,94E-01 |
| AT5G23900 | 0,57 | 0,07 | 2,77 | 0,02 | 9,81E-01 |
| AT5G23940 | 0,08 | 0,00 | 4,40 | 0,00 | 1,00E+00 |
| AT5G23990 | 0,29 | 0,04 | 4,38 | 0,01 | 9,93E-01 |
| AT5G24120 | 0,77 | 0,67 | 2,37 | 0,28 | 7,76E-01 |
| AT5G24140 | 0,08 | 0,00 | 4,40 | 0,00 | 1,00E+00 |
| AT5G24150 | 0,21 | 1,57 | 4,33 | 0,36 | 7,16E-01 |
| AT5G24160 | 0,65 | 0,74 | 2,84 | 0,26 | 7,93E-01 |
| AT5G24240 | 1,01 | 0,80 | 1,91 | 0,42 | 6,76E-01 |
| AT5G24260 | 1,06 | -1,95 | 1,97 | -0,99 | 3,23E-01 |
| AT5G24270 | 1,64 | -0,06 | 1,50 | -0,04 | 9,66E-01 |
| AT5G24280 | 2,00 | 4,75 | 3,00 | 1,58 | 1,13E-01 |
| AT5G24320 | 0,20 | 0,40 | 4,37 | 0,09 | 9,28E-01 |
| AT5G24355 | 0,08 | 0,69 | 4,36 | 0,16 | 8,75E-01 |
| AT5G24430 | 0,56 | 0,75 | 3,95 | 0,19 | 8,50E-01 |
| AT5G24470 | 1,29 | 0,05 | 2,63 | 0,02 | 9,86E-01 |
| AT5G24480 | 0,23 | 1,49 | 4,33 | 0,35 | 7,30E-01 |
| AT5G24500 | 0,24 | -0,27 | 4,35 | -0,06 | 9,51E-01 |
| AT5G24520 | 0,95 | 1,92 | 2,11 | 0,91 | 3,64E-01 |
| AT5G24590 | 1,51 | -0,06 | 1,70 | -0,03 | 9,73E-01 |
| AT5G24655 | 0,37 | 0,94 | 4,34 | 0,22 | 8,28E-01 |
| AT5G24660 | 2,17 | 3,56 | 1,73 | 2,06 | 3,96E-02 |
| AT5G24690 | 2,42 | -0,24 | 1,16 | -0,21 | 8,34E-01 |
| AT5G24710 | 4,46 | -1,25 | 0,98 | -1,28 | 2,01E-01 |
| AT5G24735 | 0,57 | -0,32 | 3,29 | -0,10 | 9,24E-01 |
| AT5G24740 | 1,40 | 1,31 | 1,71 | 0,77 | 4,44E-01 |
| AT5G24770 | 0,62 | 1,42 | 2,33 | 0,61 | 5,41E-01 |
| AT5G24780 | 0,08 | 0,00 | 4,40 | 0,00 | 1,00E+00 |
| AT5G24810 | 2,74 | 2,57 | 1,49 | 1,73 | 8,43E-02 |
| AT5G24890 | 2,39 | -0,63 | 1,24 | -0,50 | 6,15E-01 |
| AT5G24900 | 2,20 | 0,17 | 1,48 | 0,11 | 9,11E-01 |
| AT5G24940 | 0,08 | 0,44 | 4,38 | 0,10 | 9,21E-01 |
| AT5G24960 | 0,36 | 0,72 | 4,35 | 0,17 | 8,68E-01 |
| AT5G25000 | 0,16 | 0,84 | 4,35 | 0,19 | 8,46E-01 |

| | | | | | |
|---|---|---|---|---|---|
| AT5G25060 | 1,92 | -0,66 | 1,25 | -0,53 | 5,97E-01 |
| AT5G25070 | 1,26 | -0,78 | 1,54 | -0,50 | 6,16E-01 |
| AT5G25100 | 1,32 | 0,98 | 1,69 | 0,58 | 5,62E-01 |
| AT5G25200 | 0,60 | -0,62 | 2,69 | -0,23 | 8,19E-01 |
| AT5G25265 | 0,67 | 1,17 | 2,89 | 0,41 | 6,84E-01 |
| AT5G25270 | 3,67 | -0,34 | 1,08 | -0,31 | 7,55E-01 |
| AT5G25350 | 1,15 | 1,48 | 1,86 | 0,79 | 4,28E-01 |
| AT5G25370 | 0,71 | 0,00 | 3,01 | 0,00 | 1,00E+00 |
| AT5G03985 | 0,37 | 0,00 | 4,40 | 0,00 | 1,00E+00 |
| AT5G25480 | 0,27 | -1,86 | 4,40 | -0,42 | 6,72E-01 |
| AT5G25510 | 0,49 | 1,74 | 3,25 | 0,54 | 5,91E-01 |
| AT5G25530 | 1,06 | -0,36 | 3,20 | -0,11 | 9,09E-01 |
| AT5G25756 | 0,35 | 0,00 | 4,41 | 0,00 | 1,00E+00 |
| AT5G25757 | 0,87 | -1,31 | 2,03 | -0,65 | 5,19E-01 |
| AT5G25770 | 0,28 | -1,42 | 4,35 | -0,33 | 7,44E-01 |
| AT5G04085 | 0,59 | 0,00 | 4,41 | 0,00 | 1,00E+00 |
| AT5G25780 | 1,13 | -1,46 | 1,67 | -0,87 | 3,84E-01 |
| AT5G25820 | 0,43 | -2,45 | 4,36 | -0,56 | 5,75E-01 |
| AT5G25830 | 0,05 | 0,00 | 4,41 | 0,00 | 1,00E+00 |
| AT5G25920 | 0,08 | 0,44 | 4,38 | 0,10 | 9,21E-01 |
| AT5G26000 | 4,62 | -0,16 | 0,87 | -0,19 | 8,50E-01 |
| AT5G26170 | 0,17 | 1,33 | 4,33 | 0,31 | 7,59E-01 |
| AT5G26210 | 0,43 | 0,25 | 4,13 | 0,06 | 9,51E-01 |
| AT5G26230 | 0,08 | 0,00 | 4,41 | 0,00 | 1,00E+00 |
| AT5G26340 | 2,20 | -0,74 | 1,24 | -0,59 | 5,52E-01 |
| AT5G26360 | 2,70 | 0,69 | 1,18 | 0,59 | 5,58E-01 |
| AT5G26680 | 0,15 | 0,00 | 4,38 | 0,00 | 1,00E+00 |
| AT5G04245 | 1,57 | 2,82 | 2,24 | 1,26 | 2,08E-01 |
| AT5G26640 | 0,17 | 0,98 | 4,35 | 0,23 | 8,21E-01 |
| AT5G26622 | 0,46 | -2,30 | 4,38 | -0,52 | 6,00E-01 |
| AT5G04255 | 0,21 | -1,41 | 4,36 | -0,32 | 7,46E-01 |
| AT5G26760 | 0,42 | -0,95 | 4,12 | -0,23 | 8,17E-01 |
| AT5G26770 | 1,06 | -1,17 | 2,03 | -0,58 | 5,64E-01 |
| AT5G26880 | 0,63 | -2,25 | 3,06 | -0,73 | 4,64E-01 |
| AT5G26910 | 0,66 | -1,31 | 2,46 | -0,53 | 5,96E-01 |
| AT5G26960 | 0,27 | 0,11 | 4,40 | 0,03 | 9,80E-01 |
| AT5G26970 | 0,84 | -0,89 | 2,14 | -0,42 | 6,77E-01 |
| AT5G04305 | 5,44 | 1,09 | 0,87 | 1,26 | 2,09E-01 |
| AT5G27030 | 1,03 | -0,79 | 1,86 | -0,43 | 6,70E-01 |
| AT5G27100 | 0,25 | 0,38 | 4,33 | 0,09 | 9,31E-01 |
| AT5G27238 | 0,05 | 0,00 | 4,41 | 0,00 | 1,00E+00 |
| AT5G04375 | 0,24 | -0,27 | 4,35 | -0,06 | 9,51E-01 |
| AT5G27330 | 0,05 | 0,00 | 4,41 | 0,00 | 1,00E+00 |
| AT5G27380 | 0,63 | 2,53 | 3,56 | 0,71 | 4,77E-01 |
| AT5G27550 | 0,34 | 0,13 | 4,34 | 0,03 | 9,76E-01 |
| AT5G27640 | 1,34 | -0,08 | 1,65 | -0,05 | 9,59E-01 |
| AT5G27700 | 0,49 | 0,12 | 3,17 | 0,04 | 9,69E-01 |
| AT5G27715 | 0,29 | -1,78 | 4,41 | -0,40 | 6,87E-01 |
| AT5G04435 | 0,39 | 0,13 | 4,31 | 0,03 | 9,75E-01 |
| AT5G27790 | 0,75 | 1,08 | 3,54 | 0,31 | 7,60E-01 |
| AT5G27840 | 0,72 | 1,67 | 2,76 | 0,61 | 5,44E-01 |
| AT5G27870 | 1,24 | -0,31 | 1,74 | -0,18 | 8,57E-01 |

| | | | | | |
|---|---|---|---|---|---|
| AT5G27920 | 0,08 | 0,69 | 4,36 | 0,16 | 8,75E-01 |
| AT5G27960 | 0,74 | 1,75 | 2,70 | 0,65 | 5,18E-01 |
| AT5G28020 | 0,34 | 1,26 | 3,54 | 0,36 | 7,23E-01 |
| AT5G28030 | 0,66 | 1,40 | 2,87 | 0,49 | 6,27E-01 |
| AT5G28040 | 2,94 | -1,20 | 1,10 | -1,09 | 2,74E-01 |
| AT5G28050 | 0,28 | 0,08 | 4,33 | 0,02 | 9,86E-01 |
| AT5G04635 | 0,50 | 1,84 | 3,12 | 0,59 | 5,55E-01 |
| AT5G28060 | 0,65 | -0,53 | 2,24 | -0,23 | 8,14E-01 |
| AT5G04685 | 0,19 | 0,00 | 4,41 | 0,00 | 1,00E+00 |
| AT5G28190 | 0,36 | 0,90 | 4,34 | 0,21 | 8,35E-01 |
| AT5G28220 | 0,30 | 0,00 | 4,41 | 0,00 | 1,00E+00 |
| AT5G28235 | 0,34 | 0,00 | 4,41 | 0,00 | 1,00E+00 |
| AT5G28300 | 1,51 | 1,94 | 1,81 | 1,07 | 2,83E-01 |
| AT5G28310 | 0,08 | 0,44 | 4,38 | 0,10 | 9,21E-01 |
| AT5G28400 | 1,06 | -0,54 | 1,66 | -0,32 | 7,46E-01 |
| AT5G28442 | 0,36 | 1,16 | 3,40 | 0,34 | 7,32E-01 |
| AT5G28450 | 0,85 | 0,46 | 2,48 | 0,19 | 8,52E-01 |
| AT5G28530 | 0,22 | 0,48 | 4,36 | 0,11 | 9,13E-01 |
| AT5G28550 | 0,21 | 0,64 | 4,35 | 0,15 | 8,84E-01 |
| AT5G28640 | 0,08 | 0,00 | 4,40 | 0,00 | 1,00E+00 |
| AT5G28750 | 0,65 | -1,63 | 2,35 | -0,69 | 4,90E-01 |
| AT5G28770 | 0,50 | 2,45 | 3,88 | 0,63 | 5,28E-01 |
| AT5G28823 | 1,86 | 3,61 | 1,92 | 1,88 | 6,02E-02 |
| AT5G28824 | 1,47 | 2,51 | 1,81 | 1,39 | 1,65E-01 |
| AT5G28840 | 2,50 | -0,61 | 1,06 | -0,58 | 5,65E-01 |
| AT5G28850 | 0,48 | 0,36 | 3,24 | 0,11 | 9,11E-01 |
| AT5G28910 | 0,82 | -2,26 | 2,44 | -0,93 | 3,54E-01 |
| AT5G04905 | 2,64 | 0,74 | 1,21 | 0,61 | 5,41E-01 |
| AT5G28920 | 2,84 | 0,55 | 1,17 | 0,47 | 6,37E-01 |
| AT5G04925 | 0,18 | 0,00 | 4,38 | 0,00 | 1,00E+00 |
| AT5G30510 | 1,83 | 2,12 | 1,89 | 1,12 | 2,61E-01 |
| AT5G32470 | 0,85 | -0,36 | 1,99 | -0,18 | 8,56E-01 |
| AT5G32590 | 0,68 | -0,13 | 2,29 | -0,06 | 9,53E-01 |
| AT5G33210 | 0,08 | 0,00 | 4,40 | 0,00 | 1,00E+00 |
| AT5G33439 | 0,11 | 0,00 | 4,41 | 0,00 | 1,00E+00 |
| AT5G34850 | 1,64 | 0,54 | 1,43 | 0,38 | 7,04E-01 |
| AT5G35100 | 0,30 | 1,28 | 3,63 | 0,35 | 7,24E-01 |
| AT5G35200 | 0,52 | 1,22 | 3,07 | 0,40 | 6,91E-01 |
| AT5G35210 | 2,88 | -0,57 | 1,09 | -0,52 | 6,06E-01 |
| AT5G35220 | 0,25 | 0,41 | 4,33 | 0,09 | 9,25E-01 |
| AT5G35230 | 0,96 | 1,26 | 2,70 | 0,47 | 6,41E-01 |
| AT5G35370 | 0,70 | 0,45 | 2,06 | 0,22 | 8,26E-01 |
| AT5G35475 | 0,36 | 0,27 | 4,36 | 0,06 | 9,50E-01 |
| AT5G35480 | 0,65 | -0,10 | 3,12 | -0,03 | 9,74E-01 |
| AT5G35530 | 0,88 | 1,78 | 2,42 | 0,73 | 4,64E-01 |
| AT5G35560 | 0,64 | -0,92 | 2,29 | -0,40 | 6,86E-01 |
| AT5G35570 | 0,29 | 2,09 | 4,32 | 0,48 | 6,29E-01 |
| AT5G35590 | 0,12 | 0,00 | 4,39 | 0,00 | 1,00E+00 |
| AT5G35603 | 0,59 | 1,44 | 4,11 | 0,35 | 7,26E-01 |
| AT5G35604 | 0,45 | 1,88 | 3,93 | 0,48 | 6,31E-01 |
| AT5G35620 | 0,26 | 0,76 | 4,36 | 0,17 | 8,62E-01 |
| AT5G35690 | 0,37 | 0,54 | 4,30 | 0,12 | 9,01E-01 |

| | | | | | |
|---|---|---|---|---|---|
| AT5G35700 | 0,37 | 1,07 | 4,24 | 0,25 | 8,00E-01 |
| AT5G35735 | 2,19 | 0,80 | 1,34 | 0,60 | 5,48E-01 |
| AT5G35750 | 1,20 | 1,17 | 1,88 | 0,62 | 5,33E-01 |
| AT5G35753 | 0,77 | -1,64 | 2,02 | -0,81 | 4,17E-01 |
| AT5G35790 | 0,34 | -1,67 | 4,39 | -0,38 | 7,03E-01 |
| AT5G35810 | 0,15 | 0,74 | 4,35 | 0,17 | 8,65E-01 |
| AT5G35890 | 0,19 | 0,36 | 4,36 | 0,08 | 9,34E-01 |
| AT5G35910 | 0,15 | 1,01 | 4,34 | 0,23 | 8,17E-01 |
| AT5G35913 | 0,08 | 0,69 | 4,36 | 0,16 | 8,75E-01 |
| AT5G35970 | 1,60 | 0,82 | 1,54 | 0,53 | 5,97E-01 |
| AT5G35995 | 0,29 | 1,85 | 4,32 | 0,43 | 6,69E-01 |
| AT5G36002 | 0,12 | 1,23 | 4,34 | 0,28 | 7,76E-01 |
| AT5G36120 | 0,05 | 0,00 | 4,41 | 0,00 | 1,00E+00 |
| AT5G36150 | 0,29 | 0,00 | 4,41 | 0,00 | 1,00E+00 |
| AT5G36160 | 0,29 | 0,50 | 4,31 | 0,12 | 9,07E-01 |
| AT5G36200 | 1,52 | 2,80 | 1,95 | 1,44 | 1,50E-01 |
| AT5G36210 | 1,11 | 1,65 | 1,99 | 0,83 | 4,05E-01 |
| AT5G36220 | 0,80 | 1,25 | 2,19 | 0,57 | 5,68E-01 |
| AT5G36700 | 0,58 | -1,28 | 2,52 | -0,51 | 6,12E-01 |
| AT5G36790 | 0,17 | 0,18 | 4,39 | 0,04 | 9,67E-01 |
| AT5G36880 | 0,93 | -1,12 | 2,04 | -0,55 | 5,82E-01 |
| AT5G36930 | 1,17 | 1,56 | 2,22 | 0,70 | 4,83E-01 |
| AT5G37260 | 0,78 | 0,40 | 3,24 | 0,12 | 9,02E-01 |
| AT5G37310 | 0,77 | 3,35 | 3,33 | 1,01 | 3,14E-01 |
| AT5G37370 | 0,59 | 0,97 | 3,66 | 0,27 | 7,90E-01 |
| AT5G05485 | 5,35 | -0,68 | 1,28 | -0,53 | 5,95E-01 |
| AT5G37480 | 0,13 | 0,00 | 4,41 | 0,00 | 1,00E+00 |
| AT5G37590 | 0,66 | -0,31 | 3,58 | -0,09 | 9,31E-01 |
| AT5G37600 | 1,93 | -0,93 | 1,48 | -0,62 | 5,33E-01 |
| AT5G05505 | 4,00 | -0,28 | 1,36 | -0,21 | 8,35E-01 |
| AT5G37720 | 0,46 | -0,73 | 3,22 | -0,23 | 8,20E-01 |
| AT5G37770 | 1,11 | 3,17 | 2,67 | 1,19 | 2,35E-01 |
| AT5G37780 | 0,80 | 1,50 | 2,11 | 0,71 | 4,77E-01 |
| AT5G05545 | 0,31 | 0,00 | 4,41 | 0,00 | 1,00E+00 |
| AT5G37830 | 1,35 | 3,90 | 2,00 | 1,96 | 5,04E-02 |
| AT5G37890 | 0,36 | 1,83 | 4,33 | 0,42 | 6,73E-01 |
| AT5G05565 | 0,45 | 1,31 | 4,12 | 0,32 | 7,51E-01 |
| AT5G38020 | 0,42 | -1,73 | 4,40 | -0,39 | 6,94E-01 |
| AT5G38080 | 0,08 | 0,44 | 4,38 | 0,10 | 9,21E-01 |
| AT5G38140 | 0,10 | 0,00 | 4,41 | 0,00 | 1,00E+00 |
| AT5G38212 | 1,00 | 2,33 | 2,21 | 1,05 | 2,91E-01 |
| AT5G38240 | 0,41 | -2,12 | 4,32 | -0,49 | 6,23E-01 |
| AT5G38280 | 0,74 | 0,00 | 3,77 | 0,00 | 1,00E+00 |
| AT5G38420 | 5,05 | -1,37 | 0,83 | -1,66 | 9,63E-02 |
| AT5G38430 | 4,72 | -1,73 | 0,85 | -2,03 | 4,19E-02 |
| AT5G38450 | 0,46 | 0,26 | 4,20 | 0,06 | 9,50E-01 |
| AT5G05755 | 0,05 | 0,00 | 4,41 | 0,00 | 1,00E+00 |
| AT5G38510 | 0,76 | -0,34 | 2,53 | -0,13 | 8,94E-01 |
| AT5G38640 | 0,50 | 0,31 | 2,63 | 0,12 | 9,08E-01 |
| AT5G38660 | 1,59 | -0,70 | 1,52 | -0,46 | 6,46E-01 |
| AT5G38650 | 1,00 | 0,03 | 2,11 | 0,01 | 9,89E-01 |
| AT5G38830 | 0,49 | 1,18 | 4,34 | 0,27 | 7,87E-01 |

| | | | | | |
|---|---|---|---|---|---|
| AT5G39130 | 0,12 | 0,49 | 4,36 | 0,11 | 9,11E-01 |
| AT5G39160 | 0,34 | 0,00 | 4,41 | 0,00 | 1,00E+00 |
| AT5G05895 | 0,62 | 0,26 | 3,07 | 0,08 | 9,33E-01 |
| AT5G39480 | 0,08 | 0,00 | 4,40 | 0,00 | 1,00E+00 |
| AT5G39520 | 0,33 | 0,39 | 4,34 | 0,09 | 9,28E-01 |
| AT5G39530 | 0,77 | 0,68 | 2,88 | 0,24 | 8,12E-01 |
| AT5G39600 | 0,66 | 0,26 | 2,28 | 0,11 | 9,11E-01 |
| AT5G39610 | 0,52 | -1,25 | 3,85 | -0,33 | 7,45E-01 |
| AT5G05995 | 0,40 | 1,87 | 4,16 | 0,45 | 6,53E-01 |
| AT5G39680 | 0,42 | 0,95 | 4,33 | 0,22 | 8,26E-01 |
| AT5G39693 | 0,43 | 0,03 | 3,39 | 0,01 | 9,92E-01 |
| AT5G39920 | 0,25 | 0,05 | 4,38 | 0,01 | 9,90E-01 |
| AT5G39950 | 0,95 | 1,25 | 2,41 | 0,52 | 6,04E-01 |
| AT5G40170 | 0,68 | -1,21 | 2,31 | -0,52 | 6,01E-01 |
| AT5G40240 | 0,14 | 0,83 | 4,35 | 0,19 | 8,49E-01 |
| AT5G40340 | 2,24 | -0,20 | 1,36 | -0,15 | 8,84E-01 |
| AT5G40370 | 0,37 | 1,46 | 3,40 | 0,43 | 6,68E-01 |
| AT5G40384 | 2,72 | -1,50 | 1,29 | -1,17 | 2,42E-01 |
| AT5G40395 | 2,01 | -0,09 | 1,40 | -0,06 | 9,50E-01 |
| AT5G40450 | 4,44 | 0,33 | 0,98 | 0,34 | 7,34E-01 |
| AT5G40470 | 0,27 | -2,20 | 4,38 | -0,50 | 6,16E-01 |
| AT5G40580 | 0,60 | -0,95 | 4,33 | -0,22 | 8,27E-01 |
| AT5G40660 | 0,29 | 0,70 | 4,32 | 0,16 | 8,72E-01 |
| AT5G40740 | 0,85 | 1,99 | 2,32 | 0,86 | 3,92E-01 |
| AT5G40770 | 1,47 | -0,55 | 1,44 | -0,38 | 7,03E-01 |
| AT5G40820 | 0,95 | -0,51 | 2,14 | -0,24 | 8,13E-01 |
| AT5G40830 | 0,63 | 0,49 | 2,67 | 0,19 | 8,53E-01 |
| AT5G40890 | 1,74 | -1,11 | 1,37 | -0,81 | 4,18E-01 |
| AT5G40940 | 0,56 | 1,44 | 3,76 | 0,38 | 7,02E-01 |
| AT5G41020 | 0,73 | -0,43 | 2,57 | -0,17 | 8,66E-01 |
| AT5G41130 | 0,21 | 1,51 | 4,33 | 0,35 | 7,27E-01 |
| AT5G41150 | 0,33 | 0,75 | 4,32 | 0,17 | 8,62E-01 |
| AT5G41190 | 0,83 | -2,71 | 2,81 | -0,97 | 3,34E-01 |
| AT5G41330 | 0,40 | -0,54 | 3,29 | -0,16 | 8,71E-01 |
| AT5G41400 | 0,65 | -0,94 | 2,65 | -0,35 | 7,24E-01 |
| AT5G41460 | 0,17 | 0,00 | 4,41 | 0,00 | 1,00E+00 |
| AT5G41520 | 0,58 | -0,63 | 2,67 | -0,24 | 8,14E-01 |
| AT5G41670 | 0,43 | 0,21 | 2,80 | 0,07 | 9,40E-01 |
| AT5G41765 | 5,08 | 1,52 | 1,15 | 1,33 | 1,85E-01 |
| AT5G06235 | 0,77 | 3,08 | 4,29 | 0,72 | 4,72E-01 |
| AT5G41870 | 0,38 | 0,35 | 4,31 | 0,08 | 9,35E-01 |
| AT5G41990 | 1,56 | 0,79 | 1,44 | 0,55 | 5,84E-01 |
| AT5G42020 | 4,47 | -0,63 | 0,84 | -0,75 | 4,51E-01 |
| AT5G42080 | 4,02 | -0,34 | 0,92 | -0,37 | 7,10E-01 |
| AT5G42092 | 3,68 | 0,89 | 1,03 | 0,86 | 3,88E-01 |
| AT5G42100 | 1,07 | -0,91 | 1,81 | -0,50 | 6,15E-01 |
| AT5G42110 | 0,47 | -0,26 | 3,30 | -0,08 | 9,37E-01 |
| AT5G42130 | 0,21 | 1,55 | 4,33 | 0,36 | 7,20E-01 |
| AT5G42140 | 0,12 | 0,00 | 4,38 | 0,00 | 1,00E+00 |
| AT5G42190 | 0,65 | 0,93 | 2,58 | 0,36 | 7,20E-01 |
| AT5G42280 | 0,14 | 0,83 | 4,35 | 0,19 | 8,49E-01 |
| AT5G42300 | 0,48 | 1,71 | 3,90 | 0,44 | 6,60E-01 |

| | | | | | |
|---|---|---|---|---|---|
| AT5G42360 | 1,47 | -0,09 | 1,43 | -0,07 | 9,48E-01 |
| AT5G42380 | 1,28 | -1,19 | 1,95 | -0,61 | 5,41E-01 |
| AT5G42400 | 0,21 | 1,39 | 4,33 | 0,32 | 7,49E-01 |
| AT5G06305 | 1,02 | 2,02 | 1,88 | 1,08 | 2,82E-01 |
| AT5G42490 | 0,56 | 0,74 | 3,85 | 0,19 | 8,47E-01 |
| AT5G42530 | 4,10 | -0,29 | 1,01 | -0,29 | 7,72E-01 |
| AT5G42567 | 0,58 | 0,00 | 4,41 | 0,00 | 1,00E+00 |
| AT5G42570 | 0,15 | 0,67 | 4,37 | 0,15 | 8,79E-01 |
| AT5G42650 | 3,24 | -1,20 | 1,31 | -0,91 | 3,62E-01 |
| AT5G42660 | 0,12 | 0,00 | 4,38 | 0,00 | 1,00E+00 |
| AT5G42750 | 1,69 | 1,24 | 1,73 | 0,72 | 4,74E-01 |
| AT5G42790 | 0,93 | 2,09 | 2,06 | 1,01 | 3,12E-01 |
| AT5G42800 | 0,38 | 0,84 | 4,33 | 0,20 | 8,45E-01 |
| AT5G42820 | 2,08 | -0,94 | 1,23 | -0,77 | 4,44E-01 |
| AT5G42850 | 1,15 | -0,96 | 1,89 | -0,51 | 6,12E-01 |
| AT5G42870 | 0,81 | 1,80 | 2,29 | 0,79 | 4,32E-01 |
| AT5G42890 | 0,08 | 0,00 | 4,40 | 0,00 | 1,00E+00 |
| AT5G42900 | 0,25 | -0,23 | 4,35 | -0,05 | 9,58E-01 |
| AT5G42905 | 0,52 | 1,90 | 3,79 | 0,50 | 6,17E-01 |
| AT5G42930 | 2,68 | -1,22 | 1,24 | -0,98 | 3,26E-01 |
| AT5G42940 | 0,81 | -0,32 | 2,38 | -0,13 | 8,93E-01 |
| AT5G42955 | 0,65 | -0,64 | 3,85 | -0,17 | 8,67E-01 |
| AT5G42957 | 0,83 | -1,78 | 2,57 | -0,69 | 4,88E-01 |
| AT5G42990 | 0,17 | 1,52 | 4,33 | 0,35 | 7,26E-01 |
| AT5G43010 | 0,13 | 0,00 | 4,41 | 0,00 | 1,00E+00 |
| AT5G43020 | 0,41 | -0,25 | 4,14 | -0,06 | 9,51E-01 |
| AT5G43130 | 1,30 | 0,46 | 1,93 | 0,24 | 8,13E-01 |
| AT5G43190 | 1,41 | -1,43 | 1,58 | -0,91 | 3,64E-01 |
| AT5G43260 | 0,52 | -0,47 | 2,74 | -0,17 | 8,64E-01 |
| AT5G43270 | 0,76 | -1,17 | 3,08 | -0,38 | 7,05E-01 |
| AT5G43285 | 0,35 | 0,00 | 4,41 | 0,00 | 1,00E+00 |
| AT5G43310 | 2,37 | 0,78 | 1,22 | 0,64 | 5,25E-01 |
| AT5G43320 | 0,66 | -1,11 | 2,68 | -0,41 | 6,80E-01 |
| AT5G43440 | 0,40 | -0,15 | 4,35 | -0,03 | 9,73E-01 |
| AT5G43470 | 3,38 | 1,40 | 1,24 | 1,13 | 2,57E-01 |
| AT5G43670 | 0,40 | -1,66 | 4,25 | -0,39 | 6,95E-01 |
| AT5G43710 | 0,21 | 1,57 | 4,33 | 0,36 | 7,16E-01 |
| AT5G43745 | 0,36 | -1,42 | 4,35 | -0,33 | 7,45E-01 |
| AT5G43750 | 0,47 | -2,80 | 4,36 | -0,64 | 5,21E-01 |
| AT5G43780 | 0,63 | -0,64 | 2,66 | -0,24 | 8,10E-01 |
| AT5G43790 | 0,45 | -0,73 | 4,21 | -0,17 | 8,61E-01 |
| AT5G43830 | 0,91 | -0,84 | 1,86 | -0,45 | 6,52E-01 |
| AT5G43850 | 0,29 | 0,87 | 3,67 | 0,24 | 8,13E-01 |
| AT5G43880 | 0,19 | 0,00 | 4,41 | 0,00 | 1,00E+00 |
| AT5G43900 | 1,55 | 0,60 | 1,40 | 0,43 | 6,68E-01 |
| AT5G43950 | 0,65 | -0,14 | 2,53 | -0,05 | 9,57E-01 |
| AT5G43960 | 0,54 | 2,30 | 3,06 | 0,75 | 4,52E-01 |
| AT5G44000 | 0,95 | 3,03 | 2,20 | 1,38 | 1,68E-01 |
| AT5G44005 | 0,16 | 0,65 | 4,37 | 0,15 | 8,82E-01 |
| AT5G44090 | 1,96 | -0,57 | 1,61 | -0,36 | 7,22E-01 |
| AT5G44100 | 0,66 | 1,29 | 2,56 | 0,50 | 6,16E-01 |
| AT5G44250 | 0,12 | 1,16 | 4,34 | 0,27 | 7,89E-01 |

| | | | | | |
|---|---|---|---|---|---|
| AT5G44260 | 0,22 | 0,00 | 4,41 | 0,00 | 1,00E+00 |
| AT5G44283 | 0,91 | 0,44 | 2,55 | 0,17 | 8,62E-01 |
| AT5G44290 | 1,25 | 0,77 | 1,90 | 0,40 | 6,86E-01 |
| AT5G44306 | 0,93 | 0,91 | 2,91 | 0,31 | 7,54E-01 |
| AT5G44320 | 2,18 | 0,52 | 1,19 | 0,44 | 6,61E-01 |
| AT5G44495 | 1,23 | 0,23 | 1,79 | 0,13 | 8,99E-01 |
| AT5G44510 | 0,61 | 2,86 | 4,29 | 0,67 | 5,05E-01 |
| AT5G44568 | 0,48 | -0,90 | 2,91 | -0,31 | 7,56E-01 |
| AT5G06595 | 1,38 | -3,49 | 1,72 | -2,03 | 4,28E-02 |
| AT5G44670 | 0,13 | 0,73 | 4,36 | 0,17 | 8,67E-01 |
| AT5G44680 | 0,48 | 0,00 | 4,41 | 0,00 | 1,00E+00 |
| AT5G44785 | 0,10 | 0,01 | 4,38 | 0,00 | 9,99E-01 |
| AT5G44790 | 1,35 | 0,26 | 1,50 | 0,17 | 8,64E-01 |
| AT5G44800 | 4,77 | -0,03 | 0,80 | -0,03 | 9,74E-01 |
| AT5G44820 | 0,08 | 0,44 | 4,38 | 0,10 | 9,21E-01 |
| AT5G44840 | 0,14 | 0,00 | 4,41 | 0,00 | 1,00E+00 |
| AT5G45010 | 0,58 | -1,91 | 3,16 | -0,60 | 5,45E-01 |
| AT5G45040 | 0,08 | 0,69 | 4,36 | 0,16 | 8,75E-01 |
| AT5G45050 | 0,86 | 0,44 | 2,00 | 0,22 | 8,27E-01 |
| AT5G45170 | 0,11 | 0,00 | 4,41 | 0,00 | 1,00E+00 |
| AT5G45180 | 0,66 | -0,89 | 4,00 | -0,22 | 8,23E-01 |
| AT5G45230 | 0,97 | -0,73 | 1,90 | -0,38 | 7,02E-01 |
| AT5G45250 | 0,93 | -1,17 | 1,90 | -0,61 | 5,40E-01 |
| AT5G45275 | 0,48 | 1,67 | 4,33 | 0,39 | 7,00E-01 |
| AT5G45290 | 1,09 | 0,20 | 1,95 | 0,10 | 9,19E-01 |
| AT5G45340 | 0,20 | 0,15 | 4,37 | 0,04 | 9,72E-01 |
| AT5G45360 | 0,08 | 0,69 | 4,36 | 0,16 | 8,75E-01 |
| AT5G45410 | 0,46 | -0,26 | 3,99 | -0,07 | 9,48E-01 |
| AT5G45420 | 0,27 | 0,84 | 4,36 | 0,19 | 8,48E-01 |
| AT5G45440 | 0,74 | 1,87 | 3,38 | 0,55 | 5,80E-01 |
| AT5G45460 | 0,18 | -1,78 | 4,41 | -0,40 | 6,87E-01 |
| AT5G45470 | 2,49 | 2,66 | 1,86 | 1,43 | 1,53E-01 |
| AT5G45480 | 2,31 | 0,89 | 1,54 | 0,58 | 5,65E-01 |
| AT5G45500 | 2,29 | -1,38 | 1,53 | -0,90 | 3,68E-01 |
| AT5G45573 | 0,61 | 1,58 | 2,91 | 0,54 | 5,88E-01 |
| AT5G45630 | 0,05 | 0,00 | 4,41 | 0,00 | 1,00E+00 |
| AT5G45650 | 0,61 | 0,13 | 3,18 | 0,04 | 9,67E-01 |
| AT5G45775 | 0,86 | -0,35 | 1,92 | -0,18 | 8,55E-01 |
| AT5G45800 | 0,52 | 0,54 | 2,35 | 0,23 | 8,17E-01 |
| AT5G45820 | 1,25 | -0,18 | 1,78 | -0,10 | 9,21E-01 |
| AT5G46020 | 1,24 | -0,24 | 2,54 | -0,09 | 9,25E-01 |
| AT5G46040 | 0,11 | 0,48 | 4,37 | 0,11 | 9,12E-01 |
| AT5G46050 | 0,37 | -2,24 | 4,39 | -0,51 | 6,09E-01 |
| AT5G46210 | 3,09 | -0,30 | 1,07 | -0,28 | 7,77E-01 |
| AT5G46260 | 1,90 | 1,13 | 1,32 | 0,86 | 3,91E-01 |
| AT5G46270 | 2,21 | -1,26 | 1,30 | -0,97 | 3,34E-01 |
| AT5G46280 | 0,57 | 0,15 | 3,16 | 0,05 | 9,63E-01 |
| AT5G46290 | 0,63 | 0,43 | 3,11 | 0,14 | 8,89E-01 |
| AT5G46340 | 0,61 | -0,80 | 4,01 | -0,20 | 8,42E-01 |
| AT5G46395 | 0,11 | 0,00 | 4,41 | 0,00 | 1,00E+00 |
| AT5G46580 | 0,75 | 0,77 | 2,47 | 0,31 | 7,54E-01 |
| AT5G46620 | 0,64 | 0,24 | 3,13 | 0,08 | 9,39E-01 |

| | | | | | |
|---|---|---|---|---|---|
| AT5G46750 | 0,66 | -0,40 | 3,75 | -0,11 | 9,15E-01 |
| AT5G46790 | 0,53 | 1,68 | 3,02 | 0,56 | 5,79E-01 |
| AT5G46800 | 0,83 | -1,48 | 2,39 | -0,62 | 5,37E-01 |
| AT5G46860 | 0,46 | 0,00 | 4,15 | 0,00 | 1,00E+00 |
| AT5G46970 | 0,65 | 1,26 | 2,60 | 0,49 | 6,28E-01 |
| AT5G47040 | 1,34 | -0,23 | 1,57 | -0,15 | 8,83E-01 |
| AT5G47070 | 0,97 | -0,12 | 2,22 | -0,05 | 9,56E-01 |
| AT5G47080 | 0,13 | 0,73 | 4,36 | 0,17 | 8,67E-01 |
| AT5G47110 | 1,25 | -1,91 | 1,54 | -1,24 | 2,15E-01 |
| AT5G47190 | 0,93 | 0,05 | 2,16 | 0,02 | 9,83E-01 |
| AT5G47210 | 1,48 | -0,60 | 1,99 | -0,30 | 7,64E-01 |
| AT5G47220 | 0,78 | -1,20 | 2,41 | -0,50 | 6,19E-01 |
| AT5G47250 | 0,28 | 0,06 | 4,39 | 0,01 | 9,90E-01 |
| AT5G47370 | 0,28 | -1,42 | 4,36 | -0,33 | 7,44E-01 |
| AT5G47430 | 3,26 | -1,16 | 1,03 | -1,13 | 2,59E-01 |
| AT5G47460 | 0,15 | 1,16 | 4,34 | 0,27 | 7,88E-01 |
| AT5G47560 | 3,13 | -0,43 | 1,16 | -0,37 | 7,11E-01 |
| AT5G47570 | 0,30 | 1,21 | 4,34 | 0,28 | 7,80E-01 |
| AT5G47580 | 0,64 | -0,77 | 3,56 | -0,22 | 8,28E-01 |
| AT5G47650 | 0,20 | -0,31 | 4,36 | -0,07 | 9,44E-01 |
| AT5G47700 | 0,72 | 0,93 | 2,20 | 0,42 | 6,71E-01 |
| AT5G47730 | 0,16 | 0,92 | 4,35 | 0,21 | 8,33E-01 |
| AT5G47750 | 1,50 | 0,94 | 2,03 | 0,46 | 6,45E-01 |
| AT5G47760 | 0,12 | 0,00 | 4,38 | 0,00 | 1,00E+00 |
| AT5G47810 | 0,55 | 0,19 | 2,66 | 0,07 | 9,44E-01 |
| AT5G47830 | 0,54 | 1,40 | 4,35 | 0,32 | 7,47E-01 |
| AT5G47850 | 0,49 | -0,39 | 4,03 | -0,10 | 9,23E-01 |
| AT5G47880 | 0,55 | 0,88 | 3,14 | 0,28 | 7,79E-01 |
| AT5G47930 | 0,53 | -0,68 | 2,69 | -0,25 | 7,99E-01 |
| AT5G48000 | 0,12 | 0,00 | 4,39 | 0,00 | 1,00E+00 |
| AT5G07055 | 0,58 | 0,51 | 3,15 | 0,16 | 8,71E-01 |
| AT5G48120 | 1,18 | -0,62 | 1,60 | -0,39 | 6,99E-01 |
| AT5G48150 | 0,85 | -2,55 | 3,47 | -0,74 | 4,62E-01 |
| AT5G48180 | 0,98 | 1,92 | 2,03 | 0,95 | 3,44E-01 |
| AT5G48375 | 0,26 | -0,30 | 4,37 | -0,07 | 9,46E-01 |
| AT5G48380 | 2,92 | 1,13 | 1,09 | 1,03 | 3,03E-01 |
| AT5G48412 | 0,64 | -1,40 | 2,72 | -0,51 | 6,07E-01 |
| AT5G48440 | 0,43 | 1,96 | 4,18 | 0,47 | 6,40E-01 |
| AT5G48540 | 0,08 | 0,44 | 4,38 | 0,10 | 9,21E-01 |
| AT5G48580 | 0,41 | 0,79 | 4,12 | 0,19 | 8,49E-01 |
| AT5G48640 | 0,19 | 0,00 | 4,41 | 0,00 | 1,00E+00 |
| AT5G48657 | 0,40 | -0,47 | 3,35 | -0,14 | 8,88E-01 |
| AT5G07265 | 0,12 | 1,16 | 4,34 | 0,27 | 7,89E-01 |
| AT5G48770 | 0,08 | 0,00 | 4,40 | 0,00 | 1,00E+00 |
| AT5G07275 | 0,71 | 1,35 | 4,31 | 0,31 | 7,54E-01 |
| AT5G48810 | 0,80 | 0,03 | 2,26 | 0,01 | 9,88E-01 |
| AT5G48830 | 0,28 | -0,49 | 4,38 | -0,11 | 9,11E-01 |
| AT5G48860 | 3,76 | 0,94 | 1,22 | 0,77 | 4,43E-01 |
| AT5G48880 | 0,58 | -1,71 | 4,33 | -0,39 | 6,93E-01 |
| AT5G48910 | 0,76 | 1,03 | 2,22 | 0,46 | 6,43E-01 |
| AT5G48930 | 1,24 | -0,42 | 1,63 | -0,26 | 7,98E-01 |
| AT5G49030 | 1,79 | -0,84 | 1,42 | -0,60 | 5,52E-01 |

| | | | | | |
|---|---|---|---|---|---|
| AT5G49100 | 1,11 | -1,44 | 2,06 | -0,70 | 4,84E-01 |
| AT5G49300 | 0,36 | 1,57 | 4,34 | 0,36 | 7,17E-01 |
| AT5G49350 | 0,08 | 0,69 | 4,36 | 0,16 | 8,75E-01 |
| AT5G49360 | 3,80 | -0,33 | 0,95 | -0,35 | 7,26E-01 |
| AT5G49420 | 0,35 | 0,00 | 4,38 | 0,00 | 9,99E-01 |
| AT5G49430 | 1,30 | -0,67 | 1,59 | -0,42 | 6,75E-01 |
| AT5G49540 | 0,90 | 0,15 | 2,51 | 0,06 | 9,51E-01 |
| AT5G49550 | 0,27 | -0,96 | 4,39 | -0,22 | 8,27E-01 |
| AT5G49555 | 0,40 | -1,68 | 4,39 | -0,38 | 7,03E-01 |
| AT5G49570 | 0,11 | 0,65 | 4,36 | 0,15 | 8,81E-01 |
| AT5G49580 | 0,46 | 0,49 | 4,01 | 0,12 | 9,02E-01 |
| AT5G49630 | 0,19 | 0,00 | 4,41 | 0,00 | 1,00E+00 |
| AT5G49645 | 0,08 | 0,44 | 4,38 | 0,10 | 9,21E-01 |
| AT5G49660 | 0,65 | 0,70 | 2,52 | 0,28 | 7,82E-01 |
| AT5G49665 | 0,50 | -1,43 | 3,97 | -0,36 | 7,18E-01 |
| AT5G49720 | 3,89 | -0,52 | 1,05 | -0,50 | 6,20E-01 |
| AT5G49730 | 3,55 | 1,32 | 1,35 | 0,98 | 3,27E-01 |
| AT5G49740 | 2,42 | 0,41 | 1,22 | 0,34 | 7,36E-01 |
| AT5G49830 | 1,27 | 0,94 | 1,61 | 0,58 | 5,60E-01 |
| AT5G49890 | 0,22 | 0,01 | 4,38 | 0,00 | 9,98E-01 |
| AT5G49900 | 0,41 | 0,74 | 4,12 | 0,18 | 8,57E-01 |
| AT5G49930 | 0,74 | 1,10 | 2,16 | 0,51 | 6,11E-01 |
| AT5G49950 | 0,25 | -1,78 | 4,41 | -0,40 | 6,87E-01 |
| AT5G49980 | 4,39 | -0,12 | 0,97 | -0,13 | 9,00E-01 |
| AT5G50000 | 0,84 | -0,08 | 2,13 | -0,04 | 9,70E-01 |
| AT5G50150 | 0,10 | 0,21 | 4,38 | 0,05 | 9,62E-01 |
| AT5G50220 | 0,28 | -2,52 | 4,38 | -0,58 | 5,65E-01 |
| AT5G50320 | 0,77 | -0,91 | 2,89 | -0,32 | 7,52E-01 |
| AT5G50340 | 0,25 | 0,38 | 4,33 | 0,09 | 9,31E-01 |
| AT5G50350 | 0,29 | 0,00 | 4,39 | 0,00 | 1,00E+00 |
| AT5G50380 | 0,97 | 1,42 | 2,08 | 0,68 | 4,96E-01 |
| AT5G50390 | 0,17 | 1,52 | 4,33 | 0,35 | 7,26E-01 |
| AT5G50400 | 0,54 | -3,28 | 3,90 | -0,84 | 4,01E-01 |
| AT5G50460 | 0,36 | -2,50 | 4,37 | -0,57 | 5,67E-01 |
| AT5G50480 | 0,53 | 0,00 | 3,90 | 0,00 | 1,00E+00 |
| AT5G50570 | 0,31 | -1,74 | 4,40 | -0,39 | 6,93E-01 |
| AT5G50630 | 0,05 | 0,00 | 4,41 | 0,00 | 1,00E+00 |
| AT5G50640 | 0,24 | 0,24 | 4,34 | 0,06 | 9,55E-01 |
| AT5G50670 | 0,93 | -1,78 | 2,46 | -0,72 | 4,69E-01 |
| AT5G50860 | 0,49 | 0,23 | 3,83 | 0,06 | 9,52E-01 |
| AT5G50870 | 0,08 | 0,44 | 4,38 | 0,10 | 9,21E-01 |
| AT5G50900 | 0,83 | -0,05 | 2,46 | -0,02 | 9,82E-01 |
| AT5G51000 | 0,61 | 1,51 | 3,01 | 0,50 | 6,15E-01 |
| AT5G51020 | 0,37 | 1,07 | 4,24 | 0,25 | 8,00E-01 |
| AT5G51090 | 2,18 | -0,40 | 1,48 | -0,27 | 7,86E-01 |
| AT5G51110 | 0,78 | 0,53 | 2,36 | 0,22 | 8,22E-01 |
| AT5G51170 | 0,12 | 1,23 | 4,34 | 0,28 | 7,76E-01 |
| AT5G51195 | 0,57 | 1,10 | 2,54 | 0,43 | 6,67E-01 |
| AT5G51220 | 0,23 | 0,00 | 4,38 | 0,00 | 1,00E+00 |
| AT5G51290 | 0,61 | -2,86 | 2,96 | -0,97 | 3,33E-01 |
| AT5G51300 | 1,97 | -0,15 | 2,01 | -0,08 | 9,39E-01 |
| AT5G51390 | 0,29 | -1,33 | 4,35 | -0,31 | 7,60E-01 |

| | | | | | |
|---|---|---|---|---|---|
| AT5G51400 | 1,21 | -1,36 | 1,57 | -0,87 | 3,86E-01 |
| AT5G51430 | 4,74 | -1,53 | 1,08 | -1,41 | 1,58E-01 |
| AT5G51550 | 1,02 | 1,03 | 1,99 | 0,52 | 6,03E-01 |
| AT5G51640 | 0,39 | -0,77 | 3,55 | -0,22 | 8,27E-01 |
| AT5G51650 | 0,32 | 1,19 | 3,53 | 0,34 | 7,36E-01 |
| AT5G51690 | 0,72 | -0,36 | 2,37 | -0,15 | 8,78E-01 |
| AT5G51710 | 0,10 | 0,00 | 4,41 | 0,00 | 1,00E+00 |
| AT5G51820 | 0,69 | -1,39 | 2,53 | -0,55 | 5,83E-01 |
| AT5G52060 | 0,54 | 0,68 | 2,62 | 0,26 | 7,95E-01 |
| AT5G52090 | 0,73 | 0,55 | 2,23 | 0,25 | 8,04E-01 |
| AT5G52120 | 0,47 | -2,25 | 4,39 | -0,51 | 6,08E-01 |
| AT5G52180 | 0,61 | -1,44 | 2,98 | -0,48 | 6,28E-01 |
| AT5G52190 | 1,80 | 1,02 | 1,81 | 0,56 | 5,72E-01 |
| AT5G52320 | 1,77 | 1,52 | 1,66 | 0,91 | 3,61E-01 |
| AT5G52330 | 0,15 | 0,00 | 4,41 | 0,00 | 1,00E+00 |
| AT5G52370 | 0,37 | 0,40 | 4,29 | 0,09 | 9,26E-01 |
| AT5G52450 | 0,44 | -0,20 | 3,16 | -0,06 | 9,49E-01 |
| AT5G52540 | 1,60 | 0,58 | 1,56 | 0,37 | 7,09E-01 |
| AT5G52550 | 0,25 | 0,96 | 4,36 | 0,22 | 8,25E-01 |
| AT5G52650 | 0,64 | -2,97 | 3,66 | -0,81 | 4,17E-01 |
| AT5G52780 | 0,46 | -0,93 | 3,31 | -0,28 | 7,78E-01 |
| AT5G52797 | 0,48 | -0,12 | 3,21 | -0,04 | 9,71E-01 |
| AT5G52882 | 2,07 | -0,20 | 1,40 | -0,14 | 8,89E-01 |
| AT5G53000 | 0,29 | 0,24 | 4,33 | 0,06 | 9,55E-01 |
| AT5G53010 | 0,50 | -1,57 | 3,24 | -0,48 | 6,29E-01 |
| AT5G53050 | 0,49 | 0,61 | 3,18 | 0,19 | 8,48E-01 |
| AT5G53130 | 0,72 | 3,06 | 2,85 | 1,08 | 2,82E-01 |
| AT5G07955 | 0,08 | 0,44 | 4,38 | 0,10 | 9,21E-01 |
| AT5G53170 | 2,82 | -0,11 | 1,01 | -0,11 | 9,11E-01 |
| AT5G53300 | 0,89 | -0,33 | 2,53 | -0,13 | 8,95E-01 |
| AT5G53320 | 0,58 | -0,92 | 2,87 | -0,32 | 7,48E-01 |
| AT5G07975 | 0,08 | 0,44 | 4,38 | 0,10 | 9,21E-01 |
| AT5G53370 | 2,35 | 0,63 | 1,27 | 0,50 | 6,18E-01 |
| AT5G53440 | 2,08 | -0,51 | 1,24 | -0,41 | 6,84E-01 |
| AT5G53490 | 0,65 | -1,73 | 2,34 | -0,74 | 4,60E-01 |
| AT5G53500 | 0,52 | -1,73 | 3,29 | -0,53 | 5,99E-01 |
| AT5G53530 | 0,37 | -0,75 | 4,35 | -0,17 | 8,64E-01 |
| AT5G53570 | 0,72 | 1,23 | 2,38 | 0,52 | 6,05E-01 |
| AT5G53580 | 0,36 | -0,25 | 3,55 | -0,07 | 9,45E-01 |
| AT5G53730 | 0,14 | 0,09 | 4,38 | 0,02 | 9,84E-01 |
| AT5G53850 | 0,47 | -1,39 | 4,37 | -0,32 | 7,51E-01 |
| AT5G53870 | 0,19 | 0,00 | 4,41 | 0,00 | 1,00E+00 |
| AT5G53900 | 0,79 | -1,52 | 2,20 | -0,69 | 4,90E-01 |
| AT5G54030 | 0,27 | 0,00 | 4,41 | 0,00 | 1,00E+00 |
| AT5G54040 | 0,27 | 0,01 | 4,38 | 0,00 | 9,99E-01 |
| AT5G54050 | 0,38 | 1,22 | 4,33 | 0,28 | 7,78E-01 |
| AT5G54060 | 0,12 | 1,16 | 4,34 | 0,27 | 7,89E-01 |
| AT5G54090 | 1,00 | 0,37 | 2,09 | 0,18 | 8,59E-01 |
| AT5G54100 | 0,65 | 1,91 | 2,88 | 0,66 | 5,07E-01 |
| AT5G54170 | 0,64 | -0,58 | 2,17 | -0,27 | 7,88E-01 |
| AT5G54250 | 1,88 | -1,13 | 1,52 | -0,74 | 4,57E-01 |
| AT5G54280 | 1,29 | -0,79 | 1,55 | -0,51 | 6,10E-01 |

| | | | | | |
|---|---|---|---|---|---|
| AT5G54310 | 4,38 | -1,38 | 0,84 | -1,65 | 9,99E-02 |
| AT5G54380 | 1,11 | 0,96 | 1,93 | 0,50 | 6,19E-01 |
| AT5G54410 | 0,82 | 1,37 | 2,15 | 0,64 | 5,23E-01 |
| AT5G54430 | 1,12 | 0,80 | 2,48 | 0,32 | 7,47E-01 |
| AT5G54460 | 1,09 | 1,69 | 3,31 | 0,51 | 6,10E-01 |
| AT5G54470 | 0,35 | -0,87 | 3,49 | -0,25 | 8,03E-01 |
| AT5G54510 | 0,08 | 0,69 | 4,36 | 0,16 | 8,75E-01 |
| AT5G54610 | 0,49 | 1,11 | 3,83 | 0,29 | 7,73E-01 |
| AT5G54620 | 0,12 | 1,05 | 4,35 | 0,24 | 8,10E-01 |
| AT5G54630 | 0,29 | 0,38 | 4,37 | 0,09 | 9,32E-01 |
| AT5G54650 | 0,47 | -0,23 | 4,33 | -0,05 | 9,58E-01 |
| AT5G54690 | 0,39 | -1,28 | 4,38 | -0,29 | 7,70E-01 |
| AT5G54700 | 0,89 | -2,99 | 2,38 | -1,26 | 2,08E-01 |
| AT5G54710 | 0,79 | -0,69 | 2,64 | -0,26 | 7,93E-01 |
| AT5G54720 | 0,73 | -0,79 | 2,53 | -0,31 | 7,55E-01 |
| AT5G54780 | 1,16 | -1,23 | 1,62 | -0,76 | 4,48E-01 |
| AT5G54810 | 4,59 | -0,15 | 0,85 | -0,17 | 8,62E-01 |
| AT5G54930 | 0,40 | 0,78 | 4,15 | 0,19 | 8,51E-01 |
| AT5G54940 | 2,86 | 0,91 | 1,23 | 0,74 | 4,60E-01 |
| AT5G54960 | 0,98 | 0,93 | 2,29 | 0,40 | 6,86E-01 |
| AT5G55040 | 0,74 | 0,91 | 2,18 | 0,42 | 6,76E-01 |
| AT5G55055 | 0,05 | 0,00 | 4,41 | 0,00 | 1,00E+00 |
| AT5G55100 | 1,12 | -1,28 | 2,23 | -0,57 | 5,66E-01 |
| AT5G55130 | 0,31 | 1,40 | 3,59 | 0,39 | 6,95E-01 |
| AT5G55150 | 1,21 | -1,74 | 2,25 | -0,77 | 4,39E-01 |
| AT5G55160 | 0,37 | 0,70 | 4,33 | 0,16 | 8,72E-01 |
| AT5G55210 | 0,08 | 0,00 | 4,40 | 0,00 | 1,00E+00 |
| AT5G55250 | 0,40 | 1,06 | 4,35 | 0,24 | 8,08E-01 |
| AT5G55280 | 1,13 | 0,94 | 2,49 | 0,38 | 7,06E-01 |
| AT5G55290 | 0,30 | -1,78 | 4,41 | -0,40 | 6,87E-01 |
| AT5G55300 | 0,70 | -0,60 | 2,22 | -0,27 | 7,87E-01 |
| AT5G55340 | 0,19 | 0,13 | 4,41 | 0,03 | 9,76E-01 |
| AT5G55390 | 0,36 | 1,01 | 4,34 | 0,23 | 8,16E-01 |
| AT5G08275 | 0,59 | -1,87 | 3,74 | -0,50 | 6,18E-01 |
| AT5G08285 | 0,30 | -0,80 | 4,34 | -0,18 | 8,54E-01 |
| AT5G55510 | 0,05 | 0,00 | 4,41 | 0,00 | 1,00E+00 |
| AT5G55540 | 1,63 | 0,54 | 1,45 | 0,37 | 7,10E-01 |
| AT5G55550 | 0,34 | 0,58 | 4,34 | 0,13 | 8,94E-01 |
| AT5G55565 | 4,30 | 1,84 | 1,35 | 1,37 | 1,71E-01 |
| AT5G55570 | 1,30 | -2,84 | 3,08 | -0,92 | 3,57E-01 |
| AT5G55600 | 0,31 | 1,52 | 3,62 | 0,42 | 6,75E-01 |
| AT5G55670 | 0,30 | 0,85 | 4,34 | 0,20 | 8,45E-01 |
| AT5G55710 | 0,92 | 1,84 | 2,42 | 0,76 | 4,46E-01 |
| AT5G55740 | 0,27 | 1,46 | 4,33 | 0,34 | 7,37E-01 |
| AT5G55790 | 0,33 | 1,35 | 4,34 | 0,31 | 7,56E-01 |
| AT5G55835 | 0,61 | 0,72 | 2,34 | 0,31 | 7,56E-01 |
| AT5G55910 | 1,51 | -0,79 | 1,42 | -0,55 | 5,80E-01 |
| AT5G55920 | 0,81 | 0,42 | 2,39 | 0,17 | 8,61E-01 |
| AT5G55940 | 0,50 | -0,07 | 3,84 | -0,02 | 9,85E-01 |
| AT5G56000 | 1,26 | -2,58 | 2,18 | -1,18 | 2,36E-01 |
| AT5G56180 | 0,23 | -1,78 | 4,41 | -0,40 | 6,87E-01 |
| AT5G56240 | 1,23 | 0,32 | 2,06 | 0,16 | 8,76E-01 |

| | | | | | |
|---|---|---|---|---|---|
| AT5G56250 | 0,87 | 3,15 | 2,77 | 1,14 | 2,55E-01 |
| AT5G56260 | 0,25 | -1,78 | 4,41 | -0,40 | 6,87E-01 |
| AT5G56280 | 0,08 | 0,00 | 4,40 | 0,00 | 1,00E+00 |
| AT5G56350 | 0,73 | -0,57 | 1,96 | -0,29 | 7,72E-01 |
| AT5G56360 | 0,87 | 2,01 | 2,86 | 0,70 | 4,82E-01 |
| AT5G56410 | 0,30 | -1,19 | 4,35 | -0,27 | 7,84E-01 |
| AT5G56450 | 0,20 | 1,00 | 4,36 | 0,23 | 8,18E-01 |
| AT5G56710 | 0,79 | -1,04 | 3,28 | -0,32 | 7,50E-01 |
| AT5G56730 | 0,65 | 0,41 | 2,45 | 0,17 | 8,65E-01 |
| AT5G56760 | 0,72 | -1,72 | 2,79 | -0,62 | 5,37E-01 |
| AT5G56860 | 0,23 | 0,19 | 4,38 | 0,04 | 9,66E-01 |
| AT5G56890 | 0,16 | 1,33 | 4,33 | 0,31 | 7,59E-01 |
| AT5G56900 | 1,69 | -1,10 | 1,46 | -0,76 | 4,50E-01 |
| AT5G56970 | 2,67 | 0,01 | 1,32 | 0,01 | 9,94E-01 |
| AT5G56980 | 0,28 | -0,02 | 4,35 | 0,00 | 9,96E-01 |
| AT5G57040 | 0,80 | -1,34 | 1,91 | -0,70 | 4,84E-01 |
| AT5G57110 | 2,43 | -1,84 | 1,30 | -1,42 | 1,56E-01 |
| AT5G57160 | 0,81 | -1,21 | 2,36 | -0,51 | 6,08E-01 |
| AT5G57170 | 0,08 | 0,00 | 4,40 | 0,00 | 1,00E+00 |
| AT5G57250 | 0,54 | 1,49 | 3,84 | 0,39 | 6,98E-01 |
| AT5G57270 | 0,45 | -2,29 | 4,39 | -0,52 | 6,02E-01 |
| AT5G57290 | 0,69 | -0,93 | 2,62 | -0,36 | 7,22E-01 |
| AT5G57345 | 0,70 | 0,74 | 2,30 | 0,32 | 7,47E-01 |
| AT5G57350 | 4,58 | 0,56 | 0,95 | 0,59 | 5,54E-01 |
| AT5G57560 | 1,92 | -1,01 | 1,32 | -0,76 | 4,45E-01 |
| AT5G57580 | 0,90 | 1,76 | 2,38 | 0,74 | 4,61E-01 |
| AT5G57630 | 0,49 | -0,09 | 3,37 | -0,03 | 9,79E-01 |
| AT5G57670 | 0,16 | 0,09 | 4,38 | 0,02 | 9,83E-01 |
| AT5G57780 | 0,36 | -0,21 | 4,33 | -0,05 | 9,62E-01 |
| AT5G57785 | 0,56 | 1,75 | 2,64 | 0,66 | 5,06E-01 |
| AT5G57790 | 0,05 | 0,00 | 4,41 | 0,00 | 1,00E+00 |
| AT5G57850 | 0,51 | -2,06 | 4,37 | -0,47 | 6,38E-01 |
| AT5G57900 | 0,27 | 1,30 | 4,35 | 0,30 | 7,64E-01 |
| AT5G57960 | 0,12 | 0,26 | 4,37 | 0,06 | 9,52E-01 |
| AT5G57990 | 5,76 | -0,41 | 0,79 | -0,52 | 6,02E-01 |
| AT5G58030 | 0,11 | 0,65 | 4,36 | 0,15 | 8,81E-01 |
| AT5G58050 | 0,17 | 1,40 | 4,33 | 0,32 | 7,47E-01 |
| AT5G58070 | 2,01 | 0,57 | 1,56 | 0,36 | 7,16E-01 |
| AT5G58090 | 0,76 | -0,10 | 2,27 | -0,04 | 9,64E-01 |
| AT5G58130 | 0,84 | 2,23 | 2,63 | 0,85 | 3,96E-01 |
| AT5G58150 | 3,53 | 0,60 | 1,41 | 0,43 | 6,70E-01 |
| AT5G58170 | 0,90 | 2,53 | 3,15 | 0,80 | 4,23E-01 |
| AT5G58220 | 0,12 | 0,00 | 4,38 | 0,00 | 1,00E+00 |
| AT5G58260 | 0,45 | -0,78 | 3,43 | -0,23 | 8,20E-01 |
| AT5G58330 | 1,92 | -0,28 | 1,42 | -0,20 | 8,41E-01 |
| AT5G58380 | 0,93 | 0,79 | 2,12 | 0,37 | 7,08E-01 |
| AT5G58430 | 1,95 | -0,27 | 1,31 | -0,21 | 8,36E-01 |
| AT5G58440 | 0,43 | -0,22 | 2,86 | -0,08 | 9,38E-01 |
| AT5G58550 | 1,86 | 1,02 | 1,64 | 0,62 | 5,35E-01 |
| AT5G58570 | 0,61 | 2,56 | 3,87 | 0,66 | 5,09E-01 |
| AT5G08860 | 0,08 | 0,00 | 4,40 | 0,00 | 1,00E+00 |
| AT5G58600 | 0,29 | -0,93 | 4,34 | -0,21 | 8,31E-01 |

| | | | | | |
|---|---|---|---|---|---|
| AT5G58650 | 0,16 | 0,00 | 4,37 | 0,00 | 9,99E-01 |
| AT5G58670 | 0,69 | -1,01 | 2,11 | -0,48 | 6,32E-01 |
| AT5G58720 | 0,64 | -0,97 | 2,37 | -0,41 | 6,82E-01 |
| AT5G58730 | 0,50 | -0,34 | 3,95 | -0,09 | 9,32E-01 |
| AT5G58870 | 0,47 | 1,78 | 3,37 | 0,53 | 5,98E-01 |
| AT5G58900 | 0,28 | -1,42 | 4,36 | -0,33 | 7,44E-01 |
| AT5G58930 | 1,32 | 0,92 | 1,75 | 0,53 | 5,97E-01 |
| AT5G58950 | 2,42 | 1,09 | 1,35 | 0,81 | 4,19E-01 |
| AT5G59030 | 0,49 | 0,15 | 3,83 | 0,04 | 9,70E-01 |
| AT5G59130 | 0,21 | 1,72 | 4,33 | 0,40 | 6,91E-01 |
| AT5G59220 | 0,62 | -2,06 | 3,65 | -0,56 | 5,73E-01 |
| AT5G59290 | 0,24 | -0,27 | 4,35 | -0,06 | 9,51E-01 |
| AT5G08920 | 0,49 | 0,09 | 3,95 | 0,02 | 9,82E-01 |
| AT5G59480 | 0,17 | 1,50 | 4,33 | 0,35 | 7,29E-01 |
| AT5G59550 | 1,55 | 1,25 | 1,59 | 0,79 | 4,31E-01 |
| AT5G08925 | 2,86 | 1,09 | 1,75 | 0,62 | 5,33E-01 |
| AT5G59662 | 0,05 | 0,00 | 4,41 | 0,00 | 1,00E+00 |
| AT5G59700 | 0,33 | -2,48 | 4,37 | -0,57 | 5,70E-01 |
| AT5G59730 | 3,71 | -0,47 | 1,16 | -0,40 | 6,87E-01 |
| AT5G59770 | 0,46 | -1,66 | 3,25 | -0,51 | 6,09E-01 |
| AT5G59780 | 1,19 | 0,67 | 1,76 | 0,38 | 7,05E-01 |
| AT5G59840 | 0,17 | 0,02 | 4,39 | 0,00 | 9,97E-01 |
| AT5G59850 | 0,69 | -3,07 | 2,58 | -1,19 | 2,34E-01 |
| AT5G59870 | 0,19 | 0,52 | 4,38 | 0,12 | 9,05E-01 |
| AT5G59900 | 0,16 | 0,00 | 4,41 | 0,00 | 1,00E+00 |
| AT5G59960 | 0,11 | 0,65 | 4,36 | 0,15 | 8,81E-01 |
| AT5G60050 | 0,15 | 0,48 | 4,38 | 0,11 | 9,13E-01 |
| AT5G60100 | 0,30 | 0,03 | 4,39 | 0,01 | 9,94E-01 |
| AT5G60160 | 0,49 | 1,51 | 4,05 | 0,37 | 7,10E-01 |
| AT5G60180 | 0,21 | -0,49 | 4,35 | -0,11 | 9,11E-01 |
| AT5G60210 | 0,29 | 0,74 | 4,35 | 0,17 | 8,65E-01 |
| AT5G60340 | 0,36 | 0,42 | 4,38 | 0,10 | 9,23E-01 |
| AT5G60400 | 0,12 | 1,23 | 4,34 | 0,28 | 7,76E-01 |
| AT5G60460 | 0,27 | 0,29 | 4,39 | 0,07 | 9,48E-01 |
| AT5G60640 | 2,58 | -0,78 | 1,36 | -0,57 | 5,65E-01 |
| AT5G60670 | 0,64 | -1,69 | 2,16 | -0,78 | 4,33E-01 |
| AT5G60700 | 1,32 | -1,00 | 2,05 | -0,49 | 6,25E-01 |
| AT5G60710 | 1,48 | -0,62 | 1,37 | -0,45 | 6,51E-01 |
| AT5G60800 | 0,66 | -1,21 | 2,96 | -0,41 | 6,81E-01 |
| AT5G60840 | 0,12 | 0,00 | 4,41 | 0,00 | 1,00E+00 |
| AT5G60900 | 5,69 | 1,24 | 0,87 | 1,42 | 1,55E-01 |
| AT5G60920 | 1,88 | 0,75 | 1,53 | 0,49 | 6,26E-01 |
| AT5G60930 | 0,24 | -0,74 | 4,35 | -0,17 | 8,65E-01 |
| AT5G60960 | 0,29 | 0,10 | 4,32 | 0,02 | 9,82E-01 |
| AT5G60990 | 0,16 | 0,92 | 4,35 | 0,21 | 8,33E-01 |
| AT5G61020 | 2,23 | -0,44 | 1,21 | -0,36 | 7,18E-01 |
| AT5G61040 | 0,88 | -1,28 | 4,35 | -0,29 | 7,68E-01 |
| AT5G61060 | 0,37 | 0,32 | 3,42 | 0,09 | 9,25E-01 |
| AT5G61130 | 0,17 | 1,50 | 4,33 | 0,35 | 7,29E-01 |
| AT5G61150 | 0,86 | 0,36 | 2,59 | 0,14 | 8,89E-01 |
| AT5G61250 | 0,23 | -0,21 | 4,35 | -0,05 | 9,61E-01 |
| AT5G61310 | 4,76 | 0,51 | 1,01 | 0,50 | 6,17E-01 |

| | | | | | |
|---|---|---|---|---|---|
| AT5G61390 | 0,43 | -0,62 | 4,35 | -0,14 | 8,86E-01 |
| AT5G61410 | 2,49 | 0,25 | 1,18 | 0,21 | 8,35E-01 |
| AT5G61420 | 2,01 | -0,19 | 1,52 | -0,13 | 9,00E-01 |
| AT5G09000 | 2,90 | -2,89 | 1,35 | -2,15 | 3,16E-02 |
| AT5G09005 | 1,51 | 2,75 | 1,65 | 1,67 | 9,58E-02 |
| AT5G61460 | 0,21 | 1,72 | 4,33 | 0,40 | 6,91E-01 |
| AT5G61480 | 0,83 | -1,84 | 2,10 | -0,88 | 3,81E-01 |
| AT5G61500 | 0,76 | -1,96 | 2,79 | -0,70 | 4,82E-01 |
| AT5G61510 | 0,56 | -1,38 | 3,19 | -0,43 | 6,65E-01 |
| AT5G61520 | 0,33 | 0,31 | 4,32 | 0,07 | 9,42E-01 |
| AT5G61590 | 1,28 | -3,60 | 1,91 | -1,88 | 5,99E-02 |
| AT5G61600 | 0,57 | 1,19 | 2,62 | 0,45 | 6,51E-01 |
| AT5G61790 | 5,40 | -0,12 | 0,87 | -0,13 | 8,94E-01 |
| AT5G61800 | 0,12 | 1,05 | 4,35 | 0,24 | 8,10E-01 |
| AT5G61810 | 0,48 | -1,26 | 3,26 | -0,39 | 6,99E-01 |
| AT5G61820 | 1,07 | 0,38 | 2,18 | 0,17 | 8,62E-01 |
| AT5G61840 | 0,89 | 0,57 | 2,15 | 0,27 | 7,91E-01 |
| AT5G61910 | 1,60 | 1,53 | 1,58 | 0,97 | 3,32E-01 |
| AT5G61900 | 1,09 | 1,83 | 1,88 | 0,97 | 3,31E-01 |
| AT5G61960 | 0,94 | -1,09 | 2,50 | -0,43 | 6,64E-01 |
| AT5G61990 | 0,30 | -0,27 | 4,36 | -0,06 | 9,50E-01 |
| AT5G09040 | 2,31 | 0,09 | 1,57 | 0,06 | 9,54E-01 |
| AT5G62090 | 0,32 | 1,04 | 4,34 | 0,24 | 8,11E-01 |
| AT5G62110 | 0,34 | 1,50 | 4,32 | 0,35 | 7,29E-01 |
| AT5G62130 | 0,45 | 0,00 | 3,28 | 0,00 | 1,00E+00 |
| AT5G62140 | 1,15 | 2,64 | 1,81 | 1,46 | 1,44E-01 |
| AT5G62180 | 0,08 | 0,69 | 4,36 | 0,16 | 8,75E-01 |
| AT5G62190 | 2,05 | -2,66 | 1,37 | -1,93 | 5,30E-02 |
| AT5G62220 | 1,34 | -0,99 | 1,85 | -0,54 | 5,93E-01 |
| AT5G62300 | 1,07 | -2,17 | 1,78 | -1,22 | 2,23E-01 |
| AT5G62430 | 0,29 | -0,77 | 4,34 | -0,18 | 8,58E-01 |
| AT5G62470 | 0,30 | 0,00 | 4,41 | 0,00 | 1,00E+00 |
| AT5G62530 | 0,56 | 2,21 | 3,05 | 0,73 | 4,67E-01 |
| AT5G62600 | 0,50 | -0,34 | 3,95 | -0,09 | 9,32E-01 |
| AT5G62610 | 0,31 | -0,25 | 4,34 | -0,06 | 9,54E-01 |
| AT5G62630 | 0,37 | -0,45 | 4,34 | -0,10 | 9,17E-01 |
| AT5G62650 | 0,39 | -2,35 | 4,38 | -0,54 | 5,91E-01 |
| AT5G62680 | 0,33 | 0,35 | 4,32 | 0,08 | 9,35E-01 |
| AT5G62790 | 0,60 | 0,00 | 2,65 | 0,00 | 9,99E-01 |
| AT5G63080 | 0,23 | 0,66 | 4,38 | 0,15 | 8,81E-01 |
| AT5G63100 | 0,29 | 0,71 | 4,31 | 0,17 | 8,68E-01 |
| AT5G63110 | 0,58 | 1,98 | 3,11 | 0,64 | 5,24E-01 |
| AT5G63120 | 0,65 | 0,90 | 3,07 | 0,29 | 7,70E-01 |
| AT5G63160 | 0,51 | 0,83 | 3,16 | 0,26 | 7,93E-01 |
| AT5G63180 | 0,25 | 0,37 | 4,37 | 0,09 | 9,32E-01 |
| AT5G63195 | 1,93 | -1,07 | 1,57 | -0,68 | 4,94E-01 |
| AT5G63290 | 0,56 | -0,09 | 4,31 | -0,02 | 9,83E-01 |
| AT5G63310 | 0,47 | -0,10 | 4,33 | -0,02 | 9,81E-01 |
| AT5G63320 | 0,52 | 0,68 | 3,92 | 0,17 | 8,63E-01 |
| AT5G63370 | 1,26 | -0,38 | 1,49 | -0,25 | 7,99E-01 |
| AT5G63400 | 0,30 | -0,92 | 4,39 | -0,21 | 8,35E-01 |
| AT5G63410 | 0,46 | -0,17 | 2,74 | -0,06 | 9,50E-01 |

| | | | | | |
|---|---|---|---|---|---|
| AT5G63470 | 0,44 | -1,77 | 4,11 | -0,43 | 6,66E-01 |
| AT5G63610 | 0,51 | -0,35 | 3,91 | -0,09 | 9,30E-01 |
| AT5G63625 | 1,28 | 2,30 | 2,00 | 1,15 | 2,51E-01 |
| AT5G63760 | 0,90 | 0,23 | 2,26 | 0,10 | 9,18E-01 |
| AT5G63790 | 0,54 | 0,77 | 4,29 | 0,18 | 8,57E-01 |
| AT5G63860 | 0,75 | -0,25 | 2,08 | -0,12 | 9,06E-01 |
| AT5G63870 | 0,08 | 0,44 | 4,38 | 0,10 | 9,21E-01 |
| AT5G63890 | 0,39 | 0,69 | 4,30 | 0,16 | 8,72E-01 |
| AT5G63910 | 0,66 | -2,97 | 2,97 | -1,00 | 3,18E-01 |
| AT5G64020 | 0,23 | -0,21 | 4,35 | -0,05 | 9,61E-01 |
| AT5G64040 | 3,44 | -1,57 | 1,26 | -1,24 | 2,14E-01 |
| AT5G64050 | 1,12 | 0,34 | 1,70 | 0,20 | 8,41E-01 |
| AT5G64070 | 1,53 | 0,16 | 1,43 | 0,11 | 9,09E-01 |
| AT5G64130 | 0,35 | 0,58 | 3,64 | 0,16 | 8,74E-01 |
| AT5G64140 | 1,06 | -1,27 | 1,70 | -0,75 | 4,54E-01 |
| AT5G64170 | 0,64 | 0,46 | 3,19 | 0,15 | 8,84E-01 |
| AT5G64200 | 0,28 | 0,09 | 4,33 | 0,02 | 9,83E-01 |
| AT5G64280 | 1,39 | -1,40 | 1,45 | -0,96 | 3,35E-01 |
| AT5G64290 | 0,42 | 1,68 | 4,10 | 0,41 | 6,82E-01 |
| AT5G64340 | 1,19 | -0,12 | 1,74 | -0,07 | 9,43E-01 |
| AT5G64350 | 0,41 | 0,71 | 4,25 | 0,17 | 8,66E-01 |
| AT5G64360 | 0,62 | -3,75 | 3,65 | -1,03 | 3,04E-01 |
| AT5G64410 | 0,79 | 0,55 | 2,30 | 0,24 | 8,12E-01 |
| AT5G64430 | 0,96 | 1,16 | 2,22 | 0,52 | 6,02E-01 |
| AT5G64460 | 0,08 | 0,00 | 4,40 | 0,00 | 1,00E+00 |
| AT5G64550 | 0,25 | 0,38 | 4,33 | 0,09 | 9,31E-01 |
| AT5G64570 | 1,49 | -0,98 | 1,44 | -0,68 | 4,96E-01 |
| AT5G64580 | 0,77 | -1,46 | 2,50 | -0,59 | 5,58E-01 |
| AT5G64610 | 0,28 | 1,17 | 4,35 | 0,27 | 7,88E-01 |
| AT5G64660 | 0,37 | 1,16 | 3,42 | 0,34 | 7,36E-01 |
| AT5G64670 | 0,17 | 0,47 | 4,37 | 0,11 | 9,14E-01 |
| AT5G64760 | 0,83 | 1,03 | 2,97 | 0,35 | 7,28E-01 |
| AT5G64830 | 0,77 | -0,48 | 2,30 | -0,21 | 8,36E-01 |
| AT5G64840 | 5,36 | 1,16 | 0,84 | 1,38 | 1,68E-01 |
| AT5G64860 | 0,28 | 1,55 | 4,33 | 0,36 | 7,20E-01 |
| AT5G64870 | 0,67 | 1,92 | 2,83 | 0,68 | 4,96E-01 |
| AT5G64920 | 1,21 | -0,52 | 1,63 | -0,32 | 7,51E-01 |
| AT5G64930 | 0,39 | -1,40 | 4,37 | -0,32 | 7,49E-01 |
| AT5G64980 | 0,12 | 1,16 | 4,34 | 0,27 | 7,89E-01 |
| AT5G65110 | 0,46 | 1,94 | 3,17 | 0,61 | 5,42E-01 |
| AT5G65200 | 0,16 | 0,92 | 4,35 | 0,21 | 8,33E-01 |
| AT5G65240 | 0,27 | 1,17 | 3,74 | 0,31 | 7,55E-01 |
| AT5G65310 | 1,18 | -3,13 | 2,14 | -1,46 | 1,44E-01 |
| AT5G65360 | 0,52 | -1,38 | 3,14 | -0,44 | 6,61E-01 |
| AT5G65380 | 2,69 | -1,56 | 1,09 | -1,42 | 1,54E-01 |
| AT5G65430 | 0,62 | -2,31 | 2,62 | -0,88 | 3,77E-01 |
| AT5G65560 | 0,37 | 0,28 | 4,32 | 0,06 | 9,48E-01 |
| AT5G65630 | 3,70 | 0,19 | 1,00 | 0,19 | 8,48E-01 |
| AT5G65685 | 0,97 | -0,22 | 2,20 | -0,10 | 9,22E-01 |
| AT5G65840 | 0,17 | 0,35 | 4,37 | 0,08 | 9,37E-01 |
| AT5G65860 | 5,04 | 0,38 | 0,96 | 0,39 | 6,94E-01 |
| AT5G09495 | 4,86 | 0,19 | 0,97 | 0,19 | 8,47E-01 |

| | | | | | |
|---|---|---|---|---|---|
| AT5G65910 | 1,14 | 0,13 | 3,17 | 0,04 | 9,66E-01 |
| AT5G65930 | 0,99 | 2,02 | 2,08 | 0,97 | 3,32E-01 |
| AT5G09535 | 0,24 | 0,21 | 4,34 | 0,05 | 9,62E-01 |
| AT5G65970 | 0,45 | -2,08 | 4,33 | -0,48 | 6,32E-01 |
| AT5G66030 | 2,48 | -0,92 | 1,18 | -0,78 | 4,36E-01 |
| AT5G66055 | 0,63 | 1,20 | 2,51 | 0,48 | 6,33E-01 |
| AT5G66120 | 0,85 | 2,38 | 2,19 | 1,09 | 2,77E-01 |
| AT5G66190 | 3,89 | -0,51 | 1,25 | -0,41 | 6,84E-01 |
| AT5G66200 | 3,05 | 0,46 | 1,09 | 0,42 | 6,75E-01 |
| AT5G66210 | 1,05 | -0,63 | 2,11 | -0,30 | 7,64E-01 |
| AT5G66300 | 0,13 | 0,00 | 4,41 | 0,00 | 1,00E+00 |
| AT5G66310 | 0,12 | 1,16 | 4,34 | 0,27 | 7,89E-01 |
| AT5G66320 | 0,48 | -0,45 | 3,27 | -0,14 | 8,91E-01 |
| AT5G66330 | 0,11 | 0,00 | 4,41 | 0,00 | 1,00E+00 |
| AT5G66400 | 0,42 | 0,72 | 4,22 | 0,17 | 8,65E-01 |
| AT5G66440 | 0,33 | 0,35 | 4,39 | 0,08 | 9,36E-01 |
| AT5G66460 | 0,57 | -0,49 | 4,33 | -0,11 | 9,09E-01 |
| AT5G66500 | 0,15 | 1,20 | 4,34 | 0,28 | 7,81E-01 |
| AT5G66530 | 0,97 | -2,50 | 2,48 | -1,01 | 3,13E-01 |
| AT5G66540 | 0,21 | 1,42 | 4,33 | 0,33 | 7,43E-01 |
| AT5G66558 | 1,38 | -0,67 | 1,51 | -0,44 | 6,58E-01 |
| AT5G66595 | 0,74 | -1,04 | 2,47 | -0,42 | 6,72E-01 |
| AT5G66607 | 0,08 | 0,00 | 4,40 | 0,00 | 1,00E+00 |
| AT5G66650 | 0,26 | -0,30 | 4,37 | -0,07 | 9,46E-01 |
| AT5G66675 | 0,51 | -0,71 | 3,17 | -0,23 | 8,22E-01 |
| AT5G66680 | 0,76 | -0,62 | 3,61 | -0,17 | 8,64E-01 |
| AT5G66720 | 0,41 | -3,18 | 4,34 | -0,73 | 4,63E-01 |
| AT5G66730 | 0,31 | 0,70 | 4,37 | 0,16 | 8,72E-01 |
| AT5G66850 | 0,24 | 1,56 | 4,33 | 0,36 | 7,19E-01 |
| AT5G66890 | 0,75 | 0,58 | 3,43 | 0,17 | 8,66E-01 |
| AT5G66900 | 1,52 | 0,95 | 1,64 | 0,58 | 5,61E-01 |
| AT5G66910 | 0,38 | -0,15 | 3,41 | -0,04 | 9,66E-01 |
| AT5G67030 | 1,94 | -0,12 | 1,24 | -0,10 | 9,22E-01 |
| AT5G67120 | 0,79 | 2,39 | 2,73 | 0,87 | 3,82E-01 |
| AT5G67150 | 0,35 | -0,88 | 4,39 | -0,20 | 8,42E-01 |
| AT5G67190 | 0,91 | 0,36 | 2,40 | 0,15 | 8,79E-01 |
| AT5G67220 | 0,60 | 0,85 | 2,49 | 0,34 | 7,34E-01 |
| AT5G67240 | 1,06 | 0,15 | 1,94 | 0,08 | 9,36E-01 |
| AT5G67245 | 0,45 | -0,48 | 4,37 | -0,11 | 9,13E-01 |
| AT5G67250 | 1,03 | 1,41 | 2,06 | 0,69 | 4,93E-01 |
| AT5G67280 | 0,58 | -1,07 | 2,69 | -0,40 | 6,91E-01 |
| AT5G67290 | 0,82 | 0,43 | 2,37 | 0,18 | 8,57E-01 |
| AT5G09855 | 0,69 | -0,18 | 2,35 | -0,08 | 9,38E-01 |
| AT5G67310 | 0,78 | 0,97 | 2,73 | 0,35 | 7,23E-01 |
| AT5G67350 | 0,75 | 1,83 | 2,59 | 0,71 | 4,80E-01 |
| AT5G67380 | 0,30 | -0,85 | 4,36 | -0,20 | 8,45E-01 |
| AT5G67420 | 2,71 | -1,37 | 1,20 | -1,15 | 2,51E-01 |
| AT5G67460 | 0,10 | 0,01 | 4,38 | 0,00 | 9,99E-01 |
| AT5G67480 | 1,04 | 0,51 | 2,42 | 0,21 | 8,32E-01 |
| AT5G67490 | 1,16 | 0,98 | 1,85 | 0,53 | 5,95E-01 |
| AT5G67580 | 0,39 | 1,19 | 2,86 | 0,42 | 6,76E-01 |
| AT5G67610 | 0,37 | -3,12 | 4,34 | -0,72 | 4,72E-01 |

***chr11 chr17* infected by TCV relative to infected wild type.**

| |
|---|
| type |
| miRNA_precursor |
| miRNA_precursor |
| miRNA_precursor |
| miRNA_precursor |
| ncRNA |
| miRNA_precursor |
| miRNA_precursor |
| miRNA_precursor |
| protein_coding |
| protein_coding |
| miRNA_precursor |
| ncRNA |
| miRNA_precursor |
| miRNA_precursor |
| miRNA_precursor |
| miRNA_precursor |
| miRNA_precursor |
| miRNA_precursor |
| ncRNA |
| miRNA_precursor |
| protein_coding |
| miRNA_precursor |
| protein_coding |
| miRNA_precursor |
| miRNA_precursor |
| miRNA_precursor |
| miRNA_precursor |
| protein_coding |
| protein_coding |
| protein_coding |
| protein_coding |
| protein_coding |
| protein_coding |
| miRNA_precursor |
| miRNA_precursor |
| protein_coding |
| protein_coding |
| miRNA_precursor |
| protein_coding |
| ncRNA |
| protein_coding |
| protein_coding |
| miRNA_precursor |
| protein_coding |
| miRNA_precursor |
| miRNA_precursor |
| small_nucle(ol)ar_rna |
| protein_coding |
| protein_coding |

miRNA_precursor
small_nucle(ol)ar_rna
protein_coding
protein_coding
miRNA_precursor
ribosomal_rna
small_nucle(ol)ar_rna
protein_coding
ncRNA
miRNA_precursor
miRNA_precursor
protein_coding
protein_coding
miRNA_precursor
protein_coding
miRNA_precursor
miRNA_precursor
miRNA_precursor
protein_coding
miRNA_precursor
miRNA_precursor
protein_coding
ncRNA
ncRNA
protein_coding
miRNA_precursor
protein_coding
protein_coding
miRNA_precursor
miRNA_precursor
ncRNA
protein_coding
protein_coding
miRNA_precursor
protein_coding
protein_coding
small_nucle(ol)ar_rna
protein_coding
protein_coding
protein_coding
protein_coding
protein_coding
miRNA_precursor
miRNA_precursor
protein_coding
protein_coding
miRNA_precursor
protein_coding
protein_coding
miRNA_precursor
miRNA_precursor
protein_coding

protein_coding
protein_coding
protein_coding
protein_coding
ncRNA
small_nucle(ol)ar_rna
miRNA_precursor
miRNA_precursor
protein_coding
ncRNA
protein_coding
protein_coding
protein_coding
protein_coding
protein_coding
protein_coding
ncRNA
protein_coding
protein_coding
protein_coding
protein_coding
small_nucle(ol)ar_rna
protein_coding
protein_coding
protein_coding
small_nucle(ol)ar_rna
miRNA_precursor
protein_coding
ribosomal_rna
protein_coding
protein_coding
protein_coding
protein_coding
protein_coding
ncRNA
protein_coding
protein_coding
protein_coding
protein_coding
protein_coding
miRNA_precursor
ncRNA
miRNA_precursor
protein_coding
protein_coding
miRNA_precursor
ncRNA
protein_coding
small_nucle(ol)ar_rna
protein_coding
protein_coding
protein_coding

protein_coding
protein_coding
miRNA_precursor
ncRNA
miRNA_precursor
protein_coding
small_nucle(ol)ar_rna
ncRNA
protein_coding
protein_coding
protein_coding
small_nucle(ol)ar_rna
protein_coding
small_nucle(ol)ar_rna
protein_coding
protein_coding
miRNA_precursor
pre_trna
protein_coding
protein_coding
protein_coding
miRNA_precursor
miRNA_precursor
protein_coding
small_nucle(ol)ar_rna
protein_coding
protein_coding
protein_coding
small_nucle(ol)ar_rna
protein_coding
protein_coding
small_nucle(ol)ar_rna
protein_coding
protein_coding
protein_coding
miRNA_precursor
protein_coding
protein_coding
protein_coding
miRNA_precursor
ncRNA
miRNA_precursor
small_nucle(ol)ar_rna
protein_coding
protein_coding
protein_coding
small_nucle(ol)ar_rna
protein_coding
protein_coding
small_nucle(ol)ar_rna
protein_coding
protein_coding

small_nucle(ol)ar_rna
protein_coding
protein_coding
ncRNA
protein_coding
miRNA_precursor
protein_coding
protein_coding
miRNA_precursor
protein_coding
protein_coding
ribosomal_rna
small_nucle(ol)ar_rna
protein_coding
protein_coding
protein_coding
protein_coding
protein_coding
protein_coding
small_nucle(ol)ar_rna
protein_coding
protein_coding
protein_coding
protein_coding
protein_coding
miRNA_precursor
miRNA_precursor
protein_coding
protein_coding
ncRNA
protein_coding
miRNA_precursor
protein_coding
ncRNA
miRNA_precursor
protein_coding
protein_coding
protein_coding
ncRNA
protein_coding
protein_coding
small_nucle(ol)ar_rna
small_nucle(ol)ar_rna
miRNA_precursor
ncRNA
miRNA_precursor
miRNA_precursor
protein_coding
protein_coding
miRNA_precursor
protein_coding
protein_coding

miRNA_precursor
protein_coding
protein_coding
protein_coding
small_nucle(ol)ar_rna
small_nucle(ol)ar_rna
protein_coding
ncRNA
small_nucle(ol)ar_rna
protein_coding
miRNA_precursor
small_nucle(ol)ar_rna
protein_coding
ncRNA
protein_coding
protein_coding
protein_coding
protein_coding
miRNA_precursor
small_nucle(ol)ar_rna
miRNA_precursor
protein_coding
miRNA_precursor
protein_coding
protein_coding
small_nucle(ol)ar_rna
miRNA_precursor
protein_coding
protein_coding
protein_coding
small_nucle(ol)ar_rna
protein_coding
protein_coding
protein_coding
protein_coding
protein_coding
protein_coding
protein_coding
protein_coding
protein_coding
ncRNA
protein_coding
ncRNA
protein_coding
protein_coding
protein_coding
protein_coding
protein_coding
protein_coding
ncRNA
protein_coding
small_nucle(ol)ar_rna

protein_coding
protein_coding
protein_coding
protein_coding
protein_coding
protein_coding
protein_coding
protein_coding
small_nucle(ol)ar_rna
ncRNA
miRNA_precursor
protein_coding
protein_coding
protein_coding
protein_coding
small_nucle(ol)ar_rna
protein_coding
protein_coding
protein_coding
small_nucle(ol)ar_rna
small_nucle(ol)ar_rna
protein_coding
protein_coding
protein_coding
small_nucle(ol)ar_rna
ncRNA
protein_coding
protein_coding
protein_coding
small_nucle(ol)ar_rna
protein_coding
protein_coding
protein_coding
protein_coding
miRNA_precursor
protein_coding
protein_coding
protein_coding
ncRNA
protein_coding
protein_coding
protein_coding
small_nucle(ol)ar_rna
protein_coding
protein_coding
protein_coding
ncRNA
small_nucle(ol)ar_rna
protein_coding
miRNA_precursor
ncRNA
protein_coding

ncRNA
protein_coding
protein_coding
protein_coding
protein_coding
protein_coding
miRNA_precursor
protein_coding
ncRNA
protein_coding
small_nucle(ol)ar_rna
protein_coding
small_nucle(ol)ar_rna
protein_coding
protein_coding
protein_coding
protein_coding
protein_coding
protein_coding
miRNA_precursor
protein_coding
protein_coding
small_nucle(ol)ar_rna
protein_coding
protein_coding
small_nucle(ol)ar_rna
protein_coding
protein_coding
protein_coding
protein_coding
protein_coding
small_nucle(ol)ar_rna
protein_coding
protein_coding
protein_coding
protein_coding
small_nucle(ol)ar_rna
protein_coding
protein_coding
protein_coding
protein_coding
protein_coding
protein_coding
small_nucle(ol)ar_rna
small_nucle(ol)ar_rna
protein_coding
protein_coding
protein_coding
protein_coding
protein_coding
protein_coding
protein_coding

protein_coding
protein_coding
ncRNA
protein_coding
protein_coding
protein_coding
protein_coding
small_nucle(ol)ar_rna
protein_coding
protein_coding
protein_coding
miRNA_precursor
protein_coding
protein_coding
protein_coding
protein_coding
protein_coding
miRNA_precursor
miRNA_precursor
ncRNA
protein_coding
protein_coding
protein_coding
ncRNA
protein_coding
small_nucle(ol)ar_rna
small_nucle(ol)ar_rna
protein_coding
ncRNA
protein_coding
protein_coding
protein_coding
protein_coding
protein_coding
protein_coding
miRNA_precursor
protein_coding
ncRNA
protein_coding
protein_coding
protein_coding
protein_coding
protein_coding
protein_coding
protein_coding
protein_coding
protein_coding
protein_coding
protein_coding
protein_coding
protein_coding
protein_coding

protein_coding
protein_coding
protein_coding
protein_coding
protein_coding
protein_coding
protein_coding
miRNA_precursor
protein_coding
protein_coding
protein_coding
protein_coding
protein_coding
miRNA_precursor
protein_coding
small_nucle(ol)ar_rna
protein_coding
ncRNA
protein_coding
protein_coding
protein_coding
protein_coding
protein_coding
protein_coding
small_nucle(ol)ar_rna
protein_coding
protein_coding
miRNA_precursor
small_nucle(ol)ar_rna
protein_coding
protein_coding
protein_coding
protein_coding
protein_coding
small_nucle(ol)ar_rna
small_nucle(ol)ar_rna
protein_coding
ncRNA
protein_coding
protein_coding
protein_coding
ncRNA
small_nucle(ol)ar_rna
small_nucle(ol)ar_rna
protein_coding
protein_coding
protein_coding
protein_coding
protein_coding
ncRNA
protein_coding
protein_coding

small_nucle(ol)ar_rna
protein_coding
protein_coding
protein_coding
protein_coding
protein_coding
protein_coding
protein_coding
protein_coding
protein_coding
protein_coding
protein_coding
ncRNA
protein_coding
protein_coding
protein_coding
miRNA_precursor
protein_coding
protein_coding
protein_coding
miRNA_precursor
miRNA_precursor
miRNA_precursor
protein_coding
protein_coding
ncRNA
protein_coding
protein_coding
small_nucle(ol)ar_rna
protein_coding
protein_coding
protein_coding
protein_coding
protein_coding
protein_coding
protein_coding
miRNA_precursor
ncRNA
protein_coding
small_nucle(ol)ar_rna
protein_coding
miRNA_precursor
protein_coding
protein_coding
ncRNA
protein_coding
protein_coding
protein_coding
protein_coding
protein_coding
protein_coding
ncRNA

protein_coding
protein_coding
miRNA_precursor
protein_coding
miRNA_precursor
protein_coding
ncRNA
protein_coding
protein_coding
small_nucle(ol)ar_rna
ncRNA
ncRNA
protein_coding
protein_coding
protein_coding
protein_coding
protein_coding
small_nucle(ol)ar_rna
small_nucle(ol)ar_rna
protein_coding
protein_coding
protein_coding
protein_coding
protein_coding
protein_coding
ncRNA
protein_coding
protein_coding
small_nucle(ol)ar_rna
protein_coding
protein_coding
protein_coding
protein_coding
protein_coding
miRNA_precursor
protein_coding
miRNA_precursor
ncRNA
protein_coding
miRNA_precursor
protein_coding
small_nucle(ol)ar_rna
protein_coding
protein_coding
miRNA_precursor
protein_coding
protein_coding
protein_coding
protein_coding
ncRNA
miRNA_precursor
protein_coding

protein_coding
protein_coding
miRNA_precursor
protein_coding
protein_coding
protein_coding
protein_coding
protein_coding
protein_coding
protein_coding
protein_coding
protein_coding
ncRNA
protein_coding
protein_coding
protein_coding
protein_coding
protein_coding
protein_coding
protein_coding
protein_coding
protein_coding
ncRNA
protein_coding
protein_coding
protein_coding
protein_coding
protein_coding
protein_coding
protein_coding
protein_coding
protein_coding
protein_coding
ncRNA
protein_coding
protein_coding
ncRNA
protein_coding
protein_coding
protein_coding
protein_coding
protein_coding
protein_coding
protein_coding
protein_coding
protein_coding
protein_coding
protein_coding
protein_coding
protein_coding
protein_coding
protein_coding

protein_coding
protein_coding
protein_coding
ncRNA
protein_coding
protein_coding
ncRNA
protein_coding
protein_coding
protein_coding
protein_coding
protein_coding
protein_coding
protein_coding
protein_coding
protein_coding
protein_coding
small_nucle(ol)ar_rna
small_nucle(ol)ar_rna
small_nucle(ol)ar_rna
small_nucle(ol)ar_rna
protein_coding
protein_coding
protein_coding
protein_coding
protein_coding
protein_coding
protein_coding
ncRNA
protein_coding
ncRNA
protein_coding
protein_coding
ncRNA
protein_coding
protein_coding
small_nucle(ol)ar_rna
small_nucle(ol)ar_rna
protein_coding
protein_coding
protein_coding
protein_coding
protein_coding
protein_coding
protein_coding
protein_coding
protein_coding
protein_coding
protein_coding
protein_coding
protein_coding
protein_coding

protein_coding
protein_coding
protein_coding
ncRNA
protein_coding
protein_coding
protein_coding
miRNA_precursor
protein_coding
protein_coding
protein_coding
protein_coding
protein_coding
protein_coding
pre_trna
ncRNA
protein_coding
protein_coding
protein_coding
protein_coding
protein_coding
protein_coding
protein_coding
protein_coding
protein_coding
protein_coding
protein_coding
protein_coding
protein_coding
protein_coding
protein_coding
protein_coding
protein_coding
ncRNA
protein_coding
protein_coding
protein_coding
protein_coding
protein_coding
protein_coding
protein_coding
protein_coding
protein_coding
protein_coding
protein_coding
protein_coding
protein_coding
ncRNA
protein_coding
protein_coding
protein_coding
protein_coding

protein_coding
protein_coding
protein_coding
protein_coding
protein_coding
protein_coding
protein_coding
protein_coding
protein_coding
protein_coding
protein_coding
protein_coding
protein_coding
protein_coding
protein_coding
protein_coding
protein_coding
protein_coding
protein_coding
protein_coding
protein_coding
protein_coding
protein_coding
protein_coding
protein_coding
protein_coding
protein_coding
protein_coding
protein_coding
protein_coding
protein_coding
protein_coding
protein_coding
protein_coding
protein_coding
protein_coding
protein_coding
protein_coding
protein_coding
protein_coding
ncRNA
protein_coding
protein_coding
protein_coding
protein_coding
protein_coding
protein_coding
protein_coding
protein_coding
protein_coding
protein_coding
protein_coding

| |
|---|
| protein_coding |
| protein_coding |
| protein_coding |
| protein_coding |
| protein_coding |
| protein_coding |
| protein_coding |
| protein_coding |
| protein_coding |
| protein_coding |
| protein_coding |
| protein_coding |
| protein_coding |
| protein_coding |
| protein_coding |
| protein_coding |
| protein_coding |
| protein_coding |
| protein_coding |
| protein_coding |
| protein_coding |
| protein_coding |
| protein_coding |
| protein_coding |
| protein_coding |
| protein_coding |
| protein_coding |
| protein_coding |
| protein_coding |
| protein_coding |
| protein_coding |
| protein_coding |
| protein_coding |
| protein_coding |
| protein_coding |
| protein_coding |
| protein_coding |
| protein_coding |
| protein_coding |
| protein_coding |
| protein_coding |
| protein_coding |
| protein_coding |
| protein_coding |
| protein_coding |
| protein_coding |
| protein_coding |
| protein_coding |
| protein_coding |
| protein_coding |
| protein_coding |
| protein_coding |

small_nucle(ol)ar_rna
small_nucle(ol)ar_rna
protein_coding
protein_coding
protein_coding
protein_coding
protein_coding
protein_coding
protein_coding
protein_coding
protein_coding
protein_coding
protein_coding
protein_coding
protein_coding
protein_coding
protein_coding
protein_coding
protein_coding
protein_coding
protein_coding
protein_coding
protein_coding
protein_coding
protein_coding
protein_coding
protein_coding
protein_coding
protein_coding
protein_coding
protein_coding
protein_coding
protein_coding
protein_coding
protein_coding
protein_coding
pre_trna
protein_coding
protein_coding
protein_coding
protein_coding
ncRNA
protein_coding
protein_coding
protein_coding
protein_coding
protein_coding
protein_coding
protein_coding
protein_coding
protein_coding
protein_coding

protein_coding
protein_coding
protein_coding
protein_coding
protein_coding
protein_coding
protein_coding
protein_coding
protein_coding
protein_coding
protein_coding
protein_coding
protein_coding
protein_coding
protein_coding
protein_coding
protein_coding
protein_coding
protein_coding
protein_coding
pre_trna
protein_coding
ncRNA
protein_coding
protein_coding
protein_coding
ncRNA
protein_coding
protein_coding
protein_coding
protein_coding
protein_coding
protein_coding
protein_coding
protein_coding
protein_coding
protein_coding
protein_coding
protein_coding
protein_coding
protein_coding
protein_coding
protein_coding
protein_coding
protein_coding
protein_coding
protein_coding
protein_coding
protein_coding
protein_coding
protein_coding
protein_coding

protein_coding
protein_coding
protein_coding
protein_coding
protein_coding
protein_coding
protein_coding
protein_coding
protein_coding
protein_coding
protein_coding
protein_coding
protein_coding
protein_coding
ncRNA
protein_coding
protein_coding
protein_coding
protein_coding
protein_coding
protein_coding
protein_coding
protein_coding
protein_coding
protein_coding
protein_coding
protein_coding
pre_trna
protein_coding
protein_coding
protein_coding
protein_coding
protein_coding
protein_coding
protein_coding
protein_coding
protein_coding
protein_coding
protein_coding
protein_coding
protein_coding
protein_coding
protein_coding
protein_coding
protein_coding
protein_coding
protein_coding
protein_coding
protein_coding
protein_coding
protein_coding
protein_coding

protein_coding
protein_coding
protein_coding
protein_coding
protein_coding
protein_coding
pre_trna
protein_coding
protein_coding
miRNA_precursor
protein_coding
protein_coding
protein_coding
protein_coding
protein_coding
protein_coding
protein_coding
protein_coding
protein_coding
protein_coding
protein_coding
protein_coding
small_nucle(ol)ar_rna
small_nucle(ol)ar_rna
miRNA_precursor
protein_coding
protein_coding
ncRNA
protein_coding
protein_coding
protein_coding
protein_coding
protein_coding
protein_coding
protein_coding
protein_coding
protein_coding
protein_coding
ncRNA
protein_coding
protein_coding
protein_coding
protein_coding
protein_coding
protein_coding
miRNA_precursor
protein_coding
protein_coding
protein_coding
protein_coding
protein_coding
protein_coding

protein_coding
protein_coding
protein_coding
protein_coding
protein_coding
protein_coding
protein_coding
protein_coding
protein_coding
protein_coding
miRNA_precursor
protein_coding
protein_coding
protein_coding
protein_coding
protein_coding
protein_coding
protein_coding
protein_coding
protein_coding
protein_coding
protein_coding
protein_coding
protein_coding
protein_coding
protein_coding
protein_coding
protein_coding
protein_coding
protein_coding
protein_coding
protein_coding
protein_coding
protein_coding
protein_coding
protein_coding
protein_coding
protein_coding
protein_coding
protein_coding
protein_coding
protein_coding
protein_coding
protein_coding
protein_coding
protein_coding
protein_coding
ncRNA
protein_coding
protein_coding
protein_coding
protein_coding

protein_coding
protein_coding
protein_coding
protein_coding
protein_coding
protein_coding
protein_coding
protein_coding
protein_coding
protein_coding
protein_coding
protein_coding
protein_coding
protein_coding
protein_coding
protein_coding
protein_coding
ncRNA
protein_coding
protein_coding
protein_coding
protein_coding
protein_coding
protein_coding
protein_coding
protein_coding
protein_coding
protein_coding
protein_coding
protein_coding
protein_coding
protein_coding
protein_coding
protein_coding
protein_coding
protein_coding
protein_coding
protein_coding
protein_coding
protein_coding
protein_coding
protein_coding
protein_coding
protein_coding
protein_coding
protein_coding
protein_coding
ncRNA
protein_coding
protein_coding
protein_coding
protein_coding

protein_coding
protein_coding
protein_coding
protein_coding
protein_coding
protein_coding
protein_coding
protein_coding
protein_coding
protein_coding
protein_coding
protein_coding
protein_coding
protein_coding
protein_coding
protein_coding
protein_coding
protein_coding
protein_coding
small_nucle(ol)ar_rna
protein_coding
protein_coding
protein_coding
protein_coding
protein_coding
ncRNA
protein_coding
protein_coding
protein_coding
protein_coding
protein_coding
protein_coding
protein_coding
protein_coding
protein_coding
protein_coding
protein_coding
protein_coding
protein_coding
protein_coding
protein_coding
protein_coding
protein_coding
protein_coding
protein_coding
ncRNA
protein_coding
protein_coding
protein_coding
pre_trna
protein_coding
protein_coding

protein_coding
protein_coding
protein_coding
protein_coding
protein_coding
protein_coding
protein_coding
small_nucle(ol)ar_rna
small_nucle(ol)ar_rna
small_nucle(ol)ar_rna
small_nucle(ol)ar_rna
small_nucle(ol)ar_rna
small_nucle(ol)ar_rna
protein_coding
protein_coding
protein_coding
protein_coding
protein_coding
protein_coding
protein_coding
protein_coding
protein_coding
protein_coding
protein_coding
protein_coding
protein_coding
protein_coding
protein_coding
protein_coding
protein_coding
pre_trna
protein_coding
protein_coding
protein_coding
protein_coding
protein_coding
protein_coding
protein_coding
protein_coding
protein_coding
protein_coding
protein_coding
protein_coding
ncRNA
protein_coding
protein_coding
protein_coding
protein_coding
protein_coding
protein_coding
protein_coding
protein_coding

ncRNA
protein_coding
protein_coding
protein_coding
protein_coding
protein_coding
protein_coding
protein_coding
protein_coding
protein_coding
protein_coding
protein_coding
protein_coding
protein_coding
protein_coding
protein_coding
protein_coding
protein_coding
protein_coding
small_nucle(ol)ar_rna
small_nucle(ol)ar_rna
protein_coding
protein_coding
protein_coding
miRNA_precursor
protein_coding
protein_coding
protein_coding
protein_coding
miRNA_precursor
protein_coding
protein_coding
protein_coding
protein_coding
protein_coding
protein_coding
protein_coding
protein_coding
protein_coding
protein_coding
protein_coding
protein_coding
protein_coding
protein_coding
protein_coding
small_nucle(ol)ar_rna
protein_coding
ncRNA
protein_coding
protein_coding
protein_coding
protein_coding

protein_coding
protein_coding
protein_coding
protein_coding
protein_coding
protein_coding
protein_coding
protein_coding
protein_coding
protein_coding
ncRNA
protein_coding
protein_coding
protein_coding
protein_coding
protein_coding
protein_coding
protein_coding
protein_coding
protein_coding
protein_coding
ncRNA
protein_coding
protein_coding
protein_coding
protein_coding
protein_coding
protein_coding
protein_coding
protein_coding
ncRNA
protein_coding
protein_coding
protein_coding
protein_coding
ncRNA
protein_coding
protein_coding
protein_coding
protein_coding
protein_coding
protein_coding
protein_coding
protein_coding
protein_coding
protein_coding
protein_coding
protein_coding
protein_coding
protein_coding
protein_coding
protein_coding

protein_coding
ncRNA
protein_coding
protein_coding
ncRNA
protein_coding
protein_coding
protein_coding
ncRNA
protein_coding
protein_coding
protein_coding
protein_coding
protein_coding
ncRNA
protein_coding
protein_coding
protein_coding
protein_coding
protein_coding
miRNA_precursor
protein_coding
protein_coding
protein_coding
protein_coding
protein_coding
protein_coding
protein_coding
protein_coding
protein_coding
protein_coding
protein_coding
protein_coding
protein_coding
protein_coding
protein_coding
protein_coding
ncRNA
protein_coding
protein_coding
protein_coding
protein_coding
protein_coding
protein_coding
protein_coding
protein_coding
protein_coding
protein_coding
protein_coding
protein_coding
protein_coding
protein_coding

protein_coding
protein_coding
protein_coding
protein_coding
ncRNA
protein_coding
ncRNA
protein_coding
protein_coding
protein_coding
miRNA_precursor
protein_coding
protein_coding
protein_coding
protein_coding
protein_coding
protein_coding
protein_coding
protein_coding
protein_coding
protein_coding
protein_coding
protein_coding
protein_coding
protein_coding
protein_coding
protein_coding
protein_coding
ncRNA
protein_coding
protein_coding
small_nucle(ol)ar_rna
protein_coding
protein_coding
protein_coding
protein_coding
protein_coding
protein_coding
protein_coding
protein_coding
protein_coding
protein_coding
protein_coding
protein_coding
protein_coding
protein_coding
protein_coding
protein_coding
protein_coding
protein_coding
protein_coding
protein_coding

protein_coding
protein_coding
protein_coding
protein_coding
protein_coding
protein_coding
protein_coding
protein_coding
protein_coding
protein_coding
protein_coding
miRNA_precursor
protein_coding
protein_coding
protein_coding
protein_coding
protein_coding
protein_coding
protein_coding
protein_coding
protein_coding
protein_coding
protein_coding
protein_coding
protein_coding
protein_coding
protein_coding
protein_coding
protein_coding
protein_coding
protein_coding
protein_coding
protein_coding
protein_coding
protein_coding
ncRNA
protein_coding
protein_coding
protein_coding
protein_coding
protein_coding
ncRNA
protein_coding
protein_coding
protein_coding
protein_coding
protein_coding
protein_coding
protein_coding
protein_coding
ncRNA
protein_coding

protein_coding
protein_coding
protein_coding
protein_coding
protein_coding
protein_coding
protein_coding
protein_coding
protein_coding
protein_coding
protein_coding
protein_coding
protein_coding
protein_coding
protein_coding
protein_coding
protein_coding
protein_coding
protein_coding
protein_coding
protein_coding
protein_coding
protein_coding
protein_coding
protein_coding
protein_coding
ncRNA
protein_coding
protein_coding
protein_coding
protein_coding
protein_coding
protein_coding
protein_coding
ncRNA
protein_coding
ncRNA
small_nucle(ol)ar_rna
protein_coding
protein_coding
protein_coding
protein_coding
miRNA_precursor
protein_coding
protein_coding
protein_coding
protein_coding
protein_coding
protein_coding
protein_coding
protein_coding
protein_coding

protein_coding
protein_coding
protein_coding
protein_coding
protein_coding
protein_coding
protein_coding
protein_coding
small_nucle(ol)ar_rna
protein_coding
protein_coding
protein_coding
protein_coding
protein_coding
protein_coding
protein_coding
protein_coding
ncRNA
protein_coding
protein_coding
protein_coding
protein_coding
protein_coding
protein_coding
protein_coding
protein_coding
protein_coding
protein_coding
protein_coding
protein_coding
protein_coding
protein_coding
protein_coding
protein_coding
protein_coding
protein_coding
protein_coding
protein_coding
protein_coding
protein_coding
miRNA_precursor
protein_coding
protein_coding
protein_coding
ncRNA
protein_coding
protein_coding
protein_coding
protein_coding
ncRNA
protein_coding
protein_coding
protein_coding

protein_coding
protein_coding
protein_coding
protein_coding
protein_coding
protein_coding
protein_coding
protein_coding
protein_coding
protein_coding
protein_coding
protein_coding
protein_coding
protein_coding
protein_coding
protein_coding
protein_coding
protein_coding
protein_coding
protein_coding
protein_coding
protein_coding
protein_coding
protein_coding
protein_coding
protein_coding
protein_coding
protein_coding
protein_coding
protein_coding
ncRNA
protein_coding
protein_coding
protein_coding
protein_coding
protein_coding
protein_coding
protein_coding
protein_coding
protein_coding
protein_coding
protein_coding
protein_coding
protein_coding
miRNA_precursor
miRNA_precursor
miRNA_precursor
protein_coding
protein_coding
protein_coding
protein_coding
ncRNA

protein_coding
protein_coding
protein_coding
protein_coding
small_nucle(ol)ar_rna
protein_coding
protein_coding
protein_coding
protein_coding
protein_coding
protein_coding
protein_coding
protein_coding
miRNA_precursor
protein_coding
protein_coding
protein_coding
protein_coding
protein_coding
protein_coding
protein_coding
protein_coding
protein_coding
protein_coding
protein_coding
protein_coding
protein_coding
ncRNA
protein_coding
ncRNA
protein_coding
protein_coding
protein_coding
protein_coding
protein_coding
protein_coding
protein_coding
protein_coding
protein_coding
protein_coding
protein_coding
protein_coding
protein_coding
protein_coding
protein_coding
protein_coding
protein_coding
protein_coding
protein_coding
protein_coding
protein_coding
protein_coding

protein_coding
protein_coding
protein_coding
ncRNA
protein_coding
protein_coding
protein_coding
protein_coding
protein_coding
small_nucle(ol)ar_rna
protein_coding
protein_coding
protein_coding
protein_coding
protein_coding
protein_coding
protein_coding
protein_coding
pre_trna
protein_coding
ncRNA
protein_coding
protein_coding
protein_coding
protein_coding
protein_coding
protein_coding
protein_coding
protein_coding
protein_coding
protein_coding
protein_coding
ncRNA
protein_coding
protein_coding
protein_coding
protein_coding
protein_coding
protein_coding
protein_coding
protein_coding
protein_coding
protein_coding
protein_coding
protein_coding
protein_coding
protein_coding
protein_coding
protein_coding
protein_coding
protein_coding
protein_coding

protein_coding
ncRNA
protein_coding
protein_coding
protein_coding
protein_coding
protein_coding
protein_coding
protein_coding
protein_coding
protein_coding
protein_coding
protein_coding
miRNA_precursor
protein_coding
protein_coding
protein_coding
protein_coding
protein_coding
protein_coding
protein_coding
protein_coding
protein_coding
protein_coding
protein_coding
protein_coding
protein_coding
protein_coding
protein_coding
ncRNA
protein_coding
ncRNA
protein_coding
protein_coding
small_nucle(ol)ar_rna
protein_coding
protein_coding
protein_coding
protein_coding
protein_coding
protein_coding
protein_coding
protein_coding
protein_coding
ncRNA
protein_coding
protein_coding
protein_coding
ncRNA
protein_coding
protein_coding
protein_coding

protein_coding
protein_coding
protein_coding
protein_coding
protein_coding
protein_coding
protein_coding
protein_coding
protein_coding
protein_coding
protein_coding
protein_coding
protein_coding
protein_coding
protein_coding
protein_coding
protein_coding
protein_coding
protein_coding
protein_coding
protein_coding
ncRNA
protein_coding
protein_coding
protein_coding
protein_coding
protein_coding
protein_coding
protein_coding
protein_coding
protein_coding
ncRNA
protein_coding
protein_coding
protein_coding
small_nucle(ol)ar_rna
protein_coding
protein_coding
protein_coding
protein_coding
protein_coding
protein_coding
protein_coding
protein_coding
protein_coding
ncRNA
protein_coding
protein_coding
protein_coding
protein_coding
protein_coding
protein_coding

protein_coding
protein_coding
protein_coding
protein_coding
protein_coding
protein_coding
miRNA_precursor
protein_coding
protein_coding
protein_coding
protein_coding
protein_coding
protein_coding
protein_coding
protein_coding
protein_coding
protein_coding
protein_coding
protein_coding
protein_coding
protein_coding
protein_coding
protein_coding
protein_coding
protein_coding
protein_coding
protein_coding
protein_coding
protein_coding
ncRNA
protein_coding
protein_coding
protein_coding
protein_coding
protein_coding
protein_coding
protein_coding
protein_coding
protein_coding
protein_coding
protein_coding
protein_coding
protein_coding
protein_coding
protein_coding
protein_coding
protein_coding
protein_coding
protein_coding
protein_coding
protein_coding
protein_coding

protein_coding
protein_coding
protein_coding
protein_coding
protein_coding
small_nucle(ol)ar_rna
protein_coding
protein_coding
protein_coding
protein_coding
protein_coding
protein_coding
ncRNA
protein_coding
protein_coding
protein_coding
protein_coding
protein_coding
protein_coding
protein_coding
protein_coding
protein_coding
protein_coding
protein_coding
protein_coding
protein_coding
protein_coding
protein_coding
protein_coding
protein_coding
protein_coding
protein_coding
protein_coding
protein_coding
protein_coding
protein_coding
protein_coding
protein_coding
protein_coding
protein_coding
protein_coding
protein_coding
protein_coding
protein_coding
protein_coding
protein_coding
protein_coding
protein_coding
protein_coding
protein_coding
protein_coding
protein_coding

protein_coding
protein_coding
protein_coding
protein_coding
protein_coding
protein_coding
protein_coding
protein_coding
protein_coding
small_nucle(ol)ar_rna
small_nucle(ol)ar_rna
small_nucle(ol)ar_rna
protein_coding
ncRNA
protein_coding
protein_coding
protein_coding
protein_coding
protein_coding
protein_coding
protein_coding
protein_coding
protein_coding
protein_coding
protein_coding
protein_coding
protein_coding
protein_coding
protein_coding
protein_coding
protein_coding
protein_coding
protein_coding
pre_trna
protein_coding
protein_coding
miRNA_precursor
protein_coding
protein_coding
protein_coding
protein_coding
protein_coding
protein_coding
protein_coding
protein_coding
protein_coding
protein_coding
protein_coding
protein_coding
protein_coding
protein_coding
protein_coding

protein_coding
protein_coding
protein_coding
protein_coding
protein_coding
protein_coding
miRNA_precursor
protein_coding
protein_coding
protein_coding
protein_coding
protein_coding
protein_coding
miRNA_precursor
protein_coding
protein_coding
protein_coding
protein_coding
protein_coding
protein_coding
protein_coding
protein_coding
protein_coding
protein_coding
protein_coding
protein_coding
protein_coding
protein_coding
protein_coding
protein_coding
protein_coding
protein_coding
protein_coding
protein_coding
protein_coding
protein_coding
protein_coding
protein_coding
protein_coding
protein_coding
protein_coding
protein_coding
protein_coding
protein_coding
protein_coding
protein_coding
protein_coding
protein_coding
protein_coding
protein_coding
protein_coding
protein_coding

protein_coding
protein_coding
protein_coding
protein_coding
protein_coding
ncRNA
protein_coding
protein_coding
pre_trna
protein_coding
protein_coding
protein_coding
protein_coding
protein_coding
protein_coding
ncRNA
protein_coding
protein_coding
protein_coding
protein_coding
protein_coding
protein_coding
protein_coding
protein_coding
protein_coding
protein_coding
protein_coding
protein_coding
protein_coding
protein_coding
protein_coding
protein_coding
protein_coding
protein_coding
ncRNA
pre_trna
protein_coding
protein_coding
protein_coding
protein_coding
protein_coding
protein_coding
protein_coding
protein_coding
protein_coding
protein_coding
ribosomal_rna
protein_coding
protein_coding
protein_coding
protein_coding
protein_coding

protein_coding
protein_coding
ncRNA
ncRNA
protein_coding
protein_coding
protein_coding
protein_coding
protein_coding
protein_coding
protein_coding
protein_coding
protein_coding
protein_coding
protein_coding
protein_coding
protein_coding
protein_coding
protein_coding
protein_coding
protein_coding
protein_coding
protein_coding
protein_coding
protein_coding
ncRNA
protein_coding
protein_coding
protein_coding
protein_coding
ncRNA
protein_coding
ncRNA
protein_coding
small_nucle(ol)ar_rna
ncRNA
ncRNA
small_nucle(ol)ar_rna
protein_coding
protein_coding
protein_coding
protein_coding
protein_coding
protein_coding
protein_coding
protein_coding
protein_coding
protein_coding
protein_coding
protein_coding
protein_coding
protein_coding

protein_coding
protein_coding
protein_coding
protein_coding
protein_coding
protein_coding
protein_coding
protein_coding
protein_coding
protein_coding
protein_coding
protein_coding
ncRNA
protein_coding
protein_coding
ncRNA
protein_coding
protein_coding
protein_coding
protein_coding
protein_coding
protein_coding
protein_coding
protein_coding
protein_coding
protein_coding
protein_coding
protein_coding
protein_coding
protein_coding
protein_coding
protein_coding
protein_coding
protein_coding
protein_coding
ncRNA
protein_coding
protein_coding
protein_coding
protein_coding
ncRNA
miRNA_precursor
protein_coding
protein_coding
ncRNA
protein_coding
ncRNA
protein_coding
protein_coding
protein_coding
ncRNA
protein_coding

protein_coding
protein_coding
protein_coding
protein_coding
protein_coding
protein_coding
protein_coding
protein_coding
protein_coding
protein_coding
protein_coding
protein_coding
protein_coding
pre_trna
protein_coding
protein_coding
protein_coding
protein_coding
protein_coding
protein_coding
ncRNA
protein_coding
protein_coding
protein_coding
protein_coding
protein_coding
protein_coding
protein_coding
protein_coding
pre_trna
protein_coding
protein_coding
protein_coding
protein_coding
protein_coding
protein_coding
protein_coding
protein_coding
pre_trna
pre_trna
protein_coding
protein_coding
protein_coding
miRNA_precursor
protein_coding
ncRNA
protein_coding
protein_coding
protein_coding
protein_coding
protein_coding
protein_coding

protein_coding
protein_coding
protein_coding
ncRNA
protein_coding
protein_coding
ncRNA
protein_coding
protein_coding
protein_coding
protein_coding
ncRNA
protein_coding
protein_coding
protein_coding
protein_coding
protein_coding
ncRNA
protein_coding
protein_coding
protein_coding
protein_coding
protein_coding
protein_coding
protein_coding
protein_coding
protein_coding
protein_coding
protein_coding
ncRNA
protein_coding
ncRNA
protein_coding
protein_coding
protein_coding
protein_coding
protein_coding
protein_coding
protein_coding
protein_coding
protein_coding
protein_coding
protein_coding
protein_coding
protein_coding
protein_coding
protein_coding
ncRNA
protein_coding
protein_coding
protein_coding
protein_coding

protein_coding
protein_coding
protein_coding
protein_coding
protein_coding
protein_coding
protein_coding
protein_coding
protein_coding
protein_coding
protein_coding
protein_coding
protein_coding
protein_coding
protein_coding
protein_coding
protein_coding
protein_coding
protein_coding
ncRNA
protein_coding
protein_coding
protein_coding
protein_coding
protein_coding
protein_coding
protein_coding
protein_coding
protein_coding
protein_coding
protein_coding
protein_coding
protein_coding
protein_coding
protein_coding
protein_coding
protein_coding
protein_coding
protein_coding
protein_coding
protein_coding
protein_coding
protein_coding
protein_coding
protein_coding
protein_coding
protein_coding
protein_coding
protein_coding
protein_coding
protein_coding
protein_coding

protein_coding
protein_coding
protein_coding
protein_coding
protein_coding
ncRNA
protein_coding
protein_coding
protein_coding
protein_coding
protein_coding
ncRNA
ncRNA
protein_coding
protein_coding
protein_coding
protein_coding
protein_coding
protein_coding
protein_coding
protein_coding
protein_coding
protein_coding
protein_coding
miRNA_precursor
protein_coding
protein_coding
protein_coding
protein_coding
ncRNA
ncRNA
protein_coding
protein_coding
protein_coding
protein_coding
protein_coding
protein_coding
protein_coding
protein_coding
protein_coding
protein_coding
protein_coding
protein_coding
protein_coding
protein_coding
protein_coding
protein_coding
protein_coding
protein_coding
protein_coding
protein_coding
protein_coding

protein_coding
protein_coding
protein_coding
ncRNA
protein_coding
protein_coding
protein_coding
protein_coding
protein_coding
protein_coding
protein_coding
protein_coding
protein_coding
protein_coding
ncRNA
protein_coding
protein_coding
protein_coding
protein_coding
protein_coding
protein_coding
protein_coding
protein_coding
protein_coding
protein_coding
protein_coding
protein_coding
protein_coding
ncRNA
protein_coding
ncRNA
protein_coding
protein_coding
protein_coding
protein_coding
protein_coding
protein_coding
protein_coding
protein_coding
protein_coding
protein_coding
protein_coding
protein_coding
protein_coding
ncRNA
protein_coding
protein_coding
ncRNA
protein_coding
protein_coding
protein_coding
protein_coding

protein_coding
protein_coding
ncRNA
protein_coding
protein_coding
miRNA_precursor
protein_coding
protein_coding
protein_coding
miRNA_precursor
ncRNA
protein_coding
protein_coding
protein_coding
protein_coding
protein_coding
protein_coding
miRNA_precursor
protein_coding
protein_coding
protein_coding
protein_coding
miRNA_precursor
protein_coding
protein_coding
pre_trna
protein_coding
ncRNA
ncRNA
protein_coding
protein_coding
ncRNA
protein_coding
protein_coding
protein_coding
protein_coding
protein_coding
protein_coding
protein_coding
miRNA_precursor
protein_coding
protein_coding
protein_coding
protein_coding
protein_coding
protein_coding
protein_coding
protein_coding
protein_coding
protein_coding
protein_coding
protein_coding

protein_coding
protein_coding
protein_coding
protein_coding
protein_coding
protein_coding
protein_coding
protein_coding
protein_coding
protein_coding
protein_coding
protein_coding
ncRNA
protein_coding
protein_coding
protein_coding
protein_coding
protein_coding
protein_coding
protein_coding
protein_coding
ncRNA
protein_coding
protein_coding
protein_coding
protein_coding
protein_coding
protein_coding
protein_coding
protein_coding
protein_coding
protein_coding
protein_coding
protein_coding
protein_coding
protein_coding
protein_coding
protein_coding
protein_coding
protein_coding
protein_coding
protein_coding
protein_coding
protein_coding
protein_coding
protein_coding
protein_coding
protein_coding
protein_coding
protein_coding
protein_coding
protein_coding

protein_coding
protein_coding
protein_coding
protein_coding
protein_coding
protein_coding
protein_coding
protein_coding
protein_coding
protein_coding
protein_coding
protein_coding
protein_coding
protein_coding
protein_coding
protein_coding
pre_trna
protein_coding
protein_coding
protein_coding
protein_coding
protein_coding
protein_coding
protein_coding
protein_coding
protein_coding
protein_coding
protein_coding
protein_coding
protein_coding
protein_coding
protein_coding
protein_coding
protein_coding
protein_coding
protein_coding
protein_coding
protein_coding
protein_coding
protein_coding
protein_coding
protein_coding
protein_coding
pre_trna
small_nucle(ol)ar_rna
small_nucle(ol)ar_rna
protein_coding
ncRNA
protein_coding
protein_coding
protein_coding
protein_coding

protein_coding
protein_coding
protein_coding
protein_coding
protein_coding
protein_coding
protein_coding
protein_coding
protein_coding
protein_coding
protein_coding
protein_coding
protein_coding
protein_coding
protein_coding
protein_coding
protein_coding
ncRNA
protein_coding
ncRNA
protein_coding
protein_coding
protein_coding
protein_coding
protein_coding
protein_coding
protein_coding
protein_coding
protein_coding
protein_coding
protein_coding
protein_coding
protein_coding
protein_coding
protein_coding
protein_coding
protein_coding
protein_coding
miRNA_precursor
protein_coding
protein_coding
protein_coding
protein_coding
protein_coding
protein_coding
protein_coding
protein_coding
protein_coding
protein_coding
protein_coding
protein_coding
small_nucle(ol)ar_rna

protein_coding
protein_coding
protein_coding
protein_coding
protein_coding
protein_coding
pre_trna
protein_coding
protein_coding
protein_coding
protein_coding
protein_coding
pre_trna
protein_coding
protein_coding
miRNA_precursor
protein_coding
protein_coding
protein_coding
protein_coding
protein_coding
protein_coding
protein_coding
protein_coding
protein_coding
protein_coding
protein_coding
protein_coding
protein_coding
protein_coding
protein_coding
protein_coding
protein_coding
protein_coding
protein_coding
protein_coding
protein_coding
protein_coding
protein_coding
protein_coding
ncRNA
protein_coding
protein_coding
protein_coding
protein_coding
protein_coding
protein_coding
protein_coding
protein_coding
protein_coding
protein_coding
protein_coding

protein_coding
protein_coding
protein_coding
protein_coding
protein_coding
protein_coding
protein_coding
protein_coding
protein_coding
protein_coding
protein_coding
pre_trna
protein_coding
protein_coding
pre_trna
protein_coding
ncRNA
protein_coding
protein_coding
protein_coding
protein_coding
protein_coding
protein_coding
protein_coding
protein_coding
protein_coding
protein_coding
miRNA_precursor
protein_coding
protein_coding
small_nucle(ol)ar_rna
protein_coding
protein_coding
ncRNA
protein_coding
protein_coding
protein_coding
ncRNA
protein_coding
protein_coding
protein_coding
protein_coding
protein_coding
protein_coding
protein_coding
protein_coding
protein_coding
protein_coding
protein_coding
protein_coding
protein_coding
protein_coding

protein_coding
protein_coding
protein_coding
protein_coding
protein_coding
protein_coding
protein_coding
protein_coding
protein_coding
protein_coding
protein_coding
protein_coding
protein_coding
protein_coding
protein_coding
protein_coding
protein_coding
protein_coding
protein_coding
protein_coding
protein_coding
protein_coding
protein_coding
protein_coding
protein_coding
protein_coding
protein_coding
protein_coding
protein_coding
protein_coding
protein_coding
protein_coding
protein_coding
protein_coding
protein_coding
protein_coding
protein_coding
protein_coding
protein_coding
protein_coding
protein_coding
protein_coding
protein_coding
protein_coding
protein_coding
protein_coding
protein_coding
protein_coding
miRNA_precursor
protein_coding
protein_coding
protein_coding

protein_coding
protein_coding
protein_coding
protein_coding
protein_coding
protein_coding
protein_coding
protein_coding
protein_coding
protein_coding
protein_coding
protein_coding
protein_coding
protein_coding
protein_coding
protein_coding
protein_coding
protein_coding
protein_coding
ncRNA
protein_coding
protein_coding
protein_coding
protein_coding
protein_coding
protein_coding
protein_coding
protein_coding
protein_coding
protein_coding
protein_coding
protein_coding
protein_coding
protein_coding
protein_coding
protein_coding
protein_coding
protein_coding
protein_coding
protein_coding
protein_coding
protein_coding
protein_coding
protein_coding
protein_coding
protein_coding
protein_coding
protein_coding
protein_coding
protein_coding
protein_coding
protein_coding

protein_coding
protein_coding
protein_coding
protein_coding
protein_coding
small_nucle(ol)ar_rna
small_nucle(ol)ar_rna
small_nucle(ol)ar_rna
protein_coding
protein_coding
protein_coding
protein_coding
protein_coding
ncRNA
protein_coding
protein_coding
protein_coding
protein_coding
protein_coding
protein_coding
protein_coding
protein_coding
protein_coding
protein_coding
protein_coding
protein_coding
protein_coding
protein_coding
protein_coding
protein_coding
protein_coding
protein_coding
protein_coding
protein_coding
protein_coding
protein_coding
protein_coding
protein_coding
protein_coding
protein_coding
protein_coding
protein_coding
protein_coding
protein_coding
protein_coding
protein_coding
protein_coding
protein_coding
protein_coding
protein_coding
protein_coding
protein_coding

protein_coding
protein_coding
protein_coding
protein_coding
protein_coding
pre_trna
protein_coding
protein_coding
protein_coding
protein_coding
protein_coding
protein_coding
protein_coding
pre_trna
protein_coding
protein_coding
protein_coding
ncRNA
protein_coding
protein_coding
protein_coding
ncRNA
protein_coding
protein_coding
protein_coding
protein_coding
protein_coding
protein_coding
miRNA_precursor
protein_coding
protein_coding
protein_coding
protein_coding
protein_coding
protein_coding
small_nucle(ol)ar_rna
small_nucle(ol)ar_rna
small_nucle(ol)ar_rna
small_nucle(ol)ar_rna
protein_coding
protein_coding
protein_coding
protein_coding
protein_coding
protein_coding
protein_coding
protein_coding
protein_coding
protein_coding
protein_coding
protein_coding
protein_coding

protein_coding
protein_coding
protein_coding
protein_coding
protein_coding
protein_coding
protein_coding
protein_coding
protein_coding
protein_coding
protein_coding
protein_coding
protein_coding
protein_coding
protein_coding
protein_coding
protein_coding
protein_coding
protein_coding
small_nucle(ol)ar_rna
protein_coding
protein_coding
protein_coding
protein_coding
protein_coding
protein_coding
protein_coding
protein_coding
protein_coding
protein_coding
protein_coding
protein_coding
protein_coding
protein_coding
protein_coding
protein_coding
protein_coding
protein_coding
protein_coding
protein_coding
protein_coding
protein_coding
protein_coding
protein_coding
protein_coding
protein_coding
protein_coding
protein_coding
protein_coding
protein_coding
protein_coding
protein_coding

protein_coding
protein_coding
protein_coding
protein_coding
protein_coding
protein_coding
protein_coding
protein_coding
protein_coding
protein_coding
protein_coding
protein_coding
protein_coding
protein_coding
protein_coding
protein_coding
protein_coding
protein_coding
protein_coding
protein_coding
protein_coding
protein_coding
protein_coding
protein_coding
protein_coding
protein_coding
protein_coding
protein_coding
protein_coding
protein_coding
protein_coding
protein_coding
protein_coding
protein_coding
protein_coding
protein_coding
protein_coding
protein_coding
protein_coding
protein_coding
protein_coding
protein_coding
protein_coding
protein_coding
protein_coding
ncRNA
small_nucle(ol)ar_rna
protein_coding
protein_coding
protein_coding
protein_coding
protein_coding

protein_coding
protein_coding
protein_coding
protein_coding
protein_coding
protein_coding
protein_coding
protein_coding
protein_coding
protein_coding
protein_coding
protein_coding
protein_coding
protein_coding
protein_coding
protein_coding
protein_coding
protein_coding
protein_coding
protein_coding
protein_coding
protein_coding
protein_coding
protein_coding
protein_coding
protein_coding
protein_coding
protein_coding
protein_coding
protein_coding
protein_coding
protein_coding
protein_coding
protein_coding
protein_coding
protein_coding
protein_coding
protein_coding
ncRNA
ncRNA
protein_coding
protein_coding
protein_coding
protein_coding
protein_coding
protein_coding
protein_coding
protein_coding
protein_coding
protein_coding
protein_coding
protein_coding

protein_coding
protein_coding
protein_coding
protein_coding
protein_coding
protein_coding
protein_coding
protein_coding
protein_coding
protein_coding
protein_coding
protein_coding
ncRNA
ncRNA
protein_coding
protein_coding
protein_coding
protein_coding
protein_coding
protein_coding
protein_coding
protein_coding
protein_coding
protein_coding
protein_coding
protein_coding
protein_coding
protein_coding
protein_coding
protein_coding
protein_coding
protein_coding
protein_coding
protein_coding
protein_coding
protein_coding
protein_coding
protein_coding
protein_coding
protein_coding
protein_coding
protein_coding
protein_coding
protein_coding
protein_coding
protein_coding
protein_coding
protein_coding
protein_coding
protein_coding
protein_coding
protein_coding

protein_coding
protein_coding
protein_coding
protein_coding
protein_coding
protein_coding
protein_coding
protein_coding
protein_coding
small_nucle(ol)ar_rna
protein_coding
protein_coding
protein_coding
protein_coding
protein_coding
protein_coding
protein_coding
protein_coding
protein_coding
protein_coding
protein_coding
protein_coding
protein_coding
protein_coding
protein_coding
protein_coding
protein_coding
protein_coding
protein_coding
ncRNA
protein_coding
protein_coding
protein_coding
protein_coding
protein_coding
protein_coding
protein_coding
protein_coding
protein_coding
protein_coding
protein_coding
protein_coding
protein_coding
protein_coding
protein_coding
protein_coding
protein_coding
protein_coding
protein_coding
protein_coding
protein_coding
protein_coding

protein_coding
protein_coding
protein_coding
protein_coding
protein_coding
protein_coding
protein_coding
protein_coding
protein_coding
protein_coding
protein_coding
protein_coding
protein_coding
protein_coding
protein_coding
protein_coding
protein_coding
protein_coding
protein_coding
protein_coding
protein_coding
protein_coding
protein_coding
protein_coding
protein_coding
protein_coding
protein_coding
protein_coding
protein_coding
protein_coding
protein_coding
protein_coding
protein_coding
ncRNA
protein_coding
protein_coding
protein_coding
protein_coding
protein_coding
protein_coding
protein_coding
protein_coding
protein_coding
protein_coding
protein_coding
protein_coding
protein_coding
protein_coding
protein_coding
protein_coding
protein_coding
protein_coding

ncRNA
protein_coding
protein_coding
protein_coding
protein_coding
protein_coding
protein_coding
protein_coding
protein_coding
protein_coding
protein_coding
protein_coding
protein_coding
protein_coding
protein_coding
protein_coding
protein_coding
protein_coding
protein_coding
protein_coding
protein_coding
protein_coding
protein_coding
small_nucle(ol)ar_rna
small_nucle(ol)ar_rna
protein_coding
protein_coding
protein_coding
protein_coding
protein_coding
protein_coding
protein_coding
protein_coding
protein_coding
protein_coding
protein_coding
protein_coding
protein_coding
protein_coding
protein_coding
protein_coding
protein_coding
protein_coding
protein_coding
protein_coding
protein_coding
protein_coding
protein_coding
protein_coding
protein_coding
protein_coding
protein_coding

protein_coding
protein_coding
protein_coding
protein_coding
protein_coding
protein_coding
protein_coding
protein_coding
protein_coding
protein_coding
protein_coding
protein_coding
protein_coding
protein_coding
protein_coding
protein_coding
protein_coding
protein_coding
protein_coding
protein_coding
protein_coding
protein_coding
protein_coding
protein_coding
ncRNA
protein_coding
protein_coding
protein_coding
protein_coding
protein_coding
protein_coding
ncRNA
protein_coding
protein_coding
protein_coding
protein_coding
protein_coding
protein_coding
protein_coding
protein_coding
protein_coding
protein_coding
protein_coding
protein_coding
protein_coding
protein_coding
protein_coding
protein_coding
protein_coding
protein_coding
protein_coding
protein_coding

protein_coding
protein_coding
protein_coding
protein_coding
protein_coding
protein_coding
protein_coding
protein_coding
protein_coding
protein_coding
protein_coding
protein_coding
protein_coding
protein_coding
protein_coding
protein_coding
protein_coding
ncRNA
protein_coding
protein_coding
ncRNA
protein_coding
protein_coding
protein_coding
protein_coding
protein_coding
protein_coding
protein_coding
protein_coding
protein_coding
protein_coding
protein_coding
protein_coding
ncRNA
protein_coding
protein_coding
protein_coding
protein_coding
protein_coding
protein_coding
protein_coding
protein_coding
protein_coding
protein_coding
protein_coding
protein_coding
protein_coding
protein_coding
protein_coding
protein_coding
protein_coding
miRNA_precursor

protein_coding
protein_coding
protein_coding
protein_coding
protein_coding
protein_coding
protein_coding
protein_coding
protein_coding
protein_coding
protein_coding
protein_coding
miRNA_precursor
protein_coding
protein_coding
protein_coding
protein_coding
protein_coding
protein_coding
protein_coding
protein_coding
protein_coding
protein_coding
protein_coding
protein_coding
protein_coding
protein_coding
protein_coding
protein_coding
protein_coding
protein_coding
protein_coding
protein_coding
protein_coding
protein_coding
protein_coding
protein_coding
protein_coding
protein_coding
protein_coding
protein_coding
protein_coding
protein_coding
protein_coding
protein_coding
protein_coding
protein_coding
protein_coding
protein_coding
protein_coding
protein_coding
protein_coding

protein_coding
protein_coding
protein_coding
ncRNA
protein_coding
protein_coding
protein_coding
protein_coding
protein_coding
protein_coding
protein_coding
protein_coding
protein_coding
protein_coding
protein_coding
protein_coding
protein_coding
protein_coding
protein_coding
protein_coding
protein_coding
protein_coding
protein_coding
protein_coding
small_nucle(ol)ar_rna
small_nucle(ol)ar_rna
small_nucle(ol)ar_rna
protein_coding
ncRNA
protein_coding
ncRNA
protein_coding
protein_coding
protein_coding
ncRNA
protein_coding
protein_coding
protein_coding
protein_coding
protein_coding
protein_coding
protein_coding
protein_coding
protein_coding
protein_coding
protein_coding
protein_coding
protein_coding
protein_coding
protein_coding
protein_coding
protein_coding

protein_coding
protein_coding
protein_coding
protein_coding
protein_coding
ncRNA
protein_coding
protein_coding
protein_coding
protein_coding
protein_coding
protein_coding
protein_coding
protein_coding
protein_coding
protein_coding
protein_coding
protein_coding
protein_coding
protein_coding
protein_coding
protein_coding
protein_coding
protein_coding
protein_coding
protein_coding
protein_coding
protein_coding
protein_coding
protein_coding
protein_coding
protein_coding
protein_coding
protein_coding
protein_coding
protein_coding
protein_coding
protein_coding
protein_coding
protein_coding
protein_coding
protein_coding
protein_coding
protein_coding
protein_coding
protein_coding
protein_coding
protein_coding
protein_coding
protein_coding
protein_coding
protein_coding

protein_coding
protein_coding
protein_coding
protein_coding
protein_coding
protein_coding
ncRNA
protein_coding
protein_coding
protein_coding
protein_coding
protein_coding
protein_coding
protein_coding
protein_coding
protein_coding
protein_coding
protein_coding
protein_coding
protein_coding
ncRNA
small_nucle(ol)ar_rna
protein_coding
protein_coding
protein_coding
protein_coding
protein_coding
protein_coding
protein_coding
protein_coding
protein_coding
protein_coding
protein_coding
protein_coding
protein_coding
protein_coding
protein_coding
protein_coding
protein_coding
protein_coding
protein_coding
protein_coding
protein_coding
protein_coding
protein_coding
protein_coding
protein_coding
protein_coding
protein_coding
protein_coding
protein_coding
protein_coding

protein_coding
protein_coding
protein_coding
protein_coding
protein_coding
pre_trna
protein_coding
protein_coding
ncRNA
protein_coding
protein_coding
protein_coding
protein_coding
small_nucle(ol)ar_rna
small_nucle(ol)ar_rna
small_nucle(ol)ar_rna
protein_coding
protein_coding
protein_coding
protein_coding
protein_coding
protein_coding
protein_coding
protein_coding
protein_coding
protein_coding
protein_coding
protein_coding
protein_coding
protein_coding
protein_coding
protein_coding
protein_coding
protein_coding
miRNA_precursor
protein_coding
protein_coding
protein_coding
protein_coding
protein_coding
protein_coding
protein_coding
protein_coding
protein_coding
protein_coding
protein_coding
ncRNA
protein_coding
protein_coding
protein_coding
protein_coding
protein_coding

protein_coding
protein_coding
protein_coding
small_nucle(ol)ar_rna
protein_coding
protein_coding
protein_coding
protein_coding
protein_coding
protein_coding
protein_coding
protein_coding
ncRNA
protein_coding
protein_coding
protein_coding
protein_coding
protein_coding
protein_coding
protein_coding
protein_coding
protein_coding
protein_coding
protein_coding
protein_coding
protein_coding
protein_coding
protein_coding
protein_coding
protein_coding
ncRNA
protein_coding
protein_coding
protein_coding
protein_coding
protein_coding
protein_coding
protein_coding
ncRNA
protein_coding
protein_coding
protein_coding
protein_coding
protein_coding
protein_coding
ncRNA
protein_coding
ncRNA
protein_coding
protein_coding
ncRNA
protein_coding

protein_coding
protein_coding
protein_coding
protein_coding
protein_coding
protein_coding
protein_coding
protein_coding
protein_coding
protein_coding
protein_coding
protein_coding
protein_coding
protein_coding
protein_coding
protein_coding
protein_coding
protein_coding
protein_coding
protein_coding
protein_coding
protein_coding
protein_coding
protein_coding
protein_coding
ncRNA
protein_coding
protein_coding
protein_coding
protein_coding
protein_coding
protein_coding
protein_coding
protein_coding
protein_coding
protein_coding
protein_coding
protein_coding
protein_coding
protein_coding
protein_coding
protein_coding
protein_coding
pre_trna
protein_coding
protein_coding
protein_coding
protein_coding
protein_coding
protein_coding
protein_coding
protein_coding

protein_coding
protein_coding
protein_coding
protein_coding
ncRNA
miRNA_precursor
protein_coding
ncRNA
ncRNA
ncRNA
ncRNA
protein_coding
protein_coding
protein_coding
miRNA_precursor
protein_coding
protein_coding
small_nucle(ol)ar_rna
small_nucle(ol)ar_rna
ncRNA
protein_coding
protein_coding
protein_coding
protein_coding
protein_coding
protein_coding
protein_coding
protein_coding
ncRNA
ncRNA
protein_coding
protein_coding
protein_coding
protein_coding
protein_coding
protein_coding
protein_coding
ncRNA
protein_coding
protein_coding
protein_coding
protein_coding
protein_coding
protein_coding
protein_coding
protein_coding
protein_coding
protein_coding
protein_coding
protein_coding
protein_coding
protein_coding

protein_coding
protein_coding
protein_coding
protein_coding
protein_coding
ncRNA
protein_coding
protein_coding
protein_coding
protein_coding
protein_coding
protein_coding
protein_coding
protein_coding
protein_coding
protein_coding
protein_coding
protein_coding
protein_coding
protein_coding
protein_coding
protein_coding
protein_coding
protein_coding
protein_coding
protein_coding
protein_coding
protein_coding
protein_coding
protein_coding
protein_coding
protein_coding
protein_coding
protein_coding
protein_coding
protein_coding
protein_coding
protein_coding
protein_coding
protein_coding
protein_coding
protein_coding
protein_coding
protein_coding
protein_coding
protein_coding
protein_coding
protein_coding
protein_coding
protein_coding
protein_coding
protein_coding

protein_coding
protein_coding
protein_coding
protein_coding
protein_coding
protein_coding
protein_coding
protein_coding
protein_coding
protein_coding
protein_coding
protein_coding
protein_coding
protein_coding
protein_coding
protein_coding
protein_coding
protein_coding
protein_coding
protein_coding
protein_coding
protein_coding
protein_coding
protein_coding
protein_coding
protein_coding
protein_coding
protein_coding
protein_coding
protein_coding
protein_coding
protein_coding
protein_coding
protein_coding
protein_coding
protein_coding
protein_coding
protein_coding
protein_coding
protein_coding
small_nucle(ol)ar_rna
protein_coding
protein_coding
protein_coding
miRNA_precursor
ncRNA
protein_coding
protein_coding
protein_coding
protein_coding
protein_coding
protein_coding

protein_coding
protein_coding
protein_coding
protein_coding
protein_coding
protein_coding
protein_coding
protein_coding
protein_coding
protein_coding
protein_coding
protein_coding
protein_coding
protein_coding
protein_coding
protein_coding
protein_coding
protein_coding
protein_coding
protein_coding
protein_coding
protein_coding
protein_coding
protein_coding
ncRNA
protein_coding
protein_coding
protein_coding
protein_coding
protein_coding
ncRNA
protein_coding
protein_coding
protein_coding
protein_coding
protein_coding
protein_coding
protein_coding
protein_coding
protein_coding
protein_coding
protein_coding
protein_coding
protein_coding
protein_coding
protein_coding
protein_coding
protein_coding
protein_coding
protein_coding
protein_coding
ncRNA

small_nucle(ol)ar_rna
miRNA_precursor
protein_coding
protein_coding
protein_coding
protein_coding
protein_coding
protein_coding
protein_coding
protein_coding
protein_coding
protein_coding
protein_coding
protein_coding
protein_coding
protein_coding
protein_coding
protein_coding
protein_coding
protein_coding
protein_coding
ncRNA
protein_coding
ncRNA
protein_coding
protein_coding
protein_coding
protein_coding
protein_coding
protein_coding
protein_coding
protein_coding
protein_coding
protein_coding
protein_coding
ncRNA
protein_coding
protein_coding
protein_coding
protein_coding
protein_coding
protein_coding
ncRNA
protein_coding
protein_coding
protein_coding
protein_coding
protein_coding
protein_coding
protein_coding
protein_coding
protein_coding

small_nucle(ol)ar_rna
protein_coding
protein_coding
protein_coding
ncRNA
protein_coding
protein_coding
protein_coding
protein_coding
protein_coding
protein_coding
protein_coding
protein_coding
protein_coding
protein_coding
protein_coding
protein_coding
protein_coding
protein_coding
protein_coding
protein_coding
protein_coding
protein_coding
protein_coding
protein_coding
ncRNA
protein_coding
ncRNA
protein_coding
protein_coding
protein_coding
protein_coding
protein_coding
protein_coding
protein_coding
protein_coding
protein_coding
protein_coding
protein_coding
protein_coding
protein_coding
protein_coding
protein_coding
protein_coding
protein_coding
protein_coding
protein_coding
protein_coding
protein_coding
protein_coding
protein_coding
protein_coding

protein_coding
protein_coding
protein_coding
protein_coding
protein_coding
protein_coding
protein_coding
protein_coding
protein_coding
protein_coding
ncRNA
protein_coding
ncRNA
protein_coding
protein_coding
protein_coding
protein_coding
protein_coding
protein_coding
miRNA_precursor
protein_coding
ncRNA
protein_coding
protein_coding
protein_coding
protein_coding
protein_coding
protein_coding
protein_coding
protein_coding
protein_coding
protein_coding
protein_coding
protein_coding
protein_coding
protein_coding
protein_coding
protein_coding
protein_coding
protein_coding
protein_coding
protein_coding
protein_coding
protein_coding
protein_coding
protein_coding
pre_trna
protein_coding
protein_coding
protein_coding
miRNA_precursor
protein_coding

protein_coding
protein_coding
protein_coding
protein_coding
pre_trna
protein_coding
protein_coding
protein_coding
protein_coding
protein_coding
protein_coding
protein_coding
protein_coding
protein_coding
ncRNA
ncRNA
protein_coding
protein_coding
protein_coding
protein_coding
protein_coding
protein_coding
protein_coding
protein_coding
protein_coding
protein_coding
protein_coding
protein_coding
protein_coding
protein_coding
protein_coding
protein_coding
protein_coding
protein_coding
protein_coding
protein_coding
protein_coding
ncRNA
protein_coding
protein_coding
protein_coding
protein_coding
protein_coding
protein_coding
protein_coding
protein_coding
protein_coding
protein_coding
protein_coding
protein_coding
protein_coding
protein_coding

protein_coding
protein_coding
protein_coding
protein_coding
protein_coding
protein_coding
protein_coding
protein_coding
protein_coding
protein_coding
protein_coding
protein_coding
protein_coding
protein_coding
protein_coding
protein_coding
protein_coding
protein_coding
protein_coding
protein_coding
protein_coding
protein_coding
protein_coding
protein_coding
protein_coding
protein_coding
protein_coding
protein_coding
ncRNA
small_nucle(ol)ar_rna
protein_coding
protein_coding
protein_coding
protein_coding
protein_coding
protein_coding
protein_coding
protein_coding
protein_coding
protein_coding
protein_coding
protein_coding
protein_coding
protein_coding
protein_coding
protein_coding
miRNA_precursor
protein_coding
protein_coding
protein_coding
protein_coding
protein_coding

protein_coding
protein_coding
ncRNA
protein_coding
ncRNA
protein_coding
small_nucle(ol)ar_rna
small_nucle(ol)ar_rna
protein_coding
protein_coding
protein_coding
protein_coding
protein_coding
protein_coding
protein_coding
protein_coding
protein_coding
protein_coding
protein_coding
protein_coding
protein_coding
ncRNA
protein_coding
protein_coding
protein_coding
protein_coding
protein_coding
protein_coding
protein_coding
protein_coding
pre_trna
protein_coding
protein_coding
protein_coding
protein_coding
protein_coding
protein_coding
protein_coding
protein_coding
protein_coding
protein_coding
protein_coding
protein_coding
protein_coding
protein_coding
ncRNA
ncRNA
protein_coding
protein_coding
protein_coding
protein_coding
protein_coding

protein_coding
protein_coding
protein_coding
protein_coding
protein_coding
protein_coding
protein_coding
protein_coding
protein_coding
protein_coding
protein_coding
protein_coding
protein_coding
protein_coding
protein_coding
miRNA_precursor
protein_coding
protein_coding
protein_coding
protein_coding
protein_coding
protein_coding
protein_coding
protein_coding
protein_coding
protein_coding
protein_coding
protein_coding
protein_coding
protein_coding
miRNA_precursor
protein_coding
protein_coding
protein_coding
protein_coding
protein_coding
protein_coding
protein_coding
miRNA_precursor
protein_coding
protein_coding
protein_coding
protein_coding
protein_coding
protein_coding
protein_coding
protein_coding
ncRNA
protein_coding
protein_coding
protein_coding
protein_coding

protein_coding
protein_coding
protein_coding
protein_coding
protein_coding
protein_coding
protein_coding
protein_coding
ncRNA
protein_coding
protein_coding
protein_coding
protein_coding
protein_coding
protein_coding
protein_coding
protein_coding
protein_coding
protein_coding
protein_coding
protein_coding
protein_coding
protein_coding
protein_coding
protein_coding
ncRNA
protein_coding
protein_coding
protein_coding
protein_coding
protein_coding
protein_coding
protein_coding
protein_coding
protein_coding
protein_coding
protein_coding
protein_coding
protein_coding
protein_coding
protein_coding
protein_coding
protein_coding
protein_coding
protein_coding
protein_coding
protein_coding
protein_coding
protein_coding
protein_coding
protein_coding
protein_coding

protein_coding
ncRNA
protein_coding
ncRNA
protein_coding
protein_coding
protein_coding
protein_coding
protein_coding
protein_coding
protein_coding
protein_coding
protein_coding
protein_coding
protein_coding
protein_coding
ncRNA
protein_coding
protein_coding
protein_coding
protein_coding
protein_coding
protein_coding
protein_coding
protein_coding
protein_coding
ncRNA
protein_coding
protein_coding
small_nucle(ol)ar_rna
small_nucle(ol)ar_rna
small_nucle(ol)ar_rna
protein_coding
protein_coding
protein_coding
protein_coding
protein_coding
protein_coding
protein_coding
protein_coding
protein_coding
protein_coding
protein_coding
protein_coding
protein_coding
protein_coding
protein_coding
protein_coding
protein_coding
protein_coding
protein_coding
protein_coding

protein_coding
protein_coding
protein_coding
protein_coding
protein_coding
miRNA_precursor
protein_coding
ncRNA
protein_coding
protein_coding
protein_coding
protein_coding
small_nucle(ol)ar_rna
small_nucle(ol)ar_rna
miRNA_precursor
protein_coding
protein_coding
protein_coding
protein_coding
protein_coding
protein_coding
protein_coding
protein_coding
protein_coding
protein_coding
protein_coding
protein_coding
protein_coding
protein_coding
protein_coding
protein_coding
protein_coding
protein_coding
protein_coding
protein_coding
protein_coding
protein_coding
protein_coding
protein_coding
protein_coding
protein_coding
protein_coding
protein_coding
protein_coding
protein_coding
protein_coding
protein_coding
pre_trna
protein_coding
protein_coding
protein_coding
protein_coding
protein_coding

protein_coding
protein_coding
protein_coding
small_nucle(ol)ar_rna
protein_coding
protein_coding
protein_coding
protein_coding
protein_coding
protein_coding
protein_coding
protein_coding
protein_coding
protein_coding
protein_coding
protein_coding
protein_coding
protein_coding
protein_coding
protein_coding
protein_coding
protein_coding
protein_coding
protein_coding
protein_coding
protein_coding
protein_coding
protein_coding
protein_coding
protein_coding
protein_coding
protein_coding
ncRNA
ncRNA
protein_coding
protein_coding
protein_coding
protein_coding
protein_coding
protein_coding
ncRNA
protein_coding
protein_coding
protein_coding
protein_coding
protein_coding
protein_coding
protein_coding
protein_coding
protein_coding
protein_coding
protein_coding
protein_coding

protein_coding
protein_coding
protein_coding
protein_coding
protein_coding
protein_coding
protein_coding
protein_coding
protein_coding
protein_coding
protein_coding
ncRNA
protein_coding
protein_coding
protein_coding
protein_coding
pre_trna
protein_coding
protein_coding
protein_coding
protein_coding
protein_coding
protein_coding
protein_coding
protein_coding
protein_coding
miRNA_precursor
protein_coding
protein_coding
protein_coding
protein_coding
protein_coding
protein_coding
protein_coding
protein_coding
protein_coding
protein_coding
protein_coding
protein_coding
protein_coding
protein_coding
protein_coding
protein_coding
protein_coding
protein_coding
protein_coding
protein_coding
protein_coding
ncRNA
protein_coding
protein_coding
protein_coding

protein_coding
protein_coding
protein_coding
protein_coding
protein_coding
protein_coding
protein_coding
protein_coding
protein_coding
protein_coding
protein_coding
protein_coding
protein_coding
protein_coding
protein_coding
protein_coding
protein_coding
protein_coding
protein_coding
protein_coding
protein_coding
protein_coding
protein_coding
protein_coding
protein_coding
protein_coding
protein_coding
protein_coding
protein_coding
protein_coding
protein_coding
protein_coding
protein_coding
ncRNA
protein_coding
protein_coding
protein_coding
ncRNA
protein_coding
protein_coding
small_nucle(ol)ar_rna
miRNA_precursor
protein_coding
protein_coding
protein_coding
protein_coding
protein_coding
protein_coding
protein_coding
protein_coding
protein_coding
protein_coding

protein_coding
protein_coding
protein_coding
protein_coding
protein_coding
protein_coding
protein_coding
protein_coding
protein_coding
protein_coding
protein_coding
protein_coding
protein_coding
protein_coding
protein_coding
protein_coding
protein_coding
protein_coding
ncRNA
protein_coding
protein_coding
protein_coding
protein_coding
protein_coding
protein_coding
protein_coding
protein_coding
protein_coding
protein_coding
protein_coding
protein_coding
protein_coding
protein_coding
protein_coding
protein_coding
protein_coding
protein_coding
protein_coding
protein_coding
protein_coding
protein_coding
protein_coding
protein_coding
protein_coding
protein_coding
ncRNA
protein_coding
protein_coding
protein_coding
protein_coding
protein_coding
protein_coding

protein_coding
protein_coding
protein_coding
protein_coding
protein_coding
protein_coding
protein_coding
protein_coding
protein_coding
protein_coding
protein_coding
protein_coding
small_nucle(ol)ar_rna
protein_coding
protein_coding
protein_coding
protein_coding
protein_coding
protein_coding
protein_coding
protein_coding
protein_coding
protein_coding
protein_coding
protein_coding
protein_coding
protein_coding
protein_coding
ncRNA
protein_coding
protein_coding
protein_coding
protein_coding
protein_coding
ncRNA
protein_coding
protein_coding
protein_coding
protein_coding
protein_coding
protein_coding
protein_coding
protein_coding
protein_coding
protein_coding
protein_coding
protein_coding
ncRNA
protein_coding
protein_coding
protein_coding
protein_coding

protein_coding
pre_trna
protein_coding
protein_coding
protein_coding
protein_coding
protein_coding
protein_coding
protein_coding
protein_coding
protein_coding
protein_coding
protein_coding
miRNA_precursor
protein_coding
protein_coding
protein_coding
protein_coding
protein_coding
protein_coding
protein_coding
protein_coding
protein_coding
protein_coding
protein_coding
pre_trna
protein_coding
protein_coding
protein_coding
protein_coding
protein_coding
ncRNA
protein_coding
protein_coding
protein_coding
protein_coding
protein_coding
protein_coding
protein_coding
protein_coding
protein_coding
protein_coding
protein_coding
protein_coding
protein_coding
protein_coding
protein_coding
protein_coding
protein_coding
protein_coding
protein_coding
protein_coding

protein_coding
protein_coding
protein_coding
protein_coding
protein_coding
protein_coding
protein_coding
protein_coding
protein_coding
protein_coding
protein_coding
ncRNA
ncRNA
protein_coding
protein_coding
protein_coding
protein_coding
protein_coding
protein_coding
protein_coding
protein_coding
protein_coding
protein_coding
protein_coding
ncRNA
protein_coding
protein_coding
protein_coding
protein_coding
miRNA_precursor
protein_coding
protein_coding
protein_coding
protein_coding
protein_coding
protein_coding
protein_coding
small_nucle(ol)ar_rna
protein_coding
protein_coding
protein_coding
protein_coding
protein_coding
protein_coding
protein_coding
protein_coding
protein_coding
protein_coding
protein_coding
protein_coding
protein_coding
protein_coding

protein_coding
protein_coding
protein_coding
protein_coding
protein_coding
protein_coding
protein_coding
small_nucle(ol)ar_rna
small_nucle(ol)ar_rna
protein_coding
protein_coding
protein_coding
protein_coding
protein_coding
protein_coding
protein_coding
protein_coding
pre_trna
protein_coding
protein_coding
protein_coding
protein_coding
protein_coding
protein_coding
protein_coding
protein_coding
protein_coding
protein_coding
protein_coding
protein_coding
protein_coding
protein_coding
protein_coding
protein_coding
protein_coding
protein_coding
protein_coding
protein_coding
protein_coding
protein_coding
protein_coding
protein_coding
protein_coding
protein_coding
protein_coding
ncRNA
protein_coding
protein_coding
protein_coding
protein_coding
protein_coding
protein_coding

protein_coding
protein_coding
protein_coding
protein_coding
protein_coding
protein_coding
protein_coding
protein_coding
protein_coding
protein_coding
protein_coding
pre_trna
protein_coding
protein_coding
ncRNA
protein_coding
protein_coding
protein_coding
protein_coding
protein_coding
protein_coding
protein_coding
protein_coding
protein_coding
protein_coding
protein_coding
protein_coding
protein_coding
protein_coding
protein_coding
protein_coding
protein_coding
protein_coding
protein_coding
protein_coding
protein_coding
protein_coding
protein_coding
protein_coding
protein_coding
protein_coding
protein_coding
protein_coding
protein_coding
protein_coding
protein_coding
protein_coding
protein_coding
protein_coding
protein_coding
protein_coding
protein_coding

protein_coding
protein_coding
protein_coding
protein_coding
protein_coding
ncRNA
ncRNA
protein_coding
protein_coding
protein_coding
protein_coding
protein_coding
protein_coding
protein_coding
protein_coding
protein_coding
ncRNA
protein_coding
protein_coding
protein_coding
protein_coding
protein_coding
protein_coding
protein_coding
protein_coding
protein_coding
protein_coding
protein_coding
protein_coding
protein_coding
protein_coding
protein_coding
protein_coding
protein_coding
protein_coding
protein_coding
protein_coding
protein_coding
protein_coding
protein_coding
protein_coding
protein_coding
protein_coding
protein_coding
ncRNA
protein_coding
ncRNA
protein_coding
protein_coding
protein_coding
protein_coding
protein_coding

protein_coding
protein_coding
protein_coding
protein_coding
protein_coding
protein_coding
protein_coding
protein_coding
protein_coding
protein_coding
protein_coding
protein_coding
miRNA_precursor
protein_coding
protein_coding
protein_coding
protein_coding
protein_coding
protein_coding
protein_coding
protein_coding
protein_coding
protein_coding
protein_coding
protein_coding
protein_coding
protein_coding
protein_coding
protein_coding
small_nucle(ol)ar_rna
small_nucle(ol)ar_rna
small_nucle(ol)ar_rna
protein_coding
protein_coding
protein_coding
protein_coding
protein_coding
protein_coding
protein_coding
protein_coding
protein_coding
protein_coding
protein_coding
protein_coding
protein_coding
protein_coding
protein_coding
ncRNA
protein_coding
protein_coding
protein_coding
protein_coding

protein_coding
protein_coding
ncRNA
protein_coding
ncRNA
protein_coding
ncRNA
protein_coding
protein_coding
protein_coding
protein_coding
protein_coding
protein_coding
protein_coding
protein_coding
protein_coding
miRNA_precursor
protein_coding
protein_coding
protein_coding
ncRNA
protein_coding
protein_coding
protein_coding
protein_coding
protein_coding
protein_coding
protein_coding
protein_coding
protein_coding
protein_coding
protein_coding
protein_coding
protein_coding
protein_coding
protein_coding
protein_coding
protein_coding
protein_coding
protein_coding
protein_coding
protein_coding
protein_coding
protein_coding
protein_coding
protein_coding
protein_coding
protein_coding
protein_coding
protein_coding
protein_coding
protein_coding

protein_coding
protein_coding
protein_coding
protein_coding
protein_coding
protein_coding
protein_coding
protein_coding
protein_coding
protein_coding
protein_coding
protein_coding
protein_coding
protein_coding
protein_coding
ncRNA
protein_coding
protein_coding
protein_coding
protein_coding
protein_coding
protein_coding
protein_coding
protein_coding
protein_coding
protein_coding
ncRNA
protein_coding
protein_coding
protein_coding
protein_coding
protein_coding
protein_coding
protein_coding
protein_coding
protein_coding
protein_coding
protein_coding
protein_coding
protein_coding
protein_coding
protein_coding
protein_coding
protein_coding
protein_coding
protein_coding
protein_coding
protein_coding
protein_coding
protein_coding
protein_coding
protein_coding

protein_coding
protein_coding
pre_trna
protein_coding
protein_coding
protein_coding
protein_coding
ncRNA
protein_coding
protein_coding
protein_coding
protein_coding
protein_coding
ncRNA
protein_coding
protein_coding
protein_coding
protein_coding
protein_coding
protein_coding
protein_coding
protein_coding
protein_coding
protein_coding
protein_coding
protein_coding
protein_coding
protein_coding
protein_coding
protein_coding
protein_coding
ncRNA
protein_coding
protein_coding
protein_coding
protein_coding
protein_coding
protein_coding
protein_coding
protein_coding
protein_coding
protein_coding
protein_coding
protein_coding
protein_coding
protein_coding
protein_coding
protein_coding
protein_coding
protein_coding
protein_coding

protein_coding
protein_coding
protein_coding
protein_coding
protein_coding
protein_coding
protein_coding
ncRNA
protein_coding
protein_coding
protein_coding
protein_coding
protein_coding
protein_coding
protein_coding
protein_coding
protein_coding
protein_coding
protein_coding
ncRNA
protein_coding
protein_coding
protein_coding
protein_coding
protein_coding
protein_coding
protein_coding
protein_coding
protein_coding
protein_coding
protein_coding
protein_coding
protein_coding
protein_coding
protein_coding
protein_coding
protein_coding
protein_coding
small_nucle(ol)ar_rna
protein_coding
protein_coding
protein_coding
protein_coding
protein_coding
protein_coding
protein_coding
protein_coding
protein_coding
protein_coding
protein_coding
protein_coding
protein_coding

protein_coding
protein_coding
protein_coding
ncRNA
ncRNA
small_nucle(ol)ar_rna
small_nucle(ol)ar_rna
protein_coding
protein_coding
protein_coding
protein_coding
protein_coding
protein_coding
protein_coding
protein_coding
protein_coding
protein_coding
protein_coding
protein_coding
protein_coding
protein_coding
protein_coding
protein_coding
protein_coding
protein_coding
protein_coding
protein_coding
protein_coding
protein_coding
protein_coding
protein_coding
protein_coding
protein_coding
protein_coding
protein_coding
protein_coding
protein_coding
protein_coding
protein_coding
protein_coding
protein_coding
protein_coding
protein_coding
miRNA_precursor
protein_coding
protein_coding
miRNA_precursor
protein_coding
protein_coding
protein_coding
protein_coding
protein_coding

protein_coding
protein_coding
protein_coding
protein_coding
protein_coding
protein_coding
protein_coding
protein_coding
protein_coding
protein_coding
protein_coding
protein_coding
protein_coding
protein_coding
protein_coding
protein_coding
protein_coding
protein_coding
protein_coding
protein_coding
protein_coding
protein_coding
protein_coding
protein_coding
protein_coding
protein_coding
protein_coding
protein_coding
protein_coding
protein_coding
protein_coding
protein_coding
protein_coding
protein_coding
protein_coding
protein_coding
protein_coding
protein_coding
protein_coding
protein_coding
protein_coding
protein_coding
protein_coding
protein_coding
protein_coding
protein_coding
protein_coding
protein_coding
protein_coding
protein_coding
protein_coding
protein_coding

protein_coding
protein_coding
protein_coding
protein_coding
protein_coding
protein_coding
protein_coding
protein_coding
small_nucle(ol)ar_rna
small_nucle(ol)ar_rna
protein_coding
protein_coding
protein_coding
protein_coding
protein_coding
protein_coding
protein_coding
protein_coding
protein_coding
protein_coding
protein_coding
protein_coding
protein_coding
protein_coding
protein_coding
protein_coding
ncRNA
protein_coding
protein_coding
protein_coding
protein_coding
protein_coding
protein_coding
protein_coding
protein_coding
protein_coding
protein_coding
protein_coding
protein_coding
protein_coding
protein_coding
protein_coding
protein_coding
protein_coding
ncRNA
protein_coding
small_nucle(ol)ar_rna
protein_coding
protein_coding
protein_coding
protein_coding
protein_coding

protein_coding
protein_coding
protein_coding
protein_coding
protein_coding
protein_coding
protein_coding
protein_coding
protein_coding
protein_coding
protein_coding
protein_coding
protein_coding
protein_coding
protein_coding
protein_coding
protein_coding
protein_coding
protein_coding
protein_coding
protein_coding
protein_coding
protein_coding
protein_coding
protein_coding
protein_coding
protein_coding
protein_coding
pre_trna
protein_coding
protein_coding
protein_coding
protein_coding
protein_coding
ncRNA
ncRNA
protein_coding
protein_coding
protein_coding
protein_coding
protein_coding
protein_coding
protein_coding
protein_coding
protein_coding
protein_coding
protein_coding
protein_coding
protein_coding
protein_coding
protein_coding
protein_coding

protein_coding
protein_coding
protein_coding
protein_coding
protein_coding
protein_coding
protein_coding
protein_coding
ncRNA
protein_coding
protein_coding
protein_coding
protein_coding
protein_coding
protein_coding
protein_coding
protein_coding
protein_coding
protein_coding
protein_coding
protein_coding
protein_coding
protein_coding
protein_coding
protein_coding
protein_coding
protein_coding
protein_coding
protein_coding
protein_coding
protein_coding
protein_coding
protein_coding
protein_coding
protein_coding
protein_coding
protein_coding
protein_coding
protein_coding
protein_coding
miRNA_precursor
protein_coding
protein_coding
protein_coding
protein_coding
protein_coding
protein_coding
protein_coding
protein_coding
ncRNA
protein_coding
protein_coding

protein_coding
protein_coding
protein_coding
protein_coding
protein_coding
protein_coding
protein_coding
protein_coding
protein_coding
protein_coding
protein_coding
pre_trna
protein_coding
protein_coding
protein_coding
protein_coding
protein_coding
protein_coding
protein_coding
protein_coding
protein_coding
protein_coding
protein_coding
protein_coding
protein_coding
protein_coding
protein_coding
ncRNA
protein_coding
protein_coding
protein_coding
protein_coding
protein_coding
protein_coding
protein_coding
protein_coding
protein_coding
protein_coding
protein_coding
protein_coding
protein_coding
protein_coding
protein_coding
protein_coding
protein_coding
protein_coding
protein_coding
protein_coding
protein_coding
protein_coding
protein_coding
protein_coding

protein_coding
protein_coding
protein_coding
protein_coding
protein_coding
protein_coding
protein_coding
protein_coding
protein_coding
protein_coding
protein_coding
protein_coding
protein_coding
ncRNA
protein_coding
protein_coding
protein_coding
protein_coding
protein_coding
protein_coding
protein_coding
protein_coding
protein_coding
protein_coding
protein_coding
protein_coding
protein_coding
protein_coding
protein_coding
protein_coding
protein_coding
miRNA_precursor
protein_coding
protein_coding
protein_coding
protein_coding
protein_coding
miRNA_precursor
ncRNA
miRNA_precursor
protein_coding
protein_coding
protein_coding
protein_coding
protein_coding
protein_coding
protein_coding
protein_coding
protein_coding
protein_coding
protein_coding
protein_coding

protein_coding
protein_coding
protein_coding
protein_coding
protein_coding
protein_coding
protein_coding
protein_coding
protein_coding
protein_coding
protein_coding
protein_coding
protein_coding
protein_coding
protein_coding
protein_coding
protein_coding
protein_coding
protein_coding
protein_coding
protein_coding
protein_coding
protein_coding
protein_coding
protein_coding
protein_coding
protein_coding
protein_coding
protein_coding
miRNA_precursor
protein_coding
protein_coding
small_nucle(ol)ar_rna
protein_coding
protein_coding
protein_coding
protein_coding
protein_coding
protein_coding
protein_coding
protein_coding
protein_coding
protein_coding
protein_coding
ncRNA
ncRNA
protein_coding
ncRNA
protein_coding
protein_coding
protein_coding
protein_coding

protein_coding
protein_coding
protein_coding
ncRNA
protein_coding
protein_coding
protein_coding
protein_coding
protein_coding
protein_coding
miRNA_precursor
protein_coding
protein_coding
protein_coding
protein_coding
protein_coding
protein_coding
ncRNA
protein_coding
protein_coding
protein_coding
protein_coding
protein_coding
protein_coding
protein_coding
protein_coding
protein_coding
protein_coding
ncRNA
protein_coding
protein_coding
protein_coding
protein_coding
protein_coding
protein_coding
protein_coding
protein_coding
protein_coding
protein_coding
protein_coding
protein_coding
protein_coding
protein_coding
protein_coding
protein_coding
protein_coding
protein_coding
protein_coding
protein_coding
protein_coding
protein_coding

protein_coding
protein_coding
protein_coding
protein_coding
protein_coding
protein_coding
protein_coding
pre_trna
protein_coding
protein_coding
protein_coding
protein_coding
protein_coding
protein_coding
protein_coding
protein_coding
protein_coding
protein_coding
protein_coding
protein_coding
protein_coding
protein_coding
protein_coding
protein_coding
protein_coding
protein_coding
protein_coding
protein_coding
protein_coding
protein_coding
protein_coding
protein_coding
protein_coding
protein_coding
protein_coding
protein_coding
protein_coding
protein_coding
protein_coding
protein_coding
protein_coding
protein_coding
protein_coding
protein_coding
protein_coding
protein_coding
protein_coding
protein_coding
protein_coding
protein_coding
protein_coding
protein_coding

protein_coding
protein_coding
protein_coding
protein_coding
protein_coding
protein_coding
small_nucle(ol)ar_rna
protein_coding
protein_coding
protein_coding
protein_coding
protein_coding
protein_coding
protein_coding
protein_coding
small_nucle(ol)ar_rna
protein_coding
protein_coding
protein_coding
miRNA_precursor
protein_coding
protein_coding
protein_coding
protein_coding
protein_coding
protein_coding
protein_coding
protein_coding
protein_coding
protein_coding
protein_coding
protein_coding
protein_coding
protein_coding
protein_coding
protein_coding
protein_coding
protein_coding
protein_coding
protein_coding
protein_coding
protein_coding
protein_coding
protein_coding
protein_coding
protein_coding
protein_coding
protein_coding
protein_coding
protein_coding
protein_coding
protein_coding

protein_coding
protein_coding
protein_coding
ncRNA
protein_coding
protein_coding
protein_coding
protein_coding
protein_coding
protein_coding
ncRNA
protein_coding
protein_coding
protein_coding
protein_coding
protein_coding
protein_coding
protein_coding
protein_coding
protein_coding
protein_coding
protein_coding
protein_coding
protein_coding
protein_coding
protein_coding
protein_coding
protein_coding
protein_coding
protein_coding
protein_coding
protein_coding
small_nucle(ol)ar_rna
small_nucle(ol)ar_rna
protein_coding
protein_coding
protein_coding
protein_coding
miRNA_precursor
protein_coding
protein_coding
protein_coding
protein_coding
protein_coding
protein_coding
protein_coding
protein_coding
ncRNA
protein_coding
protein_coding
protein_coding
protein_coding

protein_coding
protein_coding
protein_coding
protein_coding
protein_coding
protein_coding
protein_coding
protein_coding
protein_coding
ncRNA
ncRNA
protein_coding
protein_coding
protein_coding
protein_coding
protein_coding
protein_coding
protein_coding
protein_coding
protein_coding
protein_coding
protein_coding
protein_coding
protein_coding
protein_coding
small_nucle(ol)ar_rna
protein_coding
protein_coding
protein_coding
protein_coding
protein_coding
protein_coding
small_nucle(ol)ar_rna
protein_coding
protein_coding
protein_coding
protein_coding
protein_coding
protein_coding
protein_coding
protein_coding
protein_coding
protein_coding
protein_coding
protein_coding
protein_coding
protein_coding
protein_coding
protein_coding
protein_coding
protein_coding
protein_coding

protein_coding
protein_coding
protein_coding
protein_coding
protein_coding
protein_coding
protein_coding
protein_coding
protein_coding
protein_coding
protein_coding
protein_coding
protein_coding
protein_coding
protein_coding
protein_coding
protein_coding
protein_coding
protein_coding
protein_coding
protein_coding
protein_coding
protein_coding
protein_coding
protein_coding
protein_coding
protein_coding
protein_coding
protein_coding
protein_coding
protein_coding
protein_coding
protein_coding
small_nucle(ol)ar_rna
small_nucle(ol)ar_rna
protein_coding
protein_coding
protein_coding
protein_coding
ncRNA
protein_coding
protein_coding
protein_coding
protein_coding
protein_coding
protein_coding
protein_coding
protein_coding
protein_coding
protein_coding
protein_coding
protein_coding

protein_coding
protein_coding
protein_coding
protein_coding
protein_coding
protein_coding
ncRNA
protein_coding
protein_coding
protein_coding
protein_coding
protein_coding
protein_coding
protein_coding
protein_coding
protein_coding
protein_coding
protein_coding
protein_coding
ncRNA
protein_coding
protein_coding
protein_coding
miRNA_precursor
protein_coding
protein_coding
protein_coding
protein_coding
protein_coding
protein_coding
pre_trna
protein_coding
protein_coding
protein_coding
small_nucle(ol)ar_rna
protein_coding
protein_coding
protein_coding
protein_coding
protein_coding
protein_coding
protein_coding
protein_coding
protein_coding
protein_coding
protein_coding
protein_coding
protein_coding
protein_coding
protein_coding
protein_coding
protein_coding

protein_coding
miRNA_precursor
protein_coding
protein_coding
protein_coding
protein_coding
protein_coding
protein_coding
small_nucle(ol)ar_rna
protein_coding
protein_coding
protein_coding
protein_coding
protein_coding
protein_coding
protein_coding
protein_coding
protein_coding
ncRNA
protein_coding
protein_coding
protein_coding
ncRNA
protein_coding
protein_coding
protein_coding
protein_coding
protein_coding
protein_coding
protein_coding
protein_coding
protein_coding
protein_coding
protein_coding
protein_coding
protein_coding
protein_coding
protein_coding
protein_coding
protein_coding
protein_coding
protein_coding
protein_coding
protein_coding
protein_coding
protein_coding
protein_coding
protein_coding
protein_coding
protein_coding
protein_coding
protein_coding

protein_coding
protein_coding
protein_coding
protein_coding
protein_coding
protein_coding
ncRNA
protein_coding
protein_coding
protein_coding
protein_coding
protein_coding
protein_coding
protein_coding
protein_coding
protein_coding
protein_coding
protein_coding
protein_coding
protein_coding
protein_coding
protein_coding
protein_coding
protein_coding
protein_coding
protein_coding
protein_coding
protein_coding
protein_coding
protein_coding
protein_coding
protein_coding
protein_coding
protein_coding
protein_coding
protein_coding
protein_coding
protein_coding
protein_coding
protein_coding
protein_coding
protein_coding
protein_coding
protein_coding
protein_coding
protein_coding
protein_coding
small_nucle(ol)ar_rna
small_nucle(ol)ar_rna
protein_coding
protein_coding
protein_coding

protein_coding
protein_coding
protein_coding
protein_coding
protein_coding
protein_coding
protein_coding
protein_coding
protein_coding
protein_coding
protein_coding
protein_coding
protein_coding
protein_coding
protein_coding
protein_coding
protein_coding
protein_coding
protein_coding
protein_coding
protein_coding
protein_coding
protein_coding
protein_coding
protein_coding
protein_coding
protein_coding
ncRNA
protein_coding
small_nucle(ol)ar_rna
protein_coding
protein_coding
protein_coding
protein_coding
protein_coding
protein_coding
protein_coding
protein_coding
protein_coding
protein_coding
ncRNA
protein_coding
protein_coding
protein_coding
protein_coding
protein_coding
protein_coding
protein_coding
protein_coding
protein_coding
protein_coding
protein_coding

protein_coding
protein_coding
protein_coding
miRNA_precursor
pre_trna
protein_coding
protein_coding
protein_coding
ncRNA
protein_coding
protein_coding
protein_coding
protein_coding
protein_coding
protein_coding
protein_coding
protein_coding
protein_coding
protein_coding
protein_coding
miRNA_precursor
protein_coding
protein_coding
protein_coding
protein_coding
protein_coding
ncRNA
protein_coding
protein_coding
protein_coding
protein_coding
protein_coding
protein_coding
protein_coding
protein_coding
protein_coding
protein_coding
protein_coding
protein_coding
protein_coding
protein_coding
protein_coding
protein_coding
protein_coding
protein_coding
protein_coding
protein_coding
protein_coding
protein_coding
protein_coding
protein_coding
protein_coding

protein_coding
protein_coding
protein_coding
protein_coding
protein_coding
pre_trna
protein_coding
protein_coding
protein_coding
protein_coding
protein_coding
protein_coding
protein_coding
protein_coding
protein_coding
protein_coding
protein_coding
protein_coding
protein_coding
protein_coding
protein_coding
protein_coding
protein_coding
protein_coding
protein_coding
protein_coding
protein_coding
protein_coding
protein_coding
protein_coding
protein_coding
protein_coding
ncRNA
protein_coding
protein_coding
protein_coding
protein_coding
protein_coding
protein_coding
protein_coding
protein_coding
protein_coding
protein_coding
protein_coding
protein_coding
protein_coding
protein_coding
protein_coding
protein_coding
protein_coding
protein_coding
ncRNA

protein_coding
protein_coding
pre_trna
protein_coding
protein_coding
protein_coding
protein_coding
ncRNA
protein_coding
protein_coding
protein_coding
protein_coding
protein_coding
protein_coding
protein_coding
protein_coding
protein_coding
protein_coding
protein_coding
protein_coding
protein_coding
protein_coding
protein_coding
protein_coding
protein_coding
protein_coding
protein_coding
protein_coding
protein_coding
protein_coding
protein_coding
protein_coding
protein_coding
protein_coding
protein_coding
protein_coding
protein_coding
protein_coding
protein_coding
protein_coding
pre_trna
ncRNA
protein_coding
protein_coding
protein_coding
protein_coding
ncRNA
protein_coding
protein_coding
protein_coding
protein_coding
protein_coding

pre_trna
protein_coding
protein_coding
protein_coding
protein_coding
protein_coding
protein_coding
protein_coding
protein_coding
protein_coding
protein_coding
protein_coding
protein_coding
protein_coding
protein_coding
protein_coding
protein_coding
protein_coding
protein_coding
protein_coding
protein_coding
protein_coding
protein_coding
protein_coding
protein_coding
protein_coding
protein_coding
protein_coding
protein_coding
protein_coding
protein_coding
protein_coding
protein_coding
protein_coding
protein_coding
protein_coding
small_nucle(ol)ar_rna
small_nucle(ol)ar_rna
pre_trna
protein_coding
protein_coding
protein_coding
protein_coding
protein_coding
protein_coding
protein_coding
protein_coding
protein_coding
protein_coding
protein_coding
protein_coding
protein_coding

protein_coding
protein_coding
pre_trna
protein_coding
ncRNA
protein_coding
protein_coding
protein_coding
protein_coding
protein_coding
protein_coding
ncRNA
protein_coding
protein_coding
protein_coding
protein_coding
protein_coding
protein_coding
protein_coding
protein_coding
protein_coding
protein_coding
protein_coding
protein_coding
protein_coding
protein_coding
protein_coding
protein_coding
miRNA_precursor
protein_coding
protein_coding
ncRNA
protein_coding
protein_coding
protein_coding
pre_trna
protein_coding
protein_coding
protein_coding
protein_coding
protein_coding
protein_coding
protein_coding
protein_coding
protein_coding
ncRNA
protein_coding
protein_coding
protein_coding
protein_coding
protein_coding
protein_coding

protein_coding
protein_coding
protein_coding
protein_coding
protein_coding
protein_coding
protein_coding
protein_coding
protein_coding
protein_coding
protein_coding
protein_coding
protein_coding
protein_coding
protein_coding
protein_coding
protein_coding
protein_coding
protein_coding
protein_coding
protein_coding
protein_coding
protein_coding
protein_coding
small_nucle(ol)ar_rna
small_nucle(ol)ar_rna
protein_coding
protein_coding
protein_coding
protein_coding
ncRNA
protein_coding
protein_coding
protein_coding
protein_coding
protein_coding
protein_coding
protein_coding
protein_coding
protein_coding
protein_coding
protein_coding
protein_coding
protein_coding
protein_coding
protein_coding
protein_coding
protein_coding
protein_coding
protein_coding
protein_coding
small_nucle(ol)ar_rna

protein_coding
protein_coding
protein_coding
protein_coding
protein_coding
protein_coding
protein_coding
protein_coding
protein_coding
protein_coding
protein_coding
protein_coding
protein_coding
protein_coding
protein_coding
protein_coding
protein_coding
protein_coding
protein_coding
protein_coding
protein_coding
protein_coding
protein_coding
protein_coding
protein_coding
protein_coding
protein_coding
protein_coding
protein_coding
protein_coding
protein_coding
protein_coding
protein_coding
miRNA_precursor
protein_coding
protein_coding
protein_coding
protein_coding
protein_coding
protein_coding
protein_coding
protein_coding
protein_coding
protein_coding
protein_coding
protein_coding
protein_coding
protein_coding
protein_coding
protein_coding
protein_coding
protein_coding

ncRNA
protein_coding
protein_coding
protein_coding
protein_coding
protein_coding
protein_coding
protein_coding
protein_coding
ncRNA
protein_coding
protein_coding
protein_coding
protein_coding
protein_coding
protein_coding
protein_coding
protein_coding
protein_coding
protein_coding
protein_coding
protein_coding
protein_coding
protein_coding
protein_coding
protein_coding
ncRNA
protein_coding
protein_coding
protein_coding
protein_coding
protein_coding
ncRNA
protein_coding
protein_coding
protein_coding
protein_coding
protein_coding
protein_coding
protein_coding
protein_coding
protein_coding
protein_coding
protein_coding
protein_coding
protein_coding
protein_coding
protein_coding
protein_coding
protein_coding
protein_coding
protein_coding

protein_coding
protein_coding
protein_coding
protein_coding
protein_coding
protein_coding
protein_coding
protein_coding
protein_coding
protein_coding
protein_coding
ncRNA
protein_coding
protein_coding
protein_coding
protein_coding
protein_coding
protein_coding
protein_coding
protein_coding
protein_coding
protein_coding
protein_coding
protein_coding
protein_coding
protein_coding
protein_coding
protein_coding
protein_coding
protein_coding
protein_coding
protein_coding
protein_coding
protein_coding
ncRNA
protein_coding
ncRNA
protein_coding
protein_coding
protein_coding
protein_coding
protein_coding
protein_coding
protein_coding
protein_coding
protein_coding
protein_coding
protein_coding
protein_coding
protein_coding
protein_coding
protein_coding

protein_coding
protein_coding
protein_coding
protein_coding
protein_coding
protein_coding
protein_coding
protein_coding
protein_coding
protein_coding
protein_coding
protein_coding
protein_coding
protein_coding
protein_coding
protein_coding
protein_coding
protein_coding
protein_coding
protein_coding
protein_coding
ncRNA
protein_coding
protein_coding
protein_coding
protein_coding
protein_coding
protein_coding
protein_coding
protein_coding
protein_coding
protein_coding
protein_coding
protein_coding
pre_trna
protein_coding
protein_coding
small_nucle(ol)ar_rna
protein_coding
protein_coding
protein_coding
protein_coding
protein_coding
protein_coding
protein_coding
protein_coding
protein_coding
protein_coding
protein_coding
protein_coding
pre_trna
protein_coding

protein_coding
protein_coding
protein_coding
protein_coding
protein_coding
protein_coding
protein_coding
protein_coding
protein_coding
protein_coding
protein_coding
protein_coding
protein_coding
protein_coding
protein_coding
protein_coding
protein_coding
protein_coding
protein_coding
protein_coding
protein_coding
protein_coding
protein_coding
protein_coding
protein_coding
protein_coding
protein_coding
protein_coding
protein_coding
protein_coding
protein_coding
protein_coding
protein_coding
protein_coding
protein_coding
protein_coding
protein_coding
protein_coding
protein_coding
protein_coding
protein_coding
ncRNA
protein_coding
protein_coding
protein_coding
protein_coding
protein_coding
protein_coding
protein_coding
protein_coding
protein_coding
protein_coding

protein_coding
protein_coding
protein_coding
protein_coding
protein_coding
ncRNA
small_nucle(ol)ar_rna
protein_coding
protein_coding
protein_coding
protein_coding
protein_coding
ncRNA
protein_coding
protein_coding
protein_coding
protein_coding
protein_coding
protein_coding
protein_coding
protein_coding
protein_coding
protein_coding
ncRNA
protein_coding
protein_coding
protein_coding
ncRNA
protein_coding
protein_coding
protein_coding
protein_coding
protein_coding
protein_coding
protein_coding
protein_coding
protein_coding
protein_coding
protein_coding
protein_coding
protein_coding
protein_coding
protein_coding
protein_coding
protein_coding
protein_coding
protein_coding
protein_coding
protein_coding
protein_coding
protein_coding
protein_coding
ncRNA

protein_coding
protein_coding
protein_coding
protein_coding
ncRNA
protein_coding
protein_coding
protein_coding
protein_coding
protein_coding
pre_trna
protein_coding
protein_coding
protein_coding
protein_coding
ncRNA
protein_coding
protein_coding
ncRNA
protein_coding
protein_coding
protein_coding
protein_coding
protein_coding
protein_coding
protein_coding
protein_coding
protein_coding
protein_coding
protein_coding
protein_coding
protein_coding
protein_coding
protein_coding
protein_coding
protein_coding
ncRNA
protein_coding
protein_coding
protein_coding
small_nucle(ol)ar_rna
small_nucle(ol)ar_rna
protein_coding
protein_coding
protein_coding
protein_coding
protein_coding
protein_coding
protein_coding
protein_coding
protein_coding
protein_coding

protein_coding
protein_coding
protein_coding
protein_coding
protein_coding
protein_coding
protein_coding
protein_coding
protein_coding
protein_coding
protein_coding
protein_coding
protein_coding
protein_coding
protein_coding
protein_coding
protein_coding
protein_coding
pre_trna
protein_coding
protein_coding
protein_coding
protein_coding
protein_coding
protein_coding
protein_coding
protein_coding
protein_coding
protein_coding
protein_coding
protein_coding
protein_coding
protein_coding
protein_coding
protein_coding
protein_coding
ncRNA
small_nucle(ol)ar_rna
protein_coding
protein_coding
protein_coding
protein_coding
protein_coding
protein_coding
protein_coding
protein_coding
miRNA_precursor
protein_coding
protein_coding
miRNA_precursor
protein_coding
protein_coding

| |
|---|
| protein_coding |
| protein_coding |
| protein_coding |
| protein_coding |
| protein_coding |
| protein_coding |
| protein_coding |
| protein_coding |
| protein_coding |
| protein_coding |
| protein_coding |
| protein_coding |
| protein_coding |
| protein_coding |
| protein_coding |
| protein_coding |
| protein_coding |
| protein_coding |
| protein_coding |
| protein_coding |
| protein_coding |
| protein_coding |
| protein_coding |
| protein_coding |
| protein_coding |
| protein_coding |
| ncRNA |
| protein_coding |
| ncRNA |
| protein_coding |
| protein_coding |
| protein_coding |
| protein_coding |
| protein_coding |
| ncRNA |
| protein_coding |
| protein_coding |
| protein_coding |
| protein_coding |
| protein_coding |
| protein_coding |
| protein_coding |
| protein_coding |
| protein_coding |
| protein_coding |
| protein_coding |
| protein_coding |
| protein_coding |
| protein_coding |
| protein_coding |
| protein_coding |
| protein_coding |

protein_coding
protein_coding
small_nucle(ol)ar_rna
protein_coding
protein_coding
protein_coding
protein_coding
protein_coding
protein_coding
protein_coding
protein_coding
protein_coding
protein_coding
protein_coding
protein_coding
protein_coding
protein_coding
protein_coding
ncRNA
ncRNA
small_nucle(ol)ar_rna
protein_coding
protein_coding
protein_coding
protein_coding
protein_coding
protein_coding
protein_coding
protein_coding
protein_coding
protein_coding
protein_coding
small_nucle(ol)ar_rna
protein_coding
protein_coding
protein_coding
protein_coding
protein_coding
protein_coding
protein_coding
protein_coding
protein_coding
protein_coding
protein_coding
protein_coding
protein_coding
protein_coding
protein_coding
ncRNA
protein_coding
protein_coding
protein_coding
protein_coding

protein_coding
protein_coding
protein_coding
protein_coding
protein_coding
protein_coding
protein_coding
protein_coding
protein_coding
protein_coding
protein_coding
protein_coding
protein_coding
protein_coding
protein_coding
protein_coding
protein_coding
protein_coding
protein_coding
protein_coding
protein_coding
protein_coding
protein_coding
protein_coding
protein_coding
protein_coding
ncRNA
protein_coding
protein_coding
protein_coding
small_nucle(ol)ar_rna
protein_coding
protein_coding
protein_coding
protein_coding
protein_coding
protein_coding
protein_coding
protein_coding
protein_coding
protein_coding
protein_coding
protein_coding
protein_coding
protein_coding
protein_coding
protein_coding
protein_coding
protein_coding
protein_coding
protein_coding
protein_coding

protein_coding
protein_coding
protein_coding
protein_coding
protein_coding
protein_coding
protein_coding
ncRNA
protein_coding
protein_coding
protein_coding
protein_coding
protein_coding
protein_coding
protein_coding
protein_coding
ncRNA
protein_coding
protein_coding
protein_coding
protein_coding
protein_coding
protein_coding
protein_coding
protein_coding
protein_coding
protein_coding
protein_coding
protein_coding
protein_coding
protein_coding
protein_coding
protein_coding
protein_coding
protein_coding
protein_coding
protein_coding
protein_coding
small_nucle(ol)ar_rna
protein_coding
small_nucle(ol)ar_rna
protein_coding
protein_coding
ncRNA
protein_coding
protein_coding
protein_coding
protein_coding
protein_coding
protein_coding
protein_coding
protein_coding

ncRNA
protein_coding
protein_coding
ncRNA
protein_coding
protein_coding
protein_coding
protein_coding
ncRNA
protein_coding
miRNA_precursor
protein_coding
protein_coding
protein_coding
protein_coding
protein_coding
protein_coding
protein_coding
miRNA_precursor
protein_coding
protein_coding
protein_coding
small_nucle(ol)ar_rna
protein_coding
protein_coding
protein_coding
protein_coding
protein_coding
ncRNA
protein_coding
protein_coding
protein_coding
protein_coding
protein_coding
protein_coding
protein_coding
protein_coding
protein_coding
protein_coding
protein_coding
protein_coding
protein_coding
protein_coding
protein_coding
protein_coding
protein_coding
protein_coding
protein_coding
protein_coding
protein_coding
protein_coding
protein_coding

protein_coding
protein_coding
ncRNA
ncRNA
miRNA_precursor
protein_coding
ncRNA
protein_coding
protein_coding
protein_coding
protein_coding
protein_coding
protein_coding
protein_coding
protein_coding
protein_coding
protein_coding
protein_coding
protein_coding
protein_coding
protein_coding
protein_coding
protein_coding
ncRNA
protein_coding
protein_coding
protein_coding
protein_coding
protein_coding
protein_coding
protein_coding
protein_coding
protein_coding
protein_coding
protein_coding
protein_coding
protein_coding
protein_coding
protein_coding
protein_coding
protein_coding
protein_coding
protein_coding
protein_coding
protein_coding
protein_coding
protein_coding
protein_coding
miRNA_precursor
ncRNA
protein_coding
protein_coding

protein_coding
protein_coding
protein_coding
protein_coding
protein_coding
protein_coding
protein_coding
protein_coding
protein_coding
protein_coding
protein_coding
protein_coding
protein_coding
ncRNA
protein_coding
protein_coding
ncRNA
protein_coding
protein_coding
ncRNA
protein_coding
protein_coding
protein_coding
ncRNA
small_nucle(ol)ar_rna
protein_coding
protein_coding
protein_coding
protein_coding
protein_coding
protein_coding
protein_coding
protein_coding
protein_coding
protein_coding
protein_coding
protein_coding
protein_coding
protein_coding
protein_coding
protein_coding
protein_coding
protein_coding
protein_coding
protein_coding
protein_coding
ncRNA
protein_coding
protein_coding
protein_coding
protein_coding
miRNA_precursor

protein_coding
protein_coding
protein_coding
protein_coding
protein_coding
protein_coding
protein_coding
protein_coding
protein_coding
protein_coding
protein_coding
protein_coding
protein_coding
protein_coding
protein_coding
protein_coding
protein_coding
protein_coding
protein_coding
protein_coding
protein_coding
small_nucle(ol)ar_rna
protein_coding
protein_coding
protein_coding
ncRNA
protein_coding
protein_coding
protein_coding
protein_coding
protein_coding
protein_coding
protein_coding
protein_coding
protein_coding
ncRNA
protein_coding
protein_coding
ncRNA
protein_coding
protein_coding
ncRNA
protein_coding
protein_coding
protein_coding
ncRNA
protein_coding
protein_coding
protein_coding
protein_coding
protein_coding
protein_coding

protein_coding
ncRNA
protein_coding
protein_coding
protein_coding
protein_coding
protein_coding
protein_coding
protein_coding
protein_coding
protein_coding
ncRNA
protein_coding
protein_coding
protein_coding
protein_coding
protein_coding
protein_coding
protein_coding
protein_coding
protein_coding
protein_coding
protein_coding
protein_coding
protein_coding
miRNA_precursor
protein_coding
protein_coding
protein_coding
protein_coding
protein_coding
protein_coding
protein_coding
protein_coding
protein_coding
protein_coding
protein_coding
protein_coding
protein_coding
protein_coding
protein_coding
protein_coding
protein_coding
protein_coding
protein_coding
protein_coding
protein_coding
protein_coding
protein_coding
protein_coding
protein_coding
protein_coding

protein_coding
protein_coding
protein_coding
protein_coding
protein_coding
protein_coding
protein_coding
protein_coding
protein_coding
protein_coding
protein_coding
protein_coding
protein_coding
protein_coding
protein_coding
protein_coding
protein_coding
protein_coding
protein_coding
protein_coding
protein_coding
protein_coding
protein_coding
protein_coding
protein_coding
protein_coding
protein_coding
ncRNA
protein_coding
miRNA_precursor
protein_coding
protein_coding
protein_coding
protein_coding
protein_coding
protein_coding
protein_coding
protein_coding
protein_coding
protein_coding
protein_coding
protein_coding
protein_coding
protein_coding
protein_coding
protein_coding
protein_coding
protein_coding
protein_coding
protein_coding
protein_coding
protein_coding

protein_coding
protein_coding
protein_coding
protein_coding
protein_coding
ncRNA
protein_coding
small_nucle(ol)ar_rna
small_nucle(ol)ar_rna
small_nucle(ol)ar_rna
small_nucle(ol)ar_rna
protein_coding
protein_coding
protein_coding
protein_coding
protein_coding
protein_coding
protein_coding
protein_coding
protein_coding
protein_coding
protein_coding
protein_coding
protein_coding
protein_coding
protein_coding
protein_coding
protein_coding
protein_coding
protein_coding
protein_coding
protein_coding
protein_coding
protein_coding
protein_coding
protein_coding
protein_coding
protein_coding
protein_coding
protein_coding
protein_coding
protein_coding
protein_coding
protein_coding
protein_coding
protein_coding
protein_coding
protein_coding
protein_coding
protein_coding
protein_coding
protein_coding

protein_coding
protein_coding
protein_coding
protein_coding
protein_coding
protein_coding
protein_coding
protein_coding
protein_coding
protein_coding
protein_coding
protein_coding
ncRNA
protein_coding
protein_coding
protein_coding
protein_coding
small_nucle(ol)ar_rna
protein_coding
protein_coding
protein_coding
protein_coding
protein_coding
protein_coding
protein_coding
protein_coding
protein_coding
protein_coding
protein_coding
protein_coding
protein_coding
protein_coding
protein_coding
protein_coding
protein_coding
protein_coding
protein_coding
protein_coding
protein_coding
protein_coding
protein_coding
protein_coding
protein_coding
protein_coding
protein_coding
protein_coding
ncRNA
protein_coding
protein_coding
protein_coding
protein_coding
protein_coding

protein_coding
protein_coding
miRNA_precursor
protein_coding
protein_coding
protein_coding
protein_coding
protein_coding
protein_coding
protein_coding
protein_coding
protein_coding
protein_coding
protein_coding
protein_coding
protein_coding
miRNA_precursor
protein_coding
protein_coding
protein_coding
protein_coding
protein_coding
protein_coding
protein_coding
protein_coding
protein_coding
protein_coding
protein_coding
protein_coding
protein_coding
protein_coding
protein_coding
protein_coding
ncRNA
protein_coding
protein_coding
protein_coding
protein_coding
protein_coding
protein_coding
protein_coding
protein_coding
protein_coding
protein_coding
protein_coding
protein_coding
protein_coding
protein_coding
protein_coding
protein_coding
protein_coding
protein_coding

protein_coding
protein_coding
protein_coding
protein_coding
protein_coding
protein_coding
protein_coding
protein_coding
protein_coding
protein_coding
ncRNA
protein_coding
protein_coding
protein_coding
protein_coding
protein_coding
protein_coding
protein_coding
protein_coding
protein_coding
protein_coding
protein_coding
protein_coding
protein_coding
protein_coding
ncRNA
protein_coding
protein_coding
protein_coding
protein_coding
protein_coding
protein_coding
protein_coding
protein_coding
protein_coding
protein_coding
protein_coding
protein_coding
protein_coding
protein_coding
protein_coding
protein_coding
protein_coding
protein_coding
protein_coding
ncRNA
protein_coding
protein_coding
protein_coding
protein_coding
protein_coding
protein_coding

protein_coding
protein_coding
protein_coding
protein_coding
protein_coding
protein_coding
protein_coding
protein_coding
protein_coding
protein_coding
miRNA_precursor
protein_coding
protein_coding
protein_coding
protein_coding
protein_coding
protein_coding
protein_coding
protein_coding
protein_coding
protein_coding
protein_coding
protein_coding
protein_coding
protein_coding
protein_coding
protein_coding
protein_coding
protein_coding
protein_coding
protein_coding
protein_coding
protein_coding
protein_coding
ncRNA
protein_coding
protein_coding
protein_coding
protein_coding
protein_coding
small_nucle(ol)ar_rna
protein_coding
protein_coding
protein_coding
protein_coding
protein_coding
protein_coding
protein_coding
protein_coding
protein_coding
protein_coding
protein_coding

protein_coding
protein_coding
protein_coding
protein_coding
protein_coding
protein_coding
miRNA_precursor
miRNA_precursor
protein_coding
protein_coding
protein_coding
protein_coding
protein_coding
protein_coding
protein_coding
protein_coding
protein_coding
protein_coding
protein_coding
protein_coding
protein_coding
protein_coding
protein_coding
protein_coding
protein_coding
protein_coding
protein_coding
protein_coding
protein_coding
protein_coding
protein_coding
protein_coding
protein_coding
protein_coding
protein_coding
miRNA_precursor
protein_coding
protein_coding
protein_coding
protein_coding
protein_coding
protein_coding
protein_coding
protein_coding
protein_coding
protein_coding
protein_coding
protein_coding
protein_coding
protein_coding
ncRNA
protein_coding

protein_coding
protein_coding
protein_coding
protein_coding
protein_coding
protein_coding
protein_coding
protein_coding
protein_coding
protein_coding
protein_coding
protein_coding
protein_coding
protein_coding
protein_coding
protein_coding
protein_coding
protein_coding
protein_coding
ncRNA
protein_coding
protein_coding
protein_coding
protein_coding
protein_coding
protein_coding
protein_coding
ncRNA
protein_coding
protein_coding
protein_coding
protein_coding
protein_coding
protein_coding
protein_coding
protein_coding
protein_coding
protein_coding
protein_coding
protein_coding
protein_coding
protein_coding
protein_coding
protein_coding
protein_coding
protein_coding
protein_coding
protein_coding
protein_coding
protein_coding
protein_coding
protein_coding

protein_coding
protein_coding
protein_coding
protein_coding
protein_coding
protein_coding
protein_coding
protein_coding
protein_coding
small_nucle(ol)ar_rna
small_nucle(ol)ar_rna
protein_coding
protein_coding
protein_coding
protein_coding
protein_coding
protein_coding
protein_coding
protein_coding
protein_coding
protein_coding
protein_coding
protein_coding
protein_coding
protein_coding
protein_coding
protein_coding
protein_coding
protein_coding
protein_coding
protein_coding
protein_coding
protein_coding
protein_coding
protein_coding
protein_coding
protein_coding
protein_coding
protein_coding
protein_coding
protein_coding
protein_coding
protein_coding
miRNA_precursor
protein_coding
protein_coding
protein_coding
protein_coding
protein_coding
protein_coding
protein_coding
protein_coding

ncRNA
protein_coding
protein_coding
protein_coding
protein_coding
protein_coding
protein_coding
protein_coding
protein_coding
protein_coding
ncRNA
protein_coding
protein_coding
protein_coding
protein_coding
protein_coding
protein_coding
protein_coding
protein_coding
protein_coding
protein_coding
protein_coding
protein_coding
protein_coding
protein_coding
protein_coding
protein_coding
protein_coding
pre_trna
protein_coding
protein_coding
protein_coding
protein_coding
protein_coding
protein_coding
protein_coding
protein_coding
protein_coding
protein_coding
protein_coding
protein_coding
protein_coding
protein_coding
protein_coding
protein_coding
protein_coding
protein_coding
protein_coding
protein_coding
protein_coding
protein_coding
protein_coding

protein_coding
protein_coding
ncRNA
protein_coding
protein_coding
protein_coding
protein_coding
protein_coding
protein_coding
protein_coding
protein_coding
protein_coding
protein_coding
protein_coding
protein_coding
protein_coding
protein_coding
protein_coding
protein_coding
small_nucle(ol)ar_rna
small_nucle(ol)ar_rna
protein_coding
protein_coding
protein_coding
protein_coding
protein_coding
protein_coding
protein_coding
protein_coding
protein_coding
protein_coding
protein_coding
protein_coding
protein_coding
protein_coding
protein_coding
protein_coding
protein_coding
protein_coding
pre_trna
ncRNA
protein_coding
protein_coding
protein_coding
protein_coding
protein_coding
protein_coding
protein_coding
protein_coding
protein_coding
protein_coding
protein_coding

protein_coding
protein_coding
protein_coding
protein_coding
protein_coding
protein_coding
protein_coding
protein_coding
protein_coding
protein_coding
protein_coding
protein_coding
protein_coding
protein_coding
protein_coding
protein_coding
protein_coding
protein_coding
protein_coding
protein_coding
protein_coding
protein_coding
protein_coding
protein_coding
protein_coding
protein_coding
protein_coding
protein_coding
protein_coding
protein_coding
protein_coding
protein_coding
protein_coding
protein_coding
ncRNA
protein_coding
protein_coding
protein_coding
protein_coding
protein_coding
protein_coding
protein_coding
protein_coding
protein_coding
protein_coding
protein_coding
protein_coding
protein_coding
protein_coding
protein_coding
protein_coding
protein_coding

protein_coding
protein_coding
protein_coding
protein_coding
protein_coding
ncRNA
protein_coding
protein_coding
protein_coding
protein_coding
protein_coding
protein_coding
protein_coding
protein_coding
protein_coding
protein_coding
protein_coding
protein_coding
protein_coding
pre_trna
protein_coding
protein_coding
protein_coding
protein_coding
protein_coding
protein_coding
protein_coding
protein_coding
protein_coding
protein_coding
ncRNA
ncRNA
protein_coding
protein_coding
protein_coding
protein_coding
protein_coding
protein_coding
protein_coding
protein_coding
protein_coding
protein_coding
protein_coding
protein_coding
protein_coding
protein_coding
protein_coding
protein_coding
protein_coding
protein_coding
protein_coding
protein_coding

protein_coding
protein_coding
protein_coding
ncRNA
protein_coding
small_nucle(ol)ar_rna
small_nucle(ol)ar_rna
small_nucle(ol)ar_rna
ncRNA
protein_coding
ncRNA
protein_coding
protein_coding
protein_coding
protein_coding
protein_coding
protein_coding
protein_coding
protein_coding
protein_coding
protein_coding
protein_coding
protein_coding
protein_coding
protein_coding
protein_coding
protein_coding
protein_coding
protein_coding
protein_coding
protein_coding
protein_coding
protein_coding
protein_coding
protein_coding
protein_coding
ncRNA
protein_coding
protein_coding
protein_coding
protein_coding
protein_coding
pre_trna
protein_coding
protein_coding
protein_coding
protein_coding
protein_coding
protein_coding
ncRNA
ncRNA

protein_coding
protein_coding
small_nucle(ol)ar_rna
protein_coding
protein_coding
protein_coding
protein_coding
protein_coding
ncRNA
protein_coding
ncRNA
protein_coding
protein_coding
ncRNA
protein_coding
ncRNA
protein_coding
protein_coding
protein_coding
protein_coding
protein_coding
protein_coding
protein_coding
protein_coding
protein_coding
protein_coding
protein_coding
protein_coding
protein_coding
protein_coding
protein_coding
protein_coding
protein_coding
protein_coding
protein_coding
protein_coding
protein_coding
protein_coding
protein_coding
ncRNA
protein_coding
protein_coding
ncRNA
protein_coding
protein_coding
protein_coding
ncRNA
protein_coding
protein_coding
protein_coding
protein_coding
protein_coding

ncRNA
protein_coding
protein_coding
ncRNA
protein_coding
protein_coding
protein_coding
protein_coding
protein_coding
protein_coding
protein_coding
protein_coding
protein_coding
protein_coding
protein_coding
protein_coding
protein_coding
protein_coding
protein_coding
protein_coding
protein_coding
protein_coding
protein_coding
protein_coding
protein_coding
protein_coding
protein_coding
protein_coding
protein_coding
protein_coding
protein_coding
protein_coding
protein_coding
ncRNA
protein_coding
protein_coding
protein_coding
protein_coding
protein_coding
protein_coding
protein_coding
protein_coding
protein_coding
protein_coding
protein_coding
protein_coding
protein_coding
protein_coding
protein_coding
protein_coding
protein_coding
pre_trna

ncRNA
protein_coding
protein_coding
protein_coding
protein_coding
protein_coding
protein_coding
protein_coding
protein_coding
protein_coding
ncRNA
ncRNA
protein_coding
protein_coding
protein_coding
protein_coding
protein_coding
miRNA_precursor
protein_coding
protein_coding
protein_coding
protein_coding
protein_coding
protein_coding
protein_coding
protein_coding
protein_coding
protein_coding
protein_coding
protein_coding
protein_coding
protein_coding
protein_coding
protein_coding
protein_coding
protein_coding
protein_coding
protein_coding
protein_coding
protein_coding
protein_coding
ncRNA
protein_coding
small_nucle(ol)ar_rna
small_nucle(ol)ar_rna
small_nucle(ol)ar_rna
protein_coding
protein_coding
protein_coding
protein_coding
protein_coding
protein_coding

protein_coding
protein_coding
protein_coding
protein_coding
protein_coding
protein_coding
protein_coding
protein_coding
protein_coding
ncRNA
protein_coding
protein_coding
protein_coding
protein_coding
protein_coding
protein_coding
protein_coding
protein_coding
protein_coding
protein_coding
protein_coding
protein_coding
protein_coding
protein_coding
protein_coding
protein_coding
protein_coding
protein_coding
protein_coding
protein_coding
ncRNA
protein_coding
protein_coding
protein_coding
protein_coding
protein_coding
ncRNA
protein_coding
protein_coding
protein_coding
protein_coding
protein_coding
protein_coding
protein_coding
protein_coding
protein_coding
protein_coding
protein_coding
miRNA_precursor
protein_coding
protein_coding
protein_coding

protein_coding
miRNA_precursor
protein_coding
protein_coding
protein_coding
protein_coding
protein_coding
protein_coding
protein_coding
protein_coding
protein_coding
protein_coding
protein_coding
protein_coding
protein_coding
protein_coding
protein_coding
protein_coding
protein_coding
protein_coding
protein_coding
ncRNA
ncRNA
protein_coding
protein_coding
protein_coding
protein_coding
ncRNA
protein_coding
protein_coding
protein_coding
protein_coding
miRNA_precursor
protein_coding
protein_coding
ncRNA
protein_coding
protein_coding
protein_coding
protein_coding
protein_coding
protein_coding
protein_coding
protein_coding
protein_coding
protein_coding
protein_coding
protein_coding
protein_coding
protein_coding
protein_coding
protein_coding

protein_coding
protein_coding
protein_coding
protein_coding
protein_coding
protein_coding
protein_coding
protein_coding
protein_coding
protein_coding
protein_coding
protein_coding
protein_coding
protein_coding
protein_coding
protein_coding
protein_coding
ncRNA
protein_coding
protein_coding
protein_coding
protein_coding
protein_coding
protein_coding
protein_coding
protein_coding
protein_coding
protein_coding
protein_coding
protein_coding
protein_coding
protein_coding
protein_coding
protein_coding
protein_coding
protein_coding
protein_coding
protein_coding
protein_coding
protein_coding
protein_coding
protein_coding
protein_coding
protein_coding
protein_coding
protein_coding
protein_coding
protein_coding
protein_coding
protein_coding
protein_coding
protein_coding

protein_coding
protein_coding
protein_coding
protein_coding
miRNA_precursor
protein_coding
protein_coding
protein_coding
protein_coding
protein_coding
ncRNA
protein_coding
protein_coding
protein_coding
protein_coding
protein_coding
pre_trna
protein_coding
protein_coding
protein_coding
protein_coding
protein_coding
ncRNA
protein_coding
protein_coding
protein_coding
protein_coding
protein_coding
protein_coding
protein_coding
protein_coding
protein_coding
protein_coding
protein_coding
protein_coding
protein_coding
protein_coding
protein_coding
protein_coding
protein_coding
protein_coding
small_nucle(ol)ar_rna
protein_coding
protein_coding
protein_coding
protein_coding
protein_coding
protein_coding
protein_coding
protein_coding
protein_coding
protein_coding

pre_trna
protein_coding
protein_coding
ncRNA
protein_coding
protein_coding
protein_coding
protein_coding
protein_coding
protein_coding
protein_coding
protein_coding
protein_coding
ncRNA
protein_coding
protein_coding
protein_coding
protein_coding
protein_coding
protein_coding
protein_coding
protein_coding
protein_coding
protein_coding
ncRNA
protein_coding
protein_coding
protein_coding
protein_coding
ncRNA
protein_coding
protein_coding
protein_coding
protein_coding
protein_coding
protein_coding
protein_coding
protein_coding
protein_coding
protein_coding
protein_coding
protein_coding
protein_coding
protein_coding
protein_coding
protein_coding
protein_coding
protein_coding
protein_coding
protein_coding
protein_coding
protein_coding

protein_coding
protein_coding
ncRNA
protein_coding
protein_coding
protein_coding
protein_coding
protein_coding
protein_coding
protein_coding
protein_coding
protein_coding
protein_coding
protein_coding
protein_coding
protein_coding
protein_coding
protein_coding
protein_coding
protein_coding
protein_coding
protein_coding
protein_coding
protein_coding
protein_coding
protein_coding
protein_coding
protein_coding
ncRNA
protein_coding
protein_coding
protein_coding
protein_coding
protein_coding
protein_coding
protein_coding
protein_coding
protein_coding
protein_coding
protein_coding
protein_coding
protein_coding
protein_coding
protein_coding
protein_coding
protein_coding
protein_coding
protein_coding
protein_coding
protein_coding
protein_coding
protein_coding

protein_coding
protein_coding
protein_coding
protein_coding
protein_coding
small_nucle(ol)ar_rna
protein_coding
protein_coding
protein_coding
protein_coding
protein_coding
protein_coding
protein_coding
small_nucle(ol)ar_rna
small_nucle(ol)ar_rna
small_nucle(ol)ar_rna
ncRNA
protein_coding
protein_coding
protein_coding
protein_coding
protein_coding
protein_coding
protein_coding
protein_coding
protein_coding
protein_coding
protein_coding
protein_coding
protein_coding
protein_coding
protein_coding
protein_coding
protein_coding
protein_coding
protein_coding
protein_coding
protein_coding
protein_coding
protein_coding
protein_coding
protein_coding
protein_coding
protein_coding
protein_coding
protein_coding
protein_coding
protein_coding
protein_coding
protein_coding
protein_coding
protein_coding
protein_coding

protein_coding
protein_coding
protein_coding
protein_coding
protein_coding
protein_coding
protein_coding
protein_coding
protein_coding
protein_coding
protein_coding
protein_coding
protein_coding
protein_coding
protein_coding
protein_coding
protein_coding
protein_coding
protein_coding
protein_coding
protein_coding
ncRNA
small_nucle(ol)ar_rna
small_nucle(ol)ar_rna
ncRNA
protein_coding
protein_coding
protein_coding
protein_coding
protein_coding
protein_coding
protein_coding
protein_coding
protein_coding
protein_coding
protein_coding
protein_coding
protein_coding
protein_coding
protein_coding
protein_coding
protein_coding
protein_coding
protein_coding
protein_coding
protein_coding
protein_coding
protein_coding
ncRNA
protein_coding
protein_coding
protein_coding

protein_coding
ncRNA
protein_coding
protein_coding
protein_coding
protein_coding
protein_coding
protein_coding
miRNA_precursor
protein_coding
protein_coding
ncRNA
protein_coding
protein_coding
protein_coding
protein_coding
protein_coding
protein_coding
protein_coding
protein_coding
protein_coding
protein_coding
protein_coding
protein_coding
protein_coding
protein_coding
protein_coding
protein_coding
protein_coding
protein_coding
protein_coding
protein_coding
protein_coding
protein_coding
protein_coding
protein_coding
protein_coding
protein_coding
protein_coding
protein_coding
protein_coding
protein_coding
protein_coding
ncRNA
protein_coding
protein_coding
protein_coding
protein_coding
protein_coding
protein_coding
protein_coding
protein_coding

protein_coding
protein_coding
protein_coding
miRNA_precursor
protein_coding
protein_coding
protein_coding
protein_coding
protein_coding
protein_coding
protein_coding
protein_coding
protein_coding
protein_coding
protein_coding
protein_coding
protein_coding
protein_coding
pre_trna
protein_coding
protein_coding
protein_coding
ncRNA
protein_coding
protein_coding
protein_coding
protein_coding
protein_coding
protein_coding
protein_coding
protein_coding
protein_coding
protein_coding
protein_coding
protein_coding
protein_coding
protein_coding
protein_coding
protein_coding
protein_coding
protein_coding
protein_coding
protein_coding
small_nucle(ol)ar_rna
protein_coding
protein_coding
protein_coding
protein_coding
protein_coding
protein_coding
protein_coding
protein_coding

protein_coding
protein_coding
protein_coding
protein_coding
protein_coding
protein_coding
protein_coding
protein_coding
protein_coding
ncRNA
protein_coding
protein_coding
protein_coding
protein_coding
protein_coding
protein_coding
protein_coding
protein_coding
ncRNA
ncRNA
protein_coding
protein_coding
protein_coding
ncRNA
ncRNA
protein_coding
protein_coding
protein_coding
ncRNA
protein_coding
protein_coding
protein_coding
protein_coding
protein_coding
protein_coding
protein_coding
protein_coding
protein_coding
protein_coding
protein_coding
protein_coding
protein_coding
protein_coding
protein_coding
protein_coding
protein_coding
protein_coding
protein_coding
protein_coding
protein_coding
protein_coding
protein_coding

protein_coding
protein_coding
ncRNA
protein_coding
protein_coding
protein_coding
protein_coding
protein_coding
protein_coding
protein_coding
protein_coding
protein_coding
protein_coding
protein_coding
protein_coding
protein_coding
protein_coding
protein_coding
protein_coding
protein_coding
protein_coding
protein_coding
protein_coding
protein_coding
protein_coding
protein_coding
protein_coding
protein_coding
protein_coding
protein_coding
protein_coding
protein_coding
protein_coding
protein_coding
protein_coding
small_nucle(ol)ar_rna
ncRNA
small_nucle(ol)ar_rna
protein_coding
protein_coding
protein_coding
protein_coding
protein_coding
protein_coding
protein_coding
protein_coding
protein_coding
protein_coding
protein_coding
protein_coding
protein_coding
protein_coding

protein_coding
protein_coding
protein_coding
protein_coding
protein_coding
protein_coding
protein_coding
protein_coding
protein_coding
protein_coding
protein_coding
protein_coding
protein_coding
protein_coding
protein_coding
protein_coding
protein_coding
protein_coding
ncRNA
protein_coding
protein_coding
protein_coding
protein_coding
protein_coding
protein_coding
protein_coding
protein_coding
protein_coding
protein_coding
protein_coding
protein_coding
protein_coding
protein_coding
protein_coding
protein_coding
protein_coding
protein_coding
protein_coding
protein_coding
protein_coding
protein_coding
protein_coding
protein_coding
protein_coding
protein_coding
protein_coding
protein_coding
protein_coding
protein_coding
protein_coding
protein_coding
protein_coding

protein_coding
protein_coding
protein_coding
protein_coding
protein_coding
protein_coding
protein_coding
protein_coding
protein_coding
protein_coding
miRNA_precursor
protein_coding
protein_coding
protein_coding
ncRNA
protein_coding
protein_coding
protein_coding
protein_coding
protein_coding
protein_coding
protein_coding
protein_coding
protein_coding
protein_coding
protein_coding
protein_coding
protein_coding
protein_coding
protein_coding
protein_coding
protein_coding
protein_coding
protein_coding
ncRNA
protein_coding
protein_coding
protein_coding
protein_coding
protein_coding
protein_coding
protein_coding
protein_coding
protein_coding
protein_coding
protein_coding
protein_coding
protein_coding
protein_coding
protein_coding
protein_coding
protein_coding

protein_coding
protein_coding
protein_coding
protein_coding
protein_coding
pre_trna
protein_coding
protein_coding
protein_coding
protein_coding
protein_coding
protein_coding
protein_coding
small_nucle(ol)ar_rna
protein_coding
protein_coding
protein_coding
protein_coding
protein_coding
protein_coding
pre_trna
protein_coding
protein_coding
ncRNA
protein_coding
protein_coding
protein_coding
protein_coding
protein_coding
protein_coding
protein_coding
protein_coding
protein_coding
protein_coding
protein_coding
protein_coding
protein_coding
protein_coding
protein_coding
protein_coding
protein_coding
protein_coding
protein_coding
protein_coding
protein_coding
protein_coding
protein_coding
protein_coding
protein_coding
protein_coding
protein_coding
protein_coding

protein_coding
protein_coding
protein_coding
protein_coding
protein_coding
protein_coding
protein_coding
protein_coding
protein_coding
protein_coding
protein_coding
protein_coding
protein_coding
protein_coding
protein_coding
protein_coding
protein_coding
protein_coding
protein_coding
protein_coding
protein_coding
protein_coding
protein_coding
protein_coding
protein_coding
protein_coding
protein_coding
protein_coding
protein_coding
protein_coding
protein_coding
protein_coding
protein_coding
protein_coding
protein_coding
protein_coding
protein_coding
protein_coding
protein_coding
protein_coding
protein_coding
protein_coding
protein_coding
protein_coding
protein_coding
protein_coding
protein_coding
protein_coding
protein_coding
protein_coding
protein_coding
protein_coding

protein_coding
protein_coding
protein_coding
protein_coding
protein_coding
protein_coding
protein_coding
miRNA_precursor
protein_coding
protein_coding
small_nucle(ol)ar_rna
small_nucle(ol)ar_rna
ncRNA
protein_coding
protein_coding
protein_coding
protein_coding
protein_coding
protein_coding
protein_coding
protein_coding
protein_coding
protein_coding
protein_coding
protein_coding
protein_coding
protein_coding
protein_coding
protein_coding
protein_coding
protein_coding
protein_coding
protein_coding
protein_coding
protein_coding
protein_coding
protein_coding
protein_coding
protein_coding
protein_coding
protein_coding
protein_coding
protein_coding
protein_coding
protein_coding
protein_coding
protein_coding
protein_coding
protein_coding
protein_coding
protein_coding
protein_coding

protein_coding
protein_coding
ncRNA
protein_coding
protein_coding
protein_coding
protein_coding
protein_coding
ncRNA
protein_coding
protein_coding
protein_coding
protein_coding
protein_coding
protein_coding
protein_coding
protein_coding
protein_coding
protein_coding
protein_coding
protein_coding
protein_coding
protein_coding
protein_coding
protein_coding
miRNA_precursor
protein_coding
protein_coding
protein_coding
protein_coding
protein_coding
protein_coding
protein_coding
miRNA_precursor
protein_coding
protein_coding
protein_coding
protein_coding
protein_coding
protein_coding
protein_coding
protein_coding
protein_coding
protein_coding
protein_coding
protein_coding
protein_coding
protein_coding
protein_coding
protein_coding
protein_coding
protein_coding

protein_coding
protein_coding
protein_coding
protein_coding
protein_coding
protein_coding
protein_coding
protein_coding
protein_coding
protein_coding
protein_coding
protein_coding
protein_coding
protein_coding
ncRNA
protein_coding
protein_coding
protein_coding
protein_coding
protein_coding
protein_coding
protein_coding
protein_coding
protein_coding
protein_coding
protein_coding
protein_coding
protein_coding
protein_coding
protein_coding
protein_coding
miRNA_precursor
protein_coding
protein_coding
protein_coding
protein_coding
protein_coding
protein_coding
protein_coding
protein_coding
protein_coding
protein_coding
protein_coding
protein_coding
protein_coding
protein_coding
small_nucle(ol)ar_rna
small_nucle(ol)ar_rna
ncRNA
small_nucle(ol)ar_rna
protein_coding
protein_coding

protein_coding
protein_coding
protein_coding
protein_coding
protein_coding
protein_coding
protein_coding
protein_coding
protein_coding
protein_coding
protein_coding
protein_coding
protein_coding
small_nucle(ol)ar_rna
protein_coding
protein_coding
protein_coding
protein_coding
protein_coding
protein_coding
protein_coding
protein_coding
protein_coding
protein_coding
protein_coding
protein_coding
protein_coding
protein_coding
protein_coding
protein_coding
protein_coding
protein_coding
protein_coding
protein_coding
protein_coding
protein_coding
protein_coding
protein_coding
ncRNA
protein_coding
protein_coding
protein_coding
protein_coding
protein_coding
protein_coding
protein_coding
protein_coding
protein_coding
protein_coding
protein_coding
protein_coding
protein_coding

protein_coding
protein_coding
protein_coding
protein_coding
protein_coding
protein_coding
protein_coding
protein_coding
protein_coding
protein_coding
protein_coding
protein_coding
protein_coding
protein_coding
protein_coding
protein_coding
protein_coding
protein_coding
protein_coding
protein_coding
pre_trna
protein_coding
protein_coding
ncRNA
protein_coding
protein_coding
protein_coding
protein_coding
protein_coding
protein_coding
protein_coding
protein_coding
protein_coding
protein_coding
protein_coding
protein_coding
protein_coding
protein_coding
protein_coding
protein_coding
ncRNA
protein_coding
protein_coding
protein_coding
protein_coding
protein_coding
protein_coding
protein_coding
protein_coding
protein_coding
protein_coding
protein_coding

protein_coding
protein_coding
protein_coding
protein_coding
protein_coding
protein_coding
protein_coding
protein_coding
protein_coding
protein_coding
protein_coding
protein_coding
protein_coding
protein_coding
protein_coding
protein_coding
protein_coding
protein_coding
protein_coding
protein_coding
protein_coding
protein_coding
protein_coding
protein_coding
protein_coding
protein_coding
protein_coding
protein_coding
protein_coding
protein_coding
protein_coding
protein_coding
ncRNA
protein_coding
protein_coding
protein_coding
ncRNA
protein_coding
protein_coding
protein_coding
protein_coding
protein_coding
protein_coding
protein_coding
protein_coding
protein_coding
protein_coding
protein_coding
protein_coding
protein_coding
protein_coding
protein_coding

protein_coding
protein_coding
protein_coding
protein_coding
protein_coding
protein_coding
protein_coding
protein_coding
ncRNA
small_nucle(ol)ar_rna
small_nucle(ol)ar_rna
protein_coding
protein_coding
protein_coding
protein_coding
protein_coding
protein_coding
protein_coding
protein_coding
protein_coding
protein_coding
protein_coding
protein_coding
protein_coding
protein_coding
protein_coding
protein_coding
protein_coding
protein_coding
ncRNA
protein_coding
protein_coding
protein_coding
protein_coding
protein_coding
protein_coding
protein_coding
protein_coding
protein_coding
protein_coding
protein_coding
protein_coding
protein_coding
protein_coding
protein_coding
protein_coding
protein_coding
protein_coding
protein_coding
protein_coding
protein_coding
protein_coding

protein_coding
protein_coding
protein_coding
protein_coding
protein_coding
protein_coding
protein_coding
miRNA_precursor
miRNA_precursor
protein_coding
protein_coding
protein_coding
protein_coding
protein_coding
protein_coding
protein_coding
protein_coding
protein_coding
protein_coding
protein_coding
protein_coding
protein_coding
protein_coding
ncRNA
ncRNA
protein_coding
protein_coding
protein_coding
protein_coding
protein_coding
protein_coding
protein_coding
protein_coding
protein_coding
protein_coding
ncRNA
protein_coding
protein_coding
protein_coding
protein_coding
protein_coding
pre_trna
protein_coding
protein_coding
protein_coding
protein_coding
protein_coding
protein_coding
protein_coding
protein_coding
protein_coding
protein_coding

protein_coding
protein_coding
protein_coding
ncRNA
ncRNA
protein_coding
protein_coding
protein_coding
protein_coding
protein_coding
protein_coding
protein_coding
protein_coding
protein_coding
ncRNA
protein_coding
protein_coding
ncRNA
protein_coding
protein_coding
protein_coding
protein_coding
miRNA_precursor
protein_coding
protein_coding
protein_coding
protein_coding
protein_coding
protein_coding
ncRNA
protein_coding
protein_coding
protein_coding
protein_coding
protein_coding
protein_coding
protein_coding
protein_coding
protein_coding
protein_coding
protein_coding
protein_coding
protein_coding
protein_coding
protein_coding
protein_coding
protein_coding
protein_coding
protein_coding
miRNA_precursor
protein_coding
protein_coding

protein_coding
protein_coding
protein_coding
ncRNA
ncRNA
protein_coding
protein_coding
protein_coding
protein_coding
protein_coding
protein_coding
protein_coding
protein_coding
protein_coding
protein_coding
protein_coding
protein_coding
protein_coding
protein_coding
ncRNA
miRNA_precursor
protein_coding
protein_coding
protein_coding
protein_coding
protein_coding
protein_coding
protein_coding
protein_coding
protein_coding
protein_coding
protein_coding
protein_coding
protein_coding
ncRNA
protein_coding
protein_coding
miRNA_precursor
protein_coding
protein_coding
miRNA_precursor
protein_coding
protein_coding
protein_coding
protein_coding
protein_coding
protein_coding
protein_coding
protein_coding
protein_coding
protein_coding
protein_coding

protein_coding
protein_coding
protein_coding
protein_coding
protein_coding
protein_coding
ncRNA
protein_coding
protein_coding
protein_coding
protein_coding
protein_coding
protein_coding
protein_coding
protein_coding
protein_coding
protein_coding
protein_coding
protein_coding
protein_coding
protein_coding
protein_coding
protein_coding
protein_coding
protein_coding
protein_coding
protein_coding
protein_coding
protein_coding
protein_coding
protein_coding
protein_coding
protein_coding
protein_coding
protein_coding
protein_coding
protein_coding
protein_coding
protein_coding
protein_coding
protein_coding
protein_coding
protein_coding
protein_coding
protein_coding
protein_coding
protein_coding
protein_coding
protein_coding
protein_coding
protein_coding
protein_coding

protein_coding
protein_coding
protein_coding
protein_coding
protein_coding
ncRNA
protein_coding
protein_coding
protein_coding
protein_coding
protein_coding
protein_coding
protein_coding
protein_coding
protein_coding
protein_coding
protein_coding
protein_coding
protein_coding
ncRNA
protein_coding
protein_coding
protein_coding
protein_coding
protein_coding
protein_coding
protein_coding
protein_coding
protein_coding
protein_coding
protein_coding
protein_coding
protein_coding
protein_coding
protein_coding
protein_coding
protein_coding
protein_coding
protein_coding
protein_coding
protein_coding
protein_coding
protein_coding
protein_coding
protein_coding
protein_coding
protein_coding
protein_coding
protein_coding
protein_coding
protein_coding
protein_coding

protein_coding
protein_coding
protein_coding
protein_coding
protein_coding
small_nucle(ol)ar_rna
small_nucle(ol)ar_rna
protein_coding
ncRNA
protein_coding
protein_coding
pre_trna
protein_coding
pre_trna
protein_coding
protein_coding
protein_coding
protein_coding
protein_coding
protein_coding
protein_coding
protein_coding
small_nucle(ol)ar_rna
protein_coding
protein_coding
protein_coding
protein_coding
protein_coding
protein_coding
protein_coding
ncRNA
protein_coding
protein_coding
ncRNA
ncRNA
protein_coding
protein_coding
protein_coding
protein_coding
protein_coding
protein_coding
protein_coding
protein_coding
protein_coding
protein_coding
protein_coding
protein_coding
protein_coding
protein_coding
protein_coding
protein_coding
protein_coding

miRNA_precursor
miRNA_precursor
protein_coding
protein_coding
protein_coding
protein_coding
protein_coding
protein_coding
protein_coding
protein_coding
protein_coding
protein_coding
protein_coding
protein_coding
protein_coding
protein_coding
protein_coding
protein_coding
protein_coding
protein_coding
protein_coding
protein_coding
protein_coding
protein_coding
protein_coding
protein_coding
protein_coding
protein_coding
protein_coding
protein_coding
protein_coding
protein_coding
protein_coding
protein_coding
protein_coding
protein_coding
protein_coding
protein_coding
protein_coding
protein_coding
protein_coding
protein_coding
protein_coding
protein_coding
protein_coding
protein_coding
protein_coding
protein_coding
protein_coding
protein_coding
protein_coding
protein_coding

protein_coding
protein_coding
protein_coding
protein_coding
protein_coding
protein_coding
protein_coding
protein_coding
protein_coding
protein_coding
protein_coding
protein_coding
protein_coding
protein_coding
protein_coding
protein_coding
protein_coding
protein_coding
protein_coding
protein_coding
protein_coding
pre_trna
protein_coding
protein_coding
protein_coding
protein_coding
protein_coding
protein_coding
protein_coding
protein_coding
protein_coding
protein_coding
protein_coding
protein_coding
protein_coding
protein_coding
protein_coding
protein_coding
protein_coding
protein_coding
protein_coding
protein_coding
protein_coding
protein_coding
protein_coding
protein_coding
protein_coding
protein_coding
protein_coding
protein_coding
ncRNA
protein_coding

ncRNA
protein_coding
protein_coding
protein_coding
protein_coding
protein_coding
protein_coding
protein_coding
protein_coding
ncRNA
protein_coding
protein_coding
protein_coding
protein_coding
protein_coding
protein_coding
protein_coding
protein_coding
protein_coding
protein_coding
small_nucle(ol)ar_rna
protein_coding
protein_coding
protein_coding
protein_coding
protein_coding
protein_coding
ncRNA
protein_coding
protein_coding
protein_coding
protein_coding
protein_coding
protein_coding
protein_coding
protein_coding
protein_coding
protein_coding
protein_coding
protein_coding
protein_coding
protein_coding
protein_coding
protein_coding
protein_coding
protein_coding
protein_coding
protein_coding
protein_coding
protein_coding
protein_coding
protein_coding

protein_coding
protein_coding
protein_coding
protein_coding
protein_coding
protein_coding
protein_coding
protein_coding
ncRNA
protein_coding
protein_coding
protein_coding
protein_coding
protein_coding
protein_coding
ncRNA
protein_coding
protein_coding
protein_coding
ncRNA
protein_coding
protein_coding
protein_coding
protein_coding
protein_coding
protein_coding
protein_coding
protein_coding
protein_coding
protein_coding
protein_coding
protein_coding
protein_coding
protein_coding
protein_coding
ncRNA
protein_coding
protein_coding
protein_coding
protein_coding
protein_coding
protein_coding
protein_coding
protein_coding
protein_coding
protein_coding
protein_coding
protein_coding
protein_coding
protein_coding
protein_coding
protein_coding

protein_coding
protein_coding
protein_coding
protein_coding
protein_coding
protein_coding
protein_coding
protein_coding
protein_coding
protein_coding
protein_coding
protein_coding
protein_coding
ncRNA
protein_coding
protein_coding
ncRNA
ncRNA
protein_coding
protein_coding
protein_coding
protein_coding
protein_coding
protein_coding
protein_coding
protein_coding
protein_coding
pre_trna
protein_coding
protein_coding
protein_coding
protein_coding
protein_coding
protein_coding
protein_coding
protein_coding
protein_coding
protein_coding
protein_coding
protein_coding
protein_coding
protein_coding
protein_coding
protein_coding
protein_coding
protein_coding
protein_coding
protein_coding
protein_coding
protein_coding
protein_coding
protein_coding

protein_coding
protein_coding
protein_coding
ncRNA
protein_coding
protein_coding
protein_coding
protein_coding
protein_coding
protein_coding
protein_coding
protein_coding
protein_coding
protein_coding
protein_coding
protein_coding
protein_coding
protein_coding
protein_coding
protein_coding
protein_coding
protein_coding
pre_trna
protein_coding
protein_coding
protein_coding
protein_coding
protein_coding
protein_coding
protein_coding
protein_coding
protein_coding
protein_coding
protein_coding
protein_coding
protein_coding
protein_coding
protein_coding
protein_coding
ncRNA
protein_coding
protein_coding
protein_coding
protein_coding
protein_coding
protein_coding
protein_coding
protein_coding
protein_coding
protein_coding
protein_coding
protein_coding

protein_coding
protein_coding
miRNA_precursor
protein_coding
protein_coding
protein_coding
protein_coding
protein_coding
protein_coding
protein_coding
small_nucle(ol)ar_rna
protein_coding
protein_coding
protein_coding
protein_coding
protein_coding
protein_coding
protein_coding
protein_coding
protein_coding
protein_coding
protein_coding
protein_coding
protein_coding
protein_coding
protein_coding
protein_coding
protein_coding
protein_coding
protein_coding
protein_coding
protein_coding
protein_coding
protein_coding
protein_coding
protein_coding
small_nucle(ol)ar_rna
small_nucle(ol)ar_rna
small_nucle(ol)ar_rna
small_nucle(ol)ar_rna
protein_coding
protein_coding
protein_coding
protein_coding
protein_coding
pre_trna
miRNA_precursor
protein_coding
protein_coding
protein_coding
protein_coding
protein_coding

protein_coding
protein_coding
protein_coding
ncRNA
protein_coding
ncRNA
ncRNA
protein_coding
protein_coding
protein_coding
protein_coding
protein_coding
protein_coding
protein_coding
protein_coding
protein_coding
protein_coding
protein_coding
protein_coding
protein_coding
protein_coding
protein_coding
protein_coding
protein_coding
protein_coding
protein_coding
ncRNA
protein_coding
protein_coding
protein_coding
protein_coding
protein_coding
protein_coding
protein_coding
protein_coding
protein_coding
protein_coding
protein_coding
protein_coding
ncRNA
protein_coding
protein_coding
protein_coding
miRNA_precursor
protein_coding
protein_coding
protein_coding
ncRNA
protein_coding
protein_coding
protein_coding
protein_coding

protein_coding
protein_coding
protein_coding
protein_coding
protein_coding
protein_coding
protein_coding
protein_coding
protein_coding
protein_coding
protein_coding
protein_coding
protein_coding
protein_coding
protein_coding
protein_coding
protein_coding
protein_coding
ncRNA
protein_coding
protein_coding
protein_coding
protein_coding
protein_coding
protein_coding
protein_coding
protein_coding
protein_coding
protein_coding
protein_coding
protein_coding
protein_coding
protein_coding
protein_coding
miRNA_precursor
protein_coding
protein_coding
protein_coding
protein_coding
protein_coding
protein_coding
protein_coding
protein_coding
protein_coding
protein_coding
protein_coding
protein_coding
protein_coding
miRNA_precursor
pre_trna
protein_coding
protein_coding

protein_coding
protein_coding
protein_coding
protein_coding
ncRNA
protein_coding
protein_coding
protein_coding
protein_coding
miRNA_precursor
protein_coding
protein_coding
protein_coding
protein_coding
protein_coding
protein_coding
protein_coding
miRNA_precursor
protein_coding
ncRNA
protein_coding
protein_coding
protein_coding
protein_coding
protein_coding
protein_coding
protein_coding
protein_coding
protein_coding
protein_coding
protein_coding
protein_coding
protein_coding
protein_coding
protein_coding
protein_coding
protein_coding
protein_coding
protein_coding
protein_coding
ncRNA
protein_coding
protein_coding
protein_coding
protein_coding
protein_coding
protein_coding
protein_coding
protein_coding
protein_coding
protein_coding
protein_coding

protein_coding
ncRNA
protein_coding
protein_coding
protein_coding
protein_coding
protein_coding
protein_coding
protein_coding
protein_coding
protein_coding
protein_coding
protein_coding
protein_coding
protein_coding
protein_coding
protein_coding
protein_coding
protein_coding
protein_coding
protein_coding
protein_coding
protein_coding
protein_coding
protein_coding
protein_coding
protein_coding
protein_coding
protein_coding
protein_coding
protein_coding
protein_coding
miRNA_precursor
protein_coding
protein_coding
protein_coding
protein_coding
protein_coding
protein_coding
protein_coding
protein_coding
protein_coding
protein_coding
protein_coding
ncRNA
small_nucle(ol)ar_rna
small_nucle(ol)ar_rna
small_nucle(ol)ar_rna
small_nucle(ol)ar_rna
protein_coding
protein_coding
protein_coding

protein_coding
protein_coding
ncRNA
protein_coding
protein_coding
protein_coding
protein_coding
protein_coding
protein_coding
protein_coding
protein_coding
protein_coding
protein_coding
protein_coding
protein_coding
protein_coding
protein_coding
protein_coding
protein_coding
protein_coding
protein_coding
protein_coding
protein_coding
protein_coding
protein_coding
protein_coding
protein_coding
protein_coding
protein_coding
protein_coding
protein_coding
protein_coding
protein_coding
ncRNA
protein_coding
protein_coding
protein_coding
protein_coding
protein_coding
protein_coding
protein_coding
protein_coding
protein_coding
protein_coding
protein_coding
protein_coding
protein_coding
protein_coding
protein_coding
protein_coding
small_nucle(ol)ar_rna
protein_coding

protein_coding
protein_coding
protein_coding
protein_coding
protein_coding
protein_coding
protein_coding
protein_coding
protein_coding
protein_coding
protein_coding
protein_coding
protein_coding
protein_coding
protein_coding
protein_coding
protein_coding
protein_coding
protein_coding
protein_coding
protein_coding
protein_coding
protein_coding
protein_coding
protein_coding
pre_trna
protein_coding
ncRNA
protein_coding
protein_coding
protein_coding
protein_coding
protein_coding
protein_coding
protein_coding
protein_coding
protein_coding
protein_coding
protein_coding
protein_coding
protein_coding
protein_coding
protein_coding
protein_coding
protein_coding
ncRNA
protein_coding
protein_coding
protein_coding
protein_coding
protein_coding
protein_coding

protein_coding
protein_coding
protein_coding
protein_coding
protein_coding
protein_coding
miRNA_precursor
protein_coding
protein_coding
protein_coding
protein_coding
protein_coding
protein_coding
protein_coding
protein_coding
protein_coding
protein_coding
ncRNA
protein_coding
protein_coding
ncRNA
protein_coding
protein_coding
protein_coding
protein_coding
protein_coding
protein_coding
protein_coding
protein_coding
protein_coding
protein_coding
ncRNA
protein_coding
protein_coding
protein_coding
protein_coding
protein_coding
protein_coding
protein_coding
protein_coding
protein_coding
protein_coding
protein_coding
protein_coding
protein_coding
protein_coding
protein_coding
protein_coding
protein_coding
protein_coding
protein_coding
protein_coding

protein_coding
protein_coding
protein_coding
protein_coding
pre_trna
protein_coding
protein_coding
protein_coding
protein_coding
protein_coding
protein_coding
protein_coding
protein_coding
protein_coding
protein_coding
protein_coding
protein_coding
protein_coding
protein_coding
protein_coding
protein_coding
protein_coding
protein_coding
protein_coding
protein_coding
protein_coding
ncRNA
protein_coding
protein_coding
protein_coding
protein_coding
protein_coding
protein_coding
protein_coding
protein_coding
protein_coding
protein_coding
ncRNA
protein_coding
protein_coding
ncRNA
small_nucle(ol)ar_rna
protein_coding
protein_coding
protein_coding
protein_coding
protein_coding
protein_coding
protein_coding
protein_coding
protein_coding
protein_coding

protein_coding
protein_coding
protein_coding
protein_coding
ncRNA
protein_coding
protein_coding
protein_coding
ncRNA
protein_coding
protein_coding
protein_coding
protein_coding
protein_coding
protein_coding
protein_coding
protein_coding
protein_coding
protein_coding
protein_coding
protein_coding
protein_coding
protein_coding
protein_coding
protein_coding
protein_coding
ncRNA
protein_coding
protein_coding
protein_coding
protein_coding
protein_coding
protein_coding
protein_coding
protein_coding
protein_coding
protein_coding
protein_coding
pre_trna
protein_coding
protein_coding
protein_coding
protein_coding
protein_coding
protein_coding
protein_coding
protein_coding
protein_coding
protein_coding
protein_coding
protein_coding
protein_coding

ncRNA
protein_coding
protein_coding
protein_coding
protein_coding
protein_coding
miRNA_precursor
small_nucle(ol)ar_rna
small_nucle(ol)ar_rna
protein_coding
pre_trna
protein_coding
protein_coding
protein_coding
protein_coding
protein_coding
protein_coding
protein_coding
protein_coding
protein_coding
ncRNA
protein_coding
protein_coding
protein_coding
protein_coding
protein_coding
ncRNA
protein_coding
protein_coding
protein_coding
protein_coding
protein_coding
protein_coding
protein_coding
protein_coding
protein_coding
protein_coding
protein_coding
ncRNA
protein_coding
protein_coding
protein_coding
protein_coding
protein_coding
protein_coding
protein_coding
pre_trna
protein_coding
protein_coding
protein_coding
protein_coding
protein_coding

protein_coding
protein_coding
protein_coding
protein_coding
protein_coding
protein_coding
protein_coding
protein_coding
protein_coding
protein_coding
protein_coding
protein_coding
protein_coding
protein_coding
protein_coding
protein_coding
protein_coding
protein_coding
protein_coding
protein_coding
protein_coding
protein_coding
protein_coding
small_nucle(ol)ar_rna
small_nucle(ol)ar_rna
small_nucle(ol)ar_rna
protein_coding
protein_coding
protein_coding
protein_coding
protein_coding
protein_coding
protein_coding
protein_coding
protein_coding
protein_coding
protein_coding
protein_coding
protein_coding
protein_coding
protein_coding
protein_coding
protein_coding
protein_coding
protein_coding
protein_coding
protein_coding
protein_coding
protein_coding
protein_coding
protein_coding
protein_coding

protein_coding
protein_coding
protein_coding
protein_coding
protein_coding
protein_coding
protein_coding
protein_coding
protein_coding
protein_coding
protein_coding
protein_coding
protein_coding
protein_coding
protein_coding
protein_coding
protein_coding
protein_coding
protein_coding
protein_coding
protein_coding
protein_coding
protein_coding
protein_coding
protein_coding
protein_coding
protein_coding
protein_coding
protein_coding
protein_coding
protein_coding
protein_coding
protein_coding
protein_coding
protein_coding
protein_coding
protein_coding
protein_coding
protein_coding
protein_coding
protein_coding
protein_coding
protein_coding
protein_coding
protein_coding
protein_coding
protein_coding
protein_coding
protein_coding
protein_coding
protein_coding
protein_coding

protein_coding
protein_coding
protein_coding
protein_coding
protein_coding
protein_coding
protein_coding
protein_coding
protein_coding
protein_coding
protein_coding
protein_coding
protein_coding
protein_coding
protein_coding
protein_coding
protein_coding
protein_coding
protein_coding
protein_coding
protein_coding
protein_coding
protein_coding
protein_coding
protein_coding
protein_coding
protein_coding
protein_coding
protein_coding
protein_coding
protein_coding
protein_coding
protein_coding
protein_coding
protein_coding
protein_coding
protein_coding
protein_coding
protein_coding
ncRNA
protein_coding
protein_coding
ncRNA
protein_coding
protein_coding
protein_coding
protein_coding
protein_coding
protein_coding
protein_coding
protein_coding
protein_coding

protein_coding
protein_coding
protein_coding
protein_coding
protein_coding
protein_coding
protein_coding
protein_coding
protein_coding
protein_coding
protein_coding
protein_coding
protein_coding
protein_coding
protein_coding
protein_coding
protein_coding
protein_coding
protein_coding
protein_coding
protein_coding
protein_coding
protein_coding
protein_coding
protein_coding
protein_coding
protein_coding
protein_coding
protein_coding
protein_coding
protein_coding
protein_coding
protein_coding
protein_coding
protein_coding
protein_coding
protein_coding
protein_coding
protein_coding
protein_coding
protein_coding
protein_coding
protein_coding
protein_coding
protein_coding
protein_coding
ncRNA
protein_coding
protein_coding
protein_coding
protein_coding
protein_coding

ncRNA
protein_coding
protein_coding
protein_coding
protein_coding
protein_coding
protein_coding
protein_coding
protein_coding
ncRNA
protein_coding
protein_coding
protein_coding
protein_coding
ncRNA
protein_coding
protein_coding
protein_coding
protein_coding
protein_coding
protein_coding
protein_coding
protein_coding
protein_coding
protein_coding
protein_coding
protein_coding
protein_coding
protein_coding
protein_coding
protein_coding
protein_coding
protein_coding
protein_coding
protein_coding
ncRNA
protein_coding
protein_coding
protein_coding
protein_coding
protein_coding
ncRNA
ncRNA
protein_coding
protein_coding
protein_coding
protein_coding
protein_coding
protein_coding
protein_coding
protein_coding
protein_coding

protein_coding
protein_coding
protein_coding
protein_coding
protein_coding
protein_coding
protein_coding
protein_coding
ncRNA
protein_coding
protein_coding
protein_coding
protein_coding
protein_coding
protein_coding
pre_trna
protein_coding
protein_coding
protein_coding
protein_coding
protein_coding
protein_coding
protein_coding
protein_coding
protein_coding
protein_coding
protein_coding
protein_coding
protein_coding
protein_coding
protein_coding
protein_coding
protein_coding
ncRNA
pre_trna
protein_coding
protein_coding
protein_coding
protein_coding
protein_coding
protein_coding
protein_coding
protein_coding
ncRNA
protein_coding
protein_coding
protein_coding
ncRNA
protein_coding
protein_coding
small_nucle(ol)ar_rna
small_nucle(ol)ar_rna

pre_trna
protein_coding
protein_coding
protein_coding
protein_coding
protein_coding
protein_coding
protein_coding
protein_coding
protein_coding
protein_coding
protein_coding
protein_coding
protein_coding
protein_coding
protein_coding
protein_coding
protein_coding
protein_coding
protein_coding
protein_coding
protein_coding
protein_coding
protein_coding
protein_coding
protein_coding
protein_coding
protein_coding
protein_coding
protein_coding
protein_coding
protein_coding
protein_coding
protein_coding
protein_coding
protein_coding
protein_coding
protein_coding
protein_coding
protein_coding
protein_coding
protein_coding
protein_coding
protein_coding
protein_coding
protein_coding
protein_coding
protein_coding
protein_coding
protein_coding
protein_coding
protein_coding

protein_coding
protein_coding
protein_coding
protein_coding
protein_coding
protein_coding
protein_coding
protein_coding
protein_coding
ncRNA
protein_coding
protein_coding
protein_coding
protein_coding
protein_coding
protein_coding
protein_coding
protein_coding
protein_coding
protein_coding
protein_coding
protein_coding
protein_coding
protein_coding
protein_coding
protein_coding
protein_coding
protein_coding
protein_coding
protein_coding
protein_coding
protein_coding
protein_coding
ncRNA
protein_coding
protein_coding
ncRNA
protein_coding
protein_coding
protein_coding
protein_coding
protein_coding
protein_coding
ncRNA
protein_coding
protein_coding
protein_coding
protein_coding
protein_coding
protein_coding
protein_coding
ncRNA

small_nucle(ol)ar_rna
small_nucle(ol)ar_rna
protein_coding
protein_coding
protein_coding
protein_coding
protein_coding
protein_coding
protein_coding
protein_coding
protein_coding
protein_coding
protein_coding
protein_coding
protein_coding
protein_coding
protein_coding
protein_coding
protein_coding
protein_coding
ncRNA
protein_coding
ncRNA
protein_coding
protein_coding
protein_coding
protein_coding
protein_coding
protein_coding
pre_trna
protein_coding
protein_coding
protein_coding
protein_coding
protein_coding
protein_coding
pre_trna
protein_coding
protein_coding
protein_coding
protein_coding
protein_coding
small_nucle(ol)ar_rna
small_nucle(ol)ar_rna
small_nucle(ol)ar_rna
protein_coding
protein_coding
protein_coding
protein_coding
protein_coding
protein_coding
protein_coding

protein_coding
protein_coding
protein_coding
protein_coding
protein_coding
protein_coding
protein_coding
ncRNA
protein_coding
protein_coding
protein_coding
protein_coding
protein_coding
protein_coding
protein_coding
protein_coding
protein_coding
protein_coding
protein_coding
protein_coding
protein_coding
pre_trna
protein_coding
protein_coding
protein_coding
protein_coding
protein_coding
protein_coding
protein_coding
protein_coding
protein_coding
protein_coding
protein_coding
protein_coding
protein_coding
ncRNA
protein_coding
protein_coding
protein_coding
protein_coding
protein_coding
protein_coding
protein_coding
protein_coding
protein_coding
protein_coding
ncRNA
protein_coding
protein_coding
protein_coding
protein_coding
protein_coding

protein_coding
protein_coding
protein_coding
protein_coding
protein_coding
protein_coding
protein_coding
protein_coding
protein_coding
protein_coding
protein_coding
protein_coding
protein_coding
protein_coding
protein_coding
protein_coding
protein_coding
protein_coding
protein_coding
protein_coding
protein_coding
protein_coding
protein_coding
protein_coding
protein_coding
protein_coding
protein_coding
protein_coding
protein_coding
protein_coding
protein_coding
protein_coding
protein_coding
protein_coding
protein_coding
protein_coding
protein_coding
protein_coding
ncRNA
small_nucle(ol)ar_rna
protein_coding
protein_coding
protein_coding
protein_coding
protein_coding
protein_coding
protein_coding
protein_coding
protein_coding
protein_coding
protein_coding
protein_coding

protein_coding
protein_coding
protein_coding
protein_coding
protein_coding
protein_coding
protein_coding
protein_coding
protein_coding
protein_coding
protein_coding
protein_coding
protein_coding
protein_coding
protein_coding
protein_coding
small_nucle(ol)ar_rna
protein_coding
protein_coding
protein_coding
protein_coding
protein_coding
protein_coding
protein_coding
protein_coding
protein_coding
protein_coding
protein_coding
protein_coding
protein_coding
protein_coding
protein_coding
protein_coding
protein_coding
protein_coding
protein_coding
protein_coding
protein_coding
protein_coding
protein_coding
protein_coding
protein_coding
protein_coding
protein_coding
protein_coding
protein_coding
protein_coding
protein_coding
protein_coding
protein_coding
protein_coding
protein_coding

protein_coding
protein_coding
protein_coding
protein_coding
protein_coding
protein_coding
protein_coding
protein_coding
protein_coding
protein_coding
protein_coding
protein_coding
protein_coding
protein_coding
protein_coding
protein_coding
protein_coding
protein_coding
protein_coding
protein_coding
protein_coding
protein_coding
protein_coding
protein_coding
protein_coding
protein_coding
protein_coding
protein_coding
ncRNA
small_nucle(ol)ar_rna
small_nucle(ol)ar_rna
small_nucle(ol)ar_rna
protein_coding
protein_coding
protein_coding
protein_coding
protein_coding
protein_coding
protein_coding
protein_coding
protein_coding
protein_coding
protein_coding
protein_coding
protein_coding
protein_coding
protein_coding
protein_coding
protein_coding
protein_coding
protein_coding
protein_coding

protein_coding
protein_coding
protein_coding
protein_coding
protein_coding
protein_coding
protein_coding
protein_coding
protein_coding
protein_coding
protein_coding
protein_coding
protein_coding
ncRNA
protein_coding
protein_coding
protein_coding
protein_coding
protein_coding
protein_coding
protein_coding
protein_coding
ncRNA
protein_coding
protein_coding
protein_coding
protein_coding
protein_coding
protein_coding
protein_coding
protein_coding
protein_coding
protein_coding
protein_coding
protein_coding
protein_coding
protein_coding
protein_coding
protein_coding
protein_coding
protein_coding
protein_coding
protein_coding
protein_coding
protein_coding
protein_coding
protein_coding
protein_coding
ncRNA
protein_coding
protein_coding
protein_coding

protein_coding
protein_coding
protein_coding
protein_coding
protein_coding
protein_coding
protein_coding
protein_coding
protein_coding
protein_coding
protein_coding
protein_coding
protein_coding
protein_coding
protein_coding
protein_coding
protein_coding
protein_coding
protein_coding
miRNA_precursor
protein_coding
protein_coding
protein_coding
protein_coding
protein_coding
protein_coding
protein_coding
protein_coding
protein_coding
protein_coding
protein_coding
protein_coding
protein_coding
protein_coding
protein_coding
protein_coding
protein_coding
protein_coding
protein_coding
protein_coding
protein_coding
protein_coding
miRNA_precursor
protein_coding
protein_coding
protein_coding
protein_coding
protein_coding
protein_coding
miRNA_precursor
protein_coding
protein_coding

protein_coding
protein_coding
protein_coding
protein_coding
protein_coding
protein_coding
protein_coding
protein_coding
protein_coding
protein_coding
protein_coding
protein_coding
protein_coding
protein_coding
protein_coding
protein_coding
protein_coding
protein_coding
protein_coding
protein_coding
protein_coding
protein_coding
protein_coding
protein_coding
protein_coding
protein_coding
protein_coding
protein_coding
protein_coding
protein_coding
protein_coding
protein_coding
protein_coding
protein_coding
protein_coding
protein_coding
protein_coding
protein_coding
protein_coding
protein_coding
protein_coding
protein_coding
ncRNA
protein_coding
protein_coding
protein_coding
protein_coding
protein_coding
protein_coding
protein_coding
protein_coding
protein_coding

protein_coding
protein_coding
protein_coding
protein_coding
protein_coding
protein_coding
protein_coding
protein_coding
ncRNA
protein_coding
protein_coding
protein_coding
protein_coding
protein_coding
protein_coding
small_nucle(ol)ar_rna
protein_coding
protein_coding
protein_coding
protein_coding
protein_coding
protein_coding
protein_coding
protein_coding
protein_coding
protein_coding
protein_coding
ncRNA
protein_coding
protein_coding
miRNA_precursor
protein_coding
protein_coding
protein_coding
protein_coding
protein_coding
protein_coding
small_nucle(ol)ar_rna
protein_coding
protein_coding
protein_coding
ncRNA
protein_coding
protein_coding
protein_coding
protein_coding
protein_coding
pre_trna
ncRNA
protein_coding
protein_coding
protein_coding

protein_coding
protein_coding
protein_coding
protein_coding
protein_coding
protein_coding
ncRNA
protein_coding
ncRNA
protein_coding
protein_coding
protein_coding
protein_coding
protein_coding
protein_coding
protein_coding
protein_coding
protein_coding
protein_coding
protein_coding
protein_coding
protein_coding
protein_coding
ncRNA
protein_coding
protein_coding
protein_coding
ncRNA
protein_coding
ncRNA
protein_coding
protein_coding
protein_coding
protein_coding
ncRNA
protein_coding
protein_coding
protein_coding
protein_coding
protein_coding
protein_coding
protein_coding
protein_coding
protein_coding
protein_coding
protein_coding
protein_coding
protein_coding
protein_coding
protein_coding
protein_coding
protein_coding

protein_coding
protein_coding
protein_coding
protein_coding
protein_coding
protein_coding
protein_coding
protein_coding
protein_coding
protein_coding
protein_coding
ncRNA
protein_coding
protein_coding
protein_coding
protein_coding
protein_coding
protein_coding
protein_coding
protein_coding
protein_coding
protein_coding
protein_coding
protein_coding
protein_coding
ncRNA
protein_coding
protein_coding
protein_coding
miRNA_precursor
protein_coding
protein_coding
protein_coding
ncRNA
protein_coding
protein_coding
ncRNA
protein_coding
protein_coding
protein_coding
ncRNA
protein_coding
protein_coding
protein_coding
protein_coding
protein_coding
ncRNA
protein_coding
protein_coding
protein_coding
protein_coding
protein_coding

protein_coding
protein_coding
ncRNA
protein_coding
protein_coding
protein_coding
protein_coding
protein_coding
ncRNA
protein_coding
miRNA_precursor
protein_coding
protein_coding
protein_coding
protein_coding
protein_coding
protein_coding
miRNA_precursor
small_nucle(ol)ar_rna
protein_coding
protein_coding
protein_coding
protein_coding
protein_coding
protein_coding
protein_coding
protein_coding
protein_coding
protein_coding
protein_coding
protein_coding
protein_coding
protein_coding
protein_coding
protein_coding
protein_coding
protein_coding
protein_coding
protein_coding
ncRNA
protein_coding
protein_coding
protein_coding
protein_coding
ncRNA
protein_coding
protein_coding
protein_coding
protein_coding
protein_coding
protein_coding
protein_coding

protein_coding
protein_coding
protein_coding
small_nucle(ol)ar_rna
protein_coding
protein_coding
protein_coding
protein_coding
protein_coding
protein_coding
protein_coding
protein_coding
protein_coding
protein_coding
protein_coding
protein_coding
protein_coding
protein_coding
protein_coding
protein_coding
protein_coding
protein_coding
protein_coding
protein_coding
protein_coding
protein_coding
protein_coding
protein_coding
protein_coding
protein_coding
protein_coding
protein_coding
protein_coding
protein_coding
protein_coding
protein_coding
protein_coding
protein_coding
protein_coding
protein_coding
protein_coding
protein_coding
protein_coding
protein_coding
protein_coding
protein_coding
protein_coding
protein_coding
protein_coding
protein_coding
protein_coding
protein_coding

protein_coding
pre_trna
protein_coding
protein_coding
protein_coding
protein_coding
protein_coding
protein_coding
ncRNA
protein_coding
protein_coding
protein_coding
protein_coding
protein_coding
protein_coding
protein_coding
protein_coding
protein_coding
protein_coding
protein_coding
protein_coding
protein_coding
protein_coding
protein_coding
protein_coding
protein_coding
protein_coding
protein_coding
protein_coding
protein_coding
protein_coding
protein_coding
protein_coding
protein_coding
protein_coding
protein_coding
protein_coding
protein_coding
protein_coding
protein_coding
protein_coding
protein_coding
protein_coding
protein_coding
protein_coding
protein_coding
protein_coding
protein_coding
protein_coding
protein_coding
protein_coding
protein_coding

protein_coding
protein_coding
protein_coding
protein_coding
protein_coding
protein_coding
protein_coding
protein_coding
protein_coding
protein_coding
protein_coding
protein_coding
protein_coding
protein_coding
protein_coding
protein_coding
protein_coding
protein_coding
protein_coding
protein_coding
protein_coding
protein_coding
protein_coding
protein_coding
protein_coding
protein_coding
protein_coding
protein_coding
protein_coding
protein_coding
miRNA_precursor
protein_coding
protein_coding
protein_coding
protein_coding
protein_coding
ncRNA
protein_coding
protein_coding
protein_coding
protein_coding
protein_coding
ncRNA
protein_coding
ncRNA
protein_coding
protein_coding
protein_coding
protein_coding
protein_coding
protein_coding
protein_coding

protein_coding
protein_coding
protein_coding
protein_coding
protein_coding
protein_coding
protein_coding
protein_coding
protein_coding
protein_coding
protein_coding
protein_coding
protein_coding
protein_coding
protein_coding
protein_coding
protein_coding
protein_coding
protein_coding
protein_coding
protein_coding
protein_coding
protein_coding
protein_coding
protein_coding
protein_coding
protein_coding
protein_coding
protein_coding
protein_coding
protein_coding
protein_coding
protein_coding
protein_coding
protein_coding
protein_coding
protein_coding
protein_coding
protein_coding
protein_coding
protein_coding
protein_coding
protein_coding
protein_coding
protein_coding
protein_coding
protein_coding
protein_coding
protein_coding
protein_coding
protein_coding
protein_coding

protein_coding
protein_coding
protein_coding
protein_coding
protein_coding
protein_coding
protein_coding
protein_coding
protein_coding
protein_coding
protein_coding
protein_coding
protein_coding
protein_coding
protein_coding
protein_coding
protein_coding
protein_coding
protein_coding
protein_coding
protein_coding
miRNA_precursor
protein_coding
protein_coding
protein_coding
protein_coding
protein_coding
ncRNA
protein_coding
protein_coding
protein_coding
ncRNA
protein_coding
protein_coding
protein_coding
protein_coding
protein_coding
protein_coding
protein_coding
protein_coding
protein_coding
protein_coding
protein_coding
protein_coding
protein_coding
protein_coding
protein_coding
protein_coding
protein_coding
protein_coding
protein_coding
protein_coding

protein_coding
protein_coding
protein_coding
protein_coding
protein_coding
protein_coding
protein_coding
protein_coding
protein_coding
protein_coding
protein_coding
protein_coding
protein_coding
protein_coding
protein_coding
protein_coding
protein_coding
protein_coding
protein_coding
protein_coding
protein_coding
pre_trna
protein_coding
protein_coding
protein_coding
protein_coding
protein_coding
protein_coding
protein_coding
protein_coding
protein_coding
protein_coding
protein_coding
ncRNA
ncRNA
protein_coding
protein_coding
protein_coding
protein_coding
protein_coding
protein_coding
protein_coding
protein_coding
protein_coding
protein_coding
miRNA_precursor
protein_coding
protein_coding
protein_coding
protein_coding
protein_coding
protein_coding

protein_coding
protein_coding
protein_coding
protein_coding
protein_coding
protein_coding
protein_coding
protein_coding
protein_coding
protein_coding
protein_coding
protein_coding
protein_coding
protein_coding
protein_coding
protein_coding
protein_coding
protein_coding
protein_coding
protein_coding
protein_coding
protein_coding
protein_coding
protein_coding
protein_coding
protein_coding
protein_coding
protein_coding
protein_coding
protein_coding
protein_coding
protein_coding
protein_coding
protein_coding
protein_coding
protein_coding
protein_coding
protein_coding
protein_coding
protein_coding
protein_coding
protein_coding
protein_coding
protein_coding
protein_coding
protein_coding
protein_coding
protein_coding
protein_coding
protein_coding
small_nucle(ol)ar_rna
protein_coding

protein_coding
protein_coding
protein_coding
protein_coding
protein_coding
protein_coding
protein_coding
protein_coding
protein_coding
protein_coding
protein_coding
protein_coding
ncRNA
protein_coding
protein_coding
miRNA_precursor
ncRNA
protein_coding
protein_coding
protein_coding
protein_coding
protein_coding
protein_coding
protein_coding
protein_coding
protein_coding
protein_coding
protein_coding
protein_coding
protein_coding
protein_coding
protein_coding
protein_coding
protein_coding
protein_coding
protein_coding
protein_coding
protein_coding
protein_coding
protein_coding
protein_coding
protein_coding
protein_coding
protein_coding
protein_coding
protein_coding
protein_coding
protein_coding
protein_coding
protein_coding
protein_coding
protein_coding

protein_coding
protein_coding
protein_coding
small_nucle(ol)ar_rna
small_nucle(ol)ar_rna
protein_coding
protein_coding
protein_coding
protein_coding
protein_coding
protein_coding
protein_coding
protein_coding
protein_coding
protein_coding
protein_coding
protein_coding
protein_coding
protein_coding
protein_coding
protein_coding
ncRNA
protein_coding
protein_coding
protein_coding
protein_coding
protein_coding
protein_coding
protein_coding
protein_coding
protein_coding
protein_coding
protein_coding
protein_coding
protein_coding
protein_coding
protein_coding
protein_coding
protein_coding
protein_coding
protein_coding
protein_coding
protein_coding
protein_coding
protein_coding
ncRNA
protein_coding
protein_coding
protein_coding
protein_coding
protein_coding
protein_coding

protein_coding
protein_coding
protein_coding
protein_coding
protein_coding
protein_coding
protein_coding
protein_coding
protein_coding
protein_coding
protein_coding
protein_coding
protein_coding
protein_coding
protein_coding
protein_coding
protein_coding
protein_coding
protein_coding
protein_coding
protein_coding
protein_coding
protein_coding
protein_coding
protein_coding
protein_coding
protein_coding
protein_coding
protein_coding
protein_coding
protein_coding
protein_coding
protein_coding
protein_coding
protein_coding
protein_coding
protein_coding
protein_coding
protein_coding
protein_coding
protein_coding
protein_coding
protein_coding
protein_coding
protein_coding
protein_coding
protein_coding
protein_coding
protein_coding
protein_coding
protein_coding
small_nucle(ol)ar_rna

protein_coding
protein_coding
ncRNA
protein_coding
protein_coding
protein_coding
protein_coding
protein_coding
protein_coding
protein_coding
protein_coding
protein_coding
protein_coding
protein_coding
protein_coding
protein_coding
protein_coding
protein_coding
protein_coding
protein_coding
ncRNA
protein_coding
protein_coding
protein_coding
protein_coding
protein_coding
protein_coding
protein_coding
protein_coding
protein_coding
protein_coding
protein_coding
protein_coding
protein_coding
protein_coding
protein_coding
protein_coding
protein_coding
protein_coding
protein_coding
protein_coding
protein_coding
ncRNA
protein_coding
protein_coding
protein_coding
protein_coding
protein_coding
protein_coding
protein_coding
protein_coding
protein_coding

**Dataset 1.5 Differential analysis of 21/22nt accumulation on Araport11 genes in c**

| ID | baseMean | log2FoldChange | lfcSE | stat | pvalue | padj |
|---|---|---|---|---|---|---|
| AT5G14545 | 7868,55 | 15,70 | 1,20 | 13,14 | 1,99E-39 | 4,60E-36 |
| AT5G14565 | 6609,25 | -3,46 | 0,27 | -13,00 | 1,26E-38 | 1,45E-35 |
| AT5G14550 | 7872,48 | 14,74 | 1,20 | 12,32 | 6,75E-35 | 5,19E-32 |
| AT5G13630 | 269,68 | 5,39 | 0,44 | 12,20 | 3,11E-34 | 1,79E-31 |
| AT4G13575 | 426,34 | -5,12 | 0,43 | -11,91 | 1,10E-32 | 5,09E-30 |
| AT5G38850 | 479,48 | -2,56 | 0,22 | -11,87 | 1,67E-32 | 6,42E-30 |
| AT4G06130 | 392,23 | -5,10 | 0,44 | -11,50 | 1,33E-30 | 4,38E-28 |
| AT5G23065 | 1776,68 | -2,84 | 0,26 | -10,72 | 8,05E-27 | 2,32E-24 |
| AT3G17185 | 156,12 | -2,96 | 0,28 | -10,52 | 7,04E-26 | 1,81E-23 |
| AT2G01020 | 7030,11 | -3,68 | 0,35 | -10,38 | 2,95E-25 | 6,80E-23 |
| AT3G59884 | 100,21 | 4,25 | 0,43 | 9,87 | 5,36E-23 | 1,13E-20 |
| AT1G63130 | 240,21 | -2,89 | 0,29 | -9,81 | 9,73E-23 | 1,87E-20 |
| AT5G43740 | 103,64 | -4,77 | 0,49 | -9,78 | 1,34E-22 | 2,38E-20 |
| AT5G03553 | 58,75 | -3,62 | 0,38 | -9,54 | 1,47E-21 | 2,42E-19 |
| AT5G03552 | 58,24 | -3,61 | 0,38 | -9,53 | 1,58E-21 | 2,43E-19 |
| AT1G31280 | 162,06 | 5,23 | 0,57 | 9,17 | 4,80E-20 | 6,92E-18 |
| AT1G62930 | 53,37 | -3,93 | 0,44 | -9,04 | 1,63E-19 | 2,10E-17 |
| AT1G63150 | 176,58 | -2,74 | 0,30 | -9,04 | 1,63E-19 | 2,10E-17 |
| AT2G26211 | 125,25 | 4,10 | 0,45 | 9,02 | 1,91E-19 | 2,32E-17 |
| AT4G19395 | 418,73 | 1,83 | 0,21 | 8,76 | 2,00E-18 | 2,31E-16 |
| AT3G44580 | 31,33 | -4,16 | 0,48 | -8,72 | 2,71E-18 | 2,98E-16 |
| AT1G29265 | 54,27 | 5,21 | 0,60 | 8,71 | 3,00E-18 | 3,14E-16 |
| AT4G00885 | 370,66 | -2,54 | 0,29 | -8,67 | 4,27E-18 | 4,28E-16 |
| AT4G04825 | 145,71 | -11,07 | 1,30 | -8,50 | 1,95E-17 | 1,88E-15 |
| AT5G10946 | 91,61 | -2,31 | 0,27 | -8,40 | 4,48E-17 | 4,14E-15 |
| AT5G10945 | 91,45 | -2,32 | 0,28 | -8,35 | 6,73E-17 | 5,98E-15 |
| AT1G37130 | 39,12 | 4,41 | 0,53 | 8,32 | 8,99E-17 | 7,68E-15 |
| AT2G18960 | 62,63 | 4,66 | 0,57 | 8,19 | 2,72E-16 | 2,24E-14 |
| AT1G62914 | 457,48 | -2,76 | 0,34 | -8,15 | 3,65E-16 | 2,90E-14 |
| AT1G18075 | 4243,99 | -1,43 | 0,18 | -8,07 | 6,91E-16 | 5,32E-14 |
| AT2G10606 | 1219,01 | 2,17 | 0,28 | 7,83 | 4,96E-15 | 3,70E-13 |
| AT4G06810 | 54,17 | 5,20 | 0,67 | 7,77 | 7,88E-15 | 5,68E-13 |
| AT4G06805 | 54,05 | 5,20 | 0,67 | 7,76 | 8,41E-15 | 5,88E-13 |
| AT4G04565 | 64,98 | 2,98 | 0,39 | 7,70 | 1,35E-14 | 9,17E-13 |
| AT1G48267 | 15177,13 | -2,59 | 0,34 | -7,69 | 1,49E-14 | 9,83E-13 |
| AT3G23325 | 69,65 | 4,54 | 0,59 | 7,66 | 1,81E-14 | 1,16E-12 |
| AT1G12820 | 53,20 | -2,49 | 0,33 | -7,65 | 2,04E-14 | 1,27E-12 |
| AT3G23326 | 69,32 | 4,52 | 0,60 | 7,59 | 3,32E-14 | 2,02E-12 |
| AT4G23713 | 128,07 | 2,74 | 0,36 | 7,54 | 4,74E-14 | 2,81E-12 |
| AT3G44570 | 48,95 | -8,48 | 1,13 | -7,53 | 4,99E-14 | 2,88E-12 |
| AT1G04977 | 45,78 | -2,66 | 0,36 | -7,46 | 8,38E-14 | 4,72E-12 |
| AT2G47020 | 1248,75 | 2,32 | 0,31 | 7,43 | 1,11E-13 | 6,10E-12 |
| AT2G47015 | 1248,43 | 2,32 | 0,31 | 7,42 | 1,16E-13 | 6,24E-12 |
| AT1G31173 | 488,28 | 1,81 | 0,25 | 7,35 | 1,97E-13 | 1,03E-11 |
| AT4G03039 | 93,29 | 8,50 | 1,18 | 7,20 | 6,20E-13 | 3,11E-11 |
| AT4G03038 | 93,33 | 8,51 | 1,18 | 7,20 | 6,16E-13 | 3,11E-11 |
| AT4G04365 | 93,18 | 8,50 | 1,18 | 7,19 | 6,47E-13 | 3,18E-11 |
| AT5G63715 | 27,38 | -3,56 | 0,50 | -7,19 | 6,70E-13 | 3,22E-11 |
| AT1G63080 | 109,45 | -2,17 | 0,30 | -7,12 | 1,09E-12 | 5,13E-11 |
| AT5G43603 | 44,77 | 8,70 | 1,24 | 7,04 | 1,93E-12 | 8,91E-11 |
| AT3G23690 | 28,16 | -3,53 | 0,51 | -6,93 | 4,10E-12 | 1,86E-10 |
| AT5G08185 | 1387,13 | 2,10 | 0,32 | 6,66 | 2,70E-11 | 1,20E-09 |
| AT3G25795 | 22,21 | 8,58 | 1,29 | 6,65 | 2,89E-11 | 1,26E-09 |

| | | | | | | |
|---|---|---|---|---|---|---|
| AT4G21595 | 28,61 | 8,22 | 1,24 | 6,61 | 3,79E-11 | 1,62E-09 |
| AT5G66045 | 69,41 | 8,32 | 1,27 | 6,55 | 5,89E-11 | 2,47E-09 |
| AT5G41905 | 1173,00 | -1,66 | 0,25 | -6,53 | 6,57E-11 | 2,71E-09 |
| AT2G46255 | 72,33 | -2,71 | 0,43 | -6,35 | 2,09E-10 | 8,48E-09 |
| AT1G59870 | 72,89 | 2,29 | 0,36 | 6,30 | 3,02E-10 | 1,20E-08 |
| AT1G15750 | 35,69 | 5,70 | 0,91 | 6,23 | 4,58E-10 | 1,79E-08 |
| AT1G01183 | 317,62 | -2,22 | 0,36 | -6,22 | 4,92E-10 | 1,89E-08 |
| AT1G56070 | 55,72 | 2,17 | 0,35 | 6,21 | 5,17E-10 | 1,95E-08 |
| AT4G08755 | 25,91 | 7,67 | 1,24 | 6,20 | 5,77E-10 | 2,15E-08 |
| AT5G02500 | 62,24 | 2,19 | 0,36 | 6,14 | 8,32E-10 | 3,05E-08 |
| AT1G62590 | 20,75 | -3,68 | 0,60 | -6,11 | 9,98E-10 | 3,60E-08 |
| AT4G09135 | 26,82 | 7,53 | 1,25 | 6,02 | 1,77E-09 | 6,28E-08 |
| AT2G39675 | 2246,76 | -2,04 | 0,34 | -6,01 | 1,87E-09 | 6,53E-08 |
| AT2G15530 | 40,01 | 2,40 | 0,40 | 6,00 | 1,96E-09 | 6,75E-08 |
| AT3G09840 | 22,25 | 3,67 | 0,61 | 5,99 | 2,13E-09 | 7,23E-08 |
| AT4G13493 | 160,77 | -1,94 | 0,33 | -5,98 | 2,28E-09 | 7,63E-08 |
| AT3G13080 | 13,73 | 7,46 | 1,28 | 5,81 | 6,13E-09 | 2,02E-07 |
| AT4G07070 | 70,24 | -2,86 | 0,49 | -5,80 | 6,76E-09 | 2,20E-07 |
| AT3G14210 | 12,80 | -5,41 | 0,94 | -5,74 | 9,22E-09 | 2,96E-07 |
| AT3G41768 | 91,21 | -1,74 | 0,31 | -5,68 | 1,34E-08 | 4,22E-07 |
| AT2G41310 | 487,34 | -1,56 | 0,28 | -5,64 | 1,75E-08 | 5,45E-07 |
| AT2G09250 | 487,12 | -1,55 | 0,28 | -5,62 | 1,88E-08 | 5,79E-07 |
| AT1G50055 | 182,36 | -1,68 | 0,30 | -5,60 | 2,11E-08 | 6,40E-07 |
| AT1G09797 | 19,74 | 5,12 | 0,92 | 5,58 | 2,34E-08 | 6,92E-07 |
| AT1G09793 | 19,74 | 5,12 | 0,92 | 5,58 | 2,34E-08 | 6,92E-07 |
| AT2G18320 | 12,44 | -3,73 | 0,67 | -5,58 | 2,42E-08 | 7,08E-07 |
| AT2G46685 | 3232,16 | -1,46 | 0,26 | -5,52 | 3,37E-08 | 9,72E-07 |
| AT3G02260 | 23,16 | 3,37 | 0,61 | 5,50 | 3,71E-08 | 1,06E-06 |
| AT1G62910 | 98,21 | -1,99 | 0,36 | -5,47 | 4,59E-08 | 1,29E-06 |
| AT1G70320 | 18,40 | 4,48 | 0,82 | 5,46 | 4,73E-08 | 1,31E-06 |
| AT4G23387 | 136,04 | 2,72 | 0,50 | 5,46 | 4,75E-08 | 1,31E-06 |
| AT2G47275 | 830,57 | 1,70 | 0,31 | 5,45 | 5,05E-08 | 1,37E-06 |
| AT3G47350 | 23,89 | 3,23 | 0,60 | 5,38 | 7,54E-08 | 2,02E-06 |
| AT5G35407 | 1076,14 | 1,95 | 0,37 | 5,31 | 1,10E-07 | 2,92E-06 |
| AT5G38540 | 8,36 | -5,68 | 1,07 | -5,31 | 1,11E-07 | 2,92E-06 |
| AT5G46315 | 17,01 | 6,75 | 1,27 | 5,30 | 1,18E-07 | 3,06E-06 |
| AT4G34450 | 21,43 | 4,43 | 0,85 | 5,25 | 1,55E-07 | 3,97E-06 |
| AT3G48201 | 30,66 | 3,39 | 0,65 | 5,24 | 1,56E-07 | 3,97E-06 |
| AT2G05070 | 19,17 | -3,05 | 0,59 | -5,18 | 2,24E-07 | 5,62E-06 |
| AT4G03455 | 16,01 | -4,26 | 0,83 | -5,14 | 2,72E-07 | 6,76E-06 |
| AT1G63400 | 45,16 | -2,31 | 0,45 | -5,13 | 2,85E-07 | 7,00E-06 |
| AT5G04455 | 10,87 | 6,55 | 1,29 | 5,08 | 3,73E-07 | 8,96E-06 |
| AT5G04445 | 10,87 | 6,55 | 1,29 | 5,08 | 3,73E-07 | 8,96E-06 |
| AT3G01540 | 20,21 | 4,38 | 0,86 | 5,07 | 3,92E-07 | 9,33E-06 |
| AT2G27400 | 624,49 | -1,70 | 0,34 | -5,07 | 4,08E-07 | 9,52E-06 |
| AT5G04275 | 20,42 | 6,33 | 1,25 | 5,07 | 4,05E-07 | 9,52E-06 |
| AT2G28290 | 15,57 | 5,41 | 1,07 | 5,04 | 4,77E-07 | 1,10E-05 |
| AT1G08597 | 22,78 | -3,90 | 0,78 | -5,03 | 4,90E-07 | 1,12E-05 |
| AT3G11130 | 25,04 | 3,50 | 0,70 | 5,03 | 4,97E-07 | 1,12E-05 |
| AT3G26740 | 15,30 | -3,16 | 0,63 | -5,03 | 5,03E-07 | 1,13E-05 |
| AT1G66725 | 1003,08 | 1,53 | 0,31 | 5,00 | 5,61E-07 | 1,24E-05 |
| AT1G04907 | 10,89 | 6,68 | 1,33 | 5,00 | 5,66E-07 | 1,24E-05 |
| AT1G48410 | 43,79 | 1,99 | 0,40 | 4,98 | 6,48E-07 | 1,40E-05 |
| AT1G53683 | 15,44 | -6,47 | 1,30 | -4,98 | 6,44E-07 | 1,40E-05 |
| AT2G45160 | 10,70 | -3,49 | 0,70 | -4,96 | 6,89E-07 | 1,47E-05 |
| AT2G21390 | 23,77 | 6,14 | 1,27 | 4,83 | 1,38E-06 | 2,91E-05 |

| | | | | | | |
|---|---|---|---|---|---|---|
| AT1G09620 | 17,50 | 4,57 | 0,95 | 4,80 | 1,57E-06 | 3,30E-05 |
| AT5G05170 | 36,40 | 2,71 | 0,57 | 4,78 | 1,75E-06 | 3,63E-05 |
| AT5G23110 | 11,58 | 5,82 | 1,22 | 4,76 | 1,90E-06 | 3,92E-05 |
| AT4G31570 | 11,16 | 6,48 | 1,36 | 4,76 | 1,98E-06 | 4,04E-05 |
| AT2G16030 | 25,88 | 3,08 | 0,65 | 4,71 | 2,43E-06 | 4,91E-05 |
| AT4G13495 | 869,17 | -1,59 | 0,34 | -4,69 | 2,67E-06 | 5,36E-05 |
| AT5G51174 | 65,86 | -2,06 | 0,44 | -4,67 | 2,94E-06 | 5,86E-05 |
| AT2G01021 | 3194,91 | -1,25 | 0,27 | -4,66 | 3,09E-06 | 6,10E-05 |
| AT2G23350 | 29,35 | 5,08 | 1,10 | 4,63 | 3,71E-06 | 7,19E-05 |
| AT3G26810 | 23,04 | -2,25 | 0,49 | -4,63 | 3,69E-06 | 7,19E-05 |
| AT2G43970 | 15,13 | 3,05 | 0,66 | 4,62 | 3,88E-06 | 7,46E-05 |
| AT5G52310 | 5,67 | -5,99 | 1,30 | -4,61 | 3,98E-06 | 7,60E-05 |
| AT2G41900 | 16,69 | 4,63 | 1,01 | 4,58 | 4,56E-06 | 8,60E-05 |
| AT2G43900 | 9,76 | 6,08 | 1,33 | 4,58 | 4,58E-06 | 8,60E-05 |
| AT5G42950 | 18,25 | 5,40 | 1,19 | 4,54 | 5,59E-06 | 1,04E-04 |
| AT5G64270 | 15,19 | 4,90 | 1,08 | 4,52 | 6,20E-06 | 1,14E-04 |
| AT4G16890 | 15,81 | 4,07 | 0,91 | 4,49 | 7,02E-06 | 1,29E-04 |
| AT2G37190 | 11,90 | 5,78 | 1,29 | 4,49 | 7,25E-06 | 1,32E-04 |
| AT1G62670 | 9,93 | -3,68 | 0,82 | -4,48 | 7,33E-06 | 1,32E-04 |
| AT1G70770 | 12,70 | 4,85 | 1,08 | 4,48 | 7,61E-06 | 1,36E-04 |
| AT5G08717 | 14,65 | 6,03 | 1,35 | 4,46 | 8,28E-06 | 1,47E-04 |
| AT5G53460 | 17,48 | 2,61 | 0,59 | 4,42 | 9,96E-06 | 1,75E-04 |
| AT3G23680 | 9,12 | 5,94 | 1,35 | 4,41 | 1,01E-05 | 1,77E-04 |
| AT1G53687 | 5,07 | -6,20 | 1,41 | -4,40 | 1,09E-05 | 1,89E-04 |
| AT4G07875 | 11,03 | 5,81 | 1,32 | 4,39 | 1,15E-05 | 1,98E-04 |
| AT2G42270 | 6,29 | 6,05 | 1,38 | 4,38 | 1,21E-05 | 2,07E-04 |
| AT4G02480 | 22,30 | 4,02 | 0,92 | 4,37 | 1,23E-05 | 2,09E-04 |
| AT1G51150 | 4,36 | -6,34 | 1,45 | -4,37 | 1,24E-05 | 2,10E-04 |
| AT1G29900 | 11,05 | 4,68 | 1,08 | 4,35 | 1,37E-05 | 2,26E-04 |
| AT1G73687 | 3456,65 | -1,14 | 0,26 | -4,35 | 1,37E-05 | 2,26E-04 |
| AT3G10815 | 5,65 | 5,98 | 1,38 | 4,35 | 1,36E-05 | 2,26E-04 |
| AT1G47720 | 6,21 | 5,86 | 1,35 | 4,32 | 1,53E-05 | 2,50E-04 |
| AT2G06555 | 20,57 | 2,74 | 0,64 | 4,31 | 1,64E-05 | 2,67E-04 |
| AT5G46845 | 66,07 | -1,23 | 0,29 | -4,30 | 1,68E-05 | 2,71E-04 |
| AT1G22620 | 12,70 | 3,81 | 0,89 | 4,30 | 1,72E-05 | 2,76E-04 |
| AT4G13494 | 321,70 | -2,12 | 0,50 | -4,28 | 1,87E-05 | 2,98E-04 |
| AT2G45540 | 12,39 | 4,89 | 1,15 | 4,26 | 2,07E-05 | 3,28E-04 |
| AT1G22610 | 11,17 | 4,67 | 1,10 | 4,25 | 2,13E-05 | 3,30E-04 |
| AT4G12770 | 19,67 | 3,29 | 0,77 | 4,25 | 2,13E-05 | 3,30E-04 |
| AT5G07350 | 9,52 | 5,67 | 1,33 | 4,25 | 2,11E-05 | 3,30E-04 |
| AT5G04140 | 18,61 | 2,61 | 0,62 | 4,22 | 2,46E-05 | 3,78E-04 |
| AT1G08537 | 8,63 | -5,19 | 1,24 | -4,20 | 2,71E-05 | 4,12E-04 |
| AT4G31480 | 6,09 | 5,88 | 1,40 | 4,20 | 2,70E-05 | 4,12E-04 |
| AT1G66795 | 14,00 | 2,84 | 0,68 | 4,18 | 2,94E-05 | 4,44E-04 |
| AT5G27606 | 14,40 | 4,16 | 1,00 | 4,17 | 3,05E-05 | 4,58E-04 |
| AT5G10380 | 7,30 | 5,62 | 1,35 | 4,16 | 3,20E-05 | 4,76E-04 |
| AT4G08691 | 15,19 | 5,28 | 1,27 | 4,15 | 3,36E-05 | 4,97E-04 |
| AT2G39175 | 43,76 | 1,52 | 0,37 | 4,12 | 3,81E-05 | 5,56E-04 |
| AT1G01320 | 16,50 | 2,89 | 0,70 | 4,12 | 3,81E-05 | 5,56E-04 |
| AT4G36080 | 9,35 | 4,96 | 1,21 | 4,12 | 3,83E-05 | 5,56E-04 |
| AT1G09310 | 8,77 | -3,25 | 0,79 | -4,11 | 3,89E-05 | 5,61E-04 |
| AT3G43833 | 7,44 | 5,84 | 1,42 | 4,11 | 3,94E-05 | 5,65E-04 |
| AT5G39500 | 6,69 | 5,60 | 1,36 | 4,11 | 4,00E-05 | 5,70E-04 |
| AT2G40030 | 5,36 | 5,68 | 1,39 | 4,10 | 4,21E-05 | 5,96E-04 |
| AT1G54410 | 19,25 | -2,77 | 0,68 | -4,09 | 4,33E-05 | 6,05E-04 |
| AT1G74260 | 9,63 | 4,53 | 1,11 | 4,09 | 4,31E-05 | 6,05E-04 |

| | | | | | | |
|---|---|---|---|---|---|---|
| AT5G01820 | 9,75 | 5,01 | 1,23 | 4,06 | 4,97E-05 | 6,90E-04 |
| AT1G50480 | 9,27 | 4,03 | 1,00 | 4,02 | 5,87E-05 | 8,12E-04 |
| AT5G45510 | 8,42 | 4,39 | 1,09 | 4,01 | 6,14E-05 | 8,44E-04 |
| AT3G23685 | 5,28 | 5,48 | 1,37 | 4,00 | 6,36E-05 | 8,69E-04 |
| AT2G22125 | 19,76 | 3,06 | 0,77 | 3,99 | 6,59E-05 | 8,95E-04 |
| AT1G48090 | 11,18 | 3,01 | 0,75 | 3,99 | 6,67E-05 | 9,00E-04 |
| AT4G30160 | 4,90 | 5,58 | 1,40 | 3,98 | 6,76E-05 | 9,01E-04 |
| AT5G60790 | 11,59 | 3,96 | 0,99 | 3,98 | 6,76E-05 | 9,01E-04 |
| AT5G54075 | 15,55 | 4,41 | 1,11 | 3,98 | 6,97E-05 | 9,24E-04 |
| AT5G41740 | 6,05 | 5,76 | 1,45 | 3,98 | 7,03E-05 | 9,28E-04 |
| AT4G27440 | 13,15 | -2,44 | 0,61 | -3,97 | 7,17E-05 | 9,35E-04 |
| AT5G54070 | 15,72 | 4,41 | 1,11 | 3,97 | 7,13E-05 | 9,35E-04 |
| AT3G18780 | 17,06 | 2,63 | 0,66 | 3,97 | 7,31E-05 | 9,48E-04 |
| AT4G28080 | 23,67 | 2,76 | 0,70 | 3,94 | 8,04E-05 | 1,04E-03 |
| AT3G08943 | 11,39 | 4,50 | 1,14 | 3,93 | 8,42E-05 | 1,08E-03 |
| AT4G24690 | 16,56 | 2,47 | 0,63 | 3,91 | 9,19E-05 | 1,17E-03 |
| AT5G07940 | 5,66 | 5,43 | 1,39 | 3,90 | 9,47E-05 | 1,20E-03 |
| AT1G13980 | 8,14 | 5,15 | 1,32 | 3,90 | 9,58E-05 | 1,21E-03 |
| AT4G27600 | 12,18 | 4,33 | 1,11 | 3,89 | 9,94E-05 | 1,25E-03 |
| AT3G08370 | 6,58 | 5,50 | 1,41 | 3,89 | 1,01E-04 | 1,26E-03 |
| AT1G20760 | 7,86 | 5,21 | 1,34 | 3,88 | 1,06E-04 | 1,32E-03 |
| AT5G20700 | 23,90 | 1,73 | 0,45 | 3,87 | 1,07E-04 | 1,32E-03 |
| AT1G79930 | 9,47 | 5,08 | 1,31 | 3,87 | 1,10E-04 | 1,35E-03 |
| AT1G66783 | 14,56 | 2,86 | 0,74 | 3,87 | 1,10E-04 | 1,35E-03 |
| AT1G55670 | 5,12 | -3,67 | 0,95 | -3,86 | 1,14E-04 | 1,38E-03 |
| AT4G27000 | 10,07 | 3,81 | 0,99 | 3,86 | 1,15E-04 | 1,38E-03 |
| AT5G54270 | 15,64 | -2,04 | 0,53 | -3,82 | 1,31E-04 | 1,57E-03 |
| AT1G20340 | 11,85 | -2,37 | 0,62 | -3,82 | 1,32E-04 | 1,57E-03 |
| AT1G35501 | 18,58 | -2,16 | 0,57 | -3,82 | 1,35E-04 | 1,59E-03 |
| AT1G64720 | 13,91 | 2,91 | 0,76 | 3,82 | 1,35E-04 | 1,59E-03 |
| AT1G08833 | 15,74 | -2,07 | 0,54 | -3,81 | 1,38E-04 | 1,63E-03 |
| AT5G57350 | 4,58 | 5,53 | 1,46 | 3,79 | 1,50E-04 | 1,76E-03 |
| AT3G01345 | 6,06 | 4,69 | 1,24 | 3,79 | 1,53E-04 | 1,79E-03 |
| AT1G79520 | 27,20 | -2,11 | 0,56 | -3,77 | 1,62E-04 | 1,88E-03 |
| AT4G31877 | 9,33 | -2,66 | 0,71 | -3,76 | 1,70E-04 | 1,96E-03 |
| AT3G20362 | 7,16 | -3,73 | 0,99 | -3,75 | 1,75E-04 | 2,01E-03 |
| AT1G08380 | 7,50 | -3,30 | 0,88 | -3,75 | 1,76E-04 | 2,01E-03 |
| AT3G08947 | 5,24 | 5,38 | 1,44 | 3,74 | 1,82E-04 | 2,06E-03 |
| AT1G61520 | 19,44 | -2,09 | 0,56 | -3,74 | 1,86E-04 | 2,11E-03 |
| AT2G43680 | 14,92 | 2,68 | 0,72 | 3,73 | 1,89E-04 | 2,13E-03 |
| AT3G48870 | 10,82 | 3,91 | 1,05 | 3,73 | 1,90E-04 | 2,13E-03 |
| AT5G15680 | 7,75 | 4,85 | 1,31 | 3,72 | 2,03E-04 | 2,26E-03 |
| AT4G05180 | 7,70 | -3,29 | 0,89 | -3,71 | 2,06E-04 | 2,29E-03 |
| AT2G09610 | 9,53 | -2,84 | 0,77 | -3,70 | 2,14E-04 | 2,36E-03 |
| AT4G08335 | 4,06 | 5,41 | 1,46 | 3,70 | 2,15E-04 | 2,36E-03 |
| AT1G10170 | 9,09 | 3,79 | 1,03 | 3,70 | 2,18E-04 | 2,39E-03 |
| AT3G43300 | 11,08 | 3,34 | 0,90 | 3,69 | 2,25E-04 | 2,45E-03 |
| AT3G55734 | 132,05 | 1,17 | 0,32 | 3,68 | 2,34E-04 | 2,53E-03 |
| AT2G31141 | 64,55 | 1,11 | 0,30 | 3,67 | 2,39E-04 | 2,58E-03 |
| AT5G50290 | 244,15 | -1,36 | 0,37 | -3,67 | 2,41E-04 | 2,59E-03 |
| AT4G16950 | 9,81 | 3,10 | 0,85 | 3,66 | 2,49E-04 | 2,66E-03 |
| AT2G34430 | 28,33 | -1,60 | 0,44 | -3,66 | 2,52E-04 | 2,68E-03 |
| AT1G36160 | 9,82 | 3,30 | 0,90 | 3,66 | 2,55E-04 | 2,70E-03 |
| AT2G35940 | 16,85 | 2,60 | 0,71 | 3,65 | 2,66E-04 | 2,80E-03 |
| AT4G17140 | 11,35 | 2,95 | 0,81 | 3,64 | 2,71E-04 | 2,84E-03 |
| AT1G12775 | 21,87 | -1,91 | 0,53 | -3,64 | 2,77E-04 | 2,90E-03 |

| | | | | | | |
|---|---|---|---|---|---|---|
| AT1G62020 | 12,54 | 3,19 | 0,88 | 3,63 | 2,81E-04 | 2,90E-03 |
| AT5G43725 | 9,10 | -2,79 | 0,77 | -3,63 | 2,80E-04 | 2,90E-03 |
| AT5G43730 | 9,10 | -2,79 | 0,77 | -3,63 | 2,80E-04 | 2,90E-03 |
| AT2G36910 | 8,87 | 4,24 | 1,17 | 3,62 | 2,94E-04 | 3,02E-03 |
| AT1G09877 | 23,86 | -2,01 | 0,56 | -3,61 | 3,05E-04 | 3,11E-03 |
| AT5G02490 | 4,18 | 5,14 | 1,42 | 3,61 | 3,11E-04 | 3,16E-03 |
| AT1G06680 | 10,44 | -2,61 | 0,73 | -3,60 | 3,23E-04 | 3,24E-03 |
| AT1G35710 | 7,53 | 5,19 | 1,44 | 3,60 | 3,21E-04 | 3,24E-03 |
| AT5G60900 | 5,69 | 5,04 | 1,40 | 3,60 | 3,22E-04 | 3,24E-03 |
| AT5G65950 | 6,84 | 5,42 | 1,51 | 3,59 | 3,33E-04 | 3,33E-03 |
| AT3G50370 | 9,94 | 3,54 | 0,99 | 3,59 | 3,36E-04 | 3,34E-03 |
| AT4G14910 | 7,50 | 4,81 | 1,34 | 3,58 | 3,41E-04 | 3,38E-03 |
| AT5G53902 | 6,85 | 4,99 | 1,40 | 3,58 | 3,49E-04 | 3,42E-03 |
| AT5G53905 | 6,85 | 4,99 | 1,40 | 3,58 | 3,49E-04 | 3,42E-03 |
| AT4G30960 | 5,13 | 4,75 | 1,33 | 3,57 | 3,59E-04 | 3,49E-03 |
| AT5G54390 | 4,62 | 5,72 | 1,60 | 3,57 | 3,59E-04 | 3,49E-03 |
| AT3G19820 | 10,30 | 4,04 | 1,14 | 3,55 | 3,88E-04 | 3,76E-03 |
| AT1G05423 | 13,10 | -2,03 | 0,57 | -3,54 | 3,96E-04 | 3,82E-03 |
| AT4G38600 | 8,47 | 4,30 | 1,21 | 3,54 | 3,97E-04 | 3,82E-03 |
| AT5G57870 | 8,84 | 3,97 | 1,12 | 3,54 | 4,02E-04 | 3,85E-03 |
| AT1G07810 | 6,32 | 4,13 | 1,17 | 3,54 | 4,07E-04 | 3,88E-03 |
| AT1G10290 | 9,22 | 4,15 | 1,18 | 3,52 | 4,33E-04 | 4,11E-03 |
| AT5G07970 | 3,91 | 5,18 | 1,47 | 3,52 | 4,39E-04 | 4,15E-03 |
| AT4G20850 | 8,05 | 3,62 | 1,03 | 3,51 | 4,41E-04 | 4,15E-03 |
| AT4G04570 | 4,35 | -5,05 | 1,44 | -3,51 | 4,47E-04 | 4,19E-03 |
| AT1G55860 | 12,82 | 2,65 | 0,76 | 3,51 | 4,52E-04 | 4,22E-03 |
| AT2G16018 | 9,68 | 3,32 | 0,95 | 3,50 | 4,63E-04 | 4,31E-03 |
| AT2G02750 | 20,13 | -1,77 | 0,51 | -3,49 | 4,78E-04 | 4,41E-03 |
| AT3G63500 | 4,53 | 4,59 | 1,31 | 3,49 | 4,78E-04 | 4,41E-03 |
| AT1G49040 | 3,74 | 5,19 | 1,49 | 3,48 | 5,00E-04 | 4,59E-03 |
| AT3G47470 | 25,61 | -1,74 | 0,50 | -3,48 | 5,01E-04 | 4,59E-03 |
| AT5G07745 | 20,93 | 2,13 | 0,61 | 3,48 | 5,07E-04 | 4,62E-03 |
| AT2G16500 | 6,48 | 4,64 | 1,34 | 3,48 | 5,08E-04 | 4,62E-03 |
| AT1G15500 | 7,05 | 4,59 | 1,32 | 3,47 | 5,25E-04 | 4,75E-03 |
| AT2G02741 | 19,79 | -1,76 | 0,51 | -3,45 | 5,50E-04 | 4,96E-03 |
| AT2G43160 | 6,80 | 4,57 | 1,32 | 3,45 | 5,55E-04 | 4,99E-03 |
| AT2G35945 | 15,59 | 2,49 | 0,72 | 3,45 | 5,63E-04 | 5,04E-03 |
| AT1G36730 | 3,83 | 4,90 | 1,43 | 3,44 | 5,87E-04 | 5,23E-03 |
| AT5G41750 | 8,05 | 3,84 | 1,12 | 3,44 | 5,92E-04 | 5,26E-03 |
| AT3G18290 | 9,13 | 3,98 | 1,16 | 3,43 | 5,96E-04 | 5,27E-03 |
| AT1G77300 | 4,17 | 4,90 | 1,43 | 3,42 | 6,16E-04 | 5,43E-03 |
| AT4G26850 | 7,34 | 3,44 | 1,01 | 3,42 | 6,25E-04 | 5,49E-03 |
| AT2G26780 | 3,27 | 5,01 | 1,47 | 3,42 | 6,30E-04 | 5,51E-03 |
| AT1G63330 | 7,00 | -2,76 | 0,81 | -3,41 | 6,56E-04 | 5,72E-03 |
| AT1G12294 | 124,95 | 0,91 | 0,27 | 3,40 | 6,67E-04 | 5,79E-03 |
| AT3G12630 | 8,51 | 2,90 | 0,85 | 3,40 | 6,73E-04 | 5,81E-03 |
| AT1G07977 | 4,37 | -4,04 | 1,19 | -3,40 | 6,81E-04 | 5,86E-03 |
| AT5G06320 | 5,00 | 5,17 | 1,53 | 3,39 | 7,03E-04 | 6,03E-03 |
| AT1G60072 | 5,42 | 4,63 | 1,37 | 3,38 | 7,26E-04 | 6,21E-03 |
| AT2G08770 | 10,72 | -2,45 | 0,72 | -3,38 | 7,31E-04 | 6,22E-03 |
| AT4G03580 | 13,67 | 3,03 | 0,90 | 3,38 | 7,36E-04 | 6,25E-03 |
| AT4G32020 | 12,23 | 2,36 | 0,70 | 3,37 | 7,43E-04 | 6,28E-03 |
| AT4G18600 | 3,50 | 5,11 | 1,52 | 3,37 | 7,64E-04 | 6,44E-03 |
| AT1G12290 | 131,54 | 0,88 | 0,26 | 3,36 | 7,69E-04 | 6,45E-03 |
| AT1G20960 | 9,33 | 2,63 | 0,78 | 3,36 | 7,84E-04 | 6,55E-03 |
| AT5G58140 | 8,36 | 3,36 | 1,00 | 3,35 | 8,03E-04 | 6,69E-03 |

| | | | | | | |
|---|---|---|---|---|---|---|
| AT2G28056 | 34,24 | -1,32 | 0,39 | -3,35 | 8,16E-04 | 6,78E-03 |
| AT1G21630 | 4,05 | 4,97 | 1,48 | 3,34 | 8,24E-04 | 6,81E-03 |
| AT3G19190 | 3,92 | 5,07 | 1,52 | 3,34 | 8,41E-04 | 6,93E-03 |
| AT4G12800 | 10,28 | -2,64 | 0,79 | -3,34 | 8,48E-04 | 6,96E-03 |
| AT2G20570 | 7,71 | 2,73 | 0,82 | 3,33 | 8,64E-04 | 7,07E-03 |
| AT5G62162 | 10,16 | 2,60 | 0,78 | 3,32 | 8,91E-04 | 7,27E-03 |
| AT2G42540 | 2,58 | -5,62 | 1,69 | -3,32 | 9,03E-04 | 7,33E-03 |
| AT2G01008 | 28,92 | -1,36 | 0,41 | -3,31 | 9,29E-04 | 7,44E-03 |
| AT2G39885 | 9,28 | 3,99 | 1,21 | 3,31 | 9,29E-04 | 7,44E-03 |
| AT5G60408 | 515,37 | 0,96 | 0,29 | 3,31 | 9,26E-04 | 7,44E-03 |
| AT1G13360 | 23,38 | 1,56 | 0,47 | 3,31 | 9,29E-04 | 7,44E-03 |
| AT1G18745 | 15,99 | -1,87 | 0,56 | -3,31 | 9,32E-04 | 7,44E-03 |
| AT1G80070 | 7,13 | 3,87 | 1,17 | 3,30 | 9,77E-04 | 7,78E-03 |
| AT1G06190 | 5,18 | 4,04 | 1,23 | 3,29 | 9,85E-04 | 7,81E-03 |
| AT4G16940 | 4,86 | 4,96 | 1,51 | 3,29 | 9,95E-04 | 7,87E-03 |
| AT4G05420 | 7,60 | 3,41 | 1,04 | 3,29 | 1,00E-03 | 7,87E-03 |
| AT5G04853 | 3,91 | 4,78 | 1,45 | 3,29 | 1,00E-03 | 7,87E-03 |
| AT3G22380 | 8,79 | 2,85 | 0,87 | 3,29 | 1,01E-03 | 7,88E-03 |
| AT2G42600 | 13,57 | 1,98 | 0,60 | 3,29 | 1,01E-03 | 7,88E-03 |
| AT2G48060 | 4,01 | 4,80 | 1,46 | 3,29 | 1,01E-03 | 7,88E-03 |
| AT5G46250 | 6,03 | 4,18 | 1,27 | 3,28 | 1,02E-03 | 7,91E-03 |
| AT4G01800 | 8,35 | 3,00 | 0,92 | 3,28 | 1,04E-03 | 8,02E-03 |
| AT4G39420 | 9,97 | 2,93 | 0,90 | 3,27 | 1,07E-03 | 8,24E-03 |
| AT4G13885 | 7,77 | -2,42 | 0,74 | -3,27 | 1,09E-03 | 8,38E-03 |
| AT1G48300 | 8,14 | 4,08 | 1,25 | 3,26 | 1,11E-03 | 8,48E-03 |
| AT3G49120 | 5,10 | 3,93 | 1,21 | 3,26 | 1,12E-03 | 8,55E-03 |
| AT2G26890 | 8,74 | 2,78 | 0,86 | 3,24 | 1,18E-03 | 8,99E-03 |
| AT2G06420 | 4,36 | 4,83 | 1,49 | 3,24 | 1,21E-03 | 9,12E-03 |
| AT1G02500 | 10,85 | 2,98 | 0,92 | 3,23 | 1,24E-03 | 9,32E-03 |
| AT3G61470 | 27,14 | -1,55 | 0,48 | -3,23 | 1,24E-03 | 9,32E-03 |
| AT5G54770 | 28,55 | -1,56 | 0,48 | -3,22 | 1,27E-03 | 9,48E-03 |
| AT4G00150 | 10,48 | 2,76 | 0,86 | 3,22 | 1,28E-03 | 9,56E-03 |
| AT3G23000 | 7,09 | 3,84 | 1,20 | 3,22 | 1,30E-03 | 9,68E-03 |
| AT1G75660 | 6,24 | 4,32 | 1,34 | 3,21 | 1,32E-03 | 9,74E-03 |
| AT5G58040 | 3,29 | 4,68 | 1,46 | 3,21 | 1,32E-03 | 9,74E-03 |
| AT4G27640 | 8,75 | 4,49 | 1,40 | 3,21 | 1,35E-03 | 9,93E-03 |
| AT3G01460 | 5,55 | 4,49 | 1,40 | 3,20 | 1,37E-03 | 1,01E-02 |
| AT3G07100 | 6,58 | 3,11 | 0,97 | 3,20 | 1,39E-03 | 1,02E-02 |
| AT1G58250 | 4,39 | 4,23 | 1,33 | 3,19 | 1,41E-03 | 1,03E-02 |
| AT1G31580 | 10,89 | -2,14 | 0,67 | -3,19 | 1,42E-03 | 1,03E-02 |
| AT5G25610 | 6,58 | -2,53 | 0,79 | -3,18 | 1,47E-03 | 1,07E-02 |
| AT1G09970 | 4,22 | 4,53 | 1,43 | 3,18 | 1,50E-03 | 1,08E-02 |
| AT5G26920 | 3,70 | 4,46 | 1,41 | 3,17 | 1,50E-03 | 1,08E-02 |
| AT3G52140 | 9,89 | 2,97 | 0,94 | 3,17 | 1,52E-03 | 1,10E-02 |
| AT5G06600 | 5,36 | 4,33 | 1,37 | 3,16 | 1,55E-03 | 1,11E-02 |
| AT4G38770 | 7,08 | -2,18 | 0,69 | -3,16 | 1,57E-03 | 1,12E-02 |
| AT1G80410 | 5,85 | 4,41 | 1,39 | 3,16 | 1,58E-03 | 1,12E-02 |
| AT2G24980 | 3,78 | 4,88 | 1,55 | 3,15 | 1,62E-03 | 1,15E-02 |
| AT2G32730 | 3,18 | 4,81 | 1,53 | 3,15 | 1,65E-03 | 1,16E-02 |
| AT3G41761 | 2,51 | -5,40 | 1,72 | -3,15 | 1,65E-03 | 1,16E-02 |
| AT3G23920 | 2,81 | 4,71 | 1,50 | 3,15 | 1,66E-03 | 1,17E-02 |
| AT4G13577 | 2,60 | -4,89 | 1,56 | -3,14 | 1,66E-03 | 1,17E-02 |
| AT3G17850 | 4,74 | 4,39 | 1,40 | 3,14 | 1,69E-03 | 1,18E-02 |
| AT4G00450 | 3,39 | 4,83 | 1,54 | 3,14 | 1,70E-03 | 1,19E-02 |
| AT2G02950 | 6,29 | -2,86 | 0,91 | -3,13 | 1,73E-03 | 1,20E-02 |
| AT2G30520 | 8,24 | 3,13 | 1,00 | 3,13 | 1,74E-03 | 1,21E-02 |

| | | | | | | |
|---|---|---|---|---|---|---|
| AT4G10710 | 3,86 | 4,88 | 1,56 | 3,13 | 1,78E-03 | 1,23E-02 |
| AT2G06520 | 10,63 | -2,60 | 0,83 | -3,12 | 1,78E-03 | 1,23E-02 |
| AT5G26650 | 7,57 | 3,92 | 1,26 | 3,12 | 1,82E-03 | 1,24E-02 |
| AT5G46520 | 3,26 | 4,83 | 1,55 | 3,12 | 1,82E-03 | 1,24E-02 |
| AT2G35750 | 26,17 | -1,60 | 0,51 | -3,12 | 1,82E-03 | 1,24E-02 |
| AT3G11010 | 3,23 | 5,04 | 1,62 | 3,12 | 1,82E-03 | 1,24E-02 |
| AT3G59020 | 5,79 | 3,84 | 1,23 | 3,11 | 1,88E-03 | 1,27E-02 |
| AT5G25270 | 3,67 | 4,83 | 1,56 | 3,11 | 1,90E-03 | 1,29E-02 |
| AT3G13300 | 8,42 | 3,77 | 1,22 | 3,10 | 1,94E-03 | 1,31E-02 |
| AT1G30410 | 4,36 | 4,36 | 1,41 | 3,09 | 2,00E-03 | 1,34E-02 |
| AT5G44800 | 4,77 | 4,20 | 1,36 | 3,09 | 2,00E-03 | 1,34E-02 |
| AT5G64840 | 5,36 | 3,40 | 1,11 | 3,08 | 2,08E-03 | 1,39E-02 |
| AT4G06075 | 405,29 | -1,02 | 0,33 | -3,08 | 2,10E-03 | 1,40E-02 |
| AT1G31330 | 6,03 | -2,46 | 0,80 | -3,07 | 2,11E-03 | 1,41E-02 |
| AT2G47070 | 5,92 | 3,28 | 1,07 | 3,07 | 2,12E-03 | 1,41E-02 |
| AT1G55673 | 3,49 | -3,63 | 1,18 | -3,06 | 2,18E-03 | 1,44E-02 |
| AT5G04560 | 3,61 | 4,48 | 1,46 | 3,06 | 2,20E-03 | 1,45E-02 |
| AT1G20620 | 38,72 | -1,37 | 0,45 | -3,05 | 2,25E-03 | 1,48E-02 |
| AT4G37460 | 2,98 | 4,65 | 1,52 | 3,05 | 2,26E-03 | 1,48E-02 |
| AT1G61226 | 20,72 | -1,58 | 0,52 | -3,05 | 2,28E-03 | 1,49E-02 |
| AT5G64040 | 3,44 | -3,38 | 1,11 | -3,04 | 2,33E-03 | 1,52E-02 |
| AT4G30750 | 3,95 | 4,48 | 1,47 | 3,04 | 2,36E-03 | 1,53E-02 |
| AT2G26080 | 10,37 | 2,39 | 0,79 | 3,03 | 2,41E-03 | 1,56E-02 |
| AT4G19110 | 3,30 | 4,96 | 1,64 | 3,03 | 2,45E-03 | 1,59E-02 |
| AT3G20690 | 3,62 | -3,69 | 1,22 | -3,02 | 2,49E-03 | 1,60E-02 |
| AT4G00730 | 3,19 | 4,66 | 1,54 | 3,02 | 2,49E-03 | 1,60E-02 |
| AT2G33770 | 4,11 | 4,19 | 1,39 | 3,02 | 2,52E-03 | 1,61E-02 |
| AT5G66570 | 20,06 | -1,87 | 0,62 | -3,02 | 2,56E-03 | 1,64E-02 |
| AT1G19835 | 5,86 | 4,18 | 1,39 | 3,01 | 2,59E-03 | 1,65E-02 |
| AT3G28540 | 2,73 | 4,52 | 1,50 | 3,01 | 2,61E-03 | 1,65E-02 |
| AT4G16860 | 6,11 | 3,02 | 1,00 | 3,01 | 2,61E-03 | 1,65E-02 |
| AT2G26355 | 2,39 | 4,67 | 1,55 | 3,01 | 2,64E-03 | 1,66E-02 |
| AT3G59140 | 3,91 | 5,50 | 1,83 | 3,01 | 2,63E-03 | 1,66E-02 |
| AT2G38230 | 6,88 | -2,40 | 0,80 | -3,01 | 2,65E-03 | 1,66E-02 |
| AT5G17920 | 34,31 | 1,33 | 0,44 | 3,01 | 2,65E-03 | 1,66E-02 |
| AT4G11420 | 3,20 | 4,92 | 1,64 | 3,00 | 2,70E-03 | 1,69E-02 |
| AT1G03060 | 7,63 | 2,53 | 0,84 | 3,00 | 2,72E-03 | 1,69E-02 |
| AT5G06845 | 6,24 | 3,22 | 1,07 | 3,00 | 2,72E-03 | 1,69E-02 |
| AT5G63020 | 8,08 | -2,26 | 0,76 | -2,99 | 2,77E-03 | 1,72E-02 |
| AT3G62360 | 4,39 | 4,20 | 1,40 | 2,99 | 2,80E-03 | 1,73E-02 |
| AT1G79040 | 10,12 | -2,55 | 0,85 | -2,99 | 2,83E-03 | 1,75E-02 |
| AT1G27430 | 3,11 | 4,56 | 1,53 | 2,98 | 2,92E-03 | 1,80E-02 |
| AT1G52230 | 2,95 | -3,60 | 1,21 | -2,97 | 2,94E-03 | 1,80E-02 |
| AT2G14080 | 4,62 | 4,46 | 1,50 | 2,97 | 2,95E-03 | 1,80E-02 |
| AT5G09505 | 92,76 | -0,90 | 0,30 | -2,97 | 2,95E-03 | 1,80E-02 |
| AT1G13210 | 5,46 | 3,35 | 1,13 | 2,97 | 2,97E-03 | 1,81E-02 |
| AT4G34071 | 2,35 | -4,89 | 1,65 | -2,96 | 3,04E-03 | 1,85E-02 |
| AT3G52250 | 4,06 | 4,19 | 1,41 | 2,96 | 3,05E-03 | 1,85E-02 |
| AT2G45380 | 11,33 | -1,81 | 0,61 | -2,95 | 3,14E-03 | 1,90E-02 |
| AT3G18830 | 2,46 | 4,53 | 1,53 | 2,95 | 3,17E-03 | 1,91E-02 |
| AT2G04605 | 3,46 | 4,61 | 1,56 | 2,95 | 3,18E-03 | 1,91E-02 |
| AT2G46560 | 4,43 | 3,28 | 1,12 | 2,94 | 3,28E-03 | 1,96E-02 |
| AT3G06435 | 378,12 | -0,75 | 0,25 | -2,94 | 3,28E-03 | 1,96E-02 |
| AT4G13940 | 35,08 | 1,27 | 0,43 | 2,94 | 3,31E-03 | 1,97E-02 |
| AT3G63380 | 3,14 | 4,71 | 1,60 | 2,93 | 3,34E-03 | 1,99E-02 |
| AT4G23375 | 8,41 | 3,50 | 1,19 | 2,93 | 3,36E-03 | 1,99E-02 |

| | | | | | | |
|---|---|---|---|---|---|---|
| AT5G61010 | 3,05 | 4,29 | 1,46 | 2,93 | 3,37E-03 | 1,99E-02 |
| AT1G05562 | 6,13 | -2,52 | 0,86 | -2,93 | 3,44E-03 | 2,02E-02 |
| AT3G62290 | 5,55 | 4,07 | 1,39 | 2,93 | 3,43E-03 | 2,02E-02 |
| AT5G04930 | 5,77 | 3,76 | 1,28 | 2,93 | 3,44E-03 | 2,02E-02 |
| AT1G05560 | 6,13 | -2,52 | 0,86 | -2,93 | 3,44E-03 | 2,02E-02 |
| AT3G62980 | 20,74 | 1,72 | 0,59 | 2,92 | 3,48E-03 | 2,03E-02 |
| AT1G63980 | 3,67 | 4,40 | 1,51 | 2,92 | 3,50E-03 | 2,03E-02 |
| AT4G37910 | 5,28 | 3,36 | 1,15 | 2,92 | 3,49E-03 | 2,03E-02 |
| AT1G79280 | 4,52 | 3,33 | 1,14 | 2,92 | 3,51E-03 | 2,03E-02 |
| AT1G15290 | 5,85 | 3,53 | 1,21 | 2,92 | 3,54E-03 | 2,05E-02 |
| AT4G34890 | 2,69 | 4,60 | 1,58 | 2,91 | 3,56E-03 | 2,05E-02 |
| AT5G15833 | 13,93 | 2,19 | 0,75 | 2,91 | 3,61E-03 | 2,08E-02 |
| AT3G44703 | 4,72 | -2,95 | 1,02 | -2,90 | 3,72E-03 | 2,14E-02 |
| AT1G29465 | 25,62 | -1,43 | 0,49 | -2,90 | 3,74E-03 | 2,14E-02 |
| AT2G47980 | 2,39 | 4,48 | 1,55 | 2,89 | 3,83E-03 | 2,19E-02 |
| AT5G12150 | 4,21 | 4,72 | 1,63 | 2,89 | 3,85E-03 | 2,20E-02 |
| AT2G44490 | 17,93 | -1,56 | 0,54 | -2,89 | 3,90E-03 | 2,22E-02 |
| AT2G22330 | 4,57 | 4,10 | 1,42 | 2,88 | 3,96E-03 | 2,24E-02 |
| AT4G23650 | 3,54 | 4,27 | 1,48 | 2,88 | 3,96E-03 | 2,24E-02 |
| AT1G04810 | 3,41 | 4,75 | 1,65 | 2,88 | 4,03E-03 | 2,28E-02 |
| AT1G67120 | 8,25 | 2,32 | 0,81 | 2,87 | 4,05E-03 | 2,28E-02 |
| AT5G36228 | 1,95 | -4,80 | 1,67 | -2,87 | 4,07E-03 | 2,28E-02 |
| AT2G20190 | 8,93 | 2,96 | 1,03 | 2,87 | 4,09E-03 | 2,29E-02 |
| AT2G15970 | 3,79 | -3,26 | 1,14 | -2,87 | 4,10E-03 | 2,29E-02 |
| AT3G60860 | 2,94 | 4,54 | 1,58 | 2,87 | 4,14E-03 | 2,30E-02 |
| AT3G55220 | 4,51 | 3,55 | 1,24 | 2,87 | 4,15E-03 | 2,30E-02 |
| AT4G31500 | 6,34 | 3,09 | 1,08 | 2,87 | 4,15E-03 | 2,30E-02 |
| AT1G34300 | 2,71 | 4,46 | 1,56 | 2,86 | 4,21E-03 | 2,33E-02 |
| AT3G28180 | 8,56 | -3,15 | 1,10 | -2,86 | 4,26E-03 | 2,35E-02 |
| AT1G72390 | 2,40 | 4,43 | 1,55 | 2,86 | 4,29E-03 | 2,37E-02 |
| AT3G06530 | 4,41 | 3,68 | 1,29 | 2,85 | 4,31E-03 | 2,37E-02 |
| AT5G02565 | 2,69 | -3,41 | 1,20 | -2,85 | 4,37E-03 | 2,39E-02 |
| AT2G35744 | 14,81 | -1,68 | 0,59 | -2,85 | 4,35E-03 | 2,39E-02 |
| AT3G04470 | 2,70 | 4,28 | 1,50 | 2,85 | 4,37E-03 | 2,39E-02 |
| AT2G16020 | 3,14 | 4,43 | 1,56 | 2,84 | 4,46E-03 | 2,43E-02 |
| AT5G58150 | 3,53 | 5,05 | 1,78 | 2,84 | 4,47E-03 | 2,43E-02 |
| AT3G23780 | 4,59 | 3,75 | 1,32 | 2,84 | 4,50E-03 | 2,44E-02 |
| AT5G24350 | 3,06 | 4,74 | 1,67 | 2,83 | 4,62E-03 | 2,48E-02 |
| AT2G21100 | 3,61 | -3,04 | 1,07 | -2,83 | 4,62E-03 | 2,48E-02 |
| AT4G00800 | 2,43 | 4,38 | 1,55 | 2,83 | 4,61E-03 | 2,48E-02 |
| AT1G09070 | 2,87 | 4,48 | 1,58 | 2,83 | 4,65E-03 | 2,49E-02 |
| AT4G27500 | 2,93 | 4,37 | 1,54 | 2,83 | 4,71E-03 | 2,52E-02 |
| AT3G59770 | 3,31 | 3,97 | 1,40 | 2,83 | 4,72E-03 | 2,52E-02 |
| AT4G07855 | 2,93 | 4,37 | 1,54 | 2,83 | 4,71E-03 | 2,52E-02 |
| AT5G14430 | 2,60 | 4,58 | 1,62 | 2,82 | 4,78E-03 | 2,54E-02 |
| AT5G13000 | 3,30 | 3,84 | 1,36 | 2,82 | 4,80E-03 | 2,55E-02 |
| AT4G36140 | 2,98 | 3,82 | 1,36 | 2,81 | 4,88E-03 | 2,58E-02 |
| AT4G07975 | 9,02 | 2,29 | 0,81 | 2,81 | 4,93E-03 | 2,60E-02 |
| AT1G58230 | 3,69 | 4,12 | 1,47 | 2,80 | 5,04E-03 | 2,65E-02 |
| AT1G79920 | 8,70 | 2,35 | 0,84 | 2,80 | 5,06E-03 | 2,66E-02 |
| AT3G13857 | 124,31 | 1,06 | 0,38 | 2,80 | 5,17E-03 | 2,70E-02 |
| AT3G13855 | 124,31 | 1,06 | 0,38 | 2,80 | 5,17E-03 | 2,70E-02 |
| AT4G12780 | 15,31 | 1,65 | 0,59 | 2,80 | 5,18E-03 | 2,70E-02 |
| AT5G15845 | 2,91 | 4,16 | 1,49 | 2,79 | 5,31E-03 | 2,76E-02 |
| AT1G12090 | 8,60 | -2,18 | 0,78 | -2,79 | 5,31E-03 | 2,76E-02 |
| AT5G64420 | 4,10 | 3,94 | 1,42 | 2,78 | 5,40E-03 | 2,77E-02 |

| | | | | | | |
|---|---|---|---|---|---|---|
| AT1G79570 | 2,57 | 4,27 | 1,53 | 2,78 | 5,38E-03 | 2,77E-02 |
| AT2G03820 | 4,25 | 3,90 | 1,40 | 2,78 | 5,37E-03 | 2,77E-02 |
| AT2G33830 | 1,76 | -4,68 | 1,68 | -2,78 | 5,39E-03 | 2,77E-02 |
| AT5G44320 | 2,18 | 4,37 | 1,57 | 2,78 | 5,36E-03 | 2,77E-02 |
| AT3G20760 | 5,52 | 3,71 | 1,34 | 2,77 | 5,53E-03 | 2,83E-02 |
| AT5G49980 | 4,39 | 4,03 | 1,46 | 2,77 | 5,64E-03 | 2,89E-02 |
| AT4G17788 | 20,18 | -1,43 | 0,52 | -2,76 | 5,70E-03 | 2,91E-02 |
| AT2G31880 | 6,93 | 2,80 | 1,01 | 2,76 | 5,74E-03 | 2,92E-02 |
| AT4G32940 | 4,72 | 3,31 | 1,20 | 2,76 | 5,73E-03 | 2,92E-02 |
| AT1G49240 | 7,10 | 2,66 | 0,97 | 2,76 | 5,85E-03 | 2,96E-02 |
| AT5G04720 | 2,75 | 4,97 | 1,80 | 2,76 | 5,86E-03 | 2,96E-02 |
| AT5G08712 | 123,44 | 0,95 | 0,35 | 2,74 | 6,11E-03 | 3,07E-02 |
| AT2G17930 | 7,05 | 2,43 | 0,89 | 2,74 | 6,11E-03 | 3,07E-02 |
| AT4G00570 | 3,02 | 4,27 | 1,56 | 2,74 | 6,11E-03 | 3,07E-02 |
| AT5G48160 | 6,17 | 3,02 | 1,10 | 2,74 | 6,09E-03 | 3,07E-02 |
| AT1G01960 | 2,74 | 3,92 | 1,43 | 2,74 | 6,14E-03 | 3,08E-02 |
| AT1G48953 | 8,24 | -2,30 | 0,84 | -2,74 | 6,22E-03 | 3,10E-02 |
| AT1G76550 | 2,61 | 4,23 | 1,55 | 2,74 | 6,23E-03 | 3,10E-02 |
| AT5G15850 | 2,49 | 4,19 | 1,53 | 2,73 | 6,25E-03 | 3,11E-02 |
| AT3G03710 | 4,42 | 3,78 | 1,38 | 2,73 | 6,32E-03 | 3,13E-02 |
| AT4G08620 | 2,91 | 4,56 | 1,67 | 2,73 | 6,33E-03 | 3,13E-02 |
| AT4G39080 | 4,27 | 3,48 | 1,28 | 2,72 | 6,57E-03 | 3,25E-02 |
| AT2G38440 | 3,79 | 3,80 | 1,40 | 2,72 | 6,59E-03 | 3,25E-02 |
| AT5G09000 | 2,90 | -3,51 | 1,29 | -2,71 | 6,64E-03 | 3,27E-02 |
| AT2G16586 | 20,98 | 1,87 | 0,69 | 2,71 | 6,74E-03 | 3,31E-02 |
| AT2G35387 | 52,44 | -0,99 | 0,37 | -2,71 | 6,81E-03 | 3,34E-02 |
| AT1G54090 | 3,59 | 4,39 | 1,62 | 2,70 | 6,84E-03 | 3,34E-02 |
| AT4G04970 | 2,63 | 5,23 | 1,94 | 2,70 | 6,93E-03 | 3,37E-02 |
| AT3G61198 | 7,65 | 2,87 | 1,06 | 2,70 | 6,92E-03 | 3,37E-02 |
| AT4G17330 | 4,91 | 4,11 | 1,52 | 2,70 | 6,93E-03 | 3,37E-02 |
| AT3G49500 | 2,34 | 4,54 | 1,68 | 2,70 | 7,01E-03 | 3,40E-02 |
| AT1G64140 | 5,12 | 3,33 | 1,24 | 2,69 | 7,10E-03 | 3,43E-02 |
| AT5G46260 | 1,90 | 4,32 | 1,61 | 2,69 | 7,10E-03 | 3,43E-02 |
| AT3G07870 | 5,57 | 3,93 | 1,46 | 2,69 | 7,13E-03 | 3,44E-02 |
| AT2G25660 | 3,83 | 4,06 | 1,51 | 2,69 | 7,15E-03 | 3,44E-02 |
| AT4G30100 | 3,56 | 3,84 | 1,43 | 2,69 | 7,18E-03 | 3,44E-02 |
| AT1G55591 | 5,61 | 3,81 | 1,42 | 2,69 | 7,21E-03 | 3,45E-02 |
| AT5G15970 | 2,85 | -3,98 | 1,48 | -2,69 | 7,23E-03 | 3,45E-02 |
| AT1G06273 | 2,17 | -4,76 | 1,77 | -2,68 | 7,28E-03 | 3,46E-02 |
| AT5G05505 | 4,00 | -3,02 | 1,12 | -2,68 | 7,27E-03 | 3,46E-02 |
| AT1G49700 | 4,39 | 3,24 | 1,21 | 2,68 | 7,33E-03 | 3,48E-02 |
| AT1G66100 | 8,07 | -2,25 | 0,84 | -2,68 | 7,36E-03 | 3,49E-02 |
| AT1G62310 | 4,32 | -2,96 | 1,10 | -2,68 | 7,42E-03 | 3,51E-02 |
| AT5G45430 | 3,88 | 3,03 | 1,14 | 2,67 | 7,65E-03 | 3,61E-02 |
| AT3G44300 | 1,82 | 4,27 | 1,60 | 2,67 | 7,68E-03 | 3,62E-02 |
| AT2G25800 | 4,68 | 3,76 | 1,41 | 2,66 | 7,80E-03 | 3,67E-02 |
| AT1G09770 | 3,11 | 3,76 | 1,42 | 2,66 | 7,90E-03 | 3,71E-02 |
| AT1G26985 | 10,26 | 2,09 | 0,79 | 2,65 | 7,97E-03 | 3,73E-02 |
| AT1G43320 | 2,75 | 4,77 | 1,80 | 2,65 | 8,04E-03 | 3,74E-02 |
| AT5G36230 | 2,63 | 4,27 | 1,61 | 2,65 | 8,03E-03 | 3,74E-02 |
| AT1G21580 | 3,07 | 3,86 | 1,46 | 2,65 | 8,06E-03 | 3,74E-02 |
| AT1G06227 | 343,33 | -0,98 | 0,37 | -2,65 | 8,07E-03 | 3,74E-02 |
| AT5G08670 | 5,95 | 2,63 | 0,99 | 2,65 | 8,02E-03 | 3,74E-02 |
| AT1G43910 | 2,35 | 4,37 | 1,65 | 2,65 | 8,12E-03 | 3,76E-02 |
| AT1G08623 | 2,21 | -3,71 | 1,41 | -2,64 | 8,39E-03 | 3,87E-02 |
| AT1G29910 | 27,17 | -1,33 | 0,50 | -2,63 | 8,46E-03 | 3,90E-02 |

| | | | | | | |
|---|---|---|---|---|---|---|
| AT2G43890 | 2,70 | 4,53 | 1,72 | 2,63 | 8,53E-03 | 3,91E-02 |
| AT5G09410 | 3,14 | 3,93 | 1,49 | 2,63 | 8,52E-03 | 3,91E-02 |
| AT3G07790 | 2,60 | 4,04 | 1,54 | 2,63 | 8,56E-03 | 3,92E-02 |
| AT1G10950 | 6,24 | 2,86 | 1,09 | 2,63 | 8,66E-03 | 3,93E-02 |
| AT1G18740 | 2,60 | 4,05 | 1,54 | 2,62 | 8,67E-03 | 3,93E-02 |
| AT5G07570 | 4,01 | 4,21 | 1,61 | 2,62 | 8,69E-03 | 3,93E-02 |
| AT1G27770 | 5,67 | 3,50 | 1,34 | 2,62 | 8,71E-03 | 3,93E-02 |
| AT2G48160 | 6,20 | 3,84 | 1,46 | 2,62 | 8,69E-03 | 3,93E-02 |
| AT4G03445 | 45,78 | 0,86 | 0,33 | 2,62 | 8,70E-03 | 3,93E-02 |
| AT5G12370 | 4,35 | 3,67 | 1,40 | 2,62 | 8,69E-03 | 3,93E-02 |
| AT1G80680 | 4,25 | 3,78 | 1,45 | 2,62 | 8,87E-03 | 4,00E-02 |
| AT1G06237 | 336,80 | -1,00 | 0,38 | -2,61 | 8,94E-03 | 4,02E-02 |
| AT2G03855 | 12,69 | -1,50 | 0,58 | -2,61 | 9,12E-03 | 4,09E-02 |
| AT1G77760 | 2,44 | 4,27 | 1,64 | 2,61 | 9,11E-03 | 4,09E-02 |
| AT2G46460 | 2,59 | -4,61 | 1,77 | -2,60 | 9,37E-03 | 4,19E-02 |
| AT5G13770 | 2,99 | 4,13 | 1,59 | 2,60 | 9,41E-03 | 4,20E-02 |
| AT5G62000 | 10,38 | 1,66 | 0,64 | 2,60 | 9,42E-03 | 4,20E-02 |
| AT3G06835 | 2,07 | -4,38 | 1,69 | -2,59 | 9,57E-03 | 4,26E-02 |
| AT5G46450 | 2,49 | 4,06 | 1,57 | 2,59 | 9,60E-03 | 4,26E-02 |
| AT1G59610 | 3,98 | 3,27 | 1,27 | 2,59 | 9,71E-03 | 4,30E-02 |
| AT2G01270 | 3,86 | 3,77 | 1,46 | 2,58 | 9,74E-03 | 4,31E-02 |
| AT3G48057 | 47,97 | 1,45 | 0,56 | 2,58 | 9,80E-03 | 4,32E-02 |
| AT4G04640 | 5,86 | -2,40 | 0,93 | -2,58 | 9,80E-03 | 4,32E-02 |
| AT3G14470 | 2,64 | 4,10 | 1,59 | 2,58 | 9,90E-03 | 4,35E-02 |
| AT4G16490 | 2,72 | 4,72 | 1,83 | 2,58 | 9,91E-03 | 4,35E-02 |
| AT1G15820 | 12,09 | -1,77 | 0,69 | -2,58 | 9,99E-03 | 4,38E-02 |
| AT1G27320 | 2,96 | 3,78 | 1,47 | 2,57 | 1,01E-02 | 4,41E-02 |
| AT3G03380 | 4,30 | 3,25 | 1,26 | 2,57 | 1,01E-02 | 4,41E-02 |
| AT5G24990 | 2,50 | -3,95 | 1,54 | -2,57 | 1,02E-02 | 4,42E-02 |
| AT1G63880 | 4,32 | 3,23 | 1,26 | 2,57 | 1,02E-02 | 4,42E-02 |
| AT4G06235 | 7,69 | -2,57 | 1,00 | -2,57 | 1,02E-02 | 4,42E-02 |
| AT1G09780 | 4,32 | 2,86 | 1,12 | 2,57 | 1,02E-02 | 4,43E-02 |
| AT2G45940 | 2,82 | 3,86 | 1,51 | 2,57 | 1,03E-02 | 4,43E-02 |
| AT3G01500 | 4,00 | -2,95 | 1,15 | -2,57 | 1,03E-02 | 4,43E-02 |
| AT5G57990 | 5,76 | 2,83 | 1,10 | 2,56 | 1,03E-02 | 4,45E-02 |
| AT2G28000 | 11,21 | 1,97 | 0,77 | 2,56 | 1,04E-02 | 4,48E-02 |
| AT1G52780 | 3,16 | 3,53 | 1,38 | 2,56 | 1,05E-02 | 4,52E-02 |
| AT5G53480 | 6,73 | 2,42 | 0,95 | 2,56 | 1,06E-02 | 4,53E-02 |
| AT1G13350 | 2,82 | 4,38 | 1,71 | 2,56 | 1,06E-02 | 4,53E-02 |
| AT2G05100 | 10,97 | -1,45 | 0,57 | -2,55 | 1,07E-02 | 4,58E-02 |
| AT1G15130 | 4,06 | 2,92 | 1,14 | 2,55 | 1,08E-02 | 4,60E-02 |
| AT4G21640 | 1,99 | -3,69 | 1,45 | -2,55 | 1,09E-02 | 4,63E-02 |
| AT1G12300 | 2,68 | -3,07 | 1,21 | -2,54 | 1,10E-02 | 4,68E-02 |
| AT4G06240 | 6,18 | -2,80 | 1,10 | -2,54 | 1,11E-02 | 4,70E-02 |
| AT4G02660 | 2,17 | 4,21 | 1,66 | 2,53 | 1,12E-02 | 4,76E-02 |
| AT5G20720 | 3,41 | 3,96 | 1,56 | 2,53 | 1,13E-02 | 4,76E-02 |
| AT1G13930 | 7,00 | -1,95 | 0,77 | -2,53 | 1,15E-02 | 4,77E-02 |
| AT1G32230 | 5,15 | 3,13 | 1,24 | 2,53 | 1,14E-02 | 4,77E-02 |
| AT2G40140 | 4,76 | 3,24 | 1,28 | 2,53 | 1,14E-02 | 4,77E-02 |
| AT4G12290 | 3,10 | 3,34 | 1,32 | 2,53 | 1,15E-02 | 4,77E-02 |
| AT5G11040 | 2,15 | 4,07 | 1,61 | 2,53 | 1,14E-02 | 4,77E-02 |
| AT5G47910 | 5,04 | 2,73 | 1,08 | 2,53 | 1,15E-02 | 4,77E-02 |
| AT5G61140 | 4,17 | 2,86 | 1,13 | 2,53 | 1,14E-02 | 4,77E-02 |
| AT1G17210 | 2,32 | 4,17 | 1,65 | 2,53 | 1,15E-02 | 4,77E-02 |
| AT1G64583 | 1,87 | -3,91 | 1,55 | -2,53 | 1,16E-02 | 4,77E-02 |
| AT3G09162 | 2,15 | -4,26 | 1,68 | -2,53 | 1,15E-02 | 4,77E-02 |

| | | | | | | |
|---|---|---|---|---|---|---|
| AT4G08165 | 5,10 | 4,02 | 1,59 | 2,53 | 1,16E-02 | 4,77E-02 |
| AT4G35790 | 4,32 | 3,50 | 1,38 | 2,53 | 1,14E-02 | 4,77E-02 |
| AT5G15400 | 2,82 | 3,68 | 1,45 | 2,53 | 1,14E-02 | 4,77E-02 |
| AT1G66970 | 4,60 | 2,65 | 1,05 | 2,52 | 1,16E-02 | 4,77E-02 |
| AT1G49760 | 5,88 | 2,73 | 1,08 | 2,52 | 1,16E-02 | 4,77E-02 |
| AT1G75335 | 3,97 | 3,70 | 1,47 | 2,52 | 1,17E-02 | 4,80E-02 |
| AT5G20280 | 6,18 | 2,21 | 0,88 | 2,52 | 1,19E-02 | 4,87E-02 |
| AT5G03280 | 3,28 | 3,80 | 1,51 | 2,51 | 1,19E-02 | 4,88E-02 |
| AT4G02970 | 72,42 | 0,82 | 0,33 | 2,51 | 1,21E-02 | 4,93E-02 |
| AT4G30190 | 7,61 | 2,31 | 0,92 | 2,51 | 1,22E-02 | 4,94E-02 |
| AT4G31880 | 2,70 | 3,97 | 1,58 | 2,51 | 1,22E-02 | 4,94E-02 |
| AT1G79090 | 2,69 | 4,14 | 1,65 | 2,50 | 1,23E-02 | 4,96E-02 |
| AT3G57410 | 3,70 | 3,35 | 1,34 | 2,50 | 1,22E-02 | 4,96E-02 |
| AT4G01860 | 3,28 | 4,06 | 1,62 | 2,50 | 1,24E-02 | 5,00E-02 |
| AT4G02380 | 7,09 | 2,00 | 0,80 | 2,50 | 1,24E-02 | 5,00E-02 |
| AT1G04897 | 2,68 | 3,96 | 1,59 | 2,50 | 1,25E-02 | 5,03E-02 |
| AT1G48650 | 3,21 | 3,72 | 1,49 | 2,50 | 1,26E-02 | 5,03E-02 |
| AT1G56630 | 1,84 | -5,24 | 2,10 | -2,50 | 1,25E-02 | 5,03E-02 |
| AT3G05700 | 20,03 | 1,85 | 0,74 | 2,49 | 1,26E-02 | 5,03E-02 |
| AT3G13530 | 2,08 | 4,17 | 1,67 | 2,50 | 1,25E-02 | 5,03E-02 |
| AT5G05980 | 2,96 | 4,23 | 1,70 | 2,50 | 1,26E-02 | 5,03E-02 |
| AT2G41440 | 2,60 | 3,80 | 1,52 | 2,49 | 1,27E-02 | 5,04E-02 |
| AT4G24280 | 6,53 | 2,48 | 0,99 | 2,49 | 1,28E-02 | 5,06E-02 |
| AT5G56070 | 3,26 | 4,23 | 1,70 | 2,49 | 1,28E-02 | 5,06E-02 |
| AT5G66200 | 3,05 | 3,72 | 1,49 | 2,49 | 1,27E-02 | 5,06E-02 |
| AT4G14160 | 2,71 | 4,25 | 1,71 | 2,49 | 1,29E-02 | 5,10E-02 |
| AT3G03175 | 7,33 | 2,07 | 0,83 | 2,48 | 1,30E-02 | 5,13E-02 |
| AT1G21250 | 6,46 | 2,20 | 0,89 | 2,48 | 1,30E-02 | 5,13E-02 |
| AT2G38800 | 2,71 | 3,99 | 1,62 | 2,47 | 1,34E-02 | 5,30E-02 |
| AT5G13100 | 3,43 | 3,79 | 1,54 | 2,47 | 1,35E-02 | 5,31E-02 |
| AT3G23120 | 2,24 | 3,79 | 1,53 | 2,47 | 1,35E-02 | 5,31E-02 |
| AT3G08850 | 2,35 | 3,74 | 1,52 | 2,47 | 1,36E-02 | 5,33E-02 |
| AT4G21960 | 15,77 | -1,24 | 0,50 | -2,47 | 1,36E-02 | 5,33E-02 |
| AT4G24415 | 10,68 | -1,37 | 0,56 | -2,47 | 1,37E-02 | 5,34E-02 |
| AT1G63770 | 7,56 | 1,92 | 0,78 | 2,46 | 1,37E-02 | 5,34E-02 |
| AT2G46340 | 1,90 | 4,63 | 1,88 | 2,46 | 1,37E-02 | 5,34E-02 |
| AT5G05190 | 2,00 | 4,09 | 1,66 | 2,46 | 1,39E-02 | 5,40E-02 |
| AT2G38540 | 3,59 | -2,78 | 1,13 | -2,45 | 1,44E-02 | 5,59E-02 |
| AT1G63230 | 3,31 | -2,70 | 1,10 | -2,44 | 1,46E-02 | 5,66E-02 |
| AT4G30760 | 1,76 | 4,13 | 1,70 | 2,44 | 1,48E-02 | 5,72E-02 |
| AT3G25500 | 1,85 | 3,88 | 1,59 | 2,44 | 1,49E-02 | 5,74E-02 |
| AT2G24790 | 3,54 | 3,84 | 1,58 | 2,43 | 1,50E-02 | 5,76E-02 |
| AT2G46830 | 2,43 | 3,94 | 1,62 | 2,43 | 1,50E-02 | 5,76E-02 |
| AT1G07590 | 3,38 | 2,69 | 1,10 | 2,43 | 1,50E-02 | 5,76E-02 |
| AT3G01845 | 219,09 | -0,51 | 0,21 | -2,43 | 1,50E-02 | 5,76E-02 |
| AT4G18596 | 1,78 | 4,21 | 1,74 | 2,43 | 1,53E-02 | 5,84E-02 |
| AT5G44340 | 14,19 | 1,75 | 0,72 | 2,43 | 1,53E-02 | 5,84E-02 |
| AT2G09670 | 5,26 | 3,05 | 1,26 | 2,42 | 1,53E-02 | 5,85E-02 |
| AT5G22500 | 2,19 | -3,85 | 1,59 | -2,42 | 1,54E-02 | 5,86E-02 |
| AT3G09465 | 2,48 | -2,90 | 1,20 | -2,42 | 1,55E-02 | 5,88E-02 |
| AT1G07135 | 2,46 | 3,82 | 1,58 | 2,42 | 1,56E-02 | 5,93E-02 |
| AT1G30470 | 3,87 | 3,62 | 1,50 | 2,42 | 1,57E-02 | 5,94E-02 |
| AT5G20020 | 2,46 | 4,55 | 1,88 | 2,42 | 1,57E-02 | 5,94E-02 |
| AT3G50930 | 3,98 | 3,08 | 1,28 | 2,41 | 1,58E-02 | 5,96E-02 |
| AT1G14610 | 2,42 | 3,79 | 1,57 | 2,41 | 1,59E-02 | 5,98E-02 |
| AT3G02090 | 3,68 | 3,00 | 1,24 | 2,41 | 1,60E-02 | 5,99E-02 |

| | | | | | | |
|---|---|---|---|---|---|---|
| AT4G13340 | 14,47 | 1,32 | 0,55 | 2,41 | 1,59E-02 | 5,99E-02 |
| AT1G07200 | 1,82 | 4,12 | 1,71 | 2,41 | 1,59E-02 | 5,99E-02 |
| AT5G19390 | 2,78 | 4,14 | 1,72 | 2,41 | 1,60E-02 | 6,01E-02 |
| AT1G59760 | 3,05 | 3,64 | 1,51 | 2,41 | 1,61E-02 | 6,01E-02 |
| AT1G02860 | 4,18 | 3,48 | 1,45 | 2,40 | 1,62E-02 | 6,05E-02 |
| AT1G08857 | 2,42 | 4,79 | 1,99 | 2,40 | 1,62E-02 | 6,06E-02 |
| AT1G69360 | 2,19 | 4,11 | 1,71 | 2,40 | 1,63E-02 | 6,07E-02 |
| AT1G08450 | 3,29 | 3,74 | 1,56 | 2,40 | 1,64E-02 | 6,11E-02 |
| AT3G02470 | 10,40 | 1,53 | 0,64 | 2,40 | 1,66E-02 | 6,14E-02 |
| AT5G18350 | 2,51 | 4,25 | 1,77 | 2,40 | 1,66E-02 | 6,14E-02 |
| AT2G41740 | 6,26 | 2,55 | 1,07 | 2,39 | 1,71E-02 | 6,30E-02 |
| AT4G00140 | 2,38 | 4,04 | 1,69 | 2,39 | 1,71E-02 | 6,30E-02 |
| AT2G17440 | 1,80 | 4,15 | 1,74 | 2,38 | 1,73E-02 | 6,37E-02 |
| AT5G41765 | 5,08 | 3,39 | 1,42 | 2,38 | 1,73E-02 | 6,37E-02 |
| AT1G30230 | 5,95 | 2,37 | 1,00 | 2,38 | 1,74E-02 | 6,37E-02 |
| AT3G46780 | 9,83 | 1,86 | 0,78 | 2,38 | 1,74E-02 | 6,37E-02 |
| AT5G62350 | 2,26 | -3,58 | 1,51 | -2,38 | 1,73E-02 | 6,37E-02 |
| AT5G49360 | 3,80 | 3,45 | 1,45 | 2,38 | 1,74E-02 | 6,37E-02 |
| AT3G50820 | 7,18 | -2,15 | 0,90 | -2,38 | 1,75E-02 | 6,38E-02 |
| AT3G19180 | 1,86 | 4,38 | 1,84 | 2,38 | 1,75E-02 | 6,39E-02 |
| AT4G02510 | 15,59 | 1,47 | 0,62 | 2,37 | 1,76E-02 | 6,41E-02 |
| AT5G16210 | 1,91 | 4,06 | 1,71 | 2,37 | 1,77E-02 | 6,44E-02 |
| AT3G13070 | 1,98 | 4,25 | 1,79 | 2,37 | 1,78E-02 | 6,44E-02 |
| AT1G20510 | 2,74 | 3,97 | 1,68 | 2,37 | 1,79E-02 | 6,48E-02 |
| AT1G21270 | 2,66 | 4,25 | 1,79 | 2,37 | 1,79E-02 | 6,48E-02 |
| AT3G41762 | 25,36 | -1,08 | 0,46 | -2,37 | 1,80E-02 | 6,50E-02 |
| AT5G19820 | 4,02 | 3,72 | 1,57 | 2,37 | 1,80E-02 | 6,50E-02 |
| AT1G60870 | 2,59 | 3,58 | 1,51 | 2,36 | 1,82E-02 | 6,51E-02 |
| AT1G78570 | 2,32 | 3,83 | 1,62 | 2,36 | 1,83E-02 | 6,51E-02 |
| AT3G20010 | 2,80 | 4,09 | 1,73 | 2,36 | 1,83E-02 | 6,51E-02 |
| AT3G57645 | 1,71 | 4,44 | 1,88 | 2,36 | 1,82E-02 | 6,51E-02 |
| AT5G58540 | 1,83 | 4,18 | 1,77 | 2,36 | 1,83E-02 | 6,51E-02 |
| AT5G63840 | 2,51 | 3,70 | 1,57 | 2,36 | 1,81E-02 | 6,51E-02 |
| AT1G14740 | 3,27 | 3,30 | 1,40 | 2,36 | 1,83E-02 | 6,51E-02 |
| AT1G64790 | 7,36 | 2,01 | 0,85 | 2,36 | 1,82E-02 | 6,51E-02 |
| AT3G57260 | 8,35 | 1,79 | 0,76 | 2,36 | 1,82E-02 | 6,51E-02 |
| AT1G27595 | 2,01 | 3,95 | 1,67 | 2,36 | 1,84E-02 | 6,55E-02 |
| AT4G00630 | 4,40 | 2,61 | 1,11 | 2,35 | 1,86E-02 | 6,61E-02 |
| AT4G37900 | 1,96 | 4,25 | 1,81 | 2,35 | 1,88E-02 | 6,64E-02 |
| AT5G36180 | 3,47 | -2,94 | 1,25 | -2,35 | 1,89E-02 | 6,69E-02 |
| AT1G32583 | 28,27 | -1,41 | 0,60 | -2,34 | 1,90E-02 | 6,71E-02 |
| AT4G04020 | 1,87 | -4,16 | 1,77 | -2,35 | 1,90E-02 | 6,71E-02 |
| AT2G25430 | 4,01 | 2,92 | 1,25 | 2,34 | 1,92E-02 | 6,74E-02 |
| AT3G22910 | 2,19 | 3,64 | 1,56 | 2,34 | 1,92E-02 | 6,75E-02 |
| AT1G79000 | 2,98 | 4,48 | 1,92 | 2,33 | 1,98E-02 | 6,95E-02 |
| AT3G04455 | 2,46 | -3,75 | 1,61 | -2,33 | 2,00E-02 | 7,00E-02 |
| AT4G35750 | 1,89 | 3,75 | 1,61 | 2,33 | 2,00E-02 | 7,01E-02 |
| AT1G66880 | 6,48 | 2,21 | 0,95 | 2,32 | 2,02E-02 | 7,05E-02 |
| AT3G01715 | 19,56 | 1,76 | 0,76 | 2,32 | 2,02E-02 | 7,05E-02 |
| AT2G43570 | 2,25 | 3,79 | 1,63 | 2,32 | 2,02E-02 | 7,05E-02 |
| AT2G45685 | 1,79 | 4,11 | 1,77 | 2,32 | 2,04E-02 | 7,07E-02 |
| AT2G45680 | 1,79 | 4,11 | 1,77 | 2,32 | 2,04E-02 | 7,07E-02 |
| AT1G04430 | 7,26 | 2,21 | 0,95 | 2,32 | 2,05E-02 | 7,10E-02 |
| AT4G36630 | 4,11 | 3,79 | 1,63 | 2,32 | 2,05E-02 | 7,10E-02 |
| AT1G21750 | 2,14 | 4,34 | 1,88 | 2,31 | 2,07E-02 | 7,13E-02 |
| AT2G29920 | 2,43 | 4,41 | 1,90 | 2,31 | 2,07E-02 | 7,13E-02 |

| | | | | | | |
|---|---|---|---|---|---|---|
| AT5G18360 | 2,96 | 3,15 | 1,36 | 2,31 | 2,07E-02 | 7,13E-02 |
| AT5G26360 | 2,70 | 3,42 | 1,48 | 2,31 | 2,06E-02 | 7,13E-02 |
| AT5G06640 | 3,20 | 3,70 | 1,60 | 2,31 | 2,08E-02 | 7,14E-02 |
| AT2G40805 | 280,87 | 0,94 | 0,41 | 2,31 | 2,09E-02 | 7,16E-02 |
| AT2G14610 | 3,13 | 2,98 | 1,29 | 2,31 | 2,10E-02 | 7,19E-02 |
| AT2G07671 | 2,57 | -2,73 | 1,19 | -2,31 | 2,11E-02 | 7,22E-02 |
| AT1G47128 | 7,64 | 1,59 | 0,69 | 2,30 | 2,13E-02 | 7,27E-02 |
| AT4G04835 | 19,24 | -1,30 | 0,56 | -2,30 | 2,14E-02 | 7,29E-02 |
| AT2G35382 | 3,86 | -2,49 | 1,08 | -2,30 | 2,14E-02 | 7,30E-02 |
| AT1G79990 | 2,18 | 3,94 | 1,71 | 2,30 | 2,15E-02 | 7,30E-02 |
| AT3G07855 | 4,66 | -2,03 | 0,88 | -2,30 | 2,15E-02 | 7,30E-02 |
| AT5G24660 | 2,17 | 3,98 | 1,73 | 2,30 | 2,15E-02 | 7,30E-02 |
| AT3G13724 | 5,13 | -2,13 | 0,93 | -2,30 | 2,16E-02 | 7,31E-02 |
| AT1G69550 | 1,87 | 3,94 | 1,71 | 2,30 | 2,16E-02 | 7,31E-02 |
| AT4G14504 | 4,83 | -2,49 | 1,08 | -2,29 | 2,18E-02 | 7,34E-02 |
| AT5G35630 | 4,18 | -2,38 | 1,04 | -2,29 | 2,18E-02 | 7,34E-02 |
| AT3G08580 | 9,77 | 1,65 | 0,72 | 2,29 | 2,22E-02 | 7,47E-02 |
| AT1G49050 | 1,80 | 4,37 | 1,91 | 2,29 | 2,23E-02 | 7,47E-02 |
| AT1G31540 | 3,00 | 3,34 | 1,46 | 2,29 | 2,22E-02 | 7,47E-02 |
| AT2G44410 | 2,51 | 4,31 | 1,89 | 2,28 | 2,24E-02 | 7,49E-02 |
| AT1G13640 | 2,82 | 3,38 | 1,48 | 2,28 | 2,25E-02 | 7,51E-02 |
| AT1G18260 | 1,96 | 3,81 | 1,67 | 2,28 | 2,26E-02 | 7,53E-02 |
| AT5G52882 | 2,07 | 4,01 | 1,76 | 2,28 | 2,26E-02 | 7,53E-02 |
| AT3G21805 | 21,83 | -1,67 | 0,73 | -2,28 | 2,26E-02 | 7,54E-02 |
| AT2G31660 | 3,59 | 3,31 | 1,45 | 2,28 | 2,28E-02 | 7,56E-02 |
| AT3G27690 | 1,98 | -3,68 | 1,62 | -2,28 | 2,28E-02 | 7,56E-02 |
| AT3G13090 | 2,27 | 4,72 | 2,08 | 2,27 | 2,30E-02 | 7,64E-02 |
| AT5G51430 | 4,74 | 3,50 | 1,54 | 2,27 | 2,32E-02 | 7,68E-02 |
| AT1G05630 | 1,84 | 3,90 | 1,72 | 2,27 | 2,33E-02 | 7,69E-02 |
| AT3G06510 | 3,56 | 3,18 | 1,40 | 2,27 | 2,33E-02 | 7,69E-02 |
| AT3G08530 | 12,53 | 1,37 | 0,61 | 2,27 | 2,33E-02 | 7,70E-02 |
| AT1G60950 | 5,66 | -2,10 | 0,93 | -2,25 | 2,41E-02 | 7,93E-02 |
| AT4G00360 | 2,92 | -2,68 | 1,19 | -2,25 | 2,41E-02 | 7,93E-02 |
| AT1G05570 | 10,12 | 1,63 | 0,72 | 2,25 | 2,41E-02 | 7,93E-02 |
| AT5G67300 | 3,74 | 2,91 | 1,29 | 2,25 | 2,42E-02 | 7,95E-02 |
| AT4G28760 | 2,13 | 3,67 | 1,63 | 2,25 | 2,45E-02 | 8,01E-02 |
| AT5G58410 | 3,13 | 2,74 | 1,22 | 2,25 | 2,46E-02 | 8,03E-02 |
| AT4G06225 | 6,57 | -2,33 | 1,04 | -2,25 | 2,47E-02 | 8,07E-02 |
| AT2G34208 | 2,28 | 3,63 | 1,62 | 2,24 | 2,48E-02 | 8,08E-02 |
| AT2G25450 | 4,94 | 2,22 | 0,99 | 2,24 | 2,48E-02 | 8,08E-02 |
| AT4G08470 | 1,82 | 4,50 | 2,01 | 2,24 | 2,48E-02 | 8,08E-02 |
| AT2G36350 | 2,13 | 4,02 | 1,79 | 2,24 | 2,50E-02 | 8,12E-02 |
| AT3G03870 | 3,66 | 3,10 | 1,38 | 2,24 | 2,50E-02 | 8,12E-02 |
| AT5G05485 | 5,35 | -2,63 | 1,17 | -2,24 | 2,51E-02 | 8,13E-02 |
| AT1G18160 | 2,26 | 3,71 | 1,66 | 2,24 | 2,52E-02 | 8,16E-02 |
| AT3G61580 | 5,37 | 2,16 | 0,96 | 2,24 | 2,54E-02 | 8,19E-02 |
| AT3G44006 | 2,21 | -3,27 | 1,46 | -2,23 | 2,54E-02 | 8,20E-02 |
| AT4G24520 | 3,90 | 3,70 | 1,66 | 2,23 | 2,56E-02 | 8,25E-02 |
| AT1G50440 | 6,47 | 2,28 | 1,02 | 2,23 | 2,60E-02 | 8,35E-02 |
| AT3G54050 | 3,78 | -2,42 | 1,09 | -2,22 | 2,61E-02 | 8,39E-02 |
| AT3G55800 | 2,45 | -2,73 | 1,23 | -2,22 | 2,62E-02 | 8,39E-02 |
| AT5G58550 | 1,86 | 4,16 | 1,87 | 2,22 | 2,62E-02 | 8,39E-02 |
| AT4G34670 | 4,28 | 2,77 | 1,25 | 2,22 | 2,65E-02 | 8,49E-02 |
| AT5G20230 | 2,06 | 3,81 | 1,72 | 2,22 | 2,66E-02 | 8,49E-02 |
| AT3G44330 | 1,72 | 4,43 | 2,00 | 2,22 | 2,67E-02 | 8,51E-02 |
| AT4G33050 | 2,86 | 3,27 | 1,48 | 2,21 | 2,69E-02 | 8,55E-02 |

| | | | | | | |
|---|---|---|---|---|---|---|
| AT4G28652 | 2,08 | 4,10 | 1,85 | 2,21 | 2,70E-02 | 8,56E-02 |
| AT4G28650 | 2,08 | 4,10 | 1,85 | 2,21 | 2,70E-02 | 8,56E-02 |
| AT3G57330 | 3,24 | 2,95 | 1,33 | 2,21 | 2,70E-02 | 8,56E-02 |
| AT1G11860 | 2,12 | -3,10 | 1,41 | -2,20 | 2,75E-02 | 8,67E-02 |
| AT1G06267 | 15,22 | -1,31 | 0,60 | -2,20 | 2,75E-02 | 8,67E-02 |
| AT2G08986 | 10,75 | 1,39 | 0,63 | 2,20 | 2,75E-02 | 8,67E-02 |
| AT3G10985 | 6,51 | 2,01 | 0,91 | 2,20 | 2,75E-02 | 8,67E-02 |
| AT5G44180 | 3,26 | 3,29 | 1,49 | 2,21 | 2,74E-02 | 8,67E-02 |
| AT5G39570 | 4,04 | -2,21 | 1,00 | -2,20 | 2,76E-02 | 8,69E-02 |
| AT3G22890 | 4,47 | 2,02 | 0,92 | 2,20 | 2,78E-02 | 8,74E-02 |
| AT2G39730 | 47,53 | 0,89 | 0,41 | 2,20 | 2,81E-02 | 8,81E-02 |
| AT4G03030 | 1,74 | 3,93 | 1,79 | 2,20 | 2,81E-02 | 8,81E-02 |
| AT3G03790 | 1,73 | 3,69 | 1,69 | 2,19 | 2,85E-02 | 8,84E-02 |
| AT5G01747 | 60,67 | -0,60 | 0,28 | -2,19 | 2,84E-02 | 8,84E-02 |
| AT5G10180 | 2,74 | 3,50 | 1,60 | 2,19 | 2,84E-02 | 8,84E-02 |
| AT1G11735 | 8,53 | -1,72 | 0,78 | -2,19 | 2,84E-02 | 8,84E-02 |
| AT1G62390 | 2,10 | 4,35 | 1,99 | 2,19 | 2,85E-02 | 8,84E-02 |
| AT2G47390 | 2,13 | 3,88 | 1,77 | 2,19 | 2,83E-02 | 8,84E-02 |
| AT3G13330 | 3,62 | 2,84 | 1,30 | 2,19 | 2,85E-02 | 8,84E-02 |
| AT5G65005 | 2,56 | -2,40 | 1,10 | -2,18 | 2,89E-02 | 8,96E-02 |
| AT4G27060 | 1,71 | 3,77 | 1,73 | 2,18 | 2,90E-02 | 8,97E-02 |
| AT3G52430 | 2,34 | 3,84 | 1,76 | 2,18 | 2,92E-02 | 9,00E-02 |
| AT4G21150 | 2,92 | 3,44 | 1,58 | 2,18 | 2,91E-02 | 9,00E-02 |
| AT3G11540 | 2,22 | 3,97 | 1,82 | 2,18 | 2,93E-02 | 9,02E-02 |
| AT4G35800 | 5,60 | 2,08 | 0,96 | 2,18 | 2,94E-02 | 9,06E-02 |
| AT4G21060 | 1,98 | -2,76 | 1,27 | -2,18 | 2,95E-02 | 9,07E-02 |
| AT2G28910 | 4,89 | 2,89 | 1,33 | 2,17 | 2,96E-02 | 9,09E-02 |
| AT3G15920 | 8,54 | 1,59 | 0,73 | 2,17 | 2,97E-02 | 9,09E-02 |
| AT3G25855 | 1,70 | -4,19 | 1,93 | -2,17 | 2,97E-02 | 9,09E-02 |
| AT5G18370 | 3,23 | 3,25 | 1,50 | 2,17 | 3,03E-02 | 9,26E-02 |
| AT3G45190 | 2,88 | 3,22 | 1,49 | 2,16 | 3,04E-02 | 9,29E-02 |
| AT4G32285 | 2,40 | 3,36 | 1,55 | 2,16 | 3,07E-02 | 9,33E-02 |
| AT4G34390 | 1,80 | 3,80 | 1,76 | 2,16 | 3,06E-02 | 9,33E-02 |
| AT2G45960 | 8,71 | 1,72 | 0,80 | 2,16 | 3,07E-02 | 9,33E-02 |
| AT2G29925 | 2,27 | 4,32 | 2,00 | 2,16 | 3,07E-02 | 9,33E-02 |
| AT5G46510 | 5,48 | 1,74 | 0,81 | 2,16 | 3,09E-02 | 9,37E-02 |
| AT4G38220 | 1,96 | 3,43 | 1,59 | 2,16 | 3,10E-02 | 9,40E-02 |
| AT3G44630 | 4,39 | 2,53 | 1,17 | 2,16 | 3,11E-02 | 9,40E-02 |
| AT3G54890 | 13,73 | -1,40 | 0,65 | -2,15 | 3,14E-02 | 9,50E-02 |
| AT3G08730 | 2,48 | 3,67 | 1,71 | 2,15 | 3,17E-02 | 9,56E-02 |
| AT5G66190 | 3,89 | -2,33 | 1,09 | -2,15 | 3,18E-02 | 9,57E-02 |
| AT2G33100 | 2,25 | 3,81 | 1,78 | 2,15 | 3,19E-02 | 9,59E-02 |
| AT1G28270 | 2,06 | -3,43 | 1,60 | -2,14 | 3,21E-02 | 9,62E-02 |
| AT2G16600 | 11,35 | 1,49 | 0,69 | 2,14 | 3,21E-02 | 9,62E-02 |
| AT5G51070 | 3,71 | 2,68 | 1,25 | 2,14 | 3,21E-02 | 9,62E-02 |
| AT5G61780 | 4,25 | 2,51 | 1,17 | 2,14 | 3,21E-02 | 9,62E-02 |
| AT4G21366 | 2,02 | 3,55 | 1,66 | 2,14 | 3,23E-02 | 9,66E-02 |
| AT5G53370 | 2,35 | 3,35 | 1,56 | 2,14 | 3,24E-02 | 9,68E-02 |
| AT3G48050 | 3,14 | 2,48 | 1,16 | 2,14 | 3,25E-02 | 9,69E-02 |
| AT4G34710 | 3,32 | 3,19 | 1,49 | 2,14 | 3,25E-02 | 9,69E-02 |
| AT1G74720 | 1,98 | 3,75 | 1,76 | 2,13 | 3,28E-02 | 9,76E-02 |
| AT5G20250 | 2,34 | 3,35 | 1,58 | 2,13 | 3,33E-02 | 9,90E-02 |
| AT2G47800 | 2,70 | 2,96 | 1,39 | 2,13 | 3,34E-02 | 9,91E-02 |
| AT1G07633 | 6,19 | 2,21 | 1,04 | 2,12 | 3,38E-02 | 1,00E-01 |
| AT4G00700 | 2,06 | 3,76 | 1,77 | 2,12 | 3,40E-02 | 1,01E-01 |
| AT5G02310 | 4,95 | 2,11 | 0,99 | 2,12 | 3,41E-02 | 1,01E-01 |

| | | | | | | |
|---|---|---|---|---|---|---|
| AT1G08420 | 1,72 | 3,68 | 1,74 | 2,12 | 3,42E-02 | 1,01E-01 |
| AT5G12470 | 4,58 | 3,16 | 1,49 | 2,12 | 3,44E-02 | 1,01E-01 |
| AT1G24706 | 2,66 | 3,06 | 1,45 | 2,11 | 3,46E-02 | 1,02E-01 |
| AT3G06436 | 20,96 | 0,96 | 0,45 | 2,11 | 3,46E-02 | 1,02E-01 |
| AT5G64260 | 2,41 | 3,30 | 1,56 | 2,11 | 3,45E-02 | 1,02E-01 |
| AT4G31160 | 2,16 | 3,92 | 1,86 | 2,11 | 3,48E-02 | 1,02E-01 |
| AT1G04820 | 2,61 | -2,66 | 1,26 | -2,11 | 3,49E-02 | 1,02E-01 |
| AT5G01980 | 1,71 | 3,77 | 1,79 | 2,11 | 3,51E-02 | 1,03E-01 |
| AT3G05355 | 1,75 | -3,33 | 1,58 | -2,11 | 3,52E-02 | 1,03E-01 |
| AT4G02770 | 8,37 | -1,44 | 0,68 | -2,10 | 3,54E-02 | 1,03E-01 |
| AT1G67960 | 1,96 | 3,72 | 1,77 | 2,10 | 3,55E-02 | 1,03E-01 |
| AT5G61440 | 4,44 | -1,78 | 0,85 | -2,10 | 3,61E-02 | 1,05E-01 |
| AT1G07493 | 1,97 | -3,24 | 1,55 | -2,09 | 3,62E-02 | 1,05E-01 |
| AT2G07360 | 3,99 | 2,67 | 1,28 | 2,09 | 3,62E-02 | 1,05E-01 |
| AT3G55320 | 2,94 | 2,56 | 1,22 | 2,09 | 3,62E-02 | 1,05E-01 |
| AT5G28823 | 1,86 | 4,02 | 1,92 | 2,10 | 3,60E-02 | 1,05E-01 |
| AT5G60450 | 7,78 | -1,51 | 0,72 | -2,10 | 3,61E-02 | 1,05E-01 |
| AT3G10650 | 3,29 | 3,17 | 1,51 | 2,09 | 3,63E-02 | 1,05E-01 |
| AT2G36850 | 3,05 | 3,34 | 1,60 | 2,09 | 3,64E-02 | 1,05E-01 |
| AT1G29670 | 1,73 | -3,35 | 1,61 | -2,09 | 3,68E-02 | 1,06E-01 |
| AT5G19400 | 2,38 | 3,87 | 1,85 | 2,09 | 3,68E-02 | 1,06E-01 |
| AT3G55610 | 4,81 | 2,45 | 1,18 | 2,09 | 3,68E-02 | 1,06E-01 |
| AT4G18880 | 2,25 | 3,92 | 1,88 | 2,09 | 3,69E-02 | 1,06E-01 |
| AT5G53740 | 20,74 | -0,98 | 0,47 | -2,08 | 3,72E-02 | 1,07E-01 |
| AT5G01530 | 20,05 | -1,26 | 0,61 | -2,08 | 3,74E-02 | 1,07E-01 |
| AT5G09585 | 3,20 | 2,79 | 1,35 | 2,07 | 3,85E-02 | 1,10E-01 |
| AT1G11870 | 3,10 | 2,69 | 1,30 | 2,07 | 3,86E-02 | 1,10E-01 |
| AT1G62750 | 9,17 | 1,66 | 0,80 | 2,07 | 3,86E-02 | 1,10E-01 |
| AT4G08920 | 2,64 | 3,34 | 1,62 | 2,07 | 3,87E-02 | 1,10E-01 |
| AT1G07051 | 29,68 | -1,08 | 0,52 | -2,06 | 3,90E-02 | 1,11E-01 |
| AT2G19850 | 5,66 | -2,29 | 1,11 | -2,06 | 3,92E-02 | 1,11E-01 |
| AT3G53260 | 2,56 | 3,29 | 1,59 | 2,06 | 3,92E-02 | 1,11E-01 |
| AT1G43850 | 1,82 | 3,60 | 1,75 | 2,06 | 3,94E-02 | 1,11E-01 |
| AT5G19940 | 2,37 | -2,66 | 1,29 | -2,06 | 3,94E-02 | 1,11E-01 |
| AT1G12900 | 4,70 | -1,87 | 0,91 | -2,06 | 3,94E-02 | 1,11E-01 |
| AT2G32240 | 2,12 | 3,40 | 1,65 | 2,06 | 3,95E-02 | 1,12E-01 |
| AT1G70185 | 5,33 | 2,13 | 1,03 | 2,06 | 3,96E-02 | 1,12E-01 |
| AT1G30380 | 5,72 | -2,27 | 1,10 | -2,06 | 3,97E-02 | 1,12E-01 |
| AT5G07325 | 8,05 | 1,51 | 0,73 | 2,06 | 3,97E-02 | 1,12E-01 |
| AT1G71880 | 1,96 | 3,72 | 1,81 | 2,05 | 4,01E-02 | 1,13E-01 |
| AT3G55200 | 2,62 | 3,85 | 1,88 | 2,05 | 4,01E-02 | 1,13E-01 |
| AT3G04130 | 2,12 | 4,14 | 2,02 | 2,05 | 4,05E-02 | 1,13E-01 |
| AT3G55850 | 2,66 | 3,49 | 1,70 | 2,05 | 4,05E-02 | 1,13E-01 |
| AT2G40000 | 5,19 | 2,08 | 1,01 | 2,05 | 4,07E-02 | 1,14E-01 |
| AT1G61100 | 1,97 | 3,59 | 1,76 | 2,04 | 4,11E-02 | 1,15E-01 |
| AT3G27400 | 1,80 | -2,91 | 1,43 | -2,04 | 4,14E-02 | 1,15E-01 |
| AT4G29380 | 1,74 | 3,34 | 1,64 | 2,04 | 4,14E-02 | 1,15E-01 |
| AT1G32582 | 26,60 | -1,26 | 0,62 | -2,03 | 4,19E-02 | 1,16E-01 |
| AT3G17900 | 1,92 | 3,38 | 1,66 | 2,03 | 4,20E-02 | 1,16E-01 |
| AT4G25080 | 4,26 | 2,22 | 1,09 | 2,03 | 4,22E-02 | 1,16E-01 |
| AT5G47490 | 2,21 | 3,15 | 1,55 | 2,03 | 4,19E-02 | 1,16E-01 |
| AT1G03680 | 1,73 | -3,09 | 1,52 | -2,04 | 4,18E-02 | 1,16E-01 |
| AT1G08813 | 8,42 | -1,58 | 0,78 | -2,04 | 4,17E-02 | 1,16E-01 |
| AT2G10940 | 5,77 | -2,03 | 1,00 | -2,03 | 4,22E-02 | 1,16E-01 |
| AT2G33170 | 2,19 | 3,43 | 1,69 | 2,04 | 4,18E-02 | 1,16E-01 |
| AT2G43410 | 2,84 | 2,85 | 1,40 | 2,03 | 4,21E-02 | 1,16E-01 |

| | | | | | | |
|---|---|---|---|---|---|---|
| AT3G28550 | 2,18 | 3,81 | 1,87 | 2,04 | 4,17E-02 | 1,16E-01 |
| AT3G44480 | 4,70 | 1,93 | 0,95 | 2,04 | 4,17E-02 | 1,16E-01 |
| AT4G13918 | 2,19 | 3,31 | 1,63 | 2,03 | 4,20E-02 | 1,16E-01 |
| AT5G24810 | 2,74 | 3,03 | 1,49 | 2,03 | 4,21E-02 | 1,16E-01 |
| AT1G15940 | 2,10 | 3,22 | 1,59 | 2,03 | 4,27E-02 | 1,16E-01 |
| AT2G05520 | 18,30 | -0,82 | 0,41 | -2,03 | 4,27E-02 | 1,16E-01 |
| AT4G01050 | 4,58 | 2,53 | 1,25 | 2,03 | 4,27E-02 | 1,16E-01 |
| AT5G04590 | 4,30 | 2,35 | 1,16 | 2,03 | 4,27E-02 | 1,16E-01 |
| AT5G57710 | 4,28 | 2,05 | 1,01 | 2,03 | 4,27E-02 | 1,16E-01 |
| AT1G58290 | 5,16 | 1,92 | 0,95 | 2,03 | 4,27E-02 | 1,16E-01 |
| AT4G09650 | 3,00 | -2,91 | 1,43 | -2,02 | 4,29E-02 | 1,17E-01 |
| AT4G29900 | 4,72 | 1,99 | 0,98 | 2,02 | 4,30E-02 | 1,17E-01 |
| AT2G05710 | 3,35 | 2,77 | 1,37 | 2,02 | 4,33E-02 | 1,18E-01 |
| AT1G07793 | 2,22 | -2,56 | 1,27 | -2,02 | 4,34E-02 | 1,18E-01 |
| AT4G16980 | 2,88 | -2,31 | 1,15 | -2,02 | 4,36E-02 | 1,18E-01 |
| AT4G23915 | 12,58 | 1,28 | 0,63 | 2,02 | 4,39E-02 | 1,19E-01 |
| AT4G28755 | 2,23 | -2,84 | 1,41 | -2,01 | 4,40E-02 | 1,19E-01 |
| AT3G07160 | 3,87 | 2,46 | 1,22 | 2,01 | 4,41E-02 | 1,19E-01 |
| AT5G12120 | 2,73 | 3,18 | 1,58 | 2,01 | 4,44E-02 | 1,20E-01 |
| AT5G05750 | 1,85 | 4,13 | 2,06 | 2,01 | 4,45E-02 | 1,20E-01 |
| AT4G01810 | 2,76 | 3,18 | 1,58 | 2,01 | 4,45E-02 | 1,20E-01 |
| AT3G14385 | 15,96 | 1,12 | 0,56 | 2,01 | 4,46E-02 | 1,20E-01 |
| AT4G16983 | 2,85 | -2,31 | 1,15 | -2,00 | 4,51E-02 | 1,21E-01 |
| AT5G42530 | 4,10 | -1,73 | 0,86 | -2,00 | 4,53E-02 | 1,21E-01 |
| AT5G21100 | 6,61 | 1,98 | 0,99 | 2,00 | 4,58E-02 | 1,23E-01 |
| AT2G03250 | 2,57 | -2,69 | 1,35 | -2,00 | 4,59E-02 | 1,23E-01 |
| AT4G18670 | 7,64 | -1,40 | 0,70 | -2,00 | 4,59E-02 | 1,23E-01 |
| AT5G65750 | 4,51 | 2,37 | 1,19 | 2,00 | 4,59E-02 | 1,23E-01 |
| AT3G13100 | 2,83 | 2,97 | 1,49 | 2,00 | 4,60E-02 | 1,23E-01 |
| AT3G25010 | 4,35 | 2,80 | 1,41 | 1,99 | 4,66E-02 | 1,24E-01 |
| AT4G08175 | 7,31 | -1,57 | 0,79 | -1,99 | 4,66E-02 | 1,24E-01 |
| AT3G14010 | 2,74 | 3,36 | 1,69 | 1,99 | 4,68E-02 | 1,24E-01 |
| AT1G05240 | 2,51 | -2,53 | 1,27 | -1,98 | 4,72E-02 | 1,25E-01 |
| AT3G60415 | 1,73 | 3,55 | 1,79 | 1,99 | 4,71E-02 | 1,25E-01 |
| AT5G10960 | 3,18 | 2,65 | 1,34 | 1,98 | 4,74E-02 | 1,26E-01 |
| AT3G23125 | 239,13 | -0,90 | 0,45 | -1,98 | 4,75E-02 | 1,26E-01 |
| AT1G33560 | 2,19 | 3,70 | 1,87 | 1,98 | 4,77E-02 | 1,26E-01 |
| AT5G20730 | 2,22 | 3,13 | 1,58 | 1,98 | 4,76E-02 | 1,26E-01 |
| AT5G09515 | 9,09 | -1,39 | 0,70 | -1,98 | 4,77E-02 | 1,26E-01 |
| AT1G06220 | 3,15 | 2,75 | 1,39 | 1,98 | 4,78E-02 | 1,26E-01 |
| AT5G26830 | 2,03 | 3,08 | 1,55 | 1,98 | 4,79E-02 | 1,26E-01 |
| AT4G16900 | 2,10 | 3,15 | 1,59 | 1,98 | 4,80E-02 | 1,26E-01 |
| AT2G07981 | 2,13 | 3,90 | 1,97 | 1,97 | 4,83E-02 | 1,27E-01 |
| AT4G01010 | 2,44 | -2,62 | 1,33 | -1,97 | 4,85E-02 | 1,27E-01 |
| AT1G19680 | 2,21 | 3,76 | 1,91 | 1,96 | 4,95E-02 | 1,29E-01 |
| AT2G22440 | 2,40 | -2,40 | 1,22 | -1,96 | 4,95E-02 | 1,29E-01 |
| AT2G18790 | 2,11 | 3,15 | 1,60 | 1,96 | 4,96E-02 | 1,30E-01 |
| AT1G67740 | 3,89 | -1,70 | 0,87 | -1,96 | 4,98E-02 | 1,30E-01 |
| AT1G28420 | 2,78 | 2,73 | 1,39 | 1,96 | 5,00E-02 | 1,30E-01 |
| AT3G51050 | 3,03 | 3,21 | 1,64 | 1,96 | 5,02E-02 | 1,30E-01 |
| AT5G23380 | 2,25 | 3,45 | 1,76 | 1,96 | 5,03E-02 | 1,30E-01 |
| AT5G17760 | 1,99 | 3,73 | 1,91 | 1,96 | 5,03E-02 | 1,30E-01 |
| AT5G67360 | 6,15 | 2,20 | 1,12 | 1,96 | 5,03E-02 | 1,30E-01 |
| AT5G46420 | 2,51 | 2,82 | 1,44 | 1,96 | 5,05E-02 | 1,31E-01 |
| AT3G19380 | 2,79 | 3,13 | 1,60 | 1,95 | 5,07E-02 | 1,31E-01 |
| AT5G42080 | 4,02 | 2,36 | 1,21 | 1,95 | 5,08E-02 | 1,31E-01 |

| | | | | | | |
|---|---|---|---|---|---|---|
| AT5G25280 | 3,55 | 2,86 | 1,46 | 1,95 | 5,09E-02 | 1,31E-01 |
| AT3G51860 | 2,20 | 3,14 | 1,61 | 1,95 | 5,12E-02 | 1,32E-01 |
| AT1G65720 | 3,55 | 1,93 | 0,99 | 1,95 | 5,15E-02 | 1,33E-01 |
| AT1G52110 | 2,50 | -2,47 | 1,27 | -1,95 | 5,16E-02 | 1,33E-01 |
| AT3G57480 | 3,07 | -2,39 | 1,23 | -1,95 | 5,15E-02 | 1,33E-01 |
| AT1G12013 | 2,08 | -2,83 | 1,45 | -1,95 | 5,18E-02 | 1,33E-01 |
| AT4G29780 | 2,35 | 3,64 | 1,87 | 1,95 | 5,17E-02 | 1,33E-01 |
| AT1G25440 | 6,25 | 2,17 | 1,12 | 1,94 | 5,19E-02 | 1,33E-01 |
| AT3G10745 | 164,34 | -0,67 | 0,35 | -1,94 | 5,20E-02 | 1,33E-01 |
| AT2G25320 | 3,21 | 3,01 | 1,55 | 1,94 | 5,22E-02 | 1,33E-01 |
| AT1G07000 | 3,04 | 2,66 | 1,37 | 1,94 | 5,28E-02 | 1,35E-01 |
| AT4G08850 | 2,82 | 2,95 | 1,52 | 1,94 | 5,28E-02 | 1,35E-01 |
| AT1G54115 | 2,18 | 3,26 | 1,68 | 1,94 | 5,30E-02 | 1,35E-01 |
| AT5G11740 | 4,10 | 2,03 | 1,05 | 1,93 | 5,36E-02 | 1,36E-01 |
| AT1G73805 | 1,81 | 3,48 | 1,80 | 1,93 | 5,36E-02 | 1,36E-01 |
| AT3G17240 | 1,76 | 3,43 | 1,78 | 1,93 | 5,37E-02 | 1,36E-01 |
| AT4G08630 | 1,97 | 3,43 | 1,78 | 1,93 | 5,38E-02 | 1,36E-01 |
| AT2G01600 | 1,95 | 3,29 | 1,71 | 1,93 | 5,39E-02 | 1,37E-01 |
| AT5G54310 | 4,38 | 2,42 | 1,26 | 1,93 | 5,40E-02 | 1,37E-01 |
| AT4G13920 | 2,03 | 3,30 | 1,72 | 1,92 | 5,43E-02 | 1,37E-01 |
| AT5G51630 | 1,97 | 3,18 | 1,65 | 1,92 | 5,43E-02 | 1,37E-01 |
| AT3G56150 | 2,36 | 3,15 | 1,64 | 1,92 | 5,45E-02 | 1,38E-01 |
| AT3G63070 | 1,86 | 3,78 | 1,97 | 1,92 | 5,53E-02 | 1,39E-01 |
| AT1G14320 | 3,20 | 2,55 | 1,33 | 1,91 | 5,57E-02 | 1,40E-01 |
| AT3G63460 | 4,10 | 2,44 | 1,28 | 1,91 | 5,56E-02 | 1,40E-01 |
| AT5G18525 | 1,70 | 3,47 | 1,81 | 1,91 | 5,57E-02 | 1,40E-01 |
| AT3G03050 | 2,11 | 3,42 | 1,79 | 1,91 | 5,60E-02 | 1,40E-01 |
| AT5G06630 | 1,85 | 3,44 | 1,80 | 1,91 | 5,61E-02 | 1,41E-01 |
| AT1G27340 | 2,10 | 3,27 | 1,71 | 1,91 | 5,64E-02 | 1,41E-01 |
| AT1G02890 | 1,79 | 3,13 | 1,64 | 1,91 | 5,64E-02 | 1,41E-01 |
| AT1G72440 | 1,79 | 3,17 | 1,66 | 1,91 | 5,63E-02 | 1,41E-01 |
| AT5G18490 | 1,80 | 3,48 | 1,82 | 1,91 | 5,64E-02 | 1,41E-01 |
| AT3G29360 | 2,83 | 2,55 | 1,34 | 1,91 | 5,66E-02 | 1,41E-01 |
| AT3G47910 | 2,73 | 2,88 | 1,51 | 1,90 | 5,69E-02 | 1,42E-01 |
| AT3G19420 | 2,24 | 3,33 | 1,75 | 1,90 | 5,70E-02 | 1,42E-01 |
| AT4G24680 | 3,35 | 2,28 | 1,20 | 1,90 | 5,80E-02 | 1,44E-01 |
| AT2G35384 | 13,85 | -1,34 | 0,71 | -1,90 | 5,80E-02 | 1,44E-01 |
| AT2G17200 | 1,89 | 3,11 | 1,64 | 1,89 | 5,81E-02 | 1,44E-01 |
| AT4G34830 | 2,44 | 2,74 | 1,44 | 1,89 | 5,82E-02 | 1,44E-01 |
| AT4G00231 | 1,99 | 2,68 | 1,42 | 1,89 | 5,83E-02 | 1,44E-01 |
| AT2G14560 | 27,90 | 1,01 | 0,53 | 1,89 | 5,86E-02 | 1,45E-01 |
| AT4G21280 | 3,20 | -2,26 | 1,19 | -1,89 | 5,86E-02 | 1,45E-01 |
| AT1G76630 | 2,14 | 3,12 | 1,65 | 1,89 | 5,88E-02 | 1,45E-01 |
| AT1G31163 | 6,82 | -1,31 | 0,69 | -1,89 | 5,88E-02 | 1,45E-01 |
| AT5G47480 | 1,95 | 2,91 | 1,54 | 1,88 | 5,97E-02 | 1,47E-01 |
| AT1G11700 | 23,46 | 0,87 | 0,46 | 1,88 | 5,99E-02 | 1,47E-01 |
| AT1G05960 | 4,07 | 2,69 | 1,43 | 1,88 | 5,99E-02 | 1,47E-01 |
| AT1G62180 | 2,66 | 2,56 | 1,36 | 1,88 | 6,00E-02 | 1,47E-01 |
| AT5G40910 | 2,64 | 2,50 | 1,33 | 1,88 | 6,03E-02 | 1,48E-01 |
| AT1G22530 | 2,83 | -2,78 | 1,48 | -1,88 | 6,04E-02 | 1,48E-01 |
| AT4G03295 | 3,55 | -1,88 | 1,00 | -1,87 | 6,08E-02 | 1,49E-01 |
| AT5G45500 | 2,29 | 3,54 | 1,89 | 1,87 | 6,08E-02 | 1,49E-01 |
| AT2G27150 | 1,72 | 3,88 | 2,07 | 1,87 | 6,09E-02 | 1,49E-01 |
| AT1G33680 | 2,02 | 3,49 | 1,87 | 1,87 | 6,11E-02 | 1,49E-01 |
| AT3G02170 | 3,18 | 2,87 | 1,54 | 1,87 | 6,14E-02 | 1,50E-01 |
| AT4G23140 | 2,18 | 2,80 | 1,50 | 1,87 | 6,19E-02 | 1,51E-01 |

| | | | | | | |
|---|---|---|---|---|---|---|
| AT2G25520 | 1,73 | 3,77 | 2,02 | 1,86 | 6,22E-02 | 1,51E-01 |
| AT5G13010 | 4,55 | 2,24 | 1,20 | 1,86 | 6,23E-02 | 1,51E-01 |
| AT4G28570 | 3,15 | -2,00 | 1,08 | -1,86 | 6,24E-02 | 1,51E-01 |
| AT4G39850 | 2,19 | 3,19 | 1,71 | 1,86 | 6,26E-02 | 1,52E-01 |
| AT2G04280 | 2,22 | 3,00 | 1,61 | 1,86 | 6,27E-02 | 1,52E-01 |
| AT4G20830 | 3,02 | 2,25 | 1,21 | 1,86 | 6,33E-02 | 1,53E-01 |
| AT1G63630 | 3,43 | -2,05 | 1,11 | -1,85 | 6,37E-02 | 1,54E-01 |
| AT1G63615 | 3,43 | -2,05 | 1,11 | -1,85 | 6,37E-02 | 1,54E-01 |
| AT5G02480 | 2,67 | 2,66 | 1,43 | 1,85 | 6,38E-02 | 1,54E-01 |
| AT3G15730 | 15,00 | 1,06 | 0,58 | 1,85 | 6,47E-02 | 1,55E-01 |
| AT4G12570 | 2,25 | 3,09 | 1,67 | 1,85 | 6,47E-02 | 1,55E-01 |
| AT2G41410 | 4,84 | 2,09 | 1,13 | 1,85 | 6,47E-02 | 1,55E-01 |
| AT2G28550 | 2,55 | -2,13 | 1,15 | -1,84 | 6,51E-02 | 1,56E-01 |
| AT2G09690 | 14,22 | -1,45 | 0,79 | -1,84 | 6,54E-02 | 1,57E-01 |
| AT4G21710 | 2,69 | 3,15 | 1,71 | 1,84 | 6,53E-02 | 1,57E-01 |
| AT1G61690 | 1,91 | 3,15 | 1,71 | 1,84 | 6,59E-02 | 1,57E-01 |
| AT5G46270 | 2,21 | 3,11 | 1,69 | 1,84 | 6,59E-02 | 1,57E-01 |
| AT1G23900 | 1,85 | 3,10 | 1,69 | 1,84 | 6,61E-02 | 1,57E-01 |
| AT1G64430 | 2,85 | 2,92 | 1,59 | 1,84 | 6,61E-02 | 1,57E-01 |
| AT2G03330 | 8,86 | -1,29 | 0,70 | -1,84 | 6,62E-02 | 1,57E-01 |
| AT1G11640 | 3,15 | 2,83 | 1,54 | 1,84 | 6,62E-02 | 1,57E-01 |
| AT4G18700 | 2,69 | 2,17 | 1,18 | 1,84 | 6,63E-02 | 1,57E-01 |
| AT1G63360 | 4,56 | -1,67 | 0,91 | -1,84 | 6,63E-02 | 1,58E-01 |
| AT3G14650 | 2,33 | 2,88 | 1,57 | 1,83 | 6,65E-02 | 1,58E-01 |
| AT2G07698 | 9,74 | -1,20 | 0,65 | -1,83 | 6,67E-02 | 1,58E-01 |
| AT2G07629 | 5,41 | -1,77 | 0,96 | -1,83 | 6,68E-02 | 1,58E-01 |
| AT2G20780 | 1,93 | 3,13 | 1,71 | 1,83 | 6,70E-02 | 1,58E-01 |
| AT3G06345 | 8,24 | 1,49 | 0,81 | 1,83 | 6,78E-02 | 1,60E-01 |
| AT1G74456 | 11,08 | -1,25 | 0,69 | -1,83 | 6,78E-02 | 1,60E-01 |
| AT1G50200 | 4,92 | 1,96 | 1,07 | 1,82 | 6,80E-02 | 1,60E-01 |
| AT2G25900 | 2,31 | -2,40 | 1,31 | -1,82 | 6,83E-02 | 1,61E-01 |
| AT1G04625 | 1,86 | 3,56 | 1,96 | 1,82 | 6,89E-02 | 1,62E-01 |
| AT4G38510 | 2,20 | 2,92 | 1,61 | 1,82 | 6,89E-02 | 1,62E-01 |
| AT5G08925 | 2,86 | 3,33 | 1,83 | 1,82 | 6,90E-02 | 1,62E-01 |
| AT5G05200 | 1,99 | 3,48 | 1,92 | 1,82 | 6,92E-02 | 1,62E-01 |
| AT2G36530 | 5,68 | 2,00 | 1,10 | 1,81 | 6,95E-02 | 1,63E-01 |
| AT2G44550 | 3,60 | -1,99 | 1,09 | -1,81 | 6,95E-02 | 1,63E-01 |
| AT5G14570 | 2,87 | 2,29 | 1,26 | 1,81 | 6,97E-02 | 1,63E-01 |
| AT2G04755 | 2,18 | 2,93 | 1,62 | 1,81 | 6,98E-02 | 1,63E-01 |
| AT3G17510 | 2,73 | 3,11 | 1,72 | 1,81 | 6,99E-02 | 1,63E-01 |
| AT2G46192 | 21,12 | -1,12 | 0,62 | -1,81 | 7,02E-02 | 1,63E-01 |
| AT4G16660 | 2,21 | 2,70 | 1,49 | 1,81 | 7,01E-02 | 1,63E-01 |
| AT5G19690 | 1,74 | 2,74 | 1,52 | 1,81 | 7,02E-02 | 1,63E-01 |
| AT2G47400 | 11,67 | 1,11 | 0,62 | 1,81 | 7,07E-02 | 1,64E-01 |
| AT3G04250 | 2,26 | 3,40 | 1,88 | 1,81 | 7,06E-02 | 1,64E-01 |
| AT3G24542 | 2,63 | -2,21 | 1,22 | -1,81 | 7,07E-02 | 1,64E-01 |
| AT4G30210 | 1,86 | 3,48 | 1,93 | 1,80 | 7,12E-02 | 1,65E-01 |
| AT3G45830 | 1,79 | 3,05 | 1,69 | 1,80 | 7,14E-02 | 1,65E-01 |
| AT1G24460 | 4,84 | 1,96 | 1,09 | 1,80 | 7,16E-02 | 1,65E-01 |
| AT3G07515 | 2,88 | 2,91 | 1,61 | 1,80 | 7,16E-02 | 1,65E-01 |
| AT5G11700 | 1,99 | 2,96 | 1,64 | 1,80 | 7,17E-02 | 1,65E-01 |
| AT5G11490 | 1,97 | 2,64 | 1,47 | 1,80 | 7,19E-02 | 1,66E-01 |
| AT1G73170 | 1,76 | 3,48 | 1,94 | 1,80 | 7,18E-02 | 1,66E-01 |
| AT4G26730 | 2,38 | 3,19 | 1,77 | 1,80 | 7,22E-02 | 1,66E-01 |
| AT1G29350 | 2,20 | 2,86 | 1,59 | 1,80 | 7,26E-02 | 1,66E-01 |
| AT4G33300 | 3,93 | 2,29 | 1,28 | 1,80 | 7,26E-02 | 1,66E-01 |

| | | | | | | |
|---|---|---|---|---|---|---|
| AT5G43310 | 2,37 | 2,60 | 1,45 | 1,80 | 7,25E-02 | 1,66E-01 |
| AT2G25730 | 2,88 | 2,38 | 1,33 | 1,79 | 7,30E-02 | 1,67E-01 |
| AT1G20015 | 7,84 | -1,42 | 0,79 | -1,79 | 7,32E-02 | 1,68E-01 |
| AT5G14270 | 2,74 | 2,73 | 1,52 | 1,79 | 7,33E-02 | 1,68E-01 |
| AT3G50950 | 4,46 | 1,80 | 1,01 | 1,79 | 7,39E-02 | 1,68E-01 |
| AT1G21130 | 2,17 | -2,81 | 1,57 | -1,79 | 7,37E-02 | 1,68E-01 |
| AT5G24710 | 4,46 | 2,45 | 1,37 | 1,79 | 7,39E-02 | 1,68E-01 |
| AT2G01570 | 3,41 | 2,33 | 1,31 | 1,79 | 7,41E-02 | 1,69E-01 |
| AT5G49440 | 5,06 | -1,67 | 0,94 | -1,78 | 7,44E-02 | 1,69E-01 |
| AT4G36760 | 3,13 | 2,20 | 1,23 | 1,78 | 7,49E-02 | 1,70E-01 |
| AT5G35735 | 2,19 | 2,74 | 1,54 | 1,78 | 7,49E-02 | 1,70E-01 |
| AT3G08165 | 3,95 | -2,13 | 1,20 | -1,78 | 7,52E-02 | 1,71E-01 |
| AT3G08940 | 8,84 | -1,43 | 0,80 | -1,78 | 7,54E-02 | 1,71E-01 |
| AT3G20770 | 2,68 | 2,45 | 1,38 | 1,78 | 7,54E-02 | 1,71E-01 |
| AT3G01770 | 1,96 | 3,02 | 1,70 | 1,78 | 7,55E-02 | 1,71E-01 |
| AT1G76490 | 3,44 | -1,99 | 1,12 | -1,78 | 7,57E-02 | 1,71E-01 |
| AT5G45110 | 1,72 | 2,91 | 1,64 | 1,78 | 7,57E-02 | 1,71E-01 |
| AT1G56340 | 4,49 | 1,74 | 0,98 | 1,78 | 7,58E-02 | 1,71E-01 |
| AT4G17340 | 2,05 | 3,00 | 1,69 | 1,77 | 7,61E-02 | 1,72E-01 |
| AT1G20100 | 1,77 | 3,11 | 1,76 | 1,77 | 7,63E-02 | 1,72E-01 |
| AT5G53170 | 2,82 | 2,40 | 1,35 | 1,77 | 7,63E-02 | 1,72E-01 |
| AT4G26110 | 2,20 | 2,85 | 1,61 | 1,77 | 7,64E-02 | 1,72E-01 |
| AT4G34150 | 1,90 | 2,87 | 1,62 | 1,77 | 7,71E-02 | 1,73E-01 |
| AT2G24103 | 3,17 | 2,71 | 1,53 | 1,77 | 7,76E-02 | 1,74E-01 |
| AT4G20170 | 3,89 | 2,32 | 1,32 | 1,76 | 7,80E-02 | 1,75E-01 |
| AT5G07290 | 1,99 | 3,83 | 2,18 | 1,76 | 7,85E-02 | 1,76E-01 |
| AT3G56410 | 2,56 | 2,37 | 1,35 | 1,76 | 7,86E-02 | 1,76E-01 |
| AT4G32250 | 2,07 | 3,01 | 1,71 | 1,76 | 7,86E-02 | 1,76E-01 |
| AT5G10540 | 2,27 | 2,85 | 1,62 | 1,76 | 7,88E-02 | 1,76E-01 |
| AT3G28956 | 2,37 | -2,12 | 1,21 | -1,76 | 7,87E-02 | 1,76E-01 |
| AT5G04315 | 6,08 | 1,47 | 0,84 | 1,76 | 7,92E-02 | 1,76E-01 |
| AT5G04325 | 6,08 | 1,47 | 0,84 | 1,76 | 7,92E-02 | 1,76E-01 |
| AT2G38040 | 4,80 | 1,98 | 1,14 | 1,75 | 8,05E-02 | 1,78E-01 |
| AT3G05420 | 2,70 | 2,65 | 1,52 | 1,75 | 8,05E-02 | 1,78E-01 |
| AT3G44860 | 2,37 | 2,48 | 1,42 | 1,75 | 8,03E-02 | 1,78E-01 |
| AT3G57300 | 2,34 | 2,68 | 1,53 | 1,75 | 8,05E-02 | 1,78E-01 |
| AT3G22790 | 2,45 | 2,71 | 1,55 | 1,75 | 8,04E-02 | 1,78E-01 |
| AT1G09240 | 1,84 | 3,89 | 2,23 | 1,74 | 8,18E-02 | 1,79E-01 |
| AT3G04740 | 2,26 | 2,73 | 1,56 | 1,74 | 8,13E-02 | 1,79E-01 |
| AT3G51375 | 135,65 | 0,66 | 0,38 | 1,74 | 8,17E-02 | 1,79E-01 |
| AT3G56408 | 2,59 | 2,36 | 1,36 | 1,74 | 8,17E-02 | 1,79E-01 |
| AT5G08690 | 2,64 | 2,32 | 1,33 | 1,75 | 8,08E-02 | 1,79E-01 |
| AT1G35215 | 4,81 | 1,71 | 0,98 | 1,74 | 8,18E-02 | 1,79E-01 |
| AT1G61140 | 2,12 | 2,62 | 1,50 | 1,74 | 8,14E-02 | 1,79E-01 |
| AT1G77180 | 2,45 | 2,49 | 1,43 | 1,74 | 8,17E-02 | 1,79E-01 |
| AT2G46500 | 2,79 | 2,88 | 1,65 | 1,74 | 8,13E-02 | 1,79E-01 |
| AT3G04340 | 3,02 | 2,15 | 1,23 | 1,74 | 8,14E-02 | 1,79E-01 |
| AT4G24413 | 2,10 | -2,22 | 1,28 | -1,74 | 8,18E-02 | 1,79E-01 |
| AT5G43470 | 3,38 | 2,29 | 1,31 | 1,74 | 8,16E-02 | 1,79E-01 |
| AT5G48380 | 2,92 | 2,10 | 1,20 | 1,74 | 8,15E-02 | 1,79E-01 |
| AT5G66030 | 2,48 | 2,81 | 1,61 | 1,74 | 8,15E-02 | 1,79E-01 |
| AT3G63490 | 1,82 | 3,13 | 1,80 | 1,73 | 8,31E-02 | 1,81E-01 |
| AT1G80180 | 13,61 | 1,14 | 0,66 | 1,73 | 8,33E-02 | 1,82E-01 |
| AT2G22500 | 4,35 | 2,28 | 1,32 | 1,73 | 8,41E-02 | 1,83E-01 |
| AT3G04930 | 1,93 | 2,91 | 1,69 | 1,73 | 8,45E-02 | 1,84E-01 |
| AT4G31170 | 2,81 | 3,02 | 1,75 | 1,73 | 8,44E-02 | 1,84E-01 |

| | | | | | | |
|---|---|---|---|---|---|---|
| AT5G24280 | 2,00 | 5,16 | 3,00 | 1,72 | 8,48E-02 | 1,84E-01 |
| AT1G18880 | 1,72 | 2,79 | 1,62 | 1,72 | 8,50E-02 | 1,85E-01 |
| AT5G01960 | 1,82 | 3,46 | 2,01 | 1,72 | 8,52E-02 | 1,85E-01 |
| AT2G16040 | 2,68 | 2,78 | 1,62 | 1,72 | 8,53E-02 | 1,85E-01 |
| AT3G63375 | 13,10 | -0,89 | 0,52 | -1,72 | 8,59E-02 | 1,86E-01 |
| AT4G03420 | 2,29 | 2,87 | 1,67 | 1,72 | 8,60E-02 | 1,86E-01 |
| AT3G18217 | 147,37 | 0,42 | 0,24 | 1,72 | 8,58E-02 | 1,86E-01 |
| AT5G35210 | 2,88 | 2,44 | 1,42 | 1,72 | 8,62E-02 | 1,86E-01 |
| AT1G64340 | 2,24 | 3,41 | 2,00 | 1,71 | 8,75E-02 | 1,89E-01 |
| AT5G58160 | 1,77 | 4,17 | 2,44 | 1,71 | 8,76E-02 | 1,89E-01 |
| AT5G56030 | 5,79 | 1,38 | 0,81 | 1,70 | 8,87E-02 | 1,91E-01 |
| AT4G25520 | 4,54 | -1,53 | 0,90 | -1,70 | 8,87E-02 | 1,91E-01 |
| AT5G16640 | 4,15 | -1,72 | 1,01 | -1,70 | 8,95E-02 | 1,92E-01 |
| AT3G09390 | 6,61 | 1,48 | 0,87 | 1,70 | 8,96E-02 | 1,92E-01 |
| AT1G44575 | 5,52 | -1,32 | 0,78 | -1,70 | 8,99E-02 | 1,93E-01 |
| AT3G13460 | 10,31 | 1,10 | 0,65 | 1,69 | 9,05E-02 | 1,94E-01 |
| AT5G46470 | 3,68 | 1,96 | 1,16 | 1,69 | 9,10E-02 | 1,95E-01 |
| AT5G08550 | 1,83 | 2,71 | 1,60 | 1,69 | 9,10E-02 | 1,95E-01 |
| AT4G15180 | 1,84 | 2,77 | 1,64 | 1,69 | 9,13E-02 | 1,95E-01 |
| AT5G27660 | 6,15 | 1,44 | 0,85 | 1,69 | 9,14E-02 | 1,95E-01 |
| AT4G00060 | 1,77 | 2,72 | 1,61 | 1,69 | 9,16E-02 | 1,95E-01 |
| AT2G17840 | 2,90 | -2,23 | 1,32 | -1,69 | 9,18E-02 | 1,96E-01 |
| AT4G14400 | 8,78 | -1,50 | 0,89 | -1,68 | 9,20E-02 | 1,96E-01 |
| AT5G04550 | 2,33 | 2,64 | 1,57 | 1,68 | 9,24E-02 | 1,96E-01 |
| AT5G67030 | 1,94 | 2,69 | 1,60 | 1,68 | 9,24E-02 | 1,96E-01 |
| AT4G37640 | 1,71 | 2,81 | 1,67 | 1,68 | 9,28E-02 | 1,97E-01 |
| AT3G44720 | 3,49 | 2,40 | 1,43 | 1,68 | 9,34E-02 | 1,98E-01 |
| AT3G55512 | 2,56 | 3,15 | 1,88 | 1,68 | 9,37E-02 | 1,98E-01 |
| AT2G33340 | 1,98 | 2,92 | 1,75 | 1,67 | 9,42E-02 | 1,99E-01 |
| AT2G44120 | 4,00 | 2,27 | 1,35 | 1,67 | 9,41E-02 | 1,99E-01 |
| AT1G15010 | 2,66 | 2,36 | 1,41 | 1,67 | 9,43E-02 | 1,99E-01 |
| AT3G56680 | 2,66 | 2,37 | 1,42 | 1,67 | 9,48E-02 | 2,00E-01 |
| AT5G11560 | 3,42 | 1,71 | 1,02 | 1,67 | 9,47E-02 | 2,00E-01 |
| AT2G07645 | 4,48 | -1,51 | 0,91 | -1,67 | 9,49E-02 | 2,00E-01 |
| AT3G01310 | 2,04 | 3,22 | 1,93 | 1,67 | 9,54E-02 | 2,01E-01 |
| AT2G26250 | 1,96 | -2,23 | 1,34 | -1,67 | 9,55E-02 | 2,01E-01 |
| AT4G36648 | 2,47 | 2,74 | 1,64 | 1,67 | 9,56E-02 | 2,01E-01 |
| AT3G58640 | 2,56 | 2,54 | 1,53 | 1,67 | 9,59E-02 | 2,01E-01 |
| AT3G22886 | 587,34 | -0,42 | 0,25 | -1,66 | 9,64E-02 | 2,02E-01 |
| AT1G69797 | 12,98 | 1,08 | 0,65 | 1,66 | 9,65E-02 | 2,02E-01 |
| AT2G04965 | 3,81 | -1,56 | 0,94 | -1,66 | 9,66E-02 | 2,02E-01 |
| AT3G29796 | 2,77 | 2,46 | 1,48 | 1,66 | 9,73E-02 | 2,03E-01 |
| AT3G54730 | 3,74 | -1,60 | 0,96 | -1,66 | 9,73E-02 | 2,03E-01 |
| AT4G36040 | 3,49 | 2,11 | 1,27 | 1,66 | 9,73E-02 | 2,03E-01 |
| AT2G46430 | 2,08 | 2,67 | 1,61 | 1,66 | 9,74E-02 | 2,03E-01 |
| AT1G26880 | 2,18 | -2,54 | 1,54 | -1,66 | 9,75E-02 | 2,03E-01 |
| AT2G26500 | 2,52 | 2,37 | 1,43 | 1,66 | 9,77E-02 | 2,03E-01 |
| AT2G35747 | 15,41 | -1,09 | 0,66 | -1,66 | 9,76E-02 | 2,03E-01 |
| AT3G57660 | 1,78 | 3,03 | 1,83 | 1,66 | 9,79E-02 | 2,03E-01 |
| AT4G20835 | 2,66 | 2,06 | 1,24 | 1,66 | 9,79E-02 | 2,03E-01 |
| AT2G47410 | 3,41 | 2,37 | 1,44 | 1,65 | 9,91E-02 | 2,06E-01 |
| AT5G04305 | 5,44 | 1,41 | 0,85 | 1,65 | 9,92E-02 | 2,06E-01 |
| AT1G05207 | 1,81 | -2,23 | 1,36 | -1,65 | 9,96E-02 | 2,06E-01 |
| AT1G71220 | 2,96 | 2,32 | 1,41 | 1,65 | 9,94E-02 | 2,06E-01 |
| AT1G72645 | 2,19 | -2,35 | 1,42 | -1,65 | 9,95E-02 | 2,06E-01 |
| AT4G08125 | 3,11 | -1,68 | 1,02 | -1,65 | 9,95E-02 | 2,06E-01 |

| | | | | | | |
|---|---|---|---|---|---|---|
| AT3G26450 | 2,27 | -2,38 | 1,45 | -1,64 | 1,00E-01 | 2,07E-01 |
| AT5G04415 | 4,71 | 1,57 | 0,96 | 1,64 | 1,00E-01 | 2,07E-01 |
| AT2G45180 | 2,18 | -2,21 | 1,35 | -1,64 | 1,01E-01 | 2,07E-01 |
| AT4G03550 | 3,00 | 2,37 | 1,45 | 1,64 | 1,01E-01 | 2,08E-01 |
| AT3G28899 | 2,18 | -2,47 | 1,51 | -1,64 | 1,01E-01 | 2,08E-01 |
| AT3G54580 | 1,94 | 2,87 | 1,76 | 1,63 | 1,02E-01 | 2,10E-01 |
| AT5G42390 | 8,64 | 1,31 | 0,80 | 1,63 | 1,02E-01 | 2,10E-01 |
| AT3G04240 | 3,51 | 1,96 | 1,20 | 1,63 | 1,02E-01 | 2,10E-01 |
| AT4G26090 | 1,90 | 2,59 | 1,59 | 1,63 | 1,03E-01 | 2,11E-01 |
| AT1G29470 | 3,69 | 1,94 | 1,19 | 1,63 | 1,03E-01 | 2,11E-01 |
| AT2G39800 | 6,12 | 1,50 | 0,92 | 1,63 | 1,03E-01 | 2,11E-01 |
| AT3G06650 | 3,17 | 2,57 | 1,58 | 1,63 | 1,03E-01 | 2,11E-01 |
| AT5G06530 | 2,30 | 2,22 | 1,36 | 1,63 | 1,03E-01 | 2,11E-01 |
| AT3G13440 | 3,42 | 2,00 | 1,23 | 1,63 | 1,04E-01 | 2,11E-01 |
| AT3G26440 | 2,47 | -2,39 | 1,47 | -1,63 | 1,04E-01 | 2,11E-01 |
| AT1G14580 | 3,99 | -1,83 | 1,13 | -1,63 | 1,04E-01 | 2,12E-01 |
| AT3G20950 | 1,75 | 2,77 | 1,70 | 1,62 | 1,04E-01 | 2,12E-01 |
| AT4G19520 | 3,67 | 1,58 | 0,98 | 1,62 | 1,06E-01 | 2,15E-01 |
| AT1G23720 | 2,26 | 3,30 | 2,04 | 1,62 | 1,06E-01 | 2,15E-01 |
| AT2G04435 | 3,56 | 2,04 | 1,26 | 1,62 | 1,06E-01 | 2,15E-01 |
| AT4G16340 | 3,35 | 2,14 | 1,33 | 1,62 | 1,06E-01 | 2,15E-01 |
| AT2G39770 | 2,97 | 2,24 | 1,39 | 1,61 | 1,06E-01 | 2,15E-01 |
| AT5G60410 | 2,90 | 2,04 | 1,26 | 1,61 | 1,06E-01 | 2,15E-01 |
| AT5G46760 | 2,52 | 2,36 | 1,46 | 1,61 | 1,07E-01 | 2,16E-01 |
| AT3G48115 | 1,91 | 2,30 | 1,43 | 1,61 | 1,07E-01 | 2,17E-01 |
| AT3G09200 | 3,98 | 1,90 | 1,18 | 1,61 | 1,08E-01 | 2,18E-01 |
| AT3G01290 | 2,12 | 2,56 | 1,60 | 1,61 | 1,08E-01 | 2,18E-01 |
| AT3G54590 | 2,31 | 2,87 | 1,79 | 1,60 | 1,09E-01 | 2,19E-01 |
| AT2G31400 | 2,56 | 2,34 | 1,46 | 1,60 | 1,09E-01 | 2,20E-01 |
| AT1G53310 | 3,45 | 2,09 | 1,31 | 1,60 | 1,10E-01 | 2,21E-01 |
| AT1G55540 | 2,72 | 2,24 | 1,40 | 1,60 | 1,10E-01 | 2,21E-01 |
| AT5G42020 | 4,47 | 1,70 | 1,06 | 1,60 | 1,10E-01 | 2,21E-01 |
| AT2G36480 | 1,71 | 2,92 | 1,83 | 1,60 | 1,11E-01 | 2,22E-01 |
| AT5G26742 | 9,59 | 1,24 | 0,78 | 1,59 | 1,11E-01 | 2,23E-01 |
| AT5G51300 | 1,97 | 3,56 | 2,24 | 1,59 | 1,12E-01 | 2,23E-01 |
| AT3G54920 | 2,20 | 2,46 | 1,55 | 1,59 | 1,12E-01 | 2,24E-01 |
| AT1G07243 | 1,82 | 2,62 | 1,65 | 1,59 | 1,12E-01 | 2,24E-01 |
| AT1G43170 | 3,66 | 1,61 | 1,02 | 1,59 | 1,13E-01 | 2,25E-01 |
| AT5G08530 | 3,76 | -1,78 | 1,12 | -1,59 | 1,13E-01 | 2,25E-01 |
| AT5G15020 | 2,14 | 2,37 | 1,49 | 1,58 | 1,13E-01 | 2,26E-01 |
| AT1G44100 | 2,02 | 2,59 | 1,64 | 1,58 | 1,14E-01 | 2,27E-01 |
| AT5G38990 | 2,97 | 1,96 | 1,24 | 1,58 | 1,14E-01 | 2,27E-01 |
| AT3G01835 | 72,10 | -0,48 | 0,31 | -1,58 | 1,14E-01 | 2,27E-01 |
| AT3G24480 | 4,54 | 2,03 | 1,28 | 1,58 | 1,14E-01 | 2,27E-01 |
| AT3G56825 | 56,81 | 0,57 | 0,36 | 1,58 | 1,14E-01 | 2,27E-01 |
| AT5G52170 | 2,49 | 2,45 | 1,55 | 1,58 | 1,15E-01 | 2,28E-01 |
| AT5G38410 | 6,26 | -1,51 | 0,96 | -1,58 | 1,15E-01 | 2,28E-01 |
| AT2G42490 | 2,30 | 2,53 | 1,61 | 1,57 | 1,15E-01 | 2,29E-01 |
| AT4G38550 | 2,97 | 2,24 | 1,43 | 1,57 | 1,16E-01 | 2,29E-01 |
| AT4G38552 | 2,97 | 2,24 | 1,43 | 1,57 | 1,16E-01 | 2,29E-01 |
| AT5G09065 | 4,44 | 1,61 | 1,03 | 1,57 | 1,16E-01 | 2,30E-01 |
| AT1G20450 | 3,27 | -1,65 | 1,06 | -1,57 | 1,17E-01 | 2,31E-01 |
| AT3G52590 | 1,74 | 2,78 | 1,77 | 1,57 | 1,17E-01 | 2,31E-01 |
| AT4G06320 | 7,79 | -1,05 | 0,67 | -1,57 | 1,17E-01 | 2,31E-01 |
| AT1G63950 | 1,81 | -2,98 | 1,91 | -1,56 | 1,18E-01 | 2,32E-01 |
| AT3G06720 | 1,89 | 2,85 | 1,82 | 1,56 | 1,18E-01 | 2,32E-01 |

| | | | | | | |
|---|---|---|---|---|---|---|
| AT4G22410 | 2,93 | 2,44 | 1,56 | 1,56 | 1,18E-01 | 2,32E-01 |
| AT5G54440 | 2,06 | 2,34 | 1,50 | 1,56 | 1,19E-01 | 2,33E-01 |
| AT5G42092 | 3,68 | 1,69 | 1,08 | 1,56 | 1,19E-01 | 2,33E-01 |
| AT5G01920 | 2,56 | 2,25 | 1,44 | 1,56 | 1,19E-01 | 2,33E-01 |
| AT2G46020 | 3,28 | 1,88 | 1,21 | 1,56 | 1,19E-01 | 2,33E-01 |
| AT1G13320 | 2,17 | 2,39 | 1,53 | 1,56 | 1,19E-01 | 2,33E-01 |
| AT1G18310 | 4,64 | 1,59 | 1,02 | 1,56 | 1,19E-01 | 2,33E-01 |
| AT3G19030 | 2,04 | 2,42 | 1,56 | 1,56 | 1,19E-01 | 2,33E-01 |
| AT1G18485 | 1,73 | 3,12 | 2,01 | 1,56 | 1,19E-01 | 2,33E-01 |
| AT2G46820 | 2,21 | -1,82 | 1,17 | -1,56 | 1,20E-01 | 2,34E-01 |
| AT3G13870 | 4,03 | 2,51 | 1,61 | 1,55 | 1,20E-01 | 2,35E-01 |
| AT4G38440 | 2,75 | 2,11 | 1,36 | 1,55 | 1,21E-01 | 2,35E-01 |
| AT3G57720 | 13,95 | 1,06 | 0,68 | 1,55 | 1,22E-01 | 2,38E-01 |
| AT1G72930 | 3,01 | -2,45 | 1,59 | -1,54 | 1,23E-01 | 2,39E-01 |
| AT4G14365 | 1,73 | 2,79 | 1,81 | 1,54 | 1,23E-01 | 2,39E-01 |
| AT2G44540 | 4,48 | -1,46 | 0,95 | -1,54 | 1,23E-01 | 2,40E-01 |
| AT1G07600 | 1,87 | 2,14 | 1,39 | 1,54 | 1,24E-01 | 2,41E-01 |
| AT1G23310 | 12,45 | 0,91 | 0,59 | 1,54 | 1,24E-01 | 2,41E-01 |
| AT1G72410 | 1,92 | 2,71 | 1,76 | 1,54 | 1,24E-01 | 2,41E-01 |
| AT2G21660 | 24,65 | -0,74 | 0,48 | -1,54 | 1,24E-01 | 2,41E-01 |
| AT4G26530 | 4,43 | 1,33 | 0,87 | 1,53 | 1,25E-01 | 2,42E-01 |
| AT3G62200 | 2,36 | 2,59 | 1,69 | 1,53 | 1,25E-01 | 2,42E-01 |
| AT2G04455 | 3,39 | 1,98 | 1,30 | 1,53 | 1,26E-01 | 2,42E-01 |
| AT1G58602 | 11,97 | 0,92 | 0,60 | 1,53 | 1,26E-01 | 2,43E-01 |
| AT5G59505 | 3,52 | -2,32 | 1,52 | -1,53 | 1,26E-01 | 2,43E-01 |
| AT1G65450 | 2,07 | -2,21 | 1,45 | -1,53 | 1,27E-01 | 2,44E-01 |
| AT3G08740 | 1,74 | 2,40 | 1,57 | 1,53 | 1,27E-01 | 2,44E-01 |
| AT2G47940 | 2,47 | 2,14 | 1,40 | 1,52 | 1,28E-01 | 2,47E-01 |
| AT1G22930 | 1,70 | 2,75 | 1,81 | 1,52 | 1,29E-01 | 2,48E-01 |
| AT1G05163 | 491,36 | -0,41 | 0,27 | -1,52 | 1,29E-01 | 2,48E-01 |
| AT1G15405 | 491,36 | -0,41 | 0,27 | -1,52 | 1,29E-01 | 2,48E-01 |
| AT2G44560 | 4,56 | -1,23 | 0,81 | -1,51 | 1,30E-01 | 2,49E-01 |
| AT3G61630 | 1,88 | 2,87 | 1,90 | 1,51 | 1,30E-01 | 2,49E-01 |
| AT5G27607 | 2,38 | 2,93 | 1,94 | 1,51 | 1,30E-01 | 2,49E-01 |
| AT3G07735 | 4,39 | -1,36 | 0,90 | -1,51 | 1,31E-01 | 2,49E-01 |
| AT3G48000 | 4,79 | 1,51 | 1,00 | 1,51 | 1,31E-01 | 2,49E-01 |
| AT1G61190 | 2,38 | 2,00 | 1,33 | 1,51 | 1,31E-01 | 2,51E-01 |
| AT4G13560 | 52,08 | -0,55 | 0,36 | -1,51 | 1,32E-01 | 2,51E-01 |
| AT3G11560 | 1,70 | 2,86 | 1,90 | 1,51 | 1,32E-01 | 2,51E-01 |
| AT3G45640 | 1,79 | 2,46 | 1,64 | 1,50 | 1,33E-01 | 2,53E-01 |
| AT5G06175 | 3,94 | -1,45 | 0,97 | -1,50 | 1,33E-01 | 2,53E-01 |
| AT5G17890 | 1,77 | 2,83 | 1,89 | 1,50 | 1,33E-01 | 2,53E-01 |
| AT1G77680 | 1,90 | 2,93 | 1,96 | 1,50 | 1,34E-01 | 2,54E-01 |
| AT1G77560 | 2,14 | 2,27 | 1,51 | 1,50 | 1,34E-01 | 2,54E-01 |
| AT3G55020 | 1,94 | 2,44 | 1,63 | 1,50 | 1,34E-01 | 2,54E-01 |
| AT1G04210 | 3,48 | 1,75 | 1,17 | 1,50 | 1,35E-01 | 2,55E-01 |
| AT1G09010 | 3,02 | 2,30 | 1,54 | 1,49 | 1,35E-01 | 2,55E-01 |
| AT1G21202 | 4,89 | 1,78 | 1,20 | 1,49 | 1,36E-01 | 2,56E-01 |
| AT1G64710 | 2,31 | 2,38 | 1,59 | 1,49 | 1,35E-01 | 2,56E-01 |
| AT2G18020 | 2,64 | 1,98 | 1,33 | 1,49 | 1,35E-01 | 2,56E-01 |
| AT1G05637 | 4,89 | 1,78 | 1,20 | 1,49 | 1,36E-01 | 2,56E-01 |
| AT2G43710 | 3,74 | 1,89 | 1,27 | 1,49 | 1,36E-01 | 2,56E-01 |
| AT2G02840 | 4,55 | 2,38 | 1,60 | 1,49 | 1,36E-01 | 2,56E-01 |
| AT3G16850 | 3,96 | 1,69 | 1,13 | 1,49 | 1,36E-01 | 2,56E-01 |
| AT5G21940 | 4,72 | 1,67 | 1,12 | 1,49 | 1,36E-01 | 2,56E-01 |
| AT2G21920 | 4,05 | -1,39 | 0,93 | -1,49 | 1,37E-01 | 2,57E-01 |

| | | | | | | |
|---|---|---|---|---|---|---|
| AT4G02570 | 1,99 | 2,16 | 1,45 | 1,49 | 1,37E-01 | 2,58E-01 |
| AT5G37510 | 3,25 | 1,60 | 1,08 | 1,48 | 1,38E-01 | 2,58E-01 |
| AT1G05273 | 8,42 | 0,99 | 0,67 | 1,48 | 1,39E-01 | 2,59E-01 |
| AT2G34357 | 1,72 | 2,24 | 1,51 | 1,48 | 1,39E-01 | 2,59E-01 |
| AT1G16820 | 8,42 | 0,99 | 0,67 | 1,48 | 1,39E-01 | 2,59E-01 |
| AT3G48560 | 3,89 | -1,46 | 0,98 | -1,48 | 1,38E-01 | 2,59E-01 |
| AT4G20360 | 5,64 | -1,14 | 0,77 | -1,48 | 1,39E-01 | 2,60E-01 |
| AT4G20362 | 5,64 | -1,14 | 0,77 | -1,48 | 1,39E-01 | 2,60E-01 |
| AT1G30490 | 1,77 | 2,78 | 1,88 | 1,48 | 1,40E-01 | 2,61E-01 |
| AT1G29920 | 50,33 | 0,64 | 0,44 | 1,47 | 1,40E-01 | 2,61E-01 |
| AT4G20735 | 3,19 | 2,30 | 1,56 | 1,47 | 1,41E-01 | 2,63E-01 |
| AT5G41490 | 2,24 | 2,46 | 1,68 | 1,47 | 1,41E-01 | 2,63E-01 |
| AT4G35770 | 2,85 | -1,95 | 1,33 | -1,47 | 1,42E-01 | 2,63E-01 |
| AT5G40480 | 2,70 | 1,81 | 1,23 | 1,47 | 1,42E-01 | 2,63E-01 |
| AT3G18490 | 8,11 | -1,00 | 0,68 | -1,47 | 1,42E-01 | 2,63E-01 |
| AT4G25880 | 1,93 | 2,22 | 1,51 | 1,47 | 1,42E-01 | 2,64E-01 |
| AT5G02880 | 3,00 | 1,78 | 1,21 | 1,47 | 1,42E-01 | 2,64E-01 |
| AT1G18010 | 5,80 | -1,20 | 0,82 | -1,46 | 1,43E-01 | 2,64E-01 |
| AT1G08053 | 2,32 | -1,85 | 1,26 | -1,46 | 1,43E-01 | 2,64E-01 |
| AT3G60630 | 5,65 | 1,49 | 1,02 | 1,46 | 1,43E-01 | 2,64E-01 |
| AT3G63410 | 2,71 | 1,92 | 1,31 | 1,46 | 1,43E-01 | 2,64E-01 |
| AT5G09495 | 4,86 | -1,23 | 0,84 | -1,47 | 1,43E-01 | 2,64E-01 |
| AT2G46400 | 1,95 | 2,50 | 1,71 | 1,46 | 1,43E-01 | 2,64E-01 |
| AT3G53400 | 1,89 | 2,72 | 1,86 | 1,46 | 1,44E-01 | 2,65E-01 |
| AT5G16270 | 1,71 | 2,42 | 1,66 | 1,46 | 1,45E-01 | 2,67E-01 |
| AT2G43150 | 1,94 | 2,14 | 1,47 | 1,45 | 1,46E-01 | 2,68E-01 |
| AT5G45100 | 1,72 | 2,90 | 2,00 | 1,45 | 1,46E-01 | 2,68E-01 |
| AT5G58465 | 4,24 | -1,37 | 0,94 | -1,45 | 1,46E-01 | 2,68E-01 |
| AT2G04060 | 1,72 | 3,13 | 2,16 | 1,45 | 1,46E-01 | 2,68E-01 |
| AT4G26095 | 1,81 | 2,35 | 1,62 | 1,45 | 1,46E-01 | 2,69E-01 |
| AT5G42650 | 3,24 | -1,81 | 1,25 | -1,45 | 1,48E-01 | 2,71E-01 |
| AT1G74100 | 5,98 | 1,45 | 1,00 | 1,45 | 1,48E-01 | 2,71E-01 |
| AT5G54810 | 4,59 | 1,39 | 0,96 | 1,45 | 1,48E-01 | 2,71E-01 |
| AT1G05780 | 4,77 | 1,71 | 1,18 | 1,44 | 1,49E-01 | 2,71E-01 |
| AT2G34410 | 2,89 | 1,80 | 1,24 | 1,44 | 1,49E-01 | 2,71E-01 |
| AT3G43160 | 4,62 | 1,35 | 0,94 | 1,44 | 1,49E-01 | 2,71E-01 |
| AT1G18000 | 4,70 | -1,65 | 1,14 | -1,44 | 1,49E-01 | 2,71E-01 |
| AT4G13505 | 4,72 | 1,38 | 0,95 | 1,44 | 1,49E-01 | 2,71E-01 |
| AT5G10760 | 4,94 | 1,31 | 0,91 | 1,44 | 1,49E-01 | 2,71E-01 |
| AT5G23630 | 2,35 | 2,16 | 1,50 | 1,44 | 1,49E-01 | 2,71E-01 |
| AT3G63003 | 4,44 | -1,45 | 1,01 | -1,44 | 1,50E-01 | 2,72E-01 |
| AT5G27850 | 1,94 | 2,10 | 1,46 | 1,44 | 1,50E-01 | 2,73E-01 |
| AT2G30570 | 3,97 | -1,33 | 0,93 | -1,44 | 1,50E-01 | 2,73E-01 |
| AT4G24715 | 2,06 | -1,80 | 1,25 | -1,44 | 1,50E-01 | 2,73E-01 |
| AT1G07637 | 2,62 | 2,10 | 1,46 | 1,44 | 1,51E-01 | 2,73E-01 |
| AT3G02200 | 1,75 | 2,67 | 1,86 | 1,44 | 1,51E-01 | 2,74E-01 |
| AT2G31890 | 2,01 | 2,58 | 1,80 | 1,43 | 1,52E-01 | 2,74E-01 |
| AT5G65480 | 2,00 | 2,18 | 1,52 | 1,43 | 1,52E-01 | 2,75E-01 |
| AT4G32260 | 3,86 | -1,44 | 1,01 | -1,43 | 1,52E-01 | 2,75E-01 |
| AT5G22800 | 1,71 | -1,97 | 1,38 | -1,43 | 1,53E-01 | 2,76E-01 |
| AT5G49740 | 2,42 | 2,05 | 1,44 | 1,43 | 1,53E-01 | 2,76E-01 |
| AT4G38775 | 2,66 | -1,52 | 1,07 | -1,43 | 1,54E-01 | 2,77E-01 |
| AT4G40040 | 3,01 | 1,67 | 1,17 | 1,42 | 1,54E-01 | 2,78E-01 |
| AT3G58050 | 1,93 | 2,32 | 1,63 | 1,42 | 1,54E-01 | 2,78E-01 |
| AT1G59980 | 3,55 | -1,56 | 1,10 | -1,42 | 1,55E-01 | 2,78E-01 |
| AT3G47347 | 15,63 | -0,74 | 0,52 | -1,42 | 1,55E-01 | 2,78E-01 |

| | | | | | | |
|---|---|---|---|---|---|---|
| AT1G56510 | 2,98 | 2,34 | 1,65 | 1,42 | 1,55E-01 | 2,79E-01 |
| AT4G13510 | 4,76 | 1,36 | 0,96 | 1,42 | 1,55E-01 | 2,79E-01 |
| AT1G29400 | 2,11 | 2,07 | 1,46 | 1,42 | 1,56E-01 | 2,79E-01 |
| AT2G47000 | 1,78 | 3,03 | 2,14 | 1,42 | 1,56E-01 | 2,79E-01 |
| AT5G19200 | 4,59 | 1,37 | 0,96 | 1,42 | 1,56E-01 | 2,80E-01 |
| AT1G10760 | 16,93 | 0,83 | 0,58 | 1,42 | 1,56E-01 | 2,80E-01 |
| AT5G50920 | 18,45 | 0,73 | 0,51 | 1,41 | 1,57E-01 | 2,81E-01 |
| AT4G13840 | 1,76 | -2,20 | 1,55 | -1,41 | 1,58E-01 | 2,81E-01 |
| AT1G01510 | 2,79 | 2,31 | 1,64 | 1,41 | 1,59E-01 | 2,83E-01 |
| AT3G04210 | 3,60 | 1,49 | 1,06 | 1,41 | 1,59E-01 | 2,83E-01 |
| AT5G03240 | 5,81 | 1,14 | 0,81 | 1,41 | 1,59E-01 | 2,83E-01 |
| AT1G40104 | 7,42 | 1,15 | 0,81 | 1,41 | 1,59E-01 | 2,84E-01 |
| AT5G61020 | 2,23 | 2,10 | 1,49 | 1,41 | 1,60E-01 | 2,84E-01 |
| AT2G03140 | 2,19 | 2,20 | 1,57 | 1,40 | 1,61E-01 | 2,85E-01 |
| AT3G14420 | 4,42 | -1,40 | 1,00 | -1,40 | 1,61E-01 | 2,87E-01 |
| AT1G21200 | 5,07 | 1,50 | 1,07 | 1,40 | 1,62E-01 | 2,87E-01 |
| AT5G28740 | 1,92 | 2,11 | 1,51 | 1,40 | 1,62E-01 | 2,87E-01 |
| AT5G03040 | 2,14 | 2,00 | 1,43 | 1,40 | 1,62E-01 | 2,87E-01 |
| AT3G61690 | 2,25 | 2,26 | 1,61 | 1,40 | 1,62E-01 | 2,87E-01 |
| AT3G13525 | 2,38 | -1,59 | 1,14 | -1,39 | 1,64E-01 | 2,90E-01 |
| AT5G03305 | 5,94 | 1,42 | 1,02 | 1,39 | 1,64E-01 | 2,90E-01 |
| AT1G45160 | 1,73 | 2,29 | 1,65 | 1,39 | 1,64E-01 | 2,90E-01 |
| AT3G01313 | 12,85 | 0,93 | 0,67 | 1,39 | 1,64E-01 | 2,90E-01 |
| AT3G05675 | 9,08 | 1,14 | 0,82 | 1,39 | 1,64E-01 | 2,90E-01 |
| AT1G04733 | 20,09 | 0,68 | 0,49 | 1,39 | 1,64E-01 | 2,90E-01 |
| AT1G51570 | 1,72 | 2,39 | 1,72 | 1,39 | 1,64E-01 | 2,90E-01 |
| AT1G20970 | 4,69 | 1,76 | 1,27 | 1,39 | 1,65E-01 | 2,90E-01 |
| AT4G05360 | 2,15 | 2,29 | 1,65 | 1,39 | 1,65E-01 | 2,90E-01 |
| AT4G29410 | 17,21 | -0,79 | 0,57 | -1,39 | 1,65E-01 | 2,90E-01 |
| AT4G10120 | 1,86 | 2,32 | 1,67 | 1,39 | 1,66E-01 | 2,91E-01 |
| AT4G13810 | 2,84 | 1,99 | 1,44 | 1,38 | 1,66E-01 | 2,92E-01 |
| AT4G16720 | 3,07 | 1,85 | 1,33 | 1,38 | 1,66E-01 | 2,92E-01 |
| AT5G04810 | 3,05 | 1,66 | 1,20 | 1,38 | 1,67E-01 | 2,92E-01 |
| AT1G16635 | 19,67 | -0,79 | 0,58 | -1,38 | 1,68E-01 | 2,93E-01 |
| AT3G04765 | 2,83 | -1,51 | 1,10 | -1,38 | 1,68E-01 | 2,93E-01 |
| AT5G46210 | 3,09 | 1,73 | 1,25 | 1,38 | 1,68E-01 | 2,93E-01 |
| AT3G08170 | 3,83 | 1,49 | 1,08 | 1,38 | 1,68E-01 | 2,94E-01 |
| AT2G45930 | 2,64 | 2,19 | 1,59 | 1,38 | 1,69E-01 | 2,94E-01 |
| AT1G70530 | 1,72 | 2,62 | 1,92 | 1,37 | 1,71E-01 | 2,98E-01 |
| AT5G17710 | 1,85 | 2,15 | 1,57 | 1,37 | 1,71E-01 | 2,98E-01 |
| AT4G10340 | 23,35 | -0,71 | 0,52 | -1,37 | 1,72E-01 | 2,99E-01 |
| AT3G46620 | 4,33 | 1,39 | 1,02 | 1,37 | 1,72E-01 | 2,99E-01 |
| AT4G16370 | 3,80 | 1,43 | 1,05 | 1,37 | 1,72E-01 | 2,99E-01 |
| AT2G38280 | 1,89 | 2,42 | 1,77 | 1,36 | 1,72E-01 | 2,99E-01 |
| AT1G50030 | 2,19 | 2,14 | 1,57 | 1,36 | 1,72E-01 | 2,99E-01 |
| AT4G39400 | 3,54 | 1,50 | 1,10 | 1,36 | 1,73E-01 | 3,00E-01 |
| AT1G72610 | 17,51 | -0,65 | 0,48 | -1,36 | 1,73E-01 | 3,00E-01 |
| AT4G35000 | 3,13 | -1,84 | 1,35 | -1,36 | 1,73E-01 | 3,00E-01 |
| AT1G31960 | 1,81 | -2,04 | 1,51 | -1,35 | 1,76E-01 | 3,04E-01 |
| AT2G30110 | 3,88 | 1,71 | 1,26 | 1,35 | 1,76E-01 | 3,04E-01 |
| AT3G53990 | 1,83 | -1,83 | 1,35 | -1,35 | 1,76E-01 | 3,04E-01 |
| AT3G55410 | 3,63 | 2,05 | 1,51 | 1,35 | 1,77E-01 | 3,04E-01 |
| AT4G17410 | 2,55 | 2,48 | 1,84 | 1,35 | 1,77E-01 | 3,04E-01 |
| AT1G05917 | 8,49 | -1,12 | 0,83 | -1,35 | 1,77E-01 | 3,04E-01 |
| AT4G11373 | 5,92 | 1,35 | 1,00 | 1,35 | 1,77E-01 | 3,04E-01 |
| AT1G05427 | 2,11 | -2,07 | 1,53 | -1,35 | 1,77E-01 | 3,04E-01 |

| | | | | | | |
|---|---|---|---|---|---|---|
| AT3G21870 | 1,78 | -2,22 | 1,64 | -1,35 | 1,77E-01 | 3,04E-01 |
| AT3G51770 | 2,18 | 2,06 | 1,53 | 1,35 | 1,77E-01 | 3,04E-01 |
| AT3G47250 | 1,84 | 2,20 | 1,63 | 1,35 | 1,77E-01 | 3,04E-01 |
| AT1G04425 | 1,91 | -1,91 | 1,42 | -1,35 | 1,78E-01 | 3,05E-01 |
| AT2G25510 | 7,52 | 1,18 | 0,88 | 1,35 | 1,77E-01 | 3,05E-01 |
| AT3G04400 | 2,05 | -1,71 | 1,27 | -1,34 | 1,79E-01 | 3,06E-01 |
| AT4G32760 | 1,73 | 2,36 | 1,76 | 1,34 | 1,79E-01 | 3,06E-01 |
| AT4G16180 | 1,78 | 3,09 | 2,30 | 1,34 | 1,79E-01 | 3,06E-01 |
| AT1G68520 | 2,00 | 1,74 | 1,30 | 1,34 | 1,81E-01 | 3,08E-01 |
| AT5G01400 | 1,70 | 2,74 | 2,05 | 1,34 | 1,81E-01 | 3,08E-01 |
| AT1G76135 | 2,70 | -1,81 | 1,35 | -1,34 | 1,81E-01 | 3,09E-01 |
| AT3G26730 | 1,96 | 1,96 | 1,47 | 1,34 | 1,81E-01 | 3,09E-01 |
| AT5G65610 | 2,72 | -1,85 | 1,38 | -1,34 | 1,81E-01 | 3,09E-01 |
| AT5G62165 | 7,10 | -1,08 | 0,80 | -1,34 | 1,82E-01 | 3,09E-01 |
| AT2G03150 | 2,08 | 2,18 | 1,64 | 1,33 | 1,83E-01 | 3,12E-01 |
| AT3G58030 | 2,17 | 2,04 | 1,54 | 1,33 | 1,84E-01 | 3,12E-01 |
| AT4G16143 | 2,03 | 1,75 | 1,32 | 1,33 | 1,84E-01 | 3,13E-01 |
| AT3G13405 | 1,91 | -1,95 | 1,47 | -1,33 | 1,85E-01 | 3,14E-01 |
| AT3G23450 | 2,19 | -1,93 | 1,46 | -1,32 | 1,86E-01 | 3,16E-01 |
| AT1G02080 | 3,21 | 1,90 | 1,44 | 1,32 | 1,87E-01 | 3,16E-01 |
| AT2G13360 | 3,44 | -1,31 | 0,99 | -1,32 | 1,87E-01 | 3,16E-01 |
| AT3G59765 | 1,81 | -1,86 | 1,41 | -1,32 | 1,86E-01 | 3,16E-01 |
| AT4G29190 | 1,79 | -2,05 | 1,55 | -1,32 | 1,87E-01 | 3,16E-01 |
| AT3G23810 | 2,25 | -1,76 | 1,33 | -1,32 | 1,87E-01 | 3,17E-01 |
| AT1G05247 | 19,59 | -0,77 | 0,58 | -1,32 | 1,88E-01 | 3,18E-01 |
| AT2G17510 | 1,71 | 1,82 | 1,39 | 1,32 | 1,88E-01 | 3,18E-01 |
| AT1G07233 | 2,51 | -1,57 | 1,19 | -1,31 | 1,89E-01 | 3,19E-01 |
| AT3G01695 | 8,72 | 1,10 | 0,84 | 1,31 | 1,89E-01 | 3,19E-01 |
| AT2G37220 | 3,40 | -1,38 | 1,05 | -1,31 | 1,89E-01 | 3,19E-01 |
| AT3G13920 | 8,10 | 1,16 | 0,89 | 1,31 | 1,90E-01 | 3,19E-01 |
| AT1G06230 | 2,14 | 2,06 | 1,57 | 1,31 | 1,91E-01 | 3,21E-01 |
| AT4G15560 | 2,23 | -1,66 | 1,27 | -1,31 | 1,91E-01 | 3,21E-01 |
| AT4G24190 | 5,76 | 1,09 | 0,83 | 1,31 | 1,91E-01 | 3,21E-01 |
| AT1G72600 | 18,04 | -0,61 | 0,47 | -1,30 | 1,93E-01 | 3,23E-01 |
| AT3G24615 | 9,91 | -0,84 | 0,65 | -1,30 | 1,93E-01 | 3,23E-01 |
| AT1G65960 | 3,84 | 1,51 | 1,16 | 1,30 | 1,93E-01 | 3,24E-01 |
| AT1G03080 | 1,73 | 2,02 | 1,55 | 1,30 | 1,93E-01 | 3,24E-01 |
| AT1G05035 | 3,55 | -1,50 | 1,16 | -1,30 | 1,94E-01 | 3,25E-01 |
| AT5G05690 | 2,09 | 2,10 | 1,62 | 1,30 | 1,95E-01 | 3,25E-01 |
| AT3G50480 | 1,71 | 2,20 | 1,70 | 1,29 | 1,95E-01 | 3,26E-01 |
| AT1G79750 | 4,46 | 1,41 | 1,09 | 1,30 | 1,95E-01 | 3,26E-01 |
| AT2G27310 | 2,22 | 1,75 | 1,35 | 1,30 | 1,95E-01 | 3,26E-01 |
| AT2G31610 | 1,80 | 2,24 | 1,73 | 1,29 | 1,97E-01 | 3,28E-01 |
| AT4G09735 | 2,23 | 2,36 | 1,83 | 1,29 | 1,97E-01 | 3,28E-01 |
| AT3G14050 | 1,74 | 1,97 | 1,53 | 1,29 | 1,98E-01 | 3,29E-01 |
| AT4G02130 | 3,77 | 1,45 | 1,13 | 1,29 | 1,98E-01 | 3,29E-01 |
| AT5G28560 | 3,51 | 1,42 | 1,10 | 1,29 | 1,98E-01 | 3,30E-01 |
| AT2G04885 | 2,17 | 2,06 | 1,60 | 1,28 | 1,99E-01 | 3,30E-01 |
| AT2G35260 | 3,89 | -1,27 | 0,99 | -1,28 | 1,99E-01 | 3,30E-01 |
| AT5G24300 | 2,35 | 1,81 | 1,41 | 1,29 | 1,99E-01 | 3,30E-01 |
| AT2G16485 | 2,25 | 1,75 | 1,36 | 1,28 | 1,99E-01 | 3,31E-01 |
| AT1G70290 | 2,53 | 1,83 | 1,43 | 1,28 | 2,01E-01 | 3,32E-01 |
| AT5G61455 | 11,96 | 0,78 | 0,61 | 1,28 | 2,01E-01 | 3,32E-01 |
| AT1G53900 | 2,01 | -1,74 | 1,36 | -1,28 | 2,02E-01 | 3,34E-01 |
| AT1G09800 | 20,58 | 0,62 | 0,49 | 1,28 | 2,02E-01 | 3,34E-01 |
| AT1G03130 | 1,99 | -1,60 | 1,25 | -1,28 | 2,02E-01 | 3,34E-01 |

| | | | | | | |
|---|---|---|---|---|---|---|
| AT4G00720 | 2,07 | 1,96 | 1,54 | 1,28 | 2,02E-01 | 3,34E-01 |
| AT1G77235 | 1,77 | -2,26 | 1,77 | -1,27 | 2,03E-01 | 3,34E-01 |
| AT4G30993 | 2,23 | 1,69 | 1,33 | 1,27 | 2,03E-01 | 3,34E-01 |
| AT5G64940 | 2,50 | 2,21 | 1,73 | 1,27 | 2,03E-01 | 3,34E-01 |
| AT1G01040 | 4,73 | 1,18 | 0,93 | 1,27 | 2,03E-01 | 3,35E-01 |
| AT3G14270 | 2,35 | 1,86 | 1,47 | 1,27 | 2,04E-01 | 3,35E-01 |
| AT3G23820 | 9,86 | -0,78 | 0,61 | -1,27 | 2,04E-01 | 3,35E-01 |
| AT4G01850 | 4,27 | 1,42 | 1,12 | 1,27 | 2,04E-01 | 3,35E-01 |
| AT1G42970 | 13,13 | 0,77 | 0,60 | 1,27 | 2,04E-01 | 3,36E-01 |
| AT5G42930 | 2,68 | -1,51 | 1,19 | -1,27 | 2,05E-01 | 3,36E-01 |
| AT1G78478 | 10,59 | -0,90 | 0,71 | -1,27 | 2,05E-01 | 3,36E-01 |
| AT5G04105 | 18,20 | 0,70 | 0,56 | 1,27 | 2,06E-01 | 3,37E-01 |
| AT5G47260 | 4,17 | -1,14 | 0,91 | -1,26 | 2,07E-01 | 3,38E-01 |
| AT1G07510 | 1,74 | 1,92 | 1,53 | 1,26 | 2,08E-01 | 3,40E-01 |
| AT1G16880 | 2,91 | 1,58 | 1,25 | 1,26 | 2,08E-01 | 3,40E-01 |
| AT5G26030 | 2,16 | 1,87 | 1,49 | 1,25 | 2,09E-01 | 3,42E-01 |
| AT1G09817 | 10,25 | -0,92 | 0,73 | -1,25 | 2,09E-01 | 3,42E-01 |
| AT5G65860 | 5,04 | -1,04 | 0,83 | -1,25 | 2,10E-01 | 3,42E-01 |
| AT2G47450 | 6,02 | 1,09 | 0,87 | 1,25 | 2,10E-01 | 3,42E-01 |
| AT1G52400 | 1,99 | -1,99 | 1,59 | -1,25 | 2,12E-01 | 3,45E-01 |
| AT3G05590 | 1,96 | -1,63 | 1,31 | -1,25 | 2,13E-01 | 3,46E-01 |
| AT3G07765 | 2,13 | -1,43 | 1,15 | -1,24 | 2,13E-01 | 3,47E-01 |
| AT2G01422 | 2,93 | -1,88 | 1,51 | -1,24 | 2,14E-01 | 3,47E-01 |
| AT5G42050 | 3,62 | 1,59 | 1,28 | 1,24 | 2,14E-01 | 3,48E-01 |
| AT2G24270 | 3,15 | 1,61 | 1,30 | 1,24 | 2,15E-01 | 3,49E-01 |
| AT1G78020 | 1,89 | 1,99 | 1,60 | 1,24 | 2,16E-01 | 3,50E-01 |
| AT2G36160 | 1,81 | 2,15 | 1,74 | 1,24 | 2,16E-01 | 3,50E-01 |
| AT4G13880 | 2,00 | 2,14 | 1,73 | 1,24 | 2,16E-01 | 3,50E-01 |
| AT1G62630 | 2,10 | -1,59 | 1,28 | -1,24 | 2,16E-01 | 3,50E-01 |
| AT2G23810 | 3,17 | 1,55 | 1,25 | 1,24 | 2,16E-01 | 3,50E-01 |
| AT2G15420 | 2,17 | 1,90 | 1,54 | 1,24 | 2,17E-01 | 3,50E-01 |
| AT2G36880 | 2,29 | 1,80 | 1,46 | 1,23 | 2,18E-01 | 3,51E-01 |
| AT1G80000 | 1,90 | 2,44 | 1,98 | 1,23 | 2,18E-01 | 3,52E-01 |
| AT1G10522 | 24,13 | 0,60 | 0,48 | 1,23 | 2,19E-01 | 3,53E-01 |
| AT5G02150 | 1,74 | -2,01 | 1,63 | -1,23 | 2,19E-01 | 3,53E-01 |
| AT2G19360 | 2,70 | -1,56 | 1,27 | -1,23 | 2,19E-01 | 3,53E-01 |
| AT4G39260 | 6,03 | -0,91 | 0,74 | -1,23 | 2,19E-01 | 3,53E-01 |
| AT3G16785 | 1,89 | 1,73 | 1,41 | 1,23 | 2,20E-01 | 3,53E-01 |
| AT5G64220 | 2,50 | 2,10 | 1,71 | 1,23 | 2,20E-01 | 3,54E-01 |
| AT1G62540 | 2,48 | -2,01 | 1,64 | -1,22 | 2,21E-01 | 3,55E-01 |
| AT1G13440 | 5,54 | -0,98 | 0,80 | -1,22 | 2,21E-01 | 3,55E-01 |
| AT3G24612 | 3,02 | 1,41 | 1,15 | 1,22 | 2,21E-01 | 3,55E-01 |
| AT5G28920 | 2,84 | 1,56 | 1,28 | 1,22 | 2,22E-01 | 3,56E-01 |
| AT4G30975 | 11,87 | 0,72 | 0,59 | 1,22 | 2,23E-01 | 3,57E-01 |
| AT1G16710 | 1,85 | 2,08 | 1,71 | 1,22 | 2,23E-01 | 3,57E-01 |
| AT5G49615 | 2,21 | -2,10 | 1,73 | -1,22 | 2,24E-01 | 3,58E-01 |
| AT5G19330 | 2,99 | 1,70 | 1,40 | 1,22 | 2,24E-01 | 3,58E-01 |
| AT4G11600 | 2,32 | -1,36 | 1,12 | -1,22 | 2,24E-01 | 3,58E-01 |
| AT5G40340 | 2,24 | -1,46 | 1,20 | -1,22 | 2,24E-01 | 3,58E-01 |
| AT3G55980 | 3,63 | 1,47 | 1,21 | 1,21 | 2,26E-01 | 3,61E-01 |
| AT5G46070 | 1,78 | 2,34 | 1,94 | 1,21 | 2,27E-01 | 3,62E-01 |
| AT2G44570 | 2,88 | -1,45 | 1,20 | -1,21 | 2,27E-01 | 3,62E-01 |
| AT2G06510 | 1,80 | 1,85 | 1,54 | 1,21 | 2,28E-01 | 3,63E-01 |
| AT2G35050 | 3,09 | 1,51 | 1,25 | 1,21 | 2,28E-01 | 3,63E-01 |
| AT5G65630 | 3,70 | 1,31 | 1,09 | 1,20 | 2,28E-01 | 3,63E-01 |
| AT4G31490 | 3,09 | 1,49 | 1,24 | 1,20 | 2,29E-01 | 3,63E-01 |

| | | | | | | |
|---|---|---|---|---|---|---|
| AT2G22430 | 4,62 | 1,28 | 1,06 | 1,20 | 2,29E-01 | 3,64E-01 |
| AT4G38760 | 3,63 | 1,54 | 1,29 | 1,20 | 2,30E-01 | 3,65E-01 |
| AT1G26558 | 2,72 | -1,61 | 1,35 | -1,20 | 2,31E-01 | 3,65E-01 |
| AT5G56970 | 2,67 | -1,36 | 1,13 | -1,20 | 2,31E-01 | 3,65E-01 |
| AT1G15125 | 1,84 | 2,05 | 1,71 | 1,20 | 2,31E-01 | 3,66E-01 |
| AT5G65380 | 2,69 | 1,68 | 1,41 | 1,20 | 2,32E-01 | 3,67E-01 |
| AT2G46220 | 1,83 | 1,95 | 1,63 | 1,19 | 2,33E-01 | 3,68E-01 |
| AT1G78830 | 2,12 | -1,66 | 1,39 | -1,19 | 2,33E-01 | 3,68E-01 |
| AT3G05985 | 2,35 | 1,91 | 1,60 | 1,19 | 2,33E-01 | 3,68E-01 |
| AT3G22440 | 1,87 | 1,64 | 1,38 | 1,19 | 2,34E-01 | 3,68E-01 |
| AT4G08593 | 1,89 | 2,25 | 1,89 | 1,19 | 2,34E-01 | 3,69E-01 |
| AT4G30972 | 11,83 | 0,70 | 0,59 | 1,19 | 2,34E-01 | 3,69E-01 |
| AT1G02470 | 5,92 | 1,33 | 1,12 | 1,19 | 2,35E-01 | 3,69E-01 |
| AT3G54925 | 1,76 | -2,03 | 1,71 | -1,19 | 2,34E-01 | 3,69E-01 |
| AT2G34750 | 2,36 | 1,78 | 1,50 | 1,19 | 2,35E-01 | 3,70E-01 |
| AT3G52240 | 2,27 | 1,76 | 1,49 | 1,19 | 2,36E-01 | 3,71E-01 |
| AT1G17470 | 1,97 | 1,99 | 1,68 | 1,18 | 2,37E-01 | 3,72E-01 |
| AT5G42090 | 4,37 | 1,18 | 1,00 | 1,18 | 2,38E-01 | 3,73E-01 |
| AT2G08390 | 5,34 | -1,07 | 0,90 | -1,18 | 2,38E-01 | 3,73E-01 |
| AT4G28750 | 4,56 | -1,28 | 1,09 | -1,18 | 2,38E-01 | 3,74E-01 |
| AT3G01316 | 2,03 | -1,56 | 1,33 | -1,18 | 2,39E-01 | 3,74E-01 |
| AT2G09685 | 3,15 | -1,23 | 1,05 | -1,17 | 2,43E-01 | 3,80E-01 |
| AT5G58430 | 1,95 | 1,79 | 1,54 | 1,17 | 2,44E-01 | 3,81E-01 |
| AT1G68530 | 2,97 | -1,27 | 1,09 | -1,16 | 2,44E-01 | 3,82E-01 |
| AT4G03415 | 1,93 | 2,21 | 1,90 | 1,16 | 2,44E-01 | 3,82E-01 |
| AT1G06277 | 5,95 | -0,95 | 0,82 | -1,16 | 2,45E-01 | 3,82E-01 |
| AT4G31180 | 4,29 | 1,15 | 0,99 | 1,16 | 2,45E-01 | 3,82E-01 |
| AT5G14930 | 1,80 | 2,62 | 2,25 | 1,16 | 2,45E-01 | 3,82E-01 |
| AT3G14067 | 4,84 | 1,03 | 0,88 | 1,16 | 2,45E-01 | 3,82E-01 |
| AT5G09730 | 35,07 | 0,52 | 0,45 | 1,16 | 2,46E-01 | 3,83E-01 |
| AT1G28480 | 1,91 | -1,78 | 1,53 | -1,16 | 2,46E-01 | 3,83E-01 |
| AT3G26000 | 2,02 | 1,99 | 1,72 | 1,16 | 2,46E-01 | 3,83E-01 |
| AT3G51950 | 4,09 | 1,12 | 0,97 | 1,16 | 2,47E-01 | 3,83E-01 |
| AT2G40930 | 1,74 | 2,15 | 1,87 | 1,15 | 2,49E-01 | 3,87E-01 |
| AT1G26973 | 11,41 | 0,98 | 0,85 | 1,15 | 2,50E-01 | 3,87E-01 |
| AT5G28919 | 2,64 | 1,48 | 1,29 | 1,15 | 2,50E-01 | 3,87E-01 |
| AT1G08103 | 6,40 | -1,06 | 0,92 | -1,15 | 2,50E-01 | 3,87E-01 |
| AT5G04905 | 2,64 | 1,48 | 1,29 | 1,15 | 2,50E-01 | 3,87E-01 |
| AT5G09900 | 1,74 | 1,97 | 1,72 | 1,15 | 2,51E-01 | 3,89E-01 |
| AT4G39160 | 1,99 | 1,82 | 1,59 | 1,15 | 2,51E-01 | 3,89E-01 |
| AT3G56705 | 15,48 | 0,64 | 0,56 | 1,14 | 2,52E-01 | 3,90E-01 |
| AT3G58610 | 2,68 | 1,43 | 1,25 | 1,14 | 2,53E-01 | 3,91E-01 |
| AT1G77740 | 1,74 | 1,79 | 1,57 | 1,14 | 2,55E-01 | 3,93E-01 |
| AT4G32410 | 11,90 | 0,78 | 0,69 | 1,14 | 2,55E-01 | 3,93E-01 |
| AT5G02700 | 1,99 | 1,97 | 1,73 | 1,14 | 2,55E-01 | 3,93E-01 |
| AT3G25690 | 2,72 | 1,28 | 1,13 | 1,14 | 2,56E-01 | 3,94E-01 |
| AT3G59970 | 1,71 | 1,63 | 1,44 | 1,14 | 2,56E-01 | 3,94E-01 |
| AT3G16370 | 1,93 | -1,60 | 1,41 | -1,13 | 2,57E-01 | 3,95E-01 |
| AT2G02835 | 3,94 | 1,36 | 1,20 | 1,13 | 2,58E-01 | 3,96E-01 |
| AT5G15022 | 21,95 | -0,64 | 0,57 | -1,13 | 2,58E-01 | 3,96E-01 |
| AT1G09690 | 1,78 | 1,55 | 1,38 | 1,13 | 2,58E-01 | 3,96E-01 |
| AT1G55490 | 3,72 | -1,31 | 1,16 | -1,13 | 2,58E-01 | 3,96E-01 |
| AT1G08117 | 4,98 | -1,32 | 1,17 | -1,13 | 2,60E-01 | 3,98E-01 |
| AT5G55565 | 4,30 | -1,09 | 0,97 | -1,13 | 2,60E-01 | 3,99E-01 |
| AT3G15880 | 1,94 | 1,50 | 1,34 | 1,12 | 2,62E-01 | 4,01E-01 |
| AT3G44430 | 1,70 | 1,86 | 1,66 | 1,12 | 2,64E-01 | 4,03E-01 |

| | | | | | | |
|---|---|---|---|---|---|---|
| AT5G45480 | 2,31 | 1,81 | 1,62 | 1,12 | 2,65E-01 | 4,05E-01 |
| AT1G04393 | 5,00 | 1,16 | 1,04 | 1,11 | 2,65E-01 | 4,05E-01 |
| AT3G10380 | 2,66 | 1,37 | 1,23 | 1,11 | 2,66E-01 | 4,05E-01 |
| AT1G14670 | 2,24 | 1,69 | 1,52 | 1,11 | 2,66E-01 | 4,06E-01 |
| AT1G05913 | 9,06 | -0,93 | 0,84 | -1,11 | 2,67E-01 | 4,06E-01 |
| AT1G32750 | 1,94 | 1,67 | 1,50 | 1,11 | 2,67E-01 | 4,06E-01 |
| AT1G27535 | 2,34 | -1,41 | 1,27 | -1,11 | 2,67E-01 | 4,06E-01 |
| AT1G04410 | 3,62 | -1,18 | 1,06 | -1,11 | 2,67E-01 | 4,06E-01 |
| AT3G08210 | 3,75 | -1,08 | 0,97 | -1,11 | 2,67E-01 | 4,07E-01 |
| AT1G11650 | 3,30 | 1,21 | 1,09 | 1,11 | 2,68E-01 | 4,07E-01 |
| AT4G40100 | 3,82 | -1,05 | 0,95 | -1,10 | 2,70E-01 | 4,09E-01 |
| AT5G09660 | 2,38 | -1,30 | 1,18 | -1,10 | 2,70E-01 | 4,09E-01 |
| AT3G26650 | 12,31 | 0,75 | 0,68 | 1,10 | 2,70E-01 | 4,10E-01 |
| AT3G56400 | 2,09 | 1,68 | 1,53 | 1,10 | 2,72E-01 | 4,12E-01 |
| AT5G48385 | 2,37 | 1,84 | 1,67 | 1,10 | 2,72E-01 | 4,12E-01 |
| AT1G12620 | 2,40 | -1,57 | 1,43 | -1,10 | 2,73E-01 | 4,13E-01 |
| AT2G07815 | 2,06 | -1,44 | 1,31 | -1,09 | 2,75E-01 | 4,15E-01 |
| AT2G22990 | 2,03 | -1,61 | 1,48 | -1,09 | 2,77E-01 | 4,18E-01 |
| AT3G25510 | 1,96 | 1,56 | 1,44 | 1,09 | 2,77E-01 | 4,18E-01 |
| AT4G30440 | 3,67 | 1,30 | 1,20 | 1,08 | 2,78E-01 | 4,20E-01 |
| AT2G08665 | 53,97 | -0,44 | 0,41 | -1,08 | 2,78E-01 | 4,20E-01 |
| AT2G34420 | 53,97 | -0,44 | 0,41 | -1,08 | 2,78E-01 | 4,20E-01 |
| AT5G38430 | 4,72 | -0,96 | 0,88 | -1,08 | 2,79E-01 | 4,20E-01 |
| AT3G09445 | 4,99 | -0,99 | 0,91 | -1,08 | 2,80E-01 | 4,21E-01 |
| AT3G57400 | 1,77 | 1,77 | 1,64 | 1,08 | 2,80E-01 | 4,22E-01 |
| AT2G22496 | 20,32 | 0,59 | 0,55 | 1,08 | 2,82E-01 | 4,23E-01 |
| AT3G11830 | 1,90 | 1,85 | 1,72 | 1,08 | 2,81E-01 | 4,23E-01 |
| AT5G13890 | 14,03 | 0,84 | 0,78 | 1,08 | 2,82E-01 | 4,23E-01 |
| AT5G14800 | 5,34 | 1,22 | 1,14 | 1,08 | 2,82E-01 | 4,23E-01 |
| AT3G43990 | 14,30 | 0,53 | 0,49 | 1,07 | 2,82E-01 | 4,23E-01 |
| AT5G61790 | 5,40 | 1,00 | 0,93 | 1,07 | 2,83E-01 | 4,24E-01 |
| AT5G60640 | 2,58 | -1,37 | 1,28 | -1,07 | 2,85E-01 | 4,27E-01 |
| AT2G41100 | 6,69 | 0,85 | 0,79 | 1,07 | 2,86E-01 | 4,28E-01 |
| AT2G29630 | 5,51 | 0,81 | 0,76 | 1,07 | 2,86E-01 | 4,28E-01 |
| AT4G16920 | 1,96 | 1,56 | 1,47 | 1,06 | 2,88E-01 | 4,30E-01 |
| AT1G49750 | 3,00 | -1,26 | 1,18 | -1,06 | 2,88E-01 | 4,30E-01 |
| AT4G35310 | 1,84 | 1,41 | 1,33 | 1,06 | 2,88E-01 | 4,31E-01 |
| AT3G17190 | 1,85 | 1,72 | 1,62 | 1,06 | 2,89E-01 | 4,31E-01 |
| AT1G11785 | 2,48 | -1,20 | 1,13 | -1,06 | 2,89E-01 | 4,31E-01 |
| AT1G30360 | 3,38 | 1,06 | 1,00 | 1,06 | 2,90E-01 | 4,32E-01 |
| AT4G27430 | 2,23 | 1,63 | 1,55 | 1,05 | 2,93E-01 | 4,36E-01 |
| AT5G09040 | 2,31 | -1,41 | 1,34 | -1,05 | 2,93E-01 | 4,36E-01 |
| AT4G39730 | 3,06 | -1,28 | 1,22 | -1,05 | 2,93E-01 | 4,36E-01 |
| AT2G05455 | 4,14 | -0,95 | 0,90 | -1,05 | 2,95E-01 | 4,38E-01 |
| AT5G52070 | 6,76 | -1,20 | 1,15 | -1,05 | 2,95E-01 | 4,38E-01 |
| AT2G27860 | 4,64 | 1,05 | 1,00 | 1,05 | 2,95E-01 | 4,38E-01 |
| AT3G13290 | 2,32 | 1,51 | 1,44 | 1,05 | 2,95E-01 | 4,38E-01 |
| AT1G09640 | 2,66 | 1,72 | 1,65 | 1,04 | 2,97E-01 | 4,41E-01 |
| AT1G13110 | 2,32 | -1,29 | 1,24 | -1,04 | 2,98E-01 | 4,42E-01 |
| AT3G28925 | 2,17 | 1,60 | 1,54 | 1,04 | 3,00E-01 | 4,43E-01 |
| AT4G05050 | 3,49 | 1,22 | 1,17 | 1,04 | 3,00E-01 | 4,43E-01 |
| AT5G14520 | 3,02 | 1,74 | 1,68 | 1,04 | 3,00E-01 | 4,43E-01 |
| AT1G29940 | 2,05 | 1,77 | 1,72 | 1,03 | 3,02E-01 | 4,46E-01 |
| AT3G51800 | 4,28 | -0,94 | 0,91 | -1,03 | 3,02E-01 | 4,46E-01 |
| AT4G05150 | 3,81 | 1,23 | 1,20 | 1,03 | 3,05E-01 | 4,50E-01 |
| AT1G17290 | 2,38 | 1,53 | 1,49 | 1,03 | 3,05E-01 | 4,50E-01 |

| | | | | | | |
|---|---|---|---|---|---|---|
| AT4G13430 | 3,55 | 1,16 | 1,14 | 1,02 | 3,07E-01 | 4,52E-01 |
| AT2G32690 | 2,01 | -1,29 | 1,27 | -1,02 | 3,08E-01 | 4,53E-01 |
| AT4G35830 | 9,01 | 0,83 | 0,81 | 1,02 | 3,08E-01 | 4,53E-01 |
| AT1G29370 | 3,57 | 1,04 | 1,02 | 1,02 | 3,08E-01 | 4,53E-01 |
| AT1G70940 | 2,12 | 1,74 | 1,71 | 1,02 | 3,09E-01 | 4,53E-01 |
| AT5G13530 | 2,40 | 1,27 | 1,24 | 1,02 | 3,08E-01 | 4,53E-01 |
| AT4G11830 | 1,70 | 1,84 | 1,81 | 1,02 | 3,10E-01 | 4,55E-01 |
| AT2G21030 | 1,78 | -1,54 | 1,52 | -1,01 | 3,11E-01 | 4,57E-01 |
| AT3G09630 | 1,89 | -1,37 | 1,36 | -1,01 | 3,13E-01 | 4,59E-01 |
| AT3G14310 | 2,07 | -1,64 | 1,63 | -1,01 | 3,14E-01 | 4,60E-01 |
| AT2G05315 | 4,98 | 1,06 | 1,06 | 1,00 | 3,16E-01 | 4,62E-01 |
| AT3G24180 | 1,99 | 1,37 | 1,37 | 1,00 | 3,16E-01 | 4,63E-01 |
| AT2G22300 | 2,59 | 1,63 | 1,62 | 1,00 | 3,17E-01 | 4,63E-01 |
| AT1G32060 | 5,29 | -1,03 | 1,03 | -1,00 | 3,18E-01 | 4,64E-01 |
| AT3G48860 | 1,96 | 2,26 | 2,26 | 1,00 | 3,18E-01 | 4,64E-01 |
| AT2G07890 | 2,64 | -1,20 | 1,20 | -1,00 | 3,19E-01 | 4,65E-01 |
| AT1G20330 | 2,79 | 1,39 | 1,39 | 1,00 | 3,18E-01 | 4,65E-01 |
| AT1G33140 | 1,84 | 1,56 | 1,57 | 1,00 | 3,19E-01 | 4,65E-01 |
| AT2G25170 | 3,04 | 1,33 | 1,34 | 0,99 | 3,21E-01 | 4,68E-01 |
| AT5G27807 | 30,37 | 0,41 | 0,41 | 0,99 | 3,22E-01 | 4,68E-01 |
| AT3G09440 | 5,91 | 0,80 | 0,81 | 0,99 | 3,23E-01 | 4,70E-01 |
| AT4G37260 | 2,49 | 1,28 | 1,29 | 0,99 | 3,24E-01 | 4,70E-01 |
| AT3G06055 | 2,49 | 1,25 | 1,27 | 0,99 | 3,23E-01 | 4,70E-01 |
| AT1G47660 | 2,01 | 1,72 | 1,74 | 0,99 | 3,24E-01 | 4,71E-01 |
| AT3G62420 | 1,96 | 1,96 | 1,99 | 0,98 | 3,25E-01 | 4,71E-01 |
| AT4G13570 | 36,83 | -0,40 | 0,40 | -0,98 | 3,25E-01 | 4,71E-01 |
| AT5G09670 | 2,53 | 1,22 | 1,24 | 0,98 | 3,25E-01 | 4,72E-01 |
| AT5G48620 | 2,46 | 1,41 | 1,44 | 0,98 | 3,26E-01 | 4,72E-01 |
| AT3G48430 | 1,70 | 1,88 | 1,92 | 0,98 | 3,27E-01 | 4,73E-01 |
| AT2G33730 | 2,24 | 1,18 | 1,21 | 0,98 | 3,29E-01 | 4,76E-01 |
| AT4G27870 | 2,09 | 1,38 | 1,42 | 0,98 | 3,29E-01 | 4,76E-01 |
| AT5G20950 | 3,45 | 1,13 | 1,15 | 0,98 | 3,29E-01 | 4,76E-01 |
| AT3G13750 | 2,68 | 1,19 | 1,23 | 0,97 | 3,30E-01 | 4,77E-01 |
| AT2G20990 | 4,50 | 1,03 | 1,06 | 0,97 | 3,31E-01 | 4,78E-01 |
| AT3G48350 | 1,84 | 1,99 | 2,05 | 0,97 | 3,31E-01 | 4,78E-01 |
| AT5G52190 | 1,80 | 1,83 | 1,88 | 0,97 | 3,32E-01 | 4,78E-01 |
| AT3G02540 | 1,81 | 1,67 | 1,72 | 0,97 | 3,33E-01 | 4,80E-01 |
| AT1G50920 | 2,95 | -1,01 | 1,05 | -0,97 | 3,34E-01 | 4,81E-01 |
| AT1G14920 | 1,77 | -1,37 | 1,42 | -0,96 | 3,37E-01 | 4,84E-01 |
| AT3G59710 | 6,69 | -0,81 | 0,84 | -0,96 | 3,37E-01 | 4,84E-01 |
| AT3G09225 | 6,69 | -0,81 | 0,84 | -0,96 | 3,37E-01 | 4,84E-01 |
| AT1G47265 | 3,77 | -0,99 | 1,04 | -0,96 | 3,38E-01 | 4,85E-01 |
| AT1G02780 | 1,83 | -1,28 | 1,34 | -0,96 | 3,39E-01 | 4,86E-01 |
| AT4G39800 | 2,68 | -1,23 | 1,28 | -0,96 | 3,39E-01 | 4,86E-01 |
| AT2G04030 | 4,28 | 0,93 | 0,98 | 0,95 | 3,41E-01 | 4,89E-01 |
| AT3G14840 | 1,81 | 1,52 | 1,60 | 0,95 | 3,41E-01 | 4,89E-01 |
| AT5G13887 | 13,72 | 0,85 | 0,89 | 0,95 | 3,42E-01 | 4,89E-01 |
| AT1G32270 | 1,96 | -1,38 | 1,45 | -0,95 | 3,42E-01 | 4,90E-01 |
| AT3G04775 | 14,57 | -0,48 | 0,50 | -0,95 | 3,43E-01 | 4,90E-01 |
| AT1G05010 | 2,06 | -1,55 | 1,63 | -0,95 | 3,43E-01 | 4,91E-01 |
| AT5G39770 | 12,51 | -0,57 | 0,60 | -0,94 | 3,45E-01 | 4,93E-01 |
| AT3G02020 | 19,73 | 0,53 | 0,56 | 0,94 | 3,46E-01 | 4,93E-01 |
| AT2G44310 | 1,85 | 1,37 | 1,46 | 0,94 | 3,47E-01 | 4,95E-01 |
| AT3G54280 | 2,37 | 1,28 | 1,36 | 0,94 | 3,49E-01 | 4,97E-01 |
| AT1G36180 | 2,20 | 1,48 | 1,58 | 0,94 | 3,49E-01 | 4,97E-01 |
| AT2G14910 | 2,97 | 1,12 | 1,19 | 0,94 | 3,50E-01 | 4,97E-01 |

| | | | | | | |
|---|---|---|---|---|---|---|
| AT2G21330 | 7,94 | -0,64 | 0,68 | -0,94 | 3,50E-01 | 4,97E-01 |
| AT5G17230 | 2,13 | 1,32 | 1,41 | 0,94 | 3,50E-01 | 4,97E-01 |
| AT5G58330 | 1,92 | 1,51 | 1,61 | 0,94 | 3,49E-01 | 4,97E-01 |
| AT2G18280 | 2,10 | 1,45 | 1,56 | 0,93 | 3,52E-01 | 5,00E-01 |
| AT4G19170 | 5,23 | -0,79 | 0,85 | -0,93 | 3,54E-01 | 5,01E-01 |
| AT5G47010 | 2,34 | 1,25 | 1,34 | 0,93 | 3,53E-01 | 5,01E-01 |
| AT4G00430 | 2,16 | -1,10 | 1,18 | -0,93 | 3,53E-01 | 5,01E-01 |
| AT5G07740 | 1,77 | 1,42 | 1,53 | 0,93 | 3,53E-01 | 5,01E-01 |
| AT5G25060 | 1,92 | 1,41 | 1,52 | 0,93 | 3,54E-01 | 5,01E-01 |
| AT2G39940 | 1,84 | 1,91 | 2,06 | 0,93 | 3,54E-01 | 5,01E-01 |
| AT5G11670 | 3,41 | 1,01 | 1,09 | 0,93 | 3,54E-01 | 5,01E-01 |
| AT4G26480 | 2,39 | 1,36 | 1,46 | 0,93 | 3,55E-01 | 5,01E-01 |
| AT4G32990 | 2,22 | -1,08 | 1,16 | -0,92 | 3,55E-01 | 5,01E-01 |
| AT4G14030 | 1,74 | 1,31 | 1,42 | 0,92 | 3,56E-01 | 5,02E-01 |
| AT5G19510 | 1,83 | 1,44 | 1,56 | 0,92 | 3,56E-01 | 5,03E-01 |
| AT4G03020 | 2,29 | 1,07 | 1,16 | 0,92 | 3,57E-01 | 5,03E-01 |
| AT5G18380 | 4,02 | -0,93 | 1,01 | -0,92 | 3,57E-01 | 5,03E-01 |
| AT1G79690 | 1,96 | 1,66 | 1,81 | 0,92 | 3,58E-01 | 5,04E-01 |
| AT1G63005 | 3,14 | -1,10 | 1,20 | -0,92 | 3,58E-01 | 5,04E-01 |
| AT5G48860 | 3,76 | -0,91 | 0,99 | -0,92 | 3,58E-01 | 5,04E-01 |
| AT4G25100 | 2,11 | 1,30 | 1,41 | 0,92 | 3,59E-01 | 5,05E-01 |
| AT4G11570 | 2,16 | -1,10 | 1,20 | -0,91 | 3,60E-01 | 5,06E-01 |
| AT1G31730 | 1,97 | 1,73 | 1,90 | 0,91 | 3,61E-01 | 5,07E-01 |
| AT1G29418 | 9,45 | -0,52 | 0,57 | -0,91 | 3,62E-01 | 5,07E-01 |
| AT1G78476 | 11,35 | -0,61 | 0,67 | -0,91 | 3,62E-01 | 5,08E-01 |
| AT3G53740 | 2,01 | 1,84 | 2,02 | 0,91 | 3,63E-01 | 5,08E-01 |
| AT4G35090 | 2,83 | -1,02 | 1,12 | -0,91 | 3,63E-01 | 5,08E-01 |
| AT1G06460 | 1,79 | -1,42 | 1,56 | -0,91 | 3,63E-01 | 5,09E-01 |
| AT5G54130 | 6,22 | -0,88 | 0,97 | -0,91 | 3,65E-01 | 5,09E-01 |
| AT1G31835 | 2,35 | -1,11 | 1,23 | -0,91 | 3,64E-01 | 5,09E-01 |
| AT4G19120 | 2,16 | -1,10 | 1,22 | -0,91 | 3,64E-01 | 5,09E-01 |
| AT5G49720 | 3,89 | -0,91 | 1,00 | -0,91 | 3,65E-01 | 5,09E-01 |
| AT3G57710 | 4,92 | 0,83 | 0,92 | 0,91 | 3,65E-01 | 5,09E-01 |
| AT1G74470 | 19,25 | -0,46 | 0,51 | -0,90 | 3,66E-01 | 5,10E-01 |
| AT4G36360 | 3,63 | 1,16 | 1,29 | 0,90 | 3,66E-01 | 5,10E-01 |
| AT1G26110 | 1,79 | 1,31 | 1,46 | 0,90 | 3,69E-01 | 5,13E-01 |
| AT4G23100 | 1,83 | 1,20 | 1,34 | 0,90 | 3,69E-01 | 5,13E-01 |
| AT2G29080 | 1,97 | 1,25 | 1,39 | 0,90 | 3,70E-01 | 5,15E-01 |
| AT5G14040 | 2,00 | 1,44 | 1,61 | 0,89 | 3,72E-01 | 5,17E-01 |
| AT3G22120 | 4,08 | -0,97 | 1,08 | -0,89 | 3,72E-01 | 5,17E-01 |
| AT2G41105 | 5,08 | 0,82 | 0,92 | 0,89 | 3,73E-01 | 5,18E-01 |
| AT3G01120 | 2,55 | 1,41 | 1,58 | 0,89 | 3,73E-01 | 5,18E-01 |
| AT1G80440 | 3,83 | 1,12 | 1,26 | 0,89 | 3,73E-01 | 5,18E-01 |
| AT5G04540 | 1,96 | 1,41 | 1,58 | 0,89 | 3,73E-01 | 5,18E-01 |
| AT4G03560 | 2,24 | -1,07 | 1,21 | -0,89 | 3,74E-01 | 5,18E-01 |
| AT4G03260 | 2,29 | 1,11 | 1,26 | 0,88 | 3,77E-01 | 5,22E-01 |
| AT1G78900 | 8,95 | 0,61 | 0,70 | 0,88 | 3,79E-01 | 5,24E-01 |
| AT1G67840 | 1,96 | 1,38 | 1,57 | 0,88 | 3,80E-01 | 5,25E-01 |
| AT4G23170 | 6,38 | 0,85 | 0,97 | 0,88 | 3,80E-01 | 5,25E-01 |
| AT3G52800 | 1,93 | 1,41 | 1,61 | 0,88 | 3,80E-01 | 5,25E-01 |
| AT5G51090 | 2,18 | -1,17 | 1,34 | -0,88 | 3,81E-01 | 5,25E-01 |
| AT3G23100 | 2,14 | 1,42 | 1,63 | 0,88 | 3,81E-01 | 5,25E-01 |
| AT5G39740 | 3,11 | 1,08 | 1,24 | 0,87 | 3,82E-01 | 5,27E-01 |
| AT2G19480 | 1,78 | 1,29 | 1,48 | 0,87 | 3,83E-01 | 5,28E-01 |
| AT2G18480 | 2,00 | -1,30 | 1,49 | -0,87 | 3,83E-01 | 5,28E-01 |
| AT4G06110 | 22,73 | -0,48 | 0,55 | -0,87 | 3,84E-01 | 5,29E-01 |

| | | | | | | |
|---|---|---|---|---|---|---|
| AT3G02045 | 5,72 | -0,70 | 0,81 | -0,86 | 3,88E-01 | 5,33E-01 |
| AT4G33240 | 2,23 | 1,36 | 1,57 | 0,86 | 3,88E-01 | 5,34E-01 |
| AT5G62700 | 1,86 | -1,59 | 1,85 | -0,86 | 3,89E-01 | 5,34E-01 |
| AT1G12280 | 1,78 | 1,13 | 1,31 | 0,86 | 3,91E-01 | 5,37E-01 |
| AT4G31210 | 1,90 | 1,43 | 1,67 | 0,86 | 3,92E-01 | 5,38E-01 |
| AT2G43375 | 6,65 | -0,62 | 0,72 | -0,86 | 3,92E-01 | 5,38E-01 |
| AT4G38200 | 2,17 | 1,28 | 1,50 | 0,85 | 3,93E-01 | 5,39E-01 |
| AT2G36490 | 1,90 | 2,32 | 2,73 | 0,85 | 3,96E-01 | 5,42E-01 |
| AT3G44260 | 2,79 | 1,09 | 1,29 | 0,85 | 3,96E-01 | 5,42E-01 |
| AT2G05170 | 1,80 | 1,42 | 1,68 | 0,85 | 3,97E-01 | 5,43E-01 |
| AT3G04120 | 5,88 | 0,68 | 0,80 | 0,85 | 3,97E-01 | 5,43E-01 |
| AT3G22121 | 3,96 | -0,93 | 1,10 | -0,85 | 3,98E-01 | 5,43E-01 |
| AT4G26970 | 1,79 | 1,49 | 1,76 | 0,85 | 3,97E-01 | 5,43E-01 |
| AT4G34980 | 4,56 | 0,84 | 0,99 | 0,85 | 3,97E-01 | 5,43E-01 |
| AT3G12120 | 5,50 | -0,73 | 0,86 | -0,84 | 3,98E-01 | 5,43E-01 |
| AT2G07605 | 7,80 | -0,60 | 0,72 | -0,84 | 3,99E-01 | 5,44E-01 |
| AT1G67090 | 31,55 | -0,39 | 0,46 | -0,84 | 4,00E-01 | 5,45E-01 |
| AT4G06800 | 2,19 | -1,00 | 1,19 | -0,84 | 4,00E-01 | 5,45E-01 |
| AT5G30510 | 1,83 | -1,32 | 1,57 | -0,84 | 4,02E-01 | 5,46E-01 |
| AT1G55920 | 3,12 | 1,10 | 1,31 | 0,84 | 4,03E-01 | 5,47E-01 |
| AT2G23930 | 1,70 | -1,34 | 1,60 | -0,84 | 4,03E-01 | 5,47E-01 |
| AT2G41940 | 1,76 | 1,42 | 1,70 | 0,84 | 4,03E-01 | 5,48E-01 |
| AT3G17400 | 6,94 | -0,67 | 0,80 | -0,83 | 4,04E-01 | 5,48E-01 |
| AT2G20950 | 3,17 | -0,93 | 1,12 | -0,83 | 4,05E-01 | 5,49E-01 |
| AT4G04340 | 2,36 | -1,38 | 1,66 | -0,83 | 4,05E-01 | 5,49E-01 |
| AT3G47500 | 2,57 | -0,98 | 1,18 | -0,83 | 4,05E-01 | 5,49E-01 |
| AT1G12240 | 2,62 | 1,04 | 1,25 | 0,83 | 4,06E-01 | 5,50E-01 |
| AT3G54810 | 2,87 | 1,05 | 1,27 | 0,83 | 4,06E-01 | 5,50E-01 |
| AT4G04925 | 1,79 | 1,40 | 1,69 | 0,83 | 4,07E-01 | 5,50E-01 |
| AT4G14500 | 7,04 | -0,68 | 0,81 | -0,83 | 4,06E-01 | 5,50E-01 |
| AT4G16190 | 11,22 | 0,51 | 0,62 | 0,83 | 4,07E-01 | 5,50E-01 |
| AT4G20260 | 2,30 | -1,05 | 1,27 | -0,83 | 4,07E-01 | 5,50E-01 |
| AT4G22240 | 1,83 | -1,16 | 1,40 | -0,83 | 4,07E-01 | 5,50E-01 |
| AT5G37600 | 1,93 | 1,40 | 1,69 | 0,83 | 4,07E-01 | 5,50E-01 |
| AT3G27865 | 3,38 | -0,99 | 1,20 | -0,83 | 4,08E-01 | 5,50E-01 |
| AT3G56940 | 6,51 | -0,65 | 0,79 | -0,83 | 4,08E-01 | 5,50E-01 |
| AT3G03990 | 2,37 | 1,29 | 1,57 | 0,82 | 4,10E-01 | 5,51E-01 |
| AT5G62670 | 1,85 | 1,47 | 1,78 | 0,82 | 4,10E-01 | 5,52E-01 |
| AT3G29130 | 2,05 | 1,39 | 1,70 | 0,82 | 4,13E-01 | 5,56E-01 |
| AT4G39361 | 1,80 | 1,45 | 1,78 | 0,81 | 4,15E-01 | 5,58E-01 |
| AT1G05805 | 1,94 | 1,82 | 2,25 | 0,81 | 4,17E-01 | 5,60E-01 |
| AT3G29320 | 2,29 | -0,90 | 1,11 | -0,81 | 4,17E-01 | 5,60E-01 |
| AT5G64740 | 4,40 | 0,77 | 0,95 | 0,81 | 4,17E-01 | 5,60E-01 |
| AT2G20010 | 1,89 | 1,27 | 1,56 | 0,81 | 4,17E-01 | 5,60E-01 |
| AT2G18570 | 4,57 | 0,94 | 1,16 | 0,81 | 4,18E-01 | 5,61E-01 |
| AT3G23110 | 1,95 | 1,39 | 1,72 | 0,81 | 4,19E-01 | 5,61E-01 |
| AT1G63750 | 2,64 | 1,03 | 1,27 | 0,81 | 4,19E-01 | 5,61E-01 |
| AT3G62250 | 1,91 | 1,17 | 1,45 | 0,81 | 4,19E-01 | 5,61E-01 |
| AT5G03145 | 2,30 | -1,03 | 1,28 | -0,80 | 4,21E-01 | 5,63E-01 |
| AT1G06430 | 1,82 | -1,06 | 1,32 | -0,80 | 4,22E-01 | 5,63E-01 |
| AT1G57720 | 1,72 | 1,33 | 1,66 | 0,80 | 4,23E-01 | 5,64E-01 |
| AT2G41560 | 2,99 | 1,02 | 1,28 | 0,80 | 4,23E-01 | 5,64E-01 |
| AT3G22940 | 1,72 | 1,52 | 1,89 | 0,80 | 4,24E-01 | 5,65E-01 |
| AT2G38170 | 3,76 | 0,85 | 1,06 | 0,80 | 4,25E-01 | 5,67E-01 |
| AT2G34655 | 4,12 | -0,84 | 1,05 | -0,79 | 4,27E-01 | 5,69E-01 |
| AT1G11840 | 1,70 | 1,27 | 1,61 | 0,79 | 4,28E-01 | 5,71E-01 |

| | | | | | | |
|---|---|---|---|---|---|---|
| AT2G01320 | 2,02 | 1,27 | 1,61 | 0,79 | 4,32E-01 | 5,74E-01 |
| AT3G30390 | 2,05 | 0,94 | 1,20 | 0,79 | 4,32E-01 | 5,74E-01 |
| AT1G06667 | 5,78 | -0,68 | 0,87 | -0,79 | 4,32E-01 | 5,74E-01 |
| AT3G04165 | 2,58 | 1,02 | 1,29 | 0,78 | 4,33E-01 | 5,75E-01 |
| AT4G16990 | 5,03 | 0,70 | 0,89 | 0,78 | 4,33E-01 | 5,75E-01 |
| AT1G52360 | 2,02 | 1,16 | 1,48 | 0,78 | 4,34E-01 | 5,76E-01 |
| AT1G42550 | 2,32 | 1,02 | 1,30 | 0,78 | 4,35E-01 | 5,77E-01 |
| AT2G25171 | 1,84 | 1,33 | 1,71 | 0,78 | 4,36E-01 | 5,78E-01 |
| AT5G41510 | 1,76 | 1,47 | 1,89 | 0,78 | 4,36E-01 | 5,78E-01 |
| AT1G67450 | 1,86 | 1,19 | 1,53 | 0,78 | 4,37E-01 | 5,78E-01 |
| AT2G13560 | 2,19 | -1,25 | 1,62 | -0,78 | 4,37E-01 | 5,78E-01 |
| AT1G80250 | 2,27 | 1,33 | 1,71 | 0,78 | 4,37E-01 | 5,78E-01 |
| AT3G17390 | 4,31 | -0,72 | 0,92 | -0,78 | 4,37E-01 | 5,78E-01 |
| AT1G04480 | 2,74 | 0,99 | 1,27 | 0,77 | 4,39E-01 | 5,79E-01 |
| AT3G04155 | 3,42 | -0,80 | 1,04 | -0,77 | 4,38E-01 | 5,79E-01 |
| AT4G33650 | 1,75 | 1,31 | 1,70 | 0,77 | 4,39E-01 | 5,79E-01 |
| AT1G06950 | 2,18 | 1,55 | 2,00 | 0,77 | 4,39E-01 | 5,79E-01 |
| AT3G07570 | 1,83 | -1,14 | 1,48 | -0,77 | 4,39E-01 | 5,79E-01 |
| AT3G47620 | 3,09 | 0,91 | 1,18 | 0,77 | 4,40E-01 | 5,79E-01 |
| AT5G02575 | 6,74 | -0,78 | 1,01 | -0,77 | 4,40E-01 | 5,79E-01 |
| AT3G07860 | 5,76 | -0,63 | 0,82 | -0,77 | 4,43E-01 | 5,82E-01 |
| AT5G07735 | 1,90 | -1,01 | 1,32 | -0,77 | 4,44E-01 | 5,82E-01 |
| AT1G20440 | 3,88 | -0,68 | 0,88 | -0,77 | 4,43E-01 | 5,82E-01 |
| AT2G20721 | 2,89 | 0,83 | 1,08 | 0,77 | 4,43E-01 | 5,82E-01 |
| AT5G38420 | 5,05 | -0,65 | 0,85 | -0,77 | 4,43E-01 | 5,82E-01 |
| AT3G55830 | 1,72 | 1,30 | 1,70 | 0,76 | 4,45E-01 | 5,83E-01 |
| AT3G48200 | 2,11 | -1,12 | 1,47 | -0,76 | 4,45E-01 | 5,83E-01 |
| AT3G62410 | 1,88 | -0,99 | 1,30 | -0,76 | 4,46E-01 | 5,84E-01 |
| AT1G08347 | 16,30 | -0,36 | 0,47 | -0,76 | 4,45E-01 | 5,84E-01 |
| AT4G34950 | 4,16 | -0,73 | 0,95 | -0,76 | 4,46E-01 | 5,84E-01 |
| AT1G04517 | 11,17 | 0,45 | 0,59 | 0,76 | 4,47E-01 | 5,85E-01 |
| AT1G21590 | 2,73 | 0,98 | 1,29 | 0,76 | 4,48E-01 | 5,86E-01 |
| AT3G49910 | 2,91 | 0,92 | 1,21 | 0,76 | 4,49E-01 | 5,87E-01 |
| AT4G39364 | 2,19 | 1,26 | 1,67 | 0,76 | 4,49E-01 | 5,87E-01 |
| AT1G53640 | 2,14 | -1,12 | 1,48 | -0,76 | 4,50E-01 | 5,87E-01 |
| AT4G06695 | 4,20 | -0,77 | 1,02 | -0,75 | 4,50E-01 | 5,87E-01 |
| AT5G63145 | 2,83 | -0,85 | 1,13 | -0,75 | 4,51E-01 | 5,88E-01 |
| AT1G49500 | 1,83 | -1,02 | 1,36 | -0,75 | 4,52E-01 | 5,89E-01 |
| AT1G74090 | 6,35 | 0,63 | 0,83 | 0,75 | 4,52E-01 | 5,89E-01 |
| AT1G79550 | 2,41 | 0,99 | 1,32 | 0,75 | 4,53E-01 | 5,90E-01 |
| AT4G06705 | 4,08 | -0,75 | 1,00 | -0,75 | 4,53E-01 | 5,90E-01 |
| AT1G06247 | 2,53 | -0,87 | 1,16 | -0,75 | 4,54E-01 | 5,90E-01 |
| AT3G03435 | 3,28 | 0,78 | 1,05 | 0,75 | 4,54E-01 | 5,90E-01 |
| AT5G27950 | 2,87 | -0,80 | 1,07 | -0,75 | 4,55E-01 | 5,91E-01 |
| AT5G38190 | 2,86 | 0,95 | 1,28 | 0,75 | 4,56E-01 | 5,92E-01 |
| AT2G22660 | 2,02 | 1,11 | 1,50 | 0,74 | 4,58E-01 | 5,94E-01 |
| AT1G33420 | 2,44 | -0,82 | 1,10 | -0,74 | 4,58E-01 | 5,94E-01 |
| AT5G10860 | 1,76 | 1,18 | 1,59 | 0,74 | 4,58E-01 | 5,94E-01 |
| AT2G05355 | 1,78 | -1,06 | 1,44 | -0,74 | 4,60E-01 | 5,96E-01 |
| AT2G01850 | 1,74 | -0,98 | 1,33 | -0,74 | 4,62E-01 | 5,98E-01 |
| AT5G02305 | 2,96 | -0,90 | 1,22 | -0,73 | 4,63E-01 | 5,99E-01 |
| AT3G62700 | 3,87 | 0,70 | 0,96 | 0,73 | 4,64E-01 | 6,00E-01 |
| AT3G21760 | 4,32 | -0,66 | 0,90 | -0,73 | 4,65E-01 | 6,00E-01 |
| AT2G37710 | 4,76 | 0,62 | 0,85 | 0,73 | 4,65E-01 | 6,00E-01 |
| AT3G21755 | 4,36 | -0,66 | 0,90 | -0,73 | 4,66E-01 | 6,00E-01 |
| AT3G44110 | 2,03 | 0,95 | 1,30 | 0,73 | 4,66E-01 | 6,00E-01 |

| | | | | | | |
|---|---|---|---|---|---|---|
| AT4G25730 | 1,71 | 1,37 | 1,88 | 0,73 | 4,66E-01 | 6,00E-01 |
| AT5G63800 | 1,87 | -0,89 | 1,23 | -0,73 | 4,67E-01 | 6,02E-01 |
| AT4G25530 | 5,35 | -0,66 | 0,90 | -0,73 | 4,68E-01 | 6,02E-01 |
| AT1G09027 | 2,11 | -0,94 | 1,29 | -0,73 | 4,68E-01 | 6,03E-01 |
| AT5G49910 | 2,46 | 0,97 | 1,33 | 0,72 | 4,69E-01 | 6,03E-01 |
| AT2G34250 | 2,94 | 0,79 | 1,09 | 0,72 | 4,72E-01 | 6,07E-01 |
| AT4G13564 | 27,21 | -0,30 | 0,42 | -0,72 | 4,74E-01 | 6,09E-01 |
| AT5G63190 | 2,74 | 0,91 | 1,27 | 0,71 | 4,75E-01 | 6,09E-01 |
| AT3G46530 | 2,60 | 0,89 | 1,25 | 0,72 | 4,75E-01 | 6,09E-01 |
| AT2G09700 | 2,16 | 1,10 | 1,54 | 0,71 | 4,75E-01 | 6,10E-01 |
| AT2G38195 | 4,06 | -0,90 | 1,27 | -0,71 | 4,78E-01 | 6,13E-01 |
| AT1G04080 | 1,89 | 1,10 | 1,55 | 0,71 | 4,80E-01 | 6,15E-01 |
| AT1G75040 | 3,07 | -0,71 | 1,01 | -0,71 | 4,81E-01 | 6,15E-01 |
| AT3G50590 | 4,47 | 0,80 | 1,14 | 0,71 | 4,80E-01 | 6,15E-01 |
| AT3G51550 | 4,70 | 0,68 | 0,96 | 0,71 | 4,81E-01 | 6,15E-01 |
| AT4G19410 | 1,78 | 1,11 | 1,57 | 0,71 | 4,80E-01 | 6,15E-01 |
| AT5G08610 | 1,71 | 1,10 | 1,57 | 0,70 | 4,82E-01 | 6,15E-01 |
| AT5G07875 | 4,76 | 0,91 | 1,30 | 0,70 | 4,82E-01 | 6,15E-01 |
| AT5G42820 | 2,08 | 1,00 | 1,43 | 0,70 | 4,82E-01 | 6,15E-01 |
| AT3G12780 | 5,73 | -0,54 | 0,77 | -0,70 | 4,83E-01 | 6,16E-01 |
| AT3G03105 | 2,11 | -1,02 | 1,47 | -0,70 | 4,86E-01 | 6,20E-01 |
| AT3G46740 | 3,41 | 0,82 | 1,18 | 0,70 | 4,86E-01 | 6,20E-01 |
| AT2G47470 | 1,77 | 1,16 | 1,66 | 0,70 | 4,87E-01 | 6,20E-01 |
| AT2G25490 | 3,68 | 0,75 | 1,08 | 0,69 | 4,87E-01 | 6,20E-01 |
| AT4G02555 | 3,06 | 0,81 | 1,17 | 0,69 | 4,90E-01 | 6,23E-01 |
| AT2G26975 | 1,93 | 1,06 | 1,54 | 0,69 | 4,90E-01 | 6,23E-01 |
| AT1G13470 | 1,76 | 1,20 | 1,73 | 0,69 | 4,91E-01 | 6,24E-01 |
| AT1G07890 | 4,08 | 0,69 | 1,00 | 0,69 | 4,91E-01 | 6,24E-01 |
| AT5G13180 | 1,85 | 1,11 | 1,61 | 0,69 | 4,93E-01 | 6,26E-01 |
| AT4G17090 | 4,63 | -0,61 | 0,90 | -0,68 | 4,94E-01 | 6,27E-01 |
| AT5G53440 | 2,08 | 0,96 | 1,40 | 0,68 | 4,95E-01 | 6,27E-01 |
| AT2G03890 | 2,03 | 0,92 | 1,35 | 0,68 | 4,95E-01 | 6,28E-01 |
| AT2G37270 | 1,75 | -1,09 | 1,61 | -0,68 | 4,97E-01 | 6,29E-01 |
| AT1G07920 | 7,76 | -0,50 | 0,74 | -0,68 | 4,97E-01 | 6,30E-01 |
| AT1G24510 | 1,77 | 1,03 | 1,52 | 0,68 | 4,99E-01 | 6,31E-01 |
| AT3G28900 | 3,87 | -0,75 | 1,12 | -0,67 | 5,02E-01 | 6,35E-01 |
| AT1G60140 | 2,14 | 0,95 | 1,43 | 0,67 | 5,03E-01 | 6,36E-01 |
| AT4G31980 | 2,53 | -0,91 | 1,36 | -0,67 | 5,06E-01 | 6,38E-01 |
| AT3G60190 | 1,87 | -0,83 | 1,24 | -0,67 | 5,06E-01 | 6,38E-01 |
| AT5G02960 | 2,13 | -0,94 | 1,41 | -0,67 | 5,05E-01 | 6,38E-01 |
| AT5G61310 | 4,76 | -0,58 | 0,87 | -0,67 | 5,06E-01 | 6,38E-01 |
| AT1G07930 | 3,92 | 0,76 | 1,15 | 0,66 | 5,07E-01 | 6,39E-01 |
| AT2G16676 | 2,63 | -0,76 | 1,15 | -0,66 | 5,10E-01 | 6,42E-01 |
| AT5G17165 | 3,90 | -0,76 | 1,16 | -0,66 | 5,10E-01 | 6,42E-01 |
| AT4G39350 | 3,27 | 0,70 | 1,06 | 0,66 | 5,11E-01 | 6,42E-01 |
| AT1G74290 | 1,78 | 1,09 | 1,66 | 0,66 | 5,11E-01 | 6,43E-01 |
| AT3G46600 | 2,19 | 0,80 | 1,23 | 0,66 | 5,12E-01 | 6,43E-01 |
| AT1G69830 | 1,91 | -0,82 | 1,25 | -0,66 | 5,12E-01 | 6,43E-01 |
| AT2G01910 | 2,66 | -1,09 | 1,66 | -0,65 | 5,13E-01 | 6,43E-01 |
| AT5G19120 | 2,51 | -0,81 | 1,23 | -0,66 | 5,12E-01 | 6,43E-01 |
| AT4G01840 | 3,87 | -0,72 | 1,10 | -0,65 | 5,14E-01 | 6,43E-01 |
| AT4G15470 | 1,75 | 1,13 | 1,73 | 0,65 | 5,13E-01 | 6,43E-01 |
| AT5G00760 | 4,31 | -0,58 | 0,89 | -0,65 | 5,14E-01 | 6,43E-01 |
| AT5G03350 | 2,45 | -0,76 | 1,17 | -0,65 | 5,14E-01 | 6,43E-01 |
| AT5G45472 | 4,93 | 0,69 | 1,05 | 0,65 | 5,15E-01 | 6,44E-01 |
| AT2G25250 | 2,81 | 0,86 | 1,33 | 0,65 | 5,17E-01 | 6,46E-01 |

| | | | | | | |
|---|---|---|---|---|---|---|
| AT1G65930 | 2,39 | 0,93 | 1,44 | 0,65 | 5,17E-01 | 6,46E-01 |
| AT5G02645 | 2,33 | -0,84 | 1,29 | -0,65 | 5,17E-01 | 6,46E-01 |
| AT4G02350 | 1,95 | 1,21 | 1,87 | 0,65 | 5,18E-01 | 6,47E-01 |
| AT2G39681 | 1805,59 | -0,21 | 0,32 | -0,64 | 5,19E-01 | 6,48E-01 |
| AT3G60770 | 1,96 | 0,99 | 1,54 | 0,64 | 5,20E-01 | 6,48E-01 |
| AT4G34110 | 4,15 | 0,56 | 0,87 | 0,64 | 5,20E-01 | 6,49E-01 |
| AT3G45780 | 10,09 | -0,36 | 0,56 | -0,64 | 5,21E-01 | 6,49E-01 |
| AT4G14960 | 3,57 | 0,67 | 1,04 | 0,64 | 5,22E-01 | 6,50E-01 |
| AT2G18440 | 3,48 | 0,64 | 1,01 | 0,64 | 5,25E-01 | 6,53E-01 |
| AT1G20260 | 2,71 | -0,79 | 1,25 | -0,63 | 5,26E-01 | 6,54E-01 |
| AT1G09340 | 1,73 | -0,88 | 1,39 | -0,63 | 5,26E-01 | 6,54E-01 |
| AT4G29200 | 3,91 | -0,61 | 0,96 | -0,63 | 5,29E-01 | 6,57E-01 |
| AT5G20290 | 5,09 | 0,58 | 0,92 | 0,63 | 5,28E-01 | 6,57E-01 |
| AT1G55325 | 2,49 | 0,81 | 1,29 | 0,63 | 5,30E-01 | 6,58E-01 |
| AT3G49230 | 1,94 | -1,05 | 1,67 | -0,63 | 5,30E-01 | 6,58E-01 |
| AT3G05530 | 1,78 | 0,96 | 1,53 | 0,63 | 5,31E-01 | 6,59E-01 |
| AT2G09555 | 2,08 | -0,84 | 1,35 | -0,62 | 5,33E-01 | 6,61E-01 |
| AT4G07120 | 22,20 | -0,34 | 0,55 | -0,62 | 5,34E-01 | 6,61E-01 |
| AT4G34100 | 3,45 | 0,75 | 1,20 | 0,62 | 5,34E-01 | 6,61E-01 |
| AT3G18370 | 2,55 | 1,00 | 1,60 | 0,62 | 5,34E-01 | 6,61E-01 |
| AT3G22231 | 4,08 | -0,55 | 0,89 | -0,62 | 5,37E-01 | 6,64E-01 |
| AT2G07727 | 3,34 | -0,82 | 1,34 | -0,62 | 5,38E-01 | 6,65E-01 |
| AT5G62190 | 2,05 | 1,05 | 1,72 | 0,61 | 5,39E-01 | 6,66E-01 |
| AT1G29930 | 124,68 | 0,24 | 0,39 | 0,61 | 5,40E-01 | 6,67E-01 |
| AT1G31910 | 2,79 | 0,68 | 1,11 | 0,61 | 5,41E-01 | 6,68E-01 |
| AT3G57730 | 5,17 | 0,53 | 0,86 | 0,61 | 5,42E-01 | 6,69E-01 |
| AT5G12250 | 2,28 | 0,75 | 1,24 | 0,61 | 5,45E-01 | 6,72E-01 |
| AT3G03780 | 4,16 | 0,62 | 1,02 | 0,60 | 5,46E-01 | 6,72E-01 |
| AT1G44446 | 1,72 | 0,95 | 1,57 | 0,60 | 5,46E-01 | 6,72E-01 |
| AT4G18480 | 1,72 | 0,84 | 1,39 | 0,60 | 5,46E-01 | 6,72E-01 |
| AT1G04183 | 2,59 | 0,80 | 1,33 | 0,60 | 5,48E-01 | 6,74E-01 |
| AT2G05765 | 3,87 | -0,66 | 1,10 | -0,60 | 5,48E-01 | 6,74E-01 |
| AT1G30330 | 1,89 | -0,89 | 1,48 | -0,60 | 5,50E-01 | 6,75E-01 |
| AT4G18197 | 1,87 | -0,84 | 1,41 | -0,60 | 5,50E-01 | 6,76E-01 |
| AT1G03410 | 2,63 | 0,79 | 1,33 | 0,60 | 5,51E-01 | 6,76E-01 |
| AT1G68945 | 2,15 | -0,75 | 1,25 | -0,60 | 5,52E-01 | 6,77E-01 |
| AT1G69620 | 2,14 | 0,86 | 1,45 | 0,59 | 5,55E-01 | 6,79E-01 |
| AT2G07806 | 2,71 | -0,79 | 1,34 | -0,59 | 5,54E-01 | 6,79E-01 |
| AT2G30440 | 1,74 | 0,95 | 1,61 | 0,59 | 5,55E-01 | 6,79E-01 |
| AT2G41530 | 2,42 | -0,82 | 1,39 | -0,59 | 5,54E-01 | 6,79E-01 |
| AT4G29060 | 2,00 | 0,84 | 1,43 | 0,59 | 5,55E-01 | 6,79E-01 |
| AT5G60390 | 16,82 | 0,32 | 0,55 | 0,59 | 5,56E-01 | 6,79E-01 |
| AT1G30400 | 2,52 | 0,77 | 1,32 | 0,59 | 5,57E-01 | 6,81E-01 |
| AT2G09465 | 4,22 | -0,54 | 0,93 | -0,58 | 5,59E-01 | 6,83E-01 |
| AT4G18030 | 2,74 | -0,73 | 1,24 | -0,58 | 5,59E-01 | 6,83E-01 |
| AT5G41471 | 1,79 | -1,02 | 1,74 | -0,58 | 5,60E-01 | 6,83E-01 |
| AT3G61430 | 1,94 | 0,97 | 1,66 | 0,58 | 5,61E-01 | 6,84E-01 |
| AT5G12860 | 2,32 | 0,97 | 1,67 | 0,58 | 5,63E-01 | 6,86E-01 |
| AT1G50250 | 2,35 | 0,69 | 1,19 | 0,58 | 5,64E-01 | 6,87E-01 |
| AT3G07390 | 2,06 | -0,85 | 1,48 | -0,57 | 5,67E-01 | 6,90E-01 |
| AT4G02890 | 2,39 | 0,69 | 1,20 | 0,57 | 5,68E-01 | 6,91E-01 |
| AT4G02520 | 4,48 | 0,59 | 1,04 | 0,57 | 5,68E-01 | 6,91E-01 |
| AT1G53655 | 1,78 | -0,91 | 1,59 | -0,57 | 5,69E-01 | 6,92E-01 |
| AT4G26542 | 2,99 | 0,63 | 1,11 | 0,57 | 5,69E-01 | 6,92E-01 |
| AT1G72370 | 2,39 | 0,76 | 1,33 | 0,57 | 5,71E-01 | 6,93E-01 |
| AT5G57560 | 1,92 | 0,88 | 1,56 | 0,57 | 5,71E-01 | 6,94E-01 |

| | | | | | | |
|---|---|---|---|---|---|---|
| AT1G28070 | 4,21 | -1,17 | 2,07 | -0,57 | 5,72E-01 | 6,94E-01 |
| AT1G76065 | 4,52 | 0,64 | 1,13 | 0,56 | 5,73E-01 | 6,96E-01 |
| AT1G20020 | 2,67 | -0,60 | 1,06 | -0,56 | 5,75E-01 | 6,96E-01 |
| AT5G28040 | 2,94 | 0,71 | 1,26 | 0,56 | 5,75E-01 | 6,96E-01 |
| AT5G42270 | 4,20 | 0,54 | 0,96 | 0,56 | 5,75E-01 | 6,97E-01 |
| AT4G06375 | 3,09 | 0,76 | 1,36 | 0,56 | 5,77E-01 | 6,98E-01 |
| AT3G05655 | 1,87 | 0,92 | 1,64 | 0,56 | 5,77E-01 | 6,98E-01 |
| AT1G22060 | 2,54 | 0,70 | 1,27 | 0,56 | 5,78E-01 | 6,99E-01 |
| AT2G29450 | 2,19 | 0,92 | 1,66 | 0,55 | 5,79E-01 | 6,99E-01 |
| AT4G04890 | 2,53 | -1,01 | 1,82 | -0,56 | 5,79E-01 | 6,99E-01 |
| AT1G42470 | 1,71 | 1,03 | 1,87 | 0,55 | 5,83E-01 | 7,03E-01 |
| AT5G02325 | 15,37 | -0,40 | 0,73 | -0,55 | 5,83E-01 | 7,03E-01 |
| AT5G20010 | 1,98 | 0,83 | 1,51 | 0,55 | 5,83E-01 | 7,03E-01 |
| AT1G07140 | 1,81 | 0,91 | 1,65 | 0,55 | 5,82E-01 | 7,03E-01 |
| AT1G09937 | 6,18 | 0,45 | 0,82 | 0,55 | 5,82E-01 | 7,03E-01 |
| AT5G26000 | 4,62 | -0,45 | 0,82 | -0,55 | 5,84E-01 | 7,03E-01 |
| AT1G04527 | 11,72 | -0,30 | 0,54 | -0,55 | 5,84E-01 | 7,03E-01 |
| AT3G21750 | 1,77 | 0,76 | 1,40 | 0,54 | 5,87E-01 | 7,06E-01 |
| AT4G05320 | 17,67 | 0,28 | 0,52 | 0,54 | 5,87E-01 | 7,06E-01 |
| AT4G33010 | 11,10 | 0,39 | 0,71 | 0,54 | 5,87E-01 | 7,06E-01 |
| AT1G55350 | 1,89 | -1,17 | 2,16 | -0,54 | 5,88E-01 | 7,06E-01 |
| AT5G05660 | 1,92 | 1,28 | 2,37 | 0,54 | 5,88E-01 | 7,06E-01 |
| AT5G47690 | 3,16 | 0,67 | 1,25 | 0,54 | 5,89E-01 | 7,07E-01 |
| AT3G48720 | 18,61 | -0,31 | 0,57 | -0,54 | 5,91E-01 | 7,09E-01 |
| AT1G73870 | 2,16 | -0,71 | 1,32 | -0,54 | 5,92E-01 | 7,09E-01 |
| AT1G67430 | 1,82 | 0,76 | 1,43 | 0,53 | 5,93E-01 | 7,10E-01 |
| AT4G39366 | 2,55 | -0,63 | 1,17 | -0,54 | 5,93E-01 | 7,10E-01 |
| AT5G09870 | 2,84 | 0,59 | 1,11 | 0,53 | 5,94E-01 | 7,10E-01 |
| AT1G27870 | 1,97 | -0,77 | 1,45 | -0,53 | 5,94E-01 | 7,10E-01 |
| AT3G47348 | 6,89 | -0,59 | 1,12 | -0,53 | 5,97E-01 | 7,14E-01 |
| AT2G07725 | 2,14 | -0,64 | 1,21 | -0,53 | 5,99E-01 | 7,15E-01 |
| AT4G13770 | 2,50 | 0,73 | 1,38 | 0,53 | 5,99E-01 | 7,15E-01 |
| AT3G52700 | 2,20 | -0,61 | 1,17 | -0,52 | 6,00E-01 | 7,15E-01 |
| AT5G22770 | 2,48 | 0,71 | 1,36 | 0,52 | 6,00E-01 | 7,15E-01 |
| AT5G45095 | 2,04 | 0,74 | 1,42 | 0,52 | 6,03E-01 | 7,18E-01 |
| AT4G26540 | 3,42 | 0,58 | 1,11 | 0,52 | 6,03E-01 | 7,19E-01 |
| AT3G56360 | 3,83 | -0,60 | 1,17 | -0,52 | 6,04E-01 | 7,19E-01 |
| AT5G52530 | 3,03 | 0,78 | 1,50 | 0,52 | 6,04E-01 | 7,19E-01 |
| AT5G56010 | 6,62 | 0,45 | 0,86 | 0,52 | 6,05E-01 | 7,19E-01 |
| AT1G48920 | 4,08 | -0,50 | 0,98 | -0,52 | 6,05E-01 | 7,20E-01 |
| AT5G63570 | 1,81 | -0,81 | 1,57 | -0,52 | 6,05E-01 | 7,20E-01 |
| AT5G57660 | 4,42 | 0,42 | 0,82 | 0,51 | 6,07E-01 | 7,21E-01 |
| AT2G30920 | 2,18 | -0,69 | 1,34 | -0,51 | 6,08E-01 | 7,21E-01 |
| AT3G15353 | 5,91 | 0,53 | 1,03 | 0,51 | 6,08E-01 | 7,21E-01 |
| AT1G20010 | 2,01 | -0,70 | 1,36 | -0,51 | 6,08E-01 | 7,22E-01 |
| AT3G20310 | 2,42 | 0,70 | 1,36 | 0,51 | 6,10E-01 | 7,23E-01 |
| AT1G12920 | 1,87 | 0,72 | 1,42 | 0,51 | 6,11E-01 | 7,24E-01 |
| AT2G02560 | 1,80 | 0,78 | 1,54 | 0,51 | 6,13E-01 | 7,25E-01 |
| AT5G61420 | 2,01 | 0,81 | 1,60 | 0,51 | 6,13E-01 | 7,25E-01 |
| AT3G19170 | 4,99 | 0,51 | 1,02 | 0,50 | 6,20E-01 | 7,34E-01 |
| AT1G62035 | 5,54 | -0,43 | 0,87 | -0,49 | 6,21E-01 | 7,34E-01 |
| AT1G72710 | 1,73 | 0,74 | 1,50 | 0,49 | 6,23E-01 | 7,35E-01 |
| AT5G03940 | 2,83 | 0,56 | 1,14 | 0,49 | 6,22E-01 | 7,35E-01 |
| AT4G18150 | 2,31 | -0,68 | 1,39 | -0,49 | 6,24E-01 | 7,36E-01 |
| AT3G16140 | 1,91 | -0,71 | 1,45 | -0,49 | 6,24E-01 | 7,36E-01 |
| AT5G28540 | 7,23 | 0,38 | 0,77 | 0,49 | 6,25E-01 | 7,37E-01 |

| | | | | | | |
|---|---|---|---|---|---|---|
| AT5G54940 | 2,86 | 0,55 | 1,13 | 0,49 | 6,25E-01 | 7,37E-01 |
| AT5G09590 | 2,74 | 0,70 | 1,44 | 0,49 | 6,26E-01 | 7,37E-01 |
| AT5G46430 | 2,53 | 0,67 | 1,38 | 0,49 | 6,26E-01 | 7,37E-01 |
| AT1G75950 | 1,80 | -0,59 | 1,21 | -0,49 | 6,28E-01 | 7,39E-01 |
| AT1G47497 | 2,25 | -0,74 | 1,53 | -0,48 | 6,29E-01 | 7,40E-01 |
| AT1G78206 | 14,29 | 0,26 | 0,55 | 0,48 | 6,31E-01 | 7,42E-01 |
| AT1G06040 | 2,85 | -0,60 | 1,25 | -0,48 | 6,33E-01 | 7,43E-01 |
| AT2G37160 | 3,77 | 0,48 | 1,01 | 0,48 | 6,33E-01 | 7,44E-01 |
| AT2G04955 | 3,52 | 0,53 | 1,11 | 0,48 | 6,33E-01 | 7,44E-01 |
| AT3G57520 | 3,92 | 0,56 | 1,18 | 0,48 | 6,34E-01 | 7,44E-01 |
| AT5G02810 | 1,73 | 0,85 | 1,78 | 0,48 | 6,34E-01 | 7,44E-01 |
| AT1G63740 | 1,93 | 0,66 | 1,40 | 0,47 | 6,36E-01 | 7,45E-01 |
| AT2G40840 | 4,12 | 0,43 | 0,90 | 0,47 | 6,37E-01 | 7,45E-01 |
| AT5G06165 | 4,41 | -0,46 | 0,96 | -0,47 | 6,36E-01 | 7,45E-01 |
| AT3G53430 | 2,55 | -0,54 | 1,15 | -0,47 | 6,37E-01 | 7,46E-01 |
| AT3G26520 | 6,24 | -0,40 | 0,85 | -0,47 | 6,38E-01 | 7,46E-01 |
| AT1G76030 | 2,84 | -0,60 | 1,28 | -0,47 | 6,39E-01 | 7,47E-01 |
| AT1G76062 | 4,01 | 0,57 | 1,22 | 0,47 | 6,39E-01 | 7,47E-01 |
| AT3G49470 | 2,41 | -0,64 | 1,39 | -0,47 | 6,42E-01 | 7,48E-01 |
| AT5G02870 | 2,41 | -0,63 | 1,35 | -0,47 | 6,42E-01 | 7,48E-01 |
| AT1G07670 | 1,74 | -0,66 | 1,41 | -0,47 | 6,42E-01 | 7,48E-01 |
| AT1G33240 | 1,91 | -0,77 | 1,65 | -0,47 | 6,41E-01 | 7,48E-01 |
| AT3G46510 | 1,83 | 0,66 | 1,44 | 0,46 | 6,46E-01 | 7,53E-01 |
| AT3G44444 | 1,95 | -0,70 | 1,53 | -0,46 | 6,48E-01 | 7,55E-01 |
| AT4G27520 | 5,05 | -0,45 | 1,00 | -0,46 | 6,49E-01 | 7,56E-01 |
| AT3G59940 | 11,46 | -0,30 | 0,65 | -0,45 | 6,50E-01 | 7,56E-01 |
| AT5G01410 | 2,12 | 0,63 | 1,39 | 0,45 | 6,50E-01 | 7,56E-01 |
| AT5G10840 | 2,05 | -0,62 | 1,36 | -0,45 | 6,50E-01 | 7,56E-01 |
| AT2G38220 | 2,26 | -0,57 | 1,27 | -0,45 | 6,51E-01 | 7,57E-01 |
| AT3G46970 | 7,07 | -0,32 | 0,71 | -0,45 | 6,53E-01 | 7,59E-01 |
| AT5G02380 | 2,77 | 0,62 | 1,39 | 0,45 | 6,53E-01 | 7,59E-01 |
| AT3G09275 | 11,41 | -0,29 | 0,66 | -0,45 | 6,54E-01 | 7,59E-01 |
| AT1G33415 | 2,72 | -0,48 | 1,07 | -0,45 | 6,55E-01 | 7,60E-01 |
| AT2G01010 | 3,38 | 0,55 | 1,24 | 0,45 | 6,56E-01 | 7,60E-01 |
| AT5G58950 | 2,42 | 0,55 | 1,23 | 0,44 | 6,57E-01 | 7,61E-01 |
| AT1G33950 | 1,88 | -0,76 | 1,74 | -0,44 | 6,61E-01 | 7,66E-01 |
| AT5G59732 | 3,71 | 0,53 | 1,22 | 0,43 | 6,64E-01 | 7,69E-01 |
| AT5G59730 | 3,71 | 0,53 | 1,22 | 0,43 | 6,64E-01 | 7,69E-01 |
| AT1G09210 | 5,83 | 0,35 | 0,81 | 0,43 | 6,66E-01 | 7,70E-01 |
| AT1G33590 | 1,70 | -0,59 | 1,37 | -0,43 | 6,67E-01 | 7,70E-01 |
| AT2G03445 | 309,93 | 0,11 | 0,26 | 0,43 | 6,68E-01 | 7,71E-01 |
| AT4G18210 | 1,90 | -0,68 | 1,59 | -0,43 | 6,68E-01 | 7,71E-01 |
| AT2G39010 | 4,44 | -0,46 | 1,09 | -0,43 | 6,69E-01 | 7,72E-01 |
| AT5G44870 | 1,81 | 0,60 | 1,41 | 0,43 | 6,69E-01 | 7,72E-01 |
| AT4G39330 | 1,70 | -0,64 | 1,50 | -0,42 | 6,71E-01 | 7,74E-01 |
| AT3G56070 | 2,75 | -0,45 | 1,07 | -0,42 | 6,72E-01 | 7,75E-01 |
| AT5G15650 | 3,70 | 0,47 | 1,11 | 0,42 | 6,74E-01 | 7,77E-01 |
| AT1G53210 | 1,83 | -0,68 | 1,64 | -0,42 | 6,78E-01 | 7,80E-01 |
| AT2G41650 | 5,84 | -0,31 | 0,74 | -0,42 | 6,78E-01 | 7,80E-01 |
| AT2G43110 | 4,82 | 0,42 | 1,01 | 0,41 | 6,78E-01 | 7,80E-01 |
| AT1G19410 | 3,18 | -0,41 | 0,98 | -0,41 | 6,79E-01 | 7,80E-01 |
| AT2G41840 | 3,13 | 0,51 | 1,23 | 0,41 | 6,81E-01 | 7,82E-01 |
| AT1G50010 | 2,93 | 0,52 | 1,26 | 0,41 | 6,82E-01 | 7,83E-01 |
| AT4G39090 | 7,03 | 0,31 | 0,77 | 0,41 | 6,85E-01 | 7,86E-01 |
| AT1G33070 | 1,82 | -0,68 | 1,68 | -0,40 | 6,86E-01 | 7,87E-01 |
| AT1G03743 | 2,10 | -0,56 | 1,39 | -0,40 | 6,87E-01 | 7,87E-01 |

| | | | | | | |
|---|---|---|---|---|---|---|
| AT3G00985 | 2,03 | -0,77 | 1,91 | -0,40 | 6,88E-01 | 7,88E-01 |
| AT1G01550 | 2,54 | 0,59 | 1,48 | 0,40 | 6,89E-01 | 7,89E-01 |
| AT2G41720 | 1,87 | 0,72 | 1,81 | 0,40 | 6,91E-01 | 7,91E-01 |
| AT5G45470 | 2,49 | 0,64 | 1,61 | 0,40 | 6,91E-01 | 7,91E-01 |
| AT1G15690 | 11,49 | 0,25 | 0,63 | 0,39 | 6,93E-01 | 7,93E-01 |
| AT1G26975 | 8,83 | 0,26 | 0,68 | 0,39 | 6,98E-01 | 7,98E-01 |
| AT1G07593 | 3,78 | -0,57 | 1,47 | -0,39 | 6,99E-01 | 7,99E-01 |
| AT5G18410 | 3,15 | 0,47 | 1,23 | 0,39 | 7,00E-01 | 7,99E-01 |
| AT2G04375 | 5,42 | 0,32 | 0,83 | 0,38 | 7,01E-01 | 8,00E-01 |
| AT5G40890 | 1,74 | 0,58 | 1,50 | 0,38 | 7,02E-01 | 8,01E-01 |
| AT1G28440 | 1,72 | -0,53 | 1,38 | -0,38 | 7,03E-01 | 8,01E-01 |
| AT4G04095 | 1,94 | -0,51 | 1,35 | -0,38 | 7,04E-01 | 8,02E-01 |
| AT5G01915 | 2,27 | 0,53 | 1,41 | 0,38 | 7,04E-01 | 8,02E-01 |
| AT2G29980 | 3,37 | 0,52 | 1,36 | 0,38 | 7,05E-01 | 8,02E-01 |
| AT4G37930 | 4,20 | -0,33 | 0,86 | -0,38 | 7,04E-01 | 8,02E-01 |
| AT1G68560 | 3,34 | 0,42 | 1,12 | 0,38 | 7,06E-01 | 8,03E-01 |
| AT4G02760 | 2,62 | 0,59 | 1,57 | 0,38 | 7,07E-01 | 8,03E-01 |
| AT2G31820 | 4,59 | -0,32 | 0,87 | -0,37 | 7,08E-01 | 8,04E-01 |
| AT2G08855 | 3,73 | 0,38 | 1,03 | 0,37 | 7,09E-01 | 8,04E-01 |
| AT1G76010 | 2,04 | -0,46 | 1,24 | -0,37 | 7,09E-01 | 8,04E-01 |
| AT3G00980 | 2,10 | -0,70 | 1,88 | -0,37 | 7,08E-01 | 8,04E-01 |
| AT3G52880 | 2,23 | 0,44 | 1,19 | 0,37 | 7,09E-01 | 8,04E-01 |
| AT5G11790 | 2,58 | 0,44 | 1,18 | 0,37 | 7,10E-01 | 8,04E-01 |
| AT3G59068 | 1,78 | 0,65 | 1,76 | 0,37 | 7,13E-01 | 8,08E-01 |
| AT3G59070 | 1,78 | 0,65 | 1,76 | 0,37 | 7,13E-01 | 8,08E-01 |
| AT1G74453 | 1,99 | 0,57 | 1,56 | 0,37 | 7,15E-01 | 8,09E-01 |
| AT1G08115 | 40,91 | -0,16 | 0,44 | -0,36 | 7,18E-01 | 8,13E-01 |
| AT2G41430 | 6,85 | 0,29 | 0,83 | 0,36 | 7,21E-01 | 8,15E-01 |
| AT4G31990 | 1,81 | -0,65 | 1,82 | -0,36 | 7,22E-01 | 8,16E-01 |
| AT3G05335 | 1,73 | -0,49 | 1,38 | -0,35 | 7,24E-01 | 8,17E-01 |
| AT4G22710 | 1,76 | -0,68 | 1,93 | -0,35 | 7,24E-01 | 8,17E-01 |
| AT5G47430 | 3,26 | 0,40 | 1,17 | 0,35 | 7,29E-01 | 8,23E-01 |
| AT1G65350 | 2,42 | 0,49 | 1,43 | 0,35 | 7,30E-01 | 8,23E-01 |
| AT2G26910 | 1,87 | 0,53 | 1,55 | 0,34 | 7,30E-01 | 8,23E-01 |
| AT4G23850 | 2,43 | 0,40 | 1,16 | 0,34 | 7,31E-01 | 8,23E-01 |
| AT2G07718 | 3,36 | 0,40 | 1,18 | 0,34 | 7,33E-01 | 8,25E-01 |
| AT4G12650 | 2,23 | 0,42 | 1,24 | 0,34 | 7,34E-01 | 8,26E-01 |
| AT1G19960 | 1,88 | -0,55 | 1,64 | -0,34 | 7,36E-01 | 8,28E-01 |
| AT1G23190 | 1,73 | 0,61 | 1,82 | 0,34 | 7,36E-01 | 8,28E-01 |
| AT5G28840 | 2,50 | 0,38 | 1,13 | 0,34 | 7,37E-01 | 8,28E-01 |
| AT4G01150 | 2,10 | 0,41 | 1,23 | 0,34 | 7,38E-01 | 8,29E-01 |
| AT4G11270 | 3,79 | -0,35 | 1,06 | -0,33 | 7,43E-01 | 8,33E-01 |
| AT5G49760 | 3,34 | -0,37 | 1,12 | -0,33 | 7,43E-01 | 8,33E-01 |
| AT1G05030 | 1,78 | -0,53 | 1,61 | -0,33 | 7,43E-01 | 8,33E-01 |
| AT2G42590 | 2,06 | 0,43 | 1,31 | 0,33 | 7,43E-01 | 8,33E-01 |
| AT1G50900 | 1,85 | -0,44 | 1,35 | -0,33 | 7,45E-01 | 8,35E-01 |
| AT5G58070 | 2,01 | 0,50 | 1,53 | 0,32 | 7,46E-01 | 8,36E-01 |
| AT2G30860 | 6,26 | -0,26 | 0,82 | -0,32 | 7,48E-01 | 8,37E-01 |
| AT4G03280 | 3,09 | 0,40 | 1,23 | 0,32 | 7,48E-01 | 8,37E-01 |
| AT3G11910 | 1,81 | 0,41 | 1,29 | 0,32 | 7,48E-01 | 8,37E-01 |
| AT5G13850 | 2,26 | 0,41 | 1,30 | 0,32 | 7,49E-01 | 8,38E-01 |
| AT1G19373 | 3,55 | -0,37 | 1,16 | -0,32 | 7,52E-01 | 8,38E-01 |
| AT1G69572 | 2,44 | 0,47 | 1,48 | 0,31 | 7,54E-01 | 8,38E-01 |
| AT1G78080 | 3,84 | 0,36 | 1,15 | 0,31 | 7,54E-01 | 8,38E-01 |
| AT3G14930 | 2,04 | -0,45 | 1,42 | -0,32 | 7,51E-01 | 8,38E-01 |
| AT3G20820 | 3,82 | -0,39 | 1,23 | -0,31 | 7,54E-01 | 8,38E-01 |

| | | | | | | |
|---|---|---|---|---|---|---|
| AT3G06325 | 6,42 | 0,26 | 0,82 | 0,31 | 7,54E-01 | 8,38E-01 |
| AT1G05850 | 2,93 | -0,42 | 1,32 | -0,32 | 7,52E-01 | 8,38E-01 |
| AT1G11910 | 5,12 | -0,31 | 0,98 | -0,32 | 7,52E-01 | 8,38E-01 |
| AT1G20860 | 3,03 | 0,36 | 1,14 | 0,32 | 7,52E-01 | 8,38E-01 |
| AT1G69570 | 2,44 | 0,47 | 1,48 | 0,31 | 7,54E-01 | 8,38E-01 |
| AT2G05685 | 2,44 | 0,44 | 1,40 | 0,32 | 7,52E-01 | 8,38E-01 |
| AT5G15200 | 4,33 | -0,28 | 0,89 | -0,32 | 7,52E-01 | 8,38E-01 |
| AT3G05220 | 1,84 | 0,53 | 1,71 | 0,31 | 7,57E-01 | 8,41E-01 |
| AT1G65440 | 2,14 | -0,39 | 1,26 | -0,31 | 7,57E-01 | 8,41E-01 |
| AT1G15670 | 2,03 | 0,42 | 1,40 | 0,30 | 7,63E-01 | 8,46E-01 |
| AT5G08680 | 1,77 | -0,51 | 1,69 | -0,30 | 7,63E-01 | 8,46E-01 |
| AT5G27720 | 3,01 | 0,34 | 1,13 | 0,30 | 7,63E-01 | 8,46E-01 |
| AT1G56280 | 2,59 | -0,35 | 1,15 | -0,30 | 7,63E-01 | 8,46E-01 |
| AT3G52500 | 2,36 | -0,37 | 1,23 | -0,30 | 7,62E-01 | 8,46E-01 |
| AT5G10450 | 3,14 | 0,32 | 1,07 | 0,30 | 7,64E-01 | 8,46E-01 |
| AT1G35560 | 2,43 | -0,36 | 1,20 | -0,30 | 7,64E-01 | 8,46E-01 |
| AT5G09810 | 3,96 | -0,34 | 1,16 | -0,29 | 7,69E-01 | 8,50E-01 |
| AT1G20690 | 2,85 | -0,32 | 1,11 | -0,29 | 7,70E-01 | 8,51E-01 |
| AT1G20691 | 2,85 | -0,32 | 1,11 | -0,29 | 7,70E-01 | 8,51E-01 |
| AT5G16620 | 2,01 | -0,35 | 1,20 | -0,29 | 7,73E-01 | 8,54E-01 |
| AT2G47240 | 1,70 | -0,42 | 1,46 | -0,29 | 7,74E-01 | 8,54E-01 |
| AT5G24890 | 2,39 | 0,38 | 1,35 | 0,28 | 7,76E-01 | 8,57E-01 |
| AT4G32640 | 2,41 | 0,37 | 1,31 | 0,28 | 7,77E-01 | 8,57E-01 |
| AT1G46696 | 1,91 | 0,43 | 1,53 | 0,28 | 7,77E-01 | 8,57E-01 |
| AT4G02320 | 3,77 | -0,29 | 1,01 | -0,28 | 7,78E-01 | 8,58E-01 |
| AT3G12980 | 1,82 | 0,51 | 1,83 | 0,28 | 7,79E-01 | 8,59E-01 |
| AT2G08725 | 34,05 | 0,17 | 0,60 | 0,28 | 7,80E-01 | 8,59E-01 |
| AT5G52320 | 1,77 | -0,39 | 1,42 | -0,27 | 7,84E-01 | 8,63E-01 |
| AT1G05563 | 2,75 | -0,31 | 1,15 | -0,27 | 7,86E-01 | 8,65E-01 |
| AT3G15210 | 1,70 | -0,37 | 1,36 | -0,27 | 7,89E-01 | 8,67E-01 |
| AT1G75800 | 1,71 | 0,48 | 1,82 | 0,27 | 7,90E-01 | 8,67E-01 |
| AT3G59780 | 2,03 | -0,34 | 1,27 | -0,27 | 7,90E-01 | 8,67E-01 |
| AT4G23420 | 23,63 | -0,13 | 0,50 | -0,27 | 7,90E-01 | 8,67E-01 |
| AT4G11900 | 2,31 | 0,33 | 1,25 | 0,27 | 7,90E-01 | 8,67E-01 |
| AT1G70490 | 1,85 | -0,39 | 1,49 | -0,26 | 7,93E-01 | 8,69E-01 |
| AT5G41663 | 49,62 | 0,12 | 0,48 | 0,26 | 7,93E-01 | 8,69E-01 |
| AT5G07725 | 8,72 | 0,21 | 0,79 | 0,26 | 7,94E-01 | 8,70E-01 |
| AT3G44310 | 10,52 | -0,18 | 0,69 | -0,26 | 7,95E-01 | 8,71E-01 |
| AT5G08525 | 4,34 | 0,27 | 1,06 | 0,26 | 7,97E-01 | 8,73E-01 |
| AT3G03425 | 4,95 | 0,21 | 0,82 | 0,26 | 7,97E-01 | 8,73E-01 |
| AT4G27090 | 2,08 | 0,34 | 1,33 | 0,25 | 8,01E-01 | 8,76E-01 |
| AT2G36470 | 2,06 | 0,43 | 1,72 | 0,25 | 8,01E-01 | 8,76E-01 |
| AT1G33120 | 1,97 | 0,32 | 1,28 | 0,25 | 8,03E-01 | 8,77E-01 |
| AT1G05817 | 1,94 | -0,38 | 1,51 | -0,25 | 8,03E-01 | 8,78E-01 |
| AT1G64500 | 2,64 | -0,31 | 1,24 | -0,25 | 8,05E-01 | 8,78E-01 |
| AT2G47440 | 2,19 | 0,39 | 1,57 | 0,25 | 8,04E-01 | 8,78E-01 |
| AT1G71002 | 2,81 | -0,30 | 1,23 | -0,25 | 8,05E-01 | 8,78E-01 |
| AT5G19770 | 1,75 | 0,48 | 1,95 | 0,25 | 8,05E-01 | 8,78E-01 |
| AT1G32630 | 4,04 | 0,25 | 1,04 | 0,24 | 8,08E-01 | 8,79E-01 |
| AT2G44060 | 2,21 | -0,30 | 1,24 | -0,24 | 8,08E-01 | 8,79E-01 |
| AT1G32640 | 4,04 | 0,25 | 1,04 | 0,24 | 8,08E-01 | 8,79E-01 |
| AT5G19150 | 6,35 | -0,18 | 0,74 | -0,24 | 8,08E-01 | 8,79E-01 |
| AT5G40384 | 2,72 | 0,34 | 1,43 | 0,24 | 8,11E-01 | 8,82E-01 |
| AT5G57110 | 2,43 | 0,36 | 1,49 | 0,24 | 8,11E-01 | 8,82E-01 |
| AT2G28840 | 1,76 | 0,39 | 1,62 | 0,24 | 8,11E-01 | 8,82E-01 |
| AT4G26255 | 3,93 | -0,33 | 1,41 | -0,24 | 8,13E-01 | 8,83E-01 |

| | | | | | | |
|---|---|---|---|---|---|---|
| AT5G44090 | 1,96 | 0,40 | 1,68 | 0,24 | 8,13E-01 | 8,83E-01 |
| AT1G06243 | 5,81 | -0,17 | 0,75 | -0,23 | 8,15E-01 | 8,84E-01 |
| AT5G60360 | 2,97 | -0,28 | 1,22 | -0,23 | 8,16E-01 | 8,85E-01 |
| AT4G00975 | 11,27 | 0,13 | 0,54 | 0,23 | 8,18E-01 | 8,86E-01 |
| AT5G47560 | 3,13 | 0,27 | 1,20 | 0,23 | 8,19E-01 | 8,88E-01 |
| AT5G01785 | 2,53 | 0,37 | 1,64 | 0,23 | 8,20E-01 | 8,88E-01 |
| AT1G69792 | 2,23 | -0,33 | 1,46 | -0,23 | 8,21E-01 | 8,89E-01 |
| AT1G78040 | 2,48 | 0,29 | 1,28 | 0,22 | 8,22E-01 | 8,90E-01 |
| AT1G18080 | 2,60 | 0,27 | 1,24 | 0,21 | 8,30E-01 | 8,98E-01 |
| AT1G72180 | 2,71 | 0,24 | 1,12 | 0,21 | 8,31E-01 | 8,98E-01 |
| AT1G15002 | 1,82 | -0,30 | 1,41 | -0,21 | 8,31E-01 | 8,98E-01 |
| AT2G34110 | 1,88 | 0,29 | 1,35 | 0,21 | 8,33E-01 | 9,00E-01 |
| AT4G23630 | 1,89 | -0,28 | 1,32 | -0,21 | 8,34E-01 | 9,00E-01 |
| AT5G04425 | 2,72 | 0,24 | 1,15 | 0,21 | 8,34E-01 | 9,00E-01 |
| AT4G15258 | 2,27 | 0,33 | 1,58 | 0,21 | 8,34E-01 | 9,00E-01 |
| AT3G54660 | 2,19 | -0,30 | 1,43 | -0,21 | 8,36E-01 | 9,01E-01 |
| AT2G30950 | 3,17 | 0,20 | 0,99 | 0,20 | 8,38E-01 | 9,02E-01 |
| AT2G36390 | 2,07 | -0,28 | 1,38 | -0,21 | 8,37E-01 | 9,02E-01 |
| AT3G25020 | 3,41 | 0,23 | 1,11 | 0,20 | 8,38E-01 | 9,02E-01 |
| AT5G45307 | 438,83 | 0,06 | 0,32 | 0,21 | 8,37E-01 | 9,02E-01 |
| AT1G05993 | 11,40 | 0,12 | 0,60 | 0,20 | 8,38E-01 | 9,02E-01 |
| AT5G24900 | 2,20 | -0,28 | 1,38 | -0,20 | 8,39E-01 | 9,02E-01 |
| AT3G41979 | 6596,11 | -0,07 | 0,36 | -0,20 | 8,42E-01 | 9,04E-01 |
| AT1G69940 | 3,40 | -0,20 | 1,00 | -0,20 | 8,42E-01 | 9,04E-01 |
| AT1G75380 | 2,78 | -0,21 | 1,08 | -0,20 | 8,42E-01 | 9,04E-01 |
| AT3G53460 | 6,35 | 0,16 | 0,80 | 0,20 | 8,43E-01 | 9,04E-01 |
| AT4G38970 | 12,35 | -0,10 | 0,51 | -0,20 | 8,45E-01 | 9,06E-01 |
| AT2G46765 | 2,57 | 0,25 | 1,27 | 0,19 | 8,46E-01 | 9,06E-01 |
| AT3G16830 | 1,75 | 0,29 | 1,51 | 0,19 | 8,46E-01 | 9,06E-01 |
| AT4G04205 | 2,87 | 0,24 | 1,23 | 0,19 | 8,46E-01 | 9,06E-01 |
| AT5G01155 | 4,02 | -0,19 | 1,00 | -0,19 | 8,47E-01 | 9,06E-01 |
| AT1G32720 | 2,47 | 0,25 | 1,28 | 0,19 | 8,48E-01 | 9,07E-01 |
| AT4G01290 | 1,74 | 0,30 | 1,59 | 0,19 | 8,49E-01 | 9,07E-01 |
| AT1G64385 | 1,99 | 0,36 | 1,93 | 0,19 | 8,51E-01 | 9,09E-01 |
| AT3G52850 | 1,76 | -0,26 | 1,36 | -0,19 | 8,50E-01 | 9,09E-01 |
| AT2G20722 | 3,24 | 0,22 | 1,15 | 0,19 | 8,51E-01 | 9,09E-01 |
| AT1G29070 | 1,74 | -0,27 | 1,46 | -0,19 | 8,52E-01 | 9,09E-01 |
| AT1G77230 | 3,05 | 0,21 | 1,15 | 0,19 | 8,52E-01 | 9,09E-01 |
| AT5G08180 | 2,30 | -0,22 | 1,16 | -0,19 | 8,52E-01 | 9,09E-01 |
| AT2G41640 | 6,36 | -0,13 | 0,70 | -0,18 | 8,54E-01 | 9,10E-01 |
| AT4G04465 | 2,27 | 0,23 | 1,28 | 0,18 | 8,55E-01 | 9,11E-01 |
| AT5G65700 | 1,88 | -0,25 | 1,40 | -0,18 | 8,56E-01 | 9,12E-01 |
| AT3G11940 | 3,79 | -0,19 | 1,06 | -0,18 | 8,57E-01 | 9,12E-01 |
| AT2G20580 | 2,81 | 0,29 | 1,63 | 0,18 | 8,60E-01 | 9,14E-01 |
| AT2G27720 | 1,70 | 0,31 | 1,74 | 0,18 | 8,59E-01 | 9,14E-01 |
| AT1G09570 | 2,05 | 0,26 | 1,51 | 0,17 | 8,61E-01 | 9,16E-01 |
| AT2G26530 | 1,88 | 0,33 | 1,88 | 0,17 | 8,62E-01 | 9,16E-01 |
| AT2G36460 | 1,86 | -0,25 | 1,47 | -0,17 | 8,63E-01 | 9,17E-01 |
| AT1G66730 | 1,91 | -0,21 | 1,20 | -0,17 | 8,64E-01 | 9,17E-01 |
| AT4G29090 | 2,85 | 0,21 | 1,25 | 0,17 | 8,64E-01 | 9,17E-01 |
| AT5G26340 | 2,20 | -0,21 | 1,26 | -0,17 | 8,65E-01 | 9,17E-01 |
| AT5G46110 | 7,87 | -0,12 | 0,73 | -0,17 | 8,66E-01 | 9,18E-01 |
| AT1G08327 | 2,78 | 0,26 | 1,53 | 0,17 | 8,67E-01 | 9,18E-01 |
| AT2G33150 | 2,47 | -0,19 | 1,12 | -0,17 | 8,67E-01 | 9,18E-01 |
| AT5G00765 | 3,11 | 0,17 | 1,04 | 0,17 | 8,67E-01 | 9,18E-01 |
| AT4G39363 | 6,83 | -0,13 | 0,79 | -0,17 | 8,68E-01 | 9,18E-01 |

| | | | | | | |
|---|---|---|---|---|---|---|
| AT5G01775 | 4,59 | 0,17 | 1,02 | 0,17 | 8,68E-01 | 9,18E-01 |
| AT5G52470 | 2,57 | 0,22 | 1,35 | 0,16 | 8,70E-01 | 9,19E-01 |
| AT1G32010 | 1,81 | -0,28 | 1,73 | -0,16 | 8,70E-01 | 9,19E-01 |
| AT3G52930 | 3,81 | -0,16 | 0,97 | -0,16 | 8,71E-01 | 9,19E-01 |
| AT5G61410 | 2,49 | 0,18 | 1,13 | 0,16 | 8,70E-01 | 9,19E-01 |
| AT5G40395 | 2,01 | 0,22 | 1,40 | 0,16 | 8,73E-01 | 9,21E-01 |
| AT1G47657 | 2,52 | 0,25 | 1,57 | 0,16 | 8,76E-01 | 9,22E-01 |
| AT4G11380 | 1,75 | 0,21 | 1,33 | 0,16 | 8,75E-01 | 9,22E-01 |
| AT3G08510 | 2,33 | 0,24 | 1,55 | 0,16 | 8,75E-01 | 9,22E-01 |
| AT3G11410 | 4,48 | -0,18 | 1,13 | -0,16 | 8,75E-01 | 9,22E-01 |
| AT3G01370 | 2,22 | -0,20 | 1,28 | -0,15 | 8,77E-01 | 9,23E-01 |
| AT1G67280 | 1,72 | -0,23 | 1,51 | -0,15 | 8,77E-01 | 9,23E-01 |
| AT5G04410 | 1,77 | 0,33 | 2,13 | 0,15 | 8,77E-01 | 9,23E-01 |
| AT5G67420 | 2,71 | -0,20 | 1,28 | -0,15 | 8,78E-01 | 9,23E-01 |
| AT5G41790 | 2,54 | 0,20 | 1,33 | 0,15 | 8,81E-01 | 9,26E-01 |
| AT2G41090 | 3,05 | -0,16 | 1,13 | -0,15 | 8,84E-01 | 9,28E-01 |
| AT1G55450 | 2,26 | 0,22 | 1,48 | 0,15 | 8,84E-01 | 9,28E-01 |
| AT3G06355 | 6711,54 | -0,05 | 0,36 | -0,14 | 8,85E-01 | 9,29E-01 |
| AT1G54270 | 2,59 | 0,15 | 1,06 | 0,14 | 8,86E-01 | 9,29E-01 |
| AT5G54250 | 1,88 | 0,23 | 1,62 | 0,14 | 8,86E-01 | 9,29E-01 |
| AT3G05325 | 9,46 | -0,10 | 0,67 | -0,14 | 8,87E-01 | 9,30E-01 |
| AT5G24690 | 2,42 | 0,17 | 1,17 | 0,14 | 8,88E-01 | 9,30E-01 |
| AT3G22960 | 3,17 | 0,15 | 1,11 | 0,14 | 8,90E-01 | 9,32E-01 |
| AT3G07725 | 10,80 | 0,08 | 0,61 | 0,14 | 8,92E-01 | 9,32E-01 |
| AT4G22690 | 3,37 | 0,15 | 1,10 | 0,14 | 8,92E-01 | 9,32E-01 |
| AT5G42895 | 1,80 | -0,24 | 1,74 | -0,14 | 8,92E-01 | 9,32E-01 |
| AT3G19010 | 1,81 | -0,22 | 1,59 | -0,14 | 8,91E-01 | 9,32E-01 |
| AT4G03380 | 7,25 | -0,10 | 0,71 | -0,14 | 8,91E-01 | 9,32E-01 |
| AT1G51805 | 2,63 | -0,16 | 1,15 | -0,13 | 8,93E-01 | 9,32E-01 |
| AT1G76090 | 1,72 | -0,22 | 1,64 | -0,13 | 8,93E-01 | 9,33E-01 |
| AT3G25520 | 3,31 | 0,14 | 1,09 | 0,12 | 9,01E-01 | 9,40E-01 |
| AT4G34720 | 1,83 | 0,15 | 1,21 | 0,12 | 9,01E-01 | 9,40E-01 |
| AT5G24530 | 1,77 | 0,18 | 1,44 | 0,12 | 9,02E-01 | 9,41E-01 |
| AT1G77815 | 2,65 | 0,21 | 1,74 | 0,12 | 9,03E-01 | 9,41E-01 |
| AT1G63070 | 3,60 | -0,11 | 0,93 | -0,12 | 9,06E-01 | 9,44E-01 |
| AT3G22235 | 3,04 | -0,13 | 1,10 | -0,12 | 9,07E-01 | 9,44E-01 |
| AT4G10800 | 6,52 | 0,08 | 0,71 | 0,12 | 9,08E-01 | 9,44E-01 |
| AT1G09687 | 2,04 | 0,14 | 1,26 | 0,11 | 9,08E-01 | 9,45E-01 |
| AT3G60750 | 11,99 | 0,07 | 0,60 | 0,11 | 9,10E-01 | 9,46E-01 |
| AT5G39050 | 4,52 | -0,11 | 1,00 | -0,11 | 9,11E-01 | 9,47E-01 |
| AT4G09000 | 1,74 | 0,16 | 1,47 | 0,11 | 9,12E-01 | 9,47E-01 |
| AT3G11670 | 1,77 | -0,19 | 1,70 | -0,11 | 9,12E-01 | 9,47E-01 |
| AT1G08200 | 1,82 | 0,17 | 1,63 | 0,11 | 9,16E-01 | 9,50E-01 |
| AT5G49730 | 3,55 | 0,12 | 1,18 | 0,10 | 9,17E-01 | 9,51E-01 |
| AT2G03821 | 1,77 | -0,14 | 1,37 | -0,10 | 9,17E-01 | 9,51E-01 |
| AT4G12320 | 1,93 | -0,15 | 1,46 | -0,10 | 9,20E-01 | 9,53E-01 |
| AT4G14360 | 1,93 | 0,14 | 1,44 | 0,10 | 9,20E-01 | 9,53E-01 |
| AT2G20410 | 6,12 | -0,07 | 0,74 | -0,10 | 9,21E-01 | 9,53E-01 |
| AT1G54040 | 2,21 | 0,13 | 1,36 | 0,10 | 9,21E-01 | 9,53E-01 |
| AT5G49030 | 1,79 | -0,14 | 1,46 | -0,10 | 9,22E-01 | 9,54E-01 |
| AT3G03475 | 2,58 | -0,11 | 1,15 | -0,10 | 9,24E-01 | 9,56E-01 |
| AT3G45140 | 15,78 | 0,05 | 0,53 | 0,09 | 9,24E-01 | 9,56E-01 |
| AT5G10572 | 2,06 | 0,13 | 1,35 | 0,09 | 9,26E-01 | 9,57E-01 |
| AT3G24929 | 2,38 | 0,12 | 1,32 | 0,09 | 9,28E-01 | 9,59E-01 |
| AT2G15620 | 3,38 | 0,11 | 1,22 | 0,09 | 9,30E-01 | 9,59E-01 |
| AT3G58196 | 21,50 | -0,04 | 0,51 | -0,09 | 9,30E-01 | 9,59E-01 |

| | | | | | | |
|---|---|---|---|---|---|---|
| AT5G24930 | 2,10 | 0,14 | 1,59 | 0,09 | 9,31E-01 | 9,59E-01 |
| AT1G48760 | 1,82 | -0,17 | 1,96 | -0,09 | 9,31E-01 | 9,59E-01 |
| AT5G40450 | 4,44 | 0,08 | 0,94 | 0,09 | 9,31E-01 | 9,59E-01 |
| AT5G63195 | 1,93 | 0,14 | 1,65 | 0,09 | 9,31E-01 | 9,59E-01 |
| AT3G58193 | 8,75 | 0,07 | 0,79 | 0,09 | 9,32E-01 | 9,59E-01 |
| AT3G17800 | 1,86 | 0,13 | 1,51 | 0,08 | 9,33E-01 | 9,60E-01 |
| AT2G16660 | 2,17 | 0,10 | 1,25 | 0,08 | 9,34E-01 | 9,61E-01 |
| AT3G56910 | 1,94 | -0,12 | 1,43 | -0,08 | 9,35E-01 | 9,61E-01 |
| AT3G22970 | 2,77 | -0,09 | 1,19 | -0,08 | 9,39E-01 | 9,62E-01 |
| AT5G14060 | 2,66 | -0,11 | 1,40 | -0,08 | 9,37E-01 | 9,62E-01 |
| AT5G19140 | 6,43 | -0,06 | 0,73 | -0,08 | 9,38E-01 | 9,62E-01 |
| AT5G62690 | 4,26 | 0,07 | 0,90 | 0,08 | 9,38E-01 | 9,62E-01 |
| AT1G27090 | 1,85 | -0,10 | 1,28 | -0,08 | 9,38E-01 | 9,62E-01 |
| AT4G04125 | 2,22 | 0,09 | 1,19 | 0,08 | 9,37E-01 | 9,62E-01 |
| AT4G31700 | 1,83 | -0,12 | 1,53 | -0,08 | 9,38E-01 | 9,62E-01 |
| AT1G17220 | 3,72 | 0,08 | 1,03 | 0,08 | 9,40E-01 | 9,63E-01 |
| AT4G24570 | 3,86 | -0,07 | 1,00 | -0,07 | 9,40E-01 | 9,63E-01 |
| AT1G26850 | 2,19 | -0,11 | 1,43 | -0,07 | 9,41E-01 | 9,63E-01 |
| AT3G60320 | 3,74 | 0,08 | 1,04 | 0,07 | 9,41E-01 | 9,63E-01 |
| AT1G68010 | 2,53 | 0,09 | 1,33 | 0,07 | 9,45E-01 | 9,66E-01 |
| AT5G13840 | 2,03 | 0,09 | 1,36 | 0,07 | 9,47E-01 | 9,68E-01 |
| AT3G14415 | 1,72 | -0,09 | 1,53 | -0,06 | 9,51E-01 | 9,69E-01 |
| AT4G10210 | 2,08 | -0,08 | 1,37 | -0,06 | 9,52E-01 | 9,69E-01 |
| AT5G17020 | 2,03 | -0,09 | 1,36 | -0,06 | 9,49E-01 | 9,69E-01 |
| AT5G39080 | 2,20 | -0,09 | 1,43 | -0,06 | 9,52E-01 | 9,69E-01 |
| AT5G43760 | 1,93 | 0,09 | 1,39 | 0,06 | 9,50E-01 | 9,69E-01 |
| AT1G07940 | 3,71 | 0,06 | 0,95 | 0,06 | 9,52E-01 | 9,69E-01 |
| AT1G23390 | 2,44 | 0,08 | 1,23 | 0,06 | 9,49E-01 | 9,69E-01 |
| AT1G61740 | 2,13 | -0,09 | 1,44 | -0,06 | 9,51E-01 | 9,69E-01 |
| AT4G17390 | 6,32 | -0,05 | 0,86 | -0,06 | 9,50E-01 | 9,69E-01 |
| AT2G47170 | 1,90 | -0,08 | 1,41 | -0,06 | 9,53E-01 | 9,69E-01 |
| AT2G45245 | 2,00 | 0,07 | 1,28 | 0,06 | 9,53E-01 | 9,70E-01 |
| AT1G72150 | 7,68 | 0,04 | 0,70 | 0,06 | 9,54E-01 | 9,70E-01 |
| AT2G04405 | 3,40 | -0,06 | 1,05 | -0,06 | 9,54E-01 | 9,70E-01 |
| AT5G44270 | 2,68 | -0,07 | 1,17 | -0,06 | 9,55E-01 | 9,70E-01 |
| AT5G60920 | 1,88 | 0,08 | 1,36 | 0,06 | 9,55E-01 | 9,70E-01 |
| AT5G14740 | 2,79 | -0,08 | 1,37 | -0,06 | 9,56E-01 | 9,70E-01 |
| AT4G39200 | 1,73 | -0,10 | 1,88 | -0,05 | 9,56E-01 | 9,70E-01 |
| AT5G60600 | 2,43 | 0,06 | 1,04 | 0,05 | 9,57E-01 | 9,70E-01 |
| AT1G19090 | 4,93 | 0,05 | 0,96 | 0,05 | 9,58E-01 | 9,70E-01 |
| AT2G08370 | 4,11 | 0,05 | 0,87 | 0,05 | 9,58E-01 | 9,70E-01 |
| AT1G35720 | 3,50 | -0,05 | 0,99 | -0,05 | 9,60E-01 | 9,72E-01 |
| AT3G62010 | 2,16 | -0,07 | 1,34 | -0,05 | 9,61E-01 | 9,72E-01 |
| AT5G13650 | 5,78 | 0,04 | 0,80 | 0,05 | 9,62E-01 | 9,73E-01 |
| AT2G29670 | 3,81 | 0,05 | 1,07 | 0,05 | 9,62E-01 | 9,73E-01 |
| AT3G47342 | 15,52 | -0,03 | 0,56 | -0,05 | 9,62E-01 | 9,73E-01 |
| AT1G76180 | 6,03 | 0,03 | 0,74 | 0,04 | 9,66E-01 | 9,76E-01 |
| AT2G34100 | 7,49 | -0,03 | 0,81 | -0,04 | 9,67E-01 | 9,76E-01 |
| AT3G16640 | 7,07 | -0,04 | 0,90 | -0,04 | 9,67E-01 | 9,76E-01 |
| AT5G01350 | 8,44 | -0,03 | 0,67 | -0,04 | 9,67E-01 | 9,76E-01 |
| AT2G23348 | 3,88 | 0,05 | 1,30 | 0,04 | 9,69E-01 | 9,77E-01 |
| AT4G24770 | 3,23 | -0,04 | 1,11 | -0,04 | 9,70E-01 | 9,78E-01 |
| AT3G24982 | 2,19 | -0,05 | 1,33 | -0,03 | 9,73E-01 | 9,81E-01 |
| AT2G38325 | 156,77 | -0,01 | 0,24 | -0,03 | 9,75E-01 | 9,82E-01 |
| AT4G05900 | 1,70 | 0,05 | 1,63 | 0,03 | 9,75E-01 | 9,82E-01 |
| AT1G07250 | 2,07 | 0,04 | 1,39 | 0,03 | 9,78E-01 | 9,84E-01 |

| | | | | | | |
|---|---|---|---|---|---|---|
| AT2G23347 | 3,51 | -0,03 | 1,29 | -0,03 | 9,80E-01 | 9,86E-01 |
| AT3G09105 | 30,37 | -0,01 | 0,53 | -0,02 | 9,81E-01 | 9,86E-01 |
| AT1G51980 | 1,85 | 0,03 | 1,49 | 0,02 | 9,82E-01 | 9,87E-01 |
| AT3G29190 | 2,69 | -0,03 | 1,20 | -0,02 | 9,83E-01 | 9,88E-01 |
| AT1G51690 | 2,38 | 0,02 | 1,30 | 0,02 | 9,85E-01 | 9,89E-01 |
| AT1G61275 | 11,06 | 0,01 | 0,60 | 0,02 | 9,86E-01 | 9,90E-01 |
| AT5G14780 | 2,20 | 0,02 | 1,17 | 0,02 | 9,87E-01 | 9,91E-01 |
| AT5G05010 | 2,08 | 0,02 | 1,36 | 0,01 | 9,88E-01 | 9,91E-01 |
| AT5G13390 | 2,05 | -0,02 | 1,40 | -0,01 | 9,88E-01 | 9,91E-01 |
| AT3G59410 | 1,72 | 0,02 | 1,62 | 0,01 | 9,90E-01 | 9,92E-01 |
| AT3G53420 | 2,26 | 0,02 | 1,32 | 0,01 | 9,91E-01 | 9,93E-01 |
| AT2G35110 | 1,99 | -0,01 | 1,49 | -0,01 | 9,96E-01 | 9,97E-01 |
| AT1G21680 | 4,80 | 0,00 | 0,84 | 0,00 | 9,97E-01 | 9,98E-01 |
| AT1G06148 | 2,19 | 0,00 | 1,35 | 0,00 | 9,97E-01 | 9,98E-01 |
| AT3G61897 | 2,76 | 0,00 | 2,83 | 0,00 | 1,00E+00 | 1,00E+00 |
| AT1G01046 | 1,63 | 1,95 | 1,74 | 1,12 | 2,61E-01 | |
| AT1G01160 | 0,43 | 0,41 | 3,32 | 0,12 | 9,01E-01 | |
| AT1G01180 | 0,85 | 2,40 | 2,28 | 1,05 | 2,92E-01 | |
| AT1G01220 | 0,11 | 1,07 | 4,36 | 0,25 | 8,06E-01 | |
| AT1G01225 | 0,31 | 0,42 | 4,38 | 0,09 | 9,24E-01 | |
| AT1G01230 | 0,37 | -1,23 | 4,39 | -0,28 | 7,80E-01 | |
| AT1G01240 | 0,19 | 0,78 | 4,36 | 0,18 | 8,58E-01 | |
| AT1G01260 | 0,53 | -0,56 | 2,68 | -0,21 | 8,34E-01 | |
| AT1G01300 | 1,08 | 0,26 | 1,84 | 0,14 | 8,88E-01 | |
| AT1G01350 | 0,29 | 1,56 | 4,34 | 0,36 | 7,18E-01 | |
| AT1G01448 | 0,59 | 2,35 | 4,31 | 0,55 | 5,86E-01 | |
| AT1G01480 | 0,12 | 1,65 | 4,34 | 0,38 | 7,04E-01 | |
| AT1G01520 | 0,12 | 1,65 | 4,34 | 0,38 | 7,04E-01 | |
| AT1G01540 | 0,22 | 1,61 | 4,33 | 0,37 | 7,11E-01 | |
| AT1G01560 | 1,60 | 3,73 | 1,90 | 1,97 | 4,90E-02 | |
| AT1G01630 | 0,58 | -1,36 | 3,85 | -0,35 | 7,24E-01 | |
| AT1G01710 | 0,82 | -1,31 | 3,10 | -0,42 | 6,72E-01 | |
| AT1G01720 | 1,37 | 0,87 | 1,63 | 0,54 | 5,92E-01 | |
| AT1G04047 | 0,70 | 1,62 | 2,52 | 0,64 | 5,20E-01 | |
| AT1G01730 | 0,40 | 0,42 | 4,38 | 0,09 | 9,24E-01 | |
| AT1G01790 | 0,83 | 0,07 | 1,89 | 0,04 | 9,70E-01 | |
| AT1G01810 | 0,12 | 1,58 | 4,34 | 0,36 | 7,16E-01 | |
| AT1G01950 | 0,18 | 1,06 | 4,35 | 0,24 | 8,07E-01 | |
| AT1G02010 | 0,10 | 0,62 | 4,38 | 0,14 | 8,87E-01 | |
| AT1G02100 | 0,49 | 1,68 | 3,12 | 0,54 | 5,90E-01 | |
| AT1G02120 | 0,36 | 1,12 | 3,44 | 0,32 | 7,45E-01 | |
| AT1G02150 | 1,38 | 0,40 | 1,80 | 0,22 | 8,22E-01 | |
| AT1G02205 | 1,06 | -0,57 | 1,88 | -0,30 | 7,62E-01 | |
| AT1G04093 | 0,08 | 0,00 | 4,40 | 0,00 | 1,00E+00 | |
| AT1G02300 | 0,34 | 1,50 | 4,35 | 0,35 | 7,30E-01 | |
| AT1G02305 | 0,80 | 1,02 | 2,40 | 0,42 | 6,71E-01 | |
| AT1G04103 | 0,08 | 1,10 | 4,36 | 0,25 | 8,00E-01 | |
| AT1G02390 | 0,19 | 0,79 | 4,37 | 0,18 | 8,56E-01 | |
| AT1G02420 | 0,52 | 2,73 | 3,86 | 0,71 | 4,79E-01 | |
| AT1G02560 | 1,57 | -0,50 | 1,65 | -0,30 | 7,64E-01 | |
| AT1G02690 | 0,11 | 0,00 | 4,41 | 0,00 | 1,00E+00 | |
| AT1G02750 | 0,51 | 0,48 | 4,40 | 0,11 | 9,13E-01 | |
| AT1G02816 | 0,38 | 1,78 | 3,36 | 0,53 | 5,96E-01 | |
| AT1G02850 | 0,62 | 2,19 | 2,85 | 0,77 | 4,42E-01 | |
| AT1G03160 | 0,95 | -1,23 | 2,19 | -0,56 | 5,75E-01 | |
| AT1G03190 | 0,39 | 1,99 | 3,43 | 0,58 | 5,61E-01 | |

| | | | | | |
|---|---|---|---|---|---|
| AT1G03220 | 1,11 | 2,08 | 2,15 | 0,97 | 3,33E-01 |
| AT1G03280 | 0,25 | 2,24 | 4,32 | 0,52 | 6,04E-01 |
| AT1G03290 | 0,75 | 3,17 | 2,42 | 1,31 | 1,89E-01 |
| AT1G03310 | 0,40 | 1,56 | 4,21 | 0,37 | 7,12E-01 |
| AT1G03360 | 0,35 | 1,59 | 3,49 | 0,46 | 6,48E-01 |
| AT1G03365 | 0,40 | 0,88 | 3,36 | 0,26 | 7,94E-01 |
| AT1G03370 | 0,51 | 2,38 | 3,95 | 0,60 | 5,46E-01 |
| AT1G03380 | 1,12 | 1,73 | 2,03 | 0,85 | 3,95E-01 |
| AT1G03430 | 0,37 | -0,40 | 4,36 | -0,09 | 9,26E-01 |
| AT1G03440 | 0,84 | -1,66 | 3,12 | -0,53 | 5,95E-01 |
| AT1G03545 | 0,70 | -1,33 | 2,34 | -0,57 | 5,70E-01 |
| AT1G03600 | 1,40 | 0,12 | 1,68 | 0,07 | 9,41E-01 |
| AT1G03610 | 0,72 | -0,19 | 3,12 | -0,06 | 9,51E-01 |
| AT1G03630 | 0,51 | 0,31 | 2,37 | 0,13 | 8,97E-01 |
| AT1G03687 | 0,11 | 0,00 | 4,41 | 0,00 | 1,00E+00 |
| AT1G03740 | 0,96 | 1,33 | 2,07 | 0,64 | 5,19E-01 |
| AT1G03810 | 0,77 | 1,02 | 2,28 | 0,44 | 6,57E-01 |
| AT1G03900 | 1,11 | 1,49 | 2,01 | 0,74 | 4,59E-01 |
| AT1G03905 | 0,24 | 0,00 | 4,41 | 0,00 | 1,00E+00 |
| AT1G03910 | 0,50 | -0,06 | 3,11 | -0,02 | 9,84E-01 |
| AT1G03930 | 0,53 | 1,86 | 2,70 | 0,69 | 4,92E-01 |
| AT1G04217 | 0,82 | 0,12 | 1,86 | 0,06 | 9,50E-01 |
| AT1G03935 | 0,16 | -1,36 | 4,41 | -0,31 | 7,57E-01 |
| AT1G04223 | 0,98 | -1,02 | 2,16 | -0,47 | 6,38E-01 |
| AT1G04227 | 0,32 | -0,48 | 3,61 | -0,13 | 8,93E-01 |
| AT1G03960 | 0,15 | 0,24 | 4,35 | 0,05 | 9,56E-01 |
| AT1G03970 | 0,36 | 1,63 | 4,34 | 0,38 | 7,07E-01 |
| AT1G04020 | 0,33 | -1,38 | 4,41 | -0,31 | 7,54E-01 |
| AT1G04130 | 0,50 | 1,25 | 2,70 | 0,46 | 6,44E-01 |
| AT1G04170 | 1,02 | 2,70 | 2,06 | 1,31 | 1,89E-01 |
| AT1G04247 | 0,30 | 0,00 | 4,40 | 0,00 | 1,00E+00 |
| AT1G04230 | 0,70 | -2,73 | 2,88 | -0,95 | 3,42E-01 |
| AT1G04295 | 0,43 | 1,98 | 2,75 | 0,72 | 4,72E-01 |
| AT1G04350 | 0,25 | -1,22 | 4,37 | -0,28 | 7,81E-01 |
| AT1G04420 | 1,20 | 2,60 | 1,85 | 1,41 | 1,60E-01 |
| AT1G04440 | 0,50 | 1,06 | 2,74 | 0,39 | 7,00E-01 |
| AT1G04273 | 0,37 | -0,34 | 3,50 | -0,10 | 9,22E-01 |
| AT1G04277 | 0,93 | -1,21 | 2,11 | -0,57 | 5,66E-01 |
| AT1G04510 | 0,67 | 1,81 | 2,98 | 0,61 | 5,44E-01 |
| AT1G04540 | 1,05 | 3,00 | 2,24 | 1,34 | 1,80E-01 |
| AT1G04690 | 0,77 | 1,30 | 2,38 | 0,55 | 5,83E-01 |
| AT1G04710 | 0,47 | 1,71 | 4,35 | 0,39 | 6,95E-01 |
| AT1G04760 | 0,36 | 0,84 | 4,38 | 0,19 | 8,48E-01 |
| AT1G04945 | 0,47 | -0,06 | 3,22 | -0,02 | 9,86E-01 |
| AT1G04970 | 0,34 | -0,59 | 4,31 | -0,14 | 8,91E-01 |
| AT1G05087 | 0,44 | 2,58 | 4,05 | 0,64 | 5,25E-01 |
| AT1G05090 | 0,12 | 1,65 | 4,34 | 0,38 | 7,04E-01 |
| AT1G05140 | 0,22 | 0,00 | 4,41 | 0,00 | 1,00E+00 |
| AT1G05200 | 1,12 | -1,27 | 1,81 | -0,70 | 4,83E-01 |
| AT1G05360 | 0,08 | 1,10 | 4,36 | 0,25 | 8,00E-01 |
| AT1G05380 | 1,42 | 2,68 | 2,03 | 1,32 | 1,87E-01 |
| AT1G05500 | 0,37 | 0,73 | 4,39 | 0,17 | 8,68E-01 |
| AT1G04397 | 0,08 | 1,10 | 4,36 | 0,25 | 8,00E-01 |
| AT1G05600 | 0,08 | 0,00 | 4,40 | 0,00 | 1,00E+00 |
| AT1G04423 | 1,18 | 0,49 | 2,04 | 0,24 | 8,11E-01 |
| AT1G05790 | 0,76 | 2,80 | 2,44 | 1,15 | 2,52E-01 |

| | | | | | |
|---|---|---|---|---|---|
| AT1G05830 | 0,53 | 0,73 | 3,30 | 0,22 | 8,25E-01 |
| AT1G05890 | 0,68 | 0,60 | 4,40 | 0,14 | 8,92E-01 |
| AT1G05900 | 0,21 | 0,00 | 4,38 | 0,00 | 1,00E+00 |
| AT1G05910 | 0,59 | 1,63 | 3,79 | 0,43 | 6,66E-01 |
| AT1G05980 | 0,23 | -1,21 | 4,38 | -0,28 | 7,82E-01 |
| AT1G06002 | 0,95 | -0,60 | 2,01 | -0,30 | 7,65E-01 |
| AT1G06030 | 0,96 | 1,54 | 3,34 | 0,46 | 6,45E-01 |
| AT1G06290 | 0,60 | 0,90 | 2,76 | 0,33 | 7,45E-01 |
| AT1G06380 | 0,45 | -2,21 | 4,21 | -0,53 | 5,99E-01 |
| AT1G06390 | 0,40 | 0,93 | 3,42 | 0,27 | 7,86E-01 |
| AT1G06410 | 1,61 | 2,35 | 1,78 | 1,32 | 1,88E-01 |
| AT1G06530 | 0,96 | 0,44 | 2,15 | 0,21 | 8,37E-01 |
| AT1G06640 | 1,12 | 2,38 | 1,91 | 1,25 | 2,11E-01 |
| AT1G06645 | 0,50 | 2,55 | 3,98 | 0,64 | 5,21E-01 |
| AT1G06650 | 0,69 | 2,38 | 2,98 | 0,80 | 4,24E-01 |
| AT1G06670 | 1,38 | 1,53 | 2,02 | 0,76 | 4,48E-01 |
| AT1G06750 | 0,24 | 1,23 | 4,35 | 0,28 | 7,77E-01 |
| AT1G06780 | 0,12 | 0,90 | 4,36 | 0,21 | 8,36E-01 |
| AT1G06910 | 0,23 | 1,33 | 4,34 | 0,31 | 7,60E-01 |
| AT1G06930 | 0,08 | 1,10 | 4,36 | 0,25 | 8,00E-01 |
| AT1G07010 | 0,86 | 2,07 | 2,94 | 0,71 | 4,80E-01 |
| AT1G04537 | 0,24 | 0,00 | 4,41 | 0,00 | 1,00E+00 |
| AT1G07080 | 0,86 | -1,50 | 2,01 | -0,74 | 4,57E-01 |
| AT1G07130 | 0,28 | 1,69 | 4,35 | 0,39 | 6,97E-01 |
| AT1G07170 | 0,78 | 2,05 | 2,07 | 0,99 | 3,23E-01 |
| AT1G07180 | 0,37 | -0,20 | 3,52 | -0,06 | 9,56E-01 |
| AT1G07210 | 0,17 | 1,81 | 4,33 | 0,42 | 6,76E-01 |
| AT1G07240 | 0,59 | 0,68 | 3,28 | 0,21 | 8,36E-01 |
| AT1G07280 | 1,33 | 0,42 | 1,79 | 0,23 | 8,14E-01 |
| AT1G07320 | 1,43 | -0,18 | 1,58 | -0,11 | 9,09E-01 |
| AT1G07470 | 0,61 | 1,60 | 3,73 | 0,43 | 6,68E-01 |
| AT1G07630 | 0,70 | 1,87 | 2,94 | 0,63 | 5,26E-01 |
| AT1G07705 | 0,12 | 1,39 | 4,34 | 0,32 | 7,49E-01 |
| AT1G07728 | 1,03 | 2,32 | 2,52 | 0,92 | 3,58E-01 |
| AT1G07790 | 0,25 | -0,86 | 4,36 | -0,20 | 8,44E-01 |
| AT1G07830 | 0,43 | 2,12 | 4,15 | 0,51 | 6,10E-01 |
| AT1G07990 | 0,78 | 2,55 | 3,00 | 0,85 | 3,96E-01 |
| AT1G08110 | 0,41 | 1,73 | 4,30 | 0,40 | 6,87E-01 |
| AT1G08190 | 0,34 | 0,19 | 3,59 | 0,05 | 9,58E-01 |
| AT1G08260 | 0,51 | 2,58 | 4,08 | 0,63 | 5,28E-01 |
| AT1G08360 | 1,45 | 2,98 | 1,77 | 1,68 | 9,26E-02 |
| AT1G08370 | 0,71 | 2,75 | 2,46 | 1,12 | 2,63E-01 |
| AT1G08410 | 0,87 | -0,77 | 2,51 | -0,31 | 7,58E-01 |
| AT1G08460 | 0,84 | 0,37 | 2,05 | 0,18 | 8,56E-01 |
| AT1G08465 | 0,35 | 1,54 | 4,34 | 0,35 | 7,23E-01 |
| AT1G08480 | 0,59 | 2,11 | 3,10 | 0,68 | 4,97E-01 |
| AT1G08520 | 1,05 | 0,55 | 1,78 | 0,31 | 7,59E-01 |
| AT1G08530 | 0,48 | 2,98 | 3,96 | 0,75 | 4,52E-01 |
| AT1G08550 | 0,85 | 0,40 | 2,17 | 0,19 | 8,53E-01 |
| AT1G08570 | 0,49 | 1,09 | 4,09 | 0,27 | 7,90E-01 |
| AT1G08590 | 0,74 | 3,22 | 2,84 | 1,14 | 2,56E-01 |
| AT1G08600 | 0,40 | 1,36 | 3,29 | 0,41 | 6,80E-01 |
| AT1G08620 | 1,06 | 1,90 | 1,92 | 0,99 | 3,22E-01 |
| AT1G08650 | 0,89 | -0,19 | 2,84 | -0,07 | 9,47E-01 |
| AT1G08680 | 0,17 | 0,00 | 4,41 | 0,00 | 1,00E+00 |
| AT1G08710 | 0,30 | 1,17 | 4,37 | 0,27 | 7,89E-01 |

| | | | | | |
|---|---|---|---|---|---|
| AT1G08720 | 1,46 | 2,85 | 2,05 | 1,39 | 1,64E-01 |
| AT1G08760 | 0,68 | 2,32 | 2,98 | 0,78 | 4,35E-01 |
| AT1G08770 | 0,33 | 0,43 | 3,57 | 0,12 | 9,05E-01 |
| AT1G08830 | 1,29 | 1,44 | 1,97 | 0,73 | 4,63E-01 |
| AT1G08900 | 0,34 | 1,56 | 3,47 | 0,45 | 6,54E-01 |
| AT1G08910 | 0,43 | 0,00 | 4,24 | 0,00 | 1,00E+00 |
| AT1G08920 | 0,59 | -0,07 | 3,10 | -0,02 | 9,83E-01 |
| AT1G08930 | 1,31 | 0,99 | 1,57 | 0,63 | 5,29E-01 |
| AT1G08970 | 0,82 | 1,52 | 2,58 | 0,59 | 5,56E-01 |
| AT1G08980 | 0,91 | 1,17 | 2,14 | 0,55 | 5,85E-01 |
| AT1G09040 | 0,45 | 1,65 | 3,29 | 0,50 | 6,15E-01 |
| AT1G09250 | 0,32 | 0,44 | 4,38 | 0,10 | 9,20E-01 |
| AT1G09260 | 0,62 | -3,57 | 4,32 | -0,83 | 4,08E-01 |
| AT1G09270 | 0,12 | 1,65 | 4,34 | 0,38 | 7,04E-01 |
| AT1G09300 | 0,36 | 1,27 | 4,34 | 0,29 | 7,70E-01 |
| AT1G09390 | 0,23 | 1,28 | 4,34 | 0,29 | 7,69E-01 |
| AT1G09421 | 0,17 | 1,03 | 4,37 | 0,24 | 8,13E-01 |
| AT1G09430 | 0,79 | 1,68 | 2,44 | 0,69 | 4,91E-01 |
| AT1G09450 | 0,24 | 0,65 | 4,37 | 0,15 | 8,81E-01 |
| AT1G09490 | 0,05 | 0,00 | 4,41 | 0,00 | 1,00E+00 |
| AT1G09500 | 0,08 | 1,10 | 4,36 | 0,25 | 8,00E-01 |
| AT1G09530 | 0,63 | 0,49 | 3,65 | 0,13 | 8,93E-01 |
| AT1G09560 | 0,29 | 1,89 | 4,33 | 0,44 | 6,63E-01 |
| AT1G09590 | 0,27 | -0,13 | 4,37 | -0,03 | 9,77E-01 |
| AT1G09600 | 0,94 | 3,07 | 2,79 | 1,10 | 2,71E-01 |
| AT1G09610 | 0,36 | 1,56 | 4,34 | 0,36 | 7,19E-01 |
| AT1G09645 | 0,33 | 1,15 | 4,35 | 0,26 | 7,91E-01 |
| AT1G09665 | 0,25 | 0,00 | 4,40 | 0,00 | 1,00E+00 |
| AT1G09710 | 0,91 | 1,76 | 2,90 | 0,61 | 5,44E-01 |
| AT1G09750 | 0,82 | -1,26 | 1,91 | -0,66 | 5,10E-01 |
| AT1G09795 | 0,58 | -1,69 | 4,00 | -0,42 | 6,73E-01 |
| AT1G09815 | 0,21 | -1,56 | 4,37 | -0,36 | 7,22E-01 |
| AT1G10050 | 0,78 | -0,82 | 2,14 | -0,38 | 7,01E-01 |
| AT1G10070 | 0,24 | 1,62 | 4,34 | 0,37 | 7,08E-01 |
| AT1G10130 | 1,07 | 1,70 | 1,98 | 0,86 | 3,92E-01 |
| AT1G10150 | 1,47 | 1,71 | 1,53 | 1,12 | 2,63E-01 |
| AT1G10270 | 0,58 | -0,44 | 2,62 | -0,17 | 8,68E-01 |
| AT1G10450 | 0,33 | 2,20 | 4,32 | 0,51 | 6,11E-01 |
| AT1G10490 | 0,65 | 1,61 | 2,46 | 0,65 | 5,14E-01 |
| AT1G10570 | 0,16 | 1,07 | 4,37 | 0,24 | 8,07E-01 |
| AT1G10600 | 1,27 | 3,56 | 1,94 | 1,84 | 6,64E-02 |
| AT1G10610 | 0,90 | 2,36 | 3,34 | 0,71 | 4,79E-01 |
| AT1G10660 | 0,54 | -0,65 | 4,02 | -0,16 | 8,71E-01 |
| AT1G10670 | 0,76 | -3,14 | 3,54 | -0,89 | 3,76E-01 |
| AT1G10700 | 0,45 | 0,99 | 4,37 | 0,23 | 8,22E-01 |
| AT1G10720 | 0,48 | 1,11 | 2,64 | 0,42 | 6,75E-01 |
| AT1G10745 | 1,57 | -2,36 | 1,90 | -1,25 | 2,13E-01 |
| AT1G10790 | 0,08 | 1,10 | 4,36 | 0,25 | 8,00E-01 |
| AT1G10850 | 0,85 | 2,31 | 3,49 | 0,66 | 5,09E-01 |
| AT1G10910 | 0,08 | 1,10 | 4,36 | 0,25 | 8,00E-01 |
| AT1G10960 | 0,86 | -0,64 | 2,74 | -0,24 | 8,14E-01 |
| AT1G11020 | 0,27 | 0,44 | 4,39 | 0,10 | 9,20E-01 |
| AT1G11050 | 0,22 | 0,42 | 4,38 | 0,10 | 9,24E-01 |
| AT1G11260 | 1,39 | -1,16 | 1,60 | -0,73 | 4,68E-01 |
| AT1G11300 | 0,32 | 1,36 | 4,32 | 0,31 | 7,53E-01 |
| AT1G11303 | 0,25 | 2,36 | 4,32 | 0,55 | 5,86E-01 |

| | | | | | |
|---|---|---|---|---|---|
| AT1G11320 | 0,08 | 0,85 | 4,38 | 0,20 | 8,45E-01 |
| AT1G11330 | 0,64 | 2,45 | 2,93 | 0,84 | 4,02E-01 |
| AT1G11400 | 0,26 | 0,75 | 4,36 | 0,17 | 8,63E-01 |
| AT1G11410 | 0,36 | 0,86 | 3,43 | 0,25 | 8,02E-01 |
| AT1G11430 | 0,35 | -0,66 | 3,58 | -0,18 | 8,54E-01 |
| AT1G11440 | 0,54 | 0,52 | 3,07 | 0,17 | 8,66E-01 |
| AT1G11580 | 0,31 | 1,99 | 4,32 | 0,46 | 6,45E-01 |
| AT1G11590 | 0,67 | -2,91 | 2,49 | -1,17 | 2,43E-01 |
| AT1G11593 | 0,40 | 1,05 | 4,22 | 0,25 | 8,03E-01 |
| AT1G11660 | 1,14 | 2,69 | 1,99 | 1,35 | 1,76E-01 |
| AT1G11680 | 0,83 | 1,50 | 2,71 | 0,55 | 5,79E-01 |
| AT1G11710 | 0,17 | 1,94 | 4,33 | 0,45 | 6,55E-01 |
| AT1G11750 | 0,56 | -1,82 | 3,15 | -0,58 | 5,64E-01 |
| AT1G11790 | 0,42 | -0,17 | 4,37 | -0,04 | 9,68E-01 |
| AT1G11890 | 1,50 | 0,66 | 1,85 | 0,36 | 7,19E-01 |
| AT1G11905 | 0,75 | 1,24 | 2,23 | 0,56 | 5,79E-01 |
| AT1G11930 | 0,68 | -0,13 | 2,17 | -0,06 | 9,51E-01 |
| AT1G12015 | 0,46 | -1,32 | 4,35 | -0,30 | 7,61E-01 |
| AT1G12050 | 1,22 | 0,24 | 1,87 | 0,13 | 8,98E-01 |
| AT1G12060 | 0,12 | 0,79 | 4,37 | 0,18 | 8,57E-01 |
| AT1G12140 | 0,12 | 1,65 | 4,34 | 0,38 | 7,04E-01 |
| AT1G12210 | 0,35 | -0,23 | 4,36 | -0,05 | 9,59E-01 |
| AT1G12220 | 1,65 | -1,06 | 1,41 | -0,76 | 4,49E-01 |
| AT1G12230 | 0,35 | 1,23 | 3,50 | 0,35 | 7,26E-01 |
| AT1G12250 | 0,19 | -0,55 | 4,36 | -0,13 | 9,00E-01 |
| AT1G12270 | 0,62 | -0,34 | 2,57 | -0,13 | 8,95E-01 |
| AT1G12320 | 0,82 | 2,23 | 2,83 | 0,79 | 4,31E-01 |
| AT1G12360 | 0,65 | 0,86 | 3,84 | 0,22 | 8,24E-01 |
| AT1G12410 | 0,67 | -2,74 | 3,69 | -0,74 | 4,58E-01 |
| AT1G12460 | 0,83 | -2,17 | 2,07 | -1,05 | 2,93E-01 |
| AT1G12580 | 0,43 | 1,83 | 2,80 | 0,65 | 5,13E-01 |
| AT1G12760 | 0,76 | 1,92 | 2,22 | 0,86 | 3,87E-01 |
| AT1G12810 | 0,42 | -0,47 | 2,43 | -0,19 | 8,48E-01 |
| AT1G12840 | 1,05 | 0,99 | 1,74 | 0,57 | 5,68E-01 |
| AT1G12860 | 0,23 | -0,99 | 4,33 | -0,23 | 8,20E-01 |
| AT1G12990 | 1,14 | 3,42 | 2,05 | 1,67 | 9,54E-02 |
| AT1G13040 | 0,24 | 0,99 | 4,38 | 0,23 | 8,21E-01 |
| AT1G13060 | 0,83 | -1,07 | 3,08 | -0,35 | 7,27E-01 |
| AT1G13090 | 1,29 | -0,68 | 1,88 | -0,36 | 7,19E-01 |
| AT1G13100 | 0,26 | -2,10 | 4,37 | -0,48 | 6,31E-01 |
| AT1G13180 | 0,14 | 1,18 | 4,35 | 0,27 | 7,86E-01 |
| AT1G13190 | 1,26 | 3,12 | 2,49 | 1,25 | 2,11E-01 |
| AT1G13220 | 0,56 | 2,77 | 3,78 | 0,73 | 4,63E-01 |
| AT1G13260 | 1,59 | 0,20 | 1,69 | 0,12 | 9,08E-01 |
| AT1G13270 | 0,60 | -1,76 | 2,92 | -0,60 | 5,47E-01 |
| AT1G13330 | 0,70 | 2,11 | 2,55 | 0,83 | 4,07E-01 |
| AT1G13400 | 0,23 | 0,00 | 4,41 | 0,00 | 1,00E+00 |
| AT1G13430 | 0,17 | 1,40 | 4,35 | 0,32 | 7,48E-01 |
| AT1G13460 | 0,37 | -0,04 | 4,30 | -0,01 | 9,93E-01 |
| AT1G13820 | 0,11 | 0,90 | 4,37 | 0,21 | 8,36E-01 |
| AT1G13850 | 0,13 | 1,15 | 4,36 | 0,26 | 7,92E-01 |
| AT1G13880 | 0,45 | 1,32 | 3,37 | 0,39 | 6,95E-01 |
| AT1G13990 | 0,47 | 1,90 | 3,29 | 0,58 | 5,65E-01 |
| AT1G14000 | 0,81 | 0,69 | 2,16 | 0,32 | 7,50E-01 |
| AT1G14020 | 0,41 | 0,86 | 4,22 | 0,20 | 8,38E-01 |
| AT1G05047 | 0,12 | 1,65 | 4,34 | 0,38 | 7,04E-01 |

| | | | | | |
|---|---|---|---|---|---|
| AT1G14110 | 0,52 | 1,89 | 3,83 | 0,49 | 6,22E-01 |
| AT1G14150 | 0,05 | 0,00 | 4,41 | 0,00 | 1,00E+00 |
| AT1G14185 | 0,05 | 0,00 | 4,41 | 0,00 | 1,00E+00 |
| AT1G14230 | 0,08 | 0,00 | 4,41 | 0,00 | 1,00E+00 |
| AT1G14250 | 0,32 | -2,54 | 4,35 | -0,58 | 5,59E-01 |
| AT1G14280 | 0,63 | 0,41 | 3,10 | 0,13 | 8,94E-01 |
| AT1G14310 | 0,25 | 1,63 | 4,34 | 0,38 | 7,07E-01 |
| AT1G14345 | 0,53 | 1,62 | 2,74 | 0,59 | 5,55E-01 |
| AT1G14350 | 0,37 | 0,02 | 4,32 | 0,00 | 9,96E-01 |
| AT1G14370 | 0,91 | 1,68 | 1,89 | 0,89 | 3,73E-01 |
| AT1G14470 | 0,37 | -2,46 | 4,35 | -0,57 | 5,72E-01 |
| AT1G14480 | 0,08 | 0,00 | 4,40 | 0,00 | 1,00E+00 |
| AT1G14560 | 0,21 | -0,09 | 4,37 | -0,02 | 9,83E-01 |
| AT1G14570 | 1,23 | 3,22 | 1,85 | 1,74 | 8,14E-02 |
| AT1G14650 | 1,32 | 4,07 | 2,20 | 1,85 | 6,45E-02 |
| AT1G14800 | 1,13 | 0,52 | 1,85 | 0,28 | 7,80E-01 |
| AT1G14860 | 0,08 | 0,00 | 4,40 | 0,00 | 1,00E+00 |
| AT1G15030 | 0,23 | 1,56 | 4,34 | 0,36 | 7,19E-01 |
| AT1G15120 | 0,30 | 1,61 | 4,33 | 0,37 | 7,10E-01 |
| AT1G15190 | 0,17 | 1,73 | 4,34 | 0,40 | 6,91E-01 |
| AT1G15240 | 0,58 | 1,69 | 2,62 | 0,64 | 5,19E-01 |
| AT1G15260 | 0,61 | -1,43 | 3,04 | -0,47 | 6,39E-01 |
| AT1G15390 | 0,48 | 2,27 | 4,33 | 0,53 | 5,99E-01 |
| AT1G15430 | 0,55 | 1,59 | 3,86 | 0,41 | 6,80E-01 |
| AT1G15650 | 0,36 | 0,00 | 4,41 | 0,00 | 1,00E+00 |
| AT1G05183 | 0,40 | 1,60 | 3,35 | 0,48 | 6,32E-01 |
| AT1G15720 | 0,33 | 1,56 | 4,33 | 0,36 | 7,19E-01 |
| AT1G15740 | 0,65 | 0,52 | 2,57 | 0,20 | 8,38E-01 |
| AT1G15810 | 0,77 | 1,01 | 2,04 | 0,50 | 6,20E-01 |
| AT1G15890 | 1,49 | -0,86 | 1,84 | -0,47 | 6,40E-01 |
| AT1G15910 | 0,69 | 0,00 | 3,64 | 0,00 | 1,00E+00 |
| AT1G15920 | 0,80 | 2,23 | 2,75 | 0,81 | 4,16E-01 |
| AT1G15930 | 1,08 | 0,06 | 1,83 | 0,04 | 9,72E-01 |
| AT1G15950 | 0,56 | 1,55 | 2,62 | 0,59 | 5,53E-01 |
| AT1G15980 | 1,06 | -1,53 | 1,85 | -0,83 | 4,09E-01 |
| AT1G16060 | 0,12 | 0,90 | 4,36 | 0,21 | 8,36E-01 |
| AT1G16080 | 0,31 | 0,79 | 4,33 | 0,18 | 8,54E-01 |
| AT1G16090 | 0,22 | 1,93 | 4,33 | 0,45 | 6,56E-01 |
| AT1G16110 | 0,61 | 2,72 | 3,61 | 0,75 | 4,51E-01 |
| AT1G16180 | 1,23 | 2,34 | 1,88 | 1,24 | 2,14E-01 |
| AT1G16250 | 0,81 | 0,82 | 2,23 | 0,37 | 7,12E-01 |
| AT1G16270 | 1,59 | 0,98 | 1,55 | 0,63 | 5,29E-01 |
| AT1G16350 | 0,39 | -1,73 | 4,21 | -0,41 | 6,82E-01 |
| AT1G16400 | 0,30 | -1,29 | 4,40 | -0,29 | 7,69E-01 |
| AT1G16410 | 1,19 | -2,58 | 1,67 | -1,54 | 1,24E-01 |
| AT1G16470 | 0,19 | 0,00 | 4,40 | 0,00 | 1,00E+00 |
| AT1G16500 | 0,25 | 0,44 | 4,37 | 0,10 | 9,20E-01 |
| AT1G16560 | 0,44 | 0,67 | 3,27 | 0,21 | 8,37E-01 |
| AT1G16670 | 0,35 | 0,00 | 4,41 | 0,00 | 1,00E+00 |
| AT1G16680 | 0,85 | -0,06 | 2,31 | -0,02 | 9,80E-01 |
| AT1G16690 | 0,12 | 0,79 | 4,37 | 0,18 | 8,57E-01 |
| AT1G16700 | 0,62 | 0,55 | 3,10 | 0,18 | 8,59E-01 |
| AT1G16720 | 1,64 | 1,68 | 1,57 | 1,07 | 2,85E-01 |
| AT1G16740 | 0,37 | 0,00 | 4,40 | 0,00 | 1,00E+00 |
| AT1G16810 | 0,12 | 1,65 | 4,34 | 0,38 | 7,04E-01 |
| AT1G16860 | 0,38 | -0,90 | 4,23 | -0,21 | 8,32E-01 |

| | | | | | |
|---|---|---|---|---|---|
| AT1G17050 | 0,20 | 0,32 | 4,35 | 0,07 | 9,41E-01 |
| AT1G17070 | 0,73 | 1,90 | 3,53 | 0,54 | 5,91E-01 |
| AT1G17100 | 0,86 | -0,07 | 1,77 | -0,04 | 9,67E-01 |
| AT1G17120 | 0,36 | 0,42 | 4,37 | 0,10 | 9,23E-01 |
| AT1G17150 | 0,40 | 0,69 | 3,40 | 0,20 | 8,40E-01 |
| AT1G17160 | 0,36 | 2,14 | 4,29 | 0,50 | 6,17E-01 |
| AT1G17200 | 0,19 | -1,36 | 4,41 | -0,31 | 7,58E-01 |
| AT1G17230 | 0,51 | 1,91 | 3,10 | 0,62 | 5,38E-01 |
| AT1G17255 | 1,02 | 3,77 | 2,62 | 1,44 | 1,50E-01 |
| AT1G17260 | 0,13 | 1,15 | 4,36 | 0,26 | 7,92E-01 |
| AT1G17300 | 0,61 | 2,72 | 3,61 | 0,75 | 4,51E-01 |
| AT1G17340 | 0,57 | -0,46 | 2,40 | -0,19 | 8,48E-01 |
| AT1G17360 | 1,17 | 1,07 | 2,03 | 0,53 | 5,98E-01 |
| AT1G17420 | 1,07 | 0,95 | 2,15 | 0,44 | 6,59E-01 |
| AT1G17460 | 0,31 | 0,85 | 4,38 | 0,19 | 8,46E-01 |
| AT1G17490 | 0,39 | -0,04 | 3,50 | -0,01 | 9,90E-01 |
| AT1G17500 | 0,69 | 1,59 | 2,22 | 0,72 | 4,72E-01 |
| AT1G17530 | 0,57 | 1,69 | 3,13 | 0,54 | 5,90E-01 |
| AT1G17550 | 1,05 | 1,50 | 1,88 | 0,80 | 4,24E-01 |
| AT1G17570 | 0,08 | 0,00 | 4,40 | 0,00 | 1,00E+00 |
| AT1G17580 | 1,15 | 2,93 | 2,17 | 1,35 | 1,76E-01 |
| AT1G17610 | 0,33 | 1,57 | 4,33 | 0,36 | 7,17E-01 |
| AT1G17620 | 1,43 | 1,34 | 1,79 | 0,75 | 4,54E-01 |
| AT1G17680 | 0,28 | 1,28 | 4,32 | 0,30 | 7,67E-01 |
| AT1G17745 | 0,96 | 2,41 | 2,03 | 1,18 | 2,36E-01 |
| AT1G17750 | 0,21 | 2,05 | 4,33 | 0,47 | 6,36E-01 |
| AT1G17840 | 1,51 | 0,69 | 1,63 | 0,43 | 6,70E-01 |
| AT1G17860 | 0,25 | 0,91 | 4,32 | 0,21 | 8,34E-01 |
| AT1G17970 | 0,47 | -1,57 | 4,13 | -0,38 | 7,03E-01 |
| AT1G18020 | 0,78 | 0,83 | 3,07 | 0,27 | 7,87E-01 |
| AT1G18090 | 0,88 | 3,04 | 3,50 | 0,87 | 3,86E-01 |
| AT1G18130 | 0,21 | -0,15 | 4,35 | -0,03 | 9,72E-01 |
| AT1G18170 | 0,31 | 0,61 | 4,34 | 0,14 | 8,89E-01 |
| AT1G18300 | 0,38 | 1,05 | 4,30 | 0,24 | 8,07E-01 |
| AT1G18390 | 0,88 | 1,86 | 2,09 | 0,89 | 3,73E-01 |
| AT1G18450 | 0,27 | 0,04 | 4,35 | 0,01 | 9,93E-01 |
| AT1G18460 | 0,79 | -0,39 | 1,91 | -0,21 | 8,37E-01 |
| AT1G18480 | 0,17 | 0,00 | 4,40 | 0,00 | 1,00E+00 |
| AT1G18490 | 0,64 | 2,86 | 4,31 | 0,66 | 5,07E-01 |
| AT1G18500 | 0,63 | -0,47 | 3,02 | -0,15 | 8,77E-01 |
| AT1G18570 | 0,88 | 2,91 | 2,07 | 1,40 | 1,61E-01 |
| AT1G18580 | 0,40 | 2,51 | 4,32 | 0,58 | 5,61E-01 |
| AT1G18600 | 0,74 | 1,02 | 2,43 | 0,42 | 6,75E-01 |
| AT1G18620 | 0,88 | 1,27 | 2,17 | 0,59 | 5,58E-01 |
| AT1G18700 | 1,03 | 1,90 | 2,31 | 0,82 | 4,13E-01 |
| AT1G18730 | 0,47 | -3,36 | 4,15 | -0,81 | 4,18E-01 |
| AT1G18820 | 0,52 | -2,17 | 4,33 | -0,50 | 6,16E-01 |
| AT1G18870 | 0,46 | 0,40 | 4,32 | 0,09 | 9,27E-01 |
| AT1G18879 | 1,60 | 2,80 | 1,70 | 1,65 | 9,99E-02 |
| AT1G18900 | 0,70 | 3,14 | 3,52 | 0,89 | 3,72E-01 |
| AT1G18950 | 0,83 | 2,18 | 2,20 | 0,99 | 3,21E-01 |
| AT1G19020 | 0,56 | 1,65 | 2,35 | 0,70 | 4,83E-01 |
| AT1G19025 | 0,65 | 1,15 | 3,05 | 0,38 | 7,06E-01 |
| AT1G19086 | 0,19 | 0,00 | 4,41 | 0,00 | 1,00E+00 |
| AT1G19110 | 0,80 | 0,72 | 2,20 | 0,33 | 7,43E-01 |
| AT1G19150 | 0,10 | 0,00 | 4,41 | 0,00 | 1,00E+00 |

| | | | | | |
|---|---|---|---|---|---|
| AT1G19180 | 1,37 | 1,50 | 2,04 | 0,73 | 4,63E-01 |
| AT1G19190 | 0,39 | 0,48 | 4,40 | 0,11 | 9,14E-01 |
| AT1G19290 | 0,57 | 2,37 | 3,73 | 0,64 | 5,25E-01 |
| AT1G19350 | 0,52 | 0,48 | 4,00 | 0,12 | 9,05E-01 |
| AT1G19376 | 0,56 | 2,12 | 3,04 | 0,70 | 4,86E-01 |
| AT1G19464 | 0,18 | 0,00 | 4,41 | 0,00 | 1,00E+00 |
| AT1G19580 | 0,28 | 0,48 | 4,34 | 0,11 | 9,12E-01 |
| AT1G05507 | 0,08 | 0,00 | 4,40 | 0,00 | 1,00E+00 |
| AT1G19710 | 0,37 | 1,57 | 4,28 | 0,37 | 7,14E-01 |
| AT1G19770 | 0,78 | 2,34 | 2,76 | 0,85 | 3,96E-01 |
| AT1G19800 | 0,29 | -0,05 | 4,35 | -0,01 | 9,90E-01 |
| AT1G19830 | 0,33 | 1,26 | 4,34 | 0,29 | 7,72E-01 |
| AT1G19850 | 0,68 | 3,27 | 2,87 | 1,14 | 2,55E-01 |
| AT1G19910 | 0,96 | 3,79 | 2,29 | 1,65 | 9,83E-02 |
| AT1G19990 | 0,28 | 0,65 | 4,39 | 0,15 | 8,82E-01 |
| AT1G20050 | 0,40 | 1,00 | 3,43 | 0,29 | 7,71E-01 |
| AT1G05523 | 0,59 | 1,60 | 2,43 | 0,66 | 5,10E-01 |
| AT1G20090 | 0,64 | -2,42 | 2,66 | -0,91 | 3,63E-01 |
| AT1G20110 | 0,46 | -0,24 | 2,64 | -0,09 | 9,28E-01 |
| AT1G20280 | 0,77 | -0,92 | 2,01 | -0,46 | 6,46E-01 |
| AT1G20300 | 0,15 | -1,24 | 4,39 | -0,28 | 7,77E-01 |
| AT1G20310 | 0,17 | 1,81 | 4,33 | 0,42 | 6,76E-01 |
| AT1G20375 | 1,50 | 2,34 | 1,84 | 1,27 | 2,04E-01 |
| AT1G20630 | 1,62 | 0,18 | 1,91 | 0,09 | 9,25E-01 |
| AT1G20640 | 1,05 | 1,28 | 2,19 | 0,58 | 5,59E-01 |
| AT1G20693 | 1,05 | -2,19 | 2,20 | -1,00 | 3,19E-01 |
| AT1G20696 | 0,69 | 0,27 | 2,23 | 0,12 | 9,04E-01 |
| AT1G20780 | 1,16 | -0,02 | 1,60 | -0,02 | 9,88E-01 |
| AT1G20823 | 0,81 | 1,05 | 2,29 | 0,46 | 6,46E-01 |
| AT1G20920 | 0,49 | 0,92 | 3,14 | 0,29 | 7,70E-01 |
| AT1G20980 | 1,44 | 1,00 | 1,77 | 0,56 | 5,73E-01 |
| AT1G21000 | 0,77 | 2,09 | 2,22 | 0,94 | 3,46E-01 |
| AT1G21050 | 0,58 | 0,01 | 3,04 | 0,00 | 9,99E-01 |
| AT1G21065 | 0,84 | 1,38 | 2,12 | 0,65 | 5,16E-01 |
| AT1G21140 | 0,08 | 1,10 | 4,36 | 0,25 | 8,00E-01 |
| AT1G21170 | 0,55 | -0,27 | 3,83 | -0,07 | 9,44E-01 |
| AT1G21210 | 0,63 | 2,99 | 2,90 | 1,03 | 3,03E-01 |
| AT1G21230 | 0,43 | 1,24 | 3,29 | 0,38 | 7,06E-01 |
| AT1G21240 | 0,38 | 2,01 | 4,23 | 0,48 | 6,35E-01 |
| AT1G21370 | 0,21 | 1,84 | 4,33 | 0,42 | 6,71E-01 |
| AT1G21430 | 0,53 | 1,98 | 4,32 | 0,46 | 6,47E-01 |
| AT1G21560 | 0,41 | 1,91 | 4,11 | 0,46 | 6,42E-01 |
| AT1G21640 | 0,89 | 1,52 | 1,94 | 0,78 | 4,34E-01 |
| AT1G21660 | 0,28 | 1,52 | 4,34 | 0,35 | 7,27E-01 |
| AT1G21670 | 0,86 | 0,06 | 3,33 | 0,02 | 9,86E-01 |
| AT1G21690 | 0,29 | 1,76 | 4,33 | 0,41 | 6,85E-01 |
| AT1G21720 | 0,34 | 1,50 | 4,35 | 0,35 | 7,30E-01 |
| AT1G21722 | 0,08 | 1,10 | 4,36 | 0,25 | 8,00E-01 |
| AT1G21780 | 0,75 | 1,21 | 2,53 | 0,48 | 6,33E-01 |
| AT1G21835 | 0,94 | -0,89 | 2,16 | -0,41 | 6,80E-01 |
| AT1G21840 | 0,12 | 1,65 | 4,34 | 0,38 | 7,04E-01 |
| AT1G21880 | 0,12 | 1,39 | 4,34 | 0,32 | 7,49E-01 |
| AT1G21910 | 0,33 | -2,50 | 4,36 | -0,57 | 5,66E-01 |
| AT1G21980 | 0,19 | 1,26 | 4,35 | 0,29 | 7,72E-01 |
| AT1G22040 | 0,05 | 0,00 | 4,41 | 0,00 | 1,00E+00 |
| AT1G22070 | 0,41 | 1,62 | 4,31 | 0,38 | 7,07E-01 |

| | | | | | |
|---|---|---|---|---|---|
| AT1G22190 | 1,08 | -0,51 | 1,64 | -0,31 | 7,56E-01 |
| AT1G22280 | 0,53 | 1,17 | 3,34 | 0,35 | 7,26E-01 |
| AT1G22330 | 0,11 | 0,00 | 4,41 | 0,00 | 1,00E+00 |
| AT1G22410 | 1,37 | -1,94 | 1,60 | -1,22 | 2,24E-01 |
| AT1G22500 | 0,40 | 2,57 | 4,32 | 0,60 | 5,51E-01 |
| AT1G22590 | 0,24 | 1,24 | 4,35 | 0,28 | 7,76E-01 |
| AT1G22640 | 0,56 | 0,54 | 3,81 | 0,14 | 8,88E-01 |
| AT1G22730 | 0,55 | 1,25 | 3,87 | 0,32 | 7,47E-01 |
| AT1G22750 | 0,35 | 0,79 | 3,52 | 0,23 | 8,22E-01 |
| AT1G22770 | 1,61 | -1,96 | 1,69 | -1,16 | 2,47E-01 |
| AT1G22780 | 0,54 | 1,28 | 3,27 | 0,39 | 6,96E-01 |
| AT1G22790 | 0,64 | 1,93 | 3,84 | 0,50 | 6,16E-01 |
| AT1G05767 | 0,23 | 0,69 | 4,32 | 0,16 | 8,73E-01 |
| AT1G22840 | 0,62 | 1,80 | 3,20 | 0,56 | 5,75E-01 |
| AT1G22860 | 0,86 | 2,35 | 2,09 | 1,12 | 2,62E-01 |
| AT1G22870 | 0,66 | 2,78 | 2,87 | 0,97 | 3,34E-01 |
| AT1G22880 | 1,11 | 0,54 | 2,92 | 0,18 | 8,54E-01 |
| AT1G22882 | 0,84 | 3,34 | 3,28 | 1,02 | 3,10E-01 |
| AT1G22940 | 0,55 | 0,74 | 2,69 | 0,28 | 7,82E-01 |
| AT1G23030 | 0,27 | -0,03 | 4,36 | -0,01 | 9,95E-01 |
| AT1G23040 | 0,34 | 2,01 | 4,33 | 0,46 | 6,42E-01 |
| AT1G23050 | 1,19 | -1,68 | 2,42 | -0,69 | 4,88E-01 |
| AT1G23100 | 0,08 | 1,10 | 4,36 | 0,25 | 8,00E-01 |
| AT1G23130 | 0,08 | 1,10 | 4,36 | 0,25 | 8,00E-01 |
| AT1G23230 | 1,20 | 1,82 | 1,76 | 1,03 | 3,02E-01 |
| AT1G23290 | 1,36 | -0,07 | 1,65 | -0,04 | 9,68E-01 |
| AT1G23440 | 0,65 | 1,86 | 2,95 | 0,63 | 5,29E-01 |
| AT1G23450 | 0,21 | 1,39 | 4,34 | 0,32 | 7,49E-01 |
| AT1G23490 | 1,31 | 1,23 | 1,66 | 0,74 | 4,59E-01 |
| AT1G23710 | 1,29 | 1,53 | 1,89 | 0,81 | 4,19E-01 |
| AT1G23820 | 1,10 | 2,75 | 2,93 | 0,94 | 3,48E-01 |
| AT1G05837 | 0,53 | 0,78 | 3,21 | 0,24 | 8,08E-01 |
| AT1G23870 | 0,51 | 2,78 | 3,82 | 0,73 | 4,67E-01 |
| AT1G23880 | 0,08 | 0,85 | 4,38 | 0,20 | 8,45E-01 |
| AT1G23915 | 1,01 | 3,99 | 2,48 | 1,61 | 1,08E-01 |
| AT1G24145 | 0,48 | 2,38 | 3,12 | 0,76 | 4,45E-01 |
| AT1G24147 | 0,16 | 1,07 | 4,37 | 0,24 | 8,07E-01 |
| AT1G24150 | 0,48 | 2,12 | 3,93 | 0,54 | 5,89E-01 |
| AT1G24280 | 0,44 | 1,03 | 4,31 | 0,24 | 8,11E-01 |
| AT1G24340 | 0,28 | 0,30 | 3,75 | 0,08 | 9,36E-01 |
| AT1G24360 | 0,34 | 0,46 | 4,39 | 0,11 | 9,16E-01 |
| AT1G24380 | 0,38 | 0,45 | 2,87 | 0,16 | 8,77E-01 |
| AT1G24490 | 0,45 | 1,96 | 3,22 | 0,61 | 5,42E-01 |
| AT1G24530 | 0,17 | 0,76 | 4,37 | 0,18 | 8,61E-01 |
| AT1G24560 | 0,68 | 1,24 | 2,18 | 0,57 | 5,71E-01 |
| AT1G24570 | 0,08 | 1,10 | 4,36 | 0,25 | 8,00E-01 |
| AT1G25275 | 1,34 | -1,73 | 1,68 | -1,03 | 3,03E-01 |
| AT1G25280 | 0,98 | 2,10 | 2,44 | 0,86 | 3,90E-01 |
| AT1G25480 | 0,19 | 0,00 | 4,41 | 0,00 | 1,00E+00 |
| AT1G25490 | 1,64 | -0,78 | 1,28 | -0,61 | 5,42E-01 |
| AT1G25520 | 0,81 | 1,89 | 3,52 | 0,54 | 5,91E-01 |
| AT1G25550 | 1,33 | 0,85 | 1,98 | 0,43 | 6,67E-01 |
| AT1G26130 | 0,54 | 2,83 | 3,81 | 0,74 | 4,57E-01 |
| AT1G26220 | 0,37 | 0,85 | 4,38 | 0,19 | 8,46E-01 |
| AT1G26250 | 0,44 | 1,11 | 4,11 | 0,27 | 7,87E-01 |
| AT1G26460 | 0,79 | 2,36 | 2,81 | 0,84 | 4,00E-01 |

| | | | | | |
|---|---|---|---|---|---|
| AT1G26470 | 0,34 | 1,98 | 4,34 | 0,46 | 6,48E-01 |
| AT1G26550 | 0,37 | 0,42 | 4,38 | 0,10 | 9,23E-01 |
| AT1G26560 | 0,24 | 0,33 | 4,34 | 0,08 | 9,39E-01 |
| AT1G26580 | 0,53 | 1,30 | 2,71 | 0,48 | 6,33E-01 |
| AT1G26630 | 1,29 | -0,36 | 1,69 | -0,21 | 8,32E-01 |
| AT1G26650 | 1,02 | 0,22 | 2,10 | 0,10 | 9,18E-01 |
| AT1G26665 | 0,17 | 1,94 | 4,33 | 0,45 | 6,55E-01 |
| AT1G26670 | 1,04 | 2,92 | 2,30 | 1,27 | 2,04E-01 |
| AT1G26770 | 0,19 | 0,00 | 4,41 | 0,00 | 1,00E+00 |
| AT1G26830 | 0,72 | 2,76 | 2,42 | 1,14 | 2,54E-01 |
| AT1G26920 | 0,73 | -0,76 | 2,29 | -0,33 | 7,39E-01 |
| AT1G26940 | 0,05 | 0,00 | 4,41 | 0,00 | 1,00E+00 |
| AT1G27000 | 0,38 | -0,37 | 4,32 | -0,09 | 9,31E-01 |
| AT1G27060 | 0,73 | 3,00 | 2,86 | 1,05 | 2,94E-01 |
| AT1G27130 | 0,51 | -1,08 | 3,15 | -0,34 | 7,31E-01 |
| AT1G27150 | 0,43 | 0,42 | 4,38 | 0,10 | 9,23E-01 |
| AT1G27170 | 0,35 | 1,89 | 3,43 | 0,55 | 5,81E-01 |
| AT1G27180 | 0,99 | 3,10 | 2,55 | 1,21 | 2,25E-01 |
| AT1G27290 | 1,54 | 0,87 | 1,54 | 0,56 | 5,74E-01 |
| AT1G27300 | 0,40 | 0,83 | 4,20 | 0,20 | 8,44E-01 |
| AT1G27330 | 0,50 | 0,15 | 4,35 | 0,03 | 9,72E-01 |
| AT1G27360 | 0,08 | 0,85 | 4,38 | 0,20 | 8,45E-01 |
| AT1G27400 | 0,67 | 1,13 | 2,13 | 0,53 | 5,97E-01 |
| AT1G06087 | 0,12 | 0,79 | 4,37 | 0,18 | 8,57E-01 |
| AT1G27450 | 0,37 | 0,42 | 4,30 | 0,10 | 9,22E-01 |
| AT1G27460 | 0,11 | 1,07 | 4,36 | 0,25 | 8,06E-01 |
| AT1G27480 | 0,41 | 0,46 | 4,17 | 0,11 | 9,13E-01 |
| AT1G27520 | 0,14 | 0,82 | 4,37 | 0,19 | 8,51E-01 |
| AT1G06097 | 0,36 | -1,00 | 4,35 | -0,23 | 8,18E-01 |
| AT1G27570 | 1,68 | 3,31 | 1,96 | 1,69 | 9,13E-02 |
| AT1G27630 | 0,58 | 0,42 | 4,02 | 0,10 | 9,17E-01 |
| AT1G27650 | 0,97 | 1,51 | 2,02 | 0,75 | 4,55E-01 |
| AT1G27750 | 0,63 | 1,66 | 2,67 | 0,62 | 5,36E-01 |
| AT1G27752 | 1,43 | 3,79 | 2,15 | 1,76 | 7,85E-02 |
| AT1G27760 | 0,65 | 2,51 | 2,50 | 1,01 | 3,14E-01 |
| AT1G27850 | 0,84 | 1,13 | 2,26 | 0,50 | 6,16E-01 |
| AT1G27950 | 0,55 | -1,17 | 3,80 | -0,31 | 7,58E-01 |
| AT1G28005 | 0,32 | 0,56 | 4,38 | 0,13 | 8,98E-01 |
| AT1G28060 | 0,32 | 0,66 | 4,32 | 0,15 | 8,79E-01 |
| AT1G28135 | 0,08 | 0,00 | 4,40 | 0,00 | 1,00E+00 |
| AT1G28340 | 0,87 | 2,29 | 2,62 | 0,88 | 3,81E-01 |
| AT1G28350 | 1,15 | 0,04 | 2,27 | 0,02 | 9,87E-01 |
| AT1G28380 | 1,21 | 2,38 | 2,18 | 1,09 | 2,76E-01 |
| AT1G06163 | 0,36 | -1,38 | 4,40 | -0,31 | 7,54E-01 |
| AT1G28490 | 0,17 | 1,75 | 4,33 | 0,40 | 6,86E-01 |
| AT1G28520 | 0,40 | 1,67 | 3,36 | 0,50 | 6,20E-01 |
| AT1G28680 | 0,55 | 0,53 | 3,22 | 0,17 | 8,69E-01 |
| AT1G28780 | 0,49 | 2,00 | 4,07 | 0,49 | 6,24E-01 |
| AT1G28760 | 0,10 | 0,62 | 4,38 | 0,14 | 8,87E-01 |
| AT1G29120 | 0,19 | 1,91 | 4,33 | 0,44 | 6,58E-01 |
| AT1G29150 | 0,99 | 1,35 | 2,45 | 0,55 | 5,80E-01 |
| AT1G29200 | 0,08 | 1,10 | 4,36 | 0,25 | 8,00E-01 |
| AT1G29260 | 0,92 | 1,64 | 1,86 | 0,88 | 3,78E-01 |
| AT1G29310 | 0,68 | 1,46 | 2,94 | 0,50 | 6,20E-01 |
| AT1G29340 | 0,94 | 1,66 | 2,11 | 0,79 | 4,30E-01 |
| AT1G29380 | 0,65 | 1,05 | 4,37 | 0,24 | 8,11E-01 |

| | | | | | |
|---|---|---|---|---|---|
| AT1G06233 | 0,68 | -1,16 | 2,46 | -0,47 | 6,36E-01 |
| AT1G06253 | 1,09 | -0,78 | 1,56 | -0,50 | 6,17E-01 |
| AT1G06263 | 0,52 | -2,42 | 3,99 | -0,61 | 5,44E-01 |
| AT1G06283 | 1,64 | -0,62 | 1,47 | -0,42 | 6,72E-01 |
| AT1G29620 | 0,47 | -1,17 | 3,21 | -0,37 | 7,15E-01 |
| AT1G29630 | 0,24 | 1,96 | 4,34 | 0,45 | 6,52E-01 |
| AT1G29660 | 1,40 | -3,10 | 1,64 | -1,90 | 5,80E-02 |
| AT1G29760 | 0,40 | 0,41 | 4,26 | 0,10 | 9,24E-01 |
| AT1G29770 | 0,71 | 2,65 | 4,31 | 0,62 | 5,38E-01 |
| AT1G29800 | 0,74 | 0,43 | 3,78 | 0,11 | 9,10E-01 |
| AT1G29830 | 1,13 | 0,97 | 1,97 | 0,49 | 6,25E-01 |
| AT1G29950 | 0,49 | 1,34 | 4,34 | 0,31 | 7,57E-01 |
| AT1G30010 | 0,08 | 0,00 | 4,40 | 0,00 | 1,00E+00 |
| AT1G30200 | 0,70 | 1,39 | 2,23 | 0,62 | 5,34E-01 |
| AT1G30260 | 1,16 | -1,15 | 2,33 | -0,49 | 6,21E-01 |
| AT1G30420 | 0,38 | 1,31 | 4,35 | 0,30 | 7,63E-01 |
| AT1G30430 | 0,16 | -1,36 | 4,41 | -0,31 | 7,57E-01 |
| AT1G30440 | 1,10 | -1,86 | 1,87 | -0,99 | 3,21E-01 |
| AT1G30450 | 0,86 | 0,81 | 2,05 | 0,40 | 6,92E-01 |
| AT1G30520 | 0,41 | 0,42 | 4,38 | 0,10 | 9,24E-01 |
| AT1G30530 | 0,46 | -1,24 | 3,30 | -0,37 | 7,08E-01 |
| AT1G30610 | 0,23 | 1,23 | 4,37 | 0,28 | 7,79E-01 |
| AT1G30620 | 0,56 | 1,94 | 3,26 | 0,60 | 5,51E-01 |
| AT1G30700 | 0,12 | 1,58 | 4,34 | 0,36 | 7,16E-01 |
| AT1G30720 | 0,53 | 1,98 | 4,32 | 0,46 | 6,47E-01 |
| AT1G30757 | 0,16 | 0,00 | 4,41 | 0,00 | 1,00E+00 |
| AT1G30800 | 0,20 | 0,89 | 4,37 | 0,20 | 8,39E-01 |
| AT1G30910 | 0,25 | 1,14 | 4,37 | 0,26 | 7,94E-01 |
| AT1G06393 | 0,32 | -1,60 | 4,31 | -0,37 | 7,11E-01 |
| AT1G31095 | 0,66 | -1,20 | 3,01 | -0,40 | 6,89E-01 |
| AT1G31170 | 0,22 | 0,44 | 4,33 | 0,10 | 9,19E-01 |
| AT1G31175 | 0,74 | 1,02 | 2,42 | 0,42 | 6,73E-01 |
| AT1G31190 | 0,86 | 2,47 | 2,46 | 1,00 | 3,15E-01 |
| AT1G31220 | 0,22 | 0,00 | 4,41 | 0,00 | 1,00E+00 |
| AT1G31290 | 0,15 | 0,86 | 4,37 | 0,20 | 8,43E-01 |
| AT1G31420 | 0,89 | -0,09 | 2,00 | -0,05 | 9,64E-01 |
| AT1G31480 | 0,40 | 2,57 | 4,32 | 0,60 | 5,51E-01 |
| AT1G06447 | 0,24 | 0,41 | 4,38 | 0,09 | 9,25E-01 |
| AT1G31710 | 0,82 | 2,03 | 3,52 | 0,58 | 5,65E-01 |
| AT1G31780 | 0,28 | 0,92 | 4,33 | 0,21 | 8,32E-01 |
| AT1G31800 | 0,33 | -0,07 | 3,44 | -0,02 | 9,84E-01 |
| AT1G31850 | 1,13 | 1,31 | 2,19 | 0,60 | 5,49E-01 |
| AT1G31930 | 0,12 | 1,58 | 4,34 | 0,36 | 7,16E-01 |
| AT1G31970 | 0,74 | 2,32 | 3,15 | 0,74 | 4,61E-01 |
| AT1G32050 | 0,65 | -0,13 | 2,25 | -0,06 | 9,56E-01 |
| AT1G32160 | 0,70 | 0,61 | 2,93 | 0,21 | 8,36E-01 |
| AT1G32220 | 0,21 | 0,00 | 4,41 | 0,00 | 1,00E+00 |
| AT1G32225 | 0,70 | -0,45 | 2,48 | -0,18 | 8,56E-01 |
| AT1G32320 | 0,23 | 0,00 | 4,38 | 0,00 | 1,00E+00 |
| AT1G32337 | 0,30 | -0,62 | 4,33 | -0,14 | 8,86E-01 |
| AT1G32360 | 0,41 | -0,31 | 3,31 | -0,09 | 9,26E-01 |
| AT1G06547 | 0,19 | 0,00 | 4,41 | 0,00 | 1,00E+00 |
| AT1G32385 | 0,41 | -1,51 | 2,79 | -0,54 | 5,90E-01 |
| AT1G32415 | 0,72 | 1,61 | 2,88 | 0,56 | 5,75E-01 |
| AT1G32460 | 0,41 | 0,72 | 4,32 | 0,17 | 8,67E-01 |
| AT1G32580 | 0,32 | 0,81 | 4,37 | 0,19 | 8,53E-01 |

| | | | | | |
|---|---|---|---|---|---|
| AT1G06597 | 1,64 | -2,50 | 1,39 | -1,79 | 7,32E-02 |
| AT1G32700 | 0,46 | 0,59 | 3,26 | 0,18 | 8,56E-01 |
| AT1G32810 | 0,36 | 0,95 | 3,44 | 0,28 | 7,82E-01 |
| AT1G06623 | 0,26 | 0,00 | 4,41 | 0,00 | 1,00E+00 |
| AT1G32870 | 0,59 | 2,04 | 3,78 | 0,54 | 5,89E-01 |
| AT1G32920 | 1,13 | -0,19 | 1,96 | -0,10 | 9,22E-01 |
| AT1G32928 | 0,50 | 1,10 | 4,07 | 0,27 | 7,88E-01 |
| AT1G32940 | 0,41 | 1,78 | 4,33 | 0,41 | 6,81E-01 |
| AT1G32950 | 0,62 | 2,60 | 3,64 | 0,72 | 4,74E-01 |
| AT1G32960 | 0,35 | 2,20 | 4,33 | 0,51 | 6,12E-01 |
| AT1G32990 | 0,48 | -2,48 | 4,33 | -0,57 | 5,67E-01 |
| AT1G33010 | 0,19 | 0,00 | 4,41 | 0,00 | 1,00E+00 |
| AT1G33040 | 0,24 | 0,59 | 4,34 | 0,13 | 8,93E-01 |
| AT1G33110 | 0,55 | 1,30 | 3,84 | 0,34 | 7,36E-01 |
| AT1G33170 | 0,44 | 0,73 | 3,27 | 0,22 | 8,24E-01 |
| AT1G33260 | 0,52 | 2,40 | 4,00 | 0,60 | 5,48E-01 |
| AT1G33360 | 0,47 | 1,61 | 4,07 | 0,39 | 6,93E-01 |
| AT1G33470 | 0,71 | 1,61 | 3,54 | 0,46 | 6,49E-01 |
| AT1G33600 | 1,39 | -0,54 | 1,64 | -0,33 | 7,42E-01 |
| AT1G33610 | 0,15 | 1,58 | 4,34 | 0,36 | 7,15E-01 |
| AT1G33612 | 0,17 | -1,64 | 4,37 | -0,37 | 7,08E-01 |
| AT1G33770 | 0,49 | -0,25 | 4,00 | -0,06 | 9,50E-01 |
| AT1G33811 | 0,18 | -0,43 | 4,36 | -0,10 | 9,22E-01 |
| AT1G33910 | 0,08 | 0,00 | 4,40 | 0,00 | 1,00E+00 |
| AT1G33930 | 0,36 | -2,47 | 4,35 | -0,57 | 5,70E-01 |
| AT1G33960 | 1,02 | 0,41 | 3,21 | 0,13 | 8,98E-01 |
| AT1G33980 | 0,92 | 0,27 | 4,25 | 0,06 | 9,50E-01 |
| AT1G33990 | 0,95 | -0,97 | 1,93 | -0,50 | 6,16E-01 |
| AT1G06723 | 0,59 | -1,52 | 2,59 | -0,59 | 5,57E-01 |
| AT1G34020 | 0,78 | 0,75 | 2,33 | 0,32 | 7,47E-01 |
| AT1G34070 | 0,46 | 0,63 | 4,12 | 0,15 | 8,79E-01 |
| AT1G34130 | 1,45 | 1,01 | 1,51 | 0,67 | 5,05E-01 |
| AT1G34150 | 1,09 | -0,43 | 2,56 | -0,17 | 8,67E-01 |
| AT1G34190 | 1,14 | 3,12 | 1,93 | 1,62 | 1,05E-01 |
| AT1G34200 | 0,21 | 1,97 | 4,33 | 0,46 | 6,49E-01 |
| AT1G34210 | 0,51 | 1,75 | 4,34 | 0,40 | 6,88E-01 |
| AT1G34260 | 1,42 | 3,91 | 1,99 | 1,97 | 4,93E-02 |
| AT1G34290 | 0,25 | 0,54 | 4,39 | 0,12 | 9,01E-01 |
| AT1G06747 | 0,08 | 0,00 | 4,40 | 0,00 | 1,00E+00 |
| AT1G34355 | 0,86 | 1,73 | 2,85 | 0,61 | 5,43E-01 |
| AT1G34370 | 1,49 | 3,87 | 1,98 | 1,96 | 5,05E-02 |
| AT1G34390 | 0,68 | 0,53 | 3,68 | 0,14 | 8,86E-01 |
| AT1G34410 | 0,91 | 0,70 | 3,35 | 0,21 | 8,36E-01 |
| AT1G34420 | 0,36 | 2,10 | 4,27 | 0,49 | 6,23E-01 |
| AT1G34460 | 0,31 | 0,42 | 4,38 | 0,10 | 9,24E-01 |
| AT1G34580 | 0,53 | -1,84 | 4,28 | -0,43 | 6,67E-01 |
| AT1G34770 | 0,12 | 1,65 | 4,34 | 0,38 | 7,04E-01 |
| AT1G34844 | 0,08 | 1,10 | 4,36 | 0,25 | 8,00E-01 |
| AT1G35030 | 0,42 | 0,83 | 3,33 | 0,25 | 8,04E-01 |
| AT1G35040 | 0,23 | 0,00 | 4,41 | 0,00 | 1,00E+00 |
| AT1G35140 | 0,39 | 0,00 | 4,30 | 0,00 | 1,00E+00 |
| AT1G35160 | 0,59 | -0,51 | 2,43 | -0,21 | 8,33E-01 |
| AT1G06827 | 0,56 | -0,94 | 2,68 | -0,35 | 7,24E-01 |
| AT1G35220 | 0,89 | 2,74 | 2,35 | 1,17 | 2,43E-01 |
| AT1G35230 | 1,07 | 2,09 | 2,17 | 0,96 | 3,37E-01 |
| AT1G35242 | 0,19 | 1,51 | 4,34 | 0,35 | 7,28E-01 |

| | | | | | |
|---|---|---|---|---|---|
| AT1G35420 | 0,41 | -0,91 | 3,33 | -0,27 | 7,85E-01 |
| AT1G35460 | 0,12 | 1,65 | 4,34 | 0,38 | 7,04E-01 |
| AT1G35500 | 0,14 | 0,00 | 4,41 | 0,00 | 1,00E+00 |
| AT1G35510 | 0,12 | 0,42 | 4,38 | 0,10 | 9,24E-01 |
| AT1G35515 | 0,52 | 0,96 | 2,35 | 0,41 | 6,84E-01 |
| AT1G35516 | 0,52 | 0,96 | 2,35 | 0,41 | 6,84E-01 |
| AT1G35530 | 0,08 | 1,10 | 4,36 | 0,25 | 8,00E-01 |
| AT1G35540 | 0,95 | -0,48 | 2,00 | -0,24 | 8,10E-01 |
| AT1G35614 | 1,12 | -1,07 | 2,06 | -0,52 | 6,02E-01 |
| AT1G35620 | 0,51 | 0,80 | 3,29 | 0,24 | 8,09E-01 |
| AT1G35660 | 0,28 | 2,22 | 4,33 | 0,51 | 6,07E-01 |
| AT1G35670 | 0,21 | 1,97 | 4,33 | 0,46 | 6,49E-01 |
| AT1G35680 | 0,81 | -2,10 | 2,31 | -0,91 | 3,63E-01 |
| AT1G06927 | 0,83 | 2,86 | 2,77 | 1,03 | 3,01E-01 |
| AT1G35850 | 0,23 | 0,00 | 4,41 | 0,00 | 1,00E+00 |
| AT1G36050 | 0,26 | 1,96 | 4,33 | 0,45 | 6,50E-01 |
| AT1G06963 | 0,05 | 0,00 | 4,41 | 0,00 | 1,00E+00 |
| AT1G36320 | 0,22 | -0,24 | 4,37 | -0,06 | 9,56E-01 |
| AT1G36510 | 0,10 | 0,00 | 4,41 | 0,00 | 1,00E+00 |
| AT1G06987 | 0,58 | 0,23 | 4,34 | 0,05 | 9,59E-01 |
| AT1G36675 | 0,81 | 1,10 | 2,44 | 0,45 | 6,51E-01 |
| AT1G36745 | 0,98 | 3,15 | 2,26 | 1,39 | 1,63E-01 |
| AT1G36756 | 0,94 | 2,34 | 2,50 | 0,94 | 3,50E-01 |
| AT1G36940 | 0,22 | 0,00 | 4,41 | 0,00 | 1,00E+00 |
| AT1G36980 | 0,52 | 1,53 | 3,12 | 0,49 | 6,23E-01 |
| AT1G36992 | 0,32 | 0,72 | 4,37 | 0,17 | 8,69E-01 |
| AT1G37113 | 0,08 | 0,85 | 4,38 | 0,20 | 8,45E-01 |
| AT1G42525 | 0,30 | 1,27 | 4,34 | 0,29 | 7,70E-01 |
| AT1G42540 | 1,07 | 2,37 | 2,26 | 1,05 | 2,94E-01 |
| AT1G07137 | 0,05 | 0,00 | 4,41 | 0,00 | 1,00E+00 |
| AT1G42960 | 0,66 | -3,87 | 4,29 | -0,90 | 3,66E-01 |
| AT1G42980 | 1,30 | 2,26 | 1,80 | 1,25 | 2,10E-01 |
| AT1G43040 | 0,16 | 0,00 | 4,41 | 0,00 | 1,00E+00 |
| AT1G43145 | 1,44 | 1,68 | 1,70 | 0,99 | 3,24E-01 |
| AT1G43171 | 0,72 | 0,00 | 4,41 | 0,00 | 1,00E+00 |
| AT1G43330 | 0,87 | 1,62 | 2,63 | 0,62 | 5,38E-01 |
| AT1G43670 | 0,63 | -0,88 | 2,57 | -0,34 | 7,32E-01 |
| AT1G43680 | 0,25 | 0,00 | 4,40 | 0,00 | 1,00E+00 |
| AT1G43700 | 0,72 | -1,63 | 2,04 | -0,80 | 4,25E-01 |
| AT1G43722 | 0,18 | 0,00 | 4,41 | 0,00 | 1,00E+00 |
| AT1G43770 | 1,22 | -0,96 | 1,52 | -0,63 | 5,26E-01 |
| AT1G43780 | 0,25 | -2,61 | 4,34 | -0,60 | 5,47E-01 |
| AT1G43815 | 1,28 | -0,42 | 2,25 | -0,19 | 8,52E-01 |
| AT1G43890 | 0,40 | 1,24 | 4,30 | 0,29 | 7,74E-01 |
| AT1G43970 | 0,43 | 0,00 | 4,41 | 0,00 | 1,00E+00 |
| AT1G44120 | 0,95 | 3,63 | 2,66 | 1,37 | 1,72E-01 |
| AT1G44750 | 0,33 | 2,66 | 4,32 | 0,62 | 5,37E-01 |
| AT1G44780 | 0,29 | -0,05 | 4,35 | -0,01 | 9,90E-01 |
| AT1G44810 | 0,26 | 0,66 | 3,74 | 0,18 | 8,59E-01 |
| AT1G44890 | 0,08 | 0,00 | 4,40 | 0,00 | 1,00E+00 |
| AT1G44900 | 0,12 | 1,65 | 4,34 | 0,38 | 7,04E-01 |
| AT1G44910 | 0,50 | 0,47 | 2,39 | 0,20 | 8,44E-01 |
| AT1G44960 | 0,10 | 0,62 | 4,38 | 0,14 | 8,87E-01 |
| AT1G45000 | 0,47 | -0,05 | 2,79 | -0,02 | 9,87E-01 |
| AT1G45165 | 0,70 | 2,82 | 2,48 | 1,14 | 2,54E-01 |
| AT1G45180 | 0,30 | 1,62 | 4,35 | 0,37 | 7,10E-01 |

| | | | | | |
|---|---|---|---|---|---|
| AT1G45474 | 0,41 | -2,56 | 3,33 | -0,77 | 4,42E-01 |
| AT1G45688 | 0,89 | 2,46 | 2,38 | 1,03 | 3,02E-01 |
| AT1G45976 | 0,41 | 1,11 | 4,23 | 0,26 | 7,92E-01 |
| AT1G46264 | 0,10 | 0,62 | 4,38 | 0,14 | 8,87E-01 |
| AT1G46554 | 0,28 | 1,16 | 4,35 | 0,27 | 7,90E-01 |
| AT1G47210 | 0,93 | 2,36 | 2,57 | 0,92 | 3,58E-01 |
| AT1G07327 | 1,09 | -0,16 | 2,24 | -0,07 | 9,45E-01 |
| AT1G47250 | 0,35 | 0,65 | 4,34 | 0,15 | 8,82E-01 |
| AT1G47270 | 0,13 | 1,15 | 4,36 | 0,26 | 7,92E-01 |
| AT1G07357 | 0,11 | 0,00 | 4,41 | 0,00 | 1,00E+00 |
| AT1G47480 | 1,07 | 0,55 | 1,90 | 0,29 | 7,73E-01 |
| AT1G47490 | 0,30 | 1,62 | 4,35 | 0,37 | 7,10E-01 |
| AT1G47500 | 0,37 | 1,64 | 4,28 | 0,38 | 7,03E-01 |
| AT1G47530 | 1,40 | -0,12 | 1,61 | -0,07 | 9,41E-01 |
| AT1G47550 | 0,50 | 1,67 | 4,15 | 0,40 | 6,88E-01 |
| AT1G47680 | 0,12 | 0,00 | 4,39 | 0,00 | 1,00E+00 |
| AT1G47690 | 0,26 | 2,12 | 4,32 | 0,49 | 6,23E-01 |
| AT1G47770 | 0,12 | 0,00 | 4,39 | 0,00 | 1,00E+00 |
| AT1G47890 | 1,24 | 2,39 | 2,03 | 1,18 | 2,38E-01 |
| AT1G47940 | 0,36 | 0,00 | 4,41 | 0,00 | 1,00E+00 |
| AT1G47970 | 1,63 | 1,15 | 1,59 | 0,72 | 4,70E-01 |
| AT1G48050 | 0,54 | -2,79 | 4,35 | -0,64 | 5,21E-01 |
| AT1G48100 | 0,12 | 1,39 | 4,34 | 0,32 | 7,49E-01 |
| AT1G48170 | 0,32 | 1,31 | 3,58 | 0,37 | 7,14E-01 |
| AT1G48210 | 0,24 | 1,85 | 4,34 | 0,43 | 6,69E-01 |
| AT1G48230 | 0,28 | 1,04 | 4,37 | 0,24 | 8,12E-01 |
| AT1G48315 | 0,17 | 1,92 | 4,33 | 0,44 | 6,57E-01 |
| AT1G48360 | 1,34 | 2,57 | 2,15 | 1,19 | 2,33E-01 |
| AT1G48460 | 0,51 | 1,20 | 3,11 | 0,39 | 6,99E-01 |
| AT1G07487 | 0,32 | 0,67 | 3,60 | 0,19 | 8,53E-01 |
| AT1G48480 | 0,38 | -1,58 | 4,33 | -0,36 | 7,15E-01 |
| AT1G48540 | 1,33 | 3,56 | 1,95 | 1,82 | 6,81E-02 |
| AT1G48560 | 0,14 | 0,00 | 4,41 | 0,00 | 1,00E+00 |
| AT1G48600 | 0,62 | 0,57 | 2,11 | 0,27 | 7,86E-01 |
| AT1G48640 | 0,26 | 1,08 | 4,37 | 0,25 | 8,05E-01 |
| AT1G48725 | 1,45 | -0,10 | 1,82 | -0,06 | 9,56E-01 |
| AT1G48820 | 0,24 | -1,90 | 4,37 | -0,43 | 6,64E-01 |
| AT1G49010 | 0,27 | 1,45 | 4,35 | 0,33 | 7,39E-01 |
| AT1G49032 | 0,71 | 1,78 | 2,23 | 0,80 | 4,23E-01 |
| AT1G49190 | 1,12 | -0,51 | 2,13 | -0,24 | 8,12E-01 |
| AT1G49200 | 0,08 | 0,00 | 4,40 | 0,00 | 1,00E+00 |
| AT1G49210 | 0,15 | 0,00 | 4,39 | 0,00 | 1,00E+00 |
| AT1G49230 | 0,38 | 0,51 | 3,46 | 0,15 | 8,82E-01 |
| AT1G49245 | 0,90 | 0,45 | 2,26 | 0,20 | 8,43E-01 |
| AT1G49300 | 0,19 | 1,28 | 4,34 | 0,30 | 7,68E-01 |
| AT1G49340 | 1,22 | 0,15 | 2,14 | 0,07 | 9,42E-01 |
| AT1G49390 | 0,08 | 0,85 | 4,38 | 0,20 | 8,45E-01 |
| AT1G49470 | 0,15 | 0,71 | 4,37 | 0,16 | 8,71E-01 |
| AT1G49510 | 0,25 | -2,61 | 4,34 | -0,60 | 5,47E-01 |
| AT1G49680 | 0,43 | 0,63 | 3,32 | 0,19 | 8,50E-01 |
| AT1G49720 | 0,23 | -0,34 | 4,35 | -0,08 | 9,38E-01 |
| AT1G49820 | 0,28 | 1,30 | 4,32 | 0,30 | 7,63E-01 |
| AT1G49890 | 0,23 | 0,79 | 4,37 | 0,18 | 8,56E-01 |
| AT1G49930 | 0,39 | 0,40 | 4,39 | 0,09 | 9,28E-01 |
| AT1G49975 | 1,21 | 2,52 | 2,49 | 1,01 | 3,13E-01 |
| AT1G49980 | 0,53 | 0,92 | 3,19 | 0,29 | 7,73E-01 |

| | | | | | |
|---|---|---|---|---|---|
| AT1G49990 | 0,80 | 3,57 | 4,25 | 0,84 | 4,01E-01 |
| AT1G50050 | 0,43 | -1,26 | 3,34 | -0,38 | 7,06E-01 |
| AT1G07613 | 1,47 | -2,28 | 1,58 | -1,44 | 1,49E-01 |
| AT1G50120 | 0,25 | 0,75 | 4,37 | 0,17 | 8,63E-01 |
| AT1G50170 | 0,08 | 0,85 | 4,38 | 0,20 | 8,45E-01 |
| AT1G50240 | 1,24 | 2,05 | 1,79 | 1,15 | 2,52E-01 |
| AT1G50270 | 0,45 | -0,55 | 3,36 | -0,16 | 8,71E-01 |
| AT1G50350 | 0,30 | -1,48 | 4,39 | -0,34 | 7,36E-01 |
| AT1G50360 | 0,23 | -0,37 | 4,36 | -0,08 | 9,33E-01 |
| AT1G50370 | 0,39 | 1,24 | 3,44 | 0,36 | 7,18E-01 |
| AT1G50380 | 0,57 | 2,76 | 3,15 | 0,88 | 3,81E-01 |
| AT1G50410 | 1,51 | 3,38 | 2,36 | 1,43 | 1,53E-01 |
| AT1G50560 | 0,41 | 1,58 | 4,28 | 0,37 | 7,12E-01 |
| AT1G50660 | 1,10 | 2,95 | 1,94 | 1,52 | 1,28E-01 |
| AT1G50730 | 0,12 | 1,65 | 4,34 | 0,38 | 7,04E-01 |
| AT1G50740 | 0,72 | 2,11 | 2,90 | 0,73 | 4,67E-01 |
| AT1G51060 | 0,42 | -2,74 | 4,26 | -0,64 | 5,19E-01 |
| AT1G51100 | 0,17 | 0,00 | 4,41 | 0,00 | 1,00E+00 |
| AT1G51110 | 0,52 | -1,54 | 3,15 | -0,49 | 6,24E-01 |
| AT1G51160 | 0,67 | 0,56 | 2,96 | 0,19 | 8,50E-01 |
| AT1G51172 | 1,67 | 1,73 | 1,80 | 0,96 | 3,37E-01 |
| AT1G51200 | 1,27 | 0,47 | 2,35 | 0,20 | 8,41E-01 |
| AT1G51350 | 0,44 | 2,03 | 3,24 | 0,62 | 5,32E-01 |
| AT1G51390 | 0,24 | -1,37 | 4,41 | -0,31 | 7,55E-01 |
| AT1G07737 | 0,83 | -2,37 | 2,33 | -1,02 | 3,09E-01 |
| AT1G51402 | 1,01 | -2,10 | 2,21 | -0,95 | 3,43E-01 |
| AT1G51440 | 0,14 | 0,44 | 4,39 | 0,10 | 9,20E-01 |
| AT1G51540 | 0,08 | 0,00 | 4,40 | 0,00 | 1,00E+00 |
| AT1G51580 | 0,35 | -0,77 | 4,31 | -0,18 | 8,58E-01 |
| AT1G51600 | 0,08 | 1,10 | 4,36 | 0,25 | 8,00E-01 |
| AT1G51645 | 0,08 | 1,10 | 4,36 | 0,25 | 8,00E-01 |
| AT1G51660 | 0,75 | 2,35 | 2,64 | 0,89 | 3,75E-01 |
| AT1G51670 | 0,25 | 1,75 | 4,33 | 0,40 | 6,87E-01 |
| AT1G51720 | 0,29 | 1,60 | 4,34 | 0,37 | 7,13E-01 |
| AT1G51745 | 0,21 | 1,60 | 4,35 | 0,37 | 7,13E-01 |
| AT1G51760 | 0,56 | 0,64 | 2,63 | 0,24 | 8,07E-01 |
| AT1G51950 | 0,22 | 1,24 | 4,34 | 0,29 | 7,75E-01 |
| AT1G52080 | 0,08 | 1,10 | 4,36 | 0,25 | 8,00E-01 |
| AT1G52190 | 0,40 | 1,57 | 3,35 | 0,47 | 6,39E-01 |
| AT1G52310 | 0,73 | 2,47 | 2,46 | 1,00 | 3,16E-01 |
| AT1G52320 | 0,99 | 0,67 | 1,81 | 0,37 | 7,10E-01 |
| AT1G52380 | 0,74 | 0,72 | 2,89 | 0,25 | 8,04E-01 |
| AT1G52410 | 0,68 | -1,22 | 2,47 | -0,50 | 6,20E-01 |
| AT1G52420 | 0,99 | 3,43 | 2,81 | 1,22 | 2,22E-01 |
| AT1G52500 | 0,39 | 0,13 | 4,34 | 0,03 | 9,77E-01 |
| AT1G52600 | 0,08 | 1,10 | 4,36 | 0,25 | 8,00E-01 |
| AT1G52618 | 0,68 | 1,22 | 2,91 | 0,42 | 6,75E-01 |
| AT1G52620 | 0,78 | 0,28 | 4,34 | 0,07 | 9,48E-01 |
| AT1G52720 | 0,19 | 0,00 | 4,41 | 0,00 | 1,00E+00 |
| AT1G52740 | 1,01 | 0,85 | 2,13 | 0,40 | 6,89E-01 |
| AT1G52760 | 0,33 | -0,60 | 3,58 | -0,17 | 8,67E-01 |
| AT1G52870 | 0,92 | 2,57 | 2,57 | 1,00 | 3,18E-01 |
| AT1G52905 | 0,68 | 2,48 | 2,84 | 0,88 | 3,81E-01 |
| AT1G52930 | 0,12 | 1,58 | 4,34 | 0,36 | 7,16E-01 |
| AT1G52940 | 0,76 | 1,99 | 2,49 | 0,80 | 4,25E-01 |
| AT1G52950 | 0,39 | 0,29 | 3,49 | 0,08 | 9,34E-01 |

| | | | | | |
|---|---|---|---|---|---|
| AT1G52980 | 1,44 | 2,19 | 1,83 | 1,20 | 2,31E-01 |
| AT1G53050 | 0,64 | 0,61 | 4,40 | 0,14 | 8,90E-01 |
| AT1G53090 | 0,57 | -0,75 | 4,33 | -0,17 | 8,62E-01 |
| AT1G53165 | 0,43 | 0,77 | 2,78 | 0,28 | 7,82E-01 |
| AT1G53233 | 0,99 | 3,47 | 2,22 | 1,56 | 1,18E-01 |
| AT1G53300 | 0,67 | 0,53 | 2,72 | 0,19 | 8,46E-01 |
| AT1G53330 | 0,35 | 0,00 | 4,39 | 0,00 | 1,00E+00 |
| AT1G53350 | 0,25 | 2,10 | 4,32 | 0,49 | 6,27E-01 |
| AT1G53390 | 0,38 | 0,00 | 4,40 | 0,00 | 1,00E+00 |
| AT1G53400 | 0,40 | 2,57 | 4,32 | 0,60 | 5,51E-01 |
| AT1G53430 | 1,51 | -1,88 | 1,65 | -1,14 | 2,53E-01 |
| AT1G53440 | 1,17 | -1,06 | 1,59 | -0,66 | 5,08E-01 |
| AT1G07887 | 0,84 | -0,95 | 3,01 | -0,32 | 7,52E-01 |
| AT1G53490 | 0,92 | -0,27 | 2,46 | -0,11 | 9,13E-01 |
| AT1G07893 | 0,39 | -0,57 | 4,26 | -0,13 | 8,93E-01 |
| AT1G07897 | 0,95 | 0,69 | 1,77 | 0,39 | 6,99E-01 |
| AT1G53570 | 0,96 | -0,51 | 2,41 | -0,21 | 8,34E-01 |
| AT1G53590 | 0,26 | -0,08 | 4,37 | -0,02 | 9,86E-01 |
| AT1G53670 | 0,12 | 1,58 | 4,34 | 0,36 | 7,16E-01 |
| AT1G53720 | 0,26 | -0,94 | 4,37 | -0,22 | 8,30E-01 |
| AT1G53730 | 1,50 | 0,52 | 2,23 | 0,23 | 8,16E-01 |
| AT1G53840 | 1,21 | 0,38 | 2,00 | 0,19 | 8,49E-01 |
| AT1G53870 | 0,49 | 1,87 | 3,93 | 0,48 | 6,33E-01 |
| AT1G53875 | 1,66 | -1,95 | 1,43 | -1,36 | 1,73E-01 |
| AT1G53885 | 0,24 | 0,59 | 4,34 | 0,13 | 8,93E-01 |
| AT1G53890 | 1,37 | 0,43 | 1,62 | 0,27 | 7,91E-01 |
| AT1G53910 | 0,62 | 0,60 | 2,52 | 0,24 | 8,12E-01 |
| AT1G54030 | 0,19 | 0,00 | 4,41 | 0,00 | 1,00E+00 |
| AT1G54130 | 1,27 | 0,38 | 1,82 | 0,21 | 8,36E-01 |
| AT1G54150 | 0,44 | 1,23 | 3,44 | 0,36 | 7,20E-01 |
| AT1G54200 | 0,80 | 1,59 | 2,52 | 0,63 | 5,28E-01 |
| AT1G54215 | 0,08 | 1,10 | 4,36 | 0,25 | 8,00E-01 |
| AT1G07957 | 0,34 | 1,58 | 3,47 | 0,45 | 6,49E-01 |
| AT1G54310 | 0,08 | 0,85 | 4,38 | 0,20 | 8,45E-01 |
| AT1G54340 | 0,28 | 1,26 | 4,34 | 0,29 | 7,72E-01 |
| AT1G54350 | 0,37 | 1,95 | 4,27 | 0,46 | 6,48E-01 |
| AT1G54360 | 0,57 | 0,43 | 4,39 | 0,10 | 9,22E-01 |
| AT1G54420 | 1,07 | -0,38 | 2,65 | -0,14 | 8,85E-01 |
| AT1G54440 | 0,25 | 2,24 | 4,32 | 0,52 | 6,04E-01 |
| AT1G54490 | 1,36 | 2,46 | 1,88 | 1,31 | 1,90E-01 |
| AT1G54550 | 0,56 | 2,19 | 2,94 | 0,74 | 4,57E-01 |
| AT1G54680 | 0,26 | 1,08 | 4,37 | 0,25 | 8,05E-01 |
| AT1G54720 | 0,66 | 0,69 | 3,10 | 0,22 | 8,23E-01 |
| AT1G54780 | 1,22 | -2,26 | 1,70 | -1,33 | 1,83E-01 |
| AT1G54820 | 0,40 | -0,33 | 4,31 | -0,08 | 9,39E-01 |
| AT1G54920 | 1,30 | 2,52 | 1,73 | 1,46 | 1,44E-01 |
| AT1G55000 | 0,70 | 1,53 | 2,52 | 0,61 | 5,42E-01 |
| AT1G55020 | 0,29 | 1,22 | 4,35 | 0,28 | 7,80E-01 |
| AT1G55040 | 0,41 | 1,95 | 4,18 | 0,47 | 6,40E-01 |
| AT1G55090 | 0,81 | 2,79 | 2,91 | 0,96 | 3,38E-01 |
| AT1G55120 | 1,23 | -4,02 | 1,85 | -2,17 | 2,97E-02 |
| AT1G55150 | 0,30 | 1,62 | 4,35 | 0,37 | 7,10E-01 |
| AT1G55160 | 0,99 | 2,72 | 2,41 | 1,13 | 2,59E-01 |
| AT1G55175 | 0,08 | 1,10 | 4,36 | 0,25 | 8,00E-01 |
| AT1G55320 | 0,25 | 1,03 | 4,37 | 0,24 | 8,13E-01 |
| AT1G55335 | 0,51 | -0,83 | 3,13 | -0,27 | 7,91E-01 |

| | | | | | |
|---|---|---|---|---|---|
| AT1G55370 | 0,24 | 0,00 | 4,41 | 0,00 | 1,00E+00 |
| AT1G55460 | 0,32 | 0,27 | 4,34 | 0,06 | 9,50E-01 |
| AT1G55480 | 0,48 | 1,03 | 4,11 | 0,25 | 8,02E-01 |
| AT1G55500 | 0,61 | 1,62 | 2,30 | 0,70 | 4,81E-01 |
| AT1G55510 | 0,20 | 0,75 | 4,33 | 0,17 | 8,62E-01 |
| AT1G55840 | 0,33 | 0,44 | 4,37 | 0,10 | 9,20E-01 |
| AT1G55850 | 0,50 | 1,20 | 4,29 | 0,28 | 7,80E-01 |
| AT1G08073 | 0,33 | 0,70 | 4,32 | 0,16 | 8,72E-01 |
| AT1G55890 | 0,12 | 0,79 | 4,37 | 0,18 | 8,57E-01 |
| AT1G55910 | 0,98 | 1,14 | 1,98 | 0,58 | 5,63E-01 |
| AT1G55915 | 0,81 | 2,47 | 2,98 | 0,83 | 4,08E-01 |
| AT1G08077 | 0,62 | 1,39 | 3,02 | 0,46 | 6,46E-01 |
| AT1G56045 | 0,75 | -1,53 | 2,54 | -0,60 | 5,47E-01 |
| AT1G56050 | 1,07 | 2,77 | 2,65 | 1,04 | 2,96E-01 |
| AT1G56190 | 1,11 | 1,72 | 2,12 | 0,81 | 4,18E-01 |
| AT1G56200 | 1,42 | 0,21 | 1,67 | 0,13 | 8,99E-01 |
| AT1G56210 | 0,05 | 0,00 | 4,41 | 0,00 | 1,00E+00 |
| AT1G56220 | 1,48 | -0,97 | 1,63 | -0,60 | 5,51E-01 |
| AT1G56450 | 0,83 | 1,60 | 2,15 | 0,74 | 4,57E-01 |
| AT1G56460 | 0,70 | 2,95 | 2,86 | 1,03 | 3,02E-01 |
| AT1G56500 | 0,95 | 0,28 | 1,84 | 0,15 | 8,77E-01 |
| AT1G56540 | 0,56 | 1,60 | 4,35 | 0,37 | 7,13E-01 |
| AT1G56560 | 0,52 | 2,62 | 3,86 | 0,68 | 4,98E-01 |
| AT1G56570 | 0,28 | 1,26 | 4,34 | 0,29 | 7,72E-01 |
| AT1G56600 | 0,19 | -1,23 | 4,39 | -0,28 | 7,80E-01 |
| AT1G56610 | 0,17 | -2,23 | 4,35 | -0,51 | 6,08E-01 |
| AT1G56660 | 1,02 | 1,78 | 1,83 | 0,97 | 3,31E-01 |
| AT1G56700 | 0,33 | -0,54 | 3,55 | -0,15 | 8,80E-01 |
| AT1G57620 | 0,88 | 0,99 | 3,00 | 0,33 | 7,41E-01 |
| AT1G57630 | 0,41 | 0,94 | 4,36 | 0,22 | 8,29E-01 |
| AT1G57650 | 0,46 | -1,16 | 2,71 | -0,43 | 6,69E-01 |
| AT1G57760 | 0,12 | 0,42 | 4,38 | 0,10 | 9,24E-01 |
| AT1G57765 | 0,15 | 1,16 | 4,35 | 0,27 | 7,90E-01 |
| AT1G58190 | 0,62 | 0,99 | 2,61 | 0,38 | 7,06E-01 |
| AT1G58215 | 0,57 | 2,01 | 4,32 | 0,47 | 6,42E-01 |
| AT1G58223 | 0,54 | 0,46 | 3,90 | 0,12 | 9,06E-01 |
| AT1G58260 | 0,12 | 0,90 | 4,36 | 0,21 | 8,36E-01 |
| AT1G58360 | 0,57 | -2,20 | 3,94 | -0,56 | 5,77E-01 |
| AT1G58525 | 0,11 | 1,07 | 4,36 | 0,25 | 8,06E-01 |
| AT1G58520 | 0,55 | 1,52 | 3,25 | 0,47 | 6,41E-01 |
| AT1G58643 | 0,42 | 0,00 | 4,37 | 0,00 | 1,00E+00 |
| AT1G58807 | 1,55 | 2,20 | 1,76 | 1,25 | 2,11E-01 |
| AT1G58808 | 1,22 | -2,03 | 1,99 | -1,02 | 3,09E-01 |
| AT1G58848 | 1,34 | 3,32 | 1,95 | 1,71 | 8,81E-02 |
| AT1G59124 | 1,27 | 1,11 | 1,57 | 0,71 | 4,80E-01 |
| AT1G59218 | 1,28 | 2,03 | 1,95 | 1,04 | 2,97E-01 |
| AT1G59590 | 0,59 | 1,68 | 3,19 | 0,53 | 5,98E-01 |
| AT1G59620 | 0,81 | 1,91 | 4,33 | 0,44 | 6,60E-01 |
| AT1G59650 | 0,54 | 1,38 | 3,89 | 0,35 | 7,23E-01 |
| AT1G59710 | 0,56 | -0,79 | 3,18 | -0,25 | 8,05E-01 |
| AT1G59750 | 1,67 | 3,17 | 2,37 | 1,34 | 1,82E-01 |
| AT1G59820 | 1,45 | 2,64 | 2,12 | 1,25 | 2,12E-01 |
| AT1G59890 | 0,93 | 2,89 | 2,45 | 1,18 | 2,38E-01 |
| AT1G59900 | 1,24 | 1,07 | 1,76 | 0,61 | 5,43E-01 |
| AT1G60025 | 0,05 | 0,00 | 4,41 | 0,00 | 1,00E+00 |
| AT1G60073 | 1,58 | 3,82 | 1,91 | 1,99 | 4,60E-02 |

| | | | | | |
|---|---|---|---|---|---|
| AT1G60170 | 1,29 | 3,26 | 2,37 | 1,38 | 1,69E-01 |
| AT1G60190 | 0,89 | 2,58 | 2,18 | 1,18 | 2,36E-01 |
| AT1G60220 | 0,08 | 0,85 | 4,38 | 0,20 | 8,45E-01 |
| AT1G60420 | 0,70 | 2,51 | 2,80 | 0,90 | 3,70E-01 |
| AT1G60545 | 1,07 | 2,93 | 2,98 | 0,98 | 3,27E-01 |
| AT1G60650 | 0,21 | 0,79 | 4,37 | 0,18 | 8,57E-01 |
| AT1G60720 | 1,62 | 2,12 | 2,02 | 1,05 | 2,93E-01 |
| AT1G60900 | 0,65 | 0,54 | 3,17 | 0,17 | 8,64E-01 |
| AT1G08353 | 0,98 | -0,66 | 1,93 | -0,34 | 7,32E-01 |
| AT1G61030 | 0,52 | 1,37 | 3,89 | 0,35 | 7,24E-01 |
| AT1G61040 | 0,54 | 1,38 | 3,89 | 0,35 | 7,23E-01 |
| AT1G61150 | 0,81 | -0,35 | 3,61 | -0,10 | 9,23E-01 |
| AT1G61170 | 0,33 | 2,33 | 4,32 | 0,54 | 5,89E-01 |
| AT1G61180 | 0,58 | 1,23 | 2,69 | 0,46 | 6,47E-01 |
| AT1G61210 | 1,07 | -0,48 | 1,74 | -0,28 | 7,82E-01 |
| AT1G61224 | 1,62 | 0,60 | 1,70 | 0,36 | 7,22E-01 |
| AT1G61250 | 0,18 | 0,63 | 4,37 | 0,14 | 8,86E-01 |
| AT1G61280 | 0,12 | 1,58 | 4,34 | 0,36 | 7,16E-01 |
| AT1G61350 | 0,76 | 1,26 | 2,44 | 0,52 | 6,06E-01 |
| AT1G61450 | 0,12 | 1,46 | 4,35 | 0,34 | 7,36E-01 |
| AT1G61470 | 0,18 | 0,00 | 4,40 | 0,00 | 1,00E+00 |
| AT1G61610 | 0,12 | 0,42 | 4,38 | 0,10 | 9,24E-01 |
| AT1G61620 | 0,37 | -0,55 | 4,38 | -0,12 | 9,01E-01 |
| AT1G61790 | 1,42 | -0,03 | 1,77 | -0,02 | 9,84E-01 |
| AT1G61800 | 0,56 | 2,59 | 4,32 | 0,60 | 5,49E-01 |
| AT1G61820 | 0,08 | 0,85 | 4,38 | 0,20 | 8,45E-01 |
| AT1G61850 | 0,80 | 0,74 | 2,25 | 0,33 | 7,41E-01 |
| AT1G61990 | 0,32 | 2,46 | 4,32 | 0,57 | 5,69E-01 |
| AT1G62170 | 0,36 | 0,84 | 4,38 | 0,19 | 8,48E-01 |
| AT1G08477 | 1,64 | 1,50 | 1,58 | 0,95 | 3,40E-01 |
| AT1G62270 | 0,29 | 0,00 | 4,41 | 0,00 | 1,00E+00 |
| AT1G08517 | 0,77 | -2,06 | 2,45 | -0,84 | 4,01E-01 |
| AT1G62380 | 1,16 | 0,00 | 1,60 | 0,00 | 9,99E-01 |
| AT1G62410 | 0,32 | -2,40 | 4,35 | -0,55 | 5,81E-01 |
| AT1G62480 | 0,12 | 0,42 | 4,38 | 0,10 | 9,24E-01 |
| AT1G62560 | 1,49 | -1,70 | 3,08 | -0,55 | 5,82E-01 |
| AT1G62570 | 0,39 | 0,62 | 4,32 | 0,14 | 8,86E-01 |
| AT1G62620 | 0,57 | 1,57 | 3,05 | 0,52 | 6,06E-01 |
| AT1G62660 | 1,32 | 1,18 | 1,76 | 0,67 | 5,04E-01 |
| AT1G62720 | 0,24 | -1,66 | 4,38 | -0,38 | 7,04E-01 |
| AT1G62790 | 0,51 | 0,48 | 4,40 | 0,11 | 9,13E-01 |
| AT1G62895 | 1,43 | 0,33 | 1,52 | 0,21 | 8,31E-01 |
| AT1G62920 | 0,97 | 0,42 | 2,88 | 0,14 | 8,85E-01 |
| AT1G62970 | 0,33 | 0,77 | 3,51 | 0,22 | 8,26E-01 |
| AT1G62975 | 0,08 | 0,85 | 4,38 | 0,20 | 8,45E-01 |
| AT1G63000 | 0,67 | 1,02 | 3,86 | 0,26 | 7,92E-01 |
| AT1G63110 | 0,32 | 1,91 | 4,33 | 0,44 | 6,59E-01 |
| AT1G63170 | 0,08 | 1,10 | 4,36 | 0,25 | 8,00E-01 |
| AT1G63430 | 0,74 | 3,17 | 3,85 | 0,82 | 4,09E-01 |
| AT1G63500 | 0,59 | 0,84 | 2,70 | 0,31 | 7,55E-01 |
| AT1G63540 | 0,37 | 0,00 | 4,41 | 0,00 | 1,00E+00 |
| AT1G63680 | 0,43 | 1,94 | 3,23 | 0,60 | 5,49E-01 |
| AT1G63690 | 1,03 | 0,58 | 1,91 | 0,31 | 7,60E-01 |
| AT1G63730 | 0,24 | 0,00 | 4,41 | 0,00 | 1,00E+00 |
| AT1G63830 | 0,49 | 2,00 | 4,07 | 0,49 | 6,24E-01 |
| AT1G63850 | 0,53 | 0,71 | 3,95 | 0,18 | 8,57E-01 |

| | | | | | |
|---|---|---|---|---|---|
| AT1G08643 | 0,79 | 2,80 | 2,96 | 0,95 | 3,44E-01 |
| AT1G63900 | 0,95 | 2,15 | 2,20 | 0,98 | 3,28E-01 |
| AT1G63940 | 0,81 | 1,00 | 2,03 | 0,50 | 6,20E-01 |
| AT1G64060 | 0,86 | 3,21 | 2,35 | 1,36 | 1,73E-01 |
| AT1G64065 | 0,29 | 0,42 | 4,38 | 0,10 | 9,24E-01 |
| AT1G64090 | 0,96 | 1,04 | 1,99 | 0,52 | 6,04E-01 |
| AT1G64100 | 0,61 | 1,17 | 3,69 | 0,32 | 7,51E-01 |
| AT1G64120 | 0,13 | 0,00 | 4,41 | 0,00 | 1,00E+00 |
| AT1G64130 | 0,12 | 1,58 | 4,34 | 0,36 | 7,16E-01 |
| AT1G08657 | 0,60 | -0,22 | 2,63 | -0,09 | 9,32E-01 |
| AT1G64230 | 1,39 | 0,07 | 1,55 | 0,04 | 9,64E-01 |
| AT1G64370 | 0,17 | -1,25 | 4,39 | -0,28 | 7,76E-01 |
| AT1G64390 | 1,48 | -1,32 | 2,05 | -0,64 | 5,21E-01 |
| AT1G64520 | 0,27 | 1,14 | 4,37 | 0,26 | 7,94E-01 |
| AT1G64530 | 0,54 | 1,03 | 3,07 | 0,34 | 7,36E-01 |
| AT1G64550 | 0,64 | 1,45 | 2,63 | 0,55 | 5,82E-01 |
| AT1G64570 | 0,71 | 1,94 | 2,37 | 0,82 | 4,13E-01 |
| AT1G64572 | 0,51 | 0,48 | 4,40 | 0,11 | 9,13E-01 |
| AT1G64620 | 0,45 | 0,85 | 3,29 | 0,26 | 7,95E-01 |
| AT1G64680 | 1,48 | -0,90 | 1,56 | -0,57 | 5,66E-01 |
| AT1G64740 | 0,25 | 0,54 | 4,39 | 0,12 | 9,01E-01 |
| AT1G64770 | 1,29 | -1,18 | 1,62 | -0,73 | 4,67E-01 |
| AT1G64795 | 0,74 | 0,06 | 1,96 | 0,03 | 9,75E-01 |
| AT1G64810 | 0,44 | 0,92 | 4,34 | 0,21 | 8,33E-01 |
| AT1G64820 | 0,08 | 1,10 | 4,36 | 0,25 | 8,00E-01 |
| AT1G64860 | 0,65 | -0,05 | 3,01 | -0,02 | 9,88E-01 |
| AT1G64900 | 1,03 | 1,24 | 1,87 | 0,67 | 5,05E-01 |
| AT1G64950 | 0,34 | 0,92 | 3,43 | 0,27 | 7,89E-01 |
| AT1G64960 | 0,60 | 1,46 | 3,92 | 0,37 | 7,09E-01 |
| AT1G65010 | 1,62 | -0,54 | 1,56 | -0,35 | 7,27E-01 |
| AT1G65220 | 0,59 | 1,06 | 2,70 | 0,39 | 6,95E-01 |
| AT1G65230 | 1,04 | 0,16 | 2,21 | 0,07 | 9,44E-01 |
| AT1G65260 | 0,21 | 0,00 | 4,41 | 0,00 | 1,00E+00 |
| AT1G65270 | 0,21 | 1,05 | 4,35 | 0,24 | 8,09E-01 |
| AT1G65280 | 0,39 | 1,26 | 4,29 | 0,29 | 7,68E-01 |
| AT1G65295 | 0,05 | 0,00 | 4,41 | 0,00 | 1,00E+00 |
| AT1G65360 | 0,90 | -2,65 | 2,25 | -1,18 | 2,39E-01 |
| AT1G08753 | 0,15 | 0,24 | 4,35 | 0,05 | 9,56E-01 |
| AT1G65490 | 0,49 | 0,00 | 4,40 | 0,00 | 1,00E+00 |
| AT1G65590 | 0,23 | 0,79 | 4,37 | 0,18 | 8,56E-01 |
| AT1G65681 | 0,20 | 0,00 | 4,41 | 0,00 | 1,00E+00 |
| AT1G65730 | 0,08 | 1,10 | 4,36 | 0,25 | 8,00E-01 |
| AT1G65790 | 0,93 | 1,70 | 1,92 | 0,88 | 3,76E-01 |
| AT1G65800 | 1,17 | -0,10 | 1,69 | -0,06 | 9,51E-01 |
| AT1G65820 | 0,38 | 1,23 | 2,88 | 0,43 | 6,70E-01 |
| AT1G65840 | 0,61 | 1,43 | 3,85 | 0,37 | 7,10E-01 |
| AT1G08793 | 0,15 | 0,71 | 4,37 | 0,16 | 8,71E-01 |
| AT1G66090 | 0,59 | 1,19 | 3,22 | 0,37 | 7,11E-01 |
| AT1G66150 | 1,55 | -0,49 | 1,29 | -0,38 | 7,03E-01 |
| AT1G66160 | 0,48 | 1,00 | 4,02 | 0,25 | 8,04E-01 |
| AT1G66180 | 0,53 | 0,84 | 3,21 | 0,26 | 7,95E-01 |
| AT1G66230 | 0,24 | 0,00 | 4,41 | 0,00 | 1,00E+00 |
| AT1G66340 | 0,60 | 1,62 | 2,95 | 0,55 | 5,82E-01 |
| AT1G66345 | 0,47 | 2,82 | 3,98 | 0,71 | 4,78E-01 |
| AT1G66430 | 0,50 | -3,44 | 4,31 | -0,80 | 4,24E-01 |
| AT1G66465 | 0,33 | 1,36 | 4,31 | 0,32 | 7,52E-01 |

| | | | | | |
|---|---|---|---|---|---|
| AT1G66580 | 1,55 | 1,48 | 1,58 | 0,93 | 3,50E-01 |
| AT1G66640 | 0,21 | -0,41 | 4,38 | -0,09 | 9,25E-01 |
| AT1G66750 | 0,31 | 0,79 | 4,33 | 0,18 | 8,55E-01 |
| AT1G66820 | 0,53 | -0,45 | 4,33 | -0,10 | 9,17E-01 |
| AT1G66900 | 0,20 | 1,42 | 4,36 | 0,33 | 7,44E-01 |
| AT1G67105 | 0,86 | -0,46 | 1,76 | -0,26 | 7,93E-01 |
| AT1G67190 | 0,34 | 1,67 | 4,33 | 0,39 | 6,99E-01 |
| AT1G08923 | 0,11 | 0,00 | 4,41 | 0,00 | 1,00E+00 |
| AT1G67420 | 0,13 | 1,15 | 4,36 | 0,26 | 7,92E-01 |
| AT1G08937 | 0,46 | 1,86 | 3,27 | 0,57 | 5,69E-01 |
| AT1G67470 | 0,43 | 1,28 | 3,32 | 0,39 | 6,99E-01 |
| AT1G67480 | 0,49 | 1,72 | 4,33 | 0,40 | 6,92E-01 |
| AT1G67530 | 0,54 | 0,48 | 3,96 | 0,12 | 9,04E-01 |
| AT1G67560 | 0,64 | 1,15 | 4,37 | 0,26 | 7,92E-01 |
| AT1G67620 | 0,15 | 0,42 | 4,38 | 0,09 | 9,24E-01 |
| AT1G67635 | 1,69 | 3,47 | 1,86 | 1,86 | 6,23E-02 |
| AT1G67700 | 0,46 | 0,00 | 4,03 | 0,00 | 1,00E+00 |
| AT1G67720 | 0,53 | 2,49 | 3,76 | 0,66 | 5,08E-01 |
| AT1G67792 | 0,38 | 1,31 | 4,35 | 0,30 | 7,63E-01 |
| AT1G67900 | 1,04 | -0,02 | 2,33 | -0,01 | 9,92E-01 |
| AT1G67940 | 0,66 | 1,60 | 2,94 | 0,54 | 5,86E-01 |
| AT1G08963 | 0,39 | 0,48 | 4,40 | 0,11 | 9,14E-01 |
| AT1G67980 | 0,18 | -0,69 | 4,37 | -0,16 | 8,74E-01 |
| AT1G67990 | 0,55 | -0,91 | 3,83 | -0,24 | 8,13E-01 |
| AT1G68050 | 0,24 | 0,84 | 4,33 | 0,19 | 8,46E-01 |
| AT1G68100 | 0,43 | 0,67 | 4,12 | 0,16 | 8,70E-01 |
| AT1G68110 | 0,27 | -0,03 | 4,36 | -0,01 | 9,95E-01 |
| AT1G68310 | 0,12 | 1,39 | 4,34 | 0,32 | 7,49E-01 |
| AT1G68440 | 1,32 | 0,86 | 1,83 | 0,47 | 6,38E-01 |
| AT1G68490 | 0,97 | 2,44 | 2,50 | 0,98 | 3,28E-01 |
| AT1G68570 | 0,58 | 0,96 | 2,57 | 0,38 | 7,07E-01 |
| AT1G68580 | 1,44 | 1,91 | 1,71 | 1,12 | 2,65E-01 |
| AT1G68585 | 0,27 | 0,44 | 4,39 | 0,10 | 9,20E-01 |
| AT1G68620 | 0,33 | 2,24 | 4,32 | 0,52 | 6,04E-01 |
| AT1G68650 | 0,32 | 1,25 | 4,35 | 0,29 | 7,74E-01 |
| AT1G68670 | 0,63 | 3,03 | 2,93 | 1,03 | 3,02E-01 |
| AT1G68720 | 1,50 | 1,07 | 2,01 | 0,53 | 5,94E-01 |
| AT1G68840 | 1,23 | -3,64 | 1,87 | -1,95 | 5,10E-02 |
| AT1G68890 | 1,09 | 2,31 | 1,91 | 1,21 | 2,26E-01 |
| AT1G68980 | 0,24 | 1,62 | 4,34 | 0,37 | 7,08E-01 |
| AT1G69085 | 0,08 | 0,00 | 4,41 | 0,00 | 1,00E+00 |
| AT1G69160 | 0,39 | -0,47 | 3,38 | -0,14 | 8,90E-01 |
| AT1G69170 | 0,40 | 2,57 | 4,32 | 0,60 | 5,51E-01 |
| AT1G69200 | 0,21 | 0,00 | 4,38 | 0,00 | 1,00E+00 |
| AT1G69252 | 0,90 | -2,60 | 3,03 | -0,86 | 3,90E-01 |
| AT1G69260 | 0,63 | -1,47 | 2,74 | -0,54 | 5,90E-01 |
| AT1G69350 | 0,78 | 1,85 | 2,38 | 0,78 | 4,37E-01 |
| AT1G69390 | 0,12 | 0,42 | 4,38 | 0,10 | 9,24E-01 |
| AT1G69410 | 0,27 | -0,39 | 4,31 | -0,09 | 9,27E-01 |
| AT1G69420 | 0,65 | 1,22 | 2,55 | 0,48 | 6,33E-01 |
| AT1G69510 | 0,10 | 0,00 | 4,41 | 0,00 | 1,00E+00 |
| AT1G69523 | 0,10 | 0,00 | 4,41 | 0,00 | 1,00E+00 |
| AT1G69526 | 0,08 | 1,10 | 4,36 | 0,25 | 8,00E-01 |
| AT1G69530 | 0,93 | -2,07 | 2,16 | -0,96 | 3,38E-01 |
| AT1G69580 | 0,26 | -1,10 | 4,38 | -0,25 | 8,01E-01 |
| AT1G69610 | 0,21 | 1,00 | 4,36 | 0,23 | 8,19E-01 |

| | | | | | |
|---|---|---|---|---|---|
| AT1G09087 | 0,37 | 0,34 | 3,48 | 0,10 | 9,21E-01 |
| AT1G69690 | 0,67 | -0,59 | 2,47 | -0,24 | 8,11E-01 |
| AT1G69740 | 0,99 | 2,04 | 2,00 | 1,02 | 3,07E-01 |
| AT1G69750 | 0,10 | 0,00 | 4,41 | 0,00 | 1,00E+00 |
| AT1G69870 | 0,38 | 0,69 | 4,32 | 0,16 | 8,74E-01 |
| AT1G70060 | 1,48 | 3,61 | 2,09 | 1,72 | 8,49E-02 |
| AT1G70090 | 0,99 | 0,92 | 2,19 | 0,42 | 6,75E-01 |
| AT1G70100 | 0,38 | 0,64 | 4,27 | 0,15 | 8,82E-01 |
| AT1G70160 | 1,19 | 0,95 | 1,52 | 0,63 | 5,31E-01 |
| AT1G70180 | 1,28 | 3,00 | 1,90 | 1,58 | 1,14E-01 |
| AT1G70190 | 0,40 | 1,87 | 4,35 | 0,43 | 6,67E-01 |
| AT1G70210 | 0,79 | -0,31 | 2,41 | -0,13 | 8,97E-01 |
| AT1G70230 | 0,59 | -1,30 | 2,72 | -0,48 | 6,32E-01 |
| AT1G70250 | 0,71 | 1,21 | 2,92 | 0,42 | 6,78E-01 |
| AT1G70330 | 0,33 | 0,31 | 3,58 | 0,09 | 9,30E-01 |
| AT1G70480 | 1,12 | 2,81 | 1,87 | 1,50 | 1,32E-01 |
| AT1G70510 | 0,12 | 1,46 | 4,35 | 0,34 | 7,36E-01 |
| AT1G70550 | 0,14 | 0,51 | 4,38 | 0,12 | 9,08E-01 |
| AT1G70570 | 0,36 | -0,67 | 4,36 | -0,15 | 8,78E-01 |
| AT1G70580 | 0,81 | 0,41 | 2,31 | 0,18 | 8,58E-01 |
| AT1G70620 | 1,32 | 3,65 | 2,10 | 1,73 | 8,31E-02 |
| AT1G70630 | 0,36 | 1,24 | 4,34 | 0,28 | 7,76E-01 |
| AT1G70645 | 0,05 | 0,00 | 4,41 | 0,00 | 1,00E+00 |
| AT1G70650 | 0,26 | 0,77 | 4,37 | 0,18 | 8,60E-01 |
| AT1G70660 | 0,20 | 0,89 | 4,37 | 0,20 | 8,39E-01 |
| AT1G70700 | 0,60 | -1,53 | 2,64 | -0,58 | 5,62E-01 |
| AT1G70760 | 0,35 | 1,20 | 3,44 | 0,35 | 7,28E-01 |
| AT1G70810 | 0,62 | 0,29 | 2,53 | 0,11 | 9,10E-01 |
| AT1G70820 | 0,43 | -1,44 | 2,79 | -0,51 | 6,07E-01 |
| AT1G70895 | 0,08 | 0,00 | 4,40 | 0,00 | 1,00E+00 |
| AT1G70900 | 1,03 | 1,12 | 1,99 | 0,56 | 5,74E-01 |
| AT1G70960 | 0,12 | 0,42 | 4,38 | 0,10 | 9,24E-01 |
| AT1G70980 | 0,40 | -0,17 | 4,37 | -0,04 | 9,69E-01 |
| AT1G71010 | 1,50 | 3,51 | 1,95 | 1,81 | 7,11E-02 |
| AT1G71090 | 0,35 | 1,71 | 3,52 | 0,49 | 6,27E-01 |
| AT1G71100 | 1,11 | -0,48 | 1,65 | -0,29 | 7,70E-01 |
| AT1G71110 | 0,15 | 0,86 | 4,37 | 0,20 | 8,43E-01 |
| AT1G71130 | 0,37 | 1,62 | 4,35 | 0,37 | 7,09E-01 |
| AT1G71180 | 0,42 | 2,16 | 4,24 | 0,51 | 6,11E-01 |
| AT1G71270 | 0,45 | 1,74 | 4,17 | 0,42 | 6,76E-01 |
| AT1G71300 | 0,38 | 0,82 | 4,37 | 0,19 | 8,51E-01 |
| AT1G71320 | 0,39 | 0,43 | 4,24 | 0,10 | 9,20E-01 |
| AT1G71480 | 0,80 | 2,76 | 2,84 | 0,97 | 3,31E-01 |
| AT1G09257 | 0,17 | 1,92 | 4,33 | 0,44 | 6,57E-01 |
| AT1G71500 | 0,95 | -1,26 | 2,08 | -0,60 | 5,46E-01 |
| AT1G71695 | 0,82 | 0,11 | 2,51 | 0,04 | 9,66E-01 |
| AT1G71720 | 0,53 | -2,69 | 3,76 | -0,72 | 4,75E-01 |
| AT1G71800 | 0,67 | 2,20 | 3,62 | 0,61 | 5,44E-01 |
| AT1G71820 | 1,53 | 0,53 | 1,47 | 0,36 | 7,20E-01 |
| AT1G71840 | 1,05 | 1,23 | 2,28 | 0,54 | 5,88E-01 |
| AT1G71900 | 0,51 | 0,48 | 4,40 | 0,11 | 9,13E-01 |
| AT1G71910 | 0,44 | -0,52 | 3,29 | -0,16 | 8,73E-01 |
| AT1G71920 | 0,73 | 0,42 | 2,43 | 0,18 | 8,61E-01 |
| AT1G71930 | 1,13 | 1,30 | 2,14 | 0,61 | 5,43E-01 |
| AT1G71970 | 0,14 | 1,18 | 4,35 | 0,27 | 7,86E-01 |
| AT1G71980 | 0,46 | 2,18 | 3,15 | 0,69 | 4,88E-01 |

| | | | | | |
|---|---|---|---|---|---|
| AT1G72010 | 0,42 | 0,60 | 3,29 | 0,18 | 8,55E-01 |
| AT1G72030 | 0,75 | 1,26 | 3,68 | 0,34 | 7,33E-01 |
| AT1G72050 | 0,75 | 2,33 | 3,62 | 0,64 | 5,19E-01 |
| AT1G72090 | 0,34 | 1,74 | 4,34 | 0,40 | 6,89E-01 |
| AT1G72240 | 0,60 | 1,83 | 3,85 | 0,47 | 6,35E-01 |
| AT1G72310 | 0,25 | 1,42 | 4,34 | 0,33 | 7,44E-01 |
| AT1G72330 | 0,67 | 1,80 | 2,88 | 0,62 | 5,33E-01 |
| AT1G72470 | 0,43 | 1,27 | 4,37 | 0,29 | 7,71E-01 |
| AT1G72510 | 0,57 | -1,34 | 3,15 | -0,42 | 6,71E-01 |
| AT1G72520 | 0,91 | 1,50 | 2,05 | 0,73 | 4,63E-01 |
| AT1G72530 | 0,05 | 0,00 | 4,41 | 0,00 | 1,00E+00 |
| AT1G09333 | 1,54 | -2,31 | 1,71 | -1,35 | 1,77E-01 |
| AT1G72650 | 0,79 | 3,26 | 2,33 | 1,40 | 1,62E-01 |
| AT1G72700 | 1,31 | 3,68 | 1,85 | 1,99 | 4,71E-02 |
| AT1G72750 | 0,63 | 1,60 | 2,26 | 0,71 | 4,79E-01 |
| AT1G72770 | 1,51 | 2,46 | 1,68 | 1,47 | 1,42E-01 |
| AT1G72790 | 0,71 | 1,57 | 2,99 | 0,53 | 5,98E-01 |
| AT1G72855 | 0,40 | 2,57 | 4,32 | 0,60 | 5,51E-01 |
| AT1G72900 | 0,32 | 0,98 | 3,52 | 0,28 | 7,82E-01 |
| AT1G72910 | 1,29 | 2,25 | 1,86 | 1,21 | 2,27E-01 |
| AT1G72920 | 0,44 | -0,23 | 3,41 | -0,07 | 9,47E-01 |
| AT1G72940 | 0,97 | 3,99 | 3,96 | 1,01 | 3,13E-01 |
| AT1G72970 | 0,16 | 1,33 | 4,35 | 0,31 | 7,59E-01 |
| AT1G73030 | 0,82 | 0,55 | 2,54 | 0,22 | 8,28E-01 |
| AT1G73060 | 0,24 | 0,00 | 4,41 | 0,00 | 1,00E+00 |
| AT1G73080 | 1,38 | 1,61 | 1,70 | 0,95 | 3,41E-01 |
| AT1G73100 | 0,70 | 0,98 | 2,61 | 0,37 | 7,08E-01 |
| AT1G73180 | 1,25 | 2,69 | 2,00 | 1,35 | 1,77E-01 |
| AT1G73430 | 1,05 | 1,91 | 2,23 | 0,86 | 3,91E-01 |
| AT1G73450 | 1,12 | 3,14 | 1,93 | 1,63 | 1,04E-01 |
| AT1G73460 | 0,79 | 2,47 | 2,48 | 1,00 | 3,19E-01 |
| AT1G73470 | 0,23 | 1,16 | 4,35 | 0,27 | 7,89E-01 |
| AT1G73480 | 1,37 | -1,09 | 2,02 | -0,54 | 5,91E-01 |
| AT1G73600 | 1,37 | -3,24 | 2,13 | -1,52 | 1,28E-01 |
| AT1G73630 | 0,10 | 0,42 | 4,38 | 0,10 | 9,23E-01 |
| AT1G73670 | 0,46 | 0,38 | 3,27 | 0,12 | 9,08E-01 |
| AT1G73710 | 0,22 | 1,93 | 4,33 | 0,45 | 6,56E-01 |
| AT1G73750 | 0,72 | 3,20 | 4,29 | 0,75 | 4,55E-01 |
| AT1G73770 | 0,08 | 1,10 | 4,36 | 0,25 | 8,00E-01 |
| AT1G73850 | 0,17 | 0,00 | 4,40 | 0,00 | 1,00E+00 |
| AT1G73875 | 0,39 | 1,81 | 4,21 | 0,43 | 6,67E-01 |
| AT1G73885 | 0,62 | 1,57 | 3,07 | 0,51 | 6,09E-01 |
| AT1G73920 | 0,62 | 0,54 | 2,65 | 0,20 | 8,40E-01 |
| AT1G73970 | 0,95 | -1,16 | 2,30 | -0,50 | 6,14E-01 |
| AT1G74160 | 1,08 | 0,33 | 1,79 | 0,18 | 8,54E-01 |
| AT1G74250 | 1,42 | 3,60 | 1,84 | 1,95 | 5,10E-02 |
| AT1G74360 | 1,21 | 3,48 | 2,38 | 1,46 | 1,43E-01 |
| AT1G74380 | 1,29 | 2,30 | 1,87 | 1,23 | 2,20E-01 |
| AT1G74430 | 0,38 | -1,60 | 4,38 | -0,36 | 7,16E-01 |
| AT1G74450 | 0,99 | 2,38 | 2,04 | 1,17 | 2,42E-01 |
| AT1G74580 | 0,25 | 1,14 | 4,36 | 0,26 | 7,94E-01 |
| AT1G74680 | 0,57 | 0,85 | 3,26 | 0,26 | 7,94E-01 |
| AT1G74710 | 0,60 | 0,52 | 2,55 | 0,21 | 8,37E-01 |
| AT1G74730 | 0,44 | -0,86 | 3,26 | -0,26 | 7,92E-01 |
| AT1G74750 | 0,50 | 1,40 | 4,35 | 0,32 | 7,47E-01 |
| AT1G74780 | 0,63 | 3,07 | 2,91 | 1,05 | 2,91E-01 |

| | | | | | |
|---|---|---|---|---|---|
| AT1G74790 | 0,51 | 0,82 | 3,95 | 0,21 | 8,36E-01 |
| AT1G74870 | 1,10 | 3,46 | 3,22 | 1,07 | 2,82E-01 |
| AT1G74930 | 0,81 | -0,56 | 2,82 | -0,20 | 8,43E-01 |
| AT1G74940 | 0,58 | 1,89 | 3,09 | 0,61 | 5,41E-01 |
| AT1G75080 | 0,21 | 2,05 | 4,33 | 0,47 | 6,36E-01 |
| AT1G09533 | 1,00 | 0,24 | 1,93 | 0,13 | 8,99E-01 |
| AT1G75163 | 0,94 | 1,20 | 1,97 | 0,61 | 5,43E-01 |
| AT1G75166 | 1,22 | 2,04 | 2,03 | 1,00 | 3,17E-01 |
| AT1G75170 | 0,12 | 1,39 | 4,34 | 0,32 | 7,49E-01 |
| AT1G09543 | 0,38 | 0,37 | 3,36 | 0,11 | 9,12E-01 |
| AT1G75190 | 0,22 | -1,35 | 4,41 | -0,31 | 7,59E-01 |
| AT1G75210 | 0,77 | -3,89 | 2,88 | -1,35 | 1,76E-01 |
| AT1G75280 | 0,22 | 0,00 | 4,41 | 0,00 | 1,00E+00 |
| AT1G75310 | 0,47 | -0,89 | 4,36 | -0,21 | 8,37E-01 |
| AT1G75350 | 0,83 | 2,13 | 2,10 | 1,02 | 3,09E-01 |
| AT1G75400 | 0,32 | 2,17 | 4,32 | 0,50 | 6,15E-01 |
| AT1G75420 | 0,31 | 0,85 | 4,38 | 0,19 | 8,46E-01 |
| AT1G75440 | 0,28 | 0,90 | 4,38 | 0,21 | 8,37E-01 |
| AT1G75460 | 0,84 | -0,08 | 2,13 | -0,04 | 9,69E-01 |
| AT1G75580 | 0,17 | 0,09 | 4,35 | 0,02 | 9,84E-01 |
| AT1G75710 | 0,31 | -0,08 | 4,37 | -0,02 | 9,85E-01 |
| AT1G75785 | 0,12 | 0,90 | 4,36 | 0,21 | 8,36E-01 |
| AT1G75810 | 0,60 | 0,37 | 2,63 | 0,14 | 8,87E-01 |
| AT1G75840 | 0,75 | -0,35 | 2,26 | -0,16 | 8,76E-01 |
| AT1G75900 | 0,32 | 0,78 | 4,37 | 0,18 | 8,58E-01 |
| AT1G75960 | 0,45 | 1,62 | 4,23 | 0,38 | 7,02E-01 |
| AT1G76000 | 0,58 | 3,02 | 2,96 | 1,02 | 3,08E-01 |
| AT1G76050 | 0,66 | -0,15 | 3,02 | -0,05 | 9,61E-01 |
| AT1G76140 | 0,48 | 1,45 | 4,02 | 0,36 | 7,17E-01 |
| AT1G76185 | 0,16 | 0,80 | 4,37 | 0,18 | 8,55E-01 |
| AT1G76190 | 0,11 | 0,00 | 4,41 | 0,00 | 1,00E+00 |
| AT1G76320 | 0,32 | 2,33 | 4,32 | 0,54 | 5,90E-01 |
| AT1G76350 | 0,12 | 0,79 | 4,37 | 0,18 | 8,57E-01 |
| AT1G76390 | 0,51 | 2,58 | 3,15 | 0,82 | 4,12E-01 |
| AT1G76450 | 0,08 | 0,85 | 4,38 | 0,20 | 8,45E-01 |
| AT1G76520 | 0,73 | -1,16 | 2,24 | -0,52 | 6,05E-01 |
| AT1G76580 | 0,80 | 1,03 | 3,57 | 0,29 | 7,73E-01 |
| AT1G76590 | 0,69 | -2,18 | 2,90 | -0,75 | 4,52E-01 |
| AT1G76600 | 0,49 | 2,30 | 3,20 | 0,72 | 4,72E-01 |
| AT1G76620 | 0,25 | -1,07 | 4,35 | -0,25 | 8,06E-01 |
| AT1G76680 | 0,14 | -1,28 | 4,39 | -0,29 | 7,71E-01 |
| AT1G76740 | 0,12 | 1,58 | 4,34 | 0,36 | 7,16E-01 |
| AT1G76780 | 1,47 | 0,51 | 1,73 | 0,29 | 7,70E-01 |
| AT1G76820 | 0,53 | 1,75 | 3,07 | 0,57 | 5,68E-01 |
| AT1G76850 | 1,15 | -0,11 | 1,85 | -0,06 | 9,51E-01 |
| AT1G76880 | 1,06 | -1,32 | 1,75 | -0,76 | 4,49E-01 |
| AT1G76900 | 0,91 | 0,90 | 2,69 | 0,33 | 7,38E-01 |
| AT1G76920 | 0,59 | 2,23 | 3,78 | 0,59 | 5,56E-01 |
| AT1G76950 | 0,83 | -0,31 | 2,79 | -0,11 | 9,12E-01 |
| AT1G76990 | 1,49 | -0,70 | 1,48 | -0,47 | 6,35E-01 |
| AT1G77080 | 0,41 | -0,73 | 3,35 | -0,22 | 8,28E-01 |
| AT1G77140 | 0,25 | 0,00 | 4,41 | 0,00 | 1,00E+00 |
| AT1G77320 | 0,26 | 0,90 | 4,38 | 0,21 | 8,37E-01 |
| AT1G77350 | 0,15 | 0,71 | 4,37 | 0,16 | 8,71E-01 |
| AT1G77420 | 0,12 | 1,65 | 4,34 | 0,38 | 7,04E-01 |
| AT1G77460 | 0,94 | 2,26 | 2,64 | 0,85 | 3,93E-01 |

| | | | | | |
|---|---|---|---|---|---|
| AT1G77490 | 0,64 | -1,34 | 3,09 | -0,43 | 6,64E-01 |
| AT1G77500 | 0,37 | 0,52 | 4,34 | 0,12 | 9,05E-01 |
| AT1G77510 | 1,26 | 2,02 | 1,85 | 1,09 | 2,75E-01 |
| AT1G77610 | 0,53 | 0,57 | 4,34 | 0,13 | 8,96E-01 |
| AT1G77710 | 0,32 | 0,47 | 4,39 | 0,11 | 9,15E-01 |
| AT1G77800 | 1,13 | 2,94 | 1,90 | 1,55 | 1,22E-01 |
| AT1G77840 | 0,31 | 1,61 | 3,59 | 0,45 | 6,53E-01 |
| AT1G77850 | 1,01 | 1,81 | 2,21 | 0,82 | 4,12E-01 |
| AT1G77890 | 0,98 | -0,44 | 1,93 | -0,23 | 8,20E-01 |
| AT1G77930 | 1,14 | 2,96 | 2,05 | 1,44 | 1,50E-01 |
| AT1G78010 | 0,90 | 2,35 | 2,28 | 1,03 | 3,01E-01 |
| AT1G78070 | 1,66 | -2,14 | 1,51 | -1,41 | 1,58E-01 |
| AT1G78100 | 0,64 | -0,15 | 2,19 | -0,07 | 9,44E-01 |
| AT1G78150 | 0,22 | 0,41 | 4,38 | 0,09 | 9,24E-01 |
| AT1G78200 | 0,54 | -0,36 | 2,67 | -0,14 | 8,92E-01 |
| AT1G78280 | 1,59 | 4,22 | 2,17 | 1,95 | 5,14E-02 |
| AT1G78300 | 0,54 | 1,52 | 3,15 | 0,48 | 6,29E-01 |
| AT1G78420 | 0,54 | 1,87 | 2,70 | 0,69 | 4,87E-01 |
| AT1G78490 | 0,05 | 0,00 | 4,41 | 0,00 | 1,00E+00 |
| AT1G78500 | 0,17 | 1,81 | 4,33 | 0,42 | 6,76E-01 |
| AT1G78600 | 0,20 | 0,00 | 4,41 | 0,00 | 1,00E+00 |
| AT1G78630 | 1,31 | -0,69 | 1,88 | -0,37 | 7,13E-01 |
| AT1G78670 | 0,52 | 1,30 | 3,07 | 0,42 | 6,72E-01 |
| AT1G78680 | 0,90 | 1,08 | 2,01 | 0,54 | 5,91E-01 |
| AT1G78700 | 0,37 | 1,96 | 4,32 | 0,45 | 6,50E-01 |
| AT1G78710 | 0,38 | 0,96 | 4,37 | 0,22 | 8,27E-01 |
| AT1G78770 | 0,21 | 2,05 | 4,33 | 0,47 | 6,36E-01 |
| AT1G78870 | 0,28 | 1,43 | 4,34 | 0,33 | 7,42E-01 |
| AT1G78890 | 0,36 | -0,45 | 4,36 | -0,10 | 9,18E-01 |
| AT1G78910 | 0,26 | 1,08 | 4,37 | 0,25 | 8,05E-01 |
| AT1G78920 | 1,61 | 2,09 | 1,57 | 1,33 | 1,83E-01 |
| AT1G79060 | 0,14 | 0,00 | 4,41 | 0,00 | 1,00E+00 |
| AT1G79103 | 0,05 | 0,00 | 4,41 | 0,00 | 1,00E+00 |
| AT1G79110 | 1,67 | 0,71 | 1,61 | 0,44 | 6,58E-01 |
| AT1G09847 | 0,44 | -2,53 | 3,26 | -0,78 | 4,38E-01 |
| AT1G79190 | 0,88 | 1,87 | 2,73 | 0,69 | 4,93E-01 |
| AT1G79230 | 0,71 | 1,90 | 2,90 | 0,65 | 5,13E-01 |
| AT1G79240 | 0,20 | 1,62 | 4,35 | 0,37 | 7,10E-01 |
| AT1G79270 | 1,18 | 2,17 | 1,78 | 1,22 | 2,23E-01 |
| AT1G79340 | 0,91 | -1,50 | 2,01 | -0,74 | 4,57E-01 |
| AT1G79380 | 0,91 | 3,12 | 2,77 | 1,12 | 2,61E-01 |
| AT1G79560 | 0,28 | 1,41 | 4,34 | 0,33 | 7,45E-01 |
| AT1G79590 | 0,18 | 1,66 | 4,33 | 0,38 | 7,02E-01 |
| AT1G79660 | 0,63 | 0,45 | 3,71 | 0,12 | 9,03E-01 |
| AT1G09903 | 0,08 | 1,10 | 4,36 | 0,25 | 8,00E-01 |
| AT1G79810 | 0,41 | 1,56 | 3,28 | 0,47 | 6,35E-01 |
| AT1G79840 | 0,77 | 2,68 | 2,91 | 0,92 | 3,58E-01 |
| AT1G79870 | 0,21 | 1,68 | 4,33 | 0,39 | 6,99E-01 |
| AT1G79890 | 0,21 | 0,79 | 4,37 | 0,18 | 8,57E-01 |
| AT1G79910 | 0,41 | 1,80 | 4,14 | 0,43 | 6,64E-01 |
| AT1G79940 | 1,20 | -1,28 | 1,64 | -0,78 | 4,35E-01 |
| AT1G79950 | 0,49 | -0,45 | 3,96 | -0,11 | 9,09E-01 |
| AT1G79970 | 0,27 | 0,93 | 4,36 | 0,21 | 8,30E-01 |
| AT1G79985 | 0,75 | 1,88 | 2,50 | 0,75 | 4,51E-01 |
| AT1G80010 | 1,24 | 1,64 | 1,65 | 1,00 | 3,19E-01 |
| AT1G80160 | 0,37 | 2,74 | 4,31 | 0,63 | 5,26E-01 |

| | | | | | |
|---|---|---|---|---|---|
| AT1G80260 | 0,59 | -1,22 | 3,00 | -0,40 | 6,86E-01 |
| AT1G80300 | 1,04 | 3,07 | 3,24 | 0,95 | 3,42E-01 |
| AT1G80310 | 0,64 | -0,16 | 2,59 | -0,06 | 9,52E-01 |
| AT1G80380 | 1,15 | -1,56 | 1,78 | -0,88 | 3,81E-01 |
| AT1G09963 | 0,96 | 1,23 | 2,48 | 0,50 | 6,20E-01 |
| AT1G80430 | 0,19 | 0,91 | 4,38 | 0,21 | 8,36E-01 |
| AT1G80510 | 0,61 | 1,43 | 3,85 | 0,37 | 7,10E-01 |
| AT1G80620 | 0,35 | -1,33 | 4,36 | -0,30 | 7,61E-01 |
| AT1G80770 | 0,54 | 0,54 | 3,21 | 0,17 | 8,67E-01 |
| AT1G80780 | 0,60 | 2,09 | 3,12 | 0,67 | 5,02E-01 |
| AT1G80790 | 0,69 | 1,92 | 2,77 | 0,69 | 4,89E-01 |
| AT1G80810 | 0,38 | 2,01 | 4,23 | 0,48 | 6,35E-01 |
| AT1G80840 | 1,09 | 2,40 | 2,47 | 0,97 | 3,31E-01 |
| AT1G80870 | 0,27 | 1,10 | 4,35 | 0,25 | 8,00E-01 |
| AT2G01070 | 0,81 | 2,16 | 2,47 | 0,88 | 3,82E-01 |
| AT2G01080 | 0,35 | 1,23 | 3,50 | 0,35 | 7,26E-01 |
| AT2G01100 | 0,12 | 0,42 | 4,38 | 0,10 | 9,24E-01 |
| AT2G01120 | 0,92 | 3,14 | 2,76 | 1,14 | 2,55E-01 |
| AT2G01190 | 0,50 | 1,11 | 3,10 | 0,36 | 7,21E-01 |
| AT2G01400 | 0,39 | 0,00 | 4,40 | 0,00 | 1,00E+00 |
| AT2G03975 | 0,34 | -0,49 | 3,57 | -0,14 | 8,91E-01 |
| AT2G01480 | 0,34 | 0,42 | 4,38 | 0,10 | 9,24E-01 |
| AT2G01620 | 0,60 | 1,20 | 2,57 | 0,47 | 6,41E-01 |
| AT2G01680 | 0,39 | 0,88 | 4,35 | 0,20 | 8,40E-01 |
| AT2G01750 | 0,24 | -0,42 | 4,38 | -0,10 | 9,24E-01 |
| AT2G01860 | 0,98 | 3,45 | 2,43 | 1,42 | 1,56E-01 |
| AT2G01900 | 1,22 | 0,60 | 1,66 | 0,36 | 7,18E-01 |
| AT2G01980 | 0,63 | 0,49 | 3,07 | 0,16 | 8,73E-01 |
| AT2G02040 | 0,10 | 0,42 | 4,38 | 0,10 | 9,23E-01 |
| AT2G02050 | 0,58 | -0,59 | 2,66 | -0,22 | 8,26E-01 |
| AT2G02070 | 0,84 | -0,53 | 1,95 | -0,27 | 7,86E-01 |
| AT2G02090 | 1,22 | 2,80 | 2,37 | 1,18 | 2,38E-01 |
| AT2G02100 | 0,54 | -1,34 | 3,87 | -0,35 | 7,30E-01 |
| AT2G02440 | 0,68 | -2,90 | 3,02 | -0,96 | 3,38E-01 |
| AT2G02450 | 0,66 | -0,55 | 2,58 | -0,21 | 8,30E-01 |
| AT2G02470 | 0,18 | 0,00 | 4,41 | 0,00 | 1,00E+00 |
| AT2G02550 | 1,01 | 1,23 | 2,12 | 0,58 | 5,62E-01 |
| AT2G02700 | 0,39 | 0,48 | 4,40 | 0,11 | 9,14E-01 |
| AT2G04295 | 0,12 | 0,42 | 4,38 | 0,10 | 9,24E-01 |
| AT2G02760 | 0,54 | -2,15 | 2,63 | -0,82 | 4,13E-01 |
| AT2G02790 | 0,39 | -0,60 | 4,22 | -0,14 | 8,88E-01 |
| AT2G02795 | 0,14 | 0,51 | 4,38 | 0,12 | 9,08E-01 |
| AT2G02810 | 0,62 | 2,97 | 3,00 | 0,99 | 3,22E-01 |
| AT2G04355 | 0,47 | 1,23 | 3,35 | 0,37 | 7,13E-01 |
| AT2G04365 | 0,97 | 1,40 | 2,41 | 0,58 | 5,62E-01 |
| AT2G02870 | 1,53 | 2,43 | 1,92 | 1,27 | 2,04E-01 |
| AT2G02955 | 0,36 | 0,96 | 4,35 | 0,22 | 8,25E-01 |
| AT2G03070 | 0,25 | 1,75 | 4,33 | 0,40 | 6,87E-01 |
| AT2G03120 | 0,64 | -1,68 | 2,61 | -0,64 | 5,20E-01 |
| AT2G03130 | 0,12 | 1,46 | 4,35 | 0,34 | 7,36E-01 |
| AT2G03270 | 0,39 | 1,28 | 2,86 | 0,45 | 6,54E-01 |
| AT2G03410 | 0,43 | 1,39 | 4,08 | 0,34 | 7,33E-01 |
| AT2G03450 | 0,32 | 0,00 | 4,41 | 0,00 | 1,00E+00 |
| AT2G03500 | 0,52 | -1,53 | 3,27 | -0,47 | 6,39E-01 |
| AT2G03520 | 0,08 | 1,10 | 4,36 | 0,25 | 8,00E-01 |
| AT2G03550 | 0,42 | -2,15 | 4,16 | -0,52 | 6,05E-01 |

| | | | | | |
|---|---|---|---|---|---|
| AT2G03567 | 0,47 | 0,51 | 3,32 | 0,15 | 8,79E-01 |
| AT2G03680 | 0,82 | -1,55 | 2,38 | -0,65 | 5,14E-01 |
| AT2G03710 | 0,17 | 1,94 | 4,33 | 0,45 | 6,55E-01 |
| AT2G03823 | 0,37 | 0,31 | 2,89 | 0,11 | 9,15E-01 |
| AT2G03870 | 0,41 | 1,91 | 4,11 | 0,46 | 6,42E-01 |
| AT2G03932 | 0,20 | 0,00 | 4,41 | 0,00 | 1,00E+00 |
| AT2G03931 | 0,05 | 0,00 | 4,41 | 0,00 | 1,00E+00 |
| AT2G03933 | 0,14 | 0,00 | 4,41 | 0,00 | 1,00E+00 |
| AT2G03936 | 0,18 | 0,98 | 4,36 | 0,23 | 8,22E-01 |
| AT2G03965 | 0,22 | 0,00 | 4,41 | 0,00 | 1,00E+00 |
| AT2G03980 | 0,82 | -1,74 | 2,58 | -0,67 | 5,00E-01 |
| AT2G04039 | 0,42 | 1,59 | 4,11 | 0,39 | 6,99E-01 |
| AT2G04735 | 0,15 | -1,25 | 4,39 | -0,28 | 7,76E-01 |
| AT2G04235 | 0,05 | 0,00 | 4,41 | 0,00 | 1,00E+00 |
| AT2G04270 | 1,17 | 1,26 | 2,19 | 0,57 | 5,66E-01 |
| AT2G04340 | 0,38 | 0,90 | 4,38 | 0,21 | 8,37E-01 |
| AT2G04350 | 1,50 | -0,01 | 1,41 | -0,01 | 9,96E-01 |
| AT2G04380 | 0,56 | -0,09 | 2,78 | -0,03 | 9,75E-01 |
| AT2G04390 | 0,85 | 2,76 | 2,12 | 1,30 | 1,93E-01 |
| AT2G04400 | 0,61 | 1,25 | 2,52 | 0,50 | 6,20E-01 |
| AT2G04450 | 0,21 | 2,05 | 4,33 | 0,47 | 6,36E-01 |
| AT2G04495 | 0,43 | 2,20 | 3,35 | 0,66 | 5,10E-01 |
| AT2G04550 | 0,15 | 0,24 | 4,35 | 0,05 | 9,56E-01 |
| AT2G04700 | 0,71 | 1,50 | 2,23 | 0,68 | 4,99E-01 |
| AT2G04740 | 0,38 | 1,31 | 4,35 | 0,30 | 7,63E-01 |
| AT2G04780 | 0,24 | 0,41 | 4,38 | 0,09 | 9,25E-01 |
| AT2G04795 | 0,15 | 0,71 | 4,37 | 0,16 | 8,71E-01 |
| AT2G04850 | 0,08 | 1,10 | 4,36 | 0,25 | 8,00E-01 |
| AT2G04865 | 0,25 | 2,29 | 4,32 | 0,53 | 5,96E-01 |
| AT2G04900 | 0,19 | 0,46 | 4,40 | 0,10 | 9,16E-01 |
| AT2G05060 | 0,21 | 0,79 | 4,37 | 0,18 | 8,57E-01 |
| AT2G05180 | 0,08 | 0,85 | 4,38 | 0,20 | 8,45E-01 |
| AT2G05210 | 0,12 | 0,90 | 4,36 | 0,21 | 8,36E-01 |
| AT2G05320 | 0,92 | 0,49 | 2,02 | 0,24 | 8,10E-01 |
| AT2G05350 | 0,23 | 1,63 | 4,34 | 0,37 | 7,08E-01 |
| AT2G05380 | 0,99 | -1,49 | 1,87 | -0,80 | 4,26E-01 |
| AT2G05540 | 0,30 | 0,00 | 4,41 | 0,00 | 1,00E+00 |
| AT2G04915 | 0,39 | -1,75 | 3,41 | -0,51 | 6,08E-01 |
| AT2G04935 | 1,06 | 2,30 | 1,95 | 1,18 | 2,39E-01 |
| AT2G05635 | 0,15 | 0,42 | 4,38 | 0,09 | 9,24E-01 |
| AT2G04985 | 0,95 | -0,35 | 2,15 | -0,16 | 8,72E-01 |
| AT2G05790 | 0,44 | 0,80 | 3,21 | 0,25 | 8,04E-01 |
| AT2G05812 | 1,46 | 2,14 | 1,80 | 1,19 | 2,34E-01 |
| AT2G05830 | 0,15 | 0,86 | 4,37 | 0,20 | 8,43E-01 |
| AT2G05840 | 0,17 | 0,09 | 4,35 | 0,02 | 9,84E-01 |
| AT2G05990 | 0,47 | -0,04 | 2,33 | -0,02 | 9,85E-01 |
| AT2G05995 | 0,63 | -3,16 | 3,05 | -1,03 | 3,01E-01 |
| AT2G06025 | 0,59 | -0,10 | 2,69 | -0,04 | 9,71E-01 |
| AT2G06095 | 0,70 | -3,75 | 3,45 | -1,09 | 2,78E-01 |
| AT2G06105 | 0,23 | 0,00 | 4,40 | 0,00 | 1,00E+00 |
| AT2G06845 | 0,18 | 1,23 | 4,34 | 0,28 | 7,77E-01 |
| AT2G06850 | 0,75 | -1,12 | 2,46 | -0,46 | 6,49E-01 |
| AT2G06904 | 0,38 | 1,23 | 4,35 | 0,28 | 7,77E-01 |
| AT2G06990 | 1,10 | 0,68 | 1,97 | 0,35 | 7,29E-01 |
| AT2G07050 | 0,64 | 2,40 | 2,97 | 0,81 | 4,19E-01 |
| AT2G07200 | 0,15 | 0,00 | 4,41 | 0,00 | 1,00E+00 |

| | | | | | |
|---|---|---|---|---|---|
| AT2G07565 | 0,91 | -2,84 | 2,45 | -1,16 | 2,46E-01 |
| AT2G07749 | 0,59 | -1,19 | 3,20 | -0,37 | 7,10E-01 |
| AT2G07672 | 0,25 | 0,54 | 4,39 | 0,12 | 9,01E-01 |
| AT2G07745 | 0,19 | -2,00 | 4,36 | -0,46 | 6,46E-01 |
| AT2G07674 | 0,81 | 2,79 | 3,67 | 0,76 | 4,47E-01 |
| AT2G07623 | 0,23 | 1,63 | 4,34 | 0,37 | 7,08E-01 |
| AT2G07678 | 0,32 | -1,23 | 4,39 | -0,28 | 7,80E-01 |
| AT2G07669 | 0,33 | -1,31 | 4,37 | -0,30 | 7,63E-01 |
| AT2G07681 | 0,75 | 1,58 | 2,25 | 0,71 | 4,81E-01 |
| AT2G07631 | 0,18 | -1,33 | 4,40 | -0,30 | 7,63E-01 |
| AT2G05215 | 0,68 | -0,56 | 2,47 | -0,23 | 8,21E-01 |
| AT2G07695 | 1,27 | 1,60 | 1,84 | 0,87 | 3,84E-01 |
| AT2G07798 | 1,27 | 0,36 | 1,68 | 0,21 | 8,31E-01 |
| AT2G07667 | 1,00 | 0,79 | 2,25 | 0,35 | 7,24E-01 |
| AT2G07638 | 1,46 | -1,27 | 1,50 | -0,84 | 3,98E-01 |
| AT2G07707 | 0,32 | -0,05 | 4,36 | -0,01 | 9,90E-01 |
| AT2G07708 | 0,57 | -0,62 | 4,36 | -0,14 | 8,86E-01 |
| AT2G07715 | 0,94 | -0,29 | 1,72 | -0,17 | 8,67E-01 |
| AT2G07728 | 0,11 | 0,90 | 4,37 | 0,21 | 8,36E-01 |
| AT2G07656 | 0,60 | 0,05 | 3,05 | 0,02 | 9,86E-01 |
| AT2G07827 | 0,36 | -0,03 | 3,50 | -0,01 | 9,93E-01 |
| AT2G07787 | 0,79 | 1,23 | 2,17 | 0,57 | 5,69E-01 |
| AT2G07659 | 0,28 | -0,40 | 4,38 | -0,09 | 9,27E-01 |
| AT2G07765 | 0,12 | 1,65 | 4,34 | 0,38 | 7,04E-01 |
| AT2G07741 | 0,27 | -1,36 | 4,41 | -0,31 | 7,58E-01 |
| AT2G07680 | 0,80 | 1,34 | 2,44 | 0,55 | 5,84E-01 |
| AT2G09990 | 0,53 | 0,95 | 3,06 | 0,31 | 7,55E-01 |
| AT2G05485 | 0,27 | 0,42 | 4,38 | 0,10 | 9,24E-01 |
| AT2G10545 | 0,08 | 0,00 | 4,40 | 0,00 | 1,00E+00 |
| AT2G10556 | 0,88 | 3,82 | 2,63 | 1,45 | 1,47E-01 |
| AT2G10965 | 0,86 | -0,10 | 2,08 | -0,05 | 9,62E-01 |
| AT2G11010 | 0,65 | 0,87 | 3,05 | 0,29 | 7,75E-01 |
| AT2G11035 | 0,48 | -0,07 | 4,04 | -0,02 | 9,86E-01 |
| AT2G11045 | 0,08 | 0,00 | 4,40 | 0,00 | 1,00E+00 |
| AT2G11225 | 0,91 | 2,25 | 1,93 | 1,17 | 2,42E-01 |
| AT2G11620 | 0,14 | 0,00 | 4,41 | 0,00 | 1,00E+00 |
| AT2G11851 | 0,57 | 1,57 | 3,00 | 0,52 | 6,02E-01 |
| AT2G05825 | 0,69 | 0,55 | 2,32 | 0,24 | 8,14E-01 |
| AT2G13100 | 0,26 | -1,03 | 4,37 | -0,23 | 8,14E-01 |
| AT2G13290 | 0,12 | 0,42 | 4,38 | 0,10 | 9,24E-01 |
| AT2G05885 | 0,14 | 0,00 | 4,41 | 0,00 | 1,00E+00 |
| AT2G13430 | 0,44 | 1,39 | 3,20 | 0,43 | 6,64E-01 |
| AT2G13500 | 0,11 | 0,00 | 4,41 | 0,00 | 1,00E+00 |
| AT2G13540 | 0,64 | -0,24 | 2,23 | -0,11 | 9,14E-01 |
| AT2G05925 | 0,12 | 0,42 | 4,38 | 0,10 | 9,24E-01 |
| AT2G13790 | 0,49 | 1,76 | 3,30 | 0,53 | 5,95E-01 |
| AT2G13845 | 0,67 | 2,56 | 2,26 | 1,14 | 2,56E-01 |
| AT2G13960 | 0,27 | 0,00 | 4,41 | 0,00 | 1,00E+00 |
| AT2G14070 | 0,19 | -1,36 | 4,41 | -0,31 | 7,58E-01 |
| AT2G14115 | 0,49 | 0,78 | 4,37 | 0,18 | 8,59E-01 |
| AT2G14395 | 0,63 | -1,48 | 4,31 | -0,34 | 7,31E-01 |
| AT2G14440 | 0,43 | -2,48 | 4,13 | -0,60 | 5,49E-01 |
| AT2G14530 | 0,21 | 0,00 | 4,41 | 0,00 | 1,00E+00 |
| AT2G14700 | 0,64 | 1,67 | 2,97 | 0,56 | 5,72E-01 |
| AT2G14740 | 1,25 | 3,96 | 2,41 | 1,64 | 1,01E-01 |
| AT2G14750 | 1,64 | 0,50 | 1,45 | 0,35 | 7,29E-01 |

| | | | | | |
|---|---|---|---|---|---|
| AT2G14765 | 0,40 | 0,88 | 4,36 | 0,20 | 8,41E-01 |
| AT2G14825 | 0,08 | 0,85 | 4,38 | 0,20 | 8,45E-01 |
| AT2G14835 | 0,23 | 2,14 | 4,32 | 0,50 | 6,20E-01 |
| AT2G14850 | 0,21 | 2,00 | 4,33 | 0,46 | 6,44E-01 |
| AT2G14870 | 0,17 | -2,23 | 4,35 | -0,51 | 6,08E-01 |
| AT2G14878 | 1,23 | 0,34 | 1,60 | 0,21 | 8,32E-01 |
| AT2G14890 | 0,77 | 1,25 | 2,20 | 0,57 | 5,69E-01 |
| AT2G06305 | 0,18 | 0,00 | 4,40 | 0,00 | 1,00E+00 |
| AT2G15042 | 0,11 | 0,90 | 4,37 | 0,21 | 8,36E-01 |
| AT2G15050 | 0,64 | -2,53 | 2,96 | -0,86 | 3,92E-01 |
| AT2G15080 | 0,92 | 1,89 | 2,11 | 0,90 | 3,70E-01 |
| AT2G15090 | 1,27 | 0,87 | 2,91 | 0,30 | 7,65E-01 |
| AT2G15300 | 0,31 | -0,85 | 4,35 | -0,20 | 8,45E-01 |
| AT2G15430 | 0,16 | 0,96 | 4,36 | 0,22 | 8,26E-01 |
| AT2G15450 | 0,29 | 0,96 | 4,36 | 0,22 | 8,25E-01 |
| AT2G15470 | 0,34 | 0,41 | 4,38 | 0,09 | 9,25E-01 |
| AT2G15580 | 0,29 | 0,31 | 3,69 | 0,08 | 9,34E-01 |
| AT2G15710 | 0,40 | -2,39 | 4,35 | -0,55 | 5,83E-01 |
| AT2G15730 | 0,08 | 0,00 | 4,40 | 0,00 | 1,00E+00 |
| AT2G15900 | 0,47 | 0,43 | 2,71 | 0,16 | 8,75E-01 |
| AT2G15960 | 1,09 | 2,68 | 1,91 | 1,40 | 1,61E-01 |
| AT2G06535 | 1,38 | 3,71 | 1,99 | 1,87 | 6,17E-02 |
| AT2G16019 | 1,05 | 1,29 | 2,20 | 0,59 | 5,57E-01 |
| AT2G16280 | 0,93 | 0,64 | 2,28 | 0,28 | 7,78E-01 |
| AT2G16365 | 0,96 | 1,22 | 2,18 | 0,56 | 5,76E-01 |
| AT2G16380 | 0,44 | -1,31 | 4,28 | -0,31 | 7,60E-01 |
| AT2G16390 | 0,40 | 2,51 | 4,32 | 0,58 | 5,61E-01 |
| AT2G16630 | 0,14 | 1,18 | 4,35 | 0,27 | 7,86E-01 |
| AT2G16700 | 0,28 | 1,16 | 4,32 | 0,27 | 7,88E-01 |
| AT2G16710 | 0,72 | 2,42 | 3,69 | 0,66 | 5,12E-01 |
| AT2G16800 | 0,26 | -1,90 | 4,37 | -0,43 | 6,64E-01 |
| AT2G16950 | 1,26 | 2,07 | 2,20 | 0,94 | 3,46E-01 |
| AT2G16970 | 1,25 | -3,16 | 1,75 | -1,81 | 7,02E-02 |
| AT2G17033 | 0,44 | 0,77 | 4,15 | 0,19 | 8,52E-01 |
| AT2G17040 | 1,57 | 3,02 | 1,85 | 1,63 | 1,02E-01 |
| AT2G17120 | 0,23 | 1,28 | 4,34 | 0,29 | 7,69E-01 |
| AT2G17140 | 0,42 | 0,86 | 3,39 | 0,25 | 8,00E-01 |
| AT2G17170 | 0,08 | 0,85 | 4,38 | 0,20 | 8,45E-01 |
| AT2G17240 | 0,52 | -1,67 | 3,13 | -0,53 | 5,94E-01 |
| AT2G17250 | 0,25 | 1,76 | 4,33 | 0,41 | 6,85E-01 |
| AT2G17265 | 1,56 | 2,03 | 1,55 | 1,31 | 1,92E-01 |
| AT2G17270 | 0,25 | 0,44 | 4,37 | 0,10 | 9,20E-01 |
| AT2G17290 | 0,78 | 1,29 | 2,28 | 0,57 | 5,70E-01 |
| AT2G06905 | 1,12 | -1,36 | 2,05 | -0,66 | 5,07E-01 |
| AT2G17360 | 0,51 | -0,37 | 3,27 | -0,11 | 9,10E-01 |
| AT2G17390 | 0,66 | 1,14 | 2,93 | 0,39 | 6,97E-01 |
| AT2G17410 | 1,45 | 2,29 | 1,82 | 1,26 | 2,08E-01 |
| AT2G17420 | 1,05 | 2,74 | 2,10 | 1,31 | 1,91E-01 |
| AT2G17520 | 1,17 | 1,27 | 1,81 | 0,70 | 4,84E-01 |
| AT2G17530 | 0,50 | 2,26 | 3,97 | 0,57 | 5,68E-01 |
| AT2G17570 | 0,42 | 1,76 | 4,15 | 0,42 | 6,72E-01 |
| AT2G17580 | 0,66 | 1,27 | 3,01 | 0,42 | 6,74E-01 |
| AT2G17705 | 0,42 | 0,79 | 3,32 | 0,24 | 8,12E-01 |
| AT2G17720 | 0,20 | 1,07 | 4,36 | 0,25 | 8,06E-01 |
| AT2G17760 | 0,65 | 1,60 | 2,97 | 0,54 | 5,91E-01 |
| AT2G17787 | 0,33 | 1,88 | 4,33 | 0,43 | 6,65E-01 |

| | | | | | |
|---|---|---|---|---|---|
| AT2G17790 | 0,80 | 1,43 | 2,46 | 0,58 | 5,62E-01 |
| AT2G17905 | 0,41 | -1,37 | 3,44 | -0,40 | 6,89E-01 |
| AT2G17920 | 0,15 | 0,41 | 4,38 | 0,09 | 9,25E-01 |
| AT2G17980 | 1,01 | 3,88 | 2,50 | 1,55 | 1,21E-01 |
| AT2G17990 | 0,69 | 1,48 | 2,43 | 0,61 | 5,42E-01 |
| AT2G18000 | 0,29 | 1,82 | 4,33 | 0,42 | 6,75E-01 |
| AT2G18030 | 0,68 | 3,14 | 2,94 | 1,07 | 2,86E-01 |
| AT2G18040 | 0,20 | 0,42 | 4,38 | 0,10 | 9,24E-01 |
| AT2G18090 | 1,01 | 2,18 | 1,87 | 1,17 | 2,42E-01 |
| AT2G18100 | 0,48 | 2,69 | 4,02 | 0,67 | 5,03E-01 |
| AT2G18110 | 0,62 | -1,54 | 3,03 | -0,51 | 6,12E-01 |
| AT2G18390 | 0,28 | 1,52 | 4,34 | 0,35 | 7,27E-01 |
| AT2G18420 | 0,81 | 1,99 | 3,46 | 0,58 | 5,65E-01 |
| AT2G18530 | 0,38 | 0,77 | 3,42 | 0,23 | 8,21E-01 |
| AT2G18560 | 1,54 | 0,17 | 2,35 | 0,07 | 9,42E-01 |
| AT2G18630 | 0,16 | 0,90 | 4,38 | 0,21 | 8,37E-01 |
| AT2G18670 | 1,13 | 0,11 | 1,67 | 0,07 | 9,47E-01 |
| AT2G18680 | 0,16 | 1,74 | 4,33 | 0,40 | 6,87E-01 |
| AT2G18690 | 1,29 | 3,25 | 1,93 | 1,68 | 9,33E-02 |
| AT2G18700 | 0,27 | -0,47 | 4,34 | -0,11 | 9,15E-01 |
| AT2G18730 | 0,58 | 0,15 | 2,49 | 0,06 | 9,51E-01 |
| AT2G18735 | 0,22 | -0,03 | 4,37 | -0,01 | 9,94E-01 |
| AT2G18750 | 0,67 | 0,76 | 2,10 | 0,36 | 7,17E-01 |
| AT2G18760 | 1,48 | 3,40 | 1,75 | 1,94 | 5,21E-02 |
| AT2G18850 | 0,52 | 0,00 | 4,38 | 0,00 | 1,00E+00 |
| AT2G18880 | 0,25 | 2,36 | 4,32 | 0,55 | 5,86E-01 |
| AT2G18917 | 0,08 | 0,00 | 4,40 | 0,00 | 1,00E+00 |
| AT2G07095 | 0,55 | 3,02 | 3,01 | 1,00 | 3,15E-01 |
| AT2G18938 | 0,72 | 0,43 | 2,57 | 0,17 | 8,66E-01 |
| AT2G18950 | 0,45 | 1,74 | 4,17 | 0,42 | 6,76E-01 |
| AT2G19110 | 0,15 | 1,25 | 4,35 | 0,29 | 7,75E-01 |
| AT2G19130 | 0,89 | -2,47 | 2,30 | -1,07 | 2,83E-01 |
| AT2G19160 | 0,39 | 2,27 | 4,33 | 0,53 | 5,99E-01 |
| AT2G19260 | 0,46 | 1,07 | 4,07 | 0,26 | 7,94E-01 |
| AT2G19280 | 0,36 | 1,99 | 4,34 | 0,46 | 6,46E-01 |
| AT2G19420 | 0,28 | 0,42 | 4,38 | 0,10 | 9,23E-01 |
| AT2G19430 | 0,24 | -2,17 | 4,38 | -0,49 | 6,21E-01 |
| AT2G19450 | 0,42 | -0,50 | 4,22 | -0,12 | 9,06E-01 |
| AT2G07205 | 0,75 | 1,01 | 3,68 | 0,27 | 7,83E-01 |
| AT2G19490 | 0,36 | 2,25 | 4,33 | 0,52 | 6,04E-01 |
| AT2G19520 | 0,55 | 1,16 | 4,07 | 0,28 | 7,76E-01 |
| AT2G19540 | 0,46 | 0,79 | 2,74 | 0,29 | 7,73E-01 |
| AT2G19572 | 0,52 | 1,65 | 3,07 | 0,54 | 5,92E-01 |
| AT2G19582 | 0,25 | 0,43 | 4,39 | 0,10 | 9,21E-01 |
| AT2G19600 | 0,64 | 0,32 | 3,90 | 0,08 | 9,34E-01 |
| AT2G19640 | 0,53 | 1,96 | 3,84 | 0,51 | 6,10E-01 |
| AT2G19710 | 1,44 | 2,09 | 2,17 | 0,96 | 3,35E-01 |
| AT2G19730 | 0,57 | -0,53 | 4,09 | -0,13 | 8,98E-01 |
| AT2G19740 | 0,67 | -0,05 | 2,27 | -0,02 | 9,81E-01 |
| AT2G19810 | 0,34 | 0,42 | 3,58 | 0,12 | 9,07E-01 |
| AT2G19880 | 0,75 | 1,31 | 2,45 | 0,53 | 5,93E-01 |
| AT2G20020 | 1,07 | 1,68 | 2,09 | 0,80 | 4,22E-01 |
| AT2G20100 | 0,33 | 1,77 | 4,34 | 0,41 | 6,83E-01 |
| AT2G20110 | 0,83 | 2,24 | 4,29 | 0,52 | 6,01E-01 |
| AT2G20142 | 0,28 | 1,56 | 4,34 | 0,36 | 7,18E-01 |
| AT2G20230 | 0,26 | -1,32 | 4,40 | -0,30 | 7,65E-01 |

| | | | | | |
|---|---|---|---|---|---|
| AT2G20260 | 1,52 | -1,90 | 1,63 | -1,17 | 2,42E-01 |
| AT2G20340 | 0,57 | 0,49 | 3,90 | 0,13 | 8,99E-01 |
| AT2G20360 | 0,22 | 0,44 | 4,33 | 0,10 | 9,19E-01 |
| AT2G20362 | 0,08 | 1,10 | 4,36 | 0,25 | 8,00E-01 |
| AT2G20370 | 1,01 | 1,98 | 2,54 | 0,78 | 4,35E-01 |
| AT2G20420 | 0,62 | 0,25 | 2,19 | 0,11 | 9,10E-01 |
| AT2G20450 | 0,40 | 1,26 | 4,13 | 0,31 | 7,59E-01 |
| AT2G20530 | 0,21 | -0,89 | 4,35 | -0,20 | 8,39E-01 |
| AT2G20690 | 1,05 | 0,96 | 2,38 | 0,40 | 6,86E-01 |
| AT2G20710 | 0,08 | 0,85 | 4,38 | 0,20 | 8,45E-01 |
| AT2G07335 | 0,46 | 0,19 | 4,15 | 0,04 | 9,64E-01 |
| AT2G20750 | 0,59 | -0,55 | 3,82 | -0,14 | 8,86E-01 |
| AT2G20770 | 0,17 | 1,92 | 4,33 | 0,44 | 6,57E-01 |
| AT2G20820 | 0,25 | 1,63 | 4,34 | 0,38 | 7,07E-01 |
| AT2G20860 | 1,36 | 1,48 | 1,65 | 0,90 | 3,70E-01 |
| AT2G20890 | 1,04 | -1,17 | 1,78 | -0,66 | 5,11E-01 |
| AT2G20920 | 0,65 | 2,17 | 2,48 | 0,88 | 3,80E-01 |
| AT2G20960 | 0,82 | 0,57 | 2,07 | 0,27 | 7,84E-01 |
| AT2G21160 | 0,91 | 0,92 | 3,08 | 0,30 | 7,65E-01 |
| AT2G21187 | 0,89 | 0,88 | 1,97 | 0,45 | 6,55E-01 |
| AT2G21190 | 0,23 | -1,99 | 4,37 | -0,46 | 6,49E-01 |
| AT2G21270 | 0,21 | 1,93 | 4,33 | 0,45 | 6,56E-01 |
| AT2G21380 | 1,04 | -1,65 | 2,03 | -0,81 | 4,15E-01 |
| AT2G21410 | 1,46 | 0,68 | 1,60 | 0,43 | 6,69E-01 |
| AT2G21520 | 0,84 | 1,89 | 2,10 | 0,90 | 3,68E-01 |
| AT2G21530 | 0,61 | -1,30 | 2,94 | -0,44 | 6,58E-01 |
| AT2G21550 | 0,95 | -2,20 | 2,49 | -0,88 | 3,78E-01 |
| AT2G21580 | 1,04 | 3,17 | 2,73 | 1,16 | 2,46E-01 |
| AT2G21620 | 1,16 | -4,08 | 2,11 | -1,94 | 5,30E-02 |
| AT2G21630 | 0,32 | 0,81 | 4,37 | 0,19 | 8,53E-01 |
| AT2G21770 | 0,18 | 0,00 | 4,41 | 0,00 | 1,00E+00 |
| AT2G21790 | 0,74 | 2,45 | 2,49 | 0,98 | 3,25E-01 |
| AT2G21850 | 1,02 | 1,80 | 2,62 | 0,68 | 4,94E-01 |
| AT2G07475 | 0,24 | 1,98 | 4,34 | 0,46 | 6,48E-01 |
| AT2G21960 | 0,80 | 0,59 | 2,60 | 0,23 | 8,22E-01 |
| AT2G07495 | 0,12 | 1,39 | 4,34 | 0,32 | 7,49E-01 |
| AT2G22080 | 0,65 | 1,69 | 2,55 | 0,66 | 5,07E-01 |
| AT2G22120 | 0,32 | 0,81 | 4,37 | 0,19 | 8,53E-01 |
| AT2G22425 | 0,10 | 0,00 | 4,41 | 0,00 | 1,00E+00 |
| AT2G22450 | 0,24 | 1,09 | 4,35 | 0,25 | 8,02E-01 |
| AT2G22480 | 0,23 | 0,79 | 4,37 | 0,18 | 8,56E-01 |
| AT2G22540 | 1,53 | 0,63 | 1,82 | 0,35 | 7,29E-01 |
| AT2G22670 | 0,80 | 1,23 | 2,39 | 0,52 | 6,05E-01 |
| AT2G22680 | 0,25 | 0,00 | 4,41 | 0,00 | 1,00E+00 |
| AT2G22720 | 0,99 | 2,89 | 2,71 | 1,06 | 2,87E-01 |
| AT2G07845 | 0,43 | 2,58 | 4,09 | 0,63 | 5,28E-01 |
| AT2G22790 | 0,08 | 1,10 | 4,36 | 0,25 | 8,00E-01 |
| AT2G22840 | 1,11 | 2,76 | 2,03 | 1,36 | 1,73E-01 |
| AT2G07850 | 0,72 | -1,07 | 1,89 | -0,57 | 5,72E-01 |
| AT2G22910 | 0,32 | 1,13 | 4,36 | 0,26 | 7,95E-01 |
| AT2G23080 | 0,38 | 1,84 | 4,33 | 0,43 | 6,71E-01 |
| AT2G23120 | 0,92 | 2,06 | 2,17 | 0,95 | 3,44E-01 |
| AT2G23130 | 0,41 | -0,50 | 2,77 | -0,18 | 8,57E-01 |
| AT2G23142 | 1,46 | -3,85 | 2,30 | -1,67 | 9,44E-02 |
| AT2G23200 | 1,64 | 1,55 | 1,67 | 0,93 | 3,54E-01 |
| AT2G23310 | 0,84 | 1,66 | 2,95 | 0,56 | 5,73E-01 |

| | | | | | |
|---|---|---|---|---|---|
| AT2G07895 | 0,15 | 1,62 | 4,34 | 0,37 | 7,09E-01 |
| AT2G23320 | 1,39 | 2,35 | 1,82 | 1,29 | 1,98E-01 |
| AT2G23420 | 0,47 | 1,17 | 4,37 | 0,27 | 7,88E-01 |
| AT2G23460 | 1,50 | 0,85 | 1,63 | 0,52 | 6,02E-01 |
| AT2G07915 | 1,63 | -3,88 | 2,35 | -1,65 | 9,91E-02 |
| AT2G23672 | 1,50 | -0,45 | 1,55 | -0,29 | 7,70E-01 |
| AT2G23680 | 0,31 | 0,03 | 4,36 | 0,01 | 9,94E-01 |
| AT2G23740 | 0,12 | 0,90 | 4,36 | 0,21 | 8,36E-01 |
| AT2G23890 | 0,37 | 0,76 | 3,46 | 0,22 | 8,27E-01 |
| AT2G23985 | 0,17 | -2,23 | 4,35 | -0,51 | 6,08E-01 |
| AT2G24090 | 0,67 | 0,11 | 2,54 | 0,04 | 9,65E-01 |
| AT2G24180 | 1,36 | 0,51 | 2,40 | 0,21 | 8,33E-01 |
| AT2G24240 | 0,31 | 1,24 | 4,36 | 0,28 | 7,76E-01 |
| AT2G24280 | 1,45 | -0,91 | 1,47 | -0,62 | 5,38E-01 |
| AT2G24300 | 0,15 | 0,86 | 4,37 | 0,20 | 8,43E-01 |
| AT2G08015 | 0,35 | 2,49 | 4,32 | 0,58 | 5,65E-01 |
| AT2G24350 | 0,50 | 1,24 | 3,99 | 0,31 | 7,56E-01 |
| AT2G24390 | 0,92 | -0,51 | 2,79 | -0,18 | 8,54E-01 |
| AT2G24380 | 0,65 | -1,94 | 3,72 | -0,52 | 6,02E-01 |
| AT2G24420 | 1,14 | -0,32 | 1,88 | -0,17 | 8,66E-01 |
| AT2G08045 | 0,23 | 1,68 | 4,33 | 0,39 | 6,97E-01 |
| AT2G08050 | 0,32 | 1,76 | 4,34 | 0,41 | 6,85E-01 |
| AT2G24540 | 0,88 | -0,50 | 1,83 | -0,28 | 7,83E-01 |
| AT2G24590 | 0,49 | 1,68 | 3,93 | 0,43 | 6,69E-01 |
| AT2G24592 | 0,83 | 2,62 | 2,14 | 1,22 | 2,22E-01 |
| AT2G24765 | 0,69 | 2,44 | 2,86 | 0,85 | 3,94E-01 |
| AT2G25000 | 0,52 | 3,10 | 3,80 | 0,82 | 4,15E-01 |
| AT2G25080 | 0,61 | -1,46 | 2,65 | -0,55 | 5,83E-01 |
| AT2G25110 | 0,19 | 1,57 | 4,34 | 0,36 | 7,18E-01 |
| AT2G25190 | 0,14 | 0,44 | 4,39 | 0,10 | 9,20E-01 |
| AT2G25370 | 1,47 | -3,80 | 2,10 | -1,81 | 7,00E-02 |
| AT2G25410 | 1,30 | 3,23 | 2,26 | 1,43 | 1,53E-01 |
| AT2G25480 | 0,05 | 0,00 | 4,41 | 0,00 | 1,00E+00 |
| AT2G25560 | 0,73 | 3,27 | 2,77 | 1,18 | 2,38E-01 |
| AT2G25580 | 0,08 | 0,85 | 4,38 | 0,20 | 8,45E-01 |
| AT2G25590 | 0,21 | 2,00 | 4,33 | 0,46 | 6,44E-01 |
| AT2G25625 | 0,15 | 1,58 | 4,34 | 0,36 | 7,15E-01 |
| AT2G25630 | 0,50 | 1,94 | 3,12 | 0,62 | 5,34E-01 |
| AT2G25650 | 0,70 | 2,58 | 2,87 | 0,90 | 3,70E-01 |
| AT2G25790 | 0,18 | 0,42 | 4,38 | 0,10 | 9,24E-01 |
| AT2G25840 | 0,59 | 0,67 | 3,99 | 0,17 | 8,66E-01 |
| AT2G25870 | 1,69 | 3,56 | 1,80 | 1,98 | 4,76E-02 |
| AT2G25930 | 1,26 | 2,49 | 1,73 | 1,44 | 1,49E-01 |
| AT2G26060 | 1,00 | 1,29 | 4,31 | 0,30 | 7,65E-01 |
| AT2G26110 | 0,61 | 0,49 | 3,03 | 0,16 | 8,71E-01 |
| AT2G26170 | 0,47 | 1,89 | 3,21 | 0,59 | 5,56E-01 |
| AT2G26270 | 0,21 | 0,79 | 4,37 | 0,18 | 8,57E-01 |
| AT2G26300 | 1,09 | 3,12 | 2,41 | 1,30 | 1,94E-01 |
| AT2G26340 | 0,57 | 1,57 | 3,08 | 0,51 | 6,09E-01 |
| AT2G26440 | 1,40 | 1,55 | 1,65 | 0,94 | 3,49E-01 |
| AT2G26510 | 0,24 | 1,09 | 4,35 | 0,25 | 8,02E-01 |
| AT2G26570 | 0,43 | 0,04 | 3,28 | 0,01 | 9,90E-01 |
| AT2G26600 | 0,45 | 2,16 | 3,20 | 0,68 | 4,99E-01 |
| AT2G26610 | 0,30 | 1,76 | 4,33 | 0,41 | 6,85E-01 |
| AT2G26640 | 0,71 | -0,58 | 3,55 | -0,16 | 8,71E-01 |
| AT2G26660 | 0,22 | 0,85 | 4,36 | 0,20 | 8,45E-01 |

| | | | | | |
|---|---|---|---|---|---|
| AT2G26692 | 0,08 | 1,10 | 4,36 | 0,25 | 8,00E-01 |
| AT2G26730 | 0,85 | -1,11 | 2,00 | -0,55 | 5,80E-01 |
| AT2G26920 | 0,60 | 1,53 | 3,06 | 0,50 | 6,17E-01 |
| AT2G27030 | 0,72 | 1,54 | 2,64 | 0,59 | 5,58E-01 |
| AT2G27050 | 0,99 | 2,88 | 1,89 | 1,52 | 1,28E-01 |
| AT2G27060 | 0,17 | 1,73 | 4,34 | 0,40 | 6,91E-01 |
| AT2G27080 | 0,39 | 0,83 | 4,23 | 0,20 | 8,45E-01 |
| AT2G27100 | 0,72 | 2,41 | 3,56 | 0,68 | 4,98E-01 |
| AT2G27130 | 0,30 | 0,00 | 4,41 | 0,00 | 1,00E+00 |
| AT2G27200 | 0,21 | 1,84 | 4,33 | 0,42 | 6,71E-01 |
| AT2G27210 | 0,89 | 1,95 | 2,25 | 0,87 | 3,86E-01 |
| AT2G27230 | 1,31 | 1,43 | 1,76 | 0,81 | 4,18E-01 |
| AT2G27260 | 0,40 | 2,10 | 3,31 | 0,63 | 5,27E-01 |
| AT2G27320 | 0,85 | -0,35 | 2,38 | -0,15 | 8,83E-01 |
| AT2G27380 | 0,13 | 0,00 | 4,41 | 0,00 | 1,00E+00 |
| AT2G27460 | 0,28 | -0,05 | 4,30 | -0,01 | 9,91E-01 |
| AT2G27540 | 0,95 | 0,84 | 3,19 | 0,26 | 7,93E-01 |
| AT2G27570 | 0,28 | 0,42 | 4,38 | 0,10 | 9,24E-01 |
| AT2G27600 | 1,39 | 2,21 | 1,90 | 1,17 | 2,44E-01 |
| AT2G27690 | 0,16 | 1,33 | 4,35 | 0,31 | 7,59E-01 |
| AT2G27710 | 0,57 | 0,00 | 3,23 | 0,00 | 9,99E-01 |
| AT2G27760 | 0,08 | 0,85 | 4,38 | 0,20 | 8,45E-01 |
| AT2G27810 | 0,54 | 0,29 | 2,72 | 0,11 | 9,16E-01 |
| AT2G27820 | 0,23 | 0,15 | 4,35 | 0,03 | 9,72E-01 |
| AT2G27900 | 1,32 | 2,04 | 1,71 | 1,20 | 2,31E-01 |
| AT2G28070 | 0,87 | 1,83 | 2,88 | 0,64 | 5,25E-01 |
| AT2G28120 | 0,29 | 1,97 | 4,33 | 0,46 | 6,49E-01 |
| AT2G28130 | 0,59 | 2,53 | 4,31 | 0,59 | 5,58E-01 |
| AT2G28190 | 0,42 | 0,75 | 4,33 | 0,17 | 8,62E-01 |
| AT2G28250 | 0,08 | 1,10 | 4,36 | 0,25 | 8,00E-01 |
| AT2G28310 | 0,30 | 0,41 | 4,38 | 0,09 | 9,25E-01 |
| AT2G28320 | 0,43 | 1,65 | 4,17 | 0,39 | 6,93E-01 |
| AT2G28350 | 0,96 | 3,02 | 2,04 | 1,48 | 1,38E-01 |
| AT2G28370 | 0,25 | 1,63 | 4,34 | 0,38 | 7,07E-01 |
| AT2G28460 | 0,18 | -1,25 | 4,39 | -0,28 | 7,76E-01 |
| AT2G28520 | 0,65 | 2,58 | 2,88 | 0,90 | 3,69E-01 |
| AT2G28540 | 0,87 | 1,27 | 2,30 | 0,55 | 5,81E-01 |
| AT2G28605 | 0,37 | -1,59 | 4,38 | -0,36 | 7,17E-01 |
| AT2G28660 | 0,10 | 0,00 | 4,41 | 0,00 | 1,00E+00 |
| AT2G28671 | 0,51 | 1,25 | 3,03 | 0,41 | 6,78E-01 |
| AT2G28710 | 0,24 | 1,72 | 4,34 | 0,40 | 6,92E-01 |
| AT2G28720 | 0,49 | -1,93 | 4,21 | -0,46 | 6,46E-01 |
| AT2G28730 | 1,02 | 0,67 | 1,86 | 0,36 | 7,17E-01 |
| AT2G28810 | 0,71 | 1,80 | 2,60 | 0,69 | 4,88E-01 |
| AT2G28930 | 0,12 | 0,79 | 4,37 | 0,18 | 8,57E-01 |
| AT2G29020 | 0,11 | 0,00 | 4,41 | 0,00 | 1,00E+00 |
| AT2G29065 | 0,30 | 1,45 | 4,35 | 0,33 | 7,39E-01 |
| AT2G29070 | 0,15 | 1,62 | 4,34 | 0,37 | 7,09E-01 |
| AT2G29140 | 0,94 | 2,45 | 2,48 | 0,99 | 3,24E-01 |
| AT2G29180 | 0,47 | -0,26 | 4,09 | -0,06 | 9,49E-01 |
| AT2G29210 | 1,16 | 2,80 | 2,13 | 1,31 | 1,89E-01 |
| AT2G29310 | 0,46 | -1,77 | 3,34 | -0,53 | 5,96E-01 |
| AT2G29350 | 0,65 | 3,48 | 3,82 | 0,91 | 3,62E-01 |
| AT2G29360 | 0,51 | 0,48 | 4,40 | 0,11 | 9,13E-01 |
| AT2G29510 | 0,82 | 3,34 | 3,29 | 1,01 | 3,11E-01 |
| AT2G29600 | 0,47 | 1,92 | 3,18 | 0,60 | 5,47E-01 |

| | | | | | |
|---|---|---|---|---|---|
| AT2G29640 | 0,37 | 1,57 | 4,28 | 0,37 | 7,14E-01 |
| AT2G29660 | 0,20 | 1,07 | 4,36 | 0,25 | 8,06E-01 |
| AT2G29690 | 0,14 | -1,28 | 4,39 | -0,29 | 7,71E-01 |
| AT2G29700 | 0,24 | -1,37 | 4,41 | -0,31 | 7,55E-01 |
| AT2G29970 | 1,50 | -0,35 | 1,35 | -0,26 | 7,95E-01 |
| AT2G30020 | 0,83 | 1,98 | 3,51 | 0,56 | 5,73E-01 |
| AT2G30040 | 0,38 | 1,90 | 3,33 | 0,57 | 5,69E-01 |
| AT2G30050 | 0,66 | -2,07 | 2,60 | -0,79 | 4,27E-01 |
| AT2G30140 | 0,54 | 2,87 | 3,81 | 0,75 | 4,52E-01 |
| AT2G30150 | 0,10 | 0,62 | 4,38 | 0,14 | 8,87E-01 |
| AT2G30180 | 1,06 | 0,01 | 1,79 | 0,01 | 9,96E-01 |
| AT2G08375 | 1,24 | -0,56 | 1,95 | -0,29 | 7,73E-01 |
| AT2G30230 | 0,50 | 0,77 | 3,21 | 0,24 | 8,11E-01 |
| AT2G30362 | 0,41 | 1,00 | 4,21 | 0,24 | 8,13E-01 |
| AT2G30410 | 0,72 | -2,61 | 2,51 | -1,04 | 2,98E-01 |
| AT2G30530 | 0,20 | 1,07 | 4,36 | 0,25 | 8,06E-01 |
| AT2G30550 | 1,44 | 0,06 | 1,89 | 0,03 | 9,73E-01 |
| AT2G30600 | 0,51 | -2,30 | 4,33 | -0,53 | 5,95E-01 |
| AT2G30620 | 0,85 | -0,32 | 2,22 | -0,15 | 8,84E-01 |
| AT2G30730 | 0,89 | -0,09 | 2,56 | -0,03 | 9,73E-01 |
| AT2G30780 | 0,62 | 1,34 | 3,65 | 0,37 | 7,14E-01 |
| AT2G30870 | 1,65 | 0,70 | 1,41 | 0,49 | 6,21E-01 |
| AT2G30930 | 0,49 | -0,41 | 2,71 | -0,15 | 8,79E-01 |
| AT2G30970 | 0,46 | 1,00 | 4,38 | 0,23 | 8,19E-01 |
| AT2G31010 | 0,95 | 2,11 | 2,34 | 0,90 | 3,67E-01 |
| AT2G31020 | 0,26 | 0,90 | 4,38 | 0,21 | 8,37E-01 |
| AT2G31200 | 0,16 | 0,00 | 4,39 | 0,00 | 1,00E+00 |
| AT2G31215 | 0,52 | 0,07 | 2,64 | 0,03 | 9,79E-01 |
| AT2G31280 | 1,47 | 3,82 | 2,07 | 1,85 | 6,49E-02 |
| AT2G31290 | 0,71 | 0,79 | 3,13 | 0,25 | 8,00E-01 |
| AT2G31350 | 0,38 | 2,08 | 4,28 | 0,48 | 6,28E-01 |
| AT2G31370 | 0,19 | 0,38 | 4,34 | 0,09 | 9,30E-01 |
| AT2G31380 | 1,06 | 1,60 | 1,93 | 0,83 | 4,07E-01 |
| AT2G08435 | 0,79 | 1,66 | 3,52 | 0,47 | 6,37E-01 |
| AT2G31560 | 0,67 | -0,09 | 2,56 | -0,04 | 9,72E-01 |
| AT2G31585 | 0,47 | 0,81 | 3,96 | 0,20 | 8,39E-01 |
| AT2G31750 | 0,75 | 1,27 | 2,92 | 0,44 | 6,62E-01 |
| AT2G31790 | 0,30 | 1,59 | 3,63 | 0,44 | 6,61E-01 |
| AT2G31810 | 1,10 | 0,86 | 2,15 | 0,40 | 6,89E-01 |
| AT2G31960 | 1,68 | 1,47 | 1,64 | 0,90 | 3,70E-01 |
| AT2G31970 | 0,18 | 1,66 | 4,33 | 0,38 | 7,02E-01 |
| AT2G32010 | 0,37 | 0,64 | 4,33 | 0,15 | 8,83E-01 |
| AT2G32040 | 0,08 | 0,85 | 4,38 | 0,20 | 8,45E-01 |
| AT2G32100 | 1,15 | -1,21 | 2,23 | -0,54 | 5,88E-01 |
| AT2G32150 | 1,00 | 2,53 | 2,45 | 1,03 | 3,01E-01 |
| AT2G32250 | 0,51 | 1,00 | 3,14 | 0,32 | 7,51E-01 |
| AT2G32260 | 0,44 | 2,21 | 4,08 | 0,54 | 5,88E-01 |
| AT2G32295 | 0,53 | -0,43 | 2,69 | -0,16 | 8,74E-01 |
| AT2G32320 | 0,15 | 0,86 | 4,37 | 0,20 | 8,43E-01 |
| AT2G32440 | 0,16 | 1,07 | 4,37 | 0,24 | 8,07E-01 |
| AT2G32450 | 0,56 | 2,54 | 2,57 | 0,99 | 3,23E-01 |
| AT2G32698 | 0,12 | 1,58 | 4,34 | 0,36 | 7,16E-01 |
| AT2G32700 | 1,61 | 1,16 | 1,37 | 0,85 | 3,96E-01 |
| AT2G32800 | 1,51 | 0,33 | 1,47 | 0,22 | 8,24E-01 |
| AT2G32900 | 0,29 | 1,88 | 4,33 | 0,43 | 6,65E-01 |
| AT2G32910 | 0,68 | 1,99 | 2,94 | 0,68 | 4,99E-01 |

| | | | | | |
|---|---|---|---|---|---|
| AT2G33080 | 0,74 | 3,03 | 3,44 | 0,88 | 3,78E-01 |
| AT2G33160 | 0,25 | 1,75 | 4,33 | 0,40 | 6,87E-01 |
| AT2G33220 | 0,54 | 2,16 | 3,15 | 0,69 | 4,92E-01 |
| AT2G33250 | 0,76 | 0,95 | 4,29 | 0,22 | 8,25E-01 |
| AT2G33290 | 0,29 | 0,31 | 3,69 | 0,08 | 9,34E-01 |
| AT2G33390 | 0,25 | 0,00 | 4,41 | 0,00 | 1,00E+00 |
| AT2G33420 | 0,37 | 1,57 | 4,28 | 0,37 | 7,14E-01 |
| AT2G08585 | 0,32 | 0,26 | 3,67 | 0,07 | 9,44E-01 |
| AT2G33435 | 1,31 | -0,93 | 1,67 | -0,56 | 5,77E-01 |
| AT2G33450 | 0,32 | -0,43 | 4,33 | -0,10 | 9,20E-01 |
| AT2G33490 | 0,35 | 2,11 | 4,29 | 0,49 | 6,23E-01 |
| AT2G33510 | 0,26 | 1,44 | 4,34 | 0,33 | 7,39E-01 |
| AT2G33570 | 0,17 | 0,48 | 4,40 | 0,11 | 9,12E-01 |
| AT2G33610 | 0,37 | 1,61 | 4,28 | 0,38 | 7,07E-01 |
| AT2G33650 | 0,29 | -2,03 | 4,36 | -0,47 | 6,41E-01 |
| AT2G33700 | 0,56 | 0,58 | 4,06 | 0,14 | 8,86E-01 |
| AT2G33760 | 0,64 | 1,64 | 4,35 | 0,38 | 7,07E-01 |
| AT2G33810 | 0,37 | -1,59 | 4,36 | -0,36 | 7,15E-01 |
| AT2G33840 | 0,52 | -0,50 | 3,21 | -0,16 | 8,76E-01 |
| AT2G33845 | 0,47 | 2,39 | 4,04 | 0,59 | 5,54E-01 |
| AT2G33950 | 0,39 | 0,79 | 4,31 | 0,18 | 8,55E-01 |
| AT2G34120 | 0,27 | 0,67 | 4,38 | 0,15 | 8,79E-01 |
| AT2G34240 | 0,12 | 1,46 | 4,35 | 0,34 | 7,36E-01 |
| AT2G34460 | 0,99 | 0,97 | 1,94 | 0,50 | 6,16E-01 |
| AT2G34660 | 0,69 | 0,55 | 2,58 | 0,21 | 8,30E-01 |
| AT2G34730 | 0,50 | 0,48 | 2,62 | 0,18 | 8,54E-01 |
| AT2G34770 | 0,22 | 0,85 | 4,36 | 0,20 | 8,45E-01 |
| AT2G34780 | 0,99 | 3,00 | 2,14 | 1,40 | 1,62E-01 |
| AT2G34810 | 0,16 | 1,33 | 4,35 | 0,31 | 7,59E-01 |
| AT2G34860 | 1,10 | 2,15 | 2,25 | 0,96 | 3,39E-01 |
| AT2G34970 | 0,22 | 0,41 | 4,38 | 0,09 | 9,24E-01 |
| AT2G35020 | 0,32 | 0,86 | 4,38 | 0,20 | 8,44E-01 |
| AT2G35170 | 0,10 | 0,62 | 4,38 | 0,14 | 8,87E-01 |
| AT2G35230 | 0,50 | 1,15 | 4,36 | 0,26 | 7,91E-01 |
| AT2G35330 | 0,35 | 0,00 | 4,41 | 0,00 | 1,00E+00 |
| AT2G35340 | 0,80 | 2,04 | 2,79 | 0,73 | 4,64E-01 |
| AT2G35370 | 0,43 | 0,07 | 4,29 | 0,02 | 9,88E-01 |
| AT2G35630 | 1,16 | -0,36 | 1,75 | -0,21 | 8,37E-01 |
| AT2G35635 | 0,64 | -2,54 | 4,35 | -0,58 | 5,59E-01 |
| AT2G35650 | 0,68 | 0,99 | 2,23 | 0,44 | 6,57E-01 |
| AT2G08750 | 0,08 | 0,85 | 4,38 | 0,20 | 8,45E-01 |
| AT2G35720 | 0,19 | 1,27 | 4,35 | 0,29 | 7,70E-01 |
| AT2G35760 | 0,17 | 0,00 | 4,40 | 0,00 | 1,00E+00 |
| AT2G35800 | 1,02 | 0,32 | 1,78 | 0,18 | 8,55E-01 |
| AT2G35820 | 0,71 | 2,73 | 4,31 | 0,63 | 5,26E-01 |
| AT2G35830 | 1,03 | 3,32 | 1,99 | 1,67 | 9,47E-02 |
| AT2G35840 | 0,63 | 1,54 | 2,51 | 0,61 | 5,41E-01 |
| AT2G35860 | 0,74 | -0,28 | 2,33 | -0,12 | 9,05E-01 |
| AT2G35920 | 0,77 | 1,42 | 2,40 | 0,59 | 5,55E-01 |
| AT2G35935 | 0,49 | 0,46 | 3,96 | 0,12 | 9,08E-01 |
| AT2G35960 | 0,20 | 0,57 | 4,37 | 0,13 | 8,96E-01 |
| AT2G36000 | 0,80 | 1,96 | 2,53 | 0,77 | 4,39E-01 |
| AT2G36010 | 0,59 | -1,08 | 3,07 | -0,35 | 7,24E-01 |
| AT2G36026 | 0,70 | -0,91 | 3,73 | -0,24 | 8,07E-01 |
| AT2G36090 | 0,69 | 0,94 | 3,61 | 0,26 | 7,94E-01 |
| AT2G36130 | 0,56 | 1,60 | 4,35 | 0,37 | 7,13E-01 |

| | | | | | |
|---|---|---|---|---|---|
| AT2G36320 | 1,39 | -0,16 | 1,59 | -0,10 | 9,20E-01 |
| AT2G36330 | 0,55 | 1,69 | 4,32 | 0,39 | 6,95E-01 |
| AT2G36430 | 1,44 | 0,82 | 1,96 | 0,42 | 6,74E-01 |
| AT2G36500 | 0,54 | 1,65 | 3,10 | 0,53 | 5,93E-01 |
| AT2G36510 | 0,37 | 1,64 | 4,28 | 0,38 | 7,03E-01 |
| AT2G36570 | 0,52 | 0,00 | 4,41 | 0,00 | 1,00E+00 |
| AT2G36580 | 1,14 | 1,00 | 1,66 | 0,61 | 5,45E-01 |
| AT2G36600 | 0,66 | -1,52 | 2,12 | -0,72 | 4,73E-01 |
| AT2G36720 | 1,25 | 2,91 | 2,10 | 1,39 | 1,65E-01 |
| AT2G36792 | 0,34 | 1,48 | 4,34 | 0,34 | 7,34E-01 |
| AT2G36830 | 0,86 | 0,98 | 2,24 | 0,44 | 6,62E-01 |
| AT2G36920 | 0,12 | 1,65 | 4,34 | 0,38 | 7,04E-01 |
| AT2G36950 | 0,34 | 1,96 | 4,34 | 0,45 | 6,52E-01 |
| AT2G36960 | 0,18 | 1,23 | 4,34 | 0,28 | 7,77E-01 |
| AT2G36970 | 0,19 | 1,57 | 4,34 | 0,36 | 7,18E-01 |
| AT2G36980 | 0,05 | 0,00 | 4,41 | 0,00 | 1,00E+00 |
| AT2G37025 | 0,19 | -2,00 | 4,36 | -0,46 | 6,46E-01 |
| AT2G37050 | 0,93 | 1,66 | 2,30 | 0,72 | 4,71E-01 |
| AT2G37110 | 1,16 | 2,88 | 2,32 | 1,24 | 2,15E-01 |
| AT2G37180 | 0,08 | 0,85 | 4,38 | 0,20 | 8,45E-01 |
| AT2G37250 | 0,74 | 1,39 | 2,20 | 0,63 | 5,27E-01 |
| AT2G08860 | 0,31 | 0,62 | 3,59 | 0,17 | 8,63E-01 |
| AT2G37320 | 0,08 | 1,10 | 4,36 | 0,25 | 8,00E-01 |
| AT2G37330 | 0,66 | 2,48 | 3,05 | 0,81 | 4,16E-01 |
| AT2G37362 | 0,21 | 2,17 | 4,32 | 0,50 | 6,15E-01 |
| AT2G37370 | 0,40 | 0,42 | 4,38 | 0,09 | 9,24E-01 |
| AT2G37470 | 0,26 | 2,11 | 4,32 | 0,49 | 6,26E-01 |
| AT2G37480 | 1,35 | 2,69 | 2,17 | 1,24 | 2,14E-01 |
| AT2G37555 | 0,61 | 2,43 | 2,97 | 0,82 | 4,13E-01 |
| AT2G37580 | 0,12 | 0,90 | 4,36 | 0,21 | 8,36E-01 |
| AT2G37650 | 1,11 | 2,63 | 2,09 | 1,26 | 2,09E-01 |
| AT2G37690 | 0,31 | -0,31 | 4,38 | -0,07 | 9,44E-01 |
| AT2G37770 | 0,28 | 2,33 | 4,32 | 0,54 | 5,90E-01 |
| AT2G37860 | 0,83 | 1,23 | 2,31 | 0,53 | 5,93E-01 |
| AT2G37940 | 0,73 | -1,49 | 3,09 | -0,48 | 6,31E-01 |
| AT2G37970 | 0,77 | 1,40 | 2,70 | 0,52 | 6,03E-01 |
| AT2G37990 | 0,11 | 0,90 | 4,37 | 0,21 | 8,36E-01 |
| AT2G38010 | 0,53 | 2,39 | 3,05 | 0,78 | 4,33E-01 |
| AT2G38090 | 0,20 | 0,89 | 4,37 | 0,20 | 8,39E-01 |
| AT2G38120 | 0,38 | 1,13 | 2,80 | 0,40 | 6,86E-01 |
| AT2G38140 | 0,26 | 0,34 | 3,75 | 0,09 | 9,27E-01 |
| AT2G38210 | 1,49 | -2,15 | 1,91 | -1,13 | 2,59E-01 |
| AT2G38310 | 0,71 | 1,20 | 2,98 | 0,40 | 6,87E-01 |
| AT2G38365 | 0,20 | 1,07 | 4,36 | 0,25 | 8,06E-01 |
| AT2G38420 | 0,26 | 0,90 | 4,38 | 0,21 | 8,37E-01 |
| AT2G38470 | 1,30 | 1,96 | 1,87 | 1,05 | 2,93E-01 |
| AT2G38560 | 1,56 | 3,77 | 1,96 | 1,92 | 5,44E-02 |
| AT2G38590 | 0,78 | 2,02 | 2,74 | 0,74 | 4,60E-01 |
| AT2G38670 | 0,86 | -0,60 | 2,08 | -0,29 | 7,74E-01 |
| AT2G38760 | 0,08 | 1,10 | 4,36 | 0,25 | 8,00E-01 |
| AT2G38780 | 0,82 | 0,01 | 1,87 | 0,00 | 9,96E-01 |
| AT2G38820 | 0,12 | 1,39 | 4,34 | 0,32 | 7,49E-01 |
| AT2G38840 | 0,33 | 1,39 | 4,35 | 0,32 | 7,50E-01 |
| AT2G38940 | 0,25 | 1,57 | 4,34 | 0,36 | 7,18E-01 |
| AT2G38950 | 0,65 | 0,71 | 2,62 | 0,27 | 7,86E-01 |
| AT2G39190 | 0,15 | 1,16 | 4,35 | 0,27 | 7,90E-01 |

| | | | | | |
|---|---|---|---|---|---|
| AT2G39230 | 0,19 | 0,00 | 4,40 | 0,00 | 1,00E+00 |
| AT2G39270 | 0,08 | 0,85 | 4,38 | 0,20 | 8,45E-01 |
| AT2G39340 | 0,19 | 0,00 | 4,40 | 0,00 | 1,00E+00 |
| AT2G39400 | 0,15 | 1,25 | 4,35 | 0,29 | 7,75E-01 |
| AT2G39420 | 0,56 | -0,28 | 3,19 | -0,09 | 9,31E-01 |
| AT2G39430 | 0,52 | 2,58 | 3,86 | 0,67 | 5,04E-01 |
| AT2G39460 | 0,89 | 1,14 | 1,97 | 0,58 | 5,63E-01 |
| AT2G39470 | 0,81 | 0,81 | 2,19 | 0,37 | 7,11E-01 |
| AT2G39570 | 1,60 | 1,51 | 2,87 | 0,53 | 5,98E-01 |
| AT2G39580 | 1,16 | 1,60 | 2,12 | 0,76 | 4,50E-01 |
| AT2G39670 | 1,03 | 2,87 | 2,18 | 1,32 | 1,87E-01 |
| AT2G39725 | 0,50 | 2,17 | 3,91 | 0,56 | 5,78E-01 |
| AT2G39740 | 0,63 | 1,66 | 3,07 | 0,54 | 5,88E-01 |
| AT2G39780 | 1,17 | 0,84 | 2,21 | 0,38 | 7,02E-01 |
| AT2G39810 | 1,48 | 3,05 | 2,43 | 1,25 | 2,11E-01 |
| AT2G39840 | 0,84 | 2,96 | 2,50 | 1,18 | 2,36E-01 |
| AT2G39850 | 0,54 | -1,76 | 4,36 | -0,40 | 6,87E-01 |
| AT2G39870 | 0,43 | -0,78 | 4,32 | -0,18 | 8,57E-01 |
| AT2G39930 | 1,27 | 0,22 | 1,76 | 0,12 | 9,02E-01 |
| AT2G40060 | 0,99 | 2,95 | 2,38 | 1,24 | 2,16E-01 |
| AT2G40095 | 0,64 | 2,50 | 3,07 | 0,81 | 4,16E-01 |
| AT2G40130 | 0,14 | 1,24 | 4,35 | 0,29 | 7,75E-01 |
| AT2G40150 | 0,08 | 0,85 | 4,38 | 0,20 | 8,45E-01 |
| AT2G40160 | 0,42 | 1,61 | 4,13 | 0,39 | 6,97E-01 |
| AT2G40200 | 0,38 | 1,01 | 4,36 | 0,23 | 8,17E-01 |
| AT2G09120 | 0,31 | -2,83 | 4,33 | -0,65 | 5,13E-01 |
| AT2G40310 | 0,83 | 1,72 | 2,26 | 0,76 | 4,47E-01 |
| AT2G40400 | 0,82 | 2,14 | 2,72 | 0,79 | 4,31E-01 |
| AT2G40410 | 0,54 | 0,65 | 4,10 | 0,16 | 8,73E-01 |
| AT2G40435 | 0,46 | -0,25 | 4,37 | -0,06 | 9,54E-01 |
| AT2G40460 | 0,63 | -0,39 | 2,47 | -0,16 | 8,74E-01 |
| AT2G40480 | 0,31 | 1,19 | 4,36 | 0,27 | 7,86E-01 |
| AT2G40490 | 1,23 | -1,35 | 1,96 | -0,69 | 4,89E-01 |
| AT2G40540 | 0,78 | 0,61 | 2,43 | 0,25 | 8,03E-01 |
| AT2G40550 | 0,20 | 1,07 | 4,36 | 0,25 | 8,06E-01 |
| AT2G40660 | 0,08 | 0,00 | 4,40 | 0,00 | 1,00E+00 |
| AT2G40670 | 0,10 | 0,62 | 4,38 | 0,14 | 8,87E-01 |
| AT2G40760 | 0,35 | 1,01 | 4,33 | 0,23 | 8,15E-01 |
| AT2G40765 | 0,08 | 0,85 | 4,38 | 0,20 | 8,45E-01 |
| AT2G40810 | 0,08 | 1,10 | 4,36 | 0,25 | 8,00E-01 |
| AT2G40830 | 0,33 | 2,14 | 4,32 | 0,50 | 6,20E-01 |
| AT2G40880 | 1,07 | 0,62 | 2,08 | 0,30 | 7,66E-01 |
| AT2G40955 | 1,38 | 2,59 | 1,94 | 1,34 | 1,81E-01 |
| AT2G40980 | 0,41 | 0,46 | 4,17 | 0,11 | 9,13E-01 |
| AT2G41080 | 0,59 | 1,61 | 2,64 | 0,61 | 5,42E-01 |
| AT2G41110 | 0,26 | 0,33 | 3,79 | 0,09 | 9,31E-01 |
| AT2G09230 | 0,12 | 1,65 | 4,34 | 0,38 | 7,04E-01 |
| AT2G41140 | 0,37 | 1,95 | 4,27 | 0,46 | 6,48E-01 |
| AT2G41200 | 0,84 | -0,56 | 2,29 | -0,24 | 8,08E-01 |
| AT2G41210 | 0,42 | 1,95 | 4,14 | 0,47 | 6,37E-01 |
| AT2G41220 | 0,64 | 2,42 | 2,95 | 0,82 | 4,13E-01 |
| AT2G41460 | 0,13 | 1,15 | 4,36 | 0,26 | 7,92E-01 |
| AT2G41510 | 0,08 | 0,85 | 4,38 | 0,20 | 8,45E-01 |
| AT2G41520 | 0,59 | 1,61 | 3,01 | 0,54 | 5,92E-01 |
| AT2G41590 | 1,66 | 0,38 | 1,55 | 0,25 | 8,06E-01 |
| AT2G41670 | 0,12 | 1,65 | 4,34 | 0,38 | 7,04E-01 |

| | | | | | |
|---|---|---|---|---|---|
| AT2G41730 | 0,99 | 2,75 | 2,06 | 1,34 | 1,82E-01 |
| AT2G41820 | 0,34 | 1,85 | 4,34 | 0,43 | 6,69E-01 |
| AT2G41880 | 0,26 | 0,33 | 3,79 | 0,09 | 9,31E-01 |
| AT2G41920 | 1,42 | 2,15 | 2,02 | 1,06 | 2,88E-01 |
| AT2G42130 | 0,48 | -0,90 | 3,31 | -0,27 | 7,85E-01 |
| AT2G42220 | 0,98 | -1,40 | 1,90 | -0,74 | 4,61E-01 |
| AT2G42310 | 0,61 | 1,06 | 2,52 | 0,42 | 6,76E-01 |
| AT2G42320 | 0,63 | 0,29 | 2,55 | 0,11 | 9,10E-01 |
| AT2G42430 | 0,21 | 0,00 | 4,41 | 0,00 | 1,00E+00 |
| AT2G42580 | 1,32 | 1,84 | 1,81 | 1,02 | 3,10E-01 |
| AT2G42610 | 0,29 | 0,17 | 4,35 | 0,04 | 9,68E-01 |
| AT2G42620 | 1,45 | 3,07 | 2,39 | 1,28 | 1,99E-01 |
| AT2G42700 | 0,72 | 2,60 | 3,00 | 0,87 | 3,85E-01 |
| AT2G42710 | 0,12 | 0,79 | 4,37 | 0,18 | 8,57E-01 |
| AT2G42750 | 0,57 | 2,71 | 3,74 | 0,72 | 4,69E-01 |
| AT2G42780 | 0,28 | -1,36 | 4,41 | -0,31 | 7,58E-01 |
| AT2G42890 | 0,32 | 1,24 | 4,31 | 0,29 | 7,73E-01 |
| AT2G42900 | 0,43 | 0,00 | 4,41 | 0,00 | 1,00E+00 |
| AT2G43010 | 1,28 | -0,97 | 1,78 | -0,54 | 5,88E-01 |
| AT2G43020 | 1,32 | 2,97 | 2,07 | 1,43 | 1,52E-01 |
| AT2G43030 | 0,58 | -1,07 | 3,19 | -0,33 | 7,38E-01 |
| AT2G43060 | 1,34 | -2,23 | 1,57 | -1,41 | 1,57E-01 |
| AT2G43080 | 0,63 | 2,08 | 3,08 | 0,68 | 4,98E-01 |
| AT2G43090 | 0,61 | -0,28 | 2,68 | -0,10 | 9,17E-01 |
| AT2G43100 | 0,56 | 0,74 | 3,32 | 0,22 | 8,23E-01 |
| AT2G43137 | 1,43 | -0,21 | 1,79 | -0,12 | 9,07E-01 |
| AT2G43138 | 1,21 | 2,08 | 2,11 | 0,99 | 3,22E-01 |
| AT2G43141 | 1,64 | -0,55 | 1,86 | -0,29 | 7,68E-01 |
| AT2G43210 | 0,63 | 1,86 | 2,90 | 0,64 | 5,22E-01 |
| AT2G43230 | 0,55 | 1,30 | 3,84 | 0,34 | 7,36E-01 |
| AT2G43250 | 0,31 | 1,77 | 4,33 | 0,41 | 6,83E-01 |
| AT2G43280 | 0,56 | 0,77 | 4,37 | 0,18 | 8,60E-01 |
| AT2G09440 | 0,08 | 1,10 | 4,36 | 0,25 | 8,00E-01 |
| AT2G43340 | 0,44 | 0,53 | 3,42 | 0,15 | 8,77E-01 |
| AT2G43400 | 0,37 | 2,13 | 4,30 | 0,50 | 6,20E-01 |
| AT2G43440 | 0,11 | 0,00 | 4,41 | 0,00 | 1,00E+00 |
| AT2G43540 | 1,53 | 2,16 | 1,91 | 1,13 | 2,58E-01 |
| AT2G43630 | 0,44 | 2,58 | 4,05 | 0,64 | 5,25E-01 |
| AT2G43650 | 0,83 | 1,35 | 2,43 | 0,55 | 5,79E-01 |
| AT2G43690 | 0,68 | 2,97 | 4,31 | 0,69 | 4,91E-01 |
| AT2G43790 | 0,39 | 0,16 | 2,76 | 0,06 | 9,53E-01 |
| AT2G43800 | 0,08 | 0,00 | 4,40 | 0,00 | 1,00E+00 |
| AT2G43820 | 0,45 | 1,75 | 3,23 | 0,54 | 5,88E-01 |
| AT2G44080 | 0,60 | 0,00 | 3,83 | 0,00 | 1,00E+00 |
| AT2G44100 | 0,65 | 0,75 | 2,51 | 0,30 | 7,66E-01 |
| AT2G44130 | 0,72 | -0,94 | 2,42 | -0,39 | 6,98E-01 |
| AT2G44160 | 0,80 | 1,68 | 2,44 | 0,69 | 4,93E-01 |
| AT2G44175 | 0,43 | 0,61 | 4,16 | 0,15 | 8,84E-01 |
| AT2G44200 | 0,72 | 2,80 | 2,52 | 1,11 | 2,67E-01 |
| AT2G44210 | 1,13 | -1,85 | 1,88 | -0,98 | 3,25E-01 |
| AT2G44230 | 0,76 | -2,28 | 2,69 | -0,85 | 3,97E-01 |
| AT2G44240 | 0,17 | 0,45 | 4,39 | 0,10 | 9,19E-01 |
| AT2G44260 | 0,20 | 1,68 | 4,34 | 0,39 | 6,99E-01 |
| AT2G44350 | 0,40 | -0,58 | 3,44 | -0,17 | 8,65E-01 |
| AT2G44420 | 0,20 | 1,07 | 4,36 | 0,25 | 8,06E-01 |
| AT2G44430 | 0,96 | 0,72 | 1,91 | 0,38 | 7,05E-01 |

| | | | | | |
|---|---|---|---|---|---|
| AT2G44500 | 0,91 | 3,26 | 2,85 | 1,14 | 2,53E-01 |
| AT2G44520 | 0,23 | 1,33 | 4,34 | 0,31 | 7,60E-01 |
| AT2G44530 | 0,99 | 3,33 | 2,22 | 1,50 | 1,33E-01 |
| AT2G44600 | 0,10 | 0,62 | 4,38 | 0,14 | 8,87E-01 |
| AT2G44640 | 1,50 | 2,97 | 1,78 | 1,66 | 9,60E-02 |
| AT2G44670 | 1,15 | 1,08 | 2,00 | 0,54 | 5,88E-01 |
| AT2G44710 | 1,40 | 2,75 | 1,78 | 1,54 | 1,22E-01 |
| AT2G44735 | 0,42 | 2,18 | 4,09 | 0,53 | 5,94E-01 |
| AT2G44770 | 0,15 | 1,58 | 4,34 | 0,36 | 7,15E-01 |
| AT2G44800 | 0,19 | 0,37 | 4,38 | 0,09 | 9,32E-01 |
| AT2G44870 | 0,51 | -1,70 | 2,64 | -0,65 | 5,19E-01 |
| AT2G44880 | 0,91 | 2,81 | 2,71 | 1,04 | 3,00E-01 |
| AT2G44925 | 0,92 | -0,51 | 1,89 | -0,27 | 7,86E-01 |
| AT2G44940 | 0,52 | 0,41 | 2,76 | 0,15 | 8,82E-01 |
| AT2G45000 | 0,38 | -1,25 | 3,49 | -0,36 | 7,21E-01 |
| AT2G45020 | 0,36 | -1,07 | 4,30 | -0,25 | 8,03E-01 |
| AT2G45030 | 0,91 | 2,91 | 2,75 | 1,06 | 2,89E-01 |
| AT2G45140 | 0,33 | 0,26 | 4,36 | 0,06 | 9,52E-01 |
| AT2G45240 | 0,80 | 1,65 | 3,02 | 0,55 | 5,85E-01 |
| AT2G45250 | 0,26 | 0,90 | 4,38 | 0,21 | 8,37E-01 |
| AT2G45300 | 0,10 | 0,42 | 4,38 | 0,10 | 9,23E-01 |
| AT2G45310 | 0,67 | 1,10 | 2,51 | 0,44 | 6,61E-01 |
| AT2G45390 | 0,17 | -2,23 | 4,35 | -0,51 | 6,08E-01 |
| AT2G45440 | 0,53 | 0,82 | 3,89 | 0,21 | 8,33E-01 |
| AT2G45510 | 0,84 | 0,98 | 2,15 | 0,45 | 6,50E-01 |
| AT2G45530 | 0,98 | 0,07 | 1,92 | 0,04 | 9,70E-01 |
| AT2G09650 | 0,36 | -0,09 | 4,36 | -0,02 | 9,83E-01 |
| AT2G45590 | 0,37 | 0,83 | 4,32 | 0,19 | 8,47E-01 |
| AT2G45620 | 1,35 | 0,30 | 1,77 | 0,17 | 8,65E-01 |
| AT2G45630 | 0,80 | 2,67 | 2,53 | 1,06 | 2,91E-01 |
| AT2G45710 | 0,45 | 0,00 | 4,38 | 0,00 | 1,00E+00 |
| AT2G45720 | 0,29 | -0,19 | 4,37 | -0,04 | 9,66E-01 |
| AT2G45740 | 0,35 | -0,34 | 3,52 | -0,10 | 9,23E-01 |
| AT2G45790 | 0,17 | -1,64 | 4,37 | -0,37 | 7,08E-01 |
| AT2G45810 | 0,90 | 1,47 | 2,30 | 0,64 | 5,23E-01 |
| AT2G45910 | 0,97 | 2,68 | 2,06 | 1,30 | 1,92E-01 |
| AT2G46100 | 0,37 | -1,18 | 2,86 | -0,41 | 6,79E-01 |
| AT2G46130 | 1,45 | -1,97 | 1,55 | -1,28 | 2,02E-01 |
| AT2G46170 | 0,69 | 1,22 | 3,74 | 0,33 | 7,43E-01 |
| AT2G46190 | 0,70 | -0,67 | 2,18 | -0,31 | 7,60E-01 |
| AT2G09695 | 1,56 | -1,21 | 1,48 | -0,82 | 4,13E-01 |
| AT2G09705 | 0,89 | -1,17 | 2,46 | -0,48 | 6,33E-01 |
| AT2G46225 | 0,27 | 0,42 | 4,34 | 0,10 | 9,23E-01 |
| AT2G46240 | 0,70 | 0,97 | 2,45 | 0,39 | 6,93E-01 |
| AT2G46260 | 0,08 | 0,85 | 4,38 | 0,20 | 8,45E-01 |
| AT2G46270 | 0,25 | 1,03 | 4,37 | 0,24 | 8,13E-01 |
| AT2G46370 | 0,44 | 0,71 | 3,46 | 0,21 | 8,37E-01 |
| AT2G46380 | 1,50 | 2,96 | 2,20 | 1,35 | 1,78E-01 |
| AT2G46420 | 0,84 | -0,96 | 3,00 | -0,32 | 7,49E-01 |
| AT2G46440 | 1,43 | 1,79 | 1,67 | 1,07 | 2,84E-01 |
| AT2G46450 | 1,09 | 3,13 | 2,18 | 1,44 | 1,51E-01 |
| AT2G46490 | 0,75 | 3,21 | 2,83 | 1,13 | 2,57E-01 |
| AT2G46520 | 1,16 | 3,39 | 1,82 | 1,87 | 6,19E-02 |
| AT2G46570 | 0,51 | 2,52 | 3,99 | 0,63 | 5,27E-01 |
| AT2G46590 | 0,52 | 2,15 | 3,03 | 0,71 | 4,77E-01 |
| AT2G46600 | 1,11 | 1,20 | 1,92 | 0,62 | 5,32E-01 |

| | | | | | |
|---|---|---|---|---|---|
| AT2G46630 | 0,28 | 0,10 | 4,36 | 0,02 | 9,82E-01 |
| AT2G46650 | 0,42 | 1,70 | 3,32 | 0,51 | 6,09E-01 |
| AT2G46690 | 0,43 | 1,65 | 4,17 | 0,39 | 6,93E-01 |
| AT2G46710 | 0,30 | -0,43 | 3,72 | -0,11 | 9,09E-01 |
| AT2G46790 | 0,54 | 1,03 | 2,60 | 0,39 | 6,93E-01 |
| AT2G46800 | 1,02 | 1,06 | 1,96 | 0,54 | 5,91E-01 |
| AT2G46810 | 0,59 | -1,48 | 3,15 | -0,47 | 6,38E-01 |
| AT2G46900 | 1,50 | -0,50 | 1,89 | -0,27 | 7,89E-01 |
| AT2G46910 | 0,44 | -0,14 | 4,27 | -0,03 | 9,74E-01 |
| AT2G46990 | 0,08 | 0,85 | 4,38 | 0,20 | 8,45E-01 |
| AT2G47110 | 0,66 | 0,41 | 2,29 | 0,18 | 8,59E-01 |
| AT2G09815 | 0,23 | 0,61 | 4,38 | 0,14 | 8,90E-01 |
| AT2G47190 | 0,40 | 2,51 | 4,32 | 0,58 | 5,61E-01 |
| AT2G47220 | 0,25 | 0,91 | 4,32 | 0,21 | 8,34E-01 |
| AT2G47320 | 0,28 | -0,94 | 4,37 | -0,22 | 8,30E-01 |
| AT2G47420 | 0,42 | 0,22 | 4,09 | 0,05 | 9,57E-01 |
| AT2G47510 | 1,59 | 1,21 | 1,68 | 0,72 | 4,73E-01 |
| AT2G47610 | 1,29 | 2,77 | 1,72 | 1,61 | 1,07E-01 |
| AT2G47620 | 0,26 | 0,33 | 3,79 | 0,09 | 9,31E-01 |
| AT2G47630 | 0,42 | 0,41 | 2,83 | 0,14 | 8,86E-01 |
| AT2G47640 | 0,28 | 0,00 | 4,40 | 0,00 | 1,00E+00 |
| AT2G47690 | 0,55 | -0,17 | 3,99 | -0,04 | 9,67E-01 |
| AT2G47730 | 0,80 | -0,20 | 2,39 | -0,08 | 9,34E-01 |
| AT2G47820 | 1,37 | 3,36 | 2,48 | 1,35 | 1,76E-01 |
| AT2G47850 | 0,53 | 0,65 | 3,19 | 0,21 | 8,38E-01 |
| AT2G47910 | 0,92 | 1,46 | 2,08 | 0,70 | 4,84E-01 |
| AT2G47935 | 0,72 | 0,57 | 2,75 | 0,21 | 8,37E-01 |
| AT2G47960 | 1,39 | 2,58 | 1,91 | 1,35 | 1,76E-01 |
| AT2G47970 | 0,70 | 2,66 | 3,46 | 0,77 | 4,43E-01 |
| AT2G48010 | 0,33 | 1,42 | 4,36 | 0,33 | 7,45E-01 |
| AT2G48020 | 0,56 | 1,98 | 3,24 | 0,61 | 5,41E-01 |
| AT2G48110 | 1,17 | 1,01 | 2,16 | 0,47 | 6,39E-01 |
| AT2G48120 | 0,32 | 0,52 | 4,34 | 0,12 | 9,05E-01 |
| AT3G01040 | 0,51 | 1,31 | 3,09 | 0,42 | 6,71E-01 |
| AT3G01050 | 0,18 | -0,04 | 4,35 | -0,01 | 9,92E-01 |
| AT3G01080 | 0,17 | 1,03 | 4,37 | 0,24 | 8,13E-01 |
| AT3G01085 | 1,09 | 0,73 | 1,82 | 0,40 | 6,86E-01 |
| AT3G01150 | 0,45 | 0,41 | 3,32 | 0,12 | 9,01E-01 |
| AT3G01170 | 0,42 | 1,95 | 4,14 | 0,47 | 6,37E-01 |
| AT3G01175 | 0,08 | 1,10 | 4,36 | 0,25 | 8,00E-01 |
| AT3G01210 | 0,10 | 0,42 | 4,38 | 0,10 | 9,23E-01 |
| AT3G01280 | 1,02 | -1,31 | 1,95 | -0,67 | 5,02E-01 |
| AT3G01320 | 1,14 | 3,02 | 1,86 | 1,62 | 1,05E-01 |
| AT3G01440 | 0,31 | -1,59 | 4,39 | -0,36 | 7,17E-01 |
| AT3G01475 | 1,05 | 2,59 | 1,95 | 1,33 | 1,84E-01 |
| AT3G01480 | 1,15 | -1,59 | 1,92 | -0,83 | 4,09E-01 |
| AT3G01560 | 0,68 | 1,05 | 2,52 | 0,42 | 6,77E-01 |
| AT3G01590 | 0,40 | 1,87 | 4,35 | 0,43 | 6,67E-01 |
| AT3G01610 | 0,25 | 1,17 | 4,36 | 0,27 | 7,88E-01 |
| AT3G01660 | 0,12 | 1,39 | 4,34 | 0,32 | 7,49E-01 |
| AT3G01690 | 1,08 | -0,98 | 1,95 | -0,50 | 6,14E-01 |
| AT3G01740 | 0,28 | 1,08 | 4,32 | 0,25 | 8,02E-01 |
| AT3G01750 | 0,72 | 0,94 | 2,98 | 0,31 | 7,53E-01 |
| AT3G01780 | 1,57 | 3,28 | 1,71 | 1,92 | 5,47E-02 |
| AT3G01810 | 0,37 | -1,62 | 4,33 | -0,37 | 7,08E-01 |
| AT3G01830 | 0,94 | 2,11 | 2,28 | 0,93 | 3,54E-01 |

| | | | | | |
|---|---|---|---|---|---|
| AT3G01920 | 0,37 | 0,26 | 4,38 | 0,06 | 9,53E-01 |
| AT3G02010 | 0,37 | 1,54 | 4,34 | 0,35 | 7,23E-01 |
| AT3G02030 | 0,12 | 0,90 | 4,36 | 0,21 | 8,36E-01 |
| AT3G02050 | 0,37 | 1,64 | 4,28 | 0,38 | 7,03E-01 |
| AT3G02060 | 1,23 | 1,11 | 2,03 | 0,54 | 5,86E-01 |
| AT3G02065 | 0,54 | 2,34 | 3,77 | 0,62 | 5,35E-01 |
| AT3G02070 | 0,35 | 1,07 | 3,56 | 0,30 | 7,65E-01 |
| AT3G02130 | 1,18 | 3,34 | 2,19 | 1,52 | 1,28E-01 |
| AT3G02280 | 0,25 | 0,00 | 4,41 | 0,00 | 1,00E+00 |
| AT3G02290 | 0,16 | 0,90 | 4,38 | 0,21 | 8,37E-01 |
| AT3G02300 | 0,16 | 0,00 | 4,41 | 0,00 | 1,00E+00 |
| AT3G02340 | 0,50 | -1,06 | 4,26 | -0,25 | 8,03E-01 |
| AT3G02350 | 0,84 | -0,76 | 2,17 | -0,35 | 7,25E-01 |
| AT3G02360 | 1,53 | 1,97 | 1,88 | 1,05 | 2,95E-01 |
| AT3G02370 | 0,47 | 0,10 | 3,28 | 0,03 | 9,75E-01 |
| AT3G02400 | 0,66 | 0,00 | 4,41 | 0,00 | 1,00E+00 |
| AT3G02490 | 0,33 | 1,40 | 4,35 | 0,32 | 7,47E-01 |
| AT3G02560 | 0,49 | 2,47 | 3,90 | 0,63 | 5,26E-01 |
| AT3G02610 | 0,81 | -1,57 | 2,78 | -0,57 | 5,71E-01 |
| AT3G02630 | 0,37 | 0,77 | 4,32 | 0,18 | 8,59E-01 |
| AT3G02690 | 0,66 | 1,63 | 2,58 | 0,63 | 5,28E-01 |
| AT3G02720 | 0,46 | 1,05 | 4,35 | 0,24 | 8,10E-01 |
| AT3G02830 | 1,22 | 3,10 | 2,20 | 1,41 | 1,60E-01 |
| AT3G02832 | 1,51 | 0,24 | 1,63 | 0,14 | 8,85E-01 |
| AT3G01145 | 0,57 | 0,84 | 2,58 | 0,33 | 7,44E-01 |
| AT3G02880 | 1,19 | 2,26 | 1,86 | 1,21 | 2,24E-01 |
| AT3G02900 | 0,28 | 0,00 | 4,41 | 0,00 | 1,00E+00 |
| AT3G02910 | 0,77 | 1,08 | 2,40 | 0,45 | 6,54E-01 |
| AT3G03020 | 0,43 | 2,12 | 4,15 | 0,51 | 6,10E-01 |
| AT3G03030 | 0,40 | 2,51 | 4,32 | 0,58 | 5,61E-01 |
| AT3G03040 | 0,24 | 0,65 | 4,37 | 0,15 | 8,81E-01 |
| AT3G03110 | 0,55 | 2,30 | 4,32 | 0,53 | 5,94E-01 |
| AT3G03160 | 0,47 | 2,44 | 3,98 | 0,61 | 5,40E-01 |
| AT3G03210 | 0,10 | 0,00 | 4,41 | 0,00 | 1,00E+00 |
| AT3G03320 | 0,15 | 0,00 | 4,39 | 0,00 | 1,00E+00 |
| AT3G03340 | 0,40 | 2,55 | 4,32 | 0,59 | 5,55E-01 |
| AT3G03350 | 0,24 | 1,58 | 4,34 | 0,36 | 7,16E-01 |
| AT3G03360 | 0,08 | 1,10 | 4,36 | 0,25 | 8,00E-01 |
| AT3G03610 | 0,27 | -1,54 | 4,35 | -0,35 | 7,24E-01 |
| AT3G03640 | 0,25 | -0,65 | 4,35 | -0,15 | 8,82E-01 |
| AT3G03680 | 1,60 | -0,18 | 1,58 | -0,11 | 9,09E-01 |
| AT3G03860 | 0,25 | 0,43 | 4,39 | 0,10 | 9,21E-01 |
| AT3G03880 | 0,28 | 1,28 | 4,36 | 0,29 | 7,70E-01 |
| AT3G03950 | 0,32 | -0,15 | 3,52 | -0,04 | 9,65E-01 |
| AT3G03960 | 1,45 | 1,79 | 1,53 | 1,17 | 2,43E-01 |
| AT3G04140 | 0,99 | 0,81 | 2,18 | 0,37 | 7,10E-01 |
| AT3G04490 | 0,26 | 0,00 | 4,41 | 0,00 | 1,00E+00 |
| AT3G04530 | 0,08 | 1,10 | 4,36 | 0,25 | 8,00E-01 |
| AT3G04550 | 0,64 | -1,23 | 2,05 | -0,60 | 5,50E-01 |
| AT3G04790 | 1,26 | -1,15 | 1,94 | -0,59 | 5,53E-01 |
| AT3G04810 | 0,59 | 2,35 | 4,31 | 0,55 | 5,86E-01 |
| AT3G04840 | 0,80 | 1,14 | 2,36 | 0,48 | 6,29E-01 |
| AT3G04870 | 0,49 | -2,44 | 3,16 | -0,77 | 4,41E-01 |
| AT3G04910 | 0,86 | 0,47 | 1,85 | 0,26 | 7,98E-01 |
| AT3G04920 | 0,51 | -0,85 | 4,05 | -0,21 | 8,34E-01 |
| AT3G04940 | 0,44 | 0,77 | 4,15 | 0,19 | 8,52E-01 |

| | | | | | |
|---|---|---|---|---|---|
| AT3G01555 | 0,12 | 0,79 | 4,37 | 0,18 | 8,57E-01 |
| AT3G01575 | 0,10 | 0,00 | 4,41 | 0,00 | 1,00E+00 |
| AT3G05070 | 0,17 | -0,46 | 4,37 | -0,11 | 9,16E-01 |
| AT3G05120 | 0,72 | 1,17 | 2,20 | 0,53 | 5,95E-01 |
| AT3G05200 | 0,41 | 1,07 | 4,24 | 0,25 | 8,01E-01 |
| AT3G05230 | 0,28 | 1,56 | 4,36 | 0,36 | 7,21E-01 |
| AT3G05380 | 0,92 | 1,70 | 2,25 | 0,76 | 4,50E-01 |
| AT3G05410 | 1,68 | 1,01 | 1,58 | 0,64 | 5,20E-01 |
| AT3G05430 | 0,21 | 1,99 | 4,33 | 0,46 | 6,46E-01 |
| AT3G05490 | 0,28 | -1,36 | 4,41 | -0,31 | 7,58E-01 |
| AT3G05500 | 0,55 | 1,77 | 2,67 | 0,66 | 5,08E-01 |
| AT3G05520 | 0,43 | 0,36 | 2,86 | 0,13 | 8,99E-01 |
| AT3G05545 | 0,82 | 1,57 | 2,28 | 0,69 | 4,92E-01 |
| AT3G05560 | 0,45 | -0,88 | 3,22 | -0,27 | 7,85E-01 |
| AT3G05630 | 0,21 | 2,17 | 4,32 | 0,50 | 6,15E-01 |
| AT3G05670 | 0,41 | 2,09 | 4,20 | 0,50 | 6,19E-01 |
| AT3G05710 | 0,95 | 0,95 | 2,18 | 0,43 | 6,65E-01 |
| AT3G05725 | 0,64 | -1,63 | 2,45 | -0,66 | 5,08E-01 |
| AT3G05750 | 0,61 | 3,02 | 3,64 | 0,83 | 4,08E-01 |
| AT3G05760 | 0,68 | 3,01 | 3,63 | 0,83 | 4,07E-01 |
| AT3G05830 | 0,23 | 0,00 | 4,38 | 0,00 | 1,00E+00 |
| AT3G05840 | 0,50 | 1,10 | 3,99 | 0,28 | 7,82E-01 |
| AT3G05905 | 0,51 | -1,37 | 3,20 | -0,43 | 6,67E-01 |
| AT3G06125 | 0,60 | 2,01 | 2,29 | 0,88 | 3,81E-01 |
| AT3G06170 | 0,51 | 0,95 | 3,06 | 0,31 | 7,56E-01 |
| AT3G06180 | 0,25 | 2,36 | 4,32 | 0,55 | 5,86E-01 |
| AT3G06200 | 0,41 | 1,34 | 4,36 | 0,31 | 7,58E-01 |
| AT3G06210 | 0,20 | 1,07 | 4,36 | 0,25 | 8,06E-01 |
| AT3G06240 | 0,61 | -0,03 | 2,23 | -0,01 | 9,88E-01 |
| AT3G06380 | 0,55 | 1,14 | 3,84 | 0,30 | 7,65E-01 |
| AT3G06400 | 0,95 | 2,90 | 2,57 | 1,13 | 2,59E-01 |
| AT3G06455 | 0,58 | 0,68 | 4,35 | 0,16 | 8,77E-01 |
| AT3G06470 | 0,36 | 0,82 | 4,32 | 0,19 | 8,49E-01 |
| AT3G06500 | 1,24 | 1,80 | 2,16 | 0,83 | 4,05E-01 |
| AT3G06520 | 0,50 | 2,05 | 3,90 | 0,53 | 6,00E-01 |
| AT3G06610 | 0,15 | 0,71 | 4,37 | 0,16 | 8,71E-01 |
| AT3G06660 | 0,08 | 0,00 | 4,40 | 0,00 | 1,00E+00 |
| AT3G06730 | 0,97 | 0,52 | 3,17 | 0,16 | 8,70E-01 |
| AT3G06740 | 0,50 | 0,62 | 4,15 | 0,15 | 8,82E-01 |
| AT3G06810 | 1,67 | 2,46 | 1,64 | 1,51 | 1,32E-01 |
| AT3G06830 | 0,08 | 1,10 | 4,36 | 0,25 | 8,00E-01 |
| AT3G06860 | 1,03 | 0,30 | 1,97 | 0,15 | 8,78E-01 |
| AT3G06868 | 0,29 | -1,29 | 4,40 | -0,29 | 7,70E-01 |
| AT3G06895 | 1,19 | 1,41 | 1,96 | 0,72 | 4,71E-01 |
| AT3G06920 | 0,42 | 0,42 | 4,26 | 0,10 | 9,21E-01 |
| AT3G06980 | 0,81 | 0,31 | 2,85 | 0,11 | 9,14E-01 |
| AT3G07110 | 1,39 | 1,20 | 1,74 | 0,69 | 4,90E-01 |
| AT3G07200 | 0,08 | 1,10 | 4,36 | 0,25 | 8,00E-01 |
| AT3G07230 | 0,29 | -1,36 | 4,33 | -0,31 | 7,53E-01 |
| AT3G07270 | 0,85 | 1,06 | 2,13 | 0,50 | 6,19E-01 |
| AT3G07350 | 0,42 | 0,78 | 3,28 | 0,24 | 8,12E-01 |
| AT3G07360 | 1,00 | 2,38 | 2,44 | 0,98 | 3,28E-01 |
| AT3G07400 | 0,87 | 2,16 | 2,18 | 0,99 | 3,23E-01 |
| AT3G07430 | 0,32 | -0,88 | 4,34 | -0,20 | 8,39E-01 |
| AT3G07440 | 0,08 | 1,10 | 4,36 | 0,25 | 8,00E-01 |
| AT3G07480 | 0,45 | 1,71 | 4,25 | 0,40 | 6,88E-01 |

| | | | | | |
|---|---|---|---|---|---|
| AT3G07565 | 0,19 | 0,79 | 4,37 | 0,18 | 8,56E-01 |
| AT3G07610 | 0,83 | 0,34 | 2,66 | 0,13 | 8,99E-01 |
| AT3G07640 | 0,17 | 1,94 | 4,33 | 0,45 | 6,55E-01 |
| AT3G07650 | 0,97 | 3,25 | 3,30 | 0,99 | 3,24E-01 |
| AT3G07680 | 0,59 | 1,46 | 3,24 | 0,45 | 6,52E-01 |
| AT3G07690 | 0,32 | 1,20 | 4,35 | 0,28 | 7,82E-01 |
| AT3G07770 | 1,44 | 1,45 | 1,78 | 0,81 | 4,17E-01 |
| AT3G07780 | 1,15 | 1,40 | 2,07 | 0,68 | 5,00E-01 |
| AT3G07810 | 0,90 | 0,22 | 2,05 | 0,11 | 9,15E-01 |
| AT3G07880 | 0,28 | 2,15 | 4,32 | 0,50 | 6,20E-01 |
| AT3G07950 | 0,43 | 1,49 | 4,32 | 0,35 | 7,29E-01 |
| AT3G08010 | 0,17 | -1,25 | 4,39 | -0,28 | 7,76E-01 |
| AT3G08030 | 1,37 | -1,63 | 1,60 | -1,02 | 3,08E-01 |
| AT3G08550 | 0,31 | 0,78 | 4,36 | 0,18 | 8,58E-01 |
| AT3G08560 | 1,47 | 0,20 | 1,72 | 0,12 | 9,08E-01 |
| AT3G08600 | 0,52 | 1,71 | 3,88 | 0,44 | 6,59E-01 |
| AT3G08640 | 0,20 | 0,32 | 4,35 | 0,07 | 9,41E-01 |
| AT3G08690 | 0,22 | 0,42 | 4,38 | 0,10 | 9,23E-01 |
| AT3G08920 | 0,32 | 0,43 | 4,37 | 0,10 | 9,22E-01 |
| AT3G08930 | 0,51 | 1,11 | 3,20 | 0,35 | 7,29E-01 |
| AT3G08950 | 0,44 | 0,00 | 4,27 | 0,00 | 1,00E+00 |
| AT3G08980 | 0,33 | 0,44 | 4,37 | 0,10 | 9,20E-01 |
| AT3G09020 | 0,57 | 2,15 | 3,96 | 0,54 | 5,87E-01 |
| AT3G09030 | 0,51 | 1,60 | 2,64 | 0,60 | 5,45E-01 |
| AT3G09035 | 0,19 | 1,64 | 4,33 | 0,38 | 7,05E-01 |
| AT3G09085 | 0,40 | 2,24 | 3,38 | 0,66 | 5,07E-01 |
| AT3G02105 | 0,40 | 2,51 | 4,32 | 0,58 | 5,61E-01 |
| AT3G09180 | 0,23 | 0,79 | 4,37 | 0,18 | 8,56E-01 |
| AT3G09300 | 1,07 | -0,48 | 2,44 | -0,20 | 8,43E-01 |
| AT3G09350 | 0,51 | -2,52 | 4,35 | -0,58 | 5,62E-01 |
| AT3G09370 | 0,29 | 1,89 | 4,33 | 0,44 | 6,63E-01 |
| AT3G09430 | 1,27 | 3,94 | 2,03 | 1,94 | 5,25E-02 |
| AT3G09500 | 0,81 | -0,15 | 2,13 | -0,07 | 9,42E-01 |
| AT3G09510 | 0,52 | -0,31 | 3,17 | -0,10 | 9,23E-01 |
| AT3G09570 | 0,70 | 1,47 | 2,48 | 0,60 | 5,51E-01 |
| AT3G09600 | 0,41 | 1,55 | 3,32 | 0,47 | 6,41E-01 |
| AT3G09650 | 0,58 | 2,95 | 4,31 | 0,68 | 4,94E-01 |
| AT3G09670 | 0,48 | 2,55 | 4,32 | 0,59 | 5,55E-01 |
| AT3G09690 | 0,55 | 0,91 | 4,38 | 0,21 | 8,35E-01 |
| AT3G09810 | 0,32 | 0,76 | 4,37 | 0,17 | 8,63E-01 |
| AT3G09820 | 0,77 | -0,86 | 2,25 | -0,38 | 7,01E-01 |
| AT3G09830 | 0,46 | 0,26 | 3,29 | 0,08 | 9,37E-01 |
| AT3G09880 | 0,75 | 3,02 | 2,48 | 1,22 | 2,23E-01 |
| AT3G09890 | 0,43 | 1,07 | 4,33 | 0,25 | 8,05E-01 |
| AT3G09900 | 0,19 | 1,88 | 4,33 | 0,43 | 6,64E-01 |
| AT3G10060 | 0,36 | 1,26 | 4,35 | 0,29 | 7,72E-01 |
| AT3G10120 | 0,12 | 1,46 | 4,35 | 0,34 | 7,36E-01 |
| AT3G10190 | 0,32 | 2,41 | 4,32 | 0,56 | 5,76E-01 |
| AT3G10200 | 0,08 | 1,10 | 4,36 | 0,25 | 8,00E-01 |
| AT3G10300 | 0,68 | -1,12 | 2,94 | -0,38 | 7,04E-01 |
| AT3G10310 | 0,16 | 1,07 | 4,37 | 0,24 | 8,07E-01 |
| AT3G10340 | 0,80 | -1,02 | 3,02 | -0,34 | 7,36E-01 |
| AT3G10370 | 0,28 | 1,77 | 4,33 | 0,41 | 6,83E-01 |
| AT3G10390 | 0,49 | 1,07 | 3,25 | 0,33 | 7,41E-01 |
| AT3G10420 | 1,49 | 1,69 | 1,62 | 1,05 | 2,96E-01 |
| AT3G10480 | 0,72 | 2,88 | 3,48 | 0,83 | 4,07E-01 |

| | | | | | |
|---|---|---|---|---|---|
| AT3G10490 | 0,18 | -0,04 | 4,35 | -0,01 | 9,92E-01 |
| AT3G10527 | 0,27 | 1,01 | 4,33 | 0,23 | 8,16E-01 |
| AT3G10540 | 0,56 | -0,04 | 2,65 | -0,02 | 9,88E-01 |
| AT3G10570 | 0,12 | 0,79 | 4,37 | 0,18 | 8,57E-01 |
| AT3G10760 | 0,48 | 1,09 | 3,34 | 0,33 | 7,43E-01 |
| AT3G10800 | 1,54 | 3,46 | 1,96 | 1,77 | 7,67E-02 |
| AT3G10820 | 0,24 | 0,00 | 4,41 | 0,00 | 1,00E+00 |
| AT3G10860 | 0,23 | 1,68 | 4,33 | 0,39 | 6,97E-01 |
| AT3G10920 | 1,12 | -1,69 | 1,82 | -0,93 | 3,55E-01 |
| AT3G10950 | 0,19 | 0,41 | 4,38 | 0,09 | 9,25E-01 |
| AT3G10980 | 0,79 | 2,21 | 3,13 | 0,71 | 4,79E-01 |
| AT3G11020 | 0,05 | 0,00 | 4,41 | 0,00 | 1,00E+00 |
| AT3G11080 | 0,18 | 0,63 | 4,37 | 0,14 | 8,86E-01 |
| AT3G11090 | 0,60 | 2,11 | 3,17 | 0,67 | 5,05E-01 |
| AT3G11120 | 0,39 | -1,25 | 3,43 | -0,36 | 7,16E-01 |
| AT3G11170 | 1,02 | -1,20 | 1,68 | -0,71 | 4,77E-01 |
| AT3G11230 | 0,53 | -1,97 | 4,37 | -0,45 | 6,52E-01 |
| AT3G11250 | 0,11 | 0,00 | 4,41 | 0,00 | 1,00E+00 |
| AT3G11270 | 0,10 | 0,42 | 4,38 | 0,10 | 9,23E-01 |
| AT3G11397 | 0,44 | -2,80 | 4,12 | -0,68 | 4,97E-01 |
| AT3G11400 | 0,87 | 1,53 | 2,37 | 0,65 | 5,17E-01 |
| AT3G11420 | 0,26 | -0,46 | 4,37 | -0,11 | 9,15E-01 |
| AT3G11440 | 0,65 | 2,54 | 2,96 | 0,86 | 3,91E-01 |
| AT3G02465 | 0,30 | 1,32 | 4,34 | 0,30 | 7,61E-01 |
| AT3G11630 | 0,96 | 0,64 | 1,72 | 0,37 | 7,10E-01 |
| AT3G11700 | 0,50 | 0,42 | 3,22 | 0,13 | 8,95E-01 |
| AT3G11760 | 0,23 | 1,33 | 4,34 | 0,31 | 7,60E-01 |
| AT3G11810 | 0,52 | 0,95 | 3,82 | 0,25 | 8,03E-01 |
| AT3G11900 | 0,13 | 1,15 | 4,36 | 0,26 | 7,92E-01 |
| AT3G11930 | 1,35 | -1,68 | 3,34 | -0,50 | 6,15E-01 |
| AT3G11960 | 1,04 | 2,26 | 1,96 | 1,15 | 2,50E-01 |
| AT3G12010 | 0,40 | 0,43 | 4,37 | 0,10 | 9,23E-01 |
| AT3G12020 | 0,79 | 2,20 | 2,40 | 0,92 | 3,59E-01 |
| AT3G12050 | 0,32 | 1,60 | 3,53 | 0,45 | 6,49E-01 |
| AT3G12080 | 1,08 | 2,55 | 2,51 | 1,02 | 3,08E-01 |
| AT3G12270 | 0,77 | -0,50 | 2,61 | -0,19 | 8,49E-01 |
| AT3G12290 | 0,81 | 1,19 | 2,56 | 0,46 | 6,43E-01 |
| AT3G12320 | 0,90 | 0,63 | 2,14 | 0,30 | 7,67E-01 |
| AT3G12360 | 0,71 | 0,55 | 2,52 | 0,22 | 8,29E-01 |
| AT3G12390 | 0,69 | 0,93 | 2,42 | 0,39 | 7,00E-01 |
| AT3G12480 | 0,26 | 0,95 | 4,33 | 0,22 | 8,26E-01 |
| AT3G12502 | 0,08 | 1,10 | 4,36 | 0,25 | 8,00E-01 |
| AT3G12570 | 1,24 | 1,41 | 2,56 | 0,55 | 5,81E-01 |
| AT3G12600 | 0,05 | 0,00 | 4,41 | 0,00 | 1,00E+00 |
| AT3G12640 | 0,60 | 1,07 | 2,28 | 0,47 | 6,39E-01 |
| AT3G12700 | 0,20 | 0,00 | 4,41 | 0,00 | 1,00E+00 |
| AT3G12740 | 0,52 | 1,63 | 3,32 | 0,49 | 6,23E-01 |
| AT3G12760 | 0,80 | 1,19 | 2,33 | 0,51 | 6,12E-01 |
| AT3G12810 | 1,56 | 3,54 | 1,79 | 1,98 | 4,74E-02 |
| AT3G12915 | 0,53 | 2,47 | 4,32 | 0,57 | 5,68E-01 |
| AT3G13060 | 0,87 | 1,81 | 2,27 | 0,80 | 4,26E-01 |
| AT3G13062 | 1,32 | 1,19 | 1,59 | 0,75 | 4,55E-01 |
| AT3G13065 | 0,88 | 3,15 | 3,37 | 0,93 | 3,50E-01 |
| AT3G13200 | 1,18 | 1,01 | 1,78 | 0,56 | 5,73E-01 |
| AT3G13210 | 0,13 | 1,15 | 4,36 | 0,26 | 7,92E-01 |
| AT3G13340 | 1,00 | -3,03 | 2,17 | -1,40 | 1,62E-01 |

| | | | | | |
|---|---|---|---|---|---|
| AT3G13370 | 0,63 | 0,00 | 4,41 | 0,00 | 1,00E+00 |
| AT3G13380 | 0,67 | 2,86 | 2,44 | 1,17 | 2,41E-01 |
| AT3G13437 | 0,08 | 0,85 | 4,38 | 0,20 | 8,45E-01 |
| AT3G13445 | 0,19 | -0,05 | 4,35 | -0,01 | 9,91E-01 |
| AT3G13470 | 0,61 | 2,29 | 3,89 | 0,59 | 5,55E-01 |
| AT3G02725 | 1,24 | -1,41 | 1,92 | -0,73 | 4,63E-01 |
| AT3G13662 | 0,34 | 0,56 | 4,37 | 0,13 | 8,98E-01 |
| AT3G13690 | 0,67 | -0,25 | 3,88 | -0,07 | 9,48E-01 |
| AT3G13760 | 0,08 | 0,85 | 4,38 | 0,20 | 8,45E-01 |
| AT3G13770 | 0,43 | 0,95 | 4,36 | 0,22 | 8,28E-01 |
| AT3G13810 | 0,42 | 0,47 | 4,22 | 0,11 | 9,11E-01 |
| AT3G13930 | 0,81 | -1,21 | 2,27 | -0,53 | 5,94E-01 |
| AT3G13950 | 0,55 | 1,00 | 3,98 | 0,25 | 8,01E-01 |
| AT3G14020 | 0,35 | 0,00 | 4,39 | 0,00 | 1,00E+00 |
| AT3G14100 | 0,69 | 1,69 | 2,56 | 0,66 | 5,09E-01 |
| AT3G14120 | 1,41 | 2,88 | 2,05 | 1,41 | 1,60E-01 |
| AT3G14160 | 0,19 | 1,57 | 4,34 | 0,36 | 7,18E-01 |
| AT3G14220 | 0,64 | -1,36 | 3,84 | -0,35 | 7,24E-01 |
| AT3G14290 | 0,42 | 0,85 | 4,10 | 0,21 | 8,36E-01 |
| AT3G14560 | 0,20 | 1,07 | 4,36 | 0,25 | 8,06E-01 |
| AT3G14595 | 1,01 | 0,73 | 2,03 | 0,36 | 7,20E-01 |
| AT3G14600 | 1,18 | 1,88 | 1,74 | 1,08 | 2,80E-01 |
| AT3G14610 | 0,28 | 0,42 | 4,38 | 0,10 | 9,24E-01 |
| AT3G14620 | 0,63 | 0,66 | 2,55 | 0,26 | 7,96E-01 |
| AT3G14640 | 0,33 | 1,33 | 4,35 | 0,31 | 7,59E-01 |
| AT3G14660 | 1,69 | 0,95 | 1,34 | 0,71 | 4,80E-01 |
| AT3G14670 | 1,07 | 0,71 | 1,97 | 0,36 | 7,19E-01 |
| AT3G14690 | 0,63 | 0,16 | 2,22 | 0,07 | 9,41E-01 |
| AT3G14720 | 0,35 | 0,41 | 4,38 | 0,09 | 9,25E-01 |
| AT3G14790 | 0,27 | 0,00 | 4,40 | 0,00 | 1,00E+00 |
| AT3G14830 | 0,30 | 1,76 | 4,33 | 0,41 | 6,85E-01 |
| AT3G14870 | 0,25 | 2,01 | 4,33 | 0,46 | 6,43E-01 |
| AT3G14920 | 0,30 | 1,68 | 4,34 | 0,39 | 6,99E-01 |
| AT3G14940 | 0,68 | 2,96 | 3,54 | 0,84 | 4,03E-01 |
| AT3G14990 | 0,39 | 0,93 | 2,84 | 0,33 | 7,42E-01 |
| AT3G15000 | 0,95 | 0,94 | 2,33 | 0,40 | 6,87E-01 |
| AT3G15010 | 0,55 | 2,20 | 3,05 | 0,72 | 4,71E-01 |
| AT3G15020 | 1,02 | 2,44 | 2,42 | 1,01 | 3,14E-01 |
| AT3G15030 | 0,69 | -0,39 | 3,08 | -0,13 | 8,98E-01 |
| AT3G15090 | 0,12 | 1,65 | 4,34 | 0,38 | 7,04E-01 |
| AT3G15095 | 1,44 | 0,26 | 1,46 | 0,18 | 8,56E-01 |
| AT3G15160 | 1,08 | 2,43 | 2,22 | 1,10 | 2,73E-01 |
| AT3G15180 | 0,21 | 0,00 | 4,38 | 0,00 | 1,00E+00 |
| AT3G15190 | 1,26 | 0,18 | 1,74 | 0,10 | 9,17E-01 |
| AT3G15260 | 0,39 | 1,26 | 4,27 | 0,29 | 7,69E-01 |
| AT3G15290 | 0,26 | 1,08 | 4,37 | 0,25 | 8,05E-01 |
| AT3G15350 | 1,19 | 1,23 | 2,33 | 0,53 | 5,99E-01 |
| AT3G15355 | 0,91 | 3,26 | 3,73 | 0,87 | 3,83E-01 |
| AT3G15360 | 1,20 | -1,45 | 1,74 | -0,84 | 4,03E-01 |
| AT3G15380 | 0,12 | 1,65 | 4,34 | 0,38 | 7,04E-01 |
| AT3G15450 | 0,77 | 0,58 | 2,23 | 0,26 | 7,95E-01 |
| AT3G15518 | 0,17 | 1,40 | 4,35 | 0,32 | 7,48E-01 |
| AT3G15550 | 0,17 | -0,02 | 4,36 | 0,00 | 9,97E-01 |
| AT3G03145 | 0,66 | -1,60 | 3,04 | -0,53 | 5,99E-01 |
| AT3G15635 | 0,25 | 0,44 | 4,37 | 0,10 | 9,20E-01 |
| AT3G15750 | 0,08 | 0,85 | 4,38 | 0,20 | 8,45E-01 |

| | | | | | |
|---|---|---|---|---|---|
| AT3G15760 | 0,84 | 1,39 | 2,57 | 0,54 | 5,87E-01 |
| AT3G15790 | 0,67 | 3,18 | 2,84 | 1,12 | 2,62E-01 |
| AT3G15820 | 0,32 | 2,33 | 4,32 | 0,54 | 5,90E-01 |
| AT3G15850 | 0,57 | -1,74 | 2,68 | -0,65 | 5,17E-01 |
| AT3G15930 | 0,12 | 1,58 | 4,34 | 0,36 | 7,16E-01 |
| AT3G16010 | 0,15 | 0,42 | 4,38 | 0,09 | 9,24E-01 |
| AT3G16060 | 0,47 | -0,32 | 4,09 | -0,08 | 9,37E-01 |
| AT3G16080 | 0,49 | 1,29 | 3,27 | 0,40 | 6,92E-01 |
| AT3G16130 | 0,87 | 0,08 | 2,05 | 0,04 | 9,68E-01 |
| AT3G16175 | 0,08 | 0,85 | 4,38 | 0,20 | 8,45E-01 |
| AT3G16240 | 1,39 | -3,49 | 1,81 | -1,92 | 5,43E-02 |
| AT3G16260 | 0,72 | 2,36 | 3,51 | 0,67 | 5,02E-01 |
| AT3G16270 | 0,71 | 0,07 | 2,33 | 0,03 | 9,75E-01 |
| AT3G16340 | 0,41 | 1,92 | 4,33 | 0,44 | 6,56E-01 |
| AT3G16420 | 0,28 | 0,00 | 4,41 | 0,00 | 1,00E+00 |
| AT3G16460 | 0,31 | 0,61 | 4,37 | 0,14 | 8,89E-01 |
| AT3G16525 | 0,84 | -1,92 | 2,19 | -0,88 | 3,80E-01 |
| AT3G16570 | 0,68 | 1,02 | 2,45 | 0,42 | 6,78E-01 |
| AT3G16710 | 0,43 | -0,02 | 4,22 | -0,01 | 9,95E-01 |
| AT3G16720 | 0,43 | 1,19 | 4,14 | 0,29 | 7,75E-01 |
| AT3G16740 | 0,38 | 0,77 | 3,42 | 0,23 | 8,21E-01 |
| AT3G16760 | 0,57 | 2,39 | 3,77 | 0,63 | 5,26E-01 |
| AT3G16770 | 1,33 | -0,58 | 2,29 | -0,25 | 8,00E-01 |
| AT3G16800 | 0,16 | -1,36 | 4,41 | -0,31 | 7,57E-01 |
| AT3G16857 | 0,72 | 2,40 | 3,78 | 0,64 | 5,25E-01 |
| AT3G16940 | 0,78 | 0,57 | 2,93 | 0,20 | 8,45E-01 |
| AT3G17100 | 1,17 | -1,71 | 1,92 | -0,89 | 3,74E-01 |
| AT3G17160 | 0,19 | 1,82 | 4,33 | 0,42 | 6,74E-01 |
| AT3G17205 | 0,35 | 0,44 | 4,39 | 0,10 | 9,20E-01 |
| AT3G17210 | 0,32 | 0,72 | 4,37 | 0,17 | 8,69E-01 |
| AT3G03465 | 0,80 | 1,15 | 2,28 | 0,51 | 6,13E-01 |
| AT3G17360 | 0,65 | 1,15 | 3,05 | 0,38 | 7,06E-01 |
| AT3G17380 | 0,28 | 0,65 | 4,37 | 0,15 | 8,82E-01 |
| AT3G17410 | 0,79 | 1,57 | 1,93 | 0,82 | 4,14E-01 |
| AT3G17470 | 0,76 | -0,99 | 3,83 | -0,26 | 7,96E-01 |
| AT3G17490 | 0,56 | 1,55 | 3,88 | 0,40 | 6,90E-01 |
| AT3G17626 | 0,20 | 1,07 | 4,36 | 0,25 | 8,06E-01 |
| AT3G17640 | 0,16 | -1,36 | 4,41 | -0,31 | 7,57E-01 |
| AT3G17650 | 0,88 | 2,47 | 2,04 | 1,21 | 2,25E-01 |
| AT3G17690 | 0,12 | 1,58 | 4,34 | 0,36 | 7,16E-01 |
| AT3G17700 | 0,35 | 1,65 | 4,35 | 0,38 | 7,05E-01 |
| AT3G17712 | 0,66 | 0,49 | 3,77 | 0,13 | 8,97E-01 |
| AT3G17750 | 1,26 | 1,13 | 1,68 | 0,68 | 4,99E-01 |
| AT3G17765 | 0,21 | 1,39 | 4,34 | 0,32 | 7,49E-01 |
| AT3G17790 | 0,83 | 1,07 | 2,43 | 0,44 | 6,61E-01 |
| AT3G03535 | 0,92 | -2,21 | 2,41 | -0,92 | 3,60E-01 |
| AT3G17810 | 0,53 | 1,03 | 2,78 | 0,37 | 7,12E-01 |
| AT3G17820 | 0,62 | -0,50 | 3,27 | -0,15 | 8,79E-01 |
| AT3G17860 | 0,36 | 1,58 | 4,35 | 0,36 | 7,15E-01 |
| AT3G17880 | 0,42 | 2,31 | 3,25 | 0,71 | 4,77E-01 |
| AT3G17890 | 0,10 | 0,42 | 4,38 | 0,10 | 9,23E-01 |
| AT3G17970 | 0,77 | 0,08 | 2,95 | 0,03 | 9,79E-01 |
| AT3G17980 | 0,35 | 0,00 | 4,39 | 0,00 | 1,00E+00 |
| AT3G18000 | 0,28 | 1,56 | 4,34 | 0,36 | 7,18E-01 |
| AT3G18020 | 0,41 | 0,15 | 4,22 | 0,03 | 9,72E-01 |
| AT3G18040 | 0,26 | 0,15 | 4,36 | 0,03 | 9,73E-01 |

| | | | | | |
|---|---|---|---|---|---|
| AT3G18050 | 1,52 | -0,68 | 1,44 | -0,47 | 6,36E-01 |
| AT3G18060 | 1,35 | 0,87 | 1,64 | 0,53 | 5,97E-01 |
| AT3G18070 | 0,11 | 1,07 | 4,36 | 0,25 | 8,06E-01 |
| AT3G18080 | 1,62 | 0,86 | 1,42 | 0,61 | 5,44E-01 |
| AT3G18090 | 1,26 | 3,90 | 2,93 | 1,33 | 1,84E-01 |
| AT3G18100 | 0,29 | 2,52 | 4,32 | 0,58 | 5,59E-01 |
| AT3G03625 | 0,40 | -0,58 | 3,44 | -0,17 | 8,65E-01 |
| AT3G18190 | 1,46 | -0,22 | 1,58 | -0,14 | 8,92E-01 |
| AT3G18240 | 0,43 | 0,47 | 4,21 | 0,11 | 9,11E-01 |
| AT3G18270 | 0,50 | -0,50 | 4,01 | -0,13 | 9,00E-01 |
| AT3G18340 | 0,19 | 0,00 | 4,41 | 0,00 | 1,00E+00 |
| AT3G18390 | 1,15 | -0,60 | 1,86 | -0,32 | 7,46E-01 |
| AT3G18440 | 0,08 | 1,10 | 4,36 | 0,25 | 8,00E-01 |
| AT3G18480 | 0,44 | 2,26 | 3,22 | 0,70 | 4,84E-01 |
| AT3G18740 | 0,73 | 0,52 | 2,16 | 0,24 | 8,08E-01 |
| AT3G18790 | 0,65 | 0,81 | 2,41 | 0,34 | 7,36E-01 |
| AT3G18800 | 0,33 | 1,33 | 4,35 | 0,31 | 7,59E-01 |
| AT3G18820 | 0,39 | -0,70 | 4,20 | -0,17 | 8,68E-01 |
| AT3G18827 | 0,57 | -2,25 | 3,16 | -0,71 | 4,76E-01 |
| AT3G18860 | 0,59 | 1,98 | 2,57 | 0,77 | 4,41E-01 |
| AT3G18890 | 0,86 | 0,44 | 2,25 | 0,20 | 8,45E-01 |
| AT3G18915 | 0,30 | 1,05 | 4,35 | 0,24 | 8,10E-01 |
| AT3G18960 | 0,08 | 1,10 | 4,36 | 0,25 | 8,00E-01 |
| AT3G18980 | 1,15 | 0,89 | 2,06 | 0,43 | 6,66E-01 |
| AT3G19050 | 0,62 | 1,23 | 3,01 | 0,41 | 6,84E-01 |
| AT3G19100 | 1,08 | 2,86 | 1,99 | 1,44 | 1,50E-01 |
| AT3G19240 | 1,33 | 2,00 | 1,71 | 1,17 | 2,43E-01 |
| AT3G19290 | 0,28 | 0,00 | 4,41 | 0,00 | 1,00E+00 |
| AT3G19390 | 0,35 | 0,00 | 4,39 | 0,00 | 1,00E+00 |
| AT3G19470 | 0,30 | 0,00 | 4,41 | 0,00 | 1,00E+00 |
| AT3G19480 | 0,52 | 2,32 | 3,20 | 0,72 | 4,69E-01 |
| AT3G19520 | 0,05 | 0,00 | 4,41 | 0,00 | 1,00E+00 |
| AT3G19540 | 1,20 | 2,34 | 2,12 | 1,10 | 2,70E-01 |
| AT3G19560 | 1,58 | -0,74 | 2,05 | -0,36 | 7,17E-01 |
| AT3G19640 | 0,86 | 2,12 | 2,42 | 0,87 | 3,82E-01 |
| AT3G19680 | 0,92 | -0,47 | 1,92 | -0,24 | 8,09E-01 |
| AT3G19760 | 1,20 | -0,28 | 1,74 | -0,16 | 8,74E-01 |
| AT3G19770 | 0,75 | 0,40 | 2,96 | 0,14 | 8,92E-01 |
| AT3G19780 | 1,34 | 3,72 | 2,13 | 1,75 | 8,07E-02 |
| AT3G19800 | 0,78 | 1,31 | 2,40 | 0,55 | 5,84E-01 |
| AT3G19860 | 0,48 | 0,53 | 4,01 | 0,13 | 8,95E-01 |
| AT3G19870 | 0,37 | 2,31 | 4,28 | 0,54 | 5,89E-01 |
| AT3G19910 | 1,61 | 2,54 | 1,62 | 1,56 | 1,18E-01 |
| AT3G19930 | 1,47 | 1,52 | 1,68 | 0,90 | 3,67E-01 |
| AT3G19960 | 1,65 | 2,40 | 1,75 | 1,37 | 1,71E-01 |
| AT3G19980 | 0,93 | 1,91 | 2,40 | 0,80 | 4,25E-01 |
| AT3G19990 | 0,32 | 1,25 | 4,35 | 0,29 | 7,74E-01 |
| AT3G20000 | 0,74 | 1,07 | 3,58 | 0,30 | 7,66E-01 |
| AT3G20030 | 0,79 | 1,24 | 2,42 | 0,51 | 6,08E-01 |
| AT3G20140 | 0,20 | 0,89 | 4,37 | 0,20 | 8,39E-01 |
| AT3G20150 | 0,54 | 0,00 | 4,41 | 0,00 | 1,00E+00 |
| AT3G20200 | 0,21 | 1,99 | 4,33 | 0,46 | 6,46E-01 |
| AT3G20410 | 0,37 | -1,59 | 4,38 | -0,36 | 7,17E-01 |
| AT3G20500 | 0,33 | 1,15 | 4,35 | 0,26 | 7,91E-01 |
| AT3G20520 | 0,48 | 2,55 | 4,32 | 0,59 | 5,55E-01 |
| AT3G20540 | 0,90 | 2,36 | 2,41 | 0,98 | 3,28E-01 |

| | | | | | |
|---|---|---|---|---|---|
| AT3G20600 | 0,25 | 0,00 | 4,41 | 0,00 | 1,00E+00 |
| AT3G03995 | 0,62 | 0,00 | 4,41 | 0,00 | 1,00E+00 |
| AT3G20720 | 0,42 | 1,50 | 3,42 | 0,44 | 6,61E-01 |
| AT3G20730 | 0,40 | 2,51 | 4,32 | 0,58 | 5,61E-01 |
| AT3G20790 | 0,23 | 0,00 | 4,34 | 0,00 | 9,99E-01 |
| AT3G20890 | 0,34 | 0,77 | 4,37 | 0,18 | 8,60E-01 |
| AT3G20970 | 0,63 | 1,02 | 2,96 | 0,34 | 7,31E-01 |
| AT3G21130 | 0,35 | 0,81 | 4,37 | 0,18 | 8,53E-01 |
| AT3G21170 | 1,15 | 2,21 | 2,41 | 0,92 | 3,58E-01 |
| AT3G21175 | 0,18 | 0,42 | 4,38 | 0,10 | 9,24E-01 |
| AT3G21220 | 0,51 | -1,22 | 4,10 | -0,30 | 7,66E-01 |
| AT3G21270 | 0,29 | 0,17 | 4,35 | 0,04 | 9,68E-01 |
| AT3G21295 | 0,41 | 0,06 | 3,41 | 0,02 | 9,87E-01 |
| AT3G21310 | 0,36 | 2,25 | 4,33 | 0,52 | 6,04E-01 |
| AT3G21465 | 0,14 | -1,28 | 4,39 | -0,29 | 7,71E-01 |
| AT3G21473 | 0,48 | -2,25 | 4,01 | -0,56 | 5,74E-01 |
| AT3G21560 | 0,31 | 0,85 | 4,38 | 0,19 | 8,46E-01 |
| AT3G21690 | 0,96 | 2,08 | 2,02 | 1,03 | 3,05E-01 |
| AT3G04175 | 0,99 | 0,93 | 2,08 | 0,45 | 6,55E-01 |
| AT3G21990 | 0,46 | 0,00 | 4,41 | 0,00 | 1,00E+00 |
| AT3G22072 | 0,17 | -2,23 | 4,35 | -0,51 | 6,08E-01 |
| AT3G22104 | 0,62 | 1,58 | 2,94 | 0,54 | 5,91E-01 |
| AT3G22125 | 1,40 | 0,37 | 1,66 | 0,22 | 8,25E-01 |
| AT3G22150 | 0,49 | 1,59 | 3,95 | 0,40 | 6,87E-01 |
| AT3G22200 | 0,76 | 1,15 | 2,21 | 0,52 | 6,05E-01 |
| AT3G04255 | 0,24 | -1,37 | 4,41 | -0,31 | 7,55E-01 |
| AT3G22270 | 0,79 | 0,19 | 2,59 | 0,07 | 9,43E-01 |
| AT3G22290 | 0,33 | 1,33 | 4,35 | 0,31 | 7,59E-01 |
| AT3G22330 | 0,85 | -1,03 | 2,11 | -0,49 | 6,24E-01 |
| AT3G22350 | 0,91 | 2,20 | 2,60 | 0,85 | 3,98E-01 |
| AT3G22370 | 0,72 | 1,23 | 2,63 | 0,47 | 6,39E-01 |
| AT3G22425 | 0,12 | 0,42 | 4,38 | 0,10 | 9,24E-01 |
| AT3G22430 | 0,19 | 0,43 | 4,38 | 0,10 | 9,22E-01 |
| AT3G22520 | 0,75 | 1,50 | 2,59 | 0,58 | 5,61E-01 |
| AT3G22760 | 0,51 | 0,73 | 3,21 | 0,23 | 8,19E-01 |
| AT3G22810 | 0,55 | 1,69 | 4,32 | 0,39 | 6,95E-01 |
| AT3G22845 | 0,51 | 0,48 | 4,40 | 0,11 | 9,13E-01 |
| AT3G22870 | 0,92 | 1,90 | 2,22 | 0,86 | 3,91E-01 |
| AT3G22900 | 0,24 | 1,62 | 4,34 | 0,37 | 7,08E-01 |
| AT3G22935 | 0,26 | 0,00 | 4,41 | 0,00 | 1,00E+00 |
| AT3G23010 | 0,77 | 0,86 | 3,59 | 0,24 | 8,10E-01 |
| AT3G23050 | 0,64 | 0,69 | 2,64 | 0,26 | 7,94E-01 |
| AT3G04395 | 0,16 | 1,33 | 4,35 | 0,31 | 7,59E-01 |
| AT3G23250 | 0,25 | 1,60 | 4,33 | 0,37 | 7,12E-01 |
| AT3G23280 | 0,45 | -0,17 | 4,06 | -0,04 | 9,66E-01 |
| AT3G23300 | 0,75 | 2,13 | 2,17 | 0,98 | 3,27E-01 |
| AT3G23310 | 0,63 | -0,61 | 2,26 | -0,27 | 7,87E-01 |
| AT3G23380 | 0,21 | 0,43 | 4,37 | 0,10 | 9,22E-01 |
| AT3G23390 | 0,15 | 0,42 | 4,38 | 0,09 | 9,24E-01 |
| AT3G23410 | 0,36 | -1,37 | 4,32 | -0,32 | 7,52E-01 |
| AT3G23530 | 1,09 | -0,20 | 1,97 | -0,10 | 9,20E-01 |
| AT3G23580 | 0,57 | -0,57 | 2,28 | -0,25 | 8,03E-01 |
| AT3G23590 | 1,64 | 3,10 | 1,73 | 1,79 | 7,41E-02 |
| AT3G23600 | 1,16 | 3,25 | 1,97 | 1,65 | 9,86E-02 |
| AT3G23640 | 0,83 | -0,12 | 2,59 | -0,04 | 9,64E-01 |
| AT3G23750 | 0,94 | 0,06 | 1,79 | 0,04 | 9,72E-01 |

| | | | | | |
|---|---|---|---|---|---|
| AT3G23790 | 0,38 | 0,43 | 4,37 | 0,10 | 9,22E-01 |
| AT3G23800 | 0,20 | 0,89 | 4,37 | 0,20 | 8,39E-01 |
| AT3G23890 | 0,56 | 1,74 | 3,97 | 0,44 | 6,61E-01 |
| AT3G23990 | 0,73 | 0,99 | 3,65 | 0,27 | 7,86E-01 |
| AT3G24000 | 0,34 | 1,59 | 4,35 | 0,37 | 7,14E-01 |
| AT3G24050 | 0,88 | 0,54 | 1,93 | 0,28 | 7,78E-01 |
| AT3G24070 | 0,38 | 2,04 | 4,34 | 0,47 | 6,39E-01 |
| AT3G24160 | 0,45 | -0,40 | 3,45 | -0,12 | 9,07E-01 |
| AT3G24190 | 0,59 | 1,09 | 3,04 | 0,36 | 7,20E-01 |
| AT3G24255 | 0,43 | 0,97 | 2,75 | 0,35 | 7,24E-01 |
| AT3G24490 | 0,64 | 3,15 | 3,54 | 0,89 | 3,74E-01 |
| AT3G24500 | 0,22 | 0,00 | 4,41 | 0,00 | 1,00E+00 |
| AT3G24514 | 0,29 | 0,00 | 4,41 | 0,00 | 1,00E+00 |
| AT3G24520 | 0,92 | -1,77 | 2,48 | -0,71 | 4,75E-01 |
| AT3G24550 | 0,60 | 2,90 | 2,93 | 0,99 | 3,22E-01 |
| AT3G24580 | 0,47 | 2,05 | 4,03 | 0,51 | 6,11E-01 |
| AT3G24590 | 0,21 | 0,79 | 4,37 | 0,18 | 8,57E-01 |
| AT3G24830 | 0,92 | 0,01 | 1,87 | 0,00 | 9,96E-01 |
| AT3G25012 | 0,95 | 0,95 | 1,95 | 0,48 | 6,28E-01 |
| AT3G25013 | 0,18 | 0,42 | 4,38 | 0,10 | 9,24E-01 |
| AT3G25030 | 0,21 | 0,36 | 4,35 | 0,08 | 9,35E-01 |
| AT3G25040 | 0,17 | -0,46 | 4,37 | -0,11 | 9,16E-01 |
| AT3G25070 | 0,31 | 1,82 | 3,59 | 0,51 | 6,12E-01 |
| AT3G25130 | 0,57 | -3,09 | 4,32 | -0,71 | 4,75E-01 |
| AT3G25140 | 1,51 | 1,19 | 1,96 | 0,61 | 5,43E-01 |
| AT3G25250 | 0,60 | 0,70 | 2,52 | 0,28 | 7,81E-01 |
| AT3G25290 | 0,29 | 1,15 | 4,35 | 0,26 | 7,91E-01 |
| AT3G25440 | 0,23 | 0,00 | 4,41 | 0,00 | 1,00E+00 |
| AT3G25470 | 0,67 | -0,53 | 3,71 | -0,14 | 8,87E-01 |
| AT3G25490 | 0,05 | 0,00 | 4,41 | 0,00 | 1,00E+00 |
| AT3G25585 | 0,71 | -0,46 | 2,57 | -0,18 | 8,58E-01 |
| AT3G25590 | 0,75 | 2,50 | 2,42 | 1,03 | 3,03E-01 |
| AT3G25700 | 0,66 | -0,05 | 2,19 | -0,02 | 9,82E-01 |
| AT3G04955 | 0,15 | 1,25 | 4,35 | 0,29 | 7,75E-01 |
| AT3G25720 | 0,26 | 0,41 | 4,38 | 0,09 | 9,25E-01 |
| AT3G25760 | 0,49 | -1,21 | 4,38 | -0,28 | 7,82E-01 |
| AT3G25770 | 1,09 | -2,30 | 1,82 | -1,26 | 2,06E-01 |
| AT3G25780 | 0,49 | 1,25 | 2,68 | 0,46 | 6,43E-01 |
| AT3G25860 | 0,63 | -1,43 | 3,10 | -0,46 | 6,45E-01 |
| AT3G25890 | 0,89 | 1,65 | 2,19 | 0,75 | 4,51E-01 |
| AT3G25910 | 0,36 | 2,63 | 4,32 | 0,61 | 5,43E-01 |
| AT3G26020 | 0,28 | -0,54 | 4,38 | -0,12 | 9,02E-01 |
| AT3G26030 | 1,16 | 1,82 | 1,93 | 0,94 | 3,48E-01 |
| AT3G26070 | 0,46 | -1,43 | 4,24 | -0,34 | 7,36E-01 |
| AT3G26080 | 0,26 | 1,08 | 4,37 | 0,25 | 8,05E-01 |
| AT3G26085 | 0,30 | 0,80 | 4,37 | 0,18 | 8,54E-01 |
| AT3G26090 | 0,21 | 1,00 | 4,36 | 0,23 | 8,19E-01 |
| AT3G04975 | 1,19 | -0,59 | 1,91 | -0,31 | 7,56E-01 |
| AT3G04985 | 0,58 | 0,03 | 2,59 | 0,01 | 9,92E-01 |
| AT3G26370 | 0,48 | 0,60 | 2,71 | 0,22 | 8,23E-01 |
| AT3G26395 | 0,16 | 1,74 | 4,33 | 0,40 | 6,87E-01 |
| AT3G26400 | 0,95 | -1,67 | 2,50 | -0,67 | 5,04E-01 |
| AT3G26430 | 1,60 | 1,59 | 1,51 | 1,06 | 2,90E-01 |
| AT3G26420 | 1,60 | 1,59 | 1,51 | 1,06 | 2,90E-01 |
| AT3G26512 | 0,38 | 1,57 | 3,36 | 0,47 | 6,40E-01 |
| AT3G26570 | 1,54 | 0,71 | 1,78 | 0,40 | 6,91E-01 |

| | | | | | |
|---|---|---|---|---|---|
| AT3G26580 | 0,73 | -0,08 | 2,17 | -0,04 | 9,71E-01 |
| AT3G26600 | 1,07 | 1,77 | 2,09 | 0,85 | 3,97E-01 |
| AT3G26640 | 0,37 | 0,52 | 3,45 | 0,15 | 8,81E-01 |
| AT3G26710 | 1,14 | 1,25 | 2,01 | 0,62 | 5,35E-01 |
| AT3G26720 | 0,82 | 1,23 | 2,52 | 0,49 | 6,27E-01 |
| AT3G26742 | 0,18 | 1,23 | 4,34 | 0,28 | 7,77E-01 |
| AT3G26780 | 1,24 | 3,42 | 2,02 | 1,70 | 8,99E-02 |
| AT3G26782 | 0,41 | 2,17 | 4,12 | 0,53 | 5,99E-01 |
| AT3G26800 | 0,34 | -0,42 | 3,51 | -0,12 | 9,05E-01 |
| AT3G26840 | 1,06 | 3,29 | 2,22 | 1,48 | 1,40E-01 |
| AT3G27090 | 1,24 | 1,95 | 1,96 | 0,99 | 3,20E-01 |
| AT3G27160 | 0,41 | -1,30 | 3,33 | -0,39 | 6,97E-01 |
| AT3G27170 | 0,68 | 2,07 | 2,80 | 0,74 | 4,60E-01 |
| AT3G27230 | 0,45 | 1,75 | 3,34 | 0,53 | 5,99E-01 |
| AT3G27250 | 0,85 | -1,49 | 2,55 | -0,58 | 5,60E-01 |
| AT3G27260 | 1,09 | 1,70 | 2,04 | 0,83 | 4,06E-01 |
| AT3G27270 | 0,62 | -1,35 | 3,83 | -0,35 | 7,25E-01 |
| AT3G27280 | 0,60 | 1,28 | 3,00 | 0,43 | 6,70E-01 |
| AT3G27325 | 0,56 | 1,30 | 2,70 | 0,48 | 6,29E-01 |
| AT3G27329 | 0,81 | -1,13 | 2,02 | -0,56 | 5,77E-01 |
| AT3G27331 | 0,96 | 1,35 | 3,23 | 0,42 | 6,76E-01 |
| AT3G27350 | 0,51 | 2,31 | 3,92 | 0,59 | 5,56E-01 |
| AT3G27430 | 0,35 | -0,93 | 3,55 | -0,26 | 7,95E-01 |
| AT3G27470 | 1,23 | 3,66 | 1,86 | 1,96 | 4,98E-02 |
| AT3G27540 | 0,29 | 0,37 | 3,65 | 0,10 | 9,19E-01 |
| AT3G27555 | 0,30 | 0,00 | 4,41 | 0,00 | 1,00E+00 |
| AT3G27670 | 1,19 | 2,86 | 2,27 | 1,26 | 2,07E-01 |
| AT3G05295 | 0,48 | 0,30 | 2,89 | 0,10 | 9,17E-01 |
| AT3G27820 | 1,03 | 2,57 | 1,94 | 1,33 | 1,84E-01 |
| AT3G27830 | 0,47 | 1,35 | 3,39 | 0,40 | 6,91E-01 |
| AT3G27850 | 1,55 | -2,16 | 1,59 | -1,36 | 1,75E-01 |
| AT3G27860 | 0,12 | 1,46 | 4,35 | 0,34 | 7,36E-01 |
| AT3G27870 | 0,75 | 1,85 | 2,56 | 0,72 | 4,69E-01 |
| AT3G27940 | 0,56 | 1,93 | 2,96 | 0,65 | 5,13E-01 |
| AT3G27980 | 0,34 | -1,61 | 4,38 | -0,37 | 7,13E-01 |
| AT3G27999 | 0,43 | 0,00 | 4,41 | 0,00 | 1,00E+00 |
| AT3G28040 | 1,16 | 0,07 | 2,20 | 0,03 | 9,76E-01 |
| AT3G28050 | 0,37 | 2,68 | 4,32 | 0,62 | 5,34E-01 |
| AT3G28080 | 0,70 | 0,49 | 2,33 | 0,21 | 8,34E-01 |
| AT3G28100 | 1,10 | 0,34 | 1,89 | 0,18 | 8,57E-01 |
| AT3G28170 | 0,75 | 2,30 | 3,52 | 0,65 | 5,13E-01 |
| AT3G28200 | 0,56 | -1,59 | 4,24 | -0,38 | 7,07E-01 |
| AT3G28220 | 0,69 | -3,10 | 2,90 | -1,07 | 2,84E-01 |
| AT3G28270 | 0,14 | 0,44 | 4,39 | 0,10 | 9,20E-01 |
| AT3G28290 | 1,10 | -3,77 | 2,39 | -1,58 | 1,14E-01 |
| AT3G28300 | 1,56 | -4,58 | 2,64 | -1,73 | 8,30E-02 |
| AT3G28450 | 0,78 | -0,11 | 2,52 | -0,04 | 9,65E-01 |
| AT3G28500 | 0,30 | 1,56 | 4,33 | 0,36 | 7,19E-01 |
| AT3G28510 | 0,99 | 2,07 | 2,55 | 0,81 | 4,19E-01 |
| AT3G05515 | 0,60 | -2,08 | 3,03 | -0,69 | 4,92E-01 |
| AT3G28580 | 0,77 | 3,33 | 3,41 | 0,98 | 3,29E-01 |
| AT3G28590 | 0,05 | 0,00 | 4,41 | 0,00 | 1,00E+00 |
| AT3G28650 | 0,20 | 0,00 | 4,41 | 0,00 | 1,00E+00 |
| AT3G28670 | 0,33 | 0,77 | 4,39 | 0,18 | 8,61E-01 |
| AT3G28690 | 0,39 | 0,70 | 4,21 | 0,17 | 8,67E-01 |
| AT3G28715 | 0,62 | 0,32 | 2,30 | 0,14 | 8,90E-01 |

| | | | | | |
|---|---|---|---|---|---|
| AT3G28720 | 0,91 | 2,91 | 3,18 | 0,91 | 3,61E-01 |
| AT3G28730 | 0,64 | 1,59 | 2,93 | 0,54 | 5,87E-01 |
| AT3G28870 | 0,74 | 0,93 | 3,69 | 0,25 | 8,01E-01 |
| AT3G28910 | 0,61 | 1,83 | 2,65 | 0,69 | 4,91E-01 |
| AT3G05595 | 0,35 | 0,49 | 4,35 | 0,11 | 9,10E-01 |
| AT3G28922 | 0,12 | 0,90 | 4,36 | 0,21 | 8,36E-01 |
| AT3G28958 | 0,58 | 1,42 | 3,10 | 0,46 | 6,47E-01 |
| AT3G29000 | 0,25 | 1,77 | 4,33 | 0,41 | 6,82E-01 |
| AT3G29010 | 0,38 | 0,82 | 4,37 | 0,19 | 8,51E-01 |
| AT3G29033 | 0,57 | 2,52 | 2,92 | 0,86 | 3,88E-01 |
| AT3G29080 | 1,28 | 1,10 | 1,67 | 0,66 | 5,09E-01 |
| AT3G29180 | 1,27 | 2,36 | 2,81 | 0,84 | 4,00E-01 |
| AT3G05715 | 0,40 | 0,05 | 4,28 | 0,01 | 9,91E-01 |
| AT3G29240 | 1,56 | 2,72 | 1,89 | 1,44 | 1,50E-01 |
| AT3G29270 | 0,39 | 0,40 | 4,39 | 0,09 | 9,28E-01 |
| AT3G29290 | 0,20 | 1,68 | 4,34 | 0,39 | 6,99E-01 |
| AT3G29370 | 0,05 | 0,00 | 4,41 | 0,00 | 1,00E+00 |
| AT3G29390 | 0,18 | -0,04 | 4,35 | -0,01 | 9,92E-01 |
| AT3G29450 | 0,14 | 0,00 | 4,41 | 0,00 | 1,00E+00 |
| AT3G29455 | 0,27 | 0,00 | 4,41 | 0,00 | 1,00E+00 |
| AT3G05805 | 0,31 | 1,76 | 4,33 | 0,41 | 6,84E-01 |
| AT3G29638 | 0,16 | 1,49 | 4,34 | 0,34 | 7,32E-01 |
| AT3G29670 | 0,94 | -3,32 | 2,27 | -1,46 | 1,43E-01 |
| AT3G30210 | 0,11 | 0,90 | 4,37 | 0,21 | 8,36E-01 |
| AT3G30320 | 0,40 | -0,24 | 4,36 | -0,05 | 9,56E-01 |
| AT3G30380 | 0,49 | 0,15 | 3,26 | 0,05 | 9,63E-01 |
| AT3G30383 | 0,12 | 1,46 | 4,35 | 0,34 | 7,36E-01 |
| AT3G30520 | 1,35 | 1,66 | 2,36 | 0,70 | 4,81E-01 |
| AT3G30525 | 1,14 | 2,61 | 2,71 | 0,96 | 3,37E-01 |
| AT3G30720 | 0,61 | -1,00 | 2,99 | -0,34 | 7,37E-01 |
| AT3G30770 | 0,13 | 0,00 | 4,41 | 0,00 | 1,00E+00 |
| AT3G30820 | 0,18 | 0,63 | 4,37 | 0,14 | 8,86E-01 |
| AT3G31950 | 0,59 | -0,86 | 2,25 | -0,38 | 7,04E-01 |
| AT3G32150 | 0,20 | 0,00 | 4,41 | 0,00 | 1,00E+00 |
| AT3G32280 | 0,39 | -2,17 | 4,29 | -0,51 | 6,13E-01 |
| AT3G32896 | 0,28 | 0,00 | 4,41 | 0,00 | 1,00E+00 |
| AT3G33530 | 1,04 | 1,12 | 1,96 | 0,57 | 5,67E-01 |
| AT3G42075 | 0,24 | 1,09 | 4,35 | 0,25 | 8,02E-01 |
| AT3G42170 | 0,48 | 2,12 | 3,13 | 0,67 | 5,00E-01 |
| AT3G42310 | 0,36 | -0,18 | 4,37 | -0,04 | 9,66E-01 |
| AT3G42560 | 0,10 | 0,42 | 4,38 | 0,10 | 9,23E-01 |
| AT3G42640 | 0,44 | 1,56 | 2,73 | 0,57 | 5,67E-01 |
| AT3G42870 | 1,35 | -2,39 | 1,89 | -1,26 | 2,07E-01 |
| AT3G42950 | 0,12 | 1,58 | 4,34 | 0,36 | 7,16E-01 |
| AT3G06495 | 0,51 | 0,82 | 3,13 | 0,26 | 7,95E-01 |
| AT3G43148 | 0,82 | 1,39 | 2,31 | 0,60 | 5,48E-01 |
| AT3G43150 | 0,30 | 1,23 | 3,63 | 0,34 | 7,34E-01 |
| AT3G43210 | 0,25 | 1,75 | 4,33 | 0,40 | 6,87E-01 |
| AT3G43230 | 1,49 | 4,00 | 1,71 | 2,34 | 1,94E-02 |
| AT3G43260 | 1,20 | 1,95 | 2,09 | 0,93 | 3,52E-01 |
| AT3G06565 | 0,05 | 0,00 | 4,41 | 0,00 | 1,00E+00 |
| AT3G43290 | 0,21 | 1,23 | 4,34 | 0,28 | 7,77E-01 |
| AT3G06625 | 0,13 | -1,24 | 4,39 | -0,28 | 7,77E-01 |
| AT3G43490 | 1,48 | 1,79 | 2,02 | 0,88 | 3,78E-01 |
| AT3G43503 | 0,30 | 0,84 | 3,63 | 0,23 | 8,17E-01 |
| AT3G06635 | 0,40 | 1,06 | 4,35 | 0,24 | 8,07E-01 |

| | | | | | |
|---|---|---|---|---|---|
| AT3G43670 | 0,53 | 1,21 | 2,77 | 0,44 | 6,61E-01 |
| AT3G43682 | 0,33 | 1,40 | 4,35 | 0,32 | 7,47E-01 |
| AT3G43740 | 0,28 | -0,66 | 4,37 | -0,15 | 8,80E-01 |
| AT3G43800 | 0,45 | 0,47 | 3,24 | 0,15 | 8,85E-01 |
| AT3G43837 | 0,47 | 0,82 | 4,08 | 0,20 | 8,41E-01 |
| AT3G43920 | 0,47 | 0,42 | 3,32 | 0,13 | 8,98E-01 |
| AT3G43950 | 0,37 | 0,73 | 4,39 | 0,17 | 8,68E-01 |
| AT3G44130 | 0,22 | 0,48 | 4,40 | 0,11 | 9,12E-01 |
| AT3G44150 | 0,18 | -0,69 | 4,37 | -0,16 | 8,74E-01 |
| AT3G44210 | 1,26 | -2,14 | 2,61 | -0,82 | 4,13E-01 |
| AT3G44230 | 0,25 | -0,80 | 4,37 | -0,18 | 8,55E-01 |
| AT3G44235 | 0,05 | 0,00 | 4,41 | 0,00 | 1,00E+00 |
| AT3G44320 | 0,76 | 1,32 | 2,81 | 0,47 | 6,38E-01 |
| AT3G44405 | 0,08 | 0,85 | 4,38 | 0,20 | 8,45E-01 |
| AT3G44550 | 0,99 | 2,33 | 2,41 | 0,97 | 3,34E-01 |
| AT3G44560 | 0,14 | 0,44 | 4,39 | 0,10 | 9,20E-01 |
| AT3G44620 | 0,30 | -0,39 | 3,66 | -0,11 | 9,14E-01 |
| AT3G44670 | 1,62 | 2,87 | 1,67 | 1,72 | 8,61E-02 |
| AT3G44730 | 0,52 | -0,48 | 2,36 | -0,20 | 8,39E-01 |
| AT3G44798 | 0,43 | 2,12 | 4,15 | 0,51 | 6,10E-01 |
| AT3G44880 | 0,54 | -1,78 | 4,05 | -0,44 | 6,61E-01 |
| AT3G44900 | 0,86 | 2,72 | 2,35 | 1,15 | 2,49E-01 |
| AT3G44935 | 1,28 | 2,56 | 2,10 | 1,22 | 2,22E-01 |
| AT3G44980 | 0,39 | 0,21 | 3,45 | 0,06 | 9,51E-01 |
| AT3G45010 | 0,31 | 0,43 | 4,38 | 0,10 | 9,21E-01 |
| AT3G45400 | 0,33 | 1,65 | 4,33 | 0,38 | 7,03E-01 |
| AT3G45620 | 1,31 | 2,24 | 2,41 | 0,93 | 3,54E-01 |
| AT3G45650 | 1,57 | 3,73 | 1,95 | 1,91 | 5,56E-02 |
| AT3G45660 | 1,01 | 1,62 | 2,59 | 0,62 | 5,33E-01 |
| AT3G45670 | 0,15 | 0,71 | 4,37 | 0,16 | 8,71E-01 |
| AT3G45730 | 0,15 | 1,62 | 4,34 | 0,37 | 7,09E-01 |
| AT3G45740 | 0,36 | 1,56 | 4,34 | 0,36 | 7,19E-01 |
| AT3G45870 | 0,08 | 1,10 | 4,36 | 0,25 | 8,00E-01 |
| AT3G45930 | 0,32 | 0,21 | 4,34 | 0,05 | 9,62E-01 |
| AT3G45965 | 0,21 | -1,16 | 4,33 | -0,27 | 7,90E-01 |
| AT3G45970 | 0,98 | 1,26 | 2,73 | 0,46 | 6,44E-01 |
| AT3G46040 | 0,85 | 1,84 | 2,17 | 0,85 | 3,96E-01 |
| AT3G46060 | 1,03 | 0,08 | 2,19 | 0,04 | 9,71E-01 |
| AT3G46130 | 0,12 | 1,58 | 4,34 | 0,36 | 7,16E-01 |
| AT3G46220 | 0,60 | 0,40 | 2,61 | 0,15 | 8,79E-01 |
| AT3G46290 | 1,13 | 1,70 | 1,90 | 0,89 | 3,73E-01 |
| AT3G46370 | 0,25 | -1,51 | 4,39 | -0,34 | 7,30E-01 |
| AT3G46490 | 0,32 | -1,62 | 4,37 | -0,37 | 7,11E-01 |
| AT3G46520 | 0,37 | 1,82 | 4,33 | 0,42 | 6,75E-01 |
| AT3G46560 | 0,33 | 0,44 | 4,39 | 0,10 | 9,20E-01 |
| AT3G46590 | 1,40 | 1,52 | 1,57 | 0,97 | 3,32E-01 |
| AT3G07505 | 1,14 | -1,64 | 2,03 | -0,81 | 4,20E-01 |
| AT3G46640 | 1,12 | 0,93 | 2,08 | 0,45 | 6,56E-01 |
| AT3G46658 | 0,59 | 2,22 | 2,62 | 0,85 | 3,96E-01 |
| AT3G46668 | 0,57 | -1,25 | 3,14 | -0,40 | 6,91E-01 |
| AT3G07555 | 1,02 | 2,19 | 2,12 | 1,03 | 3,03E-01 |
| AT3G07585 | 1,61 | -0,52 | 1,61 | -0,32 | 7,48E-01 |
| AT3G46930 | 0,21 | 1,99 | 4,33 | 0,46 | 6,46E-01 |
| AT3G47060 | 0,35 | 0,80 | 4,34 | 0,19 | 8,53E-01 |
| AT3G47140 | 0,11 | 0,00 | 4,41 | 0,00 | 1,00E+00 |
| AT3G07695 | 0,33 | 2,35 | 4,32 | 0,54 | 5,86E-01 |

| | | | | | |
|---|---|---|---|---|---|
| AT3G47290 | 0,08 | 1,10 | 4,36 | 0,25 | 8,00E-01 |
| AT3G47295 | 0,08 | 0,00 | 4,40 | 0,00 | 1,00E+00 |
| AT3G07715 | 0,48 | -0,12 | 2,70 | -0,05 | 9,64E-01 |
| AT3G47490 | 0,22 | 0,48 | 4,40 | 0,11 | 9,12E-01 |
| AT3G47520 | 0,29 | 0,83 | 4,33 | 0,19 | 8,47E-01 |
| AT3G47550 | 0,50 | 0,44 | 3,05 | 0,14 | 8,86E-01 |
| AT3G47570 | 0,50 | 2,20 | 3,22 | 0,68 | 4,94E-01 |
| AT3G47610 | 0,61 | 0,43 | 2,64 | 0,16 | 8,70E-01 |
| AT3G47650 | 0,65 | 2,54 | 2,96 | 0,86 | 3,91E-01 |
| AT3G47660 | 0,72 | 1,61 | 2,88 | 0,56 | 5,75E-01 |
| AT3G07795 | 0,21 | 2,05 | 4,33 | 0,47 | 6,36E-01 |
| AT3G07805 | 0,89 | 0,81 | 2,54 | 0,32 | 7,49E-01 |
| AT3G47770 | 0,17 | 1,92 | 4,33 | 0,44 | 6,57E-01 |
| AT3G47800 | 1,07 | -0,11 | 1,85 | -0,06 | 9,54E-01 |
| AT3G47820 | 1,04 | 0,67 | 1,92 | 0,35 | 7,27E-01 |
| AT3G47833 | 0,22 | -1,23 | 4,37 | -0,28 | 7,78E-01 |
| AT3G47920 | 1,60 | -3,90 | 1,62 | -2,41 | 1,59E-02 |
| AT3G47930 | 0,52 | 2,58 | 3,86 | 0,67 | 5,04E-01 |
| AT3G47950 | 0,34 | 0,66 | 4,36 | 0,15 | 8,80E-01 |
| AT3G48040 | 0,17 | 0,00 | 4,40 | 0,00 | 1,00E+00 |
| AT3G48070 | 0,40 | 0,90 | 4,29 | 0,21 | 8,34E-01 |
| AT3G48110 | 0,52 | 1,63 | 3,12 | 0,52 | 6,01E-01 |
| AT3G48140 | 0,97 | 1,81 | 2,15 | 0,84 | 4,01E-01 |
| AT3G48190 | 1,19 | 2,43 | 2,49 | 0,98 | 3,29E-01 |
| AT3G48310 | 0,40 | 0,99 | 3,33 | 0,30 | 7,65E-01 |
| AT3G48380 | 0,47 | 2,01 | 3,21 | 0,63 | 5,30E-01 |
| AT3G48420 | 0,42 | 0,49 | 4,23 | 0,12 | 9,08E-01 |
| AT3G48460 | 0,51 | -16,40 | 4,41 | -3,72 | 1,98E-04 |
| AT3G48500 | 0,57 | -0,36 | 3,88 | -0,09 | 9,26E-01 |
| AT3G48530 | 0,79 | 0,95 | 2,18 | 0,44 | 6,62E-01 |
| AT3G48620 | 0,08 | 0,85 | 4,38 | 0,20 | 8,45E-01 |
| AT3G48680 | 0,15 | 1,58 | 4,34 | 0,36 | 7,15E-01 |
| AT3G48710 | 0,27 | 1,95 | 4,33 | 0,45 | 6,53E-01 |
| AT3G48730 | 0,57 | 0,21 | 2,77 | 0,08 | 9,38E-01 |
| AT3G48750 | 0,54 | 0,65 | 2,68 | 0,24 | 8,10E-01 |
| AT3G48760 | 0,76 | 1,53 | 2,46 | 0,62 | 5,34E-01 |
| AT3G07995 | 0,83 | 1,53 | 2,34 | 0,65 | 5,15E-01 |
| AT3G48780 | 0,82 | 0,54 | 2,68 | 0,20 | 8,39E-01 |
| AT3G48810 | 0,58 | 1,48 | 3,60 | 0,41 | 6,80E-01 |
| AT3G48890 | 0,66 | -0,03 | 2,64 | -0,01 | 9,91E-01 |
| AT3G49000 | 0,27 | 1,87 | 4,33 | 0,43 | 6,65E-01 |
| AT3G49110 | 0,08 | 1,10 | 4,36 | 0,25 | 8,00E-01 |
| AT3G49130 | 1,37 | 3,31 | 2,37 | 1,40 | 1,62E-01 |
| AT3G49220 | 0,92 | -0,20 | 2,06 | -0,10 | 9,24E-01 |
| AT3G49240 | 0,54 | -2,61 | 4,10 | -0,64 | 5,25E-01 |
| AT3G49260 | 0,64 | 0,61 | 4,40 | 0,14 | 8,90E-01 |
| AT3G49430 | 0,93 | 0,67 | 2,06 | 0,33 | 7,45E-01 |
| AT3G49520 | 1,37 | 2,76 | 1,88 | 1,46 | 1,44E-01 |
| AT3G49590 | 1,44 | 2,45 | 1,86 | 1,32 | 1,86E-01 |
| AT3G49670 | 1,02 | 0,90 | 2,44 | 0,37 | 7,12E-01 |
| AT3G49680 | 0,45 | -1,28 | 3,26 | -0,39 | 6,94E-01 |
| AT3G49710 | 0,32 | 1,25 | 4,35 | 0,29 | 7,74E-01 |
| AT3G49780 | 0,25 | -0,45 | 4,38 | -0,10 | 9,19E-01 |
| AT3G50000 | 0,43 | 2,38 | 4,18 | 0,57 | 5,70E-01 |
| AT3G50080 | 0,21 | 1,21 | 4,36 | 0,28 | 7,81E-01 |
| AT3G50100 | 0,05 | 0,00 | 4,41 | 0,00 | 1,00E+00 |

| | | | | | |
|---|---|---|---|---|---|
| AT3G50270 | 0,60 | 0,87 | 3,05 | 0,28 | 7,77E-01 |
| AT3G50350 | 0,42 | 0,13 | 3,42 | 0,04 | 9,69E-01 |
| AT3G50360 | 0,31 | -0,48 | 4,36 | -0,11 | 9,12E-01 |
| AT3G50430 | 0,16 | 0,00 | 4,41 | 0,00 | 1,00E+00 |
| AT3G50530 | 0,25 | 0,66 | 4,33 | 0,15 | 8,79E-01 |
| AT3G50540 | 1,39 | -1,04 | 1,61 | -0,65 | 5,19E-01 |
| AT3G50550 | 0,51 | 0,44 | 3,26 | 0,13 | 8,93E-01 |
| AT3G50630 | 0,26 | 1,04 | 4,32 | 0,24 | 8,09E-01 |
| AT3G50700 | 0,53 | 0,83 | 2,65 | 0,31 | 7,55E-01 |
| AT3G50770 | 1,04 | 3,72 | 2,31 | 1,61 | 1,07E-01 |
| AT3G50890 | 0,25 | 0,42 | 4,38 | 0,10 | 9,24E-01 |
| AT3G50910 | 0,94 | 2,93 | 2,32 | 1,26 | 2,06E-01 |
| AT3G50970 | 0,39 | 0,00 | 4,40 | 0,00 | 1,00E+00 |
| AT3G51110 | 0,46 | 0,75 | 4,03 | 0,19 | 8,52E-01 |
| AT3G51140 | 0,97 | 0,65 | 2,23 | 0,29 | 7,71E-01 |
| AT3G51180 | 0,25 | 0,00 | 4,41 | 0,00 | 1,00E+00 |
| AT3G51260 | 0,46 | 1,68 | 3,28 | 0,51 | 6,09E-01 |
| AT3G51270 | 0,33 | 1,63 | 4,34 | 0,38 | 7,07E-01 |
| AT3G51325 | 0,36 | 1,33 | 4,36 | 0,30 | 7,61E-01 |
| AT3G51370 | 1,46 | 1,00 | 1,83 | 0,55 | 5,85E-01 |
| AT3G51420 | 0,66 | -1,18 | 2,33 | -0,51 | 6,11E-01 |
| AT3G51430 | 0,32 | 0,66 | 4,40 | 0,15 | 8,81E-01 |
| AT3G51440 | 0,80 | 2,59 | 2,18 | 1,19 | 2,36E-01 |
| AT3G51450 | 0,43 | -1,44 | 2,79 | -0,51 | 6,07E-01 |
| AT3G51460 | 0,27 | 0,59 | 4,39 | 0,14 | 8,92E-01 |
| AT3G51480 | 0,45 | 1,28 | 4,34 | 0,30 | 7,68E-01 |
| AT3G51500 | 0,44 | 2,79 | 4,31 | 0,65 | 5,16E-01 |
| AT3G51510 | 0,44 | 1,23 | 3,44 | 0,36 | 7,20E-01 |
| AT3G51540 | 0,25 | 1,75 | 4,33 | 0,40 | 6,87E-01 |
| AT3G51620 | 1,34 | 1,96 | 1,81 | 1,08 | 2,78E-01 |
| AT3G51630 | 0,87 | 3,06 | 3,21 | 0,95 | 3,40E-01 |
| AT3G51640 | 0,80 | 3,38 | 2,67 | 1,27 | 2,05E-01 |
| AT3G51650 | 1,02 | 2,64 | 2,19 | 1,21 | 2,28E-01 |
| AT3G51730 | 1,38 | -0,56 | 1,46 | -0,38 | 7,03E-01 |
| AT3G51830 | 0,25 | 1,92 | 4,33 | 0,44 | 6,57E-01 |
| AT3G51850 | 1,69 | 3,00 | 1,86 | 1,62 | 1,06E-01 |
| AT3G51870 | 1,46 | 2,69 | 2,06 | 1,31 | 1,92E-01 |
| AT3G51970 | 0,40 | 1,99 | 3,37 | 0,59 | 5,54E-01 |
| AT3G52080 | 0,33 | 1,90 | 4,33 | 0,44 | 6,60E-01 |
| AT3G52100 | 0,15 | 0,86 | 4,37 | 0,20 | 8,43E-01 |
| AT3G52105 | 0,19 | 0,78 | 4,36 | 0,18 | 8,58E-01 |
| AT3G52120 | 1,03 | 0,82 | 2,29 | 0,36 | 7,20E-01 |
| AT3G52150 | 0,83 | -3,20 | 2,83 | -1,13 | 2,59E-01 |
| AT3G52155 | 0,45 | 0,44 | 4,39 | 0,10 | 9,20E-01 |
| AT3G52200 | 0,34 | 0,54 | 4,37 | 0,12 | 9,02E-01 |
| AT3G52210 | 0,11 | 1,07 | 4,36 | 0,25 | 8,06E-01 |
| AT3G52230 | 0,83 | 1,74 | 3,06 | 0,57 | 5,69E-01 |
| AT3G52300 | 0,40 | 1,61 | 4,34 | 0,37 | 7,10E-01 |
| AT3G52340 | 0,38 | 0,51 | 4,34 | 0,12 | 9,06E-01 |
| AT3G52370 | 0,08 | 0,85 | 4,38 | 0,20 | 8,45E-01 |
| AT3G52380 | 1,54 | 0,58 | 1,86 | 0,31 | 7,55E-01 |
| AT3G52470 | 0,30 | 1,12 | 4,36 | 0,26 | 7,97E-01 |
| AT3G52570 | 0,29 | 1,85 | 4,33 | 0,43 | 6,69E-01 |
| AT3G08275 | 0,60 | 0,74 | 3,13 | 0,24 | 8,14E-01 |
| AT3G52610 | 0,30 | 1,68 | 4,34 | 0,39 | 6,99E-01 |
| AT3G52680 | 0,59 | 0,42 | 2,64 | 0,16 | 8,73E-01 |

| | | | | | |
|---|---|---|---|---|---|
| AT3G52710 | 0,88 | 1,54 | 2,00 | 0,77 | 4,43E-01 |
| AT3G08285 | 1,59 | 1,48 | 1,71 | 0,87 | 3,86E-01 |
| AT3G08295 | 0,73 | 0,46 | 3,50 | 0,13 | 8,96E-01 |
| AT3G52830 | 1,16 | 0,03 | 1,75 | 0,02 | 9,84E-01 |
| AT3G52840 | 0,82 | -2,77 | 2,58 | -1,07 | 2,83E-01 |
| AT3G52950 | 0,45 | 1,59 | 3,26 | 0,49 | 6,25E-01 |
| AT3G52960 | 0,80 | 0,34 | 1,88 | 0,18 | 8,56E-01 |
| AT3G52990 | 1,11 | -0,75 | 1,60 | -0,47 | 6,39E-01 |
| AT3G53000 | 0,31 | 1,25 | 4,36 | 0,29 | 7,74E-01 |
| AT3G53030 | 0,57 | 2,19 | 3,97 | 0,55 | 5,81E-01 |
| AT3G53050 | 0,05 | 0,00 | 4,41 | 0,00 | 1,00E+00 |
| AT3G53065 | 0,19 | 0,00 | 4,40 | 0,00 | 1,00E+00 |
| AT3G53090 | 0,36 | -1,46 | 4,40 | -0,33 | 7,40E-01 |
| AT3G53110 | 0,40 | 0,41 | 3,42 | 0,12 | 9,04E-01 |
| AT3G53180 | 0,20 | 0,43 | 4,39 | 0,10 | 9,21E-01 |
| AT3G53190 | 0,21 | 0,00 | 4,38 | 0,00 | 1,00E+00 |
| AT3G53230 | 1,14 | 2,59 | 2,75 | 0,94 | 3,47E-01 |
| AT3G53240 | 0,55 | 0,73 | 3,92 | 0,19 | 8,53E-01 |
| AT3G53305 | 0,72 | 1,28 | 2,21 | 0,58 | 5,64E-01 |
| AT3G53360 | 0,08 | 0,85 | 4,38 | 0,20 | 8,45E-01 |
| AT3G53390 | 0,27 | 0,54 | 4,37 | 0,12 | 9,01E-01 |
| AT3G53470 | 0,77 | -2,08 | 2,88 | -0,72 | 4,72E-01 |
| AT3G53520 | 0,83 | 0,96 | 2,50 | 0,38 | 7,01E-01 |
| AT3G53670 | 0,90 | 1,27 | 2,12 | 0,60 | 5,48E-01 |
| AT3G53700 | 0,94 | 0,18 | 1,83 | 0,10 | 9,22E-01 |
| AT3G53870 | 0,80 | -0,78 | 2,52 | -0,31 | 7,55E-01 |
| AT3G54220 | 0,50 | 1,08 | 3,19 | 0,34 | 7,35E-01 |
| AT3G54230 | 1,41 | 3,13 | 2,11 | 1,49 | 1,37E-01 |
| AT3G54366 | 1,11 | 2,33 | 2,13 | 1,09 | 2,74E-01 |
| AT3G54420 | 0,17 | 1,75 | 4,33 | 0,40 | 6,86E-01 |
| AT3G54540 | 1,42 | 3,64 | 1,86 | 1,96 | 4,99E-02 |
| AT3G54600 | 0,51 | -0,19 | 4,05 | -0,05 | 9,63E-01 |
| AT3G54610 | 0,47 | 1,66 | 3,93 | 0,42 | 6,72E-01 |
| AT3G54625 | 0,68 | 2,16 | 4,32 | 0,50 | 6,17E-01 |
| AT3G54680 | 0,75 | 1,04 | 2,95 | 0,35 | 7,23E-01 |
| AT3G54800 | 0,59 | 0,65 | 2,66 | 0,24 | 8,07E-01 |
| AT3G54820 | 0,15 | 1,16 | 4,35 | 0,27 | 7,90E-01 |
| AT3G54850 | 0,81 | 0,31 | 2,49 | 0,13 | 9,01E-01 |
| AT3G54980 | 0,41 | 0,86 | 4,22 | 0,20 | 8,38E-01 |
| AT3G55030 | 0,39 | 0,82 | 4,15 | 0,20 | 8,43E-01 |
| AT3G55250 | 0,63 | -0,73 | 2,28 | -0,32 | 7,48E-01 |
| AT3G55260 | 0,43 | 0,00 | 4,14 | 0,00 | 1,00E+00 |
| AT3G55270 | 1,69 | 3,48 | 1,80 | 1,94 | 5,28E-02 |
| AT3G55280 | 0,61 | -1,08 | 3,74 | -0,29 | 7,72E-01 |
| AT3G55370 | 0,12 | 0,90 | 4,36 | 0,21 | 8,36E-01 |
| AT3G55420 | 0,14 | 0,51 | 4,38 | 0,12 | 9,08E-01 |
| AT3G55440 | 1,26 | 1,58 | 1,78 | 0,89 | 3,76E-01 |
| AT3G55450 | 0,15 | 1,58 | 4,34 | 0,36 | 7,15E-01 |
| AT3G55460 | 1,02 | 0,90 | 1,83 | 0,49 | 6,22E-01 |
| AT3G55485 | 0,83 | -2,47 | 2,45 | -1,01 | 3,14E-01 |
| AT3G08485 | 0,83 | -2,47 | 2,45 | -1,01 | 3,14E-01 |
| AT3G55520 | 0,40 | 2,27 | 4,33 | 0,52 | 6,00E-01 |
| AT3G55620 | 0,54 | 1,83 | 2,64 | 0,69 | 4,88E-01 |
| AT3G55630 | 0,42 | 2,41 | 4,32 | 0,56 | 5,77E-01 |
| AT3G55677 | 0,24 | 0,00 | 4,41 | 0,00 | 1,00E+00 |
| AT3G55760 | 0,99 | -0,57 | 1,88 | -0,30 | 7,63E-01 |

| | | | | | |
|---|---|---|---|---|---|
| AT3G55770 | 0,98 | 0,64 | 1,94 | 0,33 | 7,44E-01 |
| AT3G55780 | 0,67 | 1,09 | 3,65 | 0,30 | 7,65E-01 |
| AT3G55880 | 0,30 | 1,59 | 3,63 | 0,44 | 6,61E-01 |
| AT3G55940 | 0,10 | 0,62 | 4,38 | 0,14 | 8,87E-01 |
| AT3G56130 | 0,33 | -0,25 | 4,32 | -0,06 | 9,54E-01 |
| AT3G56140 | 0,74 | 1,67 | 2,39 | 0,70 | 4,86E-01 |
| AT3G56180 | 0,21 | 0,00 | 4,38 | 0,00 | 1,00E+00 |
| AT3G56310 | 0,76 | 3,21 | 2,36 | 1,36 | 1,74E-01 |
| AT3G56330 | 0,52 | 2,57 | 3,84 | 0,67 | 5,04E-01 |
| AT3G56390 | 0,30 | 1,62 | 4,35 | 0,37 | 7,10E-01 |
| AT3G56450 | 0,39 | -0,15 | 4,37 | -0,03 | 9,73E-01 |
| AT3G08665 | 0,76 | 3,04 | 2,83 | 1,07 | 2,83E-01 |
| AT3G56470 | 0,12 | 1,58 | 4,34 | 0,36 | 7,16E-01 |
| AT3G56490 | 0,11 | 1,07 | 4,36 | 0,25 | 8,06E-01 |
| AT3G56590 | 0,57 | -0,48 | 3,06 | -0,16 | 8,77E-01 |
| AT3G56640 | 0,59 | 2,20 | 3,08 | 0,71 | 4,77E-01 |
| AT3G56650 | 0,53 | -1,50 | 3,97 | -0,38 | 7,06E-01 |
| AT3G56720 | 0,17 | -0,28 | 4,38 | -0,06 | 9,49E-01 |
| AT3G56740 | 0,49 | 2,23 | 4,01 | 0,56 | 5,78E-01 |
| AT3G56780 | 0,32 | 0,44 | 4,38 | 0,10 | 9,20E-01 |
| AT3G56840 | 0,64 | 2,53 | 2,49 | 1,02 | 3,09E-01 |
| AT3G56880 | 1,69 | 0,18 | 1,35 | 0,13 | 8,97E-01 |
| AT3G08785 | 1,51 | -0,38 | 1,69 | -0,23 | 8,20E-01 |
| AT3G56960 | 0,60 | 0,85 | 4,37 | 0,19 | 8,46E-01 |
| AT3G57040 | 1,37 | -2,04 | 1,64 | -1,24 | 2,14E-01 |
| AT3G57062 | 1,47 | 3,52 | 1,86 | 1,90 | 5,80E-02 |
| AT3G57070 | 0,24 | -1,25 | 4,39 | -0,28 | 7,76E-01 |
| AT3G57080 | 0,21 | 1,39 | 4,34 | 0,32 | 7,49E-01 |
| AT3G57090 | 0,59 | 1,11 | 2,28 | 0,49 | 6,26E-01 |
| AT3G08845 | 0,12 | 0,79 | 4,37 | 0,18 | 8,57E-01 |
| AT3G57230 | 0,38 | 0,26 | 2,89 | 0,09 | 9,29E-01 |
| AT3G57340 | 0,42 | 0,89 | 3,40 | 0,26 | 7,93E-01 |
| AT3G57390 | 0,11 | 1,07 | 4,36 | 0,25 | 8,06E-01 |
| AT3G57450 | 1,58 | 1,93 | 1,71 | 1,13 | 2,59E-01 |
| AT3G57550 | 0,50 | 2,29 | 3,18 | 0,72 | 4,72E-01 |
| AT3G57570 | 0,40 | -0,76 | 4,20 | -0,18 | 8,56E-01 |
| AT3G57587 | 0,73 | -1,43 | 2,09 | -0,68 | 4,95E-01 |
| AT3G57650 | 0,53 | 2,17 | 2,66 | 0,82 | 4,14E-01 |
| AT3G57700 | 1,50 | 1,53 | 1,53 | 1,00 | 3,18E-01 |
| AT3G08975 | 0,42 | 1,59 | 4,11 | 0,39 | 6,99E-01 |
| AT3G57760 | 0,08 | 1,10 | 4,36 | 0,25 | 8,00E-01 |
| AT3G57770 | 1,17 | 0,03 | 1,64 | 0,02 | 9,85E-01 |
| AT3G57810 | 0,21 | 0,36 | 4,35 | 0,08 | 9,35E-01 |
| AT3G57830 | 0,31 | -0,65 | 4,37 | -0,15 | 8,82E-01 |
| AT3G57890 | 1,04 | -1,18 | 1,80 | -0,66 | 5,11E-01 |
| AT3G57940 | 0,20 | 0,43 | 4,39 | 0,10 | 9,21E-01 |
| AT3G57970 | 0,17 | 1,94 | 4,33 | 0,45 | 6,55E-01 |
| AT3G58020 | 0,08 | 1,10 | 4,36 | 0,25 | 8,00E-01 |
| AT3G58110 | 1,07 | 3,54 | 3,17 | 1,12 | 2,65E-01 |
| AT3G58160 | 0,08 | 0,85 | 4,38 | 0,20 | 8,45E-01 |
| AT3G58490 | 0,30 | 0,87 | 4,38 | 0,20 | 8,43E-01 |
| AT3G58510 | 0,99 | 0,42 | 1,96 | 0,21 | 8,30E-01 |
| AT3G58530 | 0,46 | 0,41 | 4,03 | 0,10 | 9,18E-01 |
| AT3G58570 | 0,71 | 1,43 | 2,53 | 0,56 | 5,73E-01 |
| AT3G58680 | 0,61 | -1,10 | 2,64 | -0,42 | 6,77E-01 |
| AT3G58690 | 0,58 | 1,33 | 3,10 | 0,43 | 6,66E-01 |

| | | | | | |
|---|---|---|---|---|---|
| AT3G58700 | 0,20 | 0,89 | 4,37 | 0,20 | 8,39E-01 |
| AT3G58795 | 0,23 | -1,99 | 4,37 | -0,46 | 6,49E-01 |
| AT3G59030 | 0,64 | 2,58 | 3,65 | 0,71 | 4,79E-01 |
| AT3G59080 | 0,19 | 0,37 | 4,38 | 0,09 | 9,32E-01 |
| AT3G59090 | 0,80 | 1,03 | 2,52 | 0,41 | 6,82E-01 |
| AT3G59100 | 0,79 | 0,64 | 2,88 | 0,22 | 8,24E-01 |
| AT3G59110 | 0,41 | 0,30 | 4,24 | 0,07 | 9,44E-01 |
| AT3G59210 | 0,24 | 1,72 | 4,34 | 0,40 | 6,92E-01 |
| AT3G59290 | 1,04 | 0,92 | 1,92 | 0,48 | 6,32E-01 |
| AT3G59300 | 0,19 | 1,28 | 4,34 | 0,30 | 7,68E-01 |
| AT3G59350 | 0,86 | 0,85 | 2,39 | 0,36 | 7,22E-01 |
| AT3G59380 | 0,24 | 1,72 | 4,34 | 0,40 | 6,92E-01 |
| AT3G59500 | 0,47 | -1,39 | 3,42 | -0,41 | 6,83E-01 |
| AT3G59520 | 0,08 | 1,10 | 4,36 | 0,25 | 8,00E-01 |
| AT3G59530 | 0,53 | 0,43 | 4,38 | 0,10 | 9,23E-01 |
| AT3G59670 | 0,30 | -1,36 | 4,41 | -0,31 | 7,58E-01 |
| AT3G59700 | 0,10 | 0,00 | 4,41 | 0,00 | 1,00E+00 |
| AT3G59810 | 0,46 | -0,31 | 4,37 | -0,07 | 9,44E-01 |
| AT3G59920 | 0,37 | 0,87 | 4,35 | 0,20 | 8,42E-01 |
| AT3G60080 | 1,12 | 2,59 | 2,01 | 1,29 | 1,97E-01 |
| AT3G60110 | 0,48 | 0,92 | 4,38 | 0,21 | 8,33E-01 |
| AT3G60130 | 0,63 | 0,51 | 3,13 | 0,16 | 8,71E-01 |
| AT3G60140 | 0,58 | 0,54 | 3,77 | 0,14 | 8,86E-01 |
| AT3G60240 | 1,46 | 0,75 | 1,46 | 0,51 | 6,08E-01 |
| AT3G60245 | 0,58 | 0,78 | 2,39 | 0,32 | 7,46E-01 |
| AT3G60290 | 0,27 | -1,36 | 4,41 | -0,31 | 7,58E-01 |
| AT3G60300 | 0,58 | 0,49 | 2,65 | 0,18 | 8,54E-01 |
| AT3G60310 | 0,49 | 1,24 | 3,93 | 0,32 | 7,52E-01 |
| AT3G60410 | 0,24 | 1,58 | 4,34 | 0,36 | 7,16E-01 |
| AT3G60420 | 0,55 | 2,47 | 3,81 | 0,65 | 5,17E-01 |
| AT3G60440 | 0,83 | -0,22 | 3,47 | -0,06 | 9,49E-01 |
| AT3G60450 | 1,00 | 3,47 | 2,60 | 1,34 | 1,82E-01 |
| AT3G60500 | 0,28 | 1,56 | 4,34 | 0,36 | 7,18E-01 |
| AT3G60530 | 0,83 | -2,49 | 2,07 | -1,20 | 2,29E-01 |
| AT3G60600 | 1,58 | 2,88 | 1,75 | 1,64 | 1,01E-01 |
| AT3G60690 | 1,35 | 3,60 | 1,88 | 1,92 | 5,48E-02 |
| AT3G60910 | 0,08 | 0,00 | 4,41 | 0,00 | 1,00E+00 |
| AT3G60950 | 1,11 | -1,29 | 1,96 | -0,66 | 5,08E-01 |
| AT3G60960 | 1,54 | 0,10 | 1,57 | 0,06 | 9,51E-01 |
| AT3G09515 | 1,27 | 0,66 | 1,91 | 0,35 | 7,28E-01 |
| AT3G60961 | 1,33 | 3,10 | 2,01 | 1,54 | 1,23E-01 |
| AT3G09525 | 0,12 | 1,58 | 4,34 | 0,36 | 7,16E-01 |
| AT3G60970 | 0,12 | 1,58 | 4,34 | 0,36 | 7,16E-01 |
| AT3G61030 | 0,61 | -1,23 | 2,02 | -0,61 | 5,41E-01 |
| AT3G61050 | 1,51 | -1,42 | 1,59 | -0,89 | 3,74E-01 |
| AT3G61080 | 0,12 | 1,65 | 4,34 | 0,38 | 7,04E-01 |
| AT3G61110 | 1,40 | 0,01 | 1,72 | 0,01 | 9,94E-01 |
| AT3G61130 | 1,07 | -0,46 | 2,47 | -0,19 | 8,52E-01 |
| AT3G61140 | 0,73 | 0,62 | 2,08 | 0,30 | 7,66E-01 |
| AT3G61150 | 0,68 | 2,37 | 2,81 | 0,84 | 4,00E-01 |
| AT3G61180 | 0,94 | 1,30 | 2,39 | 0,54 | 5,88E-01 |
| AT3G61240 | 1,00 | 1,74 | 1,98 | 0,88 | 3,81E-01 |
| AT3G61280 | 0,41 | 2,06 | 3,35 | 0,61 | 5,39E-01 |
| AT3G61320 | 0,08 | 0,00 | 4,40 | 0,00 | 1,00E+00 |
| AT3G61440 | 1,04 | 1,98 | 2,39 | 0,83 | 4,09E-01 |
| AT3G61480 | 0,44 | 0,73 | 3,27 | 0,22 | 8,24E-01 |

| | | | | | |
|---|---|---|---|---|---|
| AT3G61490 | 0,55 | 0,91 | 4,38 | 0,21 | 8,35E-01 |
| AT3G61540 | 0,29 | 0,69 | 4,33 | 0,16 | 8,74E-01 |
| AT3G61550 | 0,37 | 0,97 | 3,48 | 0,28 | 7,81E-01 |
| AT3G61600 | 1,22 | 2,70 | 1,73 | 1,56 | 1,19E-01 |
| AT3G61670 | 0,29 | 1,62 | 4,33 | 0,37 | 7,08E-01 |
| AT3G61700 | 1,46 | -0,93 | 1,64 | -0,57 | 5,72E-01 |
| AT3G61755 | 0,16 | 0,96 | 4,36 | 0,22 | 8,26E-01 |
| AT3G61810 | 0,05 | 0,00 | 4,41 | 0,00 | 1,00E+00 |
| AT3G61850 | 0,15 | 1,16 | 4,35 | 0,27 | 7,90E-01 |
| AT3G61870 | 1,16 | -1,36 | 1,55 | -0,88 | 3,80E-01 |
| AT3G61950 | 0,11 | 1,07 | 4,36 | 0,25 | 8,06E-01 |
| AT3G62030 | 0,98 | -1,87 | 2,16 | -0,87 | 3,85E-01 |
| AT3G62130 | 0,39 | 0,47 | 4,36 | 0,11 | 9,14E-01 |
| AT3G62190 | 0,24 | 0,44 | 4,35 | 0,10 | 9,19E-01 |
| AT3G62245 | 0,42 | 0,86 | 4,13 | 0,21 | 8,36E-01 |
| AT3G62300 | 0,93 | -0,14 | 3,36 | -0,04 | 9,67E-01 |
| AT3G62370 | 0,90 | 1,29 | 2,12 | 0,61 | 5,42E-01 |
| AT3G62530 | 0,72 | 1,93 | 2,39 | 0,81 | 4,19E-01 |
| AT3G62550 | 0,70 | -1,52 | 3,65 | -0,42 | 6,77E-01 |
| AT3G62560 | 0,61 | 0,54 | 3,20 | 0,17 | 8,65E-01 |
| AT3G62580 | 0,64 | 2,01 | 2,25 | 0,89 | 3,73E-01 |
| AT3G09815 | 0,08 | 0,85 | 4,38 | 0,20 | 8,45E-01 |
| AT3G09835 | 1,22 | -3,04 | 1,81 | -1,68 | 9,33E-02 |
| AT3G62620 | 0,29 | -1,23 | 4,39 | -0,28 | 7,79E-01 |
| AT3G62650 | 1,34 | 1,78 | 1,82 | 0,98 | 3,27E-01 |
| AT3G62660 | 1,25 | -0,28 | 1,67 | -0,17 | 8,66E-01 |
| AT3G62720 | 0,40 | 2,11 | 4,10 | 0,51 | 6,08E-01 |
| AT3G62750 | 0,68 | -1,22 | 2,57 | -0,48 | 6,34E-01 |
| AT3G62820 | 0,26 | 0,41 | 4,38 | 0,09 | 9,25E-01 |
| AT3G62830 | 0,86 | 0,86 | 1,92 | 0,45 | 6,52E-01 |
| AT3G62890 | 0,20 | 1,23 | 4,36 | 0,28 | 7,77E-01 |
| AT3G63000 | 0,33 | 2,05 | 4,33 | 0,47 | 6,36E-01 |
| AT3G63010 | 0,38 | 1,57 | 3,49 | 0,45 | 6,53E-01 |
| AT3G63080 | 0,53 | 1,63 | 2,72 | 0,60 | 5,49E-01 |
| AT3G63130 | 0,43 | -0,37 | 4,11 | -0,09 | 9,28E-01 |
| AT3G63170 | 0,32 | 0,00 | 4,41 | 0,00 | 1,00E+00 |
| AT3G63215 | 0,08 | 0,00 | 4,40 | 0,00 | 1,00E+00 |
| AT3G63300 | 0,72 | 1,61 | 2,88 | 0,56 | 5,75E-01 |
| AT3G63370 | 0,57 | 1,72 | 3,13 | 0,55 | 5,81E-01 |
| AT3G63400 | 0,95 | 1,02 | 2,14 | 0,48 | 6,33E-01 |
| AT3G63420 | 0,46 | 1,91 | 4,33 | 0,44 | 6,60E-01 |
| AT3G63470 | 0,42 | 1,63 | 4,17 | 0,39 | 6,97E-01 |
| AT3G09995 | 0,45 | 1,54 | 4,00 | 0,39 | 7,00E-01 |
| AT3G63520 | 0,87 | -0,02 | 2,32 | -0,01 | 9,92E-01 |
| AT4G00030 | 0,38 | -1,02 | 4,35 | -0,24 | 8,14E-01 |
| AT4G00100 | 1,39 | -0,72 | 1,84 | -0,39 | 6,95E-01 |
| AT4G00163 | 0,65 | -2,39 | 2,67 | -0,90 | 3,70E-01 |
| AT4G00230 | 0,69 | 0,53 | 4,39 | 0,12 | 9,05E-01 |
| AT4G00300 | 1,08 | -1,71 | 1,92 | -0,89 | 3,74E-01 |
| AT4G00340 | 0,31 | 0,46 | 4,39 | 0,11 | 9,16E-01 |
| AT4G00440 | 0,33 | 1,65 | 4,34 | 0,38 | 7,04E-01 |
| AT4G00490 | 0,44 | 1,65 | 4,33 | 0,38 | 7,03E-01 |
| AT4G00580 | 0,48 | 0,42 | 4,04 | 0,10 | 9,18E-01 |
| AT4G00660 | 1,18 | 2,30 | 2,06 | 1,12 | 2,63E-01 |
| AT4G00710 | 0,48 | 0,59 | 4,32 | 0,14 | 8,92E-01 |
| AT4G00750 | 0,96 | 0,22 | 1,80 | 0,12 | 9,03E-01 |

| | | | | | |
|---|---|---|---|---|---|
| AT4G00820 | 0,05 | 0,00 | 4,41 | 0,00 | 1,00E+00 |
| AT4G00830 | 0,92 | 1,02 | 2,11 | 0,48 | 6,29E-01 |
| AT4G00889 | 0,23 | 0,00 | 4,41 | 0,00 | 1,00E+00 |
| AT4G00930 | 0,17 | 1,92 | 4,33 | 0,44 | 6,57E-01 |
| AT4G00953 | 0,67 | -0,12 | 2,54 | -0,05 | 9,64E-01 |
| AT4G00955 | 0,42 | -0,64 | 2,84 | -0,22 | 8,23E-01 |
| AT4G00960 | 1,30 | -0,76 | 2,16 | -0,35 | 7,27E-01 |
| AT4G00970 | 0,33 | 0,42 | 4,38 | 0,09 | 9,24E-01 |
| AT4G00990 | 0,61 | 1,92 | 3,93 | 0,49 | 6,26E-01 |
| AT4G01020 | 0,37 | 0,13 | 4,34 | 0,03 | 9,75E-01 |
| AT4G01026 | 0,56 | 2,13 | 2,64 | 0,81 | 4,21E-01 |
| AT4G01040 | 0,12 | 1,39 | 4,34 | 0,32 | 7,49E-01 |
| AT4G01100 | 0,48 | 0,35 | 3,19 | 0,11 | 9,12E-01 |
| AT4G01130 | 0,11 | 0,00 | 4,41 | 0,00 | 1,00E+00 |
| AT4G01280 | 0,59 | 0,67 | 3,14 | 0,21 | 8,30E-01 |
| AT4G01320 | 0,90 | 2,79 | 2,13 | 1,31 | 1,90E-01 |
| AT4G01330 | 0,80 | 1,29 | 2,51 | 0,52 | 6,07E-01 |
| AT4G01340 | 1,48 | -2,62 | 1,52 | -1,73 | 8,44E-02 |
| AT4G01360 | 0,89 | 2,27 | 2,92 | 0,78 | 4,38E-01 |
| AT4G01370 | 0,27 | 1,71 | 4,33 | 0,40 | 6,93E-01 |
| AT4G01460 | 0,10 | 0,42 | 4,38 | 0,10 | 9,23E-01 |
| AT4G01570 | 0,24 | 0,74 | 4,33 | 0,17 | 8,64E-01 |
| AT4G03905 | 0,55 | 2,30 | 3,91 | 0,59 | 5,57E-01 |
| AT4G03925 | 0,76 | -0,09 | 2,22 | -0,04 | 9,67E-01 |
| AT4G01610 | 0,62 | 2,30 | 2,49 | 0,92 | 3,55E-01 |
| AT4G01690 | 1,14 | 0,41 | 1,70 | 0,24 | 8,11E-01 |
| AT4G01720 | 0,44 | 0,48 | 4,14 | 0,12 | 9,07E-01 |
| AT4G01995 | 0,08 | 0,85 | 4,38 | 0,20 | 8,45E-01 |
| AT4G02000 | 0,69 | -0,80 | 2,10 | -0,38 | 7,03E-01 |
| AT4G02010 | 0,14 | 0,82 | 4,37 | 0,19 | 8,51E-01 |
| AT4G02020 | 0,55 | 0,53 | 3,22 | 0,17 | 8,69E-01 |
| AT4G02030 | 0,75 | 2,21 | 2,78 | 0,80 | 4,26E-01 |
| AT4G02070 | 0,12 | 1,46 | 4,35 | 0,34 | 7,36E-01 |
| AT4G02080 | 0,57 | 0,45 | 2,53 | 0,18 | 8,58E-01 |
| AT4G02100 | 0,08 | 0,00 | 4,40 | 0,00 | 1,00E+00 |
| AT4G02110 | 0,21 | 1,99 | 4,33 | 0,46 | 6,46E-01 |
| AT4G02200 | 0,55 | 2,26 | 3,84 | 0,59 | 5,57E-01 |
| AT4G02220 | 0,29 | 1,33 | 4,34 | 0,31 | 7,60E-01 |
| AT4G02260 | 0,56 | -0,07 | 3,00 | -0,02 | 9,82E-01 |
| AT4G02330 | 0,22 | -1,71 | 4,37 | -0,39 | 6,95E-01 |
| AT4G02340 | 1,02 | 0,40 | 2,33 | 0,17 | 8,62E-01 |
| AT4G02370 | 0,72 | -0,74 | 2,98 | -0,25 | 8,05E-01 |
| AT4G02390 | 0,10 | 0,42 | 4,38 | 0,10 | 9,23E-01 |
| AT4G02400 | 0,19 | 1,57 | 4,34 | 0,36 | 7,18E-01 |
| AT4G02440 | 0,90 | 0,59 | 2,03 | 0,29 | 7,73E-01 |
| AT4G04135 | 0,05 | 0,00 | 4,41 | 0,00 | 1,00E+00 |
| AT4G02500 | 1,37 | 2,62 | 2,72 | 0,96 | 3,36E-01 |
| AT4G04195 | 0,38 | 1,92 | 3,38 | 0,57 | 5,70E-01 |
| AT4G02540 | 0,29 | -1,10 | 3,68 | -0,30 | 7,65E-01 |
| AT4G02580 | 1,19 | -0,19 | 2,22 | -0,09 | 9,30E-01 |
| AT4G02600 | 0,66 | -1,07 | 2,29 | -0,47 | 6,40E-01 |
| AT4G02610 | 0,31 | 0,51 | 4,35 | 0,12 | 9,06E-01 |
| AT4G02650 | 0,24 | 1,58 | 4,34 | 0,36 | 7,16E-01 |
| AT4G02700 | 1,40 | 0,33 | 1,75 | 0,19 | 8,50E-01 |
| AT4G02725 | 0,69 | -1,53 | 2,94 | -0,52 | 6,03E-01 |
| AT4G02790 | 0,08 | 1,10 | 4,36 | 0,25 | 8,00E-01 |

| | | | | | |
|---|---|---|---|---|---|
| AT4G02880 | 0,95 | 2,04 | 2,19 | 0,93 | 3,53E-01 |
| AT4G02920 | 1,02 | 2,04 | 2,27 | 0,90 | 3,69E-01 |
| AT4G02940 | 0,12 | 0,00 | 4,39 | 0,00 | 1,00E+00 |
| AT4G02990 | 0,69 | 0,52 | 2,68 | 0,19 | 8,47E-01 |
| AT4G03000 | 0,64 | 1,06 | 2,94 | 0,36 | 7,19E-01 |
| AT4G04425 | 0,08 | 0,85 | 4,38 | 0,20 | 8,45E-01 |
| AT4G03100 | 0,38 | 0,00 | 4,39 | 0,00 | 1,00E+00 |
| AT4G03120 | 0,23 | 1,16 | 4,35 | 0,27 | 7,89E-01 |
| AT4G03180 | 0,27 | -0,02 | 4,34 | 0,00 | 9,96E-01 |
| AT4G03200 | 0,82 | 0,92 | 2,29 | 0,40 | 6,88E-01 |
| AT4G03205 | 0,48 | 1,23 | 3,21 | 0,38 | 7,01E-01 |
| AT4G03210 | 0,53 | 1,98 | 4,32 | 0,46 | 6,47E-01 |
| AT4G03285 | 0,16 | 1,33 | 4,35 | 0,31 | 7,59E-01 |
| AT4G03410 | 0,58 | -1,34 | 3,87 | -0,35 | 7,29E-01 |
| AT4G03500 | 0,47 | 15,71 | 4,40 | 3,57 | 3,54E-04 |
| AT4G03830 | 0,61 | 0,39 | 2,56 | 0,15 | 8,80E-01 |
| AT4G03930 | 0,65 | 2,25 | 2,99 | 0,75 | 4,51E-01 |
| AT4G03940 | 0,17 | 1,75 | 4,33 | 0,40 | 6,86E-01 |
| AT4G03945 | 0,92 | 2,08 | 2,34 | 0,89 | 3,73E-01 |
| AT4G04110 | 0,65 | 0,21 | 4,38 | 0,05 | 9,62E-01 |
| AT4G04180 | 0,08 | 0,00 | 4,40 | 0,00 | 1,00E+00 |
| AT4G04210 | 0,39 | 1,74 | 2,86 | 0,61 | 5,44E-01 |
| AT4G04220 | 0,89 | 2,10 | 2,15 | 0,98 | 3,29E-01 |
| AT4G04795 | 0,87 | 2,55 | 2,55 | 1,00 | 3,16E-01 |
| AT4G04315 | 0,10 | 0,42 | 4,38 | 0,10 | 9,23E-01 |
| AT4G04350 | 0,42 | -0,25 | 3,41 | -0,07 | 9,42E-01 |
| AT4G04450 | 0,17 | -1,64 | 4,37 | -0,37 | 7,08E-01 |
| AT4G04510 | 0,17 | 0,00 | 4,40 | 0,00 | 1,00E+00 |
| AT4G04540 | 0,26 | 0,15 | 4,36 | 0,03 | 9,73E-01 |
| AT4G04610 | 0,73 | -0,60 | 2,07 | -0,29 | 7,70E-01 |
| AT4G04670 | 0,48 | 2,27 | 4,33 | 0,53 | 5,99E-01 |
| AT4G04750 | 0,90 | -0,14 | 1,85 | -0,08 | 9,39E-01 |
| AT4G04760 | 1,18 | -0,96 | 2,08 | -0,46 | 6,45E-01 |
| AT4G04800 | 0,52 | 1,79 | 3,86 | 0,46 | 6,44E-01 |
| AT4G04840 | 1,00 | -1,58 | 2,09 | -0,76 | 4,49E-01 |
| AT4G04850 | 0,79 | 0,26 | 1,92 | 0,13 | 8,94E-01 |
| AT4G04860 | 0,28 | 0,67 | 3,70 | 0,18 | 8,56E-01 |
| AT4G04920 | 1,70 | 3,87 | 1,77 | 2,19 | 2,89E-02 |
| AT4G04960 | 0,39 | 1,63 | 2,86 | 0,57 | 5,69E-01 |
| AT4G05020 | 0,40 | 1,89 | 4,26 | 0,44 | 6,58E-01 |
| AT4G05080 | 0,23 | 0,00 | 4,41 | 0,00 | 1,00E+00 |
| AT4G05105 | 0,17 | 1,73 | 4,34 | 0,40 | 6,91E-01 |
| AT4G05160 | 0,46 | 2,70 | 4,05 | 0,67 | 5,05E-01 |
| AT4G05612 | 0,12 | 1,58 | 4,34 | 0,36 | 7,16E-01 |
| AT4G05632 | 0,08 | 0,00 | 4,40 | 0,00 | 1,00E+00 |
| AT4G06479 | 0,10 | 0,00 | 4,41 | 0,00 | 1,00E+00 |
| AT4G06598 | 0,74 | 0,00 | 3,57 | 0,00 | 1,00E+00 |
| AT4G06688 | 0,68 | 2,14 | 2,44 | 0,88 | 3,81E-01 |
| AT4G07390 | 0,27 | 1,59 | 3,74 | 0,42 | 6,71E-01 |
| AT4G07455 | 0,24 | 0,00 | 4,41 | 0,00 | 1,00E+00 |
| AT4G07825 | 1,05 | 0,99 | 2,67 | 0,37 | 7,11E-01 |
| AT4G07932 | 0,51 | -2,66 | 4,08 | -0,65 | 5,14E-01 |
| AT4G07965 | 0,15 | 1,58 | 4,34 | 0,36 | 7,15E-01 |
| AT4G08025 | 0,16 | 1,26 | 4,35 | 0,29 | 7,72E-01 |
| AT4G05305 | 0,12 | 0,00 | 4,41 | 0,00 | 1,00E+00 |
| AT4G08097 | 0,66 | -0,89 | 2,54 | -0,35 | 7,25E-01 |

| | | | | | |
|---|---|---|---|---|---|
| AT4G08180 | 1,01 | 0,02 | 1,94 | 0,01 | 9,92E-01 |
| AT4G08267 | 0,63 | 0,74 | 4,34 | 0,17 | 8,64E-01 |
| AT4G08270 | 0,17 | 1,03 | 4,37 | 0,24 | 8,13E-01 |
| AT4G08320 | 0,31 | 0,00 | 4,40 | 0,00 | 1,00E+00 |
| AT4G08350 | 0,65 | 1,67 | 2,57 | 0,65 | 5,16E-01 |
| AT4G08395 | 0,29 | 0,68 | 4,37 | 0,16 | 8,76E-01 |
| AT4G05455 | 0,68 | -2,17 | 2,45 | -0,88 | 3,77E-01 |
| AT4G08450 | 0,11 | 0,00 | 4,41 | 0,00 | 1,00E+00 |
| AT4G08455 | 0,63 | 2,56 | 4,32 | 0,59 | 5,53E-01 |
| AT4G08510 | 0,33 | 1,01 | 4,33 | 0,23 | 8,15E-01 |
| AT4G08545 | 1,36 | 3,02 | 1,84 | 1,64 | 1,00E-01 |
| AT4G08560 | 0,64 | -1,64 | 4,37 | -0,37 | 7,08E-01 |
| AT4G08780 | 0,43 | 0,67 | 4,17 | 0,16 | 8,72E-01 |
| AT4G05605 | 0,39 | 0,00 | 4,41 | 0,00 | 1,00E+00 |
| AT4G08930 | 0,71 | 2,25 | 2,39 | 0,94 | 3,46E-01 |
| AT4G08950 | 0,37 | 0,97 | 4,33 | 0,22 | 8,23E-01 |
| AT4G08980 | 0,65 | 1,64 | 2,91 | 0,56 | 5,72E-01 |
| AT4G08990 | 0,69 | 1,93 | 3,71 | 0,52 | 6,02E-01 |
| AT4G09020 | 1,04 | 0,12 | 1,84 | 0,06 | 9,48E-01 |
| AT4G09030 | 0,16 | 1,26 | 4,35 | 0,29 | 7,72E-01 |
| AT4G09150 | 0,62 | 2,47 | 2,98 | 0,83 | 4,06E-01 |
| AT4G09160 | 0,36 | 0,00 | 4,41 | 0,00 | 1,00E+00 |
| AT4G09250 | 0,12 | 1,39 | 4,34 | 0,32 | 7,49E-01 |
| AT4G09310 | 0,21 | 0,46 | 4,34 | 0,11 | 9,15E-01 |
| AT4G05705 | 0,10 | 0,00 | 4,41 | 0,00 | 1,00E+00 |
| AT4G09320 | 1,06 | -0,13 | 1,58 | -0,08 | 9,34E-01 |
| AT4G09360 | 0,71 | 1,27 | 2,41 | 0,53 | 5,98E-01 |
| AT4G09450 | 1,44 | -2,64 | 1,71 | -1,55 | 1,22E-01 |
| AT4G09460 | 0,37 | 1,68 | 4,29 | 0,39 | 6,95E-01 |
| AT4G09545 | 0,31 | -1,22 | 4,38 | -0,28 | 7,80E-01 |
| AT4G09570 | 0,54 | -0,46 | 2,43 | -0,19 | 8,51E-01 |
| AT4G09630 | 0,87 | -0,09 | 2,21 | -0,04 | 9,68E-01 |
| AT4G09660 | 0,30 | 0,43 | 4,38 | 0,10 | 9,23E-01 |
| AT4G09680 | 0,45 | 2,10 | 3,98 | 0,53 | 5,97E-01 |
| AT4G09690 | 1,50 | -2,23 | 1,61 | -1,39 | 1,65E-01 |
| AT4G09730 | 0,51 | -1,61 | 3,14 | -0,51 | 6,09E-01 |
| AT4G09800 | 0,41 | -1,36 | 4,31 | -0,32 | 7,52E-01 |
| AT4G09820 | 0,31 | -2,48 | 4,36 | -0,57 | 5,70E-01 |
| AT4G09830 | 0,41 | 0,96 | 3,33 | 0,29 | 7,72E-01 |
| AT4G09850 | 0,21 | 0,75 | 4,37 | 0,17 | 8,63E-01 |
| AT4G05745 | 0,45 | 1,88 | 3,16 | 0,60 | 5,51E-01 |
| AT4G09890 | 0,18 | 1,56 | 4,34 | 0,36 | 7,18E-01 |
| AT4G10060 | 0,79 | 2,41 | 2,20 | 1,09 | 2,75E-01 |
| AT4G10170 | 0,08 | 1,10 | 4,36 | 0,25 | 8,00E-01 |
| AT4G10180 | 0,21 | 2,05 | 4,33 | 0,47 | 6,36E-01 |
| AT4G10465 | 0,60 | 0,98 | 2,62 | 0,37 | 7,09E-01 |
| AT4G10610 | 1,30 | -0,12 | 2,25 | -0,05 | 9,57E-01 |
| AT4G10613 | 0,77 | 0,46 | 2,07 | 0,22 | 8,23E-01 |
| AT4G10620 | 0,24 | 0,00 | 4,41 | 0,00 | 1,00E+00 |
| AT4G10660 | 0,11 | 0,00 | 4,41 | 0,00 | 1,00E+00 |
| AT4G10840 | 1,51 | 3,36 | 2,04 | 1,64 | 1,00E-01 |
| AT4G11000 | 0,26 | 1,30 | 4,35 | 0,30 | 7,65E-01 |
| AT4G11070 | 0,37 | -1,37 | 4,37 | -0,31 | 7,54E-01 |
| AT4G11100 | 0,32 | 1,52 | 4,33 | 0,35 | 7,25E-01 |
| AT4G11110 | 0,51 | -0,80 | 2,55 | -0,31 | 7,54E-01 |
| AT4G11130 | 0,54 | 1,46 | 3,91 | 0,37 | 7,08E-01 |

| | | | | | |
|---|---|---|---|---|---|
| AT4G11150 | 1,03 | -0,89 | 1,67 | -0,53 | 5,95E-01 |
| AT4G11160 | 1,61 | 1,27 | 1,70 | 0,75 | 4,56E-01 |
| AT4G11170 | 0,50 | -2,36 | 3,13 | -0,75 | 4,51E-01 |
| AT4G11240 | 1,11 | 2,88 | 1,98 | 1,45 | 1,46E-01 |
| AT4G11280 | 0,67 | -0,37 | 3,09 | -0,12 | 9,05E-01 |
| AT4G11330 | 0,53 | 0,48 | 3,12 | 0,15 | 8,77E-01 |
| AT4G11350 | 0,19 | 1,51 | 4,34 | 0,35 | 7,28E-01 |
| AT4G11360 | 0,31 | -1,06 | 4,31 | -0,25 | 8,06E-01 |
| AT4G05885 | 0,80 | -3,21 | 2,63 | -1,22 | 2,22E-01 |
| AT4G11660 | 0,57 | 1,97 | 3,23 | 0,61 | 5,43E-01 |
| AT4G11720 | 0,60 | -1,48 | 2,98 | -0,50 | 6,19E-01 |
| AT4G11740 | 0,35 | 1,14 | 4,34 | 0,26 | 7,93E-01 |
| AT4G11880 | 0,28 | -1,36 | 4,41 | -0,31 | 7,58E-01 |
| AT4G11890 | 0,62 | 1,86 | 3,01 | 0,62 | 5,36E-01 |
| AT4G11910 | 0,12 | 1,65 | 4,34 | 0,38 | 7,04E-01 |
| AT4G12020 | 1,23 | 4,05 | 2,52 | 1,61 | 1,08E-01 |
| AT4G12030 | 0,68 | 3,49 | 3,46 | 1,01 | 3,13E-01 |
| AT4G12040 | 1,45 | 1,79 | 1,77 | 1,01 | 3,13E-01 |
| AT4G12070 | 0,19 | -1,36 | 4,41 | -0,31 | 7,58E-01 |
| AT4G12240 | 0,26 | 2,16 | 4,32 | 0,50 | 6,17E-01 |
| AT4G05985 | 0,10 | 0,42 | 4,38 | 0,10 | 9,23E-01 |
| AT4G12390 | 0,50 | 0,43 | 4,38 | 0,10 | 9,22E-01 |
| AT4G12600 | 0,92 | 1,79 | 2,91 | 0,62 | 5,38E-01 |
| AT4G12640 | 0,26 | 0,73 | 3,75 | 0,20 | 8,45E-01 |
| AT4G12680 | 0,21 | 1,60 | 4,35 | 0,37 | 7,13E-01 |
| AT4G12690 | 0,12 | 0,79 | 4,37 | 0,18 | 8,57E-01 |
| AT4G12720 | 1,54 | 3,37 | 1,79 | 1,88 | 5,97E-02 |
| AT4G12830 | 0,34 | -1,55 | 4,37 | -0,35 | 7,24E-01 |
| AT4G12900 | 0,05 | 0,00 | 4,41 | 0,00 | 1,00E+00 |
| AT4G05995 | 0,98 | 2,59 | 2,50 | 1,04 | 3,01E-01 |
| AT4G13010 | 0,63 | -2,77 | 3,63 | -0,76 | 4,45E-01 |
| AT4G13020 | 0,54 | 1,39 | 3,14 | 0,44 | 6,58E-01 |
| AT4G06010 | 0,28 | 1,69 | 4,35 | 0,39 | 6,97E-01 |
| AT4G06015 | 0,10 | 0,62 | 4,38 | 0,14 | 8,87E-01 |
| AT4G13245 | 0,48 | -1,40 | 3,30 | -0,42 | 6,72E-01 |
| AT4G13270 | 0,50 | 1,65 | 3,11 | 0,53 | 5,96E-01 |
| AT4G13390 | 1,53 | 2,53 | 1,97 | 1,28 | 2,00E-01 |
| AT4G13440 | 0,58 | 0,72 | 4,03 | 0,18 | 8,57E-01 |
| AT4G13530 | 0,95 | 0,67 | 1,91 | 0,35 | 7,28E-01 |
| AT4G13572 | 0,31 | -1,23 | 4,39 | -0,28 | 7,79E-01 |
| AT4G13610 | 0,37 | 0,76 | 2,86 | 0,27 | 7,90E-01 |
| AT4G13615 | 0,92 | 1,91 | 1,87 | 1,02 | 3,06E-01 |
| AT4G13670 | 0,92 | 3,61 | 3,14 | 1,15 | 2,50E-01 |
| AT4G13730 | 0,24 | 0,87 | 4,37 | 0,20 | 8,42E-01 |
| AT4G13750 | 0,39 | -0,16 | 3,34 | -0,05 | 9,63E-01 |
| AT4G13760 | 0,54 | 2,31 | 3,16 | 0,73 | 4,64E-01 |
| AT4G13830 | 0,70 | 0,58 | 2,92 | 0,20 | 8,42E-01 |
| AT4G13850 | 0,81 | -0,48 | 2,02 | -0,24 | 8,11E-01 |
| AT4G13860 | 0,43 | 2,62 | 4,10 | 0,64 | 5,22E-01 |
| AT4G13992 | 1,30 | -0,07 | 1,86 | -0,04 | 9,69E-01 |
| AT4G14000 | 0,12 | 0,00 | 4,41 | 0,00 | 1,00E+00 |
| AT4G14140 | 0,81 | -0,89 | 2,08 | -0,43 | 6,70E-01 |
| AT4G14220 | 0,46 | 0,00 | 4,40 | 0,00 | 1,00E+00 |
| AT4G14270 | 0,92 | 0,62 | 1,93 | 0,32 | 7,48E-01 |
| AT4G14300 | 0,77 | 0,16 | 3,50 | 0,05 | 9,62E-01 |
| AT4G14330 | 0,14 | 0,82 | 4,37 | 0,19 | 8,51E-01 |

| | | | | | |
|---|---|---|---|---|---|
| AT4G14540 | 0,71 | -1,93 | 2,49 | -0,78 | 4,37E-01 |
| AT4G14548 | 1,46 | 0,64 | 1,99 | 0,32 | 7,47E-01 |
| AT4G14560 | 0,12 | 0,68 | 4,37 | 0,16 | 8,77E-01 |
| AT4G14570 | 0,70 | 2,48 | 2,80 | 0,89 | 3,74E-01 |
| AT4G14630 | 0,43 | -1,22 | 4,09 | -0,30 | 7,65E-01 |
| AT4G14700 | 0,12 | 1,65 | 4,34 | 0,38 | 7,04E-01 |
| AT4G06245 | 1,52 | -1,86 | 1,87 | -1,00 | 3,19E-01 |
| AT4G06255 | 0,39 | 0,88 | 4,35 | 0,20 | 8,40E-01 |
| AT4G06260 | 0,42 | 0,46 | 3,34 | 0,14 | 8,90E-01 |
| AT4G14760 | 0,91 | 1,17 | 2,36 | 0,49 | 6,22E-01 |
| AT4G14800 | 0,34 | 1,77 | 4,34 | 0,41 | 6,83E-01 |
| AT4G14810 | 0,45 | -0,20 | 3,36 | -0,06 | 9,52E-01 |
| AT4G14850 | 0,15 | 0,42 | 4,38 | 0,09 | 9,24E-01 |
| AT4G14870 | 0,48 | -0,39 | 2,74 | -0,14 | 8,87E-01 |
| AT4G14890 | 0,73 | 0,70 | 2,27 | 0,31 | 7,59E-01 |
| AT4G14950 | 0,54 | 0,91 | 2,59 | 0,35 | 7,27E-01 |
| AT4G15000 | 1,11 | -2,65 | 1,78 | -1,49 | 1,36E-01 |
| AT4G15020 | 0,86 | 2,27 | 2,32 | 0,98 | 3,27E-01 |
| AT4G15030 | 0,61 | -0,11 | 2,32 | -0,05 | 9,63E-01 |
| AT4G15090 | 0,08 | 1,10 | 4,36 | 0,25 | 8,00E-01 |
| AT4G15096 | 1,15 | -1,67 | 2,24 | -0,74 | 4,57E-01 |
| AT4G15165 | 0,40 | 0,00 | 4,41 | 0,00 | 1,00E+00 |
| AT4G15233 | 0,90 | 2,45 | 2,37 | 1,03 | 3,01E-01 |
| AT4G15245 | 0,49 | 0,46 | 2,69 | 0,17 | 8,65E-01 |
| AT4G15260 | 0,57 | 0,56 | 3,07 | 0,18 | 8,56E-01 |
| AT4G15280 | 1,02 | 0,93 | 2,24 | 0,42 | 6,77E-01 |
| AT4G15415 | 0,24 | 0,92 | 4,36 | 0,21 | 8,32E-01 |
| AT4G15420 | 1,24 | 3,23 | 2,14 | 1,51 | 1,31E-01 |
| AT4G15440 | 0,99 | -1,96 | 2,46 | -0,80 | 4,25E-01 |
| AT4G15450 | 0,24 | 0,00 | 4,41 | 0,00 | 1,00E+00 |
| AT4G15475 | 0,96 | 0,00 | 4,14 | 0,00 | 1,00E+00 |
| AT4G15540 | 0,39 | 1,27 | 3,50 | 0,36 | 7,17E-01 |
| AT4G15545 | 1,17 | 0,74 | 1,61 | 0,46 | 6,45E-01 |
| AT4G15660 | 0,10 | 0,42 | 4,38 | 0,10 | 9,23E-01 |
| AT4G15700 | 0,08 | 0,00 | 4,40 | 0,00 | 1,00E+00 |
| AT4G15720 | 0,08 | 1,10 | 4,36 | 0,25 | 8,00E-01 |
| AT4G15770 | 0,18 | 0,00 | 4,41 | 0,00 | 1,00E+00 |
| AT4G15775 | 0,24 | -2,53 | 4,34 | -0,58 | 5,60E-01 |
| AT4G15800 | 0,76 | 1,19 | 2,20 | 0,54 | 5,90E-01 |
| AT4G15820 | 0,10 | 0,62 | 4,38 | 0,14 | 8,87E-01 |
| AT4G15840 | 0,73 | 0,65 | 2,07 | 0,31 | 7,54E-01 |
| AT4G15850 | 0,38 | 0,90 | 4,38 | 0,21 | 8,37E-01 |
| AT4G15880 | 0,12 | 0,90 | 4,36 | 0,21 | 8,36E-01 |
| AT4G15940 | 0,10 | 0,00 | 4,41 | 0,00 | 1,00E+00 |
| AT4G16008 | 0,08 | 1,10 | 4,36 | 0,25 | 8,00E-01 |
| AT4G16015 | 0,19 | 1,57 | 4,34 | 0,36 | 7,18E-01 |
| AT4G16130 | 1,05 | 2,04 | 2,15 | 0,95 | 3,43E-01 |
| AT4G16140 | 0,34 | 0,44 | 4,34 | 0,10 | 9,20E-01 |
| AT4G16150 | 0,50 | 2,43 | 4,09 | 0,59 | 5,53E-01 |
| AT4G16310 | 1,01 | 2,95 | 2,05 | 1,44 | 1,50E-01 |
| AT4G16360 | 0,61 | -0,95 | 2,54 | -0,37 | 7,09E-01 |
| AT4G16380 | 0,81 | 1,97 | 2,55 | 0,77 | 4,40E-01 |
| AT4G16390 | 0,67 | 1,27 | 2,29 | 0,56 | 5,79E-01 |
| AT4G16400 | 1,31 | 1,67 | 2,05 | 0,81 | 4,16E-01 |
| AT4G16430 | 0,28 | 0,00 | 4,41 | 0,00 | 1,00E+00 |
| AT4G16480 | 0,65 | 3,45 | 3,52 | 0,98 | 3,26E-01 |

| | | | | | |
|---|---|---|---|---|---|
| AT4G16510 | 0,53 | -0,62 | 3,94 | -0,16 | 8,74E-01 |
| AT4G16590 | 0,08 | 1,10 | 4,36 | 0,25 | 8,00E-01 |
| AT4G16650 | 0,41 | -0,79 | 4,20 | -0,19 | 8,51E-01 |
| AT4G16670 | 0,65 | 0,74 | 2,27 | 0,32 | 7,45E-01 |
| AT4G16710 | 0,63 | 2,66 | 2,51 | 1,06 | 2,90E-01 |
| AT4G16780 | 0,60 | 1,36 | 3,88 | 0,35 | 7,25E-01 |
| AT4G16830 | 0,78 | -2,97 | 3,11 | -0,96 | 3,39E-01 |
| AT4G16845 | 0,11 | 0,00 | 4,41 | 0,00 | 1,00E+00 |
| AT4G16850 | 0,29 | 2,02 | 4,33 | 0,47 | 6,40E-01 |
| AT4G16855 | 0,17 | 1,94 | 4,33 | 0,45 | 6,55E-01 |
| AT4G16955 | 0,78 | 1,76 | 3,06 | 0,58 | 5,64E-01 |
| AT4G17010 | 0,55 | -2,92 | 4,32 | -0,68 | 4,99E-01 |
| AT4G17080 | 0,57 | 1,93 | 4,33 | 0,45 | 6,56E-01 |
| AT4G17085 | 0,42 | 0,42 | 4,37 | 0,10 | 9,23E-01 |
| AT4G06665 | 1,22 | -0,25 | 1,72 | -0,14 | 8,86E-01 |
| AT4G17098 | 0,61 | -2,44 | 2,52 | -0,97 | 3,34E-01 |
| AT4G17245 | 0,66 | 1,73 | 2,57 | 0,67 | 5,02E-01 |
| AT4G17250 | 0,41 | 2,13 | 4,32 | 0,49 | 6,22E-01 |
| AT4G17270 | 0,64 | 0,00 | 3,13 | 0,00 | 1,00E+00 |
| AT4G17300 | 0,71 | 0,92 | 2,27 | 0,40 | 6,86E-01 |
| AT4G17380 | 0,44 | 1,61 | 2,76 | 0,58 | 5,61E-01 |
| AT4G17500 | 0,96 | 2,76 | 2,24 | 1,23 | 2,19E-01 |
| AT4G17520 | 1,02 | -2,23 | 2,66 | -0,84 | 4,03E-01 |
| AT4G17600 | 1,35 | -0,11 | 1,47 | -0,08 | 9,39E-01 |
| AT4G17616 | 0,78 | 2,55 | 3,41 | 0,75 | 4,53E-01 |
| AT4G17620 | 1,00 | 3,26 | 2,38 | 1,37 | 1,72E-01 |
| AT4G17650 | 0,32 | 0,84 | 4,33 | 0,19 | 8,46E-01 |
| AT4G17770 | 1,15 | 0,15 | 1,89 | 0,08 | 9,36E-01 |
| AT4G17810 | 0,14 | -1,60 | 4,37 | -0,37 | 7,14E-01 |
| AT4G17830 | 0,83 | 1,72 | 2,51 | 0,69 | 4,92E-01 |
| AT4G17870 | 1,03 | 1,77 | 2,32 | 0,76 | 4,45E-01 |
| AT4G17900 | 0,47 | 0,96 | 2,73 | 0,35 | 7,26E-01 |
| AT4G18070 | 1,02 | -0,69 | 2,22 | -0,31 | 7,57E-01 |
| AT4G18160 | 0,40 | 0,00 | 4,39 | 0,00 | 1,00E+00 |
| AT4G18195 | 0,22 | 0,00 | 4,41 | 0,00 | 1,00E+00 |
| AT4G18205 | 1,32 | -1,55 | 1,58 | -0,98 | 3,28E-01 |
| AT4G18220 | 1,16 | 0,46 | 1,72 | 0,27 | 7,89E-01 |
| AT4G18240 | 1,25 | 0,29 | 1,90 | 0,15 | 8,77E-01 |
| AT4G18370 | 0,87 | 1,18 | 2,32 | 0,51 | 6,12E-01 |
| AT4G18380 | 0,33 | 0,13 | 4,35 | 0,03 | 9,76E-01 |
| AT4G06795 | 0,56 | 1,78 | 3,02 | 0,59 | 5,57E-01 |
| AT4G18530 | 0,35 | 0,49 | 4,35 | 0,11 | 9,10E-01 |
| AT4G18570 | 0,37 | 0,23 | 2,89 | 0,08 | 9,38E-01 |
| AT4G18710 | 1,19 | 2,39 | 1,82 | 1,31 | 1,89E-01 |
| AT4G18725 | 0,19 | 0,00 | 4,40 | 0,00 | 1,00E+00 |
| AT4G18730 | 1,33 | 0,92 | 1,87 | 0,49 | 6,24E-01 |
| AT4G18750 | 0,50 | 1,84 | 4,06 | 0,45 | 6,50E-01 |
| AT4G18820 | 1,31 | 2,32 | 1,81 | 1,28 | 1,99E-01 |
| AT4G18830 | 0,19 | 1,26 | 4,35 | 0,29 | 7,72E-01 |
| AT4G18890 | 0,44 | 0,00 | 4,13 | 0,00 | 1,00E+00 |
| AT4G18905 | 0,35 | 1,54 | 4,34 | 0,35 | 7,23E-01 |
| AT4G18930 | 0,31 | -0,45 | 4,35 | -0,10 | 9,18E-01 |
| AT4G18950 | 1,00 | 1,03 | 2,01 | 0,51 | 6,10E-01 |
| AT4G19020 | 0,17 | 1,75 | 4,33 | 0,40 | 6,86E-01 |
| AT4G06865 | 0,10 | 0,42 | 4,38 | 0,10 | 9,23E-01 |
| AT4G19090 | 1,32 | 1,14 | 1,75 | 0,65 | 5,14E-01 |

| | | | | | |
|---|---|---|---|---|---|
| AT4G19100 | 1,38 | -0,93 | 2,22 | -0,42 | 6,76E-01 |
| AT4G19190 | 0,39 | 1,58 | 4,17 | 0,38 | 7,05E-01 |
| AT4G19200 | 0,86 | -0,24 | 2,23 | -0,11 | 9,13E-01 |
| AT4G19210 | 1,66 | 2,57 | 1,44 | 1,79 | 7,43E-02 |
| AT4G19450 | 0,65 | 2,88 | 4,30 | 0,67 | 5,03E-01 |
| AT4G19490 | 0,82 | 2,33 | 2,75 | 0,85 | 3,97E-01 |
| AT4G19500 | 0,89 | 2,55 | 2,28 | 1,12 | 2,64E-01 |
| AT4G19510 | 1,30 | 3,83 | 1,94 | 1,97 | 4,86E-02 |
| AT4G19530 | 1,16 | 0,43 | 1,98 | 0,22 | 8,27E-01 |
| AT4G19540 | 0,68 | 2,80 | 2,91 | 0,96 | 3,35E-01 |
| AT4G19580 | 0,44 | 2,58 | 4,05 | 0,64 | 5,25E-01 |
| AT4G19600 | 0,21 | -1,18 | 4,33 | -0,27 | 7,85E-01 |
| AT4G19610 | 0,37 | 0,65 | 3,43 | 0,19 | 8,49E-01 |
| AT4G19650 | 0,19 | 1,82 | 4,33 | 0,42 | 6,74E-01 |
| AT4G19710 | 1,47 | 1,38 | 1,65 | 0,83 | 4,04E-01 |
| AT4G19960 | 0,50 | 2,44 | 3,19 | 0,76 | 4,45E-01 |
| AT4G20110 | 0,83 | 2,43 | 2,25 | 1,08 | 2,81E-01 |
| AT4G20140 | 0,84 | 3,70 | 4,17 | 0,89 | 3,75E-01 |
| AT4G06950 | 0,12 | 0,42 | 4,38 | 0,10 | 9,24E-01 |
| AT4G20270 | 0,28 | 2,15 | 4,32 | 0,50 | 6,20E-01 |
| AT4G20300 | 0,68 | -1,32 | 2,48 | -0,53 | 5,94E-01 |
| AT4G20310 | 0,08 | 1,10 | 4,36 | 0,25 | 8,00E-01 |
| AT4G20400 | 1,23 | 0,52 | 2,09 | 0,25 | 8,03E-01 |
| AT4G20720 | 0,34 | 1,59 | 4,35 | 0,37 | 7,14E-01 |
| AT4G20840 | 1,22 | 2,79 | 2,46 | 1,14 | 2,55E-01 |
| AT4G20860 | 0,52 | 2,49 | 3,11 | 0,80 | 4,23E-01 |
| AT4G20890 | 0,68 | -0,99 | 2,63 | -0,38 | 7,06E-01 |
| AT4G20940 | 0,56 | 1,60 | 4,35 | 0,37 | 7,13E-01 |
| AT4G20960 | 0,48 | 1,73 | 2,74 | 0,63 | 5,27E-01 |
| AT4G20980 | 1,36 | 1,05 | 1,84 | 0,57 | 5,67E-01 |
| AT4G21105 | 0,54 | 0,79 | 4,34 | 0,18 | 8,56E-01 |
| AT4G21160 | 0,15 | -1,24 | 4,39 | -0,28 | 7,77E-01 |
| AT4G21180 | 0,08 | 0,85 | 4,38 | 0,20 | 8,45E-01 |
| AT4G21210 | 0,33 | 0,46 | 4,34 | 0,11 | 9,15E-01 |
| AT4G21300 | 0,05 | 0,00 | 4,41 | 0,00 | 1,00E+00 |
| AT4G21323 | 0,38 | 0,90 | 4,38 | 0,21 | 8,37E-01 |
| AT4G21450 | 0,15 | 1,42 | 4,34 | 0,33 | 7,43E-01 |
| AT4G21470 | 0,32 | 1,05 | 4,33 | 0,24 | 8,08E-01 |
| AT4G21515 | 0,05 | 0,00 | 4,41 | 0,00 | 1,00E+00 |
| AT4G21540 | 0,68 | 3,26 | 2,82 | 1,15 | 2,49E-01 |
| AT4G21550 | 0,37 | 0,00 | 4,40 | 0,00 | 1,00E+00 |
| AT4G21620 | 0,72 | 0,64 | 2,56 | 0,25 | 8,02E-01 |
| AT4G21670 | 1,62 | -0,23 | 1,44 | -0,16 | 8,72E-01 |
| AT4G21740 | 0,47 | -0,51 | 2,38 | -0,21 | 8,32E-01 |
| AT4G21750 | 0,81 | 1,76 | 2,75 | 0,64 | 5,21E-01 |
| AT4G21790 | 0,64 | 0,95 | 2,59 | 0,37 | 7,14E-01 |
| AT4G21800 | 0,53 | 0,00 | 3,92 | 0,00 | 1,00E+00 |
| AT4G07030 | 0,38 | 2,16 | 4,33 | 0,50 | 6,17E-01 |
| AT4G21940 | 0,15 | 1,62 | 4,34 | 0,37 | 7,09E-01 |
| AT4G22010 | 0,22 | 0,52 | 4,34 | 0,12 | 9,05E-01 |
| AT4G22050 | 0,20 | 0,57 | 4,37 | 0,13 | 8,96E-01 |
| AT4G07040 | 0,12 | 0,00 | 4,39 | 0,00 | 1,00E+00 |
| AT4G22130 | 0,44 | 1,27 | 2,81 | 0,45 | 6,50E-01 |
| AT4G22270 | 0,20 | 1,07 | 4,36 | 0,25 | 8,06E-01 |
| AT4G07075 | 1,23 | 2,06 | 1,79 | 1,15 | 2,52E-01 |
| AT4G07080 | 1,23 | 2,06 | 1,79 | 1,15 | 2,52E-01 |

| | | | | | |
|---|---|---|---|---|---|
| AT4G22330 | 0,17 | 1,94 | 4,33 | 0,45 | 6,55E-01 |
| AT4G22467 | 0,12 | 0,42 | 4,38 | 0,10 | 9,24E-01 |
| AT4G22475 | 0,49 | -1,21 | 3,96 | -0,31 | 7,59E-01 |
| AT4G22670 | 0,32 | 0,25 | 4,33 | 0,06 | 9,55E-01 |
| AT4G22720 | 0,28 | 1,56 | 4,34 | 0,36 | 7,18E-01 |
| AT4G22860 | 0,08 | 1,10 | 4,36 | 0,25 | 8,00E-01 |
| AT4G22890 | 1,50 | -2,59 | 1,70 | -1,52 | 1,28E-01 |
| AT4G22910 | 0,08 | 0,85 | 4,38 | 0,20 | 8,45E-01 |
| AT4G22960 | 0,12 | 0,42 | 4,38 | 0,10 | 9,24E-01 |
| AT4G23030 | 0,17 | 0,00 | 4,40 | 0,00 | 1,00E+00 |
| AT4G23050 | 0,25 | 1,57 | 4,34 | 0,36 | 7,18E-01 |
| AT4G23060 | 0,21 | 0,79 | 4,37 | 0,18 | 8,57E-01 |
| AT4G23150 | 0,15 | -1,25 | 4,39 | -0,28 | 7,76E-01 |
| AT4G23160 | 0,75 | 0,00 | 4,40 | 0,00 | 1,00E+00 |
| AT4G23180 | 1,57 | 1,90 | 1,67 | 1,14 | 2,54E-01 |
| AT4G23270 | 0,94 | 2,24 | 2,05 | 1,10 | 2,73E-01 |
| AT4G23400 | 0,56 | -0,11 | 2,20 | -0,05 | 9,62E-01 |
| AT4G23440 | 0,12 | 1,58 | 4,34 | 0,36 | 7,16E-01 |
| AT4G23470 | 0,84 | 2,91 | 2,07 | 1,41 | 1,60E-01 |
| AT4G23530 | 1,11 | 2,29 | 2,11 | 1,09 | 2,76E-01 |
| AT4G23570 | 0,44 | -0,30 | 3,34 | -0,09 | 9,29E-01 |
| AT4G23600 | 1,02 | -2,07 | 2,55 | -0,81 | 4,18E-01 |
| AT4G23740 | 0,40 | 2,20 | 4,33 | 0,51 | 6,11E-01 |
| AT4G23860 | 0,25 | 0,00 | 4,41 | 0,00 | 1,00E+00 |
| AT4G07190 | 0,45 | -1,12 | 3,41 | -0,33 | 7,44E-01 |
| AT4G23895 | 0,24 | 0,59 | 4,34 | 0,13 | 8,93E-01 |
| AT4G23940 | 1,06 | 3,28 | 2,20 | 1,49 | 1,35E-01 |
| AT4G23950 | 0,16 | 0,90 | 4,38 | 0,21 | 8,37E-01 |
| AT4G23980 | 1,47 | 1,18 | 1,60 | 0,74 | 4,61E-01 |
| AT4G24000 | 0,08 | 1,10 | 4,36 | 0,25 | 8,00E-01 |
| AT4G24010 | 0,33 | 0,13 | 4,35 | 0,03 | 9,76E-01 |
| AT4G24020 | 1,49 | 1,85 | 1,76 | 1,05 | 2,94E-01 |
| AT4G24060 | 0,12 | 1,58 | 4,34 | 0,36 | 7,16E-01 |
| AT4G24090 | 0,53 | -3,75 | 4,30 | -0,87 | 3,83E-01 |
| AT4G24120 | 1,68 | 4,67 | 2,10 | 2,23 | 2,57E-02 |
| AT4G24130 | 0,24 | 0,00 | 4,41 | 0,00 | 1,00E+00 |
| AT4G24160 | 0,15 | 0,86 | 4,37 | 0,20 | 8,43E-01 |
| AT4G24175 | 0,21 | 1,39 | 4,34 | 0,32 | 7,49E-01 |
| AT4G24220 | 1,52 | -0,07 | 1,44 | -0,05 | 9,60E-01 |
| AT4G24290 | 0,72 | 1,79 | 2,42 | 0,74 | 4,61E-01 |
| AT4G24350 | 0,25 | -0,45 | 4,38 | -0,10 | 9,19E-01 |
| AT4G24440 | 0,43 | 0,84 | 4,21 | 0,20 | 8,42E-01 |
| AT4G24460 | 0,18 | -0,69 | 4,37 | -0,16 | 8,74E-01 |
| AT4G24470 | 0,57 | 2,35 | 3,84 | 0,61 | 5,40E-01 |
| AT4G24550 | 0,38 | 0,71 | 3,38 | 0,21 | 8,33E-01 |
| AT4G24790 | 0,10 | 0,62 | 4,38 | 0,14 | 8,87E-01 |
| AT4G24800 | 0,98 | 2,60 | 2,62 | 0,99 | 3,21E-01 |
| AT4G24805 | 0,88 | -1,48 | 2,53 | -0,59 | 5,58E-01 |
| AT4G24880 | 0,08 | 0,00 | 4,40 | 0,00 | 1,00E+00 |
| AT4G07295 | 0,21 | 0,05 | 4,34 | 0,01 | 9,90E-01 |
| AT4G25030 | 0,40 | -0,47 | 2,77 | -0,17 | 8,64E-01 |
| AT4G25050 | 1,37 | 0,09 | 1,61 | 0,05 | 9,57E-01 |
| AT4G25070 | 0,53 | 1,69 | 4,33 | 0,39 | 6,97E-01 |
| AT4G25120 | 0,50 | -0,73 | 3,25 | -0,22 | 8,23E-01 |
| AT4G25170 | 1,05 | -1,74 | 2,06 | -0,84 | 4,00E-01 |
| AT4G25210 | 0,75 | 0,00 | 2,98 | 0,00 | 1,00E+00 |

| | | | | | |
|---|---|---|---|---|---|
| AT4G25230 | 0,77 | 1,57 | 2,21 | 0,71 | 4,76E-01 |
| AT4G25360 | 0,25 | 2,14 | 4,33 | 0,49 | 6,21E-01 |
| AT4G25370 | 0,59 | 0,69 | 3,81 | 0,18 | 8,57E-01 |
| AT4G25390 | 0,17 | 0,48 | 4,40 | 0,11 | 9,12E-01 |
| AT4G25400 | 0,31 | 0,00 | 4,41 | 0,00 | 1,00E+00 |
| AT4G25500 | 1,12 | 1,03 | 2,12 | 0,49 | 6,25E-01 |
| AT4G25515 | 0,32 | 1,20 | 4,37 | 0,28 | 7,83E-01 |
| AT4G25550 | 0,38 | 0,90 | 4,38 | 0,21 | 8,37E-01 |
| AT4G25610 | 0,54 | 1,96 | 3,15 | 0,62 | 5,34E-01 |
| AT4G25630 | 1,57 | 2,13 | 1,98 | 1,08 | 2,82E-01 |
| AT4G25631 | 1,33 | 2,70 | 2,18 | 1,24 | 2,15E-01 |
| AT4G25660 | 0,81 | -0,64 | 2,31 | -0,28 | 7,82E-01 |
| AT4G25810 | 0,12 | 1,46 | 4,35 | 0,34 | 7,36E-01 |
| AT4G25835 | 0,21 | -0,09 | 4,37 | -0,02 | 9,83E-01 |
| AT4G25900 | 0,36 | 0,22 | 4,35 | 0,05 | 9,60E-01 |
| AT4G25910 | 0,69 | 2,10 | 3,50 | 0,60 | 5,48E-01 |
| AT4G25960 | 1,53 | -3,12 | 1,67 | -1,86 | 6,22E-02 |
| AT4G25970 | 1,00 | 1,86 | 1,93 | 0,96 | 3,35E-01 |
| AT4G26000 | 0,96 | 2,84 | 2,41 | 1,18 | 2,39E-01 |
| AT4G26120 | 0,24 | 0,00 | 4,41 | 0,00 | 1,00E+00 |
| AT4G26170 | 0,08 | 0,85 | 4,38 | 0,20 | 8,45E-01 |
| AT4G26210 | 0,36 | 2,25 | 4,33 | 0,52 | 6,04E-01 |
| AT4G26230 | 0,82 | 1,93 | 2,57 | 0,75 | 4,52E-01 |
| AT4G26450 | 0,45 | -0,02 | 3,32 | -0,01 | 9,96E-01 |
| AT4G26455 | 0,54 | 2,46 | 3,76 | 0,65 | 5,13E-01 |
| AT4G26510 | 0,72 | -0,08 | 2,53 | -0,03 | 9,75E-01 |
| AT4G26520 | 1,37 | 2,40 | 2,07 | 1,16 | 2,48E-01 |
| AT4G26650 | 0,56 | 0,71 | 2,31 | 0,31 | 7,57E-01 |
| AT4G26670 | 0,33 | -0,85 | 4,31 | -0,20 | 8,44E-01 |
| AT4G26680 | 0,40 | 2,62 | 4,13 | 0,64 | 5,25E-01 |
| AT4G26700 | 0,28 | 0,46 | 4,35 | 0,11 | 9,16E-01 |
| AT4G26790 | 0,42 | 0,80 | 4,12 | 0,20 | 8,45E-01 |
| AT4G26800 | 0,12 | 0,42 | 4,38 | 0,10 | 9,24E-01 |
| AT4G26840 | 0,82 | 1,14 | 2,19 | 0,52 | 6,04E-01 |
| AT4G26870 | 0,92 | 2,50 | 2,23 | 1,12 | 2,62E-01 |
| AT4G27010 | 0,79 | 2,32 | 2,13 | 1,09 | 2,75E-01 |
| AT4G27030 | 0,63 | 0,67 | 2,44 | 0,27 | 7,83E-01 |
| AT4G07775 | 0,58 | 2,37 | 2,94 | 0,80 | 4,21E-01 |
| AT4G27080 | 1,09 | -0,62 | 2,29 | -0,27 | 7,86E-01 |
| AT4G27260 | 0,26 | 1,08 | 4,37 | 0,25 | 8,05E-01 |
| AT4G27290 | 0,45 | 0,80 | 4,37 | 0,18 | 8,55E-01 |
| AT4G27310 | 0,93 | -0,06 | 2,01 | -0,03 | 9,78E-01 |
| AT4G27380 | 0,21 | -1,16 | 4,33 | -0,27 | 7,90E-01 |
| AT4G27395 | 0,73 | -1,71 | 2,67 | -0,64 | 5,21E-01 |
| AT4G27652 | 0,57 | 0,80 | 3,75 | 0,21 | 8,31E-01 |
| AT4G27654 | 0,56 | 2,17 | 2,64 | 0,82 | 4,10E-01 |
| AT4G27657 | 0,12 | 0,42 | 4,38 | 0,10 | 9,24E-01 |
| AT4G27680 | 0,93 | 3,03 | 2,10 | 1,44 | 1,49E-01 |
| AT4G27710 | 0,52 | 0,11 | 3,19 | 0,03 | 9,73E-01 |
| AT4G27720 | 1,38 | 2,04 | 1,84 | 1,11 | 2,68E-01 |
| AT4G27740 | 0,52 | 0,47 | 3,87 | 0,12 | 9,04E-01 |
| AT4G27750 | 0,69 | -0,33 | 4,36 | -0,08 | 9,40E-01 |
| AT4G27760 | 0,28 | 0,42 | 4,38 | 0,10 | 9,24E-01 |
| AT4G27765 | 0,37 | 1,57 | 4,28 | 0,37 | 7,14E-01 |
| AT4G27780 | 0,50 | 1,32 | 3,95 | 0,33 | 7,39E-01 |
| AT4G27860 | 0,21 | 1,54 | 4,34 | 0,36 | 7,22E-01 |

| | | | | | |
|---|---|---|---|---|---|
| AT4G27880 | 0,08 | 0,85 | 4,38 | 0,20 | 8,45E-01 |
| AT4G27910 | 0,46 | 2,05 | 4,05 | 0,51 | 6,13E-01 |
| AT4G27940 | 1,14 | 2,15 | 2,25 | 0,95 | 3,40E-01 |
| AT4G28010 | 0,17 | 1,03 | 4,37 | 0,24 | 8,13E-01 |
| AT4G28210 | 0,08 | 0,00 | 4,40 | 0,00 | 1,00E+00 |
| AT4G28220 | 0,27 | 1,45 | 4,35 | 0,33 | 7,39E-01 |
| AT4G28260 | 0,80 | 3,02 | 2,86 | 1,06 | 2,91E-01 |
| AT4G28300 | 0,21 | 2,05 | 4,33 | 0,47 | 6,36E-01 |
| AT4G28362 | 0,21 | 0,79 | 4,37 | 0,18 | 8,57E-01 |
| AT4G28400 | 0,26 | 0,42 | 4,38 | 0,10 | 9,24E-01 |
| AT4G28440 | 0,16 | 0,42 | 4,37 | 0,10 | 9,23E-01 |
| AT4G28490 | 0,74 | 3,41 | 3,59 | 0,95 | 3,43E-01 |
| AT4G28600 | 1,03 | 1,99 | 2,15 | 0,93 | 3,54E-01 |
| AT4G28660 | 0,48 | -2,34 | 4,05 | -0,58 | 5,64E-01 |
| AT4G28706 | 0,39 | -2,25 | 4,33 | -0,52 | 6,03E-01 |
| AT4G28710 | 0,41 | 2,02 | 4,19 | 0,48 | 6,30E-01 |
| AT4G28740 | 0,90 | 0,61 | 2,51 | 0,24 | 8,09E-01 |
| AT4G28780 | 0,15 | 0,71 | 4,37 | 0,16 | 8,71E-01 |
| AT4G28860 | 0,76 | 0,04 | 3,06 | 0,01 | 9,90E-01 |
| AT4G28870 | 0,12 | 0,00 | 4,41 | 0,00 | 1,00E+00 |
| AT4G28880 | 1,20 | 0,80 | 1,81 | 0,44 | 6,59E-01 |
| AT4G28980 | 0,73 | 1,49 | 2,45 | 0,61 | 5,44E-01 |
| AT4G29020 | 0,17 | 0,00 | 4,40 | 0,00 | 1,00E+00 |
| AT4G29035 | 0,22 | 0,00 | 4,41 | 0,00 | 1,00E+00 |
| AT4G29040 | 1,10 | 2,97 | 2,06 | 1,44 | 1,50E-01 |
| AT4G29070 | 0,33 | 0,80 | 4,37 | 0,18 | 8,55E-01 |
| AT4G29110 | 0,63 | 0,89 | 2,98 | 0,30 | 7,65E-01 |
| AT4G29120 | 0,68 | 3,29 | 2,82 | 1,17 | 2,43E-01 |
| AT4G29160 | 0,87 | 1,66 | 1,98 | 0,84 | 4,01E-01 |
| AT4G29170 | 0,92 | 2,27 | 4,32 | 0,53 | 5,99E-01 |
| AT4G29180 | 0,96 | 1,97 | 2,64 | 0,75 | 4,55E-01 |
| AT4G29210 | 0,68 | 0,59 | 2,53 | 0,23 | 8,17E-01 |
| AT4G29350 | 0,47 | 0,49 | 4,07 | 0,12 | 9,05E-01 |
| AT4G29400 | 0,21 | 0,00 | 4,38 | 0,00 | 1,00E+00 |
| AT4G29510 | 0,71 | 1,11 | 2,30 | 0,48 | 6,30E-01 |
| AT4G29740 | 1,04 | -2,42 | 2,06 | -1,17 | 2,41E-01 |
| AT4G29750 | 0,66 | -0,19 | 2,68 | -0,07 | 9,44E-01 |
| AT4G29790 | 1,30 | 2,97 | 1,86 | 1,60 | 1,09E-01 |
| AT4G08245 | 0,54 | 1,62 | 3,16 | 0,51 | 6,08E-01 |
| AT4G08255 | 0,51 | 1,82 | 4,33 | 0,42 | 6,75E-01 |
| AT4G29950 | 0,57 | 1,91 | 2,63 | 0,73 | 4,67E-01 |
| AT4G30010 | 0,55 | 1,65 | 3,23 | 0,51 | 6,09E-01 |
| AT4G30020 | 1,12 | -1,27 | 1,85 | -0,69 | 4,92E-01 |
| AT4G30130 | 0,24 | 1,26 | 4,37 | 0,29 | 7,73E-01 |
| AT4G30150 | 1,08 | 2,31 | 2,09 | 1,10 | 2,69E-01 |
| AT4G30310 | 0,60 | 0,85 | 3,67 | 0,23 | 8,16E-01 |
| AT4G30350 | 0,36 | 1,59 | 4,35 | 0,37 | 7,14E-01 |
| AT4G30400 | 0,15 | 0,42 | 4,38 | 0,09 | 9,24E-01 |
| AT4G08285 | 0,08 | 0,85 | 4,38 | 0,20 | 8,45E-01 |
| AT4G30480 | 0,31 | -1,23 | 4,39 | -0,28 | 7,79E-01 |
| AT4G30530 | 1,06 | -1,52 | 1,72 | -0,88 | 3,77E-01 |
| AT4G30650 | 0,58 | -1,75 | 2,70 | -0,65 | 5,18E-01 |
| AT4G30660 | 0,47 | -1,60 | 2,69 | -0,59 | 5,53E-01 |
| AT4G30780 | 0,23 | 0,00 | 4,41 | 0,00 | 1,00E+00 |
| AT4G30800 | 0,17 | -2,23 | 4,35 | -0,51 | 6,08E-01 |
| AT4G30940 | 0,31 | 0,46 | 4,39 | 0,11 | 9,16E-01 |

| | | | | | |
|---|---|---|---|---|---|
| AT4G30990 | 1,34 | 2,21 | 1,84 | 1,20 | 2,31E-01 |
| AT4G08425 | 1,45 | 0,72 | 1,45 | 0,50 | 6,20E-01 |
| AT4G31080 | 0,70 | 2,73 | 2,89 | 0,95 | 3,44E-01 |
| AT4G31115 | 0,40 | -0,01 | 4,24 | 0,00 | 9,98E-01 |
| AT4G31300 | 0,37 | -1,50 | 4,39 | -0,34 | 7,33E-01 |
| AT4G31340 | 0,63 | -0,43 | 3,76 | -0,11 | 9,09E-01 |
| AT4G31390 | 0,47 | -0,05 | 2,79 | -0,02 | 9,87E-01 |
| AT4G31430 | 0,50 | 1,68 | 3,97 | 0,42 | 6,73E-01 |
| AT4G31530 | 0,77 | -0,83 | 2,52 | -0,33 | 7,42E-01 |
| AT4G31580 | 0,97 | 2,92 | 2,10 | 1,39 | 1,64E-01 |
| AT4G31710 | 0,50 | 1,32 | 3,95 | 0,33 | 7,39E-01 |
| AT4G31715 | 0,08 | 1,10 | 4,36 | 0,25 | 8,00E-01 |
| AT4G31780 | 1,36 | 2,71 | 2,85 | 0,95 | 3,42E-01 |
| AT4G31790 | 0,26 | 0,33 | 4,35 | 0,08 | 9,39E-01 |
| AT4G31800 | 1,69 | 3,10 | 1,87 | 1,65 | 9,80E-02 |
| AT4G31805 | 0,98 | 2,88 | 3,30 | 0,87 | 3,82E-01 |
| AT4G31850 | 0,54 | 1,51 | 3,29 | 0,46 | 6,46E-01 |
| AT4G31910 | 0,49 | -0,63 | 2,72 | -0,23 | 8,15E-01 |
| AT4G08615 | 0,91 | -1,48 | 2,22 | -0,66 | 5,06E-01 |
| AT4G08625 | 1,54 | -0,52 | 1,51 | -0,34 | 7,31E-01 |
| AT4G31985 | 0,37 | 1,67 | 4,33 | 0,38 | 7,01E-01 |
| AT4G32010 | 1,47 | 2,46 | 2,03 | 1,21 | 2,26E-01 |
| AT4G32030 | 0,47 | 0,15 | 4,23 | 0,03 | 9,73E-01 |
| AT4G32040 | 0,84 | 2,59 | 3,45 | 0,75 | 4,52E-01 |
| AT4G32060 | 1,61 | -0,24 | 1,39 | -0,17 | 8,65E-01 |
| AT4G32160 | 0,93 | 0,98 | 1,96 | 0,50 | 6,19E-01 |
| AT4G32190 | 0,86 | 3,40 | 2,68 | 1,27 | 2,04E-01 |
| AT4G32200 | 0,65 | 1,65 | 3,58 | 0,46 | 6,46E-01 |
| AT4G32265 | 1,11 | -2,05 | 1,62 | -1,27 | 2,05E-01 |
| AT4G32300 | 0,70 | 2,21 | 2,22 | 1,00 | 3,19E-01 |
| AT4G32330 | 0,80 | 2,01 | 2,22 | 0,90 | 3,66E-01 |
| AT4G32360 | 0,05 | 0,00 | 4,41 | 0,00 | 1,00E+00 |
| AT4G32390 | 1,34 | 2,87 | 1,76 | 1,63 | 1,03E-01 |
| AT4G32430 | 0,12 | 0,42 | 4,38 | 0,10 | 9,24E-01 |
| AT4G32551 | 0,49 | 0,64 | 2,72 | 0,24 | 8,13E-01 |
| AT4G32600 | 0,53 | 2,21 | 3,30 | 0,67 | 5,03E-01 |
| AT4G32620 | 1,61 | 4,07 | 2,08 | 1,95 | 5,07E-02 |
| AT4G32630 | 0,68 | 0,49 | 4,40 | 0,11 | 9,12E-01 |
| AT4G32700 | 0,67 | 2,78 | 3,62 | 0,77 | 4,42E-01 |
| AT4G32720 | 0,62 | 1,32 | 2,98 | 0,44 | 6,58E-01 |
| AT4G32730 | 0,45 | 2,14 | 3,98 | 0,54 | 5,90E-01 |
| AT4G32770 | 0,32 | 1,55 | 4,34 | 0,36 | 7,22E-01 |
| AT4G32800 | 0,31 | 0,82 | 4,37 | 0,19 | 8,51E-01 |
| AT4G32850 | 1,20 | -0,19 | 1,78 | -0,10 | 9,17E-01 |
| AT4G32910 | 0,98 | 0,55 | 1,88 | 0,29 | 7,70E-01 |
| AT4G32930 | 0,91 | 2,16 | 2,44 | 0,88 | 3,78E-01 |
| AT4G33000 | 0,14 | 0,00 | 4,41 | 0,00 | 1,00E+00 |
| AT4G33030 | 0,99 | 2,13 | 2,51 | 0,85 | 3,97E-01 |
| AT4G33060 | 0,15 | 1,58 | 4,34 | 0,36 | 7,15E-01 |
| AT4G33180 | 0,05 | 0,00 | 4,41 | 0,00 | 1,00E+00 |
| AT4G33220 | 0,80 | 0,00 | 3,16 | 0,00 | 9,99E-01 |
| AT4G33290 | 1,50 | 1,79 | 1,87 | 0,96 | 3,39E-01 |
| AT4G08785 | 0,69 | 1,50 | 3,77 | 0,40 | 6,91E-01 |
| AT4G33310 | 0,49 | 0,00 | 4,40 | 0,00 | 1,00E+00 |
| AT4G33380 | 0,54 | 1,47 | 3,29 | 0,45 | 6,55E-01 |
| AT4G33410 | 0,32 | 1,65 | 4,34 | 0,38 | 7,04E-01 |

| | | | | | |
|---|---|---|---|---|---|
| AT4G33510 | 1,23 | 0,37 | 1,51 | 0,25 | 8,04E-01 |
| AT4G33520 | 1,37 | 4,08 | 2,22 | 1,84 | 6,55E-02 |
| AT4G33540 | 0,41 | 1,80 | 4,25 | 0,42 | 6,71E-01 |
| AT4G33565 | 0,80 | 0,57 | 2,88 | 0,20 | 8,44E-01 |
| AT4G33580 | 1,02 | -0,96 | 3,20 | -0,30 | 7,64E-01 |
| AT4G33620 | 0,33 | 2,05 | 4,33 | 0,47 | 6,36E-01 |
| AT4G33630 | 1,15 | 0,71 | 2,09 | 0,34 | 7,35E-01 |
| AT4G33666 | 0,24 | 0,65 | 4,37 | 0,15 | 8,81E-01 |
| AT4G33690 | 0,48 | 0,00 | 4,09 | 0,00 | 1,00E+00 |
| AT4G33890 | 1,49 | 0,90 | 1,87 | 0,48 | 6,32E-01 |
| AT4G33920 | 0,65 | 1,35 | 3,64 | 0,37 | 7,11E-01 |
| AT4G33960 | 0,45 | -3,35 | 4,31 | -0,78 | 4,38E-01 |
| AT4G34030 | 0,40 | -1,38 | 3,39 | -0,41 | 6,83E-01 |
| AT4G34035 | 0,32 | 1,25 | 4,35 | 0,29 | 7,74E-01 |
| AT4G34050 | 1,48 | 0,94 | 1,76 | 0,53 | 5,94E-01 |
| AT4G34060 | 0,12 | 1,58 | 4,34 | 0,36 | 7,16E-01 |
| AT4G34120 | 1,07 | -0,78 | 1,85 | -0,42 | 6,72E-01 |
| AT4G34160 | 0,73 | 2,98 | 2,40 | 1,25 | 2,13E-01 |
| AT4G34240 | 0,30 | -1,29 | 4,40 | -0,29 | 7,69E-01 |
| AT4G34260 | 1,45 | 2,23 | 2,10 | 1,06 | 2,88E-01 |
| AT4G34280 | 0,96 | 2,84 | 2,74 | 1,04 | 3,00E-01 |
| AT4G34290 | 0,10 | 0,42 | 4,38 | 0,10 | 9,23E-01 |
| AT4G34310 | 0,63 | 0,91 | 4,38 | 0,21 | 8,36E-01 |
| AT4G34330 | 0,08 | 1,10 | 4,36 | 0,25 | 8,00E-01 |
| AT4G34370 | 0,50 | 0,50 | 3,91 | 0,13 | 8,98E-01 |
| AT4G34400 | 0,59 | 1,64 | 3,75 | 0,44 | 6,62E-01 |
| AT4G34410 | 0,77 | 1,71 | 2,83 | 0,60 | 5,47E-01 |
| AT4G34430 | 1,22 | 2,20 | 2,10 | 1,05 | 2,96E-01 |
| AT4G34440 | 0,20 | 1,07 | 4,36 | 0,25 | 8,06E-01 |
| AT4G34640 | 0,93 | 3,18 | 2,76 | 1,15 | 2,49E-01 |
| AT4G34680 | 0,14 | 0,44 | 4,39 | 0,10 | 9,20E-01 |
| AT4G34700 | 0,83 | 0,02 | 2,61 | 0,01 | 9,94E-01 |
| AT4G34870 | 1,35 | 1,89 | 1,79 | 1,06 | 2,91E-01 |
| AT4G34881 | 0,53 | -1,03 | 2,74 | -0,38 | 7,06E-01 |
| AT4G34910 | 0,61 | 3,11 | 3,01 | 1,03 | 3,02E-01 |
| AT4G34920 | 0,49 | 1,70 | 3,21 | 0,53 | 5,96E-01 |
| AT4G35040 | 0,31 | 1,99 | 4,32 | 0,46 | 6,45E-01 |
| AT4G35100 | 1,16 | -1,34 | 2,44 | -0,55 | 5,84E-01 |
| AT4G35370 | 0,13 | 1,15 | 4,36 | 0,26 | 7,92E-01 |
| AT4G35450 | 0,43 | -0,87 | 3,35 | -0,26 | 7,94E-01 |
| AT4G35460 | 0,66 | 1,60 | 2,94 | 0,54 | 5,86E-01 |
| AT4G35470 | 0,67 | 1,71 | 2,59 | 0,66 | 5,08E-01 |
| AT4G35500 | 0,79 | 2,33 | 2,75 | 0,85 | 3,98E-01 |
| AT4G35560 | 0,72 | 0,59 | 2,89 | 0,20 | 8,38E-01 |
| AT4G35600 | 1,15 | 2,52 | 1,95 | 1,29 | 1,96E-01 |
| AT4G35630 | 0,33 | 0,28 | 3,58 | 0,08 | 9,37E-01 |
| AT4G35850 | 0,77 | 0,08 | 2,12 | 0,04 | 9,69E-01 |
| AT4G35870 | 0,76 | 1,79 | 2,53 | 0,71 | 4,80E-01 |
| AT4G35880 | 0,17 | 1,40 | 4,35 | 0,32 | 7,48E-01 |
| AT4G35890 | 1,08 | 2,38 | 2,11 | 1,13 | 2,57E-01 |
| AT4G35940 | 1,33 | 0,97 | 1,86 | 0,52 | 6,01E-01 |
| AT4G36032 | 1,21 | 0,69 | 2,29 | 0,30 | 7,64E-01 |
| AT4G36052 | 0,12 | 1,65 | 4,34 | 0,38 | 7,04E-01 |
| AT4G36130 | 1,27 | 1,37 | 1,97 | 0,69 | 4,88E-01 |
| AT4G36150 | 1,48 | 2,38 | 2,41 | 0,98 | 3,25E-01 |
| AT4G36190 | 0,22 | -1,25 | 4,39 | -0,28 | 7,76E-01 |

| | | | | | |
|---|---|---|---|---|---|
| AT4G36195 | 0,86 | -0,15 | 2,18 | -0,07 | 9,44E-01 |
| AT4G36210 | 1,25 | 2,58 | 1,85 | 1,39 | 1,64E-01 |
| AT4G36250 | 0,69 | -1,13 | 2,50 | -0,45 | 6,52E-01 |
| AT4G36540 | 0,24 | 1,62 | 4,34 | 0,37 | 7,08E-01 |
| AT4G36710 | 1,27 | 0,53 | 2,36 | 0,23 | 8,21E-01 |
| AT4G36720 | 0,82 | 2,06 | 2,79 | 0,74 | 4,60E-01 |
| AT4G09375 | 0,14 | -1,28 | 4,39 | -0,29 | 7,71E-01 |
| AT4G36810 | 0,84 | 1,91 | 2,36 | 0,81 | 4,18E-01 |
| AT4G36870 | 0,25 | -1,51 | 4,39 | -0,34 | 7,30E-01 |
| AT4G36920 | 0,14 | -1,28 | 4,39 | -0,29 | 7,71E-01 |
| AT4G36925 | 1,06 | -0,79 | 1,81 | -0,43 | 6,64E-01 |
| AT4G36990 | 0,59 | 2,86 | 2,97 | 0,96 | 3,36E-01 |
| AT4G37000 | 0,23 | 0,00 | 4,41 | 0,00 | 1,00E+00 |
| AT4G37020 | 0,15 | 1,42 | 4,34 | 0,33 | 7,43E-01 |
| AT4G37080 | 0,19 | 0,48 | 4,34 | 0,11 | 9,12E-01 |
| AT4G37130 | 0,62 | 2,67 | 3,62 | 0,74 | 4,61E-01 |
| AT4G37180 | 0,47 | 0,74 | 4,08 | 0,18 | 8,57E-01 |
| AT4G37190 | 0,32 | 0,87 | 4,33 | 0,20 | 8,40E-01 |
| AT4G37210 | 0,15 | 0,24 | 4,35 | 0,05 | 9,56E-01 |
| AT4G37300 | 1,24 | 1,20 | 2,16 | 0,55 | 5,79E-01 |
| AT4G37430 | 0,86 | 2,14 | 3,30 | 0,65 | 5,18E-01 |
| AT4G37440 | 0,34 | 0,00 | 4,41 | 0,00 | 1,00E+00 |
| AT4G37550 | 0,88 | 1,07 | 2,16 | 0,50 | 6,19E-01 |
| AT4G37560 | 0,42 | 1,76 | 4,34 | 0,41 | 6,85E-01 |
| AT4G37650 | 1,47 | -0,20 | 1,97 | -0,10 | 9,19E-01 |
| AT4G37680 | 0,49 | 1,93 | 3,18 | 0,61 | 5,43E-01 |
| AT4G37730 | 0,75 | 1,25 | 2,12 | 0,59 | 5,55E-01 |
| AT4G37760 | 0,63 | -1,03 | 2,28 | -0,45 | 6,50E-01 |
| AT4G37790 | 0,15 | 0,00 | 4,39 | 0,00 | 1,00E+00 |
| AT4G37980 | 0,67 | -0,22 | 2,11 | -0,11 | 9,16E-01 |
| AT4G38040 | 0,20 | 1,07 | 4,36 | 0,25 | 8,06E-01 |
| AT4G38170 | 0,97 | 0,34 | 2,23 | 0,15 | 8,79E-01 |
| AT4G09685 | 0,46 | 1,80 | 3,99 | 0,45 | 6,51E-01 |
| AT4G38215 | 0,75 | 2,16 | 4,30 | 0,50 | 6,16E-01 |
| AT4G38225 | 0,78 | 2,22 | 2,54 | 0,88 | 3,81E-01 |
| AT4G38250 | 1,25 | 0,89 | 1,66 | 0,53 | 5,94E-01 |
| AT4G38270 | 0,51 | 1,04 | 3,03 | 0,34 | 7,31E-01 |
| AT4G38290 | 0,11 | 1,07 | 4,36 | 0,25 | 8,06E-01 |
| AT4G38320 | 0,19 | 1,50 | 4,34 | 0,34 | 7,30E-01 |
| AT4G38330 | 0,12 | 1,58 | 4,34 | 0,36 | 7,16E-01 |
| AT4G38360 | 0,15 | 1,58 | 4,34 | 0,36 | 7,15E-01 |
| AT4G38430 | 0,81 | 2,85 | 2,67 | 1,07 | 2,86E-01 |
| AT4G38460 | 1,41 | 1,55 | 1,90 | 0,82 | 4,13E-01 |
| AT4G38500 | 0,68 | 3,40 | 3,46 | 0,98 | 3,25E-01 |
| AT4G38540 | 0,10 | 0,62 | 4,38 | 0,14 | 8,87E-01 |
| AT4G38560 | 1,22 | 0,93 | 2,54 | 0,37 | 7,13E-01 |
| AT4G38580 | 0,50 | -1,67 | 4,37 | -0,38 | 7,03E-01 |
| AT4G38620 | 0,24 | -0,21 | 4,35 | -0,05 | 9,61E-01 |
| AT4G38790 | 0,69 | 0,14 | 2,91 | 0,05 | 9,62E-01 |
| AT4G38860 | 0,38 | 1,09 | 4,36 | 0,25 | 8,03E-01 |
| AT4G38880 | 0,39 | 0,40 | 4,39 | 0,09 | 9,28E-01 |
| AT4G38920 | 0,99 | 0,43 | 2,15 | 0,20 | 8,40E-01 |
| AT4G38930 | 0,41 | 0,47 | 4,39 | 0,11 | 9,14E-01 |
| AT4G39020 | 0,55 | 2,24 | 4,32 | 0,52 | 6,04E-01 |
| AT4G39030 | 0,29 | 0,42 | 4,38 | 0,10 | 9,24E-01 |
| AT4G39050 | 1,56 | 0,82 | 1,57 | 0,52 | 6,00E-01 |

| | | | | | |
|---|---|---|---|---|---|
| AT4G39210 | 1,07 | 0,04 | 2,07 | 0,02 | 9,85E-01 |
| AT4G39270 | 1,16 | 0,28 | 1,70 | 0,17 | 8,68E-01 |
| AT4G39620 | 0,18 | 1,66 | 4,33 | 0,38 | 7,02E-01 |
| AT4G39640 | 0,76 | 0,54 | 2,54 | 0,21 | 8,31E-01 |
| AT4G39670 | 0,55 | 2,36 | 3,18 | 0,74 | 4,58E-01 |
| AT4G39840 | 1,07 | 2,35 | 2,03 | 1,16 | 2,47E-01 |
| AT4G39860 | 0,51 | 0,48 | 2,40 | 0,20 | 8,41E-01 |
| AT4G39870 | 0,35 | 1,45 | 4,31 | 0,34 | 7,36E-01 |
| AT4G39940 | 0,36 | 1,27 | 4,34 | 0,29 | 7,70E-01 |
| AT4G39950 | 1,14 | -0,02 | 1,71 | -0,01 | 9,92E-01 |
| AT4G39952 | 0,28 | 0,42 | 4,38 | 0,10 | 9,24E-01 |
| AT4G39980 | 0,50 | 1,55 | 3,16 | 0,49 | 6,24E-01 |
| AT4G40063 | 0,73 | 1,72 | 2,93 | 0,59 | 5,57E-01 |
| AT5G00730 | 1,64 | -3,57 | 1,74 | -2,06 | 3,98E-02 |
| AT5G01170 | 0,10 | 0,42 | 4,38 | 0,10 | 9,23E-01 |
| AT5G01210 | 1,09 | 0,62 | 1,82 | 0,34 | 7,33E-01 |
| AT5G01240 | 0,22 | -0,28 | 4,36 | -0,06 | 9,49E-01 |
| AT5G01260 | 0,54 | 0,76 | 2,35 | 0,32 | 7,45E-01 |
| AT5G01290 | 0,28 | 2,16 | 4,32 | 0,50 | 6,18E-01 |
| AT5G01460 | 0,28 | 0,00 | 4,41 | 0,00 | 1,00E+00 |
| AT5G01542 | 0,46 | 1,79 | 4,07 | 0,44 | 6,59E-01 |
| AT5G01590 | 0,94 | 1,11 | 2,00 | 0,55 | 5,80E-01 |
| AT5G01595 | 0,81 | 3,09 | 2,45 | 1,26 | 2,06E-01 |
| AT5G01690 | 0,08 | 0,85 | 4,38 | 0,20 | 8,45E-01 |
| AT5G01732 | 0,29 | 1,97 | 4,33 | 0,46 | 6,49E-01 |
| AT5G01740 | 0,15 | 0,00 | 4,41 | 0,00 | 1,00E+00 |
| AT5G01750 | 1,69 | 0,58 | 1,83 | 0,31 | 7,54E-01 |
| AT5G01760 | 0,79 | 1,57 | 2,78 | 0,57 | 5,72E-01 |
| AT5G01800 | 0,74 | 2,51 | 2,42 | 1,04 | 3,00E-01 |
| AT5G01810 | 0,55 | 2,47 | 3,81 | 0,65 | 5,17E-01 |
| AT5G01850 | 0,24 | -0,09 | 4,36 | -0,02 | 9,83E-01 |
| AT5G02035 | 0,47 | 1,60 | 3,92 | 0,41 | 6,83E-01 |
| AT5G02040 | 0,25 | 1,57 | 4,34 | 0,36 | 7,18E-01 |
| AT5G02100 | 0,70 | 1,55 | 3,66 | 0,42 | 6,72E-01 |
| AT5G02120 | 0,39 | -2,28 | 4,34 | -0,53 | 5,99E-01 |
| AT5G00820 | 1,31 | -3,16 | 2,46 | -1,29 | 1,99E-01 |
| AT5G02160 | 1,30 | -2,81 | 1,55 | -1,82 | 6,93E-02 |
| AT5G02240 | 0,32 | 0,15 | 4,35 | 0,03 | 9,72E-01 |
| AT5G02250 | 1,33 | 1,01 | 2,27 | 0,45 | 6,56E-01 |
| AT5G02280 | 0,11 | 1,07 | 4,36 | 0,25 | 8,06E-01 |
| AT5G02420 | 0,12 | 1,39 | 4,34 | 0,32 | 7,49E-01 |
| AT5G02450 | 0,59 | 0,15 | 3,09 | 0,05 | 9,61E-01 |
| AT5G02620 | 0,32 | 0,00 | 4,39 | 0,00 | 1,00E+00 |
| AT5G02630 | 0,30 | 1,62 | 4,35 | 0,37 | 7,10E-01 |
| AT5G02650 | 0,40 | 2,57 | 4,32 | 0,60 | 5,51E-01 |
| AT5G02760 | 0,40 | -0,67 | 3,39 | -0,20 | 8,43E-01 |
| AT5G02790 | 0,39 | 0,47 | 3,52 | 0,13 | 8,93E-01 |
| AT5G02840 | 0,40 | 2,19 | 4,14 | 0,53 | 5,97E-01 |
| AT5G02850 | 0,54 | 2,07 | 3,07 | 0,68 | 5,00E-01 |
| AT5G02860 | 1,11 | 2,38 | 1,98 | 1,21 | 2,28E-01 |
| AT5G02950 | 0,61 | 0,58 | 3,75 | 0,15 | 8,77E-01 |
| AT5G02995 | 0,13 | -1,24 | 4,39 | -0,28 | 7,77E-01 |
| AT5G03020 | 0,28 | 0,42 | 4,38 | 0,10 | 9,24E-01 |
| AT5G03160 | 0,68 | 1,55 | 2,46 | 0,63 | 5,27E-01 |
| AT5G03190 | 0,92 | 2,02 | 2,06 | 0,98 | 3,28E-01 |
| AT5G03290 | 1,24 | 1,97 | 1,77 | 1,11 | 2,66E-01 |

| | | | | | |
|---|---|---|---|---|---|
| AT5G03300 | 0,81 | 0,93 | 2,38 | 0,39 | 6,97E-01 |
| AT5G03330 | 0,61 | 1,51 | 3,09 | 0,49 | 6,26E-01 |
| AT5G03355 | 0,46 | -1,24 | 3,40 | -0,37 | 7,15E-01 |
| AT5G03415 | 0,36 | 1,68 | 4,33 | 0,39 | 6,99E-01 |
| AT5G03420 | 0,63 | 2,71 | 2,55 | 1,06 | 2,87E-01 |
| AT5G03455 | 0,25 | 1,42 | 4,34 | 0,33 | 7,44E-01 |
| AT5G03460 | 0,54 | 0,80 | 3,15 | 0,25 | 7,99E-01 |
| AT5G03470 | 0,40 | 1,18 | 4,36 | 0,27 | 7,86E-01 |
| AT5G03490 | 0,11 | 0,90 | 4,37 | 0,21 | 8,36E-01 |
| AT5G03540 | 1,05 | 3,20 | 2,75 | 1,16 | 2,46E-01 |
| AT5G03650 | 0,91 | -0,15 | 1,80 | -0,08 | 9,33E-01 |
| AT5G03905 | 0,30 | 0,90 | 4,38 | 0,20 | 8,38E-01 |
| AT5G04040 | 1,27 | 1,68 | 1,89 | 0,89 | 3,74E-01 |
| AT5G04050 | 0,10 | 0,42 | 4,38 | 0,10 | 9,23E-01 |
| AT5G04060 | 0,37 | 1,62 | 4,35 | 0,37 | 7,09E-01 |
| AT5G04170 | 0,76 | 0,04 | 2,57 | 0,01 | 9,88E-01 |
| AT5G04190 | 0,13 | 0,00 | 4,41 | 0,00 | 1,00E+00 |
| AT5G04240 | 0,88 | 2,45 | 3,22 | 0,76 | 4,46E-01 |
| AT5G04280 | 1,25 | 2,79 | 1,88 | 1,48 | 1,38E-01 |
| AT5G04290 | 0,66 | 0,27 | 3,27 | 0,08 | 9,34E-01 |
| AT5G04360 | 0,45 | 0,50 | 4,13 | 0,12 | 9,03E-01 |
| AT5G00960 | 0,96 | 3,37 | 2,19 | 1,54 | 1,23E-01 |
| AT5G04460 | 0,56 | 2,62 | 3,08 | 0,85 | 3,94E-01 |
| AT5G04490 | 0,46 | 1,27 | 3,37 | 0,38 | 7,06E-01 |
| AT5G04510 | 0,58 | 0,63 | 2,63 | 0,24 | 8,12E-01 |
| AT5G04600 | 0,71 | -0,34 | 2,23 | -0,15 | 8,78E-01 |
| AT5G04620 | 0,43 | 0,89 | 4,38 | 0,20 | 8,38E-01 |
| AT5G04630 | 0,57 | 2,45 | 3,74 | 0,65 | 5,13E-01 |
| AT5G04660 | 0,24 | 0,44 | 4,35 | 0,10 | 9,19E-01 |
| AT5G00985 | 0,28 | 0,65 | 4,37 | 0,15 | 8,82E-01 |
| AT5G04800 | 0,67 | 0,23 | 2,22 | 0,10 | 9,19E-01 |
| AT5G04885 | 0,27 | 1,59 | 3,74 | 0,42 | 6,71E-01 |
| AT5G04910 | 0,42 | 0,82 | 4,14 | 0,20 | 8,43E-01 |
| AT5G05070 | 0,40 | 1,27 | 4,29 | 0,30 | 7,68E-01 |
| AT5G05090 | 0,35 | 0,88 | 3,58 | 0,24 | 8,06E-01 |
| AT5G05160 | 0,72 | 1,99 | 2,82 | 0,70 | 4,81E-01 |
| AT5G05210 | 0,28 | 0,84 | 4,37 | 0,19 | 8,47E-01 |
| AT5G05220 | 0,08 | 0,85 | 4,38 | 0,20 | 8,45E-01 |
| AT5G05230 | 0,08 | 0,85 | 4,38 | 0,20 | 8,45E-01 |
| AT5G05310 | 0,18 | 0,00 | 4,40 | 0,00 | 1,00E+00 |
| AT5G05440 | 0,16 | 0,90 | 4,38 | 0,21 | 8,37E-01 |
| AT5G05490 | 0,55 | 0,84 | 4,37 | 0,19 | 8,48E-01 |
| AT5G05560 | 1,14 | 1,81 | 1,94 | 0,93 | 3,50E-01 |
| AT5G05570 | 0,62 | 2,65 | 4,31 | 0,61 | 5,39E-01 |
| AT5G05600 | 0,31 | -2,40 | 4,35 | -0,55 | 5,81E-01 |
| AT5G05630 | 0,08 | 0,00 | 4,40 | 0,00 | 1,00E+00 |
| AT5G05680 | 1,56 | 2,92 | 1,88 | 1,55 | 1,20E-01 |
| AT5G05710 | 0,20 | 0,89 | 4,37 | 0,20 | 8,39E-01 |
| AT5G05780 | 0,96 | 1,01 | 2,36 | 0,43 | 6,69E-01 |
| AT5G05950 | 0,83 | 2,20 | 2,45 | 0,90 | 3,68E-01 |
| AT5G05970 | 0,30 | 0,98 | 4,36 | 0,23 | 8,22E-01 |
| AT5G05985 | 0,55 | 1,30 | 3,84 | 0,34 | 7,36E-01 |
| AT5G06000 | 0,23 | -1,22 | 4,38 | -0,28 | 7,81E-01 |
| AT5G06050 | 0,39 | 0,36 | 4,26 | 0,08 | 9,33E-01 |
| AT5G06100 | 0,21 | 2,05 | 4,33 | 0,47 | 6,36E-01 |
| AT5G06120 | 1,17 | 1,75 | 2,41 | 0,73 | 4,67E-01 |

| | | | | | |
|---|---|---|---|---|---|
| AT5G06170 | 0,31 | -1,47 | 4,39 | -0,34 | 7,37E-01 |
| AT5G06220 | 0,84 | 1,63 | 2,48 | 0,66 | 5,10E-01 |
| AT5G06265 | 0,26 | -0,18 | 4,37 | -0,04 | 9,66E-01 |
| AT5G06290 | 1,37 | -1,22 | 1,66 | -0,73 | 4,63E-01 |
| AT5G06310 | 0,33 | 1,81 | 4,33 | 0,42 | 6,76E-01 |
| AT5G01255 | 0,70 | 2,59 | 2,80 | 0,92 | 3,56E-01 |
| AT5G06350 | 0,36 | 0,29 | 3,46 | 0,08 | 9,32E-01 |
| AT5G06360 | 0,40 | 1,27 | 4,29 | 0,30 | 7,68E-01 |
| AT5G06380 | 0,08 | 0,85 | 4,38 | 0,20 | 8,45E-01 |
| AT5G06390 | 0,14 | 0,51 | 4,38 | 0,12 | 9,08E-01 |
| AT5G06460 | 1,60 | 3,60 | 1,91 | 1,89 | 5,90E-02 |
| AT5G06645 | 0,58 | 0,00 | 4,41 | 0,00 | 1,00E+00 |
| AT5G06660 | 0,96 | 3,30 | 3,53 | 0,94 | 3,50E-01 |
| AT5G06680 | 0,39 | 0,45 | 4,35 | 0,10 | 9,17E-01 |
| AT5G06700 | 0,91 | -1,02 | 1,74 | -0,59 | 5,57E-01 |
| AT5G06811 | 0,11 | 1,07 | 4,36 | 0,25 | 8,06E-01 |
| AT5G06820 | 0,12 | 1,58 | 4,34 | 0,36 | 7,16E-01 |
| AT5G06860 | 0,22 | -0,24 | 4,37 | -0,06 | 9,56E-01 |
| AT5G06870 | 0,67 | 0,75 | 2,89 | 0,26 | 7,95E-01 |
| AT5G06950 | 0,61 | 1,70 | 3,09 | 0,55 | 5,82E-01 |
| AT5G06980 | 0,16 | 1,74 | 4,33 | 0,40 | 6,87E-01 |
| AT5G07090 | 1,54 | -2,21 | 1,44 | -1,54 | 1,24E-01 |
| AT5G07110 | 0,85 | 2,37 | 3,35 | 0,71 | 4,79E-01 |
| AT5G01315 | 0,64 | 1,62 | 2,98 | 0,54 | 5,87E-01 |
| AT5G07130 | 0,33 | 0,70 | 4,32 | 0,16 | 8,72E-01 |
| AT5G07140 | 1,24 | 2,93 | 2,47 | 1,18 | 2,37E-01 |
| AT5G07210 | 1,09 | -1,72 | 1,88 | -0,91 | 3,61E-01 |
| AT5G07240 | 1,19 | 2,01 | 2,03 | 0,99 | 3,22E-01 |
| AT5G07270 | 0,15 | 0,42 | 4,38 | 0,09 | 9,24E-01 |
| AT5G07340 | 0,98 | 2,47 | 2,25 | 1,10 | 2,71E-01 |
| AT5G07410 | 1,62 | 1,87 | 1,92 | 0,98 | 3,29E-01 |
| AT5G07430 | 0,50 | -1,16 | 3,97 | -0,29 | 7,71E-01 |
| AT5G07440 | 0,36 | -0,38 | 4,31 | -0,09 | 9,30E-01 |
| AT5G07580 | 0,75 | 1,03 | 2,29 | 0,45 | 6,51E-01 |
| AT5G07610 | 0,48 | 2,25 | 4,05 | 0,56 | 5,79E-01 |
| AT5G07760 | 0,08 | 1,10 | 4,36 | 0,25 | 8,00E-01 |
| AT5G07770 | 0,16 | 1,07 | 4,37 | 0,24 | 8,07E-01 |
| AT5G07790 | 0,58 | 2,24 | 4,32 | 0,52 | 6,04E-01 |
| AT5G07830 | 0,86 | 0,23 | 2,60 | 0,09 | 9,31E-01 |
| AT5G07860 | 0,58 | 1,18 | 3,91 | 0,30 | 7,63E-01 |
| AT5G07870 | 0,49 | 1,64 | 4,33 | 0,38 | 7,05E-01 |
| AT5G07960 | 0,12 | 1,39 | 4,34 | 0,32 | 7,49E-01 |
| AT5G08050 | 0,48 | 1,04 | 4,15 | 0,25 | 8,02E-01 |
| AT5G08060 | 0,10 | 0,42 | 4,38 | 0,10 | 9,23E-01 |
| AT5G08070 | 0,35 | 0,00 | 4,39 | 0,00 | 1,00E+00 |
| AT5G08139 | 0,37 | 1,24 | 4,32 | 0,29 | 7,75E-01 |
| AT5G08190 | 0,29 | 2,05 | 4,33 | 0,47 | 6,36E-01 |
| AT5G08260 | 0,90 | -0,77 | 2,40 | -0,32 | 7,49E-01 |
| AT5G01545 | 0,37 | -2,46 | 4,35 | -0,57 | 5,72E-01 |
| AT5G08290 | 0,74 | 1,36 | 2,11 | 0,64 | 5,19E-01 |
| AT5G08300 | 0,94 | 2,07 | 2,44 | 0,85 | 3,96E-01 |
| AT5G08330 | 1,56 | 0,39 | 1,92 | 0,20 | 8,40E-01 |
| AT5G08335 | 0,18 | 1,23 | 4,34 | 0,28 | 7,77E-01 |
| AT5G08340 | 0,58 | 2,18 | 3,97 | 0,55 | 5,82E-01 |
| AT5G08390 | 0,26 | 2,12 | 4,32 | 0,49 | 6,23E-01 |
| AT5G08430 | 0,57 | 2,11 | 4,32 | 0,49 | 6,26E-01 |

| | | | | | |
|---|---|---|---|---|---|
| AT5G08440 | 0,59 | 1,56 | 3,89 | 0,40 | 6,89E-01 |
| AT5G08470 | 0,81 | 2,81 | 2,74 | 1,03 | 3,04E-01 |
| AT5G08510 | 0,51 | -0,33 | 3,29 | -0,10 | 9,21E-01 |
| AT5G08540 | 0,41 | 0,68 | 3,37 | 0,20 | 8,41E-01 |
| AT5G01555 | 0,36 | 0,98 | 4,38 | 0,22 | 8,23E-01 |
| AT5G08570 | 1,20 | -1,16 | 1,51 | -0,77 | 4,43E-01 |
| AT5G08590 | 0,63 | -0,26 | 2,61 | -0,10 | 9,22E-01 |
| AT5G08600 | 0,28 | 0,65 | 4,37 | 0,15 | 8,82E-01 |
| AT5G08620 | 1,12 | -0,41 | 1,73 | -0,24 | 8,14E-01 |
| AT5G08660 | 0,33 | 0,00 | 4,39 | 0,00 | 1,00E+00 |
| AT5G08695 | 0,20 | 0,89 | 4,37 | 0,20 | 8,39E-01 |
| AT5G08750 | 0,11 | 0,90 | 4,37 | 0,21 | 8,36E-01 |
| AT5G08760 | 0,55 | -2,04 | 3,93 | -0,52 | 6,04E-01 |
| AT5G08780 | 0,49 | 1,95 | 3,15 | 0,62 | 5,36E-01 |
| AT5G09220 | 0,82 | -1,80 | 3,00 | -0,60 | 5,48E-01 |
| AT5G09400 | 1,22 | 2,67 | 2,02 | 1,33 | 1,85E-01 |
| AT5G09440 | 0,77 | 2,13 | 2,37 | 0,90 | 3,68E-01 |
| AT5G09445 | 0,26 | 0,90 | 4,38 | 0,21 | 8,37E-01 |
| AT5G09450 | 1,25 | 0,25 | 2,15 | 0,12 | 9,07E-01 |
| AT5G09600 | 0,62 | 1,37 | 3,01 | 0,46 | 6,48E-01 |
| AT5G09760 | 0,14 | 0,51 | 4,38 | 0,12 | 9,08E-01 |
| AT5G10010 | 0,47 | -0,40 | 3,42 | -0,12 | 9,06E-01 |
| AT5G10020 | 0,41 | 1,93 | 4,33 | 0,45 | 6,55E-01 |
| AT5G10100 | 0,16 | 0,80 | 4,37 | 0,18 | 8,55E-01 |
| AT5G10190 | 0,41 | 1,55 | 3,32 | 0,47 | 6,41E-01 |
| AT5G10200 | 0,81 | 1,85 | 3,43 | 0,54 | 5,89E-01 |
| AT5G10340 | 1,40 | -1,93 | 2,28 | -0,85 | 3,97E-01 |
| AT5G10370 | 1,36 | 1,97 | 1,62 | 1,21 | 2,25E-01 |
| AT5G10410 | 0,05 | 0,00 | 4,41 | 0,00 | 1,00E+00 |
| AT5G10460 | 0,08 | 1,10 | 4,36 | 0,25 | 8,00E-01 |
| AT5G01765 | 1,15 | 2,01 | 1,93 | 1,04 | 2,98E-01 |
| AT5G10600 | 0,40 | 1,25 | 4,29 | 0,29 | 7,71E-01 |
| AT5G10630 | 0,28 | 2,00 | 4,33 | 0,46 | 6,44E-01 |
| AT5G10660 | 0,46 | 0,32 | 2,76 | 0,11 | 9,09E-01 |
| AT5G10695 | 0,85 | -0,03 | 2,10 | -0,01 | 9,90E-01 |
| AT5G10745 | 0,35 | 1,96 | 3,50 | 0,56 | 5,76E-01 |
| AT5G10750 | 1,12 | 0,50 | 2,22 | 0,23 | 8,21E-01 |
| AT5G10780 | 0,76 | 0,45 | 2,54 | 0,18 | 8,58E-01 |
| AT5G10790 | 0,93 | 2,37 | 2,08 | 1,14 | 2,54E-01 |
| AT5G10920 | 0,91 | 0,86 | 2,22 | 0,38 | 7,00E-01 |
| AT5G11060 | 1,02 | 0,41 | 1,73 | 0,24 | 8,13E-01 |
| AT5G11090 | 0,49 | 0,32 | 2,73 | 0,12 | 9,08E-01 |
| AT5G11110 | 0,50 | 0,99 | 4,02 | 0,25 | 8,06E-01 |
| AT5G11170 | 1,42 | 0,26 | 2,03 | 0,13 | 9,00E-01 |
| AT5G11200 | 0,33 | 2,20 | 4,32 | 0,51 | 6,11E-01 |
| AT5G11280 | 0,20 | 0,82 | 4,37 | 0,19 | 8,50E-01 |
| AT5G11340 | 0,28 | 1,64 | 4,33 | 0,38 | 7,05E-01 |
| AT5G11350 | 0,82 | 0,78 | 2,91 | 0,27 | 7,88E-01 |
| AT5G11390 | 0,30 | 1,68 | 4,34 | 0,39 | 6,99E-01 |
| AT5G11420 | 1,15 | -0,41 | 2,12 | -0,19 | 8,47E-01 |
| AT5G11430 | 0,70 | -0,11 | 2,25 | -0,05 | 9,61E-01 |
| AT5G11500 | 0,94 | 0,88 | 1,80 | 0,49 | 6,24E-01 |
| AT5G11510 | 0,81 | 2,93 | 2,35 | 1,25 | 2,13E-01 |
| AT5G11530 | 0,81 | 1,12 | 2,38 | 0,47 | 6,37E-01 |
| AT5G11580 | 0,12 | 0,79 | 4,37 | 0,18 | 8,57E-01 |
| AT5G11650 | 0,38 | 1,65 | 3,44 | 0,48 | 6,31E-01 |

| | | | | | |
|---|---|---|---|---|---|
| AT5G11680 | 0,19 | 1,57 | 4,34 | 0,36 | 7,18E-01 |
| AT5G11710 | 1,24 | 0,53 | 1,69 | 0,31 | 7,54E-01 |
| AT5G11720 | 1,68 | 3,18 | 2,08 | 1,53 | 1,26E-01 |
| AT5G11730 | 0,89 | 0,69 | 2,04 | 0,34 | 7,34E-01 |
| AT5G11760 | 0,22 | 0,51 | 4,38 | 0,12 | 9,08E-01 |
| AT5G11890 | 0,82 | 0,63 | 2,21 | 0,28 | 7,77E-01 |
| AT5G11920 | 0,12 | 0,79 | 4,37 | 0,18 | 8,57E-01 |
| AT5G11977 | 0,12 | 1,65 | 4,34 | 0,38 | 7,04E-01 |
| AT5G12090 | 0,79 | -1,64 | 2,40 | -0,69 | 4,93E-01 |
| AT5G12100 | 0,44 | 2,72 | 4,31 | 0,63 | 5,29E-01 |
| AT5G12110 | 1,52 | 1,55 | 1,60 | 0,97 | 3,34E-01 |
| AT5G12210 | 0,65 | 2,34 | 3,69 | 0,63 | 5,26E-01 |
| AT5G12260 | 0,32 | 0,15 | 4,35 | 0,03 | 9,72E-01 |
| AT5G12350 | 0,49 | 1,84 | 4,13 | 0,45 | 6,56E-01 |
| AT5G12400 | 0,93 | 3,44 | 2,84 | 1,21 | 2,25E-01 |
| AT5G12850 | 0,97 | 0,71 | 2,05 | 0,34 | 7,30E-01 |
| AT5G12950 | 0,53 | 0,96 | 2,55 | 0,38 | 7,06E-01 |
| AT5G12960 | 0,27 | -0,03 | 4,36 | -0,01 | 9,95E-01 |
| AT5G12970 | 0,24 | 0,00 | 4,41 | 0,00 | 1,00E+00 |
| AT5G13030 | 0,96 | -1,67 | 1,84 | -0,90 | 3,66E-01 |
| AT5G13200 | 1,26 | 1,29 | 1,86 | 0,69 | 4,88E-01 |
| AT5G13210 | 0,35 | 1,89 | 3,43 | 0,55 | 5,81E-01 |
| AT5G13260 | 0,92 | -1,28 | 2,06 | -0,62 | 5,34E-01 |
| AT5G13280 | 0,45 | 0,70 | 4,16 | 0,17 | 8,66E-01 |
| AT5G13310 | 0,31 | 1,94 | 4,33 | 0,45 | 6,54E-01 |
| AT5G13320 | 1,27 | 3,73 | 1,95 | 1,91 | 5,62E-02 |
| AT5G13330 | 0,37 | 2,32 | 4,32 | 0,54 | 5,91E-01 |
| AT5G13370 | 0,68 | -1,66 | 3,18 | -0,52 | 6,01E-01 |
| AT5G13380 | 0,54 | 1,34 | 4,30 | 0,31 | 7,55E-01 |
| AT5G13500 | 0,32 | 1,01 | 4,37 | 0,23 | 8,18E-01 |
| AT5G13510 | 0,48 | -0,06 | 3,24 | -0,02 | 9,86E-01 |
| AT5G13580 | 0,29 | 2,52 | 4,32 | 0,58 | 5,59E-01 |
| AT5G13640 | 0,84 | 0,48 | 3,03 | 0,16 | 8,75E-01 |
| AT5G13690 | 0,61 | 2,01 | 3,09 | 0,65 | 5,15E-01 |
| AT5G13720 | 0,21 | -0,77 | 4,37 | -0,18 | 8,60E-01 |
| AT5G13740 | 0,19 | 1,57 | 4,34 | 0,36 | 7,18E-01 |
| AT5G13750 | 0,26 | 1,08 | 4,37 | 0,25 | 8,05E-01 |
| AT5G13760 | 0,26 | 0,67 | 4,36 | 0,15 | 8,78E-01 |
| AT5G13810 | 0,93 | 0,92 | 1,93 | 0,48 | 6,34E-01 |
| AT5G13820 | 0,28 | 2,32 | 4,32 | 0,54 | 5,92E-01 |
| AT5G13930 | 1,07 | 1,94 | 2,16 | 0,90 | 3,70E-01 |
| AT5G13980 | 0,97 | 0,23 | 2,25 | 0,10 | 9,20E-01 |
| AT5G14030 | 0,98 | 3,21 | 2,44 | 1,31 | 1,89E-01 |
| AT5G14050 | 0,50 | 2,60 | 3,15 | 0,83 | 4,08E-01 |
| AT5G14120 | 1,26 | -0,52 | 1,79 | -0,29 | 7,73E-01 |
| AT5G14200 | 1,42 | 1,27 | 1,62 | 0,78 | 4,33E-01 |
| AT5G14230 | 0,39 | 1,98 | 3,42 | 0,58 | 5,62E-01 |
| AT5G14260 | 0,68 | 0,89 | 2,47 | 0,36 | 7,18E-01 |
| AT5G14280 | 0,43 | 1,05 | 4,38 | 0,24 | 8,11E-01 |
| AT5G14285 | 0,21 | 1,99 | 4,33 | 0,46 | 6,46E-01 |
| AT5G14480 | 0,25 | 0,66 | 4,33 | 0,15 | 8,79E-01 |
| AT5G14610 | 0,64 | 1,94 | 3,25 | 0,60 | 5,50E-01 |
| AT5G14670 | 0,08 | 1,10 | 4,36 | 0,25 | 8,00E-01 |
| AT5G14880 | 0,37 | 1,66 | 3,44 | 0,48 | 6,29E-01 |
| AT5G14890 | 0,99 | 3,61 | 4,01 | 0,90 | 3,69E-01 |
| AT5G14910 | 0,77 | 0,92 | 2,55 | 0,36 | 7,19E-01 |

| | | | | | |
|---|---|---|---|---|---|
| AT5G14970 | 0,18 | 1,56 | 4,34 | 0,36 | 7,18E-01 |
| AT5G14990 | 0,42 | 1,63 | 4,17 | 0,39 | 6,97E-01 |
| AT5G15010 | 0,38 | -3,08 | 4,33 | -0,71 | 4,76E-01 |
| AT5G02295 | 1,40 | -0,75 | 1,70 | -0,44 | 6,58E-01 |
| AT5G15080 | 0,92 | 0,87 | 2,88 | 0,30 | 7,62E-01 |
| AT5G15230 | 0,74 | 0,61 | 2,83 | 0,22 | 8,30E-01 |
| AT5G15240 | 0,05 | 0,00 | 4,41 | 0,00 | 1,00E+00 |
| AT5G15270 | 0,61 | 1,19 | 3,13 | 0,38 | 7,04E-01 |
| AT5G15280 | 0,54 | 1,59 | 3,84 | 0,41 | 6,79E-01 |
| AT5G15320 | 0,21 | 0,18 | 4,34 | 0,04 | 9,68E-01 |
| AT5G15330 | 0,11 | 1,07 | 4,36 | 0,25 | 8,06E-01 |
| AT5G15530 | 0,12 | 1,58 | 4,34 | 0,36 | 7,16E-01 |
| AT5G15570 | 0,44 | 2,79 | 4,31 | 0,65 | 5,16E-01 |
| AT5G15610 | 0,89 | 0,19 | 1,86 | 0,10 | 9,21E-01 |
| AT5G15685 | 0,72 | 0,40 | 2,01 | 0,20 | 8,44E-01 |
| AT5G15710 | 0,74 | 2,60 | 3,58 | 0,73 | 4,68E-01 |
| AT5G15730 | 0,59 | 2,66 | 2,58 | 1,03 | 3,03E-01 |
| AT5G15750 | 0,54 | 0,40 | 3,96 | 0,10 | 9,19E-01 |
| AT5G15820 | 0,22 | 0,85 | 4,36 | 0,20 | 8,45E-01 |
| AT5G15930 | 0,32 | 1,61 | 4,34 | 0,37 | 7,12E-01 |
| AT5G15950 | 1,23 | 3,61 | 2,07 | 1,74 | 8,13E-02 |
| AT5G15960 | 0,51 | -3,13 | 4,32 | -0,72 | 4,69E-01 |
| AT5G16010 | 0,36 | 0,42 | 4,37 | 0,10 | 9,23E-01 |
| AT5G16030 | 0,58 | -2,54 | 3,27 | -0,78 | 4,37E-01 |
| AT5G16120 | 0,30 | 1,36 | 4,35 | 0,31 | 7,55E-01 |
| AT5G16130 | 1,11 | 3,00 | 2,16 | 1,38 | 1,66E-01 |
| AT5G16150 | 1,33 | -0,63 | 1,51 | -0,42 | 6,76E-01 |
| AT5G16260 | 0,11 | 0,00 | 4,41 | 0,00 | 1,00E+00 |
| AT5G16290 | 0,43 | -0,30 | 4,28 | -0,07 | 9,44E-01 |
| AT5G16300 | 1,23 | 2,61 | 1,79 | 1,46 | 1,45E-01 |
| AT5G16440 | 0,67 | 1,23 | 2,61 | 0,47 | 6,37E-01 |
| AT5G16470 | 0,69 | -0,19 | 2,59 | -0,07 | 9,42E-01 |
| AT5G16540 | 0,32 | 1,20 | 4,35 | 0,28 | 7,82E-01 |
| AT5G16550 | 0,57 | 1,06 | 2,66 | 0,40 | 6,91E-01 |
| AT5G02555 | 0,05 | 0,00 | 4,41 | 0,00 | 1,00E+00 |
| AT5G16590 | 0,87 | 2,02 | 2,19 | 0,92 | 3,56E-01 |
| AT5G16630 | 0,92 | 3,15 | 2,60 | 1,21 | 2,27E-01 |
| AT5G16715 | 0,74 | -1,04 | 2,41 | -0,43 | 6,67E-01 |
| AT5G16730 | 1,05 | -0,12 | 1,85 | -0,07 | 9,48E-01 |
| AT5G16760 | 1,05 | 2,86 | 2,19 | 1,30 | 1,92E-01 |
| AT5G16820 | 0,94 | 2,62 | 2,29 | 1,15 | 2,51E-01 |
| AT5G16880 | 1,27 | 0,45 | 1,60 | 0,28 | 7,81E-01 |
| AT5G16900 | 0,33 | 2,12 | 4,32 | 0,49 | 6,24E-01 |
| AT5G16910 | 0,98 | 3,16 | 2,54 | 1,24 | 2,13E-01 |
| AT5G17167 | 0,20 | 0,43 | 4,39 | 0,10 | 9,21E-01 |
| AT5G17170 | 0,33 | -0,42 | 4,36 | -0,10 | 9,23E-01 |
| AT5G17190 | 0,51 | 0,00 | 3,25 | 0,00 | 1,00E+00 |
| AT5G17270 | 0,61 | 1,24 | 4,35 | 0,29 | 7,75E-01 |
| AT5G17280 | 0,29 | 1,04 | 4,33 | 0,24 | 8,10E-01 |
| AT5G17305 | 0,08 | 1,10 | 4,36 | 0,25 | 8,00E-01 |
| AT5G17440 | 0,24 | 0,83 | 4,37 | 0,19 | 8,49E-01 |
| AT5G17530 | 0,59 | -0,28 | 3,09 | -0,09 | 9,27E-01 |
| AT5G17580 | 0,17 | 1,81 | 4,33 | 0,42 | 6,76E-01 |
| AT5G17630 | 0,33 | 0,95 | 4,37 | 0,22 | 8,28E-01 |
| AT5G17670 | 0,39 | 0,48 | 4,40 | 0,11 | 9,14E-01 |
| AT5G17680 | 0,30 | 1,68 | 4,34 | 0,39 | 6,99E-01 |

| | | | | | |
|---|---|---|---|---|---|
| AT5G17850 | 0,18 | 0,00 | 4,41 | 0,00 | 1,00E+00 |
| AT5G17870 | 0,86 | -1,62 | 2,19 | -0,74 | 4,60E-01 |
| AT5G17900 | 0,30 | 0,90 | 4,36 | 0,21 | 8,37E-01 |
| AT5G17910 | 1,47 | 0,27 | 1,76 | 0,15 | 8,78E-01 |
| AT5G17980 | 0,38 | 2,23 | 4,23 | 0,53 | 5,98E-01 |
| AT5G17990 | 0,55 | 0,35 | 4,00 | 0,09 | 9,31E-01 |
| AT5G18040 | 0,71 | 2,91 | 2,43 | 1,20 | 2,32E-01 |
| AT5G18065 | 0,30 | 0,84 | 3,63 | 0,23 | 8,17E-01 |
| AT5G18070 | 0,62 | 1,36 | 2,62 | 0,52 | 6,03E-01 |
| AT5G18085 | 0,24 | 0,00 | 4,41 | 0,00 | 1,00E+00 |
| AT5G18100 | 0,33 | 0,46 | 4,34 | 0,11 | 9,15E-01 |
| AT5G18130 | 1,11 | 3,59 | 2,28 | 1,57 | 1,15E-01 |
| AT5G18140 | 1,06 | 2,75 | 2,62 | 1,05 | 2,94E-01 |
| AT5G18170 | 0,16 | 1,49 | 4,34 | 0,34 | 7,32E-01 |
| AT5G18200 | 0,08 | 0,85 | 4,38 | 0,20 | 8,45E-01 |
| AT5G18245 | 1,46 | 0,60 | 2,88 | 0,21 | 8,35E-01 |
| AT5G18255 | 1,25 | -1,13 | 1,45 | -0,78 | 4,38E-01 |
| AT5G18310 | 0,76 | -1,61 | 2,45 | -0,66 | 5,10E-01 |
| AT5G18390 | 0,33 | 1,14 | 4,36 | 0,26 | 7,94E-01 |
| AT5G18470 | 1,32 | 3,61 | 1,89 | 1,91 | 5,61E-02 |
| AT5G18500 | 0,37 | 0,13 | 4,34 | 0,03 | 9,75E-01 |
| AT5G18520 | 1,46 | 1,18 | 1,62 | 0,73 | 4,66E-01 |
| AT5G18580 | 0,90 | 0,43 | 2,09 | 0,20 | 8,38E-01 |
| AT5G18610 | 1,43 | 1,91 | 2,06 | 0,93 | 3,54E-01 |
| AT5G18630 | 0,28 | 0,00 | 4,41 | 0,00 | 1,00E+00 |
| AT5G18636 | 0,58 | -0,29 | 2,46 | -0,12 | 9,06E-01 |
| AT5G18650 | 0,68 | 0,22 | 3,08 | 0,07 | 9,43E-01 |
| AT5G18670 | 0,21 | 2,14 | 4,33 | 0,49 | 6,22E-01 |
| AT5G18750 | 0,77 | 3,59 | 3,35 | 1,07 | 2,84E-01 |
| AT5G18830 | 0,51 | 1,21 | 3,21 | 0,38 | 7,06E-01 |
| AT5G18890 | 0,98 | -1,34 | 4,28 | -0,31 | 7,54E-01 |
| AT5G18937 | 0,54 | 0,00 | 4,41 | 0,00 | 1,00E+00 |
| AT5G18940 | 1,00 | 2,05 | 2,75 | 0,75 | 4,56E-01 |
| AT5G18960 | 0,35 | 1,81 | 3,46 | 0,52 | 6,02E-01 |
| AT5G19000 | 0,44 | 0,97 | 3,40 | 0,29 | 7,74E-01 |
| AT5G19090 | 0,78 | 1,07 | 2,56 | 0,42 | 6,75E-01 |
| AT5G19180 | 0,77 | 1,60 | 2,21 | 0,72 | 4,70E-01 |
| AT5G19190 | 0,37 | 0,75 | 4,37 | 0,17 | 8,63E-01 |
| AT5G19210 | 0,49 | 0,44 | 3,39 | 0,13 | 8,97E-01 |
| AT5G19221 | 0,47 | 0,41 | 4,38 | 0,09 | 9,25E-01 |
| AT5G19240 | 0,65 | -0,14 | 2,09 | -0,06 | 9,48E-01 |
| AT5G19250 | 0,98 | -3,22 | 2,74 | -1,18 | 2,40E-01 |
| AT5G19290 | 0,41 | 1,61 | 3,36 | 0,48 | 6,31E-01 |
| AT5G19440 | 0,36 | 1,67 | 4,34 | 0,38 | 7,01E-01 |
| AT5G19500 | 1,08 | 2,91 | 2,35 | 1,24 | 2,17E-01 |
| AT5G19540 | 0,77 | -1,28 | 1,97 | -0,65 | 5,17E-01 |
| AT5G19550 | 0,90 | 1,79 | 2,02 | 0,89 | 3,75E-01 |
| AT5G19570 | 0,52 | 1,67 | 3,95 | 0,42 | 6,72E-01 |
| AT5G19620 | 0,50 | -0,19 | 4,05 | -0,05 | 9,62E-01 |
| AT5G19680 | 0,19 | -0,05 | 4,35 | -0,01 | 9,91E-01 |
| AT5G19750 | 0,55 | 1,97 | 4,28 | 0,46 | 6,45E-01 |
| AT5G19780 | 1,35 | -1,12 | 1,65 | -0,68 | 4,97E-01 |
| AT5G19990 | 0,77 | -0,54 | 2,32 | -0,23 | 8,16E-01 |
| AT5G20080 | 0,48 | -0,54 | 3,35 | -0,16 | 8,71E-01 |
| AT5G20120 | 0,55 | 1,69 | 4,32 | 0,39 | 6,95E-01 |
| AT5G20130 | 0,68 | 0,51 | 3,71 | 0,14 | 8,90E-01 |

| | | | | | |
|---|---|---|---|---|---|
| AT5G20150 | 0,56 | 2,10 | 2,61 | 0,81 | 4,20E-01 |
| AT5G20190 | 1,21 | 1,95 | 1,98 | 0,99 | 3,24E-01 |
| AT5G20200 | 1,58 | 4,01 | 1,80 | 2,23 | 2,57E-02 |
| AT5G20220 | 0,21 | 2,05 | 4,33 | 0,47 | 6,36E-01 |
| AT5G20350 | 1,36 | 1,74 | 1,67 | 1,04 | 2,98E-01 |
| AT5G20410 | 0,63 | 0,51 | 3,72 | 0,14 | 8,91E-01 |
| AT5G20480 | 0,15 | 0,42 | 4,38 | 0,09 | 9,24E-01 |
| AT5G20500 | 0,33 | 1,30 | 4,34 | 0,30 | 7,65E-01 |
| AT5G20590 | 0,85 | 0,80 | 2,13 | 0,38 | 7,07E-01 |
| AT5G20610 | 1,39 | 3,07 | 1,82 | 1,69 | 9,10E-02 |
| AT5G20660 | 1,27 | 3,09 | 1,93 | 1,60 | 1,09E-01 |
| AT5G20680 | 0,31 | -0,65 | 4,37 | -0,15 | 8,82E-01 |
| AT5G20710 | 0,12 | 1,58 | 4,34 | 0,36 | 7,16E-01 |
| AT5G20900 | 0,45 | 0,13 | 3,35 | 0,04 | 9,69E-01 |
| AT5G03255 | 0,31 | 1,77 | 4,33 | 0,41 | 6,83E-01 |
| AT5G20960 | 1,02 | 3,89 | 2,55 | 1,53 | 1,27E-01 |
| AT5G21020 | 1,50 | 0,00 | 1,49 | 0,00 | 1,00E+00 |
| AT5G21090 | 1,39 | 2,77 | 1,97 | 1,41 | 1,59E-01 |
| AT5G21105 | 1,68 | 2,50 | 1,72 | 1,46 | 1,45E-01 |
| AT5G21150 | 0,12 | 1,58 | 4,34 | 0,36 | 7,16E-01 |
| AT5G21222 | 1,01 | 2,48 | 2,72 | 0,91 | 3,61E-01 |
| AT5G21326 | 0,90 | 3,42 | 2,71 | 1,26 | 2,07E-01 |
| AT5G21482 | 0,24 | 1,24 | 4,35 | 0,28 | 7,76E-01 |
| AT5G21930 | 1,60 | 0,55 | 1,54 | 0,36 | 7,21E-01 |
| AT5G22000 | 1,27 | 1,59 | 1,88 | 0,84 | 3,99E-01 |
| AT5G22050 | 0,36 | -0,17 | 4,37 | -0,04 | 9,69E-01 |
| AT5G22070 | 0,57 | -0,27 | 3,07 | -0,09 | 9,31E-01 |
| AT5G22300 | 0,30 | 0,30 | 3,69 | 0,08 | 9,35E-01 |
| AT5G22315 | 0,20 | 0,00 | 4,41 | 0,00 | 1,00E+00 |
| AT5G22340 | 0,95 | 1,18 | 2,09 | 0,57 | 5,71E-01 |
| AT5G22420 | 0,42 | 1,76 | 4,15 | 0,42 | 6,72E-01 |
| AT5G22580 | 0,46 | -0,47 | 4,35 | -0,11 | 9,15E-01 |
| AT5G22590 | 1,03 | 0,59 | 2,73 | 0,22 | 8,29E-01 |
| AT5G22630 | 1,52 | 0,86 | 1,69 | 0,51 | 6,09E-01 |
| AT5G22640 | 1,19 | 1,55 | 1,68 | 0,92 | 3,56E-01 |
| AT5G22650 | 0,66 | -1,58 | 2,52 | -0,63 | 5,30E-01 |
| AT5G22690 | 0,12 | 1,58 | 4,34 | 0,36 | 7,16E-01 |
| AT5G22760 | 1,25 | 2,16 | 1,91 | 1,14 | 2,56E-01 |
| AT5G22794 | 0,08 | 1,10 | 4,36 | 0,25 | 8,00E-01 |
| AT5G22830 | 0,08 | 0,00 | 4,40 | 0,00 | 1,00E+00 |
| AT5G22950 | 0,12 | 0,42 | 4,38 | 0,10 | 9,24E-01 |
| AT5G23010 | 0,84 | -1,00 | 2,25 | -0,45 | 6,56E-01 |
| AT5G23070 | 0,19 | 0,79 | 4,37 | 0,18 | 8,57E-01 |
| AT5G03575 | 0,26 | -0,94 | 4,37 | -0,22 | 8,30E-01 |
| AT5G23120 | 0,65 | 0,08 | 2,14 | 0,04 | 9,71E-01 |
| AT5G23140 | 0,95 | 2,08 | 2,30 | 0,91 | 3,64E-01 |
| AT5G23150 | 1,41 | 3,59 | 2,16 | 1,66 | 9,68E-02 |
| AT5G23210 | 0,16 | 1,33 | 4,35 | 0,31 | 7,59E-01 |
| AT5G23280 | 1,01 | 1,26 | 2,59 | 0,49 | 6,25E-01 |
| AT5G23320 | 0,08 | 1,10 | 4,36 | 0,25 | 8,00E-01 |
| AT5G23330 | 0,43 | 0,89 | 4,38 | 0,20 | 8,38E-01 |
| AT5G23340 | 1,40 | 3,11 | 1,87 | 1,66 | 9,71E-02 |
| AT5G23490 | 0,49 | 1,56 | 2,68 | 0,58 | 5,60E-01 |
| AT5G23540 | 0,96 | -0,61 | 2,03 | -0,30 | 7,64E-01 |
| AT5G23570 | 0,58 | -1,22 | 2,28 | -0,53 | 5,93E-01 |
| AT5G23575 | 1,00 | 1,63 | 1,94 | 0,84 | 4,02E-01 |

| | | | | | |
|---|---|---|---|---|---|
| AT5G23600 | 0,08 | 1,10 | 4,36 | 0,25 | 8,00E-01 |
| AT5G23670 | 0,72 | 2,55 | 2,99 | 0,85 | 3,93E-01 |
| AT5G23720 | 0,51 | -0,33 | 3,29 | -0,10 | 9,21E-01 |
| AT5G23730 | 0,26 | 1,30 | 4,35 | 0,30 | 7,65E-01 |
| AT5G23890 | 0,99 | 2,68 | 2,74 | 0,98 | 3,27E-01 |
| AT5G23950 | 0,35 | 0,00 | 4,41 | 0,00 | 1,00E+00 |
| AT5G24020 | 0,16 | 1,33 | 4,35 | 0,31 | 7,59E-01 |
| AT5G24200 | 1,34 | 2,62 | 1,75 | 1,50 | 1,33E-01 |
| AT5G24210 | 0,96 | -1,15 | 2,21 | -0,52 | 6,01E-01 |
| AT5G24360 | 0,40 | 1,13 | 3,32 | 0,34 | 7,35E-01 |
| AT5G24380 | 0,41 | 0,96 | 3,33 | 0,29 | 7,72E-01 |
| AT5G24450 | 0,69 | 2,80 | 3,68 | 0,76 | 4,47E-01 |
| AT5G24460 | 0,88 | 0,68 | 1,91 | 0,36 | 7,21E-01 |
| AT5G24490 | 1,61 | -0,09 | 1,88 | -0,05 | 9,64E-01 |
| AT5G24610 | 0,55 | -0,67 | 2,57 | -0,26 | 7,94E-01 |
| AT5G24830 | 0,08 | 1,10 | 4,36 | 0,25 | 8,00E-01 |
| AT5G24980 | 0,43 | -0,54 | 3,32 | -0,16 | 8,71E-01 |
| AT5G25040 | 0,67 | 1,20 | 2,48 | 0,49 | 6,27E-01 |
| AT5G25120 | 0,51 | 0,30 | 3,24 | 0,09 | 9,26E-01 |
| AT5G03925 | 0,17 | 0,00 | 4,41 | 0,00 | 1,00E+00 |
| AT5G25130 | 1,42 | -0,26 | 1,76 | -0,15 | 8,81E-01 |
| AT5G25140 | 0,28 | 2,33 | 4,32 | 0,54 | 5,90E-01 |
| AT5G25150 | 0,27 | 0,48 | 4,38 | 0,11 | 9,13E-01 |
| AT5G25170 | 0,33 | 2,41 | 4,32 | 0,56 | 5,77E-01 |
| AT5G25210 | 0,42 | 0,83 | 3,33 | 0,25 | 8,04E-01 |
| AT5G25220 | 1,05 | 3,60 | 2,08 | 1,73 | 8,39E-02 |
| AT5G25230 | 1,66 | 3,59 | 1,74 | 2,07 | 3,88E-02 |
| AT5G25250 | 1,43 | 3,25 | 1,98 | 1,64 | 1,00E-01 |
| AT5G25260 | 1,02 | 2,30 | 2,64 | 0,87 | 3,84E-01 |
| AT5G25340 | 0,46 | 2,05 | 4,05 | 0,51 | 6,13E-01 |
| AT5G25460 | 1,53 | -2,94 | 1,96 | -1,50 | 1,33E-01 |
| AT5G25520 | 1,48 | 3,22 | 2,40 | 1,34 | 1,80E-01 |
| AT5G25560 | 0,53 | 1,23 | 3,18 | 0,39 | 6,99E-01 |
| AT5G25580 | 0,12 | 1,65 | 4,34 | 0,38 | 7,04E-01 |
| AT5G25630 | 0,47 | 1,49 | 3,15 | 0,47 | 6,36E-01 |
| AT5G25754 | 1,48 | 0,92 | 1,48 | 0,62 | 5,34E-01 |
| AT5G04145 | 0,89 | -1,23 | 2,27 | -0,54 | 5,87E-01 |
| AT5G25900 | 0,33 | 0,67 | 4,33 | 0,16 | 8,76E-01 |
| AT5G25910 | 0,60 | 2,89 | 3,67 | 0,79 | 4,31E-01 |
| AT5G25930 | 0,70 | 2,23 | 2,51 | 0,89 | 3,74E-01 |
| AT5G25980 | 1,32 | 0,41 | 1,69 | 0,24 | 8,08E-01 |
| AT5G26038 | 1,13 | 1,23 | 2,14 | 0,58 | 5,65E-01 |
| AT5G26146 | 0,89 | -0,03 | 2,15 | -0,01 | 9,89E-01 |
| AT5G26147 | 0,89 | -0,03 | 2,15 | -0,01 | 9,89E-01 |
| AT5G26200 | 0,12 | 1,58 | 4,34 | 0,36 | 7,16E-01 |
| AT5G26240 | 0,84 | 0,81 | 1,97 | 0,41 | 6,82E-01 |
| AT5G26270 | 0,26 | -0,34 | 4,36 | -0,08 | 9,38E-01 |
| AT5G26290 | 0,79 | -1,90 | 2,17 | -0,87 | 3,82E-01 |
| AT5G26570 | 1,51 | 0,75 | 1,64 | 0,46 | 6,47E-01 |
| AT5G26740 | 0,58 | 0,37 | 2,60 | 0,14 | 8,87E-01 |
| AT5G26731 | 0,38 | 0,82 | 4,38 | 0,19 | 8,51E-01 |
| AT5G26710 | 0,73 | 2,44 | 2,49 | 0,98 | 3,26E-01 |
| AT5G26670 | 0,26 | -1,63 | 4,37 | -0,37 | 7,09E-01 |
| AT5G26610 | 0,16 | 0,00 | 4,41 | 0,00 | 1,00E+00 |
| AT5G26600 | 0,82 | 0,72 | 2,53 | 0,29 | 7,74E-01 |
| AT5G26780 | 0,39 | 0,27 | 2,85 | 0,10 | 9,24E-01 |

| | | | | | |
|---|---|---|---|---|---|
| AT5G26820 | 0,42 | 1,23 | 4,36 | 0,28 | 7,78E-01 |
| AT5G26850 | 1,21 | 0,52 | 2,16 | 0,24 | 8,09E-01 |
| AT5G26860 | 1,16 | 2,51 | 1,86 | 1,35 | 1,78E-01 |
| AT5G26930 | 0,33 | 0,89 | 4,36 | 0,20 | 8,39E-01 |
| AT5G26940 | 0,66 | 1,65 | 3,72 | 0,44 | 6,58E-01 |
| AT5G26990 | 0,23 | 0,00 | 4,41 | 0,00 | 1,00E+00 |
| AT5G27120 | 0,91 | 1,06 | 3,44 | 0,31 | 7,58E-01 |
| AT5G27150 | 0,57 | 2,58 | 2,99 | 0,86 | 3,89E-01 |
| AT5G27170 | 0,25 | 0,42 | 4,38 | 0,10 | 9,24E-01 |
| AT5G27230 | 0,08 | 0,00 | 4,40 | 0,00 | 1,00E+00 |
| AT5G27240 | 0,05 | 0,00 | 4,41 | 0,00 | 1,00E+00 |
| AT5G27270 | 0,17 | -0,46 | 4,37 | -0,11 | 9,16E-01 |
| AT5G27280 | 0,34 | -0,14 | 3,57 | -0,04 | 9,68E-01 |
| AT5G27300 | 0,08 | 0,00 | 4,40 | 0,00 | 1,00E+00 |
| AT5G27320 | 0,08 | 0,00 | 4,40 | 0,00 | 1,00E+00 |
| AT5G27340 | 0,54 | 0,81 | 3,80 | 0,21 | 8,30E-01 |
| AT5G27350 | 0,64 | -0,51 | 2,51 | -0,20 | 8,38E-01 |
| AT5G27410 | 0,42 | 0,00 | 4,17 | 0,00 | 1,00E+00 |
| AT5G27420 | 1,44 | 2,83 | 1,75 | 1,62 | 1,05E-01 |
| AT5G27470 | 0,51 | 2,44 | 3,09 | 0,79 | 4,31E-01 |
| AT5G27530 | 0,32 | -1,97 | 4,38 | -0,45 | 6,53E-01 |
| AT5G27540 | 0,68 | 1,04 | 3,56 | 0,29 | 7,71E-01 |
| AT5G27600 | 0,18 | 1,06 | 4,35 | 0,24 | 8,07E-01 |
| AT5G27610 | 0,08 | 0,00 | 4,40 | 0,00 | 1,00E+00 |
| AT5G27630 | 0,17 | 0,48 | 4,40 | 0,11 | 9,12E-01 |
| AT5G27650 | 0,94 | 2,62 | 2,72 | 0,97 | 3,34E-01 |
| AT5G27710 | 0,67 | 1,73 | 2,53 | 0,69 | 4,92E-01 |
| AT5G27730 | 0,69 | 0,26 | 2,15 | 0,12 | 9,03E-01 |
| AT5G27760 | 0,27 | 0,59 | 4,39 | 0,14 | 8,92E-01 |
| AT5G27770 | 0,86 | 0,22 | 2,47 | 0,09 | 9,30E-01 |
| AT5G27800 | 0,16 | 0,51 | 4,38 | 0,12 | 9,07E-01 |
| AT5G27830 | 0,23 | 1,16 | 4,35 | 0,27 | 7,89E-01 |
| AT5G27890 | 0,40 | 1,24 | 4,34 | 0,29 | 7,75E-01 |
| AT5G27893 | 1,07 | -2,59 | 1,82 | -1,42 | 1,55E-01 |
| AT5G27970 | 1,51 | 2,76 | 2,19 | 1,26 | 2,08E-01 |
| AT5G28010 | 1,26 | -0,34 | 1,50 | -0,23 | 8,22E-01 |
| AT5G04625 | 0,10 | 0,00 | 4,41 | 0,00 | 1,00E+00 |
| AT5G04665 | 0,11 | 0,00 | 4,41 | 0,00 | 1,00E+00 |
| AT5G28090 | 0,29 | 1,47 | 4,34 | 0,34 | 7,34E-01 |
| AT5G04675 | 0,19 | 0,00 | 4,41 | 0,00 | 1,00E+00 |
| AT5G28320 | 0,89 | 3,48 | 2,62 | 1,33 | 1,84E-01 |
| AT5G28350 | 0,24 | 0,84 | 4,37 | 0,19 | 8,48E-01 |
| AT5G28410 | 0,36 | -0,78 | 4,33 | -0,18 | 8,57E-01 |
| AT5G28463 | 0,37 | -0,08 | 4,33 | -0,02 | 9,85E-01 |
| AT5G28500 | 0,70 | -1,30 | 2,60 | -0,50 | 6,17E-01 |
| AT5G04815 | 0,05 | 0,00 | 4,41 | 0,00 | 1,00E+00 |
| AT5G28810 | 0,66 | -0,46 | 2,94 | -0,16 | 8,75E-01 |
| AT5G28830 | 0,05 | 0,00 | 4,41 | 0,00 | 1,00E+00 |
| AT5G28900 | 0,40 | 0,45 | 4,28 | 0,10 | 9,17E-01 |
| AT5G29624 | 0,12 | 1,65 | 4,34 | 0,38 | 7,04E-01 |
| AT5G30341 | 0,12 | 0,00 | 4,39 | 0,00 | 1,00E+00 |
| AT5G04955 | 1,06 | 0,00 | 3,96 | 0,00 | 1,00E+00 |
| AT5G32440 | 1,33 | 2,92 | 1,92 | 1,52 | 1,29E-01 |
| AT5G32613 | 0,26 | 0,45 | 4,39 | 0,10 | 9,19E-01 |
| AT5G33290 | 0,62 | 1,01 | 3,95 | 0,26 | 7,97E-01 |
| AT5G33320 | 0,61 | 0,95 | 2,28 | 0,41 | 6,79E-01 |

| | | | | | |
|---|---|---|---|---|---|
| AT5G33390 | 0,38 | -0,29 | 4,29 | -0,07 | 9,47E-01 |
| AT5G34869 | 0,19 | 1,64 | 4,33 | 0,38 | 7,05E-01 |
| AT5G34871 | 0,12 | 1,39 | 4,34 | 0,32 | 7,49E-01 |
| AT5G34870 | 0,38 | 0,77 | 3,42 | 0,23 | 8,21E-01 |
| AT5G35160 | 1,68 | 0,96 | 1,59 | 0,60 | 5,46E-01 |
| AT5G35170 | 0,47 | -0,51 | 2,38 | -0,21 | 8,32E-01 |
| AT5G35180 | 0,52 | 0,51 | 2,63 | 0,19 | 8,48E-01 |
| AT5G35190 | 0,84 | 2,85 | 2,77 | 1,03 | 3,04E-01 |
| AT5G35300 | 0,23 | 0,43 | 4,38 | 0,10 | 9,23E-01 |
| AT5G35320 | 0,21 | 0,00 | 4,38 | 0,00 | 1,00E+00 |
| AT5G35360 | 1,57 | 3,48 | 1,72 | 2,02 | 4,38E-02 |
| AT5G35430 | 1,24 | 1,95 | 2,28 | 0,86 | 3,91E-01 |
| AT5G35450 | 0,40 | 2,27 | 4,33 | 0,52 | 6,00E-01 |
| AT5G05195 | 0,23 | 1,36 | 4,34 | 0,31 | 7,54E-01 |
| AT5G35490 | 1,40 | 2,78 | 2,33 | 1,19 | 2,32E-01 |
| AT5G35730 | 0,82 | 1,08 | 2,72 | 0,39 | 6,93E-01 |
| AT5G35840 | 1,35 | 2,73 | 2,17 | 1,26 | 2,09E-01 |
| AT5G35917 | 0,29 | 0,48 | 4,40 | 0,11 | 9,12E-01 |
| AT5G35920 | 1,48 | 2,31 | 1,81 | 1,27 | 2,03E-01 |
| AT5G35926 | 1,15 | 0,44 | 1,86 | 0,24 | 8,13E-01 |
| AT5G35980 | 1,15 | -0,50 | 1,88 | -0,27 | 7,91E-01 |
| AT5G36001 | 0,12 | 1,65 | 4,34 | 0,38 | 7,04E-01 |
| AT5G36080 | 1,49 | 3,74 | 1,88 | 1,99 | 4,63E-02 |
| AT5G36290 | 0,47 | -0,33 | 2,76 | -0,12 | 9,04E-01 |
| AT5G36300 | 0,87 | 2,00 | 2,37 | 0,85 | 3,97E-01 |
| AT5G36740 | 0,08 | 1,10 | 4,36 | 0,25 | 8,00E-01 |
| AT5G36870 | 1,07 | -3,02 | 1,96 | -1,54 | 1,24E-01 |
| AT5G36900 | 0,54 | 2,57 | 3,09 | 0,83 | 4,06E-01 |
| AT5G36940 | 0,51 | 2,90 | 3,86 | 0,75 | 4,52E-01 |
| AT5G37020 | 0,62 | -0,03 | 2,55 | -0,01 | 9,92E-01 |
| AT5G37070 | 0,43 | -1,97 | 4,37 | -0,45 | 6,52E-01 |
| AT5G37130 | 0,66 | 2,18 | 2,68 | 0,81 | 4,17E-01 |
| AT5G37150 | 0,98 | -0,38 | 2,39 | -0,16 | 8,74E-01 |
| AT5G37540 | 0,17 | 1,40 | 4,35 | 0,32 | 7,48E-01 |
| AT5G37630 | 0,94 | 3,16 | 2,21 | 1,43 | 1,53E-01 |
| AT5G37760 | 0,15 | 1,62 | 4,34 | 0,37 | 7,09E-01 |
| AT5G37793 | 0,51 | -1,19 | 2,54 | -0,47 | 6,39E-01 |
| AT5G37850 | 0,30 | 1,62 | 4,35 | 0,37 | 7,10E-01 |
| AT5G37950 | 0,11 | 0,00 | 4,41 | 0,00 | 1,00E+00 |
| AT5G38155 | 0,12 | 0,42 | 4,38 | 0,10 | 9,24E-01 |
| AT5G38210 | 1,00 | 2,75 | 2,21 | 1,24 | 2,14E-01 |
| AT5G38380 | 0,15 | -1,24 | 4,39 | -0,28 | 7,77E-01 |
| AT5G38470 | 0,73 | 1,65 | 3,69 | 0,45 | 6,55E-01 |
| AT5G38480 | 1,20 | 1,94 | 1,78 | 1,09 | 2,76E-01 |
| AT5G38560 | 0,34 | 1,23 | 3,50 | 0,35 | 7,25E-01 |
| AT5G38910 | 0,16 | 0,00 | 4,41 | 0,00 | 1,00E+00 |
| AT5G38980 | 0,57 | 0,83 | 2,72 | 0,31 | 7,59E-01 |
| AT5G39000 | 0,70 | 0,00 | 4,41 | 0,00 | 1,00E+00 |
| AT5G39030 | 0,81 | 3,18 | 2,80 | 1,14 | 2,56E-01 |
| AT5G39040 | 0,29 | 0,92 | 4,33 | 0,21 | 8,32E-01 |
| AT5G39250 | 0,34 | 0,42 | 4,37 | 0,10 | 9,24E-01 |
| AT5G39320 | 0,82 | 1,12 | 4,36 | 0,26 | 7,98E-01 |
| AT5G39350 | 0,21 | -0,31 | 4,37 | -0,07 | 9,43E-01 |
| AT5G39510 | 0,77 | 1,35 | 2,46 | 0,55 | 5,81E-01 |
| AT5G39590 | 0,82 | 2,18 | 2,40 | 0,91 | 3,64E-01 |
| AT5G39660 | 0,40 | 2,55 | 4,32 | 0,59 | 5,55E-01 |

| | | | | | |
|---|---|---|---|---|---|
| AT5G39670 | 0,40 | 2,29 | 4,16 | 0,55 | 5,82E-01 |
| AT5G39690 | 1,03 | -0,60 | 2,16 | -0,28 | 7,81E-01 |
| AT5G39710 | 0,25 | 1,75 | 4,33 | 0,40 | 6,87E-01 |
| AT5G39730 | 0,19 | 0,41 | 4,38 | 0,09 | 9,25E-01 |
| AT5G39775 | 0,53 | 0,02 | 4,19 | 0,00 | 9,97E-01 |
| AT5G39850 | 0,08 | 1,10 | 4,36 | 0,25 | 8,00E-01 |
| AT5G39960 | 0,21 | 1,39 | 4,34 | 0,32 | 7,49E-01 |
| AT5G39990 | 0,08 | 0,85 | 4,38 | 0,20 | 8,45E-01 |
| AT5G40060 | 1,33 | 2,14 | 2,50 | 0,85 | 3,93E-01 |
| AT5G40140 | 0,31 | 1,23 | 3,62 | 0,34 | 7,34E-01 |
| AT5G40200 | 0,62 | 0,00 | 2,73 | 0,00 | 9,99E-01 |
| AT5G40380 | 0,46 | -1,43 | 4,05 | -0,35 | 7,24E-01 |
| AT5G40400 | 1,55 | -0,41 | 1,50 | -0,28 | 7,82E-01 |
| AT5G40650 | 0,74 | 0,41 | 2,89 | 0,14 | 8,87E-01 |
| AT5G40670 | 1,01 | 0,54 | 2,14 | 0,25 | 7,99E-01 |
| AT5G40680 | 0,17 | 1,92 | 4,33 | 0,44 | 6,57E-01 |
| AT5G40690 | 0,62 | 2,93 | 3,66 | 0,80 | 4,23E-01 |
| AT5G40720 | 0,32 | 1,29 | 4,35 | 0,30 | 7,66E-01 |
| AT5G40760 | 1,16 | 2,73 | 2,04 | 1,34 | 1,80E-01 |
| AT5G40780 | 1,36 | 1,86 | 1,72 | 1,08 | 2,80E-01 |
| AT5G40810 | 1,32 | -0,67 | 1,62 | -0,41 | 6,80E-01 |
| AT5G40850 | 0,08 | 0,00 | 4,41 | 0,00 | 1,00E+00 |
| AT5G40870 | 0,14 | 0,51 | 4,38 | 0,12 | 9,08E-01 |
| AT5G40880 | 0,51 | 2,11 | 3,94 | 0,53 | 5,93E-01 |
| AT5G40900 | 0,15 | 0,00 | 4,39 | 0,00 | 1,00E+00 |
| AT5G41010 | 0,28 | 0,78 | 4,37 | 0,18 | 8,58E-01 |
| AT5G06115 | 0,20 | 0,42 | 4,38 | 0,10 | 9,24E-01 |
| AT5G41070 | 0,20 | 0,00 | 4,41 | 0,00 | 1,00E+00 |
| AT5G41100 | 0,28 | 0,00 | 4,41 | 0,00 | 1,00E+00 |
| AT5G41110 | 0,22 | 0,42 | 4,38 | 0,10 | 9,23E-01 |
| AT5G41140 | 0,59 | -1,19 | 3,84 | -0,31 | 7,56E-01 |
| AT5G41260 | 0,61 | 1,21 | 2,34 | 0,52 | 6,04E-01 |
| AT5G41360 | 0,66 | -0,30 | 3,72 | -0,08 | 9,36E-01 |
| AT5G41370 | 0,15 | 0,71 | 4,37 | 0,16 | 8,71E-01 |
| AT5G41410 | 1,13 | 1,96 | 1,96 | 1,00 | 3,19E-01 |
| AT5G41500 | 1,05 | 1,40 | 2,13 | 0,66 | 5,11E-01 |
| AT5G41660 | 0,71 | 0,56 | 3,00 | 0,19 | 8,51E-01 |
| AT5G41700 | 1,24 | -2,35 | 1,87 | -1,25 | 2,10E-01 |
| AT5G41761 | 0,83 | 1,24 | 2,41 | 0,52 | 6,06E-01 |
| AT5G41770 | 0,61 | 0,64 | 2,51 | 0,25 | 7,99E-01 |
| AT5G41860 | 0,77 | 3,50 | 4,29 | 0,82 | 4,14E-01 |
| AT5G41920 | 0,61 | 0,88 | 4,36 | 0,20 | 8,41E-01 |
| AT5G41940 | 0,98 | 0,66 | 1,89 | 0,35 | 7,25E-01 |
| AT5G41950 | 1,46 | 2,43 | 2,10 | 1,16 | 2,47E-01 |
| AT5G41960 | 0,15 | 0,24 | 4,35 | 0,05 | 9,56E-01 |
| AT5G41970 | 0,85 | 1,08 | 3,43 | 0,32 | 7,52E-01 |
| AT5G42010 | 0,60 | 2,69 | 3,11 | 0,86 | 3,87E-01 |
| AT5G42220 | 0,70 | 2,26 | 2,36 | 0,96 | 3,39E-01 |
| AT5G42250 | 0,33 | 2,12 | 4,32 | 0,49 | 6,24E-01 |
| AT5G42260 | 0,28 | 0,00 | 4,41 | 0,00 | 1,00E+00 |
| AT5G42290 | 0,08 | 1,10 | 4,36 | 0,25 | 8,00E-01 |
| AT5G42310 | 0,50 | -1,33 | 4,20 | -0,32 | 7,51E-01 |
| AT5G42350 | 0,37 | 1,38 | 4,34 | 0,32 | 7,50E-01 |
| AT5G42420 | 0,74 | 2,00 | 3,63 | 0,55 | 5,82E-01 |
| AT5G42460 | 0,33 | 1,53 | 4,34 | 0,35 | 7,24E-01 |
| AT5G42470 | 0,22 | -1,23 | 4,37 | -0,28 | 7,78E-01 |

| | | | | | |
|---|---|---|---|---|---|
| AT5G42480 | 0,45 | 2,11 | 4,32 | 0,49 | 6,26E-01 |
| AT5G42504 | 0,17 | -1,64 | 4,37 | -0,37 | 7,08E-01 |
| AT5G42520 | 0,71 | -0,65 | 2,31 | -0,28 | 7,77E-01 |
| AT5G42540 | 1,02 | 3,11 | 2,32 | 1,34 | 1,80E-01 |
| AT5G42620 | 0,80 | 0,52 | 2,34 | 0,22 | 8,23E-01 |
| AT5G42640 | 0,08 | 0,00 | 4,40 | 0,00 | 1,00E+00 |
| AT5G42670 | 0,16 | 0,00 | 4,41 | 0,00 | 1,00E+00 |
| AT5G42680 | 0,80 | 0,13 | 3,25 | 0,04 | 9,68E-01 |
| AT5G42720 | 0,19 | 0,37 | 4,38 | 0,09 | 9,32E-01 |
| AT5G42740 | 1,16 | 1,31 | 2,74 | 0,48 | 6,32E-01 |
| AT5G42760 | 0,12 | 0,90 | 4,36 | 0,21 | 8,36E-01 |
| AT5G06395 | 1,55 | 2,25 | 1,84 | 1,22 | 2,21E-01 |
| AT5G42825 | 0,29 | 0,21 | 4,35 | 0,05 | 9,62E-01 |
| AT5G42965 | 0,19 | 0,48 | 4,34 | 0,11 | 9,12E-01 |
| AT5G42980 | 0,59 | -0,32 | 2,64 | -0,12 | 9,02E-01 |
| AT5G43100 | 0,78 | 1,32 | 2,19 | 0,60 | 5,49E-01 |
| AT5G43150 | 0,61 | 2,12 | 3,04 | 0,70 | 4,86E-01 |
| AT5G06505 | 0,36 | -0,28 | 4,37 | -0,06 | 9,49E-01 |
| AT5G43460 | 1,01 | 1,47 | 2,02 | 0,73 | 4,68E-01 |
| AT5G43500 | 1,47 | -2,48 | 1,61 | -1,54 | 1,24E-01 |
| AT5G43525 | 0,69 | 2,16 | 2,82 | 0,77 | 4,44E-01 |
| AT5G43560 | 0,55 | -0,20 | 3,25 | -0,06 | 9,51E-01 |
| AT5G43610 | 1,12 | -1,02 | 2,17 | -0,47 | 6,38E-01 |
| AT5G43620 | 0,36 | 1,49 | 4,27 | 0,35 | 7,27E-01 |
| AT5G43700 | 0,52 | -0,07 | 3,04 | -0,02 | 9,81E-01 |
| AT5G43810 | 1,04 | 3,42 | 2,69 | 1,27 | 2,04E-01 |
| AT5G43930 | 0,73 | 0,25 | 2,67 | 0,09 | 9,26E-01 |
| AT5G43940 | 1,17 | 1,68 | 2,10 | 0,80 | 4,25E-01 |
| AT5G44020 | 1,03 | -1,13 | 2,05 | -0,55 | 5,83E-01 |
| AT5G44050 | 1,57 | -5,10 | 2,10 | -2,43 | 1,52E-02 |
| AT5G44060 | 0,47 | 0,25 | 2,79 | 0,09 | 9,28E-01 |
| AT5G44070 | 1,32 | 0,44 | 1,67 | 0,26 | 7,94E-01 |
| AT5G44130 | 0,80 | -4,02 | 2,79 | -1,44 | 1,49E-01 |
| AT5G44190 | 1,58 | -4,26 | 1,92 | -2,22 | 2,64E-02 |
| AT5G44230 | 0,12 | 1,65 | 4,34 | 0,38 | 7,04E-01 |
| AT5G44240 | 1,02 | -1,64 | 2,01 | -0,81 | 4,16E-01 |
| AT5G06565 | 0,63 | -0,83 | 2,64 | -0,31 | 7,53E-01 |
| AT5G44286 | 0,85 | -0,57 | 2,25 | -0,25 | 8,00E-01 |
| AT5G44530 | 1,17 | 1,96 | 2,53 | 0,78 | 4,37E-01 |
| AT5G44574 | 0,91 | -2,85 | 3,19 | -0,89 | 3,72E-01 |
| AT5G44580 | 0,52 | -1,38 | 4,00 | -0,34 | 7,30E-01 |
| AT5G44585 | 1,38 | -1,72 | 1,86 | -0,93 | 3,54E-01 |
| AT5G06605 | 1,31 | -1,70 | 1,86 | -0,92 | 3,59E-01 |
| AT5G44640 | 1,59 | 0,36 | 1,77 | 0,20 | 8,40E-01 |
| AT5G44650 | 0,46 | -0,01 | 3,99 | 0,00 | 9,99E-01 |
| AT5G44720 | 0,16 | 1,26 | 4,35 | 0,29 | 7,72E-01 |
| AT5G44750 | 0,74 | 1,54 | 2,20 | 0,70 | 4,85E-01 |
| AT5G44760 | 0,36 | 1,99 | 4,34 | 0,46 | 6,46E-01 |
| AT5G44770 | 0,22 | 0,90 | 4,36 | 0,21 | 8,37E-01 |
| AT5G45000 | 0,38 | -0,70 | 4,31 | -0,16 | 8,71E-01 |
| AT5G06695 | 0,59 | 1,21 | 3,04 | 0,40 | 6,90E-01 |
| AT5G45030 | 0,59 | 1,21 | 3,04 | 0,40 | 6,90E-01 |
| AT5G45060 | 1,56 | 0,60 | 1,67 | 0,36 | 7,20E-01 |
| AT5G45130 | 0,78 | 2,11 | 2,95 | 0,71 | 4,76E-01 |
| AT5G45150 | 0,08 | 0,85 | 4,38 | 0,20 | 8,45E-01 |
| AT5G45240 | 0,12 | 1,65 | 4,34 | 0,38 | 7,04E-01 |

| | | | | | |
|---|---|---|---|---|---|
| AT5G45260 | 0,75 | 2,41 | 2,88 | 0,84 | 4,02E-01 |
| AT5G45390 | 0,62 | -1,57 | 2,32 | -0,68 | 4,97E-01 |
| AT5G45400 | 0,76 | 0,98 | 2,52 | 0,39 | 6,97E-01 |
| AT5G45450 | 0,08 | 0,85 | 4,38 | 0,20 | 8,45E-01 |
| AT5G45475 | 0,74 | -0,57 | 2,19 | -0,26 | 7,94E-01 |
| AT5G45490 | 1,25 | 0,18 | 1,76 | 0,10 | 9,20E-01 |
| AT5G45570 | 0,68 | 0,87 | 2,48 | 0,35 | 7,25E-01 |
| AT5G45620 | 0,59 | 2,42 | 3,09 | 0,78 | 4,34E-01 |
| AT5G45670 | 0,10 | 0,00 | 4,41 | 0,00 | 1,00E+00 |
| AT5G45680 | 0,26 | 0,00 | 4,41 | 0,00 | 1,00E+00 |
| AT5G45710 | 0,55 | 0,41 | 3,85 | 0,11 | 9,15E-01 |
| AT5G45750 | 0,19 | 0,00 | 4,41 | 0,00 | 1,00E+00 |
| AT5G45770 | 0,12 | 1,58 | 4,34 | 0,36 | 7,16E-01 |
| AT5G45920 | 1,21 | -2,85 | 1,79 | -1,59 | 1,12E-01 |
| AT5G46330 | 0,47 | 2,44 | 3,98 | 0,61 | 5,40E-01 |
| AT5G46410 | 0,44 | 15,52 | 4,41 | 3,52 | 4,30E-04 |
| AT5G06825 | 0,67 | 0,83 | 3,11 | 0,27 | 7,89E-01 |
| AT5G46460 | 0,90 | 3,64 | 3,23 | 1,13 | 2,59E-01 |
| AT5G46490 | 1,46 | 3,25 | 1,76 | 1,85 | 6,44E-02 |
| AT5G46500 | 1,11 | 4,04 | 2,40 | 1,68 | 9,24E-02 |
| AT5G06835 | 1,13 | -3,65 | 2,40 | -1,52 | 1,29E-01 |
| AT5G46630 | 1,04 | 0,04 | 1,95 | 0,02 | 9,84E-01 |
| AT5G46640 | 1,47 | 0,65 | 2,08 | 0,31 | 7,56E-01 |
| AT5G46660 | 1,02 | 0,00 | 4,13 | 0,00 | 1,00E+00 |
| AT5G46670 | 0,74 | -0,42 | 2,84 | -0,15 | 8,83E-01 |
| AT5G46730 | 0,08 | 0,00 | 4,40 | 0,00 | 1,00E+00 |
| AT5G46780 | 0,35 | 0,27 | 4,34 | 0,06 | 9,51E-01 |
| AT5G46825 | 1,18 | -1,41 | 1,56 | -0,90 | 3,66E-01 |
| AT5G46840 | 0,62 | -0,68 | 3,09 | -0,22 | 8,27E-01 |
| AT5G46850 | 0,29 | 1,60 | 4,34 | 0,37 | 7,13E-01 |
| AT5G47020 | 1,57 | 1,91 | 1,88 | 1,01 | 3,11E-01 |
| AT5G47030 | 0,63 | 0,37 | 3,79 | 0,10 | 9,22E-01 |
| AT5G47120 | 1,06 | 3,21 | 2,09 | 1,54 | 1,24E-01 |
| AT5G47130 | 0,25 | 0,64 | 4,32 | 0,15 | 8,82E-01 |
| AT5G47140 | 0,26 | -0,08 | 4,37 | -0,02 | 9,86E-01 |
| AT5G47200 | 0,34 | 0,60 | 4,34 | 0,14 | 8,90E-01 |
| AT5G47230 | 0,27 | 0,59 | 4,37 | 0,14 | 8,92E-01 |
| AT5G47240 | 0,14 | 0,00 | 4,41 | 0,00 | 1,00E+00 |
| AT5G47280 | 1,15 | -0,23 | 1,87 | -0,12 | 9,04E-01 |
| AT5G47300 | 1,47 | 2,36 | 1,91 | 1,23 | 2,19E-01 |
| AT5G47320 | 0,19 | 0,78 | 4,36 | 0,18 | 8,58E-01 |
| AT5G47390 | 0,64 | 0,95 | 2,07 | 0,46 | 6,46E-01 |
| AT5G47420 | 0,50 | 1,24 | 3,99 | 0,31 | 7,56E-01 |
| AT5G47435 | 1,48 | 2,82 | 2,01 | 1,40 | 1,61E-01 |
| AT5G47640 | 0,70 | -1,74 | 2,20 | -0,79 | 4,31E-01 |
| AT5G47660 | 0,08 | 0,00 | 4,40 | 0,00 | 1,00E+00 |
| AT5G47720 | 0,16 | 1,74 | 4,33 | 0,40 | 6,87E-01 |
| AT5G47770 | 0,39 | -0,15 | 4,28 | -0,04 | 9,72E-01 |
| AT5G47780 | 0,52 | 1,09 | 2,67 | 0,41 | 6,83E-01 |
| AT5G47790 | 0,28 | 1,16 | 4,32 | 0,27 | 7,89E-01 |
| AT5G47800 | 0,24 | 1,58 | 4,34 | 0,36 | 7,16E-01 |
| AT5G47820 | 0,70 | 1,01 | 3,70 | 0,27 | 7,84E-01 |
| AT5G47840 | 0,45 | -0,46 | 2,78 | -0,17 | 8,68E-01 |
| AT5G47860 | 0,53 | 1,90 | 3,85 | 0,49 | 6,21E-01 |
| AT5G47890 | 1,07 | 1,59 | 1,90 | 0,84 | 4,02E-01 |
| AT5G48190 | 1,46 | -2,30 | 1,69 | -1,37 | 1,72E-01 |

| | | | | | |
|---|---|---|---|---|---|
| AT5G48220 | 0,16 | 0,90 | 4,38 | 0,21 | 8,37E-01 |
| AT5G48290 | 0,36 | 0,00 | 4,41 | 0,00 | 1,00E+00 |
| AT5G48300 | 1,23 | -1,48 | 1,89 | -0,78 | 4,34E-01 |
| AT5G48340 | 0,30 | 1,62 | 4,35 | 0,37 | 7,10E-01 |
| AT5G48360 | 0,08 | 0,00 | 4,40 | 0,00 | 1,00E+00 |
| AT5G48390 | 0,69 | 2,10 | 3,50 | 0,60 | 5,48E-01 |
| AT5G48400 | 0,56 | 2,82 | 3,76 | 0,75 | 4,54E-01 |
| AT5G48410 | 0,12 | 1,58 | 4,34 | 0,36 | 7,16E-01 |
| AT5G07095 | 1,59 | -0,15 | 1,42 | -0,10 | 9,16E-01 |
| AT5G07105 | 0,86 | -0,56 | 2,25 | -0,25 | 8,02E-01 |
| AT5G48545 | 0,30 | -0,43 | 3,72 | -0,11 | 9,09E-01 |
| AT5G48720 | 0,16 | 1,07 | 4,37 | 0,24 | 8,07E-01 |
| AT5G48730 | 0,12 | 1,58 | 4,34 | 0,36 | 7,16E-01 |
| AT5G48775 | 0,08 | 0,00 | 4,40 | 0,00 | 1,00E+00 |
| AT5G48800 | 0,90 | 3,14 | 2,87 | 1,09 | 2,74E-01 |
| AT5G48900 | 0,13 | -1,24 | 4,39 | -0,28 | 7,77E-01 |
| AT5G49010 | 1,17 | -3,60 | 2,06 | -1,75 | 7,96E-02 |
| AT5G49020 | 0,22 | -0,28 | 4,36 | -0,06 | 9,49E-01 |
| AT5G49060 | 0,17 | 1,75 | 4,33 | 0,40 | 6,86E-01 |
| AT5G49160 | 0,93 | 2,52 | 2,56 | 0,99 | 3,24E-01 |
| AT5G49215 | 0,08 | 0,00 | 4,40 | 0,00 | 1,00E+00 |
| AT5G49250 | 0,66 | 2,17 | 2,80 | 0,78 | 4,37E-01 |
| AT5G49260 | 0,08 | 0,00 | 4,40 | 0,00 | 1,00E+00 |
| AT5G49450 | 0,19 | 1,91 | 4,33 | 0,44 | 6,58E-01 |
| AT5G49460 | 0,62 | 1,61 | 3,01 | 0,53 | 5,94E-01 |
| AT5G49470 | 0,29 | 1,60 | 4,34 | 0,37 | 7,13E-01 |
| AT5G49480 | 1,15 | 1,94 | 2,33 | 0,83 | 4,05E-01 |
| AT5G49510 | 0,19 | 1,20 | 4,35 | 0,27 | 7,83E-01 |
| AT5G49520 | 0,12 | 0,90 | 4,36 | 0,21 | 8,36E-01 |
| AT5G49530 | 0,46 | 2,70 | 4,05 | 0,67 | 5,05E-01 |
| AT5G49590 | 0,86 | -2,58 | 2,84 | -0,91 | 3,63E-01 |
| AT5G49640 | 0,08 | 0,85 | 4,38 | 0,20 | 8,45E-01 |
| AT5G49650 | 0,61 | 0,05 | 2,27 | 0,02 | 9,81E-01 |
| AT5G49810 | 1,47 | -0,73 | 1,85 | -0,39 | 6,95E-01 |
| AT5G49840 | 0,10 | 0,62 | 4,38 | 0,14 | 8,87E-01 |
| AT5G49945 | 0,51 | 1,90 | 3,79 | 0,50 | 6,16E-01 |
| AT5G49970 | 0,29 | 2,17 | 4,32 | 0,50 | 6,16E-01 |
| AT5G50020 | 0,47 | 0,00 | 4,40 | 0,00 | 1,00E+00 |
| AT5G50160 | 1,06 | 2,47 | 2,29 | 1,08 | 2,80E-01 |
| AT5G50170 | 0,08 | 0,85 | 4,38 | 0,20 | 8,45E-01 |
| AT5G50200 | 0,08 | 0,00 | 4,40 | 0,00 | 1,00E+00 |
| AT5G07605 | 0,25 | 0,46 | 4,39 | 0,11 | 9,16E-01 |
| AT5G50310 | 0,21 | 2,05 | 4,33 | 0,47 | 6,36E-01 |
| AT5G50580 | 0,65 | 2,61 | 3,62 | 0,72 | 4,72E-01 |
| AT5G50680 | 1,09 | 3,75 | 2,33 | 1,61 | 1,08E-01 |
| AT5G50717 | 0,11 | 0,00 | 4,41 | 0,00 | 1,00E+00 |
| AT5G50780 | 0,59 | 0,37 | 4,31 | 0,08 | 9,32E-01 |
| AT5G50850 | 0,82 | 2,30 | 2,31 | 1,00 | 3,19E-01 |
| AT5G50950 | 1,52 | -1,07 | 1,37 | -0,78 | 4,34E-01 |
| AT5G50960 | 0,50 | 0,50 | 3,91 | 0,13 | 8,98E-01 |
| AT5G51010 | 0,08 | 1,10 | 4,36 | 0,25 | 8,00E-01 |
| AT5G51050 | 0,20 | 1,62 | 4,35 | 0,37 | 7,10E-01 |
| AT5G51055 | 0,34 | -2,13 | 3,56 | -0,60 | 5,50E-01 |
| AT5G51120 | 0,08 | 1,10 | 4,36 | 0,25 | 8,00E-01 |
| AT5G51150 | 0,28 | 0,90 | 4,38 | 0,21 | 8,37E-01 |
| AT5G51180 | 0,38 | 0,26 | 2,89 | 0,09 | 9,29E-01 |

| | | | | | |
|---|---|---|---|---|---|
| AT5G51200 | 1,04 | 1,57 | 2,14 | 0,74 | 4,61E-01 |
| AT5G51230 | 0,21 | 0,05 | 4,34 | 0,01 | 9,90E-01 |
| AT5G51280 | 0,48 | 1,63 | 2,70 | 0,60 | 5,47E-01 |
| AT5G51340 | 0,39 | 0,39 | 3,40 | 0,11 | 9,09E-01 |
| AT5G51450 | 0,53 | 0,05 | 3,88 | 0,01 | 9,91E-01 |
| AT5G51560 | 0,28 | 0,00 | 4,41 | 0,00 | 1,00E+00 |
| AT5G51570 | 0,40 | -0,31 | 3,44 | -0,09 | 9,29E-01 |
| AT5G51660 | 1,44 | 1,32 | 1,86 | 0,71 | 4,78E-01 |
| AT5G51670 | 0,12 | 1,39 | 4,34 | 0,32 | 7,49E-01 |
| AT5G51680 | 0,64 | 1,77 | 2,61 | 0,68 | 4,97E-01 |
| AT5G51750 | 0,75 | 0,57 | 3,63 | 0,16 | 8,74E-01 |
| AT5G51760 | 1,35 | 0,00 | 1,66 | 0,00 | 9,99E-01 |
| AT5G51830 | 0,27 | 1,71 | 4,33 | 0,40 | 6,93E-01 |
| AT5G51960 | 0,11 | 0,00 | 4,41 | 0,00 | 1,00E+00 |
| AT5G51970 | 1,04 | 2,36 | 1,90 | 1,24 | 2,15E-01 |
| AT5G52050 | 0,61 | 1,82 | 2,55 | 0,72 | 4,74E-01 |
| AT5G52115 | 1,12 | -3,22 | 2,64 | -1,22 | 2,22E-01 |
| AT5G52240 | 1,16 | 1,45 | 2,01 | 0,72 | 4,71E-01 |
| AT5G52280 | 0,34 | 0,42 | 4,37 | 0,10 | 9,23E-01 |
| AT5G52471 | 1,40 | -2,08 | 1,95 | -1,06 | 2,87E-01 |
| AT5G52510 | 0,69 | 0,39 | 2,94 | 0,13 | 8,95E-01 |
| AT5G52520 | 0,23 | 0,00 | 4,40 | 0,00 | 1,00E+00 |
| AT5G52580 | 0,50 | 1,80 | 3,26 | 0,55 | 5,81E-01 |
| AT5G52630 | 0,08 | 0,85 | 4,38 | 0,20 | 8,45E-01 |
| AT5G52700 | 0,14 | 0,00 | 4,41 | 0,00 | 1,00E+00 |
| AT5G52750 | 0,33 | 1,90 | 4,32 | 0,44 | 6,61E-01 |
| AT5G52760 | 0,52 | 1,66 | 2,62 | 0,63 | 5,26E-01 |
| AT5G52920 | 1,32 | 1,54 | 2,08 | 0,74 | 4,60E-01 |
| AT5G07945 | 0,52 | 2,19 | 3,96 | 0,55 | 5,81E-01 |
| AT5G52940 | 0,11 | 0,90 | 4,37 | 0,21 | 8,36E-01 |
| AT5G53020 | 0,27 | 0,42 | 4,34 | 0,10 | 9,23E-01 |
| AT5G53030 | 0,24 | -1,37 | 4,41 | -0,31 | 7,55E-01 |
| AT5G53048 | 0,25 | 0,62 | 4,34 | 0,14 | 8,86E-01 |
| AT5G53060 | 0,48 | 2,61 | 3,95 | 0,66 | 5,09E-01 |
| AT5G53080 | 0,12 | 1,58 | 4,34 | 0,36 | 7,16E-01 |
| AT5G53110 | 0,31 | -1,06 | 4,31 | -0,25 | 8,06E-01 |
| AT5G53120 | 0,82 | 2,83 | 2,56 | 1,11 | 2,68E-01 |
| AT5G53140 | 1,45 | 2,20 | 1,87 | 1,18 | 2,39E-01 |
| AT5G53150 | 0,95 | 0,57 | 2,23 | 0,26 | 7,98E-01 |
| AT5G53160 | 1,14 | 3,03 | 1,91 | 1,59 | 1,13E-01 |
| AT5G53220 | 0,48 | 2,24 | 3,10 | 0,72 | 4,70E-01 |
| AT5G53280 | 0,44 | 0,56 | 4,16 | 0,13 | 8,93E-01 |
| AT5G53350 | 0,54 | -1,76 | 3,16 | -0,56 | 5,76E-01 |
| AT5G53360 | 0,08 | 0,85 | 4,38 | 0,20 | 8,45E-01 |
| AT5G53410 | 1,13 | -0,17 | 1,63 | -0,10 | 9,19E-01 |
| AT5G53420 | 0,48 | 1,08 | 3,31 | 0,33 | 7,44E-01 |
| AT5G53430 | 1,00 | 0,40 | 1,93 | 0,21 | 8,35E-01 |
| AT5G53450 | 0,71 | -0,13 | 2,64 | -0,05 | 9,61E-01 |
| AT5G53550 | 0,99 | 3,56 | 2,29 | 1,56 | 1,20E-01 |
| AT5G53560 | 1,33 | 1,73 | 1,73 | 1,00 | 3,19E-01 |
| AT5G53590 | 0,48 | 0,64 | 3,99 | 0,16 | 8,72E-01 |
| AT5G53620 | 1,09 | 1,32 | 1,83 | 0,72 | 4,70E-01 |
| AT5G53860 | 0,25 | 1,23 | 3,80 | 0,32 | 7,46E-01 |
| AT5G53890 | 0,71 | 2,00 | 2,42 | 0,83 | 4,09E-01 |
| AT5G54035 | 0,15 | 0,00 | 4,41 | 0,00 | 1,00E+00 |
| AT5G54043 | 0,22 | 0,42 | 4,38 | 0,10 | 9,24E-01 |

| | | | | | |
|---|---|---|---|---|---|
| AT5G54080 | 0,99 | 3,22 | 2,15 | 1,50 | 1,34E-01 |
| AT5G54110 | 0,33 | 2,23 | 4,32 | 0,52 | 6,06E-01 |
| AT5G54140 | 0,24 | 1,62 | 4,34 | 0,37 | 7,08E-01 |
| AT5G54145 | 0,12 | 0,00 | 4,41 | 0,00 | 1,00E+00 |
| AT5G54160 | 0,46 | -0,23 | 4,06 | -0,06 | 9,54E-01 |
| AT5G54180 | 0,12 | 0,68 | 4,37 | 0,16 | 8,77E-01 |
| AT5G54290 | 0,66 | -0,68 | 2,71 | -0,25 | 8,02E-01 |
| AT5G54500 | 0,31 | -0,71 | 4,32 | -0,17 | 8,69E-01 |
| AT5G54540 | 0,58 | 1,54 | 2,64 | 0,58 | 5,59E-01 |
| AT5G54600 | 0,87 | 0,64 | 1,96 | 0,32 | 7,45E-01 |
| AT5G08225 | 0,36 | 1,01 | 4,37 | 0,23 | 8,17E-01 |
| AT5G54640 | 0,05 | 0,00 | 4,41 | 0,00 | 1,00E+00 |
| AT5G54680 | 0,72 | -0,83 | 2,62 | -0,32 | 7,51E-01 |
| AT5G54750 | 0,55 | 0,76 | 3,87 | 0,20 | 8,43E-01 |
| AT5G54760 | 0,49 | -1,65 | 2,74 | -0,60 | 5,47E-01 |
| AT5G54800 | 0,08 | 0,00 | 4,41 | 0,00 | 1,00E+00 |
| AT5G54830 | 0,76 | 3,46 | 2,73 | 1,27 | 2,05E-01 |
| AT5G54855 | 0,18 | -0,69 | 4,37 | -0,16 | 8,74E-01 |
| AT5G54860 | 0,26 | 1,08 | 4,37 | 0,25 | 8,05E-01 |
| AT5G54900 | 0,82 | 1,74 | 2,33 | 0,75 | 4,53E-01 |
| AT5G54920 | 0,83 | 2,24 | 4,29 | 0,52 | 6,01E-01 |
| AT5G55060 | 0,14 | 0,51 | 4,38 | 0,12 | 9,08E-01 |
| AT5G55070 | 1,01 | 2,07 | 1,94 | 1,07 | 2,85E-01 |
| AT5G55190 | 0,80 | 2,05 | 2,31 | 0,89 | 3,75E-01 |
| AT5G55220 | 0,88 | -3,35 | 2,60 | -1,29 | 1,96E-01 |
| AT5G55230 | 0,95 | 0,70 | 2,09 | 0,33 | 7,39E-01 |
| AT5G55450 | 0,62 | -0,52 | 3,05 | -0,17 | 8,64E-01 |
| AT5G55460 | 0,37 | -0,41 | 3,43 | -0,12 | 9,05E-01 |
| AT5G55500 | 0,57 | 2,85 | 3,08 | 0,93 | 3,54E-01 |
| AT5G55530 | 0,30 | 1,81 | 4,33 | 0,42 | 6,76E-01 |
| AT5G55580 | 0,70 | 0,43 | 3,86 | 0,11 | 9,12E-01 |
| AT5G55620 | 0,43 | -0,50 | 4,28 | -0,12 | 9,07E-01 |
| AT5G55630 | 0,12 | 1,39 | 4,34 | 0,32 | 7,49E-01 |
| AT5G55660 | 0,90 | -1,28 | 2,15 | -0,59 | 5,53E-01 |
| AT5G55840 | 0,22 | 1,58 | 4,33 | 0,37 | 7,15E-01 |
| AT5G55850 | 0,24 | 0,41 | 4,38 | 0,09 | 9,25E-01 |
| AT5G55900 | 0,10 | 0,00 | 4,41 | 0,00 | 1,00E+00 |
| AT5G55930 | 0,70 | 2,82 | 2,91 | 0,97 | 3,33E-01 |
| AT5G55960 | 0,59 | -1,91 | 3,04 | -0,63 | 5,31E-01 |
| AT5G55990 | 0,33 | 1,96 | 4,33 | 0,45 | 6,52E-01 |
| AT5G56020 | 0,32 | 1,20 | 4,35 | 0,28 | 7,82E-01 |
| AT5G56040 | 0,73 | 1,87 | 2,39 | 0,78 | 4,33E-01 |
| AT5G56060 | 0,67 | 1,33 | 4,33 | 0,31 | 7,60E-01 |
| AT5G56140 | 0,98 | 1,19 | 2,03 | 0,59 | 5,58E-01 |
| AT5G56160 | 0,49 | -2,71 | 4,34 | -0,62 | 5,32E-01 |
| AT5G56170 | 0,53 | 0,50 | 4,38 | 0,11 | 9,09E-01 |
| AT5G56270 | 0,76 | 1,86 | 2,83 | 0,66 | 5,12E-01 |
| AT5G56290 | 0,52 | 0,58 | 2,38 | 0,24 | 8,09E-01 |
| AT5G56340 | 0,44 | 0,73 | 3,27 | 0,22 | 8,24E-01 |
| AT5G08495 | 0,43 | -1,65 | 4,22 | -0,39 | 6,96E-01 |
| AT5G56460 | 0,74 | -1,88 | 3,86 | -0,49 | 6,26E-01 |
| AT5G56500 | 0,32 | 0,00 | 4,41 | 0,00 | 1,00E+00 |
| AT5G56590 | 0,24 | 0,41 | 4,38 | 0,09 | 9,25E-01 |
| AT5G56630 | 0,71 | 0,51 | 2,56 | 0,20 | 8,41E-01 |
| AT5G56650 | 0,64 | 1,24 | 2,37 | 0,52 | 6,00E-01 |
| AT5G56680 | 1,22 | 0,46 | 1,81 | 0,25 | 7,99E-01 |

| | | | | | |
|---|---|---|---|---|---|
| AT5G56720 | 0,12 | 0,79 | 4,37 | 0,18 | 8,57E-01 |
| AT5G56870 | 0,19 | 1,26 | 4,35 | 0,29 | 7,72E-01 |
| AT5G56910 | 0,15 | 1,58 | 4,34 | 0,36 | 7,15E-01 |
| AT5G56920 | 0,65 | 1,51 | 3,07 | 0,49 | 6,23E-01 |
| AT5G56940 | 0,08 | 0,00 | 4,41 | 0,00 | 1,00E+00 |
| AT5G56950 | 0,44 | 0,16 | 3,18 | 0,05 | 9,61E-01 |
| AT5G57015 | 0,32 | 0,76 | 4,36 | 0,18 | 8,61E-01 |
| AT5G57020 | 1,25 | 1,34 | 1,70 | 0,79 | 4,31E-01 |
| AT5G57030 | 0,86 | 1,23 | 2,24 | 0,55 | 5,82E-01 |
| AT5G57035 | 1,04 | 0,38 | 2,58 | 0,15 | 8,84E-01 |
| AT5G57070 | 0,42 | 1,13 | 4,23 | 0,27 | 7,89E-01 |
| AT5G57150 | 0,25 | 1,76 | 4,33 | 0,41 | 6,85E-01 |
| AT5G57180 | 1,05 | -2,46 | 1,85 | -1,33 | 1,83E-01 |
| AT5G57320 | 0,08 | 0,85 | 4,38 | 0,20 | 8,45E-01 |
| AT5G57330 | 0,49 | -0,21 | 2,82 | -0,07 | 9,41E-01 |
| AT5G57340 | 0,20 | 0,57 | 4,37 | 0,13 | 8,96E-01 |
| AT5G57360 | 0,96 | 2,15 | 1,89 | 1,14 | 2,54E-01 |
| AT5G57370 | 0,31 | 1,25 | 4,36 | 0,29 | 7,74E-01 |
| AT5G57460 | 1,07 | 1,89 | 1,82 | 1,04 | 3,00E-01 |
| AT5G57520 | 0,31 | 0,44 | 4,39 | 0,10 | 9,20E-01 |
| AT5G57610 | 0,89 | 1,13 | 1,86 | 0,61 | 5,43E-01 |
| AT5G57655 | 1,67 | 0,91 | 1,61 | 0,57 | 5,70E-01 |
| AT5G57735 | 0,19 | 0,00 | 4,41 | 0,00 | 1,00E+00 |
| AT5G57800 | 0,61 | -2,05 | 2,65 | -0,77 | 4,39E-01 |
| AT5G08805 | 1,03 | 2,43 | 2,42 | 1,00 | 3,16E-01 |
| AT5G57880 | 0,48 | -0,40 | 4,10 | -0,10 | 9,22E-01 |
| AT5G57930 | 1,66 | 2,56 | 1,85 | 1,38 | 1,68E-01 |
| AT5G57940 | 0,35 | 1,00 | 3,52 | 0,29 | 7,76E-01 |
| AT5G58000 | 0,08 | 1,10 | 4,36 | 0,25 | 8,00E-01 |
| AT5G58020 | 0,39 | -2,27 | 4,30 | -0,53 | 5,97E-01 |
| AT5G58060 | 0,59 | 1,57 | 2,57 | 0,61 | 5,42E-01 |
| AT5G58100 | 0,97 | 0,03 | 1,88 | 0,02 | 9,87E-01 |
| AT5G58110 | 0,27 | 0,19 | 4,34 | 0,04 | 9,65E-01 |
| AT5G58120 | 1,53 | 0,93 | 1,38 | 0,67 | 5,02E-01 |
| AT5G08820 | 0,19 | 1,50 | 4,34 | 0,34 | 7,30E-01 |
| AT5G58200 | 0,40 | -2,52 | 4,36 | -0,58 | 5,63E-01 |
| AT5G58230 | 0,77 | 2,85 | 2,44 | 1,17 | 2,43E-01 |
| AT5G58250 | 0,32 | 0,26 | 4,33 | 0,06 | 9,52E-01 |
| AT5G58270 | 0,43 | 0,64 | 4,20 | 0,15 | 8,80E-01 |
| AT5G58290 | 1,43 | 0,16 | 1,61 | 0,10 | 9,22E-01 |
| AT5G58300 | 0,99 | 2,21 | 2,28 | 0,97 | 3,34E-01 |
| AT5G58320 | 0,24 | 1,23 | 4,35 | 0,28 | 7,77E-01 |
| AT5G58350 | 1,12 | 3,67 | 1,94 | 1,89 | 5,81E-02 |
| AT5G58370 | 0,29 | 0,00 | 4,38 | 0,00 | 1,00E+00 |
| AT5G58375 | 0,48 | 0,85 | 4,38 | 0,19 | 8,46E-01 |
| AT5G58420 | 1,59 | -0,99 | 1,91 | -0,52 | 6,03E-01 |
| AT5G58450 | 0,68 | 1,82 | 2,94 | 0,62 | 5,35E-01 |
| AT5G58495 | 0,12 | 1,58 | 4,34 | 0,36 | 7,16E-01 |
| AT5G58520 | 0,39 | 0,80 | 4,37 | 0,18 | 8,56E-01 |
| AT5G58560 | 0,47 | 2,06 | 3,22 | 0,64 | 5,22E-01 |
| AT5G08855 | 0,43 | -1,25 | 4,27 | -0,29 | 7,70E-01 |
| AT5G58620 | 0,34 | 1,36 | 4,37 | 0,31 | 7,55E-01 |
| AT5G58640 | 0,71 | 0,55 | 2,97 | 0,18 | 8,54E-01 |
| AT5G58710 | 0,66 | -0,91 | 3,00 | -0,30 | 7,63E-01 |
| AT5G58990 | 0,29 | -1,23 | 4,39 | -0,28 | 7,79E-01 |
| AT5G59160 | 0,32 | 1,29 | 4,31 | 0,30 | 7,64E-01 |

| | | | | | |
|---|---|---|---|---|---|
| AT5G59170 | 0,26 | 1,08 | 4,37 | 0,25 | 8,05E-01 |
| AT5G59250 | 0,50 | -0,91 | 3,19 | -0,29 | 7,75E-01 |
| AT5G59350 | 0,35 | 1,31 | 4,35 | 0,30 | 7,64E-01 |
| AT5G59420 | 1,43 | 1,58 | 1,82 | 0,87 | 3,85E-01 |
| AT5G59430 | 0,64 | 0,41 | 2,50 | 0,16 | 8,69E-01 |
| AT5G59450 | 0,61 | 2,41 | 3,84 | 0,63 | 5,29E-01 |
| AT5G59610 | 0,51 | 2,36 | 3,96 | 0,60 | 5,50E-01 |
| AT5G59660 | 0,05 | 0,00 | 4,41 | 0,00 | 1,00E+00 |
| AT5G59670 | 0,28 | -0,34 | 3,71 | -0,09 | 9,27E-01 |
| AT5G59680 | 0,64 | 0,77 | 4,39 | 0,18 | 8,61E-01 |
| AT5G59690 | 0,64 | 2,65 | 2,92 | 0,91 | 3,65E-01 |
| AT5G59820 | 0,42 | 0,60 | 3,29 | 0,18 | 8,55E-01 |
| AT5G59830 | 0,85 | 0,59 | 2,89 | 0,20 | 8,38E-01 |
| AT5G59860 | 0,43 | 0,89 | 4,38 | 0,20 | 8,38E-01 |
| AT5G59880 | 0,87 | -0,27 | 2,51 | -0,11 | 9,13E-01 |
| AT5G59910 | 0,91 | 1,53 | 2,27 | 0,67 | 5,01E-01 |
| AT5G59920 | 0,58 | 2,54 | 3,70 | 0,69 | 4,92E-01 |
| AT5G59950 | 1,16 | 0,49 | 1,88 | 0,26 | 7,94E-01 |
| AT5G59980 | 0,39 | 0,48 | 4,40 | 0,11 | 9,14E-01 |
| AT5G60040 | 0,79 | 2,20 | 2,40 | 0,92 | 3,59E-01 |
| AT5G60110 | 0,25 | 1,75 | 4,33 | 0,40 | 6,87E-01 |
| AT5G60120 | 0,67 | 1,75 | 2,78 | 0,63 | 5,29E-01 |
| AT5G60170 | 1,68 | 1,90 | 1,69 | 1,12 | 2,62E-01 |
| AT5G60270 | 0,48 | 2,94 | 4,30 | 0,68 | 4,94E-01 |
| AT5G60280 | 0,76 | 3,39 | 3,43 | 0,99 | 3,23E-01 |
| AT5G60300 | 0,52 | 2,72 | 3,85 | 0,71 | 4,80E-01 |
| AT5G60320 | 0,25 | 1,76 | 4,33 | 0,41 | 6,85E-01 |
| AT5G60550 | 0,49 | 1,35 | 3,16 | 0,43 | 6,69E-01 |
| AT5G60560 | 0,60 | 1,75 | 3,09 | 0,57 | 5,71E-01 |
| AT5G60570 | 0,88 | 0,46 | 2,30 | 0,20 | 8,42E-01 |
| AT5G60610 | 0,72 | 1,26 | 3,60 | 0,35 | 7,27E-01 |
| AT5G60620 | 0,40 | 0,71 | 3,47 | 0,20 | 8,38E-01 |
| AT5G60680 | 1,59 | 3,64 | 1,86 | 1,96 | 5,01E-02 |
| AT5G60780 | 1,02 | 2,13 | 2,20 | 0,97 | 3,31E-01 |
| AT5G60820 | 0,45 | 1,85 | 4,10 | 0,45 | 6,52E-01 |
| AT5G60850 | 1,63 | 0,27 | 1,86 | 0,14 | 8,86E-01 |
| AT5G60890 | 0,24 | 0,00 | 4,41 | 0,00 | 1,00E+00 |
| AT5G60945 | 1,24 | -1,52 | 1,69 | -0,90 | 3,69E-01 |
| AT5G60963 | 0,50 | 1,35 | 2,62 | 0,52 | 6,06E-01 |
| AT5G60964 | 0,93 | -1,52 | 3,42 | -0,44 | 6,58E-01 |
| AT5G60980 | 1,13 | 1,48 | 2,24 | 0,66 | 5,08E-01 |
| AT5G61190 | 0,65 | 0,63 | 2,39 | 0,26 | 7,91E-01 |
| AT5G61200 | 0,25 | 0,00 | 4,40 | 0,00 | 1,00E+00 |
| AT5G61210 | 1,18 | 2,45 | 1,83 | 1,34 | 1,80E-01 |
| AT5G61230 | 1,00 | 0,23 | 1,95 | 0,12 | 9,06E-01 |
| AT5G61240 | 0,29 | -0,42 | 4,33 | -0,10 | 9,23E-01 |
| AT5G61380 | 0,78 | 1,85 | 2,37 | 0,78 | 4,36E-01 |
| AT5G61400 | 0,78 | 1,30 | 3,48 | 0,37 | 7,09E-01 |
| AT5G61470 | 0,39 | 1,23 | 3,36 | 0,37 | 7,15E-01 |
| AT5G61490 | 0,17 | 1,94 | 4,33 | 0,45 | 6,55E-01 |
| AT5G61550 | 0,85 | 3,43 | 3,29 | 1,04 | 2,97E-01 |
| AT5G61570 | 0,19 | -1,36 | 4,41 | -0,31 | 7,58E-01 |
| AT5G61580 | 0,26 | 1,30 | 4,35 | 0,30 | 7,65E-01 |
| AT5G61950 | 0,32 | 0,81 | 4,37 | 0,19 | 8,53E-01 |
| AT5G61970 | 0,49 | 0,56 | 3,39 | 0,17 | 8,69E-01 |
| AT5G61980 | 0,55 | 0,84 | 4,37 | 0,19 | 8,48E-01 |

| | | | | | |
|---|---|---|---|---|---|
| AT5G62030 | 0,41 | 0,98 | 4,16 | 0,24 | 8,13E-01 |
| AT5G62270 | 0,20 | 1,23 | 4,36 | 0,28 | 7,77E-01 |
| AT5G62290 | 0,13 | 0,00 | 4,41 | 0,00 | 1,00E+00 |
| AT5G62360 | 0,27 | 0,48 | 4,38 | 0,11 | 9,13E-01 |
| AT5G62460 | 0,44 | 1,96 | 4,15 | 0,47 | 6,36E-01 |
| AT5G62500 | 0,05 | 0,00 | 4,41 | 0,00 | 1,00E+00 |
| AT5G62520 | 0,28 | 1,30 | 4,32 | 0,30 | 7,63E-01 |
| AT5G62560 | 0,37 | -0,20 | 3,52 | -0,06 | 9,56E-01 |
| AT5G62570 | 0,48 | 1,77 | 2,68 | 0,66 | 5,09E-01 |
| AT5G62575 | 0,21 | 2,12 | 4,33 | 0,49 | 6,23E-01 |
| AT5G62620 | 0,35 | 0,93 | 4,35 | 0,21 | 8,30E-01 |
| AT5G62710 | 0,23 | 0,79 | 4,37 | 0,18 | 8,56E-01 |
| AT5G62720 | 0,31 | 2,05 | 4,33 | 0,47 | 6,37E-01 |
| AT5G62740 | 0,43 | 1,06 | 4,04 | 0,26 | 7,93E-01 |
| AT5G62770 | 0,31 | 1,29 | 4,35 | 0,30 | 7,67E-01 |
| AT5G62890 | 1,11 | 2,16 | 2,07 | 1,04 | 2,97E-01 |
| AT5G62910 | 0,36 | 1,07 | 4,33 | 0,25 | 8,06E-01 |
| AT5G62970 | 0,35 | 0,47 | 3,55 | 0,13 | 8,95E-01 |
| AT5G63010 | 0,41 | 1,95 | 3,30 | 0,59 | 5,54E-01 |
| AT5G09120 | 1,28 | -2,10 | 1,83 | -1,15 | 2,50E-01 |
| AT5G63050 | 0,55 | -0,20 | 4,05 | -0,05 | 9,60E-01 |
| AT5G63130 | 0,64 | 0,97 | 2,92 | 0,33 | 7,41E-01 |
| AT5G63200 | 0,75 | 0,00 | 3,60 | 0,00 | 1,00E+00 |
| AT5G63260 | 0,08 | 1,10 | 4,36 | 0,25 | 8,00E-01 |
| AT5G63280 | 0,12 | 1,58 | 4,34 | 0,36 | 7,16E-01 |
| AT5G09150 | 0,24 | 0,00 | 4,41 | 0,00 | 1,00E+00 |
| AT5G63300 | 0,47 | 1,64 | 4,35 | 0,38 | 7,07E-01 |
| AT5G63340 | 0,26 | 0,45 | 4,39 | 0,10 | 9,19E-01 |
| AT5G63420 | 0,69 | -1,33 | 3,08 | -0,43 | 6,65E-01 |
| AT5G63480 | 0,12 | 0,00 | 4,41 | 0,00 | 1,00E+00 |
| AT5G63510 | 0,37 | 2,01 | 4,28 | 0,47 | 6,39E-01 |
| AT5G63640 | 0,36 | 0,38 | 4,35 | 0,09 | 9,30E-01 |
| AT5G63680 | 0,63 | 1,96 | 3,14 | 0,62 | 5,32E-01 |
| AT5G63770 | 0,64 | 0,52 | 2,98 | 0,17 | 8,62E-01 |
| AT5G63780 | 0,69 | -0,55 | 2,35 | -0,23 | 8,16E-01 |
| AT5G63810 | 0,68 | -1,00 | 3,03 | -0,33 | 7,42E-01 |
| AT5G63940 | 0,43 | 0,53 | 3,38 | 0,16 | 8,76E-01 |
| AT5G63960 | 0,45 | 0,00 | 4,41 | 0,00 | 1,00E+00 |
| AT5G63980 | 0,08 | 1,10 | 4,36 | 0,25 | 8,00E-01 |
| AT5G64030 | 1,58 | -0,31 | 1,33 | -0,24 | 8,13E-01 |
| AT5G64090 | 0,72 | 2,42 | 2,44 | 0,99 | 3,21E-01 |
| AT5G64160 | 0,20 | -0,05 | 4,36 | -0,01 | 9,91E-01 |
| AT5G64190 | 0,08 | 1,10 | 4,36 | 0,25 | 8,00E-01 |
| AT5G64240 | 0,28 | 1,69 | 4,35 | 0,39 | 6,97E-01 |
| AT5G64300 | 1,43 | 0,70 | 1,61 | 0,44 | 6,61E-01 |
| AT5G64310 | 0,80 | 2,40 | 2,73 | 0,88 | 3,81E-01 |
| AT5G64330 | 1,11 | -1,87 | 1,89 | -0,99 | 3,22E-01 |
| AT5G64370 | 0,51 | 1,98 | 3,92 | 0,50 | 6,14E-01 |
| AT5G64380 | 0,10 | 0,42 | 4,38 | 0,10 | 9,23E-01 |
| AT5G64400 | 0,52 | 0,22 | 4,09 | 0,05 | 9,57E-01 |
| AT5G64440 | 0,42 | -0,55 | 3,39 | -0,16 | 8,72E-01 |
| AT5G64500 | 0,82 | -0,63 | 2,17 | -0,29 | 7,74E-01 |
| AT5G64505 | 0,26 | 1,08 | 4,37 | 0,25 | 8,05E-01 |
| AT5G64572 | 1,22 | 2,58 | 1,93 | 1,34 | 1,80E-01 |
| AT5G64690 | 0,11 | 0,00 | 4,41 | 0,00 | 1,00E+00 |
| AT5G64750 | 0,12 | 1,65 | 4,34 | 0,38 | 7,04E-01 |

| | | | | | |
|---|---|---|---|---|---|
| AT5G64813 | 0,60 | 1,92 | 2,58 | 0,74 | 4,57E-01 |
| AT5G09285 | 0,45 | 2,10 | 3,98 | 0,53 | 5,97E-01 |
| AT5G64950 | 0,48 | 2,61 | 4,32 | 0,60 | 5,46E-01 |
| AT5G64960 | 0,59 | 1,72 | 3,11 | 0,56 | 5,79E-01 |
| AT5G65010 | 1,00 | -0,82 | 1,69 | -0,49 | 6,27E-01 |
| AT5G65015 | 0,39 | 2,11 | 4,33 | 0,49 | 6,27E-01 |
| AT5G65210 | 0,26 | 1,08 | 4,37 | 0,25 | 8,05E-01 |
| AT5G65220 | 1,17 | -2,36 | 2,04 | -1,16 | 2,47E-01 |
| AT5G65230 | 0,08 | 0,85 | 4,38 | 0,20 | 8,45E-01 |
| AT5G65270 | 0,10 | 0,42 | 4,38 | 0,10 | 9,23E-01 |
| AT5G65290 | 0,47 | 0,00 | 4,40 | 0,00 | 1,00E+00 |
| AT5G65400 | 0,17 | 0,09 | 4,35 | 0,02 | 9,84E-01 |
| AT5G65440 | 0,58 | 0,41 | 3,09 | 0,13 | 8,95E-01 |
| AT5G65470 | 0,30 | 1,91 | 4,32 | 0,44 | 6,59E-01 |
| AT5G65510 | 0,12 | 0,42 | 4,38 | 0,10 | 9,24E-01 |
| AT5G65570 | 0,25 | 1,57 | 4,34 | 0,36 | 7,18E-01 |
| AT5G65620 | 1,40 | 0,60 | 1,89 | 0,31 | 7,53E-01 |
| AT5G65640 | 0,08 | 1,10 | 4,36 | 0,25 | 8,00E-01 |
| AT5G65670 | 0,91 | -0,14 | 2,06 | -0,07 | 9,44E-01 |
| AT5G65710 | 0,77 | 2,40 | 2,16 | 1,11 | 2,67E-01 |
| AT5G65720 | 1,65 | 0,83 | 1,33 | 0,62 | 5,33E-01 |
| AT5G65760 | 0,55 | 2,29 | 2,63 | 0,87 | 3,84E-01 |
| AT5G65770 | 1,30 | -0,63 | 1,93 | -0,33 | 7,43E-01 |
| AT5G65780 | 1,41 | -0,73 | 1,84 | -0,40 | 6,91E-01 |
| AT5G65890 | 0,05 | 0,00 | 4,41 | 0,00 | 1,00E+00 |
| AT5G65960 | 1,37 | 3,15 | 1,94 | 1,63 | 1,04E-01 |
| AT5G65990 | 0,70 | 0,73 | 1,97 | 0,37 | 7,11E-01 |
| AT5G66052 | 0,53 | 1,11 | 4,03 | 0,28 | 7,83E-01 |
| AT5G66070 | 0,10 | 0,42 | 4,38 | 0,10 | 9,23E-01 |
| AT5G66100 | 0,24 | 1,72 | 4,34 | 0,40 | 6,92E-01 |
| AT5G66110 | 0,30 | 1,45 | 4,35 | 0,33 | 7,39E-01 |
| AT5G66160 | 0,31 | 0,53 | 4,40 | 0,12 | 9,05E-01 |
| AT5G66170 | 0,33 | 1,53 | 4,34 | 0,35 | 7,24E-01 |
| AT5G66240 | 1,02 | 3,10 | 2,30 | 1,35 | 1,77E-01 |
| AT5G66360 | 0,60 | 2,34 | 3,78 | 0,62 | 5,36E-01 |
| AT5G66450 | 0,53 | 0,41 | 3,22 | 0,13 | 8,99E-01 |
| AT5G66560 | 1,38 | 2,14 | 1,79 | 1,20 | 2,32E-01 |
| AT5G66562 | 1,20 | 0,73 | 1,67 | 0,43 | 6,64E-01 |
| AT5G66564 | 1,37 | 0,02 | 1,60 | 0,01 | 9,88E-01 |
| AT5G66568 | 0,20 | 0,43 | 4,39 | 0,10 | 9,21E-01 |
| AT5G66760 | 1,08 | 0,46 | 1,84 | 0,25 | 8,03E-01 |
| AT5G66770 | 0,43 | 1,47 | 4,32 | 0,34 | 7,34E-01 |
| AT5G66790 | 0,74 | -0,60 | 2,09 | -0,29 | 7,74E-01 |
| AT5G66810 | 0,67 | 2,25 | 2,49 | 0,90 | 3,66E-01 |
| AT5G66930 | 0,12 | 0,79 | 4,37 | 0,18 | 8,57E-01 |
| AT5G66950 | 1,10 | -0,95 | 1,60 | -0,59 | 5,54E-01 |
| AT5G67070 | 0,51 | -1,35 | 3,98 | -0,34 | 7,35E-01 |
| AT5G67080 | 0,30 | 0,90 | 4,38 | 0,20 | 8,38E-01 |
| AT5G67170 | 0,08 | 0,85 | 4,38 | 0,20 | 8,45E-01 |
| AT5G67260 | 0,65 | 1,56 | 3,95 | 0,40 | 6,93E-01 |
| AT5G67320 | 0,96 | 2,53 | 2,53 | 1,00 | 3,18E-01 |
| AT5G67330 | 1,26 | 1,88 | 1,85 | 1,02 | 3,10E-01 |
| AT5G67340 | 0,24 | 1,96 | 4,34 | 0,45 | 6,52E-01 |
| AT5G67385 | 0,80 | 1,10 | 2,27 | 0,48 | 6,29E-01 |
| AT5G67455 | 0,10 | 0,42 | 4,38 | 0,10 | 9,23E-01 |
| AT5G67470 | 1,09 | 3,12 | 2,08 | 1,50 | 1,33E-01 |

| | | | | | |
|---|---|---|---|---|---|
| AT5G67488 | 1,20 | 1,40 | 1,86 | 0,75 | 4,52E-01 |
| AT5G67500 | 0,21 | 1,77 | 4,34 | 0,41 | 6,84E-01 |
| AT5G67530 | 0,08 | 0,85 | 4,38 | 0,20 | 8,45E-01 |
| AT5G67560 | 0,70 | 2,07 | 3,65 | 0,57 | 5,70E-01 |
| AT5G67570 | 0,48 | 0,89 | 4,17 | 0,21 | 8,31E-01 |
| AT5G67590 | 0,32 | -1,35 | 4,41 | -0,31 | 7,59E-01 |
| AT1G01020 | 0,15 | 0,42 | 4,38 | 0,09 | 9,24E-01 |
| AT1G03993 | 0,11 | 1,07 | 4,36 | 0,25 | 8,06E-01 |
| AT1G01050 | 0,66 | 1,60 | 2,94 | 0,54 | 5,86E-01 |
| AT1G01060 | 0,69 | 2,33 | 2,44 | 0,95 | 3,40E-01 |
| AT1G01090 | 1,61 | 0,43 | 1,52 | 0,28 | 7,79E-01 |
| AT1G01100 | 1,47 | -0,15 | 1,49 | -0,10 | 9,18E-01 |
| AT1G01120 | 1,43 | 0,12 | 1,57 | 0,07 | 9,41E-01 |
| AT1G01130 | 0,29 | 0,00 | 4,38 | 0,00 | 1,00E+00 |
| AT1G01140 | 0,54 | 1,57 | 2,60 | 0,60 | 5,46E-01 |
| AT1G01170 | 0,58 | 1,42 | 3,10 | 0,46 | 6,47E-01 |
| AT1G01440 | 0,35 | 1,53 | 4,34 | 0,35 | 7,24E-01 |
| AT1G01450 | 0,59 | 2,35 | 4,31 | 0,55 | 5,86E-01 |
| AT1G01470 | 1,29 | -0,62 | 1,52 | -0,41 | 6,84E-01 |
| AT1G01490 | 0,15 | -1,24 | 4,39 | -0,28 | 7,77E-01 |
| AT1G01530 | 0,08 | 0,00 | 4,41 | 0,00 | 1,00E+00 |
| AT1G01610 | 0,73 | -2,74 | 2,40 | -1,14 | 2,53E-01 |
| AT1G01620 | 1,23 | -2,00 | 2,00 | -1,00 | 3,17E-01 |
| AT1G01650 | 0,59 | 2,79 | 2,90 | 0,96 | 3,36E-01 |
| AT1G04027 | 0,46 | 2,36 | 3,12 | 0,76 | 4,50E-01 |
| AT1G01680 | 0,72 | 1,36 | 2,58 | 0,53 | 5,97E-01 |
| AT1G01725 | 1,09 | 1,19 | 1,84 | 0,65 | 5,19E-01 |
| AT1G04063 | 0,34 | -1,60 | 4,36 | -0,37 | 7,14E-01 |
| AT1G01820 | 0,86 | -0,96 | 2,24 | -0,43 | 6,67E-01 |
| AT1G01830 | 1,22 | 2,14 | 1,92 | 1,11 | 2,67E-01 |
| AT1G01880 | 1,08 | -2,64 | 2,12 | -1,24 | 2,14E-01 |
| AT1G01890 | 1,04 | -3,39 | 2,56 | -1,33 | 1,85E-01 |
| AT1G01930 | 0,16 | 0,90 | 4,38 | 0,21 | 8,37E-01 |
| AT1G01990 | 0,21 | 1,97 | 4,33 | 0,46 | 6,49E-01 |
| AT1G02020 | 0,45 | 1,23 | 4,07 | 0,30 | 7,63E-01 |
| AT1G02090 | 0,84 | 3,04 | 2,75 | 1,11 | 2,68E-01 |
| AT1G02140 | 0,21 | 0,18 | 4,34 | 0,04 | 9,68E-01 |
| AT1G02260 | 0,54 | 1,35 | 3,21 | 0,42 | 6,75E-01 |
| AT1G02280 | 0,61 | 1,60 | 3,00 | 0,53 | 5,94E-01 |
| AT1G02310 | 0,21 | 1,39 | 4,34 | 0,32 | 7,49E-01 |
| AT1G04117 | 0,17 | 1,75 | 4,33 | 0,40 | 6,86E-01 |
| AT1G02430 | 0,17 | 1,57 | 4,34 | 0,36 | 7,18E-01 |
| AT1G02450 | 0,20 | 0,82 | 4,33 | 0,19 | 8,50E-01 |
| AT1G02610 | 0,33 | 0,00 | 4,37 | 0,00 | 1,00E+00 |
| AT1G02660 | 0,36 | 0,41 | 4,38 | 0,09 | 9,25E-01 |
| AT1G02770 | 0,41 | 0,00 | 4,41 | 0,00 | 1,00E+00 |
| AT1G02870 | 0,35 | -0,48 | 4,36 | -0,11 | 9,12E-01 |
| AT1G02910 | 0,70 | 0,28 | 2,50 | 0,11 | 9,11E-01 |
| AT1G02920 | 0,41 | -0,39 | 3,41 | -0,11 | 9,10E-01 |
| AT1G02930 | 0,49 | 0,87 | 3,32 | 0,26 | 7,94E-01 |
| AT1G02960 | 0,08 | 1,10 | 4,36 | 0,25 | 8,00E-01 |
| AT1G03000 | 0,70 | 1,88 | 2,63 | 0,71 | 4,75E-01 |
| AT1G03040 | 0,62 | -0,28 | 2,69 | -0,10 | 9,18E-01 |
| AT1G03140 | 0,90 | 1,74 | 1,88 | 0,92 | 3,56E-01 |
| AT1G03240 | 1,51 | -3,77 | 2,22 | -1,70 | 8,97E-02 |
| AT1G03300 | 0,12 | 1,65 | 4,34 | 0,38 | 7,04E-01 |

| | | | | | |
|---|---|---|---|---|---|
| AT1G03400 | 0,49 | 1,60 | 2,75 | 0,58 | 5,60E-01 |
| AT1G03445 | 0,28 | 2,22 | 4,33 | 0,51 | 6,07E-01 |
| AT1G03475 | 0,69 | -1,59 | 2,26 | -0,70 | 4,82E-01 |
| AT1G03520 | 0,48 | 1,97 | 4,00 | 0,49 | 6,22E-01 |
| AT1G03530 | 0,41 | 1,46 | 4,34 | 0,34 | 7,36E-01 |
| AT1G03550 | 0,70 | -1,33 | 2,34 | -0,57 | 5,70E-01 |
| AT1G03590 | 0,41 | 0,86 | 4,22 | 0,20 | 8,38E-01 |
| AT1G03660 | 0,52 | 1,75 | 2,69 | 0,65 | 5,15E-01 |
| AT1G03730 | 1,35 | -0,66 | 2,31 | -0,29 | 7,75E-01 |
| AT1G03746 | 0,46 | -1,90 | 4,10 | -0,46 | 6,43E-01 |
| AT1G03760 | 0,08 | 0,00 | 4,40 | 0,00 | 1,00E+00 |
| AT1G03830 | 0,65 | 0,65 | 2,46 | 0,26 | 7,92E-01 |
| AT1G03870 | 0,58 | -1,01 | 3,13 | -0,32 | 7,48E-01 |
| AT1G03940 | 1,20 | -0,49 | 1,64 | -0,30 | 7,64E-01 |
| AT1G04233 | 0,08 | 0,85 | 4,38 | 0,20 | 8,45E-01 |
| AT1G03980 | 0,11 | 0,00 | 4,41 | 0,00 | 1,00E+00 |
| AT1G04000 | 0,55 | -0,24 | 3,17 | -0,08 | 9,39E-01 |
| AT1G04010 | 0,05 | 0,00 | 4,41 | 0,00 | 1,00E+00 |
| AT1G04050 | 0,08 | 0,00 | 4,40 | 0,00 | 1,00E+00 |
| AT1G04070 | 0,16 | 1,07 | 4,37 | 0,24 | 8,07E-01 |
| AT1G04110 | 0,35 | 0,00 | 4,39 | 0,00 | 1,00E+00 |
| AT1G04120 | 1,63 | 4,11 | 1,70 | 2,42 | 1,53E-02 |
| AT1G04140 | 1,54 | 3,46 | 1,73 | 2,00 | 4,58E-02 |
| AT1G04190 | 0,13 | -1,24 | 4,39 | -0,28 | 7,77E-01 |
| AT1G04220 | 0,35 | 0,45 | 4,39 | 0,10 | 9,19E-01 |
| AT1G04240 | 0,91 | -3,84 | 2,26 | -1,70 | 8,91E-02 |
| AT1G04260 | 0,47 | 1,15 | 2,68 | 0,43 | 6,69E-01 |
| AT1G04270 | 1,04 | -1,01 | 2,09 | -0,48 | 6,28E-01 |
| AT1G04280 | 0,49 | -1,13 | 3,26 | -0,35 | 7,30E-01 |
| AT1G04290 | 0,43 | 1,98 | 2,75 | 0,72 | 4,72E-01 |
| AT1G04300 | 0,83 | 2,14 | 3,02 | 0,71 | 4,79E-01 |
| AT1G04380 | 0,20 | 0,89 | 4,37 | 0,20 | 8,39E-01 |
| AT1G04390 | 0,19 | 1,57 | 4,34 | 0,36 | 7,18E-01 |
| AT1G04400 | 0,89 | -2,31 | 2,35 | -0,98 | 3,27E-01 |
| AT1G04263 | 1,35 | -0,16 | 2,15 | -0,08 | 9,40E-01 |
| AT1G04530 | 0,42 | 1,79 | 3,28 | 0,54 | 5,86E-01 |
| AT1G04570 | 0,11 | 0,90 | 4,37 | 0,21 | 8,36E-01 |
| AT1G04630 | 0,66 | 2,34 | 3,71 | 0,63 | 5,28E-01 |
| AT1G04680 | 0,74 | -1,57 | 3,00 | -0,52 | 6,00E-01 |
| AT1G04730 | 0,74 | 1,19 | 2,45 | 0,48 | 6,28E-01 |
| AT1G04750 | 0,67 | -1,86 | 2,26 | -0,82 | 4,10E-01 |
| AT1G04770 | 0,89 | 0,85 | 2,26 | 0,37 | 7,08E-01 |
| AT1G04780 | 0,86 | 0,28 | 2,03 | 0,14 | 8,89E-01 |
| AT1G04840 | 0,63 | 2,76 | 3,69 | 0,75 | 4,55E-01 |
| AT1G04850 | 0,86 | 2,20 | 2,75 | 0,80 | 4,24E-01 |
| AT1G04860 | 0,57 | 1,83 | 3,95 | 0,46 | 6,43E-01 |
| AT1G04870 | 0,84 | -1,06 | 2,98 | -0,36 | 7,22E-01 |
| AT1G04880 | 0,21 | -1,56 | 4,37 | -0,36 | 7,22E-01 |
| AT1G04910 | 0,78 | 2,84 | 2,78 | 1,02 | 3,08E-01 |
| AT1G04950 | 0,50 | 1,79 | 3,24 | 0,55 | 5,79E-01 |
| AT1G04960 | 0,36 | -1,36 | 4,41 | -0,31 | 7,58E-01 |
| AT1G04980 | 0,36 | 0,00 | 4,41 | 0,00 | 1,00E+00 |
| AT1G04985 | 0,26 | 0,75 | 4,36 | 0,17 | 8,63E-01 |
| AT1G04990 | 0,84 | 0,93 | 2,36 | 0,39 | 6,93E-01 |
| AT1G05120 | 0,53 | 1,77 | 4,33 | 0,41 | 6,84E-01 |
| AT1G05150 | 0,85 | 0,89 | 2,13 | 0,42 | 6,76E-01 |

| | | | | | |
|---|---|---|---|---|---|
| AT1G05160 | 0,59 | 2,35 | 4,31 | 0,55 | 5,86E-01 |
| AT1G05170 | 0,47 | -1,85 | 4,37 | -0,42 | 6,72E-01 |
| AT1G05190 | 0,78 | -0,20 | 2,47 | -0,08 | 9,36E-01 |
| AT1G05205 | 0,08 | 1,10 | 4,36 | 0,25 | 8,00E-01 |
| AT1G05230 | 0,45 | 1,23 | 4,07 | 0,30 | 7,63E-01 |
| AT1G05300 | 0,18 | 0,98 | 4,36 | 0,23 | 8,22E-01 |
| AT1G05350 | 0,28 | 1,56 | 4,34 | 0,36 | 7,18E-01 |
| AT1G05520 | 0,61 | 1,91 | 3,05 | 0,63 | 5,32E-01 |
| AT1G05575 | 0,16 | 0,51 | 4,38 | 0,12 | 9,07E-01 |
| AT1G05640 | 0,20 | 0,00 | 4,41 | 0,00 | 1,00E+00 |
| AT1G05670 | 0,15 | 1,42 | 4,34 | 0,33 | 7,43E-01 |
| AT1G05720 | 0,13 | -1,24 | 4,39 | -0,28 | 7,77E-01 |
| AT1G05783 | 0,96 | 1,24 | 1,82 | 0,68 | 4,96E-01 |
| AT1G05840 | 0,75 | 1,60 | 2,22 | 0,72 | 4,71E-01 |
| AT1G05870 | 0,42 | 0,43 | 4,21 | 0,10 | 9,18E-01 |
| AT1G05950 | 0,12 | 1,65 | 4,34 | 0,38 | 7,04E-01 |
| AT1G05970 | 0,08 | 0,85 | 4,38 | 0,20 | 8,45E-01 |
| AT1G06000 | 0,95 | -0,60 | 2,01 | -0,30 | 7,65E-01 |
| AT1G06130 | 0,62 | 0,37 | 3,04 | 0,12 | 9,03E-01 |
| AT1G06150 | 1,38 | 3,37 | 1,90 | 1,78 | 7,59E-02 |
| AT1G06250 | 0,72 | 0,00 | 4,41 | 0,00 | 1,00E+00 |
| AT1G06265 | 0,08 | 1,10 | 4,36 | 0,25 | 8,00E-01 |
| AT1G06360 | 0,33 | -2,50 | 4,35 | -0,58 | 5,65E-01 |
| AT1G06400 | 0,21 | 1,23 | 4,34 | 0,28 | 7,77E-01 |
| AT1G06440 | 0,58 | 1,95 | 2,68 | 0,73 | 4,67E-01 |
| AT1G06540 | 1,04 | 1,26 | 2,08 | 0,60 | 5,46E-01 |
| AT1G06570 | 0,97 | 0,45 | 2,41 | 0,19 | 8,51E-01 |
| AT1G06580 | 1,59 | -1,44 | 1,36 | -1,06 | 2,89E-01 |
| AT1G04487 | 0,39 | 1,96 | 4,18 | 0,47 | 6,40E-01 |
| AT1G06660 | 0,50 | 1,24 | 3,99 | 0,31 | 7,56E-01 |
| AT1G06690 | 0,36 | 0,38 | 4,35 | 0,09 | 9,30E-01 |
| AT1G06700 | 1,69 | 1,04 | 1,63 | 0,64 | 5,23E-01 |
| AT1G06710 | 0,26 | 0,90 | 4,38 | 0,21 | 8,37E-01 |
| AT1G06720 | 0,55 | -0,80 | 4,00 | -0,20 | 8,42E-01 |
| AT1G06760 | 1,45 | -1,04 | 1,62 | -0,64 | 5,20E-01 |
| AT1G06810 | 0,08 | 0,85 | 4,38 | 0,20 | 8,45E-01 |
| AT1G06840 | 0,17 | -0,28 | 4,38 | -0,06 | 9,49E-01 |
| AT1G06890 | 0,40 | -0,48 | 2,89 | -0,17 | 8,68E-01 |
| AT1G06900 | 1,35 | 1,14 | 1,57 | 0,72 | 4,70E-01 |
| AT1G07020 | 0,55 | -1,36 | 4,41 | -0,31 | 7,58E-01 |
| AT1G07050 | 0,42 | -0,60 | 4,04 | -0,15 | 8,82E-01 |
| AT1G07090 | 0,43 | 1,43 | 4,30 | 0,33 | 7,39E-01 |
| AT1G07110 | 1,24 | 2,64 | 2,35 | 1,13 | 2,60E-01 |
| AT1G07128 | 0,28 | 1,69 | 4,35 | 0,39 | 6,97E-01 |
| AT1G07220 | 0,25 | -2,61 | 4,34 | -0,60 | 5,47E-01 |
| AT1G07260 | 0,48 | 1,11 | 4,35 | 0,26 | 7,99E-01 |
| AT1G04573 | 0,66 | 0,66 | 4,37 | 0,15 | 8,81E-01 |
| AT1G07310 | 0,69 | -0,12 | 2,52 | -0,05 | 9,62E-01 |
| AT1G07350 | 0,45 | 1,60 | 3,27 | 0,49 | 6,25E-01 |
| AT1G07360 | 0,57 | -1,92 | 3,16 | -0,61 | 5,42E-01 |
| AT1G07430 | 0,32 | 0,84 | 4,33 | 0,19 | 8,46E-01 |
| AT1G07440 | 0,75 | 2,55 | 2,53 | 1,01 | 3,14E-01 |
| AT1G07480 | 0,80 | 0,87 | 2,29 | 0,38 | 7,06E-01 |
| AT1G07500 | 0,08 | 0,00 | 4,40 | 0,00 | 1,00E+00 |
| AT1G07520 | 1,20 | 2,02 | 1,98 | 1,02 | 3,06E-01 |
| AT1G07530 | 1,00 | 1,77 | 1,79 | 0,99 | 3,24E-01 |

| | | | | | |
|---|---|---|---|---|---|
| AT1G07640 | 0,24 | -0,46 | 4,36 | -0,11 | 9,16E-01 |
| AT1G07720 | 1,26 | -0,54 | 1,56 | -0,34 | 7,30E-01 |
| AT1G07725 | 1,03 | 2,32 | 2,52 | 0,92 | 3,58E-01 |
| AT1G07740 | 0,29 | 1,91 | 4,33 | 0,44 | 6,58E-01 |
| AT1G07730 | 0,29 | 1,91 | 4,33 | 0,44 | 6,58E-01 |
| AT1G07750 | 0,25 | -2,61 | 4,34 | -0,60 | 5,47E-01 |
| AT1G07770 | 0,87 | 0,43 | 1,77 | 0,24 | 8,09E-01 |
| AT1G07870 | 0,12 | 1,39 | 4,34 | 0,32 | 7,49E-01 |
| AT1G07970 | 0,62 | 2,36 | 3,62 | 0,65 | 5,14E-01 |
| AT1G08040 | 0,40 | 0,73 | 4,30 | 0,17 | 8,66E-01 |
| AT1G08050 | 0,33 | 2,20 | 4,32 | 0,51 | 6,11E-01 |
| AT1G08060 | 1,18 | 3,25 | 2,15 | 1,51 | 1,31E-01 |
| AT1G08125 | 0,32 | 0,44 | 4,38 | 0,10 | 9,20E-01 |
| AT1G08510 | 1,46 | -0,35 | 1,45 | -0,24 | 8,12E-01 |
| AT1G08580 | 0,24 | 0,00 | 4,41 | 0,00 | 1,00E+00 |
| AT1G08592 | 0,74 | 3,22 | 2,84 | 1,14 | 2,56E-01 |
| AT1G08610 | 0,28 | 0,87 | 4,38 | 0,20 | 8,43E-01 |
| AT1G08640 | 0,47 | 1,05 | 4,00 | 0,26 | 7,93E-01 |
| AT1G08645 | 0,64 | -0,86 | 3,73 | -0,23 | 8,17E-01 |
| AT1G08660 | 0,10 | 0,00 | 4,41 | 0,00 | 1,00E+00 |
| AT1G08700 | 1,22 | 3,41 | 2,03 | 1,67 | 9,40E-02 |
| AT1G08800 | 0,74 | 2,04 | 2,84 | 0,72 | 4,73E-01 |
| AT1G08820 | 0,89 | 0,00 | 2,79 | 0,00 | 1,00E+00 |
| AT1G08880 | 0,12 | 1,58 | 4,34 | 0,36 | 7,16E-01 |
| AT1G09020 | 0,78 | 1,42 | 2,25 | 0,63 | 5,28E-01 |
| AT1G09060 | 1,04 | 2,17 | 2,30 | 0,94 | 3,46E-01 |
| AT1G09080 | 0,40 | 1,31 | 4,31 | 0,30 | 7,61E-01 |
| AT1G09130 | 0,46 | -1,44 | 3,25 | -0,44 | 6,57E-01 |
| AT1G09160 | 0,40 | -0,96 | 3,43 | -0,28 | 7,78E-01 |
| AT1G09190 | 0,17 | 1,94 | 4,33 | 0,45 | 6,55E-01 |
| AT1G09195 | 0,70 | 2,05 | 2,92 | 0,70 | 4,82E-01 |
| AT1G09200 | 0,34 | 0,65 | 4,38 | 0,15 | 8,82E-01 |
| AT1G09230 | 0,23 | 0,00 | 4,41 | 0,00 | 1,00E+00 |
| AT1G09280 | 1,11 | -0,76 | 1,69 | -0,45 | 6,51E-01 |
| AT1G09420 | 0,88 | 2,83 | 3,32 | 0,85 | 3,94E-01 |
| AT1G09440 | 0,21 | -1,18 | 4,33 | -0,27 | 7,85E-01 |
| AT1G04707 | 0,36 | 1,56 | 4,34 | 0,36 | 7,19E-01 |
| AT1G09630 | 0,08 | 0,85 | 4,38 | 0,20 | 8,45E-01 |
| AT1G04717 | 0,54 | 2,38 | 3,79 | 0,63 | 5,31E-01 |
| AT1G09680 | 1,19 | 3,91 | 2,48 | 1,58 | 1,14E-01 |
| AT1G09730 | 0,83 | 2,57 | 2,66 | 0,97 | 3,34E-01 |
| AT1G09760 | 0,17 | 1,57 | 4,34 | 0,36 | 7,18E-01 |
| AT1G09810 | 0,42 | 0,71 | 2,79 | 0,26 | 7,98E-01 |
| AT1G09880 | 0,33 | 0,89 | 4,36 | 0,20 | 8,39E-01 |
| AT1G09910 | 0,78 | 2,15 | 2,36 | 0,91 | 3,64E-01 |
| AT1G10020 | 0,10 | 0,42 | 4,38 | 0,10 | 9,23E-01 |
| AT1G10090 | 0,57 | -0,08 | 2,57 | -0,03 | 9,75E-01 |
| AT1G10155 | 1,47 | 1,71 | 1,53 | 1,12 | 2,63E-01 |
| AT1G10180 | 0,61 | -0,47 | 2,64 | -0,18 | 8,57E-01 |
| AT1G10190 | 0,94 | -2,72 | 2,78 | -0,98 | 3,28E-01 |
| AT1G10200 | 0,57 | 2,27 | 3,83 | 0,59 | 5,54E-01 |
| AT1G10310 | 0,42 | 1,23 | 4,36 | 0,28 | 7,78E-01 |
| AT1G10320 | 0,47 | 2,82 | 3,94 | 0,72 | 4,74E-01 |
| AT1G10340 | 1,00 | 1,11 | 2,30 | 0,48 | 6,28E-01 |
| AT1G10350 | 0,52 | 2,58 | 3,86 | 0,67 | 5,04E-01 |
| AT1G10390 | 1,30 | 3,78 | 1,91 | 1,98 | 4,78E-02 |

| | | | | | |
|---|---|---|---|---|---|
| AT1G10400 | 0,08 | 1,10 | 4,36 | 0,25 | 8,00E-01 |
| AT1G10410 | 0,96 | 1,96 | 1,98 | 0,99 | 3,24E-01 |
| AT1G10430 | 0,87 | 0,96 | 2,14 | 0,45 | 6,55E-01 |
| AT1G10470 | 1,21 | -0,85 | 1,89 | -0,45 | 6,51E-01 |
| AT1G10500 | 1,39 | 1,74 | 2,26 | 0,77 | 4,41E-01 |
| AT1G10580 | 0,57 | 0,78 | 3,78 | 0,21 | 8,37E-01 |
| AT1G10586 | 1,03 | 0,36 | 2,10 | 0,17 | 8,63E-01 |
| AT1G10590 | 0,69 | 1,43 | 3,03 | 0,47 | 6,37E-01 |
| AT1G10630 | 0,65 | 0,79 | 2,63 | 0,30 | 7,63E-01 |
| AT1G10640 | 0,05 | 0,00 | 4,41 | 0,00 | 1,00E+00 |
| AT1G10680 | 0,08 | 0,85 | 4,38 | 0,20 | 8,45E-01 |
| AT1G10840 | 0,88 | 0,87 | 1,97 | 0,44 | 6,58E-01 |
| AT1G10870 | 1,11 | 0,41 | 2,03 | 0,20 | 8,40E-01 |
| AT1G10900 | 1,03 | -0,28 | 2,89 | -0,10 | 9,22E-01 |
| AT1G10920 | 0,82 | 2,96 | 4,31 | 0,69 | 4,92E-01 |
| AT1G10930 | 0,74 | 2,29 | 2,92 | 0,79 | 4,31E-01 |
| AT1G10940 | 0,48 | 1,17 | 3,33 | 0,35 | 7,26E-01 |
| AT1G04833 | 0,22 | 0,42 | 4,38 | 0,10 | 9,24E-01 |
| AT1G11100 | 0,45 | 2,40 | 3,36 | 0,71 | 4,76E-01 |
| AT1G11210 | 0,56 | 0,63 | 2,56 | 0,25 | 8,06E-01 |
| AT1G11240 | 0,05 | 0,00 | 4,41 | 0,00 | 1,00E+00 |
| AT1G11270 | 0,83 | 1,20 | 2,36 | 0,51 | 6,11E-01 |
| AT1G11280 | 0,45 | -0,50 | 2,83 | -0,18 | 8,61E-01 |
| AT1G11290 | 0,63 | 2,75 | 3,63 | 0,76 | 4,49E-01 |
| AT1G11310 | 1,46 | 2,46 | 1,68 | 1,46 | 1,43E-01 |
| AT1G11340 | 0,08 | 1,10 | 4,36 | 0,25 | 8,00E-01 |
| AT1G11350 | 0,52 | 2,32 | 3,22 | 0,72 | 4,71E-01 |
| AT1G11360 | 0,33 | -0,38 | 4,36 | -0,09 | 9,30E-01 |
| AT1G11480 | 0,70 | 0,98 | 2,23 | 0,44 | 6,61E-01 |
| AT1G11630 | 0,35 | 0,00 | 4,39 | 0,00 | 1,00E+00 |
| AT1G11820 | 0,25 | 0,64 | 4,33 | 0,15 | 8,82E-01 |
| AT1G12000 | 0,52 | 2,69 | 3,06 | 0,88 | 3,79E-01 |
| AT1G12020 | 0,63 | 0,48 | 2,28 | 0,21 | 8,32E-01 |
| AT1G12120 | 0,70 | -0,56 | 2,66 | -0,21 | 8,34E-01 |
| AT1G12211 | 0,63 | 0,74 | 3,11 | 0,24 | 8,13E-01 |
| AT1G12310 | 0,63 | 2,42 | 3,16 | 0,77 | 4,44E-01 |
| AT1G12380 | 0,49 | 2,00 | 4,07 | 0,49 | 6,24E-01 |
| AT1G12440 | 1,21 | 0,93 | 1,84 | 0,51 | 6,12E-01 |
| AT1G12500 | 0,25 | 2,10 | 4,32 | 0,49 | 6,27E-01 |
| AT1G12520 | 0,51 | -0,27 | 4,34 | -0,06 | 9,50E-01 |
| AT1G12590 | 0,78 | -1,27 | 1,79 | -0,71 | 4,79E-01 |
| AT1G12610 | 0,19 | 1,26 | 4,35 | 0,29 | 7,72E-01 |
| AT1G12615 | 1,59 | 2,39 | 2,13 | 1,12 | 2,62E-01 |
| AT1G12640 | 0,58 | 2,61 | 4,32 | 0,61 | 5,45E-01 |
| AT1G12680 | 1,43 | -0,77 | 1,75 | -0,44 | 6,61E-01 |
| AT1G12700 | 0,22 | 1,61 | 4,33 | 0,37 | 7,11E-01 |
| AT1G12780 | 0,71 | 1,83 | 2,28 | 0,80 | 4,22E-01 |
| AT1G12800 | 0,97 | -1,45 | 2,51 | -0,58 | 5,64E-01 |
| AT1G12910 | 1,07 | 2,28 | 1,99 | 1,15 | 2,52E-01 |
| AT1G13020 | 0,59 | 0,63 | 2,60 | 0,24 | 8,09E-01 |
| AT1G13120 | 0,23 | 2,01 | 4,33 | 0,46 | 6,43E-01 |
| AT1G13170 | 0,46 | 1,61 | 3,23 | 0,50 | 6,19E-01 |
| AT1G13200 | 0,66 | 1,37 | 3,64 | 0,38 | 7,06E-01 |
| AT1G13245 | 0,44 | -2,47 | 4,09 | -0,60 | 5,45E-01 |
| AT1G13310 | 0,35 | 0,00 | 4,39 | 0,00 | 1,00E+00 |
| AT1G13340 | 0,95 | 2,62 | 2,23 | 1,18 | 2,40E-01 |

| | | | | | |
|---|---|---|---|---|---|
| AT1G13390 | 0,17 | 1,92 | 4,33 | 0,44 | 6,57E-01 |
| AT1G13450 | 0,76 | 2,41 | 2,23 | 1,08 | 2,80E-01 |
| AT1G13520 | 0,54 | 1,00 | 4,37 | 0,23 | 8,20E-01 |
| AT1G13560 | 0,43 | 0,95 | 4,36 | 0,22 | 8,28E-01 |
| AT1G13825 | 0,76 | 3,24 | 3,53 | 0,92 | 3,59E-01 |
| AT1G13860 | 0,10 | 0,62 | 4,38 | 0,14 | 8,87E-01 |
| AT1G13870 | 0,12 | 0,00 | 4,39 | 0,00 | 1,00E+00 |
| AT1G13900 | 0,21 | 1,77 | 4,34 | 0,41 | 6,84E-01 |
| AT1G13940 | 0,86 | 3,16 | 2,40 | 1,32 | 1,88E-01 |
| AT1G14010 | 0,36 | 1,26 | 4,35 | 0,29 | 7,72E-01 |
| AT1G14030 | 0,62 | 1,01 | 3,02 | 0,33 | 7,38E-01 |
| AT1G14290 | 0,26 | 0,90 | 4,38 | 0,21 | 8,37E-01 |
| AT1G14300 | 0,22 | -1,55 | 4,37 | -0,36 | 7,22E-01 |
| AT1G14330 | 0,59 | 1,29 | 2,60 | 0,50 | 6,19E-01 |
| AT1G14360 | 0,18 | 0,00 | 4,41 | 0,00 | 1,00E+00 |
| AT1G14400 | 0,66 | 1,35 | 3,03 | 0,45 | 6,56E-01 |
| AT1G14450 | 0,11 | 0,00 | 4,41 | 0,00 | 1,00E+00 |
| AT1G14455 | 0,14 | -1,60 | 4,37 | -0,37 | 7,14E-01 |
| AT1G14518 | 0,84 | 1,53 | 2,39 | 0,64 | 5,21E-01 |
| AT1G14590 | 0,42 | 1,63 | 4,17 | 0,39 | 6,97E-01 |
| AT1G14690 | 0,78 | -0,54 | 2,25 | -0,24 | 8,09E-01 |
| AT1G14710 | 0,51 | 1,80 | 3,81 | 0,47 | 6,37E-01 |
| AT1G14720 | 0,26 | -1,63 | 4,37 | -0,37 | 7,09E-01 |
| AT1G14790 | 1,42 | 4,30 | 2,16 | 1,99 | 4,69E-02 |
| AT1G14810 | 1,02 | 2,53 | 2,17 | 1,17 | 2,43E-01 |
| AT1G14830 | 0,90 | 3,22 | 2,77 | 1,16 | 2,46E-01 |
| AT1G14840 | 0,51 | -0,51 | 2,28 | -0,22 | 8,23E-01 |
| AT1G14850 | 1,26 | 2,65 | 1,88 | 1,41 | 1,59E-01 |
| AT1G14880 | 1,19 | 2,10 | 1,96 | 1,07 | 2,84E-01 |
| AT1G14900 | 0,38 | -1,43 | 4,33 | -0,33 | 7,40E-01 |
| AT1G14910 | 0,80 | 1,48 | 2,44 | 0,61 | 5,43E-01 |
| AT1G14980 | 0,57 | 1,26 | 3,03 | 0,42 | 6,78E-01 |
| AT1G05137 | 0,28 | 0,92 | 4,33 | 0,21 | 8,32E-01 |
| AT1G15020 | 0,98 | 3,02 | 2,56 | 1,18 | 2,39E-01 |
| AT1G15040 | 0,21 | 1,84 | 4,33 | 0,42 | 6,71E-01 |
| AT1G15100 | 1,26 | 0,41 | 2,61 | 0,16 | 8,74E-01 |
| AT1G15140 | 0,76 | 1,59 | 2,09 | 0,76 | 4,47E-01 |
| AT1G15175 | 0,27 | 1,05 | 4,36 | 0,24 | 8,09E-01 |
| AT1G15210 | 0,17 | 1,94 | 4,33 | 0,45 | 6,55E-01 |
| AT1G15220 | 0,11 | 0,90 | 4,37 | 0,21 | 8,36E-01 |
| AT1G15340 | 0,21 | -0,43 | 4,38 | -0,10 | 9,22E-01 |
| AT1G15350 | 1,25 | 1,24 | 2,19 | 0,57 | 5,71E-01 |
| AT1G15400 | 0,43 | 1,77 | 2,79 | 0,63 | 5,27E-01 |
| AT1G15440 | 0,87 | 0,01 | 2,15 | 0,00 | 9,97E-01 |
| AT1G15520 | 0,46 | 1,63 | 3,26 | 0,50 | 6,18E-01 |
| AT1G15530 | 1,02 | 1,27 | 2,27 | 0,56 | 5,74E-01 |
| AT1G15550 | 0,24 | 0,00 | 4,41 | 0,00 | 1,00E+00 |
| AT1G15700 | 0,40 | 1,60 | 3,35 | 0,48 | 6,32E-01 |
| AT1G05187 | 0,24 | 1,62 | 4,34 | 0,37 | 7,08E-01 |
| AT1G15730 | 0,44 | 0,00 | 4,18 | 0,00 | 1,00E+00 |
| AT1G15780 | 1,01 | 0,01 | 1,90 | 0,01 | 9,94E-01 |
| AT1G15790 | 1,05 | 1,51 | 2,87 | 0,53 | 5,99E-01 |
| AT1G05203 | 0,26 | 1,08 | 4,37 | 0,25 | 8,05E-01 |
| AT1G15880 | 0,20 | 0,00 | 4,41 | 0,00 | 1,00E+00 |
| AT1G15970 | 0,11 | 0,00 | 4,41 | 0,00 | 1,00E+00 |
| AT1G16010 | 0,99 | 2,76 | 2,25 | 1,23 | 2,20E-01 |

| | | | | | |
|---|---|---|---|---|---|
| AT1G16070 | 0,40 | 2,20 | 4,33 | 0,51 | 6,11E-01 |
| AT1G16210 | 0,60 | -0,32 | 3,10 | -0,10 | 9,18E-01 |
| AT1G16240 | 0,80 | 0,90 | 2,31 | 0,39 | 6,95E-01 |
| AT1G16260 | 1,63 | 2,19 | 1,81 | 1,21 | 2,27E-01 |
| AT1G16460 | 0,57 | 0,66 | 2,35 | 0,28 | 7,80E-01 |
| AT1G16570 | 0,28 | 1,69 | 4,35 | 0,39 | 6,97E-01 |
| AT1G16610 | 1,02 | 3,03 | 3,09 | 0,98 | 3,28E-01 |
| AT1G16750 | 0,49 | 2,10 | 3,12 | 0,67 | 5,00E-01 |
| AT1G16780 | 0,12 | 1,65 | 4,34 | 0,38 | 7,04E-01 |
| AT1G16800 | 1,44 | 3,62 | 2,36 | 1,53 | 1,25E-01 |
| AT1G16870 | 0,95 | 1,81 | 3,35 | 0,54 | 5,89E-01 |
| AT1G16890 | 1,30 | -0,80 | 1,73 | -0,46 | 6,46E-01 |
| AT1G16900 | 0,65 | 2,82 | 3,67 | 0,77 | 4,42E-01 |
| AT1G16920 | 0,22 | 1,41 | 4,34 | 0,32 | 7,45E-01 |
| AT1G16970 | 0,37 | 0,68 | 3,46 | 0,20 | 8,44E-01 |
| AT1G17080 | 0,60 | 0,71 | 3,24 | 0,22 | 8,26E-01 |
| AT1G17110 | 0,57 | 0,11 | 2,54 | 0,04 | 9,65E-01 |
| AT1G17140 | 0,95 | 0,40 | 3,41 | 0,12 | 9,06E-01 |
| AT1G17147 | 0,12 | 0,00 | 4,41 | 0,00 | 1,00E+00 |
| AT1G17232 | 0,51 | 1,91 | 3,10 | 0,62 | 5,38E-01 |
| AT1G17240 | 0,05 | 0,00 | 4,41 | 0,00 | 1,00E+00 |
| AT1G17250 | 0,99 | 3,69 | 2,53 | 1,46 | 1,45E-01 |
| AT1G17370 | 0,81 | -0,04 | 2,57 | -0,02 | 9,87E-01 |
| AT1G17380 | 0,50 | 1,27 | 2,68 | 0,47 | 6,35E-01 |
| AT1G05313 | 0,32 | 1,32 | 4,35 | 0,30 | 7,61E-01 |
| AT1G17440 | 0,46 | 1,73 | 3,32 | 0,52 | 6,01E-01 |
| AT1G17450 | 1,22 | 2,65 | 1,81 | 1,46 | 1,43E-01 |
| AT1G17590 | 0,46 | 0,59 | 3,26 | 0,18 | 8,56E-01 |
| AT1G17600 | 1,00 | 1,42 | 2,14 | 0,66 | 5,08E-01 |
| AT1G17665 | 0,20 | 0,60 | 4,34 | 0,14 | 8,90E-01 |
| AT1G17670 | 0,37 | -0,16 | 4,34 | -0,04 | 9,71E-01 |
| AT1G17690 | 0,25 | 1,57 | 4,34 | 0,36 | 7,18E-01 |
| AT1G17720 | 0,66 | 1,49 | 2,27 | 0,65 | 5,13E-01 |
| AT1G17744 | 0,16 | 0,90 | 4,38 | 0,21 | 8,37E-01 |
| AT1G17790 | 0,63 | -0,63 | 3,15 | -0,20 | 8,42E-01 |
| AT1G05333 | 0,71 | 1,61 | 3,54 | 0,46 | 6,49E-01 |
| AT1G17820 | 0,71 | 1,61 | 3,54 | 0,46 | 6,49E-01 |
| AT1G17850 | 0,36 | 0,95 | 4,36 | 0,22 | 8,28E-01 |
| AT1G05347 | 0,21 | 0,65 | 4,33 | 0,15 | 8,81E-01 |
| AT1G17870 | 0,39 | 1,77 | 4,26 | 0,42 | 6,78E-01 |
| AT1G17880 | 0,59 | -0,38 | 3,88 | -0,10 | 9,22E-01 |
| AT1G17890 | 0,42 | 0,52 | 4,40 | 0,12 | 9,05E-01 |
| AT1G17930 | 0,20 | 1,07 | 4,36 | 0,25 | 8,06E-01 |
| AT1G17980 | 1,06 | -1,37 | 2,16 | -0,63 | 5,27E-01 |
| AT1G17990 | 0,53 | 0,65 | 3,19 | 0,21 | 8,38E-01 |
| AT1G18050 | 0,60 | 0,84 | 2,44 | 0,34 | 7,30E-01 |
| AT1G18060 | 0,60 | 0,62 | 2,33 | 0,27 | 7,90E-01 |
| AT1G18070 | 0,60 | 0,21 | 2,48 | 0,09 | 9,31E-01 |
| AT1G18100 | 0,36 | 2,25 | 4,33 | 0,52 | 6,04E-01 |
| AT1G18150 | 1,19 | -1,21 | 2,12 | -0,57 | 5,68E-01 |
| AT1G18210 | 0,61 | -1,09 | 2,34 | -0,47 | 6,40E-01 |
| AT1G18330 | 0,22 | 0,85 | 4,36 | 0,20 | 8,45E-01 |
| AT1G18360 | 1,25 | -1,11 | 1,70 | -0,65 | 5,14E-01 |
| AT1G18382 | 0,44 | 1,66 | 2,76 | 0,60 | 5,48E-01 |
| AT1G18470 | 0,48 | -0,12 | 2,63 | -0,05 | 9,64E-01 |
| AT1G18540 | 1,15 | 0,80 | 2,08 | 0,38 | 7,02E-01 |

| | | | | | |
|---|---|---|---|---|---|
| AT1G18720 | 0,23 | 1,68 | 4,33 | 0,39 | 6,97E-01 |
| AT1G05433 | 0,53 | 1,22 | 3,10 | 0,39 | 6,94E-01 |
| AT1G18840 | 0,43 | 1,63 | 3,35 | 0,49 | 6,26E-01 |
| AT1G18890 | 0,28 | 0,90 | 4,38 | 0,21 | 8,37E-01 |
| AT1G18990 | 0,45 | 1,81 | 4,17 | 0,43 | 6,64E-01 |
| AT1G19000 | 0,86 | -0,35 | 1,78 | -0,20 | 8,43E-01 |
| AT1G19130 | 0,49 | -0,48 | 2,71 | -0,18 | 8,59E-01 |
| AT1G19140 | 0,44 | 0,79 | 4,36 | 0,18 | 8,56E-01 |
| AT1G19210 | 0,10 | 0,00 | 4,41 | 0,00 | 1,00E+00 |
| AT1G19260 | 0,97 | 2,59 | 2,58 | 1,00 | 3,16E-01 |
| AT1G19300 | 0,88 | 2,37 | 2,44 | 0,97 | 3,32E-01 |
| AT1G19360 | 0,50 | 1,68 | 3,97 | 0,42 | 6,73E-01 |
| AT1G19370 | 0,41 | 0,64 | 4,26 | 0,15 | 8,80E-01 |
| AT1G19400 | 1,69 | 2,09 | 1,44 | 1,45 | 1,47E-01 |
| AT1G19430 | 0,39 | -0,11 | 4,25 | -0,03 | 9,80E-01 |
| AT1G19450 | 0,33 | -1,21 | 4,38 | -0,28 | 7,82E-01 |
| AT1G19570 | 0,89 | -1,18 | 2,00 | -0,59 | 5,55E-01 |
| AT1G19640 | 0,14 | -1,60 | 4,37 | -0,37 | 7,14E-01 |
| AT1G19660 | 0,55 | -0,06 | 2,70 | -0,02 | 9,82E-01 |
| AT1G19670 | 1,59 | -0,77 | 1,91 | -0,40 | 6,87E-01 |
| AT1G19700 | 0,25 | -0,85 | 4,33 | -0,20 | 8,44E-01 |
| AT1G19715 | 1,55 | 3,87 | 2,09 | 1,85 | 6,43E-02 |
| AT1G19720 | 1,66 | 2,40 | 1,71 | 1,41 | 1,60E-01 |
| AT1G19780 | 0,50 | 0,60 | 4,07 | 0,15 | 8,83E-01 |
| AT1G19860 | 0,15 | 0,86 | 4,37 | 0,20 | 8,43E-01 |
| AT1G19870 | 0,80 | 1,47 | 2,43 | 0,61 | 5,44E-01 |
| AT1G19880 | 0,41 | 0,81 | 3,45 | 0,23 | 8,15E-01 |
| AT1G19920 | 1,00 | 2,25 | 2,29 | 0,98 | 3,27E-01 |
| AT1G20040 | 0,34 | -0,44 | 4,35 | -0,10 | 9,19E-01 |
| AT1G20070 | 0,59 | 1,60 | 2,43 | 0,66 | 5,10E-01 |
| AT1G20160 | 0,33 | 2,12 | 4,32 | 0,49 | 6,24E-01 |
| AT1G20190 | 0,64 | -2,09 | 3,15 | -0,66 | 5,07E-01 |
| AT1G20200 | 0,33 | 0,90 | 3,52 | 0,26 | 7,98E-01 |
| AT1G20320 | 0,05 | 0,00 | 4,41 | 0,00 | 1,00E+00 |
| AT1G20400 | 1,62 | 0,80 | 1,77 | 0,45 | 6,51E-01 |
| AT1G20430 | 0,89 | 1,17 | 2,76 | 0,43 | 6,70E-01 |
| AT1G20490 | 0,48 | 0,59 | 4,32 | 0,14 | 8,92E-01 |
| AT1G20560 | 0,25 | 1,76 | 4,34 | 0,41 | 6,85E-01 |
| AT1G20570 | 0,08 | 1,10 | 4,36 | 0,25 | 8,00E-01 |
| AT1G20650 | 0,41 | 0,00 | 4,40 | 0,00 | 1,00E+00 |
| AT1G20670 | 0,31 | 0,87 | 4,36 | 0,20 | 8,42E-01 |
| AT1G20770 | 0,16 | 0,90 | 4,38 | 0,21 | 8,37E-01 |
| AT1G20840 | 1,09 | -0,55 | 1,86 | -0,30 | 7,68E-01 |
| AT1G20880 | 0,25 | -0,68 | 4,32 | -0,16 | 8,75E-01 |
| AT1G20890 | 0,48 | 2,44 | 3,96 | 0,62 | 5,38E-01 |
| AT1G20950 | 0,69 | 1,54 | 2,47 | 0,63 | 5,32E-01 |
| AT1G21080 | 0,61 | 0,55 | 2,68 | 0,21 | 8,36E-01 |
| AT1G05627 | 0,05 | 0,00 | 4,41 | 0,00 | 1,00E+00 |
| AT1G21160 | 0,26 | 1,59 | 4,34 | 0,37 | 7,14E-01 |
| AT1G21190 | 0,45 | -0,19 | 4,14 | -0,05 | 9,63E-01 |
| AT1G21310 | 0,44 | 0,78 | 2,75 | 0,28 | 7,76E-01 |
| AT1G21380 | 1,01 | 2,49 | 2,29 | 1,09 | 2,77E-01 |
| AT1G21390 | 0,08 | 1,10 | 4,36 | 0,25 | 8,00E-01 |
| AT1G21440 | 1,10 | -1,92 | 2,75 | -0,70 | 4,86E-01 |
| AT1G21500 | 0,51 | -1,57 | 3,18 | -0,49 | 6,21E-01 |
| AT1G21520 | 0,17 | 1,57 | 4,34 | 0,36 | 7,18E-01 |

| | | | | | |
|---|---|---|---|---|---|
| AT1G21550 | 0,17 | 1,75 | 4,33 | 0,40 | 6,86E-01 |
| AT1G21600 | 0,77 | 2,35 | 2,16 | 1,08 | 2,78E-01 |
| AT1G21650 | 0,46 | 0,78 | 3,24 | 0,24 | 8,09E-01 |
| AT1G21770 | 0,66 | 0,46 | 2,67 | 0,17 | 8,63E-01 |
| AT1G21890 | 0,36 | 0,42 | 4,38 | 0,10 | 9,24E-01 |
| AT1G21900 | 0,26 | 0,33 | 4,35 | 0,08 | 9,39E-01 |
| AT1G21920 | 0,41 | 0,89 | 3,28 | 0,27 | 7,85E-01 |
| AT1G21928 | 0,29 | 1,24 | 4,34 | 0,29 | 7,75E-01 |
| AT1G22180 | 0,30 | 1,23 | 3,63 | 0,34 | 7,34E-01 |
| AT1G22200 | 0,54 | 0,89 | 3,83 | 0,23 | 8,17E-01 |
| AT1G22300 | 0,58 | -0,58 | 2,60 | -0,22 | 8,25E-01 |
| AT1G22360 | 0,63 | -0,23 | 2,67 | -0,09 | 9,30E-01 |
| AT1G22400 | 0,23 | 1,44 | 4,34 | 0,33 | 7,40E-01 |
| AT1G22550 | 0,19 | 0,78 | 4,36 | 0,18 | 8,58E-01 |
| AT1G22570 | 0,46 | 1,37 | 3,99 | 0,34 | 7,32E-01 |
| AT1G22710 | 0,50 | 0,92 | 2,66 | 0,35 | 7,30E-01 |
| AT1G22850 | 1,38 | 0,35 | 1,59 | 0,22 | 8,24E-01 |
| AT1G23020 | 0,20 | 0,89 | 4,37 | 0,20 | 8,39E-01 |
| AT1G23052 | 1,19 | -1,68 | 2,42 | -0,69 | 4,88E-01 |
| AT1G23080 | 0,61 | 1,26 | 3,32 | 0,38 | 7,04E-01 |
| AT1G05793 | 0,08 | 0,00 | 4,40 | 0,00 | 1,00E+00 |
| AT1G23205 | 0,35 | 0,00 | 4,38 | 0,00 | 1,00E+00 |
| AT1G23260 | 0,22 | 1,92 | 4,33 | 0,44 | 6,57E-01 |
| AT1G23300 | 1,57 | -0,14 | 1,58 | -0,09 | 9,27E-01 |
| AT1G23330 | 0,10 | 0,00 | 4,41 | 0,00 | 1,00E+00 |
| AT1G23650 | 0,08 | 0,85 | 4,38 | 0,20 | 8,45E-01 |
| AT1G23670 | 0,22 | 0,00 | 4,41 | 0,00 | 1,00E+00 |
| AT1G23740 | 1,33 | -1,55 | 1,77 | -0,88 | 3,81E-01 |
| AT1G23780 | 0,31 | 1,99 | 4,32 | 0,46 | 6,45E-01 |
| AT1G23830 | 0,53 | 0,78 | 3,21 | 0,24 | 8,08E-01 |
| AT1G23840 | 0,48 | 2,10 | 3,13 | 0,67 | 5,03E-01 |
| AT1G23850 | 0,35 | 1,68 | 4,34 | 0,39 | 6,98E-01 |
| AT1G23960 | 0,23 | 0,78 | 4,37 | 0,18 | 8,59E-01 |
| AT1G24040 | 0,10 | 0,62 | 4,38 | 0,14 | 8,87E-01 |
| AT1G24100 | 1,00 | 2,21 | 2,00 | 1,10 | 2,71E-01 |
| AT1G24120 | 0,29 | -0,29 | 4,37 | -0,07 | 9,47E-01 |
| AT1G24140 | 0,74 | 1,32 | 3,00 | 0,44 | 6,61E-01 |
| AT1G24160 | 0,84 | 3,34 | 3,28 | 1,02 | 3,10E-01 |
| AT1G24190 | 0,84 | -0,88 | 2,49 | -0,35 | 7,24E-01 |
| AT1G24267 | 0,48 | 1,35 | 3,15 | 0,43 | 6,69E-01 |
| AT1G24290 | 0,17 | 1,57 | 4,34 | 0,36 | 7,18E-01 |
| AT1G24300 | 1,68 | 1,46 | 1,73 | 0,84 | 3,99E-01 |
| AT1G24350 | 0,14 | -1,28 | 4,39 | -0,29 | 7,71E-01 |
| AT1G05907 | 0,77 | 0,99 | 2,50 | 0,40 | 6,93E-01 |
| AT1G24764 | 0,53 | 0,69 | 3,24 | 0,21 | 8,32E-01 |
| AT1G24822 | 0,94 | 0,00 | 4,40 | 0,00 | 1,00E+00 |
| AT1G25220 | 0,36 | 2,25 | 4,33 | 0,52 | 6,04E-01 |
| AT1G25350 | 1,21 | 0,63 | 1,82 | 0,35 | 7,30E-01 |
| AT1G25370 | 0,12 | 1,58 | 4,34 | 0,36 | 7,16E-01 |
| AT1G25390 | 0,51 | 2,37 | 3,06 | 0,77 | 4,39E-01 |
| AT1G25400 | 0,63 | 2,71 | 2,87 | 0,94 | 3,45E-01 |
| AT1G25500 | 0,25 | 1,83 | 4,33 | 0,42 | 6,72E-01 |
| AT1G25540 | 0,94 | 3,44 | 2,29 | 1,50 | 1,32E-01 |
| AT1G25560 | 0,90 | 0,98 | 2,05 | 0,48 | 6,34E-01 |
| AT1G25570 | 0,49 | 0,00 | 4,41 | 0,00 | 1,00E+00 |
| AT1G26150 | 0,88 | 1,53 | 2,29 | 0,67 | 5,05E-01 |

| | | | | | |
|---|---|---|---|---|---|
| AT1G26160 | 0,24 | 0,65 | 4,37 | 0,15 | 8,81E-01 |
| AT1G26170 | 0,54 | 1,19 | 3,12 | 0,38 | 7,02E-01 |
| AT1G26208 | 0,43 | 1,24 | 4,36 | 0,29 | 7,76E-01 |
| AT1G26218 | 0,37 | 0,85 | 4,38 | 0,19 | 8,46E-01 |
| AT1G26233 | 0,08 | 0,00 | 4,40 | 0,00 | 1,00E+00 |
| AT1G26240 | 0,88 | 1,89 | 2,64 | 0,71 | 4,76E-01 |
| AT1G26255 | 0,44 | 1,11 | 4,11 | 0,27 | 7,87E-01 |
| AT1G26270 | 1,28 | 1,21 | 2,20 | 0,55 | 5,83E-01 |
| AT1G26380 | 0,46 | 0,00 | 4,38 | 0,00 | 1,00E+00 |
| AT1G26390 | 0,25 | 0,44 | 4,37 | 0,10 | 9,20E-01 |
| AT1G26420 | 0,40 | 0,05 | 4,28 | 0,01 | 9,91E-01 |
| AT1G26690 | 0,33 | 2,20 | 4,32 | 0,51 | 6,11E-01 |
| AT1G26761 | 0,85 | 2,81 | 4,29 | 0,66 | 5,12E-01 |
| AT1G26800 | 0,42 | -0,11 | 2,89 | -0,04 | 9,68E-01 |
| AT1G06033 | 1,64 | -3,74 | 1,71 | -2,18 | 2,91E-02 |
| AT1G26930 | 0,36 | 2,36 | 4,31 | 0,55 | 5,83E-01 |
| AT1G27008 | 0,08 | 1,10 | 4,36 | 0,25 | 8,00E-01 |
| AT1G27100 | 1,46 | 1,85 | 1,83 | 1,01 | 3,11E-01 |
| AT1G27190 | 0,77 | 2,43 | 2,18 | 1,11 | 2,66E-01 |
| AT1G27200 | 0,57 | 1,05 | 2,73 | 0,39 | 7,00E-01 |
| AT1G27210 | 0,26 | 1,24 | 4,34 | 0,29 | 7,75E-01 |
| AT1G27350 | 0,20 | 0,00 | 4,40 | 0,00 | 1,00E+00 |
| AT1G27370 | 0,51 | 1,26 | 4,10 | 0,31 | 7,59E-01 |
| AT1G27385 | 0,43 | -1,11 | 4,38 | -0,25 | 8,00E-01 |
| AT1G27440 | 0,24 | 0,83 | 4,37 | 0,19 | 8,49E-01 |
| AT1G27500 | 0,29 | 2,05 | 4,33 | 0,47 | 6,36E-01 |
| AT1G27510 | 0,44 | 2,25 | 3,22 | 0,70 | 4,85E-01 |
| AT1G06093 | 0,29 | -0,44 | 4,38 | -0,10 | 9,20E-01 |
| AT1G27600 | 1,12 | -0,98 | 1,98 | -0,49 | 6,21E-01 |
| AT1G27700 | 0,50 | 1,40 | 4,35 | 0,32 | 7,47E-01 |
| AT1G27720 | 0,50 | 2,17 | 3,91 | 0,56 | 5,78E-01 |
| AT1G27730 | 1,60 | -0,55 | 1,34 | -0,41 | 6,82E-01 |
| AT1G27900 | 0,40 | 1,62 | 3,32 | 0,49 | 6,25E-01 |
| AT1G27930 | 0,53 | 1,42 | 3,05 | 0,47 | 6,41E-01 |
| AT1G28000 | 0,35 | 0,53 | 4,38 | 0,12 | 9,04E-01 |
| AT1G28120 | 0,73 | 2,89 | 2,85 | 1,01 | 3,11E-01 |
| AT1G28140 | 1,15 | 1,49 | 1,78 | 0,84 | 4,02E-01 |
| AT1G28150 | 0,60 | 2,44 | 2,98 | 0,82 | 4,14E-01 |
| AT1G28190 | 0,15 | 1,25 | 4,35 | 0,29 | 7,75E-01 |
| AT1G28200 | 0,24 | -0,64 | 4,33 | -0,15 | 8,83E-01 |
| AT1G28230 | 0,78 | 1,24 | 2,19 | 0,56 | 5,72E-01 |
| AT1G28240 | 0,33 | -0,54 | 4,37 | -0,12 | 9,02E-01 |
| AT1G28260 | 1,27 | 2,57 | 1,88 | 1,36 | 1,73E-01 |
| AT1G28280 | 1,03 | 2,69 | 2,04 | 1,32 | 1,87E-01 |
| AT1G28290 | 0,27 | 0,00 | 4,41 | 0,00 | 1,00E+00 |
| AT1G28310 | 0,66 | -0,09 | 2,14 | -0,04 | 9,68E-01 |
| AT1G28330 | 0,60 | -1,69 | 3,15 | -0,54 | 5,91E-01 |
| AT1G28370 | 0,92 | 1,23 | 2,35 | 0,52 | 6,01E-01 |
| AT1G28390 | 0,25 | 0,00 | 4,40 | 0,00 | 1,00E+00 |
| AT1G28400 | 0,35 | -1,20 | 3,43 | -0,35 | 7,26E-01 |
| AT1G28570 | 0,12 | 1,39 | 4,34 | 0,32 | 7,49E-01 |
| AT1G28580 | 0,48 | 2,62 | 3,88 | 0,67 | 5,01E-01 |
| AT1G28600 | 1,10 | -0,29 | 1,80 | -0,16 | 8,70E-01 |
| AT1G28630 | 0,79 | -3,52 | 2,78 | -1,26 | 2,06E-01 |
| AT1G28660 | 0,08 | 1,10 | 4,36 | 0,25 | 8,00E-01 |
| AT1G28685 | 0,55 | 0,53 | 3,22 | 0,17 | 8,69E-01 |

| | | | | | |
|---|---|---|---|---|---|
| AT1G28765 | 0,17 | 1,92 | 4,33 | 0,44 | 6,57E-01 |
| AT1G28960 | 0,56 | 2,64 | 3,90 | 0,68 | 4,99E-01 |
| AT1G29050 | 0,14 | 0,00 | 4,41 | 0,00 | 1,00E+00 |
| AT1G29071 | 1,34 | 0,70 | 1,69 | 0,42 | 6,77E-01 |
| AT1G29170 | 0,29 | 1,85 | 4,33 | 0,43 | 6,69E-01 |
| AT1G29240 | 0,32 | 1,25 | 4,35 | 0,29 | 7,74E-01 |
| AT1G29300 | 0,08 | 0,00 | 4,41 | 0,00 | 1,00E+00 |
| AT1G29390 | 0,82 | 2,33 | 2,13 | 1,10 | 2,73E-01 |
| AT1G29395 | 0,39 | -0,31 | 3,30 | -0,09 | 9,25E-01 |
| AT1G29530 | 0,36 | 0,47 | 3,49 | 0,14 | 8,92E-01 |
| AT1G29690 | 0,27 | -0,51 | 4,38 | -0,12 | 9,08E-01 |
| AT1G29700 | 0,34 | -2,51 | 4,34 | -0,58 | 5,63E-01 |
| AT1G29720 | 0,71 | -1,09 | 2,55 | -0,42 | 6,71E-01 |
| AT1G29750 | 0,26 | 1,24 | 4,34 | 0,28 | 7,76E-01 |
| AT1G29880 | 1,19 | 3,35 | 2,01 | 1,67 | 9,56E-02 |
| AT1G29970 | 0,83 | 1,72 | 2,53 | 0,68 | 4,96E-01 |
| AT1G30000 | 0,84 | 2,19 | 2,65 | 0,83 | 4,08E-01 |
| AT1G30060 | 0,17 | 0,00 | 4,41 | 0,00 | 1,00E+00 |
| AT1G30070 | 0,15 | 0,00 | 4,39 | 0,00 | 1,00E+00 |
| AT1G30090 | 0,16 | 1,33 | 4,35 | 0,31 | 7,59E-01 |
| AT1G30120 | 1,11 | 2,61 | 1,97 | 1,33 | 1,85E-01 |
| AT1G30210 | 0,29 | 0,77 | 4,37 | 0,18 | 8,60E-01 |
| AT1G30250 | 0,60 | 0,09 | 2,62 | 0,04 | 9,71E-01 |
| AT1G30370 | 0,55 | 2,62 | 3,07 | 0,85 | 3,93E-01 |
| AT1G30500 | 0,64 | 2,11 | 3,81 | 0,55 | 5,80E-01 |
| AT1G06347 | 0,94 | -0,37 | 2,11 | -0,17 | 8,63E-01 |
| AT1G30540 | 0,42 | -1,22 | 4,19 | -0,29 | 7,70E-01 |
| AT1G30580 | 1,07 | -0,24 | 1,99 | -0,12 | 9,06E-01 |
| AT1G30590 | 0,66 | 0,42 | 2,96 | 0,14 | 8,86E-01 |
| AT1G30755 | 0,62 | 0,55 | 2,75 | 0,20 | 8,43E-01 |
| AT1G30780 | 0,83 | -1,31 | 2,17 | -0,60 | 5,46E-01 |
| AT1G30810 | 0,83 | -0,49 | 1,87 | -0,26 | 7,94E-01 |
| AT1G30880 | 0,08 | 0,85 | 4,38 | 0,20 | 8,45E-01 |
| AT1G30960 | 0,37 | 1,50 | 4,34 | 0,35 | 7,29E-01 |
| AT1G06407 | 0,34 | 1,24 | 3,59 | 0,34 | 7,30E-01 |
| AT1G31130 | 1,47 | 3,06 | 1,72 | 1,78 | 7,58E-02 |
| AT1G31160 | 0,42 | 1,61 | 4,13 | 0,39 | 6,97E-01 |
| AT1G31180 | 0,17 | 1,57 | 4,34 | 0,36 | 7,18E-01 |
| AT1G31230 | 1,31 | -0,44 | 1,85 | -0,24 | 8,11E-01 |
| AT1G31240 | 0,22 | -1,25 | 4,39 | -0,28 | 7,76E-01 |
| AT1G31350 | 0,45 | 1,75 | 3,23 | 0,54 | 5,88E-01 |
| AT1G31410 | 0,08 | 0,85 | 4,38 | 0,20 | 8,45E-01 |
| AT1G31470 | 0,15 | 1,62 | 4,34 | 0,37 | 7,09E-01 |
| AT1G31550 | 0,81 | 0,13 | 2,43 | 0,05 | 9,57E-01 |
| AT1G31600 | 0,81 | -0,86 | 2,82 | -0,30 | 7,61E-01 |
| AT1G31640 | 0,23 | 1,28 | 4,34 | 0,29 | 7,69E-01 |
| AT1G31650 | 0,25 | 0,00 | 4,38 | 0,00 | 1,00E+00 |
| AT1G31660 | 0,51 | -0,03 | 4,36 | -0,01 | 9,94E-01 |
| AT1G31740 | 0,22 | 1,61 | 4,33 | 0,37 | 7,11E-01 |
| AT1G31810 | 0,57 | 0,65 | 2,32 | 0,28 | 7,78E-01 |
| AT1G31812 | 0,29 | -1,23 | 4,39 | -0,28 | 7,79E-01 |
| AT1G31814 | 0,37 | 1,84 | 4,33 | 0,43 | 6,71E-01 |
| AT1G31820 | 0,94 | 3,43 | 2,74 | 1,25 | 2,10E-01 |
| AT1G31830 | 0,66 | 1,76 | 2,59 | 0,68 | 4,96E-01 |
| AT1G06453 | 1,48 | -1,57 | 1,76 | -0,89 | 3,72E-01 |
| AT1G31860 | 1,08 | 1,30 | 1,90 | 0,68 | 4,96E-01 |

| | | | | | |
|---|---|---|---|---|---|
| AT1G06463 | 0,27 | 1,56 | 4,34 | 0,36 | 7,19E-01 |
| AT1G31920 | 0,52 | 1,64 | 2,66 | 0,61 | 5,39E-01 |
| AT1G31940 | 0,35 | 0,00 | 4,41 | 0,00 | 1,00E+00 |
| AT1G06473 | 0,38 | -0,59 | 4,33 | -0,14 | 8,92E-01 |
| AT1G32000 | 0,35 | 0,44 | 4,39 | 0,10 | 9,20E-01 |
| AT1G32070 | 0,34 | 1,72 | 4,34 | 0,40 | 6,92E-01 |
| AT1G32080 | 0,85 | 0,37 | 1,93 | 0,19 | 8,48E-01 |
| AT1G32130 | 1,27 | -0,09 | 1,90 | -0,05 | 9,62E-01 |
| AT1G32200 | 0,75 | -3,20 | 2,19 | -1,46 | 1,44E-01 |
| AT1G32340 | 0,12 | 0,79 | 4,37 | 0,18 | 8,57E-01 |
| AT1G06537 | 1,54 | 3,56 | 1,82 | 1,96 | 5,00E-02 |
| AT1G32400 | 0,47 | 1,58 | 2,73 | 0,58 | 5,63E-01 |
| AT1G32450 | 0,27 | 0,96 | 4,36 | 0,22 | 8,25E-01 |
| AT1G06573 | 0,41 | 0,72 | 4,32 | 0,17 | 8,67E-01 |
| AT1G32470 | 0,50 | 0,10 | 2,64 | 0,04 | 9,71E-01 |
| AT1G32490 | 0,70 | 1,48 | 2,20 | 0,67 | 5,02E-01 |
| AT1G32500 | 0,52 | -0,47 | 2,79 | -0,17 | 8,67E-01 |
| AT1G32530 | 0,53 | 1,86 | 2,70 | 0,69 | 4,92E-01 |
| AT1G06577 | 0,16 | -1,36 | 4,41 | -0,31 | 7,57E-01 |
| AT1G32713 | 1,03 | 1,54 | 2,31 | 0,67 | 5,05E-01 |
| AT1G32790 | 1,57 | 4,30 | 1,92 | 2,24 | 2,54E-02 |
| AT1G32860 | 0,05 | 0,00 | 4,41 | 0,00 | 1,00E+00 |
| AT1G32880 | 0,26 | 1,08 | 4,37 | 0,25 | 8,05E-01 |
| AT1G32900 | 0,61 | 1,24 | 3,15 | 0,39 | 6,94E-01 |
| AT1G32970 | 0,17 | 1,94 | 4,33 | 0,45 | 6,55E-01 |
| AT1G33050 | 0,95 | 1,41 | 2,00 | 0,71 | 4,80E-01 |
| AT1G06643 | 0,36 | -0,38 | 4,31 | -0,09 | 9,30E-01 |
| AT1G33250 | 0,90 | 3,50 | 2,25 | 1,56 | 1,20E-01 |
| AT1G33270 | 0,24 | 1,58 | 4,34 | 0,36 | 7,16E-01 |
| AT1G06673 | 0,47 | 0,48 | 3,30 | 0,15 | 8,84E-01 |
| AT1G33390 | 1,05 | 0,50 | 2,43 | 0,20 | 8,38E-01 |
| AT1G33410 | 1,37 | 2,88 | 1,73 | 1,66 | 9,61E-02 |
| AT1G33430 | 0,08 | 0,85 | 4,38 | 0,20 | 8,45E-01 |
| AT1G33520 | 0,47 | 1,44 | 3,16 | 0,45 | 6,49E-01 |
| AT1G33615 | 0,17 | -1,64 | 4,37 | -0,37 | 7,08E-01 |
| AT1G33700 | 0,88 | 2,77 | 2,12 | 1,30 | 1,92E-01 |
| AT1G33720 | 1,30 | 0,88 | 1,89 | 0,46 | 6,43E-01 |
| AT1G33780 | 0,08 | 0,00 | 4,41 | 0,00 | 1,00E+00 |
| AT1G33870 | 0,60 | 0,35 | 2,49 | 0,14 | 8,89E-01 |
| AT1G33970 | 0,67 | 0,43 | 2,65 | 0,16 | 8,72E-01 |
| AT1G34030 | 0,44 | -0,59 | 2,78 | -0,21 | 8,32E-01 |
| AT1G34042 | 0,29 | 1,20 | 4,34 | 0,28 | 7,82E-01 |
| AT1G34110 | 0,91 | 1,68 | 1,89 | 0,89 | 3,73E-01 |
| AT1G34170 | 0,20 | 1,17 | 4,37 | 0,27 | 7,89E-01 |
| AT1G34220 | 1,01 | 4,01 | 2,86 | 1,40 | 1,60E-01 |
| AT1G34270 | 0,12 | 1,46 | 4,35 | 0,34 | 7,36E-01 |
| AT1G34310 | 0,97 | -0,14 | 2,73 | -0,05 | 9,60E-01 |
| AT1G34315 | 0,23 | 2,12 | 4,32 | 0,49 | 6,24E-01 |
| AT1G34380 | 0,08 | 0,85 | 4,38 | 0,20 | 8,45E-01 |
| AT1G34440 | 1,54 | -0,26 | 1,67 | -0,16 | 8,76E-01 |
| AT1G34550 | 0,63 | 0,75 | 2,47 | 0,30 | 7,62E-01 |
| AT1G34750 | 0,23 | 1,09 | 4,36 | 0,25 | 8,02E-01 |
| AT1G34780 | 0,68 | 2,93 | 2,82 | 1,04 | 2,99E-01 |
| AT1G35150 | 0,16 | 0,46 | 4,38 | 0,11 | 9,16E-01 |
| AT1G35240 | 1,26 | 2,58 | 1,87 | 1,38 | 1,67E-01 |
| AT1G35250 | 0,19 | 0,00 | 4,41 | 0,00 | 1,00E+00 |

| | | | | | |
|---|---|---|---|---|---|
| AT1G35320 | 0,59 | 1,90 | 2,96 | 0,64 | 5,20E-01 |
| AT1G35350 | 0,96 | 1,75 | 2,00 | 0,87 | 3,83E-01 |
| AT1G06883 | 0,19 | 0,00 | 4,41 | 0,00 | 1,00E+00 |
| AT1G06897 | 0,91 | -3,44 | 3,41 | -1,01 | 3,13E-01 |
| AT1G06903 | 0,10 | 0,00 | 4,41 | 0,00 | 1,00E+00 |
| AT1G35520 | 1,01 | 2,55 | 2,51 | 1,02 | 3,10E-01 |
| AT1G35545 | 0,38 | -0,48 | 3,43 | -0,14 | 8,89E-01 |
| AT1G35580 | 1,07 | 1,64 | 1,81 | 0,91 | 3,65E-01 |
| AT1G35610 | 0,22 | -0,03 | 4,37 | -0,01 | 9,94E-01 |
| AT1G35625 | 0,08 | 0,85 | 4,38 | 0,20 | 8,45E-01 |
| AT1G35730 | 0,17 | 1,92 | 4,33 | 0,44 | 6,57E-01 |
| AT1G35820 | 1,40 | 1,27 | 2,92 | 0,43 | 6,65E-01 |
| AT1G35895 | 0,55 | 1,58 | 3,12 | 0,50 | 6,14E-01 |
| AT1G36070 | 0,25 | 1,75 | 4,33 | 0,40 | 6,87E-01 |
| AT1G36085 | 0,68 | 0,00 | 4,41 | 0,00 | 1,00E+00 |
| AT1G36230 | 0,12 | 0,00 | 4,41 | 0,00 | 1,00E+00 |
| AT1G36325 | 1,41 | -1,07 | 1,62 | -0,66 | 5,11E-01 |
| AT1G36370 | 0,71 | 1,35 | 2,38 | 0,57 | 5,71E-01 |
| AT1G36390 | 0,17 | 0,09 | 4,35 | 0,02 | 9,84E-01 |
| AT1G36622 | 0,40 | 0,29 | 3,44 | 0,08 | 9,34E-01 |
| AT1G36990 | 1,02 | 1,60 | 1,96 | 0,82 | 4,14E-01 |
| AT1G07013 | 0,15 | 0,00 | 4,39 | 0,00 | 1,00E+00 |
| AT1G40087 | 0,19 | 0,00 | 4,41 | 0,00 | 1,00E+00 |
| AT1G41830 | 0,28 | 1,40 | 4,35 | 0,32 | 7,47E-01 |
| AT1G41920 | 0,51 | 0,80 | 3,22 | 0,25 | 8,03E-01 |
| AT1G42440 | 0,46 | 1,27 | 3,39 | 0,37 | 7,09E-01 |
| AT1G42560 | 0,05 | 0,00 | 4,41 | 0,00 | 1,00E+00 |
| AT1G42990 | 1,44 | 3,01 | 1,77 | 1,70 | 8,95E-02 |
| AT1G43190 | 0,38 | 1,40 | 4,36 | 0,32 | 7,48E-01 |
| AT1G43560 | 0,22 | -0,28 | 4,36 | -0,06 | 9,49E-01 |
| AT1G43580 | 0,21 | 0,18 | 4,34 | 0,04 | 9,68E-01 |
| AT1G43620 | 0,44 | 1,64 | 4,22 | 0,39 | 6,97E-01 |
| AT1G43665 | 0,88 | 2,88 | 3,20 | 0,90 | 3,68E-01 |
| AT1G43710 | 0,50 | 2,35 | 3,20 | 0,74 | 4,62E-01 |
| AT1G43730 | 0,40 | 0,27 | 4,28 | 0,06 | 9,50E-01 |
| AT1G43760 | 0,66 | -0,49 | 2,43 | -0,20 | 8,39E-01 |
| AT1G43825 | 0,81 | 0,00 | 2,89 | 0,00 | 1,00E+00 |
| AT1G43860 | 1,69 | 1,21 | 1,77 | 0,68 | 4,95E-01 |
| AT1G43950 | 0,46 | 0,97 | 4,06 | 0,24 | 8,11E-01 |
| AT1G44000 | 0,08 | 0,85 | 4,38 | 0,20 | 8,45E-01 |
| AT1G44030 | 0,25 | -0,34 | 4,38 | -0,08 | 9,38E-01 |
| AT1G44080 | 1,35 | -4,24 | 1,99 | -2,13 | 3,34E-02 |
| AT1G44085 | 0,65 | 1,26 | 2,94 | 0,43 | 6,69E-01 |
| AT1G44090 | 0,05 | 0,00 | 4,41 | 0,00 | 1,00E+00 |
| AT1G44125 | 0,91 | 3,62 | 2,64 | 1,37 | 1,71E-01 |
| AT1G44170 | 0,30 | 0,43 | 4,38 | 0,10 | 9,23E-01 |
| AT1G44350 | 0,41 | 1,11 | 4,23 | 0,26 | 7,92E-01 |
| AT1G44542 | 0,40 | 0,00 | 4,40 | 0,00 | 1,00E+00 |
| AT1G44770 | 0,41 | 1,77 | 3,33 | 0,53 | 5,95E-01 |
| AT1G44790 | 0,21 | 0,43 | 4,38 | 0,10 | 9,22E-01 |
| AT1G44920 | 0,44 | 0,33 | 3,36 | 0,10 | 9,21E-01 |
| AT1G44980 | 0,59 | -0,38 | 2,69 | -0,14 | 8,88E-01 |
| AT1G45063 | 0,43 | -2,17 | 4,30 | -0,50 | 6,14E-01 |
| AT1G45145 | 0,72 | 2,98 | 3,11 | 0,96 | 3,38E-01 |
| AT1G45150 | 0,25 | 0,00 | 4,38 | 0,00 | 1,00E+00 |
| AT1G45163 | 0,70 | 2,82 | 2,48 | 1,14 | 2,54E-01 |

| | | | | | |
|---|---|---|---|---|---|
| AT1G45207 | 0,37 | 0,00 | 4,41 | 0,00 | 1,00E+00 |
| AT1G07323 | 0,55 | 1,62 | 2,72 | 0,59 | 5,53E-01 |
| AT1G47200 | 0,16 | 0,00 | 4,41 | 0,00 | 1,00E+00 |
| AT1G47240 | 0,43 | 2,58 | 4,09 | 0,63 | 5,28E-01 |
| AT1G07333 | 0,81 | -1,46 | 3,40 | -0,43 | 6,68E-01 |
| AT1G47260 | 1,01 | 0,17 | 1,66 | 0,11 | 9,16E-01 |
| AT1G47280 | 1,31 | 0,00 | 2,27 | 0,00 | 1,00E+00 |
| AT1G07347 | 0,18 | 0,00 | 4,41 | 0,00 | 1,00E+00 |
| AT1G47380 | 0,18 | -0,04 | 4,35 | -0,01 | 9,92E-01 |
| AT1G47420 | 0,71 | 2,59 | 2,21 | 1,17 | 2,41E-01 |
| AT1G07383 | 0,55 | -1,23 | 2,70 | -0,46 | 6,48E-01 |
| AT1G07393 | 1,67 | -2,29 | 1,58 | -1,45 | 1,47E-01 |
| AT1G47640 | 0,35 | 1,23 | 3,50 | 0,35 | 7,26E-01 |
| AT1G47670 | 0,05 | 0,00 | 4,41 | 0,00 | 1,00E+00 |
| AT1G47765 | 0,55 | -0,43 | 2,45 | -0,18 | 8,60E-01 |
| AT1G47900 | 0,45 | -0,82 | 4,11 | -0,20 | 8,42E-01 |
| AT1G47960 | 1,43 | -0,93 | 1,42 | -0,66 | 5,11E-01 |
| AT1G48000 | 0,50 | -0,02 | 4,37 | 0,00 | 9,96E-01 |
| AT1G48030 | 1,26 | 4,08 | 2,13 | 1,91 | 5,58E-02 |
| AT1G48040 | 0,63 | 2,86 | 4,30 | 0,66 | 5,06E-01 |
| AT1G48110 | 0,42 | 0,85 | 4,31 | 0,20 | 8,43E-01 |
| AT1G48120 | 0,82 | 3,13 | 2,33 | 1,34 | 1,79E-01 |
| AT1G48160 | 0,15 | 0,86 | 4,37 | 0,20 | 8,43E-01 |
| AT1G48285 | 0,80 | 1,30 | 2,31 | 0,56 | 5,73E-01 |
| AT1G48320 | 0,17 | 1,92 | 4,33 | 0,44 | 6,57E-01 |
| AT1G48380 | 0,10 | 0,42 | 4,38 | 0,10 | 9,23E-01 |
| AT1G48405 | 0,82 | 3,38 | 3,28 | 1,03 | 3,03E-01 |
| AT1G48430 | 0,15 | 1,58 | 4,34 | 0,36 | 7,15E-01 |
| AT1G48450 | 1,45 | 0,31 | 1,92 | 0,16 | 8,70E-01 |
| AT1G48550 | 1,08 | 1,82 | 2,61 | 0,70 | 4,87E-01 |
| AT1G48570 | 0,26 | 0,89 | 4,32 | 0,21 | 8,37E-01 |
| AT1G48580 | 0,08 | 1,10 | 4,36 | 0,25 | 8,00E-01 |
| AT1G48620 | 0,46 | -1,23 | 3,29 | -0,37 | 7,09E-01 |
| AT1G48630 | 1,25 | 2,13 | 1,72 | 1,24 | 2,15E-01 |
| AT1G48730 | 0,75 | -1,18 | 2,38 | -0,49 | 6,22E-01 |
| AT1G48742 | 0,46 | -0,78 | 3,08 | -0,25 | 8,01E-01 |
| AT1G48790 | 0,44 | 2,30 | 3,21 | 0,72 | 4,73E-01 |
| AT1G48850 | 0,39 | 1,18 | 4,34 | 0,27 | 7,87E-01 |
| AT1G48900 | 0,23 | -0,85 | 4,35 | -0,20 | 8,45E-01 |
| AT1G48930 | 0,71 | 2,46 | 2,83 | 0,87 | 3,84E-01 |
| AT1G48970 | 1,37 | 1,19 | 1,81 | 0,66 | 5,11E-01 |
| AT1G49005 | 0,38 | -0,07 | 4,33 | -0,02 | 9,87E-01 |
| AT1G49140 | 0,12 | 0,42 | 4,38 | 0,10 | 9,24E-01 |
| AT1G49180 | 0,08 | 0,85 | 4,38 | 0,20 | 8,45E-01 |
| AT1G49205 | 0,08 | 0,00 | 4,40 | 0,00 | 1,00E+00 |
| AT1G49225 | 0,48 | 0,55 | 3,25 | 0,17 | 8,65E-01 |
| AT1G49270 | 0,10 | 0,00 | 4,41 | 0,00 | 1,00E+00 |
| AT1G49490 | 0,32 | 0,44 | 4,38 | 0,10 | 9,20E-01 |
| AT1G49540 | 0,83 | 1,19 | 2,38 | 0,50 | 6,16E-01 |
| AT1G49600 | 0,82 | 2,65 | 2,11 | 1,26 | 2,08E-01 |
| AT1G49670 | 0,26 | 0,50 | 4,38 | 0,11 | 9,08E-01 |
| AT1G49710 | 0,34 | 1,88 | 3,49 | 0,54 | 5,89E-01 |
| AT1G49780 | 1,22 | 0,90 | 1,62 | 0,55 | 5,80E-01 |
| AT1G49850 | 0,39 | 2,11 | 4,33 | 0,49 | 6,27E-01 |
| AT1G49970 | 0,66 | 0,13 | 2,16 | 0,06 | 9,53E-01 |
| AT1G07603 | 0,61 | 1,72 | 3,21 | 0,54 | 5,91E-01 |

| | | | | | |
|---|---|---|---|---|---|
| AT1G50020 | 0,58 | -2,07 | 4,02 | -0,51 | 6,07E-01 |
| AT1G50140 | 0,20 | 0,60 | 4,34 | 0,14 | 8,90E-01 |
| AT1G50190 | 0,32 | 1,61 | 4,34 | 0,37 | 7,12E-01 |
| AT1G50260 | 0,32 | 0,08 | 4,34 | 0,02 | 9,85E-01 |
| AT1G50280 | 0,61 | -1,48 | 2,36 | -0,63 | 5,29E-01 |
| AT1G50320 | 0,25 | 0,64 | 4,33 | 0,15 | 8,82E-01 |
| AT1G07627 | 0,18 | 0,00 | 4,40 | 0,00 | 1,00E+00 |
| AT1G50420 | 0,49 | 1,34 | 4,34 | 0,31 | 7,57E-01 |
| AT1G50430 | 1,02 | 2,86 | 1,92 | 1,49 | 1,37E-01 |
| AT1G50450 | 0,63 | -0,19 | 2,98 | -0,06 | 9,48E-01 |
| AT1G50500 | 0,57 | -0,28 | 3,08 | -0,09 | 9,27E-01 |
| AT1G07647 | 0,08 | 0,85 | 4,38 | 0,20 | 8,45E-01 |
| AT1G50570 | 0,65 | 0,49 | 2,72 | 0,18 | 8,57E-01 |
| AT1G50600 | 0,88 | 2,07 | 2,09 | 0,99 | 3,22E-01 |
| AT1G50640 | 1,08 | 1,26 | 1,98 | 0,64 | 5,24E-01 |
| AT1G50659 | 0,43 | 1,83 | 3,25 | 0,56 | 5,72E-01 |
| AT1G50750 | 0,08 | 0,00 | 4,40 | 0,00 | 1,00E+00 |
| AT1G50790 | 0,12 | 1,58 | 4,34 | 0,36 | 7,16E-01 |
| AT1G50820 | 0,50 | 1,10 | 3,99 | 0,28 | 7,82E-01 |
| AT1G50910 | 1,39 | -0,20 | 1,75 | -0,11 | 9,10E-01 |
| AT1G50980 | 0,39 | 0,00 | 4,40 | 0,00 | 1,00E+00 |
| AT1G51080 | 0,73 | 1,65 | 2,64 | 0,63 | 5,32E-01 |
| AT1G07733 | 0,24 | -1,37 | 4,41 | -0,31 | 7,55E-01 |
| AT1G51400 | 0,93 | -2,54 | 2,24 | -1,14 | 2,56E-01 |
| AT1G51480 | 0,33 | 0,89 | 4,36 | 0,20 | 8,39E-01 |
| AT1G51500 | 1,21 | -1,45 | 1,89 | -0,77 | 4,42E-01 |
| AT1G51510 | 0,91 | 0,23 | 2,27 | 0,10 | 9,20E-01 |
| AT1G51560 | 0,45 | 1,63 | 3,30 | 0,49 | 6,21E-01 |
| AT1G51590 | 0,26 | 0,41 | 4,38 | 0,09 | 9,25E-01 |
| AT1G51630 | 0,52 | -0,24 | 3,19 | -0,08 | 9,39E-01 |
| AT1G51680 | 0,80 | 2,35 | 2,17 | 1,08 | 2,78E-01 |
| AT1G07763 | 0,56 | -3,15 | 4,32 | -0,73 | 4,65E-01 |
| AT1G51710 | 0,45 | 0,44 | 4,10 | 0,11 | 9,14E-01 |
| AT1G51730 | 0,47 | 0,15 | 3,19 | 0,05 | 9,63E-01 |
| AT1G51790 | 0,25 | 0,00 | 4,41 | 0,00 | 1,00E+00 |
| AT1G51890 | 0,23 | 2,14 | 4,32 | 0,50 | 6,20E-01 |
| AT1G51940 | 1,08 | 0,63 | 2,50 | 0,25 | 8,02E-01 |
| AT1G52000 | 0,46 | -2,84 | 4,04 | -0,70 | 4,82E-01 |
| AT1G52040 | 0,54 | -2,76 | 3,91 | -0,71 | 4,80E-01 |
| AT1G52150 | 1,30 | 3,75 | 2,35 | 1,60 | 1,10E-01 |
| AT1G52160 | 0,61 | 1,21 | 3,17 | 0,38 | 7,02E-01 |
| AT1G52180 | 1,00 | -2,19 | 1,90 | -1,15 | 2,49E-01 |
| AT1G52185 | 0,32 | -1,65 | 4,33 | -0,38 | 7,03E-01 |
| AT1G52200 | 0,25 | 2,35 | 4,32 | 0,54 | 5,86E-01 |
| AT1G52220 | 0,76 | -0,27 | 2,10 | -0,13 | 8,98E-01 |
| AT1G52290 | 0,41 | 2,22 | 4,22 | 0,53 | 5,98E-01 |
| AT1G52300 | 0,35 | 0,87 | 4,32 | 0,20 | 8,40E-01 |
| AT1G52340 | 0,08 | 0,00 | 4,40 | 0,00 | 1,00E+00 |
| AT1G52342 | 0,45 | 2,63 | 4,32 | 0,61 | 5,43E-01 |
| AT1G52510 | 0,47 | 0,61 | 4,21 | 0,15 | 8,84E-01 |
| AT1G52570 | 1,59 | 2,75 | 2,27 | 1,21 | 2,24E-01 |
| AT1G52750 | 0,31 | 1,94 | 4,33 | 0,45 | 6,54E-01 |
| AT1G52880 | 0,40 | 1,26 | 3,38 | 0,37 | 7,10E-01 |
| AT1G52890 | 0,42 | 2,06 | 4,14 | 0,50 | 6,18E-01 |
| AT1G52970 | 1,13 | -4,01 | 1,99 | -2,02 | 4,38E-02 |
| AT1G52990 | 0,41 | 0,00 | 4,41 | 0,00 | 1,00E+00 |

| | | | | | |
|---|---|---|---|---|---|
| AT1G53000 | 0,49 | 0,84 | 4,04 | 0,21 | 8,35E-01 |
| AT1G53025 | 0,43 | 0,60 | 3,40 | 0,18 | 8,60E-01 |
| AT1G53035 | 0,12 | 1,58 | 4,34 | 0,36 | 7,16E-01 |
| AT1G53170 | 0,65 | -1,05 | 2,53 | -0,41 | 6,78E-01 |
| AT1G53200 | 0,24 | 1,58 | 4,34 | 0,36 | 7,16E-01 |
| AT1G53230 | 1,22 | 3,65 | 2,15 | 1,70 | 8,98E-02 |
| AT1G53240 | 1,16 | -0,49 | 1,77 | -0,28 | 7,80E-01 |
| AT1G53265 | 0,53 | -1,61 | 3,95 | -0,41 | 6,83E-01 |
| AT1G53320 | 0,35 | 1,54 | 4,34 | 0,35 | 7,23E-01 |
| AT1G53450 | 0,86 | -2,85 | 2,03 | -1,40 | 1,60E-01 |
| AT1G53480 | 0,84 | -0,95 | 3,01 | -0,32 | 7,52E-01 |
| AT1G53500 | 0,50 | -0,44 | 3,26 | -0,13 | 8,93E-01 |
| AT1G53510 | 0,76 | -1,11 | 2,48 | -0,45 | 6,55E-01 |
| AT1G53541 | 0,64 | 2,22 | 2,45 | 0,91 | 3,65E-01 |
| AT1G53580 | 0,12 | 0,90 | 4,36 | 0,21 | 8,36E-01 |
| AT1G53710 | 0,67 | 0,69 | 2,93 | 0,23 | 8,14E-01 |
| AT1G53750 | 1,03 | 0,55 | 2,03 | 0,27 | 7,87E-01 |
| AT1G53760 | 0,08 | 1,10 | 4,36 | 0,25 | 8,00E-01 |
| AT1G53770 | 0,33 | 1,33 | 4,35 | 0,31 | 7,59E-01 |
| AT1G53780 | 1,40 | 4,10 | 1,92 | 2,13 | 3,32E-02 |
| AT1G54050 | 0,12 | 0,90 | 4,36 | 0,21 | 8,36E-01 |
| AT1G54080 | 1,16 | 1,76 | 1,95 | 0,90 | 3,67E-01 |
| AT1G54100 | 1,08 | 0,16 | 1,99 | 0,08 | 9,35E-01 |
| AT1G54140 | 0,15 | 0,86 | 4,37 | 0,20 | 8,43E-01 |
| AT1G54170 | 0,82 | -0,98 | 2,01 | -0,49 | 6,24E-01 |
| AT1G54320 | 0,67 | -0,15 | 2,34 | -0,07 | 9,48E-01 |
| AT1G54370 | 0,38 | 2,08 | 4,28 | 0,48 | 6,28E-01 |
| AT1G54500 | 0,85 | 0,99 | 2,14 | 0,46 | 6,43E-01 |
| AT1G54590 | 0,10 | 0,42 | 4,38 | 0,10 | 9,23E-01 |
| AT1G54630 | 0,35 | -0,23 | 4,36 | -0,05 | 9,59E-01 |
| AT1G54690 | 0,48 | 1,23 | 2,77 | 0,44 | 6,57E-01 |
| AT1G54775 | 0,57 | -2,33 | 3,82 | -0,61 | 5,42E-01 |
| AT1G54840 | 0,49 | 0,00 | 4,39 | 0,00 | 1,00E+00 |
| AT1G54950 | 0,44 | -1,57 | 4,08 | -0,38 | 7,00E-01 |
| AT1G55152 | 0,78 | 2,51 | 2,49 | 1,01 | 3,13E-01 |
| AT1G55180 | 0,78 | 2,86 | 2,79 | 1,02 | 3,06E-01 |
| AT1G55200 | 0,08 | 1,10 | 4,36 | 0,25 | 8,00E-01 |
| AT1G08043 | 0,30 | 1,68 | 4,34 | 0,39 | 6,99E-01 |
| AT1G55270 | 1,00 | 2,31 | 3,17 | 0,73 | 4,67E-01 |
| AT1G55300 | 0,10 | 0,62 | 4,38 | 0,14 | 8,87E-01 |
| AT1G55330 | 0,51 | -0,83 | 3,13 | -0,27 | 7,91E-01 |
| AT1G55360 | 0,31 | 0,99 | 4,35 | 0,23 | 8,19E-01 |
| AT1G55530 | 0,83 | 0,93 | 2,14 | 0,43 | 6,64E-01 |
| AT1G55535 | 0,40 | 0,44 | 4,23 | 0,10 | 9,17E-01 |
| AT1G55590 | 0,61 | 0,22 | 2,62 | 0,08 | 9,32E-01 |
| AT1G55610 | 0,12 | 0,90 | 4,36 | 0,21 | 8,36E-01 |
| AT1G55620 | 0,57 | 1,12 | 3,14 | 0,36 | 7,22E-01 |
| AT1G55675 | 0,68 | 0,00 | 3,81 | 0,00 | 1,00E+00 |
| AT1G55680 | 1,44 | 3,34 | 2,03 | 1,65 | 9,92E-02 |
| AT1G55690 | 0,12 | 1,58 | 4,34 | 0,36 | 7,16E-01 |
| AT1G55830 | 0,23 | 1,56 | 4,34 | 0,36 | 7,19E-01 |
| AT1G55880 | 0,36 | 1,99 | 4,34 | 0,46 | 6,46E-01 |
| AT1G55950 | 0,93 | 2,89 | 2,18 | 1,32 | 1,86E-01 |
| AT1G55960 | 0,25 | -1,32 | 4,39 | -0,30 | 7,64E-01 |
| AT1G55970 | 0,17 | 1,94 | 4,33 | 0,45 | 6,55E-01 |
| AT1G56010 | 0,42 | 2,28 | 4,10 | 0,56 | 5,77E-01 |

| | | | | | |
|---|---|---|---|---|---|
| AT1G56100 | 0,86 | -2,50 | 2,83 | -0,88 | 3,77E-01 |
| AT1G56110 | 1,35 | 0,00 | 1,90 | 0,00 | 9,99E-01 |
| AT1G56120 | 0,47 | -0,58 | 4,04 | -0,14 | 8,85E-01 |
| AT1G56140 | 0,44 | 1,88 | 3,28 | 0,57 | 5,66E-01 |
| AT1G56145 | 0,80 | 2,20 | 2,74 | 0,80 | 4,23E-01 |
| AT1G56165 | 0,20 | 0,89 | 4,37 | 0,20 | 8,39E-01 |
| AT1G56230 | 0,19 | 0,00 | 4,37 | 0,00 | 9,99E-01 |
| AT1G08107 | 0,31 | -0,36 | 3,66 | -0,10 | 9,21E-01 |
| AT1G08113 | 0,55 | -1,80 | 3,17 | -0,57 | 5,69E-01 |
| AT1G08123 | 0,56 | 1,30 | 2,53 | 0,52 | 6,05E-01 |
| AT1G56300 | 0,85 | -0,60 | 2,85 | -0,21 | 8,34E-01 |
| AT1G56330 | 0,78 | -1,75 | 2,59 | -0,67 | 5,00E-01 |
| AT1G56345 | 0,19 | 0,00 | 4,40 | 0,00 | 1,00E+00 |
| AT1G56440 | 0,27 | 0,96 | 4,36 | 0,22 | 8,25E-01 |
| AT1G56520 | 1,21 | 3,16 | 2,29 | 1,38 | 1,68E-01 |
| AT1G56550 | 0,17 | 0,09 | 4,35 | 0,02 | 9,84E-01 |
| AT1G56580 | 0,24 | 0,65 | 4,37 | 0,15 | 8,81E-01 |
| AT1G56650 | 1,01 | 4,07 | 3,90 | 1,04 | 2,96E-01 |
| AT1G57680 | 1,45 | 2,00 | 1,90 | 1,05 | 2,92E-01 |
| AT1G57775 | 0,38 | -0,48 | 3,43 | -0,14 | 8,89E-01 |
| AT1G57777 | 0,31 | 1,92 | 3,61 | 0,53 | 5,94E-01 |
| AT1G57860 | 0,30 | -0,77 | 4,37 | -0,18 | 8,60E-01 |
| AT1G57870 | 0,40 | 0,48 | 4,26 | 0,11 | 9,10E-01 |
| AT1G57943 | 0,08 | 0,85 | 4,38 | 0,20 | 8,45E-01 |
| AT1G57990 | 0,77 | 2,54 | 2,53 | 1,00 | 3,15E-01 |
| AT1G58030 | 0,73 | 1,16 | 2,51 | 0,46 | 6,45E-01 |
| AT1G58060 | 0,60 | 2,47 | 3,09 | 0,80 | 4,25E-01 |
| AT1G58080 | 0,29 | 2,01 | 4,33 | 0,46 | 6,43E-01 |
| AT1G58100 | 1,20 | 0,92 | 1,98 | 0,46 | 6,44E-01 |
| AT1G58110 | 0,53 | 0,51 | 3,90 | 0,13 | 8,97E-01 |
| AT1G58200 | 0,16 | 0,00 | 4,41 | 0,00 | 1,00E+00 |
| AT1G58220 | 0,54 | 0,46 | 3,90 | 0,12 | 9,06E-01 |
| AT1G58300 | 1,33 | -3,17 | 1,75 | -1,81 | 7,05E-02 |
| AT1G58350 | 0,80 | 1,68 | 2,62 | 0,64 | 5,22E-01 |
| AT1G58390 | 0,70 | -0,68 | 3,05 | -0,22 | 8,23E-01 |
| AT1G58440 | 0,58 | -2,69 | 3,04 | -0,88 | 3,77E-01 |
| AT1G58684 | 0,49 | -2,31 | 4,35 | -0,53 | 5,96E-01 |
| AT1G58725 | 0,33 | -0,35 | 4,36 | -0,08 | 9,36E-01 |
| AT1G58766 | 0,16 | 1,33 | 4,35 | 0,31 | 7,59E-01 |
| AT1G59030 | 0,44 | 0,15 | 4,23 | 0,03 | 9,72E-01 |
| AT1G59077 | 0,16 | -1,36 | 4,41 | -0,31 | 7,57E-01 |
| AT1G59406 | 0,14 | 0,44 | 4,39 | 0,10 | 9,20E-01 |
| AT1G59453 | 0,25 | 1,02 | 4,36 | 0,23 | 8,15E-01 |
| AT1G59720 | 0,61 | 0,27 | 2,75 | 0,10 | 9,23E-01 |
| AT1G59780 | 0,52 | -0,72 | 3,20 | -0,22 | 8,23E-01 |
| AT1G59835 | 1,47 | -4,50 | 1,82 | -2,47 | 1,36E-02 |
| AT1G59910 | 1,37 | 3,03 | 2,10 | 1,45 | 1,48E-01 |
| AT1G59920 | 0,42 | -0,51 | 3,35 | -0,15 | 8,79E-01 |
| AT1G59930 | 1,11 | 3,04 | 2,39 | 1,27 | 2,03E-01 |
| AT1G59990 | 1,13 | 1,44 | 2,15 | 0,67 | 5,03E-01 |
| AT1G60000 | 0,38 | -0,01 | 3,52 | 0,00 | 9,97E-01 |
| AT1G60070 | 1,35 | 3,84 | 2,04 | 1,88 | 6,02E-02 |
| AT1G60160 | 0,43 | 1,93 | 3,21 | 0,60 | 5,48E-01 |
| AT1G60200 | 0,68 | 0,49 | 3,29 | 0,15 | 8,82E-01 |
| AT1G60230 | 0,48 | 0,00 | 4,09 | 0,00 | 1,00E+00 |
| AT1G60260 | 0,77 | -1,15 | 2,53 | -0,46 | 6,49E-01 |

| | | | | | |
|---|---|---|---|---|---|
| AT1G08317 | 0,08 | 0,85 | 4,38 | 0,20 | 8,45E-01 |
| AT1G60430 | 0,45 | 2,10 | 3,98 | 0,53 | 5,97E-01 |
| AT1G60490 | 0,25 | 0,64 | 4,33 | 0,15 | 8,82E-01 |
| AT1G60550 | 0,38 | 0,90 | 4,38 | 0,21 | 8,37E-01 |
| AT1G60560 | 0,17 | 1,81 | 4,33 | 0,42 | 6,76E-01 |
| AT1G60590 | 0,17 | -2,23 | 4,35 | -0,51 | 6,08E-01 |
| AT1G60630 | 0,08 | 0,85 | 4,38 | 0,20 | 8,45E-01 |
| AT1G60640 | 0,12 | 1,39 | 4,34 | 0,32 | 7,49E-01 |
| AT1G60700 | 0,48 | 2,55 | 4,32 | 0,59 | 5,55E-01 |
| AT1G60710 | 0,41 | 0,45 | 4,39 | 0,10 | 9,18E-01 |
| AT1G60770 | 1,20 | 2,94 | 2,11 | 1,39 | 1,64E-01 |
| AT1G60780 | 0,33 | 1,64 | 4,33 | 0,38 | 7,04E-01 |
| AT1G60800 | 0,56 | -0,70 | 2,99 | -0,23 | 8,16E-01 |
| AT1G60810 | 0,12 | 0,00 | 4,41 | 0,00 | 1,00E+00 |
| AT1G60850 | 0,11 | 1,07 | 4,36 | 0,25 | 8,06E-01 |
| AT1G60890 | 0,24 | 1,11 | 4,32 | 0,26 | 7,98E-01 |
| AT1G60930 | 0,24 | 1,23 | 4,35 | 0,28 | 7,77E-01 |
| AT1G60940 | 0,93 | -0,09 | 2,20 | -0,04 | 9,68E-01 |
| AT1G60960 | 1,57 | 1,09 | 1,80 | 0,60 | 5,47E-01 |
| AT1G60990 | 0,24 | 1,58 | 4,34 | 0,36 | 7,16E-01 |
| AT1G60995 | 0,35 | 0,44 | 4,39 | 0,10 | 9,20E-01 |
| AT1G61010 | 0,89 | 0,53 | 2,43 | 0,22 | 8,28E-01 |
| AT1G61200 | 1,13 | -1,17 | 1,59 | -0,74 | 4,60E-01 |
| AT1G61260 | 0,47 | 1,23 | 3,28 | 0,37 | 7,08E-01 |
| AT1G61300 | 1,06 | -0,13 | 2,19 | -0,06 | 9,52E-01 |
| AT1G61310 | 0,33 | 0,29 | 4,35 | 0,07 | 9,46E-01 |
| AT1G61360 | 0,15 | 0,86 | 4,37 | 0,20 | 8,43E-01 |
| AT1G61370 | 0,17 | 0,76 | 4,37 | 0,18 | 8,61E-01 |
| AT1G61380 | 0,34 | 1,97 | 4,32 | 0,45 | 6,49E-01 |
| AT1G61420 | 0,12 | 1,46 | 4,35 | 0,34 | 7,36E-01 |
| AT1G61560 | 0,37 | 1,94 | 4,33 | 0,45 | 6,54E-01 |
| AT1G08417 | 0,75 | 0,00 | 2,95 | 0,00 | 1,00E+00 |
| AT1G61660 | 0,65 | 1,13 | 2,93 | 0,39 | 6,99E-01 |
| AT1G61730 | 1,18 | 0,62 | 2,09 | 0,29 | 7,68E-01 |
| AT1G61890 | 1,56 | 3,38 | 1,85 | 1,82 | 6,84E-02 |
| AT1G61900 | 0,26 | -1,21 | 4,37 | -0,28 | 7,81E-01 |
| AT1G62030 | 1,53 | 0,06 | 1,84 | 0,03 | 9,73E-01 |
| AT1G62085 | 0,60 | 2,17 | 3,18 | 0,68 | 4,95E-01 |
| AT1G62300 | 0,27 | 0,00 | 4,41 | 0,00 | 1,00E+00 |
| AT1G62422 | 0,51 | 1,32 | 2,63 | 0,50 | 6,15E-01 |
| AT1G62610 | 0,76 | 1,78 | 2,39 | 0,74 | 4,57E-01 |
| AT1G62680 | 1,24 | 0,78 | 1,86 | 0,42 | 6,75E-01 |
| AT1G62760 | 0,29 | 1,64 | 4,33 | 0,38 | 7,06E-01 |
| AT1G62780 | 0,49 | -0,62 | 3,24 | -0,19 | 8,48E-01 |
| AT1G62810 | 0,97 | 2,32 | 2,02 | 1,15 | 2,51E-01 |
| AT1G62915 | 0,64 | -1,72 | 2,91 | -0,59 | 5,56E-01 |
| AT1G62960 | 0,36 | -1,46 | 4,40 | -0,33 | 7,40E-01 |
| AT1G63010 | 0,16 | -1,36 | 4,41 | -0,31 | 7,57E-01 |
| AT1G63020 | 0,44 | 2,79 | 4,31 | 0,65 | 5,16E-01 |
| AT1G63050 | 0,34 | 0,77 | 4,37 | 0,18 | 8,60E-01 |
| AT1G63090 | 1,34 | 0,72 | 1,61 | 0,45 | 6,55E-01 |
| AT1G63250 | 0,13 | 1,15 | 4,36 | 0,26 | 7,92E-01 |
| AT1G63290 | 0,12 | 1,39 | 4,34 | 0,32 | 7,49E-01 |
| AT1G63320 | 0,20 | 0,00 | 4,41 | 0,00 | 1,00E+00 |
| AT1G63410 | 0,62 | 0,68 | 2,56 | 0,27 | 7,89E-01 |
| AT1G63490 | 0,53 | 1,93 | 3,20 | 0,60 | 5,46E-01 |

| | | | | | |
|---|---|---|---|---|---|
| AT1G63660 | 0,92 | 0,83 | 2,22 | 0,37 | 7,10E-01 |
| AT1G63670 | 0,18 | 0,00 | 4,41 | 0,00 | 1,00E+00 |
| AT1G63700 | 1,36 | 2,75 | 1,61 | 1,71 | 8,69E-02 |
| AT1G63720 | 0,19 | 0,79 | 4,37 | 0,18 | 8,57E-01 |
| AT1G08627 | 1,18 | -3,05 | 1,96 | -1,56 | 1,19E-01 |
| AT1G63800 | 0,47 | -1,22 | 4,37 | -0,28 | 7,81E-01 |
| AT1G63810 | 0,31 | -0,60 | 4,36 | -0,14 | 8,91E-01 |
| AT1G63840 | 0,40 | 1,87 | 4,35 | 0,43 | 6,67E-01 |
| AT1G63860 | 1,09 | 1,27 | 1,98 | 0,64 | 5,21E-01 |
| AT1G63870 | 0,58 | 1,58 | 3,04 | 0,52 | 6,02E-01 |
| AT1G63970 | 0,51 | 0,38 | 3,25 | 0,12 | 9,07E-01 |
| AT1G64080 | 0,15 | 0,42 | 4,38 | 0,09 | 9,24E-01 |
| AT1G64190 | 1,11 | 0,88 | 1,78 | 0,49 | 6,23E-01 |
| AT1G64200 | 0,78 | 0,91 | 4,29 | 0,21 | 8,33E-01 |
| AT1G64280 | 0,86 | 3,14 | 2,50 | 1,26 | 2,08E-01 |
| AT1G64380 | 0,37 | -1,50 | 4,39 | -0,34 | 7,33E-01 |
| AT1G64460 | 0,72 | 2,42 | 2,44 | 0,99 | 3,21E-01 |
| AT1G64480 | 0,18 | 0,00 | 4,40 | 0,00 | 1,00E+00 |
| AT1G64510 | 0,32 | -0,55 | 4,38 | -0,13 | 9,00E-01 |
| AT1G64580 | 0,24 | -0,21 | 4,35 | -0,05 | 9,61E-01 |
| AT1G64610 | 0,62 | -0,30 | 2,71 | -0,11 | 9,12E-01 |
| AT1G64640 | 0,11 | 0,90 | 4,37 | 0,21 | 8,36E-01 |
| AT1G64650 | 0,59 | 1,90 | 2,48 | 0,76 | 4,45E-01 |
| AT1G64670 | 0,10 | 0,62 | 4,38 | 0,14 | 8,87E-01 |
| AT1G64750 | 0,36 | -0,61 | 4,37 | -0,14 | 8,88E-01 |
| AT1G64970 | 0,63 | 1,07 | 2,56 | 0,42 | 6,78E-01 |
| AT1G64980 | 1,26 | -2,44 | 1,76 | -1,38 | 1,66E-01 |
| AT1G65040 | 0,28 | -0,36 | 3,72 | -0,10 | 9,22E-01 |
| AT1G65150 | 0,20 | 0,82 | 4,33 | 0,19 | 8,50E-01 |
| AT1G65190 | 0,43 | -1,23 | 4,09 | -0,30 | 7,63E-01 |
| AT1G08747 | 0,81 | 0,01 | 2,91 | 0,00 | 9,97E-01 |
| AT1G65290 | 0,77 | 1,18 | 2,22 | 0,53 | 5,96E-01 |
| AT1G65320 | 0,39 | 0,85 | 4,29 | 0,20 | 8,43E-01 |
| AT1G65346 | 0,34 | 1,28 | 4,34 | 0,29 | 7,68E-01 |
| AT1G65420 | 0,15 | 0,24 | 4,35 | 0,05 | 9,56E-01 |
| AT1G65430 | 0,52 | 1,39 | 3,09 | 0,45 | 6,53E-01 |
| AT1G65520 | 0,29 | -0,90 | 4,34 | -0,21 | 8,36E-01 |
| AT1G65540 | 1,25 | 1,64 | 1,84 | 0,89 | 3,74E-01 |
| AT1G65550 | 0,21 | 0,66 | 4,34 | 0,15 | 8,79E-01 |
| AT1G65580 | 0,46 | -1,18 | 2,76 | -0,43 | 6,69E-01 |
| AT1G65690 | 0,12 | 1,65 | 4,34 | 0,38 | 7,04E-01 |
| AT1G65950 | 0,28 | 0,42 | 4,38 | 0,10 | 9,24E-01 |
| AT1G08797 | 0,63 | 2,87 | 4,30 | 0,67 | 5,05E-01 |
| AT1G65980 | 1,27 | 1,82 | 2,22 | 0,82 | 4,13E-01 |
| AT1G65985 | 0,33 | 0,30 | 4,33 | 0,07 | 9,44E-01 |
| AT1G65990 | 1,24 | 2,91 | 2,07 | 1,40 | 1,61E-01 |
| AT1G66200 | 1,38 | -1,11 | 1,67 | -0,66 | 5,07E-01 |
| AT1G66235 | 0,12 | 0,42 | 4,38 | 0,10 | 9,24E-01 |
| AT1G66240 | 0,12 | 1,39 | 4,34 | 0,32 | 7,49E-01 |
| AT1G66260 | 1,28 | 2,72 | 1,68 | 1,62 | 1,05E-01 |
| AT1G66290 | 0,49 | -2,31 | 3,14 | -0,74 | 4,62E-01 |
| AT1G66330 | 0,52 | 0,17 | 2,66 | 0,06 | 9,50E-01 |
| AT1G66410 | 1,68 | -0,83 | 1,68 | -0,49 | 6,24E-01 |
| AT1G66460 | 0,25 | 0,96 | 4,33 | 0,22 | 8,24E-01 |
| AT1G66490 | 0,90 | -2,56 | 2,43 | -1,05 | 2,92E-01 |
| AT1G66500 | 0,25 | 0,00 | 4,40 | 0,00 | 1,00E+00 |

| | | | | | |
|---|---|---|---|---|---|
| AT1G66570 | 1,67 | 0,24 | 1,52 | 0,16 | 8,73E-01 |
| AT1G66660 | 0,44 | 1,49 | 4,16 | 0,36 | 7,21E-01 |
| AT1G66670 | 1,37 | 1,80 | 1,88 | 0,96 | 3,38E-01 |
| AT1G66680 | 0,59 | 2,02 | 4,33 | 0,47 | 6,41E-01 |
| AT1G66700 | 0,46 | 2,54 | 4,32 | 0,59 | 5,56E-01 |
| AT1G66830 | 0,68 | -0,30 | 3,78 | -0,08 | 9,37E-01 |
| AT1G66910 | 0,65 | 1,81 | 2,26 | 0,80 | 4,24E-01 |
| AT1G66940 | 0,34 | 0,00 | 4,40 | 0,00 | 1,00E+00 |
| AT1G66980 | 1,51 | -0,35 | 1,58 | -0,22 | 8,26E-01 |
| AT1G67000 | 0,91 | -1,43 | 2,01 | -0,71 | 4,75E-01 |
| AT1G67050 | 0,28 | -0,34 | 3,71 | -0,09 | 9,27E-01 |
| AT1G67080 | 0,36 | 1,28 | 4,34 | 0,29 | 7,68E-01 |
| AT1G67140 | 0,95 | 2,88 | 2,52 | 1,14 | 2,53E-01 |
| AT1G67230 | 0,62 | -0,03 | 2,55 | -0,01 | 9,92E-01 |
| AT1G67235 | 0,29 | 0,00 | 4,41 | 0,00 | 1,00E+00 |
| AT1G67250 | 0,34 | -1,62 | 4,38 | -0,37 | 7,12E-01 |
| AT1G67300 | 0,37 | 1,85 | 4,30 | 0,43 | 6,67E-01 |
| AT1G67310 | 0,33 | 2,20 | 4,32 | 0,51 | 6,11E-01 |
| AT1G67325 | 0,61 | 2,96 | 2,96 | 1,00 | 3,18E-01 |
| AT1G67360 | 0,73 | -2,93 | 2,89 | -1,01 | 3,11E-01 |
| AT1G67481 | 1,30 | 2,59 | 1,99 | 1,31 | 1,92E-01 |
| AT1G67490 | 0,57 | 0,43 | 3,19 | 0,14 | 8,93E-01 |
| AT1G67500 | 0,52 | 1,57 | 4,00 | 0,39 | 6,95E-01 |
| AT1G67510 | 0,38 | -0,21 | 4,27 | -0,05 | 9,62E-01 |
| AT1G67580 | 1,21 | 1,89 | 2,01 | 0,94 | 3,48E-01 |
| AT1G67590 | 0,17 | 0,00 | 4,40 | 0,00 | 1,00E+00 |
| AT1G67680 | 0,43 | 2,06 | 4,32 | 0,48 | 6,33E-01 |
| AT1G67785 | 0,15 | -1,24 | 4,39 | -0,28 | 7,77E-01 |
| AT1G67790 | 0,26 | 1,08 | 4,37 | 0,25 | 8,05E-01 |
| AT1G67860 | 0,58 | 1,82 | 3,08 | 0,59 | 5,55E-01 |
| AT1G67865 | 0,24 | 0,87 | 4,33 | 0,20 | 8,40E-01 |
| AT1G67870 | 0,89 | -1,17 | 1,95 | -0,60 | 5,48E-01 |
| AT1G67930 | 1,11 | 1,95 | 1,78 | 1,10 | 2,72E-01 |
| AT1G67950 | 0,39 | 0,48 | 4,40 | 0,11 | 9,14E-01 |
| AT1G68060 | 0,56 | 0,14 | 3,18 | 0,04 | 9,65E-01 |
| AT1G68140 | 0,87 | 3,24 | 2,12 | 1,53 | 1,26E-01 |
| AT1G68325 | 0,05 | 0,00 | 4,41 | 0,00 | 1,00E+00 |
| AT1G68370 | 0,32 | 0,87 | 4,33 | 0,20 | 8,40E-01 |
| AT1G68550 | 0,87 | 2,01 | 2,01 | 1,00 | 3,19E-01 |
| AT1G08997 | 0,58 | 0,48 | 3,75 | 0,13 | 8,98E-01 |
| AT1G68590 | 1,41 | 2,82 | 1,86 | 1,51 | 1,30E-01 |
| AT1G68660 | 0,71 | -0,99 | 2,19 | -0,45 | 6,51E-01 |
| AT1G68710 | 0,46 | 2,16 | 4,33 | 0,50 | 6,18E-01 |
| AT1G68830 | 0,57 | 0,30 | 2,59 | 0,12 | 9,06E-01 |
| AT1G68910 | 0,08 | 1,10 | 4,36 | 0,25 | 8,00E-01 |
| AT1G69020 | 0,30 | 1,68 | 4,34 | 0,39 | 6,99E-01 |
| AT1G69070 | 0,46 | 1,57 | 3,30 | 0,48 | 6,34E-01 |
| AT1G69220 | 1,28 | 2,23 | 1,95 | 1,15 | 2,52E-01 |
| AT1G69270 | 0,78 | 0,73 | 3,06 | 0,24 | 8,12E-01 |
| AT1G69280 | 0,21 | 0,66 | 4,34 | 0,15 | 8,79E-01 |
| AT1G69290 | 0,15 | 1,62 | 4,34 | 0,37 | 7,09E-01 |
| AT1G69310 | 0,25 | 2,24 | 4,32 | 0,52 | 6,04E-01 |
| AT1G69450 | 0,38 | 0,00 | 4,37 | 0,00 | 1,00E+00 |
| AT1G69480 | 0,55 | 1,69 | 4,32 | 0,39 | 6,95E-01 |
| AT1G69630 | 0,59 | 0,00 | 4,41 | 0,00 | 1,00E+00 |
| AT1G69640 | 0,90 | 0,75 | 2,35 | 0,32 | 7,48E-01 |

| | | | | | |
|---|---|---|---|---|---|
| AT1G69730 | 0,73 | 2,32 | 2,37 | 0,98 | 3,27E-01 |
| AT1G69760 | 0,47 | -1,25 | 3,25 | -0,39 | 6,99E-01 |
| AT1G69770 | 1,05 | 1,96 | 2,33 | 0,84 | 4,02E-01 |
| AT1G69795 | 1,39 | 0,48 | 2,12 | 0,23 | 8,20E-01 |
| AT1G69840 | 0,65 | 1,63 | 3,05 | 0,53 | 5,93E-01 |
| AT1G69850 | 0,22 | 1,24 | 4,34 | 0,29 | 7,75E-01 |
| AT1G69890 | 1,49 | 2,89 | 1,86 | 1,56 | 1,20E-01 |
| AT1G69960 | 0,51 | 0,99 | 3,15 | 0,32 | 7,52E-01 |
| AT1G70000 | 0,98 | 2,07 | 2,02 | 1,02 | 3,06E-01 |
| AT1G70070 | 1,07 | -1,55 | 2,02 | -0,77 | 4,41E-01 |
| AT1G70200 | 0,37 | -1,21 | 4,38 | -0,28 | 7,82E-01 |
| AT1G70280 | 0,17 | 1,81 | 4,33 | 0,42 | 6,76E-01 |
| AT1G70300 | 0,38 | 1,80 | 4,27 | 0,42 | 6,74E-01 |
| AT1G70310 | 0,89 | -1,17 | 2,96 | -0,40 | 6,92E-01 |
| AT1G70340 | 0,34 | 0,12 | 3,56 | 0,03 | 9,73E-01 |
| AT1G70370 | 0,57 | -0,78 | 2,64 | -0,30 | 7,68E-01 |
| AT1G70410 | 0,70 | -2,16 | 2,69 | -0,80 | 4,22E-01 |
| AT1G70420 | 0,44 | -2,48 | 4,07 | -0,61 | 5,43E-01 |
| AT1G70520 | 0,53 | 0,26 | 4,28 | 0,06 | 9,52E-01 |
| AT1G70600 | 0,97 | 1,67 | 2,86 | 0,58 | 5,60E-01 |
| AT1G70730 | 0,91 | 0,98 | 2,27 | 0,43 | 6,65E-01 |
| AT1G70740 | 0,10 | 0,00 | 4,41 | 0,00 | 1,00E+00 |
| AT1G70750 | 0,44 | 2,45 | 4,32 | 0,57 | 5,70E-01 |
| AT1G70780 | 0,73 | 1,63 | 2,77 | 0,59 | 5,58E-01 |
| AT1G70830 | 0,27 | 1,87 | 4,33 | 0,43 | 6,65E-01 |
| AT1G70890 | 1,35 | -2,82 | 1,65 | -1,71 | 8,75E-02 |
| AT1G71020 | 0,94 | 2,46 | 2,32 | 1,06 | 2,90E-01 |
| AT1G71030 | 0,60 | -1,17 | 2,44 | -0,48 | 6,32E-01 |
| AT1G71040 | 0,83 | 2,09 | 2,10 | 1,00 | 3,19E-01 |
| AT1G71060 | 0,73 | 3,36 | 3,39 | 0,99 | 3,22E-01 |
| AT1G71070 | 0,49 | 0,79 | 3,14 | 0,25 | 8,00E-01 |
| AT1G71080 | 0,90 | 2,74 | 2,87 | 0,95 | 3,40E-01 |
| AT1G71160 | 0,69 | 2,94 | 3,53 | 0,83 | 4,05E-01 |
| AT1G71190 | 0,19 | 1,51 | 4,34 | 0,35 | 7,28E-01 |
| AT1G71210 | 0,73 | 2,79 | 2,88 | 0,97 | 3,33E-01 |
| AT1G71310 | 0,27 | 0,40 | 4,32 | 0,09 | 9,26E-01 |
| AT1G71330 | 0,30 | 1,45 | 4,35 | 0,33 | 7,39E-01 |
| AT1G71340 | 0,92 | 1,25 | 2,46 | 0,51 | 6,13E-01 |
| AT1G71350 | 0,21 | 2,17 | 4,32 | 0,50 | 6,15E-01 |
| AT1G71410 | 1,45 | 2,01 | 2,06 | 0,98 | 3,28E-01 |
| AT1G09253 | 0,32 | 1,59 | 4,34 | 0,37 | 7,14E-01 |
| AT1G71490 | 0,42 | 2,30 | 4,16 | 0,55 | 5,80E-01 |
| AT1G71710 | 0,44 | -0,49 | 2,81 | -0,18 | 8,60E-01 |
| AT1G71770 | 0,40 | 2,24 | 4,13 | 0,54 | 5,87E-01 |
| AT1G71870 | 0,88 | 2,25 | 2,81 | 0,80 | 4,23E-01 |
| AT1G71960 | 0,26 | 0,73 | 3,75 | 0,20 | 8,45E-01 |
| AT1G72020 | 0,56 | 0,15 | 2,65 | 0,06 | 9,54E-01 |
| AT1G72040 | 0,83 | 1,39 | 2,13 | 0,65 | 5,15E-01 |
| AT1G72160 | 0,64 | -1,33 | 2,32 | -0,57 | 5,66E-01 |
| AT1G72190 | 0,52 | 1,71 | 3,83 | 0,45 | 6,55E-01 |
| AT1G72270 | 0,12 | 0,79 | 4,37 | 0,18 | 8,57E-01 |
| AT1G72300 | 0,54 | 2,07 | 3,07 | 0,68 | 5,00E-01 |
| AT1G72320 | 0,39 | 1,24 | 3,44 | 0,36 | 7,18E-01 |
| AT1G72450 | 1,12 | -0,48 | 1,69 | -0,29 | 7,75E-01 |
| AT1G72480 | 0,35 | 0,80 | 4,34 | 0,19 | 8,53E-01 |
| AT1G72500 | 0,46 | 0,00 | 4,40 | 0,00 | 1,00E+00 |

| | | | | | |
|---|---|---|---|---|---|
| AT1G72550 | 0,79 | -0,08 | 2,43 | -0,03 | 9,73E-01 |
| AT1G72560 | 1,09 | -1,66 | 2,37 | -0,70 | 4,85E-01 |
| AT1G09327 | 0,20 | 0,32 | 4,35 | 0,07 | 9,41E-01 |
| AT1G72670 | 0,20 | 0,89 | 4,37 | 0,20 | 8,39E-01 |
| AT1G72720 | 0,35 | 0,00 | 4,39 | 0,00 | 1,00E+00 |
| AT1G72730 | 0,32 | 1,25 | 4,35 | 0,29 | 7,74E-01 |
| AT1G72760 | 0,65 | 3,49 | 4,29 | 0,81 | 4,16E-01 |
| AT1G72840 | 0,41 | 2,61 | 4,32 | 0,60 | 5,46E-01 |
| AT1G72860 | 0,44 | 2,79 | 4,31 | 0,65 | 5,16E-01 |
| AT1G73010 | 0,40 | 2,57 | 4,32 | 0,60 | 5,51E-01 |
| AT1G09363 | 0,41 | 0,44 | 3,38 | 0,13 | 8,98E-01 |
| AT1G73110 | 0,63 | -0,36 | 3,74 | -0,10 | 9,24E-01 |
| AT1G73200 | 0,73 | 0,81 | 2,26 | 0,36 | 7,22E-01 |
| AT1G73230 | 0,17 | 1,94 | 4,33 | 0,45 | 6,55E-01 |
| AT1G73260 | 0,63 | 2,02 | 2,97 | 0,68 | 4,98E-01 |
| AT1G73370 | 0,52 | 3,11 | 4,30 | 0,72 | 4,69E-01 |
| AT1G73390 | 0,25 | 1,33 | 4,35 | 0,31 | 7,60E-01 |
| AT1G73490 | 0,37 | 1,76 | 4,33 | 0,41 | 6,85E-01 |
| AT1G73500 | 0,92 | -0,39 | 1,99 | -0,20 | 8,43E-01 |
| AT1G73510 | 0,45 | 1,23 | 4,07 | 0,30 | 7,63E-01 |
| AT1G73530 | 0,46 | -0,57 | 4,27 | -0,13 | 8,95E-01 |
| AT1G73540 | 0,81 | 1,38 | 2,22 | 0,62 | 5,33E-01 |
| AT1G73650 | 0,96 | 1,45 | 2,35 | 0,62 | 5,38E-01 |
| AT1G73655 | 0,74 | 2,72 | 3,73 | 0,73 | 4,65E-01 |
| AT1G73660 | 1,58 | 1,62 | 1,98 | 0,82 | 4,14E-01 |
| AT1G73680 | 0,85 | 1,36 | 2,05 | 0,66 | 5,07E-01 |
| AT1G73730 | 0,17 | 1,81 | 4,33 | 0,42 | 6,76E-01 |
| AT1G73760 | 0,39 | 0,89 | 4,28 | 0,21 | 8,35E-01 |
| AT1G73930 | 0,25 | 1,42 | 4,34 | 0,33 | 7,44E-01 |
| AT1G73950 | 0,45 | 0,65 | 4,35 | 0,15 | 8,81E-01 |
| AT1G73960 | 0,29 | 1,88 | 4,33 | 0,43 | 6,65E-01 |
| AT1G73980 | 0,67 | 2,17 | 2,59 | 0,84 | 4,03E-01 |
| AT1G74050 | 0,43 | 2,16 | 3,25 | 0,67 | 5,06E-01 |
| AT1G74060 | 0,72 | -1,86 | 4,37 | -0,43 | 6,70E-01 |
| AT1G74070 | 0,42 | -2,73 | 4,23 | -0,64 | 5,19E-01 |
| AT1G74120 | 0,82 | 1,08 | 2,89 | 0,37 | 7,08E-01 |
| AT1G74170 | 0,18 | 1,56 | 4,34 | 0,36 | 7,18E-01 |
| AT1G74340 | 0,31 | 0,79 | 4,33 | 0,18 | 8,55E-01 |
| AT1G74370 | 0,53 | 0,83 | 3,95 | 0,21 | 8,33E-01 |
| AT1G74390 | 0,95 | 1,87 | 2,65 | 0,71 | 4,80E-01 |
| AT1G74420 | 0,45 | 0,80 | 4,37 | 0,18 | 8,55E-01 |
| AT1G74440 | 0,80 | 0,17 | 3,04 | 0,06 | 9,56E-01 |
| AT1G09467 | 0,50 | 0,01 | 4,06 | 0,00 | 9,97E-01 |
| AT1G74560 | 0,37 | -0,59 | 3,48 | -0,17 | 8,66E-01 |
| AT1G74690 | 0,95 | 1,69 | 2,01 | 0,84 | 4,01E-01 |
| AT1G74770 | 0,42 | 1,63 | 4,17 | 0,39 | 6,97E-01 |
| AT1G74800 | 0,56 | 2,56 | 3,14 | 0,82 | 4,15E-01 |
| AT1G74830 | 0,12 | 1,58 | 4,34 | 0,36 | 7,16E-01 |
| AT1G74850 | 0,94 | 1,31 | 2,13 | 0,62 | 5,38E-01 |
| AT1G74880 | 0,11 | 1,07 | 4,36 | 0,25 | 8,06E-01 |
| AT1G74920 | 0,86 | -1,14 | 2,63 | -0,43 | 6,66E-01 |
| AT1G74950 | 0,28 | 0,51 | 4,37 | 0,12 | 9,08E-01 |
| AT1G74960 | 1,51 | 3,26 | 2,68 | 1,22 | 2,23E-01 |
| AT1G74970 | 0,46 | -1,20 | 4,34 | -0,28 | 7,83E-01 |
| AT1G75010 | 0,20 | 0,00 | 4,41 | 0,00 | 1,00E+00 |
| AT1G75100 | 0,60 | 0,80 | 2,27 | 0,35 | 7,25E-01 |

| | | | | | |
|---|---|---|---|---|---|
| AT1G75130 | 0,12 | 1,39 | 4,34 | 0,32 | 7,49E-01 |
| AT1G75180 | 0,38 | 0,37 | 3,36 | 0,11 | 9,12E-01 |
| AT1G75200 | 0,15 | 1,42 | 4,34 | 0,33 | 7,43E-01 |
| AT1G75220 | 0,55 | 2,73 | 3,15 | 0,87 | 3,85E-01 |
| AT1G75370 | 1,12 | -0,87 | 1,61 | -0,54 | 5,87E-01 |
| AT1G75410 | 0,90 | 0,84 | 2,42 | 0,35 | 7,28E-01 |
| AT1G75450 | 0,11 | 0,90 | 4,37 | 0,21 | 8,36E-01 |
| AT1G75500 | 0,97 | -0,18 | 1,83 | -0,10 | 9,22E-01 |
| AT1G75680 | 0,99 | -1,76 | 1,71 | -1,03 | 3,02E-01 |
| AT1G75690 | 0,84 | -1,60 | 1,98 | -0,81 | 4,19E-01 |
| AT1G75750 | 1,13 | 0,38 | 2,12 | 0,18 | 8,56E-01 |
| AT1G75780 | 0,25 | 0,29 | 4,34 | 0,07 | 9,47E-01 |
| AT1G75820 | 1,17 | 2,85 | 2,26 | 1,26 | 2,06E-01 |
| AT1G75860 | 0,67 | 0,58 | 2,31 | 0,25 | 8,01E-01 |
| AT1G75945 | 0,28 | 1,36 | 4,35 | 0,31 | 7,55E-01 |
| AT1G76060 | 0,20 | 0,00 | 4,41 | 0,00 | 1,00E+00 |
| AT1G76080 | 1,03 | -0,45 | 1,62 | -0,28 | 7,83E-01 |
| AT1G76100 | 1,22 | -0,25 | 2,04 | -0,12 | 9,02E-01 |
| AT1G76120 | 0,15 | 1,25 | 4,35 | 0,29 | 7,75E-01 |
| AT1G76150 | 0,72 | 3,34 | 2,86 | 1,17 | 2,42E-01 |
| AT1G76160 | 0,88 | 1,85 | 1,91 | 0,97 | 3,33E-01 |
| AT1G76270 | 0,08 | 0,85 | 4,38 | 0,20 | 8,45E-01 |
| AT1G76280 | 0,12 | 1,39 | 4,34 | 0,32 | 7,49E-01 |
| AT1G09627 | 0,45 | 2,37 | 4,01 | 0,59 | 5,54E-01 |
| AT1G76400 | 0,57 | 2,58 | 3,03 | 0,85 | 3,95E-01 |
| AT1G76460 | 0,53 | -1,15 | 3,24 | -0,36 | 7,21E-01 |
| AT1G76640 | 0,08 | 0,85 | 4,38 | 0,20 | 8,45E-01 |
| AT1G76650 | 0,42 | -0,45 | 3,41 | -0,13 | 8,95E-01 |
| AT1G76660 | 0,13 | -1,24 | 4,39 | -0,28 | 7,77E-01 |
| AT1G76670 | 1,15 | 1,22 | 2,15 | 0,57 | 5,69E-01 |
| AT1G76700 | 0,39 | -0,15 | 4,37 | -0,03 | 9,73E-01 |
| AT1G76810 | 1,64 | -1,61 | 1,43 | -1,12 | 2,61E-01 |
| AT1G76878 | 1,06 | -1,32 | 1,75 | -0,76 | 4,49E-01 |
| AT1G76890 | 0,74 | 0,72 | 2,05 | 0,35 | 7,26E-01 |
| AT1G76955 | 0,30 | 1,45 | 4,35 | 0,33 | 7,39E-01 |
| AT1G76960 | 0,40 | 0,09 | 3,43 | 0,03 | 9,78E-01 |
| AT1G76965 | 0,25 | 1,03 | 4,37 | 0,24 | 8,13E-01 |
| AT1G76970 | 0,43 | 0,25 | 3,29 | 0,08 | 9,40E-01 |
| AT1G76980 | 0,73 | 2,63 | 3,04 | 0,87 | 3,86E-01 |
| AT1G77030 | 0,32 | 0,47 | 4,39 | 0,11 | 9,15E-01 |
| AT1G77060 | 0,12 | 0,90 | 4,36 | 0,21 | 8,36E-01 |
| AT1G77100 | 0,82 | 2,90 | 3,65 | 0,79 | 4,28E-01 |
| AT1G77145 | 0,08 | 1,10 | 4,36 | 0,25 | 8,00E-01 |
| AT1G77170 | 0,63 | 2,71 | 2,95 | 0,92 | 3,58E-01 |
| AT1G77240 | 1,26 | 3,57 | 1,92 | 1,85 | 6,38E-02 |
| AT1G77250 | 0,55 | 2,32 | 3,76 | 0,62 | 5,36E-01 |
| AT1G77260 | 0,57 | 2,53 | 3,83 | 0,66 | 5,09E-01 |
| AT1G77480 | 0,33 | -2,50 | 4,36 | -0,57 | 5,66E-01 |
| AT1G77580 | 0,18 | 1,06 | 4,35 | 0,24 | 8,07E-01 |
| AT1G77600 | 0,35 | 1,90 | 4,32 | 0,44 | 6,61E-01 |
| AT1G77660 | 0,33 | 2,05 | 4,33 | 0,47 | 6,36E-01 |
| AT1G77810 | 0,58 | 0,23 | 3,85 | 0,06 | 9,51E-01 |
| AT1G77932 | 0,47 | 1,71 | 4,35 | 0,39 | 6,95E-01 |
| AT1G77940 | 0,55 | 1,39 | 2,57 | 0,54 | 5,89E-01 |
| AT1G77960 | 1,37 | 0,06 | 1,65 | 0,04 | 9,70E-01 |
| AT1G77980 | 0,77 | -1,34 | 2,19 | -0,61 | 5,41E-01 |

| | | | | | |
|---|---|---|---|---|---|
| AT1G77990 | 0,37 | 2,16 | 4,32 | 0,50 | 6,17E-01 |
| AT1G78060 | 0,43 | -2,05 | 4,35 | -0,47 | 6,38E-01 |
| AT1G78210 | 0,32 | 0,65 | 4,39 | 0,15 | 8,82E-01 |
| AT1G78240 | 0,51 | -0,47 | 2,68 | -0,17 | 8,61E-01 |
| AT1G78370 | 1,22 | 2,43 | 2,21 | 1,10 | 2,71E-01 |
| AT1G78380 | 1,48 | -1,33 | 1,61 | -0,83 | 4,09E-01 |
| AT1G78460 | 0,60 | -1,51 | 2,56 | -0,59 | 5,55E-01 |
| AT1G78480 | 0,26 | 1,44 | 4,34 | 0,33 | 7,39E-01 |
| AT1G78510 | 0,69 | -0,68 | 3,72 | -0,18 | 8,55E-01 |
| AT1G78580 | 0,70 | 0,45 | 2,17 | 0,21 | 8,36E-01 |
| AT1G78800 | 0,21 | 1,97 | 4,33 | 0,46 | 6,49E-01 |
| AT1G78820 | 0,54 | -0,98 | 2,25 | -0,44 | 6,63E-01 |
| AT1G78850 | 0,94 | -2,95 | 1,93 | -1,53 | 1,25E-01 |
| AT1G78865 | 0,13 | 0,00 | 4,41 | 0,00 | 1,00E+00 |
| AT1G78880 | 0,88 | 1,55 | 2,72 | 0,57 | 5,70E-01 |
| AT1G78915 | 0,18 | -1,25 | 4,39 | -0,28 | 7,76E-01 |
| AT1G78990 | 0,38 | 0,96 | 4,37 | 0,22 | 8,27E-01 |
| AT1G78995 | 0,24 | 0,44 | 4,35 | 0,10 | 9,19E-01 |
| AT1G79020 | 0,25 | 0,00 | 4,40 | 0,00 | 1,00E+00 |
| AT1G79030 | 0,53 | -1,33 | 4,03 | -0,33 | 7,42E-01 |
| AT1G79050 | 0,58 | -0,47 | 3,17 | -0,15 | 8,83E-01 |
| AT1G79080 | 1,23 | 2,67 | 2,16 | 1,24 | 2,15E-01 |
| AT1G79120 | 0,10 | 0,00 | 4,41 | 0,00 | 1,00E+00 |
| AT1G79150 | 0,35 | 0,72 | 4,31 | 0,17 | 8,68E-01 |
| AT1G79350 | 0,49 | 1,60 | 2,72 | 0,59 | 5,57E-01 |
| AT1G79430 | 0,17 | 0,00 | 4,40 | 0,00 | 1,00E+00 |
| AT1G79440 | 0,39 | 0,78 | 3,50 | 0,22 | 8,24E-01 |
| AT1G09867 | 1,21 | -0,53 | 2,21 | -0,24 | 8,10E-01 |
| AT1G09883 | 0,79 | -1,36 | 3,02 | -0,45 | 6,53E-01 |
| AT1G79540 | 0,53 | 2,18 | 3,11 | 0,70 | 4,84E-01 |
| AT1G79600 | 1,43 | 2,53 | 1,88 | 1,35 | 1,77E-01 |
| AT1G79640 | 0,40 | 1,87 | 4,35 | 0,43 | 6,67E-01 |
| AT1G79670 | 0,90 | 2,66 | 2,55 | 1,04 | 2,98E-01 |
| AT1G79700 | 0,08 | 1,10 | 4,36 | 0,25 | 8,00E-01 |
| AT1G79720 | 0,47 | 0,20 | 3,23 | 0,06 | 9,50E-01 |
| AT1G79730 | 0,63 | 1,16 | 3,86 | 0,30 | 7,64E-01 |
| AT1G79740 | 0,25 | 0,00 | 4,40 | 0,00 | 1,00E+00 |
| AT1G79790 | 0,05 | 0,00 | 4,41 | 0,00 | 1,00E+00 |
| AT1G79830 | 1,06 | 0,49 | 1,79 | 0,28 | 7,83E-01 |
| AT1G79880 | 0,11 | 1,07 | 4,36 | 0,25 | 8,06E-01 |
| AT1G79900 | 0,42 | 1,63 | 4,17 | 0,39 | 6,97E-01 |
| AT1G80030 | 1,07 | 0,24 | 2,04 | 0,12 | 9,08E-01 |
| AT1G80040 | 0,61 | -0,03 | 2,60 | -0,01 | 9,90E-01 |
| AT1G80080 | 0,50 | 1,58 | 3,15 | 0,50 | 6,16E-01 |
| AT1G80325 | 1,08 | 0,72 | 1,95 | 0,37 | 7,12E-01 |
| AT1G80350 | 0,38 | 0,96 | 4,37 | 0,22 | 8,27E-01 |
| AT1G80420 | 0,34 | 0,15 | 4,34 | 0,04 | 9,72E-01 |
| AT1G80460 | 0,24 | 1,09 | 4,35 | 0,25 | 8,02E-01 |
| AT1G80480 | 0,56 | 2,37 | 3,13 | 0,76 | 4,48E-01 |
| AT1G80490 | 1,23 | 1,46 | 1,82 | 0,81 | 4,20E-01 |
| AT1G80530 | 0,68 | 0,65 | 2,88 | 0,23 | 8,21E-01 |
| AT1G80600 | 1,40 | -1,45 | 1,90 | -0,76 | 4,45E-01 |
| AT1G80610 | 0,35 | 1,57 | 3,46 | 0,45 | 6,50E-01 |
| AT1G80630 | 0,28 | 0,90 | 4,38 | 0,21 | 8,37E-01 |
| AT1G80660 | 1,14 | 2,68 | 2,04 | 1,31 | 1,89E-01 |
| AT1G80670 | 0,39 | 0,48 | 4,40 | 0,11 | 9,14E-01 |

| | | | | | |
|---|---|---|---|---|---|
| AT1G80760 | 0,62 | 1,18 | 3,75 | 0,31 | 7,54E-01 |
| AT1G80830 | 0,27 | 0,54 | 4,37 | 0,12 | 9,01E-01 |
| AT1G80850 | 0,20 | 0,00 | 4,41 | 0,00 | 1,00E+00 |
| AT1G80920 | 0,74 | 2,35 | 2,18 | 1,08 | 2,82E-01 |
| AT1G80930 | 0,69 | 0,61 | 2,46 | 0,25 | 8,05E-01 |
| AT1G80950 | 0,35 | 0,95 | 4,37 | 0,22 | 8,28E-01 |
| AT2G01023 | 0,35 | -0,49 | 2,89 | -0,17 | 8,65E-01 |
| AT2G01050 | 0,45 | 0,79 | 4,39 | 0,18 | 8,57E-01 |
| AT2G01060 | 0,57 | 2,18 | 2,60 | 0,84 | 4,03E-01 |
| AT2G01110 | 0,90 | 3,07 | 2,42 | 1,27 | 2,04E-01 |
| AT2G01130 | 1,10 | 1,57 | 1,96 | 0,80 | 4,24E-01 |
| AT2G01140 | 1,55 | 4,00 | 1,89 | 2,12 | 3,39E-02 |
| AT2G01180 | 0,86 | -0,23 | 1,91 | -0,12 | 9,06E-01 |
| AT2G01250 | 0,52 | 0,99 | 3,88 | 0,25 | 7,99E-01 |
| AT2G01275 | 0,19 | 0,41 | 4,38 | 0,09 | 9,25E-01 |
| AT2G01290 | 0,18 | 1,66 | 4,33 | 0,38 | 7,02E-01 |
| AT2G01300 | 0,08 | 0,85 | 4,38 | 0,20 | 8,45E-01 |
| AT2G01310 | 0,05 | 0,00 | 4,41 | 0,00 | 1,00E+00 |
| AT2G01350 | 0,12 | 0,00 | 4,41 | 0,00 | 1,00E+00 |
| AT2G01410 | 0,34 | -0,49 | 3,57 | -0,14 | 8,91E-01 |
| AT2G01420 | 1,24 | -2,50 | 1,79 | -1,40 | 1,61E-01 |
| AT2G01440 | 1,25 | 3,45 | 2,45 | 1,41 | 1,58E-01 |
| AT2G01450 | 1,15 | 2,67 | 1,83 | 1,46 | 1,45E-01 |
| AT2G01505 | 0,30 | 1,63 | 4,35 | 0,38 | 7,07E-01 |
| AT2G01630 | 0,87 | 2,38 | 2,79 | 0,85 | 3,95E-01 |
| AT2G01650 | 1,03 | 2,68 | 2,16 | 1,24 | 2,16E-01 |
| AT2G01670 | 0,39 | 0,98 | 3,30 | 0,30 | 7,67E-01 |
| AT2G01690 | 0,55 | 0,43 | 3,84 | 0,11 | 9,10E-01 |
| AT2G01720 | 1,57 | 4,09 | 2,19 | 1,86 | 6,26E-02 |
| AT2G01755 | 0,14 | 0,44 | 4,39 | 0,10 | 9,20E-01 |
| AT2G01820 | 1,26 | 2,23 | 2,34 | 0,95 | 3,42E-01 |
| AT2G01830 | 0,91 | 0,20 | 1,90 | 0,11 | 9,15E-01 |
| AT2G04125 | 0,12 | 0,42 | 4,38 | 0,10 | 9,24E-01 |
| AT2G01870 | 0,33 | 0,95 | 4,37 | 0,22 | 8,28E-01 |
| AT2G01918 | 1,01 | -2,03 | 2,13 | -0,95 | 3,41E-01 |
| AT2G01930 | 0,16 | 0,90 | 4,38 | 0,21 | 8,37E-01 |
| AT2G01970 | 1,49 | 0,15 | 1,67 | 0,09 | 9,26E-01 |
| AT2G01990 | 0,25 | 0,97 | 4,35 | 0,22 | 8,24E-01 |
| AT2G02080 | 1,03 | -1,00 | 2,52 | -0,40 | 6,92E-01 |
| AT2G02103 | 0,70 | -0,27 | 2,47 | -0,11 | 9,14E-01 |
| AT2G02148 | 0,42 | 1,59 | 4,11 | 0,39 | 6,99E-01 |
| AT2G02150 | 0,39 | 1,77 | 4,26 | 0,42 | 6,78E-01 |
| AT2G02160 | 0,60 | 1,57 | 2,49 | 0,63 | 5,30E-01 |
| AT2G02220 | 0,80 | 0,83 | 2,44 | 0,34 | 7,35E-01 |
| AT2G02230 | 0,17 | 1,57 | 4,34 | 0,36 | 7,18E-01 |
| AT2G02340 | 1,23 | -2,07 | 2,09 | -0,99 | 3,23E-01 |
| AT2G02360 | 0,16 | 1,33 | 4,35 | 0,31 | 7,59E-01 |
| AT2G02400 | 0,84 | 2,36 | 2,28 | 1,04 | 3,00E-01 |
| AT2G02520 | 0,08 | 0,00 | 4,41 | 0,00 | 1,00E+00 |
| AT2G02570 | 0,25 | 1,02 | 4,36 | 0,23 | 8,15E-01 |
| AT2G02635 | 0,17 | -1,18 | 4,38 | -0,27 | 7,88E-01 |
| AT2G02695 | 0,58 | 1,91 | 3,00 | 0,64 | 5,25E-01 |
| AT2G02710 | 0,66 | 1,92 | 2,49 | 0,77 | 4,41E-01 |
| AT2G02780 | 0,28 | -1,77 | 4,37 | -0,40 | 6,86E-01 |
| AT2G02800 | 0,78 | -0,56 | 2,44 | -0,23 | 8,19E-01 |
| AT2G04345 | 0,62 | 1,20 | 2,66 | 0,45 | 6,51E-01 |

| | | | | | |
|---|---|---|---|---|---|
| AT2G02930 | 0,49 | 2,30 | 3,20 | 0,72 | 4,72E-01 |
| AT2G03050 | 0,25 | 2,27 | 4,32 | 0,53 | 5,99E-01 |
| AT2G03090 | 0,08 | 1,10 | 4,36 | 0,25 | 8,00E-01 |
| AT2G03240 | 0,24 | 1,62 | 4,34 | 0,37 | 7,08E-01 |
| AT2G03340 | 0,45 | 0,82 | 3,31 | 0,25 | 8,06E-01 |
| AT2G03390 | 0,61 | -0,21 | 2,25 | -0,09 | 9,25E-01 |
| AT2G03420 | 0,43 | 1,39 | 4,08 | 0,34 | 7,33E-01 |
| AT2G03430 | 0,52 | 0,98 | 4,37 | 0,23 | 8,22E-01 |
| AT2G03440 | 0,35 | 0,42 | 4,38 | 0,10 | 9,23E-01 |
| AT2G03460 | 0,08 | 1,10 | 4,36 | 0,25 | 8,00E-01 |
| AT2G03470 | 1,22 | 2,42 | 2,39 | 1,01 | 3,11E-01 |
| AT2G03490 | 0,08 | 1,10 | 4,36 | 0,25 | 8,00E-01 |
| AT2G03620 | 0,33 | 1,53 | 4,34 | 0,35 | 7,24E-01 |
| AT2G03640 | 0,13 | 1,15 | 4,36 | 0,26 | 7,92E-01 |
| AT2G03690 | 0,23 | 0,00 | 4,41 | 0,00 | 1,00E+00 |
| AT2G03730 | 0,29 | 0,37 | 4,35 | 0,09 | 9,32E-01 |
| AT2G03750 | 1,29 | -1,29 | 1,61 | -0,80 | 4,25E-01 |
| AT2G03760 | 0,93 | 1,57 | 3,47 | 0,45 | 6,51E-01 |
| AT2G03770 | 0,15 | 1,58 | 4,34 | 0,36 | 7,15E-01 |
| AT2G03930 | 1,60 | -0,54 | 1,44 | -0,37 | 7,08E-01 |
| AT2G04655 | 0,14 | 0,00 | 4,41 | 0,00 | 1,00E+00 |
| AT2G04031 | 0,15 | 0,00 | 4,41 | 0,00 | 1,00E+00 |
| AT2G04037 | 0,16 | 0,00 | 4,39 | 0,00 | 1,00E+00 |
| AT2G04050 | 0,12 | 1,65 | 4,34 | 0,38 | 7,04E-01 |
| AT2G04062 | 0,70 | 1,72 | 2,93 | 0,59 | 5,57E-01 |
| AT2G04063 | 0,69 | 2,17 | 2,76 | 0,78 | 4,33E-01 |
| AT2G04066 | 0,42 | 1,64 | 4,34 | 0,38 | 7,06E-01 |
| AT2G04070 | 0,33 | 0,89 | 4,36 | 0,20 | 8,39E-01 |
| AT2G04080 | 0,61 | 2,01 | 2,55 | 0,79 | 4,30E-01 |
| AT2G04160 | 0,43 | 1,09 | 4,10 | 0,26 | 7,91E-01 |
| AT2G04230 | 1,09 | 1,84 | 2,04 | 0,91 | 3,65E-01 |
| AT2G04360 | 0,70 | 0,45 | 2,58 | 0,17 | 8,63E-01 |
| AT2G04378 | 0,56 | -0,09 | 2,78 | -0,03 | 9,75E-01 |
| AT2G04520 | 0,26 | -2,10 | 4,37 | -0,48 | 6,31E-01 |
| AT2G04620 | 0,17 | 1,94 | 4,33 | 0,45 | 6,55E-01 |
| AT2G04630 | 0,19 | 1,27 | 4,35 | 0,29 | 7,70E-01 |
| AT2G04842 | 0,88 | 2,08 | 3,01 | 0,69 | 4,90E-01 |
| AT2G04870 | 0,55 | -0,68 | 2,74 | -0,25 | 8,04E-01 |
| AT2G04890 | 0,11 | 0,90 | 4,37 | 0,21 | 8,36E-01 |
| AT2G05120 | 0,91 | 1,58 | 2,11 | 0,75 | 4,55E-01 |
| AT2G05220 | 0,38 | 1,86 | 4,34 | 0,43 | 6,69E-01 |
| AT2G05250 | 0,41 | 1,92 | 4,33 | 0,44 | 6,56E-01 |
| AT2G04895 | 0,17 | 0,48 | 4,40 | 0,11 | 9,12E-01 |
| AT2G05752 | 0,90 | 2,18 | 2,36 | 0,92 | 3,55E-01 |
| AT2G04995 | 1,03 | -0,28 | 2,13 | -0,13 | 8,95E-01 |
| AT2G05810 | 1,10 | 1,52 | 1,98 | 0,77 | 4,43E-01 |
| AT2G05914 | 0,51 | 0,56 | 4,37 | 0,13 | 8,98E-01 |
| AT2G05920 | 1,42 | 1,46 | 1,55 | 0,94 | 3,46E-01 |
| AT2G05940 | 0,97 | 1,79 | 2,00 | 0,89 | 3,71E-01 |
| AT2G06000 | 0,52 | 2,11 | 2,60 | 0,81 | 4,17E-01 |
| AT2G06040 | 0,24 | 0,70 | 4,34 | 0,16 | 8,72E-01 |
| AT2G06050 | 0,61 | 0,26 | 3,11 | 0,08 | 9,32E-01 |
| AT2G06530 | 0,61 | 0,17 | 3,25 | 0,05 | 9,59E-01 |
| AT2G06541 | 0,49 | 1,91 | 4,11 | 0,47 | 6,41E-01 |
| AT2G06645 | 0,15 | -1,25 | 4,39 | -0,28 | 7,76E-01 |
| AT2G06675 | 0,19 | 0,00 | 4,41 | 0,00 | 1,00E+00 |

| | | | | | |
|---|---|---|---|---|---|
| AT2G06906 | 0,12 | 1,39 | 4,34 | 0,32 | 7,49E-01 |
| AT2G06925 | 0,35 | -1,33 | 3,53 | -0,38 | 7,06E-01 |
| AT2G06960 | 0,68 | -0,33 | 2,93 | -0,11 | 9,12E-01 |
| AT2G07020 | 0,35 | 0,00 | 4,41 | 0,00 | 1,00E+00 |
| AT2G07120 | 0,12 | 0,42 | 4,38 | 0,10 | 9,24E-01 |
| AT2G07180 | 1,49 | 0,44 | 1,95 | 0,23 | 8,20E-01 |
| AT2G07240 | 1,04 | -0,19 | 1,85 | -0,10 | 9,20E-01 |
| AT2G07560 | 0,12 | 0,79 | 4,37 | 0,18 | 8,57E-01 |
| AT2G07771 | 0,58 | 0,11 | 2,71 | 0,04 | 9,68E-01 |
| AT2G07777 | 0,89 | -0,29 | 1,96 | -0,15 | 8,84E-01 |
| AT2G07779 | 0,28 | 0,42 | 4,37 | 0,10 | 9,24E-01 |
| AT2G07621 | 0,12 | 0,42 | 4,38 | 0,10 | 9,24E-01 |
| AT2G07774 | 0,25 | -1,66 | 4,38 | -0,38 | 7,04E-01 |
| AT2G07687 | 1,11 | 0,24 | 1,90 | 0,13 | 8,98E-01 |
| AT2G05205 | 0,68 | -0,56 | 2,47 | -0,23 | 8,21E-01 |
| AT2G07599 | 0,90 | -0,35 | 2,12 | -0,16 | 8,70E-01 |
| AT2G07706 | 1,10 | -2,90 | 2,31 | -1,26 | 2,08E-01 |
| AT2G07641 | 0,45 | 0,25 | 2,79 | 0,09 | 9,27E-01 |
| AT2G07719 | 0,62 | 0,54 | 2,55 | 0,21 | 8,31E-01 |
| AT2G07756 | 1,02 | -0,97 | 1,84 | -0,53 | 5,98E-01 |
| AT2G07734 | 0,71 | -1,84 | 2,94 | -0,63 | 5,31E-01 |
| AT2G07658 | 0,65 | 0,96 | 2,49 | 0,38 | 7,00E-01 |
| AT2G07662 | 1,06 | 0,34 | 2,10 | 0,16 | 8,72E-01 |
| AT2G07835 | 0,78 | 1,48 | 2,86 | 0,52 | 6,05E-01 |
| AT2G07690 | 0,71 | 2,48 | 2,49 | 1,00 | 3,19E-01 |
| AT2G07760 | 0,23 | 0,26 | 4,36 | 0,06 | 9,52E-01 |
| AT2G09795 | 0,34 | 1,59 | 4,35 | 0,37 | 7,14E-01 |
| AT2G05465 | 0,27 | 0,42 | 4,38 | 0,10 | 9,24E-01 |
| AT2G10557 | 1,21 | 4,40 | 3,63 | 1,21 | 2,25E-01 |
| AT2G10608 | 0,36 | -1,27 | 4,39 | -0,29 | 7,73E-01 |
| AT2G05565 | 0,43 | 0,00 | 4,14 | 0,00 | 1,00E+00 |
| AT2G10625 | 0,25 | 0,00 | 4,41 | 0,00 | 1,00E+00 |
| AT2G10950 | 0,47 | 0,84 | 4,00 | 0,21 | 8,35E-01 |
| AT2G10955 | 0,26 | 0,00 | 4,39 | 0,00 | 1,00E+00 |
| AT2G11000 | 0,08 | 0,00 | 4,41 | 0,00 | 1,00E+00 |
| AT2G11852 | 0,26 | 1,26 | 4,34 | 0,29 | 7,72E-01 |
| AT2G11910 | 1,43 | 1,19 | 1,57 | 0,76 | 4,49E-01 |
| AT2G12400 | 0,28 | -0,54 | 4,38 | -0,12 | 9,02E-01 |
| AT2G12405 | 0,18 | -1,25 | 4,39 | -0,28 | 7,76E-01 |
| AT2G05705 | 0,21 | 0,00 | 4,41 | 0,00 | 1,00E+00 |
| AT2G12461 | 0,12 | 1,65 | 4,34 | 0,38 | 7,04E-01 |
| AT2G05835 | 0,29 | -0,30 | 4,37 | -0,07 | 9,45E-01 |
| AT2G12945 | 0,05 | 0,00 | 4,41 | 0,00 | 1,00E+00 |
| AT2G13370 | 1,20 | 1,23 | 2,37 | 0,52 | 6,02E-01 |
| AT2G13450 | 1,51 | -2,81 | 1,65 | -1,71 | 8,80E-02 |
| AT2G13550 | 0,22 | 1,32 | 4,34 | 0,30 | 7,62E-01 |
| AT2G13640 | 0,11 | 1,07 | 4,36 | 0,25 | 8,06E-01 |
| AT2G13650 | 0,22 | 0,00 | 4,41 | 0,00 | 1,00E+00 |
| AT2G13820 | 1,49 | 0,41 | 2,90 | 0,14 | 8,87E-01 |
| AT2G14100 | 0,15 | 0,71 | 4,37 | 0,16 | 8,71E-01 |
| AT2G14120 | 0,98 | 3,10 | 2,08 | 1,49 | 1,36E-01 |
| AT2G14170 | 1,22 | 0,77 | 1,93 | 0,40 | 6,90E-01 |
| AT2G14260 | 0,26 | 2,11 | 4,32 | 0,49 | 6,26E-01 |
| AT2G14265 | 1,52 | -1,02 | 1,70 | -0,60 | 5,50E-01 |
| AT2G14288 | 0,20 | 0,00 | 4,41 | 0,00 | 1,00E+00 |
| AT2G14510 | 0,15 | 0,71 | 4,37 | 0,16 | 8,71E-01 |

| | | | | | |
|---|---|---|---|---|---|
| AT2G14540 | 0,12 | 1,39 | 4,34 | 0,32 | 7,49E-01 |
| AT2G14670 | 1,21 | -3,01 | 2,54 | -1,18 | 2,37E-01 |
| AT2G14720 | 1,05 | 2,79 | 2,34 | 1,19 | 2,33E-01 |
| AT2G14755 | 0,32 | 0,00 | 4,41 | 0,00 | 1,00E+00 |
| AT2G15000 | 0,23 | 0,00 | 4,41 | 0,00 | 1,00E+00 |
| AT2G15020 | 0,51 | -0,93 | 3,21 | -0,29 | 7,71E-01 |
| AT2G06355 | 0,23 | -0,37 | 4,36 | -0,08 | 9,33E-01 |
| AT2G15110 | 0,63 | 0,71 | 2,97 | 0,24 | 8,12E-01 |
| AT2G15290 | 0,58 | 1,37 | 3,71 | 0,37 | 7,12E-01 |
| AT2G15292 | 1,13 | 1,62 | 1,64 | 0,99 | 3,23E-01 |
| AT2G15320 | 1,25 | 0,96 | 1,65 | 0,58 | 5,59E-01 |
| AT2G15390 | 0,37 | 1,84 | 4,33 | 0,42 | 6,71E-01 |
| AT2G15555 | 1,03 | 0,24 | 1,68 | 0,14 | 8,87E-01 |
| AT2G15560 | 0,05 | 0,00 | 4,41 | 0,00 | 1,00E+00 |
| AT2G15790 | 0,41 | 1,18 | 4,19 | 0,28 | 7,78E-01 |
| AT2G15860 | 0,36 | 0,59 | 4,32 | 0,14 | 8,91E-01 |
| AT2G15880 | 1,49 | 1,16 | 1,74 | 0,67 | 5,04E-01 |
| AT2G15890 | 1,14 | 2,25 | 1,66 | 1,35 | 1,76E-01 |
| AT2G06515 | 0,69 | -3,32 | 3,66 | -0,91 | 3,64E-01 |
| AT2G15980 | 0,29 | -0,35 | 3,61 | -0,10 | 9,22E-01 |
| AT2G16016 | 1,06 | 0,87 | 2,11 | 0,41 | 6,82E-01 |
| AT2G06575 | 0,75 | 0,30 | 2,19 | 0,14 | 8,92E-01 |
| AT2G16070 | 0,64 | 0,19 | 3,81 | 0,05 | 9,60E-01 |
| AT2G16090 | 0,55 | 1,86 | 2,54 | 0,73 | 4,64E-01 |
| AT2G16370 | 0,11 | 1,07 | 4,36 | 0,25 | 8,06E-01 |
| AT2G16400 | 1,00 | 1,07 | 1,87 | 0,57 | 5,68E-01 |
| AT2G16405 | 0,20 | 0,82 | 4,33 | 0,19 | 8,50E-01 |
| AT2G16440 | 1,04 | 2,52 | 3,54 | 0,71 | 4,76E-01 |
| AT2G16450 | 0,33 | 1,53 | 4,34 | 0,35 | 7,24E-01 |
| AT2G16490 | 1,37 | -4,17 | 2,37 | -1,76 | 7,85E-02 |
| AT2G16505 | 0,85 | 0,69 | 2,20 | 0,32 | 7,52E-01 |
| AT2G16510 | 1,63 | 1,14 | 1,55 | 0,74 | 4,60E-01 |
| AT2G16620 | 0,18 | 0,63 | 4,37 | 0,14 | 8,86E-01 |
| AT2G16640 | 1,05 | 3,55 | 2,13 | 1,67 | 9,57E-02 |
| AT2G16650 | 0,37 | 2,79 | 4,31 | 0,65 | 5,18E-01 |
| AT2G16870 | 1,12 | 2,53 | 2,03 | 1,25 | 2,13E-01 |
| AT2G16880 | 0,51 | 2,02 | 4,34 | 0,46 | 6,42E-01 |
| AT2G16920 | 0,25 | 0,72 | 4,34 | 0,17 | 8,68E-01 |
| AT2G16940 | 0,88 | 2,40 | 2,13 | 1,13 | 2,59E-01 |
| AT2G17020 | 0,78 | 1,49 | 2,02 | 0,74 | 4,60E-01 |
| AT2G17110 | 0,49 | 0,40 | 2,69 | 0,15 | 8,82E-01 |
| AT2G17130 | 0,45 | 1,37 | 4,20 | 0,33 | 7,44E-01 |
| AT2G17190 | 0,94 | 1,04 | 2,37 | 0,44 | 6,60E-01 |
| AT2G17210 | 0,23 | 1,33 | 4,34 | 0,31 | 7,60E-01 |
| AT2G17220 | 0,39 | 0,00 | 4,41 | 0,00 | 1,00E+00 |
| AT2G17230 | 0,92 | 2,14 | 2,43 | 0,88 | 3,79E-01 |
| AT2G17300 | 0,17 | 1,57 | 4,34 | 0,36 | 7,18E-01 |
| AT2G06935 | 0,05 | 0,00 | 4,41 | 0,00 | 1,00E+00 |
| AT2G17340 | 0,86 | 1,60 | 2,21 | 0,72 | 4,69E-01 |
| AT2G17350 | 0,34 | -2,01 | 4,36 | -0,46 | 6,46E-01 |
| AT2G17450 | 0,63 | 1,58 | 2,32 | 0,68 | 4,95E-01 |
| AT2G17470 | 0,10 | 0,62 | 4,38 | 0,14 | 8,87E-01 |
| AT2G17550 | 0,71 | 3,13 | 2,40 | 1,30 | 1,93E-01 |
| AT2G17560 | 0,46 | 0,91 | 4,38 | 0,21 | 8,36E-01 |
| AT2G17700 | 0,43 | -0,13 | 4,10 | -0,03 | 9,74E-01 |
| AT2G17820 | 0,96 | -0,27 | 2,05 | -0,13 | 8,96E-01 |

| | | | | | |
|---|---|---|---|---|---|
| AT2G17870 | 0,45 | 0,44 | 4,15 | 0,11 | 9,15E-01 |
| AT2G18130 | 0,22 | 0,42 | 4,38 | 0,10 | 9,24E-01 |
| AT2G18160 | 1,66 | 3,00 | 1,83 | 1,64 | 1,02E-01 |
| AT2G18170 | 0,50 | 1,79 | 3,24 | 0,55 | 5,79E-01 |
| AT2G18193 | 1,29 | 2,91 | 2,14 | 1,36 | 1,74E-01 |
| AT2G18290 | 0,19 | 1,64 | 4,33 | 0,38 | 7,05E-01 |
| AT2G18300 | 1,33 | -2,44 | 2,04 | -1,19 | 2,33E-01 |
| AT2G18310 | 0,43 | 0,08 | 4,37 | 0,02 | 9,85E-01 |
| AT2G18520 | 0,25 | 0,00 | 4,40 | 0,00 | 1,00E+00 |
| AT2G18600 | 0,24 | 0,00 | 4,41 | 0,00 | 1,00E+00 |
| AT2G18660 | 0,89 | 1,44 | 2,16 | 0,67 | 5,06E-01 |
| AT2G18685 | 0,08 | 1,10 | 4,36 | 0,25 | 8,00E-01 |
| AT2G18900 | 1,01 | 2,28 | 2,00 | 1,14 | 2,55E-01 |
| AT2G18915 | 0,29 | 1,43 | 4,35 | 0,33 | 7,42E-01 |
| AT2G18940 | 0,31 | 0,80 | 3,56 | 0,22 | 8,23E-01 |
| AT2G19120 | 0,71 | -0,66 | 3,57 | -0,18 | 8,54E-01 |
| AT2G07135 | 0,16 | 0,00 | 4,41 | 0,00 | 1,00E+00 |
| AT2G07165 | 0,21 | 1,09 | 4,36 | 0,25 | 8,03E-01 |
| AT2G19290 | 0,05 | 0,00 | 4,41 | 0,00 | 1,00E+00 |
| AT2G19310 | 0,12 | 0,42 | 4,38 | 0,10 | 9,24E-01 |
| AT2G19340 | 0,43 | 1,95 | 4,32 | 0,45 | 6,52E-01 |
| AT2G19365 | 0,81 | -0,90 | 2,03 | -0,44 | 6,58E-01 |
| AT2G19390 | 0,76 | 2,80 | 2,37 | 1,18 | 2,37E-01 |
| AT2G07195 | 0,51 | 1,69 | 2,66 | 0,64 | 5,25E-01 |
| AT2G19400 | 0,15 | 0,00 | 4,39 | 0,00 | 1,00E+00 |
| AT2G19470 | 0,28 | 1,19 | 4,34 | 0,27 | 7,85E-01 |
| AT2G19560 | 0,20 | 1,68 | 4,34 | 0,39 | 6,99E-01 |
| AT2G19570 | 0,52 | 1,65 | 3,07 | 0,54 | 5,92E-01 |
| AT2G19580 | 0,25 | 0,43 | 4,39 | 0,10 | 9,21E-01 |
| AT2G19760 | 0,25 | 0,64 | 4,37 | 0,15 | 8,84E-01 |
| AT2G19780 | 0,19 | -1,36 | 4,41 | -0,31 | 7,58E-01 |
| AT2G19910 | 0,11 | 1,07 | 4,36 | 0,25 | 8,06E-01 |
| AT2G20000 | 0,44 | 1,49 | 3,28 | 0,45 | 6,50E-01 |
| AT2G20050 | 0,77 | 3,14 | 2,34 | 1,34 | 1,80E-01 |
| AT2G20120 | 0,38 | 0,00 | 4,39 | 0,00 | 1,00E+00 |
| AT2G20180 | 0,53 | -0,17 | 2,57 | -0,07 | 9,46E-01 |
| AT2G20240 | 1,31 | 2,91 | 2,52 | 1,16 | 2,47E-01 |
| AT2G20270 | 0,73 | 0,30 | 3,04 | 0,10 | 9,23E-01 |
| AT2G20280 | 0,90 | 1,42 | 2,77 | 0,51 | 6,07E-01 |
| AT2G20300 | 0,57 | 1,23 | 2,63 | 0,47 | 6,39E-01 |
| AT2G20310 | 0,15 | 1,62 | 4,34 | 0,37 | 7,09E-01 |
| AT2G20380 | 0,08 | 0,85 | 4,38 | 0,20 | 8,45E-01 |
| AT2G20562 | 0,11 | 0,00 | 4,41 | 0,00 | 1,00E+00 |
| AT2G20610 | 0,62 | 0,20 | 2,53 | 0,08 | 9,37E-01 |
| AT2G20630 | 0,57 | -1,62 | 2,60 | -0,62 | 5,33E-01 |
| AT2G20670 | 0,99 | 0,00 | 2,02 | 0,00 | 9,98E-01 |
| AT2G07325 | 0,08 | 0,85 | 4,38 | 0,20 | 8,45E-01 |
| AT2G20720 | 0,50 | 0,72 | 3,20 | 0,22 | 8,23E-01 |
| AT2G20723 | 1,21 | -0,39 | 1,70 | -0,23 | 8,20E-01 |
| AT2G20725 | 0,17 | 1,40 | 4,35 | 0,32 | 7,48E-01 |
| AT2G20760 | 1,00 | -0,22 | 2,18 | -0,10 | 9,19E-01 |
| AT2G20790 | 0,65 | -0,39 | 3,79 | -0,10 | 9,18E-01 |
| AT2G20825 | 0,24 | 1,23 | 4,35 | 0,28 | 7,77E-01 |
| AT2G20830 | 0,42 | 0,41 | 3,33 | 0,12 | 9,02E-01 |
| AT2G20850 | 0,80 | -0,75 | 3,45 | -0,22 | 8,28E-01 |
| AT2G20900 | 0,70 | 1,61 | 3,01 | 0,53 | 5,93E-01 |

| | | | | | |
|---|---|---|---|---|---|
| AT2G21060 | 0,37 | -0,29 | 4,37 | -0,07 | 9,48E-01 |
| AT2G21090 | 0,12 | 1,58 | 4,34 | 0,36 | 7,16E-01 |
| AT2G21120 | 1,14 | 2,27 | 1,91 | 1,19 | 2,35E-01 |
| AT2G21170 | 1,20 | -0,04 | 1,55 | -0,02 | 9,82E-01 |
| AT2G21185 | 0,86 | 0,88 | 1,98 | 0,44 | 6,58E-01 |
| AT2G21188 | 0,47 | 0,66 | 4,21 | 0,16 | 8,76E-01 |
| AT2G21210 | 1,12 | 2,40 | 2,16 | 1,11 | 2,66E-01 |
| AT2G21240 | 0,62 | 0,23 | 2,25 | 0,10 | 9,19E-01 |
| AT2G21280 | 0,38 | 0,96 | 4,37 | 0,22 | 8,27E-01 |
| AT2G21300 | 0,99 | 1,90 | 2,25 | 0,85 | 3,97E-01 |
| AT2G21350 | 0,05 | 0,00 | 4,41 | 0,00 | 1,00E+00 |
| AT2G21370 | 0,82 | 0,24 | 2,07 | 0,12 | 9,08E-01 |
| AT2G21440 | 0,17 | 1,92 | 4,33 | 0,44 | 6,57E-01 |
| AT2G21480 | 0,25 | 1,75 | 4,33 | 0,40 | 6,87E-01 |
| AT2G21500 | 0,51 | 1,96 | 3,06 | 0,64 | 5,22E-01 |
| AT2G21560 | 0,27 | 0,00 | 4,41 | 0,00 | 1,00E+00 |
| AT2G21870 | 1,17 | -1,62 | 2,16 | -0,75 | 4,55E-01 |
| AT2G21950 | 0,24 | 1,98 | 4,34 | 0,46 | 6,48E-01 |
| AT2G21970 | 0,61 | -1,19 | 3,78 | -0,32 | 7,52E-01 |
| AT2G22010 | 0,65 | 1,14 | 2,49 | 0,46 | 6,47E-01 |
| AT2G22090 | 0,34 | -1,47 | 4,37 | -0,34 | 7,36E-01 |
| AT2G22140 | 0,05 | 0,00 | 4,41 | 0,00 | 1,00E+00 |
| AT2G22160 | 0,35 | 1,30 | 4,35 | 0,30 | 7,65E-01 |
| AT2G22240 | 0,21 | -0,31 | 4,37 | -0,07 | 9,43E-01 |
| AT2G22250 | 0,26 | 0,90 | 4,38 | 0,21 | 8,37E-01 |
| AT2G07555 | 0,37 | -1,23 | 4,34 | -0,28 | 7,77E-01 |
| AT2G22400 | 0,55 | 0,93 | 3,83 | 0,24 | 8,07E-01 |
| AT2G22470 | 0,19 | 1,50 | 4,34 | 0,34 | 7,30E-01 |
| AT2G22490 | 1,17 | 2,18 | 2,39 | 0,91 | 3,63E-01 |
| AT2G22740 | 0,43 | 2,58 | 4,09 | 0,63 | 5,28E-01 |
| AT2G22850 | 0,76 | -1,06 | 1,88 | -0,56 | 5,74E-01 |
| AT2G22890 | 0,53 | 0,00 | 3,17 | 0,00 | 1,00E+00 |
| AT2G22940 | 0,87 | 2,41 | 2,65 | 0,91 | 3,65E-01 |
| AT2G23070 | 0,53 | 0,27 | 2,20 | 0,12 | 9,01E-01 |
| AT2G23090 | 0,40 | 1,24 | 4,30 | 0,29 | 7,74E-01 |
| AT2G23100 | 0,16 | 0,90 | 4,38 | 0,21 | 8,37E-01 |
| AT2G23140 | 0,38 | 2,28 | 4,23 | 0,54 | 5,90E-01 |
| AT2G23210 | 0,71 | 2,91 | 2,84 | 1,02 | 3,06E-01 |
| AT2G23290 | 0,64 | 0,50 | 2,61 | 0,19 | 8,47E-01 |
| AT2G23390 | 0,32 | 0,66 | 4,32 | 0,15 | 8,79E-01 |
| AT2G23430 | 0,47 | 1,54 | 3,38 | 0,46 | 6,48E-01 |
| AT2G23450 | 0,62 | -0,18 | 2,22 | -0,08 | 9,36E-01 |
| AT2G23520 | 0,43 | 0,54 | 3,41 | 0,16 | 8,74E-01 |
| AT2G07910 | 1,63 | -3,88 | 2,35 | -1,65 | 9,91E-02 |
| AT2G23550 | 1,59 | -3,88 | 2,37 | -1,64 | 1,01E-01 |
| AT2G23600 | 1,45 | -0,64 | 1,85 | -0,35 | 7,30E-01 |
| AT2G23610 | 0,63 | 0,38 | 3,09 | 0,12 | 9,02E-01 |
| AT2G23670 | 1,50 | -0,45 | 1,55 | -0,29 | 7,70E-01 |
| AT2G23760 | 0,34 | -1,36 | 4,41 | -0,31 | 7,58E-01 |
| AT2G23790 | 0,79 | 2,02 | 2,90 | 0,70 | 4,86E-01 |
| AT2G23840 | 0,70 | -1,52 | 3,02 | -0,50 | 6,14E-01 |
| AT2G23987 | 0,17 | -2,23 | 4,35 | -0,51 | 6,08E-01 |
| AT2G24120 | 0,81 | -1,22 | 2,22 | -0,55 | 5,82E-01 |
| AT2G24150 | 0,79 | 0,90 | 2,18 | 0,41 | 6,81E-01 |
| AT2G24170 | 0,17 | 1,75 | 4,33 | 0,40 | 6,86E-01 |
| AT2G24200 | 1,48 | 2,25 | 1,69 | 1,33 | 1,83E-01 |

| | | | | | |
|---|---|---|---|---|---|
| AT2G24230 | 0,59 | -0,67 | 2,66 | -0,25 | 8,02E-01 |
| AT2G24330 | 0,33 | -1,21 | 4,38 | -0,28 | 7,82E-01 |
| AT2G24340 | 1,46 | 2,05 | 1,82 | 1,13 | 2,59E-01 |
| AT2G24360 | 1,19 | 1,33 | 1,94 | 0,69 | 4,93E-01 |
| AT2G24500 | 1,13 | -0,14 | 2,12 | -0,07 | 9,46E-01 |
| AT2G08055 | 0,33 | 1,52 | 4,33 | 0,35 | 7,25E-01 |
| AT2G24545 | 1,16 | 0,28 | 1,70 | 0,17 | 8,68E-01 |
| AT2G24550 | 1,50 | 2,35 | 1,72 | 1,37 | 1,71E-01 |
| AT2G24600 | 1,31 | 3,96 | 2,24 | 1,77 | 7,72E-02 |
| AT2G24640 | 0,24 | 1,58 | 4,34 | 0,36 | 7,16E-01 |
| AT2G24710 | 0,19 | 1,91 | 4,33 | 0,44 | 6,58E-01 |
| AT2G24755 | 0,71 | 2,24 | 2,58 | 0,87 | 3,86E-01 |
| AT2G24780 | 0,92 | -1,91 | 2,29 | -0,84 | 4,03E-01 |
| AT2G24850 | 1,45 | 1,54 | 1,86 | 0,83 | 4,07E-01 |
| AT2G24870 | 1,09 | -3,30 | 2,47 | -1,33 | 1,82E-01 |
| AT2G24880 | 1,30 | -1,57 | 1,51 | -1,04 | 2,99E-01 |
| AT2G25011 | 1,24 | 0,30 | 1,77 | 0,17 | 8,65E-01 |
| AT2G25050 | 0,47 | -1,82 | 3,16 | -0,58 | 5,64E-01 |
| AT2G25070 | 0,45 | 0,34 | 2,80 | 0,12 | 9,05E-01 |
| AT2G08105 | 0,74 | -1,37 | 2,27 | -0,61 | 5,44E-01 |
| AT2G25180 | 0,76 | -0,14 | 2,94 | -0,05 | 9,61E-01 |
| AT2G25200 | 0,69 | -1,92 | 3,81 | -0,50 | 6,14E-01 |
| AT2G25300 | 0,15 | 0,24 | 4,35 | 0,05 | 9,56E-01 |
| AT2G25310 | 0,20 | 0,00 | 4,41 | 0,00 | 1,00E+00 |
| AT2G25570 | 0,45 | 0,90 | 4,37 | 0,21 | 8,37E-01 |
| AT2G25610 | 0,53 | 0,41 | 3,16 | 0,13 | 8,97E-01 |
| AT2G25670 | 0,93 | 1,33 | 2,25 | 0,59 | 5,54E-01 |
| AT2G25760 | 0,93 | 2,12 | 2,82 | 0,75 | 4,52E-01 |
| AT2G25850 | 0,16 | 1,33 | 4,35 | 0,31 | 7,59E-01 |
| AT2G25910 | 0,91 | -3,17 | 2,49 | -1,27 | 2,03E-01 |
| AT2G25970 | 1,54 | 2,37 | 1,92 | 1,24 | 2,16E-01 |
| AT2G26030 | 0,31 | 0,41 | 3,63 | 0,11 | 9,10E-01 |
| AT2G26070 | 0,39 | 0,40 | 4,39 | 0,09 | 9,28E-01 |
| AT2G26140 | 1,61 | 0,54 | 1,57 | 0,35 | 7,30E-01 |
| AT2G26190 | 1,38 | 0,37 | 1,97 | 0,19 | 8,51E-01 |
| AT2G26280 | 1,11 | 1,61 | 2,47 | 0,65 | 5,15E-01 |
| AT2G26430 | 0,38 | 1,80 | 4,27 | 0,42 | 6,74E-01 |
| AT2G26540 | 0,42 | 0,73 | 4,35 | 0,17 | 8,67E-01 |
| AT2G26560 | 1,67 | 2,32 | 1,68 | 1,38 | 1,68E-01 |
| AT2G26580 | 0,26 | -1,10 | 4,38 | -0,25 | 8,01E-01 |
| AT2G26590 | 0,25 | 0,11 | 4,36 | 0,03 | 9,80E-01 |
| AT2G26620 | 0,18 | 0,42 | 4,38 | 0,10 | 9,24E-01 |
| AT2G26680 | 0,33 | 1,53 | 4,34 | 0,35 | 7,24E-01 |
| AT2G26690 | 0,68 | 2,04 | 2,53 | 0,81 | 4,20E-01 |
| AT2G26740 | 0,33 | 1,65 | 4,33 | 0,38 | 7,03E-01 |
| AT2G08175 | 1,14 | -1,10 | 1,92 | -0,58 | 5,65E-01 |
| AT2G26790 | 0,21 | 2,12 | 4,33 | 0,49 | 6,23E-01 |
| AT2G26800 | 0,41 | 0,63 | 3,37 | 0,19 | 8,51E-01 |
| AT2G26840 | 0,08 | 1,10 | 4,36 | 0,25 | 8,00E-01 |
| AT2G26900 | 1,34 | -0,67 | 1,63 | -0,41 | 6,83E-01 |
| AT2G26930 | 0,69 | -1,53 | 2,94 | -0,52 | 6,03E-01 |
| AT2G26980 | 0,61 | -0,55 | 2,31 | -0,24 | 8,13E-01 |
| AT2G26990 | 0,56 | 1,28 | 3,10 | 0,41 | 6,79E-01 |
| AT2G27020 | 0,74 | 0,50 | 2,45 | 0,20 | 8,39E-01 |
| AT2G27040 | 0,44 | -0,46 | 3,36 | -0,14 | 8,90E-01 |
| AT2G27070 | 0,56 | -2,57 | 2,98 | -0,86 | 3,89E-01 |

| | | | | | |
|---|---|---|---|---|---|
| AT2G27090 | 0,77 | 3,66 | 3,01 | 1,22 | 2,24E-01 |
| AT2G27170 | 0,61 | -1,35 | 2,47 | -0,55 | 5,83E-01 |
| AT2G27190 | 0,39 | -2,25 | 4,33 | -0,52 | 6,03E-01 |
| AT2G27290 | 0,57 | 0,02 | 2,65 | 0,01 | 9,94E-01 |
| AT2G27330 | 0,10 | 0,00 | 4,41 | 0,00 | 1,00E+00 |
| AT2G27350 | 0,23 | 1,56 | 4,34 | 0,36 | 7,19E-01 |
| AT2G27385 | 1,03 | -2,61 | 2,02 | -1,29 | 1,96E-01 |
| AT2G27500 | 0,69 | 0,66 | 2,59 | 0,25 | 8,00E-01 |
| AT2G27510 | 0,44 | 0,44 | 4,19 | 0,10 | 9,17E-01 |
| AT2G27530 | 0,92 | 0,24 | 1,87 | 0,13 | 8,98E-01 |
| AT2G27580 | 0,43 | 2,38 | 4,18 | 0,57 | 5,70E-01 |
| AT2G27660 | 0,41 | 0,00 | 4,40 | 0,00 | 1,00E+00 |
| AT2G27680 | 0,72 | 0,25 | 1,98 | 0,13 | 8,99E-01 |
| AT2G27770 | 0,38 | 1,18 | 4,37 | 0,27 | 7,87E-01 |
| AT2G27950 | 1,06 | 2,86 | 2,02 | 1,42 | 1,56E-01 |
| AT2G28080 | 0,87 | 0,81 | 2,87 | 0,28 | 7,79E-01 |
| AT2G28170 | 0,05 | 0,00 | 4,41 | 0,00 | 1,00E+00 |
| AT2G28360 | 1,42 | -0,45 | 1,66 | -0,27 | 7,85E-01 |
| AT2G28380 | 0,74 | 2,61 | 2,87 | 0,91 | 3,63E-01 |
| AT2G28390 | 0,41 | 0,69 | 4,37 | 0,16 | 8,74E-01 |
| AT2G28400 | 0,81 | 0,35 | 2,35 | 0,15 | 8,80E-01 |
| AT2G28450 | 1,48 | 2,68 | 1,94 | 1,38 | 1,68E-01 |
| AT2G28470 | 0,87 | 0,41 | 2,81 | 0,14 | 8,85E-01 |
| AT2G28480 | 0,12 | 0,79 | 4,37 | 0,18 | 8,57E-01 |
| AT2G28620 | 1,12 | 3,20 | 2,15 | 1,49 | 1,36E-01 |
| AT2G08275 | 1,12 | 3,20 | 2,15 | 1,49 | 1,36E-01 |
| AT2G28630 | 0,37 | -0,53 | 4,30 | -0,12 | 9,02E-01 |
| AT2G28640 | 0,62 | -2,18 | 3,02 | -0,72 | 4,70E-01 |
| AT2G28670 | 0,51 | 1,25 | 3,03 | 0,41 | 6,78E-01 |
| AT2G28740 | 0,44 | 1,80 | 4,16 | 0,43 | 6,65E-01 |
| AT2G28800 | 0,62 | -0,16 | 3,03 | -0,05 | 9,58E-01 |
| AT2G08300 | 0,20 | 0,00 | 4,41 | 0,00 | 1,00E+00 |
| AT2G28880 | 0,33 | 2,00 | 4,33 | 0,46 | 6,44E-01 |
| AT2G28890 | 0,77 | 2,11 | 2,20 | 0,96 | 3,38E-01 |
| AT2G28900 | 0,81 | -3,20 | 2,59 | -1,24 | 2,16E-01 |
| AT2G28950 | 0,45 | -0,06 | 4,12 | -0,02 | 9,88E-01 |
| AT2G29030 | 0,32 | 0,00 | 4,41 | 0,00 | 1,00E+00 |
| AT2G29040 | 0,08 | 0,00 | 4,40 | 0,00 | 1,00E+00 |
| AT2G29110 | 0,41 | 1,63 | 4,12 | 0,39 | 6,93E-01 |
| AT2G29120 | 1,16 | 2,78 | 2,08 | 1,33 | 1,83E-01 |
| AT2G29170 | 0,18 | 0,98 | 4,36 | 0,23 | 8,22E-01 |
| AT2G08330 | 0,26 | 0,90 | 4,38 | 0,21 | 8,37E-01 |
| AT2G29190 | 0,61 | 2,20 | 2,63 | 0,84 | 4,03E-01 |
| AT2G29220 | 0,39 | 0,40 | 4,39 | 0,09 | 9,28E-01 |
| AT2G29390 | 0,71 | 1,40 | 2,44 | 0,57 | 5,66E-01 |
| AT2G29400 | 0,98 | 3,09 | 1,91 | 1,61 | 1,06E-01 |
| AT2G29550 | 0,75 | 1,48 | 2,36 | 0,63 | 5,29E-01 |
| AT2G29570 | 0,14 | 0,51 | 4,38 | 0,12 | 9,08E-01 |
| AT2G29605 | 0,65 | -0,42 | 2,97 | -0,14 | 8,87E-01 |
| AT2G29650 | 0,77 | -0,73 | 2,44 | -0,30 | 7,64E-01 |
| AT2G30250 | 0,71 | 1,08 | 2,30 | 0,47 | 6,41E-01 |
| AT2G30260 | 0,44 | 1,07 | 3,24 | 0,33 | 7,42E-01 |
| AT2G30270 | 0,18 | 1,52 | 4,34 | 0,35 | 7,26E-01 |
| AT2G30360 | 0,37 | 0,92 | 4,38 | 0,21 | 8,34E-01 |
| AT2G30390 | 0,49 | -0,04 | 3,31 | -0,01 | 9,92E-01 |
| AT2G30470 | 0,73 | 0,81 | 2,14 | 0,38 | 7,04E-01 |

| | | | | | |
|---|---|---|---|---|---|
| AT2G30490 | 0,91 | 0,51 | 3,07 | 0,17 | 8,68E-01 |
| AT2G30500 | 1,56 | 2,85 | 1,69 | 1,69 | 9,13E-02 |
| AT2G30505 | 0,35 | 0,74 | 4,31 | 0,17 | 8,63E-01 |
| AT2G30700 | 0,36 | 1,51 | 4,27 | 0,35 | 7,24E-01 |
| AT2G30710 | 0,35 | 1,31 | 4,35 | 0,30 | 7,64E-01 |
| AT2G30720 | 0,49 | 0,70 | 3,22 | 0,22 | 8,28E-01 |
| AT2G30790 | 0,21 | 0,00 | 4,38 | 0,00 | 1,00E+00 |
| AT2G30800 | 1,27 | 0,54 | 2,03 | 0,27 | 7,90E-01 |
| AT2G30850 | 0,41 | -1,70 | 4,35 | -0,39 | 6,96E-01 |
| AT2G30880 | 0,34 | 1,62 | 4,33 | 0,37 | 7,08E-01 |
| AT2G08420 | 0,54 | -0,82 | 3,93 | -0,21 | 8,35E-01 |
| AT2G30980 | 0,41 | 1,27 | 4,34 | 0,29 | 7,71E-01 |
| AT2G31040 | 0,19 | 0,00 | 4,37 | 0,00 | 9,99E-01 |
| AT2G31060 | 0,49 | 3,09 | 4,30 | 0,72 | 4,72E-01 |
| AT2G31070 | 0,79 | 2,68 | 2,29 | 1,17 | 2,42E-01 |
| AT2G31130 | 0,17 | 0,45 | 4,39 | 0,10 | 9,19E-01 |
| AT2G31170 | 0,45 | -0,45 | 4,33 | -0,10 | 9,17E-01 |
| AT2G31240 | 0,25 | 1,75 | 4,33 | 0,40 | 6,87E-01 |
| AT2G31250 | 0,39 | 0,48 | 4,40 | 0,11 | 9,14E-01 |
| AT2G31260 | 0,53 | 0,20 | 2,67 | 0,08 | 9,40E-01 |
| AT2G08430 | 0,08 | 1,10 | 4,36 | 0,25 | 8,00E-01 |
| AT2G31360 | 1,30 | 0,44 | 1,90 | 0,23 | 8,17E-01 |
| AT2G31390 | 0,86 | 2,07 | 2,88 | 0,72 | 4,73E-01 |
| AT2G31410 | 0,37 | 0,00 | 4,41 | 0,00 | 1,00E+00 |
| AT2G31440 | 0,44 | 2,69 | 4,31 | 0,62 | 5,32E-01 |
| AT2G31510 | 0,63 | -3,03 | 3,08 | -0,98 | 3,25E-01 |
| AT2G31570 | 0,14 | 0,82 | 4,37 | 0,19 | 8,51E-01 |
| AT2G31670 | 0,73 | 0,71 | 2,39 | 0,30 | 7,68E-01 |
| AT2G31730 | 0,53 | -1,83 | 4,35 | -0,42 | 6,74E-01 |
| AT2G31740 | 0,52 | 0,32 | 3,20 | 0,10 | 9,21E-01 |
| AT2G31751 | 0,97 | 1,51 | 2,31 | 0,65 | 5,13E-01 |
| AT2G31800 | 0,85 | 1,47 | 2,14 | 0,69 | 4,93E-01 |
| AT2G31830 | 0,34 | 1,77 | 4,34 | 0,41 | 6,83E-01 |
| AT2G31865 | 0,37 | 2,38 | 4,32 | 0,55 | 5,81E-01 |
| AT2G31870 | 0,27 | 1,65 | 3,75 | 0,44 | 6,60E-01 |
| AT2G08475 | 0,81 | 1,70 | 2,40 | 0,71 | 4,80E-01 |
| AT2G31990 | 0,31 | 1,29 | 4,34 | 0,30 | 7,66E-01 |
| AT2G32000 | 0,55 | 0,50 | 2,70 | 0,18 | 8,54E-01 |
| AT2G32030 | 0,22 | 1,61 | 4,33 | 0,37 | 7,11E-01 |
| AT2G32050 | 0,08 | 0,85 | 4,38 | 0,20 | 8,45E-01 |
| AT2G32060 | 0,34 | 1,63 | 4,34 | 0,38 | 7,07E-01 |
| AT2G32070 | 0,64 | 1,99 | 2,52 | 0,79 | 4,30E-01 |
| AT2G32080 | 0,57 | 1,40 | 2,54 | 0,55 | 5,82E-01 |
| AT2G32130 | 0,28 | 2,33 | 4,32 | 0,54 | 5,90E-01 |
| AT2G32230 | 0,70 | 1,88 | 2,86 | 0,66 | 5,09E-01 |
| AT2G32235 | 0,12 | 1,58 | 4,34 | 0,36 | 7,16E-01 |
| AT2G32390 | 0,36 | 1,99 | 4,34 | 0,46 | 6,46E-01 |
| AT2G32400 | 0,29 | 2,52 | 4,32 | 0,58 | 5,59E-01 |
| AT2G32415 | 0,42 | 1,29 | 4,36 | 0,30 | 7,67E-01 |
| AT2G32480 | 0,87 | -0,39 | 2,16 | -0,18 | 8,56E-01 |
| AT2G32520 | 0,35 | 1,37 | 4,30 | 0,32 | 7,50E-01 |
| AT2G32600 | 0,08 | 1,10 | 4,36 | 0,25 | 8,00E-01 |
| AT2G32680 | 1,55 | 2,74 | 1,78 | 1,54 | 1,23E-01 |
| AT2G32810 | 0,70 | -0,64 | 2,35 | -0,27 | 7,83E-01 |
| AT2G32920 | 1,12 | 0,36 | 1,85 | 0,19 | 8,47E-01 |
| AT2G32940 | 0,12 | 1,58 | 4,34 | 0,36 | 7,16E-01 |

| | | | | | |
|---|---|---|---|---|---|
| AT2G33040 | 0,74 | 0,96 | 2,32 | 0,41 | 6,80E-01 |
| AT2G33051 | 0,64 | 1,20 | 3,83 | 0,31 | 7,55E-01 |
| AT2G33070 | 0,08 | 0,85 | 4,38 | 0,20 | 8,45E-01 |
| AT2G33110 | 0,40 | 2,57 | 4,32 | 0,60 | 5,51E-01 |
| AT2G33120 | 0,77 | 2,08 | 2,18 | 0,96 | 3,38E-01 |
| AT2G33175 | 0,67 | 0,25 | 2,52 | 0,10 | 9,20E-01 |
| AT2G33210 | 0,70 | 2,18 | 2,47 | 0,88 | 3,79E-01 |
| AT2G33260 | 0,24 | 1,23 | 4,35 | 0,28 | 7,77E-01 |
| AT2G33320 | 0,54 | 2,30 | 2,56 | 0,90 | 3,70E-01 |
| AT2G33370 | 0,46 | 0,41 | 2,79 | 0,15 | 8,84E-01 |
| AT2G33385 | 0,08 | 0,00 | 4,40 | 0,00 | 1,00E+00 |
| AT2G33460 | 0,28 | 1,64 | 4,33 | 0,38 | 7,05E-01 |
| AT2G33470 | 0,39 | 1,91 | 3,33 | 0,57 | 5,66E-01 |
| AT2G33500 | 0,51 | -1,60 | 4,38 | -0,36 | 7,15E-01 |
| AT2G33530 | 0,99 | 1,21 | 1,98 | 0,61 | 5,41E-01 |
| AT2G33540 | 0,96 | 1,76 | 1,90 | 0,93 | 3,53E-01 |
| AT2G33550 | 0,30 | 0,00 | 4,41 | 0,00 | 1,00E+00 |
| AT2G33580 | 0,75 | 1,07 | 2,23 | 0,48 | 6,32E-01 |
| AT2G33630 | 0,50 | -0,44 | 3,27 | -0,13 | 8,94E-01 |
| AT2G33655 | 0,29 | -2,03 | 4,36 | -0,47 | 6,41E-01 |
| AT2G33800 | 0,66 | -0,68 | 2,51 | -0,27 | 7,87E-01 |
| AT2G33815 | 0,37 | -1,59 | 4,36 | -0,36 | 7,15E-01 |
| AT2G33860 | 0,43 | 1,65 | 4,17 | 0,39 | 6,93E-01 |
| AT2G33980 | 0,47 | 1,76 | 3,25 | 0,54 | 5,89E-01 |
| AT2G34040 | 1,26 | 1,06 | 1,49 | 0,71 | 4,76E-01 |
| AT2G34070 | 0,35 | 0,93 | 3,55 | 0,26 | 7,93E-01 |
| AT2G34090 | 0,11 | 0,00 | 4,41 | 0,00 | 1,00E+00 |
| AT2G34290 | 0,45 | 1,26 | 4,04 | 0,31 | 7,56E-01 |
| AT2G34300 | 0,72 | 2,53 | 4,32 | 0,58 | 5,59E-01 |
| AT2G34450 | 0,60 | 2,34 | 3,78 | 0,62 | 5,36E-01 |
| AT2G34470 | 0,95 | 2,76 | 2,09 | 1,32 | 1,86E-01 |
| AT2G34480 | 1,66 | 1,42 | 1,58 | 0,90 | 3,69E-01 |
| AT2G34490 | 0,49 | 2,46 | 4,07 | 0,60 | 5,46E-01 |
| AT2G34510 | 0,33 | -1,10 | 4,34 | -0,25 | 8,00E-01 |
| AT2G34520 | 0,36 | 0,00 | 4,41 | 0,00 | 1,00E+00 |
| AT2G34585 | 1,42 | 3,67 | 1,73 | 2,12 | 3,42E-02 |
| AT2G34680 | 1,27 | 2,14 | 2,17 | 0,99 | 3,24E-01 |
| AT2G34710 | 1,10 | 2,16 | 2,47 | 0,88 | 3,81E-01 |
| AT2G34790 | 0,43 | 0,00 | 4,19 | 0,00 | 1,00E+00 |
| AT2G34920 | 0,05 | 0,00 | 4,41 | 0,00 | 1,00E+00 |
| AT2G34930 | 0,44 | -0,46 | 3,50 | -0,13 | 8,96E-01 |
| AT2G35030 | 0,48 | 1,00 | 3,97 | 0,25 | 8,01E-01 |
| AT2G35040 | 0,75 | -0,16 | 2,89 | -0,06 | 9,56E-01 |
| AT2G35060 | 1,08 | -0,91 | 1,73 | -0,53 | 5,98E-01 |
| AT2G35120 | 0,10 | 0,00 | 4,41 | 0,00 | 1,00E+00 |
| AT2G35250 | 1,05 | -0,19 | 1,76 | -0,11 | 9,16E-01 |
| AT2G35360 | 0,45 | 0,70 | 4,16 | 0,17 | 8,66E-01 |
| AT2G35390 | 0,25 | 1,03 | 4,37 | 0,24 | 8,13E-01 |
| AT2G35420 | 0,17 | 1,57 | 4,34 | 0,36 | 7,18E-01 |
| AT2G35490 | 0,67 | 1,35 | 2,18 | 0,62 | 5,35E-01 |
| AT2G35510 | 0,57 | 1,05 | 3,25 | 0,32 | 7,48E-01 |
| AT2G35610 | 0,17 | 1,75 | 4,33 | 0,40 | 6,86E-01 |
| AT2G35680 | 0,73 | 1,56 | 2,39 | 0,65 | 5,13E-01 |
| AT2G35730 | 0,35 | 0,00 | 4,39 | 0,00 | 1,00E+00 |
| AT2G35742 | 0,34 | -0,38 | 3,55 | -0,11 | 9,16E-01 |
| AT2G08755 | 0,29 | 2,02 | 4,33 | 0,47 | 6,40E-01 |

| | | | | | |
|---|---|---|---|---|---|
| AT2G08765 | 0,51 | -2,36 | 3,22 | -0,73 | 4,63E-01 |
| AT2G35859 | 0,71 | -0,08 | 2,71 | -0,03 | 9,77E-01 |
| AT2G35880 | 0,24 | 1,62 | 4,34 | 0,37 | 7,08E-01 |
| AT2G35930 | 0,76 | 1,56 | 2,87 | 0,54 | 5,87E-01 |
| AT2G36030 | 0,62 | -2,50 | 4,32 | -0,58 | 5,63E-01 |
| AT2G36050 | 0,69 | -0,78 | 2,94 | -0,27 | 7,91E-01 |
| AT2G36060 | 0,48 | 0,13 | 2,76 | 0,05 | 9,62E-01 |
| AT2G36070 | 0,34 | 0,77 | 4,37 | 0,18 | 8,60E-01 |
| AT2G36080 | 0,43 | 0,02 | 3,33 | 0,01 | 9,96E-01 |
| AT2G36180 | 0,76 | 0,53 | 3,47 | 0,15 | 8,78E-01 |
| AT2G36200 | 0,33 | 0,77 | 4,37 | 0,18 | 8,60E-01 |
| AT2G36230 | 0,20 | 0,73 | 4,38 | 0,17 | 8,68E-01 |
| AT2G36250 | 0,55 | -2,18 | 3,24 | -0,67 | 5,01E-01 |
| AT2G36300 | 0,44 | 0,41 | 3,35 | 0,12 | 9,03E-01 |
| AT2G36360 | 0,43 | -2,18 | 3,36 | -0,65 | 5,17E-01 |
| AT2G36400 | 0,83 | 2,45 | 2,98 | 0,82 | 4,12E-01 |
| AT2G36620 | 0,25 | 1,14 | 4,36 | 0,26 | 7,94E-01 |
| AT2G36680 | 0,35 | -1,01 | 4,36 | -0,23 | 8,18E-01 |
| AT2G36710 | 0,27 | -1,36 | 4,41 | -0,31 | 7,58E-01 |
| AT2G36750 | 0,21 | 2,17 | 4,32 | 0,50 | 6,15E-01 |
| AT2G36790 | 0,34 | 1,48 | 4,34 | 0,34 | 7,34E-01 |
| AT2G36800 | 0,85 | 3,04 | 2,28 | 1,33 | 1,83E-01 |
| AT2G36810 | 1,10 | 1,41 | 2,09 | 0,67 | 5,01E-01 |
| AT2G36835 | 0,30 | 0,06 | 3,65 | 0,02 | 9,86E-01 |
| AT2G36895 | 0,35 | 0,00 | 4,39 | 0,00 | 1,00E+00 |
| AT2G36900 | 0,55 | 1,75 | 3,78 | 0,46 | 6,43E-01 |
| AT2G36930 | 0,54 | 1,38 | 3,89 | 0,35 | 7,23E-01 |
| AT2G36990 | 1,19 | 1,13 | 2,62 | 0,43 | 6,66E-01 |
| AT2G37035 | 0,78 | 1,69 | 2,35 | 0,72 | 4,72E-01 |
| AT2G37040 | 1,58 | 3,67 | 1,72 | 2,13 | 3,29E-02 |
| AT2G37080 | 0,12 | 1,46 | 4,35 | 0,34 | 7,36E-01 |
| AT2G37130 | 0,68 | 1,50 | 3,08 | 0,49 | 6,27E-01 |
| AT2G37150 | 0,75 | 2,67 | 3,41 | 0,78 | 4,34E-01 |
| AT2G37170 | 0,90 | 3,19 | 2,27 | 1,41 | 1,60E-01 |
| AT2G37200 | 0,31 | 1,61 | 3,59 | 0,45 | 6,53E-01 |
| AT2G37230 | 0,56 | 0,64 | 3,16 | 0,20 | 8,41E-01 |
| AT2G37310 | 0,77 | 3,22 | 3,47 | 0,93 | 3,53E-01 |
| AT2G37340 | 1,12 | 1,30 | 2,22 | 0,59 | 5,57E-01 |
| AT2G37360 | 0,21 | 2,17 | 4,32 | 0,50 | 6,15E-01 |
| AT2G37400 | 0,25 | 1,57 | 4,34 | 0,36 | 7,18E-01 |
| AT2G37410 | 0,19 | 1,91 | 4,33 | 0,44 | 6,58E-01 |
| AT2G37520 | 1,03 | 0,73 | 1,81 | 0,40 | 6,88E-01 |
| AT2G37540 | 0,69 | 2,82 | 2,86 | 0,98 | 3,25E-01 |
| AT2G37550 | 1,38 | 3,59 | 1,86 | 1,93 | 5,42E-02 |
| AT2G37570 | 0,98 | 0,68 | 1,86 | 0,36 | 7,16E-01 |
| AT2G37630 | 0,98 | 2,34 | 2,01 | 1,17 | 2,44E-01 |
| AT2G37660 | 1,35 | -1,67 | 1,71 | -0,98 | 3,27E-01 |
| AT2G37678 | 0,57 | -0,78 | 4,36 | -0,18 | 8,59E-01 |
| AT2G08895 | 1,00 | 2,62 | 2,26 | 1,16 | 2,46E-01 |
| AT2G08900 | 0,86 | -0,38 | 1,90 | -0,20 | 8,42E-01 |
| AT2G08910 | 1,52 | -1,93 | 1,76 | -1,10 | 2,73E-01 |
| AT2G08915 | 0,73 | 1,30 | 2,75 | 0,47 | 6,37E-01 |
| AT2G38000 | 0,32 | 0,41 | 4,38 | 0,09 | 9,25E-01 |
| AT2G38020 | 1,29 | 0,95 | 2,14 | 0,45 | 6,56E-01 |
| AT2G38160 | 0,87 | 0,47 | 3,35 | 0,14 | 8,88E-01 |
| AT2G38270 | 0,22 | -1,25 | 4,39 | -0,28 | 7,76E-01 |

| | | | | | |
|---|---|---|---|---|---|
| AT2G38290 | 0,66 | 2,54 | 2,88 | 0,88 | 3,77E-01 |
| AT2G38340 | 0,17 | 0,76 | 4,37 | 0,18 | 8,61E-01 |
| AT2G38360 | 0,39 | 0,78 | 3,37 | 0,23 | 8,18E-01 |
| AT2G38410 | 0,35 | 0,74 | 3,42 | 0,22 | 8,29E-01 |
| AT2G38480 | 0,17 | 1,75 | 4,33 | 0,40 | 6,86E-01 |
| AT2G38610 | 0,83 | 2,48 | 2,43 | 1,02 | 3,08E-01 |
| AT2G38650 | 1,16 | 0,83 | 1,98 | 0,42 | 6,74E-01 |
| AT2G38680 | 0,08 | 0,00 | 4,40 | 0,00 | 1,00E+00 |
| AT2G38750 | 0,05 | 0,00 | 4,41 | 0,00 | 1,00E+00 |
| AT2G38770 | 1,53 | 1,85 | 1,69 | 1,09 | 2,74E-01 |
| AT2G38790 | 0,55 | 0,62 | 2,40 | 0,26 | 7,96E-01 |
| AT2G38860 | 0,41 | 1,78 | 3,39 | 0,53 | 5,98E-01 |
| AT2G39000 | 1,06 | 1,97 | 2,84 | 0,69 | 4,87E-01 |
| AT2G39080 | 0,20 | 0,89 | 4,37 | 0,20 | 8,39E-01 |
| AT2G39130 | 0,08 | 0,00 | 4,40 | 0,00 | 1,00E+00 |
| AT2G39160 | 0,24 | 0,80 | 4,32 | 0,19 | 8,53E-01 |
| AT2G39170 | 0,19 | 0,46 | 4,40 | 0,10 | 9,16E-01 |
| AT2G09010 | 0,08 | 0,00 | 4,40 | 0,00 | 1,00E+00 |
| AT2G39210 | 0,91 | 2,67 | 2,72 | 0,98 | 3,27E-01 |
| AT2G39260 | 1,36 | 0,65 | 1,53 | 0,42 | 6,72E-01 |
| AT2G39350 | 0,49 | 1,96 | 3,20 | 0,61 | 5,40E-01 |
| AT2G39360 | 0,80 | 1,59 | 3,51 | 0,45 | 6,50E-01 |
| AT2G09045 | 0,52 | 2,58 | 3,86 | 0,67 | 5,04E-01 |
| AT2G39450 | 0,29 | 1,77 | 4,34 | 0,41 | 6,84E-01 |
| AT2G39480 | 1,05 | 2,38 | 2,16 | 1,10 | 2,70E-01 |
| AT2G39650 | 0,49 | 2,16 | 4,02 | 0,54 | 5,90E-01 |
| AT2G39710 | 0,60 | 1,56 | 3,21 | 0,48 | 6,28E-01 |
| AT2G39720 | 1,30 | 2,86 | 2,41 | 1,19 | 2,35E-01 |
| AT2G39750 | 1,24 | -0,21 | 1,95 | -0,11 | 9,16E-01 |
| AT2G09080 | 0,50 | 2,20 | 3,22 | 0,68 | 4,94E-01 |
| AT2G39805 | 0,30 | 1,68 | 4,34 | 0,39 | 6,99E-01 |
| AT2G39920 | 0,18 | 0,00 | 4,36 | 0,00 | 9,99E-01 |
| AT2G09090 | 0,82 | 0,17 | 2,85 | 0,06 | 9,53E-01 |
| AT2G39950 | 0,75 | 0,57 | 2,48 | 0,23 | 8,18E-01 |
| AT2G39960 | 0,25 | -0,50 | 4,31 | -0,12 | 9,07E-01 |
| AT2G39990 | 1,46 | 1,49 | 2,57 | 0,58 | 5,61E-01 |
| AT2G40070 | 0,52 | 2,73 | 3,86 | 0,71 | 4,79E-01 |
| AT2G40080 | 0,23 | 1,56 | 4,34 | 0,36 | 7,19E-01 |
| AT2G40165 | 0,38 | 0,90 | 4,38 | 0,21 | 8,37E-01 |
| AT2G40205 | 0,27 | 0,00 | 4,41 | 0,00 | 1,00E+00 |
| AT2G40270 | 1,34 | 2,71 | 1,70 | 1,60 | 1,11E-01 |
| AT2G40280 | 0,63 | -0,28 | 2,92 | -0,10 | 9,22E-01 |
| AT2G40350 | 0,12 | 1,65 | 4,34 | 0,38 | 7,04E-01 |
| AT2G40360 | 0,28 | 0,82 | 4,37 | 0,19 | 8,52E-01 |
| AT2G40430 | 0,45 | -1,32 | 4,23 | -0,31 | 7,54E-01 |
| AT2G40475 | 0,89 | 2,50 | 2,30 | 1,09 | 2,77E-01 |
| AT2G40590 | 0,50 | -1,25 | 2,67 | -0,47 | 6,41E-01 |
| AT2G40600 | 0,11 | 0,90 | 4,37 | 0,21 | 8,36E-01 |
| AT2G40610 | 0,37 | -2,44 | 4,36 | -0,56 | 5,75E-01 |
| AT2G40650 | 0,77 | 1,17 | 2,31 | 0,51 | 6,12E-01 |
| AT2G40730 | 0,34 | 0,02 | 3,50 | 0,01 | 9,95E-01 |
| AT2G40750 | 0,46 | -1,17 | 4,09 | -0,28 | 7,76E-01 |
| AT2G40770 | 0,51 | -2,06 | 4,00 | -0,52 | 6,06E-01 |
| AT2G40815 | 0,24 | 1,62 | 4,34 | 0,37 | 7,08E-01 |
| AT2G40860 | 0,30 | 1,68 | 4,34 | 0,39 | 6,99E-01 |
| AT2G40890 | 0,53 | 1,79 | 3,86 | 0,46 | 6,43E-01 |

| | | | | | |
|---|---|---|---|---|---|
| AT2G40920 | 0,35 | 0,93 | 4,33 | 0,22 | 8,29E-01 |
| AT2G40925 | 0,65 | 1,97 | 3,66 | 0,54 | 5,89E-01 |
| AT2G40950 | 0,74 | 3,26 | 3,49 | 0,93 | 3,50E-01 |
| AT2G40970 | 0,42 | 0,00 | 4,41 | 0,00 | 1,00E+00 |
| AT2G09225 | 0,24 | 0,44 | 4,35 | 0,10 | 9,19E-01 |
| AT2G41120 | 0,21 | 2,17 | 4,32 | 0,50 | 6,15E-01 |
| AT2G41160 | 0,39 | 2,11 | 4,33 | 0,49 | 6,27E-01 |
| AT2G41250 | 0,87 | 2,71 | 2,42 | 1,12 | 2,63E-01 |
| AT2G41420 | 0,11 | 0,00 | 4,41 | 0,00 | 1,00E+00 |
| AT2G41475 | 0,71 | 0,02 | 2,58 | 0,01 | 9,93E-01 |
| AT2G41500 | 0,96 | 1,16 | 2,13 | 0,55 | 5,85E-01 |
| AT2G41600 | 0,58 | 0,40 | 2,63 | 0,15 | 8,78E-01 |
| AT2G41620 | 1,68 | 2,60 | 1,84 | 1,41 | 1,57E-01 |
| AT2G41616 | 0,33 | 2,12 | 4,32 | 0,49 | 6,24E-01 |
| AT2G41630 | 0,92 | 0,67 | 2,85 | 0,24 | 8,13E-01 |
| AT2G41680 | 0,32 | 2,17 | 4,32 | 0,50 | 6,15E-01 |
| AT2G41700 | 1,62 | 2,72 | 2,03 | 1,34 | 1,82E-01 |
| AT2G41770 | 0,40 | 1,42 | 4,28 | 0,33 | 7,41E-01 |
| AT2G41790 | 0,58 | 1,06 | 2,59 | 0,41 | 6,83E-01 |
| AT2G41830 | 0,40 | 1,27 | 4,15 | 0,31 | 7,59E-01 |
| AT2G41835 | 0,70 | 2,97 | 3,67 | 0,81 | 4,18E-01 |
| AT2G41870 | 0,29 | 1,56 | 4,34 | 0,36 | 7,18E-01 |
| AT2G41890 | 0,43 | 1,17 | 4,19 | 0,28 | 7,81E-01 |
| AT2G41945 | 0,60 | 3,03 | 3,91 | 0,77 | 4,39E-01 |
| AT2G41960 | 0,92 | 1,66 | 1,90 | 0,87 | 3,82E-01 |
| AT2G42010 | 0,72 | -0,14 | 2,03 | -0,07 | 9,45E-01 |
| AT2G42020 | 0,48 | -2,42 | 4,34 | -0,56 | 5,78E-01 |
| AT2G42070 | 0,23 | 1,09 | 4,36 | 0,25 | 8,02E-01 |
| AT2G42230 | 0,42 | 2,28 | 4,10 | 0,56 | 5,77E-01 |
| AT2G42280 | 0,33 | 1,33 | 4,35 | 0,31 | 7,59E-01 |
| AT2G42365 | 0,35 | 0,00 | 4,39 | 0,00 | 1,00E+00 |
| AT2G42400 | 1,01 | 1,93 | 2,02 | 0,96 | 3,40E-01 |
| AT2G42500 | 0,82 | 0,87 | 2,34 | 0,37 | 7,10E-01 |
| AT2G42530 | 0,98 | -3,00 | 1,98 | -1,51 | 1,30E-01 |
| AT2G42690 | 1,34 | -0,01 | 1,49 | -0,01 | 9,94E-01 |
| AT2G42760 | 0,47 | 0,27 | 2,69 | 0,10 | 9,20E-01 |
| AT2G42790 | 0,53 | 0,83 | 2,64 | 0,31 | 7,55E-01 |
| AT2G42800 | 0,18 | 0,00 | 4,41 | 0,00 | 1,00E+00 |
| AT2G42810 | 0,84 | 0,61 | 1,96 | 0,31 | 7,56E-01 |
| AT2G42840 | 0,22 | 0,00 | 4,41 | 0,00 | 1,00E+00 |
| AT2G42865 | 0,08 | 1,10 | 4,36 | 0,25 | 8,00E-01 |
| AT2G42870 | 0,10 | 0,62 | 4,38 | 0,14 | 8,87E-01 |
| AT2G42880 | 0,41 | 1,88 | 3,45 | 0,55 | 5,85E-01 |
| AT2G43180 | 0,70 | 1,85 | 2,90 | 0,64 | 5,23E-01 |
| AT2G43260 | 0,20 | 1,07 | 4,36 | 0,25 | 8,06E-01 |
| AT2G43290 | 0,50 | 2,59 | 3,17 | 0,82 | 4,13E-01 |
| AT2G43320 | 0,22 | 1,92 | 4,33 | 0,44 | 6,57E-01 |
| AT2G43360 | 0,33 | 0,82 | 4,32 | 0,19 | 8,50E-01 |
| AT2G43420 | 0,11 | 0,90 | 4,37 | 0,21 | 8,36E-01 |
| AT2G43430 | 0,59 | -2,31 | 2,67 | -0,87 | 3,86E-01 |
| AT2G43460 | 0,41 | -0,58 | 3,42 | -0,17 | 8,66E-01 |
| AT2G43490 | 0,44 | 2,58 | 4,05 | 0,64 | 5,25E-01 |
| AT2G43560 | 0,25 | 0,75 | 4,37 | 0,17 | 8,63E-01 |
| AT2G43750 | 1,46 | 1,75 | 1,74 | 1,01 | 3,14E-01 |
| AT2G43865 | 0,15 | 0,71 | 4,37 | 0,16 | 8,71E-01 |
| AT2G43910 | 0,26 | 0,66 | 3,74 | 0,18 | 8,59E-01 |

| | | | | | |
|---|---|---|---|---|---|
| AT2G43940 | 0,08 | 0,85 | 4,38 | 0,20 | 8,45E-01 |
| AT2G43945 | 0,47 | -0,91 | 4,37 | -0,21 | 8,35E-01 |
| AT2G43950 | 0,24 | 1,58 | 4,34 | 0,36 | 7,16E-01 |
| AT2G44090 | 0,69 | 0,94 | 2,60 | 0,36 | 7,16E-01 |
| AT2G44140 | 0,91 | -1,22 | 2,20 | -0,55 | 5,81E-01 |
| AT2G44180 | 0,46 | 1,41 | 3,30 | 0,43 | 6,69E-01 |
| AT2G44255 | 0,20 | 1,68 | 4,34 | 0,39 | 6,99E-01 |
| AT2G44360 | 0,63 | 16,80 | 4,34 | 3,87 | 1,08E-04 |
| AT2G44610 | 0,48 | 1,04 | 4,15 | 0,25 | 8,02E-01 |
| AT2G44660 | 0,35 | 1,87 | 3,48 | 0,54 | 5,92E-01 |
| AT2G44760 | 0,32 | 1,73 | 3,58 | 0,48 | 6,29E-01 |
| AT2G44798 | 0,36 | 0,81 | 4,37 | 0,19 | 8,53E-01 |
| AT2G44950 | 0,77 | 0,66 | 2,17 | 0,30 | 7,63E-01 |
| AT2G45070 | 0,80 | 0,77 | 2,16 | 0,36 | 7,21E-01 |
| AT2G45220 | 0,54 | 2,77 | 4,32 | 0,64 | 5,21E-01 |
| AT2G45315 | 0,67 | 1,10 | 2,51 | 0,44 | 6,61E-01 |
| AT2G45460 | 0,20 | 0,51 | 4,39 | 0,12 | 9,08E-01 |
| AT2G45470 | 1,65 | -1,03 | 2,13 | -0,48 | 6,29E-01 |
| AT2G45520 | 0,22 | -0,24 | 4,37 | -0,06 | 9,56E-01 |
| AT2G09640 | 1,68 | 2,13 | 2,11 | 1,01 | 3,14E-01 |
| AT2G45560 | 0,47 | 0,00 | 4,40 | 0,00 | 1,00E+00 |
| AT2G45660 | 0,55 | 1,00 | 2,30 | 0,43 | 6,64E-01 |
| AT2G45670 | 0,95 | 2,19 | 2,24 | 0,98 | 3,28E-01 |
| AT2G45700 | 0,38 | 0,64 | 4,27 | 0,15 | 8,82E-01 |
| AT2G45820 | 0,98 | -1,78 | 1,77 | -1,01 | 3,15E-01 |
| AT2G45970 | 0,56 | 2,13 | 2,64 | 0,81 | 4,21E-01 |
| AT2G45980 | 0,45 | 2,31 | 4,09 | 0,57 | 5,72E-01 |
| AT2G46030 | 0,57 | 2,63 | 4,31 | 0,61 | 5,43E-01 |
| AT2G46040 | 0,20 | 0,89 | 4,37 | 0,20 | 8,39E-01 |
| AT2G46060 | 0,49 | -2,71 | 4,34 | -0,62 | 5,32E-01 |
| AT2G46080 | 0,51 | 0,98 | 3,26 | 0,30 | 7,65E-01 |
| AT2G46180 | 0,35 | 1,02 | 4,36 | 0,23 | 8,14E-01 |
| AT2G46280 | 1,16 | -0,65 | 1,71 | -0,38 | 7,03E-01 |
| AT2G46330 | 0,93 | -1,11 | 1,95 | -0,57 | 5,70E-01 |
| AT2G46390 | 0,46 | 1,88 | 4,07 | 0,46 | 6,45E-01 |
| AT2G46505 | 0,89 | 0,53 | 2,43 | 0,22 | 8,28E-01 |
| AT2G46510 | 1,07 | 0,14 | 2,17 | 0,06 | 9,48E-01 |
| AT2G46540 | 0,12 | 0,00 | 4,39 | 0,00 | 1,00E+00 |
| AT2G46550 | 0,57 | 0,69 | 2,73 | 0,25 | 8,00E-01 |
| AT2G46572 | 0,51 | 2,52 | 3,99 | 0,63 | 5,27E-01 |
| AT2G46610 | 0,85 | -0,45 | 2,16 | -0,21 | 8,35E-01 |
| AT2G46620 | 1,45 | 2,82 | 1,83 | 1,54 | 1,24E-01 |
| AT2G46680 | 0,94 | 1,31 | 2,29 | 0,57 | 5,69E-01 |
| AT2G46840 | 0,08 | 1,10 | 4,36 | 0,25 | 8,00E-01 |
| AT2G46920 | 0,44 | 2,53 | 4,01 | 0,63 | 5,29E-01 |
| AT2G46980 | 0,24 | 0,00 | 4,41 | 0,00 | 1,00E+00 |
| AT2G47090 | 0,25 | 2,36 | 4,32 | 0,55 | 5,86E-01 |
| AT2G47180 | 1,41 | 0,51 | 1,78 | 0,28 | 7,76E-01 |
| AT2G47250 | 1,15 | 2,37 | 2,25 | 1,05 | 2,92E-01 |
| AT2G47330 | 0,60 | 1,00 | 3,11 | 0,32 | 7,47E-01 |
| AT2G47600 | 0,61 | 2,41 | 2,52 | 0,95 | 3,40E-01 |
| AT2G47650 | 1,46 | 1,51 | 1,68 | 0,90 | 3,70E-01 |
| AT2G47680 | 0,89 | 2,19 | 2,10 | 1,04 | 2,99E-01 |
| AT2G47700 | 0,93 | 2,54 | 2,44 | 1,04 | 2,98E-01 |
| AT2G47840 | 0,52 | -1,70 | 3,29 | -0,52 | 6,05E-01 |
| AT2G47990 | 0,88 | 1,91 | 2,69 | 0,71 | 4,76E-01 |

| | | | | | |
|---|---|---|---|---|---|
| AT2G48000 | 1,00 | 3,62 | 3,32 | 1,09 | 2,76E-01 |
| AT2G48030 | 0,73 | 2,10 | 2,40 | 0,87 | 3,82E-01 |
| AT2G48130 | 0,91 | -1,27 | 2,05 | -0,62 | 5,34E-01 |
| AT3G01090 | 0,63 | 0,93 | 2,21 | 0,42 | 6,75E-01 |
| AT3G01100 | 0,90 | 1,65 | 2,23 | 0,74 | 4,59E-01 |
| AT3G01130 | 0,21 | 0,00 | 4,38 | 0,00 | 1,00E+00 |
| AT3G01160 | 0,24 | 0,80 | 4,32 | 0,19 | 8,53E-01 |
| AT3G01180 | 0,84 | 0,76 | 1,85 | 0,41 | 6,82E-01 |
| AT3G01300 | 0,08 | 1,10 | 4,36 | 0,25 | 8,00E-01 |
| AT3G01340 | 0,50 | 1,46 | 3,17 | 0,46 | 6,46E-01 |
| AT3G01380 | 1,09 | 1,54 | 1,96 | 0,78 | 4,34E-01 |
| AT3G01390 | 0,38 | -0,59 | 4,33 | -0,14 | 8,92E-01 |
| AT3G01415 | 0,40 | -1,20 | 4,39 | -0,27 | 7,85E-01 |
| AT3G01420 | 0,31 | 1,77 | 4,33 | 0,41 | 6,83E-01 |
| AT3G01430 | 0,79 | 2,02 | 2,99 | 0,68 | 4,99E-01 |
| AT3G01470 | 1,49 | 2,27 | 1,73 | 1,31 | 1,91E-01 |
| AT3G01550 | 0,10 | 0,42 | 4,38 | 0,10 | 9,23E-01 |
| AT3G01720 | 0,80 | 3,05 | 2,83 | 1,08 | 2,81E-01 |
| AT3G01820 | 0,81 | 0,27 | 3,54 | 0,08 | 9,40E-01 |
| AT3G01840 | 0,88 | 2,11 | 2,22 | 0,95 | 3,42E-01 |
| AT3G01910 | 0,46 | 1,24 | 4,05 | 0,31 | 7,59E-01 |
| AT3G01930 | 0,62 | 0,51 | 2,98 | 0,17 | 8,65E-01 |
| AT3G01980 | 0,16 | 1,74 | 4,33 | 0,40 | 6,87E-01 |
| AT3G02040 | 0,68 | 1,20 | 2,50 | 0,48 | 6,31E-01 |
| AT3G02080 | 1,20 | 0,16 | 2,12 | 0,07 | 9,40E-01 |
| AT3G02190 | 0,44 | 0,37 | 3,34 | 0,11 | 9,12E-01 |
| AT3G02210 | 0,52 | 2,23 | 4,01 | 0,56 | 5,78E-01 |
| AT3G02250 | 0,34 | -1,94 | 4,35 | -0,45 | 6,55E-01 |
| AT3G02380 | 0,91 | 1,94 | 2,32 | 0,84 | 4,02E-01 |
| AT3G02410 | 0,66 | 0,00 | 4,41 | 0,00 | 1,00E+00 |
| AT3G02450 | 0,68 | 0,37 | 2,35 | 0,16 | 8,73E-01 |
| AT3G02520 | 1,11 | -0,18 | 1,58 | -0,12 | 9,07E-01 |
| AT3G02530 | 0,67 | 1,13 | 2,65 | 0,43 | 6,69E-01 |
| AT3G02700 | 0,39 | -0,38 | 4,22 | -0,09 | 9,29E-01 |
| AT3G02730 | 0,92 | -2,18 | 2,85 | -0,77 | 4,43E-01 |
| AT3G02750 | 1,16 | 0,36 | 1,91 | 0,19 | 8,52E-01 |
| AT3G02760 | 0,94 | 2,89 | 2,71 | 1,07 | 2,86E-01 |
| AT3G01205 | 0,80 | 1,06 | 2,36 | 0,45 | 6,52E-01 |
| AT3G03090 | 0,29 | -2,22 | 4,36 | -0,51 | 6,11E-01 |
| AT3G03150 | 0,22 | -1,35 | 4,41 | -0,31 | 7,59E-01 |
| AT3G03220 | 0,16 | 0,00 | 4,41 | 0,00 | 1,00E+00 |
| AT3G03250 | 1,00 | 2,03 | 1,88 | 1,08 | 2,80E-01 |
| AT3G03300 | 0,33 | 2,63 | 4,32 | 0,61 | 5,42E-01 |
| AT3G03305 | 0,32 | 1,61 | 4,34 | 0,37 | 7,12E-01 |
| AT3G03310 | 0,29 | 1,82 | 4,33 | 0,42 | 6,75E-01 |
| AT3G03330 | 0,30 | 0,00 | 4,39 | 0,00 | 1,00E+00 |
| AT3G03405 | 0,16 | 0,00 | 4,41 | 0,00 | 1,00E+00 |
| AT3G03460 | 0,17 | 1,40 | 4,35 | 0,32 | 7,48E-01 |
| AT3G03470 | 1,68 | 4,25 | 1,83 | 2,32 | 2,04E-02 |
| AT3G03480 | 0,08 | 1,10 | 4,36 | 0,25 | 8,00E-01 |
| AT3G03550 | 0,10 | 0,42 | 4,38 | 0,10 | 9,23E-01 |
| AT3G03560 | 0,61 | 2,72 | 2,96 | 0,92 | 3,59E-01 |
| AT3G03580 | 0,26 | 1,08 | 4,37 | 0,25 | 8,05E-01 |
| AT3G03590 | 0,08 | 1,10 | 4,36 | 0,25 | 8,00E-01 |
| AT3G03600 | 0,30 | -1,59 | 4,37 | -0,36 | 7,16E-01 |
| AT3G03720 | 0,26 | 1,82 | 4,33 | 0,42 | 6,73E-01 |

| | | | | | |
|---|---|---|---|---|---|
| AT3G03770 | 0,99 | 3,98 | 3,02 | 1,32 | 1,88E-01 |
| AT3G03890 | 0,79 | 2,20 | 2,44 | 0,90 | 3,68E-01 |
| AT3G03920 | 0,42 | -2,48 | 3,32 | -0,74 | 4,56E-01 |
| AT3G03940 | 0,41 | 0,22 | 2,84 | 0,08 | 9,37E-01 |
| AT3G04030 | 0,25 | 0,00 | 4,41 | 0,00 | 1,00E+00 |
| AT3G04080 | 1,01 | 1,91 | 1,96 | 0,97 | 3,30E-01 |
| AT3G04150 | 0,26 | 1,08 | 4,37 | 0,25 | 8,05E-01 |
| AT3G04200 | 0,62 | 2,26 | 4,32 | 0,52 | 6,01E-01 |
| AT3G04220 | 0,33 | 2,27 | 4,32 | 0,53 | 5,99E-01 |
| AT3G04260 | 0,61 | -2,20 | 4,36 | -0,51 | 6,13E-01 |
| AT3G04290 | 0,84 | 0,44 | 3,01 | 0,14 | 8,85E-01 |
| AT3G04350 | 0,35 | 1,30 | 4,35 | 0,30 | 7,65E-01 |
| AT3G04480 | 0,84 | 2,95 | 2,32 | 1,27 | 2,03E-01 |
| AT3G04500 | 0,20 | 0,00 | 4,41 | 0,00 | 1,00E+00 |
| AT3G04580 | 1,03 | 1,99 | 2,02 | 0,99 | 3,23E-01 |
| AT3G04640 | 0,85 | 0,81 | 2,21 | 0,36 | 7,15E-01 |
| AT3G04690 | 0,22 | -1,66 | 4,38 | -0,38 | 7,05E-01 |
| AT3G04720 | 0,50 | 2,30 | 3,08 | 0,75 | 4,55E-01 |
| AT3G04730 | 1,02 | 0,05 | 1,67 | 0,03 | 9,77E-01 |
| AT3G04750 | 0,57 | 1,26 | 2,66 | 0,47 | 6,35E-01 |
| AT3G04760 | 0,89 | 3,23 | 2,04 | 1,59 | 1,13E-01 |
| AT3G04860 | 0,12 | 1,46 | 4,35 | 0,34 | 7,36E-01 |
| AT3G04980 | 0,46 | 2,30 | 4,33 | 0,53 | 5,95E-01 |
| AT3G05030 | 0,28 | 1,77 | 4,33 | 0,41 | 6,83E-01 |
| AT3G01585 | 0,10 | 0,00 | 4,41 | 0,00 | 1,00E+00 |
| AT3G05040 | 1,58 | 0,55 | 1,64 | 0,34 | 7,36E-01 |
| AT3G05050 | 0,38 | 2,01 | 4,33 | 0,46 | 6,42E-01 |
| AT3G05060 | 0,60 | 2,14 | 3,70 | 0,58 | 5,64E-01 |
| AT3G05090 | 0,52 | 1,76 | 2,62 | 0,67 | 5,02E-01 |
| AT3G05165 | 0,83 | 1,04 | 2,07 | 0,50 | 6,15E-01 |
| AT3G05240 | 0,13 | 1,15 | 4,36 | 0,26 | 7,92E-01 |
| AT3G05270 | 0,58 | 1,56 | 2,60 | 0,60 | 5,48E-01 |
| AT3G05280 | 0,59 | -0,55 | 3,82 | -0,14 | 8,86E-01 |
| AT3G05540 | 1,04 | -2,10 | 1,74 | -1,21 | 2,26E-01 |
| AT3G05650 | 0,98 | 3,28 | 2,27 | 1,44 | 1,49E-01 |
| AT3G05660 | 0,79 | 3,40 | 2,71 | 1,25 | 2,10E-01 |
| AT3G05680 | 1,68 | 2,02 | 1,58 | 1,28 | 2,01E-01 |
| AT3G05690 | 0,11 | 0,00 | 4,41 | 0,00 | 1,00E+00 |
| AT3G05770 | 0,46 | 1,73 | 3,32 | 0,52 | 6,01E-01 |
| AT3G05810 | 0,21 | 0,18 | 4,34 | 0,04 | 9,68E-01 |
| AT3G05850 | 0,90 | 1,59 | 2,53 | 0,63 | 5,29E-01 |
| AT3G05880 | 0,55 | -1,75 | 3,10 | -0,56 | 5,74E-01 |
| AT3G05900 | 0,51 | -1,37 | 3,20 | -0,43 | 6,67E-01 |
| AT3G05910 | 0,68 | -2,34 | 2,55 | -0,92 | 3,59E-01 |
| AT3G05932 | 0,27 | 1,70 | 4,33 | 0,39 | 6,94E-01 |
| AT3G05970 | 0,71 | -1,72 | 2,25 | -0,77 | 4,43E-01 |
| AT3G06030 | 0,29 | 2,52 | 4,32 | 0,58 | 5,59E-01 |
| AT3G06050 | 0,18 | -1,25 | 4,39 | -0,28 | 7,76E-01 |
| AT3G06060 | 0,17 | 0,00 | 4,40 | 0,00 | 1,00E+00 |
| AT3G06080 | 1,38 | -1,38 | 1,68 | -0,82 | 4,10E-01 |
| AT3G06150 | 0,98 | -1,04 | 1,58 | -0,66 | 5,09E-01 |
| AT3G06190 | 0,71 | 1,15 | 3,17 | 0,36 | 7,17E-01 |
| AT3G06250 | 0,74 | 1,80 | 2,83 | 0,64 | 5,25E-01 |
| AT3G06290 | 0,72 | 2,88 | 2,44 | 1,18 | 2,38E-01 |
| AT3G06310 | 0,08 | 0,85 | 4,38 | 0,20 | 8,45E-01 |
| AT3G06350 | 0,64 | 2,60 | 2,96 | 0,88 | 3,80E-01 |

| | | | | | |
|---|---|---|---|---|---|
| AT3G06430 | 0,12 | 1,65 | 4,34 | 0,38 | 7,04E-01 |
| AT3G06440 | 0,12 | 1,46 | 4,35 | 0,34 | 7,36E-01 |
| AT3G06450 | 0,61 | 2,30 | 2,65 | 0,87 | 3,85E-01 |
| AT3G06480 | 1,05 | 1,33 | 2,00 | 0,67 | 5,04E-01 |
| AT3G06483 | 0,45 | -0,36 | 2,83 | -0,13 | 8,98E-01 |
| AT3G06540 | 0,08 | 0,85 | 4,38 | 0,20 | 8,45E-01 |
| AT3G06550 | 0,32 | -0,21 | 4,36 | -0,05 | 9,62E-01 |
| AT3G06580 | 0,15 | 1,62 | 4,34 | 0,37 | 7,09E-01 |
| AT3G06590 | 0,40 | 0,00 | 4,40 | 0,00 | 1,00E+00 |
| AT3G06620 | 0,12 | 1,65 | 4,34 | 0,38 | 7,04E-01 |
| AT3G06665 | 0,25 | 0,00 | 4,41 | 0,00 | 1,00E+00 |
| AT3G06670 | 1,42 | 2,33 | 1,68 | 1,39 | 1,66E-01 |
| AT3G06680 | 0,28 | 0,00 | 4,41 | 0,00 | 1,00E+00 |
| AT3G06690 | 0,20 | 0,00 | 4,41 | 0,00 | 1,00E+00 |
| AT3G06700 | 0,57 | 0,03 | 2,74 | 0,01 | 9,90E-01 |
| AT3G06790 | 0,16 | 0,90 | 4,38 | 0,21 | 8,37E-01 |
| AT3G06820 | 0,08 | 1,10 | 4,36 | 0,25 | 8,00E-01 |
| AT3G06870 | 0,21 | 0,00 | 4,41 | 0,00 | 1,00E+00 |
| AT3G06900 | 1,19 | 1,41 | 1,96 | 0,72 | 4,71E-01 |
| AT3G06930 | 0,40 | 0,83 | 4,20 | 0,20 | 8,44E-01 |
| AT3G06960 | 0,89 | -2,13 | 2,51 | -0,85 | 3,95E-01 |
| AT3G07010 | 0,41 | -0,42 | 4,26 | -0,10 | 9,22E-01 |
| AT3G07020 | 0,54 | 2,46 | 3,76 | 0,65 | 5,13E-01 |
| AT3G07040 | 1,04 | 3,49 | 2,18 | 1,60 | 1,09E-01 |
| AT3G07050 | 0,53 | 0,35 | 3,25 | 0,11 | 9,14E-01 |
| AT3G07115 | 0,08 | 0,00 | 4,40 | 0,00 | 1,00E+00 |
| AT3G07180 | 0,12 | 1,65 | 4,34 | 0,38 | 7,04E-01 |
| AT3G07215 | 0,55 | -0,48 | 2,68 | -0,18 | 8,57E-01 |
| AT3G07273 | 0,08 | 1,10 | 4,36 | 0,25 | 8,00E-01 |
| AT3G07290 | 0,69 | 3,18 | 3,46 | 0,92 | 3,58E-01 |
| AT3G07300 | 0,29 | 0,92 | 4,36 | 0,21 | 8,33E-01 |
| AT3G07330 | 0,98 | 3,03 | 2,50 | 1,21 | 2,25E-01 |
| AT3G07380 | 1,22 | -0,89 | 1,62 | -0,55 | 5,80E-01 |
| AT3G07460 | 0,63 | -1,78 | 2,62 | -0,68 | 4,97E-01 |
| AT3G07470 | 0,58 | -0,11 | 3,22 | -0,03 | 9,73E-01 |
| AT3G07520 | 0,51 | 2,25 | 3,22 | 0,70 | 4,85E-01 |
| AT3G07540 | 0,08 | 1,10 | 4,36 | 0,25 | 8,00E-01 |
| AT3G07660 | 1,40 | 3,29 | 1,96 | 1,68 | 9,27E-02 |
| AT3G07700 | 0,94 | 0,65 | 1,74 | 0,38 | 7,07E-01 |
| AT3G07800 | 0,53 | 2,01 | 2,70 | 0,74 | 4,58E-01 |
| AT3G07890 | 0,34 | 1,67 | 4,33 | 0,39 | 6,99E-01 |
| AT3G08570 | 1,10 | 3,19 | 2,14 | 1,49 | 1,37E-01 |
| AT3G08590 | 1,52 | -0,30 | 1,52 | -0,20 | 8,45E-01 |
| AT3G08610 | 0,59 | -0,01 | 3,20 | 0,00 | 9,99E-01 |
| AT3G08650 | 0,24 | 1,26 | 4,37 | 0,29 | 7,73E-01 |
| AT3G08720 | 1,37 | 3,91 | 1,96 | 1,99 | 4,61E-02 |
| AT3G08760 | 0,70 | 1,24 | 2,26 | 0,55 | 5,83E-01 |
| AT3G08840 | 0,67 | 1,81 | 3,03 | 0,60 | 5,50E-01 |
| AT3G08870 | 0,75 | 2,07 | 3,38 | 0,61 | 5,41E-01 |
| AT3G08960 | 0,38 | 1,08 | 4,33 | 0,25 | 8,03E-01 |
| AT3G09040 | 0,17 | 1,92 | 4,33 | 0,44 | 6,57E-01 |
| AT3G09070 | 1,31 | 0,17 | 2,01 | 0,08 | 9,34E-01 |
| AT3G09090 | 1,29 | 2,26 | 1,88 | 1,20 | 2,28E-01 |
| AT3G09190 | 0,55 | 0,78 | 3,79 | 0,21 | 8,37E-01 |
| AT3G09210 | 0,27 | -1,63 | 4,37 | -0,37 | 7,09E-01 |
| AT3G09270 | 0,53 | 0,93 | 2,68 | 0,35 | 7,29E-01 |

| | | | | | |
|---|---|---|---|---|---|
| AT3G09320 | 0,85 | 3,37 | 2,32 | 1,45 | 1,47E-01 |
| AT3G09410 | 0,82 | 2,43 | 2,39 | 1,02 | 3,10E-01 |
| AT3G09470 | 0,82 | 3,03 | 3,44 | 0,88 | 3,78E-01 |
| AT3G09560 | 0,97 | 0,59 | 2,17 | 0,27 | 7,85E-01 |
| AT3G09580 | 0,48 | 1,26 | 3,42 | 0,37 | 7,13E-01 |
| AT3G09770 | 0,38 | 1,86 | 4,34 | 0,43 | 6,69E-01 |
| AT3G09780 | 0,23 | 0,00 | 4,41 | 0,00 | 1,00E+00 |
| AT3G09850 | 1,65 | 1,69 | 1,64 | 1,03 | 3,04E-01 |
| AT3G09870 | 0,08 | 1,10 | 4,36 | 0,25 | 8,00E-01 |
| AT3G09920 | 0,16 | 1,07 | 4,37 | 0,24 | 8,07E-01 |
| AT3G10010 | 0,35 | 0,00 | 4,41 | 0,00 | 1,00E+00 |
| AT3G10020 | 0,27 | 0,02 | 4,33 | 0,01 | 9,96E-01 |
| AT3G10030 | 0,88 | -1,06 | 2,78 | -0,38 | 7,03E-01 |
| AT3G10050 | 0,25 | 0,61 | 4,34 | 0,14 | 8,89E-01 |
| AT3G10070 | 0,46 | 1,30 | 4,27 | 0,30 | 7,61E-01 |
| AT3G10130 | 0,21 | 2,05 | 4,33 | 0,47 | 6,36E-01 |
| AT3G10160 | 0,36 | 1,56 | 3,42 | 0,46 | 6,48E-01 |
| AT3G10230 | 1,13 | 3,93 | 2,19 | 1,80 | 7,26E-02 |
| AT3G10250 | 0,08 | 1,10 | 4,36 | 0,25 | 8,00E-01 |
| AT3G10270 | 0,56 | -1,74 | 3,12 | -0,56 | 5,77E-01 |
| AT3G10320 | 0,12 | 1,65 | 4,34 | 0,38 | 7,04E-01 |
| AT3G10360 | 0,63 | 1,27 | 3,78 | 0,34 | 7,37E-01 |
| AT3G10405 | 0,68 | 0,00 | 4,41 | 0,00 | 1,00E+00 |
| AT3G10410 | 1,05 | -0,38 | 1,78 | -0,21 | 8,32E-01 |
| AT3G10520 | 0,15 | 0,00 | 4,39 | 0,00 | 1,00E+00 |
| AT3G10525 | 0,39 | 1,06 | 3,39 | 0,31 | 7,54E-01 |
| AT3G10630 | 0,99 | 3,42 | 1,99 | 1,72 | 8,59E-02 |
| AT3G10640 | 0,33 | -0,54 | 4,37 | -0,12 | 9,02E-01 |
| AT3G10660 | 0,27 | 1,05 | 4,36 | 0,24 | 8,09E-01 |
| AT3G10690 | 0,77 | 1,91 | 3,46 | 0,55 | 5,82E-01 |
| AT3G10720 | 0,36 | -2,17 | 4,38 | -0,50 | 6,20E-01 |
| AT3G10730 | 0,58 | -1,25 | 3,03 | -0,41 | 6,81E-01 |
| AT3G10770 | 0,74 | 1,60 | 2,40 | 0,67 | 5,05E-01 |
| AT3G10810 | 0,46 | 1,98 | 4,19 | 0,47 | 6,36E-01 |
| AT3G10840 | 0,41 | 0,00 | 4,41 | 0,00 | 1,00E+00 |
| AT3G10850 | 0,36 | 1,36 | 4,35 | 0,31 | 7,53E-01 |
| AT3G10900 | 0,31 | 0,64 | 4,37 | 0,15 | 8,84E-01 |
| AT3G10915 | 0,24 | 0,99 | 4,38 | 0,23 | 8,21E-01 |
| AT3G10930 | 0,61 | 1,84 | 2,52 | 0,73 | 4,66E-01 |
| AT3G10960 | 1,23 | 2,99 | 1,80 | 1,66 | 9,69E-02 |
| AT3G11030 | 0,56 | 0,42 | 3,83 | 0,11 | 9,13E-01 |
| AT3G11100 | 0,47 | 1,61 | 3,98 | 0,41 | 6,85E-01 |
| AT3G11280 | 0,31 | -0,49 | 4,37 | -0,11 | 9,11E-01 |
| AT3G11290 | 0,30 | 1,56 | 4,34 | 0,36 | 7,18E-01 |
| AT3G11310 | 0,31 | 0,99 | 4,36 | 0,23 | 8,20E-01 |
| AT3G11320 | 0,17 | 1,57 | 4,34 | 0,36 | 7,18E-01 |
| AT3G11330 | 0,97 | 3,34 | 2,57 | 1,30 | 1,95E-01 |
| AT3G11435 | 0,23 | -2,40 | 4,35 | -0,55 | 5,81E-01 |
| AT3G11450 | 1,01 | 2,33 | 2,03 | 1,15 | 2,51E-01 |
| AT3G11510 | 1,34 | -1,47 | 1,89 | -0,78 | 4,37E-01 |
| AT3G02435 | 1,34 | -1,06 | 2,11 | -0,50 | 6,15E-01 |
| AT3G02445 | 0,88 | -2,35 | 3,50 | -0,67 | 5,03E-01 |
| AT3G02455 | 0,99 | 2,48 | 2,92 | 0,85 | 3,97E-01 |
| AT3G11570 | 0,77 | 1,92 | 3,06 | 0,63 | 5,30E-01 |
| AT3G11580 | 0,26 | 1,35 | 4,35 | 0,31 | 7,56E-01 |
| AT3G11650 | 0,48 | 1,23 | 3,24 | 0,38 | 7,05E-01 |

| | | | | | |
|---|---|---|---|---|---|
| AT3G11660 | 1,33 | 3,52 | 3,36 | 1,05 | 2,96E-01 |
| AT3G11680 | 0,22 | 1,24 | 4,34 | 0,29 | 7,75E-01 |
| AT3G11710 | 1,08 | 0,20 | 2,10 | 0,10 | 9,24E-01 |
| AT3G11730 | 0,25 | 1,75 | 4,33 | 0,40 | 6,87E-01 |
| AT3G11770 | 0,28 | 0,87 | 4,38 | 0,20 | 8,43E-01 |
| AT3G11780 | 0,21 | 1,58 | 4,34 | 0,36 | 7,16E-01 |
| AT3G11820 | 1,51 | 2,36 | 1,88 | 1,26 | 2,08E-01 |
| AT3G11840 | 0,47 | 2,67 | 4,08 | 0,66 | 5,12E-01 |
| AT3G11964 | 0,92 | 0,74 | 1,91 | 0,39 | 6,99E-01 |
| AT3G12030 | 0,12 | 1,65 | 4,34 | 0,38 | 7,04E-01 |
| AT3G12130 | 0,80 | 0,44 | 2,56 | 0,17 | 8,63E-01 |
| AT3G12140 | 0,25 | -0,26 | 4,36 | -0,06 | 9,53E-01 |
| AT3G12145 | 0,15 | 0,71 | 4,37 | 0,16 | 8,71E-01 |
| AT3G12200 | 0,53 | 0,64 | 3,97 | 0,16 | 8,72E-01 |
| AT3G12260 | 0,32 | -1,30 | 4,40 | -0,29 | 7,68E-01 |
| AT3G12280 | 1,55 | 2,33 | 1,58 | 1,47 | 1,41E-01 |
| AT3G12400 | 0,71 | 1,20 | 2,02 | 0,60 | 5,52E-01 |
| AT3G12490 | 0,47 | 0,67 | 2,40 | 0,28 | 7,82E-01 |
| AT3G12500 | 0,08 | 1,10 | 4,36 | 0,25 | 8,00E-01 |
| AT3G12580 | 0,43 | 1,27 | 4,37 | 0,29 | 7,71E-01 |
| AT3G12587 | 0,28 | 0,00 | 4,41 | 0,00 | 1,00E+00 |
| AT3G12590 | 0,82 | 2,13 | 2,44 | 0,87 | 3,82E-01 |
| AT3G12610 | 0,59 | 0,23 | 2,39 | 0,09 | 9,25E-01 |
| AT3G12670 | 1,09 | 1,47 | 2,44 | 0,60 | 5,46E-01 |
| AT3G12680 | 0,32 | 0,66 | 4,40 | 0,15 | 8,81E-01 |
| AT3G12690 | 0,57 | 3,18 | 4,30 | 0,74 | 4,59E-01 |
| AT3G12750 | 0,45 | 2,14 | 3,98 | 0,54 | 5,90E-01 |
| AT3G12770 | 0,67 | 0,79 | 2,94 | 0,27 | 7,89E-01 |
| AT3G12800 | 0,96 | 0,39 | 1,88 | 0,21 | 8,36E-01 |
| AT3G12820 | 0,21 | 1,39 | 4,34 | 0,32 | 7,49E-01 |
| AT3G12920 | 1,14 | 2,03 | 2,78 | 0,73 | 4,67E-01 |
| AT3G12940 | 0,28 | 1,84 | 4,33 | 0,42 | 6,72E-01 |
| AT3G12950 | 0,48 | 2,55 | 4,32 | 0,59 | 5,55E-01 |
| AT3G12965 | 0,28 | 0,00 | 4,41 | 0,00 | 1,00E+00 |
| AT3G13000 | 0,24 | 0,00 | 4,41 | 0,00 | 1,00E+00 |
| AT3G13010 | 0,29 | 1,97 | 4,33 | 0,46 | 6,49E-01 |
| AT3G13020 | 0,33 | 0,77 | 4,37 | 0,18 | 8,61E-01 |
| AT3G13030 | 0,97 | 0,67 | 2,33 | 0,29 | 7,74E-01 |
| AT3G13040 | 0,58 | -1,02 | 2,72 | -0,37 | 7,08E-01 |
| AT3G13061 | 0,23 | 0,42 | 4,38 | 0,09 | 9,24E-01 |
| AT3G13110 | 1,06 | 1,63 | 1,92 | 0,85 | 3,95E-01 |
| AT3G13120 | 0,60 | -1,46 | 3,02 | -0,48 | 6,28E-01 |
| AT3G13150 | 0,37 | 1,58 | 4,34 | 0,36 | 7,16E-01 |
| AT3G13160 | 0,08 | 0,85 | 4,38 | 0,20 | 8,45E-01 |
| AT3G13220 | 0,37 | 2,28 | 4,32 | 0,53 | 5,98E-01 |
| AT3G13222 | 0,48 | 2,97 | 4,30 | 0,69 | 4,90E-01 |
| AT3G13225 | 0,32 | 2,47 | 4,32 | 0,57 | 5,67E-01 |
| AT3G13226 | 0,08 | 1,10 | 4,36 | 0,25 | 8,00E-01 |
| AT3G13235 | 0,44 | 0,32 | 2,82 | 0,11 | 9,09E-01 |
| AT3G13310 | 0,25 | 0,48 | 4,33 | 0,11 | 9,11E-01 |
| AT3G13390 | 0,05 | 0,00 | 4,41 | 0,00 | 1,00E+00 |
| AT3G13404 | 0,46 | -1,21 | 4,13 | -0,29 | 7,69E-01 |
| AT3G13410 | 1,64 | -0,39 | 2,12 | -0,19 | 8,53E-01 |
| AT3G13450 | 0,75 | 2,21 | 2,78 | 0,80 | 4,26E-01 |
| AT3G13480 | 0,20 | 0,82 | 4,37 | 0,19 | 8,50E-01 |
| AT3G13490 | 0,57 | 1,15 | 3,06 | 0,38 | 7,06E-01 |

| | | | | | |
|---|---|---|---|---|---|
| AT3G13570 | 0,44 | -1,01 | 4,16 | -0,24 | 8,09E-01 |
| AT3G13720 | 0,96 | -0,09 | 1,84 | -0,05 | 9,63E-01 |
| AT3G13772 | 0,72 | 3,10 | 2,76 | 1,12 | 2,61E-01 |
| AT3G13860 | 0,89 | 1,79 | 2,04 | 0,88 | 3,79E-01 |
| AT3G13990 | 1,05 | 1,05 | 2,17 | 0,49 | 6,27E-01 |
| AT3G14070 | 0,23 | 1,70 | 4,34 | 0,39 | 6,96E-01 |
| AT3G14075 | 0,26 | 0,95 | 4,33 | 0,22 | 8,26E-01 |
| AT3G14090 | 0,68 | -0,25 | 3,18 | -0,08 | 9,36E-01 |
| AT3G14110 | 0,61 | 0,45 | 2,58 | 0,17 | 8,62E-01 |
| AT3G14172 | 1,38 | 1,51 | 1,88 | 0,80 | 4,22E-01 |
| AT3G14180 | 0,40 | 2,27 | 4,33 | 0,52 | 6,00E-01 |
| AT3G14205 | 0,51 | 0,27 | 2,79 | 0,10 | 9,22E-01 |
| AT3G14230 | 0,40 | 0,83 | 4,20 | 0,20 | 8,44E-01 |
| AT3G14240 | 1,34 | -1,05 | 1,85 | -0,57 | 5,70E-01 |
| AT3G14330 | 0,56 | 1,43 | 2,99 | 0,48 | 6,33E-01 |
| AT3G14350 | 1,44 | -0,66 | 1,69 | -0,39 | 6,94E-01 |
| AT3G14400 | 1,24 | 3,63 | 3,16 | 1,15 | 2,51E-01 |
| AT3G14430 | 0,25 | 0,72 | 4,34 | 0,17 | 8,68E-01 |
| AT3G14440 | 0,33 | 2,10 | 4,32 | 0,49 | 6,26E-01 |
| AT3G14460 | 0,50 | 1,93 | 3,95 | 0,49 | 6,26E-01 |
| AT3G14590 | 0,14 | 0,51 | 4,38 | 0,12 | 9,08E-01 |
| AT3G14810 | 0,79 | 1,51 | 2,27 | 0,67 | 5,06E-01 |
| AT3G14900 | 0,17 | 1,73 | 4,34 | 0,40 | 6,91E-01 |
| AT3G15110 | 0,15 | 0,00 | 4,39 | 0,00 | 1,00E+00 |
| AT3G15120 | 0,82 | 2,40 | 2,46 | 0,97 | 3,30E-01 |
| AT3G15150 | 0,08 | 0,00 | 4,40 | 0,00 | 1,00E+00 |
| AT3G15220 | 0,77 | 0,06 | 2,50 | 0,02 | 9,81E-01 |
| AT3G15354 | 0,91 | 1,98 | 2,11 | 0,94 | 3,49E-01 |
| AT3G15356 | 0,36 | 0,59 | 4,39 | 0,13 | 8,93E-01 |
| AT3G15470 | 0,41 | 0,25 | 2,83 | 0,09 | 9,29E-01 |
| AT3G03125 | 0,17 | 1,40 | 4,35 | 0,32 | 7,48E-01 |
| AT3G15530 | 1,09 | 3,18 | 2,44 | 1,30 | 1,94E-01 |
| AT3G15570 | 0,83 | -1,64 | 2,83 | -0,58 | 5,64E-01 |
| AT3G15590 | 0,41 | 1,59 | 4,25 | 0,37 | 7,09E-01 |
| AT3G15630 | 0,46 | 1,26 | 3,25 | 0,39 | 6,99E-01 |
| AT3G15720 | 0,20 | 0,73 | 4,38 | 0,17 | 8,68E-01 |
| AT3G15810 | 0,58 | -0,43 | 2,60 | -0,17 | 8,68E-01 |
| AT3G15840 | 1,45 | -0,85 | 1,43 | -0,59 | 5,54E-01 |
| AT3G15900 | 0,29 | -1,23 | 4,39 | -0,28 | 7,79E-01 |
| AT3G15940 | 1,43 | 0,98 | 1,82 | 0,54 | 5,90E-01 |
| AT3G15970 | 0,63 | 2,44 | 2,57 | 0,95 | 3,41E-01 |
| AT3G15980 | 1,59 | 0,83 | 1,66 | 0,50 | 6,14E-01 |
| AT3G16000 | 0,51 | -1,13 | 3,20 | -0,35 | 7,24E-01 |
| AT3G16110 | 0,21 | 1,60 | 4,35 | 0,37 | 7,13E-01 |
| AT3G16190 | 0,21 | 0,79 | 4,37 | 0,18 | 8,57E-01 |
| AT3G03235 | 0,34 | 1,85 | 4,34 | 0,43 | 6,69E-01 |
| AT3G16290 | 0,84 | 0,02 | 1,96 | 0,01 | 9,93E-01 |
| AT3G16310 | 0,55 | 2,67 | 3,77 | 0,71 | 4,79E-01 |
| AT3G16500 | 0,17 | 0,76 | 4,37 | 0,18 | 8,61E-01 |
| AT3G16510 | 0,65 | 1,67 | 3,06 | 0,54 | 5,86E-01 |
| AT3G16520 | 0,76 | 0,84 | 2,14 | 0,39 | 6,95E-01 |
| AT3G16530 | 0,27 | 0,54 | 4,37 | 0,12 | 9,01E-01 |
| AT3G16560 | 0,66 | 2,60 | 2,49 | 1,04 | 2,98E-01 |
| AT3G16630 | 1,01 | 1,30 | 2,07 | 0,63 | 5,31E-01 |
| AT3G16820 | 1,66 | 0,12 | 1,60 | 0,07 | 9,40E-01 |
| AT3G16840 | 0,19 | 1,26 | 4,35 | 0,29 | 7,72E-01 |

| | | | | | |
|---|---|---|---|---|---|
| AT3G16860 | 0,47 | 0,00 | 4,08 | 0,00 | 1,00E+00 |
| AT3G16870 | 0,39 | 0,48 | 4,40 | 0,11 | 9,14E-01 |
| AT3G16890 | 0,19 | -0,45 | 4,36 | -0,10 | 9,18E-01 |
| AT3G17020 | 1,21 | 0,00 | 2,13 | 0,00 | 9,99E-01 |
| AT3G03415 | 1,34 | 0,50 | 1,76 | 0,28 | 7,76E-01 |
| AT3G03445 | 0,77 | 2,37 | 2,55 | 0,93 | 3,54E-01 |
| AT3G17170 | 0,31 | -1,70 | 4,34 | -0,39 | 6,96E-01 |
| AT3G17320 | 0,12 | 0,68 | 4,37 | 0,16 | 8,77E-01 |
| AT3G17330 | 0,20 | 1,68 | 4,34 | 0,39 | 6,99E-01 |
| AT3G17420 | 0,17 | -0,02 | 4,36 | 0,00 | 9,97E-01 |
| AT3G17440 | 0,08 | 1,10 | 4,36 | 0,25 | 8,00E-01 |
| AT3G17450 | 0,60 | 2,10 | 2,94 | 0,71 | 4,75E-01 |
| AT3G17655 | 0,19 | 1,88 | 4,33 | 0,43 | 6,64E-01 |
| AT3G17740 | 0,37 | 1,85 | 4,30 | 0,43 | 6,67E-01 |
| AT3G17760 | 0,21 | 1,39 | 4,34 | 0,32 | 7,49E-01 |
| AT3G17770 | 1,51 | 2,50 | 1,58 | 1,58 | 1,15E-01 |
| AT3G03585 | 0,05 | 0,00 | 4,41 | 0,00 | 1,00E+00 |
| AT3G17940 | 0,81 | 0,45 | 2,53 | 0,18 | 8,58E-01 |
| AT3G18030 | 0,52 | 2,23 | 4,01 | 0,56 | 5,78E-01 |
| AT3G18035 | 1,23 | -0,75 | 1,60 | -0,47 | 6,37E-01 |
| AT3G18110 | 1,16 | 3,68 | 2,44 | 1,51 | 1,32E-01 |
| AT3G18130 | 0,88 | 2,19 | 2,40 | 0,91 | 3,63E-01 |
| AT3G18140 | 0,30 | 0,00 | 4,40 | 0,00 | 1,00E+00 |
| AT3G18250 | 0,55 | 0,00 | 4,41 | 0,00 | 1,00E+00 |
| AT3G18380 | 0,25 | 1,75 | 4,33 | 0,40 | 6,87E-01 |
| AT3G18420 | 0,20 | 1,23 | 4,36 | 0,28 | 7,77E-01 |
| AT3G18535 | 0,67 | 2,02 | 2,47 | 0,82 | 4,14E-01 |
| AT3G18600 | 0,25 | 0,00 | 4,40 | 0,00 | 1,00E+00 |
| AT3G18610 | 0,55 | 1,69 | 4,32 | 0,39 | 6,95E-01 |
| AT3G18630 | 0,20 | 0,89 | 4,37 | 0,20 | 8,39E-01 |
| AT3G18700 | 1,19 | 3,75 | 2,07 | 1,81 | 7,06E-02 |
| AT3G18750 | 0,17 | 1,81 | 4,33 | 0,42 | 6,76E-01 |
| AT3G18930 | 0,90 | -0,01 | 2,36 | 0,00 | 9,97E-01 |
| AT3G18970 | 0,08 | 1,10 | 4,36 | 0,25 | 8,00E-01 |
| AT3G19000 | 0,24 | 1,59 | 4,35 | 0,37 | 7,15E-01 |
| AT3G19020 | 0,54 | 1,12 | 3,18 | 0,35 | 7,24E-01 |
| AT3G19150 | 0,16 | 0,42 | 4,37 | 0,10 | 9,23E-01 |
| AT3G03815 | 0,25 | 0,00 | 4,40 | 0,00 | 1,00E+00 |
| AT3G19300 | 0,12 | 1,39 | 4,34 | 0,32 | 7,49E-01 |
| AT3G19340 | 0,12 | 0,42 | 4,38 | 0,10 | 9,24E-01 |
| AT3G19430 | 0,25 | 0,00 | 4,40 | 0,00 | 1,00E+00 |
| AT3G19490 | 0,72 | 2,91 | 2,44 | 1,19 | 2,33E-01 |
| AT3G19510 | 0,12 | 0,90 | 4,36 | 0,21 | 8,36E-01 |
| AT3G19515 | 0,59 | 1,33 | 3,08 | 0,43 | 6,66E-01 |
| AT3G19553 | 0,42 | 0,00 | 4,18 | 0,00 | 1,00E+00 |
| AT3G19580 | 0,72 | 2,75 | 2,46 | 1,12 | 2,64E-01 |
| AT3G19620 | 0,32 | 1,20 | 4,35 | 0,28 | 7,82E-01 |
| AT3G19630 | 0,39 | 0,48 | 4,40 | 0,11 | 9,14E-01 |
| AT3G19710 | 0,69 | 2,48 | 2,48 | 1,00 | 3,18E-01 |
| AT3G19720 | 1,17 | 4,21 | 2,42 | 1,74 | 8,21E-02 |
| AT3G19740 | 0,59 | 1,24 | 3,14 | 0,39 | 6,94E-01 |
| AT3G19790 | 0,71 | 1,64 | 2,07 | 0,79 | 4,27E-01 |
| AT3G19810 | 0,70 | -1,27 | 4,36 | -0,29 | 7,71E-01 |
| AT3G19830 | 0,67 | 2,20 | 2,99 | 0,73 | 4,63E-01 |
| AT3G19850 | 0,50 | 0,51 | 3,18 | 0,16 | 8,73E-01 |
| AT3G19880 | 0,35 | 2,15 | 3,43 | 0,63 | 5,31E-01 |

| | | | | | |
|---|---|---|---|---|---|
| AT3G19895 | 0,23 | 0,61 | 4,38 | 0,14 | 8,90E-01 |
| AT3G19900 | 0,17 | 1,92 | 4,33 | 0,44 | 6,57E-01 |
| AT3G20015 | 0,62 | 0,00 | 3,18 | 0,00 | 1,00E+00 |
| AT3G20050 | 0,74 | 0,13 | 2,46 | 0,05 | 9,58E-01 |
| AT3G20060 | 0,33 | 0,00 | 4,36 | 0,00 | 9,99E-01 |
| AT3G20230 | 0,35 | 2,34 | 4,32 | 0,54 | 5,89E-01 |
| AT3G20250 | 1,61 | 2,06 | 1,89 | 1,09 | 2,76E-01 |
| AT3G20290 | 0,63 | 2,58 | 3,04 | 0,85 | 3,95E-01 |
| AT3G20320 | 0,51 | 0,67 | 2,74 | 0,25 | 8,06E-01 |
| AT3G20390 | 0,62 | -0,95 | 2,22 | -0,43 | 6,67E-01 |
| AT3G20470 | 0,38 | -0,44 | 4,37 | -0,10 | 9,21E-01 |
| AT3G20620 | 0,68 | 2,75 | 3,55 | 0,77 | 4,39E-01 |
| AT3G20630 | 1,20 | 1,77 | 1,81 | 0,98 | 3,27E-01 |
| AT3G20655 | 0,30 | 1,45 | 4,35 | 0,33 | 7,39E-01 |
| AT3G20700 | 1,05 | 1,56 | 2,51 | 0,62 | 5,34E-01 |
| AT3G03985 | 0,54 | 0,00 | 4,41 | 0,00 | 1,00E+00 |
| AT3G20830 | 0,60 | -0,45 | 3,23 | -0,14 | 8,90E-01 |
| AT3G21055 | 1,34 | -1,23 | 1,66 | -0,74 | 4,58E-01 |
| AT3G21070 | 0,23 | 1,56 | 4,34 | 0,36 | 7,19E-01 |
| AT3G21140 | 0,25 | 1,38 | 4,36 | 0,32 | 7,51E-01 |
| AT3G21160 | 0,11 | 1,07 | 4,36 | 0,25 | 8,06E-01 |
| AT3G21215 | 0,11 | 0,90 | 4,37 | 0,21 | 8,36E-01 |
| AT3G21300 | 0,80 | 0,92 | 2,35 | 0,39 | 6,94E-01 |
| AT3G21430 | 0,61 | 1,61 | 3,19 | 0,50 | 6,14E-01 |
| AT3G21540 | 0,20 | 0,32 | 4,35 | 0,07 | 9,41E-01 |
| AT3G21570 | 0,87 | -3,05 | 2,93 | -1,04 | 2,97E-01 |
| AT3G21630 | 0,12 | 0,42 | 4,38 | 0,10 | 9,24E-01 |
| AT3G21650 | 0,33 | 2,12 | 4,32 | 0,49 | 6,24E-01 |
| AT3G21670 | 1,42 | 3,97 | 2,37 | 1,68 | 9,38E-02 |
| AT3G21740 | 0,24 | 0,83 | 4,37 | 0,19 | 8,49E-01 |
| AT3G21790 | 0,96 | 2,07 | 2,13 | 0,97 | 3,30E-01 |
| AT3G21865 | 0,48 | 1,95 | 2,71 | 0,72 | 4,73E-01 |
| AT3G22070 | 0,17 | -2,23 | 4,35 | -0,51 | 6,08E-01 |
| AT3G22110 | 0,73 | 0,27 | 2,55 | 0,11 | 9,15E-01 |
| AT3G22142 | 1,40 | 0,37 | 1,66 | 0,22 | 8,25E-01 |
| AT3G22170 | 1,10 | 3,52 | 2,04 | 1,73 | 8,33E-02 |
| AT3G22190 | 0,78 | 0,90 | 2,99 | 0,30 | 7,63E-01 |
| AT3G22210 | 0,48 | -1,32 | 3,35 | -0,40 | 6,93E-01 |
| AT3G22220 | 0,36 | 0,65 | 3,46 | 0,19 | 8,51E-01 |
| AT3G22230 | 0,95 | 2,39 | 1,98 | 1,21 | 2,27E-01 |
| AT3G22240 | 1,25 | 2,18 | 1,89 | 1,16 | 2,48E-01 |
| AT3G22320 | 0,41 | 1,59 | 4,25 | 0,37 | 7,09E-01 |
| AT3G22421 | 0,96 | 3,48 | 2,67 | 1,30 | 1,93E-01 |
| AT3G22470 | 0,59 | 1,08 | 2,60 | 0,41 | 6,78E-01 |
| AT3G22530 | 0,38 | 0,10 | 4,34 | 0,02 | 9,81E-01 |
| AT3G22550 | 0,60 | 2,56 | 3,05 | 0,84 | 4,00E-01 |
| AT3G22580 | 0,15 | 0,00 | 4,39 | 0,00 | 1,00E+00 |
| AT3G22600 | 0,17 | 1,94 | 4,33 | 0,45 | 6,55E-01 |
| AT3G22630 | 0,68 | 1,78 | 2,25 | 0,79 | 4,28E-01 |
| AT3G22660 | 0,44 | 0,00 | 4,41 | 0,00 | 1,00E+00 |
| AT3G22690 | 0,29 | 1,03 | 4,33 | 0,24 | 8,12E-01 |
| AT3G22710 | 1,14 | 1,03 | 2,02 | 0,51 | 6,08E-01 |
| AT3G22720 | 0,08 | 0,00 | 4,40 | 0,00 | 1,00E+00 |
| AT3G22730 | 1,61 | -0,28 | 1,70 | -0,17 | 8,67E-01 |
| AT3G22770 | 0,68 | 0,05 | 3,19 | 0,01 | 9,88E-01 |
| AT3G22950 | 0,15 | -1,24 | 4,39 | -0,28 | 7,77E-01 |

| | | | | | |
|---|---|---|---|---|---|
| AT3G22980 | 0,74 | 1,60 | 2,44 | 0,65 | 5,14E-01 |
| AT3G23020 | 0,53 | 1,85 | 3,81 | 0,49 | 6,28E-01 |
| AT3G23030 | 0,69 | -0,65 | 2,05 | -0,32 | 7,49E-01 |
| AT3G23080 | 0,08 | 1,10 | 4,36 | 0,25 | 8,00E-01 |
| AT3G23170 | 0,50 | 1,99 | 2,65 | 0,75 | 4,53E-01 |
| AT3G23172 | 0,24 | 1,58 | 4,34 | 0,36 | 7,15E-01 |
| AT3G23400 | 0,58 | 0,49 | 3,25 | 0,15 | 8,80E-01 |
| AT3G23420 | 0,32 | 1,24 | 4,36 | 0,28 | 7,76E-01 |
| AT3G23490 | 0,26 | 0,90 | 4,37 | 0,20 | 8,38E-01 |
| AT3G23570 | 0,32 | 1,91 | 4,33 | 0,44 | 6,59E-01 |
| AT3G23633 | 0,12 | 0,00 | 4,41 | 0,00 | 1,00E+00 |
| AT3G23650 | 0,55 | 1,94 | 2,96 | 0,65 | 5,13E-01 |
| AT3G04585 | 0,72 | 2,10 | 2,84 | 0,74 | 4,60E-01 |
| AT3G23660 | 0,75 | 2,03 | 2,90 | 0,70 | 4,84E-01 |
| AT3G23700 | 0,24 | -0,36 | 4,37 | -0,08 | 9,35E-01 |
| AT3G23740 | 0,08 | 0,85 | 4,38 | 0,20 | 8,45E-01 |
| AT3G23830 | 0,27 | -0,24 | 4,36 | -0,05 | 9,57E-01 |
| AT3G23900 | 0,77 | 3,19 | 2,76 | 1,16 | 2,47E-01 |
| AT3G23970 | 0,08 | 0,85 | 4,38 | 0,20 | 8,45E-01 |
| AT3G23980 | 0,76 | 0,17 | 2,61 | 0,06 | 9,49E-01 |
| AT3G24080 | 0,22 | 1,93 | 4,33 | 0,45 | 6,56E-01 |
| AT3G24120 | 0,83 | 2,44 | 3,44 | 0,71 | 4,78E-01 |
| AT3G24140 | 0,26 | -0,44 | 4,38 | -0,10 | 9,20E-01 |
| AT3G24170 | 1,33 | 3,34 | 1,86 | 1,80 | 7,22E-02 |
| AT3G24200 | 0,11 | 1,07 | 4,36 | 0,25 | 8,06E-01 |
| AT3G24260 | 0,15 | 0,00 | 4,41 | 0,00 | 1,00E+00 |
| AT3G24270 | 0,91 | 3,32 | 2,33 | 1,43 | 1,53E-01 |
| AT3G24320 | 0,63 | 2,32 | 3,03 | 0,77 | 4,44E-01 |
| AT3G24430 | 0,54 | -1,25 | 2,69 | -0,47 | 6,42E-01 |
| AT3G24440 | 0,50 | 1,96 | 3,18 | 0,61 | 5,39E-01 |
| AT3G24465 | 0,32 | -0,15 | 3,61 | -0,04 | 9,67E-01 |
| AT3G24503 | 1,13 | -0,56 | 1,91 | -0,29 | 7,69E-01 |
| AT3G24518 | 0,96 | -1,59 | 2,48 | -0,64 | 5,22E-01 |
| AT3G24614 | 0,96 | 1,01 | 2,00 | 0,50 | 6,14E-01 |
| AT3G24670 | 0,20 | 1,62 | 4,35 | 0,37 | 7,10E-01 |
| AT3G24870 | 1,16 | -1,31 | 1,96 | -0,67 | 5,03E-01 |
| AT3G24880 | 0,67 | 0,47 | 3,17 | 0,15 | 8,82E-01 |
| AT3G24900 | 0,51 | 2,23 | 3,11 | 0,72 | 4,73E-01 |
| AT3G25090 | 0,87 | 0,29 | 2,15 | 0,13 | 8,94E-01 |
| AT3G25150 | 1,03 | 2,79 | 2,52 | 1,10 | 2,69E-01 |
| AT3G25180 | 0,15 | 0,00 | 4,39 | 0,00 | 1,00E+00 |
| AT3G25460 | 1,48 | -1,10 | 1,58 | -0,69 | 4,89E-01 |
| AT3G25530 | 0,19 | 0,43 | 4,38 | 0,10 | 9,22E-01 |
| AT3G25560 | 0,41 | 0,47 | 4,39 | 0,11 | 9,14E-01 |
| AT3G25610 | 1,05 | -1,28 | 2,36 | -0,54 | 5,88E-01 |
| AT3G25660 | 0,47 | 1,60 | 3,92 | 0,41 | 6,83E-01 |
| AT3G04935 | 0,15 | 1,25 | 4,35 | 0,29 | 7,75E-01 |
| AT3G25800 | 1,14 | -0,16 | 1,78 | -0,09 | 9,30E-01 |
| AT3G25840 | 1,32 | 0,47 | 1,77 | 0,26 | 7,92E-01 |
| AT3G25870 | 0,65 | 0,17 | 2,38 | 0,07 | 9,43E-01 |
| AT3G25880 | 1,38 | 2,54 | 2,55 | 1,00 | 3,19E-01 |
| AT3G25882 | 0,50 | 2,08 | 4,03 | 0,52 | 6,05E-01 |
| AT3G25920 | 0,58 | 0,66 | 2,29 | 0,29 | 7,74E-01 |
| AT3G25990 | 0,75 | 0,46 | 3,52 | 0,13 | 8,96E-01 |
| AT3G26170 | 1,29 | 0,28 | 1,66 | 0,17 | 8,66E-01 |
| AT3G26180 | 0,36 | 1,33 | 4,34 | 0,31 | 7,59E-01 |

| | | | | | |
|---|---|---|---|---|---|
| AT3G26210 | 1,30 | 3,31 | 2,10 | 1,58 | 1,14E-01 |
| AT3G26220 | 0,49 | 1,41 | 4,04 | 0,35 | 7,26E-01 |
| AT3G26230 | 0,79 | -0,64 | 2,56 | -0,25 | 8,04E-01 |
| AT3G26280 | 1,16 | -1,87 | 1,81 | -1,03 | 3,02E-01 |
| AT3G26290 | 0,78 | -0,71 | 1,90 | -0,37 | 7,08E-01 |
| AT3G26300 | 0,36 | -0,45 | 4,36 | -0,10 | 9,18E-01 |
| AT3G26340 | 0,62 | -1,64 | 3,82 | -0,43 | 6,69E-01 |
| AT3G26410 | 0,31 | -1,23 | 4,39 | -0,28 | 7,79E-01 |
| AT3G26500 | 1,32 | 3,45 | 1,97 | 1,75 | 8,02E-02 |
| AT3G26510 | 0,66 | 0,75 | 2,19 | 0,34 | 7,33E-01 |
| AT3G26550 | 0,20 | 1,07 | 4,36 | 0,25 | 8,06E-01 |
| AT3G26560 | 0,40 | 0,62 | 3,37 | 0,19 | 8,53E-01 |
| AT3G26590 | 0,41 | 0,00 | 4,39 | 0,00 | 1,00E+00 |
| AT3G26616 | 0,23 | 1,09 | 4,36 | 0,25 | 8,02E-01 |
| AT3G26630 | 0,69 | 1,57 | 2,47 | 0,63 | 5,26E-01 |
| AT3G26670 | 0,54 | 1,76 | 3,31 | 0,53 | 5,96E-01 |
| AT3G26744 | 0,39 | 0,36 | 4,26 | 0,08 | 9,33E-01 |
| AT3G26812 | 0,17 | 0,48 | 4,40 | 0,11 | 9,12E-01 |
| AT3G26819 | 0,05 | 0,00 | 4,41 | 0,00 | 1,00E+00 |
| AT3G26860 | 0,30 | -2,60 | 4,34 | -0,60 | 5,50E-01 |
| AT3G26870 | 1,55 | -3,50 | 1,98 | -1,77 | 7,71E-02 |
| AT3G26890 | 0,29 | 0,71 | 4,36 | 0,16 | 8,71E-01 |
| AT3G26932 | 0,90 | 1,72 | 2,35 | 0,73 | 4,65E-01 |
| AT3G26935 | 0,27 | 0,96 | 4,36 | 0,22 | 8,25E-01 |
| AT3G26980 | 0,14 | 1,24 | 4,35 | 0,29 | 7,75E-01 |
| AT3G27010 | 0,67 | 0,60 | 3,62 | 0,17 | 8,69E-01 |
| AT3G27020 | 0,51 | 1,62 | 2,39 | 0,68 | 4,97E-01 |
| AT3G27040 | 1,43 | -0,87 | 1,81 | -0,48 | 6,29E-01 |
| AT3G27060 | 0,90 | 0,65 | 2,37 | 0,28 | 7,83E-01 |
| AT3G27100 | 0,74 | 0,29 | 4,38 | 0,07 | 9,48E-01 |
| AT3G27240 | 0,37 | 0,52 | 3,45 | 0,15 | 8,81E-01 |
| AT3G27300 | 0,49 | -0,36 | 3,24 | -0,11 | 9,11E-01 |
| AT3G27380 | 0,45 | -0,45 | 4,33 | -0,10 | 9,17E-01 |
| AT3G05235 | 0,99 | -1,91 | 2,53 | -0,75 | 4,51E-01 |
| AT3G05245 | 0,33 | -1,36 | 4,41 | -0,31 | 7,58E-01 |
| AT3G27530 | 0,76 | 2,58 | 2,64 | 0,98 | 3,28E-01 |
| AT3G27600 | 1,14 | 1,92 | 2,34 | 0,82 | 4,13E-01 |
| AT3G27610 | 0,33 | 0,77 | 4,39 | 0,18 | 8,61E-01 |
| AT3G27680 | 0,86 | -2,25 | 2,42 | -0,93 | 3,52E-01 |
| AT3G27700 | 1,39 | 2,82 | 1,96 | 1,44 | 1,51E-01 |
| AT3G27740 | 0,52 | 0,98 | 3,99 | 0,25 | 8,06E-01 |
| AT3G27770 | 0,43 | 0,80 | 2,79 | 0,29 | 7,75E-01 |
| AT3G27906 | 1,35 | 0,33 | 1,80 | 0,18 | 8,55E-01 |
| AT3G27925 | 0,65 | -0,93 | 2,97 | -0,31 | 7,55E-01 |
| AT3G27930 | 0,20 | 0,73 | 4,38 | 0,17 | 8,68E-01 |
| AT3G27990 | 0,64 | -2,14 | 2,57 | -0,83 | 4,05E-01 |
| AT3G28340 | 0,22 | 0,59 | 4,37 | 0,13 | 8,94E-01 |
| AT3G28380 | 0,08 | 0,85 | 4,38 | 0,20 | 8,45E-01 |
| AT3G28410 | 0,14 | 0,44 | 4,39 | 0,10 | 9,20E-01 |
| AT3G28674 | 0,28 | 0,78 | 4,37 | 0,18 | 8,58E-01 |
| AT3G28695 | 0,43 | 0,43 | 4,22 | 0,10 | 9,20E-01 |
| AT3G28710 | 0,99 | 0,49 | 1,76 | 0,28 | 7,81E-01 |
| AT3G28780 | 0,61 | 0,46 | 3,10 | 0,15 | 8,83E-01 |
| AT3G28860 | 1,33 | 0,76 | 2,28 | 0,33 | 7,39E-01 |
| AT3G28890 | 0,43 | 0,43 | 4,33 | 0,10 | 9,22E-01 |
| AT3G28920 | 1,35 | 0,72 | 1,84 | 0,39 | 6,96E-01 |

| | | | | | |
|---|---|---|---|---|---|
| AT3G28930 | 0,77 | 1,90 | 2,94 | 0,65 | 5,18E-01 |
| AT3G28940 | 1,30 | 1,10 | 1,76 | 0,62 | 5,32E-01 |
| AT3G28950 | 0,97 | -1,38 | 2,00 | -0,69 | 4,91E-01 |
| AT3G28960 | 0,30 | 0,48 | 4,40 | 0,11 | 9,12E-01 |
| AT3G28970 | 0,52 | 2,79 | 3,98 | 0,70 | 4,83E-01 |
| AT3G29030 | 0,33 | -1,62 | 4,38 | -0,37 | 7,12E-01 |
| AT3G05665 | 0,80 | 0,42 | 2,78 | 0,15 | 8,81E-01 |
| AT3G29110 | 0,53 | -2,46 | 2,64 | -0,93 | 3,52E-01 |
| AT3G29160 | 0,59 | -17,02 | 4,35 | -3,91 | 9,28E-05 |
| AT3G29200 | 0,59 | 2,59 | 3,05 | 0,85 | 3,95E-01 |
| AT3G29350 | 0,17 | 0,00 | 4,40 | 0,00 | 1,00E+00 |
| AT3G29400 | 0,67 | 1,70 | 2,60 | 0,65 | 5,14E-01 |
| AT3G29570 | 0,38 | 0,82 | 4,38 | 0,19 | 8,51E-01 |
| AT3G05745 | 0,14 | 1,18 | 4,35 | 0,27 | 7,86E-01 |
| AT3G29633 | 0,14 | -1,60 | 4,37 | -0,37 | 7,14E-01 |
| AT3G29639 | 0,16 | 0,51 | 4,38 | 0,12 | 9,07E-01 |
| AT3G05895 | 0,08 | 0,00 | 4,40 | 0,00 | 1,00E+00 |
| AT3G29680 | 0,25 | 0,00 | 4,41 | 0,00 | 1,00E+00 |
| AT3G29720 | 0,40 | -0,17 | 4,37 | -0,04 | 9,69E-01 |
| AT3G29760 | 0,67 | 1,81 | 3,03 | 0,60 | 5,50E-01 |
| AT3G29785 | 0,22 | 0,69 | 4,37 | 0,16 | 8,74E-01 |
| AT3G30160 | 0,58 | -0,42 | 3,06 | -0,14 | 8,90E-01 |
| AT3G30200 | 0,12 | 0,90 | 4,36 | 0,21 | 8,36E-01 |
| AT3G30220 | 0,08 | 0,00 | 4,40 | 0,00 | 1,00E+00 |
| AT3G30230 | 0,54 | 2,09 | 3,91 | 0,53 | 5,93E-01 |
| AT3G30300 | 0,53 | 1,52 | 4,35 | 0,35 | 7,27E-01 |
| AT3G30840 | 0,21 | 1,97 | 4,33 | 0,46 | 6,49E-01 |
| AT3G32190 | 0,34 | 1,56 | 3,47 | 0,45 | 6,54E-01 |
| AT3G32420 | 0,10 | 0,42 | 4,38 | 0,10 | 9,23E-01 |
| AT3G32904 | 0,65 | -0,16 | 3,19 | -0,05 | 9,59E-01 |
| AT3G33528 | 0,61 | 0,53 | 2,43 | 0,22 | 8,28E-01 |
| AT3G42050 | 1,44 | 1,51 | 2,04 | 0,74 | 4,58E-01 |
| AT3G42060 | 0,32 | 0,00 | 4,36 | 0,00 | 1,00E+00 |
| AT3G42140 | 0,14 | 0,44 | 4,39 | 0,10 | 9,20E-01 |
| AT3G42155 | 0,34 | 0,00 | 4,39 | 0,00 | 1,00E+00 |
| AT3G42550 | 0,12 | 0,00 | 4,39 | 0,00 | 1,00E+00 |
| AT3G42723 | 0,12 | 0,42 | 4,38 | 0,10 | 9,24E-01 |
| AT3G42790 | 0,59 | 2,02 | 3,18 | 0,64 | 5,25E-01 |
| AT3G43153 | 0,99 | 3,71 | 2,50 | 1,48 | 1,39E-01 |
| AT3G43190 | 0,26 | 0,00 | 4,41 | 0,00 | 1,00E+00 |
| AT3G43220 | 0,79 | 0,65 | 2,85 | 0,23 | 8,20E-01 |
| AT3G43240 | 0,86 | 2,55 | 2,73 | 0,93 | 3,51E-01 |
| AT3G43270 | 0,68 | -3,57 | 2,43 | -1,47 | 1,42E-01 |
| AT3G43280 | 0,11 | 0,00 | 4,41 | 0,00 | 1,00E+00 |
| AT3G43291 | 0,65 | -0,76 | 2,57 | -0,30 | 7,67E-01 |
| AT3G06595 | 0,10 | 0,42 | 4,38 | 0,10 | 9,23E-01 |
| AT3G43432 | 0,16 | 0,46 | 4,38 | 0,11 | 9,16E-01 |
| AT3G43470 | 0,45 | 0,83 | 4,00 | 0,21 | 8,35E-01 |
| AT3G43500 | 0,45 | 0,48 | 4,04 | 0,12 | 9,05E-01 |
| AT3G43600 | 0,98 | -2,43 | 2,10 | -1,16 | 2,47E-01 |
| AT3G43610 | 0,74 | 2,21 | 2,56 | 0,87 | 3,87E-01 |
| AT3G06645 | 0,23 | 0,00 | 4,38 | 0,00 | 1,00E+00 |
| AT3G06655 | 1,06 | 1,76 | 2,14 | 0,82 | 4,11E-01 |
| AT3G43720 | 0,87 | -3,95 | 2,67 | -1,48 | 1,39E-01 |
| AT3G43810 | 0,51 | -2,48 | 3,20 | -0,78 | 4,38E-01 |
| AT3G43880 | 0,10 | 0,62 | 4,38 | 0,14 | 8,87E-01 |

| | | | | | |
|---|---|---|---|---|---|
| AT3G43930 | 0,08 | 0,85 | 4,38 | 0,20 | 8,45E-01 |
| AT3G43940 | 0,53 | 0,71 | 3,95 | 0,18 | 8,57E-01 |
| AT3G44010 | 1,00 | -0,40 | 1,99 | -0,20 | 8,41E-01 |
| AT3G44070 | 0,37 | -1,60 | 3,36 | -0,48 | 6,34E-01 |
| AT3G44160 | 0,43 | -0,37 | 4,11 | -0,09 | 9,28E-01 |
| AT3G44265 | 0,40 | 1,33 | 4,31 | 0,31 | 7,58E-01 |
| AT3G44340 | 1,56 | 2,80 | 1,73 | 1,62 | 1,05E-01 |
| AT3G44400 | 0,82 | 2,69 | 2,72 | 0,99 | 3,23E-01 |
| AT3G44435 | 0,70 | 2,39 | 3,76 | 0,63 | 5,26E-01 |
| AT3G44450 | 0,56 | -0,46 | 3,09 | -0,15 | 8,83E-01 |
| AT3G44460 | 0,11 | 0,00 | 4,41 | 0,00 | 1,00E+00 |
| AT3G44635 | 1,31 | 3,20 | 2,90 | 1,11 | 2,69E-01 |
| AT3G44680 | 0,12 | 1,39 | 4,34 | 0,32 | 7,49E-01 |
| AT3G44753 | 0,14 | 0,51 | 4,38 | 0,12 | 9,08E-01 |
| AT3G44780 | 0,10 | 0,00 | 4,41 | 0,00 | 1,00E+00 |
| AT3G44820 | 0,38 | 1,23 | 4,36 | 0,28 | 7,78E-01 |
| AT3G44990 | 0,60 | -3,79 | 3,62 | -1,05 | 2,95E-01 |
| AT3G45030 | 0,91 | -0,44 | 2,37 | -0,19 | 8,53E-01 |
| AT3G45040 | 0,29 | 0,68 | 4,37 | 0,16 | 8,76E-01 |
| AT3G45060 | 0,08 | 1,10 | 4,36 | 0,25 | 8,00E-01 |
| AT3G45090 | 1,49 | 0,98 | 1,89 | 0,52 | 6,03E-01 |
| AT3G06955 | 0,21 | 0,00 | 4,41 | 0,00 | 1,00E+00 |
| AT3G45130 | 0,82 | -1,63 | 2,44 | -0,67 | 5,04E-01 |
| AT3G45180 | 1,00 | -1,40 | 1,83 | -0,77 | 4,43E-01 |
| AT3G07065 | 0,05 | 0,00 | 4,41 | 0,00 | 1,00E+00 |
| AT3G45240 | 0,25 | 1,57 | 4,34 | 0,36 | 7,18E-01 |
| AT3G07095 | 0,20 | 0,82 | 4,33 | 0,19 | 8,50E-01 |
| AT3G45260 | 1,01 | 0,57 | 1,88 | 0,30 | 7,62E-01 |
| AT3G45300 | 0,17 | -1,64 | 4,37 | -0,37 | 7,08E-01 |
| AT3G45430 | 0,20 | 1,07 | 4,36 | 0,25 | 8,06E-01 |
| AT3G45443 | 0,08 | 0,00 | 4,40 | 0,00 | 1,00E+00 |
| AT3G45600 | 0,92 | -0,21 | 1,84 | -0,11 | 9,10E-01 |
| AT3G45630 | 0,80 | 2,32 | 2,27 | 1,02 | 3,06E-01 |
| AT3G45770 | 0,14 | 0,44 | 4,39 | 0,10 | 9,20E-01 |
| AT3G45820 | 0,59 | -2,46 | 3,05 | -0,81 | 4,20E-01 |
| AT3G45850 | 1,07 | 1,63 | 2,48 | 0,66 | 5,10E-01 |
| AT3G45860 | 0,95 | 0,49 | 2,25 | 0,22 | 8,28E-01 |
| AT3G45910 | 1,00 | 1,85 | 2,47 | 0,75 | 4,53E-01 |
| AT3G45990 | 0,46 | -0,45 | 3,31 | -0,14 | 8,92E-01 |
| AT3G46000 | 0,30 | 0,84 | 3,63 | 0,23 | 8,17E-01 |
| AT3G46030 | 0,35 | 0,43 | 4,38 | 0,10 | 9,22E-01 |
| AT3G46090 | 0,72 | 1,68 | 2,44 | 0,69 | 4,91E-01 |
| AT3G46100 | 0,51 | -3,21 | 3,97 | -0,81 | 4,18E-01 |
| AT3G46210 | 0,56 | 1,69 | 3,90 | 0,43 | 6,65E-01 |
| AT3G07375 | 0,21 | 0,46 | 4,34 | 0,11 | 9,15E-01 |
| AT3G46320 | 0,61 | 17,22 | 4,34 | 3,96 | 7,40E-05 |
| AT3G46360 | 0,82 | -4,19 | 2,79 | -1,50 | 1,34E-01 |
| AT3G46390 | 0,25 | 1,95 | 4,33 | 0,45 | 6,53E-01 |
| AT3G46440 | 0,45 | 1,28 | 4,34 | 0,30 | 7,68E-01 |
| AT3G46450 | 0,53 | 2,45 | 3,86 | 0,63 | 5,26E-01 |
| AT3G46470 | 0,38 | 1,97 | 4,25 | 0,46 | 6,42E-01 |
| AT3G46540 | 0,31 | -0,48 | 4,36 | -0,11 | 9,12E-01 |
| AT3G46550 | 1,58 | 1,63 | 1,55 | 1,06 | 2,90E-01 |
| AT3G46610 | 1,41 | 1,34 | 1,69 | 0,80 | 4,26E-01 |
| AT3G46630 | 0,81 | -0,86 | 2,11 | -0,41 | 6,85E-01 |
| AT3G46650 | 0,58 | 1,48 | 3,08 | 0,48 | 6,32E-01 |

| | | | | | |
|---|---|---|---|---|---|
| AT3G46660 | 0,59 | 2,22 | 2,62 | 0,85 | 3,96E-01 |
| AT3G46670 | 0,57 | -1,25 | 3,14 | -0,40 | 6,91E-01 |
| AT3G46690 | 0,12 | 1,65 | 4,34 | 0,38 | 7,04E-01 |
| AT3G46730 | 0,12 | 1,58 | 4,34 | 0,36 | 7,16E-01 |
| AT3G46790 | 0,53 | 1,98 | 4,32 | 0,46 | 6,47E-01 |
| AT3G46845 | 1,29 | 2,25 | 1,89 | 1,19 | 2,34E-01 |
| AT3G46920 | 0,08 | 0,85 | 4,38 | 0,20 | 8,45E-01 |
| AT3G46960 | 0,92 | 2,77 | 2,31 | 1,20 | 2,31E-01 |
| AT3G47000 | 0,33 | 0,00 | 4,39 | 0,00 | 1,00E+00 |
| AT3G47070 | 0,67 | -1,62 | 2,61 | -0,62 | 5,35E-01 |
| AT3G47090 | 0,54 | 2,31 | 3,76 | 0,62 | 5,38E-01 |
| AT3G47160 | 0,74 | -0,92 | 2,46 | -0,37 | 7,08E-01 |
| AT3G47370 | 0,35 | 0,41 | 4,38 | 0,09 | 9,25E-01 |
| AT3G47420 | 0,51 | 1,60 | 2,64 | 0,60 | 5,45E-01 |
| AT3G47480 | 0,70 | 1,60 | 3,00 | 0,53 | 5,95E-01 |
| AT3G47670 | 0,97 | 2,07 | 1,92 | 1,08 | 2,80E-01 |
| AT3G47675 | 0,21 | 0,79 | 4,37 | 0,18 | 8,57E-01 |
| AT3G47680 | 0,08 | 0,85 | 4,38 | 0,20 | 8,45E-01 |
| AT3G47700 | 0,43 | 1,98 | 2,75 | 0,72 | 4,72E-01 |
| AT3G47730 | 0,93 | 0,94 | 2,52 | 0,37 | 7,09E-01 |
| AT3G47810 | 0,27 | 1,05 | 4,36 | 0,24 | 8,09E-01 |
| AT3G47860 | 0,23 | 0,00 | 4,41 | 0,00 | 1,00E+00 |
| AT3G47890 | 1,54 | 0,38 | 1,44 | 0,26 | 7,95E-01 |
| AT3G47940 | 0,35 | 1,11 | 4,37 | 0,25 | 8,00E-01 |
| AT3G47960 | 1,01 | 1,24 | 1,96 | 0,63 | 5,26E-01 |
| AT3G47990 | 0,32 | 1,20 | 4,37 | 0,28 | 7,83E-01 |
| AT3G48060 | 1,66 | 2,79 | 1,98 | 1,41 | 1,60E-01 |
| AT3G48080 | 1,04 | 2,59 | 2,09 | 1,24 | 2,17E-01 |
| AT3G48090 | 1,08 | 3,43 | 2,69 | 1,27 | 2,02E-01 |
| AT3G48100 | 0,85 | 0,00 | 4,37 | 0,00 | 1,00E+00 |
| AT3G48131 | 0,31 | 0,89 | 4,36 | 0,20 | 8,38E-01 |
| AT3G48150 | 0,55 | 1,30 | 3,84 | 0,34 | 7,36E-01 |
| AT3G48170 | 0,90 | -0,14 | 2,53 | -0,05 | 9,57E-01 |
| AT3G48195 | 0,48 | 2,38 | 3,12 | 0,76 | 4,45E-01 |
| AT3G48360 | 1,03 | 1,09 | 2,85 | 0,38 | 7,03E-01 |
| AT3G48520 | 0,38 | 1,93 | 4,34 | 0,44 | 6,57E-01 |
| AT3G48570 | 0,28 | 0,00 | 4,41 | 0,00 | 1,00E+00 |
| AT3G48640 | 0,25 | 1,14 | 4,36 | 0,26 | 7,94E-01 |
| AT3G48690 | 0,79 | 1,27 | 2,48 | 0,51 | 6,09E-01 |
| AT3G48740 | 0,68 | 1,66 | 2,18 | 0,76 | 4,48E-01 |
| AT3G48770 | 1,49 | 1,51 | 1,75 | 0,86 | 3,90E-01 |
| AT3G08005 | 0,05 | 0,00 | 4,41 | 0,00 | 1,00E+00 |
| AT3G48840 | 0,12 | 0,79 | 4,37 | 0,18 | 8,57E-01 |
| AT3G48930 | 0,53 | 1,86 | 2,66 | 0,70 | 4,85E-01 |
| AT3G48990 | 0,58 | 1,36 | 2,97 | 0,46 | 6,47E-01 |
| AT3G49010 | 1,04 | 0,25 | 1,74 | 0,14 | 8,86E-01 |
| AT3G49055 | 0,42 | 1,08 | 4,37 | 0,25 | 8,05E-01 |
| AT3G49060 | 0,44 | -1,21 | 3,33 | -0,36 | 7,17E-01 |
| AT3G49070 | 1,57 | -4,86 | 2,95 | -1,65 | 9,98E-02 |
| AT3G49140 | 0,93 | -0,55 | 2,10 | -0,26 | 7,94E-01 |
| AT3G49160 | 0,11 | 1,07 | 4,36 | 0,25 | 8,06E-01 |
| AT3G49250 | 0,54 | -2,61 | 4,10 | -0,64 | 5,25E-01 |
| AT3G49350 | 0,36 | 1,06 | 4,31 | 0,24 | 8,06E-01 |
| AT3G49490 | 1,14 | 3,18 | 2,15 | 1,48 | 1,40E-01 |
| AT3G49530 | 1,18 | 0,16 | 2,06 | 0,08 | 9,37E-01 |
| AT3G49600 | 1,57 | 3,05 | 1,94 | 1,57 | 1,16E-01 |

| | | | | | |
|---|---|---|---|---|---|
| AT3G49601 | 0,27 | 1,19 | 4,35 | 0,27 | 7,84E-01 |
| AT3G49650 | 0,08 | 0,85 | 4,38 | 0,20 | 8,45E-01 |
| AT3G49720 | 1,09 | 0,07 | 2,43 | 0,03 | 9,76E-01 |
| AT3G49790 | 0,38 | 1,57 | 3,36 | 0,47 | 6,40E-01 |
| AT3G49800 | 1,37 | 1,99 | 1,65 | 1,21 | 2,28E-01 |
| AT3G49870 | 0,08 | 0,85 | 4,38 | 0,20 | 8,45E-01 |
| AT3G49980 | 0,21 | -0,31 | 4,37 | -0,07 | 9,43E-01 |
| AT3G49990 | 0,57 | 2,85 | 3,73 | 0,76 | 4,45E-01 |
| AT3G50060 | 0,43 | -0,54 | 2,83 | -0,19 | 8,48E-01 |
| AT3G50110 | 0,18 | 1,23 | 4,34 | 0,28 | 7,77E-01 |
| AT3G50120 | 0,78 | -2,03 | 2,08 | -0,98 | 3,28E-01 |
| AT3G50240 | 0,71 | 2,65 | 4,31 | 0,62 | 5,38E-01 |
| AT3G50320 | 1,17 | 1,07 | 2,20 | 0,49 | 6,27E-01 |
| AT3G50380 | 1,07 | 0,93 | 2,52 | 0,37 | 7,11E-01 |
| AT3G50500 | 1,35 | -0,30 | 1,90 | -0,16 | 8,75E-01 |
| AT3G50670 | 0,33 | 1,90 | 4,33 | 0,44 | 6,61E-01 |
| AT3G50690 | 0,85 | -1,27 | 1,79 | -0,71 | 4,79E-01 |
| AT3G50740 | 0,53 | -1,04 | 3,21 | -0,32 | 7,46E-01 |
| AT3G50750 | 1,00 | 0,67 | 2,22 | 0,30 | 7,64E-01 |
| AT3G50780 | 0,08 | 1,10 | 4,36 | 0,25 | 8,00E-01 |
| AT3G50790 | 0,29 | 0,46 | 4,37 | 0,10 | 9,17E-01 |
| AT3G50808 | 0,10 | 0,00 | 4,41 | 0,00 | 1,00E+00 |
| AT3G50825 | 0,41 | 0,00 | 4,41 | 0,00 | 1,00E+00 |
| AT3G50920 | 0,48 | -0,39 | 4,07 | -0,10 | 9,23E-01 |
| AT3G51000 | 0,52 | 2,08 | 3,21 | 0,65 | 5,17E-01 |
| AT3G51120 | 1,56 | 3,87 | 1,80 | 2,16 | 3,11E-02 |
| AT3G51135 | 0,51 | 1,21 | 2,73 | 0,44 | 6,57E-01 |
| AT3G51160 | 0,58 | -1,79 | 3,16 | -0,57 | 5,71E-01 |
| AT3G51265 | 0,13 | 0,00 | 4,41 | 0,00 | 1,00E+00 |
| AT3G51310 | 0,51 | 0,57 | 3,15 | 0,18 | 8,57E-01 |
| AT3G51580 | 0,25 | 2,24 | 4,32 | 0,52 | 6,04E-01 |
| AT3G51600 | 1,48 | -3,30 | 1,62 | -2,04 | 4,17E-02 |
| AT3G51660 | 0,43 | -1,50 | 4,20 | -0,36 | 7,22E-01 |
| AT3G51790 | 0,60 | 3,00 | 3,63 | 0,83 | 4,09E-01 |
| AT3G51820 | 0,59 | 2,87 | 3,82 | 0,75 | 4,52E-01 |
| AT3G51840 | 0,33 | 1,48 | 4,35 | 0,34 | 7,34E-01 |
| AT3G51880 | 0,48 | 1,23 | 3,21 | 0,38 | 7,01E-01 |
| AT3G51890 | 1,41 | 2,62 | 2,73 | 0,96 | 3,37E-01 |
| AT3G51895 | 0,82 | 2,08 | 2,66 | 0,78 | 4,34E-01 |
| AT3G51920 | 0,46 | 0,59 | 2,71 | 0,22 | 8,27E-01 |
| AT3G08215 | 0,08 | 1,10 | 4,36 | 0,25 | 8,00E-01 |
| AT3G51980 | 0,42 | -2,48 | 3,32 | -0,74 | 4,56E-01 |
| AT3G52060 | 0,46 | -0,03 | 2,73 | -0,01 | 9,91E-01 |
| AT3G52072 | 0,22 | 1,24 | 4,34 | 0,29 | 7,75E-01 |
| AT3G08230 | 0,12 | 0,90 | 4,36 | 0,21 | 8,36E-01 |
| AT3G52110 | 0,12 | 1,39 | 4,34 | 0,32 | 7,49E-01 |
| AT3G52180 | 0,66 | 1,49 | 2,68 | 0,56 | 5,79E-01 |
| AT3G52190 | 0,13 | 1,15 | 4,36 | 0,26 | 7,92E-01 |
| AT3G52220 | 0,11 | 1,07 | 4,36 | 0,25 | 8,06E-01 |
| AT3G52260 | 0,64 | 3,15 | 3,54 | 0,89 | 3,74E-01 |
| AT3G52400 | 1,57 | 1,59 | 1,59 | 1,00 | 3,16E-01 |
| AT3G52480 | 0,53 | 1,93 | 3,03 | 0,64 | 5,24E-01 |
| AT3G52560 | 0,47 | 0,19 | 3,31 | 0,06 | 9,54E-01 |
| AT3G52640 | 0,21 | 2,14 | 4,33 | 0,49 | 6,22E-01 |
| AT3G52730 | 0,69 | -0,68 | 3,72 | -0,18 | 8,55E-01 |
| AT3G52740 | 0,17 | 0,00 | 4,40 | 0,00 | 1,00E+00 |

| | | | | | |
|---|---|---|---|---|---|
| AT3G52870 | 0,19 | 0,00 | 4,41 | 0,00 | 1,00E+00 |
| AT3G53015 | 1,03 | 2,41 | 2,21 | 1,09 | 2,77E-01 |
| AT3G53016 | 0,52 | -1,77 | 4,06 | -0,44 | 6,63E-01 |
| AT3G53100 | 0,69 | -1,36 | 3,18 | -0,43 | 6,69E-01 |
| AT3G53380 | 0,40 | 0,00 | 4,27 | 0,00 | 1,00E+00 |
| AT3G53410 | 0,47 | 1,64 | 4,35 | 0,38 | 7,07E-01 |
| AT3G53500 | 0,57 | -1,16 | 2,76 | -0,42 | 6,74E-01 |
| AT3G53510 | 0,25 | 2,36 | 4,32 | 0,55 | 5,86E-01 |
| AT3G53540 | 1,08 | 1,88 | 2,23 | 0,84 | 3,98E-01 |
| AT3G53560 | 0,23 | 0,00 | 4,40 | 0,00 | 1,00E+00 |
| AT3G53570 | 0,21 | 0,79 | 4,37 | 0,18 | 8,57E-01 |
| AT3G53610 | 0,53 | 1,74 | 3,18 | 0,55 | 5,86E-01 |
| AT3G53710 | 0,37 | 1,10 | 3,41 | 0,32 | 7,48E-01 |
| AT3G53730 | 0,67 | 2,56 | 2,47 | 1,04 | 2,99E-01 |
| AT3G53810 | 0,44 | 1,72 | 4,27 | 0,40 | 6,87E-01 |
| AT3G53890 | 0,52 | 0,32 | 3,20 | 0,10 | 9,21E-01 |
| AT3G53910 | 1,01 | 2,64 | 2,20 | 1,20 | 2,31E-01 |
| AT3G53920 | 0,24 | 1,23 | 4,35 | 0,28 | 7,77E-01 |
| AT3G53960 | 0,98 | 1,92 | 2,42 | 0,79 | 4,29E-01 |
| AT3G54100 | 0,47 | 2,64 | 3,21 | 0,82 | 4,11E-01 |
| AT3G54120 | 0,20 | 0,89 | 4,37 | 0,20 | 8,39E-01 |
| AT3G54140 | 0,70 | 2,50 | 2,63 | 0,95 | 3,42E-01 |
| AT3G54210 | 1,01 | -1,18 | 1,61 | -0,73 | 4,66E-01 |
| AT3G54290 | 0,42 | 0,41 | 4,28 | 0,10 | 9,23E-01 |
| AT3G54350 | 1,02 | 3,33 | 2,00 | 1,67 | 9,48E-02 |
| AT3G54360 | 1,14 | 1,24 | 1,87 | 0,66 | 5,06E-01 |
| AT3G54400 | 0,32 | 0,71 | 4,33 | 0,16 | 8,70E-01 |
| AT3G54440 | 0,61 | -2,09 | 2,34 | -0,90 | 3,71E-01 |
| AT3G54460 | 0,97 | 3,26 | 2,70 | 1,21 | 2,28E-01 |
| AT3G54470 | 0,44 | 0,73 | 3,27 | 0,22 | 8,24E-01 |
| AT3G54500 | 1,37 | 3,08 | 2,08 | 1,48 | 1,39E-01 |
| AT3G54620 | 0,24 | 1,62 | 4,34 | 0,37 | 7,08E-01 |
| AT3G54630 | 0,68 | 2,16 | 4,32 | 0,50 | 6,17E-01 |
| AT3G54640 | 0,22 | 0,00 | 4,41 | 0,00 | 1,00E+00 |
| AT3G54650 | 0,62 | 1,81 | 3,20 | 0,57 | 5,72E-01 |
| AT3G54750 | 0,48 | 1,87 | 4,35 | 0,43 | 6,67E-01 |
| AT3G54760 | 1,04 | 1,86 | 2,04 | 0,91 | 3,62E-01 |
| AT3G54790 | 1,31 | -0,47 | 1,82 | -0,26 | 7,96E-01 |
| AT3G54860 | 0,55 | 2,38 | 3,00 | 0,79 | 4,28E-01 |
| AT3G54900 | 0,95 | 0,79 | 1,82 | 0,44 | 6,63E-01 |
| AT3G54960 | 0,90 | 0,54 | 2,00 | 0,27 | 7,89E-01 |
| AT3G55040 | 0,21 | -1,85 | 4,37 | -0,42 | 6,72E-01 |
| AT3G55050 | 0,45 | 2,29 | 3,27 | 0,70 | 4,83E-01 |
| AT3G55060 | 0,14 | 0,44 | 4,39 | 0,10 | 9,20E-01 |
| AT3G55130 | 0,51 | -2,88 | 3,14 | -0,92 | 3,58E-01 |
| AT3G55160 | 0,95 | 0,19 | 1,76 | 0,11 | 9,14E-01 |
| AT3G55252 | 0,66 | -1,09 | 2,32 | -0,47 | 6,38E-01 |
| AT3G55254 | 0,80 | 0,26 | 2,15 | 0,12 | 9,04E-01 |
| AT3G55330 | 0,44 | 0,78 | 2,75 | 0,28 | 7,76E-01 |
| AT3G55360 | 0,60 | 0,02 | 2,46 | 0,01 | 9,92E-01 |
| AT3G55400 | 0,18 | -0,76 | 4,36 | -0,17 | 8,62E-01 |
| AT3G55430 | 0,77 | -1,09 | 1,99 | -0,55 | 5,84E-01 |
| AT3G55480 | 1,64 | 2,69 | 1,72 | 1,57 | 1,17E-01 |
| AT3G55840 | 1,41 | 1,28 | 1,79 | 0,71 | 4,75E-01 |
| AT3G55860 | 0,83 | -0,89 | 2,02 | -0,44 | 6,60E-01 |
| AT3G55935 | 0,46 | 1,21 | 2,71 | 0,44 | 6,57E-01 |

| | | | | | |
|---|---|---|---|---|---|
| AT3G55950 | 0,63 | 1,27 | 2,56 | 0,50 | 6,18E-01 |
| AT3G55960 | 0,64 | 1,15 | 2,60 | 0,44 | 6,57E-01 |
| AT3G56040 | 0,77 | 0,63 | 2,16 | 0,29 | 7,72E-01 |
| AT3G56050 | 0,97 | 0,97 | 2,00 | 0,48 | 6,29E-01 |
| AT3G56085 | 0,12 | 0,42 | 4,38 | 0,10 | 9,24E-01 |
| AT3G56090 | 0,34 | -1,57 | 3,50 | -0,45 | 6,54E-01 |
| AT3G56110 | 0,41 | -0,06 | 3,36 | -0,02 | 9,86E-01 |
| AT3G56190 | 1,16 | 0,53 | 1,77 | 0,30 | 7,64E-01 |
| AT3G56210 | 0,45 | 0,00 | 4,40 | 0,00 | 1,00E+00 |
| AT3G56240 | 0,35 | 0,37 | 2,91 | 0,13 | 8,99E-01 |
| AT3G56290 | 1,19 | 2,12 | 2,24 | 0,95 | 3,43E-01 |
| AT3G56320 | 0,58 | 0,98 | 3,88 | 0,25 | 8,01E-01 |
| AT3G56340 | 0,44 | -0,46 | 3,36 | -0,14 | 8,90E-01 |
| AT3G56370 | 0,84 | 2,37 | 2,31 | 1,03 | 3,05E-01 |
| AT3G56460 | 1,07 | 3,39 | 2,01 | 1,68 | 9,24E-02 |
| AT3G56480 | 0,39 | 0,93 | 2,84 | 0,33 | 7,42E-01 |
| AT3G56670 | 0,38 | 1,50 | 4,32 | 0,35 | 7,29E-01 |
| AT3G56690 | 0,43 | -0,10 | 4,15 | -0,03 | 9,80E-01 |
| AT3G56790 | 0,33 | 0,77 | 4,39 | 0,18 | 8,61E-01 |
| AT3G56800 | 1,39 | 0,91 | 1,68 | 0,54 | 5,87E-01 |
| AT3G56820 | 0,25 | 0,00 | 4,40 | 0,00 | 1,00E+00 |
| AT3G56830 | 0,17 | 1,81 | 4,33 | 0,42 | 6,76E-01 |
| AT3G56850 | 0,48 | 1,17 | 3,33 | 0,35 | 7,26E-01 |
| AT3G56860 | 0,49 | 0,34 | 2,73 | 0,12 | 9,02E-01 |
| AT3G56950 | 0,79 | -0,90 | 2,14 | -0,42 | 6,72E-01 |
| AT3G56970 | 0,12 | 1,58 | 4,34 | 0,36 | 7,16E-01 |
| AT3G56990 | 0,46 | -1,71 | 3,16 | -0,54 | 5,88E-01 |
| AT3G57020 | 0,42 | 0,00 | 4,37 | 0,00 | 1,00E+00 |
| AT3G57030 | 0,05 | 0,00 | 4,41 | 0,00 | 1,00E+00 |
| AT3G08825 | 0,72 | -1,30 | 2,95 | -0,44 | 6,58E-01 |
| AT3G57050 | 0,52 | 1,00 | 2,60 | 0,39 | 6,99E-01 |
| AT3G08855 | 0,12 | 0,79 | 4,37 | 0,18 | 8,57E-01 |
| AT3G57140 | 0,48 | 2,54 | 3,12 | 0,81 | 4,15E-01 |
| AT3G57240 | 0,24 | 0,00 | 4,41 | 0,00 | 1,00E+00 |
| AT3G57290 | 1,20 | 1,35 | 2,28 | 0,59 | 5,53E-01 |
| AT3G57350 | 0,44 | 1,86 | 3,28 | 0,57 | 5,70E-01 |
| AT3G57420 | 0,74 | 0,37 | 2,33 | 0,16 | 8,75E-01 |
| AT3G57460 | 0,24 | 0,45 | 4,39 | 0,10 | 9,18E-01 |
| AT3G08935 | 1,61 | -4,50 | 1,89 | -2,38 | 1,75E-02 |
| AT3G57470 | 0,21 | -0,77 | 4,37 | -0,18 | 8,60E-01 |
| AT3G57530 | 1,08 | 1,12 | 2,08 | 0,54 | 5,90E-01 |
| AT3G57540 | 0,08 | 0,00 | 4,40 | 0,00 | 1,00E+00 |
| AT3G57580 | 0,45 | -0,34 | 3,37 | -0,10 | 9,20E-01 |
| AT3G57610 | 1,05 | 1,58 | 4,33 | 0,37 | 7,15E-01 |
| AT3G57640 | 0,11 | 0,00 | 4,41 | 0,00 | 1,00E+00 |
| AT3G57765 | 1,26 | 0,40 | 1,57 | 0,25 | 8,00E-01 |
| AT3G57780 | 0,43 | -1,11 | 4,38 | -0,25 | 8,00E-01 |
| AT3G57790 | 0,44 | 2,71 | 4,32 | 0,63 | 5,31E-01 |
| AT3G57800 | 0,37 | 0,75 | 4,36 | 0,17 | 8,64E-01 |
| AT3G57870 | 0,53 | 0,43 | 4,38 | 0,10 | 9,23E-01 |
| AT3G57880 | 0,23 | 1,68 | 4,33 | 0,39 | 6,97E-01 |
| AT3G57930 | 0,30 | 0,43 | 4,38 | 0,10 | 9,23E-01 |
| AT3G57965 | 0,28 | 1,30 | 4,32 | 0,30 | 7,63E-01 |
| AT3G57980 | 0,74 | 3,03 | 2,81 | 1,08 | 2,81E-01 |
| AT3G57990 | 0,40 | 1,25 | 3,35 | 0,37 | 7,09E-01 |
| AT3G58100 | 1,03 | 3,35 | 3,28 | 1,02 | 3,07E-01 |

| | | | | | |
|---|---|---|---|---|---|
| AT3G58120 | 0,15 | 0,24 | 4,35 | 0,05 | 9,56E-01 |
| AT3G58140 | 0,42 | -0,35 | 3,41 | -0,10 | 9,18E-01 |
| AT3G58170 | 0,42 | -1,36 | 4,25 | -0,32 | 7,50E-01 |
| AT3G58210 | 0,64 | 0,61 | 4,40 | 0,14 | 8,90E-01 |
| AT3G58270 | 0,99 | 2,48 | 2,57 | 0,97 | 3,33E-01 |
| AT3G58500 | 1,34 | 1,61 | 1,55 | 1,04 | 3,00E-01 |
| AT3G58560 | 0,23 | 0,96 | 4,36 | 0,22 | 8,25E-01 |
| AT3G58580 | 0,38 | 0,95 | 4,28 | 0,22 | 8,25E-01 |
| AT3G58670 | 0,12 | 0,79 | 4,37 | 0,18 | 8,57E-01 |
| AT3G58730 | 1,29 | -1,40 | 1,62 | -0,86 | 3,87E-01 |
| AT3G58750 | 0,61 | 0,02 | 3,82 | 0,01 | 9,95E-01 |
| AT3G58790 | 0,34 | -0,45 | 4,33 | -0,10 | 9,16E-01 |
| AT3G58970 | 1,53 | 3,82 | 2,34 | 1,63 | 1,02E-01 |
| AT3G58990 | 0,12 | 0,90 | 4,36 | 0,21 | 8,36E-01 |
| AT3G59040 | 1,65 | 3,36 | 1,82 | 1,85 | 6,46E-02 |
| AT3G59050 | 0,52 | 2,23 | 3,11 | 0,72 | 4,73E-01 |
| AT3G59060 | 1,47 | 1,58 | 1,72 | 0,91 | 3,61E-01 |
| AT3G59120 | 0,10 | 0,42 | 4,38 | 0,10 | 9,23E-01 |
| AT3G59280 | 0,32 | 1,13 | 4,36 | 0,26 | 7,95E-01 |
| AT3G59360 | 0,61 | 2,91 | 2,91 | 1,00 | 3,17E-01 |
| AT3G59400 | 1,33 | 0,15 | 1,62 | 0,09 | 9,25E-01 |
| AT3G59540 | 1,10 | -0,40 | 1,86 | -0,22 | 8,30E-01 |
| AT3G59570 | 0,57 | 1,01 | 2,99 | 0,34 | 7,36E-01 |
| AT3G59660 | 1,15 | 3,26 | 1,95 | 1,67 | 9,44E-02 |
| AT3G59760 | 0,20 | -1,66 | 4,38 | -0,38 | 7,04E-01 |
| AT3G59840 | 0,15 | 0,42 | 4,38 | 0,09 | 9,24E-01 |
| AT3G59910 | 0,48 | 0,73 | 4,09 | 0,18 | 8,58E-01 |
| AT3G59950 | 1,22 | -0,33 | 2,47 | -0,13 | 8,93E-01 |
| AT3G59980 | 0,35 | 1,81 | 3,46 | 0,52 | 6,02E-01 |
| AT3G60030 | 1,14 | 1,27 | 2,04 | 0,63 | 5,32E-01 |
| AT3G09335 | 0,23 | 0,15 | 4,35 | 0,03 | 9,72E-01 |
| AT3G60050 | 0,12 | 1,46 | 4,35 | 0,34 | 7,36E-01 |
| AT3G60070 | 0,35 | -0,48 | 4,36 | -0,11 | 9,12E-01 |
| AT3G60160 | 1,27 | 3,65 | 1,97 | 1,85 | 6,38E-02 |
| AT3G60176 | 0,85 | 1,65 | 2,72 | 0,61 | 5,45E-01 |
| AT3G60220 | 0,15 | 0,00 | 4,41 | 0,00 | 1,00E+00 |
| AT3G60250 | 1,02 | 0,63 | 2,08 | 0,30 | 7,61E-01 |
| AT3G60350 | 1,30 | 1,77 | 1,94 | 0,91 | 3,62E-01 |
| AT3G60380 | 0,49 | 0,43 | 3,96 | 0,11 | 9,13E-01 |
| AT3G60540 | 0,08 | 0,00 | 4,40 | 0,00 | 1,00E+00 |
| AT3G60670 | 0,12 | 0,90 | 4,36 | 0,21 | 8,36E-01 |
| AT3G60740 | 0,66 | 1,56 | 2,49 | 0,63 | 5,31E-01 |
| AT3G60760 | 0,12 | 1,65 | 4,34 | 0,38 | 7,04E-01 |
| AT3G60820 | 0,15 | 0,00 | 4,39 | 0,00 | 1,00E+00 |
| AT3G60990 | 0,73 | -0,68 | 2,59 | -0,26 | 7,93E-01 |
| AT3G61070 | 0,50 | 1,84 | 4,06 | 0,45 | 6,50E-01 |
| AT3G09565 | 0,50 | -1,36 | 3,93 | -0,35 | 7,29E-01 |
| AT3G61190 | 1,32 | 1,69 | 1,88 | 0,90 | 3,68E-01 |
| AT3G61200 | 0,69 | 1,43 | 2,39 | 0,60 | 5,51E-01 |
| AT3G61260 | 0,83 | -2,85 | 2,49 | -1,14 | 2,53E-01 |
| AT3G61310 | 0,38 | 0,90 | 4,38 | 0,21 | 8,37E-01 |
| AT3G61380 | 0,18 | 1,15 | 4,36 | 0,26 | 7,92E-01 |
| AT3G61420 | 0,37 | 0,00 | 4,40 | 0,00 | 1,00E+00 |
| AT3G61460 | 0,83 | -1,10 | 1,82 | -0,60 | 5,47E-01 |
| AT3G61530 | 0,25 | 0,91 | 4,32 | 0,21 | 8,34E-01 |
| AT3G61640 | 0,38 | 1,81 | 3,35 | 0,54 | 5,89E-01 |

| | | | | | |
|---|---|---|---|---|---|
| AT3G61740 | 0,78 | 2,73 | 2,39 | 1,14 | 2,53E-01 |
| AT3G61750 | 0,24 | 0,00 | 4,41 | 0,00 | 1,00E+00 |
| AT3G61780 | 1,45 | 3,78 | 1,76 | 2,15 | 3,17E-02 |
| AT3G61800 | 0,23 | 0,54 | 4,39 | 0,12 | 9,02E-01 |
| AT3G61820 | 1,36 | -2,08 | 1,51 | -1,38 | 1,69E-01 |
| AT3G61860 | 0,39 | 1,21 | 4,34 | 0,28 | 7,81E-01 |
| AT3G61890 | 0,44 | -0,55 | 3,50 | -0,16 | 8,74E-01 |
| AT3G61960 | 1,02 | -1,06 | 1,99 | -0,53 | 5,95E-01 |
| AT3G61990 | 0,15 | 1,16 | 4,35 | 0,27 | 7,90E-01 |
| AT3G62070 | 0,12 | 0,00 | 4,41 | 0,00 | 1,00E+00 |
| AT3G62120 | 1,00 | 0,94 | 2,05 | 0,46 | 6,46E-01 |
| AT3G62240 | 0,93 | 3,38 | 2,84 | 1,19 | 2,34E-01 |
| AT3G62260 | 0,41 | 1,79 | 4,13 | 0,43 | 6,64E-01 |
| AT3G62310 | 0,33 | -0,70 | 4,29 | -0,16 | 8,70E-01 |
| AT3G62330 | 0,73 | 2,45 | 2,91 | 0,84 | 4,00E-01 |
| AT3G62470 | 0,24 | 1,58 | 4,34 | 0,36 | 7,16E-01 |
| AT3G62540 | 0,28 | 0,42 | 4,38 | 0,10 | 9,24E-01 |
| AT3G09795 | 0,21 | 0,47 | 4,39 | 0,11 | 9,15E-01 |
| AT3G62590 | 0,64 | 2,57 | 2,94 | 0,87 | 3,83E-01 |
| AT3G62600 | 0,95 | 1,42 | 2,06 | 0,69 | 4,90E-01 |
| AT3G09825 | 1,22 | -3,04 | 1,81 | -1,68 | 9,33E-02 |
| AT3G09845 | 0,08 | 0,00 | 4,40 | 0,00 | 1,00E+00 |
| AT3G62630 | 0,91 | 1,16 | 2,39 | 0,49 | 6,26E-01 |
| AT3G62800 | 0,37 | 2,32 | 4,32 | 0,54 | 5,91E-01 |
| AT3G62840 | 0,66 | 1,71 | 3,09 | 0,55 | 5,79E-01 |
| AT3G62860 | 0,81 | 1,53 | 3,52 | 0,44 | 6,63E-01 |
| AT3G62870 | 1,22 | 1,18 | 1,82 | 0,65 | 5,18E-01 |
| AT3G62900 | 0,94 | 2,49 | 2,25 | 1,10 | 2,70E-01 |
| AT3G62910 | 0,49 | 0,00 | 4,10 | 0,00 | 1,00E+00 |
| AT3G62940 | 0,31 | -1,30 | 4,40 | -0,30 | 7,68E-01 |
| AT3G63052 | 0,42 | -0,35 | 3,41 | -0,10 | 9,18E-01 |
| AT3G63110 | 0,56 | -1,35 | 2,67 | -0,51 | 6,13E-01 |
| AT3G63140 | 0,93 | -1,72 | 2,02 | -0,85 | 3,95E-01 |
| AT3G63150 | 0,90 | 2,33 | 2,48 | 0,94 | 3,48E-01 |
| AT3G63160 | 1,15 | -2,04 | 1,89 | -1,08 | 2,79E-01 |
| AT3G63180 | 1,24 | 0,12 | 1,79 | 0,07 | 9,48E-01 |
| AT3G63190 | 0,25 | -0,34 | 4,38 | -0,08 | 9,38E-01 |
| AT3G63210 | 0,47 | -1,67 | 4,38 | -0,38 | 7,03E-01 |
| AT3G63310 | 1,42 | 2,04 | 1,98 | 1,03 | 3,02E-01 |
| AT3G63340 | 1,13 | 3,28 | 2,51 | 1,31 | 1,91E-01 |
| AT3G63390 | 0,24 | 1,29 | 4,36 | 0,30 | 7,67E-01 |
| AT3G63540 | 1,40 | -1,27 | 1,67 | -0,76 | 4,49E-01 |
| AT4G00005 | 0,19 | 0,00 | 4,37 | 0,00 | 9,99E-01 |
| AT4G00020 | 0,79 | -0,80 | 2,19 | -0,36 | 7,17E-01 |
| AT4G00070 | 0,43 | 0,47 | 4,21 | 0,11 | 9,11E-01 |
| AT4G00110 | 1,14 | -0,70 | 2,52 | -0,28 | 7,80E-01 |
| AT4G00120 | 1,32 | -1,93 | 1,89 | -1,02 | 3,06E-01 |
| AT4G00165 | 0,65 | -2,39 | 2,67 | -0,90 | 3,70E-01 |
| AT4G00180 | 0,45 | -0,45 | 4,33 | -0,10 | 9,17E-01 |
| AT4G00280 | 0,25 | 0,00 | 4,40 | 0,00 | 1,00E+00 |
| AT4G00290 | 0,50 | -1,10 | 2,75 | -0,40 | 6,89E-01 |
| AT4G00330 | 0,80 | 1,62 | 2,17 | 0,75 | 4,56E-01 |
| AT4G03585 | 1,51 | -2,30 | 1,63 | -1,42 | 1,57E-01 |
| AT4G00370 | 0,63 | -0,86 | 2,46 | -0,35 | 7,27E-01 |
| AT4G00400 | 0,42 | -0,79 | 2,79 | -0,28 | 7,78E-01 |
| AT4G00500 | 0,24 | -0,64 | 4,33 | -0,15 | 8,83E-01 |

| | | | | | |
|---|---|---|---|---|---|
| AT4G00540 | 0,54 | -1,19 | 2,64 | -0,45 | 6,51E-01 |
| AT4G00550 | 0,15 | 1,58 | 4,34 | 0,36 | 7,15E-01 |
| AT4G00560 | 0,40 | 2,74 | 4,31 | 0,64 | 5,25E-01 |
| AT4G00620 | 0,40 | -1,17 | 4,26 | -0,27 | 7,84E-01 |
| AT4G00740 | 0,31 | 0,48 | 4,40 | 0,11 | 9,14E-01 |
| AT4G00810 | 0,25 | 1,57 | 4,34 | 0,36 | 7,18E-01 |
| AT4G00890 | 0,23 | 0,00 | 4,41 | 0,00 | 1,00E+00 |
| AT4G00900 | 1,09 | 1,04 | 1,85 | 0,56 | 5,72E-01 |
| AT4G00980 | 0,33 | 1,14 | 4,36 | 0,26 | 7,94E-01 |
| AT4G00985 | 0,26 | -1,63 | 4,37 | -0,37 | 7,09E-01 |
| AT4G01000 | 0,50 | 2,87 | 3,88 | 0,74 | 4,60E-01 |
| AT4G01030 | 0,51 | 2,25 | 3,06 | 0,73 | 4,63E-01 |
| AT4G01037 | 0,21 | -2,03 | 4,35 | -0,47 | 6,41E-01 |
| AT4G01070 | 1,43 | -0,43 | 1,68 | -0,26 | 7,98E-01 |
| AT4G01090 | 0,44 | 2,29 | 3,24 | 0,71 | 4,79E-01 |
| AT4G01110 | 0,08 | 0,85 | 4,38 | 0,20 | 8,45E-01 |
| AT4G01120 | 0,18 | -0,69 | 4,37 | -0,16 | 8,74E-01 |
| AT4G01180 | 0,08 | 0,85 | 4,38 | 0,20 | 8,45E-01 |
| AT4G01190 | 0,20 | 0,89 | 4,37 | 0,20 | 8,39E-01 |
| AT4G01200 | 0,46 | 0,80 | 4,38 | 0,18 | 8,54E-01 |
| AT4G01210 | 1,50 | 3,44 | 1,91 | 1,80 | 7,22E-02 |
| AT4G01240 | 0,94 | 1,35 | 2,19 | 0,62 | 5,37E-01 |
| AT4G01310 | 0,66 | -1,26 | 3,17 | -0,40 | 6,90E-01 |
| AT4G01395 | 0,71 | -0,48 | 2,35 | -0,20 | 8,40E-01 |
| AT4G01533 | 1,62 | 1,02 | 2,06 | 0,50 | 6,20E-01 |
| AT4G01540 | 0,20 | 0,82 | 4,37 | 0,19 | 8,50E-01 |
| AT4G01550 | 0,48 | -0,37 | 4,29 | -0,09 | 9,32E-01 |
| AT4G01700 | 0,60 | -0,20 | 3,14 | -0,06 | 9,49E-01 |
| AT4G01870 | 0,45 | 0,50 | 4,13 | 0,12 | 9,03E-01 |
| AT4G01950 | 0,49 | -0,43 | 2,79 | -0,15 | 8,78E-01 |
| AT4G01985 | 0,41 | -2,31 | 4,34 | -0,53 | 5,94E-01 |
| AT4G02195 | 0,71 | -0,40 | 2,63 | -0,15 | 8,79E-01 |
| AT4G02210 | 0,65 | 2,03 | 2,89 | 0,70 | 4,83E-01 |
| AT4G02230 | 0,28 | 0,90 | 4,38 | 0,21 | 8,37E-01 |
| AT4G02405 | 0,05 | 0,00 | 4,41 | 0,00 | 1,00E+00 |
| AT4G02410 | 0,84 | 2,53 | 2,87 | 0,88 | 3,78E-01 |
| AT4G02420 | 1,44 | 1,18 | 1,81 | 0,65 | 5,13E-01 |
| AT4G02450 | 0,79 | 1,07 | 2,47 | 0,43 | 6,64E-01 |
| AT4G02460 | 0,19 | 1,50 | 4,34 | 0,34 | 7,30E-01 |
| AT4G02530 | 0,43 | -1,56 | 3,34 | -0,47 | 6,41E-01 |
| AT4G04185 | 0,38 | 1,92 | 3,38 | 0,57 | 5,70E-01 |
| AT4G02550 | 0,12 | 0,90 | 4,36 | 0,21 | 8,36E-01 |
| AT4G02560 | 0,57 | 0,97 | 2,64 | 0,37 | 7,14E-01 |
| AT4G02590 | 0,27 | -0,03 | 4,36 | -0,01 | 9,95E-01 |
| AT4G02620 | 0,23 | -0,34 | 4,35 | -0,08 | 9,38E-01 |
| AT4G02680 | 0,66 | 1,94 | 3,80 | 0,51 | 6,10E-01 |
| AT4G02710 | 0,71 | 2,84 | 2,88 | 0,99 | 3,23E-01 |
| AT4G02715 | 1,63 | 2,42 | 1,85 | 1,31 | 1,91E-01 |
| AT4G02735 | 0,41 | 0,00 | 4,41 | 0,00 | 1,00E+00 |
| AT4G02740 | 0,60 | 2,29 | 3,04 | 0,75 | 4,51E-01 |
| AT4G02750 | 0,98 | 3,22 | 2,19 | 1,47 | 1,42E-01 |
| AT4G02910 | 0,98 | 0,44 | 2,32 | 0,19 | 8,50E-01 |
| AT4G02930 | 0,46 | 1,49 | 4,35 | 0,34 | 7,33E-01 |
| AT4G03080 | 1,69 | 2,02 | 1,48 | 1,36 | 1,73E-01 |
| AT4G03090 | 0,91 | -1,00 | 1,75 | -0,57 | 5,69E-01 |
| AT4G03110 | 0,81 | 1,72 | 2,90 | 0,59 | 5,53E-01 |

| | | | | | |
|---|---|---|---|---|---|
| AT4G03130 | 0,08 | 0,85 | 4,38 | 0,20 | 8,45E-01 |
| AT4G03150 | 0,26 | 0,90 | 4,38 | 0,21 | 8,37E-01 |
| AT4G03190 | 0,84 | -0,46 | 2,24 | -0,20 | 8,38E-01 |
| AT4G03230 | 0,47 | 0,00 | 4,06 | 0,00 | 1,00E+00 |
| AT4G03270 | 0,81 | 2,45 | 2,77 | 0,88 | 3,78E-01 |
| AT4G04475 | 0,42 | -0,39 | 3,36 | -0,12 | 9,07E-01 |
| AT4G04485 | 0,66 | 1,87 | 2,42 | 0,77 | 4,40E-01 |
| AT4G03340 | 0,12 | 1,46 | 4,35 | 0,34 | 7,36E-01 |
| AT4G03390 | 0,56 | 2,04 | 2,68 | 0,76 | 4,45E-01 |
| AT4G03400 | 0,95 | 3,11 | 2,17 | 1,43 | 1,53E-01 |
| AT4G03430 | 1,44 | 3,20 | 1,69 | 1,89 | 5,82E-02 |
| AT4G03435 | 0,08 | 0,00 | 4,40 | 0,00 | 1,00E+00 |
| AT4G03438 | 0,26 | 0,00 | 4,41 | 0,00 | 1,00E+00 |
| AT4G03450 | 0,60 | 0,52 | 2,77 | 0,19 | 8,51E-01 |
| AT4G03490 | 0,42 | 0,47 | 4,34 | 0,11 | 9,14E-01 |
| AT4G03505 | 0,25 | 0,00 | 4,40 | 0,00 | 1,00E+00 |
| AT4G03520 | 1,13 | 0,14 | 1,75 | 0,08 | 9,34E-01 |
| AT4G03570 | 0,98 | -1,64 | 1,69 | -0,97 | 3,34E-01 |
| AT4G03590 | 1,16 | 0,65 | 2,33 | 0,28 | 7,80E-01 |
| AT4G03728 | 0,69 | -0,12 | 3,72 | -0,03 | 9,74E-01 |
| AT4G03740 | 1,37 | 1,85 | 2,02 | 0,91 | 3,60E-01 |
| AT4G04675 | 1,07 | -1,04 | 2,04 | -0,51 | 6,12E-01 |
| AT4G04030 | 0,82 | -2,74 | 2,46 | -1,11 | 2,65E-01 |
| AT4G04221 | 1,00 | 2,14 | 2,16 | 0,99 | 3,23E-01 |
| AT4G04805 | 1,38 | 2,75 | 2,28 | 1,21 | 2,28E-01 |
| AT4G04370 | 1,03 | 2,51 | 2,08 | 1,21 | 2,28E-01 |
| AT4G04470 | 0,08 | 0,00 | 4,40 | 0,00 | 1,00E+00 |
| AT4G04580 | 0,11 | 0,90 | 4,37 | 0,21 | 8,36E-01 |
| AT4G04614 | 0,51 | 0,43 | 4,39 | 0,10 | 9,21E-01 |
| AT4G04632 | 0,44 | 2,23 | 4,08 | 0,55 | 5,85E-01 |
| AT4G04650 | 0,82 | -0,41 | 2,47 | -0,16 | 8,69E-01 |
| AT4G04710 | 0,47 | 2,64 | 3,21 | 0,82 | 4,11E-01 |
| AT4G04720 | 1,28 | -0,77 | 1,94 | -0,40 | 6,92E-01 |
| AT4G04740 | 0,87 | 2,06 | 1,93 | 1,07 | 2,86E-01 |
| AT4G04745 | 1,61 | -2,96 | 1,68 | -1,76 | 7,79E-02 |
| AT4G04770 | 1,41 | 1,52 | 1,87 | 0,81 | 4,16E-01 |
| AT4G04910 | 0,75 | -1,43 | 2,21 | -0,65 | 5,17E-01 |
| AT4G04950 | 0,83 | -2,11 | 3,62 | -0,58 | 5,60E-01 |
| AT4G04957 | 0,91 | 2,29 | 2,64 | 0,87 | 3,85E-01 |
| AT4G04980 | 0,11 | 0,00 | 4,41 | 0,00 | 1,00E+00 |
| AT4G05070 | 1,30 | -0,95 | 1,57 | -0,61 | 5,45E-01 |
| AT4G05110 | 0,17 | 1,73 | 4,34 | 0,40 | 6,91E-01 |
| AT4G05085 | 0,05 | 0,00 | 4,41 | 0,00 | 1,00E+00 |
| AT4G05330 | 0,44 | 1,76 | 4,08 | 0,43 | 6,66E-01 |
| AT4G05410 | 0,51 | -0,16 | 4,08 | -0,04 | 9,69E-01 |
| AT4G05470 | 0,12 | 1,39 | 4,34 | 0,32 | 7,49E-01 |
| AT4G05631 | 0,41 | 0,00 | 3,42 | 0,00 | 9,99E-01 |
| AT4G06526 | 0,56 | -2,20 | 3,96 | -0,55 | 5,79E-01 |
| AT4G06639 | 0,21 | 0,43 | 4,37 | 0,10 | 9,22E-01 |
| AT4G06599 | 0,16 | 0,00 | 4,39 | 0,00 | 1,00E+00 |
| AT4G07350 | 0,08 | 0,00 | 4,40 | 0,00 | 1,00E+00 |
| AT4G07400 | 0,28 | 0,49 | 4,34 | 0,11 | 9,09E-01 |
| AT4G07410 | 1,28 | 2,33 | 2,21 | 1,05 | 2,93E-01 |
| AT4G07524 | 0,19 | 1,64 | 4,33 | 0,38 | 7,05E-01 |
| AT4G07526 | 0,83 | -2,38 | 2,35 | -1,01 | 3,11E-01 |
| AT4G05225 | 0,74 | -3,43 | 3,03 | -1,13 | 2,58E-01 |

| | | | | | |
|---|---|---|---|---|---|
| AT4G07990 | 0,20 | 0,00 | 4,41 | 0,00 | 1,00E+00 |
| AT4G08230 | 0,30 | 0,44 | 4,39 | 0,10 | 9,20E-01 |
| AT4G08310 | 0,85 | -0,37 | 2,15 | -0,17 | 8,62E-01 |
| AT4G05415 | 0,95 | 2,92 | 2,61 | 1,12 | 2,64E-01 |
| AT4G08370 | 0,46 | 1,93 | 2,69 | 0,72 | 4,72E-01 |
| AT4G08400 | 0,54 | 1,23 | 3,04 | 0,41 | 6,85E-01 |
| AT4G08410 | 1,29 | 2,71 | 2,32 | 1,17 | 2,41E-01 |
| AT4G05535 | 0,95 | -1,21 | 2,39 | -0,50 | 6,14E-01 |
| AT4G08460 | 0,62 | 0,64 | 2,58 | 0,25 | 8,02E-01 |
| AT4G08480 | 0,67 | 0,41 | 3,14 | 0,13 | 8,96E-01 |
| AT4G08485 | 0,63 | -3,93 | 3,67 | -1,07 | 2,84E-01 |
| AT4G08590 | 0,56 | 1,97 | 4,33 | 0,45 | 6,49E-01 |
| AT4G08760 | 1,19 | 0,63 | 2,34 | 0,27 | 7,87E-01 |
| AT4G08810 | 0,77 | 1,43 | 2,56 | 0,56 | 5,76E-01 |
| AT4G08870 | 1,44 | -1,86 | 1,52 | -1,23 | 2,20E-01 |
| AT4G08874 | 0,12 | 0,42 | 4,38 | 0,10 | 9,24E-01 |
| AT4G08940 | 0,46 | 2,08 | 3,33 | 0,62 | 5,32E-01 |
| AT4G09010 | 1,11 | 0,89 | 1,87 | 0,48 | 6,33E-01 |
| AT4G09012 | 0,55 | 1,49 | 3,19 | 0,47 | 6,40E-01 |
| AT4G09035 | 0,16 | 1,26 | 4,35 | 0,29 | 7,72E-01 |
| AT4G09040 | 0,47 | 0,43 | 4,38 | 0,10 | 9,22E-01 |
| AT4G09340 | 0,44 | 1,58 | 3,25 | 0,49 | 6,27E-01 |
| AT4G09350 | 0,26 | -1,10 | 4,38 | -0,25 | 8,01E-01 |
| AT4G09490 | 0,39 | -3,12 | 4,33 | -0,72 | 4,71E-01 |
| AT4G09510 | 0,38 | -0,87 | 3,34 | -0,26 | 7,95E-01 |
| AT4G09620 | 0,40 | 2,57 | 4,32 | 0,60 | 5,51E-01 |
| AT4G09640 | 0,98 | -1,68 | 1,76 | -0,96 | 3,39E-01 |
| AT4G05740 | 1,41 | -1,72 | 1,53 | -1,12 | 2,62E-01 |
| AT4G09900 | 0,13 | -1,24 | 4,39 | -0,28 | 7,77E-01 |
| AT4G09960 | 0,46 | -0,47 | 4,35 | -0,11 | 9,15E-01 |
| AT4G09980 | 0,57 | 2,05 | 2,61 | 0,79 | 4,31E-01 |
| AT4G10070 | 1,18 | 2,36 | 2,23 | 1,06 | 2,90E-01 |
| AT4G10080 | 1,04 | -1,41 | 2,07 | -0,68 | 4,95E-01 |
| AT4G10130 | 0,55 | 0,41 | 3,86 | 0,11 | 9,15E-01 |
| AT4G10320 | 1,43 | 1,37 | 1,57 | 0,87 | 3,84E-01 |
| AT4G10400 | 0,39 | 0,48 | 4,40 | 0,11 | 9,14E-01 |
| AT4G10420 | 1,07 | 1,99 | 2,68 | 0,74 | 4,58E-01 |
| AT4G10440 | 0,08 | 0,85 | 4,38 | 0,20 | 8,45E-01 |
| AT4G10480 | 0,41 | -0,13 | 4,34 | -0,03 | 9,77E-01 |
| AT4G10540 | 0,12 | 1,65 | 4,34 | 0,38 | 7,04E-01 |
| AT4G10550 | 0,52 | 2,62 | 3,86 | 0,68 | 4,98E-01 |
| AT4G10570 | 1,14 | 1,62 | 1,88 | 0,86 | 3,89E-01 |
| AT4G10590 | 1,40 | 2,94 | 2,01 | 1,46 | 1,44E-01 |
| AT4G05800 | 1,37 | 3,05 | 1,96 | 1,56 | 1,20E-01 |
| AT4G10700 | 0,54 | 0,43 | 3,82 | 0,11 | 9,10E-01 |
| AT4G10730 | 0,28 | 0,42 | 4,38 | 0,10 | 9,24E-01 |
| AT4G10740 | 1,27 | 1,92 | 2,33 | 0,83 | 4,08E-01 |
| AT4G10760 | 0,19 | 0,00 | 4,41 | 0,00 | 1,00E+00 |
| AT4G10770 | 0,10 | 0,42 | 4,38 | 0,10 | 9,23E-01 |
| AT4G10790 | 1,30 | 0,23 | 1,41 | 0,16 | 8,70E-01 |
| AT4G10920 | 0,27 | 1,65 | 3,75 | 0,44 | 6,60E-01 |
| AT4G10960 | 0,05 | 0,00 | 4,41 | 0,00 | 1,00E+00 |
| AT4G10970 | 0,10 | 0,42 | 4,38 | 0,10 | 9,23E-01 |
| AT4G11010 | 0,17 | 1,57 | 4,34 | 0,36 | 7,18E-01 |
| AT4G11050 | 0,37 | -0,41 | 3,43 | -0,12 | 9,05E-01 |
| AT4G11080 | 0,05 | 0,00 | 4,41 | 0,00 | 1,00E+00 |

| | | | | | |
|---|---|---|---|---|---|
| AT4G05855 | 1,27 | -0,43 | 2,09 | -0,20 | 8,38E-01 |
| AT4G11216 | 0,26 | 0,90 | 4,38 | 0,21 | 8,37E-01 |
| AT4G11220 | 0,29 | -1,29 | 4,40 | -0,29 | 7,69E-01 |
| AT4G11260 | 0,66 | 1,15 | 4,33 | 0,26 | 7,91E-01 |
| AT4G11300 | 0,45 | 1,82 | 4,12 | 0,44 | 6,58E-01 |
| AT4G11340 | 1,18 | -0,61 | 1,53 | -0,40 | 6,91E-01 |
| AT4G11410 | 0,24 | 1,58 | 4,34 | 0,36 | 7,16E-01 |
| AT4G05880 | 0,80 | -3,21 | 2,63 | -1,22 | 2,22E-01 |
| AT4G11650 | 0,43 | 0,89 | 4,38 | 0,20 | 8,38E-01 |
| AT4G11670 | 0,44 | 1,61 | 2,76 | 0,58 | 5,61E-01 |
| AT4G11800 | 1,18 | 1,64 | 1,73 | 0,95 | 3,43E-01 |
| AT4G05925 | 1,28 | -1,91 | 2,07 | -0,93 | 3,55E-01 |
| AT4G11820 | 0,25 | -2,61 | 4,34 | -0,60 | 5,47E-01 |
| AT4G11840 | 1,34 | 0,95 | 1,56 | 0,61 | 5,43E-01 |
| AT4G11850 | 1,56 | 0,57 | 1,61 | 0,35 | 7,23E-01 |
| AT4G11860 | 0,25 | 1,76 | 4,33 | 0,41 | 6,85E-01 |
| AT4G11920 | 0,40 | 1,89 | 4,13 | 0,46 | 6,47E-01 |
| AT4G05930 | 0,37 | 1,90 | 4,22 | 0,45 | 6,52E-01 |
| AT4G12010 | 0,59 | 3,27 | 3,64 | 0,90 | 3,69E-01 |
| AT4G05980 | 0,40 | 2,20 | 4,33 | 0,51 | 6,11E-01 |
| AT4G12300 | 1,00 | 1,95 | 2,20 | 0,89 | 3,75E-01 |
| AT4G12310 | 0,40 | 0,34 | 2,85 | 0,12 | 9,06E-01 |
| AT4G12330 | 0,81 | -4,07 | 2,63 | -1,55 | 1,22E-01 |
| AT4G12420 | 1,53 | 1,68 | 2,08 | 0,81 | 4,20E-01 |
| AT4G12590 | 0,64 | -1,18 | 2,65 | -0,44 | 6,57E-01 |
| AT4G12710 | 0,37 | 1,67 | 4,33 | 0,38 | 7,01E-01 |
| AT4G12730 | 0,13 | -1,24 | 4,39 | -0,28 | 7,77E-01 |
| AT4G12750 | 1,05 | 1,51 | 2,04 | 0,74 | 4,61E-01 |
| AT4G12790 | 0,53 | 1,93 | 3,30 | 0,59 | 5,58E-01 |
| AT4G12820 | 0,35 | -3,21 | 4,32 | -0,74 | 4,57E-01 |
| AT4G12880 | 0,35 | -3,21 | 4,32 | -0,74 | 4,57E-01 |
| AT4G12980 | 0,57 | 0,86 | 2,68 | 0,32 | 7,48E-01 |
| AT4G13150 | 0,55 | 0,93 | 4,38 | 0,21 | 8,32E-01 |
| AT4G13250 | 1,54 | 1,66 | 2,05 | 0,81 | 4,18E-01 |
| AT4G13350 | 0,30 | 1,17 | 4,35 | 0,27 | 7,88E-01 |
| AT4G13395 | 0,15 | 0,00 | 4,41 | 0,00 | 1,00E+00 |
| AT4G13420 | 0,12 | 1,58 | 4,34 | 0,36 | 7,16E-01 |
| AT4G13550 | 0,15 | 1,62 | 4,34 | 0,37 | 7,09E-01 |
| AT4G13590 | 0,56 | -0,37 | 2,72 | -0,14 | 8,91E-01 |
| AT4G13630 | 0,80 | 3,13 | 3,40 | 0,92 | 3,57E-01 |
| AT4G13640 | 0,21 | 2,17 | 4,32 | 0,50 | 6,15E-01 |
| AT4G13780 | 1,18 | 1,39 | 1,86 | 0,75 | 4,54E-01 |
| AT4G13800 | 0,45 | 1,94 | 3,22 | 0,60 | 5,47E-01 |
| AT4G13820 | 0,22 | 1,58 | 4,33 | 0,37 | 7,15E-01 |
| AT4G13870 | 0,08 | 0,85 | 4,38 | 0,20 | 8,45E-01 |
| AT4G13930 | 1,56 | 1,69 | 1,76 | 0,96 | 3,37E-01 |
| AT4G13950 | 0,08 | 1,10 | 4,36 | 0,25 | 8,00E-01 |
| AT4G13970 | 0,50 | 1,90 | 3,88 | 0,49 | 6,24E-01 |
| AT4G14010 | 0,10 | 0,00 | 4,41 | 0,00 | 1,00E+00 |
| AT4G14040 | 0,70 | 0,71 | 2,65 | 0,27 | 7,90E-01 |
| AT4G14050 | 0,14 | 1,24 | 4,35 | 0,29 | 7,75E-01 |
| AT4G14070 | 0,95 | 2,20 | 2,66 | 0,83 | 4,07E-01 |
| AT4G14090 | 0,47 | 1,23 | 2,81 | 0,44 | 6,61E-01 |
| AT4G14110 | 0,26 | 1,08 | 4,37 | 0,25 | 8,05E-01 |
| AT4G14200 | 0,21 | 1,56 | 4,34 | 0,36 | 7,19E-01 |
| AT4G14210 | 0,70 | 0,28 | 2,14 | 0,13 | 8,95E-01 |

| | | | | | |
|---|---|---|---|---|---|
| AT4G14230 | 0,73 | 2,07 | 2,94 | 0,71 | 4,81E-01 |
| AT4G14240 | 0,36 | -0,91 | 4,35 | -0,21 | 8,34E-01 |
| AT4G14340 | 0,26 | 1,08 | 4,37 | 0,25 | 8,05E-01 |
| AT4G14342 | 0,72 | 3,28 | 4,29 | 0,76 | 4,44E-01 |
| AT4G14350 | 0,16 | 0,42 | 4,37 | 0,10 | 9,23E-01 |
| AT4G14370 | 0,47 | 0,26 | 2,84 | 0,09 | 9,27E-01 |
| AT4G14385 | 0,08 | 1,10 | 4,36 | 0,25 | 8,00E-01 |
| AT4G14420 | 0,43 | 0,89 | 4,38 | 0,20 | 8,38E-01 |
| AT4G14430 | 1,12 | 0,49 | 1,59 | 0,31 | 7,58E-01 |
| AT4G14440 | 0,19 | 0,00 | 4,41 | 0,00 | 1,00E+00 |
| AT4G14510 | 1,36 | 2,33 | 1,89 | 1,23 | 2,18E-01 |
| AT4G14680 | 0,20 | 0,89 | 4,37 | 0,20 | 8,39E-01 |
| AT4G14710 | 0,27 | -0,38 | 4,37 | -0,09 | 9,31E-01 |
| AT4G14740 | 0,35 | -1,33 | 4,36 | -0,30 | 7,61E-01 |
| AT4G14746 | 0,16 | 0,96 | 4,36 | 0,22 | 8,26E-01 |
| AT4G14790 | 0,23 | 0,00 | 4,41 | 0,00 | 1,00E+00 |
| AT4G14811 | 0,45 | -0,20 | 3,36 | -0,06 | 9,52E-01 |
| AT4G14880 | 1,32 | -0,25 | 1,73 | -0,15 | 8,83E-01 |
| AT4G14900 | 1,30 | 1,68 | 1,82 | 0,92 | 3,57E-01 |
| AT4G14905 | 0,68 | -0,24 | 2,57 | -0,09 | 9,27E-01 |
| AT4G14920 | 0,44 | 1,96 | 4,15 | 0,47 | 6,36E-01 |
| AT4G14990 | 0,87 | 1,59 | 1,94 | 0,82 | 4,13E-01 |
| AT4G15050 | 0,52 | 2,10 | 3,80 | 0,55 | 5,81E-01 |
| AT4G15053 | 0,24 | 0,00 | 4,41 | 0,00 | 1,00E+00 |
| AT4G15080 | 0,43 | 1,19 | 4,14 | 0,29 | 7,74E-01 |
| AT4G15200 | 0,36 | 1,99 | 4,34 | 0,46 | 6,46E-01 |
| AT4G15210 | 0,12 | 1,58 | 4,34 | 0,36 | 7,16E-01 |
| AT4G15242 | 0,72 | 0,36 | 2,22 | 0,16 | 8,70E-01 |
| AT4G06305 | 0,44 | -2,48 | 4,07 | -0,61 | 5,43E-01 |
| AT4G06310 | 0,35 | 0,00 | 4,41 | 0,00 | 1,00E+00 |
| AT4G06315 | 1,00 | 0,00 | 3,20 | 0,00 | 1,00E+00 |
| AT4G15270 | 0,91 | 0,66 | 2,26 | 0,29 | 7,70E-01 |
| AT4G15490 | 0,92 | 3,46 | 2,04 | 1,70 | 9,00E-02 |
| AT4G15530 | 1,47 | 2,51 | 1,91 | 1,31 | 1,90E-01 |
| AT4G15550 | 0,31 | 0,51 | 4,35 | 0,12 | 9,06E-01 |
| AT4G15810 | 0,14 | 0,44 | 4,39 | 0,10 | 9,20E-01 |
| AT4G06340 | 0,10 | 0,62 | 4,38 | 0,14 | 8,87E-01 |
| AT4G15920 | 0,36 | -0,54 | 4,34 | -0,13 | 9,00E-01 |
| AT4G15955 | 0,12 | 1,65 | 4,34 | 0,38 | 7,04E-01 |
| AT4G16070 | 0,52 | 1,34 | 2,67 | 0,50 | 6,16E-01 |
| AT4G16141 | 0,51 | -1,21 | 4,04 | -0,30 | 7,65E-01 |
| AT4G16144 | 1,36 | -0,27 | 2,15 | -0,13 | 8,99E-01 |
| AT4G16155 | 0,30 | 0,41 | 4,38 | 0,09 | 9,25E-01 |
| AT4G16215 | 0,40 | 1,24 | 3,33 | 0,37 | 7,11E-01 |
| AT4G16250 | 0,73 | -1,75 | 2,89 | -0,60 | 5,46E-01 |
| AT4G16280 | 0,45 | 1,75 | 3,23 | 0,54 | 5,88E-01 |
| AT4G16330 | 0,88 | 3,55 | 2,83 | 1,26 | 2,09E-01 |
| AT4G16410 | 1,15 | 1,49 | 2,09 | 0,71 | 4,78E-01 |
| AT4G16470 | 0,11 | 1,07 | 4,36 | 0,25 | 8,06E-01 |
| AT4G16500 | 1,23 | -0,24 | 1,75 | -0,14 | 8,92E-01 |
| AT4G16515 | 0,18 | -0,69 | 4,37 | -0,16 | 8,74E-01 |
| AT4G16520 | 0,32 | 0,93 | 4,37 | 0,21 | 8,31E-01 |
| AT4G16540 | 0,20 | 0,00 | 4,41 | 0,00 | 1,00E+00 |
| AT4G16580 | 0,33 | -0,60 | 3,58 | -0,17 | 8,67E-01 |
| AT4G16630 | 0,66 | -1,36 | 3,81 | -0,36 | 7,21E-01 |
| AT4G16680 | 0,40 | 2,51 | 4,32 | 0,58 | 5,61E-01 |

| | | | | | |
|---|---|---|---|---|---|
| AT4G16690 | 0,88 | 2,09 | 2,19 | 0,96 | 3,39E-01 |
| AT4G16750 | 0,15 | -1,24 | 4,39 | -0,28 | 7,77E-01 |
| AT4G16760 | 1,69 | 1,40 | 1,52 | 0,92 | 3,59E-01 |
| AT4G16790 | 0,35 | 1,58 | 3,49 | 0,45 | 6,51E-01 |
| AT4G16880 | 1,01 | 0,86 | 2,18 | 0,39 | 6,94E-01 |
| AT4G16892 | 1,06 | 3,50 | 2,40 | 1,46 | 1,45E-01 |
| AT4G16957 | 0,81 | 3,33 | 3,37 | 0,99 | 3,23E-01 |
| AT4G16960 | 0,92 | 3,90 | 4,02 | 0,97 | 3,32E-01 |
| AT4G17040 | 0,44 | -1,01 | 4,16 | -0,24 | 8,09E-01 |
| AT4G17060 | 0,64 | 2,02 | 3,65 | 0,55 | 5,80E-01 |
| AT4G17070 | 0,40 | 2,10 | 3,31 | 0,63 | 5,27E-01 |
| AT4G17100 | 0,77 | -2,92 | 2,30 | -1,27 | 2,04E-01 |
| AT4G17190 | 0,27 | 1,45 | 4,35 | 0,33 | 7,39E-01 |
| AT4G17230 | 1,13 | 1,63 | 1,93 | 0,85 | 3,98E-01 |
| AT4G17243 | 0,66 | 1,73 | 2,57 | 0,67 | 5,02E-01 |
| AT4G17280 | 0,46 | 1,87 | 3,27 | 0,57 | 5,67E-01 |
| AT4G17430 | 0,16 | 0,00 | 4,41 | 0,00 | 1,00E+00 |
| AT4G17486 | 0,51 | 0,48 | 4,40 | 0,11 | 9,13E-01 |
| AT4G17490 | 0,05 | 0,00 | 4,41 | 0,00 | 1,00E+00 |
| AT4G17510 | 0,15 | 1,16 | 4,35 | 0,27 | 7,90E-01 |
| AT4G17550 | 0,46 | -0,11 | 4,13 | -0,03 | 9,79E-01 |
| AT4G17570 | 0,79 | 0,74 | 2,53 | 0,29 | 7,72E-01 |
| AT4G17590 | 0,76 | 3,18 | 2,77 | 1,15 | 2,50E-01 |
| AT4G17610 | 0,22 | 1,61 | 4,33 | 0,37 | 7,11E-01 |
| AT4G17670 | 0,33 | 2,00 | 4,33 | 0,46 | 6,44E-01 |
| AT4G17720 | 0,25 | -1,32 | 4,39 | -0,30 | 7,64E-01 |
| AT4G17740 | 0,24 | 0,59 | 4,34 | 0,13 | 8,93E-01 |
| AT4G17840 | 0,82 | 3,24 | 2,29 | 1,42 | 1,56E-01 |
| AT4G17880 | 0,51 | 1,76 | 3,10 | 0,57 | 5,71E-01 |
| AT4G17950 | 0,50 | -1,00 | 4,05 | -0,25 | 8,05E-01 |
| AT4G18010 | 1,46 | 3,88 | 1,93 | 2,01 | 4,46E-02 |
| AT4G18020 | 0,33 | 1,65 | 4,34 | 0,38 | 7,04E-01 |
| AT4G18040 | 0,22 | -0,28 | 4,36 | -0,06 | 9,49E-01 |
| AT4G18100 | 0,24 | -0,42 | 4,38 | -0,10 | 9,24E-01 |
| AT4G18110 | 0,16 | 0,90 | 4,38 | 0,21 | 8,37E-01 |
| AT4G18130 | 0,68 | 2,48 | 2,82 | 0,88 | 3,80E-01 |
| AT4G18140 | 0,37 | 0,22 | 4,37 | 0,05 | 9,60E-01 |
| AT4G18215 | 1,61 | 0,95 | 1,57 | 0,61 | 5,45E-01 |
| AT4G18230 | 0,15 | 1,58 | 4,34 | 0,36 | 7,15E-01 |
| AT4G18250 | 0,30 | 1,68 | 4,34 | 0,39 | 6,99E-01 |
| AT4G18253 | 0,68 | 2,14 | 2,46 | 0,87 | 3,85E-01 |
| AT4G18255 | 0,35 | 0,00 | 4,39 | 0,00 | 1,00E+00 |
| AT4G18390 | 1,03 | -1,60 | 2,01 | -0,80 | 4,26E-01 |
| AT4G18440 | 0,69 | -0,20 | 3,63 | -0,06 | 9,55E-01 |
| AT4G18520 | 0,18 | 1,66 | 4,33 | 0,38 | 7,02E-01 |
| AT4G18580 | 0,08 | 0,85 | 4,38 | 0,20 | 8,45E-01 |
| AT4G18790 | 0,08 | 1,10 | 4,36 | 0,25 | 8,00E-01 |
| AT4G18810 | 0,53 | -1,46 | 2,82 | -0,52 | 6,05E-01 |
| AT4G18840 | 0,38 | 0,36 | 3,43 | 0,10 | 9,17E-01 |
| AT4G19006 | 0,25 | 0,75 | 4,37 | 0,17 | 8,63E-01 |
| AT4G19040 | 0,87 | 2,87 | 3,23 | 0,89 | 3,73E-01 |
| AT4G19045 | 0,12 | 1,46 | 4,35 | 0,34 | 7,36E-01 |
| AT4G19050 | 0,78 | 1,67 | 2,69 | 0,62 | 5,36E-01 |
| AT4G19060 | 0,46 | 2,15 | 4,00 | 0,54 | 5,91E-01 |
| AT4G19080 | 1,21 | 1,43 | 1,77 | 0,81 | 4,18E-01 |
| AT4G19095 | 0,58 | -0,54 | 2,70 | -0,20 | 8,42E-01 |

| | | | | | |
|---|---|---|---|---|---|
| AT4G06870 | 0,16 | 0,51 | 4,38 | 0,12 | 9,07E-01 |
| AT4G19180 | 0,15 | 1,62 | 4,34 | 0,37 | 7,09E-01 |
| AT4G19220 | 0,37 | 1,64 | 4,28 | 0,38 | 7,03E-01 |
| AT4G19240 | 0,33 | 1,63 | 4,34 | 0,38 | 7,07E-01 |
| AT4G19420 | 0,68 | 1,34 | 2,48 | 0,54 | 5,89E-01 |
| AT4G19440 | 0,20 | 1,68 | 4,34 | 0,39 | 6,99E-01 |
| AT4G19640 | 0,35 | 0,41 | 3,51 | 0,12 | 9,07E-01 |
| AT4G19660 | 1,11 | 1,66 | 2,00 | 0,83 | 4,08E-01 |
| AT4G19670 | 0,77 | 2,27 | 3,52 | 0,65 | 5,18E-01 |
| AT4G06915 | 0,34 | 2,29 | 4,32 | 0,53 | 5,96E-01 |
| AT4G19700 | 1,20 | 1,65 | 2,06 | 0,80 | 4,23E-01 |
| AT4G19720 | 0,28 | -1,92 | 4,37 | -0,44 | 6,61E-01 |
| AT4G19860 | 1,15 | 1,32 | 1,82 | 0,72 | 4,69E-01 |
| AT4G19870 | 0,10 | 0,62 | 4,38 | 0,14 | 8,87E-01 |
| AT4G19880 | 0,52 | 1,10 | 2,58 | 0,43 | 6,70E-01 |
| AT4G19890 | 0,56 | 1,85 | 3,07 | 0,60 | 5,47E-01 |
| AT4G19900 | 0,17 | 1,40 | 4,35 | 0,32 | 7,48E-01 |
| AT4G19985 | 0,13 | 1,15 | 4,36 | 0,26 | 7,92E-01 |
| AT4G20095 | 0,45 | -3,35 | 4,31 | -0,78 | 4,38E-01 |
| AT4G20150 | 0,58 | -1,57 | 2,60 | -0,60 | 5,46E-01 |
| AT4G06945 | 1,00 | 3,35 | 2,82 | 1,19 | 2,35E-01 |
| AT4G20210 | 0,49 | -1,11 | 2,72 | -0,41 | 6,84E-01 |
| AT4G20330 | 0,48 | 0,76 | 2,68 | 0,28 | 7,76E-01 |
| AT4G20410 | 1,25 | -0,59 | 2,00 | -0,29 | 7,69E-01 |
| AT4G20440 | 0,64 | 1,30 | 2,23 | 0,58 | 5,59E-01 |
| AT4G20690 | 0,21 | 2,17 | 4,32 | 0,50 | 6,15E-01 |
| AT4G20880 | 0,74 | 0,30 | 2,06 | 0,15 | 8,85E-01 |
| AT4G20910 | 0,27 | 0,00 | 4,40 | 0,00 | 1,00E+00 |
| AT4G20953 | 0,51 | 2,24 | 3,24 | 0,69 | 4,90E-01 |
| AT4G21100 | 1,08 | -0,74 | 1,79 | -0,41 | 6,80E-01 |
| AT4G21140 | 0,20 | 0,89 | 4,37 | 0,20 | 8,39E-01 |
| AT4G21170 | 0,28 | -1,46 | 4,39 | -0,33 | 7,39E-01 |
| AT4G06985 | 0,08 | 0,85 | 4,38 | 0,20 | 8,45E-01 |
| AT4G21190 | 0,08 | 1,10 | 4,36 | 0,25 | 8,00E-01 |
| AT4G21350 | 0,08 | 1,10 | 4,36 | 0,25 | 8,00E-01 |
| AT4G21380 | 1,64 | 4,16 | 1,71 | 2,43 | 1,49E-02 |
| AT4G21400 | 0,16 | 1,07 | 4,37 | 0,24 | 8,07E-01 |
| AT4G21430 | 0,22 | -1,35 | 4,41 | -0,31 | 7,59E-01 |
| AT4G21510 | 0,05 | 0,00 | 4,41 | 0,00 | 1,00E+00 |
| AT4G21520 | 0,22 | 0,85 | 4,36 | 0,20 | 8,45E-01 |
| AT4G21560 | 0,38 | 0,00 | 4,41 | 0,00 | 1,00E+00 |
| AT4G21570 | 1,42 | 1,32 | 1,97 | 0,67 | 5,04E-01 |
| AT4G21660 | 1,13 | 2,67 | 2,23 | 1,20 | 2,31E-01 |
| AT4G21810 | 1,02 | 1,83 | 2,10 | 0,87 | 3,83E-01 |
| AT4G21860 | 0,50 | -0,72 | 3,28 | -0,22 | 8,26E-01 |
| AT4G21890 | 0,12 | 1,65 | 4,34 | 0,38 | 7,04E-01 |
| AT4G21910 | 0,54 | 0,94 | 4,31 | 0,22 | 8,27E-01 |
| AT4G21990 | 0,81 | 1,76 | 2,55 | 0,69 | 4,90E-01 |
| AT4G22120 | 0,12 | 1,58 | 4,34 | 0,36 | 7,16E-01 |
| AT4G22150 | 0,22 | -1,55 | 4,37 | -0,36 | 7,22E-01 |
| AT4G22190 | 0,98 | 0,44 | 1,79 | 0,25 | 8,05E-01 |
| AT4G22200 | 0,21 | 1,09 | 4,36 | 0,25 | 8,03E-01 |
| AT4G22250 | 0,16 | 1,33 | 4,35 | 0,31 | 7,59E-01 |
| AT4G22260 | 0,18 | -1,33 | 4,40 | -0,30 | 7,63E-01 |
| AT4G22265 | 0,20 | 1,07 | 4,36 | 0,25 | 8,06E-01 |
| AT4G22285 | 0,20 | -0,05 | 4,36 | -0,01 | 9,91E-01 |

| | | | | | |
|---|---|---|---|---|---|
| AT4G22290 | 0,23 | 0,61 | 4,38 | 0,14 | 8,90E-01 |
| AT4G22350 | 0,69 | 0,26 | 3,72 | 0,07 | 9,44E-01 |
| AT4G22380 | 0,51 | 1,62 | 2,67 | 0,61 | 5,45E-01 |
| AT4G22470 | 0,12 | 0,42 | 4,38 | 0,10 | 9,24E-01 |
| AT4G22485 | 0,49 | -1,21 | 3,96 | -0,31 | 7,59E-01 |
| AT4G22550 | 0,26 | 0,73 | 3,75 | 0,20 | 8,45E-01 |
| AT4G22570 | 0,24 | 0,83 | 4,37 | 0,19 | 8,49E-01 |
| AT4G22745 | 0,42 | 1,61 | 4,13 | 0,39 | 6,97E-01 |
| AT4G22750 | 1,03 | 0,21 | 2,23 | 0,09 | 9,24E-01 |
| AT4G22755 | 0,47 | 1,64 | 4,35 | 0,38 | 7,07E-01 |
| AT4G22770 | 1,29 | 0,90 | 1,47 | 0,61 | 5,43E-01 |
| AT4G22780 | 0,24 | 0,42 | 4,37 | 0,10 | 9,24E-01 |
| AT4G22970 | 0,52 | 2,75 | 3,82 | 0,72 | 4,73E-01 |
| AT4G22980 | 0,47 | 2,50 | 3,94 | 0,63 | 5,26E-01 |
| AT4G23010 | 0,83 | 1,54 | 2,37 | 0,65 | 5,15E-01 |
| AT4G23040 | 0,28 | -0,96 | 4,34 | -0,22 | 8,26E-01 |
| AT4G23130 | 0,55 | 2,82 | 3,90 | 0,72 | 4,70E-01 |
| AT4G23190 | 1,18 | 2,66 | 2,00 | 1,33 | 1,84E-01 |
| AT4G23205 | 0,08 | 1,10 | 4,36 | 0,25 | 8,00E-01 |
| AT4G23200 | 0,43 | 1,05 | 4,38 | 0,24 | 8,11E-01 |
| AT4G23210 | 0,21 | 1,97 | 4,33 | 0,46 | 6,49E-01 |
| AT4G23220 | 0,76 | 2,99 | 2,38 | 1,26 | 2,08E-01 |
| AT4G23240 | 0,37 | 2,32 | 4,32 | 0,54 | 5,91E-01 |
| AT4G23250 | 0,45 | 2,44 | 3,92 | 0,62 | 5,34E-01 |
| AT4G23260 | 0,19 | 1,50 | 4,34 | 0,34 | 7,30E-01 |
| AT4G23290 | 0,10 | 0,00 | 4,41 | 0,00 | 1,00E+00 |
| AT4G23320 | 0,08 | 1,10 | 4,36 | 0,25 | 8,00E-01 |
| AT4G23460 | 1,64 | 1,54 | 1,69 | 0,92 | 3,59E-01 |
| AT4G23490 | 0,15 | -1,24 | 4,39 | -0,28 | 7,77E-01 |
| AT4G07130 | 0,46 | 1,48 | 4,14 | 0,36 | 7,21E-01 |
| AT4G23540 | 0,67 | -0,23 | 3,12 | -0,07 | 9,42E-01 |
| AT4G23560 | 0,10 | 0,00 | 4,41 | 0,00 | 1,00E+00 |
| AT4G07135 | 1,25 | 2,94 | 2,04 | 1,44 | 1,50E-01 |
| AT4G07145 | 0,96 | 2,91 | 2,63 | 1,11 | 2,69E-01 |
| AT4G23620 | 0,31 | 1,35 | 4,34 | 0,31 | 7,57E-01 |
| AT4G23640 | 0,77 | 1,34 | 2,38 | 0,56 | 5,73E-01 |
| AT4G23670 | 1,50 | -4,43 | 1,78 | -2,49 | 1,28E-02 |
| AT4G23700 | 0,29 | 0,92 | 4,36 | 0,21 | 8,33E-01 |
| AT4G23770 | 0,22 | -0,21 | 4,37 | -0,05 | 9,61E-01 |
| AT4G23810 | 0,35 | 1,54 | 4,34 | 0,35 | 7,23E-01 |
| AT4G23820 | 0,63 | -0,52 | 2,35 | -0,22 | 8,25E-01 |
| AT4G23880 | 0,47 | -1,27 | 3,27 | -0,39 | 6,98E-01 |
| AT4G23890 | 0,66 | -1,63 | 2,67 | -0,61 | 5,42E-01 |
| AT4G24180 | 0,37 | 1,58 | 4,34 | 0,36 | 7,16E-01 |
| AT4G24200 | 0,70 | 2,08 | 2,98 | 0,70 | 4,85E-01 |
| AT4G07250 | 0,17 | -2,23 | 4,35 | -0,51 | 6,08E-01 |
| AT4G24300 | 0,45 | 0,64 | 4,37 | 0,15 | 8,83E-01 |
| AT4G24320 | 0,33 | 1,39 | 4,32 | 0,32 | 7,48E-01 |
| AT4G24330 | 0,84 | 0,37 | 1,95 | 0,19 | 8,50E-01 |
| AT4G07255 | 0,10 | 0,00 | 4,41 | 0,00 | 1,00E+00 |
| AT4G24390 | 0,29 | 1,52 | 4,34 | 0,35 | 7,27E-01 |
| AT4G24490 | 0,72 | 1,35 | 2,47 | 0,55 | 5,84E-01 |
| AT4G24530 | 0,08 | 1,10 | 4,36 | 0,25 | 8,00E-01 |
| AT4G24560 | 1,21 | 1,57 | 1,94 | 0,81 | 4,19E-01 |
| AT4G24580 | 0,46 | 0,83 | 4,14 | 0,20 | 8,41E-01 |
| AT4G24610 | 0,67 | 2,17 | 2,93 | 0,74 | 4,59E-01 |

| | | | | | |
|---|---|---|---|---|---|
| AT4G24620 | 0,42 | -1,03 | 4,21 | -0,24 | 8,07E-01 |
| AT4G24660 | 1,04 | -1,01 | 2,13 | -0,48 | 6,34E-01 |
| AT4G24670 | 0,08 | 0,00 | 4,40 | 0,00 | 1,00E+00 |
| AT4G24730 | 0,45 | 1,23 | 4,07 | 0,30 | 7,63E-01 |
| AT4G24740 | 0,63 | 1,25 | 3,11 | 0,40 | 6,88E-01 |
| AT4G24750 | 0,28 | 0,42 | 4,38 | 0,10 | 9,24E-01 |
| AT4G24780 | 1,27 | 0,82 | 2,27 | 0,36 | 7,19E-01 |
| AT4G07285 | 0,27 | -2,28 | 4,34 | -0,52 | 6,00E-01 |
| AT4G24820 | 1,43 | 2,23 | 1,79 | 1,24 | 2,14E-01 |
| AT4G24830 | 0,25 | -0,34 | 4,38 | -0,08 | 9,38E-01 |
| AT4G24840 | 0,53 | -0,49 | 3,11 | -0,16 | 8,74E-01 |
| AT4G24920 | 0,86 | 2,74 | 2,72 | 1,01 | 3,14E-01 |
| AT4G24990 | 0,31 | 0,46 | 4,39 | 0,11 | 9,16E-01 |
| AT4G25090 | 0,29 | 2,52 | 4,32 | 0,58 | 5,59E-01 |
| AT4G25270 | 0,45 | 1,23 | 4,07 | 0,30 | 7,63E-01 |
| AT4G25280 | 0,71 | 2,65 | 4,31 | 0,62 | 5,38E-01 |
| AT4G25290 | 0,52 | 1,79 | 3,86 | 0,46 | 6,44E-01 |
| AT4G25340 | 0,19 | 0,38 | 4,34 | 0,09 | 9,30E-01 |
| AT4G25435 | 0,88 | -3,26 | 2,69 | -1,21 | 2,26E-01 |
| AT4G25540 | 0,13 | 1,15 | 4,36 | 0,26 | 7,92E-01 |
| AT4G25570 | 0,43 | -1,04 | 4,15 | -0,25 | 8,02E-01 |
| AT4G25640 | 1,48 | 2,18 | 1,71 | 1,27 | 2,02E-01 |
| AT4G25650 | 1,11 | -0,17 | 1,54 | -0,11 | 9,10E-01 |
| AT4G25670 | 0,79 | 2,51 | 2,82 | 0,89 | 3,73E-01 |
| AT4G25690 | 0,56 | -0,25 | 3,30 | -0,08 | 9,39E-01 |
| AT4G25740 | 0,38 | 1,40 | 4,36 | 0,32 | 7,48E-01 |
| AT4G25870 | 0,26 | 0,90 | 4,38 | 0,21 | 8,37E-01 |
| AT4G25940 | 0,39 | 0,70 | 4,34 | 0,16 | 8,73E-01 |
| AT4G25990 | 0,26 | 0,00 | 4,41 | 0,00 | 1,00E+00 |
| AT4G26080 | 0,87 | -0,59 | 2,36 | -0,25 | 8,03E-01 |
| AT4G26100 | 0,81 | 1,15 | 2,12 | 0,55 | 5,86E-01 |
| AT4G26130 | 1,45 | -0,32 | 1,56 | -0,20 | 8,39E-01 |
| AT4G26140 | 0,21 | 0,00 | 4,38 | 0,00 | 1,00E+00 |
| AT4G26190 | 0,25 | 1,60 | 4,33 | 0,37 | 7,12E-01 |
| AT4G26300 | 0,69 | 1,17 | 2,51 | 0,47 | 6,40E-01 |
| AT4G07565 | 0,79 | -0,97 | 2,07 | -0,47 | 6,41E-01 |
| AT4G07625 | 0,79 | 0,02 | 2,53 | 0,01 | 9,93E-01 |
| AT4G07635 | 0,12 | 0,42 | 4,38 | 0,10 | 9,24E-01 |
| AT4G26370 | 0,48 | -2,50 | 4,06 | -0,62 | 5,37E-01 |
| AT4G26375 | 0,40 | 2,57 | 4,32 | 0,60 | 5,51E-01 |
| AT4G26400 | 0,60 | 1,84 | 3,92 | 0,47 | 6,38E-01 |
| AT4G26620 | 0,28 | 0,91 | 4,32 | 0,21 | 8,32E-01 |
| AT4G26630 | 0,72 | 2,86 | 2,45 | 1,17 | 2,43E-01 |
| AT4G26640 | 1,00 | 1,58 | 2,28 | 0,69 | 4,87E-01 |
| AT4G26690 | 1,51 | 1,56 | 1,72 | 0,91 | 3,65E-01 |
| AT4G26750 | 0,43 | 0,98 | 4,14 | 0,24 | 8,13E-01 |
| AT4G26760 | 0,64 | 1,62 | 3,05 | 0,53 | 5,96E-01 |
| AT4G26795 | 0,42 | 0,80 | 4,12 | 0,20 | 8,45E-01 |
| AT4G26860 | 0,37 | 0,58 | 4,33 | 0,13 | 8,93E-01 |
| AT4G26910 | 1,33 | 1,01 | 1,65 | 0,61 | 5,40E-01 |
| AT4G26940 | 0,43 | -0,13 | 4,10 | -0,03 | 9,74E-01 |
| AT4G26980 | 0,78 | 2,73 | 2,39 | 1,14 | 2,53E-01 |
| AT4G27020 | 0,48 | 1,00 | 3,37 | 0,30 | 7,66E-01 |
| AT4G07765 | 0,66 | 2,39 | 2,83 | 0,85 | 3,97E-01 |
| AT4G27130 | 0,28 | 1,23 | 3,70 | 0,33 | 7,40E-01 |
| AT4G27270 | 0,66 | 3,30 | 3,83 | 0,86 | 3,89E-01 |

| | | | | | |
|---|---|---|---|---|---|
| AT4G27280 | 0,84 | 1,38 | 2,84 | 0,49 | 6,27E-01 |
| AT4G27300 | 0,83 | 0,16 | 2,38 | 0,07 | 9,46E-01 |
| AT4G27320 | 1,19 | 0,72 | 2,20 | 0,33 | 7,42E-01 |
| AT4G27340 | 0,21 | 1,58 | 4,34 | 0,36 | 7,16E-01 |
| AT4G27410 | 0,98 | 2,67 | 2,32 | 1,15 | 2,50E-01 |
| AT4G27450 | 0,72 | 0,26 | 2,38 | 0,11 | 9,13E-01 |
| AT4G27480 | 0,65 | 3,20 | 3,54 | 0,91 | 3,65E-01 |
| AT4G27650 | 0,39 | -0,15 | 3,39 | -0,04 | 9,65E-01 |
| AT4G27700 | 0,47 | 0,49 | 3,35 | 0,15 | 8,83E-01 |
| AT4G27730 | 0,37 | 1,82 | 4,33 | 0,42 | 6,75E-01 |
| AT4G27800 | 1,09 | -0,42 | 1,89 | -0,22 | 8,24E-01 |
| AT4G27830 | 0,84 | 0,67 | 2,02 | 0,33 | 7,39E-01 |
| AT4G27840 | 0,43 | 0,00 | 4,41 | 0,00 | 1,00E+00 |
| AT4G27885 | 0,16 | 0,00 | 4,41 | 0,00 | 1,00E+00 |
| AT4G27960 | 0,38 | 0,54 | 4,34 | 0,12 | 9,01E-01 |
| AT4G28030 | 0,45 | 0,65 | 2,75 | 0,23 | 8,15E-01 |
| AT4G28050 | 0,43 | 1,28 | 4,30 | 0,30 | 7,66E-01 |
| AT4G28060 | 0,25 | 0,00 | 4,40 | 0,00 | 1,00E+00 |
| AT4G28180 | 0,69 | 1,89 | 2,58 | 0,73 | 4,64E-01 |
| AT4G28200 | 0,59 | -0,31 | 2,58 | -0,12 | 9,06E-01 |
| AT4G28230 | 0,33 | 2,08 | 4,32 | 0,48 | 6,30E-01 |
| AT4G28240 | 0,66 | -0,09 | 2,17 | -0,04 | 9,67E-01 |
| AT4G28320 | 0,20 | 0,89 | 4,37 | 0,20 | 8,39E-01 |
| AT4G28390 | 0,33 | 2,12 | 4,32 | 0,49 | 6,24E-01 |
| AT4G28450 | 0,61 | 0,00 | 4,41 | 0,00 | 1,00E+00 |
| AT4G28470 | 0,62 | 2,43 | 3,11 | 0,78 | 4,35E-01 |
| AT4G28610 | 0,93 | 2,80 | 2,61 | 1,07 | 2,83E-01 |
| AT4G28630 | 0,12 | 1,58 | 4,34 | 0,36 | 7,16E-01 |
| AT4G28770 | 0,20 | 0,42 | 4,38 | 0,10 | 9,24E-01 |
| AT4G28910 | 1,06 | 0,52 | 2,33 | 0,22 | 8,23E-01 |
| AT4G29010 | 1,63 | 0,55 | 1,61 | 0,34 | 7,31E-01 |
| AT4G29050 | 0,13 | -1,24 | 4,39 | -0,28 | 7,77E-01 |
| AT4G29080 | 0,28 | 1,19 | 4,34 | 0,27 | 7,85E-01 |
| AT4G29130 | 1,04 | -1,38 | 1,73 | -0,80 | 4,24E-01 |
| AT4G29220 | 0,63 | 0,20 | 3,06 | 0,07 | 9,47E-01 |
| AT4G29260 | 0,48 | 1,16 | 4,02 | 0,29 | 7,74E-01 |
| AT4G29300 | 1,28 | -1,96 | 1,63 | -1,20 | 2,29E-01 |
| AT4G29305 | 0,76 | -0,05 | 2,48 | -0,02 | 9,85E-01 |
| AT4G08155 | 1,40 | 0,83 | 1,79 | 0,46 | 6,45E-01 |
| AT4G29440 | 0,66 | 1,75 | 2,97 | 0,59 | 5,54E-01 |
| AT4G29520 | 0,77 | 1,04 | 2,61 | 0,40 | 6,91E-01 |
| AT4G29654 | 1,07 | 3,02 | 2,63 | 1,15 | 2,52E-01 |
| AT4G29670 | 0,65 | 0,26 | 2,34 | 0,11 | 9,13E-01 |
| AT4G29680 | 0,85 | 0,25 | 3,04 | 0,08 | 9,35E-01 |
| AT4G29720 | 0,17 | -0,02 | 4,36 | 0,00 | 9,97E-01 |
| AT4G29735 | 0,17 | 0,09 | 4,35 | 0,02 | 9,84E-01 |
| AT4G29810 | 0,27 | 1,48 | 4,35 | 0,34 | 7,34E-01 |
| AT4G29820 | 0,24 | 0,65 | 4,37 | 0,15 | 8,81E-01 |
| AT4G29840 | 0,58 | -0,57 | 2,32 | -0,24 | 8,07E-01 |
| AT4G29880 | 0,21 | 1,36 | 4,35 | 0,31 | 7,55E-01 |
| AT4G29905 | 1,17 | 0,25 | 1,85 | 0,13 | 8,94E-01 |
| AT4G29910 | 0,08 | 0,00 | 4,40 | 0,00 | 1,00E+00 |
| AT4G29920 | 0,05 | 0,00 | 4,41 | 0,00 | 1,00E+00 |
| AT4G29940 | 0,28 | 0,42 | 4,38 | 0,10 | 9,24E-01 |
| AT4G29960 | 0,50 | 2,17 | 3,91 | 0,56 | 5,78E-01 |
| AT4G29990 | 0,08 | 0,85 | 4,38 | 0,20 | 8,45E-01 |

| | | | | | |
|---|---|---|---|---|---|
| AT4G30000 | 0,38 | 0,96 | 4,37 | 0,22 | 8,27E-01 |
| AT4G30030 | 0,45 | 2,47 | 4,32 | 0,57 | 5,67E-01 |
| AT4G30060 | 0,08 | 1,10 | 4,36 | 0,25 | 8,00E-01 |
| AT4G30080 | 0,31 | 0,99 | 4,35 | 0,23 | 8,19E-01 |
| AT4G30110 | 1,05 | 3,28 | 3,13 | 1,05 | 2,94E-01 |
| AT4G30200 | 1,47 | 1,92 | 1,65 | 1,16 | 2,45E-01 |
| AT4G30240 | 0,30 | 1,56 | 4,34 | 0,36 | 7,18E-01 |
| AT4G30260 | 0,47 | 1,36 | 3,15 | 0,43 | 6,65E-01 |
| AT4G30270 | 1,68 | 2,33 | 1,49 | 1,56 | 1,18E-01 |
| AT4G30280 | 0,12 | 0,00 | 4,39 | 0,00 | 1,00E+00 |
| AT4G30360 | 0,21 | 0,00 | 4,41 | 0,00 | 1,00E+00 |
| AT4G30410 | 0,12 | 0,90 | 4,36 | 0,21 | 8,36E-01 |
| AT4G30430 | 0,08 | 0,85 | 4,38 | 0,20 | 8,45E-01 |
| AT4G30560 | 0,74 | 2,14 | 3,49 | 0,61 | 5,40E-01 |
| AT4G30580 | 0,92 | 3,18 | 3,36 | 0,95 | 3,43E-01 |
| AT4G30600 | 1,60 | 1,34 | 1,69 | 0,79 | 4,29E-01 |
| AT4G30620 | 0,23 | 0,00 | 4,41 | 0,00 | 1,00E+00 |
| AT4G30700 | 0,08 | 1,10 | 4,36 | 0,25 | 8,00E-01 |
| AT4G30710 | 0,12 | 1,58 | 4,34 | 0,36 | 7,16E-01 |
| AT4G30720 | 1,29 | 3,41 | 1,79 | 1,91 | 5,60E-02 |
| AT4G30740 | 1,03 | 2,18 | 2,45 | 0,89 | 3,72E-01 |
| AT4G30790 | 0,99 | 3,33 | 2,22 | 1,50 | 1,34E-01 |
| AT4G30890 | 0,60 | 0,85 | 3,05 | 0,28 | 7,81E-01 |
| AT4G30920 | 1,19 | -0,59 | 1,77 | -0,33 | 7,39E-01 |
| AT4G30950 | 1,64 | 0,46 | 1,43 | 0,32 | 7,49E-01 |
| AT4G31020 | 0,29 | 0,42 | 4,38 | 0,10 | 9,24E-01 |
| AT4G31120 | 0,87 | 1,23 | 2,46 | 0,50 | 6,18E-01 |
| AT4G31130 | 0,59 | -0,04 | 3,13 | -0,01 | 9,90E-01 |
| AT4G31420 | 0,28 | 0,87 | 4,38 | 0,20 | 8,43E-01 |
| AT4G31450 | 0,46 | -1,17 | 4,09 | -0,28 | 7,76E-01 |
| AT4G31540 | 0,43 | -0,84 | 4,16 | -0,20 | 8,40E-01 |
| AT4G31550 | 1,02 | 0,34 | 1,91 | 0,18 | 8,58E-01 |
| AT4G31590 | 0,63 | 2,16 | 2,62 | 0,82 | 4,10E-01 |
| AT4G31810 | 0,08 | 1,10 | 4,36 | 0,25 | 8,00E-01 |
| AT4G31820 | 0,43 | 1,08 | 2,79 | 0,39 | 6,99E-01 |
| AT4G32070 | 0,58 | 1,18 | 3,91 | 0,30 | 7,63E-01 |
| AT4G32120 | 0,15 | 0,86 | 4,37 | 0,20 | 8,43E-01 |
| AT4G32130 | 0,18 | 0,63 | 4,37 | 0,14 | 8,86E-01 |
| AT4G32150 | 1,17 | 1,34 | 1,79 | 0,75 | 4,54E-01 |
| AT4G32180 | 1,00 | 0,63 | 2,41 | 0,26 | 7,94E-01 |
| AT4G32208 | 0,37 | 1,64 | 4,28 | 0,38 | 7,03E-01 |
| AT4G32340 | 1,02 | 0,96 | 1,93 | 0,50 | 6,21E-01 |
| AT4G32420 | 0,78 | 1,33 | 2,42 | 0,55 | 5,83E-01 |
| AT4G32460 | 0,31 | 0,21 | 4,36 | 0,05 | 9,61E-01 |
| AT4G32470 | 0,29 | 0,41 | 4,38 | 0,09 | 9,25E-01 |
| AT4G32510 | 0,56 | 2,82 | 3,73 | 0,76 | 4,50E-01 |
| AT4G32520 | 1,00 | 0,61 | 1,89 | 0,33 | 7,45E-01 |
| AT4G32530 | 0,89 | 0,95 | 2,47 | 0,39 | 6,99E-01 |
| AT4G32670 | 0,15 | 1,25 | 4,35 | 0,29 | 7,75E-01 |
| AT4G32750 | 0,18 | 0,17 | 4,36 | 0,04 | 9,69E-01 |
| AT4G32820 | 0,83 | 3,19 | 2,66 | 1,20 | 2,30E-01 |
| AT4G32880 | 1,09 | 1,34 | 2,12 | 0,63 | 5,27E-01 |
| AT4G32900 | 0,12 | 1,39 | 4,34 | 0,32 | 7,49E-01 |
| AT4G32920 | 0,70 | 2,96 | 2,81 | 1,05 | 2,93E-01 |
| AT4G33040 | 0,37 | 0,53 | 4,33 | 0,12 | 9,03E-01 |
| AT4G33090 | 0,64 | 0,39 | 3,03 | 0,13 | 8,98E-01 |

| | | | | | |
|---|---|---|---|---|---|
| AT4G33110 | 0,17 | 1,23 | 4,35 | 0,28 | 7,77E-01 |
| AT4G33150 | 0,65 | 3,31 | 3,60 | 0,92 | 3,59E-01 |
| AT4G33200 | 0,29 | 1,82 | 4,33 | 0,42 | 6,75E-01 |
| AT4G33210 | 1,11 | 2,88 | 1,98 | 1,45 | 1,46E-01 |
| AT4G33250 | 0,89 | 1,10 | 2,03 | 0,54 | 5,88E-01 |
| AT4G33260 | 0,12 | 1,58 | 4,34 | 0,36 | 7,16E-01 |
| AT4G33270 | 0,12 | 0,00 | 4,39 | 0,00 | 1,00E+00 |
| AT4G33400 | 0,33 | 0,70 | 4,32 | 0,16 | 8,72E-01 |
| AT4G33430 | 1,19 | 1,97 | 2,00 | 0,99 | 3,23E-01 |
| AT4G33470 | 0,67 | 2,55 | 2,90 | 0,88 | 3,81E-01 |
| AT4G33495 | 0,29 | 1,82 | 4,33 | 0,42 | 6,75E-01 |
| AT4G33500 | 0,96 | -0,23 | 2,45 | -0,09 | 9,26E-01 |
| AT4G33530 | 1,44 | 1,09 | 1,58 | 0,69 | 4,89E-01 |
| AT4G33585 | 1,20 | 1,38 | 2,99 | 0,46 | 6,45E-01 |
| AT4G33625 | 0,16 | 1,49 | 4,34 | 0,34 | 7,32E-01 |
| AT4G33670 | 0,76 | 1,56 | 2,18 | 0,72 | 4,75E-01 |
| AT4G33680 | 0,20 | 1,42 | 4,36 | 0,33 | 7,44E-01 |
| AT4G33700 | 0,28 | -0,82 | 4,34 | -0,19 | 8,50E-01 |
| AT4G33790 | 0,16 | 1,07 | 4,37 | 0,24 | 8,07E-01 |
| AT4G33860 | 0,68 | 2,28 | 3,61 | 0,63 | 5,27E-01 |
| AT4G33865 | 0,81 | 1,19 | 2,06 | 0,58 | 5,65E-01 |
| AT4G33910 | 0,26 | 0,90 | 4,38 | 0,21 | 8,37E-01 |
| AT4G34020 | 0,41 | -1,13 | 4,31 | -0,26 | 7,94E-01 |
| AT4G34040 | 0,82 | 3,38 | 2,28 | 1,48 | 1,39E-01 |
| AT4G34070 | 0,72 | -0,97 | 2,59 | -0,38 | 7,07E-01 |
| AT4G34090 | 0,71 | 1,84 | 4,27 | 0,43 | 6,67E-01 |
| AT4G34131 | 0,52 | 2,62 | 3,86 | 0,68 | 4,98E-01 |
| AT4G34180 | 0,68 | 1,50 | 2,80 | 0,54 | 5,91E-01 |
| AT4G34190 | 1,52 | 2,43 | 1,78 | 1,36 | 1,73E-01 |
| AT4G34200 | 1,37 | 2,40 | 1,79 | 1,34 | 1,79E-01 |
| AT4G34220 | 1,59 | 0,25 | 1,63 | 0,16 | 8,76E-01 |
| AT4G34332 | 0,08 | 1,10 | 4,36 | 0,25 | 8,00E-01 |
| AT4G34340 | 0,12 | 1,58 | 4,34 | 0,36 | 7,16E-01 |
| AT4G34350 | 1,55 | 2,25 | 1,57 | 1,43 | 1,53E-01 |
| AT4G08985 | 0,45 | 1,24 | 3,25 | 0,38 | 7,03E-01 |
| AT4G34460 | 0,42 | 1,55 | 3,27 | 0,47 | 6,36E-01 |
| AT4G34480 | 1,19 | 2,16 | 1,87 | 1,15 | 2,49E-01 |
| AT4G34490 | 1,39 | 1,68 | 1,61 | 1,04 | 2,98E-01 |
| AT4G34610 | 0,37 | 0,00 | 4,39 | 0,00 | 1,00E+00 |
| AT4G34620 | 1,51 | -0,34 | 1,93 | -0,18 | 8,59E-01 |
| AT4G34630 | 0,16 | 0,00 | 4,39 | 0,00 | 1,00E+00 |
| AT4G34690 | 1,55 | 2,41 | 1,95 | 1,24 | 2,16E-01 |
| AT4G34730 | 0,55 | -0,05 | 2,65 | -0,02 | 9,85E-01 |
| AT4G34740 | 0,69 | -0,83 | 2,89 | -0,29 | 7,74E-01 |
| AT4G34860 | 0,08 | 0,85 | 4,38 | 0,20 | 8,45E-01 |
| AT4G34900 | 0,65 | 1,90 | 3,59 | 0,53 | 5,98E-01 |
| AT4G35060 | 0,12 | 1,58 | 4,34 | 0,36 | 7,16E-01 |
| AT4G35140 | 0,36 | -1,00 | 4,35 | -0,23 | 8,18E-01 |
| AT4G35180 | 0,66 | 1,38 | 3,00 | 0,46 | 6,45E-01 |
| AT4G35230 | 1,11 | 2,63 | 2,06 | 1,28 | 2,02E-01 |
| AT4G35240 | 1,02 | 0,38 | 2,21 | 0,17 | 8,62E-01 |
| AT4G35260 | 0,53 | -2,03 | 3,16 | -0,64 | 5,19E-01 |
| AT4G35300 | 1,43 | -0,90 | 2,12 | -0,43 | 6,70E-01 |
| AT4G35320 | 0,51 | 0,77 | 3,20 | 0,24 | 8,10E-01 |
| AT4G35360 | 0,54 | -2,24 | 4,03 | -0,56 | 5,79E-01 |
| AT4G35440 | 0,24 | 1,29 | 4,36 | 0,30 | 7,67E-01 |

| | | | | | |
|---|---|---|---|---|---|
| AT4G35570 | 0,29 | 1,62 | 4,33 | 0,37 | 7,08E-01 |
| AT4G35580 | 0,56 | 0,17 | 3,08 | 0,06 | 9,56E-01 |
| AT4G35780 | 0,15 | 1,62 | 4,34 | 0,37 | 7,09E-01 |
| AT4G35785 | 0,84 | 3,24 | 2,77 | 1,17 | 2,42E-01 |
| AT4G35920 | 0,70 | 0,55 | 4,39 | 0,13 | 8,99E-01 |
| AT4G36010 | 0,74 | -0,64 | 2,69 | -0,24 | 8,13E-01 |
| AT4G36030 | 1,21 | 0,69 | 2,29 | 0,30 | 7,64E-01 |
| AT4G36050 | 0,52 | 3,14 | 4,30 | 0,73 | 4,65E-01 |
| AT4G36070 | 0,08 | 0,85 | 4,38 | 0,20 | 8,45E-01 |
| AT4G36220 | 1,38 | 0,87 | 1,59 | 0,55 | 5,82E-01 |
| AT4G36270 | 0,14 | 1,18 | 4,35 | 0,27 | 7,86E-01 |
| AT4G36280 | 0,08 | 0,85 | 4,38 | 0,20 | 8,45E-01 |
| AT4G36380 | 0,24 | -2,53 | 4,34 | -0,58 | 5,60E-01 |
| AT4G36390 | 0,60 | 1,91 | 2,55 | 0,75 | 4,55E-01 |
| AT4G36460 | 0,50 | 0,43 | 3,17 | 0,13 | 8,93E-01 |
| AT4G36500 | 0,97 | -0,35 | 2,22 | -0,16 | 8,73E-01 |
| AT4G36520 | 1,04 | 2,33 | 1,76 | 1,33 | 1,84E-01 |
| AT4G36550 | 0,53 | 2,52 | 3,14 | 0,80 | 4,23E-01 |
| AT4G36640 | 1,54 | 3,50 | 2,02 | 1,73 | 8,39E-02 |
| AT4G36670 | 0,44 | 0,31 | 3,35 | 0,09 | 9,25E-01 |
| AT4G36690 | 0,56 | 0,35 | 3,29 | 0,11 | 9,14E-01 |
| AT4G36730 | 0,51 | -2,24 | 3,19 | -0,70 | 4,83E-01 |
| AT4G36770 | 0,41 | 1,49 | 4,16 | 0,36 | 7,20E-01 |
| AT4G36800 | 0,60 | 0,69 | 3,01 | 0,23 | 8,18E-01 |
| AT4G36808 | 0,88 | 1,87 | 2,36 | 0,79 | 4,28E-01 |
| AT4G36860 | 0,41 | -1,22 | 4,26 | -0,29 | 7,74E-01 |
| AT4G36890 | 0,60 | 1,32 | 3,93 | 0,34 | 7,37E-01 |
| AT4G36970 | 0,18 | 0,00 | 4,41 | 0,00 | 1,00E+00 |
| AT4G36980 | 0,30 | 1,62 | 4,35 | 0,37 | 7,10E-01 |
| AT4G37100 | 0,62 | 0,98 | 3,09 | 0,32 | 7,52E-01 |
| AT4G37270 | 0,72 | 1,54 | 2,45 | 0,63 | 5,30E-01 |
| AT4G37280 | 0,63 | 0,45 | 3,71 | 0,12 | 9,03E-01 |
| AT4G37370 | 0,86 | 2,47 | 2,57 | 0,96 | 3,35E-01 |
| AT4G37432 | 0,86 | 2,14 | 3,30 | 0,65 | 5,18E-01 |
| AT4G37470 | 1,39 | -0,98 | 1,61 | -0,61 | 5,45E-01 |
| AT4G37510 | 0,72 | 1,38 | 2,43 | 0,57 | 5,71E-01 |
| AT4G37540 | 0,41 | -1,34 | 4,41 | -0,30 | 7,61E-01 |
| AT4G37553 | 0,42 | 1,76 | 4,34 | 0,41 | 6,85E-01 |
| AT4G37590 | 0,76 | 1,57 | 2,49 | 0,63 | 5,27E-01 |
| AT4G37610 | 0,33 | 0,28 | 3,58 | 0,08 | 9,37E-01 |
| AT4G37740 | 0,08 | 0,00 | 4,40 | 0,00 | 1,00E+00 |
| AT4G37800 | 0,23 | 0,61 | 4,38 | 0,14 | 8,90E-01 |
| AT4G37830 | 0,60 | -0,35 | 2,65 | -0,13 | 8,95E-01 |
| AT4G37850 | 0,24 | 0,00 | 4,41 | 0,00 | 1,00E+00 |
| AT4G37870 | 0,70 | -0,18 | 2,11 | -0,09 | 9,31E-01 |
| AT4G37920 | 0,32 | 1,91 | 4,33 | 0,44 | 6,59E-01 |
| AT4G38010 | 0,08 | 0,85 | 4,38 | 0,20 | 8,45E-01 |
| AT4G38050 | 0,35 | 1,01 | 4,33 | 0,23 | 8,15E-01 |
| AT4G38120 | 0,89 | 2,43 | 2,27 | 1,07 | 2,84E-01 |
| AT4G38130 | 0,71 | 0,66 | 1,92 | 0,34 | 7,31E-01 |
| AT4G38180 | 0,60 | 0,95 | 2,27 | 0,42 | 6,77E-01 |
| AT4G38210 | 0,75 | 2,16 | 4,30 | 0,50 | 6,16E-01 |
| AT4G38230 | 0,40 | 0,93 | 3,42 | 0,27 | 7,86E-01 |
| AT4G38240 | 0,62 | 2,60 | 3,64 | 0,72 | 4,74E-01 |
| AT4G09695 | 1,25 | 0,89 | 1,66 | 0,53 | 5,94E-01 |
| AT4G38300 | 0,68 | 2,75 | 3,55 | 0,77 | 4,39E-01 |

| | | | | | |
|---|---|---|---|---|---|
| AT4G38350 | 1,14 | 0,84 | 1,81 | 0,46 | 6,43E-01 |
| AT4G38380 | 0,43 | 0,67 | 4,12 | 0,16 | 8,70E-01 |
| AT4G38520 | 0,55 | 1,25 | 2,38 | 0,52 | 6,00E-01 |
| AT4G38545 | 0,17 | 1,23 | 4,35 | 0,28 | 7,77E-01 |
| AT4G38570 | 0,81 | 3,18 | 2,80 | 1,14 | 2,56E-01 |
| AT4G38630 | 0,74 | -0,16 | 2,01 | -0,08 | 9,36E-01 |
| AT4G38640 | 0,43 | 1,29 | 3,35 | 0,39 | 7,00E-01 |
| AT4G38660 | 0,37 | 0,00 | 4,41 | 0,00 | 1,00E+00 |
| AT4G38670 | 0,49 | 0,81 | 3,15 | 0,26 | 7,96E-01 |
| AT4G38680 | 1,18 | 0,34 | 1,69 | 0,20 | 8,41E-01 |
| AT4G38700 | 0,34 | 0,00 | 4,40 | 0,00 | 1,00E+00 |
| AT4G38710 | 1,00 | -0,91 | 1,68 | -0,54 | 5,90E-01 |
| AT4G38730 | 0,38 | 0,90 | 4,38 | 0,21 | 8,37E-01 |
| AT4G38740 | 1,15 | -0,17 | 2,07 | -0,08 | 9,35E-01 |
| AT4G38780 | 1,10 | 4,17 | 2,47 | 1,69 | 9,16E-02 |
| AT4G38800 | 0,65 | -1,31 | 2,62 | -0,50 | 6,16E-01 |
| AT4G38810 | 0,68 | -1,44 | 2,56 | -0,56 | 5,73E-01 |
| AT4G38870 | 0,21 | 1,97 | 4,33 | 0,46 | 6,49E-01 |
| AT4G38890 | 0,48 | -0,15 | 3,36 | -0,04 | 9,65E-01 |
| AT4G38900 | 0,33 | 0,00 | 4,38 | 0,00 | 1,00E+00 |
| AT4G38932 | 0,58 | 0,70 | 2,69 | 0,26 | 7,95E-01 |
| AT4G38950 | 1,10 | 2,48 | 2,05 | 1,21 | 2,26E-01 |
| AT4G39100 | 0,33 | 2,14 | 4,33 | 0,49 | 6,21E-01 |
| AT4G39120 | 0,19 | 1,67 | 4,34 | 0,39 | 7,00E-01 |
| AT4G39140 | 0,65 | 1,77 | 3,85 | 0,46 | 6,45E-01 |
| AT4G39180 | 0,48 | 2,08 | 4,34 | 0,48 | 6,32E-01 |
| AT4G39190 | 0,69 | 1,21 | 2,50 | 0,48 | 6,29E-01 |
| AT4G39195 | 0,79 | 3,03 | 2,87 | 1,05 | 2,92E-01 |
| AT4G39240 | 1,33 | 3,71 | 2,17 | 1,71 | 8,72E-02 |
| AT4G39280 | 1,06 | -0,08 | 1,78 | -0,04 | 9,65E-01 |
| AT4G39460 | 0,85 | 1,33 | 2,38 | 0,56 | 5,76E-01 |
| AT4G39500 | 0,20 | 0,89 | 4,37 | 0,20 | 8,39E-01 |
| AT4G39510 | 0,43 | -0,85 | 4,37 | -0,20 | 8,45E-01 |
| AT4G39520 | 1,02 | 0,20 | 1,59 | 0,13 | 9,00E-01 |
| AT4G39550 | 0,17 | 1,94 | 4,33 | 0,45 | 6,55E-01 |
| AT4G39560 | 0,59 | 1,63 | 3,79 | 0,43 | 6,66E-01 |
| AT4G39570 | 0,55 | 2,11 | 3,69 | 0,57 | 5,66E-01 |
| AT4G39580 | 0,15 | 1,25 | 4,35 | 0,29 | 7,75E-01 |
| AT4G39590 | 0,08 | 0,85 | 4,38 | 0,20 | 8,45E-01 |
| AT4G39630 | 0,05 | 0,00 | 4,41 | 0,00 | 1,00E+00 |
| AT4G39680 | 0,30 | 1,12 | 4,36 | 0,26 | 7,97E-01 |
| AT4G39710 | 0,56 | 0,50 | 3,96 | 0,13 | 9,00E-01 |
| AT4G39780 | 0,21 | 0,18 | 4,34 | 0,04 | 9,68E-01 |
| AT4G39820 | 0,46 | 2,14 | 4,04 | 0,53 | 5,97E-01 |
| AT4G39838 | 1,07 | 2,35 | 2,03 | 1,16 | 2,47E-01 |
| AT4G39865 | 0,25 | 0,00 | 4,40 | 0,00 | 1,00E+00 |
| AT4G39910 | 0,28 | -0,53 | 4,37 | -0,12 | 9,03E-01 |
| AT4G40000 | 0,43 | 0,60 | 4,12 | 0,15 | 8,84E-01 |
| AT4G40020 | 0,75 | 2,17 | 2,77 | 0,78 | 4,34E-01 |
| AT4G40030 | 1,32 | 1,10 | 1,88 | 0,58 | 5,59E-01 |
| AT4G40045 | 0,12 | 1,58 | 4,34 | 0,36 | 7,16E-01 |
| AT4G40050 | 1,65 | 3,41 | 1,74 | 1,96 | 5,04E-02 |
| AT4G40060 | 1,22 | 1,32 | 1,92 | 0,69 | 4,90E-01 |
| AT4G40065 | 1,04 | 2,00 | 1,88 | 1,06 | 2,89E-01 |
| AT5G01010 | 1,03 | 2,29 | 2,32 | 0,99 | 3,23E-01 |
| AT5G01020 | 1,61 | 3,61 | 1,74 | 2,08 | 3,80E-02 |

| | | | | | |
|---|---|---|---|---|---|
| AT5G01100 | 0,27 | 0,54 | 4,37 | 0,12 | 9,01E-01 |
| AT5G01215 | 1,09 | 0,62 | 1,82 | 0,34 | 7,33E-01 |
| AT5G01220 | 0,12 | 1,65 | 4,34 | 0,38 | 7,04E-01 |
| AT5G01270 | 0,34 | 1,74 | 4,34 | 0,40 | 6,89E-01 |
| AT5G01365 | 0,66 | 1,31 | 4,29 | 0,31 | 7,60E-01 |
| AT5G01540 | 0,46 | 1,79 | 4,07 | 0,44 | 6,59E-01 |
| AT5G01560 | 1,57 | 3,54 | 1,79 | 1,97 | 4,83E-02 |
| AT5G01600 | 1,52 | 3,08 | 1,79 | 1,72 | 8,48E-02 |
| AT5G01630 | 1,03 | 3,34 | 2,65 | 1,26 | 2,08E-01 |
| AT5G01700 | 0,62 | 1,72 | 4,34 | 0,40 | 6,92E-01 |
| AT5G01710 | 1,65 | 1,05 | 1,75 | 0,60 | 5,49E-01 |
| AT5G01720 | 0,49 | 1,94 | 3,95 | 0,49 | 6,24E-01 |
| AT5G01730 | 0,29 | 1,97 | 4,33 | 0,46 | 6,49E-01 |
| AT5G01770 | 0,54 | 1,92 | 3,78 | 0,51 | 6,11E-01 |
| AT5G01780 | 0,19 | 1,28 | 4,34 | 0,30 | 7,68E-01 |
| AT5G01890 | 0,64 | 2,18 | 2,62 | 0,83 | 4,04E-01 |
| AT5G01930 | 1,04 | 1,06 | 3,15 | 0,34 | 7,37E-01 |
| AT5G01950 | 0,59 | -0,09 | 2,67 | -0,03 | 9,72E-01 |
| AT5G01970 | 1,20 | 3,76 | 2,05 | 1,83 | 6,71E-02 |
| AT5G01990 | 1,12 | 2,35 | 2,17 | 1,08 | 2,79E-01 |
| AT5G02130 | 0,08 | 0,00 | 4,40 | 0,00 | 1,00E+00 |
| AT5G02290 | 0,48 | 1,23 | 2,77 | 0,44 | 6,57E-01 |
| AT5G02502 | 0,33 | 0,00 | 4,36 | 0,00 | 9,99E-01 |
| AT5G02530 | 0,37 | 0,22 | 4,34 | 0,05 | 9,60E-01 |
| AT5G02670 | 0,44 | 2,03 | 3,24 | 0,62 | 5,32E-01 |
| AT5G02770 | 0,75 | 1,50 | 2,38 | 0,63 | 5,29E-01 |
| AT5G02800 | 0,65 | 0,98 | 3,01 | 0,33 | 7,43E-01 |
| AT5G02830 | 0,85 | 1,56 | 1,96 | 0,80 | 4,25E-01 |
| AT5G02890 | 0,18 | -1,33 | 4,40 | -0,30 | 7,63E-01 |
| AT5G02940 | 0,88 | 2,88 | 2,44 | 1,18 | 2,39E-01 |
| AT5G03140 | 0,48 | 0,99 | 2,66 | 0,37 | 7,09E-01 |
| AT5G03170 | 0,32 | -1,23 | 4,39 | -0,28 | 7,80E-01 |
| AT5G03204 | 0,29 | -1,23 | 4,39 | -0,28 | 7,79E-01 |
| AT5G03210 | 0,71 | 1,11 | 2,44 | 0,45 | 6,50E-01 |
| AT5G03340 | 1,55 | 2,56 | 1,74 | 1,47 | 1,42E-01 |
| AT5G03380 | 0,84 | 2,12 | 2,26 | 0,94 | 3,47E-01 |
| AT5G03400 | 1,62 | 2,71 | 1,70 | 1,60 | 1,09E-01 |
| AT5G03430 | 0,65 | 1,84 | 3,10 | 0,59 | 5,53E-01 |
| AT5G03450 | 0,39 | -0,24 | 4,37 | -0,05 | 9,57E-01 |
| AT5G03555 | 1,08 | 2,08 | 2,00 | 1,04 | 2,99E-01 |
| AT5G03560 | 0,43 | 1,19 | 4,14 | 0,29 | 7,75E-01 |
| AT5G03630 | 0,55 | -1,51 | 3,03 | -0,50 | 6,18E-01 |
| AT5G03700 | 0,11 | 1,07 | 4,36 | 0,25 | 8,06E-01 |
| AT5G03730 | 0,62 | -0,24 | 2,44 | -0,10 | 9,21E-01 |
| AT5G03760 | 0,44 | -2,43 | 4,10 | -0,59 | 5,53E-01 |
| AT5G03800 | 0,30 | 1,68 | 4,34 | 0,39 | 6,99E-01 |
| AT5G03850 | 0,83 | -1,56 | 3,09 | -0,50 | 6,14E-01 |
| AT5G03880 | 0,57 | 0,27 | 3,12 | 0,09 | 9,31E-01 |
| AT5G03910 | 0,83 | 0,89 | 2,27 | 0,39 | 6,96E-01 |
| AT5G03970 | 0,19 | 1,82 | 4,33 | 0,42 | 6,74E-01 |
| AT5G04020 | 0,78 | 0,54 | 1,87 | 0,29 | 7,74E-01 |
| AT5G04130 | 0,05 | 0,00 | 4,41 | 0,00 | 1,00E+00 |
| AT5G04267 | 0,16 | 0,00 | 4,41 | 0,00 | 1,00E+00 |
| AT5G04340 | 1,00 | -1,58 | 1,87 | -0,85 | 3,97E-01 |
| AT5G00955 | 1,17 | 3,38 | 1,91 | 1,77 | 7,73E-02 |
| AT5G04420 | 0,49 | 1,59 | 3,95 | 0,40 | 6,87E-01 |

| | | | | | |
|---|---|---|---|---|---|
| AT5G04430 | 1,54 | 1,92 | 1,70 | 1,13 | 2,59E-01 |
| AT5G04470 | 0,16 | 1,33 | 4,35 | 0,31 | 7,59E-01 |
| AT5G04480 | 0,41 | 1,68 | 3,29 | 0,51 | 6,09E-01 |
| AT5G04670 | 1,32 | 2,12 | 1,83 | 1,16 | 2,46E-01 |
| AT5G04700 | 0,30 | 1,36 | 4,35 | 0,31 | 7,55E-01 |
| AT5G04710 | 0,12 | 1,58 | 4,34 | 0,36 | 7,16E-01 |
| AT5G04730 | 1,06 | 3,14 | 2,15 | 1,46 | 1,45E-01 |
| AT5G04760 | 0,48 | 2,10 | 4,33 | 0,48 | 6,28E-01 |
| AT5G04830 | 0,60 | 2,17 | 3,11 | 0,70 | 4,85E-01 |
| AT5G04850 | 0,17 | 1,57 | 4,34 | 0,36 | 7,18E-01 |
| AT5G04860 | 0,46 | 0,42 | 4,22 | 0,10 | 9,21E-01 |
| AT5G04870 | 0,78 | 1,23 | 2,20 | 0,56 | 5,75E-01 |
| AT5G04940 | 0,64 | 0,96 | 3,00 | 0,32 | 7,50E-01 |
| AT5G04950 | 0,11 | 0,00 | 4,41 | 0,00 | 1,00E+00 |
| AT5G04990 | 0,74 | 0,00 | 3,54 | 0,00 | 1,00E+00 |
| AT5G05060 | 0,37 | -2,46 | 4,35 | -0,57 | 5,72E-01 |
| AT5G05080 | 0,77 | -0,24 | 4,35 | -0,06 | 9,55E-01 |
| AT5G05085 | 0,35 | 0,88 | 3,58 | 0,24 | 8,06E-01 |
| AT5G05100 | 0,72 | 0,92 | 2,92 | 0,31 | 7,54E-01 |
| AT5G01005 | 0,75 | 2,38 | 2,93 | 0,81 | 4,18E-01 |
| AT5G05260 | 0,23 | 1,63 | 4,33 | 0,38 | 7,06E-01 |
| AT5G01055 | 0,10 | 0,00 | 4,41 | 0,00 | 1,00E+00 |
| AT5G05480 | 0,31 | 1,61 | 3,59 | 0,45 | 6,53E-01 |
| AT5G05670 | 0,17 | 1,03 | 4,37 | 0,24 | 8,13E-01 |
| AT5G05700 | 0,69 | 3,01 | 2,90 | 1,04 | 3,00E-01 |
| AT5G05730 | 0,58 | -0,44 | 3,24 | -0,13 | 8,93E-01 |
| AT5G01095 | 0,95 | 3,28 | 4,29 | 0,76 | 4,44E-01 |
| AT5G05800 | 0,36 | 1,99 | 4,34 | 0,46 | 6,46E-01 |
| AT5G06043 | 0,95 | -3,21 | 2,55 | -1,26 | 2,08E-01 |
| AT5G06060 | 0,53 | -0,77 | 3,23 | -0,24 | 8,12E-01 |
| AT5G06110 | 0,69 | 1,88 | 2,92 | 0,64 | 5,20E-01 |
| AT5G06140 | 0,36 | 1,11 | 4,36 | 0,26 | 7,99E-01 |
| AT5G06160 | 0,43 | 0,60 | 2,75 | 0,22 | 8,26E-01 |
| AT5G01165 | 0,96 | 0,38 | 1,98 | 0,19 | 8,49E-01 |
| AT5G01195 | 1,58 | -0,87 | 1,47 | -0,59 | 5,53E-01 |
| AT5G06230 | 0,49 | 2,32 | 3,86 | 0,60 | 5,48E-01 |
| AT5G06260 | 0,46 | 0,99 | 3,26 | 0,30 | 7,60E-01 |
| AT5G06330 | 0,08 | 0,00 | 4,40 | 0,00 | 1,00E+00 |
| AT5G06410 | 0,25 | 1,42 | 4,34 | 0,33 | 7,44E-01 |
| AT5G06430 | 0,51 | 0,48 | 4,40 | 0,11 | 9,13E-01 |
| AT5G06440 | 0,83 | -0,25 | 3,60 | -0,07 | 9,45E-01 |
| AT5G06520 | 0,44 | 2,78 | 4,31 | 0,65 | 5,18E-01 |
| AT5G06540 | 0,48 | 2,94 | 4,30 | 0,68 | 4,94E-01 |
| AT5G06560 | 0,78 | 1,31 | 2,25 | 0,58 | 5,60E-01 |
| AT5G06670 | 0,24 | -0,36 | 4,37 | -0,08 | 9,35E-01 |
| AT5G06690 | 0,35 | 0,47 | 3,55 | 0,13 | 8,95E-01 |
| AT5G06750 | 0,11 | 0,90 | 4,37 | 0,21 | 8,36E-01 |
| AT5G06770 | 0,36 | -0,17 | 4,37 | -0,04 | 9,69E-01 |
| AT5G06865 | 0,22 | -0,24 | 4,37 | -0,06 | 9,56E-01 |
| AT5G06970 | 0,87 | 1,60 | 1,98 | 0,81 | 4,18E-01 |
| AT5G01295 | 0,28 | 0,42 | 4,38 | 0,10 | 9,24E-01 |
| AT5G07120 | 0,92 | 2,66 | 2,42 | 1,10 | 2,70E-01 |
| AT5G07220 | 0,71 | 1,54 | 2,23 | 0,69 | 4,91E-01 |
| AT5G07250 | 0,62 | -3,49 | 2,98 | -1,17 | 2,41E-01 |
| AT5G07360 | 0,50 | 0,60 | 3,24 | 0,18 | 8,54E-01 |
| AT5G07590 | 0,15 | 1,58 | 4,34 | 0,36 | 7,15E-01 |

| | | | | | |
|---|---|---|---|---|---|
| AT5G07620 | 0,13 | -1,24 | 4,39 | -0,28 | 7,77E-01 |
| AT5G07675 | 0,20 | 0,00 | 4,41 | 0,00 | 1,00E+00 |
| AT5G07810 | 0,26 | 0,90 | 4,38 | 0,21 | 8,37E-01 |
| AT5G07950 | 0,39 | 0,40 | 4,39 | 0,09 | 9,28E-01 |
| AT5G07980 | 1,38 | 2,27 | 1,94 | 1,17 | 2,42E-01 |
| AT5G08040 | 0,14 | -1,60 | 4,37 | -0,37 | 7,14E-01 |
| AT5G08075 | 0,48 | 0,42 | 4,08 | 0,10 | 9,19E-01 |
| AT5G08110 | 0,94 | 2,52 | 2,34 | 1,08 | 2,81E-01 |
| AT5G08130 | 1,09 | -0,50 | 2,14 | -0,23 | 8,16E-01 |
| AT5G08160 | 1,05 | -0,64 | 2,24 | -0,28 | 7,76E-01 |
| AT5G08170 | 0,71 | 1,10 | 3,64 | 0,30 | 7,63E-01 |
| AT5G01495 | 0,69 | 0,61 | 2,02 | 0,30 | 7,63E-01 |
| AT5G01505 | 1,02 | -1,00 | 1,75 | -0,57 | 5,66E-01 |
| AT5G01515 | 0,11 | 1,07 | 4,36 | 0,25 | 8,06E-01 |
| AT5G08230 | 0,62 | 1,82 | 3,12 | 0,58 | 5,60E-01 |
| AT5G08280 | 1,65 | -1,16 | 1,43 | -0,81 | 4,19E-01 |
| AT5G08305 | 0,42 | 2,06 | 4,14 | 0,50 | 6,18E-01 |
| AT5G08315 | 0,44 | 0,00 | 4,41 | 0,00 | 1,00E+00 |
| AT5G08350 | 0,26 | 1,08 | 4,37 | 0,25 | 8,05E-01 |
| AT5G08450 | 0,66 | 1,23 | 2,73 | 0,45 | 6,53E-01 |
| AT5G08520 | 1,35 | -3,19 | 1,63 | -1,95 | 5,08E-02 |
| AT5G08560 | 0,55 | 0,78 | 2,75 | 0,29 | 7,76E-01 |
| AT5G08650 | 1,18 | -1,02 | 1,92 | -0,53 | 5,95E-01 |
| AT5G08770 | 1,04 | -0,52 | 1,95 | -0,27 | 7,89E-01 |
| AT5G08790 | 1,51 | 3,53 | 1,66 | 2,12 | 3,38E-02 |
| AT5G09330 | 1,10 | 1,13 | 1,84 | 0,61 | 5,39E-01 |
| AT5G09443 | 0,26 | 0,90 | 4,38 | 0,21 | 8,37E-01 |
| AT5G09460 | 0,93 | 1,97 | 1,93 | 1,02 | 3,09E-01 |
| AT5G09510 | 0,57 | 0,59 | 4,02 | 0,15 | 8,83E-01 |
| AT5G09620 | 0,88 | 0,74 | 2,44 | 0,30 | 7,61E-01 |
| AT5G09650 | 0,62 | 0,68 | 2,56 | 0,27 | 7,89E-01 |
| AT5G09820 | 0,63 | 1,05 | 3,15 | 0,33 | 7,39E-01 |
| AT5G09850 | 0,67 | 2,19 | 3,08 | 0,71 | 4,76E-01 |
| AT5G09880 | 0,62 | 3,05 | 2,92 | 1,05 | 2,96E-01 |
| AT5G09890 | 0,52 | 2,93 | 3,88 | 0,75 | 4,50E-01 |
| AT5G09930 | 0,08 | 1,10 | 4,36 | 0,25 | 8,00E-01 |
| AT5G10030 | 0,52 | 2,10 | 3,93 | 0,54 | 5,92E-01 |
| AT5G10235 | 0,28 | 0,48 | 4,34 | 0,11 | 9,12E-01 |
| AT5G10290 | 0,76 | 1,97 | 2,37 | 0,83 | 4,05E-01 |
| AT5G10330 | 0,57 | 1,14 | 2,65 | 0,43 | 6,66E-01 |
| AT5G10350 | 0,66 | 1,21 | 4,34 | 0,28 | 7,80E-01 |
| AT5G10360 | 1,44 | -0,01 | 1,57 | -0,01 | 9,95E-01 |
| AT5G10390 | 0,27 | 0,00 | 4,39 | 0,00 | 1,00E+00 |
| AT5G10400 | 0,23 | 0,00 | 4,41 | 0,00 | 1,00E+00 |
| AT5G10470 | 0,57 | 1,69 | 3,23 | 0,53 | 5,99E-01 |
| AT5G10490 | 0,10 | 0,42 | 4,38 | 0,10 | 9,23E-01 |
| AT5G10550 | 0,86 | 2,12 | 2,30 | 0,92 | 3,56E-01 |
| AT5G10605 | 0,65 | 1,04 | 3,15 | 0,33 | 7,41E-01 |
| AT5G10650 | 0,25 | 1,57 | 4,34 | 0,36 | 7,18E-01 |
| AT5G10690 | 0,38 | 1,47 | 4,33 | 0,34 | 7,34E-01 |
| AT5G01815 | 0,80 | 0,01 | 2,11 | 0,00 | 9,98E-01 |
| AT5G10700 | 0,56 | 0,83 | 3,24 | 0,25 | 7,99E-01 |
| AT5G10730 | 0,23 | 0,00 | 4,38 | 0,00 | 1,00E+00 |
| AT5G10770 | 0,94 | 3,06 | 2,27 | 1,35 | 1,78E-01 |
| AT5G10800 | 0,30 | 1,62 | 4,35 | 0,37 | 7,10E-01 |
| AT5G10820 | 0,63 | 2,70 | 4,31 | 0,63 | 5,31E-01 |

| | | | | | |
|---|---|---|---|---|---|
| AT5G10930 | 1,14 | 1,72 | 1,88 | 0,92 | 3,59E-01 |
| AT5G10940 | 0,77 | 2,36 | 3,59 | 0,66 | 5,11E-01 |
| AT5G10965 | 0,75 | 1,98 | 2,40 | 0,83 | 4,09E-01 |
| AT5G10980 | 0,55 | 1,05 | 3,80 | 0,28 | 7,81E-01 |
| AT5G11000 | 0,45 | 1,39 | 4,20 | 0,33 | 7,40E-01 |
| AT5G11070 | 1,00 | 2,67 | 2,11 | 1,26 | 2,06E-01 |
| AT5G11180 | 0,81 | 0,08 | 2,16 | 0,04 | 9,69E-01 |
| AT5G11210 | 1,06 | 1,18 | 3,23 | 0,37 | 7,15E-01 |
| AT5G11250 | 1,58 | 1,32 | 1,55 | 0,85 | 3,97E-01 |
| AT5G11270 | 0,22 | 0,51 | 4,38 | 0,12 | 9,08E-01 |
| AT5G11330 | 1,01 | 1,37 | 1,88 | 0,73 | 4,67E-01 |
| AT5G11480 | 0,46 | -1,87 | 3,18 | -0,59 | 5,57E-01 |
| AT5G11520 | 1,24 | -0,07 | 1,87 | -0,04 | 9,71E-01 |
| AT5G11550 | 0,28 | 1,43 | 4,34 | 0,33 | 7,42E-01 |
| AT5G11770 | 0,54 | -0,51 | 2,67 | -0,19 | 8,50E-01 |
| AT5G11850 | 0,25 | 2,10 | 4,32 | 0,49 | 6,27E-01 |
| AT5G11860 | 0,12 | 1,58 | 4,34 | 0,36 | 7,16E-01 |
| AT5G11910 | 0,27 | 1,70 | 4,33 | 0,39 | 6,94E-01 |
| AT5G11960 | 0,36 | 2,25 | 4,33 | 0,52 | 6,04E-01 |
| AT5G12010 | 0,59 | 1,20 | 3,07 | 0,39 | 6,96E-01 |
| AT5G12080 | 0,25 | 1,75 | 4,33 | 0,40 | 6,87E-01 |
| AT5G12140 | 0,22 | 1,41 | 4,34 | 0,32 | 7,45E-01 |
| AT5G12200 | 0,44 | 0,56 | 3,20 | 0,18 | 8,61E-01 |
| AT5G12310 | 0,42 | 1,17 | 3,37 | 0,35 | 7,29E-01 |
| AT5G12390 | 0,11 | 1,07 | 4,36 | 0,25 | 8,06E-01 |
| AT5G12430 | 1,28 | 2,18 | 2,36 | 0,92 | 3,56E-01 |
| AT5G12440 | 0,88 | 2,58 | 2,68 | 0,96 | 3,35E-01 |
| AT5G12480 | 0,34 | 0,70 | 4,34 | 0,16 | 8,71E-01 |
| AT5G12890 | 0,77 | 2,22 | 3,63 | 0,61 | 5,41E-01 |
| AT5G12900 | 0,23 | 1,63 | 4,34 | 0,37 | 7,08E-01 |
| AT5G13020 | 0,81 | -0,16 | 2,18 | -0,08 | 9,40E-01 |
| AT5G13050 | 0,42 | 1,12 | 4,20 | 0,27 | 7,90E-01 |
| AT5G13090 | 0,10 | 0,62 | 4,38 | 0,14 | 8,87E-01 |
| AT5G13120 | 0,17 | -1,64 | 4,37 | -0,37 | 7,08E-01 |
| AT5G13140 | 0,23 | -1,99 | 4,37 | -0,46 | 6,49E-01 |
| AT5G13170 | 0,48 | 2,10 | 4,33 | 0,48 | 6,28E-01 |
| AT5G13190 | 0,99 | 1,87 | 1,94 | 0,97 | 3,34E-01 |
| AT5G02055 | 0,48 | -1,53 | 2,75 | -0,56 | 5,78E-01 |
| AT5G13225 | 0,48 | -1,53 | 2,75 | -0,56 | 5,78E-01 |
| AT5G13270 | 0,40 | 0,83 | 4,20 | 0,20 | 8,44E-01 |
| AT5G13300 | 0,77 | 2,63 | 2,42 | 1,09 | 2,76E-01 |
| AT5G13400 | 0,43 | 2,40 | 4,33 | 0,56 | 5,79E-01 |
| AT5G13430 | 0,54 | 2,34 | 3,77 | 0,62 | 5,35E-01 |
| AT5G13440 | 0,29 | 0,95 | 4,38 | 0,22 | 8,29E-01 |
| AT5G13450 | 0,96 | 0,51 | 2,09 | 0,24 | 8,09E-01 |
| AT5G13550 | 0,46 | 2,32 | 3,98 | 0,58 | 5,60E-01 |
| AT5G13560 | 1,05 | 0,51 | 2,07 | 0,25 | 8,04E-01 |
| AT5G13590 | 1,32 | 2,16 | 2,00 | 1,08 | 2,80E-01 |
| AT5G13680 | 1,03 | 1,53 | 1,91 | 0,80 | 4,23E-01 |
| AT5G13710 | 0,59 | 1,26 | 3,09 | 0,41 | 6,84E-01 |
| AT5G13780 | 0,62 | 2,59 | 3,07 | 0,84 | 4,00E-01 |
| AT5G13800 | 0,74 | -2,14 | 2,55 | -0,84 | 4,00E-01 |
| AT5G13825 | 0,39 | -3,07 | 4,23 | -0,73 | 4,67E-01 |
| AT5G13950 | 1,63 | 1,20 | 1,58 | 0,76 | 4,47E-01 |
| AT5G13990 | 0,12 | 0,42 | 4,38 | 0,10 | 9,24E-01 |
| AT5G14100 | 0,11 | 1,07 | 4,36 | 0,25 | 8,06E-01 |

| | | | | | |
|---|---|---|---|---|---|
| AT5G14170 | 1,07 | 2,75 | 2,20 | 1,25 | 2,12E-01 |
| AT5G14210 | 0,50 | -1,34 | 3,36 | -0,40 | 6,89E-01 |
| AT5G14240 | 0,57 | -0,47 | 2,78 | -0,17 | 8,65E-01 |
| AT5G14320 | 0,97 | -0,45 | 1,71 | -0,26 | 7,95E-01 |
| AT5G14390 | 0,52 | 1,65 | 4,11 | 0,40 | 6,88E-01 |
| AT5G14420 | 0,18 | 1,66 | 4,33 | 0,38 | 7,02E-01 |
| AT5G14530 | 0,86 | 2,46 | 2,84 | 0,87 | 3,87E-01 |
| AT5G14540 | 0,54 | 1,04 | 3,95 | 0,26 | 7,92E-01 |
| AT5G02245 | 0,10 | 0,00 | 4,41 | 0,00 | 1,00E+00 |
| AT5G14580 | 1,26 | 2,28 | 2,71 | 0,84 | 4,01E-01 |
| AT5G14590 | 0,98 | 0,42 | 1,87 | 0,23 | 8,21E-01 |
| AT5G14700 | 0,24 | 0,45 | 4,39 | 0,10 | 9,18E-01 |
| AT5G14720 | 0,96 | 3,05 | 1,89 | 1,61 | 1,07E-01 |
| AT5G14770 | 0,20 | 0,75 | 4,33 | 0,17 | 8,62E-01 |
| AT5G14820 | 0,29 | 1,62 | 4,33 | 0,37 | 7,08E-01 |
| AT5G14940 | 1,18 | 3,26 | 2,25 | 1,45 | 1,47E-01 |
| AT5G14950 | 1,52 | 1,60 | 1,56 | 1,02 | 3,07E-01 |
| AT5G15050 | 0,10 | 0,42 | 4,38 | 0,10 | 9,23E-01 |
| AT5G15210 | 0,28 | 1,36 | 4,35 | 0,31 | 7,55E-01 |
| AT5G15250 | 0,05 | 0,00 | 4,41 | 0,00 | 1,00E+00 |
| AT5G15360 | 0,52 | 0,65 | 2,61 | 0,25 | 8,03E-01 |
| AT5G15410 | 1,41 | 1,32 | 1,71 | 0,78 | 4,38E-01 |
| AT5G15420 | 1,66 | -0,75 | 1,44 | -0,52 | 6,01E-01 |
| AT5G15450 | 1,23 | 2,05 | 1,77 | 1,16 | 2,47E-01 |
| AT5G15460 | 0,08 | 1,10 | 4,36 | 0,25 | 8,00E-01 |
| AT5G15470 | 0,21 | -0,09 | 4,37 | -0,02 | 9,83E-01 |
| AT5G15490 | 0,40 | 1,31 | 3,38 | 0,39 | 6,98E-01 |
| AT5G15537 | 0,21 | 1,39 | 4,34 | 0,32 | 7,49E-01 |
| AT5G15540 | 1,01 | 2,18 | 1,99 | 1,10 | 2,73E-01 |
| AT5G15580 | 0,70 | -1,32 | 2,48 | -0,53 | 5,96E-01 |
| AT5G15690 | 0,53 | -0,43 | 2,75 | -0,16 | 8,75E-01 |
| AT5G15700 | 0,61 | 0,67 | 2,32 | 0,29 | 7,72E-01 |
| AT5G15740 | 0,28 | 0,00 | 4,41 | 0,00 | 1,00E+00 |
| AT5G15780 | 0,22 | -0,24 | 4,37 | -0,06 | 9,56E-01 |
| AT5G15800 | 0,08 | 0,00 | 4,41 | 0,00 | 1,00E+00 |
| AT5G15802 | 0,08 | 0,00 | 4,40 | 0,00 | 1,00E+00 |
| AT5G15860 | 0,19 | 1,82 | 4,33 | 0,42 | 6,74E-01 |
| AT5G16050 | 0,47 | -0,21 | 2,71 | -0,08 | 9,38E-01 |
| AT5G16070 | 0,93 | 0,19 | 2,19 | 0,08 | 9,32E-01 |
| AT5G16110 | 1,09 | 1,30 | 1,88 | 0,69 | 4,88E-01 |
| AT5G16200 | 0,36 | 0,00 | 4,41 | 0,00 | 1,00E+00 |
| AT5G16220 | 0,18 | 0,00 | 4,41 | 0,00 | 1,00E+00 |
| AT5G16280 | 0,67 | 2,39 | 2,86 | 0,83 | 4,04E-01 |
| AT5G16360 | 0,08 | 1,10 | 4,36 | 0,25 | 8,00E-01 |
| AT5G16370 | 1,57 | 2,70 | 1,59 | 1,70 | 8,96E-02 |
| AT5G16380 | 0,25 | 0,61 | 4,34 | 0,14 | 8,89E-01 |
| AT5G16390 | 0,61 | 0,26 | 3,70 | 0,07 | 9,44E-01 |
| AT5G16480 | 0,56 | 0,00 | 4,39 | 0,00 | 1,00E+00 |
| AT5G16490 | 0,08 | 0,00 | 4,40 | 0,00 | 1,00E+00 |
| AT5G16520 | 0,47 | 1,26 | 4,15 | 0,30 | 7,62E-01 |
| AT5G16560 | 0,42 | -0,89 | 4,32 | -0,21 | 8,37E-01 |
| AT5G16610 | 0,08 | 1,10 | 4,36 | 0,25 | 8,00E-01 |
| AT5G16660 | 0,23 | 1,79 | 4,33 | 0,41 | 6,79E-01 |
| AT5G16680 | 0,78 | 1,57 | 2,29 | 0,69 | 4,92E-01 |
| AT5G16740 | 0,42 | 0,52 | 4,40 | 0,12 | 9,05E-01 |
| AT5G16750 | 0,89 | 1,50 | 2,11 | 0,71 | 4,77E-01 |

| | | | | | |
|---|---|---|---|---|---|
| AT5G16780 | 0,54 | 1,12 | 3,97 | 0,28 | 7,77E-01 |
| AT5G16800 | 0,21 | 0,00 | 4,38 | 0,00 | 1,00E+00 |
| AT5G16830 | 0,10 | 0,42 | 4,38 | 0,10 | 9,23E-01 |
| AT5G16980 | 0,08 | 0,85 | 4,38 | 0,20 | 8,45E-01 |
| AT5G16990 | 0,20 | 1,42 | 4,36 | 0,33 | 7,44E-01 |
| AT5G17000 | 0,20 | 0,00 | 4,41 | 0,00 | 1,00E+00 |
| AT5G17010 | 0,24 | 1,08 | 4,37 | 0,25 | 8,05E-01 |
| AT5G02655 | 0,55 | -2,93 | 2,61 | -1,13 | 2,60E-01 |
| AT5G02665 | 1,25 | 0,26 | 2,22 | 0,12 | 9,07E-01 |
| AT5G02675 | 0,37 | 1,65 | 4,29 | 0,39 | 7,00E-01 |
| AT5G17290 | 0,65 | 1,30 | 3,06 | 0,43 | 6,70E-01 |
| AT5G17300 | 0,69 | 2,34 | 2,95 | 0,79 | 4,28E-01 |
| AT5G17320 | 0,38 | 0,00 | 4,35 | 0,00 | 1,00E+00 |
| AT5G17380 | 0,80 | 1,86 | 2,46 | 0,76 | 4,48E-01 |
| AT5G17420 | 0,40 | 2,55 | 4,32 | 0,59 | 5,55E-01 |
| AT5G17460 | 0,52 | -3,77 | 4,30 | -0,88 | 3,81E-01 |
| AT5G17520 | 0,17 | -2,23 | 4,35 | -0,51 | 6,08E-01 |
| AT5G17550 | 0,08 | 1,10 | 4,36 | 0,25 | 8,00E-01 |
| AT5G17600 | 0,90 | 2,14 | 2,22 | 0,96 | 3,35E-01 |
| AT5G17620 | 0,42 | 0,52 | 4,40 | 0,12 | 9,05E-01 |
| AT5G17640 | 0,35 | 0,00 | 4,41 | 0,00 | 1,00E+00 |
| AT5G17690 | 0,20 | 1,62 | 4,35 | 0,37 | 7,10E-01 |
| AT5G17770 | 0,49 | 1,99 | 3,20 | 0,62 | 5,34E-01 |
| AT5G17860 | 0,63 | 2,73 | 2,84 | 0,96 | 3,37E-01 |
| AT5G17880 | 1,61 | 2,22 | 2,05 | 1,08 | 2,79E-01 |
| AT5G17930 | 0,48 | 2,88 | 4,30 | 0,67 | 5,03E-01 |
| AT5G18050 | 0,24 | 0,00 | 4,41 | 0,00 | 1,00E+00 |
| AT5G18110 | 0,39 | 0,31 | 4,34 | 0,07 | 9,43E-01 |
| AT5G18190 | 0,63 | 0,76 | 2,57 | 0,29 | 7,68E-01 |
| AT5G18230 | 0,99 | 2,90 | 2,75 | 1,05 | 2,92E-01 |
| AT5G18240 | 1,40 | 0,60 | 2,92 | 0,21 | 8,37E-01 |
| AT5G18260 | 0,49 | 1,13 | 4,36 | 0,26 | 7,96E-01 |
| AT5G18280 | 0,49 | 2,39 | 3,11 | 0,77 | 4,43E-01 |
| AT5G18290 | 0,36 | 0,00 | 4,41 | 0,00 | 1,00E+00 |
| AT5G18340 | 0,58 | 1,11 | 3,99 | 0,28 | 7,81E-01 |
| AT5G18400 | 0,32 | 0,68 | 4,36 | 0,15 | 8,77E-01 |
| AT5G02895 | 0,88 | 2,99 | 2,71 | 1,10 | 2,70E-01 |
| AT5G18475 | 0,21 | 2,12 | 4,33 | 0,49 | 6,23E-01 |
| AT5G18480 | 0,62 | 0,96 | 2,92 | 0,33 | 7,41E-01 |
| AT5G18570 | 0,43 | 2,45 | 4,00 | 0,61 | 5,41E-01 |
| AT5G18590 | 0,48 | 1,68 | 3,28 | 0,51 | 6,08E-01 |
| AT5G18620 | 0,19 | 1,28 | 4,34 | 0,30 | 7,68E-01 |
| AT5G18700 | 0,89 | 1,61 | 2,77 | 0,58 | 5,61E-01 |
| AT5G02935 | 0,05 | 0,00 | 4,41 | 0,00 | 1,00E+00 |
| AT5G19010 | 0,59 | 0,97 | 2,52 | 0,39 | 7,00E-01 |
| AT5G19130 | 0,62 | 1,05 | 2,63 | 0,40 | 6,88E-01 |
| AT5G19220 | 0,51 | 0,77 | 4,37 | 0,18 | 8,60E-01 |
| AT5G19360 | 0,72 | 0,00 | 4,41 | 0,00 | 1,00E+00 |
| AT5G19410 | 0,62 | 2,20 | 2,93 | 0,75 | 4,53E-01 |
| AT5G19420 | 1,09 | 2,38 | 1,95 | 1,22 | 2,22E-01 |
| AT5G19450 | 0,87 | 0,61 | 1,95 | 0,31 | 7,55E-01 |
| AT5G19485 | 0,20 | 1,07 | 4,36 | 0,25 | 8,06E-01 |
| AT5G19590 | 0,42 | 1,80 | 3,25 | 0,55 | 5,80E-01 |
| AT5G19660 | 0,36 | 1,88 | 4,33 | 0,44 | 6,63E-01 |
| AT5G19760 | 0,90 | 0,43 | 2,55 | 0,17 | 8,65E-01 |
| AT5G19850 | 0,75 | 2,85 | 3,60 | 0,79 | 4,29E-01 |

| | | | | | |
|---|---|---|---|---|---|
| AT5G19860 | 0,51 | 2,36 | 3,96 | 0,60 | 5,50E-01 |
| AT5G19900 | 0,73 | 0,87 | 3,12 | 0,28 | 7,79E-01 |
| AT5G19920 | 0,37 | 1,22 | 4,35 | 0,28 | 7,79E-01 |
| AT5G19930 | 0,12 | 0,90 | 4,36 | 0,21 | 8,36E-01 |
| AT5G19950 | 0,27 | 0,93 | 4,36 | 0,21 | 8,30E-01 |
| AT5G19980 | 0,33 | 0,91 | 4,35 | 0,21 | 8,35E-01 |
| AT5G20040 | 0,12 | 1,65 | 4,34 | 0,38 | 7,04E-01 |
| AT5G20160 | 0,08 | 0,00 | 4,40 | 0,00 | 1,00E+00 |
| AT5G20170 | 0,19 | 0,79 | 4,37 | 0,18 | 8,56E-01 |
| AT5G20225 | 0,17 | 1,92 | 4,33 | 0,44 | 6,57E-01 |
| AT5G20260 | 0,52 | 2,39 | 4,33 | 0,55 | 5,81E-01 |
| AT5G20270 | 0,14 | 0,51 | 4,38 | 0,12 | 9,08E-01 |
| AT5G20300 | 1,26 | 2,23 | 2,53 | 0,88 | 3,79E-01 |
| AT5G20320 | 1,21 | 3,99 | 2,42 | 1,65 | 9,83E-02 |
| AT5G20360 | 0,79 | 0,37 | 2,12 | 0,17 | 8,62E-01 |
| AT5G20380 | 0,45 | -0,53 | 4,28 | -0,12 | 9,02E-01 |
| AT5G20490 | 1,59 | -0,45 | 1,73 | -0,26 | 7,97E-01 |
| AT5G20520 | 1,55 | 0,35 | 1,58 | 0,22 | 8,26E-01 |
| AT5G20570 | 0,58 | 1,27 | 3,11 | 0,41 | 6,84E-01 |
| AT5G20600 | 1,15 | 3,70 | 2,09 | 1,77 | 7,70E-02 |
| AT5G20620 | 0,96 | 2,84 | 2,41 | 1,18 | 2,39E-01 |
| AT5G20630 | 1,03 | -0,72 | 1,92 | -0,38 | 7,06E-01 |
| AT5G20650 | 0,87 | 3,43 | 2,98 | 1,15 | 2,50E-01 |
| AT5G20820 | 0,08 | 0,00 | 4,40 | 0,00 | 1,00E+00 |
| AT5G20830 | 0,97 | 0,67 | 2,45 | 0,27 | 7,85E-01 |
| AT5G20840 | 1,18 | 1,66 | 1,78 | 0,93 | 3,52E-01 |
| AT5G20890 | 1,15 | 3,09 | 2,21 | 1,40 | 1,61E-01 |
| AT5G20920 | 0,64 | 0,18 | 3,21 | 0,06 | 9,55E-01 |
| AT5G20940 | 0,27 | 1,56 | 4,34 | 0,36 | 7,19E-01 |
| AT5G03265 | 0,23 | 1,61 | 4,34 | 0,37 | 7,11E-01 |
| AT5G20980 | 0,81 | 3,47 | 2,83 | 1,22 | 2,21E-01 |
| AT5G20990 | 1,65 | 3,10 | 1,91 | 1,62 | 1,05E-01 |
| AT5G21040 | 0,32 | 1,65 | 4,34 | 0,38 | 7,04E-01 |
| AT5G21160 | 0,41 | 0,74 | 4,39 | 0,17 | 8,65E-01 |
| AT5G21274 | 0,23 | 1,79 | 4,33 | 0,41 | 6,79E-01 |
| AT5G21920 | 0,36 | 0,61 | 4,33 | 0,14 | 8,88E-01 |
| AT5G21950 | 0,08 | 0,85 | 4,38 | 0,20 | 8,45E-01 |
| AT5G22010 | 0,59 | 2,86 | 2,97 | 0,96 | 3,36E-01 |
| AT5G22020 | 0,55 | -1,50 | 3,91 | -0,38 | 7,01E-01 |
| AT5G22030 | 0,76 | 0,75 | 3,43 | 0,22 | 8,27E-01 |
| AT5G22060 | 1,34 | 3,84 | 2,25 | 1,70 | 8,83E-02 |
| AT5G22080 | 0,35 | 1,70 | 3,51 | 0,48 | 6,29E-01 |
| AT5G22090 | 0,53 | 1,16 | 3,29 | 0,35 | 7,23E-01 |
| AT5G22250 | 1,15 | 3,14 | 2,10 | 1,50 | 1,35E-01 |
| AT5G22350 | 0,24 | 1,32 | 4,35 | 0,30 | 7,62E-01 |
| AT5G22360 | 0,12 | 1,65 | 4,34 | 0,38 | 7,04E-01 |
| AT5G22390 | 0,49 | 0,25 | 3,10 | 0,08 | 9,35E-01 |
| AT5G22440 | 0,73 | 0,48 | 3,08 | 0,15 | 8,77E-01 |
| AT5G22450 | 0,87 | 1,09 | 1,98 | 0,55 | 5,82E-01 |
| AT5G22505 | 1,58 | -3,25 | 1,85 | -1,76 | 7,83E-02 |
| AT5G22510 | 0,44 | 0,00 | 4,27 | 0,00 | 1,00E+00 |
| AT5G03425 | 0,62 | -2,32 | 3,56 | -0,65 | 5,14E-01 |
| AT5G22540 | 0,22 | 0,42 | 4,38 | 0,10 | 9,23E-01 |
| AT5G22570 | 0,26 | 1,30 | 4,35 | 0,30 | 7,65E-01 |
| AT5G22700 | 0,44 | 2,74 | 4,07 | 0,67 | 5,01E-01 |
| AT5G22740 | 0,58 | 1,38 | 3,87 | 0,36 | 7,22E-01 |

| | | | | | |
|---|---|---|---|---|---|
| AT5G22750 | 0,55 | 1,69 | 4,32 | 0,39 | 6,95E-01 |
| AT5G03475 | 1,21 | -0,39 | 1,79 | -0,22 | 8,28E-01 |
| AT5G22780 | 1,47 | 1,16 | 1,75 | 0,66 | 5,07E-01 |
| AT5G22850 | 0,23 | 2,12 | 4,32 | 0,49 | 6,24E-01 |
| AT5G22860 | 0,35 | 1,00 | 3,45 | 0,29 | 7,72E-01 |
| AT5G23040 | 0,63 | -1,32 | 2,55 | -0,52 | 6,06E-01 |
| AT5G23050 | 0,45 | 1,23 | 4,07 | 0,30 | 7,63E-01 |
| AT5G23060 | 1,11 | 0,52 | 1,63 | 0,32 | 7,49E-01 |
| AT5G23080 | 1,07 | 2,97 | 2,58 | 1,15 | 2,50E-01 |
| AT5G23240 | 0,89 | 0,15 | 2,14 | 0,07 | 9,43E-01 |
| AT5G23300 | 0,32 | 1,59 | 4,34 | 0,37 | 7,14E-01 |
| AT5G23390 | 0,47 | 1,50 | 3,93 | 0,38 | 7,02E-01 |
| AT5G23405 | 0,14 | 1,24 | 4,35 | 0,29 | 7,75E-01 |
| AT5G23430 | 0,90 | 2,74 | 2,17 | 1,26 | 2,08E-01 |
| AT5G23440 | 0,16 | -1,36 | 4,41 | -0,31 | 7,57E-01 |
| AT5G23450 | 0,83 | 3,19 | 2,87 | 1,11 | 2,66E-01 |
| AT5G23580 | 0,83 | 0,48 | 2,26 | 0,21 | 8,31E-01 |
| AT5G23590 | 0,61 | 3,10 | 3,63 | 0,85 | 3,94E-01 |
| AT5G23610 | 1,31 | 2,11 | 1,86 | 1,13 | 2,57E-01 |
| AT5G23660 | 0,79 | 2,79 | 3,64 | 0,77 | 4,43E-01 |
| AT5G23740 | 0,24 | 0,45 | 4,39 | 0,10 | 9,18E-01 |
| AT5G23900 | 0,57 | -0,78 | 2,68 | -0,29 | 7,70E-01 |
| AT5G23940 | 0,08 | 0,00 | 4,40 | 0,00 | 1,00E+00 |
| AT5G23990 | 0,29 | -0,41 | 4,36 | -0,10 | 9,24E-01 |
| AT5G24120 | 0,77 | 2,85 | 2,44 | 1,17 | 2,43E-01 |
| AT5G24140 | 0,08 | 0,00 | 4,40 | 0,00 | 1,00E+00 |
| AT5G24150 | 0,21 | 1,99 | 4,33 | 0,46 | 6,46E-01 |
| AT5G24160 | 0,65 | 2,43 | 2,90 | 0,84 | 4,01E-01 |
| AT5G24240 | 1,01 | -0,68 | 1,71 | -0,40 | 6,90E-01 |
| AT5G24260 | 1,06 | 1,82 | 2,21 | 0,82 | 4,09E-01 |
| AT5G24270 | 1,64 | -0,06 | 1,44 | -0,05 | 9,64E-01 |
| AT5G24320 | 0,20 | -0,05 | 4,36 | -0,01 | 9,91E-01 |
| AT5G24355 | 0,08 | 1,10 | 4,36 | 0,25 | 8,00E-01 |
| AT5G24430 | 0,56 | -0,36 | 3,93 | -0,09 | 9,26E-01 |
| AT5G24470 | 1,29 | 0,28 | 2,61 | 0,11 | 9,16E-01 |
| AT5G24480 | 0,23 | 1,91 | 4,33 | 0,44 | 6,59E-01 |
| AT5G24500 | 0,24 | 1,09 | 4,35 | 0,25 | 8,02E-01 |
| AT5G24520 | 0,95 | 2,44 | 2,07 | 1,18 | 2,40E-01 |
| AT5G24590 | 1,51 | 2,65 | 1,88 | 1,41 | 1,58E-01 |
| AT5G24655 | 0,37 | 1,36 | 4,34 | 0,31 | 7,54E-01 |
| AT5G24735 | 0,57 | 0,13 | 3,30 | 0,04 | 9,68E-01 |
| AT5G24740 | 1,40 | 1,36 | 1,66 | 0,82 | 4,14E-01 |
| AT5G24770 | 0,62 | 0,04 | 2,27 | 0,02 | 9,86E-01 |
| AT5G24780 | 0,08 | 0,00 | 4,40 | 0,00 | 1,00E+00 |
| AT5G24940 | 0,08 | 0,85 | 4,38 | 0,20 | 8,45E-01 |
| AT5G24960 | 0,36 | 1,14 | 4,35 | 0,26 | 7,94E-01 |
| AT5G25000 | 0,16 | 1,26 | 4,35 | 0,29 | 7,72E-01 |
| AT5G25070 | 1,26 | 2,85 | 1,83 | 1,55 | 1,21E-01 |
| AT5G25100 | 1,32 | -0,27 | 1,52 | -0,18 | 8,58E-01 |
| AT5G25200 | 0,60 | -0,16 | 2,68 | -0,06 | 9,51E-01 |
| AT5G25265 | 0,67 | 1,59 | 2,89 | 0,55 | 5,81E-01 |
| AT5G25350 | 1,15 | 2,74 | 1,95 | 1,41 | 1,59E-01 |
| AT5G25370 | 0,71 | -3,71 | 2,85 | -1,30 | 1,93E-01 |
| AT5G03985 | 0,37 | -2,46 | 4,35 | -0,57 | 5,72E-01 |
| AT5G25480 | 0,27 | -1,63 | 4,37 | -0,37 | 7,09E-01 |
| AT5G25510 | 0,49 | 0,14 | 3,16 | 0,04 | 9,64E-01 |

| | | | | | |
|---|---|---|---|---|---|
| AT5G25530 | 1,06 | 2,00 | 3,24 | 0,62 | 5,37E-01 |
| AT5G25756 | 0,35 | 0,00 | 4,41 | 0,00 | 1,00E+00 |
| AT5G25757 | 0,87 | -1,76 | 1,93 | -0,92 | 3,60E-01 |
| AT5G25770 | 0,28 | 0,65 | 4,37 | 0,15 | 8,82E-01 |
| AT5G04085 | 0,59 | 0,00 | 4,41 | 0,00 | 1,00E+00 |
| AT5G25780 | 1,13 | -0,67 | 1,73 | -0,39 | 6,99E-01 |
| AT5G25820 | 0,43 | 0,00 | 4,41 | 0,00 | 1,00E+00 |
| AT5G25830 | 0,05 | 0,00 | 4,41 | 0,00 | 1,00E+00 |
| AT5G25920 | 0,08 | 0,85 | 4,38 | 0,20 | 8,45E-01 |
| AT5G26170 | 0,17 | 1,75 | 4,33 | 0,40 | 6,86E-01 |
| AT5G26210 | 0,43 | 2,07 | 4,16 | 0,50 | 6,19E-01 |
| AT5G26230 | 0,08 | 0,00 | 4,41 | 0,00 | 1,00E+00 |
| AT5G26680 | 0,15 | 0,42 | 4,38 | 0,09 | 9,24E-01 |
| AT5G04245 | 1,57 | 3,24 | 2,24 | 1,45 | 1,48E-01 |
| AT5G26640 | 0,17 | 1,40 | 4,35 | 0,32 | 7,48E-01 |
| AT5G26622 | 0,46 | 0,00 | 4,40 | 0,00 | 1,00E+00 |
| AT5G04255 | 0,21 | 0,00 | 4,38 | 0,00 | 1,00E+00 |
| AT5G26760 | 0,42 | 1,22 | 4,16 | 0,29 | 7,68E-01 |
| AT5G26770 | 1,06 | -1,76 | 1,91 | -0,92 | 3,57E-01 |
| AT5G26880 | 0,63 | -0,62 | 3,13 | -0,20 | 8,43E-01 |
| AT5G26910 | 0,66 | -0,33 | 2,54 | -0,13 | 8,98E-01 |
| AT5G26960 | 0,27 | -0,31 | 4,38 | -0,07 | 9,44E-01 |
| AT5G26970 | 0,84 | -2,26 | 1,95 | -1,16 | 2,45E-01 |
| AT5G27030 | 1,03 | 2,27 | 2,06 | 1,11 | 2,69E-01 |
| AT5G27100 | 0,25 | 1,76 | 4,33 | 0,41 | 6,85E-01 |
| AT5G27238 | 0,05 | 0,00 | 4,41 | 0,00 | 1,00E+00 |
| AT5G04375 | 0,24 | 1,09 | 4,35 | 0,25 | 8,02E-01 |
| AT5G27330 | 0,05 | 0,00 | 4,41 | 0,00 | 1,00E+00 |
| AT5G27380 | 0,63 | 1,70 | 3,52 | 0,48 | 6,29E-01 |
| AT5G27550 | 0,34 | 1,59 | 4,35 | 0,37 | 7,14E-01 |
| AT5G27640 | 1,34 | 1,90 | 1,87 | 1,01 | 3,10E-01 |
| AT5G27700 | 0,49 | 2,22 | 3,23 | 0,69 | 4,93E-01 |
| AT5G27715 | 0,29 | 0,00 | 4,41 | 0,00 | 1,00E+00 |
| AT5G04435 | 0,39 | -0,66 | 4,27 | -0,15 | 8,77E-01 |
| AT5G27790 | 0,75 | 1,50 | 3,54 | 0,42 | 6,72E-01 |
| AT5G27840 | 0,72 | 3,05 | 2,76 | 1,11 | 2,69E-01 |
| AT5G27870 | 1,24 | 3,14 | 1,99 | 1,58 | 1,15E-01 |
| AT5G27920 | 0,08 | 1,10 | 4,36 | 0,25 | 8,00E-01 |
| AT5G27960 | 0,74 | 2,16 | 2,70 | 0,80 | 4,23E-01 |
| AT5G28020 | 0,34 | 0,26 | 3,51 | 0,07 | 9,41E-01 |
| AT5G28030 | 0,66 | 2,78 | 2,87 | 0,97 | 3,34E-01 |
| AT5G28050 | 0,28 | 1,41 | 4,34 | 0,33 | 7,45E-01 |
| AT5G04635 | 0,50 | 1,30 | 3,12 | 0,42 | 6,78E-01 |
| AT5G28060 | 0,65 | -0,09 | 2,24 | -0,04 | 9,68E-01 |
| AT5G04685 | 0,19 | 0,00 | 4,41 | 0,00 | 1,00E+00 |
| AT5G28190 | 0,36 | 1,32 | 4,34 | 0,30 | 7,61E-01 |
| AT5G28220 | 0,30 | -1,48 | 4,39 | -0,34 | 7,36E-01 |
| AT5G28235 | 0,34 | 0,00 | 4,41 | 0,00 | 1,00E+00 |
| AT5G28300 | 1,51 | -1,24 | 1,50 | -0,82 | 4,10E-01 |
| AT5G28310 | 0,08 | 0,85 | 4,38 | 0,20 | 8,45E-01 |
| AT5G28400 | 1,06 | 1,76 | 1,84 | 0,95 | 3,40E-01 |
| AT5G28442 | 0,36 | 1,58 | 3,40 | 0,47 | 6,42E-01 |
| AT5G28450 | 0,85 | -1,44 | 2,34 | -0,61 | 5,39E-01 |
| AT5G28530 | 0,22 | 0,90 | 4,36 | 0,21 | 8,37E-01 |
| AT5G28550 | 0,21 | 1,05 | 4,35 | 0,24 | 8,09E-01 |
| AT5G28640 | 0,08 | 0,00 | 4,40 | 0,00 | 1,00E+00 |

| | | | | | |
|---|---|---|---|---|---|
| AT5G28750 | 0,65 | -0,73 | 2,34 | -0,31 | 7,55E-01 |
| AT5G28770 | 0,50 | 2,87 | 3,88 | 0,74 | 4,60E-01 |
| AT5G28824 | 1,47 | 3,00 | 1,82 | 1,65 | 9,85E-02 |
| AT5G28850 | 0,48 | 1,66 | 3,24 | 0,51 | 6,09E-01 |
| AT5G28910 | 0,82 | -0,70 | 2,51 | -0,28 | 7,82E-01 |
| AT5G04925 | 0,18 | -0,43 | 4,36 | -0,10 | 9,22E-01 |
| AT5G32470 | 0,85 | 1,12 | 2,13 | 0,53 | 5,98E-01 |
| AT5G32590 | 0,68 | 0,23 | 2,27 | 0,10 | 9,19E-01 |
| AT5G33210 | 0,08 | 0,00 | 4,40 | 0,00 | 1,00E+00 |
| AT5G33439 | 0,11 | 0,00 | 4,41 | 0,00 | 1,00E+00 |
| AT5G34850 | 1,64 | 0,69 | 1,42 | 0,48 | 6,29E-01 |
| AT5G35100 | 0,30 | 0,75 | 3,63 | 0,21 | 8,36E-01 |
| AT5G35200 | 0,52 | 2,60 | 3,07 | 0,84 | 3,98E-01 |
| AT5G35220 | 0,25 | 1,75 | 4,33 | 0,40 | 6,87E-01 |
| AT5G35230 | 0,96 | 1,67 | 2,70 | 0,62 | 5,35E-01 |
| AT5G35370 | 0,70 | -0,13 | 2,01 | -0,06 | 9,50E-01 |
| AT5G35475 | 0,36 | -0,25 | 4,36 | -0,06 | 9,54E-01 |
| AT5G35480 | 0,65 | 1,28 | 3,12 | 0,41 | 6,82E-01 |
| AT5G35530 | 0,88 | 2,03 | 2,37 | 0,86 | 3,90E-01 |
| AT5G35560 | 0,64 | -0,49 | 2,31 | -0,21 | 8,33E-01 |
| AT5G35570 | 0,29 | 2,50 | 4,32 | 0,58 | 5,62E-01 |
| AT5G35590 | 0,12 | 0,00 | 4,39 | 0,00 | 1,00E+00 |
| AT5G35603 | 0,59 | 0,90 | 4,11 | 0,22 | 8,27E-01 |
| AT5G35604 | 0,45 | 2,30 | 3,93 | 0,59 | 5,57E-01 |
| AT5G35620 | 0,26 | 0,27 | 4,36 | 0,06 | 9,50E-01 |
| AT5G35690 | 0,37 | 1,91 | 4,30 | 0,45 | 6,56E-01 |
| AT5G35700 | 0,37 | 2,34 | 4,26 | 0,55 | 5,83E-01 |
| AT5G35750 | 1,20 | 2,28 | 1,90 | 1,20 | 2,31E-01 |
| AT5G35753 | 0,77 | 0,73 | 2,21 | 0,33 | 7,43E-01 |
| AT5G35790 | 0,34 | -1,57 | 4,37 | -0,36 | 7,19E-01 |
| AT5G35810 | 0,15 | 1,16 | 4,35 | 0,27 | 7,90E-01 |
| AT5G35890 | 0,19 | 0,78 | 4,36 | 0,18 | 8,58E-01 |
| AT5G35910 | 0,15 | 1,42 | 4,34 | 0,33 | 7,43E-01 |
| AT5G35913 | 0,08 | 1,10 | 4,36 | 0,25 | 8,00E-01 |
| AT5G35970 | 1,60 | -0,31 | 1,31 | -0,23 | 8,15E-01 |
| AT5G35995 | 0,29 | 2,27 | 4,32 | 0,52 | 6,00E-01 |
| AT5G36002 | 0,12 | 1,65 | 4,34 | 0,38 | 7,04E-01 |
| AT5G36120 | 0,05 | 0,00 | 4,41 | 0,00 | 1,00E+00 |
| AT5G36150 | 0,29 | -2,41 | 4,35 | -0,55 | 5,79E-01 |
| AT5G36160 | 0,29 | 1,76 | 4,33 | 0,41 | 6,85E-01 |
| AT5G36200 | 1,52 | 2,30 | 1,92 | 1,20 | 2,31E-01 |
| AT5G36210 | 1,11 | -0,02 | 1,77 | -0,01 | 9,89E-01 |
| AT5G36220 | 0,80 | 1,76 | 2,16 | 0,82 | 4,13E-01 |
| AT5G36700 | 0,58 | 0,25 | 2,63 | 0,09 | 9,25E-01 |
| AT5G36790 | 0,17 | -0,28 | 4,38 | -0,06 | 9,49E-01 |
| AT5G36880 | 0,93 | 2,30 | 2,25 | 1,02 | 3,08E-01 |
| AT5G36930 | 1,17 | 1,05 | 2,11 | 0,50 | 6,19E-01 |
| AT5G37260 | 0,78 | 2,87 | 3,32 | 0,86 | 3,87E-01 |
| AT5G37310 | 0,77 | 3,77 | 3,33 | 1,13 | 2,58E-01 |
| AT5G37370 | 0,59 | 1,49 | 3,67 | 0,41 | 6,84E-01 |
| AT5G37480 | 0,13 | 0,00 | 4,41 | 0,00 | 1,00E+00 |
| AT5G37590 | 0,66 | 1,72 | 3,63 | 0,47 | 6,35E-01 |
| AT5G37720 | 0,46 | 1,68 | 3,28 | 0,51 | 6,09E-01 |
| AT5G37770 | 1,11 | 3,59 | 2,67 | 1,34 | 1,79E-01 |
| AT5G37780 | 0,80 | 1,76 | 2,08 | 0,85 | 3,97E-01 |
| AT5G05545 | 0,31 | 0,00 | 4,41 | 0,00 | 1,00E+00 |

| | | | | | |
|---|---|---|---|---|---|
| AT5G37830 | 1,35 | 3,40 | 1,98 | 1,72 | 8,51E-02 |
| AT5G37890 | 0,36 | 2,25 | 4,33 | 0,52 | 6,04E-01 |
| AT5G05565 | 0,45 | 1,73 | 4,12 | 0,42 | 6,75E-01 |
| AT5G38020 | 0,42 | -2,90 | 4,33 | -0,67 | 5,03E-01 |
| AT5G38080 | 0,08 | 0,85 | 4,38 | 0,20 | 8,45E-01 |
| AT5G38140 | 0,10 | 0,00 | 4,41 | 0,00 | 1,00E+00 |
| AT5G38212 | 1,00 | 2,75 | 2,21 | 1,24 | 2,14E-01 |
| AT5G38240 | 0,41 | 0,48 | 4,38 | 0,11 | 9,12E-01 |
| AT5G38280 | 0,74 | -3,46 | 3,65 | -0,95 | 3,43E-01 |
| AT5G38450 | 0,46 | -0,13 | 4,17 | -0,03 | 9,75E-01 |
| AT5G05755 | 0,05 | 0,00 | 4,41 | 0,00 | 1,00E+00 |
| AT5G38510 | 0,76 | -1,00 | 2,46 | -0,41 | 6,84E-01 |
| AT5G38640 | 0,50 | 0,59 | 2,68 | 0,22 | 8,25E-01 |
| AT5G38660 | 1,59 | 1,18 | 1,73 | 0,68 | 4,94E-01 |
| AT5G38650 | 1,00 | 1,90 | 2,26 | 0,84 | 4,01E-01 |
| AT5G38830 | 0,49 | -0,16 | 4,31 | -0,04 | 9,70E-01 |
| AT5G39130 | 0,12 | 0,90 | 4,36 | 0,21 | 8,36E-01 |
| AT5G39160 | 0,34 | 0,00 | 4,41 | 0,00 | 1,00E+00 |
| AT5G05895 | 0,62 | -0,39 | 3,06 | -0,13 | 9,00E-01 |
| AT5G39480 | 0,08 | 0,00 | 4,40 | 0,00 | 1,00E+00 |
| AT5G39520 | 0,33 | 1,77 | 4,34 | 0,41 | 6,83E-01 |
| AT5G39530 | 0,77 | -0,43 | 2,76 | -0,16 | 8,76E-01 |
| AT5G39600 | 0,66 | -0,88 | 2,15 | -0,41 | 6,80E-01 |
| AT5G39610 | 0,52 | 1,22 | 3,91 | 0,31 | 7,55E-01 |
| AT5G05995 | 0,40 | 2,29 | 4,16 | 0,55 | 5,82E-01 |
| AT5G39680 | 0,42 | 2,33 | 4,33 | 0,54 | 5,90E-01 |
| AT5G39693 | 0,43 | -1,60 | 3,31 | -0,48 | 6,29E-01 |
| AT5G39920 | 0,25 | -1,34 | 4,35 | -0,31 | 7,58E-01 |
| AT5G39950 | 0,95 | 1,74 | 2,37 | 0,73 | 4,63E-01 |
| AT5G40170 | 0,68 | -0,10 | 2,33 | -0,04 | 9,64E-01 |
| AT5G40240 | 0,14 | 1,24 | 4,35 | 0,29 | 7,75E-01 |
| AT5G40370 | 0,37 | 1,02 | 3,38 | 0,30 | 7,63E-01 |
| AT5G40470 | 0,27 | 0,00 | 4,40 | 0,00 | 1,00E+00 |
| AT5G40580 | 0,60 | 1,23 | 4,36 | 0,28 | 7,79E-01 |
| AT5G40660 | 0,29 | 2,05 | 4,33 | 0,47 | 6,36E-01 |
| AT5G40740 | 0,85 | 3,37 | 2,32 | 1,45 | 1,47E-01 |
| AT5G40770 | 1,47 | 1,32 | 1,61 | 0,82 | 4,11E-01 |
| AT5G40820 | 0,95 | 2,71 | 2,32 | 1,17 | 2,43E-01 |
| AT5G40830 | 0,63 | 0,91 | 2,67 | 0,34 | 7,32E-01 |
| AT5G40940 | 0,56 | 2,82 | 3,76 | 0,75 | 4,54E-01 |
| AT5G41020 | 0,73 | 1,73 | 2,62 | 0,66 | 5,08E-01 |
| AT5G41130 | 0,21 | 1,93 | 4,33 | 0,45 | 6,56E-01 |
| AT5G41150 | 0,33 | 2,12 | 4,32 | 0,49 | 6,24E-01 |
| AT5G41190 | 0,83 | 0,87 | 2,97 | 0,29 | 7,69E-01 |
| AT5G41330 | 0,40 | 1,57 | 3,35 | 0,47 | 6,39E-01 |
| AT5G41400 | 0,65 | 1,22 | 2,71 | 0,45 | 6,53E-01 |
| AT5G41460 | 0,17 | -2,23 | 4,35 | -0,51 | 6,08E-01 |
| AT5G41520 | 0,58 | 1,69 | 2,74 | 0,62 | 5,38E-01 |
| AT5G41670 | 0,43 | 1,59 | 2,80 | 0,57 | 5,71E-01 |
| AT5G06235 | 0,77 | 3,50 | 4,29 | 0,82 | 4,14E-01 |
| AT5G41870 | 0,38 | -0,40 | 4,29 | -0,09 | 9,26E-01 |
| AT5G41990 | 1,56 | 3,15 | 1,67 | 1,88 | 6,00E-02 |
| AT5G42100 | 1,07 | -1,63 | 1,69 | -0,96 | 3,35E-01 |
| AT5G42110 | 0,47 | 1,12 | 3,30 | 0,34 | 7,34E-01 |
| AT5G42130 | 0,21 | 1,97 | 4,33 | 0,46 | 6,49E-01 |
| AT5G42140 | 0,12 | 0,42 | 4,38 | 0,10 | 9,24E-01 |

| | | | | | |
|---|---|---|---|---|---|
| AT5G42190 | 0,65 | 0,93 | 2,53 | 0,37 | 7,14E-01 |
| AT5G42280 | 0,14 | 1,24 | 4,35 | 0,29 | 7,75E-01 |
| AT5G42300 | 0,48 | 2,13 | 3,90 | 0,55 | 5,84E-01 |
| AT5G42360 | 1,47 | 3,54 | 1,74 | 2,03 | 4,23E-02 |
| AT5G42380 | 1,28 | 2,23 | 2,17 | 1,03 | 3,05E-01 |
| AT5G42400 | 0,21 | 1,81 | 4,33 | 0,42 | 6,77E-01 |
| AT5G06305 | 1,02 | -0,38 | 1,64 | -0,23 | 8,17E-01 |
| AT5G42490 | 0,56 | 2,10 | 3,85 | 0,55 | 5,85E-01 |
| AT5G42567 | 0,58 | 0,00 | 4,41 | 0,00 | 1,00E+00 |
| AT5G42570 | 0,15 | 0,24 | 4,35 | 0,05 | 9,56E-01 |
| AT5G42660 | 0,12 | 0,42 | 4,38 | 0,10 | 9,24E-01 |
| AT5G42750 | 1,69 | 0,72 | 1,62 | 0,45 | 6,56E-01 |
| AT5G42790 | 0,93 | 0,50 | 1,96 | 0,26 | 7,99E-01 |
| AT5G42800 | 0,38 | 2,22 | 4,33 | 0,51 | 6,07E-01 |
| AT5G42850 | 1,15 | 1,24 | 2,08 | 0,59 | 5,52E-01 |
| AT5G42870 | 0,81 | 3,13 | 2,31 | 1,36 | 1,74E-01 |
| AT5G42890 | 0,08 | 0,00 | 4,40 | 0,00 | 1,00E+00 |
| AT5G42900 | 0,25 | 1,03 | 4,37 | 0,24 | 8,13E-01 |
| AT5G42905 | 0,52 | 2,31 | 3,79 | 0,61 | 5,42E-01 |
| AT5G42940 | 0,81 | 1,61 | 2,49 | 0,65 | 5,17E-01 |
| AT5G42955 | 0,65 | -2,05 | 3,76 | -0,55 | 5,84E-01 |
| AT5G42957 | 0,83 | -2,55 | 2,46 | -1,04 | 3,00E-01 |
| AT5G42990 | 0,17 | 1,94 | 4,33 | 0,45 | 6,55E-01 |
| AT5G43010 | 0,13 | -1,24 | 4,39 | -0,28 | 7,77E-01 |
| AT5G43020 | 0,41 | 0,98 | 4,16 | 0,24 | 8,13E-01 |
| AT5G43130 | 1,30 | 3,66 | 2,14 | 1,71 | 8,65E-02 |
| AT5G43190 | 1,41 | 0,16 | 1,73 | 0,09 | 9,27E-01 |
| AT5G43260 | 0,52 | -0,11 | 2,76 | -0,04 | 9,67E-01 |
| AT5G43270 | 0,76 | -0,58 | 3,11 | -0,19 | 8,52E-01 |
| AT5G43285 | 0,35 | -2,77 | 4,33 | -0,64 | 5,22E-01 |
| AT5G43320 | 0,66 | 0,29 | 2,72 | 0,11 | 9,14E-01 |
| AT5G43440 | 0,40 | 0,27 | 4,35 | 0,06 | 9,51E-01 |
| AT5G43670 | 0,40 | -1,22 | 4,23 | -0,29 | 7,73E-01 |
| AT5G43710 | 0,21 | 1,99 | 4,33 | 0,46 | 6,46E-01 |
| AT5G43745 | 0,36 | -0,45 | 4,36 | -0,10 | 9,18E-01 |
| AT5G43750 | 0,47 | 0,00 | 4,41 | 0,00 | 1,00E+00 |
| AT5G43780 | 0,63 | 1,51 | 2,71 | 0,56 | 5,77E-01 |
| AT5G43790 | 0,45 | 0,38 | 4,24 | 0,09 | 9,30E-01 |
| AT5G43830 | 0,91 | -0,38 | 1,88 | -0,20 | 8,40E-01 |
| AT5G43850 | 0,29 | -0,35 | 3,61 | -0,10 | 9,22E-01 |
| AT5G43880 | 0,19 | -1,25 | 4,39 | -0,28 | 7,76E-01 |
| AT5G43900 | 1,55 | 2,87 | 1,68 | 1,72 | 8,63E-02 |
| AT5G43950 | 0,65 | 1,36 | 2,60 | 0,52 | 6,02E-01 |
| AT5G43960 | 0,54 | 1,87 | 3,03 | 0,62 | 5,38E-01 |
| AT5G44000 | 0,95 | 3,45 | 2,20 | 1,57 | 1,17E-01 |
| AT5G44005 | 0,16 | 1,07 | 4,37 | 0,24 | 8,07E-01 |
| AT5G44100 | 0,66 | 1,60 | 2,54 | 0,63 | 5,29E-01 |
| AT5G44250 | 0,12 | 1,58 | 4,34 | 0,36 | 7,16E-01 |
| AT5G44260 | 0,22 | -1,66 | 4,38 | -0,38 | 7,05E-01 |
| AT5G44283 | 0,91 | -0,41 | 2,45 | -0,17 | 8,66E-01 |
| AT5G44290 | 1,25 | 1,29 | 1,89 | 0,68 | 4,95E-01 |
| AT5G44306 | 0,93 | 2,29 | 2,91 | 0,79 | 4,31E-01 |
| AT5G44495 | 1,23 | -1,49 | 1,51 | -0,99 | 3,24E-01 |
| AT5G44510 | 0,61 | 3,28 | 4,29 | 0,76 | 4,45E-01 |
| AT5G44568 | 0,48 | -1,35 | 2,85 | -0,47 | 6,35E-01 |
| AT5G06595 | 1,38 | -1,72 | 1,86 | -0,93 | 3,54E-01 |

| | | | | | |
|---|---|---|---|---|---|
| AT5G44670 | 0,13 | 1,15 | 4,36 | 0,26 | 7,92E-01 |
| AT5G44680 | 0,48 | -3,35 | 4,31 | -0,78 | 4,37E-01 |
| AT5G44785 | 0,10 | 0,42 | 4,38 | 0,10 | 9,23E-01 |
| AT5G44790 | 1,35 | 2,54 | 1,74 | 1,46 | 1,46E-01 |
| AT5G44820 | 0,08 | 0,85 | 4,38 | 0,20 | 8,45E-01 |
| AT5G44840 | 0,14 | -1,60 | 4,37 | -0,37 | 7,14E-01 |
| AT5G45010 | 0,58 | -0,56 | 3,19 | -0,17 | 8,62E-01 |
| AT5G45040 | 0,08 | 1,10 | 4,36 | 0,25 | 8,00E-01 |
| AT5G45050 | 0,86 | 1,85 | 2,11 | 0,88 | 3,81E-01 |
| AT5G45170 | 0,11 | 0,00 | 4,41 | 0,00 | 1,00E+00 |
| AT5G45180 | 0,66 | -2,83 | 3,88 | -0,73 | 4,66E-01 |
| AT5G45230 | 0,97 | -0,63 | 1,91 | -0,33 | 7,40E-01 |
| AT5G45250 | 0,93 | 2,13 | 2,12 | 1,00 | 3,15E-01 |
| AT5G45275 | 0,48 | 1,99 | 4,32 | 0,46 | 6,46E-01 |
| AT5G45290 | 1,09 | 2,20 | 2,10 | 1,05 | 2,96E-01 |
| AT5G45340 | 0,20 | 0,57 | 4,37 | 0,13 | 8,96E-01 |
| AT5G45360 | 0,08 | 1,10 | 4,36 | 0,25 | 8,00E-01 |
| AT5G45410 | 0,46 | 0,59 | 4,02 | 0,15 | 8,82E-01 |
| AT5G45420 | 0,27 | 0,42 | 4,34 | 0,10 | 9,23E-01 |
| AT5G45440 | 0,74 | 3,25 | 3,38 | 0,96 | 3,36E-01 |
| AT5G45460 | 0,18 | 0,00 | 4,41 | 0,00 | 1,00E+00 |
| AT5G45573 | 0,61 | 2,00 | 2,91 | 0,69 | 4,93E-01 |
| AT5G45630 | 0,05 | 0,00 | 4,41 | 0,00 | 1,00E+00 |
| AT5G45650 | 0,61 | 1,99 | 3,21 | 0,62 | 5,36E-01 |
| AT5G45775 | 0,86 | 0,59 | 1,98 | 0,30 | 7,64E-01 |
| AT5G45800 | 0,52 | 0,96 | 2,35 | 0,41 | 6,84E-01 |
| AT5G45820 | 1,25 | -0,37 | 1,67 | -0,22 | 8,24E-01 |
| AT5G46020 | 1,24 | -1,59 | 2,44 | -0,65 | 5,16E-01 |
| AT5G46040 | 0,11 | 0,90 | 4,37 | 0,21 | 8,36E-01 |
| AT5G46050 | 0,37 | 0,00 | 4,40 | 0,00 | 1,00E+00 |
| AT5G46280 | 0,57 | -0,25 | 3,10 | -0,08 | 9,36E-01 |
| AT5G46290 | 0,63 | -0,12 | 3,05 | -0,04 | 9,68E-01 |
| AT5G46340 | 0,61 | -2,20 | 3,91 | -0,56 | 5,73E-01 |
| AT5G46395 | 0,11 | 0,00 | 4,41 | 0,00 | 1,00E+00 |
| AT5G46580 | 0,75 | -1,53 | 2,30 | -0,67 | 5,06E-01 |
| AT5G46620 | 0,64 | -2,27 | 2,99 | -0,76 | 4,48E-01 |
| AT5G46750 | 0,66 | 1,62 | 3,81 | 0,43 | 6,70E-01 |
| AT5G46790 | 0,53 | 1,13 | 3,02 | 0,38 | 7,08E-01 |
| AT5G46800 | 0,83 | -1,92 | 2,30 | -0,84 | 4,03E-01 |
| AT5G46860 | 0,46 | -2,57 | 4,09 | -0,63 | 5,30E-01 |
| AT5G46970 | 0,65 | -0,26 | 2,52 | -0,10 | 9,19E-01 |
| AT5G47040 | 1,34 | 2,69 | 1,80 | 1,50 | 1,34E-01 |
| AT5G47070 | 0,97 | 2,29 | 2,35 | 0,98 | 3,29E-01 |
| AT5G47080 | 0,13 | 1,15 | 4,36 | 0,26 | 7,92E-01 |
| AT5G47110 | 1,25 | 0,53 | 1,82 | 0,29 | 7,70E-01 |
| AT5G47190 | 0,93 | -0,69 | 2,09 | -0,33 | 7,39E-01 |
| AT5G47210 | 1,48 | 0,69 | 2,02 | 0,34 | 7,34E-01 |
| AT5G47220 | 0,78 | 1,96 | 2,57 | 0,76 | 4,46E-01 |
| AT5G47250 | 0,28 | -0,87 | 4,35 | -0,20 | 8,42E-01 |
| AT5G47370 | 0,28 | 0,42 | 4,38 | 0,10 | 9,24E-01 |
| AT5G47460 | 0,15 | 1,58 | 4,34 | 0,36 | 7,15E-01 |
| AT5G47570 | 0,30 | 0,68 | 4,34 | 0,16 | 8,75E-01 |
| AT5G47580 | 0,64 | 1,83 | 3,63 | 0,50 | 6,14E-01 |
| AT5G47650 | 0,20 | 1,07 | 4,36 | 0,25 | 8,06E-01 |
| AT5G47700 | 0,72 | 0,16 | 2,10 | 0,08 | 9,40E-01 |
| AT5G47730 | 0,16 | 1,33 | 4,35 | 0,31 | 7,59E-01 |

| | | | | | |
|---|---|---|---|---|---|
| AT5G47750 | 1,50 | 4,08 | 2,22 | 1,84 | 6,60E-02 |
| AT5G47760 | 0,12 | 0,42 | 4,38 | 0,10 | 9,24E-01 |
| AT5G47810 | 0,55 | -0,18 | 2,62 | -0,07 | 9,45E-01 |
| AT5G47830 | 0,54 | 0,55 | 4,32 | 0,13 | 8,99E-01 |
| AT5G47850 | 0,49 | 0,99 | 4,03 | 0,25 | 8,06E-01 |
| AT5G47880 | 0,55 | 2,26 | 3,14 | 0,72 | 4,72E-01 |
| AT5G47930 | 0,53 | -0,25 | 2,69 | -0,09 | 9,27E-01 |
| AT5G48000 | 0,12 | 0,00 | 4,39 | 0,00 | 1,00E+00 |
| AT5G07055 | 0,58 | -1,37 | 3,06 | -0,45 | 6,54E-01 |
| AT5G48120 | 1,18 | 1,86 | 1,88 | 0,99 | 3,20E-01 |
| AT5G48150 | 0,85 | -0,43 | 3,56 | -0,12 | 9,03E-01 |
| AT5G48180 | 0,98 | 2,36 | 2,02 | 1,17 | 2,43E-01 |
| AT5G48375 | 0,26 | 1,08 | 4,37 | 0,25 | 8,05E-01 |
| AT5G48412 | 0,64 | -0,69 | 2,69 | -0,26 | 7,97E-01 |
| AT5G48440 | 0,43 | 2,38 | 4,18 | 0,57 | 5,70E-01 |
| AT5G48540 | 0,08 | 0,85 | 4,38 | 0,20 | 8,45E-01 |
| AT5G48580 | 0,41 | 2,17 | 4,12 | 0,53 | 5,99E-01 |
| AT5G48640 | 0,19 | -1,36 | 4,41 | -0,31 | 7,58E-01 |
| AT5G48657 | 0,40 | 0,82 | 3,37 | 0,24 | 8,07E-01 |
| AT5G07265 | 0,12 | 1,58 | 4,34 | 0,36 | 7,16E-01 |
| AT5G48770 | 0,08 | 0,00 | 4,40 | 0,00 | 1,00E+00 |
| AT5G07275 | 0,71 | 2,73 | 4,31 | 0,63 | 5,26E-01 |
| AT5G48810 | 0,80 | 1,48 | 2,35 | 0,63 | 5,29E-01 |
| AT5G48830 | 0,28 | 0,90 | 4,38 | 0,21 | 8,37E-01 |
| AT5G48880 | 0,58 | 0,34 | 4,36 | 0,08 | 9,38E-01 |
| AT5G48910 | 0,76 | 1,48 | 2,21 | 0,67 | 5,02E-01 |
| AT5G48930 | 1,24 | 1,00 | 1,72 | 0,58 | 5,60E-01 |
| AT5G49100 | 1,11 | -1,46 | 2,06 | -0,71 | 4,78E-01 |
| AT5G49300 | 0,36 | 1,99 | 4,34 | 0,46 | 6,46E-01 |
| AT5G49350 | 0,08 | 1,10 | 4,36 | 0,25 | 8,00E-01 |
| AT5G49420 | 0,35 | -0,55 | 4,38 | -0,13 | 9,00E-01 |
| AT5G49430 | 1,30 | 1,72 | 1,79 | 0,96 | 3,37E-01 |
| AT5G49540 | 0,90 | 0,26 | 2,46 | 0,11 | 9,14E-01 |
| AT5G49550 | 0,27 | 0,44 | 4,39 | 0,10 | 9,20E-01 |
| AT5G49555 | 0,40 | -1,58 | 4,37 | -0,36 | 7,18E-01 |
| AT5G49570 | 0,11 | 1,07 | 4,36 | 0,25 | 8,06E-01 |
| AT5G49580 | 0,46 | -0,67 | 3,97 | -0,17 | 8,66E-01 |
| AT5G49630 | 0,19 | 0,00 | 4,41 | 0,00 | 1,00E+00 |
| AT5G49645 | 0,08 | 0,85 | 4,38 | 0,20 | 8,45E-01 |
| AT5G49660 | 0,65 | 0,99 | 2,51 | 0,39 | 6,95E-01 |
| AT5G49665 | 0,50 | 1,05 | 4,02 | 0,26 | 7,94E-01 |
| AT5G49830 | 1,27 | 2,66 | 1,80 | 1,48 | 1,39E-01 |
| AT5G49890 | 0,22 | -0,46 | 4,37 | -0,10 | 9,17E-01 |
| AT5G49900 | 0,41 | 2,25 | 4,14 | 0,54 | 5,87E-01 |
| AT5G49930 | 0,74 | 0,69 | 2,11 | 0,33 | 7,44E-01 |
| AT5G49950 | 0,25 | -1,27 | 4,40 | -0,29 | 7,72E-01 |
| AT5G50000 | 0,84 | -1,15 | 2,00 | -0,57 | 5,66E-01 |
| AT5G50150 | 0,10 | 0,62 | 4,38 | 0,14 | 8,87E-01 |
| AT5G50220 | 0,28 | 0,00 | 4,41 | 0,00 | 1,00E+00 |
| AT5G50320 | 0,77 | 1,78 | 3,00 | 0,59 | 5,52E-01 |
| AT5G50340 | 0,25 | 1,76 | 4,33 | 0,41 | 6,85E-01 |
| AT5G50350 | 0,29 | -1,23 | 4,39 | -0,28 | 7,79E-01 |
| AT5G50380 | 0,97 | 1,85 | 2,09 | 0,88 | 3,76E-01 |
| AT5G50390 | 0,17 | 1,94 | 4,33 | 0,45 | 6,55E-01 |
| AT5G50400 | 0,54 | 0,00 | 3,99 | 0,00 | 1,00E+00 |
| AT5G50460 | 0,36 | -1,36 | 4,41 | -0,31 | 7,58E-01 |

| | | | | | |
|---|---|---|---|---|---|
| AT5G50480 | 0,53 | -3,60 | 3,78 | -0,95 | 3,41E-01 |
| AT5G50570 | 0,31 | -1,30 | 4,40 | -0,30 | 7,68E-01 |
| AT5G50630 | 0,05 | 0,00 | 4,41 | 0,00 | 1,00E+00 |
| AT5G50640 | 0,24 | 1,62 | 4,34 | 0,37 | 7,08E-01 |
| AT5G50670 | 0,93 | -0,12 | 2,51 | -0,05 | 9,60E-01 |
| AT5G50860 | 0,49 | 2,32 | 3,87 | 0,60 | 5,48E-01 |
| AT5G50870 | 0,08 | 0,85 | 4,38 | 0,20 | 8,45E-01 |
| AT5G50900 | 0,83 | 0,83 | 2,46 | 0,34 | 7,36E-01 |
| AT5G51000 | 0,61 | 0,97 | 3,01 | 0,32 | 7,48E-01 |
| AT5G51020 | 0,37 | 2,34 | 4,26 | 0,55 | 5,83E-01 |
| AT5G51110 | 0,78 | -0,58 | 2,21 | -0,26 | 7,91E-01 |
| AT5G51170 | 0,12 | 1,65 | 4,34 | 0,38 | 7,04E-01 |
| AT5G51195 | 0,57 | -0,23 | 2,48 | -0,09 | 9,26E-01 |
| AT5G51220 | 0,23 | 0,42 | 4,38 | 0,09 | 9,24E-01 |
| AT5G51290 | 0,61 | 0,46 | 3,10 | 0,15 | 8,81E-01 |
| AT5G51390 | 0,29 | 0,42 | 4,38 | 0,10 | 9,24E-01 |
| AT5G51400 | 1,21 | -0,78 | 1,60 | -0,49 | 6,27E-01 |
| AT5G51550 | 1,02 | 0,97 | 1,92 | 0,51 | 6,12E-01 |
| AT5G51640 | 0,39 | -0,23 | 3,52 | -0,07 | 9,47E-01 |
| AT5G51650 | 0,32 | 1,60 | 3,53 | 0,45 | 6,49E-01 |
| AT5G51690 | 0,72 | 2,32 | 2,49 | 0,93 | 3,52E-01 |
| AT5G51710 | 0,10 | 0,00 | 4,41 | 0,00 | 1,00E+00 |
| AT5G51820 | 0,69 | -1,27 | 2,51 | -0,51 | 6,13E-01 |
| AT5G52060 | 0,54 | 1,11 | 2,62 | 0,42 | 6,71E-01 |
| AT5G52090 | 0,73 | 1,00 | 2,22 | 0,45 | 6,53E-01 |
| AT5G52120 | 0,47 | -1,85 | 4,37 | -0,42 | 6,72E-01 |
| AT5G52180 | 0,61 | 1,51 | 3,09 | 0,49 | 6,26E-01 |
| AT5G52330 | 0,15 | 0,00 | 4,39 | 0,00 | 1,00E+00 |
| AT5G52370 | 0,37 | 1,68 | 4,29 | 0,39 | 6,95E-01 |
| AT5G52450 | 0,44 | 1,89 | 3,22 | 0,59 | 5,57E-01 |
| AT5G52540 | 1,60 | 2,12 | 1,67 | 1,27 | 2,05E-01 |
| AT5G52550 | 0,25 | 1,38 | 4,36 | 0,32 | 7,51E-01 |
| AT5G52650 | 0,64 | -1,17 | 3,70 | -0,32 | 7,52E-01 |
| AT5G52780 | 0,46 | -0,83 | 3,27 | -0,25 | 8,00E-01 |
| AT5G52797 | 0,48 | 1,23 | 3,22 | 0,38 | 7,03E-01 |
| AT5G53000 | 0,29 | 1,62 | 4,33 | 0,37 | 7,08E-01 |
| AT5G53010 | 0,50 | 0,01 | 3,32 | 0,00 | 9,97E-01 |
| AT5G53050 | 0,49 | -0,66 | 3,12 | -0,21 | 8,33E-01 |
| AT5G53130 | 0,72 | 3,48 | 2,85 | 1,22 | 2,22E-01 |
| AT5G07955 | 0,08 | 0,85 | 4,38 | 0,20 | 8,45E-01 |
| AT5G53300 | 0,89 | 2,52 | 2,66 | 0,95 | 3,42E-01 |
| AT5G53320 | 0,58 | 1,95 | 2,99 | 0,65 | 5,14E-01 |
| AT5G07975 | 0,08 | 0,85 | 4,38 | 0,20 | 8,45E-01 |
| AT5G53490 | 0,65 | -0,33 | 2,37 | -0,14 | 8,91E-01 |
| AT5G53500 | 0,52 | -1,78 | 3,24 | -0,55 | 5,84E-01 |
| AT5G53530 | 0,37 | -0,16 | 4,34 | -0,04 | 9,71E-01 |
| AT5G53570 | 0,72 | 1,52 | 2,43 | 0,63 | 5,31E-01 |
| AT5G53580 | 0,36 | 0,27 | 3,55 | 0,08 | 9,40E-01 |
| AT5G53730 | 0,14 | 0,51 | 4,38 | 0,12 | 9,08E-01 |
| AT5G53850 | 0,47 | -0,89 | 4,36 | -0,21 | 8,37E-01 |
| AT5G53870 | 0,19 | 0,00 | 4,40 | 0,00 | 1,00E+00 |
| AT5G53900 | 0,79 | 0,14 | 2,32 | 0,06 | 9,52E-01 |
| AT5G54030 | 0,27 | 0,00 | 4,41 | 0,00 | 1,00E+00 |
| AT5G54040 | 0,27 | 0,42 | 4,38 | 0,10 | 9,23E-01 |
| AT5G54050 | 0,38 | 1,64 | 4,33 | 0,38 | 7,05E-01 |
| AT5G54060 | 0,12 | 1,58 | 4,34 | 0,36 | 7,16E-01 |

| | | | | | |
|---|---|---|---|---|---|
| AT5G54090 | 1,00 | 2,28 | 2,23 | 1,03 | 3,05E-01 |
| AT5G54100 | 0,65 | 3,20 | 2,90 | 1,10 | 2,70E-01 |
| AT5G54170 | 0,64 | 0,53 | 2,28 | 0,23 | 8,15E-01 |
| AT5G54280 | 1,29 | 1,32 | 1,77 | 0,75 | 4,55E-01 |
| AT5G54380 | 1,11 | 2,14 | 1,96 | 1,09 | 2,75E-01 |
| AT5G54410 | 0,82 | -0,01 | 2,08 | 0,00 | 9,98E-01 |
| AT5G54430 | 1,12 | 1,95 | 2,52 | 0,77 | 4,41E-01 |
| AT5G54460 | 1,09 | 2,05 | 3,30 | 0,62 | 5,34E-01 |
| AT5G54470 | 0,35 | 0,41 | 3,51 | 0,12 | 9,07E-01 |
| AT5G54510 | 0,08 | 1,10 | 4,36 | 0,25 | 8,00E-01 |
| AT5G54610 | 0,49 | 2,36 | 3,86 | 0,61 | 5,41E-01 |
| AT5G54620 | 0,12 | 1,46 | 4,35 | 0,34 | 7,36E-01 |
| AT5G54630 | 0,29 | -0,84 | 4,31 | -0,20 | 8,45E-01 |
| AT5G54650 | 0,47 | 1,71 | 4,35 | 0,39 | 6,95E-01 |
| AT5G54690 | 0,39 | 0,48 | 4,40 | 0,11 | 9,14E-01 |
| AT5G54700 | 0,89 | -1,20 | 2,42 | -0,50 | 6,20E-01 |
| AT5G54710 | 0,79 | -0,17 | 2,60 | -0,06 | 9,49E-01 |
| AT5G54720 | 0,73 | -0,27 | 2,49 | -0,11 | 9,13E-01 |
| AT5G54780 | 1,16 | 1,57 | 1,88 | 0,83 | 4,05E-01 |
| AT5G54930 | 0,40 | 1,25 | 4,15 | 0,30 | 7,64E-01 |
| AT5G54960 | 0,98 | -0,70 | 2,09 | -0,33 | 7,38E-01 |
| AT5G55040 | 0,74 | 1,29 | 2,16 | 0,60 | 5,50E-01 |
| AT5G55055 | 0,05 | 0,00 | 4,41 | 0,00 | 1,00E+00 |
| AT5G55100 | 1,12 | 2,57 | 2,45 | 1,05 | 2,94E-01 |
| AT5G55130 | 0,31 | 1,82 | 3,59 | 0,51 | 6,12E-01 |
| AT5G55150 | 1,21 | 2,35 | 2,48 | 0,95 | 3,43E-01 |
| AT5G55160 | 0,37 | 2,08 | 4,33 | 0,48 | 6,32E-01 |
| AT5G55210 | 0,08 | 0,00 | 4,40 | 0,00 | 1,00E+00 |
| AT5G55250 | 0,40 | -0,05 | 4,31 | -0,01 | 9,91E-01 |
| AT5G55280 | 1,13 | -2,17 | 2,27 | -0,95 | 3,40E-01 |
| AT5G55290 | 0,30 | 0,00 | 4,41 | 0,00 | 1,00E+00 |
| AT5G55300 | 0,70 | -0,58 | 2,18 | -0,26 | 7,91E-01 |
| AT5G55340 | 0,19 | 0,46 | 4,40 | 0,10 | 9,16E-01 |
| AT5G55390 | 0,36 | 1,43 | 4,34 | 0,33 | 7,42E-01 |
| AT5G08275 | 0,59 | -0,24 | 3,81 | -0,06 | 9,49E-01 |
| AT5G08285 | 0,30 | -0,38 | 4,34 | -0,09 | 9,30E-01 |
| AT5G55510 | 0,05 | 0,00 | 4,41 | 0,00 | 1,00E+00 |
| AT5G55540 | 1,63 | 1,84 | 1,58 | 1,17 | 2,43E-01 |
| AT5G55550 | 0,34 | 1,96 | 4,34 | 0,45 | 6,52E-01 |
| AT5G55570 | 1,30 | -4,54 | 2,96 | -1,53 | 1,25E-01 |
| AT5G55600 | 0,31 | 1,00 | 3,61 | 0,28 | 7,83E-01 |
| AT5G55670 | 0,30 | 1,27 | 4,34 | 0,29 | 7,70E-01 |
| AT5G55710 | 0,92 | 2,21 | 2,40 | 0,92 | 3,58E-01 |
| AT5G55740 | 0,27 | 1,87 | 4,33 | 0,43 | 6,65E-01 |
| AT5G55790 | 0,33 | 1,77 | 4,34 | 0,41 | 6,84E-01 |
| AT5G55835 | 0,61 | 1,11 | 2,30 | 0,48 | 6,30E-01 |
| AT5G55910 | 1,51 | -0,28 | 1,44 | -0,20 | 8,45E-01 |
| AT5G55920 | 0,81 | 2,39 | 2,44 | 0,98 | 3,27E-01 |
| AT5G55940 | 0,50 | 2,17 | 3,88 | 0,56 | 5,76E-01 |
| AT5G56000 | 1,26 | -3,53 | 2,03 | -1,74 | 8,18E-02 |
| AT5G56180 | 0,23 | 0,00 | 4,41 | 0,00 | 1,00E+00 |
| AT5G56240 | 1,23 | 3,36 | 2,23 | 1,51 | 1,32E-01 |
| AT5G56250 | 0,87 | 3,57 | 2,77 | 1,29 | 1,97E-01 |
| AT5G56260 | 0,25 | -1,27 | 4,40 | -0,29 | 7,72E-01 |
| AT5G56280 | 0,08 | 0,00 | 4,40 | 0,00 | 1,00E+00 |
| AT5G56350 | 0,73 | -0,90 | 1,89 | -0,47 | 6,36E-01 |

| | | | | | |
|---|---|---|---|---|---|
| AT5G56360 | 0,87 | -0,22 | 2,72 | -0,08 | 9,37E-01 |
| AT5G56410 | 0,30 | 0,41 | 4,38 | 0,09 | 9,25E-01 |
| AT5G56450 | 0,20 | 1,42 | 4,36 | 0,33 | 7,44E-01 |
| AT5G56710 | 0,79 | 2,05 | 3,39 | 0,60 | 5,46E-01 |
| AT5G56730 | 0,65 | 1,74 | 2,46 | 0,71 | 4,80E-01 |
| AT5G56760 | 0,72 | -0,53 | 2,80 | -0,19 | 8,50E-01 |
| AT5G56860 | 0,23 | 0,61 | 4,38 | 0,14 | 8,90E-01 |
| AT5G56890 | 0,16 | 1,74 | 4,33 | 0,40 | 6,87E-01 |
| AT5G56900 | 1,69 | 2,90 | 1,79 | 1,62 | 1,06E-01 |
| AT5G56980 | 0,28 | 1,36 | 4,35 | 0,31 | 7,55E-01 |
| AT5G57040 | 0,80 | 1,89 | 2,12 | 0,89 | 3,73E-01 |
| AT5G57160 | 0,81 | 0,57 | 2,48 | 0,23 | 8,17E-01 |
| AT5G57170 | 0,08 | 0,00 | 4,40 | 0,00 | 1,00E+00 |
| AT5G57250 | 0,54 | 2,83 | 3,84 | 0,74 | 4,62E-01 |
| AT5G57270 | 0,45 | -1,47 | 4,39 | -0,34 | 7,37E-01 |
| AT5G57290 | 0,69 | -0,55 | 2,59 | -0,21 | 8,31E-01 |
| AT5G57345 | 0,70 | 1,26 | 2,26 | 0,56 | 5,76E-01 |
| AT5G57580 | 0,90 | 1,80 | 2,32 | 0,77 | 4,38E-01 |
| AT5G57630 | 0,49 | 0,55 | 3,37 | 0,16 | 8,71E-01 |
| AT5G57670 | 0,16 | 0,51 | 4,38 | 0,12 | 9,07E-01 |
| AT5G57780 | 0,36 | 1,05 | 4,35 | 0,24 | 8,09E-01 |
| AT5G57785 | 0,56 | 2,17 | 2,64 | 0,82 | 4,10E-01 |
| AT5G57790 | 0,05 | 0,00 | 4,41 | 0,00 | 1,00E+00 |
| AT5G57850 | 0,51 | -2,20 | 4,36 | -0,51 | 6,13E-01 |
| AT5G57900 | 0,27 | 0,90 | 4,33 | 0,21 | 8,35E-01 |
| AT5G57960 | 0,12 | 0,68 | 4,37 | 0,16 | 8,77E-01 |
| AT5G58030 | 0,11 | 1,07 | 4,36 | 0,25 | 8,06E-01 |
| AT5G58050 | 0,17 | 1,81 | 4,33 | 0,42 | 6,76E-01 |
| AT5G58090 | 0,76 | 2,44 | 2,38 | 1,03 | 3,05E-01 |
| AT5G58130 | 0,84 | 3,61 | 2,63 | 1,37 | 1,70E-01 |
| AT5G58170 | 0,90 | 3,89 | 3,16 | 1,23 | 2,19E-01 |
| AT5G58220 | 0,12 | 0,42 | 4,38 | 0,10 | 9,24E-01 |
| AT5G58260 | 0,45 | -1,22 | 3,37 | -0,36 | 7,17E-01 |
| AT5G58380 | 0,93 | 1,97 | 2,19 | 0,90 | 3,67E-01 |
| AT5G58440 | 0,43 | -0,54 | 2,80 | -0,19 | 8,48E-01 |
| AT5G58570 | 0,61 | 2,98 | 3,87 | 0,77 | 4,43E-01 |
| AT5G08860 | 0,08 | 0,00 | 4,40 | 0,00 | 1,00E+00 |
| AT5G58600 | 0,29 | 0,92 | 4,36 | 0,21 | 8,33E-01 |
| AT5G58650 | 0,16 | 0,42 | 4,37 | 0,10 | 9,23E-01 |
| AT5G58670 | 0,69 | -0,30 | 2,16 | -0,14 | 8,91E-01 |
| AT5G58720 | 0,64 | 1,93 | 2,52 | 0,77 | 4,43E-01 |
| AT5G58730 | 0,50 | 1,68 | 3,97 | 0,42 | 6,73E-01 |
| AT5G58870 | 0,47 | 1,14 | 3,36 | 0,34 | 7,34E-01 |
| AT5G58900 | 0,28 | 0,42 | 4,38 | 0,10 | 9,24E-01 |
| AT5G58930 | 1,32 | 2,59 | 1,90 | 1,36 | 1,74E-01 |
| AT5G59030 | 0,49 | 2,26 | 3,87 | 0,58 | 5,59E-01 |
| AT5G59130 | 0,21 | 2,14 | 4,33 | 0,49 | 6,22E-01 |
| AT5G59220 | 0,62 | 1,21 | 3,77 | 0,32 | 7,48E-01 |
| AT5G59290 | 0,24 | 1,09 | 4,35 | 0,25 | 8,02E-01 |
| AT5G08920 | 0,49 | 2,10 | 4,00 | 0,53 | 5,99E-01 |
| AT5G59480 | 0,17 | 1,92 | 4,33 | 0,44 | 6,57E-01 |
| AT5G59550 | 1,55 | 3,76 | 1,79 | 2,10 | 3,61E-02 |
| AT5G59662 | 0,05 | 0,00 | 4,41 | 0,00 | 1,00E+00 |
| AT5G59700 | 0,33 | 0,00 | 4,41 | 0,00 | 1,00E+00 |
| AT5G59770 | 0,46 | -0,51 | 3,26 | -0,16 | 8,76E-01 |
| AT5G59780 | 1,19 | 3,56 | 1,95 | 1,82 | 6,81E-02 |

| | | | | | |
|---|---|---|---|---|---|
| AT5G59840 | 0,17 | -0,46 | 4,37 | -0,11 | 9,16E-01 |
| AT5G59850 | 0,69 | -2,14 | 2,62 | -0,82 | 4,13E-01 |
| AT5G59870 | 0,19 | 0,94 | 4,38 | 0,21 | 8,30E-01 |
| AT5G59900 | 0,16 | -1,36 | 4,41 | -0,31 | 7,57E-01 |
| AT5G59960 | 0,11 | 1,07 | 4,36 | 0,25 | 8,06E-01 |
| AT5G60050 | 0,15 | 0,06 | 4,36 | 0,01 | 9,90E-01 |
| AT5G60100 | 0,30 | -0,75 | 4,35 | -0,17 | 8,63E-01 |
| AT5G60160 | 0,49 | 0,96 | 4,05 | 0,24 | 8,12E-01 |
| AT5G60180 | 0,21 | 0,79 | 4,37 | 0,18 | 8,57E-01 |
| AT5G60210 | 0,29 | 0,17 | 4,35 | 0,04 | 9,68E-01 |
| AT5G60340 | 0,36 | 0,84 | 4,38 | 0,19 | 8,48E-01 |
| AT5G60400 | 0,12 | 1,65 | 4,34 | 0,38 | 7,04E-01 |
| AT5G60460 | 0,27 | -0,13 | 4,37 | -0,03 | 9,77E-01 |
| AT5G60670 | 0,64 | 0,34 | 2,29 | 0,15 | 8,81E-01 |
| AT5G60700 | 1,32 | 2,94 | 2,29 | 1,28 | 2,00E-01 |
| AT5G60710 | 1,48 | 1,45 | 1,57 | 0,93 | 3,55E-01 |
| AT5G60800 | 0,66 | 0,29 | 3,04 | 0,10 | 9,24E-01 |
| AT5G60840 | 0,12 | 0,00 | 4,41 | 0,00 | 1,00E+00 |
| AT5G60930 | 0,24 | 0,83 | 4,37 | 0,19 | 8,49E-01 |
| AT5G60960 | 0,29 | 1,56 | 4,34 | 0,36 | 7,18E-01 |
| AT5G60990 | 0,16 | 1,33 | 4,35 | 0,31 | 7,59E-01 |
| AT5G61040 | 0,88 | 0,31 | 4,36 | 0,07 | 9,44E-01 |
| AT5G61060 | 0,37 | 1,61 | 3,44 | 0,47 | 6,40E-01 |
| AT5G61130 | 0,17 | 1,92 | 4,33 | 0,44 | 6,57E-01 |
| AT5G61150 | 0,86 | 1,40 | 2,61 | 0,54 | 5,92E-01 |
| AT5G61250 | 0,23 | 1,16 | 4,35 | 0,27 | 7,89E-01 |
| AT5G61390 | 0,43 | 1,27 | 4,37 | 0,29 | 7,71E-01 |
| AT5G09005 | 1,51 | -0,13 | 1,33 | -0,10 | 9,22E-01 |
| AT5G61460 | 0,21 | 2,14 | 4,33 | 0,49 | 6,22E-01 |
| AT5G61480 | 0,83 | 0,04 | 2,24 | 0,02 | 9,85E-01 |
| AT5G61500 | 0,76 | 0,97 | 2,91 | 0,33 | 7,38E-01 |
| AT5G61510 | 0,56 | 0,26 | 3,28 | 0,08 | 9,37E-01 |
| AT5G61520 | 0,33 | 1,57 | 4,34 | 0,36 | 7,17E-01 |
| AT5G61590 | 1,28 | -3,17 | 1,91 | -1,66 | 9,71E-02 |
| AT5G61600 | 0,57 | 0,62 | 2,61 | 0,24 | 8,12E-01 |
| AT5G61800 | 0,12 | 1,46 | 4,35 | 0,34 | 7,36E-01 |
| AT5G61810 | 0,48 | 1,17 | 3,33 | 0,35 | 7,26E-01 |
| AT5G61820 | 1,07 | 3,24 | 2,33 | 1,39 | 1,65E-01 |
| AT5G61840 | 0,89 | 2,79 | 2,22 | 1,26 | 2,08E-01 |
| AT5G61910 | 1,60 | 2,80 | 1,66 | 1,69 | 9,09E-02 |
| AT5G61900 | 1,09 | 2,19 | 1,85 | 1,18 | 2,37E-01 |
| AT5G61960 | 0,94 | 0,07 | 2,57 | 0,03 | 9,78E-01 |
| AT5G61990 | 0,30 | 1,17 | 4,37 | 0,27 | 7,89E-01 |
| AT5G62090 | 0,32 | -0,20 | 4,30 | -0,05 | 9,63E-01 |
| AT5G62110 | 0,34 | 1,91 | 4,32 | 0,44 | 6,58E-01 |
| AT5G62130 | 0,45 | -0,46 | 3,26 | -0,14 | 8,88E-01 |
| AT5G62140 | 1,15 | 0,97 | 1,65 | 0,59 | 5,56E-01 |
| AT5G62180 | 0,08 | 1,10 | 4,36 | 0,25 | 8,00E-01 |
| AT5G62220 | 1,34 | -0,77 | 1,80 | -0,42 | 6,71E-01 |
| AT5G62300 | 1,07 | -1,30 | 1,83 | -0,71 | 4,77E-01 |
| AT5G62430 | 0,29 | 0,46 | 4,37 | 0,10 | 9,17E-01 |
| AT5G62470 | 0,30 | -2,04 | 4,36 | -0,47 | 6,40E-01 |
| AT5G62530 | 0,56 | 0,56 | 2,96 | 0,19 | 8,51E-01 |
| AT5G62600 | 0,50 | 1,68 | 3,97 | 0,42 | 6,73E-01 |
| AT5G62610 | 0,31 | 1,07 | 4,35 | 0,25 | 8,05E-01 |
| AT5G62630 | 0,37 | -0,43 | 4,32 | -0,10 | 9,20E-01 |

| | | | | | |
|---|---|---|---|---|---|
| AT5G62650 | 0,39 | -2,03 | 4,35 | -0,47 | 6,41E-01 |
| AT5G62680 | 0,33 | 1,68 | 4,33 | 0,39 | 6,99E-01 |
| AT5G62790 | 0,60 | -1,56 | 2,57 | -0,61 | 5,45E-01 |
| AT5G63080 | 0,23 | 0,26 | 4,36 | 0,06 | 9,52E-01 |
| AT5G63100 | 0,29 | 2,01 | 4,33 | 0,46 | 6,43E-01 |
| AT5G63110 | 0,58 | 0,92 | 3,04 | 0,30 | 7,62E-01 |
| AT5G63120 | 0,65 | 2,08 | 3,06 | 0,68 | 4,96E-01 |
| AT5G63160 | 0,51 | 1,25 | 3,16 | 0,39 | 6,93E-01 |
| AT5G63180 | 0,25 | -0,85 | 4,33 | -0,20 | 8,44E-01 |
| AT5G63290 | 0,56 | 1,97 | 4,33 | 0,45 | 6,49E-01 |
| AT5G63310 | 0,47 | 0,22 | 4,35 | 0,05 | 9,59E-01 |
| AT5G63320 | 0,52 | 1,10 | 3,92 | 0,28 | 7,80E-01 |
| AT5G63370 | 1,26 | 2,18 | 1,79 | 1,22 | 2,22E-01 |
| AT5G63400 | 0,30 | -0,90 | 4,35 | -0,21 | 8,35E-01 |
| AT5G63410 | 0,46 | 1,21 | 2,74 | 0,44 | 6,59E-01 |
| AT5G63470 | 0,44 | 0,00 | 4,11 | 0,00 | 1,00E+00 |
| AT5G63610 | 0,51 | 1,81 | 3,95 | 0,46 | 6,46E-01 |
| AT5G63625 | 1,28 | -0,63 | 1,73 | -0,37 | 7,15E-01 |
| AT5G63760 | 0,90 | 2,72 | 2,38 | 1,14 | 2,52E-01 |
| AT5G63790 | 0,54 | 2,52 | 4,32 | 0,58 | 5,59E-01 |
| AT5G63860 | 0,75 | 1,21 | 2,20 | 0,55 | 5,81E-01 |
| AT5G63870 | 0,08 | 0,85 | 4,38 | 0,20 | 8,45E-01 |
| AT5G63890 | 0,39 | -1,20 | 4,24 | -0,28 | 7,77E-01 |
| AT5G63910 | 0,66 | -1,24 | 3,07 | -0,40 | 6,86E-01 |
| AT5G64020 | 0,23 | 1,16 | 4,35 | 0,27 | 7,89E-01 |
| AT5G64050 | 1,12 | 0,13 | 1,68 | 0,08 | 9,39E-01 |
| AT5G64070 | 1,53 | 2,37 | 1,67 | 1,42 | 1,56E-01 |
| AT5G64130 | 0,35 | -0,66 | 3,58 | -0,18 | 8,54E-01 |
| AT5G64140 | 1,06 | 0,65 | 1,88 | 0,34 | 7,31E-01 |
| AT5G64170 | 0,64 | 1,84 | 3,19 | 0,58 | 5,63E-01 |
| AT5G64200 | 0,28 | 1,43 | 4,34 | 0,33 | 7,42E-01 |
| AT5G64280 | 1,39 | 2,67 | 1,79 | 1,49 | 1,36E-01 |
| AT5G64290 | 0,42 | 1,14 | 4,10 | 0,28 | 7,81E-01 |
| AT5G64340 | 1,19 | 0,93 | 1,82 | 0,51 | 6,10E-01 |
| AT5G64350 | 0,41 | -0,60 | 4,21 | -0,14 | 8,87E-01 |
| AT5G64360 | 0,62 | 0,00 | 3,76 | 0,00 | 1,00E+00 |
| AT5G64410 | 0,79 | 0,00 | 2,21 | 0,00 | 9,99E-01 |
| AT5G64430 | 0,96 | 1,38 | 2,18 | 0,63 | 5,26E-01 |
| AT5G64460 | 0,08 | 0,00 | 4,40 | 0,00 | 1,00E+00 |
| AT5G64550 | 0,25 | 1,76 | 4,33 | 0,41 | 6,85E-01 |
| AT5G64570 | 1,49 | 1,21 | 1,65 | 0,73 | 4,63E-01 |
| AT5G64580 | 0,77 | 0,83 | 2,66 | 0,31 | 7,54E-01 |
| AT5G64610 | 0,28 | 1,58 | 4,35 | 0,36 | 7,15E-01 |
| AT5G64660 | 0,37 | 0,62 | 3,42 | 0,18 | 8,56E-01 |
| AT5G64670 | 0,17 | -0,02 | 4,36 | 0,00 | 9,97E-01 |
| AT5G64760 | 0,83 | 2,45 | 2,98 | 0,82 | 4,12E-01 |
| AT5G64830 | 0,77 | 1,51 | 2,44 | 0,62 | 5,37E-01 |
| AT5G64860 | 0,28 | 1,97 | 4,33 | 0,46 | 6,49E-01 |
| AT5G64870 | 0,67 | 2,34 | 2,83 | 0,83 | 4,07E-01 |
| AT5G64920 | 1,21 | 2,98 | 1,90 | 1,56 | 1,18E-01 |
| AT5G64930 | 0,39 | 0,40 | 4,39 | 0,09 | 9,28E-01 |
| AT5G64980 | 0,12 | 1,58 | 4,34 | 0,36 | 7,16E-01 |
| AT5G65110 | 0,46 | 0,89 | 3,14 | 0,28 | 7,77E-01 |
| AT5G65200 | 0,16 | 1,33 | 4,35 | 0,31 | 7,59E-01 |
| AT5G65240 | 0,27 | 1,59 | 3,74 | 0,42 | 6,71E-01 |
| AT5G65310 | 1,18 | -1,43 | 2,24 | -0,64 | 5,25E-01 |

| | | | | | |
|---|---|---|---|---|---|
| AT5G65360 | 0,52 | 0,81 | 3,19 | 0,25 | 7,99E-01 |
| AT5G65430 | 0,62 | 0,55 | 2,75 | 0,20 | 8,43E-01 |
| AT5G65560 | 0,37 | 1,51 | 4,35 | 0,35 | 7,28E-01 |
| AT5G65685 | 0,97 | -0,60 | 2,11 | -0,29 | 7,75E-01 |
| AT5G65840 | 0,17 | 0,76 | 4,37 | 0,18 | 8,61E-01 |
| AT5G65910 | 1,14 | 1,87 | 3,18 | 0,59 | 5,57E-01 |
| AT5G65930 | 0,99 | 2,30 | 2,14 | 1,08 | 2,81E-01 |
| AT5G09535 | 0,24 | 1,58 | 4,34 | 0,36 | 7,16E-01 |
| AT5G65970 | 0,45 | 0,44 | 4,39 | 0,10 | 9,20E-01 |
| AT5G66055 | 0,63 | 0,73 | 2,49 | 0,29 | 7,69E-01 |
| AT5G66120 | 0,85 | 1,70 | 2,15 | 0,79 | 4,28E-01 |
| AT5G66210 | 1,05 | 1,13 | 2,24 | 0,51 | 6,12E-01 |
| AT5G66300 | 0,13 | 0,00 | 4,41 | 0,00 | 1,00E+00 |
| AT5G66310 | 0,12 | 1,58 | 4,34 | 0,36 | 7,16E-01 |
| AT5G66320 | 0,48 | 1,62 | 3,33 | 0,49 | 6,26E-01 |
| AT5G66330 | 0,11 | 0,00 | 4,41 | 0,00 | 1,00E+00 |
| AT5G66400 | 0,42 | -1,50 | 4,14 | -0,36 | 7,17E-01 |
| AT5G66440 | 0,33 | 0,77 | 4,39 | 0,18 | 8,61E-01 |
| AT5G66460 | 0,57 | 1,23 | 4,36 | 0,28 | 7,79E-01 |
| AT5G66500 | 0,15 | 1,62 | 4,34 | 0,37 | 7,09E-01 |
| AT5G66530 | 0,97 | -2,09 | 2,48 | -0,84 | 3,99E-01 |
| AT5G66540 | 0,21 | 1,84 | 4,33 | 0,42 | 6,71E-01 |
| AT5G66558 | 1,38 | 2,14 | 1,79 | 1,20 | 2,32E-01 |
| AT5G66595 | 0,74 | -0,54 | 2,50 | -0,22 | 8,28E-01 |
| AT5G66607 | 0,08 | 0,00 | 4,40 | 0,00 | 1,00E+00 |
| AT5G66650 | 0,26 | 1,08 | 4,37 | 0,25 | 8,05E-01 |
| AT5G66675 | 0,51 | 0,67 | 3,17 | 0,21 | 8,33E-01 |
| AT5G66680 | 0,76 | 1,86 | 3,70 | 0,50 | 6,14E-01 |
| AT5G66720 | 0,41 | 0,00 | 4,41 | 0,00 | 1,00E+00 |
| AT5G66730 | 0,31 | 0,30 | 4,35 | 0,07 | 9,46E-01 |
| AT5G66850 | 0,24 | 1,11 | 4,32 | 0,26 | 7,98E-01 |
| AT5G66890 | 0,75 | 2,25 | 3,44 | 0,65 | 5,13E-01 |
| AT5G66900 | 1,52 | 2,73 | 1,82 | 1,50 | 1,34E-01 |
| AT5G66910 | 0,38 | 1,23 | 3,41 | 0,36 | 7,18E-01 |
| AT5G67120 | 0,79 | 2,81 | 2,73 | 1,03 | 3,04E-01 |
| AT5G67150 | 0,35 | 0,45 | 4,39 | 0,10 | 9,19E-01 |
| AT5G67190 | 0,91 | 1,17 | 2,46 | 0,47 | 6,36E-01 |
| AT5G67220 | 0,60 | 1,22 | 2,51 | 0,49 | 6,27E-01 |
| AT5G67240 | 1,06 | -0,08 | 1,90 | -0,04 | 9,66E-01 |
| AT5G67245 | 0,45 | 0,90 | 4,37 | 0,21 | 8,37E-01 |
| AT5G67250 | 1,03 | 2,79 | 2,06 | 1,36 | 1,75E-01 |
| AT5G67280 | 0,58 | -0,74 | 2,68 | -0,28 | 7,83E-01 |
| AT5G67290 | 0,82 | 2,62 | 2,44 | 1,07 | 2,83E-01 |
| AT5G09855 | 0,69 | 2,12 | 2,44 | 0,87 | 3,86E-01 |
| AT5G67310 | 0,78 | 2,24 | 2,75 | 0,82 | 4,15E-01 |
| AT5G67350 | 0,75 | 1,13 | 2,55 | 0,44 | 6,58E-01 |
| AT5G67380 | 0,30 | 0,42 | 4,38 | 0,10 | 9,24E-01 |
| AT5G67460 | 0,10 | 0,42 | 4,38 | 0,10 | 9,23E-01 |
| AT5G67480 | 1,04 | 3,01 | 2,52 | 1,19 | 2,32E-01 |
| AT5G67490 | 1,16 | 1,30 | 1,87 | 0,70 | 4,85E-01 |
| AT5G67580 | 0,39 | -0,09 | 2,80 | -0,03 | 9,75E-01 |
| AT5G67610 | 0,37 | 0,00 | 4,41 | 0,00 | 1,00E+00 |

***chr11 chr17* infected by TCV relative to uninfected wild type.**

| |
|---|
| type |
| miRNA_precursor |
| miRNA_precursor |
| protein_coding |
| protein_coding |
| protein_coding |
| protein_coding |
| miRNA_precursor |
| miRNA_precursor |
| ncRNA |
| ribosomal_rna |
| miRNA_precursor |
| protein_coding |
| protein_coding |
| protein_coding |
| miRNA_precursor |
| protein_coding |
| protein_coding |
| protein_coding |
| miRNA_precursor |
| miRNA_precursor |
| protein_coding |
| miRNA_precursor |
| miRNA_precursor |
| miRNA_precursor |
| protein_coding |
| miRNA_precursor |
| protein_coding |
| protein_coding |
| protein_coding |
| miRNA_precursor |
| miRNA_precursor |
| ncRNA |
| ncRNA |
| ncRNA |
| miRNA_precursor |
| protein_coding |
| protein_coding |
| miRNA_precursor |
| miRNA_precursor |
| protein_coding |
| ncRNA |
| protein_coding |
| miRNA_precursor |
| miRNA_precursor |
| miRNA_precursor |
| ncRNA |
| miRNA_precursor |
| miRNA_precursor |
| protein_coding |
| miRNA_precursor |
| protein_coding |
| miRNA_precursor |
| ncRNA |

miRNA_precursor
miRNA_precursor
miRNA_precursor
miRNA_precursor
protein_coding
protein_coding
miRNA_precursor
protein_coding
small_nucle(ol)ar_rna
protein_coding
protein_coding
small_nucle(ol)ar_rna
ncRNA
protein_coding
protein_coding
miRNA_precursor
protein_coding
ncRNA
protein_coding
ribosomal_rna
protein_coding
ncRNA
ncRNA
ncRNA
ncRNA
protein_coding
miRNA_precursor
protein_coding
protein_coding
protein_coding
miRNA_precursor
miRNA_precursor
protein_coding
miRNA_precursor
protein_coding
small_nucle(ol)ar_rna
protein_coding
miRNA_precursor
protein_coding
miRNA_precursor
protein_coding
small_nucle(ol)ar_rna
ncRNA
protein_coding
ncRNA
miRNA_precursor
protein_coding
miRNA_precursor
protein_coding
protein_coding
miRNA_precursor
small_nucle(ol)ar_rna
protein_coding
miRNA_precursor
protein_coding
protein_coding

protein_coding
protein_coding
protein_coding
protein_coding
protein_coding
ncRNA
small_nucle(ol)ar_rna
protein_coding
protein_coding
protein_coding
protein_coding
protein_coding
protein_coding
protein_coding
protein_coding
protein_coding
protein_coding
protein_coding
protein_coding
protein_coding
miRNA_precursor
protein_coding
protein_coding
miRNA_precursor
small_nucle(ol)ar_rna
protein_coding
protein_coding
protein_coding
protein_coding
miRNA_precursor
protein_coding
protein_coding
protein_coding
miRNA_precursor
protein_coding
miRNA_precursor
protein_coding
protein_coding
protein_coding
protein_coding
protein_coding
ncRNA
protein_coding
miRNA_precursor
protein_coding
protein_coding
protein_coding
miRNA_precursor
protein_coding
protein_coding
protein_coding
protein_coding
protein_coding
protein_coding
protein_coding
protein_coding

protein_coding
protein_coding
protein_coding
protein_coding
protein_coding
protein_coding
protein_coding
protein_coding
small_nucle(ol)ar_rna
protein_coding
protein_coding
protein_coding
protein_coding
protein_coding
protein_coding
protein_coding
protein_coding
protein_coding
protein_coding
ncRNA
protein_coding
protein_coding
protein_coding
miRNA_precursor
protein_coding
protein_coding
protein_coding
protein_coding
miRNA_precursor
protein_coding
ncRNA
protein_coding
protein_coding
protein_coding
miRNA_precursor
protein_coding
protein_coding
protein_coding
protein_coding
protein_coding
protein_coding
protein_coding
protein_coding
ncRNA
miRNA_precursor
protein_coding
protein_coding
miRNA_precursor
protein_coding
protein_coding
protein_coding
protein_coding
protein_coding
protein_coding
protein_coding
protein_coding

protein_coding
ncRNA
protein_coding
protein_coding
small_nucle(ol)ar_rna
protein_coding
protein_coding
protein_coding
protein_coding
protein_coding
protein_coding
protein_coding
small_nucle(ol)ar_rna
protein_coding
protein_coding
protein_coding
protein_coding
small_nucle(ol)ar_rna
protein_coding
protein_coding
protein_coding
protein_coding
protein_coding
protein_coding
protein_coding
protein_coding
protein_coding
protein_coding
protein_coding
protein_coding
protein_coding
ncRNA
protein_coding
protein_coding
miRNA_precursor
protein_coding
ncRNA
protein_coding
protein_coding
protein_coding
protein_coding
protein_coding
protein_coding
protein_coding
miRNA_precursor
protein_coding
ncRNA
protein_coding
miRNA_precursor
small_nucle(ol)ar_rna
protein_coding
protein_coding
protein_coding
protein_coding
protein_coding
protein_coding

miRNA_precursor
protein_coding
protein_coding
protein_coding
protein_coding
miRNA_precursor
protein_coding
protein_coding
miRNA_precursor
miRNA_precursor
protein_coding
ncRNA
protein_coding
protein_coding
protein_coding
protein_coding
protein_coding
protein_coding
protein_coding
protein_coding
protein_coding
protein_coding
protein_coding
protein_coding
protein_coding
protein_coding
protein_coding
protein_coding
protein_coding
protein_coding
protein_coding
protein_coding
protein_coding
protein_coding
protein_coding
protein_coding
protein_coding
protein_coding
protein_coding
protein_coding
protein_coding
protein_coding
protein_coding
protein_coding
protein_coding
protein_coding
protein_coding
protein_coding
protein_coding
ncRNA
protein_coding
protein_coding
protein_coding
protein_coding
protein_coding
protein_coding

protein_coding
protein_coding
protein_coding
protein_coding
protein_coding
protein_coding
protein_coding
protein_coding
protein_coding
protein_coding
protein_coding
protein_coding
miRNA_precursor
protein_coding
protein_coding
protein_coding
protein_coding
protein_coding
protein_coding
miRNA_precursor
protein_coding
protein_coding
protein_coding
protein_coding
protein_coding
protein_coding
protein_coding
protein_coding
protein_coding
protein_coding
protein_coding
ncRNA
protein_coding
protein_coding
protein_coding
protein_coding
protein_coding
ncRNA
protein_coding
protein_coding
protein_coding
protein_coding
protein_coding
protein_coding
ncRNA
protein_coding
ncRNA
protein_coding
protein_coding
protein_coding
ncRNA
protein_coding
protein_coding
protein_coding
protein_coding
miRNA_precursor

protein_coding
ncRNA
protein_coding
protein_coding
protein_coding
protein_coding
protein_coding
protein_coding
protein_coding
protein_coding
protein_coding
miRNA_precursor
protein_coding
protein_coding
protein_coding
protein_coding
protein_coding
protein_coding
protein_coding
protein_coding
protein_coding
protein_coding
protein_coding
protein_coding
protein_coding
protein_coding
protein_coding
protein_coding
protein_coding
protein_coding
protein_coding
small_nucle(ol)ar_rna
small_nucle(ol)ar_rna
protein_coding
protein_coding
protein_coding
protein_coding
protein_coding
protein_coding
protein_coding
protein_coding
protein_coding
protein_coding
ncRNA
protein_coding
protein_coding
protein_coding
ncRNA
protein_coding
protein_coding
protein_coding
small_nucle(ol)ar_rna
protein_coding
ncRNA
protein_coding
protein_coding

protein_coding
protein_coding
protein_coding
protein_coding
protein_coding
protein_coding
miRNA_precursor
protein_coding
protein_coding
protein_coding
protein_coding
miRNA_precursor
protein_coding
protein_coding
protein_coding
protein_coding
protein_coding
protein_coding
protein_coding
protein_coding
protein_coding
protein_coding
protein_coding
small_nucle(ol)ar_rna
protein_coding
small_nucle(ol)ar_rna
protein_coding
protein_coding
ncRNA
protein_coding
protein_coding
protein_coding
protein_coding
protein_coding
protein_coding
protein_coding
miRNA_precursor
protein_coding
small_nucle(ol)ar_rna
miRNA_precursor
protein_coding
protein_coding
protein_coding
protein_coding
protein_coding
protein_coding
protein_coding
miRNA_precursor
protein_coding
protein_coding
protein_coding
ncRNA
protein_coding
protein_coding
ncRNA
protein_coding

protein_coding
protein_coding
protein_coding
protein_coding
protein_coding
protein_coding
protein_coding
protein_coding
miRNA_precursor
protein_coding
protein_coding
small_nucle(ol)ar_rna
miRNA_precursor
protein_coding
protein_coding
protein_coding
protein_coding
ncRNA
protein_coding
protein_coding
protein_coding
miRNA_precursor
protein_coding
protein_coding
protein_coding
protein_coding
protein_coding
protein_coding
protein_coding
protein_coding
ncRNA
protein_coding
protein_coding
protein_coding
protein_coding
protein_coding
protein_coding
protein_coding
protein_coding
protein_coding
protein_coding
protein_coding
protein_coding
small_nucle(ol)ar_rna
protein_coding
protein_coding
protein_coding
protein_coding
protein_coding
protein_coding
protein_coding
protein_coding
protein_coding
protein_coding
protein_coding
protein_coding

small_nucle(ol)ar_rna
protein_coding
protein_coding
protein_coding
protein_coding
protein_coding
protein_coding
protein_coding
protein_coding
protein_coding
protein_coding
protein_coding
protein_coding
protein_coding
protein_coding
small_nucle(ol)ar_rna
protein_coding
protein_coding
protein_coding
protein_coding
protein_coding
protein_coding
protein_coding
protein_coding
protein_coding
protein_coding
small_nucle(ol)ar_rna
protein_coding
protein_coding
protein_coding
protein_coding
protein_coding
protein_coding
miRNA_precursor
protein_coding
protein_coding
protein_coding
protein_coding
protein_coding
protein_coding
protein_coding
protein_coding
protein_coding
protein_coding
ncRNA
protein_coding
protein_coding
miRNA_precursor
protein_coding
ncRNA
protein_coding
protein_coding
protein_coding
protein_coding
protein_coding
protein_coding

protein_coding
protein_coding
protein_coding
protein_coding
protein_coding
ncRNA
protein_coding
protein_coding
protein_coding
protein_coding
protein_coding
protein_coding
protein_coding
protein_coding
protein_coding
protein_coding
protein_coding
protein_coding
protein_coding
protein_coding
protein_coding
protein_coding
protein_coding
protein_coding
protein_coding
protein_coding
protein_coding
protein_coding
protein_coding
protein_coding
small_nucle(ol)ar_rna
protein_coding
protein_coding
protein_coding
protein_coding
protein_coding
protein_coding
protein_coding
protein_coding
protein_coding
protein_coding
protein_coding
protein_coding
protein_coding
protein_coding
ncRNA
protein_coding
protein_coding
small_nucle(ol)ar_rna
protein_coding
ncRNA
protein_coding
protein_coding
protein_coding
protein_coding
protein_coding

protein_coding
protein_coding
protein_coding
miRNA_precursor
protein_coding
protein_coding
protein_coding
miRNA_precursor
small_nucle(ol)ar_rna
protein_coding
ncRNA
protein_coding
miRNA_precursor
protein_coding
miRNA_precursor
protein_coding
protein_coding
protein_coding
protein_coding
protein_coding
protein_coding
protein_coding
protein_coding
small_nucle(ol)ar_rna
protein_coding
protein_coding
protein_coding
protein_coding
protein_coding
protein_coding
protein_coding
protein_coding
protein_coding
protein_coding
protein_coding
protein_coding
protein_coding
ncRNA
miRNA_precursor
protein_coding
protein_coding
protein_coding
protein_coding
ncRNA
protein_coding
protein_coding
protein_coding
protein_coding
protein_coding
protein_coding
protein_coding
protein_coding
protein_coding
protein_coding
protein_coding
protein_coding

ncRNA
protein_coding
protein_coding
protein_coding
small_nucle(ol)ar_rna
protein_coding
protein_coding
protein_coding
protein_coding
protein_coding
protein_coding
protein_coding
protein_coding
miRNA_precursor
protein_coding
miRNA_precursor
protein_coding
protein_coding
protein_coding
protein_coding
protein_coding
protein_coding
protein_coding
protein_coding
protein_coding
protein_coding
protein_coding
protein_coding
protein_coding
protein_coding
protein_coding
protein_coding
protein_coding
protein_coding
protein_coding
protein_coding
protein_coding
protein_coding
protein_coding
protein_coding
protein_coding
protein_coding
protein_coding
protein_coding
protein_coding
protein_coding
protein_coding
protein_coding
protein_coding
protein_coding
protein_coding
protein_coding
protein_coding
miRNA_precursor
protein_coding
protein_coding

protein_coding
protein_coding
protein_coding
protein_coding
protein_coding
protein_coding
protein_coding
protein_coding
small_nucle(ol)ar_rna
protein_coding
protein_coding
protein_coding
ncRNA
protein_coding
protein_coding
protein_coding
protein_coding
protein_coding
protein_coding
protein_coding
protein_coding
protein_coding
protein_coding
protein_coding
protein_coding
small_nucle(ol)ar_rna
protein_coding
protein_coding
protein_coding
miRNA_precursor
protein_coding
protein_coding
protein_coding
protein_coding
protein_coding
protein_coding
ncRNA
protein_coding
ncRNA
protein_coding
protein_coding
protein_coding
protein_coding
protein_coding
protein_coding
protein_coding
protein_coding
miRNA_precursor
protein_coding
protein_coding
protein_coding
protein_coding
miRNA_precursor
protein_coding
protein_coding
protein_coding

protein_coding
protein_coding
ncRNA
protein_coding
protein_coding
protein_coding
protein_coding
protein_coding
protein_coding
protein_coding
protein_coding
protein_coding
protein_coding
ncRNA
protein_coding
pre_trna
protein_coding
protein_coding
protein_coding
protein_coding
protein_coding
miRNA_precursor
protein_coding
protein_coding
protein_coding
protein_coding
protein_coding
protein_coding
protein_coding
protein_coding
small_nucle(ol)ar_rna
protein_coding
protein_coding
protein_coding
protein_coding
miRNA_precursor
protein_coding
protein_coding
small_nucle(ol)ar_rna
protein_coding
protein_coding
protein_coding
protein_coding
protein_coding
protein_coding
protein_coding
protein_coding
protein_coding
protein_coding
protein_coding
protein_coding
protein_coding
protein_coding
protein_coding
protein_coding
protein_coding

protein_coding
protein_coding
protein_coding
protein_coding
protein_coding
small_nucle(ol)ar_rna
protein_coding
protein_coding
miRNA_precursor
protein_coding
protein_coding
protein_coding
protein_coding
protein_coding
protein_coding
protein_coding
protein_coding
protein_coding
protein_coding
protein_coding
protein_coding
protein_coding
protein_coding
protein_coding
protein_coding
protein_coding
protein_coding
protein_coding
protein_coding
protein_coding
protein_coding
protein_coding
protein_coding
protein_coding
protein_coding
protein_coding
small_nucle(ol)ar_rna
protein_coding
protein_coding
protein_coding
protein_coding
protein_coding
protein_coding
protein_coding
protein_coding
protein_coding
protein_coding
protein_coding
protein_coding
protein_coding
small_nucle(ol)ar_rna
protein_coding
protein_coding
protein_coding
protein_coding
protein_coding

protein_coding
protein_coding
protein_coding
protein_coding
protein_coding
protein_coding
protein_coding
protein_coding
protein_coding
protein_coding
protein_coding
protein_coding
protein_coding
small_nucle(ol)ar_rna
protein_coding
protein_coding
protein_coding
protein_coding
protein_coding
protein_coding
pre_trna
protein_coding
protein_coding
protein_coding
protein_coding
protein_coding
protein_coding
miRNA_precursor
small_nucle(ol)ar_rna
protein_coding
protein_coding
protein_coding
protein_coding
miRNA_precursor
protein_coding
protein_coding
protein_coding
protein_coding
ncRNA
protein_coding
ncRNA
protein_coding
protein_coding
protein_coding
protein_coding
protein_coding
protein_coding
protein_coding
protein_coding
miRNA_precursor
protein_coding
protein_coding
protein_coding
protein_coding
protein_coding
protein_coding

protein_coding
protein_coding
small_nucle(ol)ar_rna
protein_coding
protein_coding
protein_coding
protein_coding
protein_coding
protein_coding
protein_coding
protein_coding
small_nucle(ol)ar_rna
protein_coding
protein_coding
protein_coding
protein_coding
protein_coding
protein_coding
protein_coding
protein_coding
protein_coding
protein_coding
protein_coding
miRNA_precursor
protein_coding
protein_coding
protein_coding
protein_coding
protein_coding
protein_coding
small_nucle(ol)ar_rna
ncRNA
protein_coding
protein_coding
protein_coding
protein_coding
protein_coding
protein_coding
protein_coding
miRNA_precursor
ncRNA
protein_coding
protein_coding
protein_coding
protein_coding
protein_coding
protein_coding
protein_coding
protein_coding
protein_coding
protein_coding
protein_coding
protein_coding
protein_coding
protein_coding
protein_coding

protein_coding
protein_coding
protein_coding
protein_coding
miRNA_precursor
protein_coding
miRNA_precursor
protein_coding
protein_coding
protein_coding
protein_coding
protein_coding
protein_coding
protein_coding
protein_coding
protein_coding
protein_coding
protein_coding
protein_coding
protein_coding
protein_coding
protein_coding
protein_coding
protein_coding
protein_coding
protein_coding
protein_coding
miRNA_precursor
protein_coding
protein_coding
protein_coding
protein_coding
protein_coding
ncRNA
protein_coding
protein_coding
ncRNA
protein_coding
miRNA_precursor
miRNA_precursor
small_nucle(ol)ar_rna
protein_coding
protein_coding
protein_coding
protein_coding
protein_coding
protein_coding
small_nucle(ol)ar_rna
protein_coding
protein_coding
protein_coding
small_nucle(ol)ar_rna
ncRNA
protein_coding
protein_coding
small_nucle(ol)ar_rna

protein_coding
miRNA_precursor
protein_coding
protein_coding
protein_coding
protein_coding
protein_coding
protein_coding
protein_coding
protein_coding
protein_coding
protein_coding
protein_coding
protein_coding
protein_coding
protein_coding
protein_coding
protein_coding
protein_coding
ncRNA
protein_coding
protein_coding
protein_coding
protein_coding
ncRNA
protein_coding
protein_coding
protein_coding
protein_coding
protein_coding
protein_coding
protein_coding
protein_coding
protein_coding
protein_coding
protein_coding
miRNA_precursor
protein_coding
protein_coding
protein_coding
protein_coding
protein_coding
ncRNA
protein_coding
small_nucle(ol)ar_rna
protein_coding
protein_coding
protein_coding
protein_coding
ncRNA
ncRNA
protein_coding
protein_coding
small_nucle(ol)ar_rna
protein_coding
protein_coding

protein_coding
protein_coding
ncRNA
protein_coding
protein_coding
protein_coding
protein_coding
protein_coding
protein_coding
protein_coding
protein_coding
protein_coding
protein_coding
protein_coding
protein_coding
protein_coding
protein_coding
protein_coding
protein_coding
protein_coding
protein_coding
protein_coding
small_nucle(ol)ar_rna
protein_coding
miRNA_precursor
protein_coding
protein_coding
protein_coding
protein_coding
ncRNA
ncRNA
protein_coding
protein_coding
protein_coding
small_nucle(ol)ar_rna
protein_coding
protein_coding
protein_coding
protein_coding
protein_coding
miRNA_precursor
protein_coding
protein_coding
pre_trna
protein_coding
protein_coding
protein_coding
miRNA_precursor
protein_coding
protein_coding
miRNA_precursor
protein_coding
protein_coding
protein_coding
protein_coding
protein_coding

protein_coding
protein_coding
small_nucle(ol)ar_rna
protein_coding
protein_coding
protein_coding
protein_coding
ncRNA
protein_coding
protein_coding
protein_coding
protein_coding
protein_coding
protein_coding
protein_coding
protein_coding
protein_coding
protein_coding
miRNA_precursor
protein_coding
protein_coding
small_nucle(ol)ar_rna
protein_coding
protein_coding
protein_coding
protein_coding
protein_coding
miRNA_precursor
protein_coding
ncRNA
protein_coding
protein_coding
protein_coding
protein_coding
protein_coding
protein_coding
protein_coding
ncRNA
protein_coding
protein_coding
pre_trna
protein_coding
protein_coding
protein_coding
ncRNA
protein_coding
protein_coding
protein_coding
protein_coding
protein_coding
protein_coding
protein_coding
protein_coding
protein_coding
protein_coding
small_nucle(ol)ar_rna

protein_coding
protein_coding
protein_coding
protein_coding
protein_coding
protein_coding
protein_coding
protein_coding
protein_coding
protein_coding
protein_coding
protein_coding
protein_coding
protein_coding
protein_coding
protein_coding
protein_coding
protein_coding
protein_coding
small_nucle(ol)ar_rna
miRNA_precursor
protein_coding
small_nucle(ol)ar_rna
protein_coding
small_nucle(ol)ar_rna
protein_coding
protein_coding
protein_coding
protein_coding
protein_coding
protein_coding
protein_coding
protein_coding
ncRNA
miRNA_precursor
protein_coding
ncRNA
protein_coding
protein_coding
protein_coding
protein_coding
protein_coding
protein_coding
protein_coding
protein_coding
protein_coding
protein_coding
protein_coding
protein_coding
protein_coding
protein_coding
protein_coding
protein_coding
small_nucle(ol)ar_rna
protein_coding
small_nucle(ol)ar_rna

protein_coding
protein_coding
protein_coding
ncRNA
protein_coding
protein_coding
protein_coding
protein_coding
protein_coding
protein_coding
miRNA_precursor
protein_coding
protein_coding
protein_coding
protein_coding
protein_coding
protein_coding
miRNA_precursor
protein_coding
protein_coding
protein_coding
ncRNA
protein_coding
protein_coding
small_nucle(ol)ar_rna
protein_coding
ncRNA
small_nucle(ol)ar_rna
protein_coding
protein_coding
protein_coding
protein_coding
protein_coding
protein_coding
small_nucle(ol)ar_rna
protein_coding
protein_coding
protein_coding
protein_coding
protein_coding
protein_coding
protein_coding
protein_coding
miRNA_precursor
protein_coding
protein_coding
protein_coding
ncRNA
protein_coding
protein_coding
protein_coding
protein_coding
small_nucle(ol)ar_rna
protein_coding
protein_coding
protein_coding

protein_coding
miRNA_precursor
protein_coding
protein_coding
protein_coding
protein_coding
protein_coding
protein_coding
protein_coding
protein_coding
miRNA_precursor
small_nucle(ol)ar_rna
protein_coding
protein_coding
protein_coding
protein_coding
miRNA_precursor
protein_coding
protein_coding
protein_coding
protein_coding
ncRNA
ncRNA
protein_coding
protein_coding
protein_coding
protein_coding
protein_coding
protein_coding
protein_coding
protein_coding
protein_coding
protein_coding
protein_coding
protein_coding
protein_coding
protein_coding
protein_coding
protein_coding
protein_coding
small_nucle(ol)ar_rna
protein_coding
ncRNA
protein_coding
ncRNA
protein_coding
protein_coding
protein_coding
protein_coding
protein_coding
protein_coding
protein_coding
protein_coding
protein_coding
protein_coding

protein_coding
protein_coding
ncRNA
protein_coding
protein_coding
protein_coding
protein_coding
protein_coding
miRNA_precursor
protein_coding
protein_coding
miRNA_precursor
protein_coding
protein_coding
protein_coding
protein_coding
protein_coding
protein_coding
small_nucle(ol)ar_rna
protein_coding
small_nucle(ol)ar_rna
small_nucle(ol)ar_rna
protein_coding
protein_coding
protein_coding
small_nucle(ol)ar_rna
protein_coding
protein_coding
protein_coding
protein_coding
protein_coding
protein_coding
protein_coding
protein_coding
miRNA_precursor
protein_coding
ncRNA
ncRNA
protein_coding
protein_coding
small_nucle(ol)ar_rna
protein_coding
protein_coding
protein_coding
protein_coding
protein_coding
protein_coding
protein_coding
protein_coding
ncRNA
protein_coding
protein_coding
small_nucle(ol)ar_rna
protein_coding
protein_coding
protein_coding

protein_coding
miRNA_precursor
protein_coding
protein_coding
ncRNA
protein_coding
protein_coding
protein_coding
small_nucle(ol)ar_rna
protein_coding
protein_coding
protein_coding
protein_coding
protein_coding
protein_coding
protein_coding
protein_coding
protein_coding
protein_coding
protein_coding
ncRNA
protein_coding
protein_coding
ncRNA
protein_coding
miRNA_precursor
protein_coding
protein_coding
protein_coding
protein_coding
protein_coding
protein_coding
protein_coding
protein_coding
protein_coding
protein_coding
protein_coding
protein_coding
protein_coding
protein_coding
protein_coding
ncRNA
protein_coding
miRNA_precursor
protein_coding
protein_coding
protein_coding
protein_coding
protein_coding
protein_coding
protein_coding
protein_coding
protein_coding
protein_coding
protein_coding
protein_coding

protein_coding
protein_coding
protein_coding
protein_coding
protein_coding
protein_coding
protein_coding
protein_coding
protein_coding
protein_coding
pre_trna
protein_coding
protein_coding
protein_coding
protein_coding
ncRNA
protein_coding
protein_coding
protein_coding
miRNA_precursor
protein_coding
protein_coding
ncRNA
protein_coding
protein_coding
protein_coding
protein_coding
protein_coding
protein_coding
protein_coding
protein_coding
protein_coding
protein_coding
protein_coding
protein_coding
protein_coding
protein_coding
protein_coding
protein_coding
protein_coding
small_nucle(ol)ar_rna
protein_coding
protein_coding
protein_coding
protein_coding
protein_coding
miRNA_precursor
protein_coding
ncRNA
protein_coding
protein_coding
protein_coding
protein_coding
protein_coding
protein_coding
protein_coding

protein_coding
protein_coding
protein_coding
protein_coding
protein_coding
protein_coding
protein_coding
protein_coding
protein_coding
protein_coding
protein_coding
protein_coding
protein_coding
protein_coding
protein_coding
protein_coding
protein_coding
protein_coding
miRNA_precursor
protein_coding
protein_coding
protein_coding
protein_coding
protein_coding
protein_coding
protein_coding
protein_coding
protein_coding
protein_coding
protein_coding
protein_coding
protein_coding
protein_coding
protein_coding
protein_coding
protein_coding
protein_coding
protein_coding
protein_coding
protein_coding
protein_coding
protein_coding
protein_coding
protein_coding
protein_coding
protein_coding
protein_coding
protein_coding
protein_coding
protein_coding
protein_coding
protein_coding
protein_coding
protein_coding
protein_coding
miRNA_precursor

small_nucle(ol)ar_rna
protein_coding
protein_coding
protein_coding
protein_coding
ncRNA
protein_coding
protein_coding
protein_coding
protein_coding
protein_coding
ncRNA
protein_coding
protein_coding
protein_coding
small_nucle(ol)ar_rna
protein_coding
ncRNA
protein_coding
protein_coding
protein_coding
protein_coding
protein_coding
protein_coding
protein_coding
protein_coding
protein_coding
protein_coding
protein_coding
protein_coding
protein_coding
protein_coding
protein_coding
protein_coding
small_nucle(ol)ar_rna
protein_coding
protein_coding
protein_coding
protein_coding
small_nucle(ol)ar_rna
protein_coding
protein_coding
protein_coding
protein_coding
protein_coding
protein_coding
protein_coding
protein_coding
miRNA_precursor
protein_coding
protein_coding
protein_coding
protein_coding
protein_coding
protein_coding
protein_coding

protein_coding
protein_coding
small_nucle(ol)ar_rna
small_nucle(ol)ar_rna
protein_coding
protein_coding
protein_coding
miRNA_precursor
protein_coding
protein_coding
protein_coding
pre_trna
protein_coding
protein_coding
small_nucle(ol)ar_rna
protein_coding
protein_coding
protein_coding
protein_coding
small_nucle(ol)ar_rna
protein_coding
small_nucle(ol)ar_rna
protein_coding
small_nucle(ol)ar_rna
protein_coding
protein_coding
protein_coding
protein_coding
small_nucle(ol)ar_rna
protein_coding
small_nucle(ol)ar_rna
protein_coding
protein_coding
small_nucle(ol)ar_rna
protein_coding
ncRNA
pre_trna
protein_coding
protein_coding
protein_coding
small_nucle(ol)ar_rna
small_nucle(ol)ar_rna
ncRNA
protein_coding
protein_coding
protein_coding
protein_coding
protein_coding
pre_trna
protein_coding
small_nucle(ol)ar_rna
protein_coding
protein_coding
protein_coding
ncRNA
protein_coding

protein_coding
protein_coding
protein_coding
small_nucle(ol)ar_rna
protein_coding
protein_coding
miRNA_precursor
protein_coding
protein_coding
small_nucle(ol)ar_rna
protein_coding
protein_coding
protein_coding
protein_coding
protein_coding
protein_coding
protein_coding
miRNA_precursor
protein_coding
protein_coding
ncRNA
protein_coding
protein_coding
protein_coding
small_nucle(ol)ar_rna
protein_coding
protein_coding
protein_coding
protein_coding
protein_coding
protein_coding
protein_coding
protein_coding
protein_coding
protein_coding
protein_coding
protein_coding
protein_coding
protein_coding
protein_coding
protein_coding
protein_coding
protein_coding
protein_coding
protein_coding
protein_coding
protein_coding
protein_coding
protein_coding
protein_coding
protein_coding
protein_coding
protein_coding
small_nucle(ol)ar_rna
protein_coding
ncRNA
protein_coding

protein_coding
ncRNA
protein_coding
ncRNA
protein_coding
protein_coding
protein_coding
protein_coding
ncRNA
protein_coding
protein_coding
protein_coding
protein_coding
protein_coding
protein_coding
protein_coding
ncRNA
small_nucle(ol)ar_rna
protein_coding
protein_coding
protein_coding
protein_coding
protein_coding
protein_coding
protein_coding
protein_coding
protein_coding
protein_coding
protein_coding
protein_coding
ncRNA
small_nucle(ol)ar_rna
protein_coding
protein_coding
protein_coding
protein_coding
protein_coding
protein_coding
protein_coding
protein_coding
protein_coding
protein_coding
protein_coding
small_nucle(ol)ar_rna
protein_coding
small_nucle(ol)ar_rna
protein_coding
protein_coding
protein_coding
protein_coding
protein_coding
protein_coding
protein_coding
ncRNA
protein_coding
protein_coding

protein_coding
protein_coding
protein_coding
protein_coding
protein_coding
small_nucle(ol)ar_rna
ncRNA
protein_coding
protein_coding
protein_coding
protein_coding
small_nucle(ol)ar_rna
protein_coding
protein_coding
ncRNA
protein_coding
small_nucle(ol)ar_rna
protein_coding
protein_coding
protein_coding
protein_coding
protein_coding
protein_coding
protein_coding
protein_coding
protein_coding
small_nucle(ol)ar_rna
protein_coding
protein_coding
small_nucle(ol)ar_rna
protein_coding
protein_coding
protein_coding
protein_coding
protein_coding
protein_coding
protein_coding
protein_coding
protein_coding
protein_coding
protein_coding
protein_coding
protein_coding
protein_coding
protein_coding
protein_coding
protein_coding
protein_coding
protein_coding
protein_coding
miRNA_precursor
protein_coding
protein_coding
protein_coding
protein_coding
protein_coding

protein_coding
protein_coding
protein_coding
protein_coding
protein_coding
miRNA_precursor
protein_coding
protein_coding
small_nucle(ol)ar_rna
protein_coding
protein_coding
protein_coding
protein_coding
ncRNA
protein_coding
protein_coding
protein_coding
miRNA_precursor
protein_coding
protein_coding
protein_coding
protein_coding
protein_coding
miRNA_precursor
protein_coding
protein_coding
protein_coding
protein_coding
protein_coding
protein_coding
protein_coding
ncRNA
ncRNA
ribosomal_rna
protein_coding
protein_coding
ncRNA
protein_coding
protein_coding
protein_coding
miRNA_precursor
protein_coding
protein_coding
protein_coding
protein_coding
protein_coding
protein_coding
protein_coding
protein_coding
protein_coding
protein_coding
protein_coding
protein_coding
protein_coding
protein_coding
small_nucle(ol)ar_rna

small_nucle(ol)ar_rna
protein_coding
protein_coding
protein_coding
protein_coding
miRNA_precursor
small_nucle(ol)ar_rna
protein_coding
small_nucle(ol)ar_rna
protein_coding
protein_coding
miRNA_precursor
miRNA_precursor
protein_coding
protein_coding
protein_coding
protein_coding
protein_coding
miRNA_precursor
protein_coding
ncRNA
protein_coding
protein_coding
ncRNA
protein_coding
small_nucle(ol)ar_rna
small_nucle(ol)ar_rna
protein_coding
protein_coding
small_nucle(ol)ar_rna
protein_coding
protein_coding
protein_coding
protein_coding
protein_coding
protein_coding
protein_coding
protein_coding
protein_coding
protein_coding
protein_coding
protein_coding
protein_coding
protein_coding
protein_coding
protein_coding
protein_coding
protein_coding
protein_coding
protein_coding
protein_coding
small_nucle(ol)ar_rna
ncRNA
protein_coding
protein_coding
protein_coding

ncRNA
protein_coding
protein_coding
protein_coding
protein_coding
ncRNA
protein_coding
protein_coding
protein_coding
protein_coding
protein_coding
protein_coding
protein_coding
protein_coding
protein_coding
protein_coding
protein_coding
protein_coding
ncRNA
protein_coding
protein_coding
protein_coding
protein_coding
protein_coding
protein_coding
protein_coding
small_nucle(ol)ar_rna
protein_coding
small_nucle(ol)ar_rna
protein_coding
protein_coding
protein_coding
protein_coding
protein_coding
protein_coding
miRNA_precursor
small_nucle(ol)ar_rna
protein_coding
small_nucle(ol)ar_rna
small_nucle(ol)ar_rna
protein_coding
protein_coding
protein_coding
ncRNA
protein_coding
protein_coding
miRNA_precursor
protein_coding
protein_coding
protein_coding
protein_coding
protein_coding
miRNA_precursor
protein_coding
protein_coding
ncRNA

protein_coding
small_nucle(ol)ar_rna
protein_coding
ncRNA
protein_coding
small_nucle(ol)ar_rna
miRNA_precursor
protein_coding
protein_coding
protein_coding
ncRNA
protein_coding
protein_coding
small_nucle(ol)ar_rna
small_nucle(ol)ar_rna
protein_coding
protein_coding
protein_coding
protein_coding
miRNA_precursor
small_nucle(ol)ar_rna
protein_coding
ribosomal_rna
protein_coding
protein_coding
protein_coding
protein_coding
protein_coding
protein_coding
small_nucle(ol)ar_rna
ncRNA
protein_coding
protein_coding
protein_coding
protein_coding
small_nucle(ol)ar_rna
protein_coding
protein_coding
protein_coding
protein_coding
small_nucle(ol)ar_rna
protein_coding
protein_coding
protein_coding
protein_coding
protein_coding
protein_coding
protein_coding
protein_coding
protein_coding
protein_coding
protein_coding
small_nucle(ol)ar_rna
protein_coding
small_nucle(ol)ar_rna
small_nucle(ol)ar_rna

ncRNA
protein_coding
protein_coding
protein_coding
protein_coding
small_nucle(ol)ar_rna
protein_coding
protein_coding
protein_coding
protein_coding
protein_coding
protein_coding
protein_coding
protein_coding
protein_coding
protein_coding
protein_coding
ncRNA
protein_coding
protein_coding
small_nucle(ol)ar_rna
protein_coding
protein_coding
small_nucle(ol)ar_rna
protein_coding
protein_coding
protein_coding
protein_coding
protein_coding
protein_coding
protein_coding
protein_coding
protein_coding
protein_coding
protein_coding
protein_coding
protein_coding
miRNA_precursor
protein_coding
protein_coding
protein_coding
protein_coding
protein_coding
protein_coding
protein_coding
protein_coding
protein_coding
protein_coding
protein_coding
protein_coding
ncRNA
protein_coding
small_nucle(ol)ar_rna
protein_coding
protein_coding
small_nucle(ol)ar_rna

protein_coding
protein_coding
protein_coding
ncRNA
small_nucle(ol)ar_rna
protein_coding
protein_coding
protein_coding
protein_coding
protein_coding
protein_coding
protein_coding
protein_coding
miRNA_precursor
protein_coding
protein_coding
protein_coding
protein_coding
protein_coding
protein_coding
protein_coding
protein_coding
protein_coding
protein_coding
protein_coding
protein_coding
protein_coding
protein_coding
protein_coding
protein_coding
protein_coding
ncRNA
protein_coding
ncRNA
protein_coding
protein_coding
protein_coding
protein_coding
protein_coding
protein_coding
small_nucle(ol)ar_rna
protein_coding
protein_coding
protein_coding
protein_coding
small_nucle(ol)ar_rna
protein_coding
protein_coding
protein_coding
protein_coding
protein_coding
protein_coding
protein_coding
miRNA_precursor
ncRNA
protein_coding

miRNA_precursor
ncRNA
protein_coding
protein_coding
protein_coding
small_nucle(ol)ar_rna
protein_coding
protein_coding
protein_coding
protein_coding
protein_coding
protein_coding
protein_coding
protein_coding
miRNA_precursor
miRNA_precursor
protein_coding
protein_coding
protein_coding
protein_coding
protein_coding
protein_coding
protein_coding
protein_coding
protein_coding
ncRNA
protein_coding
protein_coding
protein_coding
protein_coding
protein_coding
protein_coding
protein_coding
ncRNA
protein_coding
protein_coding
protein_coding
protein_coding
protein_coding
protein_coding
protein_coding
protein_coding
protein_coding
ncRNA
protein_coding
protein_coding
ncRNA
protein_coding
protein_coding
protein_coding
protein_coding
protein_coding
protein_coding
protein_coding
protein_coding
protein_coding

protein_coding
protein_coding
protein_coding
protein_coding
protein_coding
protein_coding
protein_coding
protein_coding
protein_coding
protein_coding
ncRNA
protein_coding
protein_coding
protein_coding
protein_coding
protein_coding
protein_coding
protein_coding
protein_coding
protein_coding
protein_coding
small_nucle(ol)ar_rna
small_nucle(ol)ar_rna
small_nucle(ol)ar_rna
small_nucle(ol)ar_rna
protein_coding
protein_coding
protein_coding
protein_coding
protein_coding
ncRNA
protein_coding
ncRNA
protein_coding
protein_coding
protein_coding
small_nucle(ol)ar_rna
small_nucle(ol)ar_rna
protein_coding
protein_coding
protein_coding
protein_coding
protein_coding
protein_coding
protein_coding
protein_coding
protein_coding
protein_coding
protein_coding
protein_coding
protein_coding
protein_coding
ncRNA
protein_coding
miRNA_precursor
protein_coding

protein_coding
protein_coding
protein_coding
protein_coding
pre_trna
ncRNA
protein_coding
protein_coding
protein_coding
protein_coding
protein_coding
protein_coding
protein_coding
protein_coding
protein_coding
protein_coding
protein_coding
protein_coding
protein_coding
protein_coding
protein_coding
ncRNA
protein_coding
protein_coding
protein_coding
protein_coding
protein_coding
protein_coding
protein_coding
protein_coding
protein_coding
protein_coding
protein_coding
ncRNA
protein_coding
protein_coding
protein_coding
protein_coding
protein_coding
protein_coding
protein_coding
protein_coding
protein_coding
protein_coding
protein_coding
protein_coding
protein_coding
protein_coding
protein_coding
protein_coding
protein_coding
protein_coding
protein_coding
protein_coding
protein_coding
protein_coding

protein_coding
protein_coding
protein_coding
protein_coding
protein_coding
protein_coding
protein_coding
protein_coding
protein_coding
protein_coding
protein_coding
protein_coding
protein_coding
protein_coding
protein_coding
protein_coding
ncRNA
protein_coding
protein_coding
protein_coding
protein_coding
protein_coding
protein_coding
protein_coding
protein_coding
protein_coding
protein_coding
protein_coding
protein_coding
protein_coding
protein_coding
protein_coding
protein_coding
protein_coding
protein_coding
protein_coding
protein_coding
protein_coding
protein_coding
protein_coding
protein_coding
protein_coding
protein_coding
protein_coding
protein_coding
protein_coding
protein_coding
protein_coding
protein_coding
protein_coding
protein_coding
protein_coding
protein_coding
protein_coding
protein_coding
protein_coding

protein_coding
protein_coding
protein_coding
protein_coding
protein_coding
protein_coding
protein_coding
protein_coding
protein_coding
protein_coding
protein_coding
protein_coding
protein_coding
protein_coding
protein_coding
protein_coding
protein_coding
small_nucle(ol)ar_rna
protein_coding
protein_coding
protein_coding
protein_coding
protein_coding
protein_coding
protein_coding
protein_coding
protein_coding
protein_coding
protein_coding
protein_coding
protein_coding
protein_coding
protein_coding
protein_coding
protein_coding
protein_coding
protein_coding
protein_coding
protein_coding
protein_coding
protein_coding
protein_coding
protein_coding
protein_coding
protein_coding
protein_coding
protein_coding
protein_coding
protein_coding
protein_coding
pre_trna
protein_coding
protein_coding
protein_coding
protein_coding
ncRNA

protein_coding
protein_coding
protein_coding
protein_coding
protein_coding
protein_coding
protein_coding
protein_coding
protein_coding
protein_coding
protein_coding
protein_coding
protein_coding
protein_coding
protein_coding
protein_coding
protein_coding
protein_coding
protein_coding
protein_coding
protein_coding
protein_coding
protein_coding
protein_coding
pre_trna
ncRNA
protein_coding
protein_coding
protein_coding
protein_coding
protein_coding
protein_coding
protein_coding
protein_coding
protein_coding
protein_coding
protein_coding
protein_coding
protein_coding
protein_coding
protein_coding
protein_coding
protein_coding
protein_coding
protein_coding
protein_coding
protein_coding
protein_coding
protein_coding
protein_coding
protein_coding
protein_coding
protein_coding
protein_coding
protein_coding
protein_coding

protein_coding
protein_coding
protein_coding
protein_coding
protein_coding
protein_coding
protein_coding
protein_coding
ncRNA
protein_coding
protein_coding
protein_coding
protein_coding
protein_coding
protein_coding
protein_coding
protein_coding
protein_coding
protein_coding
pre_trna
protein_coding
protein_coding
protein_coding
protein_coding
protein_coding
protein_coding
protein_coding
protein_coding
protein_coding
protein_coding
protein_coding
protein_coding
protein_coding
protein_coding
protein_coding
protein_coding
protein_coding
protein_coding
protein_coding
protein_coding
protein_coding
protein_coding
protein_coding
protein_coding
protein_coding
protein_coding
pre_trna
protein_coding
miRNA_precursor
protein_coding
protein_coding
protein_coding
protein_coding
protein_coding
protein_coding
protein_coding

protein_coding
protein_coding
protein_coding
protein_coding
small_nucle(ol)ar_rna
miRNA_precursor
protein_coding
ncRNA
protein_coding
protein_coding
protein_coding
protein_coding
protein_coding
protein_coding
protein_coding
protein_coding
ncRNA
protein_coding
protein_coding
protein_coding
protein_coding
protein_coding
miRNA_precursor
protein_coding
protein_coding
protein_coding
protein_coding
protein_coding
protein_coding
protein_coding
protein_coding
protein_coding
protein_coding
protein_coding
protein_coding
protein_coding
protein_coding
protein_coding
protein_coding
protein_coding
protein_coding
protein_coding
protein_coding
protein_coding
protein_coding
protein_coding
protein_coding
protein_coding
protein_coding
protein_coding
protein_coding
protein_coding
protein_coding
protein_coding
protein_coding
protein_coding

protein_coding
protein_coding
protein_coding
protein_coding
protein_coding
protein_coding
protein_coding
protein_coding
protein_coding
protein_coding
protein_coding
protein_coding
ncRNA
protein_coding
protein_coding
protein_coding
protein_coding
protein_coding
protein_coding
protein_coding
protein_coding
protein_coding
protein_coding
protein_coding
protein_coding
protein_coding
protein_coding
protein_coding
protein_coding
protein_coding
protein_coding
ncRNA
protein_coding
protein_coding
protein_coding
protein_coding
protein_coding
protein_coding
protein_coding
protein_coding
protein_coding
protein_coding
protein_coding
protein_coding
protein_coding
protein_coding
protein_coding
protein_coding
protein_coding
protein_coding
protein_coding
protein_coding
protein_coding
protein_coding
protein_coding
protein_coding

protein_coding
protein_coding
protein_coding
protein_coding
protein_coding
protein_coding
protein_coding
protein_coding
protein_coding
protein_coding
protein_coding
protein_coding
protein_coding
protein_coding
protein_coding
protein_coding
protein_coding
protein_coding
protein_coding
protein_coding
protein_coding
protein_coding
protein_coding
small_nucle(ol)ar_rna
protein_coding
protein_coding
protein_coding
protein_coding
ncRNA
protein_coding
protein_coding
protein_coding
protein_coding
protein_coding
protein_coding
protein_coding
protein_coding
protein_coding
protein_coding
protein_coding
protein_coding
protein_coding
protein_coding
ncRNA
protein_coding
protein_coding
protein_coding
pre_trna
protein_coding
protein_coding
protein_coding
protein_coding
protein_coding
protein_coding
protein_coding
protein_coding

small_nucle(ol)ar_rna
small_nucle(ol)ar_rna
small_nucle(ol)ar_rna
small_nucle(ol)ar_rna
protein_coding
protein_coding
protein_coding
protein_coding
protein_coding
protein_coding
protein_coding
protein_coding
protein_coding
protein_coding
protein_coding
protein_coding
pre_trna
protein_coding
protein_coding
protein_coding
protein_coding
protein_coding
protein_coding
protein_coding
protein_coding
protein_coding
protein_coding
protein_coding
ncRNA
protein_coding
protein_coding
protein_coding
protein_coding
protein_coding
protein_coding
protein_coding
protein_coding
ncRNA
protein_coding
protein_coding
protein_coding
protein_coding
protein_coding
protein_coding
protein_coding
protein_coding
protein_coding
protein_coding
protein_coding
protein_coding
protein_coding
small_nucle(ol)ar_rna
small_nucle(ol)ar_rna
protein_coding
protein_coding
protein_coding

miRNA_precursor
protein_coding
protein_coding
miRNA_precursor
protein_coding
protein_coding
protein_coding
protein_coding
protein_coding
protein_coding
protein_coding
protein_coding
protein_coding
protein_coding
protein_coding
protein_coding
protein_coding
protein_coding
protein_coding
protein_coding
protein_coding
protein_coding
protein_coding
protein_coding
protein_coding
protein_coding
protein_coding
protein_coding
ncRNA
protein_coding
protein_coding
protein_coding
protein_coding
protein_coding
protein_coding
protein_coding
protein_coding
protein_coding
ncRNA
protein_coding
protein_coding
protein_coding
protein_coding
protein_coding
protein_coding
protein_coding
protein_coding
ncRNA
protein_coding
protein_coding
protein_coding
protein_coding
ncRNA
protein_coding
protein_coding
protein_coding

protein_coding
protein_coding
protein_coding
protein_coding
protein_coding
protein_coding
protein_coding
protein_coding
protein_coding
protein_coding
protein_coding
protein_coding
protein_coding
ncRNA
protein_coding
protein_coding
ncRNA
protein_coding
protein_coding
ncRNA
protein_coding
protein_coding
protein_coding
protein_coding
protein_coding
ncRNA
protein_coding
protein_coding
protein_coding
miRNA_precursor
protein_coding
protein_coding
protein_coding
protein_coding
protein_coding
protein_coding
protein_coding
protein_coding
protein_coding
protein_coding
protein_coding
protein_coding
protein_coding
protein_coding
protein_coding
protein_coding
protein_coding
protein_coding
protein_coding
protein_coding
protein_coding
protein_coding
protein_coding
protein_coding
protein_coding
protein_coding

protein_coding
protein_coding
protein_coding
protein_coding
ncRNA
protein_coding
ncRNA
protein_coding
protein_coding
miRNA_precursor
protein_coding
protein_coding
protein_coding
protein_coding
protein_coding
protein_coding
protein_coding
protein_coding
protein_coding
protein_coding
protein_coding
protein_coding
protein_coding
protein_coding
protein_coding
protein_coding
ncRNA
protein_coding
protein_coding
small_nucle(ol)ar_rna
protein_coding
protein_coding
protein_coding
protein_coding
protein_coding
protein_coding
protein_coding
protein_coding
protein_coding
protein_coding
protein_coding
protein_coding
protein_coding
protein_coding
protein_coding
protein_coding
protein_coding
protein_coding
protein_coding
protein_coding
protein_coding
protein_coding
protein_coding
protein_coding
protein_coding
protein_coding

protein_coding
protein_coding
miRNA_precursor
protein_coding
protein_coding
protein_coding
protein_coding
protein_coding
protein_coding
protein_coding
protein_coding
protein_coding
protein_coding
protein_coding
protein_coding
protein_coding
protein_coding
protein_coding
protein_coding
protein_coding
protein_coding
protein_coding
protein_coding
protein_coding
ncRNA
protein_coding
protein_coding
protein_coding
protein_coding
protein_coding
ncRNA
protein_coding
protein_coding
protein_coding
protein_coding
protein_coding
protein_coding
protein_coding
protein_coding
protein_coding
protein_coding
protein_coding
protein_coding
protein_coding
protein_coding
protein_coding
protein_coding
protein_coding
protein_coding
protein_coding
protein_coding
protein_coding
protein_coding
protein_coding
protein_coding
protein_coding

protein_coding
protein_coding
protein_coding
protein_coding
ncRNA
protein_coding
protein_coding
protein_coding
protein_coding
protein_coding
protein_coding
protein_coding
ncRNA
protein_coding
ncRNA
small_nucle(ol)ar_rna
protein_coding
protein_coding
protein_coding
protein_coding
protein_coding
protein_coding
protein_coding
protein_coding
protein_coding
protein_coding
protein_coding
protein_coding
protein_coding
protein_coding
protein_coding
protein_coding
small_nucle(ol)ar_rna
protein_coding
protein_coding
protein_coding
protein_coding
protein_coding
protein_coding
protein_coding
protein_coding
protein_coding
protein_coding
protein_coding
protein_coding
protein_coding
protein_coding
protein_coding
protein_coding
protein_coding
protein_coding
protein_coding
protein_coding
protein_coding
protein_coding
protein_coding

protein_coding
protein_coding
protein_coding
protein_coding
protein_coding
protein_coding
protein_coding
ncRNA
protein_coding
protein_coding
protein_coding
ncRNA
protein_coding
protein_coding
protein_coding
protein_coding
protein_coding
protein_coding
protein_coding
protein_coding
protein_coding
protein_coding
protein_coding
protein_coding
protein_coding
protein_coding
protein_coding
protein_coding
protein_coding
protein_coding
protein_coding
protein_coding
protein_coding
protein_coding
protein_coding
protein_coding
protein_coding
protein_coding
protein_coding
protein_coding
protein_coding
protein_coding
ncRNA
protein_coding
protein_coding
protein_coding
protein_coding
protein_coding
protein_coding
protein_coding
protein_coding
protein_coding
protein_coding
protein_coding
miRNA_precursor
miRNA_precursor

protein_coding
protein_coding
protein_coding
protein_coding
ncRNA
protein_coding
protein_coding
protein_coding
small_nucle(ol)ar_rna
protein_coding
protein_coding
protein_coding
protein_coding
protein_coding
protein_coding
miRNA_precursor
protein_coding
protein_coding
protein_coding
protein_coding
protein_coding
protein_coding
protein_coding
protein_coding
protein_coding
protein_coding
protein_coding
protein_coding
protein_coding
ncRNA
protein_coding
ncRNA
protein_coding
protein_coding
protein_coding
protein_coding
protein_coding
protein_coding
protein_coding
protein_coding
protein_coding
protein_coding
protein_coding
protein_coding
protein_coding
protein_coding
protein_coding
protein_coding
protein_coding
protein_coding
protein_coding
protein_coding
protein_coding
protein_coding
protein_coding
protein_coding

small_nucle(ol)ar_rna
protein_coding
protein_coding
protein_coding
protein_coding
protein_coding
protein_coding
pre_trna
protein_coding
ncRNA
protein_coding
protein_coding
protein_coding
protein_coding
protein_coding
protein_coding
protein_coding
ncRNA
protein_coding
protein_coding
protein_coding
protein_coding
protein_coding
protein_coding
protein_coding
protein_coding
protein_coding
protein_coding
protein_coding
protein_coding
protein_coding
protein_coding
protein_coding
protein_coding
protein_coding
protein_coding
protein_coding
ncRNA
protein_coding
protein_coding
protein_coding
protein_coding
protein_coding
protein_coding
protein_coding
protein_coding
miRNA_precursor
protein_coding
protein_coding
protein_coding
protein_coding
protein_coding
protein_coding
protein_coding
protein_coding
protein_coding

protein_coding
protein_coding
protein_coding
protein_coding
protein_coding
ncRNA
protein_coding
ncRNA
protein_coding
small_nucle(ol)ar_rna
protein_coding
protein_coding
protein_coding
protein_coding
protein_coding
protein_coding
protein_coding
protein_coding
ncRNA
protein_coding
protein_coding
ncRNA
protein_coding
protein_coding
protein_coding
protein_coding
protein_coding
protein_coding
protein_coding
protein_coding
protein_coding
protein_coding
protein_coding
protein_coding
protein_coding
protein_coding
protein_coding
protein_coding
protein_coding
protein_coding
protein_coding
protein_coding
protein_coding
protein_coding
ncRNA
protein_coding
protein_coding
protein_coding
protein_coding
protein_coding
protein_coding
protein_coding
protein_coding
protein_coding
protein_coding
protein_coding

small_nucle(ol)ar_rna
protein_coding
protein_coding
protein_coding
protein_coding
protein_coding
protein_coding
protein_coding
protein_coding
protein_coding
protein_coding
protein_coding
protein_coding
protein_coding
protein_coding
protein_coding
protein_coding
protein_coding
protein_coding
protein_coding
protein_coding
protein_coding
miRNA_precursor
protein_coding
protein_coding
protein_coding
protein_coding
protein_coding
protein_coding
protein_coding
protein_coding
protein_coding
protein_coding
protein_coding
protein_coding
protein_coding
protein_coding
protein_coding
protein_coding
protein_coding
protein_coding
protein_coding
protein_coding
ncRNA
protein_coding
protein_coding
protein_coding
protein_coding
protein_coding
protein_coding
protein_coding
protein_coding
protein_coding
protein_coding
protein_coding
protein_coding

protein_coding
protein_coding
protein_coding
protein_coding
protein_coding
protein_coding
protein_coding
protein_coding
protein_coding
protein_coding
protein_coding
small_nucle(ol)ar_rna
protein_coding
protein_coding
protein_coding
protein_coding
protein_coding
ncRNA
protein_coding
protein_coding
protein_coding
protein_coding
protein_coding
protein_coding
protein_coding
protein_coding
protein_coding
protein_coding
protein_coding
protein_coding
protein_coding
protein_coding
protein_coding
protein_coding
protein_coding
protein_coding
protein_coding
protein_coding
protein_coding
protein_coding
protein_coding
protein_coding
protein_coding
protein_coding
protein_coding
protein_coding
protein_coding
protein_coding
protein_coding
protein_coding
protein_coding
protein_coding
protein_coding
protein_coding
protein_coding
protein_coding

protein_coding
protein_coding
protein_coding
protein_coding
protein_coding
small_nucle(ol)ar_rna
small_nucle(ol)ar_rna
small_nucle(ol)ar_rna
protein_coding
ncRNA
protein_coding
protein_coding
protein_coding
protein_coding
protein_coding
protein_coding
protein_coding
protein_coding
protein_coding
protein_coding
protein_coding
protein_coding
protein_coding
protein_coding
protein_coding
protein_coding
pre_trna
protein_coding
protein_coding
protein_coding
protein_coding
protein_coding
protein_coding
protein_coding
protein_coding
protein_coding
protein_coding
protein_coding
protein_coding
protein_coding
protein_coding
protein_coding
protein_coding
protein_coding
protein_coding
protein_coding
protein_coding
protein_coding
protein_coding
protein_coding
protein_coding
protein_coding
protein_coding
protein_coding
protein_coding
protein_coding

protein_coding
protein_coding
protein_coding
protein_coding
protein_coding
protein_coding
protein_coding
protein_coding
protein_coding
protein_coding
protein_coding
protein_coding
protein_coding
protein_coding
protein_coding
protein_coding
protein_coding
protein_coding
protein_coding
protein_coding
protein_coding
protein_coding
protein_coding
protein_coding
protein_coding
protein_coding
protein_coding
protein_coding
protein_coding
protein_coding
protein_coding
protein_coding
protein_coding
protein_coding
ncRNA
protein_coding
protein_coding
pre_trna
protein_coding
protein_coding
protein_coding
protein_coding
protein_coding
protein_coding
ncRNA
protein_coding
protein_coding
protein_coding
protein_coding
protein_coding
protein_coding
protein_coding
protein_coding
protein_coding
protein_coding
protein_coding

protein_coding
protein_coding
protein_coding
protein_coding
ncRNA
pre_trna
protein_coding
protein_coding
protein_coding
protein_coding
protein_coding
protein_coding
protein_coding
protein_coding
protein_coding
protein_coding
protein_coding
protein_coding
protein_coding
protein_coding
ncRNA
protein_coding
protein_coding
protein_coding
protein_coding
protein_coding
protein_coding
protein_coding
protein_coding
protein_coding
protein_coding
protein_coding
protein_coding
protein_coding
protein_coding
protein_coding
protein_coding
protein_coding
ncRNA
protein_coding
protein_coding
protein_coding
protein_coding
ncRNA
ncRNA
protein_coding
protein_coding
protein_coding
protein_coding
protein_coding
protein_coding
protein_coding
protein_coding
protein_coding
protein_coding
protein_coding

protein_coding
protein_coding
protein_coding
protein_coding
protein_coding
protein_coding
protein_coding
protein_coding
protein_coding
protein_coding
protein_coding
protein_coding
ncRNA
protein_coding
protein_coding
protein_coding
protein_coding
protein_coding
protein_coding
protein_coding
protein_coding
protein_coding
protein_coding
protein_coding
protein_coding
protein_coding
protein_coding
protein_coding
protein_coding
protein_coding
protein_coding
protein_coding
protein_coding
protein_coding
protein_coding
protein_coding
protein_coding
ncRNA
miRNA_precursor
protein_coding
ncRNA
protein_coding
ncRNA
protein_coding
protein_coding
protein_coding
ncRNA
protein_coding
protein_coding
protein_coding
protein_coding
protein_coding
protein_coding
protein_coding
protein_coding
protein_coding

protein_coding
protein_coding
protein_coding
pre_trna
protein_coding
protein_coding
protein_coding
protein_coding
protein_coding
protein_coding
ncRNA
protein_coding
protein_coding
protein_coding
protein_coding
protein_coding
protein_coding
protein_coding
protein_coding
protein_coding
protein_coding
protein_coding
protein_coding
pre_trna
protein_coding
protein_coding
protein_coding
ncRNA
protein_coding
protein_coding
protein_coding
protein_coding
protein_coding
protein_coding
protein_coding
protein_coding
protein_coding
ncRNA
protein_coding
protein_coding
ncRNA
protein_coding
protein_coding
protein_coding
ncRNA
protein_coding
protein_coding
protein_coding
protein_coding
ncRNA
protein_coding
protein_coding
protein_coding
protein_coding
protein_coding
protein_coding

protein_coding
protein_coding
protein_coding
protein_coding
protein_coding
ncRNA
protein_coding
ncRNA
protein_coding
protein_coding
protein_coding
protein_coding
protein_coding
protein_coding
protein_coding
protein_coding
protein_coding
protein_coding
protein_coding
protein_coding
protein_coding
ncRNA
protein_coding
protein_coding
protein_coding
protein_coding
protein_coding
protein_coding
protein_coding
protein_coding
protein_coding
protein_coding
protein_coding
protein_coding
protein_coding
protein_coding
protein_coding
protein_coding
protein_coding
protein_coding
protein_coding
protein_coding
protein_coding
ncRNA
protein_coding
protein_coding
protein_coding
protein_coding
protein_coding
protein_coding
protein_coding
protein_coding
protein_coding
protein_coding
protein_coding
protein_coding

protein_coding
protein_coding
protein_coding
protein_coding
protein_coding
protein_coding
protein_coding
protein_coding
protein_coding
protein_coding
protein_coding
protein_coding
protein_coding
protein_coding
protein_coding
protein_coding
protein_coding
protein_coding
protein_coding
protein_coding
protein_coding
ncRNA
protein_coding
protein_coding
protein_coding
protein_coding
ncRNA
ncRNA
protein_coding
protein_coding
protein_coding
protein_coding
protein_coding
protein_coding
protein_coding
protein_coding
protein_coding
protein_coding
miRNA_precursor
protein_coding
protein_coding
protein_coding
ncRNA
ncRNA
protein_coding
protein_coding
protein_coding
protein_coding
protein_coding
protein_coding
protein_coding
protein_coding
protein_coding
protein_coding
protein_coding
protein_coding

protein_coding
protein_coding
protein_coding
protein_coding
protein_coding
protein_coding
protein_coding
protein_coding
protein_coding
protein_coding
ncRNA
protein_coding
protein_coding
protein_coding
protein_coding
protein_coding
protein_coding
protein_coding
protein_coding
ncRNA
protein_coding
protein_coding
protein_coding
protein_coding
protein_coding
protein_coding
protein_coding
protein_coding
protein_coding
protein_coding
protein_coding
protein_coding
protein_coding
ncRNA
protein_coding
ncRNA
protein_coding
protein_coding
protein_coding
protein_coding
protein_coding
protein_coding
protein_coding
protein_coding
protein_coding
ncRNA
protein_coding
protein_coding
ncRNA
protein_coding
protein_coding
protein_coding
protein_coding
protein_coding
protein_coding
protein_coding

miRNA_precursor
protein_coding
protein_coding
protein_coding
miRNA_precursor
ncRNA
protein_coding
protein_coding
protein_coding
protein_coding
protein_coding
protein_coding
protein_coding
protein_coding
protein_coding
miRNA_precursor
protein_coding
protein_coding
pre_trna
protein_coding
ncRNA
ncRNA
protein_coding
protein_coding
ncRNA
protein_coding
protein_coding
protein_coding
protein_coding
protein_coding
protein_coding
protein_coding
protein_coding
protein_coding
protein_coding
protein_coding
protein_coding
protein_coding
protein_coding
protein_coding
protein_coding
protein_coding
protein_coding
protein_coding
protein_coding
protein_coding
protein_coding
protein_coding
protein_coding
protein_coding
protein_coding
protein_coding
protein_coding
protein_coding
protein_coding
protein_coding

ncRNA
protein_coding
protein_coding
protein_coding
protein_coding
protein_coding
protein_coding
protein_coding
protein_coding
protein_coding
protein_coding
protein_coding
protein_coding
protein_coding
protein_coding
protein_coding
protein_coding
protein_coding
protein_coding
protein_coding
protein_coding
protein_coding
protein_coding
protein_coding
protein_coding
protein_coding
protein_coding
protein_coding
protein_coding
protein_coding
protein_coding
protein_coding
protein_coding
protein_coding
protein_coding
protein_coding
protein_coding
protein_coding
protein_coding
protein_coding
protein_coding
protein_coding
pre_trna
protein_coding
protein_coding
protein_coding
protein_coding
protein_coding
protein_coding
protein_coding
protein_coding
protein_coding
protein_coding
protein_coding
protein_coding
protein_coding

protein_coding
protein_coding
protein_coding
protein_coding
protein_coding
protein_coding
protein_coding
protein_coding
protein_coding
protein_coding
pre_trna
small_nucle(ol)ar_rna
protein_coding
ncRNA
protein_coding
protein_coding
protein_coding
protein_coding
protein_coding
protein_coding
protein_coding
protein_coding
protein_coding
protein_coding
protein_coding
protein_coding
protein_coding
protein_coding
protein_coding
protein_coding
protein_coding
protein_coding
protein_coding
ncRNA
protein_coding
ncRNA
protein_coding
protein_coding
protein_coding
protein_coding
protein_coding
protein_coding
protein_coding
protein_coding
protein_coding
protein_coding
protein_coding
protein_coding
protein_coding
protein_coding
protein_coding
miRNA_precursor
protein_coding
protein_coding
protein_coding
protein_coding

protein_coding
protein_coding
protein_coding
protein_coding
protein_coding
protein_coding
protein_coding
small_nucle(ol)ar_rna
protein_coding
protein_coding
protein_coding
protein_coding
protein_coding
protein_coding
pre_trna
protein_coding
protein_coding
protein_coding
protein_coding
protein_coding
pre_trna
protein_coding
protein_coding
protein_coding
protein_coding
protein_coding
protein_coding
protein_coding
protein_coding
protein_coding
protein_coding
protein_coding
protein_coding
protein_coding
protein_coding
protein_coding
protein_coding
protein_coding
protein_coding
protein_coding
ncRNA
protein_coding
protein_coding
protein_coding
protein_coding
protein_coding
protein_coding
protein_coding
protein_coding
protein_coding
protein_coding
protein_coding
protein_coding
protein_coding
protein_coding
protein_coding

protein_coding
protein_coding
protein_coding
protein_coding
pre_trna
protein_coding
protein_coding
pre_trna
protein_coding
ncRNA
protein_coding
protein_coding
protein_coding
protein_coding
protein_coding
protein_coding
protein_coding
protein_coding
protein_coding
protein_coding
protein_coding
small_nucle(ol)ar_rna
protein_coding
protein_coding
ncRNA
protein_coding
protein_coding
protein_coding
ncRNA
protein_coding
protein_coding
protein_coding
protein_coding
protein_coding
protein_coding
protein_coding
protein_coding
protein_coding
protein_coding
protein_coding
protein_coding
protein_coding
protein_coding
protein_coding
protein_coding
protein_coding
protein_coding
protein_coding
protein_coding
protein_coding
protein_coding
protein_coding
protein_coding
protein_coding
protein_coding
protein_coding
protein_coding

protein_coding
protein_coding
protein_coding
protein_coding
protein_coding
protein_coding
protein_coding
protein_coding
protein_coding
protein_coding
protein_coding
protein_coding
protein_coding
protein_coding
protein_coding
protein_coding
protein_coding
protein_coding
protein_coding
protein_coding
protein_coding
protein_coding
protein_coding
protein_coding
protein_coding
miRNA_precursor
protein_coding
protein_coding
protein_coding
protein_coding
protein_coding
protein_coding
protein_coding
protein_coding
protein_coding
protein_coding
protein_coding
protein_coding
protein_coding
protein_coding
protein_coding
protein_coding
protein_coding
protein_coding
protein_coding
protein_coding
ncRNA
protein_coding
protein_coding
protein_coding
protein_coding
protein_coding
protein_coding
protein_coding
protein_coding
protein_coding

protein_coding
protein_coding
protein_coding
protein_coding
protein_coding
protein_coding
protein_coding
protein_coding
protein_coding
protein_coding
protein_coding
protein_coding
protein_coding
protein_coding
protein_coding
protein_coding
protein_coding
protein_coding
protein_coding
protein_coding
protein_coding
protein_coding
protein_coding
protein_coding
protein_coding
small_nucle(ol)ar_rna
small_nucle(ol)ar_rna
small_nucle(ol)ar_rna
protein_coding
protein_coding
protein_coding
protein_coding
ncRNA
protein_coding
protein_coding
protein_coding
protein_coding
protein_coding
protein_coding
protein_coding
protein_coding
protein_coding
protein_coding
protein_coding
protein_coding
protein_coding
protein_coding
protein_coding
protein_coding
protein_coding
protein_coding
protein_coding
protein_coding
protein_coding
protein_coding
protein_coding

protein_coding
protein_coding
protein_coding
protein_coding
protein_coding
protein_coding
protein_coding
protein_coding
protein_coding
protein_coding
protein_coding
protein_coding
protein_coding
protein_coding
protein_coding
pre_trna
protein_coding
protein_coding
protein_coding
protein_coding
protein_coding
protein_coding
pre_trna
protein_coding
protein_coding
protein_coding
ncRNA
protein_coding
protein_coding
protein_coding
protein_coding
protein_coding
protein_coding
protein_coding
protein_coding
protein_coding
protein_coding
protein_coding
protein_coding
protein_coding
small_nucle(ol)ar_rna
small_nucle(ol)ar_rna
protein_coding
protein_coding
protein_coding
protein_coding
protein_coding
protein_coding
protein_coding
protein_coding
protein_coding
protein_coding
protein_coding
protein_coding
protein_coding
protein_coding

protein_coding
protein_coding
protein_coding
protein_coding
protein_coding
protein_coding
protein_coding
protein_coding
protein_coding
protein_coding
protein_coding
small_nucle(ol)ar_rna
protein_coding
protein_coding
protein_coding
protein_coding
protein_coding
protein_coding
protein_coding
protein_coding
protein_coding
protein_coding
protein_coding
protein_coding
protein_coding
protein_coding
protein_coding
protein_coding
protein_coding
protein_coding
protein_coding
protein_coding
protein_coding
protein_coding
protein_coding
protein_coding
protein_coding
protein_coding
protein_coding
protein_coding
protein_coding
protein_coding
protein_coding
protein_coding
protein_coding
protein_coding
protein_coding
protein_coding
protein_coding
protein_coding
protein_coding
protein_coding
protein_coding
protein_coding
protein_coding
protein_coding

protein_coding
protein_coding
protein_coding
protein_coding
protein_coding
protein_coding
protein_coding
protein_coding
protein_coding
protein_coding
protein_coding
protein_coding
protein_coding
protein_coding
protein_coding
protein_coding
protein_coding
protein_coding
protein_coding
protein_coding
protein_coding
protein_coding
protein_coding
ncRNA
small_nucle(ol)ar_rna
protein_coding
protein_coding
protein_coding
protein_coding
protein_coding
protein_coding
protein_coding
protein_coding
protein_coding
protein_coding
protein_coding
protein_coding
protein_coding
protein_coding
protein_coding
protein_coding
protein_coding
protein_coding
protein_coding
protein_coding
protein_coding
protein_coding
protein_coding
protein_coding
protein_coding
protein_coding
protein_coding
protein_coding
protein_coding
protein_coding
protein_coding

ncRNA
ncRNA
protein_coding
protein_coding
protein_coding
protein_coding
protein_coding
protein_coding
protein_coding
protein_coding
protein_coding
protein_coding
protein_coding
protein_coding
protein_coding
protein_coding
protein_coding
protein_coding
protein_coding
protein_coding
protein_coding
protein_coding
ncRNA
ncRNA
protein_coding
protein_coding
protein_coding
protein_coding
protein_coding
protein_coding
protein_coding
protein_coding
protein_coding
protein_coding
protein_coding
protein_coding
protein_coding
protein_coding
protein_coding
protein_coding
protein_coding
protein_coding
protein_coding
protein_coding
protein_coding
protein_coding
protein_coding
protein_coding
protein_coding
protein_coding
protein_coding
protein_coding
protein_coding
protein_coding
protein_coding
protein_coding

protein_coding
protein_coding
protein_coding
protein_coding
protein_coding
protein_coding
protein_coding
protein_coding
protein_coding
protein_coding
protein_coding
protein_coding
protein_coding
protein_coding
protein_coding
protein_coding
protein_coding
protein_coding
protein_coding
protein_coding
protein_coding
protein_coding
protein_coding
protein_coding
protein_coding
protein_coding
ncRNA
protein_coding
protein_coding
protein_coding
protein_coding
protein_coding
protein_coding
protein_coding
protein_coding
protein_coding
protein_coding
protein_coding
protein_coding
protein_coding
protein_coding
protein_coding
protein_coding
protein_coding
protein_coding
protein_coding
protein_coding
protein_coding
protein_coding
protein_coding
protein_coding
protein_coding
protein_coding
protein_coding
protein_coding
protein_coding

protein_coding
protein_coding
protein_coding
protein_coding
protein_coding
protein_coding
protein_coding
protein_coding
protein_coding
protein_coding
protein_coding
protein_coding
protein_coding
protein_coding
protein_coding
protein_coding
protein_coding
protein_coding
protein_coding
protein_coding
protein_coding
protein_coding
protein_coding
ncRNA
protein_coding
protein_coding
protein_coding
protein_coding
protein_coding
protein_coding
protein_coding
protein_coding
protein_coding
protein_coding
protein_coding
protein_coding
protein_coding
protein_coding
protein_coding
protein_coding
protein_coding
ncRNA
protein_coding
protein_coding
protein_coding
protein_coding
protein_coding
protein_coding
protein_coding
protein_coding
protein_coding
protein_coding
protein_coding
protein_coding
protein_coding
protein_coding

protein_coding
protein_coding
protein_coding
protein_coding
protein_coding
small_nucle(ol)ar_rna
protein_coding
protein_coding
protein_coding
protein_coding
protein_coding
protein_coding
protein_coding
protein_coding
protein_coding
protein_coding
protein_coding
protein_coding
protein_coding
protein_coding
protein_coding
protein_coding
protein_coding
protein_coding
protein_coding
protein_coding
protein_coding
protein_coding
protein_coding
protein_coding
protein_coding
protein_coding
protein_coding
protein_coding
protein_coding
protein_coding
protein_coding
protein_coding
protein_coding
protein_coding
protein_coding
protein_coding
protein_coding
protein_coding
protein_coding
protein_coding
protein_coding
protein_coding
protein_coding
protein_coding
protein_coding
protein_coding
protein_coding
ncRNA
protein_coding
protein_coding

protein_coding
protein_coding
protein_coding
protein_coding
protein_coding
protein_coding
protein_coding
protein_coding
protein_coding
protein_coding
protein_coding
protein_coding
protein_coding
protein_coding
protein_coding
protein_coding
protein_coding
protein_coding
protein_coding
protein_coding
protein_coding
protein_coding
protein_coding
protein_coding
protein_coding
protein_coding
protein_coding
protein_coding
protein_coding
protein_coding
ncRNA
protein_coding
protein_coding
protein_coding
protein_coding
protein_coding
protein_coding
protein_coding
protein_coding
protein_coding
protein_coding
protein_coding
protein_coding
protein_coding
protein_coding
ncRNA
protein_coding
protein_coding
protein_coding
protein_coding
protein_coding
protein_coding
protein_coding
protein_coding
protein_coding
protein_coding

protein_coding
protein_coding
protein_coding
protein_coding
protein_coding
protein_coding
miRNA_precursor
protein_coding
protein_coding
protein_coding
protein_coding
protein_coding
protein_coding
protein_coding
protein_coding
protein_coding
protein_coding
protein_coding
miRNA_precursor
protein_coding
protein_coding
protein_coding
protein_coding
protein_coding
protein_coding
protein_coding
protein_coding
protein_coding
protein_coding
protein_coding
protein_coding
protein_coding
protein_coding
protein_coding
protein_coding
protein_coding
protein_coding
protein_coding
protein_coding
protein_coding
protein_coding
protein_coding
protein_coding
protein_coding
protein_coding
protein_coding
protein_coding
protein_coding
protein_coding
protein_coding
protein_coding
protein_coding
protein_coding
protein_coding
protein_coding
protein_coding

protein_coding
ncRNA
protein_coding
protein_coding
protein_coding
protein_coding
protein_coding
protein_coding
protein_coding
protein_coding
protein_coding
protein_coding
protein_coding
protein_coding
protein_coding
protein_coding
protein_coding
protein_coding
small_nucle(ol)ar_rna
protein_coding
ncRNA
protein_coding
protein_coding
protein_coding
protein_coding
ncRNA
protein_coding
protein_coding
protein_coding
protein_coding
protein_coding
protein_coding
protein_coding
protein_coding
protein_coding
protein_coding
protein_coding
protein_coding
protein_coding
protein_coding
protein_coding
protein_coding
ncRNA
protein_coding
protein_coding
protein_coding
protein_coding
protein_coding
protein_coding
protein_coding
protein_coding
protein_coding
protein_coding
protein_coding
protein_coding
protein_coding

protein_coding
protein_coding
protein_coding
protein_coding
protein_coding
protein_coding
protein_coding
protein_coding
protein_coding
protein_coding
protein_coding
protein_coding
protein_coding
protein_coding
protein_coding
protein_coding
protein_coding
protein_coding
protein_coding
protein_coding
protein_coding
protein_coding
protein_coding
protein_coding
protein_coding
protein_coding
protein_coding
protein_coding
protein_coding
protein_coding
protein_coding
protein_coding
protein_coding
ncRNA
protein_coding
protein_coding
protein_coding
protein_coding
protein_coding
protein_coding
protein_coding
protein_coding
protein_coding
protein_coding
protein_coding
protein_coding
protein_coding
ncRNA
small_nucle(ol)ar_rna
protein_coding
protein_coding
protein_coding
protein_coding
protein_coding
protein_coding
protein_coding

protein_coding
protein_coding
protein_coding
protein_coding
protein_coding
protein_coding
protein_coding
protein_coding
protein_coding
protein_coding
protein_coding
protein_coding
protein_coding
protein_coding
protein_coding
protein_coding
protein_coding
protein_coding
protein_coding
protein_coding
protein_coding
protein_coding
protein_coding
protein_coding
protein_coding
pre_trna
protein_coding
ncRNA
protein_coding
protein_coding
protein_coding
protein_coding
protein_coding
protein_coding
protein_coding
protein_coding
protein_coding
protein_coding
protein_coding
protein_coding
protein_coding
protein_coding
protein_coding
protein_coding
protein_coding
protein_coding
protein_coding
protein_coding
protein_coding
miRNA_precursor
protein_coding
protein_coding
protein_coding
protein_coding
protein_coding
protein_coding

protein_coding
protein_coding
protein_coding
protein_coding
ncRNA
protein_coding
protein_coding
protein_coding
protein_coding
protein_coding
protein_coding
protein_coding
small_nucle(ol)ar_rna
protein_coding
protein_coding
protein_coding
protein_coding
protein_coding
protein_coding
protein_coding
ncRNA
protein_coding
protein_coding
protein_coding
protein_coding
protein_coding
protein_coding
protein_coding
protein_coding
protein_coding
protein_coding
protein_coding
protein_coding
protein_coding
protein_coding
protein_coding
protein_coding
protein_coding
protein_coding
protein_coding
protein_coding
protein_coding
protein_coding
protein_coding
protein_coding
ncRNA
protein_coding
protein_coding
protein_coding
protein_coding
protein_coding
ncRNA
protein_coding
ncRNA
protein_coding
protein_coding
ncRNA

protein_coding
protein_coding
protein_coding
protein_coding
protein_coding
protein_coding
protein_coding
protein_coding
protein_coding
protein_coding
protein_coding
protein_coding
protein_coding
protein_coding
protein_coding
protein_coding
protein_coding
protein_coding
protein_coding
ncRNA
protein_coding
protein_coding
protein_coding
protein_coding
protein_coding
protein_coding
protein_coding
protein_coding
protein_coding
protein_coding
protein_coding
protein_coding
protein_coding
protein_coding
pre_trna
protein_coding
protein_coding
protein_coding
protein_coding
protein_coding
protein_coding
protein_coding
protein_coding
protein_coding
protein_coding
protein_coding
ncRNA
protein_coding
ncRNA
ncRNA
ncRNA
ncRNA
protein_coding
protein_coding
protein_coding
miRNA_precursor

protein_coding
protein_coding
small_nucle(ol)ar_rna
protein_coding
protein_coding
protein_coding
protein_coding
protein_coding
protein_coding
protein_coding
ncRNA
ncRNA
protein_coding
protein_coding
protein_coding
protein_coding
protein_coding
protein_coding
protein_coding
protein_coding
protein_coding
protein_coding
protein_coding
protein_coding
protein_coding
protein_coding
protein_coding
protein_coding
protein_coding
protein_coding
protein_coding
protein_coding
protein_coding
protein_coding
protein_coding
protein_coding
ncRNA
protein_coding
protein_coding
protein_coding
protein_coding
protein_coding
protein_coding
protein_coding
protein_coding
protein_coding
protein_coding
protein_coding
protein_coding
protein_coding
protein_coding
protein_coding
protein_coding
protein_coding
protein_coding
protein_coding

protein_coding
protein_coding
protein_coding
protein_coding
protein_coding
protein_coding
protein_coding
protein_coding
protein_coding
protein_coding
protein_coding
protein_coding
protein_coding
protein_coding
protein_coding
protein_coding
protein_coding
protein_coding
protein_coding
protein_coding
protein_coding
protein_coding
protein_coding
protein_coding
protein_coding
protein_coding
protein_coding
protein_coding
protein_coding
protein_coding
protein_coding
protein_coding
protein_coding
protein_coding
protein_coding
protein_coding
protein_coding
protein_coding
protein_coding
protein_coding
protein_coding
protein_coding
protein_coding
protein_coding
protein_coding
protein_coding
protein_coding
protein_coding
protein_coding
protein_coding
protein_coding
protein_coding
protein_coding
small_nucle(ol)ar_rna
protein_coding
protein_coding

protein_coding
miRNA_precursor
ncRNA
protein_coding
protein_coding
protein_coding
protein_coding
protein_coding
protein_coding
protein_coding
protein_coding
protein_coding
protein_coding
protein_coding
protein_coding
protein_coding
protein_coding
protein_coding
protein_coding
protein_coding
protein_coding
protein_coding
protein_coding
protein_coding
protein_coding
protein_coding
protein_coding
protein_coding
ncRNA
protein_coding
protein_coding
protein_coding
protein_coding
ncRNA
protein_coding
protein_coding
protein_coding
protein_coding
protein_coding
protein_coding
protein_coding
protein_coding
protein_coding
protein_coding
protein_coding
protein_coding
protein_coding
protein_coding
protein_coding
ncRNA
small_nucle(ol)ar_rna
protein_coding
protein_coding
protein_coding
protein_coding
protein_coding

protein_coding
protein_coding
protein_coding
protein_coding
protein_coding
protein_coding
protein_coding
protein_coding
protein_coding
protein_coding
protein_coding
ncRNA
protein_coding
protein_coding
protein_coding
protein_coding
protein_coding
protein_coding
protein_coding
protein_coding
protein_coding
protein_coding
ncRNA
protein_coding
protein_coding
protein_coding
protein_coding
protein_coding
protein_coding
ncRNA
protein_coding
protein_coding
protein_coding
protein_coding
protein_coding
protein_coding
protein_coding
protein_coding
protein_coding
ncRNA
protein_coding
protein_coding
protein_coding
protein_coding
protein_coding
protein_coding
protein_coding
protein_coding
protein_coding
protein_coding
protein_coding
protein_coding
protein_coding
protein_coding
protein_coding
protein_coding

protein_coding
ncRNA
protein_coding
protein_coding
protein_coding
protein_coding
protein_coding
protein_coding
protein_coding
protein_coding
protein_coding
protein_coding
protein_coding
protein_coding
protein_coding
protein_coding
protein_coding
protein_coding
protein_coding
protein_coding
protein_coding
protein_coding
protein_coding
protein_coding
protein_coding
protein_coding
protein_coding
protein_coding
protein_coding
protein_coding
protein_coding
protein_coding
protein_coding
protein_coding
protein_coding
protein_coding
protein_coding
protein_coding
protein_coding
miRNA_precursor
protein_coding
ncRNA
protein_coding
protein_coding
protein_coding
protein_coding
protein_coding
protein_coding
protein_coding
protein_coding
protein_coding
protein_coding
protein_coding
protein_coding
protein_coding
protein_coding

protein_coding
protein_coding
protein_coding
protein_coding
protein_coding
protein_coding
pre_trna
protein_coding
protein_coding
protein_coding
protein_coding
protein_coding
protein_coding
protein_coding
pre_trna
protein_coding
protein_coding
protein_coding
protein_coding
protein_coding
protein_coding
ncRNA
ncRNA
protein_coding
protein_coding
protein_coding
protein_coding
protein_coding
protein_coding
protein_coding
protein_coding
protein_coding
protein_coding
protein_coding
protein_coding
protein_coding
protein_coding
protein_coding
protein_coding
protein_coding
protein_coding
protein_coding
ncRNA
protein_coding
protein_coding
protein_coding
protein_coding
protein_coding
protein_coding
protein_coding
protein_coding
protein_coding
protein_coding
protein_coding
protein_coding
protein_coding

protein_coding
protein_coding
protein_coding
protein_coding
protein_coding
protein_coding
protein_coding
protein_coding
protein_coding
protein_coding
protein_coding
protein_coding
protein_coding
protein_coding
protein_coding
protein_coding
protein_coding
protein_coding
protein_coding
protein_coding
protein_coding
protein_coding
ncRNA
small_nucle(ol)ar_rna
protein_coding
protein_coding
protein_coding
protein_coding
protein_coding
protein_coding
protein_coding
protein_coding
protein_coding
protein_coding
protein_coding
protein_coding
protein_coding
protein_coding
protein_coding
protein_coding
protein_coding
protein_coding
protein_coding
protein_coding
protein_coding
ncRNA
protein_coding
ncRNA
protein_coding
protein_coding
protein_coding
protein_coding
protein_coding
protein_coding
protein_coding
protein_coding

protein_coding
protein_coding
protein_coding
protein_coding
protein_coding
ncRNA
protein_coding
protein_coding
protein_coding
protein_coding
protein_coding
protein_coding
pre_trna
protein_coding
protein_coding
protein_coding
protein_coding
protein_coding
protein_coding
protein_coding
protein_coding
protein_coding
protein_coding
ncRNA
ncRNA
protein_coding
protein_coding
protein_coding
protein_coding
protein_coding
protein_coding
protein_coding
protein_coding
protein_coding
protein_coding
protein_coding
protein_coding
protein_coding
protein_coding
protein_coding
protein_coding
miRNA_precursor
protein_coding
protein_coding
protein_coding
protein_coding
protein_coding
protein_coding
protein_coding
protein_coding
protein_coding
protein_coding
protein_coding
protein_coding
miRNA_precursor
protein_coding

protein_coding
protein_coding
protein_coding
protein_coding
protein_coding
protein_coding
miRNA_precursor
protein_coding
protein_coding
protein_coding
protein_coding
protein_coding
protein_coding
ncRNA
protein_coding
protein_coding
protein_coding
protein_coding
protein_coding
protein_coding
protein_coding
protein_coding
protein_coding
protein_coding
ncRNA
protein_coding
protein_coding
protein_coding
protein_coding
protein_coding
protein_coding
protein_coding
protein_coding
protein_coding
protein_coding
protein_coding
protein_coding
protein_coding
protein_coding
protein_coding
ncRNA
protein_coding
protein_coding
protein_coding
protein_coding
protein_coding
protein_coding
protein_coding
protein_coding
protein_coding
protein_coding
protein_coding
protein_coding
protein_coding
protein_coding
protein_coding

protein_coding
protein_coding
protein_coding
protein_coding
protein_coding
protein_coding
protein_coding
protein_coding
ncRNA
protein_coding
protein_coding
protein_coding
protein_coding
protein_coding
protein_coding
protein_coding
protein_coding
protein_coding
protein_coding
protein_coding
ncRNA
protein_coding
protein_coding
protein_coding
protein_coding
protein_coding
protein_coding
protein_coding
protein_coding
ncRNA
protein_coding
protein_coding
small_nucle(ol)ar_rna
small_nucle(ol)ar_rna
small_nucle(ol)ar_rna
protein_coding
protein_coding
protein_coding
protein_coding
protein_coding
protein_coding
protein_coding
protein_coding
protein_coding
protein_coding
protein_coding
protein_coding
protein_coding
protein_coding
protein_coding
protein_coding
protein_coding
protein_coding
protein_coding
protein_coding
protein_coding

protein_coding
ncRNA
protein_coding
protein_coding
protein_coding
protein_coding
small_nucle(ol)ar_rna
small_nucle(ol)ar_rna
miRNA_precursor
protein_coding
protein_coding
protein_coding
protein_coding
protein_coding
protein_coding
protein_coding
protein_coding
protein_coding
protein_coding
protein_coding
protein_coding
protein_coding
protein_coding
protein_coding
protein_coding
protein_coding
protein_coding
protein_coding
protein_coding
protein_coding
protein_coding
protein_coding
protein_coding
protein_coding
protein_coding
protein_coding
protein_coding
pre_trna
protein_coding
protein_coding
protein_coding
protein_coding
protein_coding
protein_coding
protein_coding
protein_coding
protein_coding
protein_coding
protein_coding
protein_coding
protein_coding
protein_coding
protein_coding
protein_coding
protein_coding
protein_coding

protein_coding
protein_coding
protein_coding
protein_coding
protein_coding
protein_coding
protein_coding
protein_coding
protein_coding
protein_coding
protein_coding
protein_coding
protein_coding
protein_coding
ncRNA
ncRNA
protein_coding
protein_coding
protein_coding
protein_coding
protein_coding
protein_coding
protein_coding
protein_coding
protein_coding
protein_coding
protein_coding
protein_coding
protein_coding
protein_coding
protein_coding
protein_coding
protein_coding
protein_coding
protein_coding
protein_coding
protein_coding
protein_coding
protein_coding
protein_coding
ncRNA
protein_coding
protein_coding
protein_coding
pre_trna
protein_coding
protein_coding
protein_coding
protein_coding
protein_coding
protein_coding
protein_coding
protein_coding
protein_coding
miRNA_precursor
protein_coding

protein_coding
protein_coding
protein_coding
protein_coding
protein_coding
protein_coding
protein_coding
protein_coding
protein_coding
protein_coding
protein_coding
protein_coding
protein_coding
protein_coding
protein_coding
protein_coding
protein_coding
protein_coding
ncRNA
protein_coding
protein_coding
protein_coding
protein_coding
protein_coding
protein_coding
protein_coding
protein_coding
protein_coding
protein_coding
protein_coding
protein_coding
protein_coding
protein_coding
protein_coding
protein_coding
protein_coding
protein_coding
protein_coding
protein_coding
protein_coding
protein_coding
protein_coding
protein_coding
protein_coding
protein_coding
protein_coding
protein_coding
ncRNA
protein_coding
protein_coding
protein_coding
ncRNA
protein_coding
protein_coding
small_nucle(ol)ar_rna
miRNA_precursor

protein_coding
protein_coding
protein_coding
protein_coding
protein_coding
protein_coding
protein_coding
protein_coding
protein_coding
protein_coding
protein_coding
protein_coding
protein_coding
protein_coding
protein_coding
protein_coding
protein_coding
protein_coding
protein_coding
protein_coding
protein_coding
protein_coding
protein_coding
protein_coding
ncRNA
protein_coding
protein_coding
protein_coding
protein_coding
protein_coding
protein_coding
protein_coding
protein_coding
protein_coding
protein_coding
protein_coding
protein_coding
protein_coding
protein_coding
protein_coding
protein_coding
protein_coding
protein_coding
protein_coding
protein_coding
protein_coding
protein_coding
protein_coding
protein_coding
ncRNA
protein_coding
protein_coding
protein_coding
protein_coding
protein_coding
protein_coding

protein_coding
protein_coding
protein_coding
protein_coding
protein_coding
protein_coding
protein_coding
protein_coding
protein_coding
protein_coding
small_nucle(ol)ar_rna
protein_coding
protein_coding
protein_coding
protein_coding
protein_coding
protein_coding
protein_coding
protein_coding
protein_coding
protein_coding
protein_coding
protein_coding
protein_coding
protein_coding
protein_coding
protein_coding
protein_coding
protein_coding
protein_coding
protein_coding
protein_coding
protein_coding
protein_coding
protein_coding
protein_coding
protein_coding
ncRNA
protein_coding
protein_coding
protein_coding
protein_coding
protein_coding
pre_trna
protein_coding
protein_coding
protein_coding
protein_coding
protein_coding
protein_coding
protein_coding
protein_coding
protein_coding
miRNA_precursor
protein_coding
protein_coding

protein_coding
protein_coding
protein_coding
protein_coding
protein_coding
protein_coding
protein_coding
protein_coding
pre_trna
protein_coding
protein_coding
protein_coding
protein_coding
protein_coding
protein_coding
protein_coding
protein_coding
protein_coding
protein_coding
protein_coding
protein_coding
protein_coding
protein_coding
protein_coding
protein_coding
protein_coding
protein_coding
protein_coding
protein_coding
protein_coding
protein_coding
protein_coding
protein_coding
protein_coding
protein_coding
protein_coding
protein_coding
protein_coding
ncRNA
ncRNA
protein_coding
protein_coding
protein_coding
protein_coding
protein_coding
protein_coding
protein_coding
protein_coding
ncRNA
protein_coding
protein_coding
protein_coding
protein_coding
protein_coding
protein_coding
protein_coding

protein_coding
small_nucle(ol)ar_rna
protein_coding
protein_coding
protein_coding
protein_coding
protein_coding
protein_coding
protein_coding
protein_coding
protein_coding
protein_coding
protein_coding
protein_coding
protein_coding
protein_coding
protein_coding
protein_coding
small_nucle(ol)ar_rna
small_nucle(ol)ar_rna
protein_coding
protein_coding
protein_coding
protein_coding
protein_coding
protein_coding
protein_coding
protein_coding
pre_trna
protein_coding
protein_coding
protein_coding
protein_coding
protein_coding
protein_coding
protein_coding
protein_coding
protein_coding
protein_coding
protein_coding
protein_coding
protein_coding
protein_coding
protein_coding
protein_coding
protein_coding
protein_coding
protein_coding
protein_coding
protein_coding
protein_coding
protein_coding
ncRNA
protein_coding
protein_coding
protein_coding

protein_coding
protein_coding
protein_coding
protein_coding
protein_coding
protein_coding
protein_coding
protein_coding
protein_coding
protein_coding
protein_coding
protein_coding
protein_coding
pre_trna
protein_coding
protein_coding
protein_coding
protein_coding
protein_coding
protein_coding
protein_coding
protein_coding
protein_coding
protein_coding
protein_coding
protein_coding
protein_coding
protein_coding
protein_coding
protein_coding
protein_coding
protein_coding
protein_coding
protein_coding
protein_coding
protein_coding
protein_coding
protein_coding
protein_coding
protein_coding
protein_coding
protein_coding
protein_coding
protein_coding
protein_coding
protein_coding
protein_coding
protein_coding
protein_coding
protein_coding
protein_coding
ncRNA
ncRNA
protein_coding
protein_coding
protein_coding

protein_coding
protein_coding
protein_coding
protein_coding
protein_coding
protein_coding
ncRNA
protein_coding
protein_coding
protein_coding
protein_coding
protein_coding
protein_coding
protein_coding
protein_coding
protein_coding
protein_coding
protein_coding
protein_coding
protein_coding
protein_coding
protein_coding
protein_coding
protein_coding
protein_coding
protein_coding
protein_coding
protein_coding
protein_coding
protein_coding
protein_coding
protein_coding
ncRNA
ncRNA
protein_coding
protein_coding
protein_coding
protein_coding
protein_coding
protein_coding
protein_coding
protein_coding
protein_coding
protein_coding
protein_coding
protein_coding
protein_coding
protein_coding
protein_coding
protein_coding
protein_coding
protein_coding
protein_coding
protein_coding
protein_coding
protein_coding

protein_coding
protein_coding
protein_coding
protein_coding
protein_coding
protein_coding
protein_coding
protein_coding
protein_coding
protein_coding
protein_coding
protein_coding
protein_coding
ncRNA
protein_coding
protein_coding
protein_coding
protein_coding
protein_coding
protein_coding
ncRNA
protein_coding
ncRNA
protein_coding
ncRNA
protein_coding
protein_coding
protein_coding
protein_coding
protein_coding
protein_coding
miRNA_precursor
protein_coding
protein_coding
protein_coding
ncRNA
protein_coding
protein_coding
protein_coding
protein_coding
protein_coding
protein_coding
protein_coding
protein_coding
protein_coding
protein_coding
protein_coding
protein_coding
protein_coding
protein_coding
protein_coding
protein_coding
protein_coding
protein_coding
protein_coding
protein_coding

protein_coding
protein_coding
protein_coding
protein_coding
protein_coding
protein_coding
protein_coding
protein_coding
protein_coding
protein_coding
protein_coding
protein_coding
protein_coding
protein_coding
protein_coding
protein_coding
protein_coding
protein_coding
protein_coding
protein_coding
protein_coding
ncRNA
protein_coding
protein_coding
protein_coding
protein_coding
protein_coding
protein_coding
protein_coding
ncRNA
protein_coding
protein_coding
protein_coding
protein_coding
protein_coding
protein_coding
protein_coding
protein_coding
protein_coding
protein_coding
protein_coding
protein_coding
protein_coding
protein_coding
protein_coding
protein_coding
protein_coding
protein_coding
protein_coding
protein_coding
protein_coding
pre_trna
protein_coding
protein_coding
protein_coding
protein_coding

protein_coding
protein_coding
protein_coding
protein_coding
protein_coding
ncRNA
protein_coding
protein_coding
protein_coding
protein_coding
protein_coding
protein_coding
protein_coding
protein_coding
protein_coding
protein_coding
protein_coding
protein_coding
protein_coding
protein_coding
protein_coding
protein_coding
protein_coding
ncRNA
protein_coding
protein_coding
protein_coding
protein_coding
protein_coding
protein_coding
protein_coding
protein_coding
protein_coding
protein_coding
protein_coding
protein_coding
protein_coding
protein_coding
protein_coding
protein_coding
protein_coding
protein_coding
protein_coding
protein_coding
protein_coding
protein_coding
protein_coding
protein_coding
ncRNA
protein_coding
protein_coding
protein_coding
protein_coding
protein_coding
protein_coding
protein_coding

protein_coding
protein_coding
protein_coding
protein_coding
ncRNA
protein_coding
protein_coding
protein_coding
protein_coding
protein_coding
protein_coding
protein_coding
protein_coding
protein_coding
protein_coding
protein_coding
protein_coding
protein_coding
protein_coding
protein_coding
protein_coding
protein_coding
protein_coding
protein_coding
protein_coding
protein_coding
protein_coding
protein_coding
protein_coding
protein_coding
ncRNA
protein_coding
protein_coding
protein_coding
protein_coding
protein_coding
protein_coding
protein_coding
protein_coding
protein_coding
protein_coding
protein_coding
protein_coding
protein_coding
protein_coding
protein_coding
protein_coding
protein_coding
protein_coding
protein_coding
protein_coding
protein_coding
protein_coding
protein_coding
protein_coding
protein_coding

protein_coding
protein_coding
protein_coding
protein_coding
protein_coding
protein_coding
protein_coding
miRNA_precursor
protein_coding
protein_coding
protein_coding
protein_coding
protein_coding
protein_coding
protein_coding
protein_coding
protein_coding
protein_coding
protein_coding
protein_coding
protein_coding
protein_coding
protein_coding
protein_coding
protein_coding
protein_coding
protein_coding
protein_coding
protein_coding
protein_coding
protein_coding
protein_coding
protein_coding
protein_coding
protein_coding
protein_coding
protein_coding
protein_coding
protein_coding
protein_coding
protein_coding
protein_coding
protein_coding
protein_coding
protein_coding
protein_coding
protein_coding
protein_coding
protein_coding
protein_coding
protein_coding
protein_coding
protein_coding
protein_coding
protein_coding
protein_coding

protein_coding
protein_coding
protein_coding
small_nucle(ol)ar_rna
protein_coding
protein_coding
protein_coding
protein_coding
protein_coding
protein_coding
protein_coding
protein_coding
protein_coding
protein_coding
protein_coding
protein_coding
protein_coding
protein_coding
protein_coding
protein_coding
protein_coding
protein_coding
protein_coding
protein_coding
protein_coding
protein_coding
protein_coding
protein_coding
protein_coding
protein_coding
protein_coding
protein_coding
protein_coding
protein_coding
ncRNA
protein_coding
protein_coding
protein_coding
protein_coding
protein_coding
protein_coding
protein_coding
protein_coding
protein_coding
protein_coding
protein_coding
protein_coding
protein_coding
protein_coding
protein_coding
protein_coding
protein_coding
protein_coding
protein_coding
protein_coding
protein_coding

protein_coding
protein_coding
protein_coding
protein_coding
protein_coding
protein_coding
protein_coding
protein_coding
protein_coding
pre_trna
protein_coding
protein_coding
protein_coding
protein_coding
protein_coding
ncRNA
ncRNA
protein_coding
protein_coding
protein_coding
protein_coding
protein_coding
protein_coding
protein_coding
protein_coding
protein_coding
protein_coding
protein_coding
protein_coding
protein_coding
protein_coding
protein_coding
protein_coding
protein_coding
protein_coding
protein_coding
protein_coding
protein_coding
protein_coding
ncRNA
protein_coding
protein_coding
protein_coding
protein_coding
protein_coding
protein_coding
protein_coding
protein_coding
protein_coding
protein_coding
protein_coding
protein_coding
protein_coding
protein_coding
protein_coding
protein_coding

protein_coding
protein_coding
protein_coding
protein_coding
protein_coding
protein_coding
protein_coding
protein_coding
protein_coding
protein_coding
protein_coding
protein_coding
protein_coding
protein_coding
ncRNA
protein_coding
protein_coding
protein_coding
protein_coding
protein_coding
protein_coding
protein_coding
protein_coding
protein_coding
protein_coding
protein_coding
protein_coding
protein_coding
pre_trna
protein_coding
protein_coding
protein_coding
protein_coding
protein_coding
protein_coding
protein_coding
protein_coding
protein_coding
protein_coding
protein_coding
protein_coding
protein_coding
protein_coding
ncRNA
protein_coding
protein_coding
protein_coding
protein_coding
protein_coding
protein_coding
protein_coding
protein_coding
protein_coding
protein_coding
protein_coding
protein_coding

protein_coding
protein_coding
protein_coding
protein_coding
protein_coding
protein_coding
protein_coding
protein_coding
protein_coding
protein_coding
protein_coding
protein_coding
protein_coding
protein_coding
protein_coding
protein_coding
protein_coding
protein_coding
protein_coding
ncRNA
protein_coding
protein_coding
protein_coding
protein_coding
protein_coding
protein_coding
protein_coding
protein_coding
protein_coding
protein_coding
protein_coding
protein_coding
protein_coding
protein_coding
protein_coding
protein_coding
miRNA_precursor
protein_coding
protein_coding
protein_coding
protein_coding
miRNA_precursor
ncRNA
miRNA_precursor
protein_coding
protein_coding
protein_coding
protein_coding
protein_coding
protein_coding
protein_coding
protein_coding
protein_coding
protein_coding
protein_coding
protein_coding

protein_coding
protein_coding
protein_coding
protein_coding
protein_coding
protein_coding
protein_coding
protein_coding
protein_coding
protein_coding
protein_coding
protein_coding
protein_coding
protein_coding
protein_coding
protein_coding
protein_coding
protein_coding
protein_coding
protein_coding
protein_coding
protein_coding
protein_coding
protein_coding
protein_coding
protein_coding
protein_coding
protein_coding
protein_coding
protein_coding
protein_coding
protein_coding
protein_coding
protein_coding
protein_coding
protein_coding
ncRNA
ncRNA
protein_coding
ncRNA
protein_coding
protein_coding
protein_coding
protein_coding
protein_coding
ncRNA
protein_coding
protein_coding
protein_coding
protein_coding
protein_coding
miRNA_precursor
protein_coding
protein_coding
protein_coding
protein_coding

protein_coding
protein_coding
ncRNA
protein_coding
protein_coding
protein_coding
protein_coding
protein_coding
protein_coding
protein_coding
protein_coding
protein_coding
protein_coding
ncRNA
protein_coding
protein_coding
protein_coding
protein_coding
protein_coding
protein_coding
protein_coding
protein_coding
protein_coding
protein_coding
protein_coding
protein_coding
protein_coding
protein_coding
protein_coding
protein_coding
protein_coding
protein_coding
protein_coding
protein_coding
protein_coding
protein_coding
protein_coding
protein_coding
protein_coding
pre_trna
protein_coding
protein_coding
protein_coding
protein_coding
protein_coding
protein_coding
protein_coding
protein_coding
protein_coding
protein_coding
protein_coding
protein_coding
protein_coding
protein_coding
protein_coding
protein_coding

protein_coding
protein_coding
protein_coding
protein_coding
protein_coding
protein_coding
protein_coding
protein_coding
protein_coding
protein_coding
protein_coding
protein_coding
protein_coding
protein_coding
protein_coding
protein_coding
protein_coding
protein_coding
protein_coding
protein_coding
protein_coding
protein_coding
protein_coding
protein_coding
protein_coding
protein_coding
small_nucle(ol)ar_rna
protein_coding
protein_coding
protein_coding
protein_coding
protein_coding
protein_coding
protein_coding
protein_coding
protein_coding
protein_coding
protein_coding
protein_coding
protein_coding
protein_coding
protein_coding
protein_coding
protein_coding
protein_coding
protein_coding
protein_coding
protein_coding
protein_coding
protein_coding
protein_coding
protein_coding
protein_coding
protein_coding
protein_coding
protein_coding

protein_coding
protein_coding
protein_coding
protein_coding
protein_coding
protein_coding
protein_coding
protein_coding
protein_coding
protein_coding
protein_coding
ncRNA
protein_coding
protein_coding
protein_coding
protein_coding
protein_coding
ncRNA
protein_coding
protein_coding
protein_coding
protein_coding
protein_coding
protein_coding
protein_coding
protein_coding
protein_coding
protein_coding
protein_coding
protein_coding
protein_coding
protein_coding
protein_coding
protein_coding
protein_coding
protein_coding
small_nucle(ol)ar_rna
small_nucle(ol)ar_rna
protein_coding
protein_coding
protein_coding
protein_coding
miRNA_precursor
protein_coding
protein_coding
protein_coding
protein_coding
protein_coding
protein_coding
protein_coding
ncRNA
protein_coding
protein_coding
protein_coding
protein_coding
protein_coding

protein_coding
protein_coding
protein_coding
protein_coding
ncRNA
protein_coding
protein_coding
protein_coding
protein_coding
protein_coding
protein_coding
protein_coding
protein_coding
protein_coding
protein_coding
protein_coding
small_nucle(ol)ar_rna
protein_coding
protein_coding
protein_coding
small_nucle(ol)ar_rna
protein_coding
protein_coding
protein_coding
protein_coding
protein_coding
protein_coding
protein_coding
protein_coding
protein_coding
protein_coding
protein_coding
protein_coding
protein_coding
protein_coding
protein_coding
protein_coding
protein_coding
protein_coding
protein_coding
protein_coding
protein_coding
protein_coding
protein_coding
protein_coding
protein_coding
protein_coding
protein_coding
protein_coding
protein_coding
protein_coding
protein_coding
protein_coding
protein_coding
protein_coding
protein_coding

protein_coding
protein_coding
protein_coding
protein_coding
protein_coding
protein_coding
protein_coding
protein_coding
small_nucle(ol)ar_rna
small_nucle(ol)ar_rna
protein_coding
protein_coding
protein_coding
ncRNA
protein_coding
protein_coding
protein_coding
protein_coding
protein_coding
protein_coding
protein_coding
protein_coding
protein_coding
protein_coding
protein_coding
protein_coding
protein_coding
protein_coding
protein_coding
protein_coding
protein_coding
protein_coding
protein_coding
protein_coding
protein_coding
protein_coding
protein_coding
protein_coding
protein_coding
protein_coding
protein_coding
protein_coding
ncRNA
protein_coding
protein_coding
protein_coding
miRNA_precursor
protein_coding
protein_coding
protein_coding
protein_coding
protein_coding
protein_coding
pre_trna
protein_coding
protein_coding
protein_coding

protein_coding
protein_coding
protein_coding
protein_coding
protein_coding
protein_coding
protein_coding
protein_coding
protein_coding
protein_coding
protein_coding
protein_coding
protein_coding
protein_coding
protein_coding
protein_coding
protein_coding
protein_coding
protein_coding
small_nucle(ol)ar_rna
protein_coding
protein_coding
protein_coding
protein_coding
protein_coding
protein_coding
protein_coding
protein_coding
ncRNA
protein_coding
protein_coding
protein_coding
ncRNA
protein_coding
protein_coding
protein_coding
protein_coding
protein_coding
protein_coding
protein_coding
protein_coding
protein_coding
protein_coding
protein_coding
protein_coding
protein_coding
protein_coding
protein_coding
protein_coding
protein_coding
protein_coding
protein_coding
protein_coding
protein_coding
protein_coding
protein_coding

protein_coding
protein_coding
protein_coding
protein_coding
protein_coding
protein_coding
protein_coding
protein_coding
protein_coding
protein_coding
ncRNA
protein_coding
protein_coding
protein_coding
protein_coding
protein_coding
protein_coding
protein_coding
protein_coding
protein_coding
protein_coding
protein_coding
protein_coding
protein_coding
protein_coding
protein_coding
protein_coding
protein_coding
protein_coding
protein_coding
protein_coding
protein_coding
protein_coding
protein_coding
protein_coding
protein_coding
protein_coding
protein_coding
protein_coding
protein_coding
protein_coding
protein_coding
protein_coding
protein_coding
protein_coding
protein_coding
protein_coding
protein_coding
protein_coding
small_nucle(ol)ar_rna
protein_coding
protein_coding
protein_coding
protein_coding
protein_coding
protein_coding

protein_coding
protein_coding
protein_coding
protein_coding
protein_coding
protein_coding
protein_coding
protein_coding
protein_coding
protein_coding
protein_coding
protein_coding
protein_coding
protein_coding
protein_coding
protein_coding
protein_coding
protein_coding
protein_coding
protein_coding
protein_coding
protein_coding
ncRNA
protein_coding
small_nucle(ol)ar_rna
protein_coding
protein_coding
protein_coding
protein_coding
protein_coding
protein_coding
protein_coding
protein_coding
protein_coding
ncRNA
protein_coding
protein_coding
protein_coding
protein_coding
protein_coding
protein_coding
protein_coding
protein_coding
protein_coding
protein_coding
protein_coding
protein_coding
pre_trna
protein_coding
protein_coding
ncRNA
protein_coding
protein_coding
protein_coding
protein_coding
protein_coding

protein_coding
protein_coding
protein_coding
protein_coding
protein_coding
protein_coding
protein_coding
protein_coding
protein_coding
protein_coding
protein_coding
protein_coding
protein_coding
protein_coding
protein_coding
protein_coding
protein_coding
protein_coding
protein_coding
protein_coding
protein_coding
protein_coding
protein_coding
protein_coding
protein_coding
protein_coding
protein_coding
protein_coding
protein_coding
protein_coding
protein_coding
protein_coding
protein_coding
protein_coding
protein_coding
protein_coding
protein_coding
protein_coding
pre_trna
protein_coding
protein_coding
protein_coding
protein_coding
protein_coding
protein_coding
protein_coding
protein_coding
protein_coding
protein_coding
protein_coding
protein_coding
protein_coding
protein_coding
protein_coding
protein_coding
protein_coding

protein_coding
protein_coding
protein_coding
protein_coding
protein_coding
protein_coding
protein_coding
protein_coding
protein_coding
protein_coding
protein_coding
protein_coding
protein_coding
protein_coding
protein_coding
protein_coding
protein_coding
protein_coding
protein_coding
ncRNA
protein_coding
protein_coding
protein_coding
protein_coding
protein_coding
ncRNA
protein_coding
protein_coding
protein_coding
protein_coding
protein_coding
protein_coding
protein_coding
protein_coding
protein_coding
protein_coding
protein_coding
protein_coding
protein_coding
protein_coding
protein_coding
protein_coding
protein_coding
protein_coding
protein_coding
protein_coding
protein_coding
protein_coding
protein_coding
protein_coding
protein_coding
protein_coding
pre_trna
ncRNA
protein_coding
protein_coding

protein_coding
ncRNA
protein_coding
protein_coding
protein_coding
pre_trna
protein_coding
protein_coding
protein_coding
protein_coding
protein_coding
protein_coding
protein_coding
protein_coding
protein_coding
protein_coding
protein_coding
protein_coding
protein_coding
protein_coding
protein_coding
protein_coding
protein_coding
protein_coding
protein_coding
protein_coding
protein_coding
protein_coding
protein_coding
protein_coding
protein_coding
protein_coding
protein_coding
protein_coding
protein_coding
protein_coding
protein_coding
small_nucle(ol)ar_rna
small_nucle(ol)ar_rna
pre_trna
protein_coding
protein_coding
protein_coding
protein_coding
protein_coding
protein_coding
protein_coding
protein_coding
protein_coding
protein_coding
protein_coding
protein_coding
protein_coding
protein_coding
pre_trna
protein_coding

ncRNA
protein_coding
protein_coding
protein_coding
protein_coding
protein_coding
protein_coding
ncRNA
protein_coding
protein_coding
protein_coding
protein_coding
protein_coding
protein_coding
protein_coding
protein_coding
protein_coding
protein_coding
protein_coding
protein_coding
protein_coding
protein_coding
protein_coding
protein_coding
miRNA_precursor
protein_coding
protein_coding
ncRNA
protein_coding
protein_coding
protein_coding
pre_trna
protein_coding
protein_coding
protein_coding
protein_coding
protein_coding
protein_coding
protein_coding
protein_coding
ncRNA
protein_coding
protein_coding
protein_coding
protein_coding
protein_coding
protein_coding
protein_coding
protein_coding
protein_coding
protein_coding
protein_coding
protein_coding
protein_coding
protein_coding
protein_coding

protein_coding
protein_coding
protein_coding
protein_coding
protein_coding
protein_coding
protein_coding
protein_coding
protein_coding
small_nucle(ol)ar_rna
protein_coding
protein_coding
protein_coding
protein_coding
ncRNA
protein_coding
protein_coding
protein_coding
protein_coding
protein_coding
protein_coding
protein_coding
protein_coding
protein_coding
protein_coding
protein_coding
protein_coding
protein_coding
protein_coding
protein_coding
protein_coding
protein_coding
protein_coding
protein_coding
small_nucle(ol)ar_rna
protein_coding
protein_coding
protein_coding
protein_coding
protein_coding
protein_coding
protein_coding
protein_coding
protein_coding
protein_coding
protein_coding
protein_coding
protein_coding
protein_coding
protein_coding
protein_coding
protein_coding
protein_coding
protein_coding
protein_coding
protein_coding

protein_coding
protein_coding
protein_coding
protein_coding
protein_coding
protein_coding
protein_coding
protein_coding
protein_coding
protein_coding
protein_coding
protein_coding
protein_coding
protein_coding
protein_coding
protein_coding
protein_coding
protein_coding
protein_coding
protein_coding
protein_coding
ncRNA
protein_coding
protein_coding
protein_coding
protein_coding
protein_coding
protein_coding
ncRNA
protein_coding
protein_coding
protein_coding
protein_coding
protein_coding
protein_coding
protein_coding
protein_coding
protein_coding
protein_coding
protein_coding
protein_coding
protein_coding
protein_coding
ncRNA
protein_coding
protein_coding
ncRNA
protein_coding
protein_coding
protein_coding
protein_coding
protein_coding
protein_coding
protein_coding
protein_coding
protein_coding

protein_coding
protein_coding
protein_coding
protein_coding
protein_coding
protein_coding
protein_coding
protein_coding
protein_coding
protein_coding
protein_coding
protein_coding
protein_coding
protein_coding
protein_coding
ncRNA
protein_coding
protein_coding
protein_coding
protein_coding
protein_coding
protein_coding
protein_coding
protein_coding
protein_coding
protein_coding
protein_coding
protein_coding
protein_coding
protein_coding
protein_coding
protein_coding
protein_coding
protein_coding
protein_coding
protein_coding
ncRNA
protein_coding
ncRNA
protein_coding
protein_coding
protein_coding
protein_coding
protein_coding
protein_coding
protein_coding
protein_coding
protein_coding
protein_coding
protein_coding
protein_coding
protein_coding
protein_coding
protein_coding
protein_coding
protein_coding

protein_coding
protein_coding
protein_coding
protein_coding
protein_coding
protein_coding
protein_coding
protein_coding
protein_coding
protein_coding
protein_coding
protein_coding
protein_coding
protein_coding
protein_coding
protein_coding
protein_coding
ncRNA
protein_coding
protein_coding
protein_coding
protein_coding
protein_coding
protein_coding
protein_coding
protein_coding
protein_coding
protein_coding
protein_coding
protein_coding
protein_coding
protein_coding
protein_coding
protein_coding
protein_coding
protein_coding
protein_coding
protein_coding
protein_coding
protein_coding
pre_trna
protein_coding
protein_coding
protein_coding
protein_coding
protein_coding
protein_coding
protein_coding
protein_coding
protein_coding
protein_coding
protein_coding
protein_coding
protein_coding
protein_coding
protein_coding

protein_coding
protein_coding
protein_coding
protein_coding
protein_coding
protein_coding
protein_coding
protein_coding
protein_coding
protein_coding
protein_coding
protein_coding
protein_coding
protein_coding
protein_coding
protein_coding
protein_coding
protein_coding
ncRNA
protein_coding
protein_coding
protein_coding
protein_coding
protein_coding
protein_coding
protein_coding
protein_coding
protein_coding
protein_coding
protein_coding
protein_coding
protein_coding
small_nucle(ol)ar_rna
protein_coding
protein_coding
protein_coding
protein_coding
ncRNA
protein_coding
protein_coding
protein_coding
protein_coding
protein_coding
protein_coding
protein_coding
protein_coding
protein_coding
protein_coding
ncRNA
protein_coding
protein_coding
protein_coding
ncRNA
protein_coding
protein_coding
protein_coding

protein_coding
protein_coding
protein_coding
protein_coding
protein_coding
protein_coding
protein_coding
protein_coding
protein_coding
protein_coding
protein_coding
protein_coding
protein_coding
protein_coding
protein_coding
protein_coding
protein_coding
protein_coding
protein_coding
ncRNA
protein_coding
protein_coding
protein_coding
protein_coding
ncRNA
protein_coding
protein_coding
protein_coding
protein_coding
protein_coding
pre_trna
protein_coding
protein_coding
protein_coding
protein_coding
ncRNA
protein_coding
protein_coding
ncRNA
protein_coding
protein_coding
protein_coding
protein_coding
protein_coding
protein_coding
protein_coding
protein_coding
protein_coding
protein_coding
protein_coding
protein_coding
protein_coding
protein_coding
ncRNA
protein_coding
protein_coding

protein_coding
small_nucle(ol)ar_rna
protein_coding
protein_coding
protein_coding
protein_coding
protein_coding
protein_coding
protein_coding
protein_coding
protein_coding
protein_coding
protein_coding
protein_coding
protein_coding
protein_coding
protein_coding
protein_coding
protein_coding
protein_coding
protein_coding
protein_coding
protein_coding
protein_coding
protein_coding
protein_coding
protein_coding
protein_coding
pre_trna
protein_coding
protein_coding
protein_coding
protein_coding
protein_coding
protein_coding
protein_coding
protein_coding
protein_coding
protein_coding
protein_coding
protein_coding
protein_coding
protein_coding
protein_coding
protein_coding
protein_coding
protein_coding
miRNA_precursor
protein_coding
protein_coding
protein_coding
protein_coding
protein_coding
protein_coding
protein_coding
protein_coding

protein_coding
protein_coding
protein_coding
protein_coding
protein_coding
protein_coding
protein_coding
protein_coding
protein_coding
protein_coding
protein_coding
protein_coding
protein_coding
protein_coding
protein_coding
protein_coding
protein_coding
protein_coding
ncRNA
protein_coding
ncRNA
protein_coding
protein_coding
protein_coding
protein_coding
protein_coding
protein_coding
protein_coding
protein_coding
protein_coding
protein_coding
protein_coding
protein_coding
protein_coding
protein_coding
protein_coding
protein_coding
protein_coding
protein_coding
protein_coding
protein_coding
protein_coding
protein_coding
small_nucle(ol)ar_rna
protein_coding
protein_coding
protein_coding
protein_coding
protein_coding
protein_coding
protein_coding
protein_coding
protein_coding
protein_coding
protein_coding
protein_coding

protein_coding
protein_coding
ncRNA
ncRNA
small_nucle(ol)ar_rna
protein_coding
protein_coding
protein_coding
protein_coding
protein_coding
protein_coding
protein_coding
protein_coding
protein_coding
small_nucle(ol)ar_rna
protein_coding
protein_coding
protein_coding
protein_coding
protein_coding
protein_coding
protein_coding
protein_coding
protein_coding
protein_coding
protein_coding
protein_coding
ncRNA
protein_coding
protein_coding
protein_coding
protein_coding
protein_coding
protein_coding
protein_coding
protein_coding
protein_coding
protein_coding
protein_coding
protein_coding
protein_coding
protein_coding
protein_coding
protein_coding
protein_coding
protein_coding
protein_coding
protein_coding
protein_coding
protein_coding
protein_coding
protein_coding
protein_coding
protein_coding
protein_coding
protein_coding
ncRNA

protein_coding
protein_coding
protein_coding
small_nucle(ol)ar_rna
protein_coding
protein_coding
protein_coding
protein_coding
protein_coding
protein_coding
protein_coding
protein_coding
protein_coding
protein_coding
protein_coding
protein_coding
protein_coding
protein_coding
protein_coding
protein_coding
protein_coding
protein_coding
protein_coding
protein_coding
protein_coding
ncRNA
protein_coding
protein_coding
protein_coding
protein_coding
protein_coding
protein_coding
protein_coding
protein_coding
ncRNA
protein_coding
protein_coding
protein_coding
protein_coding
protein_coding
protein_coding
protein_coding
protein_coding
protein_coding
protein_coding
protein_coding
protein_coding
protein_coding
protein_coding
protein_coding
protein_coding
protein_coding
protein_coding
protein_coding
small_nucle(ol)ar_rna
protein_coding

small_nucle(ol)ar_rna
protein_coding
protein_coding
ncRNA
protein_coding
protein_coding
protein_coding
protein_coding
protein_coding
protein_coding
ncRNA
protein_coding
protein_coding
ncRNA
protein_coding
protein_coding
protein_coding
protein_coding
ncRNA
miRNA_precursor
protein_coding
protein_coding
protein_coding
protein_coding
protein_coding
protein_coding
miRNA_precursor
protein_coding
protein_coding
small_nucle(ol)ar_rna
protein_coding
protein_coding
protein_coding
protein_coding
ncRNA
protein_coding
protein_coding
protein_coding
protein_coding
protein_coding
protein_coding
protein_coding
protein_coding
protein_coding
protein_coding
protein_coding
protein_coding
protein_coding
protein_coding
protein_coding
protein_coding
protein_coding
protein_coding
protein_coding
protein_coding
protein_coding

protein_coding
protein_coding
ncRNA
ncRNA
miRNA_precursor
protein_coding
ncRNA
protein_coding
protein_coding
protein_coding
protein_coding
protein_coding
protein_coding
protein_coding
protein_coding
protein_coding
protein_coding
protein_coding
protein_coding
protein_coding
protein_coding
ncRNA
protein_coding
protein_coding
protein_coding
protein_coding
protein_coding
protein_coding
protein_coding
protein_coding
protein_coding
protein_coding
protein_coding
protein_coding
protein_coding
protein_coding
protein_coding
protein_coding
protein_coding
protein_coding
protein_coding
protein_coding
protein_coding
protein_coding
ncRNA
protein_coding
protein_coding
protein_coding
protein_coding
protein_coding
protein_coding
protein_coding
protein_coding
protein_coding
protein_coding
protein_coding

protein_coding
ncRNA
protein_coding
protein_coding
ncRNA
protein_coding
protein_coding
ncRNA
protein_coding
protein_coding
ncRNA
small_nucle(ol)ar_rna
protein_coding
protein_coding
protein_coding
protein_coding
protein_coding
protein_coding
protein_coding
protein_coding
protein_coding
protein_coding
protein_coding
protein_coding
protein_coding
protein_coding
protein_coding
protein_coding
protein_coding
protein_coding
protein_coding
protein_coding
protein_coding
protein_coding
protein_coding
miRNA_precursor
protein_coding
protein_coding
protein_coding
protein_coding
protein_coding
protein_coding
protein_coding
protein_coding
protein_coding
protein_coding
protein_coding
protein_coding
protein_coding
protein_coding
protein_coding
protein_coding
protein_coding
protein_coding
protein_coding
ncRNA

protein_coding
protein_coding
protein_coding
protein_coding
protein_coding
protein_coding
ncRNA
protein_coding
protein_coding
protein_coding
protein_coding
ncRNA
protein_coding
protein_coding
protein_coding
ncRNA
protein_coding
protein_coding
protein_coding
protein_coding
protein_coding
protein_coding
ncRNA
protein_coding
protein_coding
protein_coding
protein_coding
protein_coding
protein_coding
protein_coding
protein_coding
ncRNA
protein_coding
protein_coding
protein_coding
protein_coding
protein_coding
protein_coding
protein_coding
protein_coding
protein_coding
protein_coding
miRNA_precursor
protein_coding
protein_coding
protein_coding
protein_coding
protein_coding
protein_coding
protein_coding
protein_coding
protein_coding
protein_coding
protein_coding
protein_coding
protein_coding

protein_coding
protein_coding
protein_coding
protein_coding
protein_coding
protein_coding
protein_coding
protein_coding
protein_coding
protein_coding
protein_coding
protein_coding
protein_coding
protein_coding
protein_coding
protein_coding
protein_coding
protein_coding
protein_coding
protein_coding
protein_coding
protein_coding
protein_coding
protein_coding
protein_coding
protein_coding
protein_coding
protein_coding
protein_coding
protein_coding
protein_coding
protein_coding
protein_coding
protein_coding
protein_coding
protein_coding
protein_coding
ncRNA
protein_coding
protein_coding
protein_coding
protein_coding
protein_coding
protein_coding
protein_coding
protein_coding
protein_coding
protein_coding
protein_coding
protein_coding
protein_coding
protein_coding
protein_coding
protein_coding
protein_coding
protein_coding

protein_coding
protein_coding
protein_coding
protein_coding
protein_coding
ncRNA
protein_coding
small_nucle(ol)ar_rna
small_nucle(ol)ar_rna
small_nucle(ol)ar_rna
protein_coding
protein_coding
protein_coding
protein_coding
protein_coding
protein_coding
protein_coding
protein_coding
protein_coding
protein_coding
protein_coding
protein_coding
protein_coding
protein_coding
protein_coding
protein_coding
protein_coding
protein_coding
protein_coding
protein_coding
protein_coding
protein_coding
protein_coding
protein_coding
protein_coding
protein_coding
protein_coding
protein_coding
protein_coding
protein_coding
protein_coding
protein_coding
protein_coding
protein_coding
protein_coding
protein_coding
protein_coding
protein_coding
protein_coding
protein_coding
protein_coding
protein_coding
protein_coding
protein_coding
protein_coding
protein_coding

ncRNA
protein_coding
protein_coding
protein_coding
protein_coding
protein_coding
protein_coding
protein_coding
protein_coding
protein_coding
protein_coding
protein_coding
protein_coding
protein_coding
protein_coding
protein_coding
protein_coding
protein_coding
protein_coding
protein_coding
protein_coding
protein_coding
protein_coding
protein_coding
protein_coding
protein_coding
protein_coding
protein_coding
protein_coding
protein_coding
protein_coding
ncRNA
protein_coding
protein_coding
protein_coding
protein_coding
protein_coding
protein_coding
protein_coding
protein_coding
protein_coding
protein_coding
protein_coding
protein_coding
protein_coding
protein_coding
protein_coding
protein_coding
protein_coding
protein_coding
protein_coding
protein_coding
protein_coding
protein_coding
protein_coding
protein_coding

protein_coding
protein_coding
protein_coding
protein_coding
ncRNA
protein_coding
protein_coding
protein_coding
protein_coding
protein_coding
protein_coding
protein_coding
protein_coding
protein_coding
protein_coding
protein_coding
protein_coding
protein_coding
protein_coding
protein_coding
protein_coding
protein_coding
protein_coding
protein_coding
protein_coding
protein_coding
protein_coding
protein_coding
protein_coding
protein_coding
ncRNA
protein_coding
protein_coding
protein_coding
protein_coding
protein_coding
protein_coding
protein_coding
protein_coding
protein_coding
protein_coding
protein_coding
ncRNA
protein_coding
protein_coding
protein_coding
protein_coding
protein_coding
protein_coding
protein_coding
protein_coding
protein_coding
protein_coding
protein_coding
protein_coding
protein_coding

protein_coding
protein_coding
protein_coding
protein_coding
protein_coding
protein_coding
protein_coding
protein_coding
protein_coding
protein_coding
protein_coding
protein_coding
protein_coding
protein_coding
protein_coding
protein_coding
protein_coding
protein_coding
protein_coding
protein_coding
miRNA_precursor
protein_coding
protein_coding
protein_coding
protein_coding
protein_coding
protein_coding
protein_coding
protein_coding
protein_coding
protein_coding
protein_coding
protein_coding
protein_coding
protein_coding
protein_coding
protein_coding
protein_coding
protein_coding
ncRNA
protein_coding
protein_coding
protein_coding
protein_coding
protein_coding
protein_coding
protein_coding
protein_coding
protein_coding
protein_coding
protein_coding
protein_coding
protein_coding
protein_coding
protein_coding
protein_coding

protein_coding
protein_coding
protein_coding
miRNA_precursor
protein_coding
protein_coding
protein_coding
protein_coding
protein_coding
protein_coding
protein_coding
protein_coding
protein_coding
protein_coding
protein_coding
protein_coding
protein_coding
protein_coding
protein_coding
protein_coding
protein_coding
protein_coding
protein_coding
protein_coding
protein_coding
protein_coding
protein_coding
protein_coding
protein_coding
protein_coding
protein_coding
protein_coding
protein_coding
protein_coding
protein_coding
protein_coding
protein_coding
protein_coding
protein_coding
protein_coding
ncRNA
protein_coding
protein_coding
protein_coding
protein_coding
protein_coding
protein_coding
protein_coding
protein_coding
protein_coding
protein_coding
protein_coding
protein_coding
protein_coding
protein_coding
protein_coding

protein_coding
protein_coding
ncRNA
protein_coding
protein_coding
protein_coding
protein_coding
protein_coding
protein_coding
protein_coding
ncRNA
protein_coding
protein_coding
protein_coding
protein_coding
protein_coding
protein_coding
protein_coding
protein_coding
protein_coding
protein_coding
protein_coding
protein_coding
protein_coding
protein_coding
protein_coding
protein_coding
protein_coding
protein_coding
protein_coding
protein_coding
protein_coding
protein_coding
protein_coding
protein_coding
protein_coding
protein_coding
protein_coding
protein_coding
protein_coding
protein_coding
protein_coding
small_nucle(ol)ar_rna
protein_coding
protein_coding
protein_coding
protein_coding
protein_coding
protein_coding
protein_coding
protein_coding
protein_coding
protein_coding
protein_coding
protein_coding
protein_coding

protein_coding
protein_coding
protein_coding
protein_coding
protein_coding
protein_coding
protein_coding
protein_coding
protein_coding
protein_coding
protein_coding
protein_coding
protein_coding
protein_coding
protein_coding
protein_coding
protein_coding
protein_coding
protein_coding
protein_coding
protein_coding
protein_coding
protein_coding
ncRNA
protein_coding
protein_coding
protein_coding
protein_coding
protein_coding
protein_coding
protein_coding
protein_coding
ncRNA
protein_coding
protein_coding
protein_coding
protein_coding
protein_coding
protein_coding
protein_coding
protein_coding
protein_coding
protein_coding
protein_coding
protein_coding
protein_coding
protein_coding
protein_coding
protein_coding
protein_coding
protein_coding
protein_coding
protein_coding
protein_coding
protein_coding
protein_coding

protein_coding
protein_coding
protein_coding
protein_coding
protein_coding
protein_coding
protein_coding
protein_coding
protein_coding
protein_coding
protein_coding
protein_coding
protein_coding
ncRNA
protein_coding
protein_coding
protein_coding
protein_coding
protein_coding
protein_coding
protein_coding
protein_coding
protein_coding
protein_coding
protein_coding
protein_coding
protein_coding
small_nucle(ol)ar_rna
small_nucle(ol)ar_rna
protein_coding
protein_coding
protein_coding
protein_coding
protein_coding
protein_coding
protein_coding
protein_coding
protein_coding
protein_coding
protein_coding
protein_coding
protein_coding
protein_coding
protein_coding
ncRNA
protein_coding
protein_coding
protein_coding
protein_coding
protein_coding
protein_coding
protein_coding
protein_coding
protein_coding
protein_coding
protein_coding

protein_coding
protein_coding
protein_coding
protein_coding
protein_coding
protein_coding
protein_coding
protein_coding
protein_coding
protein_coding
protein_coding
protein_coding
protein_coding
protein_coding
protein_coding
protein_coding
protein_coding
protein_coding
protein_coding
protein_coding
protein_coding
protein_coding
protein_coding
protein_coding
protein_coding
protein_coding
protein_coding
protein_coding
protein_coding
protein_coding
protein_coding
protein_coding
ncRNA
protein_coding
protein_coding
protein_coding
protein_coding
protein_coding
protein_coding
protein_coding
protein_coding
protein_coding
protein_coding
protein_coding
protein_coding
protein_coding
protein_coding
protein_coding
protein_coding
protein_coding
protein_coding
protein_coding
protein_coding
protein_coding
protein_coding
ncRNA

protein_coding
protein_coding
protein_coding
protein_coding
protein_coding
protein_coding
protein_coding
protein_coding
protein_coding
protein_coding
protein_coding
pre_trna
protein_coding
protein_coding
protein_coding
protein_coding
protein_coding
protein_coding
protein_coding
protein_coding
ncRNA
protein_coding
protein_coding
protein_coding
protein_coding
protein_coding
protein_coding
protein_coding
protein_coding
protein_coding
protein_coding
protein_coding
protein_coding
protein_coding
protein_coding
protein_coding
protein_coding
protein_coding
protein_coding
protein_coding
protein_coding
protein_coding
ncRNA
protein_coding
ncRNA
protein_coding
ncRNA
protein_coding
protein_coding
protein_coding
protein_coding
protein_coding
protein_coding
protein_coding
protein_coding
protein_coding

protein_coding
protein_coding
protein_coding
protein_coding
protein_coding
protein_coding
protein_coding
protein_coding
protein_coding
protein_coding
protein_coding
protein_coding
protein_coding
protein_coding
ncRNA
protein_coding
protein_coding
protein_coding
protein_coding
pre_trna
protein_coding
protein_coding
protein_coding
protein_coding
protein_coding
protein_coding
ncRNA
ncRNA
protein_coding
protein_coding
small_nucle(ol)ar_rna
protein_coding
protein_coding
protein_coding
protein_coding
ncRNA
protein_coding
protein_coding
protein_coding
ncRNA
protein_coding
ncRNA
protein_coding
protein_coding
protein_coding
protein_coding
protein_coding
protein_coding
protein_coding
protein_coding
protein_coding
protein_coding
protein_coding
protein_coding
protein_coding
protein_coding

protein_coding
protein_coding
protein_coding
protein_coding
protein_coding
protein_coding
ncRNA
protein_coding
protein_coding
ncRNA
protein_coding
protein_coding
ncRNA
protein_coding
protein_coding
protein_coding
protein_coding
protein_coding
ncRNA
protein_coding
protein_coding
ncRNA
protein_coding
protein_coding
protein_coding
protein_coding
protein_coding
protein_coding
protein_coding
protein_coding
protein_coding
protein_coding
protein_coding
protein_coding
protein_coding
protein_coding
protein_coding
protein_coding
protein_coding
protein_coding
protein_coding
protein_coding
protein_coding
protein_coding
protein_coding
protein_coding
protein_coding
ncRNA
protein_coding
protein_coding
protein_coding
protein_coding
protein_coding
protein_coding
protein_coding
protein_coding

protein_coding
protein_coding
protein_coding
protein_coding
protein_coding
protein_coding
protein_coding
pre_trna
protein_coding
protein_coding
protein_coding
protein_coding
protein_coding
protein_coding
protein_coding
protein_coding
ncRNA
ncRNA
protein_coding
protein_coding
protein_coding
protein_coding
protein_coding
miRNA_precursor
protein_coding
protein_coding
protein_coding
protein_coding
protein_coding
protein_coding
protein_coding
protein_coding
protein_coding
protein_coding
protein_coding
protein_coding
protein_coding
protein_coding
protein_coding
protein_coding
protein_coding
protein_coding
protein_coding
protein_coding
protein_coding
protein_coding
ncRNA
protein_coding
small_nucle(ol)ar_rna
protein_coding
protein_coding
protein_coding
protein_coding
protein_coding
protein_coding
protein_coding

protein_coding
protein_coding
protein_coding
protein_coding
protein_coding
ncRNA
protein_coding
protein_coding
protein_coding
protein_coding
protein_coding
protein_coding
protein_coding
protein_coding
protein_coding
protein_coding
protein_coding
protein_coding
protein_coding
protein_coding
protein_coding
protein_coding
protein_coding
protein_coding
protein_coding
ncRNA
protein_coding
protein_coding
protein_coding
protein_coding
protein_coding
protein_coding
protein_coding
protein_coding
protein_coding
protein_coding
protein_coding
protein_coding
protein_coding
protein_coding
protein_coding
protein_coding
protein_coding
miRNA_precursor
protein_coding
protein_coding
protein_coding
protein_coding
protein_coding
protein_coding
protein_coding
protein_coding
protein_coding
protein_coding
protein_coding
protein_coding

protein_coding
protein_coding
protein_coding
protein_coding
protein_coding
ncRNA
ncRNA
protein_coding
protein_coding
protein_coding
protein_coding
ncRNA
protein_coding
protein_coding
protein_coding
protein_coding
miRNA_precursor
protein_coding
protein_coding
ncRNA
protein_coding
protein_coding
protein_coding
protein_coding
protein_coding
protein_coding
protein_coding
protein_coding
protein_coding
protein_coding
protein_coding
protein_coding
protein_coding
protein_coding
protein_coding
protein_coding
protein_coding
protein_coding
protein_coding
protein_coding
protein_coding
protein_coding
protein_coding
protein_coding
protein_coding
ncRNA
protein_coding
protein_coding
protein_coding
protein_coding
protein_coding
protein_coding
protein_coding
protein_coding
protein_coding
protein_coding

protein_coding
protein_coding
protein_coding
protein_coding
protein_coding
protein_coding
protein_coding
protein_coding
protein_coding
protein_coding
protein_coding
protein_coding
protein_coding
protein_coding
protein_coding
protein_coding
protein_coding
protein_coding
protein_coding
protein_coding
protein_coding
protein_coding
protein_coding
protein_coding
protein_coding
miRNA_precursor
protein_coding
protein_coding
protein_coding
protein_coding
protein_coding
ncRNA
protein_coding
protein_coding
protein_coding
protein_coding
pre_trna
protein_coding
protein_coding
protein_coding
protein_coding
ncRNA
protein_coding
protein_coding
protein_coding
protein_coding
protein_coding
protein_coding
protein_coding
protein_coding
protein_coding
protein_coding
protein_coding
protein_coding
protein_coding
protein_coding

protein_coding
protein_coding
protein_coding
protein_coding
protein_coding
protein_coding
protein_coding
protein_coding
pre_trna
protein_coding
ncRNA
protein_coding
protein_coding
protein_coding
protein_coding
protein_coding
protein_coding
protein_coding
protein_coding
protein_coding
ncRNA
protein_coding
protein_coding
protein_coding
protein_coding
protein_coding
protein_coding
protein_coding
protein_coding
protein_coding
ncRNA
protein_coding
protein_coding
protein_coding
protein_coding
ncRNA
protein_coding
protein_coding
protein_coding
protein_coding
protein_coding
protein_coding
protein_coding
protein_coding
protein_coding
protein_coding
protein_coding
protein_coding
protein_coding
protein_coding
protein_coding
protein_coding
protein_coding
protein_coding
protein_coding
protein_coding

protein_coding
ncRNA
protein_coding
protein_coding
protein_coding
protein_coding
protein_coding
protein_coding
protein_coding
protein_coding
protein_coding
protein_coding
protein_coding
protein_coding
protein_coding
protein_coding
protein_coding
protein_coding
protein_coding
protein_coding
protein_coding
ncRNA
protein_coding
protein_coding
protein_coding
protein_coding
protein_coding
protein_coding
protein_coding
protein_coding
protein_coding
protein_coding
protein_coding
protein_coding
protein_coding
protein_coding
protein_coding
protein_coding
protein_coding
protein_coding
protein_coding
protein_coding
protein_coding
protein_coding
protein_coding
protein_coding
protein_coding
protein_coding
protein_coding
protein_coding
protein_coding
protein_coding
protein_coding
protein_coding
small_nucle(ol)ar_rna
small_nucle(ol)ar_rna

small_nucle(ol)ar_rna
ncRNA
protein_coding
protein_coding
protein_coding
protein_coding
protein_coding
protein_coding
protein_coding
protein_coding
protein_coding
protein_coding
protein_coding
protein_coding
protein_coding
protein_coding
protein_coding
protein_coding
protein_coding
protein_coding
protein_coding
protein_coding
protein_coding
protein_coding
protein_coding
protein_coding
protein_coding
protein_coding
protein_coding
protein_coding
protein_coding
protein_coding
protein_coding
protein_coding
protein_coding
protein_coding
protein_coding
protein_coding
protein_coding
protein_coding
protein_coding
protein_coding
protein_coding
protein_coding
protein_coding
protein_coding
protein_coding
protein_coding
ncRNA
small_nucle(ol)ar_rna
small_nucle(ol)ar_rna
ncRNA
protein_coding
protein_coding
protein_coding
protein_coding

protein_coding
protein_coding
protein_coding
protein_coding
protein_coding
protein_coding
protein_coding
protein_coding
protein_coding
protein_coding
protein_coding
protein_coding
protein_coding
protein_coding
protein_coding
protein_coding
protein_coding
ncRNA
protein_coding
protein_coding
protein_coding
protein_coding
ncRNA
protein_coding
protein_coding
protein_coding
protein_coding
protein_coding
protein_coding
miRNA_precursor
protein_coding
protein_coding
ncRNA
protein_coding
protein_coding
protein_coding
protein_coding
protein_coding
protein_coding
protein_coding
protein_coding
protein_coding
protein_coding
protein_coding
protein_coding
protein_coding
protein_coding
protein_coding
protein_coding
protein_coding
protein_coding
protein_coding
protein_coding
protein_coding
protein_coding
protein_coding

protein_coding
protein_coding
protein_coding
protein_coding
ncRNA
protein_coding
protein_coding
protein_coding
protein_coding
protein_coding
protein_coding
protein_coding
protein_coding
miRNA_precursor
protein_coding
protein_coding
protein_coding
protein_coding
protein_coding
protein_coding
protein_coding
protein_coding
protein_coding
protein_coding
protein_coding
protein_coding
pre_trna
protein_coding
protein_coding
protein_coding
ncRNA
protein_coding
protein_coding
protein_coding
protein_coding
protein_coding
protein_coding
protein_coding
protein_coding
protein_coding
protein_coding
protein_coding
protein_coding
protein_coding
protein_coding
protein_coding
protein_coding
protein_coding
protein_coding
protein_coding
protein_coding
protein_coding
protein_coding
protein_coding
protein_coding
protein_coding

protein_coding
protein_coding
protein_coding
protein_coding
protein_coding
protein_coding
ncRNA
protein_coding
protein_coding
protein_coding
protein_coding
ncRNA
protein_coding
protein_coding
protein_coding
ncRNA
protein_coding
protein_coding
protein_coding
ncRNA
protein_coding
protein_coding
protein_coding
protein_coding
protein_coding
protein_coding
protein_coding
protein_coding
protein_coding
protein_coding
protein_coding
protein_coding
protein_coding
protein_coding
protein_coding
protein_coding
protein_coding
protein_coding
protein_coding
ncRNA
protein_coding
protein_coding
protein_coding
protein_coding
protein_coding
protein_coding
protein_coding
protein_coding
protein_coding
protein_coding
protein_coding
protein_coding
protein_coding
protein_coding
protein_coding
protein_coding

protein_coding
protein_coding
protein_coding
protein_coding
protein_coding
protein_coding
protein_coding
protein_coding
protein_coding
protein_coding
protein_coding
protein_coding
protein_coding
protein_coding
protein_coding
protein_coding
protein_coding
protein_coding
protein_coding
protein_coding
protein_coding
protein_coding
protein_coding
protein_coding
protein_coding
protein_coding
protein_coding
protein_coding
protein_coding
protein_coding
protein_coding
protein_coding
protein_coding
protein_coding
protein_coding
protein_coding
protein_coding
ncRNA
protein_coding
protein_coding
protein_coding
protein_coding
protein_coding
protein_coding
protein_coding
protein_coding
protein_coding
protein_coding
protein_coding
protein_coding
protein_coding
protein_coding
protein_coding
protein_coding
protein_coding
protein_coding

protein_coding
protein_coding
protein_coding
protein_coding
protein_coding
protein_coding
protein_coding
protein_coding
protein_coding
protein_coding
protein_coding
protein_coding
protein_coding
protein_coding
protein_coding
protein_coding
protein_coding
protein_coding
protein_coding
protein_coding
protein_coding
protein_coding
protein_coding
protein_coding
ncRNA
protein_coding
protein_coding
protein_coding
protein_coding
protein_coding
protein_coding
protein_coding
protein_coding
protein_coding
protein_coding
protein_coding
protein_coding
protein_coding
protein_coding
protein_coding
protein_coding
protein_coding
protein_coding
protein_coding
ncRNA
protein_coding
protein_coding
protein_coding
protein_coding
protein_coding
protein_coding
protein_coding
protein_coding
protein_coding
protein_coding
protein_coding

protein_coding
protein_coding
protein_coding
protein_coding
protein_coding
protein_coding
protein_coding
protein_coding
protein_coding
protein_coding
pre_trna
protein_coding
protein_coding
protein_coding
protein_coding
protein_coding
protein_coding
protein_coding
small_nucle(ol)ar_rna
protein_coding
protein_coding
protein_coding
protein_coding
protein_coding
protein_coding
pre_trna
protein_coding
ncRNA
protein_coding
protein_coding
protein_coding
protein_coding
protein_coding
protein_coding
protein_coding
protein_coding
protein_coding
protein_coding
protein_coding
protein_coding
protein_coding
protein_coding
protein_coding
protein_coding
protein_coding
protein_coding
protein_coding
protein_coding
protein_coding
protein_coding
protein_coding
protein_coding
protein_coding
protein_coding
protein_coding
protein_coding

protein_coding
protein_coding
protein_coding
protein_coding
protein_coding
protein_coding
protein_coding
protein_coding
protein_coding
protein_coding
protein_coding
protein_coding
protein_coding
protein_coding
protein_coding
protein_coding
protein_coding
protein_coding
protein_coding
protein_coding
protein_coding
protein_coding
protein_coding
protein_coding
protein_coding
protein_coding
protein_coding
protein_coding
protein_coding
protein_coding
protein_coding
protein_coding
protein_coding
protein_coding
protein_coding
protein_coding
protein_coding
protein_coding
protein_coding
protein_coding
protein_coding
protein_coding
protein_coding
protein_coding
protein_coding
protein_coding
protein_coding
miRNA_precursor
protein_coding
protein_coding
small_nucle(ol)ar_rna
small_nucle(ol)ar_rna
ncRNA
protein_coding
protein_coding
protein_coding

protein_coding
protein_coding
protein_coding
protein_coding
protein_coding
protein_coding
protein_coding
protein_coding
protein_coding
protein_coding
protein_coding
protein_coding
protein_coding
protein_coding
protein_coding
protein_coding
protein_coding
protein_coding
protein_coding
protein_coding
protein_coding
protein_coding
protein_coding
protein_coding
protein_coding
protein_coding
protein_coding
protein_coding
protein_coding
protein_coding
protein_coding
protein_coding
protein_coding
ncRNA
protein_coding
protein_coding
protein_coding
protein_coding
protein_coding
ncRNA
protein_coding
protein_coding
protein_coding
protein_coding
protein_coding
protein_coding
protein_coding
protein_coding
protein_coding
protein_coding
protein_coding
protein_coding
protein_coding
protein_coding
protein_coding
protein_coding

protein_coding
protein_coding
protein_coding
protein_coding
protein_coding
protein_coding
protein_coding
protein_coding
protein_coding
protein_coding
protein_coding
protein_coding
protein_coding
protein_coding
protein_coding
protein_coding
protein_coding
protein_coding
protein_coding
protein_coding
protein_coding
protein_coding
protein_coding
protein_coding
protein_coding
protein_coding
protein_coding
protein_coding
protein_coding
protein_coding
ncRNA
protein_coding
protein_coding
protein_coding
protein_coding
protein_coding
protein_coding
protein_coding
protein_coding
protein_coding
protein_coding
protein_coding
protein_coding
protein_coding
protein_coding
miRNA_precursor
protein_coding
protein_coding
protein_coding
protein_coding
protein_coding
protein_coding
protein_coding
protein_coding
protein_coding
protein_coding

protein_coding
protein_coding
protein_coding
protein_coding
small_nucle(ol)ar_rna
small_nucle(ol)ar_rna
protein_coding
protein_coding
protein_coding
protein_coding
protein_coding
protein_coding
protein_coding
protein_coding
protein_coding
protein_coding
small_nucle(ol)ar_rna
protein_coding
protein_coding
protein_coding
protein_coding
protein_coding
protein_coding
protein_coding
protein_coding
protein_coding
protein_coding
protein_coding
protein_coding
protein_coding
protein_coding
protein_coding
protein_coding
protein_coding
protein_coding
protein_coding
protein_coding
ncRNA
protein_coding
protein_coding
protein_coding
protein_coding
protein_coding
protein_coding
protein_coding
protein_coding
protein_coding
protein_coding
protein_coding
protein_coding
protein_coding
protein_coding
protein_coding
protein_coding
protein_coding
protein_coding

protein_coding
protein_coding
protein_coding
protein_coding
protein_coding
protein_coding
protein_coding
protein_coding
protein_coding
protein_coding
protein_coding
protein_coding
protein_coding
pre_trna
protein_coding
ncRNA
protein_coding
protein_coding
protein_coding
protein_coding
protein_coding
protein_coding
protein_coding
protein_coding
protein_coding
protein_coding
protein_coding
protein_coding
protein_coding
protein_coding
protein_coding
protein_coding
protein_coding
protein_coding
protein_coding
protein_coding
protein_coding
protein_coding
protein_coding
protein_coding
protein_coding
protein_coding
protein_coding
protein_coding
protein_coding
protein_coding
protein_coding
protein_coding
protein_coding
protein_coding
protein_coding
protein_coding
protein_coding
protein_coding
protein_coding
protein_coding

protein_coding
protein_coding
protein_coding
protein_coding
protein_coding
protein_coding
protein_coding
protein_coding
protein_coding
protein_coding
protein_coding
protein_coding
ncRNA
protein_coding
protein_coding
protein_coding
protein_coding
protein_coding
protein_coding
protein_coding
protein_coding
protein_coding
protein_coding
protein_coding
protein_coding
protein_coding
protein_coding
protein_coding
protein_coding
protein_coding
protein_coding
protein_coding
ncRNA
small_nucle(ol)ar_rna
protein_coding
protein_coding
protein_coding
protein_coding
protein_coding
protein_coding
protein_coding
protein_coding
protein_coding
protein_coding
protein_coding
protein_coding
ncRNA
protein_coding
protein_coding
protein_coding
protein_coding
protein_coding
protein_coding
protein_coding
protein_coding
protein_coding

protein_coding
protein_coding
protein_coding
protein_coding
protein_coding
protein_coding
protein_coding
protein_coding
protein_coding
protein_coding
protein_coding
protein_coding
protein_coding
protein_coding
protein_coding
protein_coding
protein_coding
miRNA_precursor
miRNA_precursor
protein_coding
protein_coding
protein_coding
protein_coding
protein_coding
protein_coding
protein_coding
protein_coding
protein_coding
protein_coding
protein_coding
protein_coding
protein_coding
protein_coding
ncRNA
ncRNA
protein_coding
protein_coding
protein_coding
protein_coding
protein_coding
protein_coding
protein_coding
protein_coding
protein_coding
protein_coding
ncRNA
protein_coding
protein_coding
protein_coding
protein_coding
pre_trna
protein_coding
protein_coding
protein_coding
protein_coding
protein_coding

protein_coding
protein_coding
protein_coding
protein_coding
protein_coding
protein_coding
ncRNA
protein_coding
protein_coding
protein_coding
protein_coding
protein_coding
protein_coding
ncRNA
protein_coding
protein_coding
ncRNA
protein_coding
protein_coding
protein_coding
protein_coding
protein_coding
protein_coding
protein_coding
protein_coding
protein_coding
protein_coding
protein_coding
protein_coding
protein_coding
protein_coding
protein_coding
protein_coding
protein_coding
protein_coding
protein_coding
protein_coding
protein_coding
protein_coding
protein_coding
protein_coding
protein_coding
protein_coding
protein_coding
protein_coding
miRNA_precursor
protein_coding
protein_coding
protein_coding
protein_coding
protein_coding
ncRNA
ncRNA
protein_coding
protein_coding
protein_coding

protein_coding
protein_coding
protein_coding
protein_coding
protein_coding
protein_coding
protein_coding
protein_coding
protein_coding
protein_coding
protein_coding
protein_coding
protein_coding
protein_coding
protein_coding
protein_coding
protein_coding
protein_coding
protein_coding
protein_coding
protein_coding
ncRNA
protein_coding
protein_coding
miRNA_precursor
protein_coding
miRNA_precursor
protein_coding
protein_coding
protein_coding
protein_coding
protein_coding
protein_coding
protein_coding
protein_coding
protein_coding
protein_coding
protein_coding
protein_coding
protein_coding
protein_coding
protein_coding
protein_coding
protein_coding
ncRNA
protein_coding
protein_coding
protein_coding
protein_coding
protein_coding
protein_coding
protein_coding
protein_coding
protein_coding
protein_coding
protein_coding

protein_coding
protein_coding
protein_coding
protein_coding
protein_coding
protein_coding
protein_coding
protein_coding
protein_coding
protein_coding
protein_coding
protein_coding
protein_coding
protein_coding
protein_coding
protein_coding
protein_coding
protein_coding
protein_coding
protein_coding
protein_coding
protein_coding
protein_coding
protein_coding
protein_coding
protein_coding
protein_coding
protein_coding
protein_coding
protein_coding
protein_coding
protein_coding
protein_coding
protein_coding
protein_coding
protein_coding
protein_coding
protein_coding
protein_coding
protein_coding
protein_coding
ncRNA
protein_coding
protein_coding
protein_coding
protein_coding
protein_coding
protein_coding
protein_coding
protein_coding
protein_coding
protein_coding
protein_coding
protein_coding
protein_coding
protein_coding

protein_coding
protein_coding
protein_coding
protein_coding
protein_coding
protein_coding
protein_coding
protein_coding
protein_coding
protein_coding
protein_coding
protein_coding
protein_coding
protein_coding
protein_coding
protein_coding
protein_coding
protein_coding
protein_coding
protein_coding
protein_coding
protein_coding
small_nucle(ol)ar_rna
protein_coding
protein_coding
protein_coding
pre_trna
protein_coding
pre_trna
protein_coding
protein_coding
protein_coding
protein_coding
protein_coding
protein_coding
protein_coding
protein_coding
protein_coding
protein_coding
protein_coding
ncRNA
protein_coding
protein_coding
ncRNA
ncRNA
protein_coding
protein_coding
protein_coding
protein_coding
protein_coding
protein_coding
protein_coding
protein_coding
protein_coding
protein_coding
protein_coding

protein_coding
miRNA_precursor
miRNA_precursor
protein_coding
protein_coding
protein_coding
protein_coding
protein_coding
protein_coding
protein_coding
protein_coding
protein_coding
protein_coding
protein_coding
protein_coding
protein_coding
protein_coding
protein_coding
protein_coding
protein_coding
protein_coding
protein_coding
protein_coding
protein_coding
protein_coding
protein_coding
protein_coding
protein_coding
protein_coding
protein_coding
protein_coding
protein_coding
protein_coding
protein_coding
protein_coding
protein_coding
protein_coding
protein_coding
protein_coding
protein_coding
protein_coding
protein_coding
protein_coding
protein_coding
protein_coding
protein_coding
protein_coding
protein_coding
protein_coding
protein_coding
protein_coding
protein_coding
protein_coding
protein_coding
protein_coding
protein_coding

protein_coding
protein_coding
protein_coding
protein_coding
pre_trna
protein_coding
protein_coding
protein_coding
protein_coding
protein_coding
protein_coding
protein_coding
protein_coding
protein_coding
protein_coding
protein_coding
protein_coding
protein_coding
protein_coding
protein_coding
protein_coding
protein_coding
protein_coding
protein_coding
protein_coding
protein_coding
protein_coding
protein_coding
protein_coding
ncRNA
protein_coding
ncRNA
protein_coding
protein_coding
protein_coding
protein_coding
protein_coding
protein_coding
ncRNA
protein_coding
protein_coding
protein_coding
protein_coding
protein_coding
protein_coding
small_nucle(ol)ar_rna
protein_coding
protein_coding
protein_coding
protein_coding
protein_coding
protein_coding
ncRNA
protein_coding
protein_coding
protein_coding

protein_coding
protein_coding
protein_coding
protein_coding
protein_coding
protein_coding
protein_coding
protein_coding
protein_coding
protein_coding
protein_coding
protein_coding
protein_coding
protein_coding
protein_coding
protein_coding
protein_coding
protein_coding
protein_coding
protein_coding
protein_coding
protein_coding
protein_coding
protein_coding
protein_coding
protein_coding
protein_coding
protein_coding
protein_coding
protein_coding
ncRNA
protein_coding
protein_coding
protein_coding
ncRNA
protein_coding
protein_coding
protein_coding
protein_coding
protein_coding
protein_coding
protein_coding
protein_coding
protein_coding
protein_coding
protein_coding
ncRNA
protein_coding
protein_coding
protein_coding
protein_coding
protein_coding
protein_coding
protein_coding
protein_coding
protein_coding

protein_coding
protein_coding
protein_coding
protein_coding
protein_coding
protein_coding
protein_coding
protein_coding
protein_coding
protein_coding
protein_coding
protein_coding
protein_coding
protein_coding
protein_coding
protein_coding
protein_coding
ncRNA
protein_coding
protein_coding
ncRNA
ncRNA
protein_coding
protein_coding
protein_coding
protein_coding
protein_coding
protein_coding
protein_coding
protein_coding
protein_coding
protein_coding
protein_coding
protein_coding
protein_coding
protein_coding
protein_coding
protein_coding
protein_coding
protein_coding
protein_coding
protein_coding
protein_coding
protein_coding
protein_coding
protein_coding
protein_coding
protein_coding
protein_coding
protein_coding
protein_coding
protein_coding
ncRNA
protein_coding
protein_coding
protein_coding

protein_coding
protein_coding
protein_coding
protein_coding
protein_coding
protein_coding
protein_coding
protein_coding
protein_coding
pre_trna
protein_coding
protein_coding
protein_coding
protein_coding
protein_coding
protein_coding
protein_coding
protein_coding
protein_coding
protein_coding
protein_coding
protein_coding
protein_coding
protein_coding
ncRNA
protein_coding
protein_coding
protein_coding
protein_coding
protein_coding
protein_coding
protein_coding
protein_coding
protein_coding
protein_coding
protein_coding
protein_coding
protein_coding
protein_coding
protein_coding
small_nucle(ol)ar_rna
protein_coding
protein_coding
protein_coding
protein_coding
protein_coding
protein_coding
protein_coding
protein_coding
protein_coding
protein_coding
protein_coding
protein_coding
protein_coding
protein_coding
protein_coding

protein_coding
protein_coding
protein_coding
protein_coding
protein_coding
small_nucle(ol)ar_rna
small_nucle(ol)ar_rna
protein_coding
protein_coding
protein_coding
protein_coding
pre_trna
miRNA_precursor
protein_coding
protein_coding
protein_coding
protein_coding
protein_coding
protein_coding
protein_coding
protein_coding
ncRNA
protein_coding
ncRNA
ncRNA
protein_coding
protein_coding
protein_coding
protein_coding
protein_coding
protein_coding
protein_coding
protein_coding
protein_coding
protein_coding
protein_coding
protein_coding
protein_coding
protein_coding
protein_coding
protein_coding
protein_coding
ncRNA
protein_coding
protein_coding
protein_coding
protein_coding
protein_coding
protein_coding
protein_coding
protein_coding
protein_coding
protein_coding
protein_coding
protein_coding
ncRNA

protein_coding
protein_coding
protein_coding
miRNA_precursor
protein_coding
protein_coding
protein_coding
ncRNA
protein_coding
protein_coding
protein_coding
protein_coding
protein_coding
protein_coding
protein_coding
protein_coding
protein_coding
protein_coding
protein_coding
protein_coding
protein_coding
protein_coding
protein_coding
protein_coding
protein_coding
protein_coding
protein_coding
ncRNA
protein_coding
protein_coding
protein_coding
protein_coding
protein_coding
protein_coding
protein_coding
protein_coding
protein_coding
protein_coding
protein_coding
protein_coding
protein_coding
protein_coding
protein_coding
miRNA_precursor
protein_coding
protein_coding
protein_coding
protein_coding
protein_coding
protein_coding
protein_coding
protein_coding
protein_coding
protein_coding
protein_coding
protein_coding

miRNA_precursor
pre_trna
protein_coding
protein_coding
protein_coding
protein_coding
protein_coding
ncRNA
protein_coding
protein_coding
protein_coding
miRNA_precursor
protein_coding
protein_coding
protein_coding
protein_coding
protein_coding
miRNA_precursor
protein_coding
ncRNA
protein_coding
protein_coding
protein_coding
protein_coding
protein_coding
protein_coding
protein_coding
protein_coding
protein_coding
protein_coding
protein_coding
protein_coding
protein_coding
protein_coding
protein_coding
protein_coding
protein_coding
protein_coding
protein_coding
protein_coding
protein_coding
protein_coding
protein_coding
protein_coding
protein_coding
protein_coding
protein_coding
protein_coding
protein_coding
protein_coding
protein_coding
protein_coding
protein_coding
protein_coding
protein_coding
protein_coding
protein_coding

protein_coding
protein_coding
protein_coding
protein_coding
protein_coding
protein_coding
protein_coding
protein_coding
protein_coding
protein_coding
protein_coding
protein_coding
protein_coding
protein_coding
protein_coding
protein_coding
miRNA_precursor
protein_coding
protein_coding
protein_coding
protein_coding
protein_coding
protein_coding
protein_coding
protein_coding
protein_coding
protein_coding
ncRNA
small_nucle(ol)ar_rna
small_nucle(ol)ar_rna
small_nucle(ol)ar_rna
protein_coding
protein_coding
protein_coding
protein_coding
protein_coding
ncRNA
protein_coding
protein_coding
protein_coding
protein_coding
protein_coding
protein_coding
protein_coding
protein_coding
protein_coding
protein_coding
protein_coding
protein_coding
protein_coding
protein_coding
protein_coding
protein_coding
protein_coding
protein_coding
protein_coding

protein_coding
protein_coding
protein_coding
protein_coding
protein_coding
ncRNA
protein_coding
protein_coding
protein_coding
protein_coding
protein_coding
protein_coding
protein_coding
protein_coding
protein_coding
protein_coding
protein_coding
protein_coding
protein_coding
protein_coding
protein_coding
protein_coding
protein_coding
protein_coding
protein_coding
protein_coding
protein_coding
protein_coding
protein_coding
protein_coding
protein_coding
protein_coding
protein_coding
protein_coding
protein_coding
protein_coding
protein_coding
protein_coding
protein_coding
protein_coding
protein_coding
pre_trna
protein_coding
protein_coding
protein_coding
protein_coding
protein_coding
protein_coding
protein_coding
protein_coding
protein_coding
protein_coding
protein_coding
protein_coding
protein_coding
protein_coding

ncRNA
protein_coding
protein_coding
protein_coding
protein_coding
protein_coding
protein_coding
protein_coding
protein_coding
miRNA_precursor
protein_coding
protein_coding
protein_coding
protein_coding
protein_coding
protein_coding
protein_coding
protein_coding
protein_coding
protein_coding
ncRNA
protein_coding
protein_coding
protein_coding
protein_coding
protein_coding
protein_coding
protein_coding
protein_coding
protein_coding
protein_coding
protein_coding
ncRNA
protein_coding
protein_coding
protein_coding
protein_coding
protein_coding
protein_coding
protein_coding
protein_coding
protein_coding
protein_coding
protein_coding
protein_coding
protein_coding
protein_coding
protein_coding
protein_coding
protein_coding
protein_coding
protein_coding
protein_coding
protein_coding
pre_trna
protein_coding

protein_coding
protein_coding
protein_coding
protein_coding
protein_coding
protein_coding
protein_coding
protein_coding
protein_coding
protein_coding
protein_coding
protein_coding
protein_coding
protein_coding
protein_coding
protein_coding
protein_coding
protein_coding
ncRNA
protein_coding
protein_coding
protein_coding
protein_coding
protein_coding
protein_coding
protein_coding
protein_coding
protein_coding
protein_coding
ncRNA
protein_coding
protein_coding
ncRNA
small_nucle(ol)ar_rna
protein_coding
protein_coding
protein_coding
protein_coding
protein_coding
protein_coding
protein_coding
protein_coding
protein_coding
protein_coding
protein_coding
ncRNA
protein_coding
protein_coding
protein_coding
ncRNA
protein_coding
protein_coding
protein_coding
protein_coding
protein_coding
protein_coding

protein_coding
protein_coding
protein_coding
protein_coding
protein_coding
protein_coding
protein_coding
ncRNA
protein_coding
protein_coding
protein_coding
protein_coding
protein_coding
protein_coding
protein_coding
protein_coding
protein_coding
protein_coding
pre_trna
protein_coding
protein_coding
protein_coding
protein_coding
protein_coding
protein_coding
protein_coding
protein_coding
protein_coding
protein_coding
protein_coding
protein_coding
protein_coding
protein_coding
protein_coding
protein_coding
miRNA_precursor
small_nucle(ol)ar_rna
small_nucle(ol)ar_rna
protein_coding
pre_trna
protein_coding
protein_coding
protein_coding
protein_coding
protein_coding
protein_coding
protein_coding
ncRNA
protein_coding
protein_coding
protein_coding
protein_coding
protein_coding
ncRNA
protein_coding
protein_coding

protein_coding
protein_coding
protein_coding
protein_coding
protein_coding
protein_coding
protein_coding
protein_coding
protein_coding
protein_coding
protein_coding
protein_coding
protein_coding
pre_trna
protein_coding
protein_coding
protein_coding
protein_coding
protein_coding
protein_coding
protein_coding
protein_coding
protein_coding
protein_coding
protein_coding
protein_coding
protein_coding
protein_coding
protein_coding
protein_coding
protein_coding
protein_coding
protein_coding
protein_coding
protein_coding
protein_coding
protein_coding
protein_coding
small_nucle(ol)ar_rna
protein_coding
protein_coding
protein_coding
protein_coding
protein_coding
protein_coding
protein_coding
protein_coding
protein_coding
protein_coding
protein_coding
protein_coding
protein_coding
protein_coding
protein_coding
protein_coding
protein_coding

protein_coding
protein_coding
protein_coding
protein_coding
protein_coding
protein_coding
protein_coding
protein_coding
protein_coding
protein_coding
protein_coding
protein_coding
protein_coding
protein_coding
protein_coding
protein_coding
protein_coding
protein_coding
protein_coding
protein_coding
protein_coding
protein_coding
protein_coding
protein_coding
protein_coding
protein_coding
protein_coding
protein_coding
protein_coding
protein_coding
protein_coding
protein_coding
protein_coding
protein_coding
protein_coding
protein_coding
protein_coding
protein_coding
protein_coding
protein_coding
protein_coding
protein_coding
protein_coding
protein_coding
protein_coding
protein_coding
protein_coding
protein_coding
protein_coding
protein_coding
protein_coding
protein_coding
protein_coding
protein_coding
protein_coding
protein_coding

protein_coding
protein_coding
protein_coding
protein_coding
protein_coding
protein_coding
protein_coding
protein_coding
protein_coding
protein_coding
protein_coding
protein_coding
protein_coding
protein_coding
protein_coding
protein_coding
protein_coding
protein_coding
protein_coding
protein_coding
protein_coding
protein_coding
protein_coding
protein_coding
protein_coding
protein_coding
protein_coding
protein_coding
protein_coding
protein_coding
protein_coding
ncRNA
protein_coding
protein_coding
ncRNA
protein_coding
protein_coding
protein_coding
protein_coding
protein_coding
protein_coding
protein_coding
protein_coding
protein_coding
protein_coding
protein_coding
protein_coding
protein_coding
protein_coding
protein_coding
protein_coding
protein_coding
protein_coding
protein_coding
protein_coding
protein_coding

protein_coding
protein_coding
protein_coding
protein_coding
protein_coding
protein_coding
protein_coding
protein_coding
protein_coding
protein_coding
protein_coding
protein_coding
protein_coding
protein_coding
protein_coding
protein_coding
protein_coding
protein_coding
protein_coding
protein_coding
protein_coding
protein_coding
protein_coding
protein_coding
ncRNA
protein_coding
protein_coding
protein_coding
protein_coding
protein_coding
protein_coding
protein_coding
protein_coding
ncRNA
protein_coding
protein_coding
protein_coding
ncRNA
protein_coding
protein_coding
protein_coding
protein_coding
protein_coding
protein_coding
protein_coding
protein_coding
protein_coding
protein_coding
protein_coding
protein_coding
protein_coding
protein_coding
protein_coding
protein_coding
ncRNA
protein_coding

protein_coding
protein_coding
protein_coding
ncRNA
protein_coding
protein_coding
protein_coding
protein_coding
protein_coding
protein_coding
protein_coding
protein_coding
protein_coding
protein_coding
protein_coding
protein_coding
protein_coding
protein_coding
protein_coding
protein_coding
ncRNA
protein_coding
protein_coding
protein_coding
protein_coding
protein_coding
protein_coding
pre_trna
protein_coding
protein_coding
protein_coding
protein_coding
protein_coding
protein_coding
protein_coding
protein_coding
protein_coding
protein_coding
protein_coding
protein_coding
protein_coding
protein_coding
protein_coding
protein_coding
ncRNA
pre_trna
protein_coding
protein_coding
protein_coding
protein_coding
protein_coding
protein_coding
protein_coding
ncRNA
protein_coding
protein_coding

protein_coding
ncRNA
protein_coding
protein_coding
pre_trna
protein_coding
protein_coding
protein_coding
protein_coding
protein_coding
protein_coding
protein_coding
protein_coding
protein_coding
protein_coding
protein_coding
protein_coding
protein_coding
protein_coding
protein_coding
protein_coding
protein_coding
protein_coding
protein_coding
protein_coding
protein_coding
protein_coding
protein_coding
protein_coding
protein_coding
protein_coding
protein_coding
protein_coding
protein_coding
protein_coding
protein_coding
protein_coding
protein_coding
protein_coding
protein_coding
protein_coding
protein_coding
protein_coding
protein_coding
protein_coding
protein_coding
protein_coding
protein_coding
protein_coding
protein_coding
protein_coding
protein_coding
protein_coding
protein_coding
ncRNA
protein_coding

protein_coding
protein_coding
protein_coding
protein_coding
protein_coding
protein_coding
protein_coding
protein_coding
protein_coding
protein_coding
protein_coding
protein_coding
protein_coding
protein_coding
protein_coding
protein_coding
protein_coding
protein_coding
protein_coding
ncRNA
protein_coding
ncRNA
protein_coding
protein_coding
protein_coding
protein_coding
ncRNA
protein_coding
protein_coding
protein_coding
protein_coding
protein_coding
protein_coding
small_nucle(ol)ar_rna
small_nucle(ol)ar_rna
protein_coding
protein_coding
protein_coding
protein_coding
protein_coding
protein_coding
protein_coding
protein_coding
protein_coding
protein_coding
protein_coding
protein_coding
protein_coding
ncRNA
protein_coding
ncRNA
protein_coding
protein_coding
protein_coding
protein_coding
protein_coding

protein_coding
pre_trna
protein_coding
protein_coding
protein_coding
protein_coding
pre_trna
protein_coding
protein_coding
protein_coding
protein_coding
small_nucle(ol)ar_rna
small_nucle(ol)ar_rna
small_nucle(ol)ar_rna
protein_coding
protein_coding
protein_coding
protein_coding
protein_coding
protein_coding
protein_coding
protein_coding
protein_coding
protein_coding
protein_coding
protein_coding
ncRNA
protein_coding
protein_coding
protein_coding
protein_coding
protein_coding
protein_coding
protein_coding
protein_coding
protein_coding
protein_coding
pre_trna
protein_coding
protein_coding
protein_coding
protein_coding
protein_coding
protein_coding
protein_coding
protein_coding
protein_coding
protein_coding
protein_coding
protein_coding
ncRNA
protein_coding
protein_coding
protein_coding
protein_coding
protein_coding

protein_coding
protein_coding
ncRNA
protein_coding
protein_coding
protein_coding
protein_coding
protein_coding
protein_coding
protein_coding
protein_coding
protein_coding
protein_coding
protein_coding
protein_coding
protein_coding
protein_coding
protein_coding
protein_coding
protein_coding
protein_coding
protein_coding
protein_coding
protein_coding
protein_coding
protein_coding
protein_coding
protein_coding
protein_coding
protein_coding
protein_coding
protein_coding
protein_coding
protein_coding
protein_coding
protein_coding
protein_coding
ncRNA
small_nucle(ol)ar_rna
protein_coding
protein_coding
protein_coding
protein_coding
protein_coding
protein_coding
protein_coding
protein_coding
protein_coding
protein_coding
protein_coding
protein_coding
protein_coding
protein_coding
protein_coding
protein_coding
protein_coding

protein_coding
protein_coding
protein_coding
protein_coding
protein_coding
protein_coding
protein_coding
protein_coding
small_nucle(ol)ar_rna
protein_coding
protein_coding
protein_coding
protein_coding
protein_coding
protein_coding
protein_coding
protein_coding
protein_coding
protein_coding
protein_coding
protein_coding
protein_coding
protein_coding
protein_coding
protein_coding
protein_coding
protein_coding
protein_coding
protein_coding
protein_coding
protein_coding
protein_coding
protein_coding
protein_coding
protein_coding
protein_coding
protein_coding
protein_coding
protein_coding
protein_coding
protein_coding
protein_coding
protein_coding
protein_coding
protein_coding
protein_coding
protein_coding
protein_coding
protein_coding
protein_coding
protein_coding
protein_coding
protein_coding
protein_coding
protein_coding
protein_coding

protein_coding
protein_coding
protein_coding
protein_coding
protein_coding
protein_coding
protein_coding
small_nucle(ol)ar_rna
small_nucle(ol)ar_rna
small_nucle(ol)ar_rna
protein_coding
protein_coding
protein_coding
protein_coding
protein_coding
protein_coding
protein_coding
protein_coding
protein_coding
protein_coding
protein_coding
protein_coding
protein_coding
protein_coding
protein_coding
protein_coding
protein_coding
protein_coding
protein_coding
protein_coding
protein_coding
protein_coding
protein_coding
protein_coding
protein_coding
protein_coding
ncRNA
protein_coding
protein_coding
protein_coding
protein_coding
protein_coding
protein_coding
ncRNA
protein_coding
protein_coding
protein_coding
protein_coding
protein_coding
protein_coding
protein_coding
protein_coding
protein_coding
protein_coding
protein_coding
protein_coding

protein_coding
protein_coding
protein_coding
protein_coding
protein_coding
protein_coding
protein_coding
protein_coding
protein_coding
ncRNA
protein_coding
protein_coding
protein_coding
protein_coding
protein_coding
protein_coding
protein_coding
protein_coding
protein_coding
protein_coding
protein_coding
protein_coding
protein_coding
protein_coding
protein_coding
protein_coding
protein_coding
protein_coding
protein_coding
miRNA_precursor
protein_coding
protein_coding
protein_coding
protein_coding
protein_coding
protein_coding
protein_coding
protein_coding
protein_coding
protein_coding
protein_coding
protein_coding
protein_coding
protein_coding
protein_coding
protein_coding
protein_coding
protein_coding
protein_coding
protein_coding
protein_coding
miRNA_precursor
protein_coding
protein_coding
protein_coding
protein_coding

protein_coding
miRNA_precursor
protein_coding
protein_coding
protein_coding
protein_coding
protein_coding
protein_coding
protein_coding
protein_coding
protein_coding
protein_coding
protein_coding
protein_coding
protein_coding
protein_coding
protein_coding
protein_coding
protein_coding
protein_coding
protein_coding
protein_coding
protein_coding
protein_coding
protein_coding
protein_coding
protein_coding
protein_coding
protein_coding
protein_coding
protein_coding
protein_coding
protein_coding
protein_coding
protein_coding
protein_coding
protein_coding
protein_coding
protein_coding
protein_coding
ncRNA
protein_coding
protein_coding
protein_coding
protein_coding
protein_coding
protein_coding
protein_coding
protein_coding
protein_coding
protein_coding
protein_coding
protein_coding
ncRNA
protein_coding
protein_coding

protein_coding
protein_coding
protein_coding
protein_coding
small_nucle(ol)ar_rna
protein_coding
protein_coding
protein_coding
protein_coding
protein_coding
protein_coding
protein_coding
protein_coding
ncRNA
protein_coding
protein_coding
miRNA_precursor
protein_coding
protein_coding
protein_coding
protein_coding
protein_coding
protein_coding
protein_coding
protein_coding
protein_coding
ncRNA
protein_coding
protein_coding
protein_coding
protein_coding
protein_coding
pre_trna
ncRNA
protein_coding
protein_coding
protein_coding
protein_coding
protein_coding
protein_coding
protein_coding
protein_coding
ncRNA
protein_coding
ncRNA
protein_coding
protein_coding
protein_coding
protein_coding
protein_coding
protein_coding
protein_coding
protein_coding
protein_coding
protein_coding
protein_coding

protein_coding
protein_coding
ncRNA
protein_coding
protein_coding
ncRNA
protein_coding
protein_coding
protein_coding
ncRNA
protein_coding
protein_coding
protein_coding
protein_coding
protein_coding
protein_coding
protein_coding
protein_coding
protein_coding
protein_coding
protein_coding
protein_coding
protein_coding
protein_coding
protein_coding
protein_coding
protein_coding
protein_coding
protein_coding
protein_coding
protein_coding
protein_coding
protein_coding
protein_coding
protein_coding
protein_coding
ncRNA
protein_coding
protein_coding
protein_coding
protein_coding
protein_coding
protein_coding
protein_coding
protein_coding
protein_coding
protein_coding
protein_coding
protein_coding
protein_coding
protein_coding
protein_coding
protein_coding
protein_coding
protein_coding
ncRNA

protein_coding
protein_coding
ncRNA
protein_coding
protein_coding
protein_coding
ncRNA
protein_coding
protein_coding
protein_coding
ncRNA
protein_coding
protein_coding
protein_coding
protein_coding
protein_coding
protein_coding
protein_coding
ncRNA
protein_coding
protein_coding
protein_coding
protein_coding
protein_coding
ncRNA
protein_coding
miRNA_precursor
protein_coding
protein_coding
protein_coding
protein_coding
protein_coding
protein_coding
protein_coding
protein_coding
protein_coding
protein_coding
protein_coding
protein_coding
protein_coding
protein_coding
protein_coding
protein_coding
protein_coding
protein_coding
protein_coding
protein_coding
protein_coding
protein_coding
ncRNA
protein_coding
protein_coding
protein_coding
protein_coding
protein_coding
protein_coding

protein_coding
protein_coding
protein_coding
protein_coding
protein_coding
protein_coding
small_nucle(ol)ar_rna
protein_coding
protein_coding
protein_coding
protein_coding
protein_coding
protein_coding
protein_coding
protein_coding
protein_coding
protein_coding
protein_coding
protein_coding
protein_coding
protein_coding
protein_coding
protein_coding
protein_coding
protein_coding
protein_coding
protein_coding
protein_coding
protein_coding
protein_coding
protein_coding
protein_coding
protein_coding
protein_coding
protein_coding
protein_coding
protein_coding
protein_coding
protein_coding
protein_coding
protein_coding
protein_coding
protein_coding
protein_coding
protein_coding
protein_coding
protein_coding
protein_coding
protein_coding
pre_trna
protein_coding
protein_coding
protein_coding
protein_coding
protein_coding
ncRNA

protein_coding
protein_coding
protein_coding
protein_coding
protein_coding
protein_coding
protein_coding
protein_coding
protein_coding
protein_coding
protein_coding
protein_coding
protein_coding
protein_coding
protein_coding
protein_coding
protein_coding
protein_coding
protein_coding
protein_coding
protein_coding
protein_coding
protein_coding
protein_coding
protein_coding
protein_coding
protein_coding
protein_coding
protein_coding
protein_coding
protein_coding
protein_coding
protein_coding
protein_coding
protein_coding
protein_coding
protein_coding
protein_coding
protein_coding
protein_coding
protein_coding
protein_coding
protein_coding
protein_coding
protein_coding
protein_coding
protein_coding
protein_coding
protein_coding
protein_coding
protein_coding
protein_coding
protein_coding
protein_coding
protein_coding
protein_coding

protein_coding
protein_coding
protein_coding
protein_coding
protein_coding
protein_coding
protein_coding
protein_coding
miRNA_precursor
protein_coding
protein_coding
protein_coding
protein_coding
ncRNA
protein_coding
protein_coding
protein_coding
protein_coding
protein_coding
ncRNA
protein_coding
ncRNA
protein_coding
protein_coding
protein_coding
protein_coding
protein_coding
protein_coding
protein_coding
protein_coding
protein_coding
protein_coding
protein_coding
protein_coding
protein_coding
protein_coding
protein_coding
protein_coding
protein_coding
protein_coding
protein_coding
protein_coding
protein_coding
protein_coding
protein_coding
protein_coding
protein_coding
protein_coding
protein_coding
protein_coding
protein_coding
protein_coding
protein_coding
protein_coding
protein_coding
protein_coding

protein_coding
protein_coding
protein_coding
protein_coding
protein_coding
protein_coding
protein_coding
protein_coding
protein_coding
protein_coding
protein_coding
protein_coding
protein_coding
protein_coding
protein_coding
protein_coding
protein_coding
protein_coding
protein_coding
protein_coding
protein_coding
protein_coding
protein_coding
protein_coding
protein_coding
protein_coding
protein_coding
protein_coding
protein_coding
protein_coding
protein_coding
protein_coding
protein_coding
protein_coding
miRNA_precursor
protein_coding
protein_coding
protein_coding
protein_coding
ncRNA
protein_coding
protein_coding
ncRNA
protein_coding
protein_coding
protein_coding
protein_coding
protein_coding
protein_coding
protein_coding
protein_coding
protein_coding
protein_coding
protein_coding
protein_coding
protein_coding

protein_coding
protein_coding
protein_coding
protein_coding
protein_coding
protein_coding
protein_coding
protein_coding
protein_coding
protein_coding
protein_coding
protein_coding
protein_coding
protein_coding
protein_coding
protein_coding
protein_coding
protein_coding
protein_coding
protein_coding
protein_coding
protein_coding
pre_trna
protein_coding
protein_coding
protein_coding
protein_coding
protein_coding
protein_coding
protein_coding
protein_coding
protein_coding
protein_coding
protein_coding
ncRNA
ncRNA
protein_coding
protein_coding
protein_coding
protein_coding
protein_coding
protein_coding
protein_coding
protein_coding
protein_coding
miRNA_precursor
protein_coding
protein_coding
protein_coding
protein_coding
protein_coding
protein_coding
protein_coding
protein_coding
protein_coding
protein_coding

protein_coding
protein_coding
protein_coding
protein_coding
protein_coding
protein_coding
protein_coding
protein_coding
protein_coding
protein_coding
protein_coding
protein_coding
protein_coding
protein_coding
protein_coding
protein_coding
protein_coding
protein_coding
protein_coding
protein_coding
protein_coding
protein_coding
protein_coding
protein_coding
protein_coding
protein_coding
protein_coding
protein_coding
protein_coding
protein_coding
protein_coding
protein_coding
protein_coding
protein_coding
protein_coding
protein_coding
small_nucle(ol)ar_rna
protein_coding
protein_coding
protein_coding
protein_coding
protein_coding
protein_coding
protein_coding
protein_coding
protein_coding
protein_coding
protein_coding
protein_coding
ncRNA
protein_coding
protein_coding
ncRNA
protein_coding
protein_coding
protein_coding

protein_coding
protein_coding
protein_coding
protein_coding
protein_coding
protein_coding
protein_coding
protein_coding
protein_coding
protein_coding
protein_coding
protein_coding
protein_coding
protein_coding
protein_coding
protein_coding
protein_coding
protein_coding
protein_coding
protein_coding
protein_coding
protein_coding
protein_coding
protein_coding
protein_coding
protein_coding
protein_coding
small_nucle(ol)ar_rna
protein_coding
protein_coding
protein_coding
protein_coding
protein_coding
protein_coding
protein_coding
protein_coding
protein_coding
protein_coding
protein_coding
protein_coding
protein_coding
protein_coding
protein_coding
protein_coding
protein_coding
protein_coding
protein_coding
protein_coding
protein_coding
protein_coding
protein_coding
protein_coding
protein_coding
protein_coding
protein_coding
protein_coding

protein_coding
protein_coding
protein_coding
protein_coding
protein_coding
protein_coding
protein_coding
protein_coding
protein_coding
protein_coding
protein_coding
protein_coding
protein_coding
protein_coding
protein_coding
protein_coding
protein_coding
protein_coding
protein_coding
protein_coding
protein_coding
protein_coding
protein_coding
protein_coding
protein_coding
protein_coding
protein_coding
protein_coding
protein_coding
protein_coding
protein_coding
protein_coding
protein_coding
protein_coding
protein_coding
protein_coding
protein_coding
protein_coding
protein_coding
protein_coding
protein_coding
protein_coding
protein_coding
protein_coding
protein_coding
protein_coding
protein_coding
protein_coding
protein_coding
protein_coding
protein_coding
protein_coding
protein_coding
protein_coding
protein_coding
protein_coding

protein_coding
protein_coding
protein_coding
protein_coding
protein_coding
protein_coding
protein_coding
ncRNA
protein_coding
protein_coding
protein_coding
protein_coding
protein_coding
protein_coding
protein_coding
protein_coding
protein_coding
protein_coding
protein_coding
protein_coding
protein_coding
protein_coding
ncRNA
protein_coding
protein_coding
protein_coding
protein_coding
protein_coding
protein_coding
protein_coding
protein_coding
protein_coding
protein_coding
protein_coding
protein_coding
protein_coding
protein_coding
protein_coding
protein_coding
protein_coding
protein_coding
protein_coding
protein_coding
ncRNA
protein_coding
protein_coding
protein_coding
protein_coding
protein_coding
protein_coding
protein_coding
protein_coding

**Dataset 1.6 Differential analysis of 21/22nt accumulation on Araport11 genes in c**

| ID | baseMean | log2FoldChange | lfcSE | stat | pvalue | padj |
|---|---|---|---|---|---|---|
| AT2G26211 | 125,25 | -5,55 | 0,32 | -17,46 | 2,68E-68 | 8,13E-65 |
| AT5G23065 | 1776,68 | -4,02 | 0,26 | -15,27 | 1,26E-52 | 1,91E-49 |
| AT5G13630 | 269,68 | -5,95 | 0,41 | -14,67 | 9,34E-49 | 9,43E-46 |
| AT5G38850 | 479,48 | 3,25 | 0,23 | 14,02 | 1,23E-44 | 9,30E-42 |
| AT3G23325 | 69,65 | -5,15 | 0,38 | -13,57 | 6,09E-42 | 3,69E-39 |
| AT3G23326 | 69,32 | -5,22 | 0,39 | -13,44 | 3,49E-41 | 1,76E-38 |
| AT1G31280 | 162,06 | -7,31 | 0,60 | -12,14 | 6,36E-34 | 2,75E-31 |
| AT4G06810 | 54,17 | -5,20 | 0,43 | -12,02 | 2,64E-33 | 9,98E-31 |
| AT4G06805 | 54,05 | -5,19 | 0,43 | -11,98 | 4,61E-33 | 1,55E-30 |
| AT5G08185 | 1387,13 | -3,69 | 0,31 | -11,84 | 2,42E-32 | 7,32E-30 |
| AT2G10606 | 1219,01 | -3,26 | 0,28 | -11,82 | 3,20E-32 | 8,82E-30 |
| AT3G01845 | 219,09 | 2,72 | 0,23 | 11,80 | 4,05E-32 | 1,02E-29 |
| AT3G51375 | 135,65 | -4,51 | 0,40 | -11,41 | 3,87E-30 | 9,02E-28 |
| AT1G56070 | 55,72 | -3,62 | 0,33 | -11,11 | 1,15E-28 | 2,48E-26 |
| AT1G62914 | 457,48 | 3,48 | 0,34 | 10,10 | 5,45E-24 | 1,10E-21 |
| AT1G63150 | 176,58 | 3,44 | 0,34 | 10,08 | 6,67E-24 | 1,26E-21 |
| AT2G47275 | 830,57 | -3,07 | 0,31 | -9,97 | 2,12E-23 | 3,78E-21 |
| AT1G63130 | 240,21 | 3,00 | 0,31 | 9,60 | 7,99E-22 | 1,34E-19 |
| AT4G23713 | 128,07 | 6,75 | 0,71 | 9,46 | 3,13E-21 | 4,99E-19 |
| AT1G62910 | 98,21 | 4,08 | 0,43 | 9,44 | 3,62E-21 | 5,23E-19 |
| AT4G23387 | 136,04 | 4,93 | 0,52 | 9,44 | 3,60E-21 | 5,23E-19 |
| AT2G18960 | 62,63 | -5,45 | 0,59 | -9,28 | 1,71E-20 | 2,36E-18 |
| AT3G17185 | 156,12 | 2,79 | 0,30 | 9,16 | 5,01E-20 | 6,60E-18 |
| AT3G44570 | 48,95 | -4,74 | 0,52 | -9,12 | 7,84E-20 | 9,89E-18 |
| AT5G05170 | 36,40 | -5,43 | 0,60 | -9,00 | 2,28E-19 | 2,76E-17 |
| AT1G29265 | 54,27 | -3,26 | 0,36 | -8,94 | 3,91E-19 | 4,55E-17 |
| AT1G15750 | 35,69 | -6,61 | 0,74 | -8,89 | 6,27E-19 | 7,04E-17 |
| AT2G23350 | 29,35 | -6,04 | 0,68 | -8,83 | 1,01E-18 | 1,10E-16 |
| AT1G63080 | 109,45 | 3,11 | 0,35 | 8,79 | 1,50E-18 | 1,57E-16 |
| AT1G18075 | 4243,99 | 1,52 | 0,18 | 8,56 | 1,13E-17 | 1,14E-15 |
| AT5G14565 | 6609,25 | 2,26 | 0,27 | 8,49 | 1,99E-17 | 1,95E-15 |
| AT3G48201 | 30,66 | -4,31 | 0,51 | -8,37 | 5,89E-17 | 5,58E-15 |
| AT4G06075 | 405,29 | 2,66 | 0,33 | 8,07 | 7,06E-16 | 6,48E-14 |
| AT4G03039 | 93,29 | 6,33 | 0,79 | 8,01 | 1,12E-15 | 9,43E-14 |
| AT4G03038 | 93,33 | 6,33 | 0,79 | 8,01 | 1,12E-15 | 9,43E-14 |
| AT5G02500 | 62,24 | -2,57 | 0,32 | -8,02 | 1,08E-15 | 9,43E-14 |
| AT4G04365 | 93,18 | 6,33 | 0,79 | 8,01 | 1,18E-15 | 9,64E-14 |
| AT2G15530 | 40,01 | -2,65 | 0,34 | -7,79 | 6,68E-15 | 5,33E-13 |
| AT3G01540 | 20,21 | -5,15 | 0,67 | -7,71 | 1,24E-14 | 9,65E-13 |
| AT2G40805 | 280,87 | 3,19 | 0,41 | 7,70 | 1,39E-14 | 1,05E-12 |
| AT2G21390 | 23,77 | -5,38 | 0,70 | -7,66 | 1,84E-14 | 1,36E-12 |
| AT4G34450 | 21,43 | -4,98 | 0,65 | -7,63 | 2,38E-14 | 1,72E-12 |
| AT2G22125 | 19,76 | -4,88 | 0,65 | -7,49 | 7,10E-14 | 5,01E-12 |
| AT1G66725 | 1003,08 | 2,30 | 0,31 | 7,47 | 7,75E-14 | 5,34E-12 |
| AT1G09620 | 17,50 | -5,17 | 0,70 | -7,36 | 1,84E-13 | 1,24E-11 |
| AT3G11130 | 25,04 | -3,96 | 0,54 | -7,29 | 3,18E-13 | 2,09E-11 |
| AT1G12820 | 53,20 | 3,10 | 0,43 | 7,15 | 8,82E-13 | 5,69E-11 |
| AT1G12294 | 124,95 | -1,91 | 0,27 | -7,13 | 1,02E-12 | 6,44E-11 |
| AT1G37130 | 39,12 | -2,62 | 0,37 | -7,00 | 2,54E-12 | 1,57E-10 |
| AT3G44580 | 31,33 | -3,07 | 0,44 | -6,99 | 2,66E-12 | 1,61E-10 |
| AT1G01320 | 16,50 | -3,57 | 0,51 | -6,95 | 3,67E-12 | 2,18E-10 |
| AT5G46315 | 17,01 | -8,80 | 1,27 | -6,92 | 4,42E-12 | 2,58E-10 |
| AT5G60408 | 515,37 | 2,04 | 0,30 | 6,87 | 6,32E-12 | 3,61E-10 |

| | | | | | | |
|---|---|---|---|---|---|---|
| AT1G53683 | 15,44 | -8,80 | 1,29 | -6,85 | 7,64E-12 | 4,29E-10 |
| AT1G70320 | 18,40 | -3,41 | 0,50 | -6,77 | 1,29E-11 | 7,10E-10 |
| AT5G64270 | 15,19 | -5,26 | 0,78 | -6,71 | 1,90E-11 | 1,03E-09 |
| AT1G09797 | 19,74 | -3,47 | 0,52 | -6,70 | 2,02E-11 | 1,06E-09 |
| AT1G09793 | 19,74 | -3,47 | 0,52 | -6,70 | 2,02E-11 | 1,06E-09 |
| AT5G07745 | 20,93 | -3,61 | 0,54 | -6,63 | 3,36E-11 | 1,72E-09 |
| AT4G28080 | 23,67 | -4,69 | 0,71 | -6,62 | 3,67E-11 | 1,85E-09 |
| AT1G04977 | 45,78 | 3,13 | 0,48 | 6,57 | 5,00E-11 | 2,48E-09 |
| AT1G29930 | 124,68 | -2,49 | 0,38 | -6,51 | 7,75E-11 | 3,73E-09 |
| AT2G46255 | 72,33 | 3,16 | 0,49 | 6,51 | 7,66E-11 | 3,73E-09 |
| AT1G63400 | 45,16 | 8,05 | 1,24 | 6,49 | 8,42E-11 | 3,99E-09 |
| AT1G06227 | 343,33 | 2,44 | 0,38 | 6,47 | 9,51E-11 | 4,43E-09 |
| AT4G03455 | 16,01 | -8,36 | 1,31 | -6,36 | 2,04E-10 | 9,36E-09 |
| AT1G06237 | 336,80 | 2,46 | 0,39 | 6,35 | 2,16E-10 | 9,75E-09 |
| AT5G45307 | 438,83 | -1,98 | 0,31 | -6,34 | 2,35E-10 | 1,05E-08 |
| AT2G28290 | 15,57 | -4,42 | 0,70 | -6,33 | 2,51E-10 | 1,10E-08 |
| AT5G41663 | 49,62 | 3,28 | 0,52 | 6,31 | 2,73E-10 | 1,18E-08 |
| AT4G00885 | 370,66 | 1,90 | 0,30 | 6,28 | 3,40E-10 | 1,45E-08 |
| AT1G12290 | 131,54 | -1,61 | 0,26 | -6,23 | 4,76E-10 | 2,00E-08 |
| AT1G07051 | 29,68 | -3,36 | 0,54 | -6,22 | 4,99E-10 | 2,07E-08 |
| AT3G48870 | 10,82 | -8,10 | 1,32 | -6,12 | 9,44E-10 | 3,87E-08 |
| AT4G02480 | 22,30 | -4,82 | 0,79 | -6,10 | 1,07E-09 | 4,31E-08 |
| AT2G41900 | 16,69 | -4,98 | 0,82 | -6,07 | 1,26E-09 | 5,00E-08 |
| AT1G50055 | 182,36 | 1,85 | 0,30 | 6,07 | 1,31E-09 | 5,14E-08 |
| AT3G02260 | 23,16 | -3,22 | 0,53 | -6,04 | 1,57E-09 | 6,10E-08 |
| AT5G17920 | 34,31 | -2,41 | 0,40 | -6,03 | 1,69E-09 | 6,47E-08 |
| AT1G70770 | 12,70 | -4,44 | 0,74 | -6,02 | 1,80E-09 | 6,80E-08 |
| AT5G42950 | 18,25 | -4,47 | 0,76 | -5,84 | 5,13E-09 | 1,92E-07 |
| AT1G62590 | 20,75 | 7,23 | 1,24 | 5,84 | 5,35E-09 | 1,98E-07 |
| AT3G59884 | 100,21 | -1,85 | 0,32 | -5,79 | 6,95E-09 | 2,51E-07 |
| AT4G21595 | 28,61 | 7,18 | 1,24 | 5,79 | 6,92E-09 | 2,51E-07 |
| AT4G13940 | 35,08 | -2,35 | 0,41 | -5,78 | 7,34E-09 | 2,62E-07 |
| AT4G12770 | 19,67 | 4,04 | 0,71 | 5,72 | 1,04E-08 | 3,67E-07 |
| AT1G10760 | 16,93 | -3,27 | 0,58 | -5,64 | 1,68E-08 | 5,87E-07 |
| AT3G01715 | 19,56 | -3,31 | 0,59 | -5,64 | 1,71E-08 | 5,90E-07 |
| AT5G15833 | 13,93 | -3,71 | 0,66 | -5,64 | 1,75E-08 | 5,94E-07 |
| AT3G06435 | 378,12 | 1,43 | 0,25 | 5,62 | 1,94E-08 | 6,53E-07 |
| AT3G01835 | 72,10 | 1,73 | 0,31 | 5,60 | 2,19E-08 | 7,29E-07 |
| AT1G29920 | 50,33 | -2,26 | 0,40 | -5,59 | 2,23E-08 | 7,35E-07 |
| AT3G08943 | 11,39 | -5,85 | 1,05 | -5,59 | 2,32E-08 | 7,55E-07 |
| AT5G66045 | 69,41 | -2,70 | 0,48 | -5,58 | 2,37E-08 | 7,64E-07 |
| AT3G05700 | 20,03 | -3,12 | 0,56 | -5,58 | 2,40E-08 | 7,64E-07 |
| AT5G10946 | 91,61 | 1,71 | 0,31 | 5,58 | 2,42E-08 | 7,65E-07 |
| AT3G50370 | 9,94 | -3,75 | 0,67 | -5,57 | 2,57E-08 | 8,02E-07 |
| AT2G28000 | 11,21 | -3,76 | 0,68 | -5,56 | 2,77E-08 | 8,55E-07 |
| AT3G43300 | 11,08 | -4,03 | 0,73 | -5,55 | 2,82E-08 | 8,62E-07 |
| AT5G60790 | 11,59 | -4,03 | 0,73 | -5,55 | 2,85E-08 | 8,62E-07 |
| AT4G08691 | 15,19 | 7,12 | 1,28 | 5,55 | 2,92E-08 | 8,77E-07 |
| AT3G19820 | 10,30 | -7,15 | 1,29 | -5,54 | 2,99E-08 | 8,88E-07 |
| AT5G10945 | 91,45 | 1,71 | 0,31 | 5,54 | 3,05E-08 | 8,96E-07 |
| AT3G18780 | 17,06 | -3,04 | 0,55 | -5,53 | 3,23E-08 | 9,42E-07 |
| AT5G07350 | 9,52 | -4,69 | 0,85 | -5,51 | 3,68E-08 | 1,06E-06 |
| AT1G05163 | 491,36 | 1,50 | 0,27 | 5,50 | 3,91E-08 | 1,11E-06 |
| AT1G15405 | 491,36 | 1,50 | 0,27 | 5,50 | 3,91E-08 | 1,11E-06 |
| AT3G52140 | 9,89 | -3,84 | 0,70 | -5,49 | 3,96E-08 | 1,11E-06 |
| AT2G03445 | 309,93 | 1,47 | 0,27 | 5,49 | 4,06E-08 | 1,13E-06 |

| | | | | | | |
|---|---|---|---|---|---|---|
| AT5G44340 | 14,19 | -3,46 | 0,63 | -5,48 | 4,14E-08 | 1,14E-06 |
| AT2G16030 | 25,88 | 5,90 | 1,09 | 5,43 | 5,70E-08 | 1,56E-06 |
| AT5G27606 | 14,40 | 6,90 | 1,27 | 5,42 | 5,94E-08 | 1,61E-06 |
| AT1G10290 | 9,22 | -5,12 | 0,95 | -5,41 | 6,15E-08 | 1,65E-06 |
| AT2G06555 | 20,57 | 4,20 | 0,78 | 5,39 | 7,02E-08 | 1,87E-06 |
| AT2G43900 | 9,76 | -6,96 | 1,30 | -5,36 | 8,36E-08 | 2,20E-06 |
| AT1G64720 | 13,91 | -3,59 | 0,67 | -5,35 | 8,70E-08 | 2,27E-06 |
| AT4G07975 | 9,02 | -4,07 | 0,76 | -5,35 | 8,82E-08 | 2,28E-06 |
| AT5G04140 | 18,61 | -3,09 | 0,58 | -5,33 | 9,64E-08 | 2,48E-06 |
| AT4G17140 | 11,35 | -4,08 | 0,77 | -5,32 | 1,03E-07 | 2,63E-06 |
| AT4G36080 | 9,35 | -4,07 | 0,77 | -5,32 | 1,06E-07 | 2,67E-06 |
| AT3G10745 | 164,34 | -1,73 | 0,33 | -5,30 | 1,18E-07 | 2,95E-06 |
| AT1G11700 | 23,46 | 3,08 | 0,59 | 5,26 | 1,44E-07 | 3,57E-06 |
| AT4G27000 | 10,07 | -4,05 | 0,77 | -5,26 | 1,47E-07 | 3,63E-06 |
| AT5G03553 | 58,75 | 2,04 | 0,39 | 5,25 | 1,56E-07 | 3,81E-06 |
| AT2G20190 | 8,93 | -5,32 | 1,02 | -5,23 | 1,66E-07 | 4,02E-06 |
| AT2G37190 | 11,90 | -4,15 | 0,79 | -5,23 | 1,71E-07 | 4,11E-06 |
| AT5G03552 | 58,24 | 2,03 | 0,39 | 5,22 | 1,79E-07 | 4,27E-06 |
| AT1G79920 | 8,70 | -4,58 | 0,88 | -5,18 | 2,27E-07 | 5,37E-06 |
| AT5G53460 | 17,48 | -2,47 | 0,48 | -5,16 | 2,41E-07 | 5,66E-06 |
| AT1G55591 | 5,61 | -7,32 | 1,43 | -5,12 | 3,07E-07 | 7,15E-06 |
| AT3G09840 | 22,25 | -2,33 | 0,45 | -5,11 | 3,15E-07 | 7,18E-06 |
| AT2G43970 | 15,13 | -2,90 | 0,57 | -5,11 | 3,15E-07 | 7,18E-06 |
| AT4G32410 | 11,90 | -3,71 | 0,73 | -5,12 | 3,12E-07 | 7,18E-06 |
| AT1G12775 | 21,87 | 3,39 | 0,66 | 5,10 | 3,48E-07 | 7,86E-06 |
| AT1G02500 | 10,85 | -4,09 | 0,81 | -5,07 | 3,90E-07 | 8,75E-06 |
| AT4G27640 | 8,75 | -5,65 | 1,11 | -5,07 | 3,96E-07 | 8,83E-06 |
| AT1G23310 | 12,45 | -2,76 | 0,55 | -5,06 | 4,11E-07 | 9,09E-06 |
| AT1G22610 | 11,17 | -3,20 | 0,64 | -5,03 | 4,83E-07 | 1,06E-05 |
| AT5G65950 | 6,84 | -7,55 | 1,50 | -5,03 | 5,03E-07 | 1,10E-05 |
| AT2G43680 | 14,92 | -3,13 | 0,63 | -5,00 | 5,79E-07 | 1,25E-05 |
| AT4G01800 | 8,35 | -5,83 | 1,18 | -4,94 | 7,75E-07 | 1,67E-05 |
| AT5G15680 | 7,75 | -4,56 | 0,93 | -4,90 | 9,69E-07 | 2,07E-05 |
| AT5G23110 | 11,58 | -3,21 | 0,66 | -4,88 | 1,03E-06 | 2,18E-05 |
| AT1G10170 | 9,09 | -3,90 | 0,80 | -4,89 | 1,03E-06 | 2,18E-05 |
| AT1G08597 | 22,78 | 4,47 | 0,92 | 4,88 | 1,06E-06 | 2,22E-05 |
| AT3G18290 | 9,13 | -3,96 | 0,82 | -4,86 | 1,16E-06 | 2,41E-05 |
| AT3G62980 | 20,74 | -2,49 | 0,52 | -4,82 | 1,43E-06 | 2,94E-05 |
| AT5G63715 | 27,38 | 6,00 | 1,24 | 4,82 | 1,44E-06 | 2,94E-05 |
| AT4G39420 | 9,97 | -3,93 | 0,82 | -4,82 | 1,46E-06 | 2,96E-05 |
| AT4G07070 | 70,24 | 2,58 | 0,54 | 4,81 | 1,53E-06 | 3,06E-05 |
| AT1G79930 | 9,47 | -4,35 | 0,91 | -4,81 | 1,52E-06 | 3,06E-05 |
| AT2G30520 | 8,24 | -4,65 | 0,97 | -4,81 | 1,53E-06 | 3,06E-05 |
| AT2G42270 | 6,29 | -6,73 | 1,40 | -4,80 | 1,57E-06 | 3,11E-05 |
| AT5G50290 | 244,15 | 1,79 | 0,37 | 4,78 | 1,71E-06 | 3,37E-05 |
| AT4G31570 | 11,16 | -4,18 | 0,87 | -4,78 | 1,77E-06 | 3,46E-05 |
| AT1G22620 | 12,70 | -3,01 | 0,63 | -4,77 | 1,82E-06 | 3,53E-05 |
| AT3G23690 | 28,16 | 4,39 | 0,92 | 4,77 | 1,84E-06 | 3,55E-05 |
| AT4G03445 | 45,78 | 1,51 | 0,32 | 4,75 | 2,01E-06 | 3,86E-05 |
| AT5G57870 | 8,84 | -3,43 | 0,72 | -4,74 | 2,12E-06 | 4,05E-05 |
| AT1G08537 | 8,63 | 6,22 | 1,32 | 4,72 | 2,41E-06 | 4,56E-05 |
| AT1G13980 | 8,14 | -4,96 | 1,06 | -4,70 | 2,54E-06 | 4,76E-05 |
| AT1G49760 | 5,88 | -5,05 | 1,07 | -4,71 | 2,53E-06 | 4,76E-05 |
| AT5G65750 | 4,51 | -6,72 | 1,43 | -4,69 | 2,78E-06 | 5,16E-05 |
| AT5G51430 | 4,74 | -7,18 | 1,54 | -4,66 | 3,22E-06 | 5,95E-05 |
| AT1G75660 | 6,24 | -4,97 | 1,07 | -4,65 | 3,28E-06 | 6,03E-05 |

| | | | | | | |
|---|---|---|---|---|---|---|
| AT1G62930 | 53,37 | 2,03 | 0,44 | 4,65 | 3,34E-06 | 6,10E-05 |
| AT1G62020 | 12,54 | -3,52 | 0,76 | -4,63 | 3,62E-06 | 6,56E-05 |
| AT4G16940 | 4,86 | -6,94 | 1,50 | -4,63 | 3,66E-06 | 6,61E-05 |
| AT4G04835 | 19,24 | -2,32 | 0,51 | -4,60 | 4,26E-06 | 7,59E-05 |
| AT3G23680 | 9,12 | -3,78 | 0,82 | -4,60 | 4,24E-06 | 7,59E-05 |
| AT3G20760 | 5,52 | -4,77 | 1,04 | -4,59 | 4,33E-06 | 7,68E-05 |
| AT2G26890 | 8,74 | -3,70 | 0,81 | -4,59 | 4,47E-06 | 7,87E-05 |
| AT2G43160 | 6,80 | -4,93 | 1,08 | -4,56 | 5,21E-06 | 9,12E-05 |
| AT1G15500 | 7,05 | -5,05 | 1,11 | -4,54 | 5,62E-06 | 9,79E-05 |
| AT3G46780 | 9,83 | -3,34 | 0,74 | -4,52 | 6,21E-06 | 1,08E-04 |
| AT2G02741 | 19,79 | 2,61 | 0,58 | 4,49 | 7,18E-06 | 1,24E-04 |
| AT4G05420 | 7,60 | -4,01 | 0,90 | -4,45 | 8,52E-06 | 1,46E-04 |
| AT2G35945 | 15,59 | 3,25 | 0,73 | 4,44 | 9,16E-06 | 1,56E-04 |
| AT1G29900 | 11,05 | -2,77 | 0,63 | -4,41 | 1,03E-05 | 1,74E-04 |
| AT2G35940 | 16,85 | 2,99 | 0,68 | 4,41 | 1,04E-05 | 1,75E-04 |
| AT1G48300 | 8,14 | -4,65 | 1,06 | -4,40 | 1,10E-05 | 1,84E-04 |
| AT5G39500 | 6,69 | -4,40 | 1,00 | -4,39 | 1,11E-05 | 1,85E-04 |
| AT2G02750 | 20,13 | 2,48 | 0,57 | 4,39 | 1,13E-05 | 1,87E-04 |
| AT4G03580 | 13,67 | 4,13 | 0,94 | 4,39 | 1,14E-05 | 1,88E-04 |
| AT2G16586 | 20,98 | 2,57 | 0,59 | 4,38 | 1,19E-05 | 1,96E-04 |
| AT1G36160 | 9,82 | -3,22 | 0,74 | -4,37 | 1,25E-05 | 2,04E-04 |
| AT1G15290 | 5,85 | -4,12 | 0,95 | -4,35 | 1,35E-05 | 2,19E-04 |
| AT4G32020 | 12,23 | -2,48 | 0,58 | -4,29 | 1,78E-05 | 2,86E-04 |
| AT5G57990 | 5,76 | -5,21 | 1,22 | -4,29 | 1,81E-05 | 2,91E-04 |
| AT4G19395 | 418,73 | -0,89 | 0,21 | -4,28 | 1,84E-05 | 2,93E-04 |
| AT1G74260 | 9,63 | -2,95 | 0,69 | -4,28 | 1,85E-05 | 2,94E-04 |
| AT5G15022 | 21,95 | 2,60 | 0,61 | 4,27 | 1,97E-05 | 3,11E-04 |
| AT1G30410 | 4,36 | -6,20 | 1,46 | -4,24 | 2,26E-05 | 3,55E-04 |
| AT1G80180 | 13,61 | 3,25 | 0,77 | 4,23 | 2,32E-05 | 3,62E-04 |
| AT2G36910 | 8,87 | -3,38 | 0,80 | -4,23 | 2,37E-05 | 3,68E-04 |
| AT4G38600 | 8,47 | -3,89 | 0,92 | -4,22 | 2,44E-05 | 3,77E-04 |
| AT2G42600 | 13,57 | -2,40 | 0,57 | -4,22 | 2,48E-05 | 3,81E-04 |
| AT1G59870 | 72,89 | -1,43 | 0,34 | -4,20 | 2,62E-05 | 4,00E-04 |
| AT1G67120 | 8,25 | -3,12 | 0,75 | -4,18 | 2,92E-05 | 4,44E-04 |
| AT3G02020 | 19,73 | 2,32 | 0,56 | 4,18 | 2,97E-05 | 4,50E-04 |
| AT1G20960 | 9,33 | -3,04 | 0,73 | -4,17 | 3,06E-05 | 4,61E-04 |
| AT2G22496 | 20,32 | -2,03 | 0,49 | -4,17 | 3,08E-05 | 4,62E-04 |
| AT1G19835 | 5,86 | -4,46 | 1,07 | -4,17 | 3,09E-05 | 4,62E-04 |
| AT2G09670 | 5,26 | -4,37 | 1,05 | -4,16 | 3,12E-05 | 4,63E-04 |
| AT3G48720 | 18,61 | 2,45 | 0,59 | 4,16 | 3,22E-05 | 4,75E-04 |
| AT4G36630 | 4,11 | -6,76 | 1,63 | -4,15 | 3,26E-05 | 4,79E-04 |
| AT2G39730 | 47,53 | -1,67 | 0,40 | -4,15 | 3,31E-05 | 4,84E-04 |
| AT3G57520 | 3,92 | -5,03 | 1,23 | -4,11 | 4,03E-05 | 5,87E-04 |
| AT4G01860 | 3,28 | -6,55 | 1,60 | -4,10 | 4,07E-05 | 5,90E-04 |
| AT5G25270 | 3,67 | -6,39 | 1,56 | -4,10 | 4,16E-05 | 6,00E-04 |
| AT1G04430 | 7,26 | -3,47 | 0,85 | -4,08 | 4,46E-05 | 6,41E-04 |
| AT1G06220 | 3,15 | -6,20 | 1,52 | -4,07 | 4,74E-05 | 6,76E-04 |
| AT3G08370 | 6,58 | 5,66 | 1,39 | 4,07 | 4,75E-05 | 6,76E-04 |
| AT3G01313 | 12,85 | 3,37 | 0,83 | 4,06 | 4,93E-05 | 6,99E-04 |
| AT5G43725 | 9,10 | 5,13 | 1,26 | 4,05 | 5,02E-05 | 7,05E-04 |
| AT5G43730 | 9,10 | 5,13 | 1,26 | 4,05 | 5,02E-05 | 7,05E-04 |
| AT2G07360 | 3,99 | -5,01 | 1,24 | -4,05 | 5,05E-05 | 7,05E-04 |
| AT2G01008 | 28,92 | 2,00 | 0,50 | 4,04 | 5,25E-05 | 7,30E-04 |
| AT5G54310 | 4,38 | -4,18 | 1,04 | -4,04 | 5,35E-05 | 7,41E-04 |
| AT2G44120 | 4,00 | -5,82 | 1,44 | -4,04 | 5,43E-05 | 7,48E-04 |
| AT1G50480 | 9,27 | -3,11 | 0,77 | -4,03 | 5,55E-05 | 7,59E-04 |

| | | | | | | |
|---|---|---|---|---|---|---|
| AT3G57410 | 3,70 | -6,09 | 1,51 | -4,03 | 5,56E-05 | 7,59E-04 |
| AT5G14800 | 5,34 | -4,75 | 1,18 | -4,03 | 5,60E-05 | 7,61E-04 |
| AT4G06130 | 392,23 | 1,70 | 0,42 | 4,03 | 5,66E-05 | 7,66E-04 |
| AT3G55610 | 4,81 | -4,30 | 1,07 | -4,02 | 5,84E-05 | 7,86E-04 |
| AT4G23375 | 8,41 | 5,25 | 1,31 | 4,01 | 6,04E-05 | 8,09E-04 |
| AT1G49240 | 7,10 | -4,08 | 1,02 | -4,00 | 6,29E-05 | 8,35E-04 |
| AT3G23000 | 7,09 | -3,50 | 0,88 | -4,00 | 6,27E-05 | 8,35E-04 |
| AT1G20760 | 7,86 | -3,74 | 0,94 | -3,98 | 6,76E-05 | 8,91E-04 |
| AT5G18370 | 3,23 | -6,27 | 1,57 | -3,99 | 6,74E-05 | 8,91E-04 |
| AT1G26975 | 8,83 | -2,79 | 0,70 | -3,98 | 6,92E-05 | 9,07E-04 |
| AT1G26985 | 10,26 | -3,20 | 0,80 | -3,97 | 7,05E-05 | 9,10E-04 |
| AT3G13857 | 124,31 | -1,46 | 0,37 | -3,97 | 7,04E-05 | 9,10E-04 |
| AT3G13855 | 124,31 | -1,46 | 0,37 | -3,97 | 7,04E-05 | 9,10E-04 |
| AT3G13920 | 8,10 | -3,21 | 0,81 | -3,98 | 6,99E-05 | 9,10E-04 |
| AT3G01460 | 5,55 | -4,49 | 1,13 | -3,96 | 7,37E-05 | 9,47E-04 |
| AT2G31660 | 3,59 | -5,61 | 1,42 | -3,94 | 8,08E-05 | 1,03E-03 |
| AT3G14210 | 12,80 | -2,54 | 0,64 | -3,94 | 8,16E-05 | 1,04E-03 |
| AT5G53902 | 6,85 | 5,48 | 1,39 | 3,94 | 8,27E-05 | 1,04E-03 |
| AT5G53905 | 6,85 | 5,48 | 1,39 | 3,94 | 8,27E-05 | 1,04E-03 |
| AT2G22330 | 4,57 | -4,87 | 1,24 | -3,93 | 8,35E-05 | 1,05E-03 |
| AT1G64430 | 2,85 | -6,31 | 1,61 | -3,91 | 9,14E-05 | 1,14E-03 |
| AT5G12470 | 4,58 | -4,47 | 1,14 | -3,90 | 9,55E-05 | 1,19E-03 |
| AT5G06845 | 6,24 | -3,52 | 0,90 | -3,90 | 9,75E-05 | 1,21E-03 |
| AT1G05960 | 4,07 | -5,87 | 1,51 | -3,89 | 1,01E-04 | 1,25E-03 |
| AT1G07810 | 6,32 | -3,54 | 0,91 | -3,88 | 1,04E-04 | 1,29E-03 |
| AT1G21630 | 4,05 | -5,56 | 1,44 | -3,87 | 1,09E-04 | 1,34E-03 |
| AT4G39080 | 4,27 | -5,52 | 1,43 | -3,87 | 1,10E-04 | 1,34E-03 |
| AT3G08530 | 12,53 | -2,28 | 0,59 | -3,86 | 1,12E-04 | 1,36E-03 |
| AT4G13575 | 426,34 | 1,57 | 0,41 | 3,86 | 1,12E-04 | 1,36E-03 |
| AT4G14910 | 7,50 | -3,03 | 0,78 | -3,86 | 1,12E-04 | 1,36E-03 |
| AT1G62670 | 9,93 | 4,94 | 1,28 | 3,85 | 1,18E-04 | 1,42E-03 |
| AT3G12120 | 5,50 | -3,91 | 1,02 | -3,85 | 1,20E-04 | 1,44E-03 |
| AT1G73687 | 3456,65 | 1,00 | 0,26 | 3,84 | 1,24E-04 | 1,48E-03 |
| AT3G50590 | 4,47 | -4,69 | 1,23 | -3,82 | 1,33E-04 | 1,58E-03 |
| AT1G80410 | 5,85 | -4,56 | 1,19 | -3,82 | 1,35E-04 | 1,60E-03 |
| AT1G09817 | 10,25 | 2,82 | 0,74 | 3,81 | 1,41E-04 | 1,66E-03 |
| AT2G01270 | 3,86 | -5,53 | 1,46 | -3,80 | 1,44E-04 | 1,70E-03 |
| AT4G24520 | 3,90 | -5,62 | 1,48 | -3,79 | 1,50E-04 | 1,75E-03 |
| AT1G62750 | 9,17 | -2,82 | 0,75 | -3,78 | 1,54E-04 | 1,79E-03 |
| AT5G20280 | 6,18 | -3,31 | 0,88 | -3,78 | 1,54E-04 | 1,79E-03 |
| AT3G03710 | 4,42 | -5,00 | 1,32 | -3,78 | 1,55E-04 | 1,80E-03 |
| AT1G78478 | 10,59 | 2,62 | 0,70 | 3,76 | 1,70E-04 | 1,95E-03 |
| AT1G50200 | 4,92 | -5,02 | 1,34 | -3,76 | 1,70E-04 | 1,95E-03 |
| AT1G77300 | 4,17 | -4,84 | 1,29 | -3,76 | 1,71E-04 | 1,95E-03 |
| AT4G08165 | 5,10 | 5,74 | 1,53 | 3,76 | 1,71E-04 | 1,95E-03 |
| AT3G10650 | 3,29 | -5,48 | 1,46 | -3,75 | 1,74E-04 | 1,97E-03 |
| AT2G25430 | 4,01 | -4,85 | 1,30 | -3,75 | 1,80E-04 | 2,03E-03 |
| AT2G16018 | 9,68 | 3,70 | 0,99 | 3,74 | 1,87E-04 | 2,09E-03 |
| AT4G00630 | 4,40 | -4,64 | 1,24 | -3,74 | 1,86E-04 | 2,09E-03 |
| AT1G79750 | 4,46 | -4,95 | 1,33 | -3,74 | 1,87E-04 | 2,10E-03 |
| AT5G09410 | 3,14 | -5,83 | 1,56 | -3,73 | 1,89E-04 | 2,11E-03 |
| AT2G41740 | 6,26 | -3,94 | 1,06 | -3,73 | 1,95E-04 | 2,16E-03 |
| AT1G03060 | 7,63 | -3,20 | 0,86 | -3,71 | 2,07E-04 | 2,27E-03 |
| AT2G39800 | 6,12 | -3,72 | 1,00 | -3,71 | 2,07E-04 | 2,27E-03 |
| AT5G14520 | 3,02 | -5,76 | 1,55 | -3,71 | 2,07E-04 | 2,27E-03 |
| AT5G62165 | 7,10 | 5,04 | 1,36 | 3,71 | 2,09E-04 | 2,29E-03 |

| | | | | | | |
|---|---|---|---|---|---|---|
| AT3G06650 | 3,17 | -5,62 | 1,52 | -3,69 | 2,20E-04 | 2,40E-03 |
| AT4G30190 | 7,61 | -3,29 | 0,89 | -3,69 | 2,24E-04 | 2,44E-03 |
| AT4G09135 | 26,82 | -1,68 | 0,46 | -3,69 | 2,26E-04 | 2,45E-03 |
| AT2G40030 | 5,36 | -4,71 | 1,28 | -3,68 | 2,33E-04 | 2,51E-03 |
| AT4G34670 | 4,28 | -3,63 | 0,99 | -3,67 | 2,46E-04 | 2,62E-03 |
| AT1G63770 | 7,56 | -2,79 | 0,76 | -3,67 | 2,46E-04 | 2,62E-03 |
| AT4G01850 | 4,27 | -4,83 | 1,32 | -3,67 | 2,45E-04 | 2,62E-03 |
| AT4G33010 | 11,10 | -2,61 | 0,71 | -3,67 | 2,46E-04 | 2,62E-03 |
| AT1G10950 | 6,24 | -3,12 | 0,85 | -3,66 | 2,52E-04 | 2,67E-03 |
| AT2G48160 | 6,20 | -4,93 | 1,35 | -3,65 | 2,62E-04 | 2,76E-03 |
| AT4G16340 | 3,35 | -4,85 | 1,33 | -3,65 | 2,66E-04 | 2,79E-03 |
| AT3G22886 | 587,34 | 0,92 | 0,25 | 3,64 | 2,69E-04 | 2,82E-03 |
| AT1G78476 | 11,35 | 2,36 | 0,65 | 3,61 | 3,03E-04 | 3,17E-03 |
| AT5G51174 | 65,86 | 1,53 | 0,42 | 3,61 | 3,09E-04 | 3,21E-03 |
| AT1G06267 | 15,22 | 2,56 | 0,71 | 3,60 | 3,16E-04 | 3,27E-03 |
| AT3G62360 | 4,39 | -4,21 | 1,17 | -3,60 | 3,21E-04 | 3,32E-03 |
| AT5G04275 | 20,42 | 1,67 | 0,47 | 3,59 | 3,25E-04 | 3,35E-03 |
| AT3G55220 | 4,51 | -4,14 | 1,15 | -3,59 | 3,29E-04 | 3,38E-03 |
| AT5G13100 | 3,43 | -4,42 | 1,23 | -3,59 | 3,30E-04 | 3,38E-03 |
| AT5G26650 | 7,57 | 4,91 | 1,37 | 3,59 | 3,35E-04 | 3,42E-03 |
| AT3G59940 | 11,46 | 2,41 | 0,67 | 3,58 | 3,42E-04 | 3,47E-03 |
| AT5G05980 | 2,96 | -6,06 | 1,69 | -3,58 | 3,46E-04 | 3,51E-03 |
| AT5G43740 | 103,64 | 1,66 | 0,46 | 3,57 | 3,53E-04 | 3,55E-03 |
| AT1G55860 | 12,82 | -2,62 | 0,73 | -3,57 | 3,52E-04 | 3,55E-03 |
| AT4G20850 | 8,05 | -2,64 | 0,74 | -3,57 | 3,55E-04 | 3,57E-03 |
| AT2G17930 | 7,05 | -2,58 | 0,72 | -3,56 | 3,65E-04 | 3,65E-03 |
| AT1G13350 | 2,82 | -6,10 | 1,71 | -3,56 | 3,66E-04 | 3,65E-03 |
| AT5G02325 | 15,37 | 2,71 | 0,76 | 3,56 | 3,73E-04 | 3,71E-03 |
| AT1G80070 | 7,13 | -2,63 | 0,74 | -3,55 | 3,82E-04 | 3,78E-03 |
| AT5G01920 | 2,56 | -5,63 | 1,59 | -3,54 | 3,94E-04 | 3,87E-03 |
| AT2G21330 | 7,94 | -2,52 | 0,71 | -3,54 | 3,94E-04 | 3,87E-03 |
| AT3G09275 | 11,41 | 2,40 | 0,68 | 3,54 | 3,94E-04 | 3,87E-03 |
| AT1G64140 | 5,12 | -3,22 | 0,91 | -3,54 | 4,01E-04 | 3,92E-03 |
| AT4G11373 | 5,92 | 4,75 | 1,35 | 3,53 | 4,22E-04 | 4,11E-03 |
| AT5G14545 | 7868,55 | 0,72 | 0,20 | 3,52 | 4,31E-04 | 4,17E-03 |
| AT5G06600 | 5,36 | -3,45 | 0,98 | -3,52 | 4,31E-04 | 4,17E-03 |
| AT2G06420 | 4,36 | 5,13 | 1,46 | 3,51 | 4,51E-04 | 4,35E-03 |
| AT4G35800 | 5,60 | -3,09 | 0,88 | -3,50 | 4,63E-04 | 4,45E-03 |
| AT1G55540 | 2,72 | -5,14 | 1,47 | -3,49 | 4,80E-04 | 4,60E-03 |
| AT3G08947 | 5,24 | -3,78 | 1,08 | -3,49 | 4,84E-04 | 4,62E-03 |
| AT5G17165 | 3,90 | 5,05 | 1,45 | 3,49 | 4,85E-04 | 4,62E-03 |
| AT3G13300 | 8,42 | -3,95 | 1,13 | -3,49 | 4,87E-04 | 4,63E-03 |
| AT2G03820 | 4,25 | -3,93 | 1,13 | -3,49 | 4,92E-04 | 4,66E-03 |
| AT3G09105 | 30,37 | 1,90 | 0,55 | 3,48 | 4,95E-04 | 4,67E-03 |
| AT2G44540 | 4,48 | 4,82 | 1,39 | 3,48 | 5,09E-04 | 4,79E-03 |
| AT3G21805 | 21,83 | 2,80 | 0,81 | 3,47 | 5,13E-04 | 4,82E-03 |
| AT1G49040 | 3,74 | -5,05 | 1,45 | -3,47 | 5,16E-04 | 4,83E-03 |
| AT3G58196 | 21,50 | 1,89 | 0,54 | 3,47 | 5,20E-04 | 4,85E-03 |
| AT4G02320 | 3,77 | -4,09 | 1,18 | -3,46 | 5,31E-04 | 4,94E-03 |
| AT5G35407 | 1076,14 | -1,27 | 0,37 | -3,46 | 5,39E-04 | 4,99E-03 |
| AT5G12370 | 4,35 | -3,89 | 1,12 | -3,46 | 5,49E-04 | 5,07E-03 |
| AT5G58140 | 8,36 | -2,95 | 0,85 | -3,45 | 5,51E-04 | 5,07E-03 |
| AT2G25660 | 3,83 | -3,93 | 1,14 | -3,45 | 5,56E-04 | 5,10E-03 |
| AT4G37910 | 5,28 | -3,10 | 0,90 | -3,45 | 5,58E-04 | 5,11E-03 |
| AT1G30230 | 5,95 | -3,28 | 0,95 | -3,44 | 5,86E-04 | 5,31E-03 |
| AT2G26080 | 10,37 | -2,64 | 0,77 | -3,44 | 5,85E-04 | 5,31E-03 |

| | | | | | | |
|---|---|---|---|---|---|---|
| AT4G16890 | 15,81 | -2,06 | 0,60 | -3,44 | 5,84E-04 | 5,31E-03 |
| AT4G00150 | 10,48 | 2,39 | 0,70 | 3,44 | 5,89E-04 | 5,32E-03 |
| AT3G23780 | 4,59 | -3,25 | 0,95 | -3,42 | 6,17E-04 | 5,57E-03 |
| AT3G59020 | 5,79 | -3,23 | 0,94 | -3,42 | 6,20E-04 | 5,57E-03 |
| AT2G45930 | 2,64 | -5,24 | 1,53 | -3,42 | 6,24E-04 | 5,59E-03 |
| AT1G29465 | 25,62 | 1,77 | 0,52 | 3,41 | 6,39E-04 | 5,71E-03 |
| AT1G58290 | 5,16 | -2,92 | 0,86 | -3,41 | 6,42E-04 | 5,72E-03 |
| AT5G14550 | 7872,48 | 0,71 | 0,21 | 3,41 | 6,45E-04 | 5,73E-03 |
| AT2G45540 | 12,39 | -2,42 | 0,71 | -3,40 | 6,66E-04 | 5,90E-03 |
| AT5G23380 | 2,25 | -5,89 | 1,73 | -3,40 | 6,76E-04 | 5,96E-03 |
| AT4G13560 | 52,08 | -1,08 | 0,32 | -3,40 | 6,77E-04 | 5,96E-03 |
| AT3G13080 | 13,73 | -2,34 | 0,69 | -3,39 | 6,93E-04 | 6,08E-03 |
| AT4G13495 | 869,17 | 1,13 | 0,33 | 3,37 | 7,56E-04 | 6,62E-03 |
| AT3G17850 | 4,74 | -3,85 | 1,14 | -3,37 | 7,58E-04 | 6,62E-03 |
| AT5G64220 | 2,50 | -5,37 | 1,60 | -3,37 | 7,60E-04 | 6,62E-03 |
| AT2G27720 | 1,70 | -5,82 | 1,73 | -3,37 | 7,63E-04 | 6,62E-03 |
| AT3G43833 | 7,44 | 4,72 | 1,41 | 3,36 | 7,78E-04 | 6,74E-03 |
| AT1G22530 | 2,83 | -5,37 | 1,60 | -3,36 | 7,81E-04 | 6,74E-03 |
| AT4G01050 | 4,58 | -3,80 | 1,13 | -3,36 | 7,89E-04 | 6,79E-03 |
| AT3G08580 | 9,77 | -2,33 | 0,70 | -3,34 | 8,49E-04 | 7,29E-03 |
| AT1G28070 | 4,21 | -6,96 | 2,09 | -3,33 | 8,78E-04 | 7,51E-03 |
| AT5G38420 | 5,05 | -2,96 | 0,89 | -3,32 | 8,87E-04 | 7,57E-03 |
| AT4G16720 | 3,07 | -5,04 | 1,52 | -3,32 | 8,92E-04 | 7,59E-03 |
| AT2G25800 | 4,68 | -3,13 | 0,94 | -3,32 | 8,98E-04 | 7,62E-03 |
| AT4G31880 | 2,70 | -5,12 | 1,55 | -3,31 | 9,24E-04 | 7,82E-03 |
| AT3G63375 | 13,10 | -1,77 | 0,54 | -3,31 | 9,38E-04 | 7,92E-03 |
| AT1G70185 | 5,33 | -2,98 | 0,90 | -3,30 | 9,50E-04 | 7,98E-03 |
| AT5G07940 | 5,66 | -3,12 | 0,94 | -3,31 | 9,49E-04 | 7,98E-03 |
| AT2G34430 | 28,33 | -1,44 | 0,44 | -3,30 | 9,69E-04 | 8,08E-03 |
| AT3G47347 | 15,63 | 1,79 | 0,54 | 3,30 | 9,68E-04 | 8,08E-03 |
| AT4G12780 | 15,31 | 1,95 | 0,59 | 3,29 | 9,99E-04 | 8,32E-03 |
| AT5G13010 | 4,55 | -3,40 | 1,03 | -3,29 | 1,01E-03 | 8,35E-03 |
| AT2G41840 | 3,13 | -3,92 | 1,19 | -3,29 | 1,01E-03 | 8,35E-03 |
| AT1G80250 | 2,27 | -5,24 | 1,59 | -3,28 | 1,02E-03 | 8,41E-03 |
| AT2G33100 | 2,25 | -5,81 | 1,77 | -3,28 | 1,02E-03 | 8,41E-03 |
| AT1G75335 | 3,97 | -3,70 | 1,13 | -3,28 | 1,03E-03 | 8,48E-03 |
| AT3G03870 | 3,66 | -3,66 | 1,12 | -3,28 | 1,04E-03 | 8,50E-03 |
| AT4G17788 | 20,18 | -1,55 | 0,47 | -3,27 | 1,06E-03 | 8,65E-03 |
| AT1G24460 | 4,84 | -4,08 | 1,25 | -3,27 | 1,09E-03 | 8,87E-03 |
| AT1G06190 | 5,18 | -3,49 | 1,07 | -3,26 | 1,11E-03 | 9,04E-03 |
| AT3G51050 | 3,03 | -5,18 | 1,59 | -3,26 | 1,12E-03 | 9,06E-03 |
| AT1G11640 | 3,15 | -4,15 | 1,27 | -3,26 | 1,12E-03 | 9,06E-03 |
| AT1G13640 | 2,82 | -4,97 | 1,53 | -3,26 | 1,13E-03 | 9,13E-03 |
| AT3G07870 | 5,57 | -3,39 | 1,04 | -3,25 | 1,16E-03 | 9,29E-03 |
| AT5G42390 | 8,64 | -2,65 | 0,82 | -3,25 | 1,16E-03 | 9,29E-03 |
| AT5G06640 | 3,20 | 4,97 | 1,53 | 3,24 | 1,18E-03 | 9,47E-03 |
| AT3G63460 | 4,10 | -2,95 | 0,91 | -3,24 | 1,22E-03 | 9,69E-03 |
| AT5G18350 | 2,51 | -5,77 | 1,78 | -3,23 | 1,22E-03 | 9,72E-03 |
| AT5G46470 | 3,68 | -3,29 | 1,02 | -3,23 | 1,24E-03 | 9,80E-03 |
| AT1G48090 | 11,18 | -1,83 | 0,57 | -3,22 | 1,27E-03 | 1,01E-02 |
| AT1G30470 | 3,87 | -3,82 | 1,19 | -3,22 | 1,30E-03 | 1,02E-02 |
| AT1G47720 | 6,21 | -2,87 | 0,89 | -3,21 | 1,31E-03 | 1,03E-02 |
| AT5G57350 | 4,58 | -3,97 | 1,24 | -3,21 | 1,32E-03 | 1,03E-02 |
| AT3G15730 | 15,00 | -1,72 | 0,54 | -3,20 | 1,39E-03 | 1,09E-02 |
| AT1G74470 | 19,25 | -1,57 | 0,49 | -3,19 | 1,41E-03 | 1,10E-02 |
| AT4G24280 | 6,53 | -3,07 | 0,96 | -3,19 | 1,42E-03 | 1,10E-02 |

| | | | | | | |
|---|---|---|---|---|---|---|
| AT1G09010 | 3,02 | -3,71 | 1,16 | -3,19 | 1,42E-03 | 1,10E-02 |
| AT3G03780 | 4,16 | -3,47 | 1,09 | -3,19 | 1,43E-03 | 1,11E-02 |
| AT3G45780 | 10,09 | -1,85 | 0,58 | -3,19 | 1,44E-03 | 1,11E-02 |
| AT4G31480 | 6,09 | -3,18 | 1,00 | -3,19 | 1,44E-03 | 1,11E-02 |
| AT5G28560 | 3,51 | 4,56 | 1,43 | 3,19 | 1,44E-03 | 1,11E-02 |
| AT5G04105 | 18,20 | -1,54 | 0,48 | -3,19 | 1,45E-03 | 1,11E-02 |
| AT3G13870 | 4,03 | -4,78 | 1,50 | -3,18 | 1,46E-03 | 1,12E-02 |
| AT1G04810 | 3,41 | -5,18 | 1,63 | -3,18 | 1,48E-03 | 1,13E-02 |
| AT5G66030 | 2,48 | -4,95 | 1,56 | -3,18 | 1,49E-03 | 1,13E-02 |
| AT5G21100 | 6,61 | 3,27 | 1,03 | 3,17 | 1,51E-03 | 1,15E-02 |
| AT1G26973 | 11,41 | -1,79 | 0,56 | -3,16 | 1,57E-03 | 1,19E-02 |
| AT3G41768 | 91,21 | 0,96 | 0,30 | 3,16 | 1,58E-03 | 1,19E-02 |
| AT4G25530 | 5,35 | 4,18 | 1,33 | 3,15 | 1,64E-03 | 1,23E-02 |
| AT3G03380 | 4,30 | -3,36 | 1,07 | -3,14 | 1,67E-03 | 1,25E-02 |
| AT5G48160 | 6,17 | -2,93 | 0,93 | -3,14 | 1,67E-03 | 1,25E-02 |
| AT5G64420 | 4,10 | -3,27 | 1,04 | -3,14 | 1,69E-03 | 1,26E-02 |
| AT3G14470 | 2,64 | -4,99 | 1,59 | -3,14 | 1,71E-03 | 1,28E-02 |
| AT2G43890 | 2,70 | -5,51 | 1,76 | -3,13 | 1,72E-03 | 1,28E-02 |
| AT5G14740 | 2,79 | -3,91 | 1,25 | -3,13 | 1,75E-03 | 1,30E-02 |
| AT3G22790 | 2,45 | -4,94 | 1,58 | -3,13 | 1,77E-03 | 1,31E-02 |
| AT2G41650 | 5,84 | 4,08 | 1,31 | 3,12 | 1,81E-03 | 1,34E-02 |
| AT3G54730 | 3,74 | 4,35 | 1,40 | 3,12 | 1,83E-03 | 1,35E-02 |
| AT2G24980 | 3,78 | 4,72 | 1,52 | 3,11 | 1,85E-03 | 1,36E-02 |
| AT1G29910 | 27,17 | -1,54 | 0,49 | -3,11 | 1,88E-03 | 1,38E-02 |
| AT1G77680 | 1,90 | -5,87 | 1,90 | -3,10 | 1,95E-03 | 1,42E-02 |
| AT2G04605 | 3,46 | 4,74 | 1,53 | 3,10 | 1,95E-03 | 1,42E-02 |
| AT5G54810 | 4,59 | -3,12 | 1,01 | -3,10 | 1,95E-03 | 1,42E-02 |
| AT4G08175 | 7,31 | -2,75 | 0,89 | -3,10 | 1,96E-03 | 1,43E-02 |
| AT1G64710 | 2,31 | -4,34 | 1,40 | -3,09 | 1,98E-03 | 1,43E-02 |
| AT2G29980 | 3,37 | -3,66 | 1,19 | -3,09 | 2,00E-03 | 1,45E-02 |
| AT5G03305 | 5,94 | 3,21 | 1,04 | 3,09 | 2,01E-03 | 1,45E-02 |
| AT2G05070 | 19,17 | -1,58 | 0,51 | -3,08 | 2,06E-03 | 1,49E-02 |
| AT3G16850 | 3,96 | -3,43 | 1,11 | -3,08 | 2,07E-03 | 1,49E-02 |
| AT5G46250 | 6,03 | -3,31 | 1,08 | -3,08 | 2,10E-03 | 1,50E-02 |
| AT3G25510 | 1,96 | -5,16 | 1,68 | -3,08 | 2,10E-03 | 1,50E-02 |
| AT2G47070 | 5,92 | -2,48 | 0,81 | -3,06 | 2,19E-03 | 1,56E-02 |
| AT3G49500 | 2,34 | -5,28 | 1,73 | -3,06 | 2,23E-03 | 1,59E-02 |
| AT1G59760 | 3,05 | -3,75 | 1,23 | -3,05 | 2,27E-03 | 1,61E-02 |
| AT3G09200 | 3,98 | -3,15 | 1,03 | -3,05 | 2,29E-03 | 1,62E-02 |
| AT2G40930 | 1,74 | -5,61 | 1,84 | -3,05 | 2,31E-03 | 1,63E-02 |
| AT3G08730 | 2,48 | -5,06 | 1,66 | -3,05 | 2,31E-03 | 1,63E-02 |
| AT1G06247 | 2,53 | -4,90 | 1,61 | -3,04 | 2,34E-03 | 1,65E-02 |
| AT5G04810 | 3,05 | -4,08 | 1,34 | -3,04 | 2,36E-03 | 1,65E-02 |
| AT5G24710 | 4,46 | -3,17 | 1,04 | -3,04 | 2,35E-03 | 1,65E-02 |
| AT4G36360 | 3,63 | -4,42 | 1,45 | -3,04 | 2,36E-03 | 1,65E-02 |
| AT4G14504 | 4,83 | 4,15 | 1,37 | 3,04 | 2,38E-03 | 1,66E-02 |
| AT2G47020 | 1248,75 | -0,94 | 0,31 | -3,03 | 2,47E-03 | 1,72E-02 |
| AT2G47015 | 1248,43 | -0,94 | 0,31 | -3,02 | 2,50E-03 | 1,73E-02 |
| AT5G65380 | 2,69 | -3,91 | 1,30 | -3,02 | 2,56E-03 | 1,77E-02 |
| AT1G04210 | 3,48 | -3,00 | 0,99 | -3,02 | 2,57E-03 | 1,77E-02 |
| AT4G31180 | 4,29 | -2,91 | 0,97 | -3,01 | 2,60E-03 | 1,79E-02 |
| AT3G06530 | 4,41 | -3,36 | 1,12 | -3,01 | 2,60E-03 | 1,79E-02 |
| AT2G45940 | 2,82 | -4,54 | 1,51 | -3,01 | 2,62E-03 | 1,80E-02 |
| AT1G18010 | 5,80 | 3,95 | 1,32 | 3,01 | 2,65E-03 | 1,81E-02 |
| AT2G34100 | 7,49 | 2,71 | 0,90 | 3,00 | 2,73E-03 | 1,86E-02 |
| AT3G05675 | 9,08 | -2,06 | 0,69 | -3,00 | 2,73E-03 | 1,86E-02 |

| | | | | | | |
|---|---|---|---|---|---|---|
| AT1G54090 | 3,59 | -4,13 | 1,38 | -2,99 | 2,75E-03 | 1,87E-02 |
| AT1G32230 | 5,15 | -2,56 | 0,86 | -2,99 | 2,78E-03 | 1,88E-02 |
| AT2G44490 | 17,93 | -1,44 | 0,48 | -2,99 | 2,78E-03 | 1,88E-02 |
| AT4G38200 | 2,17 | -4,75 | 1,59 | -2,99 | 2,79E-03 | 1,88E-02 |
| AT3G63500 | 4,53 | -3,20 | 1,07 | -2,99 | 2,82E-03 | 1,89E-02 |
| AT4G23650 | 3,54 | -3,18 | 1,07 | -2,99 | 2,82E-03 | 1,89E-02 |
| AT3G58640 | 2,56 | -4,86 | 1,64 | -2,97 | 3,00E-03 | 2,01E-02 |
| AT5G44180 | 3,26 | -3,93 | 1,34 | -2,94 | 3,29E-03 | 2,20E-02 |
| AT2G16040 | 2,68 | 4,56 | 1,55 | 2,94 | 3,32E-03 | 2,22E-02 |
| AT3G19190 | 3,92 | -4,18 | 1,42 | -2,94 | 3,33E-03 | 2,22E-02 |
| AT5G04560 | 3,61 | -3,10 | 1,06 | -2,93 | 3,41E-03 | 2,27E-02 |
| AT1G50440 | 6,47 | 2,83 | 0,97 | 2,92 | 3,44E-03 | 2,28E-02 |
| AT1G79280 | 4,52 | -3,02 | 1,03 | -2,92 | 3,45E-03 | 2,28E-02 |
| AT3G44260 | 2,79 | 4,39 | 1,50 | 2,92 | 3,48E-03 | 2,30E-02 |
| AT1G06273 | 2,17 | -5,27 | 1,81 | -2,92 | 3,53E-03 | 2,32E-02 |
| AT2G41640 | 6,36 | 3,35 | 1,15 | 2,92 | 3,53E-03 | 2,32E-02 |
| AT1G07920 | 7,76 | -2,22 | 0,76 | -2,91 | 3,60E-03 | 2,36E-02 |
| AT1G31730 | 1,97 | -5,20 | 1,79 | -2,91 | 3,63E-03 | 2,38E-02 |
| AT3G56680 | 2,66 | -4,17 | 1,43 | -2,91 | 3,66E-03 | 2,38E-02 |
| AT4G04825 | 145,71 | 1,69 | 0,58 | 2,91 | 3,67E-03 | 2,38E-02 |
| AT4G28652 | 2,08 | -5,44 | 1,87 | -2,91 | 3,66E-03 | 2,38E-02 |
| AT5G09585 | 3,20 | 4,28 | 1,47 | 2,91 | 3,67E-03 | 2,38E-02 |
| AT4G28650 | 2,08 | -5,44 | 1,87 | -2,91 | 3,66E-03 | 2,38E-02 |
| AT3G56825 | 56,81 | 1,08 | 0,37 | 2,90 | 3,74E-03 | 2,42E-02 |
| AT1G58250 | 4,39 | -2,92 | 1,01 | -2,90 | 3,75E-03 | 2,42E-02 |
| AT4G35830 | 9,01 | -2,36 | 0,81 | -2,90 | 3,76E-03 | 2,42E-02 |
| AT5G06395 | 1,55 | -5,34 | 1,84 | -2,89 | 3,80E-03 | 2,44E-02 |
| AT5G08670 | 5,95 | -2,25 | 0,78 | -2,89 | 3,85E-03 | 2,46E-02 |
| AT5G35210 | 2,88 | -4,13 | 1,43 | -2,89 | 3,85E-03 | 2,46E-02 |
| AT5G46640 | 1,47 | -5,66 | 1,96 | -2,88 | 3,92E-03 | 2,50E-02 |
| AT2G43710 | 3,74 | -3,24 | 1,12 | -2,88 | 3,95E-03 | 2,51E-02 |
| AT1G15130 | 4,06 | -3,81 | 1,32 | -2,87 | 4,04E-03 | 2,57E-02 |
| AT3G26810 | 23,04 | 1,46 | 0,51 | 2,87 | 4,11E-03 | 2,61E-02 |
| AT3G01695 | 8,72 | -2,04 | 0,71 | -2,87 | 4,13E-03 | 2,61E-02 |
| AT3G43160 | 4,62 | 3,91 | 1,37 | 2,86 | 4,18E-03 | 2,63E-02 |
| AT4G04890 | 2,53 | -4,15 | 1,45 | -2,86 | 4,17E-03 | 2,63E-02 |
| AT3G48000 | 4,79 | -2,68 | 0,93 | -2,86 | 4,19E-03 | 2,63E-02 |
| AT1G61275 | 11,06 | 2,10 | 0,73 | 2,86 | 4,20E-03 | 2,63E-02 |
| AT5G42820 | 2,08 | -4,54 | 1,59 | -2,86 | 4,27E-03 | 2,67E-02 |
| AT5G41765 | 5,08 | 3,12 | 1,09 | 2,86 | 4,30E-03 | 2,69E-02 |
| AT3G13460 | 10,31 | -1,68 | 0,59 | -2,84 | 4,48E-03 | 2,79E-02 |
| AT3G20010 | 2,80 | -4,94 | 1,74 | -2,84 | 4,50E-03 | 2,80E-02 |
| AT5G61020 | 2,23 | -4,59 | 1,62 | -2,84 | 4,53E-03 | 2,81E-02 |
| AT5G38540 | 8,36 | -2,72 | 0,96 | -2,83 | 4,59E-03 | 2,85E-02 |
| AT3G54660 | 2,19 | -4,60 | 1,63 | -2,83 | 4,70E-03 | 2,91E-02 |
| AT1G18310 | 4,64 | 4,10 | 1,46 | 2,81 | 4,89E-03 | 3,02E-02 |
| AT1G36730 | 3,83 | -2,85 | 1,01 | -2,81 | 4,92E-03 | 3,03E-02 |
| AT5G56070 | 3,26 | 4,61 | 1,64 | 2,81 | 4,95E-03 | 3,04E-02 |
| AT5G61455 | 11,96 | -1,58 | 0,56 | -2,81 | 4,95E-03 | 3,04E-02 |
| AT2G46830 | 2,43 | -4,23 | 1,51 | -2,81 | 4,99E-03 | 3,05E-02 |
| AT3G26650 | 12,31 | -1,68 | 0,60 | -2,81 | 5,03E-03 | 3,06E-02 |
| AT3G29796 | 2,77 | 4,34 | 1,55 | 2,81 | 5,03E-03 | 3,06E-02 |
| AT3G46740 | 3,41 | -2,96 | 1,06 | -2,80 | 5,04E-03 | 3,06E-02 |
| AT5G60900 | 5,69 | -3,04 | 1,08 | -2,81 | 5,01E-03 | 3,06E-02 |
| AT3G05420 | 2,70 | -3,52 | 1,26 | -2,80 | 5,16E-03 | 3,13E-02 |
| AT4G13564 | 27,21 | -0,98 | 0,35 | -2,80 | 5,18E-03 | 3,13E-02 |

| | | | | | | |
|---|---|---|---|---|---|---|
| AT5G53480 | 6,73 | -2,58 | 0,93 | -2,79 | 5,27E-03 | 3,18E-02 |
| AT3G04340 | 3,02 | -4,21 | 1,51 | -2,79 | 5,29E-03 | 3,19E-02 |
| AT5G10380 | 7,30 | -2,10 | 0,75 | -2,79 | 5,31E-03 | 3,19E-02 |
| AT4G00570 | 3,02 | -3,80 | 1,36 | -2,79 | 5,33E-03 | 3,20E-02 |
| AT1G07633 | 6,19 | 2,71 | 0,98 | 2,77 | 5,55E-03 | 3,32E-02 |
| AT4G34710 | 3,32 | -2,98 | 1,08 | -2,77 | 5,55E-03 | 3,32E-02 |
| AT2G02950 | 6,29 | 2,64 | 0,95 | 2,77 | 5,61E-03 | 3,34E-02 |
| AT3G25520 | 3,31 | -3,00 | 1,08 | -2,77 | 5,69E-03 | 3,39E-02 |
| AT1G32720 | 2,47 | 4,21 | 1,52 | 2,76 | 5,73E-03 | 3,40E-02 |
| AT4G13570 | 36,83 | -0,96 | 0,35 | -2,76 | 5,78E-03 | 3,42E-02 |
| AT3G24480 | 4,54 | -2,25 | 0,82 | -2,76 | 5,79E-03 | 3,43E-02 |
| AT4G38440 | 2,75 | -3,15 | 1,14 | -2,76 | 5,83E-03 | 3,45E-02 |
| AT4G13340 | 14,47 | -1,29 | 0,47 | -2,75 | 5,90E-03 | 3,48E-02 |
| AT1G20970 | 4,69 | -3,13 | 1,14 | -2,75 | 5,96E-03 | 3,51E-02 |
| AT5G19200 | 4,59 | -2,99 | 1,09 | -2,75 | 5,98E-03 | 3,51E-02 |
| AT5G19330 | 2,99 | -3,71 | 1,35 | -2,75 | 5,99E-03 | 3,51E-02 |
| AT4G35790 | 4,32 | -3,64 | 1,33 | -2,75 | 6,01E-03 | 3,52E-02 |
| AT5G04853 | 3,91 | 3,91 | 1,42 | 2,74 | 6,05E-03 | 3,53E-02 |
| AT1G33680 | 2,02 | -4,93 | 1,80 | -2,74 | 6,10E-03 | 3,55E-02 |
| AT5G58040 | 3,29 | -3,38 | 1,23 | -2,74 | 6,12E-03 | 3,55E-02 |
| AT1G05993 | 11,40 | 1,88 | 0,69 | 2,74 | 6,11E-03 | 3,55E-02 |
| AT2G47800 | 2,70 | -3,78 | 1,38 | -2,74 | 6,14E-03 | 3,56E-02 |
| AT1G48650 | 3,21 | -3,55 | 1,30 | -2,74 | 6,16E-03 | 3,56E-02 |
| AT2G38040 | 4,80 | -2,75 | 1,01 | -2,73 | 6,28E-03 | 3,62E-02 |
| AT2G39885 | 9,28 | -2,08 | 0,76 | -2,73 | 6,29E-03 | 3,62E-02 |
| AT1G48920 | 4,08 | -2,30 | 0,84 | -2,73 | 6,32E-03 | 3,63E-02 |
| AT5G63020 | 8,08 | 2,78 | 1,02 | 2,73 | 6,33E-03 | 3,63E-02 |
| AT1G47657 | 2,52 | 4,46 | 1,64 | 2,73 | 6,35E-03 | 3,64E-02 |
| AT1G30490 | 1,77 | -5,16 | 1,89 | -2,73 | 6,37E-03 | 3,64E-02 |
| AT1G01510 | 2,79 | -3,81 | 1,40 | -2,73 | 6,39E-03 | 3,65E-02 |
| AT1G21680 | 4,80 | -2,49 | 0,91 | -2,73 | 6,42E-03 | 3,66E-02 |
| AT1G76630 | 2,14 | -4,84 | 1,78 | -2,72 | 6,47E-03 | 3,68E-02 |
| AT5G41490 | 2,24 | 4,38 | 1,61 | 2,72 | 6,53E-03 | 3,70E-02 |
| AT5G26742 | 9,59 | -1,94 | 0,71 | -2,72 | 6,53E-03 | 3,70E-02 |
| AT4G25080 | 4,26 | -2,65 | 0,98 | -2,71 | 6,65E-03 | 3,75E-02 |
| AT1G79000 | 2,98 | -5,15 | 1,90 | -2,71 | 6,65E-03 | 3,75E-02 |
| AT3G04130 | 2,12 | -5,49 | 2,02 | -2,71 | 6,68E-03 | 3,76E-02 |
| AT4G37900 | 1,96 | -5,02 | 1,85 | -2,71 | 6,70E-03 | 3,77E-02 |
| AT5G61010 | 3,05 | -2,95 | 1,09 | -2,71 | 6,81E-03 | 3,82E-02 |
| AT5G61780 | 4,25 | -2,58 | 0,95 | -2,71 | 6,81E-03 | 3,82E-02 |
| AT3G55512 | 2,56 | 4,81 | 1,78 | 2,70 | 6,94E-03 | 3,88E-02 |
| AT3G63380 | 3,14 | -3,87 | 1,43 | -2,70 | 7,00E-03 | 3,91E-02 |
| AT4G36760 | 3,13 | -3,31 | 1,23 | -2,69 | 7,09E-03 | 3,95E-02 |
| AT1G04625 | 1,86 | -5,26 | 1,96 | -2,69 | 7,19E-03 | 3,99E-02 |
| AT5G12120 | 2,73 | -3,60 | 1,34 | -2,69 | 7,18E-03 | 3,99E-02 |
| AT3G55410 | 3,63 | -2,77 | 1,03 | -2,69 | 7,25E-03 | 4,02E-02 |
| AT1G78900 | 8,95 | -1,81 | 0,68 | -2,68 | 7,37E-03 | 4,07E-02 |
| AT3G57480 | 3,07 | -3,32 | 1,24 | -2,68 | 7,37E-03 | 4,07E-02 |
| AT1G14610 | 2,42 | -3,52 | 1,32 | -2,68 | 7,43E-03 | 4,09E-02 |
| AT5G46210 | 3,09 | -3,48 | 1,30 | -2,68 | 7,43E-03 | 4,09E-02 |
| AT3G06436 | 20,96 | 1,29 | 0,48 | 2,68 | 7,47E-03 | 4,09E-02 |
| AT5G05660 | 1,92 | -6,08 | 2,27 | -2,68 | 7,47E-03 | 4,09E-02 |
| AT3G20770 | 2,68 | -3,42 | 1,28 | -2,67 | 7,58E-03 | 4,14E-02 |
| AT4G39200 | 1,73 | -4,87 | 1,82 | -2,67 | 7,60E-03 | 4,15E-02 |
| AT1G59610 | 3,98 | -2,93 | 1,10 | -2,67 | 7,64E-03 | 4,15E-02 |
| AT3G57300 | 2,34 | -3,42 | 1,28 | -2,67 | 7,65E-03 | 4,15E-02 |

| | | | | | | |
|---|---|---|---|---|---|---|
| AT5G03040 | 2,14 | -3,85 | 1,44 | -2,67 | 7,63E-03 | 4,15E-02 |
| AT5G45500 | 2,29 | -4,61 | 1,73 | -2,67 | 7,67E-03 | 4,16E-02 |
| AT3G23685 | 5,28 | -2,26 | 0,85 | -2,66 | 7,77E-03 | 4,19E-02 |
| AT3G52250 | 4,06 | -2,78 | 1,04 | -2,66 | 7,76E-03 | 4,19E-02 |
| AT1G51150 | 4,36 | -4,08 | 1,54 | -2,66 | 7,90E-03 | 4,26E-02 |
| AT4G24570 | 3,86 | 3,68 | 1,39 | 2,65 | 7,96E-03 | 4,28E-02 |
| AT5G47260 | 4,17 | 3,63 | 1,37 | 2,65 | 7,97E-03 | 4,28E-02 |
| AT1G13360 | 23,38 | 1,22 | 0,46 | 2,65 | 8,00E-03 | 4,29E-02 |
| AT4G01840 | 3,87 | 3,98 | 1,50 | 2,65 | 8,07E-03 | 4,32E-02 |
| AT4G18600 | 3,50 | -3,86 | 1,46 | -2,65 | 8,08E-03 | 4,32E-02 |
| AT3G61580 | 5,37 | -2,11 | 0,80 | -2,64 | 8,18E-03 | 4,36E-02 |
| AT3G52240 | 2,27 | -4,32 | 1,64 | -2,64 | 8,25E-03 | 4,39E-02 |
| AT5G24350 | 3,06 | -4,18 | 1,59 | -2,64 | 8,35E-03 | 4,44E-02 |
| AT5G60410 | 2,90 | -3,13 | 1,19 | -2,64 | 8,38E-03 | 4,45E-02 |
| AT2G31880 | 6,93 | -2,18 | 0,83 | -2,63 | 8,45E-03 | 4,48E-02 |
| AT2G18280 | 2,10 | -4,58 | 1,74 | -2,63 | 8,47E-03 | 4,48E-02 |
| AT1G35215 | 4,81 | 3,18 | 1,21 | 2,63 | 8,49E-03 | 4,48E-02 |
| AT5G19390 | 2,78 | -4,23 | 1,61 | -2,63 | 8,62E-03 | 4,54E-02 |
| AT1G80680 | 4,25 | -2,94 | 1,12 | -2,62 | 8,68E-03 | 4,56E-02 |
| AT4G00975 | 11,27 | -1,42 | 0,54 | -2,62 | 8,79E-03 | 4,62E-02 |
| AT3G17510 | 2,73 | -3,94 | 1,50 | -2,62 | 8,81E-03 | 4,62E-02 |
| AT3G17900 | 1,92 | -3,92 | 1,50 | -2,62 | 8,83E-03 | 4,62E-02 |
| AT1G25440 | 6,25 | -2,39 | 0,91 | -2,62 | 8,88E-03 | 4,64E-02 |
| AT1G79090 | 2,69 | -3,80 | 1,45 | -2,62 | 8,92E-03 | 4,65E-02 |
| AT3G18217 | 147,37 | -0,62 | 0,24 | -2,61 | 9,04E-03 | 4,70E-02 |
| AT5G19820 | 4,02 | -3,24 | 1,24 | -2,61 | 9,08E-03 | 4,72E-02 |
| AT2G35750 | 26,17 | 1,42 | 0,54 | 2,61 | 9,14E-03 | 4,74E-02 |
| AT2G41440 | 2,60 | -4,14 | 1,59 | -2,60 | 9,31E-03 | 4,82E-02 |
| AT2G38440 | 3,79 | -2,45 | 0,95 | -2,59 | 9,52E-03 | 4,91E-02 |
| AT4G30972 | 11,83 | 1,59 | 0,61 | 2,59 | 9,51E-03 | 4,91E-02 |
| AT1G72410 | 1,92 | -3,86 | 1,49 | -2,59 | 9,61E-03 | 4,95E-02 |
| AT5G07970 | 3,91 | -3,71 | 1,43 | -2,59 | 9,62E-03 | 4,95E-02 |
| AT2G04885 | 2,17 | 4,19 | 1,62 | 2,59 | 9,68E-03 | 4,96E-02 |
| AT1G52570 | 1,59 | -5,70 | 2,20 | -2,59 | 9,70E-03 | 4,96E-02 |
| AT5G20700 | 23,90 | -1,10 | 0,42 | -2,59 | 9,70E-03 | 4,96E-02 |
| AT4G30975 | 11,87 | 1,59 | 0,62 | 2,58 | 9,74E-03 | 4,98E-02 |
| AT4G00140 | 2,38 | 4,24 | 1,64 | 2,58 | 9,78E-03 | 4,98E-02 |
| AT4G01810 | 2,76 | -3,45 | 1,34 | -2,58 | 9,80E-03 | 4,98E-02 |
| AT4G11830 | 1,70 | -5,01 | 1,94 | -2,58 | 9,79E-03 | 4,98E-02 |
| AT4G21710 | 2,69 | -3,51 | 1,36 | -2,58 | 9,94E-03 | 5,04E-02 |
| AT3G06810 | 1,67 | -4,38 | 1,70 | -2,57 | 1,01E-02 | 5,12E-02 |
| AT2G15620 | 3,38 | -2,58 | 1,00 | -2,57 | 1,02E-02 | 5,14E-02 |
| AT2G20990 | 4,50 | -2,82 | 1,10 | -2,57 | 1,02E-02 | 5,15E-02 |
| AT4G24680 | 3,35 | -2,74 | 1,07 | -2,57 | 1,02E-02 | 5,15E-02 |
| AT4G14500 | 7,04 | 2,31 | 0,90 | 2,57 | 1,03E-02 | 5,18E-02 |
| AT5G57930 | 1,66 | -4,92 | 1,92 | -2,57 | 1,03E-02 | 5,18E-02 |
| AT1G48760 | 1,82 | -4,90 | 1,91 | -2,56 | 1,04E-02 | 5,21E-02 |
| AT1G09640 | 2,66 | -3,88 | 1,52 | -2,56 | 1,05E-02 | 5,25E-02 |
| AT5G39770 | 12,51 | 1,54 | 0,60 | 2,56 | 1,05E-02 | 5,25E-02 |
| AT4G22410 | 2,93 | 3,81 | 1,49 | 2,55 | 1,08E-02 | 5,37E-02 |
| AT3G25855 | 1,70 | -4,97 | 1,95 | -2,55 | 1,09E-02 | 5,44E-02 |
| AT3G45650 | 1,57 | -5,02 | 1,97 | -2,54 | 1,09E-02 | 5,44E-02 |
| AT5G24990 | 2,50 | 3,97 | 1,56 | 2,54 | 1,10E-02 | 5,44E-02 |
| AT3G19420 | 2,24 | -3,94 | 1,55 | -2,54 | 1,10E-02 | 5,45E-02 |
| AT2G35747 | 15,41 | 1,85 | 0,73 | 2,54 | 1,10E-02 | 5,45E-02 |
| AT3G44480 | 4,70 | -2,23 | 0,88 | -2,54 | 1,12E-02 | 5,51E-02 |

| | | | | | | |
|---|---|---|---|---|---|---|
| AT5G52882 | 2,07 | -4,45 | 1,75 | -2,54 | 1,12E-02 | 5,51E-02 |
| AT2G46500 | 2,79 | -3,51 | 1,39 | -2,54 | 1,12E-02 | 5,52E-02 |
| AT5G16210 | 1,91 | -4,26 | 1,68 | -2,54 | 1,12E-02 | 5,52E-02 |
| AT3G22960 | 3,17 | -3,26 | 1,29 | -2,53 | 1,13E-02 | 5,54E-02 |
| AT2G47940 | 2,47 | -4,17 | 1,65 | -2,53 | 1,15E-02 | 5,64E-02 |
| AT1G48267 | 15177,13 | 0,85 | 0,34 | 2,53 | 1,15E-02 | 5,64E-02 |
| AT1G75800 | 1,71 | -4,58 | 1,82 | -2,52 | 1,17E-02 | 5,69E-02 |
| AT2G46380 | 1,50 | -5,43 | 2,15 | -2,52 | 1,17E-02 | 5,69E-02 |
| AT4G00730 | 3,19 | -3,23 | 1,28 | -2,52 | 1,17E-02 | 5,69E-02 |
| AT2G46460 | 2,59 | 4,36 | 1,73 | 2,52 | 1,17E-02 | 5,70E-02 |
| AT3G58050 | 1,93 | -3,66 | 1,45 | -2,52 | 1,18E-02 | 5,73E-02 |
| AT5G27607 | 2,38 | 4,63 | 1,84 | 2,52 | 1,18E-02 | 5,73E-02 |
| AT2G46020 | 3,28 | -2,60 | 1,04 | -2,51 | 1,19E-02 | 5,78E-02 |
| AT5G62190 | 2,05 | -3,52 | 1,40 | -2,51 | 1,20E-02 | 5,80E-02 |
| AT5G25280 | 3,55 | -2,45 | 0,98 | -2,51 | 1,21E-02 | 5,84E-02 |
| AT1G35501 | 18,58 | 1,49 | 0,59 | 2,51 | 1,22E-02 | 5,89E-02 |
| AT4G02350 | 1,95 | -4,10 | 1,64 | -2,50 | 1,26E-02 | 6,05E-02 |
| AT1G18485 | 1,73 | -5,10 | 2,05 | -2,49 | 1,26E-02 | 6,06E-02 |
| AT1G76135 | 2,70 | 3,99 | 1,60 | 2,49 | 1,26E-02 | 6,06E-02 |
| AT3G53460 | 6,35 | -2,07 | 0,83 | -2,49 | 1,27E-02 | 6,07E-02 |
| AT3G62250 | 1,91 | -4,11 | 1,65 | -2,49 | 1,28E-02 | 6,12E-02 |
| AT3G28956 | 2,37 | 3,72 | 1,50 | 2,48 | 1,30E-02 | 6,20E-02 |
| AT5G44800 | 4,77 | -2,13 | 0,86 | -2,48 | 1,30E-02 | 6,21E-02 |
| AT4G36640 | 1,54 | -5,06 | 2,04 | -2,48 | 1,31E-02 | 6,22E-02 |
| AT5G18380 | 4,02 | -2,05 | 0,83 | -2,48 | 1,31E-02 | 6,23E-02 |
| AT3G43990 | 14,30 | 2,79 | 1,13 | 2,48 | 1,32E-02 | 6,24E-02 |
| AT5G39740 | 3,11 | -2,46 | 0,99 | -2,48 | 1,33E-02 | 6,28E-02 |
| AT2G04030 | 4,28 | -2,37 | 0,96 | -2,48 | 1,33E-02 | 6,28E-02 |
| AT4G10340 | 23,35 | -1,26 | 0,51 | -2,48 | 1,33E-02 | 6,28E-02 |
| AT1G61790 | 1,42 | -4,34 | 1,75 | -2,47 | 1,34E-02 | 6,34E-02 |
| AT4G31210 | 1,90 | -3,98 | 1,61 | -2,47 | 1,35E-02 | 6,35E-02 |
| AT5G41905 | 1173,00 | -0,62 | 0,25 | -2,47 | 1,35E-02 | 6,35E-02 |
| AT1G60140 | 2,14 | -3,27 | 1,32 | -2,47 | 1,36E-02 | 6,37E-02 |
| AT4G32640 | 2,41 | -3,58 | 1,45 | -2,46 | 1,38E-02 | 6,44E-02 |
| AT3G24929 | 2,38 | 3,89 | 1,58 | 2,46 | 1,38E-02 | 6,44E-02 |
| AT3G57660 | 1,78 | -4,06 | 1,65 | -2,46 | 1,39E-02 | 6,48E-02 |
| AT2G48060 | 4,01 | -2,44 | 1,00 | -2,45 | 1,42E-02 | 6,62E-02 |
| AT4G10710 | 3,86 | -2,93 | 1,20 | -2,45 | 1,42E-02 | 6,62E-02 |
| AT1G64790 | 7,36 | -1,85 | 0,76 | -2,45 | 1,43E-02 | 6,65E-02 |
| AT1G27340 | 2,10 | -3,64 | 1,49 | -2,45 | 1,43E-02 | 6,65E-02 |
| AT2G41100 | 6,69 | 2,59 | 1,06 | 2,45 | 1,44E-02 | 6,66E-02 |
| AT3G48350 | 1,84 | -4,52 | 1,85 | -2,45 | 1,44E-02 | 6,66E-02 |
| AT2G07645 | 4,48 | 2,98 | 1,22 | 2,44 | 1,45E-02 | 6,71E-02 |
| AT1G34300 | 2,71 | -3,18 | 1,30 | -2,44 | 1,46E-02 | 6,72E-02 |
| AT3G55270 | 1,69 | -4,38 | 1,79 | -2,44 | 1,47E-02 | 6,77E-02 |
| AT5G05680 | 1,56 | -4,41 | 1,81 | -2,44 | 1,48E-02 | 6,81E-02 |
| AT4G28760 | 2,13 | -3,46 | 1,42 | -2,43 | 1,49E-02 | 6,83E-02 |
| AT1G17210 | 2,32 | -3,86 | 1,59 | -2,43 | 1,49E-02 | 6,83E-02 |
| AT1G49700 | 4,39 | -2,11 | 0,87 | -2,43 | 1,50E-02 | 6,83E-02 |
| AT3G11940 | 3,79 | -2,33 | 0,96 | -2,43 | 1,49E-02 | 6,83E-02 |
| AT3G59040 | 1,65 | -4,39 | 1,80 | -2,43 | 1,50E-02 | 6,83E-02 |
| AT5G64740 | 4,40 | -2,00 | 0,82 | -2,43 | 1,51E-02 | 6,86E-02 |
| AT2G36160 | 1,81 | -3,80 | 1,57 | -2,43 | 1,53E-02 | 6,93E-02 |
| AT1G01183 | 317,62 | -0,85 | 0,35 | -2,43 | 1,53E-02 | 6,93E-02 |
| AT2G36530 | 5,68 | -1,80 | 0,74 | -2,43 | 1,53E-02 | 6,93E-02 |
| AT3G04250 | 2,26 | 4,35 | 1,80 | 2,42 | 1,55E-02 | 7,00E-02 |

| | | | | | | |
|---|---|---|---|---|---|---|
| AT5G17020 | 2,03 | -3,34 | 1,38 | -2,42 | 1,55E-02 | 7,00E-02 |
| AT4G08630 | 1,97 | 4,12 | 1,71 | 2,42 | 1,57E-02 | 7,07E-02 |
| AT1G27320 | 2,96 | -3,17 | 1,31 | -2,41 | 1,58E-02 | 7,10E-02 |
| AT3G27865 | 3,38 | 3,51 | 1,46 | 2,41 | 1,58E-02 | 7,11E-02 |
| AT1G04907 | 10,89 | -1,75 | 0,72 | -2,41 | 1,58E-02 | 7,11E-02 |
| AT1G54040 | 2,21 | 3,90 | 1,62 | 2,41 | 1,59E-02 | 7,16E-02 |
| AT2G26530 | 1,88 | 4,41 | 1,83 | 2,41 | 1,60E-02 | 7,16E-02 |
| AT1G29370 | 3,57 | -2,72 | 1,13 | -2,41 | 1,60E-02 | 7,18E-02 |
| AT3G06720 | 1,89 | -4,53 | 1,89 | -2,40 | 1,62E-02 | 7,25E-02 |
| AT5G61140 | 4,17 | -2,25 | 0,94 | -2,40 | 1,63E-02 | 7,26E-02 |
| AT3G18490 | 8,11 | 1,90 | 0,79 | 2,40 | 1,63E-02 | 7,26E-02 |
| AT5G09590 | 2,74 | -3,89 | 1,62 | -2,40 | 1,64E-02 | 7,29E-02 |
| AT5G47435 | 1,48 | -4,62 | 1,93 | -2,40 | 1,64E-02 | 7,30E-02 |
| AT2G33340 | 1,98 | -3,49 | 1,45 | -2,40 | 1,65E-02 | 7,32E-02 |
| AT5G24660 | 2,17 | 4,04 | 1,69 | 2,40 | 1,65E-02 | 7,32E-02 |
| AT4G31490 | 3,09 | -3,00 | 1,25 | -2,40 | 1,66E-02 | 7,33E-02 |
| AT4G38220 | 1,96 | -4,10 | 1,71 | -2,40 | 1,66E-02 | 7,33E-02 |
| AT1G10522 | 24,13 | 1,12 | 0,47 | 2,40 | 1,66E-02 | 7,33E-02 |
| AT3G08165 | 3,95 | 3,61 | 1,51 | 2,39 | 1,66E-02 | 7,33E-02 |
| AT3G60415 | 1,73 | -4,31 | 1,80 | -2,39 | 1,67E-02 | 7,34E-02 |
| AT5G26830 | 2,03 | -3,48 | 1,45 | -2,39 | 1,68E-02 | 7,35E-02 |
| AT2G46560 | 4,43 | -2,36 | 0,99 | -2,39 | 1,68E-02 | 7,35E-02 |
| AT5G38430 | 4,72 | -1,92 | 0,80 | -2,39 | 1,69E-02 | 7,39E-02 |
| AT1G17220 | 3,72 | -2,47 | 1,04 | -2,38 | 1,71E-02 | 7,46E-02 |
| AT2G01010 | 3,38 | 3,06 | 1,28 | 2,38 | 1,71E-02 | 7,46E-02 |
| AT3G03425 | 4,95 | 2,87 | 1,21 | 2,38 | 1,71E-02 | 7,46E-02 |
| AT2G40840 | 4,12 | -2,19 | 0,92 | -2,38 | 1,74E-02 | 7,52E-02 |
| AT4G29780 | 2,35 | 4,27 | 1,79 | 2,38 | 1,73E-02 | 7,52E-02 |
| AT1G69360 | 2,19 | -4,17 | 1,75 | -2,38 | 1,74E-02 | 7,52E-02 |
| AT2G27860 | 4,64 | -1,92 | 0,81 | -2,38 | 1,74E-02 | 7,52E-02 |
| AT2G41620 | 1,68 | -4,30 | 1,81 | -2,38 | 1,74E-02 | 7,52E-02 |
| AT3G53740 | 2,01 | -4,35 | 1,83 | -2,38 | 1,74E-02 | 7,52E-02 |
| AT2G34208 | 2,28 | -3,93 | 1,66 | -2,37 | 1,79E-02 | 7,73E-02 |
| AT3G58193 | 8,75 | 1,98 | 0,84 | 2,37 | 1,80E-02 | 7,77E-02 |
| AT5G46070 | 1,78 | -4,05 | 1,71 | -2,36 | 1,81E-02 | 7,80E-02 |
| AT3G07725 | 10,80 | 1,42 | 0,60 | 2,36 | 1,83E-02 | 7,82E-02 |
| AT1G07593 | 3,78 | -3,33 | 1,41 | -2,36 | 1,82E-02 | 7,82E-02 |
| AT3G49600 | 1,57 | -4,42 | 1,87 | -2,36 | 1,82E-02 | 7,82E-02 |
| AT1G79990 | 2,18 | -3,72 | 1,58 | -2,36 | 1,83E-02 | 7,83E-02 |
| AT3G01345 | 6,06 | 3,31 | 1,40 | 2,36 | 1,84E-02 | 7,85E-02 |
| AT4G38510 | 2,20 | -3,40 | 1,44 | -2,36 | 1,84E-02 | 7,85E-02 |
| AT1G24706 | 2,66 | -2,65 | 1,12 | -2,36 | 1,84E-02 | 7,85E-02 |
| AT5G41510 | 1,76 | 4,17 | 1,77 | 2,35 | 1,86E-02 | 7,89E-02 |
| AT5G54075 | 15,55 | 1,31 | 0,56 | 2,35 | 1,86E-02 | 7,89E-02 |
| AT2G08986 | 10,75 | 1,62 | 0,69 | 2,35 | 1,88E-02 | 7,97E-02 |
| AT3G02090 | 3,68 | -2,29 | 0,97 | -2,35 | 1,90E-02 | 8,03E-02 |
| AT3G25020 | 3,41 | 3,00 | 1,28 | 2,35 | 1,90E-02 | 8,03E-02 |
| AT5G18525 | 1,70 | -4,27 | 1,82 | -2,34 | 1,91E-02 | 8,06E-02 |
| AT1G64340 | 2,24 | 4,50 | 1,92 | 2,34 | 1,91E-02 | 8,07E-02 |
| AT5G54250 | 1,88 | -4,10 | 1,75 | -2,34 | 1,91E-02 | 8,07E-02 |
| AT1G29940 | 2,05 | -3,72 | 1,59 | -2,34 | 1,92E-02 | 8,09E-02 |
| AT4G34260 | 1,45 | -4,60 | 1,97 | -2,34 | 1,93E-02 | 8,09E-02 |
| AT4G32010 | 1,47 | -4,54 | 1,94 | -2,34 | 1,95E-02 | 8,18E-02 |
| AT1G32010 | 1,81 | 4,09 | 1,75 | 2,34 | 1,95E-02 | 8,18E-02 |
| AT3G57330 | 3,24 | -2,68 | 1,15 | -2,33 | 1,96E-02 | 8,19E-02 |
| AT5G15845 | 2,91 | -2,65 | 1,14 | -2,33 | 2,00E-02 | 8,34E-02 |

| | | | | | | |
|---|---|---|---|---|---|---|
| AT4G34830 | 2,44 | -3,16 | 1,36 | -2,33 | 2,00E-02 | 8,34E-02 |
| AT3G22730 | 1,61 | 3,98 | 1,71 | 2,33 | 2,00E-02 | 8,35E-02 |
| AT5G58410 | 3,13 | -3,30 | 1,42 | -2,32 | 2,02E-02 | 8,40E-02 |
| AT4G17330 | 4,91 | -2,65 | 1,14 | -2,32 | 2,02E-02 | 8,40E-02 |
| AT4G08755 | 25,91 | -0,99 | 0,42 | -2,32 | 2,03E-02 | 8,41E-02 |
| AT5G05690 | 2,09 | -3,19 | 1,38 | -2,32 | 2,03E-02 | 8,41E-02 |
| AT5G07735 | 1,90 | 3,65 | 1,57 | 2,32 | 2,05E-02 | 8,47E-02 |
| AT4G24690 | 16,56 | -1,23 | 0,53 | -2,32 | 2,05E-02 | 8,47E-02 |
| AT5G16640 | 4,15 | 2,87 | 1,24 | 2,31 | 2,06E-02 | 8,52E-02 |
| AT2G18570 | 4,57 | 2,98 | 1,29 | 2,31 | 2,08E-02 | 8,56E-02 |
| AT5G49615 | 2,21 | 4,25 | 1,84 | 2,31 | 2,08E-02 | 8,56E-02 |
| AT3G13090 | 2,27 | -4,92 | 2,13 | -2,31 | 2,09E-02 | 8,57E-02 |
| AT4G29200 | 3,91 | 2,87 | 1,24 | 2,31 | 2,09E-02 | 8,59E-02 |
| AT3G13330 | 3,62 | -2,37 | 1,03 | -2,31 | 2,10E-02 | 8,61E-02 |
| AT5G06630 | 1,85 | 4,00 | 1,73 | 2,31 | 2,10E-02 | 8,61E-02 |
| AT5G11040 | 2,15 | -3,23 | 1,40 | -2,31 | 2,11E-02 | 8,62E-02 |
| AT1G05780 | 4,77 | 2,29 | 0,99 | 2,31 | 2,11E-02 | 8,62E-02 |
| AT5G20720 | 3,41 | -2,64 | 1,15 | -2,31 | 2,12E-02 | 8,63E-02 |
| AT3G51870 | 1,46 | -4,58 | 1,99 | -2,30 | 2,16E-02 | 8,81E-02 |
| AT3G45190 | 2,88 | -2,59 | 1,13 | -2,29 | 2,18E-02 | 8,85E-02 |
| AT3G05530 | 1,78 | -3,20 | 1,40 | -2,29 | 2,18E-02 | 8,86E-02 |
| AT5G18360 | 2,96 | -2,99 | 1,30 | -2,29 | 2,18E-02 | 8,86E-02 |
| AT5G19400 | 2,38 | -3,79 | 1,65 | -2,29 | 2,19E-02 | 8,86E-02 |
| AT3G48860 | 1,96 | -5,27 | 2,30 | -2,29 | 2,19E-02 | 8,86E-02 |
| AT3G48200 | 2,11 | -3,68 | 1,61 | -2,29 | 2,22E-02 | 8,95E-02 |
| AT3G47348 | 6,89 | 3,01 | 1,32 | 2,29 | 2,22E-02 | 8,96E-02 |
| AT5G02575 | 6,74 | -1,72 | 0,75 | -2,29 | 2,22E-02 | 8,96E-02 |
| AT3G04740 | 2,26 | -2,92 | 1,28 | -2,29 | 2,23E-02 | 8,97E-02 |
| AT5G11720 | 1,68 | -4,61 | 2,02 | -2,28 | 2,23E-02 | 8,98E-02 |
| AT1G07280 | 1,33 | -4,46 | 1,96 | -2,28 | 2,26E-02 | 9,05E-02 |
| AT5G60700 | 1,32 | -5,16 | 2,26 | -2,28 | 2,26E-02 | 9,05E-02 |
| AT1G19090 | 4,93 | 2,37 | 1,04 | 2,27 | 2,29E-02 | 9,18E-02 |
| AT1G23720 | 2,26 | 4,44 | 1,95 | 2,27 | 2,31E-02 | 9,18E-02 |
| AT5G46450 | 2,49 | -2,94 | 1,29 | -2,27 | 2,31E-02 | 9,18E-02 |
| AT5G46510 | 5,48 | -2,03 | 0,89 | -2,27 | 2,31E-02 | 9,18E-02 |
| AT5G58540 | 1,83 | -4,01 | 1,76 | -2,27 | 2,31E-02 | 9,18E-02 |
| AT5G62162 | 10,16 | -1,64 | 0,72 | -2,27 | 2,31E-02 | 9,18E-02 |
| AT2G36480 | 1,71 | -3,77 | 1,66 | -2,27 | 2,30E-02 | 9,18E-02 |
| AT1G48725 | 1,45 | 3,93 | 1,73 | 2,27 | 2,33E-02 | 9,25E-02 |
| AT4G23915 | 12,58 | 1,47 | 0,65 | 2,27 | 2,34E-02 | 9,25E-02 |
| AT2G16020 | 3,14 | 3,45 | 1,52 | 2,27 | 2,35E-02 | 9,28E-02 |
| AT3G62290 | 5,55 | -1,88 | 0,83 | -2,26 | 2,35E-02 | 9,29E-02 |
| AT2G47410 | 3,41 | -3,65 | 1,61 | -2,26 | 2,37E-02 | 9,33E-02 |
| AT5G17880 | 1,61 | -4,19 | 1,85 | -2,26 | 2,37E-02 | 9,34E-02 |
| AT5G24690 | 2,42 | -3,27 | 1,45 | -2,26 | 2,37E-02 | 9,34E-02 |
| AT3G47342 | 15,52 | 1,36 | 0,60 | 2,26 | 2,39E-02 | 9,37E-02 |
| AT4G00800 | 2,43 | -3,03 | 1,34 | -2,26 | 2,39E-02 | 9,38E-02 |
| AT5G14040 | 2,00 | -3,47 | 1,54 | -2,26 | 2,39E-02 | 9,38E-02 |
| AT1G76490 | 3,44 | -2,37 | 1,05 | -2,26 | 2,40E-02 | 9,40E-02 |
| AT4G26520 | 1,37 | -4,47 | 1,98 | -2,26 | 2,40E-02 | 9,40E-02 |
| AT2G03821 | 1,77 | 3,58 | 1,59 | 2,26 | 2,41E-02 | 9,40E-02 |
| AT1G43910 | 2,35 | -3,49 | 1,55 | -2,25 | 2,43E-02 | 9,48E-02 |
| AT5G63190 | 2,74 | -2,98 | 1,32 | -2,25 | 2,46E-02 | 9,58E-02 |
| AT4G25880 | 1,93 | -3,79 | 1,69 | -2,25 | 2,47E-02 | 9,61E-02 |
| AT2G36490 | 1,90 | -5,97 | 2,66 | -2,25 | 2,47E-02 | 9,61E-02 |
| AT2G36850 | 3,05 | -2,87 | 1,28 | -2,24 | 2,49E-02 | 9,64E-02 |

| | | | | | | |
|---|---|---|---|---|---|---|
| AT2G25730 | 2,88 | -2,39 | 1,06 | -2,24 | 2,49E-02 | 9,65E-02 |
| AT2G32730 | 3,18 | -2,74 | 1,22 | -2,24 | 2,50E-02 | 9,67E-02 |
| AT3G08560 | 1,47 | -4,37 | 1,95 | -2,24 | 2,50E-02 | 9,67E-02 |
| AT5G07570 | 4,01 | 3,26 | 1,45 | 2,24 | 2,50E-02 | 9,67E-02 |
| AT1G76550 | 2,61 | -2,71 | 1,21 | -2,24 | 2,51E-02 | 9,67E-02 |
| AT3G14067 | 4,84 | -1,85 | 0,83 | -2,24 | 2,52E-02 | 9,68E-02 |
| AT4G16950 | 9,81 | -1,43 | 0,64 | -2,24 | 2,51E-02 | 9,68E-02 |
| AT3G24982 | 2,19 | 3,57 | 1,60 | 2,24 | 2,53E-02 | 9,71E-02 |
| AT4G26910 | 1,33 | -3,92 | 1,75 | -2,24 | 2,53E-02 | 9,71E-02 |
| AT4G08593 | 1,89 | 4,04 | 1,81 | 2,23 | 2,54E-02 | 9,75E-02 |
| AT3G14120 | 1,41 | -4,45 | 1,99 | -2,23 | 2,55E-02 | 9,76E-02 |
| AT3G06325 | 6,42 | 2,08 | 0,93 | 2,23 | 2,56E-02 | 9,76E-02 |
| AT1G15940 | 2,10 | -3,28 | 1,47 | -2,23 | 2,56E-02 | 9,78E-02 |
| AT3G61690 | 2,25 | -2,95 | 1,32 | -2,23 | 2,57E-02 | 9,78E-02 |
| AT1G14320 | 3,20 | -2,49 | 1,12 | -2,23 | 2,58E-02 | 9,80E-02 |
| AT1G07000 | 3,04 | -2,89 | 1,29 | -2,23 | 2,58E-02 | 9,80E-02 |
| AT4G36140 | 2,98 | -2,93 | 1,32 | -2,23 | 2,60E-02 | 9,87E-02 |
| AT3G48430 | 1,70 | -4,14 | 1,86 | -2,23 | 2,60E-02 | 9,87E-02 |
| AT5G54070 | 15,72 | 1,25 | 0,56 | 2,22 | 2,62E-02 | 9,92E-02 |
| AT3G22940 | 1,72 | 3,98 | 1,79 | 2,22 | 2,62E-02 | 9,93E-02 |
| AT5G23340 | 1,40 | -4,13 | 1,86 | -2,22 | 2,63E-02 | 9,93E-02 |
| AT3G12980 | 1,82 | -3,84 | 1,73 | -2,22 | 2,64E-02 | 9,96E-02 |
| AT5G14430 | 2,60 | -3,19 | 1,44 | -2,22 | 2,65E-02 | 9,97E-02 |
| AT5G18610 | 1,43 | -4,17 | 1,88 | -2,22 | 2,65E-02 | 9,97E-02 |
| AT2G03150 | 2,08 | -3,00 | 1,35 | -2,22 | 2,66E-02 | 1,00E-01 |
| AT2G44570 | 2,88 | 3,44 | 1,55 | 2,22 | 2,67E-02 | 1,00E-01 |
| AT3G14172 | 1,38 | -4,06 | 1,83 | -2,21 | 2,70E-02 | 1,01E-01 |
| AT5G27807 | 30,37 | 1,00 | 0,45 | 2,21 | 2,71E-02 | 1,02E-01 |
| AT4G27430 | 2,23 | -2,99 | 1,35 | -2,21 | 2,73E-02 | 1,02E-01 |
| AT3G04240 | 3,51 | -2,71 | 1,23 | -2,20 | 2,75E-02 | 1,03E-01 |
| AT4G20170 | 3,89 | 2,46 | 1,12 | 2,20 | 2,77E-02 | 1,03E-01 |
| AT1G04820 | 2,61 | -2,67 | 1,21 | -2,20 | 2,76E-02 | 1,03E-01 |
| AT5G03940 | 2,83 | -2,45 | 1,11 | -2,20 | 2,80E-02 | 1,04E-01 |
| AT5G04540 | 1,96 | -3,46 | 1,58 | -2,20 | 2,81E-02 | 1,05E-01 |
| AT4G18730 | 1,33 | -3,67 | 1,67 | -2,19 | 2,82E-02 | 1,05E-01 |
| AT5G64840 | 5,36 | -2,11 | 0,96 | -2,19 | 2,85E-02 | 1,06E-01 |
| AT5G28740 | 1,92 | -3,35 | 1,53 | -2,19 | 2,88E-02 | 1,07E-01 |
| AT3G49130 | 1,37 | -5,13 | 2,35 | -2,18 | 2,93E-02 | 1,08E-01 |
| AT1G40104 | 7,42 | 2,17 | 1,00 | 2,18 | 2,94E-02 | 1,08E-01 |
| AT3G15353 | 5,91 | 2,35 | 1,08 | 2,18 | 2,95E-02 | 1,09E-01 |
| AT3G17240 | 1,76 | -3,94 | 1,81 | -2,17 | 2,97E-02 | 1,10E-01 |
| AT1G74720 | 1,98 | -3,47 | 1,60 | -2,17 | 2,99E-02 | 1,10E-01 |
| AT2G21410 | 1,46 | -3,16 | 1,45 | -2,17 | 2,99E-02 | 1,10E-01 |
| AT5G63195 | 1,93 | -3,77 | 1,74 | -2,17 | 3,01E-02 | 1,10E-01 |
| AT1G51570 | 1,72 | -3,27 | 1,51 | -2,16 | 3,04E-02 | 1,11E-01 |
| AT3G05040 | 1,58 | -3,19 | 1,48 | -2,17 | 3,04E-02 | 1,11E-01 |
| AT3G41762 | 25,36 | 1,03 | 0,48 | 2,17 | 3,04E-02 | 1,11E-01 |
| AT5G12150 | 4,21 | -2,71 | 1,25 | -2,17 | 3,03E-02 | 1,11E-01 |
| AT5G40770 | 1,47 | -3,78 | 1,75 | -2,16 | 3,05E-02 | 1,11E-01 |
| AT3G45090 | 1,49 | -4,15 | 1,92 | -2,16 | 3,05E-02 | 1,11E-01 |
| AT4G27870 | 2,09 | -3,04 | 1,41 | -2,16 | 3,08E-02 | 1,12E-01 |
| AT1G14740 | 3,27 | -2,11 | 0,98 | -2,16 | 3,10E-02 | 1,13E-01 |
| AT2G44640 | 1,50 | -3,50 | 1,62 | -2,16 | 3,10E-02 | 1,13E-01 |
| AT3G51850 | 1,69 | -3,53 | 1,64 | -2,15 | 3,12E-02 | 1,13E-01 |
| AT2G46900 | 1,50 | -4,49 | 2,09 | -2,15 | 3,15E-02 | 1,14E-01 |
| AT4G08335 | 4,06 | 3,09 | 1,44 | 2,15 | 3,18E-02 | 1,15E-01 |

| | | | | | | |
|---|---|---|---|---|---|---|
| AT3G25880 | 1,38 | -5,36 | 2,50 | -2,15 | 3,19E-02 | 1,15E-01 |
| AT4G34050 | 1,48 | 3,77 | 1,76 | 2,15 | 3,19E-02 | 1,15E-01 |
| AT4G09735 | 2,23 | 3,56 | 1,66 | 2,14 | 3,22E-02 | 1,16E-01 |
| AT1G33560 | 2,19 | -3,74 | 1,75 | -2,14 | 3,24E-02 | 1,16E-01 |
| AT1G09570 | 2,05 | -2,51 | 1,17 | -2,14 | 3,23E-02 | 1,16E-01 |
| AT1G72390 | 2,40 | -3,22 | 1,51 | -2,13 | 3,28E-02 | 1,18E-01 |
| AT5G07410 | 1,62 | 3,91 | 1,84 | 2,13 | 3,30E-02 | 1,18E-01 |
| AT3G55020 | 1,94 | -3,08 | 1,45 | -2,13 | 3,32E-02 | 1,19E-01 |
| AT1G63980 | 3,67 | -2,21 | 1,04 | -2,13 | 3,33E-02 | 1,19E-01 |
| AT5G02480 | 2,67 | -3,12 | 1,47 | -2,13 | 3,34E-02 | 1,19E-01 |
| AT1G62030 | 1,53 | 3,84 | 1,80 | 2,13 | 3,34E-02 | 1,19E-01 |
| AT1G17470 | 1,97 | -3,22 | 1,52 | -2,12 | 3,40E-02 | 1,21E-01 |
| AT3G13070 | 1,98 | -3,92 | 1,85 | -2,12 | 3,41E-02 | 1,22E-01 |
| AT2G07981 | 2,13 | 4,06 | 1,92 | 2,12 | 3,42E-02 | 1,22E-01 |
| AT3G54590 | 2,31 | 3,56 | 1,68 | 2,11 | 3,45E-02 | 1,23E-01 |
| AT1G12620 | 2,40 | 3,35 | 1,58 | 2,11 | 3,45E-02 | 1,23E-01 |
| AT3G54580 | 1,94 | 3,61 | 1,71 | 2,11 | 3,47E-02 | 1,23E-01 |
| AT3G11540 | 2,22 | -3,41 | 1,62 | -2,11 | 3,47E-02 | 1,23E-01 |
| AT5G49980 | 4,39 | -2,14 | 1,02 | -2,11 | 3,53E-02 | 1,25E-01 |
| AT5G06530 | 2,30 | -3,52 | 1,67 | -2,10 | 3,53E-02 | 1,25E-01 |
| AT3G09850 | 1,65 | -3,50 | 1,66 | -2,10 | 3,54E-02 | 1,25E-01 |
| AT2G47000 | 1,78 | -4,31 | 2,05 | -2,10 | 3,54E-02 | 1,25E-01 |
| AT5G10960 | 3,18 | -2,29 | 1,09 | -2,10 | 3,55E-02 | 1,25E-01 |
| AT1G07637 | 2,62 | 3,09 | 1,47 | 2,10 | 3,56E-02 | 1,25E-01 |
| AT5G42080 | 4,02 | -2,03 | 0,97 | -2,10 | 3,58E-02 | 1,26E-01 |
| AT2G33730 | 2,24 | -2,80 | 1,33 | -2,10 | 3,58E-02 | 1,26E-01 |
| AT4G23420 | 23,63 | -0,97 | 0,46 | -2,10 | 3,60E-02 | 1,26E-01 |
| AT2G35050 | 3,09 | -2,66 | 1,27 | -2,09 | 3,62E-02 | 1,27E-01 |
| AT1G15010 | 2,66 | 3,14 | 1,50 | 2,09 | 3,62E-02 | 1,27E-01 |
| AT5G23630 | 2,35 | -2,61 | 1,25 | -2,09 | 3,64E-02 | 1,27E-01 |
| AT5G07725 | 8,72 | 1,79 | 0,86 | 2,09 | 3,64E-02 | 1,27E-01 |
| AT3G57260 | 8,35 | -1,18 | 0,56 | -2,09 | 3,66E-02 | 1,28E-01 |
| AT1G48360 | 1,34 | -4,31 | 2,06 | -2,09 | 3,68E-02 | 1,28E-01 |
| AT1G53310 | 3,45 | -1,92 | 0,92 | -2,08 | 3,71E-02 | 1,29E-01 |
| AT3G56940 | 6,51 | -1,62 | 0,78 | -2,08 | 3,72E-02 | 1,29E-01 |
| AT2G40000 | 5,19 | 1,99 | 0,96 | 2,08 | 3,72E-02 | 1,29E-01 |
| AT5G11560 | 3,42 | -2,23 | 1,07 | -2,08 | 3,73E-02 | 1,29E-01 |
| AT5G17710 | 1,85 | -2,66 | 1,28 | -2,08 | 3,73E-02 | 1,29E-01 |
| AT3G47910 | 2,73 | -2,21 | 1,06 | -2,08 | 3,74E-02 | 1,29E-01 |
| AT1G67635 | 1,69 | 3,74 | 1,80 | 2,08 | 3,75E-02 | 1,30E-01 |
| AT3G56360 | 3,83 | -2,18 | 1,05 | -2,08 | 3,75E-02 | 1,30E-01 |
| AT2G05455 | 4,14 | 2,91 | 1,40 | 2,08 | 3,78E-02 | 1,30E-01 |
| AT3G10985 | 6,51 | 1,74 | 0,84 | 2,08 | 3,78E-02 | 1,30E-01 |
| AT2G28450 | 1,48 | -3,74 | 1,80 | -2,08 | 3,78E-02 | 1,30E-01 |
| AT3G07100 | 6,58 | -1,57 | 0,75 | -2,08 | 3,77E-02 | 1,30E-01 |
| AT3G11910 | 1,81 | -3,49 | 1,68 | -2,08 | 3,78E-02 | 1,30E-01 |
| AT5G20290 | 5,09 | -1,78 | 0,86 | -2,08 | 3,78E-02 | 1,30E-01 |
| AT5G51070 | 3,71 | -1,96 | 0,95 | -2,08 | 3,79E-02 | 1,30E-01 |
| AT2G17440 | 1,80 | -3,72 | 1,79 | -2,07 | 3,80E-02 | 1,30E-01 |
| AT1G13320 | 2,17 | -2,71 | 1,31 | -2,07 | 3,83E-02 | 1,31E-01 |
| AT1G06243 | 5,81 | 1,75 | 0,85 | 2,07 | 3,85E-02 | 1,31E-01 |
| AT2G40140 | 4,76 | -1,96 | 0,95 | -2,07 | 3,87E-02 | 1,32E-01 |
| AT5G46110 | 7,87 | -1,50 | 0,73 | -2,06 | 3,91E-02 | 1,33E-01 |
| AT5G44640 | 1,59 | 3,77 | 1,83 | 2,06 | 3,92E-02 | 1,33E-01 |
| AT1G09770 | 3,11 | -2,48 | 1,20 | -2,06 | 3,94E-02 | 1,34E-01 |
| AT5G14270 | 2,74 | -2,19 | 1,06 | -2,06 | 3,94E-02 | 1,34E-01 |

| | | | | | | |
|---|---|---|---|---|---|---|
| AT4G09650 | 3,00 | 3,50 | 1,70 | 2,06 | 3,97E-02 | 1,34E-01 |
| AT4G27440 | 13,15 | 1,32 | 0,64 | 2,06 | 3,97E-02 | 1,34E-01 |
| AT1G12615 | 1,59 | 4,13 | 2,02 | 2,05 | 4,04E-02 | 1,37E-01 |
| AT1G27100 | 1,46 | 3,56 | 1,74 | 2,04 | 4,09E-02 | 1,38E-01 |
| AT4G29380 | 1,74 | -3,53 | 1,73 | -2,04 | 4,11E-02 | 1,39E-01 |
| AT5G06175 | 3,94 | 2,28 | 1,12 | 2,04 | 4,11E-02 | 1,39E-01 |
| AT4G37460 | 2,98 | -2,44 | 1,19 | -2,04 | 4,12E-02 | 1,39E-01 |
| AT2G03930 | 1,60 | 3,34 | 1,64 | 2,04 | 4,14E-02 | 1,39E-01 |
| AT1G79690 | 1,96 | -3,58 | 1,76 | -2,04 | 4,14E-02 | 1,39E-01 |
| AT2G24790 | 3,54 | -2,54 | 1,25 | -2,04 | 4,15E-02 | 1,39E-01 |
| AT5G45095 | 2,04 | 3,38 | 1,66 | 2,04 | 4,16E-02 | 1,39E-01 |
| AT4G11420 | 3,20 | -2,97 | 1,46 | -2,04 | 4,16E-02 | 1,39E-01 |
| AT1G73870 | 2,16 | -2,70 | 1,32 | -2,04 | 4,17E-02 | 1,40E-01 |
| AT4G29090 | 2,85 | 3,13 | 1,54 | 2,04 | 4,18E-02 | 1,40E-01 |
| AT4G06110 | 22,73 | -0,94 | 0,46 | -2,03 | 4,23E-02 | 1,41E-01 |
| AT1G02080 | 3,21 | -2,73 | 1,34 | -2,03 | 4,24E-02 | 1,41E-01 |
| AT5G53170 | 2,82 | -2,23 | 1,10 | -2,03 | 4,24E-02 | 1,41E-01 |
| AT5G50920 | 18,45 | -1,03 | 0,51 | -2,03 | 4,25E-02 | 1,41E-01 |
| AT5G64940 | 2,50 | -2,76 | 1,36 | -2,03 | 4,26E-02 | 1,42E-01 |
| AT1G36180 | 2,20 | -2,68 | 1,32 | -2,03 | 4,27E-02 | 1,42E-01 |
| AT3G54050 | 3,78 | -1,92 | 0,95 | -2,02 | 4,29E-02 | 1,42E-01 |
| AT4G30990 | 1,34 | -3,76 | 1,86 | -2,02 | 4,29E-02 | 1,42E-01 |
| AT5G04930 | 5,77 | -1,97 | 0,97 | -2,03 | 4,28E-02 | 1,42E-01 |
| AT4G16370 | 3,80 | -1,84 | 0,91 | -2,02 | 4,29E-02 | 1,42E-01 |
| AT3G17190 | 1,85 | 3,41 | 1,69 | 2,02 | 4,31E-02 | 1,42E-01 |
| AT3G16640 | 7,07 | -1,72 | 0,85 | -2,02 | 4,31E-02 | 1,42E-01 |
| AT4G05360 | 2,15 | 3,15 | 1,56 | 2,02 | 4,38E-02 | 1,44E-01 |
| AT1G47660 | 2,01 | 3,21 | 1,59 | 2,02 | 4,38E-02 | 1,44E-01 |
| AT1G67960 | 1,96 | -3,46 | 1,72 | -2,02 | 4,38E-02 | 1,44E-01 |
| AT2G20010 | 1,89 | -2,71 | 1,35 | -2,02 | 4,38E-02 | 1,44E-01 |
| AT3G04470 | 2,70 | -2,31 | 1,14 | -2,02 | 4,37E-02 | 1,44E-01 |
| AT4G39350 | 3,27 | -1,95 | 0,97 | -2,01 | 4,41E-02 | 1,44E-01 |
| AT5G17890 | 1,77 | -3,35 | 1,66 | -2,01 | 4,41E-02 | 1,44E-01 |
| AT5G13650 | 5,78 | -1,72 | 0,86 | -2,01 | 4,44E-02 | 1,45E-01 |
| AT1G48953 | 8,24 | 1,86 | 0,92 | 2,01 | 4,45E-02 | 1,45E-01 |
| AT1G19373 | 3,55 | 2,74 | 1,37 | 2,00 | 4,50E-02 | 1,46E-01 |
| AT3G61897 | 2,76 | 5,17 | 2,57 | 2,01 | 4,47E-02 | 1,46E-01 |
| AT5G15650 | 3,70 | -1,96 | 0,98 | -2,01 | 4,48E-02 | 1,46E-01 |
| AT1G34440 | 1,54 | 3,51 | 1,75 | 2,00 | 4,50E-02 | 1,46E-01 |
| AT1G63005 | 3,14 | 3,20 | 1,60 | 2,00 | 4,50E-02 | 1,46E-01 |
| AT2G34750 | 2,36 | -2,73 | 1,36 | -2,01 | 4,48E-02 | 1,46E-01 |
| AT5G13000 | 3,30 | -1,98 | 0,98 | -2,01 | 4,47E-02 | 1,46E-01 |
| AT5G04315 | 6,08 | -1,51 | 0,75 | -2,01 | 4,49E-02 | 1,46E-01 |
| AT5G04325 | 6,08 | -1,51 | 0,75 | -2,01 | 4,49E-02 | 1,46E-01 |
| AT1G51690 | 2,38 | -2,79 | 1,39 | -2,00 | 4,51E-02 | 1,46E-01 |
| AT2G05100 | 10,97 | 1,33 | 0,66 | 2,00 | 4,52E-02 | 1,46E-01 |
| AT5G46420 | 2,51 | -2,24 | 1,12 | -2,00 | 4,56E-02 | 1,47E-01 |
| AT3G58970 | 1,53 | -4,75 | 2,38 | -2,00 | 4,58E-02 | 1,47E-01 |
| AT1G42970 | 13,13 | -1,15 | 0,58 | -2,00 | 4,59E-02 | 1,48E-01 |
| AT2G41560 | 2,99 | -1,90 | 0,95 | -2,00 | 4,60E-02 | 1,48E-01 |
| AT5G20730 | 2,22 | -2,27 | 1,14 | -1,99 | 4,61E-02 | 1,48E-01 |
| AT1G04140 | 1,54 | -3,27 | 1,64 | -1,99 | 4,62E-02 | 1,48E-01 |
| AT3G11410 | 4,48 | -2,07 | 1,04 | -1,99 | 4,62E-02 | 1,48E-01 |
| AT4G02380 | 7,09 | -1,35 | 0,68 | -1,99 | 4,64E-02 | 1,48E-01 |
| AT3G54500 | 1,37 | -4,06 | 2,04 | -1,99 | 4,67E-02 | 1,49E-01 |
| AT5G38190 | 2,86 | 2,65 | 1,34 | 1,99 | 4,69E-02 | 1,50E-01 |

| | | | | | | |
|---|---|---|---|---|---|---|
| AT5G60850 | 1,63 | -3,40 | 1,71 | -1,99 | 4,69E-02 | 1,50E-01 |
| AT3G09162 | 2,15 | -2,93 | 1,48 | -1,98 | 4,74E-02 | 1,51E-01 |
| AT5G08610 | 1,71 | -3,17 | 1,60 | -1,98 | 4,75E-02 | 1,51E-01 |
| AT2G35110 | 1,99 | -2,72 | 1,37 | -1,98 | 4,75E-02 | 1,51E-01 |
| AT5G47480 | 1,95 | -2,85 | 1,44 | -1,98 | 4,76E-02 | 1,51E-01 |
| AT3G22380 | 8,79 | -1,47 | 0,74 | -1,98 | 4,78E-02 | 1,52E-01 |
| AT1G27570 | 1,68 | 3,72 | 1,88 | 1,98 | 4,79E-02 | 1,52E-01 |
| AT3G06835 | 2,07 | -2,95 | 1,49 | -1,97 | 4,83E-02 | 1,53E-01 |
| AT1G16720 | 1,64 | -3,47 | 1,76 | -1,97 | 4,86E-02 | 1,53E-01 |
| AT1G59750 | 1,67 | -4,82 | 2,45 | -1,97 | 4,87E-02 | 1,53E-01 |
| AT3G03050 | 2,11 | -3,47 | 1,76 | -1,97 | 4,87E-02 | 1,53E-01 |
| AT3G55480 | 1,64 | -3,35 | 1,70 | -1,97 | 4,86E-02 | 1,53E-01 |
| AT5G58150 | 3,53 | -3,10 | 1,57 | -1,97 | 4,86E-02 | 1,53E-01 |
| AT5G57110 | 2,43 | -2,54 | 1,29 | -1,97 | 4,90E-02 | 1,54E-01 |
| AT3G02360 | 1,53 | -3,44 | 1,75 | -1,97 | 4,91E-02 | 1,54E-01 |
| AT4G30100 | 3,56 | -2,13 | 1,08 | -1,97 | 4,94E-02 | 1,55E-01 |
| AT3G63410 | 2,71 | -2,20 | 1,12 | -1,96 | 4,94E-02 | 1,55E-01 |
| AT1G04080 | 1,89 | -2,56 | 1,30 | -1,96 | 4,96E-02 | 1,55E-01 |
| AT2G17410 | 1,45 | -3,10 | 1,58 | -1,96 | 4,97E-02 | 1,56E-01 |
| AT4G32760 | 1,73 | -3,04 | 1,55 | -1,96 | 4,98E-02 | 1,56E-01 |
| AT2G29920 | 2,43 | 3,70 | 1,89 | 1,96 | 4,99E-02 | 1,56E-01 |
| AT3G23450 | 2,19 | 3,14 | 1,60 | 1,96 | 5,00E-02 | 1,56E-01 |
| AT1G27430 | 3,11 | -2,43 | 1,24 | -1,96 | 5,05E-02 | 1,57E-01 |
| AT3G61470 | 27,14 | -0,90 | 0,46 | -1,95 | 5,07E-02 | 1,58E-01 |
| AT2G25320 | 3,21 | -2,52 | 1,29 | -1,95 | 5,10E-02 | 1,59E-01 |
| AT2G04060 | 1,72 | 4,04 | 2,07 | 1,95 | 5,12E-02 | 1,59E-01 |
| AT4G38770 | 7,08 | 1,74 | 0,90 | 1,95 | 5,14E-02 | 1,60E-01 |
| AT4G39850 | 2,19 | -2,59 | 1,33 | -1,95 | 5,16E-02 | 1,60E-01 |
| AT5G41990 | 1,56 | -3,52 | 1,81 | -1,95 | 5,16E-02 | 1,60E-01 |
| AT4G08620 | 2,91 | 3,18 | 1,63 | 1,94 | 5,18E-02 | 1,60E-01 |
| AT1G05562 | 6,13 | 1,91 | 0,98 | 1,94 | 5,23E-02 | 1,61E-01 |
| AT1G31580 | 10,89 | 1,22 | 0,63 | 1,94 | 5,24E-02 | 1,61E-01 |
| AT5G36230 | 2,63 | -2,79 | 1,44 | -1,94 | 5,20E-02 | 1,61E-01 |
| AT1G05560 | 6,13 | 1,91 | 0,98 | 1,94 | 5,23E-02 | 1,61E-01 |
| AT3G14385 | 15,96 | -0,86 | 0,44 | -1,94 | 5,23E-02 | 1,61E-01 |
| AT4G12320 | 1,93 | -3,26 | 1,68 | -1,94 | 5,21E-02 | 1,61E-01 |
| AT4G23460 | 1,64 | -2,95 | 1,52 | -1,94 | 5,23E-02 | 1,61E-01 |
| AT4G30750 | 3,95 | 2,84 | 1,46 | 1,94 | 5,21E-02 | 1,61E-01 |
| AT4G04095 | 1,94 | -2,84 | 1,47 | -1,94 | 5,26E-02 | 1,61E-01 |
| AT5G15400 | 2,82 | -2,22 | 1,15 | -1,94 | 5,27E-02 | 1,62E-01 |
| AT5G67360 | 6,15 | -1,84 | 0,95 | -1,94 | 5,30E-02 | 1,62E-01 |
| AT3G12630 | 8,51 | 1,44 | 0,74 | 1,93 | 5,34E-02 | 1,63E-01 |
| AT1G47497 | 2,25 | -3,78 | 1,96 | -1,93 | 5,39E-02 | 1,65E-01 |
| AT4G34690 | 1,55 | 3,62 | 1,88 | 1,93 | 5,42E-02 | 1,65E-01 |
| AT1G69797 | 12,98 | 1,12 | 0,58 | 1,92 | 5,44E-02 | 1,66E-01 |
| AT3G13750 | 2,68 | -2,36 | 1,23 | -1,92 | 5,44E-02 | 1,66E-01 |
| AT5G35920 | 1,48 | 3,35 | 1,74 | 1,92 | 5,46E-02 | 1,66E-01 |
| AT3G22910 | 2,19 | -3,12 | 1,62 | -1,92 | 5,46E-02 | 1,66E-01 |
| AT3G27040 | 1,43 | 3,60 | 1,88 | 1,92 | 5,52E-02 | 1,68E-01 |
| AT1G08327 | 2,78 | -2,49 | 1,30 | -1,92 | 5,54E-02 | 1,68E-01 |
| AT3G02540 | 1,81 | -2,71 | 1,42 | -1,91 | 5,57E-02 | 1,69E-01 |
| AT5G06460 | 1,60 | -3,45 | 1,80 | -1,91 | 5,59E-02 | 1,69E-01 |
| AT5G46270 | 2,21 | -2,58 | 1,35 | -1,91 | 5,60E-02 | 1,69E-01 |
| AT1G53890 | 1,37 | 3,22 | 1,69 | 1,91 | 5,62E-02 | 1,70E-01 |
| AT5G62670 | 1,85 | -3,32 | 1,74 | -1,91 | 5,64E-02 | 1,70E-01 |
| AT1G05240 | 2,51 | 2,80 | 1,47 | 1,91 | 5,66E-02 | 1,71E-01 |

| | | | | | | |
|---|---|---|---|---|---|---|
| AT2G28910 | 4,89 | -2,33 | 1,22 | -1,91 | 5,66E-02 | 1,71E-01 |
| AT5G11490 | 1,97 | -2,95 | 1,55 | -1,90 | 5,70E-02 | 1,72E-01 |
| AT5G36180 | 3,47 | 2,24 | 1,18 | 1,90 | 5,71E-02 | 1,72E-01 |
| AT5G49910 | 2,46 | -2,38 | 1,25 | -1,90 | 5,71E-02 | 1,72E-01 |
| AT1G77815 | 2,65 | 2,92 | 1,54 | 1,90 | 5,72E-02 | 1,72E-01 |
| AT1G06277 | 5,95 | 1,62 | 0,85 | 1,90 | 5,77E-02 | 1,73E-01 |
| AT4G20735 | 3,19 | 3,25 | 1,71 | 1,90 | 5,77E-02 | 1,73E-01 |
| AT5G58160 | 1,77 | -4,73 | 2,49 | -1,90 | 5,76E-02 | 1,73E-01 |
| AT5G09900 | 1,74 | -2,65 | 1,40 | -1,90 | 5,77E-02 | 1,73E-01 |
| AT5G44320 | 2,18 | -2,65 | 1,40 | -1,90 | 5,80E-02 | 1,73E-01 |
| AT4G19710 | 1,47 | -3,05 | 1,61 | -1,89 | 5,83E-02 | 1,74E-01 |
| AT2G47510 | 1,59 | -3,20 | 1,69 | -1,89 | 5,85E-02 | 1,74E-01 |
| AT3G60860 | 2,94 | -2,45 | 1,30 | -1,89 | 5,86E-02 | 1,74E-01 |
| AT5G20200 | 1,58 | -3,54 | 1,87 | -1,89 | 5,84E-02 | 1,74E-01 |
| AT1G12920 | 1,87 | -2,63 | 1,39 | -1,89 | 5,86E-02 | 1,74E-01 |
| AT2G20721 | 2,89 | 2,74 | 1,45 | 1,89 | 5,86E-02 | 1,74E-01 |
| AT1G52360 | 2,02 | -2,45 | 1,30 | -1,89 | 5,90E-02 | 1,74E-01 |
| AT1G68945 | 2,15 | 2,84 | 1,50 | 1,89 | 5,90E-02 | 1,74E-01 |
| AT2G41105 | 5,08 | 2,37 | 1,26 | 1,89 | 5,89E-02 | 1,74E-01 |
| AT4G03740 | 1,37 | 3,64 | 1,93 | 1,89 | 5,90E-02 | 1,74E-01 |
| AT4G18700 | 2,69 | -2,28 | 1,21 | -1,89 | 5,89E-02 | 1,74E-01 |
| AT5G49030 | 1,79 | -2,92 | 1,55 | -1,89 | 5,92E-02 | 1,75E-01 |
| AT1G29350 | 2,20 | -2,56 | 1,36 | -1,89 | 5,93E-02 | 1,75E-01 |
| AT2G20722 | 3,24 | 2,61 | 1,38 | 1,89 | 5,93E-02 | 1,75E-01 |
| AT2G05315 | 4,98 | 1,95 | 1,04 | 1,88 | 5,99E-02 | 1,76E-01 |
| AT1G80440 | 3,83 | 2,70 | 1,43 | 1,88 | 6,02E-02 | 1,77E-01 |
| AT1G09333 | 1,54 | 3,36 | 1,79 | 1,88 | 6,06E-02 | 1,77E-01 |
| AT2G30950 | 3,17 | -1,94 | 1,03 | -1,88 | 6,05E-02 | 1,77E-01 |
| AT5G42360 | 1,47 | -3,12 | 1,66 | -1,88 | 6,05E-02 | 1,77E-01 |
| AT3G47470 | 25,61 | -0,85 | 0,45 | -1,87 | 6,10E-02 | 1,79E-01 |
| AT1G52980 | 1,44 | -3,20 | 1,71 | -1,87 | 6,12E-02 | 1,79E-01 |
| AT4G08920 | 2,64 | -2,45 | 1,31 | -1,87 | 6,12E-02 | 1,79E-01 |
| AT5G47910 | 5,04 | -1,76 | 0,94 | -1,87 | 6,13E-02 | 1,79E-01 |
| AT3G05325 | 9,46 | 1,41 | 0,76 | 1,87 | 6,16E-02 | 1,80E-01 |
| AT3G51120 | 1,56 | -3,51 | 1,88 | -1,87 | 6,17E-02 | 1,80E-01 |
| AT2G20580 | 2,81 | -2,98 | 1,60 | -1,87 | 6,21E-02 | 1,81E-01 |
| AT1G04393 | 5,00 | 1,80 | 0,97 | 1,87 | 6,22E-02 | 1,81E-01 |
| AT5G64280 | 1,39 | -2,86 | 1,53 | -1,86 | 6,23E-02 | 1,81E-01 |
| AT3G20250 | 1,61 | -3,15 | 1,69 | -1,86 | 6,24E-02 | 1,81E-01 |
| AT2G45620 | 1,35 | -3,53 | 1,90 | -1,86 | 6,26E-02 | 1,82E-01 |
| AT1G55490 | 3,72 | -1,90 | 1,02 | -1,86 | 6,28E-02 | 1,82E-01 |
| AT1G32630 | 4,04 | 1,92 | 1,03 | 1,86 | 6,31E-02 | 1,82E-01 |
| AT1G32640 | 4,04 | 1,92 | 1,03 | 1,86 | 6,31E-02 | 1,82E-01 |
| AT1G09027 | 2,11 | 2,83 | 1,52 | 1,86 | 6,31E-02 | 1,82E-01 |
| AT2G28056 | 34,24 | 0,79 | 0,42 | 1,86 | 6,31E-02 | 1,82E-01 |
| AT2G46400 | 1,95 | -2,33 | 1,25 | -1,86 | 6,31E-02 | 1,82E-01 |
| AT4G38970 | 12,35 | -0,98 | 0,53 | -1,86 | 6,32E-02 | 1,82E-01 |
| AT1G18745 | 15,99 | 1,12 | 0,60 | 1,85 | 6,36E-02 | 1,83E-01 |
| AT5G35360 | 1,57 | -2,85 | 1,54 | -1,85 | 6,38E-02 | 1,84E-01 |
| AT2G38280 | 1,89 | -2,83 | 1,53 | -1,85 | 6,39E-02 | 1,84E-01 |
| AT5G60710 | 1,48 | -2,79 | 1,51 | -1,85 | 6,43E-02 | 1,85E-01 |
| AT1G06230 | 2,14 | -2,58 | 1,40 | -1,85 | 6,48E-02 | 1,86E-01 |
| AT1G16635 | 19,67 | 1,13 | 0,61 | 1,85 | 6,48E-02 | 1,86E-01 |
| AT1G80000 | 1,90 | -3,79 | 2,06 | -1,84 | 6,54E-02 | 1,87E-01 |
| AT2G15880 | 1,49 | -3,22 | 1,75 | -1,84 | 6,54E-02 | 1,87E-01 |
| AT1G22060 | 2,54 | -2,19 | 1,19 | -1,84 | 6,56E-02 | 1,87E-01 |

| | | | | | | |
|---|---|---|---|---|---|---|
| AT5G02490 | 4,18 | -1,79 | 0,97 | -1,84 | 6,61E-02 | 1,89E-01 |
| AT1G12300 | 2,68 | 2,70 | 1,47 | 1,84 | 6,63E-02 | 1,89E-01 |
| AT5G62000 | 10,38 | -1,07 | 0,58 | -1,84 | 6,64E-02 | 1,89E-01 |
| AT2G34655 | 4,12 | 2,41 | 1,32 | 1,83 | 6,72E-02 | 1,91E-01 |
| AT2G15420 | 2,17 | 2,92 | 1,60 | 1,83 | 6,74E-02 | 1,91E-01 |
| AT5G10450 | 3,14 | -1,89 | 1,04 | -1,83 | 6,74E-02 | 1,91E-01 |
| AT3G25795 | 22,21 | 2,50 | 1,37 | 1,83 | 6,76E-02 | 1,92E-01 |
| AT3G06670 | 1,42 | -3,02 | 1,66 | -1,83 | 6,78E-02 | 1,92E-01 |
| AT5G25520 | 1,48 | -4,48 | 2,46 | -1,82 | 6,80E-02 | 1,92E-01 |
| AT1G05247 | 19,59 | 1,12 | 0,61 | 1,82 | 6,80E-02 | 1,92E-01 |
| AT2G19850 | 5,66 | 1,97 | 1,08 | 1,82 | 6,82E-02 | 1,93E-01 |
| AT5G07290 | 1,99 | -4,06 | 2,22 | -1,82 | 6,81E-02 | 1,93E-01 |
| AT1G67090 | 31,55 | -0,82 | 0,45 | -1,82 | 6,85E-02 | 1,93E-01 |
| AT5G25230 | 1,66 | -3,07 | 1,68 | -1,82 | 6,86E-02 | 1,93E-01 |
| AT3G58610 | 2,68 | -1,97 | 1,08 | -1,82 | 6,93E-02 | 1,95E-01 |
| AT2G30110 | 3,88 | -1,66 | 0,91 | -1,82 | 6,95E-02 | 1,96E-01 |
| AT4G03030 | 1,74 | -3,04 | 1,68 | -1,81 | 7,06E-02 | 1,98E-01 |
| AT1G08420 | 1,72 | -2,84 | 1,57 | -1,81 | 7,07E-02 | 1,99E-01 |
| AT5G65610 | 2,72 | 2,90 | 1,60 | 1,81 | 7,08E-02 | 1,99E-01 |
| AT4G03550 | 3,00 | -2,52 | 1,40 | -1,80 | 7,12E-02 | 1,99E-01 |
| AT4G19100 | 1,38 | 3,88 | 2,15 | 1,80 | 7,13E-02 | 1,99E-01 |
| AT5G45430 | 3,88 | -1,82 | 1,01 | -1,80 | 7,12E-02 | 1,99E-01 |
| AT2G05520 | 18,30 | 0,84 | 0,47 | 1,80 | 7,14E-02 | 2,00E-01 |
| AT3G24180 | 1,99 | -2,45 | 1,36 | -1,80 | 7,15E-02 | 2,00E-01 |
| AT5G10840 | 2,05 | -2,37 | 1,32 | -1,80 | 7,17E-02 | 2,00E-01 |
| AT1G34370 | 1,49 | -3,59 | 2,00 | -1,80 | 7,20E-02 | 2,01E-01 |
| AT3G30520 | 1,35 | 3,99 | 2,22 | 1,80 | 7,22E-02 | 2,01E-01 |
| AT5G14930 | 1,80 | -3,67 | 2,04 | -1,80 | 7,24E-02 | 2,01E-01 |
| AT1G21590 | 2,73 | -2,26 | 1,26 | -1,80 | 7,24E-02 | 2,01E-01 |
| AT5G01980 | 1,71 | -3,02 | 1,68 | -1,80 | 7,25E-02 | 2,01E-01 |
| AT1G07613 | 1,47 | 3,06 | 1,71 | 1,79 | 7,32E-02 | 2,03E-01 |
| AT5G09870 | 2,84 | -1,81 | 1,01 | -1,79 | 7,32E-02 | 2,03E-01 |
| AT5G13770 | 2,99 | -2,65 | 1,48 | -1,79 | 7,32E-02 | 2,03E-01 |
| AT3G01316 | 2,03 | 2,89 | 1,62 | 1,79 | 7,33E-02 | 2,03E-01 |
| AT2G39681 | 1805,59 | 0,58 | 0,32 | 1,79 | 7,38E-02 | 2,04E-01 |
| AT3G62200 | 2,36 | -2,37 | 1,32 | -1,79 | 7,40E-02 | 2,05E-01 |
| AT1G33140 | 1,84 | -2,69 | 1,51 | -1,78 | 7,44E-02 | 2,05E-01 |
| AT1G29470 | 3,69 | -1,67 | 0,94 | -1,78 | 7,46E-02 | 2,06E-01 |
| AT5G26340 | 2,20 | -2,34 | 1,31 | -1,78 | 7,47E-02 | 2,06E-01 |
| AT5G03400 | 1,62 | 3,06 | 1,72 | 1,78 | 7,49E-02 | 2,06E-01 |
| AT5G04245 | 1,57 | 3,85 | 2,16 | 1,78 | 7,50E-02 | 2,06E-01 |
| AT3G04120 | 5,88 | -1,27 | 0,71 | -1,78 | 7,52E-02 | 2,06E-01 |
| AT5G63840 | 2,51 | -2,03 | 1,14 | -1,78 | 7,53E-02 | 2,07E-01 |
| AT1G61690 | 1,91 | -2,91 | 1,64 | -1,78 | 7,53E-02 | 2,07E-01 |
| AT4G16900 | 2,10 | -2,07 | 1,16 | -1,78 | 7,55E-02 | 2,07E-01 |
| AT3G55200 | 2,62 | -2,87 | 1,62 | -1,78 | 7,58E-02 | 2,07E-01 |
| AT5G64030 | 1,58 | -2,73 | 1,54 | -1,78 | 7,59E-02 | 2,07E-01 |
| AT4G03380 | 7,25 | 1,42 | 0,80 | 1,77 | 7,61E-02 | 2,08E-01 |
| AT3G11670 | 1,77 | -2,90 | 1,64 | -1,77 | 7,62E-02 | 2,08E-01 |
| AT2G46685 | 3232,16 | -0,47 | 0,26 | -1,77 | 7,68E-02 | 2,09E-01 |
| AT3G14930 | 2,04 | -2,14 | 1,21 | -1,77 | 7,68E-02 | 2,09E-01 |
| AT1G05917 | 8,49 | 1,44 | 0,81 | 1,77 | 7,71E-02 | 2,10E-01 |
| AT2G29925 | 2,27 | 3,50 | 1,98 | 1,77 | 7,71E-02 | 2,10E-01 |
| AT2G16485 | 2,25 | -2,53 | 1,43 | -1,77 | 7,72E-02 | 2,10E-01 |
| AT5G15850 | 2,49 | -2,16 | 1,23 | -1,77 | 7,72E-02 | 2,10E-01 |
| AT1G60073 | 1,58 | 3,31 | 1,87 | 1,77 | 7,75E-02 | 2,10E-01 |

| | | | | | | |
|---|---|---|---|---|---|---|
| AT2G39940 | 1,84 | -3,20 | 1,81 | -1,77 | 7,76E-02 | 2,10E-01 |
| AT4G02660 | 2,17 | -2,68 | 1,52 | -1,77 | 7,75E-02 | 2,10E-01 |
| AT1G16260 | 1,63 | -2,73 | 1,55 | -1,76 | 7,83E-02 | 2,12E-01 |
| AT5G28040 | 2,94 | -1,91 | 1,08 | -1,76 | 7,83E-02 | 2,12E-01 |
| AT5G05010 | 2,08 | -2,62 | 1,49 | -1,76 | 7,85E-02 | 2,12E-01 |
| AT4G29190 | 1,79 | -2,53 | 1,44 | -1,76 | 7,88E-02 | 2,12E-01 |
| AT5G64260 | 2,41 | 2,76 | 1,57 | 1,76 | 7,92E-02 | 2,13E-01 |
| AT4G38760 | 3,63 | 2,01 | 1,15 | 1,75 | 7,93E-02 | 2,14E-01 |
| AT1G71410 | 1,45 | -3,10 | 1,77 | -1,75 | 7,94E-02 | 2,14E-01 |
| AT5G20610 | 1,39 | -3,03 | 1,73 | -1,75 | 7,96E-02 | 2,14E-01 |
| AT3G54280 | 2,37 | -2,50 | 1,43 | -1,75 | 7,97E-02 | 2,14E-01 |
| AT5G02810 | 1,73 | -2,86 | 1,63 | -1,75 | 8,00E-02 | 2,15E-01 |
| AT3G44630 | 4,39 | -1,58 | 0,91 | -1,75 | 8,09E-02 | 2,17E-01 |
| AT1G67280 | 1,72 | -2,73 | 1,56 | -1,75 | 8,09E-02 | 2,17E-01 |
| AT4G03260 | 2,29 | -2,24 | 1,28 | -1,74 | 8,12E-02 | 2,17E-01 |
| AT4G11570 | 2,16 | -2,47 | 1,41 | -1,74 | 8,13E-02 | 2,17E-01 |
| AT4G19110 | 3,30 | -2,32 | 1,33 | -1,74 | 8,15E-02 | 2,18E-01 |
| AT1G12240 | 2,62 | -2,55 | 1,47 | -1,74 | 8,18E-02 | 2,18E-01 |
| AT5G36080 | 1,49 | 3,20 | 1,84 | 1,74 | 8,19E-02 | 2,18E-01 |
| AT1G23190 | 1,73 | -2,88 | 1,66 | -1,74 | 8,20E-02 | 2,19E-01 |
| AT2G47650 | 1,46 | -2,93 | 1,69 | -1,74 | 8,26E-02 | 2,20E-01 |
| AT3G51800 | 4,28 | 1,99 | 1,15 | 1,74 | 8,26E-02 | 2,20E-01 |
| AT1G35710 | 7,53 | -1,65 | 0,95 | -1,73 | 8,29E-02 | 2,20E-01 |
| AT1G31910 | 2,79 | -1,86 | 1,07 | -1,73 | 8,33E-02 | 2,21E-01 |
| AT4G35940 | 1,33 | -2,68 | 1,55 | -1,73 | 8,32E-02 | 2,21E-01 |
| AT4G04805 | 1,38 | 3,79 | 2,19 | 1,73 | 8,35E-02 | 2,21E-01 |
| AT5G63570 | 1,81 | -2,95 | 1,70 | -1,73 | 8,36E-02 | 2,21E-01 |
| AT1G44100 | 2,02 | 2,76 | 1,60 | 1,73 | 8,40E-02 | 2,22E-01 |
| AT5G02150 | 1,74 | -3,28 | 1,90 | -1,73 | 8,39E-02 | 2,22E-01 |
| AT5G60450 | 7,78 | 1,26 | 0,73 | 1,73 | 8,41E-02 | 2,22E-01 |
| AT5G60170 | 1,68 | -3,21 | 1,86 | -1,73 | 8,42E-02 | 2,22E-01 |
| AT3G13525 | 2,38 | 2,55 | 1,48 | 1,72 | 8,46E-02 | 2,23E-01 |
| AT5G58550 | 1,86 | -3,40 | 1,97 | -1,72 | 8,46E-02 | 2,23E-01 |
| AT3G49910 | 2,91 | -1,88 | 1,09 | -1,72 | 8,50E-02 | 2,24E-01 |
| AT3G55980 | 3,63 | 1,82 | 1,06 | 1,72 | 8,53E-02 | 2,24E-01 |
| AT4G25100 | 2,11 | 2,86 | 1,66 | 1,72 | 8,53E-02 | 2,24E-01 |
| AT1G63630 | 3,43 | 1,94 | 1,13 | 1,72 | 8,59E-02 | 2,24E-01 |
| AT1G77740 | 1,74 | -2,94 | 1,71 | -1,72 | 8,55E-02 | 2,24E-01 |
| AT1G63615 | 3,43 | 1,94 | 1,13 | 1,72 | 8,59E-02 | 2,24E-01 |
| AT3G26000 | 2,02 | -2,28 | 1,33 | -1,72 | 8,58E-02 | 2,24E-01 |
| AT3G59765 | 1,81 | -2,77 | 1,61 | -1,72 | 8,58E-02 | 2,24E-01 |
| AT5G17760 | 1,99 | -2,99 | 1,74 | -1,72 | 8,56E-02 | 2,24E-01 |
| AT5G09040 | 2,31 | 2,42 | 1,41 | 1,72 | 8,56E-02 | 2,24E-01 |
| AT5G47490 | 2,21 | -2,12 | 1,23 | -1,72 | 8,61E-02 | 2,24E-01 |
| AT3G47890 | 1,54 | -2,80 | 1,63 | -1,72 | 8,60E-02 | 2,24E-01 |
| AT4G00450 | 3,39 | -1,96 | 1,15 | -1,72 | 8,63E-02 | 2,25E-01 |
| AT2G02560 | 1,80 | -2,34 | 1,36 | -1,71 | 8,66E-02 | 2,25E-01 |
| AT3G07160 | 3,87 | -1,84 | 1,07 | -1,71 | 8,67E-02 | 2,25E-01 |
| AT3G42050 | 1,44 | -3,09 | 1,80 | -1,71 | 8,68E-02 | 2,26E-01 |
| AT5G28823 | 1,86 | 3,24 | 1,90 | 1,71 | 8,75E-02 | 2,27E-01 |
| AT5G22630 | 1,52 | -2,57 | 1,51 | -1,71 | 8,78E-02 | 2,28E-01 |
| AT4G29410 | 17,21 | 0,96 | 0,57 | 1,71 | 8,78E-02 | 2,28E-01 |
| AT1G08450 | 3,29 | -2,16 | 1,27 | -1,70 | 8,83E-02 | 2,29E-01 |
| AT1G19715 | 1,55 | -3,57 | 2,09 | -1,70 | 8,84E-02 | 2,29E-01 |
| AT5G42750 | 1,69 | 2,77 | 1,63 | 1,70 | 8,86E-02 | 2,29E-01 |
| AT5G66200 | 3,05 | -2,02 | 1,18 | -1,70 | 8,86E-02 | 2,29E-01 |

| | | | | | | |
|---|---|---|---|---|---|---|
| AT5G05190 | 2,00 | -2,51 | 1,47 | -1,70 | 8,88E-02 | 2,29E-01 |
| AT1G05805 | 1,94 | 3,80 | 2,23 | 1,70 | 8,90E-02 | 2,30E-01 |
| AT2G03855 | 12,69 | 1,11 | 0,65 | 1,70 | 8,96E-02 | 2,30E-01 |
| AT3G14010 | 2,74 | -2,36 | 1,39 | -1,70 | 8,95E-02 | 2,30E-01 |
| AT4G10120 | 1,86 | -2,24 | 1,32 | -1,70 | 8,95E-02 | 2,30E-01 |
| AT4G34110 | 4,15 | -1,42 | 0,84 | -1,70 | 8,95E-02 | 2,30E-01 |
| AT5G01400 | 1,70 | -3,24 | 1,91 | -1,70 | 9,00E-02 | 2,31E-01 |
| AT5G08925 | 2,86 | 2,82 | 1,67 | 1,69 | 9,04E-02 | 2,32E-01 |
| AT3G20820 | 3,82 | 2,10 | 1,24 | 1,69 | 9,06E-02 | 2,32E-01 |
| AT1G01100 | 1,47 | -2,46 | 1,45 | -1,69 | 9,07E-02 | 2,32E-01 |
| AT1G51172 | 1,67 | 2,88 | 1,71 | 1,69 | 9,11E-02 | 2,33E-01 |
| AT2G27150 | 1,72 | -3,41 | 2,02 | -1,69 | 9,11E-02 | 2,33E-01 |
| AT5G36200 | 1,52 | 3,20 | 1,89 | 1,69 | 9,11E-02 | 2,33E-01 |
| AT3G60600 | 1,58 | -2,93 | 1,73 | -1,69 | 9,13E-02 | 2,33E-01 |
| AT5G21940 | 4,72 | 1,42 | 0,84 | 1,69 | 9,14E-02 | 2,33E-01 |
| AT4G07120 | 22,20 | -0,85 | 0,51 | -1,68 | 9,20E-02 | 2,34E-01 |
| AT3G49590 | 1,44 | -2,58 | 1,53 | -1,68 | 9,21E-02 | 2,34E-01 |
| AT4G00720 | 2,07 | -2,47 | 1,47 | -1,68 | 9,25E-02 | 2,35E-01 |
| AT1G05913 | 9,06 | 1,37 | 0,82 | 1,68 | 9,27E-02 | 2,35E-01 |
| AT3G61198 | 7,65 | 1,29 | 0,77 | 1,68 | 9,27E-02 | 2,35E-01 |
| AT1G22850 | 1,38 | -3,40 | 2,03 | -1,68 | 9,34E-02 | 2,37E-01 |
| AT1G32750 | 1,94 | -2,59 | 1,55 | -1,68 | 9,36E-02 | 2,37E-01 |
| AT3G04930 | 1,93 | -2,48 | 1,48 | -1,67 | 9,40E-02 | 2,38E-01 |
| AT4G04205 | 2,87 | 1,93 | 1,15 | 1,67 | 9,41E-02 | 2,38E-01 |
| AT3G02170 | 3,18 | -2,60 | 1,55 | -1,67 | 9,41E-02 | 2,38E-01 |
| AT4G18670 | 7,64 | 1,38 | 0,82 | 1,68 | 9,39E-02 | 2,38E-01 |
| AT5G43310 | 2,37 | -2,39 | 1,43 | -1,67 | 9,41E-02 | 2,38E-01 |
| AT3G12810 | 1,56 | -2,66 | 1,59 | -1,67 | 9,44E-02 | 2,38E-01 |
| AT3G51770 | 2,18 | -2,52 | 1,51 | -1,67 | 9,45E-02 | 2,38E-01 |
| AT1G70940 | 2,12 | -2,11 | 1,26 | -1,67 | 9,46E-02 | 2,38E-01 |
| AT5G10540 | 2,27 | -2,15 | 1,28 | -1,67 | 9,48E-02 | 2,38E-01 |
| AT4G08870 | 1,44 | -3,06 | 1,83 | -1,67 | 9,48E-02 | 2,39E-01 |
| AT3G51370 | 1,46 | -2,85 | 1,71 | -1,67 | 9,51E-02 | 2,39E-01 |
| AT3G22125 | 1,40 | 3,01 | 1,81 | 1,66 | 9,60E-02 | 2,40E-01 |
| AT4G06225 | 6,57 | 2,03 | 1,22 | 1,67 | 9,59E-02 | 2,40E-01 |
| AT1G06700 | 1,69 | -2,64 | 1,59 | -1,66 | 9,60E-02 | 2,40E-01 |
| AT3G22142 | 1,40 | 3,01 | 1,81 | 1,66 | 9,60E-02 | 2,40E-01 |
| AT2G30550 | 1,44 | -3,04 | 1,83 | -1,66 | 9,65E-02 | 2,41E-01 |
| AT5G55565 | 4,30 | 2,17 | 1,31 | 1,66 | 9,65E-02 | 2,41E-01 |
| AT1G16270 | 1,59 | -2,51 | 1,51 | -1,66 | 9,66E-02 | 2,41E-01 |
| AT4G30440 | 3,67 | 2,02 | 1,21 | 1,66 | 9,67E-02 | 2,41E-01 |
| AT3G08850 | 2,35 | -2,40 | 1,44 | -1,66 | 9,68E-02 | 2,41E-01 |
| AT5G51660 | 1,44 | -2,66 | 1,60 | -1,66 | 9,72E-02 | 2,42E-01 |
| AT1G18879 | 1,60 | 2,87 | 1,73 | 1,66 | 9,73E-02 | 2,42E-01 |
| AT2G07629 | 5,41 | -1,41 | 0,85 | -1,66 | 9,74E-02 | 2,42E-01 |
| AT4G18596 | 1,78 | -3,13 | 1,89 | -1,66 | 9,77E-02 | 2,43E-01 |
| AT4G05050 | 3,49 | 1,85 | 1,12 | 1,65 | 9,83E-02 | 2,44E-01 |
| AT1G47530 | 1,40 | -2,63 | 1,59 | -1,65 | 9,84E-02 | 2,44E-01 |
| AT1G78206 | 14,29 | -0,83 | 0,50 | -1,65 | 9,85E-02 | 2,44E-01 |
| AT3G07515 | 2,88 | 2,64 | 1,60 | 1,65 | 9,86E-02 | 2,44E-01 |
| AT2G01600 | 1,95 | -2,69 | 1,63 | -1,65 | 9,90E-02 | 2,45E-01 |
| AT3G28540 | 2,73 | -1,95 | 1,18 | -1,65 | 9,91E-02 | 2,45E-01 |
| AT1G09210 | 5,83 | -1,27 | 0,77 | -1,65 | 9,91E-02 | 2,45E-01 |
| AT2G27400 | 624,49 | 0,55 | 0,33 | 1,65 | 9,95E-02 | 2,46E-01 |
| AT1G08347 | 16,30 | 0,82 | 0,50 | 1,65 | 9,95E-02 | 2,46E-01 |
| AT1G06283 | 1,64 | 2,64 | 1,61 | 1,64 | 1,00E-01 | 2,47E-01 |

| | | | | | | |
|---|---|---|---|---|---|---|
| AT1G16880 | 2,91 | -2,15 | 1,31 | -1,64 | 1,01E-01 | 2,47E-01 |
| AT2G45685 | 1,79 | -2,93 | 1,79 | -1,64 | 1,01E-01 | 2,47E-01 |
| AT2G45680 | 1,79 | -2,93 | 1,79 | -1,64 | 1,01E-01 | 2,47E-01 |
| AT3G26500 | 1,32 | 3,15 | 1,92 | 1,64 | 1,01E-01 | 2,48E-01 |
| AT5G25980 | 1,32 | 2,95 | 1,80 | 1,64 | 1,02E-01 | 2,49E-01 |
| AT1G50010 | 2,93 | -1,72 | 1,05 | -1,63 | 1,02E-01 | 2,50E-01 |
| AT1G13210 | 5,46 | -1,48 | 0,90 | -1,63 | 1,02E-01 | 2,50E-01 |
| AT5G40890 | 1,74 | -2,30 | 1,41 | -1,64 | 1,02E-01 | 2,50E-01 |
| AT5G56900 | 1,69 | -2,47 | 1,51 | -1,63 | 1,02E-01 | 2,50E-01 |
| AT2G16500 | 6,48 | -1,22 | 0,75 | -1,63 | 1,02E-01 | 2,50E-01 |
| AT5G66560 | 1,38 | -2,61 | 1,60 | -1,63 | 1,03E-01 | 2,50E-01 |
| AT5G66558 | 1,38 | -2,61 | 1,60 | -1,63 | 1,03E-01 | 2,50E-01 |
| AT2G23460 | 1,50 | -2,60 | 1,59 | -1,63 | 1,03E-01 | 2,51E-01 |
| AT4G06240 | 6,18 | 2,03 | 1,24 | 1,63 | 1,03E-01 | 2,51E-01 |
| AT3G14240 | 1,34 | -2,67 | 1,64 | -1,63 | 1,03E-01 | 2,51E-01 |
| AT3G53990 | 1,83 | -2,35 | 1,44 | -1,63 | 1,03E-01 | 2,51E-01 |
| AT1G60870 | 2,59 | -2,37 | 1,46 | -1,63 | 1,04E-01 | 2,51E-01 |
| AT3G26430 | 1,60 | -2,31 | 1,42 | -1,63 | 1,04E-01 | 2,51E-01 |
| AT3G26420 | 1,60 | -2,31 | 1,42 | -1,63 | 1,04E-01 | 2,51E-01 |
| AT4G31160 | 2,16 | -2,61 | 1,60 | -1,63 | 1,04E-01 | 2,51E-01 |
| AT3G00980 | 2,10 | 3,20 | 1,96 | 1,63 | 1,04E-01 | 2,51E-01 |
| AT5G04550 | 2,33 | -2,10 | 1,29 | -1,63 | 1,04E-01 | 2,51E-01 |
| AT5G19770 | 1,75 | -2,64 | 1,62 | -1,63 | 1,04E-01 | 2,51E-01 |
| AT1G09070 | 2,87 | 2,34 | 1,44 | 1,62 | 1,04E-01 | 2,52E-01 |
| AT1G60960 | 1,57 | -2,56 | 1,57 | -1,62 | 1,04E-01 | 2,52E-01 |
| AT3G44444 | 1,95 | 2,27 | 1,39 | 1,63 | 1,04E-01 | 2,52E-01 |
| AT4G34071 | 2,35 | 2,63 | 1,62 | 1,62 | 1,04E-01 | 2,52E-01 |
| AT5G05200 | 1,99 | -3,12 | 1,92 | -1,62 | 1,05E-01 | 2,52E-01 |
| AT4G10840 | 1,51 | -3,04 | 1,88 | -1,62 | 1,05E-01 | 2,53E-01 |
| AT5G01750 | 1,69 | 3,13 | 1,93 | 1,62 | 1,06E-01 | 2,54E-01 |
| AT5G02310 | 4,95 | -1,28 | 0,79 | -1,62 | 1,06E-01 | 2,54E-01 |
| AT1G64230 | 1,39 | -3,02 | 1,87 | -1,61 | 1,06E-01 | 2,55E-01 |
| AT1G67450 | 1,86 | 2,52 | 1,56 | 1,61 | 1,06E-01 | 2,55E-01 |
| AT3G02470 | 10,40 | -0,98 | 0,61 | -1,62 | 1,06E-01 | 2,55E-01 |
| AT4G26730 | 2,38 | 2,80 | 1,74 | 1,61 | 1,07E-01 | 2,55E-01 |
| AT3G52500 | 2,36 | 2,16 | 1,34 | 1,61 | 1,06E-01 | 2,55E-01 |
| AT1G01090 | 1,61 | -2,29 | 1,42 | -1,61 | 1,07E-01 | 2,55E-01 |
| AT3G05220 | 1,84 | 2,80 | 1,74 | 1,61 | 1,07E-01 | 2,56E-01 |
| AT3G55830 | 1,72 | -2,51 | 1,56 | -1,61 | 1,07E-01 | 2,56E-01 |
| AT3G02045 | 5,72 | 1,79 | 1,11 | 1,61 | 1,07E-01 | 2,56E-01 |
| AT2G03140 | 2,19 | -2,18 | 1,35 | -1,61 | 1,08E-01 | 2,56E-01 |
| AT3G51550 | 4,70 | -1,28 | 0,80 | -1,61 | 1,07E-01 | 2,56E-01 |
| AT5G51300 | 1,97 | -3,41 | 2,12 | -1,61 | 1,08E-01 | 2,56E-01 |
| AT2G31820 | 4,59 | 1,46 | 0,91 | 1,61 | 1,08E-01 | 2,57E-01 |
| AT1G05630 | 1,84 | -2,50 | 1,56 | -1,60 | 1,09E-01 | 2,58E-01 |
| AT1G60720 | 1,62 | 3,04 | 1,90 | 1,60 | 1,09E-01 | 2,58E-01 |
| AT3G03790 | 1,73 | -2,47 | 1,54 | -1,60 | 1,09E-01 | 2,58E-01 |
| AT5G20950 | 3,45 | -1,75 | 1,09 | -1,60 | 1,09E-01 | 2,58E-01 |
| AT5G55540 | 1,63 | -2,69 | 1,68 | -1,60 | 1,09E-01 | 2,58E-01 |
| AT5G55910 | 1,51 | -2,46 | 1,53 | -1,61 | 1,08E-01 | 2,58E-01 |
| AT2G17840 | 2,90 | 2,51 | 1,56 | 1,60 | 1,09E-01 | 2,58E-01 |
| AT3G41979 | 6596,11 | -0,57 | 0,36 | -1,60 | 1,09E-01 | 2,58E-01 |
| AT3G07860 | 5,76 | 1,78 | 1,11 | 1,60 | 1,10E-01 | 2,59E-01 |
| AT4G04570 | 4,35 | -1,92 | 1,20 | -1,60 | 1,10E-01 | 2,59E-01 |
| AT2G28360 | 1,42 | -2,09 | 1,31 | -1,60 | 1,10E-01 | 2,59E-01 |
| AT3G07855 | 4,66 | 1,72 | 1,08 | 1,60 | 1,11E-01 | 2,60E-01 |

| | | | | | | |
|---|---|---|---|---|---|---|
| AT3G10380 | 2,66 | -1,67 | 1,05 | -1,60 | 1,10E-01 | 2,60E-01 |
| AT1G05423 | 13,10 | 0,99 | 0,62 | 1,59 | 1,11E-01 | 2,61E-01 |
| AT4G19410 | 1,78 | -2,17 | 1,36 | -1,59 | 1,11E-01 | 2,61E-01 |
| AT1G62660 | 1,32 | -2,86 | 1,80 | -1,59 | 1,11E-01 | 2,62E-01 |
| AT2G44710 | 1,40 | -2,45 | 1,54 | -1,59 | 1,11E-01 | 2,62E-01 |
| AT5G47010 | 2,34 | -1,95 | 1,22 | -1,59 | 1,11E-01 | 2,62E-01 |
| AT2G41920 | 1,42 | 3,00 | 1,89 | 1,59 | 1,12E-01 | 2,62E-01 |
| AT1G21580 | 3,07 | -1,73 | 1,09 | -1,59 | 1,12E-01 | 2,63E-01 |
| AT3G25460 | 1,48 | 2,82 | 1,78 | 1,59 | 1,12E-01 | 2,63E-01 |
| AT1G16710 | 1,85 | -2,75 | 1,73 | -1,59 | 1,13E-01 | 2,63E-01 |
| AT3G01780 | 1,57 | -2,29 | 1,44 | -1,59 | 1,13E-01 | 2,63E-01 |
| AT1G48450 | 1,45 | -2,75 | 1,73 | -1,59 | 1,13E-01 | 2,63E-01 |
| AT1G74960 | 1,51 | -4,19 | 2,64 | -1,59 | 1,13E-01 | 2,63E-01 |
| AT5G09670 | 2,53 | -1,94 | 1,22 | -1,59 | 1,13E-01 | 2,63E-01 |
| AT5G20010 | 1,98 | -2,32 | 1,46 | -1,58 | 1,13E-01 | 2,64E-01 |
| AT3G61430 | 1,94 | -2,90 | 1,84 | -1,58 | 1,14E-01 | 2,64E-01 |
| AT4G14548 | 1,46 | 2,90 | 1,83 | 1,58 | 1,14E-01 | 2,64E-01 |
| AT5G49760 | 3,34 | -1,57 | 0,99 | -1,58 | 1,14E-01 | 2,64E-01 |
| AT3G20690 | 3,62 | 2,18 | 1,38 | 1,58 | 1,14E-01 | 2,64E-01 |
| AT3G08210 | 3,75 | 1,86 | 1,18 | 1,58 | 1,14E-01 | 2,64E-01 |
| AT1G15690 | 11,49 | -1,04 | 0,66 | -1,58 | 1,15E-01 | 2,66E-01 |
| AT3G06355 | 6711,54 | -0,56 | 0,36 | -1,58 | 1,15E-01 | 2,66E-01 |
| AT1G43145 | 1,44 | 2,75 | 1,74 | 1,57 | 1,15E-01 | 2,67E-01 |
| AT4G16490 | 2,72 | -2,62 | 1,67 | -1,57 | 1,15E-01 | 2,67E-01 |
| AT4G27060 | 1,71 | -2,40 | 1,52 | -1,57 | 1,16E-01 | 2,67E-01 |
| AT5G07980 | 1,38 | -2,93 | 1,86 | -1,57 | 1,15E-01 | 2,67E-01 |
| AT5G16370 | 1,57 | -2,57 | 1,64 | -1,57 | 1,15E-01 | 2,67E-01 |
| AT1G07320 | 1,43 | -2,36 | 1,50 | -1,57 | 1,16E-01 | 2,68E-01 |
| AT1G33420 | 2,44 | 1,99 | 1,27 | 1,57 | 1,16E-01 | 2,68E-01 |
| AT1G72645 | 2,19 | 2,20 | 1,40 | 1,57 | 1,16E-01 | 2,68E-01 |
| AT5G54440 | 2,06 | -2,11 | 1,34 | -1,57 | 1,16E-01 | 2,68E-01 |
| AT2G44410 | 2,51 | -2,75 | 1,76 | -1,57 | 1,17E-01 | 2,68E-01 |
| AT2G46620 | 1,45 | -2,46 | 1,57 | -1,57 | 1,17E-01 | 2,68E-01 |
| AT4G33890 | 1,49 | -2,75 | 1,76 | -1,57 | 1,17E-01 | 2,69E-01 |
| AT5G18245 | 1,46 | 4,22 | 2,69 | 1,57 | 1,17E-01 | 2,69E-01 |
| AT2G44550 | 3,60 | 1,82 | 1,16 | 1,57 | 1,17E-01 | 2,69E-01 |
| AT3G07770 | 1,44 | -2,34 | 1,50 | -1,56 | 1,18E-01 | 2,71E-01 |
| AT1G63880 | 4,32 | 1,46 | 0,94 | 1,56 | 1,18E-01 | 2,71E-01 |
| AT5G12110 | 1,52 | -2,39 | 1,53 | -1,56 | 1,19E-01 | 2,71E-01 |
| AT1G75040 | 3,07 | -1,54 | 0,99 | -1,56 | 1,19E-01 | 2,72E-01 |
| AT1G60070 | 1,35 | -3,26 | 2,09 | -1,56 | 1,19E-01 | 2,72E-01 |
| AT4G13390 | 1,53 | 2,92 | 1,87 | 1,56 | 1,19E-01 | 2,73E-01 |
| AT2G01910 | 2,66 | 2,50 | 1,61 | 1,56 | 1,20E-01 | 2,74E-01 |
| AT2G07915 | 1,63 | 3,58 | 2,30 | 1,55 | 1,20E-01 | 2,74E-01 |
| AT2G25900 | 2,31 | 2,42 | 1,56 | 1,55 | 1,20E-01 | 2,74E-01 |
| AT2G07910 | 1,63 | 3,58 | 2,30 | 1,55 | 1,20E-01 | 2,74E-01 |
| AT3G01120 | 2,55 | -2,29 | 1,48 | -1,55 | 1,20E-01 | 2,74E-01 |
| AT3G00985 | 2,03 | 3,11 | 2,00 | 1,55 | 1,20E-01 | 2,74E-01 |
| AT4G08470 | 1,82 | -3,26 | 2,10 | -1,55 | 1,20E-01 | 2,74E-01 |
| AT1G58230 | 3,69 | -1,80 | 1,16 | -1,55 | 1,21E-01 | 2,75E-01 |
| AT5G19510 | 1,83 | -2,19 | 1,41 | -1,55 | 1,21E-01 | 2,75E-01 |
| AT5G43900 | 1,55 | -2,52 | 1,63 | -1,55 | 1,21E-01 | 2,75E-01 |
| AT5G22505 | 1,58 | 2,86 | 1,85 | 1,55 | 1,22E-01 | 2,76E-01 |
| AT4G12570 | 2,25 | -2,25 | 1,46 | -1,55 | 1,22E-01 | 2,76E-01 |
| AT4G18030 | 2,74 | -2,02 | 1,31 | -1,55 | 1,22E-01 | 2,76E-01 |
| AT5G01775 | 4,59 | 1,59 | 1,03 | 1,54 | 1,23E-01 | 2,77E-01 |

| | | | | | | |
|---|---|---|---|---|---|---|
| AT4G40040 | 3,01 | 2,32 | 1,50 | 1,54 | 1,23E-01 | 2,77E-01 |
| AT1G06150 | 1,38 | -2,68 | 1,74 | -1,54 | 1,23E-01 | 2,77E-01 |
| AT1G77180 | 2,45 | -1,98 | 1,28 | -1,54 | 1,23E-01 | 2,77E-01 |
| AT4G12420 | 1,53 | -2,65 | 1,72 | -1,54 | 1,23E-01 | 2,78E-01 |
| AT5G35490 | 1,40 | 3,47 | 2,25 | 1,54 | 1,24E-01 | 2,79E-01 |
| AT2G22430 | 4,62 | -1,32 | 0,86 | -1,54 | 1,24E-01 | 2,79E-01 |
| AT1G08720 | 1,46 | -3,28 | 2,14 | -1,54 | 1,24E-01 | 2,80E-01 |
| AT4G04920 | 1,70 | -2,53 | 1,65 | -1,54 | 1,25E-01 | 2,80E-01 |
| AT4G19170 | 5,23 | 1,33 | 0,87 | 1,54 | 1,25E-01 | 2,80E-01 |
| AT3G63310 | 1,42 | -2,97 | 1,94 | -1,53 | 1,25E-01 | 2,81E-01 |
| AT4G36150 | 1,48 | -4,00 | 2,61 | -1,53 | 1,25E-01 | 2,81E-01 |
| AT5G42270 | 4,20 | 1,95 | 1,27 | 1,53 | 1,26E-01 | 2,81E-01 |
| AT2G21660 | 24,65 | -0,74 | 0,48 | -1,53 | 1,26E-01 | 2,81E-01 |
| AT5G11170 | 1,42 | -3,00 | 1,96 | -1,53 | 1,26E-01 | 2,81E-01 |
| AT2G01570 | 3,41 | -1,40 | 0,92 | -1,53 | 1,26E-01 | 2,81E-01 |
| AT5G01560 | 1,57 | -2,53 | 1,66 | -1,53 | 1,26E-01 | 2,81E-01 |
| AT5G17230 | 2,13 | -1,92 | 1,25 | -1,53 | 1,26E-01 | 2,81E-01 |
| AT5G01915 | 2,27 | 2,49 | 1,64 | 1,52 | 1,28E-01 | 2,86E-01 |
| AT2G39175 | 43,76 | 0,57 | 0,38 | 1,52 | 1,28E-01 | 2,86E-01 |
| AT5G15200 | 4,33 | -1,27 | 0,83 | -1,52 | 1,28E-01 | 2,86E-01 |
| AT2G23550 | 1,59 | 3,51 | 2,32 | 1,52 | 1,29E-01 | 2,88E-01 |
| AT5G18240 | 1,40 | 4,15 | 2,74 | 1,52 | 1,30E-01 | 2,88E-01 |
| AT3G62010 | 2,16 | -1,93 | 1,28 | -1,52 | 1,30E-01 | 2,88E-01 |
| AT5G02295 | 1,40 | 2,53 | 1,67 | 1,51 | 1,30E-01 | 2,89E-01 |
| AT5G41950 | 1,46 | -3,01 | 1,99 | -1,51 | 1,30E-01 | 2,89E-01 |
| AT1G71220 | 2,96 | -1,90 | 1,25 | -1,51 | 1,30E-01 | 2,89E-01 |
| AT2G06535 | 1,38 | 2,95 | 1,95 | 1,51 | 1,30E-01 | 2,89E-01 |
| AT5G40384 | 2,72 | -1,87 | 1,24 | -1,51 | 1,31E-01 | 2,90E-01 |
| AT2G38195 | 4,06 | 2,09 | 1,39 | 1,51 | 1,31E-01 | 2,90E-01 |
| AT1G54115 | 2,18 | -2,29 | 1,52 | -1,51 | 1,32E-01 | 2,92E-01 |
| AT3G15210 | 1,70 | 2,38 | 1,58 | 1,50 | 1,32E-01 | 2,92E-01 |
| AT2G35744 | 14,81 | 0,85 | 0,56 | 1,50 | 1,32E-01 | 2,92E-01 |
| AT4G31990 | 1,81 | -2,53 | 1,68 | -1,50 | 1,32E-01 | 2,92E-01 |
| AT2G20570 | 7,71 | -1,04 | 0,69 | -1,50 | 1,33E-01 | 2,93E-01 |
| AT3G19180 | 1,86 | -3,00 | 2,00 | -1,50 | 1,33E-01 | 2,94E-01 |
| AT2G31400 | 2,56 | -1,90 | 1,27 | -1,50 | 1,34E-01 | 2,94E-01 |
| AT2G46430 | 2,08 | -2,03 | 1,36 | -1,50 | 1,34E-01 | 2,94E-01 |
| AT1G20010 | 2,01 | -1,77 | 1,18 | -1,50 | 1,34E-01 | 2,94E-01 |
| AT4G06705 | 4,08 | 1,60 | 1,07 | 1,50 | 1,34E-01 | 2,94E-01 |
| AT1G35820 | 1,40 | 4,16 | 2,78 | 1,50 | 1,34E-01 | 2,95E-01 |
| AT1G26850 | 2,19 | -2,10 | 1,40 | -1,50 | 1,34E-01 | 2,95E-01 |
| AT1G06667 | 5,78 | 1,43 | 0,95 | 1,50 | 1,35E-01 | 2,95E-01 |
| AT5G49810 | 1,47 | -3,10 | 2,08 | -1,50 | 1,35E-01 | 2,95E-01 |
| AT2G22660 | 2,02 | -2,20 | 1,47 | -1,49 | 1,35E-01 | 2,95E-01 |
| AT5G47300 | 1,47 | 2,66 | 1,78 | 1,49 | 1,35E-01 | 2,95E-01 |
| AT1G07243 | 1,82 | 2,42 | 1,62 | 1,49 | 1,36E-01 | 2,97E-01 |
| AT5G26570 | 1,51 | -2,70 | 1,81 | -1,49 | 1,36E-01 | 2,97E-01 |
| AT5G13390 | 2,05 | -2,24 | 1,50 | -1,49 | 1,36E-01 | 2,97E-01 |
| AT5G30510 | 1,83 | 2,68 | 1,80 | 1,49 | 1,37E-01 | 2,99E-01 |
| AT2G26780 | 3,27 | -1,77 | 1,19 | -1,48 | 1,38E-01 | 2,99E-01 |
| AT3G52380 | 1,54 | -2,90 | 1,95 | -1,48 | 1,38E-01 | 2,99E-01 |
| AT5G08717 | 14,65 | 1,02 | 0,69 | 1,49 | 1,37E-01 | 2,99E-01 |
| AT1G45160 | 1,73 | -2,61 | 1,76 | -1,48 | 1,38E-01 | 3,00E-01 |
| AT1G35720 | 3,50 | -1,41 | 0,95 | -1,48 | 1,38E-01 | 3,00E-01 |
| AT1G63750 | 2,64 | -1,84 | 1,24 | -1,48 | 1,38E-01 | 3,00E-01 |
| AT4G16190 | 11,22 | -0,91 | 0,62 | -1,48 | 1,39E-01 | 3,01E-01 |

| | | | | | | |
|---|---|---|---|---|---|---|
| AT5G59505 | 3,52 | 2,10 | 1,42 | 1,48 | 1,39E-01 | 3,01E-01 |
| AT1G07670 | 1,74 | -2,05 | 1,38 | -1,48 | 1,39E-01 | 3,01E-01 |
| AT1G76065 | 4,52 | 1,69 | 1,14 | 1,48 | 1,39E-01 | 3,01E-01 |
| AT4G02510 | 15,59 | -0,85 | 0,57 | -1,48 | 1,39E-01 | 3,01E-01 |
| AT3G25500 | 1,85 | -1,93 | 1,31 | -1,48 | 1,40E-01 | 3,02E-01 |
| AT4G14400 | 8,78 | -1,17 | 0,79 | -1,48 | 1,40E-01 | 3,03E-01 |
| AT2G08665 | 53,97 | -0,61 | 0,41 | -1,47 | 1,41E-01 | 3,04E-01 |
| AT2G13820 | 1,49 | 4,00 | 2,72 | 1,47 | 1,41E-01 | 3,04E-01 |
| AT2G34420 | 53,97 | -0,61 | 0,41 | -1,47 | 1,41E-01 | 3,04E-01 |
| AT1G43170 | 3,66 | -1,42 | 0,97 | -1,47 | 1,41E-01 | 3,05E-01 |
| AT1G79040 | 10,12 | 1,27 | 0,86 | 1,47 | 1,42E-01 | 3,05E-01 |
| AT5G52530 | 3,03 | -2,10 | 1,43 | -1,47 | 1,42E-01 | 3,05E-01 |
| AT2G20260 | 1,52 | -2,27 | 1,55 | -1,47 | 1,42E-01 | 3,06E-01 |
| AT2G24340 | 1,46 | 2,56 | 1,75 | 1,46 | 1,43E-01 | 3,07E-01 |
| AT1G56630 | 1,84 | -3,37 | 2,30 | -1,46 | 1,44E-01 | 3,09E-01 |
| AT5G10180 | 2,74 | -1,96 | 1,34 | -1,46 | 1,44E-01 | 3,09E-01 |
| AT2G36350 | 2,13 | -2,29 | 1,57 | -1,46 | 1,44E-01 | 3,09E-01 |
| AT1G69890 | 1,49 | 2,68 | 1,84 | 1,46 | 1,45E-01 | 3,10E-01 |
| AT2G26140 | 1,61 | -2,32 | 1,59 | -1,46 | 1,45E-01 | 3,10E-01 |
| AT5G60390 | 16,82 | -0,76 | 0,52 | -1,46 | 1,45E-01 | 3,10E-01 |
| AT5G46520 | 3,26 | -1,70 | 1,17 | -1,46 | 1,45E-01 | 3,10E-01 |
| AT1G04480 | 2,74 | -1,70 | 1,17 | -1,46 | 1,45E-01 | 3,10E-01 |
| AT1G63230 | 3,31 | 1,63 | 1,12 | 1,45 | 1,46E-01 | 3,12E-01 |
| AT3G07660 | 1,40 | -2,54 | 1,75 | -1,45 | 1,46E-01 | 3,12E-01 |
| AT3G01470 | 1,49 | -2,22 | 1,53 | -1,45 | 1,47E-01 | 3,12E-01 |
| AT1G73480 | 1,37 | -3,07 | 2,11 | -1,45 | 1,47E-01 | 3,12E-01 |
| AT4G33300 | 3,93 | -1,54 | 1,06 | -1,45 | 1,47E-01 | 3,12E-01 |
| AT1G74453 | 1,99 | 2,05 | 1,42 | 1,45 | 1,47E-01 | 3,12E-01 |
| AT2G22300 | 2,59 | -1,86 | 1,29 | -1,45 | 1,48E-01 | 3,13E-01 |
| AT1G19720 | 1,66 | -2,18 | 1,51 | -1,45 | 1,48E-01 | 3,13E-01 |
| AT1G53210 | 1,83 | 2,49 | 1,72 | 1,45 | 1,48E-01 | 3,15E-01 |
| AT4G25640 | 1,48 | -2,24 | 1,55 | -1,44 | 1,48E-01 | 3,15E-01 |
| AT5G06290 | 1,37 | -2,68 | 1,86 | -1,44 | 1,49E-01 | 3,15E-01 |
| AT3G28550 | 2,18 | 2,59 | 1,79 | 1,44 | 1,49E-01 | 3,15E-01 |
| AT3G13530 | 2,08 | -2,17 | 1,50 | -1,44 | 1,49E-01 | 3,15E-01 |
| AT2G29970 | 1,50 | 2,35 | 1,63 | 1,44 | 1,50E-01 | 3,16E-01 |
| AT4G16760 | 1,69 | -2,18 | 1,51 | -1,44 | 1,50E-01 | 3,16E-01 |
| AT2G05170 | 1,80 | -2,09 | 1,45 | -1,44 | 1,50E-01 | 3,17E-01 |
| AT5G28300 | 1,51 | 2,46 | 1,71 | 1,44 | 1,50E-01 | 3,17E-01 |
| AT1G35560 | 2,43 | -1,64 | 1,14 | -1,44 | 1,51E-01 | 3,17E-01 |
| AT3G52430 | 2,34 | -2,01 | 1,41 | -1,43 | 1,52E-01 | 3,21E-01 |
| AT5G16270 | 1,71 | -2,00 | 1,40 | -1,43 | 1,52E-01 | 3,21E-01 |
| AT4G06235 | 7,69 | 1,62 | 1,13 | 1,43 | 1,53E-01 | 3,21E-01 |
| AT4G13810 | 2,84 | 2,37 | 1,66 | 1,43 | 1,53E-01 | 3,21E-01 |
| AT4G10800 | 6,52 | 1,36 | 0,96 | 1,43 | 1,54E-01 | 3,23E-01 |
| AT5G47690 | 3,16 | -1,59 | 1,11 | -1,43 | 1,54E-01 | 3,23E-01 |
| AT1G02470 | 5,92 | -1,23 | 0,86 | -1,43 | 1,54E-01 | 3,23E-01 |
| AT3G18190 | 1,46 | -2,23 | 1,56 | -1,42 | 1,54E-01 | 3,24E-01 |
| AT3G59710 | 6,69 | 1,22 | 0,86 | 1,42 | 1,55E-01 | 3,24E-01 |
| AT3G09225 | 6,69 | 1,22 | 0,86 | 1,42 | 1,55E-01 | 3,24E-01 |
| AT5G03240 | 5,81 | -1,22 | 0,86 | -1,42 | 1,55E-01 | 3,24E-01 |
| AT5G08550 | 1,83 | -2,15 | 1,51 | -1,42 | 1,55E-01 | 3,24E-01 |
| AT5G58930 | 1,32 | -2,84 | 2,00 | -1,42 | 1,55E-01 | 3,24E-01 |
| AT1G76062 | 4,01 | 1,78 | 1,25 | 1,42 | 1,56E-01 | 3,25E-01 |
| AT3G57645 | 1,71 | 2,67 | 1,88 | 1,42 | 1,56E-01 | 3,26E-01 |
| AT4G00120 | 1,32 | 2,55 | 1,80 | 1,42 | 1,56E-01 | 3,26E-01 |

| | | | | | | |
|---|---|---|---|---|---|---|
| AT3G44703 | 4,72 | -1,26 | 0,89 | -1,42 | 1,57E-01 | 3,26E-01 |
| AT5G46430 | 2,53 | 2,17 | 1,53 | 1,42 | 1,57E-01 | 3,26E-01 |
| AT2G05685 | 2,44 | 2,30 | 1,63 | 1,41 | 1,57E-01 | 3,27E-01 |
| AT5G40780 | 1,36 | -2,24 | 1,58 | -1,41 | 1,58E-01 | 3,28E-01 |
| AT5G02250 | 1,33 | -3,15 | 2,23 | -1,41 | 1,58E-01 | 3,28E-01 |
| AT4G19210 | 1,66 | -2,32 | 1,65 | -1,41 | 1,58E-01 | 3,29E-01 |
| AT1G01960 | 2,74 | -1,63 | 1,15 | -1,41 | 1,59E-01 | 3,29E-01 |
| AT3G21755 | 4,36 | 1,43 | 1,01 | 1,41 | 1,58E-01 | 3,29E-01 |
| AT1G65440 | 2,14 | -1,93 | 1,37 | -1,41 | 1,59E-01 | 3,30E-01 |
| AT3G02200 | 1,75 | -2,47 | 1,75 | -1,41 | 1,60E-01 | 3,30E-01 |
| AT3G06055 | 2,49 | 1,93 | 1,37 | 1,41 | 1,60E-01 | 3,30E-01 |
| AT3G05590 | 1,96 | -1,79 | 1,27 | -1,40 | 1,61E-01 | 3,32E-01 |
| AT1G23900 | 1,85 | -2,18 | 1,56 | -1,40 | 1,61E-01 | 3,33E-01 |
| AT1G79570 | 2,57 | -1,63 | 1,17 | -1,40 | 1,61E-01 | 3,33E-01 |
| AT4G13493 | 160,77 | -0,42 | 0,30 | -1,40 | 1,61E-01 | 3,33E-01 |
| AT4G39361 | 1,80 | 2,60 | 1,86 | 1,40 | 1,61E-01 | 3,33E-01 |
| AT5G35160 | 1,68 | -1,81 | 1,29 | -1,40 | 1,62E-01 | 3,33E-01 |
| AT1G59835 | 1,47 | -2,60 | 1,86 | -1,40 | 1,62E-01 | 3,33E-01 |
| AT1G53655 | 1,78 | 2,52 | 1,81 | 1,40 | 1,62E-01 | 3,34E-01 |
| AT1G58807 | 1,55 | -2,06 | 1,48 | -1,40 | 1,62E-01 | 3,34E-01 |
| AT2G45180 | 2,18 | 2,26 | 1,62 | 1,40 | 1,63E-01 | 3,34E-01 |
| AT3G17400 | 6,94 | 1,24 | 0,89 | 1,40 | 1,63E-01 | 3,34E-01 |
| AT4G38775 | 2,66 | 2,04 | 1,46 | 1,40 | 1,63E-01 | 3,34E-01 |
| AT3G19380 | 2,79 | 1,63 | 1,17 | 1,39 | 1,64E-01 | 3,37E-01 |
| AT1G63950 | 1,81 | 2,66 | 1,91 | 1,39 | 1,65E-01 | 3,37E-01 |
| AT2G07638 | 1,46 | -2,50 | 1,80 | -1,39 | 1,65E-01 | 3,38E-01 |
| AT3G28925 | 2,17 | 2,04 | 1,47 | 1,39 | 1,65E-01 | 3,38E-01 |
| AT2G31280 | 1,47 | -2,89 | 2,08 | -1,39 | 1,66E-01 | 3,39E-01 |
| AT3G19170 | 4,99 | 1,93 | 1,40 | 1,39 | 1,66E-01 | 3,39E-01 |
| AT2G38770 | 1,53 | -2,19 | 1,58 | -1,38 | 1,66E-01 | 3,39E-01 |
| AT5G09005 | 1,51 | 2,25 | 1,62 | 1,38 | 1,66E-01 | 3,39E-01 |
| AT2G37160 | 3,77 | 1,41 | 1,02 | 1,38 | 1,67E-01 | 3,40E-01 |
| AT1G65350 | 2,42 | 1,91 | 1,38 | 1,38 | 1,67E-01 | 3,40E-01 |
| AT5G01785 | 2,53 | 2,29 | 1,66 | 1,38 | 1,68E-01 | 3,41E-01 |
| AT2G23200 | 1,64 | -2,15 | 1,56 | -1,38 | 1,68E-01 | 3,43E-01 |
| AT1G01560 | 1,60 | -2,55 | 1,85 | -1,38 | 1,69E-01 | 3,44E-01 |
| AT3G57062 | 1,47 | -2,31 | 1,68 | -1,37 | 1,69E-01 | 3,44E-01 |
| AT1G05030 | 1,78 | 2,45 | 1,78 | 1,37 | 1,69E-01 | 3,44E-01 |
| AT1G11735 | 8,53 | -0,87 | 0,64 | -1,38 | 1,69E-01 | 3,44E-01 |
| AT1G69550 | 1,87 | -2,07 | 1,51 | -1,37 | 1,69E-01 | 3,44E-01 |
| AT2G08855 | 3,73 | 1,41 | 1,03 | 1,37 | 1,70E-01 | 3,44E-01 |
| AT3G06345 | 8,24 | 0,90 | 0,66 | 1,37 | 1,70E-01 | 3,44E-01 |
| AT3G21760 | 4,32 | 1,39 | 1,01 | 1,37 | 1,70E-01 | 3,44E-01 |
| AT4G05900 | 1,70 | 2,33 | 1,70 | 1,37 | 1,71E-01 | 3,45E-01 |
| AT4G08125 | 3,11 | 1,76 | 1,28 | 1,37 | 1,71E-01 | 3,45E-01 |
| AT5G03280 | 3,28 | -1,63 | 1,19 | -1,37 | 1,71E-01 | 3,46E-01 |
| AT1G42470 | 1,71 | -2,62 | 1,91 | -1,37 | 1,71E-01 | 3,46E-01 |
| AT4G26110 | 2,20 | -1,82 | 1,33 | -1,37 | 1,72E-01 | 3,47E-01 |
| AT4G21150 | 2,92 | -1,66 | 1,22 | -1,37 | 1,72E-01 | 3,47E-01 |
| AT1G33410 | 1,37 | -2,00 | 1,46 | -1,37 | 1,72E-01 | 3,47E-01 |
| AT2G40955 | 1,38 | 2,49 | 1,82 | 1,36 | 1,73E-01 | 3,47E-01 |
| AT5G44585 | 1,38 | -2,06 | 1,51 | -1,36 | 1,72E-01 | 3,47E-01 |
| AT5G06595 | 1,38 | -2,06 | 1,51 | -1,36 | 1,72E-01 | 3,47E-01 |
| AT1G53640 | 2,14 | -1,86 | 1,36 | -1,36 | 1,73E-01 | 3,48E-01 |
| AT5G01530 | 20,05 | -0,80 | 0,59 | -1,36 | 1,74E-01 | 3,49E-01 |
| AT1G03410 | 2,63 | 1,77 | 1,30 | 1,36 | 1,74E-01 | 3,49E-01 |

| | | | | | | |
|---|---|---|---|---|---|---|
| AT1G46696 | 1,91 | 2,31 | 1,70 | 1,36 | 1,74E-01 | 3,49E-01 |
| AT1G27595 | 2,01 | -2,00 | 1,48 | -1,35 | 1,76E-01 | 3,52E-01 |
| AT1G79600 | 1,43 | -2,31 | 1,71 | -1,35 | 1,75E-01 | 3,52E-01 |
| AT3G56705 | 15,48 | 0,77 | 0,57 | 1,35 | 1,76E-01 | 3,52E-01 |
| AT3G55840 | 1,41 | -2,33 | 1,72 | -1,35 | 1,76E-01 | 3,52E-01 |
| AT1G65450 | 2,07 | -1,79 | 1,33 | -1,35 | 1,76E-01 | 3,52E-01 |
| AT2G18790 | 2,11 | -1,85 | 1,36 | -1,35 | 1,76E-01 | 3,52E-01 |
| AT3G42870 | 1,35 | 2,64 | 1,95 | 1,35 | 1,76E-01 | 3,52E-01 |
| AT3G63070 | 1,86 | -2,40 | 1,78 | -1,35 | 1,76E-01 | 3,52E-01 |
| AT3G20950 | 1,75 | 2,23 | 1,65 | 1,35 | 1,77E-01 | 3,53E-01 |
| AT3G51620 | 1,34 | -2,64 | 1,95 | -1,35 | 1,77E-01 | 3,53E-01 |
| AT3G55320 | 2,94 | -2,15 | 1,60 | -1,34 | 1,79E-01 | 3,56E-01 |
| AT2G09690 | 14,22 | 1,03 | 0,77 | 1,34 | 1,80E-01 | 3,59E-01 |
| AT1G11650 | 3,30 | -1,36 | 1,02 | -1,34 | 1,81E-01 | 3,60E-01 |
| AT3G43230 | 1,49 | -2,33 | 1,74 | -1,34 | 1,81E-01 | 3,60E-01 |
| AT1G06537 | 1,54 | -2,44 | 1,82 | -1,34 | 1,81E-01 | 3,60E-01 |
| AT2G05765 | 3,87 | 1,50 | 1,12 | 1,34 | 1,81E-01 | 3,60E-01 |
| AT4G33240 | 2,23 | -1,87 | 1,40 | -1,34 | 1,82E-01 | 3,60E-01 |
| AT5G54130 | 6,22 | -1,01 | 0,75 | -1,34 | 1,82E-01 | 3,60E-01 |
| AT1G27870 | 1,97 | 2,25 | 1,69 | 1,34 | 1,82E-01 | 3,60E-01 |
| AT5G15020 | 2,14 | -2,15 | 1,61 | -1,33 | 1,82E-01 | 3,61E-01 |
| AT4G40050 | 1,65 | -2,04 | 1,53 | -1,33 | 1,83E-01 | 3,62E-01 |
| AT2G39810 | 1,48 | -3,05 | 2,29 | -1,33 | 1,83E-01 | 3,62E-01 |
| AT1G04183 | 2,59 | 1,73 | 1,30 | 1,33 | 1,83E-01 | 3,62E-01 |
| AT1G13110 | 2,32 | 1,98 | 1,49 | 1,33 | 1,84E-01 | 3,63E-01 |
| AT1G72600 | 18,04 | -0,62 | 0,47 | -1,33 | 1,84E-01 | 3,63E-01 |
| AT4G32620 | 1,61 | -2,76 | 2,08 | -1,33 | 1,84E-01 | 3,63E-01 |
| AT1G09780 | 4,32 | -1,16 | 0,87 | -1,33 | 1,84E-01 | 3,63E-01 |
| AT3G60320 | 3,74 | -1,12 | 0,84 | -1,33 | 1,84E-01 | 3,63E-01 |
| AT5G53440 | 2,08 | -1,68 | 1,27 | -1,33 | 1,84E-01 | 3,63E-01 |
| AT1G27535 | 2,34 | 1,82 | 1,37 | 1,33 | 1,85E-01 | 3,63E-01 |
| AT3G23100 | 2,14 | -1,85 | 1,39 | -1,33 | 1,85E-01 | 3,64E-01 |
| AT3G05355 | 1,75 | 2,16 | 1,63 | 1,33 | 1,85E-01 | 3,64E-01 |
| AT3G60770 | 1,96 | -1,82 | 1,37 | -1,33 | 1,85E-01 | 3,64E-01 |
| AT5G38660 | 1,59 | -2,07 | 1,56 | -1,32 | 1,85E-01 | 3,64E-01 |
| AT5G01350 | 8,44 | -0,85 | 0,64 | -1,32 | 1,86E-01 | 3,64E-01 |
| AT3G18050 | 1,52 | -2,11 | 1,60 | -1,32 | 1,86E-01 | 3,65E-01 |
| AT2G46192 | 21,12 | 0,81 | 0,62 | 1,32 | 1,87E-01 | 3,66E-01 |
| AT2G03330 | 8,86 | 0,99 | 0,75 | 1,32 | 1,87E-01 | 3,66E-01 |
| AT3G47350 | 23,89 | -0,63 | 0,48 | -1,32 | 1,87E-01 | 3,66E-01 |
| AT1G31960 | 1,81 | 2,17 | 1,64 | 1,32 | 1,87E-01 | 3,66E-01 |
| AT3G51950 | 4,09 | -1,17 | 0,89 | -1,32 | 1,87E-01 | 3,66E-01 |
| AT1G57720 | 1,72 | -1,90 | 1,45 | -1,32 | 1,88E-01 | 3,67E-01 |
| AT4G18197 | 1,87 | 1,95 | 1,48 | 1,32 | 1,88E-01 | 3,67E-01 |
| AT3G22235 | 3,04 | -1,61 | 1,22 | -1,32 | 1,88E-01 | 3,67E-01 |
| AT5G58330 | 1,92 | -1,93 | 1,47 | -1,32 | 1,88E-01 | 3,67E-01 |
| AT3G23110 | 1,95 | -1,98 | 1,50 | -1,31 | 1,89E-01 | 3,67E-01 |
| AT1G27730 | 1,60 | 2,05 | 1,56 | 1,31 | 1,89E-01 | 3,67E-01 |
| AT1G54490 | 1,36 | -2,08 | 1,58 | -1,31 | 1,89E-01 | 3,68E-01 |
| AT5G02305 | 2,96 | 1,75 | 1,33 | 1,31 | 1,90E-01 | 3,69E-01 |
| AT3G04455 | 2,46 | 2,23 | 1,70 | 1,31 | 1,90E-01 | 3,69E-01 |
| AT3G49070 | 1,57 | -4,00 | 3,05 | -1,31 | 1,90E-01 | 3,69E-01 |
| AT3G22970 | 2,77 | 1,56 | 1,19 | 1,31 | 1,90E-01 | 3,69E-01 |
| AT2G25250 | 2,81 | 1,65 | 1,26 | 1,31 | 1,91E-01 | 3,70E-01 |
| AT1G33415 | 2,72 | 1,52 | 1,16 | 1,31 | 1,91E-01 | 3,71E-01 |
| AT4G02570 | 1,99 | -1,86 | 1,43 | -1,30 | 1,92E-01 | 3,72E-01 |

| | | | | | | |
|---|---|---|---|---|---|---|
| AT3G61780 | 1,45 | -2,15 | 1,65 | -1,30 | 1,92E-01 | 3,72E-01 |
| AT1G07140 | 1,81 | -2,00 | 1,53 | -1,30 | 1,93E-01 | 3,73E-01 |
| AT1G76810 | 1,64 | 2,03 | 1,56 | 1,30 | 1,93E-01 | 3,73E-01 |
| AT3G01310 | 2,04 | -2,29 | 1,76 | -1,30 | 1,93E-01 | 3,73E-01 |
| AT3G02435 | 1,34 | 2,51 | 1,93 | 1,30 | 1,95E-01 | 3,75E-01 |
| AT1G61100 | 1,97 | -1,93 | 1,49 | -1,29 | 1,96E-01 | 3,78E-01 |
| AT3G49230 | 1,94 | 2,08 | 1,61 | 1,29 | 1,97E-01 | 3,79E-01 |
| AT2G32800 | 1,51 | -1,89 | 1,46 | -1,29 | 1,97E-01 | 3,79E-01 |
| AT3G18060 | 1,35 | -1,87 | 1,45 | -1,29 | 1,98E-01 | 3,80E-01 |
| AT5G26920 | 3,70 | -1,13 | 0,88 | -1,29 | 1,98E-01 | 3,81E-01 |
| AT1G06453 | 1,48 | 2,50 | 1,95 | 1,28 | 1,99E-01 | 3,84E-01 |
| AT1G05035 | 3,55 | 1,47 | 1,15 | 1,28 | 2,00E-01 | 3,84E-01 |
| AT2G01422 | 2,93 | 2,07 | 1,61 | 1,28 | 2,00E-01 | 3,84E-01 |
| AT5G24930 | 2,10 | -1,67 | 1,30 | -1,28 | 2,00E-01 | 3,84E-01 |
| AT1G07600 | 1,87 | 2,14 | 1,67 | 1,28 | 2,00E-01 | 3,84E-01 |
| AT5G22770 | 2,48 | -1,48 | 1,15 | -1,28 | 2,00E-01 | 3,84E-01 |
| AT4G20260 | 2,30 | 1,80 | 1,41 | 1,28 | 2,01E-01 | 3,85E-01 |
| AT4G27600 | 12,18 | -0,75 | 0,59 | -1,28 | 2,01E-01 | 3,86E-01 |
| AT4G04340 | 2,36 | -2,09 | 1,64 | -1,28 | 2,02E-01 | 3,86E-01 |
| AT5G13950 | 1,63 | -1,93 | 1,51 | -1,28 | 2,02E-01 | 3,86E-01 |
| AT5G26030 | 2,16 | 1,79 | 1,41 | 1,27 | 2,02E-01 | 3,86E-01 |
| AT1G08380 | 7,50 | 1,15 | 0,90 | 1,27 | 2,02E-01 | 3,86E-01 |
| AT1G50920 | 2,95 | -1,42 | 1,11 | -1,27 | 2,03E-01 | 3,87E-01 |
| AT2G23930 | 1,70 | -2,06 | 1,62 | -1,27 | 2,04E-01 | 3,87E-01 |
| AT4G06695 | 4,20 | 1,32 | 1,04 | 1,27 | 2,04E-01 | 3,87E-01 |
| AT5G37510 | 3,25 | -1,39 | 1,09 | -1,27 | 2,03E-01 | 3,87E-01 |
| AT5G66564 | 1,37 | 2,14 | 1,68 | 1,27 | 2,04E-01 | 3,87E-01 |
| AT1G06148 | 2,19 | 2,04 | 1,61 | 1,27 | 2,04E-01 | 3,87E-01 |
| AT1G06950 | 2,18 | -2,35 | 1,85 | -1,27 | 2,04E-01 | 3,87E-01 |
| AT1G07590 | 3,38 | 1,84 | 1,45 | 1,27 | 2,04E-01 | 3,87E-01 |
| AT5G20990 | 1,65 | -2,10 | 1,65 | -1,27 | 2,04E-01 | 3,87E-01 |
| AT5G09000 | 2,90 | -1,41 | 1,11 | -1,27 | 2,03E-01 | 3,87E-01 |
| AT4G08545 | 1,36 | 2,24 | 1,76 | 1,27 | 2,05E-01 | 3,88E-01 |
| AT5G25250 | 1,43 | 2,42 | 1,91 | 1,27 | 2,05E-01 | 3,89E-01 |
| AT1G29071 | 1,34 | 2,15 | 1,70 | 1,27 | 2,06E-01 | 3,89E-01 |
| AT5G11790 | 2,58 | 1,72 | 1,36 | 1,26 | 2,06E-01 | 3,90E-01 |
| AT5G28840 | 2,50 | -1,32 | 1,05 | -1,26 | 2,06E-01 | 3,90E-01 |
| AT1G63740 | 1,93 | 1,87 | 1,48 | 1,26 | 2,07E-01 | 3,91E-01 |
| AT3G08590 | 1,52 | -2,00 | 1,59 | -1,26 | 2,08E-01 | 3,92E-01 |
| AT3G44110 | 2,03 | -1,54 | 1,22 | -1,26 | 2,08E-01 | 3,93E-01 |
| AT1G69572 | 2,44 | 1,72 | 1,37 | 1,26 | 2,08E-01 | 3,93E-01 |
| AT2G38220 | 2,26 | 1,63 | 1,30 | 1,26 | 2,08E-01 | 3,93E-01 |
| AT1G69570 | 2,44 | 1,72 | 1,37 | 1,26 | 2,08E-01 | 3,93E-01 |
| AT5G12860 | 2,32 | -2,28 | 1,81 | -1,26 | 2,09E-01 | 3,93E-01 |
| AT2G04350 | 1,50 | -2,12 | 1,68 | -1,26 | 2,09E-01 | 3,93E-01 |
| AT3G08740 | 1,74 | -1,73 | 1,38 | -1,25 | 2,10E-01 | 3,94E-01 |
| AT4G30760 | 1,76 | 2,07 | 1,65 | 1,25 | 2,10E-01 | 3,94E-01 |
| AT5G09660 | 2,38 | 1,71 | 1,37 | 1,25 | 2,10E-01 | 3,94E-01 |
| AT4G07875 | 11,03 | -0,79 | 0,63 | -1,25 | 2,10E-01 | 3,95E-01 |
| AT5G47750 | 1,50 | -2,82 | 2,25 | -1,25 | 2,10E-01 | 3,95E-01 |
| AT4G29060 | 2,00 | 2,19 | 1,75 | 1,25 | 2,11E-01 | 3,95E-01 |
| AT3G44720 | 3,49 | -1,32 | 1,05 | -1,25 | 2,11E-01 | 3,95E-01 |
| AT4G09450 | 1,44 | -2,51 | 2,01 | -1,25 | 2,11E-01 | 3,95E-01 |
| AT3G15940 | 1,43 | -2,54 | 2,03 | -1,25 | 2,11E-01 | 3,96E-01 |
| AT1G18080 | 2,60 | -1,54 | 1,23 | -1,25 | 2,12E-01 | 3,97E-01 |
| AT5G07090 | 1,54 | -1,65 | 1,33 | -1,25 | 2,12E-01 | 3,97E-01 |

| | | | | | | |
|---|---|---|---|---|---|---|
| AT3G51890 | 1,41 | -3,28 | 2,63 | -1,25 | 2,12E-01 | 3,97E-01 |
| AT3G48560 | 3,89 | -1,17 | 0,94 | -1,25 | 2,13E-01 | 3,97E-01 |
| AT5G42020 | 4,47 | -1,01 | 0,81 | -1,25 | 2,13E-01 | 3,97E-01 |
| AT5G61310 | 4,76 | 1,18 | 0,95 | 1,24 | 2,14E-01 | 3,98E-01 |
| AT1G55350 | 1,89 | -2,61 | 2,10 | -1,24 | 2,14E-01 | 3,98E-01 |
| AT3G55734 | 132,05 | 0,37 | 0,30 | 1,24 | 2,14E-01 | 3,98E-01 |
| AT4G14030 | 1,74 | -1,94 | 1,56 | -1,24 | 2,14E-01 | 3,98E-01 |
| AT2G37270 | 1,75 | -1,99 | 1,60 | -1,24 | 2,14E-01 | 3,98E-01 |
| AT1G50410 | 1,51 | -2,94 | 2,37 | -1,24 | 2,14E-01 | 3,99E-01 |
| AT4G25730 | 1,71 | -2,17 | 1,75 | -1,24 | 2,15E-01 | 3,99E-01 |
| AT1G29070 | 1,74 | -1,72 | 1,39 | -1,24 | 2,16E-01 | 4,00E-01 |
| AT4G03295 | 3,55 | 1,41 | 1,14 | 1,24 | 2,15E-01 | 4,00E-01 |
| AT4G30160 | 4,90 | -1,17 | 0,95 | -1,24 | 2,16E-01 | 4,01E-01 |
| AT3G08935 | 1,61 | -2,28 | 1,85 | -1,23 | 2,17E-01 | 4,03E-01 |
| AT3G15880 | 1,94 | -1,57 | 1,27 | -1,23 | 2,18E-01 | 4,03E-01 |
| AT2G36470 | 2,06 | 2,24 | 1,82 | 1,23 | 2,18E-01 | 4,03E-01 |
| AT2G26500 | 2,52 | -1,44 | 1,17 | -1,23 | 2,19E-01 | 4,05E-01 |
| AT5G44790 | 1,35 | -1,99 | 1,62 | -1,23 | 2,20E-01 | 4,06E-01 |
| AT3G10815 | 5,65 | 1,25 | 1,02 | 1,23 | 2,20E-01 | 4,07E-01 |
| AT1G09970 | 4,22 | -1,31 | 1,07 | -1,22 | 2,22E-01 | 4,09E-01 |
| AT5G62350 | 2,26 | 1,72 | 1,41 | 1,22 | 2,22E-01 | 4,09E-01 |
| AT1G20400 | 1,62 | 1,86 | 1,53 | 1,22 | 2,22E-01 | 4,09E-01 |
| AT1G33590 | 1,70 | 1,92 | 1,58 | 1,22 | 2,22E-01 | 4,10E-01 |
| AT3G18370 | 2,55 | -1,73 | 1,42 | -1,22 | 2,23E-01 | 4,10E-01 |
| AT4G32285 | 2,40 | -1,52 | 1,25 | -1,22 | 2,23E-01 | 4,10E-01 |
| AT5G01960 | 1,82 | -2,49 | 2,04 | -1,22 | 2,23E-01 | 4,11E-01 |
| AT1G12680 | 1,43 | 2,26 | 1,86 | 1,22 | 2,24E-01 | 4,11E-01 |
| AT2G37040 | 1,58 | -1,93 | 1,59 | -1,22 | 2,24E-01 | 4,11E-01 |
| AT2G41940 | 1,76 | 1,78 | 1,46 | 1,22 | 2,24E-01 | 4,12E-01 |
| AT3G63490 | 1,82 | -2,11 | 1,73 | -1,21 | 2,25E-01 | 4,12E-01 |
| AT2G14265 | 1,52 | 2,20 | 1,81 | 1,21 | 2,25E-01 | 4,12E-01 |
| AT1G20375 | 1,50 | -1,74 | 1,43 | -1,21 | 2,25E-01 | 4,12E-01 |
| AT1G06460 | 1,79 | -1,72 | 1,42 | -1,21 | 2,25E-01 | 4,12E-01 |
| AT3G07790 | 2,60 | -1,34 | 1,11 | -1,21 | 2,25E-01 | 4,12E-01 |
| AT1G68520 | 2,00 | -1,47 | 1,22 | -1,21 | 2,25E-01 | 4,12E-01 |
| AT4G02420 | 1,44 | 2,21 | 1,82 | 1,21 | 2,25E-01 | 4,12E-01 |
| AT3G59140 | 3,91 | -2,32 | 1,92 | -1,21 | 2,26E-01 | 4,13E-01 |
| AT2G38560 | 1,56 | -2,25 | 1,86 | -1,21 | 2,26E-01 | 4,13E-01 |
| AT3G09465 | 2,48 | 1,60 | 1,33 | 1,21 | 2,27E-01 | 4,13E-01 |
| AT5G01020 | 1,61 | -1,82 | 1,50 | -1,21 | 2,27E-01 | 4,13E-01 |
| AT2G30500 | 1,56 | -2,45 | 2,03 | -1,21 | 2,27E-01 | 4,14E-01 |
| AT3G07390 | 2,06 | 1,83 | 1,52 | 1,21 | 2,28E-01 | 4,15E-01 |
| AT2G05920 | 1,42 | 2,09 | 1,73 | 1,21 | 2,28E-01 | 4,15E-01 |
| AT4G19120 | 2,16 | -1,38 | 1,15 | -1,21 | 2,28E-01 | 4,15E-01 |
| AT5G52170 | 2,49 | -1,66 | 1,38 | -1,20 | 2,28E-01 | 4,15E-01 |
| AT4G33650 | 1,75 | -1,93 | 1,60 | -1,20 | 2,29E-01 | 4,15E-01 |
| AT3G14840 | 1,81 | -1,66 | 1,38 | -1,20 | 2,29E-01 | 4,16E-01 |
| AT4G02760 | 2,62 | -1,39 | 1,16 | -1,20 | 2,30E-01 | 4,17E-01 |
| AT4G13770 | 2,50 | 1,99 | 1,66 | 1,20 | 2,31E-01 | 4,19E-01 |
| AT2G20860 | 1,36 | -1,96 | 1,64 | -1,20 | 2,31E-01 | 4,19E-01 |
| AT1G26110 | 1,79 | -1,73 | 1,44 | -1,20 | 2,31E-01 | 4,19E-01 |
| AT1G78080 | 3,84 | -1,17 | 0,98 | -1,20 | 2,31E-01 | 4,19E-01 |
| AT2G35387 | 52,44 | 0,44 | 0,37 | 1,20 | 2,32E-01 | 4,19E-01 |
| AT3G23125 | 239,13 | 0,53 | 0,44 | 1,20 | 2,32E-01 | 4,20E-01 |
| AT3G05335 | 1,73 | 1,91 | 1,60 | 1,19 | 2,32E-01 | 4,20E-01 |
| AT5G51630 | 1,97 | -1,92 | 1,61 | -1,19 | 2,32E-01 | 4,20E-01 |

| | | | | | | |
|---|---|---|---|---|---|---|
| AT2G27900 | 1,32 | -1,69 | 1,41 | -1,19 | 2,33E-01 | 4,20E-01 |
| AT5G09515 | 9,09 | 0,92 | 0,77 | 1,19 | 2,32E-01 | 4,20E-01 |
| AT5G27850 | 1,94 | -1,73 | 1,45 | -1,19 | 2,33E-01 | 4,20E-01 |
| AT5G53140 | 1,45 | -1,96 | 1,64 | -1,19 | 2,33E-01 | 4,20E-01 |
| AT3G22890 | 4,47 | -1,03 | 0,86 | -1,19 | 2,33E-01 | 4,20E-01 |
| AT1G50900 | 1,85 | -1,70 | 1,42 | -1,19 | 2,34E-01 | 4,21E-01 |
| AT5G04415 | 4,71 | -0,92 | 0,77 | -1,19 | 2,34E-01 | 4,21E-01 |
| AT5G66570 | 20,06 | -0,73 | 0,61 | -1,19 | 2,35E-01 | 4,22E-01 |
| AT2G35382 | 3,86 | 1,20 | 1,01 | 1,19 | 2,35E-01 | 4,22E-01 |
| AT2G08770 | 10,72 | 0,86 | 0,72 | 1,19 | 2,35E-01 | 4,22E-01 |
| AT2G21030 | 1,78 | 2,02 | 1,70 | 1,19 | 2,35E-01 | 4,22E-01 |
| AT2G43750 | 1,46 | -1,70 | 1,43 | -1,19 | 2,36E-01 | 4,22E-01 |
| AT2G30870 | 1,65 | 1,91 | 1,61 | 1,18 | 2,36E-01 | 4,23E-01 |
| AT3G20310 | 2,42 | 1,48 | 1,25 | 1,18 | 2,37E-01 | 4,24E-01 |
| AT4G20835 | 2,66 | 1,58 | 1,34 | 1,18 | 2,36E-01 | 4,24E-01 |
| AT1G18880 | 1,72 | 1,93 | 1,63 | 1,18 | 2,37E-01 | 4,24E-01 |
| AT1G53875 | 1,66 | 1,89 | 1,60 | 1,18 | 2,37E-01 | 4,24E-01 |
| AT2G25370 | 1,47 | 2,40 | 2,03 | 1,18 | 2,37E-01 | 4,24E-01 |
| AT4G16990 | 5,03 | -1,01 | 0,86 | -1,18 | 2,37E-01 | 4,24E-01 |
| AT4G14360 | 1,93 | -1,48 | 1,25 | -1,18 | 2,38E-01 | 4,25E-01 |
| AT2G47820 | 1,37 | -2,83 | 2,40 | -1,18 | 2,38E-01 | 4,25E-01 |
| AT5G65960 | 1,37 | -2,44 | 2,07 | -1,18 | 2,39E-01 | 4,25E-01 |
| AT3G29360 | 2,83 | -1,35 | 1,15 | -1,18 | 2,39E-01 | 4,25E-01 |
| AT3G54890 | 13,73 | -0,73 | 0,62 | -1,18 | 2,39E-01 | 4,25E-01 |
| AT4G39800 | 2,68 | -1,28 | 1,09 | -1,18 | 2,39E-01 | 4,25E-01 |
| AT5G27640 | 1,34 | -2,05 | 1,74 | -1,18 | 2,39E-01 | 4,25E-01 |
| AT1G76010 | 2,04 | -1,61 | 1,37 | -1,18 | 2,39E-01 | 4,25E-01 |
| AT1G02890 | 1,79 | -1,75 | 1,49 | -1,17 | 2,40E-01 | 4,27E-01 |
| AT2G03250 | 2,57 | 1,67 | 1,42 | 1,17 | 2,40E-01 | 4,27E-01 |
| AT2G41430 | 6,85 | 0,96 | 0,82 | 1,17 | 2,41E-01 | 4,27E-01 |
| AT1G09340 | 1,73 | -1,54 | 1,32 | -1,17 | 2,42E-01 | 4,28E-01 |
| AT2G39010 | 4,44 | -1,24 | 1,06 | -1,17 | 2,42E-01 | 4,28E-01 |
| AT5G61420 | 2,01 | -1,84 | 1,57 | -1,17 | 2,42E-01 | 4,28E-01 |
| AT5G64570 | 1,49 | -1,69 | 1,44 | -1,17 | 2,41E-01 | 4,28E-01 |
| AT5G04455 | 10,87 | 0,74 | 0,63 | 1,17 | 2,42E-01 | 4,29E-01 |
| AT5G04445 | 10,87 | 0,74 | 0,63 | 1,17 | 2,42E-01 | 4,29E-01 |
| AT5G22500 | 2,19 | 1,90 | 1,63 | 1,17 | 2,44E-01 | 4,30E-01 |
| AT1G80600 | 1,40 | -2,24 | 1,92 | -1,17 | 2,44E-01 | 4,30E-01 |
| AT5G06165 | 4,41 | 1,17 | 1,00 | 1,17 | 2,44E-01 | 4,30E-01 |
| AT1G07200 | 1,82 | -1,82 | 1,56 | -1,17 | 2,44E-01 | 4,30E-01 |
| AT4G24715 | 2,06 | -1,35 | 1,16 | -1,16 | 2,44E-01 | 4,31E-01 |
| AT2G43150 | 1,94 | -1,70 | 1,46 | -1,16 | 2,45E-01 | 4,32E-01 |
| AT1G72930 | 3,01 | 1,79 | 1,54 | 1,16 | 2,45E-01 | 4,32E-01 |
| AT4G32060 | 1,61 | 1,88 | 1,62 | 1,16 | 2,46E-01 | 4,32E-01 |
| AT4G03415 | 1,93 | -1,79 | 1,55 | -1,16 | 2,47E-01 | 4,34E-01 |
| AT5G02880 | 3,00 | -1,34 | 1,16 | -1,16 | 2,47E-01 | 4,34E-01 |
| AT4G26090 | 1,90 | -1,47 | 1,27 | -1,16 | 2,48E-01 | 4,36E-01 |
| AT2G47180 | 1,41 | -1,84 | 1,60 | -1,16 | 2,48E-01 | 4,36E-01 |
| AT3G19780 | 1,34 | -2,51 | 2,17 | -1,15 | 2,49E-01 | 4,37E-01 |
| AT2G46220 | 1,83 | -1,77 | 1,54 | -1,15 | 2,50E-01 | 4,37E-01 |
| AT3G59970 | 1,71 | 1,85 | 1,61 | 1,15 | 2,50E-01 | 4,37E-01 |
| AT5G48385 | 2,37 | -1,63 | 1,42 | -1,15 | 2,50E-01 | 4,37E-01 |
| AT2G29670 | 3,81 | -1,19 | 1,04 | -1,15 | 2,50E-01 | 4,37E-01 |
| AT4G22240 | 1,83 | -1,44 | 1,25 | -1,15 | 2,50E-01 | 4,37E-01 |
| AT4G26095 | 1,81 | -1,49 | 1,29 | -1,15 | 2,49E-01 | 4,37E-01 |
| AT3G61630 | 1,88 | -1,78 | 1,55 | -1,15 | 2,51E-01 | 4,39E-01 |

| | | | | | | |
|---|---|---|---|---|---|---|
| AT4G11160 | 1,61 | -1,88 | 1,64 | -1,15 | 2,51E-01 | 4,39E-01 |
| AT1G33950 | 1,88 | 1,86 | 1,62 | 1,15 | 2,51E-01 | 4,39E-01 |
| AT2G25970 | 1,54 | -1,92 | 1,68 | -1,15 | 2,51E-01 | 4,39E-01 |
| AT3G46620 | 4,33 | -1,05 | 0,92 | -1,15 | 2,52E-01 | 4,39E-01 |
| AT1G72610 | 17,51 | -0,55 | 0,48 | -1,15 | 2,52E-01 | 4,39E-01 |
| AT3G44330 | 1,72 | -2,53 | 2,21 | -1,14 | 2,52E-01 | 4,39E-01 |
| AT1G47970 | 1,63 | -1,62 | 1,42 | -1,14 | 2,53E-01 | 4,40E-01 |
| AT4G15180 | 1,84 | -1,54 | 1,35 | -1,14 | 2,53E-01 | 4,40E-01 |
| AT5G57660 | 4,42 | -0,92 | 0,80 | -1,14 | 2,53E-01 | 4,40E-01 |
| AT5G08530 | 3,76 | 1,46 | 1,28 | 1,14 | 2,54E-01 | 4,42E-01 |
| AT2G47980 | 2,39 | -1,47 | 1,29 | -1,14 | 2,55E-01 | 4,42E-01 |
| AT3G62650 | 1,34 | -1,89 | 1,66 | -1,14 | 2,54E-01 | 4,42E-01 |
| AT5G54940 | 2,86 | 1,34 | 1,18 | 1,14 | 2,55E-01 | 4,42E-01 |
| AT5G08330 | 1,56 | -1,66 | 1,46 | -1,14 | 2,56E-01 | 4,45E-01 |
| AT3G25140 | 1,51 | -2,48 | 2,19 | -1,13 | 2,58E-01 | 4,46E-01 |
| AT3G50480 | 1,71 | 1,71 | 1,52 | 1,13 | 2,59E-01 | 4,48E-01 |
| AT1G06670 | 1,38 | -2,03 | 1,79 | -1,13 | 2,59E-01 | 4,48E-01 |
| AT3G11830 | 1,90 | -1,53 | 1,36 | -1,13 | 2,60E-01 | 4,49E-01 |
| AT4G19520 | 3,67 | -1,05 | 0,93 | -1,13 | 2,60E-01 | 4,50E-01 |
| AT1G43320 | 2,75 | 1,93 | 1,72 | 1,12 | 2,61E-01 | 4,52E-01 |
| AT1G64583 | 1,87 | 1,80 | 1,61 | 1,12 | 2,62E-01 | 4,52E-01 |
| AT4G01210 | 1,50 | -1,90 | 1,69 | -1,12 | 2,62E-01 | 4,52E-01 |
| AT5G67030 | 1,94 | -1,42 | 1,26 | -1,12 | 2,62E-01 | 4,52E-01 |
| AT2G41530 | 2,42 | -1,35 | 1,20 | -1,12 | 2,62E-01 | 4,52E-01 |
| AT4G28750 | 4,56 | -1,17 | 1,04 | -1,12 | 2,62E-01 | 4,52E-01 |
| AT4G13930 | 1,56 | -1,62 | 1,45 | -1,12 | 2,63E-01 | 4,52E-01 |
| AT3G54540 | 1,42 | -2,13 | 1,90 | -1,12 | 2,63E-01 | 4,53E-01 |
| AT5G04305 | 5,44 | 0,94 | 0,84 | 1,12 | 2,64E-01 | 4,54E-01 |
| AT1G21270 | 2,66 | -1,68 | 1,51 | -1,11 | 2,65E-01 | 4,54E-01 |
| AT1G54270 | 2,59 | 1,47 | 1,32 | 1,12 | 2,64E-01 | 4,54E-01 |
| AT2G47960 | 1,39 | -1,84 | 1,65 | -1,11 | 2,65E-01 | 4,54E-01 |
| AT3G53400 | 1,89 | -1,77 | 1,59 | -1,11 | 2,65E-01 | 4,54E-01 |
| AT4G04925 | 1,79 | 1,78 | 1,60 | 1,12 | 2,65E-01 | 4,54E-01 |
| AT1G68590 | 1,41 | -1,74 | 1,56 | -1,11 | 2,65E-01 | 4,54E-01 |
| AT1G78830 | 2,12 | -1,44 | 1,30 | -1,11 | 2,65E-01 | 4,54E-01 |
| AT5G46260 | 1,90 | -1,61 | 1,44 | -1,11 | 2,65E-01 | 4,54E-01 |
| AT1G62390 | 2,10 | -1,98 | 1,78 | -1,11 | 2,67E-01 | 4,56E-01 |
| AT4G18010 | 1,46 | 2,16 | 1,94 | 1,11 | 2,67E-01 | 4,56E-01 |
| AT5G61790 | 5,40 | -0,95 | 0,86 | -1,11 | 2,68E-01 | 4,58E-01 |
| AT4G34890 | 2,69 | -1,46 | 1,32 | -1,11 | 2,68E-01 | 4,58E-01 |
| AT5G25060 | 1,92 | -1,36 | 1,23 | -1,11 | 2,68E-01 | 4,58E-01 |
| AT4G06245 | 1,52 | -2,58 | 2,33 | -1,11 | 2,69E-01 | 4,59E-01 |
| AT2G44560 | 4,56 | 0,99 | 0,90 | 1,10 | 2,69E-01 | 4,59E-01 |
| AT2G08725 | 34,05 | 0,66 | 0,60 | 1,10 | 2,70E-01 | 4,61E-01 |
| AT1G07940 | 3,71 | -1,10 | 1,00 | -1,10 | 2,70E-01 | 4,61E-01 |
| AT3G50820 | 7,18 | 1,06 | 0,96 | 1,10 | 2,71E-01 | 4,61E-01 |
| AT4G23980 | 1,47 | -1,65 | 1,50 | -1,10 | 2,71E-01 | 4,62E-01 |
| AT1G29418 | 9,45 | 0,68 | 0,62 | 1,10 | 2,72E-01 | 4,63E-01 |
| AT1G62540 | 2,48 | 1,72 | 1,57 | 1,10 | 2,73E-01 | 4,63E-01 |
| AT1G01120 | 1,43 | 1,75 | 1,60 | 1,10 | 2,73E-01 | 4,63E-01 |
| AT3G14050 | 1,74 | 1,77 | 1,61 | 1,10 | 2,73E-01 | 4,63E-01 |
| AT5G11700 | 1,99 | -1,49 | 1,36 | -1,10 | 2,73E-01 | 4,64E-01 |
| AT1G08103 | 6,40 | 1,01 | 0,92 | 1,09 | 2,74E-01 | 4,65E-01 |
| AT1G53730 | 1,50 | -2,22 | 2,03 | -1,09 | 2,74E-01 | 4,65E-01 |
| AT3G04400 | 2,05 | -1,39 | 1,27 | -1,09 | 2,75E-01 | 4,65E-01 |
| AT3G44670 | 1,62 | -1,68 | 1,54 | -1,09 | 2,75E-01 | 4,65E-01 |

| | | | | | | |
|---|---|---|---|---|---|---|
| AT3G21670 | 1,42 | -2,62 | 2,40 | -1,09 | 2,75E-01 | 4,65E-01 |
| AT1G04425 | 1,91 | 1,62 | 1,48 | 1,09 | 2,75E-01 | 4,65E-01 |
| AT5G44270 | 2,68 | 1,33 | 1,22 | 1,09 | 2,77E-01 | 4,68E-01 |
| AT2G47390 | 2,13 | -1,71 | 1,57 | -1,09 | 2,77E-01 | 4,68E-01 |
| AT1G61190 | 2,38 | -1,48 | 1,36 | -1,09 | 2,77E-01 | 4,68E-01 |
| AT1G25550 | 1,33 | 2,13 | 1,97 | 1,09 | 2,77E-01 | 4,68E-01 |
| AT1G01550 | 2,54 | -1,08 | 1,00 | -1,08 | 2,79E-01 | 4,70E-01 |
| AT1G56280 | 2,59 | -1,21 | 1,12 | -1,08 | 2,79E-01 | 4,70E-01 |
| AT2G38325 | 156,77 | -0,26 | 0,24 | -1,08 | 2,79E-01 | 4,70E-01 |
| AT2G04455 | 3,39 | 1,23 | 1,14 | 1,08 | 2,80E-01 | 4,71E-01 |
| AT3G54920 | 2,20 | -1,19 | 1,10 | -1,08 | 2,80E-01 | 4,71E-01 |
| AT2G25870 | 1,69 | -1,64 | 1,52 | -1,08 | 2,80E-01 | 4,71E-01 |
| AT3G25690 | 2,72 | -1,25 | 1,16 | -1,08 | 2,81E-01 | 4,72E-01 |
| AT5G62700 | 1,86 | -1,55 | 1,44 | -1,08 | 2,81E-01 | 4,72E-01 |
| AT2G37550 | 1,38 | -2,06 | 1,91 | -1,08 | 2,81E-01 | 4,72E-01 |
| AT3G22231 | 4,08 | 1,11 | 1,04 | 1,07 | 2,83E-01 | 4,74E-01 |
| AT4G16920 | 1,96 | -1,42 | 1,32 | -1,07 | 2,83E-01 | 4,74E-01 |
| AT3G28300 | 1,56 | -3,05 | 2,84 | -1,07 | 2,83E-01 | 4,75E-01 |
| AT4G04465 | 2,27 | 1,53 | 1,42 | 1,07 | 2,84E-01 | 4,75E-01 |
| AT5G23150 | 1,41 | -2,18 | 2,04 | -1,07 | 2,85E-01 | 4,77E-01 |
| AT1G62035 | 5,54 | 0,79 | 0,74 | 1,07 | 2,85E-01 | 4,77E-01 |
| AT4G17090 | 4,63 | -1,02 | 0,96 | -1,07 | 2,85E-01 | 4,77E-01 |
| AT3G28860 | 1,33 | -2,50 | 2,34 | -1,07 | 2,86E-01 | 4,77E-01 |
| AT1G12280 | 1,78 | -1,38 | 1,29 | -1,07 | 2,86E-01 | 4,78E-01 |
| AT2G47400 | 11,67 | -0,58 | 0,54 | -1,06 | 2,87E-01 | 4,79E-01 |
| AT2G23600 | 1,45 | -1,85 | 1,74 | -1,06 | 2,88E-01 | 4,80E-01 |
| AT1G14580 | 3,99 | 1,19 | 1,12 | 1,06 | 2,88E-01 | 4,81E-01 |
| AT3G14650 | 2,33 | -1,25 | 1,18 | -1,06 | 2,90E-01 | 4,83E-01 |
| AT4G24020 | 1,49 | -1,51 | 1,42 | -1,06 | 2,90E-01 | 4,83E-01 |
| AT4G26255 | 3,93 | -1,17 | 1,11 | -1,06 | 2,90E-01 | 4,83E-01 |
| AT1G69940 | 3,40 | 1,12 | 1,06 | 1,06 | 2,90E-01 | 4,83E-01 |
| AT2G31141 | 64,55 | 0,33 | 0,31 | 1,06 | 2,90E-01 | 4,83E-01 |
| AT4G03020 | 2,29 | -1,32 | 1,24 | -1,06 | 2,90E-01 | 4,83E-01 |
| AT2G16600 | 11,35 | 0,71 | 0,67 | 1,05 | 2,92E-01 | 4,85E-01 |
| AT3G08785 | 1,51 | -2,06 | 1,96 | -1,05 | 2,92E-01 | 4,85E-01 |
| AT1G73660 | 1,58 | -1,88 | 1,78 | -1,05 | 2,92E-01 | 4,85E-01 |
| AT2G34410 | 2,89 | -1,39 | 1,32 | -1,05 | 2,93E-01 | 4,86E-01 |
| AT2G46765 | 2,57 | 1,16 | 1,11 | 1,05 | 2,93E-01 | 4,86E-01 |
| AT1G11870 | 3,10 | 1,19 | 1,14 | 1,05 | 2,93E-01 | 4,86E-01 |
| AT2G28840 | 1,76 | -1,62 | 1,54 | -1,05 | 2,93E-01 | 4,86E-01 |
| AT1G18260 | 1,96 | -1,45 | 1,38 | -1,05 | 2,94E-01 | 4,86E-01 |
| AT2G30920 | 2,18 | 1,52 | 1,45 | 1,05 | 2,94E-01 | 4,87E-01 |
| AT3G01770 | 1,96 | -1,51 | 1,44 | -1,05 | 2,95E-01 | 4,87E-01 |
| AT1G58848 | 1,34 | -1,94 | 1,85 | -1,05 | 2,95E-01 | 4,87E-01 |
| AT3G56910 | 1,94 | -1,75 | 1,67 | -1,05 | 2,95E-01 | 4,87E-01 |
| AT2G25171 | 1,84 | 1,80 | 1,72 | 1,05 | 2,96E-01 | 4,88E-01 |
| AT3G10800 | 1,54 | -1,91 | 1,83 | -1,04 | 2,97E-01 | 4,90E-01 |
| AT5G08680 | 1,77 | -1,66 | 1,59 | -1,04 | 2,97E-01 | 4,90E-01 |
| AT5G39050 | 4,52 | 1,27 | 1,22 | 1,04 | 2,97E-01 | 4,90E-01 |
| AT3G04775 | 14,57 | 0,52 | 0,50 | 1,04 | 2,97E-01 | 4,90E-01 |
| AT2G07605 | 7,80 | 0,80 | 0,77 | 1,04 | 2,98E-01 | 4,91E-01 |
| AT2G44310 | 1,85 | 1,47 | 1,42 | 1,04 | 3,00E-01 | 4,94E-01 |
| AT5G01710 | 1,65 | -1,52 | 1,47 | -1,04 | 3,00E-01 | 4,94E-01 |
| AT3G63003 | 4,44 | 1,41 | 1,36 | 1,03 | 3,01E-01 | 4,95E-01 |
| AT4G33520 | 1,37 | -2,31 | 2,24 | -1,03 | 3,02E-01 | 4,96E-01 |
| AT4G32990 | 2,22 | 1,57 | 1,52 | 1,03 | 3,02E-01 | 4,97E-01 |

| | | | | | | |
|---|---|---|---|---|---|---|
| AT1G30330 | 1,89 | 1,53 | 1,49 | 1,03 | 3,03E-01 | 4,98E-01 |
| AT1G08360 | 1,45 | -1,78 | 1,74 | -1,03 | 3,04E-01 | 4,99E-01 |
| AT1G53780 | 1,40 | -2,06 | 2,00 | -1,03 | 3,05E-01 | 5,00E-01 |
| AT1G10500 | 1,39 | -2,60 | 2,54 | -1,03 | 3,05E-01 | 5,00E-01 |
| AT3G24170 | 1,33 | -1,84 | 1,79 | -1,03 | 3,05E-01 | 5,00E-01 |
| AT3G16820 | 1,66 | 1,38 | 1,35 | 1,02 | 3,06E-01 | 5,01E-01 |
| AT4G30950 | 1,64 | -1,67 | 1,64 | -1,02 | 3,06E-01 | 5,01E-01 |
| AT1G70060 | 1,48 | -2,02 | 1,98 | -1,02 | 3,08E-01 | 5,03E-01 |
| AT2G09685 | 3,15 | 1,19 | 1,16 | 1,02 | 3,08E-01 | 5,03E-01 |
| AT3G07110 | 1,39 | -1,81 | 1,78 | -1,02 | 3,09E-01 | 5,04E-01 |
| AT4G31780 | 1,36 | -2,73 | 2,68 | -1,02 | 3,09E-01 | 5,04E-01 |
| AT1G66783 | 14,56 | 0,67 | 0,65 | 1,02 | 3,09E-01 | 5,04E-01 |
| AT4G39240 | 1,33 | -2,37 | 2,33 | -1,02 | 3,09E-01 | 5,04E-01 |
| AT1G19680 | 2,21 | -1,86 | 1,83 | -1,02 | 3,10E-01 | 5,05E-01 |
| AT2G07727 | 3,34 | -1,28 | 1,26 | -1,01 | 3,10E-01 | 5,05E-01 |
| AT2G07806 | 2,71 | -1,24 | 1,22 | -1,01 | 3,10E-01 | 5,05E-01 |
| AT5G65720 | 1,65 | -1,70 | 1,67 | -1,02 | 3,10E-01 | 5,05E-01 |
| AT1G04263 | 1,35 | -2,60 | 2,57 | -1,01 | 3,10E-01 | 5,05E-01 |
| AT2G07815 | 2,06 | 1,47 | 1,45 | 1,01 | 3,11E-01 | 5,05E-01 |
| AT4G31170 | 2,81 | -1,37 | 1,36 | -1,01 | 3,11E-01 | 5,06E-01 |
| AT5G20520 | 1,55 | -1,60 | 1,58 | -1,01 | 3,11E-01 | 5,06E-01 |
| AT4G10590 | 1,40 | -2,09 | 2,07 | -1,01 | 3,13E-01 | 5,08E-01 |
| AT5G04590 | 4,30 | -1,01 | 1,00 | -1,01 | 3,13E-01 | 5,08E-01 |
| AT2G26250 | 1,96 | 1,42 | 1,41 | 1,01 | 3,13E-01 | 5,08E-01 |
| AT3G22120 | 4,08 | -0,94 | 0,94 | -1,01 | 3,13E-01 | 5,08E-01 |
| AT1G15125 | 1,84 | -1,45 | 1,44 | -1,01 | 3,14E-01 | 5,08E-01 |
| AT1G20690 | 2,85 | 1,24 | 1,23 | 1,00 | 3,15E-01 | 5,08E-01 |
| AT1G62630 | 2,10 | 1,42 | 1,41 | 1,00 | 3,15E-01 | 5,08E-01 |
| AT4G39364 | 2,19 | 1,55 | 1,55 | 1,01 | 3,15E-01 | 5,08E-01 |
| AT1G06900 | 1,35 | -1,70 | 1,69 | -1,01 | 3,15E-01 | 5,08E-01 |
| AT1G20691 | 2,85 | 1,24 | 1,23 | 1,00 | 3,15E-01 | 5,08E-01 |
| AT1G69830 | 1,91 | -1,24 | 1,23 | -1,01 | 3,15E-01 | 5,08E-01 |
| AT4G13250 | 1,54 | -1,86 | 1,85 | -1,01 | 3,15E-01 | 5,08E-01 |
| AT5G02700 | 1,99 | 1,42 | 1,41 | 1,01 | 3,14E-01 | 5,08E-01 |
| AT5G02960 | 2,13 | -1,34 | 1,33 | -1,01 | 3,14E-01 | 5,08E-01 |
| AT1G64390 | 1,48 | -2,11 | 2,10 | -1,00 | 3,16E-01 | 5,09E-01 |
| AT4G06800 | 2,19 | 1,25 | 1,25 | 1,00 | 3,16E-01 | 5,09E-01 |
| AT5G17910 | 1,47 | -1,82 | 1,82 | -1,00 | 3,16E-01 | 5,09E-01 |
| AT5G13850 | 2,26 | -1,27 | 1,27 | -1,00 | 3,17E-01 | 5,09E-01 |
| AT5G65480 | 2,00 | -1,35 | 1,35 | -1,00 | 3,17E-01 | 5,09E-01 |
| AT1G55450 | 2,26 | 1,57 | 1,57 | 1,00 | 3,17E-01 | 5,10E-01 |
| AT2G23142 | 1,46 | -2,60 | 2,61 | -1,00 | 3,18E-01 | 5,10E-01 |
| AT4G25050 | 1,37 | -1,58 | 1,58 | -1,00 | 3,18E-01 | 5,10E-01 |
| AT5G48860 | 3,76 | 1,17 | 1,18 | 1,00 | 3,18E-01 | 5,10E-01 |
| AT5G15410 | 1,41 | -1,94 | 1,95 | -1,00 | 3,20E-01 | 5,12E-01 |
| AT1G05380 | 1,42 | -2,03 | 2,04 | -0,99 | 3,21E-01 | 5,14E-01 |
| AT3G48060 | 1,66 | -1,78 | 1,79 | -0,99 | 3,21E-01 | 5,14E-01 |
| AT3G52930 | 3,81 | -0,98 | 0,99 | -0,99 | 3,21E-01 | 5,14E-01 |
| AT1G31330 | 6,03 | 0,79 | 0,79 | 0,99 | 3,22E-01 | 5,15E-01 |
| AT3G62700 | 3,87 | -1,00 | 1,01 | -0,99 | 3,22E-01 | 5,15E-01 |
| AT5G26360 | 2,70 | -1,22 | 1,24 | -0,99 | 3,23E-01 | 5,16E-01 |
| AT5G45060 | 1,56 | -1,68 | 1,70 | -0,99 | 3,24E-01 | 5,18E-01 |
| AT2G43540 | 1,53 | 1,76 | 1,78 | 0,99 | 3,24E-01 | 5,18E-01 |
| AT4G16180 | 1,78 | -2,19 | 2,22 | -0,98 | 3,25E-01 | 5,18E-01 |
| AT1G55680 | 1,44 | -2,12 | 2,15 | -0,98 | 3,25E-01 | 5,19E-01 |
| AT4G03420 | 2,29 | -1,59 | 1,62 | -0,98 | 3,26E-01 | 5,19E-01 |

| | | | | | | |
|---|---|---|---|---|---|---|
| AT2G29080 | 1,97 | -1,39 | 1,42 | -0,98 | 3,26E-01 | 5,20E-01 |
| AT1G09690 | 1,78 | -1,36 | 1,39 | -0,98 | 3,27E-01 | 5,20E-01 |
| AT4G36040 | 3,49 | -0,92 | 0,94 | -0,98 | 3,27E-01 | 5,20E-01 |
| AT4G00100 | 1,39 | -1,92 | 1,96 | -0,98 | 3,28E-01 | 5,21E-01 |
| AT5G04425 | 2,72 | 1,21 | 1,23 | 0,98 | 3,28E-01 | 5,21E-01 |
| AT1G12220 | 1,65 | 1,43 | 1,47 | 0,98 | 3,28E-01 | 5,21E-01 |
| AT2G36880 | 2,29 | -1,12 | 1,14 | -0,98 | 3,28E-01 | 5,22E-01 |
| AT2G41700 | 1,62 | 2,06 | 2,11 | 0,98 | 3,29E-01 | 5,22E-01 |
| AT3G51860 | 2,20 | -1,29 | 1,33 | -0,98 | 3,29E-01 | 5,22E-01 |
| AT1G68580 | 1,44 | -1,44 | 1,48 | -0,97 | 3,30E-01 | 5,22E-01 |
| AT4G28755 | 2,23 | -1,23 | 1,27 | -0,97 | 3,30E-01 | 5,22E-01 |
| AT5G21020 | 1,50 | 1,60 | 1,64 | 0,97 | 3,30E-01 | 5,22E-01 |
| AT1G16800 | 1,44 | -2,22 | 2,28 | -0,97 | 3,30E-01 | 5,22E-01 |
| AT3G14350 | 1,44 | -1,31 | 1,34 | -0,97 | 3,30E-01 | 5,22E-01 |
| AT4G26850 | 7,34 | -0,65 | 0,66 | -0,97 | 3,30E-01 | 5,22E-01 |
| AT1G30400 | 2,52 | -1,31 | 1,35 | -0,97 | 3,31E-01 | 5,23E-01 |
| AT2G25490 | 3,68 | 1,12 | 1,16 | 0,97 | 3,32E-01 | 5,24E-01 |
| AT5G65620 | 1,40 | -1,46 | 1,51 | -0,97 | 3,33E-01 | 5,25E-01 |
| AT2G42540 | 2,58 | -1,82 | 1,88 | -0,97 | 3,33E-01 | 5,26E-01 |
| AT3G07570 | 1,83 | -1,33 | 1,37 | -0,97 | 3,34E-01 | 5,26E-01 |
| AT5G53370 | 2,35 | -1,28 | 1,32 | -0,97 | 3,34E-01 | 5,26E-01 |
| AT5G60360 | 2,97 | -1,28 | 1,32 | -0,97 | 3,34E-01 | 5,26E-01 |
| AT5G02870 | 2,41 | -1,26 | 1,31 | -0,96 | 3,35E-01 | 5,27E-01 |
| AT1G24300 | 1,68 | -1,55 | 1,60 | -0,96 | 3,35E-01 | 5,27E-01 |
| AT2G14910 | 2,97 | -1,12 | 1,16 | -0,96 | 3,35E-01 | 5,27E-01 |
| AT3G48050 | 3,14 | -0,95 | 0,98 | -0,96 | 3,35E-01 | 5,27E-01 |
| AT1G21200 | 5,07 | 0,92 | 0,95 | 0,96 | 3,36E-01 | 5,28E-01 |
| AT5G47020 | 1,57 | -1,54 | 1,61 | -0,96 | 3,37E-01 | 5,28E-01 |
| AT5G56010 | 6,62 | -0,77 | 0,80 | -0,96 | 3,37E-01 | 5,28E-01 |
| AT3G25010 | 4,35 | -0,76 | 0,79 | -0,96 | 3,37E-01 | 5,28E-01 |
| AT5G51090 | 2,18 | 1,30 | 1,35 | 0,96 | 3,36E-01 | 5,28E-01 |
| AT3G08940 | 8,84 | -0,74 | 0,77 | -0,96 | 3,38E-01 | 5,29E-01 |
| AT3G53260 | 2,56 | -1,26 | 1,32 | -0,96 | 3,39E-01 | 5,30E-01 |
| AT1G58602 | 11,97 | -0,54 | 0,56 | -0,96 | 3,39E-01 | 5,32E-01 |
| AT5G11250 | 1,58 | -1,36 | 1,42 | -0,95 | 3,40E-01 | 5,32E-01 |
| AT4G06320 | 7,79 | 0,70 | 0,73 | 0,95 | 3,40E-01 | 5,32E-01 |
| AT3G56150 | 2,36 | -1,23 | 1,29 | -0,95 | 3,41E-01 | 5,33E-01 |
| AT3G26520 | 6,24 | 0,89 | 0,94 | 0,95 | 3,42E-01 | 5,34E-01 |
| AT4G17410 | 2,55 | -1,36 | 1,44 | -0,95 | 3,42E-01 | 5,35E-01 |
| AT5G41740 | 6,05 | -0,89 | 0,94 | -0,95 | 3,43E-01 | 5,35E-01 |
| AT1G31540 | 3,00 | -1,18 | 1,24 | -0,95 | 3,43E-01 | 5,35E-01 |
| AT4G20830 | 3,02 | 1,11 | 1,18 | 0,95 | 3,45E-01 | 5,37E-01 |
| AT2G42590 | 2,06 | 1,54 | 1,64 | 0,94 | 3,47E-01 | 5,41E-01 |
| AT5G05750 | 1,85 | -2,17 | 2,31 | -0,94 | 3,48E-01 | 5,41E-01 |
| AT1G72440 | 1,79 | -1,73 | 1,85 | -0,94 | 3,49E-01 | 5,43E-01 |
| AT4G34200 | 1,37 | -1,60 | 1,71 | -0,94 | 3,49E-01 | 5,43E-01 |
| AT5G24900 | 2,20 | 1,30 | 1,39 | 0,94 | 3,49E-01 | 5,43E-01 |
| AT1G61140 | 2,12 | -1,13 | 1,20 | -0,94 | 3,50E-01 | 5,44E-01 |
| AT1G11310 | 1,46 | -1,41 | 1,51 | -0,93 | 3,50E-01 | 5,44E-01 |
| AT5G54770 | 28,55 | 0,47 | 0,51 | 0,93 | 3,50E-01 | 5,44E-01 |
| AT3G26740 | 15,30 | 0,53 | 0,57 | 0,93 | 3,51E-01 | 5,45E-01 |
| AT4G00700 | 2,06 | -1,41 | 1,51 | -0,93 | 3,51E-01 | 5,45E-01 |
| AT2G07180 | 1,49 | -1,72 | 1,85 | -0,93 | 3,51E-01 | 5,45E-01 |
| AT3G54230 | 1,41 | -1,79 | 1,93 | -0,93 | 3,54E-01 | 5,48E-01 |
| AT1G73080 | 1,38 | -1,32 | 1,42 | -0,93 | 3,54E-01 | 5,48E-01 |
| AT5G65005 | 2,56 | -1,05 | 1,14 | -0,92 | 3,56E-01 | 5,51E-01 |

| | | | | | | |
|---|---|---|---|---|---|---|
| AT2G24270 | 3,15 | -0,98 | 1,06 | -0,92 | 3,57E-01 | 5,53E-01 |
| AT1G04733 | 20,09 | 0,46 | 0,50 | 0,92 | 3,58E-01 | 5,53E-01 |
| AT1G54410 | 19,25 | -0,59 | 0,64 | -0,92 | 3,58E-01 | 5,53E-01 |
| AT3G43490 | 1,48 | 1,84 | 2,00 | 0,92 | 3,58E-01 | 5,54E-01 |
| AT1G49050 | 1,80 | -1,71 | 1,86 | -0,92 | 3,60E-01 | 5,55E-01 |
| AT1G76030 | 2,84 | -1,11 | 1,21 | -0,92 | 3,60E-01 | 5,55E-01 |
| AT1G70530 | 1,72 | -1,71 | 1,87 | -0,91 | 3,61E-01 | 5,57E-01 |
| AT1G31835 | 2,35 | 1,28 | 1,41 | 0,91 | 3,61E-01 | 5,57E-01 |
| AT2G36430 | 1,44 | 1,70 | 1,86 | 0,91 | 3,62E-01 | 5,58E-01 |
| AT2G31960 | 1,68 | -1,30 | 1,43 | -0,91 | 3,62E-01 | 5,58E-01 |
| AT1G59820 | 1,45 | -1,69 | 1,86 | -0,91 | 3,64E-01 | 5,60E-01 |
| AT1G43860 | 1,69 | -1,51 | 1,66 | -0,91 | 3,64E-01 | 5,61E-01 |
| AT4G12650 | 2,23 | -1,09 | 1,21 | -0,91 | 3,65E-01 | 5,61E-01 |
| AT4G26480 | 2,39 | -1,28 | 1,41 | -0,91 | 3,65E-01 | 5,61E-01 |
| AT5G63800 | 1,87 | 1,42 | 1,57 | 0,91 | 3,65E-01 | 5,61E-01 |
| AT1G20510 | 2,74 | 1,23 | 1,36 | 0,90 | 3,66E-01 | 5,61E-01 |
| AT5G62220 | 1,34 | 1,51 | 1,67 | 0,90 | 3,66E-01 | 5,61E-01 |
| AT4G39366 | 2,55 | 1,25 | 1,39 | 0,90 | 3,67E-01 | 5,62E-01 |
| AT3G57710 | 4,92 | 0,72 | 0,79 | 0,90 | 3,67E-01 | 5,62E-01 |
| AT3G46550 | 1,58 | 1,34 | 1,49 | 0,90 | 3,67E-01 | 5,62E-01 |
| AT3G56070 | 2,75 | 1,22 | 1,35 | 0,90 | 3,67E-01 | 5,62E-01 |
| AT2G39770 | 2,97 | -0,97 | 1,08 | -0,90 | 3,69E-01 | 5,63E-01 |
| AT4G27500 | 2,93 | -0,99 | 1,10 | -0,90 | 3,69E-01 | 5,63E-01 |
| AT5G58465 | 4,24 | 0,90 | 1,00 | 0,90 | 3,69E-01 | 5,63E-01 |
| AT5G65700 | 1,88 | -1,11 | 1,23 | -0,90 | 3,68E-01 | 5,63E-01 |
| AT4G07855 | 2,93 | -0,99 | 1,10 | -0,90 | 3,69E-01 | 5,63E-01 |
| AT5G40910 | 2,64 | -0,95 | 1,06 | -0,90 | 3,69E-01 | 5,64E-01 |
| AT2G09555 | 2,08 | 1,18 | 1,31 | 0,90 | 3,70E-01 | 5,64E-01 |
| AT5G01195 | 1,58 | 1,38 | 1,54 | 0,90 | 3,70E-01 | 5,65E-01 |
| AT4G34490 | 1,39 | -1,47 | 1,64 | -0,90 | 3,70E-01 | 5,65E-01 |
| AT3G62420 | 1,96 | -1,50 | 1,68 | -0,89 | 3,72E-01 | 5,66E-01 |
| AT2G42490 | 2,30 | -1,04 | 1,17 | -0,89 | 3,72E-01 | 5,67E-01 |
| AT2G01320 | 2,02 | -1,17 | 1,31 | -0,89 | 3,73E-01 | 5,67E-01 |
| AT2G04755 | 2,18 | -1,44 | 1,62 | -0,89 | 3,73E-01 | 5,67E-01 |
| AT4G02770 | 8,37 | 0,64 | 0,71 | 0,89 | 3,73E-01 | 5,67E-01 |
| AT5G27660 | 6,15 | -0,66 | 0,74 | -0,89 | 3,74E-01 | 5,69E-01 |
| AT1G04897 | 2,68 | 1,12 | 1,26 | 0,89 | 3,75E-01 | 5,69E-01 |
| AT1G72370 | 2,39 | -1,01 | 1,13 | -0,89 | 3,75E-01 | 5,69E-01 |
| AT1G50030 | 2,19 | -1,24 | 1,41 | -0,89 | 3,76E-01 | 5,70E-01 |
| AT3G59770 | 3,31 | -1,11 | 1,26 | -0,89 | 3,76E-01 | 5,70E-01 |
| AT5G52320 | 1,77 | 1,41 | 1,60 | 0,88 | 3,78E-01 | 5,73E-01 |
| AT1G72150 | 7,68 | 0,66 | 0,75 | 0,88 | 3,80E-01 | 5,75E-01 |
| AT2G19710 | 1,44 | -1,62 | 1,84 | -0,88 | 3,80E-01 | 5,75E-01 |
| AT3G10420 | 1,49 | -1,43 | 1,63 | -0,88 | 3,79E-01 | 5,75E-01 |
| AT3G50950 | 4,46 | -0,76 | 0,87 | -0,88 | 3,80E-01 | 5,75E-01 |
| AT3G29130 | 2,05 | 1,44 | 1,64 | 0,88 | 3,80E-01 | 5,75E-01 |
| AT5G37600 | 1,93 | -1,30 | 1,49 | -0,88 | 3,80E-01 | 5,75E-01 |
| AT1G21202 | 4,89 | 0,91 | 1,04 | 0,87 | 3,84E-01 | 5,76E-01 |
| AT1G65010 | 1,62 | -1,47 | 1,69 | -0,87 | 3,84E-01 | 5,76E-01 |
| AT1G77230 | 3,05 | -0,95 | 1,09 | -0,87 | 3,85E-01 | 5,76E-01 |
| AT2G04435 | 3,56 | 0,93 | 1,07 | 0,88 | 3,81E-01 | 5,76E-01 |
| AT2G14750 | 1,64 | -1,21 | 1,38 | -0,87 | 3,82E-01 | 5,76E-01 |
| AT2G34357 | 1,72 | -1,58 | 1,80 | -0,87 | 3,82E-01 | 5,76E-01 |
| AT3G04165 | 2,58 | 1,07 | 1,23 | 0,87 | 3,84E-01 | 5,76E-01 |
| AT3G52800 | 1,93 | 1,38 | 1,58 | 0,88 | 3,81E-01 | 5,76E-01 |
| AT4G02970 | 72,42 | -0,27 | 0,32 | -0,87 | 3,85E-01 | 5,76E-01 |

| | | | | | | |
|---|---|---|---|---|---|---|
| AT4G21366 | 2,02 | 1,20 | 1,37 | 0,87 | 3,83E-01 | 5,76E-01 |
| AT4G31980 | 2,53 | 1,24 | 1,43 | 0,87 | 3,84E-01 | 5,76E-01 |
| AT5G01155 | 4,02 | 0,89 | 1,02 | 0,87 | 3,83E-01 | 5,76E-01 |
| AT5G14060 | 2,66 | 1,08 | 1,23 | 0,87 | 3,83E-01 | 5,76E-01 |
| AT1G11785 | 2,48 | 1,03 | 1,17 | 0,88 | 3,81E-01 | 5,76E-01 |
| AT1G05637 | 4,89 | 0,91 | 1,04 | 0,87 | 3,84E-01 | 5,76E-01 |
| AT1G66670 | 1,37 | 1,60 | 1,83 | 0,87 | 3,82E-01 | 5,76E-01 |
| AT2G01720 | 1,57 | -2,02 | 2,32 | -0,87 | 3,84E-01 | 5,76E-01 |
| AT2G04280 | 2,22 | -1,41 | 1,61 | -0,87 | 3,83E-01 | 5,76E-01 |
| AT2G38170 | 3,76 | -0,94 | 1,08 | -0,87 | 3,85E-01 | 5,76E-01 |
| AT4G13577 | 2,60 | -1,04 | 1,19 | -0,87 | 3,85E-01 | 5,76E-01 |
| AT5G07740 | 1,77 | -1,15 | 1,32 | -0,87 | 3,85E-01 | 5,76E-01 |
| AT5G14950 | 1,52 | -1,24 | 1,43 | -0,87 | 3,84E-01 | 5,76E-01 |
| AT5G35735 | 2,19 | 1,11 | 1,28 | 0,87 | 3,84E-01 | 5,76E-01 |
| AT3G56800 | 1,39 | 1,35 | 1,57 | 0,86 | 3,88E-01 | 5,79E-01 |
| AT3G12280 | 1,55 | -1,37 | 1,59 | -0,86 | 3,89E-01 | 5,81E-01 |
| AT1G53900 | 2,01 | 1,17 | 1,36 | 0,86 | 3,90E-01 | 5,81E-01 |
| AT5G54390 | 4,62 | -1,09 | 1,27 | -0,86 | 3,90E-01 | 5,81E-01 |
| AT5G00765 | 3,11 | -0,90 | 1,04 | -0,86 | 3,90E-01 | 5,81E-01 |
| AT1G61890 | 1,56 | 1,46 | 1,70 | 0,86 | 3,90E-01 | 5,81E-01 |
| AT2G31610 | 1,80 | -1,19 | 1,39 | -0,86 | 3,92E-01 | 5,82E-01 |
| AT3G19930 | 1,47 | -1,14 | 1,33 | -0,86 | 3,92E-01 | 5,82E-01 |
| AT5G00730 | 1,64 | 1,59 | 1,85 | 0,86 | 3,92E-01 | 5,82E-01 |
| AT2G36460 | 1,86 | -1,26 | 1,48 | -0,86 | 3,92E-01 | 5,82E-01 |
| AT2G43410 | 2,84 | -1,06 | 1,24 | -0,86 | 3,92E-01 | 5,82E-01 |
| AT2G05355 | 1,78 | -1,17 | 1,37 | -0,85 | 3,93E-01 | 5,84E-01 |
| AT2G24103 | 3,17 | -0,87 | 1,01 | -0,85 | 3,94E-01 | 5,85E-01 |
| AT3G22121 | 3,96 | -0,82 | 0,96 | -0,85 | 3,94E-01 | 5,85E-01 |
| AT1G27770 | 5,67 | -0,82 | 0,97 | -0,85 | 3,96E-01 | 5,88E-01 |
| AT2G31890 | 2,01 | -1,25 | 1,47 | -0,85 | 3,96E-01 | 5,88E-01 |
| AT1G07250 | 2,07 | -0,95 | 1,12 | -0,84 | 3,98E-01 | 5,91E-01 |
| AT1G71002 | 2,81 | -0,93 | 1,10 | -0,84 | 3,99E-01 | 5,91E-01 |
| AT1G78920 | 1,61 | -1,42 | 1,68 | -0,84 | 3,99E-01 | 5,91E-01 |
| AT2G17040 | 1,57 | -1,28 | 1,52 | -0,84 | 4,00E-01 | 5,92E-01 |
| AT4G29010 | 1,63 | 1,48 | 1,76 | 0,84 | 4,00E-01 | 5,92E-01 |
| AT1G55325 | 2,49 | -0,86 | 1,02 | -0,84 | 4,01E-01 | 5,93E-01 |
| AT4G18210 | 1,90 | 1,52 | 1,81 | 0,84 | 4,01E-01 | 5,93E-01 |
| AT2G24200 | 1,48 | 1,49 | 1,78 | 0,84 | 4,02E-01 | 5,94E-01 |
| AT5G49360 | 3,80 | -0,77 | 0,92 | -0,84 | 4,03E-01 | 5,95E-01 |
| AT3G45830 | 1,79 | -1,26 | 1,51 | -0,83 | 4,04E-01 | 5,97E-01 |
| AT1G05563 | 2,75 | 1,06 | 1,27 | 0,83 | 4,05E-01 | 5,97E-01 |
| AT4G14960 | 3,57 | -0,83 | 1,00 | -0,83 | 4,06E-01 | 5,98E-01 |
| AT1G17620 | 1,43 | 1,44 | 1,73 | 0,83 | 4,06E-01 | 5,99E-01 |
| AT4G19090 | 1,32 | -1,36 | 1,64 | -0,83 | 4,06E-01 | 5,99E-01 |
| AT1G20020 | 2,67 | -0,83 | 1,00 | -0,83 | 4,07E-01 | 6,00E-01 |
| AT1G03600 | 1,40 | 1,47 | 1,77 | 0,83 | 4,09E-01 | 6,01E-01 |
| AT1G55920 | 3,12 | 1,01 | 1,23 | 0,83 | 4,09E-01 | 6,01E-01 |
| AT2G32700 | 1,61 | -1,22 | 1,47 | -0,83 | 4,09E-01 | 6,01E-01 |
| AT4G18150 | 2,31 | 1,22 | 1,48 | 0,82 | 4,10E-01 | 6,01E-01 |
| AT5G44190 | 1,58 | 1,56 | 1,90 | 0,82 | 4,10E-01 | 6,01E-01 |
| AT1G50910 | 1,39 | -1,23 | 1,49 | -0,82 | 4,10E-01 | 6,01E-01 |
| AT1G73170 | 1,76 | -1,40 | 1,70 | -0,82 | 4,09E-01 | 6,01E-01 |
| AT3G05655 | 1,87 | 1,28 | 1,55 | 0,82 | 4,10E-01 | 6,01E-01 |
| AT1G62895 | 1,43 | -1,53 | 1,86 | -0,82 | 4,11E-01 | 6,02E-01 |
| AT1G66580 | 1,55 | -1,30 | 1,58 | -0,82 | 4,11E-01 | 6,02E-01 |
| AT5G49440 | 5,06 | -0,72 | 0,87 | -0,82 | 4,11E-01 | 6,02E-01 |

| | | | | | | |
|---|---|---|---|---|---|---|
| AT5G46490 | 1,46 | -1,21 | 1,47 | -0,82 | 4,12E-01 | 6,03E-01 |
| AT3G11660 | 1,33 | 2,74 | 3,34 | 0,82 | 4,12E-01 | 6,03E-01 |
| AT4G05800 | 1,37 | 1,49 | 1,82 | 0,82 | 4,12E-01 | 6,03E-01 |
| AT1G53687 | 5,07 | 0,84 | 1,02 | 0,82 | 4,14E-01 | 6,04E-01 |
| AT3G07585 | 1,61 | 1,34 | 1,64 | 0,82 | 4,13E-01 | 6,04E-01 |
| AT4G14160 | 2,71 | -1,09 | 1,33 | -0,82 | 4,13E-01 | 6,04E-01 |
| AT1G14920 | 1,77 | -1,25 | 1,52 | -0,82 | 4,14E-01 | 6,04E-01 |
| AT3G44860 | 2,37 | 1,22 | 1,50 | 0,82 | 4,14E-01 | 6,04E-01 |
| AT1G20100 | 1,77 | -1,41 | 1,73 | -0,81 | 4,16E-01 | 6,05E-01 |
| AT4G03080 | 1,69 | -1,19 | 1,46 | -0,81 | 4,15E-01 | 6,05E-01 |
| AT5G10860 | 1,76 | -1,21 | 1,48 | -0,81 | 4,16E-01 | 6,05E-01 |
| AT1G63700 | 1,36 | -1,23 | 1,51 | -0,81 | 4,16E-01 | 6,06E-01 |
| AT5G43190 | 1,41 | -1,26 | 1,55 | -0,81 | 4,17E-01 | 6,06E-01 |
| AT5G13840 | 2,03 | -1,11 | 1,37 | -0,81 | 4,17E-01 | 6,06E-01 |
| AT4G34950 | 4,16 | -0,70 | 0,87 | -0,81 | 4,18E-01 | 6,08E-01 |
| AT5G46845 | 66,07 | 0,25 | 0,31 | 0,81 | 4,18E-01 | 6,08E-01 |
| AT3G16370 | 1,93 | -1,03 | 1,27 | -0,81 | 4,20E-01 | 6,10E-01 |
| AT3G08510 | 2,33 | -1,24 | 1,54 | -0,81 | 4,20E-01 | 6,10E-01 |
| AT2G24550 | 1,50 | 1,41 | 1,75 | 0,81 | 4,21E-01 | 6,10E-01 |
| AT3G05680 | 1,68 | -1,29 | 1,61 | -0,80 | 4,21E-01 | 6,10E-01 |
| AT3G44430 | 1,70 | -1,11 | 1,38 | -0,80 | 4,22E-01 | 6,12E-01 |
| AT4G21280 | 3,20 | -0,88 | 1,10 | -0,80 | 4,22E-01 | 6,12E-01 |
| AT1G66980 | 1,51 | -1,19 | 1,48 | -0,80 | 4,22E-01 | 6,12E-01 |
| AT5G09810 | 3,96 | -0,88 | 1,09 | -0,80 | 4,23E-01 | 6,13E-01 |
| AT1G20620 | 38,72 | -0,35 | 0,43 | -0,80 | 4,24E-01 | 6,13E-01 |
| AT2G18440 | 3,48 | -0,78 | 0,98 | -0,80 | 4,26E-01 | 6,16E-01 |
| AT3G17770 | 1,51 | -1,16 | 1,46 | -0,80 | 4,26E-01 | 6,16E-01 |
| AT1G01040 | 4,73 | 0,72 | 0,91 | 0,79 | 4,27E-01 | 6,16E-01 |
| AT2G26910 | 1,87 | -0,93 | 1,17 | -0,79 | 4,28E-01 | 6,18E-01 |
| AT3G17390 | 4,31 | 0,78 | 0,98 | 0,79 | 4,28E-01 | 6,18E-01 |
| AT1G53430 | 1,51 | -1,30 | 1,65 | -0,79 | 4,30E-01 | 6,20E-01 |
| AT1G56200 | 1,42 | -1,38 | 1,75 | -0,79 | 4,30E-01 | 6,20E-01 |
| AT3G02832 | 1,51 | 1,26 | 1,60 | 0,79 | 4,30E-01 | 6,20E-01 |
| AT5G19780 | 1,35 | -1,45 | 1,84 | -0,79 | 4,30E-01 | 6,20E-01 |
| AT1G56110 | 1,35 | -1,41 | 1,80 | -0,79 | 4,32E-01 | 6,21E-01 |
| AT4G31700 | 1,83 | 1,23 | 1,57 | 0,79 | 4,32E-01 | 6,21E-01 |
| AT4G22690 | 3,37 | -0,69 | 0,88 | -0,79 | 4,32E-01 | 6,22E-01 |
| AT3G09630 | 1,89 | -1,19 | 1,52 | -0,78 | 4,33E-01 | 6,22E-01 |
| AT3G60750 | 11,99 | -0,45 | 0,58 | -0,78 | 4,33E-01 | 6,22E-01 |
| AT4G34100 | 3,45 | -0,84 | 1,08 | -0,78 | 4,33E-01 | 6,22E-01 |
| AT1G77560 | 2,14 | -1,08 | 1,38 | -0,78 | 4,33E-01 | 6,22E-01 |
| AT5G08520 | 1,35 | -1,22 | 1,56 | -0,78 | 4,34E-01 | 6,22E-01 |
| AT5G58420 | 1,59 | -1,38 | 1,77 | -0,78 | 4,34E-01 | 6,23E-01 |
| AT2G19360 | 2,70 | 0,97 | 1,24 | 0,78 | 4,35E-01 | 6,23E-01 |
| AT2G38540 | 3,59 | -0,80 | 1,02 | -0,78 | 4,35E-01 | 6,23E-01 |
| AT1G79520 | 27,20 | 0,41 | 0,52 | 0,78 | 4,35E-01 | 6,23E-01 |
| AT2G22540 | 1,53 | -1,04 | 1,33 | -0,78 | 4,36E-01 | 6,24E-01 |
| AT5G62690 | 4,26 | -0,73 | 0,93 | -0,78 | 4,36E-01 | 6,24E-01 |
| AT4G04745 | 1,61 | 1,24 | 1,59 | 0,78 | 4,36E-01 | 6,24E-01 |
| AT5G40395 | 2,01 | 0,99 | 1,27 | 0,78 | 4,36E-01 | 6,24E-01 |
| AT3G46590 | 1,40 | -1,46 | 1,88 | -0,78 | 4,37E-01 | 6,24E-01 |
| AT3G46530 | 2,60 | -0,86 | 1,10 | -0,78 | 4,37E-01 | 6,24E-01 |
| AT1G65930 | 2,39 | -1,08 | 1,39 | -0,78 | 4,38E-01 | 6,25E-01 |
| AT1G32060 | 5,29 | 0,86 | 1,11 | 0,78 | 4,38E-01 | 6,25E-01 |
| AT1G78570 | 2,32 | -0,96 | 1,24 | -0,77 | 4,39E-01 | 6,26E-01 |
| AT2G20950 | 3,17 | 0,94 | 1,22 | 0,77 | 4,39E-01 | 6,26E-01 |

| | | | | | | |
|---|---|---|---|---|---|---|
| AT2G34250 | 2,94 | -0,85 | 1,10 | -0,77 | 4,40E-01 | 6,26E-01 |
| AT1G29400 | 2,11 | -0,94 | 1,21 | -0,77 | 4,40E-01 | 6,26E-01 |
| AT1G31130 | 1,47 | 1,37 | 1,78 | 0,77 | 4,39E-01 | 6,26E-01 |
| AT2G24850 | 1,45 | -1,21 | 1,56 | -0,77 | 4,39E-01 | 6,26E-01 |
| AT5G10360 | 1,44 | -1,09 | 1,41 | -0,77 | 4,40E-01 | 6,26E-01 |
| AT2G43060 | 1,34 | 1,29 | 1,68 | 0,77 | 4,41E-01 | 6,26E-01 |
| AT4G30960 | 5,13 | 0,72 | 0,94 | 0,77 | 4,41E-01 | 6,26E-01 |
| AT2G21100 | 3,61 | 0,91 | 1,18 | 0,77 | 4,41E-01 | 6,26E-01 |
| AT2G14610 | 3,13 | 0,88 | 1,14 | 0,77 | 4,41E-01 | 6,27E-01 |
| AT5G39080 | 2,20 | 1,17 | 1,53 | 0,77 | 4,43E-01 | 6,29E-01 |
| AT4G01533 | 1,62 | 1,37 | 1,78 | 0,77 | 4,44E-01 | 6,29E-01 |
| AT4G23100 | 1,83 | -1,00 | 1,30 | -0,77 | 4,44E-01 | 6,29E-01 |
| AT1G14650 | 1,32 | -1,74 | 2,28 | -0,76 | 4,46E-01 | 6,30E-01 |
| AT1G14670 | 2,24 | -0,90 | 1,18 | -0,76 | 4,45E-01 | 6,30E-01 |
| AT1G20630 | 1,62 | -1,32 | 1,73 | -0,76 | 4,46E-01 | 6,30E-01 |
| AT2G07698 | 9,74 | 0,50 | 0,66 | 0,76 | 4,46E-01 | 6,30E-01 |
| AT4G22710 | 1,76 | 1,30 | 1,70 | 0,76 | 4,45E-01 | 6,30E-01 |
| AT5G27420 | 1,44 | -1,28 | 1,68 | -0,76 | 4,46E-01 | 6,30E-01 |
| AT3G21055 | 1,34 | -1,28 | 1,68 | -0,76 | 4,46E-01 | 6,30E-01 |
| AT3G46970 | 7,07 | -0,54 | 0,70 | -0,76 | 4,45E-01 | 6,30E-01 |
| AT4G23670 | 1,50 | 1,36 | 1,78 | 0,76 | 4,45E-01 | 6,30E-01 |
| AT5G27720 | 3,01 | 0,88 | 1,15 | 0,76 | 4,46E-01 | 6,30E-01 |
| AT5G40810 | 1,32 | -1,39 | 1,83 | -0,76 | 4,47E-01 | 6,31E-01 |
| AT1G09877 | 23,86 | 0,39 | 0,52 | 0,76 | 4,47E-01 | 6,31E-01 |
| AT5G04720 | 2,75 | -1,20 | 1,59 | -0,76 | 4,48E-01 | 6,31E-01 |
| AT5G56970 | 2,67 | 0,92 | 1,21 | 0,76 | 4,48E-01 | 6,32E-01 |
| AT4G16660 | 2,21 | -1,03 | 1,37 | -0,76 | 4,50E-01 | 6,33E-01 |
| AT2G18760 | 1,48 | -1,14 | 1,51 | -0,75 | 4,51E-01 | 6,34E-01 |
| AT5G45472 | 4,93 | 0,85 | 1,13 | 0,75 | 4,51E-01 | 6,35E-01 |
| AT4G37650 | 1,47 | 1,33 | 1,76 | 0,75 | 4,52E-01 | 6,35E-01 |
| AT2G09465 | 4,22 | 0,71 | 0,94 | 0,75 | 4,52E-01 | 6,35E-01 |
| AT1G03240 | 1,51 | 1,58 | 2,11 | 0,75 | 4,52E-01 | 6,35E-01 |
| AT1G12090 | 8,60 | -0,53 | 0,70 | -0,75 | 4,52E-01 | 6,35E-01 |
| AT2G01850 | 1,74 | 1,21 | 1,62 | 0,75 | 4,53E-01 | 6,36E-01 |
| AT3G27400 | 1,80 | 1,10 | 1,47 | 0,75 | 4,53E-01 | 6,36E-01 |
| AT2G04965 | 3,81 | 0,68 | 0,91 | 0,75 | 4,53E-01 | 6,36E-01 |
| AT1G31163 | 6,82 | 0,55 | 0,73 | 0,75 | 4,54E-01 | 6,36E-01 |
| AT5G09505 | 92,76 | 0,23 | 0,31 | 0,75 | 4,55E-01 | 6,38E-01 |
| AT3G09440 | 5,91 | -0,58 | 0,78 | -0,75 | 4,56E-01 | 6,38E-01 |
| AT4G11900 | 2,31 | 0,88 | 1,18 | 0,75 | 4,56E-01 | 6,39E-01 |
| AT1G33120 | 1,97 | -1,10 | 1,48 | -0,74 | 4,57E-01 | 6,39E-01 |
| AT2G05710 | 3,35 | -0,78 | 1,05 | -0,74 | 4,58E-01 | 6,39E-01 |
| AT2G18320 | 12,44 | 0,42 | 0,56 | 0,74 | 4,58E-01 | 6,39E-01 |
| AT3G13290 | 2,32 | -1,10 | 1,48 | -0,74 | 4,58E-01 | 6,39E-01 |
| AT4G15560 | 2,23 | -0,95 | 1,27 | -0,74 | 4,58E-01 | 6,39E-01 |
| AT3G60190 | 1,87 | -1,05 | 1,41 | -0,74 | 4,57E-01 | 6,39E-01 |
| AT2G09695 | 1,56 | -1,10 | 1,49 | -0,74 | 4,59E-01 | 6,41E-01 |
| AT1G09800 | 20,58 | 0,37 | 0,50 | 0,74 | 4,60E-01 | 6,42E-01 |
| AT4G08425 | 1,45 | 1,20 | 1,62 | 0,74 | 4,61E-01 | 6,43E-01 |
| AT1G74290 | 1,78 | 1,21 | 1,65 | 0,73 | 4,63E-01 | 6,45E-01 |
| AT5G13530 | 2,40 | -0,88 | 1,20 | -0,73 | 4,65E-01 | 6,48E-01 |
| AT2G18300 | 1,33 | -1,49 | 2,04 | -0,73 | 4,65E-01 | 6,48E-01 |
| AT5G07095 | 1,59 | -1,00 | 1,37 | -0,73 | 4,66E-01 | 6,49E-01 |
| AT4G09690 | 1,50 | 1,31 | 1,81 | 0,73 | 4,67E-01 | 6,49E-01 |
| AT1G19670 | 1,59 | 1,38 | 1,90 | 0,73 | 4,67E-01 | 6,50E-01 |
| AT4G23140 | 2,18 | -0,98 | 1,35 | -0,73 | 4,68E-01 | 6,50E-01 |

| | | | | | | |
|---|---|---|---|---|---|---|
| AT2G04375 | 5,42 | -0,59 | 0,82 | -0,73 | 4,68E-01 | 6,50E-01 |
| AT2G45960 | 8,71 | -0,52 | 0,71 | -0,72 | 4,69E-01 | 6,50E-01 |
| AT4G35770 | 2,85 | 0,89 | 1,22 | 0,73 | 4,68E-01 | 6,50E-01 |
| AT4G30200 | 1,47 | -1,22 | 1,69 | -0,72 | 4,69E-01 | 6,50E-01 |
| AT5G10760 | 4,94 | -0,61 | 0,85 | -0,72 | 4,69E-01 | 6,50E-01 |
| AT1G21250 | 6,46 | -0,50 | 0,69 | -0,72 | 4,69E-01 | 6,50E-01 |
| AT2G41310 | 487,34 | -0,20 | 0,28 | -0,72 | 4,71E-01 | 6,52E-01 |
| AT2G43375 | 6,65 | 0,52 | 0,72 | 0,72 | 4,71E-01 | 6,52E-01 |
| AT4G37470 | 1,39 | 1,24 | 1,72 | 0,72 | 4,71E-01 | 6,52E-01 |
| AT5G45470 | 2,49 | 1,35 | 1,87 | 0,72 | 4,71E-01 | 6,52E-01 |
| AT1G02560 | 1,57 | -1,09 | 1,52 | -0,72 | 4,72E-01 | 6,52E-01 |
| AT3G03990 | 2,37 | -0,90 | 1,25 | -0,72 | 4,72E-01 | 6,52E-01 |
| AT3G60961 | 1,33 | -1,66 | 2,31 | -0,72 | 4,72E-01 | 6,52E-01 |
| AT1G15820 | 12,09 | -0,47 | 0,66 | -0,72 | 4,72E-01 | 6,52E-01 |
| AT1G20330 | 2,79 | -1,00 | 1,39 | -0,72 | 4,73E-01 | 6,52E-01 |
| AT2G23810 | 3,17 | 0,95 | 1,32 | 0,72 | 4,72E-01 | 6,52E-01 |
| AT2G09250 | 487,12 | -0,20 | 0,28 | -0,72 | 4,72E-01 | 6,52E-01 |
| AT1G72180 | 2,71 | -0,77 | 1,07 | -0,72 | 4,73E-01 | 6,52E-01 |
| AT5G65630 | 3,70 | -0,71 | 0,99 | -0,72 | 4,73E-01 | 6,52E-01 |
| AT3G07765 | 2,13 | 1,11 | 1,56 | 0,72 | 4,74E-01 | 6,53E-01 |
| AT3G53430 | 2,55 | -0,75 | 1,05 | -0,72 | 4,74E-01 | 6,53E-01 |
| AT1G10745 | 1,57 | 1,19 | 1,66 | 0,71 | 4,75E-01 | 6,53E-01 |
| AT5G65780 | 1,41 | -1,28 | 1,79 | -0,71 | 4,75E-01 | 6,53E-01 |
| AT4G34720 | 1,83 | -0,96 | 1,35 | -0,71 | 4,75E-01 | 6,53E-01 |
| AT5G42092 | 3,68 | -0,76 | 1,06 | -0,71 | 4,76E-01 | 6,54E-01 |
| AT3G16830 | 1,75 | -1,04 | 1,47 | -0,71 | 4,76E-01 | 6,54E-01 |
| AT5G40060 | 1,33 | -1,81 | 2,54 | -0,71 | 4,77E-01 | 6,54E-01 |
| AT5G40480 | 2,70 | -0,79 | 1,11 | -0,71 | 4,77E-01 | 6,54E-01 |
| AT2G46440 | 1,43 | -0,93 | 1,32 | -0,71 | 4,79E-01 | 6,56E-01 |
| AT4G13920 | 2,03 | 1,15 | 1,63 | 0,71 | 4,79E-01 | 6,56E-01 |
| AT1G06760 | 1,45 | -0,98 | 1,39 | -0,71 | 4,79E-01 | 6,56E-01 |
| AT3G13724 | 5,13 | 0,66 | 0,93 | 0,71 | 4,80E-01 | 6,57E-01 |
| AT1G07493 | 1,97 | -1,14 | 1,62 | -0,70 | 4,81E-01 | 6,58E-01 |
| AT3G11820 | 1,51 | -1,18 | 1,67 | -0,70 | 4,81E-01 | 6,58E-01 |
| AT4G30270 | 1,68 | 1,15 | 1,63 | 0,70 | 4,81E-01 | 6,58E-01 |
| AT1G28420 | 2,78 | -1,08 | 1,53 | -0,70 | 4,82E-01 | 6,59E-01 |
| AT5G45510 | 8,42 | -0,61 | 0,86 | -0,70 | 4,82E-01 | 6,59E-01 |
| AT1G19410 | 3,18 | 0,75 | 1,06 | 0,70 | 4,82E-01 | 6,59E-01 |
| AT4G00231 | 1,99 | -1,01 | 1,44 | -0,70 | 4,83E-01 | 6,59E-01 |
| AT4G13885 | 7,77 | 0,51 | 0,72 | 0,70 | 4,83E-01 | 6,60E-01 |
| AT1G30360 | 3,38 | -0,64 | 0,91 | -0,70 | 4,84E-01 | 6,60E-01 |
| AT5G45480 | 2,31 | 1,04 | 1,49 | 0,70 | 4,84E-01 | 6,60E-01 |
| AT1G05207 | 1,81 | -1,04 | 1,49 | -0,70 | 4,85E-01 | 6,60E-01 |
| AT1G75950 | 1,80 | -0,96 | 1,37 | -0,70 | 4,85E-01 | 6,60E-01 |
| AT3G44006 | 2,21 | -0,96 | 1,38 | -0,70 | 4,85E-01 | 6,60E-01 |
| AT2G47170 | 1,90 | 1,03 | 1,47 | 0,70 | 4,86E-01 | 6,61E-01 |
| AT3G04765 | 2,83 | 0,83 | 1,19 | 0,70 | 4,86E-01 | 6,61E-01 |
| AT3G17800 | 1,86 | -1,01 | 1,46 | -0,70 | 4,86E-01 | 6,61E-01 |
| AT5G58290 | 1,43 | 1,28 | 1,84 | 0,69 | 4,88E-01 | 6,63E-01 |
| AT1G61520 | 19,44 | -0,38 | 0,55 | -0,69 | 4,88E-01 | 6,63E-01 |
| AT4G24120 | 1,68 | -1,64 | 2,37 | -0,69 | 4,89E-01 | 6,63E-01 |
| AT3G23920 | 2,81 | -0,86 | 1,25 | -0,69 | 4,89E-01 | 6,63E-01 |
| AT1G04120 | 1,63 | -1,07 | 1,55 | -0,69 | 4,90E-01 | 6,64E-01 |
| AT2G20780 | 1,93 | -1,02 | 1,48 | -0,69 | 4,90E-01 | 6,64E-01 |
| AT2G33150 | 2,47 | -0,91 | 1,32 | -0,69 | 4,91E-01 | 6,66E-01 |
| AT4G16143 | 2,03 | -0,87 | 1,26 | -0,69 | 4,93E-01 | 6,67E-01 |

| | | | | | | |
|---|---|---|---|---|---|---|
| AT1G77760 | 2,44 | -0,89 | 1,30 | -0,68 | 4,94E-01 | 6,69E-01 |
| AT3G44300 | 1,82 | -1,00 | 1,46 | -0,68 | 4,96E-01 | 6,70E-01 |
| AT3G49470 | 2,41 | 1,06 | 1,56 | 0,68 | 4,95E-01 | 6,70E-01 |
| AT3G47250 | 1,84 | -0,90 | 1,33 | -0,68 | 4,95E-01 | 6,70E-01 |
| AT3G53420 | 2,26 | 1,01 | 1,49 | 0,68 | 4,96E-01 | 6,70E-01 |
| AT4G16144 | 1,36 | 1,55 | 2,28 | 0,68 | 4,96E-01 | 6,70E-01 |
| AT3G24615 | 9,91 | 0,44 | 0,65 | 0,68 | 4,96E-01 | 6,70E-01 |
| AT3G23120 | 2,24 | -0,98 | 1,45 | -0,68 | 4,97E-01 | 6,71E-01 |
| AT3G52880 | 2,23 | 0,97 | 1,43 | 0,68 | 4,97E-01 | 6,71E-01 |
| AT2G26560 | 1,67 | 1,08 | 1,59 | 0,68 | 4,98E-01 | 6,71E-01 |
| AT4G16980 | 2,88 | 0,75 | 1,11 | 0,68 | 4,98E-01 | 6,72E-01 |
| AT5G08690 | 2,64 | -0,79 | 1,17 | -0,68 | 4,99E-01 | 6,72E-01 |
| AT1G68530 | 2,97 | 0,72 | 1,07 | 0,67 | 5,00E-01 | 6,73E-01 |
| AT1G66100 | 8,07 | 0,47 | 0,70 | 0,67 | 5,01E-01 | 6,74E-01 |
| AT1G70490 | 1,85 | -0,86 | 1,28 | -0,67 | 5,00E-01 | 6,74E-01 |
| AT5G24590 | 1,51 | 1,08 | 1,60 | 0,67 | 5,01E-01 | 6,74E-01 |
| AT1G08623 | 2,21 | -1,03 | 1,53 | -0,67 | 5,02E-01 | 6,75E-01 |
| AT5G47040 | 1,34 | -1,04 | 1,56 | -0,67 | 5,03E-01 | 6,75E-01 |
| AT1G60950 | 5,66 | 0,62 | 0,93 | 0,66 | 5,07E-01 | 6,81E-01 |
| AT3G19030 | 2,04 | -0,82 | 1,24 | -0,66 | 5,08E-01 | 6,82E-01 |
| AT3G29190 | 2,69 | 0,97 | 1,47 | 0,66 | 5,09E-01 | 6,83E-01 |
| AT3G16785 | 1,89 | -0,77 | 1,17 | -0,66 | 5,09E-01 | 6,83E-01 |
| AT2G29450 | 2,19 | 1,06 | 1,61 | 0,66 | 5,10E-01 | 6,83E-01 |
| AT3G57450 | 1,58 | -0,88 | 1,34 | -0,66 | 5,11E-01 | 6,85E-01 |
| AT3G20362 | 7,16 | -0,55 | 0,84 | -0,66 | 5,12E-01 | 6,86E-01 |
| AT2G08910 | 1,52 | 0,94 | 1,43 | 0,65 | 5,13E-01 | 6,86E-01 |
| AT4G24413 | 2,10 | 0,93 | 1,42 | 0,66 | 5,12E-01 | 6,86E-01 |
| AT2G09700 | 2,16 | 1,04 | 1,58 | 0,65 | 5,13E-01 | 6,87E-01 |
| AT3G07735 | 4,39 | 0,60 | 0,92 | 0,65 | 5,13E-01 | 6,87E-01 |
| AT2G33170 | 2,19 | -0,97 | 1,49 | -0,65 | 5,15E-01 | 6,88E-01 |
| AT1G27290 | 1,54 | 0,90 | 1,39 | 0,65 | 5,16E-01 | 6,89E-01 |
| AT2G14560 | 27,90 | -0,31 | 0,48 | -0,65 | 5,16E-01 | 6,89E-01 |
| AT5G01820 | 9,75 | -0,57 | 0,88 | -0,65 | 5,16E-01 | 6,89E-01 |
| AT1G15890 | 1,49 | -1,08 | 1,67 | -0,65 | 5,16E-01 | 6,89E-01 |
| AT2G38210 | 1,49 | 1,29 | 1,99 | 0,65 | 5,17E-01 | 6,89E-01 |
| AT5G24300 | 2,35 | -0,95 | 1,47 | -0,65 | 5,17E-01 | 6,89E-01 |
| AT2G17265 | 1,56 | -1,01 | 1,57 | -0,65 | 5,18E-01 | 6,90E-01 |
| AT2G42690 | 1,34 | 1,08 | 1,67 | 0,65 | 5,18E-01 | 6,90E-01 |
| AT1G64385 | 1,99 | -1,12 | 1,75 | -0,64 | 5,20E-01 | 6,92E-01 |
| AT5G52310 | 5,67 | -0,83 | 1,29 | -0,64 | 5,20E-01 | 6,92E-01 |
| AT1G79550 | 2,41 | -0,73 | 1,13 | -0,64 | 5,20E-01 | 6,92E-01 |
| AT3G54925 | 1,76 | 1,13 | 1,76 | 0,64 | 5,20E-01 | 6,92E-01 |
| AT5G04430 | 1,54 | -1,18 | 1,83 | -0,64 | 5,20E-01 | 6,92E-01 |
| AT1G42550 | 2,32 | -0,79 | 1,24 | -0,64 | 5,23E-01 | 6,93E-01 |
| AT3G59068 | 1,78 | 1,01 | 1,58 | 0,64 | 5,23E-01 | 6,93E-01 |
| AT5G16620 | 2,01 | -0,77 | 1,21 | -0,64 | 5,23E-01 | 6,93E-01 |
| AT5G46760 | 2,52 | -0,82 | 1,28 | -0,64 | 5,23E-01 | 6,93E-01 |
| AT1G06040 | 2,85 | -0,82 | 1,28 | -0,64 | 5,23E-01 | 6,93E-01 |
| AT1G05427 | 2,11 | 0,99 | 1,55 | 0,64 | 5,22E-01 | 6,93E-01 |
| AT3G59070 | 1,78 | 1,01 | 1,58 | 0,64 | 5,23E-01 | 6,93E-01 |
| AT3G16140 | 1,91 | -0,83 | 1,30 | -0,64 | 5,24E-01 | 6,94E-01 |
| AT4G24220 | 1,52 | -0,97 | 1,52 | -0,64 | 5,25E-01 | 6,95E-01 |
| AT3G41761 | 2,51 | 1,12 | 1,77 | 0,63 | 5,26E-01 | 6,96E-01 |
| AT1G20860 | 3,03 | 0,72 | 1,14 | 0,63 | 5,26E-01 | 6,96E-01 |
| AT3G11930 | 1,35 | -2,17 | 3,45 | -0,63 | 5,28E-01 | 6,98E-01 |
| AT1G61740 | 2,13 | 0,83 | 1,32 | 0,63 | 5,28E-01 | 6,98E-01 |

| | | | | | | |
|---|---|---|---|---|---|---|
| AT5G08790 | 1,51 | -0,94 | 1,50 | -0,63 | 5,28E-01 | 6,98E-01 |
| AT1G63330 | 7,00 | -0,42 | 0,67 | -0,63 | 5,29E-01 | 6,99E-01 |
| AT4G23170 | 6,38 | 0,55 | 0,88 | 0,63 | 5,29E-01 | 6,99E-01 |
| AT5G64070 | 1,53 | -0,90 | 1,44 | -0,63 | 5,30E-01 | 6,99E-01 |
| AT3G14270 | 2,35 | -0,74 | 1,19 | -0,63 | 5,31E-01 | 6,99E-01 |
| AT1G03680 | 1,73 | 0,98 | 1,56 | 0,63 | 5,31E-01 | 6,99E-01 |
| AT1G20440 | 3,88 | -0,57 | 0,91 | -0,63 | 5,31E-01 | 6,99E-01 |
| AT2G35384 | 13,85 | 0,40 | 0,64 | 0,63 | 5,31E-01 | 6,99E-01 |
| AT4G15530 | 1,47 | -1,07 | 1,71 | -0,63 | 5,31E-01 | 6,99E-01 |
| AT5G49740 | 2,42 | -0,77 | 1,23 | -0,63 | 5,31E-01 | 6,99E-01 |
| AT4G16983 | 2,85 | 0,70 | 1,12 | 0,62 | 5,33E-01 | 7,02E-01 |
| AT2G37660 | 1,35 | 1,08 | 1,74 | 0,62 | 5,35E-01 | 7,03E-01 |
| AT5G66900 | 1,52 | -1,06 | 1,71 | -0,62 | 5,36E-01 | 7,04E-01 |
| AT5G51760 | 1,35 | -1,02 | 1,65 | -0,62 | 5,36E-01 | 7,04E-01 |
| AT1G76180 | 6,03 | 0,46 | 0,74 | 0,62 | 5,38E-01 | 7,06E-01 |
| AT1G66880 | 6,48 | -0,55 | 0,89 | -0,62 | 5,38E-01 | 7,06E-01 |
| AT1G28440 | 1,72 | -0,79 | 1,28 | -0,61 | 5,40E-01 | 7,08E-01 |
| AT5G41790 | 2,54 | -0,84 | 1,38 | -0,61 | 5,40E-01 | 7,08E-01 |
| AT1G05570 | 10,12 | -0,35 | 0,58 | -0,61 | 5,40E-01 | 7,08E-01 |
| AT1G08813 | 8,42 | 0,41 | 0,67 | 0,61 | 5,40E-01 | 7,08E-01 |
| AT5G20350 | 1,36 | -1,09 | 1,78 | -0,61 | 5,41E-01 | 7,08E-01 |
| AT1G48540 | 1,33 | -1,11 | 1,82 | -0,61 | 5,41E-01 | 7,09E-01 |
| AT3G15095 | 1,44 | -0,90 | 1,48 | -0,61 | 5,42E-01 | 7,09E-01 |
| AT2G30570 | 3,97 | 0,60 | 0,98 | 0,61 | 5,42E-01 | 7,09E-01 |
| AT2G39570 | 1,60 | -1,70 | 2,79 | -0,61 | 5,43E-01 | 7,10E-01 |
| AT4G33530 | 1,44 | -1,10 | 1,81 | -0,61 | 5,44E-01 | 7,11E-01 |
| AT4G13494 | 321,70 | 0,29 | 0,48 | 0,61 | 5,45E-01 | 7,12E-01 |
| AT1G23740 | 1,33 | -1,15 | 1,91 | -0,60 | 5,46E-01 | 7,12E-01 |
| AT4G02130 | 3,77 | -0,53 | 0,88 | -0,60 | 5,45E-01 | 7,12E-01 |
| AT4G37640 | 1,71 | -0,89 | 1,47 | -0,60 | 5,45E-01 | 7,12E-01 |
| AT1G01046 | 1,63 | 0,88 | 1,46 | 0,60 | 5,47E-01 | 7,14E-01 |
| AT4G32260 | 3,86 | 0,69 | 1,14 | 0,60 | 5,48E-01 | 7,14E-01 |
| AT3G23590 | 1,64 | -1,16 | 1,94 | -0,60 | 5,48E-01 | 7,14E-01 |
| AT3G52700 | 2,20 | 0,74 | 1,24 | 0,60 | 5,48E-01 | 7,14E-01 |
| AT3G58030 | 2,17 | -0,92 | 1,53 | -0,60 | 5,49E-01 | 7,14E-01 |
| AT3G59780 | 2,03 | -0,97 | 1,62 | -0,60 | 5,50E-01 | 7,16E-01 |
| AT2G22440 | 2,40 | 0,75 | 1,26 | 0,60 | 5,52E-01 | 7,16E-01 |
| AT2G09610 | 9,53 | 0,38 | 0,64 | 0,60 | 5,51E-01 | 7,16E-01 |
| AT3G28899 | 2,18 | 0,79 | 1,32 | 0,60 | 5,51E-01 | 7,16E-01 |
| AT4G08625 | 1,54 | 0,94 | 1,58 | 0,60 | 5,52E-01 | 7,16E-01 |
| AT2G28550 | 2,55 | 0,74 | 1,24 | 0,60 | 5,51E-01 | 7,16E-01 |
| AT4G03430 | 1,44 | -0,84 | 1,40 | -0,60 | 5,51E-01 | 7,16E-01 |
| AT5G43470 | 3,38 | -0,77 | 1,29 | -0,59 | 5,52E-01 | 7,16E-01 |
| AT4G04565 | 64,98 | -0,18 | 0,31 | -0,59 | 5,53E-01 | 7,17E-01 |
| AT1G06580 | 1,59 | 0,99 | 1,67 | 0,59 | 5,53E-01 | 7,17E-01 |
| AT1G04517 | 11,17 | 0,35 | 0,59 | 0,59 | 5,53E-01 | 7,17E-01 |
| AT1G08053 | 2,32 | 0,87 | 1,47 | 0,59 | 5,55E-01 | 7,18E-01 |
| AT1G06680 | 10,44 | -0,39 | 0,65 | -0,59 | 5,55E-01 | 7,19E-01 |
| AT3G21870 | 1,78 | 1,08 | 1,84 | 0,59 | 5,56E-01 | 7,19E-01 |
| AT5G44090 | 1,96 | -0,93 | 1,58 | -0,59 | 5,57E-01 | 7,20E-01 |
| AT3G28180 | 8,56 | -0,60 | 1,03 | -0,59 | 5,57E-01 | 7,20E-01 |
| AT3G48770 | 1,49 | 0,90 | 1,53 | 0,59 | 5,57E-01 | 7,20E-01 |
| AT1G61226 | 20,72 | -0,29 | 0,49 | -0,59 | 5,58E-01 | 7,20E-01 |
| AT3G57400 | 1,77 | 0,88 | 1,51 | 0,59 | 5,58E-01 | 7,20E-01 |
| AT2G32240 | 2,12 | 0,90 | 1,54 | 0,59 | 5,59E-01 | 7,20E-01 |
| AT4G34190 | 1,52 | -0,96 | 1,63 | -0,59 | 5,58E-01 | 7,20E-01 |

| | | | | | | |
|---|---|---|---|---|---|---|
| AT3G08170 | 3,83 | -0,57 | 0,97 | -0,58 | 5,59E-01 | 7,21E-01 |
| AT2G47440 | 2,19 | -0,83 | 1,43 | -0,58 | 5,59E-01 | 7,21E-01 |
| AT5G25130 | 1,42 | 0,87 | 1,50 | 0,58 | 5,61E-01 | 7,22E-01 |
| AT2G47450 | 6,02 | -0,46 | 0,78 | -0,58 | 5,61E-01 | 7,23E-01 |
| AT3G18080 | 1,62 | -0,84 | 1,45 | -0,58 | 5,62E-01 | 7,23E-01 |
| AT3G56408 | 2,59 | 0,87 | 1,50 | 0,58 | 5,62E-01 | 7,23E-01 |
| AT2G32690 | 2,01 | 0,82 | 1,40 | 0,58 | 5,61E-01 | 7,23E-01 |
| AT3G49800 | 1,37 | -0,86 | 1,48 | -0,58 | 5,62E-01 | 7,23E-01 |
| AT4G21640 | 1,99 | 0,96 | 1,65 | 0,58 | 5,62E-01 | 7,23E-01 |
| AT5G20020 | 2,46 | -0,95 | 1,64 | -0,58 | 5,63E-01 | 7,23E-01 |
| AT5G52920 | 1,32 | -1,20 | 2,09 | -0,57 | 5,67E-01 | 7,28E-01 |
| AT1G09937 | 6,18 | -0,40 | 0,70 | -0,57 | 5,67E-01 | 7,28E-01 |
| AT4G26530 | 4,43 | -0,50 | 0,88 | -0,57 | 5,68E-01 | 7,29E-01 |
| AT5G07325 | 8,05 | -0,37 | 0,65 | -0,57 | 5,69E-01 | 7,29E-01 |
| AT4G05320 | 17,67 | -0,31 | 0,54 | -0,57 | 5,70E-01 | 7,30E-01 |
| AT4G15470 | 1,75 | -0,74 | 1,30 | -0,57 | 5,70E-01 | 7,30E-01 |
| AT5G35630 | 4,18 | -0,56 | 1,00 | -0,57 | 5,71E-01 | 7,31E-01 |
| AT1G19400 | 1,69 | 0,82 | 1,46 | 0,57 | 5,72E-01 | 7,32E-01 |
| AT1G17840 | 1,51 | 1,00 | 1,78 | 0,56 | 5,75E-01 | 7,35E-01 |
| AT4G18480 | 1,72 | -0,87 | 1,55 | -0,56 | 5,75E-01 | 7,35E-01 |
| AT4G35750 | 1,89 | -0,71 | 1,27 | -0,56 | 5,77E-01 | 7,38E-01 |
| AT1G20980 | 1,44 | -1,03 | 1,85 | -0,56 | 5,78E-01 | 7,38E-01 |
| AT5G22800 | 1,71 | -0,83 | 1,50 | -0,55 | 5,79E-01 | 7,40E-01 |
| AT2G36390 | 2,07 | -0,84 | 1,53 | -0,55 | 5,80E-01 | 7,41E-01 |
| AT3G01290 | 2,12 | 0,68 | 1,23 | 0,55 | 5,81E-01 | 7,41E-01 |
| AT5G42090 | 4,37 | -0,55 | 1,00 | -0,55 | 5,81E-01 | 7,41E-01 |
| AT1G68560 | 3,34 | 0,68 | 1,24 | 0,55 | 5,84E-01 | 7,45E-01 |
| AT3G03960 | 1,45 | -0,93 | 1,70 | -0,55 | 5,85E-01 | 7,45E-01 |
| AT3G61700 | 1,46 | 0,84 | 1,53 | 0,55 | 5,85E-01 | 7,45E-01 |
| AT4G24820 | 1,43 | -0,89 | 1,63 | -0,55 | 5,85E-01 | 7,45E-01 |
| AT5G47560 | 3,13 | 0,58 | 1,06 | 0,55 | 5,86E-01 | 7,46E-01 |
| AT5G50950 | 1,52 | 0,80 | 1,48 | 0,54 | 5,87E-01 | 7,47E-01 |
| AT3G59060 | 1,47 | -0,75 | 1,38 | -0,54 | 5,87E-01 | 7,47E-01 |
| AT4G34150 | 1,90 | 0,82 | 1,52 | 0,54 | 5,87E-01 | 7,47E-01 |
| AT3G26870 | 1,55 | 1,15 | 2,12 | 0,54 | 5,88E-01 | 7,47E-01 |
| AT4G26540 | 3,42 | -0,56 | 1,04 | -0,54 | 5,88E-01 | 7,47E-01 |
| AT3G13062 | 1,32 | -0,80 | 1,50 | -0,54 | 5,91E-01 | 7,49E-01 |
| AT4G06375 | 3,09 | -0,69 | 1,29 | -0,54 | 5,91E-01 | 7,49E-01 |
| AT5G14780 | 2,20 | 0,76 | 1,41 | 0,54 | 5,91E-01 | 7,49E-01 |
| AT5G48620 | 2,46 | -0,78 | 1,45 | -0,54 | 5,90E-01 | 7,49E-01 |
| AT3G11510 | 1,34 | -0,98 | 1,82 | -0,54 | 5,91E-01 | 7,49E-01 |
| AT4G25520 | 4,54 | -0,49 | 0,91 | -0,54 | 5,90E-01 | 7,49E-01 |
| AT5G52470 | 2,57 | -0,63 | 1,17 | -0,54 | 5,92E-01 | 7,50E-01 |
| AT4G40100 | 3,82 | 0,59 | 1,10 | 0,53 | 5,93E-01 | 7,50E-01 |
| AT5G10572 | 2,06 | 0,72 | 1,35 | 0,53 | 5,93E-01 | 7,50E-01 |
| AT1G33070 | 1,82 | 0,94 | 1,76 | 0,53 | 5,94E-01 | 7,52E-01 |
| AT4G33290 | 1,50 | 0,79 | 1,49 | 0,53 | 5,95E-01 | 7,53E-01 |
| AT1G63360 | 4,56 | 0,49 | 0,92 | 0,53 | 5,96E-01 | 7,53E-01 |
| AT1G71880 | 1,96 | 0,81 | 1,54 | 0,53 | 5,96E-01 | 7,53E-01 |
| AT4G26690 | 1,51 | 0,91 | 1,72 | 0,53 | 5,97E-01 | 7,54E-01 |
| AT3G21750 | 1,77 | 0,76 | 1,44 | 0,53 | 5,98E-01 | 7,55E-01 |
| AT3G19960 | 1,65 | -0,79 | 1,50 | -0,53 | 5,99E-01 | 7,56E-01 |
| AT4G35000 | 3,13 | -0,69 | 1,32 | -0,53 | 5,99E-01 | 7,56E-01 |
| AT3G57730 | 5,17 | -0,41 | 0,78 | -0,53 | 6,00E-01 | 7,56E-01 |
| AT5G58430 | 1,95 | -0,67 | 1,27 | -0,52 | 6,00E-01 | 7,56E-01 |
| AT3G56410 | 2,56 | 0,78 | 1,49 | 0,52 | 6,00E-01 | 7,57E-01 |

| | | | | | | |
|---|---|---|---|---|---|---|
| AT3G22440 | 1,87 | 0,77 | 1,47 | 0,52 | 6,01E-01 | 7,57E-01 |
| AT5G52070 | 6,76 | 0,55 | 1,05 | 0,52 | 6,01E-01 | 7,57E-01 |
| AT5G42530 | 4,10 | 0,49 | 0,93 | 0,52 | 6,02E-01 | 7,57E-01 |
| AT1G18160 | 2,26 | -0,81 | 1,55 | -0,52 | 6,02E-01 | 7,58E-01 |
| AT1G48410 | 43,79 | -0,19 | 0,36 | -0,52 | 6,03E-01 | 7,58E-01 |
| AT4G39330 | 1,70 | -0,66 | 1,28 | -0,52 | 6,04E-01 | 7,59E-01 |
| AT5G34850 | 1,64 | -0,77 | 1,48 | -0,52 | 6,04E-01 | 7,59E-01 |
| AT3G26570 | 1,54 | -0,93 | 1,81 | -0,51 | 6,07E-01 | 7,60E-01 |
| AT1G08510 | 1,46 | 0,89 | 1,72 | 0,52 | 6,06E-01 | 7,60E-01 |
| AT1G70290 | 2,53 | -0,59 | 1,15 | -0,51 | 6,07E-01 | 7,60E-01 |
| AT1G74456 | 11,08 | 0,32 | 0,62 | 0,52 | 6,06E-01 | 7,60E-01 |
| AT2G25450 | 4,94 | -0,46 | 0,90 | -0,52 | 6,05E-01 | 7,60E-01 |
| AT2G39675 | 2246,76 | 0,17 | 0,34 | 0,52 | 6,06E-01 | 7,60E-01 |
| AT4G02890 | 2,39 | -0,65 | 1,26 | -0,51 | 6,07E-01 | 7,60E-01 |
| AT5G19940 | 2,37 | -0,64 | 1,25 | -0,51 | 6,07E-01 | 7,60E-01 |
| AT3G29240 | 1,56 | -0,81 | 1,57 | -0,51 | 6,08E-01 | 7,60E-01 |
| AT4G39160 | 1,99 | -0,65 | 1,27 | -0,51 | 6,07E-01 | 7,60E-01 |
| AT3G14660 | 1,69 | 0,74 | 1,44 | 0,51 | 6,09E-01 | 7,61E-01 |
| AT1G18000 | 4,70 | 0,55 | 1,08 | 0,51 | 6,09E-01 | 7,61E-01 |
| AT1G44446 | 1,72 | -0,79 | 1,55 | -0,51 | 6,09E-01 | 7,61E-01 |
| AT3G30390 | 2,05 | -0,62 | 1,20 | -0,51 | 6,09E-01 | 7,61E-01 |
| AT5G00760 | 4,31 | -0,42 | 0,83 | -0,51 | 6,10E-01 | 7,61E-01 |
| AT5G06320 | 5,00 | 0,54 | 1,07 | 0,51 | 6,12E-01 | 7,63E-01 |
| AT3G55800 | 2,45 | -0,65 | 1,28 | -0,51 | 6,12E-01 | 7,64E-01 |
| AT3G15840 | 1,45 | -0,66 | 1,31 | -0,51 | 6,13E-01 | 7,65E-01 |
| AT2G22500 | 4,35 | 0,49 | 0,97 | 0,50 | 6,14E-01 | 7,65E-01 |
| AT5G38410 | 6,26 | 0,48 | 0,96 | 0,50 | 6,14E-01 | 7,65E-01 |
| AT2G01970 | 1,49 | -0,81 | 1,61 | -0,50 | 6,15E-01 | 7,66E-01 |
| AT1G66150 | 1,55 | -0,69 | 1,37 | -0,50 | 6,15E-01 | 7,66E-01 |
| AT1G72710 | 1,73 | -0,73 | 1,45 | -0,50 | 6,15E-01 | 7,66E-01 |
| AT3G16770 | 1,33 | 1,25 | 2,49 | 0,50 | 6,16E-01 | 7,66E-01 |
| AT5G38990 | 2,97 | -0,58 | 1,15 | -0,50 | 6,16E-01 | 7,66E-01 |
| AT2G32680 | 1,55 | 0,86 | 1,72 | 0,50 | 6,16E-01 | 7,66E-01 |
| AT1G43850 | 1,82 | -0,73 | 1,46 | -0,50 | 6,18E-01 | 7,68E-01 |
| AT4G24190 | 5,76 | 0,39 | 0,79 | 0,50 | 6,20E-01 | 7,69E-01 |
| AT1G76990 | 1,49 | 0,74 | 1,50 | 0,50 | 6,21E-01 | 7,70E-01 |
| AT5G21105 | 1,68 | 0,85 | 1,72 | 0,49 | 6,21E-01 | 7,70E-01 |
| AT5G21930 | 1,60 | -0,77 | 1,56 | -0,49 | 6,22E-01 | 7,71E-01 |
| AT5G42050 | 3,62 | 0,48 | 0,97 | 0,49 | 6,22E-01 | 7,71E-01 |
| AT3G23810 | 2,25 | -0,68 | 1,38 | -0,49 | 6,22E-01 | 7,71E-01 |
| AT4G21570 | 1,42 | 1,00 | 2,03 | 0,49 | 6,22E-01 | 7,71E-01 |
| AT5G67420 | 2,71 | -0,53 | 1,08 | -0,49 | 6,24E-01 | 7,73E-01 |
| AT1G19180 | 1,37 | 0,84 | 1,71 | 0,49 | 6,24E-01 | 7,73E-01 |
| AT3G27850 | 1,55 | -0,85 | 1,73 | -0,49 | 6,25E-01 | 7,73E-01 |
| AT5G40400 | 1,55 | 0,67 | 1,36 | 0,49 | 6,25E-01 | 7,73E-01 |
| AT1G49750 | 3,00 | 0,62 | 1,28 | 0,49 | 6,27E-01 | 7,75E-01 |
| AT4G02520 | 4,48 | 0,43 | 0,89 | 0,49 | 6,27E-01 | 7,75E-01 |
| AT5G32440 | 1,33 | -0,80 | 1,65 | -0,49 | 6,27E-01 | 7,75E-01 |
| AT5G40450 | 4,44 | -0,47 | 0,97 | -0,49 | 6,28E-01 | 7,75E-01 |
| AT5G52540 | 1,60 | -0,76 | 1,59 | -0,48 | 6,31E-01 | 7,79E-01 |
| AT1G30380 | 5,72 | 0,53 | 1,11 | 0,48 | 6,31E-01 | 7,79E-01 |
| AT4G31877 | 9,33 | 0,33 | 0,68 | 0,48 | 6,32E-01 | 7,79E-01 |
| AT1G27090 | 1,85 | -0,61 | 1,27 | -0,48 | 6,33E-01 | 7,80E-01 |
| AT5G24200 | 1,34 | -0,67 | 1,40 | -0,48 | 6,34E-01 | 7,81E-01 |
| AT3G46600 | 2,19 | -0,55 | 1,16 | -0,48 | 6,35E-01 | 7,81E-01 |
| AT3G48057 | 47,97 | 0,26 | 0,55 | 0,47 | 6,35E-01 | 7,81E-01 |

| | | | | | | |
|---|---|---|---|---|---|---|
| AT4G20360 | 5,64 | 0,40 | 0,84 | 0,47 | 6,35E-01 | 7,81E-01 |
| AT4G35310 | 1,84 | 0,79 | 1,66 | 0,47 | 6,35E-01 | 7,81E-01 |
| AT4G20362 | 5,64 | 0,40 | 0,84 | 0,47 | 6,35E-01 | 7,81E-01 |
| AT4G08155 | 1,40 | 0,68 | 1,45 | 0,47 | 6,37E-01 | 7,82E-01 |
| AT2G07890 | 2,64 | 0,56 | 1,18 | 0,47 | 6,37E-01 | 7,83E-01 |
| AT3G52850 | 1,76 | 0,80 | 1,70 | 0,47 | 6,37E-01 | 7,83E-01 |
| AT2G04955 | 3,52 | 0,44 | 0,94 | 0,47 | 6,37E-01 | 7,83E-01 |
| AT1G60072 | 5,42 | -0,41 | 0,87 | -0,47 | 6,38E-01 | 7,83E-01 |
| AT1G08117 | 4,98 | 0,52 | 1,10 | 0,47 | 6,38E-01 | 7,83E-01 |
| AT2G39990 | 1,46 | -1,24 | 2,64 | -0,47 | 6,39E-01 | 7,83E-01 |
| AT5G49730 | 3,55 | 0,63 | 1,33 | 0,47 | 6,38E-01 | 7,83E-01 |
| AT2G15970 | 3,79 | -0,56 | 1,20 | -0,47 | 6,40E-01 | 7,83E-01 |
| AT2G37480 | 1,35 | -0,98 | 2,09 | -0,47 | 6,39E-01 | 7,83E-01 |
| AT4G02715 | 1,63 | -1,01 | 2,16 | -0,47 | 6,40E-01 | 7,83E-01 |
| AT5G28824 | 1,47 | 0,85 | 1,83 | 0,47 | 6,40E-01 | 7,84E-01 |
| AT4G11380 | 1,75 | -0,87 | 1,87 | -0,47 | 6,41E-01 | 7,84E-01 |
| AT5G60680 | 1,59 | -0,78 | 1,67 | -0,47 | 6,41E-01 | 7,84E-01 |
| AT1G03743 | 2,10 | 0,65 | 1,40 | 0,47 | 6,41E-01 | 7,84E-01 |
| AT5G56030 | 5,79 | -0,34 | 0,74 | -0,46 | 6,43E-01 | 7,86E-01 |
| AT1G66730 | 1,91 | 0,66 | 1,42 | 0,46 | 6,45E-01 | 7,88E-01 |
| AT1G10150 | 1,47 | -0,66 | 1,44 | -0,46 | 6,46E-01 | 7,88E-01 |
| AT5G52471 | 1,40 | 0,82 | 1,79 | 0,46 | 6,46E-01 | 7,88E-01 |
| AT1G10155 | 1,47 | -0,66 | 1,44 | -0,46 | 6,46E-01 | 7,88E-01 |
| AT1G25490 | 1,64 | -0,64 | 1,41 | -0,46 | 6,47E-01 | 7,89E-01 |
| AT2G30440 | 1,74 | 0,64 | 1,40 | 0,46 | 6,48E-01 | 7,89E-01 |
| AT1G49500 | 1,83 | 0,61 | 1,33 | 0,46 | 6,49E-01 | 7,91E-01 |
| AT1G26880 | 2,18 | -0,66 | 1,46 | -0,45 | 6,50E-01 | 7,91E-01 |
| AT1G47128 | 7,64 | -0,30 | 0,66 | -0,45 | 6,50E-01 | 7,91E-01 |
| AT5G60920 | 1,88 | 0,67 | 1,47 | 0,45 | 6,51E-01 | 7,92E-01 |
| AT3G49520 | 1,37 | 0,74 | 1,63 | 0,45 | 6,52E-01 | 7,93E-01 |
| AT1G03730 | 1,35 | 1,14 | 2,53 | 0,45 | 6,52E-01 | 7,93E-01 |
| AT4G16860 | 6,11 | -0,37 | 0,83 | -0,45 | 6,52E-01 | 7,93E-01 |
| AT4G10320 | 1,43 | 0,68 | 1,52 | 0,45 | 6,53E-01 | 7,94E-01 |
| AT3G19240 | 1,33 | -0,62 | 1,39 | -0,45 | 6,55E-01 | 7,95E-01 |
| AT3G49120 | 5,10 | 0,41 | 0,92 | 0,45 | 6,55E-01 | 7,95E-01 |
| AT5G48190 | 1,46 | 0,85 | 1,90 | 0,45 | 6,55E-01 | 7,95E-01 |
| AT1G09687 | 2,04 | 0,58 | 1,31 | 0,45 | 6,56E-01 | 7,95E-01 |
| AT3G51730 | 1,38 | -0,74 | 1,67 | -0,44 | 6,56E-01 | 7,96E-01 |
| AT4G27520 | 5,05 | 0,46 | 1,02 | 0,44 | 6,56E-01 | 7,96E-01 |
| AT4G09000 | 1,74 | 0,58 | 1,31 | 0,44 | 6,57E-01 | 7,96E-01 |
| AT4G34620 | 1,51 | -0,86 | 1,93 | -0,44 | 6,57E-01 | 7,96E-01 |
| AT5G53560 | 1,33 | 0,77 | 1,73 | 0,44 | 6,57E-01 | 7,96E-01 |
| AT1G06430 | 1,82 | -0,73 | 1,65 | -0,44 | 6,58E-01 | 7,96E-01 |
| AT5G04670 | 1,32 | -0,91 | 2,05 | -0,44 | 6,58E-01 | 7,96E-01 |
| AT1G02150 | 1,38 | -0,74 | 1,67 | -0,44 | 6,60E-01 | 7,97E-01 |
| AT5G43760 | 1,93 | -0,66 | 1,51 | -0,44 | 6,60E-01 | 7,97E-01 |
| AT4G26130 | 1,45 | 0,85 | 1,94 | 0,44 | 6,61E-01 | 7,98E-01 |
| AT5G14570 | 2,87 | 0,53 | 1,21 | 0,44 | 6,61E-01 | 7,98E-01 |
| AT4G39363 | 6,83 | 0,33 | 0,75 | 0,44 | 6,62E-01 | 7,99E-01 |
| AT5G15970 | 2,85 | -0,65 | 1,49 | -0,44 | 6,62E-01 | 7,99E-01 |
| AT1G21750 | 2,14 | -0,70 | 1,62 | -0,44 | 6,63E-01 | 8,00E-01 |
| AT1G44080 | 1,35 | -0,87 | 2,00 | -0,44 | 6,63E-01 | 8,00E-01 |
| AT1G34260 | 1,42 | -0,87 | 2,02 | -0,43 | 6,65E-01 | 8,01E-01 |
| AT1G07510 | 1,74 | -0,73 | 1,69 | -0,43 | 6,66E-01 | 8,03E-01 |
| AT3G19010 | 1,81 | -0,65 | 1,52 | -0,43 | 6,69E-01 | 8,05E-01 |
| AT3G08285 | 1,59 | 0,62 | 1,45 | 0,43 | 6,69E-01 | 8,05E-01 |

| | | | | | | |
|---|---|---|---|---|---|---|
| AT1G72770 | 1,51 | -0,64 | 1,50 | -0,42 | 6,71E-01 | 8,07E-01 |
| AT3G05410 | 1,68 | 0,59 | 1,39 | 0,42 | 6,71E-01 | 8,07E-01 |
| AT1G04527 | 11,72 | -0,23 | 0,55 | -0,42 | 6,71E-01 | 8,07E-01 |
| AT3G56400 | 2,09 | 0,57 | 1,35 | 0,42 | 6,72E-01 | 8,07E-01 |
| AT1G07890 | 4,08 | -0,37 | 0,87 | -0,42 | 6,73E-01 | 8,09E-01 |
| AT1G76780 | 1,47 | -0,79 | 1,87 | -0,42 | 6,73E-01 | 8,09E-01 |
| AT1G55673 | 3,49 | -0,40 | 0,95 | -0,42 | 6,74E-01 | 8,09E-01 |
| AT3G50540 | 1,39 | -0,74 | 1,76 | -0,42 | 6,74E-01 | 8,09E-01 |
| AT4G36648 | 2,47 | -0,64 | 1,52 | -0,42 | 6,74E-01 | 8,09E-01 |
| AT1G18740 | 2,60 | 0,48 | 1,15 | 0,42 | 6,75E-01 | 8,09E-01 |
| AT1G07135 | 2,46 | 0,52 | 1,25 | 0,42 | 6,75E-01 | 8,09E-01 |
| AT1G74100 | 5,98 | 0,36 | 0,87 | 0,42 | 6,75E-01 | 8,09E-01 |
| AT5G24740 | 1,40 | -0,74 | 1,78 | -0,42 | 6,75E-01 | 8,09E-01 |
| AT5G67300 | 3,74 | 0,42 | 1,00 | 0,42 | 6,77E-01 | 8,10E-01 |
| AT4G04020 | 1,87 | 0,83 | 1,99 | 0,41 | 6,78E-01 | 8,12E-01 |
| AT1G47265 | 3,77 | 0,39 | 0,95 | 0,41 | 6,79E-01 | 8,12E-01 |
| AT1G64500 | 2,64 | 0,47 | 1,14 | 0,41 | 6,79E-01 | 8,12E-01 |
| AT5G10370 | 1,36 | -0,82 | 1,98 | -0,41 | 6,80E-01 | 8,12E-01 |
| AT1G36325 | 1,41 | -0,62 | 1,49 | -0,41 | 6,80E-01 | 8,12E-01 |
| AT5G09730 | 35,07 | 0,19 | 0,47 | 0,41 | 6,80E-01 | 8,12E-01 |
| AT5G11740 | 4,10 | -0,36 | 0,86 | -0,41 | 6,81E-01 | 8,13E-01 |
| AT3G09390 | 6,61 | -0,33 | 0,80 | -0,41 | 6,81E-01 | 8,13E-01 |
| AT1G33600 | 1,39 | 0,64 | 1,58 | 0,41 | 6,83E-01 | 8,15E-01 |
| AT5G52190 | 1,80 | -0,76 | 1,85 | -0,41 | 6,83E-01 | 8,15E-01 |
| AT4G12800 | 10,28 | 0,31 | 0,76 | 0,41 | 6,84E-01 | 8,15E-01 |
| AT1G74090 | 6,35 | -0,30 | 0,74 | -0,41 | 6,85E-01 | 8,16E-01 |
| AT4G38550 | 2,97 | 0,47 | 1,15 | 0,41 | 6,85E-01 | 8,16E-01 |
| AT4G38552 | 2,97 | 0,47 | 1,15 | 0,41 | 6,85E-01 | 8,16E-01 |
| AT2G36320 | 1,39 | 0,74 | 1,83 | 0,40 | 6,86E-01 | 8,17E-01 |
| AT2G47240 | 1,70 | 0,61 | 1,52 | 0,40 | 6,87E-01 | 8,17E-01 |
| AT4G05150 | 3,81 | 0,48 | 1,20 | 0,40 | 6,87E-01 | 8,17E-01 |
| AT5G01747 | 60,67 | 0,11 | 0,28 | 0,40 | 6,87E-01 | 8,17E-01 |
| AT2G18480 | 2,00 | 0,64 | 1,59 | 0,40 | 6,89E-01 | 8,18E-01 |
| AT4G13430 | 3,55 | -0,37 | 0,93 | -0,40 | 6,89E-01 | 8,19E-01 |
| AT5G64300 | 1,43 | 0,69 | 1,72 | 0,40 | 6,90E-01 | 8,19E-01 |
| AT3G50500 | 1,35 | 0,84 | 2,11 | 0,40 | 6,91E-01 | 8,20E-01 |
| AT1G78280 | 1,59 | -0,87 | 2,19 | -0,40 | 6,92E-01 | 8,21E-01 |
| AT1G11840 | 1,70 | 0,56 | 1,42 | 0,40 | 6,92E-01 | 8,21E-01 |
| AT1G02860 | 4,18 | -0,54 | 1,36 | -0,39 | 6,95E-01 | 8,23E-01 |
| AT2G11910 | 1,43 | -0,70 | 1,77 | -0,39 | 6,95E-01 | 8,23E-01 |
| AT1G23290 | 1,36 | 0,72 | 1,84 | 0,39 | 6,96E-01 | 8,24E-01 |
| AT1G34130 | 1,45 | -0,66 | 1,70 | -0,39 | 6,96E-01 | 8,24E-01 |
| AT4G00060 | 1,77 | -0,66 | 1,69 | -0,39 | 6,96E-01 | 8,24E-01 |
| AT1G29660 | 1,40 | 0,60 | 1,55 | 0,39 | 6,98E-01 | 8,25E-01 |
| AT4G13510 | 4,76 | 0,34 | 0,87 | 0,39 | 6,98E-01 | 8,25E-01 |
| AT2G21920 | 4,05 | -0,33 | 0,84 | -0,39 | 6,98E-01 | 8,25E-01 |
| AT2G02840 | 4,55 | 0,41 | 1,05 | 0,39 | 6,99E-01 | 8,26E-01 |
| AT1G07930 | 3,92 | -0,37 | 0,95 | -0,39 | 7,00E-01 | 8,26E-01 |
| AT4G30993 | 2,23 | 0,55 | 1,42 | 0,39 | 7,00E-01 | 8,26E-01 |
| AT5G08712 | 123,44 | -0,13 | 0,34 | -0,39 | 7,00E-01 | 8,26E-01 |
| AT2G13450 | 1,51 | -0,64 | 1,65 | -0,38 | 7,00E-01 | 8,26E-01 |
| AT5G25100 | 1,32 | -0,68 | 1,76 | -0,38 | 7,01E-01 | 8,26E-01 |
| AT4G23180 | 1,57 | -0,51 | 1,33 | -0,38 | 7,01E-01 | 8,26E-01 |
| AT5G61440 | 4,44 | -0,33 | 0,87 | -0,38 | 7,02E-01 | 8,27E-01 |
| AT3G13440 | 3,42 | 0,42 | 1,09 | 0,38 | 7,02E-01 | 8,28E-01 |
| AT4G34390 | 1,80 | -0,56 | 1,47 | -0,38 | 7,03E-01 | 8,28E-01 |

| | | | | | | |
|---|---|---|---|---|---|---|
| AT4G40030 | 1,32 | -0,65 | 1,71 | -0,38 | 7,03E-01 | 8,28E-01 |
| AT3G52590 | 1,74 | -0,52 | 1,38 | -0,38 | 7,05E-01 | 8,30E-01 |
| AT3G27906 | 1,35 | 0,61 | 1,64 | 0,38 | 7,07E-01 | 8,32E-01 |
| AT3G28900 | 3,87 | -0,35 | 0,94 | -0,38 | 7,08E-01 | 8,32E-01 |
| AT4G02700 | 1,40 | 0,65 | 1,73 | 0,37 | 7,08E-01 | 8,32E-01 |
| AT1G11890 | 1,50 | -0,65 | 1,75 | -0,37 | 7,09E-01 | 8,33E-01 |
| AT4G34220 | 1,59 | 0,58 | 1,55 | 0,37 | 7,10E-01 | 8,34E-01 |
| AT1G08200 | 1,82 | -0,55 | 1,49 | -0,37 | 7,11E-01 | 8,34E-01 |
| AT1G56340 | 4,49 | -0,32 | 0,87 | -0,37 | 7,11E-01 | 8,34E-01 |
| AT5G24890 | 2,39 | 0,43 | 1,17 | 0,37 | 7,11E-01 | 8,34E-01 |
| AT1G28270 | 2,06 | 0,52 | 1,41 | 0,37 | 7,12E-01 | 8,34E-01 |
| AT1G69620 | 2,14 | 0,50 | 1,35 | 0,37 | 7,12E-01 | 8,34E-01 |
| AT5G28540 | 7,23 | -0,26 | 0,70 | -0,37 | 7,12E-01 | 8,34E-01 |
| AT1G32790 | 1,57 | -0,79 | 2,15 | -0,37 | 7,14E-01 | 8,36E-01 |
| AT1G71010 | 1,50 | -0,62 | 1,70 | -0,37 | 7,14E-01 | 8,36E-01 |
| AT1G13470 | 1,76 | 0,56 | 1,53 | 0,37 | 7,14E-01 | 8,36E-01 |
| AT4G00430 | 2,16 | -0,49 | 1,34 | -0,36 | 7,17E-01 | 8,38E-01 |
| AT1G11260 | 1,39 | -0,63 | 1,75 | -0,36 | 7,17E-01 | 8,38E-01 |
| AT4G00360 | 2,92 | -0,40 | 1,11 | -0,36 | 7,17E-01 | 8,38E-01 |
| AT3G04210 | 3,60 | 0,39 | 1,07 | 0,36 | 7,17E-01 | 8,38E-01 |
| AT2G44060 | 2,21 | -0,44 | 1,21 | -0,36 | 7,20E-01 | 8,40E-01 |
| AT3G61110 | 1,40 | -0,66 | 1,83 | -0,36 | 7,19E-01 | 8,40E-01 |
| AT5G54270 | 15,64 | -0,18 | 0,49 | -0,36 | 7,19E-01 | 8,40E-01 |
| AT1G77960 | 1,37 | -0,56 | 1,56 | -0,36 | 7,20E-01 | 8,40E-01 |
| AT3G51600 | 1,48 | -0,56 | 1,56 | -0,36 | 7,20E-01 | 8,40E-01 |
| AT1G24510 | 1,77 | 0,51 | 1,43 | 0,36 | 7,21E-01 | 8,40E-01 |
| AT1G08115 | 40,91 | -0,15 | 0,41 | -0,36 | 7,21E-01 | 8,40E-01 |
| AT4G24415 | 10,68 | 0,23 | 0,65 | 0,36 | 7,22E-01 | 8,41E-01 |
| AT3G01500 | 4,00 | 0,41 | 1,15 | 0,36 | 7,22E-01 | 8,41E-01 |
| AT3G55850 | 2,66 | -0,44 | 1,25 | -0,36 | 7,22E-01 | 8,41E-01 |
| AT5G09495 | 4,86 | -0,34 | 0,95 | -0,36 | 7,22E-01 | 8,41E-01 |
| AT3G15920 | 8,54 | 0,24 | 0,67 | 0,35 | 7,23E-01 | 8,41E-01 |
| AT5G53740 | 20,74 | 0,22 | 0,62 | 0,35 | 7,24E-01 | 8,42E-01 |
| AT5G03350 | 2,45 | 0,45 | 1,27 | 0,35 | 7,24E-01 | 8,42E-01 |
| AT3G59410 | 1,72 | -0,51 | 1,45 | -0,35 | 7,25E-01 | 8,43E-01 |
| AT1G13930 | 7,00 | 0,28 | 0,80 | 0,35 | 7,29E-01 | 8,43E-01 |
| AT1G56510 | 2,98 | -0,43 | 1,25 | -0,35 | 7,30E-01 | 8,43E-01 |
| AT2G25520 | 1,73 | -0,62 | 1,79 | -0,34 | 7,30E-01 | 8,43E-01 |
| AT3G03475 | 2,58 | 0,45 | 1,29 | 0,35 | 7,30E-01 | 8,43E-01 |
| AT3G47620 | 3,09 | -0,37 | 1,07 | -0,35 | 7,27E-01 | 8,43E-01 |
| AT3G47920 | 1,60 | 0,60 | 1,74 | 0,34 | 7,30E-01 | 8,43E-01 |
| AT4G20980 | 1,36 | -0,69 | 2,00 | -0,34 | 7,30E-01 | 8,43E-01 |
| AT5G14200 | 1,42 | 0,54 | 1,57 | 0,35 | 7,29E-01 | 8,43E-01 |
| AT5G25460 | 1,53 | 0,71 | 2,03 | 0,35 | 7,27E-01 | 8,43E-01 |
| AT5G57655 | 1,67 | 0,59 | 1,69 | 0,35 | 7,28E-01 | 8,43E-01 |
| AT1G20450 | 3,27 | -0,41 | 1,17 | -0,35 | 7,27E-01 | 8,43E-01 |
| AT1G55670 | 5,12 | 0,29 | 0,83 | 0,35 | 7,30E-01 | 8,43E-01 |
| AT2G26190 | 1,38 | -0,56 | 1,61 | -0,35 | 7,26E-01 | 8,43E-01 |
| AT2G34585 | 1,42 | 0,57 | 1,63 | 0,35 | 7,29E-01 | 8,43E-01 |
| AT2G37710 | 4,76 | 0,34 | 0,98 | 0,35 | 7,29E-01 | 8,43E-01 |
| AT5G05485 | 5,35 | 0,41 | 1,18 | 0,35 | 7,28E-01 | 8,43E-01 |
| AT5G60640 | 2,58 | -0,44 | 1,26 | -0,35 | 7,27E-01 | 8,43E-01 |
| AT1G67840 | 1,96 | 0,55 | 1,61 | 0,34 | 7,32E-01 | 8,44E-01 |
| AT2G29630 | 5,51 | -0,26 | 0,77 | -0,34 | 7,32E-01 | 8,44E-01 |
| AT3G08720 | 1,37 | -0,67 | 1,97 | -0,34 | 7,32E-01 | 8,44E-01 |
| AT4G01290 | 1,74 | -0,53 | 1,55 | -0,34 | 7,32E-01 | 8,44E-01 |

| | | | | | | |
|---|---|---|---|---|---|---|
| AT3G24612 | 3,02 | 0,37 | 1,08 | 0,34 | 7,33E-01 | 8,44E-01 |
| AT4G15258 | 2,27 | 0,41 | 1,20 | 0,34 | 7,33E-01 | 8,44E-01 |
| AT2G23320 | 1,39 | -0,55 | 1,61 | -0,34 | 7,34E-01 | 8,45E-01 |
| AT5G39570 | 4,04 | 0,37 | 1,08 | 0,34 | 7,34E-01 | 8,45E-01 |
| AT1G07977 | 4,37 | -0,44 | 1,31 | -0,34 | 7,35E-01 | 8,46E-01 |
| AT2G26975 | 1,93 | 0,43 | 1,27 | 0,34 | 7,35E-01 | 8,46E-01 |
| AT5G44870 | 1,81 | 0,46 | 1,38 | 0,34 | 7,38E-01 | 8,47E-01 |
| AT1G20340 | 11,85 | -0,20 | 0,59 | -0,34 | 7,37E-01 | 8,47E-01 |
| AT1G78040 | 2,48 | 0,44 | 1,32 | 0,33 | 7,39E-01 | 8,49E-01 |
| AT1G08477 | 1,64 | 0,53 | 1,59 | 0,33 | 7,40E-01 | 8,50E-01 |
| AT4G05740 | 1,41 | 0,52 | 1,58 | 0,33 | 7,41E-01 | 8,50E-01 |
| AT3G58500 | 1,34 | 0,58 | 1,76 | 0,33 | 7,42E-01 | 8,51E-01 |
| AT3G13405 | 1,91 | 0,48 | 1,46 | 0,33 | 7,43E-01 | 8,52E-01 |
| AT3G08030 | 1,37 | -0,49 | 1,51 | -0,32 | 7,45E-01 | 8,53E-01 |
| AT3G03175 | 7,33 | 0,22 | 0,69 | 0,33 | 7,45E-01 | 8,53E-01 |
| AT1G13440 | 5,54 | -0,26 | 0,79 | -0,32 | 7,45E-01 | 8,53E-01 |
| AT1G66200 | 1,38 | -0,45 | 1,37 | -0,32 | 7,45E-01 | 8,53E-01 |
| AT4G13918 | 2,19 | 0,45 | 1,39 | 0,33 | 7,45E-01 | 8,53E-01 |
| AT1G08833 | 15,74 | 0,18 | 0,55 | 0,32 | 7,47E-01 | 8,54E-01 |
| AT5G47430 | 3,26 | -0,30 | 0,94 | -0,32 | 7,47E-01 | 8,55E-01 |
| AT4G13505 | 4,72 | 0,28 | 0,87 | 0,32 | 7,48E-01 | 8,55E-01 |
| AT4G17340 | 2,05 | 0,53 | 1,66 | 0,32 | 7,50E-01 | 8,56E-01 |
| AT4G21960 | 15,77 | -0,16 | 0,49 | -0,32 | 7,50E-01 | 8,56E-01 |
| AT1G32270 | 1,96 | 0,52 | 1,64 | 0,32 | 7,50E-01 | 8,57E-01 |
| AT5G01600 | 1,52 | -0,47 | 1,47 | -0,32 | 7,51E-01 | 8,58E-01 |
| AT2G33830 | 1,76 | -0,48 | 1,51 | -0,32 | 7,52E-01 | 8,59E-01 |
| AT3G50930 | 3,98 | 0,29 | 0,93 | 0,31 | 7,54E-01 | 8,60E-01 |
| AT2G01020 | 7030,11 | 0,11 | 0,35 | 0,31 | 7,55E-01 | 8,60E-01 |
| AT4G32940 | 4,72 | 0,25 | 0,81 | 0,31 | 7,55E-01 | 8,60E-01 |
| AT2G24280 | 1,45 | -0,51 | 1,65 | -0,31 | 7,55E-01 | 8,60E-01 |
| AT4G02500 | 1,37 | -0,90 | 2,89 | -0,31 | 7,55E-01 | 8,60E-01 |
| AT4G04125 | 2,22 | -0,40 | 1,29 | -0,31 | 7,57E-01 | 8,62E-01 |
| AT4G11840 | 1,34 | -0,53 | 1,72 | -0,31 | 7,59E-01 | 8,63E-01 |
| AT4G37930 | 4,20 | -0,29 | 0,95 | -0,31 | 7,58E-01 | 8,63E-01 |
| AT5G20490 | 1,59 | 0,59 | 1,91 | 0,31 | 7,59E-01 | 8,63E-01 |
| AT1G22930 | 1,70 | -0,49 | 1,61 | -0,30 | 7,62E-01 | 8,65E-01 |
| AT2G03890 | 2,03 | 0,39 | 1,29 | 0,30 | 7,61E-01 | 8,65E-01 |
| AT3G14420 | 4,42 | 0,31 | 1,02 | 0,30 | 7,62E-01 | 8,65E-01 |
| AT3G04155 | 3,42 | 0,32 | 1,04 | 0,30 | 7,62E-01 | 8,65E-01 |
| AT4G11600 | 2,32 | 0,42 | 1,39 | 0,30 | 7,61E-01 | 8,65E-01 |
| AT1G03130 | 1,99 | 0,39 | 1,29 | 0,30 | 7,63E-01 | 8,66E-01 |
| AT3G06510 | 3,56 | -0,34 | 1,13 | -0,30 | 7,65E-01 | 8,68E-01 |
| AT4G04640 | 5,86 | 0,24 | 0,81 | 0,29 | 7,70E-01 | 8,73E-01 |
| AT1G12013 | 2,08 | 0,39 | 1,36 | 0,29 | 7,71E-01 | 8,73E-01 |
| AT1G20015 | 7,84 | 0,23 | 0,78 | 0,29 | 7,71E-01 | 8,73E-01 |
| AT1G67430 | 1,82 | 0,40 | 1,38 | 0,29 | 7,72E-01 | 8,73E-01 |
| AT5G43603 | 44,77 | -0,12 | 0,41 | -0,29 | 7,70E-01 | 8,73E-01 |
| AT2G18160 | 1,66 | -0,43 | 1,46 | -0,29 | 7,71E-01 | 8,73E-01 |
| AT5G19150 | 6,35 | 0,20 | 0,71 | 0,29 | 7,72E-01 | 8,73E-01 |
| AT5G19140 | 6,43 | 0,20 | 0,70 | 0,29 | 7,72E-01 | 8,73E-01 |
| AT1G31173 | 488,28 | 0,07 | 0,24 | 0,29 | 7,74E-01 | 8,74E-01 |
| AT5G37830 | 1,35 | 0,62 | 2,14 | 0,29 | 7,74E-01 | 8,74E-01 |
| AT4G28570 | 3,15 | 0,32 | 1,12 | 0,29 | 7,74E-01 | 8,75E-01 |
| AT1G61224 | 1,62 | -0,45 | 1,57 | -0,29 | 7,75E-01 | 8,75E-01 |
| AT1G64680 | 1,48 | -0,52 | 1,82 | -0,29 | 7,75E-01 | 8,75E-01 |
| AT4G08850 | 2,82 | -0,37 | 1,28 | -0,29 | 7,75E-01 | 8,75E-01 |

| | | | | | | |
|---|---|---|---|---|---|---|
| AT4G34350 | 1,55 | -0,49 | 1,70 | -0,29 | 7,75E-01 | 8,75E-01 |
| AT3G23820 | 9,86 | -0,16 | 0,55 | -0,28 | 7,77E-01 | 8,76E-01 |
| AT3G57700 | 1,50 | 0,42 | 1,51 | 0,28 | 7,80E-01 | 8,79E-01 |
| AT1G05273 | 8,42 | 0,17 | 0,63 | 0,28 | 7,82E-01 | 8,79E-01 |
| AT2G07718 | 3,36 | 0,28 | 1,01 | 0,28 | 7,81E-01 | 8,79E-01 |
| AT2G38230 | 6,88 | 0,19 | 0,70 | 0,28 | 7,81E-01 | 8,79E-01 |
| AT1G16820 | 8,42 | 0,17 | 0,63 | 0,28 | 7,82E-01 | 8,79E-01 |
| AT3G52400 | 1,57 | 0,39 | 1,42 | 0,28 | 7,82E-01 | 8,79E-01 |
| AT3G03105 | 2,11 | 0,34 | 1,24 | 0,28 | 7,82E-01 | 8,80E-01 |
| AT1G20260 | 2,71 | -0,34 | 1,25 | -0,28 | 7,83E-01 | 8,80E-01 |
| AT1G68720 | 1,50 | -0,62 | 2,24 | -0,28 | 7,83E-01 | 8,80E-01 |
| AT1G63070 | 3,60 | -0,40 | 1,46 | -0,27 | 7,84E-01 | 8,80E-01 |
| AT5G01410 | 2,12 | 0,42 | 1,52 | 0,27 | 7,84E-01 | 8,81E-01 |
| AT5G15420 | 1,66 | 0,45 | 1,65 | 0,27 | 7,85E-01 | 8,81E-01 |
| AT2G26355 | 2,39 | -0,35 | 1,30 | -0,27 | 7,85E-01 | 8,81E-01 |
| AT4G26970 | 1,79 | -0,48 | 1,75 | -0,27 | 7,86E-01 | 8,81E-01 |
| AT2G46340 | 1,90 | -0,51 | 1,87 | -0,27 | 7,86E-01 | 8,81E-01 |
| AT3G61820 | 1,36 | -0,45 | 1,68 | -0,27 | 7,86E-01 | 8,81E-01 |
| AT5G59420 | 1,43 | -0,51 | 1,89 | -0,27 | 7,88E-01 | 8,83E-01 |
| AT1G25275 | 1,34 | 0,46 | 1,74 | 0,27 | 7,89E-01 | 8,84E-01 |
| AT2G41090 | 3,05 | 0,29 | 1,10 | 0,27 | 7,90E-01 | 8,84E-01 |
| AT4G35300 | 1,43 | -0,55 | 2,07 | -0,27 | 7,90E-01 | 8,84E-01 |
| AT5G64040 | 3,44 | 0,29 | 1,10 | 0,27 | 7,90E-01 | 8,84E-01 |
| AT3G14415 | 1,72 | 0,39 | 1,47 | 0,27 | 7,91E-01 | 8,84E-01 |
| AT4G04970 | 2,63 | 0,53 | 1,99 | 0,26 | 7,91E-01 | 8,84E-01 |
| AT2G09640 | 1,68 | -0,56 | 2,09 | -0,27 | 7,91E-01 | 8,84E-01 |
| AT5G22780 | 1,47 | -0,38 | 1,44 | -0,26 | 7,91E-01 | 8,84E-01 |
| AT1G15670 | 2,03 | -0,32 | 1,20 | -0,26 | 7,92E-01 | 8,84E-01 |
| AT5G26000 | 4,62 | -0,22 | 0,83 | -0,26 | 7,93E-01 | 8,85E-01 |
| AT1G33240 | 1,91 | -0,37 | 1,46 | -0,26 | 7,97E-01 | 8,89E-01 |
| AT3G27700 | 1,39 | 0,52 | 2,04 | 0,26 | 7,98E-01 | 8,90E-01 |
| AT3G60240 | 1,46 | 0,39 | 1,52 | 0,25 | 7,99E-01 | 8,91E-01 |
| AT1G22770 | 1,61 | -0,32 | 1,27 | -0,25 | 8,01E-01 | 8,91E-01 |
| AT1G71820 | 1,53 | 0,39 | 1,55 | 0,25 | 8,01E-01 | 8,91E-01 |
| AT3G60960 | 1,54 | -0,39 | 1,55 | -0,25 | 8,01E-01 | 8,91E-01 |
| AT4G31800 | 1,69 | 0,39 | 1,55 | 0,25 | 8,00E-01 | 8,91E-01 |
| AT5G02565 | 2,69 | 0,35 | 1,38 | 0,25 | 8,01E-01 | 8,91E-01 |
| AT1G62310 | 4,32 | 0,22 | 0,86 | 0,25 | 8,01E-01 | 8,91E-01 |
| AT3G03435 | 3,28 | 0,29 | 1,13 | 0,25 | 8,01E-01 | 8,91E-01 |
| AT3G59400 | 1,33 | -0,38 | 1,49 | -0,25 | 7,99E-01 | 8,91E-01 |
| AT5G08280 | 1,65 | 0,47 | 1,87 | 0,25 | 8,02E-01 | 8,91E-01 |
| AT3G15980 | 1,59 | -0,34 | 1,35 | -0,25 | 8,02E-01 | 8,92E-01 |
| AT4G22890 | 1,50 | -0,42 | 1,70 | -0,25 | 8,03E-01 | 8,92E-01 |
| AT1G14790 | 1,42 | -0,55 | 2,21 | -0,25 | 8,03E-01 | 8,92E-01 |
| AT2G02835 | 3,94 | 0,22 | 0,88 | 0,25 | 8,05E-01 | 8,94E-01 |
| AT3G45640 | 1,79 | 0,33 | 1,35 | 0,24 | 8,07E-01 | 8,94E-01 |
| AT4G25630 | 1,57 | 0,42 | 1,73 | 0,25 | 8,06E-01 | 8,94E-01 |
| AT1G11910 | 5,12 | -0,25 | 1,04 | -0,24 | 8,08E-01 | 8,94E-01 |
| AT1G42990 | 1,44 | 0,37 | 1,52 | 0,24 | 8,07E-01 | 8,94E-01 |
| AT1G52110 | 2,50 | -0,33 | 1,37 | -0,24 | 8,08E-01 | 8,94E-01 |
| AT2G45380 | 11,33 | 0,13 | 0,55 | 0,24 | 8,07E-01 | 8,94E-01 |
| AT3G12780 | 5,73 | -0,19 | 0,79 | -0,25 | 8,06E-01 | 8,94E-01 |
| AT5G35970 | 1,60 | 0,37 | 1,51 | 0,24 | 8,07E-01 | 8,94E-01 |
| AT5G45100 | 1,72 | -0,46 | 1,90 | -0,24 | 8,10E-01 | 8,96E-01 |
| AT5G65860 | 5,04 | -0,23 | 0,94 | -0,24 | 8,10E-01 | 8,96E-01 |
| AT1G66410 | 1,68 | -0,36 | 1,50 | -0,24 | 8,11E-01 | 8,97E-01 |

| | | | | | | |
|---|---|---|---|---|---|---|
| AT3G01370 | 2,22 | 0,31 | 1,32 | 0,24 | 8,12E-01 | 8,97E-01 |
| AT3G48115 | 1,91 | 0,35 | 1,46 | 0,24 | 8,12E-01 | 8,97E-01 |
| AT5G66190 | 3,89 | 0,27 | 1,13 | 0,24 | 8,13E-01 | 8,97E-01 |
| AT4G27720 | 1,38 | 0,41 | 1,74 | 0,23 | 8,15E-01 | 8,99E-01 |
| AT2G23348 | 3,88 | -0,20 | 0,87 | -0,23 | 8,16E-01 | 9,00E-01 |
| AT5G59732 | 3,71 | 0,25 | 1,08 | 0,23 | 8,18E-01 | 9,02E-01 |
| AT5G63145 | 2,83 | 0,27 | 1,19 | 0,23 | 8,18E-01 | 9,02E-01 |
| AT1G50250 | 2,35 | -0,28 | 1,21 | -0,23 | 8,18E-01 | 9,02E-01 |
| AT5G59730 | 3,71 | 0,25 | 1,08 | 0,23 | 8,18E-01 | 9,02E-01 |
| AT3G56880 | 1,69 | -0,32 | 1,39 | -0,23 | 8,21E-01 | 9,03E-01 |
| AT4G11270 | 3,79 | -0,24 | 1,06 | -0,23 | 8,20E-01 | 9,03E-01 |
| AT5G27950 | 2,87 | 0,32 | 1,43 | 0,23 | 8,20E-01 | 9,03E-01 |
| AT5G44050 | 1,57 | 0,51 | 2,27 | 0,23 | 8,20E-01 | 9,03E-01 |
| AT1G66970 | 4,60 | -0,20 | 0,89 | -0,23 | 8,20E-01 | 9,03E-01 |
| AT1G59980 | 3,55 | -0,23 | 1,01 | -0,23 | 8,21E-01 | 9,03E-01 |
| AT1G62560 | 1,49 | 0,67 | 2,98 | 0,23 | 8,22E-01 | 9,03E-01 |
| AT1G74250 | 1,42 | 0,39 | 1,75 | 0,22 | 8,22E-01 | 9,03E-01 |
| AT1G03080 | 1,73 | -0,33 | 1,45 | -0,22 | 8,22E-01 | 9,03E-01 |
| AT2G45245 | 2,00 | -0,26 | 1,16 | -0,22 | 8,22E-01 | 9,03E-01 |
| AT2G24180 | 1,36 | -0,59 | 2,68 | -0,22 | 8,25E-01 | 9,05E-01 |
| AT5G24490 | 1,61 | 0,47 | 2,13 | 0,22 | 8,25E-01 | 9,05E-01 |
| AT1G21130 | 2,17 | 0,31 | 1,39 | 0,22 | 8,26E-01 | 9,06E-01 |
| AT2G01021 | 3194,91 | 0,06 | 0,27 | 0,22 | 8,27E-01 | 9,07E-01 |
| AT1G26558 | 2,72 | 0,26 | 1,21 | 0,22 | 8,29E-01 | 9,09E-01 |
| AT1G63090 | 1,34 | 0,35 | 1,66 | 0,21 | 8,31E-01 | 9,10E-01 |
| AT2G39260 | 1,36 | -0,34 | 1,60 | -0,21 | 8,31E-01 | 9,10E-01 |
| AT4G01070 | 1,43 | 0,37 | 1,74 | 0,21 | 8,31E-01 | 9,10E-01 |
| AT5G02380 | 2,77 | 0,31 | 1,46 | 0,21 | 8,31E-01 | 9,10E-01 |
| AT4G31500 | 6,34 | -0,15 | 0,72 | -0,21 | 8,32E-01 | 9,10E-01 |
| AT3G09445 | 4,99 | -0,18 | 0,85 | -0,21 | 8,33E-01 | 9,11E-01 |
| AT1G09240 | 1,84 | 0,45 | 2,13 | 0,21 | 8,34E-01 | 9,11E-01 |
| AT4G21670 | 1,62 | 0,29 | 1,41 | 0,21 | 8,34E-01 | 9,11E-01 |
| AT1G12900 | 4,70 | -0,18 | 0,83 | -0,21 | 8,34E-01 | 9,11E-01 |
| AT1G66795 | 14,00 | 0,12 | 0,59 | 0,21 | 8,35E-01 | 9,11E-01 |
| AT3G24542 | 2,63 | -0,25 | 1,21 | -0,21 | 8,35E-01 | 9,11E-01 |
| AT5G60600 | 2,43 | 0,25 | 1,20 | 0,21 | 8,35E-01 | 9,11E-01 |
| AT4G11850 | 1,56 | -0,34 | 1,62 | -0,21 | 8,36E-01 | 9,11E-01 |
| AT4G18215 | 1,61 | 0,33 | 1,59 | 0,21 | 8,36E-01 | 9,11E-01 |
| AT3G45140 | 15,78 | -0,11 | 0,53 | -0,21 | 8,37E-01 | 9,12E-01 |
| AT4G27090 | 2,08 | -0,32 | 1,56 | -0,21 | 8,37E-01 | 9,12E-01 |
| AT3G46610 | 1,41 | -0,33 | 1,61 | -0,20 | 8,38E-01 | 9,12E-01 |
| AT4G34980 | 4,56 | -0,16 | 0,77 | -0,20 | 8,40E-01 | 9,15E-01 |
| AT4G39050 | 1,56 | 0,33 | 1,66 | 0,20 | 8,41E-01 | 9,15E-01 |
| AT3G61190 | 1,32 | -0,34 | 1,69 | -0,20 | 8,42E-01 | 9,16E-01 |
| AT4G12040 | 1,45 | -0,37 | 1,85 | -0,20 | 8,43E-01 | 9,16E-01 |
| AT5G07875 | 4,76 | 0,18 | 0,91 | 0,20 | 8,43E-01 | 9,16E-01 |
| AT1G05850 | 2,93 | 0,28 | 1,40 | 0,20 | 8,43E-01 | 9,16E-01 |
| AT2G05812 | 1,46 | 0,30 | 1,54 | 0,19 | 8,46E-01 | 9,19E-01 |
| AT2G19480 | 1,78 | -0,32 | 1,66 | -0,19 | 8,46E-01 | 9,19E-01 |
| AT5G08180 | 2,30 | 0,20 | 1,05 | 0,19 | 8,46E-01 | 9,19E-01 |
| AT3G54810 | 2,87 | 0,23 | 1,18 | 0,19 | 8,47E-01 | 9,19E-01 |
| AT2G16490 | 1,37 | 0,44 | 2,29 | 0,19 | 8,48E-01 | 9,19E-01 |
| AT5G09065 | 4,44 | 0,16 | 0,85 | 0,19 | 8,49E-01 | 9,20E-01 |
| AT2G17200 | 1,89 | -0,31 | 1,61 | -0,19 | 8,49E-01 | 9,20E-01 |
| AT4G39400 | 3,54 | -0,19 | 1,02 | -0,19 | 8,50E-01 | 9,21E-01 |
| AT3G57040 | 1,37 | 0,31 | 1,67 | 0,19 | 8,51E-01 | 9,21E-01 |

| | | | | | | |
|---|---|---|---|---|---|---|
| AT4G21060 | 1,98 | -0,25 | 1,36 | -0,19 | 8,53E-01 | 9,23E-01 |
| AT1G70890 | 1,35 | 0,33 | 1,78 | 0,19 | 8,53E-01 | 9,23E-01 |
| AT2G10940 | 5,77 | 0,20 | 1,07 | 0,18 | 8,54E-01 | 9,24E-01 |
| AT3G44340 | 1,56 | -0,30 | 1,60 | -0,18 | 8,54E-01 | 9,24E-01 |
| AT2G18020 | 2,64 | -0,25 | 1,41 | -0,18 | 8,57E-01 | 9,26E-01 |
| AT2G23672 | 1,50 | -0,32 | 1,75 | -0,18 | 8,57E-01 | 9,26E-01 |
| AT2G41590 | 1,66 | -0,28 | 1,54 | -0,18 | 8,57E-01 | 9,26E-01 |
| AT2G23670 | 1,50 | -0,32 | 1,75 | -0,18 | 8,57E-01 | 9,26E-01 |
| AT5G40340 | 2,24 | 0,23 | 1,28 | 0,18 | 8,57E-01 | 9,26E-01 |
| AT2G06510 | 1,80 | 0,26 | 1,45 | 0,18 | 8,59E-01 | 9,26E-01 |
| AT5G13180 | 1,85 | -0,25 | 1,41 | -0,18 | 8,58E-01 | 9,26E-01 |
| AT4G39090 | 7,03 | -0,14 | 0,81 | -0,18 | 8,58E-01 | 9,26E-01 |
| AT2G08390 | 5,34 | 0,16 | 0,93 | 0,18 | 8,60E-01 | 9,27E-01 |
| AT3G13100 | 2,83 | -0,28 | 1,59 | -0,18 | 8,60E-01 | 9,27E-01 |
| AT5G36228 | 1,95 | -0,27 | 1,54 | -0,18 | 8,61E-01 | 9,28E-01 |
| AT4G25960 | 1,53 | 0,28 | 1,60 | 0,17 | 8,62E-01 | 9,28E-01 |
| AT5G27970 | 1,51 | -0,39 | 2,25 | -0,17 | 8,63E-01 | 9,29E-01 |
| AT1G78020 | 1,89 | 0,21 | 1,21 | 0,17 | 8,64E-01 | 9,30E-01 |
| AT5G19120 | 2,51 | -0,18 | 1,06 | -0,17 | 8,64E-01 | 9,30E-01 |
| AT2G07725 | 2,14 | -0,21 | 1,24 | -0,17 | 8,65E-01 | 9,31E-01 |
| AT1G58300 | 1,33 | -0,29 | 1,75 | -0,17 | 8,66E-01 | 9,31E-01 |
| AT5G02645 | 2,33 | -0,19 | 1,14 | -0,17 | 8,66E-01 | 9,31E-01 |
| AT1G52780 | 3,16 | 0,22 | 1,32 | 0,17 | 8,68E-01 | 9,32E-01 |
| AT4G34870 | 1,35 | -0,25 | 1,52 | -0,17 | 8,68E-01 | 9,32E-01 |
| AT3G03470 | 1,68 | 0,30 | 1,78 | 0,17 | 8,68E-01 | 9,32E-01 |
| AT2G45470 | 1,65 | -0,30 | 1,82 | -0,17 | 8,69E-01 | 9,32E-01 |
| AT4G30210 | 1,86 | 0,27 | 1,62 | 0,16 | 8,69E-01 | 9,33E-01 |
| AT1G04410 | 3,62 | 0,19 | 1,15 | 0,16 | 8,70E-01 | 9,33E-01 |
| AT2G25170 | 3,04 | 0,20 | 1,26 | 0,16 | 8,71E-01 | 9,34E-01 |
| AT2G46130 | 1,45 | -0,26 | 1,65 | -0,16 | 8,73E-01 | 9,35E-01 |
| AT5G22060 | 1,34 | 0,37 | 2,29 | 0,16 | 8,73E-01 | 9,35E-01 |
| AT3G19910 | 1,61 | 0,25 | 1,58 | 0,16 | 8,73E-01 | 9,36E-01 |
| AT1G05010 | 2,06 | 0,25 | 1,57 | 0,16 | 8,74E-01 | 9,36E-01 |
| AT1G73600 | 1,37 | 0,32 | 2,07 | 0,16 | 8,75E-01 | 9,36E-01 |
| AT2G14080 | 4,62 | -0,18 | 1,12 | -0,16 | 8,75E-01 | 9,36E-01 |
| AT4G01150 | 2,10 | -0,19 | 1,21 | -0,16 | 8,77E-01 | 9,36E-01 |
| AT2G41410 | 4,84 | -0,16 | 1,02 | -0,16 | 8,76E-01 | 9,36E-01 |
| AT3G47500 | 2,57 | 0,19 | 1,20 | 0,16 | 8,76E-01 | 9,36E-01 |
| AT4G21380 | 1,64 | -0,26 | 1,69 | -0,16 | 8,76E-01 | 9,36E-01 |
| AT5G24810 | 2,74 | 0,24 | 1,55 | 0,16 | 8,77E-01 | 9,36E-01 |
| AT1G44575 | 5,52 | 0,12 | 0,77 | 0,15 | 8,79E-01 | 9,37E-01 |
| AT1G52400 | 1,99 | 0,27 | 1,75 | 0,15 | 8,79E-01 | 9,37E-01 |
| AT1G02780 | 1,83 | -0,19 | 1,24 | -0,15 | 8,79E-01 | 9,37E-01 |
| AT1G67740 | 3,89 | -0,16 | 1,04 | -0,15 | 8,79E-01 | 9,37E-01 |
| AT2G45160 | 10,70 | 0,10 | 0,67 | 0,15 | 8,79E-01 | 9,37E-01 |
| AT5G03145 | 2,30 | 0,23 | 1,54 | 0,15 | 8,80E-01 | 9,37E-01 |
| AT1G15002 | 1,82 | 0,21 | 1,37 | 0,15 | 8,80E-01 | 9,37E-01 |
| AT5G28919 | 2,64 | 0,18 | 1,19 | 0,15 | 8,82E-01 | 9,39E-01 |
| AT5G04905 | 2,64 | 0,18 | 1,19 | 0,15 | 8,82E-01 | 9,39E-01 |
| AT3G16240 | 1,39 | 0,22 | 1,51 | 0,15 | 8,82E-01 | 9,39E-01 |
| AT1G27752 | 1,43 | 0,33 | 2,21 | 0,15 | 8,82E-01 | 9,39E-01 |
| AT2G42620 | 1,45 | -0,35 | 2,44 | -0,14 | 8,86E-01 | 9,40E-01 |
| AT3G61050 | 1,51 | -0,24 | 1,69 | -0,14 | 8,86E-01 | 9,40E-01 |
| AT4G02555 | 3,06 | 0,16 | 1,14 | 0,14 | 8,86E-01 | 9,40E-01 |
| AT1G28480 | 1,91 | 0,29 | 1,99 | 0,14 | 8,85E-01 | 9,40E-01 |
| AT1G65720 | 3,55 | -0,14 | 0,99 | -0,15 | 8,84E-01 | 9,40E-01 |

| | | | | | | |
|---|---|---|---|---|---|---|
| AT1G73805 | 1,81 | -0,22 | 1,52 | -0,14 | 8,85E-01 | 9,40E-01 |
| AT3G60630 | 5,65 | 0,11 | 0,76 | 0,15 | 8,85E-01 | 9,40E-01 |
| AT4G24770 | 3,23 | -0,17 | 1,13 | -0,15 | 8,84E-01 | 9,40E-01 |
| AT4G39260 | 6,03 | 0,11 | 0,77 | 0,14 | 8,87E-01 | 9,40E-01 |
| AT5G42930 | 2,68 | 0,16 | 1,12 | 0,15 | 8,85E-01 | 9,40E-01 |
| AT5G59550 | 1,55 | -0,23 | 1,61 | -0,14 | 8,87E-01 | 9,40E-01 |
| AT4G30600 | 1,60 | -0,21 | 1,50 | -0,14 | 8,87E-01 | 9,40E-01 |
| AT2G38800 | 2,71 | -0,20 | 1,44 | -0,14 | 8,88E-01 | 9,40E-01 |
| AT1G06033 | 1,64 | -0,24 | 1,70 | -0,14 | 8,89E-01 | 9,41E-01 |
| AT1G76090 | 1,72 | -0,20 | 1,44 | -0,14 | 8,89E-01 | 9,41E-01 |
| AT4G36220 | 1,38 | -0,21 | 1,50 | -0,14 | 8,90E-01 | 9,42E-01 |
| AT5G43500 | 1,47 | 0,26 | 1,90 | 0,14 | 8,91E-01 | 9,42E-01 |
| AT1G09310 | 8,77 | -0,10 | 0,74 | -0,13 | 8,93E-01 | 9,44E-01 |
| AT5G48380 | 2,92 | -0,15 | 1,10 | -0,13 | 8,93E-01 | 9,44E-01 |
| AT5G61410 | 2,49 | 0,15 | 1,13 | 0,13 | 8,94E-01 | 9,44E-01 |
| AT1G06597 | 1,64 | -0,21 | 1,58 | -0,13 | 8,95E-01 | 9,45E-01 |
| AT5G20250 | 2,34 | 0,18 | 1,36 | 0,13 | 8,95E-01 | 9,45E-01 |
| AT3G60690 | 1,35 | 0,23 | 1,75 | 0,13 | 8,96E-01 | 9,45E-01 |
| AT1G29670 | 1,73 | 0,21 | 1,58 | 0,13 | 8,97E-01 | 9,46E-01 |
| AT2G22990 | 2,03 | 0,15 | 1,15 | 0,13 | 8,97E-01 | 9,46E-01 |
| AT1G05817 | 1,94 | -0,15 | 1,18 | -0,13 | 8,97E-01 | 9,46E-01 |
| AT1G17290 | 2,38 | 0,16 | 1,22 | 0,13 | 8,99E-01 | 9,47E-01 |
| AT1G77235 | 1,77 | 0,22 | 1,70 | 0,13 | 8,99E-01 | 9,47E-01 |
| AT2G18560 | 1,54 | 0,30 | 2,44 | 0,12 | 9,01E-01 | 9,48E-01 |
| AT4G17600 | 1,35 | 0,22 | 1,77 | 0,12 | 9,01E-01 | 9,48E-01 |
| AT1G23300 | 1,57 | 0,21 | 1,66 | 0,12 | 9,02E-01 | 9,48E-01 |
| AT3G11010 | 3,23 | 0,18 | 1,42 | 0,12 | 9,02E-01 | 9,48E-01 |
| AT3G13410 | 1,64 | 0,26 | 2,08 | 0,12 | 9,02E-01 | 9,48E-01 |
| AT3G14310 | 2,07 | 0,22 | 1,78 | 0,13 | 9,00E-01 | 9,48E-01 |
| AT5G25610 | 6,58 | -0,10 | 0,82 | -0,12 | 9,02E-01 | 9,48E-01 |
| AT1G13260 | 1,59 | -0,21 | 1,71 | -0,12 | 9,04E-01 | 9,50E-01 |
| AT1G56220 | 1,48 | 0,19 | 1,56 | 0,12 | 9,05E-01 | 9,50E-01 |
| AT3G06080 | 1,38 | 0,18 | 1,50 | 0,12 | 9,05E-01 | 9,50E-01 |
| AT5G24530 | 1,77 | -0,16 | 1,32 | -0,12 | 9,06E-01 | 9,51E-01 |
| AT4G23630 | 1,89 | -0,15 | 1,26 | -0,12 | 9,08E-01 | 9,52E-01 |
| AT3G26450 | 2,27 | 0,15 | 1,32 | 0,11 | 9,09E-01 | 9,53E-01 |
| AT1G78070 | 1,66 | 0,18 | 1,60 | 0,11 | 9,10E-01 | 9,53E-01 |
| AT3G26730 | 1,96 | 0,18 | 1,55 | 0,11 | 9,09E-01 | 9,53E-01 |
| AT4G14510 | 1,36 | -0,23 | 2,02 | -0,11 | 9,10E-01 | 9,53E-01 |
| AT5G49720 | 3,89 | -0,11 | 0,98 | -0,11 | 9,10E-01 | 9,53E-01 |
| AT1G07793 | 2,22 | 0,16 | 1,46 | 0,11 | 9,16E-01 | 9,56E-01 |
| AT3G03680 | 1,60 | -0,17 | 1,62 | -0,11 | 9,15E-01 | 9,56E-01 |
| AT5G04410 | 1,77 | -0,23 | 2,20 | -0,11 | 9,16E-01 | 9,56E-01 |
| AT1G47960 | 1,43 | 0,17 | 1,55 | 0,11 | 9,14E-01 | 9,56E-01 |
| AT1G75380 | 2,78 | -0,12 | 1,08 | -0,11 | 9,14E-01 | 9,56E-01 |
| AT2G16510 | 1,63 | -0,20 | 1,86 | -0,11 | 9,15E-01 | 9,56E-01 |
| AT2G27310 | 2,22 | 0,13 | 1,22 | 0,11 | 9,14E-01 | 9,56E-01 |
| AT2G37220 | 3,40 | 0,12 | 1,12 | 0,11 | 9,15E-01 | 9,56E-01 |
| AT3G05985 | 2,35 | -0,14 | 1,34 | -0,11 | 9,16E-01 | 9,56E-01 |
| AT5G13887 | 13,72 | 0,09 | 0,86 | 0,11 | 9,16E-01 | 9,56E-01 |
| AT5G61910 | 1,60 | 0,17 | 1,60 | 0,11 | 9,14E-01 | 9,56E-01 |
| AT2G43141 | 1,64 | -0,18 | 1,77 | -0,10 | 9,18E-01 | 9,57E-01 |
| AT5G35840 | 1,35 | -0,24 | 2,31 | -0,10 | 9,18E-01 | 9,57E-01 |
| AT5G28920 | 2,84 | -0,12 | 1,16 | -0,10 | 9,18E-01 | 9,57E-01 |
| AT4G13880 | 2,00 | -0,15 | 1,49 | -0,10 | 9,19E-01 | 9,57E-01 |
| AT5G20230 | 2,06 | -0,15 | 1,45 | -0,10 | 9,19E-01 | 9,57E-01 |

| | | | | | | |
|---|---|---|---|---|---|---|
| AT5G41750 | 8,05 | -0,07 | 0,64 | -0,10 | 9,19E-01 | 9,57E-01 |
| AT1G51980 | 1,85 | 0,15 | 1,49 | 0,10 | 9,20E-01 | 9,57E-01 |
| AT2G43137 | 1,43 | 0,19 | 1,88 | 0,10 | 9,20E-01 | 9,57E-01 |
| AT5G58950 | 2,42 | -0,14 | 1,36 | -0,10 | 9,20E-01 | 9,57E-01 |
| AT5G41471 | 1,79 | 0,17 | 1,77 | 0,10 | 9,22E-01 | 9,58E-01 |
| AT2G34110 | 1,88 | 0,13 | 1,40 | 0,09 | 9,25E-01 | 9,61E-01 |
| AT2G35260 | 3,89 | -0,09 | 1,00 | -0,09 | 9,25E-01 | 9,61E-01 |
| AT5G10340 | 1,40 | -0,19 | 1,98 | -0,09 | 9,25E-01 | 9,61E-01 |
| AT2G40270 | 1,34 | 0,15 | 1,64 | 0,09 | 9,29E-01 | 9,65E-01 |
| AT1G68010 | 2,53 | 0,12 | 1,34 | 0,09 | 9,30E-01 | 9,65E-01 |
| AT5G05505 | 4,00 | -0,11 | 1,28 | -0,08 | 9,32E-01 | 9,68E-01 |
| AT4G12720 | 1,54 | 0,13 | 1,54 | 0,08 | 9,33E-01 | 9,68E-01 |
| AT4G32250 | 2,07 | -0,12 | 1,46 | -0,08 | 9,33E-01 | 9,68E-01 |
| AT3G28920 | 1,35 | 0,15 | 1,83 | 0,08 | 9,34E-01 | 9,68E-01 |
| AT5G18490 | 1,80 | -0,15 | 1,83 | -0,08 | 9,34E-01 | 9,68E-01 |
| AT1G68440 | 1,32 | -0,12 | 1,52 | -0,08 | 9,35E-01 | 9,69E-01 |
| AT2G23347 | 3,51 | 0,07 | 0,90 | 0,08 | 9,35E-01 | 9,69E-01 |
| AT2G07671 | 2,57 | 0,11 | 1,41 | 0,08 | 9,36E-01 | 9,69E-01 |
| AT2G13560 | 2,19 | -0,13 | 1,63 | -0,08 | 9,37E-01 | 9,70E-01 |
| AT2G43110 | 4,82 | 0,07 | 0,92 | 0,08 | 9,38E-01 | 9,70E-01 |
| AT3G46510 | 1,83 | 0,10 | 1,39 | 0,07 | 9,40E-01 | 9,73E-01 |
| AT1G51805 | 2,63 | 0,10 | 1,32 | 0,07 | 9,42E-01 | 9,74E-01 |
| AT1G78380 | 1,48 | 0,12 | 1,68 | 0,07 | 9,42E-01 | 9,74E-01 |
| AT2G26440 | 1,40 | -0,15 | 2,07 | -0,07 | 9,43E-01 | 9,74E-01 |
| AT5G45110 | 1,72 | 0,10 | 1,42 | 0,07 | 9,44E-01 | 9,75E-01 |
| AT5G13890 | 14,03 | 0,05 | 0,75 | 0,07 | 9,44E-01 | 9,75E-01 |
| AT1G06410 | 1,61 | -0,10 | 1,53 | -0,07 | 9,46E-01 | 9,76E-01 |
| AT3G27690 | 1,98 | 0,10 | 1,43 | 0,07 | 9,46E-01 | 9,76E-01 |
| AT3G44310 | 10,52 | -0,05 | 0,69 | -0,07 | 9,47E-01 | 9,76E-01 |
| AT4G25631 | 1,33 | 0,13 | 1,94 | 0,07 | 9,47E-01 | 9,76E-01 |
| AT5G25754 | 1,48 | 0,10 | 1,46 | 0,07 | 9,47E-01 | 9,76E-01 |
| AT4G14365 | 1,73 | 0,11 | 1,60 | 0,07 | 9,46E-01 | 9,76E-01 |
| AT1G11860 | 2,12 | 0,09 | 1,37 | 0,07 | 9,48E-01 | 9,76E-01 |
| AT5G24270 | 1,64 | 0,09 | 1,41 | 0,06 | 9,49E-01 | 9,77E-01 |
| AT3G29320 | 2,29 | 0,09 | 1,42 | 0,06 | 9,51E-01 | 9,77E-01 |
| AT5G21090 | 1,39 | 0,11 | 1,72 | 0,06 | 9,50E-01 | 9,77E-01 |
| AT3G03415 | 1,34 | -0,11 | 1,71 | -0,06 | 9,51E-01 | 9,77E-01 |
| AT4G03585 | 1,51 | 0,10 | 1,58 | 0,06 | 9,51E-01 | 9,77E-01 |
| AT4G29900 | 4,72 | 0,05 | 0,82 | 0,06 | 9,52E-01 | 9,78E-01 |
| AT1G32582 | 26,60 | 0,03 | 0,57 | 0,06 | 9,53E-01 | 9,79E-01 |
| AT2G01140 | 1,55 | 0,10 | 1,78 | 0,06 | 9,53E-01 | 9,79E-01 |
| AT2G41720 | 1,87 | 0,10 | 1,64 | 0,06 | 9,54E-01 | 9,79E-01 |
| AT4G13840 | 1,76 | 0,10 | 1,64 | 0,06 | 9,53E-01 | 9,79E-01 |
| AT1G01720 | 1,37 | -0,08 | 1,44 | -0,06 | 9,55E-01 | 9,79E-01 |
| AT2G27600 | 1,39 | -0,10 | 1,86 | -0,06 | 9,55E-01 | 9,79E-01 |
| AT4G01340 | 1,48 | -0,10 | 1,85 | -0,06 | 9,55E-01 | 9,79E-01 |
| AT3G62410 | 1,88 | -0,07 | 1,35 | -0,06 | 9,56E-01 | 9,79E-01 |
| AT5G16150 | 1,33 | -0,09 | 1,64 | -0,06 | 9,56E-01 | 9,79E-01 |
| AT1G66570 | 1,67 | -0,08 | 1,40 | -0,06 | 9,56E-01 | 9,79E-01 |
| AT2G16660 | 2,17 | -0,06 | 1,15 | -0,05 | 9,57E-01 | 9,79E-01 |
| AT2G08370 | 4,11 | -0,05 | 0,87 | -0,05 | 9,58E-01 | 9,80E-01 |
| AT4G10210 | 2,08 | -0,07 | 1,36 | -0,05 | 9,58E-01 | 9,80E-01 |
| AT5G47210 | 1,48 | -0,10 | 1,89 | -0,05 | 9,58E-01 | 9,80E-01 |
| AT4G35090 | 2,83 | 0,06 | 1,07 | 0,05 | 9,59E-01 | 9,81E-01 |
| AT5G03340 | 1,55 | -0,10 | 1,86 | -0,05 | 9,59E-01 | 9,81E-01 |
| AT4G38460 | 1,41 | -0,08 | 1,55 | -0,05 | 9,60E-01 | 9,81E-01 |

| | | | | | | |
|---|---|---|---|---|---|---|
| AT3G19560 | 1,58 | 0,09 | 1,90 | 0,05 | 9,60E-01 | 9,81E-01 |
| AT4G33050 | 2,86 | -0,05 | 1,07 | -0,05 | 9,60E-01 | 9,81E-01 |
| AT5G57560 | 1,92 | 0,06 | 1,21 | 0,05 | 9,61E-01 | 9,81E-01 |
| AT3G63540 | 1,40 | 0,08 | 1,65 | 0,05 | 9,63E-01 | 9,82E-01 |
| AT5G18410 | 3,15 | -0,05 | 1,00 | -0,05 | 9,63E-01 | 9,82E-01 |
| AT4G26542 | 2,99 | -0,05 | 1,11 | -0,04 | 9,66E-01 | 9,85E-01 |
| AT1G32583 | 28,27 | 0,02 | 0,55 | 0,04 | 9,66E-01 | 9,85E-01 |
| AT5G42650 | 3,24 | 0,05 | 1,16 | 0,04 | 9,67E-01 | 9,86E-01 |
| AT2G06520 | 10,63 | 0,03 | 0,81 | 0,04 | 9,68E-01 | 9,86E-01 |
| AT2G46820 | 2,21 | 0,05 | 1,32 | 0,04 | 9,68E-01 | 9,86E-01 |
| AT2G30860 | 6,26 | 0,03 | 0,75 | 0,04 | 9,69E-01 | 9,86E-01 |
| AT4G05180 | 7,70 | -0,03 | 0,84 | -0,04 | 9,69E-01 | 9,86E-01 |
| AT1G79110 | 1,67 | 0,06 | 1,52 | 0,04 | 9,70E-01 | 9,87E-01 |
| AT2G02870 | 1,53 | 0,07 | 1,80 | 0,04 | 9,70E-01 | 9,87E-01 |
| AT5G08525 | 4,34 | 0,04 | 1,09 | 0,04 | 9,71E-01 | 9,87E-01 |
| AT1G07233 | 2,51 | 0,04 | 1,07 | 0,03 | 9,73E-01 | 9,89E-01 |
| AT4G37260 | 2,49 | -0,04 | 1,27 | -0,03 | 9,73E-01 | 9,89E-01 |
| AT4G01010 | 2,44 | 0,04 | 1,31 | 0,03 | 9,73E-01 | 9,89E-01 |
| AT1G59910 | 1,37 | 0,06 | 1,91 | 0,03 | 9,75E-01 | 9,90E-01 |
| AT1G22410 | 1,37 | -0,04 | 1,45 | -0,03 | 9,80E-01 | 9,90E-01 |
| AT2G04405 | 3,40 | 0,03 | 1,11 | 0,03 | 9,77E-01 | 9,90E-01 |
| AT3G11560 | 1,70 | 0,05 | 1,87 | 0,03 | 9,78E-01 | 9,90E-01 |
| AT3G26440 | 2,47 | -0,03 | 1,29 | -0,03 | 9,79E-01 | 9,90E-01 |
| AT4G03280 | 3,09 | 0,03 | 1,12 | 0,03 | 9,80E-01 | 9,90E-01 |
| AT4G03560 | 2,24 | -0,04 | 1,47 | -0,03 | 9,77E-01 | 9,90E-01 |
| AT4G12290 | 3,10 | 0,03 | 1,24 | 0,03 | 9,80E-01 | 9,90E-01 |
| AT4G39730 | 3,06 | 0,04 | 1,33 | 0,03 | 9,78E-01 | 9,90E-01 |
| AT5G19690 | 1,74 | 0,04 | 1,41 | 0,03 | 9,79E-01 | 9,90E-01 |
| AT5G42895 | 1,80 | 0,04 | 1,43 | 0,03 | 9,80E-01 | 9,90E-01 |
| AT1G23390 | 2,44 | -0,04 | 1,27 | -0,03 | 9,78E-01 | 9,90E-01 |
| AT1G07393 | 1,67 | -0,04 | 1,68 | -0,02 | 9,81E-01 | 9,90E-01 |
| AT1G48970 | 1,37 | 0,05 | 1,99 | 0,03 | 9,80E-01 | 9,90E-01 |
| AT2G26900 | 1,34 | -0,05 | 1,78 | -0,03 | 9,79E-01 | 9,90E-01 |
| AT2G33770 | 4,11 | 0,02 | 0,91 | 0,03 | 9,79E-01 | 9,90E-01 |
| AT4G14880 | 1,32 | 0,04 | 1,50 | 0,02 | 9,80E-01 | 9,90E-01 |
| AT5G58070 | 2,01 | -0,04 | 1,53 | -0,03 | 9,77E-01 | 9,90E-01 |
| AT1G19960 | 1,88 | -0,05 | 1,94 | -0,02 | 9,81E-01 | 9,90E-01 |
| AT1G62180 | 2,66 | 0,03 | 1,20 | 0,02 | 9,82E-01 | 9,90E-01 |
| AT2G17510 | 1,71 | -0,03 | 1,54 | -0,02 | 9,82E-01 | 9,91E-01 |
| AT5G58120 | 1,53 | 0,03 | 1,49 | 0,02 | 9,84E-01 | 9,92E-01 |
| AT1G57680 | 1,45 | 0,04 | 2,01 | 0,02 | 9,84E-01 | 9,92E-01 |
| AT2G43570 | 2,25 | -0,03 | 1,61 | -0,02 | 9,86E-01 | 9,93E-01 |
| AT2G13360 | 3,44 | 0,02 | 0,91 | 0,02 | 9,86E-01 | 9,94E-01 |
| AT1G65960 | 3,84 | -0,02 | 1,24 | -0,02 | 9,87E-01 | 9,94E-01 |
| AT4G04770 | 1,41 | -0,03 | 1,87 | -0,02 | 9,88E-01 | 9,94E-01 |
| AT4G17390 | 6,32 | 0,01 | 0,82 | 0,01 | 9,89E-01 | 9,95E-01 |
| AT2G20410 | 6,12 | 0,01 | 0,89 | 0,01 | 9,90E-01 | 9,96E-01 |
| AT5G12250 | 2,28 | 0,01 | 1,25 | 0,01 | 9,91E-01 | 9,96E-01 |
| AT3G57720 | 13,95 | -0,01 | 0,62 | -0,01 | 9,92E-01 | 9,97E-01 |
| AT2G47470 | 1,77 | -0,02 | 1,77 | -0,01 | 9,93E-01 | 9,98E-01 |
| AT2G16676 | 2,63 | -0,01 | 1,23 | -0,01 | 9,93E-01 | 9,98E-01 |
| AT3G18830 | 2,46 | -0,01 | 1,32 | -0,01 | 9,93E-01 | 9,98E-01 |
| AT4G18880 | 2,25 | 0,01 | 1,55 | 0,01 | 9,94E-01 | 9,98E-01 |
| AT5G11670 | 3,41 | -0,01 | 0,99 | -0,01 | 9,94E-01 | 9,98E-01 |
| AT4G23850 | 2,43 | -0,01 | 1,13 | -0,01 | 9,95E-01 | 9,98E-01 |
| AT4G32390 | 1,34 | -0,01 | 1,93 | -0,01 | 9,96E-01 | 9,99E-01 |

| | | | | | | |
|---|---|---|---|---|---|---|
| AT2G25510 | 7,52 | 0,00 | 0,78 | 0,00 | 9,97E-01 | 9,99E-01 |
| AT5G18520 | 1,46 | -0,01 | 1,61 | 0,00 | 9,97E-01 | 9,99E-01 |
| AT1G52230 | 2,95 | 0,00 | 1,03 | 0,00 | 9,99E-01 | 1,00E+00 |
| AT5G57710 | 4,28 | 0,00 | 0,88 | 0,00 | 1,00E+00 | 1,00E+00 |
| AT1G08857 | 2,42 | 0,00 | 1,86 | 0,00 | 9,99E-01 | 1,00E+00 |
| AT1G69792 | 2,23 | 0,00 | 2,11 | 0,00 | 1,00E+00 | 1,00E+00 |
| AT1G69795 | 1,39 | 0,00 | 2,56 | 0,00 | 1,00E+00 | 1,00E+00 |
| AT2G34480 | 1,66 | 0,00 | 1,35 | 0,00 | 9,99E-01 | 1,00E+00 |
| AT5G24280 | 2,00 | 0,00 | 3,20 | 0,00 | 1,00E+00 | 1,00E+00 |
| AT1G01160 | 0,43 | 0,64 | 3,20 | 0,20 | 8,43E-01 | |
| AT1G01180 | 0,85 | 1,13 | 2,36 | 0,48 | 6,34E-01 | |
| AT1G01220 | 0,11 | -0,68 | 4,41 | -0,15 | 8,77E-01 | |
| AT1G01225 | 0,31 | -2,15 | 4,35 | -0,49 | 6,21E-01 | |
| AT1G01230 | 0,37 | -2,13 | 4,37 | -0,49 | 6,26E-01 | |
| AT1G01240 | 0,19 | 0,75 | 4,34 | 0,17 | 8,62E-01 | |
| AT1G01260 | 0,53 | -1,00 | 2,75 | -0,36 | 7,16E-01 | |
| AT1G01300 | 1,08 | 2,65 | 1,93 | 1,37 | 1,70E-01 | |
| AT1G01350 | 0,29 | -1,77 | 4,39 | -0,40 | 6,87E-01 | |
| AT1G01448 | 0,59 | 0,00 | 4,41 | 0,00 | 1,00E+00 | |
| AT1G01480 | 0,12 | 0,00 | 4,41 | 0,00 | 1,00E+00 | |
| AT1G01520 | 0,12 | 0,00 | 4,41 | 0,00 | 1,00E+00 | |
| AT1G01540 | 0,22 | 0,38 | 4,35 | 0,09 | 9,31E-01 | |
| AT1G01630 | 0,58 | -2,89 | 3,77 | -0,77 | 4,44E-01 | |
| AT1G01710 | 0,82 | -1,16 | 3,07 | -0,38 | 7,07E-01 | |
| AT1G04047 | 0,70 | 1,86 | 2,49 | 0,75 | 4,56E-01 | |
| AT1G01730 | 0,40 | -3,36 | 4,38 | -0,77 | 4,43E-01 | |
| AT1G01790 | 0,83 | -0,99 | 1,87 | -0,53 | 5,96E-01 | |
| AT1G01810 | 0,12 | 0,00 | 4,41 | 0,00 | 1,00E+00 | |
| AT1G01950 | 0,18 | 0,38 | 4,35 | 0,09 | 9,30E-01 | |
| AT1G02010 | 0,10 | -0,18 | 4,38 | -0,04 | 9,67E-01 | |
| AT1G02100 | 0,49 | 0,48 | 3,19 | 0,15 | 8,79E-01 | |
| AT1G02120 | 0,36 | 0,07 | 3,50 | 0,02 | 9,84E-01 | |
| AT1G02205 | 1,06 | 1,91 | 2,00 | 0,96 | 3,38E-01 | |
| AT1G04093 | 0,08 | -0,21 | 4,39 | -0,05 | 9,62E-01 | |
| AT1G02300 | 0,34 | -1,00 | 4,38 | -0,23 | 8,20E-01 | |
| AT1G02305 | 0,80 | -1,96 | 2,54 | -0,77 | 4,41E-01 | |
| AT1G04103 | 0,08 | 0,00 | 4,41 | 0,00 | 1,00E+00 | |
| AT1G02390 | 0,19 | 0,39 | 4,35 | 0,09 | 9,29E-01 | |
| AT1G02420 | 0,52 | -2,59 | 3,96 | -0,65 | 5,13E-01 | |
| AT1G02690 | 0,11 | 0,46 | 4,35 | 0,11 | 9,15E-01 | |
| AT1G02750 | 0,51 | -4,07 | 4,33 | -0,94 | 3,48E-01 | |
| AT1G02816 | 0,38 | -0,52 | 3,37 | -0,16 | 8,77E-01 | |
| AT1G02850 | 0,62 | -0,71 | 3,03 | -0,24 | 8,14E-01 | |
| AT1G03160 | 0,95 | -0,80 | 2,17 | -0,37 | 7,10E-01 | |
| AT1G03190 | 0,39 | -0,73 | 3,47 | -0,21 | 8,33E-01 | |
| AT1G03220 | 1,11 | -0,33 | 2,02 | -0,16 | 8,71E-01 | |
| AT1G03280 | 0,25 | -0,73 | 4,40 | -0,16 | 8,69E-01 | |
| AT1G03290 | 0,75 | 1,46 | 2,42 | 0,60 | 5,46E-01 | |
| AT1G03310 | 0,40 | -1,89 | 4,24 | -0,44 | 6,56E-01 | |
| AT1G03360 | 0,35 | -0,42 | 3,49 | -0,12 | 9,05E-01 | |
| AT1G03365 | 0,40 | -1,69 | 3,45 | -0,49 | 6,25E-01 | |
| AT1G03370 | 0,51 | 1,34 | 3,95 | 0,34 | 7,35E-01 | |
| AT1G03380 | 1,12 | -0,68 | 2,02 | -0,34 | 7,37E-01 | |
| AT1G03430 | 0,37 | -1,21 | 4,36 | -0,28 | 7,81E-01 | |
| AT1G03440 | 0,84 | -1,17 | 3,09 | -0,38 | 7,06E-01 | |
| AT1G03545 | 0,70 | -1,63 | 2,24 | -0,73 | 4,68E-01 | |

| | | | | | |
|---|---|---|---|---|---|
| AT1G03610 | 0,72 | 0,77 | 3,20 | 0,24 | 8,10E-01 |
| AT1G03630 | 0,51 | 0,72 | 2,28 | 0,32 | 7,51E-01 |
| AT1G03687 | 0,11 | 0,46 | 4,35 | 0,11 | 9,15E-01 |
| AT1G03740 | 0,96 | 1,41 | 1,96 | 0,72 | 4,71E-01 |
| AT1G03810 | 0,77 | -0,55 | 1,98 | -0,28 | 7,81E-01 |
| AT1G03900 | 1,11 | 0,25 | 2,02 | 0,12 | 9,02E-01 |
| AT1G03905 | 0,24 | -3,18 | 4,39 | -0,73 | 4,68E-01 |
| AT1G03910 | 0,50 | -0,79 | 3,09 | -0,26 | 7,97E-01 |
| AT1G03930 | 0,53 | -2,01 | 2,68 | -0,75 | 4,53E-01 |
| AT1G04217 | 0,82 | -0,69 | 1,98 | -0,35 | 7,29E-01 |
| AT1G03935 | 0,16 | -0,01 | 4,37 | 0,00 | 9,98E-01 |
| AT1G04223 | 0,98 | 1,07 | 1,87 | 0,57 | 5,67E-01 |
| AT1G04227 | 0,32 | -0,02 | 3,67 | -0,01 | 9,95E-01 |
| AT1G03960 | 0,15 | 0,00 | 4,41 | 0,00 | 1,00E+00 |
| AT1G03970 | 0,36 | 1,54 | 4,30 | 0,36 | 7,20E-01 |
| AT1G04020 | 0,33 | -3,14 | 4,39 | -0,72 | 4,74E-01 |
| AT1G04130 | 0,50 | 0,75 | 2,60 | 0,29 | 7,71E-01 |
| AT1G04170 | 1,02 | -3,58 | 2,05 | -1,75 | 8,07E-02 |
| AT1G04247 | 0,30 | -0,40 | 4,34 | -0,09 | 9,26E-01 |
| AT1G04230 | 0,70 | -0,11 | 3,04 | -0,04 | 9,72E-01 |
| AT1G04295 | 0,43 | -0,02 | 2,75 | -0,01 | 9,96E-01 |
| AT1G04350 | 0,25 | 0,44 | 4,36 | 0,10 | 9,20E-01 |
| AT1G04420 | 1,20 | -1,65 | 1,72 | -0,96 | 3,39E-01 |
| AT1G04440 | 0,50 | -0,56 | 2,63 | -0,21 | 8,31E-01 |
| AT1G04273 | 0,37 | -0,41 | 3,45 | -0,12 | 9,06E-01 |
| AT1G04277 | 0,93 | -1,22 | 2,05 | -0,60 | 5,51E-01 |
| AT1G04510 | 0,67 | -2,82 | 2,99 | -0,94 | 3,46E-01 |
| AT1G04540 | 1,05 | -2,94 | 2,16 | -1,36 | 1,74E-01 |
| AT1G04690 | 0,77 | 0,02 | 2,35 | 0,01 | 9,92E-01 |
| AT1G04710 | 0,47 | -3,16 | 4,39 | -0,72 | 4,72E-01 |
| AT1G04760 | 0,36 | 0,11 | 4,37 | 0,02 | 9,80E-01 |
| AT1G04945 | 0,47 | -2,46 | 3,22 | -0,77 | 4,44E-01 |
| AT1G04970 | 0,34 | 0,00 | 4,41 | 0,00 | 1,00E+00 |
| AT1G05087 | 0,44 | -0,71 | 4,14 | -0,17 | 8,63E-01 |
| AT1G05090 | 0,12 | 0,00 | 4,41 | 0,00 | 1,00E+00 |
| AT1G05140 | 0,22 | -0,56 | 4,35 | -0,13 | 8,98E-01 |
| AT1G05200 | 1,12 | -1,68 | 1,97 | -0,85 | 3,94E-01 |
| AT1G05360 | 0,08 | 0,00 | 4,41 | 0,00 | 1,00E+00 |
| AT1G05500 | 0,37 | 0,67 | 4,34 | 0,15 | 8,78E-01 |
| AT1G04397 | 0,08 | 0,00 | 4,41 | 0,00 | 1,00E+00 |
| AT1G05600 | 0,08 | -0,21 | 4,39 | -0,05 | 9,62E-01 |
| AT1G04423 | 1,18 | 1,18 | 1,83 | 0,64 | 5,20E-01 |
| AT1G05790 | 0,76 | -2,47 | 2,56 | -0,97 | 3,34E-01 |
| AT1G05830 | 0,53 | -0,34 | 3,31 | -0,10 | 9,18E-01 |
| AT1G05890 | 0,68 | -4,21 | 4,32 | -0,97 | 3,30E-01 |
| AT1G05900 | 0,21 | -1,17 | 4,36 | -0,27 | 7,88E-01 |
| AT1G05910 | 0,59 | -4,08 | 3,77 | -1,08 | 2,79E-01 |
| AT1G05980 | 0,23 | 0,56 | 4,35 | 0,13 | 8,98E-01 |
| AT1G06002 | 0,95 | -1,20 | 1,91 | -0,63 | 5,28E-01 |
| AT1G06030 | 0,96 | -3,96 | 3,36 | -1,18 | 2,38E-01 |
| AT1G06290 | 0,60 | -1,21 | 2,73 | -0,44 | 6,57E-01 |
| AT1G06380 | 0,45 | -0,28 | 4,17 | -0,07 | 9,47E-01 |
| AT1G06390 | 0,40 | -1,71 | 3,38 | -0,51 | 6,12E-01 |
| AT1G06530 | 0,96 | -3,77 | 2,07 | -1,82 | 6,85E-02 |
| AT1G06640 | 1,12 | 0,66 | 1,71 | 0,39 | 7,00E-01 |
| AT1G06645 | 0,50 | -1,21 | 4,03 | -0,30 | 7,63E-01 |

| | | | | | |
|---|---|---|---|---|---|
| AT1G06650 | 0,69 | -1,54 | 2,95 | -0,52 | 6,02E-01 |
| AT1G06750 | 0,24 | -2,54 | 4,40 | -0,58 | 5,64E-01 |
| AT1G06780 | 0,12 | -0,14 | 4,38 | -0,03 | 9,74E-01 |
| AT1G06910 | 0,23 | -1,73 | 4,39 | -0,39 | 6,93E-01 |
| AT1G06930 | 0,08 | 0,00 | 4,41 | 0,00 | 1,00E+00 |
| AT1G07010 | 0,86 | -1,69 | 2,83 | -0,60 | 5,50E-01 |
| AT1G04537 | 0,24 | -3,18 | 4,39 | -0,73 | 4,68E-01 |
| AT1G07080 | 0,86 | -0,12 | 2,13 | -0,06 | 9,54E-01 |
| AT1G07130 | 0,28 | 0,18 | 4,37 | 0,04 | 9,67E-01 |
| AT1G07170 | 0,78 | -0,73 | 2,26 | -0,32 | 7,46E-01 |
| AT1G07180 | 0,37 | -0,10 | 3,47 | -0,03 | 9,78E-01 |
| AT1G07210 | 0,17 | 0,00 | 4,41 | 0,00 | 1,00E+00 |
| AT1G07240 | 0,59 | -1,52 | 3,23 | -0,47 | 6,37E-01 |
| AT1G07470 | 0,61 | -4,16 | 3,73 | -1,11 | 2,65E-01 |
| AT1G07630 | 0,70 | -0,92 | 2,85 | -0,32 | 7,46E-01 |
| AT1G07705 | 0,12 | -0,68 | 4,41 | -0,15 | 8,77E-01 |
| AT1G07728 | 1,03 | -5,02 | 2,48 | -2,02 | 4,29E-02 |
| AT1G07790 | 0,25 | 0,16 | 4,37 | 0,04 | 9,70E-01 |
| AT1G07830 | 0,43 | 0,07 | 4,19 | 0,02 | 9,86E-01 |
| AT1G07990 | 0,78 | -0,56 | 3,00 | -0,19 | 8,51E-01 |
| AT1G08110 | 0,41 | -2,29 | 4,32 | -0,53 | 5,96E-01 |
| AT1G08190 | 0,34 | 1,19 | 3,55 | 0,34 | 7,36E-01 |
| AT1G08260 | 0,51 | 0,82 | 4,09 | 0,20 | 8,42E-01 |
| AT1G08370 | 0,71 | -2,45 | 2,56 | -0,96 | 3,37E-01 |
| AT1G08410 | 0,87 | -2,21 | 2,50 | -0,88 | 3,77E-01 |
| AT1G08460 | 0,84 | 0,26 | 2,08 | 0,12 | 9,01E-01 |
| AT1G08465 | 0,35 | -0,99 | 4,38 | -0,23 | 8,21E-01 |
| AT1G08480 | 0,59 | -0,11 | 3,24 | -0,03 | 9,73E-01 |
| AT1G08520 | 1,05 | -2,13 | 1,71 | -1,24 | 2,13E-01 |
| AT1G08530 | 0,48 | -0,68 | 4,07 | -0,17 | 8,67E-01 |
| AT1G08550 | 0,85 | -0,95 | 2,41 | -0,39 | 6,94E-01 |
| AT1G08570 | 0,49 | -0,72 | 4,10 | -0,17 | 8,61E-01 |
| AT1G08590 | 0,74 | -1,64 | 3,01 | -0,54 | 5,86E-01 |
| AT1G08600 | 0,40 | 0,00 | 3,35 | 0,00 | 9,99E-01 |
| AT1G08620 | 1,06 | -2,07 | 1,79 | -1,16 | 2,47E-01 |
| AT1G08650 | 0,89 | -1,09 | 2,97 | -0,37 | 7,15E-01 |
| AT1G08680 | 0,17 | 1,16 | 4,31 | 0,27 | 7,88E-01 |
| AT1G08710 | 0,30 | -2,67 | 4,40 | -0,61 | 5,45E-01 |
| AT1G08760 | 0,68 | -1,01 | 3,05 | -0,33 | 7,40E-01 |
| AT1G08770 | 0,33 | 0,24 | 3,48 | 0,07 | 9,44E-01 |
| AT1G08830 | 1,29 | -1,84 | 2,22 | -0,83 | 4,08E-01 |
| AT1G08900 | 0,34 | -0,54 | 3,48 | -0,15 | 8,78E-01 |
| AT1G08910 | 0,43 | -1,02 | 4,13 | -0,25 | 8,05E-01 |
| AT1G08920 | 0,59 | -3,68 | 3,19 | -1,15 | 2,48E-01 |
| AT1G08930 | 1,31 | -0,68 | 1,55 | -0,44 | 6,61E-01 |
| AT1G08970 | 0,82 | -0,61 | 2,68 | -0,23 | 8,20E-01 |
| AT1G08980 | 0,91 | -2,96 | 2,07 | -1,43 | 1,52E-01 |
| AT1G09040 | 0,45 | -2,44 | 3,33 | -0,73 | 4,65E-01 |
| AT1G09250 | 0,32 | -2,68 | 4,36 | -0,61 | 5,39E-01 |
| AT1G09260 | 0,62 | 0,00 | 4,41 | 0,00 | 1,00E+00 |
| AT1G09270 | 0,12 | 0,00 | 4,41 | 0,00 | 1,00E+00 |
| AT1G09300 | 0,36 | -2,00 | 4,37 | -0,46 | 6,47E-01 |
| AT1G09390 | 0,23 | 0,68 | 4,34 | 0,16 | 8,76E-01 |
| AT1G09421 | 0,17 | -0,80 | 4,39 | -0,18 | 8,56E-01 |
| AT1G09430 | 0,79 | -0,24 | 2,32 | -0,10 | 9,17E-01 |
| AT1G09450 | 0,24 | -0,79 | 4,36 | -0,18 | 8,56E-01 |

| | | | | | |
|---|---|---|---|---|---|
| AT1G09490 | 0,05 | -0,24 | 4,38 | -0,05 | 9,57E-01 |
| AT1G09500 | 0,08 | 0,00 | 4,41 | 0,00 | 1,00E+00 |
| AT1G09530 | 0,63 | -2,71 | 3,72 | -0,73 | 4,67E-01 |
| AT1G09560 | 0,29 | -2,60 | 4,41 | -0,59 | 5,55E-01 |
| AT1G09590 | 0,27 | 0,80 | 4,33 | 0,18 | 8,54E-01 |
| AT1G09600 | 0,94 | -4,12 | 2,84 | -1,45 | 1,47E-01 |
| AT1G09610 | 0,36 | -3,08 | 4,39 | -0,70 | 4,83E-01 |
| AT1G09645 | 0,33 | -3,35 | 4,37 | -0,77 | 4,44E-01 |
| AT1G09665 | 0,25 | -3,06 | 4,38 | -0,70 | 4,85E-01 |
| AT1G09710 | 0,91 | -2,88 | 2,82 | -1,02 | 3,07E-01 |
| AT1G09750 | 0,82 | 0,75 | 1,96 | 0,38 | 7,03E-01 |
| AT1G09795 | 0,58 | -0,28 | 3,90 | -0,07 | 9,44E-01 |
| AT1G09815 | 0,21 | 0,41 | 4,36 | 0,09 | 9,24E-01 |
| AT1G10050 | 0,78 | -2,30 | 2,36 | -0,97 | 3,30E-01 |
| AT1G10070 | 0,24 | -2,60 | 4,41 | -0,59 | 5,55E-01 |
| AT1G10130 | 1,07 | -0,56 | 1,83 | -0,31 | 7,60E-01 |
| AT1G10270 | 0,58 | -0,81 | 2,70 | -0,30 | 7,65E-01 |
| AT1G10450 | 0,33 | -1,64 | 4,38 | -0,37 | 7,08E-01 |
| AT1G10490 | 0,65 | -2,46 | 2,53 | -0,97 | 3,32E-01 |
| AT1G10570 | 0,16 | -0,62 | 4,40 | -0,14 | 8,87E-01 |
| AT1G10600 | 1,27 | -2,26 | 1,96 | -1,15 | 2,51E-01 |
| AT1G10610 | 0,90 | -4,59 | 3,32 | -1,38 | 1,67E-01 |
| AT1G10660 | 0,54 | -2,87 | 3,99 | -0,72 | 4,72E-01 |
| AT1G10670 | 0,76 | 0,89 | 3,58 | 0,25 | 8,04E-01 |
| AT1G10700 | 0,45 | -1,25 | 4,37 | -0,29 | 7,75E-01 |
| AT1G10720 | 0,48 | -0,92 | 2,68 | -0,34 | 7,33E-01 |
| AT1G10790 | 0,08 | 0,00 | 4,41 | 0,00 | 1,00E+00 |
| AT1G10850 | 0,85 | -4,44 | 3,48 | -1,28 | 2,01E-01 |
| AT1G10910 | 0,08 | 0,00 | 4,41 | 0,00 | 1,00E+00 |
| AT1G10960 | 0,86 | -3,29 | 2,93 | -1,12 | 2,61E-01 |
| AT1G11020 | 0,27 | -1,27 | 4,38 | -0,29 | 7,72E-01 |
| AT1G11050 | 0,22 | 1,04 | 4,32 | 0,24 | 8,10E-01 |
| AT1G11300 | 0,32 | -0,68 | 4,41 | -0,15 | 8,77E-01 |
| AT1G11303 | 0,25 | 0,00 | 4,41 | 0,00 | 1,00E+00 |
| AT1G11320 | 0,08 | 0,00 | 4,41 | 0,00 | 1,00E+00 |
| AT1G11330 | 0,64 | -0,87 | 2,91 | -0,30 | 7,65E-01 |
| AT1G11400 | 0,26 | -0,53 | 4,34 | -0,12 | 9,03E-01 |
| AT1G11410 | 0,36 | 0,43 | 3,45 | 0,12 | 9,01E-01 |
| AT1G11430 | 0,35 | 0,59 | 3,61 | 0,16 | 8,71E-01 |
| AT1G11440 | 0,54 | -0,34 | 3,07 | -0,11 | 9,11E-01 |
| AT1G11580 | 0,31 | -1,72 | 4,39 | -0,39 | 6,95E-01 |
| AT1G11590 | 0,67 | 2,21 | 2,41 | 0,92 | 3,58E-01 |
| AT1G11593 | 0,40 | 2,03 | 4,14 | 0,49 | 6,23E-01 |
| AT1G11660 | 1,14 | -3,85 | 1,97 | -1,96 | 5,01E-02 |
| AT1G11680 | 0,83 | -2,50 | 2,80 | -0,89 | 3,71E-01 |
| AT1G11710 | 0,17 | 0,00 | 4,41 | 0,00 | 1,00E+00 |
| AT1G11750 | 0,56 | 0,01 | 3,26 | 0,00 | 9,99E-01 |
| AT1G11790 | 0,42 | -2,98 | 4,38 | -0,68 | 4,96E-01 |
| AT1G11905 | 0,75 | -2,24 | 2,18 | -1,03 | 3,05E-01 |
| AT1G11930 | 0,68 | -2,95 | 2,25 | -1,31 | 1,90E-01 |
| AT1G12015 | 0,46 | 0,31 | 4,35 | 0,07 | 9,43E-01 |
| AT1G12050 | 1,22 | -0,88 | 1,41 | -0,63 | 5,31E-01 |
| AT1G12060 | 0,12 | -0,78 | 4,39 | -0,18 | 8,59E-01 |
| AT1G12140 | 0,12 | 0,00 | 4,41 | 0,00 | 1,00E+00 |
| AT1G12210 | 0,35 | -1,23 | 4,38 | -0,28 | 7,80E-01 |
| AT1G12230 | 0,35 | -0,48 | 3,50 | -0,14 | 8,90E-01 |

| | | | | | |
|---|---|---|---|---|---|
| AT1G12250 | 0,19 | -0,02 | 4,37 | 0,00 | 9,96E-01 |
| AT1G12270 | 0,62 | -0,91 | 2,72 | -0,34 | 7,38E-01 |
| AT1G12320 | 0,82 | 2,64 | 2,75 | 0,96 | 3,36E-01 |
| AT1G12360 | 0,65 | -2,16 | 3,79 | -0,57 | 5,67E-01 |
| AT1G12410 | 0,67 | -2,60 | 3,83 | -0,68 | 4,97E-01 |
| AT1G12460 | 0,83 | 0,50 | 2,08 | 0,24 | 8,09E-01 |
| AT1G12580 | 0,43 | 0,02 | 2,79 | 0,01 | 9,94E-01 |
| AT1G12760 | 0,76 | -1,40 | 2,05 | -0,68 | 4,96E-01 |
| AT1G12810 | 0,42 | 1,24 | 2,43 | 0,51 | 6,10E-01 |
| AT1G12840 | 1,05 | -1,69 | 1,88 | -0,90 | 3,67E-01 |
| AT1G12860 | 0,23 | -0,83 | 4,38 | -0,19 | 8,50E-01 |
| AT1G12990 | 1,14 | -0,89 | 2,08 | -0,43 | 6,68E-01 |
| AT1G13040 | 0,24 | 0,20 | 4,36 | 0,04 | 9,64E-01 |
| AT1G13060 | 0,83 | -1,74 | 3,11 | -0,56 | 5,75E-01 |
| AT1G13090 | 1,29 | -1,50 | 1,65 | -0,91 | 3,64E-01 |
| AT1G13100 | 0,26 | 0,64 | 4,34 | 0,15 | 8,83E-01 |
| AT1G13180 | 0,14 | -0,11 | 4,37 | -0,03 | 9,80E-01 |
| AT1G13190 | 1,26 | 2,69 | 2,45 | 1,10 | 2,73E-01 |
| AT1G13220 | 0,56 | 0,03 | 3,84 | 0,01 | 9,94E-01 |
| AT1G13270 | 0,60 | 0,00 | 3,15 | 0,00 | 1,00E+00 |
| AT1G13330 | 0,70 | -1,24 | 2,53 | -0,49 | 6,22E-01 |
| AT1G13400 | 0,23 | -1,14 | 4,39 | -0,26 | 7,96E-01 |
| AT1G13430 | 0,17 | 0,00 | 4,41 | 0,00 | 1,00E+00 |
| AT1G13460 | 0,37 | -2,49 | 4,38 | -0,57 | 5,70E-01 |
| AT1G13820 | 0,11 | -0,80 | 4,39 | -0,18 | 8,56E-01 |
| AT1G13850 | 0,13 | 0,00 | 4,41 | 0,00 | 1,00E+00 |
| AT1G13880 | 0,45 | 1,00 | 3,39 | 0,30 | 7,67E-01 |
| AT1G13990 | 0,47 | -1,61 | 3,28 | -0,49 | 6,23E-01 |
| AT1G14000 | 0,81 | -0,84 | 2,38 | -0,35 | 7,24E-01 |
| AT1G14020 | 0,41 | -2,34 | 4,21 | -0,56 | 5,78E-01 |
| AT1G05047 | 0,12 | 0,00 | 4,41 | 0,00 | 1,00E+00 |
| AT1G14110 | 0,52 | 1,92 | 3,80 | 0,51 | 6,13E-01 |
| AT1G14150 | 0,05 | -0,24 | 4,38 | -0,05 | 9,57E-01 |
| AT1G14185 | 0,05 | -0,24 | 4,38 | -0,05 | 9,57E-01 |
| AT1G14230 | 0,08 | 0,05 | 4,38 | 0,01 | 9,91E-01 |
| AT1G14250 | 0,32 | -0,26 | 4,38 | -0,06 | 9,53E-01 |
| AT1G14280 | 0,63 | 0,09 | 2,92 | 0,03 | 9,76E-01 |
| AT1G14310 | 0,25 | -1,64 | 4,38 | -0,37 | 7,09E-01 |
| AT1G14345 | 0,53 | -0,63 | 2,66 | -0,24 | 8,14E-01 |
| AT1G14350 | 0,37 | -1,74 | 4,35 | -0,40 | 6,89E-01 |
| AT1G14370 | 0,91 | 0,64 | 1,91 | 0,34 | 7,36E-01 |
| AT1G14470 | 0,37 | 0,00 | 4,41 | 0,00 | 1,00E+00 |
| AT1G14480 | 0,08 | -0,21 | 4,39 | -0,05 | 9,62E-01 |
| AT1G14560 | 0,21 | -0,75 | 4,40 | -0,17 | 8,65E-01 |
| AT1G14570 | 1,23 | -1,76 | 1,64 | -1,07 | 2,84E-01 |
| AT1G14800 | 1,13 | -0,29 | 1,72 | -0,17 | 8,67E-01 |
| AT1G14860 | 0,08 | -0,21 | 4,39 | -0,05 | 9,62E-01 |
| AT1G15030 | 0,23 | 0,17 | 4,37 | 0,04 | 9,70E-01 |
| AT1G15120 | 0,30 | -1,15 | 4,37 | -0,26 | 7,93E-01 |
| AT1G15190 | 0,17 | 0,00 | 4,41 | 0,00 | 1,00E+00 |
| AT1G15240 | 0,58 | 0,68 | 2,68 | 0,26 | 7,98E-01 |
| AT1G15260 | 0,61 | -1,69 | 3,18 | -0,53 | 5,96E-01 |
| AT1G15390 | 0,48 | -2,60 | 4,41 | -0,59 | 5,55E-01 |
| AT1G15430 | 0,55 | -2,06 | 3,91 | -0,53 | 5,99E-01 |
| AT1G15650 | 0,36 | -3,60 | 4,36 | -0,82 | 4,10E-01 |
| AT1G05183 | 0,40 | -2,44 | 3,38 | -0,72 | 4,71E-01 |

| | | | | | |
|---|---|---|---|---|---|
| AT1G15720 | 0,33 | -1,28 | 4,35 | -0,30 | 7,68E-01 |
| AT1G15740 | 0,65 | 0,50 | 2,35 | 0,21 | 8,32E-01 |
| AT1G15810 | 0,77 | -0,52 | 1,95 | -0,26 | 7,91E-01 |
| AT1G15910 | 0,69 | -3,81 | 3,51 | -1,08 | 2,78E-01 |
| AT1G15920 | 0,80 | -4,53 | 2,72 | -1,67 | 9,58E-02 |
| AT1G15930 | 1,08 | -2,54 | 2,10 | -1,21 | 2,27E-01 |
| AT1G15950 | 0,56 | -2,15 | 2,55 | -0,85 | 3,97E-01 |
| AT1G15980 | 1,06 | 1,10 | 1,95 | 0,57 | 5,72E-01 |
| AT1G16060 | 0,12 | -0,14 | 4,38 | -0,03 | 9,74E-01 |
| AT1G16080 | 0,31 | 0,00 | 4,41 | 0,00 | 1,00E+00 |
| AT1G16090 | 0,22 | -0,21 | 4,38 | -0,05 | 9,61E-01 |
| AT1G16110 | 0,61 | -2,58 | 3,73 | -0,69 | 4,89E-01 |
| AT1G16180 | 1,23 | -0,81 | 1,82 | -0,44 | 6,57E-01 |
| AT1G16250 | 0,81 | -0,15 | 2,25 | -0,07 | 9,46E-01 |
| AT1G16350 | 0,39 | -0,68 | 4,28 | -0,16 | 8,74E-01 |
| AT1G16400 | 0,30 | -0,30 | 4,34 | -0,07 | 9,45E-01 |
| AT1G16410 | 1,19 | 0,64 | 1,58 | 0,41 | 6,84E-01 |
| AT1G16470 | 0,19 | 0,64 | 4,34 | 0,15 | 8,83E-01 |
| AT1G16500 | 0,25 | -1,21 | 4,35 | -0,28 | 7,80E-01 |
| AT1G16560 | 0,44 | -3,45 | 3,32 | -1,04 | 2,99E-01 |
| AT1G16670 | 0,35 | -1,29 | 4,31 | -0,30 | 7,65E-01 |
| AT1G16680 | 0,85 | -1,14 | 2,51 | -0,45 | 6,51E-01 |
| AT1G16690 | 0,12 | -0,78 | 4,39 | -0,18 | 8,59E-01 |
| AT1G16700 | 0,62 | -1,05 | 3,06 | -0,34 | 7,33E-01 |
| AT1G16740 | 0,37 | -2,11 | 4,38 | -0,48 | 6,31E-01 |
| AT1G16810 | 0,12 | 0,00 | 4,41 | 0,00 | 1,00E+00 |
| AT1G16860 | 0,38 | -0,80 | 4,29 | -0,19 | 8,51E-01 |
| AT1G17050 | 0,20 | 0,00 | 4,41 | 0,00 | 1,00E+00 |
| AT1G17070 | 0,73 | -0,68 | 3,69 | -0,18 | 8,54E-01 |
| AT1G17100 | 0,86 | -0,60 | 1,83 | -0,33 | 7,42E-01 |
| AT1G17120 | 0,36 | -1,18 | 4,33 | -0,27 | 7,85E-01 |
| AT1G17150 | 0,40 | 1,05 | 3,40 | 0,31 | 7,59E-01 |
| AT1G17160 | 0,36 | -1,69 | 4,36 | -0,39 | 6,99E-01 |
| AT1G17200 | 0,19 | 0,52 | 4,35 | 0,12 | 9,04E-01 |
| AT1G17230 | 0,51 | 2,17 | 3,04 | 0,71 | 4,76E-01 |
| AT1G17255 | 1,02 | -0,97 | 2,76 | -0,35 | 7,27E-01 |
| AT1G17260 | 0,13 | 0,00 | 4,41 | 0,00 | 1,00E+00 |
| AT1G17300 | 0,61 | -2,58 | 3,73 | -0,69 | 4,89E-01 |
| AT1G17340 | 0,57 | -0,37 | 2,18 | -0,17 | 8,66E-01 |
| AT1G17360 | 1,17 | -2,42 | 2,24 | -1,08 | 2,80E-01 |
| AT1G17420 | 1,07 | 0,46 | 2,05 | 0,22 | 8,22E-01 |
| AT1G17460 | 0,31 | -1,42 | 4,37 | -0,32 | 7,46E-01 |
| AT1G17490 | 0,39 | 1,36 | 3,44 | 0,40 | 6,92E-01 |
| AT1G17500 | 0,69 | -1,74 | 2,26 | -0,77 | 4,42E-01 |
| AT1G17530 | 0,57 | -0,32 | 3,12 | -0,10 | 9,19E-01 |
| AT1G17550 | 1,05 | -2,04 | 1,70 | -1,20 | 2,30E-01 |
| AT1G17570 | 0,08 | -0,21 | 4,39 | -0,05 | 9,62E-01 |
| AT1G17580 | 1,15 | -2,51 | 2,08 | -1,21 | 2,27E-01 |
| AT1G17610 | 0,33 | 1,08 | 4,33 | 0,25 | 8,03E-01 |
| AT1G17680 | 0,28 | 0,00 | 4,41 | 0,00 | 1,00E+00 |
| AT1G17745 | 0,96 | -0,32 | 1,83 | -0,17 | 8,61E-01 |
| AT1G17750 | 0,21 | -0,70 | 4,40 | -0,16 | 8,73E-01 |
| AT1G17860 | 0,25 | 0,00 | 4,41 | 0,00 | 1,00E+00 |
| AT1G17970 | 0,47 | -0,45 | 4,08 | -0,11 | 9,12E-01 |
| AT1G18020 | 0,78 | 0,48 | 2,90 | 0,17 | 8,69E-01 |
| AT1G18090 | 0,88 | -3,58 | 3,58 | -1,00 | 3,18E-01 |

| | | | | | |
|---|---|---|---|---|---|
| AT1G18130 | 0,21 | -0,35 | 4,38 | -0,08 | 9,36E-01 |
| AT1G18170 | 0,31 | 0,00 | 4,41 | 0,00 | 1,00E+00 |
| AT1G18300 | 0,38 | -0,16 | 4,27 | -0,04 | 9,69E-01 |
| AT1G18390 | 0,88 | -0,96 | 1,88 | -0,51 | 6,08E-01 |
| AT1G18450 | 0,27 | 0,05 | 4,37 | 0,01 | 9,91E-01 |
| AT1G18460 | 0,79 | -0,76 | 2,15 | -0,35 | 7,23E-01 |
| AT1G18480 | 0,17 | -1,25 | 4,37 | -0,29 | 7,74E-01 |
| AT1G18490 | 0,64 | -2,58 | 4,40 | -0,58 | 5,59E-01 |
| AT1G18500 | 0,63 | -0,88 | 3,03 | -0,29 | 7,71E-01 |
| AT1G18570 | 0,88 | -0,47 | 1,97 | -0,24 | 8,12E-01 |
| AT1G18580 | 0,40 | 0,00 | 4,41 | 0,00 | 1,00E+00 |
| AT1G18600 | 0,74 | -2,13 | 2,47 | -0,86 | 3,88E-01 |
| AT1G18620 | 0,88 | 0,41 | 2,11 | 0,19 | 8,47E-01 |
| AT1G18700 | 1,03 | -4,26 | 2,22 | -1,92 | 5,50E-02 |
| AT1G18730 | 0,47 | 0,22 | 4,21 | 0,05 | 9,58E-01 |
| AT1G18820 | 0,52 | 0,40 | 4,36 | 0,09 | 9,26E-01 |
| AT1G18870 | 0,46 | 0,00 | 4,41 | 0,00 | 1,00E+00 |
| AT1G18900 | 0,70 | -0,71 | 3,65 | -0,19 | 8,46E-01 |
| AT1G18950 | 0,83 | 0,95 | 2,10 | 0,45 | 6,51E-01 |
| AT1G19020 | 0,56 | -0,93 | 2,27 | -0,41 | 6,81E-01 |
| AT1G19025 | 0,65 | -3,56 | 2,99 | -1,19 | 2,34E-01 |
| AT1G19086 | 0,19 | 1,38 | 4,31 | 0,32 | 7,48E-01 |
| AT1G19110 | 0,80 | -3,13 | 2,22 | -1,41 | 1,58E-01 |
| AT1G19150 | 0,10 | 0,45 | 4,35 | 0,10 | 9,19E-01 |
| AT1G19190 | 0,39 | -3,07 | 4,38 | -0,70 | 4,84E-01 |
| AT1G19290 | 0,57 | -2,56 | 3,83 | -0,67 | 5,04E-01 |
| AT1G19350 | 0,52 | -0,71 | 3,90 | -0,18 | 8,55E-01 |
| AT1G19376 | 0,56 | 2,17 | 2,99 | 0,73 | 4,68E-01 |
| AT1G19464 | 0,18 | 1,18 | 4,31 | 0,27 | 7,85E-01 |
| AT1G19580 | 0,28 | -2,60 | 4,41 | -0,59 | 5,55E-01 |
| AT1G05507 | 0,08 | -0,21 | 4,39 | -0,05 | 9,62E-01 |
| AT1G19710 | 0,37 | -3,35 | 4,32 | -0,78 | 4,38E-01 |
| AT1G19770 | 0,78 | 2,56 | 2,70 | 0,95 | 3,43E-01 |
| AT1G19800 | 0,29 | -2,41 | 4,38 | -0,55 | 5,82E-01 |
| AT1G19830 | 0,33 | 1,47 | 4,31 | 0,34 | 7,33E-01 |
| AT1G19850 | 0,68 | 0,92 | 2,92 | 0,32 | 7,51E-01 |
| AT1G19910 | 0,96 | -0,18 | 2,34 | -0,08 | 9,37E-01 |
| AT1G19990 | 0,28 | 1,35 | 4,30 | 0,31 | 7,54E-01 |
| AT1G20050 | 0,40 | -1,14 | 3,47 | -0,33 | 7,43E-01 |
| AT1G05523 | 0,59 | -0,52 | 2,49 | -0,21 | 8,35E-01 |
| AT1G20090 | 0,64 | -1,03 | 2,58 | -0,40 | 6,90E-01 |
| AT1G20110 | 0,46 | 0,06 | 2,79 | 0,02 | 9,83E-01 |
| AT1G20280 | 0,77 | 1,14 | 2,01 | 0,57 | 5,72E-01 |
| AT1G20300 | 0,15 | -0,14 | 4,39 | -0,03 | 9,74E-01 |
| AT1G20310 | 0,17 | 0,00 | 4,41 | 0,00 | 1,00E+00 |
| AT1G20640 | 1,05 | -1,34 | 1,83 | -0,73 | 4,64E-01 |
| AT1G20693 | 1,05 | 1,50 | 2,18 | 0,69 | 4,91E-01 |
| AT1G20696 | 0,69 | -0,66 | 2,27 | -0,29 | 7,70E-01 |
| AT1G20780 | 1,16 | -1,65 | 2,08 | -0,79 | 4,30E-01 |
| AT1G20823 | 0,81 | 2,96 | 2,19 | 1,35 | 1,76E-01 |
| AT1G20920 | 0,49 | -3,33 | 3,21 | -1,04 | 2,99E-01 |
| AT1G21000 | 0,77 | 0,85 | 2,13 | 0,40 | 6,91E-01 |
| AT1G21050 | 0,58 | 2,50 | 2,94 | 0,85 | 3,95E-01 |
| AT1G21065 | 0,84 | 1,68 | 2,09 | 0,81 | 4,20E-01 |
| AT1G21140 | 0,08 | 0,00 | 4,41 | 0,00 | 1,00E+00 |
| AT1G21170 | 0,55 | 0,82 | 3,79 | 0,22 | 8,29E-01 |

| | | | | | |
|---|---|---|---|---|---|
| AT1G21210 | 0,63 | -1,71 | 3,01 | -0,57 | 5,69E-01 |
| AT1G21230 | 0,43 | 2,06 | 3,19 | 0,65 | 5,18E-01 |
| AT1G21240 | 0,38 | -1,69 | 4,30 | -0,39 | 6,95E-01 |
| AT1G21370 | 0,21 | -0,03 | 4,36 | -0,01 | 9,95E-01 |
| AT1G21430 | 0,53 | -2,54 | 4,40 | -0,58 | 5,64E-01 |
| AT1G21560 | 0,41 | -3,33 | 4,16 | -0,80 | 4,23E-01 |
| AT1G21640 | 0,89 | -0,53 | 1,84 | -0,29 | 7,74E-01 |
| AT1G21660 | 0,28 | -1,75 | 4,39 | -0,40 | 6,91E-01 |
| AT1G21670 | 0,86 | -4,33 | 3,30 | -1,31 | 1,89E-01 |
| AT1G21690 | 0,29 | -1,17 | 4,36 | -0,27 | 7,88E-01 |
| AT1G21720 | 0,34 | -1,00 | 4,38 | -0,23 | 8,20E-01 |
| AT1G21722 | 0,08 | 0,00 | 4,41 | 0,00 | 1,00E+00 |
| AT1G21780 | 0,75 | -0,14 | 2,47 | -0,06 | 9,54E-01 |
| AT1G21835 | 0,94 | 0,59 | 2,02 | 0,29 | 7,72E-01 |
| AT1G21840 | 0,12 | 0,00 | 4,41 | 0,00 | 1,00E+00 |
| AT1G21880 | 0,12 | -0,68 | 4,41 | -0,15 | 8,77E-01 |
| AT1G21910 | 0,33 | 0,00 | 4,41 | 0,00 | 1,00E+00 |
| AT1G21980 | 0,19 | 0,42 | 4,35 | 0,10 | 9,23E-01 |
| AT1G22040 | 0,05 | -0,24 | 4,38 | -0,05 | 9,57E-01 |
| AT1G22070 | 0,41 | -0,56 | 4,32 | -0,13 | 8,98E-01 |
| AT1G22190 | 1,08 | -1,69 | 1,74 | -0,97 | 3,32E-01 |
| AT1G22280 | 0,53 | -0,96 | 3,25 | -0,30 | 7,67E-01 |
| AT1G22330 | 0,11 | 0,46 | 4,35 | 0,11 | 9,15E-01 |
| AT1G22500 | 0,40 | 0,00 | 4,41 | 0,00 | 1,00E+00 |
| AT1G22590 | 0,24 | 0,92 | 4,33 | 0,21 | 8,32E-01 |
| AT1G22640 | 0,56 | 1,82 | 3,69 | 0,49 | 6,23E-01 |
| AT1G22730 | 0,55 | -3,97 | 3,84 | -1,03 | 3,01E-01 |
| AT1G22750 | 0,35 | 0,09 | 3,44 | 0,03 | 9,80E-01 |
| AT1G22780 | 0,54 | -1,16 | 3,20 | -0,36 | 7,16E-01 |
| AT1G22790 | 0,64 | -3,07 | 3,84 | -0,80 | 4,25E-01 |
| AT1G05767 | 0,23 | -0,81 | 4,39 | -0,19 | 8,53E-01 |
| AT1G22840 | 0,62 | 1,07 | 3,23 | 0,33 | 7,39E-01 |
| AT1G22860 | 0,86 | -2,81 | 1,98 | -1,42 | 1,56E-01 |
| AT1G22870 | 0,66 | -1,64 | 3,03 | -0,54 | 5,88E-01 |
| AT1G22880 | 1,11 | -4,70 | 2,81 | -1,68 | 9,37E-02 |
| AT1G22882 | 0,84 | -2,60 | 3,44 | -0,76 | 4,49E-01 |
| AT1G22940 | 0,55 | -2,54 | 2,75 | -0,92 | 3,55E-01 |
| AT1G23030 | 0,27 | -2,58 | 4,40 | -0,59 | 5,58E-01 |
| AT1G23040 | 0,34 | 0,54 | 4,35 | 0,12 | 9,02E-01 |
| AT1G23050 | 1,19 | 1,61 | 2,37 | 0,68 | 4,95E-01 |
| AT1G23100 | 0,08 | 0,00 | 4,41 | 0,00 | 1,00E+00 |
| AT1G23130 | 0,08 | 0,00 | 4,41 | 0,00 | 1,00E+00 |
| AT1G23230 | 1,20 | 0,00 | 1,71 | 0,00 | 9,99E-01 |
| AT1G23440 | 0,65 | -2,60 | 3,10 | -0,84 | 4,00E-01 |
| AT1G23450 | 0,21 | -2,60 | 4,41 | -0,59 | 5,55E-01 |
| AT1G23490 | 1,31 | 0,85 | 1,58 | 0,54 | 5,90E-01 |
| AT1G23710 | 1,29 | 2,23 | 1,80 | 1,24 | 2,15E-01 |
| AT1G23820 | 1,10 | -2,70 | 2,88 | -0,94 | 3,49E-01 |
| AT1G05837 | 0,53 | -0,58 | 3,20 | -0,18 | 8,58E-01 |
| AT1G23870 | 0,51 | 0,40 | 3,85 | 0,10 | 9,17E-01 |
| AT1G23880 | 0,08 | 0,00 | 4,41 | 0,00 | 1,00E+00 |
| AT1G23915 | 1,01 | 0,45 | 2,58 | 0,18 | 8,61E-01 |
| AT1G24145 | 0,48 | -0,50 | 3,13 | -0,16 | 8,74E-01 |
| AT1G24147 | 0,16 | -0,62 | 4,40 | -0,14 | 8,87E-01 |
| AT1G24150 | 0,48 | -2,16 | 3,96 | -0,55 | 5,85E-01 |
| AT1G24280 | 0,44 | 0,00 | 4,41 | 0,00 | 1,00E+00 |

| | | | | | |
|---|---|---|---|---|---|
| AT1G24340 | 0,28 | 0,79 | 3,73 | 0,21 | 8,32E-01 |
| AT1G24360 | 0,34 | -0,13 | 4,33 | -0,03 | 9,77E-01 |
| AT1G24380 | 0,38 | 1,21 | 2,82 | 0,43 | 6,68E-01 |
| AT1G24490 | 0,45 | -0,51 | 3,24 | -0,16 | 8,76E-01 |
| AT1G24530 | 0,17 | -0,35 | 4,38 | -0,08 | 9,37E-01 |
| AT1G24560 | 0,68 | 0,53 | 2,15 | 0,25 | 8,05E-01 |
| AT1G24570 | 0,08 | 0,00 | 4,41 | 0,00 | 1,00E+00 |
| AT1G25280 | 0,98 | -2,92 | 2,34 | -1,25 | 2,11E-01 |
| AT1G25480 | 0,19 | 1,18 | 4,31 | 0,27 | 7,84E-01 |
| AT1G25520 | 0,81 | -1,48 | 3,60 | -0,41 | 6,81E-01 |
| AT1G26130 | 0,54 | -1,61 | 3,93 | -0,41 | 6,82E-01 |
| AT1G26220 | 0,37 | 1,46 | 4,31 | 0,34 | 7,35E-01 |
| AT1G26250 | 0,44 | 2,25 | 4,03 | 0,56 | 5,77E-01 |
| AT1G26460 | 0,79 | -2,26 | 2,89 | -0,78 | 4,36E-01 |
| AT1G26470 | 0,34 | -2,60 | 4,41 | -0,59 | 5,55E-01 |
| AT1G26550 | 0,37 | 0,62 | 4,29 | 0,14 | 8,86E-01 |
| AT1G26560 | 0,24 | 0,05 | 4,38 | 0,01 | 9,90E-01 |
| AT1G26580 | 0,53 | -1,05 | 2,64 | -0,40 | 6,90E-01 |
| AT1G26630 | 1,29 | 0,42 | 1,68 | 0,25 | 8,04E-01 |
| AT1G26650 | 1,02 | -2,97 | 2,21 | -1,34 | 1,80E-01 |
| AT1G26665 | 0,17 | 0,00 | 4,41 | 0,00 | 1,00E+00 |
| AT1G26670 | 1,04 | -2,38 | 2,19 | -1,08 | 2,78E-01 |
| AT1G26770 | 0,19 | 1,18 | 4,31 | 0,27 | 7,84E-01 |
| AT1G26830 | 0,72 | 0,69 | 2,38 | 0,29 | 7,71E-01 |
| AT1G26920 | 0,73 | 0,92 | 2,20 | 0,42 | 6,76E-01 |
| AT1G26940 | 0,05 | -0,24 | 4,38 | -0,05 | 9,57E-01 |
| AT1G27000 | 0,38 | -1,69 | 4,40 | -0,38 | 7,01E-01 |
| AT1G27060 | 0,73 | -3,39 | 2,96 | -1,14 | 2,53E-01 |
| AT1G27130 | 0,51 | 1,18 | 3,15 | 0,37 | 7,09E-01 |
| AT1G27150 | 0,43 | -2,45 | 4,35 | -0,56 | 5,73E-01 |
| AT1G27170 | 0,35 | -0,91 | 3,46 | -0,26 | 7,92E-01 |
| AT1G27180 | 0,99 | -3,92 | 2,63 | -1,49 | 1,37E-01 |
| AT1G27300 | 0,40 | -2,52 | 4,18 | -0,60 | 5,46E-01 |
| AT1G27330 | 0,50 | -3,04 | 4,37 | -0,70 | 4,87E-01 |
| AT1G27360 | 0,08 | 0,00 | 4,41 | 0,00 | 1,00E+00 |
| AT1G27400 | 0,67 | -0,93 | 2,06 | -0,45 | 6,52E-01 |
| AT1G06087 | 0,12 | -0,78 | 4,39 | -0,18 | 8,59E-01 |
| AT1G27450 | 0,37 | -0,35 | 4,38 | -0,08 | 9,36E-01 |
| AT1G27460 | 0,11 | -0,68 | 4,41 | -0,15 | 8,77E-01 |
| AT1G27480 | 0,41 | -2,97 | 4,22 | -0,70 | 4,81E-01 |
| AT1G27520 | 0,14 | -0,38 | 4,38 | -0,09 | 9,31E-01 |
| AT1G06097 | 0,36 | -1,18 | 4,38 | -0,27 | 7,88E-01 |
| AT1G27630 | 0,58 | -0,65 | 3,89 | -0,17 | 8,67E-01 |
| AT1G27650 | 0,97 | -2,13 | 2,12 | -1,00 | 3,15E-01 |
| AT1G27750 | 0,63 | -0,42 | 2,54 | -0,16 | 8,69E-01 |
| AT1G27760 | 0,65 | -1,62 | 2,54 | -0,64 | 5,23E-01 |
| AT1G27850 | 0,84 | -2,12 | 2,15 | -0,99 | 3,25E-01 |
| AT1G27950 | 0,55 | 1,56 | 3,78 | 0,41 | 6,79E-01 |
| AT1G28005 | 0,32 | 1,91 | 4,29 | 0,45 | 6,56E-01 |
| AT1G28060 | 0,32 | -0,83 | 4,38 | -0,19 | 8,50E-01 |
| AT1G28135 | 0,08 | -0,21 | 4,39 | -0,05 | 9,62E-01 |
| AT1G28340 | 0,87 | -0,68 | 2,85 | -0,24 | 8,12E-01 |
| AT1G28350 | 1,15 | -2,19 | 1,93 | -1,14 | 2,55E-01 |
| AT1G28380 | 1,21 | -0,76 | 2,08 | -0,36 | 7,16E-01 |
| AT1G06163 | 0,36 | 0,22 | 4,32 | 0,05 | 9,60E-01 |
| AT1G28490 | 0,17 | -0,71 | 4,40 | -0,16 | 8,72E-01 |

| | | | | | |
|---|---|---|---|---|---|
| AT1G28520 | 0,40 | -0,88 | 3,39 | -0,26 | 7,95E-01 |
| AT1G28680 | 0,55 | -0,59 | 3,22 | -0,18 | 8,55E-01 |
| AT1G28780 | 0,49 | 1,26 | 4,05 | 0,31 | 7,55E-01 |
| AT1G28760 | 0,10 | -0,18 | 4,38 | -0,04 | 9,67E-01 |
| AT1G29120 | 0,19 | -0,76 | 4,40 | -0,17 | 8,63E-01 |
| AT1G29150 | 0,99 | -2,27 | 2,26 | -1,00 | 3,15E-01 |
| AT1G29200 | 0,08 | 0,00 | 4,41 | 0,00 | 1,00E+00 |
| AT1G29260 | 0,92 | -1,23 | 1,68 | -0,73 | 4,63E-01 |
| AT1G29310 | 0,68 | -3,08 | 3,05 | -1,01 | 3,13E-01 |
| AT1G29340 | 0,94 | 1,11 | 2,12 | 0,52 | 6,01E-01 |
| AT1G29380 | 0,65 | -3,54 | 4,35 | -0,81 | 4,16E-01 |
| AT1G06233 | 0,68 | -0,46 | 2,52 | -0,18 | 8,56E-01 |
| AT1G06253 | 1,09 | 0,70 | 1,77 | 0,39 | 6,94E-01 |
| AT1G06263 | 0,52 | -0,76 | 4,11 | -0,19 | 8,53E-01 |
| AT1G29620 | 0,47 | 0,95 | 3,22 | 0,30 | 7,68E-01 |
| AT1G29630 | 0,24 | -0,68 | 4,41 | -0,15 | 8,77E-01 |
| AT1G29760 | 0,40 | -1,72 | 4,30 | -0,40 | 6,89E-01 |
| AT1G29770 | 0,71 | -2,60 | 4,41 | -0,59 | 5,55E-01 |
| AT1G29800 | 0,74 | -4,17 | 3,66 | -1,14 | 2,55E-01 |
| AT1G29830 | 1,13 | -1,94 | 1,79 | -1,08 | 2,80E-01 |
| AT1G29950 | 0,49 | -3,94 | 4,34 | -0,91 | 3,65E-01 |
| AT1G30010 | 0,08 | -0,21 | 4,39 | -0,05 | 9,62E-01 |
| AT1G30200 | 0,70 | -1,96 | 2,22 | -0,88 | 3,78E-01 |
| AT1G30260 | 1,16 | 0,18 | 2,28 | 0,08 | 9,37E-01 |
| AT1G30420 | 0,38 | -3,44 | 4,37 | -0,79 | 4,31E-01 |
| AT1G30430 | 0,16 | -0,01 | 4,37 | 0,00 | 9,98E-01 |
| AT1G30440 | 1,10 | -0,53 | 1,76 | -0,30 | 7,64E-01 |
| AT1G30450 | 0,86 | -1,13 | 2,18 | -0,52 | 6,03E-01 |
| AT1G30520 | 0,41 | -3,81 | 4,35 | -0,88 | 3,81E-01 |
| AT1G30530 | 0,46 | 0,93 | 3,30 | 0,28 | 7,79E-01 |
| AT1G30610 | 0,23 | 0,14 | 4,37 | 0,03 | 9,75E-01 |
| AT1G30620 | 0,56 | -0,88 | 3,20 | -0,28 | 7,82E-01 |
| AT1G30700 | 0,12 | 0,00 | 4,41 | 0,00 | 1,00E+00 |
| AT1G30720 | 0,53 | -2,54 | 4,40 | -0,58 | 5,64E-01 |
| AT1G30757 | 0,16 | 0,84 | 4,33 | 0,20 | 8,45E-01 |
| AT1G30800 | 0,20 | -2,55 | 4,40 | -0,58 | 5,62E-01 |
| AT1G30910 | 0,25 | 0,42 | 4,36 | 0,10 | 9,23E-01 |
| AT1G06393 | 0,32 | -0,36 | 4,38 | -0,08 | 9,34E-01 |
| AT1G31095 | 0,66 | 0,63 | 2,83 | 0,22 | 8,23E-01 |
| AT1G31170 | 0,22 | -0,39 | 4,38 | -0,09 | 9,29E-01 |
| AT1G31175 | 0,74 | -2,78 | 2,49 | -1,12 | 2,64E-01 |
| AT1G31190 | 0,86 | 1,24 | 2,39 | 0,52 | 6,04E-01 |
| AT1G31220 | 0,22 | -0,56 | 4,35 | -0,13 | 8,98E-01 |
| AT1G31290 | 0,15 | 0,05 | 4,37 | 0,01 | 9,91E-01 |
| AT1G31420 | 0,89 | -2,48 | 2,26 | -1,10 | 2,72E-01 |
| AT1G31480 | 0,40 | 0,00 | 4,41 | 0,00 | 1,00E+00 |
| AT1G06447 | 0,24 | -1,46 | 4,36 | -0,34 | 7,37E-01 |
| AT1G31710 | 0,82 | 3,05 | 3,45 | 0,88 | 3,77E-01 |
| AT1G31780 | 0,28 | -0,17 | 4,39 | -0,04 | 9,70E-01 |
| AT1G31800 | 0,33 | -0,36 | 3,54 | -0,10 | 9,20E-01 |
| AT1G31850 | 1,13 | -3,30 | 2,05 | -1,61 | 1,07E-01 |
| AT1G31930 | 0,12 | 0,00 | 4,41 | 0,00 | 1,00E+00 |
| AT1G31970 | 0,74 | -1,98 | 3,08 | -0,64 | 5,21E-01 |
| AT1G32050 | 0,65 | -0,47 | 2,37 | -0,20 | 8,42E-01 |
| AT1G32160 | 0,70 | 0,05 | 2,99 | 0,02 | 9,87E-01 |
| AT1G32220 | 0,21 | 1,00 | 4,32 | 0,23 | 8,16E-01 |

| | | | | | |
|---|---|---|---|---|---|
| AT1G32225 | 0,70 | 1,71 | 2,31 | 0,74 | 4,61E-01 |
| AT1G32320 | 0,23 | 0,11 | 4,38 | 0,03 | 9,80E-01 |
| AT1G32337 | 0,30 | 0,36 | 4,35 | 0,08 | 9,35E-01 |
| AT1G32360 | 0,41 | -0,68 | 3,45 | -0,20 | 8,44E-01 |
| AT1G06547 | 0,19 | 1,05 | 4,32 | 0,24 | 8,08E-01 |
| AT1G32385 | 0,41 | 0,87 | 2,79 | 0,31 | 7,56E-01 |
| AT1G32415 | 0,72 | -4,52 | 2,82 | -1,60 | 1,10E-01 |
| AT1G32460 | 0,41 | 1,11 | 4,32 | 0,26 | 7,97E-01 |
| AT1G32580 | 0,32 | -3,01 | 4,38 | -0,69 | 4,92E-01 |
| AT1G32700 | 0,46 | -1,28 | 3,30 | -0,39 | 6,98E-01 |
| AT1G32810 | 0,36 | -1,68 | 3,51 | -0,48 | 6,32E-01 |
| AT1G06623 | 0,26 | 1,54 | 4,30 | 0,36 | 7,20E-01 |
| AT1G32870 | 0,59 | -2,06 | 3,87 | -0,53 | 5,94E-01 |
| AT1G32920 | 1,13 | 1,72 | 1,82 | 0,95 | 3,43E-01 |
| AT1G32928 | 0,50 | -1,51 | 4,05 | -0,37 | 7,09E-01 |
| AT1G32940 | 0,41 | -3,05 | 4,38 | -0,70 | 4,87E-01 |
| AT1G32950 | 0,62 | -2,58 | 3,75 | -0,69 | 4,92E-01 |
| AT1G32960 | 0,35 | 0,14 | 4,37 | 0,03 | 9,75E-01 |
| AT1G32990 | 0,48 | -0,78 | 4,39 | -0,18 | 8,60E-01 |
| AT1G33010 | 0,19 | 1,18 | 4,31 | 0,27 | 7,84E-01 |
| AT1G33040 | 0,24 | -0,04 | 4,38 | -0,01 | 9,93E-01 |
| AT1G33110 | 0,55 | -3,82 | 3,83 | -1,00 | 3,19E-01 |
| AT1G33170 | 0,44 | -3,42 | 3,33 | -1,03 | 3,04E-01 |
| AT1G33260 | 0,52 | -1,16 | 4,03 | -0,29 | 7,73E-01 |
| AT1G33360 | 0,47 | 0,84 | 4,05 | 0,21 | 8,35E-01 |
| AT1G33470 | 0,71 | -4,36 | 3,51 | -1,24 | 2,13E-01 |
| AT1G33610 | 0,15 | -0,77 | 4,39 | -0,18 | 8,61E-01 |
| AT1G33612 | 0,17 | -0,68 | 4,41 | -0,15 | 8,77E-01 |
| AT1G33770 | 0,49 | -1,25 | 4,05 | -0,31 | 7,57E-01 |
| AT1G33811 | 0,18 | -0,42 | 4,37 | -0,10 | 9,24E-01 |
| AT1G33910 | 0,08 | -0,21 | 4,39 | -0,05 | 9,62E-01 |
| AT1G33930 | 0,36 | 0,95 | 4,32 | 0,22 | 8,27E-01 |
| AT1G33960 | 1,02 | -3,68 | 3,11 | -1,18 | 2,37E-01 |
| AT1G33980 | 0,92 | -3,87 | 4,23 | -0,92 | 3,60E-01 |
| AT1G33990 | 0,95 | -1,50 | 2,01 | -0,75 | 4,55E-01 |
| AT1G06723 | 0,59 | -2,71 | 2,63 | -1,03 | 3,04E-01 |
| AT1G34020 | 0,78 | -1,67 | 2,56 | -0,65 | 5,14E-01 |
| AT1G34070 | 0,46 | 2,33 | 4,00 | 0,58 | 5,60E-01 |
| AT1G34150 | 1,09 | -0,36 | 2,41 | -0,15 | 8,80E-01 |
| AT1G34190 | 1,14 | -1,51 | 1,85 | -0,81 | 4,15E-01 |
| AT1G34200 | 0,21 | -0,73 | 4,40 | -0,17 | 8,67E-01 |
| AT1G34210 | 0,51 | -2,30 | 4,39 | -0,52 | 6,00E-01 |
| AT1G34290 | 0,25 | 1,50 | 4,30 | 0,35 | 7,28E-01 |
| AT1G06747 | 0,08 | -0,21 | 4,39 | -0,05 | 9,62E-01 |
| AT1G34355 | 0,86 | -3,82 | 2,78 | -1,38 | 1,69E-01 |
| AT1G34390 | 0,68 | 2,38 | 3,65 | 0,65 | 5,14E-01 |
| AT1G34410 | 0,91 | 3,42 | 3,20 | 1,07 | 2,85E-01 |
| AT1G34420 | 0,36 | 0,80 | 4,29 | 0,19 | 8,52E-01 |
| AT1G34460 | 0,31 | 1,74 | 4,30 | 0,40 | 6,86E-01 |
| AT1G34580 | 0,53 | 0,00 | 4,41 | 0,00 | 1,00E+00 |
| AT1G34770 | 0,12 | 0,00 | 4,41 | 0,00 | 1,00E+00 |
| AT1G34844 | 0,08 | 0,00 | 4,41 | 0,00 | 1,00E+00 |
| AT1G35030 | 0,42 | 0,37 | 3,35 | 0,11 | 9,13E-01 |
| AT1G35040 | 0,23 | 1,08 | 4,31 | 0,25 | 8,03E-01 |
| AT1G35140 | 0,39 | 0,63 | 4,18 | 0,15 | 8,81E-01 |
| AT1G35160 | 0,59 | -0,76 | 2,67 | -0,29 | 7,75E-01 |

| | | | | | |
|---|---|---|---|---|---|
| AT1G06827 | 0,56 | -0,07 | 2,67 | -0,03 | 9,80E-01 |
| AT1G35220 | 0,89 | -2,99 | 2,40 | -1,24 | 2,13E-01 |
| AT1G35230 | 1,07 | -2,39 | 2,07 | -1,16 | 2,47E-01 |
| AT1G35242 | 0,19 | 0,05 | 4,36 | 0,01 | 9,90E-01 |
| AT1G35420 | 0,41 | 0,12 | 3,38 | 0,04 | 9,71E-01 |
| AT1G35460 | 0,12 | 0,00 | 4,41 | 0,00 | 1,00E+00 |
| AT1G35500 | 0,14 | 0,87 | 4,33 | 0,20 | 8,40E-01 |
| AT1G35510 | 0,12 | -0,31 | 4,38 | -0,07 | 9,44E-01 |
| AT1G35515 | 0,52 | -0,07 | 2,36 | -0,03 | 9,76E-01 |
| AT1G35516 | 0,52 | -0,07 | 2,36 | -0,03 | 9,76E-01 |
| AT1G35530 | 0,08 | 0,00 | 4,41 | 0,00 | 1,00E+00 |
| AT1G35540 | 0,95 | 1,45 | 1,88 | 0,77 | 4,39E-01 |
| AT1G35614 | 1,12 | 0,88 | 2,11 | 0,42 | 6,75E-01 |
| AT1G35620 | 0,51 | 1,16 | 3,27 | 0,35 | 7,23E-01 |
| AT1G35660 | 0,28 | -0,68 | 4,41 | -0,15 | 8,77E-01 |
| AT1G35670 | 0,21 | -0,73 | 4,40 | -0,17 | 8,67E-01 |
| AT1G35680 | 0,81 | -2,59 | 2,11 | -1,22 | 2,21E-01 |
| AT1G06927 | 0,83 | 2,49 | 2,71 | 0,92 | 3,59E-01 |
| AT1G35850 | 0,23 | 1,44 | 4,30 | 0,33 | 7,38E-01 |
| AT1G36050 | 0,26 | -0,37 | 4,38 | -0,08 | 9,33E-01 |
| AT1G06963 | 0,05 | -0,24 | 4,38 | -0,05 | 9,57E-01 |
| AT1G36320 | 0,22 | 0,62 | 4,34 | 0,14 | 8,86E-01 |
| AT1G36510 | 0,10 | 0,45 | 4,35 | 0,10 | 9,19E-01 |
| AT1G06987 | 0,58 | -0,07 | 4,37 | -0,02 | 9,87E-01 |
| AT1G36675 | 0,81 | 0,15 | 2,49 | 0,06 | 9,52E-01 |
| AT1G36745 | 0,98 | 2,35 | 2,22 | 1,06 | 2,90E-01 |
| AT1G36756 | 0,94 | 3,17 | 2,41 | 1,31 | 1,89E-01 |
| AT1G36940 | 0,22 | 1,22 | 4,31 | 0,28 | 7,78E-01 |
| AT1G36980 | 0,52 | -0,68 | 3,26 | -0,21 | 8,35E-01 |
| AT1G36992 | 0,32 | -2,30 | 4,37 | -0,53 | 5,99E-01 |
| AT1G37113 | 0,08 | 0,00 | 4,41 | 0,00 | 1,00E+00 |
| AT1G42525 | 0,30 | 1,17 | 4,32 | 0,27 | 7,87E-01 |
| AT1G42540 | 1,07 | -3,20 | 2,27 | -1,41 | 1,59E-01 |
| AT1G07137 | 0,05 | -0,24 | 4,38 | -0,05 | 9,57E-01 |
| AT1G42960 | 0,66 | -0,69 | 4,41 | -0,16 | 8,76E-01 |
| AT1G42980 | 1,30 | -2,07 | 1,91 | -1,08 | 2,78E-01 |
| AT1G43040 | 0,16 | 0,84 | 4,33 | 0,20 | 8,45E-01 |
| AT1G43171 | 0,72 | -4,76 | 4,30 | -1,11 | 2,68E-01 |
| AT1G43330 | 0,87 | 3,30 | 2,51 | 1,31 | 1,89E-01 |
| AT1G43670 | 0,63 | -0,98 | 2,59 | -0,38 | 7,05E-01 |
| AT1G43680 | 0,25 | -3,06 | 4,38 | -0,70 | 4,85E-01 |
| AT1G43700 | 0,72 | -0,32 | 1,91 | -0,17 | 8,68E-01 |
| AT1G43722 | 0,18 | 1,18 | 4,31 | 0,27 | 7,85E-01 |
| AT1G43770 | 1,22 | -1,31 | 1,75 | -0,75 | 4,56E-01 |
| AT1G43780 | 0,25 | 0,00 | 4,41 | 0,00 | 1,00E+00 |
| AT1G43815 | 1,28 | 2,81 | 2,01 | 1,40 | 1,62E-01 |
| AT1G43890 | 0,40 | -2,59 | 4,31 | -0,60 | 5,48E-01 |
| AT1G43970 | 0,43 | 2,58 | 4,27 | 0,60 | 5,45E-01 |
| AT1G44120 | 0,95 | 0,83 | 2,73 | 0,30 | 7,63E-01 |
| AT1G44750 | 0,33 | 0,00 | 4,41 | 0,00 | 1,00E+00 |
| AT1G44780 | 0,29 | -2,41 | 4,38 | -0,55 | 5,82E-01 |
| AT1G44810 | 0,26 | 0,06 | 3,78 | 0,01 | 9,88E-01 |
| AT1G44890 | 0,08 | -0,21 | 4,39 | -0,05 | 9,62E-01 |
| AT1G44900 | 0,12 | 0,00 | 4,41 | 0,00 | 1,00E+00 |
| AT1G44910 | 0,50 | 0,25 | 2,33 | 0,11 | 9,14E-01 |
| AT1G44960 | 0,10 | -0,18 | 4,38 | -0,04 | 9,67E-01 |

| | | | | | |
|---|---|---|---|---|---|
| AT1G45000 | 0,47 | -1,71 | 2,75 | -0,62 | 5,32E-01 |
| AT1G45165 | 0,70 | -0,35 | 2,51 | -0,14 | 8,90E-01 |
| AT1G45180 | 0,30 | -2,60 | 4,41 | -0,59 | 5,55E-01 |
| AT1G45474 | 0,41 | 1,14 | 3,30 | 0,35 | 7,30E-01 |
| AT1G45688 | 0,89 | 1,30 | 2,30 | 0,57 | 5,71E-01 |
| AT1G45976 | 0,41 | -1,02 | 4,24 | -0,24 | 8,09E-01 |
| AT1G46264 | 0,10 | -0,18 | 4,38 | -0,04 | 9,67E-01 |
| AT1G46554 | 0,28 | -1,32 | 4,37 | -0,30 | 7,63E-01 |
| AT1G47210 | 0,93 | -3,48 | 2,55 | -1,37 | 1,72E-01 |
| AT1G07327 | 1,09 | 3,13 | 2,25 | 1,39 | 1,64E-01 |
| AT1G47250 | 0,35 | -0,81 | 4,39 | -0,18 | 8,54E-01 |
| AT1G47270 | 0,13 | 0,00 | 4,41 | 0,00 | 1,00E+00 |
| AT1G07357 | 0,11 | 0,46 | 4,35 | 0,11 | 9,15E-01 |
| AT1G47480 | 1,07 | 1,56 | 2,06 | 0,76 | 4,50E-01 |
| AT1G47490 | 0,30 | -2,60 | 4,41 | -0,59 | 5,55E-01 |
| AT1G47500 | 0,37 | -3,33 | 4,32 | -0,77 | 4,41E-01 |
| AT1G47550 | 0,50 | -0,78 | 4,12 | -0,19 | 8,51E-01 |
| AT1G47680 | 0,12 | 0,07 | 4,38 | 0,02 | 9,87E-01 |
| AT1G47690 | 0,26 | -0,17 | 4,37 | -0,04 | 9,69E-01 |
| AT1G47770 | 0,12 | 0,07 | 4,38 | 0,02 | 9,87E-01 |
| AT1G47890 | 1,24 | -3,47 | 1,90 | -1,82 | 6,84E-02 |
| AT1G47940 | 0,36 | 2,19 | 4,28 | 0,51 | 6,08E-01 |
| AT1G48050 | 0,54 | -2,56 | 4,40 | -0,58 | 5,61E-01 |
| AT1G48100 | 0,12 | -0,68 | 4,41 | -0,15 | 8,77E-01 |
| AT1G48170 | 0,32 | -0,49 | 3,57 | -0,14 | 8,91E-01 |
| AT1G48210 | 0,24 | -0,68 | 4,41 | -0,15 | 8,77E-01 |
| AT1G48230 | 0,28 | -1,64 | 4,41 | -0,37 | 7,10E-01 |
| AT1G48315 | 0,17 | 0,00 | 4,41 | 0,00 | 1,00E+00 |
| AT1G48460 | 0,51 | 1,56 | 3,08 | 0,51 | 6,12E-01 |
| AT1G07487 | 0,32 | 0,79 | 3,59 | 0,22 | 8,26E-01 |
| AT1G48480 | 0,38 | -0,38 | 4,29 | -0,09 | 9,30E-01 |
| AT1G48560 | 0,14 | 0,87 | 4,33 | 0,20 | 8,40E-01 |
| AT1G48600 | 0,62 | -0,50 | 2,24 | -0,22 | 8,25E-01 |
| AT1G48640 | 0,26 | -2,60 | 4,41 | -0,59 | 5,55E-01 |
| AT1G48820 | 0,24 | -0,68 | 4,41 | -0,15 | 8,77E-01 |
| AT1G49010 | 0,27 | 0,92 | 4,33 | 0,21 | 8,32E-01 |
| AT1G49032 | 0,71 | -0,88 | 2,08 | -0,42 | 6,72E-01 |
| AT1G49190 | 1,12 | 3,32 | 2,13 | 1,56 | 1,18E-01 |
| AT1G49200 | 0,08 | -0,21 | 4,39 | -0,05 | 9,62E-01 |
| AT1G49210 | 0,15 | -0,17 | 4,39 | -0,04 | 9,68E-01 |
| AT1G49230 | 0,38 | -1,26 | 3,37 | -0,37 | 7,09E-01 |
| AT1G49245 | 0,90 | -0,25 | 2,12 | -0,12 | 9,06E-01 |
| AT1G49300 | 0,19 | 0,49 | 4,34 | 0,11 | 9,10E-01 |
| AT1G49340 | 1,22 | -1,69 | 1,85 | -0,91 | 3,62E-01 |
| AT1G49390 | 0,08 | 0,00 | 4,41 | 0,00 | 1,00E+00 |
| AT1G49470 | 0,15 | 0,48 | 4,35 | 0,11 | 9,12E-01 |
| AT1G49510 | 0,25 | 0,00 | 4,41 | 0,00 | 1,00E+00 |
| AT1G49680 | 0,43 | 1,65 | 3,27 | 0,50 | 6,14E-01 |
| AT1G49720 | 0,23 | 0,00 | 4,41 | 0,00 | 1,00E+00 |
| AT1G49820 | 0,28 | 0,00 | 4,41 | 0,00 | 1,00E+00 |
| AT1G49890 | 0,23 | -1,68 | 4,36 | -0,39 | 7,00E-01 |
| AT1G49930 | 0,39 | -3,10 | 4,38 | -0,71 | 4,79E-01 |
| AT1G49975 | 1,21 | 1,27 | 2,36 | 0,54 | 5,92E-01 |
| AT1G49980 | 0,53 | -3,29 | 3,11 | -1,06 | 2,89E-01 |
| AT1G49990 | 0,80 | 0,00 | 4,37 | 0,00 | 1,00E+00 |
| AT1G50050 | 0,43 | 0,77 | 3,21 | 0,24 | 8,09E-01 |

| | | | | | |
|---|---|---|---|---|---|
| AT1G50120 | 0,25 | -1,64 | 4,36 | -0,38 | 7,07E-01 |
| AT1G50170 | 0,08 | 0,00 | 4,41 | 0,00 | 1,00E+00 |
| AT1G50240 | 1,24 | -3,20 | 1,94 | -1,65 | 9,95E-02 |
| AT1G50270 | 0,45 | -1,90 | 3,29 | -0,58 | 5,63E-01 |
| AT1G50350 | 0,30 | 1,51 | 4,30 | 0,35 | 7,26E-01 |
| AT1G50360 | 0,23 | 0,59 | 4,34 | 0,14 | 8,92E-01 |
| AT1G50370 | 0,39 | -1,43 | 3,42 | -0,42 | 6,76E-01 |
| AT1G50380 | 0,57 | -0,08 | 3,16 | -0,02 | 9,80E-01 |
| AT1G50560 | 0,41 | -2,40 | 4,30 | -0,56 | 5,76E-01 |
| AT1G50660 | 1,10 | 0,66 | 1,79 | 0,37 | 7,10E-01 |
| AT1G50730 | 0,12 | 0,00 | 4,41 | 0,00 | 1,00E+00 |
| AT1G50740 | 0,72 | -2,46 | 2,91 | -0,85 | 3,97E-01 |
| AT1G51060 | 0,42 | 0,64 | 4,25 | 0,15 | 8,81E-01 |
| AT1G51100 | 0,17 | 1,16 | 4,31 | 0,27 | 7,88E-01 |
| AT1G51110 | 0,52 | -0,98 | 3,20 | -0,31 | 7,59E-01 |
| AT1G51160 | 0,67 | -3,91 | 2,99 | -1,31 | 1,91E-01 |
| AT1G51200 | 1,27 | 0,10 | 2,27 | 0,04 | 9,66E-01 |
| AT1G51350 | 0,44 | -0,86 | 3,30 | -0,26 | 7,95E-01 |
| AT1G51390 | 0,24 | -0,11 | 4,38 | -0,03 | 9,80E-01 |
| AT1G07737 | 0,83 | 2,37 | 2,30 | 1,03 | 3,02E-01 |
| AT1G51402 | 1,01 | 2,18 | 2,18 | 1,00 | 3,17E-01 |
| AT1G51440 | 0,14 | 0,48 | 4,35 | 0,11 | 9,13E-01 |
| AT1G51540 | 0,08 | -0,21 | 4,39 | -0,05 | 9,62E-01 |
| AT1G51580 | 0,35 | 0,18 | 4,37 | 0,04 | 9,67E-01 |
| AT1G51600 | 0,08 | 0,00 | 4,41 | 0,00 | 1,00E+00 |
| AT1G51645 | 0,08 | 0,00 | 4,41 | 0,00 | 1,00E+00 |
| AT1G51660 | 0,75 | -2,20 | 2,56 | -0,86 | 3,89E-01 |
| AT1G51670 | 0,25 | -2,57 | 4,40 | -0,58 | 5,60E-01 |
| AT1G51720 | 0,29 | -1,64 | 4,37 | -0,38 | 7,08E-01 |
| AT1G51745 | 0,21 | 0,00 | 4,41 | 0,00 | 1,00E+00 |
| AT1G51760 | 0,56 | 0,23 | 2,61 | 0,09 | 9,30E-01 |
| AT1G51950 | 0,22 | 0,75 | 4,34 | 0,17 | 8,63E-01 |
| AT1G52080 | 0,08 | 0,00 | 4,41 | 0,00 | 1,00E+00 |
| AT1G52190 | 0,40 | -2,48 | 3,37 | -0,74 | 4,62E-01 |
| AT1G52310 | 0,73 | -1,70 | 2,49 | -0,68 | 4,95E-01 |
| AT1G52320 | 0,99 | 1,16 | 1,86 | 0,63 | 5,30E-01 |
| AT1G52380 | 0,74 | -2,66 | 2,94 | -0,90 | 3,66E-01 |
| AT1G52410 | 0,68 | -1,64 | 2,66 | -0,62 | 5,38E-01 |
| AT1G52420 | 0,99 | -1,03 | 2,78 | -0,37 | 7,11E-01 |
| AT1G52500 | 0,39 | -2,60 | 4,41 | -0,59 | 5,55E-01 |
| AT1G52600 | 0,08 | 0,00 | 4,41 | 0,00 | 1,00E+00 |
| AT1G52618 | 0,68 | 2,87 | 2,80 | 1,02 | 3,06E-01 |
| AT1G52620 | 0,78 | -0,53 | 4,34 | -0,12 | 9,03E-01 |
| AT1G52720 | 0,19 | 1,18 | 4,31 | 0,27 | 7,84E-01 |
| AT1G52740 | 1,01 | 1,00 | 2,07 | 0,48 | 6,31E-01 |
| AT1G52760 | 0,33 | -0,47 | 3,57 | -0,13 | 8,94E-01 |
| AT1G52870 | 0,92 | 2,93 | 2,50 | 1,17 | 2,40E-01 |
| AT1G52905 | 0,68 | 2,36 | 2,78 | 0,85 | 3,95E-01 |
| AT1G52930 | 0,12 | 0,00 | 4,41 | 0,00 | 1,00E+00 |
| AT1G52940 | 0,76 | -0,05 | 2,37 | -0,02 | 9,82E-01 |
| AT1G52950 | 0,39 | 1,58 | 3,44 | 0,46 | 6,46E-01 |
| AT1G53050 | 0,64 | -4,30 | 4,32 | -1,00 | 3,19E-01 |
| AT1G53090 | 0,57 | -2,33 | 4,36 | -0,54 | 5,93E-01 |
| AT1G53165 | 0,43 | -0,68 | 2,83 | -0,24 | 8,10E-01 |
| AT1G53233 | 0,99 | 0,31 | 2,24 | 0,14 | 8,89E-01 |
| AT1G53300 | 0,67 | 0,40 | 2,48 | 0,16 | 8,73E-01 |

| | | | | | |
|---|---|---|---|---|---|
| AT1G53330 | 0,35 | -2,99 | 4,38 | -0,68 | 4,95E-01 |
| AT1G53350 | 0,25 | -0,13 | 4,37 | -0,03 | 9,77E-01 |
| AT1G53390 | 0,38 | -0,47 | 4,32 | -0,11 | 9,14E-01 |
| AT1G53400 | 0,40 | 0,00 | 4,41 | 0,00 | 1,00E+00 |
| AT1G53440 | 1,17 | 0,22 | 1,62 | 0,13 | 8,94E-01 |
| AT1G07887 | 0,84 | 1,38 | 2,99 | 0,46 | 6,45E-01 |
| AT1G53490 | 0,92 | 1,44 | 2,49 | 0,58 | 5,64E-01 |
| AT1G07893 | 0,39 | -0,54 | 4,37 | -0,12 | 9,01E-01 |
| AT1G07897 | 0,95 | -0,11 | 1,74 | -0,06 | 9,48E-01 |
| AT1G53570 | 0,96 | -3,91 | 2,24 | -1,75 | 8,01E-02 |
| AT1G53590 | 0,26 | 0,47 | 4,36 | 0,11 | 9,14E-01 |
| AT1G53670 | 0,12 | 0,00 | 4,41 | 0,00 | 1,00E+00 |
| AT1G53720 | 0,26 | 0,00 | 4,41 | 0,00 | 1,00E+00 |
| AT1G53840 | 1,21 | 1,88 | 2,00 | 0,94 | 3,49E-01 |
| AT1G53870 | 0,49 | 1,80 | 3,90 | 0,46 | 6,44E-01 |
| AT1G53885 | 0,24 | -0,04 | 4,38 | -0,01 | 9,93E-01 |
| AT1G53910 | 0,62 | 2,12 | 2,43 | 0,87 | 3,83E-01 |
| AT1G54030 | 0,19 | 1,18 | 4,31 | 0,27 | 7,84E-01 |
| AT1G54130 | 1,27 | -3,02 | 1,97 | -1,53 | 1,26E-01 |
| AT1G54150 | 0,44 | -0,91 | 3,41 | -0,27 | 7,89E-01 |
| AT1G54200 | 0,80 | 0,23 | 2,35 | 0,10 | 9,22E-01 |
| AT1G54215 | 0,08 | 0,00 | 4,41 | 0,00 | 1,00E+00 |
| AT1G07957 | 0,34 | 1,47 | 3,42 | 0,43 | 6,68E-01 |
| AT1G54310 | 0,08 | 0,00 | 4,41 | 0,00 | 1,00E+00 |
| AT1G54340 | 0,28 | -1,10 | 4,35 | -0,25 | 8,00E-01 |
| AT1G54350 | 0,37 | -1,68 | 4,34 | -0,39 | 6,99E-01 |
| AT1G54360 | 0,57 | -4,07 | 4,33 | -0,94 | 3,47E-01 |
| AT1G54420 | 1,07 | 3,44 | 2,54 | 1,36 | 1,75E-01 |
| AT1G54440 | 0,25 | -0,73 | 4,40 | -0,16 | 8,69E-01 |
| AT1G54550 | 0,56 | 2,23 | 2,88 | 0,77 | 4,39E-01 |
| AT1G54680 | 0,26 | -2,60 | 4,41 | -0,59 | 5,55E-01 |
| AT1G54720 | 0,66 | 2,69 | 3,01 | 0,89 | 3,71E-01 |
| AT1G54780 | 1,22 | 1,53 | 1,79 | 0,85 | 3,93E-01 |
| AT1G54820 | 0,40 | -1,17 | 4,33 | -0,27 | 7,86E-01 |
| AT1G54920 | 1,30 | -3,49 | 1,84 | -1,89 | 5,82E-02 |
| AT1G55000 | 0,70 | 0,12 | 2,67 | 0,04 | 9,65E-01 |
| AT1G55020 | 0,29 | -0,89 | 4,35 | -0,20 | 8,38E-01 |
| AT1G55040 | 0,41 | -2,09 | 4,20 | -0,50 | 6,19E-01 |
| AT1G55090 | 0,81 | -1,30 | 2,88 | -0,45 | 6,52E-01 |
| AT1G55120 | 1,23 | -0,12 | 1,76 | -0,07 | 9,47E-01 |
| AT1G55150 | 0,30 | -2,60 | 4,41 | -0,59 | 5,55E-01 |
| AT1G55160 | 0,99 | 0,13 | 2,33 | 0,06 | 9,55E-01 |
| AT1G55175 | 0,08 | 0,00 | 4,41 | 0,00 | 1,00E+00 |
| AT1G55320 | 0,25 | -2,48 | 4,39 | -0,57 | 5,71E-01 |
| AT1G55335 | 0,51 | -1,98 | 3,23 | -0,61 | 5,40E-01 |
| AT1G55370 | 0,24 | -3,18 | 4,39 | -0,73 | 4,68E-01 |
| AT1G55460 | 0,32 | -0,10 | 4,37 | -0,02 | 9,82E-01 |
| AT1G55480 | 0,48 | -1,62 | 4,08 | -0,40 | 6,92E-01 |
| AT1G55500 | 0,61 | -2,42 | 2,21 | -1,10 | 2,73E-01 |
| AT1G55510 | 0,20 | 0,00 | 4,41 | 0,00 | 1,00E+00 |
| AT1G55840 | 0,33 | -2,06 | 4,35 | -0,47 | 6,36E-01 |
| AT1G55850 | 0,50 | 0,00 | 4,41 | 0,00 | 1,00E+00 |
| AT1G08073 | 0,33 | -2,52 | 4,40 | -0,57 | 5,66E-01 |
| AT1G55890 | 0,12 | -0,78 | 4,39 | -0,18 | 8,59E-01 |
| AT1G55910 | 0,98 | -1,69 | 1,74 | -0,97 | 3,32E-01 |
| AT1G55915 | 0,81 | -0,69 | 2,91 | -0,24 | 8,12E-01 |

| | | | | | |
|---|---|---|---|---|---|
| AT1G08077 | 0,62 | 1,70 | 2,91 | 0,58 | 5,60E-01 |
| AT1G56045 | 0,75 | -0,97 | 2,73 | -0,35 | 7,23E-01 |
| AT1G56050 | 1,07 | -4,46 | 2,67 | -1,67 | 9,48E-02 |
| AT1G56190 | 1,11 | -0,62 | 2,11 | -0,30 | 7,68E-01 |
| AT1G56210 | 0,05 | -0,24 | 4,38 | -0,05 | 9,57E-01 |
| AT1G56450 | 0,83 | -0,16 | 2,24 | -0,07 | 9,43E-01 |
| AT1G56460 | 0,70 | 0,60 | 2,94 | 0,21 | 8,37E-01 |
| AT1G56500 | 0,95 | -0,80 | 1,95 | -0,41 | 6,83E-01 |
| AT1G56540 | 0,56 | -3,21 | 4,38 | -0,73 | 4,64E-01 |
| AT1G56560 | 0,52 | -2,60 | 3,96 | -0,66 | 5,11E-01 |
| AT1G56570 | 0,28 | -1,10 | 4,35 | -0,25 | 8,00E-01 |
| AT1G56600 | 0,19 | 0,20 | 4,37 | 0,05 | 9,63E-01 |
| AT1G56610 | 0,17 | 0,00 | 4,41 | 0,00 | 1,00E+00 |
| AT1G56660 | 1,02 | 1,39 | 1,80 | 0,77 | 4,41E-01 |
| AT1G56700 | 0,33 | 0,44 | 3,59 | 0,12 | 9,02E-01 |
| AT1G57620 | 0,88 | -1,33 | 3,11 | -0,43 | 6,69E-01 |
| AT1G57630 | 0,41 | -0,82 | 4,33 | -0,19 | 8,50E-01 |
| AT1G57650 | 0,46 | 0,96 | 2,76 | 0,35 | 7,28E-01 |
| AT1G57760 | 0,12 | -0,06 | 4,37 | -0,01 | 9,88E-01 |
| AT1G57765 | 0,15 | 0,05 | 4,37 | 0,01 | 9,91E-01 |
| AT1G58190 | 0,62 | -1,85 | 2,45 | -0,75 | 4,51E-01 |
| AT1G58215 | 0,57 | -2,54 | 4,40 | -0,58 | 5,64E-01 |
| AT1G58223 | 0,54 | -2,87 | 3,85 | -0,75 | 4,56E-01 |
| AT1G58260 | 0,12 | -0,14 | 4,38 | -0,03 | 9,74E-01 |
| AT1G58360 | 0,57 | -1,40 | 3,90 | -0,36 | 7,19E-01 |
| AT1G58525 | 0,11 | -0,68 | 4,41 | -0,15 | 8,77E-01 |
| AT1G58520 | 0,55 | -0,56 | 3,29 | -0,17 | 8,65E-01 |
| AT1G58643 | 0,42 | -2,02 | 4,36 | -0,46 | 6,43E-01 |
| AT1G58808 | 1,22 | -2,85 | 1,88 | -1,52 | 1,29E-01 |
| AT1G59124 | 1,27 | -2,27 | 1,56 | -1,45 | 1,46E-01 |
| AT1G59218 | 1,28 | -0,25 | 1,78 | -0,14 | 8,89E-01 |
| AT1G59590 | 0,59 | -1,53 | 3,13 | -0,49 | 6,24E-01 |
| AT1G59620 | 0,81 | -3,59 | 4,34 | -0,83 | 4,08E-01 |
| AT1G59650 | 0,54 | -1,06 | 3,90 | -0,27 | 7,86E-01 |
| AT1G59710 | 0,56 | -1,64 | 3,33 | -0,49 | 6,22E-01 |
| AT1G59890 | 0,93 | 0,19 | 2,48 | 0,08 | 9,40E-01 |
| AT1G59900 | 1,24 | 2,09 | 1,89 | 1,11 | 2,68E-01 |
| AT1G60025 | 0,05 | -0,24 | 4,38 | -0,05 | 9,57E-01 |
| AT1G60170 | 1,29 | -0,72 | 2,27 | -0,31 | 7,53E-01 |
| AT1G60190 | 0,89 | -2,13 | 2,16 | -0,99 | 3,23E-01 |
| AT1G60220 | 0,08 | 0,00 | 4,41 | 0,00 | 1,00E+00 |
| AT1G60420 | 0,70 | -0,33 | 2,93 | -0,11 | 9,10E-01 |
| AT1G60545 | 1,07 | -4,33 | 3,01 | -1,44 | 1,51E-01 |
| AT1G60650 | 0,21 | -2,50 | 4,39 | -0,57 | 5,70E-01 |
| AT1G60900 | 0,65 | -0,76 | 3,24 | -0,23 | 8,16E-01 |
| AT1G08353 | 0,98 | -2,27 | 1,97 | -1,15 | 2,48E-01 |
| AT1G61030 | 0,52 | -2,55 | 4,00 | -0,64 | 5,23E-01 |
| AT1G61040 | 0,54 | -1,06 | 3,90 | -0,27 | 7,86E-01 |
| AT1G61150 | 0,81 | -2,76 | 3,62 | -0,76 | 4,46E-01 |
| AT1G61170 | 0,33 | 0,02 | 4,36 | 0,00 | 9,97E-01 |
| AT1G61180 | 0,58 | -1,50 | 2,74 | -0,55 | 5,84E-01 |
| AT1G61210 | 1,07 | -2,48 | 1,76 | -1,41 | 1,58E-01 |
| AT1G61250 | 0,18 | 0,78 | 4,34 | 0,18 | 8,57E-01 |
| AT1G61280 | 0,12 | 0,00 | 4,41 | 0,00 | 1,00E+00 |
| AT1G61350 | 0,76 | 1,76 | 2,28 | 0,77 | 4,40E-01 |
| AT1G61450 | 0,12 | 0,00 | 4,41 | 0,00 | 1,00E+00 |

| | | | | | |
|---|---|---|---|---|---|
| AT1G61470 | 0,18 | 0,57 | 4,34 | 0,13 | 8,95E-01 |
| AT1G61610 | 0,12 | -0,06 | 4,37 | -0,01 | 9,88E-01 |
| AT1G61620 | 0,37 | -3,03 | 4,38 | -0,69 | 4,90E-01 |
| AT1G61800 | 0,56 | -2,59 | 4,41 | -0,59 | 5,56E-01 |
| AT1G61820 | 0,08 | 0,00 | 4,41 | 0,00 | 1,00E+00 |
| AT1G61850 | 0,80 | -2,14 | 2,34 | -0,91 | 3,60E-01 |
| AT1G61990 | 0,32 | -0,74 | 4,40 | -0,17 | 8,67E-01 |
| AT1G62170 | 0,36 | 0,11 | 4,37 | 0,02 | 9,80E-01 |
| AT1G62270 | 0,29 | 1,83 | 4,29 | 0,43 | 6,70E-01 |
| AT1G08517 | 0,77 | 0,42 | 2,45 | 0,17 | 8,65E-01 |
| AT1G62380 | 1,16 | 1,75 | 1,81 | 0,96 | 3,36E-01 |
| AT1G62410 | 0,32 | 0,60 | 4,34 | 0,14 | 8,90E-01 |
| AT1G62480 | 0,12 | -0,06 | 4,37 | -0,01 | 9,88E-01 |
| AT1G62570 | 0,39 | 0,37 | 4,33 | 0,08 | 9,33E-01 |
| AT1G62620 | 0,57 | -2,95 | 3,03 | -0,97 | 3,31E-01 |
| AT1G62720 | 0,24 | 0,78 | 4,33 | 0,18 | 8,58E-01 |
| AT1G62790 | 0,51 | -4,07 | 4,33 | -0,94 | 3,48E-01 |
| AT1G62920 | 0,97 | 2,07 | 2,93 | 0,71 | 4,80E-01 |
| AT1G62970 | 0,33 | 0,59 | 3,53 | 0,17 | 8,67E-01 |
| AT1G62975 | 0,08 | 0,00 | 4,41 | 0,00 | 1,00E+00 |
| AT1G63000 | 0,67 | 0,70 | 3,78 | 0,19 | 8,53E-01 |
| AT1G63110 | 0,32 | -1,65 | 4,40 | -0,38 | 7,08E-01 |
| AT1G63170 | 0,08 | 0,00 | 4,41 | 0,00 | 1,00E+00 |
| AT1G63430 | 0,74 | -0,79 | 3,92 | -0,20 | 8,39E-01 |
| AT1G63500 | 0,59 | 0,21 | 2,69 | 0,08 | 9,37E-01 |
| AT1G63540 | 0,37 | 2,18 | 4,28 | 0,51 | 6,11E-01 |
| AT1G63680 | 0,43 | -0,51 | 3,26 | -0,16 | 8,76E-01 |
| AT1G63690 | 1,03 | -1,79 | 1,62 | -1,11 | 2,67E-01 |
| AT1G63730 | 0,24 | 1,71 | 4,29 | 0,40 | 6,91E-01 |
| AT1G63830 | 0,49 | 1,26 | 4,05 | 0,31 | 7,55E-01 |
| AT1G63850 | 0,53 | -2,15 | 3,96 | -0,54 | 5,87E-01 |
| AT1G08643 | 0,79 | 2,06 | 2,93 | 0,70 | 4,81E-01 |
| AT1G63900 | 0,95 | -1,58 | 1,98 | -0,80 | 4,24E-01 |
| AT1G63940 | 0,81 | -1,43 | 1,91 | -0,75 | 4,53E-01 |
| AT1G64060 | 0,86 | -0,01 | 2,41 | 0,00 | 9,98E-01 |
| AT1G64065 | 0,29 | -0,47 | 4,35 | -0,11 | 9,15E-01 |
| AT1G64090 | 0,96 | -4,00 | 2,15 | -1,86 | 6,23E-02 |
| AT1G64100 | 0,61 | -3,16 | 3,75 | -0,84 | 3,99E-01 |
| AT1G64120 | 0,13 | 0,52 | 4,35 | 0,12 | 9,04E-01 |
| AT1G64130 | 0,12 | 0,00 | 4,41 | 0,00 | 1,00E+00 |
| AT1G08657 | 0,60 | -0,51 | 2,67 | -0,19 | 8,49E-01 |
| AT1G64370 | 0,17 | 0,66 | 4,34 | 0,15 | 8,80E-01 |
| AT1G64520 | 0,27 | 0,89 | 4,33 | 0,21 | 8,37E-01 |
| AT1G64530 | 0,54 | -3,43 | 3,15 | -1,09 | 2,76E-01 |
| AT1G64550 | 0,64 | -1,01 | 2,63 | -0,38 | 7,00E-01 |
| AT1G64570 | 0,71 | -1,72 | 2,59 | -0,66 | 5,07E-01 |
| AT1G64572 | 0,51 | -3,74 | 4,35 | -0,86 | 3,90E-01 |
| AT1G64620 | 0,45 | 0,74 | 3,18 | 0,23 | 8,16E-01 |
| AT1G64740 | 0,25 | 1,50 | 4,30 | 0,35 | 7,28E-01 |
| AT1G64770 | 1,29 | -0,62 | 1,58 | -0,39 | 6,95E-01 |
| AT1G64795 | 0,74 | 0,87 | 1,95 | 0,45 | 6,56E-01 |
| AT1G64810 | 0,44 | -0,57 | 4,34 | -0,13 | 8,96E-01 |
| AT1G64820 | 0,08 | 0,00 | 4,41 | 0,00 | 1,00E+00 |
| AT1G64860 | 0,65 | -1,72 | 3,22 | -0,53 | 5,93E-01 |
| AT1G64900 | 1,03 | -1,87 | 2,03 | -0,92 | 3,57E-01 |
| AT1G64950 | 0,34 | 0,36 | 3,48 | 0,10 | 9,18E-01 |

| | | | | | |
|---|---|---|---|---|---|
| AT1G64960 | 0,60 | -2,04 | 3,96 | -0,51 | 6,07E-01 |
| AT1G65220 | 0,59 | -0,27 | 2,68 | -0,10 | 9,19E-01 |
| AT1G65230 | 1,04 | -2,18 | 1,94 | -1,12 | 2,62E-01 |
| AT1G65260 | 0,21 | 1,49 | 4,30 | 0,35 | 7,29E-01 |
| AT1G65270 | 0,21 | 0,76 | 4,34 | 0,18 | 8,61E-01 |
| AT1G65280 | 0,39 | -0,65 | 4,27 | -0,15 | 8,79E-01 |
| AT1G65295 | 0,05 | -0,24 | 4,38 | -0,05 | 9,57E-01 |
| AT1G65360 | 0,90 | -3,89 | 2,37 | -1,64 | 1,01E-01 |
| AT1G08753 | 0,15 | 0,00 | 4,41 | 0,00 | 1,00E+00 |
| AT1G65490 | 0,49 | -2,10 | 4,29 | -0,49 | 6,24E-01 |
| AT1G65590 | 0,23 | -1,68 | 4,36 | -0,39 | 7,00E-01 |
| AT1G65681 | 0,20 | 1,40 | 4,30 | 0,33 | 7,44E-01 |
| AT1G65730 | 0,08 | 0,00 | 4,41 | 0,00 | 1,00E+00 |
| AT1G65790 | 0,93 | -1,38 | 1,82 | -0,76 | 4,47E-01 |
| AT1G65800 | 1,17 | 0,82 | 1,74 | 0,47 | 6,36E-01 |
| AT1G65820 | 0,38 | 0,20 | 2,84 | 0,07 | 9,44E-01 |
| AT1G65840 | 0,61 | -0,84 | 3,81 | -0,22 | 8,25E-01 |
| AT1G08793 | 0,15 | 0,48 | 4,35 | 0,11 | 9,12E-01 |
| AT1G66090 | 0,59 | -1,13 | 3,21 | -0,35 | 7,24E-01 |
| AT1G66160 | 0,48 | -0,68 | 4,11 | -0,17 | 8,69E-01 |
| AT1G66180 | 0,53 | 0,25 | 3,09 | 0,08 | 9,35E-01 |
| AT1G66230 | 0,24 | -3,18 | 4,39 | -0,73 | 4,68E-01 |
| AT1G66340 | 0,60 | -1,64 | 3,01 | -0,55 | 5,86E-01 |
| AT1G66345 | 0,47 | -0,70 | 4,08 | -0,17 | 8,65E-01 |
| AT1G66430 | 0,50 | -0,68 | 4,41 | -0,15 | 8,77E-01 |
| AT1G66465 | 0,33 | 0,00 | 4,41 | 0,00 | 1,00E+00 |
| AT1G66640 | 0,21 | 0,51 | 4,35 | 0,12 | 9,06E-01 |
| AT1G66750 | 0,31 | 0,00 | 4,41 | 0,00 | 1,00E+00 |
| AT1G66820 | 0,53 | -2,47 | 4,39 | -0,56 | 5,73E-01 |
| AT1G66900 | 0,20 | -0,57 | 4,39 | -0,13 | 8,97E-01 |
| AT1G67105 | 0,86 | 1,56 | 1,92 | 0,81 | 4,17E-01 |
| AT1G67190 | 0,34 | -0,57 | 4,34 | -0,13 | 8,96E-01 |
| AT1G08923 | 0,11 | 0,46 | 4,35 | 0,11 | 9,15E-01 |
| AT1G67420 | 0,13 | 0,00 | 4,41 | 0,00 | 1,00E+00 |
| AT1G08937 | 0,46 | 1,70 | 3,21 | 0,53 | 5,95E-01 |
| AT1G67470 | 0,43 | -2,06 | 3,31 | -0,62 | 5,33E-01 |
| AT1G67480 | 0,49 | -1,53 | 4,34 | -0,35 | 7,24E-01 |
| AT1G67530 | 0,54 | -3,20 | 3,88 | -0,83 | 4,09E-01 |
| AT1G67560 | 0,64 | -3,69 | 4,35 | -0,85 | 3,95E-01 |
| AT1G67620 | 0,15 | 0,41 | 4,35 | 0,09 | 9,26E-01 |
| AT1G67700 | 0,46 | 0,91 | 3,90 | 0,23 | 8,16E-01 |
| AT1G67720 | 0,53 | -3,32 | 3,82 | -0,87 | 3,85E-01 |
| AT1G67792 | 0,38 | -3,44 | 4,37 | -0,79 | 4,31E-01 |
| AT1G67900 | 1,04 | -0,80 | 2,32 | -0,34 | 7,31E-01 |
| AT1G67940 | 0,66 | -4,52 | 2,90 | -1,56 | 1,19E-01 |
| AT1G08963 | 0,39 | -3,07 | 4,38 | -0,70 | 4,84E-01 |
| AT1G67980 | 0,18 | 0,00 | 4,41 | 0,00 | 1,00E+00 |
| AT1G67990 | 0,55 | -3,40 | 3,84 | -0,89 | 3,75E-01 |
| AT1G68050 | 0,24 | -0,68 | 4,41 | -0,15 | 8,77E-01 |
| AT1G68100 | 0,43 | -2,60 | 4,19 | -0,62 | 5,35E-01 |
| AT1G68110 | 0,27 | -2,58 | 4,40 | -0,59 | 5,58E-01 |
| AT1G68310 | 0,12 | -0,68 | 4,41 | -0,15 | 8,77E-01 |
| AT1G68490 | 0,97 | -1,76 | 2,40 | -0,73 | 4,62E-01 |
| AT1G68570 | 0,58 | 1,17 | 2,55 | 0,46 | 6,47E-01 |
| AT1G68585 | 0,27 | -1,27 | 4,38 | -0,29 | 7,72E-01 |
| AT1G68620 | 0,33 | -2,55 | 4,40 | -0,58 | 5,62E-01 |

| | | | | | |
|---|---|---|---|---|---|
| AT1G68650 | 0,32 | -3,12 | 4,39 | -0,71 | 4,77E-01 |
| AT1G68670 | 0,63 | 0,73 | 2,97 | 0,25 | 8,05E-01 |
| AT1G68840 | 1,23 | -2,30 | 1,74 | -1,32 | 1,86E-01 |
| AT1G68890 | 1,09 | -2,88 | 2,08 | -1,38 | 1,66E-01 |
| AT1G68980 | 0,24 | -2,60 | 4,41 | -0,59 | 5,55E-01 |
| AT1G69085 | 0,08 | 0,05 | 4,38 | 0,01 | 9,91E-01 |
| AT1G69160 | 0,39 | 1,82 | 3,28 | 0,56 | 5,78E-01 |
| AT1G69170 | 0,40 | 0,00 | 4,41 | 0,00 | 1,00E+00 |
| AT1G69200 | 0,21 | -1,17 | 4,36 | -0,27 | 7,88E-01 |
| AT1G69252 | 0,90 | -0,07 | 2,90 | -0,03 | 9,80E-01 |
| AT1G69260 | 0,63 | 0,71 | 2,65 | 0,27 | 7,89E-01 |
| AT1G69350 | 0,78 | -2,21 | 2,46 | -0,90 | 3,67E-01 |
| AT1G69390 | 0,12 | -0,31 | 4,38 | -0,07 | 9,44E-01 |
| AT1G69410 | 0,27 | -0,72 | 4,40 | -0,16 | 8,70E-01 |
| AT1G69420 | 0,65 | -3,55 | 2,47 | -1,44 | 1,50E-01 |
| AT1G69510 | 0,10 | 0,45 | 4,35 | 0,10 | 9,19E-01 |
| AT1G69523 | 0,10 | 0,45 | 4,35 | 0,10 | 9,19E-01 |
| AT1G69526 | 0,08 | 0,00 | 4,41 | 0,00 | 1,00E+00 |
| AT1G69530 | 0,93 | 0,12 | 1,82 | 0,06 | 9,48E-01 |
| AT1G69580 | 0,26 | -0,35 | 4,38 | -0,08 | 9,36E-01 |
| AT1G69610 | 0,21 | -0,37 | 4,38 | -0,08 | 9,33E-01 |
| AT1G09087 | 0,37 | -1,09 | 3,50 | -0,31 | 7,56E-01 |
| AT1G69690 | 0,67 | 1,72 | 2,50 | 0,69 | 4,92E-01 |
| AT1G69740 | 0,99 | 0,47 | 1,81 | 0,26 | 7,93E-01 |
| AT1G69750 | 0,10 | 0,45 | 4,35 | 0,10 | 9,19E-01 |
| AT1G69870 | 0,38 | -0,79 | 4,39 | -0,18 | 8,57E-01 |
| AT1G70090 | 0,99 | 0,45 | 2,00 | 0,22 | 8,23E-01 |
| AT1G70100 | 0,38 | -0,68 | 4,36 | -0,16 | 8,76E-01 |
| AT1G70160 | 1,19 | -0,93 | 1,68 | -0,55 | 5,79E-01 |
| AT1G70180 | 1,28 | -3,82 | 1,89 | -2,02 | 4,35E-02 |
| AT1G70190 | 0,40 | -0,74 | 4,40 | -0,17 | 8,67E-01 |
| AT1G70210 | 0,79 | 2,59 | 2,32 | 1,11 | 2,66E-01 |
| AT1G70230 | 0,59 | 0,25 | 2,65 | 0,10 | 9,24E-01 |
| AT1G70250 | 0,71 | -4,08 | 2,93 | -1,39 | 1,64E-01 |
| AT1G70330 | 0,33 | 1,21 | 3,53 | 0,34 | 7,31E-01 |
| AT1G70480 | 1,12 | -2,24 | 1,72 | -1,30 | 1,93E-01 |
| AT1G70510 | 0,12 | 0,00 | 4,41 | 0,00 | 1,00E+00 |
| AT1G70550 | 0,14 | 0,40 | 4,35 | 0,09 | 9,27E-01 |
| AT1G70570 | 0,36 | -2,50 | 4,39 | -0,57 | 5,69E-01 |
| AT1G70580 | 0,81 | -3,53 | 2,50 | -1,41 | 1,58E-01 |
| AT1G70620 | 1,32 | -2,46 | 2,11 | -1,17 | 2,44E-01 |
| AT1G70630 | 0,36 | -2,20 | 4,35 | -0,51 | 6,12E-01 |
| AT1G70645 | 0,05 | -0,24 | 4,38 | -0,05 | 9,57E-01 |
| AT1G70650 | 0,26 | -1,28 | 4,35 | -0,29 | 7,69E-01 |
| AT1G70660 | 0,20 | -2,55 | 4,40 | -0,58 | 5,62E-01 |
| AT1G70700 | 0,60 | 1,87 | 2,58 | 0,73 | 4,67E-01 |
| AT1G70760 | 0,35 | 0,19 | 3,51 | 0,05 | 9,57E-01 |
| AT1G70810 | 0,62 | -3,54 | 2,66 | -1,33 | 1,84E-01 |
| AT1G70820 | 0,43 | -0,46 | 2,79 | -0,16 | 8,69E-01 |
| AT1G70895 | 0,08 | -0,21 | 4,39 | -0,05 | 9,62E-01 |
| AT1G70900 | 1,03 | -2,02 | 1,96 | -1,03 | 3,02E-01 |
| AT1G70960 | 0,12 | -0,06 | 4,37 | -0,01 | 9,88E-01 |
| AT1G70980 | 0,40 | 0,01 | 4,37 | 0,00 | 9,98E-01 |
| AT1G71090 | 0,35 | -0,79 | 3,56 | -0,22 | 8,25E-01 |
| AT1G71100 | 1,11 | 0,43 | 1,86 | 0,23 | 8,18E-01 |
| AT1G71110 | 0,15 | 0,05 | 4,37 | 0,01 | 9,91E-01 |

| | | | | | |
|---|---|---|---|---|---|
| AT1G71130 | 0,37 | -0,27 | 4,38 | -0,06 | 9,50E-01 |
| AT1G71180 | 0,42 | -1,16 | 4,29 | -0,27 | 7,87E-01 |
| AT1G71270 | 0,45 | 0,60 | 4,16 | 0,14 | 8,85E-01 |
| AT1G71300 | 0,38 | -3,32 | 4,36 | -0,76 | 4,47E-01 |
| AT1G71320 | 0,39 | 2,21 | 4,13 | 0,54 | 5,92E-01 |
| AT1G71480 | 0,80 | -2,09 | 2,91 | -0,72 | 4,72E-01 |
| AT1G09257 | 0,17 | 0,00 | 4,41 | 0,00 | 1,00E+00 |
| AT1G71500 | 0,95 | -1,34 | 1,84 | -0,73 | 4,68E-01 |
| AT1G71695 | 0,82 | -1,78 | 2,45 | -0,73 | 4,68E-01 |
| AT1G71720 | 0,53 | 0,00 | 3,89 | 0,00 | 1,00E+00 |
| AT1G71800 | 0,67 | -4,11 | 3,62 | -1,14 | 2,56E-01 |
| AT1G71840 | 1,05 | 2,49 | 2,10 | 1,19 | 2,35E-01 |
| AT1G71900 | 0,51 | -3,74 | 4,35 | -0,86 | 3,90E-01 |
| AT1G71910 | 0,44 | 2,06 | 3,18 | 0,65 | 5,16E-01 |
| AT1G71920 | 0,73 | 2,54 | 2,36 | 1,08 | 2,82E-01 |
| AT1G71930 | 1,13 | 2,56 | 2,05 | 1,25 | 2,11E-01 |
| AT1G71970 | 0,14 | -0,11 | 4,37 | -0,03 | 9,80E-01 |
| AT1G71980 | 0,46 | 1,69 | 3,12 | 0,54 | 5,87E-01 |
| AT1G72010 | 0,42 | 0,36 | 3,30 | 0,11 | 9,13E-01 |
| AT1G72030 | 0,75 | -1,55 | 3,64 | -0,43 | 6,70E-01 |
| AT1G72050 | 0,75 | -4,27 | 3,62 | -1,18 | 2,38E-01 |
| AT1G72090 | 0,34 | -1,64 | 4,41 | -0,37 | 7,10E-01 |
| AT1G72240 | 0,60 | -2,10 | 3,90 | -0,54 | 5,89E-01 |
| AT1G72310 | 0,25 | -2,61 | 4,41 | -0,59 | 5,53E-01 |
| AT1G72330 | 0,67 | -2,41 | 2,85 | -0,85 | 3,96E-01 |
| AT1G72470 | 0,43 | -3,13 | 4,39 | -0,71 | 4,76E-01 |
| AT1G72510 | 0,57 | 1,11 | 3,22 | 0,34 | 7,30E-01 |
| AT1G72520 | 0,91 | 1,06 | 2,00 | 0,53 | 5,95E-01 |
| AT1G72530 | 0,05 | -0,24 | 4,38 | -0,05 | 9,57E-01 |
| AT1G72650 | 0,79 | -0,42 | 2,39 | -0,17 | 8,62E-01 |
| AT1G72700 | 1,31 | 0,81 | 1,88 | 0,43 | 6,65E-01 |
| AT1G72750 | 0,63 | -0,27 | 2,11 | -0,13 | 9,00E-01 |
| AT1G72790 | 0,71 | 1,31 | 2,98 | 0,44 | 6,60E-01 |
| AT1G72855 | 0,40 | 0,00 | 4,41 | 0,00 | 1,00E+00 |
| AT1G72900 | 0,32 | 0,02 | 3,57 | 0,01 | 9,95E-01 |
| AT1G72910 | 1,29 | 1,05 | 1,76 | 0,60 | 5,50E-01 |
| AT1G72920 | 0,44 | -1,58 | 3,36 | -0,47 | 6,38E-01 |
| AT1G72940 | 0,97 | 0,00 | 4,10 | 0,00 | 1,00E+00 |
| AT1G72970 | 0,16 | -0,26 | 4,38 | -0,06 | 9,52E-01 |
| AT1G73030 | 0,82 | -0,80 | 2,20 | -0,36 | 7,18E-01 |
| AT1G73060 | 0,24 | -0,68 | 4,34 | -0,16 | 8,75E-01 |
| AT1G73100 | 0,70 | -1,52 | 2,55 | -0,60 | 5,51E-01 |
| AT1G73180 | 1,25 | -4,01 | 1,92 | -2,09 | 3,70E-02 |
| AT1G73430 | 1,05 | -1,22 | 2,23 | -0,55 | 5,83E-01 |
| AT1G73450 | 1,12 | -1,51 | 1,81 | -0,83 | 4,04E-01 |
| AT1G73460 | 0,79 | -1,50 | 2,43 | -0,62 | 5,38E-01 |
| AT1G73470 | 0,23 | -1,72 | 4,40 | -0,39 | 6,95E-01 |
| AT1G73630 | 0,10 | -0,34 | 4,38 | -0,08 | 9,38E-01 |
| AT1G73670 | 0,46 | -1,61 | 3,39 | -0,48 | 6,34E-01 |
| AT1G73710 | 0,22 | -0,21 | 4,38 | -0,05 | 9,61E-01 |
| AT1G73750 | 0,72 | 0,00 | 4,41 | 0,00 | 1,00E+00 |
| AT1G73770 | 0,08 | 0,00 | 4,41 | 0,00 | 1,00E+00 |
| AT1G73850 | 0,17 | -1,25 | 4,37 | -0,29 | 7,74E-01 |
| AT1G73875 | 0,39 | -1,32 | 4,25 | -0,31 | 7,56E-01 |
| AT1G73885 | 0,62 | 0,87 | 3,10 | 0,28 | 7,78E-01 |
| AT1G73920 | 0,62 | -1,37 | 2,42 | -0,57 | 5,71E-01 |

| | | | | | |
|---|---|---|---|---|---|
| AT1G73970 | 0,95 | 1,75 | 2,45 | 0,71 | 4,75E-01 |
| AT1G74160 | 1,08 | -3,48 | 1,77 | -1,96 | 4,95E-02 |
| AT1G74360 | 1,21 | -2,20 | 2,57 | -0,86 | 3,92E-01 |
| AT1G74380 | 1,29 | -0,77 | 1,54 | -0,50 | 6,18E-01 |
| AT1G74430 | 0,38 | -0,46 | 4,35 | -0,10 | 9,17E-01 |
| AT1G74450 | 0,99 | 1,65 | 1,94 | 0,85 | 3,98E-01 |
| AT1G74580 | 0,25 | -1,17 | 4,36 | -0,27 | 7,88E-01 |
| AT1G74680 | 0,57 | -2,40 | 3,13 | -0,77 | 4,44E-01 |
| AT1G74710 | 0,60 | -0,17 | 2,64 | -0,06 | 9,49E-01 |
| AT1G74730 | 0,44 | -1,29 | 3,30 | -0,39 | 6,96E-01 |
| AT1G74750 | 0,50 | -3,94 | 4,34 | -0,91 | 3,63E-01 |
| AT1G74780 | 0,63 | -1,72 | 3,03 | -0,57 | 5,71E-01 |
| AT1G74790 | 0,51 | -1,41 | 3,91 | -0,36 | 7,18E-01 |
| AT1G74870 | 1,10 | -1,72 | 3,35 | -0,51 | 6,09E-01 |
| AT1G74930 | 0,81 | 3,14 | 2,66 | 1,18 | 2,38E-01 |
| AT1G74940 | 0,58 | 1,20 | 3,05 | 0,39 | 6,94E-01 |
| AT1G75080 | 0,21 | -0,70 | 4,40 | -0,16 | 8,73E-01 |
| AT1G09533 | 1,00 | 0,21 | 1,85 | 0,11 | 9,10E-01 |
| AT1G75163 | 0,94 | 1,93 | 2,04 | 0,95 | 3,44E-01 |
| AT1G75166 | 1,22 | 1,20 | 1,98 | 0,61 | 5,42E-01 |
| AT1G75170 | 0,12 | -0,68 | 4,41 | -0,15 | 8,77E-01 |
| AT1G09543 | 0,38 | 1,23 | 3,36 | 0,37 | 7,14E-01 |
| AT1G75190 | 0,22 | 0,80 | 4,34 | 0,18 | 8,54E-01 |
| AT1G75210 | 0,77 | -1,58 | 2,99 | -0,53 | 5,98E-01 |
| AT1G75280 | 0,22 | 1,22 | 4,31 | 0,28 | 7,78E-01 |
| AT1G75310 | 0,47 | -3,10 | 4,38 | -0,71 | 4,79E-01 |
| AT1G75350 | 0,83 | -0,34 | 2,19 | -0,15 | 8,78E-01 |
| AT1G75400 | 0,32 | 0,46 | 4,34 | 0,11 | 9,16E-01 |
| AT1G75420 | 0,31 | -1,42 | 4,37 | -0,32 | 7,46E-01 |
| AT1G75440 | 0,28 | -1,75 | 4,39 | -0,40 | 6,90E-01 |
| AT1G75460 | 0,84 | -3,17 | 2,23 | -1,42 | 1,55E-01 |
| AT1G75580 | 0,17 | 0,00 | 4,41 | 0,00 | 1,00E+00 |
| AT1G75710 | 0,31 | 1,14 | 4,31 | 0,27 | 7,91E-01 |
| AT1G75785 | 0,12 | -0,14 | 4,38 | -0,03 | 9,74E-01 |
| AT1G75810 | 0,60 | -2,98 | 2,62 | -1,14 | 2,55E-01 |
| AT1G75840 | 0,75 | -1,89 | 2,22 | -0,85 | 3,95E-01 |
| AT1G75900 | 0,32 | -0,09 | 4,33 | -0,02 | 9,83E-01 |
| AT1G75960 | 0,45 | -1,49 | 4,21 | -0,35 | 7,24E-01 |
| AT1G76000 | 0,58 | 0,45 | 3,02 | 0,15 | 8,82E-01 |
| AT1G76050 | 0,66 | -2,44 | 2,97 | -0,82 | 4,10E-01 |
| AT1G76140 | 0,48 | 0,67 | 4,00 | 0,17 | 8,67E-01 |
| AT1G76185 | 0,16 | -0,11 | 4,37 | -0,03 | 9,80E-01 |
| AT1G76190 | 0,11 | 0,46 | 4,35 | 0,11 | 9,15E-01 |
| AT1G76320 | 0,32 | 0,01 | 4,36 | 0,00 | 9,98E-01 |
| AT1G76350 | 0,12 | -0,78 | 4,39 | -0,18 | 8,59E-01 |
| AT1G76390 | 0,51 | -0,71 | 3,21 | -0,22 | 8,25E-01 |
| AT1G76450 | 0,08 | 0,00 | 4,41 | 0,00 | 1,00E+00 |
| AT1G76520 | 0,73 | -2,06 | 2,18 | -0,94 | 3,45E-01 |
| AT1G76580 | 0,80 | -1,80 | 3,48 | -0,52 | 6,05E-01 |
| AT1G76590 | 0,69 | -0,02 | 2,90 | -0,01 | 9,94E-01 |
| AT1G76600 | 0,49 | 0,92 | 3,21 | 0,29 | 7,75E-01 |
| AT1G76620 | 0,25 | -0,40 | 4,38 | -0,09 | 9,27E-01 |
| AT1G76680 | 0,14 | -0,35 | 4,38 | -0,08 | 9,37E-01 |
| AT1G76740 | 0,12 | 0,00 | 4,41 | 0,00 | 1,00E+00 |
| AT1G76820 | 0,53 | -1,57 | 3,23 | -0,49 | 6,27E-01 |
| AT1G76850 | 1,15 | -2,50 | 1,78 | -1,40 | 1,62E-01 |

| | | | | | |
|---|---|---|---|---|---|
| AT1G76880 | 1,06 | 0,31 | 1,81 | 0,17 | 8,63E-01 |
| AT1G76900 | 0,91 | -1,62 | 2,76 | -0,59 | 5,57E-01 |
| AT1G76920 | 0,59 | -2,58 | 3,81 | -0,68 | 4,99E-01 |
| AT1G76950 | 0,83 | -2,99 | 2,74 | -1,09 | 2,74E-01 |
| AT1G77080 | 0,41 | 1,16 | 3,28 | 0,35 | 7,23E-01 |
| AT1G77140 | 0,25 | -1,15 | 4,38 | -0,26 | 7,93E-01 |
| AT1G77320 | 0,26 | -2,60 | 4,41 | -0,59 | 5,55E-01 |
| AT1G77350 | 0,15 | 0,48 | 4,35 | 0,11 | 9,12E-01 |
| AT1G77420 | 0,12 | 0,00 | 4,41 | 0,00 | 1,00E+00 |
| AT1G77460 | 0,94 | -2,53 | 2,55 | -0,99 | 3,20E-01 |
| AT1G77490 | 0,64 | -0,43 | 2,92 | -0,15 | 8,83E-01 |
| AT1G77500 | 0,37 | -1,64 | 4,41 | -0,37 | 7,10E-01 |
| AT1G77510 | 1,26 | -2,64 | 1,67 | -1,58 | 1,13E-01 |
| AT1G77610 | 0,53 | -1,73 | 4,32 | -0,40 | 6,89E-01 |
| AT1G77710 | 0,32 | 0,19 | 4,32 | 0,04 | 9,65E-01 |
| AT1G77800 | 1,13 | -2,72 | 1,83 | -1,49 | 1,37E-01 |
| AT1G77840 | 0,31 | -0,95 | 3,62 | -0,26 | 7,94E-01 |
| AT1G77850 | 1,01 | 1,34 | 2,11 | 0,63 | 5,26E-01 |
| AT1G77890 | 0,98 | -3,94 | 1,89 | -2,08 | 3,73E-02 |
| AT1G77930 | 1,14 | -2,09 | 2,31 | -0,91 | 3,65E-01 |
| AT1G78010 | 0,90 | -1,83 | 2,12 | -0,86 | 3,90E-01 |
| AT1G78100 | 0,64 | 0,60 | 2,22 | 0,27 | 7,86E-01 |
| AT1G78150 | 0,22 | -1,27 | 4,36 | -0,29 | 7,70E-01 |
| AT1G78200 | 0,54 | -1,26 | 2,56 | -0,49 | 6,23E-01 |
| AT1G78300 | 0,54 | 1,08 | 3,18 | 0,34 | 7,33E-01 |
| AT1G78420 | 0,54 | -1,54 | 2,69 | -0,57 | 5,67E-01 |
| AT1G78490 | 0,05 | -0,24 | 4,38 | -0,05 | 9,57E-01 |
| AT1G78500 | 0,17 | 0,00 | 4,41 | 0,00 | 1,00E+00 |
| AT1G78600 | 0,20 | -1,00 | 4,37 | -0,23 | 8,19E-01 |
| AT1G78630 | 1,31 | -1,23 | 1,66 | -0,74 | 4,58E-01 |
| AT1G78670 | 0,52 | -1,47 | 3,13 | -0,47 | 6,38E-01 |
| AT1G78680 | 0,90 | 1,06 | 1,92 | 0,55 | 5,81E-01 |
| AT1G78700 | 0,37 | -1,89 | 4,37 | -0,43 | 6,65E-01 |
| AT1G78710 | 0,38 | -3,70 | 4,35 | -0,85 | 3,95E-01 |
| AT1G78770 | 0,21 | -0,70 | 4,40 | -0,16 | 8,73E-01 |
| AT1G78870 | 0,28 | -1,31 | 4,37 | -0,30 | 7,65E-01 |
| AT1G78890 | 0,36 | -3,05 | 4,38 | -0,70 | 4,85E-01 |
| AT1G78910 | 0,26 | -2,60 | 4,41 | -0,59 | 5,55E-01 |
| AT1G79060 | 0,14 | 0,87 | 4,33 | 0,20 | 8,40E-01 |
| AT1G79103 | 0,05 | -0,24 | 4,38 | -0,05 | 9,57E-01 |
| AT1G09847 | 0,44 | 1,35 | 3,22 | 0,42 | 6,76E-01 |
| AT1G79190 | 0,88 | -4,91 | 2,67 | -1,84 | 6,57E-02 |
| AT1G79230 | 0,71 | -2,78 | 2,90 | -0,96 | 3,38E-01 |
| AT1G79240 | 0,20 | -0,68 | 4,41 | -0,15 | 8,77E-01 |
| AT1G79270 | 1,18 | -1,85 | 1,68 | -1,10 | 2,70E-01 |
| AT1G79340 | 0,91 | 2,05 | 2,07 | 0,99 | 3,22E-01 |
| AT1G79380 | 0,91 | -1,85 | 2,85 | -0,65 | 5,16E-01 |
| AT1G79560 | 0,28 | -0,92 | 4,35 | -0,21 | 8,33E-01 |
| AT1G79590 | 0,18 | -0,27 | 4,38 | -0,06 | 9,51E-01 |
| AT1G79660 | 0,63 | -4,50 | 3,61 | -1,25 | 2,13E-01 |
| AT1G09903 | 0,08 | 0,00 | 4,41 | 0,00 | 1,00E+00 |
| AT1G79810 | 0,41 | 0,06 | 3,27 | 0,02 | 9,85E-01 |
| AT1G79840 | 0,77 | -2,42 | 3,00 | -0,81 | 4,20E-01 |
| AT1G79870 | 0,21 | 0,09 | 4,36 | 0,02 | 9,84E-01 |
| AT1G79890 | 0,21 | -2,50 | 4,39 | -0,57 | 5,70E-01 |
| AT1G79910 | 0,41 | -3,36 | 4,19 | -0,80 | 4,23E-01 |

| | | | | | |
|---|---|---|---|---|---|
| AT1G79940 | 1,20 | -1,20 | 1,61 | -0,74 | 4,57E-01 |
| AT1G79950 | 0,49 | -3,71 | 3,95 | -0,94 | 3,47E-01 |
| AT1G79970 | 0,27 | -0,50 | 4,33 | -0,11 | 9,09E-01 |
| AT1G79985 | 0,75 | 0,28 | 2,47 | 0,11 | 9,09E-01 |
| AT1G80010 | 1,24 | -2,21 | 1,74 | -1,27 | 2,04E-01 |
| AT1G80160 | 0,37 | -0,68 | 4,40 | -0,15 | 8,77E-01 |
| AT1G80260 | 0,59 | -2,06 | 3,04 | -0,68 | 4,98E-01 |
| AT1G80300 | 1,04 | -4,63 | 3,23 | -1,43 | 1,52E-01 |
| AT1G80310 | 0,64 | -0,74 | 2,60 | -0,29 | 7,75E-01 |
| AT1G80380 | 1,15 | -1,57 | 1,79 | -0,88 | 3,81E-01 |
| AT1G09963 | 0,96 | -3,53 | 2,47 | -1,43 | 1,53E-01 |
| AT1G80430 | 0,19 | 0,12 | 4,37 | 0,03 | 9,79E-01 |
| AT1G80510 | 0,61 | -0,84 | 3,81 | -0,22 | 8,25E-01 |
| AT1G80620 | 0,35 | -0,14 | 4,39 | -0,03 | 9,75E-01 |
| AT1G80770 | 0,54 | -0,70 | 3,17 | -0,22 | 8,25E-01 |
| AT1G80780 | 0,60 | 0,08 | 3,11 | 0,03 | 9,79E-01 |
| AT1G80790 | 0,69 | -2,53 | 2,97 | -0,85 | 3,94E-01 |
| AT1G80810 | 0,38 | -1,69 | 4,30 | -0,39 | 6,95E-01 |
| AT1G80840 | 1,09 | 3,41 | 2,38 | 1,43 | 1,52E-01 |
| AT1G80870 | 0,27 | -1,00 | 4,36 | -0,23 | 8,18E-01 |
| AT2G01070 | 0,81 | -2,96 | 2,49 | -1,19 | 2,35E-01 |
| AT2G01080 | 0,35 | -0,48 | 3,50 | -0,14 | 8,90E-01 |
| AT2G01100 | 0,12 | -0,06 | 4,37 | -0,01 | 9,88E-01 |
| AT2G01120 | 0,92 | -3,56 | 2,87 | -1,24 | 2,14E-01 |
| AT2G01190 | 0,50 | 1,16 | 3,11 | 0,37 | 7,11E-01 |
| AT2G01400 | 0,39 | -2,16 | 4,38 | -0,49 | 6,21E-01 |
| AT2G03975 | 0,34 | -0,47 | 3,53 | -0,13 | 8,93E-01 |
| AT2G01480 | 0,34 | -2,12 | 4,34 | -0,49 | 6,25E-01 |
| AT2G01620 | 0,60 | -1,26 | 2,44 | -0,52 | 6,06E-01 |
| AT2G01680 | 0,39 | -0,95 | 4,32 | -0,22 | 8,25E-01 |
| AT2G01750 | 0,24 | 0,99 | 4,32 | 0,23 | 8,19E-01 |
| AT2G01860 | 0,98 | 0,38 | 2,47 | 0,15 | 8,77E-01 |
| AT2G01900 | 1,22 | 2,12 | 1,80 | 1,18 | 2,39E-01 |
| AT2G01980 | 0,63 | -1,42 | 3,12 | -0,46 | 6,48E-01 |
| AT2G02040 | 0,10 | -0,34 | 4,38 | -0,08 | 9,38E-01 |
| AT2G02050 | 0,58 | 0,97 | 2,51 | 0,39 | 6,98E-01 |
| AT2G02070 | 0,84 | -0,33 | 1,92 | -0,17 | 8,62E-01 |
| AT2G02090 | 1,22 | -2,02 | 2,21 | -0,91 | 3,61E-01 |
| AT2G02100 | 0,54 | -0,76 | 3,97 | -0,19 | 8,49E-01 |
| AT2G02440 | 0,68 | -0,75 | 3,07 | -0,25 | 8,06E-01 |
| AT2G02450 | 0,66 | 2,05 | 2,56 | 0,80 | 4,22E-01 |
| AT2G02470 | 0,18 | -1,07 | 4,38 | -0,25 | 8,06E-01 |
| AT2G02550 | 1,01 | 3,02 | 2,02 | 1,50 | 1,35E-01 |
| AT2G02700 | 0,39 | -3,07 | 4,38 | -0,70 | 4,84E-01 |
| AT2G04295 | 0,12 | -0,31 | 4,38 | -0,07 | 9,44E-01 |
| AT2G02760 | 0,54 | 0,76 | 2,69 | 0,28 | 7,79E-01 |
| AT2G02790 | 0,39 | -1,64 | 4,25 | -0,39 | 7,00E-01 |
| AT2G02795 | 0,14 | 0,40 | 4,35 | 0,09 | 9,27E-01 |
| AT2G02810 | 0,62 | -0,75 | 3,10 | -0,24 | 8,08E-01 |
| AT2G04355 | 0,47 | -0,85 | 3,28 | -0,26 | 7,96E-01 |
| AT2G04365 | 0,97 | 2,75 | 2,33 | 1,18 | 2,39E-01 |
| AT2G02955 | 0,36 | -1,70 | 4,35 | -0,39 | 6,96E-01 |
| AT2G03070 | 0,25 | -2,57 | 4,40 | -0,58 | 5,60E-01 |
| AT2G03120 | 0,64 | -1,40 | 2,66 | -0,53 | 5,99E-01 |
| AT2G03130 | 0,12 | 0,00 | 4,41 | 0,00 | 1,00E+00 |
| AT2G03270 | 0,39 | 0,27 | 2,80 | 0,09 | 9,24E-01 |

| | | | | | |
|---|---|---|---|---|---|
| AT2G03410 | 0,43 | -0,80 | 4,15 | -0,19 | 8,48E-01 |
| AT2G03450 | 0,32 | -2,39 | 4,36 | -0,55 | 5,84E-01 |
| AT2G03500 | 0,52 | 0,35 | 3,16 | 0,11 | 9,12E-01 |
| AT2G03520 | 0,08 | 0,00 | 4,41 | 0,00 | 1,00E+00 |
| AT2G03550 | 0,42 | -0,80 | 4,24 | -0,19 | 8,51E-01 |
| AT2G03567 | 0,47 | 1,07 | 3,32 | 0,32 | 7,47E-01 |
| AT2G03680 | 0,82 | -1,50 | 2,21 | -0,68 | 4,97E-01 |
| AT2G03710 | 0,17 | 0,00 | 4,41 | 0,00 | 1,00E+00 |
| AT2G03823 | 0,37 | 1,35 | 2,84 | 0,48 | 6,34E-01 |
| AT2G03870 | 0,41 | -3,33 | 4,16 | -0,80 | 4,23E-01 |
| AT2G03932 | 0,20 | 1,40 | 4,30 | 0,33 | 7,44E-01 |
| AT2G03931 | 0,05 | -0,24 | 4,38 | -0,05 | 9,57E-01 |
| AT2G03933 | 0,14 | 0,87 | 4,33 | 0,20 | 8,40E-01 |
| AT2G03936 | 0,18 | 0,48 | 4,35 | 0,11 | 9,11E-01 |
| AT2G03965 | 0,22 | 1,41 | 4,30 | 0,33 | 7,44E-01 |
| AT2G03980 | 0,82 | -1,27 | 2,55 | -0,50 | 6,19E-01 |
| AT2G04039 | 0,42 | -3,35 | 4,14 | -0,81 | 4,18E-01 |
| AT2G04735 | 0,15 | 0,16 | 4,37 | 0,04 | 9,72E-01 |
| AT2G04235 | 0,05 | -0,24 | 4,38 | -0,05 | 9,57E-01 |
| AT2G04270 | 1,17 | -0,28 | 2,29 | -0,12 | 9,02E-01 |
| AT2G04340 | 0,38 | -3,33 | 4,37 | -0,76 | 4,46E-01 |
| AT2G04380 | 0,56 | -0,73 | 2,65 | -0,28 | 7,82E-01 |
| AT2G04390 | 0,85 | -1,18 | 2,12 | -0,56 | 5,78E-01 |
| AT2G04400 | 0,61 | -0,38 | 2,52 | -0,15 | 8,79E-01 |
| AT2G04450 | 0,21 | -0,70 | 4,40 | -0,16 | 8,73E-01 |
| AT2G04495 | 0,43 | -0,69 | 3,40 | -0,20 | 8,38E-01 |
| AT2G04550 | 0,15 | 0,00 | 4,41 | 0,00 | 1,00E+00 |
| AT2G04700 | 0,71 | -0,29 | 2,29 | -0,13 | 9,00E-01 |
| AT2G04740 | 0,38 | -3,44 | 4,37 | -0,79 | 4,31E-01 |
| AT2G04780 | 0,24 | -1,46 | 4,36 | -0,34 | 7,37E-01 |
| AT2G04795 | 0,15 | 0,48 | 4,35 | 0,11 | 9,12E-01 |
| AT2G04850 | 0,08 | 0,00 | 4,41 | 0,00 | 1,00E+00 |
| AT2G04865 | 0,25 | 0,00 | 4,41 | 0,00 | 1,00E+00 |
| AT2G04900 | 0,19 | 0,04 | 4,37 | 0,01 | 9,92E-01 |
| AT2G05060 | 0,21 | -2,50 | 4,39 | -0,57 | 5,70E-01 |
| AT2G05180 | 0,08 | 0,00 | 4,41 | 0,00 | 1,00E+00 |
| AT2G05210 | 0,12 | -0,14 | 4,38 | -0,03 | 9,74E-01 |
| AT2G05320 | 0,92 | 1,79 | 2,12 | 0,85 | 3,97E-01 |
| AT2G05350 | 0,23 | 0,29 | 4,36 | 0,07 | 9,46E-01 |
| AT2G05380 | 0,99 | -3,03 | 2,01 | -1,51 | 1,31E-01 |
| AT2G05540 | 0,30 | 0,27 | 4,31 | 0,06 | 9,50E-01 |
| AT2G04915 | 0,39 | 0,50 | 3,45 | 0,14 | 8,86E-01 |
| AT2G04935 | 1,06 | 1,15 | 1,75 | 0,66 | 5,10E-01 |
| AT2G05635 | 0,15 | 0,41 | 4,35 | 0,09 | 9,26E-01 |
| AT2G04985 | 0,95 | 2,22 | 2,14 | 1,04 | 3,00E-01 |
| AT2G05790 | 0,44 | -0,84 | 3,24 | -0,26 | 7,95E-01 |
| AT2G05830 | 0,15 | 0,05 | 4,37 | 0,01 | 9,91E-01 |
| AT2G05840 | 0,17 | 0,00 | 4,41 | 0,00 | 1,00E+00 |
| AT2G05990 | 0,47 | -0,89 | 2,46 | -0,36 | 7,18E-01 |
| AT2G05995 | 0,63 | -0,33 | 3,08 | -0,11 | 9,14E-01 |
| AT2G06025 | 0,59 | -0,08 | 2,56 | -0,03 | 9,75E-01 |
| AT2G06095 | 0,70 | -0,70 | 3,59 | -0,19 | 8,46E-01 |
| AT2G06105 | 0,23 | 1,25 | 4,31 | 0,29 | 7,72E-01 |
| AT2G06845 | 0,18 | 0,40 | 4,35 | 0,09 | 9,27E-01 |
| AT2G06850 | 0,75 | -0,49 | 2,61 | -0,19 | 8,51E-01 |
| AT2G06904 | 0,38 | 1,94 | 4,29 | 0,45 | 6,50E-01 |

| | | | | | |
|---|---|---|---|---|---|
| AT2G06990 | 1,10 | 1,50 | 1,99 | 0,75 | 4,50E-01 |
| AT2G07050 | 0,64 | 2,02 | 2,93 | 0,69 | 4,90E-01 |
| AT2G07200 | 0,15 | 0,79 | 4,33 | 0,18 | 8,56E-01 |
| AT2G07565 | 0,91 | 1,52 | 2,36 | 0,64 | 5,20E-01 |
| AT2G07749 | 0,59 | 1,80 | 3,17 | 0,57 | 5,70E-01 |
| AT2G07672 | 0,25 | 1,50 | 4,30 | 0,35 | 7,28E-01 |
| AT2G07745 | 0,19 | -0,80 | 4,39 | -0,18 | 8,56E-01 |
| AT2G07674 | 0,81 | 0,10 | 3,68 | 0,03 | 9,79E-01 |
| AT2G07623 | 0,23 | 0,29 | 4,36 | 0,07 | 9,46E-01 |
| AT2G07678 | 0,32 | -3,05 | 4,38 | -0,70 | 4,86E-01 |
| AT2G07669 | 0,33 | -0,47 | 4,35 | -0,11 | 9,14E-01 |
| AT2G07681 | 0,75 | 0,38 | 2,07 | 0,19 | 8,53E-01 |
| AT2G07631 | 0,18 | 0,40 | 4,35 | 0,09 | 9,28E-01 |
| AT2G05215 | 0,68 | 1,43 | 2,56 | 0,56 | 5,76E-01 |
| AT2G07695 | 1,27 | 2,19 | 1,94 | 1,13 | 2,59E-01 |
| AT2G07798 | 1,27 | 1,31 | 1,81 | 0,72 | 4,71E-01 |
| AT2G07667 | 1,00 | 1,51 | 2,27 | 0,66 | 5,07E-01 |
| AT2G07707 | 0,32 | 0,20 | 4,35 | 0,05 | 9,63E-01 |
| AT2G07708 | 0,57 | -0,68 | 4,35 | -0,16 | 8,76E-01 |
| AT2G07715 | 0,94 | 0,19 | 1,70 | 0,11 | 9,11E-01 |
| AT2G07728 | 0,11 | -0,80 | 4,39 | -0,18 | 8,56E-01 |
| AT2G07656 | 0,60 | 2,46 | 2,93 | 0,84 | 4,01E-01 |
| AT2G07827 | 0,36 | -0,86 | 3,49 | -0,25 | 8,06E-01 |
| AT2G07787 | 0,79 | 0,73 | 2,12 | 0,34 | 7,32E-01 |
| AT2G07659 | 0,28 | 1,29 | 4,31 | 0,30 | 7,64E-01 |
| AT2G07765 | 0,12 | 0,00 | 4,41 | 0,00 | 1,00E+00 |
| AT2G07741 | 0,27 | 1,09 | 4,32 | 0,25 | 8,01E-01 |
| AT2G07680 | 0,80 | 2,75 | 2,36 | 1,17 | 2,43E-01 |
| AT2G09990 | 0,53 | 0,45 | 3,16 | 0,14 | 8,86E-01 |
| AT2G05485 | 0,27 | 1,02 | 4,33 | 0,24 | 8,13E-01 |
| AT2G10545 | 0,08 | -0,21 | 4,39 | -0,05 | 9,62E-01 |
| AT2G10556 | 0,88 | 0,48 | 2,73 | 0,18 | 8,60E-01 |
| AT2G10965 | 0,86 | 2,42 | 2,00 | 1,21 | 2,27E-01 |
| AT2G11010 | 0,65 | 0,79 | 2,92 | 0,27 | 7,86E-01 |
| AT2G11035 | 0,48 | -1,38 | 4,03 | -0,34 | 7,32E-01 |
| AT2G11045 | 0,08 | -0,21 | 4,39 | -0,05 | 9,62E-01 |
| AT2G11225 | 0,91 | -0,61 | 1,67 | -0,37 | 7,13E-01 |
| AT2G11620 | 0,14 | 0,87 | 4,33 | 0,20 | 8,40E-01 |
| AT2G11851 | 0,57 | 2,50 | 2,92 | 0,86 | 3,91E-01 |
| AT2G05825 | 0,69 | -0,33 | 2,16 | -0,15 | 8,79E-01 |
| AT2G13100 | 0,26 | -1,28 | 4,35 | -0,29 | 7,69E-01 |
| AT2G13290 | 0,12 | -0,31 | 4,38 | -0,07 | 9,44E-01 |
| AT2G05885 | 0,14 | 0,87 | 4,33 | 0,20 | 8,40E-01 |
| AT2G13430 | 0,44 | 0,70 | 3,23 | 0,22 | 8,28E-01 |
| AT2G13500 | 0,11 | 0,46 | 4,35 | 0,11 | 9,15E-01 |
| AT2G13540 | 0,64 | -0,53 | 2,40 | -0,22 | 8,26E-01 |
| AT2G05925 | 0,12 | -0,06 | 4,37 | -0,01 | 9,88E-01 |
| AT2G13790 | 0,49 | 0,07 | 3,28 | 0,02 | 9,83E-01 |
| AT2G13845 | 0,67 | -0,15 | 2,28 | -0,06 | 9,49E-01 |
| AT2G13960 | 0,27 | -0,14 | 4,34 | -0,03 | 9,73E-01 |
| AT2G14070 | 0,19 | 0,52 | 4,35 | 0,12 | 9,04E-01 |
| AT2G14115 | 0,49 | -0,93 | 4,31 | -0,22 | 8,30E-01 |
| AT2G14395 | 0,63 | -0,95 | 4,36 | -0,22 | 8,28E-01 |
| AT2G14440 | 0,43 | 0,60 | 4,11 | 0,15 | 8,85E-01 |
| AT2G14530 | 0,21 | 1,00 | 4,32 | 0,23 | 8,16E-01 |
| AT2G14700 | 0,64 | 2,60 | 2,88 | 0,90 | 3,68E-01 |

| | | | | | |
|---|---|---|---|---|---|
| AT2G14740 | 1,25 | -1,21 | 2,56 | -0,47 | 6,37E-01 |
| AT2G14765 | 0,40 | 1,82 | 4,30 | 0,42 | 6,71E-01 |
| AT2G14825 | 0,08 | 0,00 | 4,41 | 0,00 | 1,00E+00 |
| AT2G14835 | 0,23 | -0,75 | 4,40 | -0,17 | 8,65E-01 |
| AT2G14850 | 0,21 | -0,60 | 4,40 | -0,14 | 8,91E-01 |
| AT2G14870 | 0,17 | 0,00 | 4,41 | 0,00 | 1,00E+00 |
| AT2G14878 | 1,23 | 0,57 | 1,58 | 0,36 | 7,16E-01 |
| AT2G14890 | 0,77 | -0,31 | 2,09 | -0,15 | 8,81E-01 |
| AT2G06305 | 0,18 | 0,57 | 4,34 | 0,13 | 8,95E-01 |
| AT2G15042 | 0,11 | -0,80 | 4,39 | -0,18 | 8,56E-01 |
| AT2G15050 | 0,64 | 2,06 | 2,84 | 0,72 | 4,69E-01 |
| AT2G15080 | 0,92 | 0,12 | 2,06 | 0,06 | 9,52E-01 |
| AT2G15090 | 1,27 | 2,32 | 2,99 | 0,78 | 4,38E-01 |
| AT2G15300 | 0,31 | -2,57 | 4,40 | -0,58 | 5,59E-01 |
| AT2G15430 | 0,16 | 0,21 | 4,36 | 0,05 | 9,61E-01 |
| AT2G15450 | 0,29 | 0,00 | 4,36 | 0,00 | 9,99E-01 |
| AT2G15470 | 0,34 | 1,56 | 4,30 | 0,36 | 7,18E-01 |
| AT2G15580 | 0,29 | 0,91 | 3,66 | 0,25 | 8,03E-01 |
| AT2G15710 | 0,40 | 1,45 | 4,30 | 0,34 | 7,36E-01 |
| AT2G15730 | 0,08 | -0,21 | 4,39 | -0,05 | 9,62E-01 |
| AT2G15900 | 0,47 | -0,07 | 2,73 | -0,03 | 9,79E-01 |
| AT2G15960 | 1,09 | -2,67 | 1,83 | -1,46 | 1,44E-01 |
| AT2G16019 | 1,05 | 3,34 | 2,10 | 1,59 | 1,11E-01 |
| AT2G16280 | 0,93 | 3,03 | 2,21 | 1,37 | 1,70E-01 |
| AT2G16365 | 0,96 | -0,03 | 2,33 | -0,01 | 9,90E-01 |
| AT2G16380 | 0,44 | -0,66 | 4,21 | -0,16 | 8,75E-01 |
| AT2G16390 | 0,40 | 0,00 | 4,41 | 0,00 | 1,00E+00 |
| AT2G16630 | 0,14 | -0,11 | 4,37 | -0,03 | 9,80E-01 |
| AT2G16700 | 0,28 | -0,68 | 4,41 | -0,15 | 8,77E-01 |
| AT2G16710 | 0,72 | 0,54 | 3,74 | 0,14 | 8,86E-01 |
| AT2G16800 | 0,26 | 0,39 | 4,36 | 0,09 | 9,28E-01 |
| AT2G16950 | 1,26 | -1,85 | 2,34 | -0,79 | 4,29E-01 |
| AT2G16970 | 1,25 | -3,13 | 1,91 | -1,64 | 1,01E-01 |
| AT2G17033 | 0,44 | -3,22 | 4,21 | -0,77 | 4,44E-01 |
| AT2G17120 | 0,23 | 0,68 | 4,34 | 0,16 | 8,76E-01 |
| AT2G17140 | 0,42 | 0,24 | 3,43 | 0,07 | 9,45E-01 |
| AT2G17170 | 0,08 | 0,00 | 4,41 | 0,00 | 1,00E+00 |
| AT2G17240 | 0,52 | -1,46 | 3,21 | -0,46 | 6,49E-01 |
| AT2G17250 | 0,25 | -2,60 | 4,41 | -0,59 | 5,55E-01 |
| AT2G17270 | 0,25 | -1,21 | 4,35 | -0,28 | 7,80E-01 |
| AT2G17290 | 0,78 | -1,55 | 2,30 | -0,67 | 5,00E-01 |
| AT2G06905 | 1,12 | -4,10 | 1,94 | -2,11 | 3,45E-02 |
| AT2G17360 | 0,51 | 0,27 | 3,25 | 0,08 | 9,33E-01 |
| AT2G17390 | 0,66 | -3,80 | 2,99 | -1,27 | 2,03E-01 |
| AT2G17420 | 1,05 | 0,01 | 1,90 | 0,01 | 9,94E-01 |
| AT2G17520 | 1,17 | 0,57 | 1,86 | 0,30 | 7,61E-01 |
| AT2G17530 | 0,50 | -3,48 | 4,02 | -0,87 | 3,87E-01 |
| AT2G17570 | 0,42 | -3,38 | 4,19 | -0,81 | 4,21E-01 |
| AT2G17580 | 0,66 | -3,89 | 3,05 | -1,28 | 2,02E-01 |
| AT2G17705 | 0,42 | 0,70 | 3,21 | 0,22 | 8,27E-01 |
| AT2G17720 | 0,20 | -2,60 | 4,41 | -0,59 | 5,55E-01 |
| AT2G17760 | 0,65 | -2,61 | 2,91 | -0,90 | 3,70E-01 |
| AT2G17787 | 0,33 | -1,43 | 4,39 | -0,33 | 7,44E-01 |
| AT2G17790 | 0,80 | -0,11 | 2,50 | -0,05 | 9,64E-01 |
| AT2G17905 | 0,41 | 0,43 | 3,31 | 0,13 | 8,95E-01 |
| AT2G17920 | 0,15 | 0,14 | 4,37 | 0,03 | 9,75E-01 |

| | | | | | |
|---|---|---|---|---|---|
| AT2G17980 | 1,01 | -1,59 | 2,71 | -0,59 | 5,57E-01 |
| AT2G17990 | 0,69 | -1,34 | 2,53 | -0,53 | 5,98E-01 |
| AT2G18000 | 0,29 | -2,57 | 4,40 | -0,58 | 5,59E-01 |
| AT2G18030 | 0,68 | -1,55 | 3,07 | -0,50 | 6,15E-01 |
| AT2G18040 | 0,20 | 0,47 | 4,36 | 0,11 | 9,14E-01 |
| AT2G18090 | 1,01 | -0,57 | 1,87 | -0,31 | 7,60E-01 |
| AT2G18100 | 0,48 | 0,45 | 4,06 | 0,11 | 9,12E-01 |
| AT2G18110 | 0,62 | 0,13 | 3,06 | 0,04 | 9,66E-01 |
| AT2G18390 | 0,28 | -1,75 | 4,39 | -0,40 | 6,91E-01 |
| AT2G18420 | 0,81 | -4,40 | 3,43 | -1,28 | 2,00E-01 |
| AT2G18530 | 0,38 | 0,00 | 3,46 | 0,00 | 9,99E-01 |
| AT2G18630 | 0,16 | -0,68 | 4,41 | -0,15 | 8,77E-01 |
| AT2G18670 | 1,13 | 1,60 | 1,78 | 0,90 | 3,70E-01 |
| AT2G18680 | 0,16 | -0,64 | 4,40 | -0,15 | 8,85E-01 |
| AT2G18690 | 1,29 | 0,94 | 1,80 | 0,52 | 6,03E-01 |
| AT2G18700 | 0,27 | -0,74 | 4,40 | -0,17 | 8,67E-01 |
| AT2G18730 | 0,58 | 0,66 | 2,57 | 0,26 | 7,97E-01 |
| AT2G18735 | 0,22 | 0,09 | 4,38 | 0,02 | 9,83E-01 |
| AT2G18750 | 0,67 | -0,68 | 2,01 | -0,34 | 7,33E-01 |
| AT2G18850 | 0,52 | -3,26 | 4,37 | -0,75 | 4,56E-01 |
| AT2G18880 | 0,25 | 0,00 | 4,41 | 0,00 | 1,00E+00 |
| AT2G18917 | 0,08 | -0,21 | 4,39 | -0,05 | 9,62E-01 |
| AT2G07095 | 0,55 | 0,05 | 3,10 | 0,02 | 9,86E-01 |
| AT2G18938 | 0,72 | -1,47 | 2,44 | -0,60 | 5,46E-01 |
| AT2G18950 | 0,45 | 0,60 | 4,16 | 0,14 | 8,85E-01 |
| AT2G19110 | 0,15 | -0,81 | 4,39 | -0,18 | 8,54E-01 |
| AT2G19130 | 0,89 | -0,77 | 2,05 | -0,38 | 7,06E-01 |
| AT2G19160 | 0,39 | -0,68 | 4,41 | -0,15 | 8,77E-01 |
| AT2G19260 | 0,46 | -0,75 | 4,03 | -0,19 | 8,52E-01 |
| AT2G19280 | 0,36 | 0,00 | 4,41 | 0,00 | 1,00E+00 |
| AT2G19420 | 0,28 | 0,99 | 4,32 | 0,23 | 8,18E-01 |
| AT2G19430 | 0,24 | -0,01 | 4,38 | 0,00 | 9,98E-01 |
| AT2G19450 | 0,42 | -0,64 | 4,32 | -0,15 | 8,81E-01 |
| AT2G07205 | 0,75 | 1,19 | 3,58 | 0,33 | 7,39E-01 |
| AT2G19490 | 0,36 | 0,00 | 4,41 | 0,00 | 1,00E+00 |
| AT2G19520 | 0,55 | -0,12 | 4,00 | -0,03 | 9,75E-01 |
| AT2G19540 | 0,46 | -0,29 | 2,76 | -0,10 | 9,17E-01 |
| AT2G19572 | 0,52 | 2,29 | 2,99 | 0,77 | 4,43E-01 |
| AT2G19582 | 0,25 | 1,48 | 4,30 | 0,34 | 7,30E-01 |
| AT2G19600 | 0,64 | 0,03 | 3,88 | 0,01 | 9,93E-01 |
| AT2G19640 | 0,53 | -3,73 | 3,87 | -0,96 | 3,35E-01 |
| AT2G19730 | 0,57 | 1,83 | 4,09 | 0,45 | 6,54E-01 |
| AT2G19740 | 0,67 | 0,76 | 2,26 | 0,34 | 7,37E-01 |
| AT2G19810 | 0,34 | 0,43 | 3,47 | 0,12 | 9,01E-01 |
| AT2G19880 | 0,75 | -2,57 | 2,52 | -1,02 | 3,08E-01 |
| AT2G20020 | 1,07 | -1,42 | 2,15 | -0,66 | 5,09E-01 |
| AT2G20100 | 0,33 | -1,64 | 4,41 | -0,37 | 7,10E-01 |
| AT2G20110 | 0,83 | 0,00 | 4,41 | 0,00 | 1,00E+00 |
| AT2G20142 | 0,28 | -1,56 | 4,39 | -0,35 | 7,23E-01 |
| AT2G20230 | 0,26 | -1,30 | 4,38 | -0,30 | 7,67E-01 |
| AT2G20340 | 0,57 | -0,54 | 4,02 | -0,13 | 8,94E-01 |
| AT2G20360 | 0,22 | -0,39 | 4,38 | -0,09 | 9,29E-01 |
| AT2G20362 | 0,08 | 0,00 | 4,41 | 0,00 | 1,00E+00 |
| AT2G20370 | 1,01 | -1,83 | 2,35 | -0,78 | 4,38E-01 |
| AT2G20420 | 0,62 | 0,02 | 2,27 | 0,01 | 9,93E-01 |
| AT2G20450 | 0,40 | -2,57 | 4,21 | -0,61 | 5,43E-01 |

| | | | | | |
|---|---|---|---|---|---|
| AT2G20530 | 0,21 | -0,74 | 4,40 | -0,17 | 8,67E-01 |
| AT2G20690 | 1,05 | 1,07 | 2,53 | 0,42 | 6,73E-01 |
| AT2G20710 | 0,08 | 0,00 | 4,41 | 0,00 | 1,00E+00 |
| AT2G07335 | 0,46 | -3,51 | 4,18 | -0,84 | 4,01E-01 |
| AT2G20750 | 0,59 | -4,05 | 3,78 | -1,07 | 2,84E-01 |
| AT2G20770 | 0,17 | 0,00 | 4,41 | 0,00 | 1,00E+00 |
| AT2G20820 | 0,25 | -1,64 | 4,38 | -0,37 | 7,09E-01 |
| AT2G20890 | 1,04 | -1,07 | 1,96 | -0,54 | 5,86E-01 |
| AT2G20920 | 0,65 | -3,20 | 2,48 | -1,29 | 1,97E-01 |
| AT2G20960 | 0,82 | -1,47 | 2,03 | -0,73 | 4,68E-01 |
| AT2G21160 | 0,91 | -0,58 | 3,10 | -0,19 | 8,50E-01 |
| AT2G21187 | 0,89 | 0,67 | 1,78 | 0,38 | 7,05E-01 |
| AT2G21190 | 0,23 | 0,09 | 4,37 | 0,02 | 9,83E-01 |
| AT2G21270 | 0,21 | 0,00 | 4,41 | 0,00 | 1,00E+00 |
| AT2G21380 | 1,04 | -0,97 | 1,84 | -0,52 | 6,01E-01 |
| AT2G21520 | 0,84 | -0,16 | 2,12 | -0,07 | 9,41E-01 |
| AT2G21530 | 0,61 | 0,94 | 3,02 | 0,31 | 7,55E-01 |
| AT2G21550 | 0,95 | 0,88 | 2,67 | 0,33 | 7,42E-01 |
| AT2G21580 | 1,04 | -4,10 | 2,78 | -1,47 | 1,41E-01 |
| AT2G21620 | 1,16 | -0,96 | 2,13 | -0,45 | 6,51E-01 |
| AT2G21630 | 0,32 | -3,01 | 4,38 | -0,69 | 4,92E-01 |
| AT2G21770 | 0,18 | -1,07 | 4,38 | -0,25 | 8,06E-01 |
| AT2G21790 | 0,74 | -2,33 | 2,45 | -0,95 | 3,41E-01 |
| AT2G21850 | 1,02 | -3,88 | 2,53 | -1,54 | 1,24E-01 |
| AT2G07475 | 0,24 | -0,68 | 4,41 | -0,15 | 8,77E-01 |
| AT2G21960 | 0,80 | -3,28 | 2,50 | -1,31 | 1,90E-01 |
| AT2G07495 | 0,12 | -0,68 | 4,41 | -0,15 | 8,77E-01 |
| AT2G22080 | 0,65 | -1,64 | 2,68 | -0,61 | 5,42E-01 |
| AT2G22120 | 0,32 | -3,01 | 4,38 | -0,69 | 4,92E-01 |
| AT2G22425 | 0,10 | 0,45 | 4,35 | 0,10 | 9,19E-01 |
| AT2G22450 | 0,24 | -1,38 | 4,38 | -0,31 | 7,53E-01 |
| AT2G22480 | 0,23 | -1,68 | 4,36 | -0,39 | 7,00E-01 |
| AT2G22670 | 0,80 | 2,04 | 2,23 | 0,91 | 3,62E-01 |
| AT2G22680 | 0,25 | -1,15 | 4,38 | -0,26 | 7,93E-01 |
| AT2G22720 | 0,99 | -3,67 | 2,69 | -1,36 | 1,72E-01 |
| AT2G07845 | 0,43 | -0,73 | 4,18 | -0,17 | 8,61E-01 |
| AT2G22790 | 0,08 | 0,00 | 4,41 | 0,00 | 1,00E+00 |
| AT2G22840 | 1,11 | -1,60 | 2,33 | -0,69 | 4,93E-01 |
| AT2G07850 | 0,72 | 1,20 | 2,04 | 0,59 | 5,58E-01 |
| AT2G22910 | 0,32 | -0,86 | 4,38 | -0,20 | 8,44E-01 |
| AT2G23080 | 0,38 | -0,55 | 4,34 | -0,13 | 8,98E-01 |
| AT2G23120 | 0,92 | -1,01 | 2,23 | -0,45 | 6,50E-01 |
| AT2G23130 | 0,41 | 1,17 | 2,78 | 0,42 | 6,74E-01 |
| AT2G23310 | 0,84 | -3,28 | 2,86 | -1,15 | 2,51E-01 |
| AT2G07895 | 0,15 | -0,73 | 4,40 | -0,17 | 8,68E-01 |
| AT2G23420 | 0,47 | -3,32 | 4,36 | -0,76 | 4,47E-01 |
| AT2G23680 | 0,31 | 0,00 | 4,41 | 0,00 | 1,00E+00 |
| AT2G23740 | 0,12 | -0,14 | 4,38 | -0,03 | 9,74E-01 |
| AT2G23890 | 0,37 | -1,62 | 3,53 | -0,46 | 6,45E-01 |
| AT2G23985 | 0,17 | 0,00 | 4,41 | 0,00 | 1,00E+00 |
| AT2G24090 | 0,67 | -0,75 | 2,62 | -0,28 | 7,76E-01 |
| AT2G24240 | 0,31 | 0,59 | 4,34 | 0,14 | 8,92E-01 |
| AT2G24300 | 0,15 | 0,05 | 4,37 | 0,01 | 9,91E-01 |
| AT2G08015 | 0,35 | -0,21 | 4,37 | -0,05 | 9,62E-01 |
| AT2G24350 | 0,50 | -3,97 | 3,97 | -1,00 | 3,17E-01 |
| AT2G24390 | 0,92 | -3,34 | 2,96 | -1,13 | 2,60E-01 |

| | | | | | |
|---|---|---|---|---|---|
| AT2G24380 | 0,65 | -3,07 | 3,83 | -0,80 | 4,23E-01 |
| AT2G24420 | 1,14 | -0,70 | 2,21 | -0,31 | 7,53E-01 |
| AT2G08045 | 0,23 | 0,43 | 4,34 | 0,10 | 9,20E-01 |
| AT2G08050 | 0,32 | 1,13 | 4,32 | 0,26 | 7,94E-01 |
| AT2G24540 | 0,88 | -0,95 | 1,60 | -0,59 | 5,53E-01 |
| AT2G24590 | 0,49 | -3,73 | 3,96 | -0,94 | 3,47E-01 |
| AT2G24592 | 0,83 | -1,22 | 2,14 | -0,57 | 5,69E-01 |
| AT2G24765 | 0,69 | -3,37 | 2,95 | -1,14 | 2,53E-01 |
| AT2G25000 | 0,52 | -0,70 | 3,92 | -0,18 | 8,59E-01 |
| AT2G25080 | 0,61 | 0,30 | 2,63 | 0,12 | 9,08E-01 |
| AT2G25110 | 0,19 | 0,01 | 4,36 | 0,00 | 9,98E-01 |
| AT2G25190 | 0,14 | 0,48 | 4,35 | 0,11 | 9,13E-01 |
| AT2G25410 | 1,30 | 3,32 | 2,19 | 1,52 | 1,30E-01 |
| AT2G25480 | 0,05 | -0,24 | 4,38 | -0,05 | 9,57E-01 |
| AT2G25560 | 0,73 | -1,64 | 2,94 | -0,56 | 5,76E-01 |
| AT2G25580 | 0,08 | 0,00 | 4,41 | 0,00 | 1,00E+00 |
| AT2G25590 | 0,21 | -0,60 | 4,40 | -0,14 | 8,91E-01 |
| AT2G25625 | 0,15 | -0,77 | 4,39 | -0,18 | 8,61E-01 |
| AT2G25630 | 0,50 | 2,10 | 3,05 | 0,69 | 4,91E-01 |
| AT2G25650 | 0,70 | -3,35 | 2,96 | -1,13 | 2,58E-01 |
| AT2G25790 | 0,18 | 0,78 | 4,34 | 0,18 | 8,57E-01 |
| AT2G25840 | 0,59 | -0,35 | 3,87 | -0,09 | 9,29E-01 |
| AT2G25930 | 1,26 | -1,23 | 1,62 | -0,76 | 4,47E-01 |
| AT2G26060 | 1,00 | -2,45 | 4,38 | -0,56 | 5,75E-01 |
| AT2G26110 | 0,61 | 1,29 | 2,85 | 0,45 | 6,51E-01 |
| AT2G26170 | 0,47 | -1,46 | 3,21 | -0,45 | 6,50E-01 |
| AT2G26270 | 0,21 | -2,50 | 4,39 | -0,57 | 5,70E-01 |
| AT2G26300 | 1,09 | -2,59 | 2,65 | -0,98 | 3,28E-01 |
| AT2G26340 | 0,57 | -2,94 | 3,08 | -0,95 | 3,40E-01 |
| AT2G26510 | 0,24 | -1,38 | 4,38 | -0,31 | 7,53E-01 |
| AT2G26570 | 0,43 | -1,60 | 3,37 | -0,48 | 6,35E-01 |
| AT2G26600 | 0,45 | -0,91 | 3,26 | -0,28 | 7,79E-01 |
| AT2G26610 | 0,30 | -1,26 | 4,36 | -0,29 | 7,73E-01 |
| AT2G26640 | 0,71 | 0,00 | 3,68 | 0,00 | 1,00E+00 |
| AT2G26660 | 0,22 | -1,25 | 4,37 | -0,29 | 7,75E-01 |
| AT2G26692 | 0,08 | 0,00 | 4,41 | 0,00 | 1,00E+00 |
| AT2G26730 | 0,85 | 1,67 | 2,16 | 0,78 | 4,37E-01 |
| AT2G26920 | 0,60 | -1,83 | 3,14 | -0,58 | 5,61E-01 |
| AT2G27030 | 0,72 | 0,07 | 2,46 | 0,03 | 9,78E-01 |
| AT2G27050 | 0,99 | -0,35 | 1,80 | -0,20 | 8,45E-01 |
| AT2G27060 | 0,17 | 0,00 | 4,41 | 0,00 | 1,00E+00 |
| AT2G27080 | 0,39 | 1,73 | 4,15 | 0,42 | 6,77E-01 |
| AT2G27100 | 0,72 | -4,09 | 3,56 | -1,15 | 2,50E-01 |
| AT2G27130 | 0,30 | 0,27 | 4,31 | 0,06 | 9,50E-01 |
| AT2G27200 | 0,21 | -0,03 | 4,36 | -0,01 | 9,95E-01 |
| AT2G27210 | 0,89 | 1,09 | 2,36 | 0,46 | 6,44E-01 |
| AT2G27230 | 1,31 | -2,02 | 1,43 | -1,41 | 1,59E-01 |
| AT2G27260 | 0,40 | -0,89 | 3,35 | -0,26 | 7,92E-01 |
| AT2G27320 | 0,85 | 0,96 | 2,39 | 0,40 | 6,89E-01 |
| AT2G27380 | 0,13 | 0,52 | 4,35 | 0,12 | 9,04E-01 |
| AT2G27460 | 0,28 | 0,00 | 4,41 | 0,00 | 1,00E+00 |
| AT2G27540 | 0,95 | 3,44 | 3,04 | 1,13 | 2,58E-01 |
| AT2G27570 | 0,28 | -3,07 | 4,38 | -0,70 | 4,84E-01 |
| AT2G27690 | 0,16 | -0,26 | 4,38 | -0,06 | 9,52E-01 |
| AT2G27710 | 0,57 | -1,56 | 3,17 | -0,49 | 6,22E-01 |
| AT2G27760 | 0,08 | 0,00 | 4,41 | 0,00 | 1,00E+00 |

| | | | | | |
|---|---|---|---|---|---|
| AT2G27810 | 0,54 | -0,52 | 2,66 | -0,20 | 8,44E-01 |
| AT2G27820 | 0,23 | 0,01 | 4,36 | 0,00 | 9,99E-01 |
| AT2G28070 | 0,87 | -3,24 | 2,85 | -1,13 | 2,57E-01 |
| AT2G28120 | 0,29 | -2,54 | 4,40 | -0,58 | 5,63E-01 |
| AT2G28130 | 0,59 | 0,00 | 4,41 | 0,00 | 1,00E+00 |
| AT2G28190 | 0,42 | -0,63 | 4,27 | -0,15 | 8,82E-01 |
| AT2G28250 | 0,08 | 0,00 | 4,41 | 0,00 | 1,00E+00 |
| AT2G28310 | 0,30 | -1,46 | 4,36 | -0,34 | 7,37E-01 |
| AT2G28320 | 0,43 | 0,52 | 4,17 | 0,12 | 9,01E-01 |
| AT2G28350 | 0,96 | -2,17 | 2,01 | -1,08 | 2,80E-01 |
| AT2G28370 | 0,25 | -1,64 | 4,38 | -0,37 | 7,09E-01 |
| AT2G28460 | 0,18 | 0,68 | 4,34 | 0,16 | 8,75E-01 |
| AT2G28520 | 0,65 | -1,29 | 2,89 | -0,45 | 6,54E-01 |
| AT2G28540 | 0,87 | -4,41 | 2,33 | -1,89 | 5,84E-02 |
| AT2G28605 | 0,37 | -0,21 | 4,33 | -0,05 | 9,61E-01 |
| AT2G28660 | 0,10 | 0,45 | 4,35 | 0,10 | 9,19E-01 |
| AT2G28671 | 0,51 | 1,22 | 3,05 | 0,40 | 6,90E-01 |
| AT2G28710 | 0,24 | 0,38 | 4,36 | 0,09 | 9,31E-01 |
| AT2G28720 | 0,49 | 0,72 | 4,14 | 0,17 | 8,63E-01 |
| AT2G28730 | 1,02 | -0,88 | 1,50 | -0,59 | 5,58E-01 |
| AT2G28810 | 0,71 | -0,74 | 2,54 | -0,29 | 7,70E-01 |
| AT2G28930 | 0,12 | -0,78 | 4,39 | -0,18 | 8,59E-01 |
| AT2G29020 | 0,11 | 0,46 | 4,35 | 0,11 | 9,15E-01 |
| AT2G29065 | 0,30 | -2,60 | 4,41 | -0,59 | 5,55E-01 |
| AT2G29070 | 0,15 | -0,73 | 4,40 | -0,17 | 8,68E-01 |
| AT2G29140 | 0,94 | -0,17 | 2,50 | -0,07 | 9,47E-01 |
| AT2G29180 | 0,47 | -1,65 | 4,12 | -0,40 | 6,88E-01 |
| AT2G29210 | 1,16 | -3,11 | 2,01 | -1,54 | 1,23E-01 |
| AT2G29310 | 0,46 | -1,83 | 3,28 | -0,56 | 5,77E-01 |
| AT2G29350 | 0,65 | 0,19 | 3,89 | 0,05 | 9,60E-01 |
| AT2G29360 | 0,51 | -3,74 | 4,35 | -0,86 | 3,90E-01 |
| AT2G29510 | 0,82 | -3,04 | 3,40 | -0,89 | 3,72E-01 |
| AT2G29600 | 0,47 | 1,95 | 3,13 | 0,62 | 5,34E-01 |
| AT2G29640 | 0,37 | -3,35 | 4,32 | -0,78 | 4,38E-01 |
| AT2G29660 | 0,20 | -2,60 | 4,41 | -0,59 | 5,55E-01 |
| AT2G29690 | 0,14 | -0,35 | 4,38 | -0,08 | 9,37E-01 |
| AT2G29700 | 0,24 | -0,11 | 4,38 | -0,03 | 9,80E-01 |
| AT2G30020 | 0,83 | 2,96 | 3,44 | 0,86 | 3,91E-01 |
| AT2G30040 | 0,38 | -0,55 | 3,35 | -0,17 | 8,68E-01 |
| AT2G30050 | 0,66 | -1,09 | 2,50 | -0,43 | 6,64E-01 |
| AT2G30140 | 0,54 | -1,61 | 3,94 | -0,41 | 6,82E-01 |
| AT2G30150 | 0,10 | -0,18 | 4,38 | -0,04 | 9,67E-01 |
| AT2G30180 | 1,06 | -0,73 | 1,72 | -0,42 | 6,71E-01 |
| AT2G08375 | 1,24 | 1,69 | 1,83 | 0,92 | 3,58E-01 |
| AT2G30230 | 0,50 | 0,44 | 3,26 | 0,13 | 8,93E-01 |
| AT2G30362 | 0,41 | 1,86 | 4,13 | 0,45 | 6,53E-01 |
| AT2G30410 | 0,72 | -1,04 | 2,39 | -0,44 | 6,63E-01 |
| AT2G30530 | 0,20 | -2,60 | 4,41 | -0,59 | 5,55E-01 |
| AT2G30600 | 0,51 | -1,84 | 4,34 | -0,42 | 6,72E-01 |
| AT2G30620 | 0,85 | 1,26 | 2,37 | 0,53 | 5,96E-01 |
| AT2G30730 | 0,89 | 0,26 | 2,46 | 0,11 | 9,15E-01 |
| AT2G30780 | 0,62 | -0,30 | 3,76 | -0,08 | 9,36E-01 |
| AT2G30930 | 0,49 | -1,14 | 2,62 | -0,43 | 6,64E-01 |
| AT2G30970 | 0,46 | -2,30 | 4,38 | -0,52 | 6,00E-01 |
| AT2G31010 | 0,95 | -4,43 | 2,31 | -1,92 | 5,52E-02 |
| AT2G31020 | 0,26 | -2,60 | 4,41 | -0,59 | 5,55E-01 |

| | | | | | |
|---|---|---|---|---|---|
| AT2G31200 | 0,16 | 0,32 | 4,36 | 0,07 | 9,42E-01 |
| AT2G31215 | 0,52 | 1,44 | 2,58 | 0,56 | 5,78E-01 |
| AT2G31290 | 0,71 | -2,92 | 3,16 | -0,92 | 3,55E-01 |
| AT2G31350 | 0,38 | -1,64 | 4,36 | -0,38 | 7,07E-01 |
| AT2G31370 | 0,19 | -0,77 | 4,39 | -0,18 | 8,61E-01 |
| AT2G31380 | 1,06 | -3,91 | 1,90 | -2,05 | 3,99E-02 |
| AT2G08435 | 0,79 | -3,71 | 3,54 | -1,05 | 2,95E-01 |
| AT2G31560 | 0,67 | 1,01 | 2,42 | 0,42 | 6,76E-01 |
| AT2G31585 | 0,47 | -2,47 | 4,01 | -0,61 | 5,39E-01 |
| AT2G31750 | 0,75 | 0,42 | 3,05 | 0,14 | 8,90E-01 |
| AT2G31790 | 0,30 | 1,20 | 3,60 | 0,33 | 7,39E-01 |
| AT2G31810 | 1,10 | -2,20 | 1,97 | -1,12 | 2,64E-01 |
| AT2G31970 | 0,18 | -0,27 | 4,38 | -0,06 | 9,51E-01 |
| AT2G32010 | 0,37 | 0,83 | 4,34 | 0,19 | 8,48E-01 |
| AT2G32040 | 0,08 | 0,00 | 4,41 | 0,00 | 1,00E+00 |
| AT2G32100 | 1,15 | 3,73 | 2,07 | 1,80 | 7,17E-02 |
| AT2G32150 | 1,00 | 0,93 | 2,31 | 0,40 | 6,86E-01 |
| AT2G32250 | 0,51 | -2,16 | 3,22 | -0,67 | 5,03E-01 |
| AT2G32260 | 0,44 | -1,19 | 4,14 | -0,29 | 7,74E-01 |
| AT2G32295 | 0,53 | -0,36 | 2,74 | -0,13 | 8,95E-01 |
| AT2G32320 | 0,15 | 0,05 | 4,37 | 0,01 | 9,91E-01 |
| AT2G32440 | 0,16 | -0,62 | 4,40 | -0,14 | 8,87E-01 |
| AT2G32450 | 0,56 | -0,04 | 2,60 | -0,01 | 9,89E-01 |
| AT2G32698 | 0,12 | 0,00 | 4,41 | 0,00 | 1,00E+00 |
| AT2G32900 | 0,29 | -1,64 | 4,38 | -0,37 | 7,08E-01 |
| AT2G32910 | 0,68 | -0,86 | 3,00 | -0,29 | 7,75E-01 |
| AT2G33080 | 0,74 | -2,60 | 3,58 | -0,73 | 4,68E-01 |
| AT2G33160 | 0,25 | -2,57 | 4,40 | -0,58 | 5,60E-01 |
| AT2G33220 | 0,54 | -2,53 | 3,18 | -0,80 | 4,26E-01 |
| AT2G33250 | 0,76 | -0,68 | 4,41 | -0,15 | 8,77E-01 |
| AT2G33290 | 0,29 | 0,91 | 3,66 | 0,25 | 8,03E-01 |
| AT2G33390 | 0,25 | -1,15 | 4,38 | -0,26 | 7,93E-01 |
| AT2G33420 | 0,37 | -3,35 | 4,32 | -0,78 | 4,38E-01 |
| AT2G08585 | 0,32 | 0,97 | 3,64 | 0,27 | 7,90E-01 |
| AT2G33435 | 1,31 | -2,11 | 1,63 | -1,29 | 1,96E-01 |
| AT2G33450 | 0,32 | 0,17 | 4,36 | 0,04 | 9,69E-01 |
| AT2G33490 | 0,35 | -1,70 | 4,34 | -0,39 | 6,96E-01 |
| AT2G33510 | 0,26 | -1,25 | 4,37 | -0,29 | 7,75E-01 |
| AT2G33570 | 0,17 | 0,78 | 4,33 | 0,18 | 8,58E-01 |
| AT2G33610 | 0,37 | -1,59 | 4,34 | -0,37 | 7,13E-01 |
| AT2G33650 | 0,29 | 0,00 | 4,41 | 0,00 | 1,00E+00 |
| AT2G33700 | 0,56 | 0,00 | 3,93 | 0,00 | 1,00E+00 |
| AT2G33760 | 0,64 | -3,59 | 4,36 | -0,82 | 4,10E-01 |
| AT2G33810 | 0,37 | -1,13 | 4,34 | -0,26 | 7,95E-01 |
| AT2G33840 | 0,52 | -1,94 | 3,18 | -0,61 | 5,42E-01 |
| AT2G33845 | 0,47 | 0,07 | 4,10 | 0,02 | 9,87E-01 |
| AT2G33950 | 0,39 | -2,00 | 4,29 | -0,47 | 6,40E-01 |
| AT2G34120 | 0,27 | 1,51 | 4,30 | 0,35 | 7,25E-01 |
| AT2G34240 | 0,12 | 0,00 | 4,41 | 0,00 | 1,00E+00 |
| AT2G34460 | 0,99 | 0,30 | 1,76 | 0,17 | 8,66E-01 |
| AT2G34660 | 0,69 | -0,14 | 2,64 | -0,05 | 9,58E-01 |
| AT2G34730 | 0,50 | -0,87 | 2,72 | -0,32 | 7,49E-01 |
| AT2G34770 | 0,22 | -1,25 | 4,37 | -0,29 | 7,75E-01 |
| AT2G34780 | 0,99 | -1,37 | 2,01 | -0,69 | 4,93E-01 |
| AT2G34810 | 0,16 | -0,26 | 4,38 | -0,06 | 9,52E-01 |
| AT2G34860 | 1,10 | -3,73 | 2,12 | -1,76 | 7,87E-02 |

| | | | | | |
|---|---|---|---|---|---|
| AT2G34970 | 0,22 | -1,27 | 4,36 | -0,29 | 7,70E-01 |
| AT2G35020 | 0,32 | -0,94 | 4,37 | -0,21 | 8,30E-01 |
| AT2G35170 | 0,10 | -0,18 | 4,38 | -0,04 | 9,67E-01 |
| AT2G35230 | 0,50 | -1,04 | 4,33 | -0,24 | 8,09E-01 |
| AT2G35330 | 0,35 | -1,04 | 4,31 | -0,24 | 8,09E-01 |
| AT2G35340 | 0,80 | -4,56 | 2,76 | -1,65 | 9,86E-02 |
| AT2G35370 | 0,43 | -1,50 | 4,29 | -0,35 | 7,27E-01 |
| AT2G35630 | 1,16 | -2,01 | 1,77 | -1,14 | 2,56E-01 |
| AT2G35635 | 0,64 | -3,92 | 4,34 | -0,90 | 3,67E-01 |
| AT2G35650 | 0,68 | -3,20 | 2,22 | -1,44 | 1,50E-01 |
| AT2G08750 | 0,08 | 0,00 | 4,41 | 0,00 | 1,00E+00 |
| AT2G35720 | 0,19 | 0,25 | 4,36 | 0,06 | 9,54E-01 |
| AT2G35760 | 0,17 | -1,25 | 4,37 | -0,29 | 7,74E-01 |
| AT2G35800 | 1,02 | -1,75 | 1,75 | -1,00 | 3,15E-01 |
| AT2G35820 | 0,71 | -2,60 | 4,41 | -0,59 | 5,55E-01 |
| AT2G35830 | 1,03 | -0,62 | 1,93 | -0,32 | 7,47E-01 |
| AT2G35840 | 0,63 | -1,41 | 2,39 | -0,59 | 5,55E-01 |
| AT2G35860 | 0,74 | 0,52 | 2,30 | 0,22 | 8,23E-01 |
| AT2G35920 | 0,77 | -2,56 | 2,43 | -1,05 | 2,92E-01 |
| AT2G35935 | 0,49 | 2,57 | 3,83 | 0,67 | 5,02E-01 |
| AT2G35960 | 0,20 | 1,02 | 4,33 | 0,24 | 8,14E-01 |
| AT2G36000 | 0,80 | -0,13 | 2,58 | -0,05 | 9,60E-01 |
| AT2G36010 | 0,59 | -2,64 | 3,07 | -0,86 | 3,90E-01 |
| AT2G36026 | 0,70 | 0,35 | 3,79 | 0,09 | 9,27E-01 |
| AT2G36090 | 0,69 | 3,08 | 3,49 | 0,88 | 3,76E-01 |
| AT2G36130 | 0,56 | -3,21 | 4,38 | -0,73 | 4,64E-01 |
| AT2G36330 | 0,55 | 0,00 | 4,41 | 0,00 | 1,00E+00 |
| AT2G36500 | 0,54 | -4,09 | 3,08 | -1,33 | 1,85E-01 |
| AT2G36510 | 0,37 | -3,33 | 4,32 | -0,77 | 4,41E-01 |
| AT2G36570 | 0,52 | -3,29 | 4,32 | -0,76 | 4,46E-01 |
| AT2G36580 | 1,14 | -0,57 | 1,56 | -0,37 | 7,15E-01 |
| AT2G36600 | 0,66 | -1,64 | 2,36 | -0,69 | 4,88E-01 |
| AT2G36720 | 1,25 | -4,12 | 2,04 | -2,02 | 4,38E-02 |
| AT2G36792 | 0,34 | 1,52 | 4,30 | 0,35 | 7,23E-01 |
| AT2G36830 | 0,86 | -2,44 | 2,09 | -1,16 | 2,44E-01 |
| AT2G36920 | 0,12 | 0,00 | 4,41 | 0,00 | 1,00E+00 |
| AT2G36950 | 0,34 | -2,60 | 4,41 | -0,59 | 5,55E-01 |
| AT2G36960 | 0,18 | 0,40 | 4,35 | 0,09 | 9,27E-01 |
| AT2G36970 | 0,19 | 0,01 | 4,36 | 0,00 | 9,98E-01 |
| AT2G36980 | 0,05 | -0,24 | 4,38 | -0,05 | 9,57E-01 |
| AT2G37025 | 0,19 | -0,80 | 4,39 | -0,18 | 8,56E-01 |
| AT2G37050 | 0,93 | -3,42 | 2,31 | -1,48 | 1,38E-01 |
| AT2G37110 | 1,16 | 1,46 | 2,25 | 0,65 | 5,16E-01 |
| AT2G37180 | 0,08 | 0,00 | 4,41 | 0,00 | 1,00E+00 |
| AT2G37250 | 0,74 | -0,39 | 2,28 | -0,17 | 8,65E-01 |
| AT2G08860 | 0,31 | 0,14 | 3,65 | 0,04 | 9,70E-01 |
| AT2G37320 | 0,08 | 0,00 | 4,41 | 0,00 | 1,00E+00 |
| AT2G37330 | 0,66 | -2,67 | 3,10 | -0,86 | 3,90E-01 |
| AT2G37362 | 0,21 | 0,00 | 4,41 | 0,00 | 1,00E+00 |
| AT2G37370 | 0,40 | 2,01 | 4,29 | 0,47 | 6,40E-01 |
| AT2G37470 | 0,26 | -0,33 | 4,38 | -0,08 | 9,40E-01 |
| AT2G37555 | 0,61 | -1,64 | 3,09 | -0,53 | 5,96E-01 |
| AT2G37580 | 0,12 | -0,14 | 4,38 | -0,03 | 9,74E-01 |
| AT2G37650 | 1,11 | -1,63 | 1,91 | -0,85 | 3,95E-01 |
| AT2G37690 | 0,31 | -0,76 | 4,37 | -0,17 | 8,62E-01 |
| AT2G37770 | 0,28 | -0,76 | 4,40 | -0,17 | 8,62E-01 |

| | | | | | |
|---|---|---|---|---|---|
| AT2G37860 | 0,83 | -1,11 | 2,03 | -0,54 | 5,86E-01 |
| AT2G37940 | 0,73 | -1,03 | 2,85 | -0,36 | 7,17E-01 |
| AT2G37970 | 0,77 | 1,27 | 2,69 | 0,47 | 6,38E-01 |
| AT2G37990 | 0,11 | -0,80 | 4,39 | -0,18 | 8,56E-01 |
| AT2G38010 | 0,53 | -0,53 | 3,09 | -0,17 | 8,64E-01 |
| AT2G38090 | 0,20 | -2,55 | 4,40 | -0,58 | 5,62E-01 |
| AT2G38120 | 0,38 | 0,81 | 2,81 | 0,29 | 7,72E-01 |
| AT2G38140 | 0,26 | 0,36 | 3,77 | 0,10 | 9,24E-01 |
| AT2G38310 | 0,71 | -1,69 | 2,94 | -0,57 | 5,67E-01 |
| AT2G38365 | 0,20 | -2,60 | 4,41 | -0,59 | 5,55E-01 |
| AT2G38420 | 0,26 | -2,60 | 4,41 | -0,59 | 5,55E-01 |
| AT2G38470 | 1,30 | -0,58 | 1,63 | -0,35 | 7,24E-01 |
| AT2G38590 | 0,78 | 2,95 | 2,65 | 1,11 | 2,65E-01 |
| AT2G38670 | 0,86 | -2,43 | 2,24 | -1,09 | 2,77E-01 |
| AT2G38760 | 0,08 | 0,00 | 4,41 | 0,00 | 1,00E+00 |
| AT2G38780 | 0,82 | -0,24 | 1,78 | -0,13 | 8,94E-01 |
| AT2G38820 | 0,12 | -0,68 | 4,41 | -0,15 | 8,77E-01 |
| AT2G38840 | 0,33 | -2,03 | 4,36 | -0,46 | 6,42E-01 |
| AT2G38940 | 0,25 | -2,53 | 4,40 | -0,57 | 5,65E-01 |
| AT2G38950 | 0,65 | -3,28 | 2,60 | -1,26 | 2,07E-01 |
| AT2G39190 | 0,15 | 0,05 | 4,37 | 0,01 | 9,91E-01 |
| AT2G39230 | 0,19 | 0,64 | 4,34 | 0,15 | 8,83E-01 |
| AT2G39270 | 0,08 | 0,00 | 4,41 | 0,00 | 1,00E+00 |
| AT2G39340 | 0,19 | 0,64 | 4,34 | 0,15 | 8,83E-01 |
| AT2G39400 | 0,15 | -0,81 | 4,39 | -0,18 | 8,54E-01 |
| AT2G39420 | 0,56 | -1,85 | 3,13 | -0,59 | 5,53E-01 |
| AT2G39430 | 0,52 | -2,60 | 3,96 | -0,66 | 5,11E-01 |
| AT2G39460 | 0,89 | 1,01 | 1,93 | 0,53 | 5,99E-01 |
| AT2G39470 | 0,81 | -1,69 | 2,35 | -0,72 | 4,72E-01 |
| AT2G39580 | 1,16 | -5,08 | 2,08 | -2,44 | 1,46E-02 |
| AT2G39670 | 1,03 | 0,08 | 2,34 | 0,03 | 9,74E-01 |
| AT2G39725 | 0,50 | -3,56 | 3,95 | -0,90 | 3,68E-01 |
| AT2G39740 | 0,63 | -1,17 | 3,00 | -0,39 | 6,95E-01 |
| AT2G39780 | 1,17 | -2,36 | 2,14 | -1,10 | 2,70E-01 |
| AT2G39840 | 0,84 | -0,51 | 2,45 | -0,21 | 8,34E-01 |
| AT2G39850 | 0,54 | 0,98 | 4,33 | 0,23 | 8,20E-01 |
| AT2G39870 | 0,43 | -0,21 | 4,29 | -0,05 | 9,60E-01 |
| AT2G39930 | 1,27 | -2,56 | 1,63 | -1,57 | 1,16E-01 |
| AT2G40060 | 0,99 | 0,28 | 2,40 | 0,12 | 9,06E-01 |
| AT2G40095 | 0,64 | -2,44 | 3,15 | -0,78 | 4,38E-01 |
| AT2G40130 | 0,14 | -0,29 | 4,38 | -0,07 | 9,47E-01 |
| AT2G40150 | 0,08 | 0,00 | 4,41 | 0,00 | 1,00E+00 |
| AT2G40160 | 0,42 | -3,39 | 4,16 | -0,81 | 4,15E-01 |
| AT2G40200 | 0,38 | 0,02 | 4,32 | 0,00 | 9,96E-01 |
| AT2G09120 | 0,31 | 0,00 | 4,41 | 0,00 | 1,00E+00 |
| AT2G40310 | 0,83 | 2,60 | 2,19 | 1,19 | 2,35E-01 |
| AT2G40400 | 0,82 | -2,05 | 2,84 | -0,72 | 4,70E-01 |
| AT2G40410 | 0,54 | -2,36 | 4,00 | -0,59 | 5,55E-01 |
| AT2G40435 | 0,46 | -2,97 | 4,38 | -0,68 | 4,97E-01 |
| AT2G40460 | 0,63 | 1,34 | 2,54 | 0,53 | 5,99E-01 |
| AT2G40480 | 0,31 | -1,04 | 4,38 | -0,24 | 8,13E-01 |
| AT2G40490 | 1,23 | -0,18 | 1,96 | -0,09 | 9,27E-01 |
| AT2G40540 | 0,78 | 0,12 | 2,48 | 0,05 | 9,61E-01 |
| AT2G40550 | 0,20 | -2,60 | 4,41 | -0,59 | 5,55E-01 |
| AT2G40660 | 0,08 | -0,21 | 4,39 | -0,05 | 9,62E-01 |
| AT2G40670 | 0,10 | -0,18 | 4,38 | -0,04 | 9,67E-01 |

| | | | | | |
|---|---|---|---|---|---|
| AT2G40760 | 0,35 | 0,00 | 4,41 | 0,00 | 1,00E+00 |
| AT2G40765 | 0,08 | 0,00 | 4,41 | 0,00 | 1,00E+00 |
| AT2G40810 | 0,08 | 0,00 | 4,41 | 0,00 | 1,00E+00 |
| AT2G40830 | 0,33 | -2,60 | 4,41 | -0,59 | 5,55E-01 |
| AT2G40880 | 1,07 | -1,77 | 2,20 | -0,81 | 4,21E-01 |
| AT2G40980 | 0,41 | -2,97 | 4,22 | -0,70 | 4,81E-01 |
| AT2G41080 | 0,59 | -0,83 | 2,53 | -0,33 | 7,44E-01 |
| AT2G41110 | 0,26 | 0,48 | 3,79 | 0,13 | 8,99E-01 |
| AT2G09230 | 0,12 | 0,00 | 4,41 | 0,00 | 1,00E+00 |
| AT2G41140 | 0,37 | -1,68 | 4,34 | -0,39 | 6,99E-01 |
| AT2G41200 | 0,84 | -0,51 | 2,12 | -0,24 | 8,10E-01 |
| AT2G41210 | 0,42 | -1,89 | 4,18 | -0,45 | 6,51E-01 |
| AT2G41220 | 0,64 | 0,37 | 2,94 | 0,13 | 8,99E-01 |
| AT2G41460 | 0,13 | 0,00 | 4,41 | 0,00 | 1,00E+00 |
| AT2G41510 | 0,08 | 0,00 | 4,41 | 0,00 | 1,00E+00 |
| AT2G41520 | 0,59 | -2,85 | 2,99 | -0,95 | 3,41E-01 |
| AT2G41670 | 0,12 | 0,00 | 4,41 | 0,00 | 1,00E+00 |
| AT2G41730 | 0,99 | -2,93 | 1,96 | -1,50 | 1,35E-01 |
| AT2G41820 | 0,34 | -2,60 | 4,41 | -0,59 | 5,55E-01 |
| AT2G41880 | 0,26 | 0,48 | 3,79 | 0,13 | 8,99E-01 |
| AT2G42130 | 0,48 | -0,41 | 3,29 | -0,12 | 9,02E-01 |
| AT2G42220 | 0,98 | -1,04 | 2,13 | -0,49 | 6,27E-01 |
| AT2G42310 | 0,61 | -0,65 | 2,54 | -0,26 | 7,98E-01 |
| AT2G42320 | 0,63 | -1,43 | 2,69 | -0,53 | 5,94E-01 |
| AT2G42430 | 0,21 | 1,49 | 4,30 | 0,35 | 7,29E-01 |
| AT2G42580 | 1,32 | -2,13 | 1,92 | -1,11 | 2,67E-01 |
| AT2G42610 | 0,29 | -0,76 | 4,40 | -0,17 | 8,62E-01 |
| AT2G42700 | 0,72 | -2,45 | 3,08 | -0,80 | 4,26E-01 |
| AT2G42710 | 0,12 | -0,78 | 4,39 | -0,18 | 8,59E-01 |
| AT2G42750 | 0,57 | -2,55 | 3,84 | -0,66 | 5,06E-01 |
| AT2G42780 | 0,28 | 0,49 | 4,35 | 0,11 | 9,11E-01 |
| AT2G42890 | 0,32 | -0,82 | 4,38 | -0,19 | 8,53E-01 |
| AT2G42900 | 0,43 | -1,71 | 4,34 | -0,39 | 6,94E-01 |
| AT2G43010 | 1,28 | -2,38 | 1,75 | -1,36 | 1,75E-01 |
| AT2G43020 | 1,32 | 2,29 | 2,00 | 1,14 | 2,52E-01 |
| AT2G43030 | 0,58 | 0,51 | 3,13 | 0,16 | 8,70E-01 |
| AT2G43080 | 0,63 | -0,26 | 3,08 | -0,08 | 9,33E-01 |
| AT2G43090 | 0,61 | 0,24 | 2,66 | 0,09 | 9,27E-01 |
| AT2G43100 | 0,56 | 0,20 | 3,32 | 0,06 | 9,53E-01 |
| AT2G43138 | 1,21 | 2,06 | 1,97 | 1,04 | 2,97E-01 |
| AT2G43210 | 0,63 | -2,60 | 3,08 | -0,85 | 3,97E-01 |
| AT2G43230 | 0,55 | -3,82 | 3,83 | -1,00 | 3,19E-01 |
| AT2G43250 | 0,31 | -1,69 | 4,37 | -0,39 | 6,99E-01 |
| AT2G43280 | 0,56 | -3,38 | 4,36 | -0,78 | 4,38E-01 |
| AT2G09440 | 0,08 | 0,00 | 4,41 | 0,00 | 1,00E+00 |
| AT2G43340 | 0,44 | -1,23 | 3,32 | -0,37 | 7,10E-01 |
| AT2G43400 | 0,37 | -1,51 | 4,36 | -0,35 | 7,29E-01 |
| AT2G43440 | 0,11 | 0,27 | 4,37 | 0,06 | 9,52E-01 |
| AT2G43630 | 0,44 | -0,71 | 4,14 | -0,17 | 8,63E-01 |
| AT2G43650 | 0,83 | -0,44 | 2,55 | -0,17 | 8,61E-01 |
| AT2G43690 | 0,68 | -2,57 | 4,40 | -0,58 | 5,60E-01 |
| AT2G43790 | 0,39 | 0,04 | 2,88 | 0,01 | 9,89E-01 |
| AT2G43800 | 0,08 | -0,21 | 4,39 | -0,05 | 9,62E-01 |
| AT2G43820 | 0,45 | -2,53 | 3,26 | -0,78 | 4,38E-01 |
| AT2G44080 | 0,60 | -3,09 | 3,69 | -0,84 | 4,01E-01 |
| AT2G44100 | 0,65 | 0,42 | 2,62 | 0,16 | 8,72E-01 |

| | | | | | |
|---|---|---|---|---|---|
| AT2G44130 | 0,72 | 1,40 | 2,49 | 0,56 | 5,72E-01 |
| AT2G44160 | 0,80 | -0,60 | 2,31 | -0,26 | 7,95E-01 |
| AT2G44175 | 0,43 | 2,36 | 4,06 | 0,58 | 5,61E-01 |
| AT2G44200 | 0,72 | -0,27 | 2,57 | -0,11 | 9,15E-01 |
| AT2G44210 | 1,13 | -0,43 | 1,62 | -0,27 | 7,90E-01 |
| AT2G44230 | 0,76 | 1,01 | 2,77 | 0,36 | 7,17E-01 |
| AT2G44240 | 0,17 | 0,42 | 4,36 | 0,10 | 9,23E-01 |
| AT2G44260 | 0,20 | -0,68 | 4,41 | -0,15 | 8,77E-01 |
| AT2G44350 | 0,40 | 1,33 | 3,35 | 0,40 | 6,92E-01 |
| AT2G44420 | 0,20 | -2,60 | 4,41 | -0,59 | 5,55E-01 |
| AT2G44430 | 0,96 | 1,76 | 1,88 | 0,93 | 3,50E-01 |
| AT2G44500 | 0,91 | 1,78 | 2,84 | 0,63 | 5,32E-01 |
| AT2G44520 | 0,23 | -1,73 | 4,39 | -0,39 | 6,93E-01 |
| AT2G44530 | 0,99 | -0,71 | 2,37 | -0,30 | 7,66E-01 |
| AT2G44600 | 0,10 | -0,18 | 4,38 | -0,04 | 9,67E-01 |
| AT2G44670 | 1,15 | -0,01 | 1,92 | -0,01 | 9,96E-01 |
| AT2G44735 | 0,42 | -1,76 | 4,16 | -0,42 | 6,73E-01 |
| AT2G44770 | 0,15 | -0,77 | 4,39 | -0,18 | 8,61E-01 |
| AT2G44800 | 0,19 | -0,03 | 4,38 | -0,01 | 9,94E-01 |
| AT2G44870 | 0,51 | -0,17 | 2,69 | -0,06 | 9,50E-01 |
| AT2G44880 | 0,91 | -3,82 | 2,78 | -1,38 | 1,69E-01 |
| AT2G44925 | 0,92 | 0,20 | 1,71 | 0,12 | 9,06E-01 |
| AT2G44940 | 0,52 | -1,72 | 2,76 | -0,62 | 5,34E-01 |
| AT2G45000 | 0,38 | 0,31 | 3,38 | 0,09 | 9,26E-01 |
| AT2G45020 | 0,36 | -0,44 | 4,38 | -0,10 | 9,21E-01 |
| AT2G45030 | 0,91 | -1,47 | 2,78 | -0,53 | 5,97E-01 |
| AT2G45140 | 0,33 | -2,55 | 4,40 | -0,58 | 5,63E-01 |
| AT2G45240 | 0,80 | -1,39 | 3,11 | -0,45 | 6,55E-01 |
| AT2G45250 | 0,26 | -2,60 | 4,41 | -0,59 | 5,55E-01 |
| AT2G45300 | 0,10 | -0,34 | 4,38 | -0,08 | 9,38E-01 |
| AT2G45310 | 0,67 | 0,31 | 2,49 | 0,12 | 9,01E-01 |
| AT2G45390 | 0,17 | 0,00 | 4,41 | 0,00 | 1,00E+00 |
| AT2G45440 | 0,53 | -2,60 | 3,85 | -0,68 | 4,99E-01 |
| AT2G45510 | 0,84 | -0,97 | 2,02 | -0,48 | 6,30E-01 |
| AT2G45530 | 0,98 | -0,85 | 2,02 | -0,42 | 6,76E-01 |
| AT2G09650 | 0,36 | 0,73 | 4,33 | 0,17 | 8,66E-01 |
| AT2G45590 | 0,37 | -0,93 | 4,31 | -0,22 | 8,29E-01 |
| AT2G45630 | 0,80 | -1,07 | 2,55 | -0,42 | 6,75E-01 |
| AT2G45710 | 0,45 | -1,23 | 4,34 | -0,28 | 7,76E-01 |
| AT2G45720 | 0,29 | 0,04 | 4,37 | 0,01 | 9,93E-01 |
| AT2G45740 | 0,35 | 0,14 | 3,60 | 0,04 | 9,70E-01 |
| AT2G45790 | 0,17 | -0,68 | 4,41 | -0,15 | 8,77E-01 |
| AT2G45810 | 0,90 | 1,10 | 2,43 | 0,45 | 6,52E-01 |
| AT2G45910 | 0,97 | -0,62 | 1,88 | -0,33 | 7,42E-01 |
| AT2G46100 | 0,37 | 1,14 | 2,84 | 0,40 | 6,88E-01 |
| AT2G46170 | 0,69 | -2,95 | 3,67 | -0,80 | 4,21E-01 |
| AT2G46190 | 0,70 | 2,16 | 2,16 | 1,00 | 3,18E-01 |
| AT2G09705 | 0,89 | 2,09 | 2,54 | 0,82 | 4,10E-01 |
| AT2G46225 | 0,27 | 0,36 | 4,36 | 0,08 | 9,35E-01 |
| AT2G46240 | 0,70 | -0,72 | 2,66 | -0,27 | 7,88E-01 |
| AT2G46260 | 0,08 | 0,00 | 4,41 | 0,00 | 1,00E+00 |
| AT2G46270 | 0,25 | -2,48 | 4,39 | -0,57 | 5,71E-01 |
| AT2G46370 | 0,44 | -0,81 | 3,36 | -0,24 | 8,09E-01 |
| AT2G46420 | 0,84 | 0,22 | 2,86 | 0,08 | 9,40E-01 |
| AT2G46450 | 1,09 | -2,27 | 2,10 | -1,08 | 2,80E-01 |
| AT2G46490 | 0,75 | 0,80 | 2,89 | 0,28 | 7,82E-01 |

| | | | | | |
|---|---|---|---|---|---|
| AT2G46520 | 1,16 | -0,25 | 1,75 | -0,15 | 8,85E-01 |
| AT2G46570 | 0,51 | -1,73 | 4,07 | -0,42 | 6,71E-01 |
| AT2G46590 | 0,52 | 2,00 | 2,98 | 0,67 | 5,03E-01 |
| AT2G46600 | 1,11 | -2,74 | 1,70 | -1,61 | 1,07E-01 |
| AT2G46630 | 0,28 | 0,97 | 4,32 | 0,22 | 8,23E-01 |
| AT2G46650 | 0,42 | 0,04 | 3,30 | 0,01 | 9,90E-01 |
| AT2G46690 | 0,43 | 0,52 | 4,17 | 0,12 | 9,01E-01 |
| AT2G46710 | 0,30 | 1,17 | 3,64 | 0,32 | 7,49E-01 |
| AT2G46790 | 0,54 | -0,81 | 2,61 | -0,31 | 7,56E-01 |
| AT2G46800 | 1,02 | -0,81 | 1,81 | -0,45 | 6,55E-01 |
| AT2G46810 | 0,59 | 1,44 | 3,15 | 0,46 | 6,48E-01 |
| AT2G46910 | 0,44 | 1,70 | 4,23 | 0,40 | 6,87E-01 |
| AT2G46990 | 0,08 | 0,00 | 4,41 | 0,00 | 1,00E+00 |
| AT2G47110 | 0,66 | 1,10 | 2,16 | 0,51 | 6,10E-01 |
| AT2G09815 | 0,23 | 1,25 | 4,31 | 0,29 | 7,72E-01 |
| AT2G47190 | 0,40 | 0,00 | 4,41 | 0,00 | 1,00E+00 |
| AT2G47220 | 0,25 | 0,00 | 4,41 | 0,00 | 1,00E+00 |
| AT2G47320 | 0,28 | -0,08 | 4,39 | -0,02 | 9,86E-01 |
| AT2G47420 | 0,42 | 0,00 | 4,20 | 0,00 | 1,00E+00 |
| AT2G47610 | 1,29 | -1,58 | 1,91 | -0,83 | 4,08E-01 |
| AT2G47620 | 0,26 | 0,48 | 3,79 | 0,13 | 8,99E-01 |
| AT2G47630 | 0,42 | -0,27 | 2,82 | -0,10 | 9,23E-01 |
| AT2G47640 | 0,28 | -0,43 | 4,35 | -0,10 | 9,21E-01 |
| AT2G47690 | 0,55 | -2,27 | 4,01 | -0,57 | 5,71E-01 |
| AT2G47730 | 0,80 | 1,48 | 2,20 | 0,67 | 5,01E-01 |
| AT2G47850 | 0,53 | -1,46 | 3,07 | -0,48 | 6,34E-01 |
| AT2G47910 | 0,92 | 1,28 | 2,02 | 0,64 | 5,24E-01 |
| AT2G47935 | 0,72 | -2,32 | 2,80 | -0,83 | 4,06E-01 |
| AT2G47970 | 0,70 | -3,71 | 3,51 | -1,06 | 2,91E-01 |
| AT2G48010 | 0,33 | 0,14 | 4,37 | 0,03 | 9,74E-01 |
| AT2G48020 | 0,56 | -1,27 | 3,20 | -0,40 | 6,91E-01 |
| AT2G48110 | 1,17 | -3,06 | 2,00 | -1,53 | 1,26E-01 |
| AT2G48120 | 0,32 | -2,60 | 4,41 | -0,59 | 5,55E-01 |
| AT3G01040 | 0,51 | -1,70 | 3,20 | -0,53 | 5,95E-01 |
| AT3G01050 | 0,18 | -0,34 | 4,38 | -0,08 | 9,38E-01 |
| AT3G01080 | 0,17 | -0,80 | 4,39 | -0,18 | 8,56E-01 |
| AT3G01085 | 1,09 | 1,15 | 1,89 | 0,61 | 5,43E-01 |
| AT3G01150 | 0,45 | -1,30 | 3,24 | -0,40 | 6,89E-01 |
| AT3G01170 | 0,42 | -1,89 | 4,18 | -0,45 | 6,51E-01 |
| AT3G01175 | 0,08 | 0,00 | 4,41 | 0,00 | 1,00E+00 |
| AT3G01210 | 0,10 | -0,34 | 4,38 | -0,08 | 9,38E-01 |
| AT3G01280 | 1,02 | -1,85 | 1,62 | -1,14 | 2,56E-01 |
| AT3G01320 | 1,14 | -1,76 | 1,65 | -1,07 | 2,86E-01 |
| AT3G01440 | 0,31 | 0,78 | 4,33 | 0,18 | 8,57E-01 |
| AT3G01475 | 1,05 | -2,79 | 1,85 | -1,51 | 1,32E-01 |
| AT3G01480 | 1,15 | -1,51 | 1,84 | -0,82 | 4,13E-01 |
| AT3G01560 | 0,68 | -1,70 | 2,52 | -0,68 | 4,99E-01 |
| AT3G01590 | 0,40 | -0,74 | 4,40 | -0,17 | 8,67E-01 |
| AT3G01610 | 0,25 | -0,40 | 4,37 | -0,09 | 9,27E-01 |
| AT3G01660 | 0,12 | -0,68 | 4,41 | -0,15 | 8,77E-01 |
| AT3G01690 | 1,08 | 0,57 | 1,86 | 0,31 | 7,59E-01 |
| AT3G01740 | 0,28 | -0,79 | 4,39 | -0,18 | 8,57E-01 |
| AT3G01750 | 0,72 | -4,09 | 3,00 | -1,36 | 1,72E-01 |
| AT3G01810 | 0,37 | 0,00 | 4,41 | 0,00 | 1,00E+00 |
| AT3G01830 | 0,94 | -1,24 | 2,10 | -0,59 | 5,54E-01 |
| AT3G01920 | 0,37 | -1,07 | 4,36 | -0,24 | 8,07E-01 |

| | | | | | |
|---|---|---|---|---|---|
| AT3G02010 | 0,37 | -3,40 | 4,37 | -0,78 | 4,36E-01 |
| AT3G02030 | 0,12 | -0,14 | 4,38 | -0,03 | 9,74E-01 |
| AT3G02050 | 0,37 | -3,33 | 4,32 | -0,77 | 4,41E-01 |
| AT3G02060 | 1,23 | -0,41 | 2,00 | -0,21 | 8,36E-01 |
| AT3G02065 | 0,54 | -3,44 | 3,83 | -0,90 | 3,68E-01 |
| AT3G02070 | 0,35 | -0,08 | 3,53 | -0,02 | 9,81E-01 |
| AT3G02130 | 1,18 | -2,26 | 2,17 | -1,04 | 2,98E-01 |
| AT3G02280 | 0,25 | -1,15 | 4,38 | -0,26 | 7,93E-01 |
| AT3G02290 | 0,16 | -0,68 | 4,41 | -0,15 | 8,77E-01 |
| AT3G02300 | 0,16 | 0,61 | 4,34 | 0,14 | 8,89E-01 |
| AT3G02340 | 0,50 | 0,22 | 4,35 | 0,05 | 9,59E-01 |
| AT3G02350 | 0,84 | 0,14 | 2,12 | 0,07 | 9,47E-01 |
| AT3G02370 | 0,47 | -0,93 | 3,39 | -0,27 | 7,84E-01 |
| AT3G02400 | 0,66 | -4,85 | 4,30 | -1,13 | 2,58E-01 |
| AT3G02490 | 0,33 | -1,64 | 4,41 | -0,37 | 7,10E-01 |
| AT3G02560 | 0,49 | -1,72 | 3,98 | -0,43 | 6,65E-01 |
| AT3G02610 | 0,81 | 0,57 | 2,88 | 0,20 | 8,42E-01 |
| AT3G02630 | 0,37 | -2,59 | 4,31 | -0,60 | 5,47E-01 |
| AT3G02690 | 0,66 | -2,39 | 2,50 | -0,96 | 3,38E-01 |
| AT3G02720 | 0,46 | -2,07 | 4,34 | -0,48 | 6,34E-01 |
| AT3G02830 | 1,22 | -0,82 | 2,09 | -0,39 | 6,95E-01 |
| AT3G01145 | 0,57 | 2,21 | 2,50 | 0,89 | 3,76E-01 |
| AT3G02880 | 1,19 | -2,65 | 1,59 | -1,67 | 9,59E-02 |
| AT3G02900 | 0,28 | 1,74 | 4,29 | 0,41 | 6,85E-01 |
| AT3G02910 | 0,77 | 0,04 | 2,34 | 0,02 | 9,85E-01 |
| AT3G03020 | 0,43 | 0,07 | 4,19 | 0,02 | 9,86E-01 |
| AT3G03030 | 0,40 | 0,00 | 4,41 | 0,00 | 1,00E+00 |
| AT3G03040 | 0,24 | -0,79 | 4,36 | -0,18 | 8,56E-01 |
| AT3G03110 | 0,55 | -3,19 | 4,39 | -0,73 | 4,67E-01 |
| AT3G03160 | 0,47 | -0,98 | 4,02 | -0,24 | 8,08E-01 |
| AT3G03210 | 0,10 | 0,45 | 4,35 | 0,10 | 9,19E-01 |
| AT3G03320 | 0,15 | -0,17 | 4,39 | -0,04 | 9,68E-01 |
| AT3G03340 | 0,40 | -0,68 | 4,41 | -0,15 | 8,77E-01 |
| AT3G03350 | 0,24 | -2,59 | 4,41 | -0,59 | 5,56E-01 |
| AT3G03360 | 0,08 | 0,00 | 4,41 | 0,00 | 1,00E+00 |
| AT3G03610 | 0,27 | 0,00 | 4,41 | 0,00 | 1,00E+00 |
| AT3G03640 | 0,25 | 0,34 | 4,36 | 0,08 | 9,37E-01 |
| AT3G03860 | 0,25 | 1,48 | 4,30 | 0,34 | 7,30E-01 |
| AT3G03880 | 0,28 | 0,87 | 4,32 | 0,20 | 8,40E-01 |
| AT3G03950 | 0,32 | -0,68 | 3,64 | -0,19 | 8,52E-01 |
| AT3G04140 | 0,99 | -3,84 | 2,35 | -1,63 | 1,02E-01 |
| AT3G04490 | 0,26 | -0,77 | 4,37 | -0,18 | 8,59E-01 |
| AT3G04530 | 0,08 | 0,00 | 4,41 | 0,00 | 1,00E+00 |
| AT3G04550 | 0,64 | 0,41 | 2,27 | 0,18 | 8,57E-01 |
| AT3G04790 | 1,26 | -2,26 | 2,11 | -1,07 | 2,84E-01 |
| AT3G04810 | 0,59 | 0,00 | 4,41 | 0,00 | 1,00E+00 |
| AT3G04840 | 0,80 | -1,82 | 2,25 | -0,81 | 4,18E-01 |
| AT3G04870 | 0,49 | 0,06 | 3,13 | 0,02 | 9,85E-01 |
| AT3G04910 | 0,86 | 1,34 | 1,86 | 0,72 | 4,71E-01 |
| AT3G04920 | 0,51 | -2,32 | 4,06 | -0,57 | 5,67E-01 |
| AT3G04940 | 0,44 | -3,22 | 4,21 | -0,77 | 4,44E-01 |
| AT3G01555 | 0,12 | -0,78 | 4,39 | -0,18 | 8,59E-01 |
| AT3G01575 | 0,10 | 0,45 | 4,35 | 0,10 | 9,19E-01 |
| AT3G05070 | 0,17 | -0,29 | 4,38 | -0,07 | 9,46E-01 |
| AT3G05120 | 0,72 | 0,25 | 2,23 | 0,11 | 9,12E-01 |
| AT3G05200 | 0,41 | -0,76 | 4,21 | -0,18 | 8,57E-01 |

| | | | | | |
|---|---|---|---|---|---|
| AT3G05230 | 0,28 | -0,55 | 4,39 | -0,13 | 9,00E-01 |
| AT3G05380 | 0,92 | -3,30 | 2,26 | -1,46 | 1,44E-01 |
| AT3G05430 | 0,21 | 0,00 | 4,41 | 0,00 | 1,00E+00 |
| AT3G05490 | 0,28 | 0,49 | 4,35 | 0,11 | 9,11E-01 |
| AT3G05500 | 0,55 | -1,19 | 2,64 | -0,45 | 6,51E-01 |
| AT3G05520 | 0,43 | -0,11 | 2,83 | -0,04 | 9,68E-01 |
| AT3G05545 | 0,82 | 0,82 | 2,22 | 0,37 | 7,10E-01 |
| AT3G05560 | 0,45 | 0,46 | 3,32 | 0,14 | 8,89E-01 |
| AT3G05630 | 0,21 | 0,00 | 4,41 | 0,00 | 1,00E+00 |
| AT3G05670 | 0,41 | -1,23 | 4,22 | -0,29 | 7,71E-01 |
| AT3G05710 | 0,95 | -2,19 | 2,01 | -1,09 | 2,77E-01 |
| AT3G05725 | 0,64 | 2,31 | 2,40 | 0,96 | 3,36E-01 |
| AT3G05750 | 0,61 | -0,75 | 3,75 | -0,20 | 8,41E-01 |
| AT3G05760 | 0,68 | -0,68 | 3,76 | -0,18 | 8,56E-01 |
| AT3G05830 | 0,23 | 0,11 | 4,38 | 0,03 | 9,80E-01 |
| AT3G05840 | 0,50 | -4,09 | 3,95 | -1,04 | 3,00E-01 |
| AT3G05905 | 0,51 | -0,62 | 3,25 | -0,19 | 8,47E-01 |
| AT3G06125 | 0,60 | -1,01 | 2,22 | -0,46 | 6,48E-01 |
| AT3G06170 | 0,51 | 0,99 | 3,06 | 0,32 | 7,48E-01 |
| AT3G06180 | 0,25 | 0,00 | 4,41 | 0,00 | 1,00E+00 |
| AT3G06200 | 0,41 | -0,07 | 4,32 | -0,02 | 9,87E-01 |
| AT3G06210 | 0,20 | -2,60 | 4,41 | -0,59 | 5,55E-01 |
| AT3G06240 | 0,61 | -2,40 | 2,36 | -1,02 | 3,09E-01 |
| AT3G06380 | 0,55 | -2,64 | 3,93 | -0,67 | 5,01E-01 |
| AT3G06400 | 0,95 | 0,93 | 2,53 | 0,37 | 7,13E-01 |
| AT3G06455 | 0,58 | -1,53 | 4,35 | -0,35 | 7,26E-01 |
| AT3G06470 | 0,36 | -0,83 | 4,38 | -0,19 | 8,50E-01 |
| AT3G06500 | 1,24 | 0,27 | 1,99 | 0,14 | 8,91E-01 |
| AT3G06520 | 0,50 | -3,38 | 3,95 | -0,86 | 3,92E-01 |
| AT3G06610 | 0,15 | 0,48 | 4,35 | 0,11 | 9,12E-01 |
| AT3G06660 | 0,08 | -0,21 | 4,39 | -0,05 | 9,62E-01 |
| AT3G06730 | 0,97 | -2,17 | 3,01 | -0,72 | 4,71E-01 |
| AT3G06740 | 0,50 | -0,73 | 4,06 | -0,18 | 8,57E-01 |
| AT3G06830 | 0,08 | 0,00 | 4,41 | 0,00 | 1,00E+00 |
| AT3G06860 | 1,03 | -1,20 | 1,90 | -0,63 | 5,29E-01 |
| AT3G06868 | 0,29 | -1,07 | 4,36 | -0,25 | 8,05E-01 |
| AT3G06895 | 1,19 | 0,32 | 1,77 | 0,18 | 8,55E-01 |
| AT3G06920 | 0,42 | -1,45 | 4,32 | -0,34 | 7,36E-01 |
| AT3G06980 | 0,81 | -3,05 | 3,05 | -1,00 | 3,17E-01 |
| AT3G07200 | 0,08 | 0,00 | 4,41 | 0,00 | 1,00E+00 |
| AT3G07230 | 0,29 | -0,70 | 4,40 | -0,16 | 8,74E-01 |
| AT3G07270 | 0,85 | -3,57 | 2,09 | -1,71 | 8,76E-02 |
| AT3G07350 | 0,42 | -1,16 | 3,35 | -0,35 | 7,28E-01 |
| AT3G07360 | 1,00 | 0,42 | 2,35 | 0,18 | 8,59E-01 |
| AT3G07400 | 0,87 | -2,89 | 2,06 | -1,40 | 1,61E-01 |
| AT3G07430 | 0,32 | 0,48 | 4,34 | 0,11 | 9,12E-01 |
| AT3G07440 | 0,08 | 0,00 | 4,41 | 0,00 | 1,00E+00 |
| AT3G07480 | 0,45 | -1,14 | 4,25 | -0,27 | 7,88E-01 |
| AT3G07565 | 0,19 | 0,39 | 4,35 | 0,09 | 9,29E-01 |
| AT3G07610 | 0,83 | -1,36 | 2,42 | -0,56 | 5,74E-01 |
| AT3G07640 | 0,17 | 0,00 | 4,41 | 0,00 | 1,00E+00 |
| AT3G07650 | 0,97 | -2,39 | 3,44 | -0,69 | 4,88E-01 |
| AT3G07680 | 0,59 | -1,91 | 3,15 | -0,60 | 5,46E-01 |
| AT3G07690 | 0,32 | -3,36 | 4,37 | -0,77 | 4,42E-01 |
| AT3G07780 | 1,15 | -0,80 | 2,15 | -0,37 | 7,11E-01 |
| AT3G07810 | 0,90 | -1,26 | 1,81 | -0,70 | 4,87E-01 |

| | | | | | |
|---|---|---|---|---|---|
| AT3G07880 | 0,28 | 0,01 | 4,36 | 0,00 | 9,99E-01 |
| AT3G07950 | 0,43 | 0,00 | 4,41 | 0,00 | 1,00E+00 |
| AT3G08010 | 0,17 | 0,66 | 4,34 | 0,15 | 8,80E-01 |
| AT3G08550 | 0,31 | -0,13 | 4,33 | -0,03 | 9,77E-01 |
| AT3G08600 | 0,52 | 0,42 | 3,87 | 0,11 | 9,14E-01 |
| AT3G08640 | 0,20 | 0,00 | 4,41 | 0,00 | 1,00E+00 |
| AT3G08690 | 0,22 | -0,80 | 4,35 | -0,18 | 8,54E-01 |
| AT3G08920 | 0,32 | -0,24 | 4,32 | -0,05 | 9,57E-01 |
| AT3G08930 | 0,51 | -0,51 | 3,25 | -0,16 | 8,76E-01 |
| AT3G08950 | 0,44 | -1,03 | 4,13 | -0,25 | 8,02E-01 |
| AT3G08980 | 0,33 | -2,06 | 4,35 | -0,47 | 6,36E-01 |
| AT3G09020 | 0,57 | -0,06 | 4,04 | -0,01 | 9,89E-01 |
| AT3G09030 | 0,51 | -0,93 | 2,56 | -0,36 | 7,17E-01 |
| AT3G09035 | 0,19 | 0,02 | 4,36 | 0,00 | 9,97E-01 |
| AT3G09085 | 0,40 | -0,91 | 3,42 | -0,27 | 7,90E-01 |
| AT3G02105 | 0,40 | 0,00 | 4,41 | 0,00 | 1,00E+00 |
| AT3G09180 | 0,23 | -1,68 | 4,36 | -0,39 | 7,00E-01 |
| AT3G09300 | 1,07 | -3,54 | 2,47 | -1,43 | 1,53E-01 |
| AT3G09350 | 0,51 | -3,16 | 4,39 | -0,72 | 4,72E-01 |
| AT3G09370 | 0,29 | -2,60 | 4,41 | -0,59 | 5,55E-01 |
| AT3G09430 | 1,27 | -4,09 | 2,14 | -1,91 | 5,63E-02 |
| AT3G09500 | 0,81 | -3,17 | 2,16 | -1,47 | 1,42E-01 |
| AT3G09510 | 0,52 | 1,98 | 3,06 | 0,65 | 5,18E-01 |
| AT3G09570 | 0,70 | -2,39 | 2,59 | -0,92 | 3,55E-01 |
| AT3G09600 | 0,41 | -2,51 | 3,35 | -0,75 | 4,54E-01 |
| AT3G09650 | 0,58 | -2,58 | 4,40 | -0,59 | 5,58E-01 |
| AT3G09670 | 0,48 | -2,60 | 4,41 | -0,59 | 5,55E-01 |
| AT3G09690 | 0,55 | -4,20 | 4,33 | -0,97 | 3,32E-01 |
| AT3G09810 | 0,32 | -0,44 | 4,33 | -0,10 | 9,18E-01 |
| AT3G09820 | 0,77 | -0,02 | 2,48 | -0,01 | 9,94E-01 |
| AT3G09830 | 0,46 | 1,40 | 3,23 | 0,43 | 6,64E-01 |
| AT3G09880 | 0,75 | -1,51 | 2,54 | -0,59 | 5,52E-01 |
| AT3G09890 | 0,43 | -2,51 | 4,39 | -0,57 | 5,68E-01 |
| AT3G09900 | 0,19 | -0,72 | 4,40 | -0,16 | 8,70E-01 |
| AT3G10060 | 0,36 | -0,37 | 4,35 | -0,08 | 9,32E-01 |
| AT3G10120 | 0,12 | 0,00 | 4,41 | 0,00 | 1,00E+00 |
| AT3G10190 | 0,32 | -0,79 | 4,39 | -0,18 | 8,57E-01 |
| AT3G10200 | 0,08 | 0,00 | 4,41 | 0,00 | 1,00E+00 |
| AT3G10300 | 0,68 | 2,16 | 2,84 | 0,76 | 4,48E-01 |
| AT3G10310 | 0,16 | -0,62 | 4,40 | -0,14 | 8,87E-01 |
| AT3G10340 | 0,80 | 2,44 | 2,96 | 0,83 | 4,08E-01 |
| AT3G10370 | 0,28 | 0,38 | 4,35 | 0,09 | 9,30E-01 |
| AT3G10390 | 0,49 | -1,67 | 3,20 | -0,52 | 6,01E-01 |
| AT3G10480 | 0,72 | -3,11 | 3,57 | -0,87 | 3,83E-01 |
| AT3G10490 | 0,18 | -0,34 | 4,38 | -0,08 | 9,38E-01 |
| AT3G10527 | 0,27 | -0,68 | 4,41 | -0,15 | 8,77E-01 |
| AT3G10540 | 0,56 | -1,10 | 2,56 | -0,43 | 6,68E-01 |
| AT3G10570 | 0,12 | -0,78 | 4,39 | -0,18 | 8,59E-01 |
| AT3G10760 | 0,48 | -1,24 | 3,27 | -0,38 | 7,05E-01 |
| AT3G10820 | 0,24 | -3,18 | 4,39 | -0,73 | 4,68E-01 |
| AT3G10860 | 0,23 | 0,43 | 4,34 | 0,10 | 9,20E-01 |
| AT3G10920 | 1,12 | -1,28 | 2,01 | -0,64 | 5,24E-01 |
| AT3G10950 | 0,19 | 0,65 | 4,34 | 0,15 | 8,82E-01 |
| AT3G10980 | 0,79 | -2,13 | 3,03 | -0,70 | 4,83E-01 |
| AT3G11020 | 0,05 | -0,24 | 4,38 | -0,05 | 9,57E-01 |
| AT3G11080 | 0,18 | 0,78 | 4,34 | 0,18 | 8,57E-01 |

| | | | | | |
|---|---|---|---|---|---|
| AT3G11090 | 0,60 | -0,49 | 3,20 | -0,15 | 8,78E-01 |
| AT3G11120 | 0,39 | 0,40 | 3,33 | 0,12 | 9,04E-01 |
| AT3G11170 | 1,02 | 0,10 | 1,71 | 0,06 | 9,55E-01 |
| AT3G11230 | 0,53 | -3,00 | 4,38 | -0,68 | 4,94E-01 |
| AT3G11250 | 0,11 | 0,46 | 4,35 | 0,11 | 9,15E-01 |
| AT3G11270 | 0,10 | -0,34 | 4,38 | -0,08 | 9,38E-01 |
| AT3G11397 | 0,44 | -1,71 | 4,18 | -0,41 | 6,82E-01 |
| AT3G11400 | 0,87 | -3,09 | 2,22 | -1,40 | 1,63E-01 |
| AT3G11420 | 0,26 | 0,84 | 4,33 | 0,19 | 8,46E-01 |
| AT3G11440 | 0,65 | -3,45 | 3,04 | -1,14 | 2,56E-01 |
| AT3G02465 | 0,30 | -0,55 | 4,34 | -0,13 | 8,98E-01 |
| AT3G11630 | 0,96 | 1,81 | 1,84 | 0,99 | 3,24E-01 |
| AT3G11700 | 0,50 | 0,91 | 3,20 | 0,28 | 7,76E-01 |
| AT3G11760 | 0,23 | -1,73 | 4,39 | -0,39 | 6,93E-01 |
| AT3G11810 | 0,52 | -0,62 | 3,77 | -0,16 | 8,69E-01 |
| AT3G11900 | 0,13 | 0,00 | 4,41 | 0,00 | 1,00E+00 |
| AT3G11960 | 1,04 | -2,96 | 2,08 | -1,42 | 1,56E-01 |
| AT3G12010 | 0,40 | -0,68 | 4,31 | -0,16 | 8,74E-01 |
| AT3G12020 | 0,79 | -3,54 | 2,37 | -1,50 | 1,35E-01 |
| AT3G12050 | 0,32 | 1,37 | 3,49 | 0,39 | 6,94E-01 |
| AT3G12080 | 1,08 | -3,50 | 2,48 | -1,41 | 1,57E-01 |
| AT3G12270 | 0,77 | -0,56 | 2,69 | -0,21 | 8,34E-01 |
| AT3G12290 | 0,81 | -0,60 | 2,33 | -0,26 | 7,96E-01 |
| AT3G12320 | 0,90 | -3,85 | 2,06 | -1,87 | 6,20E-02 |
| AT3G12360 | 0,71 | -0,59 | 2,47 | -0,24 | 8,12E-01 |
| AT3G12390 | 0,69 | -2,93 | 2,44 | -1,20 | 2,30E-01 |
| AT3G12480 | 0,26 | -0,76 | 4,40 | -0,17 | 8,63E-01 |
| AT3G12502 | 0,08 | 0,00 | 4,41 | 0,00 | 1,00E+00 |
| AT3G12570 | 1,24 | -0,68 | 2,86 | -0,24 | 8,12E-01 |
| AT3G12600 | 0,05 | -0,24 | 4,38 | -0,05 | 9,57E-01 |
| AT3G12640 | 0,60 | -1,68 | 2,25 | -0,75 | 4,55E-01 |
| AT3G12700 | 0,20 | -1,00 | 4,37 | -0,23 | 8,19E-01 |
| AT3G12740 | 0,52 | 0,65 | 3,26 | 0,20 | 8,42E-01 |
| AT3G12760 | 0,80 | -2,85 | 2,34 | -1,22 | 2,24E-01 |
| AT3G12915 | 0,53 | -3,07 | 4,38 | -0,70 | 4,83E-01 |
| AT3G13060 | 0,87 | -0,33 | 2,09 | -0,16 | 8,75E-01 |
| AT3G13065 | 0,88 | -4,04 | 3,41 | -1,18 | 2,37E-01 |
| AT3G13200 | 1,18 | 0,77 | 1,73 | 0,44 | 6,58E-01 |
| AT3G13210 | 0,13 | 0,00 | 4,41 | 0,00 | 1,00E+00 |
| AT3G13340 | 1,00 | 0,41 | 2,00 | 0,21 | 8,36E-01 |
| AT3G13370 | 0,63 | 3,00 | 4,27 | 0,70 | 4,83E-01 |
| AT3G13380 | 0,67 | 0,27 | 2,45 | 0,11 | 9,12E-01 |
| AT3G13437 | 0,08 | 0,00 | 4,41 | 0,00 | 1,00E+00 |
| AT3G13445 | 0,19 | -0,83 | 4,38 | -0,19 | 8,50E-01 |
| AT3G13470 | 0,61 | -1,13 | 3,85 | -0,29 | 7,68E-01 |
| AT3G02725 | 1,24 | -1,05 | 1,77 | -0,59 | 5,54E-01 |
| AT3G13662 | 0,34 | 0,18 | 4,32 | 0,04 | 9,67E-01 |
| AT3G13690 | 0,67 | -1,51 | 3,83 | -0,39 | 6,93E-01 |
| AT3G13760 | 0,08 | 0,00 | 4,41 | 0,00 | 1,00E+00 |
| AT3G13770 | 0,43 | -2,54 | 4,35 | -0,58 | 5,60E-01 |
| AT3G13810 | 0,42 | 0,19 | 4,15 | 0,05 | 9,64E-01 |
| AT3G13930 | 0,81 | -1,96 | 1,98 | -0,99 | 3,21E-01 |
| AT3G13950 | 0,55 | 0,03 | 3,93 | 0,01 | 9,95E-01 |
| AT3G14020 | 0,35 | -2,99 | 4,38 | -0,68 | 4,95E-01 |
| AT3G14100 | 0,69 | 1,63 | 2,54 | 0,64 | 5,20E-01 |
| AT3G14160 | 0,19 | 0,01 | 4,36 | 0,00 | 9,98E-01 |

| | | | | | |
|---|---|---|---|---|---|
| AT3G14220 | 0,64 | -3,57 | 3,74 | -0,95 | 3,40E-01 |
| AT3G14290 | 0,42 | -2,60 | 4,19 | -0,62 | 5,35E-01 |
| AT3G14560 | 0,20 | -2,60 | 4,41 | -0,59 | 5,55E-01 |
| AT3G14595 | 1,01 | -0,90 | 2,10 | -0,43 | 6,67E-01 |
| AT3G14600 | 1,18 | -1,51 | 1,71 | -0,88 | 3,78E-01 |
| AT3G14610 | 0,28 | -3,07 | 4,38 | -0,70 | 4,84E-01 |
| AT3G14620 | 0,63 | 0,47 | 2,50 | 0,19 | 8,50E-01 |
| AT3G14640 | 0,33 | -3,01 | 4,38 | -0,69 | 4,92E-01 |
| AT3G14670 | 1,07 | 1,72 | 1,91 | 0,90 | 3,70E-01 |
| AT3G14690 | 0,63 | 1,46 | 2,26 | 0,65 | 5,17E-01 |
| AT3G14720 | 0,35 | -0,30 | 4,31 | -0,07 | 9,45E-01 |
| AT3G14790 | 0,27 | -2,23 | 4,36 | -0,51 | 6,10E-01 |
| AT3G14830 | 0,30 | -1,26 | 4,36 | -0,29 | 7,73E-01 |
| AT3G14870 | 0,25 | -0,25 | 4,38 | -0,06 | 9,54E-01 |
| AT3G14920 | 0,30 | -2,60 | 4,41 | -0,59 | 5,55E-01 |
| AT3G14940 | 0,68 | -1,35 | 3,59 | -0,38 | 7,07E-01 |
| AT3G14990 | 0,39 | 0,90 | 2,85 | 0,32 | 7,51E-01 |
| AT3G15000 | 0,95 | -0,67 | 2,27 | -0,29 | 7,68E-01 |
| AT3G15010 | 0,55 | 0,17 | 3,18 | 0,05 | 9,58E-01 |
| AT3G15020 | 1,02 | -3,63 | 2,41 | -1,51 | 1,32E-01 |
| AT3G15030 | 0,69 | -3,24 | 2,98 | -1,09 | 2,76E-01 |
| AT3G15090 | 0,12 | 0,00 | 4,41 | 0,00 | 1,00E+00 |
| AT3G15160 | 1,08 | -2,25 | 2,16 | -1,04 | 2,98E-01 |
| AT3G15180 | 0,21 | -1,17 | 4,36 | -0,27 | 7,88E-01 |
| AT3G15190 | 1,26 | -2,29 | 1,60 | -1,43 | 1,54E-01 |
| AT3G15260 | 0,39 | -0,79 | 4,34 | -0,18 | 8,56E-01 |
| AT3G15290 | 0,26 | -2,60 | 4,41 | -0,59 | 5,55E-01 |
| AT3G15350 | 1,19 | -4,34 | 2,25 | -1,93 | 5,33E-02 |
| AT3G15355 | 0,91 | -1,70 | 3,77 | -0,45 | 6,53E-01 |
| AT3G15360 | 1,20 | -0,29 | 1,71 | -0,17 | 8,63E-01 |
| AT3G15380 | 0,12 | 0,00 | 4,41 | 0,00 | 1,00E+00 |
| AT3G15450 | 0,77 | -3,32 | 2,17 | -1,53 | 1,26E-01 |
| AT3G15518 | 0,17 | 0,00 | 4,41 | 0,00 | 1,00E+00 |
| AT3G15550 | 0,17 | 0,00 | 4,41 | 0,00 | 1,00E+00 |
| AT3G03145 | 0,66 | 0,63 | 2,90 | 0,22 | 8,28E-01 |
| AT3G15635 | 0,25 | 0,32 | 4,36 | 0,07 | 9,42E-01 |
| AT3G15750 | 0,08 | 0,00 | 4,41 | 0,00 | 1,00E+00 |
| AT3G15760 | 0,84 | -0,18 | 2,61 | -0,07 | 9,45E-01 |
| AT3G15790 | 0,67 | -1,71 | 2,96 | -0,58 | 5,63E-01 |
| AT3G15820 | 0,32 | 0,01 | 4,36 | 0,00 | 9,98E-01 |
| AT3G15850 | 0,57 | 1,58 | 2,64 | 0,60 | 5,50E-01 |
| AT3G15930 | 0,12 | 0,00 | 4,41 | 0,00 | 1,00E+00 |
| AT3G16010 | 0,15 | 0,41 | 4,35 | 0,09 | 9,26E-01 |
| AT3G16060 | 0,47 | -1,12 | 4,09 | -0,27 | 7,84E-01 |
| AT3G16080 | 0,49 | -1,63 | 3,26 | -0,50 | 6,16E-01 |
| AT3G16130 | 0,87 | -0,38 | 1,99 | -0,19 | 8,50E-01 |
| AT3G16175 | 0,08 | 0,00 | 4,41 | 0,00 | 1,00E+00 |
| AT3G16260 | 0,72 | -3,90 | 3,53 | -1,10 | 2,69E-01 |
| AT3G16270 | 0,71 | 0,46 | 2,53 | 0,18 | 8,56E-01 |
| AT3G16340 | 0,41 | -3,11 | 4,39 | -0,71 | 4,78E-01 |
| AT3G16420 | 0,28 | -2,28 | 4,37 | -0,52 | 6,01E-01 |
| AT3G16460 | 0,31 | -0,05 | 4,33 | -0,01 | 9,90E-01 |
| AT3G16525 | 0,84 | 1,28 | 2,13 | 0,60 | 5,47E-01 |
| AT3G16570 | 0,68 | -0,22 | 2,49 | -0,09 | 9,31E-01 |
| AT3G16710 | 0,43 | -1,39 | 4,27 | -0,32 | 7,45E-01 |
| AT3G16720 | 0,43 | -0,63 | 4,12 | -0,15 | 8,79E-01 |

| | | | | | |
|---|---|---|---|---|---|
| AT3G16740 | 0,38 | 0,00 | 3,46 | 0,00 | 9,99E-01 |
| AT3G16760 | 0,57 | 0,40 | 3,81 | 0,11 | 9,16E-01 |
| AT3G16800 | 0,16 | -0,01 | 4,37 | 0,00 | 9,98E-01 |
| AT3G16857 | 0,72 | -3,16 | 3,78 | -0,83 | 4,04E-01 |
| AT3G16940 | 0,78 | -1,75 | 2,92 | -0,60 | 5,49E-01 |
| AT3G17100 | 1,17 | -1,61 | 1,80 | -0,90 | 3,71E-01 |
| AT3G17160 | 0,19 | -0,79 | 4,39 | -0,18 | 8,58E-01 |
| AT3G17205 | 0,35 | 0,08 | 4,32 | 0,02 | 9,85E-01 |
| AT3G17210 | 0,32 | -2,30 | 4,37 | -0,53 | 5,99E-01 |
| AT3G03465 | 0,80 | 2,25 | 2,29 | 0,98 | 3,27E-01 |
| AT3G17360 | 0,65 | -3,56 | 2,99 | -1,19 | 2,34E-01 |
| AT3G17380 | 0,28 | -3,29 | 4,38 | -0,75 | 4,53E-01 |
| AT3G17410 | 0,79 | -0,59 | 1,87 | -0,32 | 7,51E-01 |
| AT3G17470 | 0,76 | -3,07 | 3,71 | -0,83 | 4,08E-01 |
| AT3G17490 | 0,56 | 2,57 | 3,81 | 0,67 | 5,00E-01 |
| AT3G17626 | 0,20 | -2,60 | 4,41 | -0,59 | 5,55E-01 |
| AT3G17640 | 0,16 | -0,01 | 4,37 | 0,00 | 9,98E-01 |
| AT3G17650 | 0,88 | 0,84 | 2,14 | 0,39 | 6,95E-01 |
| AT3G17690 | 0,12 | 0,00 | 4,41 | 0,00 | 1,00E+00 |
| AT3G17700 | 0,35 | 0,71 | 4,33 | 0,16 | 8,69E-01 |
| AT3G17712 | 0,66 | -3,67 | 3,66 | -1,00 | 3,15E-01 |
| AT3G17750 | 1,26 | -1,43 | 1,74 | -0,82 | 4,11E-01 |
| AT3G17765 | 0,21 | -2,60 | 4,41 | -0,59 | 5,55E-01 |
| AT3G17790 | 0,83 | -2,57 | 2,52 | -1,02 | 3,07E-01 |
| AT3G03535 | 0,92 | -1,56 | 2,18 | -0,72 | 4,73E-01 |
| AT3G17810 | 0,53 | 0,02 | 2,79 | 0,01 | 9,94E-01 |
| AT3G17820 | 0,62 | -2,66 | 3,13 | -0,85 | 3,95E-01 |
| AT3G17860 | 0,36 | 1,17 | 4,32 | 0,27 | 7,87E-01 |
| AT3G17880 | 0,42 | 1,22 | 3,25 | 0,37 | 7,08E-01 |
| AT3G17890 | 0,10 | -0,34 | 4,38 | -0,08 | 9,38E-01 |
| AT3G17970 | 0,77 | -1,03 | 3,01 | -0,34 | 7,33E-01 |
| AT3G17980 | 0,35 | -2,99 | 4,38 | -0,68 | 4,95E-01 |
| AT3G18000 | 0,28 | -1,56 | 4,39 | -0,35 | 7,23E-01 |
| AT3G18020 | 0,41 | -2,04 | 4,24 | -0,48 | 6,31E-01 |
| AT3G18040 | 0,26 | 0,30 | 4,36 | 0,07 | 9,46E-01 |
| AT3G18070 | 0,11 | -0,68 | 4,41 | -0,15 | 8,77E-01 |
| AT3G18090 | 1,26 | -3,37 | 3,06 | -1,10 | 2,71E-01 |
| AT3G18100 | 0,29 | 0,00 | 4,41 | 0,00 | 1,00E+00 |
| AT3G03625 | 0,40 | 1,33 | 3,35 | 0,40 | 6,92E-01 |
| AT3G18240 | 0,43 | -1,25 | 4,20 | -0,30 | 7,67E-01 |
| AT3G18270 | 0,50 | -1,49 | 3,98 | -0,37 | 7,08E-01 |
| AT3G18340 | 0,19 | 1,18 | 4,31 | 0,27 | 7,84E-01 |
| AT3G18390 | 1,15 | -1,04 | 1,87 | -0,56 | 5,76E-01 |
| AT3G18440 | 0,08 | 0,00 | 4,41 | 0,00 | 1,00E+00 |
| AT3G18480 | 0,44 | -0,87 | 3,27 | -0,26 | 7,91E-01 |
| AT3G18740 | 0,73 | 0,05 | 2,34 | 0,02 | 9,83E-01 |
| AT3G18790 | 0,65 | -2,50 | 2,49 | -1,00 | 3,16E-01 |
| AT3G18800 | 0,33 | -3,01 | 4,38 | -0,69 | 4,92E-01 |
| AT3G18820 | 0,39 | 0,76 | 4,18 | 0,18 | 8,55E-01 |
| AT3G18827 | 0,57 | -1,49 | 3,10 | -0,48 | 6,30E-01 |
| AT3G18860 | 0,59 | 0,55 | 2,51 | 0,22 | 8,27E-01 |
| AT3G18890 | 0,86 | -2,45 | 2,13 | -1,15 | 2,50E-01 |
| AT3G18915 | 0,30 | 1,31 | 4,32 | 0,30 | 7,62E-01 |
| AT3G18960 | 0,08 | 0,00 | 4,41 | 0,00 | 1,00E+00 |
| AT3G18980 | 1,15 | -3,08 | 1,94 | -1,59 | 1,13E-01 |
| AT3G19050 | 0,62 | -4,49 | 2,95 | -1,52 | 1,28E-01 |

| | | | | | |
|---|---|---|---|---|---|
| AT3G19100 | 1,08 | -2,98 | 1,93 | -1,55 | 1,21E-01 |
| AT3G19290 | 0,28 | -2,15 | 4,38 | -0,49 | 6,25E-01 |
| AT3G19390 | 0,35 | -2,99 | 4,38 | -0,68 | 4,95E-01 |
| AT3G19470 | 0,30 | -1,47 | 4,34 | -0,34 | 7,35E-01 |
| AT3G19480 | 0,52 | -0,31 | 3,21 | -0,10 | 9,24E-01 |
| AT3G19520 | 0,05 | -0,24 | 4,38 | -0,05 | 9,57E-01 |
| AT3G19540 | 1,20 | -1,71 | 1,85 | -0,93 | 3,54E-01 |
| AT3G19640 | 0,86 | -2,92 | 2,44 | -1,20 | 2,32E-01 |
| AT3G19680 | 0,92 | -1,74 | 2,10 | -0,83 | 4,06E-01 |
| AT3G19760 | 1,20 | -0,55 | 1,75 | -0,32 | 7,52E-01 |
| AT3G19770 | 0,75 | -2,48 | 3,05 | -0,81 | 4,17E-01 |
| AT3G19800 | 0,78 | -3,35 | 2,54 | -1,32 | 1,87E-01 |
| AT3G19860 | 0,48 | -3,49 | 4,04 | -0,87 | 3,87E-01 |
| AT3G19870 | 0,37 | 0,27 | 4,32 | 0,06 | 9,50E-01 |
| AT3G19980 | 0,93 | -2,39 | 2,28 | -1,05 | 2,95E-01 |
| AT3G19990 | 0,32 | -3,12 | 4,39 | -0,71 | 4,77E-01 |
| AT3G20000 | 0,74 | -3,46 | 3,64 | -0,95 | 3,43E-01 |
| AT3G20030 | 0,79 | 2,58 | 2,37 | 1,09 | 2,78E-01 |
| AT3G20140 | 0,20 | -2,55 | 4,40 | -0,58 | 5,62E-01 |
| AT3G20150 | 0,54 | -4,47 | 4,31 | -1,04 | 3,00E-01 |
| AT3G20200 | 0,21 | 0,00 | 4,41 | 0,00 | 1,00E+00 |
| AT3G20410 | 0,37 | -0,21 | 4,33 | -0,05 | 9,61E-01 |
| AT3G20500 | 0,33 | -3,35 | 4,37 | -0,77 | 4,44E-01 |
| AT3G20520 | 0,48 | -2,60 | 4,41 | -0,59 | 5,55E-01 |
| AT3G20540 | 0,90 | -3,00 | 2,30 | -1,30 | 1,93E-01 |
| AT3G20600 | 0,25 | -1,15 | 4,38 | -0,26 | 7,93E-01 |
| AT3G03995 | 0,62 | 2,70 | 4,27 | 0,63 | 5,27E-01 |
| AT3G20720 | 0,42 | 0,25 | 3,39 | 0,07 | 9,41E-01 |
| AT3G20730 | 0,40 | 0,00 | 4,41 | 0,00 | 1,00E+00 |
| AT3G20790 | 0,23 | 0,00 | 4,41 | 0,00 | 1,00E+00 |
| AT3G20890 | 0,34 | -1,60 | 4,35 | -0,37 | 7,13E-01 |
| AT3G20970 | 0,63 | -1,56 | 3,07 | -0,51 | 6,12E-01 |
| AT3G21130 | 0,35 | 1,27 | 4,32 | 0,29 | 7,69E-01 |
| AT3G21170 | 1,15 | 3,60 | 2,30 | 1,56 | 1,18E-01 |
| AT3G21175 | 0,18 | 0,78 | 4,34 | 0,18 | 8,57E-01 |
| AT3G21220 | 0,51 | -0,19 | 3,99 | -0,05 | 9,61E-01 |
| AT3G21270 | 0,29 | -0,76 | 4,40 | -0,17 | 8,62E-01 |
| AT3G21295 | 0,41 | -0,91 | 3,43 | -0,27 | 7,91E-01 |
| AT3G21310 | 0,36 | 0,00 | 4,41 | 0,00 | 1,00E+00 |
| AT3G21465 | 0,14 | -0,35 | 4,38 | -0,08 | 9,37E-01 |
| AT3G21473 | 0,48 | 1,20 | 3,97 | 0,30 | 7,63E-01 |
| AT3G21560 | 0,31 | -1,42 | 4,37 | -0,32 | 7,46E-01 |
| AT3G21690 | 0,96 | 2,22 | 1,99 | 1,12 | 2,63E-01 |
| AT3G04175 | 0,99 | 0,56 | 1,94 | 0,29 | 7,72E-01 |
| AT3G21990 | 0,46 | 2,31 | 4,28 | 0,54 | 5,89E-01 |
| AT3G22072 | 0,17 | 0,00 | 4,41 | 0,00 | 1,00E+00 |
| AT3G22104 | 0,62 | -2,19 | 2,90 | -0,75 | 4,51E-01 |
| AT3G22150 | 0,49 | -3,80 | 3,96 | -0,96 | 3,37E-01 |
| AT3G22200 | 0,76 | -0,50 | 2,26 | -0,22 | 8,25E-01 |
| AT3G04255 | 0,24 | -0,11 | 4,38 | -0,03 | 9,80E-01 |
| AT3G22270 | 0,79 | -1,52 | 2,50 | -0,61 | 5,43E-01 |
| AT3G22290 | 0,33 | -3,01 | 4,38 | -0,69 | 4,92E-01 |
| AT3G22330 | 0,85 | -0,63 | 2,19 | -0,29 | 7,72E-01 |
| AT3G22350 | 0,91 | 3,08 | 2,52 | 1,22 | 2,21E-01 |
| AT3G22370 | 0,72 | -1,23 | 2,42 | -0,51 | 6,12E-01 |
| AT3G22425 | 0,12 | -0,31 | 4,38 | -0,07 | 9,44E-01 |

| | | | | | |
|---|---|---|---|---|---|
| AT3G22430 | 0,19 | 0,80 | 4,34 | 0,18 | 8,53E-01 |
| AT3G22520 | 0,75 | -1,52 | 2,56 | -0,59 | 5,53E-01 |
| AT3G22760 | 0,51 | -0,88 | 3,23 | -0,27 | 7,86E-01 |
| AT3G22810 | 0,55 | 0,00 | 4,41 | 0,00 | 1,00E+00 |
| AT3G22845 | 0,51 | -4,07 | 4,33 | -0,94 | 3,48E-01 |
| AT3G22870 | 0,92 | 2,08 | 2,11 | 0,99 | 3,24E-01 |
| AT3G22900 | 0,24 | -2,60 | 4,41 | -0,59 | 5,55E-01 |
| AT3G22935 | 0,26 | 1,81 | 4,29 | 0,42 | 6,73E-01 |
| AT3G23010 | 0,77 | -3,80 | 3,49 | -1,09 | 2,76E-01 |
| AT3G23050 | 0,64 | 0,20 | 2,61 | 0,08 | 9,38E-01 |
| AT3G04395 | 0,16 | -0,26 | 4,38 | -0,06 | 9,52E-01 |
| AT3G23250 | 0,25 | 0,67 | 4,34 | 0,16 | 8,77E-01 |
| AT3G23280 | 0,45 | 1,16 | 4,07 | 0,28 | 7,76E-01 |
| AT3G23300 | 0,75 | -1,75 | 2,04 | -0,86 | 3,91E-01 |
| AT3G23310 | 0,63 | 0,25 | 2,26 | 0,11 | 9,10E-01 |
| AT3G23380 | 0,21 | 0,17 | 4,37 | 0,04 | 9,68E-01 |
| AT3G23390 | 0,15 | 0,41 | 4,35 | 0,09 | 9,26E-01 |
| AT3G23410 | 0,36 | -0,15 | 4,37 | -0,03 | 9,73E-01 |
| AT3G23530 | 1,09 | -2,81 | 1,91 | -1,47 | 1,42E-01 |
| AT3G23580 | 0,57 | -0,46 | 2,31 | -0,20 | 8,42E-01 |
| AT3G23600 | 1,16 | -0,65 | 1,80 | -0,36 | 7,16E-01 |
| AT3G23640 | 0,83 | -2,69 | 2,46 | -1,09 | 2,74E-01 |
| AT3G23750 | 0,94 | -0,91 | 1,67 | -0,54 | 5,86E-01 |
| AT3G23790 | 0,38 | -1,13 | 4,34 | -0,26 | 7,94E-01 |
| AT3G23800 | 0,20 | -2,55 | 4,40 | -0,58 | 5,62E-01 |
| AT3G23890 | 0,56 | -2,45 | 3,97 | -0,62 | 5,37E-01 |
| AT3G23990 | 0,73 | -4,03 | 3,68 | -1,10 | 2,73E-01 |
| AT3G24000 | 0,34 | -1,76 | 4,39 | -0,40 | 6,88E-01 |
| AT3G24050 | 0,88 | 0,11 | 2,11 | 0,05 | 9,58E-01 |
| AT3G24070 | 0,38 | -2,60 | 4,41 | -0,59 | 5,55E-01 |
| AT3G24160 | 0,45 | 1,42 | 3,45 | 0,41 | 6,80E-01 |
| AT3G24190 | 0,59 | -1,64 | 3,16 | -0,52 | 6,04E-01 |
| AT3G24255 | 0,43 | -0,90 | 2,81 | -0,32 | 7,48E-01 |
| AT3G24490 | 0,64 | -2,60 | 3,67 | -0,71 | 4,78E-01 |
| AT3G24500 | 0,22 | 1,22 | 4,31 | 0,28 | 7,78E-01 |
| AT3G24514 | 0,29 | 1,81 | 4,29 | 0,42 | 6,73E-01 |
| AT3G24520 | 0,92 | 0,22 | 2,42 | 0,09 | 9,27E-01 |
| AT3G24550 | 0,60 | 1,22 | 2,94 | 0,41 | 6,79E-01 |
| AT3G24580 | 0,47 | 0,46 | 4,05 | 0,11 | 9,09E-01 |
| AT3G24590 | 0,21 | -2,50 | 4,39 | -0,57 | 5,70E-01 |
| AT3G24830 | 0,92 | 0,59 | 2,14 | 0,28 | 7,82E-01 |
| AT3G25012 | 0,95 | -1,59 | 1,74 | -0,91 | 3,62E-01 |
| AT3G25013 | 0,18 | 0,78 | 4,34 | 0,18 | 8,57E-01 |
| AT3G25030 | 0,21 | -0,23 | 4,38 | -0,05 | 9,58E-01 |
| AT3G25040 | 0,17 | -0,29 | 4,38 | -0,07 | 9,46E-01 |
| AT3G25070 | 0,31 | 0,88 | 3,58 | 0,24 | 8,07E-01 |
| AT3G25130 | 0,57 | -2,61 | 4,41 | -0,59 | 5,53E-01 |
| AT3G25250 | 0,60 | -0,68 | 2,72 | -0,25 | 8,03E-01 |
| AT3G25290 | 0,29 | -1,98 | 4,37 | -0,45 | 6,50E-01 |
| AT3G25440 | 0,23 | -1,14 | 4,39 | -0,26 | 7,96E-01 |
| AT3G25470 | 0,67 | -3,29 | 3,76 | -0,88 | 3,81E-01 |
| AT3G25490 | 0,05 | -0,24 | 4,38 | -0,05 | 9,57E-01 |
| AT3G25585 | 0,71 | -2,58 | 2,67 | -0,97 | 3,34E-01 |
| AT3G25590 | 0,75 | 1,17 | 2,36 | 0,50 | 6,18E-01 |
| AT3G25700 | 0,66 | -0,39 | 2,24 | -0,17 | 8,62E-01 |
| AT3G04955 | 0,15 | -0,81 | 4,39 | -0,18 | 8,54E-01 |

| | | | | | |
|---|---|---|---|---|---|
| AT3G25720 | 0,26 | 1,25 | 4,32 | 0,29 | 7,73E-01 |
| AT3G25760 | 0,49 | -3,79 | 4,35 | -0,87 | 3,83E-01 |
| AT3G25770 | 1,09 | 0,59 | 1,98 | 0,30 | 7,66E-01 |
| AT3G25780 | 0,49 | 0,96 | 2,58 | 0,37 | 7,10E-01 |
| AT3G25860 | 0,63 | 0,43 | 3,24 | 0,13 | 8,93E-01 |
| AT3G25890 | 0,89 | -0,47 | 2,15 | -0,22 | 8,28E-01 |
| AT3G25910 | 0,36 | -0,77 | 4,39 | -0,18 | 8,61E-01 |
| AT3G26020 | 0,28 | 0,13 | 4,36 | 0,03 | 9,77E-01 |
| AT3G26030 | 1,16 | -2,47 | 1,85 | -1,33 | 1,82E-01 |
| AT3G26070 | 0,46 | -1,76 | 4,16 | -0,42 | 6,72E-01 |
| AT3G26080 | 0,26 | -2,60 | 4,41 | -0,59 | 5,55E-01 |
| AT3G26085 | 0,30 | -1,92 | 4,37 | -0,44 | 6,60E-01 |
| AT3G26090 | 0,21 | -0,37 | 4,38 | -0,08 | 9,33E-01 |
| AT3G04975 | 1,19 | 2,90 | 2,01 | 1,44 | 1,49E-01 |
| AT3G04985 | 0,58 | 1,95 | 2,59 | 0,75 | 4,52E-01 |
| AT3G26370 | 0,48 | -2,50 | 2,75 | -0,91 | 3,65E-01 |
| AT3G26395 | 0,16 | -0,64 | 4,40 | -0,15 | 8,85E-01 |
| AT3G26400 | 0,95 | -2,08 | 2,26 | -0,92 | 3,59E-01 |
| AT3G26512 | 0,38 | 1,67 | 3,30 | 0,51 | 6,13E-01 |
| AT3G26580 | 0,73 | 0,80 | 2,25 | 0,36 | 7,21E-01 |
| AT3G26600 | 1,07 | -0,25 | 1,98 | -0,13 | 9,00E-01 |
| AT3G26640 | 0,37 | 0,51 | 3,35 | 0,15 | 8,79E-01 |
| AT3G26710 | 1,14 | -4,81 | 2,00 | -2,41 | 1,62E-02 |
| AT3G26720 | 0,82 | -1,24 | 2,53 | -0,49 | 6,24E-01 |
| AT3G26742 | 0,18 | 0,40 | 4,35 | 0,09 | 9,27E-01 |
| AT3G26780 | 1,24 | -1,27 | 1,95 | -0,65 | 5,14E-01 |
| AT3G26782 | 0,41 | -1,70 | 4,20 | -0,41 | 6,85E-01 |
| AT3G26800 | 0,34 | -0,43 | 3,49 | -0,12 | 9,02E-01 |
| AT3G26840 | 1,06 | -3,04 | 2,32 | -1,31 | 1,90E-01 |
| AT3G27090 | 1,24 | -2,05 | 1,85 | -1,11 | 2,68E-01 |
| AT3G27160 | 0,41 | 0,44 | 3,36 | 0,13 | 8,97E-01 |
| AT3G27170 | 0,68 | -2,60 | 2,98 | -0,87 | 3,83E-01 |
| AT3G27230 | 0,45 | -1,85 | 3,32 | -0,56 | 5,77E-01 |
| AT3G27250 | 0,85 | -1,13 | 2,44 | -0,46 | 6,44E-01 |
| AT3G27260 | 1,09 | 0,24 | 1,94 | 0,12 | 9,02E-01 |
| AT3G27270 | 0,62 | -2,18 | 3,87 | -0,56 | 5,73E-01 |
| AT3G27280 | 0,60 | -1,98 | 3,05 | -0,65 | 5,16E-01 |
| AT3G27325 | 0,56 | -1,10 | 2,60 | -0,42 | 6,73E-01 |
| AT3G27329 | 0,81 | 1,15 | 2,15 | 0,53 | 5,94E-01 |
| AT3G27331 | 0,96 | 3,38 | 3,12 | 1,08 | 2,79E-01 |
| AT3G27350 | 0,51 | 0,44 | 3,96 | 0,11 | 9,12E-01 |
| AT3G27430 | 0,35 | 0,84 | 3,55 | 0,24 | 8,12E-01 |
| AT3G27470 | 1,23 | 1,14 | 1,87 | 0,61 | 5,42E-01 |
| AT3G27540 | 0,29 | 0,59 | 3,66 | 0,16 | 8,72E-01 |
| AT3G27555 | 0,30 | 0,06 | 4,32 | 0,01 | 9,88E-01 |
| AT3G27670 | 1,19 | -1,86 | 2,12 | -0,88 | 3,80E-01 |
| AT3G05295 | 0,48 | 0,37 | 2,84 | 0,13 | 8,97E-01 |
| AT3G27820 | 1,03 | 1,70 | 1,95 | 0,87 | 3,85E-01 |
| AT3G27830 | 0,47 | 0,85 | 3,33 | 0,26 | 7,98E-01 |
| AT3G27860 | 0,12 | 0,00 | 4,41 | 0,00 | 1,00E+00 |
| AT3G27870 | 0,75 | -2,16 | 2,51 | -0,86 | 3,90E-01 |
| AT3G27940 | 0,56 | 2,40 | 2,88 | 0,83 | 4,05E-01 |
| AT3G27980 | 0,34 | 1,65 | 4,29 | 0,38 | 7,01E-01 |
| AT3G27999 | 0,43 | 2,23 | 4,28 | 0,52 | 6,02E-01 |
| AT3G28040 | 1,16 | -2,32 | 2,08 | -1,11 | 2,65E-01 |
| AT3G28050 | 0,37 | -0,75 | 4,40 | -0,17 | 8,65E-01 |

| | | | | | |
|---|---|---|---|---|---|
| AT3G28080 | 0,70 | 0,00 | 2,17 | 0,00 | 9,98E-01 |
| AT3G28100 | 1,10 | 2,11 | 1,83 | 1,15 | 2,49E-01 |
| AT3G28170 | 0,75 | -4,17 | 3,53 | -1,18 | 2,37E-01 |
| AT3G28200 | 0,56 | 0,00 | 4,41 | 0,00 | 1,00E+00 |
| AT3G28220 | 0,69 | -0,27 | 3,05 | -0,09 | 9,28E-01 |
| AT3G28270 | 0,14 | 0,48 | 4,35 | 0,11 | 9,13E-01 |
| AT3G28290 | 1,10 | -0,68 | 2,64 | -0,26 | 7,97E-01 |
| AT3G28450 | 0,78 | 1,95 | 2,52 | 0,77 | 4,40E-01 |
| AT3G28500 | 0,30 | 0,96 | 4,33 | 0,22 | 8,25E-01 |
| AT3G28510 | 0,99 | -1,78 | 2,52 | -0,71 | 4,80E-01 |
| AT3G05515 | 0,60 | 2,68 | 2,89 | 0,93 | 3,55E-01 |
| AT3G28580 | 0,77 | 0,71 | 3,45 | 0,21 | 8,37E-01 |
| AT3G28590 | 0,05 | -0,24 | 4,38 | -0,05 | 9,57E-01 |
| AT3G28650 | 0,20 | -1,00 | 4,37 | -0,23 | 8,19E-01 |
| AT3G28670 | 0,33 | -0,20 | 4,39 | -0,05 | 9,63E-01 |
| AT3G28690 | 0,39 | -0,45 | 4,24 | -0,11 | 9,16E-01 |
| AT3G28715 | 0,62 | -1,77 | 2,32 | -0,76 | 4,46E-01 |
| AT3G28720 | 0,91 | -2,02 | 3,21 | -0,63 | 5,29E-01 |
| AT3G28730 | 0,64 | -3,21 | 2,92 | -1,10 | 2,71E-01 |
| AT3G28870 | 0,74 | 0,02 | 3,73 | 0,01 | 9,96E-01 |
| AT3G28910 | 0,61 | -0,84 | 2,59 | -0,32 | 7,47E-01 |
| AT3G05595 | 0,35 | -0,39 | 4,38 | -0,09 | 9,30E-01 |
| AT3G28922 | 0,12 | -0,14 | 4,38 | -0,03 | 9,74E-01 |
| AT3G28958 | 0,58 | 1,59 | 3,01 | 0,53 | 5,97E-01 |
| AT3G29000 | 0,25 | -0,08 | 4,37 | -0,02 | 9,85E-01 |
| AT3G29010 | 0,38 | -3,32 | 4,36 | -0,76 | 4,47E-01 |
| AT3G29033 | 0,57 | 1,86 | 2,89 | 0,64 | 5,21E-01 |
| AT3G29080 | 1,28 | 1,85 | 1,70 | 1,09 | 2,76E-01 |
| AT3G29180 | 1,27 | -3,35 | 2,73 | -1,23 | 2,21E-01 |
| AT3G05715 | 0,40 | 0,65 | 4,28 | 0,15 | 8,80E-01 |
| AT3G29270 | 0,39 | -3,10 | 4,38 | -0,71 | 4,79E-01 |
| AT3G29290 | 0,20 | -0,68 | 4,41 | -0,15 | 8,77E-01 |
| AT3G29370 | 0,05 | -0,24 | 4,38 | -0,05 | 9,57E-01 |
| AT3G29390 | 0,18 | -0,34 | 4,38 | -0,08 | 9,38E-01 |
| AT3G29450 | 0,14 | 0,87 | 4,33 | 0,20 | 8,40E-01 |
| AT3G29455 | 0,27 | 1,85 | 4,29 | 0,43 | 6,67E-01 |
| AT3G05805 | 0,31 | 0,99 | 4,33 | 0,23 | 8,20E-01 |
| AT3G29638 | 0,16 | -0,07 | 4,37 | -0,02 | 9,87E-01 |
| AT3G29670 | 0,94 | 2,70 | 2,20 | 1,23 | 2,19E-01 |
| AT3G30210 | 0,11 | -0,80 | 4,39 | -0,18 | 8,56E-01 |
| AT3G30320 | 0,40 | 1,51 | 4,31 | 0,35 | 7,26E-01 |
| AT3G30380 | 0,49 | -2,58 | 3,28 | -0,79 | 4,32E-01 |
| AT3G30383 | 0,12 | 0,00 | 4,41 | 0,00 | 1,00E+00 |
| AT3G30525 | 1,14 | 3,29 | 2,63 | 1,25 | 2,11E-01 |
| AT3G30720 | 0,61 | -0,33 | 3,14 | -0,10 | 9,17E-01 |
| AT3G30770 | 0,13 | 0,52 | 4,35 | 0,12 | 9,04E-01 |
| AT3G30820 | 0,18 | 0,78 | 4,34 | 0,18 | 8,57E-01 |
| AT3G31950 | 0,59 | 1,15 | 2,27 | 0,50 | 6,14E-01 |
| AT3G32150 | 0,20 | 1,44 | 4,30 | 0,33 | 7,38E-01 |
| AT3G32280 | 0,39 | 0,99 | 4,24 | 0,23 | 8,16E-01 |
| AT3G32896 | 0,28 | 0,07 | 4,33 | 0,02 | 9,87E-01 |
| AT3G33530 | 1,04 | -0,58 | 1,76 | -0,33 | 7,41E-01 |
| AT3G42075 | 0,24 | -1,38 | 4,38 | -0,31 | 7,53E-01 |
| AT3G42170 | 0,48 | -2,44 | 3,18 | -0,77 | 4,42E-01 |
| AT3G42310 | 0,36 | 1,61 | 4,30 | 0,38 | 7,07E-01 |
| AT3G42560 | 0,10 | -0,34 | 4,38 | -0,08 | 9,38E-01 |

| | | | | | |
|---|---|---|---|---|---|
| AT3G42640 | 0,44 | 0,43 | 2,67 | 0,16 | 8,73E-01 |
| AT3G42950 | 0,12 | 0,00 | 4,41 | 0,00 | 1,00E+00 |
| AT3G06495 | 0,51 | 1,53 | 3,14 | 0,49 | 6,27E-01 |
| AT3G43148 | 0,82 | 2,64 | 2,22 | 1,19 | 2,34E-01 |
| AT3G43150 | 0,30 | 1,47 | 3,57 | 0,41 | 6,81E-01 |
| AT3G43210 | 0,25 | -2,57 | 4,40 | -0,58 | 5,60E-01 |
| AT3G43260 | 1,20 | 3,24 | 2,04 | 1,59 | 1,12E-01 |
| AT3G06565 | 0,05 | -0,24 | 4,38 | -0,05 | 9,57E-01 |
| AT3G43290 | 0,21 | 0,65 | 4,34 | 0,15 | 8,80E-01 |
| AT3G06625 | 0,13 | -0,18 | 4,39 | -0,04 | 9,67E-01 |
| AT3G43503 | 0,30 | 0,15 | 3,68 | 0,04 | 9,67E-01 |
| AT3G06635 | 0,40 | 1,82 | 4,30 | 0,42 | 6,72E-01 |
| AT3G43670 | 0,53 | -0,91 | 2,66 | -0,34 | 7,31E-01 |
| AT3G43682 | 0,33 | 0,71 | 4,34 | 0,16 | 8,70E-01 |
| AT3G43740 | 0,28 | 0,72 | 4,34 | 0,17 | 8,68E-01 |
| AT3G43800 | 0,45 | -1,71 | 3,33 | -0,51 | 6,07E-01 |
| AT3G43837 | 0,47 | 0,31 | 3,99 | 0,08 | 9,39E-01 |
| AT3G43920 | 0,47 | -0,75 | 3,20 | -0,23 | 8,16E-01 |
| AT3G43950 | 0,37 | 0,67 | 4,34 | 0,15 | 8,78E-01 |
| AT3G44130 | 0,22 | 0,58 | 4,35 | 0,13 | 8,94E-01 |
| AT3G44150 | 0,18 | 0,00 | 4,41 | 0,00 | 1,00E+00 |
| AT3G44210 | 1,26 | 3,78 | 2,42 | 1,57 | 1,18E-01 |
| AT3G44230 | 0,25 | 0,18 | 4,37 | 0,04 | 9,68E-01 |
| AT3G44235 | 0,05 | -0,24 | 4,38 | -0,05 | 9,57E-01 |
| AT3G44320 | 0,76 | -3,33 | 2,91 | -1,15 | 2,52E-01 |
| AT3G44405 | 0,08 | 0,00 | 4,41 | 0,00 | 1,00E+00 |
| AT3G44550 | 0,99 | 1,71 | 2,42 | 0,70 | 4,81E-01 |
| AT3G44560 | 0,14 | 0,48 | 4,35 | 0,11 | 9,13E-01 |
| AT3G44620 | 0,30 | 1,15 | 3,59 | 0,32 | 7,48E-01 |
| AT3G44730 | 0,52 | 1,14 | 2,34 | 0,49 | 6,26E-01 |
| AT3G44798 | 0,43 | 0,07 | 4,19 | 0,02 | 9,86E-01 |
| AT3G44880 | 0,54 | -0,53 | 3,97 | -0,13 | 8,93E-01 |
| AT3G44900 | 0,86 | -4,09 | 2,39 | -1,71 | 8,78E-02 |
| AT3G44935 | 1,28 | 3,21 | 1,99 | 1,61 | 1,08E-01 |
| AT3G44980 | 0,39 | -0,74 | 3,45 | -0,21 | 8,30E-01 |
| AT3G45010 | 0,31 | -0,43 | 4,32 | -0,10 | 9,20E-01 |
| AT3G45400 | 0,33 | -0,94 | 4,35 | -0,22 | 8,29E-01 |
| AT3G45620 | 1,31 | -0,83 | 2,50 | -0,33 | 7,38E-01 |
| AT3G45660 | 1,01 | -5,04 | 2,51 | -2,01 | 4,44E-02 |
| AT3G45670 | 0,15 | 0,48 | 4,35 | 0,11 | 9,12E-01 |
| AT3G45730 | 0,15 | -0,73 | 4,40 | -0,17 | 8,68E-01 |
| AT3G45740 | 0,36 | -3,08 | 4,39 | -0,70 | 4,83E-01 |
| AT3G45870 | 0,08 | 0,00 | 4,41 | 0,00 | 1,00E+00 |
| AT3G45930 | 0,32 | -2,55 | 4,40 | -0,58 | 5,62E-01 |
| AT3G45965 | 0,21 | 0,00 | 4,41 | 0,00 | 1,00E+00 |
| AT3G45970 | 0,98 | -3,06 | 2,68 | -1,14 | 2,53E-01 |
| AT3G46040 | 0,85 | -1,76 | 2,25 | -0,78 | 4,34E-01 |
| AT3G46060 | 1,03 | -0,78 | 1,95 | -0,40 | 6,88E-01 |
| AT3G46130 | 0,12 | 0,00 | 4,41 | 0,00 | 1,00E+00 |
| AT3G46220 | 0,60 | -2,92 | 2,60 | -1,12 | 2,62E-01 |
| AT3G46290 | 1,13 | -1,90 | 1,91 | -0,99 | 3,20E-01 |
| AT3G46370 | 0,25 | 1,14 | 4,32 | 0,26 | 7,93E-01 |
| AT3G46490 | 0,32 | 1,11 | 4,32 | 0,26 | 7,97E-01 |
| AT3G46520 | 0,37 | -3,03 | 4,38 | -0,69 | 4,90E-01 |
| AT3G46560 | 0,33 | 1,84 | 4,29 | 0,43 | 6,68E-01 |
| AT3G07505 | 1,14 | 0,64 | 1,85 | 0,35 | 7,27E-01 |

| | | | | | |
|---|---|---|---|---|---|
| AT3G46640 | 1,12 | -3,32 | 2,22 | -1,50 | 1,34E-01 |
| AT3G46658 | 0,59 | -0,41 | 2,66 | -0,15 | 8,78E-01 |
| AT3G46668 | 0,57 | -1,58 | 3,31 | -0,48 | 6,34E-01 |
| AT3G07555 | 1,02 | 0,95 | 2,16 | 0,44 | 6,61E-01 |
| AT3G46930 | 0,21 | 0,00 | 4,41 | 0,00 | 1,00E+00 |
| AT3G47060 | 0,35 | 0,00 | 4,41 | 0,00 | 1,00E+00 |
| AT3G47140 | 0,11 | 0,46 | 4,35 | 0,11 | 9,15E-01 |
| AT3G07695 | 0,33 | 0,28 | 4,35 | 0,06 | 9,50E-01 |
| AT3G47290 | 0,08 | 0,00 | 4,41 | 0,00 | 1,00E+00 |
| AT3G47295 | 0,08 | -0,21 | 4,39 | -0,05 | 9,62E-01 |
| AT3G07715 | 0,48 | -0,24 | 2,72 | -0,09 | 9,30E-01 |
| AT3G47490 | 0,22 | 0,58 | 4,35 | 0,13 | 8,94E-01 |
| AT3G47520 | 0,29 | -0,73 | 4,40 | -0,17 | 8,68E-01 |
| AT3G47550 | 0,50 | 0,97 | 3,11 | 0,31 | 7,54E-01 |
| AT3G47570 | 0,50 | -2,50 | 3,28 | -0,76 | 4,46E-01 |
| AT3G47610 | 0,61 | -0,82 | 2,41 | -0,34 | 7,33E-01 |
| AT3G47650 | 0,65 | -3,45 | 3,04 | -1,14 | 2,56E-01 |
| AT3G47660 | 0,72 | -4,52 | 2,82 | -1,60 | 1,10E-01 |
| AT3G07795 | 0,21 | -0,70 | 4,40 | -0,16 | 8,73E-01 |
| AT3G07805 | 0,89 | 0,00 | 2,85 | 0,00 | 1,00E+00 |
| AT3G47770 | 0,17 | 0,00 | 4,41 | 0,00 | 1,00E+00 |
| AT3G47800 | 1,07 | -0,63 | 1,86 | -0,34 | 7,33E-01 |
| AT3G47820 | 1,04 | 0,10 | 1,76 | 0,06 | 9,54E-01 |
| AT3G47833 | 0,22 | -0,26 | 4,38 | -0,06 | 9,53E-01 |
| AT3G47930 | 0,52 | -2,60 | 3,96 | -0,66 | 5,11E-01 |
| AT3G47950 | 0,34 | -1,87 | 4,33 | -0,43 | 6,65E-01 |
| AT3G48040 | 0,17 | -1,25 | 4,37 | -0,29 | 7,74E-01 |
| AT3G48070 | 0,40 | -1,64 | 4,37 | -0,38 | 7,07E-01 |
| AT3G48110 | 0,52 | -2,87 | 3,12 | -0,92 | 3,58E-01 |
| AT3G48140 | 0,97 | -0,19 | 2,12 | -0,09 | 9,30E-01 |
| AT3G48190 | 1,19 | -2,90 | 2,35 | -1,23 | 2,17E-01 |
| AT3G48310 | 0,40 | -1,64 | 3,44 | -0,48 | 6,33E-01 |
| AT3G48380 | 0,47 | -0,39 | 3,22 | -0,12 | 9,04E-01 |
| AT3G48420 | 0,42 | -1,53 | 4,28 | -0,36 | 7,21E-01 |
| AT3G48460 | 0,51 | -2,18 | 4,31 | -0,51 | 6,12E-01 |
| AT3G48500 | 0,57 | -0,51 | 3,98 | -0,13 | 8,97E-01 |
| AT3G48530 | 0,79 | -0,25 | 2,12 | -0,12 | 9,04E-01 |
| AT3G48620 | 0,08 | 0,00 | 4,41 | 0,00 | 1,00E+00 |
| AT3G48680 | 0,15 | -0,77 | 4,39 | -0,18 | 8,61E-01 |
| AT3G48710 | 0,27 | 0,24 | 4,37 | 0,06 | 9,55E-01 |
| AT3G48730 | 0,57 | -1,47 | 2,68 | -0,55 | 5,84E-01 |
| AT3G48750 | 0,54 | -2,40 | 2,71 | -0,89 | 3,76E-01 |
| AT3G48760 | 0,76 | -2,35 | 2,56 | -0,92 | 3,60E-01 |
| AT3G07995 | 0,83 | 1,89 | 2,22 | 0,85 | 3,93E-01 |
| AT3G48780 | 0,82 | 1,29 | 2,72 | 0,47 | 6,36E-01 |
| AT3G48810 | 0,58 | 0,00 | 3,77 | 0,00 | 1,00E+00 |
| AT3G48890 | 0,66 | -2,45 | 2,77 | -0,88 | 3,77E-01 |
| AT3G49000 | 0,27 | 0,36 | 4,36 | 0,08 | 9,33E-01 |
| AT3G49110 | 0,08 | 0,00 | 4,41 | 0,00 | 1,00E+00 |
| AT3G49220 | 0,92 | -2,40 | 1,97 | -1,22 | 2,22E-01 |
| AT3G49240 | 0,54 | 0,10 | 4,16 | 0,02 | 9,80E-01 |
| AT3G49260 | 0,64 | -4,30 | 4,32 | -1,00 | 3,19E-01 |
| AT3G49430 | 0,93 | -2,14 | 2,17 | -0,99 | 3,24E-01 |
| AT3G49670 | 1,02 | -3,88 | 2,33 | -1,66 | 9,61E-02 |
| AT3G49680 | 0,45 | 0,69 | 3,27 | 0,21 | 8,33E-01 |
| AT3G49710 | 0,32 | -3,12 | 4,39 | -0,71 | 4,77E-01 |

| | | | | | |
|---|---|---|---|---|---|
| AT3G49780 | 0,25 | 1,02 | 4,32 | 0,24 | 8,13E-01 |
| AT3G50000 | 0,43 | 0,07 | 4,23 | 0,02 | 9,87E-01 |
| AT3G50080 | 0,21 | -0,80 | 4,39 | -0,18 | 8,55E-01 |
| AT3G50100 | 0,05 | -0,24 | 4,38 | -0,05 | 9,57E-01 |
| AT3G50270 | 0,60 | -3,40 | 3,12 | -1,09 | 2,76E-01 |
| AT3G50350 | 0,42 | -0,11 | 3,38 | -0,03 | 9,75E-01 |
| AT3G50360 | 0,31 | -0,47 | 4,37 | -0,11 | 9,13E-01 |
| AT3G50430 | 0,16 | 1,16 | 4,31 | 0,27 | 7,88E-01 |
| AT3G50530 | 0,25 | -0,53 | 4,39 | -0,12 | 9,03E-01 |
| AT3G50550 | 0,51 | -2,19 | 3,15 | -0,70 | 4,87E-01 |
| AT3G50630 | 0,26 | -0,72 | 4,40 | -0,16 | 8,70E-01 |
| AT3G50700 | 0,53 | 0,24 | 2,65 | 0,09 | 9,27E-01 |
| AT3G50770 | 1,04 | -1,46 | 2,42 | -0,60 | 5,48E-01 |
| AT3G50890 | 0,25 | 1,33 | 4,31 | 0,31 | 7,57E-01 |
| AT3G50910 | 0,94 | -3,12 | 2,36 | -1,32 | 1,87E-01 |
| AT3G50970 | 0,39 | -2,16 | 4,38 | -0,49 | 6,21E-01 |
| AT3G51110 | 0,46 | -2,13 | 3,99 | -0,54 | 5,93E-01 |
| AT3G51140 | 0,97 | -1,87 | 1,99 | -0,94 | 3,48E-01 |
| AT3G51180 | 0,25 | -1,15 | 4,38 | -0,26 | 7,93E-01 |
| AT3G51260 | 0,46 | -2,67 | 3,31 | -0,81 | 4,20E-01 |
| AT3G51270 | 0,33 | -1,21 | 4,35 | -0,28 | 7,81E-01 |
| AT3G51325 | 0,36 | -0,62 | 4,36 | -0,14 | 8,87E-01 |
| AT3G51420 | 0,66 | -0,05 | 2,25 | -0,02 | 9,81E-01 |
| AT3G51430 | 0,32 | -0,84 | 4,37 | -0,19 | 8,48E-01 |
| AT3G51440 | 0,80 | -1,69 | 2,21 | -0,76 | 4,45E-01 |
| AT3G51450 | 0,43 | -0,46 | 2,79 | -0,16 | 8,69E-01 |
| AT3G51460 | 0,27 | -1,08 | 4,39 | -0,25 | 8,06E-01 |
| AT3G51480 | 0,45 | -3,76 | 4,35 | -0,86 | 3,88E-01 |
| AT3G51500 | 0,44 | 0,00 | 4,41 | 0,00 | 1,00E+00 |
| AT3G51510 | 0,44 | -0,91 | 3,41 | -0,27 | 7,89E-01 |
| AT3G51540 | 0,25 | -2,57 | 4,40 | -0,58 | 5,60E-01 |
| AT3G51630 | 0,87 | -3,69 | 3,26 | -1,13 | 2,58E-01 |
| AT3G51640 | 0,80 | -1,64 | 2,86 | -0,57 | 5,66E-01 |
| AT3G51650 | 1,02 | -3,80 | 2,14 | -1,78 | 7,55E-02 |
| AT3G51830 | 0,25 | -0,78 | 4,39 | -0,18 | 8,60E-01 |
| AT3G51970 | 0,40 | -0,82 | 3,40 | -0,24 | 8,09E-01 |
| AT3G52080 | 0,33 | -1,61 | 4,39 | -0,37 | 7,14E-01 |
| AT3G52100 | 0,15 | 0,05 | 4,37 | 0,01 | 9,91E-01 |
| AT3G52105 | 0,19 | 0,75 | 4,34 | 0,17 | 8,62E-01 |
| AT3G52120 | 1,03 | -2,89 | 2,03 | -1,42 | 1,55E-01 |
| AT3G52150 | 0,83 | 2,35 | 2,78 | 0,85 | 3,97E-01 |
| AT3G52155 | 0,45 | -3,74 | 4,35 | -0,86 | 3,90E-01 |
| AT3G52200 | 0,34 | -0,07 | 4,32 | -0,02 | 9,87E-01 |
| AT3G52210 | 0,11 | -0,68 | 4,41 | -0,15 | 8,77E-01 |
| AT3G52230 | 0,83 | -2,83 | 2,92 | -0,97 | 3,34E-01 |
| AT3G52300 | 0,40 | -1,65 | 4,35 | -0,38 | 7,04E-01 |
| AT3G52340 | 0,38 | 0,31 | 4,36 | 0,07 | 9,43E-01 |
| AT3G52370 | 0,08 | 0,00 | 4,41 | 0,00 | 1,00E+00 |
| AT3G52470 | 0,30 | 1,51 | 4,30 | 0,35 | 7,25E-01 |
| AT3G52570 | 0,29 | 0,69 | 4,34 | 0,16 | 8,74E-01 |
| AT3G08275 | 0,60 | 0,40 | 3,02 | 0,13 | 8,95E-01 |
| AT3G52610 | 0,30 | -2,60 | 4,41 | -0,59 | 5,55E-01 |
| AT3G52680 | 0,59 | 0,01 | 2,41 | 0,00 | 9,97E-01 |
| AT3G52710 | 0,88 | 0,29 | 1,99 | 0,15 | 8,84E-01 |
| AT3G08295 | 0,73 | 2,99 | 3,34 | 0,89 | 3,71E-01 |
| AT3G52830 | 1,16 | 2,77 | 1,86 | 1,49 | 1,35E-01 |

| | | | | | |
|---|---|---|---|---|---|
| AT3G52840 | 0,82 | -0,76 | 2,71 | -0,28 | 7,78E-01 |
| AT3G52950 | 0,45 | -2,49 | 3,30 | -0,75 | 4,50E-01 |
| AT3G52960 | 0,80 | -0,63 | 1,85 | -0,34 | 7,33E-01 |
| AT3G52990 | 1,11 | 0,61 | 1,76 | 0,34 | 7,31E-01 |
| AT3G53000 | 0,31 | -1,19 | 4,38 | -0,27 | 7,86E-01 |
| AT3G53030 | 0,57 | -1,19 | 3,96 | -0,30 | 7,65E-01 |
| AT3G53050 | 0,05 | -0,24 | 4,38 | -0,05 | 9,57E-01 |
| AT3G53065 | 0,19 | 0,64 | 4,34 | 0,15 | 8,83E-01 |
| AT3G53090 | 0,36 | -1,24 | 4,38 | -0,28 | 7,77E-01 |
| AT3G53110 | 0,40 | -1,36 | 3,33 | -0,41 | 6,82E-01 |
| AT3G53180 | 0,20 | 0,68 | 4,35 | 0,16 | 8,75E-01 |
| AT3G53190 | 0,21 | -1,17 | 4,36 | -0,27 | 7,88E-01 |
| AT3G53230 | 1,14 | -3,84 | 2,71 | -1,42 | 1,56E-01 |
| AT3G53240 | 0,55 | -3,21 | 3,87 | -0,83 | 4,08E-01 |
| AT3G53305 | 0,72 | 1,99 | 2,16 | 0,92 | 3,57E-01 |
| AT3G53360 | 0,08 | 0,00 | 4,41 | 0,00 | 1,00E+00 |
| AT3G53390 | 0,27 | -0,35 | 4,34 | -0,08 | 9,37E-01 |
| AT3G53470 | 0,77 | -2,83 | 2,85 | -0,99 | 3,21E-01 |
| AT3G53520 | 0,83 | -2,93 | 2,53 | -1,16 | 2,45E-01 |
| AT3G53670 | 0,90 | -1,12 | 1,99 | -0,56 | 5,74E-01 |
| AT3G53700 | 0,94 | 0,05 | 1,96 | 0,02 | 9,82E-01 |
| AT3G53870 | 0,80 | -0,06 | 2,64 | -0,02 | 9,82E-01 |
| AT3G54220 | 0,50 | -1,49 | 3,26 | -0,46 | 6,47E-01 |
| AT3G54366 | 1,11 | 2,90 | 2,09 | 1,39 | 1,65E-01 |
| AT3G54420 | 0,17 | -0,71 | 4,40 | -0,16 | 8,72E-01 |
| AT3G54600 | 0,51 | 1,04 | 4,01 | 0,26 | 7,96E-01 |
| AT3G54610 | 0,47 | 0,00 | 4,06 | 0,00 | 1,00E+00 |
| AT3G54625 | 0,68 | -2,69 | 4,40 | -0,61 | 5,41E-01 |
| AT3G54680 | 0,75 | -1,56 | 2,81 | -0,55 | 5,79E-01 |
| AT3G54800 | 0,59 | 2,36 | 2,57 | 0,92 | 3,59E-01 |
| AT3G54820 | 0,15 | 0,05 | 4,37 | 0,01 | 9,91E-01 |
| AT3G54850 | 0,81 | -3,05 | 2,44 | -1,25 | 2,11E-01 |
| AT3G54980 | 0,41 | -2,34 | 4,21 | -0,56 | 5,78E-01 |
| AT3G55030 | 0,39 | -0,69 | 4,26 | -0,16 | 8,72E-01 |
| AT3G55250 | 0,63 | -0,53 | 2,34 | -0,22 | 8,23E-01 |
| AT3G55260 | 0,43 | 1,14 | 4,00 | 0,29 | 7,75E-01 |
| AT3G55280 | 0,61 | 1,26 | 3,70 | 0,34 | 7,34E-01 |
| AT3G55370 | 0,12 | -0,14 | 4,38 | -0,03 | 9,74E-01 |
| AT3G55420 | 0,14 | 0,40 | 4,35 | 0,09 | 9,27E-01 |
| AT3G55440 | 1,26 | 1,18 | 1,80 | 0,66 | 5,12E-01 |
| AT3G55450 | 0,15 | -0,77 | 4,39 | -0,18 | 8,61E-01 |
| AT3G55460 | 1,02 | -0,21 | 2,03 | -0,10 | 9,19E-01 |
| AT3G55485 | 0,83 | 1,26 | 2,30 | 0,55 | 5,85E-01 |
| AT3G08485 | 0,83 | 1,26 | 2,30 | 0,55 | 5,85E-01 |
| AT3G55520 | 0,40 | 0,00 | 4,41 | 0,00 | 1,00E+00 |
| AT3G55620 | 0,54 | -1,11 | 2,60 | -0,43 | 6,69E-01 |
| AT3G55630 | 0,42 | -2,60 | 4,41 | -0,59 | 5,55E-01 |
| AT3G55677 | 0,24 | -0,68 | 4,34 | -0,16 | 8,75E-01 |
| AT3G55760 | 0,99 | -4,21 | 2,07 | -2,03 | 4,21E-02 |
| AT3G55770 | 0,98 | -1,90 | 1,86 | -1,02 | 3,07E-01 |
| AT3G55780 | 0,67 | -4,49 | 3,59 | -1,25 | 2,11E-01 |
| AT3G55880 | 0,30 | 1,20 | 3,60 | 0,33 | 7,39E-01 |
| AT3G55940 | 0,10 | -0,18 | 4,38 | -0,04 | 9,67E-01 |
| AT3G56130 | 0,33 | 0,15 | 4,35 | 0,04 | 9,72E-01 |
| AT3G56140 | 0,74 | -0,99 | 2,47 | -0,40 | 6,88E-01 |
| AT3G56180 | 0,21 | -1,17 | 4,36 | -0,27 | 7,88E-01 |

| | | | | | |
|---|---|---|---|---|---|
| AT3G56310 | 0,76 | -0,46 | 2,41 | -0,19 | 8,50E-01 |
| AT3G56330 | 0,52 | -2,56 | 3,93 | -0,65 | 5,15E-01 |
| AT3G56390 | 0,30 | -2,60 | 4,41 | -0,59 | 5,55E-01 |
| AT3G56450 | 0,39 | -2,82 | 4,39 | -0,64 | 5,20E-01 |
| AT3G08665 | 0,76 | -1,50 | 2,97 | -0,51 | 6,13E-01 |
| AT3G56470 | 0,12 | 0,00 | 4,41 | 0,00 | 1,00E+00 |
| AT3G56490 | 0,11 | -0,68 | 4,41 | -0,15 | 8,77E-01 |
| AT3G56590 | 0,57 | 0,34 | 2,97 | 0,11 | 9,09E-01 |
| AT3G56640 | 0,59 | -1,10 | 3,16 | -0,35 | 7,28E-01 |
| AT3G56650 | 0,53 | -2,10 | 4,00 | -0,53 | 5,99E-01 |
| AT3G56720 | 0,17 | -0,13 | 4,39 | -0,03 | 9,77E-01 |
| AT3G56740 | 0,49 | -1,01 | 4,01 | -0,25 | 8,00E-01 |
| AT3G56780 | 0,32 | -1,82 | 4,36 | -0,42 | 6,75E-01 |
| AT3G56840 | 0,64 | -1,48 | 2,49 | -0,59 | 5,54E-01 |
| AT3G56960 | 0,60 | -3,60 | 4,35 | -0,83 | 4,07E-01 |
| AT3G57070 | 0,24 | 1,41 | 4,30 | 0,33 | 7,42E-01 |
| AT3G57080 | 0,21 | -2,60 | 4,41 | -0,59 | 5,55E-01 |
| AT3G57090 | 0,59 | 0,46 | 2,23 | 0,20 | 8,38E-01 |
| AT3G08845 | 0,12 | -0,78 | 4,39 | -0,18 | 8,59E-01 |
| AT3G57230 | 0,38 | -0,41 | 2,88 | -0,14 | 8,87E-01 |
| AT3G57340 | 0,42 | -0,55 | 3,43 | -0,16 | 8,72E-01 |
| AT3G57390 | 0,11 | -0,68 | 4,41 | -0,15 | 8,77E-01 |
| AT3G57550 | 0,50 | -0,34 | 3,19 | -0,11 | 9,15E-01 |
| AT3G57570 | 0,40 | -1,57 | 4,24 | -0,37 | 7,12E-01 |
| AT3G57587 | 0,73 | 0,50 | 2,19 | 0,23 | 8,18E-01 |
| AT3G57650 | 0,53 | 1,16 | 2,63 | 0,44 | 6,58E-01 |
| AT3G08975 | 0,42 | -3,35 | 4,14 | -0,81 | 4,18E-01 |
| AT3G57760 | 0,08 | 0,00 | 4,41 | 0,00 | 1,00E+00 |
| AT3G57770 | 1,17 | 0,82 | 1,60 | 0,51 | 6,11E-01 |
| AT3G57810 | 0,21 | -0,23 | 4,38 | -0,05 | 9,58E-01 |
| AT3G57830 | 0,31 | 0,01 | 4,37 | 0,00 | 9,99E-01 |
| AT3G57890 | 1,04 | -1,78 | 1,91 | -0,93 | 3,52E-01 |
| AT3G57940 | 0,20 | 0,68 | 4,35 | 0,16 | 8,75E-01 |
| AT3G57970 | 0,17 | 0,00 | 4,41 | 0,00 | 1,00E+00 |
| AT3G58020 | 0,08 | 0,00 | 4,41 | 0,00 | 1,00E+00 |
| AT3G58110 | 1,07 | -2,60 | 3,34 | -0,78 | 4,36E-01 |
| AT3G58160 | 0,08 | 0,00 | 4,41 | 0,00 | 1,00E+00 |
| AT3G58490 | 0,30 | 1,33 | 4,31 | 0,31 | 7,57E-01 |
| AT3G58510 | 0,99 | -0,62 | 1,92 | -0,32 | 7,49E-01 |
| AT3G58530 | 0,46 | -2,47 | 3,97 | -0,62 | 5,34E-01 |
| AT3G58570 | 0,71 | -0,26 | 2,53 | -0,10 | 9,17E-01 |
| AT3G58680 | 0,61 | -0,64 | 2,56 | -0,25 | 8,04E-01 |
| AT3G58690 | 0,58 | -2,81 | 3,07 | -0,91 | 3,60E-01 |
| AT3G58700 | 0,20 | -2,55 | 4,40 | -0,58 | 5,62E-01 |
| AT3G58795 | 0,23 | 0,09 | 4,37 | 0,02 | 9,83E-01 |
| AT3G59030 | 0,64 | -3,15 | 3,73 | -0,85 | 3,97E-01 |
| AT3G59080 | 0,19 | -0,03 | 4,38 | -0,01 | 9,94E-01 |
| AT3G59090 | 0,80 | -0,54 | 2,60 | -0,21 | 8,35E-01 |
| AT3G59100 | 0,79 | -3,90 | 2,74 | -1,42 | 1,55E-01 |
| AT3G59110 | 0,41 | 1,00 | 4,25 | 0,24 | 8,13E-01 |
| AT3G59210 | 0,24 | 0,38 | 4,36 | 0,09 | 9,31E-01 |
| AT3G59290 | 1,04 | -0,54 | 1,84 | -0,29 | 7,69E-01 |
| AT3G59300 | 0,19 | 0,49 | 4,34 | 0,11 | 9,10E-01 |
| AT3G59350 | 0,86 | -3,55 | 2,41 | -1,47 | 1,42E-01 |
| AT3G59380 | 0,24 | 0,38 | 4,36 | 0,09 | 9,31E-01 |
| AT3G59500 | 0,47 | 0,85 | 3,43 | 0,25 | 8,04E-01 |

| | | | | | |
|---|---|---|---|---|---|
| AT3G59520 | 0,08 | 0,00 | 4,41 | 0,00 | 1,00E+00 |
| AT3G59530 | 0,53 | -4,36 | 4,32 | -1,01 | 3,13E-01 |
| AT3G59670 | 0,30 | -0,57 | 4,35 | -0,13 | 8,96E-01 |
| AT3G59700 | 0,10 | 0,45 | 4,35 | 0,10 | 9,19E-01 |
| AT3G59810 | 0,46 | -2,97 | 4,38 | -0,68 | 4,98E-01 |
| AT3G59920 | 0,37 | 0,27 | 4,31 | 0,06 | 9,49E-01 |
| AT3G60080 | 1,12 | 0,03 | 1,78 | 0,02 | 9,86E-01 |
| AT3G60110 | 0,48 | -1,08 | 4,39 | -0,25 | 8,05E-01 |
| AT3G60130 | 0,63 | -1,64 | 3,28 | -0,50 | 6,17E-01 |
| AT3G60140 | 0,58 | -0,68 | 3,91 | -0,17 | 8,62E-01 |
| AT3G60245 | 0,58 | -0,83 | 2,35 | -0,35 | 7,23E-01 |
| AT3G60290 | 0,27 | 0,27 | 4,36 | 0,06 | 9,51E-01 |
| AT3G60300 | 0,58 | -0,65 | 2,71 | -0,24 | 8,10E-01 |
| AT3G60310 | 0,49 | -0,21 | 3,99 | -0,05 | 9,57E-01 |
| AT3G60410 | 0,24 | -2,59 | 4,41 | -0,59 | 5,56E-01 |
| AT3G60420 | 0,55 | -3,44 | 3,88 | -0,89 | 3,74E-01 |
| AT3G60440 | 0,83 | -4,49 | 3,41 | -1,31 | 1,89E-01 |
| AT3G60450 | 1,00 | -3,50 | 2,73 | -1,28 | 2,00E-01 |
| AT3G60500 | 0,28 | -1,56 | 4,39 | -0,35 | 7,23E-01 |
| AT3G60530 | 0,83 | 2,16 | 2,06 | 1,05 | 2,95E-01 |
| AT3G60910 | 0,08 | 0,05 | 4,38 | 0,01 | 9,91E-01 |
| AT3G60950 | 1,11 | 2,85 | 2,04 | 1,40 | 1,62E-01 |
| AT3G09515 | 1,27 | -0,61 | 1,76 | -0,35 | 7,29E-01 |
| AT3G09525 | 0,12 | 0,00 | 4,41 | 0,00 | 1,00E+00 |
| AT3G60970 | 0,12 | 0,00 | 4,41 | 0,00 | 1,00E+00 |
| AT3G61030 | 0,61 | 1,17 | 2,10 | 0,56 | 5,78E-01 |
| AT3G61080 | 0,12 | 0,00 | 4,41 | 0,00 | 1,00E+00 |
| AT3G61130 | 1,07 | -2,74 | 2,21 | -1,24 | 2,16E-01 |
| AT3G61140 | 0,73 | -1,93 | 1,94 | -1,00 | 3,19E-01 |
| AT3G61150 | 0,68 | -2,86 | 2,82 | -1,01 | 3,11E-01 |
| AT3G61180 | 0,94 | -0,93 | 2,51 | -0,37 | 7,12E-01 |
| AT3G61240 | 1,00 | -0,87 | 1,79 | -0,49 | 6,28E-01 |
| AT3G61280 | 0,41 | -0,68 | 3,38 | -0,20 | 8,42E-01 |
| AT3G61320 | 0,08 | -0,21 | 4,39 | -0,05 | 9,62E-01 |
| AT3G61440 | 1,04 | -1,47 | 2,31 | -0,64 | 5,25E-01 |
| AT3G61480 | 0,44 | -3,42 | 3,33 | -1,03 | 3,04E-01 |
| AT3G61490 | 0,55 | -4,20 | 4,33 | -0,97 | 3,32E-01 |
| AT3G61540 | 0,29 | 0,09 | 4,35 | 0,02 | 9,84E-01 |
| AT3G61550 | 0,37 | 0,54 | 3,50 | 0,15 | 8,77E-01 |
| AT3G61600 | 1,22 | -3,13 | 1,92 | -1,63 | 1,03E-01 |
| AT3G61670 | 0,29 | -1,50 | 4,39 | -0,34 | 7,33E-01 |
| AT3G61755 | 0,16 | 0,21 | 4,36 | 0,05 | 9,61E-01 |
| AT3G61810 | 0,05 | -0,24 | 4,38 | -0,05 | 9,57E-01 |
| AT3G61850 | 0,15 | 0,05 | 4,37 | 0,01 | 9,91E-01 |
| AT3G61870 | 1,16 | -0,20 | 1,62 | -0,12 | 9,04E-01 |
| AT3G61950 | 0,11 | -0,68 | 4,41 | -0,15 | 8,77E-01 |
| AT3G62030 | 0,98 | -0,75 | 2,12 | -0,35 | 7,24E-01 |
| AT3G62130 | 0,39 | -0,73 | 4,30 | -0,17 | 8,65E-01 |
| AT3G62190 | 0,24 | -0,20 | 4,39 | -0,05 | 9,64E-01 |
| AT3G62245 | 0,42 | 1,84 | 4,04 | 0,46 | 6,49E-01 |
| AT3G62300 | 0,93 | -2,84 | 3,44 | -0,83 | 4,09E-01 |
| AT3G62370 | 0,90 | 0,49 | 2,27 | 0,22 | 8,29E-01 |
| AT3G62530 | 0,72 | 0,83 | 2,47 | 0,34 | 7,37E-01 |
| AT3G62550 | 0,70 | -0,78 | 3,79 | -0,21 | 8,36E-01 |
| AT3G62560 | 0,61 | -1,97 | 3,03 | -0,65 | 5,15E-01 |
| AT3G62580 | 0,64 | -0,61 | 2,16 | -0,28 | 7,76E-01 |

| | | | | | |
|---|---|---|---|---|---|
| AT3G09815 | 0,08 | 0,00 | 4,41 | 0,00 | 1,00E+00 |
| AT3G09835 | 1,22 | 2,45 | 1,84 | 1,33 | 1,82E-01 |
| AT3G62620 | 0,29 | 0,39 | 4,35 | 0,09 | 9,28E-01 |
| AT3G62660 | 1,25 | 1,31 | 1,76 | 0,75 | 4,56E-01 |
| AT3G62720 | 0,40 | 1,10 | 4,11 | 0,27 | 7,89E-01 |
| AT3G62750 | 0,68 | -0,97 | 2,33 | -0,42 | 6,77E-01 |
| AT3G62820 | 0,26 | -0,32 | 4,33 | -0,07 | 9,41E-01 |
| AT3G62830 | 0,86 | -0,14 | 1,95 | -0,07 | 9,43E-01 |
| AT3G62890 | 0,20 | 0,46 | 4,35 | 0,10 | 9,16E-01 |
| AT3G63000 | 0,33 | -1,64 | 4,37 | -0,37 | 7,08E-01 |
| AT3G63010 | 0,38 | 0,04 | 3,47 | 0,01 | 9,90E-01 |
| AT3G63080 | 0,53 | -1,44 | 2,65 | -0,54 | 5,87E-01 |
| AT3G63130 | 0,43 | -1,28 | 4,10 | -0,31 | 7,55E-01 |
| AT3G63170 | 0,32 | -2,39 | 4,36 | -0,55 | 5,84E-01 |
| AT3G63215 | 0,08 | -0,21 | 4,39 | -0,05 | 9,62E-01 |
| AT3G63300 | 0,72 | -4,52 | 2,82 | -1,60 | 1,10E-01 |
| AT3G63370 | 0,57 | -3,06 | 3,11 | -0,98 | 3,25E-01 |
| AT3G63400 | 0,95 | -1,33 | 2,20 | -0,60 | 5,46E-01 |
| AT3G63420 | 0,46 | -2,94 | 4,38 | -0,67 | 5,02E-01 |
| AT3G63470 | 0,42 | -3,55 | 4,20 | -0,85 | 3,98E-01 |
| AT3G09995 | 0,45 | -2,59 | 4,12 | -0,63 | 5,30E-01 |
| AT3G63520 | 0,87 | -1,82 | 2,23 | -0,81 | 4,16E-01 |
| AT4G00030 | 0,38 | 0,12 | 4,36 | 0,03 | 9,78E-01 |
| AT4G00163 | 0,65 | -0,64 | 2,60 | -0,25 | 8,05E-01 |
| AT4G00230 | 0,69 | -3,92 | 4,35 | -0,90 | 3,67E-01 |
| AT4G00300 | 1,08 | 0,79 | 1,75 | 0,45 | 6,54E-01 |
| AT4G00340 | 0,31 | -0,32 | 4,34 | -0,07 | 9,41E-01 |
| AT4G00440 | 0,33 | -1,65 | 4,41 | -0,37 | 7,09E-01 |
| AT4G00490 | 0,44 | -0,68 | 4,41 | -0,15 | 8,77E-01 |
| AT4G00580 | 0,48 | 0,32 | 3,95 | 0,08 | 9,35E-01 |
| AT4G00660 | 1,18 | -2,41 | 2,03 | -1,19 | 2,35E-01 |
| AT4G00710 | 0,48 | -2,60 | 4,41 | -0,59 | 5,55E-01 |
| AT4G00750 | 0,96 | -0,27 | 1,76 | -0,16 | 8,77E-01 |
| AT4G00820 | 0,05 | -0,24 | 4,38 | -0,05 | 9,57E-01 |
| AT4G00830 | 0,92 | -0,50 | 2,08 | -0,24 | 8,11E-01 |
| AT4G00889 | 0,23 | 1,59 | 4,30 | 0,37 | 7,11E-01 |
| AT4G00930 | 0,17 | 0,00 | 4,41 | 0,00 | 1,00E+00 |
| AT4G00953 | 0,67 | -0,13 | 2,61 | -0,05 | 9,61E-01 |
| AT4G00955 | 0,42 | 1,40 | 2,83 | 0,49 | 6,21E-01 |
| AT4G00960 | 1,30 | 2,54 | 2,03 | 1,25 | 2,10E-01 |
| AT4G00970 | 0,33 | 1,68 | 4,30 | 0,39 | 6,96E-01 |
| AT4G00990 | 0,61 | -3,21 | 3,92 | -0,82 | 4,14E-01 |
| AT4G01020 | 0,37 | -1,08 | 4,34 | -0,25 | 8,03E-01 |
| AT4G01026 | 0,56 | -1,21 | 2,59 | -0,47 | 6,41E-01 |
| AT4G01040 | 0,12 | -0,68 | 4,41 | -0,15 | 8,77E-01 |
| AT4G01100 | 0,48 | 1,11 | 3,17 | 0,35 | 7,26E-01 |
| AT4G01130 | 0,11 | 0,46 | 4,35 | 0,11 | 9,15E-01 |
| AT4G01280 | 0,59 | -0,13 | 3,18 | -0,04 | 9,67E-01 |
| AT4G01320 | 0,90 | -3,13 | 2,18 | -1,44 | 1,51E-01 |
| AT4G01330 | 0,80 | 1,14 | 2,47 | 0,46 | 6,43E-01 |
| AT4G01360 | 0,89 | 0,61 | 2,87 | 0,21 | 8,32E-01 |
| AT4G01370 | 0,27 | 0,78 | 4,33 | 0,18 | 8,57E-01 |
| AT4G01460 | 0,10 | -0,34 | 4,38 | -0,08 | 9,38E-01 |
| AT4G01570 | 0,24 | -0,77 | 4,39 | -0,18 | 8,61E-01 |
| AT4G03905 | 0,55 | 1,39 | 3,89 | 0,36 | 7,21E-01 |
| AT4G03925 | 0,76 | 0,31 | 2,24 | 0,14 | 8,91E-01 |

| | | | | | |
|---|---|---|---|---|---|
| AT4G01610 | 0,62 | 0,83 | 2,45 | 0,34 | 7,34E-01 |
| AT4G01690 | 1,14 | -1,64 | 1,62 | -1,01 | 3,12E-01 |
| AT4G01720 | 0,44 | -2,74 | 4,11 | -0,67 | 5,06E-01 |
| AT4G01995 | 0,08 | 0,00 | 4,41 | 0,00 | 1,00E+00 |
| AT4G02000 | 0,69 | 1,69 | 2,17 | 0,78 | 4,35E-01 |
| AT4G02010 | 0,14 | -0,38 | 4,38 | -0,09 | 9,31E-01 |
| AT4G02020 | 0,55 | -0,59 | 3,22 | -0,18 | 8,55E-01 |
| AT4G02030 | 0,75 | -4,44 | 2,77 | -1,60 | 1,09E-01 |
| AT4G02070 | 0,12 | 0,00 | 4,41 | 0,00 | 1,00E+00 |
| AT4G02080 | 0,57 | -1,59 | 2,73 | -0,58 | 5,61E-01 |
| AT4G02100 | 0,08 | -0,21 | 4,39 | -0,05 | 9,62E-01 |
| AT4G02110 | 0,21 | 0,00 | 4,41 | 0,00 | 1,00E+00 |
| AT4G02200 | 0,55 | 0,74 | 3,85 | 0,19 | 8,48E-01 |
| AT4G02220 | 0,29 | -0,98 | 4,35 | -0,23 | 8,22E-01 |
| AT4G02260 | 0,56 | -0,49 | 2,97 | -0,16 | 8,70E-01 |
| AT4G02330 | 0,22 | 0,25 | 4,36 | 0,06 | 9,54E-01 |
| AT4G02340 | 1,02 | -0,93 | 2,15 | -0,43 | 6,67E-01 |
| AT4G02370 | 0,72 | 0,85 | 3,08 | 0,28 | 7,82E-01 |
| AT4G02390 | 0,10 | -0,34 | 4,38 | -0,08 | 9,38E-01 |
| AT4G02400 | 0,19 | 0,01 | 4,36 | 0,00 | 9,98E-01 |
| AT4G02440 | 0,90 | 0,38 | 1,83 | 0,20 | 8,38E-01 |
| AT4G04135 | 0,05 | -0,24 | 4,38 | -0,05 | 9,57E-01 |
| AT4G04195 | 0,38 | 1,49 | 3,35 | 0,44 | 6,56E-01 |
| AT4G02540 | 0,29 | 0,45 | 3,69 | 0,12 | 9,04E-01 |
| AT4G02580 | 1,19 | -1,99 | 2,12 | -0,94 | 3,48E-01 |
| AT4G02600 | 0,66 | -1,24 | 2,22 | -0,56 | 5,77E-01 |
| AT4G02610 | 0,31 | 0,10 | 4,37 | 0,02 | 9,82E-01 |
| AT4G02650 | 0,24 | -2,59 | 4,41 | -0,59 | 5,56E-01 |
| AT4G02725 | 0,69 | -2,06 | 3,04 | -0,68 | 4,97E-01 |
| AT4G02790 | 0,08 | 0,00 | 4,41 | 0,00 | 1,00E+00 |
| AT4G02880 | 0,95 | -1,96 | 2,02 | -0,97 | 3,32E-01 |
| AT4G02920 | 1,02 | -0,67 | 2,39 | -0,28 | 7,79E-01 |
| AT4G02940 | 0,12 | 0,07 | 4,38 | 0,02 | 9,87E-01 |
| AT4G02990 | 0,69 | -0,93 | 2,40 | -0,39 | 6,97E-01 |
| AT4G03000 | 0,64 | -1,64 | 2,99 | -0,55 | 5,85E-01 |
| AT4G04425 | 0,08 | 0,00 | 4,41 | 0,00 | 1,00E+00 |
| AT4G03100 | 0,38 | -1,96 | 4,34 | -0,45 | 6,52E-01 |
| AT4G03120 | 0,23 | -1,72 | 4,40 | -0,39 | 6,95E-01 |
| AT4G03180 | 0,27 | -0,68 | 4,41 | -0,15 | 8,77E-01 |
| AT4G03200 | 0,82 | -2,60 | 2,55 | -1,02 | 3,06E-01 |
| AT4G03205 | 0,48 | -2,88 | 3,19 | -0,90 | 3,68E-01 |
| AT4G03210 | 0,53 | -2,54 | 4,40 | -0,58 | 5,64E-01 |
| AT4G03285 | 0,16 | -0,26 | 4,38 | -0,06 | 9,52E-01 |
| AT4G03410 | 0,58 | 0,50 | 3,74 | 0,13 | 8,93E-01 |
| AT4G03500 | 0,47 | -1,01 | 4,34 | -0,23 | 8,16E-01 |
| AT4G03830 | 0,61 | -0,11 | 2,50 | -0,04 | 9,66E-01 |
| AT4G03930 | 0,65 | -0,11 | 3,14 | -0,04 | 9,71E-01 |
| AT4G03940 | 0,17 | -0,71 | 4,40 | -0,16 | 8,72E-01 |
| AT4G03945 | 0,92 | 2,36 | 2,30 | 1,03 | 3,05E-01 |
| AT4G04110 | 0,65 | -1,74 | 4,35 | -0,40 | 6,89E-01 |
| AT4G04180 | 0,08 | -0,21 | 4,39 | -0,05 | 9,62E-01 |
| AT4G04210 | 0,39 | -0,05 | 2,85 | -0,02 | 9,87E-01 |
| AT4G04220 | 0,89 | -2,53 | 1,99 | -1,28 | 2,02E-01 |
| AT4G04795 | 0,87 | 2,96 | 2,47 | 1,20 | 2,31E-01 |
| AT4G04315 | 0,10 | -0,34 | 4,38 | -0,08 | 9,38E-01 |
| AT4G04350 | 0,42 | -0,18 | 3,34 | -0,05 | 9,58E-01 |

| | | | | | |
|---|---|---|---|---|---|
| AT4G04450 | 0,17 | -0,68 | 4,41 | -0,15 | 8,77E-01 |
| AT4G04510 | 0,17 | -1,25 | 4,37 | -0,29 | 7,74E-01 |
| AT4G04540 | 0,26 | 0,30 | 4,36 | 0,07 | 9,46E-01 |
| AT4G04610 | 0,73 | -0,23 | 2,15 | -0,11 | 9,14E-01 |
| AT4G04670 | 0,48 | -2,60 | 4,41 | -0,59 | 5,55E-01 |
| AT4G04750 | 0,90 | 2,11 | 1,92 | 1,10 | 2,70E-01 |
| AT4G04760 | 1,18 | 1,30 | 2,03 | 0,64 | 5,23E-01 |
| AT4G04800 | 0,52 | -1,64 | 3,88 | -0,42 | 6,73E-01 |
| AT4G04840 | 1,00 | 0,84 | 2,01 | 0,42 | 6,76E-01 |
| AT4G04850 | 0,79 | -2,07 | 1,97 | -1,05 | 2,92E-01 |
| AT4G04860 | 0,28 | 0,05 | 3,74 | 0,01 | 9,90E-01 |
| AT4G04960 | 0,39 | -0,07 | 2,85 | -0,02 | 9,82E-01 |
| AT4G05020 | 0,40 | -0,43 | 4,28 | -0,10 | 9,21E-01 |
| AT4G05080 | 0,23 | 1,44 | 4,30 | 0,33 | 7,38E-01 |
| AT4G05105 | 0,17 | 0,00 | 4,41 | 0,00 | 1,00E+00 |
| AT4G05160 | 0,46 | -0,23 | 4,12 | -0,06 | 9,56E-01 |
| AT4G05612 | 0,12 | 0,00 | 4,41 | 0,00 | 1,00E+00 |
| AT4G05632 | 0,08 | -0,21 | 4,39 | -0,05 | 9,62E-01 |
| AT4G06479 | 0,10 | 0,45 | 4,35 | 0,10 | 9,19E-01 |
| AT4G06598 | 0,74 | -4,02 | 3,43 | -1,17 | 2,41E-01 |
| AT4G06688 | 0,68 | -0,73 | 2,37 | -0,31 | 7,57E-01 |
| AT4G07390 | 0,27 | 0,86 | 3,72 | 0,23 | 8,17E-01 |
| AT4G07455 | 0,24 | -3,18 | 4,39 | -0,73 | 4,68E-01 |
| AT4G07825 | 1,05 | -2,05 | 2,58 | -0,79 | 4,27E-01 |
| AT4G07932 | 0,51 | 1,75 | 4,03 | 0,43 | 6,64E-01 |
| AT4G07965 | 0,15 | -0,77 | 4,39 | -0,18 | 8,61E-01 |
| AT4G08025 | 0,16 | -0,01 | 4,37 | 0,00 | 9,99E-01 |
| AT4G05305 | 0,12 | 0,75 | 4,34 | 0,17 | 8,63E-01 |
| AT4G08097 | 0,66 | 2,07 | 2,52 | 0,82 | 4,12E-01 |
| AT4G08180 | 1,01 | -2,54 | 2,24 | -1,13 | 2,57E-01 |
| AT4G08267 | 0,63 | -0,94 | 4,37 | -0,22 | 8,29E-01 |
| AT4G08270 | 0,17 | -0,80 | 4,39 | -0,18 | 8,56E-01 |
| AT4G08320 | 0,31 | 1,39 | 4,30 | 0,32 | 7,47E-01 |
| AT4G08350 | 0,65 | 1,41 | 2,57 | 0,55 | 5,84E-01 |
| AT4G08395 | 0,29 | 1,53 | 4,30 | 0,36 | 7,22E-01 |
| AT4G05455 | 0,68 | 1,36 | 2,53 | 0,54 | 5,91E-01 |
| AT4G08450 | 0,11 | 0,27 | 4,37 | 0,06 | 9,52E-01 |
| AT4G08455 | 0,63 | -0,69 | 4,41 | -0,16 | 8,75E-01 |
| AT4G08510 | 0,33 | -0,74 | 4,40 | -0,17 | 8,66E-01 |
| AT4G08560 | 0,64 | -4,65 | 4,31 | -1,08 | 2,81E-01 |
| AT4G08780 | 0,43 | -2,88 | 4,14 | -0,69 | 4,87E-01 |
| AT4G05605 | 0,39 | 0,33 | 4,31 | 0,08 | 9,39E-01 |
| AT4G08930 | 0,71 | -1,70 | 2,53 | -0,67 | 5,02E-01 |
| AT4G08950 | 0,37 | 0,12 | 4,30 | 0,03 | 9,77E-01 |
| AT4G08980 | 0,65 | 0,93 | 2,93 | 0,32 | 7,52E-01 |
| AT4G08990 | 0,69 | 2,66 | 3,64 | 0,73 | 4,64E-01 |
| AT4G09020 | 1,04 | 1,64 | 2,02 | 0,81 | 4,17E-01 |
| AT4G09030 | 0,16 | -0,01 | 4,37 | 0,00 | 9,99E-01 |
| AT4G09150 | 0,62 | -1,93 | 2,98 | -0,65 | 5,17E-01 |
| AT4G09160 | 0,36 | -4,02 | 4,33 | -0,93 | 3,54E-01 |
| AT4G09250 | 0,12 | -0,68 | 4,41 | -0,15 | 8,77E-01 |
| AT4G09310 | 0,21 | 0,00 | 4,41 | 0,00 | 1,00E+00 |
| AT4G05705 | 0,10 | 0,45 | 4,35 | 0,10 | 9,19E-01 |
| AT4G09320 | 1,06 | 1,14 | 1,72 | 0,66 | 5,06E-01 |
| AT4G09360 | 0,71 | 2,30 | 2,39 | 0,97 | 3,34E-01 |
| AT4G09460 | 0,37 | -1,60 | 4,35 | -0,37 | 7,14E-01 |

| | | | | | |
|---|---|---|---|---|---|
| AT4G09545 | 0,31 | 1,48 | 4,30 | 0,34 | 7,31E-01 |
| AT4G09570 | 0,54 | -0,72 | 2,24 | -0,32 | 7,49E-01 |
| AT4G09630 | 0,87 | 0,64 | 1,98 | 0,32 | 7,47E-01 |
| AT4G09660 | 0,30 | 1,68 | 4,29 | 0,39 | 6,96E-01 |
| AT4G09680 | 0,45 | -3,33 | 4,03 | -0,83 | 4,08E-01 |
| AT4G09730 | 0,51 | -1,48 | 3,21 | -0,46 | 6,44E-01 |
| AT4G09800 | 0,41 | -1,68 | 4,22 | -0,40 | 6,91E-01 |
| AT4G09820 | 0,31 | -0,02 | 4,39 | -0,01 | 9,96E-01 |
| AT4G09830 | 0,41 | -1,64 | 3,44 | -0,48 | 6,33E-01 |
| AT4G09850 | 0,21 | -0,07 | 4,37 | -0,02 | 9,87E-01 |
| AT4G05745 | 0,45 | 0,03 | 3,15 | 0,01 | 9,92E-01 |
| AT4G09890 | 0,18 | -0,33 | 4,38 | -0,08 | 9,40E-01 |
| AT4G10060 | 0,79 | -0,65 | 2,16 | -0,30 | 7,64E-01 |
| AT4G10170 | 0,08 | 0,00 | 4,41 | 0,00 | 1,00E+00 |
| AT4G10180 | 0,21 | -0,70 | 4,40 | -0,16 | 8,73E-01 |
| AT4G10465 | 0,60 | 2,15 | 2,55 | 0,84 | 4,00E-01 |
| AT4G10610 | 1,30 | -1,86 | 2,05 | -0,90 | 3,66E-01 |
| AT4G10613 | 0,77 | 2,36 | 2,07 | 1,14 | 2,53E-01 |
| AT4G10620 | 0,24 | -3,18 | 4,39 | -0,73 | 4,68E-01 |
| AT4G10660 | 0,11 | 0,46 | 4,35 | 0,11 | 9,15E-01 |
| AT4G11000 | 0,26 | -1,26 | 4,36 | -0,29 | 7,72E-01 |
| AT4G11070 | 0,37 | -1,77 | 4,35 | -0,41 | 6,84E-01 |
| AT4G11100 | 0,32 | -0,50 | 4,33 | -0,12 | 9,08E-01 |
| AT4G11110 | 0,51 | 0,09 | 2,72 | 0,03 | 9,72E-01 |
| AT4G11130 | 0,54 | -4,10 | 3,89 | -1,06 | 2,91E-01 |
| AT4G11150 | 1,03 | 2,47 | 1,80 | 1,37 | 1,71E-01 |
| AT4G11170 | 0,50 | -0,30 | 3,13 | -0,09 | 9,25E-01 |
| AT4G11240 | 1,11 | -3,53 | 2,00 | -1,77 | 7,73E-02 |
| AT4G11280 | 0,67 | -0,42 | 3,02 | -0,14 | 8,89E-01 |
| AT4G11330 | 0,53 | -2,14 | 3,18 | -0,67 | 5,00E-01 |
| AT4G11350 | 0,19 | 0,05 | 4,36 | 0,01 | 9,90E-01 |
| AT4G11360 | 0,31 | 0,00 | 4,41 | 0,00 | 1,00E+00 |
| AT4G05885 | 0,80 | -0,79 | 2,82 | -0,28 | 7,80E-01 |
| AT4G11660 | 0,57 | -0,36 | 3,18 | -0,11 | 9,09E-01 |
| AT4G11720 | 0,60 | -1,73 | 3,00 | -0,58 | 5,64E-01 |
| AT4G11740 | 0,35 | 0,00 | 4,32 | 0,00 | 1,00E+00 |
| AT4G11880 | 0,28 | 0,49 | 4,35 | 0,11 | 9,11E-01 |
| AT4G11890 | 0,62 | -0,76 | 3,13 | -0,24 | 8,07E-01 |
| AT4G11910 | 0,12 | 0,00 | 4,41 | 0,00 | 1,00E+00 |
| AT4G12020 | 1,23 | -1,66 | 2,61 | -0,64 | 5,24E-01 |
| AT4G12030 | 0,68 | -0,74 | 3,59 | -0,21 | 8,36E-01 |
| AT4G12070 | 0,19 | 0,52 | 4,35 | 0,12 | 9,04E-01 |
| AT4G12240 | 0,26 | -0,29 | 4,38 | -0,07 | 9,47E-01 |
| AT4G05985 | 0,10 | -0,34 | 4,38 | -0,08 | 9,38E-01 |
| AT4G12390 | 0,50 | -0,77 | 4,35 | -0,18 | 8,59E-01 |
| AT4G12600 | 0,92 | -1,66 | 3,09 | -0,54 | 5,91E-01 |
| AT4G12640 | 0,26 | 0,06 | 3,79 | 0,02 | 9,87E-01 |
| AT4G12680 | 0,21 | 0,00 | 4,41 | 0,00 | 1,00E+00 |
| AT4G12690 | 0,12 | -0,78 | 4,39 | -0,18 | 8,59E-01 |
| AT4G12830 | 0,34 | 0,42 | 4,36 | 0,10 | 9,22E-01 |
| AT4G12900 | 0,05 | -0,24 | 4,38 | -0,05 | 9,57E-01 |
| AT4G05995 | 0,98 | 3,17 | 2,42 | 1,31 | 1,89E-01 |
| AT4G13010 | 0,63 | 0,46 | 3,64 | 0,13 | 8,99E-01 |
| AT4G13020 | 0,54 | 0,19 | 3,22 | 0,06 | 9,54E-01 |
| AT4G06010 | 0,28 | 0,18 | 4,37 | 0,04 | 9,67E-01 |
| AT4G06015 | 0,10 | -0,18 | 4,38 | -0,04 | 9,67E-01 |

| | | | | | |
|---|---|---|---|---|---|
| AT4G13245 | 0,48 | 0,21 | 3,19 | 0,07 | 9,48E-01 |
| AT4G13270 | 0,50 | -0,68 | 3,26 | -0,21 | 8,35E-01 |
| AT4G13440 | 0,58 | -0,88 | 4,00 | -0,22 | 8,26E-01 |
| AT4G13530 | 0,95 | -0,64 | 2,13 | -0,30 | 7,62E-01 |
| AT4G13572 | 0,31 | -3,42 | 4,37 | -0,78 | 4,34E-01 |
| AT4G13610 | 0,37 | 1,18 | 2,83 | 0,42 | 6,76E-01 |
| AT4G13615 | 0,92 | -1,56 | 2,01 | -0,78 | 4,36E-01 |
| AT4G13670 | 0,92 | -2,60 | 3,31 | -0,79 | 4,31E-01 |
| AT4G13730 | 0,24 | 0,11 | 4,37 | 0,03 | 9,80E-01 |
| AT4G13750 | 0,39 | 0,78 | 3,37 | 0,23 | 8,17E-01 |
| AT4G13760 | 0,54 | 1,21 | 3,14 | 0,38 | 7,01E-01 |
| AT4G13830 | 0,70 | 1,04 | 2,89 | 0,36 | 7,18E-01 |
| AT4G13850 | 0,81 | 0,20 | 2,20 | 0,09 | 9,28E-01 |
| AT4G13860 | 0,43 | -0,70 | 4,19 | -0,17 | 8,67E-01 |
| AT4G13992 | 1,30 | 2,30 | 1,98 | 1,17 | 2,44E-01 |
| AT4G14000 | 0,12 | 0,75 | 4,34 | 0,17 | 8,63E-01 |
| AT4G14140 | 0,81 | 1,37 | 2,05 | 0,67 | 5,05E-01 |
| AT4G14220 | 0,46 | -2,79 | 4,34 | -0,64 | 5,21E-01 |
| AT4G14270 | 0,92 | 0,04 | 1,71 | 0,02 | 9,82E-01 |
| AT4G14300 | 0,77 | -3,69 | 3,53 | -1,04 | 2,96E-01 |
| AT4G14330 | 0,14 | -0,38 | 4,38 | -0,09 | 9,31E-01 |
| AT4G14540 | 0,71 | 1,99 | 2,48 | 0,80 | 4,22E-01 |
| AT4G14560 | 0,12 | 0,18 | 4,36 | 0,04 | 9,68E-01 |
| AT4G14570 | 0,70 | -2,82 | 2,82 | -1,00 | 3,18E-01 |
| AT4G14630 | 0,43 | 2,23 | 3,98 | 0,56 | 5,75E-01 |
| AT4G14700 | 0,12 | 0,00 | 4,41 | 0,00 | 1,00E+00 |
| AT4G06255 | 0,39 | -0,95 | 4,32 | -0,22 | 8,25E-01 |
| AT4G06260 | 0,42 | -0,96 | 3,23 | -0,30 | 7,66E-01 |
| AT4G14760 | 0,91 | -3,52 | 2,35 | -1,50 | 1,35E-01 |
| AT4G14800 | 0,34 | -1,64 | 4,41 | -0,37 | 7,10E-01 |
| AT4G14810 | 0,45 | -0,62 | 3,33 | -0,19 | 8,52E-01 |
| AT4G14850 | 0,15 | 0,41 | 4,35 | 0,09 | 9,26E-01 |
| AT4G14870 | 0,48 | 0,55 | 2,63 | 0,21 | 8,35E-01 |
| AT4G14890 | 0,73 | 0,10 | 2,20 | 0,04 | 9,65E-01 |
| AT4G14950 | 0,54 | -0,44 | 2,61 | -0,17 | 8,66E-01 |
| AT4G15000 | 1,11 | -1,76 | 1,80 | -0,98 | 3,27E-01 |
| AT4G15020 | 0,86 | -1,55 | 2,48 | -0,62 | 5,32E-01 |
| AT4G15030 | 0,61 | -0,12 | 2,40 | -0,05 | 9,59E-01 |
| AT4G15090 | 0,08 | 0,00 | 4,41 | 0,00 | 1,00E+00 |
| AT4G15096 | 1,15 | 3,64 | 2,13 | 1,71 | 8,67E-02 |
| AT4G15165 | 0,40 | -2,68 | 4,33 | -0,62 | 5,37E-01 |
| AT4G15233 | 0,90 | -3,00 | 2,41 | -1,24 | 2,14E-01 |
| AT4G15245 | 0,49 | 1,79 | 2,61 | 0,69 | 4,93E-01 |
| AT4G15260 | 0,57 | -1,10 | 2,93 | -0,38 | 7,06E-01 |
| AT4G15280 | 1,02 | -0,27 | 2,11 | -0,13 | 8,99E-01 |
| AT4G15415 | 0,24 | -0,96 | 4,36 | -0,22 | 8,26E-01 |
| AT4G15420 | 1,24 | -3,99 | 2,10 | -1,90 | 5,77E-02 |
| AT4G15440 | 0,99 | -0,40 | 2,22 | -0,18 | 8,57E-01 |
| AT4G15450 | 0,24 | -0,68 | 4,34 | -0,16 | 8,75E-01 |
| AT4G15475 | 0,96 | -5,17 | 4,01 | -1,29 | 1,98E-01 |
| AT4G15540 | 0,39 | 0,10 | 3,45 | 0,03 | 9,77E-01 |
| AT4G15545 | 1,17 | -0,28 | 1,68 | -0,17 | 8,68E-01 |
| AT4G15660 | 0,10 | -0,34 | 4,38 | -0,08 | 9,38E-01 |
| AT4G15700 | 0,08 | -0,21 | 4,39 | -0,05 | 9,62E-01 |
| AT4G15720 | 0,08 | 0,00 | 4,41 | 0,00 | 1,00E+00 |
| AT4G15770 | 0,18 | -1,07 | 4,38 | -0,25 | 8,06E-01 |

| | | | | | |
|---|---|---|---|---|---|
| AT4G15775 | 0,24 | -0,68 | 4,41 | -0,15 | 8,77E-01 |
| AT4G15800 | 0,76 | -1,96 | 1,98 | -0,99 | 3,21E-01 |
| AT4G15820 | 0,10 | -0,18 | 4,38 | -0,04 | 9,67E-01 |
| AT4G15840 | 0,73 | -2,37 | 1,99 | -1,19 | 2,34E-01 |
| AT4G15850 | 0,38 | -3,33 | 4,37 | -0,76 | 4,46E-01 |
| AT4G15880 | 0,12 | -0,14 | 4,38 | -0,03 | 9,74E-01 |
| AT4G15940 | 0,10 | 0,45 | 4,35 | 0,10 | 9,19E-01 |
| AT4G16008 | 0,08 | 0,00 | 4,41 | 0,00 | 1,00E+00 |
| AT4G16015 | 0,19 | 0,01 | 4,36 | 0,00 | 9,98E-01 |
| AT4G16130 | 1,05 | -2,30 | 2,11 | -1,09 | 2,76E-01 |
| AT4G16140 | 0,34 | 1,23 | 4,31 | 0,29 | 7,76E-01 |
| AT4G16150 | 0,50 | -0,80 | 4,11 | -0,19 | 8,45E-01 |
| AT4G16310 | 1,01 | 0,57 | 2,02 | 0,28 | 7,79E-01 |
| AT4G16360 | 0,61 | -1,64 | 2,72 | -0,60 | 5,46E-01 |
| AT4G16380 | 0,81 | -0,56 | 2,37 | -0,23 | 8,14E-01 |
| AT4G16390 | 0,67 | -1,42 | 2,26 | -0,63 | 5,31E-01 |
| AT4G16400 | 1,31 | -2,87 | 1,88 | -1,53 | 1,27E-01 |
| AT4G16430 | 0,28 | -2,15 | 4,38 | -0,49 | 6,25E-01 |
| AT4G16480 | 0,65 | -0,71 | 3,65 | -0,20 | 8,45E-01 |
| AT4G16510 | 0,53 | -2,58 | 4,07 | -0,63 | 5,26E-01 |
| AT4G16590 | 0,08 | 0,00 | 4,41 | 0,00 | 1,00E+00 |
| AT4G16650 | 0,41 | -2,58 | 4,26 | -0,61 | 5,44E-01 |
| AT4G16670 | 0,65 | -3,24 | 2,25 | -1,44 | 1,49E-01 |
| AT4G16710 | 0,63 | -0,83 | 2,58 | -0,32 | 7,48E-01 |
| AT4G16780 | 0,60 | -0,97 | 3,95 | -0,25 | 8,06E-01 |
| AT4G16830 | 0,78 | -1,06 | 3,09 | -0,34 | 7,31E-01 |
| AT4G16845 | 0,11 | 0,46 | 4,35 | 0,11 | 9,15E-01 |
| AT4G16850 | 0,29 | -2,60 | 4,41 | -0,59 | 5,55E-01 |
| AT4G16855 | 0,17 | 0,00 | 4,41 | 0,00 | 1,00E+00 |
| AT4G16955 | 0,78 | -3,35 | 2,96 | -1,13 | 2,57E-01 |
| AT4G17010 | 0,55 | 0,00 | 4,41 | 0,00 | 1,00E+00 |
| AT4G17080 | 0,57 | -3,26 | 4,37 | -0,75 | 4,56E-01 |
| AT4G17085 | 0,42 | -3,24 | 4,37 | -0,74 | 4,58E-01 |
| AT4G06665 | 1,22 | -0,68 | 1,89 | -0,36 | 7,21E-01 |
| AT4G17098 | 0,61 | 0,74 | 2,60 | 0,29 | 7,74E-01 |
| AT4G17245 | 0,66 | -1,29 | 2,54 | -0,51 | 6,12E-01 |
| AT4G17250 | 0,41 | -2,10 | 4,36 | -0,48 | 6,30E-01 |
| AT4G17270 | 0,64 | -1,06 | 2,86 | -0,37 | 7,12E-01 |
| AT4G17300 | 0,71 | 0,23 | 2,19 | 0,10 | 9,17E-01 |
| AT4G17380 | 0,44 | -1,67 | 2,73 | -0,61 | 5,40E-01 |
| AT4G17500 | 0,96 | 1,59 | 2,18 | 0,73 | 4,67E-01 |
| AT4G17520 | 1,02 | -1,11 | 2,61 | -0,43 | 6,70E-01 |
| AT4G17616 | 0,78 | -3,99 | 3,44 | -1,16 | 2,46E-01 |
| AT4G17620 | 1,00 | -1,97 | 2,46 | -0,80 | 4,23E-01 |
| AT4G17650 | 0,32 | -2,60 | 4,41 | -0,59 | 5,55E-01 |
| AT4G17770 | 1,15 | -0,88 | 1,88 | -0,47 | 6,39E-01 |
| AT4G17810 | 0,14 | 0,00 | 4,41 | 0,00 | 1,00E+00 |
| AT4G17830 | 0,83 | -0,27 | 2,53 | -0,11 | 9,14E-01 |
| AT4G17870 | 1,03 | 2,59 | 2,27 | 1,14 | 2,54E-01 |
| AT4G17900 | 0,47 | -0,82 | 2,81 | -0,29 | 7,70E-01 |
| AT4G18070 | 1,02 | 0,30 | 2,29 | 0,13 | 8,95E-01 |
| AT4G18160 | 0,40 | -2,42 | 4,33 | -0,56 | 5,76E-01 |
| AT4G18195 | 0,22 | -0,45 | 4,34 | -0,10 | 9,17E-01 |
| AT4G18205 | 1,32 | 1,98 | 1,72 | 1,15 | 2,49E-01 |
| AT4G18220 | 1,16 | -0,21 | 1,67 | -0,13 | 9,00E-01 |
| AT4G18240 | 1,25 | -0,57 | 1,90 | -0,30 | 7,65E-01 |

| | | | | | |
|---|---|---|---|---|---|
| AT4G18370 | 0,87 | -2,75 | 2,33 | -1,18 | 2,39E-01 |
| AT4G18380 | 0,33 | -2,54 | 4,40 | -0,58 | 5,64E-01 |
| AT4G06795 | 0,56 | -2,87 | 3,02 | -0,95 | 3,42E-01 |
| AT4G18530 | 0,35 | -0,39 | 4,38 | -0,09 | 9,30E-01 |
| AT4G18570 | 0,37 | -0,53 | 2,90 | -0,18 | 8,55E-01 |
| AT4G18710 | 1,19 | -1,58 | 1,98 | -0,80 | 4,25E-01 |
| AT4G18725 | 0,19 | 0,64 | 4,34 | 0,15 | 8,83E-01 |
| AT4G18750 | 0,50 | -0,95 | 4,10 | -0,23 | 8,17E-01 |
| AT4G18820 | 1,31 | -2,77 | 2,08 | -1,33 | 1,83E-01 |
| AT4G18830 | 0,19 | 0,42 | 4,35 | 0,10 | 9,23E-01 |
| AT4G18890 | 0,44 | 1,04 | 4,00 | 0,26 | 7,95E-01 |
| AT4G18905 | 0,35 | -0,99 | 4,38 | -0,23 | 8,21E-01 |
| AT4G18930 | 0,31 | 0,36 | 4,36 | 0,08 | 9,33E-01 |
| AT4G18950 | 1,00 | 0,77 | 2,07 | 0,37 | 7,09E-01 |
| AT4G19020 | 0,17 | -0,71 | 4,40 | -0,16 | 8,72E-01 |
| AT4G06865 | 0,10 | -0,34 | 4,38 | -0,08 | 9,38E-01 |
| AT4G19190 | 0,39 | -0,69 | 4,27 | -0,16 | 8,71E-01 |
| AT4G19200 | 0,86 | -0,49 | 2,13 | -0,23 | 8,17E-01 |
| AT4G19450 | 0,65 | -0,19 | 4,39 | -0,04 | 9,65E-01 |
| AT4G19490 | 0,82 | -2,61 | 2,69 | -0,97 | 3,32E-01 |
| AT4G19500 | 0,89 | -2,44 | 2,20 | -1,11 | 2,68E-01 |
| AT4G19510 | 1,30 | -2,37 | 1,94 | -1,22 | 2,22E-01 |
| AT4G19530 | 1,16 | -0,79 | 1,72 | -0,46 | 6,46E-01 |
| AT4G19540 | 0,68 | -1,54 | 3,04 | -0,51 | 6,11E-01 |
| AT4G19580 | 0,44 | -0,71 | 4,14 | -0,17 | 8,63E-01 |
| AT4G19600 | 0,21 | 0,00 | 4,41 | 0,00 | 1,00E+00 |
| AT4G19610 | 0,37 | -1,16 | 3,49 | -0,33 | 7,40E-01 |
| AT4G19650 | 0,19 | -0,79 | 4,39 | -0,18 | 8,58E-01 |
| AT4G19960 | 0,50 | -0,26 | 3,22 | -0,08 | 9,36E-01 |
| AT4G20110 | 0,83 | -1,19 | 2,18 | -0,54 | 5,87E-01 |
| AT4G20140 | 0,84 | 0,00 | 4,30 | 0,00 | 1,00E+00 |
| AT4G06950 | 0,12 | -0,31 | 4,38 | -0,07 | 9,44E-01 |
| AT4G20270 | 0,28 | 0,01 | 4,36 | 0,00 | 9,99E-01 |
| AT4G20300 | 0,68 | -0,35 | 2,42 | -0,14 | 8,87E-01 |
| AT4G20310 | 0,08 | 0,00 | 4,41 | 0,00 | 1,00E+00 |
| AT4G20400 | 1,23 | -3,55 | 2,18 | -1,63 | 1,03E-01 |
| AT4G20720 | 0,34 | -1,76 | 4,39 | -0,40 | 6,88E-01 |
| AT4G20840 | 1,22 | -5,11 | 2,42 | -2,11 | 3,46E-02 |
| AT4G20860 | 0,52 | -0,23 | 3,11 | -0,07 | 9,41E-01 |
| AT4G20890 | 0,68 | -0,31 | 2,60 | -0,12 | 9,05E-01 |
| AT4G20940 | 0,56 | -3,21 | 4,38 | -0,73 | 4,64E-01 |
| AT4G20960 | 0,48 | 0,25 | 2,69 | 0,09 | 9,25E-01 |
| AT4G21105 | 0,54 | -1,97 | 4,32 | -0,45 | 6,49E-01 |
| AT4G21160 | 0,15 | -0,14 | 4,39 | -0,03 | 9,74E-01 |
| AT4G21180 | 0,08 | 0,00 | 4,41 | 0,00 | 1,00E+00 |
| AT4G21210 | 0,33 | 0,00 | 4,36 | 0,00 | 1,00E+00 |
| AT4G21300 | 0,05 | -0,24 | 4,38 | -0,05 | 9,57E-01 |
| AT4G21323 | 0,38 | -3,33 | 4,37 | -0,76 | 4,46E-01 |
| AT4G21450 | 0,15 | -0,68 | 4,41 | -0,15 | 8,77E-01 |
| AT4G21470 | 0,32 | -0,75 | 4,40 | -0,17 | 8,65E-01 |
| AT4G21515 | 0,05 | -0,24 | 4,38 | -0,05 | 9,57E-01 |
| AT4G21540 | 0,68 | 0,18 | 2,95 | 0,06 | 9,50E-01 |
| AT4G21550 | 0,37 | -2,11 | 4,38 | -0,48 | 6,31E-01 |
| AT4G21620 | 0,72 | -1,06 | 2,52 | -0,42 | 6,74E-01 |
| AT4G21740 | 0,47 | -0,34 | 2,45 | -0,14 | 8,89E-01 |
| AT4G21750 | 0,81 | -3,67 | 2,82 | -1,30 | 1,93E-01 |

| | | | | | |
|---|---|---|---|---|---|
| AT4G21790 | 0,64 | -0,47 | 2,53 | -0,19 | 8,52E-01 |
| AT4G21800 | 0,53 | -1,61 | 3,81 | -0,42 | 6,73E-01 |
| AT4G07030 | 0,38 | -2,60 | 4,41 | -0,59 | 5,55E-01 |
| AT4G21940 | 0,15 | -0,73 | 4,40 | -0,17 | 8,68E-01 |
| AT4G22010 | 0,22 | -0,68 | 4,41 | -0,15 | 8,77E-01 |
| AT4G22050 | 0,20 | 1,02 | 4,33 | 0,24 | 8,14E-01 |
| AT4G07040 | 0,12 | 0,07 | 4,38 | 0,02 | 9,87E-01 |
| AT4G22130 | 0,44 | 0,22 | 2,73 | 0,08 | 9,35E-01 |
| AT4G22270 | 0,20 | -2,60 | 4,41 | -0,59 | 5,55E-01 |
| AT4G07075 | 1,23 | 1,31 | 1,78 | 0,74 | 4,61E-01 |
| AT4G07080 | 1,23 | 1,31 | 1,78 | 0,74 | 4,61E-01 |
| AT4G22330 | 0,17 | 0,00 | 4,41 | 0,00 | 1,00E+00 |
| AT4G22467 | 0,12 | -0,31 | 4,38 | -0,07 | 9,44E-01 |
| AT4G22475 | 0,49 | 2,14 | 3,85 | 0,56 | 5,78E-01 |
| AT4G22670 | 0,32 | 0,91 | 4,33 | 0,21 | 8,33E-01 |
| AT4G22720 | 0,28 | -1,56 | 4,39 | -0,35 | 7,23E-01 |
| AT4G22860 | 0,08 | 0,00 | 4,41 | 0,00 | 1,00E+00 |
| AT4G22910 | 0,08 | 0,00 | 4,41 | 0,00 | 1,00E+00 |
| AT4G22960 | 0,12 | -0,31 | 4,38 | -0,07 | 9,44E-01 |
| AT4G23030 | 0,17 | -1,25 | 4,37 | -0,29 | 7,74E-01 |
| AT4G23050 | 0,25 | -2,53 | 4,40 | -0,57 | 5,65E-01 |
| AT4G23060 | 0,21 | -2,50 | 4,39 | -0,57 | 5,70E-01 |
| AT4G23150 | 0,15 | 0,16 | 4,37 | 0,04 | 9,72E-01 |
| AT4G23160 | 0,75 | -1,65 | 4,22 | -0,39 | 6,97E-01 |
| AT4G23270 | 0,94 | -0,95 | 2,16 | -0,44 | 6,62E-01 |
| AT4G23400 | 0,56 | 1,16 | 2,28 | 0,51 | 6,12E-01 |
| AT4G23440 | 0,12 | 0,00 | 4,41 | 0,00 | 1,00E+00 |
| AT4G23470 | 0,84 | 0,71 | 2,05 | 0,34 | 7,30E-01 |
| AT4G23530 | 1,11 | -2,86 | 1,93 | -1,48 | 1,38E-01 |
| AT4G23570 | 0,44 | -0,92 | 3,37 | -0,27 | 7,83E-01 |
| AT4G23600 | 1,02 | -0,85 | 2,79 | -0,31 | 7,60E-01 |
| AT4G23740 | 0,40 | -0,19 | 4,39 | -0,04 | 9,65E-01 |
| AT4G23860 | 0,25 | -1,15 | 4,38 | -0,26 | 7,93E-01 |
| AT4G07190 | 0,45 | -0,67 | 3,43 | -0,20 | 8,45E-01 |
| AT4G23895 | 0,24 | -0,04 | 4,38 | -0,01 | 9,93E-01 |
| AT4G23940 | 1,06 | -3,01 | 2,28 | -1,32 | 1,86E-01 |
| AT4G23950 | 0,16 | -0,68 | 4,41 | -0,15 | 8,77E-01 |
| AT4G24000 | 0,08 | 0,00 | 4,41 | 0,00 | 1,00E+00 |
| AT4G24010 | 0,33 | -2,54 | 4,40 | -0,58 | 5,64E-01 |
| AT4G24060 | 0,12 | 0,00 | 4,41 | 0,00 | 1,00E+00 |
| AT4G24090 | 0,53 | 0,00 | 4,41 | 0,00 | 1,00E+00 |
| AT4G24130 | 0,24 | -3,18 | 4,39 | -0,73 | 4,68E-01 |
| AT4G24160 | 0,15 | 0,05 | 4,37 | 0,01 | 9,91E-01 |
| AT4G24175 | 0,21 | -2,60 | 4,41 | -0,59 | 5,55E-01 |
| AT4G24290 | 0,72 | -1,40 | 2,46 | -0,57 | 5,69E-01 |
| AT4G24350 | 0,25 | 1,02 | 4,32 | 0,24 | 8,13E-01 |
| AT4G24440 | 0,43 | -2,14 | 4,25 | -0,50 | 6,15E-01 |
| AT4G24460 | 0,18 | 0,00 | 4,41 | 0,00 | 1,00E+00 |
| AT4G24470 | 0,57 | -1,64 | 3,94 | -0,42 | 6,77E-01 |
| AT4G24550 | 0,38 | -1,70 | 3,46 | -0,49 | 6,22E-01 |
| AT4G24790 | 0,10 | -0,18 | 4,38 | -0,04 | 9,67E-01 |
| AT4G24800 | 0,98 | -3,24 | 2,59 | -1,25 | 2,10E-01 |
| AT4G24805 | 0,88 | 0,19 | 2,42 | 0,08 | 9,38E-01 |
| AT4G24880 | 0,08 | -0,21 | 4,39 | -0,05 | 9,62E-01 |
| AT4G07295 | 0,21 | -0,07 | 4,37 | -0,02 | 9,87E-01 |
| AT4G25030 | 0,40 | 1,10 | 2,80 | 0,39 | 6,95E-01 |

| | | | | | |
|---|---|---|---|---|---|
| AT4G25070 | 0,53 | -2,52 | 4,40 | -0,57 | 5,66E-01 |
| AT4G25120 | 0,50 | -2,66 | 3,29 | -0,81 | 4,18E-01 |
| AT4G25170 | 1,05 | -1,20 | 1,92 | -0,62 | 5,33E-01 |
| AT4G25210 | 0,75 | -0,11 | 2,70 | -0,04 | 9,66E-01 |
| AT4G25230 | 0,77 | 1,15 | 2,23 | 0,51 | 6,07E-01 |
| AT4G25360 | 0,25 | -0,75 | 4,40 | -0,17 | 8,65E-01 |
| AT4G25370 | 0,59 | -3,32 | 3,89 | -0,85 | 3,93E-01 |
| AT4G25390 | 0,17 | 0,78 | 4,33 | 0,18 | 8,58E-01 |
| AT4G25400 | 0,31 | 1,80 | 4,29 | 0,42 | 6,74E-01 |
| AT4G25500 | 1,12 | -0,46 | 2,01 | -0,23 | 8,17E-01 |
| AT4G25515 | 0,32 | -1,83 | 4,38 | -0,42 | 6,77E-01 |
| AT4G25550 | 0,38 | -3,33 | 4,37 | -0,76 | 4,46E-01 |
| AT4G25610 | 0,54 | -2,56 | 3,19 | -0,80 | 4,23E-01 |
| AT4G25660 | 0,81 | -2,15 | 2,30 | -0,93 | 3,50E-01 |
| AT4G25810 | 0,12 | 0,00 | 4,41 | 0,00 | 1,00E+00 |
| AT4G25835 | 0,21 | -0,75 | 4,40 | -0,17 | 8,65E-01 |
| AT4G25900 | 0,36 | -0,68 | 4,36 | -0,16 | 8,76E-01 |
| AT4G25910 | 0,69 | -4,02 | 3,52 | -1,14 | 2,53E-01 |
| AT4G25970 | 1,00 | -0,73 | 1,79 | -0,40 | 6,85E-01 |
| AT4G26000 | 0,96 | -2,17 | 2,40 | -0,91 | 3,65E-01 |
| AT4G26120 | 0,24 | -3,18 | 4,39 | -0,73 | 4,68E-01 |
| AT4G26170 | 0,08 | 0,00 | 4,41 | 0,00 | 1,00E+00 |
| AT4G26210 | 0,36 | 0,00 | 4,41 | 0,00 | 1,00E+00 |
| AT4G26230 | 0,82 | -2,59 | 2,52 | -1,03 | 3,03E-01 |
| AT4G26450 | 0,45 | -0,56 | 3,29 | -0,17 | 8,64E-01 |
| AT4G26455 | 0,54 | -3,33 | 3,82 | -0,87 | 3,84E-01 |
| AT4G26510 | 0,72 | -1,57 | 2,48 | -0,64 | 5,25E-01 |
| AT4G26650 | 0,56 | -1,62 | 2,32 | -0,70 | 4,85E-01 |
| AT4G26670 | 0,33 | -0,35 | 4,38 | -0,08 | 9,36E-01 |
| AT4G26680 | 0,40 | 0,00 | 4,18 | 0,00 | 9,99E-01 |
| AT4G26700 | 0,28 | 0,10 | 4,38 | 0,02 | 9,81E-01 |
| AT4G26790 | 0,42 | 1,93 | 4,03 | 0,48 | 6,31E-01 |
| AT4G26800 | 0,12 | -0,06 | 4,37 | -0,01 | 9,88E-01 |
| AT4G26840 | 0,82 | -1,16 | 2,11 | -0,55 | 5,83E-01 |
| AT4G26870 | 0,92 | -2,09 | 2,12 | -0,99 | 3,23E-01 |
| AT4G27010 | 0,79 | 0,25 | 2,12 | 0,12 | 9,06E-01 |
| AT4G27030 | 0,63 | -0,68 | 2,69 | -0,25 | 8,01E-01 |
| AT4G07775 | 0,58 | 1,98 | 2,90 | 0,68 | 4,95E-01 |
| AT4G27080 | 1,09 | -1,87 | 2,10 | -0,89 | 3,72E-01 |
| AT4G27260 | 0,26 | -2,60 | 4,41 | -0,59 | 5,55E-01 |
| AT4G27290 | 0,45 | -3,77 | 4,36 | -0,87 | 3,87E-01 |
| AT4G27310 | 0,93 | 1,16 | 2,06 | 0,56 | 5,74E-01 |
| AT4G27380 | 0,21 | 0,00 | 4,41 | 0,00 | 1,00E+00 |
| AT4G27395 | 0,73 | 0,00 | 2,95 | 0,00 | 1,00E+00 |
| AT4G27652 | 0,57 | 2,41 | 3,64 | 0,66 | 5,08E-01 |
| AT4G27654 | 0,56 | 1,41 | 2,59 | 0,54 | 5,86E-01 |
| AT4G27657 | 0,12 | -0,31 | 4,38 | -0,07 | 9,44E-01 |
| AT4G27680 | 0,93 | -1,40 | 2,09 | -0,67 | 5,04E-01 |
| AT4G27710 | 0,52 | 0,05 | 3,30 | 0,02 | 9,88E-01 |
| AT4G27740 | 0,52 | 2,55 | 3,74 | 0,68 | 4,95E-01 |
| AT4G27750 | 0,69 | -2,83 | 4,37 | -0,65 | 5,17E-01 |
| AT4G27760 | 0,28 | -3,07 | 4,38 | -0,70 | 4,84E-01 |
| AT4G27765 | 0,37 | -3,35 | 4,32 | -0,78 | 4,38E-01 |
| AT4G27780 | 0,50 | -3,74 | 3,95 | -0,95 | 3,44E-01 |
| AT4G27860 | 0,21 | 0,28 | 4,35 | 0,07 | 9,48E-01 |
| AT4G27880 | 0,08 | 0,00 | 4,41 | 0,00 | 1,00E+00 |

| | | | | | |
|---|---|---|---|---|---|
| AT4G27910 | 0,46 | -3,43 | 4,10 | -0,84 | 4,03E-01 |
| AT4G27940 | 1,14 | -3,87 | 2,16 | -1,79 | 7,35E-02 |
| AT4G28010 | 0,17 | -0,80 | 4,39 | -0,18 | 8,56E-01 |
| AT4G28210 | 0,08 | -0,21 | 4,39 | -0,05 | 9,62E-01 |
| AT4G28220 | 0,27 | 0,92 | 4,33 | 0,21 | 8,32E-01 |
| AT4G28260 | 0,80 | -1,64 | 3,03 | -0,54 | 5,88E-01 |
| AT4G28300 | 0,21 | -0,70 | 4,40 | -0,16 | 8,73E-01 |
| AT4G28362 | 0,21 | -2,50 | 4,39 | -0,57 | 5,70E-01 |
| AT4G28400 | 0,26 | -0,50 | 4,34 | -0,11 | 9,09E-01 |
| AT4G28440 | 0,16 | -0,03 | 4,37 | -0,01 | 9,95E-01 |
| AT4G28490 | 0,74 | 0,96 | 3,63 | 0,26 | 7,92E-01 |
| AT4G28600 | 1,03 | -1,67 | 2,17 | -0,77 | 4,42E-01 |
| AT4G28660 | 0,48 | -0,93 | 4,08 | -0,23 | 8,20E-01 |
| AT4G28706 | 0,39 | 0,72 | 4,30 | 0,17 | 8,66E-01 |
| AT4G28710 | 0,41 | -1,03 | 4,21 | -0,25 | 8,06E-01 |
| AT4G28740 | 0,90 | 0,37 | 2,52 | 0,14 | 8,85E-01 |
| AT4G28780 | 0,15 | 0,48 | 4,35 | 0,11 | 9,12E-01 |
| AT4G28860 | 0,76 | -3,31 | 3,06 | -1,08 | 2,79E-01 |
| AT4G28870 | 0,12 | 0,75 | 4,34 | 0,17 | 8,63E-01 |
| AT4G28880 | 1,20 | -2,27 | 1,46 | -1,55 | 1,20E-01 |
| AT4G28980 | 0,73 | -1,72 | 2,60 | -0,66 | 5,09E-01 |
| AT4G29020 | 0,17 | -1,25 | 4,37 | -0,29 | 7,74E-01 |
| AT4G29035 | 0,22 | -0,45 | 4,34 | -0,10 | 9,17E-01 |
| AT4G29040 | 1,10 | 1,06 | 2,03 | 0,52 | 6,02E-01 |
| AT4G29070 | 0,33 | -0,43 | 4,35 | -0,10 | 9,22E-01 |
| AT4G29110 | 0,63 | 1,12 | 2,83 | 0,40 | 6,91E-01 |
| AT4G29120 | 0,68 | 0,46 | 2,89 | 0,16 | 8,74E-01 |
| AT4G29160 | 0,87 | -1,50 | 1,86 | -0,81 | 4,19E-01 |
| AT4G29170 | 0,92 | -4,28 | 4,32 | -0,99 | 3,22E-01 |
| AT4G29180 | 0,96 | -4,85 | 2,59 | -1,87 | 6,13E-02 |
| AT4G29210 | 0,68 | -2,51 | 2,62 | -0,96 | 3,38E-01 |
| AT4G29350 | 0,47 | 1,15 | 4,06 | 0,28 | 7,78E-01 |
| AT4G29400 | 0,21 | -1,17 | 4,36 | -0,27 | 7,88E-01 |
| AT4G29510 | 0,71 | -1,44 | 2,28 | -0,63 | 5,29E-01 |
| AT4G29740 | 1,04 | -0,59 | 1,99 | -0,30 | 7,66E-01 |
| AT4G29750 | 0,66 | -0,74 | 2,59 | -0,29 | 7,75E-01 |
| AT4G29790 | 1,30 | -4,09 | 2,01 | -2,03 | 4,22E-02 |
| AT4G08245 | 0,54 | 0,78 | 3,10 | 0,25 | 8,01E-01 |
| AT4G08255 | 0,51 | 1,88 | 4,30 | 0,44 | 6,62E-01 |
| AT4G29950 | 0,57 | -0,68 | 2,57 | -0,27 | 7,90E-01 |
| AT4G30010 | 0,55 | 1,10 | 3,24 | 0,34 | 7,34E-01 |
| AT4G30020 | 1,12 | -3,12 | 2,02 | -1,55 | 1,22E-01 |
| AT4G30130 | 0,24 | 0,06 | 4,38 | 0,01 | 9,90E-01 |
| AT4G30150 | 1,08 | -2,37 | 2,08 | -1,14 | 2,54E-01 |
| AT4G30310 | 0,60 | -2,81 | 3,71 | -0,76 | 4,49E-01 |
| AT4G30350 | 0,36 | 1,22 | 4,31 | 0,28 | 7,77E-01 |
| AT4G30400 | 0,15 | 0,41 | 4,35 | 0,09 | 9,26E-01 |
| AT4G08285 | 0,08 | 0,00 | 4,41 | 0,00 | 1,00E+00 |
| AT4G30480 | 0,31 | -3,42 | 4,37 | -0,78 | 4,34E-01 |
| AT4G30530 | 1,06 | 2,45 | 1,85 | 1,32 | 1,85E-01 |
| AT4G30650 | 0,58 | 1,12 | 2,72 | 0,41 | 6,81E-01 |
| AT4G30660 | 0,47 | -0,73 | 2,75 | -0,27 | 7,90E-01 |
| AT4G30780 | 0,23 | -1,14 | 4,39 | -0,26 | 7,96E-01 |
| AT4G30800 | 0,17 | 0,00 | 4,41 | 0,00 | 1,00E+00 |
| AT4G30940 | 0,31 | -0,32 | 4,34 | -0,07 | 9,41E-01 |
| AT4G31080 | 0,70 | -1,56 | 2,87 | -0,54 | 5,87E-01 |

| | | | | | |
|---|---|---|---|---|---|
| AT4G31115 | 0,40 | -3,56 | 4,24 | -0,84 | 4,01E-01 |
| AT4G31300 | 0,37 | 1,37 | 4,30 | 0,32 | 7,50E-01 |
| AT4G31340 | 0,63 | -3,12 | 3,82 | -0,82 | 4,13E-01 |
| AT4G31390 | 0,47 | -1,71 | 2,75 | -0,62 | 5,32E-01 |
| AT4G31430 | 0,50 | -2,28 | 4,02 | -0,57 | 5,72E-01 |
| AT4G31530 | 0,77 | -0,63 | 2,44 | -0,26 | 7,97E-01 |
| AT4G31580 | 0,97 | -0,40 | 2,01 | -0,20 | 8,41E-01 |
| AT4G31710 | 0,50 | -3,74 | 3,95 | -0,95 | 3,44E-01 |
| AT4G31715 | 0,08 | 0,00 | 4,41 | 0,00 | 1,00E+00 |
| AT4G31790 | 0,26 | 0,58 | 4,35 | 0,13 | 8,94E-01 |
| AT4G31805 | 0,98 | -4,36 | 3,31 | -1,32 | 1,88E-01 |
| AT4G31850 | 0,54 | -1,33 | 3,22 | -0,41 | 6,80E-01 |
| AT4G31910 | 0,49 | 1,88 | 2,68 | 0,70 | 4,83E-01 |
| AT4G08615 | 0,91 | 1,04 | 2,27 | 0,46 | 6,46E-01 |
| AT4G31985 | 0,37 | -3,19 | 4,38 | -0,73 | 4,66E-01 |
| AT4G32030 | 0,47 | -1,26 | 4,20 | -0,30 | 7,64E-01 |
| AT4G32040 | 0,84 | -1,58 | 3,58 | -0,44 | 6,58E-01 |
| AT4G32160 | 0,93 | 1,07 | 2,08 | 0,51 | 6,08E-01 |
| AT4G32190 | 0,86 | -1,72 | 2,84 | -0,61 | 5,45E-01 |
| AT4G32200 | 0,65 | -2,60 | 3,70 | -0,70 | 4,82E-01 |
| AT4G32265 | 1,11 | -0,55 | 1,90 | -0,29 | 7,71E-01 |
| AT4G32300 | 0,70 | -0,35 | 2,10 | -0,17 | 8,67E-01 |
| AT4G32330 | 0,80 | 0,12 | 2,04 | 0,06 | 9,55E-01 |
| AT4G32360 | 0,05 | -0,24 | 4,38 | -0,05 | 9,57E-01 |
| AT4G32430 | 0,12 | -0,31 | 4,38 | -0,07 | 9,44E-01 |
| AT4G32551 | 0,49 | -0,38 | 2,74 | -0,14 | 8,90E-01 |
| AT4G32600 | 0,53 | 0,29 | 3,29 | 0,09 | 9,30E-01 |
| AT4G32630 | 0,68 | -4,07 | 4,33 | -0,94 | 3,47E-01 |
| AT4G32700 | 0,67 | -3,64 | 3,67 | -0,99 | 3,21E-01 |
| AT4G32720 | 0,62 | -2,27 | 3,08 | -0,74 | 4,62E-01 |
| AT4G32730 | 0,45 | -3,32 | 4,03 | -0,82 | 4,10E-01 |
| AT4G32770 | 0,32 | -0,61 | 4,36 | -0,14 | 8,88E-01 |
| AT4G32800 | 0,31 | -0,71 | 4,36 | -0,16 | 8,70E-01 |
| AT4G32850 | 1,20 | -2,19 | 1,85 | -1,19 | 2,35E-01 |
| AT4G32910 | 0,98 | -2,60 | 2,27 | -1,15 | 2,52E-01 |
| AT4G32930 | 0,91 | -3,16 | 2,35 | -1,34 | 1,79E-01 |
| AT4G33000 | 0,14 | 0,87 | 4,33 | 0,20 | 8,40E-01 |
| AT4G33030 | 0,99 | -3,60 | 2,60 | -1,38 | 1,67E-01 |
| AT4G33060 | 0,15 | -0,77 | 4,39 | -0,18 | 8,61E-01 |
| AT4G33180 | 0,05 | -0,24 | 4,38 | -0,05 | 9,57E-01 |
| AT4G33220 | 0,80 | 1,83 | 3,22 | 0,57 | 5,70E-01 |
| AT4G08785 | 0,69 | -2,12 | 3,70 | -0,57 | 5,66E-01 |
| AT4G33310 | 0,49 | -2,10 | 4,29 | -0,49 | 6,24E-01 |
| AT4G33380 | 0,54 | -0,91 | 3,32 | -0,27 | 7,83E-01 |
| AT4G33410 | 0,32 | -1,63 | 4,41 | -0,37 | 7,11E-01 |
| AT4G33510 | 1,23 | -0,06 | 1,56 | -0,04 | 9,71E-01 |
| AT4G33540 | 0,41 | 0,55 | 4,26 | 0,13 | 8,98E-01 |
| AT4G33565 | 0,80 | 1,28 | 2,67 | 0,48 | 6,31E-01 |
| AT4G33580 | 1,02 | -0,86 | 3,05 | -0,28 | 7,79E-01 |
| AT4G33620 | 0,33 | -1,64 | 4,37 | -0,37 | 7,08E-01 |
| AT4G33630 | 1,15 | -2,84 | 1,92 | -1,48 | 1,38E-01 |
| AT4G33666 | 0,24 | -0,79 | 4,36 | -0,18 | 8,56E-01 |
| AT4G33690 | 0,48 | -0,27 | 3,94 | -0,07 | 9,45E-01 |
| AT4G33920 | 0,65 | 2,50 | 3,54 | 0,71 | 4,80E-01 |
| AT4G33960 | 0,45 | 0,00 | 4,41 | 0,00 | 1,00E+00 |
| AT4G34030 | 0,40 | 0,45 | 3,40 | 0,13 | 8,95E-01 |

| | | | | | |
|---|---|---|---|---|---|
| AT4G34035 | 0,32 | -3,12 | 4,39 | -0,71 | 4,77E-01 |
| AT4G34060 | 0,12 | 0,00 | 4,41 | 0,00 | 1,00E+00 |
| AT4G34120 | 1,07 | -1,61 | 1,78 | -0,91 | 3,63E-01 |
| AT4G34160 | 0,73 | -0,17 | 2,44 | -0,07 | 9,44E-01 |
| AT4G34240 | 0,30 | -0,30 | 4,34 | -0,07 | 9,45E-01 |
| AT4G34280 | 0,96 | -4,51 | 2,74 | -1,65 | 9,94E-02 |
| AT4G34290 | 0,10 | -0,34 | 4,38 | -0,08 | 9,38E-01 |
| AT4G34310 | 0,63 | -2,85 | 4,33 | -0,66 | 5,10E-01 |
| AT4G34330 | 0,08 | 0,00 | 4,41 | 0,00 | 1,00E+00 |
| AT4G34370 | 0,50 | -3,03 | 3,96 | -0,76 | 4,44E-01 |
| AT4G34400 | 0,59 | -3,94 | 3,74 | -1,05 | 2,92E-01 |
| AT4G34410 | 0,77 | 2,92 | 2,74 | 1,07 | 2,86E-01 |
| AT4G34430 | 1,22 | -3,85 | 2,26 | -1,71 | 8,81E-02 |
| AT4G34440 | 0,20 | -2,60 | 4,41 | -0,59 | 5,55E-01 |
| AT4G34640 | 0,93 | -3,99 | 2,81 | -1,42 | 1,56E-01 |
| AT4G34680 | 0,14 | 0,48 | 4,35 | 0,11 | 9,13E-01 |
| AT4G34700 | 0,83 | -0,41 | 2,45 | -0,17 | 8,67E-01 |
| AT4G34881 | 0,53 | -0,08 | 2,71 | -0,03 | 9,75E-01 |
| AT4G34910 | 0,61 | -1,64 | 3,12 | -0,52 | 6,00E-01 |
| AT4G34920 | 0,49 | -0,85 | 3,18 | -0,27 | 7,89E-01 |
| AT4G35040 | 0,31 | -1,72 | 4,39 | -0,39 | 6,95E-01 |
| AT4G35100 | 1,16 | -1,98 | 2,52 | -0,78 | 4,32E-01 |
| AT4G35370 | 0,13 | 0,00 | 4,41 | 0,00 | 1,00E+00 |
| AT4G35450 | 0,43 | -0,56 | 3,35 | -0,17 | 8,67E-01 |
| AT4G35460 | 0,66 | -4,52 | 2,90 | -1,56 | 1,19E-01 |
| AT4G35470 | 0,67 | 0,04 | 2,62 | 0,02 | 9,86E-01 |
| AT4G35500 | 0,79 | -4,52 | 2,74 | -1,65 | 9,92E-02 |
| AT4G35560 | 0,72 | -2,46 | 2,97 | -0,83 | 4,08E-01 |
| AT4G35600 | 1,15 | -3,30 | 1,79 | -1,84 | 6,62E-02 |
| AT4G35630 | 0,33 | -1,07 | 3,61 | -0,29 | 7,68E-01 |
| AT4G35850 | 0,77 | -2,50 | 2,27 | -1,10 | 2,71E-01 |
| AT4G35870 | 0,76 | -1,39 | 2,49 | -0,56 | 5,78E-01 |
| AT4G35880 | 0,17 | 0,00 | 4,41 | 0,00 | 1,00E+00 |
| AT4G35890 | 1,08 | -1,69 | 1,96 | -0,86 | 3,88E-01 |
| AT4G36032 | 1,21 | 0,62 | 2,03 | 0,31 | 7,59E-01 |
| AT4G36052 | 0,12 | 0,00 | 4,41 | 0,00 | 1,00E+00 |
| AT4G36130 | 1,27 | -5,02 | 1,94 | -2,59 | 9,61E-03 |
| AT4G36190 | 0,22 | 1,03 | 4,32 | 0,24 | 8,11E-01 |
| AT4G36195 | 0,86 | 1,01 | 2,22 | 0,45 | 6,49E-01 |
| AT4G36210 | 1,25 | -1,22 | 1,54 | -0,80 | 4,26E-01 |
| AT4G36250 | 0,69 | 0,48 | 2,46 | 0,20 | 8,45E-01 |
| AT4G36540 | 0,24 | -2,60 | 4,41 | -0,59 | 5,55E-01 |
| AT4G36710 | 1,27 | -0,72 | 2,60 | -0,28 | 7,82E-01 |
| AT4G36720 | 0,82 | -3,81 | 2,83 | -1,35 | 1,78E-01 |
| AT4G09375 | 0,14 | -0,35 | 4,38 | -0,08 | 9,37E-01 |
| AT4G36810 | 0,84 | -0,33 | 2,49 | -0,13 | 8,95E-01 |
| AT4G36870 | 0,25 | 1,14 | 4,32 | 0,26 | 7,93E-01 |
| AT4G36920 | 0,14 | -0,35 | 4,38 | -0,08 | 9,37E-01 |
| AT4G36925 | 1,06 | -0,68 | 1,77 | -0,38 | 7,01E-01 |
| AT4G36990 | 0,59 | -1,72 | 3,08 | -0,56 | 5,78E-01 |
| AT4G37000 | 0,23 | -1,14 | 4,39 | -0,26 | 7,96E-01 |
| AT4G37020 | 0,15 | -0,68 | 4,41 | -0,15 | 8,77E-01 |
| AT4G37080 | 0,19 | -0,68 | 4,41 | -0,15 | 8,77E-01 |
| AT4G37130 | 0,62 | -3,49 | 3,68 | -0,95 | 3,44E-01 |
| AT4G37180 | 0,47 | 0,93 | 4,08 | 0,23 | 8,20E-01 |
| AT4G37190 | 0,32 | -2,60 | 4,41 | -0,59 | 5,55E-01 |

| | | | | | |
|---|---|---|---|---|---|
| AT4G37210 | 0,15 | 0,00 | 4,41 | 0,00 | 1,00E+00 |
| AT4G37300 | 1,24 | -2,26 | 1,95 | -1,16 | 2,47E-01 |
| AT4G37430 | 0,86 | -0,73 | 3,49 | -0,21 | 8,35E-01 |
| AT4G37440 | 0,34 | -1,45 | 4,34 | -0,33 | 7,38E-01 |
| AT4G37550 | 0,88 | -0,73 | 2,03 | -0,36 | 7,20E-01 |
| AT4G37560 | 0,42 | -2,95 | 4,38 | -0,67 | 5,00E-01 |
| AT4G37680 | 0,49 | 0,01 | 3,18 | 0,00 | 9,98E-01 |
| AT4G37730 | 0,75 | 0,87 | 2,09 | 0,42 | 6,75E-01 |
| AT4G37760 | 0,63 | 0,34 | 2,32 | 0,15 | 8,82E-01 |
| AT4G37790 | 0,15 | -0,17 | 4,39 | -0,04 | 9,68E-01 |
| AT4G37980 | 0,67 | 1,90 | 2,14 | 0,89 | 3,74E-01 |
| AT4G38040 | 0,20 | -2,60 | 4,41 | -0,59 | 5,55E-01 |
| AT4G38170 | 0,97 | -1,56 | 2,11 | -0,74 | 4,61E-01 |
| AT4G09685 | 0,46 | -3,36 | 4,02 | -0,83 | 4,04E-01 |
| AT4G38215 | 0,75 | -3,01 | 4,38 | -0,69 | 4,92E-01 |
| AT4G38225 | 0,78 | -0,42 | 2,40 | -0,18 | 8,60E-01 |
| AT4G38250 | 1,25 | -1,79 | 1,61 | -1,12 | 2,65E-01 |
| AT4G38270 | 0,51 | -0,75 | 3,20 | -0,23 | 8,15E-01 |
| AT4G38290 | 0,11 | -0,68 | 4,41 | -0,15 | 8,77E-01 |
| AT4G38320 | 0,19 | -0,81 | 4,39 | -0,19 | 8,53E-01 |
| AT4G38330 | 0,12 | 0,00 | 4,41 | 0,00 | 1,00E+00 |
| AT4G38360 | 0,15 | -0,77 | 4,39 | -0,18 | 8,61E-01 |
| AT4G38430 | 0,81 | -2,86 | 2,69 | -1,06 | 2,88E-01 |
| AT4G38500 | 0,68 | -0,68 | 3,61 | -0,19 | 8,51E-01 |
| AT4G38540 | 0,10 | -0,18 | 4,38 | -0,04 | 9,67E-01 |
| AT4G38560 | 1,22 | -4,15 | 2,40 | -1,73 | 8,43E-02 |
| AT4G38580 | 0,50 | -3,93 | 4,34 | -0,90 | 3,66E-01 |
| AT4G38620 | 0,24 | -0,68 | 4,41 | -0,15 | 8,77E-01 |
| AT4G38790 | 0,69 | -2,52 | 3,07 | -0,82 | 4,11E-01 |
| AT4G38860 | 0,38 | 1,30 | 4,32 | 0,30 | 7,63E-01 |
| AT4G38880 | 0,39 | -3,10 | 4,38 | -0,71 | 4,79E-01 |
| AT4G38920 | 0,99 | -0,97 | 2,12 | -0,46 | 6,48E-01 |
| AT4G38930 | 0,41 | -2,31 | 4,37 | -0,53 | 5,96E-01 |
| AT4G39020 | 0,55 | 1,10 | 4,33 | 0,25 | 7,99E-01 |
| AT4G39030 | 0,29 | -2,97 | 4,38 | -0,68 | 4,98E-01 |
| AT4G39210 | 1,07 | -3,67 | 2,28 | -1,61 | 1,07E-01 |
| AT4G39270 | 1,16 | -0,91 | 1,51 | -0,61 | 5,44E-01 |
| AT4G39620 | 0,18 | -0,27 | 4,38 | -0,06 | 9,51E-01 |
| AT4G39640 | 0,76 | -0,36 | 2,51 | -0,14 | 8,85E-01 |
| AT4G39670 | 0,55 | 1,06 | 3,17 | 0,33 | 7,39E-01 |
| AT4G39840 | 1,07 | -0,16 | 1,77 | -0,09 | 9,27E-01 |
| AT4G39860 | 0,51 | 0,27 | 2,33 | 0,12 | 9,06E-01 |
| AT4G39870 | 0,35 | -0,73 | 4,40 | -0,17 | 8,68E-01 |
| AT4G39940 | 0,36 | -2,00 | 4,37 | -0,46 | 6,47E-01 |
| AT4G39950 | 1,14 | -0,99 | 1,86 | -0,53 | 5,95E-01 |
| AT4G39952 | 0,28 | -3,07 | 4,38 | -0,70 | 4,84E-01 |
| AT4G39980 | 0,50 | -0,65 | 3,12 | -0,21 | 8,35E-01 |
| AT4G40063 | 0,73 | -3,17 | 3,04 | -1,05 | 2,96E-01 |
| AT5G01170 | 0,10 | -0,34 | 4,38 | -0,08 | 9,38E-01 |
| AT5G01210 | 1,09 | 1,51 | 1,78 | 0,85 | 3,97E-01 |
| AT5G01240 | 0,22 | 0,38 | 4,35 | 0,09 | 9,31E-01 |
| AT5G01260 | 0,54 | -1,75 | 2,37 | -0,74 | 4,58E-01 |
| AT5G01290 | 0,28 | -0,80 | 4,39 | -0,18 | 8,55E-01 |
| AT5G01460 | 0,28 | -2,28 | 4,37 | -0,52 | 6,01E-01 |
| AT5G01542 | 0,46 | -1,29 | 4,07 | -0,32 | 7,51E-01 |
| AT5G01590 | 0,94 | -1,58 | 2,26 | -0,70 | 4,85E-01 |

| | | | | | |
|---|---|---|---|---|---|
| AT5G01595 | 0,81 | -0,02 | 2,47 | -0,01 | 9,94E-01 |
| AT5G01690 | 0,08 | 0,00 | 4,41 | 0,00 | 1,00E+00 |
| AT5G01732 | 0,29 | -2,54 | 4,40 | -0,58 | 5,63E-01 |
| AT5G01740 | 0,15 | 0,79 | 4,33 | 0,18 | 8,56E-01 |
| AT5G01760 | 0,79 | -4,64 | 2,72 | -1,71 | 8,81E-02 |
| AT5G01800 | 0,74 | -3,19 | 2,44 | -1,31 | 1,91E-01 |
| AT5G01810 | 0,55 | -3,44 | 3,88 | -0,89 | 3,74E-01 |
| AT5G01850 | 0,24 | -0,30 | 4,38 | -0,07 | 9,46E-01 |
| AT5G02035 | 0,47 | 0,00 | 4,05 | 0,00 | 1,00E+00 |
| AT5G02040 | 0,25 | -2,53 | 4,40 | -0,57 | 5,65E-01 |
| AT5G02100 | 0,70 | -0,68 | 3,82 | -0,18 | 8,59E-01 |
| AT5G02120 | 0,39 | 0,08 | 4,37 | 0,02 | 9,86E-01 |
| AT5G00820 | 1,31 | -3,32 | 2,68 | -1,24 | 2,16E-01 |
| AT5G02160 | 1,30 | 0,12 | 1,81 | 0,06 | 9,49E-01 |
| AT5G02240 | 0,32 | -1,65 | 4,40 | -0,38 | 7,07E-01 |
| AT5G02280 | 0,11 | -0,68 | 4,41 | -0,15 | 8,77E-01 |
| AT5G02420 | 0,12 | -0,68 | 4,41 | -0,15 | 8,77E-01 |
| AT5G02450 | 0,59 | -2,01 | 3,20 | -0,63 | 5,31E-01 |
| AT5G02620 | 0,32 | -1,30 | 4,34 | -0,30 | 7,64E-01 |
| AT5G02630 | 0,30 | -2,60 | 4,41 | -0,59 | 5,55E-01 |
| AT5G02650 | 0,40 | 0,00 | 4,41 | 0,00 | 1,00E+00 |
| AT5G02760 | 0,40 | -1,64 | 3,50 | -0,47 | 6,39E-01 |
| AT5G02790 | 0,39 | 0,72 | 3,41 | 0,21 | 8,33E-01 |
| AT5G02840 | 0,40 | -1,58 | 4,22 | -0,37 | 7,09E-01 |
| AT5G02850 | 0,54 | -2,39 | 3,12 | -0,77 | 4,44E-01 |
| AT5G02860 | 1,11 | -1,26 | 2,00 | -0,63 | 5,28E-01 |
| AT5G02950 | 0,61 | 0,00 | 3,89 | 0,00 | 1,00E+00 |
| AT5G02995 | 0,13 | -0,18 | 4,39 | -0,04 | 9,67E-01 |
| AT5G03020 | 0,28 | -3,07 | 4,38 | -0,70 | 4,84E-01 |
| AT5G03160 | 0,68 | -0,97 | 2,31 | -0,42 | 6,75E-01 |
| AT5G03190 | 0,92 | -1,64 | 1,88 | -0,87 | 3,84E-01 |
| AT5G03290 | 1,24 | -1,10 | 1,49 | -0,74 | 4,62E-01 |
| AT5G03300 | 0,81 | 1,11 | 2,46 | 0,45 | 6,53E-01 |
| AT5G03330 | 0,61 | -3,34 | 3,04 | -1,10 | 2,71E-01 |
| AT5G03355 | 0,46 | -0,90 | 3,27 | -0,27 | 7,85E-01 |
| AT5G03415 | 0,36 | -1,09 | 4,34 | -0,25 | 8,02E-01 |
| AT5G03420 | 0,63 | -0,46 | 2,59 | -0,18 | 8,60E-01 |
| AT5G03455 | 0,25 | -2,61 | 4,41 | -0,59 | 5,53E-01 |
| AT5G03460 | 0,54 | -3,37 | 3,07 | -1,10 | 2,72E-01 |
| AT5G03470 | 0,40 | -2,58 | 4,36 | -0,59 | 5,54E-01 |
| AT5G03490 | 0,11 | -0,80 | 4,39 | -0,18 | 8,56E-01 |
| AT5G03540 | 1,05 | -2,90 | 2,76 | -1,05 | 2,93E-01 |
| AT5G03650 | 0,91 | -0,40 | 1,76 | -0,23 | 8,19E-01 |
| AT5G03905 | 0,30 | -1,09 | 4,39 | -0,25 | 8,04E-01 |
| AT5G04040 | 1,27 | -0,75 | 2,16 | -0,35 | 7,29E-01 |
| AT5G04050 | 0,10 | -0,34 | 4,38 | -0,08 | 9,38E-01 |
| AT5G04060 | 0,37 | -0,27 | 4,38 | -0,06 | 9,50E-01 |
| AT5G04170 | 0,76 | -1,72 | 2,46 | -0,70 | 4,84E-01 |
| AT5G04190 | 0,13 | 0,52 | 4,35 | 0,12 | 9,04E-01 |
| AT5G04240 | 0,88 | 0,00 | 3,41 | 0,00 | 1,00E+00 |
| AT5G04280 | 1,25 | 0,12 | 2,02 | 0,06 | 9,52E-01 |
| AT5G04290 | 0,66 | -0,91 | 3,18 | -0,29 | 7,75E-01 |
| AT5G04360 | 0,45 | -3,39 | 4,17 | -0,81 | 4,16E-01 |
| AT5G00960 | 0,96 | 2,20 | 2,16 | 1,02 | 3,09E-01 |
| AT5G04460 | 0,56 | -1,64 | 3,20 | -0,51 | 6,10E-01 |
| AT5G04490 | 0,46 | 0,30 | 3,31 | 0,09 | 9,27E-01 |

| | | | | | |
|---|---|---|---|---|---|
| AT5G04510 | 0,58 | -2,37 | 2,71 | -0,87 | 3,83E-01 |
| AT5G04600 | 0,71 | -1,63 | 2,20 | -0,74 | 4,59E-01 |
| AT5G04620 | 0,43 | -3,24 | 4,37 | -0,74 | 4,58E-01 |
| AT5G04630 | 0,57 | -2,57 | 3,84 | -0,67 | 5,04E-01 |
| AT5G04660 | 0,24 | -0,20 | 4,39 | -0,05 | 9,64E-01 |
| AT5G00985 | 0,28 | -3,29 | 4,38 | -0,75 | 4,53E-01 |
| AT5G04800 | 0,67 | -1,36 | 2,07 | -0,66 | 5,11E-01 |
| AT5G04885 | 0,27 | 0,86 | 3,72 | 0,23 | 8,17E-01 |
| AT5G04910 | 0,42 | -0,89 | 4,10 | -0,22 | 8,28E-01 |
| AT5G05070 | 0,40 | -1,99 | 4,30 | -0,46 | 6,43E-01 |
| AT5G05090 | 0,35 | 0,09 | 3,52 | 0,03 | 9,79E-01 |
| AT5G05160 | 0,72 | -3,27 | 2,81 | -1,16 | 2,45E-01 |
| AT5G05210 | 0,28 | 0,63 | 4,34 | 0,15 | 8,84E-01 |
| AT5G05220 | 0,08 | 0,00 | 4,41 | 0,00 | 1,00E+00 |
| AT5G05230 | 0,08 | 0,00 | 4,41 | 0,00 | 1,00E+00 |
| AT5G05310 | 0,18 | 0,57 | 4,34 | 0,13 | 8,95E-01 |
| AT5G05440 | 0,16 | -0,68 | 4,41 | -0,15 | 8,77E-01 |
| AT5G05490 | 0,55 | -3,77 | 4,34 | -0,87 | 3,86E-01 |
| AT5G05560 | 1,14 | -0,83 | 1,81 | -0,46 | 6,45E-01 |
| AT5G05570 | 0,62 | -0,70 | 4,40 | -0,16 | 8,74E-01 |
| AT5G05600 | 0,31 | -0,15 | 4,39 | -0,03 | 9,73E-01 |
| AT5G05630 | 0,08 | -0,21 | 4,39 | -0,05 | 9,62E-01 |
| AT5G05710 | 0,20 | -2,55 | 4,40 | -0,58 | 5,62E-01 |
| AT5G05780 | 0,96 | -0,75 | 2,65 | -0,28 | 7,77E-01 |
| AT5G05950 | 0,83 | -1,73 | 2,33 | -0,74 | 4,59E-01 |
| AT5G05970 | 0,30 | -1,29 | 4,35 | -0,30 | 7,67E-01 |
| AT5G05985 | 0,55 | -3,82 | 3,83 | -1,00 | 3,19E-01 |
| AT5G06000 | 0,23 | 0,82 | 4,33 | 0,19 | 8,50E-01 |
| AT5G06050 | 0,39 | -3,20 | 4,30 | -0,74 | 4,57E-01 |
| AT5G06100 | 0,21 | -0,70 | 4,40 | -0,16 | 8,73E-01 |
| AT5G06120 | 1,17 | -4,36 | 2,45 | -1,78 | 7,52E-02 |
| AT5G06170 | 0,31 | 1,55 | 4,30 | 0,36 | 7,18E-01 |
| AT5G06220 | 0,84 | -3,07 | 2,50 | -1,23 | 2,19E-01 |
| AT5G06265 | 0,26 | 0,99 | 4,32 | 0,23 | 8,19E-01 |
| AT5G06310 | 0,33 | 0,97 | 4,33 | 0,22 | 8,23E-01 |
| AT5G01255 | 0,70 | 0,48 | 2,79 | 0,17 | 8,65E-01 |
| AT5G06350 | 0,36 | 0,46 | 3,46 | 0,13 | 8,95E-01 |
| AT5G06360 | 0,40 | -1,99 | 4,30 | -0,46 | 6,43E-01 |
| AT5G06380 | 0,08 | 0,00 | 4,41 | 0,00 | 1,00E+00 |
| AT5G06390 | 0,14 | 0,40 | 4,35 | 0,09 | 9,27E-01 |
| AT5G06645 | 0,58 | 2,92 | 4,27 | 0,68 | 4,94E-01 |
| AT5G06660 | 0,96 | 0,66 | 3,59 | 0,19 | 8,53E-01 |
| AT5G06680 | 0,39 | -1,68 | 4,30 | -0,39 | 6,96E-01 |
| AT5G06700 | 0,91 | -0,71 | 1,80 | -0,39 | 6,94E-01 |
| AT5G06811 | 0,11 | -0,68 | 4,41 | -0,15 | 8,77E-01 |
| AT5G06820 | 0,12 | 0,00 | 4,41 | 0,00 | 1,00E+00 |
| AT5G06860 | 0,22 | 0,62 | 4,34 | 0,14 | 8,86E-01 |
| AT5G06870 | 0,67 | 0,66 | 2,72 | 0,24 | 8,09E-01 |
| AT5G06950 | 0,61 | -3,09 | 3,07 | -1,01 | 3,15E-01 |
| AT5G06980 | 0,16 | -0,64 | 4,40 | -0,15 | 8,85E-01 |
| AT5G07110 | 0,85 | -3,74 | 3,38 | -1,11 | 2,69E-01 |
| AT5G01315 | 0,64 | -3,00 | 2,96 | -1,01 | 3,11E-01 |
| AT5G07130 | 0,33 | -2,52 | 4,40 | -0,57 | 5,66E-01 |
| AT5G07140 | 1,24 | -4,68 | 2,48 | -1,89 | 5,93E-02 |
| AT5G07210 | 1,09 | 1,09 | 2,00 | 0,54 | 5,86E-01 |
| AT5G07240 | 1,19 | -2,63 | 2,21 | -1,19 | 2,35E-01 |

| | | | | | |
|---|---|---|---|---|---|
| AT5G07270 | 0,15 | 0,41 | 4,35 | 0,09 | 9,26E-01 |
| AT5G07340 | 0,98 | 0,68 | 2,37 | 0,29 | 7,73E-01 |
| AT5G07430 | 0,50 | 0,51 | 3,88 | 0,13 | 8,96E-01 |
| AT5G07440 | 0,36 | -2,60 | 4,41 | -0,59 | 5,55E-01 |
| AT5G07580 | 0,75 | -0,26 | 2,15 | -0,12 | 9,05E-01 |
| AT5G07610 | 0,48 | 1,66 | 4,02 | 0,41 | 6,79E-01 |
| AT5G07760 | 0,08 | 0,00 | 4,41 | 0,00 | 1,00E+00 |
| AT5G07770 | 0,16 | -0,62 | 4,40 | -0,14 | 8,87E-01 |
| AT5G07790 | 0,58 | -0,68 | 4,41 | -0,15 | 8,77E-01 |
| AT5G07830 | 0,86 | -0,42 | 2,55 | -0,16 | 8,70E-01 |
| AT5G07860 | 0,58 | -3,14 | 3,87 | -0,81 | 4,18E-01 |
| AT5G07870 | 0,49 | -3,76 | 4,35 | -0,86 | 3,87E-01 |
| AT5G07960 | 0,12 | -0,68 | 4,41 | -0,15 | 8,77E-01 |
| AT5G08050 | 0,48 | 0,04 | 4,20 | 0,01 | 9,93E-01 |
| AT5G08060 | 0,10 | -0,34 | 4,38 | -0,08 | 9,38E-01 |
| AT5G08070 | 0,35 | -2,99 | 4,38 | -0,68 | 4,95E-01 |
| AT5G08139 | 0,37 | 1,71 | 4,28 | 0,40 | 6,89E-01 |
| AT5G08190 | 0,29 | -2,57 | 4,40 | -0,58 | 5,59E-01 |
| AT5G08260 | 0,90 | -3,92 | 2,31 | -1,70 | 8,97E-02 |
| AT5G01545 | 0,37 | 0,00 | 4,41 | 0,00 | 1,00E+00 |
| AT5G08290 | 0,74 | 0,23 | 2,19 | 0,10 | 9,17E-01 |
| AT5G08300 | 0,94 | -0,45 | 2,25 | -0,20 | 8,41E-01 |
| AT5G08335 | 0,18 | 0,40 | 4,35 | 0,09 | 9,27E-01 |
| AT5G08340 | 0,58 | -2,54 | 4,01 | -0,63 | 5,27E-01 |
| AT5G08390 | 0,26 | -0,17 | 4,37 | -0,04 | 9,69E-01 |
| AT5G08430 | 0,57 | -3,43 | 4,37 | -0,78 | 4,33E-01 |
| AT5G08440 | 0,59 | -2,99 | 3,88 | -0,77 | 4,40E-01 |
| AT5G08470 | 0,81 | -2,74 | 2,77 | -0,99 | 3,21E-01 |
| AT5G08510 | 0,51 | -3,06 | 3,23 | -0,95 | 3,43E-01 |
| AT5G08540 | 0,41 | 0,35 | 3,40 | 0,10 | 9,17E-01 |
| AT5G01555 | 0,36 | -0,66 | 4,36 | -0,15 | 8,79E-01 |
| AT5G08570 | 1,20 | -1,32 | 1,64 | -0,80 | 4,23E-01 |
| AT5G08590 | 0,63 | -0,57 | 2,72 | -0,21 | 8,35E-01 |
| AT5G08600 | 0,28 | -3,29 | 4,38 | -0,75 | 4,53E-01 |
| AT5G08620 | 1,12 | -2,27 | 1,79 | -1,27 | 2,04E-01 |
| AT5G08660 | 0,33 | -2,04 | 4,35 | -0,47 | 6,39E-01 |
| AT5G08695 | 0,20 | -2,55 | 4,40 | -0,58 | 5,62E-01 |
| AT5G08750 | 0,11 | -0,80 | 4,39 | -0,18 | 8,56E-01 |
| AT5G08760 | 0,55 | -0,43 | 3,89 | -0,11 | 9,13E-01 |
| AT5G08780 | 0,49 | -2,44 | 3,22 | -0,76 | 4,48E-01 |
| AT5G09220 | 0,82 | 1,26 | 2,99 | 0,42 | 6,73E-01 |
| AT5G09400 | 1,22 | -1,92 | 1,98 | -0,97 | 3,34E-01 |
| AT5G09440 | 0,77 | 0,06 | 2,27 | 0,02 | 9,80E-01 |
| AT5G09445 | 0,26 | -2,60 | 4,41 | -0,59 | 5,55E-01 |
| AT5G09450 | 1,25 | -3,14 | 2,03 | -1,54 | 1,23E-01 |
| AT5G09600 | 0,62 | -1,64 | 3,14 | -0,52 | 6,01E-01 |
| AT5G09760 | 0,14 | 0,40 | 4,35 | 0,09 | 9,27E-01 |
| AT5G10010 | 0,47 | -1,03 | 3,35 | -0,31 | 7,59E-01 |
| AT5G10020 | 0,41 | -3,25 | 4,37 | -0,74 | 4,57E-01 |
| AT5G10100 | 0,16 | -0,11 | 4,37 | -0,03 | 9,80E-01 |
| AT5G10190 | 0,41 | -2,51 | 3,35 | -0,75 | 4,54E-01 |
| AT5G10200 | 0,81 | -4,18 | 3,41 | -1,23 | 2,20E-01 |
| AT5G10410 | 0,05 | -0,24 | 4,38 | -0,05 | 9,57E-01 |
| AT5G10460 | 0,08 | 0,00 | 4,41 | 0,00 | 1,00E+00 |
| AT5G01765 | 1,15 | -1,28 | 1,86 | -0,69 | 4,91E-01 |
| AT5G10600 | 0,40 | -0,61 | 4,26 | -0,14 | 8,86E-01 |

| | | | | | |
|---|---|---|---|---|---|
| AT5G10630 | 0,28 | 0,45 | 4,35 | 0,10 | 9,18E-01 |
| AT5G10660 | 0,46 | -0,33 | 2,75 | -0,12 | 9,05E-01 |
| AT5G10695 | 0,85 | 2,59 | 2,07 | 1,25 | 2,12E-01 |
| AT5G10745 | 0,35 | 1,26 | 3,48 | 0,36 | 7,18E-01 |
| AT5G10750 | 1,12 | -1,49 | 2,14 | -0,69 | 4,88E-01 |
| AT5G10780 | 0,76 | -1,33 | 2,56 | -0,52 | 6,02E-01 |
| AT5G10790 | 0,93 | -2,53 | 1,93 | -1,31 | 1,90E-01 |
| AT5G10920 | 0,91 | -1,74 | 2,17 | -0,80 | 4,22E-01 |
| AT5G11060 | 1,02 | 0,58 | 1,63 | 0,35 | 7,23E-01 |
| AT5G11090 | 0,49 | -0,32 | 2,69 | -0,12 | 9,07E-01 |
| AT5G11110 | 0,50 | -0,34 | 4,00 | -0,09 | 9,32E-01 |
| AT5G11200 | 0,33 | -1,64 | 4,38 | -0,37 | 7,08E-01 |
| AT5G11280 | 0,20 | 0,11 | 4,37 | 0,03 | 9,80E-01 |
| AT5G11340 | 0,28 | -1,70 | 4,40 | -0,39 | 7,00E-01 |
| AT5G11350 | 0,82 | -4,35 | 2,90 | -1,50 | 1,33E-01 |
| AT5G11390 | 0,30 | -2,60 | 4,41 | -0,59 | 5,55E-01 |
| AT5G11420 | 1,15 | 0,18 | 1,89 | 0,09 | 9,26E-01 |
| AT5G11430 | 0,70 | -1,42 | 2,22 | -0,64 | 5,22E-01 |
| AT5G11500 | 0,94 | -0,86 | 2,10 | -0,41 | 6,84E-01 |
| AT5G11510 | 0,81 | -2,51 | 2,48 | -1,01 | 3,11E-01 |
| AT5G11530 | 0,81 | -2,17 | 2,36 | -0,92 | 3,57E-01 |
| AT5G11580 | 0,12 | -0,78 | 4,39 | -0,18 | 8,59E-01 |
| AT5G11650 | 0,38 | -0,13 | 3,44 | -0,04 | 9,70E-01 |
| AT5G11680 | 0,19 | 0,01 | 4,36 | 0,00 | 9,98E-01 |
| AT5G11710 | 1,24 | -1,11 | 1,56 | -0,71 | 4,77E-01 |
| AT5G11730 | 0,89 | -0,97 | 1,81 | -0,53 | 5,93E-01 |
| AT5G11760 | 0,22 | -1,25 | 4,38 | -0,29 | 7,75E-01 |
| AT5G11890 | 0,82 | -0,65 | 2,00 | -0,32 | 7,47E-01 |
| AT5G11920 | 0,12 | -0,78 | 4,39 | -0,18 | 8,59E-01 |
| AT5G11977 | 0,12 | 0,00 | 4,41 | 0,00 | 1,00E+00 |
| AT5G12090 | 0,79 | 2,86 | 2,31 | 1,23 | 2,17E-01 |
| AT5G12100 | 0,44 | 0,00 | 4,41 | 0,00 | 1,00E+00 |
| AT5G12210 | 0,65 | -2,18 | 3,71 | -0,59 | 5,56E-01 |
| AT5G12260 | 0,32 | -1,65 | 4,40 | -0,38 | 7,07E-01 |
| AT5G12350 | 0,49 | -2,69 | 4,16 | -0,65 | 5,18E-01 |
| AT5G12400 | 0,93 | -1,45 | 2,85 | -0,51 | 6,11E-01 |
| AT5G12850 | 0,97 | -0,57 | 1,77 | -0,32 | 7,49E-01 |
| AT5G12950 | 0,53 | 1,88 | 2,51 | 0,75 | 4,54E-01 |
| AT5G12960 | 0,27 | -2,58 | 4,40 | -0,59 | 5,58E-01 |
| AT5G12970 | 0,24 | -3,18 | 4,39 | -0,73 | 4,68E-01 |
| AT5G13030 | 0,96 | 1,54 | 2,03 | 0,76 | 4,50E-01 |
| AT5G13200 | 1,26 | 0,92 | 1,65 | 0,56 | 5,78E-01 |
| AT5G13210 | 0,35 | -0,91 | 3,46 | -0,26 | 7,92E-01 |
| AT5G13260 | 0,92 | -2,87 | 2,18 | -1,32 | 1,88E-01 |
| AT5G13280 | 0,45 | -2,73 | 4,13 | -0,66 | 5,09E-01 |
| AT5G13310 | 0,31 | -1,71 | 4,38 | -0,39 | 6,97E-01 |
| AT5G13320 | 1,27 | -0,18 | 1,88 | -0,10 | 9,23E-01 |
| AT5G13330 | 0,37 | -2,60 | 4,41 | -0,59 | 5,55E-01 |
| AT5G13370 | 0,68 | -0,82 | 3,22 | -0,26 | 7,98E-01 |
| AT5G13380 | 0,54 | 0,00 | 4,41 | 0,00 | 1,00E+00 |
| AT5G13500 | 0,32 | -0,23 | 4,33 | -0,05 | 9,57E-01 |
| AT5G13510 | 0,48 | -1,64 | 3,38 | -0,49 | 6,28E-01 |
| AT5G13580 | 0,29 | 0,00 | 4,41 | 0,00 | 1,00E+00 |
| AT5G13640 | 0,84 | -3,37 | 2,96 | -1,14 | 2,54E-01 |
| AT5G13690 | 0,61 | -3,11 | 3,09 | -1,01 | 3,13E-01 |
| AT5G13720 | 0,21 | -0,68 | 4,41 | -0,15 | 8,77E-01 |

| | | | | | |
|---|---|---|---|---|---|
| AT5G13740 | 0,19 | 0,01 | 4,36 | 0,00 | 9,98E-01 |
| AT5G13750 | 0,26 | -2,60 | 4,41 | -0,59 | 5,55E-01 |
| AT5G13760 | 0,26 | -0,58 | 4,35 | -0,13 | 8,93E-01 |
| AT5G13810 | 0,93 | -0,66 | 1,75 | -0,38 | 7,04E-01 |
| AT5G13820 | 0,28 | -0,74 | 4,40 | -0,17 | 8,66E-01 |
| AT5G13930 | 1,07 | -3,84 | 2,32 | -1,65 | 9,81E-02 |
| AT5G13980 | 0,97 | -2,39 | 2,29 | -1,04 | 2,97E-01 |
| AT5G14030 | 0,98 | -2,12 | 2,46 | -0,86 | 3,89E-01 |
| AT5G14050 | 0,50 | 0,45 | 3,20 | 0,14 | 8,87E-01 |
| AT5G14120 | 1,26 | -0,95 | 1,44 | -0,66 | 5,10E-01 |
| AT5G14230 | 0,39 | 1,51 | 3,39 | 0,44 | 6,57E-01 |
| AT5G14260 | 0,68 | -1,47 | 2,49 | -0,59 | 5,54E-01 |
| AT5G14280 | 0,43 | -3,12 | 4,39 | -0,71 | 4,77E-01 |
| AT5G14285 | 0,21 | 0,00 | 4,41 | 0,00 | 1,00E+00 |
| AT5G14480 | 0,25 | -0,53 | 4,39 | -0,12 | 9,03E-01 |
| AT5G14610 | 0,64 | -0,75 | 3,17 | -0,24 | 8,12E-01 |
| AT5G14670 | 0,08 | 0,00 | 4,41 | 0,00 | 1,00E+00 |
| AT5G14880 | 0,37 | -0,90 | 3,48 | -0,26 | 7,97E-01 |
| AT5G14890 | 0,99 | 0,00 | 4,15 | 0,00 | 1,00E+00 |
| AT5G14910 | 0,77 | -0,70 | 2,64 | -0,27 | 7,90E-01 |
| AT5G14970 | 0,18 | -0,33 | 4,38 | -0,08 | 9,40E-01 |
| AT5G14990 | 0,42 | -3,55 | 4,20 | -0,85 | 3,98E-01 |
| AT5G15010 | 0,38 | -0,75 | 4,40 | -0,17 | 8,65E-01 |
| AT5G15080 | 0,92 | -4,51 | 2,86 | -1,58 | 1,15E-01 |
| AT5G15230 | 0,74 | -3,75 | 2,90 | -1,29 | 1,96E-01 |
| AT5G15240 | 0,05 | -0,24 | 4,38 | -0,05 | 9,57E-01 |
| AT5G15270 | 0,61 | 0,43 | 3,17 | 0,14 | 8,92E-01 |
| AT5G15280 | 0,54 | -3,89 | 3,84 | -1,01 | 3,11E-01 |
| AT5G15320 | 0,21 | -0,44 | 4,38 | -0,10 | 9,20E-01 |
| AT5G15330 | 0,11 | -0,68 | 4,41 | -0,15 | 8,77E-01 |
| AT5G15530 | 0,12 | 0,00 | 4,41 | 0,00 | 1,00E+00 |
| AT5G15570 | 0,44 | 0,00 | 4,41 | 0,00 | 1,00E+00 |
| AT5G15610 | 0,89 | -1,48 | 1,77 | -0,83 | 4,04E-01 |
| AT5G15685 | 0,72 | 0,65 | 2,04 | 0,32 | 7,51E-01 |
| AT5G15710 | 0,74 | 0,09 | 3,68 | 0,03 | 9,80E-01 |
| AT5G15730 | 0,59 | -0,88 | 2,66 | -0,33 | 7,41E-01 |
| AT5G15750 | 0,54 | 0,12 | 3,87 | 0,03 | 9,76E-01 |
| AT5G15820 | 0,22 | -1,25 | 4,37 | -0,29 | 7,75E-01 |
| AT5G15930 | 0,32 | -1,70 | 4,40 | -0,39 | 6,99E-01 |
| AT5G15950 | 1,23 | -4,45 | 2,13 | -2,09 | 3,65E-02 |
| AT5G15960 | 0,51 | 0,00 | 4,41 | 0,00 | 1,00E+00 |
| AT5G16010 | 0,36 | -1,18 | 4,33 | -0,27 | 7,85E-01 |
| AT5G16030 | 0,58 | 0,78 | 3,32 | 0,23 | 8,15E-01 |
| AT5G16120 | 0,30 | -1,64 | 4,41 | -0,37 | 7,10E-01 |
| AT5G16130 | 1,11 | -2,50 | 2,08 | -1,20 | 2,29E-01 |
| AT5G16260 | 0,11 | 0,46 | 4,35 | 0,11 | 9,15E-01 |
| AT5G16290 | 0,43 | -0,10 | 4,34 | -0,02 | 9,81E-01 |
| AT5G16300 | 1,23 | -0,56 | 1,74 | -0,32 | 7,47E-01 |
| AT5G16440 | 0,67 | -2,44 | 2,49 | -0,98 | 3,28E-01 |
| AT5G16470 | 0,69 | -1,63 | 2,45 | -0,66 | 5,06E-01 |
| AT5G16540 | 0,32 | -3,36 | 4,37 | -0,77 | 4,42E-01 |
| AT5G16550 | 0,57 | -1,88 | 2,68 | -0,70 | 4,84E-01 |
| AT5G02555 | 0,05 | -0,24 | 4,38 | -0,05 | 9,57E-01 |
| AT5G16590 | 0,87 | -1,00 | 2,23 | -0,45 | 6,53E-01 |
| AT5G16630 | 0,92 | -1,62 | 2,79 | -0,58 | 5,62E-01 |
| AT5G16715 | 0,74 | -3,04 | 2,47 | -1,23 | 2,19E-01 |

| | | | | | |
|---|---|---|---|---|---|
| AT5G16730 | 1,05 | 0,40 | 2,02 | 0,20 | 8,42E-01 |
| AT5G16760 | 1,05 | -0,56 | 2,15 | -0,26 | 7,96E-01 |
| AT5G16820 | 0,94 | -2,91 | 2,35 | -1,24 | 2,16E-01 |
| AT5G16880 | 1,27 | -1,74 | 1,65 | -1,05 | 2,93E-01 |
| AT5G16900 | 0,33 | -2,59 | 4,41 | -0,59 | 5,56E-01 |
| AT5G16910 | 0,98 | -3,86 | 2,62 | -1,47 | 1,41E-01 |
| AT5G17167 | 0,20 | 0,68 | 4,35 | 0,16 | 8,75E-01 |
| AT5G17170 | 0,33 | -1,38 | 4,34 | -0,32 | 7,51E-01 |
| AT5G17190 | 0,51 | -0,23 | 3,05 | -0,08 | 9,39E-01 |
| AT5G17270 | 0,61 | -4,28 | 4,32 | -0,99 | 3,22E-01 |
| AT5G17280 | 0,29 | -0,19 | 4,39 | -0,04 | 9,65E-01 |
| AT5G17305 | 0,08 | 0,00 | 4,41 | 0,00 | 1,00E+00 |
| AT5G17440 | 0,24 | -1,92 | 4,38 | -0,44 | 6,61E-01 |
| AT5G17530 | 0,59 | -2,55 | 3,12 | -0,82 | 4,14E-01 |
| AT5G17580 | 0,17 | 0,00 | 4,41 | 0,00 | 1,00E+00 |
| AT5G17630 | 0,33 | -1,56 | 4,40 | -0,36 | 7,22E-01 |
| AT5G17670 | 0,39 | -3,07 | 4,38 | -0,70 | 4,84E-01 |
| AT5G17680 | 0,30 | -2,60 | 4,41 | -0,59 | 5,55E-01 |
| AT5G17850 | 0,18 | 1,20 | 4,31 | 0,28 | 7,81E-01 |
| AT5G17870 | 0,86 | -0,07 | 2,30 | -0,03 | 9,77E-01 |
| AT5G17900 | 0,30 | -0,48 | 4,34 | -0,11 | 9,13E-01 |
| AT5G17980 | 0,38 | -1,26 | 4,27 | -0,30 | 7,68E-01 |
| AT5G17990 | 0,55 | -2,33 | 4,00 | -0,58 | 5,61E-01 |
| AT5G18040 | 0,71 | -0,64 | 2,53 | -0,25 | 8,00E-01 |
| AT5G18065 | 0,30 | 0,15 | 3,68 | 0,04 | 9,67E-01 |
| AT5G18070 | 0,62 | -1,97 | 2,53 | -0,78 | 4,37E-01 |
| AT5G18085 | 0,24 | -3,18 | 4,39 | -0,73 | 4,68E-01 |
| AT5G18100 | 0,33 | 0,00 | 4,36 | 0,00 | 1,00E+00 |
| AT5G18130 | 1,11 | -1,55 | 2,26 | -0,69 | 4,92E-01 |
| AT5G18140 | 1,06 | -2,69 | 2,54 | -1,06 | 2,88E-01 |
| AT5G18170 | 0,16 | -0,07 | 4,37 | -0,02 | 9,87E-01 |
| AT5G18200 | 0,08 | 0,00 | 4,41 | 0,00 | 1,00E+00 |
| AT5G18255 | 1,25 | -2,12 | 1,66 | -1,28 | 2,02E-01 |
| AT5G18310 | 0,76 | 2,37 | 2,36 | 1,00 | 3,16E-01 |
| AT5G18390 | 0,33 | -2,10 | 4,36 | -0,48 | 6,30E-01 |
| AT5G18470 | 1,32 | -1,16 | 1,76 | -0,66 | 5,10E-01 |
| AT5G18500 | 0,37 | -1,08 | 4,34 | -0,25 | 8,03E-01 |
| AT5G18580 | 0,90 | -0,87 | 2,15 | -0,40 | 6,87E-01 |
| AT5G18630 | 0,28 | -0,07 | 4,32 | -0,02 | 9,87E-01 |
| AT5G18636 | 0,58 | -0,31 | 2,25 | -0,14 | 8,89E-01 |
| AT5G18650 | 0,68 | -1,32 | 3,01 | -0,44 | 6,60E-01 |
| AT5G18670 | 0,21 | 0,00 | 4,41 | 0,00 | 1,00E+00 |
| AT5G18750 | 0,77 | 0,00 | 3,45 | 0,00 | 9,99E-01 |
| AT5G18830 | 0,51 | 0,11 | 3,28 | 0,03 | 9,73E-01 |
| AT5G18890 | 0,98 | -3,12 | 4,29 | -0,73 | 4,67E-01 |
| AT5G18937 | 0,54 | 2,90 | 4,27 | 0,68 | 4,97E-01 |
| AT5G18940 | 1,00 | -3,77 | 2,69 | -1,40 | 1,61E-01 |
| AT5G18960 | 0,35 | -0,84 | 3,49 | -0,24 | 8,09E-01 |
| AT5G19000 | 0,44 | -2,06 | 3,34 | -0,62 | 5,36E-01 |
| AT5G19090 | 0,78 | 0,45 | 2,33 | 0,19 | 8,48E-01 |
| AT5G19180 | 0,77 | 0,25 | 2,25 | 0,11 | 9,11E-01 |
| AT5G19190 | 0,37 | -1,20 | 4,32 | -0,28 | 7,81E-01 |
| AT5G19210 | 0,49 | -0,45 | 3,25 | -0,14 | 8,90E-01 |
| AT5G19221 | 0,47 | -2,13 | 4,33 | -0,49 | 6,22E-01 |
| AT5G19240 | 0,65 | 1,74 | 2,14 | 0,81 | 4,16E-01 |
| AT5G19250 | 0,98 | 0,29 | 2,73 | 0,11 | 9,15E-01 |

| | | | | | |
|---|---|---|---|---|---|
| AT5G19290 | 0,41 | -0,28 | 3,34 | -0,08 | 9,33E-01 |
| AT5G19440 | 0,36 | -1,64 | 4,41 | -0,37 | 7,10E-01 |
| AT5G19500 | 1,08 | -2,95 | 2,27 | -1,30 | 1,93E-01 |
| AT5G19540 | 0,77 | -1,20 | 2,03 | -0,59 | 5,56E-01 |
| AT5G19550 | 0,90 | 0,54 | 2,10 | 0,26 | 7,96E-01 |
| AT5G19570 | 0,52 | -1,26 | 3,98 | -0,32 | 7,52E-01 |
| AT5G19620 | 0,50 | 1,05 | 4,01 | 0,26 | 7,94E-01 |
| AT5G19680 | 0,19 | -0,83 | 4,38 | -0,19 | 8,50E-01 |
| AT5G19750 | 0,55 | 0,00 | 4,41 | 0,00 | 1,00E+00 |
| AT5G19990 | 0,77 | -2,88 | 2,12 | -1,36 | 1,75E-01 |
| AT5G20080 | 0,48 | -0,95 | 3,28 | -0,29 | 7,73E-01 |
| AT5G20120 | 0,55 | 0,00 | 4,41 | 0,00 | 1,00E+00 |
| AT5G20130 | 0,68 | -3,76 | 3,75 | -1,00 | 3,16E-01 |
| AT5G20150 | 0,56 | -1,70 | 2,59 | -0,66 | 5,11E-01 |
| AT5G20190 | 1,21 | -0,13 | 1,90 | -0,07 | 9,46E-01 |
| AT5G20220 | 0,21 | -0,70 | 4,40 | -0,16 | 8,73E-01 |
| AT5G20410 | 0,63 | -4,46 | 3,61 | -1,23 | 2,17E-01 |
| AT5G20480 | 0,15 | 0,41 | 4,35 | 0,09 | 9,26E-01 |
| AT5G20500 | 0,33 | -0,33 | 4,34 | -0,08 | 9,40E-01 |
| AT5G20590 | 0,85 | -1,76 | 2,20 | -0,80 | 4,25E-01 |
| AT5G20660 | 1,27 | -2,79 | 1,75 | -1,60 | 1,11E-01 |
| AT5G20680 | 0,31 | 0,01 | 4,37 | 0,00 | 9,99E-01 |
| AT5G20710 | 0,12 | 0,00 | 4,41 | 0,00 | 1,00E+00 |
| AT5G20900 | 0,45 | 0,01 | 3,31 | 0,00 | 9,96E-01 |
| AT5G03255 | 0,31 | -1,69 | 4,37 | -0,39 | 6,99E-01 |
| AT5G20960 | 1,02 | 1,15 | 2,62 | 0,44 | 6,60E-01 |
| AT5G21150 | 0,12 | 0,00 | 4,41 | 0,00 | 1,00E+00 |
| AT5G21222 | 1,01 | -3,18 | 2,65 | -1,20 | 2,31E-01 |
| AT5G21326 | 0,90 | -1,11 | 2,86 | -0,39 | 6,98E-01 |
| AT5G21482 | 0,24 | 0,92 | 4,33 | 0,21 | 8,32E-01 |
| AT5G22000 | 1,27 | -2,83 | 1,67 | -1,69 | 9,07E-02 |
| AT5G22050 | 0,36 | 0,60 | 4,34 | 0,14 | 8,90E-01 |
| AT5G22070 | 0,57 | 1,75 | 2,99 | 0,58 | 5,59E-01 |
| AT5G22300 | 0,30 | 1,04 | 3,66 | 0,28 | 7,76E-01 |
| AT5G22315 | 0,20 | -1,00 | 4,37 | -0,23 | 8,19E-01 |
| AT5G22340 | 0,95 | -2,50 | 2,26 | -1,11 | 2,68E-01 |
| AT5G22420 | 0,42 | -3,38 | 4,19 | -0,81 | 4,21E-01 |
| AT5G22580 | 0,46 | -0,62 | 4,40 | -0,14 | 8,88E-01 |
| AT5G22590 | 1,03 | 2,57 | 2,52 | 1,02 | 3,07E-01 |
| AT5G22640 | 1,19 | -1,64 | 1,86 | -0,88 | 3,79E-01 |
| AT5G22650 | 0,66 | 1,80 | 2,43 | 0,74 | 4,59E-01 |
| AT5G22690 | 0,12 | 0,00 | 4,41 | 0,00 | 1,00E+00 |
| AT5G22760 | 1,25 | -3,26 | 1,83 | -1,78 | 7,56E-02 |
| AT5G22794 | 0,08 | 0,00 | 4,41 | 0,00 | 1,00E+00 |
| AT5G22830 | 0,08 | -0,21 | 4,39 | -0,05 | 9,62E-01 |
| AT5G22950 | 0,12 | -0,31 | 4,38 | -0,07 | 9,44E-01 |
| AT5G23010 | 0,84 | 0,71 | 2,09 | 0,34 | 7,35E-01 |
| AT5G23070 | 0,19 | 0,09 | 4,37 | 0,02 | 9,84E-01 |
| AT5G03575 | 0,26 | 0,00 | 4,41 | 0,00 | 1,00E+00 |
| AT5G23120 | 0,65 | -0,57 | 2,23 | -0,26 | 7,98E-01 |
| AT5G23140 | 0,95 | -3,97 | 2,22 | -1,79 | 7,40E-02 |
| AT5G23210 | 0,16 | -0,26 | 4,38 | -0,06 | 9,52E-01 |
| AT5G23280 | 1,01 | -2,26 | 2,32 | -0,97 | 3,31E-01 |
| AT5G23320 | 0,08 | 0,00 | 4,41 | 0,00 | 1,00E+00 |
| AT5G23330 | 0,43 | -3,24 | 4,37 | -0,74 | 4,58E-01 |
| AT5G23490 | 0,49 | -1,00 | 2,61 | -0,38 | 7,03E-01 |

| | | | | | |
|---|---|---|---|---|---|
| AT5G23540 | 0,96 | -0,26 | 1,82 | -0,14 | 8,85E-01 |
| AT5G23570 | 0,58 | 0,70 | 2,39 | 0,29 | 7,71E-01 |
| AT5G23575 | 1,00 | -1,67 | 1,61 | -1,04 | 3,00E-01 |
| AT5G23600 | 0,08 | 0,00 | 4,41 | 0,00 | 1,00E+00 |
| AT5G23670 | 0,72 | -2,42 | 3,06 | -0,79 | 4,30E-01 |
| AT5G23720 | 0,51 | -3,06 | 3,23 | -0,95 | 3,43E-01 |
| AT5G23730 | 0,26 | -1,26 | 4,36 | -0,29 | 7,72E-01 |
| AT5G23890 | 0,99 | -3,26 | 2,71 | -1,20 | 2,29E-01 |
| AT5G23950 | 0,35 | -1,29 | 4,31 | -0,30 | 7,65E-01 |
| AT5G24020 | 0,16 | -0,26 | 4,38 | -0,06 | 9,52E-01 |
| AT5G24210 | 0,96 | -2,07 | 2,05 | -1,01 | 3,13E-01 |
| AT5G24360 | 0,40 | -1,64 | 3,40 | -0,48 | 6,30E-01 |
| AT5G24380 | 0,41 | -1,64 | 3,44 | -0,48 | 6,33E-01 |
| AT5G24450 | 0,69 | 1,42 | 3,67 | 0,39 | 6,99E-01 |
| AT5G24460 | 0,88 | -1,58 | 1,99 | -0,79 | 4,27E-01 |
| AT5G24610 | 0,55 | -1,23 | 2,73 | -0,45 | 6,52E-01 |
| AT5G24830 | 0,08 | 0,00 | 4,41 | 0,00 | 1,00E+00 |
| AT5G24980 | 0,43 | -0,24 | 3,41 | -0,07 | 9,43E-01 |
| AT5G25040 | 0,67 | 1,66 | 2,35 | 0,71 | 4,80E-01 |
| AT5G25120 | 0,51 | 2,18 | 3,15 | 0,69 | 4,88E-01 |
| AT5G03925 | 0,17 | 1,16 | 4,31 | 0,27 | 7,88E-01 |
| AT5G25140 | 0,28 | -0,76 | 4,40 | -0,17 | 8,62E-01 |
| AT5G25150 | 0,27 | -0,62 | 4,34 | -0,14 | 8,86E-01 |
| AT5G25170 | 0,33 | -0,24 | 4,37 | -0,05 | 9,57E-01 |
| AT5G25210 | 0,42 | 0,37 | 3,35 | 0,11 | 9,13E-01 |
| AT5G25220 | 1,05 | 2,20 | 2,07 | 1,06 | 2,89E-01 |
| AT5G25260 | 1,02 | -2,60 | 2,86 | -0,91 | 3,62E-01 |
| AT5G25340 | 0,46 | -3,43 | 4,10 | -0,84 | 4,03E-01 |
| AT5G25560 | 0,53 | -3,02 | 3,16 | -0,95 | 3,40E-01 |
| AT5G25580 | 0,12 | 0,00 | 4,41 | 0,00 | 1,00E+00 |
| AT5G25630 | 0,47 | -1,27 | 3,22 | -0,40 | 6,92E-01 |
| AT5G04145 | 0,89 | -2,87 | 2,23 | -1,29 | 1,97E-01 |
| AT5G25900 | 0,33 | -2,60 | 4,41 | -0,59 | 5,55E-01 |
| AT5G25910 | 0,60 | -2,60 | 3,79 | -0,69 | 4,92E-01 |
| AT5G25930 | 0,70 | 0,81 | 2,45 | 0,33 | 7,40E-01 |
| AT5G26038 | 1,13 | 1,71 | 2,07 | 0,83 | 4,09E-01 |
| AT5G26146 | 0,89 | 1,13 | 2,30 | 0,49 | 6,24E-01 |
| AT5G26147 | 0,89 | 1,13 | 2,30 | 0,49 | 6,24E-01 |
| AT5G26200 | 0,12 | 0,00 | 4,41 | 0,00 | 1,00E+00 |
| AT5G26240 | 0,84 | -2,92 | 2,10 | -1,39 | 1,64E-01 |
| AT5G26270 | 0,26 | 0,59 | 4,35 | 0,14 | 8,92E-01 |
| AT5G26290 | 0,79 | -0,60 | 2,12 | -0,29 | 7,75E-01 |
| AT5G26740 | 0,58 | 0,11 | 2,74 | 0,04 | 9,69E-01 |
| AT5G26731 | 0,38 | 0,66 | 4,34 | 0,15 | 8,80E-01 |
| AT5G26710 | 0,73 | -2,65 | 2,56 | -1,04 | 3,00E-01 |
| AT5G26670 | 0,26 | -2,69 | 4,40 | -0,61 | 5,41E-01 |
| AT5G26610 | 0,16 | 1,16 | 4,31 | 0,27 | 7,88E-01 |
| AT5G26600 | 0,82 | -1,05 | 2,59 | -0,40 | 6,86E-01 |
| AT5G26780 | 0,39 | -0,56 | 2,86 | -0,20 | 8,44E-01 |
| AT5G26820 | 0,42 | -3,34 | 4,37 | -0,76 | 4,45E-01 |
| AT5G26850 | 1,21 | -4,76 | 2,23 | -2,13 | 3,32E-02 |
| AT5G26860 | 1,16 | -0,93 | 1,86 | -0,50 | 6,18E-01 |
| AT5G26930 | 0,33 | -3,41 | 4,38 | -0,78 | 4,35E-01 |
| AT5G26940 | 0,66 | -4,42 | 3,69 | -1,20 | 2,31E-01 |
| AT5G26990 | 0,23 | -1,14 | 4,39 | -0,26 | 7,96E-01 |
| AT5G27120 | 0,91 | -3,83 | 3,49 | -1,10 | 2,73E-01 |

| | | | | | |
|---|---|---|---|---|---|
| AT5G27150 | 0,57 | -0,50 | 3,03 | -0,16 | 8,69E-01 |
| AT5G27170 | 0,25 | 1,14 | 4,32 | 0,26 | 7,92E-01 |
| AT5G27230 | 0,08 | -0,21 | 4,39 | -0,05 | 9,62E-01 |
| AT5G27240 | 0,05 | -0,24 | 4,38 | -0,05 | 9,57E-01 |
| AT5G27270 | 0,17 | -0,29 | 4,38 | -0,07 | 9,46E-01 |
| AT5G27280 | 0,34 | -0,77 | 3,59 | -0,21 | 8,30E-01 |
| AT5G27300 | 0,08 | -0,21 | 4,39 | -0,05 | 9,62E-01 |
| AT5G27320 | 0,08 | -0,21 | 4,39 | -0,05 | 9,62E-01 |
| AT5G27340 | 0,54 | 2,34 | 3,69 | 0,63 | 5,25E-01 |
| AT5G27350 | 0,64 | 1,92 | 2,54 | 0,75 | 4,51E-01 |
| AT5G27410 | 0,42 | -1,26 | 4,07 | -0,31 | 7,56E-01 |
| AT5G27470 | 0,51 | -0,76 | 3,14 | -0,24 | 8,08E-01 |
| AT5G27530 | 0,32 | 0,89 | 4,33 | 0,21 | 8,37E-01 |
| AT5G27540 | 0,68 | -3,09 | 3,50 | -0,88 | 3,77E-01 |
| AT5G27600 | 0,18 | 0,38 | 4,35 | 0,09 | 9,30E-01 |
| AT5G27610 | 0,08 | -0,21 | 4,39 | -0,05 | 9,62E-01 |
| AT5G27630 | 0,17 | 0,78 | 4,33 | 0,18 | 8,58E-01 |
| AT5G27650 | 0,94 | -2,93 | 2,69 | -1,09 | 2,76E-01 |
| AT5G27710 | 0,67 | 0,24 | 2,45 | 0,10 | 9,21E-01 |
| AT5G27730 | 0,69 | 2,09 | 2,13 | 0,98 | 3,27E-01 |
| AT5G27760 | 0,27 | -1,08 | 4,39 | -0,25 | 8,06E-01 |
| AT5G27770 | 0,86 | -3,14 | 2,34 | -1,34 | 1,80E-01 |
| AT5G27800 | 0,16 | 0,75 | 4,34 | 0,17 | 8,63E-01 |
| AT5G27830 | 0,23 | -1,72 | 4,40 | -0,39 | 6,95E-01 |
| AT5G27890 | 0,40 | 1,64 | 4,30 | 0,38 | 7,03E-01 |
| AT5G27893 | 1,07 | -0,35 | 1,80 | -0,19 | 8,47E-01 |
| AT5G28010 | 1,26 | 0,36 | 1,61 | 0,22 | 8,24E-01 |
| AT5G04625 | 0,10 | 0,45 | 4,35 | 0,10 | 9,19E-01 |
| AT5G04665 | 0,11 | 0,46 | 4,35 | 0,11 | 9,15E-01 |
| AT5G28090 | 0,29 | 0,96 | 4,33 | 0,22 | 8,24E-01 |
| AT5G04675 | 0,19 | 1,18 | 4,31 | 0,27 | 7,84E-01 |
| AT5G28320 | 0,89 | -3,37 | 2,75 | -1,22 | 2,21E-01 |
| AT5G28350 | 0,24 | -1,38 | 4,38 | -0,32 | 7,53E-01 |
| AT5G28410 | 0,36 | -1,64 | 4,35 | -0,38 | 7,06E-01 |
| AT5G28463 | 0,37 | 1,44 | 4,26 | 0,34 | 7,35E-01 |
| AT5G28500 | 0,70 | 0,44 | 2,58 | 0,17 | 8,64E-01 |
| AT5G04815 | 0,05 | -0,24 | 4,38 | -0,05 | 9,57E-01 |
| AT5G28810 | 0,66 | 2,78 | 2,77 | 1,01 | 3,15E-01 |
| AT5G28830 | 0,05 | -0,24 | 4,38 | -0,05 | 9,57E-01 |
| AT5G28900 | 0,40 | -1,91 | 4,31 | -0,44 | 6,58E-01 |
| AT5G29624 | 0,12 | 0,00 | 4,41 | 0,00 | 1,00E+00 |
| AT5G30341 | 0,12 | 0,07 | 4,38 | 0,02 | 9,87E-01 |
| AT5G04955 | 1,06 | 3,87 | 3,80 | 1,02 | 3,07E-01 |
| AT5G32613 | 0,26 | 1,28 | 4,31 | 0,30 | 7,66E-01 |
| AT5G33290 | 0,62 | -3,44 | 3,85 | -0,89 | 3,71E-01 |
| AT5G33320 | 0,61 | -1,29 | 2,27 | -0,57 | 5,70E-01 |
| AT5G33390 | 0,38 | 0,00 | 4,41 | 0,00 | 1,00E+00 |
| AT5G34869 | 0,19 | 0,02 | 4,36 | 0,00 | 9,97E-01 |
| AT5G34871 | 0,12 | -0,68 | 4,41 | -0,15 | 8,77E-01 |
| AT5G34870 | 0,38 | 0,00 | 3,46 | 0,00 | 9,99E-01 |
| AT5G35170 | 0,47 | -0,34 | 2,45 | -0,14 | 8,89E-01 |
| AT5G35180 | 0,52 | -0,57 | 2,67 | -0,21 | 8,31E-01 |
| AT5G35190 | 0,84 | 2,21 | 2,73 | 0,81 | 4,19E-01 |
| AT5G35300 | 0,23 | 1,14 | 4,31 | 0,26 | 7,91E-01 |
| AT5G35320 | 0,21 | -1,17 | 4,36 | -0,27 | 7,88E-01 |
| AT5G35430 | 1,24 | -4,26 | 2,23 | -1,91 | 5,59E-02 |

| | | | | | |
|---|---|---|---|---|---|
| AT5G35450 | 0,40 | 0,00 | 4,41 | 0,00 | 1,00E+00 |
| AT5G05195 | 0,23 | 0,77 | 4,34 | 0,18 | 8,59E-01 |
| AT5G35730 | 0,82 | -4,23 | 2,73 | -1,55 | 1,22E-01 |
| AT5G35917 | 0,29 | 1,54 | 4,30 | 0,36 | 7,21E-01 |
| AT5G35926 | 1,15 | 0,36 | 1,80 | 0,20 | 8,43E-01 |
| AT5G35980 | 1,15 | 0,24 | 1,83 | 0,13 | 8,98E-01 |
| AT5G36001 | 0,12 | 0,00 | 4,41 | 0,00 | 1,00E+00 |
| AT5G36290 | 0,47 | 0,29 | 2,66 | 0,11 | 9,12E-01 |
| AT5G36300 | 0,87 | 2,20 | 2,34 | 0,94 | 3,46E-01 |
| AT5G36740 | 0,08 | 0,00 | 4,41 | 0,00 | 1,00E+00 |
| AT5G36870 | 1,07 | 0,87 | 2,04 | 0,43 | 6,69E-01 |
| AT5G36900 | 0,54 | 0,89 | 3,11 | 0,29 | 7,75E-01 |
| AT5G36940 | 0,51 | -0,76 | 3,96 | -0,19 | 8,48E-01 |
| AT5G37020 | 0,62 | 1,07 | 2,61 | 0,41 | 6,82E-01 |
| AT5G37070 | 0,43 | -3,00 | 4,38 | -0,68 | 4,94E-01 |
| AT5G37130 | 0,66 | -0,82 | 2,61 | -0,31 | 7,55E-01 |
| AT5G37150 | 0,98 | 2,30 | 2,18 | 1,05 | 2,92E-01 |
| AT5G37540 | 0,17 | 0,00 | 4,41 | 0,00 | 1,00E+00 |
| AT5G37630 | 0,94 | -2,37 | 2,36 | -1,00 | 3,16E-01 |
| AT5G37760 | 0,15 | -0,73 | 4,40 | -0,17 | 8,68E-01 |
| AT5G37793 | 0,51 | 0,69 | 2,66 | 0,26 | 7,95E-01 |
| AT5G37850 | 0,30 | -2,60 | 4,41 | -0,59 | 5,55E-01 |
| AT5G37950 | 0,11 | 0,46 | 4,35 | 0,11 | 9,15E-01 |
| AT5G38155 | 0,12 | -0,31 | 4,38 | -0,07 | 9,44E-01 |
| AT5G38210 | 1,00 | 2,76 | 2,13 | 1,29 | 1,96E-01 |
| AT5G38380 | 0,15 | -0,14 | 4,39 | -0,03 | 9,74E-01 |
| AT5G38470 | 0,73 | -2,58 | 3,64 | -0,71 | 4,78E-01 |
| AT5G38480 | 1,20 | -2,17 | 1,64 | -1,33 | 1,84E-01 |
| AT5G38560 | 0,34 | -0,18 | 3,48 | -0,05 | 9,60E-01 |
| AT5G38910 | 0,16 | 0,61 | 4,34 | 0,14 | 8,89E-01 |
| AT5G38980 | 0,57 | 0,74 | 2,66 | 0,28 | 7,81E-01 |
| AT5G39000 | 0,70 | -2,87 | 4,30 | -0,67 | 5,05E-01 |
| AT5G39030 | 0,81 | -2,28 | 2,88 | -0,79 | 4,30E-01 |
| AT5G39040 | 0,29 | 0,00 | 4,41 | 0,00 | 1,00E+00 |
| AT5G39250 | 0,34 | -1,27 | 4,34 | -0,29 | 7,70E-01 |
| AT5G39320 | 0,82 | -3,96 | 4,34 | -0,91 | 3,61E-01 |
| AT5G39350 | 0,21 | 0,17 | 4,36 | 0,04 | 9,68E-01 |
| AT5G39510 | 0,77 | 1,08 | 2,30 | 0,47 | 6,38E-01 |
| AT5G39590 | 0,82 | 0,56 | 2,37 | 0,23 | 8,15E-01 |
| AT5G39660 | 0,40 | -0,68 | 4,41 | -0,15 | 8,77E-01 |
| AT5G39670 | 0,40 | 0,86 | 4,17 | 0,21 | 8,36E-01 |
| AT5G39690 | 1,03 | 0,45 | 2,06 | 0,22 | 8,28E-01 |
| AT5G39710 | 0,25 | -2,57 | 4,40 | -0,58 | 5,60E-01 |
| AT5G39730 | 0,19 | 0,65 | 4,34 | 0,15 | 8,82E-01 |
| AT5G39775 | 0,53 | 0,48 | 4,13 | 0,12 | 9,07E-01 |
| AT5G39850 | 0,08 | 0,00 | 4,41 | 0,00 | 1,00E+00 |
| AT5G39960 | 0,21 | -2,60 | 4,41 | -0,59 | 5,55E-01 |
| AT5G39990 | 0,08 | 0,00 | 4,41 | 0,00 | 1,00E+00 |
| AT5G40140 | 0,31 | -0,47 | 3,62 | -0,13 | 8,96E-01 |
| AT5G40200 | 0,62 | -0,05 | 2,76 | -0,02 | 9,86E-01 |
| AT5G40380 | 0,46 | 1,91 | 3,92 | 0,49 | 6,27E-01 |
| AT5G40650 | 0,74 | -3,99 | 2,75 | -1,45 | 1,47E-01 |
| AT5G40670 | 1,01 | -1,95 | 2,22 | -0,87 | 3,82E-01 |
| AT5G40680 | 0,17 | 0,00 | 4,41 | 0,00 | 1,00E+00 |
| AT5G40690 | 0,62 | -1,46 | 3,74 | -0,39 | 6,97E-01 |
| AT5G40720 | 0,32 | 1,55 | 4,30 | 0,36 | 7,18E-01 |

| | | | | | |
|---|---|---|---|---|---|
| AT5G40760 | 1,16 | -2,74 | 1,94 | -1,41 | 1,58E-01 |
| AT5G40850 | 0,08 | 0,05 | 4,38 | 0,01 | 9,91E-01 |
| AT5G40870 | 0,14 | 0,40 | 4,35 | 0,09 | 9,27E-01 |
| AT5G40880 | 0,51 | -1,62 | 4,03 | -0,40 | 6,87E-01 |
| AT5G40900 | 0,15 | -0,17 | 4,39 | -0,04 | 9,68E-01 |
| AT5G41010 | 0,28 | 0,70 | 4,34 | 0,16 | 8,72E-01 |
| AT5G06115 | 0,20 | 0,47 | 4,36 | 0,11 | 9,14E-01 |
| AT5G41070 | 0,20 | -1,00 | 4,37 | -0,23 | 8,19E-01 |
| AT5G41100 | 0,28 | -0,25 | 4,33 | -0,06 | 9,54E-01 |
| AT5G41110 | 0,22 | -0,80 | 4,35 | -0,18 | 8,54E-01 |
| AT5G41140 | 0,59 | -3,20 | 3,76 | -0,85 | 3,94E-01 |
| AT5G41260 | 0,61 | -2,00 | 2,18 | -0,92 | 3,60E-01 |
| AT5G41360 | 0,66 | -2,42 | 3,70 | -0,65 | 5,14E-01 |
| AT5G41370 | 0,15 | 0,48 | 4,35 | 0,11 | 9,12E-01 |
| AT5G41410 | 1,13 | -0,38 | 1,79 | -0,21 | 8,34E-01 |
| AT5G41500 | 1,05 | 3,26 | 2,04 | 1,59 | 1,11E-01 |
| AT5G41660 | 0,71 | 2,85 | 2,90 | 0,98 | 3,27E-01 |
| AT5G41700 | 1,24 | -1,97 | 1,93 | -1,02 | 3,08E-01 |
| AT5G41761 | 0,83 | 1,89 | 2,25 | 0,84 | 4,00E-01 |
| AT5G41770 | 0,61 | 0,32 | 2,65 | 0,12 | 9,05E-01 |
| AT5G41860 | 0,77 | -0,19 | 4,39 | -0,04 | 9,65E-01 |
| AT5G41920 | 0,61 | -0,27 | 4,32 | -0,06 | 9,50E-01 |
| AT5G41940 | 0,98 | -2,50 | 2,11 | -1,19 | 2,36E-01 |
| AT5G41960 | 0,15 | 0,00 | 4,41 | 0,00 | 1,00E+00 |
| AT5G41970 | 0,85 | -4,76 | 3,34 | -1,42 | 1,54E-01 |
| AT5G42010 | 0,60 | -0,95 | 3,18 | -0,30 | 7,66E-01 |
| AT5G42220 | 0,70 | -0,49 | 2,26 | -0,21 | 8,30E-01 |
| AT5G42250 | 0,33 | -2,59 | 4,41 | -0,59 | 5,56E-01 |
| AT5G42260 | 0,28 | 1,92 | 4,29 | 0,45 | 6,54E-01 |
| AT5G42290 | 0,08 | 0,00 | 4,41 | 0,00 | 1,00E+00 |
| AT5G42310 | 0,50 | -0,38 | 4,06 | -0,09 | 9,26E-01 |
| AT5G42350 | 0,37 | -2,34 | 4,36 | -0,54 | 5,91E-01 |
| AT5G42420 | 0,74 | -3,78 | 3,64 | -1,04 | 2,99E-01 |
| AT5G42460 | 0,33 | -3,08 | 4,39 | -0,70 | 4,83E-01 |
| AT5G42470 | 0,22 | -0,26 | 4,38 | -0,06 | 9,53E-01 |
| AT5G42480 | 0,45 | -3,07 | 4,39 | -0,70 | 4,83E-01 |
| AT5G42504 | 0,17 | -0,68 | 4,41 | -0,15 | 8,77E-01 |
| AT5G42520 | 0,71 | -1,89 | 2,33 | -0,81 | 4,18E-01 |
| AT5G42540 | 1,02 | -3,03 | 2,40 | -1,26 | 2,06E-01 |
| AT5G42620 | 0,80 | -1,42 | 2,36 | -0,60 | 5,48E-01 |
| AT5G42640 | 0,08 | -0,21 | 4,39 | -0,05 | 9,62E-01 |
| AT5G42670 | 0,16 | 0,84 | 4,33 | 0,20 | 8,45E-01 |
| AT5G42680 | 0,80 | 0,89 | 3,19 | 0,28 | 7,79E-01 |
| AT5G42720 | 0,19 | -0,03 | 4,38 | -0,01 | 9,94E-01 |
| AT5G42740 | 1,16 | 0,11 | 2,93 | 0,04 | 9,71E-01 |
| AT5G42760 | 0,12 | -0,14 | 4,38 | -0,03 | 9,74E-01 |
| AT5G42825 | 0,29 | -2,60 | 4,41 | -0,59 | 5,55E-01 |
| AT5G42965 | 0,19 | -0,68 | 4,41 | -0,15 | 8,77E-01 |
| AT5G42980 | 0,59 | -0,09 | 2,71 | -0,03 | 9,75E-01 |
| AT5G43100 | 0,78 | -1,33 | 2,16 | -0,61 | 5,39E-01 |
| AT5G43150 | 0,61 | -2,66 | 3,04 | -0,87 | 3,82E-01 |
| AT5G06505 | 0,36 | -0,38 | 4,35 | -0,09 | 9,30E-01 |
| AT5G43460 | 1,01 | -3,83 | 1,98 | -1,93 | 5,35E-02 |
| AT5G43525 | 0,69 | 2,67 | 2,74 | 0,97 | 3,30E-01 |
| AT5G43560 | 0,55 | -0,15 | 3,14 | -0,05 | 9,62E-01 |
| AT5G43610 | 1,12 | 0,03 | 2,17 | 0,01 | 9,91E-01 |

| | | | | | |
|---|---|---|---|---|---|
| AT5G43620 | 0,36 | -0,80 | 4,35 | -0,18 | 8,54E-01 |
| AT5G43700 | 0,52 | -3,05 | 3,17 | -0,96 | 3,36E-01 |
| AT5G43810 | 1,04 | -1,73 | 2,70 | -0,64 | 5,22E-01 |
| AT5G43930 | 0,73 | -1,32 | 2,51 | -0,52 | 6,00E-01 |
| AT5G43940 | 1,17 | 0,49 | 1,94 | 0,25 | 8,01E-01 |
| AT5G44020 | 1,03 | 0,40 | 2,05 | 0,20 | 8,44E-01 |
| AT5G44060 | 0,47 | 0,33 | 2,74 | 0,12 | 9,03E-01 |
| AT5G44070 | 1,32 | 0,53 | 1,68 | 0,31 | 7,54E-01 |
| AT5G44130 | 0,80 | -1,64 | 2,97 | -0,55 | 5,80E-01 |
| AT5G44230 | 0,12 | 0,00 | 4,41 | 0,00 | 1,00E+00 |
| AT5G44240 | 1,02 | -2,98 | 1,88 | -1,59 | 1,12E-01 |
| AT5G06565 | 0,63 | 1,46 | 2,66 | 0,55 | 5,83E-01 |
| AT5G44286 | 0,85 | 0,55 | 2,15 | 0,26 | 7,98E-01 |
| AT5G44530 | 1,17 | -3,53 | 2,72 | -1,30 | 1,95E-01 |
| AT5G44574 | 0,91 | 0,51 | 3,08 | 0,16 | 8,70E-01 |
| AT5G44580 | 0,52 | -2,28 | 3,91 | -0,58 | 5,60E-01 |
| AT5G06605 | 1,31 | -2,62 | 1,57 | -1,66 | 9,60E-02 |
| AT5G44650 | 0,46 | -2,60 | 4,11 | -0,63 | 5,26E-01 |
| AT5G44720 | 0,16 | -0,01 | 4,37 | 0,00 | 9,99E-01 |
| AT5G44750 | 0,74 | -2,60 | 2,30 | -1,13 | 2,59E-01 |
| AT5G44760 | 0,36 | 0,00 | 4,41 | 0,00 | 1,00E+00 |
| AT5G44770 | 0,22 | 0,78 | 4,34 | 0,18 | 8,58E-01 |
| AT5G45000 | 0,38 | -2,17 | 4,33 | -0,50 | 6,16E-01 |
| AT5G06695 | 0,59 | -3,60 | 3,09 | -1,17 | 2,44E-01 |
| AT5G45030 | 0,59 | -3,60 | 3,09 | -1,17 | 2,44E-01 |
| AT5G45130 | 0,78 | -1,57 | 3,08 | -0,51 | 6,11E-01 |
| AT5G45150 | 0,08 | 0,00 | 4,41 | 0,00 | 1,00E+00 |
| AT5G45240 | 0,12 | 0,00 | 4,41 | 0,00 | 1,00E+00 |
| AT5G45260 | 0,75 | -3,23 | 2,88 | -1,12 | 2,62E-01 |
| AT5G45390 | 0,62 | 0,28 | 2,27 | 0,12 | 9,01E-01 |
| AT5G45400 | 0,76 | -3,20 | 2,50 | -1,28 | 2,00E-01 |
| AT5G45450 | 0,08 | 0,00 | 4,41 | 0,00 | 1,00E+00 |
| AT5G45475 | 0,74 | 0,42 | 2,26 | 0,19 | 8,51E-01 |
| AT5G45490 | 1,25 | 0,82 | 1,77 | 0,47 | 6,41E-01 |
| AT5G45570 | 0,68 | 2,55 | 2,38 | 1,07 | 2,85E-01 |
| AT5G45620 | 0,59 | 0,83 | 3,10 | 0,27 | 7,89E-01 |
| AT5G45670 | 0,10 | 0,45 | 4,35 | 0,10 | 9,19E-01 |
| AT5G45680 | 0,26 | -0,77 | 4,37 | -0,18 | 8,59E-01 |
| AT5G45710 | 0,55 | -2,93 | 3,79 | -0,77 | 4,39E-01 |
| AT5G45750 | 0,19 | 1,18 | 4,31 | 0,27 | 7,84E-01 |
| AT5G45770 | 0,12 | 0,00 | 4,41 | 0,00 | 1,00E+00 |
| AT5G45920 | 1,21 | 0,22 | 2,11 | 0,10 | 9,18E-01 |
| AT5G46330 | 0,47 | -0,98 | 4,02 | -0,24 | 8,08E-01 |
| AT5G46410 | 0,44 | -1,33 | 4,36 | -0,30 | 7,61E-01 |
| AT5G06825 | 0,67 | 2,03 | 2,95 | 0,69 | 4,90E-01 |
| AT5G46460 | 0,90 | -3,05 | 3,35 | -0,91 | 3,62E-01 |
| AT5G46500 | 1,11 | -1,71 | 2,59 | -0,66 | 5,08E-01 |
| AT5G06835 | 1,13 | -2,60 | 2,64 | -0,99 | 3,24E-01 |
| AT5G46630 | 1,04 | -2,70 | 2,04 | -1,32 | 1,87E-01 |
| AT5G46660 | 1,02 | 3,66 | 3,97 | 0,92 | 3,57E-01 |
| AT5G46670 | 0,74 | 3,15 | 2,67 | 1,18 | 2,38E-01 |
| AT5G46730 | 0,08 | -0,21 | 4,39 | -0,05 | 9,62E-01 |
| AT5G46780 | 0,35 | 0,27 | 4,35 | 0,06 | 9,51E-01 |
| AT5G46825 | 1,18 | 1,61 | 1,68 | 0,96 | 3,37E-01 |
| AT5G46840 | 0,62 | -1,94 | 3,11 | -0,62 | 5,33E-01 |
| AT5G46850 | 0,29 | -1,64 | 4,37 | -0,38 | 7,08E-01 |

| | | | | | |
|---|---|---|---|---|---|
| AT5G47030 | 0,63 | -2,60 | 3,81 | -0,68 | 4,94E-01 |
| AT5G47120 | 1,06 | 2,70 | 2,04 | 1,32 | 1,86E-01 |
| AT5G47130 | 0,25 | -0,06 | 4,37 | -0,01 | 9,89E-01 |
| AT5G47140 | 0,26 | 0,47 | 4,36 | 0,11 | 9,14E-01 |
| AT5G47200 | 0,34 | -0,62 | 4,40 | -0,14 | 8,87E-01 |
| AT5G47230 | 0,27 | 1,50 | 4,31 | 0,35 | 7,27E-01 |
| AT5G47240 | 0,14 | 0,87 | 4,33 | 0,20 | 8,40E-01 |
| AT5G47280 | 1,15 | 3,05 | 1,90 | 1,61 | 1,08E-01 |
| AT5G47320 | 0,19 | 0,75 | 4,34 | 0,17 | 8,62E-01 |
| AT5G47390 | 0,64 | -0,87 | 2,00 | -0,43 | 6,64E-01 |
| AT5G47420 | 0,50 | -3,97 | 3,97 | -1,00 | 3,17E-01 |
| AT5G47640 | 0,70 | -1,69 | 2,22 | -0,76 | 4,46E-01 |
| AT5G47660 | 0,08 | -0,21 | 4,39 | -0,05 | 9,62E-01 |
| AT5G47720 | 0,16 | -0,64 | 4,40 | -0,15 | 8,85E-01 |
| AT5G47770 | 0,39 | 0,97 | 4,25 | 0,23 | 8,20E-01 |
| AT5G47780 | 0,52 | -0,21 | 2,69 | -0,08 | 9,39E-01 |
| AT5G47790 | 0,28 | 0,00 | 4,41 | 0,00 | 1,00E+00 |
| AT5G47800 | 0,24 | -2,59 | 4,41 | -0,59 | 5,56E-01 |
| AT5G47820 | 0,70 | -3,29 | 3,65 | -0,90 | 3,68E-01 |
| AT5G47840 | 0,45 | 0,62 | 2,65 | 0,24 | 8,14E-01 |
| AT5G47860 | 0,53 | -0,23 | 3,92 | -0,06 | 9,53E-01 |
| AT5G47890 | 1,07 | -4,09 | 2,08 | -1,97 | 4,86E-02 |
| AT5G48220 | 0,16 | -0,68 | 4,41 | -0,15 | 8,77E-01 |
| AT5G48290 | 0,36 | -3,60 | 4,36 | -0,82 | 4,10E-01 |
| AT5G48300 | 1,23 | 0,43 | 2,00 | 0,21 | 8,31E-01 |
| AT5G48340 | 0,30 | -2,60 | 4,41 | -0,59 | 5,55E-01 |
| AT5G48360 | 0,08 | -0,21 | 4,39 | -0,05 | 9,62E-01 |
| AT5G48390 | 0,69 | -4,02 | 3,52 | -1,14 | 2,53E-01 |
| AT5G48400 | 0,56 | -2,60 | 3,88 | -0,67 | 5,02E-01 |
| AT5G48410 | 0,12 | 0,00 | 4,41 | 0,00 | 1,00E+00 |
| AT5G07105 | 0,86 | -0,19 | 1,92 | -0,10 | 9,22E-01 |
| AT5G48545 | 0,30 | 1,17 | 3,64 | 0,32 | 7,49E-01 |
| AT5G48720 | 0,16 | -0,62 | 4,40 | -0,14 | 8,87E-01 |
| AT5G48730 | 0,12 | 0,00 | 4,41 | 0,00 | 1,00E+00 |
| AT5G48775 | 0,08 | -0,21 | 4,39 | -0,05 | 9,62E-01 |
| AT5G48800 | 0,90 | -2,26 | 2,96 | -0,76 | 4,45E-01 |
| AT5G48900 | 0,13 | -0,18 | 4,39 | -0,04 | 9,67E-01 |
| AT5G49010 | 1,17 | -0,50 | 1,85 | -0,27 | 7,88E-01 |
| AT5G49020 | 0,22 | 0,38 | 4,35 | 0,09 | 9,31E-01 |
| AT5G49060 | 0,17 | -0,71 | 4,40 | -0,16 | 8,72E-01 |
| AT5G49160 | 0,93 | -1,71 | 2,45 | -0,70 | 4,86E-01 |
| AT5G49215 | 0,08 | -0,21 | 4,39 | -0,05 | 9,62E-01 |
| AT5G49250 | 0,66 | 2,58 | 2,73 | 0,95 | 3,44E-01 |
| AT5G49260 | 0,08 | -0,21 | 4,39 | -0,05 | 9,62E-01 |
| AT5G49450 | 0,19 | -0,76 | 4,40 | -0,17 | 8,63E-01 |
| AT5G49460 | 0,62 | -3,06 | 2,98 | -1,03 | 3,04E-01 |
| AT5G49470 | 0,29 | -1,64 | 4,37 | -0,38 | 7,08E-01 |
| AT5G49480 | 1,15 | -3,51 | 2,17 | -1,62 | 1,06E-01 |
| AT5G49510 | 0,19 | 0,05 | 4,37 | 0,01 | 9,91E-01 |
| AT5G49520 | 0,12 | -0,14 | 4,38 | -0,03 | 9,74E-01 |
| AT5G49530 | 0,46 | -0,23 | 4,12 | -0,06 | 9,56E-01 |
| AT5G49590 | 0,86 | 1,05 | 2,74 | 0,38 | 7,01E-01 |
| AT5G49640 | 0,08 | 0,00 | 4,41 | 0,00 | 1,00E+00 |
| AT5G49650 | 0,61 | -0,46 | 2,33 | -0,20 | 8,44E-01 |
| AT5G49840 | 0,10 | -0,18 | 4,38 | -0,04 | 9,67E-01 |
| AT5G49945 | 0,51 | 0,00 | 3,93 | 0,00 | 1,00E+00 |

| | | | | | |
|---|---|---|---|---|---|
| AT5G49970 | 0,29 | 0,07 | 4,36 | 0,02 | 9,87E-01 |
| AT5G50020 | 0,47 | -3,81 | 4,35 | -0,88 | 3,81E-01 |
| AT5G50160 | 1,06 | 0,74 | 2,30 | 0,32 | 7,46E-01 |
| AT5G50170 | 0,08 | 0,00 | 4,41 | 0,00 | 1,00E+00 |
| AT5G50200 | 0,08 | -0,21 | 4,39 | -0,05 | 9,62E-01 |
| AT5G07605 | 0,25 | 0,93 | 4,33 | 0,21 | 8,30E-01 |
| AT5G50310 | 0,21 | -0,70 | 4,40 | -0,16 | 8,73E-01 |
| AT5G50580 | 0,65 | -3,04 | 3,69 | -0,82 | 4,11E-01 |
| AT5G50680 | 1,09 | 0,73 | 2,34 | 0,31 | 7,53E-01 |
| AT5G50717 | 0,11 | 0,46 | 4,35 | 0,11 | 9,15E-01 |
| AT5G50780 | 0,59 | -3,01 | 4,38 | -0,69 | 4,92E-01 |
| AT5G50850 | 0,82 | 0,05 | 2,42 | 0,02 | 9,85E-01 |
| AT5G50960 | 0,50 | -3,03 | 3,96 | -0,76 | 4,44E-01 |
| AT5G51010 | 0,08 | 0,00 | 4,41 | 0,00 | 1,00E+00 |
| AT5G51050 | 0,20 | -0,68 | 4,41 | -0,15 | 8,77E-01 |
| AT5G51055 | 0,34 | 1,27 | 3,49 | 0,36 | 7,16E-01 |
| AT5G51120 | 0,08 | 0,00 | 4,41 | 0,00 | 1,00E+00 |
| AT5G51150 | 0,28 | -1,75 | 4,39 | -0,40 | 6,90E-01 |
| AT5G51180 | 0,38 | -0,41 | 2,88 | -0,14 | 8,87E-01 |
| AT5G51200 | 1,04 | -0,45 | 2,21 | -0,20 | 8,38E-01 |
| AT5G51230 | 0,21 | -0,07 | 4,37 | -0,02 | 9,87E-01 |
| AT5G51280 | 0,48 | 0,20 | 2,64 | 0,07 | 9,40E-01 |
| AT5G51340 | 0,39 | -1,39 | 3,44 | -0,41 | 6,85E-01 |
| AT5G51450 | 0,53 | -2,66 | 4,00 | -0,66 | 5,06E-01 |
| AT5G51560 | 0,28 | -2,28 | 4,37 | -0,52 | 6,01E-01 |
| AT5G51570 | 0,40 | 1,19 | 3,37 | 0,35 | 7,24E-01 |
| AT5G51670 | 0,12 | -0,68 | 4,41 | -0,15 | 8,77E-01 |
| AT5G51680 | 0,64 | -2,14 | 2,56 | -0,84 | 4,03E-01 |
| AT5G51750 | 0,75 | 0,55 | 3,63 | 0,15 | 8,79E-01 |
| AT5G51830 | 0,27 | 0,78 | 4,33 | 0,18 | 8,57E-01 |
| AT5G51960 | 0,11 | 0,46 | 4,35 | 0,11 | 9,15E-01 |
| AT5G51970 | 1,04 | 0,38 | 1,73 | 0,22 | 8,25E-01 |
| AT5G52050 | 0,61 | 1,17 | 2,48 | 0,47 | 6,35E-01 |
| AT5G52115 | 1,12 | 3,15 | 2,54 | 1,24 | 2,15E-01 |
| AT5G52240 | 1,16 | -1,02 | 2,11 | -0,49 | 6,27E-01 |
| AT5G52280 | 0,34 | -1,73 | 4,34 | -0,40 | 6,90E-01 |
| AT5G52510 | 0,69 | -0,68 | 3,12 | -0,22 | 8,28E-01 |
| AT5G52520 | 0,23 | -0,48 | 4,34 | -0,11 | 9,12E-01 |
| AT5G52580 | 0,50 | 1,22 | 3,22 | 0,38 | 7,06E-01 |
| AT5G52630 | 0,08 | 0,00 | 4,41 | 0,00 | 1,00E+00 |
| AT5G52700 | 0,14 | 0,87 | 4,33 | 0,20 | 8,40E-01 |
| AT5G52750 | 0,33 | 0,95 | 4,33 | 0,22 | 8,27E-01 |
| AT5G52760 | 0,52 | 0,90 | 2,56 | 0,35 | 7,24E-01 |
| AT5G07945 | 0,52 | 1,69 | 3,92 | 0,43 | 6,66E-01 |
| AT5G52940 | 0,11 | -0,80 | 4,39 | -0,18 | 8,56E-01 |
| AT5G53020 | 0,27 | 0,36 | 4,36 | 0,08 | 9,35E-01 |
| AT5G53030 | 0,24 | -0,11 | 4,38 | -0,03 | 9,80E-01 |
| AT5G53048 | 0,25 | -0,03 | 4,38 | -0,01 | 9,94E-01 |
| AT5G53060 | 0,48 | 0,02 | 4,01 | 0,00 | 9,97E-01 |
| AT5G53080 | 0,12 | 0,00 | 4,41 | 0,00 | 1,00E+00 |
| AT5G53110 | 0,31 | 0,00 | 4,41 | 0,00 | 1,00E+00 |
| AT5G53120 | 0,82 | -0,82 | 2,50 | -0,33 | 7,43E-01 |
| AT5G53150 | 0,95 | -2,77 | 2,10 | -1,32 | 1,88E-01 |
| AT5G53160 | 1,14 | 1,52 | 1,86 | 0,82 | 4,12E-01 |
| AT5G53220 | 0,48 | -0,49 | 3,11 | -0,16 | 8,75E-01 |
| AT5G53280 | 0,44 | -2,96 | 4,12 | -0,72 | 4,72E-01 |

| | | | | | |
|---|---|---|---|---|---|
| AT5G53350 | 0,54 | -1,28 | 3,24 | -0,40 | 6,92E-01 |
| AT5G53360 | 0,08 | 0,00 | 4,41 | 0,00 | 1,00E+00 |
| AT5G53410 | 1,13 | 1,25 | 1,88 | 0,67 | 5,06E-01 |
| AT5G53420 | 0,48 | -1,85 | 3,26 | -0,57 | 5,72E-01 |
| AT5G53430 | 1,00 | -1,67 | 1,88 | -0,89 | 3,75E-01 |
| AT5G53450 | 0,71 | -2,38 | 2,52 | -0,94 | 3,45E-01 |
| AT5G53550 | 0,99 | -1,15 | 2,31 | -0,50 | 6,18E-01 |
| AT5G53590 | 0,48 | -2,60 | 4,08 | -0,64 | 5,24E-01 |
| AT5G53620 | 1,09 | -2,72 | 1,73 | -1,57 | 1,17E-01 |
| AT5G53860 | 0,25 | 1,17 | 3,76 | 0,31 | 7,56E-01 |
| AT5G53890 | 0,71 | 0,58 | 2,51 | 0,23 | 8,16E-01 |
| AT5G54035 | 0,15 | 1,00 | 4,32 | 0,23 | 8,17E-01 |
| AT5G54043 | 0,22 | 1,04 | 4,32 | 0,24 | 8,10E-01 |
| AT5G54080 | 0,99 | -1,71 | 2,23 | -0,77 | 4,43E-01 |
| AT5G54110 | 0,33 | -2,60 | 4,41 | -0,59 | 5,55E-01 |
| AT5G54140 | 0,24 | -2,60 | 4,41 | -0,59 | 5,55E-01 |
| AT5G54145 | 0,12 | 0,75 | 4,34 | 0,17 | 8,63E-01 |
| AT5G54160 | 0,46 | -0,03 | 4,16 | -0,01 | 9,94E-01 |
| AT5G54180 | 0,12 | 0,18 | 4,36 | 0,04 | 9,68E-01 |
| AT5G54290 | 0,66 | -1,03 | 2,58 | -0,40 | 6,90E-01 |
| AT5G54500 | 0,31 | 0,21 | 4,37 | 0,05 | 9,62E-01 |
| AT5G54540 | 0,58 | -0,54 | 2,69 | -0,20 | 8,40E-01 |
| AT5G54600 | 0,87 | -1,67 | 2,05 | -0,81 | 4,16E-01 |
| AT5G08225 | 0,36 | -0,20 | 4,39 | -0,05 | 9,64E-01 |
| AT5G54640 | 0,05 | -0,24 | 4,38 | -0,05 | 9,57E-01 |
| AT5G54680 | 0,72 | -0,85 | 2,45 | -0,35 | 7,29E-01 |
| AT5G54750 | 0,55 | -0,18 | 3,94 | -0,05 | 9,63E-01 |
| AT5G54760 | 0,49 | 1,15 | 2,72 | 0,42 | 6,71E-01 |
| AT5G54800 | 0,08 | 0,05 | 4,38 | 0,01 | 9,91E-01 |
| AT5G54830 | 0,76 | -1,72 | 2,88 | -0,60 | 5,51E-01 |
| AT5G54855 | 0,18 | 0,00 | 4,41 | 0,00 | 1,00E+00 |
| AT5G54860 | 0,26 | -2,60 | 4,41 | -0,59 | 5,55E-01 |
| AT5G54900 | 0,82 | -2,79 | 2,47 | -1,13 | 2,59E-01 |
| AT5G54920 | 0,83 | 0,00 | 4,41 | 0,00 | 1,00E+00 |
| AT5G55060 | 0,14 | 0,40 | 4,35 | 0,09 | 9,27E-01 |
| AT5G55070 | 1,01 | -1,30 | 1,92 | -0,68 | 4,98E-01 |
| AT5G55190 | 0,80 | -1,25 | 2,38 | -0,53 | 5,99E-01 |
| AT5G55220 | 0,88 | -0,87 | 2,56 | -0,34 | 7,34E-01 |
| AT5G55230 | 0,95 | -1,07 | 1,85 | -0,58 | 5,63E-01 |
| AT5G55450 | 0,62 | -2,87 | 3,02 | -0,95 | 3,42E-01 |
| AT5G55460 | 0,37 | -0,24 | 3,40 | -0,07 | 9,43E-01 |
| AT5G55500 | 0,57 | -1,64 | 3,17 | -0,52 | 6,06E-01 |
| AT5G55530 | 0,30 | -1,25 | 4,37 | -0,29 | 7,75E-01 |
| AT5G55580 | 0,70 | -1,77 | 3,90 | -0,45 | 6,50E-01 |
| AT5G55620 | 0,43 | 1,51 | 4,21 | 0,36 | 7,20E-01 |
| AT5G55630 | 0,12 | -0,68 | 4,41 | -0,15 | 8,77E-01 |
| AT5G55660 | 0,90 | -1,67 | 2,10 | -0,80 | 4,26E-01 |
| AT5G55840 | 0,22 | 0,41 | 4,35 | 0,09 | 9,25E-01 |
| AT5G55850 | 0,24 | 0,77 | 4,34 | 0,18 | 8,60E-01 |
| AT5G55900 | 0,10 | 0,45 | 4,35 | 0,10 | 9,19E-01 |
| AT5G55930 | 0,70 | -1,76 | 2,96 | -0,60 | 5,50E-01 |
| AT5G55960 | 0,59 | 0,86 | 3,07 | 0,28 | 7,79E-01 |
| AT5G55990 | 0,33 | -1,19 | 4,36 | -0,27 | 7,85E-01 |
| AT5G56020 | 0,32 | -3,36 | 4,37 | -0,77 | 4,42E-01 |
| AT5G56040 | 0,73 | -0,55 | 2,46 | -0,22 | 8,24E-01 |
| AT5G56060 | 0,67 | 0,07 | 4,37 | 0,02 | 9,88E-01 |

| | | | | | |
|---|---|---|---|---|---|
| AT5G56140 | 0,98 | -1,20 | 1,86 | -0,64 | 5,20E-01 |
| AT5G56160 | 0,49 | -2,60 | 4,41 | -0,59 | 5,55E-01 |
| AT5G56170 | 0,53 | -0,41 | 4,33 | -0,09 | 9,25E-01 |
| AT5G56270 | 0,76 | -4,54 | 2,79 | -1,63 | 1,04E-01 |
| AT5G56290 | 0,52 | -0,11 | 2,39 | -0,05 | 9,62E-01 |
| AT5G56340 | 0,44 | -3,42 | 3,33 | -1,03 | 3,04E-01 |
| AT5G08495 | 0,43 | 0,08 | 4,15 | 0,02 | 9,85E-01 |
| AT5G56460 | 0,74 | 0,33 | 3,99 | 0,08 | 9,35E-01 |
| AT5G56500 | 0,32 | -2,39 | 4,36 | -0,55 | 5,84E-01 |
| AT5G56590 | 0,24 | -1,46 | 4,36 | -0,34 | 7,37E-01 |
| AT5G56630 | 0,71 | -2,32 | 2,51 | -0,93 | 3,55E-01 |
| AT5G56650 | 0,64 | -1,49 | 2,18 | -0,68 | 4,94E-01 |
| AT5G56680 | 1,22 | -2,72 | 1,85 | -1,47 | 1,41E-01 |
| AT5G56720 | 0,12 | -0,78 | 4,39 | -0,18 | 8,59E-01 |
| AT5G56870 | 0,19 | 0,42 | 4,35 | 0,10 | 9,23E-01 |
| AT5G56910 | 0,15 | -0,77 | 4,39 | -0,18 | 8,61E-01 |
| AT5G56920 | 0,65 | 1,35 | 2,97 | 0,45 | 6,51E-01 |
| AT5G56940 | 0,08 | 0,05 | 4,38 | 0,01 | 9,91E-01 |
| AT5G56950 | 0,44 | -2,54 | 3,33 | -0,76 | 4,45E-01 |
| AT5G57015 | 0,32 | -0,24 | 4,33 | -0,06 | 9,56E-01 |
| AT5G57020 | 1,25 | -0,68 | 1,74 | -0,39 | 6,96E-01 |
| AT5G57030 | 0,86 | -2,54 | 2,48 | -1,03 | 3,05E-01 |
| AT5G57035 | 1,04 | -3,39 | 2,54 | -1,33 | 1,83E-01 |
| AT5G57070 | 0,42 | -2,40 | 4,24 | -0,57 | 5,71E-01 |
| AT5G57150 | 0,25 | -2,60 | 4,41 | -0,59 | 5,55E-01 |
| AT5G57180 | 1,05 | -0,74 | 1,86 | -0,40 | 6,91E-01 |
| AT5G57320 | 0,08 | 0,00 | 4,41 | 0,00 | 1,00E+00 |
| AT5G57330 | 0,49 | -0,12 | 2,85 | -0,04 | 9,65E-01 |
| AT5G57340 | 0,20 | 1,02 | 4,33 | 0,24 | 8,14E-01 |
| AT5G57360 | 0,96 | -0,69 | 1,85 | -0,37 | 7,08E-01 |
| AT5G57370 | 0,31 | -1,19 | 4,38 | -0,27 | 7,86E-01 |
| AT5G57460 | 1,07 | -1,75 | 1,74 | -1,01 | 3,14E-01 |
| AT5G57520 | 0,31 | 1,91 | 4,29 | 0,45 | 6,56E-01 |
| AT5G57610 | 0,89 | -0,87 | 1,88 | -0,46 | 6,44E-01 |
| AT5G57735 | 0,19 | 0,88 | 4,33 | 0,20 | 8,39E-01 |
| AT5G57800 | 0,61 | -0,33 | 2,50 | -0,13 | 8,95E-01 |
| AT5G08805 | 1,03 | 0,81 | 2,35 | 0,34 | 7,31E-01 |
| AT5G57880 | 0,48 | -4,25 | 4,04 | -1,05 | 2,92E-01 |
| AT5G57940 | 0,35 | 0,23 | 3,57 | 0,06 | 9,49E-01 |
| AT5G58000 | 0,08 | 0,00 | 4,41 | 0,00 | 1,00E+00 |
| AT5G58020 | 0,39 | 0,00 | 4,41 | 0,00 | 1,00E+00 |
| AT5G58060 | 0,59 | -1,74 | 2,47 | -0,70 | 4,82E-01 |
| AT5G58100 | 0,97 | -0,73 | 1,71 | -0,43 | 6,71E-01 |
| AT5G58110 | 0,27 | 0,61 | 4,34 | 0,14 | 8,89E-01 |
| AT5G08820 | 0,19 | -0,81 | 4,39 | -0,19 | 8,53E-01 |
| AT5G58200 | 0,40 | -0,71 | 4,40 | -0,16 | 8,72E-01 |
| AT5G58230 | 0,77 | -1,94 | 2,49 | -0,78 | 4,36E-01 |
| AT5G58250 | 0,32 | 0,67 | 4,35 | 0,15 | 8,77E-01 |
| AT5G58270 | 0,43 | -1,35 | 4,23 | -0,32 | 7,49E-01 |
| AT5G58300 | 0,99 | -3,99 | 2,21 | -1,80 | 7,11E-02 |
| AT5G58320 | 0,24 | -2,54 | 4,40 | -0,58 | 5,64E-01 |
| AT5G58350 | 1,12 | -1,32 | 1,98 | -0,67 | 5,04E-01 |
| AT5G58370 | 0,29 | -2,10 | 4,36 | -0,48 | 6,30E-01 |
| AT5G58375 | 0,48 | -2,15 | 4,33 | -0,50 | 6,20E-01 |
| AT5G58450 | 0,68 | -2,45 | 2,94 | -0,83 | 4,05E-01 |
| AT5G58495 | 0,12 | 0,00 | 4,41 | 0,00 | 1,00E+00 |

| | | | | | |
|---|---|---|---|---|---|
| AT5G58520 | 0,39 | -1,07 | 4,34 | -0,25 | 8,05E-01 |
| AT5G58560 | 0,47 | -0,33 | 3,21 | -0,10 | 9,18E-01 |
| AT5G08855 | 0,43 | 0,51 | 4,17 | 0,12 | 9,02E-01 |
| AT5G58620 | 0,34 | -2,72 | 4,39 | -0,62 | 5,37E-01 |
| AT5G58640 | 0,71 | -2,58 | 2,98 | -0,86 | 3,87E-01 |
| AT5G58710 | 0,66 | 0,28 | 3,14 | 0,09 | 9,30E-01 |
| AT5G58990 | 0,29 | 0,39 | 4,35 | 0,09 | 9,28E-01 |
| AT5G59160 | 0,32 | -0,79 | 4,39 | -0,18 | 8,57E-01 |
| AT5G59170 | 0,26 | -2,60 | 4,41 | -0,59 | 5,55E-01 |
| AT5G59250 | 0,50 | -2,00 | 3,23 | -0,62 | 5,35E-01 |
| AT5G59350 | 0,35 | -2,19 | 4,36 | -0,50 | 6,16E-01 |
| AT5G59430 | 0,64 | 0,71 | 2,58 | 0,28 | 7,83E-01 |
| AT5G59450 | 0,61 | -1,36 | 3,82 | -0,36 | 7,21E-01 |
| AT5G59610 | 0,51 | -1,66 | 4,05 | -0,41 | 6,82E-01 |
| AT5G59660 | 0,05 | -0,24 | 4,38 | -0,05 | 9,57E-01 |
| AT5G59670 | 0,28 | 1,14 | 3,64 | 0,31 | 7,53E-01 |
| AT5G59680 | 0,64 | -4,19 | 4,33 | -0,97 | 3,33E-01 |
| AT5G59690 | 0,64 | -0,42 | 2,92 | -0,14 | 8,85E-01 |
| AT5G59820 | 0,42 | 0,36 | 3,30 | 0,11 | 9,13E-01 |
| AT5G59830 | 0,85 | -4,38 | 2,88 | -1,52 | 1,28E-01 |
| AT5G59860 | 0,43 | -3,24 | 4,37 | -0,74 | 4,58E-01 |
| AT5G59880 | 0,87 | -1,04 | 2,58 | -0,40 | 6,87E-01 |
| AT5G59910 | 0,91 | 1,40 | 2,08 | 0,67 | 5,01E-01 |
| AT5G59920 | 0,58 | -3,50 | 3,76 | -0,93 | 3,51E-01 |
| AT5G59950 | 1,16 | -2,25 | 1,81 | -1,24 | 2,14E-01 |
| AT5G59980 | 0,39 | -3,07 | 4,38 | -0,70 | 4,84E-01 |
| AT5G60040 | 0,79 | -3,54 | 2,37 | -1,50 | 1,35E-01 |
| AT5G60110 | 0,25 | -2,57 | 4,40 | -0,58 | 5,60E-01 |
| AT5G60120 | 0,67 | -1,72 | 2,84 | -0,61 | 5,45E-01 |
| AT5G60270 | 0,48 | 0,00 | 4,41 | 0,00 | 1,00E+00 |
| AT5G60280 | 0,76 | -0,01 | 3,57 | 0,00 | 9,99E-01 |
| AT5G60300 | 0,52 | -0,68 | 3,96 | -0,17 | 8,64E-01 |
| AT5G60320 | 0,25 | -2,60 | 4,41 | -0,59 | 5,55E-01 |
| AT5G60550 | 0,49 | -1,37 | 3,24 | -0,42 | 6,72E-01 |
| AT5G60560 | 0,60 | 1,97 | 3,01 | 0,65 | 5,14E-01 |
| AT5G60570 | 0,88 | -1,20 | 1,89 | -0,64 | 5,25E-01 |
| AT5G60610 | 0,72 | -4,31 | 3,55 | -1,21 | 2,24E-01 |
| AT5G60620 | 0,40 | -0,87 | 3,51 | -0,25 | 8,05E-01 |
| AT5G60780 | 1,02 | -4,05 | 2,13 | -1,90 | 5,76E-02 |
| AT5G60820 | 0,45 | -2,05 | 4,15 | -0,50 | 6,20E-01 |
| AT5G60890 | 0,24 | -3,18 | 4,39 | -0,73 | 4,68E-01 |
| AT5G60945 | 1,24 | 0,14 | 1,54 | 0,09 | 9,28E-01 |
| AT5G60963 | 0,50 | -0,84 | 2,72 | -0,31 | 7,57E-01 |
| AT5G60964 | 0,93 | 1,56 | 3,30 | 0,47 | 6,37E-01 |
| AT5G60980 | 1,13 | -2,18 | 2,05 | -1,06 | 2,88E-01 |
| AT5G61190 | 0,65 | -3,37 | 2,56 | -1,32 | 1,88E-01 |
| AT5G61200 | 0,25 | -3,06 | 4,38 | -0,70 | 4,85E-01 |
| AT5G61210 | 1,18 | -0,42 | 1,81 | -0,23 | 8,18E-01 |
| AT5G61230 | 1,00 | 0,04 | 2,23 | 0,02 | 9,84E-01 |
| AT5G61240 | 0,29 | -0,31 | 4,38 | -0,07 | 9,43E-01 |
| AT5G61380 | 0,78 | -2,31 | 2,51 | -0,92 | 3,56E-01 |
| AT5G61400 | 0,78 | -3,69 | 3,48 | -1,06 | 2,88E-01 |
| AT5G61470 | 0,39 | 2,02 | 3,28 | 0,62 | 5,38E-01 |
| AT5G61490 | 0,17 | 0,00 | 4,41 | 0,00 | 1,00E+00 |
| AT5G61550 | 0,85 | -3,16 | 3,41 | -0,93 | 3,54E-01 |
| AT5G61570 | 0,19 | 0,52 | 4,35 | 0,12 | 9,04E-01 |

| | | | | | |
|---|---|---|---|---|---|
| AT5G61580 | 0,26 | -1,26 | 4,36 | -0,29 | 7,72E-01 |
| AT5G61950 | 0,32 | -3,01 | 4,38 | -0,69 | 4,92E-01 |
| AT5G61970 | 0,49 | -0,03 | 3,36 | -0,01 | 9,92E-01 |
| AT5G61980 | 0,55 | -3,77 | 4,34 | -0,87 | 3,86E-01 |
| AT5G62030 | 0,41 | -0,38 | 4,13 | -0,09 | 9,27E-01 |
| AT5G62270 | 0,20 | 0,46 | 4,35 | 0,10 | 9,16E-01 |
| AT5G62290 | 0,13 | 0,52 | 4,35 | 0,12 | 9,04E-01 |
| AT5G62360 | 0,27 | -0,62 | 4,34 | -0,14 | 8,86E-01 |
| AT5G62460 | 0,44 | -2,27 | 4,19 | -0,54 | 5,88E-01 |
| AT5G62500 | 0,05 | -0,24 | 4,38 | -0,05 | 9,57E-01 |
| AT5G62520 | 0,28 | 0,00 | 4,41 | 0,00 | 1,00E+00 |
| AT5G62560 | 0,37 | -0,10 | 3,47 | -0,03 | 9,78E-01 |
| AT5G62570 | 0,48 | 0,30 | 2,62 | 0,11 | 9,10E-01 |
| AT5G62575 | 0,21 | 0,00 | 4,41 | 0,00 | 1,00E+00 |
| AT5G62620 | 0,35 | -0,54 | 4,31 | -0,13 | 9,00E-01 |
| AT5G62710 | 0,23 | -1,68 | 4,36 | -0,39 | 7,00E-01 |
| AT5G62720 | 0,31 | 0,14 | 4,38 | 0,03 | 9,75E-01 |
| AT5G62740 | 0,43 | -0,68 | 4,16 | -0,16 | 8,70E-01 |
| AT5G62770 | 0,31 | -1,66 | 4,36 | -0,38 | 7,02E-01 |
| AT5G62890 | 1,11 | 0,04 | 2,05 | 0,02 | 9,86E-01 |
| AT5G62910 | 0,36 | -1,63 | 4,40 | -0,37 | 7,10E-01 |
| AT5G62970 | 0,35 | 0,96 | 3,52 | 0,27 | 7,86E-01 |
| AT5G63010 | 0,41 | -0,91 | 3,35 | -0,27 | 7,86E-01 |
| AT5G09120 | 1,28 | 2,89 | 1,90 | 1,52 | 1,28E-01 |
| AT5G63050 | 0,55 | -3,14 | 4,00 | -0,78 | 4,33E-01 |
| AT5G63130 | 0,64 | -1,61 | 3,11 | -0,52 | 6,05E-01 |
| AT5G63200 | 0,75 | -2,26 | 3,50 | -0,65 | 5,18E-01 |
| AT5G63260 | 0,08 | 0,00 | 4,41 | 0,00 | 1,00E+00 |
| AT5G63280 | 0,12 | 0,00 | 4,41 | 0,00 | 1,00E+00 |
| AT5G09150 | 0,24 | -3,18 | 4,39 | -0,73 | 4,68E-01 |
| AT5G63300 | 0,47 | -3,21 | 4,38 | -0,73 | 4,64E-01 |
| AT5G63340 | 0,26 | 1,28 | 4,31 | 0,30 | 7,66E-01 |
| AT5G63420 | 0,69 | -3,20 | 3,09 | -1,03 | 3,01E-01 |
| AT5G63480 | 0,12 | 0,75 | 4,34 | 0,17 | 8,63E-01 |
| AT5G63510 | 0,37 | -1,61 | 4,35 | -0,37 | 7,12E-01 |
| AT5G63640 | 0,36 | -2,50 | 4,39 | -0,57 | 5,70E-01 |
| AT5G63680 | 0,63 | -1,50 | 3,16 | -0,47 | 6,36E-01 |
| AT5G63770 | 0,64 | -0,53 | 3,03 | -0,18 | 8,61E-01 |
| AT5G63780 | 0,69 | -0,66 | 2,16 | -0,31 | 7,59E-01 |
| AT5G63810 | 0,68 | -2,29 | 3,07 | -0,75 | 4,56E-01 |
| AT5G63940 | 0,43 | 0,59 | 3,25 | 0,18 | 8,56E-01 |
| AT5G63960 | 0,45 | -3,25 | 4,32 | -0,75 | 4,52E-01 |
| AT5G63980 | 0,08 | 0,00 | 4,41 | 0,00 | 1,00E+00 |
| AT5G64090 | 0,72 | -2,46 | 2,51 | -0,98 | 3,26E-01 |
| AT5G64160 | 0,20 | -0,75 | 4,40 | -0,17 | 8,64E-01 |
| AT5G64190 | 0,08 | 0,00 | 4,41 | 0,00 | 1,00E+00 |
| AT5G64240 | 0,28 | 0,18 | 4,37 | 0,04 | 9,67E-01 |
| AT5G64310 | 0,80 | 1,08 | 2,69 | 0,40 | 6,88E-01 |
| AT5G64330 | 1,11 | -2,07 | 1,91 | -1,08 | 2,78E-01 |
| AT5G64370 | 0,51 | -1,71 | 3,97 | -0,43 | 6,67E-01 |
| AT5G64380 | 0,10 | -0,34 | 4,38 | -0,08 | 9,38E-01 |
| AT5G64400 | 0,52 | 0,11 | 4,20 | 0,03 | 9,79E-01 |
| AT5G64440 | 0,42 | 1,61 | 3,29 | 0,49 | 6,23E-01 |
| AT5G64500 | 0,82 | -3,35 | 2,14 | -1,57 | 1,17E-01 |
| AT5G64505 | 0,26 | -2,60 | 4,41 | -0,59 | 5,55E-01 |
| AT5G64572 | 1,22 | -1,77 | 1,63 | -1,08 | 2,80E-01 |

| | | | | | |
|---|---|---|---|---|---|
| AT5G64690 | 0,11 | 0,46 | 4,35 | 0,11 | 9,15E-01 |
| AT5G64750 | 0,12 | 0,00 | 4,41 | 0,00 | 1,00E+00 |
| AT5G64813 | 0,60 | -2,09 | 2,52 | -0,83 | 4,06E-01 |
| AT5G09285 | 0,45 | -3,33 | 4,03 | -0,83 | 4,08E-01 |
| AT5G64950 | 0,48 | 0,00 | 4,41 | 0,00 | 1,00E+00 |
| AT5G64960 | 0,59 | -1,64 | 3,24 | -0,51 | 6,14E-01 |
| AT5G65010 | 1,00 | -2,26 | 1,75 | -1,29 | 1,96E-01 |
| AT5G65015 | 0,39 | -0,75 | 4,40 | -0,17 | 8,65E-01 |
| AT5G65210 | 0,26 | -2,60 | 4,41 | -0,59 | 5,55E-01 |
| AT5G65220 | 1,17 | 0,31 | 2,19 | 0,14 | 8,86E-01 |
| AT5G65230 | 0,08 | 0,00 | 4,41 | 0,00 | 1,00E+00 |
| AT5G65270 | 0,10 | -0,34 | 4,38 | -0,08 | 9,38E-01 |
| AT5G65290 | 0,47 | -3,70 | 4,35 | -0,85 | 3,95E-01 |
| AT5G65400 | 0,17 | 0,00 | 4,41 | 0,00 | 1,00E+00 |
| AT5G65440 | 0,58 | -1,94 | 2,98 | -0,65 | 5,15E-01 |
| AT5G65470 | 0,30 | 0,74 | 4,33 | 0,17 | 8,65E-01 |
| AT5G65510 | 0,12 | -0,31 | 4,38 | -0,07 | 9,44E-01 |
| AT5G65570 | 0,25 | -2,53 | 4,40 | -0,57 | 5,65E-01 |
| AT5G65640 | 0,08 | 0,00 | 4,41 | 0,00 | 1,00E+00 |
| AT5G65670 | 0,91 | -0,12 | 2,09 | -0,06 | 9,56E-01 |
| AT5G65710 | 0,77 | 0,55 | 2,12 | 0,26 | 7,95E-01 |
| AT5G65760 | 0,55 | 0,51 | 2,59 | 0,20 | 8,45E-01 |
| AT5G65770 | 1,30 | -1,18 | 1,83 | -0,64 | 5,20E-01 |
| AT5G65890 | 0,05 | -0,24 | 4,38 | -0,05 | 9,57E-01 |
| AT5G65990 | 0,70 | 0,82 | 2,10 | 0,39 | 6,97E-01 |
| AT5G66052 | 0,53 | -3,33 | 3,99 | -0,84 | 4,03E-01 |
| AT5G66070 | 0,10 | -0,34 | 4,38 | -0,08 | 9,38E-01 |
| AT5G66100 | 0,24 | 0,38 | 4,36 | 0,09 | 9,31E-01 |
| AT5G66110 | 0,30 | -2,60 | 4,41 | -0,59 | 5,55E-01 |
| AT5G66160 | 0,31 | -0,91 | 4,37 | -0,21 | 8,35E-01 |
| AT5G66170 | 0,33 | -3,08 | 4,39 | -0,70 | 4,83E-01 |
| AT5G66240 | 1,02 | -2,06 | 2,21 | -0,93 | 3,51E-01 |
| AT5G66360 | 0,60 | -3,19 | 3,85 | -0,83 | 4,07E-01 |
| AT5G66450 | 0,53 | -0,81 | 3,10 | -0,26 | 7,93E-01 |
| AT5G66562 | 1,20 | 0,28 | 1,58 | 0,17 | 8,62E-01 |
| AT5G66568 | 0,20 | 0,68 | 4,35 | 0,16 | 8,75E-01 |
| AT5G66760 | 1,08 | -3,57 | 1,91 | -1,87 | 6,16E-02 |
| AT5G66770 | 0,43 | -0,32 | 4,37 | -0,07 | 9,42E-01 |
| AT5G66790 | 0,74 | -3,00 | 2,21 | -1,36 | 1,74E-01 |
| AT5G66810 | 0,67 | -3,19 | 2,50 | -1,28 | 2,02E-01 |
| AT5G66930 | 0,12 | -0,78 | 4,39 | -0,18 | 8,59E-01 |
| AT5G66950 | 1,10 | 0,73 | 1,79 | 0,41 | 6,82E-01 |
| AT5G67070 | 0,51 | 0,13 | 3,88 | 0,03 | 9,74E-01 |
| AT5G67080 | 0,30 | -1,09 | 4,39 | -0,25 | 8,04E-01 |
| AT5G67170 | 0,08 | 0,00 | 4,41 | 0,00 | 1,00E+00 |
| AT5G67260 | 0,65 | -0,36 | 3,86 | -0,09 | 9,25E-01 |
| AT5G67320 | 0,96 | -0,70 | 2,76 | -0,25 | 8,00E-01 |
| AT5G67330 | 1,26 | -2,29 | 1,71 | -1,34 | 1,80E-01 |
| AT5G67340 | 0,24 | -0,68 | 4,41 | -0,15 | 8,77E-01 |
| AT5G67385 | 0,80 | 1,04 | 2,23 | 0,47 | 6,41E-01 |
| AT5G67455 | 0,10 | -0,34 | 4,38 | -0,08 | 9,38E-01 |
| AT5G67470 | 1,09 | -1,13 | 1,92 | -0,59 | 5,57E-01 |
| AT5G67488 | 1,20 | 0,28 | 1,82 | 0,16 | 8,76E-01 |
| AT5G67500 | 0,21 | -0,77 | 4,40 | -0,17 | 8,62E-01 |
| AT5G67530 | 0,08 | 0,00 | 4,41 | 0,00 | 1,00E+00 |
| AT5G67560 | 0,70 | 0,04 | 3,73 | 0,01 | 9,92E-01 |

| | | | | | |
|---|---|---|---|---|---|
| AT5G67570 | 0,48 | -1,79 | 4,10 | -0,44 | 6,62E-01 |
| AT5G67590 | 0,32 | -0,50 | 4,34 | -0,11 | 9,08E-01 |
| AT1G01020 | 0,15 | 0,41 | 4,35 | 0,09 | 9,26E-01 |
| AT1G03993 | 0,11 | -0,68 | 4,41 | -0,15 | 8,77E-01 |
| AT1G01050 | 0,66 | -4,52 | 2,90 | -1,56 | 1,19E-01 |
| AT1G01060 | 0,69 | -3,25 | 2,43 | -1,34 | 1,81E-01 |
| AT1G01130 | 0,29 | -0,69 | 4,34 | -0,16 | 8,73E-01 |
| AT1G01140 | 0,54 | 1,00 | 2,50 | 0,40 | 6,88E-01 |
| AT1G01170 | 0,58 | 1,59 | 3,01 | 0,53 | 5,97E-01 |
| AT1G01440 | 0,35 | -0,74 | 4,37 | -0,17 | 8,66E-01 |
| AT1G01450 | 0,59 | 0,00 | 4,41 | 0,00 | 1,00E+00 |
| AT1G01470 | 1,29 | 0,54 | 1,58 | 0,34 | 7,32E-01 |
| AT1G01490 | 0,15 | -0,14 | 4,39 | -0,03 | 9,74E-01 |
| AT1G01530 | 0,08 | 0,05 | 4,38 | 0,01 | 9,91E-01 |
| AT1G01610 | 0,73 | 1,01 | 2,48 | 0,41 | 6,83E-01 |
| AT1G01620 | 1,23 | 0,48 | 2,02 | 0,24 | 8,11E-01 |
| AT1G01650 | 0,59 | 1,48 | 2,89 | 0,51 | 6,09E-01 |
| AT1G04027 | 0,46 | 1,48 | 3,10 | 0,48 | 6,34E-01 |
| AT1G01680 | 0,72 | -1,03 | 2,46 | -0,42 | 6,75E-01 |
| AT1G01725 | 1,09 | 1,89 | 1,99 | 0,95 | 3,41E-01 |
| AT1G04063 | 0,34 | 0,89 | 4,34 | 0,20 | 8,38E-01 |
| AT1G01820 | 0,86 | -2,26 | 2,17 | -1,04 | 2,98E-01 |
| AT1G01830 | 1,22 | 0,05 | 2,06 | 0,03 | 9,79E-01 |
| AT1G01880 | 1,08 | -0,71 | 2,47 | -0,29 | 7,74E-01 |
| AT1G01890 | 1,04 | -0,65 | 2,79 | -0,23 | 8,17E-01 |
| AT1G01930 | 0,16 | -0,68 | 4,41 | -0,15 | 8,77E-01 |
| AT1G01990 | 0,21 | -0,73 | 4,40 | -0,17 | 8,67E-01 |
| AT1G02020 | 0,45 | -3,79 | 4,08 | -0,93 | 3,53E-01 |
| AT1G02090 | 0,84 | -2,14 | 2,84 | -0,75 | 4,52E-01 |
| AT1G02140 | 0,21 | -0,44 | 4,38 | -0,10 | 9,20E-01 |
| AT1G02260 | 0,54 | -3,14 | 3,19 | -0,99 | 3,25E-01 |
| AT1G02280 | 0,61 | -0,68 | 3,14 | -0,22 | 8,29E-01 |
| AT1G02310 | 0,21 | -2,60 | 4,41 | -0,59 | 5,55E-01 |
| AT1G04117 | 0,17 | -0,71 | 4,40 | -0,16 | 8,72E-01 |
| AT1G02430 | 0,17 | -0,75 | 4,40 | -0,17 | 8,65E-01 |
| AT1G02450 | 0,20 | 0,00 | 4,41 | 0,00 | 1,00E+00 |
| AT1G02610 | 0,33 | -1,12 | 4,35 | -0,26 | 7,97E-01 |
| AT1G02660 | 0,36 | -2,24 | 4,32 | -0,52 | 6,03E-01 |
| AT1G02770 | 0,41 | -3,93 | 4,34 | -0,91 | 3,65E-01 |
| AT1G02870 | 0,35 | -3,17 | 4,38 | -0,72 | 4,69E-01 |
| AT1G02910 | 0,70 | -4,43 | 2,45 | -1,81 | 7,06E-02 |
| AT1G02920 | 0,41 | 2,15 | 3,28 | 0,66 | 5,12E-01 |
| AT1G02930 | 0,49 | -1,80 | 3,22 | -0,56 | 5,77E-01 |
| AT1G02960 | 0,08 | 0,00 | 4,41 | 0,00 | 1,00E+00 |
| AT1G03000 | 0,70 | -0,42 | 2,70 | -0,15 | 8,77E-01 |
| AT1G03040 | 0,62 | -1,15 | 2,65 | -0,44 | 6,63E-01 |
| AT1G03140 | 0,90 | 0,00 | 1,79 | 0,00 | 1,00E+00 |
| AT1G03300 | 0,12 | 0,00 | 4,41 | 0,00 | 1,00E+00 |
| AT1G03400 | 0,49 | -1,95 | 2,72 | -0,72 | 4,73E-01 |
| AT1G03445 | 0,28 | -0,68 | 4,41 | -0,15 | 8,77E-01 |
| AT1G03475 | 0,69 | -0,08 | 2,23 | -0,04 | 9,70E-01 |
| AT1G03520 | 0,48 | -1,95 | 4,03 | -0,48 | 6,28E-01 |
| AT1G03530 | 0,41 | -3,42 | 4,37 | -0,78 | 4,34E-01 |
| AT1G03550 | 0,70 | -1,63 | 2,24 | -0,73 | 4,68E-01 |
| AT1G03590 | 0,41 | -2,34 | 4,21 | -0,56 | 5,78E-01 |
| AT1G03660 | 0,52 | -1,62 | 2,70 | -0,60 | 5,48E-01 |

| | | | | | |
|---|---|---|---|---|---|
| AT1G03746 | 0,46 | -0,11 | 4,06 | -0,03 | 9,79E-01 |
| AT1G03760 | 0,08 | -0,21 | 4,39 | -0,05 | 9,62E-01 |
| AT1G03830 | 0,65 | -2,58 | 2,55 | -1,01 | 3,12E-01 |
| AT1G03870 | 0,58 | -2,24 | 3,20 | -0,70 | 4,83E-01 |
| AT1G03940 | 1,20 | -0,32 | 1,94 | -0,17 | 8,67E-01 |
| AT1G04233 | 0,08 | 0,00 | 4,41 | 0,00 | 1,00E+00 |
| AT1G03980 | 0,11 | 0,46 | 4,35 | 0,11 | 9,15E-01 |
| AT1G04000 | 0,55 | 1,56 | 3,07 | 0,51 | 6,10E-01 |
| AT1G04010 | 0,05 | -0,24 | 4,38 | -0,05 | 9,57E-01 |
| AT1G04050 | 0,08 | -0,21 | 4,39 | -0,05 | 9,62E-01 |
| AT1G04070 | 0,16 | -0,62 | 4,40 | -0,14 | 8,87E-01 |
| AT1G04110 | 0,35 | -2,99 | 4,38 | -0,68 | 4,95E-01 |
| AT1G04190 | 0,13 | -0,18 | 4,39 | -0,04 | 9,67E-01 |
| AT1G04220 | 0,35 | -0,52 | 4,35 | -0,12 | 9,05E-01 |
| AT1G04240 | 0,91 | -0,46 | 2,34 | -0,20 | 8,45E-01 |
| AT1G04260 | 0,47 | -0,89 | 2,76 | -0,32 | 7,46E-01 |
| AT1G04270 | 1,04 | 0,25 | 2,19 | 0,11 | 9,10E-01 |
| AT1G04280 | 0,49 | 0,84 | 3,32 | 0,25 | 8,01E-01 |
| AT1G04290 | 0,43 | -0,02 | 2,75 | -0,01 | 9,96E-01 |
| AT1G04300 | 0,83 | -2,99 | 2,94 | -1,02 | 3,08E-01 |
| AT1G04380 | 0,20 | -2,55 | 4,40 | -0,58 | 5,62E-01 |
| AT1G04390 | 0,19 | 0,01 | 4,36 | 0,00 | 9,98E-01 |
| AT1G04400 | 0,89 | -2,03 | 2,22 | -0,92 | 3,60E-01 |
| AT1G04530 | 0,42 | -0,42 | 3,30 | -0,13 | 8,99E-01 |
| AT1G04570 | 0,11 | -0,80 | 4,39 | -0,18 | 8,56E-01 |
| AT1G04630 | 0,66 | -3,36 | 3,77 | -0,89 | 3,73E-01 |
| AT1G04680 | 0,74 | 1,32 | 3,10 | 0,43 | 6,70E-01 |
| AT1G04730 | 0,74 | -3,12 | 2,48 | -1,26 | 2,09E-01 |
| AT1G04750 | 0,67 | -0,93 | 2,23 | -0,42 | 6,77E-01 |
| AT1G04770 | 0,89 | -0,02 | 2,17 | -0,01 | 9,93E-01 |
| AT1G04780 | 0,86 | -0,10 | 2,03 | -0,05 | 9,61E-01 |
| AT1G04840 | 0,63 | -3,51 | 3,75 | -0,94 | 3,49E-01 |
| AT1G04850 | 0,86 | -1,26 | 2,74 | -0,46 | 6,44E-01 |
| AT1G04860 | 0,57 | -2,84 | 3,94 | -0,72 | 4,72E-01 |
| AT1G04870 | 0,84 | -2,75 | 3,06 | -0,90 | 3,68E-01 |
| AT1G04880 | 0,21 | 0,41 | 4,36 | 0,09 | 9,24E-01 |
| AT1G04910 | 0,78 | -1,33 | 2,86 | -0,47 | 6,41E-01 |
| AT1G04950 | 0,50 | -2,91 | 3,25 | -0,90 | 3,70E-01 |
| AT1G04960 | 0,36 | -2,43 | 4,37 | -0,56 | 5,78E-01 |
| AT1G04980 | 0,36 | -3,60 | 4,36 | -0,82 | 4,10E-01 |
| AT1G04985 | 0,26 | -0,53 | 4,34 | -0,12 | 9,03E-01 |
| AT1G04990 | 0,84 | -2,57 | 2,47 | -1,04 | 2,99E-01 |
| AT1G05120 | 0,53 | -3,32 | 4,37 | -0,76 | 4,48E-01 |
| AT1G05150 | 0,85 | -2,42 | 2,08 | -1,16 | 2,46E-01 |
| AT1G05160 | 0,59 | 0,00 | 4,41 | 0,00 | 1,00E+00 |
| AT1G05170 | 0,47 | -2,12 | 4,39 | -0,48 | 6,30E-01 |
| AT1G05190 | 0,78 | -0,44 | 2,31 | -0,19 | 8,49E-01 |
| AT1G05205 | 0,08 | 0,00 | 4,41 | 0,00 | 1,00E+00 |
| AT1G05230 | 0,45 | -3,79 | 4,08 | -0,93 | 3,53E-01 |
| AT1G05300 | 0,18 | 0,48 | 4,35 | 0,11 | 9,11E-01 |
| AT1G05350 | 0,28 | -1,56 | 4,39 | -0,35 | 7,23E-01 |
| AT1G05520 | 0,61 | -2,30 | 3,00 | -0,77 | 4,43E-01 |
| AT1G05575 | 0,16 | 0,75 | 4,34 | 0,17 | 8,63E-01 |
| AT1G05640 | 0,20 | -1,00 | 4,37 | -0,23 | 8,19E-01 |
| AT1G05670 | 0,15 | -0,68 | 4,41 | -0,15 | 8,77E-01 |
| AT1G05720 | 0,13 | -0,18 | 4,39 | -0,04 | 9,67E-01 |

| | | | | | |
|---|---|---|---|---|---|
| AT1G05783 | 0,96 | 0,21 | 1,97 | 0,11 | 9,14E-01 |
| AT1G05840 | 0,75 | -1,16 | 2,02 | -0,58 | 5,65E-01 |
| AT1G05870 | 0,42 | -0,50 | 4,14 | -0,12 | 9,04E-01 |
| AT1G05950 | 0,12 | 0,00 | 4,41 | 0,00 | 1,00E+00 |
| AT1G05970 | 0,08 | 0,00 | 4,41 | 0,00 | 1,00E+00 |
| AT1G06000 | 0,95 | -1,20 | 1,91 | -0,63 | 5,28E-01 |
| AT1G06130 | 0,62 | -4,16 | 3,03 | -1,37 | 1,69E-01 |
| AT1G06250 | 0,72 | -4,76 | 4,30 | -1,11 | 2,68E-01 |
| AT1G06265 | 0,08 | 0,00 | 4,41 | 0,00 | 1,00E+00 |
| AT1G06360 | 0,33 | 0,23 | 4,36 | 0,05 | 9,57E-01 |
| AT1G06400 | 0,21 | 0,65 | 4,34 | 0,15 | 8,80E-01 |
| AT1G06440 | 0,58 | -1,32 | 2,66 | -0,50 | 6,19E-01 |
| AT1G06540 | 1,04 | -3,79 | 2,06 | -1,84 | 6,57E-02 |
| AT1G06570 | 0,97 | -3,40 | 2,56 | -1,32 | 1,85E-01 |
| AT1G04487 | 0,39 | -1,21 | 4,24 | -0,28 | 7,76E-01 |
| AT1G06660 | 0,50 | -3,97 | 3,97 | -1,00 | 3,17E-01 |
| AT1G06690 | 0,36 | -2,50 | 4,39 | -0,57 | 5,70E-01 |
| AT1G06710 | 0,26 | -2,60 | 4,41 | -0,59 | 5,55E-01 |
| AT1G06720 | 0,55 | -1,97 | 3,98 | -0,50 | 6,20E-01 |
| AT1G06810 | 0,08 | 0,00 | 4,41 | 0,00 | 1,00E+00 |
| AT1G06840 | 0,17 | -0,13 | 4,39 | -0,03 | 9,77E-01 |
| AT1G06890 | 0,40 | 0,85 | 2,90 | 0,29 | 7,69E-01 |
| AT1G07020 | 0,55 | -4,01 | 4,34 | -0,92 | 3,55E-01 |
| AT1G07050 | 0,42 | 0,00 | 4,18 | 0,00 | 1,00E+00 |
| AT1G07090 | 0,43 | -1,34 | 4,31 | -0,31 | 7,56E-01 |
| AT1G07110 | 1,24 | -5,11 | 2,31 | -2,21 | 2,71E-02 |
| AT1G07128 | 0,28 | 0,18 | 4,37 | 0,04 | 9,67E-01 |
| AT1G07220 | 0,25 | 0,00 | 4,41 | 0,00 | 1,00E+00 |
| AT1G07260 | 0,48 | -1,77 | 4,33 | -0,41 | 6,82E-01 |
| AT1G04573 | 0,66 | -4,57 | 4,31 | -1,06 | 2,88E-01 |
| AT1G07310 | 0,69 | -2,01 | 2,66 | -0,76 | 4,50E-01 |
| AT1G07350 | 0,45 | -2,49 | 3,31 | -0,75 | 4,52E-01 |
| AT1G07360 | 0,57 | 0,90 | 3,15 | 0,29 | 7,75E-01 |
| AT1G07430 | 0,32 | -2,60 | 4,41 | -0,59 | 5,55E-01 |
| AT1G07440 | 0,75 | -1,14 | 2,51 | -0,45 | 6,49E-01 |
| AT1G07480 | 0,80 | -0,94 | 2,24 | -0,42 | 6,75E-01 |
| AT1G07500 | 0,08 | -0,21 | 4,39 | -0,05 | 9,62E-01 |
| AT1G07520 | 1,20 | -3,34 | 2,23 | -1,50 | 1,34E-01 |
| AT1G07530 | 1,00 | -2,29 | 1,80 | -1,27 | 2,05E-01 |
| AT1G07640 | 0,24 | 0,03 | 4,37 | 0,01 | 9,94E-01 |
| AT1G07720 | 1,26 | 2,56 | 1,75 | 1,46 | 1,43E-01 |
| AT1G07725 | 1,03 | -5,02 | 2,48 | -2,02 | 4,29E-02 |
| AT1G07740 | 0,29 | -2,60 | 4,41 | -0,59 | 5,55E-01 |
| AT1G07730 | 0,29 | -2,60 | 4,41 | -0,59 | 5,55E-01 |
| AT1G07750 | 0,25 | 0,00 | 4,41 | 0,00 | 1,00E+00 |
| AT1G07770 | 0,87 | -0,10 | 1,74 | -0,06 | 9,52E-01 |
| AT1G07870 | 0,12 | -0,68 | 4,41 | -0,15 | 8,77E-01 |
| AT1G07970 | 0,62 | -3,72 | 3,66 | -1,01 | 3,10E-01 |
| AT1G08040 | 0,40 | -1,55 | 4,35 | -0,36 | 7,21E-01 |
| AT1G08050 | 0,33 | -1,64 | 4,38 | -0,37 | 7,08E-01 |
| AT1G08060 | 1,18 | -2,45 | 2,07 | -1,18 | 2,36E-01 |
| AT1G08125 | 0,32 | -1,82 | 4,36 | -0,42 | 6,75E-01 |
| AT1G08580 | 0,24 | -3,18 | 4,39 | -0,73 | 4,68E-01 |
| AT1G08592 | 0,74 | -1,64 | 3,01 | -0,54 | 5,86E-01 |
| AT1G08610 | 0,28 | -1,32 | 4,38 | -0,30 | 7,63E-01 |
| AT1G08640 | 0,47 | -2,17 | 3,96 | -0,55 | 5,84E-01 |

| | | | | | |
|---|---|---|---|---|---|
| AT1G08645 | 0,64 | -1,83 | 3,73 | -0,49 | 6,25E-01 |
| AT1G08660 | 0,10 | 0,45 | 4,35 | 0,10 | 9,19E-01 |
| AT1G08700 | 1,22 | -1,64 | 2,35 | -0,70 | 4,85E-01 |
| AT1G08800 | 0,74 | -2,21 | 3,01 | -0,73 | 4,62E-01 |
| AT1G08820 | 0,89 | -0,52 | 2,50 | -0,21 | 8,36E-01 |
| AT1G08880 | 0,12 | 0,00 | 4,41 | 0,00 | 1,00E+00 |
| AT1G09020 | 0,78 | -1,28 | 2,19 | -0,58 | 5,59E-01 |
| AT1G09060 | 1,04 | -1,99 | 2,31 | -0,86 | 3,90E-01 |
| AT1G09080 | 0,40 | -0,61 | 4,31 | -0,14 | 8,87E-01 |
| AT1G09130 | 0,46 | 0,63 | 3,25 | 0,19 | 8,47E-01 |
| AT1G09160 | 0,40 | 0,44 | 3,45 | 0,13 | 8,98E-01 |
| AT1G09190 | 0,17 | 0,00 | 4,41 | 0,00 | 1,00E+00 |
| AT1G09195 | 0,70 | -2,61 | 2,91 | -0,90 | 3,70E-01 |
| AT1G09200 | 0,34 | 1,93 | 4,29 | 0,45 | 6,53E-01 |
| AT1G09230 | 0,23 | -1,14 | 4,39 | -0,26 | 7,96E-01 |
| AT1G09280 | 1,11 | -1,13 | 1,82 | -0,62 | 5,35E-01 |
| AT1G09420 | 0,88 | 0,05 | 3,42 | 0,01 | 9,89E-01 |
| AT1G09440 | 0,21 | 0,00 | 4,41 | 0,00 | 1,00E+00 |
| AT1G04707 | 0,36 | -3,08 | 4,39 | -0,70 | 4,83E-01 |
| AT1G09630 | 0,08 | 0,00 | 4,41 | 0,00 | 1,00E+00 |
| AT1G04717 | 0,54 | -3,53 | 3,85 | -0,92 | 3,59E-01 |
| AT1G09680 | 1,19 | -2,31 | 2,67 | -0,87 | 3,86E-01 |
| AT1G09730 | 0,83 | -0,77 | 2,82 | -0,27 | 7,84E-01 |
| AT1G09760 | 0,17 | -0,75 | 4,40 | -0,17 | 8,65E-01 |
| AT1G09810 | 0,42 | -0,49 | 2,81 | -0,17 | 8,63E-01 |
| AT1G09880 | 0,33 | -3,41 | 4,38 | -0,78 | 4,35E-01 |
| AT1G09910 | 0,78 | -3,53 | 2,34 | -1,51 | 1,32E-01 |
| AT1G10020 | 0,10 | -0,34 | 4,38 | -0,08 | 9,38E-01 |
| AT1G10090 | 0,57 | -1,24 | 2,69 | -0,46 | 6,46E-01 |
| AT1G10180 | 0,61 | -2,76 | 2,53 | -1,09 | 2,76E-01 |
| AT1G10190 | 0,94 | 2,91 | 2,66 | 1,10 | 2,74E-01 |
| AT1G10200 | 0,57 | -1,65 | 3,86 | -0,43 | 6,69E-01 |
| AT1G10310 | 0,42 | -3,34 | 4,37 | -0,76 | 4,45E-01 |
| AT1G10320 | 0,47 | -0,72 | 4,04 | -0,18 | 8,58E-01 |
| AT1G10340 | 1,00 | -2,69 | 2,08 | -1,29 | 1,96E-01 |
| AT1G10350 | 0,52 | -2,60 | 3,96 | -0,66 | 5,11E-01 |
| AT1G10390 | 1,30 | -0,51 | 1,93 | -0,27 | 7,91E-01 |
| AT1G10400 | 0,08 | 0,00 | 4,41 | 0,00 | 1,00E+00 |
| AT1G10410 | 0,96 | 0,27 | 1,96 | 0,14 | 8,93E-01 |
| AT1G10430 | 0,87 | 0,74 | 2,14 | 0,35 | 7,29E-01 |
| AT1G10470 | 1,21 | -0,42 | 1,71 | -0,24 | 8,07E-01 |
| AT1G10580 | 0,57 | 0,00 | 3,94 | 0,00 | 1,00E+00 |
| AT1G10586 | 1,03 | 2,88 | 2,07 | 1,39 | 1,63E-01 |
| AT1G10590 | 0,69 | -1,92 | 2,94 | -0,65 | 5,13E-01 |
| AT1G10630 | 0,65 | -3,12 | 2,63 | -1,19 | 2,35E-01 |
| AT1G10640 | 0,05 | -0,24 | 4,38 | -0,05 | 9,57E-01 |
| AT1G10680 | 0,08 | 0,00 | 4,41 | 0,00 | 1,00E+00 |
| AT1G10840 | 0,88 | -1,52 | 1,86 | -0,82 | 4,14E-01 |
| AT1G10870 | 1,11 | -2,85 | 1,87 | -1,53 | 1,27E-01 |
| AT1G10900 | 1,03 | -3,94 | 2,79 | -1,41 | 1,58E-01 |
| AT1G10920 | 0,82 | -3,53 | 4,36 | -0,81 | 4,17E-01 |
| AT1G10930 | 0,74 | -3,11 | 2,92 | -1,07 | 2,87E-01 |
| AT1G10940 | 0,48 | -1,59 | 3,27 | -0,49 | 6,27E-01 |
| AT1G04833 | 0,22 | 1,04 | 4,32 | 0,24 | 8,10E-01 |
| AT1G11100 | 0,45 | -0,58 | 3,39 | -0,17 | 8,64E-01 |
| AT1G11210 | 0,56 | 2,15 | 2,49 | 0,86 | 3,87E-01 |

| | | | | | |
|---|---|---|---|---|---|
| AT1G11240 | 0,05 | -0,24 | 4,38 | -0,05 | 9,57E-01 |
| AT1G11270 | 0,83 | 2,08 | 2,21 | 0,94 | 3,47E-01 |
| AT1G11280 | 0,45 | -1,24 | 2,72 | -0,46 | 6,49E-01 |
| AT1G11290 | 0,63 | -3,41 | 3,70 | -0,92 | 3,57E-01 |
| AT1G11340 | 0,08 | 0,00 | 4,41 | 0,00 | 1,00E+00 |
| AT1G11350 | 0,52 | -1,39 | 3,23 | -0,43 | 6,66E-01 |
| AT1G11360 | 0,33 | -0,43 | 4,34 | -0,10 | 9,20E-01 |
| AT1G11480 | 0,70 | -1,16 | 2,14 | -0,54 | 5,87E-01 |
| AT1G11630 | 0,35 | -2,99 | 4,38 | -0,68 | 4,95E-01 |
| AT1G11820 | 0,25 | -0,28 | 4,38 | -0,06 | 9,49E-01 |
| AT1G12000 | 0,52 | 1,23 | 3,06 | 0,40 | 6,88E-01 |
| AT1G12020 | 0,63 | 0,98 | 2,17 | 0,45 | 6,52E-01 |
| AT1G12120 | 0,70 | -1,57 | 2,51 | -0,63 | 5,32E-01 |
| AT1G12211 | 0,63 | -1,17 | 3,22 | -0,36 | 7,16E-01 |
| AT1G12310 | 0,63 | 1,70 | 3,12 | 0,55 | 5,85E-01 |
| AT1G12380 | 0,49 | 1,26 | 4,05 | 0,31 | 7,55E-01 |
| AT1G12440 | 1,21 | -1,31 | 1,59 | -0,83 | 4,09E-01 |
| AT1G12500 | 0,25 | -0,13 | 4,37 | -0,03 | 9,77E-01 |
| AT1G12520 | 0,51 | -3,38 | 4,36 | -0,77 | 4,39E-01 |
| AT1G12590 | 0,78 | 0,68 | 1,87 | 0,37 | 7,15E-01 |
| AT1G12610 | 0,19 | 0,42 | 4,35 | 0,10 | 9,23E-01 |
| AT1G12640 | 0,58 | -2,96 | 4,38 | -0,68 | 4,99E-01 |
| AT1G12700 | 0,22 | 0,38 | 4,35 | 0,09 | 9,31E-01 |
| AT1G12780 | 0,71 | 0,36 | 2,29 | 0,16 | 8,75E-01 |
| AT1G12800 | 0,97 | -3,56 | 2,43 | -1,47 | 1,43E-01 |
| AT1G12910 | 1,07 | -2,47 | 2,17 | -1,14 | 2,56E-01 |
| AT1G13020 | 0,59 | -2,91 | 2,61 | -1,12 | 2,64E-01 |
| AT1G13120 | 0,23 | -0,79 | 4,39 | -0,18 | 8,56E-01 |
| AT1G13170 | 0,46 | -2,51 | 3,24 | -0,77 | 4,39E-01 |
| AT1G13200 | 0,66 | -4,42 | 3,60 | -1,23 | 2,20E-01 |
| AT1G13245 | 0,44 | -1,58 | 4,13 | -0,38 | 7,03E-01 |
| AT1G13310 | 0,35 | -2,99 | 4,38 | -0,68 | 4,95E-01 |
| AT1G13340 | 0,95 | -0,68 | 2,13 | -0,32 | 7,48E-01 |
| AT1G13390 | 0,17 | 0,00 | 4,41 | 0,00 | 1,00E+00 |
| AT1G13450 | 0,76 | -1,70 | 2,24 | -0,76 | 4,47E-01 |
| AT1G13520 | 0,54 | -2,26 | 4,36 | -0,52 | 6,05E-01 |
| AT1G13560 | 0,43 | -2,54 | 4,35 | -0,58 | 5,60E-01 |
| AT1G13825 | 0,76 | 0,06 | 3,63 | 0,02 | 9,87E-01 |
| AT1G13860 | 0,10 | -0,18 | 4,38 | -0,04 | 9,67E-01 |
| AT1G13870 | 0,12 | 0,07 | 4,38 | 0,02 | 9,87E-01 |
| AT1G13900 | 0,21 | -0,77 | 4,40 | -0,17 | 8,62E-01 |
| AT1G13940 | 0,86 | -0,84 | 2,43 | -0,35 | 7,29E-01 |
| AT1G14010 | 0,36 | -0,37 | 4,35 | -0,08 | 9,32E-01 |
| AT1G14030 | 0,62 | -1,73 | 3,20 | -0,54 | 5,90E-01 |
| AT1G14290 | 0,26 | -2,60 | 4,41 | -0,59 | 5,55E-01 |
| AT1G14300 | 0,22 | -0,75 | 4,40 | -0,17 | 8,65E-01 |
| AT1G14330 | 0,59 | -0,77 | 2,70 | -0,29 | 7,74E-01 |
| AT1G14360 | 0,18 | -1,07 | 4,38 | -0,25 | 8,06E-01 |
| AT1G14400 | 0,66 | -0,89 | 3,18 | -0,28 | 7,80E-01 |
| AT1G14450 | 0,11 | 0,27 | 4,37 | 0,06 | 9,52E-01 |
| AT1G14455 | 0,14 | 0,00 | 4,41 | 0,00 | 1,00E+00 |
| AT1G14518 | 0,84 | 1,69 | 2,27 | 0,75 | 4,55E-01 |
| AT1G14590 | 0,42 | -3,55 | 4,20 | -0,85 | 3,98E-01 |
| AT1G14690 | 0,78 | -3,19 | 2,27 | -1,40 | 1,60E-01 |
| AT1G14710 | 0,51 | 0,00 | 3,95 | 0,00 | 1,00E+00 |
| AT1G14720 | 0,26 | -2,69 | 4,40 | -0,61 | 5,41E-01 |

| | | | | | |
|---|---|---|---|---|---|
| AT1G14810 | 1,02 | -2,92 | 2,08 | -1,40 | 1,61E-01 |
| AT1G14830 | 0,90 | -0,51 | 2,77 | -0,18 | 8,53E-01 |
| AT1G14840 | 0,51 | 0,88 | 2,38 | 0,37 | 7,12E-01 |
| AT1G14850 | 1,26 | -2,58 | 1,68 | -1,54 | 1,24E-01 |
| AT1G14880 | 1,19 | 2,00 | 1,88 | 1,07 | 2,87E-01 |
| AT1G14900 | 0,38 | -2,60 | 4,41 | -0,59 | 5,55E-01 |
| AT1G14910 | 0,80 | -0,55 | 2,64 | -0,21 | 8,36E-01 |
| AT1G14980 | 0,57 | -3,35 | 3,11 | -1,08 | 2,81E-01 |
| AT1G05137 | 0,28 | -0,17 | 4,39 | -0,04 | 9,70E-01 |
| AT1G15020 | 0,98 | -3,78 | 2,65 | -1,43 | 1,54E-01 |
| AT1G15040 | 0,21 | -0,03 | 4,36 | -0,01 | 9,95E-01 |
| AT1G15100 | 1,26 | -0,81 | 2,39 | -0,34 | 7,35E-01 |
| AT1G15140 | 0,76 | -0,78 | 2,23 | -0,35 | 7,27E-01 |
| AT1G15175 | 0,27 | -1,67 | 4,36 | -0,38 | 7,02E-01 |
| AT1G15210 | 0,17 | 0,00 | 4,41 | 0,00 | 1,00E+00 |
| AT1G15220 | 0,11 | -0,80 | 4,39 | -0,18 | 8,56E-01 |
| AT1G15340 | 0,21 | 0,62 | 4,34 | 0,14 | 8,87E-01 |
| AT1G15350 | 1,25 | -2,63 | 1,98 | -1,33 | 1,84E-01 |
| AT1G15400 | 0,43 | 0,27 | 2,76 | 0,10 | 9,21E-01 |
| AT1G15440 | 0,87 | -3,39 | 2,14 | -1,59 | 1,12E-01 |
| AT1G15520 | 0,46 | -0,18 | 3,24 | -0,06 | 9,55E-01 |
| AT1G15530 | 1,02 | -4,28 | 2,31 | -1,85 | 6,44E-02 |
| AT1G15550 | 0,24 | -3,18 | 4,39 | -0,73 | 4,68E-01 |
| AT1G15700 | 0,40 | -2,44 | 3,38 | -0,72 | 4,71E-01 |
| AT1G05187 | 0,24 | -2,60 | 4,41 | -0,59 | 5,55E-01 |
| AT1G15730 | 0,44 | 0,78 | 4,05 | 0,19 | 8,48E-01 |
| AT1G15780 | 1,01 | 0,29 | 2,06 | 0,14 | 8,87E-01 |
| AT1G15790 | 1,05 | -2,19 | 2,72 | -0,80 | 4,21E-01 |
| AT1G05203 | 0,26 | -2,60 | 4,41 | -0,59 | 5,55E-01 |
| AT1G15880 | 0,20 | -1,00 | 4,37 | -0,23 | 8,19E-01 |
| AT1G15970 | 0,11 | 0,46 | 4,35 | 0,11 | 9,15E-01 |
| AT1G16010 | 0,99 | -1,70 | 2,44 | -0,70 | 4,85E-01 |
| AT1G16070 | 0,40 | -0,19 | 4,39 | -0,04 | 9,65E-01 |
| AT1G16210 | 0,60 | -2,85 | 3,05 | -0,93 | 3,50E-01 |
| AT1G16240 | 0,80 | -0,28 | 2,53 | -0,11 | 9,12E-01 |
| AT1G16460 | 0,57 | -0,07 | 2,35 | -0,03 | 9,75E-01 |
| AT1G16570 | 0,28 | 0,18 | 4,37 | 0,04 | 9,67E-01 |
| AT1G16610 | 1,02 | -1,80 | 3,01 | -0,60 | 5,50E-01 |
| AT1G16750 | 0,49 | -2,49 | 3,16 | -0,79 | 4,32E-01 |
| AT1G16780 | 0,12 | 0,00 | 4,41 | 0,00 | 1,00E+00 |
| AT1G16870 | 0,95 | -5,03 | 3,30 | -1,53 | 1,27E-01 |
| AT1G16890 | 1,30 | -0,40 | 1,58 | -0,25 | 8,01E-01 |
| AT1G16900 | 0,65 | -3,07 | 3,76 | -0,82 | 4,14E-01 |
| AT1G16920 | 0,22 | 0,37 | 4,35 | 0,09 | 9,32E-01 |
| AT1G16970 | 0,37 | -1,25 | 3,52 | -0,35 | 7,23E-01 |
| AT1G17080 | 0,60 | -0,64 | 3,24 | -0,20 | 8,42E-01 |
| AT1G17110 | 0,57 | -1,52 | 2,71 | -0,56 | 5,76E-01 |
| AT1G17140 | 0,95 | -4,36 | 3,40 | -1,28 | 2,00E-01 |
| AT1G17147 | 0,12 | 0,75 | 4,34 | 0,17 | 8,63E-01 |
| AT1G17232 | 0,51 | 2,17 | 3,04 | 0,71 | 4,76E-01 |
| AT1G17240 | 0,05 | -0,24 | 4,38 | -0,05 | 9,57E-01 |
| AT1G17250 | 0,99 | -1,64 | 2,74 | -0,60 | 5,48E-01 |
| AT1G17370 | 0,81 | -0,12 | 2,70 | -0,05 | 9,64E-01 |
| AT1G17380 | 0,50 | 0,92 | 2,57 | 0,36 | 7,21E-01 |
| AT1G05313 | 0,32 | 1,34 | 4,31 | 0,31 | 7,57E-01 |
| AT1G17440 | 0,46 | 0,53 | 3,28 | 0,16 | 8,73E-01 |

| | | | | | |
|---|---|---|---|---|---|
| AT1G17450 | 1,22 | -1,38 | 1,94 | -0,71 | 4,79E-01 |
| AT1G17590 | 0,46 | -1,28 | 3,30 | -0,39 | 6,98E-01 |
| AT1G17600 | 1,00 | 0,32 | 2,01 | 0,16 | 8,74E-01 |
| AT1G17665 | 0,20 | 0,00 | 4,41 | 0,00 | 1,00E+00 |
| AT1G17670 | 0,37 | -1,96 | 4,36 | -0,45 | 6,53E-01 |
| AT1G17690 | 0,25 | -2,53 | 4,40 | -0,57 | 5,65E-01 |
| AT1G17720 | 0,66 | 0,34 | 2,27 | 0,15 | 8,81E-01 |
| AT1G17744 | 0,16 | -0,68 | 4,41 | -0,15 | 8,77E-01 |
| AT1G17790 | 0,63 | -3,23 | 3,08 | -1,05 | 2,95E-01 |
| AT1G05333 | 0,71 | -4,36 | 3,51 | -1,24 | 2,13E-01 |
| AT1G17820 | 0,71 | -4,36 | 3,51 | -1,24 | 2,13E-01 |
| AT1G17850 | 0,36 | -2,77 | 4,35 | -0,64 | 5,24E-01 |
| AT1G05347 | 0,21 | 0,00 | 4,41 | 0,00 | 1,00E+00 |
| AT1G17870 | 0,39 | -2,20 | 4,28 | -0,51 | 6,07E-01 |
| AT1G17880 | 0,59 | -4,39 | 3,85 | -1,14 | 2,53E-01 |
| AT1G17890 | 0,42 | -2,18 | 4,39 | -0,50 | 6,19E-01 |
| AT1G17930 | 0,20 | -2,60 | 4,41 | -0,59 | 5,55E-01 |
| AT1G17980 | 1,06 | 0,76 | 2,26 | 0,34 | 7,37E-01 |
| AT1G17990 | 0,53 | -1,46 | 3,07 | -0,48 | 6,34E-01 |
| AT1G18050 | 0,60 | 0,86 | 2,54 | 0,34 | 7,37E-01 |
| AT1G18060 | 0,60 | -1,49 | 2,25 | -0,66 | 5,09E-01 |
| AT1G18070 | 0,60 | -1,64 | 2,61 | -0,63 | 5,30E-01 |
| AT1G18100 | 0,36 | 0,00 | 4,41 | 0,00 | 1,00E+00 |
| AT1G18150 | 1,19 | -4,11 | 2,21 | -1,86 | 6,36E-02 |
| AT1G18210 | 0,61 | 0,95 | 2,24 | 0,42 | 6,73E-01 |
| AT1G18330 | 0,22 | -1,25 | 4,37 | -0,29 | 7,75E-01 |
| AT1G18360 | 1,25 | -1,67 | 1,64 | -1,02 | 3,07E-01 |
| AT1G18382 | 0,44 | 0,41 | 2,72 | 0,15 | 8,79E-01 |
| AT1G18470 | 0,48 | 1,38 | 2,66 | 0,52 | 6,04E-01 |
| AT1G18540 | 1,15 | -0,51 | 2,04 | -0,25 | 8,03E-01 |
| AT1G18720 | 0,23 | 0,43 | 4,34 | 0,10 | 9,20E-01 |
| AT1G05433 | 0,53 | 2,51 | 3,00 | 0,84 | 4,03E-01 |
| AT1G18840 | 0,43 | -1,54 | 3,34 | -0,46 | 6,44E-01 |
| AT1G18890 | 0,28 | -1,75 | 4,39 | -0,40 | 6,90E-01 |
| AT1G18990 | 0,45 | -2,52 | 4,19 | -0,60 | 5,47E-01 |
| AT1G19000 | 0,86 | -0,30 | 1,75 | -0,17 | 8,62E-01 |
| AT1G19130 | 0,49 | 0,32 | 2,83 | 0,11 | 9,09E-01 |
| AT1G19140 | 0,44 | -2,61 | 4,35 | -0,60 | 5,49E-01 |
| AT1G19210 | 0,10 | 0,45 | 4,35 | 0,10 | 9,19E-01 |
| AT1G19260 | 0,97 | 2,94 | 2,51 | 1,17 | 2,41E-01 |
| AT1G19300 | 0,88 | 1,33 | 2,47 | 0,54 | 5,91E-01 |
| AT1G19360 | 0,50 | -2,28 | 4,02 | -0,57 | 5,72E-01 |
| AT1G19370 | 0,41 | -1,68 | 4,33 | -0,39 | 6,98E-01 |
| AT1G19430 | 0,39 | -1,67 | 4,26 | -0,39 | 6,94E-01 |
| AT1G19450 | 0,33 | -0,83 | 4,36 | -0,19 | 8,50E-01 |
| AT1G19570 | 0,89 | -0,39 | 1,95 | -0,20 | 8,41E-01 |
| AT1G19640 | 0,14 | 0,00 | 4,41 | 0,00 | 1,00E+00 |
| AT1G19660 | 0,55 | 0,62 | 2,61 | 0,24 | 8,11E-01 |
| AT1G19700 | 0,25 | -0,27 | 4,38 | -0,06 | 9,51E-01 |
| AT1G19780 | 0,50 | -0,62 | 3,98 | -0,16 | 8,76E-01 |
| AT1G19860 | 0,15 | 0,05 | 4,37 | 0,01 | 9,91E-01 |
| AT1G19870 | 0,80 | -0,87 | 2,60 | -0,33 | 7,39E-01 |
| AT1G19880 | 0,41 | 0,80 | 3,34 | 0,24 | 8,10E-01 |
| AT1G19920 | 1,00 | -3,94 | 2,24 | -1,76 | 7,88E-02 |
| AT1G20040 | 0,34 | -0,51 | 4,39 | -0,12 | 9,07E-01 |
| AT1G20070 | 0,59 | -0,52 | 2,49 | -0,21 | 8,35E-01 |

| | | | | | |
|---|---|---|---|---|---|
| AT1G20160 | 0,33 | -2,59 | 4,41 | -0,59 | 5,56E-01 |
| AT1G20190 | 0,64 | 0,15 | 3,30 | 0,05 | 9,64E-01 |
| AT1G20200 | 0,33 | 0,48 | 3,54 | 0,14 | 8,91E-01 |
| AT1G20320 | 0,05 | -0,24 | 4,38 | -0,05 | 9,57E-01 |
| AT1G20430 | 0,89 | -2,52 | 2,96 | -0,85 | 3,94E-01 |
| AT1G20490 | 0,48 | -2,60 | 4,41 | -0,59 | 5,55E-01 |
| AT1G20560 | 0,25 | -0,32 | 4,38 | -0,07 | 9,42E-01 |
| AT1G20570 | 0,08 | 0,00 | 4,41 | 0,00 | 1,00E+00 |
| AT1G20650 | 0,41 | -1,13 | 4,34 | -0,26 | 7,95E-01 |
| AT1G20670 | 0,31 | -2,23 | 4,36 | -0,51 | 6,09E-01 |
| AT1G20770 | 0,16 | -0,68 | 4,41 | -0,15 | 8,77E-01 |
| AT1G20840 | 1,09 | 0,11 | 1,89 | 0,06 | 9,55E-01 |
| AT1G20880 | 0,25 | 0,00 | 4,41 | 0,00 | 1,00E+00 |
| AT1G20890 | 0,48 | -0,68 | 4,07 | -0,17 | 8,67E-01 |
| AT1G20950 | 0,69 | -2,21 | 2,53 | -0,87 | 3,83E-01 |
| AT1G21080 | 0,61 | -0,10 | 2,43 | -0,04 | 9,68E-01 |
| AT1G05627 | 0,05 | -0,24 | 4,38 | -0,05 | 9,57E-01 |
| AT1G21160 | 0,26 | 0,58 | 4,35 | 0,13 | 8,94E-01 |
| AT1G21190 | 0,45 | 0,52 | 4,13 | 0,13 | 8,99E-01 |
| AT1G21310 | 0,44 | -0,82 | 2,82 | -0,29 | 7,71E-01 |
| AT1G21380 | 1,01 | -1,22 | 2,11 | -0,58 | 5,64E-01 |
| AT1G21390 | 0,08 | 0,00 | 4,41 | 0,00 | 1,00E+00 |
| AT1G21440 | 1,10 | -1,31 | 2,56 | -0,51 | 6,09E-01 |
| AT1G21500 | 0,51 | 0,75 | 3,06 | 0,24 | 8,07E-01 |
| AT1G21520 | 0,17 | -0,75 | 4,40 | -0,17 | 8,65E-01 |
| AT1G21550 | 0,17 | -0,71 | 4,40 | -0,16 | 8,72E-01 |
| AT1G21600 | 0,77 | 0,18 | 2,04 | 0,09 | 9,30E-01 |
| AT1G21650 | 0,46 | -1,44 | 3,30 | -0,44 | 6,62E-01 |
| AT1G21770 | 0,66 | 0,02 | 2,56 | 0,01 | 9,93E-01 |
| AT1G21890 | 0,36 | 1,88 | 4,29 | 0,44 | 6,61E-01 |
| AT1G21900 | 0,26 | 0,58 | 4,35 | 0,13 | 8,94E-01 |
| AT1G21920 | 0,41 | -1,24 | 3,36 | -0,37 | 7,11E-01 |
| AT1G21928 | 0,29 | -0,69 | 4,35 | -0,16 | 8,74E-01 |
| AT1G22180 | 0,30 | 1,47 | 3,57 | 0,41 | 6,81E-01 |
| AT1G22200 | 0,54 | -3,05 | 3,90 | -0,78 | 4,34E-01 |
| AT1G22300 | 0,58 | 1,55 | 2,59 | 0,60 | 5,51E-01 |
| AT1G22360 | 0,63 | -0,79 | 2,76 | -0,29 | 7,74E-01 |
| AT1G22400 | 0,23 | 0,05 | 4,36 | 0,01 | 9,90E-01 |
| AT1G22550 | 0,19 | 0,75 | 4,34 | 0,17 | 8,62E-01 |
| AT1G22570 | 0,46 | -0,75 | 4,08 | -0,18 | 8,54E-01 |
| AT1G22710 | 0,50 | 1,79 | 2,62 | 0,68 | 4,95E-01 |
| AT1G23020 | 0,20 | -2,55 | 4,40 | -0,58 | 5,62E-01 |
| AT1G23052 | 1,19 | 1,61 | 2,37 | 0,68 | 4,95E-01 |
| AT1G23080 | 0,61 | -0,57 | 3,20 | -0,18 | 8,60E-01 |
| AT1G05793 | 0,08 | -0,21 | 4,39 | -0,05 | 9,62E-01 |
| AT1G23205 | 0,35 | -0,42 | 4,34 | -0,10 | 9,23E-01 |
| AT1G23260 | 0,22 | -0,31 | 4,38 | -0,07 | 9,44E-01 |
| AT1G23330 | 0,10 | 0,45 | 4,35 | 0,10 | 9,19E-01 |
| AT1G23650 | 0,08 | 0,00 | 4,41 | 0,00 | 1,00E+00 |
| AT1G23670 | 0,22 | 1,56 | 4,30 | 0,36 | 7,16E-01 |
| AT1G23780 | 0,31 | -1,72 | 4,39 | -0,39 | 6,95E-01 |
| AT1G23830 | 0,53 | -0,58 | 3,20 | -0,18 | 8,58E-01 |
| AT1G23840 | 0,48 | -2,47 | 3,17 | -0,78 | 4,36E-01 |
| AT1G23850 | 0,35 | -1,16 | 4,39 | -0,27 | 7,91E-01 |
| AT1G23960 | 0,23 | 0,62 | 4,35 | 0,14 | 8,87E-01 |
| AT1G24040 | 0,10 | -0,18 | 4,38 | -0,04 | 9,67E-01 |

| | | | | | |
|---|---|---|---|---|---|
| AT1G24100 | 1,00 | 2,22 | 1,98 | 1,12 | 2,61E-01 |
| AT1G24120 | 0,29 | 0,68 | 4,34 | 0,16 | 8,76E-01 |
| AT1G24140 | 0,74 | -3,75 | 2,92 | -1,28 | 1,99E-01 |
| AT1G24160 | 0,84 | -2,60 | 3,44 | -0,76 | 4,49E-01 |
| AT1G24190 | 0,84 | -0,90 | 2,72 | -0,33 | 7,42E-01 |
| AT1G24267 | 0,48 | -1,67 | 3,27 | -0,51 | 6,09E-01 |
| AT1G24290 | 0,17 | -0,75 | 4,40 | -0,17 | 8,65E-01 |
| AT1G24350 | 0,14 | -0,35 | 4,38 | -0,08 | 9,37E-01 |
| AT1G05907 | 0,77 | 1,34 | 2,43 | 0,55 | 5,81E-01 |
| AT1G24764 | 0,53 | 0,44 | 3,30 | 0,13 | 8,94E-01 |
| AT1G24822 | 0,94 | -4,56 | 4,31 | -1,06 | 2,90E-01 |
| AT1G25220 | 0,36 | 0,00 | 4,41 | 0,00 | 1,00E+00 |
| AT1G25350 | 1,21 | -2,17 | 1,74 | -1,25 | 2,12E-01 |
| AT1G25370 | 0,12 | 0,00 | 4,41 | 0,00 | 1,00E+00 |
| AT1G25390 | 0,51 | -0,53 | 3,08 | -0,17 | 8,64E-01 |
| AT1G25400 | 0,63 | -0,23 | 2,88 | -0,08 | 9,38E-01 |
| AT1G25500 | 0,25 | 0,18 | 4,36 | 0,04 | 9,68E-01 |
| AT1G25540 | 0,94 | 0,16 | 2,34 | 0,07 | 9,45E-01 |
| AT1G25560 | 0,90 | 1,53 | 2,02 | 0,76 | 4,49E-01 |
| AT1G25570 | 0,49 | -4,43 | 4,31 | -1,03 | 3,04E-01 |
| AT1G26150 | 0,88 | -0,62 | 2,24 | -0,27 | 7,83E-01 |
| AT1G26160 | 0,24 | -0,79 | 4,36 | -0,18 | 8,56E-01 |
| AT1G26170 | 0,54 | 0,43 | 3,17 | 0,13 | 8,93E-01 |
| AT1G26208 | 0,43 | 1,17 | 4,33 | 0,27 | 7,87E-01 |
| AT1G26218 | 0,37 | 1,46 | 4,31 | 0,34 | 7,35E-01 |
| AT1G26233 | 0,08 | -0,21 | 4,39 | -0,05 | 9,62E-01 |
| AT1G26240 | 0,88 | 3,20 | 2,55 | 1,26 | 2,09E-01 |
| AT1G26255 | 0,44 | 2,25 | 4,03 | 0,56 | 5,77E-01 |
| AT1G26270 | 1,28 | -3,72 | 1,97 | -1,89 | 5,84E-02 |
| AT1G26380 | 0,46 | -1,21 | 4,34 | -0,28 | 7,80E-01 |
| AT1G26390 | 0,25 | -1,21 | 4,35 | -0,28 | 7,80E-01 |
| AT1G26420 | 0,40 | 0,65 | 4,28 | 0,15 | 8,80E-01 |
| AT1G26690 | 0,33 | -1,64 | 4,38 | -0,37 | 7,08E-01 |
| AT1G26761 | 0,85 | -0,68 | 4,41 | -0,15 | 8,77E-01 |
| AT1G26800 | 0,42 | 0,20 | 2,80 | 0,07 | 9,42E-01 |
| AT1G26930 | 0,36 | 0,43 | 4,33 | 0,10 | 9,21E-01 |
| AT1G27008 | 0,08 | 0,00 | 4,41 | 0,00 | 1,00E+00 |
| AT1G27190 | 0,77 | -0,96 | 2,14 | -0,45 | 6,53E-01 |
| AT1G27200 | 0,57 | 0,13 | 2,71 | 0,05 | 9,63E-01 |
| AT1G27210 | 0,26 | 1,00 | 4,33 | 0,23 | 8,18E-01 |
| AT1G27350 | 0,20 | -0,96 | 4,36 | -0,22 | 8,26E-01 |
| AT1G27370 | 0,51 | -2,06 | 4,06 | -0,51 | 6,12E-01 |
| AT1G27385 | 0,43 | -2,92 | 4,38 | -0,67 | 5,04E-01 |
| AT1G27440 | 0,24 | -1,92 | 4,38 | -0,44 | 6,61E-01 |
| AT1G27500 | 0,29 | -2,57 | 4,40 | -0,58 | 5,59E-01 |
| AT1G27510 | 0,44 | -0,50 | 3,23 | -0,15 | 8,78E-01 |
| AT1G06093 | 0,29 | -1,19 | 4,38 | -0,27 | 7,86E-01 |
| AT1G27600 | 1,12 | -4,78 | 2,07 | -2,31 | 2,09E-02 |
| AT1G27700 | 0,50 | -3,94 | 4,34 | -0,91 | 3,63E-01 |
| AT1G27720 | 0,50 | -3,56 | 3,95 | -0,90 | 3,68E-01 |
| AT1G27900 | 0,40 | 1,78 | 3,26 | 0,55 | 5,85E-01 |
| AT1G27930 | 0,53 | -1,87 | 3,14 | -0,59 | 5,52E-01 |
| AT1G28000 | 0,35 | 2,05 | 4,28 | 0,48 | 6,33E-01 |
| AT1G28120 | 0,73 | -3,42 | 2,95 | -1,16 | 2,46E-01 |
| AT1G28140 | 1,15 | -2,64 | 2,01 | -1,31 | 1,89E-01 |
| AT1G28150 | 0,60 | -1,65 | 3,11 | -0,53 | 5,95E-01 |

| | | | | | |
|---|---|---|---|---|---|
| AT1G28190 | 0,15 | -0,81 | 4,39 | -0,18 | 8,54E-01 |
| AT1G28200 | 0,24 | -0,80 | 4,39 | -0,18 | 8,55E-01 |
| AT1G28230 | 0,78 | 1,55 | 2,21 | 0,70 | 4,84E-01 |
| AT1G28240 | 0,33 | 0,38 | 4,35 | 0,09 | 9,31E-01 |
| AT1G28260 | 1,27 | -0,82 | 2,13 | -0,39 | 7,00E-01 |
| AT1G28280 | 1,03 | 0,69 | 1,92 | 0,36 | 7,21E-01 |
| AT1G28290 | 0,27 | -0,14 | 4,34 | -0,03 | 9,73E-01 |
| AT1G28310 | 0,66 | 2,22 | 2,14 | 1,04 | 3,00E-01 |
| AT1G28330 | 0,60 | -0,73 | 3,18 | -0,23 | 8,18E-01 |
| AT1G28370 | 0,92 | 2,54 | 2,26 | 1,12 | 2,62E-01 |
| AT1G28390 | 0,25 | -3,06 | 4,38 | -0,70 | 4,85E-01 |
| AT1G28400 | 0,35 | 0,61 | 3,46 | 0,18 | 8,60E-01 |
| AT1G28570 | 0,12 | -0,68 | 4,41 | -0,15 | 8,77E-01 |
| AT1G28580 | 0,48 | -1,63 | 3,97 | -0,41 | 6,82E-01 |
| AT1G28600 | 1,10 | -2,36 | 1,70 | -1,39 | 1,64E-01 |
| AT1G28630 | 0,79 | 1,79 | 2,77 | 0,65 | 5,17E-01 |
| AT1G28660 | 0,08 | 0,00 | 4,41 | 0,00 | 1,00E+00 |
| AT1G28685 | 0,55 | -0,59 | 3,22 | -0,18 | 8,55E-01 |
| AT1G28765 | 0,17 | 0,00 | 4,41 | 0,00 | 1,00E+00 |
| AT1G28960 | 0,56 | -1,56 | 3,99 | -0,39 | 6,96E-01 |
| AT1G29050 | 0,14 | 0,87 | 4,33 | 0,20 | 8,40E-01 |
| AT1G29170 | 0,29 | 0,44 | 4,35 | 0,10 | 9,20E-01 |
| AT1G29240 | 0,32 | -3,12 | 4,39 | -0,71 | 4,77E-01 |
| AT1G29300 | 0,08 | 0,05 | 4,38 | 0,01 | 9,91E-01 |
| AT1G29390 | 0,82 | -1,56 | 2,26 | -0,69 | 4,89E-01 |
| AT1G29395 | 0,39 | -0,79 | 3,45 | -0,23 | 8,20E-01 |
| AT1G29530 | 0,36 | -1,15 | 3,50 | -0,33 | 7,43E-01 |
| AT1G29690 | 0,27 | 0,70 | 4,34 | 0,16 | 8,72E-01 |
| AT1G29700 | 0,34 | -2,55 | 4,40 | -0,58 | 5,62E-01 |
| AT1G29720 | 0,71 | -2,74 | 2,54 | -1,08 | 2,80E-01 |
| AT1G29750 | 0,26 | -1,06 | 4,37 | -0,24 | 8,08E-01 |
| AT1G29880 | 1,19 | 0,07 | 1,89 | 0,04 | 9,71E-01 |
| AT1G29970 | 0,83 | -2,19 | 2,46 | -0,89 | 3,73E-01 |
| AT1G30000 | 0,84 | -1,75 | 2,57 | -0,68 | 4,98E-01 |
| AT1G30060 | 0,17 | 1,16 | 4,31 | 0,27 | 7,88E-01 |
| AT1G30070 | 0,15 | -0,17 | 4,39 | -0,04 | 9,68E-01 |
| AT1G30090 | 0,16 | -0,26 | 4,38 | -0,06 | 9,52E-01 |
| AT1G30120 | 1,11 | -0,51 | 1,91 | -0,27 | 7,88E-01 |
| AT1G30210 | 0,29 | -1,60 | 4,35 | -0,37 | 7,13E-01 |
| AT1G30250 | 0,60 | 1,10 | 2,50 | 0,44 | 6,60E-01 |
| AT1G30370 | 0,55 | -1,64 | 3,20 | -0,51 | 6,07E-01 |
| AT1G30500 | 0,64 | -1,24 | 3,88 | -0,32 | 7,50E-01 |
| AT1G06347 | 0,94 | -0,03 | 2,10 | -0,01 | 9,89E-01 |
| AT1G30540 | 0,42 | -1,90 | 4,15 | -0,46 | 6,48E-01 |
| AT1G30580 | 1,07 | -1,65 | 2,07 | -0,80 | 4,26E-01 |
| AT1G30590 | 0,66 | 1,82 | 2,78 | 0,66 | 5,12E-01 |
| AT1G30755 | 0,62 | -1,38 | 2,50 | -0,55 | 5,81E-01 |
| AT1G30780 | 0,83 | 0,89 | 2,21 | 0,40 | 6,88E-01 |
| AT1G30810 | 0,83 | -2,78 | 1,92 | -1,45 | 1,47E-01 |
| AT1G30880 | 0,08 | 0,00 | 4,41 | 0,00 | 1,00E+00 |
| AT1G30960 | 0,37 | -3,34 | 4,37 | -0,76 | 4,45E-01 |
| AT1G06407 | 0,34 | -0,08 | 3,57 | -0,02 | 9,83E-01 |
| AT1G31160 | 0,42 | -3,39 | 4,16 | -0,81 | 4,15E-01 |
| AT1G31180 | 0,17 | -0,75 | 4,40 | -0,17 | 8,65E-01 |
| AT1G31230 | 1,31 | 0,66 | 1,71 | 0,39 | 6,97E-01 |
| AT1G31240 | 0,22 | 1,03 | 4,32 | 0,24 | 8,11E-01 |

| | | | | | |
|---|---|---|---|---|---|
| AT1G31350 | 0,45 | -2,53 | 3,26 | -0,78 | 4,38E-01 |
| AT1G31410 | 0,08 | 0,00 | 4,41 | 0,00 | 1,00E+00 |
| AT1G31470 | 0,15 | -0,73 | 4,40 | -0,17 | 8,68E-01 |
| AT1G31550 | 0,81 | -1,64 | 2,70 | -0,61 | 5,45E-01 |
| AT1G31600 | 0,81 | -1,15 | 2,91 | -0,40 | 6,92E-01 |
| AT1G31640 | 0,23 | 0,68 | 4,34 | 0,16 | 8,76E-01 |
| AT1G31650 | 0,25 | -0,89 | 4,35 | -0,21 | 8,37E-01 |
| AT1G31660 | 0,51 | -3,68 | 4,36 | -0,85 | 3,98E-01 |
| AT1G31740 | 0,22 | 0,38 | 4,35 | 0,09 | 9,31E-01 |
| AT1G31810 | 0,57 | -1,70 | 2,32 | -0,73 | 4,63E-01 |
| AT1G31812 | 0,29 | 0,39 | 4,35 | 0,09 | 9,28E-01 |
| AT1G31814 | 0,37 | -2,49 | 4,39 | -0,57 | 5,70E-01 |
| AT1G31820 | 0,94 | -2,10 | 2,78 | -0,75 | 4,51E-01 |
| AT1G31830 | 0,66 | -0,87 | 2,70 | -0,32 | 7,48E-01 |
| AT1G31860 | 1,08 | 0,40 | 1,71 | 0,23 | 8,16E-01 |
| AT1G06463 | 0,27 | 0,65 | 4,34 | 0,15 | 8,82E-01 |
| AT1G31920 | 0,52 | -1,61 | 2,64 | -0,61 | 5,42E-01 |
| AT1G31940 | 0,35 | -2,02 | 4,35 | -0,47 | 6,42E-01 |
| AT1G06473 | 0,38 | -0,28 | 4,38 | -0,06 | 9,50E-01 |
| AT1G32000 | 0,35 | 0,08 | 4,32 | 0,02 | 9,85E-01 |
| AT1G32070 | 0,34 | -2,60 | 4,41 | -0,59 | 5,55E-01 |
| AT1G32080 | 0,85 | -0,35 | 1,93 | -0,18 | 8,55E-01 |
| AT1G32130 | 1,27 | -2,77 | 1,96 | -1,41 | 1,57E-01 |
| AT1G32200 | 0,75 | 0,28 | 2,16 | 0,13 | 8,98E-01 |
| AT1G32340 | 0,12 | -0,78 | 4,39 | -0,18 | 8,59E-01 |
| AT1G32400 | 0,47 | 0,52 | 2,68 | 0,20 | 8,45E-01 |
| AT1G32450 | 0,27 | -0,77 | 4,35 | -0,18 | 8,60E-01 |
| AT1G06573 | 0,41 | 1,11 | 4,32 | 0,26 | 7,97E-01 |
| AT1G32470 | 0,50 | 1,51 | 2,65 | 0,57 | 5,69E-01 |
| AT1G32490 | 0,70 | -0,52 | 2,27 | -0,23 | 8,18E-01 |
| AT1G32500 | 0,52 | -1,68 | 2,66 | -0,63 | 5,28E-01 |
| AT1G32530 | 0,53 | -2,01 | 2,68 | -0,75 | 4,53E-01 |
| AT1G06577 | 0,16 | -0,01 | 4,37 | 0,00 | 9,98E-01 |
| AT1G32713 | 1,03 | 2,69 | 2,26 | 1,19 | 2,33E-01 |
| AT1G32860 | 0,05 | -0,24 | 4,38 | -0,05 | 9,57E-01 |
| AT1G32880 | 0,26 | -2,60 | 4,41 | -0,59 | 5,55E-01 |
| AT1G32900 | 0,61 | 0,54 | 3,01 | 0,18 | 8,58E-01 |
| AT1G32970 | 0,17 | 0,00 | 4,41 | 0,00 | 1,00E+00 |
| AT1G33050 | 0,95 | -2,30 | 1,93 | -1,19 | 2,34E-01 |
| AT1G06643 | 0,36 | -2,60 | 4,41 | -0,59 | 5,55E-01 |
| AT1G33250 | 0,90 | -0,17 | 2,30 | -0,07 | 9,40E-01 |
| AT1G33270 | 0,24 | -2,59 | 4,41 | -0,59 | 5,56E-01 |
| AT1G06673 | 0,47 | 1,60 | 3,24 | 0,49 | 6,21E-01 |
| AT1G33390 | 1,05 | -2,98 | 2,37 | -1,26 | 2,08E-01 |
| AT1G33430 | 0,08 | 0,00 | 4,41 | 0,00 | 1,00E+00 |
| AT1G33520 | 0,47 | -1,72 | 3,27 | -0,53 | 5,98E-01 |
| AT1G33615 | 0,17 | -0,68 | 4,41 | -0,15 | 8,77E-01 |
| AT1G33700 | 0,88 | -1,43 | 2,16 | -0,66 | 5,07E-01 |
| AT1G33720 | 1,30 | -0,56 | 1,84 | -0,30 | 7,62E-01 |
| AT1G33780 | 0,08 | 0,05 | 4,38 | 0,01 | 9,91E-01 |
| AT1G33870 | 0,60 | 0,43 | 2,62 | 0,16 | 8,70E-01 |
| AT1G33970 | 0,67 | -1,34 | 2,54 | -0,53 | 5,97E-01 |
| AT1G34030 | 0,44 | -1,11 | 2,86 | -0,39 | 6,99E-01 |
| AT1G34042 | 0,29 | 1,25 | 4,32 | 0,29 | 7,73E-01 |
| AT1G34110 | 0,91 | 0,64 | 1,91 | 0,34 | 7,36E-01 |
| AT1G34170 | 0,20 | -0,63 | 4,40 | -0,14 | 8,86E-01 |

| | | | | | |
|---|---|---|---|---|---|
| AT1G34220 | 1,01 | -0,83 | 2,95 | -0,28 | 7,78E-01 |
| AT1G34270 | 0,12 | 0,00 | 4,41 | 0,00 | 1,00E+00 |
| AT1G34310 | 0,97 | 3,14 | 2,57 | 1,22 | 2,21E-01 |
| AT1G34315 | 0,23 | -0,77 | 4,39 | -0,18 | 8,60E-01 |
| AT1G34380 | 0,08 | 0,00 | 4,41 | 0,00 | 1,00E+00 |
| AT1G34550 | 0,63 | 0,41 | 2,42 | 0,17 | 8,64E-01 |
| AT1G34750 | 0,23 | 0,89 | 4,33 | 0,21 | 8,37E-01 |
| AT1G34780 | 0,68 | -1,64 | 2,99 | -0,55 | 5,83E-01 |
| AT1G35150 | 0,16 | 0,63 | 4,35 | 0,14 | 8,85E-01 |
| AT1G35240 | 1,26 | 2,50 | 1,88 | 1,33 | 1,84E-01 |
| AT1G35250 | 0,19 | 1,05 | 4,32 | 0,24 | 8,08E-01 |
| AT1G35320 | 0,59 | 2,42 | 2,88 | 0,84 | 4,01E-01 |
| AT1G35350 | 0,96 | 1,68 | 1,93 | 0,87 | 3,85E-01 |
| AT1G06883 | 0,19 | 1,38 | 4,31 | 0,32 | 7,48E-01 |
| AT1G06897 | 0,91 | 1,73 | 3,41 | 0,51 | 6,11E-01 |
| AT1G06903 | 0,10 | 0,45 | 4,35 | 0,10 | 9,19E-01 |
| AT1G35520 | 1,01 | 3,24 | 2,43 | 1,33 | 1,82E-01 |
| AT1G35545 | 0,38 | 1,69 | 3,32 | 0,51 | 6,11E-01 |
| AT1G35580 | 1,07 | -0,04 | 1,70 | -0,02 | 9,84E-01 |
| AT1G35610 | 0,22 | 0,09 | 4,38 | 0,02 | 9,83E-01 |
| AT1G35625 | 0,08 | 0,00 | 4,41 | 0,00 | 1,00E+00 |
| AT1G35730 | 0,17 | 0,00 | 4,41 | 0,00 | 1,00E+00 |
| AT1G35895 | 0,55 | 0,26 | 3,26 | 0,08 | 9,37E-01 |
| AT1G36070 | 0,25 | -2,57 | 4,40 | -0,58 | 5,60E-01 |
| AT1G36085 | 0,68 | 3,22 | 4,26 | 0,76 | 4,50E-01 |
| AT1G36230 | 0,12 | 0,75 | 4,34 | 0,17 | 8,63E-01 |
| AT1G36370 | 0,71 | -0,25 | 2,37 | -0,11 | 9,15E-01 |
| AT1G36390 | 0,17 | 0,00 | 4,41 | 0,00 | 1,00E+00 |
| AT1G36622 | 0,40 | -0,73 | 3,46 | -0,21 | 8,33E-01 |
| AT1G36990 | 1,02 | -1,71 | 1,89 | -0,90 | 3,65E-01 |
| AT1G07013 | 0,15 | -0,17 | 4,39 | -0,04 | 9,68E-01 |
| AT1G40087 | 0,19 | 1,18 | 4,31 | 0,27 | 7,84E-01 |
| AT1G41830 | 0,28 | 1,25 | 4,31 | 0,29 | 7,72E-01 |
| AT1G41920 | 0,51 | 0,51 | 3,25 | 0,16 | 8,74E-01 |
| AT1G42440 | 0,46 | -0,44 | 3,33 | -0,13 | 8,96E-01 |
| AT1G42560 | 0,05 | -0,24 | 4,38 | -0,05 | 9,57E-01 |
| AT1G43190 | 0,38 | -1,12 | 4,35 | -0,26 | 7,97E-01 |
| AT1G43560 | 0,22 | 0,38 | 4,35 | 0,09 | 9,31E-01 |
| AT1G43580 | 0,21 | -0,44 | 4,38 | -0,10 | 9,20E-01 |
| AT1G43620 | 0,44 | -0,97 | 4,20 | -0,23 | 8,18E-01 |
| AT1G43665 | 0,88 | 2,06 | 3,18 | 0,65 | 5,18E-01 |
| AT1G43710 | 0,50 | -0,72 | 3,26 | -0,22 | 8,24E-01 |
| AT1G43730 | 0,40 | 1,53 | 4,23 | 0,36 | 7,18E-01 |
| AT1G43760 | 0,66 | 1,13 | 2,55 | 0,44 | 6,59E-01 |
| AT1G43825 | 0,81 | -0,04 | 2,58 | -0,02 | 9,87E-01 |
| AT1G43950 | 0,46 | 0,67 | 4,00 | 0,17 | 8,68E-01 |
| AT1G44000 | 0,08 | 0,00 | 4,41 | 0,00 | 1,00E+00 |
| AT1G44030 | 0,25 | 1,02 | 4,32 | 0,24 | 8,13E-01 |
| AT1G44085 | 0,65 | 2,72 | 2,81 | 0,97 | 3,33E-01 |
| AT1G44090 | 0,05 | -0,24 | 4,38 | -0,05 | 9,57E-01 |
| AT1G44125 | 0,91 | 0,51 | 2,75 | 0,18 | 8,54E-01 |
| AT1G44170 | 0,30 | 1,68 | 4,29 | 0,39 | 6,96E-01 |
| AT1G44350 | 0,41 | -1,02 | 4,24 | -0,24 | 8,09E-01 |
| AT1G44542 | 0,40 | -0,37 | 4,30 | -0,09 | 9,31E-01 |
| AT1G44770 | 0,41 | -0,08 | 3,32 | -0,02 | 9,80E-01 |
| AT1G44790 | 0,21 | 1,09 | 4,32 | 0,25 | 8,01E-01 |

| | | | | | |
|---|---|---|---|---|---|
| AT1G44920 | 0,44 | 0,12 | 3,33 | 0,04 | 9,71E-01 |
| AT1G44980 | 0,59 | 1,53 | 2,71 | 0,56 | 5,72E-01 |
| AT1G45063 | 0,43 | 0,00 | 4,41 | 0,00 | 1,00E+00 |
| AT1G45145 | 0,72 | -0,10 | 3,14 | -0,03 | 9,74E-01 |
| AT1G45150 | 0,25 | -0,89 | 4,35 | -0,21 | 8,37E-01 |
| AT1G45163 | 0,70 | -0,35 | 2,51 | -0,14 | 8,90E-01 |
| AT1G45207 | 0,37 | -3,94 | 4,34 | -0,91 | 3,63E-01 |
| AT1G07323 | 0,55 | -0,35 | 2,63 | -0,13 | 8,93E-01 |
| AT1G47200 | 0,16 | 0,61 | 4,34 | 0,14 | 8,89E-01 |
| AT1G47240 | 0,43 | -0,73 | 4,18 | -0,17 | 8,61E-01 |
| AT1G07333 | 0,81 | 3,09 | 3,23 | 0,95 | 3,40E-01 |
| AT1G47260 | 1,01 | 0,17 | 1,67 | 0,10 | 9,17E-01 |
| AT1G47280 | 1,31 | -0,60 | 1,72 | -0,35 | 7,27E-01 |
| AT1G07347 | 0,18 | 1,18 | 4,31 | 0,27 | 7,85E-01 |
| AT1G47380 | 0,18 | -0,34 | 4,38 | -0,08 | 9,38E-01 |
| AT1G47420 | 0,71 | -0,10 | 2,22 | -0,05 | 9,64E-01 |
| AT1G07383 | 0,55 | 0,53 | 2,63 | 0,20 | 8,41E-01 |
| AT1G47640 | 0,35 | -0,48 | 3,50 | -0,14 | 8,90E-01 |
| AT1G47670 | 0,05 | -0,24 | 4,38 | -0,05 | 9,57E-01 |
| AT1G47765 | 0,55 | -0,39 | 2,27 | -0,17 | 8,64E-01 |
| AT1G47900 | 0,45 | -3,33 | 4,15 | -0,80 | 4,23E-01 |
| AT1G48000 | 0,50 | -0,29 | 4,35 | -0,07 | 9,47E-01 |
| AT1G48030 | 1,26 | 0,15 | 2,22 | 0,07 | 9,47E-01 |
| AT1G48040 | 0,63 | 0,00 | 4,41 | 0,00 | 1,00E+00 |
| AT1G48110 | 0,42 | -0,80 | 4,39 | -0,18 | 8,56E-01 |
| AT1G48120 | 0,82 | -1,97 | 2,39 | -0,82 | 4,10E-01 |
| AT1G48160 | 0,15 | 0,05 | 4,37 | 0,01 | 9,91E-01 |
| AT1G48285 | 0,80 | -0,54 | 2,18 | -0,25 | 8,03E-01 |
| AT1G48320 | 0,17 | 0,00 | 4,41 | 0,00 | 1,00E+00 |
| AT1G48380 | 0,10 | -0,34 | 4,38 | -0,08 | 9,38E-01 |
| AT1G48405 | 0,82 | -2,60 | 3,43 | -0,76 | 4,48E-01 |
| AT1G48430 | 0,15 | -0,77 | 4,39 | -0,18 | 8,61E-01 |
| AT1G48550 | 1,08 | -1,82 | 2,72 | -0,67 | 5,02E-01 |
| AT1G48570 | 0,26 | -0,38 | 4,38 | -0,09 | 9,32E-01 |
| AT1G48580 | 0,08 | 0,00 | 4,41 | 0,00 | 1,00E+00 |
| AT1G48620 | 0,46 | 0,49 | 3,17 | 0,16 | 8,76E-01 |
| AT1G48630 | 1,25 | -3,12 | 1,91 | -1,63 | 1,03E-01 |
| AT1G48730 | 0,75 | 2,89 | 2,26 | 1,28 | 2,00E-01 |
| AT1G48742 | 0,46 | -0,80 | 3,24 | -0,25 | 8,06E-01 |
| AT1G48790 | 0,44 | -0,91 | 3,26 | -0,28 | 7,80E-01 |
| AT1G48850 | 0,39 | -2,01 | 4,33 | -0,46 | 6,43E-01 |
| AT1G48900 | 0,23 | -0,70 | 4,40 | -0,16 | 8,73E-01 |
| AT1G48930 | 0,71 | -4,13 | 2,84 | -1,45 | 1,47E-01 |
| AT1G49005 | 0,38 | 1,10 | 4,31 | 0,26 | 7,98E-01 |
| AT1G49140 | 0,12 | -0,31 | 4,38 | -0,07 | 9,44E-01 |
| AT1G49180 | 0,08 | 0,00 | 4,41 | 0,00 | 1,00E+00 |
| AT1G49205 | 0,08 | -0,21 | 4,39 | -0,05 | 9,62E-01 |
| AT1G49225 | 0,48 | -0,50 | 3,13 | -0,16 | 8,72E-01 |
| AT1G49270 | 0,10 | 0,45 | 4,35 | 0,10 | 9,19E-01 |
| AT1G49490 | 0,32 | -2,68 | 4,36 | -0,61 | 5,39E-01 |
| AT1G49540 | 0,83 | -0,17 | 2,58 | -0,07 | 9,47E-01 |
| AT1G49600 | 0,82 | -2,46 | 2,05 | -1,20 | 2,29E-01 |
| AT1G49670 | 0,26 | 1,51 | 4,30 | 0,35 | 7,26E-01 |
| AT1G49710 | 0,34 | 1,16 | 3,47 | 0,34 | 7,37E-01 |
| AT1G49780 | 1,22 | 0,97 | 1,79 | 0,54 | 5,89E-01 |
| AT1G49850 | 0,39 | -0,75 | 4,40 | -0,17 | 8,65E-01 |

| | | | | | |
|---|---|---|---|---|---|
| AT1G49970 | 0,66 | 0,45 | 2,21 | 0,20 | 8,40E-01 |
| AT1G07603 | 0,61 | -2,39 | 3,15 | -0,76 | 4,48E-01 |
| AT1G50020 | 0,58 | -2,51 | 3,96 | -0,63 | 5,26E-01 |
| AT1G50140 | 0,20 | 0,00 | 4,41 | 0,00 | 1,00E+00 |
| AT1G50190 | 0,32 | -1,70 | 4,40 | -0,39 | 6,99E-01 |
| AT1G50260 | 0,32 | -1,96 | 4,37 | -0,45 | 6,53E-01 |
| AT1G50280 | 0,61 | 0,20 | 2,31 | 0,09 | 9,31E-01 |
| AT1G50320 | 0,25 | -0,28 | 4,38 | -0,06 | 9,49E-01 |
| AT1G07627 | 0,18 | 0,57 | 4,34 | 0,13 | 8,95E-01 |
| AT1G50420 | 0,49 | -3,94 | 4,34 | -0,91 | 3,65E-01 |
| AT1G50430 | 1,02 | -1,19 | 1,77 | -0,67 | 5,02E-01 |
| AT1G50450 | 0,63 | -2,85 | 2,94 | -0,97 | 3,33E-01 |
| AT1G50500 | 0,57 | -1,97 | 3,19 | -0,62 | 5,37E-01 |
| AT1G07647 | 0,08 | 0,00 | 4,41 | 0,00 | 1,00E+00 |
| AT1G50570 | 0,65 | -0,44 | 2,61 | -0,17 | 8,67E-01 |
| AT1G50600 | 0,88 | 0,36 | 2,09 | 0,17 | 8,63E-01 |
| AT1G50640 | 1,08 | -2,08 | 1,80 | -1,15 | 2,49E-01 |
| AT1G50659 | 0,43 | 1,86 | 3,20 | 0,58 | 5,61E-01 |
| AT1G50750 | 0,08 | -0,21 | 4,39 | -0,05 | 9,62E-01 |
| AT1G50790 | 0,12 | 0,00 | 4,41 | 0,00 | 1,00E+00 |
| AT1G50820 | 0,50 | -4,09 | 3,95 | -1,04 | 3,00E-01 |
| AT1G50980 | 0,39 | -2,16 | 4,38 | -0,49 | 6,21E-01 |
| AT1G51080 | 0,73 | -1,21 | 2,47 | -0,49 | 6,23E-01 |
| AT1G07733 | 0,24 | -0,11 | 4,38 | -0,03 | 9,80E-01 |
| AT1G51400 | 0,93 | 2,50 | 2,21 | 1,13 | 2,58E-01 |
| AT1G51480 | 0,33 | -3,41 | 4,38 | -0,78 | 4,35E-01 |
| AT1G51500 | 1,21 | 0,99 | 2,07 | 0,48 | 6,34E-01 |
| AT1G51510 | 0,91 | 0,04 | 2,08 | 0,02 | 9,83E-01 |
| AT1G51560 | 0,45 | -2,66 | 3,32 | -0,80 | 4,24E-01 |
| AT1G51590 | 0,26 | 1,25 | 4,32 | 0,29 | 7,73E-01 |
| AT1G51630 | 0,52 | 1,28 | 3,10 | 0,41 | 6,81E-01 |
| AT1G51680 | 0,80 | 0,08 | 2,04 | 0,04 | 9,71E-01 |
| AT1G07763 | 0,56 | -2,60 | 4,41 | -0,59 | 5,55E-01 |
| AT1G51710 | 0,45 | -3,37 | 4,14 | -0,81 | 4,15E-01 |
| AT1G51730 | 0,47 | -1,14 | 3,24 | -0,35 | 7,24E-01 |
| AT1G51790 | 0,25 | -1,15 | 4,38 | -0,26 | 7,93E-01 |
| AT1G51890 | 0,23 | -0,75 | 4,40 | -0,17 | 8,65E-01 |
| AT1G51940 | 1,08 | -0,08 | 2,64 | -0,03 | 9,76E-01 |
| AT1G52000 | 0,46 | -1,63 | 4,11 | -0,40 | 6,91E-01 |
| AT1G52040 | 0,54 | -3,42 | 3,95 | -0,87 | 3,86E-01 |
| AT1G52150 | 1,30 | -2,80 | 2,36 | -1,19 | 2,36E-01 |
| AT1G52160 | 0,61 | -0,21 | 3,06 | -0,07 | 9,45E-01 |
| AT1G52180 | 1,00 | 0,53 | 2,01 | 0,27 | 7,91E-01 |
| AT1G52185 | 0,32 | -0,52 | 4,39 | -0,12 | 9,05E-01 |
| AT1G52200 | 0,25 | 0,00 | 4,41 | 0,00 | 1,00E+00 |
| AT1G52220 | 0,76 | -0,33 | 2,25 | -0,15 | 8,84E-01 |
| AT1G52290 | 0,41 | -1,33 | 4,28 | -0,31 | 7,55E-01 |
| AT1G52300 | 0,35 | 0,37 | 4,34 | 0,09 | 9,31E-01 |
| AT1G52340 | 0,08 | -0,21 | 4,39 | -0,05 | 9,62E-01 |
| AT1G52342 | 0,45 | -2,60 | 4,41 | -0,59 | 5,56E-01 |
| AT1G52510 | 0,47 | -1,39 | 4,22 | -0,33 | 7,41E-01 |
| AT1G52750 | 0,31 | -1,71 | 4,38 | -0,39 | 6,97E-01 |
| AT1G52880 | 0,40 | -1,64 | 3,47 | -0,47 | 6,37E-01 |
| AT1G52890 | 0,42 | -1,61 | 4,21 | -0,38 | 7,03E-01 |
| AT1G52970 | 1,13 | -1,13 | 2,00 | -0,56 | 5,72E-01 |
| AT1G52990 | 0,41 | 2,42 | 4,28 | 0,57 | 5,71E-01 |

| | | | | | |
|---|---|---|---|---|---|
| AT1G53000 | 0,49 | -3,19 | 4,00 | -0,80 | 4,24E-01 |
| AT1G53025 | 0,43 | -1,47 | 3,31 | -0,44 | 6,57E-01 |
| AT1G53035 | 0,12 | 0,00 | 4,41 | 0,00 | 1,00E+00 |
| AT1G53170 | 0,65 | 1,64 | 2,56 | 0,64 | 5,22E-01 |
| AT1G53200 | 0,24 | -2,59 | 4,41 | -0,59 | 5,56E-01 |
| AT1G53230 | 1,22 | -0,16 | 2,08 | -0,08 | 9,37E-01 |
| AT1G53240 | 1,16 | 1,99 | 1,92 | 1,04 | 2,99E-01 |
| AT1G53265 | 0,53 | -0,75 | 4,08 | -0,18 | 8,54E-01 |
| AT1G53320 | 0,35 | -0,99 | 4,38 | -0,23 | 8,21E-01 |
| AT1G53450 | 0,86 | -0,19 | 1,89 | -0,10 | 9,19E-01 |
| AT1G53480 | 0,84 | 1,38 | 2,99 | 0,46 | 6,45E-01 |
| AT1G53500 | 0,50 | 0,74 | 3,35 | 0,22 | 8,25E-01 |
| AT1G53510 | 0,76 | -1,42 | 2,40 | -0,59 | 5,54E-01 |
| AT1G53541 | 0,64 | -0,87 | 2,38 | -0,36 | 7,16E-01 |
| AT1G53580 | 0,12 | -0,14 | 4,38 | -0,03 | 9,74E-01 |
| AT1G53710 | 0,67 | -3,82 | 2,95 | -1,29 | 1,96E-01 |
| AT1G53750 | 1,03 | -1,71 | 1,82 | -0,94 | 3,49E-01 |
| AT1G53760 | 0,08 | 0,00 | 4,41 | 0,00 | 1,00E+00 |
| AT1G53770 | 0,33 | -3,01 | 4,38 | -0,69 | 4,92E-01 |
| AT1G54050 | 0,12 | -0,14 | 4,38 | -0,03 | 9,74E-01 |
| AT1G54080 | 1,16 | -2,63 | 2,15 | -1,22 | 2,21E-01 |
| AT1G54100 | 1,08 | -0,05 | 1,95 | -0,03 | 9,79E-01 |
| AT1G54140 | 0,15 | 0,05 | 4,37 | 0,01 | 9,91E-01 |
| AT1G54170 | 0,82 | -2,18 | 2,26 | -0,96 | 3,35E-01 |
| AT1G54320 | 0,67 | 0,40 | 2,49 | 0,16 | 8,73E-01 |
| AT1G54370 | 0,38 | -1,64 | 4,36 | -0,38 | 7,07E-01 |
| AT1G54500 | 0,85 | 0,80 | 1,99 | 0,40 | 6,89E-01 |
| AT1G54590 | 0,10 | -0,34 | 4,38 | -0,08 | 9,38E-01 |
| AT1G54630 | 0,35 | -1,23 | 4,38 | -0,28 | 7,80E-01 |
| AT1G54690 | 0,48 | -1,36 | 2,70 | -0,50 | 6,15E-01 |
| AT1G54775 | 0,57 | 0,93 | 3,81 | 0,24 | 8,08E-01 |
| AT1G54840 | 0,49 | -2,99 | 4,34 | -0,69 | 4,92E-01 |
| AT1G54950 | 0,44 | 1,36 | 4,01 | 0,34 | 7,34E-01 |
| AT1G55152 | 0,78 | 0,72 | 2,45 | 0,29 | 7,69E-01 |
| AT1G55180 | 0,78 | -2,77 | 2,82 | -0,98 | 3,26E-01 |
| AT1G55200 | 0,08 | 0,00 | 4,41 | 0,00 | 1,00E+00 |
| AT1G08043 | 0,30 | -2,60 | 4,41 | -0,59 | 5,55E-01 |
| AT1G55270 | 1,00 | -4,55 | 3,15 | -1,45 | 1,48E-01 |
| AT1G55300 | 0,10 | -0,18 | 4,38 | -0,04 | 9,67E-01 |
| AT1G55330 | 0,51 | -1,98 | 3,23 | -0,61 | 5,40E-01 |
| AT1G55360 | 0,31 | -0,72 | 4,33 | -0,17 | 8,67E-01 |
| AT1G55530 | 0,83 | -1,73 | 2,22 | -0,78 | 4,36E-01 |
| AT1G55535 | 0,40 | -3,08 | 4,26 | -0,72 | 4,70E-01 |
| AT1G55590 | 0,61 | -2,09 | 2,62 | -0,80 | 4,24E-01 |
| AT1G55610 | 0,12 | -0,14 | 4,38 | -0,03 | 9,74E-01 |
| AT1G55620 | 0,57 | 1,19 | 3,13 | 0,38 | 7,03E-01 |
| AT1G55675 | 0,68 | -2,34 | 3,62 | -0,65 | 5,18E-01 |
| AT1G55690 | 0,12 | 0,00 | 4,41 | 0,00 | 1,00E+00 |
| AT1G55830 | 0,23 | 0,39 | 4,35 | 0,09 | 9,28E-01 |
| AT1G55880 | 0,36 | 0,00 | 4,41 | 0,00 | 1,00E+00 |
| AT1G55950 | 0,93 | 2,37 | 2,13 | 1,11 | 2,66E-01 |
| AT1G55960 | 0,25 | -1,24 | 4,38 | -0,28 | 7,76E-01 |
| AT1G55970 | 0,17 | 0,00 | 4,41 | 0,00 | 1,00E+00 |
| AT1G56010 | 0,42 | -1,65 | 4,18 | -0,39 | 6,93E-01 |
| AT1G56100 | 0,86 | -0,64 | 2,89 | -0,22 | 8,24E-01 |
| AT1G56120 | 0,47 | -1,01 | 4,05 | -0,25 | 8,03E-01 |

| | | | | | |
|---|---|---|---|---|---|
| AT1G56140 | 0,44 | -0,27 | 3,28 | -0,08 | 9,34E-01 |
| AT1G56145 | 0,80 | -4,54 | 2,72 | -1,67 | 9,48E-02 |
| AT1G56165 | 0,20 | -2,55 | 4,40 | -0,58 | 5,62E-01 |
| AT1G56230 | 0,19 | -0,11 | 4,39 | -0,02 | 9,80E-01 |
| AT1G08107 | 0,31 | 1,52 | 3,56 | 0,43 | 6,69E-01 |
| AT1G08113 | 0,55 | 0,56 | 3,23 | 0,17 | 8,63E-01 |
| AT1G08123 | 0,56 | 1,90 | 2,48 | 0,77 | 4,43E-01 |
| AT1G56300 | 0,85 | -0,65 | 2,77 | -0,24 | 8,14E-01 |
| AT1G56330 | 0,78 | -0,73 | 2,30 | -0,32 | 7,51E-01 |
| AT1G56345 | 0,19 | 0,64 | 4,34 | 0,15 | 8,83E-01 |
| AT1G56440 | 0,27 | -0,77 | 4,35 | -0,18 | 8,60E-01 |
| AT1G56520 | 1,21 | -1,17 | 2,11 | -0,56 | 5,79E-01 |
| AT1G56550 | 0,17 | 0,00 | 4,41 | 0,00 | 1,00E+00 |
| AT1G56580 | 0,24 | -0,79 | 4,36 | -0,18 | 8,56E-01 |
| AT1G56650 | 1,01 | 0,00 | 4,04 | 0,00 | 1,00E+00 |
| AT1G57775 | 0,38 | 1,69 | 3,32 | 0,51 | 6,11E-01 |
| AT1G57777 | 0,31 | 0,89 | 3,61 | 0,25 | 8,04E-01 |
| AT1G57860 | 0,30 | -2,60 | 4,41 | -0,59 | 5,55E-01 |
| AT1G57870 | 0,40 | 0,71 | 4,26 | 0,17 | 8,67E-01 |
| AT1G57943 | 0,08 | 0,00 | 4,41 | 0,00 | 1,00E+00 |
| AT1G57990 | 0,77 | -0,96 | 2,44 | -0,39 | 6,93E-01 |
| AT1G58030 | 0,73 | -1,38 | 2,51 | -0,55 | 5,84E-01 |
| AT1G58060 | 0,60 | -1,00 | 3,10 | -0,32 | 7,46E-01 |
| AT1G58080 | 0,29 | -1,64 | 4,38 | -0,37 | 7,09E-01 |
| AT1G58100 | 1,20 | -3,26 | 1,85 | -1,76 | 7,76E-02 |
| AT1G58110 | 0,53 | -3,19 | 3,94 | -0,81 | 4,19E-01 |
| AT1G58200 | 0,16 | 0,61 | 4,34 | 0,14 | 8,89E-01 |
| AT1G58220 | 0,54 | -2,87 | 3,85 | -0,75 | 4,56E-01 |
| AT1G58350 | 0,80 | -1,34 | 2,42 | -0,55 | 5,80E-01 |
| AT1G58390 | 0,70 | 2,07 | 2,93 | 0,71 | 4,78E-01 |
| AT1G58440 | 0,58 | 1,18 | 2,98 | 0,40 | 6,93E-01 |
| AT1G58684 | 0,49 | 0,84 | 4,33 | 0,19 | 8,47E-01 |
| AT1G58725 | 0,33 | 1,17 | 4,32 | 0,27 | 7,86E-01 |
| AT1G58766 | 0,16 | -0,26 | 4,38 | -0,06 | 9,52E-01 |
| AT1G59030 | 0,44 | 1,04 | 4,26 | 0,24 | 8,07E-01 |
| AT1G59077 | 0,16 | -0,01 | 4,37 | 0,00 | 9,98E-01 |
| AT1G59406 | 0,14 | 0,48 | 4,35 | 0,11 | 9,13E-01 |
| AT1G59453 | 0,25 | 1,15 | 4,32 | 0,27 | 7,91E-01 |
| AT1G59720 | 0,61 | -1,12 | 2,65 | -0,42 | 6,73E-01 |
| AT1G59780 | 0,52 | -0,21 | 3,12 | -0,07 | 9,46E-01 |
| AT1G59920 | 0,42 | 1,86 | 3,22 | 0,58 | 5,64E-01 |
| AT1G59930 | 1,11 | 2,83 | 2,34 | 1,21 | 2,27E-01 |
| AT1G59990 | 1,13 | -2,01 | 2,13 | -0,94 | 3,45E-01 |
| AT1G60000 | 0,38 | -0,25 | 3,52 | -0,07 | 9,43E-01 |
| AT1G60160 | 0,43 | -0,14 | 3,20 | -0,04 | 9,66E-01 |
| AT1G60200 | 0,68 | -0,92 | 3,05 | -0,30 | 7,63E-01 |
| AT1G60230 | 0,48 | -2,26 | 3,97 | -0,57 | 5,69E-01 |
| AT1G60260 | 0,77 | -0,62 | 2,43 | -0,26 | 7,98E-01 |
| AT1G08317 | 0,08 | 0,00 | 4,41 | 0,00 | 1,00E+00 |
| AT1G60430 | 0,45 | -3,33 | 4,03 | -0,83 | 4,08E-01 |
| AT1G60490 | 0,25 | -0,28 | 4,38 | -0,06 | 9,49E-01 |
| AT1G60550 | 0,38 | -3,33 | 4,37 | -0,76 | 4,46E-01 |
| AT1G60560 | 0,17 | 0,00 | 4,41 | 0,00 | 1,00E+00 |
| AT1G60590 | 0,17 | 0,00 | 4,41 | 0,00 | 1,00E+00 |
| AT1G60630 | 0,08 | 0,00 | 4,41 | 0,00 | 1,00E+00 |
| AT1G60640 | 0,12 | -0,68 | 4,41 | -0,15 | 8,77E-01 |

| | | | | | |
|---|---|---|---|---|---|
| AT1G60700 | 0,48 | -2,60 | 4,41 | -0,59 | 5,55E-01 |
| AT1G60710 | 0,41 | -4,00 | 4,34 | -0,92 | 3,56E-01 |
| AT1G60770 | 1,20 | -4,78 | 2,11 | -2,26 | 2,36E-02 |
| AT1G60780 | 0,33 | 1,17 | 4,32 | 0,27 | 7,87E-01 |
| AT1G60800 | 0,56 | -3,04 | 3,13 | -0,97 | 3,32E-01 |
| AT1G60810 | 0,12 | 0,75 | 4,34 | 0,17 | 8,63E-01 |
| AT1G60850 | 0,11 | -0,68 | 4,41 | -0,15 | 8,77E-01 |
| AT1G60890 | 0,24 | 0,00 | 4,41 | 0,00 | 1,00E+00 |
| AT1G60930 | 0,24 | -2,54 | 4,40 | -0,58 | 5,64E-01 |
| AT1G60940 | 0,93 | -1,71 | 2,11 | -0,81 | 4,19E-01 |
| AT1G60990 | 0,24 | -2,59 | 4,41 | -0,59 | 5,56E-01 |
| AT1G60995 | 0,35 | 0,08 | 4,32 | 0,02 | 9,85E-01 |
| AT1G61010 | 0,89 | -3,28 | 2,19 | -1,49 | 1,35E-01 |
| AT1G61200 | 1,13 | 1,31 | 1,76 | 0,75 | 4,56E-01 |
| AT1G61260 | 0,47 | 0,64 | 3,21 | 0,20 | 8,41E-01 |
| AT1G61300 | 1,06 | -1,17 | 2,26 | -0,52 | 6,06E-01 |
| AT1G61310 | 0,33 | 1,30 | 4,31 | 0,30 | 7,63E-01 |
| AT1G61360 | 0,15 | 0,05 | 4,37 | 0,01 | 9,91E-01 |
| AT1G61370 | 0,17 | -0,35 | 4,38 | -0,08 | 9,37E-01 |
| AT1G61380 | 0,34 | 0,73 | 4,33 | 0,17 | 8,66E-01 |
| AT1G61420 | 0,12 | 0,00 | 4,41 | 0,00 | 1,00E+00 |
| AT1G61560 | 0,37 | -0,19 | 4,39 | -0,04 | 9,65E-01 |
| AT1G08417 | 0,75 | -0,76 | 2,64 | -0,29 | 7,74E-01 |
| AT1G61660 | 0,65 | 1,45 | 2,94 | 0,49 | 6,23E-01 |
| AT1G61730 | 1,18 | -4,12 | 2,21 | -1,86 | 6,25E-02 |
| AT1G61900 | 0,26 | 0,48 | 4,35 | 0,11 | 9,12E-01 |
| AT1G62085 | 0,60 | -0,43 | 3,19 | -0,13 | 8,94E-01 |
| AT1G62300 | 0,27 | -0,27 | 4,34 | -0,06 | 9,50E-01 |
| AT1G62422 | 0,51 | -0,47 | 2,66 | -0,18 | 8,60E-01 |
| AT1G62610 | 0,76 | -3,10 | 2,30 | -1,35 | 1,77E-01 |
| AT1G62680 | 1,24 | 0,25 | 1,98 | 0,13 | 8,99E-01 |
| AT1G62760 | 0,29 | 0,78 | 4,33 | 0,18 | 8,58E-01 |
| AT1G62780 | 0,49 | 0,20 | 3,18 | 0,06 | 9,49E-01 |
| AT1G62810 | 0,97 | -2,10 | 2,11 | -0,99 | 3,21E-01 |
| AT1G62915 | 0,64 | 1,35 | 2,93 | 0,46 | 6,46E-01 |
| AT1G62960 | 0,36 | -1,24 | 4,38 | -0,28 | 7,77E-01 |
| AT1G63010 | 0,16 | -0,01 | 4,37 | 0,00 | 9,98E-01 |
| AT1G63020 | 0,44 | 0,00 | 4,41 | 0,00 | 1,00E+00 |
| AT1G63050 | 0,34 | -1,60 | 4,35 | -0,37 | 7,13E-01 |
| AT1G63250 | 0,13 | 0,00 | 4,41 | 0,00 | 1,00E+00 |
| AT1G63290 | 0,12 | -0,68 | 4,41 | -0,15 | 8,77E-01 |
| AT1G63320 | 0,20 | 1,44 | 4,30 | 0,33 | 7,38E-01 |
| AT1G63410 | 0,62 | -0,44 | 2,60 | -0,17 | 8,66E-01 |
| AT1G63490 | 0,53 | -0,53 | 3,23 | -0,16 | 8,70E-01 |
| AT1G63660 | 0,92 | -1,86 | 2,08 | -0,90 | 3,70E-01 |
| AT1G63670 | 0,18 | -1,07 | 4,38 | -0,25 | 8,06E-01 |
| AT1G63720 | 0,19 | 0,09 | 4,37 | 0,02 | 9,84E-01 |
| AT1G08627 | 1,18 | -1,76 | 2,12 | -0,83 | 4,07E-01 |
| AT1G63800 | 0,47 | -1,18 | 4,33 | -0,27 | 7,85E-01 |
| AT1G63810 | 0,31 | -2,60 | 4,41 | -0,59 | 5,55E-01 |
| AT1G63840 | 0,40 | -0,74 | 4,40 | -0,17 | 8,67E-01 |
| AT1G63860 | 1,09 | -2,20 | 1,75 | -1,26 | 2,09E-01 |
| AT1G63870 | 0,58 | -0,82 | 3,13 | -0,26 | 7,93E-01 |
| AT1G63970 | 0,51 | -0,92 | 3,28 | -0,28 | 7,80E-01 |
| AT1G64080 | 0,15 | 0,41 | 4,35 | 0,09 | 9,26E-01 |
| AT1G64190 | 1,11 | -1,32 | 1,70 | -0,78 | 4,37E-01 |

| | | | | | |
|---|---|---|---|---|---|
| AT1G64200 | 0,78 | 0,00 | 4,41 | 0,00 | 1,00E+00 |
| AT1G64280 | 0,86 | -1,64 | 2,56 | -0,64 | 5,22E-01 |
| AT1G64380 | 0,37 | 1,37 | 4,30 | 0,32 | 7,50E-01 |
| AT1G64460 | 0,72 | -2,46 | 2,51 | -0,98 | 3,26E-01 |
| AT1G64480 | 0,18 | 0,57 | 4,34 | 0,13 | 8,95E-01 |
| AT1G64510 | 0,32 | 0,56 | 4,34 | 0,13 | 8,97E-01 |
| AT1G64580 | 0,24 | -0,68 | 4,41 | -0,15 | 8,77E-01 |
| AT1G64610 | 0,62 | -1,84 | 2,53 | -0,73 | 4,68E-01 |
| AT1G64640 | 0,11 | -0,80 | 4,39 | -0,18 | 8,56E-01 |
| AT1G64650 | 0,59 | 1,05 | 2,42 | 0,43 | 6,64E-01 |
| AT1G64670 | 0,10 | -0,18 | 4,38 | -0,04 | 9,67E-01 |
| AT1G64750 | 0,36 | -2,50 | 4,39 | -0,57 | 5,69E-01 |
| AT1G64970 | 0,63 | -0,77 | 2,63 | -0,29 | 7,71E-01 |
| AT1G64980 | 1,26 | -0,60 | 1,60 | -0,37 | 7,08E-01 |
| AT1G65040 | 0,28 | 1,23 | 3,64 | 0,34 | 7,35E-01 |
| AT1G65150 | 0,20 | 0,00 | 4,41 | 0,00 | 1,00E+00 |
| AT1G65190 | 0,43 | -0,37 | 4,15 | -0,09 | 9,30E-01 |
| AT1G08747 | 0,81 | -2,77 | 2,86 | -0,97 | 3,33E-01 |
| AT1G65290 | 0,77 | -0,55 | 2,25 | -0,24 | 8,07E-01 |
| AT1G65320 | 0,39 | -0,39 | 4,38 | -0,09 | 9,29E-01 |
| AT1G65346 | 0,34 | 1,35 | 4,31 | 0,31 | 7,54E-01 |
| AT1G65420 | 0,15 | 0,00 | 4,41 | 0,00 | 1,00E+00 |
| AT1G65430 | 0,52 | -1,38 | 3,16 | -0,44 | 6,63E-01 |
| AT1G65520 | 0,29 | 0,34 | 4,35 | 0,08 | 9,38E-01 |
| AT1G65540 | 1,25 | -1,62 | 2,22 | -0,73 | 4,65E-01 |
| AT1G65550 | 0,21 | 0,00 | 4,41 | 0,00 | 1,00E+00 |
| AT1G65580 | 0,46 | -0,33 | 2,77 | -0,12 | 9,04E-01 |
| AT1G65690 | 0,12 | 0,00 | 4,41 | 0,00 | 1,00E+00 |
| AT1G65950 | 0,28 | -3,07 | 4,38 | -0,70 | 4,84E-01 |
| AT1G08797 | 0,63 | -0,68 | 4,41 | -0,15 | 8,77E-01 |
| AT1G65980 | 1,27 | -4,23 | 2,13 | -1,98 | 4,72E-02 |
| AT1G65985 | 0,33 | -1,01 | 4,35 | -0,23 | 8,15E-01 |
| AT1G65990 | 1,24 | 3,33 | 1,99 | 1,68 | 9,36E-02 |
| AT1G66235 | 0,12 | -0,31 | 4,38 | -0,07 | 9,44E-01 |
| AT1G66240 | 0,12 | -0,68 | 4,41 | -0,15 | 8,77E-01 |
| AT1G66260 | 1,28 | 0,50 | 1,79 | 0,28 | 7,79E-01 |
| AT1G66290 | 0,49 | -0,71 | 3,30 | -0,21 | 8,30E-01 |
| AT1G66330 | 0,52 | -0,77 | 2,69 | -0,29 | 7,74E-01 |
| AT1G66460 | 0,25 | 0,00 | 4,41 | 0,00 | 1,00E+00 |
| AT1G66490 | 0,90 | -3,32 | 2,41 | -1,38 | 1,67E-01 |
| AT1G66500 | 0,25 | -3,06 | 4,38 | -0,70 | 4,85E-01 |
| AT1G66660 | 0,44 | -0,91 | 4,15 | -0,22 | 8,28E-01 |
| AT1G66680 | 0,59 | -4,04 | 4,33 | -0,93 | 3,51E-01 |
| AT1G66700 | 0,46 | -0,35 | 4,38 | -0,08 | 9,37E-01 |
| AT1G66830 | 0,68 | -1,50 | 3,79 | -0,40 | 6,92E-01 |
| AT1G66910 | 0,65 | -2,45 | 2,18 | -1,12 | 2,61E-01 |
| AT1G66940 | 0,34 | 0,28 | 4,30 | 0,07 | 9,48E-01 |
| AT1G67000 | 0,91 | 2,67 | 2,00 | 1,33 | 1,83E-01 |
| AT1G67050 | 0,28 | 1,14 | 3,64 | 0,31 | 7,53E-01 |
| AT1G67080 | 0,36 | -2,39 | 4,36 | -0,55 | 5,84E-01 |
| AT1G67140 | 0,95 | -2,64 | 2,46 | -1,07 | 2,83E-01 |
| AT1G67230 | 0,62 | 1,07 | 2,61 | 0,41 | 6,82E-01 |
| AT1G67235 | 0,29 | 1,90 | 4,29 | 0,44 | 6,58E-01 |
| AT1G67250 | 0,34 | -0,97 | 4,39 | -0,22 | 8,25E-01 |
| AT1G67300 | 0,37 | -1,72 | 4,36 | -0,39 | 6,94E-01 |
| AT1G67310 | 0,33 | -1,64 | 4,38 | -0,37 | 7,08E-01 |

| | | | | | |
|---|---|---|---|---|---|
| AT1G67325 | 0,61 | -1,64 | 3,06 | -0,54 | 5,92E-01 |
| AT1G67360 | 0,73 | 0,17 | 2,87 | 0,06 | 9,52E-01 |
| AT1G67481 | 1,30 | -1,28 | 1,70 | -0,75 | 4,51E-01 |
| AT1G67490 | 0,57 | -1,63 | 3,07 | -0,53 | 5,95E-01 |
| AT1G67500 | 0,52 | -2,88 | 4,01 | -0,72 | 4,73E-01 |
| AT1G67510 | 0,38 | 0,37 | 4,26 | 0,09 | 9,31E-01 |
| AT1G67580 | 1,21 | -1,56 | 1,83 | -0,85 | 3,95E-01 |
| AT1G67590 | 0,17 | -1,25 | 4,37 | -0,29 | 7,74E-01 |
| AT1G67680 | 0,43 | -2,29 | 4,36 | -0,53 | 5,99E-01 |
| AT1G67785 | 0,15 | -0,14 | 4,39 | -0,03 | 9,74E-01 |
| AT1G67790 | 0,26 | -2,60 | 4,41 | -0,59 | 5,55E-01 |
| AT1G67860 | 0,58 | -0,87 | 3,10 | -0,28 | 7,79E-01 |
| AT1G67865 | 0,24 | -0,68 | 4,41 | -0,15 | 8,77E-01 |
| AT1G67870 | 0,89 | 2,61 | 1,95 | 1,34 | 1,80E-01 |
| AT1G67930 | 1,11 | -1,36 | 1,64 | -0,83 | 4,06E-01 |
| AT1G67950 | 0,39 | -3,07 | 4,38 | -0,70 | 4,84E-01 |
| AT1G68060 | 0,56 | 0,02 | 3,25 | 0,01 | 9,96E-01 |
| AT1G68140 | 0,87 | 0,46 | 2,12 | 0,22 | 8,28E-01 |
| AT1G68325 | 0,05 | -0,24 | 4,38 | -0,05 | 9,57E-01 |
| AT1G68370 | 0,32 | -2,60 | 4,41 | -0,59 | 5,55E-01 |
| AT1G68550 | 0,87 | 2,00 | 2,03 | 0,99 | 3,24E-01 |
| AT1G08997 | 0,58 | -0,11 | 3,80 | -0,03 | 9,76E-01 |
| AT1G68660 | 0,71 | -0,79 | 2,31 | -0,34 | 7,33E-01 |
| AT1G68710 | 0,46 | -2,98 | 4,38 | -0,68 | 4,97E-01 |
| AT1G68830 | 0,57 | -0,91 | 2,72 | -0,34 | 7,37E-01 |
| AT1G68910 | 0,08 | 0,00 | 4,41 | 0,00 | 1,00E+00 |
| AT1G69020 | 0,30 | -2,60 | 4,41 | -0,59 | 5,55E-01 |
| AT1G69070 | 0,46 | -2,83 | 3,31 | -0,86 | 3,92E-01 |
| AT1G69220 | 1,28 | -4,00 | 2,00 | -2,00 | 4,51E-02 |
| AT1G69270 | 0,78 | -2,95 | 3,06 | -0,96 | 3,35E-01 |
| AT1G69280 | 0,21 | 0,00 | 4,41 | 0,00 | 1,00E+00 |
| AT1G69290 | 0,15 | -0,73 | 4,40 | -0,17 | 8,68E-01 |
| AT1G69310 | 0,25 | -0,73 | 4,40 | -0,16 | 8,69E-01 |
| AT1G69450 | 0,38 | -0,61 | 4,28 | -0,14 | 8,86E-01 |
| AT1G69480 | 0,55 | 0,00 | 4,41 | 0,00 | 1,00E+00 |
| AT1G69630 | 0,59 | -4,32 | 4,32 | -1,00 | 3,17E-01 |
| AT1G69640 | 0,90 | 0,87 | 2,43 | 0,36 | 7,21E-01 |
| AT1G69730 | 0,73 | 1,35 | 2,30 | 0,59 | 5,56E-01 |
| AT1G69760 | 0,47 | 0,92 | 3,10 | 0,30 | 7,66E-01 |
| AT1G69770 | 1,05 | -3,94 | 2,25 | -1,76 | 7,91E-02 |
| AT1G69840 | 0,65 | 0,60 | 2,98 | 0,20 | 8,40E-01 |
| AT1G69850 | 0,22 | 0,75 | 4,34 | 0,17 | 8,63E-01 |
| AT1G69960 | 0,51 | -1,96 | 3,21 | -0,61 | 5,40E-01 |
| AT1G70000 | 0,98 | -2,43 | 2,15 | -1,13 | 2,57E-01 |
| AT1G70070 | 1,07 | -0,83 | 2,23 | -0,37 | 7,09E-01 |
| AT1G70200 | 0,37 | -0,44 | 4,34 | -0,10 | 9,20E-01 |
| AT1G70280 | 0,17 | 0,00 | 4,41 | 0,00 | 1,00E+00 |
| AT1G70300 | 0,38 | -1,84 | 4,31 | -0,43 | 6,70E-01 |
| AT1G70310 | 0,89 | -2,59 | 3,12 | -0,83 | 4,07E-01 |
| AT1G70340 | 0,34 | -0,91 | 3,58 | -0,26 | 7,98E-01 |
| AT1G70370 | 0,57 | 1,69 | 2,66 | 0,64 | 5,25E-01 |
| AT1G70410 | 0,70 | -1,16 | 2,52 | -0,46 | 6,44E-01 |
| AT1G70420 | 0,44 | 0,04 | 4,08 | 0,01 | 9,92E-01 |
| AT1G70520 | 0,53 | 0,00 | 4,41 | 0,00 | 1,00E+00 |
| AT1G70600 | 0,97 | -2,77 | 2,94 | -0,94 | 3,45E-01 |
| AT1G70730 | 0,91 | -2,36 | 2,47 | -0,96 | 3,40E-01 |

| | | | | | |
|---|---|---|---|---|---|
| AT1G70740 | 0,10 | 0,45 | 4,35 | 0,10 | 9,19E-01 |
| AT1G70750 | 0,44 | 0,00 | 4,41 | 0,00 | 1,00E+00 |
| AT1G70780 | 0,73 | 1,22 | 2,79 | 0,44 | 6,60E-01 |
| AT1G70830 | 0,27 | 0,36 | 4,36 | 0,08 | 9,33E-01 |
| AT1G71020 | 0,94 | -4,49 | 2,31 | -1,95 | 5,12E-02 |
| AT1G71030 | 0,60 | -0,25 | 2,50 | -0,10 | 9,21E-01 |
| AT1G71040 | 0,83 | -1,20 | 1,92 | -0,62 | 5,32E-01 |
| AT1G71060 | 0,73 | -2,60 | 3,54 | -0,74 | 4,62E-01 |
| AT1G71070 | 0,49 | -3,33 | 3,19 | -1,04 | 2,97E-01 |
| AT1G71080 | 0,90 | -2,99 | 2,89 | -1,03 | 3,01E-01 |
| AT1G71160 | 0,69 | -3,05 | 3,62 | -0,84 | 3,99E-01 |
| AT1G71190 | 0,19 | 0,05 | 4,36 | 0,01 | 9,90E-01 |
| AT1G71210 | 0,73 | -2,15 | 2,89 | -0,74 | 4,58E-01 |
| AT1G71310 | 0,27 | 0,00 | 4,41 | 0,00 | 1,00E+00 |
| AT1G71330 | 0,30 | -2,60 | 4,41 | -0,59 | 5,55E-01 |
| AT1G71340 | 0,92 | 0,52 | 2,28 | 0,23 | 8,19E-01 |
| AT1G71350 | 0,21 | 0,00 | 4,41 | 0,00 | 1,00E+00 |
| AT1G09253 | 0,32 | -1,48 | 4,39 | -0,34 | 7,36E-01 |
| AT1G71490 | 0,42 | -1,64 | 4,25 | -0,39 | 6,99E-01 |
| AT1G71710 | 0,44 | -0,77 | 2,89 | -0,27 | 7,90E-01 |
| AT1G71770 | 0,40 | 0,63 | 4,14 | 0,15 | 8,79E-01 |
| AT1G71870 | 0,88 | 1,82 | 2,78 | 0,65 | 5,13E-01 |
| AT1G71960 | 0,26 | 0,06 | 3,79 | 0,02 | 9,87E-01 |
| AT1G72020 | 0,56 | -0,42 | 2,61 | -0,16 | 8,72E-01 |
| AT1G72040 | 0,83 | -0,24 | 2,19 | -0,11 | 9,13E-01 |
| AT1G72160 | 0,64 | 0,32 | 2,21 | 0,15 | 8,84E-01 |
| AT1G72190 | 0,52 | 0,00 | 3,95 | 0,00 | 1,00E+00 |
| AT1G72270 | 0,12 | -0,78 | 4,39 | -0,18 | 8,59E-01 |
| AT1G72300 | 0,54 | -2,39 | 3,12 | -0,77 | 4,44E-01 |
| AT1G72320 | 0,39 | -1,43 | 3,42 | -0,42 | 6,76E-01 |
| AT1G72450 | 1,12 | 0,24 | 1,70 | 0,14 | 8,88E-01 |
| AT1G72480 | 0,35 | 0,00 | 4,41 | 0,00 | 1,00E+00 |
| AT1G72500 | 0,46 | -2,79 | 4,34 | -0,64 | 5,21E-01 |
| AT1G72550 | 0,79 | -3,05 | 2,50 | -1,22 | 2,23E-01 |
| AT1G72560 | 1,09 | -0,72 | 2,30 | -0,31 | 7,54E-01 |
| AT1G09327 | 0,20 | 0,00 | 4,41 | 0,00 | 1,00E+00 |
| AT1G72670 | 0,20 | -2,55 | 4,40 | -0,58 | 5,62E-01 |
| AT1G72720 | 0,35 | -2,99 | 4,38 | -0,68 | 4,95E-01 |
| AT1G72730 | 0,32 | -3,12 | 4,39 | -0,71 | 4,77E-01 |
| AT1G72760 | 0,65 | 0,00 | 4,41 | 0,00 | 1,00E+00 |
| AT1G72840 | 0,41 | -0,68 | 4,41 | -0,15 | 8,77E-01 |
| AT1G72860 | 0,44 | 0,00 | 4,41 | 0,00 | 1,00E+00 |
| AT1G73010 | 0,40 | 0,00 | 4,41 | 0,00 | 1,00E+00 |
| AT1G09363 | 0,41 | 0,66 | 3,26 | 0,20 | 8,39E-01 |
| AT1G73110 | 0,63 | 0,00 | 3,89 | 0,00 | 1,00E+00 |
| AT1G73200 | 0,73 | -2,28 | 2,20 | -1,03 | 3,01E-01 |
| AT1G73230 | 0,17 | 0,00 | 4,41 | 0,00 | 1,00E+00 |
| AT1G73260 | 0,63 | -4,13 | 2,98 | -1,39 | 1,65E-01 |
| AT1G73370 | 0,52 | 0,00 | 4,41 | 0,00 | 1,00E+00 |
| AT1G73390 | 0,25 | -0,15 | 4,37 | -0,03 | 9,72E-01 |
| AT1G73490 | 0,37 | -2,10 | 4,36 | -0,48 | 6,30E-01 |
| AT1G73500 | 0,92 | -1,24 | 1,92 | -0,65 | 5,18E-01 |
| AT1G73510 | 0,45 | -3,79 | 4,08 | -0,93 | 3,53E-01 |
| AT1G73530 | 0,46 | 1,82 | 4,20 | 0,43 | 6,64E-01 |
| AT1G73540 | 0,81 | 1,34 | 2,14 | 0,63 | 5,31E-01 |
| AT1G73650 | 0,96 | -1,68 | 2,55 | -0,66 | 5,09E-01 |

| | | | | | |
|---|---|---|---|---|---|
| AT1G73655 | 0,74 | -0,85 | 3,80 | -0,22 | 8,24E-01 |
| AT1G73680 | 0,85 | 1,33 | 2,16 | 0,62 | 5,38E-01 |
| AT1G73730 | 0,17 | 0,00 | 4,41 | 0,00 | 1,00E+00 |
| AT1G73760 | 0,39 | -0,41 | 4,23 | -0,10 | 9,23E-01 |
| AT1G73930 | 0,25 | -2,61 | 4,41 | -0,59 | 5,53E-01 |
| AT1G73950 | 0,45 | -2,48 | 4,39 | -0,57 | 5,72E-01 |
| AT1G73960 | 0,29 | -1,64 | 4,38 | -0,37 | 7,08E-01 |
| AT1G73980 | 0,67 | 0,21 | 2,56 | 0,08 | 9,36E-01 |
| AT1G74050 | 0,43 | -0,79 | 3,30 | -0,24 | 8,10E-01 |
| AT1G74060 | 0,72 | -1,19 | 4,29 | -0,28 | 7,81E-01 |
| AT1G74070 | 0,42 | 0,91 | 4,22 | 0,21 | 8,30E-01 |
| AT1G74120 | 0,82 | -0,82 | 2,83 | -0,29 | 7,71E-01 |
| AT1G74170 | 0,18 | -0,33 | 4,38 | -0,08 | 9,40E-01 |
| AT1G74340 | 0,31 | 0,00 | 4,41 | 0,00 | 1,00E+00 |
| AT1G74370 | 0,53 | -0,61 | 3,92 | -0,15 | 8,77E-01 |
| AT1G74390 | 0,95 | -4,35 | 2,69 | -1,62 | 1,06E-01 |
| AT1G74420 | 0,45 | -3,77 | 4,36 | -0,87 | 3,87E-01 |
| AT1G74440 | 0,80 | 0,60 | 2,94 | 0,20 | 8,39E-01 |
| AT1G09467 | 0,50 | -0,68 | 4,19 | -0,16 | 8,71E-01 |
| AT1G74560 | 0,37 | 1,58 | 3,40 | 0,47 | 6,42E-01 |
| AT1G74690 | 0,95 | -2,57 | 2,22 | -1,16 | 2,46E-01 |
| AT1G74770 | 0,42 | -3,55 | 4,20 | -0,85 | 3,98E-01 |
| AT1G74800 | 0,56 | 0,92 | 3,15 | 0,29 | 7,70E-01 |
| AT1G74830 | 0,12 | 0,00 | 4,41 | 0,00 | 1,00E+00 |
| AT1G74850 | 0,94 | -2,58 | 2,02 | -1,28 | 2,01E-01 |
| AT1G74880 | 0,11 | -0,68 | 4,41 | -0,15 | 8,77E-01 |
| AT1G74920 | 0,86 | -0,74 | 2,57 | -0,29 | 7,73E-01 |
| AT1G74950 | 0,28 | -0,17 | 4,33 | -0,04 | 9,68E-01 |
| AT1G74970 | 0,46 | -1,70 | 4,33 | -0,39 | 6,95E-01 |
| AT1G75010 | 0,20 | 1,44 | 4,30 | 0,33 | 7,38E-01 |
| AT1G75100 | 0,60 | 0,50 | 2,21 | 0,23 | 8,21E-01 |
| AT1G75130 | 0,12 | -0,68 | 4,41 | -0,15 | 8,77E-01 |
| AT1G75180 | 0,38 | 1,23 | 3,36 | 0,37 | 7,14E-01 |
| AT1G75200 | 0,15 | -0,68 | 4,41 | -0,15 | 8,77E-01 |
| AT1G75220 | 0,55 | 0,79 | 3,17 | 0,25 | 8,03E-01 |
| AT1G75370 | 1,12 | -1,55 | 1,55 | -1,00 | 3,17E-01 |
| AT1G75410 | 0,90 | -1,07 | 2,55 | -0,42 | 6,74E-01 |
| AT1G75450 | 0,11 | -0,80 | 4,39 | -0,18 | 8,56E-01 |
| AT1G75500 | 0,97 | -0,25 | 1,84 | -0,14 | 8,92E-01 |
| AT1G75680 | 0,99 | 1,03 | 1,79 | 0,57 | 5,66E-01 |
| AT1G75690 | 0,84 | -1,22 | 1,90 | -0,64 | 5,20E-01 |
| AT1G75750 | 1,13 | -4,35 | 2,28 | -1,91 | 5,67E-02 |
| AT1G75780 | 0,25 | -0,01 | 4,36 | 0,00 | 9,98E-01 |
| AT1G75820 | 1,17 | -2,66 | 2,25 | -1,18 | 2,38E-01 |
| AT1G75860 | 0,67 | -2,08 | 2,26 | -0,92 | 3,56E-01 |
| AT1G75945 | 0,28 | -1,64 | 4,41 | -0,37 | 7,10E-01 |
| AT1G76060 | 0,20 | -1,00 | 4,37 | -0,23 | 8,19E-01 |
| AT1G76080 | 1,03 | 2,14 | 1,81 | 1,18 | 2,37E-01 |
| AT1G76100 | 1,22 | -0,53 | 1,87 | -0,28 | 7,77E-01 |
| AT1G76120 | 0,15 | -0,81 | 4,39 | -0,18 | 8,54E-01 |
| AT1G76150 | 0,72 | 0,71 | 2,92 | 0,24 | 8,08E-01 |
| AT1G76160 | 0,88 | -1,55 | 2,01 | -0,77 | 4,41E-01 |
| AT1G76270 | 0,08 | 0,00 | 4,41 | 0,00 | 1,00E+00 |
| AT1G76280 | 0,12 | -0,68 | 4,41 | -0,15 | 8,77E-01 |
| AT1G09627 | 0,45 | -1,60 | 4,10 | -0,39 | 6,96E-01 |
| AT1G76400 | 0,57 | -0,45 | 3,08 | -0,15 | 8,83E-01 |

| | | | | | |
|---|---|---|---|---|---|
| AT1G76460 | 0,53 | -2,23 | 3,25 | -0,69 | 4,93E-01 |
| AT1G76640 | 0,08 | 0,00 | 4,41 | 0,00 | 1,00E+00 |
| AT1G76650 | 0,42 | 2,04 | 3,29 | 0,62 | 5,35E-01 |
| AT1G76660 | 0,13 | -0,18 | 4,39 | -0,04 | 9,67E-01 |
| AT1G76670 | 1,15 | -2,88 | 2,05 | -1,40 | 1,61E-01 |
| AT1G76700 | 0,39 | -2,82 | 4,39 | -0,64 | 5,20E-01 |
| AT1G76878 | 1,06 | 0,31 | 1,81 | 0,17 | 8,63E-01 |
| AT1G76890 | 0,74 | 0,07 | 1,88 | 0,04 | 9,70E-01 |
| AT1G76955 | 0,30 | -2,60 | 4,41 | -0,59 | 5,55E-01 |
| AT1G76960 | 0,40 | -0,13 | 3,41 | -0,04 | 9,70E-01 |
| AT1G76965 | 0,25 | -2,48 | 4,39 | -0,57 | 5,71E-01 |
| AT1G76970 | 0,43 | 0,74 | 3,33 | 0,22 | 8,24E-01 |
| AT1G76980 | 0,73 | -0,34 | 3,04 | -0,11 | 9,10E-01 |
| AT1G77030 | 0,32 | 0,19 | 4,32 | 0,04 | 9,65E-01 |
| AT1G77060 | 0,12 | -0,14 | 4,38 | -0,03 | 9,74E-01 |
| AT1G77100 | 0,82 | 1,86 | 3,63 | 0,51 | 6,10E-01 |
| AT1G77145 | 0,08 | 0,00 | 4,41 | 0,00 | 1,00E+00 |
| AT1G77170 | 0,63 | -2,51 | 3,02 | -0,83 | 4,06E-01 |
| AT1G77240 | 1,26 | -3,54 | 1,96 | -1,81 | 7,10E-02 |
| AT1G77250 | 0,55 | -3,38 | 3,82 | -0,89 | 3,76E-01 |
| AT1G77260 | 0,57 | -1,46 | 3,86 | -0,38 | 7,05E-01 |
| AT1G77480 | 0,33 | 0,00 | 4,41 | 0,00 | 1,00E+00 |
| AT1G77580 | 0,18 | 0,38 | 4,35 | 0,09 | 9,30E-01 |
| AT1G77600 | 0,35 | -1,04 | 4,36 | -0,24 | 8,11E-01 |
| AT1G77660 | 0,33 | -1,64 | 4,37 | -0,37 | 7,08E-01 |
| AT1G77810 | 0,58 | -0,25 | 3,78 | -0,06 | 9,48E-01 |
| AT1G77932 | 0,47 | -3,16 | 4,39 | -0,72 | 4,72E-01 |
| AT1G77940 | 0,55 | -0,44 | 2,61 | -0,17 | 8,67E-01 |
| AT1G77980 | 0,77 | 1,44 | 2,04 | 0,70 | 4,81E-01 |
| AT1G77990 | 0,37 | 1,04 | 4,32 | 0,24 | 8,10E-01 |
| AT1G78060 | 0,43 | 0,08 | 4,37 | 0,02 | 9,85E-01 |
| AT1G78210 | 0,32 | -0,32 | 4,34 | -0,07 | 9,42E-01 |
| AT1G78240 | 0,51 | -2,46 | 2,77 | -0,89 | 3,74E-01 |
| AT1G78370 | 1,22 | -0,83 | 2,02 | -0,41 | 6,82E-01 |
| AT1G78460 | 0,60 | -2,33 | 2,62 | -0,89 | 3,73E-01 |
| AT1G78480 | 0,26 | -1,25 | 4,37 | -0,29 | 7,75E-01 |
| AT1G78510 | 0,69 | -2,17 | 3,74 | -0,58 | 5,62E-01 |
| AT1G78580 | 0,70 | -2,32 | 2,32 | -1,00 | 3,17E-01 |
| AT1G78800 | 0,21 | -0,73 | 4,40 | -0,17 | 8,67E-01 |
| AT1G78820 | 0,54 | -0,54 | 2,37 | -0,23 | 8,19E-01 |
| AT1G78850 | 0,94 | -0,41 | 2,08 | -0,20 | 8,45E-01 |
| AT1G78865 | 0,13 | 0,52 | 4,35 | 0,12 | 9,04E-01 |
| AT1G78880 | 0,88 | -1,48 | 2,86 | -0,52 | 6,05E-01 |
| AT1G78915 | 0,18 | 0,68 | 4,34 | 0,16 | 8,75E-01 |
| AT1G78990 | 0,38 | -3,70 | 4,35 | -0,85 | 3,95E-01 |
| AT1G78995 | 0,24 | -0,20 | 4,39 | -0,05 | 9,64E-01 |
| AT1G79020 | 0,25 | -3,06 | 4,38 | -0,70 | 4,85E-01 |
| AT1G79030 | 0,53 | -1,45 | 3,91 | -0,37 | 7,11E-01 |
| AT1G79050 | 0,58 | -2,57 | 3,10 | -0,83 | 4,07E-01 |
| AT1G79080 | 1,23 | -3,43 | 2,34 | -1,47 | 1,43E-01 |
| AT1G79120 | 0,10 | 0,45 | 4,35 | 0,10 | 9,19E-01 |
| AT1G79150 | 0,35 | -0,68 | 4,41 | -0,15 | 8,77E-01 |
| AT1G79350 | 0,49 | -1,81 | 2,71 | -0,67 | 5,03E-01 |
| AT1G79430 | 0,17 | -1,25 | 4,37 | -0,29 | 7,74E-01 |
| AT1G79440 | 0,39 | -0,93 | 3,54 | -0,26 | 7,94E-01 |
| AT1G09867 | 1,21 | 3,07 | 2,29 | 1,34 | 1,81E-01 |

| | | | | | |
|---|---|---|---|---|---|
| AT1G09883 | 0,79 | 0,35 | 2,77 | 0,12 | 9,01E-01 |
| AT1G79540 | 0,53 | -2,68 | 3,16 | -0,85 | 3,97E-01 |
| AT1G79640 | 0,40 | -0,74 | 4,40 | -0,17 | 8,67E-01 |
| AT1G79670 | 0,90 | -0,97 | 2,64 | -0,37 | 7,12E-01 |
| AT1G79700 | 0,08 | 0,00 | 4,41 | 0,00 | 1,00E+00 |
| AT1G79720 | 0,47 | -0,71 | 3,37 | -0,21 | 8,33E-01 |
| AT1G79730 | 0,63 | -0,98 | 3,83 | -0,26 | 7,98E-01 |
| AT1G79740 | 0,25 | -3,06 | 4,38 | -0,70 | 4,85E-01 |
| AT1G79790 | 0,05 | -0,24 | 4,38 | -0,05 | 9,57E-01 |
| AT1G79830 | 1,06 | 0,37 | 1,80 | 0,20 | 8,39E-01 |
| AT1G79880 | 0,11 | -0,68 | 4,41 | -0,15 | 8,77E-01 |
| AT1G79900 | 0,42 | -3,55 | 4,20 | -0,85 | 3,98E-01 |
| AT1G80030 | 1,07 | -2,85 | 2,25 | -1,27 | 2,05E-01 |
| AT1G80040 | 0,61 | -3,38 | 2,52 | -1,34 | 1,81E-01 |
| AT1G80080 | 0,50 | -1,83 | 3,13 | -0,58 | 5,60E-01 |
| AT1G80325 | 1,08 | -1,70 | 2,02 | -0,84 | 4,01E-01 |
| AT1G80350 | 0,38 | -3,70 | 4,35 | -0,85 | 3,95E-01 |
| AT1G80420 | 0,34 | 0,44 | 4,36 | 0,10 | 9,20E-01 |
| AT1G80460 | 0,24 | -1,38 | 4,38 | -0,31 | 7,53E-01 |
| AT1G80480 | 0,56 | -0,24 | 3,15 | -0,08 | 9,39E-01 |
| AT1G80490 | 1,23 | -2,68 | 1,65 | -1,62 | 1,05E-01 |
| AT1G80530 | 0,68 | -0,82 | 3,07 | -0,27 | 7,90E-01 |
| AT1G80610 | 0,35 | 1,40 | 3,41 | 0,41 | 6,82E-01 |
| AT1G80630 | 0,28 | -1,75 | 4,39 | -0,40 | 6,90E-01 |
| AT1G80660 | 1,14 | -3,88 | 2,01 | -1,93 | 5,38E-02 |
| AT1G80670 | 0,39 | -3,07 | 4,38 | -0,70 | 4,84E-01 |
| AT1G80760 | 0,62 | -2,82 | 3,71 | -0,76 | 4,47E-01 |
| AT1G80830 | 0,27 | -0,35 | 4,34 | -0,08 | 9,37E-01 |
| AT1G80850 | 0,20 | 1,40 | 4,30 | 0,33 | 7,44E-01 |
| AT1G80920 | 0,74 | -0,02 | 2,07 | -0,01 | 9,92E-01 |
| AT1G80930 | 0,69 | 0,92 | 2,41 | 0,38 | 7,04E-01 |
| AT1G80950 | 0,35 | -0,35 | 4,34 | -0,08 | 9,35E-01 |
| AT2G01023 | 0,35 | 0,54 | 2,94 | 0,18 | 8,55E-01 |
| AT2G01050 | 0,45 | 1,42 | 4,30 | 0,33 | 7,42E-01 |
| AT2G01060 | 0,57 | 1,50 | 2,55 | 0,59 | 5,58E-01 |
| AT2G01110 | 0,90 | -2,39 | 2,44 | -0,98 | 3,26E-01 |
| AT2G01130 | 1,10 | -3,47 | 2,05 | -1,69 | 9,01E-02 |
| AT2G01180 | 0,86 | 1,24 | 1,90 | 0,65 | 5,14E-01 |
| AT2G01250 | 0,52 | -2,60 | 3,98 | -0,65 | 5,14E-01 |
| AT2G01275 | 0,19 | 0,65 | 4,34 | 0,15 | 8,82E-01 |
| AT2G01290 | 0,18 | -0,27 | 4,38 | -0,06 | 9,51E-01 |
| AT2G01300 | 0,08 | 0,00 | 4,41 | 0,00 | 1,00E+00 |
| AT2G01310 | 0,05 | -0,24 | 4,38 | -0,05 | 9,57E-01 |
| AT2G01350 | 0,12 | 0,75 | 4,34 | 0,17 | 8,63E-01 |
| AT2G01410 | 0,34 | -0,47 | 3,53 | -0,13 | 8,93E-01 |
| AT2G01420 | 1,24 | 1,56 | 1,76 | 0,89 | 3,74E-01 |
| AT2G01440 | 1,25 | -2,62 | 2,41 | -1,09 | 2,77E-01 |
| AT2G01450 | 1,15 | -0,13 | 1,56 | -0,08 | 9,35E-01 |
| AT2G01505 | 0,30 | 0,82 | 4,33 | 0,19 | 8,51E-01 |
| AT2G01630 | 0,87 | -3,71 | 2,76 | -1,35 | 1,79E-01 |
| AT2G01650 | 1,03 | -1,46 | 2,03 | -0,72 | 4,71E-01 |
| AT2G01670 | 0,39 | 0,75 | 3,32 | 0,22 | 8,22E-01 |
| AT2G01690 | 0,55 | -0,70 | 3,76 | -0,19 | 8,52E-01 |
| AT2G01755 | 0,14 | 0,48 | 4,35 | 0,11 | 9,13E-01 |
| AT2G01820 | 1,26 | -3,88 | 2,41 | -1,61 | 1,07E-01 |
| AT2G01830 | 0,91 | 0,18 | 2,05 | 0,09 | 9,30E-01 |

| | | | | | |
|---|---|---|---|---|---|
| AT2G04125 | 0,12 | -0,31 | 4,38 | -0,07 | 9,44E-01 |
| AT2G01870 | 0,33 | -1,56 | 4,40 | -0,36 | 7,22E-01 |
| AT2G01918 | 1,01 | 1,34 | 2,10 | 0,64 | 5,23E-01 |
| AT2G01930 | 0,16 | -0,68 | 4,41 | -0,15 | 8,77E-01 |
| AT2G01990 | 0,25 | 1,04 | 4,33 | 0,24 | 8,11E-01 |
| AT2G02080 | 1,03 | 0,27 | 2,59 | 0,10 | 9,17E-01 |
| AT2G02103 | 0,70 | 2,09 | 2,49 | 0,84 | 4,02E-01 |
| AT2G02148 | 0,42 | -3,35 | 4,14 | -0,81 | 4,18E-01 |
| AT2G02150 | 0,39 | -2,20 | 4,28 | -0,51 | 6,07E-01 |
| AT2G02160 | 0,60 | -0,39 | 2,54 | -0,15 | 8,79E-01 |
| AT2G02220 | 0,80 | -0,34 | 2,56 | -0,13 | 8,94E-01 |
| AT2G02230 | 0,17 | -0,75 | 4,40 | -0,17 | 8,65E-01 |
| AT2G02340 | 1,23 | -0,91 | 2,01 | -0,45 | 6,53E-01 |
| AT2G02360 | 0,16 | -0,26 | 4,38 | -0,06 | 9,52E-01 |
| AT2G02400 | 0,84 | 1,62 | 2,21 | 0,73 | 4,64E-01 |
| AT2G02520 | 0,08 | 0,05 | 4,38 | 0,01 | 9,91E-01 |
| AT2G02570 | 0,25 | 1,15 | 4,32 | 0,27 | 7,91E-01 |
| AT2G02635 | 0,17 | -0,08 | 4,37 | -0,02 | 9,85E-01 |
| AT2G02695 | 0,58 | 2,40 | 2,93 | 0,82 | 4,13E-01 |
| AT2G02710 | 0,66 | -1,09 | 2,43 | -0,45 | 6,55E-01 |
| AT2G02780 | 0,28 | 0,84 | 4,33 | 0,19 | 8,46E-01 |
| AT2G02800 | 0,78 | 0,55 | 2,50 | 0,22 | 8,24E-01 |
| AT2G04345 | 0,62 | 0,36 | 2,52 | 0,14 | 8,87E-01 |
| AT2G02930 | 0,49 | 0,92 | 3,21 | 0,29 | 7,75E-01 |
| AT2G03050 | 0,25 | -0,69 | 4,40 | -0,16 | 8,76E-01 |
| AT2G03090 | 0,08 | 0,00 | 4,41 | 0,00 | 1,00E+00 |
| AT2G03240 | 0,24 | -2,60 | 4,41 | -0,59 | 5,55E-01 |
| AT2G03340 | 0,45 | -1,69 | 3,26 | -0,52 | 6,03E-01 |
| AT2G03390 | 0,61 | -0,92 | 2,42 | -0,38 | 7,03E-01 |
| AT2G03420 | 0,43 | -0,80 | 4,15 | -0,19 | 8,48E-01 |
| AT2G03430 | 0,52 | -3,19 | 4,37 | -0,73 | 4,66E-01 |
| AT2G03440 | 0,35 | 0,19 | 4,31 | 0,04 | 9,65E-01 |
| AT2G03460 | 0,08 | 0,00 | 4,41 | 0,00 | 1,00E+00 |
| AT2G03470 | 1,22 | -1,82 | 2,27 | -0,80 | 4,23E-01 |
| AT2G03490 | 0,08 | 0,00 | 4,41 | 0,00 | 1,00E+00 |
| AT2G03620 | 0,33 | -3,08 | 4,39 | -0,70 | 4,83E-01 |
| AT2G03640 | 0,13 | 0,00 | 4,41 | 0,00 | 1,00E+00 |
| AT2G03690 | 0,23 | -1,14 | 4,39 | -0,26 | 7,96E-01 |
| AT2G03730 | 0,29 | 0,58 | 4,35 | 0,13 | 8,94E-01 |
| AT2G03750 | 1,29 | 1,76 | 1,70 | 1,04 | 2,99E-01 |
| AT2G03760 | 0,93 | -0,26 | 3,39 | -0,08 | 9,39E-01 |
| AT2G03770 | 0,15 | -0,77 | 4,39 | -0,18 | 8,61E-01 |
| AT2G04655 | 0,14 | 0,87 | 4,33 | 0,20 | 8,40E-01 |
| AT2G04031 | 0,15 | 0,79 | 4,33 | 0,18 | 8,56E-01 |
| AT2G04037 | 0,16 | 0,32 | 4,36 | 0,07 | 9,42E-01 |
| AT2G04050 | 0,12 | 0,00 | 4,41 | 0,00 | 1,00E+00 |
| AT2G04062 | 0,70 | 2,74 | 2,84 | 0,96 | 3,35E-01 |
| AT2G04063 | 0,69 | 2,66 | 2,69 | 0,99 | 3,21E-01 |
| AT2G04066 | 0,42 | -0,54 | 4,34 | -0,12 | 9,01E-01 |
| AT2G04070 | 0,33 | -3,41 | 4,38 | -0,78 | 4,35E-01 |
| AT2G04080 | 0,61 | -1,06 | 2,50 | -0,42 | 6,73E-01 |
| AT2G04160 | 0,43 | 1,73 | 4,02 | 0,43 | 6,66E-01 |
| AT2G04230 | 1,09 | -2,42 | 2,24 | -1,08 | 2,80E-01 |
| AT2G04360 | 0,70 | -2,82 | 2,38 | -1,18 | 2,36E-01 |
| AT2G04378 | 0,56 | -0,73 | 2,65 | -0,28 | 7,82E-01 |
| AT2G04520 | 0,26 | 0,64 | 4,34 | 0,15 | 8,83E-01 |

| | | | | | |
|---|---|---|---|---|---|
| AT2G04620 | 0,17 | 0,00 | 4,41 | 0,00 | 1,00E+00 |
| AT2G04630 | 0,19 | 0,25 | 4,36 | 0,06 | 9,54E-01 |
| AT2G04842 | 0,88 | 0,49 | 3,10 | 0,16 | 8,76E-01 |
| AT2G04870 | 0,55 | 0,55 | 2,62 | 0,21 | 8,34E-01 |
| AT2G04890 | 0,11 | -0,80 | 4,39 | -0,18 | 8,56E-01 |
| AT2G05120 | 0,91 | -2,51 | 2,17 | -1,16 | 2,47E-01 |
| AT2G05220 | 0,38 | -1,25 | 4,37 | -0,29 | 7,75E-01 |
| AT2G05250 | 0,41 | -3,11 | 4,39 | -0,71 | 4,78E-01 |
| AT2G04895 | 0,17 | 0,78 | 4,33 | 0,18 | 8,58E-01 |
| AT2G05752 | 0,90 | 1,41 | 2,27 | 0,62 | 5,34E-01 |
| AT2G04995 | 1,03 | 2,44 | 2,11 | 1,15 | 2,48E-01 |
| AT2G05810 | 1,10 | 0,09 | 1,77 | 0,05 | 9,59E-01 |
| AT2G05914 | 0,51 | 0,90 | 4,33 | 0,21 | 8,36E-01 |
| AT2G05940 | 0,97 | 1,64 | 1,95 | 0,84 | 3,99E-01 |
| AT2G06000 | 0,52 | -1,67 | 2,58 | -0,65 | 5,17E-01 |
| AT2G06040 | 0,24 | -0,71 | 4,40 | -0,16 | 8,72E-01 |
| AT2G06050 | 0,61 | 2,69 | 3,00 | 0,90 | 3,69E-01 |
| AT2G06530 | 0,61 | 1,07 | 3,30 | 0,33 | 7,45E-01 |
| AT2G06541 | 0,49 | 1,33 | 4,08 | 0,33 | 7,44E-01 |
| AT2G06645 | 0,15 | 0,16 | 4,37 | 0,04 | 9,72E-01 |
| AT2G06675 | 0,19 | 1,18 | 4,31 | 0,27 | 7,84E-01 |
| AT2G06906 | 0,12 | -0,68 | 4,41 | -0,15 | 8,77E-01 |
| AT2G06925 | 0,35 | 0,64 | 3,51 | 0,18 | 8,55E-01 |
| AT2G06960 | 0,68 | 2,93 | 2,75 | 1,06 | 2,87E-01 |
| AT2G07020 | 0,35 | 1,97 | 4,28 | 0,46 | 6,45E-01 |
| AT2G07120 | 0,12 | -0,06 | 4,37 | -0,01 | 9,88E-01 |
| AT2G07240 | 1,04 | 2,61 | 1,96 | 1,33 | 1,82E-01 |
| AT2G07560 | 0,12 | -0,78 | 4,39 | -0,18 | 8,59E-01 |
| AT2G07771 | 0,58 | -0,56 | 2,58 | -0,22 | 8,29E-01 |
| AT2G07777 | 0,89 | -0,73 | 2,09 | -0,35 | 7,26E-01 |
| AT2G07779 | 0,28 | -1,20 | 4,34 | -0,28 | 7,83E-01 |
| AT2G07621 | 0,12 | -0,31 | 4,38 | -0,07 | 9,44E-01 |
| AT2G07774 | 0,25 | 0,80 | 4,33 | 0,19 | 8,53E-01 |
| AT2G07687 | 1,11 | -0,34 | 2,04 | -0,17 | 8,68E-01 |
| AT2G05205 | 0,68 | 1,43 | 2,56 | 0,56 | 5,76E-01 |
| AT2G07599 | 0,90 | 0,79 | 2,20 | 0,36 | 7,19E-01 |
| AT2G07706 | 1,10 | 2,18 | 2,35 | 0,93 | 3,54E-01 |
| AT2G07641 | 0,45 | -0,41 | 2,79 | -0,15 | 8,83E-01 |
| AT2G07719 | 0,62 | -0,87 | 2,69 | -0,32 | 7,46E-01 |
| AT2G07756 | 1,02 | 1,08 | 1,91 | 0,56 | 5,72E-01 |
| AT2G07734 | 0,71 | -0,82 | 2,81 | -0,29 | 7,70E-01 |
| AT2G07658 | 0,65 | -0,18 | 2,50 | -0,07 | 9,42E-01 |
| AT2G07662 | 1,06 | 1,68 | 2,26 | 0,74 | 4,58E-01 |
| AT2G07835 | 0,78 | 0,11 | 3,00 | 0,04 | 9,71E-01 |
| AT2G07690 | 0,71 | -1,42 | 2,53 | -0,56 | 5,74E-01 |
| AT2G07760 | 0,23 | -0,68 | 4,41 | -0,15 | 8,77E-01 |
| AT2G09795 | 0,34 | -1,76 | 4,39 | -0,40 | 6,88E-01 |
| AT2G05465 | 0,27 | 1,02 | 4,33 | 0,24 | 8,13E-01 |
| AT2G10557 | 1,21 | 0,00 | 3,79 | 0,00 | 1,00E+00 |
| AT2G10608 | 0,36 | -1,79 | 4,36 | -0,41 | 6,81E-01 |
| AT2G05565 | 0,43 | 1,14 | 4,00 | 0,29 | 7,75E-01 |
| AT2G10625 | 0,25 | 1,52 | 4,30 | 0,35 | 7,23E-01 |
| AT2G10950 | 0,47 | 2,32 | 3,90 | 0,59 | 5,53E-01 |
| AT2G10955 | 0,26 | 0,92 | 4,32 | 0,21 | 8,32E-01 |
| AT2G11000 | 0,08 | 0,05 | 4,38 | 0,01 | 9,91E-01 |
| AT2G11852 | 0,26 | 0,98 | 4,33 | 0,23 | 8,22E-01 |

| | | | | | |
|---|---|---|---|---|---|
| AT2G12400 | 0,28 | 0,13 | 4,36 | 0,03 | 9,77E-01 |
| AT2G12405 | 0,18 | 0,68 | 4,34 | 0,16 | 8,75E-01 |
| AT2G05705 | 0,21 | -0,69 | 4,36 | -0,16 | 8,74E-01 |
| AT2G12461 | 0,12 | 0,00 | 4,41 | 0,00 | 1,00E+00 |
| AT2G05835 | 0,29 | 0,47 | 4,35 | 0,11 | 9,14E-01 |
| AT2G12945 | 0,05 | -0,24 | 4,38 | -0,05 | 9,57E-01 |
| AT2G13370 | 1,20 | -1,73 | 2,27 | -0,76 | 4,47E-01 |
| AT2G13550 | 0,22 | 0,49 | 4,34 | 0,11 | 9,10E-01 |
| AT2G13640 | 0,11 | -0,68 | 4,41 | -0,15 | 8,77E-01 |
| AT2G13650 | 0,22 | -0,56 | 4,35 | -0,13 | 8,98E-01 |
| AT2G14100 | 0,15 | 0,48 | 4,35 | 0,11 | 9,12E-01 |
| AT2G14120 | 0,98 | -0,20 | 2,01 | -0,10 | 9,21E-01 |
| AT2G14170 | 1,22 | 1,42 | 2,13 | 0,67 | 5,04E-01 |
| AT2G14260 | 0,26 | -0,33 | 4,38 | -0,08 | 9,40E-01 |
| AT2G14288 | 0,20 | 0,94 | 4,32 | 0,22 | 8,28E-01 |
| AT2G14510 | 0,15 | 0,48 | 4,35 | 0,11 | 9,12E-01 |
| AT2G14540 | 0,12 | -0,68 | 4,41 | -0,15 | 8,77E-01 |
| AT2G14670 | 1,21 | 3,47 | 2,41 | 1,44 | 1,50E-01 |
| AT2G14720 | 1,05 | -1,06 | 2,20 | -0,48 | 6,30E-01 |
| AT2G14755 | 0,32 | 2,00 | 4,28 | 0,47 | 6,40E-01 |
| AT2G15000 | 0,23 | -1,14 | 4,39 | -0,26 | 7,96E-01 |
| AT2G15020 | 0,51 | -1,97 | 3,27 | -0,60 | 5,47E-01 |
| AT2G06355 | 0,23 | 0,59 | 4,34 | 0,14 | 8,92E-01 |
| AT2G15110 | 0,63 | 1,77 | 2,94 | 0,60 | 5,47E-01 |
| AT2G15290 | 0,58 | -3,06 | 3,81 | -0,80 | 4,22E-01 |
| AT2G15292 | 1,13 | -0,52 | 1,76 | -0,29 | 7,69E-01 |
| AT2G15320 | 1,25 | 1,81 | 1,69 | 1,07 | 2,86E-01 |
| AT2G15390 | 0,37 | -0,95 | 4,34 | -0,22 | 8,28E-01 |
| AT2G15555 | 1,03 | -0,59 | 1,77 | -0,33 | 7,40E-01 |
| AT2G15560 | 0,05 | -0,24 | 4,38 | -0,05 | 9,57E-01 |
| AT2G15790 | 0,41 | -0,65 | 4,17 | -0,16 | 8,75E-01 |
| AT2G15860 | 0,36 | 0,93 | 4,33 | 0,22 | 8,30E-01 |
| AT2G15890 | 1,14 | -0,28 | 1,66 | -0,17 | 8,67E-01 |
| AT2G06515 | 0,69 | -1,75 | 3,74 | -0,47 | 6,41E-01 |
| AT2G15980 | 0,29 | -0,33 | 3,70 | -0,09 | 9,30E-01 |
| AT2G16016 | 1,06 | 0,75 | 2,01 | 0,37 | 7,09E-01 |
| AT2G06575 | 0,75 | 1,72 | 2,04 | 0,84 | 3,99E-01 |
| AT2G16070 | 0,64 | -1,87 | 3,82 | -0,49 | 6,25E-01 |
| AT2G16090 | 0,55 | -0,94 | 2,46 | -0,38 | 7,04E-01 |
| AT2G16370 | 0,11 | -0,68 | 4,41 | -0,15 | 8,77E-01 |
| AT2G16400 | 1,00 | 0,36 | 2,01 | 0,18 | 8,56E-01 |
| AT2G16405 | 0,20 | 0,00 | 4,41 | 0,00 | 1,00E+00 |
| AT2G16440 | 1,04 | -2,92 | 3,49 | -0,84 | 4,02E-01 |
| AT2G16450 | 0,33 | -3,08 | 4,39 | -0,70 | 4,83E-01 |
| AT2G16505 | 0,85 | 2,11 | 2,21 | 0,95 | 3,41E-01 |
| AT2G16620 | 0,18 | 0,78 | 4,34 | 0,18 | 8,57E-01 |
| AT2G16640 | 1,05 | -0,89 | 2,14 | -0,42 | 6,77E-01 |
| AT2G16650 | 0,37 | 0,00 | 4,41 | 0,00 | 1,00E+00 |
| AT2G16870 | 1,12 | -2,84 | 1,92 | -1,48 | 1,40E-01 |
| AT2G16880 | 0,51 | -3,20 | 4,39 | -0,73 | 4,66E-01 |
| AT2G16920 | 0,25 | 0,00 | 4,41 | 0,00 | 1,00E+00 |
| AT2G16940 | 0,88 | -1,11 | 2,13 | -0,52 | 6,02E-01 |
| AT2G17020 | 0,78 | -1,73 | 2,08 | -0,83 | 4,06E-01 |
| AT2G17110 | 0,49 | 0,09 | 2,71 | 0,03 | 9,74E-01 |
| AT2G17130 | 0,45 | -2,89 | 4,18 | -0,69 | 4,90E-01 |
| AT2G17190 | 0,94 | 0,74 | 2,44 | 0,31 | 7,60E-01 |

| | | | | | |
|---|---|---|---|---|---|
| AT2G17210 | 0,23 | -1,73 | 4,39 | -0,39 | 6,93E-01 |
| AT2G17220 | 0,39 | -2,96 | 4,33 | -0,68 | 4,95E-01 |
| AT2G17230 | 0,92 | 1,51 | 2,47 | 0,61 | 5,42E-01 |
| AT2G17300 | 0,17 | -0,75 | 4,40 | -0,17 | 8,65E-01 |
| AT2G06935 | 0,05 | -0,24 | 4,38 | -0,05 | 9,57E-01 |
| AT2G17340 | 0,86 | -1,82 | 2,29 | -0,79 | 4,27E-01 |
| AT2G17350 | 0,34 | -0,16 | 4,39 | -0,04 | 9,71E-01 |
| AT2G17450 | 0,63 | -1,37 | 2,18 | -0,63 | 5,31E-01 |
| AT2G17470 | 0,10 | -0,18 | 4,38 | -0,04 | 9,67E-01 |
| AT2G17550 | 0,71 | -0,91 | 2,51 | -0,36 | 7,16E-01 |
| AT2G17560 | 0,46 | -2,47 | 4,35 | -0,57 | 5,71E-01 |
| AT2G17700 | 0,43 | -1,64 | 4,09 | -0,40 | 6,88E-01 |
| AT2G17820 | 0,96 | -0,99 | 2,15 | -0,46 | 6,46E-01 |
| AT2G17870 | 0,45 | -1,67 | 4,10 | -0,41 | 6,83E-01 |
| AT2G18130 | 0,22 | 1,04 | 4,32 | 0,24 | 8,10E-01 |
| AT2G18170 | 0,50 | -2,91 | 3,25 | -0,90 | 3,70E-01 |
| AT2G18193 | 1,29 | -4,24 | 2,09 | -2,03 | 4,22E-02 |
| AT2G18290 | 0,19 | 0,02 | 4,36 | 0,00 | 9,97E-01 |
| AT2G18310 | 0,43 | -0,44 | 4,37 | -0,10 | 9,19E-01 |
| AT2G18520 | 0,25 | -3,06 | 4,38 | -0,70 | 4,85E-01 |
| AT2G18600 | 0,24 | -3,18 | 4,39 | -0,73 | 4,68E-01 |
| AT2G18660 | 0,89 | 0,82 | 2,13 | 0,38 | 7,01E-01 |
| AT2G18685 | 0,08 | 0,00 | 4,41 | 0,00 | 1,00E+00 |
| AT2G18900 | 1,01 | -2,65 | 1,82 | -1,46 | 1,45E-01 |
| AT2G18915 | 0,29 | -1,19 | 4,35 | -0,27 | 7,84E-01 |
| AT2G18940 | 0,31 | 0,10 | 3,61 | 0,03 | 9,78E-01 |
| AT2G19120 | 0,71 | -4,08 | 3,56 | -1,15 | 2,52E-01 |
| AT2G07135 | 0,16 | 1,16 | 4,31 | 0,27 | 7,88E-01 |
| AT2G07165 | 0,21 | -0,14 | 4,37 | -0,03 | 9,75E-01 |
| AT2G19290 | 0,05 | -0,24 | 4,38 | -0,05 | 9,57E-01 |
| AT2G19310 | 0,12 | -0,06 | 4,37 | -0,01 | 9,88E-01 |
| AT2G19340 | 0,43 | -0,68 | 4,41 | -0,15 | 8,77E-01 |
| AT2G19365 | 0,81 | 0,76 | 2,12 | 0,36 | 7,20E-01 |
| AT2G19390 | 0,76 | 0,75 | 2,34 | 0,32 | 7,47E-01 |
| AT2G07195 | 0,51 | 0,53 | 2,59 | 0,21 | 8,38E-01 |
| AT2G19400 | 0,15 | -0,17 | 4,39 | -0,04 | 9,68E-01 |
| AT2G19470 | 0,28 | -0,71 | 4,35 | -0,16 | 8,70E-01 |
| AT2G19560 | 0,20 | -0,68 | 4,41 | -0,15 | 8,77E-01 |
| AT2G19570 | 0,52 | 2,29 | 2,99 | 0,77 | 4,43E-01 |
| AT2G19580 | 0,25 | 1,48 | 4,30 | 0,34 | 7,30E-01 |
| AT2G19760 | 0,25 | 1,31 | 4,31 | 0,30 | 7,61E-01 |
| AT2G19780 | 0,19 | 0,52 | 4,35 | 0,12 | 9,04E-01 |
| AT2G19910 | 0,11 | -0,68 | 4,41 | -0,15 | 8,77E-01 |
| AT2G20000 | 0,44 | -1,69 | 3,37 | -0,50 | 6,16E-01 |
| AT2G20050 | 0,77 | -0,26 | 2,39 | -0,11 | 9,14E-01 |
| AT2G20120 | 0,38 | -1,96 | 4,34 | -0,45 | 6,52E-01 |
| AT2G20180 | 0,53 | 0,67 | 2,63 | 0,26 | 7,98E-01 |
| AT2G20240 | 1,31 | -1,49 | 2,63 | -0,57 | 5,71E-01 |
| AT2G20270 | 0,73 | -0,77 | 3,11 | -0,25 | 8,05E-01 |
| AT2G20280 | 0,90 | -3,05 | 2,70 | -1,13 | 2,58E-01 |
| AT2G20300 | 0,57 | 0,83 | 2,67 | 0,31 | 7,56E-01 |
| AT2G20310 | 0,15 | -0,73 | 4,40 | -0,17 | 8,68E-01 |
| AT2G20380 | 0,08 | 0,00 | 4,41 | 0,00 | 1,00E+00 |
| AT2G20562 | 0,11 | 0,46 | 4,35 | 0,11 | 9,15E-01 |
| AT2G20610 | 0,62 | 0,01 | 2,46 | 0,01 | 9,96E-01 |
| AT2G20630 | 0,57 | -0,19 | 2,62 | -0,07 | 9,42E-01 |

| | | | | | |
|---|---|---|---|---|---|
| AT2G20670 | 0,99 | -4,84 | 2,03 | -2,38 | 1,73E-02 |
| AT2G07325 | 0,08 | 0,00 | 4,41 | 0,00 | 1,00E+00 |
| AT2G20720 | 0,50 | -3,45 | 3,26 | -1,06 | 2,90E-01 |
| AT2G20723 | 1,21 | 1,73 | 1,79 | 0,97 | 3,34E-01 |
| AT2G20725 | 0,17 | 0,00 | 4,41 | 0,00 | 1,00E+00 |
| AT2G20760 | 1,00 | -0,48 | 2,20 | -0,22 | 8,28E-01 |
| AT2G20790 | 0,65 | -1,86 | 3,76 | -0,49 | 6,21E-01 |
| AT2G20825 | 0,24 | -2,54 | 4,40 | -0,58 | 5,64E-01 |
| AT2G20830 | 0,42 | 0,82 | 3,20 | 0,26 | 7,98E-01 |
| AT2G20850 | 0,80 | -2,21 | 3,45 | -0,64 | 5,21E-01 |
| AT2G20900 | 0,70 | -3,36 | 2,96 | -1,14 | 2,56E-01 |
| AT2G21060 | 0,37 | 1,01 | 4,32 | 0,23 | 8,16E-01 |
| AT2G21090 | 0,12 | 0,00 | 4,41 | 0,00 | 1,00E+00 |
| AT2G21120 | 1,14 | -1,32 | 1,58 | -0,84 | 4,03E-01 |
| AT2G21170 | 1,20 | -1,55 | 1,64 | -0,94 | 3,45E-01 |
| AT2G21185 | 0,86 | 0,56 | 1,79 | 0,31 | 7,55E-01 |
| AT2G21188 | 0,47 | -2,46 | 4,25 | -0,58 | 5,62E-01 |
| AT2G21210 | 1,12 | -1,65 | 1,95 | -0,84 | 3,98E-01 |
| AT2G21240 | 0,62 | 0,90 | 2,37 | 0,38 | 7,03E-01 |
| AT2G21280 | 0,38 | -3,70 | 4,35 | -0,85 | 3,95E-01 |
| AT2G21300 | 0,99 | -1,70 | 2,31 | -0,74 | 4,62E-01 |
| AT2G21350 | 0,05 | -0,24 | 4,38 | -0,05 | 9,57E-01 |
| AT2G21370 | 0,82 | -0,97 | 1,88 | -0,52 | 6,05E-01 |
| AT2G21440 | 0,17 | 0,00 | 4,41 | 0,00 | 1,00E+00 |
| AT2G21480 | 0,25 | -2,57 | 4,40 | -0,58 | 5,60E-01 |
| AT2G21500 | 0,51 | 0,26 | 3,04 | 0,08 | 9,32E-01 |
| AT2G21560 | 0,27 | 1,88 | 4,29 | 0,44 | 6,61E-01 |
| AT2G21870 | 1,17 | -2,75 | 2,11 | -1,30 | 1,92E-01 |
| AT2G21950 | 0,24 | -0,68 | 4,41 | -0,15 | 8,77E-01 |
| AT2G21970 | 0,61 | -2,56 | 3,94 | -0,65 | 5,16E-01 |
| AT2G22010 | 0,65 | 0,29 | 2,48 | 0,12 | 9,07E-01 |
| AT2G22090 | 0,34 | -1,04 | 4,36 | -0,24 | 8,12E-01 |
| AT2G22140 | 0,05 | -0,24 | 4,38 | -0,05 | 9,57E-01 |
| AT2G22160 | 0,35 | 1,75 | 4,29 | 0,41 | 6,83E-01 |
| AT2G22240 | 0,21 | 0,17 | 4,36 | 0,04 | 9,68E-01 |
| AT2G22250 | 0,26 | -2,60 | 4,41 | -0,59 | 5,55E-01 |
| AT2G07555 | 0,37 | 2,00 | 4,23 | 0,47 | 6,37E-01 |
| AT2G22400 | 0,55 | -1,63 | 3,78 | -0,43 | 6,67E-01 |
| AT2G22470 | 0,19 | -0,81 | 4,39 | -0,19 | 8,53E-01 |
| AT2G22490 | 1,17 | -5,28 | 2,33 | -2,27 | 2,33E-02 |
| AT2G22740 | 0,43 | -0,73 | 4,18 | -0,17 | 8,61E-01 |
| AT2G22850 | 0,76 | 1,46 | 2,02 | 0,72 | 4,70E-01 |
| AT2G22890 | 0,53 | 0,14 | 2,96 | 0,05 | 9,64E-01 |
| AT2G22940 | 0,87 | 2,96 | 2,58 | 1,15 | 2,51E-01 |
| AT2G23070 | 0,53 | 1,48 | 2,24 | 0,66 | 5,08E-01 |
| AT2G23090 | 0,40 | -2,59 | 4,31 | -0,60 | 5,48E-01 |
| AT2G23100 | 0,16 | -0,68 | 4,41 | -0,15 | 8,77E-01 |
| AT2G23140 | 0,38 | 0,74 | 4,24 | 0,17 | 8,61E-01 |
| AT2G23210 | 0,71 | -2,24 | 2,91 | -0,77 | 4,40E-01 |
| AT2G23290 | 0,64 | 0,30 | 2,61 | 0,12 | 9,08E-01 |
| AT2G23390 | 0,32 | -0,83 | 4,38 | -0,19 | 8,50E-01 |
| AT2G23430 | 0,47 | 0,60 | 3,32 | 0,18 | 8,56E-01 |
| AT2G23450 | 0,62 | -1,11 | 2,18 | -0,51 | 6,10E-01 |
| AT2G23520 | 0,43 | -1,49 | 3,31 | -0,45 | 6,52E-01 |
| AT2G23610 | 0,63 | 0,47 | 2,96 | 0,16 | 8,73E-01 |
| AT2G23760 | 0,34 | -0,20 | 4,33 | -0,05 | 9,63E-01 |

| | | | | | |
|---|---|---|---|---|---|
| AT2G23790 | 0,79 | -0,05 | 3,05 | -0,02 | 9,86E-01 |
| AT2G23840 | 0,70 | -1,69 | 3,05 | -0,55 | 5,80E-01 |
| AT2G23987 | 0,17 | 0,00 | 4,41 | 0,00 | 1,00E+00 |
| AT2G24120 | 0,81 | 1,09 | 2,15 | 0,51 | 6,13E-01 |
| AT2G24150 | 0,79 | -2,54 | 2,10 | -1,21 | 2,25E-01 |
| AT2G24170 | 0,17 | -0,71 | 4,40 | -0,16 | 8,72E-01 |
| AT2G24230 | 0,59 | 0,84 | 2,52 | 0,33 | 7,38E-01 |
| AT2G24330 | 0,33 | -0,83 | 4,36 | -0,19 | 8,50E-01 |
| AT2G24360 | 1,19 | -0,79 | 1,78 | -0,44 | 6,57E-01 |
| AT2G24500 | 1,13 | -1,21 | 2,16 | -0,56 | 5,77E-01 |
| AT2G08055 | 0,33 | -0,65 | 4,35 | -0,15 | 8,81E-01 |
| AT2G24545 | 1,16 | -0,91 | 1,51 | -0,61 | 5,44E-01 |
| AT2G24600 | 1,31 | 0,24 | 2,31 | 0,10 | 9,18E-01 |
| AT2G24640 | 0,24 | -2,59 | 4,41 | -0,59 | 5,56E-01 |
| AT2G24710 | 0,19 | -0,76 | 4,40 | -0,17 | 8,63E-01 |
| AT2G24755 | 0,71 | -1,06 | 2,56 | -0,41 | 6,80E-01 |
| AT2G24780 | 0,92 | -0,86 | 2,33 | -0,37 | 7,11E-01 |
| AT2G24870 | 1,09 | 0,95 | 2,36 | 0,40 | 6,87E-01 |
| AT2G24880 | 1,30 | -0,46 | 1,68 | -0,27 | 7,85E-01 |
| AT2G25011 | 1,24 | 1,45 | 1,88 | 0,77 | 4,41E-01 |
| AT2G25050 | 0,47 | -0,36 | 3,28 | -0,11 | 9,13E-01 |
| AT2G25070 | 0,45 | 0,19 | 2,75 | 0,07 | 9,46E-01 |
| AT2G08105 | 0,74 | -1,76 | 2,53 | -0,69 | 4,87E-01 |
| AT2G25180 | 0,76 | -0,23 | 3,14 | -0,07 | 9,41E-01 |
| AT2G25200 | 0,69 | 0,71 | 3,85 | 0,18 | 8,54E-01 |
| AT2G25300 | 0,15 | 0,00 | 4,41 | 0,00 | 1,00E+00 |
| AT2G25310 | 0,20 | -1,00 | 4,37 | -0,23 | 8,19E-01 |
| AT2G25570 | 0,45 | -1,07 | 4,39 | -0,24 | 8,07E-01 |
| AT2G25610 | 0,53 | -1,95 | 3,08 | -0,63 | 5,26E-01 |
| AT2G25670 | 0,93 | 0,10 | 2,37 | 0,04 | 9,67E-01 |
| AT2G25760 | 0,93 | -2,35 | 2,85 | -0,82 | 4,10E-01 |
| AT2G25850 | 0,16 | -0,26 | 4,38 | -0,06 | 9,52E-01 |
| AT2G25910 | 0,91 | 1,09 | 2,42 | 0,45 | 6,53E-01 |
| AT2G26030 | 0,31 | 0,86 | 3,61 | 0,24 | 8,12E-01 |
| AT2G26070 | 0,39 | -3,10 | 4,38 | -0,71 | 4,79E-01 |
| AT2G26280 | 1,11 | -1,88 | 2,21 | -0,85 | 3,97E-01 |
| AT2G26430 | 0,38 | -1,84 | 4,31 | -0,43 | 6,70E-01 |
| AT2G26540 | 0,42 | -1,97 | 4,30 | -0,46 | 6,47E-01 |
| AT2G26580 | 0,26 | -0,35 | 4,38 | -0,08 | 9,36E-01 |
| AT2G26590 | 0,25 | 0,44 | 4,36 | 0,10 | 9,19E-01 |
| AT2G26620 | 0,18 | 0,78 | 4,34 | 0,18 | 8,57E-01 |
| AT2G26680 | 0,33 | -3,08 | 4,39 | -0,70 | 4,83E-01 |
| AT2G26690 | 0,68 | -0,93 | 2,62 | -0,36 | 7,21E-01 |
| AT2G26740 | 0,33 | -0,94 | 4,35 | -0,22 | 8,29E-01 |
| AT2G08175 | 1,14 | 0,69 | 1,80 | 0,38 | 7,01E-01 |
| AT2G26790 | 0,21 | 0,00 | 4,41 | 0,00 | 1,00E+00 |
| AT2G26800 | 0,41 | -1,61 | 3,46 | -0,46 | 6,42E-01 |
| AT2G26840 | 0,08 | 0,00 | 4,41 | 0,00 | 1,00E+00 |
| AT2G26930 | 0,69 | -2,06 | 3,04 | -0,68 | 4,97E-01 |
| AT2G26980 | 0,61 | -0,32 | 2,36 | -0,14 | 8,91E-01 |
| AT2G26990 | 0,56 | -1,85 | 3,18 | -0,58 | 5,61E-01 |
| AT2G27020 | 0,74 | 0,21 | 2,64 | 0,08 | 9,37E-01 |
| AT2G27040 | 0,44 | -0,65 | 3,31 | -0,20 | 8,45E-01 |
| AT2G27070 | 0,56 | -0,69 | 3,16 | -0,22 | 8,27E-01 |
| AT2G27090 | 0,77 | -0,84 | 3,08 | -0,27 | 7,84E-01 |
| AT2G27170 | 0,61 | 0,36 | 2,62 | 0,14 | 8,90E-01 |

| | | | | | |
|---|---|---|---|---|---|
| AT2G27190 | 0,39 | 0,72 | 4,30 | 0,17 | 8,66E-01 |
| AT2G27290 | 0,57 | 0,61 | 2,72 | 0,23 | 8,22E-01 |
| AT2G27330 | 0,10 | 0,45 | 4,35 | 0,10 | 9,19E-01 |
| AT2G27350 | 0,23 | 0,39 | 4,35 | 0,09 | 9,28E-01 |
| AT2G27385 | 1,03 | 0,68 | 2,09 | 0,32 | 7,47E-01 |
| AT2G27500 | 0,69 | 0,59 | 2,38 | 0,25 | 8,04E-01 |
| AT2G27510 | 0,44 | -0,93 | 4,20 | -0,22 | 8,25E-01 |
| AT2G27530 | 0,92 | -1,03 | 1,80 | -0,57 | 5,67E-01 |
| AT2G27580 | 0,43 | 0,07 | 4,23 | 0,02 | 9,87E-01 |
| AT2G27660 | 0,41 | -2,91 | 4,34 | -0,67 | 5,03E-01 |
| AT2G27680 | 0,72 | -1,26 | 2,02 | -0,62 | 5,33E-01 |
| AT2G27770 | 0,38 | 1,43 | 4,30 | 0,33 | 7,40E-01 |
| AT2G27950 | 1,06 | -2,53 | 1,87 | -1,35 | 1,77E-01 |
| AT2G28080 | 0,87 | 1,85 | 2,69 | 0,69 | 4,92E-01 |
| AT2G28170 | 0,05 | -0,24 | 4,38 | -0,05 | 9,57E-01 |
| AT2G28380 | 0,74 | 0,07 | 2,99 | 0,03 | 9,80E-01 |
| AT2G28390 | 0,41 | -3,82 | 4,35 | -0,88 | 3,79E-01 |
| AT2G28400 | 0,81 | -2,32 | 2,53 | -0,92 | 3,60E-01 |
| AT2G28470 | 0,87 | 3,31 | 2,68 | 1,24 | 2,16E-01 |
| AT2G28480 | 0,12 | -0,78 | 4,39 | -0,18 | 8,59E-01 |
| AT2G28620 | 1,12 | 2,41 | 2,12 | 1,14 | 2,56E-01 |
| AT2G08275 | 1,12 | 2,41 | 2,12 | 1,14 | 2,56E-01 |
| AT2G28630 | 0,37 | 0,59 | 4,28 | 0,14 | 8,91E-01 |
| AT2G28640 | 0,62 | 0,42 | 2,93 | 0,14 | 8,85E-01 |
| AT2G28670 | 0,51 | 1,22 | 3,05 | 0,40 | 6,90E-01 |
| AT2G28740 | 0,44 | 0,81 | 4,15 | 0,19 | 8,46E-01 |
| AT2G28800 | 0,62 | 2,46 | 2,94 | 0,84 | 4,03E-01 |
| AT2G08300 | 0,20 | -1,00 | 4,37 | -0,23 | 8,19E-01 |
| AT2G28880 | 0,33 | -2,56 | 4,40 | -0,58 | 5,61E-01 |
| AT2G28890 | 0,77 | -0,63 | 2,12 | -0,30 | 7,67E-01 |
| AT2G28900 | 0,81 | 0,93 | 2,54 | 0,37 | 7,14E-01 |
| AT2G28950 | 0,45 | -1,17 | 4,11 | -0,28 | 7,76E-01 |
| AT2G29030 | 0,32 | 0,10 | 4,32 | 0,02 | 9,81E-01 |
| AT2G29040 | 0,08 | -0,21 | 4,39 | -0,05 | 9,62E-01 |
| AT2G29110 | 0,41 | -2,07 | 4,13 | -0,50 | 6,17E-01 |
| AT2G29120 | 1,16 | -1,64 | 2,35 | -0,70 | 4,85E-01 |
| AT2G29170 | 0,18 | 0,48 | 4,35 | 0,11 | 9,11E-01 |
| AT2G08330 | 0,26 | -2,60 | 4,41 | -0,59 | 5,55E-01 |
| AT2G29190 | 0,61 | -1,75 | 2,64 | -0,67 | 5,05E-01 |
| AT2G29220 | 0,39 | -3,10 | 4,38 | -0,71 | 4,79E-01 |
| AT2G29390 | 0,71 | -1,27 | 2,58 | -0,49 | 6,21E-01 |
| AT2G29400 | 0,98 | -1,63 | 1,80 | -0,91 | 3,64E-01 |
| AT2G29550 | 0,75 | 2,09 | 2,30 | 0,91 | 3,64E-01 |
| AT2G29570 | 0,14 | 0,40 | 4,35 | 0,09 | 9,27E-01 |
| AT2G29605 | 0,65 | 2,77 | 2,80 | 0,99 | 3,23E-01 |
| AT2G29650 | 0,77 | -2,58 | 2,58 | -1,00 | 3,17E-01 |
| AT2G30250 | 0,71 | -2,19 | 2,25 | -0,98 | 3,28E-01 |
| AT2G30260 | 0,44 | -1,69 | 3,35 | -0,50 | 6,14E-01 |
| AT2G30270 | 0,18 | -0,07 | 4,37 | -0,02 | 9,88E-01 |
| AT2G30360 | 0,37 | 1,28 | 4,31 | 0,30 | 7,66E-01 |
| AT2G30390 | 0,49 | -3,07 | 3,27 | -0,94 | 3,47E-01 |
| AT2G30470 | 0,73 | -1,09 | 2,36 | -0,46 | 6,43E-01 |
| AT2G30490 | 0,91 | -0,77 | 3,06 | -0,25 | 8,02E-01 |
| AT2G30505 | 0,35 | -0,82 | 4,38 | -0,19 | 8,52E-01 |
| AT2G30700 | 0,36 | -0,70 | 4,36 | -0,16 | 8,73E-01 |
| AT2G30710 | 0,35 | -2,19 | 4,36 | -0,50 | 6,16E-01 |

| | | | | | |
|---|---|---|---|---|---|
| AT2G30720 | 0,49 | -1,62 | 3,26 | -0,50 | 6,20E-01 |
| AT2G30790 | 0,21 | -1,17 | 4,36 | -0,27 | 7,88E-01 |
| AT2G30800 | 1,27 | -3,18 | 1,84 | -1,72 | 8,46E-02 |
| AT2G30850 | 0,41 | -2,96 | 4,38 | -0,68 | 4,99E-01 |
| AT2G30880 | 0,34 | -1,48 | 4,36 | -0,34 | 7,34E-01 |
| AT2G08420 | 0,54 | -2,60 | 4,06 | -0,64 | 5,21E-01 |
| AT2G30980 | 0,41 | -2,83 | 4,34 | -0,65 | 5,15E-01 |
| AT2G31040 | 0,19 | -0,11 | 4,39 | -0,02 | 9,80E-01 |
| AT2G31060 | 0,49 | -0,68 | 4,41 | -0,15 | 8,77E-01 |
| AT2G31070 | 0,79 | 1,19 | 2,24 | 0,53 | 5,96E-01 |
| AT2G31130 | 0,17 | 0,42 | 4,36 | 0,10 | 9,23E-01 |
| AT2G31170 | 0,45 | -0,81 | 4,39 | -0,18 | 8,54E-01 |
| AT2G31240 | 0,25 | -2,57 | 4,40 | -0,58 | 5,60E-01 |
| AT2G31250 | 0,39 | -3,07 | 4,38 | -0,70 | 4,84E-01 |
| AT2G31260 | 0,53 | 2,18 | 2,57 | 0,85 | 3,97E-01 |
| AT2G08430 | 0,08 | 0,00 | 4,41 | 0,00 | 1,00E+00 |
| AT2G31360 | 1,30 | -0,85 | 1,95 | -0,43 | 6,65E-01 |
| AT2G31390 | 0,86 | -3,03 | 2,88 | -1,05 | 2,92E-01 |
| AT2G31410 | 0,37 | -3,94 | 4,34 | -0,91 | 3,63E-01 |
| AT2G31440 | 0,44 | 0,00 | 4,41 | 0,00 | 1,00E+00 |
| AT2G31510 | 0,63 | -0,93 | 3,15 | -0,29 | 7,68E-01 |
| AT2G31570 | 0,14 | -0,38 | 4,38 | -0,09 | 9,31E-01 |
| AT2G31670 | 0,73 | -0,75 | 2,63 | -0,28 | 7,77E-01 |
| AT2G31730 | 0,53 | -2,10 | 4,36 | -0,48 | 6,30E-01 |
| AT2G31740 | 0,52 | -4,03 | 3,18 | -1,27 | 2,04E-01 |
| AT2G31751 | 0,97 | -0,56 | 2,48 | -0,23 | 8,21E-01 |
| AT2G31800 | 0,85 | -0,22 | 2,01 | -0,11 | 9,12E-01 |
| AT2G31830 | 0,34 | -1,64 | 4,41 | -0,37 | 7,10E-01 |
| AT2G31865 | 0,37 | -2,54 | 4,40 | -0,58 | 5,64E-01 |
| AT2G31870 | 0,27 | 0,87 | 3,74 | 0,23 | 8,15E-01 |
| AT2G08475 | 0,81 | -0,61 | 2,37 | -0,26 | 7,97E-01 |
| AT2G31990 | 0,31 | -0,74 | 4,35 | -0,17 | 8,64E-01 |
| AT2G32000 | 0,55 | -2,03 | 2,51 | -0,81 | 4,19E-01 |
| AT2G32030 | 0,22 | 0,38 | 4,35 | 0,09 | 9,31E-01 |
| AT2G32050 | 0,08 | 0,00 | 4,41 | 0,00 | 1,00E+00 |
| AT2G32060 | 0,34 | 1,17 | 4,32 | 0,27 | 7,87E-01 |
| AT2G32070 | 0,64 | 0,05 | 2,62 | 0,02 | 9,86E-01 |
| AT2G32080 | 0,57 | -0,90 | 2,58 | -0,35 | 7,27E-01 |
| AT2G32130 | 0,28 | -0,76 | 4,40 | -0,17 | 8,62E-01 |
| AT2G32230 | 0,70 | -4,44 | 2,83 | -1,57 | 1,17E-01 |
| AT2G32235 | 0,12 | 0,00 | 4,41 | 0,00 | 1,00E+00 |
| AT2G32390 | 0,36 | 0,00 | 4,41 | 0,00 | 1,00E+00 |
| AT2G32400 | 0,29 | 0,00 | 4,41 | 0,00 | 1,00E+00 |
| AT2G32415 | 0,42 | -3,76 | 4,35 | -0,86 | 3,87E-01 |
| AT2G32480 | 0,87 | -1,96 | 2,28 | -0,86 | 3,91E-01 |
| AT2G32520 | 0,35 | -0,39 | 4,38 | -0,09 | 9,29E-01 |
| AT2G32600 | 0,08 | 0,00 | 4,41 | 0,00 | 1,00E+00 |
| AT2G32810 | 0,70 | 1,32 | 2,49 | 0,53 | 5,94E-01 |
| AT2G32920 | 1,12 | -1,12 | 1,71 | -0,66 | 5,10E-01 |
| AT2G32940 | 0,12 | 0,00 | 4,41 | 0,00 | 1,00E+00 |
| AT2G33040 | 0,74 | 2,00 | 2,35 | 0,85 | 3,95E-01 |
| AT2G33051 | 0,64 | -3,35 | 3,88 | -0,86 | 3,88E-01 |
| AT2G33070 | 0,08 | 0,00 | 4,41 | 0,00 | 1,00E+00 |
| AT2G33110 | 0,40 | 0,00 | 4,41 | 0,00 | 1,00E+00 |
| AT2G33120 | 0,77 | 0,02 | 2,02 | 0,01 | 9,94E-01 |
| AT2G33175 | 0,67 | 1,24 | 2,38 | 0,52 | 6,02E-01 |

| | | | | | |
|---|---|---|---|---|---|
| AT2G33210 | 0,70 | -0,87 | 2,42 | -0,36 | 7,18E-01 |
| AT2G33260 | 0,24 | -2,54 | 4,40 | -0,58 | 5,64E-01 |
| AT2G33320 | 0,54 | 0,26 | 2,55 | 0,10 | 9,20E-01 |
| AT2G33370 | 0,46 | 0,18 | 2,74 | 0,07 | 9,48E-01 |
| AT2G33385 | 0,08 | -0,21 | 4,39 | -0,05 | 9,62E-01 |
| AT2G33460 | 0,28 | -1,70 | 4,40 | -0,39 | 7,00E-01 |
| AT2G33470 | 0,39 | -0,46 | 3,34 | -0,14 | 8,92E-01 |
| AT2G33500 | 0,51 | -3,91 | 4,34 | -0,90 | 3,68E-01 |
| AT2G33530 | 0,99 | -2,45 | 1,79 | -1,37 | 1,71E-01 |
| AT2G33540 | 0,96 | 0,20 | 1,75 | 0,11 | 9,10E-01 |
| AT2G33550 | 0,30 | 0,06 | 4,32 | 0,01 | 9,88E-01 |
| AT2G33580 | 0,75 | 0,46 | 2,15 | 0,22 | 8,29E-01 |
| AT2G33630 | 0,50 | -2,43 | 3,20 | -0,76 | 4,48E-01 |
| AT2G33655 | 0,29 | 0,00 | 4,41 | 0,00 | 1,00E+00 |
| AT2G33800 | 0,66 | -1,26 | 2,61 | -0,48 | 6,29E-01 |
| AT2G33815 | 0,37 | -1,13 | 4,34 | -0,26 | 7,95E-01 |
| AT2G33860 | 0,43 | 0,52 | 4,17 | 0,12 | 9,01E-01 |
| AT2G33980 | 0,47 | -2,08 | 3,27 | -0,64 | 5,25E-01 |
| AT2G34040 | 1,26 | -0,17 | 1,81 | -0,10 | 9,24E-01 |
| AT2G34070 | 0,35 | 0,75 | 3,56 | 0,21 | 8,34E-01 |
| AT2G34090 | 0,11 | 0,46 | 4,35 | 0,11 | 9,15E-01 |
| AT2G34290 | 0,45 | 1,97 | 3,97 | 0,50 | 6,20E-01 |
| AT2G34300 | 0,72 | -1,64 | 4,38 | -0,37 | 7,09E-01 |
| AT2G34450 | 0,60 | -3,19 | 3,85 | -0,83 | 4,07E-01 |
| AT2G34470 | 0,95 | -2,68 | 2,06 | -1,30 | 1,92E-01 |
| AT2G34490 | 0,49 | 1,41 | 4,06 | 0,35 | 7,28E-01 |
| AT2G34510 | 0,33 | -2,62 | 4,41 | -0,59 | 5,53E-01 |
| AT2G34520 | 0,36 | -4,02 | 4,33 | -0,93 | 3,54E-01 |
| AT2G34680 | 1,27 | -0,09 | 2,15 | -0,04 | 9,66E-01 |
| AT2G34710 | 1,10 | -3,26 | 2,43 | -1,34 | 1,80E-01 |
| AT2G34790 | 0,43 | -0,70 | 4,06 | -0,17 | 8,64E-01 |
| AT2G34920 | 0,05 | -0,24 | 4,38 | -0,05 | 9,57E-01 |
| AT2G34930 | 0,44 | 0,81 | 3,40 | 0,24 | 8,12E-01 |
| AT2G35030 | 0,48 | -3,17 | 4,04 | -0,78 | 4,33E-01 |
| AT2G35040 | 0,75 | -1,47 | 3,07 | -0,48 | 6,31E-01 |
| AT2G35060 | 1,08 | 1,07 | 1,80 | 0,60 | 5,50E-01 |
| AT2G35120 | 0,10 | 0,45 | 4,35 | 0,10 | 9,19E-01 |
| AT2G35250 | 1,05 | 1,13 | 1,99 | 0,57 | 5,72E-01 |
| AT2G35360 | 0,45 | -2,73 | 4,13 | -0,66 | 5,09E-01 |
| AT2G35390 | 0,25 | -2,48 | 4,39 | -0,57 | 5,71E-01 |
| AT2G35420 | 0,17 | -0,75 | 4,40 | -0,17 | 8,65E-01 |
| AT2G35490 | 0,67 | 0,58 | 2,16 | 0,27 | 7,88E-01 |
| AT2G35510 | 0,57 | -2,60 | 3,15 | -0,83 | 4,09E-01 |
| AT2G35610 | 0,17 | -0,71 | 4,40 | -0,16 | 8,72E-01 |
| AT2G35680 | 0,73 | -0,01 | 2,37 | 0,00 | 9,97E-01 |
| AT2G35730 | 0,35 | -2,99 | 4,38 | -0,68 | 4,95E-01 |
| AT2G35742 | 0,34 | 1,55 | 3,46 | 0,45 | 6,55E-01 |
| AT2G08755 | 0,29 | 0,11 | 4,36 | 0,02 | 9,80E-01 |
| AT2G08765 | 0,51 | -0,12 | 3,18 | -0,04 | 9,70E-01 |
| AT2G35859 | 0,71 | 0,01 | 2,73 | 0,00 | 9,98E-01 |
| AT2G35880 | 0,24 | -2,60 | 4,41 | -0,59 | 5,55E-01 |
| AT2G35930 | 0,76 | 2,52 | 2,75 | 0,92 | 3,58E-01 |
| AT2G36030 | 0,62 | 0,10 | 4,37 | 0,02 | 9,82E-01 |
| AT2G36050 | 0,69 | 0,68 | 2,88 | 0,24 | 8,14E-01 |
| AT2G36060 | 0,48 | -1,63 | 2,73 | -0,60 | 5,50E-01 |
| AT2G36070 | 0,34 | -1,60 | 4,35 | -0,37 | 7,13E-01 |

| | | | | | |
|---|---|---|---|---|---|
| AT2G36080 | 0,43 | -0,47 | 3,31 | -0,14 | 8,88E-01 |
| AT2G36180 | 0,76 | 2,68 | 3,30 | 0,81 | 4,17E-01 |
| AT2G36200 | 0,33 | -0,69 | 4,35 | -0,16 | 8,74E-01 |
| AT2G36230 | 0,20 | 0,84 | 4,33 | 0,19 | 8,47E-01 |
| AT2G36250 | 0,55 | 0,27 | 3,16 | 0,08 | 9,33E-01 |
| AT2G36300 | 0,44 | -0,39 | 3,25 | -0,12 | 9,05E-01 |
| AT2G36360 | 0,43 | 1,24 | 3,28 | 0,38 | 7,06E-01 |
| AT2G36400 | 0,83 | -0,73 | 3,05 | -0,24 | 8,11E-01 |
| AT2G36620 | 0,25 | -1,17 | 4,36 | -0,27 | 7,88E-01 |
| AT2G36680 | 0,35 | -2,13 | 4,35 | -0,49 | 6,25E-01 |
| AT2G36710 | 0,27 | 1,09 | 4,32 | 0,25 | 8,01E-01 |
| AT2G36750 | 0,21 | 0,00 | 4,41 | 0,00 | 1,00E+00 |
| AT2G36790 | 0,34 | 1,52 | 4,30 | 0,35 | 7,23E-01 |
| AT2G36800 | 0,85 | -2,44 | 2,43 | -1,00 | 3,15E-01 |
| AT2G36810 | 1,10 | -1,90 | 2,06 | -0,92 | 3,57E-01 |
| AT2G36835 | 0,30 | 0,83 | 3,61 | 0,23 | 8,18E-01 |
| AT2G36895 | 0,35 | -2,99 | 4,38 | -0,68 | 4,95E-01 |
| AT2G36900 | 0,55 | -2,43 | 3,78 | -0,64 | 5,21E-01 |
| AT2G36930 | 0,54 | -1,06 | 3,90 | -0,27 | 7,86E-01 |
| AT2G36990 | 1,19 | -4,72 | 2,46 | -1,92 | 5,53E-02 |
| AT2G37035 | 0,78 | -0,29 | 2,48 | -0,12 | 9,07E-01 |
| AT2G37080 | 0,12 | 0,00 | 4,41 | 0,00 | 1,00E+00 |
| AT2G37130 | 0,68 | 0,24 | 3,01 | 0,08 | 9,37E-01 |
| AT2G37150 | 0,75 | -3,79 | 3,46 | -1,10 | 2,73E-01 |
| AT2G37170 | 0,90 | -2,41 | 2,43 | -0,99 | 3,20E-01 |
| AT2G37200 | 0,31 | -0,95 | 3,62 | -0,26 | 7,94E-01 |
| AT2G37230 | 0,56 | -0,58 | 3,18 | -0,18 | 8,56E-01 |
| AT2G37310 | 0,77 | -3,19 | 3,59 | -0,89 | 3,74E-01 |
| AT2G37340 | 1,12 | -5,03 | 2,20 | -2,29 | 2,21E-02 |
| AT2G37360 | 0,21 | 0,00 | 4,41 | 0,00 | 1,00E+00 |
| AT2G37400 | 0,25 | -2,53 | 4,40 | -0,57 | 5,65E-01 |
| AT2G37410 | 0,19 | -0,76 | 4,40 | -0,17 | 8,63E-01 |
| AT2G37520 | 1,03 | 0,26 | 1,91 | 0,14 | 8,92E-01 |
| AT2G37540 | 0,69 | -3,41 | 2,96 | -1,15 | 2,49E-01 |
| AT2G37570 | 0,98 | -1,11 | 1,84 | -0,60 | 5,47E-01 |
| AT2G37630 | 0,98 | -3,09 | 1,87 | -1,65 | 9,86E-02 |
| AT2G37678 | 0,57 | -1,03 | 4,36 | -0,24 | 8,14E-01 |
| AT2G08895 | 1,00 | 2,00 | 2,24 | 0,89 | 3,71E-01 |
| AT2G08900 | 0,86 | 1,78 | 2,05 | 0,87 | 3,85E-01 |
| AT2G08915 | 0,73 | 1,29 | 2,75 | 0,47 | 6,40E-01 |
| AT2G38000 | 0,32 | -0,26 | 4,34 | -0,06 | 9,53E-01 |
| AT2G38020 | 1,29 | -3,69 | 2,19 | -1,68 | 9,26E-02 |
| AT2G38160 | 0,87 | -5,00 | 3,22 | -1,55 | 1,20E-01 |
| AT2G38270 | 0,22 | 1,03 | 4,32 | 0,24 | 8,11E-01 |
| AT2G38290 | 0,66 | -2,53 | 2,92 | -0,87 | 3,86E-01 |
| AT2G38340 | 0,17 | -0,35 | 4,38 | -0,08 | 9,37E-01 |
| AT2G38360 | 0,39 | 0,32 | 3,28 | 0,10 | 9,22E-01 |
| AT2G38410 | 0,35 | -1,67 | 3,47 | -0,48 | 6,30E-01 |
| AT2G38480 | 0,17 | -0,71 | 4,40 | -0,16 | 8,72E-01 |
| AT2G38610 | 0,83 | -1,64 | 2,53 | -0,65 | 5,18E-01 |
| AT2G38650 | 1,16 | 1,02 | 2,18 | 0,47 | 6,41E-01 |
| AT2G38680 | 0,08 | -0,21 | 4,39 | -0,05 | 9,62E-01 |
| AT2G38750 | 0,05 | -0,24 | 4,38 | -0,05 | 9,57E-01 |
| AT2G38790 | 0,55 | 0,49 | 2,34 | 0,21 | 8,35E-01 |
| AT2G38860 | 0,41 | -0,74 | 3,42 | -0,22 | 8,29E-01 |
| AT2G39000 | 1,06 | -3,49 | 2,97 | -1,17 | 2,41E-01 |

| | | | | | |
|---|---|---|---|---|---|
| AT2G39080 | 0,20 | -2,55 | 4,40 | -0,58 | 5,62E-01 |
| AT2G39130 | 0,08 | -0,21 | 4,39 | -0,05 | 9,62E-01 |
| AT2G39160 | 0,24 | -0,80 | 4,39 | -0,18 | 8,56E-01 |
| AT2G39170 | 0,19 | 0,04 | 4,37 | 0,01 | 9,92E-01 |
| AT2G09010 | 0,08 | -0,21 | 4,39 | -0,05 | 9,62E-01 |
| AT2G39210 | 0,91 | -3,96 | 2,77 | -1,43 | 1,53E-01 |
| AT2G39350 | 0,49 | -1,92 | 3,19 | -0,60 | 5,47E-01 |
| AT2G39360 | 0,80 | -4,63 | 3,45 | -1,34 | 1,81E-01 |
| AT2G09045 | 0,52 | -2,60 | 3,96 | -0,66 | 5,11E-01 |
| AT2G39450 | 0,29 | -2,51 | 4,39 | -0,57 | 5,67E-01 |
| AT2G39480 | 1,05 | -0,67 | 1,92 | -0,35 | 7,28E-01 |
| AT2G39650 | 0,49 | 1,42 | 4,00 | 0,35 | 7,23E-01 |
| AT2G39710 | 0,60 | -0,24 | 3,25 | -0,08 | 9,40E-01 |
| AT2G39720 | 1,30 | -3,52 | 2,32 | -1,51 | 1,30E-01 |
| AT2G39750 | 1,24 | -2,50 | 1,86 | -1,34 | 1,79E-01 |
| AT2G09080 | 0,50 | -2,50 | 3,28 | -0,76 | 4,46E-01 |
| AT2G39805 | 0,30 | -2,60 | 4,41 | -0,59 | 5,55E-01 |
| AT2G39920 | 0,18 | -0,78 | 4,39 | -0,18 | 8,59E-01 |
| AT2G09090 | 0,82 | -3,80 | 2,87 | -1,32 | 1,85E-01 |
| AT2G39950 | 0,75 | -2,33 | 2,65 | -0,88 | 3,80E-01 |
| AT2G39960 | 0,25 | 0,00 | 4,41 | 0,00 | 1,00E+00 |
| AT2G40070 | 0,52 | -2,59 | 3,96 | -0,65 | 5,13E-01 |
| AT2G40080 | 0,23 | 0,39 | 4,35 | 0,09 | 9,28E-01 |
| AT2G40165 | 0,38 | -3,33 | 4,37 | -0,76 | 4,46E-01 |
| AT2G40205 | 0,27 | -0,27 | 4,34 | -0,06 | 9,50E-01 |
| AT2G40280 | 0,63 | -3,09 | 3,06 | -1,01 | 3,13E-01 |
| AT2G40350 | 0,12 | 0,00 | 4,41 | 0,00 | 1,00E+00 |
| AT2G40360 | 0,28 | 1,38 | 4,30 | 0,32 | 7,48E-01 |
| AT2G40430 | 0,45 | -1,31 | 4,13 | -0,32 | 7,51E-01 |
| AT2G40475 | 0,89 | 2,72 | 2,21 | 1,23 | 2,18E-01 |
| AT2G40590 | 0,50 | -2,13 | 2,69 | -0,79 | 4,29E-01 |
| AT2G40600 | 0,11 | -0,80 | 4,39 | -0,18 | 8,56E-01 |
| AT2G40610 | 0,37 | -0,71 | 4,40 | -0,16 | 8,71E-01 |
| AT2G40650 | 0,77 | 2,62 | 2,24 | 1,17 | 2,41E-01 |
| AT2G40730 | 0,34 | 1,14 | 3,44 | 0,33 | 7,41E-01 |
| AT2G40750 | 0,46 | -2,60 | 4,21 | -0,62 | 5,37E-01 |
| AT2G40770 | 0,51 | -1,30 | 4,01 | -0,32 | 7,46E-01 |
| AT2G40815 | 0,24 | -2,60 | 4,41 | -0,59 | 5,55E-01 |
| AT2G40860 | 0,30 | -2,60 | 4,41 | -0,59 | 5,55E-01 |
| AT2G40890 | 0,53 | 0,06 | 3,93 | 0,02 | 9,88E-01 |
| AT2G40920 | 0,35 | -0,79 | 4,39 | -0,18 | 8,57E-01 |
| AT2G40925 | 0,65 | -3,34 | 3,69 | -0,90 | 3,67E-01 |
| AT2G40950 | 0,74 | -1,43 | 3,60 | -0,40 | 6,90E-01 |
| AT2G40970 | 0,42 | -3,74 | 4,35 | -0,86 | 3,90E-01 |
| AT2G09225 | 0,24 | -0,20 | 4,39 | -0,05 | 9,64E-01 |
| AT2G41120 | 0,21 | 0,00 | 4,41 | 0,00 | 1,00E+00 |
| AT2G41160 | 0,39 | -0,75 | 4,40 | -0,17 | 8,65E-01 |
| AT2G41250 | 0,87 | 1,03 | 2,37 | 0,43 | 6,64E-01 |
| AT2G41420 | 0,11 | 0,46 | 4,35 | 0,11 | 9,15E-01 |
| AT2G41475 | 0,71 | -1,19 | 2,50 | -0,47 | 6,36E-01 |
| AT2G41500 | 0,96 | -3,57 | 2,11 | -1,69 | 9,08E-02 |
| AT2G41600 | 0,58 | -1,21 | 2,59 | -0,47 | 6,41E-01 |
| AT2G41616 | 0,33 | -2,59 | 4,41 | -0,59 | 5,56E-01 |
| AT2G41630 | 0,92 | -2,89 | 2,68 | -1,08 | 2,79E-01 |
| AT2G41680 | 0,32 | 0,42 | 4,34 | 0,10 | 9,23E-01 |
| AT2G41770 | 0,40 | -0,92 | 4,28 | -0,21 | 8,30E-01 |

| | | | | | |
|---|---|---|---|---|---|
| AT2G41790 | 0,58 | -0,23 | 2,63 | -0,09 | 9,31E-01 |
| AT2G41830 | 0,40 | -2,60 | 4,24 | -0,61 | 5,39E-01 |
| AT2G41835 | 0,70 | 0,09 | 3,76 | 0,03 | 9,80E-01 |
| AT2G41870 | 0,29 | -1,77 | 4,39 | -0,40 | 6,87E-01 |
| AT2G41890 | 0,43 | -0,14 | 4,14 | -0,03 | 9,73E-01 |
| AT2G41945 | 0,60 | -0,95 | 3,96 | -0,24 | 8,11E-01 |
| AT2G41960 | 0,92 | -0,88 | 2,12 | -0,42 | 6,77E-01 |
| AT2G42010 | 0,72 | 0,36 | 2,06 | 0,17 | 8,61E-01 |
| AT2G42020 | 0,48 | 0,08 | 4,37 | 0,02 | 9,86E-01 |
| AT2G42070 | 0,23 | 0,89 | 4,33 | 0,21 | 8,37E-01 |
| AT2G42230 | 0,42 | -1,65 | 4,18 | -0,39 | 6,93E-01 |
| AT2G42280 | 0,33 | -3,01 | 4,38 | -0,69 | 4,92E-01 |
| AT2G42365 | 0,35 | -2,99 | 4,38 | -0,68 | 4,95E-01 |
| AT2G42400 | 1,01 | -2,89 | 2,17 | -1,33 | 1,83E-01 |
| AT2G42500 | 0,82 | -0,60 | 2,24 | -0,27 | 7,90E-01 |
| AT2G42530 | 0,98 | -0,41 | 2,18 | -0,19 | 8,49E-01 |
| AT2G42760 | 0,47 | 2,01 | 2,61 | 0,77 | 4,41E-01 |
| AT2G42790 | 0,53 | -2,54 | 2,71 | -0,94 | 3,49E-01 |
| AT2G42800 | 0,18 | 1,18 | 4,31 | 0,27 | 7,85E-01 |
| AT2G42810 | 0,84 | -2,34 | 1,95 | -1,20 | 2,30E-01 |
| AT2G42840 | 0,22 | 1,22 | 4,31 | 0,28 | 7,78E-01 |
| AT2G42865 | 0,08 | 0,00 | 4,41 | 0,00 | 1,00E+00 |
| AT2G42870 | 0,10 | -0,18 | 4,38 | -0,04 | 9,67E-01 |
| AT2G42880 | 0,41 | -0,17 | 3,44 | -0,05 | 9,60E-01 |
| AT2G43180 | 0,70 | -4,54 | 2,87 | -1,58 | 1,13E-01 |
| AT2G43260 | 0,20 | -2,60 | 4,41 | -0,59 | 5,55E-01 |
| AT2G43290 | 0,50 | 0,48 | 3,21 | 0,15 | 8,82E-01 |
| AT2G43320 | 0,22 | -0,31 | 4,38 | -0,07 | 9,44E-01 |
| AT2G43360 | 0,33 | -2,56 | 4,40 | -0,58 | 5,61E-01 |
| AT2G43420 | 0,11 | -0,80 | 4,39 | -0,18 | 8,56E-01 |
| AT2G43430 | 0,59 | -1,24 | 2,57 | -0,48 | 6,30E-01 |
| AT2G43460 | 0,41 | -0,45 | 3,38 | -0,13 | 8,93E-01 |
| AT2G43490 | 0,44 | -0,71 | 4,14 | -0,17 | 8,63E-01 |
| AT2G43560 | 0,25 | -1,64 | 4,36 | -0,38 | 7,07E-01 |
| AT2G43865 | 0,15 | 0,48 | 4,35 | 0,11 | 9,12E-01 |
| AT2G43910 | 0,26 | 0,06 | 3,78 | 0,01 | 9,88E-01 |
| AT2G43940 | 0,08 | 0,00 | 4,41 | 0,00 | 1,00E+00 |
| AT2G43945 | 0,47 | -3,08 | 4,39 | -0,70 | 4,83E-01 |
| AT2G43950 | 0,24 | -2,59 | 4,41 | -0,59 | 5,56E-01 |
| AT2G44090 | 0,69 | -3,34 | 2,59 | -1,29 | 1,96E-01 |
| AT2G44140 | 0,91 | 1,89 | 2,29 | 0,83 | 4,08E-01 |
| AT2G44180 | 0,46 | 0,36 | 3,24 | 0,11 | 9,12E-01 |
| AT2G44255 | 0,20 | -0,68 | 4,41 | -0,15 | 8,77E-01 |
| AT2G44360 | 0,63 | 15,76 | 4,34 | 3,63 | 2,82E-04 |
| AT2G44610 | 0,48 | 0,04 | 4,20 | 0,01 | 9,93E-01 |
| AT2G44660 | 0,35 | -0,98 | 3,51 | -0,28 | 7,81E-01 |
| AT2G44760 | 0,32 | -0,89 | 3,60 | -0,25 | 8,05E-01 |
| AT2G44798 | 0,36 | -0,52 | 4,34 | -0,12 | 9,04E-01 |
| AT2G44950 | 0,77 | -3,13 | 2,23 | -1,40 | 1,61E-01 |
| AT2G45070 | 0,80 | 2,18 | 2,13 | 1,02 | 3,07E-01 |
| AT2G45220 | 0,54 | -2,50 | 4,39 | -0,57 | 5,69E-01 |
| AT2G45315 | 0,67 | 0,31 | 2,49 | 0,12 | 9,01E-01 |
| AT2G45460 | 0,20 | 1,10 | 4,31 | 0,25 | 7,99E-01 |
| AT2G45520 | 0,22 | 0,62 | 4,34 | 0,14 | 8,86E-01 |
| AT2G45560 | 0,47 | -3,81 | 4,35 | -0,88 | 3,81E-01 |
| AT2G45660 | 0,55 | 0,43 | 2,29 | 0,19 | 8,51E-01 |

| | | | | | |
|---|---|---|---|---|---|
| AT2G45670 | 0,95 | -3,48 | 2,41 | -1,44 | 1,49E-01 |
| AT2G45700 | 0,38 | -0,68 | 4,36 | -0,16 | 8,76E-01 |
| AT2G45820 | 0,98 | 0,49 | 1,89 | 0,26 | 7,94E-01 |
| AT2G45970 | 0,56 | -1,21 | 2,59 | -0,47 | 6,41E-01 |
| AT2G45980 | 0,45 | -1,12 | 4,12 | -0,27 | 7,86E-01 |
| AT2G46030 | 0,57 | -3,12 | 4,38 | -0,71 | 4,76E-01 |
| AT2G46040 | 0,20 | -2,55 | 4,40 | -0,58 | 5,62E-01 |
| AT2G46060 | 0,49 | -2,60 | 4,41 | -0,59 | 5,55E-01 |
| AT2G46080 | 0,51 | -1,58 | 3,17 | -0,50 | 6,17E-01 |
| AT2G46180 | 0,35 | -0,50 | 4,36 | -0,12 | 9,08E-01 |
| AT2G46280 | 1,16 | -3,12 | 1,97 | -1,59 | 1,12E-01 |
| AT2G46330 | 0,93 | 1,38 | 2,13 | 0,65 | 5,18E-01 |
| AT2G46390 | 0,46 | -3,62 | 4,10 | -0,88 | 3,78E-01 |
| AT2G46505 | 0,89 | -3,28 | 2,19 | -1,49 | 1,35E-01 |
| AT2G46510 | 1,07 | 0,01 | 1,88 | 0,00 | 9,96E-01 |
| AT2G46540 | 0,12 | 0,07 | 4,38 | 0,02 | 9,87E-01 |
| AT2G46550 | 0,57 | -1,56 | 2,74 | -0,57 | 5,68E-01 |
| AT2G46572 | 0,51 | -1,73 | 4,07 | -0,42 | 6,71E-01 |
| AT2G46610 | 0,85 | -1,66 | 1,79 | -0,93 | 3,54E-01 |
| AT2G46680 | 0,94 | -0,17 | 2,28 | -0,07 | 9,41E-01 |
| AT2G46840 | 0,08 | 0,00 | 4,41 | 0,00 | 1,00E+00 |
| AT2G46920 | 0,44 | -1,71 | 4,08 | -0,42 | 6,76E-01 |
| AT2G46980 | 0,24 | -3,18 | 4,39 | -0,73 | 4,68E-01 |
| AT2G47090 | 0,25 | 0,00 | 4,41 | 0,00 | 1,00E+00 |
| AT2G47250 | 1,15 | -3,46 | 2,42 | -1,43 | 1,52E-01 |
| AT2G47330 | 0,60 | 0,50 | 3,14 | 0,16 | 8,73E-01 |
| AT2G47600 | 0,61 | -1,79 | 2,53 | -0,71 | 4,79E-01 |
| AT2G47680 | 0,89 | -2,50 | 1,94 | -1,29 | 1,97E-01 |
| AT2G47700 | 0,93 | -1,50 | 2,28 | -0,66 | 5,10E-01 |
| AT2G47840 | 0,52 | -0,46 | 3,34 | -0,14 | 8,90E-01 |
| AT2G47990 | 0,88 | -4,81 | 2,63 | -1,83 | 6,74E-02 |
| AT2G48000 | 1,00 | -3,25 | 3,46 | -0,94 | 3,47E-01 |
| AT2G48030 | 0,73 | 0,05 | 2,50 | 0,02 | 9,85E-01 |
| AT2G48130 | 0,91 | 0,98 | 2,01 | 0,49 | 6,24E-01 |
| AT3G01090 | 0,63 | 0,74 | 2,16 | 0,34 | 7,32E-01 |
| AT3G01100 | 0,90 | -2,72 | 2,05 | -1,33 | 1,84E-01 |
| AT3G01130 | 0,21 | -1,17 | 4,36 | -0,27 | 7,88E-01 |
| AT3G01160 | 0,24 | -0,80 | 4,39 | -0,18 | 8,56E-01 |
| AT3G01180 | 0,84 | -1,10 | 1,89 | -0,58 | 5,59E-01 |
| AT3G01300 | 0,08 | 0,00 | 4,41 | 0,00 | 1,00E+00 |
| AT3G01340 | 0,50 | -0,33 | 3,26 | -0,10 | 9,19E-01 |
| AT3G01380 | 1,09 | -3,25 | 2,02 | -1,61 | 1,08E-01 |
| AT3G01390 | 0,38 | -0,28 | 4,38 | -0,06 | 9,50E-01 |
| AT3G01415 | 0,40 | 1,20 | 4,32 | 0,28 | 7,82E-01 |
| AT3G01420 | 0,31 | -1,69 | 4,37 | -0,39 | 6,99E-01 |
| AT3G01430 | 0,79 | -1,85 | 2,89 | -0,64 | 5,22E-01 |
| AT3G01550 | 0,10 | -0,34 | 4,38 | -0,08 | 9,38E-01 |
| AT3G01720 | 0,80 | -0,74 | 2,93 | -0,25 | 8,01E-01 |
| AT3G01820 | 0,81 | -3,55 | 3,62 | -0,98 | 3,27E-01 |
| AT3G01840 | 0,88 | -2,02 | 2,16 | -0,94 | 3,49E-01 |
| AT3G01910 | 0,46 | -0,78 | 4,14 | -0,19 | 8,51E-01 |
| AT3G01930 | 0,62 | -3,74 | 2,99 | -1,25 | 2,11E-01 |
| AT3G01980 | 0,16 | -0,64 | 4,40 | -0,15 | 8,85E-01 |
| AT3G02040 | 0,68 | -2,06 | 2,35 | -0,87 | 3,83E-01 |
| AT3G02080 | 1,20 | -2,59 | 2,04 | -1,27 | 2,05E-01 |
| AT3G02190 | 0,44 | -0,78 | 3,36 | -0,23 | 8,17E-01 |

| | | | | | |
|---|---|---|---|---|---|
| AT3G02210 | 0,52 | -0,95 | 4,04 | -0,23 | 8,14E-01 |
| AT3G02250 | 0,34 | -0,29 | 4,38 | -0,07 | 9,48E-01 |
| AT3G02380 | 0,91 | -2,98 | 2,20 | -1,35 | 1,76E-01 |
| AT3G02410 | 0,66 | -4,85 | 4,30 | -1,13 | 2,58E-01 |
| AT3G02450 | 0,68 | -1,67 | 2,20 | -0,76 | 4,48E-01 |
| AT3G02520 | 1,11 | -0,64 | 1,74 | -0,37 | 7,11E-01 |
| AT3G02530 | 0,67 | 0,05 | 2,69 | 0,02 | 9,84E-01 |
| AT3G02700 | 0,39 | -2,06 | 4,22 | -0,49 | 6,25E-01 |
| AT3G02730 | 0,92 | -3,77 | 2,74 | -1,38 | 1,68E-01 |
| AT3G02750 | 1,16 | -2,55 | 1,80 | -1,42 | 1,56E-01 |
| AT3G02760 | 0,94 | -3,14 | 2,72 | -1,15 | 2,50E-01 |
| AT3G01205 | 0,80 | 0,29 | 2,29 | 0,13 | 8,99E-01 |
| AT3G03090 | 0,29 | -2,60 | 4,41 | -0,59 | 5,55E-01 |
| AT3G03150 | 0,22 | 0,80 | 4,34 | 0,18 | 8,54E-01 |
| AT3G03220 | 0,16 | 1,16 | 4,31 | 0,27 | 7,88E-01 |
| AT3G03250 | 1,00 | 1,13 | 1,86 | 0,61 | 5,42E-01 |
| AT3G03300 | 0,33 | 0,00 | 4,41 | 0,00 | 1,00E+00 |
| AT3G03305 | 0,32 | -1,70 | 4,40 | -0,39 | 6,99E-01 |
| AT3G03310 | 0,29 | -2,57 | 4,40 | -0,58 | 5,59E-01 |
| AT3G03330 | 0,30 | -1,76 | 4,35 | -0,40 | 6,86E-01 |
| AT3G03405 | 0,16 | 1,16 | 4,31 | 0,27 | 7,88E-01 |
| AT3G03460 | 0,17 | 0,00 | 4,41 | 0,00 | 1,00E+00 |
| AT3G03480 | 0,08 | 0,00 | 4,41 | 0,00 | 1,00E+00 |
| AT3G03550 | 0,10 | -0,34 | 4,38 | -0,08 | 9,38E-01 |
| AT3G03560 | 0,61 | -1,64 | 3,04 | -0,54 | 5,90E-01 |
| AT3G03580 | 0,26 | -2,60 | 4,41 | -0,59 | 5,55E-01 |
| AT3G03590 | 0,08 | 0,00 | 4,41 | 0,00 | 1,00E+00 |
| AT3G03600 | 0,30 | 0,55 | 4,35 | 0,13 | 9,00E-01 |
| AT3G03720 | 0,26 | 0,42 | 4,34 | 0,10 | 9,23E-01 |
| AT3G03770 | 0,99 | -0,73 | 3,20 | -0,23 | 8,19E-01 |
| AT3G03890 | 0,79 | -2,00 | 2,35 | -0,85 | 3,95E-01 |
| AT3G03920 | 0,42 | -0,88 | 3,35 | -0,26 | 7,92E-01 |
| AT3G03940 | 0,41 | 1,61 | 2,77 | 0,58 | 5,61E-01 |
| AT3G04030 | 0,25 | 1,72 | 4,29 | 0,40 | 6,89E-01 |
| AT3G04080 | 1,01 | -2,85 | 2,11 | -1,35 | 1,78E-01 |
| AT3G04150 | 0,26 | -2,60 | 4,41 | -0,59 | 5,55E-01 |
| AT3G04200 | 0,62 | -2,55 | 4,40 | -0,58 | 5,62E-01 |
| AT3G04220 | 0,33 | -0,31 | 4,38 | -0,07 | 9,43E-01 |
| AT3G04260 | 0,61 | -2,02 | 4,36 | -0,46 | 6,43E-01 |
| AT3G04290 | 0,84 | -1,34 | 3,13 | -0,43 | 6,68E-01 |
| AT3G04350 | 0,35 | 1,75 | 4,29 | 0,41 | 6,83E-01 |
| AT3G04480 | 0,84 | -0,18 | 2,33 | -0,08 | 9,39E-01 |
| AT3G04500 | 0,20 | -1,00 | 4,37 | -0,23 | 8,19E-01 |
| AT3G04580 | 1,03 | -2,48 | 2,17 | -1,14 | 2,53E-01 |
| AT3G04640 | 0,85 | 1,73 | 2,11 | 0,82 | 4,12E-01 |
| AT3G04690 | 0,22 | -0,06 | 4,38 | -0,01 | 9,89E-01 |
| AT3G04720 | 0,50 | 1,67 | 3,05 | 0,55 | 5,84E-01 |
| AT3G04730 | 1,02 | -0,74 | 1,65 | -0,45 | 6,54E-01 |
| AT3G04750 | 0,57 | -2,16 | 2,59 | -0,83 | 4,04E-01 |
| AT3G04760 | 0,89 | 0,47 | 2,05 | 0,23 | 8,18E-01 |
| AT3G04860 | 0,12 | 0,00 | 4,41 | 0,00 | 1,00E+00 |
| AT3G04980 | 0,46 | -2,64 | 4,40 | -0,60 | 5,48E-01 |
| AT3G05030 | 0,28 | 0,38 | 4,35 | 0,09 | 9,30E-01 |
| AT3G01585 | 0,10 | 0,45 | 4,35 | 0,10 | 9,19E-01 |
| AT3G05050 | 0,38 | -1,64 | 4,41 | -0,37 | 7,10E-01 |
| AT3G05060 | 0,60 | -2,46 | 3,72 | -0,66 | 5,09E-01 |

| | | | | | |
|---|---|---|---|---|---|
| AT3G05090 | 0,52 | 0,49 | 2,55 | 0,19 | 8,46E-01 |
| AT3G05165 | 0,83 | -0,73 | 1,95 | -0,38 | 7,07E-01 |
| AT3G05240 | 0,13 | 0,00 | 4,41 | 0,00 | 1,00E+00 |
| AT3G05270 | 0,58 | -3,24 | 2,57 | -1,26 | 2,06E-01 |
| AT3G05280 | 0,59 | -4,05 | 3,78 | -1,07 | 2,84E-01 |
| AT3G05540 | 1,04 | 0,53 | 1,90 | 0,28 | 7,81E-01 |
| AT3G05650 | 0,98 | -0,71 | 2,22 | -0,32 | 7,49E-01 |
| AT3G05660 | 0,79 | -1,11 | 2,83 | -0,39 | 6,95E-01 |
| AT3G05690 | 0,11 | 0,46 | 4,35 | 0,11 | 9,15E-01 |
| AT3G05770 | 0,46 | 0,53 | 3,28 | 0,16 | 8,73E-01 |
| AT3G05810 | 0,21 | -0,44 | 4,38 | -0,10 | 9,20E-01 |
| AT3G05850 | 0,90 | -2,66 | 2,36 | -1,13 | 2,59E-01 |
| AT3G05880 | 0,55 | -1,60 | 3,22 | -0,50 | 6,19E-01 |
| AT3G05900 | 0,51 | -0,62 | 3,25 | -0,19 | 8,47E-01 |
| AT3G05910 | 0,68 | 0,24 | 2,70 | 0,09 | 9,29E-01 |
| AT3G05932 | 0,27 | -1,72 | 4,38 | -0,39 | 6,95E-01 |
| AT3G05970 | 0,71 | 0,42 | 2,27 | 0,18 | 8,53E-01 |
| AT3G06030 | 0,29 | 0,00 | 4,41 | 0,00 | 1,00E+00 |
| AT3G06050 | 0,18 | 0,68 | 4,34 | 0,16 | 8,75E-01 |
| AT3G06060 | 0,17 | -1,25 | 4,37 | -0,29 | 7,74E-01 |
| AT3G06150 | 0,98 | -0,63 | 1,71 | -0,37 | 7,15E-01 |
| AT3G06190 | 0,71 | -1,22 | 3,00 | -0,41 | 6,85E-01 |
| AT3G06250 | 0,74 | -1,61 | 2,90 | -0,56 | 5,78E-01 |
| AT3G06290 | 0,72 | -0,18 | 2,46 | -0,07 | 9,42E-01 |
| AT3G06310 | 0,08 | 0,00 | 4,41 | 0,00 | 1,00E+00 |
| AT3G06350 | 0,64 | -3,44 | 3,05 | -1,13 | 2,60E-01 |
| AT3G06430 | 0,12 | 0,00 | 4,41 | 0,00 | 1,00E+00 |
| AT3G06440 | 0,12 | 0,00 | 4,41 | 0,00 | 1,00E+00 |
| AT3G06450 | 0,61 | -1,41 | 2,62 | -0,54 | 5,91E-01 |
| AT3G06480 | 1,05 | -2,53 | 1,81 | -1,39 | 1,64E-01 |
| AT3G06483 | 0,45 | 0,33 | 2,73 | 0,12 | 9,03E-01 |
| AT3G06540 | 0,08 | 0,00 | 4,41 | 0,00 | 1,00E+00 |
| AT3G06550 | 0,32 | 0,42 | 4,35 | 0,10 | 9,24E-01 |
| AT3G06580 | 0,15 | -0,73 | 4,40 | -0,17 | 8,68E-01 |
| AT3G06590 | 0,40 | -1,50 | 4,35 | -0,34 | 7,31E-01 |
| AT3G06620 | 0,12 | 0,00 | 4,41 | 0,00 | 1,00E+00 |
| AT3G06665 | 0,25 | 1,72 | 4,29 | 0,40 | 6,89E-01 |
| AT3G06680 | 0,28 | -0,27 | 4,34 | -0,06 | 9,51E-01 |
| AT3G06690 | 0,20 | -1,00 | 4,37 | -0,23 | 8,19E-01 |
| AT3G06700 | 0,57 | -0,99 | 2,67 | -0,37 | 7,11E-01 |
| AT3G06790 | 0,16 | -0,68 | 4,41 | -0,15 | 8,77E-01 |
| AT3G06820 | 0,08 | 0,00 | 4,41 | 0,00 | 1,00E+00 |
| AT3G06870 | 0,21 | -0,69 | 4,36 | -0,16 | 8,74E-01 |
| AT3G06900 | 1,19 | 0,32 | 1,77 | 0,18 | 8,55E-01 |
| AT3G06930 | 0,40 | -2,52 | 4,18 | -0,60 | 5,46E-01 |
| AT3G06960 | 0,89 | 1,70 | 2,51 | 0,68 | 4,97E-01 |
| AT3G07010 | 0,41 | -1,49 | 4,24 | -0,35 | 7,25E-01 |
| AT3G07020 | 0,54 | -3,33 | 3,82 | -0,87 | 3,84E-01 |
| AT3G07040 | 1,04 | -2,92 | 2,24 | -1,30 | 1,93E-01 |
| AT3G07050 | 0,53 | -0,02 | 3,20 | -0,01 | 9,94E-01 |
| AT3G07115 | 0,08 | -0,21 | 4,39 | -0,05 | 9,62E-01 |
| AT3G07180 | 0,12 | 0,00 | 4,41 | 0,00 | 1,00E+00 |
| AT3G07215 | 0,55 | 1,26 | 2,51 | 0,50 | 6,16E-01 |
| AT3G07273 | 0,08 | 0,00 | 4,41 | 0,00 | 1,00E+00 |
| AT3G07290 | 0,69 | -2,54 | 3,58 | -0,71 | 4,78E-01 |
| AT3G07300 | 0,29 | -3,07 | 4,39 | -0,70 | 4,83E-01 |

| | | | | | |
|---|---|---|---|---|---|
| AT3G07330 | 0,98 | 0,59 | 2,50 | 0,24 | 8,13E-01 |
| AT3G07380 | 1,22 | 1,16 | 1,59 | 0,73 | 4,66E-01 |
| AT3G07460 | 0,63 | 0,41 | 2,57 | 0,16 | 8,75E-01 |
| AT3G07470 | 0,58 | -1,71 | 3,21 | -0,53 | 5,95E-01 |
| AT3G07520 | 0,51 | 0,05 | 3,21 | 0,02 | 9,87E-01 |
| AT3G07540 | 0,08 | 0,00 | 4,41 | 0,00 | 1,00E+00 |
| AT3G07700 | 0,94 | -1,07 | 1,77 | -0,61 | 5,44E-01 |
| AT3G07800 | 0,53 | -1,86 | 2,70 | -0,69 | 4,91E-01 |
| AT3G07890 | 0,34 | -0,57 | 4,34 | -0,13 | 8,96E-01 |
| AT3G08570 | 1,10 | -4,46 | 2,18 | -2,05 | 4,03E-02 |
| AT3G08610 | 0,59 | -2,76 | 3,17 | -0,87 | 3,83E-01 |
| AT3G08650 | 0,24 | 0,06 | 4,38 | 0,01 | 9,90E-01 |
| AT3G08760 | 0,70 | -0,04 | 2,31 | -0,02 | 9,88E-01 |
| AT3G08840 | 0,67 | -3,44 | 2,99 | -1,15 | 2,50E-01 |
| AT3G08870 | 0,75 | 0,00 | 3,56 | 0,00 | 1,00E+00 |
| AT3G08960 | 0,38 | 0,22 | 4,30 | 0,05 | 9,60E-01 |
| AT3G09040 | 0,17 | 0,00 | 4,41 | 0,00 | 1,00E+00 |
| AT3G09070 | 1,31 | 2,17 | 2,10 | 1,03 | 3,01E-01 |
| AT3G09090 | 1,29 | -2,30 | 2,04 | -1,12 | 2,61E-01 |
| AT3G09190 | 0,55 | -2,46 | 3,72 | -0,66 | 5,08E-01 |
| AT3G09210 | 0,27 | -2,69 | 4,40 | -0,61 | 5,41E-01 |
| AT3G09270 | 0,53 | -0,36 | 2,71 | -0,13 | 8,94E-01 |
| AT3G09320 | 0,85 | -0,59 | 2,45 | -0,24 | 8,11E-01 |
| AT3G09410 | 0,82 | -4,10 | 2,41 | -1,70 | 8,95E-02 |
| AT3G09470 | 0,82 | -1,71 | 3,54 | -0,48 | 6,28E-01 |
| AT3G09560 | 0,97 | -1,80 | 2,26 | -0,80 | 4,26E-01 |
| AT3G09580 | 0,48 | -1,59 | 3,35 | -0,48 | 6,34E-01 |
| AT3G09770 | 0,38 | -1,25 | 4,37 | -0,29 | 7,75E-01 |
| AT3G09780 | 0,23 | -1,14 | 4,39 | -0,26 | 7,96E-01 |
| AT3G09870 | 0,08 | 0,00 | 4,41 | 0,00 | 1,00E+00 |
| AT3G09920 | 0,16 | -0,62 | 4,40 | -0,14 | 8,87E-01 |
| AT3G10010 | 0,35 | -2,02 | 4,35 | -0,47 | 6,42E-01 |
| AT3G10020 | 0,27 | -0,68 | 4,41 | -0,15 | 8,77E-01 |
| AT3G10030 | 0,88 | -2,62 | 2,96 | -0,88 | 3,77E-01 |
| AT3G10050 | 0,25 | -0,70 | 4,40 | -0,16 | 8,73E-01 |
| AT3G10070 | 0,46 | -1,87 | 4,23 | -0,44 | 6,58E-01 |
| AT3G10130 | 0,21 | -0,70 | 4,40 | -0,16 | 8,73E-01 |
| AT3G10160 | 0,36 | -0,48 | 3,41 | -0,14 | 8,87E-01 |
| AT3G10230 | 1,13 | -0,23 | 2,29 | -0,10 | 9,20E-01 |
| AT3G10250 | 0,08 | 0,00 | 4,41 | 0,00 | 1,00E+00 |
| AT3G10270 | 0,56 | -1,06 | 3,18 | -0,33 | 7,39E-01 |
| AT3G10320 | 0,12 | 0,00 | 4,41 | 0,00 | 1,00E+00 |
| AT3G10360 | 0,63 | -2,70 | 3,77 | -0,71 | 4,75E-01 |
| AT3G10405 | 0,68 | -3,45 | 4,32 | -0,80 | 4,25E-01 |
| AT3G10410 | 1,05 | -1,06 | 1,88 | -0,56 | 5,74E-01 |
| AT3G10520 | 0,15 | -0,17 | 4,39 | -0,04 | 9,68E-01 |
| AT3G10525 | 0,39 | -1,72 | 3,48 | -0,49 | 6,22E-01 |
| AT3G10630 | 0,99 | -1,27 | 1,98 | -0,64 | 5,23E-01 |
| AT3G10640 | 0,33 | 0,38 | 4,35 | 0,09 | 9,31E-01 |
| AT3G10660 | 0,27 | -1,67 | 4,36 | -0,38 | 7,02E-01 |
| AT3G10690 | 0,77 | -3,77 | 3,46 | -1,09 | 2,76E-01 |
| AT3G10720 | 0,36 | 1,08 | 4,32 | 0,25 | 8,02E-01 |
| AT3G10730 | 0,58 | -2,42 | 3,07 | -0,79 | 4,31E-01 |
| AT3G10770 | 0,74 | -1,64 | 2,61 | -0,63 | 5,30E-01 |
| AT3G10810 | 0,46 | -2,47 | 4,23 | -0,58 | 5,59E-01 |
| AT3G10840 | 0,41 | -3,93 | 4,34 | -0,91 | 3,65E-01 |

| | | | | | |
|---|---|---|---|---|---|
| AT3G10850 | 0,36 | 0,01 | 4,33 | 0,00 | 9,99E-01 |
| AT3G10900 | 0,31 | 1,71 | 4,30 | 0,40 | 6,90E-01 |
| AT3G10915 | 0,24 | 0,20 | 4,36 | 0,04 | 9,64E-01 |
| AT3G10930 | 0,61 | 1,02 | 2,46 | 0,42 | 6,77E-01 |
| AT3G10960 | 1,23 | 0,73 | 1,65 | 0,45 | 6,56E-01 |
| AT3G11030 | 0,56 | -2,53 | 3,75 | -0,67 | 5,01E-01 |
| AT3G11100 | 0,47 | -0,79 | 4,06 | -0,19 | 8,45E-01 |
| AT3G11280 | 0,31 | -1,29 | 4,37 | -0,30 | 7,68E-01 |
| AT3G11290 | 0,30 | -1,27 | 4,36 | -0,29 | 7,71E-01 |
| AT3G11310 | 0,31 | 1,73 | 4,29 | 0,40 | 6,87E-01 |
| AT3G11320 | 0,17 | -0,75 | 4,40 | -0,17 | 8,65E-01 |
| AT3G11330 | 0,97 | -3,86 | 2,65 | -1,45 | 1,46E-01 |
| AT3G11435 | 0,23 | 0,00 | 4,41 | 0,00 | 1,00E+00 |
| AT3G11450 | 1,01 | -1,58 | 1,77 | -0,89 | 3,72E-01 |
| AT3G02445 | 0,88 | 2,91 | 3,40 | 0,86 | 3,92E-01 |
| AT3G02455 | 0,99 | 1,16 | 2,96 | 0,39 | 6,94E-01 |
| AT3G11570 | 0,77 | -2,33 | 2,96 | -0,79 | 4,31E-01 |
| AT3G11580 | 0,26 | 0,94 | 4,32 | 0,22 | 8,28E-01 |
| AT3G11650 | 0,48 | 0,87 | 3,16 | 0,28 | 7,83E-01 |
| AT3G11680 | 0,22 | 0,75 | 4,34 | 0,17 | 8,63E-01 |
| AT3G11710 | 1,08 | -0,48 | 2,01 | -0,24 | 8,12E-01 |
| AT3G11730 | 0,25 | -2,57 | 4,40 | -0,58 | 5,60E-01 |
| AT3G11770 | 0,28 | -1,32 | 4,38 | -0,30 | 7,63E-01 |
| AT3G11780 | 0,21 | -0,30 | 4,38 | -0,07 | 9,46E-01 |
| AT3G11840 | 0,47 | 0,06 | 4,14 | 0,02 | 9,88E-01 |
| AT3G11964 | 0,92 | -1,32 | 1,84 | -0,72 | 4,71E-01 |
| AT3G12030 | 0,12 | 0,00 | 4,41 | 0,00 | 1,00E+00 |
| AT3G12130 | 0,80 | -3,07 | 2,49 | -1,23 | 2,18E-01 |
| AT3G12140 | 0,25 | 0,54 | 4,34 | 0,12 | 9,01E-01 |
| AT3G12145 | 0,15 | 0,48 | 4,35 | 0,11 | 9,12E-01 |
| AT3G12200 | 0,53 | -0,23 | 3,90 | -0,06 | 9,54E-01 |
| AT3G12260 | 0,32 | 0,78 | 4,34 | 0,18 | 8,57E-01 |
| AT3G12400 | 0,71 | -0,57 | 1,94 | -0,29 | 7,68E-01 |
| AT3G12490 | 0,47 | 0,08 | 2,40 | 0,03 | 9,75E-01 |
| AT3G12500 | 0,08 | 0,00 | 4,41 | 0,00 | 1,00E+00 |
| AT3G12580 | 0,43 | -3,13 | 4,39 | -0,71 | 4,76E-01 |
| AT3G12587 | 0,28 | -2,28 | 4,37 | -0,52 | 6,01E-01 |
| AT3G12590 | 0,82 | -3,62 | 2,40 | -1,51 | 1,32E-01 |
| AT3G12610 | 0,59 | -1,10 | 2,26 | -0,49 | 6,26E-01 |
| AT3G12670 | 1,09 | -0,84 | 2,59 | -0,33 | 7,44E-01 |
| AT3G12680 | 0,32 | -0,84 | 4,37 | -0,19 | 8,48E-01 |
| AT3G12690 | 0,57 | 0,00 | 4,41 | 0,00 | 1,00E+00 |
| AT3G12750 | 0,45 | -3,32 | 4,03 | -0,82 | 4,10E-01 |
| AT3G12770 | 0,67 | -3,60 | 2,84 | -1,27 | 2,05E-01 |
| AT3G12800 | 0,96 | 1,35 | 1,90 | 0,71 | 4,75E-01 |
| AT3G12820 | 0,21 | -2,60 | 4,41 | -0,59 | 5,55E-01 |
| AT3G12920 | 1,14 | 0,90 | 2,65 | 0,34 | 7,34E-01 |
| AT3G12940 | 0,28 | 0,57 | 4,35 | 0,13 | 8,96E-01 |
| AT3G12950 | 0,48 | -2,60 | 4,41 | -0,59 | 5,55E-01 |
| AT3G12965 | 0,28 | -0,27 | 4,34 | -0,06 | 9,51E-01 |
| AT3G13000 | 0,24 | -3,18 | 4,39 | -0,73 | 4,68E-01 |
| AT3G13010 | 0,29 | -2,54 | 4,40 | -0,58 | 5,63E-01 |
| AT3G13020 | 0,33 | -2,93 | 4,38 | -0,67 | 5,03E-01 |
| AT3G13030 | 0,97 | -4,78 | 2,28 | -2,10 | 3,59E-02 |
| AT3G13040 | 0,58 | 0,58 | 2,66 | 0,22 | 8,26E-01 |
| AT3G13061 | 0,23 | 0,83 | 4,34 | 0,19 | 8,48E-01 |

| | | | | | |
|---|---|---|---|---|---|
| AT3G13110 | 1,06 | -2,76 | 2,11 | -1,31 | 1,90E-01 |
| AT3G13120 | 0,60 | -2,07 | 3,05 | -0,68 | 4,99E-01 |
| AT3G13150 | 0,37 | -2,98 | 4,38 | -0,68 | 4,97E-01 |
| AT3G13160 | 0,08 | 0,00 | 4,41 | 0,00 | 1,00E+00 |
| AT3G13220 | 0,37 | -2,58 | 4,40 | -0,58 | 5,59E-01 |
| AT3G13222 | 0,48 | 0,00 | 4,41 | 0,00 | 1,00E+00 |
| AT3G13225 | 0,32 | -0,78 | 4,39 | -0,18 | 8,60E-01 |
| AT3G13226 | 0,08 | 0,00 | 4,41 | 0,00 | 1,00E+00 |
| AT3G13235 | 0,44 | -0,01 | 2,81 | 0,00 | 9,98E-01 |
| AT3G13310 | 0,25 | -0,13 | 4,37 | -0,03 | 9,77E-01 |
| AT3G13390 | 0,05 | -0,24 | 4,38 | -0,05 | 9,57E-01 |
| AT3G13404 | 0,46 | 0,50 | 4,04 | 0,12 | 9,02E-01 |
| AT3G13450 | 0,75 | -4,44 | 2,77 | -1,60 | 1,09E-01 |
| AT3G13480 | 0,20 | 0,11 | 4,37 | 0,03 | 9,80E-01 |
| AT3G13490 | 0,57 | -2,35 | 3,11 | -0,76 | 4,50E-01 |
| AT3G13570 | 0,44 | -3,25 | 4,21 | -0,77 | 4,40E-01 |
| AT3G13720 | 0,96 | 1,93 | 1,80 | 1,07 | 2,84E-01 |
| AT3G13772 | 0,72 | -1,65 | 2,93 | -0,56 | 5,74E-01 |
| AT3G13860 | 0,89 | -2,11 | 2,00 | -1,05 | 2,92E-01 |
| AT3G13990 | 1,05 | -4,83 | 2,18 | -2,22 | 2,67E-02 |
| AT3G14070 | 0,23 | -0,82 | 4,38 | -0,19 | 8,52E-01 |
| AT3G14075 | 0,26 | -0,76 | 4,40 | -0,17 | 8,63E-01 |
| AT3G14090 | 0,68 | -1,30 | 3,10 | -0,42 | 6,76E-01 |
| AT3G14110 | 0,61 | -2,93 | 2,58 | -1,14 | 2,55E-01 |
| AT3G14180 | 0,40 | 0,00 | 4,41 | 0,00 | 1,00E+00 |
| AT3G14205 | 0,51 | -1,18 | 2,73 | -0,43 | 6,66E-01 |
| AT3G14230 | 0,40 | -2,52 | 4,18 | -0,60 | 5,46E-01 |
| AT3G14330 | 0,56 | -3,38 | 3,07 | -1,10 | 2,71E-01 |
| AT3G14400 | 1,24 | -3,99 | 3,23 | -1,24 | 2,17E-01 |
| AT3G14430 | 0,25 | 0,00 | 4,41 | 0,00 | 1,00E+00 |
| AT3G14440 | 0,33 | 0,62 | 4,34 | 0,14 | 8,87E-01 |
| AT3G14460 | 0,50 | -1,34 | 3,95 | -0,34 | 7,34E-01 |
| AT3G14590 | 0,14 | 0,40 | 4,35 | 0,09 | 9,27E-01 |
| AT3G14810 | 0,79 | -1,64 | 2,52 | -0,65 | 5,14E-01 |
| AT3G14900 | 0,17 | 0,00 | 4,41 | 0,00 | 1,00E+00 |
| AT3G15110 | 0,15 | -0,17 | 4,39 | -0,04 | 9,68E-01 |
| AT3G15120 | 0,82 | -2,18 | 2,35 | -0,93 | 3,52E-01 |
| AT3G15150 | 0,08 | -0,21 | 4,39 | -0,05 | 9,62E-01 |
| AT3G15220 | 0,77 | -3,50 | 2,41 | -1,45 | 1,46E-01 |
| AT3G15354 | 0,91 | -0,86 | 2,28 | -0,38 | 7,06E-01 |
| AT3G15356 | 0,36 | -2,56 | 4,37 | -0,59 | 5,58E-01 |
| AT3G15470 | 0,41 | -0,10 | 2,80 | -0,03 | 9,72E-01 |
| AT3G03125 | 0,17 | 0,00 | 4,41 | 0,00 | 1,00E+00 |
| AT3G15530 | 1,09 | 1,30 | 2,41 | 0,54 | 5,89E-01 |
| AT3G15570 | 0,83 | 1,33 | 2,67 | 0,50 | 6,18E-01 |
| AT3G15590 | 0,41 | -2,00 | 4,27 | -0,47 | 6,40E-01 |
| AT3G15630 | 0,46 | 2,11 | 3,16 | 0,67 | 5,04E-01 |
| AT3G15720 | 0,20 | 0,84 | 4,33 | 0,19 | 8,47E-01 |
| AT3G15810 | 0,58 | 1,19 | 2,45 | 0,48 | 6,28E-01 |
| AT3G15900 | 0,29 | 0,39 | 4,35 | 0,09 | 9,28E-01 |
| AT3G15970 | 0,63 | -0,65 | 2,64 | -0,25 | 8,06E-01 |
| AT3G16000 | 0,51 | 1,48 | 3,20 | 0,46 | 6,45E-01 |
| AT3G16110 | 0,21 | 0,00 | 4,41 | 0,00 | 1,00E+00 |
| AT3G16190 | 0,21 | -2,50 | 4,39 | -0,57 | 5,70E-01 |
| AT3G03235 | 0,34 | -2,60 | 4,41 | -0,59 | 5,55E-01 |
| AT3G16290 | 0,84 | 1,47 | 2,02 | 0,73 | 4,65E-01 |

| | | | | | |
|---|---|---|---|---|---|
| AT3G16310 | 0,55 | -1,80 | 3,85 | -0,47 | 6,40E-01 |
| AT3G16500 | 0,17 | -0,35 | 4,38 | -0,08 | 9,37E-01 |
| AT3G16510 | 0,65 | 0,13 | 3,01 | 0,04 | 9,66E-01 |
| AT3G16520 | 0,76 | 0,88 | 2,12 | 0,41 | 6,80E-01 |
| AT3G16530 | 0,27 | -0,35 | 4,34 | -0,08 | 9,37E-01 |
| AT3G16560 | 0,66 | -1,83 | 2,50 | -0,73 | 4,65E-01 |
| AT3G16630 | 1,01 | 0,07 | 2,01 | 0,03 | 9,74E-01 |
| AT3G16840 | 0,19 | 0,42 | 4,35 | 0,10 | 9,23E-01 |
| AT3G16860 | 0,47 | 0,51 | 3,97 | 0,13 | 8,97E-01 |
| AT3G16870 | 0,39 | -3,07 | 4,38 | -0,70 | 4,84E-01 |
| AT3G16890 | 0,19 | -0,25 | 4,38 | -0,06 | 9,54E-01 |
| AT3G17020 | 1,21 | -1,17 | 2,06 | -0,57 | 5,70E-01 |
| AT3G03445 | 0,77 | -1,59 | 2,59 | -0,62 | 5,38E-01 |
| AT3G17170 | 0,31 | -0,68 | 4,41 | -0,15 | 8,77E-01 |
| AT3G17320 | 0,12 | 0,18 | 4,36 | 0,04 | 9,68E-01 |
| AT3G17330 | 0,20 | -0,68 | 4,41 | -0,15 | 8,77E-01 |
| AT3G17420 | 0,17 | 0,00 | 4,41 | 0,00 | 1,00E+00 |
| AT3G17440 | 0,08 | 0,00 | 4,41 | 0,00 | 1,00E+00 |
| AT3G17450 | 0,60 | -2,89 | 2,96 | -0,98 | 3,29E-01 |
| AT3G17655 | 0,19 | -0,72 | 4,40 | -0,16 | 8,70E-01 |
| AT3G17740 | 0,37 | -1,72 | 4,36 | -0,39 | 6,94E-01 |
| AT3G17760 | 0,21 | -2,60 | 4,41 | -0,59 | 5,55E-01 |
| AT3G03585 | 0,05 | -0,24 | 4,38 | -0,05 | 9,57E-01 |
| AT3G17940 | 0,81 | -2,28 | 2,51 | -0,91 | 3,63E-01 |
| AT3G18030 | 0,52 | -0,95 | 4,04 | -0,23 | 8,14E-01 |
| AT3G18035 | 1,23 | 0,38 | 1,65 | 0,23 | 8,16E-01 |
| AT3G18110 | 1,16 | -3,87 | 2,55 | -1,52 | 1,30E-01 |
| AT3G18130 | 0,88 | -2,05 | 2,37 | -0,87 | 3,86E-01 |
| AT3G18140 | 0,30 | 0,14 | 4,31 | 0,03 | 9,74E-01 |
| AT3G18250 | 0,55 | -2,75 | 4,30 | -0,64 | 5,23E-01 |
| AT3G18380 | 0,25 | -2,57 | 4,40 | -0,58 | 5,60E-01 |
| AT3G18420 | 0,20 | 0,46 | 4,35 | 0,10 | 9,16E-01 |
| AT3G18535 | 0,67 | 1,09 | 2,39 | 0,46 | 6,48E-01 |
| AT3G18600 | 0,25 | -3,06 | 4,38 | -0,70 | 4,85E-01 |
| AT3G18610 | 0,55 | 0,00 | 4,41 | 0,00 | 1,00E+00 |
| AT3G18630 | 0,20 | -2,55 | 4,40 | -0,58 | 5,62E-01 |
| AT3G18700 | 1,19 | 2,09 | 2,08 | 1,01 | 3,14E-01 |
| AT3G18750 | 0,17 | 0,00 | 4,41 | 0,00 | 1,00E+00 |
| AT3G18930 | 0,90 | 2,19 | 2,20 | 0,99 | 3,20E-01 |
| AT3G18970 | 0,08 | 0,00 | 4,41 | 0,00 | 1,00E+00 |
| AT3G19000 | 0,24 | 0,23 | 4,37 | 0,05 | 9,58E-01 |
| AT3G19020 | 0,54 | -1,75 | 3,08 | -0,57 | 5,70E-01 |
| AT3G19150 | 0,16 | -0,03 | 4,37 | -0,01 | 9,95E-01 |
| AT3G03815 | 0,25 | -3,06 | 4,38 | -0,70 | 4,85E-01 |
| AT3G19300 | 0,12 | -0,68 | 4,41 | -0,15 | 8,77E-01 |
| AT3G19340 | 0,12 | -0,31 | 4,38 | -0,07 | 9,44E-01 |
| AT3G19430 | 0,25 | -3,06 | 4,38 | -0,70 | 4,85E-01 |
| AT3G19490 | 0,72 | -0,84 | 2,58 | -0,33 | 7,44E-01 |
| AT3G19510 | 0,12 | -0,14 | 4,38 | -0,03 | 9,74E-01 |
| AT3G19515 | 0,59 | -1,64 | 3,21 | -0,51 | 6,10E-01 |
| AT3G19553 | 0,42 | -0,71 | 4,06 | -0,18 | 8,61E-01 |
| AT3G19580 | 0,72 | 0,67 | 2,45 | 0,27 | 7,85E-01 |
| AT3G19620 | 0,32 | -3,36 | 4,37 | -0,77 | 4,42E-01 |
| AT3G19630 | 0,39 | -3,07 | 4,38 | -0,70 | 4,84E-01 |
| AT3G19710 | 0,69 | 0,25 | 2,47 | 0,10 | 9,19E-01 |
| AT3G19720 | 1,17 | 0,75 | 2,51 | 0,30 | 7,67E-01 |

| | | | | | |
|---|---|---|---|---|---|
| AT3G19740 | 0,59 | -0,56 | 3,17 | -0,18 | 8,61E-01 |
| AT3G19790 | 0,71 | -1,31 | 1,85 | -0,71 | 4,79E-01 |
| AT3G19810 | 0,70 | -2,87 | 4,37 | -0,66 | 5,11E-01 |
| AT3G19830 | 0,67 | 2,10 | 2,93 | 0,72 | 4,74E-01 |
| AT3G19850 | 0,50 | -1,70 | 3,07 | -0,56 | 5,79E-01 |
| AT3G19880 | 0,35 | 0,88 | 3,44 | 0,26 | 7,97E-01 |
| AT3G19895 | 0,23 | 1,25 | 4,31 | 0,29 | 7,72E-01 |
| AT3G19900 | 0,17 | 0,00 | 4,41 | 0,00 | 1,00E+00 |
| AT3G20015 | 0,62 | -1,56 | 2,94 | -0,53 | 5,96E-01 |
| AT3G20050 | 0,74 | -1,52 | 2,50 | -0,61 | 5,43E-01 |
| AT3G20060 | 0,33 | 0,37 | 4,35 | 0,09 | 9,31E-01 |
| AT3G20230 | 0,35 | 0,36 | 4,36 | 0,08 | 9,35E-01 |
| AT3G20290 | 0,63 | 1,15 | 3,04 | 0,38 | 7,04E-01 |
| AT3G20320 | 0,51 | -0,01 | 2,74 | 0,00 | 9,98E-01 |
| AT3G20390 | 0,62 | 1,49 | 2,22 | 0,67 | 5,03E-01 |
| AT3G20470 | 0,38 | -0,75 | 4,37 | -0,17 | 8,63E-01 |
| AT3G20620 | 0,68 | -3,13 | 3,64 | -0,86 | 3,90E-01 |
| AT3G20630 | 1,20 | -2,23 | 1,59 | -1,40 | 1,60E-01 |
| AT3G20655 | 0,30 | -2,60 | 4,41 | -0,59 | 5,55E-01 |
| AT3G20700 | 1,05 | 3,54 | 2,38 | 1,49 | 1,37E-01 |
| AT3G03985 | 0,54 | 2,78 | 4,27 | 0,65 | 5,15E-01 |
| AT3G20830 | 0,60 | 0,50 | 3,31 | 0,15 | 8,81E-01 |
| AT3G21070 | 0,23 | 0,39 | 4,35 | 0,09 | 9,28E-01 |
| AT3G21140 | 0,25 | -0,81 | 4,39 | -0,18 | 8,54E-01 |
| AT3G21160 | 0,11 | -0,68 | 4,41 | -0,15 | 8,77E-01 |
| AT3G21215 | 0,11 | -0,80 | 4,39 | -0,18 | 8,56E-01 |
| AT3G21300 | 0,80 | -1,27 | 2,20 | -0,58 | 5,63E-01 |
| AT3G21430 | 0,61 | 0,25 | 3,13 | 0,08 | 9,37E-01 |
| AT3G21540 | 0,20 | 0,00 | 4,41 | 0,00 | 1,00E+00 |
| AT3G21570 | 0,87 | 1,17 | 2,86 | 0,41 | 6,81E-01 |
| AT3G21630 | 0,12 | -0,31 | 4,38 | -0,07 | 9,44E-01 |
| AT3G21650 | 0,33 | -2,59 | 4,41 | -0,59 | 5,56E-01 |
| AT3G21740 | 0,24 | -1,92 | 4,38 | -0,44 | 6,61E-01 |
| AT3G21790 | 0,96 | 0,56 | 2,15 | 0,26 | 7,95E-01 |
| AT3G21865 | 0,48 | 0,06 | 2,69 | 0,02 | 9,83E-01 |
| AT3G22070 | 0,17 | 0,00 | 4,41 | 0,00 | 1,00E+00 |
| AT3G22110 | 0,73 | -0,87 | 2,40 | -0,36 | 7,17E-01 |
| AT3G22170 | 1,10 | -1,07 | 2,04 | -0,53 | 5,99E-01 |
| AT3G22190 | 0,78 | -2,70 | 2,89 | -0,94 | 3,49E-01 |
| AT3G22210 | 0,48 | -0,54 | 3,34 | -0,16 | 8,71E-01 |
| AT3G22220 | 0,36 | -1,64 | 3,55 | -0,46 | 6,43E-01 |
| AT3G22230 | 0,95 | -1,46 | 1,81 | -0,81 | 4,20E-01 |
| AT3G22240 | 1,25 | 1,65 | 1,84 | 0,89 | 3,71E-01 |
| AT3G22320 | 0,41 | -2,00 | 4,27 | -0,47 | 6,40E-01 |
| AT3G22421 | 0,96 | 2,02 | 2,66 | 0,76 | 4,46E-01 |
| AT3G22470 | 0,59 | -0,94 | 2,66 | -0,35 | 7,24E-01 |
| AT3G22530 | 0,38 | -2,66 | 4,40 | -0,60 | 5,46E-01 |
| AT3G22550 | 0,60 | -1,14 | 3,11 | -0,37 | 7,14E-01 |
| AT3G22580 | 0,15 | -0,17 | 4,39 | -0,04 | 9,68E-01 |
| AT3G22600 | 0,17 | 0,00 | 4,41 | 0,00 | 1,00E+00 |
| AT3G22630 | 0,68 | -1,25 | 2,13 | -0,59 | 5,57E-01 |
| AT3G22660 | 0,44 | -2,71 | 4,33 | -0,63 | 5,32E-01 |
| AT3G22690 | 0,29 | -0,25 | 4,38 | -0,06 | 9,55E-01 |
| AT3G22710 | 1,14 | -1,26 | 1,75 | -0,72 | 4,71E-01 |
| AT3G22720 | 0,08 | -0,21 | 4,39 | -0,05 | 9,62E-01 |
| AT3G22770 | 0,68 | 0,96 | 3,28 | 0,29 | 7,69E-01 |

| | | | | | |
|---|---|---|---|---|---|
| AT3G22950 | 0,15 | -0,14 | 4,39 | -0,03 | 9,74E-01 |
| AT3G22980 | 0,74 | -1,22 | 2,56 | -0,48 | 6,33E-01 |
| AT3G23020 | 0,53 | -3,73 | 3,84 | -0,97 | 3,31E-01 |
| AT3G23030 | 0,69 | -1,29 | 2,04 | -0,64 | 5,25E-01 |
| AT3G23080 | 0,08 | 0,00 | 4,41 | 0,00 | 1,00E+00 |
| AT3G23170 | 0,50 | 0,28 | 2,63 | 0,11 | 9,16E-01 |
| AT3G23172 | 0,24 | 0,52 | 4,35 | 0,12 | 9,04E-01 |
| AT3G23400 | 0,58 | -0,26 | 3,21 | -0,08 | 9,35E-01 |
| AT3G23420 | 0,32 | 1,42 | 4,30 | 0,33 | 7,41E-01 |
| AT3G23490 | 0,26 | 1,27 | 4,31 | 0,29 | 7,69E-01 |
| AT3G23570 | 0,32 | -1,65 | 4,40 | -0,38 | 7,08E-01 |
| AT3G23633 | 0,12 | 0,75 | 4,34 | 0,17 | 8,63E-01 |
| AT3G23650 | 0,55 | 2,34 | 2,89 | 0,81 | 4,17E-01 |
| AT3G04585 | 0,72 | 2,67 | 2,77 | 0,96 | 3,35E-01 |
| AT3G23660 | 0,75 | -3,02 | 2,90 | -1,04 | 2,96E-01 |
| AT3G23700 | 0,24 | 0,41 | 4,35 | 0,09 | 9,25E-01 |
| AT3G23740 | 0,08 | 0,00 | 4,41 | 0,00 | 1,00E+00 |
| AT3G23830 | 0,27 | 0,00 | 4,41 | 0,00 | 1,00E+00 |
| AT3G23900 | 0,77 | -0,99 | 2,81 | -0,35 | 7,23E-01 |
| AT3G23970 | 0,08 | 0,00 | 4,41 | 0,00 | 1,00E+00 |
| AT3G23980 | 0,76 | -0,94 | 2,42 | -0,39 | 6,98E-01 |
| AT3G24080 | 0,22 | -0,21 | 4,38 | -0,05 | 9,61E-01 |
| AT3G24120 | 0,83 | -1,58 | 3,56 | -0,44 | 6,57E-01 |
| AT3G24140 | 0,26 | 1,04 | 4,32 | 0,24 | 8,09E-01 |
| AT3G24200 | 0,11 | -0,68 | 4,41 | -0,15 | 8,77E-01 |
| AT3G24260 | 0,15 | 0,79 | 4,33 | 0,18 | 8,56E-01 |
| AT3G24270 | 0,91 | 2,03 | 2,31 | 0,88 | 3,81E-01 |
| AT3G24320 | 0,63 | -2,16 | 3,11 | -0,69 | 4,88E-01 |
| AT3G24430 | 0,54 | -0,69 | 2,67 | -0,26 | 7,95E-01 |
| AT3G24440 | 0,50 | -2,75 | 3,22 | -0,86 | 3,92E-01 |
| AT3G24465 | 0,32 | 0,58 | 3,62 | 0,16 | 8,72E-01 |
| AT3G24503 | 1,13 | -0,03 | 1,73 | -0,02 | 9,87E-01 |
| AT3G24518 | 0,96 | 0,35 | 2,40 | 0,15 | 8,84E-01 |
| AT3G24614 | 0,96 | -0,09 | 1,74 | -0,05 | 9,59E-01 |
| AT3G24670 | 0,20 | -0,68 | 4,41 | -0,15 | 8,77E-01 |
| AT3G24870 | 1,16 | 0,28 | 2,13 | 0,13 | 8,96E-01 |
| AT3G24880 | 0,67 | -1,28 | 2,93 | -0,44 | 6,63E-01 |
| AT3G24900 | 0,51 | -0,43 | 3,14 | -0,14 | 8,91E-01 |
| AT3G25090 | 0,87 | 2,03 | 1,98 | 1,03 | 3,03E-01 |
| AT3G25150 | 1,03 | -3,25 | 2,52 | -1,29 | 1,97E-01 |
| AT3G25180 | 0,15 | -0,17 | 4,39 | -0,04 | 9,68E-01 |
| AT3G25530 | 0,19 | 0,80 | 4,34 | 0,18 | 8,53E-01 |
| AT3G25560 | 0,41 | -2,31 | 4,37 | -0,53 | 5,96E-01 |
| AT3G25610 | 1,05 | -3,37 | 2,33 | -1,45 | 1,48E-01 |
| AT3G25660 | 0,47 | 0,00 | 4,05 | 0,00 | 1,00E+00 |
| AT3G04935 | 0,15 | -0,81 | 4,39 | -0,18 | 8,54E-01 |
| AT3G25800 | 1,14 | -2,44 | 2,15 | -1,14 | 2,56E-01 |
| AT3G25840 | 1,32 | -2,05 | 1,57 | -1,30 | 1,94E-01 |
| AT3G25870 | 0,65 | 0,94 | 2,46 | 0,38 | 7,03E-01 |
| AT3G25882 | 0,50 | -1,35 | 4,03 | -0,34 | 7,38E-01 |
| AT3G25920 | 0,58 | -1,22 | 2,27 | -0,54 | 5,91E-01 |
| AT3G25990 | 0,75 | -4,91 | 3,40 | -1,44 | 1,49E-01 |
| AT3G26170 | 1,29 | 1,02 | 1,83 | 0,56 | 5,75E-01 |
| AT3G26180 | 0,36 | -0,02 | 4,32 | -0,01 | 9,95E-01 |
| AT3G26210 | 1,30 | 0,59 | 1,98 | 0,30 | 7,66E-01 |
| AT3G26220 | 0,49 | 0,59 | 4,01 | 0,15 | 8,82E-01 |

| | | | | | |
|---|---|---|---|---|---|
| AT3G26230 | 0,79 | -0,52 | 2,40 | -0,22 | 8,30E-01 |
| AT3G26280 | 1,16 | -0,96 | 1,62 | -0,59 | 5,53E-01 |
| AT3G26290 | 0,78 | 0,50 | 1,97 | 0,26 | 7,99E-01 |
| AT3G26300 | 0,36 | -3,05 | 4,38 | -0,70 | 4,85E-01 |
| AT3G26340 | 0,62 | -3,39 | 3,75 | -0,91 | 3,65E-01 |
| AT3G26410 | 0,31 | -3,42 | 4,37 | -0,78 | 4,34E-01 |
| AT3G26510 | 0,66 | 1,19 | 2,12 | 0,56 | 5,76E-01 |
| AT3G26550 | 0,20 | -2,60 | 4,41 | -0,59 | 5,55E-01 |
| AT3G26560 | 0,40 | 1,44 | 3,32 | 0,43 | 6,65E-01 |
| AT3G26590 | 0,41 | -1,46 | 4,31 | -0,34 | 7,35E-01 |
| AT3G26616 | 0,23 | 0,89 | 4,33 | 0,21 | 8,37E-01 |
| AT3G26630 | 0,69 | -3,56 | 2,40 | -1,48 | 1,38E-01 |
| AT3G26670 | 0,54 | -1,86 | 3,26 | -0,57 | 5,68E-01 |
| AT3G26744 | 0,39 | -3,20 | 4,30 | -0,74 | 4,57E-01 |
| AT3G26812 | 0,17 | 0,78 | 4,33 | 0,18 | 8,58E-01 |
| AT3G26819 | 0,05 | -0,24 | 4,38 | -0,05 | 9,57E-01 |
| AT3G26860 | 0,30 | 0,00 | 4,41 | 0,00 | 1,00E+00 |
| AT3G26890 | 0,29 | -0,28 | 4,34 | -0,06 | 9,49E-01 |
| AT3G26932 | 0,90 | 2,39 | 2,33 | 1,03 | 3,05E-01 |
| AT3G26935 | 0,27 | -0,77 | 4,35 | -0,18 | 8,60E-01 |
| AT3G26980 | 0,14 | -0,29 | 4,38 | -0,07 | 9,47E-01 |
| AT3G27010 | 0,67 | -2,48 | 3,68 | -0,67 | 5,01E-01 |
| AT3G27020 | 0,51 | -0,84 | 2,32 | -0,36 | 7,16E-01 |
| AT3G27060 | 0,90 | 1,99 | 2,39 | 0,83 | 4,05E-01 |
| AT3G27100 | 0,74 | -3,72 | 4,36 | -0,85 | 3,94E-01 |
| AT3G27240 | 0,37 | 0,51 | 3,35 | 0,15 | 8,79E-01 |
| AT3G27300 | 0,49 | -2,24 | 3,32 | -0,68 | 5,00E-01 |
| AT3G27380 | 0,45 | -0,81 | 4,39 | -0,18 | 8,54E-01 |
| AT3G05235 | 0,99 | 2,01 | 2,42 | 0,83 | 4,06E-01 |
| AT3G05245 | 0,33 | 1,02 | 4,33 | 0,24 | 8,14E-01 |
| AT3G27530 | 0,76 | -1,52 | 2,64 | -0,58 | 5,65E-01 |
| AT3G27600 | 1,14 | 0,97 | 2,13 | 0,46 | 6,49E-01 |
| AT3G27610 | 0,33 | -0,20 | 4,39 | -0,05 | 9,63E-01 |
| AT3G27680 | 0,86 | 1,61 | 2,25 | 0,72 | 4,73E-01 |
| AT3G27740 | 0,52 | -3,40 | 4,05 | -0,84 | 4,01E-01 |
| AT3G27770 | 0,43 | -0,85 | 2,86 | -0,30 | 7,68E-01 |
| AT3G27925 | 0,65 | 0,00 | 3,10 | 0,00 | 1,00E+00 |
| AT3G27930 | 0,20 | 0,84 | 4,33 | 0,19 | 8,47E-01 |
| AT3G27990 | 0,64 | 1,08 | 2,43 | 0,44 | 6,57E-01 |
| AT3G28340 | 0,22 | 1,18 | 4,32 | 0,27 | 7,84E-01 |
| AT3G28380 | 0,08 | 0,00 | 4,41 | 0,00 | 1,00E+00 |
| AT3G28410 | 0,14 | 0,48 | 4,35 | 0,11 | 9,13E-01 |
| AT3G28674 | 0,28 | 0,70 | 4,34 | 0,16 | 8,72E-01 |
| AT3G28695 | 0,43 | 0,48 | 4,12 | 0,12 | 9,08E-01 |
| AT3G28710 | 0,99 | 0,07 | 1,70 | 0,04 | 9,66E-01 |
| AT3G28780 | 0,61 | 1,84 | 3,11 | 0,59 | 5,55E-01 |
| AT3G28890 | 0,43 | -2,58 | 4,40 | -0,59 | 5,58E-01 |
| AT3G28930 | 0,77 | 1,36 | 2,93 | 0,46 | 6,42E-01 |
| AT3G28940 | 1,30 | 0,74 | 1,70 | 0,44 | 6,61E-01 |
| AT3G28950 | 0,97 | -0,36 | 1,91 | -0,19 | 8,50E-01 |
| AT3G28960 | 0,30 | 1,21 | 4,31 | 0,28 | 7,79E-01 |
| AT3G28970 | 0,52 | -1,64 | 4,05 | -0,40 | 6,86E-01 |
| AT3G29030 | 0,33 | 1,52 | 4,30 | 0,35 | 7,24E-01 |
| AT3G05665 | 0,80 | 2,23 | 2,58 | 0,86 | 3,88E-01 |
| AT3G29110 | 0,53 | 0,22 | 2,60 | 0,09 | 9,31E-01 |
| AT3G29160 | 0,59 | -1,22 | 4,35 | -0,28 | 7,79E-01 |

| | | | | | |
|---|---|---|---|---|---|
| AT3G29200 | 0,59 | -2,50 | 3,12 | -0,80 | 4,22E-01 |
| AT3G29350 | 0,17 | -1,25 | 4,37 | -0,29 | 7,74E-01 |
| AT3G29400 | 0,67 | -0,16 | 2,48 | -0,06 | 9,49E-01 |
| AT3G29570 | 0,38 | 0,66 | 4,34 | 0,15 | 8,80E-01 |
| AT3G05745 | 0,14 | -0,11 | 4,37 | -0,03 | 9,80E-01 |
| AT3G29633 | 0,14 | 0,00 | 4,41 | 0,00 | 1,00E+00 |
| AT3G29639 | 0,16 | 0,75 | 4,34 | 0,17 | 8,63E-01 |
| AT3G05895 | 0,08 | -0,21 | 4,39 | -0,05 | 9,62E-01 |
| AT3G29680 | 0,25 | -0,18 | 4,34 | -0,04 | 9,67E-01 |
| AT3G29720 | 0,40 | 0,01 | 4,37 | 0,00 | 9,98E-01 |
| AT3G29760 | 0,67 | -3,44 | 2,99 | -1,15 | 2,50E-01 |
| AT3G29785 | 0,22 | 1,07 | 4,32 | 0,25 | 8,05E-01 |
| AT3G30160 | 0,58 | 2,57 | 2,90 | 0,89 | 3,75E-01 |
| AT3G30200 | 0,12 | -0,14 | 4,38 | -0,03 | 9,74E-01 |
| AT3G30220 | 0,08 | -0,21 | 4,39 | -0,05 | 9,62E-01 |
| AT3G30230 | 0,54 | 1,22 | 3,88 | 0,31 | 7,54E-01 |
| AT3G30300 | 0,53 | -1,69 | 4,35 | -0,39 | 6,97E-01 |
| AT3G30840 | 0,21 | -0,73 | 4,40 | -0,17 | 8,67E-01 |
| AT3G32190 | 0,34 | -0,54 | 3,48 | -0,15 | 8,78E-01 |
| AT3G32420 | 0,10 | -0,34 | 4,38 | -0,08 | 9,38E-01 |
| AT3G32904 | 0,65 | 0,49 | 3,09 | 0,16 | 8,73E-01 |
| AT3G33528 | 0,61 | 0,41 | 2,54 | 0,16 | 8,71E-01 |
| AT3G42060 | 0,32 | 1,32 | 4,31 | 0,31 | 7,59E-01 |
| AT3G42140 | 0,14 | 0,48 | 4,35 | 0,11 | 9,13E-01 |
| AT3G42155 | 0,34 | 0,35 | 4,30 | 0,08 | 9,35E-01 |
| AT3G42550 | 0,12 | 0,07 | 4,38 | 0,02 | 9,87E-01 |
| AT3G42723 | 0,12 | -0,31 | 4,38 | -0,07 | 9,44E-01 |
| AT3G42790 | 0,59 | -0,18 | 3,12 | -0,06 | 9,55E-01 |
| AT3G43153 | 0,99 | 1,25 | 2,55 | 0,49 | 6,23E-01 |
| AT3G43190 | 0,26 | -0,77 | 4,37 | -0,18 | 8,59E-01 |
| AT3G43220 | 0,79 | -2,10 | 2,94 | -0,71 | 4,75E-01 |
| AT3G43240 | 0,86 | -3,18 | 2,72 | -1,17 | 2,42E-01 |
| AT3G43270 | 0,68 | 1,21 | 2,45 | 0,49 | 6,22E-01 |
| AT3G43280 | 0,11 | 0,46 | 4,35 | 0,11 | 9,15E-01 |
| AT3G43291 | 0,65 | 1,16 | 2,42 | 0,48 | 6,32E-01 |
| AT3G06595 | 0,10 | -0,34 | 4,38 | -0,08 | 9,38E-01 |
| AT3G43432 | 0,16 | 0,63 | 4,35 | 0,14 | 8,85E-01 |
| AT3G43470 | 0,45 | 2,14 | 3,90 | 0,55 | 5,82E-01 |
| AT3G43500 | 0,45 | 2,44 | 3,91 | 0,62 | 5,33E-01 |
| AT3G43600 | 0,98 | -4,09 | 2,20 | -1,86 | 6,30E-02 |
| AT3G43610 | 0,74 | -2,11 | 2,47 | -0,85 | 3,94E-01 |
| AT3G06645 | 0,23 | 0,11 | 4,38 | 0,03 | 9,80E-01 |
| AT3G06655 | 1,06 | 3,19 | 2,06 | 1,55 | 1,22E-01 |
| AT3G43720 | 0,87 | -3,38 | 2,81 | -1,20 | 2,29E-01 |
| AT3G43810 | 0,51 | 0,31 | 3,14 | 0,10 | 9,22E-01 |
| AT3G43880 | 0,10 | -0,18 | 4,38 | -0,04 | 9,67E-01 |
| AT3G43930 | 0,08 | 0,00 | 4,41 | 0,00 | 1,00E+00 |
| AT3G43940 | 0,53 | 1,80 | 3,83 | 0,47 | 6,38E-01 |
| AT3G44010 | 1,00 | 0,01 | 1,90 | 0,01 | 9,95E-01 |
| AT3G44070 | 0,37 | 0,44 | 3,42 | 0,13 | 8,98E-01 |
| AT3G44160 | 0,43 | -1,28 | 4,10 | -0,31 | 7,55E-01 |
| AT3G44265 | 0,40 | 1,57 | 4,26 | 0,37 | 7,12E-01 |
| AT3G44400 | 0,82 | -0,41 | 2,65 | -0,15 | 8,78E-01 |
| AT3G44435 | 0,70 | -0,12 | 3,72 | -0,03 | 9,75E-01 |
| AT3G44450 | 0,56 | 0,63 | 3,02 | 0,21 | 8,35E-01 |
| AT3G44460 | 0,11 | 0,46 | 4,35 | 0,11 | 9,15E-01 |

| | | | | | |
|---|---|---|---|---|---|
| AT3G44635 | 1,31 | -4,46 | 2,92 | -1,52 | 1,27E-01 |
| AT3G44680 | 0,12 | -0,68 | 4,41 | -0,15 | 8,77E-01 |
| AT3G44753 | 0,14 | 0,40 | 4,35 | 0,09 | 9,27E-01 |
| AT3G44780 | 0,10 | 0,45 | 4,35 | 0,10 | 9,19E-01 |
| AT3G44820 | 0,38 | -2,07 | 4,36 | -0,47 | 6,36E-01 |
| AT3G44990 | 0,60 | -0,74 | 3,74 | -0,20 | 8,43E-01 |
| AT3G45030 | 0,91 | -1,15 | 2,49 | -0,46 | 6,45E-01 |
| AT3G45040 | 0,29 | 1,53 | 4,30 | 0,36 | 7,22E-01 |
| AT3G45060 | 0,08 | 0,00 | 4,41 | 0,00 | 1,00E+00 |
| AT3G06955 | 0,21 | 1,49 | 4,30 | 0,35 | 7,29E-01 |
| AT3G45130 | 0,82 | 2,95 | 2,31 | 1,28 | 2,02E-01 |
| AT3G45180 | 1,00 | -0,61 | 1,84 | -0,33 | 7,39E-01 |
| AT3G07065 | 0,05 | -0,24 | 4,38 | -0,05 | 9,57E-01 |
| AT3G45240 | 0,25 | -2,53 | 4,40 | -0,57 | 5,65E-01 |
| AT3G07095 | 0,20 | 0,00 | 4,41 | 0,00 | 1,00E+00 |
| AT3G45260 | 1,01 | -1,98 | 1,77 | -1,12 | 2,63E-01 |
| AT3G45300 | 0,17 | -0,68 | 4,41 | -0,15 | 8,77E-01 |
| AT3G45430 | 0,20 | -2,60 | 4,41 | -0,59 | 5,55E-01 |
| AT3G45443 | 0,08 | -0,21 | 4,39 | -0,05 | 9,62E-01 |
| AT3G45600 | 0,92 | -1,63 | 2,02 | -0,81 | 4,19E-01 |
| AT3G45630 | 0,80 | -1,62 | 2,14 | -0,76 | 4,48E-01 |
| AT3G45770 | 0,14 | 0,48 | 4,35 | 0,11 | 9,13E-01 |
| AT3G45820 | 0,59 | 1,59 | 2,96 | 0,54 | 5,90E-01 |
| AT3G45850 | 1,07 | -2,45 | 2,35 | -1,04 | 2,98E-01 |
| AT3G45860 | 0,95 | -2,28 | 1,90 | -1,20 | 2,29E-01 |
| AT3G45910 | 1,00 | 2,06 | 2,47 | 0,83 | 4,05E-01 |
| AT3G45990 | 0,46 | 0,30 | 3,40 | 0,09 | 9,29E-01 |
| AT3G46000 | 0,30 | 0,15 | 3,68 | 0,04 | 9,67E-01 |
| AT3G46030 | 0,35 | 0,26 | 4,32 | 0,06 | 9,52E-01 |
| AT3G46090 | 0,72 | 1,42 | 2,31 | 0,61 | 5,40E-01 |
| AT3G46100 | 0,51 | 0,41 | 4,00 | 0,10 | 9,18E-01 |
| AT3G46210 | 0,56 | 1,76 | 3,84 | 0,46 | 6,47E-01 |
| AT3G07375 | 0,21 | 0,00 | 4,41 | 0,00 | 1,00E+00 |
| AT3G46320 | 0,61 | 14,66 | 4,34 | 3,38 | 7,38E-04 |
| AT3G46360 | 0,82 | 0,57 | 2,88 | 0,20 | 8,42E-01 |
| AT3G46390 | 0,25 | 0,15 | 4,36 | 0,03 | 9,73E-01 |
| AT3G46440 | 0,45 | -3,76 | 4,35 | -0,86 | 3,88E-01 |
| AT3G46450 | 0,53 | -1,43 | 3,89 | -0,37 | 7,12E-01 |
| AT3G46470 | 0,38 | 1,17 | 4,25 | 0,27 | 7,84E-01 |
| AT3G46540 | 0,31 | -0,47 | 4,37 | -0,11 | 9,13E-01 |
| AT3G46630 | 0,81 | 0,08 | 2,29 | 0,04 | 9,72E-01 |
| AT3G46650 | 0,58 | -3,40 | 3,18 | -1,07 | 2,85E-01 |
| AT3G46660 | 0,59 | -0,41 | 2,66 | -0,15 | 8,78E-01 |
| AT3G46670 | 0,57 | -1,58 | 3,31 | -0,48 | 6,34E-01 |
| AT3G46690 | 0,12 | 0,00 | 4,41 | 0,00 | 1,00E+00 |
| AT3G46730 | 0,12 | 0,00 | 4,41 | 0,00 | 1,00E+00 |
| AT3G46790 | 0,53 | -2,54 | 4,40 | -0,58 | 5,64E-01 |
| AT3G46845 | 1,29 | 0,45 | 1,81 | 0,25 | 8,04E-01 |
| AT3G46920 | 0,08 | 0,00 | 4,41 | 0,00 | 1,00E+00 |
| AT3G46960 | 0,92 | -1,73 | 2,20 | -0,79 | 4,32E-01 |
| AT3G47000 | 0,33 | -2,04 | 4,35 | -0,47 | 6,39E-01 |
| AT3G47070 | 0,67 | -0,19 | 2,58 | -0,07 | 9,43E-01 |
| AT3G47090 | 0,54 | -2,07 | 3,79 | -0,55 | 5,85E-01 |
| AT3G47160 | 0,74 | 2,06 | 2,51 | 0,82 | 4,12E-01 |
| AT3G47370 | 0,35 | -1,12 | 4,33 | -0,26 | 7,96E-01 |
| AT3G47420 | 0,51 | -0,93 | 2,56 | -0,36 | 7,17E-01 |

| | | | | | |
|---|---|---|---|---|---|
| AT3G47480 | 0,70 | 0,99 | 2,87 | 0,34 | 7,31E-01 |
| AT3G47670 | 0,97 | -0,15 | 2,00 | -0,07 | 9,42E-01 |
| AT3G47675 | 0,21 | -2,50 | 4,39 | -0,57 | 5,70E-01 |
| AT3G47680 | 0,08 | 0,00 | 4,41 | 0,00 | 1,00E+00 |
| AT3G47700 | 0,43 | -0,02 | 2,75 | -0,01 | 9,96E-01 |
| AT3G47730 | 0,93 | 0,00 | 2,83 | 0,00 | 1,00E+00 |
| AT3G47810 | 0,27 | -1,67 | 4,36 | -0,38 | 7,02E-01 |
| AT3G47860 | 0,23 | -0,44 | 4,35 | -0,10 | 9,19E-01 |
| AT3G47940 | 0,35 | -1,51 | 4,36 | -0,35 | 7,29E-01 |
| AT3G47960 | 1,01 | 0,34 | 1,80 | 0,19 | 8,51E-01 |
| AT3G47990 | 0,32 | -1,83 | 4,38 | -0,42 | 6,77E-01 |
| AT3G48080 | 1,04 | -2,50 | 2,03 | -1,24 | 2,16E-01 |
| AT3G48090 | 1,08 | -1,73 | 2,73 | -0,63 | 5,28E-01 |
| AT3G48100 | 0,85 | -3,83 | 4,25 | -0,90 | 3,67E-01 |
| AT3G48131 | 0,31 | 1,34 | 4,31 | 0,31 | 7,56E-01 |
| AT3G48150 | 0,55 | -3,82 | 3,83 | -1,00 | 3,19E-01 |
| AT3G48170 | 0,90 | -2,09 | 2,58 | -0,81 | 4,17E-01 |
| AT3G48195 | 0,48 | -0,50 | 3,13 | -0,16 | 8,74E-01 |
| AT3G48360 | 1,03 | -4,58 | 2,82 | -1,62 | 1,04E-01 |
| AT3G48520 | 0,38 | -2,60 | 4,41 | -0,59 | 5,55E-01 |
| AT3G48570 | 0,28 | -0,25 | 4,33 | -0,06 | 9,54E-01 |
| AT3G48640 | 0,25 | -1,17 | 4,36 | -0,27 | 7,88E-01 |
| AT3G48690 | 0,79 | -3,39 | 2,46 | -1,38 | 1,68E-01 |
| AT3G48740 | 0,68 | 0,43 | 2,17 | 0,20 | 8,43E-01 |
| AT3G08005 | 0,05 | -0,24 | 4,38 | -0,05 | 9,57E-01 |
| AT3G48840 | 0,12 | -0,78 | 4,39 | -0,18 | 8,59E-01 |
| AT3G48930 | 0,53 | -1,78 | 2,66 | -0,67 | 5,03E-01 |
| AT3G48990 | 0,58 | -3,31 | 3,05 | -1,08 | 2,78E-01 |
| AT3G49010 | 1,04 | -0,51 | 1,80 | -0,28 | 7,77E-01 |
| AT3G49055 | 0,42 | -3,33 | 4,36 | -0,76 | 4,46E-01 |
| AT3G49060 | 0,44 | 0,68 | 3,21 | 0,21 | 8,33E-01 |
| AT3G49140 | 0,93 | -2,13 | 2,10 | -1,02 | 3,08E-01 |
| AT3G49160 | 0,11 | -0,68 | 4,41 | -0,15 | 8,77E-01 |
| AT3G49250 | 0,54 | 0,10 | 4,16 | 0,02 | 9,80E-01 |
| AT3G49350 | 0,36 | -1,69 | 4,37 | -0,39 | 6,98E-01 |
| AT3G49490 | 1,14 | -2,80 | 2,08 | -1,34 | 1,79E-01 |
| AT3G49530 | 1,18 | -1,43 | 1,97 | -0,73 | 4,66E-01 |
| AT3G49601 | 0,27 | 1,29 | 4,31 | 0,30 | 7,65E-01 |
| AT3G49650 | 0,08 | 0,00 | 4,41 | 0,00 | 1,00E+00 |
| AT3G49720 | 1,09 | 2,70 | 2,21 | 1,22 | 2,22E-01 |
| AT3G49790 | 0,38 | 1,67 | 3,30 | 0,51 | 6,13E-01 |
| AT3G49870 | 0,08 | 0,00 | 4,41 | 0,00 | 1,00E+00 |
| AT3G49980 | 0,21 | 0,17 | 4,36 | 0,04 | 9,68E-01 |
| AT3G49990 | 0,57 | -0,27 | 3,80 | -0,07 | 9,43E-01 |
| AT3G50060 | 0,43 | 0,26 | 2,72 | 0,10 | 9,23E-01 |
| AT3G50110 | 0,18 | 0,40 | 4,35 | 0,09 | 9,27E-01 |
| AT3G50120 | 0,78 | 0,25 | 2,32 | 0,11 | 9,15E-01 |
| AT3G50240 | 0,71 | -2,60 | 4,41 | -0,59 | 5,55E-01 |
| AT3G50320 | 1,17 | 3,62 | 2,08 | 1,74 | 8,22E-02 |
| AT3G50380 | 1,07 | -2,29 | 2,37 | -0,97 | 3,33E-01 |
| AT3G50670 | 0,33 | -0,06 | 4,37 | -0,01 | 9,89E-01 |
| AT3G50690 | 0,85 | 0,51 | 1,95 | 0,26 | 7,94E-01 |
| AT3G50740 | 0,53 | 0,74 | 3,20 | 0,23 | 8,18E-01 |
| AT3G50750 | 1,00 | 0,50 | 1,98 | 0,25 | 8,02E-01 |
| AT3G50780 | 0,08 | 0,00 | 4,41 | 0,00 | 1,00E+00 |
| AT3G50790 | 0,29 | -0,95 | 4,34 | -0,22 | 8,27E-01 |

| | | | | | |
|---|---|---|---|---|---|
| AT3G50808 | 0,10 | 0,45 | 4,35 | 0,10 | 9,19E-01 |
| AT3G50825 | 0,41 | 2,32 | 4,28 | 0,54 | 5,88E-01 |
| AT3G50920 | 0,48 | -4,07 | 4,02 | -1,01 | 3,11E-01 |
| AT3G51000 | 0,52 | 0,14 | 3,20 | 0,04 | 9,65E-01 |
| AT3G51135 | 0,51 | -1,12 | 2,63 | -0,43 | 6,70E-01 |
| AT3G51160 | 0,58 | -2,33 | 3,04 | -0,77 | 4,44E-01 |
| AT3G51265 | 0,13 | 0,52 | 4,35 | 0,12 | 9,04E-01 |
| AT3G51310 | 0,51 | -1,64 | 3,26 | -0,50 | 6,14E-01 |
| AT3G51580 | 0,25 | -0,73 | 4,40 | -0,16 | 8,69E-01 |
| AT3G51660 | 0,43 | -0,34 | 4,15 | -0,08 | 9,35E-01 |
| AT3G51790 | 0,60 | -2,60 | 3,76 | -0,69 | 4,88E-01 |
| AT3G51820 | 0,59 | -1,68 | 3,92 | -0,43 | 6,67E-01 |
| AT3G51840 | 0,33 | -0,92 | 4,34 | -0,21 | 8,32E-01 |
| AT3G51880 | 0,48 | -2,88 | 3,19 | -0,90 | 3,68E-01 |
| AT3G51895 | 0,82 | -2,60 | 2,89 | -0,90 | 3,68E-01 |
| AT3G51920 | 0,46 | -0,06 | 2,69 | -0,02 | 9,83E-01 |
| AT3G08215 | 0,08 | 0,00 | 4,41 | 0,00 | 1,00E+00 |
| AT3G51980 | 0,42 | -0,88 | 3,35 | -0,26 | 7,92E-01 |
| AT3G52060 | 0,46 | -0,85 | 2,85 | -0,30 | 7,65E-01 |
| AT3G52072 | 0,22 | 0,75 | 4,34 | 0,17 | 8,63E-01 |
| AT3G08230 | 0,12 | -0,14 | 4,38 | -0,03 | 9,74E-01 |
| AT3G52110 | 0,12 | -0,68 | 4,41 | -0,15 | 8,77E-01 |
| AT3G52180 | 0,66 | 0,38 | 2,67 | 0,14 | 8,86E-01 |
| AT3G52190 | 0,13 | 0,00 | 4,41 | 0,00 | 1,00E+00 |
| AT3G52220 | 0,11 | -0,68 | 4,41 | -0,15 | 8,77E-01 |
| AT3G52260 | 0,64 | -2,60 | 3,67 | -0,71 | 4,78E-01 |
| AT3G52480 | 0,53 | 2,14 | 2,97 | 0,72 | 4,71E-01 |
| AT3G52560 | 0,47 | -0,85 | 3,39 | -0,25 | 8,02E-01 |
| AT3G52640 | 0,21 | 0,00 | 4,41 | 0,00 | 1,00E+00 |
| AT3G52730 | 0,69 | -2,17 | 3,74 | -0,58 | 5,62E-01 |
| AT3G52740 | 0,17 | -1,25 | 4,37 | -0,29 | 7,74E-01 |
| AT3G52870 | 0,19 | 1,18 | 4,31 | 0,27 | 7,84E-01 |
| AT3G53015 | 1,03 | -1,74 | 2,34 | -0,74 | 4,56E-01 |
| AT3G53016 | 0,52 | -0,49 | 4,02 | -0,12 | 9,03E-01 |
| AT3G53100 | 0,69 | -1,30 | 2,97 | -0,44 | 6,61E-01 |
| AT3G53380 | 0,40 | 0,57 | 4,16 | 0,14 | 8,91E-01 |
| AT3G53410 | 0,47 | -3,21 | 4,38 | -0,73 | 4,64E-01 |
| AT3G53500 | 0,57 | -1,26 | 2,72 | -0,46 | 6,44E-01 |
| AT3G53510 | 0,25 | 0,00 | 4,41 | 0,00 | 1,00E+00 |
| AT3G53540 | 1,08 | -3,78 | 2,32 | -1,63 | 1,03E-01 |
| AT3G53560 | 0,23 | -0,48 | 4,34 | -0,11 | 9,12E-01 |
| AT3G53570 | 0,21 | -2,50 | 4,39 | -0,57 | 5,70E-01 |
| AT3G53610 | 0,53 | 1,52 | 3,12 | 0,49 | 6,27E-01 |
| AT3G53710 | 0,37 | 0,42 | 3,45 | 0,12 | 9,04E-01 |
| AT3G53730 | 0,67 | 0,81 | 2,43 | 0,33 | 7,39E-01 |
| AT3G53810 | 0,44 | 0,06 | 4,26 | 0,01 | 9,89E-01 |
| AT3G53890 | 0,52 | -4,03 | 3,18 | -1,27 | 2,04E-01 |
| AT3G53910 | 1,01 | 2,81 | 2,12 | 1,33 | 1,85E-01 |
| AT3G53920 | 0,24 | -2,54 | 4,40 | -0,58 | 5,64E-01 |
| AT3G53960 | 0,98 | -2,71 | 2,56 | -1,06 | 2,90E-01 |
| AT3G54100 | 0,47 | 0,06 | 3,28 | 0,02 | 9,85E-01 |
| AT3G54120 | 0,20 | -2,55 | 4,40 | -0,58 | 5,62E-01 |
| AT3G54140 | 0,70 | -0,45 | 2,57 | -0,18 | 8,61E-01 |
| AT3G54210 | 1,01 | 2,18 | 1,80 | 1,21 | 2,25E-01 |
| AT3G54290 | 0,42 | -2,60 | 4,22 | -0,62 | 5,37E-01 |
| AT3G54350 | 1,02 | -0,14 | 2,09 | -0,07 | 9,45E-01 |

| | | | | | |
|---|---|---|---|---|---|
| AT3G54360 | 1,14 | -2,12 | 2,05 | -1,04 | 3,00E-01 |
| AT3G54400 | 0,32 | -2,60 | 4,41 | -0,59 | 5,55E-01 |
| AT3G54440 | 0,61 | -0,33 | 2,14 | -0,15 | 8,79E-01 |
| AT3G54460 | 0,97 | -3,15 | 2,75 | -1,15 | 2,51E-01 |
| AT3G54470 | 0,44 | -3,42 | 3,33 | -1,03 | 3,04E-01 |
| AT3G54620 | 0,24 | -2,60 | 4,41 | -0,59 | 5,55E-01 |
| AT3G54630 | 0,68 | -2,69 | 4,40 | -0,61 | 5,41E-01 |
| AT3G54640 | 0,22 | -0,56 | 4,35 | -0,13 | 8,98E-01 |
| AT3G54650 | 0,62 | -1,02 | 3,15 | -0,32 | 7,46E-01 |
| AT3G54750 | 0,48 | -2,54 | 4,40 | -0,58 | 5,64E-01 |
| AT3G54760 | 1,04 | -0,87 | 1,90 | -0,46 | 6,47E-01 |
| AT3G54790 | 1,31 | -2,89 | 1,70 | -1,70 | 8,98E-02 |
| AT3G54860 | 0,55 | -0,12 | 3,01 | -0,04 | 9,68E-01 |
| AT3G54900 | 0,95 | -0,89 | 2,10 | -0,42 | 6,72E-01 |
| AT3G54960 | 0,90 | 0,85 | 1,99 | 0,43 | 6,69E-01 |
| AT3G55040 | 0,21 | 0,00 | 4,41 | 0,00 | 1,00E+00 |
| AT3G55050 | 0,45 | 0,47 | 3,31 | 0,14 | 8,87E-01 |
| AT3G55060 | 0,14 | 0,48 | 4,35 | 0,11 | 9,13E-01 |
| AT3G55130 | 0,51 | 1,18 | 3,12 | 0,38 | 7,04E-01 |
| AT3G55160 | 0,95 | -2,18 | 1,89 | -1,15 | 2,51E-01 |
| AT3G55252 | 0,66 | -0,20 | 2,19 | -0,09 | 9,26E-01 |
| AT3G55254 | 0,80 | 1,15 | 2,26 | 0,51 | 6,10E-01 |
| AT3G55330 | 0,44 | -0,82 | 2,82 | -0,29 | 7,71E-01 |
| AT3G55360 | 0,60 | 0,97 | 2,53 | 0,38 | 7,02E-01 |
| AT3G55400 | 0,18 | 0,00 | 4,41 | 0,00 | 1,00E+00 |
| AT3G55430 | 0,77 | -0,78 | 2,04 | -0,38 | 7,00E-01 |
| AT3G55860 | 0,83 | 2,07 | 2,11 | 0,98 | 3,28E-01 |
| AT3G55935 | 0,46 | -0,93 | 2,62 | -0,36 | 7,21E-01 |
| AT3G55950 | 0,63 | 1,22 | 2,42 | 0,50 | 6,14E-01 |
| AT3G55960 | 0,64 | -0,26 | 2,64 | -0,10 | 9,23E-01 |
| AT3G56040 | 0,77 | -1,24 | 2,15 | -0,58 | 5,63E-01 |
| AT3G56050 | 0,97 | -0,37 | 2,01 | -0,19 | 8,53E-01 |
| AT3G56085 | 0,12 | -0,31 | 4,38 | -0,07 | 9,44E-01 |
| AT3G56090 | 0,34 | 0,13 | 3,56 | 0,04 | 9,71E-01 |
| AT3G56110 | 0,41 | 0,07 | 3,33 | 0,02 | 9,84E-01 |
| AT3G56190 | 1,16 | 1,08 | 1,86 | 0,58 | 5,62E-01 |
| AT3G56210 | 0,45 | -2,14 | 4,32 | -0,49 | 6,21E-01 |
| AT3G56240 | 0,35 | 1,41 | 2,85 | 0,49 | 6,21E-01 |
| AT3G56290 | 1,19 | -4,13 | 2,13 | -1,94 | 5,20E-02 |
| AT3G56320 | 0,58 | -3,06 | 3,86 | -0,79 | 4,27E-01 |
| AT3G56340 | 0,44 | -0,65 | 3,31 | -0,20 | 8,45E-01 |
| AT3G56370 | 0,84 | -3,07 | 2,25 | -1,36 | 1,73E-01 |
| AT3G56460 | 1,07 | -0,63 | 2,03 | -0,31 | 7,55E-01 |
| AT3G56480 | 0,39 | 0,90 | 2,85 | 0,32 | 7,51E-01 |
| AT3G56670 | 0,38 | 0,06 | 4,30 | 0,01 | 9,90E-01 |
| AT3G56690 | 0,43 | 0,16 | 4,17 | 0,04 | 9,69E-01 |
| AT3G56790 | 0,33 | -0,20 | 4,39 | -0,05 | 9,63E-01 |
| AT3G56820 | 0,25 | -3,06 | 4,38 | -0,70 | 4,85E-01 |
| AT3G56830 | 0,17 | 0,00 | 4,41 | 0,00 | 1,00E+00 |
| AT3G56850 | 0,48 | -1,59 | 3,27 | -0,49 | 6,27E-01 |
| AT3G56860 | 0,49 | 1,36 | 2,67 | 0,51 | 6,09E-01 |
| AT3G56950 | 0,79 | -0,61 | 2,30 | -0,27 | 7,91E-01 |
| AT3G56970 | 0,12 | 0,00 | 4,41 | 0,00 | 1,00E+00 |
| AT3G56990 | 0,46 | -0,77 | 3,30 | -0,23 | 8,15E-01 |
| AT3G57020 | 0,42 | -2,02 | 4,36 | -0,46 | 6,43E-01 |
| AT3G57030 | 0,05 | -0,24 | 4,38 | -0,05 | 9,57E-01 |

| | | | | | |
|---|---|---|---|---|---|
| AT3G08825 | 0,72 | 1,00 | 3,01 | 0,33 | 7,39E-01 |
| AT3G57050 | 0,52 | -0,56 | 2,64 | -0,21 | 8,31E-01 |
| AT3G08855 | 0,12 | -0,78 | 4,39 | -0,18 | 8,59E-01 |
| AT3G57140 | 0,48 | -0,88 | 3,16 | -0,28 | 7,79E-01 |
| AT3G57240 | 0,24 | -3,18 | 4,39 | -0,73 | 4,68E-01 |
| AT3G57290 | 1,20 | 0,83 | 2,27 | 0,37 | 7,15E-01 |
| AT3G57350 | 0,44 | -2,43 | 3,32 | -0,73 | 4,64E-01 |
| AT3G57420 | 0,74 | -0,64 | 2,21 | -0,29 | 7,73E-01 |
| AT3G57460 | 0,24 | 1,49 | 4,30 | 0,35 | 7,30E-01 |
| AT3G57470 | 0,21 | -0,68 | 4,41 | -0,15 | 8,77E-01 |
| AT3G57530 | 1,08 | 0,63 | 1,93 | 0,33 | 7,44E-01 |
| AT3G57540 | 0,08 | -0,21 | 4,39 | -0,05 | 9,62E-01 |
| AT3G57580 | 0,45 | 0,64 | 3,27 | 0,19 | 8,45E-01 |
| AT3G57610 | 1,05 | -4,37 | 4,28 | -1,02 | 3,06E-01 |
| AT3G57640 | 0,11 | 0,46 | 4,35 | 0,11 | 9,15E-01 |
| AT3G57765 | 1,26 | 0,80 | 1,55 | 0,51 | 6,08E-01 |
| AT3G57780 | 0,43 | -2,92 | 4,38 | -0,67 | 5,04E-01 |
| AT3G57790 | 0,44 | -0,19 | 4,39 | -0,04 | 9,65E-01 |
| AT3G57800 | 0,37 | 1,86 | 4,29 | 0,43 | 6,65E-01 |
| AT3G57870 | 0,53 | -4,36 | 4,32 | -1,01 | 3,13E-01 |
| AT3G57880 | 0,23 | 0,43 | 4,34 | 0,10 | 9,20E-01 |
| AT3G57930 | 0,30 | -1,06 | 4,36 | -0,24 | 8,08E-01 |
| AT3G57965 | 0,28 | 0,00 | 4,41 | 0,00 | 1,00E+00 |
| AT3G57980 | 0,74 | 1,74 | 2,79 | 0,62 | 5,33E-01 |
| AT3G57990 | 0,40 | 0,30 | 3,31 | 0,09 | 9,28E-01 |
| AT3G58100 | 1,03 | -2,60 | 3,44 | -0,76 | 4,49E-01 |
| AT3G58120 | 0,15 | 0,00 | 4,41 | 0,00 | 1,00E+00 |
| AT3G58140 | 0,42 | 0,31 | 3,32 | 0,09 | 9,25E-01 |
| AT3G58170 | 0,42 | -1,52 | 4,31 | -0,35 | 7,25E-01 |
| AT3G58210 | 0,64 | -4,30 | 4,32 | -1,00 | 3,19E-01 |
| AT3G58270 | 0,99 | -0,15 | 2,63 | -0,06 | 9,55E-01 |
| AT3G58560 | 0,23 | 0,77 | 4,34 | 0,18 | 8,59E-01 |
| AT3G58580 | 0,38 | -0,75 | 4,36 | -0,17 | 8,63E-01 |
| AT3G58670 | 0,12 | -0,78 | 4,39 | -0,18 | 8,59E-01 |
| AT3G58730 | 1,29 | 0,08 | 1,78 | 0,04 | 9,66E-01 |
| AT3G58750 | 0,61 | -3,27 | 3,86 | -0,85 | 3,96E-01 |
| AT3G58790 | 0,34 | 0,41 | 4,36 | 0,09 | 9,25E-01 |
| AT3G58990 | 0,12 | -0,14 | 4,38 | -0,03 | 9,74E-01 |
| AT3G59050 | 0,52 | -0,41 | 3,11 | -0,13 | 8,95E-01 |
| AT3G59120 | 0,10 | -0,34 | 4,38 | -0,08 | 9,38E-01 |
| AT3G59280 | 0,32 | -0,86 | 4,38 | -0,20 | 8,44E-01 |
| AT3G59360 | 0,61 | 0,54 | 2,98 | 0,18 | 8,57E-01 |
| AT3G59540 | 1,10 | -2,78 | 1,93 | -1,44 | 1,49E-01 |
| AT3G59570 | 0,57 | 0,90 | 3,04 | 0,30 | 7,66E-01 |
| AT3G59660 | 1,15 | -0,66 | 1,90 | -0,35 | 7,28E-01 |
| AT3G59760 | 0,20 | -0,11 | 4,38 | -0,03 | 9,80E-01 |
| AT3G59840 | 0,15 | 0,41 | 4,35 | 0,09 | 9,26E-01 |
| AT3G59910 | 0,48 | -3,04 | 4,04 | -0,75 | 4,51E-01 |
| AT3G59950 | 1,22 | 0,37 | 2,35 | 0,16 | 8,73E-01 |
| AT3G59980 | 0,35 | -0,84 | 3,49 | -0,24 | 8,09E-01 |
| AT3G60030 | 1,14 | -3,64 | 2,11 | -1,72 | 8,52E-02 |
| AT3G09335 | 0,23 | 0,01 | 4,36 | 0,00 | 9,99E-01 |
| AT3G60050 | 0,12 | 0,00 | 4,41 | 0,00 | 1,00E+00 |
| AT3G60070 | 0,35 | -3,17 | 4,38 | -0,72 | 4,69E-01 |
| AT3G60160 | 1,27 | -0,69 | 1,96 | -0,35 | 7,24E-01 |
| AT3G60176 | 0,85 | 2,40 | 2,68 | 0,90 | 3,71E-01 |

| | | | | | |
|---|---|---|---|---|---|
| AT3G60220 | 0,15 | 0,79 | 4,33 | 0,18 | 8,56E-01 |
| AT3G60250 | 1,02 | -4,09 | 1,97 | -2,07 | 3,81E-02 |
| AT3G60350 | 1,30 | -5,13 | 1,94 | -2,64 | 8,26E-03 |
| AT3G60380 | 0,49 | -2,48 | 3,91 | -0,63 | 5,26E-01 |
| AT3G60540 | 0,08 | -0,21 | 4,39 | -0,05 | 9,62E-01 |
| AT3G60670 | 0,12 | -0,14 | 4,38 | -0,03 | 9,74E-01 |
| AT3G60740 | 0,66 | 0,54 | 2,35 | 0,23 | 8,17E-01 |
| AT3G60760 | 0,12 | 0,00 | 4,41 | 0,00 | 1,00E+00 |
| AT3G60820 | 0,15 | -0,17 | 4,39 | -0,04 | 9,68E-01 |
| AT3G60990 | 0,73 | -0,22 | 2,71 | -0,08 | 9,36E-01 |
| AT3G61070 | 0,50 | -0,95 | 4,10 | -0,23 | 8,17E-01 |
| AT3G09565 | 0,50 | -0,83 | 3,91 | -0,21 | 8,32E-01 |
| AT3G61200 | 0,69 | 0,19 | 2,57 | 0,08 | 9,40E-01 |
| AT3G61260 | 0,83 | 0,36 | 2,55 | 0,14 | 8,89E-01 |
| AT3G61310 | 0,38 | -3,33 | 4,37 | -0,76 | 4,46E-01 |
| AT3G61380 | 0,18 | -0,40 | 4,37 | -0,09 | 9,28E-01 |
| AT3G61420 | 0,37 | -2,11 | 4,38 | -0,48 | 6,31E-01 |
| AT3G61460 | 0,83 | 1,89 | 1,94 | 0,97 | 3,31E-01 |
| AT3G61530 | 0,25 | 0,00 | 4,41 | 0,00 | 1,00E+00 |
| AT3G61640 | 0,38 | 1,47 | 3,32 | 0,44 | 6,57E-01 |
| AT3G61740 | 0,78 | -2,61 | 2,49 | -1,05 | 2,95E-01 |
| AT3G61750 | 0,24 | -3,18 | 4,39 | -0,73 | 4,68E-01 |
| AT3G61800 | 0,23 | 1,33 | 4,31 | 0,31 | 7,57E-01 |
| AT3G61860 | 0,39 | -1,69 | 4,31 | -0,39 | 6,95E-01 |
| AT3G61890 | 0,44 | 1,17 | 3,51 | 0,33 | 7,38E-01 |
| AT3G61960 | 1,02 | -1,63 | 1,83 | -0,89 | 3,75E-01 |
| AT3G61990 | 0,15 | 0,05 | 4,37 | 0,01 | 9,91E-01 |
| AT3G62070 | 0,12 | 0,75 | 4,34 | 0,17 | 8,63E-01 |
| AT3G62120 | 1,00 | -0,75 | 2,15 | -0,35 | 7,29E-01 |
| AT3G62240 | 0,93 | -0,89 | 2,89 | -0,31 | 7,57E-01 |
| AT3G62260 | 0,41 | -0,95 | 4,13 | -0,23 | 8,18E-01 |
| AT3G62310 | 0,33 | 0,00 | 4,41 | 0,00 | 1,00E+00 |
| AT3G62330 | 0,73 | -2,21 | 2,97 | -0,74 | 4,57E-01 |
| AT3G62470 | 0,24 | -2,59 | 4,41 | -0,59 | 5,56E-01 |
| AT3G62540 | 0,28 | -3,07 | 4,38 | -0,70 | 4,84E-01 |
| AT3G09795 | 0,21 | 1,14 | 4,32 | 0,26 | 7,92E-01 |
| AT3G62590 | 0,64 | 0,63 | 2,99 | 0,21 | 8,34E-01 |
| AT3G62600 | 0,95 | -0,62 | 1,85 | -0,34 | 7,36E-01 |
| AT3G09825 | 1,22 | 2,45 | 1,84 | 1,33 | 1,82E-01 |
| AT3G09845 | 0,08 | -0,21 | 4,39 | -0,05 | 9,62E-01 |
| AT3G62630 | 0,91 | -1,29 | 2,14 | -0,60 | 5,47E-01 |
| AT3G62800 | 0,37 | -2,60 | 4,41 | -0,59 | 5,55E-01 |
| AT3G62840 | 0,66 | -3,03 | 3,05 | -1,00 | 3,19E-01 |
| AT3G62860 | 0,81 | -4,35 | 3,48 | -1,25 | 2,11E-01 |
| AT3G62870 | 1,22 | -3,61 | 1,90 | -1,90 | 5,73E-02 |
| AT3G62900 | 0,94 | 0,71 | 2,38 | 0,30 | 7,65E-01 |
| AT3G62910 | 0,49 | -2,26 | 3,98 | -0,57 | 5,70E-01 |
| AT3G62940 | 0,31 | -0,30 | 4,34 | -0,07 | 9,44E-01 |
| AT3G63052 | 0,42 | 0,31 | 3,32 | 0,09 | 9,25E-01 |
| AT3G63110 | 0,56 | 1,70 | 2,62 | 0,65 | 5,16E-01 |
| AT3G63140 | 0,93 | 1,00 | 2,05 | 0,49 | 6,25E-01 |
| AT3G63150 | 0,90 | -1,23 | 2,37 | -0,52 | 6,03E-01 |
| AT3G63160 | 1,15 | 2,48 | 2,01 | 1,23 | 2,18E-01 |
| AT3G63180 | 1,24 | -4,00 | 1,79 | -2,23 | 2,57E-02 |
| AT3G63190 | 0,25 | 1,02 | 4,32 | 0,24 | 8,13E-01 |
| AT3G63210 | 0,47 | 2,37 | 4,27 | 0,56 | 5,79E-01 |

| | | | | | |
|---|---|---|---|---|---|
| AT3G63340 | 1,13 | -3,20 | 2,53 | -1,27 | 2,05E-01 |
| AT3G63390 | 0,24 | 0,71 | 4,34 | 0,16 | 8,70E-01 |
| AT4G00005 | 0,19 | -0,11 | 4,39 | -0,02 | 9,80E-01 |
| AT4G00020 | 0,79 | 0,38 | 2,26 | 0,17 | 8,66E-01 |
| AT4G00070 | 0,43 | -1,25 | 4,20 | -0,30 | 7,67E-01 |
| AT4G00110 | 1,14 | -3,12 | 2,30 | -1,35 | 1,76E-01 |
| AT4G00165 | 0,65 | -0,64 | 2,60 | -0,25 | 8,05E-01 |
| AT4G00180 | 0,45 | -0,81 | 4,39 | -0,18 | 8,54E-01 |
| AT4G00280 | 0,25 | -3,06 | 4,38 | -0,70 | 4,85E-01 |
| AT4G00290 | 0,50 | -0,40 | 2,74 | -0,15 | 8,84E-01 |
| AT4G00330 | 0,80 | 0,48 | 2,15 | 0,22 | 8,22E-01 |
| AT4G00370 | 0,63 | -2,60 | 2,70 | -0,97 | 3,34E-01 |
| AT4G00400 | 0,42 | 1,16 | 2,75 | 0,42 | 6,74E-01 |
| AT4G00500 | 0,24 | -0,80 | 4,39 | -0,18 | 8,55E-01 |
| AT4G00540 | 0,54 | 2,32 | 2,54 | 0,91 | 3,60E-01 |
| AT4G00550 | 0,15 | -0,77 | 4,39 | -0,18 | 8,61E-01 |
| AT4G00560 | 0,40 | -0,78 | 4,39 | -0,18 | 8,59E-01 |
| AT4G00620 | 0,40 | -0,26 | 4,19 | -0,06 | 9,51E-01 |
| AT4G00740 | 0,31 | -0,43 | 4,35 | -0,10 | 9,22E-01 |
| AT4G00810 | 0,25 | -2,53 | 4,40 | -0,57 | 5,65E-01 |
| AT4G00890 | 0,23 | 1,59 | 4,30 | 0,37 | 7,11E-01 |
| AT4G00900 | 1,09 | -1,64 | 2,25 | -0,73 | 4,66E-01 |
| AT4G00980 | 0,33 | -2,10 | 4,36 | -0,48 | 6,30E-01 |
| AT4G00985 | 0,26 | -2,69 | 4,40 | -0,61 | 5,41E-01 |
| AT4G01000 | 0,50 | -0,25 | 3,96 | -0,06 | 9,49E-01 |
| AT4G01030 | 0,51 | -0,14 | 3,07 | -0,04 | 9,65E-01 |
| AT4G01037 | 0,21 | -0,29 | 4,38 | -0,07 | 9,47E-01 |
| AT4G01090 | 0,44 | -0,97 | 3,29 | -0,29 | 7,69E-01 |
| AT4G01110 | 0,08 | 0,00 | 4,41 | 0,00 | 1,00E+00 |
| AT4G01120 | 0,18 | 0,00 | 4,41 | 0,00 | 1,00E+00 |
| AT4G01180 | 0,08 | 0,00 | 4,41 | 0,00 | 1,00E+00 |
| AT4G01190 | 0,20 | -2,55 | 4,40 | -0,58 | 5,62E-01 |
| AT4G01200 | 0,46 | -3,86 | 4,34 | -0,89 | 3,75E-01 |
| AT4G01240 | 0,94 | -0,15 | 1,87 | -0,08 | 9,36E-01 |
| AT4G01310 | 0,66 | -0,90 | 2,93 | -0,31 | 7,59E-01 |
| AT4G01395 | 0,71 | -1,17 | 2,06 | -0,57 | 5,71E-01 |
| AT4G01540 | 0,20 | 0,11 | 4,37 | 0,03 | 9,80E-01 |
| AT4G01550 | 0,48 | 0,00 | 4,41 | 0,00 | 1,00E+00 |
| AT4G01700 | 0,60 | -1,18 | 3,18 | -0,37 | 7,11E-01 |
| AT4G01870 | 0,45 | -3,39 | 4,17 | -0,81 | 4,16E-01 |
| AT4G01950 | 0,49 | 0,90 | 2,64 | 0,34 | 7,33E-01 |
| AT4G01985 | 0,41 | -0,25 | 4,38 | -0,06 | 9,54E-01 |
| AT4G02195 | 0,71 | -2,39 | 2,48 | -0,96 | 3,36E-01 |
| AT4G02210 | 0,65 | 0,70 | 2,84 | 0,24 | 8,07E-01 |
| AT4G02230 | 0,28 | -1,75 | 4,39 | -0,40 | 6,90E-01 |
| AT4G02405 | 0,05 | -0,24 | 4,38 | -0,05 | 9,57E-01 |
| AT4G02410 | 0,84 | -2,65 | 2,81 | -0,94 | 3,45E-01 |
| AT4G02450 | 0,79 | 2,15 | 2,42 | 0,89 | 3,74E-01 |
| AT4G02460 | 0,19 | -0,81 | 4,39 | -0,19 | 8,53E-01 |
| AT4G02530 | 0,43 | 1,27 | 3,30 | 0,38 | 7,01E-01 |
| AT4G04185 | 0,38 | 1,49 | 3,35 | 0,44 | 6,56E-01 |
| AT4G02550 | 0,12 | -0,14 | 4,38 | -0,03 | 9,74E-01 |
| AT4G02560 | 0,57 | -2,74 | 2,69 | -1,02 | 3,08E-01 |
| AT4G02590 | 0,27 | -2,58 | 4,40 | -0,59 | 5,58E-01 |
| AT4G02620 | 0,23 | 0,00 | 4,41 | 0,00 | 1,00E+00 |
| AT4G02680 | 0,66 | -3,05 | 3,81 | -0,80 | 4,23E-01 |

| | | | | | |
|---|---|---|---|---|---|
| AT4G02710 | 0,71 | -2,74 | 2,94 | -0,93 | 3,50E-01 |
| AT4G02735 | 0,41 | -3,93 | 4,34 | -0,91 | 3,65E-01 |
| AT4G02740 | 0,60 | -0,78 | 3,03 | -0,26 | 7,96E-01 |
| AT4G02750 | 0,98 | -4,09 | 2,26 | -1,81 | 7,10E-02 |
| AT4G02910 | 0,98 | 0,33 | 2,34 | 0,14 | 8,88E-01 |
| AT4G02930 | 0,46 | -3,35 | 4,37 | -0,77 | 4,44E-01 |
| AT4G03090 | 0,91 | 1,63 | 1,93 | 0,85 | 3,98E-01 |
| AT4G03110 | 0,81 | -2,81 | 2,91 | -0,97 | 3,33E-01 |
| AT4G03130 | 0,08 | 0,00 | 4,41 | 0,00 | 1,00E+00 |
| AT4G03150 | 0,26 | -2,60 | 4,41 | -0,59 | 5,55E-01 |
| AT4G03190 | 0,84 | -3,26 | 2,23 | -1,46 | 1,45E-01 |
| AT4G03230 | 0,47 | -2,70 | 3,95 | -0,68 | 4,94E-01 |
| AT4G03270 | 0,81 | -3,87 | 2,81 | -1,38 | 1,69E-01 |
| AT4G04475 | 0,42 | 0,35 | 3,30 | 0,11 | 9,15E-01 |
| AT4G04485 | 0,66 | -0,48 | 2,48 | -0,19 | 8,47E-01 |
| AT4G03340 | 0,12 | 0,00 | 4,41 | 0,00 | 1,00E+00 |
| AT4G03390 | 0,56 | -1,63 | 2,69 | -0,61 | 5,45E-01 |
| AT4G03400 | 0,95 | 1,15 | 2,14 | 0,54 | 5,92E-01 |
| AT4G03435 | 0,08 | -0,21 | 4,39 | -0,05 | 9,62E-01 |
| AT4G03438 | 0,26 | -0,21 | 4,34 | -0,05 | 9,62E-01 |
| AT4G03450 | 0,60 | -1,67 | 2,52 | -0,66 | 5,08E-01 |
| AT4G03490 | 0,42 | -1,38 | 4,27 | -0,32 | 7,46E-01 |
| AT4G03505 | 0,25 | -3,06 | 4,38 | -0,70 | 4,85E-01 |
| AT4G03520 | 1,13 | -0,50 | 2,05 | -0,24 | 8,09E-01 |
| AT4G03570 | 0,98 | 1,36 | 1,77 | 0,77 | 4,43E-01 |
| AT4G03590 | 1,16 | 3,32 | 2,20 | 1,51 | 1,32E-01 |
| AT4G03728 | 0,69 | 1,83 | 3,64 | 0,50 | 6,14E-01 |
| AT4G04675 | 1,07 | -0,65 | 2,01 | -0,32 | 7,47E-01 |
| AT4G04030 | 0,82 | 1,87 | 2,46 | 0,76 | 4,47E-01 |
| AT4G04221 | 1,00 | -1,62 | 1,92 | -0,85 | 3,97E-01 |
| AT4G04370 | 1,03 | -3,51 | 2,07 | -1,70 | 9,00E-02 |
| AT4G04470 | 0,08 | -0,21 | 4,39 | -0,05 | 9,62E-01 |
| AT4G04580 | 0,11 | -0,80 | 4,39 | -0,18 | 8,56E-01 |
| AT4G04614 | 0,51 | -3,74 | 4,35 | -0,86 | 3,90E-01 |
| AT4G04632 | 0,44 | 1,05 | 4,09 | 0,26 | 7,97E-01 |
| AT4G04650 | 0,82 | 2,05 | 2,26 | 0,91 | 3,63E-01 |
| AT4G04710 | 0,47 | 0,06 | 3,28 | 0,02 | 9,85E-01 |
| AT4G04720 | 1,28 | -2,55 | 2,16 | -1,18 | 2,39E-01 |
| AT4G04740 | 0,87 | -0,08 | 2,01 | -0,04 | 9,70E-01 |
| AT4G04910 | 0,75 | -0,90 | 2,17 | -0,42 | 6,78E-01 |
| AT4G04950 | 0,83 | -3,39 | 3,67 | -0,92 | 3,56E-01 |
| AT4G04957 | 0,91 | 3,16 | 2,55 | 1,24 | 2,16E-01 |
| AT4G04980 | 0,11 | 0,46 | 4,35 | 0,11 | 9,15E-01 |
| AT4G05070 | 1,30 | -1,09 | 1,64 | -0,67 | 5,06E-01 |
| AT4G05110 | 0,17 | 0,00 | 4,41 | 0,00 | 1,00E+00 |
| AT4G05085 | 0,05 | -0,24 | 4,38 | -0,05 | 9,57E-01 |
| AT4G05330 | 0,44 | -1,81 | 4,11 | -0,44 | 6,59E-01 |
| AT4G05410 | 0,51 | -2,50 | 4,07 | -0,62 | 5,38E-01 |
| AT4G05470 | 0,12 | -0,68 | 4,41 | -0,15 | 8,77E-01 |
| AT4G05631 | 0,41 | -0,90 | 3,43 | -0,26 | 7,93E-01 |
| AT4G06526 | 0,56 | 0,11 | 3,90 | 0,03 | 9,78E-01 |
| AT4G06639 | 0,21 | 0,17 | 4,37 | 0,04 | 9,68E-01 |
| AT4G06599 | 0,16 | 0,32 | 4,36 | 0,07 | 9,42E-01 |
| AT4G07350 | 0,08 | -0,21 | 4,39 | -0,05 | 9,62E-01 |
| AT4G07400 | 0,28 | 0,00 | 4,36 | 0,00 | 1,00E+00 |
| AT4G07410 | 1,28 | -2,51 | 2,00 | -1,26 | 2,08E-01 |

| | | | | | |
|---|---|---|---|---|---|
| AT4G07524 | 0,19 | 0,02 | 4,36 | 0,00 | 9,97E-01 |
| AT4G07526 | 0,83 | -2,80 | 2,42 | -1,16 | 2,47E-01 |
| AT4G05225 | 0,74 | 1,69 | 3,01 | 0,56 | 5,75E-01 |
| AT4G07990 | 0,20 | -1,00 | 4,37 | -0,23 | 8,19E-01 |
| AT4G08230 | 0,30 | 1,64 | 4,30 | 0,38 | 7,03E-01 |
| AT4G08310 | 0,85 | -1,50 | 1,74 | -0,86 | 3,88E-01 |
| AT4G05415 | 0,95 | -3,93 | 2,68 | -1,47 | 1,42E-01 |
| AT4G08370 | 0,46 | 0,22 | 2,67 | 0,08 | 9,34E-01 |
| AT4G08400 | 0,54 | 2,56 | 2,93 | 0,87 | 3,82E-01 |
| AT4G08410 | 1,29 | 3,59 | 2,22 | 1,62 | 1,06E-01 |
| AT4G05535 | 0,95 | -2,19 | 2,41 | -0,91 | 3,63E-01 |
| AT4G08460 | 0,62 | -2,41 | 2,58 | -0,93 | 3,51E-01 |
| AT4G08480 | 0,67 | -0,91 | 3,01 | -0,30 | 7,63E-01 |
| AT4G08485 | 0,63 | -0,68 | 3,80 | -0,18 | 8,58E-01 |
| AT4G08590 | 0,56 | -3,29 | 4,38 | -0,75 | 4,54E-01 |
| AT4G08760 | 1,19 | 3,72 | 2,20 | 1,69 | 9,11E-02 |
| AT4G08810 | 0,77 | -3,25 | 2,57 | -1,27 | 2,05E-01 |
| AT4G08874 | 0,12 | -0,31 | 4,38 | -0,07 | 9,44E-01 |
| AT4G08940 | 0,46 | -0,67 | 3,36 | -0,20 | 8,42E-01 |
| AT4G09010 | 1,11 | -0,99 | 1,83 | -0,54 | 5,88E-01 |
| AT4G09012 | 0,55 | 0,22 | 3,26 | 0,07 | 9,45E-01 |
| AT4G09035 | 0,16 | -0,01 | 4,37 | 0,00 | 9,99E-01 |
| AT4G09040 | 0,47 | -1,76 | 4,32 | -0,41 | 6,85E-01 |
| AT4G09340 | 0,44 | -2,53 | 3,27 | -0,77 | 4,39E-01 |
| AT4G09350 | 0,26 | -0,35 | 4,38 | -0,08 | 9,36E-01 |
| AT4G09490 | 0,39 | 0,00 | 4,41 | 0,00 | 1,00E+00 |
| AT4G09510 | 0,38 | -1,67 | 3,41 | -0,49 | 6,24E-01 |
| AT4G09620 | 0,40 | 0,00 | 4,41 | 0,00 | 1,00E+00 |
| AT4G09640 | 0,98 | -0,07 | 1,82 | -0,04 | 9,68E-01 |
| AT4G09900 | 0,13 | -0,18 | 4,39 | -0,04 | 9,67E-01 |
| AT4G09960 | 0,46 | -0,62 | 4,40 | -0,14 | 8,88E-01 |
| AT4G09980 | 0,57 | -1,75 | 2,62 | -0,67 | 5,03E-01 |
| AT4G10070 | 1,18 | -3,88 | 2,15 | -1,80 | 7,16E-02 |
| AT4G10080 | 1,04 | -1,22 | 2,03 | -0,60 | 5,47E-01 |
| AT4G10130 | 0,55 | -3,18 | 3,79 | -0,84 | 4,01E-01 |
| AT4G10400 | 0,39 | -3,07 | 4,38 | -0,70 | 4,84E-01 |
| AT4G10420 | 1,07 | -4,90 | 2,61 | -1,88 | 6,06E-02 |
| AT4G10440 | 0,08 | 0,00 | 4,41 | 0,00 | 1,00E+00 |
| AT4G10480 | 0,41 | -0,32 | 4,38 | -0,07 | 9,42E-01 |
| AT4G10540 | 0,12 | 0,00 | 4,41 | 0,00 | 1,00E+00 |
| AT4G10550 | 0,52 | -2,60 | 3,96 | -0,66 | 5,11E-01 |
| AT4G10570 | 1,14 | -0,36 | 1,71 | -0,21 | 8,32E-01 |
| AT4G10700 | 0,54 | 2,71 | 3,69 | 0,73 | 4,63E-01 |
| AT4G10730 | 0,28 | -3,07 | 4,38 | -0,70 | 4,84E-01 |
| AT4G10740 | 1,27 | 3,77 | 2,21 | 1,71 | 8,76E-02 |
| AT4G10760 | 0,19 | 1,18 | 4,31 | 0,27 | 7,84E-01 |
| AT4G10770 | 0,10 | -0,34 | 4,38 | -0,08 | 9,38E-01 |
| AT4G10790 | 1,30 | 0,80 | 1,81 | 0,44 | 6,58E-01 |
| AT4G10920 | 0,27 | 0,87 | 3,74 | 0,23 | 8,15E-01 |
| AT4G10960 | 0,05 | -0,24 | 4,38 | -0,05 | 9,57E-01 |
| AT4G10970 | 0,10 | -0,34 | 4,38 | -0,08 | 9,38E-01 |
| AT4G11010 | 0,17 | -0,75 | 4,40 | -0,17 | 8,65E-01 |
| AT4G11050 | 0,37 | -0,24 | 3,40 | -0,07 | 9,43E-01 |
| AT4G11080 | 0,05 | -0,24 | 4,38 | -0,05 | 9,57E-01 |
| AT4G05855 | 1,27 | -3,64 | 2,12 | -1,72 | 8,57E-02 |
| AT4G11216 | 0,26 | -2,60 | 4,41 | -0,59 | 5,55E-01 |

| | | | | | |
|---|---|---|---|---|---|
| AT4G11220 | 0,29 | -0,34 | 4,34 | -0,08 | 9,37E-01 |
| AT4G11260 | 0,66 | -1,61 | 4,32 | -0,37 | 7,09E-01 |
| AT4G11300 | 0,45 | -1,87 | 4,14 | -0,45 | 6,52E-01 |
| AT4G11340 | 1,18 | 0,66 | 1,74 | 0,38 | 7,06E-01 |
| AT4G11410 | 0,24 | -2,59 | 4,41 | -0,59 | 5,56E-01 |
| AT4G05880 | 0,80 | -0,79 | 2,82 | -0,28 | 7,80E-01 |
| AT4G11650 | 0,43 | -3,24 | 4,37 | -0,74 | 4,58E-01 |
| AT4G11670 | 0,44 | -1,67 | 2,73 | -0,61 | 5,40E-01 |
| AT4G11800 | 1,18 | 1,26 | 1,83 | 0,69 | 4,90E-01 |
| AT4G05925 | 1,28 | 0,72 | 1,89 | 0,38 | 7,03E-01 |
| AT4G11820 | 0,25 | 0,00 | 4,41 | 0,00 | 1,00E+00 |
| AT4G11860 | 0,25 | -2,60 | 4,41 | -0,59 | 5,55E-01 |
| AT4G11920 | 0,40 | 1,32 | 4,13 | 0,32 | 7,49E-01 |
| AT4G05930 | 0,37 | 1,16 | 4,22 | 0,27 | 7,83E-01 |
| AT4G12010 | 0,59 | -0,71 | 3,76 | -0,19 | 8,51E-01 |
| AT4G05980 | 0,40 | -0,19 | 4,39 | -0,04 | 9,65E-01 |
| AT4G12300 | 1,00 | 0,56 | 2,23 | 0,25 | 8,01E-01 |
| AT4G12310 | 0,40 | -0,32 | 2,85 | -0,11 | 9,10E-01 |
| AT4G12330 | 0,81 | 0,08 | 2,77 | 0,03 | 9,77E-01 |
| AT4G12590 | 0,64 | 1,20 | 2,54 | 0,47 | 6,37E-01 |
| AT4G12710 | 0,37 | -3,19 | 4,38 | -0,73 | 4,66E-01 |
| AT4G12730 | 0,13 | -0,18 | 4,39 | -0,04 | 9,67E-01 |
| AT4G12750 | 1,05 | -3,51 | 2,03 | -1,73 | 8,42E-02 |
| AT4G12790 | 0,53 | 0,49 | 3,25 | 0,15 | 8,81E-01 |
| AT4G12820 | 0,35 | 0,00 | 4,41 | 0,00 | 1,00E+00 |
| AT4G12880 | 0,35 | 0,00 | 4,41 | 0,00 | 1,00E+00 |
| AT4G12980 | 0,57 | 1,27 | 2,65 | 0,48 | 6,30E-01 |
| AT4G13150 | 0,55 | -3,76 | 4,35 | -0,86 | 3,87E-01 |
| AT4G13350 | 0,30 | 0,97 | 4,33 | 0,22 | 8,23E-01 |
| AT4G13395 | 0,15 | 0,79 | 4,33 | 0,18 | 8,56E-01 |
| AT4G13420 | 0,12 | 0,00 | 4,41 | 0,00 | 1,00E+00 |
| AT4G13550 | 0,15 | -0,73 | 4,40 | -0,17 | 8,68E-01 |
| AT4G13590 | 0,56 | -0,33 | 2,55 | -0,13 | 8,96E-01 |
| AT4G13630 | 0,80 | -2,58 | 3,54 | -0,73 | 4,65E-01 |
| AT4G13640 | 0,21 | 0,00 | 4,41 | 0,00 | 1,00E+00 |
| AT4G13780 | 1,18 | -2,65 | 1,85 | -1,43 | 1,52E-01 |
| AT4G13800 | 0,45 | -2,47 | 3,26 | -0,76 | 4,49E-01 |
| AT4G13820 | 0,22 | 0,41 | 4,35 | 0,09 | 9,25E-01 |
| AT4G13870 | 0,08 | 0,00 | 4,41 | 0,00 | 1,00E+00 |
| AT4G13950 | 0,08 | 0,00 | 4,41 | 0,00 | 1,00E+00 |
| AT4G13970 | 0,50 | -2,05 | 3,89 | -0,53 | 5,98E-01 |
| AT4G14010 | 0,10 | 0,45 | 4,35 | 0,10 | 9,19E-01 |
| AT4G14040 | 0,70 | -2,55 | 2,45 | -1,04 | 2,98E-01 |
| AT4G14050 | 0,14 | -0,29 | 4,38 | -0,07 | 9,47E-01 |
| AT4G14070 | 0,95 | -2,79 | 2,61 | -1,07 | 2,85E-01 |
| AT4G14090 | 0,47 | -0,57 | 2,75 | -0,21 | 8,34E-01 |
| AT4G14110 | 0,26 | -2,60 | 4,41 | -0,59 | 5,55E-01 |
| AT4G14200 | 0,21 | -0,05 | 4,37 | -0,01 | 9,90E-01 |
| AT4G14210 | 0,70 | -1,16 | 1,91 | -0,60 | 5,46E-01 |
| AT4G14230 | 0,73 | -2,44 | 3,04 | -0,80 | 4,22E-01 |
| AT4G14240 | 0,36 | -2,01 | 4,36 | -0,46 | 6,45E-01 |
| AT4G14340 | 0,26 | -2,60 | 4,41 | -0,59 | 5,55E-01 |
| AT4G14342 | 0,72 | 0,00 | 4,41 | 0,00 | 1,00E+00 |
| AT4G14350 | 0,16 | -0,03 | 4,37 | -0,01 | 9,95E-01 |
| AT4G14370 | 0,47 | 0,18 | 2,79 | 0,06 | 9,49E-01 |
| AT4G14385 | 0,08 | 0,00 | 4,41 | 0,00 | 1,00E+00 |

| | | | | | |
|---|---|---|---|---|---|
| AT4G14420 | 0,43 | -3,24 | 4,37 | -0,74 | 4,58E-01 |
| AT4G14430 | 1,12 | 0,49 | 1,94 | 0,25 | 8,02E-01 |
| AT4G14440 | 0,19 | 1,05 | 4,32 | 0,24 | 8,08E-01 |
| AT4G14680 | 0,20 | -2,55 | 4,40 | -0,58 | 5,62E-01 |
| AT4G14710 | 0,27 | 0,65 | 4,34 | 0,15 | 8,80E-01 |
| AT4G14740 | 0,35 | -0,14 | 4,39 | -0,03 | 9,75E-01 |
| AT4G14746 | 0,16 | 0,21 | 4,36 | 0,05 | 9,61E-01 |
| AT4G14790 | 0,23 | -1,14 | 4,39 | -0,26 | 7,96E-01 |
| AT4G14811 | 0,45 | -0,62 | 3,33 | -0,19 | 8,52E-01 |
| AT4G14900 | 1,30 | -1,42 | 1,59 | -0,89 | 3,72E-01 |
| AT4G14905 | 0,68 | -2,05 | 2,47 | -0,83 | 4,08E-01 |
| AT4G14920 | 0,44 | -2,27 | 4,19 | -0,54 | 5,88E-01 |
| AT4G14990 | 0,87 | -1,97 | 1,85 | -1,06 | 2,88E-01 |
| AT4G15050 | 0,52 | 1,71 | 3,79 | 0,45 | 6,52E-01 |
| AT4G15053 | 0,24 | 1,19 | 4,31 | 0,28 | 7,83E-01 |
| AT4G15080 | 0,43 | -1,13 | 4,13 | -0,27 | 7,84E-01 |
| AT4G15200 | 0,36 | 0,00 | 4,41 | 0,00 | 1,00E+00 |
| AT4G15210 | 0,12 | 0,00 | 4,41 | 0,00 | 1,00E+00 |
| AT4G15242 | 0,72 | 1,49 | 2,08 | 0,71 | 4,75E-01 |
| AT4G06305 | 0,44 | 0,04 | 4,08 | 0,01 | 9,92E-01 |
| AT4G06310 | 0,35 | -1,98 | 4,34 | -0,46 | 6,48E-01 |
| AT4G06315 | 1,00 | 2,68 | 2,99 | 0,90 | 3,71E-01 |
| AT4G15270 | 0,91 | -2,42 | 2,13 | -1,14 | 2,56E-01 |
| AT4G15490 | 0,92 | 0,21 | 2,08 | 0,10 | 9,19E-01 |
| AT4G15550 | 0,31 | 0,10 | 4,37 | 0,02 | 9,82E-01 |
| AT4G15810 | 0,14 | 0,48 | 4,35 | 0,11 | 9,13E-01 |
| AT4G06340 | 0,10 | -0,18 | 4,38 | -0,04 | 9,67E-01 |
| AT4G15920 | 0,36 | 0,81 | 4,30 | 0,19 | 8,51E-01 |
| AT4G15955 | 0,12 | 0,00 | 4,41 | 0,00 | 1,00E+00 |
| AT4G16070 | 0,52 | -0,53 | 2,74 | -0,19 | 8,46E-01 |
| AT4G16141 | 0,51 | -3,59 | 3,97 | -0,90 | 3,66E-01 |
| AT4G16155 | 0,30 | -1,46 | 4,36 | -0,34 | 7,37E-01 |
| AT4G16215 | 0,40 | 2,03 | 3,25 | 0,62 | 5,33E-01 |
| AT4G16250 | 0,73 | -3,37 | 3,01 | -1,12 | 2,63E-01 |
| AT4G16280 | 0,45 | -2,53 | 3,26 | -0,78 | 4,38E-01 |
| AT4G16330 | 0,88 | 0,87 | 2,89 | 0,30 | 7,63E-01 |
| AT4G16410 | 1,15 | -2,67 | 1,95 | -1,37 | 1,72E-01 |
| AT4G16470 | 0,11 | -0,68 | 4,41 | -0,15 | 8,77E-01 |
| AT4G16500 | 1,23 | -0,50 | 1,49 | -0,34 | 7,37E-01 |
| AT4G16515 | 0,18 | 0,00 | 4,41 | 0,00 | 1,00E+00 |
| AT4G16520 | 0,32 | -0,20 | 4,34 | -0,05 | 9,64E-01 |
| AT4G16540 | 0,20 | -1,00 | 4,37 | -0,23 | 8,19E-01 |
| AT4G16580 | 0,33 | -0,47 | 3,57 | -0,13 | 8,94E-01 |
| AT4G16630 | 0,66 | -3,12 | 3,69 | -0,85 | 3,97E-01 |
| AT4G16680 | 0,40 | 0,00 | 4,41 | 0,00 | 1,00E+00 |
| AT4G16690 | 0,88 | 0,99 | 2,09 | 0,47 | 6,37E-01 |
| AT4G16750 | 0,15 | -0,14 | 4,39 | -0,03 | 9,74E-01 |
| AT4G16790 | 0,35 | -0,79 | 3,52 | -0,22 | 8,23E-01 |
| AT4G16880 | 1,01 | 0,20 | 2,02 | 0,10 | 9,22E-01 |
| AT4G16892 | 1,06 | -0,54 | 2,41 | -0,22 | 8,23E-01 |
| AT4G16957 | 0,81 | -3,17 | 3,49 | -0,91 | 3,64E-01 |
| AT4G16960 | 0,92 | 0,00 | 4,16 | 0,00 | 1,00E+00 |
| AT4G17040 | 0,44 | -3,25 | 4,21 | -0,77 | 4,40E-01 |
| AT4G17060 | 0,64 | -3,91 | 3,65 | -1,07 | 2,84E-01 |
| AT4G17070 | 0,40 | -0,89 | 3,35 | -0,26 | 7,92E-01 |
| AT4G17100 | 0,77 | 0,69 | 2,42 | 0,28 | 7,76E-01 |

| | | | | | |
|---|---|---|---|---|---|
| AT4G17190 | 0,27 | 0,92 | 4,33 | 0,21 | 8,32E-01 |
| AT4G17230 | 1,13 | 2,08 | 1,84 | 1,13 | 2,59E-01 |
| AT4G17243 | 0,66 | -1,29 | 2,54 | -0,51 | 6,12E-01 |
| AT4G17280 | 0,46 | -2,49 | 3,31 | -0,75 | 4,52E-01 |
| AT4G17430 | 0,16 | 0,84 | 4,33 | 0,20 | 8,45E-01 |
| AT4G17486 | 0,51 | -4,07 | 4,33 | -0,94 | 3,48E-01 |
| AT4G17490 | 0,05 | -0,24 | 4,38 | -0,05 | 9,57E-01 |
| AT4G17510 | 0,15 | 0,05 | 4,37 | 0,01 | 9,91E-01 |
| AT4G17550 | 0,46 | -1,50 | 4,18 | -0,36 | 7,20E-01 |
| AT4G17570 | 0,79 | -0,78 | 2,28 | -0,34 | 7,31E-01 |
| AT4G17590 | 0,76 | 1,56 | 2,76 | 0,57 | 5,72E-01 |
| AT4G17610 | 0,22 | 0,38 | 4,35 | 0,09 | 9,31E-01 |
| AT4G17670 | 0,33 | -2,56 | 4,40 | -0,58 | 5,61E-01 |
| AT4G17720 | 0,25 | -1,24 | 4,38 | -0,28 | 7,76E-01 |
| AT4G17740 | 0,24 | -0,04 | 4,38 | -0,01 | 9,93E-01 |
| AT4G17840 | 0,82 | -0,12 | 2,34 | -0,05 | 9,60E-01 |
| AT4G17880 | 0,51 | -0,68 | 3,26 | -0,21 | 8,35E-01 |
| AT4G17950 | 0,50 | -3,37 | 4,10 | -0,82 | 4,12E-01 |
| AT4G18020 | 0,33 | -1,65 | 4,41 | -0,37 | 7,09E-01 |
| AT4G18040 | 0,22 | 0,38 | 4,35 | 0,09 | 9,31E-01 |
| AT4G18100 | 0,24 | 0,99 | 4,32 | 0,23 | 8,19E-01 |
| AT4G18110 | 0,16 | -0,68 | 4,41 | -0,15 | 8,77E-01 |
| AT4G18130 | 0,68 | -2,87 | 2,84 | -1,01 | 3,13E-01 |
| AT4G18140 | 0,37 | -1,07 | 4,36 | -0,24 | 8,07E-01 |
| AT4G18230 | 0,15 | -0,77 | 4,39 | -0,18 | 8,61E-01 |
| AT4G18250 | 0,30 | -2,60 | 4,41 | -0,59 | 5,55E-01 |
| AT4G18253 | 0,68 | -2,77 | 2,41 | -1,15 | 2,52E-01 |
| AT4G18255 | 0,35 | -2,99 | 4,38 | -0,68 | 4,95E-01 |
| AT4G18390 | 1,03 | -0,49 | 2,28 | -0,22 | 8,29E-01 |
| AT4G18440 | 0,69 | 2,13 | 3,53 | 0,60 | 5,46E-01 |
| AT4G18520 | 0,18 | -0,27 | 4,38 | -0,06 | 9,51E-01 |
| AT4G18580 | 0,08 | 0,00 | 4,41 | 0,00 | 1,00E+00 |
| AT4G18790 | 0,08 | 0,00 | 4,41 | 0,00 | 1,00E+00 |
| AT4G18810 | 0,53 | -0,15 | 2,80 | -0,05 | 9,58E-01 |
| AT4G18840 | 0,38 | 0,37 | 3,44 | 0,11 | 9,15E-01 |
| AT4G19006 | 0,25 | -1,64 | 4,36 | -0,38 | 7,07E-01 |
| AT4G19040 | 0,87 | -0,16 | 3,38 | -0,05 | 9,63E-01 |
| AT4G19045 | 0,12 | 0,00 | 4,41 | 0,00 | 1,00E+00 |
| AT4G19050 | 0,78 | -3,74 | 2,76 | -1,35 | 1,76E-01 |
| AT4G19060 | 0,46 | -1,02 | 4,04 | -0,25 | 8,00E-01 |
| AT4G19080 | 1,21 | -1,61 | 1,96 | -0,82 | 4,11E-01 |
| AT4G19095 | 0,58 | -0,28 | 2,54 | -0,11 | 9,12E-01 |
| AT4G06870 | 0,16 | 0,75 | 4,34 | 0,17 | 8,63E-01 |
| AT4G19180 | 0,15 | -0,73 | 4,40 | -0,17 | 8,68E-01 |
| AT4G19220 | 0,37 | -3,33 | 4,32 | -0,77 | 4,41E-01 |
| AT4G19240 | 0,33 | -1,21 | 4,35 | -0,28 | 7,81E-01 |
| AT4G19420 | 0,68 | 0,82 | 2,44 | 0,34 | 7,37E-01 |
| AT4G19440 | 0,20 | -0,68 | 4,41 | -0,15 | 8,77E-01 |
| AT4G19640 | 0,35 | 0,29 | 3,41 | 0,08 | 9,33E-01 |
| AT4G19660 | 1,11 | -1,46 | 1,79 | -0,81 | 4,17E-01 |
| AT4G19670 | 0,77 | -3,95 | 3,53 | -1,12 | 2,63E-01 |
| AT4G06915 | 0,34 | 0,36 | 4,34 | 0,08 | 9,34E-01 |
| AT4G19700 | 1,20 | 1,44 | 1,98 | 0,73 | 4,66E-01 |
| AT4G19720 | 0,28 | 0,33 | 4,36 | 0,08 | 9,40E-01 |
| AT4G19860 | 1,15 | -1,58 | 1,74 | -0,91 | 3,65E-01 |
| AT4G19870 | 0,10 | -0,18 | 4,38 | -0,04 | 9,67E-01 |

| | | | | | |
|---|---|---|---|---|---|
| AT4G19880 | 0,52 | -0,50 | 2,59 | -0,19 | 8,48E-01 |
| AT4G19890 | 0,56 | -2,93 | 3,07 | -0,95 | 3,40E-01 |
| AT4G19900 | 0,17 | 0,00 | 4,41 | 0,00 | 1,00E+00 |
| AT4G19985 | 0,13 | 0,00 | 4,41 | 0,00 | 1,00E+00 |
| AT4G20095 | 0,45 | 0,00 | 4,41 | 0,00 | 1,00E+00 |
| AT4G20150 | 0,58 | -0,93 | 2,70 | -0,34 | 7,31E-01 |
| AT4G06945 | 1,00 | -1,79 | 2,77 | -0,65 | 5,19E-01 |
| AT4G20210 | 0,49 | 2,01 | 2,63 | 0,76 | 4,45E-01 |
| AT4G20330 | 0,48 | -2,48 | 2,72 | -0,91 | 3,62E-01 |
| AT4G20410 | 1,25 | -2,80 | 1,83 | -1,53 | 1,26E-01 |
| AT4G20440 | 0,64 | 0,20 | 2,23 | 0,09 | 9,30E-01 |
| AT4G20690 | 0,21 | 0,00 | 4,41 | 0,00 | 1,00E+00 |
| AT4G20880 | 0,74 | 0,33 | 1,84 | 0,18 | 8,58E-01 |
| AT4G20910 | 0,27 | -2,23 | 4,36 | -0,51 | 6,10E-01 |
| AT4G20953 | 0,51 | 1,89 | 3,21 | 0,59 | 5,56E-01 |
| AT4G21100 | 1,08 | -2,29 | 1,75 | -1,31 | 1,89E-01 |
| AT4G21140 | 0,20 | -2,55 | 4,40 | -0,58 | 5,62E-01 |
| AT4G21170 | 0,28 | 1,34 | 4,31 | 0,31 | 7,56E-01 |
| AT4G06985 | 0,08 | 0,00 | 4,41 | 0,00 | 1,00E+00 |
| AT4G21190 | 0,08 | 0,00 | 4,41 | 0,00 | 1,00E+00 |
| AT4G21350 | 0,08 | 0,00 | 4,41 | 0,00 | 1,00E+00 |
| AT4G21400 | 0,16 | -0,62 | 4,40 | -0,14 | 8,87E-01 |
| AT4G21430 | 0,22 | 0,80 | 4,34 | 0,18 | 8,54E-01 |
| AT4G21510 | 0,05 | -0,24 | 4,38 | -0,05 | 9,57E-01 |
| AT4G21520 | 0,22 | -1,25 | 4,37 | -0,29 | 7,75E-01 |
| AT4G21560 | 0,38 | -1,70 | 4,34 | -0,39 | 6,95E-01 |
| AT4G21660 | 1,13 | -1,73 | 2,05 | -0,84 | 4,01E-01 |
| AT4G21810 | 1,02 | -1,51 | 1,99 | -0,76 | 4,47E-01 |
| AT4G21860 | 0,50 | -1,58 | 3,41 | -0,46 | 6,44E-01 |
| AT4G21890 | 0,12 | 0,00 | 4,41 | 0,00 | 1,00E+00 |
| AT4G21910 | 0,54 | 0,43 | 4,35 | 0,10 | 9,21E-01 |
| AT4G21990 | 0,81 | -2,22 | 2,41 | -0,92 | 3,56E-01 |
| AT4G22120 | 0,12 | 0,00 | 4,41 | 0,00 | 1,00E+00 |
| AT4G22150 | 0,22 | -0,75 | 4,40 | -0,17 | 8,65E-01 |
| AT4G22190 | 0,98 | -2,30 | 1,85 | -1,24 | 2,14E-01 |
| AT4G22200 | 0,21 | -0,14 | 4,37 | -0,03 | 9,75E-01 |
| AT4G22250 | 0,16 | -0,26 | 4,38 | -0,06 | 9,52E-01 |
| AT4G22260 | 0,18 | 0,40 | 4,35 | 0,09 | 9,28E-01 |
| AT4G22265 | 0,20 | -2,60 | 4,41 | -0,59 | 5,55E-01 |
| AT4G22285 | 0,20 | -0,75 | 4,40 | -0,17 | 8,64E-01 |
| AT4G22290 | 0,23 | 1,25 | 4,31 | 0,29 | 7,72E-01 |
| AT4G22350 | 0,69 | -0,67 | 3,65 | -0,18 | 8,54E-01 |
| AT4G22380 | 0,51 | 0,75 | 2,60 | 0,29 | 7,75E-01 |
| AT4G22470 | 0,12 | -0,31 | 4,38 | -0,07 | 9,44E-01 |
| AT4G22485 | 0,49 | 2,14 | 3,85 | 0,56 | 5,78E-01 |
| AT4G22550 | 0,26 | 0,06 | 3,79 | 0,02 | 9,87E-01 |
| AT4G22570 | 0,24 | -1,92 | 4,38 | -0,44 | 6,61E-01 |
| AT4G22745 | 0,42 | -3,39 | 4,16 | -0,81 | 4,15E-01 |
| AT4G22750 | 1,03 | -1,84 | 2,31 | -0,80 | 4,25E-01 |
| AT4G22755 | 0,47 | -3,21 | 4,38 | -0,73 | 4,64E-01 |
| AT4G22770 | 1,29 | 0,58 | 1,71 | 0,34 | 7,36E-01 |
| AT4G22780 | 0,24 | -1,00 | 4,35 | -0,23 | 8,18E-01 |
| AT4G22970 | 0,52 | -0,26 | 3,90 | -0,07 | 9,47E-01 |
| AT4G22980 | 0,47 | 0,97 | 3,95 | 0,25 | 8,05E-01 |
| AT4G23010 | 0,83 | -3,90 | 2,28 | -1,71 | 8,76E-02 |
| AT4G23040 | 0,28 | -0,72 | 4,40 | -0,16 | 8,70E-01 |

| | | | | | |
|---|---|---|---|---|---|
| AT4G23130 | 0,55 | -1,73 | 3,99 | -0,43 | 6,65E-01 |
| AT4G23190 | 1,18 | -0,60 | 1,77 | -0,34 | 7,35E-01 |
| AT4G23205 | 0,08 | 0,00 | 4,41 | 0,00 | 1,00E+00 |
| AT4G23200 | 0,43 | -3,12 | 4,39 | -0,71 | 4,77E-01 |
| AT4G23210 | 0,21 | -0,73 | 4,40 | -0,17 | 8,67E-01 |
| AT4G23220 | 0,76 | -0,18 | 2,40 | -0,08 | 9,40E-01 |
| AT4G23240 | 0,37 | -2,60 | 4,41 | -0,59 | 5,55E-01 |
| AT4G23250 | 0,45 | 0,94 | 3,94 | 0,24 | 8,11E-01 |
| AT4G23260 | 0,19 | -0,81 | 4,39 | -0,19 | 8,53E-01 |
| AT4G23290 | 0,10 | 0,45 | 4,35 | 0,10 | 9,19E-01 |
| AT4G23320 | 0,08 | 0,00 | 4,41 | 0,00 | 1,00E+00 |
| AT4G23490 | 0,15 | -0,14 | 4,39 | -0,03 | 9,74E-01 |
| AT4G07130 | 0,46 | -2,34 | 4,15 | -0,56 | 5,72E-01 |
| AT4G23540 | 0,67 | -3,30 | 3,08 | -1,07 | 2,83E-01 |
| AT4G23560 | 0,10 | 0,45 | 4,35 | 0,10 | 9,19E-01 |
| AT4G07135 | 1,25 | -2,83 | 1,86 | -1,52 | 1,28E-01 |
| AT4G07145 | 0,96 | -4,57 | 2,63 | -1,74 | 8,23E-02 |
| AT4G23620 | 0,31 | -0,78 | 4,35 | -0,18 | 8,58E-01 |
| AT4G23640 | 0,77 | -0,27 | 2,34 | -0,11 | 9,10E-01 |
| AT4G23700 | 0,29 | -3,07 | 4,39 | -0,70 | 4,83E-01 |
| AT4G23770 | 0,22 | 0,55 | 4,35 | 0,13 | 8,99E-01 |
| AT4G23810 | 0,35 | -0,99 | 4,38 | -0,23 | 8,21E-01 |
| AT4G23820 | 0,63 | -0,14 | 2,15 | -0,06 | 9,48E-01 |
| AT4G23880 | 0,47 | 1,03 | 3,12 | 0,33 | 7,41E-01 |
| AT4G23890 | 0,66 | -1,72 | 2,70 | -0,64 | 5,24E-01 |
| AT4G24180 | 0,37 | -2,98 | 4,38 | -0,68 | 4,97E-01 |
| AT4G24200 | 0,70 | -2,03 | 2,91 | -0,70 | 4,85E-01 |
| AT4G07250 | 0,17 | 0,00 | 4,41 | 0,00 | 1,00E+00 |
| AT4G24300 | 0,45 | 2,28 | 4,28 | 0,53 | 5,94E-01 |
| AT4G24320 | 0,33 | 0,00 | 4,41 | 0,00 | 1,00E+00 |
| AT4G24330 | 0,84 | 1,08 | 1,93 | 0,56 | 5,78E-01 |
| AT4G07255 | 0,10 | 0,45 | 4,35 | 0,10 | 9,19E-01 |
| AT4G24390 | 0,29 | -0,95 | 4,36 | -0,22 | 8,28E-01 |
| AT4G24490 | 0,72 | -1,70 | 2,48 | -0,69 | 4,92E-01 |
| AT4G24530 | 0,08 | 0,00 | 4,41 | 0,00 | 1,00E+00 |
| AT4G24560 | 1,21 | -1,84 | 2,11 | -0,87 | 3,83E-01 |
| AT4G24580 | 0,46 | -2,32 | 4,08 | -0,57 | 5,68E-01 |
| AT4G24610 | 0,67 | -4,14 | 2,93 | -1,41 | 1,58E-01 |
| AT4G24620 | 0,42 | -0,44 | 4,16 | -0,11 | 9,16E-01 |
| AT4G24660 | 1,04 | 1,42 | 2,20 | 0,64 | 5,21E-01 |
| AT4G24670 | 0,08 | -0,21 | 4,39 | -0,05 | 9,62E-01 |
| AT4G24730 | 0,45 | -3,79 | 4,08 | -0,93 | 3,53E-01 |
| AT4G24740 | 0,63 | -2,06 | 3,05 | -0,68 | 4,98E-01 |
| AT4G24750 | 0,28 | -3,07 | 4,38 | -0,70 | 4,84E-01 |
| AT4G24780 | 1,27 | 2,20 | 2,09 | 1,06 | 2,91E-01 |
| AT4G07285 | 0,27 | 0,07 | 4,37 | 0,02 | 9,87E-01 |
| AT4G24830 | 0,25 | 1,02 | 4,32 | 0,24 | 8,13E-01 |
| AT4G24840 | 0,53 | 0,37 | 3,09 | 0,12 | 9,04E-01 |
| AT4G24920 | 0,86 | -3,94 | 2,77 | -1,42 | 1,55E-01 |
| AT4G24990 | 0,31 | -0,32 | 4,34 | -0,07 | 9,41E-01 |
| AT4G25090 | 0,29 | 0,00 | 4,41 | 0,00 | 1,00E+00 |
| AT4G25270 | 0,45 | -3,79 | 4,08 | -0,93 | 3,53E-01 |
| AT4G25280 | 0,71 | -2,60 | 4,41 | -0,59 | 5,55E-01 |
| AT4G25290 | 0,52 | -1,64 | 3,88 | -0,42 | 6,73E-01 |
| AT4G25340 | 0,19 | -0,77 | 4,39 | -0,18 | 8,61E-01 |
| AT4G25435 | 0,88 | -3,72 | 2,73 | -1,36 | 1,73E-01 |

| | | | | | |
|---|---|---|---|---|---|
| AT4G25540 | 0,13 | 0,00 | 4,41 | 0,00 | 1,00E+00 |
| AT4G25570 | 0,43 | -0,89 | 4,17 | -0,21 | 8,31E-01 |
| AT4G25650 | 1,11 | -1,23 | 1,78 | -0,69 | 4,89E-01 |
| AT4G25670 | 0,79 | -3,15 | 2,84 | -1,11 | 2,67E-01 |
| AT4G25690 | 0,56 | -2,26 | 3,20 | -0,71 | 4,79E-01 |
| AT4G25740 | 0,38 | -1,12 | 4,35 | -0,26 | 7,97E-01 |
| AT4G25870 | 0,26 | -2,60 | 4,41 | -0,59 | 5,55E-01 |
| AT4G25940 | 0,39 | -2,51 | 4,39 | -0,57 | 5,68E-01 |
| AT4G25990 | 0,26 | -0,16 | 4,34 | -0,04 | 9,70E-01 |
| AT4G26080 | 0,87 | -1,05 | 2,62 | -0,40 | 6,88E-01 |
| AT4G26100 | 0,81 | -0,73 | 2,00 | -0,36 | 7,17E-01 |
| AT4G26140 | 0,21 | -1,17 | 4,36 | -0,27 | 7,88E-01 |
| AT4G26190 | 0,25 | 0,67 | 4,34 | 0,16 | 8,77E-01 |
| AT4G26300 | 0,69 | -2,40 | 2,65 | -0,91 | 3,65E-01 |
| AT4G07565 | 0,79 | 2,64 | 2,05 | 1,29 | 1,97E-01 |
| AT4G07625 | 0,79 | -0,52 | 2,68 | -0,19 | 8,47E-01 |
| AT4G07635 | 0,12 | -0,31 | 4,38 | -0,07 | 9,44E-01 |
| AT4G26370 | 0,48 | -0,72 | 4,06 | -0,18 | 8,60E-01 |
| AT4G26375 | 0,40 | 0,00 | 4,41 | 0,00 | 1,00E+00 |
| AT4G26400 | 0,60 | -2,61 | 3,95 | -0,66 | 5,10E-01 |
| AT4G26620 | 0,28 | -0,82 | 4,38 | -0,19 | 8,51E-01 |
| AT4G26630 | 0,72 | 0,43 | 2,45 | 0,17 | 8,61E-01 |
| AT4G26640 | 1,00 | 0,05 | 2,44 | 0,02 | 9,83E-01 |
| AT4G26750 | 0,43 | -0,68 | 4,24 | -0,16 | 8,73E-01 |
| AT4G26760 | 0,64 | -1,58 | 3,17 | -0,50 | 6,18E-01 |
| AT4G26795 | 0,42 | 1,93 | 4,03 | 0,48 | 6,31E-01 |
| AT4G26860 | 0,37 | 0,37 | 4,34 | 0,09 | 9,32E-01 |
| AT4G26940 | 0,43 | -1,64 | 4,09 | -0,40 | 6,88E-01 |
| AT4G26980 | 0,78 | -2,61 | 2,49 | -1,05 | 2,95E-01 |
| AT4G27020 | 0,48 | -0,41 | 3,39 | -0,12 | 9,03E-01 |
| AT4G07765 | 0,66 | 2,35 | 2,77 | 0,85 | 3,97E-01 |
| AT4G27130 | 0,28 | 1,39 | 3,65 | 0,38 | 7,02E-01 |
| AT4G27270 | 0,66 | -1,24 | 3,91 | -0,32 | 7,51E-01 |
| AT4G27280 | 0,84 | 1,73 | 2,72 | 0,64 | 5,24E-01 |
| AT4G27300 | 0,83 | -0,26 | 2,38 | -0,11 | 9,12E-01 |
| AT4G27320 | 1,19 | -2,55 | 2,13 | -1,20 | 2,32E-01 |
| AT4G27340 | 0,21 | -0,30 | 4,38 | -0,07 | 9,46E-01 |
| AT4G27410 | 0,98 | -4,55 | 2,31 | -1,96 | 4,94E-02 |
| AT4G27450 | 0,72 | -1,45 | 2,27 | -0,64 | 5,23E-01 |
| AT4G27480 | 0,65 | -2,57 | 3,66 | -0,70 | 4,83E-01 |
| AT4G27650 | 0,39 | -1,64 | 3,48 | -0,47 | 6,37E-01 |
| AT4G27700 | 0,47 | -1,41 | 3,23 | -0,44 | 6,61E-01 |
| AT4G27730 | 0,37 | -3,03 | 4,38 | -0,69 | 4,90E-01 |
| AT4G27800 | 1,09 | -2,23 | 2,00 | -1,12 | 2,65E-01 |
| AT4G27830 | 0,84 | -0,25 | 1,84 | -0,14 | 8,92E-01 |
| AT4G27840 | 0,43 | -1,77 | 4,32 | -0,41 | 6,81E-01 |
| AT4G27885 | 0,16 | 0,84 | 4,33 | 0,20 | 8,45E-01 |
| AT4G27960 | 0,38 | 1,93 | 4,22 | 0,46 | 6,48E-01 |
| AT4G28030 | 0,45 | -0,88 | 2,81 | -0,31 | 7,54E-01 |
| AT4G28050 | 0,43 | 0,00 | 4,41 | 0,00 | 1,00E+00 |
| AT4G28060 | 0,25 | -3,06 | 4,38 | -0,70 | 4,85E-01 |
| AT4G28180 | 0,69 | 0,74 | 2,51 | 0,30 | 7,67E-01 |
| AT4G28200 | 0,59 | -1,62 | 2,78 | -0,58 | 5,61E-01 |
| AT4G28230 | 0,33 | -2,60 | 4,41 | -0,59 | 5,55E-01 |
| AT4G28240 | 0,66 | 0,70 | 2,19 | 0,32 | 7,49E-01 |
| AT4G28320 | 0,20 | -2,55 | 4,40 | -0,58 | 5,62E-01 |

| | | | | | |
|---|---|---|---|---|---|
| AT4G28390 | 0,33 | -2,59 | 4,41 | -0,59 | 5,56E-01 |
| AT4G28450 | 0,61 | -4,76 | 4,30 | -1,11 | 2,68E-01 |
| AT4G28470 | 0,62 | -2,13 | 3,10 | -0,69 | 4,91E-01 |
| AT4G28610 | 0,93 | -1,53 | 2,53 | -0,60 | 5,46E-01 |
| AT4G28630 | 0,12 | 0,00 | 4,41 | 0,00 | 1,00E+00 |
| AT4G28770 | 0,20 | 0,47 | 4,36 | 0,11 | 9,14E-01 |
| AT4G28910 | 1,06 | -4,23 | 2,40 | -1,76 | 7,77E-02 |
| AT4G29050 | 0,13 | -0,18 | 4,39 | -0,04 | 9,67E-01 |
| AT4G29080 | 0,28 | -0,71 | 4,35 | -0,16 | 8,70E-01 |
| AT4G29130 | 1,04 | -2,76 | 1,95 | -1,41 | 1,57E-01 |
| AT4G29220 | 0,63 | -1,25 | 3,12 | -0,40 | 6,89E-01 |
| AT4G29260 | 0,48 | 2,41 | 3,94 | 0,61 | 5,41E-01 |
| AT4G29300 | 1,28 | 1,25 | 1,70 | 0,74 | 4,61E-01 |
| AT4G29305 | 0,76 | 1,93 | 2,29 | 0,84 | 3,99E-01 |
| AT4G29440 | 0,66 | -0,95 | 2,90 | -0,33 | 7,43E-01 |
| AT4G29520 | 0,77 | -1,46 | 2,53 | -0,58 | 5,63E-01 |
| AT4G29654 | 1,07 | 2,92 | 2,57 | 1,14 | 2,56E-01 |
| AT4G29670 | 0,65 | -0,80 | 2,22 | -0,36 | 7,17E-01 |
| AT4G29680 | 0,85 | -2,28 | 2,99 | -0,76 | 4,46E-01 |
| AT4G29720 | 0,17 | 0,00 | 4,41 | 0,00 | 1,00E+00 |
| AT4G29735 | 0,17 | 0,00 | 4,41 | 0,00 | 1,00E+00 |
| AT4G29810 | 0,27 | 0,15 | 4,37 | 0,03 | 9,73E-01 |
| AT4G29820 | 0,24 | -0,79 | 4,36 | -0,18 | 8,56E-01 |
| AT4G29840 | 0,58 | -1,13 | 2,30 | -0,49 | 6,24E-01 |
| AT4G29880 | 0,21 | 0,08 | 4,37 | 0,02 | 9,85E-01 |
| AT4G29905 | 1,17 | 2,05 | 1,82 | 1,12 | 2,61E-01 |
| AT4G29910 | 0,08 | -0,21 | 4,39 | -0,05 | 9,62E-01 |
| AT4G29920 | 0,05 | -0,24 | 4,38 | -0,05 | 9,57E-01 |
| AT4G29940 | 0,28 | -3,07 | 4,38 | -0,70 | 4,84E-01 |
| AT4G29960 | 0,50 | -3,56 | 3,95 | -0,90 | 3,68E-01 |
| AT4G29990 | 0,08 | 0,00 | 4,41 | 0,00 | 1,00E+00 |
| AT4G30000 | 0,38 | -3,70 | 4,35 | -0,85 | 3,95E-01 |
| AT4G30030 | 0,45 | -0,52 | 4,32 | -0,12 | 9,04E-01 |
| AT4G30060 | 0,08 | 0,00 | 4,41 | 0,00 | 1,00E+00 |
| AT4G30080 | 0,31 | -0,72 | 4,33 | -0,17 | 8,67E-01 |
| AT4G30110 | 1,05 | -3,33 | 3,22 | -1,03 | 3,02E-01 |
| AT4G30240 | 0,30 | -1,27 | 4,36 | -0,29 | 7,71E-01 |
| AT4G30260 | 0,47 | -1,72 | 3,26 | -0,53 | 5,98E-01 |
| AT4G30280 | 0,12 | 0,07 | 4,38 | 0,02 | 9,87E-01 |
| AT4G30360 | 0,21 | 1,49 | 4,30 | 0,35 | 7,29E-01 |
| AT4G30410 | 0,12 | -0,14 | 4,38 | -0,03 | 9,74E-01 |
| AT4G30430 | 0,08 | 0,00 | 4,41 | 0,00 | 1,00E+00 |
| AT4G30560 | 0,74 | -4,28 | 3,49 | -1,23 | 2,20E-01 |
| AT4G30580 | 0,92 | -3,25 | 3,47 | -0,94 | 3,49E-01 |
| AT4G30620 | 0,23 | -1,14 | 4,39 | -0,26 | 7,96E-01 |
| AT4G30700 | 0,08 | 0,00 | 4,41 | 0,00 | 1,00E+00 |
| AT4G30710 | 0,12 | 0,00 | 4,41 | 0,00 | 1,00E+00 |
| AT4G30720 | 1,29 | -2,29 | 1,66 | -1,38 | 1,68E-01 |
| AT4G30740 | 1,03 | 3,42 | 2,34 | 1,46 | 1,44E-01 |
| AT4G30790 | 0,99 | 0,59 | 2,24 | 0,26 | 7,92E-01 |
| AT4G30890 | 0,60 | -1,95 | 2,95 | -0,66 | 5,08E-01 |
| AT4G30920 | 1,19 | -1,28 | 1,92 | -0,67 | 5,05E-01 |
| AT4G31020 | 0,29 | -2,97 | 4,38 | -0,68 | 4,98E-01 |
| AT4G31120 | 0,87 | -3,28 | 2,43 | -1,35 | 1,77E-01 |
| AT4G31130 | 0,59 | 0,45 | 3,19 | 0,14 | 8,87E-01 |
| AT4G31420 | 0,28 | -1,32 | 4,38 | -0,30 | 7,63E-01 |

| | | | | | |
|---|---|---|---|---|---|
| AT4G31450 | 0,46 | -2,60 | 4,21 | -0,62 | 5,37E-01 |
| AT4G31540 | 0,43 | 1,21 | 4,15 | 0,29 | 7,71E-01 |
| AT4G31550 | 1,02 | -1,56 | 1,71 | -0,91 | 3,63E-01 |
| AT4G31590 | 0,63 | -0,24 | 2,62 | -0,09 | 9,26E-01 |
| AT4G31810 | 0,08 | 0,00 | 4,41 | 0,00 | 1,00E+00 |
| AT4G31820 | 0,43 | -0,91 | 2,85 | -0,32 | 7,48E-01 |
| AT4G32070 | 0,58 | -3,14 | 3,87 | -0,81 | 4,18E-01 |
| AT4G32120 | 0,15 | 0,05 | 4,37 | 0,01 | 9,91E-01 |
| AT4G32130 | 0,18 | 0,78 | 4,34 | 0,18 | 8,57E-01 |
| AT4G32150 | 1,17 | -1,01 | 1,57 | -0,64 | 5,19E-01 |
| AT4G32180 | 1,00 | -3,42 | 2,63 | -1,30 | 1,92E-01 |
| AT4G32208 | 0,37 | -3,33 | 4,32 | -0,77 | 4,41E-01 |
| AT4G32340 | 1,02 | 0,33 | 1,71 | 0,19 | 8,46E-01 |
| AT4G32420 | 0,78 | -1,64 | 2,63 | -0,63 | 5,32E-01 |
| AT4G32460 | 0,31 | -0,35 | 4,38 | -0,08 | 9,37E-01 |
| AT4G32470 | 0,29 | -0,02 | 4,32 | 0,00 | 9,96E-01 |
| AT4G32510 | 0,56 | -2,60 | 3,84 | -0,68 | 4,98E-01 |
| AT4G32520 | 1,00 | -1,18 | 1,96 | -0,60 | 5,46E-01 |
| AT4G32530 | 0,89 | -2,52 | 2,45 | -1,03 | 3,04E-01 |
| AT4G32670 | 0,15 | -0,81 | 4,39 | -0,18 | 8,54E-01 |
| AT4G32750 | 0,18 | -0,68 | 4,41 | -0,15 | 8,77E-01 |
| AT4G32820 | 0,83 | -2,22 | 2,74 | -0,81 | 4,18E-01 |
| AT4G32880 | 1,09 | -2,47 | 1,89 | -1,30 | 1,93E-01 |
| AT4G32900 | 0,12 | -0,68 | 4,41 | -0,15 | 8,77E-01 |
| AT4G32920 | 0,70 | -0,55 | 2,86 | -0,19 | 8,47E-01 |
| AT4G33040 | 0,37 | -1,66 | 4,38 | -0,38 | 7,04E-01 |
| AT4G33090 | 0,64 | 1,53 | 3,03 | 0,51 | 6,13E-01 |
| AT4G33110 | 0,17 | 0,25 | 4,36 | 0,06 | 9,55E-01 |
| AT4G33150 | 0,65 | -2,60 | 3,73 | -0,70 | 4,86E-01 |
| AT4G33200 | 0,29 | -2,57 | 4,40 | -0,58 | 5,59E-01 |
| AT4G33210 | 1,11 | -3,53 | 2,00 | -1,77 | 7,73E-02 |
| AT4G33250 | 0,89 | -1,76 | 1,92 | -0,91 | 3,61E-01 |
| AT4G33260 | 0,12 | 0,00 | 4,41 | 0,00 | 1,00E+00 |
| AT4G33270 | 0,12 | 0,07 | 4,38 | 0,02 | 9,87E-01 |
| AT4G33400 | 0,33 | -2,52 | 4,40 | -0,57 | 5,66E-01 |
| AT4G33430 | 1,19 | 0,00 | 1,92 | 0,00 | 9,99E-01 |
| AT4G33470 | 0,67 | -2,23 | 2,96 | -0,75 | 4,51E-01 |
| AT4G33495 | 0,29 | -2,57 | 4,40 | -0,58 | 5,59E-01 |
| AT4G33500 | 0,96 | 0,06 | 2,66 | 0,02 | 9,81E-01 |
| AT4G33585 | 1,20 | -4,08 | 2,87 | -1,42 | 1,55E-01 |
| AT4G33625 | 0,16 | -0,07 | 4,37 | -0,02 | 9,87E-01 |
| AT4G33670 | 0,76 | -2,38 | 2,00 | -1,19 | 2,36E-01 |
| AT4G33680 | 0,20 | -0,57 | 4,39 | -0,13 | 8,97E-01 |
| AT4G33700 | 0,28 | 0,03 | 4,36 | 0,01 | 9,94E-01 |
| AT4G33790 | 0,16 | -0,62 | 4,40 | -0,14 | 8,87E-01 |
| AT4G33860 | 0,68 | -3,98 | 3,62 | -1,10 | 2,71E-01 |
| AT4G33865 | 0,81 | -0,34 | 2,22 | -0,15 | 8,78E-01 |
| AT4G33910 | 0,26 | -2,60 | 4,41 | -0,59 | 5,55E-01 |
| AT4G34020 | 0,41 | 0,00 | 4,41 | 0,00 | 1,00E+00 |
| AT4G34040 | 0,82 | -0,53 | 2,37 | -0,22 | 8,24E-01 |
| AT4G34070 | 0,72 | -2,41 | 2,73 | -0,88 | 3,78E-01 |
| AT4G34090 | 0,71 | 0,00 | 4,41 | 0,00 | 1,00E+00 |
| AT4G34131 | 0,52 | -2,60 | 3,96 | -0,66 | 5,11E-01 |
| AT4G34180 | 0,68 | -0,17 | 2,99 | -0,06 | 9,53E-01 |
| AT4G34332 | 0,08 | 0,00 | 4,41 | 0,00 | 1,00E+00 |
| AT4G34340 | 0,12 | 0,00 | 4,41 | 0,00 | 1,00E+00 |

| | | | | | |
|---|---|---|---|---|---|
| AT4G08985 | 0,45 | 2,16 | 3,16 | 0,68 | 4,95E-01 |
| AT4G34460 | 0,42 | 0,03 | 3,24 | 0,01 | 9,94E-01 |
| AT4G34480 | 1,19 | -1,18 | 1,76 | -0,67 | 5,02E-01 |
| AT4G34610 | 0,37 | -1,70 | 4,33 | -0,39 | 6,94E-01 |
| AT4G34630 | 0,16 | 0,32 | 4,36 | 0,07 | 9,42E-01 |
| AT4G34730 | 0,55 | -0,19 | 2,58 | -0,07 | 9,42E-01 |
| AT4G34740 | 0,69 | -2,41 | 2,90 | -0,83 | 4,06E-01 |
| AT4G34860 | 0,08 | 0,00 | 4,41 | 0,00 | 1,00E+00 |
| AT4G34900 | 0,65 | -4,11 | 3,60 | -1,14 | 2,54E-01 |
| AT4G35060 | 0,12 | 0,00 | 4,41 | 0,00 | 1,00E+00 |
| AT4G35140 | 0,36 | -1,18 | 4,38 | -0,27 | 7,88E-01 |
| AT4G35180 | 0,66 | -1,96 | 2,94 | -0,67 | 5,05E-01 |
| AT4G35230 | 1,11 | -0,37 | 1,80 | -0,21 | 8,37E-01 |
| AT4G35240 | 1,02 | -2,98 | 2,10 | -1,42 | 1,55E-01 |
| AT4G35260 | 0,53 | -0,47 | 3,09 | -0,15 | 8,79E-01 |
| AT4G35320 | 0,51 | 1,22 | 3,08 | 0,40 | 6,92E-01 |
| AT4G35360 | 0,54 | -2,79 | 4,01 | -0,70 | 4,86E-01 |
| AT4G35440 | 0,24 | 0,71 | 4,34 | 0,16 | 8,70E-01 |
| AT4G35570 | 0,29 | -1,50 | 4,39 | -0,34 | 7,33E-01 |
| AT4G35580 | 0,56 | -1,23 | 3,05 | -0,40 | 6,87E-01 |
| AT4G35780 | 0,15 | -0,73 | 4,40 | -0,17 | 8,68E-01 |
| AT4G35785 | 0,84 | -1,64 | 2,94 | -0,56 | 5,77E-01 |
| AT4G35920 | 0,70 | -4,82 | 4,30 | -1,12 | 2,61E-01 |
| AT4G36010 | 0,74 | 0,14 | 2,73 | 0,05 | 9,60E-01 |
| AT4G36030 | 1,21 | 0,62 | 2,03 | 0,31 | 7,59E-01 |
| AT4G36050 | 0,52 | 0,00 | 4,41 | 0,00 | 1,00E+00 |
| AT4G36070 | 0,08 | 0,00 | 4,41 | 0,00 | 1,00E+00 |
| AT4G36270 | 0,14 | -0,11 | 4,37 | -0,03 | 9,80E-01 |
| AT4G36280 | 0,08 | 0,00 | 4,41 | 0,00 | 1,00E+00 |
| AT4G36380 | 0,24 | -0,68 | 4,41 | -0,15 | 8,77E-01 |
| AT4G36390 | 0,60 | -1,16 | 2,48 | -0,47 | 6,40E-01 |
| AT4G36460 | 0,50 | 1,81 | 3,10 | 0,58 | 5,59E-01 |
| AT4G36500 | 0,97 | -0,67 | 2,16 | -0,31 | 7,56E-01 |
| AT4G36520 | 1,04 | 0,81 | 1,82 | 0,44 | 6,57E-01 |
| AT4G36550 | 0,53 | -0,38 | 3,18 | -0,12 | 9,05E-01 |
| AT4G36670 | 0,44 | 1,19 | 3,31 | 0,36 | 7,19E-01 |
| AT4G36690 | 0,56 | -0,02 | 3,25 | -0,01 | 9,94E-01 |
| AT4G36730 | 0,51 | -0,30 | 3,15 | -0,10 | 9,23E-01 |
| AT4G36770 | 0,41 | -1,64 | 4,16 | -0,39 | 6,94E-01 |
| AT4G36800 | 0,60 | -0,83 | 3,09 | -0,27 | 7,87E-01 |
| AT4G36808 | 0,88 | -0,01 | 2,45 | 0,00 | 9,97E-01 |
| AT4G36860 | 0,41 | -1,94 | 4,31 | -0,45 | 6,53E-01 |
| AT4G36890 | 0,60 | -3,11 | 3,86 | -0,80 | 4,21E-01 |
| AT4G36970 | 0,18 | -1,07 | 4,38 | -0,25 | 8,06E-01 |
| AT4G36980 | 0,30 | -2,60 | 4,41 | -0,59 | 5,55E-01 |
| AT4G37100 | 0,62 | -2,21 | 2,99 | -0,74 | 4,60E-01 |
| AT4G37270 | 0,72 | -0,86 | 2,58 | -0,34 | 7,37E-01 |
| AT4G37280 | 0,63 | -4,50 | 3,61 | -1,25 | 2,13E-01 |
| AT4G37370 | 0,86 | -2,56 | 2,54 | -1,01 | 3,13E-01 |
| AT4G37432 | 0,86 | -0,73 | 3,49 | -0,21 | 8,35E-01 |
| AT4G37510 | 0,72 | -1,06 | 2,39 | -0,44 | 6,57E-01 |
| AT4G37540 | 0,41 | 0,20 | 4,33 | 0,05 | 9,63E-01 |
| AT4G37553 | 0,42 | -2,95 | 4,38 | -0,67 | 5,00E-01 |
| AT4G37590 | 0,76 | 1,23 | 2,49 | 0,49 | 6,22E-01 |
| AT4G37610 | 0,33 | -1,07 | 3,61 | -0,29 | 7,68E-01 |
| AT4G37740 | 0,08 | -0,21 | 4,39 | -0,05 | 9,62E-01 |

| | | | | | |
|---|---|---|---|---|---|
| AT4G37800 | 0,23 | 1,25 | 4,31 | 0,29 | 7,72E-01 |
| AT4G37830 | 0,60 | -1,47 | 2,53 | -0,58 | 5,62E-01 |
| AT4G37850 | 0,24 | -3,18 | 4,39 | -0,73 | 4,68E-01 |
| AT4G37870 | 0,70 | 0,05 | 2,22 | 0,02 | 9,83E-01 |
| AT4G37920 | 0,32 | -1,71 | 4,40 | -0,39 | 6,97E-01 |
| AT4G38010 | 0,08 | 0,00 | 4,41 | 0,00 | 1,00E+00 |
| AT4G38050 | 0,35 | 0,00 | 4,41 | 0,00 | 1,00E+00 |
| AT4G38120 | 0,89 | -2,30 | 2,19 | -1,05 | 2,95E-01 |
| AT4G38130 | 0,71 | 0,17 | 2,01 | 0,08 | 9,33E-01 |
| AT4G38180 | 0,60 | -1,62 | 2,29 | -0,71 | 4,80E-01 |
| AT4G38210 | 0,75 | -3,01 | 4,38 | -0,69 | 4,92E-01 |
| AT4G38230 | 0,40 | -1,71 | 3,38 | -0,51 | 6,12E-01 |
| AT4G38240 | 0,62 | -2,58 | 3,75 | -0,69 | 4,92E-01 |
| AT4G09695 | 1,25 | -1,79 | 1,61 | -1,12 | 2,65E-01 |
| AT4G38300 | 0,68 | -3,13 | 3,64 | -0,86 | 3,90E-01 |
| AT4G38350 | 1,14 | -1,50 | 2,02 | -0,74 | 4,58E-01 |
| AT4G38380 | 0,43 | -2,60 | 4,19 | -0,62 | 5,35E-01 |
| AT4G38520 | 0,55 | -0,65 | 2,24 | -0,29 | 7,70E-01 |
| AT4G38545 | 0,17 | 0,25 | 4,36 | 0,06 | 9,55E-01 |
| AT4G38570 | 0,81 | -2,28 | 2,88 | -0,79 | 4,30E-01 |
| AT4G38630 | 0,74 | -1,63 | 2,06 | -0,79 | 4,27E-01 |
| AT4G38640 | 0,43 | -1,31 | 3,34 | -0,39 | 6,94E-01 |
| AT4G38660 | 0,37 | -1,05 | 4,33 | -0,24 | 8,09E-01 |
| AT4G38670 | 0,49 | -3,37 | 3,21 | -1,05 | 2,94E-01 |
| AT4G38680 | 1,18 | 1,58 | 1,73 | 0,91 | 3,61E-01 |
| AT4G38700 | 0,34 | 0,28 | 4,30 | 0,07 | 9,48E-01 |
| AT4G38710 | 1,00 | 1,35 | 1,70 | 0,79 | 4,29E-01 |
| AT4G38730 | 0,38 | -3,33 | 4,37 | -0,76 | 4,46E-01 |
| AT4G38740 | 1,15 | 0,68 | 1,76 | 0,38 | 7,02E-01 |
| AT4G38780 | 1,10 | 0,24 | 2,63 | 0,09 | 9,27E-01 |
| AT4G38800 | 0,65 | 0,34 | 2,53 | 0,13 | 8,94E-01 |
| AT4G38810 | 0,68 | -0,64 | 2,61 | -0,25 | 8,06E-01 |
| AT4G38870 | 0,21 | -0,73 | 4,40 | -0,17 | 8,67E-01 |
| AT4G38890 | 0,48 | 0,26 | 3,27 | 0,08 | 9,37E-01 |
| AT4G38900 | 0,33 | -1,63 | 4,34 | -0,38 | 7,07E-01 |
| AT4G38932 | 0,58 | -1,84 | 2,68 | -0,69 | 4,92E-01 |
| AT4G38950 | 1,10 | -3,85 | 2,02 | -1,91 | 5,65E-02 |
| AT4G39100 | 0,33 | -2,53 | 4,40 | -0,57 | 5,65E-01 |
| AT4G39120 | 0,19 | -0,65 | 4,40 | -0,15 | 8,83E-01 |
| AT4G39140 | 0,65 | -1,42 | 3,81 | -0,37 | 7,09E-01 |
| AT4G39180 | 0,48 | -0,30 | 4,38 | -0,07 | 9,45E-01 |
| AT4G39190 | 0,69 | 1,79 | 2,37 | 0,75 | 4,51E-01 |
| AT4G39195 | 0,79 | -1,64 | 3,03 | -0,54 | 5,89E-01 |
| AT4G39280 | 1,06 | 1,88 | 1,80 | 1,04 | 2,98E-01 |
| AT4G39460 | 0,85 | -3,82 | 2,45 | -1,56 | 1,19E-01 |
| AT4G39500 | 0,20 | -2,55 | 4,40 | -0,58 | 5,62E-01 |
| AT4G39510 | 0,43 | -2,49 | 4,39 | -0,57 | 5,71E-01 |
| AT4G39520 | 1,02 | -1,32 | 1,55 | -0,85 | 3,96E-01 |
| AT4G39550 | 0,17 | 0,00 | 4,41 | 0,00 | 1,00E+00 |
| AT4G39560 | 0,59 | -4,08 | 3,77 | -1,08 | 2,79E-01 |
| AT4G39570 | 0,55 | 0,00 | 3,84 | 0,00 | 1,00E+00 |
| AT4G39580 | 0,15 | -0,81 | 4,39 | -0,18 | 8,54E-01 |
| AT4G39590 | 0,08 | 0,00 | 4,41 | 0,00 | 1,00E+00 |
| AT4G39630 | 0,05 | -0,24 | 4,38 | -0,05 | 9,57E-01 |
| AT4G39680 | 0,30 | 1,51 | 4,30 | 0,35 | 7,25E-01 |
| AT4G39710 | 0,56 | 0,12 | 3,88 | 0,03 | 9,76E-01 |

| | | | | | |
|---|---|---|---|---|---|
| AT4G39780 | 0,21 | -0,44 | 4,38 | -0,10 | 9,20E-01 |
| AT4G39820 | 0,46 | -1,61 | 4,12 | -0,39 | 6,97E-01 |
| AT4G39838 | 1,07 | -0,16 | 1,77 | -0,09 | 9,27E-01 |
| AT4G39865 | 0,25 | -3,06 | 4,38 | -0,70 | 4,85E-01 |
| AT4G39910 | 0,28 | -0,29 | 4,38 | -0,07 | 9,48E-01 |
| AT4G40000 | 0,43 | -2,70 | 4,18 | -0,65 | 5,19E-01 |
| AT4G40020 | 0,75 | -4,46 | 2,76 | -1,62 | 1,05E-01 |
| AT4G40045 | 0,12 | 0,00 | 4,41 | 0,00 | 1,00E+00 |
| AT4G40060 | 1,22 | 2,14 | 1,82 | 1,17 | 2,41E-01 |
| AT4G40065 | 1,04 | -2,18 | 1,86 | -1,17 | 2,41E-01 |
| AT5G01010 | 1,03 | -2,66 | 2,40 | -1,11 | 2,67E-01 |
| AT5G01100 | 0,27 | -0,35 | 4,34 | -0,08 | 9,37E-01 |
| AT5G01215 | 1,09 | 1,51 | 1,78 | 0,85 | 3,97E-01 |
| AT5G01220 | 0,12 | 0,00 | 4,41 | 0,00 | 1,00E+00 |
| AT5G01270 | 0,34 | -1,64 | 4,41 | -0,37 | 7,10E-01 |
| AT5G01365 | 0,66 | 0,00 | 4,41 | 0,00 | 1,00E+00 |
| AT5G01540 | 0,46 | -1,29 | 4,07 | -0,32 | 7,51E-01 |
| AT5G01630 | 1,03 | -2,17 | 2,63 | -0,82 | 4,10E-01 |
| AT5G01700 | 0,62 | -3,72 | 4,36 | -0,85 | 3,94E-01 |
| AT5G01720 | 0,49 | 0,41 | 3,96 | 0,10 | 9,18E-01 |
| AT5G01730 | 0,29 | -2,54 | 4,40 | -0,58 | 5,63E-01 |
| AT5G01770 | 0,54 | -3,71 | 3,81 | -0,97 | 3,30E-01 |
| AT5G01780 | 0,19 | 0,49 | 4,34 | 0,11 | 9,10E-01 |
| AT5G01890 | 0,64 | -0,23 | 2,62 | -0,09 | 9,31E-01 |
| AT5G01930 | 1,04 | -5,28 | 3,04 | -1,73 | 8,29E-02 |
| AT5G01950 | 0,59 | -1,72 | 2,59 | -0,66 | 5,07E-01 |
| AT5G01970 | 1,20 | -2,47 | 2,22 | -1,11 | 2,66E-01 |
| AT5G01990 | 1,12 | -4,11 | 2,13 | -1,93 | 5,33E-02 |
| AT5G02130 | 0,08 | -0,21 | 4,39 | -0,05 | 9,62E-01 |
| AT5G02290 | 0,48 | -1,36 | 2,70 | -0,50 | 6,15E-01 |
| AT5G02502 | 0,33 | 0,37 | 4,35 | 0,09 | 9,31E-01 |
| AT5G02530 | 0,37 | -1,68 | 4,40 | -0,38 | 7,03E-01 |
| AT5G02670 | 0,44 | -0,86 | 3,30 | -0,26 | 7,95E-01 |
| AT5G02770 | 0,75 | -0,54 | 2,22 | -0,24 | 8,09E-01 |
| AT5G02800 | 0,65 | -3,43 | 2,93 | -1,17 | 2,42E-01 |
| AT5G02830 | 0,85 | 0,63 | 1,96 | 0,32 | 7,48E-01 |
| AT5G02890 | 0,18 | 0,40 | 4,35 | 0,09 | 9,28E-01 |
| AT5G02940 | 0,88 | 0,83 | 2,42 | 0,34 | 7,32E-01 |
| AT5G03140 | 0,48 | -0,34 | 2,69 | -0,13 | 8,99E-01 |
| AT5G03170 | 0,32 | -3,05 | 4,38 | -0,70 | 4,86E-01 |
| AT5G03204 | 0,29 | 0,39 | 4,35 | 0,09 | 9,28E-01 |
| AT5G03210 | 0,71 | 1,68 | 2,41 | 0,69 | 4,88E-01 |
| AT5G03380 | 0,84 | 1,24 | 2,35 | 0,53 | 5,97E-01 |
| AT5G03430 | 0,65 | -0,90 | 3,02 | -0,30 | 7,66E-01 |
| AT5G03450 | 0,39 | -2,31 | 4,39 | -0,53 | 5,98E-01 |
| AT5G03555 | 1,08 | -2,21 | 2,08 | -1,06 | 2,87E-01 |
| AT5G03560 | 0,43 | -0,63 | 4,12 | -0,15 | 8,79E-01 |
| AT5G03630 | 0,55 | 0,13 | 3,13 | 0,04 | 9,67E-01 |
| AT5G03700 | 0,11 | -0,68 | 4,41 | -0,15 | 8,77E-01 |
| AT5G03730 | 0,62 | -1,63 | 2,68 | -0,61 | 5,42E-01 |
| AT5G03760 | 0,44 | -1,05 | 4,12 | -0,26 | 7,99E-01 |
| AT5G03800 | 0,30 | -2,60 | 4,41 | -0,59 | 5,55E-01 |
| AT5G03850 | 0,83 | -1,89 | 2,87 | -0,66 | 5,11E-01 |
| AT5G03880 | 0,57 | 1,12 | 3,18 | 0,35 | 7,25E-01 |
| AT5G03910 | 0,83 | -1,50 | 2,47 | -0,61 | 5,45E-01 |
| AT5G03970 | 0,19 | -0,79 | 4,39 | -0,18 | 8,58E-01 |

| | | | | | |
|---|---|---|---|---|---|
| AT5G04020 | 0,78 | -1,04 | 1,95 | -0,53 | 5,94E-01 |
| AT5G04130 | 0,05 | -0,24 | 4,38 | -0,05 | 9,57E-01 |
| AT5G04267 | 0,16 | 0,69 | 4,35 | 0,16 | 8,75E-01 |
| AT5G04340 | 1,00 | -0,94 | 1,96 | -0,48 | 6,32E-01 |
| AT5G00955 | 1,17 | 1,45 | 1,87 | 0,78 | 4,37E-01 |
| AT5G04420 | 0,49 | -3,80 | 3,96 | -0,96 | 3,37E-01 |
| AT5G04470 | 0,16 | -0,26 | 4,38 | -0,06 | 9,52E-01 |
| AT5G04480 | 0,41 | 0,08 | 3,27 | 0,02 | 9,82E-01 |
| AT5G04700 | 0,30 | -1,64 | 4,41 | -0,37 | 7,10E-01 |
| AT5G04710 | 0,12 | 0,00 | 4,41 | 0,00 | 1,00E+00 |
| AT5G04730 | 1,06 | -3,57 | 2,14 | -1,67 | 9,46E-02 |
| AT5G04760 | 0,48 | -2,60 | 4,41 | -0,59 | 5,55E-01 |
| AT5G04830 | 0,60 | -0,74 | 3,17 | -0,23 | 8,15E-01 |
| AT5G04850 | 0,17 | -0,75 | 4,40 | -0,17 | 8,65E-01 |
| AT5G04860 | 0,46 | -2,41 | 4,14 | -0,58 | 5,60E-01 |
| AT5G04870 | 0,78 | -0,90 | 2,21 | -0,41 | 6,85E-01 |
| AT5G04940 | 0,64 | -2,48 | 2,90 | -0,86 | 3,92E-01 |
| AT5G04950 | 0,11 | 0,46 | 4,35 | 0,11 | 9,15E-01 |
| AT5G04990 | 0,74 | -3,85 | 3,40 | -1,13 | 2,56E-01 |
| AT5G05060 | 0,37 | 0,00 | 4,41 | 0,00 | 1,00E+00 |
| AT5G05080 | 0,77 | -1,60 | 4,33 | -0,37 | 7,12E-01 |
| AT5G05085 | 0,35 | 0,09 | 3,52 | 0,03 | 9,79E-01 |
| AT5G05100 | 0,72 | -4,11 | 2,94 | -1,40 | 1,62E-01 |
| AT5G01005 | 0,75 | -2,56 | 2,97 | -0,86 | 3,87E-01 |
| AT5G05260 | 0,23 | 0,50 | 4,34 | 0,12 | 9,08E-01 |
| AT5G01055 | 0,10 | 0,45 | 4,35 | 0,10 | 9,19E-01 |
| AT5G05480 | 0,31 | -0,95 | 3,62 | -0,26 | 7,94E-01 |
| AT5G05670 | 0,17 | -0,80 | 4,39 | -0,18 | 8,56E-01 |
| AT5G05700 | 0,69 | -1,72 | 3,03 | -0,57 | 5,71E-01 |
| AT5G05730 | 0,58 | -3,18 | 3,16 | -1,00 | 3,15E-01 |
| AT5G01095 | 0,95 | 0,00 | 4,41 | 0,00 | 1,00E+00 |
| AT5G05800 | 0,36 | 0,00 | 4,41 | 0,00 | 1,00E+00 |
| AT5G06043 | 0,95 | -0,05 | 2,65 | -0,02 | 9,84E-01 |
| AT5G06060 | 0,53 | -0,43 | 3,20 | -0,13 | 8,94E-01 |
| AT5G06110 | 0,69 | -3,11 | 2,90 | -1,07 | 2,83E-01 |
| AT5G06140 | 0,36 | -2,93 | 4,37 | -0,67 | 5,02E-01 |
| AT5G06160 | 0,43 | -0,45 | 2,75 | -0,16 | 8,70E-01 |
| AT5G01165 | 0,96 | 1,22 | 1,97 | 0,62 | 5,35E-01 |
| AT5G06230 | 0,49 | -3,32 | 3,92 | -0,85 | 3,96E-01 |
| AT5G06260 | 0,46 | -1,05 | 3,29 | -0,32 | 7,49E-01 |
| AT5G06330 | 0,08 | -0,21 | 4,39 | -0,05 | 9,62E-01 |
| AT5G06410 | 0,25 | -2,61 | 4,41 | -0,59 | 5,53E-01 |
| AT5G06430 | 0,51 | -3,74 | 4,35 | -0,86 | 3,90E-01 |
| AT5G06440 | 0,83 | -4,90 | 3,50 | -1,40 | 1,62E-01 |
| AT5G06520 | 0,44 | 0,00 | 4,41 | 0,00 | 1,00E+00 |
| AT5G06540 | 0,48 | 0,00 | 4,41 | 0,00 | 1,00E+00 |
| AT5G06560 | 0,78 | -1,73 | 2,18 | -0,79 | 4,28E-01 |
| AT5G06670 | 0,24 | 0,41 | 4,35 | 0,09 | 9,25E-01 |
| AT5G06690 | 0,35 | 0,96 | 3,52 | 0,27 | 7,86E-01 |
| AT5G06750 | 0,11 | -0,80 | 4,39 | -0,18 | 8,56E-01 |
| AT5G06770 | 0,36 | 0,60 | 4,34 | 0,14 | 8,90E-01 |
| AT5G06865 | 0,22 | 0,62 | 4,34 | 0,14 | 8,86E-01 |
| AT5G06970 | 0,87 | -1,24 | 1,98 | -0,63 | 5,30E-01 |
| AT5G01295 | 0,28 | -3,07 | 4,38 | -0,70 | 4,84E-01 |
| AT5G07120 | 0,92 | -1,36 | 2,32 | -0,59 | 5,57E-01 |
| AT5G07220 | 0,71 | 1,53 | 2,22 | 0,69 | 4,91E-01 |

| | | | | | |
|---|---|---|---|---|---|
| AT5G07250 | 0,62 | -1,64 | 3,12 | -0,53 | 5,98E-01 |
| AT5G07360 | 0,50 | 0,77 | 3,23 | 0,24 | 8,12E-01 |
| AT5G07590 | 0,15 | -0,77 | 4,39 | -0,18 | 8,61E-01 |
| AT5G07620 | 0,13 | -0,18 | 4,39 | -0,04 | 9,67E-01 |
| AT5G07675 | 0,20 | 0,94 | 4,32 | 0,22 | 8,28E-01 |
| AT5G07810 | 0,26 | -2,60 | 4,41 | -0,59 | 5,55E-01 |
| AT5G07950 | 0,39 | -3,10 | 4,38 | -0,71 | 4,79E-01 |
| AT5G08040 | 0,14 | 0,00 | 4,41 | 0,00 | 1,00E+00 |
| AT5G08075 | 0,48 | -0,66 | 4,01 | -0,17 | 8,69E-01 |
| AT5G08110 | 0,94 | -4,47 | 2,33 | -1,92 | 5,46E-02 |
| AT5G08130 | 1,09 | 1,46 | 2,02 | 0,72 | 4,69E-01 |
| AT5G08160 | 1,05 | -2,46 | 1,96 | -1,25 | 2,10E-01 |
| AT5G08170 | 0,71 | -3,35 | 3,57 | -0,94 | 3,47E-01 |
| AT5G01495 | 0,69 | -0,07 | 1,89 | -0,04 | 9,72E-01 |
| AT5G01505 | 1,02 | 0,81 | 1,63 | 0,49 | 6,21E-01 |
| AT5G01515 | 0,11 | -0,68 | 4,41 | -0,15 | 8,77E-01 |
| AT5G08230 | 0,62 | -1,43 | 3,07 | -0,47 | 6,42E-01 |
| AT5G08305 | 0,42 | -1,61 | 4,21 | -0,38 | 7,03E-01 |
| AT5G08315 | 0,44 | -3,34 | 4,32 | -0,77 | 4,39E-01 |
| AT5G08350 | 0,26 | -2,60 | 4,41 | -0,59 | 5,55E-01 |
| AT5G08450 | 0,66 | 0,93 | 2,81 | 0,33 | 7,42E-01 |
| AT5G08560 | 0,55 | 0,40 | 2,71 | 0,15 | 8,83E-01 |
| AT5G08650 | 1,18 | -1,52 | 1,87 | -0,81 | 4,16E-01 |
| AT5G08770 | 1,04 | 0,88 | 2,09 | 0,42 | 6,75E-01 |
| AT5G09330 | 1,10 | -0,97 | 1,53 | -0,63 | 5,27E-01 |
| AT5G09443 | 0,26 | -2,60 | 4,41 | -0,59 | 5,55E-01 |
| AT5G09460 | 0,93 | -2,38 | 1,89 | -1,26 | 2,08E-01 |
| AT5G09510 | 0,57 | -0,54 | 4,02 | -0,14 | 8,92E-01 |
| AT5G09620 | 0,88 | 2,09 | 2,34 | 0,89 | 3,73E-01 |
| AT5G09650 | 0,62 | -0,44 | 2,60 | -0,17 | 8,66E-01 |
| AT5G09820 | 0,63 | -1,87 | 3,00 | -0,62 | 5,33E-01 |
| AT5G09850 | 0,67 | 0,23 | 3,05 | 0,08 | 9,39E-01 |
| AT5G09880 | 0,62 | 0,84 | 2,96 | 0,28 | 7,77E-01 |
| AT5G09890 | 0,52 | 0,35 | 3,93 | 0,09 | 9,30E-01 |
| AT5G09930 | 0,08 | 0,00 | 4,41 | 0,00 | 1,00E+00 |
| AT5G10030 | 0,52 | -1,67 | 4,01 | -0,42 | 6,77E-01 |
| AT5G10235 | 0,28 | -2,60 | 4,41 | -0,59 | 5,55E-01 |
| AT5G10290 | 0,76 | 0,53 | 2,44 | 0,22 | 8,29E-01 |
| AT5G10330 | 0,57 | -0,37 | 2,69 | -0,14 | 8,91E-01 |
| AT5G10350 | 0,66 | -2,93 | 4,38 | -0,67 | 5,03E-01 |
| AT5G10390 | 0,27 | -0,23 | 4,33 | -0,05 | 9,58E-01 |
| AT5G10400 | 0,23 | 1,59 | 4,30 | 0,37 | 7,11E-01 |
| AT5G10470 | 0,57 | -1,91 | 3,20 | -0,60 | 5,50E-01 |
| AT5G10490 | 0,10 | -0,34 | 4,38 | -0,08 | 9,38E-01 |
| AT5G10550 | 0,86 | 0,67 | 2,36 | 0,28 | 7,76E-01 |
| AT5G10605 | 0,65 | -1,45 | 3,01 | -0,48 | 6,30E-01 |
| AT5G10650 | 0,25 | -2,53 | 4,40 | -0,57 | 5,65E-01 |
| AT5G10690 | 0,38 | -0,29 | 4,34 | -0,07 | 9,47E-01 |
| AT5G01815 | 0,80 | 2,36 | 2,07 | 1,14 | 2,55E-01 |
| AT5G10700 | 0,56 | -0,41 | 3,11 | -0,13 | 8,94E-01 |
| AT5G10730 | 0,23 | 0,11 | 4,38 | 0,03 | 9,80E-01 |
| AT5G10770 | 0,94 | 0,92 | 2,24 | 0,41 | 6,80E-01 |
| AT5G10800 | 0,30 | -2,60 | 4,41 | -0,59 | 5,55E-01 |
| AT5G10820 | 0,63 | 0,00 | 4,41 | 0,00 | 1,00E+00 |
| AT5G10930 | 1,14 | -2,19 | 1,67 | -1,31 | 1,91E-01 |
| AT5G10940 | 0,77 | -2,98 | 3,60 | -0,83 | 4,08E-01 |

| | | | | | |
|---|---|---|---|---|---|
| AT5G10965 | 0,75 | -3,51 | 2,37 | -1,48 | 1,38E-01 |
| AT5G10980 | 0,55 | -0,19 | 3,82 | -0,05 | 9,60E-01 |
| AT5G11000 | 0,45 | -0,91 | 4,17 | -0,22 | 8,27E-01 |
| AT5G11070 | 1,00 | 0,35 | 1,95 | 0,18 | 8,55E-01 |
| AT5G11180 | 0,81 | -0,36 | 2,33 | -0,15 | 8,78E-01 |
| AT5G11210 | 1,06 | -5,06 | 3,13 | -1,62 | 1,06E-01 |
| AT5G11270 | 0,22 | -1,25 | 4,38 | -0,29 | 7,75E-01 |
| AT5G11330 | 1,01 | -0,68 | 1,71 | -0,40 | 6,92E-01 |
| AT5G11480 | 0,46 | 0,44 | 3,25 | 0,14 | 8,92E-01 |
| AT5G11520 | 1,24 | -0,78 | 1,90 | -0,41 | 6,83E-01 |
| AT5G11550 | 0,28 | -1,31 | 4,37 | -0,30 | 7,65E-01 |
| AT5G11770 | 0,54 | 1,24 | 2,73 | 0,45 | 6,50E-01 |
| AT5G11850 | 0,25 | -0,13 | 4,37 | -0,03 | 9,77E-01 |
| AT5G11860 | 0,12 | 0,00 | 4,41 | 0,00 | 1,00E+00 |
| AT5G11910 | 0,27 | -1,72 | 4,38 | -0,39 | 6,95E-01 |
| AT5G11960 | 0,36 | 0,00 | 4,41 | 0,00 | 1,00E+00 |
| AT5G12010 | 0,59 | -2,07 | 3,08 | -0,67 | 5,01E-01 |
| AT5G12080 | 0,25 | -2,57 | 4,40 | -0,58 | 5,60E-01 |
| AT5G12140 | 0,22 | 0,37 | 4,35 | 0,09 | 9,32E-01 |
| AT5G12200 | 0,44 | -0,64 | 3,36 | -0,19 | 8,48E-01 |
| AT5G12310 | 0,42 | 0,33 | 3,30 | 0,10 | 9,19E-01 |
| AT5G12390 | 0,11 | -0,68 | 4,41 | -0,15 | 8,77E-01 |
| AT5G12430 | 1,28 | -5,50 | 2,28 | -2,41 | 1,61E-02 |
| AT5G12440 | 0,88 | -4,56 | 2,67 | -1,71 | 8,75E-02 |
| AT5G12480 | 0,34 | 0,77 | 4,34 | 0,18 | 8,59E-01 |
| AT5G12890 | 0,77 | -4,10 | 3,63 | -1,13 | 2,58E-01 |
| AT5G12900 | 0,23 | 0,29 | 4,36 | 0,07 | 9,46E-01 |
| AT5G13020 | 0,81 | -3,16 | 2,21 | -1,43 | 1,53E-01 |
| AT5G13050 | 0,42 | -0,99 | 4,17 | -0,24 | 8,12E-01 |
| AT5G13090 | 0,10 | -0,18 | 4,38 | -0,04 | 9,67E-01 |
| AT5G13120 | 0,17 | -0,68 | 4,41 | -0,15 | 8,77E-01 |
| AT5G13140 | 0,23 | 0,09 | 4,37 | 0,02 | 9,83E-01 |
| AT5G13170 | 0,48 | -2,60 | 4,41 | -0,59 | 5,55E-01 |
| AT5G13190 | 0,99 | 0,24 | 1,81 | 0,13 | 8,94E-01 |
| AT5G02055 | 0,48 | 1,15 | 2,74 | 0,42 | 6,75E-01 |
| AT5G13225 | 0,48 | 1,15 | 2,74 | 0,42 | 6,75E-01 |
| AT5G13270 | 0,40 | -2,52 | 4,18 | -0,60 | 5,46E-01 |
| AT5G13300 | 0,77 | -1,99 | 2,46 | -0,81 | 4,18E-01 |
| AT5G13400 | 0,43 | -0,76 | 4,40 | -0,17 | 8,63E-01 |
| AT5G13430 | 0,54 | -3,44 | 3,83 | -0,90 | 3,68E-01 |
| AT5G13440 | 0,29 | 1,40 | 4,30 | 0,32 | 7,45E-01 |
| AT5G13450 | 0,96 | 0,87 | 2,07 | 0,42 | 6,75E-01 |
| AT5G13550 | 0,46 | -1,81 | 4,05 | -0,45 | 6,55E-01 |
| AT5G13560 | 1,05 | -3,18 | 2,17 | -1,47 | 1,42E-01 |
| AT5G13590 | 1,32 | -3,02 | 1,94 | -1,56 | 1,19E-01 |
| AT5G13680 | 1,03 | -1,27 | 1,73 | -0,73 | 4,62E-01 |
| AT5G13710 | 0,59 | -1,54 | 3,01 | -0,51 | 6,10E-01 |
| AT5G13780 | 0,62 | -1,76 | 3,06 | -0,57 | 5,66E-01 |
| AT5G13800 | 0,74 | -0,01 | 2,39 | -0,01 | 9,96E-01 |
| AT5G13825 | 0,39 | -0,23 | 4,28 | -0,05 | 9,58E-01 |
| AT5G13990 | 0,12 | -0,31 | 4,38 | -0,07 | 9,44E-01 |
| AT5G14100 | 0,11 | -0,68 | 4,41 | -0,15 | 8,77E-01 |
| AT5G14170 | 1,07 | -3,97 | 2,16 | -1,84 | 6,54E-02 |
| AT5G14210 | 0,50 | -0,14 | 3,33 | -0,04 | 9,67E-01 |
| AT5G14240 | 0,57 | -1,26 | 2,62 | -0,48 | 6,32E-01 |
| AT5G14320 | 0,97 | -0,27 | 1,72 | -0,16 | 8,75E-01 |

| | | | | | |
|---|---|---|---|---|---|
| AT5G14390 | 0,52 | -1,52 | 4,07 | -0,37 | 7,09E-01 |
| AT5G14420 | 0,18 | -0,27 | 4,38 | -0,06 | 9,51E-01 |
| AT5G14530 | 0,86 | -1,00 | 2,90 | -0,35 | 7,30E-01 |
| AT5G14540 | 0,54 | -2,77 | 3,91 | -0,71 | 4,78E-01 |
| AT5G02245 | 0,10 | 0,45 | 4,35 | 0,10 | 9,19E-01 |
| AT5G14580 | 1,26 | -3,33 | 2,63 | -1,27 | 2,05E-01 |
| AT5G14590 | 0,98 | -1,40 | 1,98 | -0,71 | 4,78E-01 |
| AT5G14700 | 0,24 | 1,49 | 4,30 | 0,35 | 7,30E-01 |
| AT5G14720 | 0,96 | -0,96 | 1,89 | -0,51 | 6,13E-01 |
| AT5G14770 | 0,20 | 0,00 | 4,41 | 0,00 | 1,00E+00 |
| AT5G14820 | 0,29 | -1,50 | 4,39 | -0,34 | 7,33E-01 |
| AT5G14940 | 1,18 | -0,79 | 2,40 | -0,33 | 7,41E-01 |
| AT5G15050 | 0,10 | -0,34 | 4,38 | -0,08 | 9,38E-01 |
| AT5G15210 | 0,28 | -1,64 | 4,41 | -0,37 | 7,10E-01 |
| AT5G15250 | 0,05 | -0,24 | 4,38 | -0,05 | 9,57E-01 |
| AT5G15360 | 0,52 | 2,07 | 2,53 | 0,82 | 4,14E-01 |
| AT5G15450 | 1,23 | 0,14 | 1,96 | 0,07 | 9,43E-01 |
| AT5G15460 | 0,08 | 0,00 | 4,41 | 0,00 | 1,00E+00 |
| AT5G15470 | 0,21 | -0,75 | 4,40 | -0,17 | 8,65E-01 |
| AT5G15490 | 0,40 | -1,31 | 3,37 | -0,39 | 6,98E-01 |
| AT5G15537 | 0,21 | -2,60 | 4,41 | -0,59 | 5,55E-01 |
| AT5G15540 | 1,01 | -1,97 | 1,99 | -0,99 | 3,22E-01 |
| AT5G15580 | 0,70 | -2,88 | 2,47 | -1,16 | 2,45E-01 |
| AT5G15690 | 0,53 | -0,81 | 2,63 | -0,31 | 7,59E-01 |
| AT5G15700 | 0,61 | 0,26 | 2,29 | 0,11 | 9,11E-01 |
| AT5G15740 | 0,28 | 0,07 | 4,33 | 0,02 | 9,87E-01 |
| AT5G15780 | 0,22 | 0,62 | 4,34 | 0,14 | 8,86E-01 |
| AT5G15800 | 0,08 | 0,05 | 4,38 | 0,01 | 9,91E-01 |
| AT5G15802 | 0,08 | -0,21 | 4,39 | -0,05 | 9,62E-01 |
| AT5G15860 | 0,19 | -0,79 | 4,39 | -0,18 | 8,58E-01 |
| AT5G16050 | 0,47 | 1,03 | 2,75 | 0,37 | 7,09E-01 |
| AT5G16070 | 0,93 | -3,50 | 2,14 | -1,64 | 1,02E-01 |
| AT5G16110 | 1,09 | -2,63 | 1,69 | -1,55 | 1,21E-01 |
| AT5G16200 | 0,36 | -4,02 | 4,33 | -0,93 | 3,54E-01 |
| AT5G16220 | 0,18 | 1,18 | 4,31 | 0,27 | 7,85E-01 |
| AT5G16280 | 0,67 | -4,11 | 2,88 | -1,43 | 1,54E-01 |
| AT5G16360 | 0,08 | 0,00 | 4,41 | 0,00 | 1,00E+00 |
| AT5G16380 | 0,25 | -0,70 | 4,40 | -0,16 | 8,73E-01 |
| AT5G16390 | 0,61 | -3,70 | 3,71 | -1,00 | 3,18E-01 |
| AT5G16480 | 0,56 | -2,52 | 4,36 | -0,58 | 5,63E-01 |
| AT5G16490 | 0,08 | -0,21 | 4,39 | -0,05 | 9,62E-01 |
| AT5G16520 | 0,47 | -0,10 | 4,22 | -0,02 | 9,80E-01 |
| AT5G16560 | 0,42 | 0,55 | 4,35 | 0,13 | 8,99E-01 |
| AT5G16610 | 0,08 | 0,00 | 4,41 | 0,00 | 1,00E+00 |
| AT5G16660 | 0,23 | 0,01 | 4,36 | 0,00 | 9,98E-01 |
| AT5G16680 | 0,78 | -0,59 | 2,15 | -0,28 | 7,82E-01 |
| AT5G16740 | 0,42 | -2,18 | 4,39 | -0,50 | 6,19E-01 |
| AT5G16750 | 0,89 | -1,31 | 2,00 | -0,66 | 5,12E-01 |
| AT5G16780 | 0,54 | 0,57 | 3,87 | 0,15 | 8,84E-01 |
| AT5G16800 | 0,21 | -1,17 | 4,36 | -0,27 | 7,88E-01 |
| AT5G16830 | 0,10 | -0,34 | 4,38 | -0,08 | 9,38E-01 |
| AT5G16980 | 0,08 | 0,00 | 4,41 | 0,00 | 1,00E+00 |
| AT5G16990 | 0,20 | -0,57 | 4,39 | -0,13 | 8,97E-01 |
| AT5G17000 | 0,20 | -1,00 | 4,37 | -0,23 | 8,19E-01 |
| AT5G17010 | 0,24 | 0,12 | 4,37 | 0,03 | 9,77E-01 |
| AT5G02655 | 0,55 | 1,49 | 2,56 | 0,58 | 5,61E-01 |

| | | | | | |
|---|---|---|---|---|---|
| AT5G02665 | 1,25 | -1,56 | 2,01 | -0,78 | 4,37E-01 |
| AT5G02675 | 0,37 | -0,94 | 4,29 | -0,22 | 8,27E-01 |
| AT5G17290 | 0,65 | -0,98 | 3,06 | -0,32 | 7,50E-01 |
| AT5G17300 | 0,69 | -2,98 | 2,96 | -1,01 | 3,14E-01 |
| AT5G17320 | 0,38 | -1,51 | 4,39 | -0,34 | 7,31E-01 |
| AT5G17380 | 0,80 | -0,29 | 2,58 | -0,11 | 9,11E-01 |
| AT5G17420 | 0,40 | -0,68 | 4,41 | -0,15 | 8,77E-01 |
| AT5G17460 | 0,52 | 0,00 | 4,41 | 0,00 | 1,00E+00 |
| AT5G17520 | 0,17 | 0,00 | 4,41 | 0,00 | 1,00E+00 |
| AT5G17550 | 0,08 | 0,00 | 4,41 | 0,00 | 1,00E+00 |
| AT5G17600 | 0,90 | 0,06 | 2,07 | 0,03 | 9,77E-01 |
| AT5G17620 | 0,42 | -2,18 | 4,39 | -0,50 | 6,19E-01 |
| AT5G17640 | 0,35 | -1,10 | 4,33 | -0,25 | 8,00E-01 |
| AT5G17690 | 0,20 | -0,68 | 4,41 | -0,15 | 8,77E-01 |
| AT5G17770 | 0,49 | 0,28 | 3,18 | 0,09 | 9,31E-01 |
| AT5G17860 | 0,63 | 1,67 | 2,82 | 0,59 | 5,55E-01 |
| AT5G17930 | 0,48 | 0,00 | 4,41 | 0,00 | 1,00E+00 |
| AT5G18050 | 0,24 | -3,18 | 4,39 | -0,73 | 4,68E-01 |
| AT5G18110 | 0,39 | 0,62 | 4,34 | 0,14 | 8,87E-01 |
| AT5G18190 | 0,63 | -2,41 | 2,64 | -0,92 | 3,60E-01 |
| AT5G18230 | 0,99 | 0,27 | 2,91 | 0,09 | 9,27E-01 |
| AT5G18260 | 0,49 | -3,73 | 4,36 | -0,86 | 3,92E-01 |
| AT5G18280 | 0,49 | -0,92 | 3,18 | -0,29 | 7,73E-01 |
| AT5G18290 | 0,36 | -3,60 | 4,36 | -0,82 | 4,10E-01 |
| AT5G18340 | 0,58 | -3,74 | 3,92 | -0,95 | 3,40E-01 |
| AT5G18400 | 0,32 | -0,09 | 4,33 | -0,02 | 9,84E-01 |
| AT5G02895 | 0,88 | -0,92 | 2,66 | -0,35 | 7,29E-01 |
| AT5G18475 | 0,21 | 0,00 | 4,41 | 0,00 | 1,00E+00 |
| AT5G18480 | 0,62 | -1,89 | 3,06 | -0,62 | 5,37E-01 |
| AT5G18570 | 0,43 | 0,72 | 4,02 | 0,18 | 8,59E-01 |
| AT5G18590 | 0,48 | -0,24 | 3,26 | -0,07 | 9,41E-01 |
| AT5G18620 | 0,19 | 0,49 | 4,34 | 0,11 | 9,10E-01 |
| AT5G18700 | 0,89 | -3,59 | 2,71 | -1,33 | 1,84E-01 |
| AT5G02935 | 0,05 | -0,24 | 4,38 | -0,05 | 9,57E-01 |
| AT5G19010 | 0,59 | 0,40 | 2,52 | 0,16 | 8,73E-01 |
| AT5G19130 | 0,62 | -1,67 | 2,62 | -0,64 | 5,23E-01 |
| AT5G19220 | 0,51 | -2,28 | 4,32 | -0,53 | 5,98E-01 |
| AT5G19360 | 0,72 | -4,76 | 4,30 | -1,11 | 2,68E-01 |
| AT5G19410 | 0,62 | -4,10 | 2,94 | -1,40 | 1,63E-01 |
| AT5G19420 | 1,09 | -2,23 | 1,77 | -1,26 | 2,08E-01 |
| AT5G19450 | 0,87 | -1,64 | 1,75 | -0,94 | 3,47E-01 |
| AT5G19485 | 0,20 | -2,60 | 4,41 | -0,59 | 5,55E-01 |
| AT5G19590 | 0,42 | 1,66 | 3,20 | 0,52 | 6,04E-01 |
| AT5G19660 | 0,36 | -0,50 | 4,33 | -0,12 | 9,08E-01 |
| AT5G19760 | 0,90 | -2,84 | 2,40 | -1,18 | 2,37E-01 |
| AT5G19850 | 0,75 | 0,13 | 3,70 | 0,03 | 9,73E-01 |
| AT5G19860 | 0,51 | -1,66 | 4,05 | -0,41 | 6,82E-01 |
| AT5G19900 | 0,73 | -1,58 | 2,90 | -0,54 | 5,86E-01 |
| AT5G19920 | 0,37 | -3,42 | 4,38 | -0,78 | 4,35E-01 |
| AT5G19930 | 0,12 | -0,14 | 4,38 | -0,03 | 9,74E-01 |
| AT5G19950 | 0,27 | -0,50 | 4,33 | -0,11 | 9,09E-01 |
| AT5G19980 | 0,33 | -1,04 | 4,34 | -0,24 | 8,11E-01 |
| AT5G20040 | 0,12 | 0,00 | 4,41 | 0,00 | 1,00E+00 |
| AT5G20160 | 0,08 | -0,21 | 4,39 | -0,05 | 9,62E-01 |
| AT5G20170 | 0,19 | 0,39 | 4,35 | 0,09 | 9,29E-01 |
| AT5G20225 | 0,17 | 0,00 | 4,41 | 0,00 | 1,00E+00 |

| | | | | | |
|---|---|---|---|---|---|
| AT5G20260 | 0,52 | -2,60 | 4,41 | -0,59 | 5,55E-01 |
| AT5G20270 | 0,14 | 0,40 | 4,35 | 0,09 | 9,27E-01 |
| AT5G20300 | 1,26 | -1,12 | 2,46 | -0,46 | 6,47E-01 |
| AT5G20320 | 1,21 | -3,42 | 2,59 | -1,32 | 1,87E-01 |
| AT5G20360 | 0,79 | -0,63 | 2,18 | -0,29 | 7,74E-01 |
| AT5G20380 | 0,45 | 0,00 | 4,41 | 0,00 | 1,00E+00 |
| AT5G20570 | 0,58 | -3,04 | 3,07 | -0,99 | 3,21E-01 |
| AT5G20600 | 1,15 | -4,09 | 2,19 | -1,87 | 6,17E-02 |
| AT5G20620 | 0,96 | -2,17 | 2,40 | -0,91 | 3,65E-01 |
| AT5G20630 | 1,03 | -2,02 | 1,73 | -1,17 | 2,41E-01 |
| AT5G20650 | 0,87 | -0,45 | 3,06 | -0,15 | 8,82E-01 |
| AT5G20820 | 0,08 | -0,21 | 4,39 | -0,05 | 9,62E-01 |
| AT5G20830 | 0,97 | -2,70 | 2,35 | -1,15 | 2,50E-01 |
| AT5G20840 | 1,18 | -0,01 | 1,50 | 0,00 | 9,97E-01 |
| AT5G20890 | 1,15 | -1,86 | 2,11 | -0,88 | 3,78E-01 |
| AT5G20920 | 0,64 | -1,26 | 3,22 | -0,39 | 6,95E-01 |
| AT5G20940 | 0,27 | -1,70 | 4,37 | -0,39 | 6,98E-01 |
| AT5G03265 | 0,23 | 0,15 | 4,38 | 0,03 | 9,74E-01 |
| AT5G20980 | 0,81 | -0,95 | 2,92 | -0,33 | 7,44E-01 |
| AT5G21040 | 0,32 | -1,63 | 4,41 | -0,37 | 7,11E-01 |
| AT5G21160 | 0,41 | 1,13 | 4,31 | 0,26 | 7,94E-01 |
| AT5G21274 | 0,23 | 0,01 | 4,36 | 0,00 | 9,98E-01 |
| AT5G21920 | 0,36 | -0,20 | 4,37 | -0,04 | 9,64E-01 |
| AT5G21950 | 0,08 | 0,00 | 4,41 | 0,00 | 1,00E+00 |
| AT5G22010 | 0,59 | -1,72 | 3,08 | -0,56 | 5,78E-01 |
| AT5G22020 | 0,55 | -2,80 | 3,86 | -0,73 | 4,68E-01 |
| AT5G22030 | 0,76 | -2,85 | 3,35 | -0,85 | 3,95E-01 |
| AT5G22080 | 0,35 | 1,47 | 3,47 | 0,42 | 6,71E-01 |
| AT5G22090 | 0,53 | -0,54 | 3,34 | -0,16 | 8,71E-01 |
| AT5G22250 | 1,15 | 2,00 | 2,03 | 0,99 | 3,24E-01 |
| AT5G22350 | 0,24 | 0,94 | 4,32 | 0,22 | 8,29E-01 |
| AT5G22360 | 0,12 | 0,00 | 4,41 | 0,00 | 1,00E+00 |
| AT5G22390 | 0,49 | -2,60 | 3,26 | -0,80 | 4,25E-01 |
| AT5G22440 | 0,73 | -2,79 | 2,89 | -0,96 | 3,35E-01 |
| AT5G22450 | 0,87 | 0,07 | 1,82 | 0,04 | 9,67E-01 |
| AT5G22510 | 0,44 | -1,03 | 4,13 | -0,25 | 8,02E-01 |
| AT5G03425 | 0,62 | -2,53 | 3,69 | -0,69 | 4,92E-01 |
| AT5G22540 | 0,22 | -0,80 | 4,35 | -0,18 | 8,54E-01 |
| AT5G22570 | 0,26 | -1,26 | 4,36 | -0,29 | 7,72E-01 |
| AT5G22700 | 0,44 | -0,69 | 4,17 | -0,17 | 8,69E-01 |
| AT5G22740 | 0,58 | -2,70 | 3,84 | -0,70 | 4,83E-01 |
| AT5G22750 | 0,55 | 0,00 | 4,41 | 0,00 | 1,00E+00 |
| AT5G03475 | 1,21 | -0,03 | 1,73 | -0,02 | 9,86E-01 |
| AT5G22850 | 0,23 | -0,77 | 4,39 | -0,18 | 8,60E-01 |
| AT5G22860 | 0,35 | 0,47 | 3,48 | 0,14 | 8,92E-01 |
| AT5G23040 | 0,63 | 1,76 | 2,47 | 0,71 | 4,78E-01 |
| AT5G23050 | 0,45 | -3,79 | 4,08 | -0,93 | 3,53E-01 |
| AT5G23060 | 1,11 | 0,28 | 1,83 | 0,15 | 8,77E-01 |
| AT5G23080 | 1,07 | -3,10 | 2,58 | -1,20 | 2,30E-01 |
| AT5G23240 | 0,89 | 0,88 | 2,20 | 0,40 | 6,87E-01 |
| AT5G23300 | 0,32 | -1,48 | 4,39 | -0,34 | 7,36E-01 |
| AT5G23390 | 0,47 | -0,80 | 4,01 | -0,20 | 8,41E-01 |
| AT5G23405 | 0,14 | -0,29 | 4,38 | -0,07 | 9,47E-01 |
| AT5G23430 | 0,90 | -0,21 | 2,10 | -0,10 | 9,21E-01 |
| AT5G23440 | 0,16 | -0,01 | 4,37 | 0,00 | 9,98E-01 |
| AT5G23450 | 0,83 | -1,69 | 2,89 | -0,59 | 5,58E-01 |

| | | | | | |
|---|---|---|---|---|---|
| AT5G23580 | 0,83 | -2,32 | 2,19 | -1,06 | 2,88E-01 |
| AT5G23590 | 0,61 | -2,58 | 3,75 | -0,69 | 4,92E-01 |
| AT5G23610 | 1,31 | -2,44 | 2,14 | -1,14 | 2,55E-01 |
| AT5G23660 | 0,79 | 2,26 | 3,60 | 0,63 | 5,30E-01 |
| AT5G23740 | 0,24 | 1,49 | 4,30 | 0,35 | 7,30E-01 |
| AT5G23900 | 0,57 | -0,85 | 2,81 | -0,30 | 7,63E-01 |
| AT5G23940 | 0,08 | -0,21 | 4,39 | -0,05 | 9,62E-01 |
| AT5G23990 | 0,29 | 0,99 | 4,33 | 0,23 | 8,19E-01 |
| AT5G24120 | 0,77 | -1,94 | 2,49 | -0,78 | 4,36E-01 |
| AT5G24140 | 0,08 | -0,21 | 4,39 | -0,05 | 9,62E-01 |
| AT5G24150 | 0,21 | 0,00 | 4,41 | 0,00 | 1,00E+00 |
| AT5G24160 | 0,65 | -0,70 | 2,87 | -0,25 | 8,06E-01 |
| AT5G24240 | 1,01 | 0,84 | 1,86 | 0,45 | 6,53E-01 |
| AT5G24260 | 1,06 | -2,51 | 1,96 | -1,28 | 2,00E-01 |
| AT5G24320 | 0,20 | -0,75 | 4,40 | -0,17 | 8,64E-01 |
| AT5G24355 | 0,08 | 0,00 | 4,41 | 0,00 | 1,00E+00 |
| AT5G24430 | 0,56 | 0,56 | 3,94 | 0,14 | 8,87E-01 |
| AT5G24470 | 1,29 | -0,70 | 2,61 | -0,27 | 7,90E-01 |
| AT5G24480 | 0,23 | 0,01 | 4,36 | 0,00 | 9,99E-01 |
| AT5G24500 | 0,24 | -1,38 | 4,38 | -0,31 | 7,53E-01 |
| AT5G24520 | 0,95 | 0,74 | 2,13 | 0,35 | 7,28E-01 |
| AT5G24655 | 0,37 | 1,39 | 4,32 | 0,32 | 7,47E-01 |
| AT5G24735 | 0,57 | 0,47 | 3,24 | 0,14 | 8,85E-01 |
| AT5G24770 | 0,62 | 1,54 | 2,27 | 0,68 | 4,97E-01 |
| AT5G24780 | 0,08 | -0,21 | 4,39 | -0,05 | 9,62E-01 |
| AT5G24940 | 0,08 | 0,00 | 4,41 | 0,00 | 1,00E+00 |
| AT5G24960 | 0,36 | 1,91 | 4,29 | 0,45 | 6,56E-01 |
| AT5G25000 | 0,16 | -0,01 | 4,37 | 0,00 | 9,99E-01 |
| AT5G25070 | 1,26 | -3,09 | 1,73 | -1,79 | 7,34E-02 |
| AT5G25200 | 0,60 | 1,25 | 2,53 | 0,49 | 6,22E-01 |
| AT5G25265 | 0,67 | -1,59 | 3,01 | -0,53 | 5,99E-01 |
| AT5G25350 | 1,15 | -2,49 | 2,14 | -1,16 | 2,44E-01 |
| AT5G25370 | 0,71 | 1,08 | 2,88 | 0,37 | 7,08E-01 |
| AT5G03985 | 0,37 | 0,00 | 4,41 | 0,00 | 1,00E+00 |
| AT5G25480 | 0,27 | -2,69 | 4,40 | -0,61 | 5,41E-01 |
| AT5G25510 | 0,49 | -0,05 | 3,31 | -0,02 | 9,87E-01 |
| AT5G25530 | 1,06 | -3,79 | 3,32 | -1,14 | 2,54E-01 |
| AT5G25756 | 0,35 | 2,26 | 4,28 | 0,53 | 5,98E-01 |
| AT5G25757 | 0,87 | -1,92 | 2,06 | -0,93 | 3,51E-01 |
| AT5G25770 | 0,28 | -3,29 | 4,38 | -0,75 | 4,53E-01 |
| AT5G04085 | 0,59 | 2,78 | 4,27 | 0,65 | 5,15E-01 |
| AT5G25780 | 1,13 | -3,49 | 1,84 | -1,89 | 5,82E-02 |
| AT5G25820 | 0,43 | -1,71 | 4,34 | -0,39 | 6,94E-01 |
| AT5G25830 | 0,05 | -0,24 | 4,38 | -0,05 | 9,57E-01 |
| AT5G25920 | 0,08 | 0,00 | 4,41 | 0,00 | 1,00E+00 |
| AT5G26170 | 0,17 | -0,71 | 4,40 | -0,16 | 8,72E-01 |
| AT5G26210 | 0,43 | -2,21 | 4,19 | -0,53 | 5,98E-01 |
| AT5G26230 | 0,08 | 0,05 | 4,38 | 0,01 | 9,91E-01 |
| AT5G26680 | 0,15 | 0,41 | 4,35 | 0,09 | 9,26E-01 |
| AT5G26640 | 0,17 | 0,00 | 4,41 | 0,00 | 1,00E+00 |
| AT5G26622 | 0,46 | -0,62 | 4,30 | -0,14 | 8,86E-01 |
| AT5G04255 | 0,21 | -1,17 | 4,36 | -0,27 | 7,88E-01 |
| AT5G26760 | 0,42 | -2,18 | 4,14 | -0,52 | 6,00E-01 |
| AT5G26770 | 1,06 | 0,12 | 1,89 | 0,06 | 9,49E-01 |
| AT5G26880 | 0,63 | -2,84 | 3,06 | -0,93 | 3,54E-01 |
| AT5G26910 | 0,66 | -2,68 | 2,54 | -1,05 | 2,92E-01 |

| | | | | | |
|---|---|---|---|---|---|
| AT5G26960 | 0,27 | 1,08 | 4,31 | 0,25 | 8,02E-01 |
| AT5G26970 | 0,84 | 0,57 | 1,93 | 0,29 | 7,68E-01 |
| AT5G27030 | 1,03 | -1,29 | 1,82 | -0,71 | 4,80E-01 |
| AT5G27100 | 0,25 | -2,60 | 4,41 | -0,59 | 5,55E-01 |
| AT5G27238 | 0,05 | -0,24 | 4,38 | -0,05 | 9,57E-01 |
| AT5G04375 | 0,24 | -1,38 | 4,38 | -0,31 | 7,53E-01 |
| AT5G27330 | 0,05 | -0,24 | 4,38 | -0,05 | 9,57E-01 |
| AT5G27380 | 0,63 | 0,00 | 3,69 | 0,00 | 1,00E+00 |
| AT5G27550 | 0,34 | -1,76 | 4,39 | -0,40 | 6,88E-01 |
| AT5G27700 | 0,49 | -2,32 | 3,26 | -0,71 | 4,77E-01 |
| AT5G27715 | 0,29 | 0,18 | 4,32 | 0,04 | 9,67E-01 |
| AT5G04435 | 0,39 | 1,12 | 4,21 | 0,26 | 7,91E-01 |
| AT5G27790 | 0,75 | 3,07 | 3,45 | 0,89 | 3,73E-01 |
| AT5G27840 | 0,72 | -1,67 | 2,93 | -0,57 | 5,68E-01 |
| AT5G27870 | 1,24 | -3,24 | 1,92 | -1,68 | 9,24E-02 |
| AT5G27920 | 0,08 | 0,00 | 4,41 | 0,00 | 1,00E+00 |
| AT5G27960 | 0,74 | 2,80 | 2,62 | 1,07 | 2,84E-01 |
| AT5G28020 | 0,34 | 0,09 | 3,59 | 0,02 | 9,81E-01 |
| AT5G28030 | 0,66 | -1,64 | 3,03 | -0,54 | 5,88E-01 |
| AT5G28050 | 0,28 | -0,92 | 4,35 | -0,21 | 8,33E-01 |
| AT5G04635 | 0,50 | 0,36 | 3,15 | 0,11 | 9,10E-01 |
| AT5G28060 | 0,65 | -0,86 | 2,24 | -0,38 | 7,02E-01 |
| AT5G04685 | 0,19 | 1,18 | 4,31 | 0,27 | 7,84E-01 |
| AT5G28190 | 0,36 | 1,18 | 4,32 | 0,27 | 7,85E-01 |
| AT5G28220 | 0,30 | 1,51 | 4,30 | 0,35 | 7,26E-01 |
| AT5G28235 | 0,34 | 2,16 | 4,28 | 0,50 | 6,14E-01 |
| AT5G28310 | 0,08 | 0,00 | 4,41 | 0,00 | 1,00E+00 |
| AT5G28400 | 1,06 | -2,74 | 1,79 | -1,53 | 1,26E-01 |
| AT5G28442 | 0,36 | 1,68 | 3,34 | 0,50 | 6,15E-01 |
| AT5G28450 | 0,85 | 2,68 | 2,32 | 1,16 | 2,47E-01 |
| AT5G28530 | 0,22 | 0,78 | 4,34 | 0,18 | 8,58E-01 |
| AT5G28550 | 0,21 | 0,76 | 4,34 | 0,18 | 8,61E-01 |
| AT5G28640 | 0,08 | -0,21 | 4,39 | -0,05 | 9,62E-01 |
| AT5G28750 | 0,65 | -0,57 | 2,16 | -0,26 | 7,93E-01 |
| AT5G28770 | 0,50 | -0,25 | 3,96 | -0,06 | 9,49E-01 |
| AT5G28850 | 0,48 | -0,36 | 3,24 | -0,11 | 9,12E-01 |
| AT5G28910 | 0,82 | -1,27 | 2,35 | -0,54 | 5,91E-01 |
| AT5G04925 | 0,18 | -0,42 | 4,37 | -0,10 | 9,24E-01 |
| AT5G32470 | 0,85 | -0,98 | 2,01 | -0,49 | 6,25E-01 |
| AT5G32590 | 0,68 | 0,76 | 2,17 | 0,35 | 7,25E-01 |
| AT5G33210 | 0,08 | -0,21 | 4,39 | -0,05 | 9,62E-01 |
| AT5G33439 | 0,11 | 0,46 | 4,35 | 0,11 | 9,15E-01 |
| AT5G35100 | 0,30 | 0,50 | 3,65 | 0,14 | 8,91E-01 |
| AT5G35200 | 0,52 | -0,96 | 3,13 | -0,31 | 7,60E-01 |
| AT5G35220 | 0,25 | -2,57 | 4,40 | -0,58 | 5,60E-01 |
| AT5G35230 | 0,96 | 3,10 | 2,57 | 1,21 | 2,27E-01 |
| AT5G35370 | 0,70 | -0,03 | 2,06 | -0,01 | 9,88E-01 |
| AT5G35475 | 0,36 | 1,31 | 4,32 | 0,30 | 7,61E-01 |
| AT5G35480 | 0,65 | 1,20 | 3,01 | 0,40 | 6,90E-01 |
| AT5G35530 | 0,88 | -0,62 | 2,54 | -0,25 | 8,06E-01 |
| AT5G35560 | 0,64 | -1,62 | 2,32 | -0,70 | 4,84E-01 |
| AT5G35570 | 0,29 | 0,00 | 4,41 | 0,00 | 1,00E+00 |
| AT5G35590 | 0,12 | 0,07 | 4,38 | 0,02 | 9,87E-01 |
| AT5G35603 | 0,59 | 2,06 | 4,06 | 0,51 | 6,12E-01 |
| AT5G35604 | 0,45 | 1,15 | 3,93 | 0,29 | 7,70E-01 |
| AT5G35620 | 0,26 | 0,10 | 4,37 | 0,02 | 9,82E-01 |

| | | | | | |
|---|---|---|---|---|---|
| AT5G35690 | 0,37 | -1,64 | 4,38 | -0,38 | 7,08E-01 |
| AT5G35700 | 0,37 | -1,64 | 4,31 | -0,38 | 7,04E-01 |
| AT5G35750 | 1,20 | -2,47 | 2,08 | -1,19 | 2,34E-01 |
| AT5G35753 | 0,77 | -3,58 | 2,16 | -1,66 | 9,67E-02 |
| AT5G35790 | 0,34 | -1,48 | 4,39 | -0,34 | 7,35E-01 |
| AT5G35810 | 0,15 | 0,05 | 4,37 | 0,01 | 9,91E-01 |
| AT5G35890 | 0,19 | 0,75 | 4,34 | 0,17 | 8,62E-01 |
| AT5G35910 | 0,15 | -0,68 | 4,41 | -0,15 | 8,77E-01 |
| AT5G35913 | 0,08 | 0,00 | 4,41 | 0,00 | 1,00E+00 |
| AT5G35995 | 0,29 | -0,76 | 4,40 | -0,17 | 8,62E-01 |
| AT5G36002 | 0,12 | 0,00 | 4,41 | 0,00 | 1,00E+00 |
| AT5G36120 | 0,05 | -0,24 | 4,38 | -0,05 | 9,57E-01 |
| AT5G36150 | 0,29 | 0,14 | 4,38 | 0,03 | 9,74E-01 |
| AT5G36160 | 0,29 | -1,17 | 4,36 | -0,27 | 7,88E-01 |
| AT5G36210 | 1,11 | -1,49 | 2,15 | -0,69 | 4,89E-01 |
| AT5G36220 | 0,80 | -0,19 | 2,25 | -0,08 | 9,32E-01 |
| AT5G36700 | 0,58 | -2,76 | 2,61 | -1,06 | 2,90E-01 |
| AT5G36790 | 0,17 | -0,13 | 4,39 | -0,03 | 9,77E-01 |
| AT5G36880 | 0,93 | -3,80 | 2,19 | -1,74 | 8,23E-02 |
| AT5G36930 | 1,17 | 0,77 | 2,21 | 0,35 | 7,27E-01 |
| AT5G37260 | 0,78 | -3,69 | 3,37 | -1,10 | 2,73E-01 |
| AT5G37310 | 0,77 | -0,71 | 3,48 | -0,20 | 8,38E-01 |
| AT5G37370 | 0,59 | -2,71 | 3,77 | -0,72 | 4,72E-01 |
| AT5G37480 | 0,13 | 0,52 | 4,35 | 0,12 | 9,04E-01 |
| AT5G37590 | 0,66 | -3,29 | 3,64 | -0,90 | 3,66E-01 |
| AT5G37720 | 0,46 | -2,67 | 3,31 | -0,81 | 4,20E-01 |
| AT5G37770 | 1,11 | 2,30 | 2,66 | 0,86 | 3,87E-01 |
| AT5G37780 | 0,80 | 0,41 | 2,11 | 0,19 | 8,47E-01 |
| AT5G05545 | 0,31 | 1,80 | 4,29 | 0,42 | 6,74E-01 |
| AT5G37890 | 0,36 | 0,00 | 4,41 | 0,00 | 1,00E+00 |
| AT5G05565 | 0,45 | 0,99 | 4,10 | 0,24 | 8,10E-01 |
| AT5G38020 | 0,42 | -2,56 | 4,40 | -0,58 | 5,61E-01 |
| AT5G38080 | 0,08 | 0,00 | 4,41 | 0,00 | 1,00E+00 |
| AT5G38140 | 0,10 | 0,45 | 4,35 | 0,10 | 9,19E-01 |
| AT5G38212 | 1,00 | 2,76 | 2,13 | 1,29 | 1,96E-01 |
| AT5G38240 | 0,41 | -3,83 | 4,35 | -0,88 | 3,78E-01 |
| AT5G38280 | 0,74 | 0,88 | 3,68 | 0,24 | 8,12E-01 |
| AT5G38450 | 0,46 | 0,93 | 4,13 | 0,22 | 8,23E-01 |
| AT5G05755 | 0,05 | -0,24 | 4,38 | -0,05 | 9,57E-01 |
| AT5G38510 | 0,76 | -1,19 | 2,56 | -0,47 | 6,42E-01 |
| AT5G38640 | 0,50 | -0,77 | 2,67 | -0,29 | 7,73E-01 |
| AT5G38650 | 1,00 | -1,95 | 2,19 | -0,89 | 3,73E-01 |
| AT5G38830 | 0,49 | 0,00 | 4,41 | 0,00 | 1,00E+00 |
| AT5G39130 | 0,12 | -0,14 | 4,38 | -0,03 | 9,74E-01 |
| AT5G39160 | 0,34 | 1,95 | 4,28 | 0,46 | 6,49E-01 |
| AT5G05895 | 0,62 | 2,29 | 2,92 | 0,78 | 4,34E-01 |
| AT5G39480 | 0,08 | -0,21 | 4,39 | -0,05 | 9,62E-01 |
| AT5G39520 | 0,33 | -1,64 | 4,41 | -0,37 | 7,10E-01 |
| AT5G39530 | 0,77 | -2,67 | 3,01 | -0,89 | 3,75E-01 |
| AT5G39600 | 0,66 | -1,58 | 2,39 | -0,66 | 5,09E-01 |
| AT5G39610 | 0,52 | -2,88 | 3,89 | -0,74 | 4,59E-01 |
| AT5G05995 | 0,40 | 0,86 | 4,17 | 0,21 | 8,36E-01 |
| AT5G39680 | 0,42 | -2,60 | 4,41 | -0,59 | 5,55E-01 |
| AT5G39693 | 0,43 | 1,31 | 3,29 | 0,40 | 6,90E-01 |
| AT5G39920 | 0,25 | -0,68 | 4,41 | -0,15 | 8,77E-01 |
| AT5G39950 | 0,95 | -0,43 | 2,49 | -0,17 | 8,64E-01 |

| | | | | | |
|---|---|---|---|---|---|
| AT5G40170 | 0,68 | -0,29 | 2,19 | -0,13 | 8,94E-01 |
| AT5G40240 | 0,14 | -0,29 | 4,38 | -0,07 | 9,47E-01 |
| AT5G40370 | 0,37 | 0,44 | 3,43 | 0,13 | 8,97E-01 |
| AT5G40470 | 0,27 | -2,23 | 4,36 | -0,51 | 6,10E-01 |
| AT5G40580 | 0,60 | -3,47 | 4,37 | -0,79 | 4,28E-01 |
| AT5G40660 | 0,29 | -2,57 | 4,40 | -0,58 | 5,59E-01 |
| AT5G40740 | 0,85 | -0,59 | 2,45 | -0,24 | 8,11E-01 |
| AT5G40820 | 0,95 | -2,68 | 2,23 | -1,20 | 2,30E-01 |
| AT5G40830 | 0,63 | 0,19 | 2,66 | 0,07 | 9,43E-01 |
| AT5G40940 | 0,56 | -2,60 | 3,88 | -0,67 | 5,02E-01 |
| AT5G41020 | 0,73 | -0,81 | 2,54 | -0,32 | 7,50E-01 |
| AT5G41130 | 0,21 | 0,00 | 4,41 | 0,00 | 1,00E+00 |
| AT5G41150 | 0,33 | -2,59 | 4,41 | -0,59 | 5,56E-01 |
| AT5G41190 | 0,83 | -2,89 | 2,79 | -1,03 | 3,01E-01 |
| AT5G41330 | 0,40 | -2,48 | 3,37 | -0,74 | 4,62E-01 |
| AT5G41400 | 0,65 | 0,18 | 2,54 | 0,07 | 9,43E-01 |
| AT5G41460 | 0,17 | 0,00 | 4,41 | 0,00 | 1,00E+00 |
| AT5G41520 | 0,58 | -0,98 | 2,65 | -0,37 | 7,10E-01 |
| AT5G41670 | 0,43 | 0,36 | 2,77 | 0,13 | 8,95E-01 |
| AT5G06235 | 0,77 | -0,19 | 4,39 | -0,04 | 9,65E-01 |
| AT5G41870 | 0,38 | 0,15 | 4,31 | 0,03 | 9,72E-01 |
| AT5G42100 | 1,07 | 0,42 | 1,69 | 0,25 | 8,02E-01 |
| AT5G42110 | 0,47 | 0,59 | 3,21 | 0,18 | 8,55E-01 |
| AT5G42130 | 0,21 | -0,73 | 4,40 | -0,17 | 8,67E-01 |
| AT5G42140 | 0,12 | -0,31 | 4,38 | -0,07 | 9,44E-01 |
| AT5G42190 | 0,65 | 0,09 | 2,57 | 0,04 | 9,72E-01 |
| AT5G42280 | 0,14 | -0,29 | 4,38 | -0,07 | 9,47E-01 |
| AT5G42300 | 0,48 | 1,49 | 3,89 | 0,38 | 7,01E-01 |
| AT5G42380 | 1,28 | -1,08 | 1,86 | -0,58 | 5,63E-01 |
| AT5G42400 | 0,21 | -0,19 | 4,37 | -0,04 | 9,66E-01 |
| AT5G06305 | 1,02 | 2,17 | 1,82 | 1,19 | 2,33E-01 |
| AT5G42490 | 0,56 | -1,62 | 3,93 | -0,41 | 6,80E-01 |
| AT5G42567 | 0,58 | 2,98 | 4,27 | 0,70 | 4,84E-01 |
| AT5G42570 | 0,15 | 0,00 | 4,41 | 0,00 | 1,00E+00 |
| AT5G42660 | 0,12 | -0,31 | 4,38 | -0,07 | 9,44E-01 |
| AT5G42790 | 0,93 | 1,64 | 2,03 | 0,81 | 4,20E-01 |
| AT5G42800 | 0,38 | -2,60 | 4,41 | -0,59 | 5,55E-01 |
| AT5G42850 | 1,15 | -3,54 | 2,06 | -1,72 | 8,57E-02 |
| AT5G42870 | 0,81 | 0,17 | 2,33 | 0,07 | 9,41E-01 |
| AT5G42890 | 0,08 | -0,21 | 4,39 | -0,05 | 9,62E-01 |
| AT5G42900 | 0,25 | -2,48 | 4,39 | -0,57 | 5,71E-01 |
| AT5G42905 | 0,52 | 1,48 | 3,79 | 0,39 | 6,97E-01 |
| AT5G42940 | 0,81 | -3,07 | 2,54 | -1,21 | 2,27E-01 |
| AT5G42955 | 0,65 | -2,60 | 3,87 | -0,67 | 5,02E-01 |
| AT5G42957 | 0,83 | 1,52 | 2,33 | 0,65 | 5,12E-01 |
| AT5G42990 | 0,17 | 0,00 | 4,41 | 0,00 | 1,00E+00 |
| AT5G43010 | 0,13 | -0,18 | 4,39 | -0,04 | 9,67E-01 |
| AT5G43020 | 0,41 | -0,38 | 4,13 | -0,09 | 9,27E-01 |
| AT5G43130 | 1,30 | -3,43 | 2,18 | -1,57 | 1,15E-01 |
| AT5G43260 | 0,52 | 0,38 | 2,66 | 0,14 | 8,87E-01 |
| AT5G43270 | 0,76 | 0,14 | 3,03 | 0,05 | 9,63E-01 |
| AT5G43285 | 0,35 | -0,79 | 4,39 | -0,18 | 8,57E-01 |
| AT5G43320 | 0,66 | -0,95 | 2,63 | -0,36 | 7,19E-01 |
| AT5G43440 | 0,40 | -0,30 | 4,34 | -0,07 | 9,44E-01 |
| AT5G43670 | 0,40 | -0,28 | 4,18 | -0,07 | 9,47E-01 |
| AT5G43710 | 0,21 | 0,00 | 4,41 | 0,00 | 1,00E+00 |

| | | | | | |
|---|---|---|---|---|---|
| AT5G43745 | 0,36 | -3,05 | 4,38 | -0,70 | 4,85E-01 |
| AT5G43750 | 0,47 | -1,41 | 4,29 | -0,33 | 7,43E-01 |
| AT5G43780 | 0,63 | 0,09 | 2,58 | 0,03 | 9,73E-01 |
| AT5G43790 | 0,45 | -2,07 | 4,25 | -0,49 | 6,25E-01 |
| AT5G43830 | 0,91 | -1,70 | 1,90 | -0,90 | 3,70E-01 |
| AT5G43850 | 0,29 | -0,33 | 3,70 | -0,09 | 9,30E-01 |
| AT5G43880 | 0,19 | 0,99 | 4,32 | 0,23 | 8,18E-01 |
| AT5G43950 | 0,65 | -1,97 | 2,61 | -0,75 | 4,52E-01 |
| AT5G43960 | 0,54 | -0,68 | 3,20 | -0,21 | 8,32E-01 |
| AT5G44000 | 0,95 | 0,80 | 2,29 | 0,35 | 7,26E-01 |
| AT5G44005 | 0,16 | -0,62 | 4,40 | -0,14 | 8,87E-01 |
| AT5G44100 | 0,66 | -0,55 | 2,64 | -0,21 | 8,34E-01 |
| AT5G44250 | 0,12 | 0,00 | 4,41 | 0,00 | 1,00E+00 |
| AT5G44260 | 0,22 | -0,06 | 4,38 | -0,01 | 9,89E-01 |
| AT5G44283 | 0,91 | 0,33 | 2,54 | 0,13 | 8,97E-01 |
| AT5G44290 | 1,25 | -0,54 | 1,92 | -0,28 | 7,78E-01 |
| AT5G44306 | 0,93 | 0,74 | 2,86 | 0,26 | 7,96E-01 |
| AT5G44495 | 1,23 | 1,73 | 1,66 | 1,05 | 2,96E-01 |
| AT5G44510 | 0,61 | 0,00 | 4,41 | 0,00 | 1,00E+00 |
| AT5G44568 | 0,48 | -0,07 | 2,82 | -0,03 | 9,79E-01 |
| AT5G44670 | 0,13 | 0,00 | 4,41 | 0,00 | 1,00E+00 |
| AT5G44680 | 0,48 | -0,69 | 4,41 | -0,16 | 8,76E-01 |
| AT5G44785 | 0,10 | -0,34 | 4,38 | -0,08 | 9,38E-01 |
| AT5G44820 | 0,08 | 0,00 | 4,41 | 0,00 | 1,00E+00 |
| AT5G44840 | 0,14 | 0,00 | 4,41 | 0,00 | 1,00E+00 |
| AT5G45010 | 0,58 | -1,82 | 3,14 | -0,58 | 5,62E-01 |
| AT5G45040 | 0,08 | 0,00 | 4,41 | 0,00 | 1,00E+00 |
| AT5G45050 | 0,86 | -1,41 | 2,07 | -0,68 | 4,95E-01 |
| AT5G45170 | 0,11 | 0,46 | 4,35 | 0,11 | 9,15E-01 |
| AT5G45180 | 0,66 | -1,64 | 4,02 | -0,41 | 6,83E-01 |
| AT5G45230 | 0,97 | 0,80 | 1,78 | 0,45 | 6,52E-01 |
| AT5G45250 | 0,93 | -2,38 | 1,96 | -1,21 | 2,25E-01 |
| AT5G45275 | 0,48 | -0,68 | 4,41 | -0,15 | 8,77E-01 |
| AT5G45290 | 1,09 | -2,30 | 2,11 | -1,09 | 2,77E-01 |
| AT5G45340 | 0,20 | 1,02 | 4,33 | 0,24 | 8,14E-01 |
| AT5G45360 | 0,08 | 0,00 | 4,41 | 0,00 | 1,00E+00 |
| AT5G45410 | 0,46 | -3,04 | 4,07 | -0,75 | 4,55E-01 |
| AT5G45420 | 0,27 | 0,36 | 4,36 | 0,08 | 9,35E-01 |
| AT5G45440 | 0,74 | -2,60 | 3,53 | -0,74 | 4,61E-01 |
| AT5G45460 | 0,18 | -1,07 | 4,38 | -0,25 | 8,06E-01 |
| AT5G45573 | 0,61 | 2,34 | 2,84 | 0,83 | 4,09E-01 |
| AT5G45630 | 0,05 | -0,24 | 4,38 | -0,05 | 9,57E-01 |
| AT5G45650 | 0,61 | -0,12 | 3,17 | -0,04 | 9,69E-01 |
| AT5G45775 | 0,86 | -1,60 | 1,98 | -0,81 | 4,19E-01 |
| AT5G45800 | 0,52 | -0,07 | 2,36 | -0,03 | 9,76E-01 |
| AT5G45820 | 1,25 | 0,59 | 1,65 | 0,36 | 7,22E-01 |
| AT5G46020 | 1,24 | -0,31 | 2,46 | -0,13 | 9,00E-01 |
| AT5G46040 | 0,11 | -0,80 | 4,39 | -0,18 | 8,56E-01 |
| AT5G46050 | 0,37 | -2,11 | 4,38 | -0,48 | 6,31E-01 |
| AT5G46280 | 0,57 | 1,13 | 3,03 | 0,37 | 7,08E-01 |
| AT5G46290 | 0,63 | -1,57 | 3,19 | -0,49 | 6,21E-01 |
| AT5G46340 | 0,61 | -1,64 | 4,03 | -0,41 | 6,84E-01 |
| AT5G46395 | 0,11 | 0,46 | 4,35 | 0,11 | 9,15E-01 |
| AT5G46580 | 0,75 | 0,73 | 2,46 | 0,30 | 7,67E-01 |
| AT5G46620 | 0,64 | 0,01 | 3,13 | 0,00 | 9,97E-01 |
| AT5G46750 | 0,66 | -1,59 | 3,75 | -0,42 | 6,71E-01 |

| | | | | | |
|---|---|---|---|---|---|
| AT5G46790 | 0,53 | 0,74 | 3,04 | 0,24 | 8,08E-01 |
| AT5G46800 | 0,83 | -0,93 | 2,27 | -0,41 | 6,83E-01 |
| AT5G46860 | 0,46 | 1,08 | 4,05 | 0,27 | 7,90E-01 |
| AT5G46970 | 0,65 | 1,68 | 2,55 | 0,66 | 5,10E-01 |
| AT5G47070 | 0,97 | 0,24 | 2,18 | 0,11 | 9,13E-01 |
| AT5G47080 | 0,13 | 0,00 | 4,41 | 0,00 | 1,00E+00 |
| AT5G47110 | 1,25 | -0,92 | 1,42 | -0,65 | 5,18E-01 |
| AT5G47190 | 0,93 | -0,82 | 2,14 | -0,38 | 7,01E-01 |
| AT5G47220 | 0,78 | -1,91 | 2,42 | -0,79 | 4,30E-01 |
| AT5G47250 | 0,28 | 0,18 | 4,36 | 0,04 | 9,67E-01 |
| AT5G47370 | 0,28 | -3,07 | 4,38 | -0,70 | 4,84E-01 |
| AT5G47460 | 0,15 | -0,77 | 4,39 | -0,18 | 8,61E-01 |
| AT5G47570 | 0,30 | 0,56 | 4,35 | 0,13 | 8,97E-01 |
| AT5G47580 | 0,64 | -3,83 | 3,63 | -1,05 | 2,93E-01 |
| AT5G47650 | 0,20 | -2,60 | 4,41 | -0,59 | 5,55E-01 |
| AT5G47700 | 0,72 | -0,55 | 2,26 | -0,24 | 8,10E-01 |
| AT5G47730 | 0,16 | -0,26 | 4,38 | -0,06 | 9,52E-01 |
| AT5G47760 | 0,12 | -0,31 | 4,38 | -0,07 | 9,44E-01 |
| AT5G47810 | 0,55 | -1,58 | 2,75 | -0,57 | 5,66E-01 |
| AT5G47830 | 0,54 | 0,00 | 4,41 | 0,00 | 1,00E+00 |
| AT5G47850 | 0,49 | -0,50 | 4,02 | -0,12 | 9,02E-01 |
| AT5G47880 | 0,55 | -0,91 | 3,21 | -0,28 | 7,77E-01 |
| AT5G47930 | 0,53 | 0,95 | 2,57 | 0,37 | 7,11E-01 |
| AT5G48000 | 0,12 | 0,07 | 4,38 | 0,02 | 9,87E-01 |
| AT5G07055 | 0,58 | 1,35 | 3,10 | 0,44 | 6,62E-01 |
| AT5G48120 | 1,18 | -2,52 | 1,71 | -1,48 | 1,39E-01 |
| AT5G48150 | 0,85 | -4,52 | 3,49 | -1,29 | 1,96E-01 |
| AT5G48180 | 0,98 | 0,56 | 2,05 | 0,28 | 7,83E-01 |
| AT5G48375 | 0,26 | -2,60 | 4,41 | -0,59 | 5,55E-01 |
| AT5G48412 | 0,64 | -0,35 | 2,55 | -0,14 | 8,92E-01 |
| AT5G48440 | 0,43 | 0,07 | 4,23 | 0,02 | 9,87E-01 |
| AT5G48540 | 0,08 | 0,00 | 4,41 | 0,00 | 1,00E+00 |
| AT5G48580 | 0,41 | -1,70 | 4,20 | -0,41 | 6,85E-01 |
| AT5G48640 | 0,19 | 0,52 | 4,35 | 0,12 | 9,04E-01 |
| AT5G48657 | 0,40 | 0,47 | 3,27 | 0,14 | 8,87E-01 |
| AT5G07265 | 0,12 | 0,00 | 4,41 | 0,00 | 1,00E+00 |
| AT5G48770 | 0,08 | -0,21 | 4,39 | -0,05 | 9,62E-01 |
| AT5G07275 | 0,71 | -2,60 | 4,41 | -0,59 | 5,55E-01 |
| AT5G48810 | 0,80 | -1,13 | 2,28 | -0,50 | 6,20E-01 |
| AT5G48830 | 0,28 | -1,75 | 4,39 | -0,40 | 6,90E-01 |
| AT5G48880 | 0,58 | -2,22 | 4,33 | -0,51 | 6,08E-01 |
| AT5G48910 | 0,76 | 0,41 | 2,20 | 0,19 | 8,53E-01 |
| AT5G48930 | 1,24 | -1,98 | 1,70 | -1,16 | 2,45E-01 |
| AT5G49100 | 1,11 | 1,55 | 1,85 | 0,84 | 4,03E-01 |
| AT5G49300 | 0,36 | 0,00 | 4,41 | 0,00 | 1,00E+00 |
| AT5G49350 | 0,08 | 0,00 | 4,41 | 0,00 | 1,00E+00 |
| AT5G49420 | 0,35 | 0,87 | 4,33 | 0,20 | 8,41E-01 |
| AT5G49430 | 1,30 | -1,35 | 1,56 | -0,86 | 3,88E-01 |
| AT5G49540 | 0,90 | -2,34 | 2,62 | -0,89 | 3,73E-01 |
| AT5G49550 | 0,27 | -1,27 | 4,38 | -0,29 | 7,72E-01 |
| AT5G49555 | 0,40 | -0,70 | 4,35 | -0,16 | 8,73E-01 |
| AT5G49570 | 0,11 | -0,68 | 4,41 | -0,15 | 8,77E-01 |
| AT5G49580 | 0,46 | -0,47 | 4,03 | -0,12 | 9,07E-01 |
| AT5G49630 | 0,19 | 1,18 | 4,31 | 0,27 | 7,84E-01 |
| AT5G49645 | 0,08 | 0,00 | 4,41 | 0,00 | 1,00E+00 |
| AT5G49660 | 0,65 | -0,23 | 2,54 | -0,09 | 9,29E-01 |

| | | | | | |
|---|---|---|---|---|---|
| AT5G49665 | 0,50 | -2,82 | 4,00 | -0,70 | 4,81E-01 |
| AT5G49830 | 1,27 | -1,77 | 1,76 | -1,01 | 3,14E-01 |
| AT5G49890 | 0,22 | 0,53 | 4,35 | 0,12 | 9,04E-01 |
| AT5G49900 | 0,41 | -1,81 | 4,20 | -0,43 | 6,67E-01 |
| AT5G49930 | 0,74 | -0,89 | 2,27 | -0,39 | 6,94E-01 |
| AT5G49950 | 0,25 | -1,07 | 4,39 | -0,24 | 8,07E-01 |
| AT5G50000 | 0,84 | 0,69 | 2,03 | 0,34 | 7,33E-01 |
| AT5G50150 | 0,10 | -0,18 | 4,38 | -0,04 | 9,67E-01 |
| AT5G50220 | 0,28 | -2,28 | 4,37 | -0,52 | 6,01E-01 |
| AT5G50320 | 0,77 | -2,17 | 2,93 | -0,74 | 4,59E-01 |
| AT5G50340 | 0,25 | -2,60 | 4,41 | -0,59 | 5,55E-01 |
| AT5G50350 | 0,29 | 0,39 | 4,35 | 0,09 | 9,28E-01 |
| AT5G50380 | 0,97 | 0,78 | 2,08 | 0,37 | 7,08E-01 |
| AT5G50390 | 0,17 | 0,00 | 4,41 | 0,00 | 1,00E+00 |
| AT5G50400 | 0,54 | -2,52 | 3,88 | -0,65 | 5,17E-01 |
| AT5G50460 | 0,36 | -2,43 | 4,37 | -0,56 | 5,78E-01 |
| AT5G50480 | 0,53 | -0,68 | 3,90 | -0,17 | 8,62E-01 |
| AT5G50570 | 0,31 | -0,30 | 4,34 | -0,07 | 9,44E-01 |
| AT5G50630 | 0,05 | -0,24 | 4,38 | -0,05 | 9,57E-01 |
| AT5G50640 | 0,24 | -2,60 | 4,41 | -0,59 | 5,55E-01 |
| AT5G50670 | 0,93 | -1,04 | 2,33 | -0,45 | 6,55E-01 |
| AT5G50860 | 0,49 | -3,32 | 3,93 | -0,84 | 3,98E-01 |
| AT5G50870 | 0,08 | 0,00 | 4,41 | 0,00 | 1,00E+00 |
| AT5G50900 | 0,83 | -1,06 | 2,48 | -0,43 | 6,67E-01 |
| AT5G51000 | 0,61 | 2,22 | 2,94 | 0,76 | 4,50E-01 |
| AT5G51020 | 0,37 | -1,64 | 4,31 | -0,38 | 7,04E-01 |
| AT5G51110 | 0,78 | -0,35 | 2,39 | -0,15 | 8,83E-01 |
| AT5G51170 | 0,12 | 0,00 | 4,41 | 0,00 | 1,00E+00 |
| AT5G51195 | 0,57 | 0,95 | 2,54 | 0,38 | 7,07E-01 |
| AT5G51220 | 0,23 | 0,83 | 4,34 | 0,19 | 8,48E-01 |
| AT5G51290 | 0,61 | -3,66 | 2,97 | -1,23 | 2,17E-01 |
| AT5G51390 | 0,29 | -2,97 | 4,38 | -0,68 | 4,98E-01 |
| AT5G51400 | 1,21 | -1,66 | 1,57 | -1,06 | 2,89E-01 |
| AT5G51550 | 1,02 | 1,42 | 1,90 | 0,75 | 4,55E-01 |
| AT5G51640 | 0,39 | -0,06 | 3,45 | -0,02 | 9,85E-01 |
| AT5G51650 | 0,32 | 1,37 | 3,49 | 0,39 | 6,94E-01 |
| AT5G51690 | 0,72 | -2,15 | 2,46 | -0,87 | 3,83E-01 |
| AT5G51710 | 0,10 | 0,45 | 4,35 | 0,10 | 9,19E-01 |
| AT5G51820 | 0,69 | -2,54 | 2,57 | -0,99 | 3,23E-01 |
| AT5G52060 | 0,54 | -0,18 | 2,64 | -0,07 | 9,47E-01 |
| AT5G52090 | 0,73 | 1,36 | 2,13 | 0,64 | 5,25E-01 |
| AT5G52120 | 0,47 | -2,12 | 4,39 | -0,48 | 6,30E-01 |
| AT5G52180 | 0,61 | -3,34 | 3,04 | -1,10 | 2,71E-01 |
| AT5G52330 | 0,15 | -0,17 | 4,39 | -0,04 | 9,68E-01 |
| AT5G52370 | 0,37 | -1,60 | 4,35 | -0,37 | 7,14E-01 |
| AT5G52450 | 0,44 | -2,45 | 3,25 | -0,75 | 4,51E-01 |
| AT5G52550 | 0,25 | -0,81 | 4,39 | -0,18 | 8,54E-01 |
| AT5G52650 | 0,64 | -3,11 | 3,64 | -0,85 | 3,93E-01 |
| AT5G52780 | 0,46 | -0,37 | 3,24 | -0,11 | 9,10E-01 |
| AT5G52797 | 0,48 | 0,62 | 3,18 | 0,20 | 8,45E-01 |
| AT5G53000 | 0,29 | -1,50 | 4,39 | -0,34 | 7,33E-01 |
| AT5G53010 | 0,50 | -2,83 | 3,30 | -0,86 | 3,92E-01 |
| AT5G53050 | 0,49 | 0,43 | 3,17 | 0,14 | 8,92E-01 |
| AT5G53130 | 0,72 | 0,49 | 2,92 | 0,17 | 8,66E-01 |
| AT5G07955 | 0,08 | 0,00 | 4,41 | 0,00 | 1,00E+00 |
| AT5G53300 | 0,89 | -3,26 | 2,64 | -1,24 | 2,16E-01 |

| | | | | | |
|---|---|---|---|---|---|
| AT5G53320 | 0,58 | -4,10 | 2,99 | -1,37 | 1,71E-01 |
| AT5G07975 | 0,08 | 0,00 | 4,41 | 0,00 | 1,00E+00 |
| AT5G53490 | 0,65 | -0,08 | 2,12 | -0,04 | 9,71E-01 |
| AT5G53500 | 0,52 | 0,68 | 3,13 | 0,22 | 8,28E-01 |
| AT5G53530 | 0,37 | -1,96 | 4,36 | -0,45 | 6,53E-01 |
| AT5G53570 | 0,72 | -0,71 | 2,46 | -0,29 | 7,72E-01 |
| AT5G53580 | 0,36 | -0,99 | 3,57 | -0,28 | 7,81E-01 |
| AT5G53730 | 0,14 | 0,40 | 4,35 | 0,09 | 9,27E-01 |
| AT5G53850 | 0,47 | -3,10 | 4,38 | -0,71 | 4,79E-01 |
| AT5G53870 | 0,19 | 0,64 | 4,34 | 0,15 | 8,83E-01 |
| AT5G53900 | 0,79 | -1,10 | 2,13 | -0,52 | 6,06E-01 |
| AT5G54030 | 0,27 | 1,85 | 4,29 | 0,43 | 6,67E-01 |
| AT5G54040 | 0,27 | 1,56 | 4,30 | 0,36 | 7,17E-01 |
| AT5G54050 | 0,38 | 1,22 | 4,33 | 0,28 | 7,77E-01 |
| AT5G54060 | 0,12 | 0,00 | 4,41 | 0,00 | 1,00E+00 |
| AT5G54090 | 1,00 | -4,04 | 2,32 | -1,74 | 8,14E-02 |
| AT5G54100 | 0,65 | -1,64 | 3,01 | -0,54 | 5,87E-01 |
| AT5G54170 | 0,64 | -0,72 | 2,16 | -0,33 | 7,40E-01 |
| AT5G54280 | 1,29 | -0,60 | 1,50 | -0,40 | 6,87E-01 |
| AT5G54380 | 1,11 | -0,14 | 1,93 | -0,07 | 9,42E-01 |
| AT5G54410 | 0,82 | 2,34 | 2,08 | 1,12 | 2,61E-01 |
| AT5G54430 | 1,12 | -0,67 | 2,51 | -0,27 | 7,90E-01 |
| AT5G54460 | 1,09 | 2,99 | 3,23 | 0,93 | 3,54E-01 |
| AT5G54470 | 0,35 | 0,29 | 3,41 | 0,08 | 9,33E-01 |
| AT5G54510 | 0,08 | 0,00 | 4,41 | 0,00 | 1,00E+00 |
| AT5G54610 | 0,49 | -1,01 | 3,89 | -0,26 | 7,96E-01 |
| AT5G54620 | 0,12 | 0,00 | 4,41 | 0,00 | 1,00E+00 |
| AT5G54630 | 0,29 | -0,41 | 4,37 | -0,09 | 9,25E-01 |
| AT5G54650 | 0,47 | -3,16 | 4,39 | -0,72 | 4,72E-01 |
| AT5G54690 | 0,39 | -3,07 | 4,38 | -0,70 | 4,84E-01 |
| AT5G54700 | 0,89 | 0,12 | 2,12 | 0,06 | 9,53E-01 |
| AT5G54710 | 0,79 | 1,70 | 2,43 | 0,70 | 4,83E-01 |
| AT5G54720 | 0,73 | 1,66 | 2,29 | 0,72 | 4,69E-01 |
| AT5G54780 | 1,16 | -3,28 | 1,75 | -1,87 | 6,18E-02 |
| AT5G54930 | 0,40 | -2,60 | 4,24 | -0,61 | 5,39E-01 |
| AT5G54960 | 0,98 | 0,48 | 2,28 | 0,21 | 8,32E-01 |
| AT5G55040 | 0,74 | 0,21 | 2,16 | 0,10 | 9,21E-01 |
| AT5G55055 | 0,05 | -0,24 | 4,38 | -0,05 | 9,57E-01 |
| AT5G55100 | 1,12 | -5,08 | 2,41 | -2,11 | 3,52E-02 |
| AT5G55130 | 0,31 | 0,88 | 3,58 | 0,24 | 8,07E-01 |
| AT5G55150 | 1,21 | -5,32 | 2,42 | -2,19 | 2,83E-02 |
| AT5G55160 | 0,37 | -1,64 | 4,41 | -0,37 | 7,09E-01 |
| AT5G55210 | 0,08 | -0,21 | 4,39 | -0,05 | 9,62E-01 |
| AT5G55250 | 0,40 | 1,06 | 4,32 | 0,25 | 8,06E-01 |
| AT5G55280 | 1,13 | 1,88 | 2,39 | 0,79 | 4,32E-01 |
| AT5G55290 | 0,30 | 0,06 | 4,32 | 0,01 | 9,88E-01 |
| AT5G55300 | 0,70 | -2,54 | 2,36 | -1,08 | 2,82E-01 |
| AT5G55340 | 0,19 | 0,04 | 4,37 | 0,01 | 9,92E-01 |
| AT5G55390 | 0,36 | 1,68 | 4,30 | 0,39 | 6,96E-01 |
| AT5G08275 | 0,59 | -3,81 | 3,80 | -1,00 | 3,16E-01 |
| AT5G08285 | 0,30 | -1,29 | 4,34 | -0,30 | 7,66E-01 |
| AT5G55510 | 0,05 | -0,24 | 4,38 | -0,05 | 9,57E-01 |
| AT5G55550 | 0,34 | -2,60 | 4,41 | -0,59 | 5,55E-01 |
| AT5G55570 | 1,30 | -3,66 | 3,08 | -1,19 | 2,35E-01 |
| AT5G55600 | 0,31 | 0,06 | 3,66 | 0,02 | 9,86E-01 |
| AT5G55670 | 0,30 | 1,17 | 4,32 | 0,27 | 7,87E-01 |

| | | | | | |
|---|---|---|---|---|---|
| AT5G55710 | 0,92 | -1,55 | 2,59 | -0,60 | 5,51E-01 |
| AT5G55740 | 0,27 | 0,36 | 4,36 | 0,08 | 9,33E-01 |
| AT5G55790 | 0,33 | 0,21 | 4,36 | 0,05 | 9,61E-01 |
| AT5G55835 | 0,61 | -0,09 | 2,34 | -0,04 | 9,70E-01 |
| AT5G55920 | 0,81 | -1,30 | 2,48 | -0,52 | 6,01E-01 |
| AT5G55940 | 0,50 | -3,46 | 3,93 | -0,88 | 3,78E-01 |
| AT5G56000 | 1,26 | 0,72 | 1,89 | 0,38 | 7,05E-01 |
| AT5G56180 | 0,23 | -1,14 | 4,39 | -0,26 | 7,96E-01 |
| AT5G56240 | 1,23 | -2,15 | 2,19 | -0,98 | 3,25E-01 |
| AT5G56250 | 0,87 | 0,51 | 2,85 | 0,18 | 8,58E-01 |
| AT5G56260 | 0,25 | -1,07 | 4,39 | -0,24 | 8,07E-01 |
| AT5G56280 | 0,08 | -0,21 | 4,39 | -0,05 | 9,62E-01 |
| AT5G56350 | 0,73 | -2,44 | 2,12 | -1,15 | 2,49E-01 |
| AT5G56360 | 0,87 | -0,79 | 2,96 | -0,27 | 7,89E-01 |
| AT5G56410 | 0,30 | -1,46 | 4,36 | -0,34 | 7,37E-01 |
| AT5G56450 | 0,20 | -0,57 | 4,39 | -0,13 | 8,97E-01 |
| AT5G56710 | 0,79 | -4,32 | 3,38 | -1,28 | 2,01E-01 |
| AT5G56730 | 0,65 | -2,41 | 2,62 | -0,92 | 3,58E-01 |
| AT5G56760 | 0,72 | -2,53 | 2,79 | -0,91 | 3,65E-01 |
| AT5G56860 | 0,23 | 1,25 | 4,31 | 0,29 | 7,72E-01 |
| AT5G56890 | 0,16 | -0,64 | 4,40 | -0,15 | 8,85E-01 |
| AT5G56980 | 0,28 | -1,64 | 4,41 | -0,37 | 7,10E-01 |
| AT5G57040 | 0,80 | -2,39 | 1,96 | -1,22 | 2,24E-01 |
| AT5G57160 | 0,81 | -3,02 | 2,47 | -1,23 | 2,20E-01 |
| AT5G57170 | 0,08 | -0,21 | 4,39 | -0,05 | 9,62E-01 |
| AT5G57250 | 0,54 | -1,61 | 3,96 | -0,41 | 6,84E-01 |
| AT5G57270 | 0,45 | -2,26 | 4,37 | -0,52 | 6,04E-01 |
| AT5G57290 | 0,69 | -0,29 | 2,50 | -0,12 | 9,07E-01 |
| AT5G57345 | 0,70 | -0,06 | 2,29 | -0,03 | 9,80E-01 |
| AT5G57580 | 0,90 | -1,54 | 2,57 | -0,60 | 5,49E-01 |
| AT5G57630 | 0,49 | -0,52 | 3,35 | -0,16 | 8,76E-01 |
| AT5G57670 | 0,16 | 0,75 | 4,34 | 0,17 | 8,63E-01 |
| AT5G57780 | 0,36 | -0,04 | 4,32 | -0,01 | 9,93E-01 |
| AT5G57785 | 0,56 | 1,41 | 2,59 | 0,54 | 5,86E-01 |
| AT5G57790 | 0,05 | -0,24 | 4,38 | -0,05 | 9,57E-01 |
| AT5G57850 | 0,51 | -2,02 | 4,36 | -0,46 | 6,43E-01 |
| AT5G57900 | 0,27 | -0,68 | 4,41 | -0,15 | 8,77E-01 |
| AT5G57960 | 0,12 | 0,18 | 4,36 | 0,04 | 9,68E-01 |
| AT5G58030 | 0,11 | -0,68 | 4,41 | -0,15 | 8,77E-01 |
| AT5G58050 | 0,17 | 0,00 | 4,41 | 0,00 | 1,00E+00 |
| AT5G58090 | 0,76 | -1,98 | 2,36 | -0,84 | 4,01E-01 |
| AT5G58130 | 0,84 | -1,67 | 2,81 | -0,60 | 5,51E-01 |
| AT5G58170 | 0,90 | -2,58 | 3,32 | -0,78 | 4,38E-01 |
| AT5G58220 | 0,12 | -0,06 | 4,37 | -0,01 | 9,88E-01 |
| AT5G58260 | 0,45 | -0,88 | 3,42 | -0,26 | 7,96E-01 |
| AT5G58380 | 0,93 | -0,79 | 2,16 | -0,37 | 7,14E-01 |
| AT5G58440 | 0,43 | -0,91 | 2,89 | -0,32 | 7,53E-01 |
| AT5G58570 | 0,61 | 0,79 | 3,90 | 0,20 | 8,40E-01 |
| AT5G08860 | 0,08 | -0,21 | 4,39 | -0,05 | 9,62E-01 |
| AT5G58600 | 0,29 | -3,07 | 4,39 | -0,70 | 4,83E-01 |
| AT5G58650 | 0,16 | -0,03 | 4,37 | -0,01 | 9,95E-01 |
| AT5G58670 | 0,69 | -2,86 | 2,25 | -1,27 | 2,04E-01 |
| AT5G58720 | 0,64 | -2,73 | 2,47 | -1,11 | 2,69E-01 |
| AT5G58730 | 0,50 | -2,28 | 4,02 | -0,57 | 5,72E-01 |
| AT5G58870 | 0,47 | 1,04 | 3,37 | 0,31 | 7,57E-01 |
| AT5G58900 | 0,28 | -3,07 | 4,38 | -0,70 | 4,84E-01 |

| | | | | | |
|---|---|---|---|---|---|
| AT5G59030 | 0,49 | -3,34 | 3,93 | -0,85 | 3,95E-01 |
| AT5G59130 | 0,21 | 0,00 | 4,41 | 0,00 | 1,00E+00 |
| AT5G59220 | 0,62 | -4,49 | 3,72 | -1,21 | 2,27E-01 |
| AT5G59290 | 0,24 | -1,38 | 4,38 | -0,31 | 7,53E-01 |
| AT5G08920 | 0,49 | -2,06 | 4,02 | -0,51 | 6,08E-01 |
| AT5G59480 | 0,17 | 0,00 | 4,41 | 0,00 | 1,00E+00 |
| AT5G59662 | 0,05 | -0,24 | 4,38 | -0,05 | 9,57E-01 |
| AT5G59700 | 0,33 | -2,34 | 4,37 | -0,54 | 5,92E-01 |
| AT5G59770 | 0,46 | -1,99 | 3,24 | -0,61 | 5,39E-01 |
| AT5G59780 | 1,19 | -2,59 | 1,97 | -1,32 | 1,88E-01 |
| AT5G59840 | 0,17 | -0,29 | 4,38 | -0,07 | 9,46E-01 |
| AT5G59850 | 0,69 | -2,03 | 2,51 | -0,81 | 4,17E-01 |
| AT5G59870 | 0,19 | 0,40 | 4,36 | 0,09 | 9,27E-01 |
| AT5G59900 | 0,16 | -0,01 | 4,37 | 0,00 | 9,98E-01 |
| AT5G59960 | 0,11 | -0,68 | 4,41 | -0,15 | 8,77E-01 |
| AT5G60050 | 0,15 | 0,00 | 4,41 | 0,00 | 1,00E+00 |
| AT5G60100 | 0,30 | 0,04 | 4,37 | 0,01 | 9,93E-01 |
| AT5G60160 | 0,49 | 1,03 | 4,05 | 0,25 | 7,99E-01 |
| AT5G60180 | 0,21 | -2,50 | 4,39 | -0,57 | 5,70E-01 |
| AT5G60210 | 0,29 | -0,76 | 4,40 | -0,17 | 8,62E-01 |
| AT5G60340 | 0,36 | 0,11 | 4,37 | 0,02 | 9,80E-01 |
| AT5G60400 | 0,12 | 0,00 | 4,41 | 0,00 | 1,00E+00 |
| AT5G60460 | 0,27 | 0,80 | 4,33 | 0,18 | 8,54E-01 |
| AT5G60670 | 0,64 | -2,81 | 2,21 | -1,27 | 2,04E-01 |
| AT5G60800 | 0,66 | -2,82 | 3,04 | -0,93 | 3,53E-01 |
| AT5G60840 | 0,12 | 0,75 | 4,34 | 0,17 | 8,63E-01 |
| AT5G60930 | 0,24 | -1,92 | 4,38 | -0,44 | 6,61E-01 |
| AT5G60960 | 0,29 | -1,77 | 4,39 | -0,40 | 6,87E-01 |
| AT5G60990 | 0,16 | -0,26 | 4,38 | -0,06 | 9,52E-01 |
| AT5G61040 | 0,88 | -1,22 | 4,31 | -0,28 | 7,77E-01 |
| AT5G61060 | 0,37 | -0,74 | 3,45 | -0,21 | 8,31E-01 |
| AT5G61130 | 0,17 | 0,00 | 4,41 | 0,00 | 1,00E+00 |
| AT5G61150 | 0,86 | -0,31 | 2,58 | -0,12 | 9,05E-01 |
| AT5G61250 | 0,23 | -1,72 | 4,40 | -0,39 | 6,95E-01 |
| AT5G61390 | 0,43 | -3,13 | 4,39 | -0,71 | 4,76E-01 |
| AT5G61460 | 0,21 | 0,00 | 4,41 | 0,00 | 1,00E+00 |
| AT5G61480 | 0,83 | -1,60 | 2,07 | -0,77 | 4,39E-01 |
| AT5G61500 | 0,76 | -2,44 | 2,81 | -0,87 | 3,84E-01 |
| AT5G61510 | 0,56 | -2,76 | 3,25 | -0,85 | 3,97E-01 |
| AT5G61520 | 0,33 | -1,00 | 4,35 | -0,23 | 8,19E-01 |
| AT5G61590 | 1,28 | -1,03 | 1,63 | -0,63 | 5,26E-01 |
| AT5G61600 | 0,57 | 2,22 | 2,52 | 0,88 | 3,78E-01 |
| AT5G61800 | 0,12 | 0,00 | 4,41 | 0,00 | 1,00E+00 |
| AT5G61810 | 0,48 | -1,59 | 3,27 | -0,49 | 6,27E-01 |
| AT5G61820 | 1,07 | -2,06 | 2,28 | -0,90 | 3,67E-01 |
| AT5G61840 | 0,89 | -0,71 | 2,16 | -0,33 | 7,43E-01 |
| AT5G61900 | 1,09 | 0,75 | 1,89 | 0,40 | 6,89E-01 |
| AT5G61960 | 0,94 | -1,68 | 2,51 | -0,67 | 5,03E-01 |
| AT5G61990 | 0,30 | -2,67 | 4,40 | -0,61 | 5,45E-01 |
| AT5G62090 | 0,32 | -0,50 | 4,39 | -0,11 | 9,10E-01 |
| AT5G62110 | 0,34 | 0,96 | 4,32 | 0,22 | 8,25E-01 |
| AT5G62130 | 0,45 | 2,01 | 3,15 | 0,64 | 5,23E-01 |
| AT5G62140 | 1,15 | 2,41 | 1,77 | 1,36 | 1,73E-01 |
| AT5G62180 | 0,08 | 0,00 | 4,41 | 0,00 | 1,00E+00 |
| AT5G62300 | 1,07 | -3,58 | 1,91 | -1,87 | 6,09E-02 |
| AT5G62430 | 0,29 | -0,95 | 4,34 | -0,22 | 8,27E-01 |

| | | | | | |
|---|---|---|---|---|---|
| AT5G62470 | 0,30 | 1,03 | 4,32 | 0,24 | 8,12E-01 |
| AT5G62530 | 0,56 | -0,78 | 3,15 | -0,25 | 8,04E-01 |
| AT5G62600 | 0,50 | -2,28 | 4,02 | -0,57 | 5,72E-01 |
| AT5G62610 | 0,31 | -0,30 | 4,33 | -0,07 | 9,46E-01 |
| AT5G62630 | 0,37 | -2,70 | 4,40 | -0,61 | 5,40E-01 |
| AT5G62650 | 0,39 | -3,18 | 4,38 | -0,73 | 4,68E-01 |
| AT5G62680 | 0,33 | -1,26 | 4,37 | -0,29 | 7,73E-01 |
| AT5G62790 | 0,60 | 1,81 | 2,51 | 0,72 | 4,72E-01 |
| AT5G63080 | 0,23 | -0,68 | 4,41 | -0,15 | 8,77E-01 |
| AT5G63100 | 0,29 | -1,64 | 4,38 | -0,37 | 7,09E-01 |
| AT5G63110 | 0,58 | -0,37 | 3,17 | -0,12 | 9,06E-01 |
| AT5G63120 | 0,65 | -0,71 | 3,14 | -0,23 | 8,20E-01 |
| AT5G63160 | 0,51 | 2,27 | 3,05 | 0,74 | 4,57E-01 |
| AT5G63180 | 0,25 | -0,27 | 4,38 | -0,06 | 9,51E-01 |
| AT5G63290 | 0,56 | -3,29 | 4,38 | -0,75 | 4,54E-01 |
| AT5G63310 | 0,47 | -2,35 | 4,37 | -0,54 | 5,91E-01 |
| AT5G63320 | 0,52 | 2,48 | 3,82 | 0,65 | 5,16E-01 |
| AT5G63370 | 1,26 | -3,06 | 1,67 | -1,84 | 6,65E-02 |
| AT5G63400 | 0,30 | -2,60 | 4,41 | -0,59 | 5,55E-01 |
| AT5G63410 | 0,46 | 0,77 | 2,66 | 0,29 | 7,72E-01 |
| AT5G63470 | 0,44 | 1,09 | 3,98 | 0,27 | 7,84E-01 |
| AT5G63610 | 0,51 | -2,42 | 3,99 | -0,61 | 5,43E-01 |
| AT5G63625 | 1,28 | 2,42 | 1,94 | 1,25 | 2,12E-01 |
| AT5G63760 | 0,90 | -1,00 | 2,29 | -0,44 | 6,63E-01 |
| AT5G63790 | 0,54 | -2,97 | 4,38 | -0,68 | 4,97E-01 |
| AT5G63860 | 0,75 | -1,98 | 2,17 | -0,91 | 3,61E-01 |
| AT5G63870 | 0,08 | 0,00 | 4,41 | 0,00 | 1,00E+00 |
| AT5G63890 | 0,39 | -0,75 | 4,33 | -0,17 | 8,63E-01 |
| AT5G63910 | 0,66 | -1,90 | 2,89 | -0,66 | 5,12E-01 |
| AT5G64020 | 0,23 | -1,72 | 4,40 | -0,39 | 6,95E-01 |
| AT5G64050 | 1,12 | -0,96 | 1,74 | -0,55 | 5,81E-01 |
| AT5G64130 | 0,35 | 0,59 | 3,61 | 0,16 | 8,71E-01 |
| AT5G64140 | 1,06 | -0,01 | 1,54 | -0,01 | 9,94E-01 |
| AT5G64170 | 0,64 | 0,22 | 3,14 | 0,07 | 9,43E-01 |
| AT5G64200 | 0,28 | -1,31 | 4,37 | -0,30 | 7,65E-01 |
| AT5G64290 | 0,42 | -0,65 | 4,18 | -0,16 | 8,76E-01 |
| AT5G64340 | 1,19 | -0,49 | 1,72 | -0,29 | 7,75E-01 |
| AT5G64350 | 0,41 | -0,53 | 4,28 | -0,12 | 9,02E-01 |
| AT5G64360 | 0,62 | -3,61 | 3,65 | -0,99 | 3,22E-01 |
| AT5G64410 | 0,79 | 0,69 | 2,25 | 0,31 | 7,58E-01 |
| AT5G64430 | 0,96 | -0,81 | 2,29 | -0,35 | 7,23E-01 |
| AT5G64460 | 0,08 | -0,21 | 4,39 | -0,05 | 9,62E-01 |
| AT5G64550 | 0,25 | -2,60 | 4,41 | -0,59 | 5,55E-01 |
| AT5G64580 | 0,77 | -2,60 | 2,54 | -1,02 | 3,07E-01 |
| AT5G64610 | 0,28 | 0,66 | 4,34 | 0,15 | 8,79E-01 |
| AT5G64660 | 0,37 | 1,16 | 3,39 | 0,34 | 7,31E-01 |
| AT5G64670 | 0,17 | 0,00 | 4,41 | 0,00 | 1,00E+00 |
| AT5G64760 | 0,83 | -0,73 | 3,05 | -0,24 | 8,11E-01 |
| AT5G64830 | 0,77 | -3,18 | 2,46 | -1,29 | 1,97E-01 |
| AT5G64860 | 0,28 | 0,01 | 4,36 | 0,00 | 9,98E-01 |
| AT5G64870 | 0,67 | 2,38 | 2,77 | 0,86 | 3,90E-01 |
| AT5G64920 | 1,21 | -2,66 | 1,73 | -1,54 | 1,25E-01 |
| AT5G64930 | 0,39 | -3,10 | 4,38 | -0,71 | 4,79E-01 |
| AT5G64980 | 0,12 | 0,00 | 4,41 | 0,00 | 1,00E+00 |
| AT5G65110 | 0,46 | 0,07 | 3,24 | 0,02 | 9,83E-01 |
| AT5G65200 | 0,16 | -0,26 | 4,38 | -0,06 | 9,52E-01 |

| | | | | | |
|---|---|---|---|---|---|
| AT5G65240 | 0,27 | 0,86 | 3,72 | 0,23 | 8,17E-01 |
| AT5G65310 | 1,18 | -1,98 | 1,93 | -1,02 | 3,06E-01 |
| AT5G65360 | 0,52 | -0,80 | 3,08 | -0,26 | 7,95E-01 |
| AT5G65430 | 0,62 | -1,38 | 2,50 | -0,55 | 5,81E-01 |
| AT5G65560 | 0,37 | -0,72 | 4,34 | -0,17 | 8,68E-01 |
| AT5G65685 | 0,97 | 0,10 | 2,14 | 0,05 | 9,61E-01 |
| AT5G65840 | 0,17 | -0,35 | 4,38 | -0,08 | 9,37E-01 |
| AT5G65910 | 1,14 | -3,77 | 3,31 | -1,14 | 2,55E-01 |
| AT5G65930 | 0,99 | -0,07 | 2,17 | -0,03 | 9,74E-01 |
| AT5G09535 | 0,24 | -2,59 | 4,41 | -0,59 | 5,56E-01 |
| AT5G65970 | 0,45 | -3,74 | 4,35 | -0,86 | 3,90E-01 |
| AT5G66055 | 0,63 | -1,64 | 2,63 | -0,62 | 5,33E-01 |
| AT5G66120 | 0,85 | 1,26 | 2,20 | 0,57 | 5,67E-01 |
| AT5G66210 | 1,05 | 0,13 | 2,05 | 0,06 | 9,50E-01 |
| AT5G66300 | 0,13 | 0,52 | 4,35 | 0,12 | 9,04E-01 |
| AT5G66310 | 0,12 | 0,00 | 4,41 | 0,00 | 1,00E+00 |
| AT5G66320 | 0,48 | -1,19 | 3,28 | -0,36 | 7,18E-01 |
| AT5G66330 | 0,11 | 0,27 | 4,37 | 0,06 | 9,52E-01 |
| AT5G66400 | 0,42 | 0,00 | 4,27 | 0,00 | 1,00E+00 |
| AT5G66440 | 0,33 | -0,20 | 4,39 | -0,05 | 9,63E-01 |
| AT5G66460 | 0,57 | -2,10 | 4,36 | -0,48 | 6,30E-01 |
| AT5G66500 | 0,15 | -0,73 | 4,40 | -0,17 | 8,68E-01 |
| AT5G66530 | 0,97 | -3,22 | 2,51 | -1,28 | 2,00E-01 |
| AT5G66540 | 0,21 | -0,03 | 4,36 | -0,01 | 9,95E-01 |
| AT5G66595 | 0,74 | 1,30 | 2,29 | 0,57 | 5,70E-01 |
| AT5G66607 | 0,08 | -0,21 | 4,39 | -0,05 | 9,62E-01 |
| AT5G66650 | 0,26 | -2,60 | 4,41 | -0,59 | 5,55E-01 |
| AT5G66675 | 0,51 | 1,23 | 3,03 | 0,40 | 6,86E-01 |
| AT5G66680 | 0,76 | -0,80 | 3,61 | -0,22 | 8,24E-01 |
| AT5G66720 | 0,41 | -3,06 | 4,33 | -0,71 | 4,81E-01 |
| AT5G66730 | 0,31 | 0,06 | 4,37 | 0,01 | 9,90E-01 |
| AT5G66850 | 0,24 | 0,00 | 4,41 | 0,00 | 1,00E+00 |
| AT5G66890 | 0,75 | -3,03 | 3,55 | -0,85 | 3,93E-01 |
| AT5G66910 | 0,38 | 0,07 | 3,39 | 0,02 | 9,84E-01 |
| AT5G67120 | 0,79 | 2,40 | 2,68 | 0,89 | 3,71E-01 |
| AT5G67150 | 0,35 | -0,52 | 4,35 | -0,12 | 9,05E-01 |
| AT5G67190 | 0,91 | -0,18 | 2,40 | -0,07 | 9,42E-01 |
| AT5G67220 | 0,60 | -0,40 | 2,52 | -0,16 | 8,73E-01 |
| AT5G67240 | 1,06 | -1,01 | 1,98 | -0,51 | 6,08E-01 |
| AT5G67245 | 0,45 | -1,07 | 4,39 | -0,24 | 8,07E-01 |
| AT5G67250 | 1,03 | 1,50 | 2,00 | 0,75 | 4,51E-01 |
| AT5G67280 | 0,58 | 0,07 | 2,59 | 0,03 | 9,79E-01 |
| AT5G67290 | 0,82 | -0,63 | 2,37 | -0,26 | 7,91E-01 |
| AT5G09855 | 0,69 | -0,66 | 2,35 | -0,28 | 7,79E-01 |
| AT5G67310 | 0,78 | -3,37 | 2,90 | -1,16 | 2,45E-01 |
| AT5G67350 | 0,75 | 1,89 | 2,52 | 0,75 | 4,54E-01 |
| AT5G67380 | 0,30 | -0,05 | 4,32 | -0,01 | 9,91E-01 |
| AT5G67460 | 0,10 | -0,34 | 4,38 | -0,08 | 9,38E-01 |
| AT5G67480 | 1,04 | -2,15 | 2,51 | -0,86 | 3,92E-01 |
| AT5G67490 | 1,16 | 0,33 | 1,83 | 0,18 | 8,58E-01 |
| AT5G67580 | 0,39 | 0,56 | 2,87 | 0,20 | 8,44E-01 |
| AT5G67610 | 0,37 | -3,94 | 4,34 | -0,91 | 3,63E-01 |

**chr11 chr17 uninfected by TCV relative to infected wild type.**

| |
|---|
| type |
| miRNA_precursor |
| miRNA_precursor |
| protein_coding |
| protein_coding |
| protein_coding |
| miRNA_precursor |
| protein_coding |
| ncRNA |
| ncRNA |
| miRNA_precursor |
| miRNA_precursor |
| ncRNA |
| miRNA_precursor |
| protein_coding |
| protein_coding |
| protein_coding |
| miRNA_precursor |
| protein_coding |
| miRNA_precursor |
| protein_coding |
| miRNA_precursor |
| protein_coding |
| ncRNA |
| protein_coding |
| protein_coding |
| miRNA_precursor |
| protein_coding |
| protein_coding |
| protein_coding |
| miRNA_precursor |
| miRNA_precursor |
| miRNA_precursor |
| miRNA_precursor |
| miRNA_precursor |
| ncRNA |
| protein_coding |
| miRNA_precursor |
| protein_coding |
| protein_coding |
| miRNA_precursor |
| protein_coding |
| protein_coding |
| protein_coding |
| miRNA_precursor |
| protein_coding |
| protein_coding |
| protein_coding |
| miRNA_precursor |
| protein_coding |
| protein_coding |
| protein_coding |
| small_nucle(ol)ar_rna |
| miRNA_precursor |

miRNA_precursor
protein_coding
protein_coding
ncRNA
ncRNA
ncRNA
protein_coding
ncRNA
protein_coding
miRNA_precursor
protein_coding
ncRNA
miRNA_precursor
small_nucle(ol)ar_rna
miRNA_precursor
protein_coding
miRNA_precursor
miRNA_precursor
protein_coding
miRNA_precursor
protein_coding
protein_coding
protein_coding
ncRNA
protein_coding
protein_coding
protein_coding
protein_coding
protein_coding
miRNA_precursor
miRNA_precursor
protein_coding
protein_coding
protein_coding
small_nucle(ol)ar_rna
miRNA_precursor
protein_coding
ncRNA
protein_coding
protein_coding
miRNA_precursor
protein_coding
protein_coding
protein_coding
protein_coding
protein_coding
protein_coding
protein_coding
protein_coding
miRNA_precursor
protein_coding
protein_coding
ncRNA
ncRNA
protein_coding
miRNA_precursor

protein_coding
protein_coding
protein_coding
protein_coding
protein_coding
protein_coding
protein_coding
ncRNA
protein_coding
protein_coding
protein_coding
miRNA_precursor
protein_coding
protein_coding
protein_coding
protein_coding
protein_coding
miRNA_precursor
protein_coding
protein_coding
miRNA_precursor
protein_coding
protein_coding
protein_coding
protein_coding
protein_coding
protein_coding
protein_coding
protein_coding
protein_coding
protein_coding
protein_coding
protein_coding
protein_coding
protein_coding
miRNA_precursor
protein_coding
protein_coding
miRNA_precursor
protein_coding
ncRNA
protein_coding
protein_coding
protein_coding
protein_coding
protein_coding
protein_coding
protein_coding
miRNA_precursor
protein_coding
ncRNA
protein_coding
protein_coding
protein_coding
protein_coding
protein_coding

protein_coding
protein_coding
protein_coding
miRNA_precursor
protein_coding
protein_coding
protein_coding
protein_coding
protein_coding
protein_coding
miRNA_precursor
protein_coding
ncRNA
protein_coding
protein_coding
protein_coding
protein_coding
protein_coding
protein_coding
protein_coding
protein_coding
protein_coding
protein_coding
protein_coding
miRNA_precursor
protein_coding
ncRNA
protein_coding
protein_coding
protein_coding
protein_coding
protein_coding
protein_coding
protein_coding
protein_coding
protein_coding
miRNA_precursor
protein_coding
miRNA_precursor
protein_coding
protein_coding
protein_coding
protein_coding
protein_coding
protein_coding
protein_coding
protein_coding
ncRNA
small_nucle(ol)ar_rna
ncRNA
protein_coding
protein_coding
protein_coding
protein_coding
protein_coding
protein_coding

protein_coding
protein_coding
miRNA_precursor
protein_coding
miRNA_precursor
protein_coding
protein_coding
protein_coding
protein_coding
miRNA_precursor
miRNA_precursor
protein_coding
small_nucle(ol)ar_rna
protein_coding
protein_coding
protein_coding
protein_coding
small_nucle(ol)ar_rna
protein_coding
protein_coding
protein_coding
protein_coding
ncRNA
protein_coding
protein_coding
protein_coding
protein_coding
protein_coding
protein_coding
protein_coding
protein_coding
protein_coding
miRNA_precursor
protein_coding
protein_coding
miRNA_precursor
protein_coding
protein_coding
protein_coding
protein_coding
protein_coding
miRNA_precursor
protein_coding
protein_coding
small_nucle(ol)ar_rna
protein_coding
protein_coding
protein_coding
protein_coding
protein_coding
protein_coding
protein_coding
protein_coding
protein_coding
protein_coding
protein_coding

protein_coding
protein_coding
small_nucle(ol)ar_rna
protein_coding
protein_coding
protein_coding
protein_coding
protein_coding
protein_coding
protein_coding
protein_coding
miRNA_precursor
protein_coding
small_nucle(ol)ar_rna
small_nucle(ol)ar_rna
protein_coding
miRNA_precursor
protein_coding
protein_coding
protein_coding
protein_coding
protein_coding
protein_coding
protein_coding
protein_coding
protein_coding
protein_coding
small_nucle(ol)ar_rna
protein_coding
protein_coding
protein_coding
ncRNA
protein_coding
protein_coding
miRNA_precursor
protein_coding
protein_coding
protein_coding
protein_coding
protein_coding
protein_coding
protein_coding
protein_coding
ncRNA
protein_coding
small_nucle(ol)ar_rna
protein_coding
small_nucle(ol)ar_rna
protein_coding
miRNA_precursor
protein_coding
protein_coding
protein_coding
protein_coding
protein_coding
protein_coding

protein_coding
protein_coding
protein_coding
protein_coding
protein_coding
protein_coding
protein_coding
protein_coding
protein_coding
protein_coding
protein_coding
protein_coding
ncRNA
protein_coding
protein_coding
protein_coding
protein_coding
protein_coding
protein_coding
protein_coding
protein_coding
protein_coding
protein_coding
protein_coding
protein_coding
miRNA_precursor
ncRNA
protein_coding
protein_coding
small_nucle(ol)ar_rna
protein_coding
protein_coding
protein_coding
pre_trna
protein_coding
protein_coding
protein_coding
miRNA_precursor
protein_coding
protein_coding
protein_coding
pre_trna
protein_coding
protein_coding
protein_coding
protein_coding
protein_coding
protein_coding
protein_coding
protein_coding
protein_coding
protein_coding
protein_coding
protein_coding
protein_coding
protein_coding

protein_coding
protein_coding
protein_coding
protein_coding
protein_coding
small_nucle(ol)ar_rna
protein_coding
protein_coding
protein_coding
protein_coding
miRNA_precursor
ribosomal_rna
protein_coding
protein_coding
protein_coding
protein_coding
protein_coding
protein_coding
protein_coding
protein_coding
protein_coding
protein_coding
protein_coding
protein_coding
protein_coding
ncRNA
protein_coding
small_nucle(ol)ar_rna
protein_coding
protein_coding
miRNA_precursor
protein_coding
protein_coding
protein_coding
protein_coding
protein_coding
protein_coding
protein_coding
protein_coding
protein_coding
protein_coding
small_nucle(ol)ar_rna
protein_coding
protein_coding
protein_coding
miRNA_precursor
protein_coding
miRNA_precursor
protein_coding
protein_coding
protein_coding
protein_coding
protein_coding
protein_coding
protein_coding
protein_coding

protein_coding
protein_coding
protein_coding
protein_coding
protein_coding
protein_coding
protein_coding
protein_coding
protein_coding
protein_coding
protein_coding
protein_coding
protein_coding
protein_coding
small_nucle(ol)ar_rna
protein_coding
protein_coding
protein_coding
protein_coding
miRNA_precursor
ncRNA
small_nucle(ol)ar_rna
protein_coding
small_nucle(ol)ar_rna
protein_coding
protein_coding
ncRNA
protein_coding
protein_coding
protein_coding
protein_coding
protein_coding
protein_coding
small_nucle(ol)ar_rna
protein_coding
protein_coding
protein_coding
small_nucle(ol)ar_rna
protein_coding
protein_coding
protein_coding
protein_coding
protein_coding
protein_coding
protein_coding
protein_coding
protein_coding
protein_coding
small_nucle(ol)ar_rna
protein_coding
protein_coding
protein_coding
protein_coding
protein_coding
protein_coding
miRNA_precursor

protein_coding
protein_coding
protein_coding
protein_coding
miRNA_precursor
protein_coding
protein_coding
protein_coding
protein_coding
protein_coding
protein_coding
protein_coding
protein_coding
protein_coding
protein_coding
protein_coding
protein_coding
protein_coding
protein_coding
protein_coding
small_nucle(ol)ar_rna
protein_coding
protein_coding
protein_coding
miRNA_precursor
protein_coding
protein_coding
protein_coding
protein_coding
protein_coding
protein_coding
protein_coding
protein_coding
protein_coding
protein_coding
protein_coding
protein_coding
protein_coding
protein_coding
protein_coding
miRNA_precursor
protein_coding
protein_coding
protein_coding
protein_coding
protein_coding
protein_coding
protein_coding
protein_coding
protein_coding
protein_coding
protein_coding
protein_coding
protein_coding
protein_coding
protein_coding

protein_coding
protein_coding
protein_coding
protein_coding
protein_coding
protein_coding
protein_coding
protein_coding
protein_coding
protein_coding
protein_coding
protein_coding
protein_coding
protein_coding
protein_coding
protein_coding
protein_coding
protein_coding
protein_coding
ncRNA
protein_coding
protein_coding
protein_coding
protein_coding
miRNA_precursor
protein_coding
protein_coding
protein_coding
protein_coding
miRNA_precursor
protein_coding
protein_coding
ncRNA
protein_coding
protein_coding
ncRNA
protein_coding
protein_coding
protein_coding
protein_coding
protein_coding
protein_coding
protein_coding
protein_coding
protein_coding
protein_coding
protein_coding
protein_coding
protein_coding
protein_coding
protein_coding
protein_coding
protein_coding
protein_coding
small_nucle(ol)ar_rna
protein_coding

protein_coding
protein_coding
protein_coding
protein_coding
protein_coding
miRNA_precursor
protein_coding
protein_coding
protein_coding
protein_coding
protein_coding
protein_coding
protein_coding
protein_coding
protein_coding
miRNA_precursor
protein_coding
protein_coding
miRNA_precursor
protein_coding
protein_coding
protein_coding
protein_coding
protein_coding
protein_coding
protein_coding
protein_coding
protein_coding
protein_coding
protein_coding
protein_coding
miRNA_precursor
protein_coding
protein_coding
protein_coding
protein_coding
protein_coding
protein_coding
protein_coding
protein_coding
protein_coding
protein_coding
ncRNA
protein_coding
protein_coding
protein_coding
protein_coding
protein_coding
protein_coding
protein_coding
protein_coding
protein_coding
protein_coding
miRNA_precursor
protein_coding
protein_coding

protein_coding
protein_coding
protein_coding
small_nucle(ol)ar_rna
small_nucle(ol)ar_rna
protein_coding
protein_coding
protein_coding
protein_coding
protein_coding
protein_coding
protein_coding
protein_coding
protein_coding
protein_coding
protein_coding
protein_coding
protein_coding
small_nucle(ol)ar_rna
protein_coding
protein_coding
protein_coding
protein_coding
protein_coding
ribosomal_rna
small_nucle(ol)ar_rna
protein_coding
protein_coding
protein_coding
protein_coding
protein_coding
protein_coding
miRNA_precursor
small_nucle(ol)ar_rna
protein_coding
small_nucle(ol)ar_rna
small_nucle(ol)ar_rna
protein_coding
protein_coding
protein_coding
protein_coding
protein_coding
protein_coding
small_nucle(ol)ar_rna
protein_coding
protein_coding
protein_coding
protein_coding
protein_coding
protein_coding
protein_coding
protein_coding
protein_coding
protein_coding
protein_coding
ncRNA

protein_coding
protein_coding
protein_coding
protein_coding
small_nucle(ol)ar_rna
protein_coding
small_nucle(ol)ar_rna
protein_coding
protein_coding
protein_coding
ncRNA
protein_coding
protein_coding
protein_coding
protein_coding
protein_coding
protein_coding
protein_coding
protein_coding
protein_coding
protein_coding
protein_coding
protein_coding
protein_coding
protein_coding
small_nucle(ol)ar_rna
small_nucle(ol)ar_rna
protein_coding
protein_coding
protein_coding
protein_coding
protein_coding
protein_coding
protein_coding
protein_coding
protein_coding
miRNA_precursor
protein_coding
protein_coding
pre_trna
protein_coding
protein_coding
protein_coding
protein_coding
protein_coding
small_nucle(ol)ar_rna
protein_coding
protein_coding
protein_coding
protein_coding
protein_coding
protein_coding
protein_coding
protein_coding
protein_coding
protein_coding

protein_coding
protein_coding
protein_coding
protein_coding
protein_coding
protein_coding
protein_coding
protein_coding
protein_coding
protein_coding
protein_coding
ncRNA
protein_coding
protein_coding
protein_coding
protein_coding
protein_coding
protein_coding
protein_coding
protein_coding
protein_coding
protein_coding
protein_coding
protein_coding
protein_coding
protein_coding
protein_coding
miRNA_precursor
protein_coding
protein_coding
protein_coding
protein_coding
protein_coding
protein_coding
protein_coding
protein_coding
protein_coding
protein_coding
protein_coding
protein_coding
protein_coding
protein_coding
protein_coding
ncRNA
protein_coding
protein_coding
protein_coding
protein_coding
protein_coding
protein_coding
protein_coding
protein_coding
protein_coding
protein_coding
protein_coding
miRNA_precursor

protein_coding
protein_coding
miRNA_precursor
protein_coding
protein_coding
protein_coding
protein_coding
protein_coding
protein_coding
protein_coding
protein_coding
protein_coding
protein_coding
protein_coding
protein_coding
protein_coding
protein_coding
protein_coding
protein_coding
protein_coding
protein_coding
protein_coding
protein_coding
ncRNA
protein_coding
protein_coding
protein_coding
protein_coding
protein_coding
protein_coding
small_nucle(ol)ar_rna
protein_coding
protein_coding
protein_coding
protein_coding
protein_coding
protein_coding
protein_coding
protein_coding
protein_coding
protein_coding
miRNA_precursor
protein_coding
protein_coding
protein_coding
protein_coding
protein_coding
protein_coding
protein_coding
protein_coding
small_nucle(ol)ar_rna
protein_coding
protein_coding
protein_coding
protein_coding
protein_coding

protein_coding
protein_coding
protein_coding
protein_coding
protein_coding
miRNA_precursor
protein_coding
protein_coding
protein_coding
protein_coding
protein_coding
protein_coding
protein_coding
protein_coding
miRNA_precursor
protein_coding
protein_coding
protein_coding
protein_coding
protein_coding
protein_coding
protein_coding
protein_coding
protein_coding
protein_coding
protein_coding
protein_coding
protein_coding
protein_coding
protein_coding
protein_coding
protein_coding
protein_coding
protein_coding
protein_coding
small_nucle(ol)ar_rna
miRNA_precursor
protein_coding
protein_coding
miRNA_precursor
protein_coding
protein_coding
small_nucle(ol)ar_rna
ncRNA
protein_coding
protein_coding
protein_coding
protein_coding
protein_coding
protein_coding
protein_coding
protein_coding
protein_coding
protein_coding
protein_coding
protein_coding

protein_coding
protein_coding
protein_coding
protein_coding
protein_coding
protein_coding
protein_coding
ncRNA
protein_coding
protein_coding
protein_coding
protein_coding
protein_coding
protein_coding
protein_coding
protein_coding
protein_coding
protein_coding
protein_coding
protein_coding
protein_coding
protein_coding
protein_coding
protein_coding
protein_coding
protein_coding
protein_coding
protein_coding
protein_coding
protein_coding
ncRNA
protein_coding
protein_coding
protein_coding
miRNA_precursor
protein_coding
protein_coding
protein_coding
miRNA_precursor
protein_coding
protein_coding
protein_coding
protein_coding
protein_coding
miRNA_precursor
protein_coding
protein_coding
protein_coding
protein_coding
small_nucle(ol)ar_rna
protein_coding
protein_coding
protein_coding
protein_coding
protein_coding
protein_coding

protein_coding
protein_coding
protein_coding
protein_coding
protein_coding
small_nucle(ol)ar_rna
protein_coding
protein_coding
protein_coding
protein_coding
protein_coding
protein_coding
protein_coding
protein_coding
protein_coding
small_nucle(ol)ar_rna
protein_coding
protein_coding
protein_coding
protein_coding
protein_coding
protein_coding
protein_coding
small_nucle(ol)ar_rna
pre_trna
protein_coding
small_nucle(ol)ar_rna
protein_coding
protein_coding
protein_coding
protein_coding
protein_coding
protein_coding
small_nucle(ol)ar_rna
protein_coding
protein_coding
miRNA_precursor
protein_coding
protein_coding
protein_coding
protein_coding
protein_coding
protein_coding
small_nucle(ol)ar_rna
miRNA_precursor
protein_coding
protein_coding
ncRNA
protein_coding
protein_coding
protein_coding
protein_coding
ncRNA
protein_coding
protein_coding
protein_coding

protein_coding
protein_coding
protein_coding
protein_coding
protein_coding
protein_coding
ncRNA
protein_coding
protein_coding
small_nucle(ol)ar_rna
protein_coding
protein_coding
protein_coding
protein_coding
protein_coding
protein_coding
protein_coding
protein_coding
protein_coding
protein_coding
protein_coding
protein_coding
protein_coding
protein_coding
protein_coding
protein_coding
protein_coding
protein_coding
protein_coding
protein_coding
miRNA_precursor
protein_coding
protein_coding
small_nucle(ol)ar_rna
ncRNA
protein_coding
protein_coding
protein_coding
protein_coding
protein_coding
ncRNA
protein_coding
protein_coding
protein_coding
protein_coding
protein_coding
protein_coding
protein_coding
protein_coding
miRNA_precursor
protein_coding
small_nucle(ol)ar_rna
protein_coding
protein_coding
protein_coding
miRNA_precursor

protein_coding
protein_coding
protein_coding
protein_coding
protein_coding
protein_coding
protein_coding
protein_coding
protein_coding
protein_coding
protein_coding
protein_coding
protein_coding
protein_coding
protein_coding
protein_coding
protein_coding
protein_coding
protein_coding
protein_coding
protein_coding
protein_coding
protein_coding
protein_coding
protein_coding
ncRNA
protein_coding
protein_coding
protein_coding
protein_coding
protein_coding
small_nucle(ol)ar_rna
protein_coding
protein_coding
protein_coding
protein_coding
protein_coding
protein_coding
protein_coding
protein_coding
ncRNA
protein_coding
ncRNA
protein_coding
protein_coding
protein_coding
protein_coding
protein_coding
protein_coding
protein_coding
protein_coding
protein_coding
protein_coding
protein_coding
protein_coding
protein_coding

protein_coding
protein_coding
miRNA_precursor
protein_coding
protein_coding
protein_coding
protein_coding
miRNA_precursor
protein_coding
protein_coding
protein_coding
protein_coding
protein_coding
protein_coding
protein_coding
small_nucle(ol)ar_rna
protein_coding
protein_coding
ncRNA
ncRNA
protein_coding
protein_coding
protein_coding
small_nucle(ol)ar_rna
protein_coding
protein_coding
protein_coding
protein_coding
protein_coding
protein_coding
protein_coding
protein_coding
protein_coding
protein_coding
ncRNA
protein_coding
protein_coding
protein_coding
protein_coding
protein_coding
protein_coding
protein_coding
protein_coding
miRNA_precursor
protein_coding
protein_coding
protein_coding
protein_coding
miRNA_precursor
miRNA_precursor
protein_coding
protein_coding
protein_coding
ncRNA
small_nucle(ol)ar_rna
small_nucle(ol)ar_rna

protein_coding
ncRNA
protein_coding
protein_coding
protein_coding
protein_coding
protein_coding
protein_coding
protein_coding
protein_coding
protein_coding
ncRNA
protein_coding
small_nucle(ol)ar_rna
protein_coding
protein_coding
protein_coding
protein_coding
protein_coding
protein_coding
ncRNA
protein_coding
protein_coding
protein_coding
protein_coding
miRNA_precursor
ncRNA
protein_coding
protein_coding
protein_coding
protein_coding
protein_coding
protein_coding
protein_coding
protein_coding
protein_coding
protein_coding
protein_coding
protein_coding
small_nucle(ol)ar_rna
protein_coding
protein_coding
protein_coding
protein_coding
protein_coding
protein_coding
protein_coding
protein_coding
protein_coding
protein_coding
protein_coding
ribosomal_rna
protein_coding
protein_coding
protein_coding
ncRNA

protein_coding
small_nucle(ol)ar_rna
protein_coding
protein_coding
protein_coding
protein_coding
protein_coding
protein_coding
protein_coding
protein_coding
protein_coding
protein_coding
protein_coding
protein_coding
protein_coding
protein_coding
ncRNA
protein_coding
protein_coding
small_nucle(ol)ar_rna
protein_coding
ncRNA
protein_coding
protein_coding
protein_coding
protein_coding
protein_coding
protein_coding
protein_coding
protein_coding
protein_coding
protein_coding
protein_coding
protein_coding
ncRNA
protein_coding
protein_coding
protein_coding
protein_coding
protein_coding
protein_coding
protein_coding
protein_coding
miRNA_precursor
protein_coding
miRNA_precursor
protein_coding
small_nucle(ol)ar_rna
protein_coding
protein_coding
protein_coding
protein_coding
protein_coding
protein_coding
protein_coding
ncRNA

protein_coding
protein_coding
protein_coding
protein_coding
protein_coding
protein_coding
protein_coding
protein_coding
protein_coding
protein_coding
protein_coding
protein_coding
protein_coding
protein_coding
protein_coding
protein_coding
protein_coding
miRNA_precursor
miRNA_precursor
protein_coding
protein_coding
protein_coding
protein_coding
small_nucle(ol)ar_rna
protein_coding
protein_coding
ncRNA
miRNA_precursor
protein_coding
protein_coding
protein_coding
small_nucle(ol)ar_rna
protein_coding
protein_coding
protein_coding
protein_coding
protein_coding
protein_coding
small_nucle(ol)ar_rna
protein_coding
protein_coding
small_nucle(ol)ar_rna
protein_coding
protein_coding
protein_coding
miRNA_precursor
protein_coding
protein_coding
protein_coding
protein_coding
protein_coding
miRNA_precursor
protein_coding
protein_coding
protein_coding
protein_coding

miRNA_precursor
protein_coding
protein_coding
protein_coding
protein_coding
protein_coding
ncRNA
protein_coding
protein_coding
protein_coding
protein_coding
protein_coding
protein_coding
protein_coding
protein_coding
protein_coding
protein_coding
protein_coding
protein_coding
protein_coding
protein_coding
protein_coding
protein_coding
protein_coding
protein_coding
protein_coding
protein_coding
small_nucle(ol)ar_rna
protein_coding
protein_coding
protein_coding
protein_coding
protein_coding
protein_coding
protein_coding
protein_coding
protein_coding
protein_coding
protein_coding
protein_coding
protein_coding
protein_coding
ncRNA
protein_coding
protein_coding
protein_coding
protein_coding
protein_coding
protein_coding
small_nucle(ol)ar_rna
protein_coding
protein_coding
protein_coding
miRNA_precursor
small_nucle(ol)ar_rna
protein_coding

protein_coding
protein_coding
ncRNA
protein_coding
protein_coding
protein_coding
protein_coding
ncRNA
protein_coding
protein_coding
ncRNA
protein_coding
protein_coding
protein_coding
miRNA_precursor
small_nucle(ol)ar_rna
protein_coding
protein_coding
protein_coding
protein_coding
protein_coding
protein_coding
protein_coding
protein_coding
protein_coding
protein_coding
protein_coding
protein_coding
protein_coding
protein_coding
small_nucle(ol)ar_rna
protein_coding
protein_coding
small_nucle(ol)ar_rna
protein_coding
protein_coding
protein_coding
protein_coding
miRNA_precursor
protein_coding
miRNA_precursor
miRNA_precursor
protein_coding
ncRNA
small_nucle(ol)ar_rna
protein_coding
protein_coding
protein_coding
protein_coding
protein_coding
protein_coding
protein_coding
ncRNA
protein_coding
protein_coding
protein_coding

protein_coding
protein_coding
protein_coding
small_nucle(ol)ar_rna
protein_coding
protein_coding
protein_coding
protein_coding
protein_coding
protein_coding
protein_coding
protein_coding
small_nucle(ol)ar_rna
protein_coding
protein_coding
ncRNA
small_nucle(ol)ar_rna
protein_coding
protein_coding
protein_coding
protein_coding
protein_coding
protein_coding
ncRNA
protein_coding
protein_coding
protein_coding
protein_coding
protein_coding
protein_coding
protein_coding
protein_coding
small_nucle(ol)ar_rna
protein_coding
protein_coding
protein_coding
protein_coding
ncRNA
protein_coding
protein_coding
protein_coding
protein_coding
protein_coding
protein_coding
protein_coding
protein_coding
protein_coding
protein_coding
protein_coding
small_nucle(ol)ar_rna
ncRNA
protein_coding
protein_coding
protein_coding
ncRNA
protein_coding

protein_coding
protein_coding
protein_coding
protein_coding
small_nucle(ol)ar_rna
protein_coding
protein_coding
protein_coding
protein_coding
protein_coding
small_nucle(ol)ar_rna
protein_coding
ncRNA
protein_coding
protein_coding
protein_coding
protein_coding
protein_coding
protein_coding
protein_coding
protein_coding
protein_coding
protein_coding
protein_coding
ncRNA
protein_coding
small_nucle(ol)ar_rna
protein_coding
protein_coding
protein_coding
protein_coding
small_nucle(ol)ar_rna
protein_coding
protein_coding
small_nucle(ol)ar_rna
protein_coding
protein_coding
protein_coding
protein_coding
protein_coding
ncRNA
protein_coding
protein_coding
protein_coding
protein_coding
protein_coding
protein_coding
protein_coding
small_nucle(ol)ar_rna
protein_coding
protein_coding
protein_coding
protein_coding
protein_coding
protein_coding
protein_coding

protein_coding
protein_coding
protein_coding
protein_coding
protein_coding
miRNA_precursor
protein_coding
protein_coding
protein_coding
protein_coding
protein_coding
small_nucle(ol)ar_rna
protein_coding
ncRNA
protein_coding
protein_coding
protein_coding
protein_coding
protein_coding
protein_coding
protein_coding
protein_coding
protein_coding
protein_coding
protein_coding
protein_coding
protein_coding
protein_coding
protein_coding
protein_coding
protein_coding
miRNA_precursor
protein_coding
protein_coding
protein_coding
protein_coding
protein_coding
protein_coding
ncRNA
protein_coding
protein_coding
protein_coding
protein_coding
protein_coding
protein_coding
protein_coding
protein_coding
protein_coding
protein_coding
protein_coding
protein_coding
protein_coding
small_nucle(ol)ar_rna
miRNA_precursor
small_nucle(ol)ar_rna
protein_coding

protein_coding
small_nucle(ol)ar_rna
protein_coding
protein_coding
protein_coding
protein_coding
miRNA_precursor
protein_coding
small_nucle(ol)ar_rna
small_nucle(ol)ar_rna
protein_coding
protein_coding
protein_coding
protein_coding
protein_coding
protein_coding
protein_coding
protein_coding
protein_coding
protein_coding
protein_coding
protein_coding
protein_coding
protein_coding
protein_coding
protein_coding
protein_coding
protein_coding
protein_coding
protein_coding
protein_coding
protein_coding
protein_coding
protein_coding
small_nucle(ol)ar_rna
ncRNA
protein_coding
protein_coding
ncRNA
protein_coding
protein_coding
protein_coding
protein_coding
protein_coding
protein_coding
protein_coding
protein_coding
protein_coding
protein_coding
protein_coding
protein_coding
protein_coding
protein_coding
protein_coding
ncRNA
protein_coding

protein_coding
protein_coding
protein_coding
protein_coding
protein_coding
protein_coding
protein_coding
protein_coding
protein_coding
protein_coding
protein_coding
protein_coding
protein_coding
protein_coding
protein_coding
protein_coding
protein_coding
protein_coding
protein_coding
protein_coding
protein_coding
protein_coding
protein_coding
protein_coding
protein_coding
protein_coding
protein_coding
small_nucle(ol)ar_rna
protein_coding
protein_coding
protein_coding
protein_coding
protein_coding
protein_coding
protein_coding
protein_coding
protein_coding
protein_coding
protein_coding
protein_coding
protein_coding
small_nucle(ol)ar_rna
protein_coding
small_nucle(ol)ar_rna
protein_coding
protein_coding
protein_coding
protein_coding
protein_coding
protein_coding
protein_coding
protein_coding
ncRNA
protein_coding
protein_coding
protein_coding

protein_coding
ncRNA
protein_coding
protein_coding
protein_coding
protein_coding
protein_coding
protein_coding
miRNA_precursor
small_nucle(ol)ar_rna
protein_coding
protein_coding
protein_coding
protein_coding
protein_coding
protein_coding
protein_coding
protein_coding
small_nucle(ol)ar_rna
protein_coding
miRNA_precursor
protein_coding
protein_coding
protein_coding
protein_coding
protein_coding
protein_coding
protein_coding
protein_coding
ncRNA
protein_coding
protein_coding
protein_coding
protein_coding
ncRNA
protein_coding
protein_coding
protein_coding
protein_coding
protein_coding
protein_coding
protein_coding
protein_coding
protein_coding
protein_coding
miRNA_precursor
protein_coding
protein_coding
protein_coding
ncRNA
small_nucle(ol)ar_rna
protein_coding
protein_coding
pre_trna
protein_coding
protein_coding

protein_coding
protein_coding
protein_coding
protein_coding
protein_coding
protein_coding
protein_coding
protein_coding
small_nucle(ol)ar_rna
protein_coding
protein_coding
miRNA_precursor
protein_coding
protein_coding
protein_coding
protein_coding
protein_coding
small_nucle(ol)ar_rna
protein_coding
protein_coding
protein_coding
protein_coding
protein_coding
protein_coding
protein_coding
protein_coding
protein_coding
protein_coding
small_nucle(ol)ar_rna
protein_coding
ncRNA
protein_coding
protein_coding
protein_coding
protein_coding
protein_coding
ncRNA
protein_coding
protein_coding
protein_coding
protein_coding
protein_coding
protein_coding
protein_coding
protein_coding
protein_coding
protein_coding
protein_coding
protein_coding
protein_coding
protein_coding
protein_coding
protein_coding
protein_coding
protein_coding
protein_coding

protein_coding
protein_coding
protein_coding
protein_coding
small_nucle(ol)ar_rna
protein_coding
protein_coding
protein_coding
protein_coding
protein_coding
protein_coding
protein_coding
protein_coding
protein_coding
protein_coding
protein_coding
protein_coding
protein_coding
protein_coding
protein_coding
protein_coding
protein_coding
protein_coding
protein_coding
protein_coding
protein_coding
protein_coding
protein_coding
protein_coding
protein_coding
protein_coding
protein_coding
protein_coding
protein_coding
protein_coding
small_nucle(ol)ar_rna
protein_coding
protein_coding
protein_coding
protein_coding
protein_coding
protein_coding
protein_coding
protein_coding
protein_coding
protein_coding
protein_coding
protein_coding
protein_coding
protein_coding
protein_coding
protein_coding
protein_coding
protein_coding
protein_coding
protein_coding

protein_coding
small_nucle(ol)ar_rna
protein_coding
protein_coding
protein_coding
protein_coding
protein_coding
protein_coding
protein_coding
protein_coding
protein_coding
protein_coding
protein_coding
protein_coding
protein_coding
protein_coding
protein_coding
small_nucle(ol)ar_rna
protein_coding
protein_coding
protein_coding
protein_coding
protein_coding
miRNA_precursor
protein_coding
ncRNA
protein_coding
ncRNA
small_nucle(ol)ar_rna
protein_coding
protein_coding
protein_coding
protein_coding
ncRNA
protein_coding
protein_coding
small_nucle(ol)ar_rna
protein_coding
protein_coding
protein_coding
protein_coding
protein_coding
protein_coding
protein_coding
protein_coding
protein_coding
protein_coding
miRNA_precursor
protein_coding
protein_coding
ncRNA
protein_coding
protein_coding
small_nucle(ol)ar_rna
protein_coding
protein_coding

protein_coding
protein_coding
ncRNA
protein_coding
protein_coding
miRNA_precursor
protein_coding
protein_coding
protein_coding
protein_coding
protein_coding
protein_coding
protein_coding
protein_coding
protein_coding
protein_coding
protein_coding
protein_coding
small_nucle(ol)ar_rna
protein_coding
protein_coding
protein_coding
ncRNA
protein_coding
protein_coding
pre_trna
miRNA_precursor
ncRNA
protein_coding
protein_coding
protein_coding
miRNA_precursor
protein_coding
protein_coding
protein_coding
protein_coding
protein_coding
protein_coding
protein_coding
protein_coding
small_nucle(ol)ar_rna
protein_coding
protein_coding
protein_coding
protein_coding
protein_coding
protein_coding
protein_coding
protein_coding
protein_coding
protein_coding
protein_coding
ncRNA
protein_coding
protein_coding
protein_coding

protein_coding
protein_coding
miRNA_precursor
miRNA_precursor
ncRNA
protein_coding
protein_coding
protein_coding
protein_coding
protein_coding
protein_coding
protein_coding
protein_coding
protein_coding
protein_coding
miRNA_precursor
protein_coding
protein_coding
protein_coding
protein_coding
protein_coding
protein_coding
protein_coding
protein_coding
protein_coding
ncRNA
protein_coding
protein_coding
protein_coding
protein_coding
protein_coding
protein_coding
ncRNA
protein_coding
protein_coding
protein_coding
protein_coding
protein_coding
protein_coding
protein_coding
pre_trna
protein_coding
protein_coding
protein_coding
protein_coding
protein_coding
protein_coding
protein_coding
protein_coding
small_nucle(ol)ar_rna
protein_coding
protein_coding
protein_coding
protein_coding
protein_coding
protein_coding

protein_coding
protein_coding
protein_coding
protein_coding
protein_coding
protein_coding
protein_coding
protein_coding
protein_coding
protein_coding
ncRNA
protein_coding
protein_coding
protein_coding
protein_coding
protein_coding
protein_coding
protein_coding
protein_coding
protein_coding
protein_coding
protein_coding
protein_coding
small_nucle(ol)ar_rna
protein_coding
protein_coding
protein_coding
protein_coding
ncRNA
protein_coding
small_nucle(ol)ar_rna
protein_coding
protein_coding
protein_coding
protein_coding
small_nucle(ol)ar_rna
protein_coding
ncRNA
protein_coding
protein_coding
protein_coding
protein_coding
protein_coding
protein_coding
protein_coding
protein_coding
small_nucle(ol)ar_rna
protein_coding
small_nucle(ol)ar_rna
protein_coding
protein_coding
protein_coding
small_nucle(ol)ar_rna
protein_coding
protein_coding
protein_coding

small_nucle(ol)ar_rna
protein_coding
protein_coding
protein_coding
protein_coding
protein_coding
protein_coding
ncRNA
protein_coding
protein_coding
protein_coding
protein_coding
protein_coding
protein_coding
protein_coding
protein_coding
ncRNA
protein_coding
protein_coding
ncRNA
protein_coding
protein_coding
protein_coding
protein_coding
ncRNA
protein_coding
protein_coding
protein_coding
protein_coding
protein_coding
protein_coding
miRNA_precursor
ncRNA
protein_coding
protein_coding
protein_coding
protein_coding
protein_coding
protein_coding
protein_coding
protein_coding
protein_coding
ncRNA
protein_coding
protein_coding
protein_coding
miRNA_precursor
protein_coding
protein_coding
protein_coding
protein_coding
protein_coding
protein_coding
protein_coding
protein_coding
protein_coding

protein_coding
protein_coding
protein_coding
protein_coding
protein_coding
protein_coding
small_nucle(ol)ar_rna
protein_coding
protein_coding
protein_coding
protein_coding
protein_coding
protein_coding
protein_coding
protein_coding
protein_coding
ncRNA
protein_coding
protein_coding
protein_coding
protein_coding
protein_coding
protein_coding
protein_coding
protein_coding
small_nucle(ol)ar_rna
protein_coding
small_nucle(ol)ar_rna
small_nucle(ol)ar_rna
protein_coding
protein_coding
protein_coding
protein_coding
protein_coding
protein_coding
protein_coding
protein_coding
protein_coding
protein_coding
protein_coding
protein_coding
protein_coding
protein_coding
protein_coding
ncRNA
protein_coding
protein_coding
protein_coding
small_nucle(ol)ar_rna
protein_coding
protein_coding
protein_coding
ncRNA
protein_coding
protein_coding
protein_coding

protein_coding
protein_coding
protein_coding
protein_coding
protein_coding
protein_coding
protein_coding
small_nucle(ol)ar_rna
protein_coding
protein_coding
protein_coding
protein_coding
protein_coding
protein_coding
protein_coding
protein_coding
protein_coding
protein_coding
protein_coding
miRNA_precursor
protein_coding
protein_coding
protein_coding
protein_coding
protein_coding
protein_coding
miRNA_precursor
protein_coding
protein_coding
protein_coding
miRNA_precursor
protein_coding
protein_coding
protein_coding
protein_coding
protein_coding
protein_coding
ncRNA
protein_coding
small_nucle(ol)ar_rna
protein_coding
protein_coding
protein_coding
ncRNA
protein_coding
small_nucle(ol)ar_rna
miRNA_precursor
protein_coding
protein_coding
protein_coding
protein_coding
protein_coding
miRNA_precursor
protein_coding
protein_coding
protein_coding

ncRNA
protein_coding
protein_coding
protein_coding
protein_coding
ncRNA
protein_coding
protein_coding
protein_coding
protein_coding
protein_coding
ncRNA
protein_coding
ncRNA
protein_coding
protein_coding
protein_coding
protein_coding
protein_coding
protein_coding
protein_coding
protein_coding
protein_coding
protein_coding
protein_coding
protein_coding
protein_coding
protein_coding
protein_coding
protein_coding
protein_coding
protein_coding
protein_coding
protein_coding
protein_coding
protein_coding
protein_coding
small_nucle(ol)ar_rna
protein_coding
protein_coding
protein_coding
protein_coding
protein_coding
protein_coding
small_nucle(ol)ar_rna
protein_coding
protein_coding
protein_coding
protein_coding
protein_coding
protein_coding
protein_coding
protein_coding
protein_coding
protein_coding
protein_coding

protein_coding
protein_coding
protein_coding
protein_coding
protein_coding
protein_coding
protein_coding
protein_coding
protein_coding
protein_coding
small_nucle(ol)ar_rna
protein_coding
ncRNA
protein_coding
protein_coding
protein_coding
protein_coding
protein_coding
protein_coding
protein_coding
protein_coding
small_nucle(ol)ar_rna
protein_coding
protein_coding
protein_coding
protein_coding
protein_coding
protein_coding
protein_coding
protein_coding
protein_coding
protein_coding
protein_coding
protein_coding
protein_coding
protein_coding
protein_coding
protein_coding
protein_coding
protein_coding
protein_coding
protein_coding
protein_coding
protein_coding
protein_coding
protein_coding
protein_coding
protein_coding
protein_coding
protein_coding
protein_coding
miRNA_precursor
protein_coding
protein_coding
protein_coding
miRNA_precursor

protein_coding
protein_coding
ncRNA
small_nucle(ol)ar_rna
ncRNA
protein_coding
small_nucle(ol)ar_rna
miRNA_precursor
small_nucle(ol)ar_rna
protein_coding
protein_coding
protein_coding
protein_coding
protein_coding
ncRNA
protein_coding
protein_coding
small_nucle(ol)ar_rna
protein_coding
protein_coding
protein_coding
small_nucle(ol)ar_rna
protein_coding
protein_coding
protein_coding
protein_coding
protein_coding
protein_coding
protein_coding
protein_coding
protein_coding
protein_coding
protein_coding
protein_coding
protein_coding
protein_coding
miRNA_precursor
protein_coding
protein_coding
protein_coding
protein_coding
protein_coding
protein_coding
protein_coding
protein_coding
protein_coding
protein_coding
protein_coding
small_nucle(ol)ar_rna
protein_coding
protein_coding
protein_coding
protein_coding
protein_coding
protein_coding
miRNA_precursor

protein_coding
protein_coding
small_nucle(ol)ar_rna
protein_coding
protein_coding
protein_coding
protein_coding
protein_coding
ncRNA
protein_coding
protein_coding
protein_coding
protein_coding
protein_coding
protein_coding
protein_coding
protein_coding
protein_coding
protein_coding
protein_coding
protein_coding
protein_coding
protein_coding
protein_coding
protein_coding
protein_coding
protein_coding
ncRNA
protein_coding
protein_coding
protein_coding
miRNA_precursor
protein_coding
protein_coding
protein_coding
protein_coding
protein_coding
protein_coding
protein_coding
protein_coding
protein_coding
protein_coding
protein_coding
protein_coding
protein_coding
protein_coding
protein_coding
protein_coding
protein_coding
miRNA_precursor
protein_coding
protein_coding
protein_coding
protein_coding
protein_coding
protein_coding

protein_coding
protein_coding
protein_coding
protein_coding
protein_coding
protein_coding
protein_coding
protein_coding
protein_coding
protein_coding
protein_coding
protein_coding
protein_coding
protein_coding
protein_coding
protein_coding
protein_coding
protein_coding
protein_coding
protein_coding
protein_coding
protein_coding
protein_coding
protein_coding
protein_coding
protein_coding
small_nucle(ol)ar_rna
miRNA_precursor
protein_coding
protein_coding
small_nucle(ol)ar_rna
protein_coding
protein_coding
protein_coding
protein_coding
protein_coding
protein_coding
protein_coding
ncRNA
protein_coding
protein_coding
protein_coding
protein_coding
protein_coding
protein_coding
protein_coding
protein_coding
protein_coding
protein_coding
protein_coding
ncRNA
protein_coding
protein_coding
protein_coding
protein_coding
protein_coding

small_nucle(ol)ar_rna
small_nucle(ol)ar_rna
protein_coding
protein_coding
ncRNA
protein_coding
protein_coding
protein_coding
protein_coding
ncRNA
ncRNA
protein_coding
miRNA_precursor
protein_coding
small_nucle(ol)ar_rna
protein_coding
protein_coding
ncRNA
ncRNA
protein_coding
ncRNA
protein_coding
protein_coding
protein_coding
protein_coding
protein_coding
protein_coding
ribosomal_rna
protein_coding
protein_coding
protein_coding
miRNA_precursor
protein_coding
protein_coding
protein_coding
protein_coding
protein_coding
protein_coding
small_nucle(ol)ar_rna
protein_coding
protein_coding
protein_coding
protein_coding
small_nucle(ol)ar_rna
small_nucle(ol)ar_rna
protein_coding
miRNA_precursor
protein_coding
protein_coding
protein_coding
miRNA_precursor
protein_coding
protein_coding
miRNA_precursor
protein_coding
protein_coding

protein_coding
protein_coding
protein_coding
small_nucle(ol)ar_rna
protein_coding
protein_coding
protein_coding
protein_coding
ncRNA
protein_coding
protein_coding
protein_coding
protein_coding
protein_coding
ncRNA
protein_coding
protein_coding
protein_coding
protein_coding
protein_coding
protein_coding
protein_coding
protein_coding
protein_coding
protein_coding
ncRNA
protein_coding
protein_coding
protein_coding
protein_coding
protein_coding
protein_coding
protein_coding
protein_coding
protein_coding
protein_coding
small_nucle(ol)ar_rna
protein_coding
ncRNA
protein_coding
protein_coding
protein_coding
protein_coding
protein_coding
protein_coding
protein_coding
protein_coding
protein_coding
protein_coding
protein_coding
protein_coding
protein_coding
protein_coding
protein_coding
protein_coding
protein_coding

protein_coding
ncRNA
protein_coding
protein_coding
protein_coding
ncRNA
pre_trna
protein_coding
protein_coding
protein_coding
protein_coding
protein_coding
protein_coding
protein_coding
protein_coding
protein_coding
protein_coding
protein_coding
ncRNA
protein_coding
protein_coding
protein_coding
protein_coding
ncRNA
protein_coding
protein_coding
protein_coding
protein_coding
protein_coding
ncRNA
protein_coding
protein_coding
protein_coding
miRNA_precursor
protein_coding
protein_coding
protein_coding
protein_coding
protein_coding
protein_coding
protein_coding
protein_coding
protein_coding
protein_coding
protein_coding
miRNA_precursor
protein_coding
ncRNA
protein_coding
protein_coding
protein_coding
protein_coding
ncRNA
protein_coding
protein_coding
protein_coding

protein_coding
protein_coding
protein_coding
protein_coding
protein_coding
ncRNA
protein_coding
protein_coding
protein_coding
protein_coding
protein_coding
protein_coding
small_nucle(ol)ar_rna
protein_coding
protein_coding
protein_coding
protein_coding
protein_coding
protein_coding
protein_coding
protein_coding
ncRNA
protein_coding
protein_coding
protein_coding
protein_coding
protein_coding
protein_coding
protein_coding
protein_coding
protein_coding
protein_coding
protein_coding
protein_coding
protein_coding
protein_coding
protein_coding
protein_coding
protein_coding
protein_coding
protein_coding
protein_coding
protein_coding
protein_coding
protein_coding
miRNA_precursor
ncRNA
protein_coding
ncRNA
protein_coding
protein_coding
protein_coding
protein_coding
small_nucle(ol)ar_rna
protein_coding
protein_coding

protein_coding
protein_coding
protein_coding
protein_coding
protein_coding
protein_coding
protein_coding
protein_coding
small_nucle(ol)ar_rna
protein_coding
protein_coding
protein_coding
protein_coding
protein_coding
protein_coding
miRNA_precursor
protein_coding
protein_coding
protein_coding
protein_coding
ncRNA
protein_coding
miRNA_precursor
protein_coding
protein_coding
protein_coding
protein_coding
protein_coding
protein_coding
protein_coding
protein_coding
protein_coding
protein_coding
protein_coding
protein_coding
protein_coding
protein_coding
protein_coding
protein_coding
protein_coding
ncRNA
protein_coding
protein_coding
protein_coding
protein_coding
protein_coding
protein_coding
protein_coding
miRNA_precursor
miRNA_precursor
protein_coding
small_nucle(ol)ar_rna
protein_coding
protein_coding
protein_coding
protein_coding

protein_coding
protein_coding
small_nucle(ol)ar_rna
protein_coding
small_nucle(ol)ar_rna
protein_coding
protein_coding
protein_coding
protein_coding
protein_coding
miRNA_precursor
protein_coding
protein_coding
protein_coding
protein_coding
protein_coding
miRNA_precursor
protein_coding
protein_coding
protein_coding
protein_coding
protein_coding
protein_coding
protein_coding
protein_coding
protein_coding
protein_coding
protein_coding
protein_coding
small_nucle(ol)ar_rna
protein_coding
protein_coding
protein_coding
protein_coding
protein_coding
protein_coding
small_nucle(ol)ar_rna
ncRNA
protein_coding
miRNA_precursor
protein_coding
protein_coding
protein_coding
protein_coding
protein_coding
protein_coding
protein_coding
protein_coding
protein_coding
protein_coding
small_nucle(ol)ar_rna
protein_coding
protein_coding
protein_coding
protein_coding
protein_coding

protein_coding
protein_coding
protein_coding
protein_coding
protein_coding
ncRNA
protein_coding
protein_coding
protein_coding
protein_coding
protein_coding
protein_coding
protein_coding
protein_coding
small_nucle(ol)ar_rna
ncRNA
protein_coding
protein_coding
protein_coding
protein_coding
ncRNA
protein_coding
protein_coding
protein_coding
protein_coding
protein_coding
protein_coding
protein_coding
protein_coding
protein_coding
small_nucle(ol)ar_rna
protein_coding
protein_coding
protein_coding
protein_coding
protein_coding
protein_coding
protein_coding
protein_coding
protein_coding
protein_coding
protein_coding
protein_coding
protein_coding
protein_coding
protein_coding
protein_coding
protein_coding
protein_coding
protein_coding
protein_coding
protein_coding
protein_coding
protein_coding
protein_coding
protein_coding

protein_coding
protein_coding
protein_coding
protein_coding
ncRNA
miRNA_precursor
miRNA_precursor
protein_coding
protein_coding
protein_coding
protein_coding
protein_coding
protein_coding
protein_coding
protein_coding
protein_coding
protein_coding
protein_coding
ncRNA
protein_coding
protein_coding
protein_coding
protein_coding
protein_coding
ncRNA
protein_coding
protein_coding
protein_coding
protein_coding
protein_coding
protein_coding
protein_coding
protein_coding
ncRNA
protein_coding
protein_coding
ncRNA
protein_coding
protein_coding
protein_coding
protein_coding
protein_coding
protein_coding
protein_coding
protein_coding
protein_coding
protein_coding
protein_coding
protein_coding
protein_coding
protein_coding
protein_coding
protein_coding
protein_coding
protein_coding
ncRNA

protein_coding
protein_coding
protein_coding
protein_coding
protein_coding
protein_coding
protein_coding
protein_coding
protein_coding
small_nucle(ol)ar_rna
small_nucle(ol)ar_rna
small_nucle(ol)ar_rna
small_nucle(ol)ar_rna
protein_coding
protein_coding
protein_coding
protein_coding
protein_coding
ncRNA
protein_coding
ncRNA
protein_coding
protein_coding
protein_coding
small_nucle(ol)ar_rna
small_nucle(ol)ar_rna
protein_coding
protein_coding
protein_coding
protein_coding
protein_coding
protein_coding
protein_coding
protein_coding
protein_coding
protein_coding
protein_coding
protein_coding
protein_coding
ncRNA
protein_coding
miRNA_precursor
protein_coding
protein_coding
protein_coding
protein_coding
protein_coding
pre_trna
ncRNA
protein_coding
protein_coding
protein_coding
protein_coding
protein_coding
protein_coding
protein_coding

protein_coding
protein_coding
protein_coding
protein_coding
protein_coding
protein_coding
ncRNA
protein_coding
protein_coding
protein_coding
protein_coding
protein_coding
protein_coding
protein_coding
protein_coding
protein_coding
ncRNA
protein_coding
protein_coding
protein_coding
protein_coding
protein_coding
protein_coding
protein_coding
protein_coding
protein_coding
protein_coding
protein_coding
protein_coding
protein_coding
protein_coding
protein_coding
protein_coding
protein_coding
protein_coding
protein_coding
protein_coding
protein_coding
protein_coding
protein_coding
protein_coding
protein_coding
protein_coding
protein_coding
protein_coding
protein_coding
protein_coding
protein_coding
protein_coding
protein_coding
protein_coding
protein_coding
protein_coding
ncRNA
protein_coding
protein_coding

protein_coding
protein_coding
protein_coding
protein_coding
protein_coding
protein_coding
protein_coding
protein_coding
protein_coding
protein_coding
protein_coding
protein_coding
protein_coding
protein_coding
protein_coding
protein_coding
protein_coding
protein_coding
protein_coding
protein_coding
protein_coding
protein_coding
protein_coding
protein_coding
protein_coding
protein_coding
protein_coding
protein_coding
protein_coding
protein_coding
protein_coding
protein_coding
protein_coding
protein_coding
protein_coding
protein_coding
protein_coding
protein_coding
protein_coding
protein_coding
protein_coding
protein_coding
protein_coding
protein_coding
protein_coding
protein_coding
protein_coding
protein_coding
protein_coding
protein_coding
small_nucle(ol)ar_rna
protein_coding
protein_coding
protein_coding
protein_coding
protein_coding

protein_coding
protein_coding
protein_coding
protein_coding
protein_coding
protein_coding
protein_coding
protein_coding
protein_coding
protein_coding
protein_coding
protein_coding
protein_coding
protein_coding
protein_coding
protein_coding
protein_coding
protein_coding
protein_coding
protein_coding
protein_coding
protein_coding
protein_coding
protein_coding
protein_coding
pre_trna
protein_coding
protein_coding
protein_coding
protein_coding
ncRNA
protein_coding
protein_coding
protein_coding
protein_coding
protein_coding
protein_coding
protein_coding
protein_coding
protein_coding
protein_coding
protein_coding
protein_coding
protein_coding
protein_coding
protein_coding
protein_coding
protein_coding
protein_coding
protein_coding
protein_coding
protein_coding
protein_coding
protein_coding
protein_coding
pre_trna
ncRNA

protein_coding
protein_coding
protein_coding
protein_coding
protein_coding
protein_coding
protein_coding
protein_coding
protein_coding
protein_coding
protein_coding
protein_coding
protein_coding
protein_coding
protein_coding
protein_coding
protein_coding
protein_coding
protein_coding
protein_coding
protein_coding
protein_coding
protein_coding
protein_coding
protein_coding
protein_coding
protein_coding
protein_coding
protein_coding
protein_coding
protein_coding
protein_coding
protein_coding
protein_coding
protein_coding
ncRNA
protein_coding
protein_coding
protein_coding
protein_coding
protein_coding
protein_coding
protein_coding
protein_coding
protein_coding
protein_coding
pre_trna
protein_coding
protein_coding
protein_coding
protein_coding
protein_coding
protein_coding
protein_coding
protein_coding
protein_coding

protein_coding
protein_coding
protein_coding
protein_coding
protein_coding
protein_coding
protein_coding
protein_coding
protein_coding
protein_coding
protein_coding
protein_coding
protein_coding
protein_coding
protein_coding
pre_trna
protein_coding
protein_coding
protein_coding
protein_coding
protein_coding
protein_coding
protein_coding
protein_coding
protein_coding
protein_coding
protein_coding
small_nucle(ol)ar_rna
miRNA_precursor
protein_coding
ncRNA
protein_coding
protein_coding
protein_coding
protein_coding
protein_coding
protein_coding
protein_coding
protein_coding
ncRNA
protein_coding
protein_coding
protein_coding
protein_coding
protein_coding
protein_coding
protein_coding
protein_coding
protein_coding
protein_coding
protein_coding
protein_coding
protein_coding
protein_coding
protein_coding
protein_coding

protein_coding
protein_coding
protein_coding
protein_coding
protein_coding
protein_coding
protein_coding
protein_coding
protein_coding
protein_coding
protein_coding
protein_coding
protein_coding
protein_coding
protein_coding
protein_coding
protein_coding
protein_coding
protein_coding
protein_coding
protein_coding
protein_coding
protein_coding
protein_coding
protein_coding
protein_coding
protein_coding
protein_coding
protein_coding
protein_coding
ncRNA
protein_coding
protein_coding
protein_coding
protein_coding
protein_coding
protein_coding
protein_coding
protein_coding
protein_coding
protein_coding
protein_coding
protein_coding
protein_coding
protein_coding
protein_coding
protein_coding
protein_coding
ncRNA
protein_coding
protein_coding
protein_coding
protein_coding
protein_coding
protein_coding
protein_coding

protein_coding
protein_coding
protein_coding
protein_coding
protein_coding
protein_coding
protein_coding
protein_coding
protein_coding
protein_coding
protein_coding
protein_coding
protein_coding
protein_coding
protein_coding
protein_coding
protein_coding
protein_coding
protein_coding
protein_coding
protein_coding
protein_coding
protein_coding
protein_coding
protein_coding
protein_coding
protein_coding
protein_coding
protein_coding
protein_coding
protein_coding
protein_coding
protein_coding
protein_coding
protein_coding
protein_coding
small_nucle(ol)ar_rna
protein_coding
protein_coding
protein_coding
protein_coding
ncRNA
protein_coding
protein_coding
protein_coding
protein_coding
protein_coding
protein_coding
protein_coding
protein_coding
protein_coding
protein_coding
protein_coding
protein_coding
ncRNA
protein_coding

protein_coding
protein_coding
pre_trna
protein_coding
protein_coding
protein_coding
protein_coding
protein_coding
protein_coding
protein_coding
protein_coding
small_nucle(ol)ar_rna
small_nucle(ol)ar_rna
small_nucle(ol)ar_rna
protein_coding
protein_coding
protein_coding
protein_coding
protein_coding
protein_coding
protein_coding
protein_coding
protein_coding
protein_coding
protein_coding
pre_trna
protein_coding
protein_coding
protein_coding
protein_coding
protein_coding
protein_coding
protein_coding
protein_coding
protein_coding
protein_coding
protein_coding
ncRNA
protein_coding
protein_coding
protein_coding
protein_coding
protein_coding
protein_coding
protein_coding
protein_coding
ncRNA
protein_coding
protein_coding
protein_coding
protein_coding
protein_coding
protein_coding
protein_coding
protein_coding
protein_coding

protein_coding
protein_coding
protein_coding
protein_coding
small_nucle(ol)ar_rna
small_nucle(ol)ar_rna
protein_coding
protein_coding
protein_coding
protein_coding
protein_coding
miRNA_precursor
protein_coding
protein_coding
protein_coding
protein_coding
protein_coding
protein_coding
protein_coding
protein_coding
protein_coding
protein_coding
protein_coding
protein_coding
protein_coding
protein_coding
protein_coding
protein_coding
protein_coding
protein_coding
protein_coding
protein_coding
protein_coding
protein_coding
protein_coding
ncRNA
protein_coding
protein_coding
protein_coding
protein_coding
protein_coding
protein_coding
protein_coding
ncRNA
protein_coding
protein_coding
protein_coding
protein_coding
protein_coding
protein_coding
protein_coding
ncRNA
protein_coding
protein_coding
protein_coding
protein_coding

ncRNA
protein_coding
protein_coding
protein_coding
protein_coding
protein_coding
protein_coding
protein_coding
protein_coding
protein_coding
protein_coding
protein_coding
protein_coding
protein_coding
protein_coding
protein_coding
protein_coding
ncRNA
protein_coding
protein_coding
ncRNA
protein_coding
protein_coding
ncRNA
protein_coding
protein_coding
protein_coding
protein_coding
protein_coding
ncRNA
protein_coding
protein_coding
protein_coding
miRNA_precursor
protein_coding
protein_coding
protein_coding
protein_coding
protein_coding
protein_coding
protein_coding
protein_coding
protein_coding
protein_coding
protein_coding
protein_coding
protein_coding
protein_coding
protein_coding
protein_coding
protein_coding
protein_coding
protein_coding
protein_coding
protein_coding
protein_coding

protein_coding
protein_coding
protein_coding
protein_coding
protein_coding
protein_coding
protein_coding
ncRNA
protein_coding
ncRNA
protein_coding
protein_coding
miRNA_precursor
protein_coding
protein_coding
protein_coding
protein_coding
protein_coding
protein_coding
protein_coding
protein_coding
protein_coding
protein_coding
protein_coding
protein_coding
protein_coding
protein_coding
ncRNA
protein_coding
small_nucle(ol)ar_rna
protein_coding
protein_coding
protein_coding
protein_coding
protein_coding
protein_coding
protein_coding
protein_coding
protein_coding
protein_coding
protein_coding
protein_coding
protein_coding
protein_coding
protein_coding
protein_coding
protein_coding
protein_coding
protein_coding
protein_coding
protein_coding
protein_coding
protein_coding
protein_coding
protein_coding
protein_coding

protein_coding
protein_coding
protein_coding
protein_coding
protein_coding
protein_coding
protein_coding
protein_coding
protein_coding
protein_coding
protein_coding
protein_coding
protein_coding
protein_coding
protein_coding
protein_coding
protein_coding
protein_coding
protein_coding
ncRNA
protein_coding
protein_coding
protein_coding
protein_coding
protein_coding
ncRNA
protein_coding
protein_coding
protein_coding
protein_coding
protein_coding
protein_coding
protein_coding
protein_coding
protein_coding
protein_coding
protein_coding
protein_coding
protein_coding
protein_coding
protein_coding
protein_coding
protein_coding
protein_coding
protein_coding
protein_coding
protein_coding
protein_coding
protein_coding
protein_coding
protein_coding
protein_coding
protein_coding
protein_coding
ncRNA
protein_coding

protein_coding
protein_coding
protein_coding
protein_coding
protein_coding
ncRNA
protein_coding
ncRNA
small_nucle(ol)ar_rna
protein_coding
protein_coding
protein_coding
protein_coding
protein_coding
protein_coding
protein_coding
protein_coding
protein_coding
protein_coding
protein_coding
protein_coding
protein_coding
small_nucle(ol)ar_rna
protein_coding
protein_coding
protein_coding
protein_coding
protein_coding
protein_coding
protein_coding
protein_coding
protein_coding
protein_coding
protein_coding
protein_coding
protein_coding
protein_coding
protein_coding
protein_coding
protein_coding
protein_coding
protein_coding
protein_coding
protein_coding
protein_coding
protein_coding
protein_coding
protein_coding
protein_coding
protein_coding
protein_coding
protein_coding
ncRNA
protein_coding
protein_coding
protein_coding

ncRNA
protein_coding
protein_coding
protein_coding
protein_coding
protein_coding
protein_coding
protein_coding
protein_coding
protein_coding
protein_coding
protein_coding
protein_coding
protein_coding
protein_coding
protein_coding
protein_coding
protein_coding
protein_coding
protein_coding
protein_coding
protein_coding
protein_coding
protein_coding
protein_coding
protein_coding
protein_coding
protein_coding
ncRNA
protein_coding
protein_coding
protein_coding
protein_coding
protein_coding
protein_coding
protein_coding
protein_coding
miRNA_precursor
protein_coding
protein_coding
protein_coding
protein_coding
ncRNA
protein_coding
protein_coding
small_nucle(ol)ar_rna
protein_coding
protein_coding
protein_coding
protein_coding
protein_coding
protein_coding
protein_coding
protein_coding
protein_coding
protein_coding

protein_coding
protein_coding
protein_coding
protein_coding
protein_coding
protein_coding
protein_coding
protein_coding
protein_coding
ncRNA
protein_coding
protein_coding
protein_coding
protein_coding
protein_coding
protein_coding
protein_coding
protein_coding
protein_coding
protein_coding
protein_coding
protein_coding
protein_coding
protein_coding
protein_coding
protein_coding
protein_coding
protein_coding
protein_coding
protein_coding
protein_coding
small_nucle(ol)ar_rna
protein_coding
protein_coding
protein_coding
protein_coding
protein_coding
protein_coding
pre_trna
protein_coding
ncRNA
protein_coding
protein_coding
protein_coding
protein_coding
protein_coding
ncRNA
protein_coding
protein_coding
protein_coding
protein_coding
protein_coding
protein_coding
protein_coding
protein_coding
protein_coding

protein_coding
protein_coding
protein_coding
protein_coding
protein_coding
protein_coding
protein_coding
protein_coding
ncRNA
protein_coding
protein_coding
protein_coding
protein_coding
protein_coding
protein_coding
protein_coding
protein_coding
miRNA_precursor
protein_coding
protein_coding
protein_coding
protein_coding
protein_coding
protein_coding
protein_coding
protein_coding
protein_coding
protein_coding
protein_coding
protein_coding
ncRNA
protein_coding
ncRNA
protein_coding
small_nucle(ol)ar_rna
protein_coding
protein_coding
protein_coding
protein_coding
protein_coding
protein_coding
protein_coding
ncRNA
protein_coding
protein_coding
ncRNA
protein_coding
protein_coding
protein_coding
protein_coding
protein_coding
protein_coding
protein_coding
protein_coding
protein_coding
protein_coding

protein_coding
protein_coding
protein_coding
protein_coding
protein_coding
protein_coding
protein_coding
protein_coding
protein_coding
ncRNA
protein_coding
protein_coding
protein_coding
protein_coding
protein_coding
protein_coding
protein_coding
protein_coding
protein_coding
protein_coding
protein_coding
small_nucle(ol)ar_rna
protein_coding
protein_coding
protein_coding
protein_coding
protein_coding
protein_coding
protein_coding
protein_coding
protein_coding
protein_coding
protein_coding
protein_coding
protein_coding
protein_coding
protein_coding
protein_coding
protein_coding
protein_coding
protein_coding
protein_coding
miRNA_precursor
protein_coding
protein_coding
protein_coding
protein_coding
protein_coding
protein_coding
protein_coding
protein_coding
protein_coding
protein_coding
protein_coding
protein_coding
protein_coding

protein_coding
protein_coding
protein_coding
protein_coding
protein_coding
protein_coding
ncRNA
protein_coding
protein_coding
protein_coding
protein_coding
protein_coding
protein_coding
protein_coding
protein_coding
protein_coding
protein_coding
protein_coding
protein_coding
protein_coding
protein_coding
protein_coding
protein_coding
protein_coding
protein_coding
protein_coding
protein_coding
protein_coding
protein_coding
protein_coding
protein_coding
protein_coding
protein_coding
ncRNA
protein_coding
protein_coding
protein_coding
protein_coding
protein_coding
protein_coding
protein_coding
protein_coding
protein_coding
protein_coding
protein_coding
protein_coding
protein_coding
protein_coding
protein_coding
protein_coding
protein_coding
protein_coding
protein_coding
protein_coding
protein_coding
protein_coding

protein_coding
protein_coding
protein_coding
protein_coding
protein_coding
protein_coding
protein_coding
protein_coding
protein_coding
protein_coding
protein_coding
protein_coding
protein_coding
protein_coding
protein_coding
protein_coding
protein_coding
small_nucle(ol)ar_rna
small_nucle(ol)ar_rna
small_nucle(ol)ar_rna
protein_coding
ncRNA
protein_coding
protein_coding
protein_coding
protein_coding
protein_coding
protein_coding
protein_coding
protein_coding
protein_coding
protein_coding
protein_coding
protein_coding
protein_coding
protein_coding
protein_coding
protein_coding
pre_trna
protein_coding
protein_coding
protein_coding
protein_coding
protein_coding
protein_coding
protein_coding
protein_coding
protein_coding
protein_coding
protein_coding
protein_coding
protein_coding
protein_coding
protein_coding
protein_coding
protein_coding

protein_coding
protein_coding
protein_coding
protein_coding
protein_coding
protein_coding
protein_coding
protein_coding
protein_coding
protein_coding
protein_coding
protein_coding
protein_coding
protein_coding
protein_coding
protein_coding
protein_coding
protein_coding
protein_coding
protein_coding
protein_coding
protein_coding
protein_coding
protein_coding
protein_coding
protein_coding
protein_coding
protein_coding
protein_coding
protein_coding
protein_coding
protein_coding
protein_coding
protein_coding
protein_coding
protein_coding
protein_coding
protein_coding
protein_coding
protein_coding
ncRNA
protein_coding
protein_coding
pre_trna
protein_coding
protein_coding
protein_coding
protein_coding
protein_coding
protein_coding
ncRNA
protein_coding
protein_coding
protein_coding
protein_coding
protein_coding

| |
|---|
| protein_coding |
| protein_coding |
| protein_coding |
| protein_coding |
| protein_coding |
| protein_coding |
| protein_coding |
| protein_coding |
| protein_coding |
| protein_coding |
| ncRNA |
| pre_trna |
| protein_coding |
| protein_coding |
| protein_coding |
| protein_coding |
| protein_coding |
| protein_coding |
| protein_coding |
| protein_coding |
| protein_coding |
| protein_coding |
| protein_coding |
| protein_coding |
| protein_coding |
| protein_coding |
| ncRNA |
| protein_coding |
| protein_coding |
| protein_coding |
| protein_coding |
| protein_coding |
| protein_coding |
| protein_coding |
| protein_coding |
| protein_coding |
| protein_coding |
| protein_coding |
| protein_coding |
| protein_coding |
| protein_coding |
| protein_coding |
| protein_coding |
| protein_coding |
| ncRNA |
| protein_coding |
| protein_coding |
| protein_coding |
| protein_coding |
| ncRNA |
| ncRNA |
| protein_coding |
| protein_coding |
| protein_coding |
| protein_coding |
| protein_coding |

protein_coding
protein_coding
protein_coding
protein_coding
protein_coding
protein_coding
protein_coding
protein_coding
protein_coding
protein_coding
protein_coding
protein_coding
protein_coding
protein_coding
protein_coding
protein_coding
protein_coding
ncRNA
protein_coding
protein_coding
protein_coding
protein_coding
protein_coding
protein_coding
protein_coding
protein_coding
protein_coding
protein_coding
protein_coding
protein_coding
protein_coding
protein_coding
protein_coding
protein_coding
protein_coding
protein_coding
protein_coding
protein_coding
protein_coding
protein_coding
protein_coding
ncRNA
miRNA_precursor
protein_coding
ncRNA
protein_coding
protein_coding
protein_coding
protein_coding
ncRNA
protein_coding
protein_coding
protein_coding
protein_coding
protein_coding
protein_coding

protein_coding
protein_coding
protein_coding
protein_coding
protein_coding
protein_coding
pre_trna
protein_coding
protein_coding
protein_coding
protein_coding
protein_coding
protein_coding
ncRNA
protein_coding
protein_coding
protein_coding
protein_coding
protein_coding
protein_coding
protein_coding
protein_coding
protein_coding
protein_coding
protein_coding
pre_trna
protein_coding
protein_coding
protein_coding
ncRNA
protein_coding
protein_coding
protein_coding
protein_coding
protein_coding
protein_coding
protein_coding
protein_coding
protein_coding
ncRNA
protein_coding
protein_coding
ncRNA
protein_coding
protein_coding
protein_coding
ncRNA
protein_coding
protein_coding
protein_coding
protein_coding
ncRNA
protein_coding
protein_coding
protein_coding
protein_coding

protein_coding
protein_coding
protein_coding
protein_coding
protein_coding
protein_coding
ncRNA
protein_coding
ncRNA
protein_coding
protein_coding
protein_coding
protein_coding
protein_coding
protein_coding
protein_coding
protein_coding
protein_coding
protein_coding
protein_coding
protein_coding
protein_coding
protein_coding
protein_coding
protein_coding
protein_coding
protein_coding
protein_coding
protein_coding
protein_coding
protein_coding
protein_coding
protein_coding
protein_coding
protein_coding
protein_coding
protein_coding
protein_coding
protein_coding
protein_coding
protein_coding
ncRNA
protein_coding
protein_coding
protein_coding
protein_coding
protein_coding
protein_coding
protein_coding
protein_coding
protein_coding
protein_coding
protein_coding
protein_coding
protein_coding
protein_coding

protein_coding
protein_coding
protein_coding
protein_coding
protein_coding
protein_coding
protein_coding
protein_coding
protein_coding
protein_coding
protein_coding
protein_coding
protein_coding
protein_coding
protein_coding
protein_coding
protein_coding
ncRNA
protein_coding
protein_coding
protein_coding
ncRNA
ncRNA
protein_coding
protein_coding
protein_coding
protein_coding
protein_coding
protein_coding
protein_coding
protein_coding
protein_coding
protein_coding
miRNA_precursor
protein_coding
protein_coding
protein_coding
ncRNA
ncRNA
protein_coding
protein_coding
protein_coding
protein_coding
protein_coding
protein_coding
protein_coding
protein_coding
protein_coding
protein_coding
protein_coding
protein_coding
protein_coding
protein_coding
protein_coding
protein_coding
protein_coding

protein_coding
protein_coding
protein_coding
ncRNA
protein_coding
protein_coding
protein_coding
protein_coding
protein_coding
protein_coding
protein_coding
ncRNA
protein_coding
protein_coding
protein_coding
protein_coding
protein_coding
protein_coding
protein_coding
protein_coding
protein_coding
protein_coding
protein_coding
protein_coding
ncRNA
protein_coding
ncRNA
protein_coding
protein_coding
protein_coding
protein_coding
protein_coding
protein_coding
protein_coding
protein_coding
ncRNA
protein_coding
protein_coding
ncRNA
protein_coding
protein_coding
protein_coding
protein_coding
protein_coding
miRNA_precursor
protein_coding
protein_coding
protein_coding
protein_coding
protein_coding
protein_coding
protein_coding
protein_coding
miRNA_precursor
protein_coding
protein_coding

pre_trna
protein_coding
ncRNA
ncRNA
protein_coding
protein_coding
ncRNA
protein_coding
protein_coding
protein_coding
protein_coding
protein_coding
protein_coding
protein_coding
protein_coding
protein_coding
protein_coding
protein_coding
protein_coding
protein_coding
protein_coding
protein_coding
protein_coding
protein_coding
protein_coding
protein_coding
protein_coding
protein_coding
protein_coding
protein_coding
protein_coding
protein_coding
protein_coding
protein_coding
protein_coding
ncRNA
protein_coding
protein_coding
protein_coding
protein_coding
protein_coding
protein_coding
protein_coding
protein_coding
protein_coding
protein_coding
protein_coding
protein_coding
protein_coding
protein_coding
protein_coding
protein_coding
protein_coding
protein_coding
protein_coding
protein_coding

protein_coding
protein_coding
protein_coding
protein_coding
protein_coding
protein_coding
protein_coding
protein_coding
protein_coding
protein_coding
protein_coding
protein_coding
protein_coding
protein_coding
protein_coding
protein_coding
protein_coding
protein_coding
protein_coding
pre_trna
protein_coding
protein_coding
protein_coding
protein_coding
protein_coding
protein_coding
protein_coding
protein_coding
protein_coding
protein_coding
protein_coding
protein_coding
protein_coding
protein_coding
protein_coding
protein_coding
protein_coding
protein_coding
protein_coding
protein_coding
protein_coding
protein_coding
pre_trna
small_nucle(ol)ar_rna
protein_coding
ncRNA
protein_coding
protein_coding
protein_coding
protein_coding
protein_coding
protein_coding
protein_coding
protein_coding
protein_coding
protein_coding

protein_coding
protein_coding
protein_coding
protein_coding
protein_coding
protein_coding
ncRNA
protein_coding
ncRNA
protein_coding
protein_coding
protein_coding
protein_coding
protein_coding
protein_coding
protein_coding
protein_coding
protein_coding
protein_coding
protein_coding
protein_coding
protein_coding
protein_coding
miRNA_precursor
protein_coding
protein_coding
protein_coding
protein_coding
protein_coding
protein_coding
protein_coding
protein_coding
protein_coding
small_nucle(ol)ar_rna
protein_coding
protein_coding
protein_coding
protein_coding
protein_coding
protein_coding
pre_trna
protein_coding
protein_coding
protein_coding
protein_coding
protein_coding
pre_trna
protein_coding
protein_coding
protein_coding
protein_coding
protein_coding
protein_coding
protein_coding
protein_coding
protein_coding

protein_coding
protein_coding
protein_coding
protein_coding
protein_coding
protein_coding
protein_coding
protein_coding
protein_coding
protein_coding
ncRNA
protein_coding
protein_coding
protein_coding
protein_coding
protein_coding
protein_coding
protein_coding
protein_coding
protein_coding
protein_coding
protein_coding
protein_coding
protein_coding
protein_coding
protein_coding
protein_coding
protein_coding
pre_trna
protein_coding
protein_coding
pre_trna
protein_coding
ncRNA
protein_coding
protein_coding
protein_coding
protein_coding
protein_coding
protein_coding
protein_coding
protein_coding
protein_coding
protein_coding
protein_coding
small_nucle(ol)ar_rna
protein_coding
protein_coding
ncRNA
protein_coding
protein_coding
ncRNA
protein_coding
protein_coding
protein_coding
protein_coding

protein_coding
protein_coding
protein_coding
protein_coding
protein_coding
protein_coding
protein_coding
protein_coding
protein_coding
protein_coding
protein_coding
protein_coding
protein_coding
protein_coding
protein_coding
protein_coding
protein_coding
protein_coding
protein_coding
protein_coding
protein_coding
protein_coding
protein_coding
protein_coding
protein_coding
protein_coding
protein_coding
protein_coding
protein_coding
protein_coding
protein_coding
protein_coding
protein_coding
protein_coding
protein_coding
protein_coding
protein_coding
protein_coding
protein_coding
protein_coding
protein_coding
protein_coding
protein_coding
protein_coding
miRNA_precursor
protein_coding
protein_coding
protein_coding
protein_coding
protein_coding
protein_coding
protein_coding
protein_coding
protein_coding
protein_coding
protein_coding

protein_coding
protein_coding
protein_coding
protein_coding
protein_coding
protein_coding
protein_coding
protein_coding
ncRNA
protein_coding
protein_coding
protein_coding
protein_coding
protein_coding
protein_coding
protein_coding
protein_coding
protein_coding
protein_coding
protein_coding
protein_coding
protein_coding
protein_coding
protein_coding
protein_coding
protein_coding
protein_coding
protein_coding
protein_coding
protein_coding
protein_coding
protein_coding
protein_coding
protein_coding
protein_coding
protein_coding
protein_coding
protein_coding
protein_coding
small_nucle(ol)ar_rna
protein_coding
protein_coding
protein_coding
protein_coding
ncRNA
protein_coding
protein_coding
protein_coding
protein_coding
protein_coding
protein_coding
protein_coding
protein_coding
protein_coding
protein_coding
protein_coding

protein_coding
protein_coding
protein_coding
protein_coding
protein_coding
protein_coding
protein_coding
protein_coding
protein_coding
protein_coding
protein_coding
protein_coding
protein_coding
protein_coding
protein_coding
protein_coding
protein_coding
protein_coding
protein_coding
protein_coding
protein_coding
protein_coding
protein_coding
protein_coding
pre_trna
protein_coding
protein_coding
protein_coding
protein_coding
protein_coding
protein_coding
pre_trna
protein_coding
protein_coding
protein_coding
ncRNA
protein_coding
protein_coding
protein_coding
protein_coding
protein_coding
protein_coding
protein_coding
protein_coding
protein_coding
protein_coding
protein_coding
small_nucle(ol)ar_rna
protein_coding
protein_coding
protein_coding
protein_coding
protein_coding
protein_coding
protein_coding
protein_coding

protein_coding
protein_coding
protein_coding
protein_coding
protein_coding
protein_coding
protein_coding
protein_coding
protein_coding
protein_coding
protein_coding
protein_coding
protein_coding
protein_coding
small_nucle(ol)ar_rna
protein_coding
protein_coding
protein_coding
protein_coding
protein_coding
protein_coding
protein_coding
protein_coding
protein_coding
protein_coding
protein_coding
protein_coding
protein_coding
protein_coding
protein_coding
protein_coding
protein_coding
protein_coding
protein_coding
protein_coding
protein_coding
protein_coding
protein_coding
protein_coding
protein_coding
protein_coding
protein_coding
protein_coding
protein_coding
protein_coding
protein_coding
protein_coding
protein_coding
protein_coding
protein_coding
protein_coding
protein_coding
protein_coding
protein_coding
protein_coding
protein_coding

protein_coding
protein_coding
protein_coding
protein_coding
protein_coding
protein_coding
protein_coding
protein_coding
protein_coding
protein_coding
protein_coding
protein_coding
protein_coding
protein_coding
protein_coding
protein_coding
protein_coding
protein_coding
protein_coding
protein_coding
protein_coding
small_nucle(ol)ar_rna
protein_coding
protein_coding
protein_coding
protein_coding
protein_coding
protein_coding
protein_coding
protein_coding
protein_coding
protein_coding
protein_coding
protein_coding
protein_coding
protein_coding
protein_coding
protein_coding
protein_coding
protein_coding
protein_coding
protein_coding
protein_coding
protein_coding
protein_coding
protein_coding
protein_coding
protein_coding
protein_coding
protein_coding
protein_coding
ncRNA
ncRNA
protein_coding
protein_coding
protein_coding

| |
|---|
| protein_coding |
| protein_coding |
| protein_coding |
| protein_coding |
| protein_coding |
| protein_coding |
| protein_coding |
| protein_coding |
| protein_coding |
| protein_coding |
| protein_coding |
| protein_coding |
| protein_coding |
| protein_coding |
| protein_coding |
| protein_coding |
| ncRNA |
| ncRNA |
| protein_coding |
| protein_coding |
| protein_coding |
| protein_coding |
| protein_coding |
| protein_coding |
| protein_coding |
| protein_coding |
| protein_coding |
| protein_coding |
| protein_coding |
| protein_coding |
| protein_coding |
| protein_coding |
| protein_coding |
| protein_coding |
| protein_coding |
| protein_coding |
| protein_coding |
| protein_coding |
| protein_coding |
| protein_coding |
| protein_coding |
| protein_coding |
| protein_coding |
| protein_coding |
| protein_coding |
| protein_coding |
| protein_coding |
| protein_coding |
| protein_coding |
| protein_coding |
| protein_coding |
| protein_coding |
| protein_coding |
| protein_coding |
| protein_coding |
| protein_coding |

protein_coding
protein_coding
protein_coding
protein_coding
protein_coding
protein_coding
protein_coding
protein_coding
protein_coding
protein_coding
protein_coding
protein_coding
protein_coding
protein_coding
protein_coding
ncRNA
protein_coding
protein_coding
protein_coding
protein_coding
protein_coding
protein_coding
protein_coding
protein_coding
protein_coding
protein_coding
protein_coding
protein_coding
protein_coding
protein_coding
protein_coding
protein_coding
protein_coding
protein_coding
protein_coding
protein_coding
protein_coding
protein_coding
protein_coding
protein_coding
protein_coding
protein_coding
protein_coding
protein_coding
protein_coding
protein_coding
protein_coding
protein_coding
protein_coding
protein_coding
protein_coding
protein_coding
protein_coding
protein_coding
protein_coding
protein_coding

protein_coding
protein_coding
protein_coding
protein_coding
protein_coding
protein_coding
protein_coding
protein_coding
protein_coding
protein_coding
ncRNA
protein_coding
protein_coding
protein_coding
protein_coding
protein_coding
protein_coding
protein_coding
protein_coding
protein_coding
protein_coding
protein_coding
protein_coding
protein_coding
protein_coding
protein_coding
protein_coding
ncRNA
protein_coding
protein_coding
protein_coding
protein_coding
protein_coding
protein_coding
protein_coding
protein_coding
protein_coding
protein_coding
protein_coding
protein_coding
protein_coding
protein_coding
protein_coding
protein_coding
protein_coding
small_nucle(ol)ar_rna
protein_coding
protein_coding
protein_coding
protein_coding
protein_coding
protein_coding
protein_coding
protein_coding
protein_coding
protein_coding

protein_coding
protein_coding
protein_coding
protein_coding
protein_coding
protein_coding
protein_coding
protein_coding
protein_coding
protein_coding
protein_coding
protein_coding
protein_coding
protein_coding
protein_coding
protein_coding
protein_coding
protein_coding
protein_coding
protein_coding
protein_coding
protein_coding
protein_coding
protein_coding
protein_coding
protein_coding
protein_coding
protein_coding
protein_coding
protein_coding
protein_coding
protein_coding
protein_coding
protein_coding
ncRNA
protein_coding
protein_coding
protein_coding
protein_coding
protein_coding
protein_coding
protein_coding
protein_coding
protein_coding
protein_coding
protein_coding
protein_coding
protein_coding
protein_coding
protein_coding
protein_coding
protein_coding
protein_coding
protein_coding
protein_coding
protein_coding

protein_coding
protein_coding
protein_coding
protein_coding
protein_coding
protein_coding
protein_coding
protein_coding
protein_coding
ncRNA
protein_coding
protein_coding
protein_coding
protein_coding
protein_coding
protein_coding
protein_coding
protein_coding
protein_coding
protein_coding
protein_coding
protein_coding
protein_coding
protein_coding
ncRNA
protein_coding
protein_coding
protein_coding
protein_coding
protein_coding
protein_coding
protein_coding
protein_coding
protein_coding
protein_coding
protein_coding
protein_coding
protein_coding
miRNA_precursor
protein_coding
protein_coding
protein_coding
protein_coding
protein_coding
protein_coding
protein_coding
protein_coding
protein_coding
protein_coding
miRNA_precursor
protein_coding
protein_coding
protein_coding
protein_coding
protein_coding
protein_coding

protein_coding
protein_coding
protein_coding
protein_coding
protein_coding
protein_coding
protein_coding
protein_coding
protein_coding
protein_coding
protein_coding
protein_coding
protein_coding
protein_coding
protein_coding
protein_coding
protein_coding
protein_coding
protein_coding
protein_coding
protein_coding
protein_coding
protein_coding
protein_coding
protein_coding
protein_coding
ncRNA
protein_coding
protein_coding
protein_coding
protein_coding
protein_coding
protein_coding
protein_coding
protein_coding
protein_coding
protein_coding
protein_coding
protein_coding
protein_coding
protein_coding
protein_coding
protein_coding
small_nucle(ol)ar_rna
protein_coding
ncRNA
protein_coding
protein_coding
protein_coding
ncRNA
protein_coding
protein_coding
protein_coding
protein_coding
protein_coding
protein_coding

protein_coding
protein_coding
protein_coding
protein_coding
protein_coding
protein_coding
protein_coding
protein_coding
protein_coding
protein_coding
ncRNA
protein_coding
protein_coding
protein_coding
protein_coding
protein_coding
protein_coding
protein_coding
protein_coding
protein_coding
protein_coding
protein_coding
protein_coding
protein_coding
protein_coding
protein_coding
protein_coding
protein_coding
protein_coding
protein_coding
protein_coding
protein_coding
protein_coding
protein_coding
protein_coding
protein_coding
protein_coding
protein_coding
protein_coding
protein_coding
protein_coding
protein_coding
protein_coding
protein_coding
protein_coding
protein_coding
protein_coding
protein_coding
protein_coding
protein_coding
protein_coding
protein_coding
protein_coding
protein_coding
protein_coding
ncRNA

protein_coding
protein_coding
protein_coding
protein_coding
protein_coding
protein_coding
protein_coding
protein_coding
protein_coding
protein_coding
protein_coding
protein_coding
protein_coding
ncRNA
small_nucle(ol)ar_rna
protein_coding
protein_coding
protein_coding
protein_coding
protein_coding
protein_coding
protein_coding
protein_coding
protein_coding
protein_coding
protein_coding
protein_coding
protein_coding
protein_coding
protein_coding
protein_coding
protein_coding
protein_coding
protein_coding
protein_coding
protein_coding
protein_coding
protein_coding
protein_coding
protein_coding
protein_coding
protein_coding
protein_coding
protein_coding
pre_trna
protein_coding
ncRNA
protein_coding
protein_coding
protein_coding
protein_coding
protein_coding
protein_coding
protein_coding
protein_coding
protein_coding

protein_coding
protein_coding
protein_coding
protein_coding
protein_coding
protein_coding
protein_coding
protein_coding
protein_coding
protein_coding
miRNA_precursor
protein_coding
protein_coding
protein_coding
protein_coding
protein_coding
protein_coding
protein_coding
protein_coding
protein_coding
protein_coding
ncRNA
protein_coding
protein_coding
protein_coding
protein_coding
protein_coding
protein_coding
protein_coding
small_nucle(ol)ar_rna
protein_coding
protein_coding
protein_coding
protein_coding
protein_coding
protein_coding
ncRNA
protein_coding
protein_coding
protein_coding
protein_coding
protein_coding
protein_coding
protein_coding
protein_coding
protein_coding
protein_coding
protein_coding
protein_coding
protein_coding
protein_coding
protein_coding
protein_coding
protein_coding
protein_coding
protein_coding

protein_coding
protein_coding
ncRNA
protein_coding
protein_coding
protein_coding
protein_coding
ncRNA
protein_coding
ncRNA
protein_coding
ncRNA
protein_coding
protein_coding
protein_coding
protein_coding
protein_coding
protein_coding
protein_coding
protein_coding
protein_coding
protein_coding
protein_coding
protein_coding
protein_coding
protein_coding
protein_coding
protein_coding
protein_coding
protein_coding
ncRNA
protein_coding
protein_coding
protein_coding
protein_coding
protein_coding
protein_coding
protein_coding
protein_coding
protein_coding
protein_coding
protein_coding
protein_coding
protein_coding
pre_trna
protein_coding
protein_coding
protein_coding
protein_coding
protein_coding
protein_coding
protein_coding
protein_coding
protein_coding
protein_coding
ncRNA

protein_coding
ncRNA
ncRNA
ncRNA
protein_coding
protein_coding
protein_coding
miRNA_precursor
protein_coding
protein_coding
small_nucle(ol)ar_rna
protein_coding
protein_coding
protein_coding
protein_coding
protein_coding
protein_coding
protein_coding
ncRNA
ncRNA
protein_coding
protein_coding
protein_coding
protein_coding
protein_coding
protein_coding
protein_coding
protein_coding
protein_coding
protein_coding
protein_coding
protein_coding
protein_coding
protein_coding
protein_coding
protein_coding
protein_coding
protein_coding
protein_coding
protein_coding
protein_coding
protein_coding
protein_coding
ncRNA
protein_coding
protein_coding
protein_coding
protein_coding
protein_coding
protein_coding
protein_coding
protein_coding
protein_coding
protein_coding
protein_coding
protein_coding

protein_coding
protein_coding
protein_coding
protein_coding
protein_coding
protein_coding
protein_coding
protein_coding
protein_coding
protein_coding
protein_coding
protein_coding
protein_coding
protein_coding
protein_coding
protein_coding
protein_coding
protein_coding
protein_coding
protein_coding
protein_coding
protein_coding
protein_coding
protein_coding
protein_coding
protein_coding
protein_coding
protein_coding
protein_coding
protein_coding
protein_coding
protein_coding
protein_coding
protein_coding
protein_coding
protein_coding
protein_coding
protein_coding
protein_coding
protein_coding
protein_coding
protein_coding
protein_coding
protein_coding
protein_coding
protein_coding
protein_coding
protein_coding
protein_coding
protein_coding
small_nucle(ol)ar_rna
protein_coding
protein_coding
protein_coding
ncRNA
protein_coding

protein_coding
protein_coding
protein_coding
protein_coding
protein_coding
protein_coding
protein_coding
protein_coding
protein_coding
protein_coding
protein_coding
protein_coding
protein_coding
protein_coding
protein_coding
protein_coding
protein_coding
protein_coding
protein_coding
protein_coding
protein_coding
protein_coding
protein_coding
ncRNA
protein_coding
protein_coding
protein_coding
ncRNA
protein_coding
protein_coding
protein_coding
protein_coding
protein_coding
protein_coding
protein_coding
protein_coding
protein_coding
protein_coding
protein_coding
protein_coding
protein_coding
protein_coding
ncRNA
small_nucle(ol)ar_rna
protein_coding
protein_coding
protein_coding
protein_coding
protein_coding
protein_coding
protein_coding
protein_coding
protein_coding
protein_coding
protein_coding
protein_coding

protein_coding
protein_coding
protein_coding
protein_coding
ncRNA
protein_coding
protein_coding
protein_coding
protein_coding
protein_coding
protein_coding
protein_coding
protein_coding
protein_coding
protein_coding
protein_coding
protein_coding
protein_coding
ncRNA
protein_coding
protein_coding
protein_coding
protein_coding
protein_coding
protein_coding
protein_coding
ncRNA
protein_coding
protein_coding
protein_coding
protein_coding
protein_coding
protein_coding
protein_coding
protein_coding
protein_coding
protein_coding
protein_coding
protein_coding
protein_coding
protein_coding
protein_coding
protein_coding
protein_coding
ncRNA
protein_coding
protein_coding
protein_coding
protein_coding
protein_coding
protein_coding
protein_coding
protein_coding
protein_coding
protein_coding
protein_coding

protein_coding
protein_coding
protein_coding
protein_coding
protein_coding
protein_coding
protein_coding
protein_coding
protein_coding
protein_coding
protein_coding
protein_coding
protein_coding
protein_coding
protein_coding
protein_coding
protein_coding
protein_coding
protein_coding
protein_coding
protein_coding
protein_coding
miRNA_precursor
ncRNA
protein_coding
protein_coding
protein_coding
protein_coding
protein_coding
protein_coding
protein_coding
protein_coding
protein_coding
protein_coding
protein_coding
protein_coding
protein_coding
protein_coding
protein_coding
protein_coding
protein_coding
pre_trna
protein_coding
protein_coding
protein_coding
protein_coding
protein_coding
protein_coding
protein_coding
pre_trna
protein_coding
protein_coding
protein_coding
protein_coding
protein_coding
protein_coding

ncRNA
ncRNA
protein_coding
protein_coding
protein_coding
protein_coding
protein_coding
protein_coding
protein_coding
protein_coding
protein_coding
protein_coding
protein_coding
protein_coding
protein_coding
protein_coding
protein_coding
protein_coding
protein_coding
protein_coding
ncRNA
protein_coding
protein_coding
protein_coding
protein_coding
protein_coding
protein_coding
protein_coding
protein_coding
protein_coding
protein_coding
protein_coding
protein_coding
protein_coding
protein_coding
protein_coding
protein_coding
protein_coding
protein_coding
protein_coding
protein_coding
protein_coding
protein_coding
protein_coding
protein_coding
protein_coding
protein_coding
protein_coding
protein_coding
protein_coding
protein_coding
protein_coding
protein_coding
protein_coding
ncRNA
small_nucle(ol)ar_rna

protein_coding
protein_coding
protein_coding
protein_coding
protein_coding
protein_coding
protein_coding
protein_coding
protein_coding
protein_coding
protein_coding
protein_coding
protein_coding
protein_coding
protein_coding
protein_coding
protein_coding
protein_coding
protein_coding
protein_coding
protein_coding
ncRNA
ncRNA
protein_coding
protein_coding
protein_coding
protein_coding
protein_coding
protein_coding
protein_coding
protein_coding
protein_coding
protein_coding
protein_coding
protein_coding
ncRNA
protein_coding
protein_coding
protein_coding
protein_coding
protein_coding
protein_coding
pre_trna
protein_coding
protein_coding
protein_coding
protein_coding
protein_coding
protein_coding
protein_coding
protein_coding
protein_coding
protein_coding
ncRNA
ncRNA
protein_coding

protein_coding
protein_coding
protein_coding
protein_coding
protein_coding
protein_coding
protein_coding
protein_coding
protein_coding
protein_coding
protein_coding
protein_coding
protein_coding
protein_coding
miRNA_precursor
protein_coding
protein_coding
protein_coding
protein_coding
protein_coding
protein_coding
protein_coding
protein_coding
protein_coding
protein_coding
protein_coding
protein_coding
miRNA_precursor
protein_coding
protein_coding
protein_coding
protein_coding
protein_coding
protein_coding
protein_coding
miRNA_precursor
protein_coding
protein_coding
protein_coding
protein_coding
protein_coding
ncRNA
protein_coding
protein_coding
protein_coding
protein_coding
protein_coding
protein_coding
protein_coding
protein_coding
protein_coding
protein_coding
ncRNA
protein_coding
protein_coding
protein_coding

protein_coding
protein_coding
protein_coding
protein_coding
protein_coding
protein_coding
protein_coding
protein_coding
protein_coding
protein_coding
ncRNA
protein_coding
protein_coding
protein_coding
protein_coding
protein_coding
protein_coding
protein_coding
protein_coding
protein_coding
protein_coding
protein_coding
protein_coding
protein_coding
protein_coding
protein_coding
protein_coding
protein_coding
protein_coding
protein_coding
protein_coding
protein_coding
ncRNA
protein_coding
protein_coding
protein_coding
protein_coding
protein_coding
protein_coding
protein_coding
protein_coding
protein_coding
protein_coding
ncRNA
protein_coding
protein_coding
protein_coding
protein_coding
protein_coding
protein_coding
protein_coding
ncRNA
protein_coding
protein_coding
small_nucle(ol)ar_rna
small_nucle(ol)ar_rna

small_nucle(ol)ar_rna
protein_coding
protein_coding
protein_coding
protein_coding
protein_coding
protein_coding
protein_coding
protein_coding
protein_coding
protein_coding
protein_coding
protein_coding
protein_coding
protein_coding
protein_coding
protein_coding
protein_coding
protein_coding
protein_coding
protein_coding
protein_coding
protein_coding
protein_coding
protein_coding
protein_coding
small_nucle(ol)ar_rna
miRNA_precursor
protein_coding
protein_coding
protein_coding
protein_coding
protein_coding
protein_coding
protein_coding
protein_coding
protein_coding
protein_coding
protein_coding
protein_coding
protein_coding
protein_coding
protein_coding
protein_coding
protein_coding
protein_coding
protein_coding
protein_coding
protein_coding
protein_coding
protein_coding
protein_coding
protein_coding
protein_coding
protein_coding
protein_coding

pre_trna
protein_coding
protein_coding
protein_coding
protein_coding
protein_coding
protein_coding
protein_coding
protein_coding
protein_coding
protein_coding
protein_coding
protein_coding
protein_coding
protein_coding
protein_coding
protein_coding
protein_coding
protein_coding
protein_coding
protein_coding
protein_coding
protein_coding
protein_coding
protein_coding
protein_coding
protein_coding
protein_coding
protein_coding
protein_coding
protein_coding
protein_coding
protein_coding
ncRNA
ncRNA
protein_coding
protein_coding
protein_coding
protein_coding
protein_coding
protein_coding
protein_coding
protein_coding
protein_coding
protein_coding
protein_coding
protein_coding
protein_coding
protein_coding
protein_coding
protein_coding
protein_coding
protein_coding
protein_coding
protein_coding
protein_coding

protein_coding
protein_coding
ncRNA
protein_coding
protein_coding
protein_coding
pre_trna
protein_coding
protein_coding
protein_coding
protein_coding
protein_coding
protein_coding
protein_coding
protein_coding
miRNA_precursor
protein_coding
protein_coding
protein_coding
protein_coding
protein_coding
protein_coding
protein_coding
protein_coding
protein_coding
protein_coding
protein_coding
protein_coding
protein_coding
protein_coding
protein_coding
ncRNA
protein_coding
protein_coding
protein_coding
protein_coding
protein_coding
protein_coding
protein_coding
protein_coding
protein_coding
protein_coding
protein_coding
protein_coding
protein_coding
protein_coding
protein_coding
protein_coding
protein_coding
protein_coding
protein_coding
protein_coding
protein_coding
protein_coding
protein_coding
protein_coding

protein_coding
protein_coding
ncRNA
protein_coding
protein_coding
protein_coding
ncRNA
protein_coding
protein_coding
small_nucle(ol)ar_rna
miRNA_precursor
protein_coding
protein_coding
protein_coding
protein_coding
protein_coding
protein_coding
protein_coding
protein_coding
protein_coding
protein_coding
protein_coding
protein_coding
protein_coding
protein_coding
protein_coding
protein_coding
protein_coding
protein_coding
protein_coding
protein_coding
protein_coding
protein_coding
ncRNA
protein_coding
protein_coding
protein_coding
protein_coding
protein_coding
protein_coding
protein_coding
protein_coding
protein_coding
protein_coding
protein_coding
protein_coding
protein_coding
protein_coding
protein_coding
protein_coding
protein_coding
protein_coding
protein_coding
protein_coding
ncRNA
protein_coding

protein_coding
protein_coding
protein_coding
protein_coding
protein_coding
protein_coding
protein_coding
protein_coding
protein_coding
protein_coding
protein_coding
protein_coding
protein_coding
protein_coding
protein_coding
protein_coding
protein_coding
protein_coding
protein_coding
protein_coding
protein_coding
protein_coding
protein_coding
protein_coding
protein_coding
protein_coding
protein_coding
protein_coding
protein_coding
protein_coding
protein_coding
protein_coding
protein_coding
protein_coding
protein_coding
protein_coding
protein_coding
ncRNA
protein_coding
protein_coding
protein_coding
protein_coding
protein_coding
pre_trna
protein_coding
protein_coding
protein_coding
protein_coding
protein_coding
protein_coding
protein_coding
protein_coding
miRNA_precursor
protein_coding
protein_coding
protein_coding

protein_coding
protein_coding
protein_coding
protein_coding
protein_coding
protein_coding
protein_coding
pre_trna
protein_coding
protein_coding
protein_coding
protein_coding
protein_coding
protein_coding
protein_coding
protein_coding
protein_coding
protein_coding
protein_coding
protein_coding
protein_coding
protein_coding
protein_coding
protein_coding
protein_coding
protein_coding
protein_coding
protein_coding
protein_coding
protein_coding
protein_coding
protein_coding
protein_coding
protein_coding
protein_coding
protein_coding
protein_coding
ncRNA
ncRNA
protein_coding
protein_coding
protein_coding
protein_coding
protein_coding
protein_coding
protein_coding
protein_coding
ncRNA
protein_coding
protein_coding
protein_coding
protein_coding
protein_coding
protein_coding
protein_coding
protein_coding

protein_coding
protein_coding
protein_coding
protein_coding
protein_coding
protein_coding
protein_coding
protein_coding
protein_coding
protein_coding
protein_coding
protein_coding
protein_coding
small_nucle(ol)ar_rna
protein_coding
protein_coding
protein_coding
protein_coding
protein_coding
protein_coding
pre_trna
protein_coding
protein_coding
protein_coding
protein_coding
protein_coding
protein_coding
protein_coding
protein_coding
protein_coding
protein_coding
protein_coding
protein_coding
protein_coding
protein_coding
protein_coding
protein_coding
protein_coding
protein_coding
protein_coding
protein_coding
ncRNA
protein_coding
protein_coding
protein_coding
protein_coding
protein_coding
protein_coding
protein_coding
protein_coding
protein_coding
protein_coding
protein_coding
protein_coding
protein_coding
protein_coding

pre_trna
protein_coding
protein_coding
protein_coding
protein_coding
protein_coding
protein_coding
protein_coding
protein_coding
protein_coding
protein_coding
protein_coding
protein_coding
protein_coding
protein_coding
protein_coding
protein_coding
protein_coding
protein_coding
protein_coding
protein_coding
protein_coding
protein_coding
protein_coding
protein_coding
protein_coding
protein_coding
protein_coding
protein_coding
protein_coding
protein_coding
protein_coding
protein_coding
protein_coding
ncRNA
ncRNA
protein_coding
protein_coding
protein_coding
protein_coding
protein_coding
protein_coding
protein_coding
protein_coding
ncRNA
protein_coding
protein_coding
protein_coding
protein_coding
protein_coding
protein_coding
protein_coding
protein_coding
protein_coding
protein_coding
protein_coding

protein_coding
protein_coding
protein_coding
protein_coding
protein_coding
protein_coding
protein_coding
protein_coding
protein_coding
protein_coding
protein_coding
protein_coding
protein_coding
ncRNA
ncRNA
protein_coding
protein_coding
protein_coding
protein_coding
protein_coding
protein_coding
protein_coding
protein_coding
protein_coding
protein_coding
protein_coding
protein_coding
protein_coding
protein_coding
protein_coding
protein_coding
protein_coding
protein_coding
protein_coding
protein_coding
protein_coding
protein_coding
protein_coding
protein_coding
protein_coding
protein_coding
protein_coding
protein_coding
protein_coding
protein_coding
protein_coding
protein_coding
protein_coding
protein_coding
protein_coding
protein_coding
protein_coding
protein_coding
protein_coding
ncRNA
protein_coding

ncRNA
protein_coding
ncRNA
protein_coding
protein_coding
protein_coding
protein_coding
protein_coding
miRNA_precursor
protein_coding
protein_coding
protein_coding
ncRNA
protein_coding
protein_coding
protein_coding
protein_coding
protein_coding
protein_coding
protein_coding
protein_coding
protein_coding
protein_coding
protein_coding
protein_coding
protein_coding
protein_coding
protein_coding
protein_coding
protein_coding
protein_coding
protein_coding
protein_coding
protein_coding
protein_coding
protein_coding
protein_coding
protein_coding
protein_coding
protein_coding
protein_coding
protein_coding
protein_coding
protein_coding
protein_coding
protein_coding
protein_coding
protein_coding
protein_coding
protein_coding
protein_coding
protein_coding
protein_coding
ncRNA
protein_coding
protein_coding

protein_coding
protein_coding
protein_coding
protein_coding
protein_coding
ncRNA
protein_coding
protein_coding
protein_coding
protein_coding
protein_coding
protein_coding
protein_coding
protein_coding
protein_coding
protein_coding
protein_coding
protein_coding
protein_coding
protein_coding
protein_coding
protein_coding
protein_coding
protein_coding
protein_coding
protein_coding
pre_trna
protein_coding
protein_coding
protein_coding
protein_coding
protein_coding
protein_coding
protein_coding
protein_coding
ncRNA
protein_coding
protein_coding
protein_coding
protein_coding
protein_coding
protein_coding
protein_coding
protein_coding
protein_coding
protein_coding
protein_coding
protein_coding
protein_coding
protein_coding
protein_coding
ncRNA
protein_coding
protein_coding
protein_coding
protein_coding

protein_coding
protein_coding
protein_coding
protein_coding
protein_coding
protein_coding
protein_coding
protein_coding
protein_coding
protein_coding
protein_coding
protein_coding
protein_coding
protein_coding
protein_coding
protein_coding
protein_coding
protein_coding
protein_coding
ncRNA
protein_coding
protein_coding
protein_coding
protein_coding
protein_coding
protein_coding
protein_coding
protein_coding
protein_coding
protein_coding
ncRNA
protein_coding
protein_coding
protein_coding
protein_coding
protein_coding
protein_coding
protein_coding
protein_coding
protein_coding
protein_coding
protein_coding
protein_coding
protein_coding
protein_coding
protein_coding
protein_coding
protein_coding
protein_coding
protein_coding
protein_coding
protein_coding
protein_coding
protein_coding
ncRNA
protein_coding

protein_coding
protein_coding
protein_coding
protein_coding
protein_coding
protein_coding
protein_coding
protein_coding
protein_coding
protein_coding
protein_coding
protein_coding
protein_coding
protein_coding
protein_coding
protein_coding
protein_coding
protein_coding
protein_coding
protein_coding
protein_coding
protein_coding
protein_coding
protein_coding
protein_coding
protein_coding
protein_coding
protein_coding
protein_coding
miRNA_precursor
protein_coding
protein_coding
protein_coding
protein_coding
protein_coding
protein_coding
protein_coding
protein_coding
protein_coding
protein_coding
protein_coding
protein_coding
protein_coding
protein_coding
protein_coding
protein_coding
protein_coding
protein_coding
protein_coding
protein_coding
protein_coding
protein_coding
protein_coding
protein_coding
protein_coding
protein_coding

protein_coding
protein_coding
protein_coding
protein_coding
protein_coding
protein_coding
protein_coding
protein_coding
protein_coding
protein_coding
protein_coding
protein_coding
protein_coding
protein_coding
protein_coding
protein_coding
protein_coding
protein_coding
protein_coding
protein_coding
protein_coding
protein_coding
protein_coding
protein_coding
protein_coding
protein_coding
protein_coding
protein_coding
protein_coding
protein_coding
protein_coding
protein_coding
protein_coding
protein_coding
protein_coding
protein_coding
protein_coding
protein_coding
protein_coding
protein_coding
protein_coding
protein_coding
protein_coding
protein_coding
protein_coding
protein_coding
protein_coding
protein_coding
protein_coding
protein_coding
protein_coding
protein_coding
ncRNA
protein_coding
protein_coding
protein_coding

protein_coding
protein_coding
protein_coding
protein_coding
protein_coding
protein_coding
protein_coding
protein_coding
protein_coding
protein_coding
protein_coding
protein_coding
protein_coding
protein_coding
protein_coding
protein_coding
protein_coding
protein_coding
protein_coding
protein_coding
protein_coding
protein_coding
protein_coding
protein_coding
protein_coding
protein_coding
pre_trna
protein_coding
protein_coding
protein_coding
protein_coding
protein_coding
ncRNA
protein_coding
protein_coding
protein_coding
protein_coding
protein_coding
protein_coding
protein_coding
protein_coding
protein_coding
protein_coding
protein_coding
protein_coding
protein_coding
protein_coding
protein_coding
protein_coding
protein_coding
protein_coding
protein_coding
protein_coding
ncRNA
protein_coding
protein_coding

protein_coding
protein_coding
protein_coding
protein_coding
protein_coding
protein_coding
protein_coding
protein_coding
protein_coding
protein_coding
protein_coding
protein_coding
protein_coding
protein_coding
protein_coding
protein_coding
protein_coding
protein_coding
protein_coding
protein_coding
protein_coding
protein_coding
protein_coding
protein_coding
ncRNA
protein_coding
protein_coding
protein_coding
protein_coding
protein_coding
protein_coding
protein_coding
protein_coding
protein_coding
pre_trna
protein_coding
protein_coding
protein_coding
protein_coding
protein_coding
protein_coding
protein_coding
protein_coding
protein_coding
protein_coding
protein_coding
protein_coding
protein_coding
ncRNA
protein_coding
protein_coding
protein_coding
protein_coding
protein_coding
protein_coding
protein_coding

protein_coding
protein_coding
protein_coding
protein_coding
protein_coding
protein_coding
protein_coding
protein_coding
protein_coding
protein_coding
protein_coding
protein_coding
protein_coding
protein_coding
protein_coding
protein_coding
protein_coding
protein_coding
protein_coding
protein_coding
ncRNA
protein_coding
protein_coding
protein_coding
protein_coding
protein_coding
protein_coding
protein_coding
protein_coding
protein_coding
protein_coding
miRNA_precursor
protein_coding
protein_coding
protein_coding
miRNA_precursor
ncRNA
miRNA_precursor
protein_coding
protein_coding
protein_coding
protein_coding
protein_coding
protein_coding
protein_coding
protein_coding
protein_coding
protein_coding
protein_coding
protein_coding
protein_coding
protein_coding
protein_coding
protein_coding
protein_coding
protein_coding

protein_coding
protein_coding
protein_coding
protein_coding
protein_coding
protein_coding
protein_coding
protein_coding
protein_coding
protein_coding
protein_coding
protein_coding
protein_coding
protein_coding
protein_coding
protein_coding
protein_coding
protein_coding
protein_coding
protein_coding
protein_coding
protein_coding
protein_coding
protein_coding
protein_coding
protein_coding
protein_coding
ncRNA
ncRNA
protein_coding
ncRNA
protein_coding
protein_coding
protein_coding
protein_coding
protein_coding
ncRNA
protein_coding
protein_coding
protein_coding
protein_coding
protein_coding
miRNA_precursor
protein_coding
protein_coding
protein_coding
protein_coding
protein_coding
ncRNA
protein_coding
protein_coding
protein_coding
protein_coding
protein_coding
protein_coding
protein_coding

protein_coding
ncRNA
protein_coding
protein_coding
protein_coding
protein_coding
protein_coding
protein_coding
protein_coding
protein_coding
protein_coding
protein_coding
protein_coding
protein_coding
protein_coding
protein_coding
protein_coding
protein_coding
protein_coding
protein_coding
protein_coding
protein_coding
protein_coding
pre_trna
protein_coding
protein_coding
protein_coding
protein_coding
protein_coding
protein_coding
protein_coding
protein_coding
protein_coding
protein_coding
protein_coding
protein_coding
protein_coding
protein_coding
protein_coding
protein_coding
protein_coding
protein_coding
protein_coding
protein_coding
protein_coding
protein_coding
protein_coding
protein_coding
protein_coding
protein_coding
protein_coding
protein_coding
protein_coding
protein_coding
protein_coding
protein_coding

protein_coding
protein_coding
protein_coding
protein_coding
protein_coding
protein_coding
small_nucle(ol)ar_rna
protein_coding
protein_coding
protein_coding
protein_coding
protein_coding
protein_coding
protein_coding
protein_coding
protein_coding
protein_coding
protein_coding
protein_coding
protein_coding
protein_coding
protein_coding
protein_coding
protein_coding
protein_coding
protein_coding
protein_coding
protein_coding
protein_coding
protein_coding
protein_coding
protein_coding
protein_coding
protein_coding
protein_coding
protein_coding
protein_coding
protein_coding
protein_coding
protein_coding
protein_coding
protein_coding
protein_coding
protein_coding
protein_coding
protein_coding
protein_coding
protein_coding
protein_coding
protein_coding
protein_coding
ncRNA
protein_coding
protein_coding
protein_coding
protein_coding

protein_coding
protein_coding
protein_coding
protein_coding
protein_coding
protein_coding
protein_coding
protein_coding
protein_coding
protein_coding
protein_coding
small_nucle(ol)ar_rna
small_nucle(ol)ar_rna
protein_coding
protein_coding
protein_coding
miRNA_precursor
protein_coding
protein_coding
protein_coding
protein_coding
protein_coding
protein_coding
ncRNA
protein_coding
protein_coding
protein_coding
protein_coding
protein_coding
protein_coding
protein_coding
protein_coding
ncRNA
protein_coding
protein_coding
protein_coding
protein_coding
protein_coding
protein_coding
protein_coding
protein_coding
protein_coding
protein_coding
protein_coding
small_nucle(ol)ar_rna
protein_coding
protein_coding
small_nucle(ol)ar_rna
protein_coding
protein_coding
protein_coding
protein_coding
protein_coding
protein_coding
protein_coding
protein_coding

protein_coding
protein_coding
protein_coding
protein_coding
protein_coding
protein_coding
protein_coding
protein_coding
protein_coding
protein_coding
protein_coding
protein_coding
protein_coding
protein_coding
protein_coding
protein_coding
protein_coding
protein_coding
protein_coding
protein_coding
protein_coding
protein_coding
protein_coding
protein_coding
protein_coding
protein_coding
protein_coding
protein_coding
protein_coding
protein_coding
small_nucle(ol)ar_rna
protein_coding
protein_coding
protein_coding
ncRNA
protein_coding
protein_coding
protein_coding
protein_coding
protein_coding
protein_coding
protein_coding
protein_coding
protein_coding
protein_coding
protein_coding
protein_coding
protein_coding
protein_coding
protein_coding
protein_coding
protein_coding
protein_coding
protein_coding
protein_coding
protein_coding

protein_coding
protein_coding
protein_coding
protein_coding
protein_coding
ncRNA
protein_coding
protein_coding
protein_coding
miRNA_precursor
protein_coding
protein_coding
protein_coding
protein_coding
protein_coding
pre_trna
protein_coding
protein_coding
protein_coding
protein_coding
protein_coding
protein_coding
protein_coding
protein_coding
protein_coding
protein_coding
protein_coding
protein_coding
protein_coding
protein_coding
protein_coding
protein_coding
protein_coding
protein_coding
protein_coding
protein_coding
protein_coding
protein_coding
protein_coding
protein_coding
protein_coding
protein_coding
protein_coding
ncRNA
protein_coding
protein_coding
protein_coding
ncRNA
protein_coding
protein_coding
protein_coding
protein_coding
protein_coding
protein_coding
protein_coding
protein_coding

protein_coding
protein_coding
protein_coding
protein_coding
protein_coding
protein_coding
protein_coding
protein_coding
protein_coding
protein_coding
protein_coding
protein_coding
protein_coding
protein_coding
protein_coding
protein_coding
protein_coding
protein_coding
protein_coding
protein_coding
protein_coding
protein_coding
protein_coding
ncRNA
protein_coding
protein_coding
protein_coding
protein_coding
protein_coding
protein_coding
protein_coding
protein_coding
protein_coding
protein_coding
protein_coding
protein_coding
protein_coding
protein_coding
protein_coding
protein_coding
protein_coding
protein_coding
protein_coding
protein_coding
protein_coding
protein_coding
protein_coding
protein_coding
protein_coding
protein_coding
protein_coding
protein_coding
protein_coding
protein_coding
protein_coding
protein_coding

protein_coding
protein_coding
protein_coding
protein_coding
protein_coding
protein_coding
small_nucle(ol)ar_rna
protein_coding
protein_coding
protein_coding
protein_coding
protein_coding
protein_coding
protein_coding
protein_coding
protein_coding
protein_coding
protein_coding
protein_coding
protein_coding
protein_coding
protein_coding
protein_coding
protein_coding
protein_coding
protein_coding
protein_coding
protein_coding
protein_coding
protein_coding
protein_coding
protein_coding
protein_coding
protein_coding
ncRNA
protein_coding
small_nucle(ol)ar_rna
protein_coding
protein_coding
protein_coding
protein_coding
protein_coding
protein_coding
protein_coding
ncRNA
protein_coding
protein_coding
protein_coding
protein_coding
protein_coding
protein_coding
protein_coding
protein_coding
protein_coding
protein_coding
pre_trna

protein_coding
protein_coding
ncRNA
protein_coding
protein_coding
protein_coding
protein_coding
protein_coding
protein_coding
protein_coding
protein_coding
protein_coding
protein_coding
protein_coding
protein_coding
protein_coding
protein_coding
protein_coding
protein_coding
protein_coding
protein_coding
protein_coding
protein_coding
protein_coding
protein_coding
protein_coding
protein_coding
protein_coding
protein_coding
protein_coding
protein_coding
protein_coding
protein_coding
protein_coding
protein_coding
protein_coding
protein_coding
protein_coding
protein_coding
protein_coding
protein_coding
protein_coding
pre_trna
protein_coding
protein_coding
protein_coding
protein_coding
protein_coding
protein_coding
protein_coding
protein_coding
protein_coding
protein_coding
protein_coding
protein_coding
protein_coding

protein_coding
protein_coding
protein_coding
protein_coding
protein_coding
protein_coding
protein_coding
protein_coding
protein_coding
protein_coding
protein_coding
protein_coding
protein_coding
protein_coding
protein_coding
protein_coding
protein_coding
protein_coding
protein_coding
protein_coding
protein_coding
protein_coding
protein_coding
ncRNA
protein_coding
protein_coding
protein_coding
protein_coding
protein_coding
ncRNA
protein_coding
protein_coding
protein_coding
protein_coding
protein_coding
protein_coding
protein_coding
protein_coding
protein_coding
protein_coding
protein_coding
protein_coding
protein_coding
protein_coding
protein_coding
protein_coding
protein_coding
protein_coding
protein_coding
protein_coding
protein_coding
protein_coding
protein_coding
protein_coding
pre_trna
ncRNA

protein_coding
protein_coding
protein_coding
ncRNA
protein_coding
protein_coding
protein_coding
pre_trna
protein_coding
protein_coding
protein_coding
protein_coding
protein_coding
protein_coding
protein_coding
protein_coding
protein_coding
protein_coding
protein_coding
protein_coding
protein_coding
protein_coding
protein_coding
protein_coding
protein_coding
protein_coding
protein_coding
protein_coding
protein_coding
protein_coding
protein_coding
protein_coding
protein_coding
protein_coding
small_nucle(ol)ar_rna
pre_trna
protein_coding
protein_coding
protein_coding
protein_coding
protein_coding
protein_coding
protein_coding
protein_coding
protein_coding
protein_coding
protein_coding
protein_coding
protein_coding
protein_coding
pre_trna
protein_coding
ncRNA
protein_coding
protein_coding
protein_coding

protein_coding
protein_coding
protein_coding
ncRNA
protein_coding
protein_coding
protein_coding
protein_coding
protein_coding
protein_coding
protein_coding
protein_coding
protein_coding
protein_coding
protein_coding
protein_coding
protein_coding
miRNA_precursor
protein_coding
protein_coding
ncRNA
protein_coding
protein_coding
protein_coding
pre_trna
protein_coding
protein_coding
protein_coding
protein_coding
protein_coding
protein_coding
protein_coding
protein_coding
ncRNA
protein_coding
protein_coding
protein_coding
protein_coding
protein_coding
protein_coding
protein_coding
protein_coding
protein_coding
protein_coding
protein_coding
protein_coding
protein_coding
protein_coding
protein_coding
protein_coding
protein_coding
protein_coding
protein_coding
protein_coding
protein_coding
protein_coding

small_nucle(ol)ar_rna
protein_coding
protein_coding
protein_coding
protein_coding
ncRNA
protein_coding
protein_coding
protein_coding
protein_coding
protein_coding
protein_coding
protein_coding
protein_coding
protein_coding
protein_coding
protein_coding
protein_coding
protein_coding
protein_coding
protein_coding
protein_coding
protein_coding
protein_coding
protein_coding
protein_coding
protein_coding
protein_coding
protein_coding
protein_coding
protein_coding
protein_coding
protein_coding
protein_coding
protein_coding
protein_coding
protein_coding
protein_coding
protein_coding
protein_coding
protein_coding
protein_coding
protein_coding
protein_coding
protein_coding
protein_coding
protein_coding
protein_coding
protein_coding
protein_coding
protein_coding
protein_coding
protein_coding
protein_coding
protein_coding
protein_coding

protein_coding
protein_coding
protein_coding
protein_coding
protein_coding
protein_coding
protein_coding
protein_coding
ncRNA
protein_coding
protein_coding
protein_coding
protein_coding
protein_coding
ncRNA
protein_coding
protein_coding
protein_coding
protein_coding
protein_coding
protein_coding
protein_coding
protein_coding
protein_coding
protein_coding
protein_coding
ncRNA
protein_coding
protein_coding
ncRNA
protein_coding
protein_coding
protein_coding
protein_coding
protein_coding
protein_coding
protein_coding
protein_coding
protein_coding
protein_coding
protein_coding
protein_coding
protein_coding
protein_coding
protein_coding
protein_coding
protein_coding
protein_coding
protein_coding
protein_coding
protein_coding
protein_coding
protein_coding
ncRNA
protein_coding
protein_coding

protein_coding
protein_coding
protein_coding
protein_coding
protein_coding
protein_coding
protein_coding
protein_coding
protein_coding
protein_coding
protein_coding
protein_coding
protein_coding
protein_coding
protein_coding
protein_coding
protein_coding
protein_coding
ncRNA
protein_coding
ncRNA
protein_coding
protein_coding
protein_coding
protein_coding
protein_coding
protein_coding
protein_coding
protein_coding
protein_coding
protein_coding
protein_coding
protein_coding
protein_coding
protein_coding
protein_coding
protein_coding
protein_coding
protein_coding
protein_coding
protein_coding
protein_coding
protein_coding
protein_coding
protein_coding
protein_coding
protein_coding
protein_coding
protein_coding
protein_coding
protein_coding
protein_coding
protein_coding
ncRNA
protein_coding
protein_coding

protein_coding
protein_coding
protein_coding
protein_coding
protein_coding
protein_coding
protein_coding
protein_coding
protein_coding
protein_coding
protein_coding
protein_coding
protein_coding
protein_coding
protein_coding
protein_coding
protein_coding
protein_coding
protein_coding
pre_trna
protein_coding
protein_coding
protein_coding
protein_coding
protein_coding
protein_coding
protein_coding
protein_coding
protein_coding
protein_coding
protein_coding
protein_coding
protein_coding
protein_coding
protein_coding
protein_coding
protein_coding
protein_coding
protein_coding
protein_coding
protein_coding
protein_coding
protein_coding
protein_coding
protein_coding
protein_coding
protein_coding
protein_coding
protein_coding
protein_coding
protein_coding
ncRNA
protein_coding
protein_coding
protein_coding
protein_coding

protein_coding
protein_coding
protein_coding
protein_coding
protein_coding
protein_coding
protein_coding
protein_coding
small_nucle(ol)ar_rna
protein_coding
protein_coding
protein_coding
protein_coding
ncRNA
protein_coding
protein_coding
protein_coding
protein_coding
protein_coding
protein_coding
protein_coding
protein_coding
protein_coding
protein_coding
ncRNA
protein_coding
protein_coding
protein_coding
ncRNA
protein_coding
protein_coding
protein_coding
protein_coding
protein_coding
protein_coding
protein_coding
protein_coding
protein_coding
protein_coding
protein_coding
protein_coding
protein_coding
protein_coding
protein_coding
protein_coding
protein_coding
protein_coding
protein_coding
protein_coding
ncRNA
protein_coding
protein_coding
protein_coding
protein_coding
ncRNA
protein_coding

protein_coding
protein_coding
protein_coding
protein_coding
pre_trna
protein_coding
protein_coding
protein_coding
protein_coding
ncRNA
protein_coding
protein_coding
ncRNA
protein_coding
protein_coding
protein_coding
protein_coding
protein_coding
protein_coding
protein_coding
protein_coding
protein_coding
protein_coding
protein_coding
protein_coding
protein_coding
protein_coding
ncRNA
protein_coding
protein_coding
protein_coding
small_nucle(ol)ar_rna
protein_coding
protein_coding
protein_coding
protein_coding
protein_coding
protein_coding
protein_coding
protein_coding
protein_coding
protein_coding
protein_coding
protein_coding
protein_coding
protein_coding
protein_coding
protein_coding
protein_coding
protein_coding
protein_coding
protein_coding
protein_coding
protein_coding
pre_trna
protein_coding

protein_coding
protein_coding
protein_coding
protein_coding
protein_coding
protein_coding
protein_coding
protein_coding
protein_coding
protein_coding
protein_coding
protein_coding
protein_coding
protein_coding
protein_coding
protein_coding
miRNA_precursor
protein_coding
protein_coding
protein_coding
protein_coding
protein_coding
protein_coding
protein_coding
protein_coding
protein_coding
protein_coding
protein_coding
protein_coding
protein_coding
protein_coding
protein_coding
protein_coding
protein_coding
protein_coding
protein_coding
protein_coding
protein_coding
protein_coding
protein_coding
protein_coding
protein_coding
ncRNA
protein_coding
ncRNA
protein_coding
protein_coding
protein_coding
protein_coding
protein_coding
protein_coding
protein_coding
protein_coding
protein_coding
protein_coding
protein_coding

protein_coding
protein_coding
protein_coding
protein_coding
protein_coding
protein_coding
protein_coding
protein_coding
small_nucle(ol)ar_rna
protein_coding
protein_coding
protein_coding
protein_coding
protein_coding
protein_coding
protein_coding
protein_coding
protein_coding
protein_coding
protein_coding
protein_coding
protein_coding
protein_coding
ncRNA
ncRNA
small_nucle(ol)ar_rna
protein_coding
protein_coding
protein_coding
protein_coding
protein_coding
protein_coding
protein_coding
protein_coding
protein_coding
protein_coding
protein_coding
protein_coding
protein_coding
protein_coding
protein_coding
protein_coding
protein_coding
protein_coding
protein_coding
protein_coding
ncRNA
protein_coding
protein_coding
protein_coding
protein_coding
protein_coding
protein_coding
protein_coding
protein_coding
protein_coding

protein_coding
protein_coding
protein_coding
protein_coding
protein_coding
protein_coding
protein_coding
protein_coding
protein_coding
protein_coding
protein_coding
protein_coding
protein_coding
protein_coding
protein_coding
protein_coding
protein_coding
ncRNA
protein_coding
protein_coding
protein_coding
protein_coding
protein_coding
protein_coding
protein_coding
protein_coding
protein_coding
protein_coding
protein_coding
protein_coding
protein_coding
protein_coding
protein_coding
protein_coding
protein_coding
protein_coding
protein_coding
protein_coding
protein_coding
protein_coding
protein_coding
protein_coding
ncRNA
protein_coding
protein_coding
protein_coding
protein_coding
protein_coding
protein_coding
protein_coding
protein_coding
ncRNA
protein_coding
protein_coding
protein_coding
protein_coding

protein_coding
protein_coding
protein_coding
protein_coding
protein_coding
protein_coding
protein_coding
protein_coding
protein_coding
protein_coding
protein_coding
protein_coding
protein_coding
protein_coding
protein_coding
small_nucle(ol)ar_rna
protein_coding
protein_coding
ncRNA
protein_coding
protein_coding
protein_coding
protein_coding
protein_coding
protein_coding
protein_coding
protein_coding
ncRNA
protein_coding
protein_coding
protein_coding
protein_coding
ncRNA
miRNA_precursor
protein_coding
protein_coding
protein_coding
protein_coding
protein_coding
miRNA_precursor
protein_coding
protein_coding
small_nucle(ol)ar_rna
protein_coding
protein_coding
protein_coding
ncRNA
protein_coding
protein_coding
protein_coding
protein_coding
protein_coding
protein_coding
protein_coding
protein_coding
protein_coding

protein_coding
protein_coding
protein_coding
protein_coding
protein_coding
protein_coding
protein_coding
protein_coding
protein_coding
protein_coding
protein_coding
protein_coding
protein_coding
ncRNA
ncRNA
miRNA_precursor
protein_coding
ncRNA
protein_coding
protein_coding
protein_coding
protein_coding
protein_coding
protein_coding
protein_coding
protein_coding
protein_coding
protein_coding
protein_coding
protein_coding
ncRNA
protein_coding
protein_coding
protein_coding
protein_coding
protein_coding
protein_coding
protein_coding
protein_coding
protein_coding
protein_coding
protein_coding
protein_coding
protein_coding
protein_coding
protein_coding
protein_coding
protein_coding
protein_coding
protein_coding
ncRNA
protein_coding
protein_coding
protein_coding
protein_coding
protein_coding

protein_coding
protein_coding
protein_coding
protein_coding
protein_coding
protein_coding
protein_coding
ncRNA
protein_coding
protein_coding
ncRNA
protein_coding
protein_coding
ncRNA
protein_coding
protein_coding
ncRNA
protein_coding
protein_coding
protein_coding
protein_coding
protein_coding
protein_coding
protein_coding
protein_coding
protein_coding
protein_coding
protein_coding
protein_coding
protein_coding
protein_coding
protein_coding
protein_coding
protein_coding
protein_coding
protein_coding
protein_coding
protein_coding
miRNA_precursor
protein_coding
protein_coding
protein_coding
protein_coding
protein_coding
protein_coding
protein_coding
protein_coding
protein_coding
protein_coding
protein_coding
protein_coding
protein_coding
protein_coding
protein_coding
protein_coding
protein_coding

protein_coding
ncRNA
protein_coding
protein_coding
protein_coding
protein_coding
protein_coding
protein_coding
ncRNA
protein_coding
protein_coding
protein_coding
protein_coding
ncRNA
protein_coding
protein_coding
protein_coding
ncRNA
protein_coding
protein_coding
protein_coding
protein_coding
protein_coding
ncRNA
protein_coding
protein_coding
protein_coding
protein_coding
protein_coding
protein_coding
protein_coding
protein_coding
ncRNA
protein_coding
protein_coding
protein_coding
protein_coding
protein_coding
protein_coding
protein_coding
protein_coding
protein_coding
protein_coding
miRNA_precursor
protein_coding
protein_coding
protein_coding
protein_coding
protein_coding
protein_coding
protein_coding
protein_coding
protein_coding
protein_coding
protein_coding
protein_coding

protein_coding
protein_coding
protein_coding
protein_coding
protein_coding
protein_coding
protein_coding
protein_coding
protein_coding
protein_coding
protein_coding
protein_coding
protein_coding
protein_coding
protein_coding
protein_coding
protein_coding
protein_coding
protein_coding
protein_coding
protein_coding
protein_coding
protein_coding
protein_coding
protein_coding
protein_coding
protein_coding
protein_coding
protein_coding
protein_coding
protein_coding
protein_coding
protein_coding
protein_coding
protein_coding
protein_coding
ncRNA
protein_coding
protein_coding
protein_coding
protein_coding
protein_coding
protein_coding
protein_coding
protein_coding
protein_coding
protein_coding
protein_coding
protein_coding
protein_coding
protein_coding
protein_coding
protein_coding
protein_coding
protein_coding
protein_coding

protein_coding
protein_coding
ncRNA
protein_coding
small_nucle(ol)ar_rna
small_nucle(ol)ar_rna
small_nucle(ol)ar_rna
protein_coding
protein_coding
protein_coding
protein_coding
protein_coding
protein_coding
protein_coding
protein_coding
protein_coding
protein_coding
protein_coding
protein_coding
protein_coding
protein_coding
protein_coding
protein_coding
protein_coding
protein_coding
protein_coding
protein_coding
protein_coding
protein_coding
protein_coding
protein_coding
protein_coding
protein_coding
protein_coding
protein_coding
protein_coding
protein_coding
protein_coding
protein_coding
protein_coding
protein_coding
protein_coding
protein_coding
protein_coding
protein_coding
protein_coding
protein_coding
protein_coding
ncRNA
protein_coding
protein_coding
protein_coding
protein_coding
protein_coding
protein_coding
protein_coding

protein_coding
protein_coding
protein_coding
protein_coding
protein_coding
protein_coding
protein_coding
protein_coding
protein_coding
protein_coding
protein_coding
protein_coding
protein_coding
protein_coding
protein_coding
protein_coding
protein_coding
protein_coding
protein_coding
protein_coding
protein_coding
protein_coding
ncRNA
protein_coding
protein_coding
protein_coding
protein_coding
protein_coding
protein_coding
protein_coding
protein_coding
protein_coding
protein_coding
protein_coding
protein_coding
protein_coding
protein_coding
protein_coding
protein_coding
protein_coding
protein_coding
protein_coding
protein_coding
protein_coding
protein_coding
protein_coding
protein_coding
ncRNA
protein_coding
protein_coding
protein_coding
protein_coding
protein_coding
protein_coding
protein_coding
protein_coding

protein_coding
protein_coding
protein_coding
protein_coding
protein_coding
protein_coding
protein_coding
protein_coding
protein_coding
protein_coding
protein_coding
protein_coding
protein_coding
protein_coding
protein_coding
protein_coding
protein_coding
ncRNA
protein_coding
protein_coding
protein_coding
protein_coding
protein_coding
protein_coding
protein_coding
protein_coding
protein_coding
protein_coding
protein_coding
ncRNA
protein_coding
protein_coding
protein_coding
protein_coding
protein_coding
protein_coding
protein_coding
protein_coding
protein_coding
protein_coding
protein_coding
protein_coding
protein_coding
protein_coding
protein_coding
protein_coding
protein_coding
protein_coding
protein_coding
protein_coding
protein_coding
protein_coding
protein_coding
protein_coding
protein_coding
protein_coding

protein_coding
protein_coding
miRNA_precursor
protein_coding
protein_coding
protein_coding
protein_coding
protein_coding
protein_coding
protein_coding
protein_coding
protein_coding
protein_coding
protein_coding
protein_coding
protein_coding
protein_coding
protein_coding
protein_coding
protein_coding
protein_coding
ncRNA
protein_coding
protein_coding
protein_coding
protein_coding
protein_coding
protein_coding
protein_coding
protein_coding
protein_coding
protein_coding
protein_coding
protein_coding
protein_coding
protein_coding
protein_coding
protein_coding
protein_coding
protein_coding
protein_coding
protein_coding
protein_coding
protein_coding
protein_coding
protein_coding
protein_coding
protein_coding
protein_coding
protein_coding
protein_coding
protein_coding
protein_coding
protein_coding
protein_coding
protein_coding

protein_coding
protein_coding
protein_coding
protein_coding
protein_coding
protein_coding
protein_coding
protein_coding
protein_coding
protein_coding
protein_coding
protein_coding
protein_coding
protein_coding
protein_coding
protein_coding
protein_coding
ncRNA
protein_coding
protein_coding
protein_coding
protein_coding
protein_coding
protein_coding
protein_coding
protein_coding
protein_coding
protein_coding
protein_coding
protein_coding
protein_coding
protein_coding
protein_coding
protein_coding
protein_coding
ncRNA
protein_coding
protein_coding
protein_coding
protein_coding
protein_coding
protein_coding
protein_coding
ncRNA
protein_coding
protein_coding
protein_coding
protein_coding
protein_coding
protein_coding
protein_coding
protein_coding
protein_coding
protein_coding
protein_coding
protein_coding

protein_coding
protein_coding
protein_coding
protein_coding
protein_coding
protein_coding
protein_coding
protein_coding
protein_coding
protein_coding
protein_coding
protein_coding
protein_coding
protein_coding
protein_coding
protein_coding
protein_coding
protein_coding
small_nucle(ol)ar_rna
protein_coding
protein_coding
protein_coding
protein_coding
protein_coding
protein_coding
protein_coding
protein_coding
protein_coding
protein_coding
protein_coding
protein_coding
protein_coding
protein_coding
protein_coding
protein_coding
protein_coding
protein_coding
protein_coding
protein_coding
protein_coding
protein_coding
protein_coding
protein_coding
protein_coding
protein_coding
protein_coding
protein_coding
protein_coding
protein_coding
protein_coding
protein_coding
protein_coding
protein_coding
protein_coding
ncRNA
protein_coding

protein_coding
protein_coding
protein_coding
protein_coding
protein_coding
protein_coding
ncRNA
protein_coding
protein_coding
protein_coding
protein_coding
protein_coding
protein_coding
protein_coding
protein_coding
protein_coding
protein_coding
protein_coding
protein_coding
protein_coding
protein_coding
protein_coding
protein_coding
protein_coding
protein_coding
protein_coding
protein_coding
protein_coding
protein_coding
protein_coding
protein_coding
protein_coding
protein_coding
protein_coding
protein_coding
protein_coding
protein_coding
protein_coding
protein_coding
protein_coding
protein_coding
ncRNA
protein_coding
protein_coding
protein_coding
protein_coding
protein_coding
protein_coding
protein_coding
protein_coding
protein_coding
protein_coding
protein_coding
protein_coding
protein_coding
small_nucle(ol)ar_rna

small_nucle(ol)ar_rna
protein_coding
protein_coding
protein_coding
protein_coding
protein_coding
protein_coding
protein_coding
protein_coding
protein_coding
protein_coding
protein_coding
protein_coding
protein_coding
protein_coding
ncRNA
protein_coding
protein_coding
protein_coding
protein_coding
protein_coding
protein_coding
protein_coding
protein_coding
protein_coding
protein_coding
protein_coding
protein_coding
protein_coding
protein_coding
protein_coding
protein_coding
protein_coding
protein_coding
protein_coding
protein_coding
protein_coding
protein_coding
protein_coding
protein_coding
protein_coding
protein_coding
protein_coding
protein_coding
protein_coding
protein_coding
protein_coding
protein_coding
protein_coding
protein_coding
protein_coding
protein_coding
protein_coding
protein_coding
protein_coding
protein_coding

ncRNA
protein_coding
protein_coding
protein_coding
protein_coding
protein_coding
protein_coding
protein_coding
protein_coding
protein_coding
protein_coding
protein_coding
protein_coding
protein_coding
protein_coding
protein_coding
protein_coding
protein_coding
protein_coding
protein_coding
protein_coding
protein_coding
ncRNA
protein_coding
protein_coding
protein_coding
protein_coding
protein_coding
protein_coding
protein_coding
protein_coding
protein_coding
protein_coding
protein_coding
pre_trna
protein_coding
protein_coding
protein_coding
protein_coding
protein_coding
protein_coding
protein_coding
ncRNA
protein_coding
protein_coding
protein_coding
protein_coding
protein_coding
protein_coding
protein_coding
protein_coding
protein_coding
protein_coding
protein_coding
protein_coding
protein_coding

protein_coding
protein_coding
protein_coding
protein_coding
protein_coding
protein_coding
protein_coding
protein_coding
ncRNA
protein_coding
ncRNA
protein_coding
ncRNA
protein_coding
protein_coding
protein_coding
protein_coding
protein_coding
protein_coding
protein_coding
protein_coding
protein_coding
protein_coding
protein_coding
protein_coding
protein_coding
protein_coding
protein_coding
protein_coding
protein_coding
protein_coding
protein_coding
protein_coding
protein_coding
ncRNA
protein_coding
protein_coding
protein_coding
protein_coding
pre_trna
protein_coding
protein_coding
protein_coding
protein_coding
protein_coding
protein_coding
ncRNA
ncRNA
protein_coding
protein_coding
small_nucle(ol)ar_rna
protein_coding
protein_coding
protein_coding
protein_coding
ncRNA

protein_coding
protein_coding
ncRNA
protein_coding
ncRNA
protein_coding
protein_coding
protein_coding
protein_coding
protein_coding
protein_coding
protein_coding
protein_coding
protein_coding
protein_coding
protein_coding
protein_coding
protein_coding
protein_coding
protein_coding
protein_coding
protein_coding
ncRNA
protein_coding
protein_coding
ncRNA
protein_coding
protein_coding
ncRNA
protein_coding
protein_coding
protein_coding
protein_coding
ncRNA
protein_coding
protein_coding
ncRNA
protein_coding
protein_coding
protein_coding
protein_coding
protein_coding
protein_coding
protein_coding
protein_coding
protein_coding
protein_coding
protein_coding
protein_coding
protein_coding
protein_coding
protein_coding
protein_coding
protein_coding
protein_coding
protein_coding
protein_coding

protein_coding
protein_coding
protein_coding
protein_coding
ncRNA
protein_coding
protein_coding
protein_coding
protein_coding
protein_coding
protein_coding
protein_coding
protein_coding
protein_coding
protein_coding
protein_coding
protein_coding
protein_coding
pre_trna
protein_coding
protein_coding
protein_coding
protein_coding
protein_coding
protein_coding
protein_coding
protein_coding
ncRNA
ncRNA
protein_coding
protein_coding
protein_coding
protein_coding
protein_coding
miRNA_precursor
protein_coding
protein_coding
protein_coding
protein_coding
protein_coding
protein_coding
protein_coding
protein_coding
protein_coding
protein_coding
protein_coding
protein_coding
protein_coding
protein_coding
protein_coding
protein_coding
protein_coding
protein_coding
protein_coding
protein_coding

protein_coding
ncRNA
protein_coding
small_nucle(ol)ar_rna
protein_coding
protein_coding
protein_coding
protein_coding
protein_coding
protein_coding
protein_coding
protein_coding
protein_coding
protein_coding
protein_coding
protein_coding
ncRNA
protein_coding
protein_coding
protein_coding
protein_coding
protein_coding
protein_coding
protein_coding
protein_coding
protein_coding
protein_coding
protein_coding
protein_coding
protein_coding
protein_coding
protein_coding
protein_coding
protein_coding
protein_coding
protein_coding
ncRNA
protein_coding
protein_coding
protein_coding
protein_coding
protein_coding
protein_coding
protein_coding
protein_coding
protein_coding
protein_coding
protein_coding
protein_coding
protein_coding
protein_coding
protein_coding
protein_coding
protein_coding
protein_coding
protein_coding

protein_coding
protein_coding
protein_coding
protein_coding
protein_coding
protein_coding
protein_coding
protein_coding
protein_coding
protein_coding
ncRNA
ncRNA
protein_coding
protein_coding
protein_coding
ncRNA
protein_coding
protein_coding
protein_coding
miRNA_precursor
protein_coding
protein_coding
ncRNA
protein_coding
protein_coding
protein_coding
protein_coding
protein_coding
protein_coding
protein_coding
protein_coding
protein_coding
protein_coding
protein_coding
protein_coding
protein_coding
protein_coding
protein_coding
protein_coding
protein_coding
protein_coding
protein_coding
protein_coding
protein_coding
ncRNA
protein_coding
protein_coding
protein_coding
protein_coding
protein_coding
protein_coding
protein_coding
protein_coding
protein_coding
protein_coding
protein_coding

protein_coding
protein_coding
protein_coding
protein_coding
protein_coding
protein_coding
protein_coding
protein_coding
protein_coding
protein_coding
protein_coding
protein_coding
protein_coding
protein_coding
protein_coding
protein_coding
protein_coding
protein_coding
protein_coding
protein_coding
protein_coding
miRNA_precursor
protein_coding
protein_coding
protein_coding
protein_coding
protein_coding
ncRNA
protein_coding
protein_coding
protein_coding
protein_coding
pre_trna
protein_coding
protein_coding
protein_coding
protein_coding
ncRNA
protein_coding
protein_coding
protein_coding
protein_coding
protein_coding
protein_coding
protein_coding
protein_coding
protein_coding
protein_coding
protein_coding
protein_coding
protein_coding
protein_coding
protein_coding
protein_coding
protein_coding
protein_coding

protein_coding
protein_coding
protein_coding
pre_trna
protein_coding
ncRNA
protein_coding
protein_coding
protein_coding
protein_coding
protein_coding
protein_coding
protein_coding
protein_coding
protein_coding
ncRNA
protein_coding
protein_coding
protein_coding
protein_coding
protein_coding
protein_coding
protein_coding
protein_coding
protein_coding
ncRNA
protein_coding
protein_coding
protein_coding
protein_coding
ncRNA
protein_coding
protein_coding
protein_coding
protein_coding
protein_coding
protein_coding
protein_coding
protein_coding
protein_coding
protein_coding
protein_coding
protein_coding
protein_coding
protein_coding
protein_coding
protein_coding
protein_coding
protein_coding
protein_coding
protein_coding
ncRNA
protein_coding
protein_coding
protein_coding
protein_coding

protein_coding
protein_coding
protein_coding
protein_coding
protein_coding
protein_coding
protein_coding
protein_coding
protein_coding
protein_coding
protein_coding
protein_coding
protein_coding
protein_coding
protein_coding
ncRNA
protein_coding
protein_coding
protein_coding
protein_coding
protein_coding
protein_coding
protein_coding
protein_coding
protein_coding
protein_coding
protein_coding
protein_coding
protein_coding
protein_coding
protein_coding
protein_coding
protein_coding
protein_coding
protein_coding
protein_coding
protein_coding
protein_coding
protein_coding
protein_coding
protein_coding
protein_coding
protein_coding
protein_coding
protein_coding
protein_coding
small_nucle(ol)ar_rna
small_nucle(ol)ar_rna
small_nucle(ol)ar_rna
ncRNA
protein_coding
protein_coding
protein_coding
protein_coding
protein_coding
protein_coding

protein_coding
protein_coding
protein_coding
protein_coding
protein_coding
protein_coding
protein_coding
protein_coding
protein_coding
protein_coding
protein_coding
protein_coding
protein_coding
protein_coding
protein_coding
protein_coding
protein_coding
protein_coding
protein_coding
protein_coding
protein_coding
protein_coding
protein_coding
protein_coding
protein_coding
protein_coding
protein_coding
protein_coding
protein_coding
protein_coding
protein_coding
protein_coding
protein_coding
protein_coding
protein_coding
protein_coding
protein_coding
ncRNA
small_nucle(ol)ar_rna
ncRNA
protein_coding
protein_coding
protein_coding
protein_coding
protein_coding
protein_coding
protein_coding
protein_coding
protein_coding
protein_coding
protein_coding
protein_coding
protein_coding
protein_coding
protein_coding
protein_coding

protein_coding
protein_coding
protein_coding
protein_coding
ncRNA
protein_coding
protein_coding
protein_coding
ncRNA
protein_coding
protein_coding
protein_coding
protein_coding
protein_coding
protein_coding
miRNA_precursor
protein_coding
protein_coding
ncRNA
protein_coding
protein_coding
protein_coding
protein_coding
protein_coding
protein_coding
protein_coding
protein_coding
protein_coding
protein_coding
protein_coding
protein_coding
protein_coding
protein_coding
protein_coding
protein_coding
protein_coding
protein_coding
protein_coding
protein_coding
protein_coding
protein_coding
protein_coding
protein_coding
protein_coding
ncRNA
protein_coding
protein_coding
protein_coding
protein_coding
protein_coding
protein_coding
protein_coding
miRNA_precursor
protein_coding
protein_coding
protein_coding

protein_coding
protein_coding
protein_coding
protein_coding
protein_coding
protein_coding
protein_coding
protein_coding
pre_trna
protein_coding
protein_coding
protein_coding
ncRNA
protein_coding
protein_coding
protein_coding
protein_coding
protein_coding
protein_coding
protein_coding
protein_coding
protein_coding
protein_coding
protein_coding
protein_coding
protein_coding
protein_coding
protein_coding
protein_coding
protein_coding
protein_coding
protein_coding
protein_coding
protein_coding
protein_coding
protein_coding
protein_coding
protein_coding
protein_coding
protein_coding
protein_coding
protein_coding
ncRNA
protein_coding
protein_coding
protein_coding
protein_coding
ncRNA
protein_coding
protein_coding
protein_coding
ncRNA
protein_coding
protein_coding
protein_coding
protein_coding

protein_coding
protein_coding
protein_coding
protein_coding
protein_coding
protein_coding
protein_coding
protein_coding
protein_coding
protein_coding
protein_coding
protein_coding
protein_coding
protein_coding
protein_coding
protein_coding
protein_coding
ncRNA
protein_coding
protein_coding
protein_coding
protein_coding
protein_coding
protein_coding
protein_coding
protein_coding
protein_coding
protein_coding
protein_coding
protein_coding
protein_coding
protein_coding
protein_coding
protein_coding
protein_coding
protein_coding
protein_coding
protein_coding
protein_coding
protein_coding
protein_coding
protein_coding
protein_coding
protein_coding
protein_coding
protein_coding
protein_coding
protein_coding
protein_coding
protein_coding
protein_coding
protein_coding
protein_coding
protein_coding
protein_coding
protein_coding

protein_coding
protein_coding
protein_coding
protein_coding
protein_coding
protein_coding
protein_coding
protein_coding
protein_coding
protein_coding
protein_coding
ncRNA
protein_coding
protein_coding
protein_coding
protein_coding
protein_coding
protein_coding
protein_coding
protein_coding
protein_coding
protein_coding
protein_coding
protein_coding
protein_coding
protein_coding
protein_coding
protein_coding
protein_coding
protein_coding
protein_coding
protein_coding
protein_coding
protein_coding
protein_coding
protein_coding
protein_coding
protein_coding
protein_coding
protein_coding
protein_coding
protein_coding
protein_coding
protein_coding
protein_coding
protein_coding
protein_coding
protein_coding
protein_coding
protein_coding
protein_coding
protein_coding
protein_coding
ncRNA
protein_coding
protein_coding

protein_coding
protein_coding
protein_coding
protein_coding
protein_coding
protein_coding
protein_coding
protein_coding
protein_coding
protein_coding
protein_coding
protein_coding
protein_coding
protein_coding
protein_coding
ncRNA
protein_coding
protein_coding
protein_coding
protein_coding
protein_coding
protein_coding
protein_coding
protein_coding
protein_coding
protein_coding
protein_coding
protein_coding
protein_coding
protein_coding
protein_coding
protein_coding
protein_coding
protein_coding
protein_coding
protein_coding
pre_trna
protein_coding
protein_coding
protein_coding
protein_coding
protein_coding
protein_coding
small_nucle(ol)ar_rna
protein_coding
protein_coding
protein_coding
protein_coding
protein_coding
protein_coding
pre_trna
protein_coding
ncRNA
protein_coding
protein_coding
protein_coding

protein_coding
protein_coding
protein_coding
protein_coding
protein_coding
protein_coding
protein_coding
protein_coding
protein_coding
protein_coding
protein_coding
protein_coding
protein_coding
protein_coding
protein_coding
protein_coding
protein_coding
protein_coding
protein_coding
protein_coding
protein_coding
protein_coding
protein_coding
protein_coding
protein_coding
protein_coding
protein_coding
protein_coding
protein_coding
protein_coding
protein_coding
protein_coding
protein_coding
protein_coding
protein_coding
protein_coding
protein_coding
protein_coding
protein_coding
protein_coding
protein_coding
protein_coding
protein_coding
protein_coding
protein_coding
protein_coding
protein_coding
protein_coding
protein_coding
protein_coding
protein_coding
protein_coding
protein_coding
protein_coding
protein_coding
protein_coding

protein_coding
protein_coding
protein_coding
protein_coding
protein_coding
protein_coding
protein_coding
protein_coding
protein_coding
protein_coding
protein_coding
protein_coding
miRNA_precursor
protein_coding
small_nucle(ol)ar_rna
ncRNA
protein_coding
protein_coding
protein_coding
protein_coding
protein_coding
protein_coding
protein_coding
protein_coding
protein_coding
protein_coding
protein_coding
protein_coding
protein_coding
protein_coding
protein_coding
protein_coding
protein_coding
protein_coding
protein_coding
protein_coding
protein_coding
protein_coding
protein_coding
protein_coding
protein_coding
protein_coding
protein_coding
protein_coding
protein_coding
protein_coding
protein_coding
protein_coding
protein_coding
ncRNA
protein_coding
protein_coding
protein_coding
protein_coding
protein_coding
ncRNA

protein_coding
protein_coding
protein_coding
protein_coding
protein_coding
protein_coding
protein_coding
protein_coding
protein_coding
protein_coding
protein_coding
protein_coding
protein_coding
protein_coding
protein_coding
protein_coding
protein_coding
protein_coding
protein_coding
protein_coding
protein_coding
protein_coding
protein_coding
protein_coding
protein_coding
protein_coding
protein_coding
protein_coding
protein_coding
protein_coding
protein_coding
protein_coding
protein_coding
protein_coding
protein_coding
protein_coding
protein_coding
protein_coding
protein_coding
protein_coding
protein_coding
protein_coding
ncRNA
protein_coding
protein_coding
protein_coding
protein_coding
protein_coding
protein_coding
protein_coding
protein_coding
protein_coding
protein_coding
protein_coding
miRNA_precursor
protein_coding

protein_coding
protein_coding
protein_coding
protein_coding
protein_coding
protein_coding
protein_coding
protein_coding
protein_coding
protein_coding
protein_coding
protein_coding
small_nucle(ol)ar_rna
protein_coding
protein_coding
protein_coding
protein_coding
protein_coding
protein_coding
protein_coding
protein_coding
protein_coding
small_nucle(ol)ar_rna
protein_coding
protein_coding
protein_coding
protein_coding
protein_coding
protein_coding
protein_coding
protein_coding
protein_coding
protein_coding
protein_coding
protein_coding
protein_coding
protein_coding
protein_coding
protein_coding
protein_coding
protein_coding
protein_coding
protein_coding
ncRNA
protein_coding
protein_coding
protein_coding
protein_coding
protein_coding
protein_coding
protein_coding
protein_coding
protein_coding
protein_coding
protein_coding
protein_coding

protein_coding
protein_coding
protein_coding
protein_coding
protein_coding
protein_coding
protein_coding
protein_coding
protein_coding
protein_coding
protein_coding
protein_coding
protein_coding
protein_coding
protein_coding
protein_coding
protein_coding
protein_coding
pre_trna
protein_coding
ncRNA
protein_coding
protein_coding
protein_coding
protein_coding
protein_coding
protein_coding
protein_coding
protein_coding
protein_coding
protein_coding
protein_coding
protein_coding
protein_coding
protein_coding
protein_coding
protein_coding
protein_coding
protein_coding
protein_coding
protein_coding
protein_coding
protein_coding
protein_coding
protein_coding
protein_coding
protein_coding
protein_coding
protein_coding
protein_coding
protein_coding
protein_coding
protein_coding
protein_coding
protein_coding
protein_coding

protein_coding
protein_coding
protein_coding
protein_coding
protein_coding
protein_coding
protein_coding
protein_coding
protein_coding
protein_coding
protein_coding
protein_coding
protein_coding
ncRNA
protein_coding
protein_coding
protein_coding
protein_coding
protein_coding
protein_coding
protein_coding
protein_coding
protein_coding
protein_coding
protein_coding
protein_coding
protein_coding
protein_coding
protein_coding
protein_coding
protein_coding
protein_coding
ncRNA
small_nucle(ol)ar_rna
protein_coding
protein_coding
protein_coding
protein_coding
protein_coding
protein_coding
protein_coding
protein_coding
protein_coding
protein_coding
protein_coding
ncRNA
protein_coding
protein_coding
protein_coding
protein_coding
protein_coding
protein_coding
protein_coding
protein_coding
protein_coding
protein_coding

protein_coding
protein_coding
protein_coding
protein_coding
protein_coding
protein_coding
protein_coding
protein_coding
protein_coding
protein_coding
protein_coding
protein_coding
protein_coding
protein_coding
miRNA_precursor
miRNA_precursor
protein_coding
protein_coding
protein_coding
protein_coding
protein_coding
protein_coding
protein_coding
protein_coding
protein_coding
protein_coding
protein_coding
protein_coding
ncRNA
ncRNA
protein_coding
protein_coding
protein_coding
protein_coding
protein_coding
protein_coding
protein_coding
protein_coding
ncRNA
protein_coding
protein_coding
protein_coding
protein_coding
pre_trna
protein_coding
protein_coding
protein_coding
protein_coding
protein_coding
protein_coding
protein_coding
protein_coding
protein_coding
ncRNA
protein_coding
protein_coding

protein_coding
protein_coding
protein_coding
protein_coding
ncRNA
protein_coding
protein_coding
ncRNA
protein_coding
protein_coding
protein_coding
protein_coding
protein_coding
protein_coding
protein_coding
protein_coding
protein_coding
protein_coding
protein_coding
protein_coding
protein_coding
protein_coding
protein_coding
protein_coding
protein_coding
protein_coding
protein_coding
protein_coding
protein_coding
protein_coding
protein_coding
protein_coding
protein_coding
protein_coding
protein_coding
miRNA_precursor
protein_coding
protein_coding
protein_coding
protein_coding
protein_coding
ncRNA
ncRNA
protein_coding
protein_coding
protein_coding
protein_coding
protein_coding
protein_coding
protein_coding
protein_coding
protein_coding
protein_coding
protein_coding
protein_coding
protein_coding

protein_coding
protein_coding
protein_coding
protein_coding
protein_coding
protein_coding
protein_coding
protein_coding
protein_coding
ncRNA
protein_coding
protein_coding
miRNA_precursor
protein_coding
miRNA_precursor
protein_coding
protein_coding
protein_coding
protein_coding
protein_coding
protein_coding
protein_coding
protein_coding
protein_coding
protein_coding
protein_coding
protein_coding
protein_coding
protein_coding
protein_coding
protein_coding
protein_coding
ncRNA
protein_coding
protein_coding
protein_coding
protein_coding
protein_coding
protein_coding
protein_coding
protein_coding
protein_coding
protein_coding
protein_coding
protein_coding
protein_coding
protein_coding
protein_coding
protein_coding
protein_coding
protein_coding
protein_coding
protein_coding
protein_coding
protein_coding
protein_coding

protein_coding
protein_coding
protein_coding
protein_coding
protein_coding
protein_coding
protein_coding
protein_coding
protein_coding
protein_coding
protein_coding
protein_coding
protein_coding
protein_coding
protein_coding
protein_coding
protein_coding
protein_coding
protein_coding
protein_coding
protein_coding
protein_coding
protein_coding
protein_coding
ncRNA
protein_coding
protein_coding
protein_coding
protein_coding
protein_coding
protein_coding
protein_coding
protein_coding
protein_coding
protein_coding
protein_coding
protein_coding
protein_coding
protein_coding
protein_coding
protein_coding
protein_coding
protein_coding
protein_coding
protein_coding
protein_coding
protein_coding
protein_coding
protein_coding
protein_coding
protein_coding
protein_coding
protein_coding
protein_coding
protein_coding
protein_coding

protein_coding
small_nucle(ol)ar_rna
protein_coding
protein_coding
pre_trna
protein_coding
pre_trna
protein_coding
protein_coding
protein_coding
protein_coding
protein_coding
protein_coding
protein_coding
protein_coding
protein_coding
ncRNA
protein_coding
protein_coding
ncRNA
ncRNA
protein_coding
protein_coding
protein_coding
protein_coding
protein_coding
protein_coding
protein_coding
protein_coding
protein_coding
protein_coding
protein_coding
miRNA_precursor
miRNA_precursor
protein_coding
protein_coding
protein_coding
protein_coding
protein_coding
protein_coding
protein_coding
protein_coding
protein_coding
protein_coding
protein_coding
protein_coding
protein_coding
protein_coding
protein_coding
protein_coding
protein_coding
protein_coding
protein_coding
protein_coding
protein_coding
protein_coding

protein_coding
protein_coding
protein_coding
protein_coding
protein_coding
protein_coding
protein_coding
protein_coding
protein_coding
protein_coding
protein_coding
protein_coding
protein_coding
protein_coding
protein_coding
protein_coding
protein_coding
protein_coding
protein_coding
protein_coding
protein_coding
protein_coding
protein_coding
protein_coding
protein_coding
protein_coding
protein_coding
protein_coding
protein_coding
protein_coding
protein_coding
protein_coding
pre_trna
protein_coding
protein_coding
protein_coding
protein_coding
protein_coding
protein_coding
protein_coding
protein_coding
protein_coding
protein_coding
protein_coding
protein_coding
protein_coding
protein_coding
protein_coding
protein_coding
protein_coding
protein_coding
protein_coding
protein_coding
protein_coding
protein_coding
protein_coding

ncRNA
protein_coding
ncRNA
protein_coding
protein_coding
protein_coding
protein_coding
protein_coding
protein_coding
protein_coding
protein_coding
protein_coding
protein_coding
protein_coding
protein_coding
small_nucle(ol)ar_rna
protein_coding
protein_coding
protein_coding
protein_coding
protein_coding
protein_coding
ncRNA
protein_coding
protein_coding
protein_coding
protein_coding
protein_coding
protein_coding
protein_coding
protein_coding
protein_coding
protein_coding
protein_coding
protein_coding
protein_coding
protein_coding
protein_coding
protein_coding
protein_coding
protein_coding
protein_coding
protein_coding
protein_coding
protein_coding
protein_coding
protein_coding
protein_coding
protein_coding
protein_coding
protein_coding
ncRNA
protein_coding
protein_coding
protein_coding
ncRNA

protein_coding
protein_coding
protein_coding
protein_coding
protein_coding
protein_coding
protein_coding
protein_coding
protein_coding
protein_coding
protein_coding
ncRNA
protein_coding
protein_coding
protein_coding
protein_coding
protein_coding
protein_coding
protein_coding
protein_coding
protein_coding
protein_coding
protein_coding
protein_coding
protein_coding
protein_coding
protein_coding
protein_coding
protein_coding
protein_coding
protein_coding
protein_coding
protein_coding
protein_coding
protein_coding
ncRNA
protein_coding
protein_coding
ncRNA
ncRNA
protein_coding
protein_coding
protein_coding
protein_coding
protein_coding
protein_coding
protein_coding
protein_coding
protein_coding
protein_coding
protein_coding
protein_coding
protein_coding
protein_coding
protein_coding
protein_coding

protein_coding
protein_coding
protein_coding
protein_coding
protein_coding
protein_coding
protein_coding
protein_coding
protein_coding
protein_coding
protein_coding
protein_coding
protein_coding
protein_coding
protein_coding
protein_coding
protein_coding
protein_coding
protein_coding
protein_coding
protein_coding
protein_coding
protein_coding
pre_trna
protein_coding
protein_coding
protein_coding
protein_coding
protein_coding
protein_coding
protein_coding
protein_coding
protein_coding
protein_coding
protein_coding
protein_coding
protein_coding
protein_coding
protein_coding
protein_coding
protein_coding
protein_coding
protein_coding
protein_coding
protein_coding
protein_coding
protein_coding
protein_coding
protein_coding
protein_coding
small_nucle(ol)ar_rna
protein_coding
protein_coding
protein_coding
protein_coding
protein_coding

protein_coding
protein_coding
protein_coding
protein_coding
protein_coding
protein_coding
protein_coding
protein_coding
protein_coding
protein_coding
protein_coding
protein_coding
protein_coding
small_nucle(ol)ar_rna
small_nucle(ol)ar_rna
protein_coding
protein_coding
protein_coding
pre_trna
miRNA_precursor
protein_coding
protein_coding
protein_coding
protein_coding
protein_coding
protein_coding
protein_coding
ncRNA
protein_coding
ncRNA
protein_coding
protein_coding
protein_coding
protein_coding
protein_coding
protein_coding
protein_coding
protein_coding
protein_coding
protein_coding
protein_coding
protein_coding
protein_coding
protein_coding
protein_coding
ncRNA
protein_coding
protein_coding
protein_coding
protein_coding
protein_coding
protein_coding
protein_coding
protein_coding
protein_coding
protein_coding

protein_coding
protein_coding
ncRNA
protein_coding
protein_coding
protein_coding
miRNA_precursor
protein_coding
protein_coding
protein_coding
ncRNA
protein_coding
protein_coding
protein_coding
protein_coding
protein_coding
protein_coding
protein_coding
protein_coding
protein_coding
protein_coding
protein_coding
protein_coding
protein_coding
protein_coding
protein_coding
protein_coding
protein_coding
protein_coding
protein_coding
protein_coding
protein_coding
protein_coding
protein_coding
protein_coding
protein_coding
protein_coding
protein_coding
protein_coding
protein_coding
protein_coding
protein_coding
protein_coding
protein_coding
protein_coding
protein_coding
protein_coding
protein_coding
protein_coding
protein_coding
protein_coding
protein_coding
protein_coding
protein_coding
protein_coding
miRNA_precursor
pre_trna

protein_coding
protein_coding
protein_coding
protein_coding
protein_coding
ncRNA
protein_coding
protein_coding
protein_coding
miRNA_precursor
protein_coding
protein_coding
protein_coding
miRNA_precursor
protein_coding
ncRNA
protein_coding
protein_coding
protein_coding
protein_coding
protein_coding
protein_coding
protein_coding
protein_coding
protein_coding
protein_coding
protein_coding
protein_coding
protein_coding
protein_coding
protein_coding
protein_coding
protein_coding
protein_coding
protein_coding
protein_coding
protein_coding
protein_coding
protein_coding
protein_coding
protein_coding
protein_coding
protein_coding
protein_coding
protein_coding
protein_coding
protein_coding
protein_coding
protein_coding
protein_coding
protein_coding
protein_coding
protein_coding
protein_coding
protein_coding
protein_coding

protein_coding
protein_coding
protein_coding
protein_coding
protein_coding
protein_coding
protein_coding
protein_coding
miRNA_precursor
protein_coding
protein_coding
protein_coding
protein_coding
protein_coding
protein_coding
protein_coding
protein_coding
protein_coding
ncRNA
small_nucle(ol)ar_rna
small_nucle(ol)ar_rna
small_nucle(ol)ar_rna
protein_coding
protein_coding
protein_coding
protein_coding
ncRNA
protein_coding
protein_coding
protein_coding
protein_coding
protein_coding
protein_coding
protein_coding
protein_coding
protein_coding
protein_coding
protein_coding
protein_coding
protein_coding
protein_coding
protein_coding
protein_coding
protein_coding
protein_coding
protein_coding
protein_coding
protein_coding
protein_coding
ncRNA
protein_coding
protein_coding
protein_coding
protein_coding
protein_coding
protein_coding

protein_coding
protein_coding
protein_coding
protein_coding
protein_coding
protein_coding
protein_coding
protein_coding
protein_coding
protein_coding
protein_coding
protein_coding
protein_coding
protein_coding
protein_coding
protein_coding
protein_coding
protein_coding
protein_coding
protein_coding
protein_coding
protein_coding
protein_coding
protein_coding
protein_coding
protein_coding
protein_coding
pre_trna
protein_coding
protein_coding
protein_coding
protein_coding
protein_coding
protein_coding
protein_coding
protein_coding
protein_coding
protein_coding
protein_coding
protein_coding
protein_coding
protein_coding
ncRNA
protein_coding
protein_coding
protein_coding
protein_coding
protein_coding
protein_coding
protein_coding
protein_coding
miRNA_precursor
protein_coding
protein_coding
protein_coding
protein_coding

protein_coding
protein_coding
protein_coding
protein_coding
protein_coding
protein_coding
ncRNA
protein_coding
protein_coding
protein_coding
protein_coding
protein_coding
protein_coding
protein_coding
protein_coding
protein_coding
protein_coding
protein_coding
ncRNA
protein_coding
protein_coding
protein_coding
protein_coding
protein_coding
protein_coding
protein_coding
protein_coding
protein_coding
protein_coding
protein_coding
protein_coding
protein_coding
protein_coding
protein_coding
protein_coding
protein_coding
protein_coding
protein_coding
pre_trna
protein_coding
protein_coding
protein_coding
protein_coding
protein_coding
protein_coding
protein_coding
protein_coding
protein_coding
protein_coding
protein_coding
protein_coding
protein_coding
protein_coding
protein_coding
protein_coding
protein_coding

protein_coding
protein_coding
ncRNA
protein_coding
protein_coding
protein_coding
protein_coding
protein_coding
protein_coding
protein_coding
protein_coding
protein_coding
ncRNA
protein_coding
protein_coding
ncRNA
small_nucle(ol)ar_rna
protein_coding
protein_coding
protein_coding
protein_coding
protein_coding
protein_coding
protein_coding
protein_coding
protein_coding
protein_coding
ncRNA
protein_coding
protein_coding
protein_coding
ncRNA
protein_coding
protein_coding
protein_coding
protein_coding
protein_coding
protein_coding
protein_coding
protein_coding
protein_coding
protein_coding
protein_coding
protein_coding
protein_coding
ncRNA
protein_coding
protein_coding
protein_coding
protein_coding
protein_coding
protein_coding
protein_coding
protein_coding
protein_coding
pre_trna

protein_coding
protein_coding
protein_coding
protein_coding
protein_coding
protein_coding
protein_coding
protein_coding
protein_coding
protein_coding
protein_coding
protein_coding
protein_coding
protein_coding
miRNA_precursor
small_nucle(ol)ar_rna
small_nucle(ol)ar_rna
protein_coding
pre_trna
protein_coding
protein_coding
protein_coding
protein_coding
protein_coding
protein_coding
ncRNA
protein_coding
protein_coding
protein_coding
protein_coding
ncRNA
protein_coding
protein_coding
protein_coding
protein_coding
protein_coding
protein_coding
protein_coding
protein_coding
protein_coding
protein_coding
protein_coding
protein_coding
protein_coding
protein_coding
protein_coding
pre_trna
protein_coding
protein_coding
protein_coding
protein_coding
protein_coding
protein_coding
protein_coding
protein_coding
protein_coding

protein_coding
protein_coding
protein_coding
protein_coding
protein_coding
protein_coding
protein_coding
protein_coding
protein_coding
protein_coding
protein_coding
protein_coding
protein_coding
protein_coding
protein_coding
protein_coding
protein_coding
protein_coding
protein_coding
protein_coding
protein_coding
protein_coding
protein_coding
protein_coding
protein_coding
protein_coding
protein_coding
protein_coding
protein_coding
protein_coding
protein_coding
protein_coding
protein_coding
protein_coding
protein_coding
protein_coding
protein_coding
protein_coding
protein_coding
protein_coding
protein_coding
protein_coding
protein_coding
protein_coding
protein_coding
protein_coding
protein_coding
protein_coding
protein_coding
protein_coding
protein_coding
protein_coding
protein_coding
protein_coding
protein_coding
protein_coding

protein_coding
protein_coding
protein_coding
protein_coding
protein_coding
protein_coding
protein_coding
protein_coding
protein_coding
protein_coding
protein_coding
protein_coding
protein_coding
protein_coding
protein_coding
protein_coding
protein_coding
protein_coding
protein_coding
protein_coding
protein_coding
protein_coding
protein_coding
protein_coding
protein_coding
protein_coding
protein_coding
protein_coding
protein_coding
protein_coding
protein_coding
protein_coding
protein_coding
protein_coding
protein_coding
protein_coding
protein_coding
protein_coding
protein_coding
protein_coding
protein_coding
protein_coding
protein_coding
protein_coding
protein_coding
protein_coding
protein_coding
protein_coding
protein_coding
protein_coding
protein_coding
protein_coding
protein_coding
protein_coding
ncRNA
protein_coding

ncRNA
protein_coding
protein_coding
protein_coding
protein_coding
protein_coding
protein_coding
protein_coding
protein_coding
protein_coding
protein_coding
protein_coding
protein_coding
protein_coding
protein_coding
protein_coding
protein_coding
protein_coding
protein_coding
protein_coding
protein_coding
protein_coding
protein_coding
protein_coding
protein_coding
protein_coding
protein_coding
protein_coding
protein_coding
protein_coding
protein_coding
protein_coding
protein_coding
protein_coding
protein_coding
protein_coding
protein_coding
protein_coding
protein_coding
protein_coding
ncRNA
protein_coding
protein_coding
protein_coding
protein_coding
protein_coding
protein_coding
protein_coding
protein_coding
ncRNA
protein_coding
protein_coding
ncRNA
protein_coding
protein_coding
protein_coding

protein_coding
protein_coding
protein_coding
protein_coding
protein_coding
protein_coding
protein_coding
protein_coding
protein_coding
protein_coding
protein_coding
protein_coding
protein_coding
ncRNA
protein_coding
protein_coding
protein_coding
protein_coding
ncRNA
protein_coding
protein_coding
protein_coding
protein_coding
protein_coding
protein_coding
protein_coding
protein_coding
protein_coding
protein_coding
protein_coding
protein_coding
protein_coding
protein_coding
protein_coding
protein_coding
ncRNA
protein_coding
protein_coding
protein_coding
protein_coding
protein_coding
protein_coding
pre_trna
protein_coding
protein_coding
protein_coding
protein_coding
protein_coding
protein_coding
protein_coding
protein_coding
protein_coding
protein_coding
protein_coding
protein_coding
protein_coding

protein_coding
protein_coding
ncRNA
pre_trna
protein_coding
protein_coding
protein_coding
protein_coding
protein_coding
ncRNA
protein_coding
protein_coding
ncRNA
protein_coding
protein_coding
pre_trna
protein_coding
protein_coding
protein_coding
protein_coding
protein_coding
protein_coding
protein_coding
protein_coding
protein_coding
protein_coding
protein_coding
protein_coding
protein_coding
protein_coding
protein_coding
protein_coding
protein_coding
protein_coding
protein_coding
protein_coding
protein_coding
protein_coding
protein_coding
protein_coding
protein_coding
protein_coding
protein_coding
protein_coding
protein_coding
protein_coding
protein_coding
protein_coding
protein_coding
protein_coding
protein_coding
protein_coding
protein_coding
protein_coding
protein_coding
protein_coding
protein_coding

protein_coding
protein_coding
protein_coding
protein_coding
ncRNA
protein_coding
protein_coding
protein_coding
protein_coding
protein_coding
protein_coding
protein_coding
protein_coding
protein_coding
protein_coding
protein_coding
protein_coding
protein_coding
protein_coding
protein_coding
protein_coding
protein_coding
protein_coding
ncRNA
protein_coding
ncRNA
protein_coding
protein_coding
protein_coding
protein_coding
ncRNA
protein_coding
protein_coding
protein_coding
protein_coding
protein_coding
protein_coding
small_nucle(ol)ar_rna
protein_coding
protein_coding
protein_coding
protein_coding
protein_coding
protein_coding
protein_coding
protein_coding
protein_coding
protein_coding
protein_coding
protein_coding
protein_coding
ncRNA
protein_coding
ncRNA
protein_coding
protein_coding

protein_coding
protein_coding
protein_coding
protein_coding
pre_trna
protein_coding
protein_coding
protein_coding
pre_trna
protein_coding
protein_coding
protein_coding
protein_coding
small_nucle(ol)ar_rna
small_nucle(ol)ar_rna
small_nucle(ol)ar_rna
protein_coding
protein_coding
protein_coding
protein_coding
protein_coding
protein_coding
protein_coding
protein_coding
protein_coding
ncRNA
protein_coding
protein_coding
protein_coding
protein_coding
protein_coding
protein_coding
protein_coding
protein_coding
protein_coding
protein_coding
pre_trna
protein_coding
protein_coding
protein_coding
protein_coding
protein_coding
protein_coding
protein_coding
protein_coding
protein_coding
protein_coding
protein_coding
ncRNA
protein_coding
protein_coding
protein_coding
protein_coding
protein_coding
protein_coding
protein_coding

ncRNA
protein_coding
protein_coding
protein_coding
protein_coding
protein_coding
protein_coding
protein_coding
protein_coding
protein_coding
protein_coding
protein_coding
protein_coding
protein_coding
protein_coding
protein_coding
protein_coding
protein_coding
protein_coding
protein_coding
protein_coding
protein_coding
protein_coding
protein_coding
protein_coding
protein_coding
protein_coding
protein_coding
protein_coding
protein_coding
protein_coding
protein_coding
protein_coding
protein_coding
ncRNA
small_nucle(ol)ar_rna
protein_coding
protein_coding
protein_coding
protein_coding
protein_coding
protein_coding
protein_coding
protein_coding
protein_coding
protein_coding
protein_coding
protein_coding
protein_coding
protein_coding
protein_coding
protein_coding
protein_coding
protein_coding
protein_coding
protein_coding

protein_coding
protein_coding
protein_coding
protein_coding
small_nucle(ol)ar_rna
protein_coding
protein_coding
protein_coding
protein_coding
protein_coding
protein_coding
protein_coding
protein_coding
protein_coding
protein_coding
protein_coding
protein_coding
protein_coding
protein_coding
protein_coding
protein_coding
protein_coding
protein_coding
protein_coding
protein_coding
protein_coding
protein_coding
protein_coding
protein_coding
protein_coding
protein_coding
protein_coding
protein_coding
protein_coding
protein_coding
protein_coding
protein_coding
protein_coding
protein_coding
protein_coding
protein_coding
protein_coding
protein_coding
protein_coding
protein_coding
protein_coding
protein_coding
protein_coding
protein_coding
protein_coding
protein_coding
protein_coding
protein_coding
protein_coding
protein_coding
small_nucle(ol)ar_rna

small_nucle(ol)ar_rna
small_nucle(ol)ar_rna
protein_coding
protein_coding
protein_coding
protein_coding
protein_coding
protein_coding
protein_coding
protein_coding
protein_coding
protein_coding
protein_coding
protein_coding
protein_coding
protein_coding
protein_coding
protein_coding
protein_coding
protein_coding
protein_coding
protein_coding
protein_coding
protein_coding
protein_coding
protein_coding
ncRNA
protein_coding
protein_coding
protein_coding
protein_coding
protein_coding
protein_coding
ncRNA
protein_coding
protein_coding
protein_coding
protein_coding
protein_coding
protein_coding
protein_coding
protein_coding
protein_coding
protein_coding
protein_coding
protein_coding
protein_coding
protein_coding
protein_coding
protein_coding
protein_coding
protein_coding
protein_coding
protein_coding
protein_coding
ncRNA

protein_coding
protein_coding
protein_coding
protein_coding
protein_coding
protein_coding
protein_coding
protein_coding
protein_coding
protein_coding
protein_coding
protein_coding
protein_coding
protein_coding
protein_coding
protein_coding
protein_coding
miRNA_precursor
protein_coding
protein_coding
protein_coding
protein_coding
protein_coding
protein_coding
protein_coding
protein_coding
protein_coding
protein_coding
protein_coding
protein_coding
protein_coding
protein_coding
protein_coding
protein_coding
protein_coding
protein_coding
miRNA_precursor
protein_coding
protein_coding
protein_coding
protein_coding
protein_coding
miRNA_precursor
protein_coding
protein_coding
protein_coding
protein_coding
protein_coding
protein_coding
protein_coding
protein_coding
protein_coding
protein_coding
protein_coding
protein_coding
protein_coding

protein_coding
protein_coding
protein_coding
protein_coding
protein_coding
protein_coding
protein_coding
protein_coding
protein_coding
protein_coding
protein_coding
protein_coding
protein_coding
protein_coding
protein_coding
protein_coding
protein_coding
protein_coding
protein_coding
protein_coding
protein_coding
protein_coding
ncRNA
protein_coding
protein_coding
protein_coding
protein_coding
protein_coding
protein_coding
protein_coding
protein_coding
protein_coding
protein_coding
ncRNA
protein_coding
protein_coding
protein_coding
protein_coding
protein_coding
protein_coding
small_nucle(ol)ar_rna
protein_coding
protein_coding
protein_coding
protein_coding
protein_coding
protein_coding
protein_coding
protein_coding
protein_coding
protein_coding
miRNA_precursor
protein_coding
protein_coding
protein_coding
protein_coding

protein_coding
protein_coding
protein_coding
protein_coding
protein_coding
ncRNA
protein_coding
protein_coding
protein_coding
protein_coding
pre_trna
ncRNA
protein_coding
protein_coding
protein_coding
protein_coding
protein_coding
protein_coding
protein_coding
protein_coding
ncRNA
protein_coding
ncRNA
protein_coding
protein_coding
protein_coding
protein_coding
protein_coding
protein_coding
protein_coding
protein_coding
protein_coding
protein_coding
protein_coding
protein_coding
protein_coding
protein_coding
ncRNA
protein_coding
protein_coding
protein_coding
ncRNA
protein_coding
protein_coding
protein_coding
protein_coding
protein_coding
protein_coding
protein_coding
protein_coding
protein_coding
protein_coding
protein_coding
protein_coding
protein_coding
protein_coding

protein_coding
protein_coding
protein_coding
protein_coding
protein_coding
protein_coding
protein_coding
protein_coding
protein_coding
protein_coding
ncRNA
protein_coding
protein_coding
protein_coding
protein_coding
protein_coding
protein_coding
protein_coding
protein_coding
protein_coding
protein_coding
protein_coding
protein_coding
protein_coding
protein_coding
protein_coding
protein_coding
protein_coding
ncRNA
protein_coding
ncRNA
protein_coding
protein_coding
protein_coding
ncRNA
protein_coding
protein_coding
protein_coding
ncRNA
protein_coding
protein_coding
protein_coding
protein_coding
protein_coding
protein_coding
ncRNA
protein_coding
protein_coding
protein_coding
protein_coding
protein_coding
ncRNA
protein_coding
miRNA_precursor
protein_coding
protein_coding

protein_coding
protein_coding
protein_coding
protein_coding
protein_coding
protein_coding
protein_coding
protein_coding
protein_coding
protein_coding
protein_coding
protein_coding
protein_coding
protein_coding
protein_coding
protein_coding
protein_coding
protein_coding
protein_coding
ncRNA
protein_coding
protein_coding
protein_coding
protein_coding
protein_coding
protein_coding
protein_coding
protein_coding
protein_coding
protein_coding
small_nucle(ol)ar_rna
protein_coding
protein_coding
protein_coding
protein_coding
protein_coding
protein_coding
protein_coding
protein_coding
protein_coding
protein_coding
protein_coding
protein_coding
protein_coding
protein_coding
protein_coding
protein_coding
protein_coding
protein_coding
protein_coding
protein_coding
protein_coding
protein_coding
protein_coding
protein_coding
protein_coding

protein_coding
protein_coding
protein_coding
protein_coding
protein_coding
protein_coding
protein_coding
protein_coding
protein_coding
protein_coding
protein_coding
protein_coding
protein_coding
protein_coding
pre_trna
protein_coding
protein_coding
protein_coding
protein_coding
protein_coding
protein_coding
protein_coding
protein_coding
protein_coding
protein_coding
protein_coding
protein_coding
protein_coding
protein_coding
protein_coding
protein_coding
protein_coding
protein_coding
protein_coding
protein_coding
protein_coding
protein_coding
protein_coding
protein_coding
protein_coding
protein_coding
protein_coding
protein_coding
protein_coding
protein_coding
protein_coding
protein_coding
protein_coding
protein_coding
protein_coding
protein_coding
protein_coding
protein_coding
protein_coding
protein_coding
protein_coding

protein_coding
protein_coding
protein_coding
protein_coding
protein_coding
protein_coding
protein_coding
protein_coding
protein_coding
protein_coding
protein_coding
protein_coding
protein_coding
protein_coding
protein_coding
protein_coding
protein_coding
protein_coding
protein_coding
protein_coding
protein_coding
protein_coding
protein_coding
protein_coding
miRNA_precursor
protein_coding
protein_coding
protein_coding
protein_coding
ncRNA
protein_coding
protein_coding
protein_coding
protein_coding
protein_coding
ncRNA
protein_coding
ncRNA
protein_coding
protein_coding
protein_coding
protein_coding
protein_coding
protein_coding
protein_coding
protein_coding
protein_coding
protein_coding
protein_coding
protein_coding
protein_coding
protein_coding
protein_coding
protein_coding
protein_coding
protein_coding

protein_coding
protein_coding
protein_coding
protein_coding
protein_coding
protein_coding
protein_coding
protein_coding
protein_coding
protein_coding
protein_coding
protein_coding
protein_coding
protein_coding
protein_coding
protein_coding
protein_coding
protein_coding
protein_coding
protein_coding
protein_coding
protein_coding
protein_coding
protein_coding
protein_coding
protein_coding
protein_coding
protein_coding
protein_coding
protein_coding
protein_coding
protein_coding
protein_coding
protein_coding
protein_coding
protein_coding
protein_coding
protein_coding
protein_coding
protein_coding
protein_coding
protein_coding
protein_coding
protein_coding
protein_coding
protein_coding
protein_coding
protein_coding
protein_coding
miRNA_precursor
protein_coding
protein_coding
protein_coding
protein_coding
ncRNA
protein_coding

protein_coding
ncRNA
protein_coding
protein_coding
protein_coding
protein_coding
protein_coding
protein_coding
protein_coding
protein_coding
protein_coding
protein_coding
protein_coding
protein_coding
protein_coding
protein_coding
protein_coding
protein_coding
protein_coding
protein_coding
protein_coding
protein_coding
protein_coding
protein_coding
protein_coding
protein_coding
protein_coding
protein_coding
protein_coding
protein_coding
protein_coding
protein_coding
protein_coding
protein_coding
protein_coding
protein_coding
protein_coding
pre_trna
protein_coding
protein_coding
protein_coding
protein_coding
protein_coding
protein_coding
protein_coding
protein_coding
protein_coding
protein_coding
protein_coding
ncRNA
ncRNA
protein_coding
protein_coding
protein_coding
protein_coding
protein_coding

protein_coding
protein_coding
protein_coding
miRNA_precursor
protein_coding
protein_coding
protein_coding
protein_coding
protein_coding
protein_coding
protein_coding
protein_coding
protein_coding
protein_coding
protein_coding
protein_coding
protein_coding
protein_coding
protein_coding
protein_coding
protein_coding
protein_coding
protein_coding
protein_coding
protein_coding
protein_coding
protein_coding
protein_coding
protein_coding
protein_coding
protein_coding
protein_coding
protein_coding
protein_coding
protein_coding
protein_coding
protein_coding
protein_coding
protein_coding
protein_coding
protein_coding
protein_coding
protein_coding
protein_coding
protein_coding
protein_coding
protein_coding
protein_coding
small_nucle(ol)ar_rna
protein_coding
protein_coding
protein_coding
protein_coding
protein_coding
protein_coding
protein_coding

protein_coding
protein_coding
protein_coding
protein_coding
ncRNA
protein_coding
ncRNA
protein_coding
protein_coding
protein_coding
protein_coding
protein_coding
protein_coding
protein_coding
protein_coding
protein_coding
protein_coding
protein_coding
protein_coding
protein_coding
protein_coding
protein_coding
protein_coding
protein_coding
protein_coding
protein_coding
protein_coding
protein_coding
protein_coding
protein_coding
protein_coding
protein_coding
protein_coding
protein_coding
protein_coding
protein_coding
protein_coding
protein_coding
protein_coding
protein_coding
protein_coding
protein_coding
protein_coding
protein_coding
protein_coding
protein_coding
protein_coding
protein_coding
protein_coding
protein_coding
protein_coding
protein_coding
protein_coding
protein_coding
protein_coding
protein_coding

protein_coding
protein_coding
protein_coding
protein_coding
protein_coding
protein_coding
protein_coding
protein_coding
protein_coding
protein_coding
protein_coding
protein_coding
protein_coding
protein_coding
protein_coding
protein_coding
protein_coding
protein_coding
protein_coding
protein_coding
protein_coding
protein_coding
protein_coding
protein_coding
protein_coding
protein_coding
protein_coding
protein_coding
protein_coding
protein_coding
protein_coding
protein_coding
protein_coding
protein_coding
protein_coding
protein_coding
protein_coding
protein_coding
protein_coding
protein_coding
protein_coding
protein_coding
protein_coding
protein_coding
protein_coding
protein_coding
protein_coding
protein_coding
protein_coding
protein_coding
protein_coding
protein_coding
protein_coding
protein_coding
protein_coding
protein_coding

protein_coding
protein_coding
protein_coding
protein_coding
protein_coding
protein_coding
protein_coding
protein_coding
protein_coding
ncRNA
protein_coding
protein_coding
protein_coding
protein_coding
protein_coding
protein_coding
protein_coding
protein_coding
protein_coding
protein_coding
protein_coding
protein_coding
protein_coding
protein_coding
protein_coding
protein_coding
protein_coding
protein_coding
protein_coding
protein_coding
protein_coding
protein_coding
protein_coding
protein_coding
protein_coding
protein_coding
protein_coding
protein_coding
protein_coding
protein_coding
protein_coding
protein_coding
protein_coding
ncRNA
protein_coding
protein_coding
protein_coding
protein_coding
protein_coding
protein_coding
protein_coding
protein_coding
